# Differential CpG methylation at *Nnat* in the early establishment of beta cell heterogeneity

**DOI:** 10.1101/2023.02.04.527050

**Authors:** Vanessa Yu, Fiona Yong, Angellica Marta, Sanjay Khadayate, Adrien Osakwe, Supriyo Bhattacharya, Sneha S. Varghese, Pauline Chabosseau, Sayed M. Tabibi, Keran Chen, Eleni Georgiadou, Nazia Parveen, Mara Suleiman, Zoe Stamoulis, Lorella Marselli, Carmela De Luca, Marta Tesi, Giada Ostinelli, Luis Delgadillo-Silva, Xiwei Wu, Yuki Hatanaka, Alex Montoya, James Elliott, Bhavik Patel, Nikita Demchenko, Chad Whilding, Petra Hajkova, Pavel Shliaha, Holger Kramer, Yusuf Ali, Piero Marchetti, Robert Sladek, Sangeeta Dhawan, Dominic J. Withers, Guy A. Rutter, Steven J. Millership

## Abstract

**Aims/hypothesis:** Beta cells within the pancreatic islet represent a heterogenous population wherein individual sub-groups of cells make distinct contributions to the overall control of insulin secretion. These include a subpopulation of highly-connected ‘hub’ cells, important for the propagation of intercellular Ca^2+^ waves. Functional subpopulations have also been demonstrated in human beta cells, with an altered subtype distribution apparent in type 2 diabetes. At present, the molecular mechanisms through which beta cell hierarchy is established are poorly understood. Changes at the level of the epigenome provide one such possibility which we explore here by focussing on the imprinted gene neuronatin (*Nnat*), which is required for normal insulin synthesis and secretion.

**Methods:** Single cell RNA-seq datasets were examined using Seurat 4.0 and ClusterProfiler running under R. Transgenic mice expressing eGFP under the control of the *Nnat* enhancer/promoter regions were generated for fluorescence-activated cell (FAC) sorting of beta cells and downstream analysis of CpG methylation by bisulphite and RNA sequencing, respectively. Animals deleted for the de novo methyltransferase, DNMT3A from the pancreatic progenitor stage were used to explore control of promoter methylation. Proteomics was performed using affinity purification mass spectrometry and Ca^2+^ dynamics explored by rapid confocal imaging of Cal-520 and Cal-590. Insulin secretion was measured using Homogeneous Time Resolved Fluorescence Imaging.

**Results:** *Nnat* mRNA was differentially expressed in a discrete beta cell population in a developmental stage- and DNA methylation (DNMT3A)-dependent manner. Thus, pseudo-time analysis of embryonic data sets demonstrated the early establishment of *Nnat*-positive and negative subpopulations during embryogenesis. NNAT expression is also restricted to a subset of beta cells across the human islet that is maintained throughout adult life. NNAT^+^ beta cells also displayed a discrete transcriptome at adult stages, representing a sub-population specialised for insulin production, reminiscent of recently-described “β_HI_” cells and were diminished in *db/db* mice. ‘Hub’ cells were less abundant in the NNAT^+^ population, consistent with epigenetic control of this functional specialization.

**Conclusions/interpretation:** These findings demonstrate that differential DNA methylation at *Nnat* represents a novel means through which beta cell heterogeneity is established during development. We therefore hypothesise that changes in methylation at this locus may thus contribute to a loss of beta cell hierarchy and connectivity, potentially contributing to defective insulin secretion in some forms of diabetes.

**Research in context:** What is already known about this subject?

- Neuronatin (*Nnat*/*NNAT*) is an imprinted gene in humans and mice and is required for glucose-stimulated insulin secretion *in vivo*

- Pancreatic beta cells are functionally heterogeneous with specific highly-connected subpopulations known to coordinate islet wide Ca^2+^ dynamics

- Functional subpopulations have been described in human beta cells and their distribution is altered in type 2 diabetes

What is the key question?

- Does NNAT mark a discrete subpopulation of functional beta cells and which epigenetic pathways coordinate its formation and maintenance?

What are the new findings?

- A subpopulation of NNAT^+^ beta cells is established prior to the first week of postnatal life in mice via de novo DNA methylation at the *Nnat* promoter

- NNAT^+^ beta cells are transcriptionally highly differentiated and appear to be functionally specialised for insulin production, possibly corresponding to recently-described “β_HI_” and “CD63^hi^” beta cells. NNAT is expressed in a subset of beta cells across the human islet, and its deficiency in human beta cells diminishes glucose-stimulated insulin secretion

- NNAT^+^ cells are likelier to belong to the population of ‘follower’, rather than ‘hub’ cells, consistent with a role in insulin production rather than glucose detection

How might this impact on clinical practice in the foreseeable future?

- Epigenome-modifying compounds may provide a way of enhancing beta cell function and the ensemble behaviour of the islet to stimulate insulin secretion

## Introduction

Insulin-producing beta cells are central to the modulation of glucose homeostasis and their impaired function, loss of identity, or lowered numbers result in type 2 diabetes [1, 2]. Previous studies have provided an understanding of the transcriptional machinery that orchestrates beta cell development from early pancreatic and endocrine precursors [3, 4]. To bolster these transcriptional programmes *in vivo*, chronic regulation via the epigenome appears to step in to maintain beta cell identity in the long term [5–11].

Early reports [12–14], and more recent studies based on single cell transcriptomic profiling [15–17], electrophysiology [18] and functional imaging [19–22] have demonstrated functional heterogeneity amongst individual beta cells (reviewed in [23, 24]). Identified subpopulations have been associated with known markers or maturation states (*Flattop*/*Cfap126* [21], PSA-NCAM [25], CD81 [26], CD24 [11, 27], TH [10], NPY [28], CD63 [29]), are ‘virgin’ beta cells [30] or are defined by their roles in coordinating islet-wide Ca^2+^ dynamics (e.g. ‘hubs’ [19], ‘leaders’ [20] [31] and ‘first responders’ [32]). Furthermore, loss of beta cell heterogeneity or intercellular connectivity may contribute to the development of type 2 diabetes [19, 20, 33]. Importantly, functional subpopulations have also been demonstrated in human beta cells [16, 17, 34, 35], and the distribution of antigenically-defined sub-groups (based on CD9 and ST8SIA1 positivity) is altered in type 2 diabetes [15]. The features underlying beta cell heterogeneity include pathways governing glucose sensing and metabolism [19–21, 36] insulin content and secretory competence [9, 18, 30, 37, 38] and cilia activity and localisation within the islet [31]. Recently, two discrete populations of “CD63^hi^” and “CD63^lo^” cells have been described [29], with “CD63^hi^” cells enriched for CD63 and for insulin content and glucose-stimulated insulin secretion (GSIS). Epigenomically-defined (by histone methylation, H3K27me3) CD24-positive “β_HI_” beta cells with enhanced insulin content and GSIS [11] may partly overlap the “CD63^hi^” population [29].

Imprinted genes are expressed from a single allele in a parent of origin-specific manner and their expression is also controlled via epigenetic modifications, notably DNA methylation. Imprinted genes often play key physiological roles, particularly in early (fetal and postnatal) growth and development, controlling a wide range of cellular processes. Thus, human imprinting disorders involving altered expression from specific imprinted *loci* are associated with severe childhood developmental and metabolic complications (reviewed in [39, 40]).

Imprinted genes play key functional roles in pancreatic beta cells by modulating insulin secretory machinery or beta cell mass [41]. Correspondingly, imprinted gene expression is dysregulated both in beta cells with diminished GSIS, in pancreatic islets from subjects with type 2 diabetes, and single nucleotide polymorphisms (SNPs) at imprinted *loci* are associated with type 2 diabetes risk [23]. Monoallelic expression of imprinted genes is maintained trans-generationally by differential methylation between parental alleles at imprinted *loci* in the germline [42, 43], with additional ‘somatic’ or ‘secondary’ differentially-methylated regions (DMRs) also established post-fertilisation [44].

We have previously shown [45] that the paternally expressed, imprinted gene *Neuronatin* (*Nnat*) is nutrient-regulated in pancreatic beta cells, and controls insulin content and GSIS by modulating early insulin precursor processing at the signal peptidase complex [45]. At extra-pancreatic sites, changes in *Nnat* expression also modulate appetite and metabolism [46–48]. Here, we show that differential methylation of the *Nnat* gene contributes to the functional heterogeneity of embryonic and adult pancreatic beta cells, marking an insulin mRNA and protein-enriched, mature beta cell subpopulation.

## Methods

For details, please refer to ESM Methods.

### Bioinformatic clustering analysis

Single cell RNA-seq embryonic and adult islet datasets [9, 49–52], were assessed for hypervariability of gene expression and unbiased clustering using the scran and RaceID package, respectively, as described in the ESM methods.

### Pseudo-time analysis

scRNA-seq datasets [49] were processed with Seurat v4.3] and clustered and processed for pseudo-time analysis, followed by integration with a dataset from adult mouse islets [51, 52], according to ESM methods.

### Animal models

Transgenic mice lines expressing *Cre* recombinase under the control of the rat insulin promoter (RIP) [53] and with a tdTomato reporter downstream of a stop codon flanked by *loxP* sites [54] have been described previously. Mice with *Nnat*-driven *Egfp* expression were purchased from the Mutant Mouse Resource and Research Centre (MMRRC, USA) repository (Tg(Nnat-EGFP)EA106Gsat/Mmucd, #010611-UCD). Mice with global deletion of *Nnat* [45] and conditional *Dnmt3a* null mice generated by crossing to *Pdx1*-*Cre* are described in the ESM methods. *db/db* and control C57BLKS/J mice on the BKS background were obtained from the Jackson Laboratory (JAX 000642 and JAX 000662, respectively).

### Primary islet isolation and FACS

To isolate primary islets, the pancreas was inflated with Liberase TM (Roche), digested at 37°C, purified using a Histopaque 1119/1083/1077 gradient (Sigma-Aldrich) and hand-picked. For FACS-based experiments, cells from purified islets were isolated using Accutase (Sigma-Aldrich) sorted using an FACSAria III flow cytometer (BD, UK). Total insulin content was assessed using ultra-sensitive insulin homogeneous time-resolved fluorescence (HTRF) assay kits (Cisbio).

### Intracellular calcium imaging

Pancreatic islets from reporter mice expressing *Nnat*-eGFP were isolated as above, with Ca^2+^ imaging of whole islets performed after loading the cytosol with 2 μM Cal-590 AM (Stratech). Images were captured on an Axiovert microscope (Zeiss, Germany) equipped with a 10x 0.3–0.5 NA objective, a Hamamatsu ImagEM camera coupled to a Nipkow spinning-disk head (CSU-10, Yokogawa UK Ltd) and illuminated at 490 nm or 530 nm.

For experiments involving global *Nnat* null mice [45], islets were incubated with 4.5 μM Cal-520 AM (Stratech), and imaging performed on a Nikon Eclipse Ti microscope equipped with a 40x/1.2 NA oil objective and an ibidi heating system. Cal-520 AM was excited with a 491 nm laser line and emitted light filtered at 525/50 nm. Images were acquired with an ORCA-Flash 4.0 camera (Hamamatsu) and Metamorph software (Molecular Device). Pearson-based connectivity and correlation analyses in an imaged islet were performed with Ca^2+^ signals smoothed, binarised and analysed as described in ESM methods.

### Histological techniques and immunofluorescence

Dissected tissues were fixed, cryoprotected and embedded in optimal cutting temperature (OCT) and stored at -80°C. Sections (10 μm) were cut using a CM1950 Cryostat (Leica) and immunostained using primary and secondary antibodies (listed in ESM Methods) and imaged using a TCS SP5 confocal microscope (Leica, Germany). Immunostaining of paraffin-embedded pancreas from mice with beta cell-selective deletion of *Dnmt3a* was performed according to [10, 28] as described in ESM methods. Human pancreata were processed, immunostained and imaged as described in ESM methods with the approval of the Ethics Committee of the University of Pisa, upon written consent of donors’ next-of-kin.

### Cell culture, RNA silencing and GSIS

Human EndoC-βH1 and rat INS1E beta cells were cultured as previously described [45] and incubated with lentiviruses expressing an *NNAT*-targeting shRNA (Sigma-Aldrich) or Silencer Select siRNAs (Ambion), respectively, as described in ESM methods. GSIS assays were performed using ultra-sensitive insulin HTRF assay kits (Cisbio) for insulin quantification.

### Immunoprecipitation and mass spectrometry

Cells were processed for immunoprecipitation and incubated with antibodies against NNAT (ab27266, Abcam, 1:500) prior to mass spectrometry analysis [45] and as described in ESM methods.

### Western immunoblotting, RT-PCR, RNA sequencing

For Western blotting, EndoC-βH1 cells were processed as previously described [45] using primary antibodies against NNAT (ab27266, Abcam, 1:2000) and β-tubulin (clone 9F3, #2128, Cell Signalling, 1:5000). For RT-PCR, mRNA was purified using an Allprep or RNeasy kits (both Qiagen), reverse transcribed, and cDNA assessed by quantitative RT-PCR using Taqman reagents (all Applied Biosystems) on a QuantStudio 7 Real Time PCR cycler with *Hprt*/*HPRT1* as an internal mRNA control.

For RNA sequencing, RNA from FACS-purified cells was quantified and assessed for integrity using a Bioanalyser 2100 and a RNA 6000 pico assay (Agilent Technologies). RNA was processed using a NEBNext Ultra II Directional RNA library Prep Kit for Illumina paired with a Poly(A) mRNA magnetic isolation module and AMPure XP SPRIselect beads (Beckman Coulter). Libraries were sequenced using a NextSeq 500 High Output sequencer (Illumina, USA), with 2 x 75 bp length at 50 million reads per sample.

### Bisulphite sequencing

Genomic DNA was extracted from FACS-purified cells using Allprep DNA/RNA/protein mini kits from Qiagen. Sperm cells were lysed using differential extraction. Bisulphite sequencing experiments were performed using an EZ DNA Methylation-Gold kit from Zymo Research, an EpiTaq hot start kit (TaKaRa Bio), a CloneJET PCR cloning kit (Thermo Scientific) and DNA purification using a Wizard SV 96 plasmid system (Promega) before Sanger sequencing (Genewiz, now Azenta Life Sciences) and analysed using BISMA software.

### Statistical analysis

Data are shown as mean±SEM in all panels. Data were assessed using GraphPad Prism 9.0 with <5% error probability was considered significant (i.e., P<0.05) and further statistical information, such as *n* numbers and P values, are provided in the figure legends.

## Results

### Neuronatin expression is heterogeneous across the islet and represents a highly differentiated beta cell subtype

We have previously shown that *Nnat* is crucial for insulin storage and GSIS in the mouse [45]. NNAT expression is diminished in rodent models of obesity and diabetes including the Zucker diabetic fatty (ZDF) rat model (85% decrease, P=0.0023, n=5, [55]) and obese *ob/ob* mice (ESM Fig. 1a). Pancreatic sections of *db/db* mice displayed a loss of NNAT immunoreactivity across islets, and a significant reduction of NNAT^+^ beta cells (ESM Fig. 1b, c).

Reanalysis of published single cell RNA-sequencing datasets (scRNA-seq) from primary mouse beta cells at embryonic (E17.5) [49, 50] and adult [9, 51, 52] stages identified *Nnat* as amongst the most highly variable genes between individual cells (ESM Table 1). Pseudo-time analysis of embryonic (E12.5 to E17.5) mouse beta cells [49] clearly demarcated the differentiation of beta cells through embryonic cell states, with late embryonic (E17.5) beta cells overlapping with high levels of *Nnat* expression (Fig. 1a-c, ESM Fig. 2a-k). Analysis of the proportions of different beta cell progenitors revealed that *Nnat*^+^ cells become more prominent at later stages of development (Fig. 1d). Integrating cells from the embryonic beta cell trajectory with adult beta cells [51, 52] allowed us to evaluate the progressive change in beta cell marker and *Nnat* expression from development to maturity (Fig. 1e-f, ESM Fig. 3a-i), and revealed large increases in *Ins1* expression alongside *Nnat* downregulation in most beta cells. Nevertheless, a considerable number of adult beta cells (with high expression of *Ins1*/*Ins2*) also expressed *Nnat* (Fig. 1f). Thus, *Nnat* appears to mark late-stage beta cell differentiation, peaking in expression around E17.5 and then is gradually downregulated across most beta cells in adulthood.

**Fig. 1.**
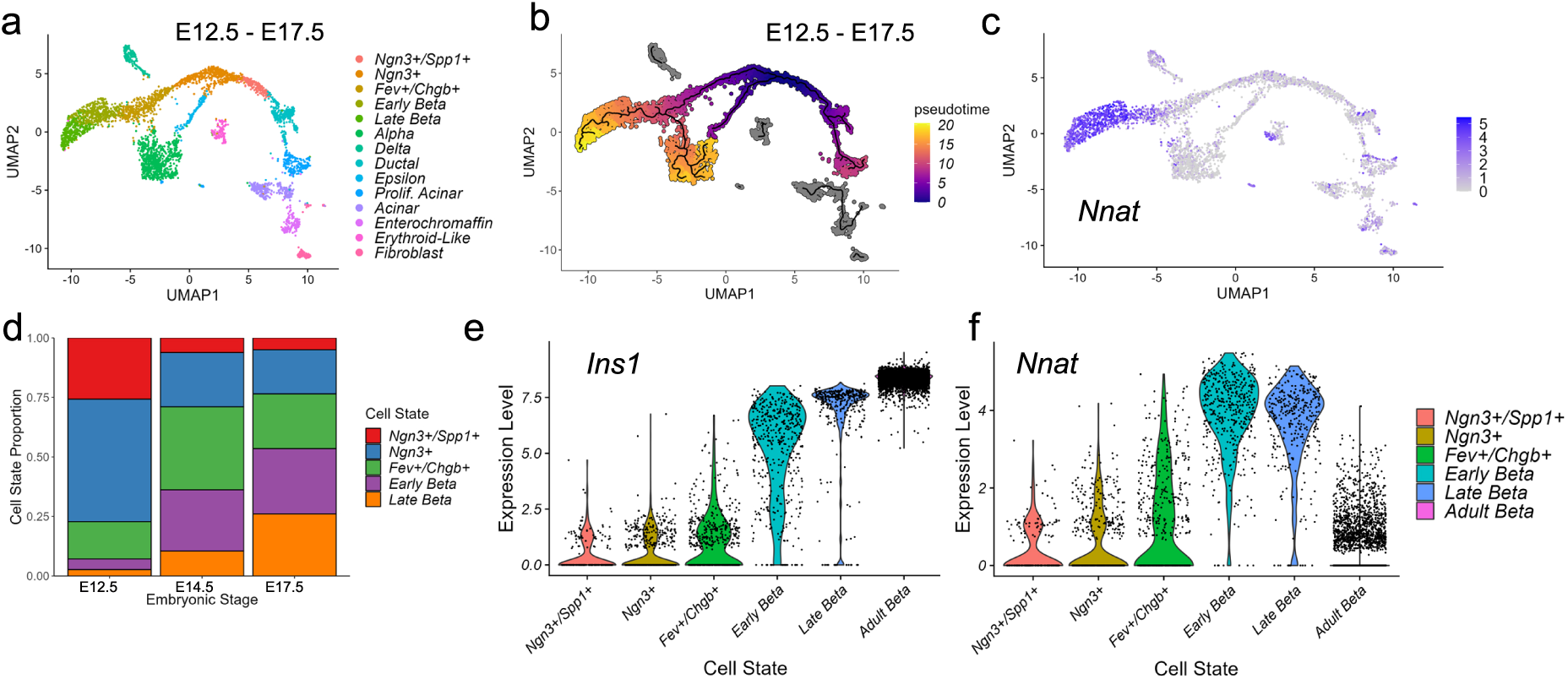
*Nnat* expression is a marker for late-stage beta cell differentiation during islet development. (a-c) UMAP projection of scRNA-seq data from embryonic mouse islets (E12.5-E17.5), with cells labelled by their cellular state (a), pseudo-time (b), and corresponding *Nnat* expression (c). (d) Changes in the distribution of beta cell precursors from E12.5 to E17.5. (e, f) Violin plots showing the fluctuations in *Ins1* (e) and *Nnat* (f) expression from E12.5 to adulthood in the beta cell development trajectory.

In light of the abundance of *Nnat*^+^ beta cells during late embryogenesis, we further evaluated the E17.5 dataset [49] and again identified two beta cell populations with distinct levels of *Nnat* expression (Fig. 2a-c, ESM Fig. 4a-c). Embryonic *Nnat^+^* beta cells were enriched for *Ins1*, *Ins2*, *Nkx6.1*, *Pdx1*, *Ucn3*, *Slc2a2*, *Iapp*, *Ero1b*, *G6pc2*, *Dlk1* and *Npy* expression compared to *Nnat*^-^ beta cells that had higher *Neurog3*, *Pax4*, *Gcg*, *Arx*, and *Ghrl* expression, indicating that the *Nnat^+^* beta cells were more fully differentiated (Fig. 2d, ESM Table 2). *Nnat^+^* beta cells were also de-enriched for *Cd24a* (Fig. 2d, ESM Table 2). Geneset enrichment analysis (GSEA) revealed upregulation of processes related to protein synthesis, transport from ER, ER stress, as well as oxidative phosphorylation and carbohydrate metabolism, with downregulated processes included mRNA processing, splicing, and histone methylation (Fig. 2e). Indeed, a separate cell clustering analysis [56] identified *Nnat* as a highly differentially-expressed gene between two beta cell clusters at both the late embryonic [49, 50] and adult [9] stages (ESM Fig. 5a-o, ESM Table 3).

**Fig. 2.**
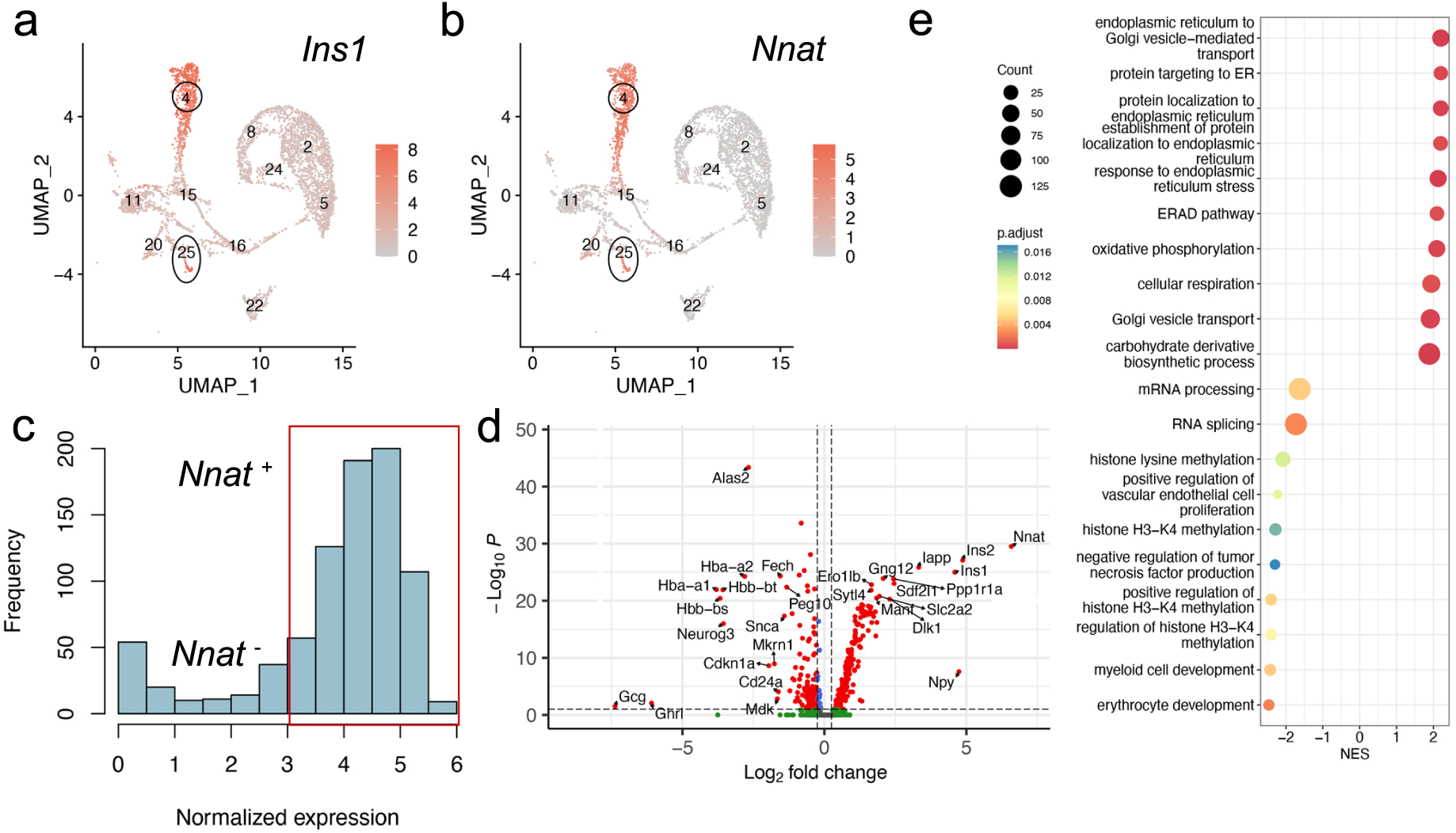
Transcriptomic analysis of islet cells at E17.5 reveal *Nnat*^+^ cells to be a more differentiated beta cell subcluster. (a, b) Islet cells plotted in the UMAP space and coloured by *Ins1* (a) and *Nnat* (b) expression, respectively. Individual cellular clusters are labelled. Beta cell clusters are highlighted with black circles. (c) Distribution of *Nnat* expression among beta cell clusters, along with the expression range used in defining *Nnat*^+^ beta cells (highlighted by the red rectangle). (d) Volcano plot showing the log_2_ fold change in gene expression between *Nnat*^+^ and *Nnat*^-^ beta cells against FDR-corrected P values. Genes with FDR < 0.1 are coloured red. Top 10 upregulated and downregulated genes are labelled by gene name. (e) Geneset enrichment analysis (GSEA) of differentially expressed genes between *Nnat*^+^ and *Nnat*^-^ beta cells (using GO-BP terms), showing the top 10 upregulated and downregulated processes. The normalized enrichment score (NES) is plotted along the X-axis. Circle diameters are proportional to the number of genes in each process, with the colours defining statistical significance.

We confirmed some of these findings at the protein level using immunofluorescence in postnatal and adult mice (Fig. 3a, b). We noted that NNAT protein expression is highly dynamic in beta cells across the mouse islet throughout the postnatal stage, transitioning from expression throughout the majority (90.4±3.2%, n=15 mice) of beta cells at late embryogenesis to a subset (24.4±3.0%, n=5 mice) of beta cells by P14 (Fig. 3a, b). Specific staining for endogenous NNAT was confirmed by the absence of reactivity in pancreatic cryosections from mice with constitutive deletion of the paternal *Nnat* allele (Fig. 3a). NNAT^+^ beta cells persisted throughout the islet into adulthood (Fig. 3a, b). NNAT expression was not apparent in other mouse islet cell types, including alpha and delta cells at P14. However, NNAT expression was detectable in a small number of alpha and delta cells at E17.5 (Fig. 3c).

**Fig. 3.**
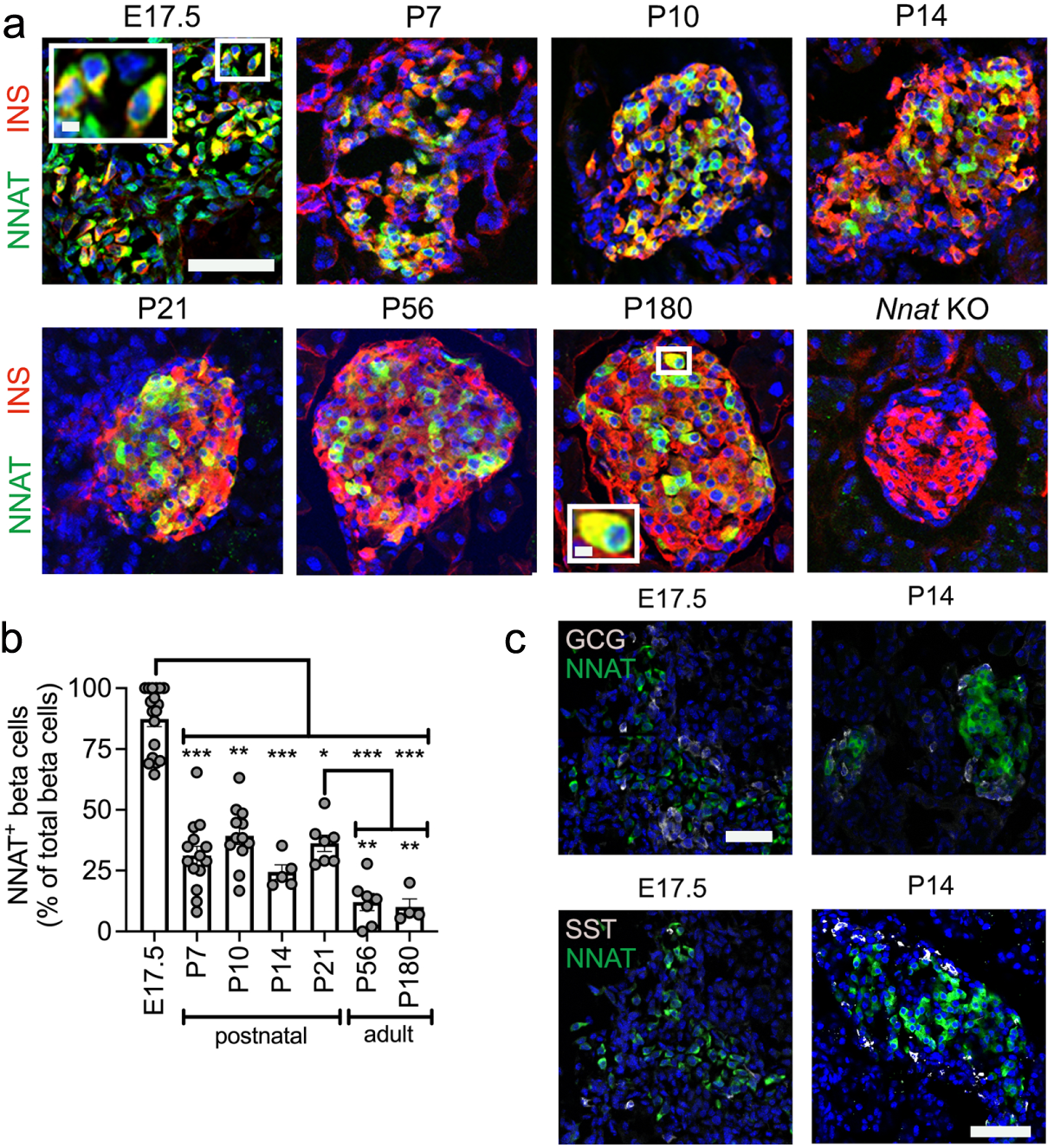
A subpopulation of NNAT^+^ beta cells develop during the early postnatal period in mice. (a) Representative confocal microscopy of pancreatic cryosections from wild type mice on a C57BL/6J background of developmental stages from embryonic day 17.5 (E17.5) through the postnatal (P) period into adulthood. Sections were immunostained with antibodies against endogenous neuronatin (NNAT, green) and insulin (INS, red). Nuclei are visualised with DAPI and sections from P56 mice with constituent deletion of *Nnat* were used as an immunostaining control. Scale bar = 100μm (inset = 10μm) (n = 4-15 mice per timepoint, Kruskal-Wallace test with Dunn’s multiple comparisons). (b) Quantification of NNAT^+^ beta cells from images shown in a, expressed as NNAT/INS co-positive cells as a percentage of total INS-positive cells. (c) Representative confocal microscopy of pancreatic cryosections as in a from E17.5 and postnatal (P) day 14 mice (n = 18 and n = 5 mice, respectively) immunostained with antibodies against endogenous neuronatin (NNAT, green) and GCG or SST (both grey). Scale bar = 100μm. Representative images from three independent experiments and breeding pairs. * P < 0.05, ** P < 0.01, *** P < 0.001.

### NNAT immunoreactivity is detected in a subset of beta cells in human islets and NNAT deficiency in human beta cells blunts GSIS

Whilst the role of NNAT has previously been examined in mouse beta cells [45], no data currently exist for human beta cells. We therefore next compared the functionality of human NNAT^+^ and NNAT^-^ beta cells. Transient silencing of *NNAT* in human EndoC-βH1 beta cells completely abrogated glucose-stimulated insulin release (Fig. 4a-c). As in mouse beta cells [45], NNAT interacted *in cellulo* with subunits of the signal peptidase complex (SPC), SEC11A, SPCS1, SPCS2 and SPCS3 (ESM Fig. 6a), likely via SPCS1 (ESM Fig. 6b).

**Fig. 4.**
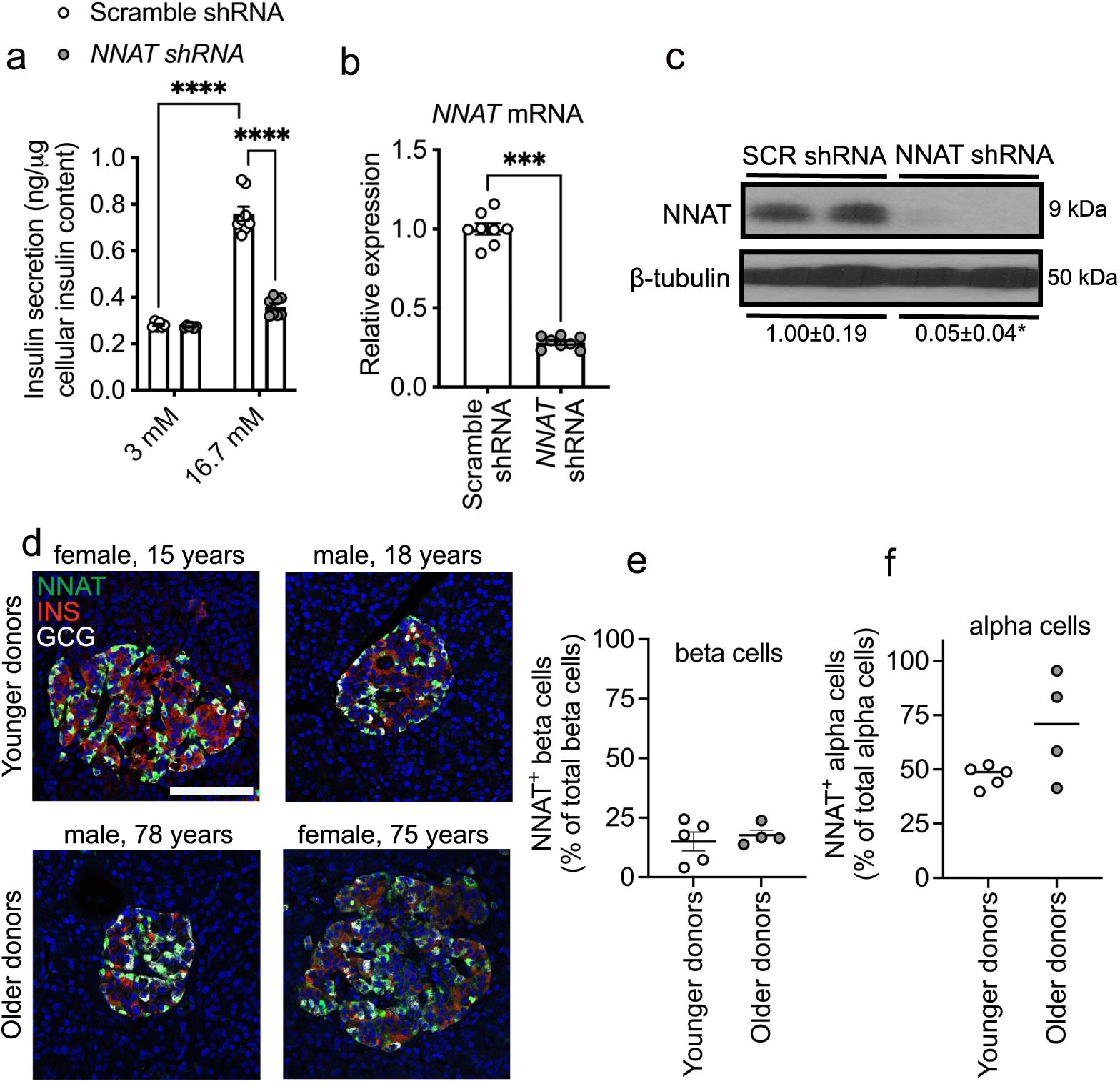
NNAT is expressed in a subset of human beta cells and NNAT deficiency in human beta cells reduces insulin secretion. (a) GSIS analysis at low (3 mM) and high (16.7 mM) glucose in human EndoC-βH1 beta cells following 72h lentiviral-mediated shRNA knockdown of neuronatin with data expressed as of insulin secreted into culture media as a percentage of total cellular insulin content (n = 8 independent cultures per group, two-way ANOVA with Sidak’s multiple corrections test). (b, c) RT-PCR (n = 8, unpaired Student’s t test) and Western blot (n = 4, Mann-Whitney U test) analysis of neuronatin expression in EndoC-βH1 beta cells after transient *NNAT* silencing as in a. Western blotting analysis of NNAT protein levels shown via a representative blot of two independent experiments. β-tubulin was used as a loading control. Mean values for band intensities in multiple experiments quantified by densitometry are shown below the panel, expressed relative to scramble (SCR) shRNA controls. (d) Representative confocal microscopy of human pancreatic cryosections from younger (15.6 ± 0.9 years, n = 5) and older (71.0 ± 3.9 years, n = 4) donors. Sections were immunostained with antibodies against endogenous neuronatin (NNAT, green), insulin (INS, red) and glucagon (GCG, grey). Donor sex and age are indicated in the text. Nuclei are visualised with DAPI. Scale bar = 100μm. (e, f) Quantification of NNAT^+^ beta (e) and alpha (f) cells from images shown in d, expressed as NNAT/INS or NNAT/GCG co-positive cells as a percentage of total INS-positive or GCG-positive cells. * P < 0.05, *** P < 0.001, **** P < 0.0001).

Heterogeneous expression of NNAT in human islet cells was also observed in whole pancreatic sections from multiple donors. Thus, NNAT was expressed in a subset of beta cells in both younger (15.6±0.9 years, 15.0±4.0%, n=5) and older (71.0±3.9 years, 17.7±2.1%, n= 4) donors (Fig. 4d, e, ESM Fig. 7a, ESM Table 4). Interestingly, NNAT was also expressed in a large fraction of human alpha cells (but not in delta cells, Fig. 4d, f, ESM Fig. 7b) in both younger (47.0±2.2%) and older donors (69.6±12.2%).

### Heterogeneous NNAT expression in beta cells during postnatal development is associated with altered CpG methylation

To extend the findings above using an orthogonal approach, and to facilitate subsequent functional studies, we utilised a Bacterial Artificial Chromosome (BAC) reporter mouse line expressing eGFP under the control of the *Nnat* promoter and upstream enhancers (ESM results and ESM Fig. 8a-e). Aligning with studies of the endogenous gene (Fig. 1-4) adult reporter mice expressed eGFP in a subpopulation (∼15% of total) of beta cells (Fig. 5a).

**Fig. 5.**
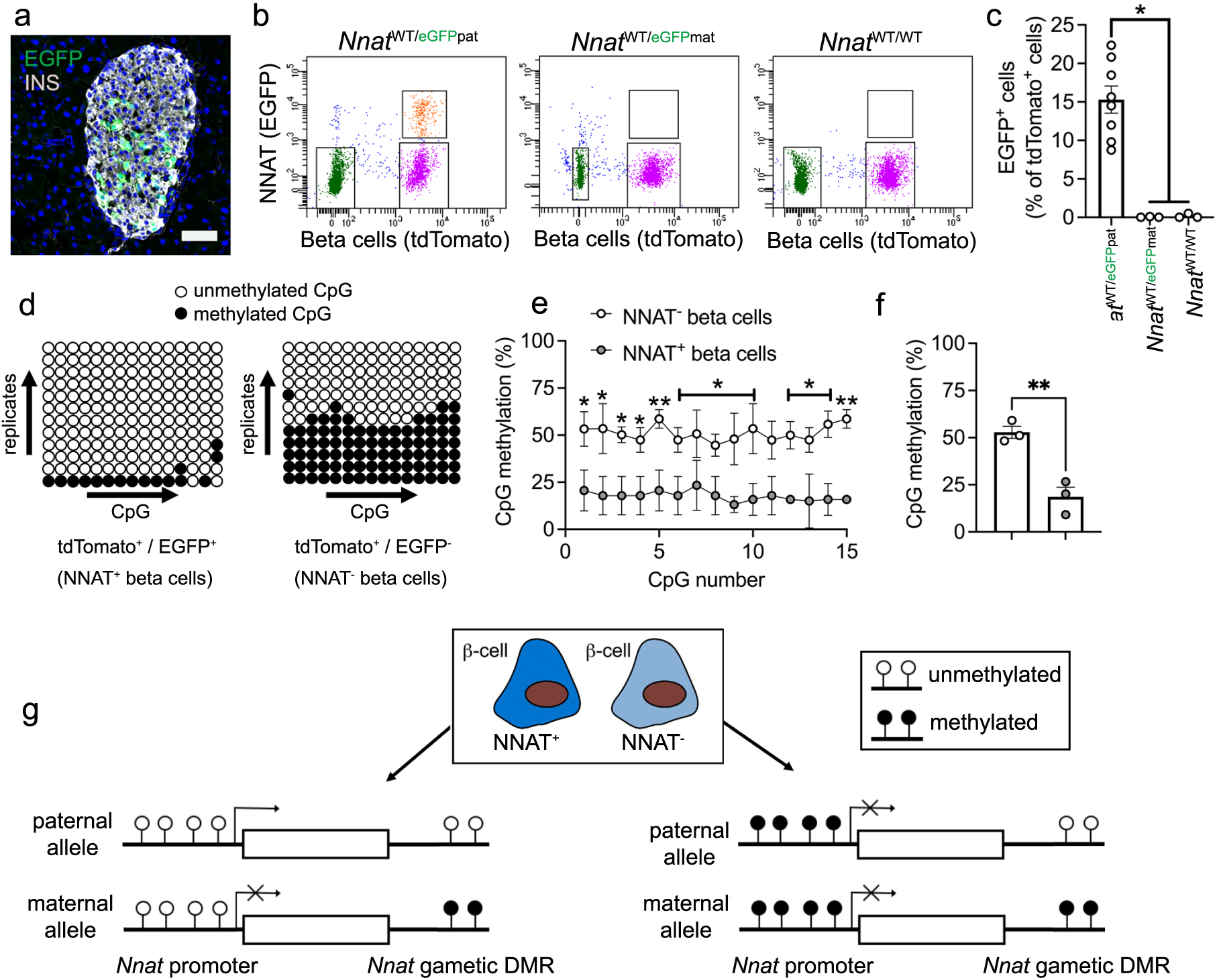
Beta cell heterogeneity of NNAT expression is associated with changes in CpG methylation at the *Nnat* promoter. (a) Representative confocal microscopy of pancreatic cryosections from P56 mice with *Nnat*-driven EGFP expression from the paternal allele (*Nnat*^WT/eGFPpat^) (n = 7 mice). Sections were immunostained with antibodies against EGFP (green) and insulin (INS, grey). Nuclei are visualised with DAPI. Scale bar = 50μm. (b) Separation of dispersed primary islet cells by FACS from reporter mice with insulin-driven expression of tdTomato (to label beta cells) and *Nnat*-driven EGFP expression from the paternal (*Nnat*^WT/eGFPpat^), or maternal *(Nnat*^WT/eGFPmat^) allele or wild type (*Nnat*^WT/WT^) at this locus (representative image of one dispersed islet preparation, each from a single mouse) (n = 8, 3 and 3 mice, respectively, Kruskal-Wallace test with Dunn’s multiple comparisons). (c) Quantification of data in b expressed as percentage of EGFP/tdTomato co-positive primary islet cells. (d) Representative bisulphite analysis of CpG methylation at the *Nnat* promoter in FACS-purified islet cell populations from b (n = 3 *Nnat*^WT/eGFPpat^ mice with paternally expressed *Nnat*-driven EGFP, n = >12 clones each). Closed circles = methylated CpG, open circles = unmethylated CpG. (e, f) Quantification of data in d expressed as percentage CpG methylation across the *Nnat* promoter at individual CpGs (e) and across the entire *Nnat* promoter (f) (paired Student’s *t* test). (g) Schematic summarising level of CpG methylation at the *Nnat* promoter and gametic DMR in NNAT^+^ vs NNAT^-^ beta cells. Representative images from three independent experiments and breeding pairs. * P < 0.05, ** P < 0.01.

To isolate and purify beta cells based on NNAT levels, we crossed *Nnat*-eGFP reporter mice to animals expressing *Cre* recombinase under the rat insulin promoter [53] and to transgenic mice expressing a tdTomato reporter downstream of a *lox*P-flanked stop codon [54]. Dispersion of primary adult islets into single cells, and subsequent FACS analysis, verified the presence of NNAT^+^ and NNAT^-^ beta cells, with the NNAT^+^ fraction comprising 15.3±1.8% (n=8 mice) of the total beta cell compartment (Fig. 5b, c, ESM Fig. 9a, b and ESM Fig. 10a-f). Collection of FACS-purified NNAT^+^ and NNAT^-^ beta cells and subsequent bisulphite sequencing analysis revealed that CpG methylation at the gametic DMR (known to control monoallelic *Nnat* expression; introduction) was unchanged (ESM Fig. 9c-e). Nevertheless, CpG methylation at the *Nnat* promoter was significantly altered across this genomic region, with minimal CpG methylation observed in NNAT^+^ beta cells (Fig. 5d-f).

Recent whole-genome methylation analysis in mouse germ cells has revealed that the *Nnat* promoter region is unmethylated in sperm, and this was confirmed by targeted bisulphite sequencing analysis (ESM Fig. 9f-h), whereas it is fully methylated in oocytes [57]. Thus, the ‘classical’ imprinting of *Nnat* is unaltered between beta cell subtypes with near binary (‘on/off’) expression of neuronatin. However, and overlaying this control, a second DMR at the *Nnat* promoter within the non-imprinted allele dictates beta cell subtype specificity of expression (Fig. 5g).

### The *de novo* DNA methyltransferase DNMT3A establishes NNAT beta cell subtype specificity

The above data indicate that beta cells transition from a state where the majority (>90%) are NNAT-positive at late embryogenesis to 31.2±3.9% (n=14 mice) NNAT-positive by postnatal day 7. This is associated with differential CpG methylation at the *Nnat* promoter (Fig. 5d-g). To ask whether this apparent transition in the early postnatal period is driven by *de novo* CpG methylation, we assessed endogenous NNAT beta cell immunoreactivity in postnatal mice conditionally deleted for the methyltransferase DNMT3A at the pancreatic progenitor stage (using *Pdx1*-*Cre*) [10]. NNAT staining in control mice at postnatal (P) day 6 demonstrated expression in a subpopulation of beta cells whereas deletion of DNMT3A resulted in a loss of this heterogenous expression across the islet (Fig. 6a, b, ESM Fig. 11a). These findings demonstrate that *de novo* methylation is likely to drive NNAT restriction across the beta cell complement in the first few days of postnatal life. Interestingly, a high number of NNAT^-^ beta cells were highly enriched for the repressive histone modification H3K27me3 (Fig. 6c), suggesting that other epigenetic pathways may play an additional role in silencing this gene in the early postnatal period.

**Fig. 6.**
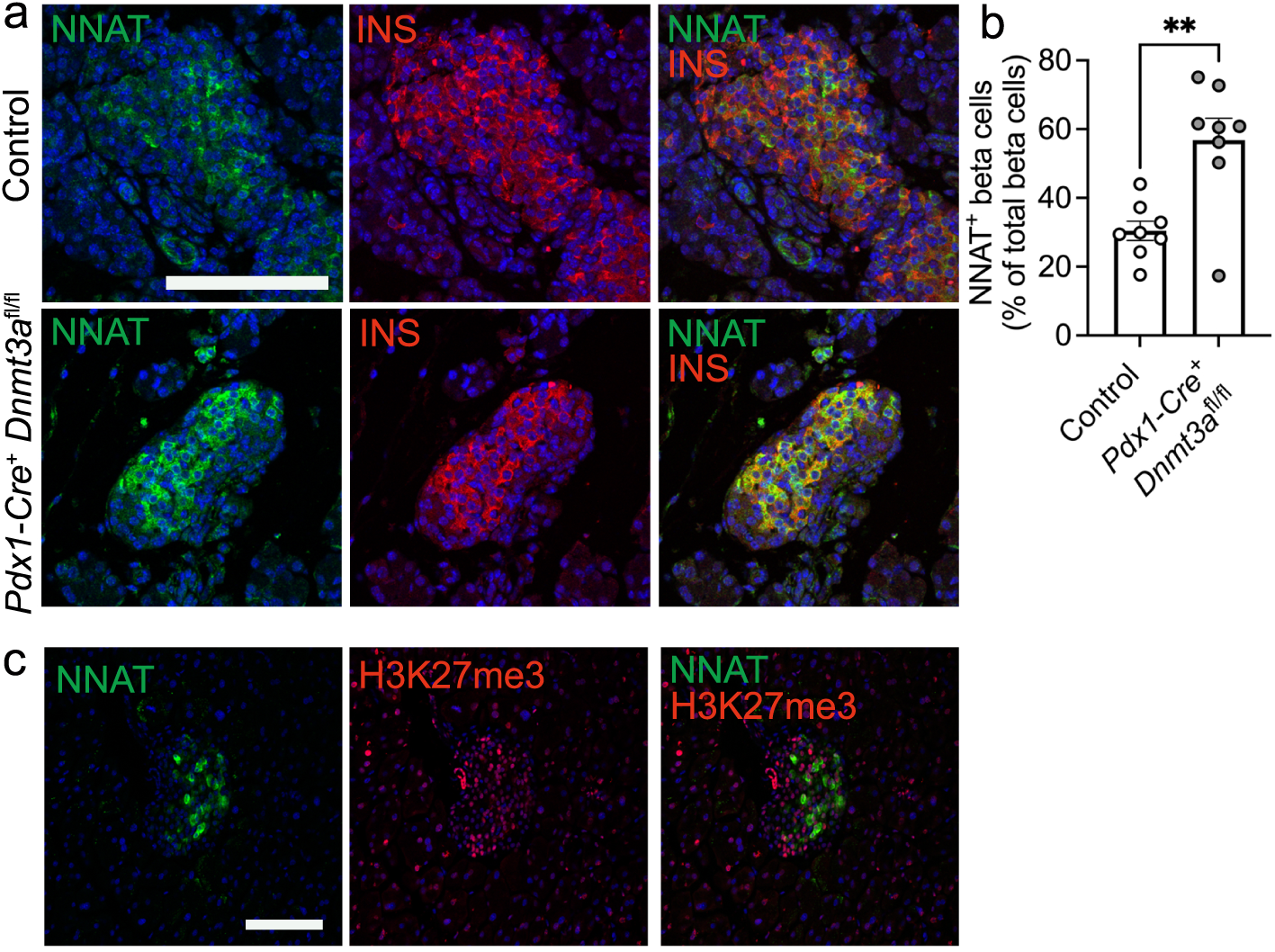
Postnatal restriction of NNAT in a subset of beta cells is at least partially driven by the *de novo* methyltransferase, DNMT3A. (a) Representative confocal microscopy of pancreatic cryosections from mice with conditional deletion of DNMT3A under the control of the *Pdx1* promoter (*Pdx1*-*Cre^+^ Dnmt3a*^fl/fl^) vs control (*Pdx1*-*Cre^-^ Dnmt3a*^fl/fl^) mice at postnatal (P) day 6. Sections were immunostained with antibodies against endogenous neuronatin (NNAT, green) and insulin (INS, red). (b) Quantification of NNAT^+^ beta cells from images shown in a, expressed as NNAT/INS co-positive cells as a percentage of total INS-positive cells. Scale bar = 50μm (n = 8 mice per genotype, unpaired Students t test, ** P < 0.01). (c) Representative confocal microscopy of pancreatic cryosections from P56 (8 week old) wild type mice on a C57BL/6J background immunostained with antibodies against endogenous neuronatin (NNAT, green) and H3K27me3 (red). Scale bar = 100μm, n = 3 mice. Nuclei are visualised with DAPI. Representative images from three independent experiments and breeding pairs.

### NNAT^+^ and NNAT^-^ beta cells have distinct transcriptional signatures

In contrast to the embryonic populations described above, RNA-seq revealed that FACS-sorted adult (8 weeks) NNAT^+^ and NNAT^-^ beta cells displayed high transcriptional overlap and were similar to each other when compared with non-beta endocrine cells (Fig. 7a). Major beta cell identity markers such as *Ins1*, *Ins2*, *Mafa*, *Slc2a2* (*Glut2*) and *Nkx6.1* were not differentially expressed (ESM Table 5).

**Fig. 7.**
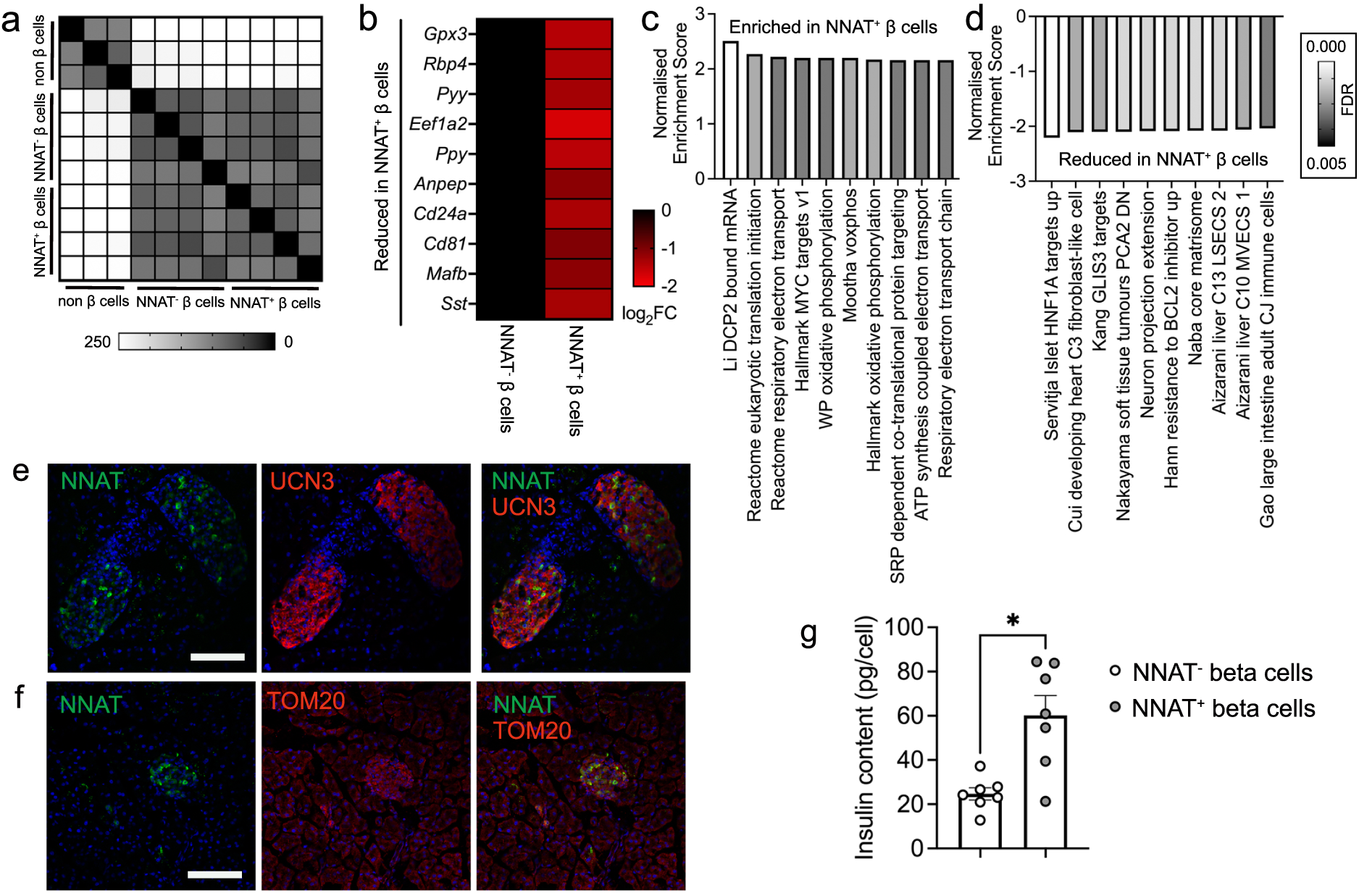
NNAT^+^ adult beta cells are transcriptionally distinct and have significantly higher insulin content. (a) Correlation matrix of differentially expressed genes between NNAT^+^ and NNAT^-^ beta cells as assessed by RNA-sequencing analysis (n = 4 FACS-purified populations from individual mouse islet preparations). (b) Heatmap of top 10 most differentially expressed genes reduced in NNAT^+^ beta cells compared with NNAT^-^ beta cells. (c, d) Gene set enrichment analysis (GSEA) showing categories significantly enriched (c) and reduced (d) in NNAT^+^ (vs NNAT^-^) beta cells. (e, f) Representative confocal microscopy of pancreatic cryosections from P56 (8 week old) wild type mice on a C57BL/6J background immunostained with antibodies against endogenous neuronatin (NNAT, green) and UCN3 (red, e) or TOM20 (red, f). Scale bar = 100μm, n = 3 mice. Nuclei are visualised with DAPI. Representative images from three independent experiments and breeding pairs. (g) Insulin content assessed in NNAT^+^ and NNAT^-^ beta cells (* P < 0.05, n = 7 FACS-purified populations from individual mouse islet preparations, Wilcoxon matched-pairs signed rank test).

Nevertheless, we identified 241 (1.8%) and 79 (0.6%) genes that were significantly down- and up-regulated in NNAT^+^ vs NNAT^-^ beta cells, respectively (>2-fold, FDR < 0.01). Differentially-expressed genes included several markers of non-beta cell islet lineages (*Pyy*, *Ppy*, *Mafb*, *Sst*), that were lower in the NNAT^+^ beta cell fraction; genes linked with beta cell heterogeneity and plasticity (*Gpx3*, *Rbp4*), the beta cell immaturity marker *Cd81* [26] and the *Cd24a* antigen [27] (Fig. 7b). Interestingly, *Npy* was enriched in NNAT^+^ beta cells (ESM Table 5), consistent our observations at late embryogenesis (E17.5, Fig. 2d). We did not observe any clear differences between NNAT^+^ and NNAT^-^ beta cells in the expression of other imprinted genes known to be functional in the beta cell (*Plagl1*/*ZAC*, *Dlk1*, *Rasgrf1*, *Cdkn1c*, *Grb10* and *Gtl2*/*MEG3*), nor changes in ‘disallowed’ genes whose expression are known to be restricted in mature functional beta cells (*Hk1*, *Mct1* (*Slc16a1*) and *Ldha*) [58] (ESM Table 5). Likewise, no differences were apparent in the levels of transcripts encoding proteins previously demonstrated by others to mark specific beta cell subpopulations such as *Flattop*/*Cfap126* [21], *CD9* and *ST8SIA1* [15], *Cd63* [29], ‘virgin’ beta cells (via *Ucn3*) [30] or those differentially-expressed in beta cells implicated in the control of Ca^2+^ dynamics (‘hubs’ [19], and ‘leaders’ [20, 32]).

GSEA did, however, reveal enrichment of pathways in NNAT^+^ beta cells including translation initiation, the electron transport chain (ETC), oxidative phosphorylation and signal recognition particle (SRP)-dependent co-translational protein targeting (Fig. 7c). Pathways involving genes associated with hepatic nuclear factor 1 alpha (HNF1A) and GLI-similar zinc finger protein family 3 (GLIS3) targets, were reduced in the NNAT^+^ beta cell fraction (Fig. 7d). NNAT^+^ beta cells displayed lower levels of the late maturation marker UCN3 (Fig. 7e) but no differences in mitochondrial volume (Fig. 7f). Furthermore, FACS-sorted primary NNAT^+^ beta cells had a significantly higher insulin content than NNAT^-^ cells (Fig. 7g).

### The NNAT^+^ beta cell population shows impaired glucose-stimulated Ca^2+^ dynamics and is de-enriched for highly connected ‘hub’ cells

To determine whether NNAT may influence beta cell connectivity and membership of the ‘hub’ cell subgroup [19], we studied glucose-induced Ca^2+^ dynamics in *Nnat* deficient (*Nnat*^+/-p^) islets [45] by high-speed confocal imaging of the Ca^2+^ probe Cal-520 and off-line analysis using Pearson’s R-based correlation analyses [20]. The mean Pearson’s coefficient of correlation and the identification of highly-connected ‘hub’ cells (displaying coordinated Ca^2+^ responses with at least 50% of all cells) was not significantly different between wild type vs *Nnat* deficient islets under low or high (11 mM) glucose conditions (ESM Fig. 12a-d). Interestingly, while there were no significant differences in the average connectivity between the highly-connected wild type vs *Nnat* deficient ‘hub’ cells in low glucose (P=0.16), wild type ‘hub’ cells were significantly more connected than *Nnat* deficient ‘hub’ cells in high (11 mM) glucose (P<0.001) (ESM Fig. 12e). The proportions of highly-connected ‘hub’ cells and ‘follower’ cells were not significantly different between the wild type and *Nnat* deficient groups in both low and high glucose conditions (ESM Fig. 12f). Thus, NNAT is not involved in setting beta cell connectivity but likely reflects a marker of less well-connected cells.

We further explored this question using islets from reporter mice with paternally expressed *Nnat*-eGFP. In this case, intracellular Ca^2+^ changes were monitored using the red-shifted calcium probe Cal-590 [59]. *Nnat*-GFP^+^ and *Nnat*-GFP^-^ cells responded similarly to challenge with high (11 mM) glucose (Fig. 8a). Beta cell-beta cell connectivity (Fig. 8b) was similarly not different between NNAT^+^ and NNAT^-^ populations. Thus, considered across the whole population, the mean Pearson’s coefficient of correlation was 0.87±0.01 (Fig. 8c), with 14.9% beta cells being significantly connected.

**Fig. 8.**
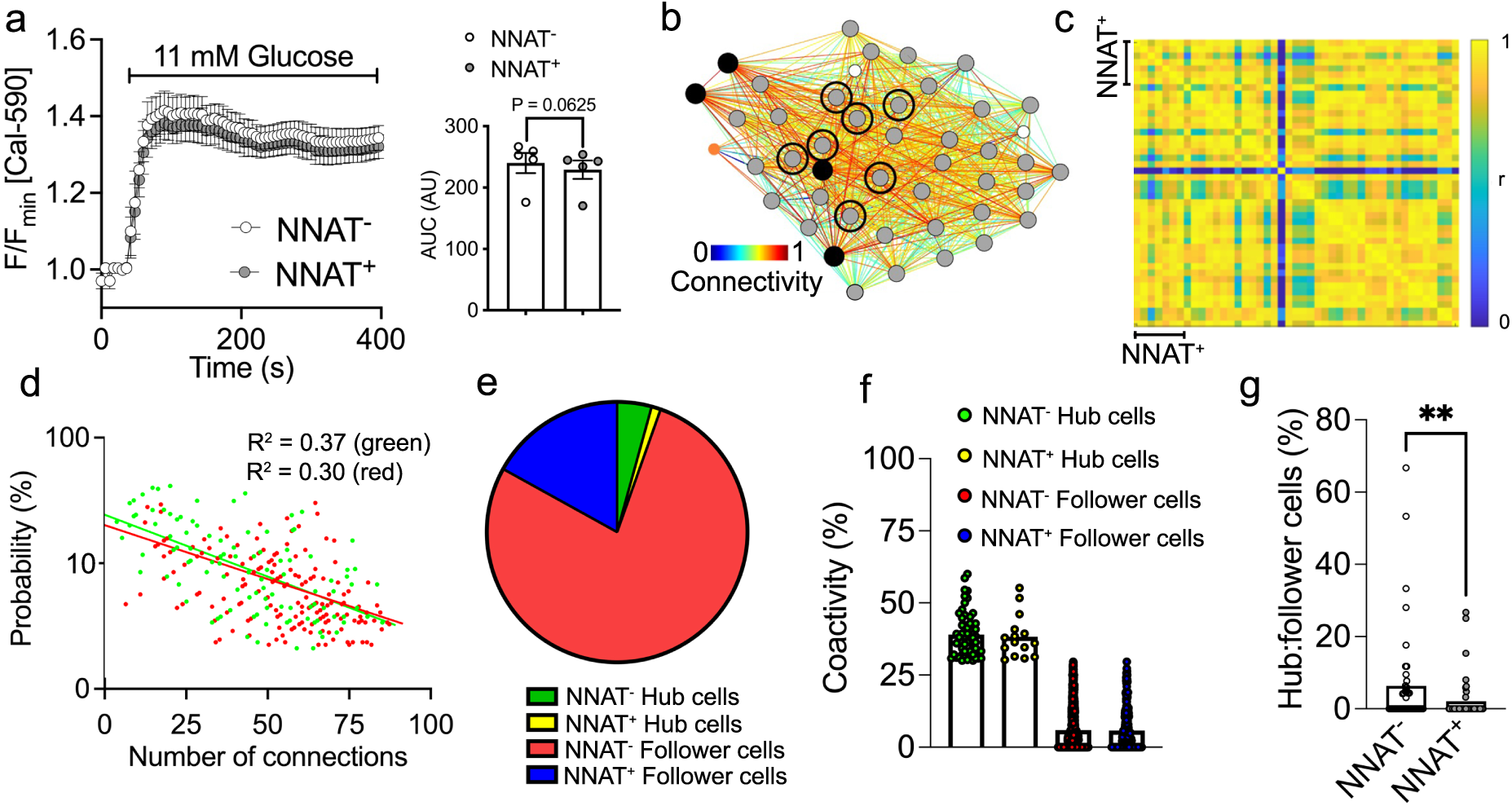
NNAT^+^ beta cells show unaltered glucose-induced Ca^2+^ dynamics but are de-enriched for highly connected ‘hub’ cells within individual islets. (a) Ca^2+^ bound Cal-590 fluorescence in response to high glucose (11 mM) in NNAT^-^ and NNAT^+^ cells from primary islets from *Nnat*^WT/eGFPpat^ reporter mice expressed as normalised intensity over time (F / F_min_) (n = 61 islets total from five mice per genotype, inset shows quantification of area under the curve (AUC), P = 0.063, Wilcoxon matched-pairs signed rank test). (b) Representative cartesian map of beta cells with colour coded lines connecting cells according of the strength of coactivation (colour coded R values from 0 to 1, blue to red). Beta cells are represented by differently coloured nodes depending on their coactivity with the other beta cells, where black nodes indicate coactivity with ≥80% of the remaining beta cells, while grey and white nodes represent coactivity with ≥60% and ≥40%, respectively. Nodes circled with a solid black line indicate NNAT^+^ cells. (c) Representative heatmaps depicting connectivity strength (r) of all cell pairs (colour coded r values from 0 to 1, blue to yellow). (d) Log-log graphs of beta cell-beta cell connectivity distribution. NNAT^+^ cells are represented by green circles while NNAT^-^ cells are represented by red circles (45 islets total using primary islet preparations, each from an individual *Nnat*^WT/eGFPpat^ reporter mouse). (e) Categorisation of beta cells based on data from d. (f) Percentage coactivity of beta cells between all cells in identified ‘hub’ and ‘follower’ cells. (g) The proportion of cells designated as ‘hub’ vs ‘follower’ cells in both the NNAT^-^ and NNAT^+^ populations assessed in each of 45 islets (** P < 0.01, Wilcoxon matched-pairs signed rank test). Representative images from three independent experiments and breeding pairs.

Pooled data over 45 islets uncovered a scale-free network topography (Fig. 8d) where a small proportion of beta cells (4.4%) were identified as ‘hub’ cells, defined as those displaying coordinated Ca^2+^ responses with at least 30% of all beta cells (R^2^=0.31). Of the 45 islets analysed, 34 (75.6%) displayed obedience to a power law, and this adherence pertained for both NNAT^+^ (Fig. 8d; R^2^ = 0.38) and NNAT^-^ populations (R^2^=0.30). Cells could be categorised as belonging to one of four distinct groups (Fig. 8e). Of all cells examined, 4.27% were NNAT^-^ ‘hub’ cells that displayed coordinated Ca^2+^ responses to an average of 39.0% of all beta cells (Fig. 8e, f). Similarly, the fraction of these cells (1.15%, Fig. 8e) that were identified as NNAT^+^ ‘hub’ cells had an average of 38.3% coactivity with the remaining cells (Fig. 8f). There were no significant differences in connectivity between these two groups (Fig. 8f; P=0.76) nor in connectivity between NNAT^+^ and NNAT^-^ ’follower’ cells (P=0.72). However, when considered within each individual islet, a significantly lower ratio of ‘hubs’ to ‘followers’ was observed in NNAT^+^ *versus* NNAT^-^ cells (Fig. 8g). Overlap with ‘leader’ cells [20, 31] was not explored in the absence of readily identifiable Ca^2+^ waves.

## Discussion

We describe here a novel subgroup of beta cells characterized by transient methylation of a second DMR (but not the gametic DMR) in the *Nnat* locus. This is consistent with previous findings that methylation at gametic DMRs is thought to be relatively stable [60, 61], whereas ‘secondary’ imprinting regions have been shown to be more sensitive to nutrient-or physiology-based changes [10, 62]. We have previously described the importance of *Nnat* for the normal secretory function of beta cells [45] and therefore hypothesise that NNAT^+^ and NNAT^-^ cells display differences in secretory function. Correspondingly, transcriptomic analysis of purified adult NNAT^+^ and NNAT^-^ beta cells demonstrated that the NNAT^+^ fraction is enriched for functional pathways including translation initiation, ETC/oxidative phosphorylation pathways and co-translational protein ER membrane targeting.

Extending these studies to man, NNAT deficiency in human EndoC-βH1 beta cells severely blunted GSIS, likely via enhancement of signal peptidase-mediated processing, consistent with our previous work in rodents [45]. We show here that E17.5 *Nnat*^+^ beta cells are enriched for expression of the SPC and translocon apparatus (*Spcs1*, *Spcs2*, *Sec11a*, *Sec11c*, *Sec61*) and that the SPC/NNAT interaction is via SPCS1, and likely to be how NNAT exerts its effect on cellular insulin content. Importantly, we demonstrate that heterogeneous expression of NNAT is also a feature of the adult human pancreatic islet. Finally, and in line with our previous studies, NNAT^+^ and NNAT^-^ cells did not differ in their glucose-induced Ca^2+^ dynamics, although their alignment with a highly-connected group of ‘hub’ cells differed compared to that of NNAT^-^ cells. Thus, NNAT^+^ cells are likelier to belong to the population of ‘follower’ cells consistent with a role in insulin production rather than glucose detection [19]. Interestingly, Ca^2+^ responses to 11 mM glucose were significantly higher in *Nnat* deficient versus wild type mouse islets, in contrast to previous findings where differences were not observed in response to 16.7 mM glucose [45]. A similar strong tendency was also seen when comparing NNAT^+^ versus NNAT^-^ cells in the same islet in *Nnat*-eGFP reporter mice, and suggests that NNAT^+^ cells are less responsive to metabolic stimulation by glucose, in line with their enrichment in a ‘follower’ subset of cells [19, 20]. The molecular underpinnings of the weaker Ca^2+^ responses are unclear, but do not appear to involve differences in the levels of transcripts encoding *Slc2a2* (*Glut2*) or *Gck*. Nevertheless, the transcriptome of NNAT^+^ cells does not show enrichment for genes imputed to be enriched in ‘hub’ cells [20] nor in ‘leader’ cells [31] which appear to be discrete populations.

Interestingly, bimodal expression of *Nnat* was already clearly evident at embryonic stages, with *Nnat*^+^ beta cells enriched for markers of late-stage beta cell differentiation, as well as *Ins1* and *Ins2* mRNAs. These differences were, however, less marked in the adult islet, though CD24a [11] was more weakly expressed in the NNAT^+^ population. De-enrichment of markers of other islet cell types (and the immaturity marker *Cd81*) was also a common feature of both embryonic and adult NNAT^+^ beta cells. Interestingly, both embryonic and adult NNAT^+^ beta cells were enriched for *Npy* and adult NNAT^+^ beta cells had lower UCN3 protein, suggesting that these cells were not fully matured. NNAT^+^ beta cells however had significantly higher insulin content, suggesting a possible, at least partial, overlap with the recently-described “CD63^hi^” population recently defined by [29], and the “β_HI_” population described by [11].

Our work also provides evidence that the epigenome controls the fate of specific beta cell subpopulations of beta cells. CpG methylation plays a crucial role in early beta cell developmental maturation, including the silencing of ‘disallowed’ genes such as *Hk1*, *Mct1* (*Slc16a1*) and *Ldha* via DNMT3A [8, 58]. Here we show that islets transition from a state where the majority of beta cells are NNAT^+^ in late embryogenesis to comprising a restricted subpopulation of NNAT^+^ beta cells by postnatal (P) day 7. Whether this represents changes at the level of individual beta cells, or the turnover of the NNAT^+^ population and replacement with a largely NNAT^-^ population, was not determined, and would require investigation using other approaches, including fate mapping (lineage tracing).

Adult NNAT^+^ beta cells are virtually unmethylated at the *Nnat* promoter whereas robust promoter methylation was apparent in NNAT^-^ cells. These findings, and the fact that the *Nnat* promoter is differentially methylated between sperm and oocytes (see results), indicates that promoter methylation during this transition is likely to be acquired on the paternal allele (from which *Nnat* is selectively expressed via genomic imprinting). Our studies also show that specific deletion of the *de novo* methyltransferase DNMT3A at the pancreatic progenitor stage resulted in partial loss of NNAT-based beta cell heterogeneity. Thus, DNA methylation modulates restricted NNAT expression in specific beta cells during early maturation. We also observe heterogeneity between beta cells at the level of the histone modification, H3K27me3, as recently documented [11], and find that NNAT^-^ beta cells in the adult are more likely to be marked by this repressive epigenetic modification. This again suggests there may be at least a partial overlap with the “β_HI_” population described in [11].

Might these findings be relevant for the pathogenesis of type 2 diabetes? Whether demethylation of the second DMR at the *Nnat* locus may occur in type 2 diabetes is an interesting possibility which remains to be explored. In rodents, even mild hyperglycaemia deregulates *Nnat* expression alongside that of several other critical beta cell identity genes [63] and we show in our present work a significant reduction of NNAT^+^ beta cells in *db/db* mice. Moreover, altered CpG methylation is a common observation in islets from patients with diabetes at imprinted and non-imprinted *loci* (reviewed in [23, 41]).

Our previous findings [45] and the current studies have shown that neuronatin is crucial for GSIS in a cell autonomous manner. We now demonstrate postnatal acquisition of beta cell heterogeneity as conferred by ‘on/off’ expression of NNAT, likely from a common NNAT^+^ beta cell precursor at late embryogenesis. Since *Nnat* deletion lowers insulin content and GSIS [45], and NNAT^+^ beta cells are retained well into adulthood, we suggest that NNAT^+^ beta cells may play a distinct functional role in the mature islet, contributing to a highly differentiated, INS high population. Interestingly these appear to be enriched in the “follower” population, which appear to respond to upstream signals from ‘leader’ and ‘hub’ cells (see Introduction and references therein).

In conclusion, the present work demonstrates how an effector gene in pancreatic beta cells, neuronatin, is controlled at the level of the epigenome (DNA methylome), contributing to a functional hierarchy between cells. Chemical modification of the epigenome [64] may in future provide an attractive therapeutic angle not only for beta cell replacement or regeneration [65] but also to modulate beta cell function and cell-cell connectivity in type 2 diabetes.

## Acknowledgements

We thank the LMS/NIHR Imperial Biomedical Research Centre Flow Cytometry Facility and the MRC LMS Genomics Facility for the support.

## Data availability

RNA-seq and proteomics data will be deposited to GEO and ProteomeXchange (PRIDE) repositories, respectively. ImageJ scripts and CellProfiler pipelines can be made available upon reasonable request. Pseudo-time analysis scripts will be made available via GitHub. Transcriptomic data for *ob/ob* mice can be obtained from a publicly available source from the Alan Attie laboratory (http://diabetes.wisc.edu/search).

## Authors relationships and activities

G.A.R. has received grant funding from and is a consultant for Sun Pharmaceuticals Inc. Other authors declare no conflicts of interest.

## Author contributions

SJM and GAR designed the research. VY, FY, AM, SK, AO, SB, SSV, PC, SMT, KC, EG, NP, MS, ZS, LM, CDL, MT, GO, LDS, XW, YH, AM, JE, BP, ND, CW, PS, HK and YA performed experiments and analyses. PH, PM, RS, SD and DJW contributed new reagents/analytical tools. SJM and GAR drafted and/or wrote the manuscript. SJM, SD, DJW and GAR provided funding. SJM and GAR supervised the work. SJM serves as a guarantor of this work.

