## Supplemental data for "Differential CpG methylation at *Nnat* in the early establishment of beta cell heterogeneity"

**Electronic supplementary material (ESM)**

**ESM Methods**

*Bioinformatic clustering analysis*

Single cell RNA-seq islet data were downloaded from Gene Expression Omnibus database (accession number GSE101099 [1], E17.5 timepoint, batches '1SW', '2SW' and 'Batch 2'; GSE155742 [2], E16.5 newly synthesised beta cells batches 'Green 1' and 'Green 2'; GSE128565 [3, 4], adult islets, batch 'No STZ', and GSE110648 [5], adult islets, 'B6_28wks_1' and 'B6_28wks_2' (chow fed mice)), filtered according to the respective published work and hypervariability of gene expression was assessed by variance analysis using the scran package [6]. This method avoids artefacts resulting from technical dropouts or of prioritising low abundance genes with large variances due to technical noise. The R package RaceID [7] was used for further analysis. Mitochondrial genes were removed from the expression matrix prior to downstream analysis and cells were filtered to have at least 5000 transcripts with an expression value of >2. A distance matrix was then constructed using Pearson correlation as the distance metric. A K-nearest neighbour graph layout utilizing the Fruchterman-Rheingold algorithm was performed to cluster cells into different groups, with a value of k = 3. This algorithm also identifies outlier populations, and so a total of six clusters were identified using the clustering method, with differential expression analysis performed comparing individual clusters to all other cells. Further analysis of E17.5 mouse scRNA-seq data in the form of Seurat R object was downloaded from https://doi.org/10.6084/m9.figshare.c.4158458 [1]. Normalization was performed using the sctransform function [8] in Seurat v4 [9], followed by principal component analysis (PCA) and clustering in the top 30 PC space using the 'leiden' algorithm. The clusters were assigned their appropriate cell-types based on the expression of known cell-type specific marker genes. For visualization purpose, cells were mapped to the UMAP space using Seurat's built-in function, with extraction of islet clusters identifying two distinct subpopulations of beta cells. Based on this distribution, cells with *Nnat* expression >3 were designated as *Nnat*+. For geneset enrichment analysis (GSEA), differentially expressed genes were ranked according to their fold change. GSEA was performed for the Gene Ontology biological process terms [10, 11] and (org.Mm.eg.db: Genome wide annotation for Mouse. R package version 3.8.2) using the R package ClusterProfiler [12]. The enrichment and pathway module plots were prepared using the DOSE [13] and enrichplot (package version 1.14.2. 2022) R packages respectively. Volcano plots were prepared using the R package EnhancedVolcano (R package version 1.12.0, https://github.com/kevinblighe/EnhancedVolcano).

*Pseudo-time analysis*

scRNA-seq datasets collected from islets during mouse embryogenesis (E12.5-E17.5) were obtained from [1] with Seurat v4.3 used for quality control and batch correction [14]. The dataset was filtered to select for islet cells by visualizing the expression of *Cdh1* and *Chga* as well as the lack of expression of *Vim* and *Col3a1* [1], which resulted in 4556 cells for downstream analyses. Following filtering, clustering was achieved by running the Louvain community detection algorithm on a shared nearest neighbours (SNN) graph generated from the single cell expression profiles [14]. Using the developmental and cell type markers identified in [1] we labelled each cluster as a known islet cell type or precursor of which 818 cells were identified as ‘early’ or ‘late’ beta cells. This also allowed us to evaluate the change in beta cell precursor/state proportion across embryonic stages. After labelling, we used the Monocle3 package to identify cell fate trajectories using a pseudo-time analysis [15]. After defining the development trajectory of beta cells during embryogenesis, we integrated our data with a dataset from adult mouse islets [3, 4].

*Study approval*

All animal procedures were in accordance with the UK Animals (Scientific Procedures) Act 1986 and approved by the UK Home Office (PPL: PP4712156). All animal studies have been approved by the Imperial College Animal Welfare and Ethical Review Body. Animals were euthanised by cervical dislocation. Animal work was designed and reported according to the Animal Research: Reporting of In Vivo Experiments (ARRIVE) guidelines. Power calculations were based on known effect sizes. Where possible, blinding for experimental mice and tissues was used and all studies were replicated in at least three independent cohorts or using litters from three different breeding cages.

*Animal models*

Transgenic mice lines expressing *Cre* recombinase under the control of the rat insulin promoter (RIP) [16] and with a tdTomato reporter downstream of a stop codon flanked by *loxP* sites [17] have been described previously. Mice with *Nnat*-driven *Egfp* expression were purchased from the Mutant Mouse Resource and Research Centre (MMRRC, USA) repository (Tg(Nnat-EGFP)EA106Gsat/Mmucd, #010611-UCD) with the presence of the *Egfp* allele assessed by standard PCR using DNA from ear biopsy (primers 5'-CATCACCCCTCCTTCTCAAC-3' and 5'-GAACTTCAGGGTCAGCTTGC-3'). Mice with global deletion of *Nnat* were used as previously described [18]. Conditional *Dnmt3a* null mice [19], generated by crossing to *Pdx1-Cre* [20] have been described elsewhere [21-23]. Mice containing the *Dnmt3a* floxed allele but lacking *Pdx1-Cre* were used as controls. Islet transcriptomic data from 7 week old Zucker diabetic fatty (ZDF) rats vs ZDF/+ controls was according to [24]. Similarly, islet transcriptomic data in the mouse *ob/ob* strain vs *+/ob* control mice (4 and 10 weeks old on a C57BL/6 background) was obtained from a publicly available source from the Alan Attie laboratory (http://diabetes.wisc.edu/search). 6-8 week old *db/db* and control C57BLKS/J mice on the BKS background were obtained from the Jackson Laboratory (JAX 000642 and JAX 000662, respectively). All other mouse strains were on a C57BL/6J background and experimental cohorts were group-housed and maintained on a 12-hour light/dark cycle. Animals were fed standard mouse chow (RM3, Special Diet Services) with access to drinking water *ad libitum* in specific pathogen-free barrier facilities. Male mice were 10-12 weeks old at the time of experiments unless otherwise stated. For experiments involving the *Nnat*-eGFP reporter line, mice were hemizygous for the reporter via paternal transmission, unless otherwise stated. Sperm was isolated from young adult male C57BL/6J mice.

*Primary islet isolation and FACS*

To isolate primary islets, the pancreas was inflated with Liberase TM (Roche), digested at 37°C, purified using a Histopaque 1119/1083/1077 gradient (Sigma-Aldrich) and hand-picked. For FACS-based experiments, cells from purified islets were isolated immediately following isolation using Accutase (Sigma-Aldrich) for 5 min. at 37°C followed by gentle disaggregation by pipetting. Islet cells were centrifuged and resuspended in FACS buffer (25 mM HEPES, 5 mM EDTA, 1% fetal bovine serum (FBS) in PBS, pH 7.4) prior to cell sorting using an FACSAria III flow cytometer (BD, UK) equipped with 488 nm and 561 nm lasers for excitation of EGFP and tdTomato, respectively. Emission from EGFP and tdTomato expressing cells was collected using a 530/30 and 580/30 bandpass filter, respectively. DAPI (0.2 $\mu$g/mL) was used to exclude dead cells and was excited using a 405 nm laser with emission collected using a 450/50 bandpass filter. Forward scatter area *vs* forward scatter height was used to offset and remove doublets. Total insulin content in FACS-sorted populations was assessed using ultra-sensitive insulin homogeneous time-resolved fluorescence (HTRF) assay kits (Cisbio).

*Intracellular calcium imaging*

Pancreatic islets were isolated as above and left to settle overnight in RPMI medium (Gibco) supplemented with 10% (v/v) FBS, (Sigma-Aldrich), 2 mM L-glutamine and 100 U/mL penicillin/100 $\mu$g/mL streptomycin (both Gibco) at 37°C and 5% $CO_2$. For experiments involving reporter mice expressing *Nnat*-eGFP, $Ca^{2+}$ imaging of whole islets was performed after loading the cytosol with 2 $\mu$M Cal-590 AM (Stratech) in Krebs-Ringer bicarbonate buffer (140 mM NaCl, 3.6 mM KCl, 0.5 mM $NaH_2PO_4$, 25 mM $NaHCO_3$ (saturated with $CO_2$), 1.5 mM $CaCl_2$, 0.5 mM $MgSO_4$, 10 mM HEPES, pH 7.4) containing 3 mM or 11 mM glucose. Images were captured at 0.5 Hz on an Axiovert microscope (Zeiss, Germany) equipped with a 10x 0.3–0.5 NA objective, a Hamamatsu ImagEM camera coupled to a Nipkow spinning-disk head (CSU-10, Yokogawa UK Ltd) and illuminated at 490 nm or 530 nm. Data were analysed using ImageJ and changes in fluorescence intensity over time were measured after drawing a region of interest around individual cells for each islet. Pearson-based connectivity and correlation analyses were performed as previously described [25]. The first 50 datapoints were removed to account for the initial spike in response to the addition of 11 mM glucose. Signal binarisation-based connectivity and correlation analyses were performed as previously described [26]. Connectivity between cells was explored using Pearson's R-based correlation analyses in MATLAB as described [22]. We note that although the $Ca^{2+}$ dye used here was not selectively targeted towards beta cells, at the concentration of glucose used (11 mM), pancreatic alpha cells are expected to be largely inactive. We do not exclude a contribution of delta cells, although these represent <5% of the overall endocrine cell population.

For experiments involving global *Nnat* null mice [18], islets were incubated at 37 °C for 40 min in Krebs-Ringer bicarbonate buffer containing 4.5 μM Cal-520 AM (Stratech) and either 3 mM or 11 mM glucose, and then transferred into an imaging chamber. Imaging was performed on a Nikon Eclipse Ti microscope equipped with a 40x/1.2 NA oil objective and an ibidi heating system. Cal-520 AM was excited with a 491 nm laser line and emitted light filtered at 525/50 nm. Images were acquired with an ORCA-Flash 4.0 camera (Hamamatsu) and Metamorph software (Molecular Device) was used for data capture, in streaming mode and with an interval of 230 ms.

Pearson's R-based connectivity and correlation analyses in an imaged islet were performed as previously described [25]. The first 50 datapoints were removed to account for the initial spike in response to the addition of 11 mM glucose. $Ca^{2+}$ signals were first smoothed by fitting them to a smoothing spline [27]. The smoothing spline *s* was constructed and minimised thus:

$$p\sum_i w_i\big(y_i - s(x_i)\big)^2 + (1-p)\int\left(\frac{d^2s}{dx^2}\right)^2 dx$$

where $p$ = specified smoothing parameter between 0 and 1, $w$ = specified weight

Once the data was fitted to a smoothing spline model, a baseline trend was then set according to the means of each individual fluorescence trace. A 20% threshold above this baseline was then imposed to minimise false positives from any residual fluctuations in baseline fluorescence. Cell signals with deflection above the de-trended baseline were represented as '1' and inactivity represented as '0', thus binarising the signal at each time point. The coactivity of every cell pair was then measured as:

$$C_{ij} = \frac{T_{ij}}{\sqrt{T_i T_j}}$$

where $T_{ij}$ = total coactivity time, $T_i$ and $T_j$ = total activity time for two cells

The significance at P < 0.001 of each coactivity measured against chance was assessed by subjecting the activity events of each cell to a simulation with 10,000 iterations obtained through shuffling the existing data. Synchronised $Ca^{2+}$-spiking behaviour was assessed by calculating the percentage of coactivity using the binarised cell activity dataset. A topographic representation of the connectivity was plotted in MATLAB (available from: https://www.mathworks.com/matlabcentral/fileexchange/24035-wgplot-weighted-graph-plot-a-better-version-of-gplot) with the edge colours representing the strength of the coactivity between any two cells. A 95% threshold was imposed to determine the probability of the data, which was then plotted as a function of the number of connections for each cell to determine if the dataset obeyed a power-law relationship [28] (available from: https://www.mathworks.com/matlabcentral/fileexchange/29545-power-law-exponential-and-logarithmic-fit). Finally, the data and figures were written from MATLAB to Microsoft Excel files (available from: https://www.github.com/michellehirsch/xlswritefig) for easy visualisation and dissemination.

*Histological techniques and immunofluorescence*

Dissected tissues were fixed in 4% paraformaldehyde at 4°C, washed with PBS and cryoprotected overnight at 4°C using 30% (w/v) sucrose in PBS before embedding in optimal cutting temperature (OCT) and stored at -80°C. Sections (10 μm) were cut using a CM1950 Cryostat (Leica) and permeabilised with 0.2% Triton X-100 in PBS, followed by blocking in 2% (w/v) BSA and 5% (v/v) chicken serum in PBS containing 0.1% Tween-20. Sections were immunostained using primary antibodies diluted in blocking buffer against NNAT (clone EPR13554(B), ab181353 from Abcam, 1:1000), GFP (GFP-1010, Aves Labs, 1:3000) INS (clone K36AC10, I2018, Sigma-Aldrich, 1:5000), somatostatin (SST) (13-2366, American Research Products, 1:1000), glucagon (GCG) (clone K79bB10, G2654, Sigma-Aldrich, 1:3000) urocortin 3 (UCN3) (a kind gift from Joan Vaughan and Paul E. Sawchenko, The Salk Institute for Biological Studies, PBL #7218 [29] 1:1000), translocase of outer mitochondrial membrane (TOM20) (FL-145, sc-11415, Santa Cruz Biotechnology, 1:250) and H3K27me3 (clone C36B11, #9733, Cell Signalling, 1:1000), DNMT1 (clone 60B1220.1, NB100-56519, Novus Biologicals, 1:500) and DNMT3A (clone 64B1446, NB120-13888, Novus Biologicals, 1:500) with overnight incubation at 4°C followed by incubation with secondary antibody (Alexa Fluor, Thermo Fisher Scientific, 1:500) and counterstaining with DAPI (0.1 μg/mL). Sections were imaged using a TCS SP5 confocal microscope (Leica, Germany), capturing multiple islets from at least three regions of tissue >100μm apart. Primary antibodies were first assessed individually by comparative staining *vs* IgG controls. For immunostaining of paraffin-embedded sections from *db/db* mice and mice with beta cell-selective deletion of *Dnmt3a*, slides were immunostained according to [23, 30, 31] using antigen retrieval with Tris-EDTA buffer (10 mM Tris base, 1 mM EDTA, 0.05% Tween-20, pH 9.0) and antibodies against NNAT (ab27266, Abcam, 1:200) and INS (ab195956, Abcam, 1:800).

Human pancreata were obtained from 9 non-diabetic (ND) heart-beating organ donors (mean glycemia during intensive care unit stay: 147±18 mg/dL (mean±SEM, see ESM Table 4 for details)), received before November 2021 and processed as previously described to assess pancreatic islet cell features [32, 33], with the approval of the Ethics Committee of the University of Pisa, upon written consent of donors' next-of-kin. Tissue samples were obtained from the body of the pancreas close to the head. Specimens (approximately 4-5 $mm^3$) were cleaned of any residual fat tissue, wiped and fixed for 24h either in 4% paraformaldehyde or 10% buffered formalin at 4°C. Samples were washed three times in PBS and stored in 70% ethanol solution at 4°C until further processing steps and paraffin embedding. Sequential sections (2 µm thickness) were prepared, and H&E staining performed on one section from each tissue block to observe the overall morphology of the pancreatic tissue and evaluate the quality (adequate and uniform fixation, preservation of tissue structures, absence of artifacts such as shrinkage, tearing or crumbling) before proceeding with immunofluorescence staining. H&E staining was performed using the automated stainer Dako CoverStainer (Agilent). Sections were pre-heated at 60°C for 15 min, deparaffinized with Histoclear (Sigma Aldrich) and hydrated through passages in a decreasing alcohol series. The staining was performed by incubation of the sections with Hematoxylin (Agilent) for 3.5 min followed by rinsing in distilled water for 1 min, treatment with Bluing Buffer (Agilent) for 1 min, and rinsing in tap water for 1 min and ethanol 95% for 1 min, followed by incubation with Eosin (Agilent) for 4 min. After dehydration through passages in increasing alcohol series, sections were mounted using mounting medium (Agilent), covered with a coverslip and allowed to dry. Immunosections were deparaffinised with xylene, rehydrated through a graded alcohol series and antigen retrieval carried out using a citrate-based unmasking solution (pH 6.0, Vector Laboratories). Sections were then permeabilised with 0.4% Triton X-100 in PBS, followed by blocking in 2% (w/v) BSA and 5% (v/v) chicken serum in PBS containing 0.1% Tween-20. Sections were immunostained using primary antibodies diluted in blocking buffer against neuronatin (NNAT) (PA5-115646 from Thermo Fisher Scientific, 1:1000), insulin (INS) (IR002,

Dako, 1:100), somatostatin (SST) (13-2366, American Research Products, 1:1000) and glucagon (GCG) (clone K79bB10, G2654, Sigma-Aldrich, 1:3000) with overnight incubation at 4°C followed by incubation with secondary antibody (Alexa Fluor, Thermo Fisher Scientific, 1:500) and counterstaining with DAPI (0.1 mg/mL). Experiments to validate primary antibodies targeting NNAT (PA5-115646 from Thermo Fisher Scientific) used for experiments involving human pancreatic sections were carried out in HEK293T cells (cultured in DMEM, 10% FBS, 2 mM L-glutamine, 37 °C, 5% $CO_2$ on poly-L-lysine coated coverslips). Cells were transfected with plasmids expressing human *NNAT* (Origene, RC200328) using Lipofectamine 2000. Transfected cells were fixed with 4% paraformaldehyde and immunostained as above using primary antibodies targeting NNAT (PA5-115646 from Thermo Fisher Scientific, 1:500). Transfected cells were visualised using co-transfection with a GFP-expressing plasmid (a kind gift from Connie Cepko, Addgene #11150 [34]). Sections were imaged using a TCS SP5 confocal microscope (Leica, Germany).

Cell classification was carried out with a custom-written Fiji script [35] and CellProfiler pipeline (software version 4.1.3). Nuclei segmentation was batch processed in Fiji using the Stardist plugin, with subsequently generated nuclei label images imported into CellProfiler for cell type classification following measurement of fluorescence intensity in acquired fluorescent channels.

*Cell culture, RNA silencing and GSIS*

Human EndoC-βH1 [36] and rat INS1E [37] beta cells were mycoplasma free and cultured as previously described [18]. EndoC-βH1 beta cells were incubated with lentiviruses expressing an *NNAT*-targeting shRNA (clone ID: TRCN0000163961) *vs* scrambled shRNA control (multiplicity of infections (MOI) = 2, MISSION transduction particles, Sigma-Aldrich) for 72h in complete media without BSA. INS1E cells were transfected with Silencer Select siRNAs (30 nM; *Nnat*, s137247; *Sec11a*, s80747; *Spcs1*, s226238; *Spcs2*, s18919; *Spcs3*, s184336; all

Ambion) using Lipofectamine 2000 (Invitrogen) in OptiMEM (Gibco) for 48h. GSIS assays were performed as previously described [36]. Briefly, EndoC-βH1 cells were cultured in low (2.8 mM) glucose complete media for 16h prior to GSIS assay. Cells were incubated in Krebs-Ringer bicarbonate buffer (115 mM NaCl, 5 mM KCl, 24 mM $NaHCO_3$ (saturated with $CO_2$), 1 mM $CaCl_2$, 1 mM $MgSO_4$, 10 mM HEPES, 0.2% BSA, pH 7.4) containing 0.5 mM glucose for 1h prior to incubation with low (3 mM) or high (16.7 mM) glucose for 1h. Cells were collected in lysis buffer (20 mM Tris-HCl pH 7.5, 137 mM NaCl, 0.1% Triton X-100, 2 mM EGTA, 1% glycerol with protease inhibitors (Complete Mini from Roche)), centrifuged for 5 min at 3,000 rpm and 4°C, with supernatants and collected media stored at -80°C until insulin quantification with ultra-sensitive insulin HTRF assay kits (Cisbio).

*Immunoprecipitation and mass spectrometry*

Cells were homogenised in protein lysis buffer (50 mM Tris-HCl pH 7.5, 150 mM NaCl, 1% Triton X-100, 1 mM EDTA with protease inhibitors, Complete Mini from Roche), solubilised for 30 min on ice, clarified and normalised for protein content. Supernatants were incubated with protein A-agarose beads (Calbiochem) and antibodies against NNAT (ab27266, Abcam, 1:500) or rabbit IgG overnight at 4°C. Agarose beads were washed seven times with protein lysis buffer, with the final three washes not containing Triton X-100. For mass spectrometry, beads were sedimented and resuspended in 100 μL of 20 mM HEPES (pH 8.0), 1M urea, 1 mM dithiothreitol (DTT) and 1.5 µg of trypsin gold (Promega), transferred to protein lo-bind tubes, and digested sequentially for 30 min and overnight at 37 °C (EndoC-βH1) or overnight at 37 °C only (INS1E). Digests were subsequently acidified with trifluoroacetic acid, followed by de-salting using reversed-phase spin tips (Glygen Corp) according to the manufacturer's recommendations. Eluents were subsequently dried to completion using vacuum centrifugation. Dried peptides were re-suspended in 0.1% trifluoroacetic acid (TFA) and analysed by LC-MS/MS analysis using an Ultimate 3000 nano-HPLC coupled to a Q-Exactive mass spectrometer (Thermo Scientific, UK) via an EASY-Spray source. Peptides were loaded onto a trap column (Acclaim PepMap 100 C18, 100μm × 2cm) at 8 μL/min in 2% acetonitrile, 0.1% TFA. Peptides were then eluted on-line to an analytical column (Acclaim Pepmap RSLC C18, 75 μm × 75 cm). A stepped gradient separation was used with 4-25% of 75% acetonitrile, 0.1% formic acid for 75-90 min, followed by 25-45% for 10-30 min, with subsequent column conditioning and equilibration. Eluted peptides were analysed by the Q-Exactive operating in positive polarity and data-dependent acquisition mode. Ions for fragmentation were determined from an initial MS1 survey scan at 70000 (EndoC-βH1) or 120,000 resolution (INS1E), followed by higher-energy collisional dissociation of the top 12 (EndoC-βH1) or 20 (INS1E) most abundant ions at a resolution of 17,500 (EndoC-βH1) or 15,000 (INS1E). MS1 and MS2 scan AGC targets set to 3e6 and 5e4 for a maximum injection time of 50ms and 50ms (EndoC-βH1) and 25ms and 110ms (INS1E), respectively. A survey scan m/z range of 400–1800 m/z (EndoC-βH1) or 350-1750 (INS1E) was used, with a normalised collision energy set to 27%, minimum AGC of 1e3, charge state exclusion enabled for unassigned and +1 ions and a dynamic exclusion of 30 (EndoC-βH1) or 50 (INS1E) seconds. Data was processed using the MaxQuant [38] software platform (v1.6.1.0) for EndoC-βH1 and (v1.6.10.43) for INS1E with database searches carried out by the in-built Andromeda search engine against the Swissprot Homo Sapiens database (downloaded 4$^{th}$ January 2018, entries: 20,256) or Ratttus Norvegicus (Downloaded 31$^{st}$ May 2022, entries: 22,831). A reverse decoy database approach was used at a 1% FDR for peptide spectrum matches and protein identifications. Search parameters included: maximum missed cleavages set to 2, variable modifications of methionine oxidation, protein N-terminal acetylation, asparagine deamidation and cyclisation of glutamine to pyroglutamate. Label-free quantification was enabled with an label-free quantification (LFQ) minimum ratio count of 2 (EndoC-βH1) or 1 (INS1E). For post processing of raw protein intensities for experiments with EndoC-βH1, technical intensities per sample were averaged, with the matrix containing an intensity column per replicate for each digest and condition. Intensity per digest was then generated by averaging raw intensities, with the matrix containing an intensity per digest for each condition (four columns). Intensities per digest for each condition were then summed, with the final matrix containing an intensity per condition.

*Western immunoblotting, RT-PCR, RNA sequencing*

For Western blotting, EndoC-βH1 cells were homogenised in protein lysis buffer as above, clarified and normalised for protein content using the BCA method (Bio-Rad) as previously described [18]. Briefly, proteins were transferred from SDS-PAGE to PVDF and immunoblotted using antibodies against NNAT (ab27266, Abcam, 1:2000) and β-tubulin (clone 9F3, #2128, Cell Signalling, 1:5000), HRP-conjugated secondary antibodies (Cell Signalling, 1:3000) and Immobilon Forte Western HRP substrate (Millipore). Scanned blots were assessed by densitometry using Image J. For RT-PCR, mRNA was purified using an Allprep DNA/RNA/protein mini kit or by direct homogenisation into Trizol reagent with purification using chloroform and RNeasy kits (both Qiagen). Normalised quantities of mRNA were reverse transcribed, and cDNA assessed by quantitative RT-PCR using Taqman reagents on a QuantStudio 7 Real Time PCR cycler with *Hprt*/*HPRT1* as an internal mRNA control (all Applied Biosystems, USA, *NNAT* Hs00193590_m1; *HPRT1* Hs02800695_m1; *Egfp* Mr04097229_mr; *Mafa* Mm00845209_s1; *Neurod1* Mm01280117_m1; *Nkx6.1* Mm00454962_m1; *Pdx1* Mm0435565_m1; *Nnat* Mm00440480_m1; *Hprt* Mm00446968_m1; *Ins2* Mm00731595_gH; *Slc2a2* (*Glut2*) Mm00446229_m1; *Sst* Mm00436671_m1; *Gcg* Mm00801714_m1).

For RNA sequencing, RNA from FACS-purified cells (four biological replicates each from an individual mouse) was quantified and assessed for integrity (RNA integrity number > 9) using a Bioanalyser 2100 and a RNA 6000 pico assay (Agilent Technologies). and 20 ng of RNA was processed for RNA sequencing using a NEBNext Ultra II Directional RNA library Prep Kit for Illumina paired with a Poly(A) mRNA magnetic isolation module and AMPure XP SPRIselect beads (Beckman Coulter), all according to manufacturer's instructions. Libraries were assessed on a Bioanalyser 2100 using a High-Sensitivity DNA assay (Agilent Technologies), with subsequent cDNA quantification using a Quant-iT dsDNA High Sensitivity Assay kit (Thermo Fisher Scientific). Libraries were sequenced using a NextSeq 500 High Output sequencer (Illumina, USA), with 2 x 75 bp length at 50 million reads per sample. Quality assessment of paired end reads was performed using the FASTQC tool (version 0.11.2, http://www.bioinformatics.babraham.ac.uk/projects/fastqc/) and raw reads were aligned to the mouse genome (mm10) using STAR aligner (version 2.7.1a) [39]. Gene counts were obtained using STAR and gencode gene annotations (vM25) and differential expression analysis was performed using the DESeq2 Bioconductor package (version 1.32.0) [40]. Differentially expressed genes were obtained by comparing $NNAT^+$ beta cells, $NNAT^-$ beta cells and the non-beta cell fraction. Gene set enrichment analysis was performed using GSEA (version 4.1.0) [41] and the Molecular Signatures Database v7.2.0, with Wald statistical analysis from the differential expression data used to create preranked files as an input to the GSEA preranked analysis.

*Bisulphite sequencing*

Genomic DNA was extracted from FACS-purified cells using Allprep DNA/RNA/protein mini kits from Qiagen. Sperm cells were lysed using differential extraction, first by incubation in 100 mM Tris-HCl pH 7.5, 200 mM NaCl, 5 mM EDTA, 0.2% SDS, 0.1 mg/mL proteinase K at 55°C for 2h to lyse epithelial cells, followed by pelleting and resuspension of sperm cells in the above lysis buffer plus 10 mM DTT and subsequent DNA extraction using phenol/chloroform/isoamyl alcohol (25:24:1) and standard isopropanol precipitation. Bisulphite conversion and desulphonation was performed using an EZ DNA Methylation-Gold kit from Zymo Research. Eluted DNA was used for amplification of PCR products with primers spanning the *Nnat* promoter (5’-TTTAGGTGGTAAGAGGGTATTTAAGGTA-3’ and 5’-AATACATACTCACCTACAACA-3’) and gametic DMR (5’-TTGATTGGTGGATAAGTTGTGTTT-3’ and 5’-CCACCCTTAAAAAAATACCCATAAT-3’), designed by BiSearch [42] and an EpiTaq hot start kit (TaKaRa Bio). PCR fragments were gel purified (Qiagen), ligated into pJET1.2/blunt using a CloneJET PCR cloning kit (Thermo Scientific) and antibiotic-resistant plasmids prepared in 96 well format using a Wizard SV 96 plasmid DNA purification system (Promega) before Sanger sequencing (Genewiz, now Azenta Life Sciences). Sequencing data was quantified as percentage of methylated CpGs across >12 clones, using Bisulphite Sequencing DNA Methylation Analysis (BISMA) software [43].

*Statistical analysis*

Data are shown as mean ± SEM in all panels. Data were assessed for normal distribution using the D'Agostino-Pearson normality test with statistical significance between biological groups determined using GraphPad Prism 9 with ANOVA, 2-tailed Student's *t* test (parametric) or Mann-Whitney *U*/Kruskal-Wallis tests (nonparametric) used where appropriate. Sidak's (or Dunn's as nonparametric equivalent) post hoc tests were utilised to correct for multiple corrections. Paired analyses were assessed using paired Student's *t* tests or the Wilcoxon matched-pairs signed rank test. <5% error probability was considered significant (i.e., $P < 0.05$) and further statistical information, such as *n* numbers and P values, are provided in the figure legends.

**ESM results**

*Validation of Nnat-eGFP reporter mice*

To extend the findings in Fig. 1-4 using an orthogonal approach, and to facilitate subsequent functional studies, we utilised a Bacterial Artificial Chromosome (BAC) reporter mouse line expressing eGFP under the control of the *Nnat* promoter and upstream enhancers (ESM results and ESM Fig. 8a-e). Aligning with studies of the endogenous gene (Fig. 1-4), adult reporter mice expressed eGFP in a subpopulation (~15% of total) of beta cells (Fig. 5a). eGFP expression was allele-specific with expression from the paternal, but not maternal allele, indicating the preservation of *Nnat* imprinting in the reporter (ESM Fig. 8a, b). Furthermore, eGFP expression was restricted to tissues known to express *Nnat* endogenously (pancreatic islets, hypothalamus) with no evidence of expression in *Nnat*-deficient tissue systems (liver, kidney, muscle) (ESM Fig. 8a-c). Possession of the transgene did not affect endogenous *Nnat* expression (ESM Fig. 8b) nor the expression of key beta cell functional genes, and eGFP staining overlapped with endogenous NNAT reactivity (ESM Fig. 8d, e). Furthermore, through multiple (>6) generations of mice, eGFP reporter expression was silenced via maternal inheritance and re-established through paternal inheritance. Thus, *Nnat* imprinting status for the transgene was restored transgenerationally via the germline. Taken together, these data demonstrate that the *Nnat*-eGFP reporter mouse line mimics endogenous *Nnat* expression.

*Assessment of beta cell connectivity and proportion of 'hub' cells in Nnat deficient islets*

To determine whether NNAT may influence beta cell connectivity and membership of the 'hub' cell subgroup [44], we studied glucose-induced $Ca^{2+}$ dynamics in *Nnat* deficient (*Nnat*$^{+/-p}$) islets [18] by high-speed confocal imaging of the $Ca^{2+}$ probe Cal-520 and off-line analysis using Pearson's R-based correlation analyses [45]. The mean Pearson's coefficient of correlation in wild type islets under low glucose conditions was 0.72±0.05, a correlation not significantly different when compared with high glucose conditions (0.79±0.03) (ESM Fig. 12a, b). Similarly, the mean Pearson's coefficient of correlation in *Nnat* deficient islets under low glucose conditions was 0.70±0.039, with this correlation increased significantly under high glucose conditions (0.80±0.020; $P = 0.03$) (ESM Fig. 12a, b). Connectivity was further analysed using signal binarisation-based connectivity and correlation analyses. Surprisingly, the pooled data over 31 wild type islets under low glucose conditions did not conform to a scale-free network, with 53.3% of all beta cells identified to be highly connected ($R^2 = 0.17$) (ESM Fig. 12c, d). However, these same 31 wild type islets displayed an obedience to a power-law distribution only when in under high glucose conditions, where a small proportion of beta cells (15.4%) were identified to be highly-connected 'hub' cells, as assessed by counting the number of beta cells displaying coordinated $Ca^{2+}$ responses with at least 50% of all beta cells ($R^2 = 0.003$), and this behaviour was similar in the *Nnat* deficient islets (ESM Fig. 12c, d). Pooled data over 38 *Nnat* deficient islets revealed an adherence to a power law only in the high glucose condition, where 9.9% of beta cells were identified as 'hub' cells ($R^2 = 0.11$). This is in contrast to the 51.7% of identified highly-connected beta cells in the low glucose condition ($R^2 = 0.15$) (ESM Fig. 12c, d).

The average connectivity of highly-connected wild type beta cells in low glucose was significantly higher than 'follower' cells (70.2±0.6% vs 14.1±0.9%; $P < 0.001$) (ESM Fig. 12e). This was true also under high glucose conditions (66.7±0.9% vs 13.4±0.5%; $P < 0.001$) (ESM Fig. 12e). When looking at *Nnat* deficient islets, beta cell 'hubs' were also significantly more connected than followers in both low glucose (69.1±0.5% vs 15.3±0.8%; $P < 0.001$) and high glucose (59.7±0.6% vs 12.2±0.5%; $P < 0.001$) (ESM Fig. 12e). Interestingly, while there were no significant differences in the average connectivity between the highly-connected wild type vs *Nnat* deficient 'hub' cells in low glucose ($P = 0.16$), the wild type 'hub' cells were significantly more connected than the *Nnat* deficient' hub' cells in high (11 mM) glucose ($P < 0.001$) (ESM Fig. 12e). The proportion of highly-connected 'hubs' to 'followers' were not significantly different between the wild type and *Nnat* deficient groups (ESM Fig. 12f). However, this proportion decreased significantly in both conditions under high glucose (ESM Fig. 12f). In wild type islets, the 53.5% of highly-connected beta cells decreased significantly to only 13.6% when in high glucose ($P < 0.001$) (ESM Fig. 12f). Similarly, in the *Nnat* deficient islets, the 53.1% of highly-connected beta cells also decreased significantly to 9.3% under high glucose conditions ($P < 0.001$) (ESM Fig. 12f). Interestingly however, there were no significant differences in the proportion of ‘hub’/’follower’ cells between the wild type and *Nnat* deficient islets under high glucose conditions ($P = 0.36$) (ESM Fig. 12f). Thus, NNAT is not involved in setting beta cell connectivity but likely reflects a marker of less well-connected cells.

**ESM Tables**

| Unique identifier | ID | Sex | Age | Body mass index (BMI, KG/m$^2$) |
|---|---|---|---|---|
| 18/40 | HP 25/08/2012 | M | 17 | 24.5 |
| 26/88 | HP 11/08/2020 | F | 15 | 24.7 |
| 27/119 | HP 22/11/2021 | M | 18 | 20.4 |
| 20/46 | HP 25/08/2014B | M | 15 | 19.5 |
| 20/60 | HP 27/09/2014 | M | 13 | 25.4 |
| 18/29 | HP 20/06/2012 | F | 60 | 20.8 |
| 17/71 | HP 18/11/2011 | M | 71 | 25.4 |
| 17/17 | HP 18/03/2011 | M | 78 | 24.2 |
| 16/58 | HP 20/12/2010 | F | 75 | 23.4 |

ESM Table 4: Donor characteristics for human pancreatic sections

**ESM Figures and Figure Legends**

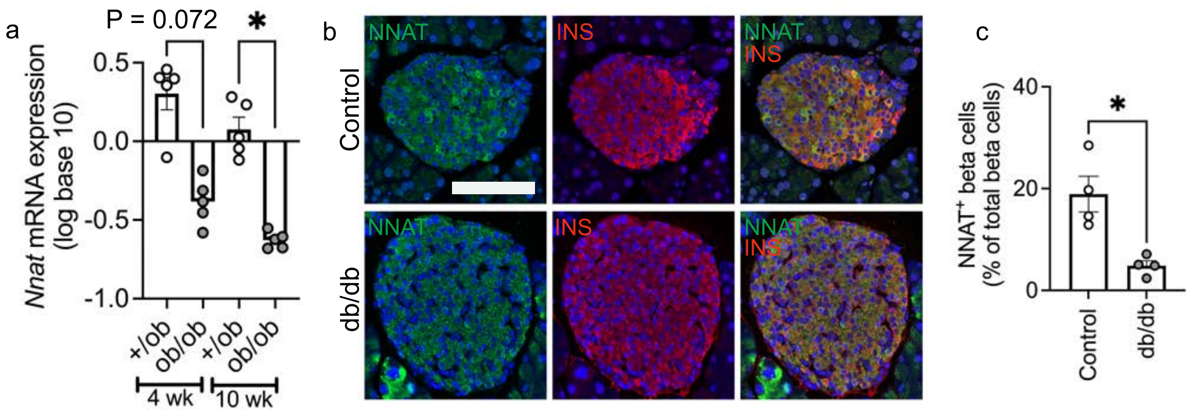


**ESM Fig. 1.** (a) Meta-analysis of *Nnat* mRNA expression in isolated islets of 4 week and 10 week old *ob/ob* vs *+/ob* control mice (n = 5 mice per genotype and age, Kruskal-Wallace test with Dunn's multiple comparisons). (b) Representative confocal microscopy of pancreatic sections from *db/db* vs control mice immunostained with antibodies against NNAT (green) and insulin (INS, red). Nuclei are visualised with DAPI. Scale bar = 100µm (n = 4 mice per genotype). (c) Quantification of $NNAT^{+}$ beta cells from images shown in b, expressed as NNAT/INS co-positive cells as a percentage of total INS-positive cells. * P < 0.05.

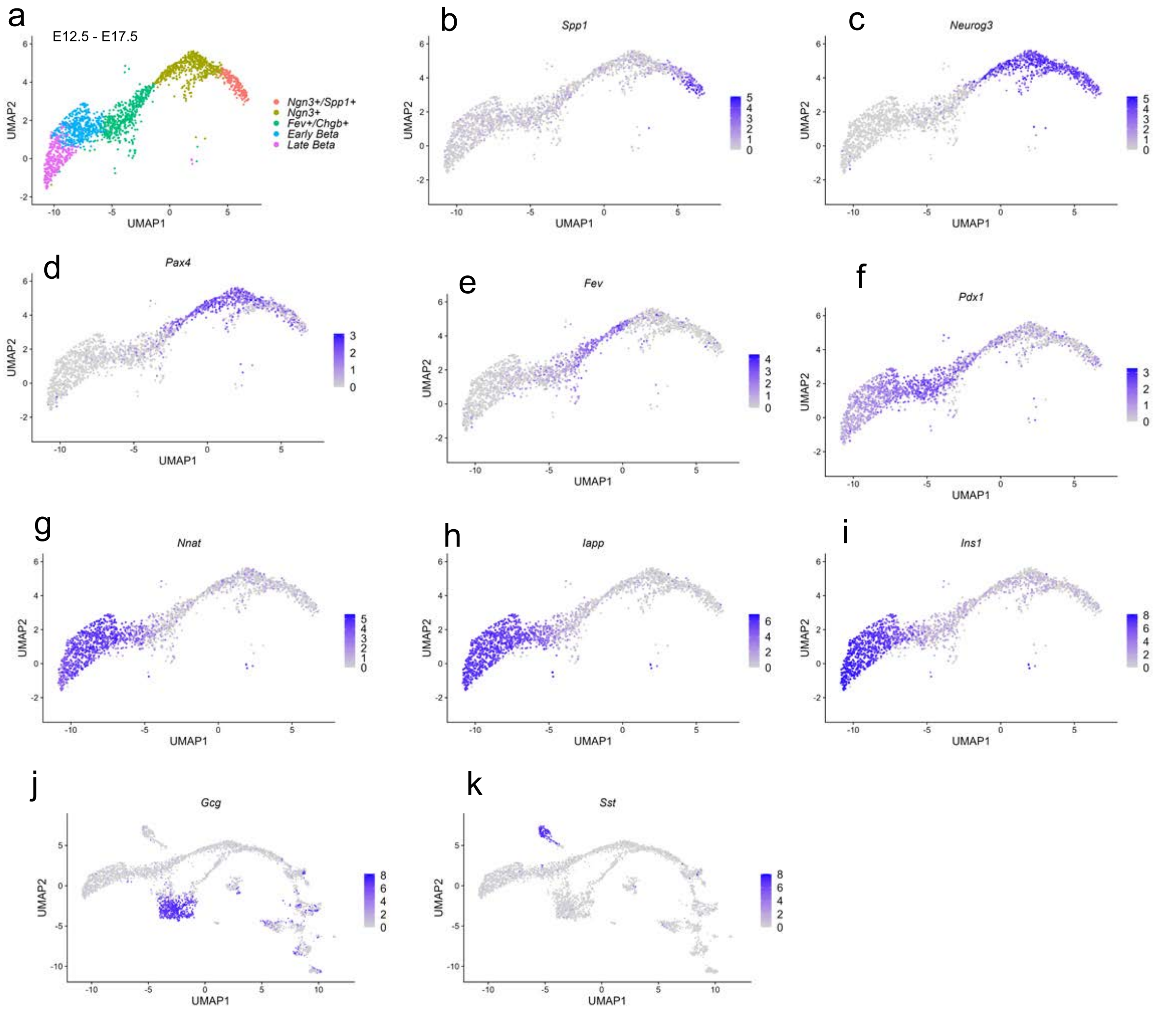


**ESM Fig. 2.** UMAP projections of beta cell precursors and identity markers, and other islet cell markers, from pseudo-time analysis of embryonic scRNA-seq islet data. (a) Plot representing the five beta cell states found in the embryonic data and (b-i) plots representing the shift in expression of markers across precursor states from E12.5 to E17.5 (b-i) and markers of other islet cell types (j, k).

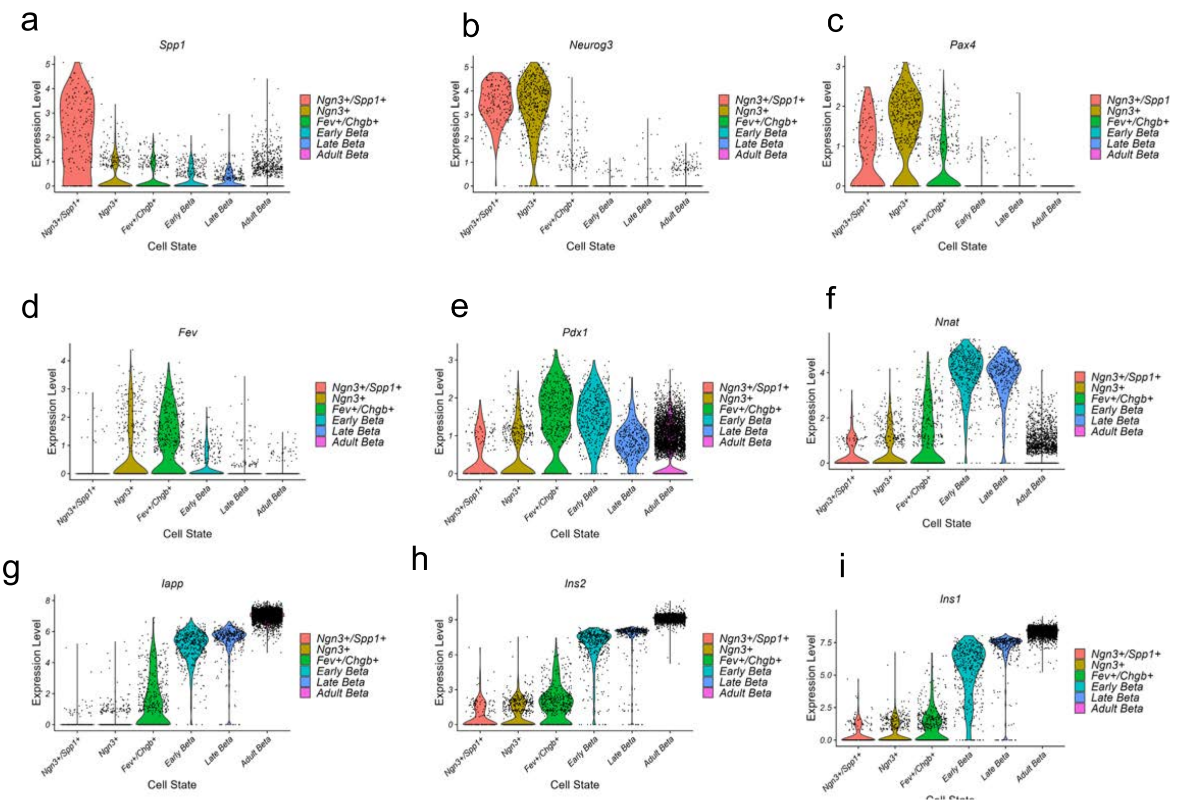


**ESM Fig. 3.** (a-i) Violin plots representing the progressive change in marker expression from early beta cell precursors during embryogenesis (E12.5 – E17.5) to mature beta cells in adulthood.

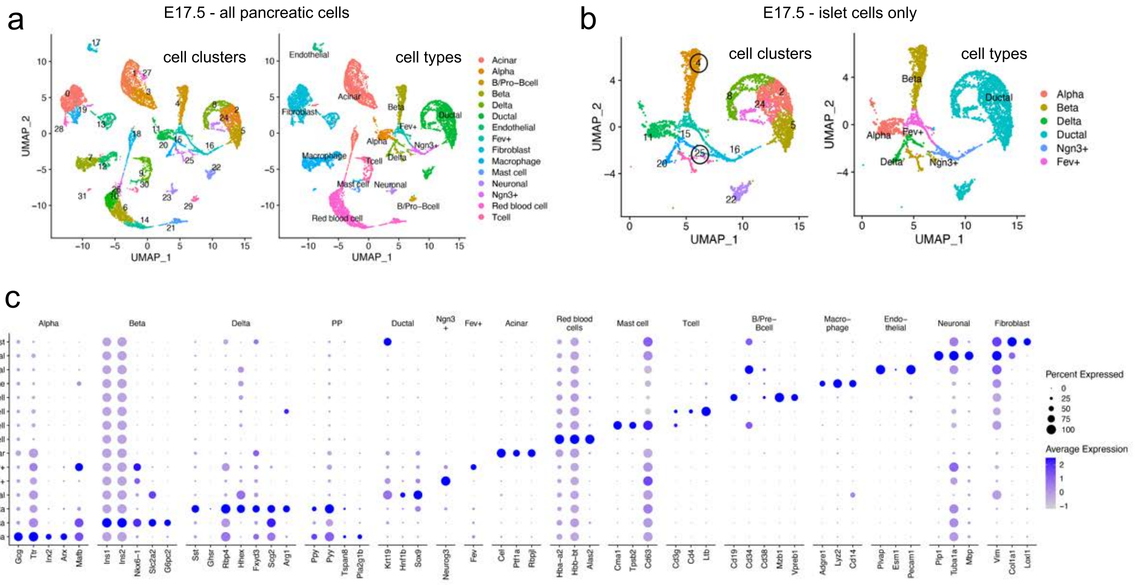


**ESM Fig. 4.** (a, b) Cell clusters and assigned cell types within the E17.5 pancreas for all cells (a) and islet cells only (b), displayed in the UMAP space. In panel b, the beta cell clusters are labelled using black circles. (c) Dot plot showing the expressions of known marker genes within each assigned cell type. Each dot is coloured according to the average gene expressions, while the dot diameters are proportional to the percentage of cells expressing the gene.

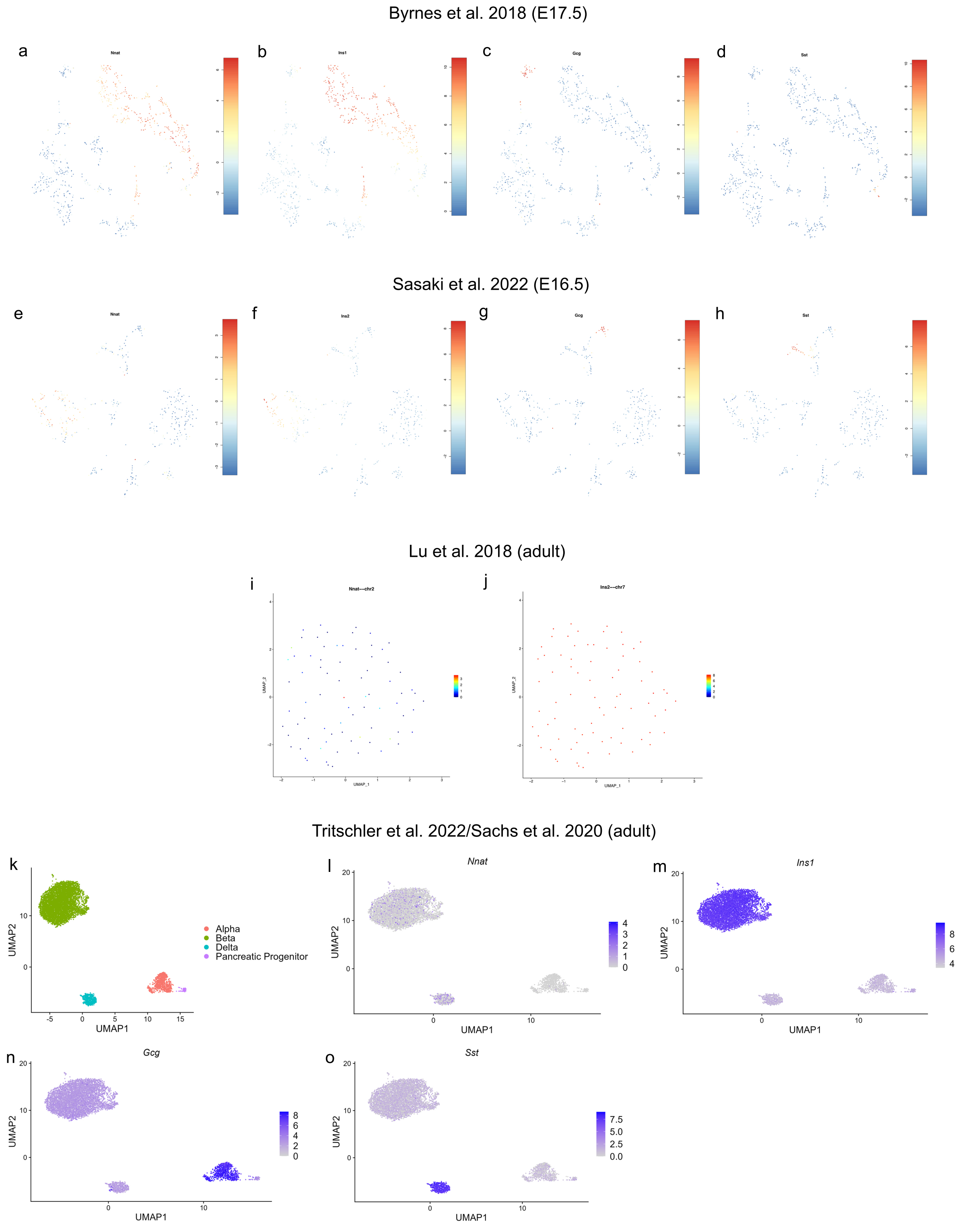


**ESM Fig. 5.** UMAP visualisation of gene expressions of *Nnat* and key islet cell markers in multiple islet scRNA-seq datasets at the embryonic stage (E17.5, a-d and E16.5, e-h) and adult stage (beta cells only, i, j and all islet cells, k-o).

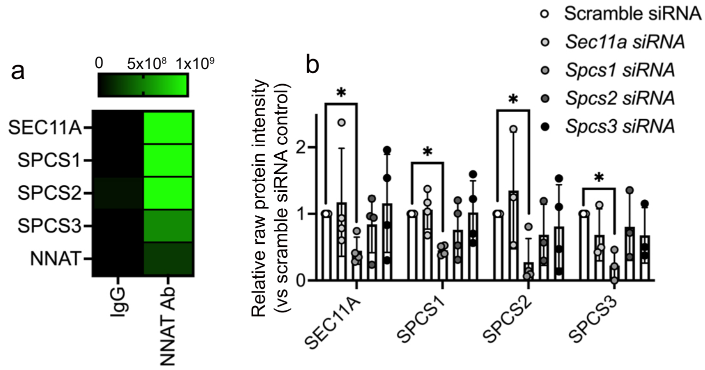


**ESM Fig. 6.** (a) Heatmap from affinity purification/mass spectrometry (AP/MS) analysis showing co-immunoprecipitation of signal peptidase complex subunits with endogenous NNAT in human EndoC-βH1 beta cells using antibodies against NNAT (NNAT Ab) vs control IPs with rabbit immunoglobulins (IgG). (b) INS1E cells transiently transfected with siRNA targeting subunits of the signal peptidase complex followed by NNAT immunoprecipitation as above and detection of signal peptidase complex components by affinity purification/mass spectrometry analysis, with relative protein abundance measured vs scramble siRNA controls (n = 4 independent cultures per group, Kruskal-Wallace test with Dunn's multiple comparisons).

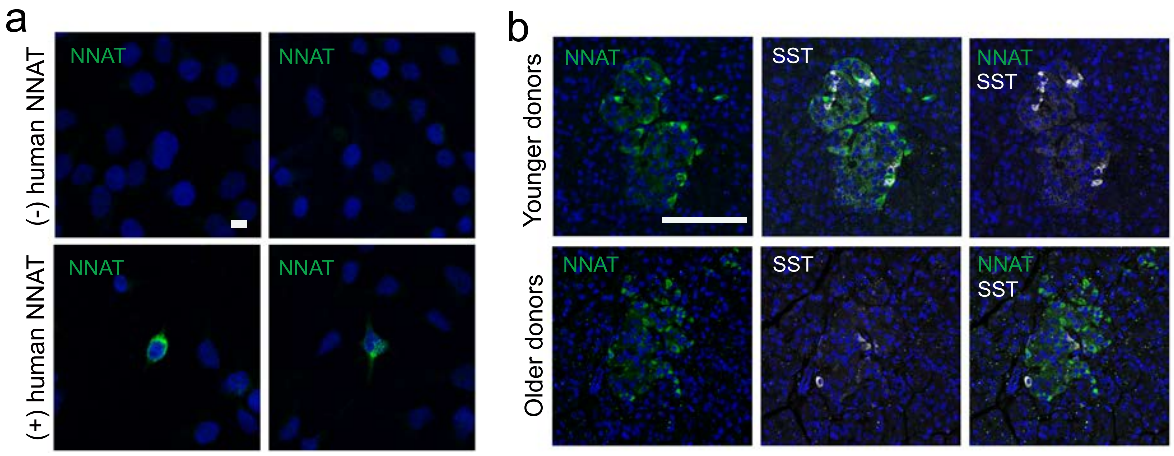


**ESM Fig. 7.** (a) Representative confocal microscopy of HEK293T cells transfected with a plasmid expressing human NNAT and immunostained with antibodies against NNAT (green). Cells transfected with empty vector were used as a control and nuclei are visualised with DAPI. Scale bar = 10μm (three independent experiments, each in triplicate per group). (b) Representative confocal microscopy of human pancreatic cryosections from younger (15.6±0.9 years, n = 5) and older (71.0±3.9 years, n = 4) donors. Sections were immunostained with antibodies against endogenous neuronatin (NNAT, green) and somatostatin (SST, grey). Nuclei are visualised with DAPI. Scale bar = 100μm.

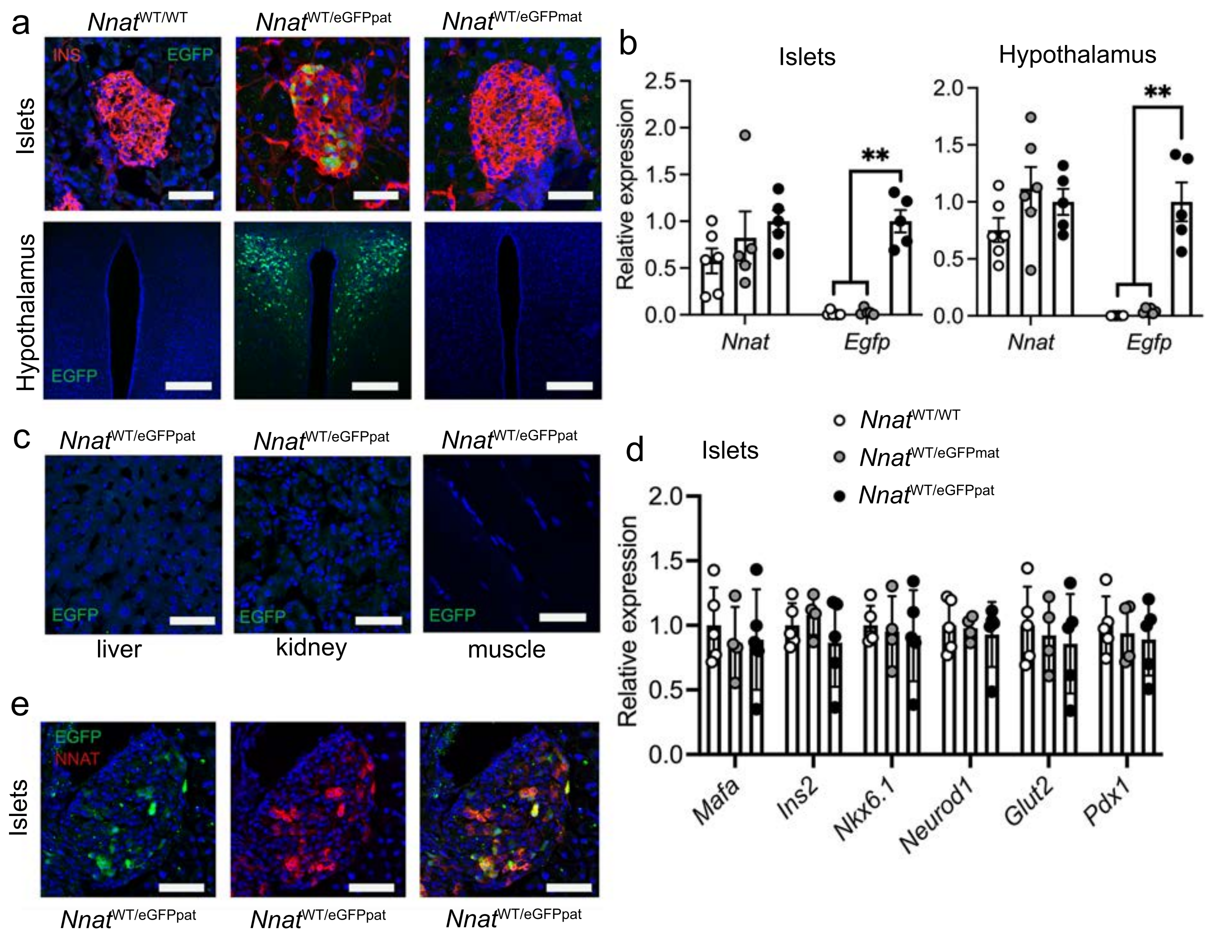


**ESM Fig. 8.** (a) Representative confocal microscopy of pancreatic and hypothalamic cryosections from P56 mice with *Nnat*-driven EGFP expression from the paternal (*Nnat*$^{WT/eGFPpat}$), or maternal (*Nnat*$^{WT/eGFPmat}$) allele or wild type (*Nnat*$^{WT/WT}$) at this locus. Sections were immunostained with antibodies against EGFP (green) and insulin (INS, red). Nuclei are visualised with DAPI. Scale bar = 100μm (n = 3 mice per genotype). (b) RT-PCR analysis of *Nnat* and *Egfp* mRNA expression in isolated islets and hypothalamus of *Nnat*$^{WT/eGFPpat}$, *Nnat*$^{WT/eGFPmat}$ or *Nnat*$^{WT/WT}$ mice. *Hprt* mRNA was used as an internal control and data are expressed relative to *Nnat*$^{WT/eGFPpat}$ mice (n = 5-6 mice per genotype, one way ANOVA with Bonferroni test for multiple comparisons, ** $P < 0.01$). (c) Representative confocal microscopy of cryosections using multiple tissues from P56 *Nnat*$^{WT/eGFPpat}$ mice (n = 4 mice per genotype). Sections were immunostained with antibodies against EGFP (green) and nuclei are visualised with DAPI. Scale bar = 50μm. (d) RT-PCR analysis of mRNAs encoding key beta cell genes in isolated islets of *Nnat*$^{WT/eGFPpat}$, *Nnat*$^{WT/eGFPmat}$ or *Nnat*$^{WT/WT}$ mice. *Hprt* mRNA was used as an internal control and data are expressed relative to *Nnat*$^{WT/WT}$ mice (n = 4-5 mice per genotype). (e) Representative confocal microscopy of pancreatic cryosections from P56 *Nnat*$^{WT/eGFPpat}$ mice (n = 3 mice per genotype). Sections were immunostained with antibodies against EGFP (green) and endogenous NNAT (red). Nuclei are visualised with DAPI. Scale bar = 100μm. Representative images from three independent experiments and breeding pairs.

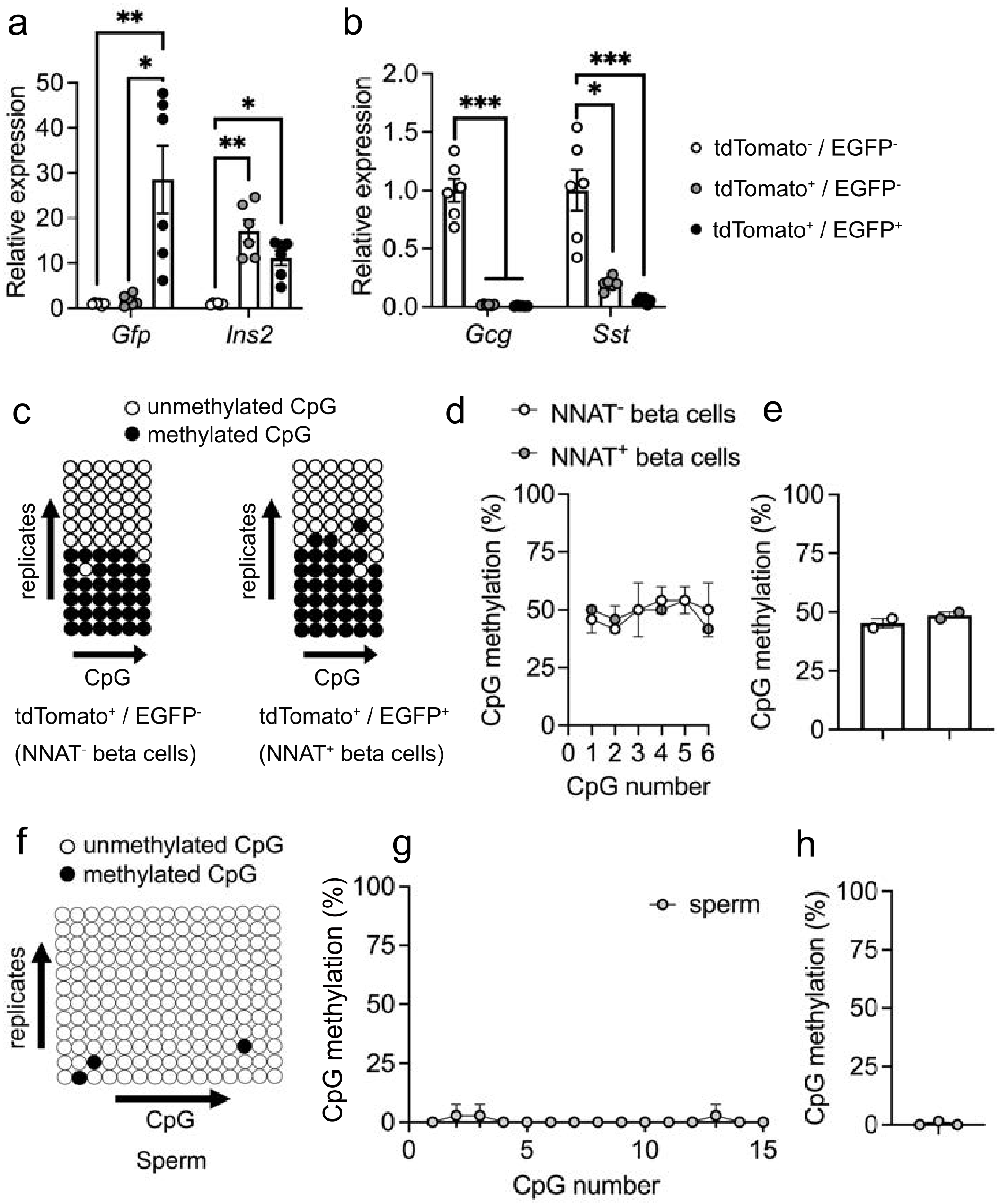


**ESM Fig. 9.** (a, b) RT-PCR analysis of *Egfp*, *Ins2*, *Gcg*, and *Sst* mRNA expression in FACS-sorted primary islet cells from reporter mice with insulin-driven expression of tdTomato (to label beta cells) and *Nnat*-driven EGFP expression from the paternal (*Nnat*$^{WT/eGFPpat}$) (n = 6 mice, * P < 0.05, ** P < 0.01, *** P < 0.001, one way ANOVA with Bonferroni test for multiple comparisons). *Hprt* mRNA was used as an internal control and data are expressed relative to tdTomato$^{-}$/*Egfp*$^{-}$ cells. (c) Representative bisulphite analysis of CpG methylation at the *Nnat* gametic DMR in FACS-purified islet cell populations (n = 2 *Nnat*$^{WT/eGFPpat}$ mice with paternally expressed *Nnat*-driven EGFP, n = 12 clones each, two independent experiments and breeding pairs). (d, e) Quantification of data in C expressed as percentage CpG methylation across the *Nnat* gametic DMR at individual CpGs (d) and across the entire *Nnat* gametic DMR (e). (f) Representative bisulphite analysis of CpG methylation at the *Nnat* promoter in purified sperm cells from male C57BL/6J mice (n = 3, n = 12 clones each, three independent experiments and breeding pairs). (g, h) Quantification of data in f expressed as percentage CpG methylation across the *Nnat* promoter at individual CpGs (g) and across the entire *Nnat* promoter (h). Closed circles = methylated CpG, open circles = unmethylated CpG.

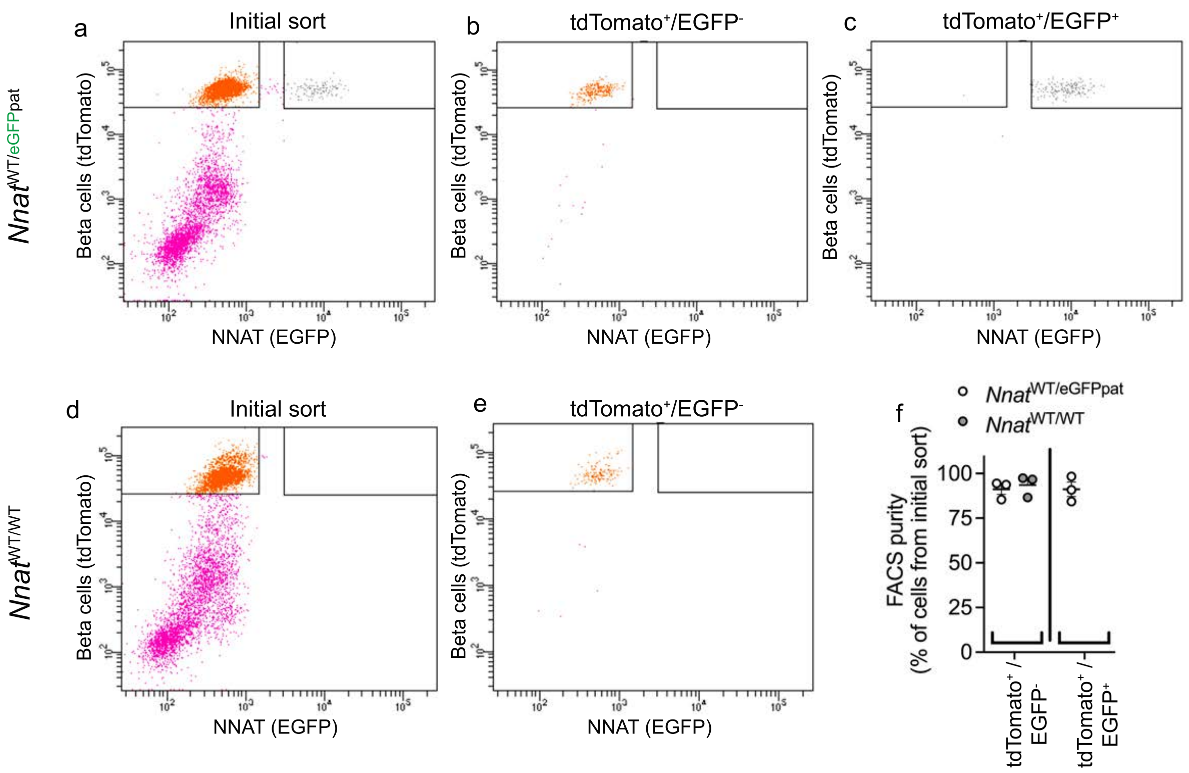


**ESM Fig. 10.** Separation of dispersed primary islet cells by FACS from reporter mice with insulin-driven expression of tdTomato (to label beta cells) and *Nnat*-driven EGFP expression from the paternal (*Nnat*$^{WT/eGFPpat}$) allele or wild type (*Nnat*$^{WT/WT}$) at this locus (representative image of one dispersed islet preparation). The initial sort (a, d) collecting separate populations of tdTomato$^{+}$/EGFP$^{-}$ (b, e) and tdTomato$^{+}$/EGFP$^{+}$ (c) cells were then re-sorted to assess purity (three independent experiments). (f) Quantification of data in a-e to expressed as purity of specific cell populations as a percentage of total cells.

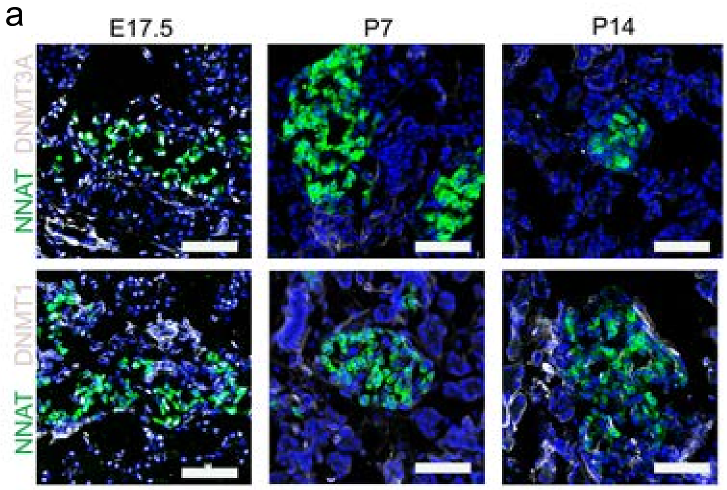


**ESM Fig. 11.** (a) Representative confocal microscopy of pancreatic cryosections from wild type mice on a C57BL/6J background at embryonic (E) day 17.5 and postnatal (P) day 7 and 14 (n = 3 mice per developmental stage). Sections were immunostained with antibodies against endogenous neuronatin (NNAT, green) and DNMT3A or DNMT1 (both grey). Nuclei are visualised with DAPI. Scale bar = 50μm.

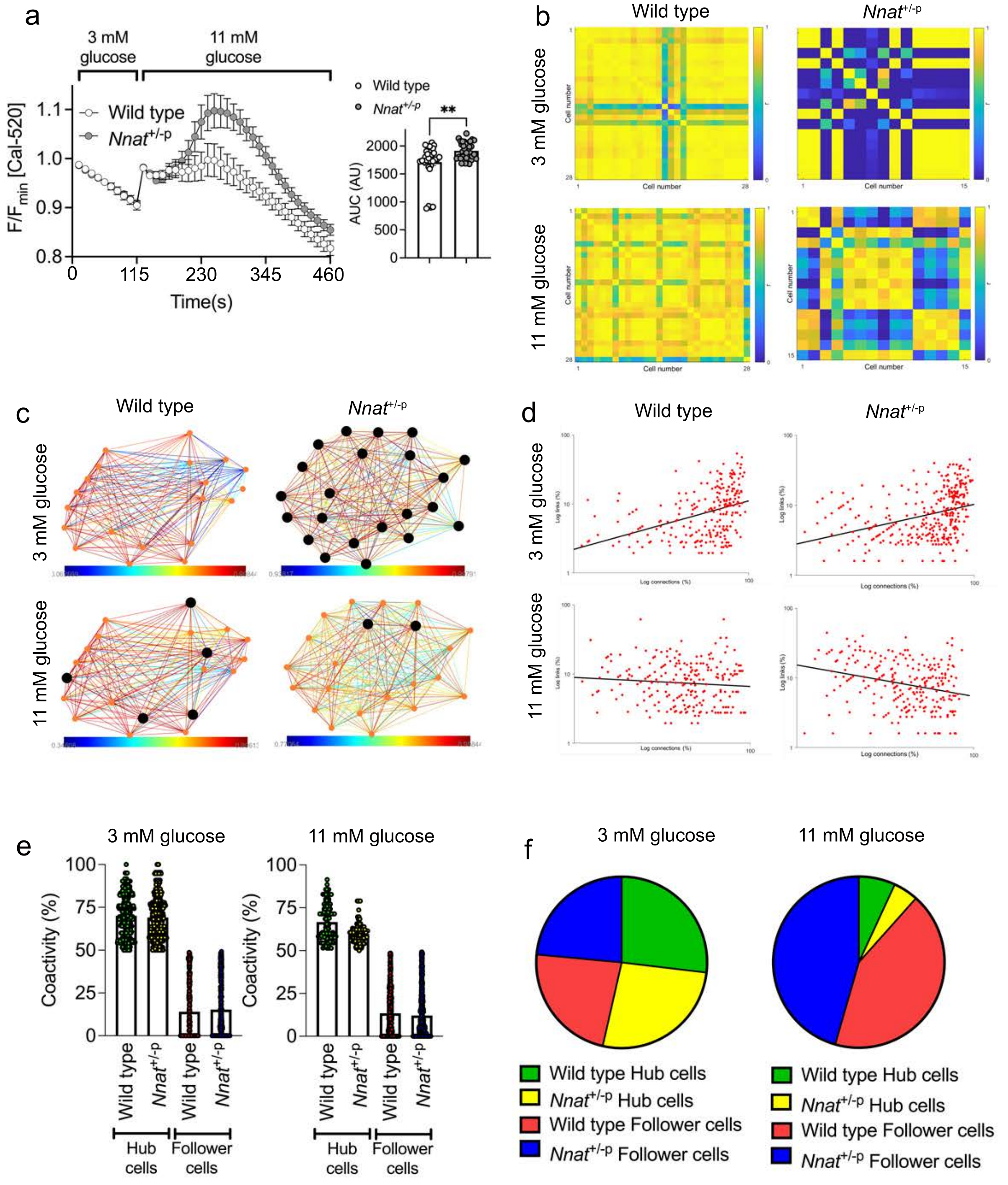


**ESM Fig. 12.** (a) $Ca^{2+}$ bound Cal-520 fluorescence in response to high (11 mM) vs low (3 mM) glucose in primary islets from wild type and *Nnat*$^{+/-p}$ mice expressed as normalised intensity over time (F / $F_{min}$) (n = 31 wild type and 38 *Nnat* deficient islets total from 9 mice per genotype, inset shows quantification of area under the curve (AUC), ** $P < 0.01$, Mann-Whitney U test). (b) Representative heatmaps depicting connectivity strength (r) of all cell pairs (colour coded r values from 0 to 1, blue to yellow). (c) Representative cartesian map of cells with colour coded lines connecting cells according of the strength of coactivation (colour coded R values from 0 to 1, blue to red). Beta cells are represented by differently coloured nodes depending on their coactivity with the other beta cells, where black nodes indicate coactivity with ≥80% of the remaining beta cells, while grey and white nodes represent coactivity with ≥60% and ≥40%, respectively. (d) Log-log graphs of beta cell-beta cell connectivity distribution. (e) Percentage coactivity of beta cells between all cells for identified ‘hub’ and ‘follower’ cells. (f) The proportion of cells designated as ‘hub’ cells vs ‘follower’ cells. Representative images from three independent experiments and breeding pairs.

**ESM References**

[11] Gene Ontology C (2021) The Gene Ontology resource: enriching a GOld mine. Nucleic Acids Res 49(D1): D325-D334. 10.1093/nar/gkaa1113
