## Supplemental Table 1 for "Differential CpG methylation at *Nnat* in the early establishment of beta cell heterogeneity"

| | mean | total | tech | bio | p.value | FDR |
|---|---|---|---|---|---|---|
| tdTomato | 4.9067814 | 13.8906671 | 0.55312303 | 13.3375441 | 0 | 0 |
| pTimer | 4.74201973 | 13.0745206 | 0.57485182 | 12.4996688 | 0 | 0 |
| Iapp | 3.43887049 | 8.02127344 | 0.78002771 | 7.24124574 | 0 | 0 |
| eGFP | 2.18740845 | 7.77509563 | 0.92107106 | 6.85402457 | 2.75E-240 | 8.37E-237 |
| Pyy | 3.06147803 | 7.12220513 | 0.82873288 | 6.29347225 | 3.77E-250 | 1.43E-246 |
| Col1a1 | 2.52793167 | 7.06264049 | 0.89518871 | 6.16745179 | 2.57E-206 | 4.35E-203 |
| Sparc | 2.81380131 | 6.88217292 | 0.86330078 | 6.01887213 | 3.20E-211 | 6.08E-208 |
| Ins2 | 3.29370114 | 6.5360137 | 0.80060476 | 5.73540894 | 7.10E-223 | 1.80E-219 |
| Lgals1 | 2.6701039 | 6.4342541 | 0.881112 | 5.5531421 | 4.89E-173 | 7.43E-170 |
| Itm2a | 2.41395137 | 6.38148793 | 0.90487845 | 5.47660948 | 9.29E-160 | 1.18E-156 |
| Col3a1 | 2.31752395 | 6.31525371 | 0.91275048 | 5.40250323 | 6.77E-153 | 7.91E-150 |
| Rbp4 | 2.84645428 | 6.11154947 | 0.85832146 | 5.25322802 | 2.56E-163 | 3.54E-160 |
| Dcn | 2.14739932 | 6.17541631 | 0.92291459 | 5.25250172 | 1.56E-141 | 1.48E-138 |
| Ptn | 2.22134983 | 6.00155831 | 0.91924144 | 5.08231687 | 1.06E-133 | 9.49E-131 |
| Col1a2 | 2.21927045 | 5.98381704 | 0.91936384 | 5.0644532 | 9.61E-133 | 8.11E-130 |
| Dlk1 | 3.89379542 | 5.67836784 | 0.7017544 | 4.97661344 | 1.87E-218 | 4.05E-215 |
| Cdkn1c | 2.79123684 | 5.80290115 | 0.86689976 | 4.93600139 | 1.16E-141 | 1.17E-138 |
| Chga | 2.59922515 | 5.5456777 | 0.88836264 | 4.65731506 | 1.91E-120 | 1.53E-117 |
| H19 | 2.40274047 | 5.52076998 | 0.90583364 | 4.61493635 | 7.14E-114 | 5.43E-111 |
| Pcsk2 | 2.35101872 | 5.40259163 | 0.91013437 | 4.49245725 | 4.98E-107 | 3.60E-104 |
| Ins1 | 1.97259384 | 5.3070931 | 0.92732899 | 4.3797641 | 3.53E-98 | 2.44E-95 |
| Chgb | 2.31415816 | 5.14672834 | 0.91300781 | 4.23372052 | 1.01E-94 | 6.68E-92 |
| Sst | 1.76559687 | 4.87853204 | 0.9224897 | 3.95604234 | 2.48E-81 | 1.51E-78 |
| Ttr | 2.30598148 | 4.84631153 | 0.91362822 | 3.93268331 | 6.33E-82 | 4.01E-79 |
| Gcg | 1.58739261 | 4.39809348 | 0.90650599 | 3.49158749 | 4.99E-66 | 2.71E-63 |
| Mest | 2.11310761 | 4.25734498 | 0.92426843 | 3.33307655 | 3.84E-58 | 1.82E-55 |
| Meg3 | 5.12076979 | 3.81070924 | 0.55268172 | 3.25802752 | 1.12E-151 | 1.21E-148 |
| S100a6 | 1.5898619 | 4.13585065 | 0.90679477 | 3.22905589 | 9.57E-57 | 4.41E-54 |

mean variance for all transcripts = 0.286

**mean**: A numeric vector of mean log-CP
**total**: A numeric vector of the variances
**bio**: A numeric vector containing the bio
**tech**: A numeric vector containing the te

| | | | | | | |
|---|---|---|---|---|---|---|
| Vim | 2.21684069 | 4.04546426 | 0.91950624 | 3.12595802 | 6.62E-52 | 2.72E-49 |
| Serpinh1 | 2.44457609 | 3.92941075 | 0.9022743 | 3.02713644 | 1.32E-50 | 5.26E-48 |
| Nnat | 1.78801745 | **3.9480564** | 0.92365053 | 3.02440587 | 2.65E-48 | **1.01E-45** |
| Lum | 1.45003474 | 3.90680448 | 0.88439227 | 3.02241221 | 1.97E-52 | 8.32E-50 |
| Xist | 2.40006129 | 3.92078712 | 0.90606027 | 3.01472685 | 8.27E-50 | 3.22E-47 |
| Pcsk1n | 2.01093901 | 3.82535305 | 0.92700893 | 2.89834412 | 3.19E-44 | 1.13E-41 |
| Igf2 | 1.74605533 | 3.71356303 | 0.92134686 | 2.79221617 | 1.13E-41 | 3.82E-39 |
| Crip1 | 2.02865512 | 3.58491929 | 0.92675021 | 2.65816908 | 1.54E-37 | 4.49E-35 |
| Gsn | 1.47530568 | 3.3997702 | 0.88906513 | 2.51070507 | 1.88E-36 | 5.30E-34 |
| Tmsb4x | 3.86894713 | 3.20768555 | 0.7060061 | 2.50167945 | 3.29E-56 | 1.43E-53 |
| Col6a1 | 1.47338509 | 3.27900631 | 0.88872669 | 2.39027962 | 2.99E-33 | 7.99E-31 |
| Gpc3 | 1.62139515 | 3.28793287 | 0.91046752 | 2.37746535 | 1.86E-31 | 4.57E-29 |
| Fos | 1.79344239 | 3.28344589 | 0.92391482 | 2.35953108 | 3.57E-30 | 8.61E-28 |
| Igfbp4 | 1.60290951 | 3.18222292 | 0.90831845 | 2.27390447 | 4.52E-29 | 1.06E-26 |
| Sparcl1 | 1.39560168 | 3.06679826 | 0.87341709 | 2.19338117 | 3.06E-29 | 7.26E-27 |
| Anxa2 | 1.65488408 | 2.95288861 | 0.91412249 | 2.03876612 | 1.79E-23 | 3.28E-21 |
| Tmsb15b2 | 1.15644699 | 2.80932289 | 0.80863545 | 2.00068744 | 1.93E-28 | 4.38E-26 |
| Ppy | 0.85477692 | 2.63994203 | 0.6814571 | 1.95848492 | 1.11E-37 | 3.30E-35 |
| Rcn3 | 1.48501868 | 2.83399096 | 0.89076148 | 1.94322948 | 1.53E-22 | 2.65E-20 |
| Fstl1 | 1.51121026 | 2.79426898 | 0.89521749 | 1.89905149 | 2.04E-21 | 3.33E-19 |
| Mafb | 1.4336376 | 2.7258978 | 0.88122093 | 1.84467687 | 6.63E-21 | 1.01E-18 |
| Col6a2 | 1.37774285 | 2.70211464 | 0.86948777 | 1.83262688 | 3.63E-21 | 5.64E-19 |
| Mdk | 1.88521374 | 2.75086123 | 0.92679787 | 1.82406336 | 1.07E-18 | 1.42E-16 |
| Slc38a5 | 1.06296061 | 2.56470452 | 0.77565599 | 1.78904852 | 5.69E-25 | 1.14E-22 |
| Cd24a | 1.11894842 | 2.55650963 | 0.79590821 | 1.76060143 | 4.02E-23 | 7.19E-21 |
| S100a10 | 1.78683366 | 2.68176933 | 0.92359208 | 1.75817725 | 1.31E-17 | 1.65E-15 |
| Mfap4 | 0.9837919 | 2.49751004 | 0.74395354 | 1.75355651 | 5.41E-26 | 1.14E-23 |
| Aplp1 | 1.73511692 | 2.65170458 | 0.92064157 | 1.73106301 | 3.16E-17 | 3.88E-15 |
| Mfap2 | 1.65684955 | 2.62544068 | 0.91433498 | 1.71110569 | 4.41E-17 | 5.32E-15 |
| Ifitm2 | 2.55497419 | 2.59436575 | 0.89261303 | 1.70175272 | 1.17E-17 | 1.50E-15 |

| | | | | | | |
|---|---|---|---|---|---|---|
| Dbpht2 | 1.40152469 | 2.56100926 | 0.87467158 | 1.68633768 | 5.13E-18 | 6.72E-16 |
| 1700086L19F | 1.53483146 | 2.5803748 | 0.89908586 | 1.68128895 | 4.65E-17 | 5.57E-15 |
| Tmem27 | 1.11957338 | 2.43319851 | 0.79612824 | 1.63707027 | 3.08E-20 | 4.42E-18 |
| Pax6 | 1.51431963 | 2.51510875 | 0.89573523 | 1.61937351 | 4.60E-16 | 5.10E-14 |
| Cxcl12 | 1.14753185 | 2.41143256 | 0.80574508 | 1.60568748 | 4.03E-19 | 5.42E-17 |
| Ghrl | 0.80193198 | 2.25763693 | 0.65330777 | 1.60432916 | 4.78E-28 | 1.07E-25 |
| Stmn1 | 1.54541067 | 2.48283111 | 0.90077112 | 1.58206 | 2.91E-15 | 3.09E-13 |
| Spp1 | 0.65548563 | 2.14909208 | 0.56845677 | 1.5806353 | 2.12E-35 | 5.74E-33 |
| Apoe | 0.80764393 | 2.23113637 | 0.65641065 | 1.57472572 | 7.42E-27 | 1.63E-24 |
| Bgn | 1.14966128 | 2.34991955 | 0.80643807 | 1.54348148 | 8.80E-18 | 1.13E-15 |
| Cpe | 2.94092037 | 2.38159084 | 0.844903 | 1.53668784 | 3.09E-16 | 3.51E-14 |
| Isl1 | 1.39139802 | 2.39880612 | 0.87250462 | 1.5263015 | 3.72E-15 | 3.92E-13 |
| Spint2 | 1.84019812 | 2.4508766 | 0.92562643 | 1.52525017 | 1.19E-13 | 1.15E-11 |
| Fam183b | 1.47116552 | 2.410471 | 0.8883343 | 1.5221367 | 1.29E-14 | 1.29E-12 |
| Cryba2 | 1.29387295 | 2.36475341 | 0.84923084 | 1.51552257 | 1.06E-15 | 1.16E-13 |
| Krt8 | 1.69948342 | 2.39844657 | 0.9181646 | 1.48028197 | 3.82E-13 | 3.59E-11 |
| Bex2 | 1.65827515 | 2.3849457 | 0.91448856 | 1.47045714 | 4.39E-13 | 4.10E-11 |
| Hmgn3 | 1.90137542 | 2.38280149 | 0.92700618 | 1.45579531 | 1.46E-12 | 1.22E-10 |
| Eln | 1.02472626 | 2.18926411 | 0.76076391 | 1.42850019 | 3.48E-17 | 4.24E-15 |
| Ifitm3 | 1.18763655 | 2.2338497 | 0.8186443 | 1.41520541 | 7.64E-15 | 7.90E-13 |
| Gnas | 4.62956392 | 1.99225817 | 0.58934448 | 1.40291369 | 1.79E-26 | 3.84E-24 |
| Postn | 1.05995632 | 2.17491909 | 0.7745129 | 1.40040619 | 4.56E-16 | 5.10E-14 |
| Meis2 | 2.16261266 | 2.32200686 | 0.92227493 | 1.39973193 | 7.54E-12 | 5.91E-10 |
| Mt1 | 0.69515781 | 1.97422876 | 0.59249939 | 1.38172937 | 1.75E-25 | 3.59E-23 |
| Scg2 | 1.01133899 | 2.08640729 | 0.75536471 | 1.33104258 | 2.37E-15 | 2.54E-13 |
| Hmgb2 | 1.51070887 | 2.21930975 | 0.89513376 | 1.32417599 | 2.41E-11 | 1.79E-09 |
| Epcam | 1.52361318 | 2.17812822 | 0.89726781 | 1.2808604 | 1.10E-10 | 7.59E-09 |
| Pcolce | 1.10270275 | 2.04414481 | 0.79020473 | 1.25394008 | 8.66E-13 | 7.44E-11 |
| Akap12 | 1.0636845 | 2.01520836 | 0.77593091 | 1.23927745 | 6.22E-13 | 5.57E-11 |
| Slc25a5 | 2.61248347 | 2.12234957 | 0.88704651 | 1.23530306 | 2.99E-10 | 1.98E-08 |

| | | | | | | |
|---|---|---|---|---|---|---|
| Neurod1 | 1.2714865 | 2.05626534 | 0.84322801 | 1.21303734 | 8.01E-11 | 5.61E-09 |
| Ndn | 1.93067712 | 2.13977998 | 0.92723475 | 1.21254523 | 3.06E-09 | 1.77E-07 |
| Csrp2 | 1.37517775 | 2.05831152 | 0.86891497 | 1.18939654 | 5.81E-10 | 3.70E-08 |
| Scg3 | 1.26993893 | 2.02122088 | 0.84280712 | 1.17841376 | 2.55E-10 | 1.71E-08 |
| Neurog3 | 0.42749758 | 1.58670442 | 0.41222456 | 1.17447986 | 4.56E-37 | 1.31E-34 |
| Actb | 4.71519502 | 1.74908343 | 0.57869656 | 1.17038687 | 1.22E-19 | 1.72E-17 |
| Gpx3 | 1.40858589 | 2.04403463 | 0.87615393 | 1.1678807 | 1.55E-09 | 9.33E-08 |
| Tuba1a | 3.47367555 | 1.94184649 | 0.77447291 | 1.16737358 | 1.04E-11 | 7.95E-10 |
| Scgn | 1.31137744 | 2.01550617 | 0.85365172 | 1.16185445 | 7.21E-10 | 4.55E-08 |
| Fabp5 | 1.3650649 | 2.02782744 | 0.8666252 | 1.16120224 | 1.29E-09 | 7.91E-08 |
| Fbln1 | 1.03249138 | 1.9241793 | 0.76385775 | 1.16032155 | 7.24E-12 | 5.70E-10 |
| Ifitm1 | 0.85454828 | 1.83948462 | 0.68133795 | 1.15814667 | 2.07E-14 | 2.05E-12 |
| Rarres2 | 0.95178106 | 1.88576814 | 0.7297007 | 1.15606744 | 9.39E-13 | 7.98E-11 |
| Cald1 | 1.16497248 | 1.95817962 | 0.81137283 | 1.14680679 | 1.66E-10 | 1.12E-08 |
| Ccdc80 | 0.96910197 | 1.86516997 | 0.73762281 | 1.12754716 | 5.39E-12 | 4.36E-10 |
| Ppic | 1.29016827 | 1.96510484 | 0.84824841 | 1.11685643 | 2.41E-09 | 1.41E-07 |
| Krt7 | 1.48672305 | 1.98587427 | 0.89105653 | 1.09481775 | 2.35E-08 | 1.28E-06 |
| Lrpprc | 0.99986618 | 1.83164814 | 0.75066663 | 1.08098151 | 7.68E-11 | 5.40E-09 |
| Emp3 | 1.0566326 | 1.83274374 | 0.77324427 | 1.05949947 | 5.60E-10 | 3.58E-08 |
| Krt18 | 1.32286576 | 1.88463741 | 0.85651344 | 1.02812397 | 4.74E-08 | 2.47E-06 |
| Ldha | 1.25222057 | 1.85215119 | 0.83780551 | 1.01434568 | 3.68E-08 | 1.95E-06 |
| Plagl1 | 1.12592347 | 1.81042796 | 0.79835599 | 1.01207197 | 8.72E-09 | 4.93E-07 |
| Nrp1 | 1.0100885 | 1.76622638 | 0.75485518 | 1.0113712 | 1.29E-09 | 7.91E-08 |
| Junb | 1.19247516 | 1.82543894 | 0.82017245 | 1.00526649 | 2.53E-08 | 1.37E-06 |
| Maged2 | 1.96492264 | 1.92388375 | 0.92737801 | 0.99650574 | 8.90E-07 | 3.85E-05 |
| Ebf1 | 0.87085772 | 1.68547298 | 0.68977643 | 0.99569655 | 6.94E-11 | 4.90E-09 |
| Cd34 | 0.921335 | 1.69602021 | 0.71509821 | 0.980922 | 5.36E-10 | 3.44E-08 |
| Itih5 | 0.81818995 | 1.63761272 | 0.66210091 | 0.97551181 | 2.88E-11 | 2.11E-09 |
| Clu | 0.74750135 | 1.59745905 | 0.62299182 | 0.97446722 | 1.78E-12 | 1.48E-10 |
| Tm4sf4 | 1.17733717 | 1.77490389 | 0.81535682 | 0.95954706 | 8.39E-08 | 4.14E-06 |

| | | | | | | |
|---|---|---|---|---|---|---|
| Gng12 | 1.72906975 | 1.87818956 | 0.92024086 | 0.9579487 | 1.85E-06 | 7.59E-05 |
| Pmp22 | 0.9662323 | 1.68866369 | 0.73637592 | 0.95228776 | 4.49E-09 | 2.57E-07 |
| Igf1 | 0.74693769 | 1.57099457 | 0.62267065 | 0.94832392 | 6.41E-12 | 5.09E-10 |
| B2m | 1.16606084 | 1.75856008 | 0.81172042 | 0.94683966 | 1.08E-07 | 5.23E-06 |
| Btg2 | 1.25392801 | 1.77496434 | 0.8382923 | 0.93667204 | 3.39E-07 | 1.54E-05 |
| Pdx1 | 1.09429039 | 1.72266862 | 0.78721565 | 0.93545297 | 6.36E-08 | 3.24E-06 |
| Cav1 | 0.46797763 | 1.37713207 | 0.44246235 | 0.93466972 | 2.97E-21 | 4.80E-19 |
| Cthrc1 | 0.67907559 | 1.51229743 | 0.58285253 | 0.9294449 | 6.75E-13 | 5.96E-11 |
| Prkcdbp | 1.04136453 | 1.6866051 | 0.76736204 | 0.91924306 | 5.03E-08 | 2.61E-06 |
| Tubb3 | 0.92575095 | 1.62027129 | 0.71724921 | 0.90302208 | 1.09E-08 | 6.03E-07 |
| Hhex | 0.50191764 | 1.36628807 | 0.46692097 | 0.8993671 | 5.50E-18 | 7.15E-16 |
| Tmsb10 | 4.90326963 | 1.44275517 | 0.55350045 | 0.88925472 | 4.59E-13 | 4.25E-11 |
| Tuba1b | 2.63316317 | 1.76534959 | 0.88495985 | 0.88038974 | 4.87E-06 | 0.00018434 |
| Syt13 | 1.35313792 | 1.73738851 | 0.86386329 | 0.87352523 | 3.47E-06 | 0.00013468 |
| Gch1 | 1.16289515 | 1.66595688 | 0.81070823 | 0.85524865 | 1.37E-06 | 5.74E-05 |
| Laptm4a | 2.91097365 | 1.7035516 | 0.84923282 | 0.85431878 | 3.87E-06 | 0.00014782 |
| Jun | 1.61023857 | 1.76100677 | 0.90918029 | 0.85182648 | 1.56E-05 | 0.00054237 |
| Dbi | 1.47968842 | 1.73995792 | 0.88983372 | 0.85012421 | 1.08E-05 | 0.00038526 |
| Mt2 | 0.38069 | 1.22538316 | 0.37572812 | 0.84965504 | 4.45E-24 | 8.67E-22 |
| Map1b | 1.96255317 | 1.76946403 | 0.92737343 | 0.8420906 | 2.71E-05 | 0.00088574 |
| S100a16 | 0.91643833 | 1.55308127 | 0.71270428 | 0.84037699 | 7.94E-08 | 3.93E-06 |
| Rbp1 | 0.99178728 | 1.58761253 | 0.74732666 | 0.84028587 | 2.89E-07 | 1.31E-05 |
| Ccnd2 | 1.18604467 | 1.65352529 | 0.81813856 | 0.83538674 | 2.82E-06 | 0.0001117 |
| Col4a1 | 0.98246514 | 1.57347099 | 0.74338532 | 0.83008567 | 3.45E-07 | 1.56E-05 |
| Ctrb1 | 0.36668054 | 1.18067556 | 0.3644691 | 0.81620645 | 1.19E-23 | 2.25E-21 |
| Igfbp6 | 0.36463806 | 1.17763329 | 0.36281398 | 0.81481931 | 8.95E-24 | 1.72E-21 |
| Igfbp7 | 1.19681484 | 1.62927427 | 0.82148231 | 0.80779196 | 6.17E-06 | 0.00022873 |
| Egr1 | 1.22479979 | 1.63526434 | 0.82981275 | 0.80545159 | 7.98E-06 | 0.00029008 |
| Acta2 | 0.5433453 | 1.28724267 | 0.49573422 | 0.79150845 | 6.33E-13 | 5.62E-11 |
| Clps | 0.38821126 | 1.17091131 | 0.38170799 | 0.78920333 | 1.94E-20 | 2.83E-18 |

| | | | | | | |
|---|---|---|---|---|---|---|
| Pam | 1.53040848 | 1.6776 | 0.89837289 | 0.77922711 | 5.76E-05 | 0.00179135 |
| Tspan7 | 1.29129421 | 1.62348023 | 0.84854745 | 0.77493278 | 2.45E-05 | 0.0008058 |
| Agtr2 | 0.52059138 | 1.25087694 | 0.48004725 | 0.7708297 | 4.72E-13 | 4.35E-11 |
| Hadh | 1.58492326 | 1.67124956 | 0.9062157 | 0.76503386 | 8.74E-05 | 0.00261412 |
| Ahnak | 0.95802702 | 1.49094321 | 0.73263162 | 0.75831159 | 2.10E-06 | 8.50E-05 |
| Col14a1 | 0.69242389 | 1.3372808 | 0.59086928 | 0.74641152 | 9.78E-09 | 5.48E-07 |
| Cystm1 | 1.34873468 | 1.60608575 | 0.86283287 | 0.74325289 | 6.42E-05 | 0.00197566 |
| Islr | 0.74637706 | 1.36238177 | 0.62235106 | 0.7400307 | 6.24E-08 | 3.20E-06 |
| Cdkn1a | 0.59333402 | 1.26409616 | 0.52906236 | 0.73503379 | 3.28E-10 | 2.17E-08 |
| Tubb4b | 1.84503549 | 1.6551226 | 0.9257721 | 0.7293505 | 0.00023056 | 0.0063839 |
| Id3 | 0.55150057 | 1.23046936 | 0.50127544 | 0.72919393 | 5.00E-11 | 3.57E-09 |
| H2afz | 2.69330128 | 1.60371826 | 0.87858924 | 0.72512902 | 0.0001217 | 0.00353045 |
| Sec61b | 3.18155059 | 1.53949346 | 0.81473598 | 0.72475748 | 3.83E-05 | 0.00122183 |
| Ier2 | 1.28446941 | 1.5709585 | 0.8467287 | 0.7242298 | 7.16E-05 | 0.00216492 |
| Krt19 | 0.47571523 | 1.16476484 | 0.44810793 | 0.71665691 | 5.81E-13 | 5.22E-11 |
| Mid1ip1 | 1.39894015 | 1.58119208 | 0.87412542 | 0.70706666 | 0.00016158 | 0.00461701 |
| Cd63 | 3.23589526 | 1.51039463 | 0.80814756 | 0.70224707 | 5.60E-05 | 0.00174407 |
| Scg5 | 1.04009831 | 1.46795487 | 0.76686585 | 0.70108902 | 2.41E-05 | 0.0007926 |
| Rpl36a | 3.87098599 | 1.40016428 | 0.70565679 | 0.69450749 | 6.06E-06 | 0.00022578 |
| Zeb2 | 0.72207397 | 1.30031755 | 0.60834869 | 0.69196886 | 2.13E-07 | 9.89E-06 |
| Nfib | 1.33579947 | 1.55089624 | 0.85971376 | 0.69118248 | 0.0001757 | 0.00497358 |
| Hba-a1 | 0.49252657 | 1.15022491 | 0.46023279 | 0.68999211 | 1.32E-11 | 1.00E-09 |
| Cd81 | 3.23422982 | 1.49573831 | 0.80834485 | 0.68739345 | 7.83E-05 | 0.00235639 |
| Igfbp5 | 0.67037374 | 1.26490684 | 0.57757709 | 0.68732976 | 6.10E-08 | 3.13E-06 |
| Mfap5 | 0.47098936 | 1.12300559 | 0.44466497 | 0.67834063 | 5.93E-12 | 4.77E-10 |
| Sfrp1 | 0.66587715 | 1.25281418 | 0.57483561 | 0.67797858 | 7.89E-08 | 3.92E-06 |
| Peg10 | 0.67610027 | 1.25893773 | 0.58105314 | 0.67788459 | 1.07E-07 | 5.22E-06 |
| Mgp | 0.47481597 | 1.12413546 | 0.44745397 | 0.6766815 | 8.90E-12 | 6.90E-10 |
| Manf | 2.28011675 | 1.59169931 | 0.9154895 | 0.67620981 | 0.00051233 | 0.01328997 |
| Tppp3 | 0.66155977 | 1.24767387 | 0.57219226 | 0.67548161 | 7.68E-08 | 3.85E-06 |

| | | | | | | |
|---|---|---|---|---|---|---|
| mt-Co3 | 5.90367246 | 1.38462343 | 0.71027395 | 0.67434949 | 1.22E-05 | 0.00042938 |
| Hbb-bs | 0.89178772 | 1.37081071 | 0.70043009 | 0.67038062 | 1.05E-05 | 0.00037317 |
| Snhg18 | 0.8900876 | 1.36916189 | 0.69957421 | 0.66958768 | 1.05E-05 | 0.00037317 |
| Atp1b1 | 1.26675996 | 1.50788245 | 0.84192064 | 0.66596181 | 0.00021865 | 0.00606509 |
| Pcsk1 | 0.90718375 | 1.36130915 | 0.70812927 | 0.65317988 | 2.06E-05 | 0.00068976 |
| Hist1h2bc | 0.93579242 | 1.37339947 | 0.72211271 | 0.65128676 | 3.04E-05 | 0.00098504 |
| Gpx8 | 0.88943284 | 1.34216187 | 0.69924429 | 0.64291758 | 2.18E-05 | 0.0007268 |
| Cplx2 | 1.1282149 | 1.43968282 | 0.79915631 | 0.64052651 | 0.0001832 | 0.00515715 |
| Hba-a2 | 0.16237332 | 0.81855514 | 0.18003014 | 0.638525 | 2.60E-56 | 1.16E-53 |
| Rpl39 | 4.72130359 | 1.21623254 | 0.57781946 | 0.63841308 | 4.51E-07 | 2.01E-05 |
| Crip2 | 0.83789744 | 1.28896273 | 0.67260197 | 0.61636076 | 2.31E-05 | 0.00076397 |
| Col5a2 | 0.76717675 | 1.24368965 | 0.63410994 | 0.60957971 | 9.62E-06 | 0.00034637 |
| Peg3 | 2.75584471 | 1.47872251 | 0.87117825 | 0.60754425 | 0.00096667 | 0.02314468 |
| Ccnd3 | 1.45257547 | 1.49227163 | 0.88487317 | 0.60739845 | 0.00113823 | 0.02645604 |
| Gast | 0.29294265 | 0.90805169 | 0.30247181 | 0.60557988 | 2.78E-19 | 3.83E-17 |
| Ramp2 | 0.99266039 | 1.35192617 | 0.74768889 | 0.60423728 | 0.00016365 | 0.00466731 |
| S100a11 | 3.23753845 | 1.41003118 | 0.80795285 | 0.60207833 | 0.0004618 | 0.01208232 |
| Myl9 | 0.63944212 | 1.15738123 | 0.55849544 | 0.59888579 | 9.34E-07 | 4.04E-05 |
| Rrbp1 | 2.0729308 | 1.52350957 | 0.92564171 | 0.59786786 | 0.00204331 | 0.04362404 |
| Top2a | 0.33987728 | 0.93592314 | 0.34247596 | 0.59344718 | 6.62E-15 | 6.90E-13 |
| Resp18 | 0.49478266 | 1.05509011 | 0.46184522 | 0.59324489 | 5.63E-09 | 3.20E-07 |
| Rap1b | 2.11411876 | 1.5064193 | 0.92422998 | 0.58218932 | 0.00255221 | 0.05271225 |
| Zbtb20 | 1.39701077 | 1.45482376 | 0.87371645 | 0.5811073 | 0.00155442 | 0.03434419 |
| mt-Co2 | 4.66166135 | 1.1610546 | 0.58548599 | 0.57556861 | 6.20E-06 | 0.00022889 |
| Col5a1 | 0.759704 | 1.20096013 | 0.62990864 | 0.57105148 | 2.79E-05 | 0.00090693 |
| Gas6 | 0.63892449 | 1.12847033 | 0.55817161 | 0.57029872 | 2.78E-06 | 0.00011048 |
| Gng11 | 0.57734528 | 1.08836535 | 0.51856669 | 0.56979866 | 5.18E-07 | 2.29E-05 |
| Sepp1 | 1.37829562 | 1.43628899 | 0.86961095 | 0.56667804 | 0.00188399 | 0.04062211 |
| Stmn2 | 0.2451328 | 0.82362279 | 0.25963943 | 0.56398335 | 2.30E-22 | 3.93E-20 |
| Nrk | 0.62687248 | 1.11424495 | 0.55059065 | 0.5636543 | 2.67E-06 | 0.00010647 |

| | | | | | | |
|---|---|---|---|---|---|---|
| Emp1 | 0.75870037 | 1.19226626 | 0.62934235 | 0.56292391 | 3.50E-05 | 0.00112174 |
| Mmp14 | 1.12815566 | 1.36177794 | 0.79913565 | 0.56264229 | 0.00087406 | 0.0211908 |
| Lmna | 1.46516932 | 1.44705009 | 0.88723074 | 0.55981935 | 0.00251535 | 0.05209246 |
| 2810417H13 | 0.39599921 | 0.94570524 | 0.38785322 | 0.55785202 | 8.07E-11 | 5.63E-09 |
| Nid1 | 0.79383589 | 1.20513119 | 0.64888538 | 0.55624581 | 6.92E-05 | 0.00210127 |
| Hist3h2ba | 0.9378243 | 1.27710663 | 0.72309219 | 0.55401443 | 0.00032948 | 0.00883322 |
| Emb | 0.99884291 | 1.30367137 | 0.75024505 | 0.55342632 | 0.00052023 | 0.01347202 |
| mt-Co1 | 5.40613774 | 1.14408726 | 0.59529687 | 0.54879039 | 2.08E-05 | 0.00069523 |
| mt-Cytb | 5.03201416 | 1.09241501 | 0.54391516 | 0.54849986 | 3.68E-06 | 0.00014193 |
| Slc30a8 | 0.60198695 | 1.07859536 | 0.53467828 | 0.54391708 | 3.06E-06 | 0.00012007 |
| Hspa5 | 3.23498863 | 1.35087183 | 0.80825497 | 0.54261685 | 0.00142007 | 0.03169829 |
| Tubb6 | 0.61936079 | 1.08244563 | 0.54582535 | 0.53662028 | 6.19E-06 | 0.00022889 |
| Gm42418 | 2.55242775 | 1.42896411 | 0.89285264 | 0.53611147 | 0.00379986 | 0.07433926 |
| H2-D1 | 1.2504839 | 1.37320498 | 0.83730941 | 0.53589556 | 0.00221861 | 0.0465174 |
| Cks2 | 0.51426054 | 1.01145157 | 0.47562287 | 0.5358287 | 2.75E-07 | 1.26E-05 |
| Igfbp2 | 0.39553129 | 0.9215332 | 0.38748547 | 0.53404773 | 4.47E-10 | 2.92E-08 |
| Osr1 | 0.5297419 | 1.01968616 | 0.48639494 | 0.53329122 | 5.46E-07 | 2.41E-05 |
| Bst2 | 0.36991253 | 0.89885043 | 0.36708127 | 0.53176917 | 5.97E-11 | 4.24E-09 |
| Tuba4a | 0.98487995 | 1.27545658 | 0.744419 | 0.53103759 | 0.00075876 | 0.01875441 |
| Matn2 | 0.68780679 | 1.11848787 | 0.58810681 | 0.53038105 | 3.05E-05 | 0.00098504 |
| Ppp1r1a | 0.49970752 | 0.99457417 | 0.4653523 | 0.52922186 | 2.14E-07 | 9.90E-06 |
| Tpm2 | 0.65069443 | 1.08944118 | 0.56549653 | 0.52394465 | 1.90E-05 | 0.00064571 |
| Fbn1 | 0.6606757 | 1.09480126 | 0.57164978 | 0.52315149 | 2.37E-05 | 0.0007803 |
| Vcan | 0.64159842 | 1.08205996 | 0.55984288 | 0.52221708 | 1.69E-05 | 0.00058233 |
| Prdx4 | 1.27614711 | 1.36564454 | 0.84449088 | 0.52115365 | 0.00303959 | 0.06144249 |
| Abcc8 | 1.00849199 | 1.27370202 | 0.75420376 | 0.51949825 | 0.00109877 | 0.02577527 |
| Hbb-bt | 0.24438967 | 0.77737533 | 0.25895615 | 0.51841919 | 2.79E-19 | 3.83E-17 |
| Fbn2 | 0.71586653 | 1.1231037 | 0.60472641 | 0.5183773 | 6.93E-05 | 0.00210127 |
| Sdpr | 0.54267271 | 1.01269241 | 0.49527516 | 0.51741725 | 1.71E-06 | 7.05E-05 |
| Selm | 1.81029075 | 1.44198945 | 0.92466584 | 0.51732362 | 0.00643788 | 0.11553978 |

| | | | | | | |
|---|---|---|---|---|---|---|
| Olfml3 | 0.67952112 | 1.0985515 | 0.58312158 | 0.51542992 | 4.26E-05 | 0.00134765 |
| Pnliprp1 | 0.17468259 | 0.70600432 | 0.1923391 | 0.51366522 | 8.22E-33 | 2.15E-30 |
| Il11ra1 | 0.97856701 | 1.25386526 | 0.74171177 | 0.51215349 | 0.00107137 | 0.02524942 |
| Nkain4 | 0.55468357 | 1.01090981 | 0.50342712 | 0.50748269 | 3.71E-06 | 0.00014267 |
| Arpc1b | 1.58381453 | 1.41311854 | 0.90608487 | 0.50703366 | 0.00642851 | 0.11553978 |
| Ubqln2 | 1.21521847 | 1.33353601 | 0.8269648 | 0.50657121 | 0.00323234 | 0.06482167 |
| Tm4sf1 | 0.33663549 | 0.84617053 | 0.33977468 | 0.50639585 | 1.73E-11 | 1.29E-09 |
| Ube2c | 0.28353173 | 0.80018499 | 0.29421123 | 0.50597377 | 1.04E-14 | 1.07E-12 |
| Nbl1 | 0.5686315 | 1.01856533 | 0.51278327 | 0.50578207 | 5.80E-06 | 0.00021669 |
| Plvap | 0.22264254 | 0.74251686 | 0.23871751 | 0.50379936 | 3.23E-21 | 5.17E-19 |
| Cyba | 1.01624678 | 1.25587938 | 0.75735846 | 0.49852092 | 0.00171532 | 0.03735614 |
| Ngfrap1 | 2.54907823 | 1.39072091 | 0.89316712 | 0.49755379 | 0.00663388 | 0.11835875 |
| Ambp | 0.5434936 | 0.99245895 | 0.49583535 | 0.4966236 | 4.24E-06 | 0.00016155 |
| Dpysl3 | 0.61764524 | 1.03795163 | 0.54473264 | 0.49321899 | 2.85E-05 | 0.00092452 |
| Pcbd1 | 1.10025286 | 1.28225904 | 0.78933696 | 0.49292207 | 0.00275013 | 0.05618915 |
| Kdelr2 | 1.53594205 | 1.39052733 | 0.8992641 | 0.49126323 | 0.00757839 | 0.13302432 |
| Pak3 | 0.83082871 | 1.1594033 | 0.66885423 | 0.49054907 | 0.00055611 | 0.01423142 |
| Serpine2 | 0.87353554 | 1.18162494 | 0.6911499 | 0.49047504 | 0.00080296 | 0.01965517 |
| Txndc5 | 1.32778794 | 1.34704928 | 0.8577374 | 0.48931189 | 0.00560511 | 0.10377978 |
| mt-Nd1 | 4.54276216 | 1.08929916 | 0.60052917 | 0.48876999 | 0.00014829 | 0.00426109 |
| Calu | 1.00979605 | 1.23921168 | 0.75473593 | 0.48447574 | 0.00216068 | 0.04542806 |
| Ndufa4l2 | 0.498088 | 0.94488758 | 0.46420078 | 0.48068679 | 2.08E-06 | 8.44E-05 |
| Mgst1 | 0.62779636 | 1.03097368 | 0.5511746 | 0.47979907 | 5.44E-05 | 0.00169917 |
| Egfl7 | 0.53834376 | 0.9714541 | 0.49231387 | 0.47914023 | 7.57E-06 | 0.00027594 |
| Smoc2 | 0.44271456 | 0.89936125 | 0.42373154 | 0.47562972 | 3.02E-07 | 1.37E-05 |
| Tagln2 | 1.48104548 | 1.36355785 | 0.89007065 | 0.47348721 | 0.00901721 | 0.15383899 |
| Mlxipl | 0.81883668 | 1.13380162 | 0.66244812 | 0.47135349 | 0.00078024 | 0.01916065 |
| Col6a3 | 0.64696217 | 1.03452674 | 0.56318209 | 0.47134465 | 9.93E-05 | 0.00294386 |
| Upk3b | 0.19429438 | 0.67677608 | 0.21160201 | 0.46517407 | 7.35E-23 | 1.30E-20 |
| Kdelr3 | 0.6536508 | 1.03205452 | 0.56732456 | 0.46472996 | 0.00013544 | 0.00390529 |

| | | | | | | |
|---|---|---|---|---|---|---|
| Ptrf | 0.69648555 | 1.05775413 | 0.5932897 | 0.46446443 | 0.00025043 | 0.00683435 |
| Vwa5b2 | 0.91857933 | 1.17699708 | 0.71375213 | 0.46324495 | 0.00195507 | 0.04185772 |
| Id2 | 0.69102778 | 1.05172995 | 0.5900354 | 0.46169455 | 0.000252 | 0.00686486 |
| Clec3b | 0.3379338 | 0.80110243 | 0.34085759 | 0.46024485 | 9.70E-10 | 6.04E-08 |
| Cxcl13 | 0.21230751 | 0.68752239 | 0.22892907 | 0.45859332 | 2.66E-19 | 3.71E-17 |
| Ddah2 | 1.25706223 | 1.29684916 | 0.83918341 | 0.45766575 | 0.0076647 | 0.13407494 |
| Fscn1 | 0.80055295 | 1.10985013 | 0.65255652 | 0.45729361 | 0.00091861 | 0.02216469 |
| Mxra8 | 0.64606518 | 1.01984126 | 0.56262474 | 0.45721652 | 0.0001515 | 0.00434525 |
| Kcnq1ot1 | 1.73595658 | 1.37660572 | 0.92069661 | 0.45590912 | 0.01385497 | 0.21802223 |
| Hsp90aa1 | 3.00876974 | 1.29122763 | 0.83553281 | 0.45569482 | 0.00766246 | 0.13407494 |
| Ppib | 2.72066698 | 1.3307583 | 0.8754815 | 0.45527679 | 0.01039299 | 0.17284887 |
| Gabarapl2 | 1.87073791 | 1.38145158 | 0.92646631 | 0.45498528 | 0.01450958 | 0.22575246 |
| Hes1 | 0.54294744 | 0.94619691 | 0.49546271 | 0.4507342 | 2.62E-05 | 0.00085973 |
| Htra3 | 0.55693297 | 0.95411173 | 0.50494398 | 0.44916775 | 3.83E-05 | 0.00122183 |
| Des | 0.51243348 | 0.92240998 | 0.47434101 | 0.44806897 | 1.34E-05 | 0.00047089 |
| Cryab | 0.42728598 | 0.8478073 | 0.41206336 | 0.43574393 | 1.29E-06 | 5.46E-05 |
| Penk | 0.35651038 | 0.78803423 | 0.35619429 | 0.43183994 | 3.53E-08 | 1.88E-06 |
| Zcchc18 | 0.6995897 | 1.02504936 | 0.59513396 | 0.4299154 | 0.00066037 | 0.0166196 |
| Aebp1 | 0.5979057 | 0.95957089 | 0.53203481 | 0.42753608 | 0.00017684 | 0.00499661 |
| Birc5 | 0.31816915 | 0.75090571 | 0.32421569 | 0.42669003 | 2.45E-09 | 1.43E-07 |
| H1f0 | 1.6977024 | 1.3440277 | 0.91803259 | 0.42599511 | 0.01956121 | 0.28373093 |
| Tshz2 | 1.64462839 | 1.33900094 | 0.91301924 | 0.4259817 | 0.01903228 | 0.27875656 |
| Pecam1 | 0.21308362 | 0.65448793 | 0.22966795 | 0.42481998 | 9.92E-17 | 1.16E-14 |
| Ap1s2 | 1.14375151 | 1.22931358 | 0.80451078 | 0.42480281 | 0.00945255 | 0.15929963 |
| Tubb2a | 1.47563291 | 1.31316875 | 0.8891227 | 0.42404606 | 0.01699346 | 0.25620715 |
| Serf2 | 3.28789495 | 1.22473265 | 0.80137396 | 0.42335869 | 0.00942273 | 0.15897319 |
| Copz2 | 0.78458405 | 1.06523866 | 0.64379456 | 0.4214441 | 0.00180595 | 0.03916165 |
| Calr | 3.13148749 | 1.24096311 | 0.82058959 | 0.42037353 | 0.01138101 | 0.18602448 |
| Tubb5 | 3.17728045 | 1.23505919 | 0.81526183 | 0.41979736 | 0.01103542 | 0.18154698 |
| Gpx7 | 0.70083399 | 1.01541135 | 0.59587188 | 0.41953947 | 0.00087389 | 0.0211908 |

| | | | | | | |
|---|---|---|---|---|---|---|
| Cenpa | 0.34148406 | 0.76299729 | 0.34381157 | 0.41918573 | 2.98E-08 | 1.60E-06 |
| Cd9 | 0.95565864 | 1.14845031 | 0.73152198 | 0.41692833 | 0.00564234 | 0.10411931 |
| Pdia6 | 2.64719639 | 1.29937773 | 0.88350845 | 0.41586928 | 0.01819439 | 0.27088439 |
| Akr1c19 | 0.92815642 | 1.1324149 | 0.71841777 | 0.41399713 | 0.00520603 | 0.09733929 |
| Igf2r | 0.8063605 | 1.06742705 | 0.65571469 | 0.41171235 | 0.00262466 | 0.05391545 |
| Sdf2l1 | 0.87405383 | 1.1001266 | 0.69141541 | 0.40871119 | 0.00429572 | 0.08244845 |
| Auts2 | 0.9709154 | 1.14649774 | 0.73840903 | 0.40808871 | 0.00700696 | 0.12416293 |
| Lpl | 0.48939287 | 0.86384107 | 0.45798707 | 0.405854 | 4.08E-05 | 0.00129528 |
| Prss23 | 0.5235203 | 0.88444041 | 0.48208482 | 0.40235559 | 0.00010348 | 0.00304246 |
| Hist1h1c | 0.61666191 | 0.94520145 | 0.54410548 | 0.40109597 | 0.0005243 | 0.01353134 |
| Spats2l | 0.59886152 | 0.93365034 | 0.53265476 | 0.40099559 | 0.00040881 | 0.01082627 |
| Mmp2 | 0.56396702 | 0.90965318 | 0.50966756 | 0.39998563 | 0.00024251 | 0.00665919 |
| Cpa1 | 0.18527279 | 0.60253397 | 0.2027928 | 0.39974117 | 9.50E-19 | 1.27E-16 |
| Cldn5 | 0.17406222 | 0.58962447 | 0.1917229 | 0.39790157 | 1.40E-20 | 2.07E-18 |
| Rcn1 | 1.3098367 | 1.24758802 | 0.85326651 | 0.39432151 | 0.0199658 | 0.28849829 |
| Acly | 1.40935662 | 1.2698017 | 0.87631486 | 0.39348684 | 0.02295683 | 0.32133221 |
| P4ha2 | 0.62124067 | 0.93973733 | 0.54702085 | 0.39271648 | 0.00070748 | 0.017659 |
| Eno1 | 1.74797451 | 1.31128895 | 0.92146802 | 0.38982093 | 0.03000744 | 0.39942481 |
| Dek | 0.97651657 | 1.12844413 | 0.74082903 | 0.3876151 | 0.01000871 | 0.1671894 |
| Cmtm3 | 0.68483196 | 0.97364524 | 0.58632097 | 0.38732427 | 0.00165824 | 0.03637354 |
| Plac8 | 0.39783245 | 0.77070897 | 0.38929239 | 0.38141658 | 6.63E-06 | 0.00024296 |
| Tpm4 | 1.19938351 | 1.20363207 | 0.82225454 | 0.38137754 | 0.01960586 | 0.28410746 |
| Fxyd1 | 0.60325317 | 0.91336224 | 0.53549649 | 0.37786575 | 0.00085349 | 0.02075814 |
| Maf | 0.45312434 | 0.80879413 | 0.43150446 | 0.37728967 | 5.07E-05 | 0.00159012 |
| Tsc22d1 | 2.15330306 | 1.29915093 | 0.92267383 | 0.3764771 | 0.03484398 | 0.44842781 |
| Tpd52 | 1.08730163 | 1.16061717 | 0.78468671 | 0.37593047 | 0.01659337 | 0.25048249 |
| Rps19 | 5.13634185 | 0.92984047 | 0.55495411 | 0.37488636 | 0.00133611 | 0.03017858 |
| Myl12a | 2.18605294 | 1.29505993 | 0.9211401 | 0.37391983 | 0.03556772 | 0.45510514 |
| Col18a1 | 0.50753124 | 0.84458049 | 0.47089092 | 0.37368957 | 0.0002094 | 0.00584054 |
| Zfp36l1 | 0.77685825 | 1.0121092 | 0.63951311 | 0.37259609 | 0.00479713 | 0.09086026 |

| | | | | | | |
|---|---|---|---|---|---|---|
| Fbln2 | 0.58729111 | 0.89207771 | 0.52511321 | 0.3669645 | 0.00094588 | 0.02271451 |
| Fxyd3 | 0.65862537 | 0.93671912 | 0.57039008 | 0.36632904 | 0.00215076 | 0.04534495 |
| Nid2 | 0.51121227 | 0.83883694 | 0.473483 | 0.36535394 | 0.00030138 | 0.00813738 |
| Tmem176b | 1.73633658 | 1.28438307 | 0.92072146 | 0.36366161 | 0.0395526 | 0.49281891 |
| Adamts5 | 0.45584255 | 0.79567203 | 0.43352133 | 0.3621507 | 0.00010211 | 0.00301408 |
| Cldn4 | 0.75421071 | 0.98877265 | 0.62680343 | 0.36196922 | 0.00512516 | 0.09594528 |
| Hmgb1 | 3.83854356 | 1.07315579 | 0.7112051 | 0.36195069 | 0.01183375 | 0.19197958 |
| Fkbp7 | 0.63923028 | 0.91817289 | 0.55836294 | 0.35980996 | 0.00208665 | 0.04417722 |
| Ffar2 | 0.74007889 | 0.97779051 | 0.61875049 | 0.35904002 | 0.00494524 | 0.0929203 |
| Col4a2 | 0.58133466 | 0.8797376 | 0.52119953 | 0.35853808 | 0.00111345 | 0.02595941 |
| Timp2 | 0.66275134 | 0.9297204 | 0.57292278 | 0.35679762 | 0.00281507 | 0.05720839 |
| BC048546 | 0.59600858 | 0.88705843 | 0.53080279 | 0.35625564 | 0.00142379 | 0.03173462 |
| Eng | 0.38854217 | 0.73819941 | 0.38197007 | 0.35622935 | 1.69E-05 | 0.00058306 |
| Ddx5 | 3.19962725 | 1.1653234 | 0.81250508 | 0.35281833 | 0.02677624 | 0.3643918 |
| Ccl21a | 0.07797758 | 0.44302464 | 0.09089794 | 0.3521267 | 9.13E-67 | 5.14E-64 |
| Pf4 | 0.08221892 | 0.44712533 | 0.09558011 | 0.35154522 | 2.14E-60 | 1.12E-57 |
| Tpm1 | 1.33725794 | 1.21045516 | 0.86007146 | 0.3503837 | 0.03506335 | 0.45016727 |
| C1qa | 0.22973995 | 0.59549187 | 0.24537473 | 0.35011713 | 1.12E-10 | 7.67E-09 |
| Lgals7 | 0.14927186 | 0.5167302 | 0.16674072 | 0.34998947 | 5.23E-21 | 8.03E-19 |
| Oat | 0.95074271 | 1.07915159 | 0.72921203 | 0.34993956 | 0.01644638 | 0.2489047 |
| Gap43 | 0.3077887 | 0.66447792 | 0.31533708 | 0.34914084 | 4.28E-07 | 1.92E-05 |
| Mif | 2.27985283 | 1.26425775 | 0.91550798 | 0.34874976 | 0.04518106 | 0.54819953 |
| Ociad2 | 0.66529061 | 0.92274134 | 0.57447706 | 0.34826428 | 0.00351888 | 0.06973982 |
| Serpinf1 | 0.53319015 | 0.83468297 | 0.48877325 | 0.34590972 | 0.00082703 | 0.0202118 |
| Rps26 | 4.38353013 | 0.96756017 | 0.6221591 | 0.34540107 | 0.00679265 | 0.12090751 |
| Gadd45a | 0.49123801 | 0.80413255 | 0.45931016 | 0.3448224 | 0.00042269 | 0.01115502 |
| 6330403K07Rik | 0.51741297 | 0.8217602 | 0.47782956 | 0.34393064 | 0.00068751 | 0.01724558 |
| Btg1 | 1.69994615 | 1.26105293 | 0.91819878 | 0.34285415 | 0.04845891 | 0.57824787 |
| Glul | 1.25593838 | 1.18108368 | 0.83886425 | 0.34221943 | 0.03486887 | 0.44842781 |
| Rhoc | 1.16188914 | 1.152266 | 0.81038582 | 0.34188018 | 0.03036316 | 0.40204743 |

| | | | | | | |
|---|---|---|---|---|---|---|
| Htra1 | 0.46138656 | 0.77944207 | 0.43761883 | 0.34182324 | 0.00025786 | 0.00701207 |
| C1qb | 0.07972844 | 0.42918064 | 0.09283388 | 0.33634675 | 1.14E-58 | 5.78E-56 |
| mt-Atp6 | 5.79330892 | 1.01446065 | 0.67872224 | 0.33573841 | 0.01393631 | 0.21862316 |
| Anxa4 | 1.27297104 | 1.17933206 | 0.84363103 | 0.33570103 | 0.03844582 | 0.48219046 |
| Ogn | 0.44079447 | 0.75792059 | 0.42228881 | 0.33563178 | 0.00020518 | 0.00574385 |
| Cdo1 | 0.53774201 | 0.82732011 | 0.49190131 | 0.3354188 | 0.00121719 | 0.02794945 |
| Lrrc17 | 0.46722215 | 0.77536188 | 0.44190871 | 0.33345317 | 0.00039748 | 0.01056314 |
| Dpt | 0.34461232 | 0.67844973 | 0.34640568 | 0.33204406 | 1.02E-05 | 0.00036519 |
| Sfrp2 | 0.25882492 | 0.60385546 | 0.27212959 | 0.33172587 | 2.99E-08 | 1.61E-06 |
| Mfge8 | 0.64821169 | 0.89517183 | 0.56395777 | 0.33121407 | 0.00451498 | 0.08622137 |
| Loxl1 | 0.55489222 | 0.83476774 | 0.50356795 | 0.33119979 | 0.00172836 | 0.0375863 |
| Cdk1 | 0.3306063 | 0.66412904 | 0.33472714 | 0.3294019 | 6.07E-06 | 0.00022578 |
| Hist1h2al | 0.72178072 | 0.93749893 | 0.60817799 | 0.32932094 | 0.00803677 | 0.13961935 |
| Atp5l | 3.75932542 | 1.05414404 | 0.72533843 | 0.32880561 | 0.02193908 | 0.31095399 |
| Cdh5 | 0.18737405 | 0.53363677 | 0.20485242 | 0.32878435 | 4.84E-13 | 4.40E-11 |
| Hsp90b1 | 2.85561327 | 1.18550305 | 0.85699141 | 0.32851164 | 0.04417746 | 0.53852572 |
| Osr2 | 0.42920501 | 0.74181872 | 0.4135241 | 0.32829462 | 0.0002083 | 0.00582042 |
| Smim22 | 0.82509094 | 0.99167139 | 0.66579673 | 0.32587466 | 0.01478118 | 0.2290405 |
| Nkx6-1 | 0.66566701 | 0.89915973 | 0.57470717 | 0.32445256 | 0.00604033 | 0.1097001 |
| Hist1h2ap | 0.27059035 | 0.60657663 | 0.28271801 | 0.32385862 | 1.77E-07 | 8.29E-06 |
| Cyb5r3 | 1.66086215 | 1.23826485 | 0.91476608 | 0.32349877 | 0.05795805 | 0.65406109 |
| Cnn2 | 0.58908583 | 0.84973848 | 0.52628838 | 0.32345009 | 0.00314553 | 0.06324898 |
| Capn6 | 0.48391993 | 0.77720687 | 0.4540492 | 0.32315767 | 0.00077797 | 0.01913569 |
| Tcf4 | 0.91187046 | 1.03352885 | 0.71045473 | 0.32307413 | 0.02160847 | 0.30784482 |
| Runx1t1 | 0.93551366 | 1.04465436 | 0.72197821 | 0.32267615 | 0.0234673 | 0.32667262 |
| Ypel3 | 1.41064728 | 1.1990519 | 0.87658398 | 0.32246792 | 0.05098261 | 0.60204511 |
| Emilin1 | 0.54027759 | 0.81569845 | 0.49363818 | 0.32206027 | 0.00186337 | 0.04023457 |
| Fkbp11 | 0.55060896 | 0.82257102 | 0.50067161 | 0.32189941 | 0.00213001 | 0.0450324 |
| Npepl1 | 0.86282146 | 1.00721088 | 0.68563583 | 0.32157505 | 0.01853165 | 0.27376052 |
| Msn | 0.4807062 | 0.77308696 | 0.45172748 | 0.32135947 | 0.00078181 | 0.01916819 |

| | | | | | | |
|---|---|---|---|---|---|---|
| Cst3 | 2.67415122 | 1.20133569 | 0.88068524 | 0.32065045 | 0.05276563 | 0.61699263 |
| Cldn7 | 0.86530881 | 1.00715592 | 0.68692115 | 0.32023477 | 0.01910985 | 0.27931618 |
| Lmo4 | 0.60359458 | 0.85592885 | 0.53571695 | 0.3202119 | 0.003939 | 0.07666679 |
| Gadd45g | 0.56904405 | 0.83275339 | 0.5130581 | 0.31969529 | 0.00280167 | 0.05708864 |
| Ptgis | 0.48145252 | 0.77143856 | 0.45226727 | 0.31917128 | 0.00085248 | 0.02075814 |
| Sep-11 | 0.76610127 | 0.94937197 | 0.6335071 | 0.31586487 | 0.01332635 | 0.21145496 |
| Slc9a3r1 | 0.42288452 | 0.72452194 | 0.40870283 | 0.3158191 | 0.00029597 | 0.0080198 |
| Tinagl1 | 0.33000657 | 0.64628055 | 0.33422336 | 0.31205719 | 1.66E-05 | 0.000575 |
| Efemp2 | 0.9210865 | 1.02674533 | 0.71497694 | 0.31176839 | 0.0262797 | 0.35988984 |
| Gm13305 | 0.620052 | 0.85740792 | 0.54626515 | 0.31114277 | 0.00566848 | 0.10411931 |
| Immp1l | 1.43535792 | 1.19261306 | 0.88155817 | 0.31105489 | 0.05836855 | 0.65722983 |
| Prrx1 | 0.47822737 | 0.76052219 | 0.44993191 | 0.31059029 | 0.00107451 | 0.02528433 |
| Apoc1 | 0.20846458 | 0.53547231 | 0.22526129 | 0.31021102 | 4.62E-10 | 3.00E-08 |
| Anxa5 | 1.89480512 | 1.23461273 | 0.92693971 | 0.30767303 | 0.07002552 | 0.74657023 |
| Hn1 | 1.59551355 | 1.21241632 | 0.90745005 | 0.30496627 | 0.06758354 | 0.73016166 |
| Csrp1 | 0.65790416 | 0.87263107 | 0.56994645 | 0.30268463 | 0.00911402 | 0.15531645 |
| Cenpf | 0.23169059 | 0.548395 | 0.24719547 | 0.30119953 | 3.03E-08 | 1.62E-06 |
| Cdh11 | 0.50343804 | 0.76864961 | 0.46799824 | 0.30065137 | 0.00214544 | 0.04529563 |
| Nusap1 | 0.24073872 | 0.55560817 | 0.25559133 | 0.30001683 | 9.03E-08 | 4.41E-06 |
| Arf4 | 2.16124912 | 1.22144462 | 0.92233885 | 0.29910577 | 0.07469801 | 0.77879592 |
| Fabp4 | 0.178635 | 0.49518685 | 0.19625495 | 0.2989319 | 6.37E-12 | 5.09E-10 |
| Serping1 | 0.78007723 | 0.9398733 | 0.64130023 | 0.29857307 | 0.01923736 | 0.28090978 |
| Uqcc2 | 2.12318281 | 1.22218957 | 0.92387925 | 0.29831032 | 0.07558133 | 0.78584943 |
| Figf | 0.40869269 | 0.69473206 | 0.39776545 | 0.29696661 | 0.00045165 | 0.01185754 |
| G0s2 | 0.34641199 | 0.64468724 | 0.34789414 | 0.2967931 | 7.45E-05 | 0.0022481 |
| Phlda3 | 0.57285141 | 0.81054623 | 0.51558969 | 0.29495654 | 0.00549172 | 0.10192875 |
| Txnip | 1.25408544 | 1.13244453 | 0.83833713 | 0.2941074 | 0.05942509 | 0.66469518 |
| Cox17 | 1.61864194 | 1.20165255 | 0.91015261 | 0.29149994 | 0.07724921 | 0.79719302 |
| Fcgrt | 0.61440745 | 0.83262965 | 0.54266508 | 0.28996457 | 0.00876402 | 0.14985587 |
| Ltbp4 | 0.52767843 | 0.77469161 | 0.48496815 | 0.28972346 | 0.00395559 | 0.07669505 |

| | | | | | | |
|---|---|---|---|---|---|---|
| Ifi27 | 0.56656813 | 0.79904244 | 0.51140675 | 0.28763569 | 0.00620288 | 0.11211645 |
| Aldh2 | 0.79878831 | 0.93876484 | 0.65159399 | 0.28717085 | 0.02504059 | 0.34478439 |
| Gja4 | 0.11238716 | 0.41514815 | 0.12823721 | 0.28691094 | 1.31E-23 | 2.42E-21 |
| 0610011F06I | 1.11952735 | 1.08212442 | 0.79611204 | 0.28601238 | 0.0551178 | 0.63521283 |
| Rps20 | 4.10017548 | 0.95304785 | 0.66780504 | 0.28524281 | 0.02879056 | 0.38593062 |
| Gatsl2 | 0.67257617 | 0.86361047 | 0.57891598 | 0.28469449 | 0.01439963 | 0.22450139 |
| 2700094K13I | 1.0904801 | 1.07038438 | 0.78584743 | 0.28453695 | 0.05373715 | 0.62276574 |
| Ly6h | 0.46096682 | 0.72126653 | 0.43730936 | 0.28395718 | 0.0019468 | 0.04173942 |
| Ckb | 0.63415332 | 0.8389671 | 0.55517991 | 0.28378719 | 0.01153035 | 0.18806099 |
| Cirbp | 1.37245928 | 1.15187642 | 0.86830581 | 0.28357061 | 0.0732727 | 0.77027545 |
| Lox | 0.47839062 | 0.73303914 | 0.45005028 | 0.28298886 | 0.00259202 | 0.05338923 |
| Reep5 | 1.31833385 | 1.13790257 | 0.85538174 | 0.28252083 | 0.0710083 | 0.75390782 |
| Cd248 | 0.54146822 | 0.77663864 | 0.49445238 | 0.28218626 | 0.0055883 | 0.10359485 |
| Tubb2b | 0.50243266 | 0.74916697 | 0.46728605 | 0.28188092 | 0.00366221 | 0.0720171 |
| Lyz2 | 0.10814632 | 0.40543296 | 0.12370417 | 0.28172879 | 2.15E-24 | 4.24E-22 |
| Hnrnpa1 | 2.5162952 | 1.17756669 | 0.89629209 | 0.2812746 | 0.08148964 | 0.82790075 |
| Rps26-ps1 | 1.51653532 | 1.17709624 | 0.89610264 | 0.2809936 | 0.08165546 | 0.82804847 |
| Rph3al | 0.79695419 | 0.93157548 | 0.65059211 | 0.28098337 | 0.0274285 | 0.371274 |
| Sh3bgrl3 | 1.59294449 | 1.18594836 | 0.90715315 | 0.27879521 | 0.08592753 | 0.85989757 |
| Ncam1 | 0.82115788 | 0.94209854 | 0.66369286 | 0.27840568 | 0.0311018 | 0.40827162 |
| Nfia | 0.70221192 | 0.8750737 | 0.59668816 | 0.27838554 | 0.01903489 | 0.27875656 |
| Plxdc2 | 0.5065338 | 0.74824246 | 0.47018703 | 0.27805543 | 0.00428173 | 0.08228392 |
| Cldn3 | 0.46597159 | 0.71771219 | 0.44099137 | 0.27672081 | 0.00263862 | 0.05412906 |
| Grb10 | 1.85096135 | 1.20201111 | 0.92594459 | 0.27606652 | 0.0925128 | 0.90084241 |
| Nkx2-2 | 0.72319104 | 0.88452697 | 0.60899857 | 0.2755284 | 0.02214585 | 0.3131526 |
| Tyrobp | 0.0723122 | 0.35858298 | 0.08460354 | 0.27397944 | 2.72E-47 | 1.01E-44 |
| Tagln | 0.25029329 | 0.53779219 | 0.26436869 | 0.2734235 | 2.13E-06 | 8.63E-05 |
| Alcam | 0.89470149 | 0.9736728 | 0.7018943 | 0.2717785 | 0.0425918 | 0.52297082 |
| Tmed3 | 1.7264357 | 1.19163958 | 0.92006396 | 0.27157561 | 0.09472444 | 0.90650187 |
| Fam159b | 0.38362259 | 0.64844335 | 0.37806498 | 0.27037837 | 0.00073806 | 0.01836219 |

| | | | | | | |
|---|---|---|---|---|---|---|
| Marcksl1 | 2.41931081 | 1.17327268 | 0.90441824 | 0.26885444 | 0.0931614 | 0.90318385 |
| mt-Nd2 | 3.09297441 | 1.09095887 | 0.82509454 | 0.26586433 | 0.07600382 | 0.78862391 |
| Klf2 | 0.30033559 | 0.57472543 | 0.30890362 | 0.26582181 | 6.52E-05 | 0.00199916 |
| Cks1b | 0.94751688 | 0.99333671 | 0.72769127 | 0.26564544 | 0.05230761 | 0.61257938 |
| Timp3 | 0.41778988 | 0.66956386 | 0.40479444 | 0.26476942 | 0.00181991 | 0.03940815 |
| Igfbp3 | 0.30436048 | 0.57693548 | 0.31238401 | 0.26455147 | 8.33E-05 | 0.00249785 |
| Pdlim2 | 0.4815407 | 0.71679122 | 0.45233103 | 0.26446019 | 0.00467245 | 0.08889359 |
| Sytl4 | 0.44796585 | 0.69143258 | 0.42766258 | 0.26377 | 0.00305404 | 0.06165266 |
| Cldn6 | 0.64012696 | 0.8215343 | 0.55892367 | 0.26261063 | 0.01836417 | 0.27287764 |
| Ccnb2 | 0.24111777 | 0.51841051 | 0.25594129 | 0.26246922 | 2.57E-06 | 0.00010308 |
| Clec14a | 0.19258651 | 0.47218066 | 0.20994106 | 0.26223961 | 1.40E-08 | 7.73E-07 |
| H2afv | 1.68503052 | 1.1784478 | 0.91707305 | 0.26137474 | 0.1025703 | 0.90650187 |
| Dusp1 | 0.60924523 | 0.80050561 | 0.53935616 | 0.26114945 | 0.01567874 | 0.23953021 |
| Slit2 | 0.44964985 | 0.68967579 | 0.42891885 | 0.26075694 | 0.00343933 | 0.06861061 |
| Arl6ip1 | 1.34142953 | 1.12130389 | 0.86109102 | 0.26021287 | 0.08956855 | 0.88411139 |
| C1qc | 0.07639138 | 0.34766486 | 0.08914028 | 0.25852458 | 2.46E-38 | 7.48E-36 |
| Gpx1 | 1.22305896 | 1.0875194 | 0.82929582 | 0.25822358 | 0.08313954 | 0.83917943 |
| Fkbp10 | 0.57577709 | 0.77527248 | 0.51752918 | 0.25774329 | 0.01341335 | 0.21259066 |
| Mgst3 | 0.83783702 | 0.93021162 | 0.67257003 | 0.2576416 | 0.04428593 | 0.53941543 |
| Mef2c | 0.3316995 | 0.59287039 | 0.33564464 | 0.25722575 | 0.00032847 | 0.00882158 |
| Hepacam2 | 0.72498143 | 0.86723827 | 0.61003889 | 0.25719938 | 0.03044327 | 0.40263779 |
| Esam | 0.2001943 | 0.47432204 | 0.21731555 | 0.25700649 | 7.30E-08 | 3.69E-06 |
| Zfos1 | 1.22000934 | 1.08332343 | 0.82838781 | 0.25493561 | 0.08563548 | 0.85918035 |
| Mageh1 | 1.26124321 | 1.09511087 | 0.84036721 | 0.25474366 | 0.08889139 | 0.87871497 |
| Fcer1g | 0.07705123 | 0.34239121 | 0.0898719 | 0.25251932 | 4.17E-36 | 1.15E-33 |
| Sepw1 | 2.59706941 | 1.14018456 | 0.88857554 | 0.25160902 | 0.10404809 | 0.90650187 |
| Hic1 | 0.45054189 | 0.68042932 | 0.42958349 | 0.25084583 | 0.00471698 | 0.08962858 |
| Marcks | 1.79229207 | 1.17381717 | 0.92385926 | 0.24995791 | 0.11452759 | 0.90650187 |
| H2afx | 0.88132296 | 0.94466472 | 0.69512789 | 0.24953682 | 0.05525751 | 0.63634042 |
| Kcnj8 | 0.36434409 | 0.61208648 | 0.36257549 | 0.24951098 | 0.00110927 | 0.02594157 |

| | | | | | | |
|---|---|---|---|---|---|---|
| Serpinb6a | 0.71800896 | 0.85494397 | 0.60597871 | 0.24896526 | 0.0338898 | 0.43768805 |
| Icam2 | 0.15316586 | 0.41949632 | 0.17071111 | 0.24878521 | 4.62E-11 | 3.31E-09 |
| Ehd2 | 0.51122751 | 0.71974699 | 0.47349372 | 0.24625327 | 0.01038723 | 0.17284887 |
| Spc25 | 0.33497905 | 0.58439949 | 0.338391 | 0.24600848 | 0.00061478 | 0.01560147 |
| Tpi1 | 1.90480259 | 1.17227303 | 0.92703786 | 0.24523517 | 0.11979258 | 0.90650187 |
| Sat1 | 0.47186469 | 0.690402 | 0.44530383 | 0.24509817 | 0.00720472 | 0.12705213 |
| Rplp1 | 4.52678401 | 0.84630248 | 0.60245439 | 0.24384809 | 0.03597821 | 0.45842817 |
| Filip1l | 0.43644006 | 0.66270359 | 0.41900712 | 0.24369647 | 0.00486069 | 0.09161836 |
| Gars | 1.40373464 | 1.11882222 | 0.87513705 | 0.24368517 | 0.10787738 | 0.90650187 |
| Nr2f2 | 0.51310943 | 0.71825223 | 0.47481551 | 0.24343672 | 0.01132607 | 0.18532566 |
| Fxyd6 | 1.18889858 | 1.06245121 | 0.81904463 | 0.24340659 | 0.09322383 | 0.90318385 |
| Ece1 | 0.80204305 | 0.89580326 | 0.65336824 | 0.24243502 | 0.04951639 | 0.58758681 |
| Mrfap1 | 3.21433485 | 1.05282938 | 0.81069652 | 0.24213285 | 0.09212525 | 0.8999974 |
| Gstm1 | 0.52854152 | 0.72768809 | 0.48556526 | 0.24212282 | 0.01331967 | 0.21145496 |
| Ifi27l2a | 0.22018048 | 0.47622612 | 0.23639559 | 0.23983053 | 3.24E-06 | 0.0001259 |
| Tuba1c | 0.27152132 | 0.52280389 | 0.28355027 | 0.23925363 | 8.80E-05 | 0.0026289 |
| Ccnb1 | 0.19242505 | 0.44856873 | 0.20978388 | 0.23878486 | 2.09E-07 | 9.74E-06 |
| Ccnd1 | 0.44252985 | 0.66160624 | 0.42359286 | 0.23801338 | 0.00624569 | 0.11275623 |
| Sox4 | 0.92910694 | 0.95681683 | 0.71887892 | 0.23793791 | 0.07058776 | 0.7508779 |
| Spint1 | 0.73833063 | 0.85526624 | 0.61774725 | 0.23751899 | 0.04369761 | 0.53353204 |
| Ntrk2 | 0.35529998 | 0.59189532 | 0.35520386 | 0.23669146 | 0.00152652 | 0.0338024 |
| Btbd17 | 0.29142326 | 0.53706775 | 0.3011436 | 0.23592415 | 0.00024809 | 0.00678277 |
| Nenf | 2.01115436 | 1.16290817 | 0.9270068 | 0.23590136 | 0.12896401 | 0.90650187 |
| Ccna2 | 0.19205135 | 0.44503447 | 0.20941996 | 0.23561451 | 2.84E-07 | 1.30E-05 |
| Cdc20 | 0.22453395 | 0.47577277 | 0.24049667 | 0.2352761 | 6.83E-06 | 0.00024975 |
| Basp1 | 0.50563033 | 0.70306957 | 0.4695489 | 0.23352067 | 0.01352025 | 0.21363965 |
| Aplp2 | 1.07928209 | 1.01525435 | 0.7817413 | 0.23351305 | 0.09209864 | 0.8999974 |
| Prnp | 1.26117984 | 1.07297981 | 0.84034931 | 0.23263051 | 0.10922517 | 0.90650187 |
| Fbln5 | 0.24350689 | 0.49037535 | 0.25814376 | 0.23223159 | 3.18E-05 | 0.00102491 |
| Colec12 | 0.43705575 | 0.65155768 | 0.41947196 | 0.23208572 | 0.00695328 | 0.12347759 |

| | | | | | | |
|---|---|---|---|---|---|---|
| Aqp1 | 0.26677899 | 0.5111418 | 0.2793023 | 0.2318395 | 0.00011206 | 0.00326957 |
| Plat | 0.4307725 | 0.64631069 | 0.41471524 | 0.23159545 | 0.00652092 | 0.11661715 |
| Ly6e | 1.27350604 | 1.07466689 | 0.84377609 | 0.2308908 | 0.11189816 | 0.90650187 |
| Rps8 | 5.08411867 | 0.7763585 | 0.54817066 | 0.22818784 | 0.03211504 | 0.42043551 |
| Id1 | 0.24948829 | 0.49142447 | 0.26363277 | 0.22779171 | 6.12E-05 | 0.00189121 |
| Vimp | 1.24550966 | 1.06339739 | 0.83588306 | 0.22751433 | 0.11313471 | 0.90650187 |
| Rps4x | 4.64177548 | 0.81537024 | 0.5879031 | 0.22746714 | 0.04271065 | 0.52400696 |
| Stard10 | 0.7102419 | 0.82834125 | 0.60142637 | 0.22691488 | 0.04674163 | 0.56167548 |
| Aplnr | 0.11787777 | 0.36015718 | 0.13407313 | 0.22608405 | 3.28E-14 | 3.23E-12 |
| Lockd | 0.24795941 | 0.48787114 | 0.26223336 | 0.22563778 | 6.53E-05 | 0.00199916 |
| Ccl6 | 0.05270724 | 0.28793032 | 0.0624603 | 0.22547002 | 2.96E-58 | 1.45E-55 |
| Hes6 | 0.76143932 | 0.85594574 | 0.63088669 | 0.22505906 | 0.05638086 | 0.64487995 |
| Nupr1 | 0.31479091 | 0.54539631 | 0.32133657 | 0.22405975 | 0.00096823 | 0.02314468 |
| Rab3a | 0.74924172 | 0.84758762 | 0.62398254 | 0.22360507 | 0.05557004 | 0.63800617 |
| Gpm6a | 0.14552651 | 0.38643123 | 0.1629047 | 0.22352654 | 5.30E-10 | 3.43E-08 |
| Blvrb | 0.45358348 | 0.65482488 | 0.4318455 | 0.22297938 | 0.01085375 | 0.17894566 |
| Ubc | 2.92492328 | 1.0689415 | 0.84716142 | 0.22178008 | 0.12224764 | 0.90650187 |
| Ecscr | 0.19005182 | 0.42924415 | 0.20747024 | 0.22177391 | 1.01E-06 | 4.31E-05 |
| Fosb | 0.51713306 | 0.69830166 | 0.47763388 | 0.22066777 | 0.01999372 | 0.28862725 |
| Svbp | 0.7309935 | 0.83198544 | 0.61352103 | 0.21846442 | 0.05671017 | 0.64678604 |
| Grn | 0.74190351 | 0.83675663 | 0.61979559 | 0.21696104 | 0.05983076 | 0.6677587 |
| Gnao1 | 0.73967883 | 0.83437431 | 0.61852104 | 0.21585327 | 0.06039806 | 0.67260871 |
| Rgcc | 0.1781158 | 0.41153926 | 0.19574154 | 0.21579772 | 4.76E-07 | 2.12E-05 |
| Hspb1 | 0.24377953 | 0.47259209 | 0.25839474 | 0.21419735 | 0.00011426 | 0.00332637 |
| S100a1 | 0.55338198 | 0.71654337 | 0.502548 | 0.21399537 | 0.02917684 | 0.39007667 |
| Gm8730 | 4.32912695 | 0.84382595 | 0.6299561 | 0.21386985 | 0.06561342 | 0.7130802 |
| Pdia4 | 1.5162456 | 1.10908523 | 0.89605467 | 0.21303056 | 0.14527611 | 0.90650187 |
| Trf | 0.1505738 | 0.38056926 | 0.16807014 | 0.21249911 | 9.50E-09 | 5.35E-07 |
| Enpp2 | 0.55302337 | 0.71475119 | 0.50230561 | 0.21244558 | 0.03003846 | 0.39948788 |
| Cldn10 | 0.19025111 | 0.41984766 | 0.20766476 | 0.21218291 | 2.78E-06 | 0.00011048 |

| | | | | | | |
|---|---|---|---|---|---|---|
| Cdca8 | 0.22985939 | 0.45606262 | 0.24548633 | 0.2105763 | 6.85E-05 | 0.00208769 |
| Axl | 0.47565645 | 0.65829772 | 0.4480652 | 0.21023252 | 0.01849467 | 0.27348002 |
| Fkbp4 | 1.91575535 | 1.13732444 | 0.92712656 | 0.21019788 | 0.15674964 | 0.90650187 |
| Prc1 | 0.19646874 | 0.42340376 | 0.21371214 | 0.20969162 | 6.44E-06 | 0.00023663 |
| Pim2 | 0.51089695 | 0.68290247 | 0.47326131 | 0.20964117 | 0.02446071 | 0.33833232 |
| Bambi | 0.88631033 | 0.90703062 | 0.69766292 | 0.2093677 | 0.09108219 | 0.89440591 |
| Ranbp1 | 2.16039426 | 1.13167695 | 0.92237881 | 0.20929814 | 0.15654532 | 0.90650187 |
| Emcn | 0.1378566 | 0.36427066 | 0.15499803 | 0.20927263 | 9.73E-10 | 6.04E-08 |
| Thbd | 0.30042335 | 0.51800335 | 0.30897967 | 0.20902368 | 0.0013173 | 0.02979808 |
| Slc9a3r2 | 0.41745975 | 0.61351433 | 0.40454045 | 0.20897388 | 0.01082447 | 0.1786566 |
| Arg1 | 0.22669718 | 0.4512019 | 0.24252696 | 0.20867494 | 6.54E-05 | 0.00199916 |
| Lhfp | 0.42876893 | 0.62112175 | 0.41319239 | 0.20792936 | 0.01263837 | 0.20308233 |
| Rgs5 | 0.29065503 | 0.50833764 | 0.30047124 | 0.2078664 | 0.00105083 | 0.02488117 |
| Ctsz | 1.36321312 | 1.07379075 | 0.86619916 | 0.20759159 | 0.14334176 | 0.90650187 |
| Pgrmc1 | 1.63560176 | 1.11885129 | 0.9120452 | 0.2068061 | 0.15671751 | 0.90650187 |
| Galk1 | 0.5580306 | 0.71164749 | 0.50568304 | 0.20596445 | 0.03509417 | 0.45018271 |
| Tmsb15l | 0.41663959 | 0.60972082 | 0.40390908 | 0.20581174 | 0.01174845 | 0.19120241 |
| Gfra3 | 0.41321737 | 0.6070709 | 0.40126922 | 0.20580168 | 0.01130241 | 0.18513783 |
| 8430408G22 | 0.11883592 | 0.3408149 | 0.13508775 | 0.20572715 | 6.43E-12 | 5.09E-10 |
| Rbms3 | 0.74776851 | 0.82731494 | 0.623144 | 0.20417094 | 0.07261517 | 0.76707662 |
| Hmgn2 | 2.92400445 | 1.0513844 | 0.84729089 | 0.20409351 | 0.14211898 | 0.90650187 |
| Sox18 | 0.13941755 | 0.36035842 | 0.15661275 | 0.20374567 | 3.66E-09 | 2.11E-07 |
| Nefm | 0.28692102 | 0.49944349 | 0.29719555 | 0.20224794 | 0.00124181 | 0.02842881 |
| Phgdh | 0.38115701 | 0.57777445 | 0.37610072 | 0.20167373 | 0.00856664 | 0.14730942 |
| Celf3 | 0.56410933 | 0.71136775 | 0.50976282 | 0.20160493 | 0.03935821 | 0.49201005 |
| Ctsl | 1.82099184 | 1.12632185 | 0.92502558 | 0.20129627 | 0.16666821 | 0.90650187 |
| Eif4ebp1 | 0.72887166 | 0.81316139 | 0.61229405 | 0.20086734 | 0.07236348 | 0.76601483 |
| Abi3bp | 0.38562873 | 0.58050401 | 0.37965971 | 0.2008443 | 0.00934201 | 0.15778647 |
| mt-Nd4 | 4.18087853 | 0.85518147 | 0.65445771 | 0.20072376 | 0.08636593 | 0.86189704 |
| Plp2 | 0.89110212 | 0.90062913 | 0.70008508 | 0.20054405 | 0.10142501 | 0.90650187 |

| | | | | | | |
|---|---|---|---|---|---|---|
| Atp6v1g1 | 1.92615683 | 1.12771147 | 0.92720597 | 0.20050551 | 0.16818835 | 0.90650187 |
| Ddost | 2.16688408 | 1.12237286 | 0.92207319 | 0.20029968 | 0.16709514 | 0.90650187 |
| Spon1 | 0.28775988 | 0.49816018 | 0.29793255 | 0.20022764 | 0.00140542 | 0.0315101 |
| Smarca1 | 0.6646184 | 0.77417802 | 0.57406593 | 0.20011209 | 0.06060895 | 0.6744631 |
| Ndufa1 | 1.65639679 | 1.11386717 | 0.91428611 | 0.19958106 | 0.16591388 | 0.90650187 |
| Insm1 | 0.62711981 | 0.75020137 | 0.55074703 | 0.19945434 | 0.05370036 | 0.62276574 |
| Mmp23 | 0.39096886 | 0.58327235 | 0.38388939 | 0.19938296 | 0.01047341 | 0.17399602 |
| Lgals3 | 0.13096745 | 0.34708937 | 0.14783708 | 0.19925229 | 1.04E-09 | 6.42E-08 |
| Tspan13 | 0.71061619 | 0.80034897 | 0.60164645 | 0.19870251 | 0.07102138 | 0.75390782 |
| Smc4 | 0.52447359 | 0.68065594 | 0.48274682 | 0.19790912 | 0.03418644 | 0.44077021 |
| Plk2 | 0.3585022 | 0.55556982 | 0.35782153 | 0.19774829 | 0.00700821 | 0.12416293 |
| Vkorc1 | 1.16858029 | 1.00995472 | 0.81253289 | 0.19742183 | 0.14003706 | 0.90650187 |
| Lpar4 | 0.36552567 | 0.56089436 | 0.36353367 | 0.19736069 | 0.0079001 | 0.13755945 |
| Ndrg2 | 0.47875875 | 0.64680021 | 0.45031716 | 0.19648304 | 0.02620726 | 0.35922146 |
| Cdc42ep2 | 0.45566934 | 0.62971336 | 0.43339297 | 0.19632039 | 0.0220157 | 0.31160212 |
| Serpinb1a | 0.32789501 | 0.52857976 | 0.3324472 | 0.19613256 | 0.00436108 | 0.08359735 |
| Dynll2 | 1.1777953 | 1.0115019 | 0.81550383 | 0.19599807 | 0.14265735 | 0.90650187 |
| Glud1 | 1.32560175 | 1.05301559 | 0.85719469 | 0.19582091 | 0.15491669 | 0.90650187 |
| Rhoj | 0.48310412 | 0.64913055 | 0.45346049 | 0.19567005 | 0.02753666 | 0.37240641 |
| F13a1 | 0.06272922 | 0.26940631 | 0.07385062 | 0.19555569 | 2.72E-32 | 6.79E-30 |
| Aldoa | 2.27613199 | 1.11096192 | 0.91576784 | 0.19519407 | 0.17167368 | 0.90650187 |
| Creld2 | 0.84710671 | 0.87263143 | 0.67744823 | 0.1951832 | 0.10012368 | 0.90650187 |
| Pros1 | 0.43567285 | 0.61356459 | 0.4184275 | 0.19513709 | 0.01907475 | 0.27907146 |
| Tcf7l2 | 0.75789839 | 0.82390504 | 0.62888951 | 0.19501553 | 0.08401514 | 0.84633148 |
| Tmem100 | 0.19201189 | 0.40368012 | 0.20938152 | 0.1942986 | 1.85E-05 | 0.00063079 |
| Cygb | 0.33360069 | 0.53073537 | 0.33723785 | 0.19349753 | 0.00537437 | 0.09987259 |
| Ldhb | 0.45739234 | 0.62781767 | 0.43466893 | 0.19314874 | 0.02410901 | 0.33407568 |
| Fkbp2 | 1.31889361 | 1.04805269 | 0.85552029 | 0.1925324 | 0.15854131 | 0.90650187 |
| Atp2a2 | 0.9545975 | 0.92336663 | 0.73102413 | 0.19234251 | 0.12106082 | 0.90650187 |
| Smpd3 | 0.64781549 | 0.75563901 | 0.56371191 | 0.1919271 | 0.06506411 | 0.70962289 |

| | | | | | | |
|---|---|---|---|---|---|---|
| Rpl23 | 4.62605306 | 0.78161914 | 0.58975921 | 0.19185993 | 0.07405486 | 0.77635035 |
| Selenbp1 | 0.43019653 | 0.6060146 | 0.41427776 | 0.19173683 | 0.01981792 | 0.2866918 |
| Cxxc4 | 0.68516049 | 0.77820154 | 0.58651841 | 0.19168312 | 0.07312692 | 0.76980766 |
| Sox17 | 0.08958651 | 0.29506332 | 0.10365292 | 0.19141039 | 1.11E-16 | 1.29E-14 |
| Rarres1 | 0.10509726 | 0.3115394 | 0.12043119 | 0.19110821 | 8.66E-13 | 7.44E-11 |
| Tyms | 0.42372593 | 0.59975424 | 0.40934636 | 0.19040788 | 0.01932573 | 0.28165907 |
| Plod2 | 0.50663639 | 0.66064464 | 0.47025946 | 0.19038518 | 0.03594534 | 0.45839356 |
| Cck | 0.14647617 | 0.35419216 | 0.1638789 | 0.19031327 | 1.22E-07 | 5.84E-06 |
| Sfr1 | 2.10532049 | 1.11454904 | 0.92455995 | 0.18998909 | 0.18048095 | 0.90650187 |
| Atp6v1e1 | 1.36440555 | 1.05615803 | 0.86647362 | 0.18968441 | 0.16522406 | 0.90650187 |
| Adamts1 | 0.48089168 | 0.64144474 | 0.45186167 | 0.18958307 | 0.0310768 | 0.40827162 |
| Fxyd5 | 0.18296963 | 0.3898518 | 0.20052976 | 0.18932204 | 1.35E-05 | 0.00047447 |
| Cyr61 | 0.26412607 | 0.46480147 | 0.27691667 | 0.1878848 | 0.00127943 | 0.02915831 |
| Glipr2 | 0.39225799 | 0.57255432 | 0.38490713 | 0.18764719 | 0.01510636 | 0.23203229 |
| Smc2 | 0.34754037 | 0.53562499 | 0.34882578 | 0.18679921 | 0.00864082 | 0.14793535 |
| Rplp0 | 5.08711266 | 0.73519098 | 0.5485382 | 0.18665279 | 0.06517537 | 0.71020133 |
| Pkm | 2.39753334 | 1.0927369 | 0.90627352 | 0.18646338 | 0.18018199 | 0.90650187 |
| Rps2 | 4.74405855 | 0.76096074 | 0.57455214 | 0.1864086 | 0.0746033 | 0.77834237 |
| Loxl2 | 0.45614429 | 0.61920053 | 0.4337449 | 0.18545564 | 0.02866404 | 0.38525378 |
| Ndufv3 | 1.93056399 | 1.11247474 | 0.92723407 | 0.18524067 | 0.18723462 | 0.90650187 |
| Ednrb | 0.14458437 | 0.34690146 | 0.16193718 | 0.18496428 | 1.91E-07 | 8.93E-06 |
| Cyb5a | 1.14705357 | 0.9903504 | 0.8055892 | 0.18476119 | 0.15396005 | 0.90650187 |
| Gas1 | 0.42259576 | 0.59297241 | 0.40848186 | 0.18449055 | 0.02232935 | 0.31488391 |
| Plpp3 | 0.39423594 | 0.57032238 | 0.38646619 | 0.18385619 | 0.01721759 | 0.25913331 |
| Foxa2 | 0.56678443 | 0.69505479 | 0.51155119 | 0.1835036 | 0.05538659 | 0.63691881 |
| Fdps | 0.84721309 | 0.86084395 | 0.677504 | 0.18333995 | 0.11448247 | 0.90650187 |
| Etv1 | 0.30734825 | 0.49806608 | 0.31495826 | 0.18310782 | 0.00487531 | 0.09171982 |
| Eva1b | 1.38245107 | 1.05345005 | 0.8705341 | 0.18291595 | 0.17512426 | 0.90650187 |
| Gstz1 | 0.81134172 | 0.84110465 | 0.65841187 | 0.18269278 | 0.10868638 | 0.90650187 |
| 1110008P14I | 0.90158936 | 0.88790402 | 0.70534234 | 0.18256168 | 0.12493952 | 0.90650187 |

| | | | | | | |
|---|---|---|---|---|---|---|
| Ywhaq | 2.65918295 | 1.06395641 | 0.88225864 | 0.18169777 | 0.17995003 | 0.90650187 |
| Cela2a | 0.09518867 | 0.29070294 | 0.10974044 | 0.1809625 | 1.14E-13 | 1.11E-11 |
| S100a13 | 0.77462957 | 0.81919429 | 0.63827306 | 0.18092124 | 0.10381192 | 0.90650187 |
| Slbp | 0.95278874 | 0.91093591 | 0.73017455 | 0.18076136 | 0.1355462 | 0.90650187 |
| Kdm6b | 0.74136238 | 0.79995743 | 0.61948584 | 0.18047159 | 0.09764022 | 0.90650187 |
| Sez6l2 | 0.60476226 | 0.71685883 | 0.53647046 | 0.18038837 | 0.06747854 | 0.73016166 |
| Rab3b | 0.43275876 | 0.59640365 | 0.41622204 | 0.18018162 | 0.02714639 | 0.36811082 |
| Krtcap2 | 2.12697688 | 1.1038745 | 0.92372922 | 0.18014528 | 0.19297641 | 0.90650187 |
| Arap1 | 0.51325052 | 0.65492296 | 0.47491451 | 0.18000845 | 0.04599108 | 0.55397026 |
| Cldn11 | 0.36069819 | 0.53951948 | 0.35961181 | 0.17990766 | 0.01307217 | 0.20850997 |
| Sulf1 | 0.24290237 | 0.43719357 | 0.25758699 | 0.17960658 | 0.00096836 | 0.02314468 |
| Mcm6 | 0.46986446 | 0.62302958 | 0.44384318 | 0.1791864 | 0.03634594 | 0.46195203 |
| Ddc | 0.36091212 | 0.53778091 | 0.35978601 | 0.1779949 | 0.01392653 | 0.21862316 |
| Rps5 | 4.88183483 | 0.73376258 | 0.55580716 | 0.17795542 | 0.07731433 | 0.79732372 |
| Msln | 0.09713513 | 0.28966815 | 0.11184535 | 0.1778228 | 7.86E-13 | 6.82E-11 |
| Col15a1 | 0.28765165 | 0.47532601 | 0.2978375 | 0.17748852 | 0.00403361 | 0.07800875 |
| Aspn | 0.20206292 | 0.39638443 | 0.21911713 | 0.1772673 | 0.00016125 | 0.00461604 |
| Idh2 | 1.70469314 | 1.09569035 | 0.91854672 | 0.17714363 | 0.1956273 | 0.90650187 |
| Sox9 | 0.46380319 | 0.61651795 | 0.43939822 | 0.17711973 | 0.03656594 | 0.46397228 |
| Lmnb1 | 0.70368712 | 0.7744937 | 0.59756101 | 0.17693269 | 0.0940357 | 0.90650187 |
| Hmgn1 | 2.80837575 | 1.04102707 | 0.86416769 | 0.17685938 | 0.18145563 | 0.90650187 |
| Zim1 | 0.53547967 | 0.66712825 | 0.49034823 | 0.17678002 | 0.05449747 | 0.62997415 |
| Pi15 | 0.28605795 | 0.47281777 | 0.2964366 | 0.17638118 | 0.00408265 | 0.07885695 |
| Sgk1 | 0.23714038 | 0.42860692 | 0.25226214 | 0.17634478 | 0.00094269 | 0.0226738 |
| Rps12 | 3.65185469 | 0.92080684 | 0.74482888 | 0.17597795 | 0.14677933 | 0.90650187 |
| Scd2 | 0.98839506 | 0.92175106 | 0.74591637 | 0.17583469 | 0.14732811 | 0.90650187 |
| Papss2 | 0.52132077 | 0.6563749 | 0.48055517 | 0.17581973 | 0.05192115 | 0.6091644 |
| Ptgds | 0.18773521 | 0.38084743 | 0.20520595 | 0.17564147 | 7.09E-05 | 0.00214595 |
| Mrps33 | 1.78159253 | 1.09877571 | 0.92332993 | 0.17544578 | 0.1991313 | 0.90650187 |
| Frzb | 0.32668592 | 0.50682608 | 0.33142844 | 0.17539764 | 0.00931905 | 0.15774927 |

| | | | | | | |
|---|---|---|---|---|---|---|
| Cpxm1 | 0.3730332 | 0.54427091 | 0.3695947 | 0.17467621 | 0.01781773 | 0.26605824 |
| Dab2 | 0.31789434 | 0.4986343 | 0.32398185 | 0.17465244 | 0.00827531 | 0.14309708 |
| Gnb1 | 1.27104234 | 1.01747804 | 0.84310729 | 0.17437075 | 0.17893363 | 0.90650187 |
| Tspo | 0.54193783 | 0.66883601 | 0.49477328 | 0.17406273 | 0.0589114 | 0.66187153 |
| Igfbpl1 | 0.2437418 | 0.43239332 | 0.25836001 | 0.17403331 | 0.00137405 | 0.03094356 |
| Cd164 | 1.28356513 | 1.02050335 | 0.84648661 | 0.17401674 | 0.18038289 | 0.90650187 |
| Fcna | 0.05027624 | 0.23368776 | 0.05967475 | 0.174013 | 9.85E-39 | 3.06E-36 |
| Wls | 0.86640938 | 0.86078762 | 0.68748845 | 0.17329916 | 0.1312228 | 0.90650187 |
| Wfdc2 | 0.21535469 | 0.4046247 | 0.2318265 | 0.1727982 | 0.00046043 | 0.01206716 |
| Serinc3 | 1.04348196 | 0.94085153 | 0.76818319 | 0.17266834 | 0.15883415 | 0.90650187 |
| Pdgfra | 0.31541963 | 0.49427466 | 0.32187315 | 0.17240152 | 0.00862927 | 0.14793535 |
| Prps1 | 0.76033772 | 0.80128829 | 0.63026598 | 0.17102231 | 0.11384847 | 0.90650187 |
| Pdia3 | 2.8754312 | 1.02519496 | 0.85432966 | 0.1708653 | 0.18696984 | 0.90650187 |
| Ssr3 | 1.54360329 | 1.07020437 | 0.90048519 | 0.16971918 | 0.20104714 | 0.90650187 |
| Lrp1 | 0.41555177 | 0.57260128 | 0.4030709 | 0.16953038 | 0.03075521 | 0.40582465 |
| Pcp4 | 0.3278261 | 0.5016975 | 0.33238917 | 0.16930833 | 0.01177327 | 0.19120241 |
| Tpst1 | 0.7301421 | 0.78229 | 0.61302895 | 0.16926105 | 0.10982504 | 0.90650187 |
| Lsr | 0.78318792 | 0.81224879 | 0.64302283 | 0.16922596 | 0.12100821 | 0.90650187 |
| Hap1 | 0.53324717 | 0.65794401 | 0.48881251 | 0.1691315 | 0.0619997 | 0.6859224 |
| Thra | 0.91683392 | 0.88170715 | 0.71289802 | 0.16880913 | 0.14624209 | 0.90650187 |
| Dpysl2 | 1.00855679 | 0.92303057 | 0.75423023 | 0.16880035 | 0.15988077 | 0.90650187 |
| Rpl22l1 | 3.87939428 | 0.87285818 | 0.70421706 | 0.16864111 | 0.14352768 | 0.90650187 |
| Ccdc34 | 0.71263861 | 0.77103793 | 0.60283444 | 0.16820349 | 0.10741047 | 0.90650187 |
| Rhob | 0.82577474 | 0.83380355 | 0.66616183 | 0.16764172 | 0.13162323 | 0.90650187 |
| Nkx2-3 | 0.29845098 | 0.47467544 | 0.30726899 | 0.16740645 | 0.00771645 | 0.13482509 |
| Mki67 | 0.21822291 | 0.40169448 | 0.23454503 | 0.16714946 | 0.00076698 | 0.01889617 |
| Bloc1s1 | 0.64015806 | 0.72593735 | 0.5589431 | 0.16699424 | 0.0920553 | 0.8999974 |
| Lamb1 | 0.77565486 | 0.80558314 | 0.63884382 | 0.16673932 | 0.12296114 | 0.90650187 |
| Nsg1 | 0.35141329 | 0.51845204 | 0.35201536 | 0.16643668 | 0.01777967 | 0.26601254 |
| Rpl23a-ps3 | 3.15254667 | 0.98450793 | 0.81813631 | 0.16637162 | 0.18298878 | 0.90650187 |

| | | | | | | |
|---|---|---|---|---|---|---|
| Gpihbp1 | 0.1013862 | 0.28267825 | 0.11643188 | 0.16624638 | 1.09E-10 | 7.55E-09 |
| Mien1 | 1.29153725 | 1.01461849 | 0.84861195 | 0.16600654 | 0.19224445 | 0.90650187 |
| Cd47 | 1.37206079 | 1.03296142 | 0.86821634 | 0.16474508 | 0.19945792 | 0.90650187 |
| Ffar4 | 0.5844132 | 0.68783304 | 0.52322486 | 0.16460818 | 0.08096623 | 0.82601856 |
| Tcf21 | 0.30180905 | 0.47442911 | 0.31017942 | 0.16424968 | 0.00928413 | 0.15744466 |
| Col6a6 | 0.29266072 | 0.46646062 | 0.30222551 | 0.1642351 | 0.00784975 | 0.13683959 |
| Taldo1 | 1.57844846 | 1.06964639 | 0.90544734 | 0.16419905 | 0.2100661 | 0.90650187 |
| Krtdap | 0.13480099 | 0.31554899 | 0.15182882 | 0.16372017 | 8.18E-07 | 3.55E-05 |
| Cela3b | 0.101235 | 0.27994434 | 0.11626857 | 0.16367577 | 1.95E-10 | 1.31E-08 |
| Ubl5 | 2.39011078 | 1.06998036 | 0.90689637 | 0.16308399 | 0.2120183 | 0.90650187 |
| Slc39a1 | 0.49673442 | 0.62607978 | 0.46323703 | 0.16284276 | 0.05905288 | 0.66199323 |
| Sema3c | 0.27578497 | 0.44972027 | 0.28735033 | 0.16236993 | 0.00600194 | 0.10913328 |
| Trip6 | 0.45536096 | 0.59545514 | 0.43316438 | 0.16229076 | 0.04789604 | 0.57238027 |
| Ptbp3 | 1.02417603 | 0.92247877 | 0.76054379 | 0.16193498 | 0.17193191 | 0.90650187 |
| Acaa2 | 0.53180384 | 0.64972334 | 0.48781799 | 0.16190535 | 0.07004128 | 0.74657023 |
| Usp9x | 0.91662075 | 0.87467761 | 0.71279363 | 0.16188398 | 0.15633138 | 0.90650187 |
| Lyve1 | 0.12070613 | 0.29886636 | 0.13706497 | 0.16180139 | 7.69E-08 | 3.85E-06 |
| Fam162a | 0.8270002 | 0.82842926 | 0.66681566 | 0.16161359 | 0.14063633 | 0.90650187 |
| Ssr4 | 2.19854227 | 1.0820573 | 0.9204956 | 0.1615617 | 0.2176145 | 0.90650187 |
| Rpl12 | 3.12798545 | 0.98246777 | 0.82102974 | 0.16143803 | 0.19102267 | 0.90650187 |
| Lama4 | 0.40772479 | 0.55841649 | 0.39701394 | 0.16140255 | 0.03535601 | 0.4531591 |
| F2r | 0.40124795 | 0.55294849 | 0.39196682 | 0.16098167 | 0.03393863 | 0.43794666 |
| Entpd2 | 0.30191125 | 0.47117879 | 0.31026784 | 0.16091095 | 0.01056711 | 0.17497889 |
| Gstm5 | 0.75522527 | 0.78797803 | 0.62737798 | 0.16060005 | 0.12755743 | 0.90650187 |
| Aldh1a2 | 0.10543845 | 0.28130859 | 0.12079802 | 0.16051057 | 1.74E-09 | 1.03E-07 |
| Arhgap31 | 0.30391913 | 0.47216254 | 0.31200307 | 0.16015947 | 0.01124325 | 0.18456665 |
| Cmtm7 | 0.35448085 | 0.51460965 | 0.35453291 | 0.16007674 | 0.02236122 | 0.31488391 |
| Slc6a6 | 0.72734897 | 0.77148037 | 0.61141222 | 0.16006815 | 0.12223979 | 0.90650187 |
| Mia | 0.28227947 | 0.45299509 | 0.29310593 | 0.15988916 | 0.00765258 | 0.13407494 |
| Ctsb | 1.54871409 | 1.06056027 | 0.9012907 | 0.15926958 | 0.21605227 | 0.90650187 |

| | | | | | | |
|---|---|---|---|---|---|---|
| Alas2 | 0.03284707 | 0.19867722 | 0.03943965 | 0.15923757 | 2.44E-72 | 1.42E-69 |
| Sox11 | 0.26828827 | 0.43983484 | 0.28065654 | 0.1591783 | 0.00584464 | 0.1065174 |
| Tmed10 | 2.33386388 | 1.07032691 | 0.91148523 | 0.15884168 | 0.21925199 | 0.90650187 |
| Pid1 | 0.28823418 | 0.45672928 | 0.29834898 | 0.15838031 | 0.00913805 | 0.15555157 |
| Slc24a5 | 0.59705852 | 0.68951535 | 0.53148489 | 0.15803046 | 0.09310971 | 0.90318385 |
| Dmkn | 0.08589902 | 0.25759191 | 0.09962203 | 0.15796987 | 8.99E-13 | 7.67E-11 |
| Rnase4 | 0.97797899 | 0.89907475 | 0.7414588 | 0.15761595 | 0.17232258 | 0.90650187 |
| Pebp1 | 2.89683887 | 1.00893286 | 0.85133576 | 0.15759711 | 0.20526556 | 0.90650187 |
| Ankrd44 | 0.52869841 | 0.64325998 | 0.48567375 | 0.15758623 | 0.07458592 | 0.77834237 |
| Klf4 | 0.32657456 | 0.48847273 | 0.33133455 | 0.15713818 | 0.01749967 | 0.26285818 |
| Foxp1 | 0.89017186 | 0.85668664 | 0.69961666 | 0.15706998 | 0.15912101 | 0.90650187 |
| Tgfb1i1 | 0.42113417 | 0.56403288 | 0.40736246 | 0.15667041 | 0.04365446 | 0.53343368 |
| Lxn | 0.39680714 | 0.54515568 | 0.38848779 | 0.15666788 | 0.03650187 | 0.46354636 |
| Slc43a3 | 0.36231453 | 0.51724241 | 0.36092703 | 0.15631538 | 0.02709179 | 0.36769854 |
| Cpa2 | 0.08970007 | 0.25990653 | 0.10377676 | 0.15612978 | 1.13E-11 | 8.58E-10 |
| Mapre3 | 0.63160951 | 0.7093712 | 0.55357979 | 0.15579141 | 0.10544781 | 0.90650187 |
| Rnf130 | 0.71449783 | 0.75958413 | 0.6039248 | 0.15565933 | 0.12593105 | 0.90650187 |
| Prrg2 | 0.65167478 | 0.72168614 | 0.56610323 | 0.15558291 | 0.11089315 | 0.90650187 |
| Col5a3 | 0.22821551 | 0.39930873 | 0.24394915 | 0.15535957 | 0.00231873 | 0.04848274 |
| Tfpi | 0.46723748 | 0.59708944 | 0.44191994 | 0.1551695 | 0.05926575 | 0.6638899 |
| Dnajb9 | 0.7076624 | 0.75505356 | 0.59990776 | 0.1551458 | 0.12513263 | 0.90650187 |
| Fap | 0.3215387 | 0.48167491 | 0.32707742 | 0.15459749 | 0.01780842 | 0.26605824 |
| Dtymk | 0.84824772 | 0.83246601 | 0.67804612 | 0.15441989 | 0.15566068 | 0.90650187 |
| Pdpn | 0.21122028 | 0.38215202 | 0.22789295 | 0.15425907 | 0.00130969 | 0.02971432 |
| Fam174b | 0.65031021 | 0.71950946 | 0.56525861 | 0.15425085 | 0.1125379 | 0.90650187 |
| Fn1 | 0.31930819 | 0.47915637 | 0.3251842 | 0.15397217 | 0.01764741 | 0.26481566 |
| Cav2 | 0.16456838 | 0.33578898 | 0.18223761 | 0.15355138 | 8.99E-05 | 0.00267965 |
| Cotl1 | 1.17040435 | 0.96630453 | 0.81312332 | 0.15318121 | 0.20115835 | 0.90650187 |
| Tmem119 | 0.31586111 | 0.47526096 | 0.32224972 | 0.15301124 | 0.01739168 | 0.26149455 |
| Efemp1 | 0.17728381 | 0.34788041 | 0.1949182 | 0.15296221 | 0.00024269 | 0.00665919 |

| | | | | | | |
|---|---|---|---|---|---|---|
| Ass1 | 0.28079822 | 0.44461491 | 0.29179664 | 0.15281827 | 0.00994993 | 0.1665737 |
| Tmem176a | 0.97844589 | 0.89410669 | 0.74165968 | 0.15244701 | 0.18041424 | 0.90650187 |
| Cbx3 | 1.97416294 | 1.079467 | 0.9273178 | 0.1521492 | 0.23287526 | 0.90650187 |
| Slc2a2 | 0.23617747 | 0.40348531 | 0.25136907 | 0.15211624 | 0.00356979 | 0.07065667 |
| Ubl3 | 0.79317026 | 0.8003695 | 0.64852051 | 0.15184899 | 0.14895531 | 0.90650187 |
| Fabp7 | 0.06472588 | 0.22785088 | 0.07610211 | 0.15174877 | 3.84E-19 | 5.21E-17 |
| Lman1 | 1.38374102 | 1.02244315 | 0.87081964 | 0.1516235 | 0.21945046 | 0.90650187 |
| Nucb2 | 0.737561 | 0.7683773 | 0.61730513 | 0.15107217 | 0.13830492 | 0.90650187 |
| Rbms1 | 0.8475749 | 0.82810307 | 0.67769363 | 0.15040944 | 0.16190179 | 0.90650187 |
| Celf2 | 0.85265907 | 0.8306623 | 0.68035261 | 0.15030969 | 0.16300957 | 0.90650187 |
| Psap | 1.9395317 | 1.07652446 | 0.92728116 | 0.1492433 | 0.23714764 | 0.90650187 |
| Cadm1 | 0.28674652 | 0.4457183 | 0.29704215 | 0.14867615 | 0.01303646 | 0.20837769 |
| Gjd2 | 0.45374586 | 0.5801759 | 0.43196608 | 0.14820982 | 0.0635908 | 0.698442 |
| PISD | 0.84952291 | 0.82633124 | 0.67871368 | 0.14761757 | 0.16679652 | 0.90650187 |
| Cd200 | 0.56185551 | 0.65580389 | 0.50825275 | 0.14755114 | 0.09842008 | 0.90650187 |
| Plpp1 | 0.41681083 | 0.55138267 | 0.40404094 | 0.14734172 | 0.05248947 | 0.61423585 |
| Lrp11 | 0.61951651 | 0.69284611 | 0.54592445 | 0.14692165 | 0.11576603 | 0.90650187 |
| Socs3 | 0.26444203 | 0.4239714 | 0.27720114 | 0.14677026 | 0.0092907 | 0.15744466 |
| Fkbp9 | 1.13545502 | 0.94822612 | 0.80167278 | 0.14655334 | 0.2081954 | 0.90650187 |
| P3h4 | 0.69371446 | 0.73774314 | 0.59163925 | 0.14610389 | 0.13613897 | 0.90650187 |
| Nme1 | 2.01697647 | 1.07304816 | 0.92694595 | 0.14610221 | 0.2417445 | 0.90650187 |
| Fndc3a | 1.04206017 | 0.91339619 | 0.76763201 | 0.14576418 | 0.19928784 | 0.90650187 |
| Flt1 | 0.12635104 | 0.28821736 | 0.14300685 | 0.14521052 | 3.18E-06 | 0.00012391 |
| Gpm6b | 0.28785812 | 0.4431195 | 0.29801882 | 0.14510068 | 0.01521316 | 0.2333554 |
| Nfkbia | 0.70099061 | 0.74082958 | 0.59596471 | 0.14486487 | 0.13993279 | 0.90650187 |
| Alox5ap | 0.10473403 | 0.264904 | 0.12004051 | 0.14486349 | 4.05E-08 | 2.14E-06 |
| Uqcrq | 2.14800003 | 1.06761359 | 0.92289029 | 0.1447233 | 0.24285791 | 0.90650187 |
| Hspg2 | 0.37654172 | 0.51711548 | 0.37241095 | 0.14470453 | 0.04204872 | 0.51797615 |
| S100b | 0.19800619 | 0.35917051 | 0.21520114 | 0.14396937 | 0.00146921 | 0.03265137 |
| Add3 | 0.53231062 | 0.63167791 | 0.48816733 | 0.14351059 | 0.09562135 | 0.90650187 |

| | | | | | | |
|---|---|---|---|---|---|---|
| Pttg1 | 0.30320421 | 0.4548835 | 0.31138564 | 0.14349786 | 0.02024585 | 0.29116103 |
| Anxa1 | 0.24243928 | 0.40049422 | 0.25716024 | 0.14333398 | 0.0066087 | 0.11804792 |
| Bicc1 | 0.60809356 | 0.68193237 | 0.53861591 | 0.14331646 | 0.11842445 | 0.90650187 |
| Rac2 | 0.04506818 | 0.19691068 | 0.05367702 | 0.14323366 | 9.23E-33 | 2.38E-30 |
| Trim47 | 0.41316947 | 0.54389345 | 0.4012322 | 0.14266125 | 0.05697674 | 0.64823856 |
| Dpy30 | 1.24153596 | 0.97719711 | 0.83473131 | 0.1424658 | 0.22400303 | 0.90650187 |
| Ctla2a | 0.10778858 | 0.2654107 | 0.12332075 | 0.14208995 | 1.51E-07 | 7.17E-06 |
| Srm | 0.84009291 | 0.81577024 | 0.67376121 | 0.14200903 | 0.17437853 | 0.90650187 |
| Cd302 | 0.30969394 | 0.45889412 | 0.31697376 | 0.14192037 | 0.0232704 | 0.32452608 |
| Epha3 | 0.36474407 | 0.50475608 | 0.36289998 | 0.14185611 | 0.04112574 | 0.50784112 |
| Rpl35 | 4.5960936 | 0.73501712 | 0.59357138 | 0.14144574 | 0.14471551 | 0.90650187 |
| Akr1c13 | 0.48407463 | 0.59559604 | 0.45416079 | 0.14143525 | 0.08310951 | 0.83917943 |
| Vcam1 | 0.32567566 | 0.47196959 | 0.33057626 | 0.14139333 | 0.02862058 | 0.38501019 |
| Gas7 | 0.18615811 | 0.34487937 | 0.20366116 | 0.14121821 | 0.00102609 | 0.02437133 |
| Hmgb3 | 0.53667124 | 0.63236273 | 0.49116663 | 0.1411961 | 0.10062783 | 0.90650187 |
| Gm9844 | 1.01648048 | 0.89854523 | 0.75745269 | 0.14109254 | 0.20380869 | 0.90650187 |
| Sulf2 | 0.47330435 | 0.58725564 | 0.44635343 | 0.14090221 | 0.08025436 | 0.82093036 |
| Kdr | 0.13523504 | 0.29307275 | 0.15227968 | 0.14079308 | 1.98E-05 | 0.00066453 |
| Phip | 1.17991912 | 0.95657677 | 0.81618439 | 0.14039239 | 0.22222763 | 0.90650187 |
| Myl12b | 1.39569667 | 1.01378611 | 0.87343729 | 0.14034882 | 0.23750692 | 0.90650187 |
| Rnaset2a | 0.81298901 | 0.79963827 | 0.65930147 | 0.1403368 | 0.17200353 | 0.90650187 |
| Spc24 | 0.30074055 | 0.44958852 | 0.30925445 | 0.14033406 | 0.02183025 | 0.31013243 |
| Atf3 | 0.2342737 | 0.38982449 | 0.24960067 | 0.14022382 | 0.00625347 | 0.1127627 |
| Ero1lb | 0.4525505 | 0.57101162 | 0.43107801 | 0.13993361 | 0.07449581 | 0.77834237 |
| Cdca3 | 0.18854373 | 0.34583347 | 0.20599687 | 0.1398366 | 0.00127306 | 0.02905676 |
| Lmo2 | 0.1278019 | 0.28426652 | 0.14452767 | 0.13973885 | 8.61E-06 | 0.00031171 |
| Gck | 0.58164654 | 0.66087455 | 0.52140496 | 0.13946959 | 0.11719101 | 0.90650187 |
| Fhl1 | 0.25556341 | 0.40833517 | 0.26917101 | 0.13916417 | 0.01076883 | 0.17793155 |
| Ldb2 | 0.33660482 | 0.47884549 | 0.33974908 | 0.1390964 | 0.03437401 | 0.44281295 |
| Aprt | 1.27890278 | 0.9842953 | 0.84523431 | 0.13906098 | 0.23226526 | 0.90650187 |

| | | | | | | |
|---|---|---|---|---|---|---|
| Dkk2 | 0.196686 | 0.35204094 | 0.21392271 | 0.13811823 | 0.00205051 | 0.04365509 |
| Ostc | 2.23330123 | 1.05647703 | 0.9185283 | 0.13794873 | 0.25217493 | 0.90650187 |
| Klf6 | 0.57908071 | 0.6574851 | 0.51971316 | 0.13777194 | 0.11929949 | 0.90650187 |
| C1s1 | 0.24522047 | 0.39738498 | 0.25972001 | 0.13766497 | 0.00922651 | 0.1567064 |
| Ehd4 | 0.24080751 | 0.39328627 | 0.25565486 | 0.13763141 | 0.00834912 | 0.14405785 |
| Rpl9 | 4.53227465 | 0.73939952 | 0.60179249 | 0.13760704 | 0.15468592 | 0.90650187 |
| Lst1 | 0.10473419 | 0.25764235 | 0.12004068 | 0.13760167 | 1.73E-07 | 8.16E-06 |
| Gmfg | 0.10122156 | 0.25365387 | 0.11625405 | 0.13739982 | 7.43E-08 | 3.74E-06 |
| Pdcd4 | 1.01099653 | 0.89256852 | 0.75522523 | 0.13734329 | 0.20941108 | 0.90650187 |
| Anp32b | 1.96163865 | 1.06419412 | 0.92737142 | 0.1368227 | 0.25594529 | 0.90650187 |
| Uchl1 | 0.39139027 | 0.52100803 | 0.38422223 | 0.13678581 | 0.05674869 | 0.64678604 |
| Rgs17 | 0.47902626 | 0.58722885 | 0.45051104 | 0.1367178 | 0.08864919 | 0.87809516 |
| Tsc22d3 | 0.39325702 | 0.52220424 | 0.38569497 | 0.13650927 | 0.05780891 | 0.65286276 |
| Mapt | 0.28573351 | 0.43257687 | 0.29615112 | 0.13642575 | 0.02028369 | 0.29142951 |
| Spink4 | 0.1806732 | 0.33468531 | 0.19826758 | 0.13641772 | 0.00111115 | 0.02594569 |
| Cycs | 1.64646211 | 1.04959111 | 0.91321481 | 0.13637631 | 0.25337994 | 0.90650187 |
| Col12a1 | 0.35623272 | 0.49181495 | 0.35596719 | 0.13584776 | 0.04488835 | 0.54587824 |
| Plp1 | 0.1127645 | 0.26390052 | 0.12863946 | 0.13526106 | 1.47E-06 | 6.17E-05 |
| Ccl9 | 0.03785539 | 0.18051287 | 0.04530216 | 0.13521071 | 1.76E-40 | 5.70E-38 |
| Higd1a | 1.11867186 | 0.93085268 | 0.79581079 | 0.13504189 | 0.2253108 | 0.90650187 |
| Siva1 | 0.99627738 | 0.88422115 | 0.74918621 | 0.13503494 | 0.21148313 | 0.90650187 |
| Mylk | 0.17537531 | 0.32743604 | 0.19302666 | 0.13440938 | 0.00098211 | 0.02343655 |
| Rasgrp2 | 0.21711199 | 0.36789205 | 0.23349309 | 0.13439896 | 0.00525023 | 0.09804523 |
| Csf1r | 0.04845593 | 0.1916125 | 0.05758312 | 0.13402939 | 2.15E-25 | 4.35E-23 |
| Chd7 | 0.62057178 | 0.68011951 | 0.54659569 | 0.13352381 | 0.13874299 | 0.90650187 |
| Pgf | 0.39364578 | 0.51917424 | 0.38600133 | 0.13317292 | 0.06254405 | 0.68993619 |
| Ptprn2 | 0.66416017 | 0.70687328 | 0.57378554 | 0.13308774 | 0.15123657 | 0.90650187 |
| Fam46a | 0.39963742 | 0.52334217 | 0.39070684 | 0.13263533 | 0.06562721 | 0.7130802 |
| Rab11a | 1.48978805 | 1.02349953 | 0.89158516 | 0.13191437 | 0.25534967 | 0.90650187 |
| Anxa3 | 0.28513246 | 0.42744339 | 0.29562201 | 0.13182138 | 0.02372011 | 0.32958806 |

| | | | | | | |
|---|---|---|---|---|---|---|
| P3h1 | 0.52457015 | 0.61452251 | 0.48281384 | 0.13170867 | 0.11261572 | 0.90650187 |
| Lig1 | 0.34134207 | 0.47471759 | 0.34369362 | 0.13102396 | 0.04505972 | 0.54752423 |
| Ndufa12 | 1.87534362 | 1.05701942 | 0.92657588 | 0.13044354 | 0.26570557 | 0.90650187 |
| Srgn | 0.0747871 | 0.21759822 | 0.08735891 | 0.13023932 | 1.70E-11 | 1.28E-09 |
| Ptpn18 | 0.14684286 | 0.29428795 | 0.16425478 | 0.13003317 | 0.00021627 | 0.00601014 |
| Rps11 | 4.47881597 | 0.73944417 | 0.60950719 | 0.12993698 | 0.17163329 | 0.90650187 |
| Fam111a | 0.2631319 | 0.40588767 | 0.27602095 | 0.12986673 | 0.01823538 | 0.27122902 |
| Fgfbp1 | 0.06107434 | 0.20146817 | 0.07198007 | 0.1294881 | 6.35E-16 | 7.00E-14 |
| Ets1 | 0.1969102 | 0.34351482 | 0.21413994 | 0.12937488 | 0.00361718 | 0.0712237 |
| Rpl22 | 2.8890383 | 0.98183466 | 0.85249368 | 0.12934099 | 0.24999968 | 0.90650187 |
| Sdc1 | 0.43803088 | 0.54913954 | 0.42020762 | 0.12893191 | 0.08627729 | 0.86189704 |
| Vtn | 0.10922189 | 0.25341325 | 0.12485595 | 0.1285573 | 2.35E-06 | 9.47E-05 |
| Psip1 | 1.08542656 | 0.91245899 | 0.78400019 | 0.1284588 | 0.23317989 | 0.90650187 |
| Tspan4 | 0.42330552 | 0.53740429 | 0.40902489 | 0.12837941 | 0.08145832 | 0.82790075 |
| Sgce | 0.5581538 | 0.63379693 | 0.50576594 | 0.12803098 | 0.1302155 | 0.90650187 |
| Rps18-ps3 | 1.57720096 | 1.0330445 | 0.9052981 | 0.1277464 | 0.26522525 | 0.90650187 |
| Ptma | 4.785946 | 0.69613516 | 0.56846443 | 0.12767073 | 0.15903464 | 0.90650187 |
| Phactr2 | 0.38723721 | 0.50856904 | 0.38093605 | 0.127633 | 0.06817725 | 0.73292957 |
| Pls3 | 0.49066947 | 0.58647875 | 0.45890272 | 0.12757603 | 0.10824993 | 0.90650187 |
| Agl | 0.30500671 | 0.44043912 | 0.31294147 | 0.12749765 | 0.03505377 | 0.45016727 |
| Ffar1 | 0.26793816 | 0.40774687 | 0.28034258 | 0.12740428 | 0.0216737 | 0.30848499 |
| Myadm | 0.75461358 | 0.75434735 | 0.62703163 | 0.12731572 | 0.18335353 | 0.90650187 |
| Amotl2 | 0.35304175 | 0.4806259 | 0.3533528 | 0.12727309 | 0.05466082 | 0.63138125 |
| Phlda1 | 0.45107939 | 0.55716189 | 0.4299837 | 0.12717819 | 0.09427231 | 0.90650187 |
| Gja1 | 0.22708202 | 0.37000771 | 0.24288765 | 0.12712006 | 0.00999072 | 0.1670726 |
| Zeb1 | 0.3839858 | 0.50541717 | 0.37835394 | 0.12706323 | 0.06772182 | 0.73113595 |
| Ran | 2.39709423 | 1.03328286 | 0.90631051 | 0.12697235 | 0.26670112 | 0.90650187 |
| Rpsa | 4.12620759 | 0.79041554 | 0.66363507 | 0.12678047 | 0.19786105 | 0.90650187 |
| Pdap1 | 2.38222099 | 1.03400566 | 0.90756386 | 0.1264418 | 0.26783899 | 0.90650187 |
| Tgfb1 | 0.16677394 | 0.31068321 | 0.18445018 | 0.12623304 | 0.00117326 | 0.02702237 |

| | | | | | | |
|---|---|---|---|---|---|---|
| Ctss | 0.03948861 | 0.17291379 | 0.04720553 | 0.12570826 | 1.23E-32 | 3.12E-30 |
| Myh9 | 0.542492 | 0.62081773 | 0.49515178 | 0.12566595 | 0.12960446 | 0.90650187 |
| H2-K1 | 0.53136764 | 0.61315053 | 0.48751717 | 0.12563337 | 0.1259732 | 0.90650187 |
| Sep-07 | 1.95282085 | 1.05280178 | 0.92734504 | 0.12545673 | 0.27377749 | 0.90650187 |
| Lurap1l | 0.37821945 | 0.49898829 | 0.37375418 | 0.12523411 | 0.06816567 | 0.73292957 |
| Car2 | 0.06861603 | 0.20567595 | 0.08047199 | 0.12520396 | 2.31E-12 | 1.88E-10 |
| Pdgfrb | 0.26302503 | 0.40100288 | 0.2759246 | 0.12507828 | 0.02194079 | 0.31095399 |
| Nfix | 0.54597422 | 0.62255794 | 0.49752495 | 0.12503299 | 0.131949 | 0.90650187 |
| Prkar1a | 2.22404245 | 1.04401145 | 0.91908219 | 0.12492925 | 0.27282756 | 0.90650187 |
| Madcam1 | 0.08167428 | 0.21951691 | 0.09498029 | 0.12453661 | 2.79E-09 | 1.62E-07 |
| Wt1 | 0.18539497 | 0.32713741 | 0.20291269 | 0.12422472 | 0.003248 | 0.06504981 |
| Rpl18a | 4.95565215 | 0.6711726 | 0.54724694 | 0.12392566 | 0.15703456 | 0.90650187 |
| Rogdi | 0.7240083 | 0.73281313 | 0.60947364 | 0.12333949 | 0.18415071 | 0.90650187 |
| Tk1 | 0.18643892 | 0.32686718 | 0.20393642 | 0.12293077 | 0.00368369 | 0.07234588 |
| Ptger3 | 0.23605646 | 0.37408725 | 0.25125678 | 0.12283047 | 0.01487894 | 0.2300154 |
| Tgm2 | 0.24692506 | 0.38393732 | 0.26128514 | 0.12265218 | 0.01845078 | 0.27348002 |
| Hcfc1r1 | 1.24581256 | 0.95847525 | 0.83597015 | 0.12250511 | 0.25737157 | 0.90650187 |
| Dhrs7 | 0.51581294 | 0.59914138 | 0.47671034 | 0.12243104 | 0.12678065 | 0.90650187 |
| Cela1 | 0.07886815 | 0.21414874 | 0.0918832 | 0.12226553 | 1.65E-09 | 9.86E-08 |
| Tmem256 | 1.85562203 | 1.04820435 | 0.92607569 | 0.12212867 | 0.27884502 | 0.90650187 |
| Msmo1 | 0.49346464 | 0.58298131 | 0.46090378 | 0.12207753 | 0.1195004 | 0.90650187 |
| Alad | 0.72533875 | 0.73205656 | 0.61024634 | 0.12181022 | 0.18743721 | 0.90650187 |
| Pcdh17 | 0.11021477 | 0.24765853 | 0.12591792 | 0.12174062 | 8.61E-06 | 0.00031171 |
| Tpx2 | 0.15698581 | 0.29613927 | 0.17458908 | 0.12155019 | 0.00098385 | 0.02344132 |
| Stbd1 | 0.37898823 | 0.49535056 | 0.37436895 | 0.12098161 | 0.07540894 | 0.78459363 |
| Tmem45a | 0.31026371 | 0.43839009 | 0.31746259 | 0.1209275 | 0.04518821 | 0.54819953 |
| Calml4 | 0.41572439 | 0.52400103 | 0.40320396 | 0.12079707 | 0.09145106 | 0.89629118 |
| Gip | 0.08850441 | 0.22290899 | 0.10247202 | 0.12043696 | 8.71E-08 | 4.27E-06 |
| Ctsf | 0.78085728 | 0.76213425 | 0.64173261 | 0.12040165 | 0.20211708 | 0.90650187 |
| Maged1 | 2.75942294 | 0.99097795 | 0.87073601 | 0.12024194 | 0.26963938 | 0.90650187 |

| | | | | | | |
|---|---|---|---|---|---|---|
| Fibin | 0.16876168 | 0.30660134 | 0.18643956 | 0.12016179 | 0.00208354 | 0.04417626 |
| Atp8b1 | 0.61120165 | 0.66070321 | 0.54061196 | 0.12009125 | 0.16168787 | 0.90650187 |
| Cyp20a1 | 0.51094581 | 0.59268542 | 0.47329567 | 0.11938975 | 0.13105608 | 0.90650187 |
| Rrm2 | 0.17628107 | 0.3132677 | 0.19392487 | 0.11934283 | 0.00311088 | 0.06271674 |
| Tspan6 | 0.58408812 | 0.64222205 | 0.52301126 | 0.1192108 | 0.15545983 | 0.90650187 |
| Ube2l6 | 0.19165869 | 0.32814085 | 0.20903742 | 0.11910343 | 0.00565492 | 0.10411931 |
| Angptl2 | 0.35183741 | 0.47138711 | 0.3523639 | 0.11902321 | 0.06659263 | 0.7225372 |
| Hmgcr | 0.69540883 | 0.71157324 | 0.59264887 | 0.11892437 | 0.1861747 | 0.90650187 |
| Lifr | 0.38768443 | 0.5001412 | 0.38129056 | 0.11885064 | 0.08291572 | 0.83803318 |
| Mcpt8 | 0.03230309 | 0.15759216 | 0.03880053 | 0.11879163 | 1.73E-42 | 5.98E-40 |
| Pax4 | 0.19424949 | 0.33034541 | 0.2115584 | 0.11878702 | 0.00627757 | 0.11306327 |
| Rps12-ps3 | 2.37759723 | 1.02671554 | 0.90795242 | 0.11876312 | 0.28045184 | 0.90650187 |
| Hspe1 | 2.19897868 | 1.03918064 | 0.92047274 | 0.1187079 | 0.28321287 | 0.90650187 |
| Rab34 | 0.78429452 | 0.7622358 | 0.64363459 | 0.11860122 | 0.20634109 | 0.90650187 |
| Sdc4 | 0.25906829 | 0.39063696 | 0.27234995 | 0.11828701 | 0.02675294 | 0.3643918 |
| Dstn | 2.30758011 | 1.03160195 | 0.91350745 | 0.1180945 | 0.2827438 | 0.90650187 |
| Srpx | 0.31190552 | 0.43674782 | 0.31886959 | 0.11787824 | 0.05014687 | 0.59321604 |
| Lsp1 | 0.20792793 | 0.34262318 | 0.22474787 | 0.11787531 | 0.00986027 | 0.16543711 |
| Emilin2 | 0.24265722 | 0.37515294 | 0.25736111 | 0.11779184 | 0.02093985 | 0.29944183 |
| Sod2 | 0.94096416 | 0.84229631 | 0.72458994 | 0.11770637 | 0.23509538 | 0.90650187 |
| Oxct1 | 0.69668998 | 0.71096638 | 0.59341131 | 0.11755507 | 0.1892464 | 0.90650187 |
| Tacc3 | 0.2242578 | 0.35766535 | 0.24023714 | 0.1174282 | 0.01488949 | 0.2300154 |
| 1810037I17R | 1.58228262 | 1.02332595 | 0.9059036 | 0.11742235 | 0.2822281 | 0.90650187 |
| P4ha1 | 0.59552312 | 0.64789859 | 0.53048719 | 0.1174114 | 0.16257375 | 0.90650187 |
| Ucp2 | 0.7321224 | 0.73142983 | 0.61417294 | 0.11725689 | 0.1980108 | 0.90650187 |
| Rasip1 | 0.14896359 | 0.28326249 | 0.16642561 | 0.11683687 | 0.00090126 | 0.0217806 |
| Ghr | 0.79458502 | 0.76589309 | 0.64929579 | 0.11659729 | 0.21234248 | 0.90650187 |
| Mgll | 0.15152961 | 0.28550677 | 0.16904489 | 0.11646188 | 0.00109654 | 0.02576265 |
| 5330417C22I | 0.3976683 | 0.50555673 | 0.38916363 | 0.1163931 | 0.09182253 | 0.89877283 |
| Poldip2 | 0.6178778 | 0.66103571 | 0.54488086 | 0.11615485 | 0.17164337 | 0.90650187 |

| | | | | | | |
|---|---|---|---|---|---|---|
| Pycr2 | 1.02011265 | 0.87505717 | 0.75891455 | 0.11614262 | 0.24814274 | 0.90650187 |
| Thy1 | 0.17350325 | 0.30708353 | 0.19116727 | 0.11591626 | 0.00351268 | 0.06973982 |
| Vamp5 | 0.14739885 | 0.28070556 | 0.16482441 | 0.11588115 | 0.00088746 | 0.02148122 |
| Ppp1r14a | 0.24272886 | 0.37322269 | 0.25742713 | 0.11579556 | 0.0227654 | 0.31940792 |
| Rnase1 | 0.07517034 | 0.20337948 | 0.08778479 | 0.11559469 | 2.40E-09 | 1.41E-07 |
| Sema3d | 0.16809827 | 0.30136127 | 0.18577609 | 0.11558518 | 0.00283792 | 0.05759567 |
| Tmem167 | 1.55101626 | 1.01696665 | 0.90162853 | 0.11533813 | 0.28478297 | 0.90650187 |
| Elk3 | 0.30605826 | 0.42910253 | 0.31384778 | 0.11525475 | 0.05128015 | 0.60427093 |
| Rpl13-ps3 | 1.70766501 | 1.03386092 | 0.91876016 | 0.11510076 | 0.28878481 | 0.90650187 |
| Fam198b | 0.15124306 | 0.28377852 | 0.16875278 | 0.11502574 | 0.00122185 | 0.02801413 |
| Slc29a1 | 0.57608229 | 0.63254929 | 0.51773122 | 0.11481807 | 0.16208885 | 0.90650187 |
| Rest | 0.3790229 | 0.48920206 | 0.37439666 | 0.1148054 | 0.08640886 | 0.86189704 |
| Ssr2 | 2.83792844 | 0.97436164 | 0.85956962 | 0.11479201 | 0.27635888 | 0.90650187 |
| Vstm4 | 0.26364895 | 0.39102122 | 0.27648691 | 0.11453431 | 0.03276786 | 0.42536655 |
| Fam212a | 0.30872556 | 0.43060099 | 0.31614228 | 0.11445871 | 0.05375098 | 0.62276574 |
| Tmem108 | 0.48326374 | 0.56799229 | 0.45357572 | 0.11441657 | 0.13105355 | 0.90650187 |
| Cel | 0.0511893 | 0.1750275 | 0.06072202 | 0.11430548 | 2.92E-17 | 3.60E-15 |
| Tie1 | 0.11199617 | 0.24202451 | 0.12782021 | 0.11420431 | 3.56E-05 | 0.00114022 |
| Minos1 | 2.59049841 | 1.00335484 | 0.88922254 | 0.1141323 | 0.28413659 | 0.90650187 |
| Crtap | 0.48541355 | 0.56881052 | 0.45512588 | 0.11368464 | 0.13340134 | 0.90650187 |
| Rtn1 | 0.0906536 | 0.21848377 | 0.10481583 | 0.11366794 | 7.14E-07 | 3.12E-05 |
| Rsu1 | 0.78909159 | 0.75993379 | 0.64628023 | 0.11365356 | 0.21716737 | 0.90650187 |
| Pim1 | 0.21485965 | 0.34485692 | 0.23135644 | 0.11350048 | 0.0145933 | 0.22659495 |
| Fam114a1 | 0.34751939 | 0.46203694 | 0.34880847 | 0.11322848 | 0.07449464 | 0.77834237 |
| Map1lc3a | 1.44850939 | 0.9972022 | 0.8841027 | 0.1130995 | 0.28477732 | 0.90650187 |
| Creb3l1 | 0.45037615 | 0.54239853 | 0.42946005 | 0.11293848 | 0.12118247 | 0.90650187 |
| Cnih2 | 0.53277215 | 0.60125901 | 0.48848534 | 0.11277367 | 0.15236794 | 0.90650187 |
| Raph1 | 0.43689333 | 0.53201572 | 0.41934936 | 0.11266636 | 0.11616105 | 0.90650187 |
| St3gal4 | 0.41390507 | 0.51435644 | 0.40180041 | 0.11255604 | 0.10650261 | 0.90650187 |
| Tnfaip8 | 0.39010735 | 0.4956665 | 0.38320852 | 0.11245798 | 0.09601007 | 0.90650187 |

| | | | | | | |
|---|---|---|---|---|---|---|
| Tapbp | 0.54436683 | 0.60887021 | 0.49643056 | 0.11243965 | 0.15698827 | 0.90650187 |
| Gprasp2 | 0.4456672 | 0.53835548 | 0.42594454 | 0.11241094 | 0.12034996 | 0.90650187 |
| Pbk | 0.12810268 | 0.25721491 | 0.14484264 | 0.11237227 | 0.00028133 | 0.00763657 |
| Gltscr2 | 1.61425835 | 1.02195383 | 0.90964751 | 0.11230633 | 0.29155054 | 0.90650187 |
| Thbs2 | 0.3337766 | 0.44963868 | 0.33738511 | 0.11225357 | 0.06955328 | 0.74403902 |
| Hpgd | 0.17136733 | 0.30093663 | 0.18904069 | 0.11189594 | 0.00425145 | 0.08180549 |
| Snca | 0.03914125 | 0.15863676 | 0.04680106 | 0.1118357 | 1.16E-26 | 2.52E-24 |
| Exoc7 | 0.67542349 | 0.69192691 | 0.58064321 | 0.1112837 | 0.19709963 | 0.90650187 |
| 1500011B03 | 0.74393934 | 0.73217226 | 0.6209597 | 0.11121256 | 0.21295831 | 0.90650187 |
| Atp6v1b2 | 0.87611191 | 0.80362916 | 0.69246864 | 0.11116051 | 0.23772376 | 0.90650187 |
| Ube2b | 1.75084518 | 1.03278175 | 0.92164782 | 0.11113393 | 0.29595885 | 0.90650187 |
| Tead2 | 0.5294097 | 0.59729149 | 0.48616542 | 0.11112607 | 0.15477614 | 0.90650187 |
| Mthfd2 | 0.34208076 | 0.45539889 | 0.34430701 | 0.11109188 | 0.0757304 | 0.78632364 |
| Prss35 | 0.20320027 | 0.33127118 | 0.22021188 | 0.1110593 | 0.01247909 | 0.2011608 |
| Arx | 0.15755583 | 0.28615904 | 0.17516633 | 0.11099271 | 0.00242431 | 0.05048219 |
| Gstm2 | 0.36306355 | 0.47246627 | 0.36153579 | 0.11093048 | 0.08627576 | 0.86189704 |
| Gsta3 | 0.12114345 | 0.24821057 | 0.1375267 | 0.11068387 | 0.00017318 | 0.00491146 |
| Rps15a | 4.64432507 | 0.6982094 | 0.58760239 | 0.11060702 | 0.20134792 | 0.90650187 |
| Bcam | 0.32853484 | 0.443553 | 0.3329858 | 0.11056719 | 0.06995132 | 0.74657023 |
| Syt7 | 0.57031378 | 0.62436901 | 0.51390332 | 0.11046569 | 0.16963672 | 0.90650187 |
| Bub3 | 0.98123183 | 0.85316357 | 0.74285649 | 0.11030708 | 0.25458521 | 0.90650187 |
| Hmcn1 | 0.20265855 | 0.3299943 | 0.21969062 | 0.11030368 | 0.01280488 | 0.20554071 |
| Tmem88 | 0.13716013 | 0.26455125 | 0.15427663 | 0.11027462 | 0.00074236 | 0.01843848 |
| Anpep | 0.32433725 | 0.4397008 | 0.32944592 | 0.11025489 | 0.06840175 | 0.73378613 |
| Pdzk1ip1 | 0.50263519 | 0.57765488 | 0.46742958 | 0.11022531 | 0.14724435 | 0.90650187 |
| Smim1 | 0.44660929 | 0.53655216 | 0.42664913 | 0.10990303 | 0.12606945 | 0.90650187 |
| Nudt19 | 0.37518264 | 0.48104244 | 0.37132119 | 0.10972124 | 0.09448592 | 0.90650187 |
| Aurkb | 0.13476581 | 0.26149327 | 0.15179227 | 0.109701 | 0.0006571 | 0.01656478 |
| Snhg6 | 0.72845171 | 0.72155171 | 0.61205096 | 0.10950075 | 0.21320378 | 0.90650187 |
| Dhrs4 | 0.7467168 | 0.73200773 | 0.62254475 | 0.10946298 | 0.21720202 | 0.90650187 |

| | | | | | | |
|---|---|---|---|---|---|---|
| Qpct | 0.25672683 | 0.37965252 | 0.27022754 | 0.10942497 | 0.03591549 | 0.45839356 |
| Atox1 | 1.64159813 | 1.02192513 | 0.91269436 | 0.10923077 | 0.29734645 | 0.90650187 |
| Ckap4 | 0.4384921 | 0.52964003 | 0.42055533 | 0.10908469 | 0.12443211 | 0.90650187 |
| Try10 | 0.04299848 | 0.16028438 | 0.05128204 | 0.10900234 | 1.70E-21 | 2.82E-19 |
| Zfp36l2 | 0.4380435 | 0.52921309 | 0.42021714 | 0.10899595 | 0.12443432 | 0.90650187 |
| Pvrl2 | 0.67658554 | 0.69032268 | 0.58134693 | 0.10897575 | 0.20232474 | 0.90650187 |
| Arhgdib | 0.12516383 | 0.2502755 | 0.14176051 | 0.10851499 | 0.00033322 | 0.00890201 |
| Pdlim7 | 0.53279442 | 0.59676442 | 0.48850068 | 0.10826374 | 0.1622493 | 0.90650187 |
| Sema5a | 0.37545009 | 0.47972555 | 0.37153576 | 0.10818979 | 0.09773887 | 0.90650187 |
| Col9a2 | 0.55405834 | 0.61117085 | 0.50300496 | 0.10816589 | 0.16954102 | 0.90650187 |
| Gdi1 | 1.20939823 | 0.93333608 | 0.82524351 | 0.10809257 | 0.28018325 | 0.90650187 |
| Rangap1 | 0.85284923 | 0.78836551 | 0.68045186 | 0.10791366 | 0.24039457 | 0.90650187 |
| Ptprf | 0.82183105 | 0.77184275 | 0.66405342 | 0.10778933 | 0.23526653 | 0.90650187 |
| Meis1 | 0.54910511 | 0.60736908 | 0.49965205 | 0.10771703 | 0.16893004 | 0.90650187 |
| Lima1 | 0.47027779 | 0.55173324 | 0.44414523 | 0.107588 | 0.14076553 | 0.90650187 |
| Rpl23a | 4.97251708 | 0.65316271 | 0.54570597 | 0.10745674 | 0.19067847 | 0.90650187 |
| Rnaset2b | 0.75616985 | 0.73451115 | 0.62791248 | 0.10659867 | 0.22520964 | 0.90650187 |
| Ier5 | 0.35782684 | 0.46376748 | 0.35727015 | 0.10649733 | 0.09255668 | 0.90084241 |
| Laptm5 | 0.05986665 | 0.17700293 | 0.07061244 | 0.10639048 | 1.06E-11 | 8.06E-10 |
| Fkbp14 | 0.42248746 | 0.51454099 | 0.40839897 | 0.10614202 | 0.1239626 | 0.90650187 |
| Bex4 | 0.59092125 | 0.63337076 | 0.52748828 | 0.10588248 | 0.1860988 | 0.90650187 |
| BC028528 | 0.27186256 | 0.38965381 | 0.28385512 | 0.10579869 | 0.04876281 | 0.58103391 |
| Cnot6l | 0.66219899 | 0.67832002 | 0.57258424 | 0.10573578 | 0.20583905 | 0.90650187 |
| Emp2 | 0.22811384 | 0.34952427 | 0.24385399 | 0.10567028 | 0.02702515 | 0.36712178 |
| Folr2 | 0.0404909 | 0.1539551 | 0.04837156 | 0.10558354 | 1.45E-22 | 2.54E-20 |
| Rpn2 | 1.91904416 | 1.03272602 | 0.92715366 | 0.10557236 | 0.30635511 | 0.90650187 |
| Ptp4a2 | 0.92578393 | 0.82283709 | 0.71726525 | 0.10557184 | 0.25644841 | 0.90650187 |
| Plin3 | 0.5901861 | 0.63196612 | 0.52700791 | 0.10495821 | 0.18797555 | 0.90650187 |
| Nrros | 0.09875489 | 0.21853952 | 0.11359298 | 0.10494655 | 2.00E-05 | 0.00067189 |
| Cxcl2 | 0.02884214 | 0.1392905 | 0.03472346 | 0.10456704 | 3.57E-41 | 1.18E-38 |

| | | | | | | |
|---|---|---|---|---|---|---|
| Zdbf2 | 0.54198771 | 0.59937411 | 0.49480735 | 0.10456676 | 0.17374164 | 0.90650187 |
| Rgs2 | 0.1826816 | 0.30475007 | 0.20024635 | 0.10450372 | 0.01016897 | 0.16967999 |
| Myct1 | 0.09022518 | 0.20884226 | 0.10434913 | 0.10449313 | 4.26E-06 | 0.00016185 |
| Actg1 | 4.87967324 | 0.66039928 | 0.55604008 | 0.1043592 | 0.20203789 | 0.90650187 |
| Eef2 | 3.80201969 | 0.82179349 | 0.7176852 | 0.10410829 | 0.25950084 | 0.90650187 |
| Man2a1 | 0.50319444 | 0.57163036 | 0.46782574 | 0.10380462 | 0.16196257 | 0.90650187 |
| Cdc42ep5 | 0.42544869 | 0.51435463 | 0.41066232 | 0.10369231 | 0.13082147 | 0.90650187 |
| Tmed9 | 1.79496152 | 1.02740801 | 0.92398778 | 0.10342023 | 0.30938708 | 0.90650187 |
| Tle6 | 0.29984738 | 0.41184226 | 0.30848048 | 0.10336178 | 0.06816759 | 0.73292957 |
| Wbp5 | 3.18031454 | 0.91812172 | 0.81488824 | 0.10323348 | 0.28665337 | 0.90650187 |
| Arpc5 | 1.76575215 | 1.02572319 | 0.92249812 | 0.10322507 | 0.30943537 | 0.90650187 |
| Gimap6 | 0.12471131 | 0.24440751 | 0.141285 | 0.10312251 | 0.00058766 | 0.01496314 |
| Dusp10 | 0.36568957 | 0.46665478 | 0.36366649 | 0.10298829 | 0.10402048 | 0.90650187 |
| Rbms2 | 0.39777802 | 0.4920896 | 0.3892497 | 0.10283991 | 0.12009125 | 0.90650187 |
| Gm12840 | 0.12177208 | 0.24009137 | 0.13819 | 0.10190136 | 0.00052237 | 0.01350437 |
| Shisa5 | 0.84597836 | 0.77875506 | 0.67685645 | 0.1018986 | 0.25166099 | 0.90650187 |
| Bax | 1.49122622 | 0.99360916 | 0.89183234 | 0.10177681 | 0.30595915 | 0.90650187 |
| Ccl12 | 0.02607918 | 0.13317949 | 0.03145509 | 0.1017244 | 3.61E-47 | 1.31E-44 |
| Pclo | 0.49754581 | 0.5655192 | 0.46381489 | 0.10170431 | 0.16482408 | 0.90650187 |
| Dpm3 | 1.59325131 | 1.00820673 | 0.90718869 | 0.10101803 | 0.31028917 | 0.90650187 |
| Creb5 | 0.22451851 | 0.34149411 | 0.24048216 | 0.10101195 | 0.03092781 | 0.40774819 |
| Stmn4 | 0.10227395 | 0.21838015 | 0.11739016 | 0.10098999 | 6.55E-05 | 0.00199969 |
| Fev | 0.304383 | 0.41339236 | 0.31240344 | 0.10098892 | 0.07534387 | 0.78445357 |
| Gm2a | 0.36769285 | 0.46598489 | 0.36528818 | 0.10069671 | 0.11019484 | 0.90650187 |
| Rspo1 | 0.1176461 | 0.23443261 | 0.13382764 | 0.10060497 | 0.00041591 | 0.01099512 |
| Fcgr3 | 0.04442635 | 0.15341854 | 0.05293502 | 0.10048352 | 1.60E-17 | 2.01E-15 |
| Ugdh | 0.55756809 | 0.60573832 | 0.5053717 | 0.10036662 | 0.18864633 | 0.90650187 |
| Tnfrsf1a | 0.44933754 | 0.52900239 | 0.42868602 | 0.10031637 | 0.14909755 | 0.90650187 |
| Vat1 | 0.6231537 | 0.64854553 | 0.54823541 | 0.10031012 | 0.20799197 | 0.90650187 |
| Clic4 | 0.41252925 | 0.50081167 | 0.40073735 | 0.10007432 | 0.13346051 | 0.90650187 |

| | | | | | | |
|---|---|---|---|---|---|---|
| Ppm1b | 0.79498707 | 0.74955857 | 0.64951595 | 0.10004262 | 0.24675373 | 0.90650187 |
| Zyx | 0.44237683 | 0.52333593 | 0.42347796 | 0.09985796 | 0.1472517 | 0.90650187 |
| Flna | 0.52023016 | 0.57947404 | 0.47979557 | 0.09967847 | 0.17785163 | 0.90650187 |
| Tmem97 | 0.3621912 | 0.46044235 | 0.36082675 | 0.0996156 | 0.10985021 | 0.90650187 |
| Trappc2l | 1.29814757 | 0.94988674 | 0.85031965 | 0.0995671 | 0.30133923 | 0.90650187 |
| Atp6v0b | 1.65490045 | 1.01354968 | 0.91412426 | 0.09942542 | 0.31435983 | 0.90650187 |
| Ube2s | 1.55028784 | 1.00094351 | 0.90152178 | 0.09942173 | 0.31197111 | 0.90650187 |
| 2010107G23 | 0.42452748 | 0.50840593 | 0.40995891 | 0.09844703 | 0.14285897 | 0.90650187 |
| Ralb | 0.57297169 | 0.61407324 | 0.51566952 | 0.09840371 | 0.19812373 | 0.90650187 |
| Cd52 | 0.0381489 | 0.14390711 | 0.04564452 | 0.09826259 | 5.32E-22 | 8.99E-20 |
| Kif21a | 0.52492718 | 0.58118218 | 0.4830616 | 0.09812058 | 0.18326249 | 0.90650187 |
| Ly6c1 | 0.07243538 | 0.18277506 | 0.08474089 | 0.09803418 | 1.35E-07 | 6.44E-06 |
| Gucy1a3 | 0.13777472 | 0.25291622 | 0.15491325 | 0.09800297 | 0.00245844 | 0.05105289 |
| Ptp4a3 | 0.43181358 | 0.51350758 | 0.41550537 | 0.09800221 | 0.14719203 | 0.90650187 |
| St3gal6 | 0.34353501 | 0.44343061 | 0.34551323 | 0.09791738 | 0.10385794 | 0.90650187 |
| Epha4 | 0.21719726 | 0.33134184 | 0.23357388 | 0.09776796 | 0.03138513 | 0.41163528 |
| Cdh1 | 0.57742132 | 0.61593074 | 0.51861696 | 0.09731378 | 0.20209039 | 0.90650187 |
| Stxbp5l | 0.31399502 | 0.41793867 | 0.32065681 | 0.09728186 | 0.08871316 | 0.87809516 |
| Sec61g | 3.6207496 | 0.84765351 | 0.75039243 | 0.09726108 | 0.28223623 | 0.90650187 |
| Rem1 | 0.24289402 | 0.35429048 | 0.25757931 | 0.09671118 | 0.04754332 | 0.56905986 |
| Ndc80 | 0.10715592 | 0.21919693 | 0.12264231 | 0.09655462 | 0.00023266 | 0.00642551 |
| Cenpm | 0.21083308 | 0.32407553 | 0.22752365 | 0.09655188 | 0.02961136 | 0.39519077 |
| Hmmr | 0.14173783 | 0.25550707 | 0.15900768 | 0.09649939 | 0.00348816 | 0.06940258 |
| Try5 | 0.04417276 | 0.14889273 | 0.05264168 | 0.09625105 | 2.17E-16 | 2.49E-14 |
| Tmc4 | 0.66710606 | 0.67163979 | 0.57558597 | 0.09605382 | 0.2290791 | 0.90650187 |
| Mn1 | 0.17452932 | 0.28815513 | 0.1921869 | 0.09596822 | 0.01321256 | 0.21023232 |
| Amer1 | 0.39119155 | 0.47993759 | 0.38406529 | 0.0958723 | 0.13355673 | 0.90650187 |
| Gltp | 0.44561278 | 0.52169134 | 0.42590382 | 0.09578752 | 0.15869515 | 0.90650187 |
| Dlc1 | 0.45931943 | 0.53186174 | 0.43609356 | 0.09576818 | 0.16446389 | 0.90650187 |
| Vwa1 | 0.16330582 | 0.27670516 | 0.18096857 | 0.09573659 | 0.00934068 | 0.15778647 |

| | | | | | | |
|---|---|---|---|---|---|---|
| Cenpe | 0.14009077 | 0.25267721 | 0.15730829 | 0.09536893 | 0.00351763 | 0.06973982 |
| Gpd2 | 0.54610437 | 0.59276476 | 0.49761349 | 0.09515127 | 0.19764377 | 0.90650187 |
| F11r | 0.56885478 | 0.60796416 | 0.51293203 | 0.09503213 | 0.20506979 | 0.90650187 |
| Cnrip1 | 0.29004849 | 0.39479993 | 0.29994001 | 0.09485993 | 0.07986491 | 0.81752624 |
| Nme4 | 0.36272044 | 0.45601786 | 0.36125699 | 0.09476088 | 0.12178266 | 0.90650187 |
| Ctsk | 0.17348056 | 0.28586427 | 0.1911447 | 0.09471956 | 0.01379846 | 0.21735794 |
| Dut | 0.8481777 | 0.77255758 | 0.67800944 | 0.09454813 | 0.26764932 | 0.90650187 |
| Myt1 | 0.29321509 | 0.39724438 | 0.30270975 | 0.09453463 | 0.08251691 | 0.83622636 |
| Gm5617 | 0.61217655 | 0.63565529 | 0.54123693 | 0.09441836 | 0.21901251 | 0.90650187 |
| Gpx2 | 0.31813699 | 0.41843875 | 0.32418832 | 0.09425043 | 0.09809902 | 0.90650187 |
| Sqle | 0.4845259 | 0.54871833 | 0.45448621 | 0.09423213 | 0.17833162 | 0.90650187 |
| Hagh | 0.69690492 | 0.68774822 | 0.59353914 | 0.09420907 | 0.24021048 | 0.90650187 |
| Chmp5 | 1.50568446 | 0.98849548 | 0.89429116 | 0.09420431 | 0.31978554 | 0.90650187 |
| Rgs3 | 0.45447029 | 0.52657636 | 0.43250379 | 0.09407257 | 0.16678447 | 0.90650187 |
| Nasp | 0.74772205 | 0.71715965 | 0.62311754 | 0.09404211 | 0.25112952 | 0.90650187 |
| Gapdh | 2.85455919 | 0.95114303 | 0.85713257 | 0.09401046 | 0.3129184 | 0.90650187 |
| Cd93 | 0.10269634 | 0.21175915 | 0.11784577 | 0.09391339 | 0.00019796 | 0.00555213 |
| Gimap4 | 0.09756735 | 0.20615978 | 0.11231204 | 0.09384774 | 0.00010171 | 0.00300785 |
| Tmem258 | 2.34221647 | 1.00464345 | 0.91083105 | 0.09381239 | 0.32351868 | 0.90650187 |
| Cd44 | 0.22140565 | 0.33133477 | 0.23755179 | 0.09378297 | 0.03962289 | 0.49329043 |
| Dnmt3a | 1.10195912 | 0.88367938 | 0.78994157 | 0.09373782 | 0.29891123 | 0.90650187 |
| Creg1 | 0.61891784 | 0.6389118 | 0.54554337 | 0.09336843 | 0.22337191 | 0.90650187 |
| Hspa8 | 4.67720048 | 0.67621065 | 0.58345731 | 0.09275334 | 0.23986819 | 0.90650187 |
| Spry2 | 0.23184767 | 0.34005599 | 0.24734191 | 0.09271408 | 0.047817 | 0.57188529 |
| Pea15a | 0.77580964 | 0.73143881 | 0.63892994 | 0.09250887 | 0.25989574 | 0.90650187 |
| Cox7a2l | 2.29560827 | 1.00672308 | 0.91439153 | 0.09233155 | 0.32675214 | 0.90650187 |
| H3f3b | 4.34136506 | 0.7207656 | 0.62851299 | 0.09225261 | 0.25703212 | 0.90650187 |
| Sbspon | 0.13185369 | 0.24100369 | 0.14876145 | 0.09224224 | 0.00292047 | 0.05911336 |
| Tmed6 | 0.17714532 | 0.28688111 | 0.19478107 | 0.09210003 | 0.01777424 | 0.26601254 |
| Bex1 | 0.57927917 | 0.61170641 | 0.51984415 | 0.09186226 | 0.21605438 | 0.90650187 |

| | | | | | | |
|---|---|---|---|---|---|---|
| Ace2 | 0.20019753 | 0.30901434 | 0.21731867 | 0.09169567 | 0.03034248 | 0.40204743 |
| Tspan12 | 0.22556569 | 0.33299783 | 0.2414656 | 0.09153224 | 0.0459756 | 0.55397026 |
| Cartpt | 0.05814431 | 0.16012255 | 0.06865826 | 0.09146429 | 1.59E-09 | 9.50E-08 |
| Elovl1 | 0.68425517 | 0.67738962 | 0.58597418 | 0.09141543 | 0.2439851 | 0.90650187 |
| Coro1a | 0.04792549 | 0.14835562 | 0.05697267 | 0.09138295 | 4.99E-13 | 4.52E-11 |
| Entpd3 | 0.41515143 | 0.49414506 | 0.4027622 | 0.09138285 | 0.15656763 | 0.90650187 |
| Srek1ip1 | 0.55150624 | 0.59264964 | 0.50127928 | 0.09137036 | 0.20887812 | 0.90650187 |
| Ank | 0.42864614 | 0.50426588 | 0.41309897 | 0.09116691 | 0.16327125 | 0.90650187 |
| Prss2 | 0.04116145 | 0.14031673 | 0.04915079 | 0.09116593 | 8.20E-17 | 9.67E-15 |
| Svil | 0.52175695 | 0.5719869 | 0.48085875 | 0.09112815 | 0.19975617 | 0.90650187 |
| Hoxb4 | 0.32264614 | 0.41872025 | 0.3280155 | 0.09070475 | 0.109475 | 0.90650187 |
| Prph | 0.04787833 | 0.14762288 | 0.05691837 | 0.09070451 | 6.98E-13 | 6.13E-11 |
| Gata6 | 0.53766239 | 0.58253198 | 0.49184671 | 0.09068527 | 0.20620269 | 0.90650187 |
| Nrarp | 0.11605895 | 0.22275912 | 0.13214402 | 0.0906151 | 0.00114999 | 0.02660735 |
| Fth1 | 3.67385029 | 0.83120987 | 0.74082258 | 0.09038728 | 0.29376958 | 0.90650187 |
| Acot1 | 0.36152075 | 0.45054943 | 0.3602814 | 0.09026804 | 0.13267372 | 0.90650187 |
| Tmpo | 1.03977081 | 0.85688459 | 0.7667368 | 0.09014779 | 0.30059685 | 0.90650187 |
| Efna1 | 0.18041874 | 0.28791442 | 0.19801656 | 0.08989785 | 0.02178169 | 0.30973193 |
| Clmp | 0.25346488 | 0.35696257 | 0.26726197 | 0.0897006 | 0.06784041 | 0.73189638 |
| Itgb1 | 1.71496891 | 1.00894128 | 0.91927654 | 0.08966474 | 0.33228322 | 0.90650187 |
| Klc3 | 0.50318157 | 0.55745044 | 0.46781662 | 0.08963382 | 0.19716821 | 0.90650187 |
| Adam15 | 0.21880415 | 0.32458157 | 0.2350949 | 0.08948667 | 0.04530598 | 0.54876195 |
| Kpna2 | 0.25373443 | 0.35690918 | 0.26750742 | 0.08940176 | 0.06867467 | 0.7361944 |
| Abcg2 | 0.29726821 | 0.39559208 | 0.30624148 | 0.0893506 | 0.097303 | 0.90650187 |
| Knstrn | 0.1776201 | 0.28456626 | 0.19525108 | 0.08931518 | 0.02099661 | 0.29997134 |
| Hoxa5 | 0.24446649 | 0.34829997 | 0.25902681 | 0.08927316 | 0.06274014 | 0.69159738 |
| Slfn2 | 0.04674394 | 0.14487817 | 0.05561136 | 0.08926681 | 4.80E-13 | 4.40E-11 |
| Cd59a | 0.17961994 | 0.28637716 | 0.19722811 | 0.08914905 | 0.0222443 | 0.31425236 |
| Myef2 | 0.82056376 | 0.75247367 | 0.66337448 | 0.08909918 | 0.27522082 | 0.90650187 |
| Dnajb11 | 1.12040259 | 0.88549749 | 0.79641997 | 0.08907752 | 0.30951358 | 0.90650187 |

| | | | | | | |
|---|---|---|---|---|---|---|
| Col16a1 | 0.31138641 | 0.40742291 | 0.31842498 | 0.08899793 | 0.10702217 | 0.90650187 |
| Prss1 | 0.05843123 | 0.15777324 | 0.0689841 | 0.08878913 | 5.27E-09 | 3.00E-07 |
| Rpa2 | 0.32293867 | 0.41701601 | 0.3282631 | 0.08875291 | 0.1146887 | 0.90650187 |
| Litaf | 0.54828542 | 0.58778767 | 0.49909574 | 0.08869193 | 0.21476168 | 0.90650187 |
| Cd55 | 0.19396767 | 0.29994873 | 0.21128452 | 0.08866421 | 0.03105073 | 0.40827162 |
| Chodl | 0.10803911 | 0.2119161 | 0.12358928 | 0.08832682 | 0.00074356 | 0.01843848 |
| Tram1 | 1.44054888 | 0.97039828 | 0.88257077 | 0.0878275 | 0.32910104 | 0.90650187 |
| Snap25 | 0.43593284 | 0.50643017 | 0.41862397 | 0.0878062 | 0.17554838 | 0.90650187 |
| Hand2 | 0.31280798 | 0.40736403 | 0.31964196 | 0.08772207 | 0.11122377 | 0.90650187 |
| Podxl | 0.13081734 | 0.23539873 | 0.14768042 | 0.08771831 | 0.00413829 | 0.07983008 |
| Tmem263 | 0.58912337 | 0.61390645 | 0.52631295 | 0.0875935 | 0.22968767 | 0.90650187 |
| Tax1bp3 | 0.73115576 | 0.7010457 | 0.61361476 | 0.08743094 | 0.26322563 | 0.90650187 |
| Actn4 | 0.87852972 | 0.78110819 | 0.69370382 | 0.08740437 | 0.28769484 | 0.90650187 |
| Napa | 1.34023835 | 0.94811289 | 0.86080042 | 0.08731246 | 0.32602163 | 0.90650187 |
| Kctd12 | 0.542862 | 0.58269449 | 0.49540439 | 0.0872901 | 0.21672124 | 0.90650187 |
| Ramp1 | 0.32642439 | 0.4183981 | 0.33120792 | 0.08719018 | 0.12093977 | 0.90650187 |
| Npy | 0.03253778 | 0.12615915 | 0.03907633 | 0.08708282 | 1.94E-23 | 3.51E-21 |
| Aldoc | 0.34001546 | 0.42959591 | 0.34259091 | 0.087005 | 0.12944593 | 0.90650187 |
| Ost4 | 1.84186921 | 1.01266805 | 0.92567724 | 0.08699081 | 0.33805499 | 0.90650187 |
| Ptgr1 | 0.20601962 | 0.30978921 | 0.22291974 | 0.08686947 | 0.04160067 | 0.51287251 |
| Prox1 | 0.46501195 | 0.52711124 | 0.44028671 | 0.08682453 | 0.19033201 | 0.90650187 |
| Fzd2 | 0.29637719 | 0.39213333 | 0.3054665 | 0.08666683 | 0.10359939 | 0.90650187 |
| Hsd11b2 | 0.14111396 | 0.24495795 | 0.15836436 | 0.08659359 | 0.00753159 | 0.13235574 |
| Cenpw | 0.21619437 | 0.31918503 | 0.23262323 | 0.0865618 | 0.04903718 | 0.58372295 |
| Gm10073 | 1.8688603 | 1.0128106 | 0.92642054 | 0.08639006 | 0.33923229 | 0.90650187 |
| Fam43a | 0.32157881 | 0.413499 | 0.32711142 | 0.08638758 | 0.12018749 | 0.90650187 |
| Cd151 | 0.54061861 | 0.58017967 | 0.49387147 | 0.0863082 | 0.21860678 | 0.90650187 |
| Atp1a1 | 1.49699936 | 0.97908645 | 0.89281906 | 0.0862674 | 0.33376132 | 0.90650187 |
| Rbm47 | 0.3918296 | 0.47079435 | 0.38456907 | 0.08622528 | 0.15944015 | 0.90650187 |
| B3gnt9 | 0.27412733 | 0.37206137 | 0.28587501 | 0.08618636 | 0.09007827 | 0.88741401 |

| | | | | | | |
|---|---|---|---|---|---|---|
| P3h3 | 0.21922822 | 0.32164893 | 0.23549587 | 0.08615306 | 0.05193586 | 0.6091644 |
| Pcna | 0.53215968 | 0.57404228 | 0.4880633 | 0.08597898 | 0.21676837 | 0.90650187 |
| Inhba | 0.06267475 | 0.15966685 | 0.07378911 | 0.08587774 | 1.15E-07 | 5.55E-06 |
| Ctnna1 | 0.91323629 | 0.79698203 | 0.71113083 | 0.0858512 | 0.29573902 | 0.90650187 |
| Ttll7 | 0.45582683 | 0.51927686 | 0.43350968 | 0.08576718 | 0.18955552 | 0.90650187 |
| Sec11c | 0.57511211 | 0.6027553 | 0.5170888 | 0.0856665 | 0.23071111 | 0.90650187 |
| Sycn | 0.05719639 | 0.15268223 | 0.06758085 | 0.08510138 | 1.08E-08 | 6.01E-07 |
| Ccl11 | 0.12232747 | 0.22365182 | 0.13877563 | 0.08487619 | 0.00327432 | 0.06549076 |
| Vasp | 0.97147503 | 0.82337028 | 0.73865139 | 0.0847189 | 0.30506621 | 0.90650187 |
| Cdkn2c | 0.12362772 | 0.2248349 | 0.14014521 | 0.08468969 | 0.00361048 | 0.07118415 |
| Paip1 | 0.41988377 | 0.49101577 | 0.40640355 | 0.08461222 | 0.17733622 | 0.90650187 |
| Mecom | 0.12475595 | 0.22593726 | 0.14133192 | 0.08460534 | 0.00389235 | 0.07598603 |
| Dap | 1.00092761 | 0.83560055 | 0.7511035 | 0.08449705 | 0.30849588 | 0.90650187 |
| Mbnl1 | 0.335007 | 0.42289667 | 0.33841438 | 0.0844823 | 0.13354085 | 0.90650187 |
| Snap23 | 0.55875148 | 0.59061989 | 0.50616802 | 0.08445187 | 0.22912561 | 0.90650187 |
| Metrn | 0.22985035 | 0.32986217 | 0.24547788 | 0.08438429 | 0.06323142 | 0.69600353 |
| Acvrl1 | 0.23170255 | 0.33152333 | 0.24720662 | 0.08431672 | 0.06472199 | 0.70728895 |
| Fgfr1 | 0.32987964 | 0.4183424 | 0.3341167 | 0.0842257 | 0.13121433 | 0.90650187 |
| Parm1 | 0.46403979 | 0.52379282 | 0.43957221 | 0.08422061 | 0.19717262 | 0.90650187 |
| Stmn3 | 0.10737281 | 0.20693043 | 0.12287495 | 0.08405548 | 0.0011785 | 0.02710184 |
| Armcx2 | 0.84892705 | 0.76238423 | 0.67840183 | 0.08398241 | 0.29104257 | 0.90650187 |
| Fkbp3 | 1.45403351 | 0.96910821 | 0.88514834 | 0.08395986 | 0.336628 | 0.90650187 |
| Elavl4 | 0.1542737 | 0.25576901 | 0.1718375 | 0.08393151 | 0.0149508 | 0.23049408 |
| Klhl7 | 0.96819088 | 0.82103483 | 0.7372273 | 0.08380753 | 0.30664819 | 0.90650187 |
| Ms4a7 | 0.03112191 | 0.12118551 | 0.0374112 | 0.08377431 | 1.20E-23 | 2.25E-21 |
| Sin3b | 1.35381961 | 0.94775852 | 0.86402229 | 0.08373624 | 0.3332911 | 0.90650187 |
| Hells | 0.22110906 | 0.32069508 | 0.23727204 | 0.08342304 | 0.05902145 | 0.66199323 |
| Gpc1 | 0.28377736 | 0.37778717 | 0.29442786 | 0.08335931 | 0.10407775 | 0.90650187 |
| Lmcd1 | 0.14991689 | 0.25067001 | 0.16739961 | 0.0832704 | 0.0135041 | 0.21360641 |
| Notch2 | 0.22283608 | 0.32208121 | 0.23889977 | 0.08318144 | 0.0608237 | 0.67586334 |

| | | | | | | |
|---|---|---|---|---|---|---|
| Kmt2d | 0.3990766 | 0.47336916 | 0.39026762 | 0.08310153 | 0.17191521 | 0.90650187 |
| Pdgfa | 0.40418952 | 0.47730883 | 0.39426301 | 0.08304583 | 0.17453287 | 0.90650187 |
| Sapcd1 | 0.18264314 | 0.28320047 | 0.20020849 | 0.08299198 | 0.03267712 | 0.42491438 |
| Actg2 | 0.08076568 | 0.17692859 | 0.09397872 | 0.08294987 | 4.36E-05 | 0.00137677 |
| Airn | 0.3870119 | 0.463561 | 0.38075738 | 0.08280362 | 0.16682461 | 0.90650187 |
| Isg15 | 0.04602065 | 0.13748776 | 0.05477699 | 0.08271077 | 9.56E-12 | 7.37E-10 |
| Lgals9 | 0.2688664 | 0.36381251 | 0.28117471 | 0.0826378 | 0.09567897 | 0.90650187 |
| Ctsd | 1.37491595 | 0.95148325 | 0.8688564 | 0.08262685 | 0.33623153 | 0.90650187 |
| Nrep | 0.80233892 | 0.73605424 | 0.6535293 | 0.08252495 | 0.28727178 | 0.90650187 |
| Rrm1 | 0.37684384 | 0.45513368 | 0.372653 | 0.08248068 | 0.16256694 | 0.90650187 |
| Tpst2 | 0.71670675 | 0.68764886 | 0.60521781 | 0.08243105 | 0.27242527 | 0.90650187 |
| Dctpp1 | 0.625159 | 0.63187239 | 0.54950639 | 0.08236599 | 0.25259185 | 0.90650187 |
| Ccne1 | 0.17509852 | 0.27510827 | 0.192752 | 0.08235627 | 0.0287524 | 0.38575923 |
| Uqcrfs1 | 1.54546908 | 0.98313529 | 0.90078034 | 0.08235494 | 0.34220783 | 0.90650187 |
| Abhd2 | 0.33866748 | 0.42381246 | 0.34146891 | 0.08234355 | 0.14185191 | 0.90650187 |
| Ezr | 0.43221759 | 0.49801702 | 0.41581179 | 0.08220523 | 0.18973092 | 0.90650187 |
| Adamts2 | 0.21483027 | 0.3134647 | 0.23132854 | 0.08213617 | 0.05722888 | 0.64823856 |
| Cpne3 | 0.37224434 | 0.45104195 | 0.36896015 | 0.08208181 | 0.16133037 | 0.90650187 |
| Ddit4 | 0.27501157 | 0.36872983 | 0.28666231 | 0.08206752 | 0.10155932 | 0.90650187 |
| Sipa1 | 0.35461215 | 0.43657814 | 0.3546405 | 0.08193764 | 0.15217922 | 0.90650187 |
| Vps8 | 0.26392156 | 0.35860982 | 0.27673248 | 0.08187734 | 0.09419958 | 0.90650187 |
| Rnh1 | 1.00250458 | 0.83362709 | 0.75175171 | 0.08187538 | 0.31412786 | 0.90650187 |
| Twist2 | 0.24998895 | 0.34594113 | 0.26409054 | 0.08185059 | 0.08412576 | 0.84688451 |
| Gmnn | 0.23466563 | 0.33180036 | 0.24996504 | 0.08183532 | 0.07277457 | 0.7682266 |
| Cox6b2 | 0.35427393 | 0.43617632 | 0.35436333 | 0.08181298 | 0.15235832 | 0.90650187 |
| Spi1 | 0.04155623 | 0.13141985 | 0.04960924 | 0.08181061 | 1.14E-13 | 1.11E-11 |
| Eif3e | 2.14619074 | 1.00463963 | 0.92296332 | 0.08167631 | 0.34700926 | 0.90650187 |
| Flrt2 | 0.33905444 | 0.4234545 | 0.34179115 | 0.08166335 | 0.14407738 | 0.90650187 |
| Sncg | 0.06106866 | 0.15353624 | 0.07197364 | 0.0815626 | 2.35E-07 | 1.08E-05 |
| Snrk | 0.41915627 | 0.48733554 | 0.40584485 | 0.08149069 | 0.18602354 | 0.90650187 |

| | | | | | | |
|---|---|---|---|---|---|---|
| Mcm7 | 0.37874577 | 0.45547161 | 0.37417511 | 0.0812965 | 0.16704946 | 0.90650187 |
| Usmg5 | 2.89640006 | 0.93257833 | 0.85140094 | 0.08117739 | 0.33583057 | 0.90650187 |
| Gngt2 | 0.20799221 | 0.30588131 | 0.22480938 | 0.08107193 | 0.05444551 | 0.62985251 |
| Spock2 | 0.49601279 | 0.54370883 | 0.46272272 | 0.08098611 | 0.21826279 | 0.90650187 |
| Slitrk6 | 0.26760228 | 0.36084046 | 0.28004128 | 0.08079917 | 0.09980337 | 0.90650187 |
| Impdh2 | 1.15537985 | 0.88904152 | 0.80829098 | 0.08075053 | 0.32847483 | 0.90650187 |
| Atp6v1f | 1.89439846 | 1.00756539 | 0.92693534 | 0.08063005 | 0.34948773 | 0.90650187 |
| Nop10 | 1.54584435 | 0.98144662 | 0.9008396 | 0.08060702 | 0.34539164 | 0.90650187 |
| Sphkap | 0.37443332 | 0.45129489 | 0.37071974 | 0.08057515 | 0.16696127 | 0.90650187 |
| Mllt11 | 0.41352507 | 0.48188133 | 0.40150693 | 0.08037439 | 0.18675184 | 0.90650187 |
| Fgd2 | 0.48795948 | 0.5371076 | 0.45695767 | 0.08014993 | 0.21776754 | 0.90650187 |
| Mbnl2 | 1.28597686 | 0.92717131 | 0.84713169 | 0.08003962 | 0.33723008 | 0.90650187 |
| Adamtsl1 | 0.17468692 | 0.2722879 | 0.19234341 | 0.07994449 | 0.03231956 | 0.422433 |
| Hebp1 | 0.28845967 | 0.37840893 | 0.29854688 | 0.07986205 | 0.11717833 | 0.90650187 |
| Col27a1 | 0.3109282 | 0.39775366 | 0.31803233 | 0.07972133 | 0.13255756 | 0.90650187 |
| Timm8b | 1.25938157 | 0.91956054 | 0.8398408 | 0.07971974 | 0.33651696 | 0.90650187 |
| Phyh | 0.50823759 | 0.55109041 | 0.47138899 | 0.07970142 | 0.22612977 | 0.90650187 |
| Cd53 | 0.03222941 | 0.11833035 | 0.03871394 | 0.07961641 | 3.05E-20 | 4.42E-18 |
| Wnk3 | 0.24693256 | 0.34090739 | 0.26129202 | 0.07961536 | 0.08777716 | 0.87266231 |
| Tkt | 1.04091694 | 0.84679863 | 0.76718824 | 0.07961039 | 0.32228583 | 0.90650187 |
| Nkd1 | 0.30242324 | 0.39020743 | 0.31071065 | 0.07949678 | 0.12767944 | 0.90650187 |
| Prr5 | 0.39840555 | 0.4692299 | 0.38974177 | 0.07948814 | 0.18228672 | 0.90650187 |
| Lgmn | 0.43160153 | 0.49480869 | 0.4153445 | 0.07946419 | 0.19751287 | 0.90650187 |
| Kcnmb2 | 0.49280596 | 0.53988043 | 0.4604327 | 0.07944773 | 0.22151399 | 0.90650187 |
| Tgfbi | 0.11662126 | 0.21205249 | 0.13274086 | 0.07931163 | 0.00395116 | 0.07669505 |
| Ms4a6c | 0.04216589 | 0.12959481 | 0.05031674 | 0.07927807 | 1.24E-12 | 1.04E-10 |
| Pmepa1 | 0.31743038 | 0.40279262 | 0.32358692 | 0.0792057 | 0.13826017 | 0.90650187 |
| Iscu | 0.95377518 | 0.80973718 | 0.73063803 | 0.07909914 | 0.31515769 | 0.90650187 |
| Cat | 0.48090538 | 0.53090637 | 0.45187158 | 0.07903479 | 0.21841403 | 0.90650187 |
| Itm2b | 3.43023673 | 0.86012831 | 0.78139504 | 0.07873328 | 0.32709868 | 0.90650187 |

| | | | | | | |
|---|---|---|---|---|---|---|
| Ifi30 | 0.56115624 | 0.5864139 | 0.50778361 | 0.07863029 | 0.2455998 | 0.90650187 |
| Mcm5 | 0.25621118 | 0.34834573 | 0.26975943 | 0.0785863 | 0.09764393 | 0.90650187 |
| Plk1 | 0.10100983 | 0.19459558 | 0.11602531 | 0.07857027 | 0.00130415 | 0.02963291 |
| Akr1c12 | 0.40490595 | 0.4731663 | 0.39482125 | 0.07834505 | 0.18884807 | 0.90650187 |
| Tmem255a | 0.12101882 | 0.21563521 | 0.13739514 | 0.07824007 | 0.00567776 | 0.10411931 |
| Capsl | 0.3698214 | 0.44514997 | 0.36700773 | 0.07814224 | 0.17193512 | 0.90650187 |
| Hmgcs1 | 0.59471282 | 0.60809589 | 0.5299601 | 0.07813579 | 0.25608988 | 0.90650187 |
| Asf1b | 0.10747536 | 0.20107984 | 0.12298493 | 0.07809491 | 0.00237926 | 0.04961208 |
| Slc7a2 | 0.40428235 | 0.47239787 | 0.39433536 | 0.0780625 | 0.18941589 | 0.90650187 |
| Gjb5 | 0.10570113 | 0.19914257 | 0.12108033 | 0.07806223 | 0.00207751 | 0.04416816 |
| Agrn | 0.23217796 | 0.32551204 | 0.24764978 | 0.07786225 | 0.08109866 | 0.82626054 |
| Ly6a | 0.1000663 | 0.19254919 | 0.11500528 | 0.07754391 | 0.00136097 | 0.03069444 |
| Sod1 | 0.78283006 | 0.7203316 | 0.64282488 | 0.07750672 | 0.29597377 | 0.90650187 |
| Gm26522 | 0.03100525 | 0.11476923 | 0.03727385 | 0.07749538 | 1.20E-20 | 1.81E-18 |
| Cib1 | 0.72949793 | 0.69001543 | 0.61265642 | 0.07735901 | 0.28728336 | 0.90650187 |
| Sox10 | 0.04791838 | 0.13429941 | 0.05696448 | 0.07733493 | 7.93E-10 | 4.96E-08 |
| Abhd17a | 0.53688643 | 0.56847534 | 0.49131434 | 0.07716099 | 0.24253132 | 0.90650187 |
| Sub1 | 3.11141664 | 0.900249 | 0.82310783 | 0.07714117 | 0.33847103 | 0.90650187 |
| Bace2 | 0.09852802 | 0.19045317 | 0.11334842 | 0.07710475 | 0.00124685 | 0.02850141 |
| Ubald2 | 0.94468016 | 0.80344907 | 0.72635069 | 0.07709838 | 0.31850688 | 0.90650187 |
| Rab27a | 0.41194181 | 0.47737038 | 0.40028302 | 0.07708736 | 0.19595895 | 0.90650187 |
| Sptssa | 1.27632391 | 0.92160821 | 0.84453866 | 0.07706955 | 0.3424852 | 0.90650187 |
| Cdk2ap2 | 0.67105927 | 0.65492119 | 0.5779941 | 0.07692709 | 0.27703244 | 0.90650187 |
| Fam213a | 0.52259119 | 0.55813916 | 0.48143905 | 0.07670011 | 0.23939548 | 0.90650187 |
| Hyou1 | 0.36192653 | 0.43730712 | 0.36061151 | 0.07669561 | 0.17220161 | 0.90650187 |
| Tenm4 | 0.45464048 | 0.50922493 | 0.43263006 | 0.07659487 | 0.21562009 | 0.90650187 |
| Heg1 | 0.33929076 | 0.41853184 | 0.34198789 | 0.07654395 | 0.15986363 | 0.90650187 |
| Snx3 | 1.50469999 | 0.97066027 | 0.89412529 | 0.07653497 | 0.35177483 | 0.90650187 |
| Dnajc9 | 0.50186989 | 0.54330782 | 0.46688711 | 0.07642071 | 0.23341001 | 0.90650187 |
| Rnd3 | 0.30199753 | 0.3867004 | 0.31034248 | 0.07635791 | 0.13701809 | 0.90650187 |

| | | | | | | |
|---|---|---|---|---|---|---|
| Gm27033 | 0.32507993 | 0.40640419 | 0.33007332 | 0.07633087 | 0.15195952 | 0.90650187 |
| Pabpc4 | 0.55688815 | 0.58122049 | 0.50491378 | 0.07630671 | 0.25083752 | 0.90650187 |
| Hotairm1 | 0.1666632 | 0.26064398 | 0.18433921 | 0.07630477 | 0.03286946 | 0.42632142 |
| Acat1 | 1.23009127 | 0.9073587 | 0.83137784 | 0.07598086 | 0.34226456 | 0.90650187 |
| Mmp16 | 0.22545158 | 0.31705631 | 0.24135849 | 0.07569783 | 0.08161564 | 0.82804847 |
| Ogt | 1.18831732 | 0.89453006 | 0.81886031 | 0.07566975 | 0.34060508 | 0.90650187 |
| Unc93b1 | 0.1469904 | 0.23997118 | 0.16440597 | 0.07556521 | 0.02051064 | 0.29413421 |
| Skap2 | 0.36650359 | 0.4397953 | 0.36432584 | 0.07546945 | 0.17855145 | 0.90650187 |
| Xbp1 | 1.20681731 | 0.89991157 | 0.82447652 | 0.07543506 | 0.34209636 | 0.90650187 |
| Tff2 | 0.07653561 | 0.16439775 | 0.08930024 | 0.0750975 | 9.25E-05 | 0.00275285 |
| Smo | 0.36803186 | 0.44065569 | 0.36556229 | 0.0750934 | 0.18056565 | 0.90650187 |
| Taf1d | 0.67751365 | 0.65672979 | 0.58190848 | 0.07482131 | 0.28379229 | 0.90650187 |
| 1110008F13l | 0.96342683 | 0.80985669 | 0.73515368 | 0.07470301 | 0.32572724 | 0.90650187 |
| Pnmal2 | 0.64116321 | 0.63425568 | 0.55957113 | 0.07468455 | 0.27647553 | 0.90650187 |
| Mpzl1 | 1.19281107 | 0.89493808 | 0.82027407 | 0.07466401 | 0.34286611 | 0.90650187 |
| St6galnac3 | 0.19615239 | 0.28801173 | 0.21340545 | 0.07460628 | 0.06007092 | 0.669456 |
| Cbfa2t3 | 0.41794954 | 0.47951797 | 0.40491725 | 0.07460073 | 0.20638036 | 0.90650187 |
| Pdlim4 | 0.32820067 | 0.40728076 | 0.33270455 | 0.07457621 | 0.15950626 | 0.90650187 |
| Psmg2 | 0.39593388 | 0.46235698 | 0.38780189 | 0.07455509 | 0.19636697 | 0.90650187 |
| Arhgef25 | 0.32607383 | 0.40534279 | 0.33091223 | 0.07443056 | 0.15867228 | 0.90650187 |
| Ech1 | 1.02623357 | 0.83568206 | 0.76136631 | 0.07431575 | 0.33217017 | 0.90650187 |
| Sh3bgrl | 0.80568641 | 0.72966163 | 0.65534887 | 0.07431276 | 0.30709346 | 0.90650187 |
| Ltc4s | 0.02095183 | 0.09966981 | 0.02535771 | 0.0743121 | 4.24E-39 | 1.34E-36 |
| Nt5dc2 | 1.10000072 | 0.86345899 | 0.78924753 | 0.07421147 | 0.33796916 | 0.90650187 |
| Scand1 | 1.58043726 | 0.97970662 | 0.90568446 | 0.07402216 | 0.35817463 | 0.90650187 |
| Syce2 | 0.527539 | 0.55888632 | 0.48487164 | 0.07401468 | 0.24869159 | 0.90650187 |
| Ngfr | 0.14611093 | 0.23732157 | 0.16350434 | 0.07381723 | 0.02237186 | 0.31488391 |
| Cacna2d1 | 0.53611694 | 0.56460206 | 0.49078604 | 0.07381602 | 0.25186402 | 0.90650187 |
| Npr3 | 0.1735204 | 0.26480315 | 0.19118432 | 0.07361883 | 0.04346144 | 0.53150233 |
| Rps27a | 5.0313333 | 0.61733894 | 0.54386287 | 0.07347607 | 0.27405221 | 0.90650187 |

| | | | | | | |
|---|---|---|---|---|---|---|
| Lhfpl2 | 0.27566396 | 0.36069154 | 0.28724272 | 0.07344882 | 0.12782039 | 0.90650187 |
| Gria2 | 0.16573125 | 0.25679782 | 0.18340485 | 0.07339297 | 0.03762033 | 0.47484348 |
| Efna5 | 0.19083574 | 0.28162127 | 0.20823514 | 0.07338614 | 0.05859023 | 0.65923772 |
| Fam151a | 0.21696502 | 0.30673667 | 0.23335383 | 0.07338284 | 0.08105488 | 0.82626054 |
| Lrpap1 | 1.30135253 | 0.92440926 | 0.85113192 | 0.07327734 | 0.3509563 | 0.90650187 |
| Lbh | 0.30077174 | 0.38250424 | 0.30928147 | 0.07322277 | 0.14628445 | 0.90650187 |
| Racgap1 | 0.18617578 | 0.27668346 | 0.20367849 | 0.07300497 | 0.05552972 | 0.63800617 |
| Ccl3 | 0.0354309 | 0.1153932 | 0.04246904 | 0.07292416 | 1.14E-14 | 1.16E-12 |
| Rspo3 | 0.15097435 | 0.24138241 | 0.16847877 | 0.07290364 | 0.02719674 | 0.36846491 |
| Efnb1 | 0.49085103 | 0.53190117 | 0.45903286 | 0.07286831 | 0.24018443 | 0.90650187 |
| Kif12 | 0.53624745 | 0.56365667 | 0.49087567 | 0.072781 | 0.25490301 | 0.90650187 |
| Cd14 | 0.06760818 | 0.1520798 | 0.07934198 | 0.07273783 | 2.30E-05 | 0.00076008 |
| Egfl6 | 0.10913932 | 0.19745501 | 0.12476759 | 0.07268742 | 0.00479974 | 0.09086026 |
| Plin2 | 0.37891012 | 0.44692408 | 0.37430651 | 0.07261757 | 0.19421374 | 0.90650187 |
| Mbp | 0.23751933 | 0.32522975 | 0.25261334 | 0.07261641 | 0.10063581 | 0.90650187 |
| Cd82 | 0.33822184 | 0.4136407 | 0.34109764 | 0.07254306 | 0.17220927 | 0.90650187 |
| Hmgcs2 | 0.10289529 | 0.19059066 | 0.11806028 | 0.07253038 | 0.00315559 | 0.06336608 |
| Bpnt1 | 0.56221507 | 0.58099514 | 0.50849386 | 0.07250128 | 0.26308722 | 0.90650187 |
| Mcm3 | 0.25200863 | 0.33825725 | 0.26593471 | 0.07232254 | 0.11332964 | 0.90650187 |
| Rgs10 | 0.20982779 | 0.29881851 | 0.22656412 | 0.07225439 | 0.07812976 | 0.80409648 |
| Eef1b2 | 2.72714851 | 0.94696907 | 0.87472927 | 0.07223981 | 0.35675659 | 0.90650187 |
| Mrpl32 | 0.83831346 | 0.745051 | 0.67282189 | 0.07222911 | 0.31659315 | 0.90650187 |
| Tesc | 0.18617714 | 0.27586554 | 0.20367982 | 0.07218572 | 0.05756388 | 0.65057885 |
| Sumf2 | 0.37926989 | 0.44674209 | 0.37459406 | 0.07214803 | 0.19593301 | 0.90650187 |
| Cntfr | 0.29565882 | 0.37698243 | 0.30484101 | 0.07214142 | 0.14638503 | 0.90650187 |
| Dnajb6 | 1.56897051 | 0.97631986 | 0.90421764 | 0.07210222 | 0.36148503 | 0.90650187 |
| Arhgap18 | 0.14790964 | 0.23739399 | 0.16534742 | 0.07204657 | 0.02636796 | 0.36044904 |
| Prkacb | 0.51583296 | 0.54866182 | 0.47672435 | 0.07193747 | 0.2511613 | 0.90650187 |
| Ocrl | 0.59845314 | 0.604145 | 0.53238995 | 0.07175505 | 0.27452811 | 0.90650187 |
| Gamt | 0.27485517 | 0.35821609 | 0.28652312 | 0.07169297 | 0.13299029 | 0.90650187 |

| | | | | | | |
|---|---|---|---|---|---|---|
| Mrgprf | 0.18820936 | 0.27726019 | 0.20566987 | 0.07159033 | 0.06087822 | 0.67597503 |
| Coa3 | 1.73012751 | 0.99189976 | 0.92031148 | 0.07158828 | 0.3647426 | 0.90650187 |
| Avp | 0.04989925 | 0.1306986 | 0.05924198 | 0.07145662 | 4.11E-08 | 2.15E-06 |
| Hk2 | 0.15115919 | 0.2401196 | 0.16866726 | 0.07145233 | 0.02983071 | 0.39776898 |
| Limd2 | 0.78748327 | 0.71681351 | 0.64539436 | 0.07141915 | 0.31137752 | 0.90650187 |
| Rnaseh2c | 0.71269951 | 0.67409707 | 0.60287018 | 0.07122689 | 0.29971095 | 0.90650187 |
| Upk1b | 0.05480093 | 0.13606858 | 0.06485222 | 0.07121636 | 5.25E-07 | 2.32E-05 |
| Nefl | 0.08125609 | 0.16571391 | 0.09451946 | 0.07119445 | 0.00040619 | 0.01077568 |
| Gm2000 | 1.2443864 | 0.90671704 | 0.83555985 | 0.07115719 | 0.35249518 | 0.90650187 |
| Tspan15 | 0.35597331 | 0.42688506 | 0.35575497 | 0.07113009 | 0.18703916 | 0.90650187 |
| St18 | 0.42121025 | 0.4783365 | 0.40742077 | 0.07091573 | 0.21952364 | 0.90650187 |
| Wfdc15b | 0.14070631 | 0.2288194 | 0.15794375 | 0.07087565 | 0.02302577 | 0.32170478 |
| C1qtnf7 | 0.13596709 | 0.22386677 | 0.15303957 | 0.0708272 | 0.01982192 | 0.2866918 |
| Klf13 | 0.31794868 | 0.3948356 | 0.3240281 | 0.0708075 | 0.16565815 | 0.90650187 |
| Idi1 | 0.23361762 | 0.31973309 | 0.24899041 | 0.07074268 | 0.10328072 | 0.90650187 |
| 1700011H14 | 0.12428364 | 0.21139373 | 0.14083533 | 0.07055841 | 0.01296542 | 0.20746037 |
| Cbx5 | 0.8120574 | 0.72934309 | 0.6587985 | 0.07054459 | 0.31702346 | 0.90650187 |
| Olfml2b | 0.27208774 | 0.35456228 | 0.28405623 | 0.07050606 | 0.13491564 | 0.90650187 |
| Ttyh1 | 0.18081261 | 0.26890442 | 0.19840508 | 0.07049933 | 0.05709254 | 0.64823856 |
| Hist1h2ae | 0.13952111 | 0.22720751 | 0.15671978 | 0.07048772 | 0.02277733 | 0.31940792 |
| H2-Ke6 | 0.81232896 | 0.72924644 | 0.65894516 | 0.07030128 | 0.31764617 | 0.90650187 |
| Chic1 | 0.44386546 | 0.49475726 | 0.42459504 | 0.07016221 | 0.23128722 | 0.90650187 |
| Nrm | 0.28783587 | 0.36813524 | 0.29799928 | 0.07013596 | 0.14771111 | 0.90650187 |
| Neat1 | 0.14196098 | 0.22933014 | 0.15923768 | 0.07009247 | 0.02518282 | 0.34642905 |
| Kif22 | 0.15941822 | 0.24705109 | 0.17704975 | 0.07000134 | 0.03940022 | 0.49213045 |
| Plod1 | 0.39561948 | 0.45755124 | 0.38755479 | 0.06999645 | 0.21100961 | 0.90650187 |
| Plxnd1 | 0.12660194 | 0.21323357 | 0.14327003 | 0.06996354 | 0.01496793 | 0.23052438 |
| Hsd17b10 | 0.99020496 | 0.81660055 | 0.7466694 | 0.06993114 | 0.3385718 | 0.90650187 |
| Adgre5 | 0.13850896 | 0.22554976 | 0.15567321 | 0.06987655 | 0.02299497 | 0.32156992 |
| Gnai1 | 0.49466403 | 0.53136734 | 0.46176054 | 0.0696068 | 0.25138412 | 0.90650187 |

| | | | | | | |
|---|---|---|---|---|---|---|
| Acads | 0.51969643 | 0.54898922 | 0.47942355 | 0.06956566 | 0.25944105 | 0.90650187 |
| Ndrg1 | 0.54696928 | 0.56764793 | 0.49820163 | 0.0694463 | 0.26773073 | 0.90650187 |
| Adgrg1 | 0.41580577 | 0.47266205 | 0.40326669 | 0.06939536 | 0.22213176 | 0.90650187 |
| Fam64a | 0.10406494 | 0.18837348 | 0.11932042 | 0.06905306 | 0.00504457 | 0.09455303 |
| Rfk | 0.48119677 | 0.52111959 | 0.45208234 | 0.06903725 | 0.24860502 | 0.90650187 |
| Man2b1 | 0.31296266 | 0.38866952 | 0.31977427 | 0.06889525 | 0.16908075 | 0.90650187 |
| Tmem184a | 0.44592964 | 0.49500416 | 0.42614088 | 0.06886328 | 0.23625576 | 0.90650187 |
| Nop56 | 0.89318071 | 0.76990605 | 0.70113051 | 0.06877554 | 0.33138934 | 0.90650187 |
| Rbpj | 0.39069422 | 0.45240067 | 0.3836724 | 0.06872827 | 0.21291331 | 0.90650187 |
| Col9a3 | 0.21706212 | 0.30210355 | 0.23344584 | 0.06865771 | 0.09552532 | 0.90650187 |
| Akap13 | 0.58106531 | 0.58960194 | 0.52102206 | 0.06857988 | 0.27922143 | 0.90650187 |
| Rpl9-ps6 | 1.32575903 | 0.92562282 | 0.85723378 | 0.06838904 | 0.36142033 | 0.90650187 |
| Baiap3 | 0.52424178 | 0.55094804 | 0.48258589 | 0.06836215 | 0.26442721 | 0.90650187 |
| Xlr3a | 0.14473565 | 0.23042347 | 0.16209261 | 0.06833086 | 0.03046072 | 0.40263779 |
| Lpar1 | 0.22071431 | 0.30518234 | 0.23689956 | 0.06828279 | 0.10003009 | 0.90650187 |
| Ikbip | 0.50605295 | 0.53812643 | 0.46984746 | 0.06827897 | 0.25912593 | 0.90650187 |
| Medag | 0.21304781 | 0.29789062 | 0.22963388 | 0.06825674 | 0.09318082 | 0.90318385 |
| Pltp | 0.19983864 | 0.28515508 | 0.21697224 | 0.06818285 | 0.0812041 | 0.82678063 |
| Klhdc8b | 0.84277246 | 0.74322267 | 0.67517221 | 0.06805046 | 0.32705073 | 0.90650187 |
| Mdm1 | 0.28529934 | 0.3637489 | 0.29576895 | 0.06797995 | 0.15344148 | 0.90650187 |
| Ethe1 | 0.33855219 | 0.40923549 | 0.34137287 | 0.06786262 | 0.18841385 | 0.90650187 |
| Barx1 | 0.12877376 | 0.21323176 | 0.14554498 | 0.06768678 | 0.01934529 | 0.28167412 |
| Kif20a | 0.11059273 | 0.1939669 | 0.12632186 | 0.06764504 | 0.00864197 | 0.14793535 |
| F3 | 0.42558092 | 0.47831905 | 0.41076323 | 0.06755581 | 0.23234599 | 0.90650187 |
| Lfng | 0.24021237 | 0.32245592 | 0.25510515 | 0.06735077 | 0.12025908 | 0.90650187 |
| Pole4 | 0.51952225 | 0.54646801 | 0.47930211 | 0.0671659 | 0.26665054 | 0.90650187 |
| Scara3 | 0.27587902 | 0.35454499 | 0.28743397 | 0.06711102 | 0.14964091 | 0.90650187 |
| Rap2b | 0.25473806 | 0.33552765 | 0.26842069 | 0.06710696 | 0.13319141 | 0.90650187 |
| Fam69a | 0.19774973 | 0.28203922 | 0.21495293 | 0.06708629 | 0.08265032 | 0.83684877 |
| Grcc10 | 1.5663791 | 0.97090321 | 0.9038489 | 0.06705431 | 0.37077232 | 0.90650187 |

| | | | | | | |
|---|---|---|---|---|---|---|
| Lamp2 | 1.09496622 | 0.85446757 | 0.78745676 | 0.06701081 | 0.35259972 | 0.90650187 |
| Bola3 | 0.73628087 | 0.68357433 | 0.61656915 | 0.06700518 | 0.31450425 | 0.90650187 |
| Arl4a | 0.68773122 | 0.65504359 | 0.58806151 | 0.06698208 | 0.30629891 | 0.90650187 |
| Srsf3 | 2.23670732 | 0.98529036 | 0.91832209 | 0.06696827 | 0.37289555 | 0.90650187 |
| Wipi1 | 0.67694882 | 0.64846121 | 0.58156678 | 0.06689443 | 0.30455171 | 0.90650187 |
| Praf2 | 0.38177416 | 0.44346598 | 0.37659283 | 0.06687315 | 0.21493191 | 0.90650187 |
| Stom | 0.2255686 | 0.30833604 | 0.24146832 | 0.06686772 | 0.10914591 | 0.90650187 |
| Ptprz1 | 0.1193594 | 0.20246525 | 0.13564162 | 0.06682363 | 0.01425773 | 0.22251726 |
| Rfc1 | 0.65380744 | 0.63401589 | 0.56742129 | 0.0665946 | 0.30092138 | 0.90650187 |
| Cnn3 | 1.7965275 | 0.99061864 | 0.92406251 | 0.06655613 | 0.37440938 | 0.90650187 |
| Mboat4 | 0.10684978 | 0.1888292 | 0.12231384 | 0.06651537 | 0.00781246 | 0.13634581 |
| Ppp1r14b | 0.7401062 | 0.68522754 | 0.61876615 | 0.06646139 | 0.31650286 | 0.90650187 |
| Surf4 | 1.16207249 | 0.87689445 | 0.81044461 | 0.06644984 | 0.35774105 | 0.90650187 |
| Pcsk2os1 | 0.43018175 | 0.4806786 | 0.41426654 | 0.06641207 | 0.23802008 | 0.90650187 |
| Clic1 | 2.18414164 | 0.98753852 | 0.92123515 | 0.06630337 | 0.37449921 | 0.90650187 |
| Rab13 | 0.33149869 | 0.40176457 | 0.33547618 | 0.06628838 | 0.18985543 | 0.90650187 |
| Hdlbp | 1.20308223 | 0.88957969 | 0.82336248 | 0.06621722 | 0.36034804 | 0.90650187 |
| Ppp1r14c | 0.20294537 | 0.28612458 | 0.21996665 | 0.06615793 | 0.09059891 | 0.89081117 |
| G6pc2 | 0.14851525 | 0.23212375 | 0.16596712 | 0.06615664 | 0.03819096 | 0.47978578 |
| Stip1 | 1.02700095 | 0.82777976 | 0.76167265 | 0.06610711 | 0.3498064 | 0.90650187 |
| Ak1 | 0.37867193 | 0.44011465 | 0.37411607 | 0.06599858 | 0.21644421 | 0.90650187 |
| Qdpr | 0.81931034 | 0.72865506 | 0.66270231 | 0.06595275 | 0.32908872 | 0.90650187 |
| Sep-04 | 0.150628 | 0.23394092 | 0.16812545 | 0.06581547 | 0.04090268 | 0.50559243 |
| Ttc3 | 1.7497375 | 0.98734898 | 0.92157865 | 0.06577034 | 0.37551946 | 0.90650187 |
| Mmd | 0.22093368 | 0.30285503 | 0.23710658 | 0.06574845 | 0.10883563 | 0.90650187 |
| Hs6st2 | 0.13224089 | 0.21490599 | 0.14916502 | 0.06574097 | 0.02503904 | 0.34478439 |
| Idh1 | 0.42476461 | 0.47586906 | 0.41014004 | 0.06572903 | 0.23809224 | 0.90650187 |
| Tnnt1 | 0.10057378 | 0.18121063 | 0.11555404 | 0.06565659 | 0.00576935 | 0.10540855 |
| Kitl | 0.23886081 | 0.3193256 | 0.25385548 | 0.06547012 | 0.12578679 | 0.90650187 |
| Lhx1 | 0.21236316 | 0.29442103 | 0.22898208 | 0.06543895 | 0.10195788 | 0.90650187 |

| | | | | | | |
|---|---|---|---|---|---|---|
| Rpl17 | 4.8064935 | 0.63095753 | 0.56554461 | 0.06541292 | 0.30355791 | 0.90650187 |
| Kif5b | 1.65068172 | 0.97907462 | 0.91366521 | 0.06540941 | 0.37514328 | 0.90650187 |
| Gcsh | 0.79029653 | 0.71233608 | 0.646943 | 0.06539308 | 0.32658511 | 0.90650187 |
| Matn4 | 0.07300363 | 0.15074751 | 0.08537421 | 0.0653733 | 0.00033189 | 0.00888214 |
| Tssc4 | 0.72952588 | 0.67802982 | 0.61267259 | 0.06535723 | 0.31766517 | 0.90650187 |
| Rpl37 | 4.99397367 | 0.60867392 | 0.54374808 | 0.06492584 | 0.29777024 | 0.90650187 |
| Gnb4 | 0.2751374 | 0.35165411 | 0.28677429 | 0.06487982 | 0.15726214 | 0.90650187 |
| Cfl2 | 1.09285195 | 0.85156066 | 0.78670188 | 0.06485877 | 0.35699111 | 0.90650187 |
| Pla2g1b | 0.1315012 | 0.21324875 | 0.1483939 | 0.06485485 | 0.02601224 | 0.35719246 |
| Rgs4 | 0.10097494 | 0.18083771 | 0.11598761 | 0.06485009 | 0.00646679 | 0.11592181 |
| Procr | 0.15904526 | 0.24151147 | 0.17667289 | 0.06483857 | 0.05138949 | 0.60504342 |
| Camk1 | 0.48592679 | 0.52030525 | 0.45549551 | 0.06480974 | 0.26351743 | 0.90650187 |
| Cdc25b | 0.11453759 | 0.19528486 | 0.1305273 | 0.06475757 | 0.01370718 | 0.21614402 |
| Ccl4 | 0.02159117 | 0.0908535 | 0.0261203 | 0.0647332 | 1.57E-28 | 3.63E-26 |
| Odc1 | 0.66766534 | 0.64065567 | 0.57592715 | 0.06472851 | 0.30866353 | 0.90650187 |
| Esco2 | 0.07535081 | 0.15261572 | 0.08798526 | 0.06463047 | 0.00054623 | 0.01400207 |
| Shisa4 | 0.29820282 | 0.3716319 | 0.30705352 | 0.06457838 | 0.17489831 | 0.90650187 |
| Cct7 | 2.06824051 | 0.99014992 | 0.92578426 | 0.06436566 | 0.37862911 | 0.90650187 |
| Cisd1 | 1.00964 | 0.81903449 | 0.75467229 | 0.06436221 | 0.35229068 | 0.90650187 |
| Fmod | 0.11543025 | 0.19578692 | 0.13147626 | 0.06431066 | 0.01483245 | 0.22960081 |
| Slc16a3 | 0.14361807 | 0.22512083 | 0.16094378 | 0.06417705 | 0.03813876 | 0.47967968 |
| Nhp2 | 0.63714833 | 0.62123491 | 0.55705933 | 0.06417557 | 0.30427214 | 0.90650187 |
| Rpl36-ps3 | 1.32284975 | 0.92067667 | 0.85650944 | 0.06416722 | 0.3695475 | 0.90650187 |
| Ppp1r17 | 0.05154238 | 0.12527825 | 0.06112666 | 0.06415159 | 1.54E-06 | 6.43E-05 |
| Skp2 | 0.26912321 | 0.34542092 | 0.28140479 | 0.06401613 | 0.15593145 | 0.90650187 |
| 2610524H06 | 0.66847978 | 0.64039266 | 0.5764237 | 0.06396896 | 0.31088131 | 0.90650187 |
| Arpc3 | 1.58221513 | 0.96978966 | 0.9058956 | 0.06389406 | 0.37692964 | 0.90650187 |
| Arhgap10 | 0.25115943 | 0.32894994 | 0.26515979 | 0.06379015 | 0.14242433 | 0.90650187 |
| Hey1 | 0.06187342 | 0.13659458 | 0.0728838 | 0.06371078 | 5.09E-05 | 0.00159316 |
| Jund | 0.88627807 | 0.76133365 | 0.69764655 | 0.0636871 | 0.34243277 | 0.90650187 |

| | | | | | | |
|---|---|---|---|---|---|---|
| Ftl1 | 4.88940643 | 0.61862808 | 0.55499175 | 0.06363634 | 0.30511643 | 0.90650187 |
| Epas1 | 0.07936802 | 0.1560107 | 0.09243573 | 0.06357497 | 0.00111567 | 0.02597148 |
| Cyp51 | 0.34824154 | 0.4129692 | 0.34940415 | 0.06356505 | 0.2093257 | 0.90650187 |
| Incenp | 0.14415328 | 0.22503989 | 0.16149414 | 0.06354576 | 0.04012081 | 0.49826502 |
| Jag1 | 0.15513228 | 0.23606563 | 0.17270948 | 0.06335615 | 0.05146511 | 0.60504342 |
| Rasd1 | 0.21930276 | 0.29885191 | 0.23556633 | 0.06328558 | 0.11617543 | 0.90650187 |
| Stab1 | 0.07527465 | 0.1511585 | 0.08790067 | 0.06325783 | 0.00068884 | 0.01725053 |
| Nrgn | 0.07351743 | 0.14913287 | 0.08594646 | 0.06318641 | 0.00054091 | 0.01390299 |
| Lgi2 | 0.23159503 | 0.31026647 | 0.24710635 | 0.06316012 | 0.12791714 | 0.90650187 |
| 1700025G04 | 0.31377021 | 0.38357948 | 0.32046471 | 0.06311477 | 0.19063689 | 0.90650187 |
| Myh10 | 0.41922418 | 0.46872986 | 0.40589702 | 0.06283284 | 0.24566982 | 0.90650187 |
| Rab25 | 0.49315305 | 0.52348275 | 0.46068097 | 0.06280178 | 0.27224343 | 0.90650187 |
| Ccr2 | 0.02424256 | 0.09202654 | 0.02927582 | 0.06275072 | 7.95E-22 | 1.33E-19 |
| Bcl2 | 0.31335469 | 0.38279532 | 0.32010953 | 0.0626858 | 0.19199565 | 0.90650187 |
| Mxd3 | 0.1008463 | 0.17844982 | 0.1158486 | 0.06260122 | 0.00814669 | 0.14136738 |
| Stxbp6 | 0.29353231 | 0.36558675 | 0.30298672 | 0.06260003 | 0.17917702 | 0.90650187 |
| Ctsh | 0.19109584 | 0.27101679 | 0.20848878 | 0.06252801 | 0.09121919 | 0.89511343 |
| Golga4 | 0.79185423 | 0.71024299 | 0.64779854 | 0.06244445 | 0.33413134 | 0.90650187 |
| Slc16a10 | 0.26129778 | 0.33680993 | 0.27436598 | 0.06244395 | 0.15581884 | 0.90650187 |
| Gsto1 | 0.41414321 | 0.46437204 | 0.40198426 | 0.06238778 | 0.24511088 | 0.90650187 |
| Metrnl | 0.20911404 | 0.28822376 | 0.22588223 | 0.06234153 | 0.10992093 | 0.90650187 |
| Sec23a | 0.39520225 | 0.44945762 | 0.38722676 | 0.06223087 | 0.23747466 | 0.90650187 |
| Elp4 | 0.4613837 | 0.49981156 | 0.43761672 | 0.06219484 | 0.26375316 | 0.90650187 |
| Hyal2 | 0.39492747 | 0.44911629 | 0.38701055 | 0.06210574 | 0.23779567 | 0.90650187 |
| Gpr83 | 0.07911086 | 0.15419198 | 0.09215152 | 0.06204046 | 0.00138135 | 0.03106195 |
| Rfx3 | 0.37814969 | 0.43561706 | 0.37369837 | 0.06191869 | 0.23068291 | 0.90650187 |
| Hpcal1 | 0.61517181 | 0.60506387 | 0.54315376 | 0.06191011 | 0.30617494 | 0.90650187 |
| Cox14 | 1.55422539 | 0.96395761 | 0.90209722 | 0.06186038 | 0.38023912 | 0.90650187 |
| Camk2n1 | 0.53788668 | 0.55380587 | 0.49200052 | 0.06180535 | 0.28826541 | 0.90650187 |
| Hlx | 0.14550663 | 0.22467872 | 0.16288429 | 0.06179443 | 0.04584424 | 0.55307801 |

| | | | | | | |
|---|---|---|---|---|---|---|
| Enho | 0.41108651 | 0.46135553 | 0.39962106 | 0.06173447 | 0.24611432 | 0.90650187 |
| Ms4a6b | 0.02458216 | 0.09134687 | 0.02967918 | 0.06166769 | 1.27E-20 | 1.89E-18 |
| BC023829 | 0.47760044 | 0.51110124 | 0.44947712 | 0.06162413 | 0.27109631 | 0.90650187 |
| Pla2g2f | 0.26801452 | 0.34191777 | 0.28041107 | 0.0615067 | 0.16475024 | 0.90650187 |
| C3ar1 | 0.0297963 | 0.09728928 | 0.03584936 | 0.06143991 | 1.28E-14 | 1.29E-12 |
| Nmt2 | 0.63316343 | 0.61584723 | 0.55455766 | 0.06128956 | 0.31159752 | 0.90650187 |
| Trim35 | 1.27381651 | 0.90498987 | 0.84386024 | 0.06112963 | 0.37371051 | 0.90650187 |
| Wisp1 | 0.17603103 | 0.25480088 | 0.19367701 | 0.06112387 | 0.08030547 | 0.82093036 |
| Abcd3 | 0.78982402 | 0.70770704 | 0.64668327 | 0.06102377 | 0.33742311 | 0.90650187 |
| Dhrs3 | 0.09271871 | 0.16807707 | 0.10706187 | 0.0610152 | 0.00564523 | 0.10411931 |
| Ank2 | 0.28049265 | 0.35246517 | 0.29152629 | 0.06093888 | 0.17637138 | 0.90650187 |
| Isyna1 | 0.85233741 | 0.74110594 | 0.6801847 | 0.06092124 | 0.34525097 | 0.90650187 |
| Akirin1 | 0.73286249 | 0.67551152 | 0.6146 | 0.06091152 | 0.32975373 | 0.90650187 |
| Ppp2r2a | 0.71079631 | 0.66264841 | 0.60175234 | 0.06089607 | 0.32639631 | 0.90650187 |
| Dynlt1c | 0.81841383 | 0.72310477 | 0.66222112 | 0.06088364 | 0.34137164 | 0.90650187 |
| Dok4 | 0.14806626 | 0.22637017 | 0.16550773 | 0.06086245 | 0.05104648 | 0.60204511 |
| Fli1 | 0.07525143 | 0.14864666 | 0.08787487 | 0.06077179 | 0.00105443 | 0.0249276 |
| Zc3h15 | 1.20539812 | 0.88474725 | 0.82405379 | 0.06069347 | 0.37167165 | 0.90650187 |
| Mpst | 0.44726449 | 0.4877967 | 0.42713878 | 0.06065793 | 0.26391565 | 0.90650187 |
| Smco4 | 0.31276968 | 0.38018791 | 0.3196092 | 0.06057871 | 0.19972052 | 0.90650187 |
| Tcea1 | 0.9054696 | 0.76781905 | 0.70727663 | 0.06054242 | 0.35177192 | 0.90650187 |
| Sep-09 | 0.36146641 | 0.42077313 | 0.36023718 | 0.06053595 | 0.22751349 | 0.90650187 |
| Ppp1r1b | 0.0797213 | 0.15323052 | 0.092826 | 0.06040453 | 0.00190862 | 0.04109476 |
| Prmt1 | 1.89135803 | 0.98729182 | 0.92690179 | 0.06039003 | 0.38604482 | 0.90650187 |
| Arsi | 0.18253336 | 0.26031438 | 0.20010045 | 0.06021394 | 0.09048646 | 0.89028133 |
| Fam216a | 0.74391055 | 0.68108705 | 0.62094325 | 0.0601438 | 0.3333811 | 0.90650187 |
| Spock3 | 0.14598764 | 0.2234553 | 0.16337788 | 0.06007742 | 0.05105165 | 0.60204511 |
| Rundc3a | 0.45972827 | 0.49638782 | 0.43639546 | 0.05999236 | 0.27055065 | 0.90650187 |
| Tbxa2r | 0.20793402 | 0.28473507 | 0.2247537 | 0.05998137 | 0.1177271 | 0.90650187 |
| C1qtnf2 | 0.17320808 | 0.25077933 | 0.19087369 | 0.05990564 | 0.08146993 | 0.82790075 |

| | | | | | | |
|---|---|---|---|---|---|---|
| Neurod2 | 0.07940031 | 0.15236987 | 0.0924714 | 0.05989846 | 0.0019906 | 0.04255862 |
| Gpsm3 | 0.11206426 | 0.18773213 | 0.12789284 | 0.05983929 | 0.01876105 | 0.27607614 |
| Ckap2l | 0.11932603 | 0.19541099 | 0.13560632 | 0.05980467 | 0.0249636 | 0.34434814 |
| Rpl10a | 4.27291525 | 0.69885178 | 0.63910148 | 0.0597503 | 0.33884236 | 0.90650187 |
| Ywhah | 1.88859087 | 0.98655768 | 0.92686984 | 0.05968784 | 0.38733004 | 0.90650187 |
| Ube2e3 | 1.47336111 | 0.94840152 | 0.88872245 | 0.05967907 | 0.38264963 | 0.90650187 |
| Rassf1 | 0.53172629 | 0.54743184 | 0.48776451 | 0.05966734 | 0.29328169 | 0.90650187 |
| Chd4 | 1.19435854 | 0.88039006 | 0.82074172 | 0.05964833 | 0.37331374 | 0.90650187 |
| Chst12 | 0.37481453 | 0.43066841 | 0.37102578 | 0.05964263 | 0.23741761 | 0.90650187 |
| Cpb1 | 0.02760272 | 0.09287715 | 0.03325882 | 0.05961833 | 8.00E-16 | 8.75E-14 |
| Tmem5 | 0.55833124 | 0.5654594 | 0.50588534 | 0.05957406 | 0.30030502 | 0.90650187 |
| Palmd | 0.1031101 | 0.17782565 | 0.11829184 | 0.05953382 | 0.01263072 | 0.20308233 |
| Wnt5a | 0.14975588 | 0.22676767 | 0.16723519 | 0.05953248 | 0.05676209 | 0.64678604 |
| Ica1 | 0.43776278 | 0.47950402 | 0.42000543 | 0.05949859 | 0.26442216 | 0.90650187 |
| Lamc1 | 0.52597394 | 0.54320466 | 0.48378755 | 0.05941712 | 0.29253502 | 0.90650187 |
| Tspan18 | 0.18990543 | 0.26671149 | 0.20732732 | 0.05938417 | 0.10144831 | 0.90650187 |
| Nt5c3 | 0.70381561 | 0.65697471 | 0.59763698 | 0.05933773 | 0.32946461 | 0.90650187 |
| Tshz1 | 0.6267682 | 0.60982991 | 0.55052471 | 0.0593052 | 0.31600407 | 0.90650187 |
| Tln1 | 0.82184733 | 0.72326782 | 0.66406213 | 0.05920568 | 0.34592113 | 0.90650187 |
| Msrb1 | 0.49360304 | 0.52015627 | 0.46100272 | 0.05915354 | 0.2841904 | 0.90650187 |
| Mif4gd | 0.20076157 | 0.2769611 | 0.21786286 | 0.05909825 | 0.11392207 | 0.90650187 |
| Bmp7 | 0.1226894 | 0.19822545 | 0.13915706 | 0.05906839 | 0.02957753 | 0.39508621 |
| C1ra | 0.13830192 | 0.21452268 | 0.15545898 | 0.0590637 | 0.0456074 | 0.55065774 |
| Deb1 | 0.77522702 | 0.69765912 | 0.6386057 | 0.05905342 | 0.34050106 | 0.90650187 |
| Snf8 | 0.9532482 | 0.78937857 | 0.73039048 | 0.0589881 | 0.35978337 | 0.90650187 |
| Casp8 | 0.25403592 | 0.32674776 | 0.26778187 | 0.05896589 | 0.16380782 | 0.90650187 |
| Bpgm | 0.2456874 | 0.31910091 | 0.26014902 | 0.05895189 | 0.15686873 | 0.90650187 |
| Galnt6 | 0.02313178 | 0.08687411 | 0.0279552 | 0.05891891 | 3.64E-21 | 5.64E-19 |
| Hspa1a | 0.14783629 | 0.22416504 | 0.16527233 | 0.05889272 | 0.05658139 | 0.64620111 |
| Gimap1 | 0.08028007 | 0.1522651 | 0.09344293 | 0.05882217 | 0.00256701 | 0.05294594 |

| | | | | | | |
|---|---|---|---|---|---|---|
| Ralgds | 0.2753124 | 0.34563511 | 0.28693 | 0.0587051 | 0.18152755 | 0.90650187 |
| Fermt2 | 0.75768225 | 0.68746572 | 0.62876742 | 0.05869831 | 0.3390645 | 0.90650187 |
| Snrpn | 0.38619981 | 0.43877872 | 0.38011309 | 0.05866563 | 0.24631782 | 0.90650187 |
| Vps36 | 0.62435538 | 0.60751851 | 0.54899731 | 0.05852119 | 0.31779057 | 0.90650187 |
| Tmem109 | 0.70533137 | 0.65704476 | 0.59853262 | 0.05851214 | 0.3319266 | 0.90650187 |
| Rbm15 | 0.24032811 | 0.31370398 | 0.25521208 | 0.0584919 | 0.15412861 | 0.90650187 |
| Pdlim1 | 0.24373892 | 0.31683586 | 0.25835736 | 0.0584785 | 0.15714737 | 0.90650187 |
| Dnajc3 | 0.97540109 | 0.79880243 | 0.7403481 | 0.05845433 | 0.36279278 | 0.90650187 |
| Msra | 0.15564786 | 0.23167413 | 0.1732327 | 0.05844143 | 0.0668382 | 0.7246844 |
| Clec4a2 | 0.0321534 | 0.09704134 | 0.03862459 | 0.05841675 | 8.86E-12 | 6.90E-10 |
| Slc40a1 | 0.25168346 | 0.32400038 | 0.26563807 | 0.05836231 | 0.16435218 | 0.90650187 |
| Ppp1r18 | 0.17361109 | 0.24962092 | 0.1912745 | 0.05834642 | 0.08753544 | 0.87082866 |
| Zfp771 | 0.53005389 | 0.54490765 | 0.48661043 | 0.05829723 | 0.29715681 | 0.90650187 |
| Tnxb | 0.1889684 | 0.26470309 | 0.20641201 | 0.05829108 | 0.10465838 | 0.90650187 |
| Hjurp | 0.44348751 | 0.48242684 | 0.42431158 | 0.05811526 | 0.27129984 | 0.90650187 |
| Cebpzos | 0.61070757 | 0.59840521 | 0.54029502 | 0.0581102 | 0.31627634 | 0.90650187 |
| Gm10260 | 3.05966819 | 0.88705838 | 0.82896449 | 0.05809389 | 0.37769188 | 0.90650187 |
| Slc35c1 | 0.11839847 | 0.19267027 | 0.13462466 | 0.05804561 | 0.02763197 | 0.37303156 |
| Xpnpep1 | 0.59462196 | 0.5878444 | 0.52990098 | 0.05794342 | 0.31344226 | 0.90650187 |
| Hist1h1b | 0.08778629 | 0.15961768 | 0.10168744 | 0.05793024 | 0.00566078 | 0.10411931 |
| Emid1 | 0.21105707 | 0.28557536 | 0.22773731 | 0.05783806 | 0.12943983 | 0.90650187 |
| Gm1673 | 1.15056545 | 0.86456099 | 0.80673182 | 0.05782917 | 0.37498628 | 0.90650187 |
| Smarcd2 | 0.65637165 | 0.62672129 | 0.56900287 | 0.05771843 | 0.32601181 | 0.90650187 |
| Hacd4 | 0.15510633 | 0.23034449 | 0.17268314 | 0.05766135 | 0.06884464 | 0.7374964 |
| Tgfb2 | 0.20629926 | 0.28082517 | 0.22318787 | 0.05763729 | 0.12547222 | 0.90650187 |
| Amd1 | 0.67096893 | 0.6354313 | 0.57793916 | 0.05749214 | 0.32915802 | 0.90650187 |
| Mrps36 | 1.07635017 | 0.83813629 | 0.78065844 | 0.05747786 | 0.37171351 | 0.90650187 |
| Rpp25 | 0.10671722 | 0.17926473 | 0.12217157 | 0.05709317 | 0.01887685 | 0.27751155 |
| Amotl1 | 0.52197241 | 0.53801175 | 0.48100867 | 0.05700308 | 0.29915334 | 0.90650187 |
| Mfng | 0.28005164 | 0.34804868 | 0.29113596 | 0.05691271 | 0.19241007 | 0.90650187 |

| | | | | | | |
|---|---|---|---|---|---|---|
| Ebpl | 0.50157677 | 0.52358424 | 0.46667924 | 0.056905 | 0.29388189 | 0.90650187 |
| Pole3 | 0.38579327 | 0.43663522 | 0.37979037 | 0.05684485 | 0.25289959 | 0.90650187 |
| Necap2 | 0.30679569 | 0.37131171 | 0.31448277 | 0.05682894 | 0.2108868 | 0.90650187 |
| Swap70 | 0.09833648 | 0.16988214 | 0.11314188 | 0.05674026 | 0.01289178 | 0.20649943 |
| Ppp1r15a | 0.4897759 | 0.51485945 | 0.45826191 | 0.05659753 | 0.29148433 | 0.90650187 |
| Rnft1 | 0.40844097 | 0.45415957 | 0.39757007 | 0.05658949 | 0.26343849 | 0.90650187 |
| Bok | 0.15831023 | 0.23233573 | 0.17592972 | 0.05640602 | 0.07703007 | 0.79547148 |
| Cdk4 | 2.10343813 | 0.98099796 | 0.9246292 | 0.05636876 | 0.39318769 | 0.90650187 |
| Tmed5 | 0.46209023 | 0.49448057 | 0.43813737 | 0.05634321 | 0.28376526 | 0.90650187 |
| Igsf3 | 0.30384688 | 0.36804282 | 0.3119407 | 0.05610212 | 0.21198971 | 0.90650187 |
| Upp1 | 0.19511086 | 0.26845865 | 0.21239496 | 0.05606369 | 0.1203056 | 0.90650187 |
| Ttll6 | 0.06164571 | 0.12867285 | 0.07262636 | 0.05604649 | 0.00030099 | 0.00813738 |
| Ctsc | 0.11813704 | 0.1901297 | 0.13434779 | 0.05578191 | 0.03245778 | 0.42387524 |
| Tcea3 | 0.13243836 | 0.20514419 | 0.14937076 | 0.05577343 | 0.04846311 | 0.57824787 |
| Pofut2 | 1.09430745 | 0.84298266 | 0.78722174 | 0.05576093 | 0.3764212 | 0.90650187 |
| Cept1 | 0.39013906 | 0.43889177 | 0.3832336 | 0.05565817 | 0.25925339 | 0.90650187 |
| Utrn | 0.33256015 | 0.39202311 | 0.33636625 | 0.05565686 | 0.23098934 | 0.90650187 |
| Kank3 | 0.1358372 | 0.20851803 | 0.15290479 | 0.05561323 | 0.05294797 | 0.61864883 |
| Mdfi | 0.32013233 | 0.38145474 | 0.32588426 | 0.05557049 | 0.22420322 | 0.90650187 |
| Ggta1 | 0.20279915 | 0.27538671 | 0.21982594 | 0.05556077 | 0.13058664 | 0.90650187 |
| Kif23 | 0.1160618 | 0.18762669 | 0.13214705 | 0.05547964 | 0.03099197 | 0.40824004 |
| Ush1c | 0.29262663 | 0.35765499 | 0.30219573 | 0.05545926 | 0.20728952 | 0.90650187 |
| Riok3 | 0.60139881 | 0.58974346 | 0.53429792 | 0.05544554 | 0.32227999 | 0.90650187 |
| Aacs | 0.40787647 | 0.45252993 | 0.39713176 | 0.05539817 | 0.26758184 | 0.90650187 |
| Fyb | 0.09254331 | 0.16225207 | 0.10687133 | 0.05538074 | 0.01061936 | 0.17565269 |
| Gatm | 0.12452418 | 0.19646391 | 0.14108828 | 0.05537564 | 0.04050524 | 0.50180941 |
| Klhl4 | 0.06425927 | 0.13092727 | 0.07557648 | 0.05535079 | 0.00056521 | 0.01443993 |
| Cnpy3 | 0.50862546 | 0.52698451 | 0.47166236 | 0.05532215 | 0.30103245 | 0.90650187 |
| Casz1 | 0.35064906 | 0.40665284 | 0.35138696 | 0.05526588 | 0.24221299 | 0.90650187 |
| Nfic | 0.63645247 | 0.61168239 | 0.5566231 | 0.05505929 | 0.33006109 | 0.90650187 |

| | | | | | | |
|---|---|---|---|---|---|---|
| Apex1 | 0.69728634 | 0.64882306 | 0.59376594 | 0.05505712 | 0.34008902 | 0.90650187 |
| Dusp3 | 0.33863667 | 0.39647071 | 0.34144324 | 0.05502746 | 0.23685367 | 0.90650187 |
| Tbrg1 | 1.40410605 | 0.93022209 | 0.87521514 | 0.05500695 | 0.38996713 | 0.90650187 |
| Prex2 | 0.1015194 | 0.17154356 | 0.11657572 | 0.05496784 | 0.01803248 | 0.26900066 |
| Colgalt1 | 0.53603777 | 0.5456888 | 0.49073166 | 0.05495714 | 0.30928983 | 0.90650187 |
| 1810008I18R | 0.20796601 | 0.27973119 | 0.22478431 | 0.05494688 | 0.13858559 | 0.90650187 |
| Gm266 | 0.17588533 | 0.24845469 | 0.19353254 | 0.05492216 | 0.10354485 | 0.90650187 |
| Trib1 | 0.26036802 | 0.32842081 | 0.27352581 | 0.054895 | 0.18614108 | 0.90650187 |
| Arl13b | 0.28764291 | 0.35258629 | 0.29782982 | 0.05475647 | 0.20686951 | 0.90650187 |
| Narf | 0.25742961 | 0.32554916 | 0.27086513 | 0.05468403 | 0.18472389 | 0.90650187 |
| Pard6g | 0.2231419 | 0.29385649 | 0.23918765 | 0.05466883 | 0.15479333 | 0.90650187 |
| Flnc | 0.17770511 | 0.24996419 | 0.19533521 | 0.05462898 | 0.10687989 | 0.90650187 |
| Ehbp1l1 | 0.19131258 | 0.26317869 | 0.20870009 | 0.0544786 | 0.12292855 | 0.90650187 |
| Zcchc5 | 0.20830299 | 0.27954723 | 0.22510672 | 0.05444051 | 0.14115621 | 0.90650187 |
| Metap2 | 1.83958632 | 0.97997454 | 0.92560769 | 0.05436685 | 0.39700043 | 0.90650187 |
| Dok2 | 0.03955694 | 0.10158075 | 0.04728507 | 0.05429568 | 1.66E-07 | 7.84E-06 |
| Spcs2 | 2.29956915 | 0.96836024 | 0.91410679 | 0.05425345 | 0.39594675 | 0.90650187 |
| Nucks1 | 0.9935581 | 0.80224495 | 0.74806101 | 0.05418394 | 0.37372391 | 0.90650187 |
| Ctnnal1 | 0.25773614 | 0.32530746 | 0.27114307 | 0.05416438 | 0.18725205 | 0.90650187 |
| Npnt | 0.21456881 | 0.28516206 | 0.23108016 | 0.05408191 | 0.14906592 | 0.90650187 |
| Phb | 0.74326623 | 0.67463735 | 0.62057503 | 0.05406232 | 0.34927238 | 0.90650187 |
| Lsm4 | 1.23287734 | 0.88622101 | 0.83219817 | 0.05402283 | 0.38644749 | 0.90650187 |
| Rps23 | 4.89747676 | 0.60809431 | 0.55412334 | 0.05397098 | 0.33250848 | 0.90650187 |
| Rarb | 0.09469314 | 0.16312845 | 0.10920374 | 0.05392471 | 0.01407331 | 0.22054475 |
| Rrs1 | 0.36667593 | 0.41838134 | 0.36446537 | 0.05391597 | 0.25538283 | 0.90650187 |
| Fcrls | 0.02570263 | 0.08485405 | 0.03100872 | 0.05384533 | 5.83E-15 | 6.12E-13 |
| Asb4 | 0.21375788 | 0.28410164 | 0.23030936 | 0.05379229 | 0.14955598 | 0.90650187 |
| Bmp2k | 0.11003772 | 0.17947056 | 0.12572864 | 0.05374192 | 0.02870039 | 0.38540161 |
| Nav2 | 0.66842159 | 0.63008151 | 0.57638823 | 0.05369328 | 0.33938976 | 0.90650187 |
| Fam132a | 0.30238424 | 0.36436306 | 0.31067693 | 0.05368612 | 0.22117907 | 0.90650187 |

| | | | | | | |
|---|---|---|---|---|---|---|
| Fam49b | 0.58243255 | 0.57559684 | 0.52192245 | 0.05367439 | 0.32376896 | 0.90650187 |
| Nudc | 1.64200957 | 0.96637345 | 0.91273859 | 0.05363485 | 0.39695557 | 0.90650187 |
| Pbdc1 | 0.74122593 | 0.67300372 | 0.61940771 | 0.05359601 | 0.35024144 | 0.90650187 |
| Pou2f2 | 0.02999453 | 0.08967157 | 0.03608309 | 0.05358848 | 2.02E-11 | 1.51E-09 |
| Dbn1 | 0.99676959 | 0.80295812 | 0.74938956 | 0.05356856 | 0.37532404 | 0.90650187 |
| Psrc1 | 0.09247505 | 0.16019124 | 0.10679718 | 0.05339407 | 0.01312089 | 0.20906769 |
| Srpx2 | 0.19968856 | 0.27019178 | 0.21682732 | 0.05336446 | 0.13694886 | 0.90650187 |
| Zfyve21 | 0.68927853 | 0.64226925 | 0.58898867 | 0.05328058 | 0.34378514 | 0.90650187 |
| Crot | 0.39035006 | 0.43665858 | 0.3834004 | 0.05325819 | 0.26843944 | 0.90650187 |
| Cpt2 | 0.53667391 | 0.54436369 | 0.49116847 | 0.05319522 | 0.31508972 | 0.90650187 |
| Rac1 | 0.71401404 | 0.65683284 | 0.60364124 | 0.0531916 | 0.34762609 | 0.90650187 |
| Serp1 | 1.4974411 | 0.94598363 | 0.89289441 | 0.05308922 | 0.39576498 | 0.90650187 |
| Vwf | 0.04379133 | 0.10521286 | 0.05220027 | 0.05301259 | 3.17E-06 | 0.00012384 |
| Nap1l5 | 0.11871056 | 0.18779271 | 0.13495507 | 0.05283764 | 0.0408817 | 0.50559243 |
| Gadd45b | 0.27743559 | 0.34163298 | 0.28881687 | 0.05281611 | 0.20811744 | 0.90650187 |
| Adra2a | 0.2255137 | 0.29420811 | 0.24141679 | 0.05279132 | 0.16549199 | 0.90650187 |
| Snd1 | 0.9224855 | 0.76825545 | 0.71565933 | 0.05259611 | 0.37193909 | 0.90650187 |
| Nme2 | 0.48596093 | 0.50804727 | 0.45552009 | 0.05252718 | 0.30410361 | 0.90650187 |
| Scn3a | 0.15826602 | 0.22840158 | 0.175885 | 0.05251658 | 0.09218981 | 0.90003543 |
| Afap1l1 | 0.07714317 | 0.14245881 | 0.08997378 | 0.05248503 | 0.00475318 | 0.09020358 |
| Pbx1 | 1.20118806 | 0.87517123 | 0.82279568 | 0.05237556 | 0.38859311 | 0.90650187 |
| Efcab1 | 0.54956707 | 0.55217768 | 0.49996539 | 0.05221229 | 0.32122976 | 0.90650187 |
| Fmo2 | 0.11063304 | 0.17839084 | 0.12636492 | 0.05202592 | 0.03360193 | 0.43470892 |
| Ccdc85b | 0.23803424 | 0.30497001 | 0.25309033 | 0.05187968 | 0.18107346 | 0.90650187 |
| Cd36 | 0.06426641 | 0.12745388 | 0.07558453 | 0.05186935 | 0.00114129 | 0.02648666 |
| Wfdc1 | 0.10741917 | 0.17466799 | 0.12292467 | 0.05174331 | 0.03065093 | 0.40479999 |
| Limch1 | 0.20146013 | 0.27025197 | 0.21853637 | 0.0517156 | 0.14639251 | 0.90650187 |
| Arhgap29 | 0.40403866 | 0.44573438 | 0.3941454 | 0.05158898 | 0.28032464 | 0.90650187 |
| Cdc42ep1 | 0.18147946 | 0.25057737 | 0.19906248 | 0.05151489 | 0.12497566 | 0.90650187 |
| Ptprb | 0.05724854 | 0.11913872 | 0.06764017 | 0.05149855 | 0.00035627 | 0.00948449 |

| | | | | | | |
|---|---|---|---|---|---|---|
| D630033O11 | 0.11277449 | 0.18013698 | 0.12865011 | 0.05148687 | 0.03760597 | 0.47484348 |
| Cdca2 | 0.08547759 | 0.15064012 | 0.09916014 | 0.05147998 | 0.01050009 | 0.17424873 |
| Aldh1b1 | 0.10012592 | 0.16654751 | 0.11506976 | 0.05147774 | 0.0233621 | 0.32550623 |
| Ncl | 2.49231633 | 0.94978174 | 0.89832156 | 0.05146018 | 0.39949065 | 0.90650187 |
| Pamr1 | 0.23153299 | 0.29844898 | 0.24704851 | 0.05140048 | 0.17749725 | 0.90650187 |
| Taf1b | 0.20847813 | 0.27666329 | 0.22527424 | 0.05138905 | 0.15526236 | 0.90650187 |
| Fam192a | 0.50168656 | 0.51814591 | 0.46675711 | 0.05138879 | 0.31226151 | 0.90650187 |
| Iah1 | 0.39688804 | 0.43992736 | 0.38855131 | 0.05137605 | 0.27832702 | 0.90650187 |
| Tmem203 | 0.25705248 | 0.32186699 | 0.27052304 | 0.05134395 | 0.19940337 | 0.90650187 |
| Lsm2 | 0.73754607 | 0.668503 | 0.61729655 | 0.05120645 | 0.35614761 | 0.90650187 |
| Tln2 | 0.22625087 | 0.29326363 | 0.24210846 | 0.05115517 | 0.1737849 | 0.90650187 |
| Smagp | 0.09643706 | 0.16222471 | 0.11109106 | 0.05113365 | 0.02036589 | 0.29233414 |
| Stt3a | 1.15261086 | 0.85844461 | 0.80739525 | 0.05104936 | 0.38932324 | 0.90650187 |
| Fyn | 0.30380996 | 0.36290142 | 0.31190882 | 0.0509926 | 0.23367649 | 0.90650187 |
| Ly86 | 0.02220853 | 0.07775091 | 0.02685604 | 0.05089487 | 1.81E-17 | 2.25E-15 |
| Tnfrsf12a | 0.30187139 | 0.36112221 | 0.31023336 | 0.05088885 | 0.23293018 | 0.90650187 |
| Mrps28 | 0.75064585 | 0.6756448 | 0.62478083 | 0.05086397 | 0.35870561 | 0.90650187 |
| Pxdn | 0.28996924 | 0.35073267 | 0.29987058 | 0.05086209 | 0.2254142 | 0.90650187 |
| Dnajc25 | 0.30069925 | 0.36007119 | 0.30921868 | 0.0508525 | 0.23235861 | 0.90650187 |
| Abracl | 0.88228185 | 0.74637938 | 0.69561605 | 0.05076334 | 0.37280891 | 0.90650187 |
| Lgals3bp | 0.41632876 | 0.45442548 | 0.40366967 | 0.05075582 | 0.28809013 | 0.90650187 |
| Lrrn4 | 0.07456216 | 0.13776729 | 0.08710883 | 0.05065845 | 0.00486389 | 0.09161836 |
| Cd79a | 0.16011804 | 0.22841114 | 0.17775645 | 0.0506547 | 0.10260438 | 0.90650187 |
| Acp5 | 0.13174887 | 0.19927501 | 0.14865216 | 0.05062284 | 0.06502157 | 0.70962289 |
| Cxx1b | 0.57618208 | 0.56837916 | 0.51779726 | 0.05058189 | 0.33204383 | 0.90650187 |
| Dse | 0.2078709 | 0.27526032 | 0.22469329 | 0.05056704 | 0.15853914 | 0.90650187 |
| Cmbl | 0.1326686 | 0.20017202 | 0.14961059 | 0.05056143 | 0.06649584 | 0.72200237 |
| Aig1 | 0.48103854 | 0.50246446 | 0.4519679 | 0.05049656 | 0.30970393 | 0.90650187 |
| Dusp6 | 0.21407832 | 0.28105439 | 0.23061403 | 0.05044036 | 0.16543775 | 0.90650187 |
| Tmem132a | 0.30847946 | 0.36632806 | 0.31593084 | 0.05039722 | 0.23911193 | 0.90650187 |

| | | | | | | |
|---|---|---|---|---|---|---|
| Rftn2 | 0.12009476 | 0.18678401 | 0.13641909 | 0.05036491 | 0.05036916 | 0.59538226 |
| Dnajc10 | 0.86889203 | 0.73910643 | 0.68876637 | 0.05034007 | 0.37262183 | 0.90650187 |
| Gdap1l1 | 0.39523133 | 0.43758471 | 0.38724963 | 0.05033507 | 0.28168468 | 0.90650187 |
| Fgl2 | 0.04244949 | 0.10095366 | 0.05064567 | 0.05030799 | 5.03E-06 | 0.00018918 |
| Edem2 | 0.59210805 | 0.57846756 | 0.52826311 | 0.05020446 | 0.33633137 | 0.90650187 |
| Stra13 | 0.87951993 | 0.74421295 | 0.69420901 | 0.05000395 | 0.37440174 | 0.90650187 |
| Gstt1 | 0.22628132 | 0.2920626 | 0.24213702 | 0.04992558 | 0.17966893 | 0.90650187 |
| Canx | 2.02438139 | 0.97665171 | 0.92682422 | 0.04982749 | 0.40555177 | 0.90650187 |
| Fhl2 | 0.20862244 | 0.27523249 | 0.22541225 | 0.04982024 | 0.16291189 | 0.90650187 |
| Gm8186 | 1.07735706 | 0.83084374 | 0.78103068 | 0.04981306 | 0.38838352 | 0.90650187 |
| Mpp1 | 0.25151873 | 0.31529039 | 0.26548775 | 0.04980264 | 0.2021552 | 0.90650187 |
| Mrpl1 | 0.64318371 | 0.6106338 | 0.56083141 | 0.04980238 | 0.34650476 | 0.90650187 |
| Ano1 | 0.16368209 | 0.23105683 | 0.18134696 | 0.04970987 | 0.11149685 | 0.90650187 |
| Ppp3ca | 0.99531584 | 0.79840641 | 0.74878868 | 0.04961773 | 0.38415597 | 0.90650187 |
| Plpp2 | 0.1928192 | 0.2597769 | 0.21016753 | 0.04960936 | 0.14700371 | 0.90650187 |
| Ahsa1 | 1.30792766 | 0.90239337 | 0.85278817 | 0.04960519 | 0.39797459 | 0.90650187 |
| Vta1 | 0.56142692 | 0.55751254 | 0.50796524 | 0.0495473 | 0.33227943 | 0.90650187 |
| Smtnl2 | 0.20936365 | 0.27561309 | 0.22612075 | 0.04949234 | 0.16526788 | 0.90650187 |
| Fech | 0.27529681 | 0.33628111 | 0.28691613 | 0.04936498 | 0.22217063 | 0.90650187 |
| Cmas | 0.50652266 | 0.51951779 | 0.47017917 | 0.04933862 | 0.32042768 | 0.90650187 |
| Dad1 | 2.52600523 | 0.94464281 | 0.89537207 | 0.04927074 | 0.40336997 | 0.90650187 |
| Cpz | 0.15862086 | 0.22550378 | 0.17624386 | 0.04925992 | 0.10701853 | 0.90650187 |
| Gpc6 | 0.20484758 | 0.27101958 | 0.22179506 | 0.04922452 | 0.1619085 | 0.90650187 |
| Znrd1 | 0.62381655 | 0.59781829 | 0.54865578 | 0.04916251 | 0.34518602 | 0.90650187 |
| Rarg | 0.14889576 | 0.21551441 | 0.16635627 | 0.04915814 | 0.09447813 | 0.90650187 |
| Arpc4 | 1.23753922 | 0.88266105 | 0.83356509 | 0.04909597 | 0.3967221 | 0.90650187 |
| Pld4 | 0.02603365 | 0.08043054 | 0.03140113 | 0.04902942 | 1.94E-12 | 1.60E-10 |
| Pcyt1b | 0.20301033 | 0.26901915 | 0.22002914 | 0.04899001 | 0.16112991 | 0.90650187 |
| Banf1 | 1.02655133 | 0.81043382 | 0.76149319 | 0.04894063 | 0.38754783 | 0.90650187 |
| Mmrn2 | 0.08756673 | 0.15037621 | 0.10144741 | 0.0489288 | 0.01601054 | 0.24313303 |

| | | | | | | |
|---|---|---|---|---|---|---|
| Fam20c | 0.49978892 | 0.51431671 | 0.46541014 | 0.04890657 | 0.32019396 | 0.90650187 |
| Bcl7c | 0.68980027 | 0.63816201 | 0.58930104 | 0.04886097 | 0.35621277 | 0.90650187 |
| Cnn1 | 0.03044178 | 0.08541807 | 0.03661021 | 0.04880786 | 1.54E-09 | 9.32E-08 |
| Tmem47 | 0.11684019 | 0.18175784 | 0.13297313 | 0.04878471 | 0.05144638 | 0.60504342 |
| Slc38a4 | 0.47287023 | 0.49475219 | 0.44603708 | 0.04871511 | 0.31364717 | 0.90650187 |
| Mmrn1 | 0.03066279 | 0.0855556 | 0.03687057 | 0.04868503 | 2.18E-09 | 1.28E-07 |
| Coq3 | 0.09421858 | 0.1572391 | 0.10868942 | 0.04854967 | 0.02352917 | 0.32723407 |
| Rab3d | 0.44577792 | 0.47454967 | 0.42602738 | 0.04852229 | 0.30631203 | 0.90650187 |
| Tcp11l2 | 0.31208895 | 0.3675331 | 0.31902663 | 0.04850647 | 0.24954157 | 0.90650187 |
| Dkc1 | 0.41320786 | 0.44973529 | 0.40126187 | 0.04847342 | 0.29561993 | 0.90650187 |
| Trim30a | 0.03833452 | 0.09431524 | 0.04586097 | 0.04845427 | 1.32E-06 | 5.56E-05 |
| Col23a1 | 0.10269426 | 0.16627776 | 0.11784352 | 0.04843425 | 0.03383746 | 0.43738367 |
| Cyth3 | 0.35215101 | 0.40101296 | 0.35262152 | 0.04839144 | 0.27090256 | 0.90650187 |
| Rab26os | 0.3214441 | 0.37524522 | 0.32699724 | 0.04824798 | 0.25593043 | 0.90650187 |
| Mapk3 | 1.1812078 | 0.86480367 | 0.81659657 | 0.0482071 | 0.3964901 | 0.90650187 |
| Bsg | 3.50180263 | 0.81812224 | 0.76997511 | 0.04814713 | 0.39051111 | 0.90650187 |
| Dcaf12 | 0.28526139 | 0.34378961 | 0.29573554 | 0.04805407 | 0.23503457 | 0.90650187 |
| Nemf | 0.51073505 | 0.52117728 | 0.47314745 | 0.04802982 | 0.3258941 | 0.90650187 |
| Jam3 | 0.63141746 | 0.6014484 | 0.55345885 | 0.04798955 | 0.34994404 | 0.90650187 |
| Ezh2 | 0.41258143 | 0.4487049 | 0.40077769 | 0.04792721 | 0.29749109 | 0.90650187 |
| Tmem37 | 0.12159829 | 0.18591823 | 0.13800668 | 0.04791155 | 0.06137005 | 0.68093876 |
| Tceal3 | 0.0719084 | 0.13200597 | 0.08415314 | 0.04785283 | 0.00573655 | 0.10493536 |
| Dnajc24 | 0.53400506 | 0.53717162 | 0.48933421 | 0.04783741 | 0.33192518 | 0.90650187 |
| Mtap | 0.26180129 | 0.32262363 | 0.27482063 | 0.047803 | 0.21967834 | 0.90650187 |
| Fam110b | 0.34825938 | 0.39719321 | 0.34941886 | 0.04777435 | 0.27165136 | 0.90650187 |
| Ap2a2 | 0.94542769 | 0.77441473 | 0.72670426 | 0.04771047 | 0.38519364 | 0.90650187 |
| C030010L15F | 0.01938342 | 0.07117471 | 0.0234842 | 0.04769051 | 8.75E-20 | 1.24E-17 |
| Mrpl57 | 1.07230913 | 0.82683941 | 0.7791606 | 0.04767881 | 0.39279645 | 0.90650187 |
| Slc3a2 | 0.7936263 | 0.6964452 | 0.64877052 | 0.04767468 | 0.3719535 | 0.90650187 |
| Tff3 | 0.05621446 | 0.11413479 | 0.06646339 | 0.0476714 | 0.0007147 | 0.01781012 |

| | | | | | | |
|---|---|---|---|---|---|---|
| Rbpms | 0.78704494 | 0.6928018 | 0.64515273 | 0.04764907 | 0.37132755 | 0.90650187 |
| Elof1 | 1.49212098 | 0.93959035 | 0.89198585 | 0.0476045 | 0.40622829 | 0.90650187 |
| Acvr1c | 0.25969676 | 0.32050899 | 0.27291873 | 0.04759027 | 0.21910968 | 0.90650187 |
| Gpc4 | 0.51473733 | 0.52343971 | 0.47595703 | 0.04748268 | 0.32870027 | 0.90650187 |
| 2210013O21 | 1.00709116 | 0.80110621 | 0.75363136 | 0.04747485 | 0.38971955 | 0.90650187 |
| Foxa3 | 0.26896575 | 0.32869663 | 0.28126373 | 0.0474329 | 0.22671279 | 0.90650187 |
| Rbbp7 | 1.54410082 | 0.9479842 | 0.90056398 | 0.04742021 | 0.40745817 | 0.90650187 |
| Slc4a7 | 0.32264047 | 0.37540837 | 0.32801069 | 0.04739767 | 0.2603094 | 0.90650187 |
| Maob | 0.15727999 | 0.22227097 | 0.17488703 | 0.04738394 | 0.11419941 | 0.90650187 |
| Pla2g16 | 0.38337272 | 0.42520249 | 0.37786613 | 0.04733635 | 0.28879323 | 0.90650187 |
| Ttyh2 | 0.26921576 | 0.32881949 | 0.28148769 | 0.0473318 | 0.22737394 | 0.90650187 |
| Herpud1 | 0.68891525 | 0.63606317 | 0.5887711 | 0.04729207 | 0.36051369 | 0.90650187 |
| Chmp2a | 1.65098487 | 0.96098202 | 0.91369833 | 0.0472837 | 0.40902289 | 0.90650187 |
| Bcl2l1 | 0.31391213 | 0.3677785 | 0.32058599 | 0.04719251 | 0.25641916 | 0.90650187 |
| Tcf19 | 0.17138714 | 0.23613862 | 0.18906043 | 0.04707819 | 0.13414558 | 0.90650187 |
| Desi1 | 0.53521682 | 0.53723923 | 0.49016759 | 0.04707164 | 0.33471905 | 0.90650187 |
| Ddx21 | 0.60162855 | 0.58146215 | 0.53444652 | 0.04701563 | 0.34786792 | 0.90650187 |
| Epb41l2 | 0.31335603 | 0.36709836 | 0.32011067 | 0.04698769 | 0.25702264 | 0.90650187 |
| Hmgn5 | 0.4067686 | 0.44323913 | 0.39627082 | 0.04696831 | 0.29912494 | 0.90650187 |
| Crabp1 | 0.09774928 | 0.15947577 | 0.11250841 | 0.04696736 | 0.03173769 | 0.4159005 |
| Becn1 | 0.55362291 | 0.54966091 | 0.50271081 | 0.0469501 | 0.33900055 | 0.90650187 |
| Etnk1 | 0.4272844 | 0.45894937 | 0.41206216 | 0.04688721 | 0.30648059 | 0.90650187 |
| Ctxn1 | 0.62800156 | 0.59818817 | 0.55130424 | 0.04688393 | 0.35269205 | 0.90650187 |
| Cpq | 0.46744588 | 0.48893565 | 0.4420727 | 0.04686295 | 0.31872507 | 0.90650187 |
| Tes | 0.47684243 | 0.49578531 | 0.44892688 | 0.04685843 | 0.32131424 | 0.90650187 |
| Vit | 0.05129135 | 0.10767415 | 0.06083899 | 0.04683516 | 0.00031044 | 0.00836709 |
| Tbc1d15 | 0.44855162 | 0.47493429 | 0.4280998 | 0.0468345 | 0.31335836 | 0.90650187 |
| Prelp | 0.15103742 | 0.21530508 | 0.16854308 | 0.046762 | 0.10870841 | 0.90650187 |
| Ndufb4 | 1.90383029 | 0.97377659 | 0.92702908 | 0.04674752 | 0.41130876 | 0.90650187 |
| 0610040J01F | 0.45776196 | 0.48166617 | 0.43494238 | 0.0467238 | 0.316478 | 0.90650187 |

| | | | | | | |
|---|---|---|---|---|---|---|
| Acot9 | 0.24589647 | 0.30703809 | 0.26034104 | 0.04669704 | 0.21260857 | 0.90650187 |
| Lin7c | 0.58994508 | 0.57354321 | 0.52685036 | 0.04669285 | 0.34679123 | 0.90650187 |
| Vamp3 | 0.77018949 | 0.68248626 | 0.63579588 | 0.04669038 | 0.37203511 | 0.90650187 |
| Orai1 | 0.26160147 | 0.32129647 | 0.27464023 | 0.04665624 | 0.22505687 | 0.90650187 |
| Gng5 | 2.61280527 | 0.93355538 | 0.88701442 | 0.04654096 | 0.40778071 | 0.90650187 |
| Stx16 | 1.00585718 | 0.79962879 | 0.75312648 | 0.04650231 | 0.39185134 | 0.90650187 |
| Bbip1 | 0.87921032 | 0.74054867 | 0.69405109 | 0.04649758 | 0.38291608 | 0.90650187 |
| Fblim1 | 0.16586282 | 0.22993153 | 0.18353682 | 0.0463947 | 0.130556 | 0.90650187 |
| Gm43861 | 0.28181197 | 0.33895251 | 0.29269292 | 0.04625959 | 0.24114626 | 0.90650187 |
| Vars | 0.40708899 | 0.44276113 | 0.3965199 | 0.04624123 | 0.30207695 | 0.90650187 |
| Klf3 | 0.56042512 | 0.55351926 | 0.50729279 | 0.04622647 | 0.34270173 | 0.90650187 |
| Cmip | 0.59501442 | 0.57636509 | 0.53015633 | 0.04620875 | 0.34919978 | 0.90650187 |
| Fxyd7 | 0.06216268 | 0.11941528 | 0.0732107 | 0.04620459 | 0.00251036 | 0.0520599 |
| Timeless | 0.12096526 | 0.18352486 | 0.13733859 | 0.04618627 | 0.06745266 | 0.73016166 |
| Amigo2 | 0.2903671 | 0.34639675 | 0.3002191 | 0.04617766 | 0.24705298 | 0.90650187 |
| Rab3ip | 0.39348693 | 0.43199041 | 0.38587616 | 0.04611425 | 0.2976146 | 0.90650187 |
| Tmem121 | 0.12080048 | 0.18326861 | 0.1371646 | 0.04610401 | 0.06755319 | 0.73016166 |
| Il33 | 0.06276626 | 0.1199652 | 0.07389244 | 0.04607275 | 0.00278664 | 0.05685867 |
| Slc7a14 | 0.39160038 | 0.4304052 | 0.38438812 | 0.04601707 | 0.29729154 | 0.90650187 |
| Ckap2 | 0.11186358 | 0.17365622 | 0.12767875 | 0.04597748 | 0.05470226 | 0.63138125 |
| Rnf2 | 0.61684016 | 0.59016591 | 0.54421925 | 0.04594666 | 0.35370826 | 0.90650187 |
| Atad2 | 0.18416161 | 0.24762361 | 0.2017017 | 0.04592191 | 0.15573512 | 0.90650187 |
| Lrrc8c | 0.15851174 | 0.22205133 | 0.17613352 | 0.04591781 | 0.12323533 | 0.90650187 |
| Erlec1 | 0.77421774 | 0.68395011 | 0.63804367 | 0.04590644 | 0.37453883 | 0.90650187 |
| Ggps1 | 0.15150654 | 0.21479971 | 0.16902137 | 0.04577833 | 0.11428254 | 0.90650187 |
| Birc6 | 0.74314451 | 0.66626963 | 0.62050544 | 0.04576419 | 0.37150227 | 0.90650187 |
| 1810011O10 | 0.09396524 | 0.15417297 | 0.10841473 | 0.04575824 | 0.03030417 | 0.40204743 |
| Npl | 0.09937992 | 0.16000919 | 0.11426639 | 0.0457428 | 0.03756639 | 0.47484348 |
| Adm | 0.07037046 | 0.12816738 | 0.08243555 | 0.04573183 | 0.00682703 | 0.12137734 |
| Cpeb1 | 0.05416293 | 0.1098419 | 0.06412404 | 0.04571786 | 0.00076341 | 0.01883852 |

| | | | | | | |
|---|---|---|---|---|---|---|
| Clec11a | 0.25544925 | 0.31472548 | 0.26906726 | 0.04565822 | 0.22531108 | 0.90650187 |
| Hspa4 | 0.89860495 | 0.74949082 | 0.70385063 | 0.04564019 | 0.38657041 | 0.90650187 |
| Pgm5 | 0.13266085 | 0.19523788 | 0.14960252 | 0.04563536 | 0.08753274 | 0.87082866 |
| Fn3krp | 0.07212157 | 0.12991598 | 0.08439094 | 0.04552505 | 0.00823791 | 0.14262465 |
| Ppp1r9a | 0.44866908 | 0.47367922 | 0.42818744 | 0.04549178 | 0.31835167 | 0.90650187 |
| Cltb | 1.0622364 | 0.82085752 | 0.77538076 | 0.04547676 | 0.39714701 | 0.90650187 |
| Acot7 | 0.19236231 | 0.25514182 | 0.20972278 | 0.04541903 | 0.16783106 | 0.90650187 |
| Mrpl4 | 0.6123174 | 0.58669045 | 0.54132718 | 0.04536327 | 0.3547446 | 0.90650187 |
| Nov | 0.09465203 | 0.1544575 | 0.10915919 | 0.0452983 | 0.03253228 | 0.42448348 |
| Serpina6 | 0.0255265 | 0.07608593 | 0.03079986 | 0.04528608 | 3.15E-11 | 2.30E-09 |
| Osgep | 0.55436546 | 0.54845975 | 0.50321236 | 0.04524739 | 0.3446755 | 0.90650187 |
| Cited4 | 0.09260171 | 0.15217515 | 0.10693477 | 0.04524038 | 0.03000085 | 0.39942481 |
| Chaf1a | 0.12803674 | 0.18994495 | 0.1447736 | 0.04517136 | 0.08270709 | 0.83684877 |
| C5ar1 | 0.02622331 | 0.07679647 | 0.03162587 | 0.0451706 | 1.08E-10 | 7.49E-09 |
| Pop5 | 0.87849371 | 0.73885107 | 0.69368544 | 0.04516563 | 0.38611776 | 0.90650187 |
| Etv5 | 0.11584567 | 0.17700873 | 0.13191755 | 0.04509118 | 0.06430984 | 0.70481173 |
| Mitd1 | 0.30581934 | 0.35872422 | 0.31364194 | 0.04508228 | 0.26141056 | 0.90650187 |
| Efhd2 | 0.45020726 | 0.47437345 | 0.42933423 | 0.04503922 | 0.32047699 | 0.90650187 |
| Adgra2 | 0.20297913 | 0.2649737 | 0.21999913 | 0.04497457 | 0.18172299 | 0.90650187 |
| Gbp7 | 0.07701225 | 0.13477722 | 0.08982869 | 0.04494853 | 0.01305758 | 0.20849614 |
| Irak4 | 0.08946643 | 0.14845351 | 0.10352196 | 0.04493156 | 0.02683297 | 0.36483716 |
| Cnih4 | 0.58818001 | 0.57058677 | 0.52569549 | 0.04489127 | 0.35211065 | 0.90650187 |
| Wsb1 | 0.60326314 | 0.58023795 | 0.53550293 | 0.04473502 | 0.3551778 | 0.90650187 |
| Zeb2os | 0.0846089 | 0.1429398 | 0.09820724 | 0.04473256 | 0.02143699 | 0.30568827 |
| mt-Nd3 | 2.25099264 | 0.96214627 | 0.91743994 | 0.04470633 | 0.41424761 | 0.90650187 |
| Insig1 | 0.23889987 | 0.29857887 | 0.25389162 | 0.04468725 | 0.21696995 | 0.90650187 |
| Leprot | 0.65233968 | 0.61119322 | 0.56651442 | 0.0446788 | 0.3629413 | 0.90650187 |
| Kctd12b | 0.24414906 | 0.30341344 | 0.2587348 | 0.04467864 | 0.22134085 | 0.90650187 |
| Lrrc8a | 0.29586552 | 0.34969672 | 0.30502104 | 0.04467568 | 0.25747902 | 0.90650187 |
| Ncf1 | 0.02716294 | 0.07736011 | 0.03273854 | 0.04462157 | 6.84E-10 | 4.33E-08 |

| | | | | | | |
|---|---|---|---|---|---|---|
| Shisa3 | 0.06481503 | 0.12082206 | 0.07620251 | 0.04461955 | 0.00461957 | 0.08799756 |
| Gng2 | 0.19103122 | 0.25301627 | 0.20842577 | 0.0445905 | 0.17077918 | 0.90650187 |
| Sdcbp | 1.23493786 | 0.87737524 | 0.83280322 | 0.04457202 | 0.40596732 | 0.90650187 |
| Snx15 | 0.51183409 | 0.51847313 | 0.47392 | 0.04455313 | 0.33799893 | 0.90650187 |
| Ddr2 | 0.21650636 | 0.27747073 | 0.23291908 | 0.04455165 | 0.19756962 | 0.90650187 |
| Cx3cr1 | 0.02148106 | 0.07052115 | 0.02598901 | 0.04453214 | 1.29E-14 | 1.29E-12 |
| Polr2k | 1.32668013 | 0.90181935 | 0.85746257 | 0.04435678 | 0.40905659 | 0.90650187 |
| Epha2 | 0.0399263 | 0.09202265 | 0.04771491 | 0.04430774 | 1.83E-05 | 0.00062457 |
| Skap1 | 0.06456101 | 0.12011324 | 0.07591643 | 0.04419681 | 0.00482492 | 0.09122337 |
| Scamp3 | 0.63421696 | 0.59940041 | 0.55521989 | 0.04418052 | 0.36176302 | 0.90650187 |
| Ccdc88a | 0.7362637 | 0.66073429 | 0.61655927 | 0.04417503 | 0.3750463 | 0.90650187 |
| Ctps | 0.31972518 | 0.36964397 | 0.32553848 | 0.04410549 | 0.27348335 | 0.90650187 |
| Ccdc102a | 0.2769629 | 0.33247998 | 0.28839715 | 0.04408283 | 0.24840064 | 0.90650187 |
| Psmb9 | 0.06588708 | 0.12148906 | 0.07740883 | 0.04408023 | 0.00567818 | 0.10411931 |
| Grap | 0.06071688 | 0.11556476 | 0.07157549 | 0.04398926 | 0.0031456 | 0.06324898 |
| Eif4a2 | 2.39300379 | 0.9506147 | 0.9066542 | 0.0439605 | 0.41466851 | 0.90650187 |
| Tspan14 | 0.21480876 | 0.27525965 | 0.23130811 | 0.04395155 | 0.19913277 | 0.90650187 |
| Pear1 | 0.24466186 | 0.30309831 | 0.25920648 | 0.04389183 | 0.22579067 | 0.90650187 |
| Bcl10 | 0.567873 | 0.55615783 | 0.51227771 | 0.04388012 | 0.35167699 | 0.90650187 |
| Hprt | 0.92215217 | 0.75936698 | 0.71549681 | 0.04387017 | 0.39258838 | 0.90650187 |
| Nfatc4 | 0.19941518 | 0.26038625 | 0.21656329 | 0.04382297 | 0.1841674 | 0.90650187 |
| Muc1 | 0.21782338 | 0.27794421 | 0.23416686 | 0.04377735 | 0.20295761 | 0.90650187 |
| Klk8 | 0.12856764 | 0.18910459 | 0.14532931 | 0.04377528 | 0.09027224 | 0.88874891 |
| Dcdc2a | 0.1249087 | 0.18525724 | 0.14149245 | 0.0437648 | 0.08455593 | 0.85008909 |
| Prss50 | 0.18973632 | 0.25091377 | 0.2071622 | 0.04375157 | 0.17389395 | 0.90650187 |
| Ctse | 0.02469231 | 0.07355968 | 0.02980997 | 0.04374971 | 3.41E-11 | 2.48E-09 |
| Ahnak2 | 0.1502774 | 0.21143556 | 0.16776767 | 0.0436679 | 0.12360806 | 0.90650187 |
| Lipa | 0.27113559 | 0.3268507 | 0.28320554 | 0.04364516 | 0.24663476 | 0.90650187 |
| Fam92a | 0.74096101 | 0.66288917 | 0.619256 | 0.04363317 | 0.37704906 | 0.90650187 |
| Cfp | 0.02871996 | 0.07821176 | 0.03457919 | 0.04363257 | 1.01E-08 | 5.67E-07 |

| | | | | | | |
|---|---|---|---|---|---|---|
| Bag2 | 0.27073706 | 0.32635172 | 0.28284922 | 0.0435025 | 0.24707007 | 0.90650187 |
| Lpar6 | 0.34123408 | 0.38701648 | 0.34360392 | 0.04341256 | 0.28716733 | 0.90650187 |
| Lama2 | 0.10892262 | 0.16792901 | 0.12453562 | 0.04339338 | 0.06068678 | 0.67483526 |
| 1110012L19F | 0.37931272 | 0.41798742 | 0.37462828 | 0.04335914 | 0.30344059 | 0.90650187 |
| Sema6a | 0.25137377 | 0.30851837 | 0.26535545 | 0.04316292 | 0.2348013 | 0.90650187 |
| Ncaph | 0.09246263 | 0.14993251 | 0.10678368 | 0.04314883 | 0.03621807 | 0.46071206 |
| Ensa | 0.50966191 | 0.51551091 | 0.47239235 | 0.04311856 | 0.34245153 | 0.90650187 |
| Ninj1 | 0.58109585 | 0.56405358 | 0.52104218 | 0.0430114 | 0.35681719 | 0.90650187 |
| Cfap57 | 0.03435311 | 0.08420693 | 0.04120665 | 0.04300028 | 1.75E-06 | 7.21E-05 |
| Rrad | 0.06195595 | 0.11590376 | 0.07297707 | 0.04292669 | 0.00446135 | 0.08530447 |
| Pygb | 0.29720257 | 0.3490827 | 0.30618442 | 0.04289828 | 0.26668953 | 0.90650187 |
| Plcxd3 | 0.28651287 | 0.33970237 | 0.29683672 | 0.04286565 | 0.26044246 | 0.90650187 |
| Hspd1 | 1.83966475 | 0.96842058 | 0.9256101 | 0.04281049 | 0.41854614 | 0.90650187 |
| Apbb1 | 0.44163818 | 0.46568675 | 0.42292309 | 0.04276367 | 0.32652996 | 0.90650187 |
| Raly | 1.54275683 | 0.94302124 | 0.90035101 | 0.04267023 | 0.41656455 | 0.90650187 |
| Atp2b1 | 1.46928326 | 0.93061766 | 0.88799151 | 0.04262615 | 0.41550675 | 0.90650187 |
| Chchd2 | 3.49854333 | 0.81299357 | 0.77047779 | 0.04251577 | 0.40310696 | 0.90650187 |
| Armcx1 | 0.50682716 | 0.51280914 | 0.47039412 | 0.04241501 | 0.34426296 | 0.90650187 |
| Gpr180 | 0.45946491 | 0.4785883 | 0.436201 | 0.0423873 | 0.33287202 | 0.90650187 |
| Robo4 | 0.0573012 | 0.11005627 | 0.06770005 | 0.04235622 | 0.00270647 | 0.05537394 |
| Cracr2b | 0.09462775 | 0.15139551 | 0.10913289 | 0.04226262 | 0.04257118 | 0.52297082 |
| Bin1 | 0.48099963 | 0.49417449 | 0.45193975 | 0.04223474 | 0.33890574 | 0.90650187 |
| Clk4 | 0.49965601 | 0.50751664 | 0.46531571 | 0.04220094 | 0.34340572 | 0.90650187 |
| Vgll3 | 0.09899228 | 0.15604251 | 0.11384881 | 0.0421937 | 0.04971802 | 0.58951921 |
| Tmem222 | 0.51042254 | 0.51510988 | 0.47292764 | 0.04218225 | 0.34586042 | 0.90650187 |
| Snai2 | 0.18420655 | 0.24388147 | 0.20174585 | 0.04213562 | 0.17657752 | 0.90650187 |
| Arf3 | 0.70115353 | 0.6381682 | 0.59606126 | 0.04210694 | 0.37674292 | 0.90650187 |
| Prdx3 | 0.82204644 | 0.70622789 | 0.66416874 | 0.04205915 | 0.38915485 | 0.90650187 |
| Gm21541 | 0.01639563 | 0.06188596 | 0.01990425 | 0.04198171 | 3.41E-21 | 5.39E-19 |
| Car9 | 0.07748643 | 0.1323311 | 0.09035408 | 0.04197702 | 0.01944469 | 0.28258006 |

| | | | | | | |
|---|---|---|---|---|---|---|
| B4galt1 | 0.38721017 | 0.42288524 | 0.38091461 | 0.04197063 | 0.31212571 | 0.90650187 |
| Specc1 | 0.090147 | 0.14621238 | 0.10426394 | 0.04194844 | 0.03683982 | 0.46705764 |
| Dnm3os | 0.15539615 | 0.2149234 | 0.1729773 | 0.0419461 | 0.14050875 | 0.90650187 |
| Coro1c | 0.42303642 | 0.45070721 | 0.40881905 | 0.04188816 | 0.32437372 | 0.90650187 |
| Nab2 | 0.2880585 | 0.34002478 | 0.29819476 | 0.04183003 | 0.266439 | 0.90650187 |
| Ostf1 | 0.70857635 | 0.64222965 | 0.6004462 | 0.04178345 | 0.37852465 | 0.90650187 |
| Ccl2 | 0.02883654 | 0.07644931 | 0.03471685 | 0.04173246 | 4.55E-08 | 2.37E-06 |
| C1qbp | 0.87645323 | 0.73436122 | 0.69264315 | 0.04171806 | 0.39444206 | 0.90650187 |
| Atp5g1 | 2.84197468 | 0.90068813 | 0.85897758 | 0.04171054 | 0.41454401 | 0.90650187 |
| Sephs2 | 0.53190426 | 0.52956602 | 0.48788722 | 0.0416788 | 0.3520561 | 0.90650187 |
| Rassf2 | 0.12741103 | 0.18576145 | 0.1441182 | 0.04164325 | 0.0994718 | 0.90650187 |
| Sdc2 | 0.46631942 | 0.48284146 | 0.44124663 | 0.04159483 | 0.33758166 | 0.90650187 |
| Ebf2 | 0.12075118 | 0.17867313 | 0.13711254 | 0.04156059 | 0.08890623 | 0.87871497 |
| Smpdl3a | 0.26951832 | 0.32329452 | 0.28175865 | 0.04153588 | 0.25611936 | 0.90650187 |
| Ndufa6 | 2.30874742 | 0.95493271 | 0.91341911 | 0.0415136 | 0.41994019 | 0.90650187 |
| Sult5a1 | 0.14908626 | 0.20802127 | 0.16655103 | 0.04147024 | 0.13416207 | 0.90650187 |
| Syne2 | 0.41769127 | 0.44617259 | 0.40471858 | 0.04145401 | 0.32442908 | 0.90650187 |
| Spred1 | 0.23893264 | 0.29533602 | 0.25392194 | 0.04141408 | 0.23420514 | 0.90650187 |
| Nxt1 | 0.38255017 | 0.41861803 | 0.3772112 | 0.04140683 | 0.31277549 | 0.90650187 |
| Ackr3 | 0.12914468 | 0.18731966 | 0.14593294 | 0.04138672 | 0.1036944 | 0.90650187 |
| Ppp1cb | 1.77374337 | 0.96430253 | 0.92292457 | 0.04137796 | 0.42100927 | 0.90650187 |
| Lbp | 0.12458913 | 0.1825244 | 0.14115657 | 0.04136783 | 0.09631387 | 0.90650187 |
| Bnip1 | 0.28022233 | 0.33262293 | 0.29128706 | 0.04133587 | 0.26406416 | 0.90650187 |
| Glrx | 0.44748431 | 0.46863615 | 0.42730298 | 0.04133316 | 0.33358872 | 0.90650187 |
| Nmi | 0.17907378 | 0.23796449 | 0.19668861 | 0.04127587 | 0.1754286 | 0.90650187 |
| Ipo11 | 0.14197534 | 0.20050953 | 0.15925247 | 0.04125706 | 0.12472015 | 0.90650187 |
| Psmd9 | 0.77792127 | 0.68135442 | 0.6401038 | 0.04125063 | 0.387251 | 0.90650187 |
| Chd2 | 0.45705506 | 0.47565268 | 0.43441932 | 0.04123336 | 0.33652736 | 0.90650187 |
| Uqcrb | 2.81212168 | 0.90478385 | 0.86356926 | 0.04121459 | 0.41598705 | 0.90650187 |
| Rnf121 | 0.39736822 | 0.43013139 | 0.38892819 | 0.0412032 | 0.31883137 | 0.90650187 |

| | | | | | | |
|---|---|---|---|---|---|---|
| Blcap | 0.42152218 | 0.44885878 | 0.40765979 | 0.04119899 | 0.32661387 | 0.90650187 |
| Gyltl1b | 0.07859851 | 0.1327657 | 0.09158501 | 0.04118069 | 0.0228073 | 0.31953347 |
| Tekt2 | 0.36155653 | 0.40147533 | 0.36031051 | 0.04116481 | 0.30576091 | 0.90650187 |
| Cd74 | 0.02637932 | 0.07293775 | 0.03181072 | 0.04112703 | 4.53E-09 | 2.59E-07 |
| Prdx2 | 2.8506446 | 0.89881068 | 0.85770707 | 0.0411036 | 0.41564583 | 0.90650187 |
| Myo1f | 0.02463654 | 0.07084106 | 0.02974375 | 0.04109732 | 4.06E-10 | 2.66E-08 |
| Rhog | 0.2720548 | 0.32511572 | 0.28402681 | 0.0410889 | 0.2600711 | 0.90650187 |
| Kif20b | 0.10177831 | 0.15793505 | 0.11685526 | 0.04107979 | 0.05904663 | 0.66199323 |
| Rras2 | 0.2449669 | 0.30055353 | 0.25948694 | 0.04106658 | 0.24085194 | 0.90650187 |
| AI607873 | 0.02067081 | 0.06608418 | 0.02502231 | 0.04106187 | 1.49E-13 | 1.41E-11 |
| Klhl13 | 0.31110701 | 0.35924621 | 0.31818557 | 0.04106064 | 0.28308926 | 0.90650187 |
| Sema5b | 0.13186195 | 0.18982446 | 0.14877006 | 0.0410544 | 0.10994814 | 0.90650187 |
| Degs1 | 1.08470466 | 0.82475544 | 0.78373553 | 0.0410199 | 0.40800557 | 0.90650187 |
| Vldlr | 0.22734286 | 0.28415066 | 0.24313204 | 0.04101862 | 0.22662289 | 0.90650187 |
| Slc25a4 | 2.77234157 | 0.91019042 | 0.86918408 | 0.04100634 | 0.41693737 | 0.90650187 |
| Mpeg1 | 0.02325213 | 0.06909476 | 0.02809838 | 0.04099638 | 4.40E-11 | 3.18E-09 |
| Nuf2 | 0.10837254 | 0.16492872 | 0.12394654 | 0.04098218 | 0.07079265 | 0.7525308 |
| Pde10a | 0.13606933 | 0.19411842 | 0.15314565 | 0.04097277 | 0.11714454 | 0.90650187 |
| Sbsn | 0.19390998 | 0.25218182 | 0.21122844 | 0.04095338 | 0.19436503 | 0.90650187 |
| Fermt3 | 0.02880048 | 0.07557941 | 0.03467427 | 0.04090514 | 7.84E-08 | 3.91E-06 |
| Nrip1 | 0.41238599 | 0.44143639 | 0.40062658 | 0.04080981 | 0.32532783 | 0.90650187 |
| Tac1 | 0.02685126 | 0.07315114 | 0.03236962 | 0.04078152 | 1.07E-08 | 5.94E-07 |
| Syp | 0.36239184 | 0.40171454 | 0.36098989 | 0.04072465 | 0.30800054 | 0.90650187 |
| Acer2 | 0.3649378 | 0.40378058 | 0.36305709 | 0.04072349 | 0.30901085 | 0.90650187 |
| Npc2 | 1.77589861 | 0.96367414 | 0.92303736 | 0.04063678 | 0.4224154 | 0.90650187 |
| Pou3f4 | 0.13879398 | 0.19654593 | 0.15596804 | 0.04057789 | 0.12371698 | 0.90650187 |
| Pld3 | 0.74609073 | 0.66276186 | 0.62218778 | 0.04057408 | 0.38594381 | 0.90650187 |
| Fam46c | 0.1836381 | 0.24172635 | 0.20118718 | 0.04053917 | 0.18518284 | 0.90650187 |
| Adgrf5 | 0.12483844 | 0.18187067 | 0.14141861 | 0.04045206 | 0.10175006 | 0.90650187 |
| Plekhg2 | 0.18070343 | 0.23872197 | 0.1982974 | 0.04042457 | 0.18239546 | 0.90650187 |

| | | | | | | |
|---|---|---|---|---|---|---|
| Sesn3 | 0.56903202 | 0.55345776 | 0.51305008 | 0.04040768 | 0.36311881 | 0.90650187 |
| Fcgr1 | 0.02260636 | 0.0677086 | 0.02732984 | 0.04037875 | 2.55E-11 | 1.88E-09 |
| Notch1 | 0.24803345 | 0.30267696 | 0.26230119 | 0.04037577 | 0.24689037 | 0.90650187 |
| Tipin | 0.53594692 | 0.53090389 | 0.49066926 | 0.04023463 | 0.35772835 | 0.90650187 |
| Sesn2 | 0.27200624 | 0.32410731 | 0.28398345 | 0.04012386 | 0.26496358 | 0.90650187 |
| Rai2 | 0.34492151 | 0.3867615 | 0.34666163 | 0.04009987 | 0.30354114 | 0.90650187 |
| Mrps6 | 0.41553662 | 0.4431377 | 0.40305921 | 0.04007849 | 0.32922573 | 0.90650187 |
| Paics | 0.82594574 | 0.70631755 | 0.66625311 | 0.04006445 | 0.39460678 | 0.90650187 |
| Ssbp1 | 0.94804163 | 0.76797998 | 0.72793893 | 0.04004105 | 0.40340814 | 0.90650187 |
| C2 | 0.09104434 | 0.14524964 | 0.10524127 | 0.04000838 | 0.04551056 | 0.54992531 |
| Acadm | 0.49834294 | 0.50437671 | 0.46438217 | 0.03999454 | 0.3509065 | 0.90650187 |
| Ftsj1 | 0.22197711 | 0.27804921 | 0.23809057 | 0.03995864 | 0.2278022 | 0.90650187 |
| Znhit1 | 1.0607222 | 0.81470964 | 0.77480464 | 0.03990501 | 0.40944866 | 0.90650187 |
| Klf14 | 0.11870747 | 0.17483362 | 0.1349518 | 0.03988183 | 0.09445772 | 0.90650187 |
| Snai1 | 0.11404364 | 0.16986529 | 0.13000177 | 0.03986352 | 0.08641083 | 0.86189704 |
| Uqcr11 | 2.62756692 | 0.92539056 | 0.8855352 | 0.03985537 | 0.42070751 | 0.90650187 |
| Ebp | 0.4403905 | 0.46180775 | 0.42198493 | 0.03982282 | 0.33741334 | 0.90650187 |
| Nphp1 | 0.33680475 | 0.37971969 | 0.33991594 | 0.03980374 | 0.30133139 | 0.90650187 |
| Tmem163 | 0.33208023 | 0.37575909 | 0.33596393 | 0.03979516 | 0.29924086 | 0.90650187 |
| Mlec | 0.87215928 | 0.73021001 | 0.69044436 | 0.03976564 | 0.39895941 | 0.90650187 |
| Sod3 | 0.12077987 | 0.17684955 | 0.13714284 | 0.03970672 | 0.09902478 | 0.90650187 |
| Rbp7 | 0.03605541 | 0.08285957 | 0.04319968 | 0.03965989 | 2.24E-05 | 0.00074472 |
| Crb3 | 0.44496769 | 0.46507609 | 0.42542096 | 0.03965513 | 0.33929346 | 0.90650187 |
| Spink2 | 0.09842869 | 0.15284318 | 0.11324131 | 0.03960186 | 0.06001085 | 0.66927728 |
| Glrx5 | 1.09028217 | 0.82528107 | 0.78577526 | 0.03950581 | 0.41156997 | 0.90650187 |
| Acyp2 | 0.28911355 | 0.33859514 | 0.29912051 | 0.03947463 | 0.27870859 | 0.90650187 |
| Tmem151a | 0.10041171 | 0.15483526 | 0.11537882 | 0.03945644 | 0.06422065 | 0.70434212 |
| Psmg4 | 0.60046324 | 0.57313243 | 0.53369247 | 0.03943996 | 0.3712551 | 0.90650187 |
| Gm10036 | 1.63529353 | 0.95137664 | 0.9120116 | 0.03936504 | 0.4239161 | 0.90650187 |
| Ramp3 | 0.10505894 | 0.15974517 | 0.12038998 | 0.03935519 | 0.07307612 | 0.76980604 |

| | | | | | | |
|---|---|---|---|---|---|---|
| Ren1 | 0.03442347 | 0.08055416 | 0.04128913 | 0.03926503 | 1.18E-05 | 0.00041736 |
| Tmem165 | 0.81544129 | 0.69984614 | 0.66062321 | 0.03922293 | 0.39591027 | 0.90650187 |
| Egfl8 | 0.06591849 | 0.11664219 | 0.07744415 | 0.03919804 | 0.01222048 | 0.19741073 |
| Mybbp1a | 0.43161395 | 0.45450751 | 0.41535392 | 0.03915358 | 0.33758332 | 0.90650187 |
| Nek8 | 0.10793906 | 0.16260564 | 0.12348205 | 0.0391236 | 0.07948718 | 0.81530674 |
| Xaf1 | 0.10647888 | 0.16103803 | 0.12191571 | 0.03912233 | 0.07685056 | 0.79500368 |
| Mfsd10 | 0.4796409 | 0.49007604 | 0.45095633 | 0.03911971 | 0.34987834 | 0.90650187 |
| Ehd1 | 0.55116673 | 0.54013752 | 0.50104941 | 0.03908811 | 0.36436598 | 0.90650187 |
| Eif2s3y | 0.27333195 | 0.32421459 | 0.2851662 | 0.03904839 | 0.27134641 | 0.90650187 |
| Irf8 | 0.03024997 | 0.0753847 | 0.03638418 | 0.03900051 | 9.43E-07 | 4.06E-05 |
| Hdc | 0.0227036 | 0.06639297 | 0.02744561 | 0.03894736 | 1.41E-10 | 9.55E-09 |
| Tox3 | 0.44429589 | 0.46383113 | 0.42491775 | 0.03891338 | 0.341959 | 0.90650187 |
| Tnnt2 | 0.03464171 | 0.08032833 | 0.04154486 | 0.03878347 | 1.66E-05 | 0.00057533 |
| Fmo1 | 0.1835647 | 0.23986216 | 0.20111502 | 0.03874714 | 0.19586128 | 0.90650187 |
| Dcaf12l2 | 0.0270379 | 0.07133503 | 0.03259054 | 0.03874449 | 6.28E-08 | 3.21E-06 |
| Gnl3 | 0.55544119 | 0.54267717 | 0.50393836 | 0.03873881 | 0.36627208 | 0.90650187 |
| Gsr | 0.53207665 | 0.52659966 | 0.48800607 | 0.03859359 | 0.3625776 | 0.90650187 |
| Calcrl | 0.05791317 | 0.10695102 | 0.06839566 | 0.03855536 | 0.00610467 | 0.1104728 |
| Marf1 | 0.30881952 | 0.35477691 | 0.31622299 | 0.03855391 | 0.29390636 | 0.90650187 |
| Btbd3 | 0.29969863 | 0.34680588 | 0.30835151 | 0.03845437 | 0.28964935 | 0.90650187 |
| Commd8 | 0.5820703 | 0.56010871 | 0.521684 | 0.03842472 | 0.37166687 | 0.90650187 |
| Fzd1 | 0.20010198 | 0.25565082 | 0.21722645 | 0.03842437 | 0.21582638 | 0.90650187 |
| Trip10 | 0.17967423 | 0.23568089 | 0.19728172 | 0.03839917 | 0.19343746 | 0.90650187 |
| Med28 | 1.35796171 | 0.9033435 | 0.86498544 | 0.03835806 | 0.42185826 | 0.90650187 |
| Cdc42ep3 | 0.19798512 | 0.2535089 | 0.21518075 | 0.03832815 | 0.21422274 | 0.90650187 |
| Gm11808 | 2.21872294 | 0.95765102 | 0.91939599 | 0.03825503 | 0.42662367 | 0.90650187 |
| Cldn15 | 0.0600732 | 0.10903234 | 0.0708465 | 0.03818585 | 0.00828402 | 0.14309708 |
| Pfkl | 0.64679305 | 0.60125998 | 0.56307704 | 0.03818295 | 0.3815311 | 0.90650187 |
| Tomm6 | 0.55628819 | 0.5426309 | 0.50450949 | 0.0381214 | 0.36846715 | 0.90650187 |
| Ncf2 | 0.03144487 | 0.07586063 | 0.03779128 | 0.03806935 | 3.76E-06 | 0.0001444 |

| | | | | | | |
|---|---|---|---|---|---|---|
| Vcl | 0.2309292 | 0.28452974 | 0.24648524 | 0.0380445 | 0.24630282 | 0.90650187 |
| Slc28a3 | 0.12553357 | 0.18019113 | 0.14214885 | 0.03804229 | 0.11707311 | 0.90650187 |
| Rabl6 | 0.63515506 | 0.59384207 | 0.55580906 | 0.03803301 | 0.38048593 | 0.90650187 |
| Ifi203 | 0.04038801 | 0.08623692 | 0.04825194 | 0.03798499 | 0.00023291 | 0.00642551 |
| Trp53 | 0.87233956 | 0.7285162 | 0.69053683 | 0.03797937 | 0.40341895 | 0.90650187 |
| Ddt | 0.53959623 | 0.53114042 | 0.49317184 | 0.03796858 | 0.36607734 | 0.90650187 |
| Neu1 | 0.35695041 | 0.3944196 | 0.35655405 | 0.03786555 | 0.31842155 | 0.90650187 |
| Sf3b5 | 1.26546994 | 0.87942078 | 0.84155826 | 0.03786252 | 0.42073554 | 0.90650187 |
| Chst5 | 0.01679056 | 0.05820586 | 0.02037827 | 0.03782759 | 7.77E-17 | 9.23E-15 |
| Tmx1 | 1.03589406 | 0.80298528 | 0.76520607 | 0.03777921 | 0.41313601 | 0.90650187 |
| Morc3 | 0.31635386 | 0.36041583 | 0.32266983 | 0.037746 | 0.30151498 | 0.90650187 |
| Stx6 | 0.53070955 | 0.52479609 | 0.48706309 | 0.037733 | 0.36527155 | 0.90650187 |
| Hacd1 | 0.52177801 | 0.51859429 | 0.48087341 | 0.03772088 | 0.36364821 | 0.90650187 |
| Atf5 | 0.30297695 | 0.34888529 | 0.31118928 | 0.03769602 | 0.29510906 | 0.90650187 |
| Ssbp2 | 0.77476929 | 0.67603495 | 0.63835086 | 0.03768409 | 0.39649123 | 0.90650187 |
| Capg | 0.17788861 | 0.23319838 | 0.19551678 | 0.0376816 | 0.19577944 | 0.90650187 |
| Hoxb2 | 0.22563441 | 0.27914766 | 0.2415301 | 0.03761756 | 0.24434643 | 0.90650187 |
| Dst | 0.74308819 | 0.658083 | 0.62047324 | 0.03760976 | 0.39378448 | 0.90650187 |
| Rrp15 | 0.37187985 | 0.40626309 | 0.36866679 | 0.0375963 | 0.32514517 | 0.90650187 |
| Cpn1 | 0.25998101 | 0.31076083 | 0.27317585 | 0.03758498 | 0.27038467 | 0.90650187 |
| Megf6 | 0.12970836 | 0.18409665 | 0.14652221 | 0.03757445 | 0.12713479 | 0.90650187 |
| P4ha3 | 0.19897735 | 0.25368365 | 0.21614028 | 0.03754338 | 0.21999861 | 0.90650187 |
| Stxbp1 | 0.32782243 | 0.36992893 | 0.33238608 | 0.03754285 | 0.30778836 | 0.90650187 |
| Isx | 0.01683842 | 0.05797689 | 0.02043571 | 0.03754118 | 1.58E-16 | 1.82E-14 |
| Stc1 | 0.11183689 | 0.16514521 | 0.12765028 | 0.03749493 | 0.0958077 | 0.90650187 |
| Slc11a1 | 0.0354645 | 0.07995133 | 0.04250837 | 0.03744296 | 4.50E-05 | 0.00141756 |
| Itpkb | 0.21788105 | 0.27165651 | 0.23422146 | 0.03743505 | 0.23868758 | 0.90650187 |
| Plscr4 | 0.17765076 | 0.23268557 | 0.19528142 | 0.03740415 | 0.19724301 | 0.90650187 |
| Aoc3 | 0.13474518 | 0.18917012 | 0.15177083 | 0.0373993 | 0.13665228 | 0.90650187 |
| Smim19 | 0.21703045 | 0.27080762 | 0.23341582 | 0.03739179 | 0.23818315 | 0.90650187 |

| | | | | | | |
|---|---|---|---|---|---|---|
| Tmtc3 | 0.1773976 | 0.23241722 | 0.19503085 | 0.03738638 | 0.19705181 | 0.90650187 |
| Sep-08 | 0.26146654 | 0.31178853 | 0.27451839 | 0.03727013 | 0.27306808 | 0.90650187 |
| Mapk15 | 0.33784504 | 0.3780189 | 0.34078359 | 0.03723531 | 0.31357442 | 0.90650187 |
| Fam92b | 0.29862443 | 0.34462882 | 0.30741958 | 0.03720924 | 0.29525942 | 0.90650187 |
| Slc1a5 | 0.24188338 | 0.2938246 | 0.2566477 | 0.0371769 | 0.25979686 | 0.90650187 |
| Borcs8 | 0.46838555 | 0.47992092 | 0.44276113 | 0.03715979 | 0.35453398 | 0.90650187 |
| Zfp503 | 0.28433508 | 0.33206848 | 0.29491954 | 0.03714894 | 0.28774594 | 0.90650187 |
| Ppp2r2b | 0.35394247 | 0.39122376 | 0.35409162 | 0.03713214 | 0.32053882 | 0.90650187 |
| Dlgap5 | 0.09039465 | 0.14165166 | 0.10453377 | 0.03711789 | 0.05721995 | 0.64823856 |
| Snta1 | 0.13468175 | 0.18880093 | 0.15170492 | 0.037096 | 0.13850242 | 0.90650187 |
| 3110002H16 | 0.17036016 | 0.2250787 | 0.18803616 | 0.03704254 | 0.19057737 | 0.90650187 |
| Hip1r | 0.28405267 | 0.33170231 | 0.29467061 | 0.0370317 | 0.28818798 | 0.90650187 |
| Adrbk1 | 0.18294675 | 0.23753393 | 0.20050726 | 0.03702668 | 0.20583795 | 0.90650187 |
| Ophn1 | 0.12535233 | 0.17895954 | 0.1419585 | 0.03700104 | 0.12328246 | 0.90650187 |
| Gabarapl1 | 0.55196396 | 0.53857453 | 0.50158907 | 0.03698546 | 0.37153001 | 0.90650187 |
| Ptbp1 | 0.85638744 | 0.71919435 | 0.68229579 | 0.03689856 | 0.40500296 | 0.90650187 |
| Brd9 | 0.42282257 | 0.44545333 | 0.40865543 | 0.0367979 | 0.34446398 | 0.90650187 |
| Zfp65 | 0.11243078 | 0.16507633 | 0.12828371 | 0.03679262 | 0.10114944 | 0.90650187 |
| Paqr4 | 0.10040577 | 0.15213487 | 0.1153724 | 0.03676247 | 0.07830639 | 0.80536905 |
| Gata2 | 0.04285582 | 0.08784915 | 0.05111671 | 0.03673244 | 0.0007001 | 0.01750365 |
| Gpr119 | 0.09915809 | 0.15061319 | 0.11402744 | 0.03658574 | 0.07688017 | 0.79500368 |
| Tcf15 | 0.06280595 | 0.11051867 | 0.07393725 | 0.03658142 | 0.01392041 | 0.21862316 |
| Ptprc | 0.02203512 | 0.06320919 | 0.02664944 | 0.03655975 | 5.34E-10 | 3.44E-08 |
| Col26a1 | 0.11925635 | 0.17208025 | 0.13553261 | 0.03654764 | 0.11530299 | 0.90650187 |
| Chka | 0.70565722 | 0.63526718 | 0.59872501 | 0.03654218 | 0.39306856 | 0.90650187 |
| Eif3j1 | 1.06094372 | 0.81137926 | 0.77488898 | 0.03649029 | 0.41708829 | 0.90650187 |
| Ctnnb1 | 1.61031361 | 0.94565924 | 0.90918905 | 0.0364702 | 0.42923344 | 0.90650187 |
| Tgfbr2 | 0.21243382 | 0.2654915 | 0.22904937 | 0.03644213 | 0.23968901 | 0.90650187 |
| Spcs3 | 0.71955334 | 0.64331013 | 0.60688006 | 0.03643007 | 0.39478737 | 0.90650187 |
| Pcdh1 | 0.21205111 | 0.26507409 | 0.22868483 | 0.03638925 | 0.2396581 | 0.90650187 |

| | | | | | | |
|---|---|---|---|---|---|---|
| Cxxc5 | 0.67035424 | 0.61393664 | 0.57756522 | 0.03637142 | 0.38975551 | 0.90650187 |
| Myo1c | 0.20427245 | 0.25755986 | 0.22124264 | 0.03631722 | 0.23277117 | 0.90650187 |
| Atp6v1c1 | 0.58453139 | 0.55960757 | 0.5233025 | 0.03630506 | 0.37888061 | 0.90650187 |
| Zhx3 | 0.1488663 | 0.20259324 | 0.16632614 | 0.0362671 | 0.16618378 | 0.90650187 |
| Pik3r1 | 0.77036488 | 0.6721598 | 0.6358939 | 0.0362659 | 0.39992585 | 0.90650187 |
| Mrc1 | 0.01918889 | 0.05950144 | 0.02325156 | 0.03624989 | 2.09E-12 | 1.71E-10 |
| Ms4a6d | 0.01893322 | 0.0591866 | 0.02294569 | 0.03624091 | 1.10E-12 | 9.27E-11 |
| Scn9a | 0.15995325 | 0.21381151 | 0.17759009 | 0.03622143 | 0.18227489 | 0.90650187 |
| Baz2b | 0.71232484 | 0.63885119 | 0.60265026 | 0.03620092 | 0.39471705 | 0.90650187 |
| Psmc5 | 1.31705474 | 0.89124541 | 0.85506476 | 0.03618065 | 0.42539581 | 0.90650187 |
| Ifnar2 | 0.39885093 | 0.42619911 | 0.39009082 | 0.03610829 | 0.34035215 | 0.90650187 |
| Lrrc58 | 0.95921554 | 0.76927048 | 0.73318767 | 0.03608282 | 0.41340902 | 0.90650187 |
| Adamts9 | 0.19484237 | 0.24821169 | 0.21213429 | 0.0360774 | 0.22480707 | 0.90650187 |
| Klhl17 | 0.08704761 | 0.1369209 | 0.10087962 | 0.03604127 | 0.05611127 | 0.64256775 |
| Pdcd5 | 1.47252146 | 0.9245962 | 0.88857417 | 0.03602203 | 0.42848952 | 0.90650187 |
| Lbr | 0.33236421 | 0.372215 | 0.33620202 | 0.03601298 | 0.31696581 | 0.90650187 |
| Fndc4 | 0.22536991 | 0.27722827 | 0.24128182 | 0.03594645 | 0.25388574 | 0.90650187 |
| Smim20 | 0.52633828 | 0.51998194 | 0.48404006 | 0.03594188 | 0.37066079 | 0.90650187 |
| Fam178a | 0.30992941 | 0.35311387 | 0.31717581 | 0.03593806 | 0.30723056 | 0.90650187 |
| Cped1 | 0.12808828 | 0.18073108 | 0.14482755 | 0.03590353 | 0.13521135 | 0.90650187 |
| Sult1a1 | 0.11332439 | 0.16508288 | 0.129236 | 0.03584688 | 0.10876892 | 0.90650187 |
| Hsph1 | 0.53036493 | 0.52264539 | 0.4868252 | 0.03582019 | 0.3717946 | 0.90650187 |
| Gpr183 | 0.04468016 | 0.08901954 | 0.05322852 | 0.03579102 | 0.00139837 | 0.03139815 |
| Slc2a1 | 0.2540174 | 0.30353044 | 0.26776501 | 0.03576542 | 0.27632278 | 0.90650187 |
| Depdc1a | 0.07549277 | 0.12390473 | 0.08814292 | 0.03576181 | 0.03563958 | 0.45564114 |
| Csgalnact1 | 0.09397977 | 0.14417035 | 0.10843049 | 0.03573985 | 0.07141688 | 0.7575771 |
| 4933434E20I | 0.70186044 | 0.63219748 | 0.59648003 | 0.03571745 | 0.39504084 | 0.90650187 |
| Unc5c | 0.16794114 | 0.22133046 | 0.18561887 | 0.03571159 | 0.19619273 | 0.90650187 |
| S1pr2 | 0.18832522 | 0.2414788 | 0.20578319 | 0.03569562 | 0.22031051 | 0.90650187 |
| Ets2 | 0.33850037 | 0.37701896 | 0.3413297 | 0.03568926 | 0.3210265 | 0.90650187 |

| | | | | | | |
|---|---|---|---|---|---|---|
| Abcc9 | 0.14092502 | 0.19384773 | 0.15816944 | 0.03567829 | 0.15797973 | 0.90650187 |
| Tlcd2 | 0.13117799 | 0.18369608 | 0.14805677 | 0.03563932 | 0.14228286 | 0.90650187 |
| Parp1 | 0.54795949 | 0.53445779 | 0.49887442 | 0.03558337 | 0.37558648 | 0.90650187 |
| Ift20 | 1.34424727 | 0.89732545 | 0.86177673 | 0.03554872 | 0.42724857 | 0.90650187 |
| Gm2115 | 0.09274401 | 0.14263534 | 0.10708934 | 0.035546 | 0.07002254 | 0.74657023 |
| Flrt3 | 0.13871454 | 0.1914211 | 0.15588588 | 0.03553522 | 0.15543328 | 0.90650187 |
| Tcp11l1 | 0.16359547 | 0.21678183 | 0.18125987 | 0.03552196 | 0.19181789 | 0.90650187 |
| Chrac1 | 0.68499004 | 0.62193092 | 0.58641598 | 0.03551494 | 0.39387327 | 0.90650187 |
| Vip | 0.02437583 | 0.06494854 | 0.02943413 | 0.03551441 | 4.07E-08 | 2.14E-06 |
| Dnajc15 | 0.66953442 | 0.61251768 | 0.57706619 | 0.03545149 | 0.39238378 | 0.90650187 |
| Slf1 | 0.20432681 | 0.25672217 | 0.22129487 | 0.0354273 | 0.23832489 | 0.90650187 |
| Rnf138rt1 | 0.28729894 | 0.33293293 | 0.29752765 | 0.03540528 | 0.29839597 | 0.90650187 |
| Lyn | 0.08517557 | 0.13415882 | 0.09882897 | 0.03532985 | 0.05600349 | 0.6420129 |
| Aifm2 | 0.15369315 | 0.2065622 | 0.17124741 | 0.03531479 | 0.17962982 | 0.90650187 |
| Cdca7 | 0.21709331 | 0.26877011 | 0.23347539 | 0.03529471 | 0.25077695 | 0.90650187 |
| Adam12 | 0.19049332 | 0.24313536 | 0.20790111 | 0.03523426 | 0.22559766 | 0.90650187 |
| Thbs1 | 0.16466765 | 0.21747715 | 0.18233731 | 0.03513984 | 0.19579088 | 0.90650187 |
| Gdf10 | 0.06142926 | 0.10746299 | 0.07238159 | 0.0350814 | 0.01559402 | 0.23847555 |
| Fgf13 | 0.04745003 | 0.09144936 | 0.05642512 | 0.03502424 | 0.00289468 | 0.05866928 |
| Aif1 | 0.02657233 | 0.06706151 | 0.03203933 | 0.03502219 | 5.88E-07 | 2.59E-05 |
| B4gat1 | 0.3772098 | 0.40789881 | 0.3729461 | 0.03495272 | 0.33846914 | 0.90650187 |
| 9130401M01 | 0.45168073 | 0.46537982 | 0.43043121 | 0.03494861 | 0.35906527 | 0.90650187 |
| Dll4 | 0.12524908 | 0.17677789 | 0.14185006 | 0.03492783 | 0.13683619 | 0.90650187 |
| Tsen54 | 0.20032555 | 0.25236489 | 0.21744222 | 0.03492267 | 0.23761543 | 0.90650187 |
| Aard | 0.08512614 | 0.13369402 | 0.09877475 | 0.03491927 | 0.0580182 | 0.65425424 |
| Rad51 | 0.09971686 | 0.14951472 | 0.11462919 | 0.03488554 | 0.08803608 | 0.87466439 |
| Itgb5 | 0.16793538 | 0.22049862 | 0.18561311 | 0.03488551 | 0.20170673 | 0.90650187 |
| Cyth4 | 0.09016265 | 0.13906767 | 0.104281 | 0.03478667 | 0.06903831 | 0.73905027 |
| Dpp4 | 0.18130134 | 0.23363958 | 0.19888693 | 0.03475266 | 0.21863652 | 0.90650187 |
| Baz1b | 0.74551451 | 0.65659694 | 0.62185907 | 0.03473788 | 0.40193664 | 0.90650187 |

| | | | | | | |
|---|---|---|---|---|---|---|
| Pmm1 | 0.32072301 | 0.36112121 | 0.32638564 | 0.03473557 | 0.31806259 | 0.90650187 |
| Fez2 | 0.25744551 | 0.3055779 | 0.27087954 | 0.03469836 | 0.28452177 | 0.90650187 |
| Nudt7 | 0.30761916 | 0.34986049 | 0.31519128 | 0.03466921 | 0.3124241 | 0.90650187 |
| Scpep1 | 0.68636803 | 0.62190119 | 0.58724366 | 0.03465753 | 0.39651901 | 0.90650187 |
| Svip | 0.37941829 | 0.40935391 | 0.37471264 | 0.03464127 | 0.3405416 | 0.90650187 |
| Sct | 0.17265557 | 0.22489936 | 0.1903239 | 0.03457545 | 0.20965549 | 0.90650187 |
| Nfkb1 | 0.22396526 | 0.2745136 | 0.23996214 | 0.03455147 | 0.26105046 | 0.90650187 |
| Tspan8 | 0.07626474 | 0.12354779 | 0.08899979 | 0.03454801 | 0.0422008 | 0.51942868 |
| Pqlc1 | 0.29267636 | 0.33677387 | 0.30223917 | 0.03453469 | 0.3057381 | 0.90650187 |
| Uap1l1 | 0.14988286 | 0.20185067 | 0.16736486 | 0.0344858 | 0.17982703 | 0.90650187 |
| Stat1 | 0.10457096 | 0.15427706 | 0.11986507 | 0.03441199 | 0.10092716 | 0.90650187 |
| Adap2os | 0.02880625 | 0.06901511 | 0.03468108 | 0.03433403 | 5.38E-06 | 0.00020161 |
| Sox6 | 0.2561614 | 0.30402145 | 0.26971422 | 0.03430723 | 0.28587565 | 0.90650187 |
| Card19 | 0.54155928 | 0.52881445 | 0.49451462 | 0.03429983 | 0.37890804 | 0.90650187 |
| Ddit3 | 0.24265765 | 0.29164488 | 0.25736151 | 0.03428337 | 0.27685716 | 0.90650187 |
| Cdc42ep4 | 0.31554237 | 0.35625507 | 0.32197786 | 0.03427721 | 0.31800972 | 0.90650187 |
| Nsdhl | 0.21854316 | 0.26909803 | 0.23484804 | 0.03424999 | 0.25838171 | 0.90650187 |
| Prss3 | 0.02625544 | 0.0659132 | 0.03166395 | 0.03424925 | 7.60E-07 | 3.31E-05 |
| Stard5 | 0.28067612 | 0.32592475 | 0.29168863 | 0.03423612 | 0.30090814 | 0.90650187 |
| Slc15a3 | 0.02706244 | 0.06683737 | 0.03261959 | 0.03421778 | 1.56E-06 | 6.48E-05 |
| Fuca1 | 0.8171771 | 0.69575571 | 0.66155677 | 0.03419895 | 0.40911813 | 0.90650187 |
| Jam2 | 0.06291356 | 0.10823938 | 0.07405873 | 0.03418065 | 0.02009455 | 0.28925871 |
| Kcne3 | 0.0253548 | 0.06475093 | 0.0305962 | 0.03415473 | 3.48E-07 | 1.56E-05 |
| Pitx2 | 0.04082238 | 0.08290463 | 0.04875685 | 0.03414777 | 0.00092421 | 0.02226462 |
| Pth1r | 0.18167681 | 0.23336907 | 0.19925694 | 0.03411213 | 0.2233067 | 0.90650187 |
| C530008M17 | 0.10934207 | 0.15905216 | 0.12498456 | 0.0340676 | 0.1128024 | 0.90650187 |
| Txn1 | 2.15949553 | 0.956477 | 0.92241655 | 0.03406045 | 0.43480454 | 0.90650187 |
| Atg3 | 0.7048111 | 0.63226177 | 0.59822533 | 0.03403644 | 0.40015891 | 0.90650187 |
| Gfra2 | 0.10800417 | 0.15757968 | 0.12355183 | 0.03402785 | 0.11040443 | 0.90650187 |
| Spink1 | 0.02717945 | 0.06675777 | 0.03275807 | 0.03399969 | 1.97E-06 | 8.04E-05 |

| | | | | | | |
|---|---|---|---|---|---|---|
| Asrgl1 | 0.16148334 | 0.21297918 | 0.17913359 | 0.0338456 | 0.20046671 | 0.90650187 |
| Hoxb5 | 0.11186067 | 0.16147027 | 0.12767565 | 0.03379462 | 0.11965505 | 0.90650187 |
| Man1a | 0.17301052 | 0.22445327 | 0.19067715 | 0.03377612 | 0.21549908 | 0.90650187 |
| Hmha1 | 0.08797135 | 0.13562862 | 0.1018897 | 0.03373892 | 0.07049941 | 0.75046326 |
| Dynll1 | 3.47475419 | 0.8080179 | 0.77429961 | 0.03371828 | 0.42324768 | 0.90650187 |
| Rcsd1 | 0.19651468 | 0.24742744 | 0.21375666 | 0.03367077 | 0.24187989 | 0.90650187 |
| Ahr | 0.16162406 | 0.21294528 | 0.17927541 | 0.03366987 | 0.20187766 | 0.90650187 |
| Dync1li1 | 0.52968979 | 0.52002385 | 0.48635894 | 0.03366491 | 0.37914882 | 0.90650187 |
| Crabp2 | 0.01521471 | 0.05205857 | 0.0184853 | 0.03357327 | 3.40E-16 | 3.82E-14 |
| 2010107G12 | 0.0156793 | 0.05261396 | 0.01904381 | 0.03357015 | 2.31E-15 | 2.49E-13 |
| Zfhx3 | 0.43730924 | 0.45319296 | 0.41966327 | 0.03352969 | 0.361224 | 0.90650187 |
| Mgat2 | 0.92746644 | 0.75159982 | 0.71808281 | 0.03351701 | 0.41780902 | 0.90650187 |
| Kifc3 | 0.28310273 | 0.32730625 | 0.29383273 | 0.03347352 | 0.30627218 | 0.90650187 |
| Dynlt3 | 0.55389796 | 0.53631845 | 0.50289663 | 0.03342182 | 0.38382549 | 0.90650187 |
| Gria3 | 0.24089403 | 0.28914242 | 0.25573473 | 0.03340769 | 0.28070517 | 0.90650187 |
| Haus8 | 0.16289774 | 0.21393777 | 0.18055802 | 0.03337976 | 0.20557854 | 0.90650187 |
| Kif3a | 0.45519197 | 0.46641318 | 0.43303909 | 0.0333741 | 0.36594197 | 0.90650187 |
| Serpini2 | 0.02315333 | 0.06132167 | 0.02798084 | 0.03334084 | 5.88E-08 | 3.03E-06 |
| Inpp1 | 0.14825645 | 0.19903349 | 0.16570235 | 0.03333114 | 0.18559764 | 0.90650187 |
| Dnajc12 | 0.33490895 | 0.37164215 | 0.3383324 | 0.03330975 | 0.33080835 | 0.90650187 |
| Trp53i11 | 0.4147383 | 0.43572868 | 0.40244352 | 0.03328516 | 0.35655382 | 0.90650187 |
| Itfg2 | 0.30581 | 0.34691678 | 0.3136339 | 0.03328287 | 0.31854606 | 0.90650187 |
| Gm26523 | 0.14783706 | 0.19853548 | 0.16527312 | 0.03326236 | 0.18547135 | 0.90650187 |
| Timp1 | 0.19656802 | 0.24704878 | 0.21380836 | 0.03324042 | 0.24473541 | 0.90650187 |
| Camk2b | 0.39381651 | 0.41937407 | 0.38613584 | 0.03323823 | 0.35098081 | 0.90650187 |
| Ugt2b34 | 0.05877013 | 0.10260494 | 0.06936883 | 0.03323611 | 0.01658638 | 0.25048249 |
| Gm16141 | 0.20603819 | 0.25616868 | 0.22293755 | 0.03323113 | 0.25377314 | 0.90650187 |
| Tmem206 | 0.33476696 | 0.37143493 | 0.33821367 | 0.03322126 | 0.33117439 | 0.90650187 |
| Snrpb2 | 1.05642792 | 0.80637329 | 0.77316601 | 0.03320728 | 0.42428721 | 0.90650187 |
| Pycr1 | 0.19750105 | 0.24789939 | 0.21471219 | 0.0331872 | 0.2459974 | 0.90650187 |

| | | | | | | |
|---|---|---|---|---|---|---|
| Tril | 0.14163749 | 0.19206226 | 0.15890424 | 0.03315802 | 0.17679462 | 0.90650187 |
| Mnda | 0.01661504 | 0.05329502 | 0.02016763 | 0.03312739 | 1.41E-13 | 1.34E-11 |
| Mkrn1 | 0.78769926 | 0.67862333 | 0.64551339 | 0.03310993 | 0.40981308 | 0.90650187 |
| Adam33 | 0.10508366 | 0.15345128 | 0.12041657 | 0.03303472 | 0.11130935 | 0.90650187 |
| Dtx1 | 0.06310799 | 0.10730265 | 0.07427819 | 0.03302447 | 0.02404685 | 0.33351834 |
| Wasf2 | 0.77618276 | 0.67213195 | 0.6391375 | 0.03299445 | 0.40924119 | 0.90650187 |
| Prss53 | 0.18708775 | 0.23749813 | 0.20457208 | 0.03292605 | 0.23714265 | 0.90650187 |
| Aldh1a3 | 0.04702342 | 0.08885796 | 0.05593355 | 0.03292441 | 0.00443744 | 0.08495408 |
| Ddx3y | 0.27466456 | 0.31925282 | 0.28635343 | 0.03289939 | 0.30475975 | 0.90650187 |
| Cckar | 0.02032166 | 0.05748742 | 0.02460543 | 0.03288199 | 1.42E-09 | 8.65E-08 |
| A230083N12 | 0.01154633 | 0.04690386 | 0.01406309 | 0.03284077 | 1.50E-25 | 3.13E-23 |
| C3 | 0.05044939 | 0.09270107 | 0.05987345 | 0.03282762 | 0.00739558 | 0.13011596 |
| Chid1 | 0.57461517 | 0.54958266 | 0.51675953 | 0.03282313 | 0.38882892 | 0.90650187 |
| Polr2c | 1.17492883 | 0.84736163 | 0.81458283 | 0.0327788 | 0.4290116 | 0.90650187 |
| Lrrc59 | 0.82933831 | 0.70083767 | 0.66806137 | 0.03277631 | 0.41367189 | 0.90650187 |
| Aurka | 0.10130508 | 0.14909758 | 0.11634427 | 0.03275331 | 0.10537061 | 0.90650187 |
| Ell | 0.1555639 | 0.20589456 | 0.17314752 | 0.03274704 | 0.20023322 | 0.90650187 |
| Lrrc42 | 0.63591549 | 0.58901404 | 0.55628629 | 0.03272774 | 0.3968356 | 0.90650187 |
| Nrbp1 | 0.65638156 | 0.60173633 | 0.56900897 | 0.03272735 | 0.39909306 | 0.90650187 |
| Rab2b | 0.2488332 | 0.29575939 | 0.26303343 | 0.03272596 | 0.29009362 | 0.90650187 |
| Vps37a | 0.56032111 | 0.53992745 | 0.50722294 | 0.03270451 | 0.38719324 | 0.90650187 |
| Por | 0.78560274 | 0.67705887 | 0.6443571 | 0.03270177 | 0.4107489 | 0.90650187 |
| Rock1 | 1.12178097 | 0.82956461 | 0.79690435 | 0.03266025 | 0.42771355 | 0.90650187 |
| Calm1 | 3.67412231 | 0.77342952 | 0.74077309 | 0.03265643 | 0.42231243 | 0.90650187 |
| Spon2 | 0.08168384 | 0.12761922 | 0.09499083 | 0.03262838 | 0.06337797 | 0.69711178 |
| Ddrgk1 | 0.76673511 | 0.66646144 | 0.63386245 | 0.03259899 | 0.40957684 | 0.90650187 |
| Rab7b | 0.13989063 | 0.18964561 | 0.15710156 | 0.03254404 | 0.17854605 | 0.90650187 |
| Fam210b | 0.1725975 | 0.22280446 | 0.1902661 | 0.03253836 | 0.22354794 | 0.90650187 |
| Mcee | 0.69578665 | 0.62533019 | 0.5928738 | 0.03245639 | 0.40385898 | 0.90650187 |
| Flt3l | 0.04180147 | 0.08232394 | 0.04989391 | 0.03243003 | 0.00192894 | 0.04141506 |

| | | | | | | |
|---|---|---|---|---|---|---|
| Tbc1d20 | 0.32741978 | 0.36444206 | 0.33204693 | 0.03239513 | 0.33224525 | 0.90650187 |
| Nrsn1 | 0.05211397 | 0.09414461 | 0.06178133 | 0.03236328 | 0.00993567 | 0.16651835 |
| Rbmx | 0.53487609 | 0.52229148 | 0.48993334 | 0.03235814 | 0.38452625 | 0.90650187 |
| Chpf | 0.74942367 | 0.65635677 | 0.62408604 | 0.03227073 | 0.4090937 | 0.90650187 |
| Atp1b3 | 0.47438156 | 0.47939911 | 0.44713786 | 0.03226126 | 0.37419875 | 0.90650187 |
| C1s2 | 0.09827261 | 0.14532644 | 0.113073 | 0.03225344 | 0.1023839 | 0.90650187 |
| Snapc5 | 0.46782404 | 0.47459084 | 0.44234983 | 0.03224101 | 0.37296089 | 0.90650187 |
| Tmod3 | 0.46409538 | 0.47183876 | 0.43961308 | 0.03222568 | 0.3722562 | 0.90650187 |
| Bend5 | 0.16234842 | 0.21217057 | 0.18000507 | 0.0321655 | 0.21348271 | 0.90650187 |
| Gm10110 | 0.09772395 | 0.14464487 | 0.11248107 | 0.0321638 | 0.10182614 | 0.90650187 |
| Eif3j2 | 0.36252475 | 0.3932443 | 0.36109794 | 0.03214636 | 0.34613906 | 0.90650187 |
| Pitpna | 0.46586858 | 0.47306025 | 0.44091576 | 0.03214449 | 0.37293036 | 0.90650187 |
| Kcnk1 | 0.29224099 | 0.33399 | 0.3018587 | 0.03213131 | 0.31803099 | 0.90650187 |
| Rap1gap2 | 0.42368215 | 0.44142929 | 0.40931289 | 0.03211641 | 0.36361188 | 0.90650187 |
| Ifi204 | 0.02863306 | 0.0665364 | 0.03447656 | 0.03205984 | 1.78E-05 | 0.00061308 |
| Mndal | 0.05153457 | 0.09317291 | 0.06111772 | 0.0320552 | 0.0098597 | 0.16543711 |
| Mar-05 | 0.86405195 | 0.71832095 | 0.68627269 | 0.03204826 | 0.41776853 | 0.90650187 |
| 2310030G06 | 0.2502946 | 0.29639547 | 0.26436988 | 0.03202559 | 0.29510292 | 0.90650187 |
| Mis18bp1 | 0.09708467 | 0.14381023 | 0.11179085 | 0.03201938 | 0.10145219 | 0.90650187 |
| Dera | 0.36553427 | 0.39546802 | 0.36354064 | 0.03192738 | 0.3481099 | 0.90650187 |
| Rpa1 | 0.27930508 | 0.32236924 | 0.2904748 | 0.03189445 | 0.31272803 | 0.90650187 |
| Gjc1 | 0.2411726 | 0.28788439 | 0.2559919 | 0.03189249 | 0.2898403 | 0.90650187 |
| Fam133b | 0.75287024 | 0.65792921 | 0.62604357 | 0.03188564 | 0.41043602 | 0.90650187 |
| Camk4 | 0.07947923 | 0.12444415 | 0.09255859 | 0.03188556 | 0.06282674 | 0.69205016 |
| Lcp1 | 0.03884379 | 0.07832691 | 0.04645454 | 0.03187237 | 0.00114374 | 0.0265031 |
| Egflam | 0.19581489 | 0.24490066 | 0.21307814 | 0.03182253 | 0.25336536 | 0.90650187 |
| Adgre1 | 0.02162364 | 0.05793595 | 0.02615901 | 0.03177694 | 3.33E-08 | 1.77E-06 |
| Selplg | 0.02423741 | 0.06102744 | 0.0292697 | 0.03175774 | 7.05E-07 | 3.09E-05 |
| Gnaz | 0.41652873 | 0.43552724 | 0.4038237 | 0.03170355 | 0.36353813 | 0.90650187 |
| Lyar | 0.58402185 | 0.55465774 | 0.52296771 | 0.03169003 | 0.39381543 | 0.90650187 |

| | | | | | | |
|---|---|---|---|---|---|---|
| Endod1 | 0.12869192 | 0.17713688 | 0.14545935 | 0.03167753 | 0.16648587 | 0.90650187 |
| Msi2 | 0.5666751 | 0.5431009 | 0.51147819 | 0.03162271 | 0.39171397 | 0.90650187 |
| Mphosph10 | 0.37091328 | 0.39947057 | 0.36788832 | 0.03158225 | 0.35136284 | 0.90650187 |
| Ctxn2 | 0.10254313 | 0.14926031 | 0.11768054 | 0.03157977 | 0.11643745 | 0.90650187 |
| C8b | 0.02820098 | 0.06553733 | 0.03396609 | 0.03157125 | 1.80E-05 | 0.00061656 |
| Zfp36 | 0.24567021 | 0.29166164 | 0.26013324 | 0.0315284 | 0.29500835 | 0.90650187 |
| Ier3 | 0.12408364 | 0.17214376 | 0.14062495 | 0.03151881 | 0.15952526 | 0.90650187 |
| Slc16a12 | 0.31577834 | 0.35366874 | 0.32217914 | 0.0314896 | 0.3319587 | 0.90650187 |
| Ssu2 | 0.03505118 | 0.07351173 | 0.0420245 | 0.03148724 | 0.00043281 | 0.01138264 |
| Josd2 | 0.8222874 | 0.69569964 | 0.66429773 | 0.0314019 | 0.41677628 | 0.90650187 |
| Prdx5 | 1.92655005 | 0.95860554 | 0.92720861 | 0.03139693 | 0.44017082 | 0.90650187 |
| Reck | 0.19277713 | 0.24149553 | 0.21012659 | 0.03136894 | 0.25345193 | 0.90650187 |
| Rorc | 0.30965727 | 0.34825415 | 0.31694229 | 0.03131186 | 0.33025928 | 0.90650187 |
| Cxx1a | 0.82240826 | 0.69565213 | 0.66436242 | 0.03128971 | 0.41707726 | 0.90650187 |
| Hs3st6 | 0.0644347 | 0.10705525 | 0.07577414 | 0.03128111 | 0.03323532 | 0.43033228 |
| Acot13 | 0.54539134 | 0.52840456 | 0.49712828 | 0.03127628 | 0.3898575 | 0.90650187 |
| Slc16a2 | 0.15805384 | 0.20691434 | 0.17567035 | 0.03124399 | 0.21456626 | 0.90650187 |
| Alg2 | 0.67283926 | 0.61031538 | 0.57907576 | 0.03123963 | 0.40523147 | 0.90650187 |
| Mettl1 | 0.60532118 | 0.56804543 | 0.53683085 | 0.03121458 | 0.39801272 | 0.90650187 |
| Gtf2f2 | 0.37660407 | 0.40365488 | 0.37246091 | 0.03119397 | 0.35482584 | 0.90650187 |
| Igsf6 | 0.0216253 | 0.0573471 | 0.026161 | 0.0311861 | 5.80E-08 | 3.00E-06 |
| Elmo1 | 0.09066755 | 0.13601425 | 0.10483102 | 0.03118323 | 0.09301776 | 0.90318385 |
| 1700056E22l | 0.17073415 | 0.21958508 | 0.1884093 | 0.03117578 | 0.2309849 | 0.90650187 |
| Myo6 | 0.50508707 | 0.50030915 | 0.46916493 | 0.03114422 | 0.38395519 | 0.90650187 |
| Gm11627 | 0.12527486 | 0.17300983 | 0.14187714 | 0.03113269 | 0.16465174 | 0.90650187 |
| Upf3b | 0.58563275 | 0.55514617 | 0.52402566 | 0.03112052 | 0.39588496 | 0.90650187 |
| Mad2l1 | 0.15024827 | 0.19885453 | 0.16773793 | 0.0311166 | 0.20477319 | 0.90650187 |
| Pnrc1 | 0.86872216 | 0.71976226 | 0.68867901 | 0.03108326 | 0.42048586 | 0.90650187 |
| Tnfaip6 | 0.0661768 | 0.10880053 | 0.07773456 | 0.03106598 | 0.03781292 | 0.47621722 |
| Cdh2 | 0.25885886 | 0.30319747 | 0.27216032 | 0.03103715 | 0.3060856 | 0.90650187 |

| | | | | | | |
|---|---|---|---|---|---|---|
| Abca1 | 0.2351497 | 0.28145049 | 0.25041483 | 0.03103566 | 0.29082538 | 0.90650187 |
| Pcsk5 | 0.06716261 | 0.10986892 | 0.07884192 | 0.031027 | 0.04010245 | 0.49826502 |
| Fat4 | 0.22862027 | 0.27535078 | 0.2443279 | 0.03102289 | 0.28621747 | 0.90650187 |
| Chchd10 | 0.48107008 | 0.48300568 | 0.45199071 | 0.03101497 | 0.38016354 | 0.90650187 |
| Tmem141 | 0.19845714 | 0.24664621 | 0.2156374 | 0.03100881 | 0.26131998 | 0.90650187 |
| Pnpo | 0.12295179 | 0.17037349 | 0.13943349 | 0.03094 | 0.16195109 | 0.90650187 |
| Rhbdd2 | 0.30953008 | 0.34774004 | 0.31683312 | 0.03090692 | 0.33226522 | 0.90650187 |
| Fam101b | 0.30218154 | 0.34133614 | 0.31050164 | 0.0308345 | 0.32943536 | 0.90650187 |
| Ccdc8 | 0.64722087 | 0.59417285 | 0.56334276 | 0.03083009 | 0.40388845 | 0.90650187 |
| Uba7 | 0.1371222 | 0.18503719 | 0.15423733 | 0.03079986 | 0.18733788 | 0.90650187 |
| Cmpk1 | 1.16118526 | 0.84095978 | 0.81016003 | 0.03079976 | 0.43289514 | 0.90650187 |
| Cdkn2d | 0.45171814 | 0.46118936 | 0.43045904 | 0.03073033 | 0.37548131 | 0.90650187 |
| Kctd10 | 0.50208369 | 0.49769446 | 0.4670387 | 0.03065576 | 0.38521851 | 0.90650187 |
| Def8 | 0.22298564 | 0.26969625 | 0.23904059 | 0.03065566 | 0.28429649 | 0.90650187 |
| Gm16174 | 0.03832938 | 0.07647001 | 0.04585498 | 0.03061503 | 0.00149821 | 0.03324718 |
| Dcp2 | 0.24604424 | 0.2910911 | 0.26047674 | 0.03061436 | 0.30066073 | 0.90650187 |
| Sh3glb1 | 0.98025008 | 0.77300892 | 0.74243509 | 0.03057383 | 0.4273707 | 0.90650187 |
| Cdkn3 | 0.07332136 | 0.11628308 | 0.08572813 | 0.03055496 | 0.0565412 | 0.64620111 |
| Ciao1 | 0.30580843 | 0.34414352 | 0.31363254 | 0.03051098 | 0.33269631 | 0.90650187 |
| Isg20 | 0.19734255 | 0.24501907 | 0.21455872 | 0.03046035 | 0.26397735 | 0.90650187 |
| Fbxo5 | 0.12303657 | 0.16997844 | 0.13952279 | 0.03045564 | 0.165922 | 0.90650187 |
| Ddx6 | 0.76582203 | 0.66380284 | 0.63335048 | 0.03045235 | 0.4153707 | 0.90650187 |
| Lrrk1 | 0.13573285 | 0.18320424 | 0.15279649 | 0.03040775 | 0.18815597 | 0.90650187 |
| Tbx3os1 | 0.05040577 | 0.09021227 | 0.05982339 | 0.03038888 | 0.0119646 | 0.19389539 |
| Vwa5a | 0.25899096 | 0.30259372 | 0.27227994 | 0.03031378 | 0.31032006 | 0.90650187 |
| Runx1 | 0.07286488 | 0.11550553 | 0.08521962 | 0.03028591 | 0.05706384 | 0.64823856 |
| Ttyh3 | 0.35966885 | 0.38902681 | 0.35877313 | 0.03025368 | 0.35387579 | 0.90650187 |
| Pqbp1 | 0.60756146 | 0.56852724 | 0.53827364 | 0.03025359 | 0.401346 | 0.90650187 |
| Asph | 0.56095091 | 0.53787031 | 0.5076458 | 0.03022451 | 0.39562605 | 0.90650187 |
| Ilk | 0.95613406 | 0.76191774 | 0.73174489 | 0.03017285 | 0.42727716 | 0.90650187 |

| | | | | | | |
|---|---|---|---|---|---|---|
| Cxcl10 | 0.02187957 | 0.05663246 | 0.02646408 | 0.03016838 | 2.01E-07 | 9.38E-06 |
| Otub1 | 0.67843953 | 0.61258751 | 0.58246824 | 0.03011927 | 0.4090921 | 0.90650187 |
| Grina | 0.59402191 | 0.55962148 | 0.52951039 | 0.03011109 | 0.40021004 | 0.90650187 |
| Mrc2 | 0.12748923 | 0.1743009 | 0.14420013 | 0.03010077 | 0.1767066 | 0.90650187 |
| Fign | 0.09343043 | 0.13792144 | 0.10783457 | 0.03008688 | 0.1074199 | 0.90650187 |
| Pla1a | 0.03809295 | 0.07565367 | 0.04557927 | 0.0300744 | 0.00167675 | 0.03672663 |
| Slc1a3 | 0.1582067 | 0.20586045 | 0.17582499 | 0.03003545 | 0.22379935 | 0.90650187 |
| Slc25a24 | 0.21625666 | 0.26268846 | 0.23268231 | 0.03000616 | 0.28322254 | 0.90650187 |
| Mcts1 | 0.79993712 | 0.68222507 | 0.65222076 | 0.0300043 | 0.4189765 | 0.90650187 |
| Prickle1 | 0.13975048 | 0.18693015 | 0.15695678 | 0.02997337 | 0.19795218 | 0.90650187 |
| Ccne2 | 0.13534147 | 0.18235942 | 0.1523902 | 0.02996922 | 0.19098352 | 0.90650187 |
| Disp2 | 0.26796565 | 0.31030248 | 0.28036724 | 0.02993524 | 0.31751315 | 0.90650187 |
| Cd7 | 0.01578497 | 0.04909563 | 0.01917078 | 0.02992484 | 1.97E-12 | 1.62E-10 |
| Gls | 0.3630305 | 0.39142845 | 0.36150894 | 0.02991951 | 0.35646231 | 0.90650187 |
| Mesdc1 | 0.40633216 | 0.42583941 | 0.39593141 | 0.02990799 | 0.3685057 | 0.90650187 |
| Thsd7a | 0.05939956 | 0.09996858 | 0.07008291 | 0.02988568 | 0.02899629 | 0.38800405 |
| Utp14a | 0.22548441 | 0.27127261 | 0.2413893 | 0.0298833 | 0.29103854 | 0.90650187 |
| Tlr2 | 0.05803973 | 0.09841318 | 0.06853946 | 0.02987372 | 0.02633178 | 0.36027851 |
| Dock11 | 0.19473797 | 0.24190044 | 0.21203291 | 0.02986752 | 0.26558532 | 0.90650187 |
| Erp27 | 0.02426552 | 0.05916462 | 0.02930309 | 0.02986153 | 2.94E-06 | 0.00011596 |
| Mar-02 | 0.63957437 | 0.58843634 | 0.55857816 | 0.02985818 | 0.4060821 | 0.90650187 |
| Eml4 | 0.4608359 | 0.4670516 | 0.43721281 | 0.02983879 | 0.38079153 | 0.90650187 |
| Trafd1 | 0.40335848 | 0.42344929 | 0.39361496 | 0.02983433 | 0.36807432 | 0.90650187 |
| Rsad2 | 0.02954981 | 0.06539174 | 0.03555864 | 0.0298331 | 9.58E-05 | 0.00284484 |
| Samsn1 | 0.02690082 | 0.06218786 | 0.03242829 | 0.02975957 | 2.25E-05 | 0.00074817 |
| Gsta4 | 0.1198696 | 0.16593396 | 0.1361811 | 0.02975286 | 0.16570526 | 0.90650187 |
| Copz1 | 1.10295207 | 0.82004566 | 0.79029291 | 0.02975275 | 0.43354062 | 0.90650187 |
| Col13a1 | 0.106542 | 0.15169367 | 0.12198348 | 0.02971019 | 0.13945575 | 0.90650187 |
| Rpl32 | 5.07863481 | 0.5772038 | 0.5474979 | 0.0297059 | 0.40469705 | 0.90650187 |
| Prmt5 | 0.36435955 | 0.39226369 | 0.36258804 | 0.02967566 | 0.35798643 | 0.90650187 |

| | | | | | | |
|---|---|---|---|---|---|---|
| Arap3 | 0.05006349 | 0.08910352 | 0.05943055 | 0.02967298 | 0.01322164 | 0.21023232 |
| Slc35b2 | 0.5473259 | 0.52811159 | 0.49844401 | 0.02966759 | 0.39565716 | 0.90650187 |
| Vsir | 0.03464909 | 0.07120884 | 0.04155351 | 0.02965534 | 0.0007552 | 0.01869664 |
| Pacsin2 | 0.30222423 | 0.34014348 | 0.31053856 | 0.02960492 | 0.33584931 | 0.90650187 |
| Fam122a | 0.59529889 | 0.55992637 | 0.53034137 | 0.029585 | 0.40206822 | 0.90650187 |
| Antxr2 | 0.20638754 | 0.25283317 | 0.2232725 | 0.02956067 | 0.27806944 | 0.90650187 |
| Gm648 | 0.03598891 | 0.07264723 | 0.04312191 | 0.02952532 | 0.00116774 | 0.02696394 |
| Clcc1 | 0.40573291 | 0.42497245 | 0.39546515 | 0.0295073 | 0.37005594 | 0.90650187 |
| Homer3 | 0.18188097 | 0.22896215 | 0.19945806 | 0.02950409 | 0.25539769 | 0.90650187 |
| Sncb | 0.02589724 | 0.06068353 | 0.03123944 | 0.02944409 | 1.39E-05 | 0.00048828 |
| Hmx2 | 0.01741542 | 0.05056817 | 0.02112777 | 0.0294404 | 2.92E-10 | 1.95E-08 |
| Cxadr | 0.26900164 | 0.31072686 | 0.28129589 | 0.02943097 | 0.32091998 | 0.90650187 |
| Arhgap20 | 0.10524912 | 0.1500227 | 0.12059448 | 0.02942821 | 0.13899528 | 0.90650187 |
| Alkbh8 | 0.16191139 | 0.2089842 | 0.17956491 | 0.02941928 | 0.23319893 | 0.90650187 |
| B4galt3 | 0.56486428 | 0.53966356 | 0.51026796 | 0.0293956 | 0.39893557 | 0.90650187 |
| Lipt2 | 0.15203171 | 0.19894484 | 0.16955652 | 0.02938832 | 0.22049188 | 0.90650187 |
| Gm21092 | 0.72992975 | 0.64228833 | 0.61290617 | 0.02938216 | 0.41561733 | 0.90650187 |
| Traf5 | 0.11731429 | 0.16283381 | 0.13347592 | 0.02935789 | 0.16408503 | 0.90650187 |
| Tmem171 | 0.04047618 | 0.07771208 | 0.04835444 | 0.02935764 | 0.00347658 | 0.06926271 |
| Crmp1 | 0.07016223 | 0.11145871 | 0.08220273 | 0.02925598 | 0.0568028 | 0.64678604 |
| Il18 | 0.05052878 | 0.08919615 | 0.05996453 | 0.02923162 | 0.01511163 | 0.23203229 |
| Rimbp2 | 0.45820924 | 0.46449143 | 0.43527315 | 0.02921828 | 0.38269239 | 0.90650187 |
| Chchd7 | 0.73962004 | 0.64770522 | 0.61848731 | 0.02921791 | 0.41682817 | 0.90650187 |
| Wfdc10 | 0.05474072 | 0.09396807 | 0.06478353 | 0.02918453 | 0.02260394 | 0.31785621 |
| Cmtm8 | 0.1773405 | 0.22415813 | 0.19497432 | 0.02918381 | 0.25289108 | 0.90650187 |
| Psmc1 | 1.07299263 | 0.80859589 | 0.77941438 | 0.02918151 | 0.43390345 | 0.90650187 |
| Sp3os | 0.36385937 | 0.39135307 | 0.36218209 | 0.02917098 | 0.36014944 | 0.90650187 |
| Tnc | 0.05669797 | 0.09617566 | 0.06701383 | 0.02916184 | 0.02652236 | 0.3622339 |
| Nudt11 | 0.13123905 | 0.17720333 | 0.14812047 | 0.02908287 | 0.19136501 | 0.90650187 |
| Man1b1 | 0.39113042 | 0.41307135 | 0.38401701 | 0.02905434 | 0.36830345 | 0.90650187 |

| | | | | | | |
|---|---|---|---|---|---|---|
| Cd38 | 0.0498679 | 0.08825159 | 0.05920598 | 0.0290456 | 0.01459354 | 0.22659495 |
| Safb | 0.86057747 | 0.71349904 | 0.68447283 | 0.02902621 | 0.42523321 | 0.90650187 |
| Tgs1 | 0.35388575 | 0.38306495 | 0.35404512 | 0.02901983 | 0.35778314 | 0.90650187 |
| Dnase1l1 | 0.08758438 | 0.13045846 | 0.1014667 | 0.02899175 | 0.10200158 | 0.90650187 |
| Vegfc | 0.07144413 | 0.11259318 | 0.08363499 | 0.02895819 | 0.06186938 | 0.6854785 |
| Rnf114 | 0.39648974 | 0.4171612 | 0.38823856 | 0.02892265 | 0.37025246 | 0.90650187 |
| Cox4i2 | 0.0627489 | 0.10279011 | 0.07387284 | 0.02891728 | 0.04091038 | 0.50559243 |
| Arhgef1 | 0.59354408 | 0.55810197 | 0.5291992 | 0.02890277 | 0.40407962 | 0.90650187 |
| Chst2 | 0.07632978 | 0.11797353 | 0.08907194 | 0.02890159 | 0.07458259 | 0.77834237 |
| Ptms | 0.96857391 | 0.76628823 | 0.73739361 | 0.02889461 | 0.43085408 | 0.90650187 |
| Rabif | 0.30716452 | 0.34368163 | 0.31480018 | 0.02888145 | 0.34168689 | 0.90650187 |
| Alkbh5 | 0.49554328 | 0.4912471 | 0.46238792 | 0.02885918 | 0.39071146 | 0.90650187 |
| 1810055G02 | 0.17973266 | 0.22611921 | 0.19733941 | 0.0287798 | 0.25838153 | 0.90650187 |
| Sh3pxd2b | 0.1875236 | 0.23377666 | 0.20499884 | 0.02877782 | 0.2662888 | 0.90650187 |
| Utp11l | 0.8195972 | 0.69162885 | 0.6628562 | 0.02877265 | 0.42349092 | 0.90650187 |
| Rara | 0.24978377 | 0.29264929 | 0.26390297 | 0.02874632 | 0.31410449 | 0.90650187 |
| Hcls1 | 0.03279601 | 0.06811733 | 0.03937968 | 0.02873765 | 0.00058887 | 0.01496881 |
| Arl4c | 0.17504505 | 0.22142488 | 0.19269892 | 0.02872595 | 0.25375686 | 0.90650187 |
| Mprip | 0.51842367 | 0.50723952 | 0.47853569 | 0.02870383 | 0.39486581 | 0.90650187 |
| Gpr17 | 0.02910328 | 0.06372524 | 0.03503174 | 0.0286935 | 0.00013565 | 0.00390529 |
| Haus4 | 0.13097562 | 0.17653851 | 0.14784561 | 0.0286929 | 0.19413085 | 0.90650187 |
| Clec1b | 0.02239985 | 0.05577182 | 0.02708393 | 0.02868789 | 1.25E-06 | 5.27E-05 |
| Msr1 | 0.02037107 | 0.05335172 | 0.02466444 | 0.02868728 | 1.17E-07 | 5.63E-06 |
| Cybb | 0.01768247 | 0.05010151 | 0.02144789 | 0.02865362 | 1.43E-09 | 8.71E-08 |
| Chd1l | 0.10268912 | 0.14646323 | 0.11783797 | 0.02862526 | 0.14008689 | 0.90650187 |
| Tars2 | 0.15550808 | 0.20170804 | 0.17309089 | 0.02861715 | 0.23117192 | 0.90650187 |
| Zc3hc1 | 0.24119383 | 0.28462198 | 0.25601149 | 0.02861049 | 0.30965891 | 0.90650187 |
| Naga | 0.34974847 | 0.37923878 | 0.3506458 | 0.02859298 | 0.35848493 | 0.90650187 |
| Ggh | 0.46985143 | 0.47239946 | 0.44383366 | 0.0285658 | 0.38739085 | 0.90650187 |
| Pmf1 | 0.32912969 | 0.36203185 | 0.33348622 | 0.02854563 | 0.35177478 | 0.90650187 |

| | | | | | | |
|---|---|---|---|---|---|---|
| Nrxn2 | 0.10319726 | 0.14692961 | 0.11838577 | 0.02854383 | 0.1418883 | 0.90650187 |
| Ncaph2 | 0.56257758 | 0.53726591 | 0.50873687 | 0.02852904 | 0.40156376 | 0.90650187 |
| Rbm46 | 0.0802943 | 0.12198438 | 0.09345863 | 0.02852575 | 0.08740607 | 0.87068126 |
| 1700023F06I | 0.30467473 | 0.34117512 | 0.31265513 | 0.02851998 | 0.34254684 | 0.90650187 |
| Tspan32 | 0.02631135 | 0.06021985 | 0.03173019 | 0.02848966 | 3.28E-05 | 0.00105687 |
| Eps8 | 0.44962197 | 0.45737521 | 0.42889807 | 0.02847714 | 0.38393172 | 0.90650187 |
| Ssna1 | 0.88997254 | 0.7279683 | 0.69951625 | 0.02845205 | 0.42825439 | 0.90650187 |
| Trim2 | 0.13109035 | 0.1764 | 0.14796533 | 0.02843468 | 0.19646452 | 0.90650187 |
| Plaur | 0.04214793 | 0.07867919 | 0.05029592 | 0.02838327 | 0.00605764 | 0.10975223 |
| Rcan2 | 0.16848295 | 0.21453123 | 0.18616086 | 0.02837037 | 0.24904518 | 0.90650187 |
| Ece2 | 0.23691189 | 0.28041839 | 0.2520503 | 0.02836809 | 0.30841433 | 0.90650187 |
| Tcn2 | 0.37927407 | 0.40294395 | 0.3745974 | 0.02834655 | 0.36828158 | 0.90650187 |
| Prkcd | 0.20646932 | 0.25169717 | 0.22335089 | 0.02834628 | 0.28630623 | 0.90650187 |
| Gbp9 | 0.01889815 | 0.05124347 | 0.02290373 | 0.02833974 | 1.89E-08 | 1.03E-06 |
| Cenpc1 | 0.25327226 | 0.29540286 | 0.26708653 | 0.02831633 | 0.31870609 | 0.90650187 |
| Abi3 | 0.06889473 | 0.10907373 | 0.08078421 | 0.02828952 | 0.05976013 | 0.66746046 |
| AI464131 | 0.09530996 | 0.13816051 | 0.10987176 | 0.02828875 | 0.12618471 | 0.90650187 |
| Srpk2 | 0.74044314 | 0.64721905 | 0.61895933 | 0.02825972 | 0.41957834 | 0.90650187 |
| Myzap | 0.04369614 | 0.08030248 | 0.05209008 | 0.02821241 | 0.008025 | 0.13957447 |
| Arf6 | 0.40135137 | 0.42021842 | 0.39204766 | 0.02817076 | 0.37469603 | 0.90650187 |
| Upf1 | 0.29235921 | 0.33008315 | 0.30196203 | 0.02812112 | 0.33943323 | 0.90650187 |
| Rab5a | 0.84156031 | 0.70265393 | 0.67453428 | 0.02811965 | 0.42648659 | 0.90650187 |
| 2810004N23 | 0.75470357 | 0.65513536 | 0.6270826 | 0.02805277 | 0.42117995 | 0.90650187 |
| Atpif1 | 3.21746345 | 0.83836014 | 0.81032734 | 0.0280328 | 0.43888641 | 0.90650187 |
| Ttc4 | 0.49049794 | 0.48680731 | 0.45877975 | 0.02802756 | 0.39296906 | 0.90650187 |
| Dfna5 | 0.23117733 | 0.27473065 | 0.24671676 | 0.02801389 | 0.30685517 | 0.90650187 |
| Tst | 0.15718529 | 0.20279746 | 0.17479113 | 0.02800633 | 0.23813706 | 0.90650187 |
| Evi5 | 0.5556144 | 0.53205709 | 0.50405518 | 0.02800191 | 0.40246652 | 0.90650187 |
| Dynlt1f | 0.92284135 | 0.74383011 | 0.71583279 | 0.02799732 | 0.43098204 | 0.90650187 |
| Nfkbiz | 0.13005566 | 0.17487949 | 0.14688508 | 0.02799441 | 0.1984208 | 0.90650187 |

| | | | | | | |
|---|---|---|---|---|---|---|
| Fam174a | 0.67430441 | 0.60788371 | 0.57996486 | 0.02791885 | 0.415271 | 0.90650187 |
| Mier1 | 0.66525737 | 0.60235898 | 0.57445674 | 0.02790224 | 0.41452119 | 0.90650187 |
| Has2 | 0.06004095 | 0.09866398 | 0.07080996 | 0.02785402 | 0.0401682 | 0.49844643 |
| Palld | 0.11727337 | 0.16128422 | 0.13343253 | 0.02785169 | 0.1767188 | 0.90650187 |
| Dio3 | 0.09597464 | 0.1384254 | 0.11059102 | 0.02783438 | 0.13159065 | 0.90650187 |
| Nkiras2 | 0.71077174 | 0.62956291 | 0.60173789 | 0.02782502 | 0.41856353 | 0.90650187 |
| Myo10 | 0.22024266 | 0.264267 | 0.2364543 | 0.0278127 | 0.30051846 | 0.90650187 |
| 0610009B22 | 0.80449657 | 0.68251256 | 0.65470267 | 0.02780989 | 0.42511046 | 0.90650187 |
| Ppa2 | 0.57394722 | 0.54409453 | 0.51631672 | 0.02777781 | 0.40548558 | 0.90650187 |
| Scn7a | 0.0597693 | 0.09827117 | 0.0705021 | 0.02776906 | 0.03997156 | 0.49722398 |
| Wdr1 | 0.73640243 | 0.64440209 | 0.61663907 | 0.02776302 | 0.42067963 | 0.90650187 |
| Slc44a3 | 0.42825359 | 0.44050079 | 0.41280023 | 0.02770056 | 0.38273018 | 0.90650187 |
| Nosip | 0.26055936 | 0.30138573 | 0.27369878 | 0.02768695 | 0.32645953 | 0.90650187 |
| Nupl2 | 0.09160652 | 0.13353056 | 0.10585298 | 0.02767758 | 0.12253626 | 0.90650187 |
| Socs2 | 0.40542114 | 0.42289888 | 0.39522246 | 0.02767643 | 0.37778076 | 0.90650187 |
| 4833439L19 | 0.6447518 | 0.58941746 | 0.5618079 | 0.02760957 | 0.41352932 | 0.90650187 |
| 2810474O19 | 0.40737073 | 0.42434699 | 0.39673886 | 0.02760813 | 0.3785241 | 0.90650187 |
| Hpgds | 0.06630734 | 0.10548498 | 0.07788127 | 0.02760371 | 0.05755091 | 0.65057885 |
| Ehf | 0.09905399 | 0.14150271 | 0.1139153 | 0.02758741 | 0.14082643 | 0.90650187 |
| B3gnt8 | 0.17596217 | 0.22116756 | 0.19360873 | 0.02755883 | 0.26343194 | 0.90650187 |
| Hk1 | 0.13064312 | 0.17500048 | 0.14749856 | 0.02750191 | 0.20357834 | 0.90650187 |
| Abi1 | 0.35782739 | 0.38472899 | 0.35727059 | 0.02745839 | 0.36629903 | 0.90650187 |
| 1810034E14 | 0.03300117 | 0.0670737 | 0.03962061 | 0.02745309 | 0.00103376 | 0.02451512 |
| Tgfb3 | 0.11046267 | 0.15360039 | 0.12618287 | 0.02741751 | 0.16703238 | 0.90650187 |
| Nkd2 | 0.05002621 | 0.08680066 | 0.05938775 | 0.02741291 | 0.02008209 | 0.28925871 |
| 1810058I24R | 1.46760204 | 0.91504188 | 0.88768117 | 0.02736072 | 0.44550531 | 0.90650187 |
| Faim | 0.40634459 | 0.42325524 | 0.39594108 | 0.02731417 | 0.37954269 | 0.90650187 |
| Slc2a5 | 0.27780899 | 0.31643717 | 0.28914827 | 0.0272889 | 0.33740264 | 0.90650187 |
| Klhl32 | 0.29480925 | 0.33136466 | 0.30410068 | 0.02726397 | 0.34510566 | 0.90650187 |
| Nsl1 | 0.05325952 | 0.09030083 | 0.06309188 | 0.02720895 | 0.02760576 | 0.37300897 |

| | | | | | | |
|---|---|---|---|---|---|---|
| Myo18a | 0.31348246 | 0.34738749 | 0.32021875 | 0.02716873 | 0.35301831 | 0.90650187 |
| Prkd1 | 0.02532651 | 0.05768501 | 0.03056264 | 0.02712236 | 3.99E-05 | 0.00126788 |
| Crybb1 | 0.07388194 | 0.1134458 | 0.08635219 | 0.02709361 | 0.08153197 | 0.82790075 |
| Wee1 | 0.07305411 | 0.11251662 | 0.08543045 | 0.02708617 | 0.07934328 | 0.81530674 |
| Hoxc4 | 0.1472974 | 0.19179676 | 0.16472049 | 0.02707626 | 0.23246424 | 0.90650187 |
| Enox1 | 0.04854462 | 0.08475918 | 0.05768514 | 0.02707404 | 0.0184663 | 0.27348002 |
| Mafg | 0.42638429 | 0.43844373 | 0.41137608 | 0.02706765 | 0.3849479 | 0.90650187 |
| Zfp395 | 0.38930271 | 0.40961629 | 0.38257208 | 0.02704421 | 0.37666099 | 0.90650187 |
| Hexb | 0.29560171 | 0.33179573 | 0.30479126 | 0.02700446 | 0.34683478 | 0.90650187 |
| Iqsec1 | 0.13153761 | 0.17542911 | 0.14843188 | 0.02699723 | 0.20937838 | 0.90650187 |
| Timd4 | 0.01349697 | 0.04338794 | 0.0164173 | 0.02697064 | 1.40E-13 | 1.34E-11 |
| Ggcx | 0.24846969 | 0.28965547 | 0.26270068 | 0.02695479 | 0.3241418 | 0.90650187 |
| Mettl23 | 0.39371918 | 0.41300628 | 0.38605916 | 0.02694712 | 0.3781643 | 0.90650187 |
| BC002059 | 0.0934753 | 0.13482706 | 0.10788326 | 0.0269438 | 0.13343738 | 0.90650187 |
| Fgb | 0.01323703 | 0.04304402 | 0.01610394 | 0.02694007 | 5.15E-14 | 5.05E-12 |
| Sugt1 | 0.7920607 | 0.67483434 | 0.64791186 | 0.02692248 | 0.42672162 | 0.90650187 |
| Klk1 | 0.02087337 | 0.05210046 | 0.02526408 | 0.02683638 | 1.17E-06 | 4.97E-05 |
| Rpa3 | 0.68276148 | 0.61190317 | 0.58507536 | 0.02682781 | 0.41923652 | 0.90650187 |
| Cir1 | 0.56078948 | 0.53435416 | 0.50753743 | 0.02681672 | 0.4071462 | 0.90650187 |
| Lat2 | 0.02234587 | 0.05383085 | 0.02701964 | 0.02681121 | 5.14E-06 | 0.00019282 |
| Nsa2 | 1.39904783 | 0.90090373 | 0.87414822 | 0.02675551 | 0.44588337 | 0.90650187 |
| Fam83d | 0.0564769 | 0.09350717 | 0.0667622 | 0.02674497 | 0.03746341 | 0.47417256 |
| Exoc3 | 0.53139738 | 0.51426976 | 0.48753767 | 0.02673209 | 0.40370981 | 0.90650187 |
| Rhoq | 0.13014631 | 0.17368024 | 0.14697977 | 0.02670047 | 0.20966254 | 0.90650187 |
| Nde1 | 0.17856605 | 0.22287672 | 0.19618679 | 0.02668993 | 0.2726574 | 0.90650187 |
| Slc4a3 | 0.14227745 | 0.18623888 | 0.15956375 | 0.02667513 | 0.22868153 | 0.90650187 |
| Pop4 | 0.30378005 | 0.33855363 | 0.31188299 | 0.02667063 | 0.35191131 | 0.90650187 |
| Rtn4 | 0.64854929 | 0.59081263 | 0.5641672 | 0.02664543 | 0.41684769 | 0.90650187 |
| Mapre2 | 0.43490978 | 0.4444907 | 0.41785058 | 0.02664012 | 0.38842353 | 0.90650187 |
| Elf4 | 0.12121848 | 0.16423742 | 0.13760589 | 0.02663153 | 0.19478993 | 0.90650187 |

| | | | | | | |
|---|---|---|---|---|---|---|
| Naalad2 | 0.11001289 | 0.1523276 | 0.12570209 | 0.02662551 | 0.17318767 | 0.90650187 |
| Spry4 | 0.10880448 | 0.15102753 | 0.12440913 | 0.02661839 | 0.17075769 | 0.90650187 |
| Fndc1 | 0.13559843 | 0.17921431 | 0.15265697 | 0.02655734 | 0.2196456 | 0.90650187 |
| Cdon | 0.2358092 | 0.27757436 | 0.25102727 | 0.02654709 | 0.31912755 | 0.90650187 |
| Ankrd37 | 0.27250058 | 0.31095697 | 0.28442469 | 0.02653228 | 0.33917926 | 0.90650187 |
| Plk4 | 0.15119801 | 0.19522514 | 0.16870684 | 0.0265183 | 0.24234298 | 0.90650187 |
| Magt1 | 0.79096689 | 0.67380005 | 0.64731132 | 0.02648874 | 0.42782299 | 0.90650187 |
| Notch4 | 0.05396239 | 0.0903738 | 0.06389503 | 0.02647877 | 0.03271521 | 0.42504607 |
| Tgoln1 | 0.5163973 | 0.50357233 | 0.47711929 | 0.02645305 | 0.40265578 | 0.90650187 |
| Arsj | 0.08498211 | 0.12503478 | 0.09861676 | 0.02641802 | 0.11684363 | 0.90650187 |
| Sigmar1 | 0.64610937 | 0.58906808 | 0.56265221 | 0.02641587 | 0.41733496 | 0.90650187 |
| Clcn3 | 0.54435311 | 0.52281684 | 0.49642121 | 0.02639563 | 0.40656849 | 0.90650187 |
| Esco1 | 0.23602505 | 0.27760475 | 0.25122762 | 0.02637714 | 0.32033672 | 0.90650187 |
| Abcc3 | 0.10054513 | 0.1418712 | 0.11552307 | 0.02634813 | 0.1553058 | 0.90650187 |
| Fxyd2 | 0.11888656 | 0.16148795 | 0.13514135 | 0.0263466 | 0.19305402 | 0.90650187 |
| Hspb2 | 0.04824108 | 0.08365312 | 0.05733591 | 0.02631721 | 0.0206485 | 0.29555349 |
| Tmem204 | 0.10259778 | 0.14405647 | 0.11773947 | 0.02631699 | 0.1601901 | 0.90650187 |
| Glt8d2 | 0.07657266 | 0.11564253 | 0.08934133 | 0.0263012 | 0.09531086 | 0.90650187 |
| Vps26a | 0.29793683 | 0.33311956 | 0.30682249 | 0.02629707 | 0.35159309 | 0.90650187 |
| Aspm | 0.06207761 | 0.09940009 | 0.07311457 | 0.02628551 | 0.05499417 | 0.63426882 |
| Clstn1 | 0.22050238 | 0.26293903 | 0.23669952 | 0.02623951 | 0.31106946 | 0.90650187 |
| Mgarp | 0.01477642 | 0.04413092 | 0.0179581 | 0.02617283 | 4.61E-11 | 3.31E-09 |
| Mrpl18 | 0.89908693 | 0.73023264 | 0.70409177 | 0.02614087 | 0.43445139 | 0.90650187 |
| Renbp | 0.12756315 | 0.17041162 | 0.14427757 | 0.02613404 | 0.21033341 | 0.90650187 |
| Wdr86 | 0.12844815 | 0.17132448 | 0.14520426 | 0.02612022 | 0.2119416 | 0.90650187 |
| S1pr1 | 0.05695992 | 0.09342164 | 0.06731188 | 0.02610976 | 0.04231631 | 0.52042905 |
| Nrp2 | 0.13812717 | 0.1813344 | 0.15527812 | 0.02605628 | 0.22783665 | 0.90650187 |
| Ranbp3 | 0.4986318 | 0.49062713 | 0.46458763 | 0.0260395 | 0.4016146 | 0.90650187 |
| Gypc | 0.13395713 | 0.17696563 | 0.15095163 | 0.026014 | 0.22180064 | 0.90650187 |
| Gng13 | 0.08080106 | 0.12000195 | 0.09401773 | 0.02598422 | 0.10960007 | 0.90650187 |

| | | | | | | |
|---|---|---|---|---|---|---|
| Cd164l2 | 0.37432986 | 0.39659965 | 0.37063666 | 0.025963 | 0.37774315 | 0.90650187 |
| Dqx1 | 0.01712686 | 0.04671786 | 0.02078173 | 0.02593613 | 1.44E-08 | 7.92E-07 |
| Dtx2 | 0.179008 | 0.22253167 | 0.19662361 | 0.02590806 | 0.27901366 | 0.90650187 |
| Col11a1 | 0.08849697 | 0.1283424 | 0.10246391 | 0.02587849 | 0.13076308 | 0.90650187 |
| Efs | 0.20129656 | 0.24424733 | 0.21837871 | 0.02586862 | 0.2992302 | 0.90650187 |
| Tmem173 | 0.09960434 | 0.14034581 | 0.11450804 | 0.02583776 | 0.15790314 | 0.90650187 |
| Ube2j1 | 0.41535161 | 0.42874716 | 0.40291657 | 0.02583058 | 0.38782076 | 0.90650187 |
| Ubtd2 | 0.12348418 | 0.16582122 | 0.13999411 | 0.02582711 | 0.20606326 | 0.90650187 |
| Desi2 | 0.58487111 | 0.54933529 | 0.52352564 | 0.02580965 | 0.41325998 | 0.90650187 |
| Sar1a | 1.32851836 | 0.88371905 | 0.85791839 | 0.02580065 | 0.44682181 | 0.90650187 |
| Mapre1 | 1.20484538 | 0.8496477 | 0.82388896 | 0.02575874 | 0.44472861 | 0.90650187 |
| Slc45a3 | 0.0148454 | 0.04378974 | 0.01804109 | 0.02574865 | 1.11E-10 | 7.62E-09 |
| Efna4 | 0.15155764 | 0.19481974 | 0.16907346 | 0.02574627 | 0.24921225 | 0.90650187 |
| Pdgfrl | 0.15796247 | 0.20130398 | 0.17557789 | 0.02572608 | 0.25740041 | 0.90650187 |
| Ly6g6e | 0.03584656 | 0.06866835 | 0.04295541 | 0.02571294 | 0.00389403 | 0.07598603 |
| Usp54 | 0.04856114 | 0.08341454 | 0.05770414 | 0.0257104 | 0.02380886 | 0.33051919 |
| Heyl | 0.09083627 | 0.13063667 | 0.10501475 | 0.02562192 | 0.13903681 | 0.90650187 |
| Maff | 0.18970695 | 0.2327218 | 0.20713352 | 0.02558828 | 0.29143794 | 0.90650187 |
| Ceacam1 | 0.04375029 | 0.07770734 | 0.05215277 | 0.02555457 | 0.01469102 | 0.22787574 |
| Smad7 | 0.17444406 | 0.2176508 | 0.19210222 | 0.02554857 | 0.27717916 | 0.90650187 |
| Otud1 | 0.03212211 | 0.06412987 | 0.0385878 | 0.02554207 | 0.00162714 | 0.03574293 |
| Gm7808 | 0.04216162 | 0.07584753 | 0.05031179 | 0.02553574 | 0.01202385 | 0.19464811 |
| Acsl1 | 0.05968352 | 0.09592486 | 0.07040487 | 0.02552 | 0.05354403 | 0.62226513 |
| Crim1 | 0.08672791 | 0.12602942 | 0.10052978 | 0.02549964 | 0.12973605 | 0.90650187 |
| Tsku | 0.08521277 | 0.12436902 | 0.09886976 | 0.02549926 | 0.12578263 | 0.90650187 |
| Fam185a | 0.05669836 | 0.09251351 | 0.06701426 | 0.02549925 | 0.04536355 | 0.5490217 |
| Trip13 | 0.08106582 | 0.11980507 | 0.0943097 | 0.02549537 | 0.11471846 | 0.90650187 |
| Txnrd2 | 0.19483873 | 0.23760228 | 0.21213075 | 0.02547153 | 0.29673835 | 0.90650187 |
| Amt | 0.10878513 | 0.14984987 | 0.12438842 | 0.02546145 | 0.18141524 | 0.90650187 |
| Ppp6c | 0.66667525 | 0.60077635 | 0.57532305 | 0.02545331 | 0.4220384 | 0.90650187 |

| | | | | | | |
|---|---|---|---|---|---|---|
| Sqrdl | 0.21609625 | 0.25797936 | 0.23253017 | 0.0254492 | 0.3132893 | 0.90650187 |
| Cercam | 0.1804166 | 0.22338202 | 0.19801445 | 0.02536757 | 0.28449988 | 0.90650187 |
| Lrwd1 | 0.17798072 | 0.22097527 | 0.19560791 | 0.02536736 | 0.28212938 | 0.90650187 |
| Mast4 | 0.62582447 | 0.57522125 | 0.54992768 | 0.02529356 | 0.41899204 | 0.90650187 |
| Pbxip1 | 0.47335995 | 0.47168039 | 0.44639394 | 0.02528644 | 0.4005878 | 0.90650187 |
| Pip5k1a | 0.50178966 | 0.49209923 | 0.46683023 | 0.025269 | 0.40491873 | 0.90650187 |
| Itgb1bp1 | 0.39155781 | 0.40960585 | 0.38435451 | 0.02525134 | 0.38511744 | 0.90650187 |
| Gm5786 | 0.08376838 | 0.12252135 | 0.09728425 | 0.0252371 | 0.12440148 | 0.90650187 |
| Msl3 | 0.27428981 | 0.31123837 | 0.28601973 | 0.02521864 | 0.34753891 | 0.90650187 |
| Slc39a13 | 0.20222289 | 0.24445076 | 0.21927119 | 0.02517957 | 0.30484979 | 0.90650187 |
| Galnt12 | 0.28815053 | 0.32344787 | 0.29827555 | 0.02517232 | 0.35376429 | 0.90650187 |
| Pcdh11x | 0.07673814 | 0.11468396 | 0.08952483 | 0.02515913 | 0.10576905 | 0.90650187 |
| Rcc1 | 0.14678399 | 0.18934343 | 0.16419444 | 0.02514899 | 0.24796255 | 0.90650187 |
| Sgol2a | 0.07743816 | 0.11544918 | 0.09030062 | 0.02514856 | 0.10784068 | 0.90650187 |
| Lsm3 | 0.81781354 | 0.68704359 | 0.66189874 | 0.02514485 | 0.43294393 | 0.90650187 |
| Tm6sf1 | 0.07333121 | 0.11087321 | 0.08573909 | 0.02513412 | 0.09625087 | 0.90650187 |
| Ubash3b | 0.11181386 | 0.15275697 | 0.1276257 | 0.02513127 | 0.19067775 | 0.90650187 |
| Eif4b | 1.07221331 | 0.80421604 | 0.77912501 | 0.02509103 | 0.44307923 | 0.90650187 |
| Shcbp1 | 0.12213409 | 0.1636258 | 0.13857177 | 0.02505403 | 0.21076379 | 0.90650187 |
| Ccdc85a | 0.03161158 | 0.06303721 | 0.03798742 | 0.02504979 | 0.00168645 | 0.03683298 |
| Ccdc14 | 0.02828088 | 0.05910175 | 0.03406051 | 0.02504124 | 0.00054078 | 0.01390299 |
| Atg16l2 | 0.11961149 | 0.16090146 | 0.13590822 | 0.02499324 | 0.20681059 | 0.90650187 |
| Trim30d | 0.0203182 | 0.04959434 | 0.0246013 | 0.02499304 | 3.15E-06 | 0.00012322 |
| Nsrp1 | 0.66147321 | 0.59711311 | 0.57213916 | 0.02497395 | 0.42306786 | 0.90650187 |
| P4hb | 1.55678815 | 0.92743559 | 0.90246966 | 0.02496594 | 0.45106015 | 0.90650187 |
| Suclg1 | 1.21429082 | 0.85165192 | 0.82669123 | 0.02496069 | 0.44661101 | 0.90650187 |
| 0610007P14I | 0.71201094 | 0.62741292 | 0.60246595 | 0.02494697 | 0.42697374 | 0.90650187 |
| Smap2 | 0.35432771 | 0.3793372 | 0.35440741 | 0.02492979 | 0.37724914 | 0.90650187 |
| Mtss1 | 0.42628401 | 0.43622594 | 0.41129961 | 0.02492632 | 0.39380299 | 0.90650187 |
| Nedd9 | 0.07229112 | 0.10948725 | 0.08458003 | 0.02490722 | 0.09524227 | 0.90650187 |

| | | | | | | |
|---|---|---|---|---|---|---|
| Mcm4 | 0.27132467 | 0.30828044 | 0.28337454 | 0.0249059 | 0.34799987 | 0.90650187 |
| Htra2 | 0.38825608 | 0.40664643 | 0.38174349 | 0.02490294 | 0.38590505 | 0.90650187 |
| Zfp623 | 0.10797148 | 0.14839695 | 0.1235168 | 0.02488015 | 0.18526414 | 0.90650187 |
| Cdc16 | 0.81386654 | 0.68463829 | 0.6597749 | 0.02486339 | 0.43347607 | 0.90650187 |
| Dtd1 | 0.44353257 | 0.44920287 | 0.42434538 | 0.02485749 | 0.39727119 | 0.90650187 |
| Adat2 | 0.10337018 | 0.14342694 | 0.11857212 | 0.02485482 | 0.17569982 | 0.90650187 |
| Tspan5 | 0.44984286 | 0.45389354 | 0.42906271 | 0.02483084 | 0.39848233 | 0.90650187 |
| Zmat1 | 0.07392985 | 0.11121176 | 0.0864055 | 0.02480625 | 0.10092578 | 0.90650187 |
| Pex1 | 0.09144816 | 0.13046209 | 0.10568071 | 0.02478138 | 0.14859844 | 0.90650187 |
| 4930452B06I | 0.036022 | 0.06792593 | 0.04316061 | 0.02476533 | 0.0053728 | 0.09987259 |
| Arfip1 | 0.38738231 | 0.40579896 | 0.38105109 | 0.02474787 | 0.38639559 | 0.90650187 |
| Llgl2 | 0.34710041 | 0.37320649 | 0.34846265 | 0.02474383 | 0.37612417 | 0.90650187 |
| Anln | 0.08532983 | 0.12373905 | 0.09899813 | 0.02474092 | 0.13328096 | 0.90650187 |
| Slc7a10 | 0.07413266 | 0.11130653 | 0.08663115 | 0.02467538 | 0.10271113 | 0.90650187 |
| Ncf4 | 0.02050312 | 0.04949737 | 0.02482212 | 0.02467525 | 4.95E-06 | 0.00018675 |
| Ccl19 | 0.01728392 | 0.04562601 | 0.02097009 | 0.02465592 | 8.61E-08 | 4.24E-06 |
| Uaca | 0.29499177 | 0.32891164 | 0.30425979 | 0.02465185 | 0.35935119 | 0.90650187 |
| Rufy3 | 0.45871802 | 0.46029341 | 0.43564924 | 0.02464418 | 0.40072041 | 0.90650187 |
| Tns1 | 0.28790957 | 0.32269954 | 0.29806399 | 0.02463555 | 0.35664626 | 0.90650187 |
| Mybl2 | 0.05902854 | 0.09424781 | 0.06966207 | 0.02458575 | 0.05832602 | 0.65722983 |
| Habp2 | 0.09889746 | 0.13833017 | 0.11374663 | 0.02458354 | 0.16832419 | 0.90650187 |
| Melk | 0.06905819 | 0.10553533 | 0.08096728 | 0.02456805 | 0.08867836 | 0.87809516 |
| Rap2a | 0.18062335 | 0.22277924 | 0.19821841 | 0.02456082 | 0.29086973 | 0.90650187 |
| Scin | 0.0863461 | 0.12467187 | 0.10011176 | 0.02456011 | 0.13771853 | 0.90650187 |
| Gale | 0.26150596 | 0.29911143 | 0.27455399 | 0.02455744 | 0.34544905 | 0.90650187 |
| Sde2 | 0.5450976 | 0.52145988 | 0.4969283 | 0.02453158 | 0.41314441 | 0.90650187 |
| Rit1 | 0.39062151 | 0.40813228 | 0.38361494 | 0.02451734 | 0.38815744 | 0.90650187 |
| Bag5 | 0.40548411 | 0.41977644 | 0.39527147 | 0.02450497 | 0.3914251 | 0.90650187 |
| Rbm28 | 0.41459487 | 0.42682483 | 0.40233286 | 0.02449198 | 0.39333939 | 0.90650187 |
| Olfm1 | 0.28741114 | 0.32206966 | 0.29762623 | 0.02444342 | 0.35751524 | 0.90650187 |

| | | | | | | |
|---|---|---|---|---|---|---|
| AW551984 | 0.20639117 | 0.24765712 | 0.22327598 | 0.02438114 | 0.31367917 | 0.90650187 |
| P2ry1 | 0.11629218 | 0.15674738 | 0.13239163 | 0.02435576 | 0.20672227 | 0.90650187 |
| Supt6 | 0.38483567 | 0.4033848 | 0.37902967 | 0.02435513 | 0.38756957 | 0.90650187 |
| Stat5a | 0.06230028 | 0.09771086 | 0.07336616 | 0.02434469 | 0.07008458 | 0.74657023 |
| Mab21l2 | 0.04942845 | 0.08304368 | 0.05870122 | 0.02434246 | 0.03262641 | 0.42461821 |
| Epm2a | 0.03685936 | 0.06846669 | 0.04413936 | 0.02432732 | 0.0071391 | 0.12611093 |
| Layn | 0.06007097 | 0.09515304 | 0.07084397 | 0.02430907 | 0.06357409 | 0.698442 |
| Tfdp2 | 0.23962549 | 0.27883267 | 0.25456274 | 0.02426994 | 0.33584022 | 0.90650187 |
| Rasgrp3 | 0.06915761 | 0.1053469 | 0.0810786 | 0.0242683 | 0.09165201 | 0.89768186 |
| Pcdh18 | 0.20526396 | 0.24645509 | 0.22219478 | 0.02426031 | 0.31369883 | 0.90650187 |
| Vsnl1 | 0.03516295 | 0.0663936 | 0.04215536 | 0.02423824 | 0.00529251 | 0.09871345 |
| Sike1 | 0.32123811 | 0.3510487 | 0.32682263 | 0.02422607 | 0.37087559 | 0.90650187 |
| Rwdd4a | 0.56389527 | 0.53382306 | 0.50961953 | 0.02420353 | 0.4163903 | 0.90650187 |
| Cela3a | 0.01462874 | 0.04194688 | 0.01778039 | 0.02416649 | 7.60E-10 | 4.77E-08 |
| Gdpd3 | 0.06248589 | 0.09769776 | 0.07357583 | 0.02412194 | 0.07248947 | 0.76681446 |
| Rnf26 | 0.33916518 | 0.36599997 | 0.34188335 | 0.02411662 | 0.37691425 | 0.90650187 |
| Fam149b | 0.14864525 | 0.19020608 | 0.16610009 | 0.02410599 | 0.2594028 | 0.90650187 |
| Gm8797 | 3.11563479 | 0.84667166 | 0.82257947 | 0.02409219 | 0.44820191 | 0.90650187 |
| Qars | 0.54079791 | 0.51804785 | 0.4939941 | 0.02405375 | 0.41431181 | 0.90650187 |
| Arhgef40 | 0.29495709 | 0.3282663 | 0.30422956 | 0.02403674 | 0.36270378 | 0.90650187 |
| Gm5771 | 0.02086697 | 0.04928788 | 0.02525645 | 0.02403143 | 1.17E-05 | 0.00041406 |
| Mrpl48 | 0.75292226 | 0.65009519 | 0.62607308 | 0.02402211 | 0.4322787 | 0.90650187 |
| Lrrfip2 | 0.35765717 | 0.38115168 | 0.35713156 | 0.02402011 | 0.38246829 | 0.90650187 |
| Spin2c | 0.06282206 | 0.09796887 | 0.07395543 | 0.02401344 | 0.07444054 | 0.77834237 |
| Adamts15 | 0.09529752 | 0.13385216 | 0.10985829 | 0.02399387 | 0.16578573 | 0.90650187 |
| Riok1 | 0.28743478 | 0.32162293 | 0.297647 | 0.02397593 | 0.36013403 | 0.90650187 |
| Tpmt | 0.11398864 | 0.15390629 | 0.12994324 | 0.02396305 | 0.20615884 | 0.90650187 |
| Gpatch11 | 0.41479655 | 0.42644015 | 0.40248847 | 0.02395168 | 0.39567671 | 0.90650187 |
| Crem | 0.24899751 | 0.28713385 | 0.2631838 | 0.02395006 | 0.34290204 | 0.90650187 |
| Derl2 | 0.74737172 | 0.64683836 | 0.62291797 | 0.02392039 | 0.43222445 | 0.90650187 |

| | | | | | | |
|---|---|---|---|---|---|---|
| Dhrs11 | 0.0661409 | 0.10158622 | 0.07769419 | 0.02389203 | 0.08579934 | 0.85918035 |
| Lars | 0.56070121 | 0.53136306 | 0.50747818 | 0.02388488 | 0.41713194 | 0.90650187 |
| Tnfrsf11b | 0.05153465 | 0.08499692 | 0.0611178 | 0.02387912 | 0.04120002 | 0.50834544 |
| Klk14 | 0.01855065 | 0.04636245 | 0.02248782 | 0.02387463 | 1.18E-06 | 5.01E-05 |
| Tmem181a | 0.3920991 | 0.40864089 | 0.38478176 | 0.02385913 | 0.39140523 | 0.90650187 |
| Cdt1 | 0.12517306 | 0.16561791 | 0.1417702 | 0.02384772 | 0.22728656 | 0.90650187 |
| Cpox | 0.39908247 | 0.41407914 | 0.39027222 | 0.02380692 | 0.39312417 | 0.90650187 |
| Pald1 | 0.06916863 | 0.10485811 | 0.08109094 | 0.02376716 | 0.09629169 | 0.90650187 |
| Dhcr7 | 0.21353987 | 0.2538619 | 0.23010202 | 0.02375988 | 0.32310107 | 0.90650187 |
| Pde3a | 0.13226254 | 0.1729307 | 0.14918758 | 0.02374313 | 0.23962296 | 0.90650187 |
| Ephb4 | 0.25581622 | 0.2931189 | 0.2694007 | 0.0237182 | 0.34775309 | 0.90650187 |
| Slc39a6 | 0.43599886 | 0.4423641 | 0.41867385 | 0.02369025 | 0.40069435 | 0.90650187 |
| Sardh | 0.1637099 | 0.20504761 | 0.18137492 | 0.02367269 | 0.28087921 | 0.90650187 |
| Chchd4 | 0.48945486 | 0.48169973 | 0.45803156 | 0.02366817 | 0.40915436 | 0.90650187 |
| Nrn1 | 0.05376155 | 0.0873049 | 0.06366561 | 0.02363929 | 0.0494029 | 0.5871567 |
| Cenpk | 0.11381749 | 0.15339526 | 0.12976106 | 0.0236342 | 0.20905447 | 0.90650187 |
| Mcm2 | 0.29895508 | 0.33133901 | 0.30770655 | 0.02363247 | 0.36638947 | 0.90650187 |
| Foxm1 | 0.05212778 | 0.08542263 | 0.06179714 | 0.0236255 | 0.04460424 | 0.54285755 |
| Soat1 | 0.52414343 | 0.50610506 | 0.48251761 | 0.02358745 | 0.41397964 | 0.90650187 |
| Tmem214 | 0.58620966 | 0.54795625 | 0.52440418 | 0.02355207 | 0.42087272 | 0.90650187 |
| Celf4 | 0.29543301 | 0.32814372 | 0.3046443 | 0.02349942 | 0.36582867 | 0.90650187 |
| Rpl13a-ps1 | 0.13572516 | 0.17625685 | 0.15278851 | 0.02346834 | 0.24735217 | 0.90650187 |
| Snx7 | 0.25906229 | 0.2958004 | 0.27234452 | 0.02345588 | 0.3509039 | 0.90650187 |
| Il10rb | 0.33713087 | 0.36363038 | 0.34018804 | 0.02344234 | 0.37967042 | 0.90650187 |
| Cep120 | 0.12711645 | 0.16720648 | 0.14380948 | 0.023397 | 0.23475536 | 0.90650187 |
| Nudt9 | 0.31044996 | 0.34101747 | 0.31762233 | 0.02339515 | 0.37166356 | 0.90650187 |
| Sptbn1 | 1.19346852 | 0.84385436 | 0.82047286 | 0.0233815 | 0.44959325 | 0.90650187 |
| Wnt4 | 0.19516453 | 0.2358045 | 0.21244706 | 0.02335744 | 0.31250191 | 0.90650187 |
| Epb42 | 0.0302051 | 0.0596564 | 0.0363313 | 0.0233251 | 0.00215775 | 0.04542806 |
| AI662270 | 0.01619894 | 0.04298514 | 0.01966807 | 0.02331707 | 6.81E-08 | 3.45E-06 |

| | | | | | | |
|---|---|---|---|---|---|---|
| Pgm1 | 0.2419476 | 0.28000422 | 0.25670692 | 0.0232973 | 0.34330542 | 0.90650187 |
| Tsg101 | 1.13006841 | 0.82308075 | 0.79980232 | 0.02327843 | 0.44852437 | 0.90650187 |
| Sdk2 | 0.18709867 | 0.22780988 | 0.20458278 | 0.0232271 | 0.30687512 | 0.90650187 |
| Pxdc1 | 0.12993528 | 0.16997821 | 0.14675932 | 0.02321889 | 0.24092125 | 0.90650187 |
| Rab37 | 0.13556459 | 0.17583868 | 0.15262184 | 0.02321684 | 0.24943593 | 0.90650187 |
| 1110008L16F | 0.1435908 | 0.18409721 | 0.16091573 | 0.02318148 | 0.26094557 | 0.90650187 |
| Pttg1ip | 0.6938597 | 0.61484995 | 0.59172585 | 0.0231241 | 0.43103877 | 0.90650187 |
| Gm8113 | 0.0189387 | 0.04604954 | 0.02295225 | 0.02309729 | 3.84E-06 | 0.00014718 |
| Tc2n | 0.02582917 | 0.05423663 | 0.03115875 | 0.02307788 | 0.00049622 | 0.01289414 |
| Tmem59 | 2.59755539 | 0.91157632 | 0.88852757 | 0.02304875 | 0.45409545 | 0.90650187 |
| Pla2g12a | 0.32755161 | 0.35519856 | 0.33215799 | 0.02304057 | 0.37889839 | 0.90650187 |
| Ccr1 | 0.01632697 | 0.04284866 | 0.01982181 | 0.02302684 | 1.21E-07 | 5.80E-06 |
| Tmem70 | 0.60881831 | 0.56207012 | 0.53908184 | 0.02298828 | 0.42482079 | 0.90650187 |
| Gid4 | 0.21549945 | 0.25494522 | 0.23196391 | 0.02298131 | 0.32980977 | 0.90650187 |
| Sp110 | 0.01773305 | 0.0444333 | 0.02150851 | 0.02292479 | 1.08E-06 | 4.60E-05 |
| Mospd1 | 0.188881 | 0.22922875 | 0.20632659 | 0.02290217 | 0.31084363 | 0.90650187 |
| Hnrnph3 | 0.91048159 | 0.73266554 | 0.70976647 | 0.02289907 | 0.44297617 | 0.90650187 |
| Tmem120a | 0.24655896 | 0.28384527 | 0.26094924 | 0.02289603 | 0.34824485 | 0.90650187 |
| Pds5b | 0.37672732 | 0.39545449 | 0.37255965 | 0.02289484 | 0.39235166 | 0.90650187 |
| Rpp21 | 0.83100874 | 0.69183318 | 0.66894994 | 0.02288324 | 0.43956431 | 0.90650187 |
| Cnpy4 | 0.29591976 | 0.32787307 | 0.30506827 | 0.02280481 | 0.36982269 | 0.90650187 |
| Ddit4l | 0.05665242 | 0.08973891 | 0.06696198 | 0.02277692 | 0.06524657 | 0.71046786 |
| Mesp2 | 0.03558858 | 0.06541675 | 0.04265357 | 0.02276317 | 0.0088338 | 0.15087931 |
| Vamp2 | 0.8044115 | 0.67736012 | 0.65465645 | 0.02270366 | 0.43873594 | 0.90650187 |
| Slc38a2 | 0.65704964 | 0.59211912 | 0.56942046 | 0.02269865 | 0.42967015 | 0.90650187 |
| Ces1d | 0.05383389 | 0.08639117 | 0.06374825 | 0.02264292 | 0.05716254 | 0.64823856 |
| Cntln | 0.36935087 | 0.3892395 | 0.36662793 | 0.02261157 | 0.39197303 | 0.90650187 |
| Szrd1 | 0.54091348 | 0.51668176 | 0.49407314 | 0.02260863 | 0.41939967 | 0.90650187 |
| Dennd1c | 0.08108458 | 0.1169308 | 0.09433038 | 0.02260042 | 0.14341224 | 0.90650187 |
| Tbpl1 | 0.57983667 | 0.54280618 | 0.520212 | 0.02259417 | 0.42344645 | 0.90650187 |

| | | | | | | |
|---|---|---|---|---|---|---|
| Sptan1 | 0.65026275 | 0.58782118 | 0.56522922 | 0.02259196 | 0.42948364 | 0.90650187 |
| Trappc2 | 0.35486015 | 0.37743435 | 0.35484366 | 0.02259068 | 0.38857921 | 0.90650187 |
| Ubac2 | 0.33956536 | 0.36479691 | 0.34221643 | 0.02258048 | 0.38463319 | 0.90650187 |
| Nmnat3 | 0.06497008 | 0.09895221 | 0.07637709 | 0.02257512 | 0.0944218 | 0.90650187 |
| Snx1 | 0.5456081 | 0.51983454 | 0.49727581 | 0.02255872 | 0.4200861 | 0.90650187 |
| As3mt | 0.37562555 | 0.39420969 | 0.37167649 | 0.0225332 | 0.3937653 | 0.90650187 |
| Jagn1 | 0.87498767 | 0.71438059 | 0.69189352 | 0.02248707 | 0.4425585 | 0.90650187 |
| Txndc16 | 0.1083988 | 0.14644963 | 0.12397466 | 0.02247497 | 0.21014168 | 0.90650187 |
| Zc4h2 | 0.09965043 | 0.13702778 | 0.11455767 | 0.02247011 | 0.19160659 | 0.90650187 |
| Tlx2 | 0.01635371 | 0.04232023 | 0.01985392 | 0.02246631 | 2.45E-07 | 1.12E-05 |
| Pcnxl4 | 0.15147883 | 0.19145182 | 0.16899313 | 0.02245869 | 0.27732455 | 0.90650187 |
| Efna3 | 0.06041097 | 0.09368382 | 0.07122911 | 0.0224547 | 0.08053825 | 0.82275666 |
| Arhgap30 | 0.01611946 | 0.04202554 | 0.01957261 | 0.02245293 | 1.70E-07 | 8.02E-06 |
| Romo1 | 2.23378012 | 0.94092634 | 0.91849939 | 0.02242695 | 0.45678045 | 0.90650187 |
| Cyp2j6 | 0.11255249 | 0.15079949 | 0.12841347 | 0.02238601 | 0.2191722 | 0.90650187 |
| Btbd10 | 0.30206194 | 0.33277246 | 0.3103982 | 0.02237426 | 0.37431356 | 0.90650187 |
| Lsm1 | 0.65961605 | 0.59335532 | 0.57099901 | 0.02235631 | 0.43090966 | 0.90650187 |
| Lrrtm1 | 0.02929308 | 0.05758413 | 0.03525574 | 0.02232839 | 0.00243492 | 0.05063374 |
| Mcam | 0.11733173 | 0.15580556 | 0.13349441 | 0.02231115 | 0.22874029 | 0.90650187 |
| Nmral1 | 0.24605372 | 0.28274886 | 0.26048545 | 0.02226341 | 0.35198712 | 0.90650187 |
| Tjap1 | 0.29063428 | 0.32271459 | 0.30045307 | 0.02226152 | 0.37093082 | 0.90650187 |
| E330009J07F | 0.16122392 | 0.20112971 | 0.17887208 | 0.02225764 | 0.29006963 | 0.90650187 |
| Rnls | 0.07572708 | 0.11064175 | 0.0884031 | 0.02223865 | 0.13171195 | 0.90650187 |
| Rapgef2 | 0.09418782 | 0.13087289 | 0.10865608 | 0.0222168 | 0.181678 | 0.90650187 |
| Aen | 0.25144398 | 0.28763206 | 0.26541953 | 0.02221253 | 0.35492943 | 0.90650187 |
| Samhd1 | 0.18825242 | 0.22791547 | 0.20571198 | 0.02220349 | 0.31567207 | 0.90650187 |
| Cast | 0.18511389 | 0.22483698 | 0.20263686 | 0.02220012 | 0.31311369 | 0.90650187 |
| Gfra1 | 0.04715862 | 0.07825954 | 0.05608937 | 0.02217017 | 0.03944246 | 0.4922536 |
| Hspb11 | 0.50041912 | 0.48799628 | 0.46585773 | 0.02213855 | 0.41634038 | 0.90650187 |
| Rec114 | 0.0182573 | 0.04426394 | 0.02213657 | 0.02212737 | 4.42E-06 | 0.00016755 |

| | | | | | | |
|---|---|---|---|---|---|---|
| Cxcl16 | 0.0513188 | 0.08296342 | 0.06087045 | 0.02209297 | 0.05331403 | 0.62090631 |
| Sgcb | 0.36096779 | 0.38189794 | 0.35983133 | 0.02206661 | 0.39257029 | 0.90650187 |
| Tro | 0.18786721 | 0.22735529 | 0.20533513 | 0.02202017 | 0.31677098 | 0.90650187 |
| Ell2 | 0.17103897 | 0.21073302 | 0.18871331 | 0.0220197 | 0.30197511 | 0.90650187 |
| Fuca2 | 0.22561207 | 0.26345841 | 0.24150913 | 0.02194928 | 0.3430939 | 0.90650187 |
| Glipr1 | 0.02087335 | 0.04715762 | 0.02526406 | 0.02189355 | 5.85E-05 | 0.00181345 |
| Sep-06 | 0.08318879 | 0.11853431 | 0.09664719 | 0.02188712 | 0.1570226 | 0.90650187 |
| Serpine1 | 0.07677861 | 0.11145605 | 0.0895697 | 0.02188635 | 0.13867679 | 0.90650187 |
| Cmss1 | 0.09253281 | 0.12871039 | 0.10685993 | 0.02185046 | 0.181668 | 0.90650187 |
| Nol7 | 1.08833075 | 0.80688195 | 0.78506294 | 0.02181901 | 0.45083366 | 0.90650187 |
| Stt3b | 0.75557205 | 0.64939001 | 0.62757426 | 0.02181576 | 0.43859266 | 0.90650187 |
| Rasl11b | 0.11471801 | 0.15251739 | 0.13071918 | 0.02179821 | 0.2292462 | 0.90650187 |
| Angpt2 | 0.06585028 | 0.0991638 | 0.07736745 | 0.02179635 | 0.10520499 | 0.90650187 |
| Rpgrip1 | 0.24830405 | 0.28434296 | 0.26254901 | 0.02179394 | 0.35605434 | 0.90650187 |
| Pla2g2d | 0.15649032 | 0.19587077 | 0.17408701 | 0.02178376 | 0.28900799 | 0.90650187 |
| Gbp2 | 0.08577456 | 0.12126573 | 0.09948564 | 0.02178009 | 0.16521124 | 0.90650187 |
| 4930430F08I | 0.11958503 | 0.15764643 | 0.13588024 | 0.02176618 | 0.2381933 | 0.90650187 |
| Stk24 | 0.17115 | 0.21058871 | 0.18882402 | 0.02176469 | 0.30417846 | 0.90650187 |
| Gstk1 | 0.10633017 | 0.14350684 | 0.12175605 | 0.02175079 | 0.21354712 | 0.90650187 |
| Dact2 | 0.24293545 | 0.27935327 | 0.25761747 | 0.02173579 | 0.35379814 | 0.90650187 |
| Obsl1 | 0.34018069 | 0.3644516 | 0.34272833 | 0.02172327 | 0.3890571 | 0.90650187 |
| Magel2 | 0.10020919 | 0.13688287 | 0.11515982 | 0.02172304 | 0.20084916 | 0.90650187 |
| Tmem26 | 0.09063356 | 0.1264912 | 0.10479401 | 0.02169719 | 0.17867001 | 0.90650187 |
| Map2k1 | 0.56873263 | 0.53454627 | 0.51285065 | 0.02169562 | 0.42541214 | 0.90650187 |
| Arid4b | 1.03478341 | 0.78646136 | 0.76476647 | 0.02169489 | 0.44982126 | 0.90650187 |
| Pla2g4a | 0.12798009 | 0.16640805 | 0.14471428 | 0.02169377 | 0.25256806 | 0.90650187 |
| Scarf1 | 0.0459248 | 0.07635029 | 0.05466636 | 0.02168393 | 0.0389172 | 0.4872985 |
| Rnf128 | 0.14992245 | 0.18908619 | 0.16740529 | 0.0216809 | 0.28238977 | 0.90650187 |
| Ccdc186 | 0.36215375 | 0.382475 | 0.3607963 | 0.0216787 | 0.39468939 | 0.90650187 |
| Ndor1 | 0.05913232 | 0.09137867 | 0.0697798 | 0.02159886 | 0.08440407 | 0.84912391 |

| | | | | | | |
|---|---|---|---|---|---|---|
| Oasl2 | 0.03347317 | 0.0617727 | 0.04017467 | 0.02159803 | 0.00842483 | 0.14519943 |
| Fgf10 | 0.06054536 | 0.09297325 | 0.0713813 | 0.02159194 | 0.08935462 | 0.88257281 |
| Tbx3 | 0.07434082 | 0.10844559 | 0.0868627 | 0.0215829 | 0.13466517 | 0.90650187 |
| Apba3 | 0.1542135 | 0.19335532 | 0.17177633 | 0.02157899 | 0.28826217 | 0.90650187 |
| Casc5 | 0.07669741 | 0.11104035 | 0.08947966 | 0.02156069 | 0.14204043 | 0.90650187 |
| A330009N23 | 0.05917829 | 0.09137426 | 0.06983195 | 0.02154231 | 0.08512273 | 0.85522185 |
| Fkbp1b | 0.11271262 | 0.15011481 | 0.12858417 | 0.02153064 | 0.22832033 | 0.90650187 |
| Slit3 | 0.0872718 | 0.12265381 | 0.10112488 | 0.02152894 | 0.17196074 | 0.90650187 |
| Ttc23 | 0.08513458 | 0.12030735 | 0.098784 | 0.02152334 | 0.16636701 | 0.90650187 |
| Cryz | 0.14178128 | 0.18054877 | 0.15905247 | 0.0214963 | 0.27397532 | 0.90650187 |
| Plscr2 | 0.03980425 | 0.06905275 | 0.0475729 | 0.02147985 | 0.02236123 | 0.31488391 |
| 3632451O06 | 0.14823097 | 0.18714817 | 0.16567627 | 0.02147189 | 0.28225444 | 0.90650187 |
| Cct4 | 1.61381337 | 0.93106671 | 0.90959598 | 0.02147073 | 0.45821284 | 0.90650187 |
| Exosc8 | 0.34433103 | 0.36764329 | 0.34617274 | 0.02147054 | 0.39137842 | 0.90650187 |
| Gm9493 | 2.17020745 | 0.94335916 | 0.92191465 | 0.02144451 | 0.45881937 | 0.90650187 |
| Lcn2 | 0.0190345 | 0.04450089 | 0.02306687 | 0.02143402 | 1.81E-05 | 0.00061853 |
| Oaf | 0.28313597 | 0.31529377 | 0.29386206 | 0.02143171 | 0.37288447 | 0.90650187 |
| Cacng8 | 0.01689135 | 0.04192151 | 0.02049921 | 0.0214223 | 1.69E-06 | 7.02E-05 |
| Syn1 | 0.05895772 | 0.09098537 | 0.06958171 | 0.02140367 | 0.08573593 | 0.85918035 |
| Mpp6 | 0.21493815 | 0.25283345 | 0.231431 | 0.02140245 | 0.34049075 | 0.90650187 |
| Hist1h2bg | 0.02323592 | 0.04947265 | 0.02807909 | 0.02139355 | 0.00035315 | 0.00941795 |
| Meox1 | 0.07986189 | 0.11437445 | 0.09298127 | 0.02139318 | 0.15318998 | 0.90650187 |
| Chchd3 | 0.77877106 | 0.66196844 | 0.64057563 | 0.02139281 | 0.44098733 | 0.90650187 |
| Elavl2 | 0.07636428 | 0.11049329 | 0.08911022 | 0.02138307 | 0.14303634 | 0.90650187 |
| Them6 | 0.05551086 | 0.08704081 | 0.06566179 | 0.02137902 | 0.07388431 | 0.7756322 |
| Irs4 | 0.02587398 | 0.05258027 | 0.03121187 | 0.02136839 | 0.00116895 | 0.02696394 |
| Pfkp | 0.09874524 | 0.13491662 | 0.11358258 | 0.02133404 | 0.20185566 | 0.90650187 |
| Rwdd1 | 1.34014712 | 0.88207513 | 0.86077815 | 0.02129698 | 0.45620819 | 0.90650187 |
| Hbp1 | 0.61250998 | 0.56269595 | 0.54145056 | 0.0212454 | 0.43076127 | 0.90650187 |
| Ifngr2 | 0.43879908 | 0.442026 | 0.42078668 | 0.02123931 | 0.41122572 | 0.90650187 |

| | | | | | | |
|---|---|---|---|---|---|---|
| Wbp2 | 0.6793648 | 0.60424881 | 0.58302719 | 0.02122162 | 0.43572548 | 0.90650187 |
| Adgrl4 | 0.03272502 | 0.06051713 | 0.03929629 | 0.02122084 | 0.00818137 | 0.14180727 |
| Pik3r4 | 0.10249855 | 0.13883087 | 0.11763244 | 0.02119842 | 0.21152584 | 0.90650187 |
| Rfc2 | 0.74315178 | 0.64169771 | 0.6205096 | 0.02118812 | 0.43967192 | 0.90650187 |
| Unc5b | 0.08661892 | 0.12158653 | 0.10041047 | 0.02117606 | 0.17423632 | 0.90650187 |
| Nedd4 | 2.27821592 | 0.93679344 | 0.91562248 | 0.02117096 | 0.45906443 | 0.90650187 |
| Map3k7cl | 0.02967154 | 0.05683945 | 0.03570222 | 0.02113722 | 0.00424436 | 0.08177261 |
| Otud6b | 0.32728098 | 0.35306279 | 0.33192999 | 0.0211328 | 0.38857463 | 0.90650187 |
| Hgs | 0.4454341 | 0.44690245 | 0.4257701 | 0.02113235 | 0.41268254 | 0.90650187 |
| Il6st | 0.34600365 | 0.36864322 | 0.34755674 | 0.02108647 | 0.39368877 | 0.90650187 |
| Rad21 | 0.8632823 | 0.70693306 | 0.68587442 | 0.02105865 | 0.44571498 | 0.90650187 |
| Gpat4 | 0.57450506 | 0.53773287 | 0.51668655 | 0.02104632 | 0.42815083 | 0.90650187 |
| Cenpq | 0.14226901 | 0.18059536 | 0.15955506 | 0.0210403 | 0.2788585 | 0.90650187 |
| Ppm1f | 0.07983003 | 0.11398223 | 0.09294608 | 0.02103615 | 0.15716973 | 0.90650187 |
| Ddx41 | 0.66270504 | 0.59392565 | 0.5728944 | 0.02103124 | 0.4351803 | 0.90650187 |
| Adamtsl5 | 0.13546883 | 0.17354725 | 0.15252243 | 0.02102482 | 0.26999922 | 0.90650187 |
| Mrps7 | 0.83573748 | 0.69245614 | 0.67145901 | 0.02099713 | 0.44471791 | 0.90650187 |
| Tshz3 | 0.18493005 | 0.22345024 | 0.20245639 | 0.02099385 | 0.32240288 | 0.90650187 |
| Tmco3 | 0.31762965 | 0.34473828 | 0.32375656 | 0.02098173 | 0.38663247 | 0.90650187 |
| Mycl | 0.19363621 | 0.23191228 | 0.21096229 | 0.02094999 | 0.32943311 | 0.90650187 |
| Kdelc2 | 0.4021277 | 0.41360038 | 0.39265424 | 0.02094614 | 0.40626986 | 0.90650187 |
| Hnf4a | 0.13865072 | 0.17674607 | 0.15581987 | 0.0209262 | 0.27524257 | 0.90650187 |
| Armcx3 | 0.99247637 | 0.76852659 | 0.74761257 | 0.02091402 | 0.45051385 | 0.90650187 |
| Pon2 | 0.26688869 | 0.30031195 | 0.27940079 | 0.02091116 | 0.36967208 | 0.90650187 |
| Rgs19 | 0.26138593 | 0.29535587 | 0.27444559 | 0.02091028 | 0.36741219 | 0.90650187 |
| Scara5 | 0.0438396 | 0.07315996 | 0.05225615 | 0.02090381 | 0.0376726 | 0.47484348 |
| Sema3g | 0.02582376 | 0.05202851 | 0.03115233 | 0.02087618 | 0.00144527 | 0.03216631 |
| Lypd2 | 0.04433846 | 0.07370241 | 0.05283336 | 0.02086905 | 0.03954387 | 0.49281891 |
| Inppl1 | 0.29219659 | 0.32264331 | 0.30181989 | 0.02082342 | 0.3795301 | 0.90650187 |
| Snx4 | 0.73882729 | 0.63884478 | 0.6180324 | 0.02081238 | 0.44049798 | 0.90650187 |

| | | | | | | |
|---|---|---|---|---|---|---|
| Elovl6 | 0.17663148 | 0.21507242 | 0.19427212 | 0.0208003 | 0.31704378 | 0.90650187 |
| Hspa9 | 1.17552209 | 0.83557257 | 0.81477368 | 0.02079889 | 0.45482348 | 0.90650187 |
| Anapc16 | 1.0173755 | 0.77860745 | 0.7578134 | 0.02079405 | 0.45145531 | 0.90650187 |
| B3gat3 | 0.30327553 | 0.33224116 | 0.31144726 | 0.0207939 | 0.38330478 | 0.90650187 |
| Pkib | 0.09528971 | 0.13064318 | 0.10984983 | 0.02079334 | 0.20003297 | 0.90650187 |
| Nnmt | 0.12357629 | 0.16085327 | 0.14009108 | 0.02076219 | 0.25499256 | 0.90650187 |
| Phf11d | 0.05894833 | 0.09032085 | 0.06957106 | 0.0207498 | 0.09243366 | 0.90084241 |
| Nup54 | 0.23937324 | 0.27507522 | 0.2543295 | 0.02074572 | 0.35844095 | 0.90650187 |
| Exosc1 | 0.47553951 | 0.46871575 | 0.44798019 | 0.02073556 | 0.4184842 | 0.90650187 |
| Tagln3 | 0.0534817 | 0.08407014 | 0.06334584 | 0.02072429 | 0.07291365 | 0.76862723 |
| Atp2a3 | 0.37005328 | 0.38790047 | 0.36719483 | 0.02070564 | 0.40103001 | 0.90650187 |
| Bcl2l13 | 0.21294311 | 0.25022453 | 0.22953422 | 0.02069031 | 0.34430993 | 0.90650187 |
| Dnaic1 | 0.04492916 | 0.07418655 | 0.05351636 | 0.0206702 | 0.0429828 | 0.52685717 |
| Eid1 | 1.13342262 | 0.82162993 | 0.80096826 | 0.02066167 | 0.45435007 | 0.90650187 |
| Gm9008 | 0.10675703 | 0.14285017 | 0.12221429 | 0.02063587 | 0.22643458 | 0.90650187 |
| Gm26737 | 0.02680942 | 0.0529107 | 0.03232007 | 0.02059063 | 0.00231124 | 0.04839283 |
| Poli | 0.06201133 | 0.09361167 | 0.07303967 | 0.020572 | 0.10526161 | 0.90650187 |
| Tecpr1 | 0.35411978 | 0.37477008 | 0.35423698 | 0.0205331 | 0.39832443 | 0.90650187 |
| Fbxo27 | 0.04864491 | 0.07833193 | 0.05780049 | 0.02053144 | 0.05715275 | 0.64823856 |
| 1500009L16I | 0.54265399 | 0.51579033 | 0.49526238 | 0.02052794 | 0.42690308 | 0.90650187 |
| Scarb2 | 0.38875173 | 0.40266267 | 0.38213599 | 0.02052668 | 0.40563088 | 0.90650187 |
| Foxd3 | 0.01056648 | 0.03338148 | 0.01287819 | 0.02050329 | 7.32E-13 | 6.40E-11 |
| Zfp57 | 0.17499242 | 0.21314299 | 0.19264669 | 0.02049629 | 0.31811296 | 0.90650187 |
| A330069E16 | 0.07392057 | 0.10688857 | 0.08639518 | 0.02049338 | 0.14582228 | 0.90650187 |
| Rpl24 | 4.47454808 | 0.63063719 | 0.61015574 | 0.02048145 | 0.44068676 | 0.90650187 |
| Tsr1 | 0.23824027 | 0.27375336 | 0.25328112 | 0.02047225 | 0.35967405 | 0.90650187 |
| Naaa | 0.1719235 | 0.21006271 | 0.18959491 | 0.02046781 | 0.31563948 | 0.90650187 |
| Zfp560 | 0.38742378 | 0.40154909 | 0.38108397 | 0.02046513 | 0.40565368 | 0.90650187 |
| Fam167b | 0.04238778 | 0.07103416 | 0.0505741 | 0.02046006 | 0.03604915 | 0.45894732 |
| Klhdc3 | 0.42304808 | 0.42928112 | 0.40882797 | 0.02045316 | 0.41199808 | 0.90650187 |

| | | | | | | |
|---|---|---|---|---|---|---|
| Zfp729a | 0.08151718 | 0.11525942 | 0.09480721 | 0.02045221 | 0.16877312 | 0.90650187 |
| Uxs1 | 0.25735242 | 0.29123711 | 0.27079512 | 0.02044199 | 0.36858877 | 0.90650187 |
| Sfxn2 | 0.41445855 | 0.42266921 | 0.40222766 | 0.02044155 | 0.41062814 | 0.90650187 |
| Oas1a | 0.02059428 | 0.0453712 | 0.02493096 | 0.02044023 | 0.00013378 | 0.0038662 |
| Nav1 | 0.46271634 | 0.4590094 | 0.43859846 | 0.02041094 | 0.41805023 | 0.90650187 |
| Hsd17b7 | 0.10462426 | 0.14033022 | 0.11992241 | 0.02040781 | 0.22466532 | 0.90650187 |
| Icam1 | 0.07917943 | 0.1126312 | 0.09222731 | 0.02040389 | 0.16267505 | 0.90650187 |
| Mocs3 | 0.03532982 | 0.0627248 | 0.04235073 | 0.02037408 | 0.01623034 | 0.24597939 |
| Fbl | 1.14397726 | 0.82495575 | 0.80458463 | 0.02037112 | 0.45519071 | 0.90650187 |
| Tmem126a | 0.66477732 | 0.59447644 | 0.57416315 | 0.02031329 | 0.43751175 | 0.90650187 |
| Lrrfip1 | 0.43357363 | 0.43713913 | 0.41683937 | 0.02029976 | 0.41429993 | 0.90650187 |
| Lyl1 | 0.05690337 | 0.08754719 | 0.06724754 | 0.02029964 | 0.08980278 | 0.88584823 |
| 5033428I22R | 0.09873939 | 0.13386659 | 0.11357627 | 0.02029033 | 0.21353813 | 0.90650187 |
| Aco2 | 1.21711971 | 0.84779307 | 0.82752457 | 0.0202685 | 0.4566465 | 0.90650187 |
| Ntng1 | 0.02083859 | 0.04547864 | 0.02522258 | 0.02025606 | 0.00017833 | 0.00502935 |
| Vsig8 | 0.01250452 | 0.03546435 | 0.0152203 | 0.02024404 | 1.68E-09 | 9.98E-08 |
| Stk32a | 0.09632892 | 0.13121543 | 0.11097415 | 0.02024128 | 0.20872237 | 0.90650187 |
| Agpat2 | 0.32382501 | 0.34924446 | 0.3290129 | 0.02023157 | 0.3922852 | 0.90650187 |
| Carhsp1 | 0.73503682 | 0.63607998 | 0.61585316 | 0.02022683 | 0.44195688 | 0.90650187 |
| Gm6133 | 1.60082171 | 0.92829569 | 0.90807056 | 0.02022512 | 0.46056303 | 0.90650187 |
| Sdhaf4 | 0.79876308 | 0.67179163 | 0.65158021 | 0.02021142 | 0.44516028 | 0.90650187 |
| Tfam | 0.41398418 | 0.42205936 | 0.40186148 | 0.02019788 | 0.41159664 | 0.90650187 |
| Gdpd1 | 0.27847034 | 0.30992897 | 0.28973492 | 0.02019405 | 0.37833697 | 0.90650187 |
| Efcab10 | 0.25372731 | 0.2876774 | 0.26750093 | 0.02017647 | 0.36869436 | 0.90650187 |
| Cnot7 | 0.85744566 | 0.70300189 | 0.68284629 | 0.0201556 | 0.44780019 | 0.90650187 |
| C1qtnf4 | 0.41593793 | 0.42351203 | 0.40336854 | 0.02014349 | 0.41215481 | 0.90650187 |
| Smtn | 0.21332813 | 0.25001456 | 0.2299006 | 0.02011395 | 0.34865836 | 0.90650187 |
| Cadm3 | 0.07756382 | 0.11055353 | 0.0904398 | 0.02011374 | 0.16140517 | 0.90650187 |
| Gse1 | 0.37322916 | 0.38984771 | 0.36975226 | 0.02009545 | 0.40454054 | 0.90650187 |
| Sec23b | 0.58864718 | 0.54609606 | 0.52600133 | 0.02009473 | 0.43257008 | 0.90650187 |

| | | | | | | |
|---|---|---|---|---|---|---|
| Mrps18b | 0.38814641 | 0.40174704 | 0.38165661 | 0.02009043 | 0.40748589 | 0.90650187 |
| BC017643 | 0.2651978 | 0.29796025 | 0.27788122 | 0.02007903 | 0.37401846 | 0.90650187 |
| Sntb2 | 0.35648589 | 0.37622583 | 0.35617426 | 0.02005157 | 0.40118747 | 0.90650187 |
| Jak1 | 0.7041797 | 0.61789475 | 0.59785222 | 0.02004252 | 0.44076283 | 0.90650187 |
| Oip5 | 0.04640233 | 0.07523975 | 0.05521739 | 0.02002236 | 0.05347879 | 0.62198243 |
| Csk | 0.45096549 | 0.44991255 | 0.42989892 | 0.02001363 | 0.41801978 | 0.90650187 |
| Krt15 | 0.02924972 | 0.05519804 | 0.03520458 | 0.01999346 | 0.00578888 | 0.10563846 |
| E2f5 | 0.23695398 | 0.27207434 | 0.25208932 | 0.01998501 | 0.36225543 | 0.90650187 |
| Timmdc1 | 0.28779547 | 0.31794602 | 0.2979638 | 0.01998222 | 0.3828005 | 0.90650187 |
| Gm17634 | 0.01839861 | 0.04226637 | 0.02230579 | 0.01996057 | 3.47E-05 | 0.0011158 |
| Sash1 | 0.2354447 | 0.27064438 | 0.25068884 | 0.01995554 | 0.36171334 | 0.90650187 |
| Ilvbl | 0.30199866 | 0.33029682 | 0.31034346 | 0.01995336 | 0.38750503 | 0.90650187 |
| Tmem189 | 0.09498706 | 0.12941492 | 0.10952212 | 0.0198928 | 0.20969856 | 0.90650187 |
| Gm11837 | 0.11075368 | 0.14638138 | 0.12649381 | 0.01988756 | 0.24229305 | 0.90650187 |
| Psmb8 | 0.07286536 | 0.10507284 | 0.08522015 | 0.01985269 | 0.15018543 | 0.90650187 |
| Oplah | 0.07241142 | 0.1045625 | 0.08471417 | 0.01984833 | 0.14879978 | 0.90650187 |
| Prune2 | 0.07464112 | 0.10700576 | 0.08719662 | 0.01980914 | 0.15626127 | 0.90650187 |
| Lrrc32 | 0.08687492 | 0.12048268 | 0.10069067 | 0.01979201 | 0.19110277 | 0.90650187 |
| Ntan1 | 0.73841398 | 0.63757637 | 0.61779511 | 0.01978126 | 0.44340397 | 0.90650187 |
| Stim2 | 0.26182579 | 0.29462209 | 0.27484274 | 0.01977935 | 0.37450964 | 0.90650187 |
| 4931406P16I | 0.35563945 | 0.37523934 | 0.35548175 | 0.01975759 | 0.40242093 | 0.90650187 |
| Ankrd12 | 0.72817543 | 0.63163699 | 0.61189099 | 0.01974601 | 0.44296275 | 0.90650187 |
| Slc39a7 | 1.02334559 | 0.77995682 | 0.76021135 | 0.01974547 | 0.45403683 | 0.90650187 |
| Spry1 | 0.13141984 | 0.16805061 | 0.14830905 | 0.01974156 | 0.27700603 | 0.90650187 |
| Gldc | 0.03630328 | 0.06320194 | 0.0434895 | 0.01971244 | 0.02194945 | 0.31095399 |
| Slc37a4 | 0.36527007 | 0.38303884 | 0.36332649 | 0.01971235 | 0.40470111 | 0.90650187 |
| Ints12 | 0.19018627 | 0.22726318 | 0.20760147 | 0.01966171 | 0.33686347 | 0.90650187 |
| Lysmd2 | 0.64078831 | 0.57898884 | 0.55933695 | 0.01965189 | 0.43794042 | 0.90650187 |
| Aatf | 0.29591466 | 0.32465331 | 0.30506383 | 0.01958949 | 0.38764133 | 0.90650187 |
| Prr13 | 0.82895799 | 0.68742448 | 0.6678589 | 0.01956558 | 0.4481889 | 0.90650187 |

| | | | | | | |
|---|---|---|---|---|---|---|
| Pmvk | 0.54273477 | 0.51487807 | 0.49531753 | 0.01956054 | 0.43031942 | 0.90650187 |
| Ntpcr | 0.2454476 | 0.27948383 | 0.25992872 | 0.01955511 | 0.36901835 | 0.90650187 |
| Dync1li2 | 0.32407969 | 0.34878269 | 0.32922822 | 0.01955447 | 0.39587213 | 0.90650187 |
| Coasy | 0.32371086 | 0.34844394 | 0.32891637 | 0.01952756 | 0.39591579 | 0.90650187 |
| Prkar1b | 0.30491292 | 0.33235889 | 0.31286059 | 0.0194983 | 0.39086629 | 0.90650187 |
| Klf10 | 0.43689882 | 0.43885045 | 0.41935351 | 0.01949694 | 0.41812642 | 0.90650187 |
| G3bp2 | 0.99373216 | 0.767597 | 0.74813312 | 0.01946388 | 0.45396118 | 0.90650187 |
| Gem | 0.12288381 | 0.15880934 | 0.13936189 | 0.01944745 | 0.26750766 | 0.90650187 |
| Spryd3 | 0.28496711 | 0.31491316 | 0.29547639 | 0.01943677 | 0.38497625 | 0.90650187 |
| Lsamp | 0.05408692 | 0.08346373 | 0.06403724 | 0.01942649 | 0.08872799 | 0.87809516 |
| Nhsl2 | 0.17856605 | 0.21559839 | 0.19618679 | 0.0194116 | 0.33001628 | 0.90650187 |
| Cryl1 | 0.19558234 | 0.23225724 | 0.21285253 | 0.01940471 | 0.34263442 | 0.90650187 |
| Cdkal1 | 0.22321357 | 0.25865903 | 0.2392551 | 0.01940394 | 0.35921976 | 0.90650187 |
| Elavl3 | 0.05806368 | 0.08792734 | 0.06856666 | 0.01936068 | 0.10468968 | 0.90650187 |
| Kank2 | 0.22936416 | 0.26435779 | 0.24502353 | 0.01933427 | 0.36287183 | 0.90650187 |
| Wbscr16 | 0.17115265 | 0.20815117 | 0.18882666 | 0.01932451 | 0.32456793 | 0.90650187 |
| Srp14 | 2.00782104 | 0.94633281 | 0.92703895 | 0.01929386 | 0.46314106 | 0.90650187 |
| Pde1c | 0.23505856 | 0.26962333 | 0.25033016 | 0.01929317 | 0.36593962 | 0.90650187 |
| Cct8 | 1.79136892 | 0.94310337 | 0.92381449 | 0.01928887 | 0.4630223 | 0.90650187 |
| Kif11 | 0.07227145 | 0.10382958 | 0.0845581 | 0.01927149 | 0.15548462 | 0.90650187 |
| Adamts8 | 0.09115121 | 0.12462573 | 0.10535758 | 0.01926815 | 0.20810113 | 0.90650187 |
| Fam13c | 0.05476229 | 0.08407027 | 0.06480813 | 0.01926214 | 0.09319841 | 0.90318385 |
| Dennd5b | 0.16917128 | 0.20610294 | 0.18684894 | 0.01925399 | 0.32343996 | 0.90650187 |
| Clic3 | 0.0171314 | 0.04002596 | 0.02078718 | 0.01923878 | 1.94E-05 | 0.00065548 |
| 2610001J05F | 1.12886678 | 0.81859563 | 0.79938366 | 0.01921197 | 0.45745661 | 0.90650187 |
| S1pr3 | 0.09276895 | 0.12632712 | 0.10711643 | 0.01921068 | 0.21264055 | 0.90650187 |
| Meox2 | 0.05877734 | 0.08856335 | 0.06937701 | 0.01918633 | 0.10945341 | 0.90650187 |
| Sowahc | 0.19226671 | 0.22878346 | 0.2096297 | 0.01915376 | 0.34230046 | 0.90650187 |
| A430033K04 | 0.12811645 | 0.16400941 | 0.14485705 | 0.01915236 | 0.27834054 | 0.90650187 |
| Slitrk4 | 0.05547081 | 0.08474784 | 0.06561614 | 0.0191317 | 0.09745238 | 0.90650187 |

| | | | | | | |
|---|---|---|---|---|---|---|
| Cpb2 | 0.02650162 | 0.05104713 | 0.03195559 | 0.01909155 | 0.00395388 | 0.07669505 |
| Pcdh7 | 0.11798683 | 0.15326756 | 0.13418868 | 0.01907888 | 0.26366922 | 0.90650187 |
| Tmem229b | 0.1060825 | 0.14055992 | 0.12149006 | 0.01906987 | 0.24264763 | 0.90650187 |
| Nabp1 | 0.08501124 | 0.11771798 | 0.09864872 | 0.01906926 | 0.1950719 | 0.90650187 |
| Poc1a | 0.0922543 | 0.12561938 | 0.10655729 | 0.01906209 | 0.21322617 | 0.90650187 |
| Guca1a | 0.09174742 | 0.12505636 | 0.10600622 | 0.01905014 | 0.21217135 | 0.90650187 |
| Tbc1d8b | 0.07966751 | 0.11178864 | 0.09276659 | 0.01902205 | 0.18099392 | 0.90650187 |
| Tob2 | 0.3483705 | 0.36852813 | 0.34951048 | 0.01901765 | 0.40443061 | 0.90650187 |
| Entpd1 | 0.03828986 | 0.06480586 | 0.0458089 | 0.01899696 | 0.03262115 | 0.42461821 |
| Krtap17-1 | 0.04453728 | 0.07205849 | 0.0530633 | 0.01899519 | 0.05575942 | 0.63969729 |
| Cdyl | 0.08062178 | 0.11281217 | 0.09381998 | 0.01899219 | 0.18407769 | 0.90650187 |
| Gcnt1 | 0.06402476 | 0.0942943 | 0.07531218 | 0.01898212 | 0.13125116 | 0.90650187 |
| Zfp930 | 0.02579005 | 0.05008595 | 0.03111237 | 0.01897358 | 0.00335291 | 0.06697454 |
| Elf3 | 0.17648137 | 0.21309221 | 0.19412338 | 0.01896883 | 0.33199754 | 0.90650187 |
| Ccnjl | 0.07508357 | 0.10664954 | 0.08768838 | 0.01896116 | 0.16820355 | 0.90650187 |
| Cit | 0.17579996 | 0.21240541 | 0.19344788 | 0.01895753 | 0.33154135 | 0.90650187 |
| Myo9b | 0.13926031 | 0.17540293 | 0.15645023 | 0.0189527 | 0.29509872 | 0.90650187 |
| Tceal8 | 1.58522198 | 0.92520201 | 0.9062509 | 0.01895111 | 0.46296587 | 0.90650187 |
| Csf2ra | 0.25005022 | 0.28309678 | 0.26414654 | 0.01895024 | 0.3748883 | 0.90650187 |
| Tm4sf5 | 0.04198413 | 0.06904446 | 0.05010587 | 0.01893859 | 0.04644826 | 0.55859172 |
| Clec1a | 0.04282016 | 0.06999834 | 0.05107539 | 0.01892295 | 0.04977339 | 0.58971569 |
| Cenph | 0.0982707 | 0.13198938 | 0.11307094 | 0.01891844 | 0.22849403 | 0.90650187 |
| Tgif1 | 0.15089104 | 0.18729735 | 0.16839379 | 0.01890357 | 0.3088702 | 0.90650187 |
| Pde2a | 0.05868331 | 0.08815726 | 0.06927029 | 0.01888697 | 0.11273282 | 0.90650187 |
| 4930550C14I | 0.06112713 | 0.09091984 | 0.07203981 | 0.01888003 | 0.12199038 | 0.90650187 |
| Hpn | 0.16557732 | 0.20212591 | 0.18325043 | 0.01887548 | 0.32350707 | 0.90650187 |
| Sostdc1 | 0.02497363 | 0.04900323 | 0.03014392 | 0.01885931 | 0.00270659 | 0.05537394 |
| Commd4 | 0.506189 | 0.48878936 | 0.46994356 | 0.0188458 | 0.42925198 | 0.90650187 |
| Mrpl47 | 0.28380851 | 0.31328431 | 0.29445533 | 0.01882898 | 0.38809988 | 0.90650187 |
| Pacrg | 0.2198665 | 0.2549267 | 0.23609904 | 0.01882766 | 0.36147692 | 0.90650187 |

| | | | | | | |
|---|---|---|---|---|---|---|
| Grrp1 | 0.05752563 | 0.08677588 | 0.06795522 | 0.01882066 | 0.10911644 | 0.90650187 |
| Hmga2 | 0.16130674 | 0.19776653 | 0.17895557 | 0.01881096 | 0.3201423 | 0.90650187 |
| Itm2c | 1.5299496 | 0.91709503 | 0.89829864 | 0.01879639 | 0.46294312 | 0.90650187 |
| Sema7a | 0.01332327 | 0.03500362 | 0.01620792 | 0.01879571 | 1.27E-07 | 6.05E-06 |
| Golt1b | 0.53958308 | 0.51195487 | 0.49316284 | 0.01879203 | 0.43274086 | 0.90650187 |
| Bag3 | 0.1190871 | 0.15412493 | 0.13535355 | 0.01877138 | 0.2687708 | 0.90650187 |
| Kazald1 | 0.07471254 | 0.10603922 | 0.08727603 | 0.01876319 | 0.16960011 | 0.90650187 |
| Smarcd3 | 0.13344739 | 0.1691809 | 0.15042136 | 0.01875954 | 0.28964359 | 0.90650187 |
| Gas2l3 | 0.0622985 | 0.09211621 | 0.07336416 | 0.01875205 | 0.12791395 | 0.90650187 |
| Cpne2 | 0.09973836 | 0.13339015 | 0.11465233 | 0.01873782 | 0.23375001 | 0.90650187 |
| Cyth2 | 0.97031824 | 0.75687842 | 0.73815027 | 0.01872815 | 0.45509745 | 0.90650187 |
| Gar1 | 0.55560281 | 0.52277331 | 0.50404737 | 0.01872594 | 0.43440926 | 0.90650187 |
| Slu7 | 0.42573562 | 0.42957618 | 0.41088128 | 0.0186949 | 0.41985168 | 0.90650187 |
| Slco2b1 | 0.02894766 | 0.05352887 | 0.03484804 | 0.01868083 | 0.00858291 | 0.14742239 |
| Smc1a | 0.96007575 | 0.75226645 | 0.73358979 | 0.01867666 | 0.45494317 | 0.90650187 |
| Phyhipl | 0.24428954 | 0.27748987 | 0.25886404 | 0.01862583 | 0.37453306 | 0.90650187 |
| Susd6 | 0.20029556 | 0.23601851 | 0.21741328 | 0.01860523 | 0.35181139 | 0.90650187 |
| Gm11273 | 0.31993726 | 0.34426185 | 0.32571861 | 0.01854324 | 0.40009954 | 0.90650187 |
| Nebl | 0.02550358 | 0.04931479 | 0.03077268 | 0.01854212 | 0.00369536 | 0.0724815 |
| Krt14 | 0.01844 | 0.04088483 | 0.02235535 | 0.01852948 | 0.00011445 | 0.00332637 |
| Itgb2 | 0.02108037 | 0.04403714 | 0.02551109 | 0.01852605 | 0.00062247 | 0.01574394 |
| Pja2 | 0.94126278 | 0.74325456 | 0.72473163 | 0.01852293 | 0.45476862 | 0.90650187 |
| Dpm2 | 0.57466513 | 0.53529694 | 0.51679264 | 0.0185043 | 0.43676369 | 0.90650187 |
| Nop9 | 0.21762629 | 0.2524831 | 0.23398024 | 0.01850286 | 0.3625868 | 0.90650187 |
| Pcid2 | 0.24650095 | 0.27938229 | 0.260896 | 0.01848629 | 0.37638006 | 0.90650187 |
| Kcna5 | 0.06032008 | 0.08960333 | 0.07112617 | 0.01847716 | 0.12406989 | 0.90650187 |
| Nfam1 | 0.02160354 | 0.0445998 | 0.02613505 | 0.01846475 | 0.00084221 | 0.02054968 |
| Tnfsf11 | 0.01473531 | 0.03634905 | 0.01790863 | 0.01844042 | 2.35E-06 | 9.47E-05 |
| Hsf2 | 0.23636107 | 0.2699648 | 0.25153942 | 0.01842538 | 0.37234734 | 0.90650187 |
| Celf1 | 0.71671562 | 0.62363937 | 0.60522299 | 0.01841638 | 0.44619694 | 0.90650187 |

| | | | | | | |
|---|---|---|---|---|---|---|
| Sh3gl1 | 0.60782849 | 0.55685415 | 0.53844542 | 0.01840873 | 0.43959768 | 0.90650187 |
| Prcp | 0.14819971 | 0.18404568 | 0.16564429 | 0.01840139 | 0.31070198 | 0.90650187 |
| Chst4 | 0.03857623 | 0.06454085 | 0.04614275 | 0.0183981 | 0.03815096 | 0.47967968 |
| Tmem106b | 0.42284701 | 0.4270721 | 0.40867413 | 0.01839798 | 0.42068735 | 0.90650187 |
| Far1 | 0.41907597 | 0.42417998 | 0.40578316 | 0.01839682 | 0.42013481 | 0.90650187 |
| Vrk2 | 0.10699894 | 0.14086252 | 0.12247389 | 0.01838863 | 0.25223343 | 0.90650187 |
| Orc2 | 0.2108014 | 0.2458803 | 0.22749343 | 0.01838687 | 0.35968137 | 0.90650187 |
| Aldh9a1 | 0.29792244 | 0.32519282 | 0.30680999 | 0.01838282 | 0.39497988 | 0.90650187 |
| Dock1 | 0.16876322 | 0.20479463 | 0.1864411 | 0.01835353 | 0.33082656 | 0.90650187 |
| Nuak2 | 0.0830082 | 0.11478086 | 0.09644861 | 0.01833225 | 0.19905859 | 0.90650187 |
| Gm15261 | 0.05268502 | 0.08075549 | 0.06243488 | 0.01832061 | 0.09603174 | 0.90650187 |
| Cpsf2 | 0.34880888 | 0.36819079 | 0.34987184 | 0.01831895 | 0.40797092 | 0.90650187 |
| Gucy1b3 | 0.10662134 | 0.1403703 | 0.12206864 | 0.01830166 | 0.25253736 | 0.90650187 |
| Thbs3 | 0.28918627 | 0.31747343 | 0.29918427 | 0.01828916 | 0.39290317 | 0.90650187 |
| Smarcc2 | 0.50985128 | 0.49081324 | 0.47252565 | 0.01828759 | 0.43169795 | 0.90650187 |
| Zdhhc16 | 0.34624477 | 0.36604059 | 0.34775599 | 0.01828459 | 0.40759161 | 0.90650187 |
| Klhl22 | 0.33164528 | 0.35388244 | 0.33559916 | 0.01828328 | 0.40431472 | 0.90650187 |
| Ptcd1 | 0.13124311 | 0.16635321 | 0.1481247 | 0.01822851 | 0.29216021 | 0.90650187 |
| Adcy2 | 0.03807375 | 0.06378242 | 0.04555688 | 0.01822554 | 0.03765967 | 0.47484348 |
| Vegfa | 0.52469939 | 0.5011089 | 0.48290354 | 0.01820536 | 0.4334496 | 0.90650187 |
| Zfp260 | 0.63674994 | 0.574998 | 0.55680962 | 0.01818839 | 0.44226965 | 0.90650187 |
| Vps4b | 0.60112403 | 0.55230775 | 0.53412015 | 0.0181876 | 0.43983814 | 0.90650187 |
| Yes1 | 0.40814045 | 0.41551923 | 0.39733676 | 0.01818247 | 0.41939763 | 0.90650187 |
| Dbp | 0.15228231 | 0.18798578 | 0.16981177 | 0.01817402 | 0.31711246 | 0.90650187 |
| Sae1 | 0.78474816 | 0.66202066 | 0.64388521 | 0.01813545 | 0.4501773 | 0.90650187 |
| Mnat1 | 0.46782917 | 0.46048559 | 0.44235358 | 0.018132 | 0.42770321 | 0.90650187 |
| Ing1 | 0.51151237 | 0.49181373 | 0.47369393 | 0.0181198 | 0.43248389 | 0.90650187 |
| Rad51ap1 | 0.08587582 | 0.117704 | 0.09959661 | 0.01810739 | 0.20947519 | 0.90650187 |
| Itpk1 | 0.07650555 | 0.10732855 | 0.08926691 | 0.01806164 | 0.18419514 | 0.90650187 |
| Malat1 | 7.71186343 | 3.30783338 | 3.28977844 | 0.01805495 | 0.49026743 | 0.90650187 |

| | | | | | | |
|---|---|---|---|---|---|---|
| 9430020K01I | 0.06755935 | 0.09731671 | 0.07928719 | 0.01802952 | 0.15602997 | 0.90650187 |
| Clec2d | 0.07063693 | 0.10075697 | 0.08273339 | 0.01802358 | 0.16640198 | 0.90650187 |
| Fmn2 | 0.19853283 | 0.233705 | 0.21571058 | 0.01799441 | 0.35537495 | 0.90650187 |
| Aff2 | 0.02702698 | 0.05056673 | 0.03257763 | 0.0179891 | 0.0070476 | 0.12471545 |
| Nespas | 0.24426529 | 0.27680136 | 0.25884174 | 0.01795962 | 0.37886749 | 0.90650187 |
| Oxr1 | 0.31190061 | 0.33682171 | 0.31886538 | 0.01795634 | 0.40115973 | 0.90650187 |
| Taf9b | 0.19953558 | 0.23463099 | 0.21667958 | 0.01795141 | 0.3563217 | 0.90650187 |
| Adgrd1 | 0.09118823 | 0.12334607 | 0.10539787 | 0.0179482 | 0.22451225 | 0.90650187 |
| Gpn1 | 0.24941928 | 0.28150623 | 0.26356965 | 0.01793658 | 0.38112223 | 0.90650187 |
| Trmt12 | 0.10813974 | 0.14161146 | 0.12369712 | 0.01791434 | 0.2598423 | 0.90650187 |
| Prss8 | 0.09206583 | 0.12425353 | 0.10635244 | 0.0179011 | 0.2271461 | 0.90650187 |
| AU020206 | 0.04590109 | 0.0725209 | 0.05463899 | 0.01788191 | 0.07284423 | 0.76842826 |
| Gchfr | 0.1142356 | 0.14808598 | 0.13020603 | 0.01787994 | 0.27077444 | 0.90650187 |
| Mis18a | 0.12682619 | 0.16137161 | 0.14350519 | 0.01786642 | 0.28996785 | 0.90650187 |
| Ift81 | 0.47451561 | 0.465094 | 0.44723542 | 0.01785859 | 0.42955084 | 0.90650187 |
| Ktn1 | 1.11344114 | 0.81181333 | 0.79398207 | 0.01783126 | 0.46023588 | 0.90650187 |
| Yars | 0.30407792 | 0.3299096 | 0.31214015 | 0.01776945 | 0.40010374 | 0.90650187 |
| Capn2 | 0.68604574 | 0.60481705 | 0.58705016 | 0.01776688 | 0.44648588 | 0.90650187 |
| Endou | 0.01049875 | 0.03055692 | 0.01279623 | 0.01776069 | 3.41E-10 | 2.24E-08 |
| Bet1 | 0.66463815 | 0.59181709 | 0.57407801 | 0.01773908 | 0.44536926 | 0.90650187 |
| Galnt16 | 0.15190304 | 0.18716282 | 0.16942543 | 0.01773739 | 0.32081647 | 0.90650187 |
| Cfb | 0.01750698 | 0.0389741 | 0.02123754 | 0.01773657 | 0.00010251 | 0.00301979 |
| Zxdb | 0.0999819 | 0.13264758 | 0.11491445 | 0.01773312 | 0.24634789 | 0.90650187 |
| Fundc2 | 1.20934744 | 0.84295593 | 0.82522844 | 0.0177275 | 0.46195877 | 0.90650187 |
| St3gal5 | 0.29500783 | 0.32199947 | 0.30427379 | 0.01772568 | 0.39782459 | 0.90650187 |
| Atp1a2 | 0.10611974 | 0.13923787 | 0.12153006 | 0.01770781 | 0.25857104 | 0.90650187 |
| Il17ra | 0.08264682 | 0.11375517 | 0.09605106 | 0.0177041 | 0.20627547 | 0.90650187 |
| Ppan | 0.36029303 | 0.37697838 | 0.3592818 | 0.01769657 | 0.41333664 | 0.90650187 |
| Nle1 | 0.15348966 | 0.1887278 | 0.17104048 | 0.01768732 | 0.32285832 | 0.90650187 |
| Fam162b | 0.03642063 | 0.06130718 | 0.04362668 | 0.0176805 | 0.03579926 | 0.45729797 |

| | | | | | | |
|---|---|---|---|---|---|---|
| Ap1b1 | 0.36051841 | 0.37712683 | 0.35946539 | 0.01766143 | 0.41354949 | 0.90650187 |
| Vmp1 | 0.67179429 | 0.59608615 | 0.57844095 | 0.0176452 | 0.44606396 | 0.90650187 |
| Nol12 | 0.5312863 | 0.50509893 | 0.48746105 | 0.01763788 | 0.43610338 | 0.90650187 |
| Stag1 | 0.52607416 | 0.50146751 | 0.48385702 | 0.01761049 | 0.43573066 | 0.90650187 |
| Il3ra | 0.16710642 | 0.20238755 | 0.18478323 | 0.01760432 | 0.33595304 | 0.90650187 |
| Trim27 | 0.76833182 | 0.65235405 | 0.63475681 | 0.01759724 | 0.45095668 | 0.90650187 |
| Cyp11a1 | 0.01081352 | 0.03076987 | 0.01317707 | 0.01759279 | 1.47E-09 | 8.88E-08 |
| 1500011K16I | 0.55891086 | 0.52386654 | 0.50627521 | 0.01759133 | 0.43861972 | 0.90650187 |
| Zfp943 | 0.18222794 | 0.21738521 | 0.19979976 | 0.01758545 | 0.3477945 | 0.90650187 |
| Azin2 | 0.14305025 | 0.17789539 | 0.16035952 | 0.01753587 | 0.31343312 | 0.90650187 |
| Aga | 0.51151123 | 0.4911609 | 0.47369313 | 0.01746776 | 0.43489122 | 0.90650187 |
| Trem2 | 0.0150815 | 0.03577609 | 0.01832511 | 0.01745099 | 1.15E-05 | 0.0004085 |
| Rab5c | 0.80659329 | 0.67328389 | 0.65584098 | 0.01744291 | 0.45294014 | 0.90650187 |
| Ggt5 | 0.06082033 | 0.08909446 | 0.0716926 | 0.01740186 | 0.14027676 | 0.90650187 |
| Ccdc103 | 0.06853025 | 0.0977666 | 0.08037587 | 0.01739074 | 0.16805363 | 0.90650187 |
| Ripk3 | 0.06201105 | 0.09042292 | 0.07303936 | 0.01738357 | 0.14501255 | 0.90650187 |
| Armc2 | 0.05187379 | 0.07888765 | 0.0615063 | 0.01738135 | 0.1045028 | 0.90650187 |
| Cdc42se1 | 0.62458689 | 0.5665031 | 0.54914401 | 0.01735909 | 0.44412029 | 0.90650187 |
| Gjb1 | 0.05770872 | 0.08549306 | 0.06816334 | 0.01732972 | 0.12918698 | 0.90650187 |
| Optn | 0.08781422 | 0.11902969 | 0.10171796 | 0.01731172 | 0.22464097 | 0.90650187 |
| Htatsf1 | 0.44277993 | 0.44105169 | 0.42378061 | 0.01727108 | 0.42811328 | 0.90650187 |
| Itgav | 0.23492131 | 0.2674638 | 0.25020264 | 0.01726116 | 0.37953715 | 0.90650187 |
| Cyb5d2 | 0.12216744 | 0.15586075 | 0.13860693 | 0.01725382 | 0.28999816 | 0.90650187 |
| Tmem150a | 0.18332446 | 0.21812792 | 0.20087878 | 0.01724913 | 0.35132824 | 0.90650187 |
| 2700081O15 | 0.22312509 | 0.2563985 | 0.23917183 | 0.01722667 | 0.37440811 | 0.90650187 |
| Loxl3 | 0.11118826 | 0.14417522 | 0.12695796 | 0.01721726 | 0.27329227 | 0.90650187 |
| Nav3 | 0.05206557 | 0.0789388 | 0.06172591 | 0.01721289 | 0.10754288 | 0.90650187 |
| Cope | 1.94518503 | 0.9444945 | 0.92731191 | 0.01718259 | 0.46717438 | 0.90650187 |
| Gins2 | 0.23259769 | 0.2652154 | 0.24804085 | 0.01717455 | 0.37911068 | 0.90650187 |
| Ctbs | 0.17066896 | 0.2054965 | 0.18834427 | 0.01715224 | 0.34279211 | 0.90650187 |

| | | | | | | |
|---|---|---|---|---|---|---|
| Exosc2 | 0.28998105 | 0.31703206 | 0.29988092 | 0.01715114 | 0.39964799 | 0.90650187 |
| Mtrf1l | 0.18986182 | 0.22442788 | 0.20728475 | 0.01714313 | 0.35656105 | 0.90650187 |
| Cetn2 | 0.94155479 | 0.74200099 | 0.72487015 | 0.01713084 | 0.45816293 | 0.90650187 |
| Tmem106a | 0.04272862 | 0.06809924 | 0.05096928 | 0.01712996 | 0.06757516 | 0.73016166 |
| Piezo1 | 0.1316551 | 0.16567971 | 0.14855439 | 0.01712531 | 0.30415463 | 0.90650187 |
| Phox2a | 0.01523096 | 0.03562621 | 0.01850484 | 0.01712137 | 1.95E-05 | 0.00065749 |
| Malsu1 | 0.40119381 | 0.40902537 | 0.39192449 | 0.01710088 | 0.42309752 | 0.90650187 |
| Itpr1 | 0.40433153 | 0.41144605 | 0.39437369 | 0.01707236 | 0.42369509 | 0.90650187 |
| Adgrg6 | 0.09279269 | 0.12420603 | 0.10714222 | 0.01706382 | 0.23946591 | 0.90650187 |
| Prickle2 | 0.06128433 | 0.08928112 | 0.07221766 | 0.01706347 | 0.14676701 | 0.90650187 |
| Miat | 0.15880491 | 0.19349159 | 0.17642994 | 0.01706164 | 0.33362982 | 0.90650187 |
| Pithd1 | 0.40194364 | 0.40957164 | 0.39251047 | 0.01706118 | 0.42338695 | 0.90650187 |
| Susd2 | 0.05999096 | 0.08779804 | 0.07075332 | 0.01704472 | 0.14209323 | 0.90650187 |
| Inpp5d | 0.01547437 | 0.0358411 | 0.01879749 | 0.01704361 | 2.78E-05 | 0.00090657 |
| Nup50 | 0.38579695 | 0.39681261 | 0.37979329 | 0.01701933 | 0.42104655 | 0.90650187 |
| Ecel1 | 0.03147545 | 0.05484288 | 0.03782727 | 0.01701561 | 0.02276421 | 0.31940792 |
| Tenm3 | 0.25242001 | 0.28331853 | 0.26630987 | 0.01700866 | 0.38823134 | 0.90650187 |
| Ncoa3 | 0.17605969 | 0.21071401 | 0.19370542 | 0.01700859 | 0.34813778 | 0.90650187 |
| Top1 | 1.62573728 | 0.92796664 | 0.910959 | 0.01700764 | 0.46692591 | 0.90650187 |
| Rabgef1 | 0.14074464 | 0.17498317 | 0.15798331 | 0.01699986 | 0.31619292 | 0.90650187 |
| Tmem215 | 0.0938048 | 0.12522859 | 0.10824074 | 0.01698786 | 0.2426773 | 0.90650187 |
| Yif1b | 0.71040759 | 0.61850973 | 0.6015238 | 0.01698593 | 0.4500497 | 0.90650187 |
| Gm5611 | 0.1813504 | 0.21588223 | 0.19893528 | 0.01694695 | 0.35245033 | 0.90650187 |
| Dcps | 0.71194995 | 0.61936573 | 0.60243014 | 0.0169356 | 0.45027148 | 0.90650187 |
| Rbp2 | 0.10162681 | 0.13362451 | 0.1166917 | 0.01693281 | 0.2594345 | 0.90650187 |
| Cux2 | 0.04827084 | 0.07429778 | 0.05737015 | 0.01692763 | 0.09480729 | 0.90650187 |
| Avpi1 | 0.17829443 | 0.21284231 | 0.19591821 | 0.01692409 | 0.35047926 | 0.90650187 |
| Zfp655 | 0.27990046 | 0.30790921 | 0.29100212 | 0.01690709 | 0.39809243 | 0.90650187 |
| Ptgs1 | 0.09798432 | 0.12965961 | 0.11276203 | 0.01689758 | 0.25264747 | 0.90650187 |
| Hfe | 0.06778846 | 0.09642476 | 0.07954422 | 0.01688054 | 0.17273102 | 0.90650187 |

| | | | | | | |
|---|---|---|---|---|---|---|
| Afap1l2 | 0.09477145 | 0.12613706 | 0.10928858 | 0.01684849 | 0.24655918 | 0.90650187 |
| Ctnnbip1 | 0.84724785 | 0.69434979 | 0.67752222 | 0.01682758 | 0.45604003 | 0.90650187 |
| Cdh19 | 0.02066231 | 0.04180987 | 0.02501217 | 0.01679771 | 0.00141518 | 0.03163897 |
| Fam81a | 0.08234593 | 0.1125122 | 0.09571993 | 0.01679227 | 0.21772538 | 0.90650187 |
| Notch3 | 0.15782111 | 0.19222277 | 0.17543485 | 0.01678793 | 0.33526727 | 0.90650187 |
| Inpp5a | 0.0750247 | 0.10440136 | 0.08762297 | 0.0167784 | 0.19731177 | 0.90650187 |
| Nhlrc1 | 0.0210092 | 0.04216555 | 0.02542617 | 0.01673938 | 0.00171248 | 0.03734777 |
| Nup107 | 0.11504082 | 0.14779367 | 0.13106238 | 0.01673128 | 0.28518002 | 0.90650187 |
| Tmem107 | 0.56722044 | 0.52852863 | 0.51184227 | 0.01668636 | 0.44238332 | 0.90650187 |
| Acot2 | 0.17196263 | 0.20631819 | 0.18963389 | 0.0166843 | 0.34784992 | 0.90650187 |
| Gdf6 | 0.02981924 | 0.05253278 | 0.03587641 | 0.01665637 | 0.01951052 | 0.28326589 |
| Zfand5 | 0.71725652 | 0.62218324 | 0.60553915 | 0.0166441 | 0.45137279 | 0.90650187 |
| Armcx6 | 0.33377976 | 0.35403173 | 0.33738775 | 0.01664398 | 0.41320419 | 0.90650187 |
| Serpina1b | 0.03413108 | 0.05758319 | 0.04094637 | 0.01663682 | 0.03543703 | 0.45381491 |
| Saxo2 | 0.10185796 | 0.13357469 | 0.11694124 | 0.01663345 | 0.2635848 | 0.90650187 |
| Dpysl4 | 0.11851202 | 0.15135159 | 0.13474489 | 0.0166067 | 0.29187995 | 0.90650187 |
| Gpr153 | 0.12045475 | 0.15339082 | 0.13679946 | 0.01659136 | 0.2948835 | 0.90650187 |
| Arsb | 0.22508433 | 0.2575615 | 0.24101368 | 0.01654782 | 0.3800947 | 0.90650187 |
| Pcyox1 | 0.30300535 | 0.32774609 | 0.31121382 | 0.01653227 | 0.40665461 | 0.90650187 |
| Efr3a | 0.41514114 | 0.41928018 | 0.40275427 | 0.01652591 | 0.42762923 | 0.90650187 |
| Kctd8 | 0.13482096 | 0.16836816 | 0.15184956 | 0.01651859 | 0.31433322 | 0.90650187 |
| Cyyr1 | 0.0377489 | 0.06167266 | 0.04517792 | 0.01649474 | 0.05228206 | 0.61257938 |
| Bnc2 | 0.15717117 | 0.1912475 | 0.17477683 | 0.01647066 | 0.33762773 | 0.90650187 |
| Hes5 | 0.01403854 | 0.03352375 | 0.01706982 | 0.01645393 | 9.13E-06 | 0.0003296 |
| Mapk7 | 0.24743403 | 0.27820364 | 0.26175193 | 0.01645172 | 0.38996253 | 0.90650187 |
| Stx4a | 0.56459159 | 0.52652901 | 0.51008554 | 0.01644348 | 0.44302201 | 0.90650187 |
| G430095P16 | 0.03974841 | 0.0639363 | 0.04750792 | 0.01642837 | 0.06210957 | 0.68663823 |
| Tom1l2 | 0.09247817 | 0.12321872 | 0.10680056 | 0.01641816 | 0.24717355 | 0.90650187 |
| Tle1 | 0.41089285 | 0.41588128 | 0.39947111 | 0.01641017 | 0.4275464 | 0.90650187 |
| Cd2ap | 0.63057127 | 0.56932724 | 0.55292569 | 0.01640155 | 0.4475432 | 0.90650187 |

| | | | | | | |
|---|---|---|---|---|---|---|
| Srebf2 | 0.36976269 | 0.3833552 | 0.36696035 | 0.01639485 | 0.42128067 | 0.90650187 |
| Arf5 | 2.89038821 | 0.86868686 | 0.85229343 | 0.01639342 | 0.46592839 | 0.90650187 |
| Ifih1 | 0.05555328 | 0.0820981 | 0.06571013 | 0.01638797 | 0.13377443 | 0.90650187 |
| Nptn | 1.09995976 | 0.80560815 | 0.78923299 | 0.01637516 | 0.46325436 | 0.90650187 |
| Hdgfrp3 | 0.16078324 | 0.19479802 | 0.17842768 | 0.01637034 | 0.34168296 | 0.90650187 |
| Tbc1d4 | 0.07535942 | 0.10434504 | 0.08799483 | 0.01635021 | 0.20439281 | 0.90650187 |
| Mcur1 | 0.1763341 | 0.21031605 | 0.19397743 | 0.01633862 | 0.35403433 | 0.90650187 |
| Ahsa2 | 0.52911901 | 0.50229146 | 0.48596452 | 0.01632693 | 0.44063523 | 0.90650187 |
| Ptpn1 | 0.5156342 | 0.49290644 | 0.47658521 | 0.01632123 | 0.43949681 | 0.90650187 |
| Ten1 | 0.74396046 | 0.6372917 | 0.62097177 | 0.01631994 | 0.4534949 | 0.90650187 |
| Zfp354c | 0.15143539 | 0.18526879 | 0.16894886 | 0.01631993 | 0.33380454 | 0.90650187 |
| Arhgap36 | 0.05224369 | 0.07824835 | 0.06192983 | 0.01631853 | 0.12071531 | 0.90650187 |
| Sgms1 | 0.41882799 | 0.42190523 | 0.40559261 | 0.01631262 | 0.42904783 | 0.90650187 |
| Rac3 | 0.18792661 | 0.22169807 | 0.20539325 | 0.01630482 | 0.36207894 | 0.90650187 |
| Dsel | 0.13792128 | 0.17134858 | 0.15506499 | 0.01628359 | 0.32030747 | 0.90650187 |
| Cdk5rap3 | 0.61664156 | 0.56034989 | 0.54409249 | 0.01625739 | 0.44716234 | 0.90650187 |
| Ifit3 | 0.02631363 | 0.04797777 | 0.03173289 | 0.01624488 | 0.0114284 | 0.18659838 |
| Cebpg | 0.34025097 | 0.35901709 | 0.34278678 | 0.01623031 | 0.41664218 | 0.90650187 |
| Rab3il1 | 0.09374028 | 0.12438967 | 0.10817075 | 0.01621893 | 0.25252464 | 0.90650187 |
| Nkap | 0.39315604 | 0.40183164 | 0.38561537 | 0.01621627 | 0.42584954 | 0.90650187 |
| Cdk2ap1 | 0.3840842 | 0.39462182 | 0.3784322 | 0.01618962 | 0.42458153 | 0.90650187 |
| Plac9b | 0.11925529 | 0.15170921 | 0.13553149 | 0.01617772 | 0.29783065 | 0.90650187 |
| Zbtb8a | 0.19217248 | 0.22571396 | 0.20953793 | 0.01617603 | 0.36572606 | 0.90650187 |
| Gpr137 | 0.21520917 | 0.24785442 | 0.23168835 | 0.01616607 | 0.37820731 | 0.90650187 |
| Pip4k2b | 0.11548268 | 0.14768666 | 0.13153197 | 0.0161547 | 0.29253044 | 0.90650187 |
| Lmf2 | 0.2285182 | 0.26036318 | 0.2442324 | 0.01613079 | 0.38452475 | 0.90650187 |
| Cbs | 0.05649473 | 0.08290413 | 0.0667825 | 0.01612164 | 0.14159244 | 0.90650187 |
| Impa2 | 0.13531569 | 0.16846204 | 0.15236343 | 0.01609861 | 0.31927764 | 0.90650187 |
| Mar-02 | 0.43407573 | 0.4332982 | 0.41721952 | 0.01607868 | 0.43198474 | 0.90650187 |
| Lix1 | 0.03164089 | 0.05406369 | 0.0380219 | 0.01604179 | 0.03035223 | 0.40204743 |

| | | | | | | |
|---|---|---|---|---|---|---|
| Gm42637 | 0.03023439 | 0.05240689 | 0.03636583 | 0.01604106 | 0.02494138 | 0.34434814 |
| Adal | 0.09358317 | 0.12403612 | 0.1080003 | 0.01603582 | 0.25460097 | 0.90650187 |
| Kcnab2 | 0.02005272 | 0.04030069 | 0.02428419 | 0.0160165 | 0.00168353 | 0.03682198 |
| Carm1 | 0.17422115 | 0.20789176 | 0.19188081 | 0.01601095 | 0.35533697 | 0.90650187 |
| Bin3 | 0.31282068 | 0.33566318 | 0.31965283 | 0.01601035 | 0.41189788 | 0.90650187 |
| Tssc1 | 0.13556018 | 0.16862611 | 0.15261726 | 0.01600885 | 0.32049188 | 0.90650187 |
| Zbp1 | 0.01420028 | 0.03325483 | 0.0172646 | 0.01599022 | 1.92E-05 | 0.00064846 |
| Kcne4 | 0.07843478 | 0.10739342 | 0.0914039 | 0.01598952 | 0.21837848 | 0.90650187 |
| Yipf5 | 0.91895261 | 0.72991805 | 0.71393464 | 0.01598341 | 0.46035983 | 0.90650187 |
| Irf5 | 0.02744763 | 0.04903301 | 0.03307536 | 0.01595764 | 0.01598277 | 0.24313303 |
| Mal | 0.02811058 | 0.04981287 | 0.03385926 | 0.01595361 | 0.01810082 | 0.26975545 |
| Lrrc1 | 0.1181024 | 0.15024794 | 0.1343111 | 0.01593684 | 0.29892373 | 0.90650187 |
| Ift74 | 0.46224674 | 0.45418094 | 0.43825266 | 0.01592828 | 0.4358201 | 0.90650187 |
| Sh2d4a | 0.05563198 | 0.08172721 | 0.06579983 | 0.01592738 | 0.14094244 | 0.90650187 |
| Baz2a | 0.44728758 | 0.4430811 | 0.42715603 | 0.01592508 | 0.43418098 | 0.90650187 |
| St3gal2 | 0.11878683 | 0.15096026 | 0.1350358 | 0.01592446 | 0.30004879 | 0.90650187 |
| Hcst | 0.01485101 | 0.03396936 | 0.01804784 | 0.01592151 | 4.39E-05 | 0.00138567 |
| Upp2 | 0.11504813 | 0.1469907 | 0.13107016 | 0.01592054 | 0.29460225 | 0.90650187 |
| Pot1b | 0.04077692 | 0.06461046 | 0.04870403 | 0.01590643 | 0.07326373 | 0.77027545 |
| Net1 | 0.1156968 | 0.14765535 | 0.13175943 | 0.01589592 | 0.29586405 | 0.90650187 |
| Npr1 | 0.0784419 | 0.10730656 | 0.09141177 | 0.01589478 | 0.21975909 | 0.90650187 |
| Galnt9 | 0.01178284 | 0.03021837 | 0.01434887 | 0.0158695 | 4.40E-07 | 1.97E-05 |
| Efcab7 | 0.06974397 | 0.09759092 | 0.08173489 | 0.01585602 | 0.1942288 | 0.90650187 |
| Prdm16 | 0.19270421 | 0.22589951 | 0.21005563 | 0.01584388 | 0.36869219 | 0.90650187 |
| Scarf2 | 0.18156549 | 0.21497506 | 0.19914725 | 0.01582781 | 0.36192146 | 0.90650187 |
| Ddr1 | 0.71015802 | 0.6172009 | 0.60137704 | 0.01582386 | 0.45343956 | 0.90650187 |
| Lmbr1 | 0.16853263 | 0.20201505 | 0.18621054 | 0.01580451 | 0.35296855 | 0.90650187 |
| Snx6 | 1.18780928 | 0.83450281 | 0.81869912 | 0.01580369 | 0.46580659 | 0.90650187 |
| Dnajb2 | 0.49038919 | 0.47450224 | 0.45870178 | 0.01580046 | 0.43914648 | 0.90650187 |
| Gm9780 | 0.08188566 | 0.11097566 | 0.09521314 | 0.01576252 | 0.23087466 | 0.90650187 |

| | | | | | | |
|---|---|---|---|---|---|---|
| Nt5dc1 | 0.09511278 | 0.12539475 | 0.10965827 | 0.01573648 | 0.26174799 | 0.90650187 |
| Lmbrd1 | 0.32590525 | 0.34650574 | 0.33077 | 0.01573574 | 0.41625206 | 0.90650187 |
| Prom1 | 0.11406064 | 0.14574315 | 0.13001986 | 0.01572329 | 0.29542425 | 0.90650187 |
| Kti12 | 0.35078306 | 0.36719877 | 0.35149718 | 0.01570159 | 0.42129255 | 0.90650187 |
| Apold1 | 0.05493847 | 0.08070097 | 0.06500913 | 0.01569184 | 0.14161841 | 0.90650187 |
| Cgn | 0.16734661 | 0.20069071 | 0.18502376 | 0.01566695 | 0.35329744 | 0.90650187 |
| Hs3st1 | 0.10569302 | 0.13673355 | 0.12107162 | 0.01566193 | 0.28261611 | 0.90650187 |
| Ppp1r3c | 0.11922963 | 0.15115286 | 0.13550435 | 0.01564851 | 0.30383793 | 0.90650187 |
| Sppl2a | 0.3385259 | 0.35698937 | 0.34135097 | 0.01563839 | 0.41930662 | 0.90650187 |
| Slc20a2 | 0.11799228 | 0.14982983 | 0.13419445 | 0.01563538 | 0.30223948 | 0.90650187 |
| Tgm7 | 0.01042326 | 0.02833875 | 0.01270486 | 0.01563388 | 2.24E-08 | 1.22E-06 |
| St8sia6 | 0.01735597 | 0.03668356 | 0.02105649 | 0.01562707 | 0.00048462 | 0.01265759 |
| Gcc2 | 0.64256005 | 0.57604896 | 0.56044269 | 0.01560627 | 0.45073927 | 0.90650187 |
| AB124611 | 0.01225742 | 0.03052704 | 0.01492202 | 0.01560502 | 1.67E-06 | 6.92E-05 |
| Slc9a9 | 0.0494196 | 0.07428522 | 0.05869104 | 0.01559418 | 0.11876262 | 0.90650187 |
| Foxs1 | 0.10012768 | 0.13065931 | 0.11507167 | 0.01558764 | 0.27351949 | 0.90650187 |
| Khk | 0.26203102 | 0.29061475 | 0.27502799 | 0.01558676 | 0.40054083 | 0.90650187 |
| Tmsb15b1 | 0.14649988 | 0.17946269 | 0.1639032 | 0.01555949 | 0.33650309 | 0.90650187 |
| Rbak | 0.09849371 | 0.12885059 | 0.11331143 | 0.01553917 | 0.27104478 | 0.90650187 |
| Sec24d | 0.39458906 | 0.40224787 | 0.38674421 | 0.01550366 | 0.42927752 | 0.90650187 |
| Ccbl1 | 0.06926858 | 0.09669854 | 0.08120285 | 0.01549569 | 0.1981239 | 0.90650187 |
| Efnb2 | 0.07626709 | 0.10449018 | 0.08900239 | 0.01548779 | 0.21958248 | 0.90650187 |
| Mars2 | 0.04981628 | 0.07462917 | 0.0591467 | 0.01548247 | 0.12227295 | 0.90650187 |
| Ncapd2 | 0.15492007 | 0.18797409 | 0.17249404 | 0.01548005 | 0.34496135 | 0.90650187 |
| Tmcc3 | 0.20769052 | 0.23999116 | 0.22452064 | 0.01547051 | 0.3796794 | 0.90650187 |
| Slc35g2 | 0.26551552 | 0.29363304 | 0.27816695 | 0.01546609 | 0.40238607 | 0.90650187 |
| Mex3b | 0.16229243 | 0.19540743 | 0.1799487 | 0.01545873 | 0.35126564 | 0.90650187 |
| Hopx | 0.14920439 | 0.18210062 | 0.16667178 | 0.01542884 | 0.34034173 | 0.90650187 |
| N4bp3 | 0.09558302 | 0.1255726 | 0.11016732 | 0.01540528 | 0.26708509 | 0.90650187 |
| Bfar | 0.59304218 | 0.5442697 | 0.52887221 | 0.0153975 | 0.44850924 | 0.90650187 |

| | | | | | | |
|---|---|---|---|---|---|---|
| Gm43194 | 0.14445165 | 0.17717948 | 0.1618008 | 0.01537868 | 0.33631486 | 0.90650187 |
| Hgf | 0.050515 | 0.07531765 | 0.05994872 | 0.01536894 | 0.12720328 | 0.90650187 |
| Hspa12b | 0.16975284 | 0.2027915 | 0.18742989 | 0.01536161 | 0.35779513 | 0.90650187 |
| Frmd8 | 0.09174426 | 0.12135542 | 0.10600279 | 0.01535263 | 0.25983069 | 0.90650187 |
| Trmt10b | 0.16357327 | 0.19654871 | 0.18123754 | 0.01531117 | 0.35361821 | 0.90650187 |
| 9130008F23l | 0.05166256 | 0.07657115 | 0.06126435 | 0.0153068 | 0.13334282 | 0.90650187 |
| Lrrc49 | 0.16049332 | 0.1934413 | 0.17813519 | 0.01530611 | 0.35123634 | 0.90650187 |
| Kctd1 | 0.11013143 | 0.14113018 | 0.12582883 | 0.01530135 | 0.29438976 | 0.90650187 |
| Akr1b7 | 0.01735736 | 0.03635145 | 0.02105816 | 0.01529329 | 0.00062203 | 0.01574394 |
| Map3k15 | 0.144537 | 0.1771721 | 0.1618885 | 0.0152836 | 0.33735171 | 0.90650187 |
| Hsbp1 | 1.92267157 | 0.94242219 | 0.92718141 | 0.01524078 | 0.4708729 | 0.90650187 |
| Mrpl23 | 1.27975886 | 0.86069429 | 0.84546477 | 0.01522952 | 0.46808699 | 0.90650187 |
| Oser1 | 0.69581782 | 0.60810173 | 0.59289236 | 0.01520937 | 0.45460208 | 0.90650187 |
| Dtx4 | 0.06328487 | 0.08967039 | 0.07447777 | 0.01519261 | 0.18224235 | 0.90650187 |
| Aldh5a1 | 0.14516524 | 0.17769398 | 0.16253382 | 0.01516015 | 0.33919584 | 0.90650187 |
| Ubap2 | 0.39271253 | 0.40042463 | 0.38526567 | 0.01515896 | 0.43057116 | 0.90650187 |
| Msc | 0.02484234 | 0.04514444 | 0.02998808 | 0.01515636 | 0.01232431 | 0.19887672 |
| Ttc7 | 0.06112235 | 0.08718271 | 0.0720344 | 0.0151483 | 0.1749255 | 0.90650187 |
| Zfp536 | 0.0225064 | 0.04235906 | 0.02721081 | 0.01514825 | 0.00666426 | 0.11876141 |
| Pinx1 | 0.17196868 | 0.20478787 | 0.18963991 | 0.01514795 | 0.36125595 | 0.90650187 |
| Rgl1 | 0.15421629 | 0.18688997 | 0.17177916 | 0.01511081 | 0.34787484 | 0.90650187 |
| Nfe2l3 | 0.06345071 | 0.08975656 | 0.07466487 | 0.01509169 | 0.18444053 | 0.90650187 |
| 1500015O10 | 0.04466884 | 0.06828864 | 0.05321543 | 0.01507321 | 0.10397674 | 0.90650187 |
| Map4 | 0.35584172 | 0.37068223 | 0.35564729 | 0.01503494 | 0.42546288 | 0.90650187 |
| Evl | 0.26640123 | 0.29398853 | 0.278963 | 0.01502553 | 0.40537849 | 0.90650187 |
| Lilrb4a | 0.01406312 | 0.03211023 | 0.01709943 | 0.0150108 | 4.76E-05 | 0.00149446 |
| Lgi4 | 0.04308923 | 0.06638671 | 0.05138719 | 0.01499952 | 0.09720587 | 0.90650187 |
| Pglyrp1 | 0.04209619 | 0.06523161 | 0.05023588 | 0.01499572 | 0.09224756 | 0.90003543 |
| Plek | 0.02258618 | 0.04228769 | 0.02730581 | 0.01498188 | 0.00736049 | 0.12964872 |
| Ptprn | 0.1345499 | 0.16654589 | 0.1515679 | 0.01497799 | 0.33021617 | 0.90650187 |

| | | | | | | |
|---|---|---|---|---|---|---|
| Spata6 | 0.15980802 | 0.19241097 | 0.17744345 | 0.01496752 | 0.35383356 | 0.90650187 |
| Iqck | 0.02565439 | 0.04591378 | 0.03095153 | 0.01496225 | 0.01581537 | 0.24113279 |
| Sertad1 | 0.22362405 | 0.25459495 | 0.23964126 | 0.01495369 | 0.39073374 | 0.90650187 |
| Psmb10 | 0.35118572 | 0.36677732 | 0.35182829 | 0.01494903 | 0.42508877 | 0.90650187 |
| Nudt2 | 0.29799885 | 0.32182475 | 0.30687636 | 0.01494839 | 0.41427878 | 0.90650187 |
| Bik | 0.10576918 | 0.13607129 | 0.12115346 | 0.01491783 | 0.29205365 | 0.90650187 |
| Gipr | 0.26362528 | 0.29138124 | 0.27646558 | 0.01491566 | 0.40522483 | 0.90650187 |
| Nova1 | 0.04306417 | 0.06626812 | 0.05135815 | 0.01490998 | 0.09841765 | 0.90650187 |
| Sec13 | 1.01997945 | 0.77377055 | 0.75886103 | 0.01490952 | 0.4651991 | 0.90650187 |
| Gm16010 | 0.01533053 | 0.03353164 | 0.01862456 | 0.01490707 | 0.00018666 | 0.00524483 |
| Slc43a2 | 0.26333523 | 0.29107057 | 0.27620422 | 0.01486635 | 0.40544452 | 0.90650187 |
| Pcdh9 | 0.10945337 | 0.1399576 | 0.12510365 | 0.01485395 | 0.29880498 | 0.90650187 |
| B3gnt5 | 0.03316503 | 0.05465122 | 0.039813 | 0.01483822 | 0.04877306 | 0.58103391 |
| Fam107b | 0.22041883 | 0.25145761 | 0.23662064 | 0.01483697 | 0.39021617 | 0.90650187 |
| Cul1 | 0.71757662 | 0.62055442 | 0.60572618 | 0.01482824 | 0.4566691 | 0.90650187 |
| Pkp4 | 0.33596636 | 0.35402028 | 0.33921602 | 0.01480427 | 0.42308098 | 0.90650187 |
| Fes | 0.0987004 | 0.12833829 | 0.11353424 | 0.01480405 | 0.28106683 | 0.90650187 |
| Rnf6 | 0.6159434 | 0.55844923 | 0.54364672 | 0.01480251 | 0.4518274 | 0.90650187 |
| Scp2 | 1.44562516 | 0.89834078 | 0.88355341 | 0.01478738 | 0.47034494 | 0.90650187 |
| Tex264 | 0.62163552 | 0.56204305 | 0.5472717 | 0.01477135 | 0.45224518 | 0.90650187 |
| Slc46a1 | 0.04773132 | 0.07151961 | 0.0567491 | 0.01477051 | 0.1236176 | 0.90650187 |
| Synpo | 0.06991969 | 0.09668291 | 0.08193147 | 0.01475144 | 0.21173547 | 0.90650187 |
| Mlx | 0.47887796 | 0.46514525 | 0.45040357 | 0.01474168 | 0.44215633 | 0.90650187 |
| Gmppb | 0.37314113 | 0.3843984 | 0.36968149 | 0.01471692 | 0.4297627 | 0.90650187 |
| Trappc1 | 1.00984139 | 0.76947054 | 0.75475442 | 0.01471612 | 0.46546296 | 0.90650187 |
| Fam155a | 0.08212594 | 0.11014648 | 0.09547775 | 0.01466873 | 0.24730303 | 0.90650187 |
| Gxylt2 | 0.09190042 | 0.12083996 | 0.1061726 | 0.01466735 | 0.26956006 | 0.90650187 |
| Mllt3 | 0.21177427 | 0.24307946 | 0.22842105 | 0.01465841 | 0.38771217 | 0.90650187 |
| Sep-01 | 0.13340966 | 0.16503778 | 0.1503821 | 0.01465568 | 0.33241572 | 0.90650187 |
| Cryba4 | 0.01155584 | 0.02872293 | 0.01407459 | 0.01464835 | 1.86E-06 | 7.61E-05 |

| | | | | | | |
|---|---|---|---|---|---|---|
| Gm28875 | 0.02824423 | 0.0486481 | 0.0340172 | 0.0146309 | 0.02793368 | 0.37677004 |
| Thg1l | 0.07235574 | 0.09927951 | 0.08465208 | 0.01462743 | 0.22119094 | 0.90650187 |
| Rbm4b | 0.95753811 | 0.74702826 | 0.73240272 | 0.01462554 | 0.46463018 | 0.90650187 |
| Gt(ROSA)26S | 0.27207005 | 0.29866489 | 0.28404043 | 0.01462446 | 0.40947642 | 0.90650187 |
| Rab32 | 0.05572435 | 0.08052548 | 0.06590509 | 0.01462038 | 0.1620143 | 0.90650187 |
| Cbx1 | 1.57530269 | 0.91968944 | 0.90507025 | 0.01461918 | 0.47137742 | 0.90650187 |
| Stard9 | 0.07790662 | 0.10543599 | 0.09081939 | 0.0146166 | 0.2371556 | 0.90650187 |
| Zfp251 | 0.09160692 | 0.12044963 | 0.10585341 | 0.01459622 | 0.2699353 | 0.90650187 |
| Dffb | 0.07752186 | 0.10498493 | 0.09039333 | 0.0145916 | 0.23649423 | 0.90650187 |
| Avpr1b | 0.02366347 | 0.04316534 | 0.02858758 | 0.01457776 | 0.01169627 | 0.19056267 |
| Scamp2 | 0.50893171 | 0.48645188 | 0.47187813 | 0.01457375 | 0.44539659 | 0.90650187 |
| Snx2 | 0.92397961 | 0.73092891 | 0.71638729 | 0.01454162 | 0.46404851 | 0.90650187 |
| Rnpep | 0.5107934 | 0.48772339 | 0.47318849 | 0.0145349 | 0.44569112 | 0.90650187 |
| Rab30 | 0.11546072 | 0.1460374 | 0.13150863 | 0.01452877 | 0.31166362 | 0.90650187 |
| Tspan11 | 0.06190434 | 0.08742355 | 0.07291875 | 0.01450481 | 0.18826486 | 0.90650187 |
| Kif2c | 0.06345208 | 0.08915983 | 0.07466641 | 0.01449341 | 0.19408751 | 0.90650187 |
| Cdc25c | 0.07098539 | 0.09758334 | 0.08312272 | 0.01446062 | 0.21964604 | 0.90650187 |
| P2ry6 | 0.01817144 | 0.03647532 | 0.02203375 | 0.01444157 | 0.0017854 | 0.03877115 |
| Dnah2 | 0.02449267 | 0.04400204 | 0.0295729 | 0.01442914 | 0.0150379 | 0.23136755 |
| Gm10334 | 0.02664849 | 0.04653904 | 0.03212953 | 0.01440951 | 0.02308838 | 0.32228332 |
| Wscd1 | 0.07953271 | 0.10702275 | 0.09261768 | 0.01440507 | 0.24464559 | 0.90650187 |
| Slfn5 | 0.02399163 | 0.04337768 | 0.02897766 | 0.01440002 | 0.01358114 | 0.21437892 |
| Gnptab | 0.07203166 | 0.09867855 | 0.08429065 | 0.01438791 | 0.22397463 | 0.90650187 |
| Rcbtb2 | 0.39010204 | 0.39757762 | 0.38320433 | 0.01437329 | 0.43378475 | 0.90650187 |
| Oasl1 | 0.01101709 | 0.0277943 | 0.0134233 | 0.014371 | 9.71E-07 | 4.17E-05 |
| Dusp7 | 0.1226715 | 0.15350225 | 0.1391382 | 0.01436405 | 0.32313645 | 0.90650187 |
| Gm17056 | 0.02043673 | 0.03909602 | 0.02474285 | 0.01435316 | 0.00495625 | 0.0930123 |
| Cyp2d22 | 0.08827142 | 0.11655818 | 0.10221755 | 0.01434063 | 0.26641331 | 0.90650187 |
| Pkn3 | 0.01982006 | 0.03833899 | 0.02400618 | 0.01433281 | 0.00397443 | 0.07696215 |
| Cdca7l | 0.09930569 | 0.12849885 | 0.11418643 | 0.01431241 | 0.28868741 | 0.90650187 |

| | | | | | | |
|---|---|---|---|---|---|---|
| Tmem254c | 0.0135989 | 0.03085044 | 0.01654015 | 0.01431029 | 6.00E-05 | 0.00185646 |
| Zbtb8os | 0.3362332 | 0.35373597 | 0.33943885 | 0.01429712 | 0.42573294 | 0.90650187 |
| Tsix | 0.32765336 | 0.34654029 | 0.33224369 | 0.0142966 | 0.42414664 | 0.90650187 |
| Stard4 | 0.15738724 | 0.18928867 | 0.17499563 | 0.01429304 | 0.35826446 | 0.90650187 |
| Fhl4 | 0.02501987 | 0.04448734 | 0.0301988 | 0.01428854 | 0.01771388 | 0.26555099 |
| Dync2h1 | 0.34894535 | 0.36424849 | 0.3499843 | 0.0142642 | 0.42810996 | 0.90650187 |
| Zcchc12 | 0.03137806 | 0.05196883 | 0.03771267 | 0.01425616 | 0.04642712 | 0.55859172 |
| Dclk1 | 0.05792622 | 0.08266602 | 0.06841049 | 0.01425553 | 0.17712289 | 0.90650187 |
| Gulp1 | 0.12491548 | 0.15575427 | 0.14149957 | 0.0142547 | 0.32713024 | 0.90650187 |
| Gm5239 | 0.03265863 | 0.05347164 | 0.03921831 | 0.01425333 | 0.0530805 | 0.61929297 |
| H2-T22 | 0.25295804 | 0.28103251 | 0.26680026 | 0.01423225 | 0.40627123 | 0.90650187 |
| Homer2 | 0.11222118 | 0.14229101 | 0.1280602 | 0.01423081 | 0.31064529 | 0.90650187 |
| Pigs | 0.48524321 | 0.46923333 | 0.45500317 | 0.01423016 | 0.44471097 | 0.90650187 |
| Sfxn3 | 0.21043846 | 0.24134954 | 0.22714712 | 0.01420242 | 0.39052058 | 0.90650187 |
| Hivep2 | 0.13877222 | 0.17013448 | 0.15594554 | 0.01418895 | 0.34292609 | 0.90650187 |
| Capn15 | 0.14032995 | 0.17173768 | 0.15755526 | 0.01418242 | 0.34451433 | 0.90650187 |
| Zdhhc20 | 0.64402768 | 0.57553451 | 0.56135714 | 0.01417738 | 0.45530222 | 0.90650187 |
| Gkap1 | 0.36548496 | 0.37767361 | 0.36350067 | 0.01417294 | 0.43119415 | 0.90650187 |
| C1rb | 0.04969989 | 0.0731751 | 0.05901303 | 0.01416207 | 0.14301611 | 0.90650187 |
| Cwc27 | 0.27219297 | 0.29828357 | 0.28415016 | 0.01413341 | 0.41249908 | 0.90650187 |
| Zfhx4 | 0.16444326 | 0.19624497 | 0.18211193 | 0.01413304 | 0.3650445 | 0.90650187 |
| Thsd1 | 0.05366108 | 0.07767888 | 0.06355082 | 0.01412806 | 0.1615009 | 0.90650187 |
| Hist1h1e | 0.1999214 | 0.23116948 | 0.21705213 | 0.01411735 | 0.38623407 | 0.90650187 |
| Brip1 | 0.02691801 | 0.04656371 | 0.03244863 | 0.01411508 | 0.02656712 | 0.36251953 |
| Slc29a3 | 0.05920108 | 0.08396977 | 0.0698578 | 0.01411197 | 0.18457775 | 0.90650187 |
| S100a4 | 0.05649679 | 0.08089205 | 0.06678484 | 0.01410722 | 0.17384937 | 0.90650187 |
| Rragd | 0.06763805 | 0.09343276 | 0.07937549 | 0.01405726 | 0.21555041 | 0.90650187 |
| Med11 | 0.21594887 | 0.24642817 | 0.23239037 | 0.0140378 | 0.39414112 | 0.90650187 |
| Zfp229 | 0.05064325 | 0.07412685 | 0.06009585 | 0.014031 | 0.14964737 | 0.90650187 |
| Hps6 | 0.09203181 | 0.12032237 | 0.10631545 | 0.01400692 | 0.27903766 | 0.90650187 |

| | | | | | | |
|---|---|---|---|---|---|---|
| Ahi1 | 0.36696628 | 0.37869607 | 0.36470038 | 0.01399569 | 0.43226757 | 0.90650187 |
| Gm6710 | 0.1551821 | 0.18675068 | 0.17276006 | 0.01399062 | 0.35941668 | 0.90650187 |
| Slc7a7 | 0.05831706 | 0.08284295 | 0.06885446 | 0.01398849 | 0.18321771 | 0.90650187 |
| Bmx | 0.01734606 | 0.03501192 | 0.0210446 | 0.01396731 | 0.00158604 | 0.03499177 |
| Qprt | 0.04467928 | 0.06719074 | 0.0532275 | 0.01396324 | 0.1217624 | 0.90650187 |
| Terf2ip | 0.34644105 | 0.36186911 | 0.34791815 | 0.01395096 | 0.42925874 | 0.90650187 |
| Sertad2 | 0.35336838 | 0.36754543 | 0.3536208 | 0.01392463 | 0.43051784 | 0.90650187 |
| Rnf13 | 0.40025359 | 0.40511055 | 0.39118913 | 0.01392142 | 0.43714655 | 0.90650187 |
| Aspa | 0.01219001 | 0.0287571 | 0.01484064 | 0.01391646 | 1.53E-05 | 0.0005351 |
| Itga2 | 0.01232435 | 0.02891664 | 0.01500282 | 0.01391382 | 1.87E-05 | 0.00063611 |
| Etohi1 | 0.02691893 | 0.04635067 | 0.03244973 | 0.01390095 | 0.02842728 | 0.38274854 |
| Alg14 | 0.25447582 | 0.28207586 | 0.26818216 | 0.0138937 | 0.40892417 | 0.90650187 |
| Tox | 0.43471265 | 0.43158702 | 0.41770148 | 0.01388554 | 0.44125658 | 0.90650187 |
| Acaca | 0.17198008 | 0.20353364 | 0.18965127 | 0.01388237 | 0.37243313 | 0.90650187 |
| Serpina1a | 0.02392988 | 0.04277352 | 0.02890427 | 0.01386925 | 0.0164561 | 0.2489047 |
| Ppt1 | 0.51405612 | 0.48934833 | 0.47547956 | 0.01386878 | 0.44841402 | 0.90650187 |
| Mrps25 | 0.40366999 | 0.40771919 | 0.39385794 | 0.01386125 | 0.43783658 | 0.90650187 |
| Mutyh | 0.0333402 | 0.05386684 | 0.04001861 | 0.01384823 | 0.0619781 | 0.6859224 |
| Azin1 | 0.60698915 | 0.55175173 | 0.53790533 | 0.01384641 | 0.45444614 | 0.90650187 |
| Tlr4 | 0.03284828 | 0.05328565 | 0.03944107 | 0.01384459 | 0.05932145 | 0.66402454 |
| Zdhhc9 | 0.10242956 | 0.13138982 | 0.11755803 | 0.01383179 | 0.30046398 | 0.90650187 |
| BC022687 | 0.23703864 | 0.26599337 | 0.25216782 | 0.01382555 | 0.40371675 | 0.90650187 |
| Flot1 | 0.29725191 | 0.32004926 | 0.30622731 | 0.01382195 | 0.42048301 | 0.90650187 |
| Adamts10 | 0.22704567 | 0.25665543 | 0.24285359 | 0.01380184 | 0.4002684 | 0.90650187 |
| Akr1b8 | 0.07941187 | 0.10628321 | 0.09248418 | 0.01379904 | 0.25356806 | 0.90650187 |
| Slc7a8 | 0.15964561 | 0.19106194 | 0.17727944 | 0.0137825 | 0.36481362 | 0.90650187 |
| Spata1 | 0.04623071 | 0.06877079 | 0.05501939 | 0.01375139 | 0.13325783 | 0.90650187 |
| Ifit2 | 0.05005243 | 0.07315937 | 0.05941785 | 0.01374151 | 0.151944 | 0.90650187 |
| 1700034P13I | 0.0117798 | 0.02808498 | 0.0143452 | 0.01373978 | 1.03E-05 | 0.00036977 |
| Emg1 | 1.15348557 | 0.82140498 | 0.80767851 | 0.01372647 | 0.46988738 | 0.90650187 |

| | | | | | | |
|---|---|---|---|---|---|---|
| Slc33a1 | 0.32617768 | 0.3447141 | 0.33099984 | 0.01371426 | 0.42693061 | 0.90650187 |
| Zbtb49 | 0.04226061 | 0.06413745 | 0.05042661 | 0.01371084 | 0.11338017 | 0.90650187 |
| Tbx2 | 0.08861758 | 0.1162987 | 0.1025956 | 0.0137031 | 0.2763317 | 0.90650187 |
| mt-Nd6 | 0.15950496 | 0.19080837 | 0.17713737 | 0.013671 | 0.3657615 | 0.90650187 |
| Dbf4 | 0.13158467 | 0.16214148 | 0.14848096 | 0.01366052 | 0.34126842 | 0.90650187 |
| Tns2 | 0.12382367 | 0.15401183 | 0.14035142 | 0.0136604 | 0.33261969 | 0.90650187 |
| H2-T23 | 0.14705285 | 0.17811204 | 0.16446996 | 0.01364208 | 0.35615927 | 0.90650187 |
| Cwf19l2 | 0.36575929 | 0.3773632 | 0.36372298 | 0.01364022 | 0.43379608 | 0.90650187 |
| Tmem160 | 0.70491857 | 0.6118839 | 0.59828881 | 0.01359509 | 0.45976778 | 0.90650187 |
| Lsm5 | 0.98376368 | 0.75753551 | 0.74394146 | 0.01359405 | 0.46762769 | 0.90650187 |
| Clcn5 | 0.27893104 | 0.30372728 | 0.29014334 | 0.01358393 | 0.41756199 | 0.90650187 |
| Mfsd1 | 0.26860137 | 0.29450477 | 0.28093721 | 0.01356756 | 0.41500229 | 0.90650187 |
| Tmlhe | 0.01528792 | 0.03213552 | 0.01857334 | 0.01356219 | 0.00058485 | 0.01491667 |
| Smc6 | 1.14087054 | 0.81710362 | 0.80354292 | 0.0135607 | 0.47009754 | 0.90650187 |
| Pccb | 0.30238618 | 0.32422588 | 0.31067861 | 0.01354727 | 0.4231457 | 0.90650187 |
| Foxo3 | 0.25196485 | 0.27943831 | 0.26589478 | 0.01354353 | 0.41042965 | 0.90650187 |
| Tnfaip8l1 | 0.07801508 | 0.10444042 | 0.09093945 | 0.01350097 | 0.254627 | 0.90650187 |
| Slc31a2 | 0.29540593 | 0.31811547 | 0.30462071 | 0.01349477 | 0.42193671 | 0.90650187 |
| Gimap9 | 0.07883397 | 0.10532731 | 0.0918454 | 0.01348191 | 0.25701779 | 0.90650187 |
| Ercc1 | 0.30093351 | 0.32290169 | 0.30942157 | 0.01348012 | 0.42321507 | 0.90650187 |
| Cgnl1 | 0.16344884 | 0.19458731 | 0.18111242 | 0.01347489 | 0.37041431 | 0.90650187 |
| Angptl6 | 0.09426177 | 0.12220024 | 0.10873625 | 0.013464 | 0.29099982 | 0.90650187 |
| Hoxd3os1 | 0.02160099 | 0.03958173 | 0.026132 | 0.01344972 | 0.01106636 | 0.18185916 |
| Il6ra | 0.15647005 | 0.18748579 | 0.17406646 | 0.01341932 | 0.36590247 | 0.90650187 |
| Fam177a | 0.43947943 | 0.43470414 | 0.42129917 | 0.01340496 | 0.44375683 | 0.90650187 |
| Ruvbl2 | 0.51362683 | 0.48858133 | 0.4751785 | 0.01340283 | 0.4501065 | 0.90650187 |
| Gm13470 | 0.03717686 | 0.05787539 | 0.0445102 | 0.0133652 | 0.09095545 | 0.89373869 |
| Fbrs | 0.17578265 | 0.20678384 | 0.19343072 | 0.01335312 | 0.37946204 | 0.90650187 |
| Abhd5 | 0.21520187 | 0.24502532 | 0.23168142 | 0.0133439 | 0.39895658 | 0.90650187 |
| Tubg1 | 0.57341946 | 0.52930798 | 0.51596666 | 0.01334133 | 0.45424244 | 0.90650187 |

| | | | | | | |
|---|---|---|---|---|---|---|
| Plekhg4 | 0.01067686 | 0.02634604 | 0.01301175 | 0.0133343 | 2.61E-06 | 0.00010438 |
| Ppp1r11 | 0.9140499 | 0.72486495 | 0.71153324 | 0.01333171 | 0.46680827 | 0.90650187 |
| Smim15 | 0.56487501 | 0.52359559 | 0.51027513 | 0.01332046 | 0.45380637 | 0.90650187 |
| Lgr4 | 0.13791334 | 0.16837543 | 0.15505677 | 0.01331866 | 0.35128374 | 0.90650187 |
| Ctrc | 0.01289595 | 0.02900536 | 0.0156926 | 0.01331275 | 8.12E-05 | 0.00243827 |
| Morn5 | 0.01702093 | 0.03396665 | 0.02065467 | 0.01331198 | 0.0020837 | 0.04417626 |
| Mxra7 | 0.29137412 | 0.31440174 | 0.30110061 | 0.01330113 | 0.42215451 | 0.90650187 |
| Bvht | 0.05181201 | 0.07471994 | 0.06143554 | 0.0132844 | 0.16820381 | 0.90650187 |
| Ccdc184 | 0.12874593 | 0.15875513 | 0.14551585 | 0.01323928 | 0.34293401 | 0.90650187 |
| Elfn1 | 0.02570115 | 0.04423924 | 0.03100696 | 0.01323228 | 0.0289027 | 0.38709249 |
| Nr0b2 | 0.03389205 | 0.05389702 | 0.04066608 | 0.01323094 | 0.07403204 | 0.77635035 |
| Irgm1 | 0.08077691 | 0.10721956 | 0.0939911 | 0.01322846 | 0.26576181 | 0.90650187 |
| Ska3 | 0.04915171 | 0.07157983 | 0.05838319 | 0.01319664 | 0.15748152 | 0.90650187 |
| Wwc2 | 0.20334795 | 0.23354993 | 0.22035393 | 0.013196 | 0.39503206 | 0.90650187 |
| Gja5 | 0.01367901 | 0.02982221 | 0.01663669 | 0.01318552 | 0.00021303 | 0.00593082 |
| St6galnac4 | 0.20798037 | 0.23797749 | 0.22479805 | 0.01317945 | 0.39718633 | 0.90650187 |
| Abhd12 | 0.32039081 | 0.33927625 | 0.3261037 | 0.01317255 | 0.42874321 | 0.90650187 |
| Ncs1 | 0.1835814 | 0.21430107 | 0.20113144 | 0.01316963 | 0.38549192 | 0.90650187 |
| Dlst | 0.41770281 | 0.41789228 | 0.40472746 | 0.01316482 | 0.44251138 | 0.90650187 |
| Mir143hg | 0.05937537 | 0.08321939 | 0.07005547 | 0.01316391 | 0.20175741 | 0.90650187 |
| C1qtnf9 | 0.02183983 | 0.03957078 | 0.02641672 | 0.01315406 | 0.0134259 | 0.21259066 |
| Enah | 0.61696752 | 0.55745376 | 0.54430052 | 0.01315324 | 0.45722399 | 0.90650187 |
| Rhbdf1 | 0.21732158 | 0.24682434 | 0.23369165 | 0.01313269 | 0.40136009 | 0.90650187 |
| Zfp235 | 0.11835024 | 0.14769161 | 0.13457359 | 0.01311802 | 0.33238003 | 0.90650187 |
| Papss1 | 0.85777495 | 0.69612175 | 0.6830175 | 0.01310425 | 0.46601438 | 0.90650187 |
| Bub1b | 0.07061829 | 0.09580463 | 0.08271256 | 0.01309207 | 0.24081976 | 0.90650187 |
| Asah2 | 0.03499941 | 0.05505491 | 0.04196386 | 0.01309105 | 0.08274348 | 0.83684877 |
| Slc8b1 | 0.0650174 | 0.08951355 | 0.07643036 | 0.01308319 | 0.22333156 | 0.90650187 |
| Abcg1 | 0.16922872 | 0.19998615 | 0.18690634 | 0.01307981 | 0.37786 | 0.90650187 |
| Dpagt1 | 0.19637808 | 0.22669345 | 0.21362426 | 0.01306919 | 0.39282061 | 0.90650187 |

| | | | | | | |
|---|---|---|---|---|---|---|
| Ddx55 | 0.2690027 | 0.29435484 | 0.28129683 | 0.01305801 | 0.41825159 | 0.90650187 |
| Lrrc41 | 0.46375342 | 0.45238249 | 0.43936161 | 0.01302089 | 0.44759116 | 0.90650187 |
| Mdh1 | 1.75303715 | 0.93480208 | 0.92178396 | 0.01301813 | 0.47496918 | 0.90650187 |
| Bnip2 | 0.55764948 | 0.51843732 | 0.5054265 | 0.01301082 | 0.45444451 | 0.90650187 |
| Gm10146 | 0.18116208 | 0.21174677 | 0.19874965 | 0.01299711 | 0.38563362 | 0.90650187 |
| Slc25a53 | 0.28142636 | 0.3053473 | 0.29235211 | 0.0129952 | 0.42167534 | 0.90650187 |
| Mboat1 | 0.08672672 | 0.11352062 | 0.10052847 | 0.01299215 | 0.28280004 | 0.90650187 |
| Svep1 | 0.1118269 | 0.14063017 | 0.12763962 | 0.01299055 | 0.32547144 | 0.90650187 |
| Zswim7 | 0.19167739 | 0.22202996 | 0.20905564 | 0.01297432 | 0.39131193 | 0.90650187 |
| Eya2 | 0.06579223 | 0.09027215 | 0.07730217 | 0.01296997 | 0.22786505 | 0.90650187 |
| Acbd7 | 0.0290262 | 0.04790846 | 0.03494076 | 0.0129677 | 0.04948011 | 0.58758681 |
| Irf2bp2 | 0.52078778 | 0.49315028 | 0.48018405 | 0.01296624 | 0.4522245 | 0.90650187 |
| Nfil3 | 0.13197861 | 0.16185699 | 0.14889167 | 0.01296532 | 0.34933427 | 0.90650187 |
| Abca9 | 0.03168619 | 0.05103788 | 0.03807519 | 0.01296269 | 0.06507561 | 0.70962289 |
| Zfp266 | 0.37021385 | 0.38025061 | 0.36732437 | 0.01292623 | 0.43784234 | 0.90650187 |
| Cnp | 0.29566994 | 0.31777563 | 0.3048507 | 0.01292493 | 0.42524904 | 0.90650187 |
| Gclm | 0.47920301 | 0.46355868 | 0.45063912 | 0.01291955 | 0.44929099 | 0.90650187 |
| Ccl24 | 0.01707579 | 0.03362526 | 0.02072048 | 0.01290478 | 0.0028137 | 0.05720839 |
| Slc35a2 | 0.42947626 | 0.42662583 | 0.41373035 | 0.01289548 | 0.44489725 | 0.90650187 |
| Prss12 | 0.07146058 | 0.09654817 | 0.08365336 | 0.01289481 | 0.24658635 | 0.90650187 |
| Sh2d3c | 0.32168154 | 0.3400607 | 0.32719848 | 0.01286222 | 0.43063512 | 0.90650187 |
| Tyk2 | 0.05216685 | 0.074701 | 0.06184186 | 0.01285914 | 0.17763882 | 0.90650187 |
| Igsf8 | 0.60160745 | 0.54726474 | 0.53443287 | 0.01283187 | 0.45749756 | 0.90650187 |
| Zcchc9 | 0.6934374 | 0.6042975 | 0.59147402 | 0.01282347 | 0.46160815 | 0.90650187 |
| Tbxas1 | 0.01029634 | 0.02534281 | 0.01255124 | 0.01279157 | 2.94E-06 | 0.00011596 |
| Tmod2 | 0.27084631 | 0.29571966 | 0.28294691 | 0.01277275 | 0.42047324 | 0.90650187 |
| Deptor | 0.2866073 | 0.30968612 | 0.29691976 | 0.01276636 | 0.42420667 | 0.90650187 |
| Numbl | 0.21245928 | 0.24183765 | 0.22907361 | 0.01276404 | 0.4021793 | 0.90650187 |
| Sptlc2 | 0.38946221 | 0.3954582 | 0.38269828 | 0.01275992 | 0.44108249 | 0.90650187 |
| Fam72a | 0.03817552 | 0.05842443 | 0.04567557 | 0.01274886 | 0.10733097 | 0.90650187 |

| | | | | | | |
|---|---|---|---|---|---|---|
| L1cam | 0.03164768 | 0.05077637 | 0.03802989 | 0.01274648 | 0.06810768 | 0.73292957 |
| Fam129a | 0.05452841 | 0.07726936 | 0.06454125 | 0.01272811 | 0.19032124 | 0.90650187 |
| Itpripl2 | 0.10191937 | 0.12972554 | 0.11700753 | 0.01271801 | 0.31447315 | 0.90650187 |
| Tll1 | 0.01061249 | 0.02564059 | 0.01293387 | 0.01270672 | 6.28E-06 | 0.00023125 |
| Fmnl3 | 0.09642696 | 0.12378599 | 0.11108014 | 0.01270585 | 0.30554828 | 0.90650187 |
| Thnsl2 | 0.04321074 | 0.06422249 | 0.05152796 | 0.01269453 | 0.13670835 | 0.90650187 |
| Isl2 | 0.02239224 | 0.03974452 | 0.02707486 | 0.01266966 | 0.01874821 | 0.27607614 |
| Epb41l4aos | 0.34234014 | 0.35718785 | 0.34452228 | 0.01266558 | 0.43508892 | 0.90650187 |
| Tmem41a | 0.15056635 | 0.1807254 | 0.16806255 | 0.01266285 | 0.36882807 | 0.90650187 |
| CAAA011473 | 0.02049407 | 0.03745059 | 0.02481132 | 0.01263927 | 0.01176674 | 0.19120241 |
| Bdnf | 0.09914656 | 0.12665234 | 0.11401502 | 0.01263732 | 0.31109573 | 0.90650187 |
| Il16 | 0.02359054 | 0.04109738 | 0.02850086 | 0.01259651 | 0.0247174 | 0.34157204 |
| Srxn1 | 0.09956791 | 0.12706337 | 0.11446882 | 0.01259456 | 0.3123739 | 0.90650187 |
| Zbtb7a | 0.74405315 | 0.63360706 | 0.62102472 | 0.01258234 | 0.46411552 | 0.90650187 |
| Prim1 | 0.20442851 | 0.23397138 | 0.22139257 | 0.01257881 | 0.40029448 | 0.90650187 |
| Rnf170 | 0.21682509 | 0.24578907 | 0.23322121 | 0.01256786 | 0.40533358 | 0.90650187 |
| Msx1 | 0.05622808 | 0.07904048 | 0.0664789 | 0.01256157 | 0.20044774 | 0.90650187 |
| D830030K20 | 0.01997857 | 0.03675292 | 0.02419559 | 0.01255733 | 0.01052083 | 0.17440259 |
| Lpgat1 | 0.30566681 | 0.326054 | 0.31351051 | 0.01254349 | 0.4294134 | 0.90650187 |
| Ssx2ip | 0.16121196 | 0.19139936 | 0.17886003 | 0.01253934 | 0.3776464 | 0.90650187 |
| Tatdn3 | 0.09244073 | 0.11928023 | 0.10675988 | 0.01252034 | 0.30105741 | 0.90650187 |
| St3gal1 | 0.1390456 | 0.16873969 | 0.15622825 | 0.01251144 | 0.36091151 | 0.90650187 |
| Plekhb2 | 0.14559379 | 0.17548396 | 0.16297375 | 0.01251021 | 0.36645614 | 0.90650187 |
| Rexo2 | 0.80804368 | 0.66913453 | 0.65662728 | 0.01250725 | 0.46625843 | 0.90650187 |
| 1810043G02 | 0.44714615 | 0.43955644 | 0.42705036 | 0.01250608 | 0.44820859 | 0.90650187 |
| Lnx1 | 0.08502354 | 0.11116544 | 0.09866221 | 0.01250323 | 0.28658773 | 0.90650187 |
| Ptprs | 0.50414591 | 0.48097049 | 0.46849927 | 0.01247121 | 0.45289922 | 0.90650187 |
| Cmtm5 | 0.01866855 | 0.03509984 | 0.02262895 | 0.01247089 | 0.00714305 | 0.12611093 |
| Slc37a3 | 0.31578205 | 0.33463469 | 0.3221823 | 0.01245239 | 0.43178821 | 0.90650187 |
| E2f2 | 0.03371883 | 0.052909 | 0.04046289 | 0.01244611 | 0.08574404 | 0.85918035 |

| | | | | | | |
|---|---|---|---|---|---|---|
| Clip3 | 0.3367453 | 0.35230781 | 0.33986633 | 0.01244149 | 0.43536145 | 0.90650187 |
| Plcg2 | 0.01509617 | 0.0307753 | 0.01834274 | 0.01243255 | 0.00129257 | 0.0294137 |
| Rfc3 | 0.35013252 | 0.36339386 | 0.35096194 | 0.01243191 | 0.43743563 | 0.90650187 |
| Sgol1 | 0.08242616 | 0.10821797 | 0.09580824 | 0.01240973 | 0.28236676 | 0.90650187 |
| Slc16a1 | 0.09197046 | 0.1186405 | 0.10624875 | 0.01239175 | 0.30205848 | 0.90650187 |
| Gm26772 | 0.0135757 | 0.02888773 | 0.01651219 | 0.01237553 | 0.0004313 | 0.0113625 |
| Nrxn3 | 0.04256657 | 0.06315316 | 0.05078142 | 0.01237175 | 0.13938834 | 0.90650187 |
| Cpsf3l | 0.40338896 | 0.40601012 | 0.39363874 | 0.01237138 | 0.44444155 | 0.90650187 |
| Itgam | 0.01588227 | 0.03165667 | 0.01928769 | 0.01236899 | 0.00217965 | 0.04576364 |
| Mrpl41 | 0.93672758 | 0.7349202 | 0.72256371 | 0.01235649 | 0.46970004 | 0.90650187 |
| Ung | 0.14317797 | 0.17284292 | 0.16049097 | 0.01235195 | 0.36611919 | 0.90650187 |
| St6gal1 | 0.15006604 | 0.17990059 | 0.1675519 | 0.01234868 | 0.37159041 | 0.90650187 |
| Prkd3 | 0.23099332 | 0.25889293 | 0.24654507 | 0.01234786 | 0.4119032 | 0.90650187 |
| Mtmr11 | 0.04067438 | 0.06090472 | 0.04858485 | 0.01231987 | 0.12980958 | 0.90650187 |
| Fbxo6 | 0.22611625 | 0.25428904 | 0.2419822 | 0.01230685 | 0.41056308 | 0.90650187 |
| Angptl4 | 0.11776246 | 0.14624352 | 0.13395096 | 0.01229257 | 0.341648 | 0.90650187 |
| Hes7 | 0.02946753 | 0.0477114 | 0.03546158 | 0.01224983 | 0.06230749 | 0.68782581 |
| Dhcr24 | 0.14819572 | 0.17787851 | 0.16564021 | 0.0122383 | 0.37128083 | 0.90650187 |
| Bcl2a1b | 0.01368526 | 0.02887971 | 0.01664422 | 0.01223549 | 0.00054145 | 0.01390299 |
| Tcaf1 | 0.57950704 | 0.53222258 | 0.51999453 | 0.01222805 | 0.45836975 | 0.90650187 |
| Edaradd | 0.01958973 | 0.03595704 | 0.02373087 | 0.01222618 | 0.01099958 | 0.18115337 |
| Ifngr1 | 0.34633671 | 0.36004991 | 0.34783195 | 0.01221796 | 0.4379548 | 0.90650187 |
| Gm4631 | 0.12495102 | 0.15375366 | 0.14153691 | 0.01221674 | 0.35059223 | 0.90650187 |
| Ntrk1 | 0.12081304 | 0.14938702 | 0.13717787 | 0.01220915 | 0.34617446 | 0.90650187 |
| Taf13 | 0.3107157 | 0.33004502 | 0.31785018 | 0.01219484 | 0.4322837 | 0.90650187 |
| Ptk2 | 0.1830377 | 0.21277145 | 0.20059673 | 0.01217472 | 0.39365124 | 0.90650187 |
| Dapk1 | 0.36724301 | 0.37709797 | 0.3649243 | 0.01217367 | 0.44105187 | 0.90650187 |
| Kdm7a | 0.49745282 | 0.47592208 | 0.46374868 | 0.0121734 | 0.45355002 | 0.90650187 |
| 2310010G23 | 0.0260396 | 0.04353702 | 0.03140818 | 0.01212884 | 0.04301229 | 0.52685717 |
| Tmbim1 | 0.39584131 | 0.39985661 | 0.38772914 | 0.01212747 | 0.4447051 | 0.90650187 |

| | | | | | | |
|---|---|---|---|---|---|---|
| Gng3 | 0.0435059 | 0.0639875 | 0.05186981 | 0.01211768 | 0.14950227 | 0.90650187 |
| Aktip | 0.41151145 | 0.41205294 | 0.39995001 | 0.01210293 | 0.44649217 | 0.90650187 |
| Tufm | 0.41475922 | 0.41455207 | 0.40245967 | 0.0120924 | 0.44686982 | 0.90650187 |
| Ccng1 | 0.28174516 | 0.30470019 | 0.29263388 | 0.01206632 | 0.42727823 | 0.90650187 |
| Sec14l1 | 0.37677564 | 0.38466015 | 0.37259837 | 0.01206179 | 0.44278437 | 0.90650187 |
| Lepr | 0.08051747 | 0.1057637 | 0.0937049 | 0.0120588 | 0.28362675 | 0.90650187 |
| Magi3 | 0.12076499 | 0.14918113 | 0.13712713 | 0.012054 | 0.34797774 | 0.90650187 |
| Smg6 | 0.40042988 | 0.40337859 | 0.39132706 | 0.01205153 | 0.44555121 | 0.90650187 |
| Cacna1a | 0.42633453 | 0.42337467 | 0.41133814 | 0.01203654 | 0.44824884 | 0.90650187 |
| Plcd1 | 0.12708349 | 0.15579193 | 0.14377494 | 0.01201699 | 0.35510557 | 0.90650187 |
| Ch25h | 0.02502378 | 0.04221636 | 0.03020344 | 0.01201292 | 0.03851651 | 0.48267882 |
| Fam13a | 0.07323996 | 0.09764398 | 0.08563747 | 0.01200651 | 0.26654997 | 0.90650187 |
| AU019823 | 0.36961828 | 0.3788471 | 0.36684379 | 0.0120033 | 0.44217292 | 0.90650187 |
| Fance | 0.22474778 | 0.25269594 | 0.24069757 | 0.01199837 | 0.41231033 | 0.90650187 |
| Dand5 | 0.07827634 | 0.10319952 | 0.0912286 | 0.01197092 | 0.27982956 | 0.90650187 |
| Pik3r6 | 0.02767144 | 0.0452995 | 0.03334008 | 0.01195942 | 0.05539153 | 0.63691881 |
| Qsox1 | 0.47188102 | 0.45727389 | 0.44531575 | 0.01195814 | 0.45248771 | 0.90650187 |
| Trappc6b | 0.91291459 | 0.72292293 | 0.71097165 | 0.01195128 | 0.47021489 | 0.90650187 |
| Snrnp25 | 0.25808542 | 0.28341032 | 0.27145967 | 0.01195066 | 0.42241779 | 0.90650187 |
| Hoxb7 | 0.06671201 | 0.09028027 | 0.07833594 | 0.01194433 | 0.2489343 | 0.90650187 |
| Rassf8 | 0.12934037 | 0.15807661 | 0.14613756 | 0.01193905 | 0.35822987 | 0.90650187 |
| Vangl1 | 0.18767517 | 0.21708546 | 0.2051472 | 0.01193827 | 0.39793098 | 0.90650187 |
| Rem2 | 0.256038 | 0.28151322 | 0.26960215 | 0.01191106 | 0.4221456 | 0.90650187 |
| Abcf2 | 0.4203962 | 0.41870616 | 0.40679666 | 0.0119095 | 0.44822354 | 0.90650187 |
| Ip6k2 | 0.73238181 | 0.62623045 | 0.61432266 | 0.01190779 | 0.46566488 | 0.90650187 |
| Cdca5 | 0.03535851 | 0.05427787 | 0.0423843 | 0.01189356 | 0.10610846 | 0.90650187 |
| 1810062O18 | 0.04156673 | 0.06151437 | 0.04962143 | 0.01189294 | 0.1433263 | 0.90650187 |
| Smarcc1 | 0.87918419 | 0.7059298 | 0.69403776 | 0.01189204 | 0.46964051 | 0.90650187 |
| Prr15l | 0.24609107 | 0.27240816 | 0.26051974 | 0.01188842 | 0.4196189 | 0.90650187 |
| Bri3 | 1.20423236 | 0.8355867 | 0.82370603 | 0.01188067 | 0.47443697 | 0.90650187 |

| | | | | | | |
|---|---|---|---|---|---|---|
| Sstr5 | 0.01171756 | 0.02613865 | 0.01427001 | 0.01186865 | 0.00010887 | 0.00318861 |
| Anp32e | 0.88057526 | 0.70661086 | 0.69474699 | 0.01186387 | 0.46974315 | 0.90650187 |
| Lix1l | 0.1772193 | 0.20671577 | 0.19485433 | 0.01186144 | 0.39334201 | 0.90650187 |
| Ptpre | 0.01984996 | 0.03589056 | 0.02404191 | 0.01184865 | 0.01422796 | 0.22228087 |
| Muc16 | 0.02938158 | 0.04720452 | 0.03536016 | 0.01184435 | 0.0682286 | 0.73296321 |
| Zc3hav1 | 0.08981013 | 0.11571924 | 0.10389675 | 0.01182249 | 0.30647434 | 0.90650187 |
| Ly96 | 0.14300433 | 0.17213201 | 0.16031226 | 0.01181975 | 0.37154187 | 0.90650187 |
| Cyp39a1 | 0.10195968 | 0.12885431 | 0.11705104 | 0.01180328 | 0.32697219 | 0.90650187 |
| Dusp18 | 0.2926667 | 0.31402274 | 0.30223074 | 0.011792 | 0.43114799 | 0.90650187 |
| Bmp4 | 0.08326401 | 0.10850819 | 0.0967299 | 0.01177829 | 0.29414423 | 0.90650187 |
| Cep70 | 0.29053673 | 0.31214576 | 0.30036765 | 0.01177811 | 0.43080603 | 0.90650187 |
| Rnf31 | 0.10547694 | 0.1326118 | 0.12083939 | 0.01177241 | 0.33247119 | 0.90650187 |
| Shprh | 0.21233446 | 0.24072459 | 0.22895474 | 0.01176985 | 0.40961525 | 0.90650187 |
| Gm10941 | 0.21994294 | 0.24793854 | 0.23617124 | 0.0117673 | 0.41235019 | 0.90650187 |
| Dcaf7 | 0.4451641 | 0.43732907 | 0.42556801 | 0.01176106 | 0.4511092 | 0.90650187 |
| Evi2a | 0.01987781 | 0.03583564 | 0.02407519 | 0.01176045 | 0.01494194 | 0.23049408 |
| Iqgap3 | 0.03929052 | 0.05872365 | 0.04697489 | 0.01174876 | 0.1330953 | 0.90650187 |
| Nudcd2 | 0.75865588 | 0.64106043 | 0.62931724 | 0.01174318 | 0.4669433 | 0.90650187 |
| Cyb5rl | 0.14098222 | 0.16996044 | 0.15822845 | 0.011732 | 0.37084219 | 0.90650187 |
| Nip7 | 0.43785318 | 0.43179415 | 0.42007362 | 0.01172054 | 0.45064279 | 0.90650187 |
| Sp100 | 0.02959007 | 0.04732629 | 0.03560613 | 0.01172016 | 0.07168964 | 0.75994015 |
| Ino80b | 0.51964436 | 0.49109287 | 0.47938726 | 0.01170561 | 0.45677883 | 0.90650187 |
| Dock10 | 0.07775768 | 0.10234747 | 0.09065448 | 0.01169298 | 0.28318275 | 0.90650187 |
| Mxd4 | 0.64641355 | 0.57453403 | 0.56284125 | 0.01169278 | 0.46320779 | 0.90650187 |
| Slc46a3 | 0.02023488 | 0.03618566 | 0.02450179 | 0.01168387 | 0.01700632 | 0.25620715 |
| Ptprm | 0.04899662 | 0.06988829 | 0.05820492 | 0.01168338 | 0.18610024 | 0.90650187 |
| Spred2 | 0.10754677 | 0.13472445 | 0.12306151 | 0.01166294 | 0.33675898 | 0.90650187 |
| Tab3 | 0.05823564 | 0.08042159 | 0.06876199 | 0.0116596 | 0.22547947 | 0.90650187 |
| Ephb6 | 0.039211 | 0.05854177 | 0.04688229 | 0.01165948 | 0.13444796 | 0.90650187 |
| Rfx6 | 0.30675105 | 0.32609725 | 0.31444434 | 0.0116529 | 0.43457099 | 0.90650187 |

| | | | | | | |
|---|---|---|---|---|---|---|
| Neil3 | 0.05980255 | 0.0821883 | 0.07053979 | 0.0116485 | 0.23143789 | 0.90650187 |
| Slc22a21 | 0.01789034 | 0.03333544 | 0.02169699 | 0.01163844 | 0.00854748 | 0.14714629 |
| Ttc12 | 0.04237095 | 0.06218656 | 0.05055459 | 0.01163196 | 0.15318298 | 0.90650187 |
| Gm4890 | 0.07697483 | 0.10141704 | 0.08978721 | 0.01162983 | 0.28236706 | 0.90650187 |
| Pappa | 0.07934076 | 0.10402667 | 0.0924056 | 0.01162107 | 0.2880515 | 0.90650187 |
| Cdkl1 | 0.08461067 | 0.1098224 | 0.09820919 | 0.01161321 | 0.29955127 | 0.90650187 |
| Lmntd1 | 0.04188006 | 0.06159372 | 0.04998511 | 0.01160861 | 0.1509299 | 0.90650187 |
| Psat1 | 0.18545787 | 0.21457276 | 0.2029744 | 0.01159836 | 0.39973592 | 0.90650187 |
| Ccdc67 | 0.01181402 | 0.02597538 | 0.01438654 | 0.01158884 | 0.00017109 | 0.00486106 |
| Nfe2l2 | 0.66997421 | 0.58891979 | 0.57733394 | 0.01158586 | 0.46445599 | 0.90650187 |
| Gnpnat1 | 0.49141481 | 0.47102205 | 0.45943682 | 0.01158523 | 0.45537162 | 0.90650187 |
| Zg16 | 0.01284842 | 0.02721577 | 0.01563526 | 0.01158051 | 0.00049613 | 0.01289414 |
| Sult1d1 | 0.07624389 | 0.10054652 | 0.08897665 | 0.01156986 | 0.28160721 | 0.90650187 |
| A830018L16I | 0.02922273 | 0.04673757 | 0.03517272 | 0.01156485 | 0.0719086 | 0.76173001 |
| Exoc1 | 0.26385851 | 0.28822095 | 0.2766757 | 0.01154525 | 0.42641519 | 0.90650187 |
| Ripply3 | 0.20507313 | 0.23355594 | 0.2220116 | 0.01154433 | 0.40859288 | 0.90650187 |
| Arid5a | 0.07058091 | 0.09421041 | 0.08267078 | 0.01153963 | 0.26745057 | 0.90650187 |
| A930033H14 | 0.043198 | 0.06303835 | 0.0515132 | 0.01152514 | 0.15995966 | 0.90650187 |
| 4930404N11 | 0.02531307 | 0.04206466 | 0.0305467 | 0.01151796 | 0.04684334 | 0.56245304 |
| Rad50 | 0.22987912 | 0.25701096 | 0.24550476 | 0.0115062 | 0.41747608 | 0.90650187 |
| Ngb | 0.05198463 | 0.07313868 | 0.06163323 | 0.01150545 | 0.20330105 | 0.90650187 |
| Itsn1 | 0.17244272 | 0.2016166 | 0.19011201 | 0.01150459 | 0.39395519 | 0.90650187 |
| Mvb12a | 0.50392283 | 0.47983786 | 0.46834141 | 0.01149644 | 0.45655091 | 0.90650187 |
| Hat1 | 0.3471399 | 0.35999146 | 0.34849525 | 0.0114962 | 0.44170328 | 0.90650187 |
| Cers6 | 0.30403587 | 0.32359875 | 0.31210385 | 0.0114949 | 0.43497056 | 0.90650187 |
| Tmed1 | 0.69048707 | 0.60120171 | 0.58971201 | 0.0114897 | 0.46548824 | 0.90650187 |
| Cd68 | 0.02291117 | 0.03917966 | 0.02769268 | 0.01148699 | 0.03258809 | 0.42461821 |
| Klhl24 | 0.21363244 | 0.24167034 | 0.23019007 | 0.01148026 | 0.41226763 | 0.90650187 |
| Hck | 0.02529047 | 0.04199208 | 0.03051988 | 0.0114722 | 0.04735389 | 0.56723915 |
| Rasl11a | 0.07974657 | 0.10432197 | 0.09285391 | 0.01146806 | 0.29148173 | 0.90650187 |

| | | | | | | |
|---|---|---|---|---|---|---|
| Gm33370 | 0.0188283 | 0.03426677 | 0.02282015 | 0.01144662 | 0.0128763 | 0.20646902 |
| Zfp423 | 0.10394698 | 0.13063626 | 0.11919342 | 0.01144284 | 0.33476671 | 0.90650187 |
| Arid5b | 0.46065244 | 0.44851435 | 0.43707748 | 0.01143687 | 0.45369678 | 0.90650187 |
| Zfp462 | 0.12558844 | 0.15363781 | 0.14220646 | 0.01143135 | 0.36041015 | 0.90650187 |
| E430025E21I | 0.14240577 | 0.17109871 | 0.15969594 | 0.01140277 | 0.37545883 | 0.90650187 |
| Asxl3 | 0.15249608 | 0.18142624 | 0.17002943 | 0.01139681 | 0.38285843 | 0.90650187 |
| Cd84 | 0.01345463 | 0.02775659 | 0.01636628 | 0.01139032 | 0.00098746 | 0.02349052 |
| Mfsd2a | 0.0202222 | 0.03587687 | 0.02448664 | 0.01139023 | 0.01932362 | 0.28165907 |
| Stat2 | 0.0473471 | 0.06769571 | 0.05630655 | 0.01138915 | 0.18426907 | 0.90650187 |
| Zcwpw1 | 0.09481124 | 0.12071933 | 0.10933168 | 0.01138764 | 0.32166771 | 0.90650187 |
| Borcs7 | 0.48797572 | 0.46834373 | 0.45696933 | 0.0113744 | 0.45594483 | 0.90650187 |
| Kras | 0.73118095 | 0.62499281 | 0.61362931 | 0.0113635 | 0.46719374 | 0.90650187 |
| Csf1 | 0.12679292 | 0.15482613 | 0.1434703 | 0.01135583 | 0.36246618 | 0.90650187 |
| Ackr4 | 0.05113621 | 0.07199691 | 0.06066116 | 0.01133575 | 0.20305731 | 0.90650187 |
| Syt11 | 0.15311812 | 0.18197128 | 0.17066254 | 0.01130874 | 0.38415644 | 0.90650187 |
| Higd1b | 0.0154658 | 0.03009239 | 0.01878719 | 0.01130519 | 0.00373487 | 0.07316198 |
| Ccdc120 | 0.05178211 | 0.07270216 | 0.06140129 | 0.01130086 | 0.20661844 | 0.90650187 |
| Ttll1 | 0.24108857 | 0.26720146 | 0.25591433 | 0.01128713 | 0.42227614 | 0.90650187 |
| Tceal5 | 0.0672996 | 0.09027159 | 0.0789957 | 0.01127588 | 0.26285514 | 0.90650187 |
| Ncald | 0.23124011 | 0.25803041 | 0.24677532 | 0.01125509 | 0.41966205 | 0.90650187 |
| Zcchc24 | 0.13363111 | 0.16185723 | 0.15061252 | 0.01124471 | 0.36997924 | 0.90650187 |
| Cfh | 0.01273643 | 0.02672524 | 0.01550015 | 0.01122509 | 0.00064213 | 0.0162143 |
| Apln | 0.04464313 | 0.06440982 | 0.0531857 | 0.01122412 | 0.17407474 | 0.90650187 |
| Sned1 | 0.10404441 | 0.1304719 | 0.11929832 | 0.01117358 | 0.3385663 | 0.90650187 |
| Gcat | 0.32083349 | 0.33764706 | 0.32647939 | 0.01116766 | 0.43956669 | 0.90650187 |
| Ttpa | 0.01994043 | 0.03530496 | 0.02415002 | 0.01115493 | 0.02001532 | 0.28866497 |
| Stam2 | 0.31185872 | 0.32997653 | 0.31882951 | 0.01114702 | 0.43824169 | 0.90650187 |
| Edn1 | 0.03003556 | 0.04726973 | 0.03613146 | 0.01113827 | 0.08527322 | 0.85616789 |
| Rnase6 | 0.01324709 | 0.02725198 | 0.01611607 | 0.0111359 | 0.00106375 | 0.02510886 |
| AF251705 | 0.01574344 | 0.0302488 | 0.01912089 | 0.01112792 | 0.00483739 | 0.09134554 |

| | | | | | | |
|---|---|---|---|---|---|---|
| Th | 0.10396787 | 0.13034279 | 0.11921591 | 0.01112688 | 0.33909817 | 0.90650187 |
| Tnfaip2 | 0.03826605 | 0.05690463 | 0.04578114 | 0.01112348 | 0.14003712 | 0.90650187 |
| Fam45a | 0.30578345 | 0.32472399 | 0.31361102 | 0.01111297 | 0.43741266 | 0.90650187 |
| Pbld2 | 0.0753831 | 0.09911171 | 0.08802112 | 0.01109059 | 0.28769109 | 0.90650187 |
| Senp8 | 0.16048361 | 0.18921055 | 0.1781254 | 0.01108515 | 0.39102049 | 0.90650187 |
| Slc26a7 | 0.05240938 | 0.07319932 | 0.06211948 | 0.01107984 | 0.21390823 | 0.90650187 |
| BC049730 | 0.04103467 | 0.0600605 | 0.04900353 | 0.01105697 | 0.15790878 | 0.90650187 |
| Cacybp | 1.60235365 | 0.91928926 | 0.90825255 | 0.01103671 | 0.47845914 | 0.90650187 |
| Ocel1 | 0.21185665 | 0.23953533 | 0.22849956 | 0.01103577 | 0.41499747 | 0.90650187 |
| Mul1 | 0.20289455 | 0.23093825 | 0.21991774 | 0.01102051 | 0.41185412 | 0.90650187 |
| Eps15 | 0.22665692 | 0.25350063 | 0.24248922 | 0.01101142 | 0.42000737 | 0.90650187 |
| Hsdl1 | 0.23740681 | 0.26351518 | 0.25250908 | 0.01100611 | 0.42317779 | 0.90650187 |
| Psd | 0.12424888 | 0.15179407 | 0.14079877 | 0.0109953 | 0.3642327 | 0.90650187 |
| Mga | 0.51725654 | 0.48868655 | 0.47772021 | 0.01096634 | 0.45935793 | 0.90650187 |
| Reg1 | 0.01734063 | 0.03198706 | 0.0210381 | 0.01094896 | 0.01034349 | 0.17240278 |
| Nckap1l | 0.01578303 | 0.03007575 | 0.01916846 | 0.0109073 | 0.00570877 | 0.10455304 |
| Ccdc138 | 0.03099479 | 0.04814557 | 0.03726154 | 0.01088403 | 0.09704849 | 0.90650187 |
| Hook1 | 0.25717591 | 0.28150009 | 0.27063502 | 0.01086507 | 0.42917462 | 0.90650187 |
| Fdx1l | 0.41233038 | 0.41143866 | 0.40058357 | 0.0108551 | 0.45205623 | 0.90650187 |
| C030039L03I | 0.04717141 | 0.06694948 | 0.05610411 | 0.01084537 | 0.19506798 | 0.90650187 |
| Mtus1 | 0.10224243 | 0.12817519 | 0.11735616 | 0.01081903 | 0.34096195 | 0.90650187 |
| Batf | 0.01592231 | 0.03015422 | 0.01933579 | 0.01081843 | 0.00643532 | 0.11553978 |
| Alkbh4 | 0.13176816 | 0.15945644 | 0.14867228 | 0.01078415 | 0.37354895 | 0.90650187 |
| Sema4b | 0.20816592 | 0.23575163 | 0.22497559 | 0.01077605 | 0.41568731 | 0.90650187 |
| Yaf2 | 0.4271161 | 0.42269893 | 0.41193392 | 0.01076501 | 0.45375638 | 0.90650187 |
| Gsap | 0.03866176 | 0.05700622 | 0.04624242 | 0.0107638 | 0.15038142 | 0.90650187 |
| Plscr1 | 0.16238791 | 0.1907867 | 0.18004483 | 0.01074186 | 0.39541442 | 0.90650187 |
| Hoxa3 | 0.06113143 | 0.08276829 | 0.07204467 | 0.01072362 | 0.25407721 | 0.90650187 |
| Heph | 0.11056322 | 0.1370132 | 0.12629032 | 0.01072288 | 0.35291541 | 0.90650187 |
| Snhg11 | 0.03213736 | 0.04932149 | 0.03860574 | 0.01071576 | 0.1086084 | 0.90650187 |

| | | | | | | |
|---|---|---|---|---|---|---|
| Mpz | 0.05789971 | 0.07908884 | 0.06838038 | 0.01070846 | 0.24315559 | 0.90650187 |
| Ildr2 | 0.03738568 | 0.05546132 | 0.044754 | 0.01070732 | 0.14375416 | 0.90650187 |
| Slpi | 0.03063367 | 0.04751841 | 0.03683627 | 0.01068214 | 0.09866767 | 0.90650187 |
| Uhrf1 | 0.15331156 | 0.18154039 | 0.17085933 | 0.01068106 | 0.3905401 | 0.90650187 |
| Plxna4 | 0.03771284 | 0.05580615 | 0.04513585 | 0.01067031 | 0.1466384 | 0.90650187 |
| Kcmf1 | 0.30028298 | 0.31952133 | 0.30885804 | 0.01066329 | 0.43900828 | 0.90650187 |
| Cwc22 | 0.35749807 | 0.36765634 | 0.3570016 | 0.01065474 | 0.44722346 | 0.90650187 |
| 1700037H04 | 0.32703848 | 0.34236976 | 0.33172563 | 0.01064413 | 0.44328456 | 0.90650187 |
| Wwtr1 | 0.17452846 | 0.20280864 | 0.19218605 | 0.01062259 | 0.40294972 | 0.90650187 |
| Ncapg | 0.0770631 | 0.10050389 | 0.08988504 | 0.01061884 | 0.29972376 | 0.90650187 |
| Myc | 0.14172969 | 0.16960981 | 0.15899929 | 0.01061052 | 0.38335948 | 0.90650187 |
| Cdx2 | 0.01371848 | 0.02728129 | 0.01668425 | 0.01059703 | 0.00237412 | 0.0495728 |
| Parvb | 0.07193813 | 0.09477261 | 0.0841863 | 0.01058631 | 0.28807098 | 0.90650187 |
| Map2 | 0.40432256 | 0.40494353 | 0.3943667 | 0.01057683 | 0.45254648 | 0.90650187 |
| Gins4 | 0.33888855 | 0.35221244 | 0.34165302 | 0.01055942 | 0.44535741 | 0.90650187 |
| 2310022B05 | 0.75581515 | 0.63826678 | 0.62771181 | 0.01055497 | 0.47020568 | 0.90650187 |
| Ergic1 | 0.49393374 | 0.47177692 | 0.46123906 | 0.01053785 | 0.45954978 | 0.90650187 |
| Rab27b | 0.06289234 | 0.08456303 | 0.07403478 | 0.01052825 | 0.26362953 | 0.90650187 |
| Susd4 | 0.04234591 | 0.06103644 | 0.05052555 | 0.01051088 | 0.17752868 | 0.90650187 |
| Slc41a3 | 0.1154246 | 0.14198035 | 0.13147025 | 0.0105101 | 0.36114714 | 0.90650187 |
| Nyap1 | 0.12078898 | 0.14764148 | 0.13715246 | 0.01048902 | 0.36693316 | 0.90650187 |
| 1810064F22I | 0.02703631 | 0.04307269 | 0.03258866 | 0.01048402 | 0.07633184 | 0.79148727 |
| Mcfd2 | 0.59140155 | 0.53826928 | 0.52780195 | 0.01046733 | 0.4648728 | 0.90650187 |
| Pirb | 0.01297598 | 0.02624532 | 0.01578913 | 0.01045619 | 0.00161971 | 0.03563117 |
| Ikzf1 | 0.01297598 | 0.02624532 | 0.01578913 | 0.01045619 | 0.00161971 | 0.03563117 |
| Hilpda | 0.28912311 | 0.30958499 | 0.29912889 | 0.0104561 | 0.43825419 | 0.90650187 |
| Usf3 | 0.11077984 | 0.13692987 | 0.12652176 | 0.01040811 | 0.35729053 | 0.90650187 |
| Kctd18 | 0.10398614 | 0.12964249 | 0.11923558 | 0.01040691 | 0.34900294 | 0.90650187 |
| Galk2 | 0.35471352 | 0.36512689 | 0.35472354 | 0.01040335 | 0.44813254 | 0.90650187 |
| Brip1os | 0.26986786 | 0.29246761 | 0.28207157 | 0.01039603 | 0.43492568 | 0.90650187 |

| | | | | | | |
|---|---|---|---|---|---|---|
| Slc18a1 | 0.03748408 | 0.05526077 | 0.04486886 | 0.0103919 | 0.15159194 | 0.90650187 |
| Tmem39a | 0.44604696 | 0.43661106 | 0.42622863 | 0.01038243 | 0.45688293 | 0.90650187 |
| Gng8 | 0.03953826 | 0.05764219 | 0.04726333 | 0.01037887 | 0.16447276 | 0.90650187 |
| Rab4a | 0.34566753 | 0.35765566 | 0.34727886 | 0.01037679 | 0.44716153 | 0.90650187 |
| Trh | 0.01053615 | 0.02321582 | 0.01284148 | 0.01037434 | 0.00016439 | 0.00467954 |
| Phox2b | 0.01148296 | 0.0243589 | 0.01398651 | 0.01037239 | 0.00048886 | 0.01274648 |
| Gm13139 | 0.038757 | 0.056717 | 0.04635341 | 0.01036359 | 0.16012649 | 0.90650187 |
| Gm14964 | 0.13176411 | 0.15903056 | 0.14866806 | 0.0103625 | 0.37833037 | 0.90650187 |
| Chsy1 | 0.14315595 | 0.17082229 | 0.16046832 | 0.01035397 | 0.38711515 | 0.90650187 |
| Dpf1 | 0.02471789 | 0.04019216 | 0.02984035 | 0.01035181 | 0.06151131 | 0.68200837 |
| Tjp1 | 0.33191558 | 0.34616897 | 0.33582587 | 0.0103431 | 0.44554693 | 0.90650187 |
| Ncoa7 | 0.37280406 | 0.37975024 | 0.36941043 | 0.01033981 | 0.45048636 | 0.90650187 |
| Bhlhe40 | 0.04917077 | 0.06874305 | 0.0584051 | 0.01033795 | 0.21567276 | 0.90650187 |
| Fez1 | 0.12701933 | 0.15403886 | 0.14370768 | 0.01033118 | 0.37463738 | 0.90650187 |
| Lypd1 | 0.02699697 | 0.04285754 | 0.0325421 | 0.01031544 | 0.07938778 | 0.81530674 |
| Podn | 0.04657097 | 0.06572612 | 0.0554119 | 0.01031422 | 0.20397879 | 0.90650187 |
| Gstp1 | 0.54367227 | 0.50626966 | 0.49595717 | 0.01031249 | 0.46317505 | 0.90650187 |
| Cdh13 | 0.02237693 | 0.03736075 | 0.02705663 | 0.01030412 | 0.04522349 | 0.54819953 |
| Atg4a | 0.1201651 | 0.14679137 | 0.13649342 | 0.01029795 | 0.36865961 | 0.90650187 |
| Exoc3l | 0.03132606 | 0.04792591 | 0.03765147 | 0.01027444 | 0.11254017 | 0.90650187 |
| Ift43 | 0.32907714 | 0.34371137 | 0.33344203 | 0.01026934 | 0.44554873 | 0.90650187 |
| Capns1 | 1.89477646 | 0.93720721 | 0.9269394 | 0.01026781 | 0.48036224 | 0.90650187 |
| Zfp511 | 0.25284274 | 0.27693774 | 0.2666952 | 0.01024254 | 0.43221592 | 0.90650187 |
| Sox8 | 0.02409694 | 0.03930445 | 0.0291028 | 0.01020165 | 0.05957452 | 0.66587668 |
| Rars2 | 0.33782161 | 0.35096059 | 0.34076406 | 0.01019653 | 0.44708726 | 0.90650187 |
| Cerk | 0.11267667 | 0.13868992 | 0.12854584 | 0.01014408 | 0.36286136 | 0.90650187 |
| Agr2 | 0.01000558 | 0.02232699 | 0.01219919 | 0.0101278 | 0.0001118 | 0.0032681 |
| Ctgf | 0.04714048 | 0.06618752 | 0.05606847 | 0.01011905 | 0.21118151 | 0.90650187 |
| Baiap2l2 | 0.03037585 | 0.04664836 | 0.03653252 | 0.01011584 | 0.10916406 | 0.90650187 |
| Bace1 | 0.55656422 | 0.51481081 | 0.50469552 | 0.01011529 | 0.46450104 | 0.90650187 |

| | | | | | | |
|---|---|---|---|---|---|---|
| Stoml2 | 0.79358801 | 0.65884275 | 0.64874953 | 0.01009323 | 0.47242929 | 0.90650187 |
| Dcx | 0.23281042 | 0.25832394 | 0.24823899 | 0.01008494 | 0.42833806 | 0.90650187 |
| Cytip | 0.11510209 | 0.14118192 | 0.13112751 | 0.0100544 | 0.36659935 | 0.90650187 |
| Mar-03 | 0.00954988 | 0.02169273 | 0.01164718 | 0.01004556 | 6.30E-05 | 0.00194144 |
| Gm7879 | 0.04443737 | 0.06298901 | 0.05294776 | 0.01004125 | 0.19959053 | 0.90650187 |
| Tspan9 | 0.28603424 | 0.30645376 | 0.29641574 | 0.01003802 | 0.44016577 | 0.90650187 |
| Casp3 | 0.11678652 | 0.14294546 | 0.13291619 | 0.01002926 | 0.36864446 | 0.90650187 |
| Ndnl2 | 0.66244427 | 0.58276008 | 0.57273459 | 0.01002549 | 0.4689862 | 0.90650187 |
| Sdad1 | 0.23187214 | 0.2573891 | 0.24736473 | 0.01002437 | 0.42851478 | 0.90650187 |
| Rab6b | 0.03240347 | 0.04893888 | 0.03891851 | 0.01002037 | 0.1261848 | 0.90650187 |
| Cdk17 | 0.17093722 | 0.19862023 | 0.18861184 | 0.01000839 | 0.40675546 | 0.90650187 |
| St8sia1 | 0.02872974 | 0.04458142 | 0.03459073 | 0.00999068 | 0.09957021 | 0.90650187 |
| Rtl1 | 0.02957313 | 0.04555947 | 0.03558615 | 0.00997331 | 0.10639728 | 0.90650187 |
| Cog8 | 0.48349237 | 0.46370731 | 0.45374072 | 0.00996659 | 0.46110534 | 0.90650187 |
| Tox2 | 0.04189333 | 0.05996051 | 0.05000051 | 0.00996 | 0.1879285 | 0.90650187 |
| Cttnbp2 | 0.09712663 | 0.12179597 | 0.11183617 | 0.0099598 | 0.34608471 | 0.90650187 |
| Ppil2 | 0.68510753 | 0.59641045 | 0.58648659 | 0.00992387 | 0.47001842 | 0.90650187 |
| Vnn1 | 0.04872332 | 0.06780462 | 0.05789067 | 0.00991394 | 0.22323215 | 0.90650187 |
| Adamts12 | 0.02054144 | 0.03477719 | 0.02486787 | 0.00990932 | 0.03824045 | 0.48001077 |
| Hoxd9 | 0.01295443 | 0.02566809 | 0.01576314 | 0.00990496 | 0.00260745 | 0.05363445 |
| Mvd | 0.23697901 | 0.26201325 | 0.25211254 | 0.00990071 | 0.43070351 | 0.90650187 |
| Zfp467 | 0.27042179 | 0.29244899 | 0.28256724 | 0.00988175 | 0.43822593 | 0.90650187 |
| Uchl4 | 0.03804224 | 0.05539339 | 0.04552014 | 0.00987325 | 0.16746137 | 0.90650187 |
| Hoxb6 | 0.06969349 | 0.09154646 | 0.08167842 | 0.00986804 | 0.29559944 | 0.90650187 |
| Fstl3 | 0.11423354 | 0.14006752 | 0.13020384 | 0.00986367 | 0.3681415 | 0.90650187 |
| Aamdc | 0.28015782 | 0.30108383 | 0.29122996 | 0.00985387 | 0.44021715 | 0.90650187 |
| Galns | 0.16824172 | 0.19576943 | 0.1859196 | 0.00984983 | 0.40690117 | 0.90650187 |
| Ect2 | 0.05952303 | 0.08006312 | 0.07022291 | 0.00984021 | 0.26665712 | 0.90650187 |
| Lyrm9 | 0.2652754 | 0.28779089 | 0.27795102 | 0.00983987 | 0.43747231 | 0.90650187 |
| Gm12688 | 0.01282213 | 0.02543748 | 0.01560355 | 0.00983393 | 0.00254105 | 0.05255305 |

| | | | | | | |
|---|---|---|---|---|---|---|
| Pipox | 0.02050081 | 0.03465063 | 0.02481936 | 0.00983127 | 0.03912232 | 0.48946365 |
| Ctsa | 0.66383674 | 0.58341106 | 0.57358756 | 0.00982351 | 0.46965491 | 0.90650187 |
| Hhatl | 0.01406454 | 0.02691703 | 0.01710114 | 0.00981589 | 0.00535939 | 0.09983827 |
| Rbm18 | 0.58899404 | 0.53603956 | 0.52622833 | 0.00981124 | 0.46697115 | 0.90650187 |
| Chpf2 | 0.17042049 | 0.19789913 | 0.18809636 | 0.00980276 | 0.40839108 | 0.90650187 |
| Klf9 | 0.12088571 | 0.14705595 | 0.13725459 | 0.00980135 | 0.37544697 | 0.90650187 |
| Pan3 | 0.34338084 | 0.35517762 | 0.34538544 | 0.00979218 | 0.44985036 | 0.90650187 |
| C130074G19 | 0.04218843 | 0.06013493 | 0.0503429 | 0.00979203 | 0.1936016 | 0.90650187 |
| Xiap | 0.47143867 | 0.45477963 | 0.44499296 | 0.00978666 | 0.46105685 | 0.90650187 |
| Hip1 | 0.16365809 | 0.19109585 | 0.18132284 | 0.00977301 | 0.40531591 | 0.90650187 |
| Kxd1 | 0.39982212 | 0.40061634 | 0.39085144 | 0.0097649 | 0.45578139 | 0.90650187 |
| Map2k3os | 0.03005602 | 0.04591088 | 0.03615558 | 0.0097553 | 0.11516902 | 0.90650187 |
| Zfp105 | 0.11477779 | 0.14053675 | 0.13078274 | 0.00975401 | 0.37011025 | 0.90650187 |
| Jade3 | 0.17768805 | 0.20504784 | 0.19531833 | 0.00972951 | 0.41237039 | 0.90650187 |
| Katnal2 | 0.04724755 | 0.06590859 | 0.05619185 | 0.00971674 | 0.22102448 | 0.90650187 |
| Bmf | 0.07882721 | 0.10155364 | 0.09183793 | 0.00971571 | 0.31906701 | 0.90650187 |
| Cdkn1b | 0.388663 | 0.39177859 | 0.38206574 | 0.00971285 | 0.45500896 | 0.90650187 |
| Clip2 | 0.10280107 | 0.12766463 | 0.11795869 | 0.00970594 | 0.3572587 | 0.90650187 |
| Coro2b | 0.07963835 | 0.10242625 | 0.09273438 | 0.00969187 | 0.32110184 | 0.90650187 |
| A630001G21 | 0.01218462 | 0.02451961 | 0.01483412 | 0.00968548 | 0.00185026 | 0.04000829 |
| Zmym6 | 0.05693024 | 0.07695258 | 0.06727812 | 0.00967446 | 0.26132384 | 0.90650187 |
| Kcne1l | 0.15512155 | 0.18236739 | 0.17269859 | 0.00966881 | 0.40172125 | 0.90650187 |
| Sez6l | 0.09923378 | 0.12376795 | 0.11410899 | 0.00965896 | 0.35334452 | 0.90650187 |
| Sirpa | 0.15525148 | 0.18248009 | 0.17283048 | 0.00964961 | 0.40198568 | 0.90650187 |
| Fam229b | 0.19054289 | 0.21759181 | 0.20794947 | 0.00964234 | 0.41834204 | 0.90650187 |
| Hoxd4 | 0.02984077 | 0.04554347 | 0.0359018 | 0.00964167 | 0.11625903 | 0.90650187 |
| Smad6 | 0.02618461 | 0.0412198 | 0.03158002 | 0.00963979 | 0.08738755 | 0.87068126 |
| Pira2 | 0.00963715 | 0.02139229 | 0.01175292 | 0.00963937 | 0.0001331 | 0.00385385 |
| Robo2 | 0.20405556 | 0.23067183 | 0.22103423 | 0.00963761 | 0.42315107 | 0.90650187 |
| Slc2a12 | 0.02487542 | 0.03963675 | 0.03002735 | 0.0096094 | 0.07741302 | 0.79780022 |

| | | | | | | |
|---|---|---|---|---|---|---|
| Shq1 | 0.12271371 | 0.14878412 | 0.13918267 | 0.00960145 | 0.37954424 | 0.90650187 |
| Smarca2 | 1.01213464 | 0.76528052 | 0.75568858 | 0.00959194 | 0.47750044 | 0.90650187 |
| Sorbs2 | 0.39948912 | 0.40016876 | 0.39059072 | 0.00957804 | 0.45659536 | 0.90650187 |
| Fam3c | 0.49823424 | 0.47388272 | 0.46430484 | 0.00957789 | 0.46346583 | 0.90650187 |
| Mob3a | 0.1803857 | 0.20755467 | 0.19798396 | 0.0095707 | 0.41492104 | 0.90650187 |
| Eci1 | 0.7367846 | 0.62642905 | 0.61685885 | 0.0095702 | 0.47250638 | 0.90650187 |
| Zadh2 | 0.17343201 | 0.20065411 | 0.19109642 | 0.00955769 | 0.41202185 | 0.90650187 |
| Ddx25 | 0.0668887 | 0.0880843 | 0.07853439 | 0.00954991 | 0.294394 | 0.90650187 |
| Camk2d | 0.30243449 | 0.32026614 | 0.31072038 | 0.00954576 | 0.44568331 | 0.90650187 |
| Sh3pxd2a | 0.19803408 | 0.22477371 | 0.21522813 | 0.00954558 | 0.42184841 | 0.90650187 |
| Phf8 | 0.1908892 | 0.21783166 | 0.20828728 | 0.00954437 | 0.41928945 | 0.90650187 |
| Gm5475 | 0.00773117 | 0.0189789 | 0.00944066 | 0.00953824 | 3.54E-06 | 0.00013674 |
| Morn1 | 0.04114205 | 0.05866621 | 0.04912827 | 0.00953795 | 0.19404488 | 0.90650187 |
| Cby1 | 0.49847958 | 0.47401703 | 0.46447937 | 0.00953766 | 0.46363249 | 0.90650187 |
| Nme6 | 0.1871944 | 0.21420083 | 0.20467652 | 0.00952431 | 0.41805591 | 0.90650187 |
| Dlx5 | 0.01082724 | 0.02271472 | 0.01319367 | 0.00952105 | 0.00066804 | 0.01678481 |
| Apbb1ip | 0.0275031 | 0.04265959 | 0.03314098 | 0.00951861 | 0.10082791 | 0.90650187 |
| Ndufaf2 | 0.46682431 | 0.45112536 | 0.44161699 | 0.00950837 | 0.46187248 | 0.90650187 |
| Myd88 | 0.14781678 | 0.17476072 | 0.16525235 | 0.00950837 | 0.39905519 | 0.90650187 |
| Was | 0.01372414 | 0.02619573 | 0.01669107 | 0.00950467 | 0.00567824 | 0.10411931 |
| Rab7 | 1.39500701 | 0.88279065 | 0.87329058 | 0.00950007 | 0.4807141 | 0.90650187 |
| Atp6v0e | 1.82343344 | 0.93459625 | 0.92510456 | 0.0094917 | 0.48180955 | 0.90650187 |
| Manbal | 0.61449047 | 0.5521885 | 0.54271817 | 0.00947032 | 0.46908309 | 0.90650187 |
| Bcl3 | 0.05322483 | 0.07250796 | 0.06305222 | 0.00945574 | 0.25248409 | 0.90650187 |
| H2-Q2 | 0.11529371 | 0.14078034 | 0.13133117 | 0.00944917 | 0.37453809 | 0.90650187 |
| Sec24c | 0.33845615 | 0.35073476 | 0.34129286 | 0.0094419 | 0.45105825 | 0.90650187 |
| Asap3 | 0.04787252 | 0.06634165 | 0.05691168 | 0.00942997 | 0.23067859 | 0.90650187 |
| Mrps15 | 0.69631062 | 0.60261113 | 0.59318563 | 0.00942551 | 0.4718425 | 0.90650187 |
| 9330158H04 | 0.01784167 | 0.03104777 | 0.02163868 | 0.00940909 | 0.02661371 | 0.36282959 |
| Bri3bp | 0.33996911 | 0.3519512 | 0.34255235 | 0.00939885 | 0.45145852 | 0.90650187 |

| | | | | | | |
|---|---|---|---|---|---|---|
| Lrmp | 0.02677347 | 0.04167526 | 0.03227751 | 0.00939775 | 0.09777071 | 0.90650187 |
| Rbpjl | 0.01933461 | 0.03281599 | 0.02342583 | 0.00939015 | 0.03737434 | 0.47343941 |
| Piezo2 | 0.06650913 | 0.08747842 | 0.07810802 | 0.0093704 | 0.29690368 | 0.90650187 |
| Arhgef28 | 0.09951804 | 0.12376919 | 0.11441512 | 0.00935407 | 0.35813324 | 0.90650187 |
| Pnliprp2 | 0.01519066 | 0.02780327 | 0.01845638 | 0.00934689 | 0.01218029 | 0.19697076 |
| Crlf3 | 0.20617295 | 0.23241245 | 0.22306677 | 0.00934567 | 0.4261226 | 0.90650187 |
| Ube2g2 | 0.42389961 | 0.41881702 | 0.40947912 | 0.0093379 | 0.45962478 | 0.90650187 |
| Pthlh | 0.02160322 | 0.0354705 | 0.02613467 | 0.00933583 | 0.05613644 | 0.64256775 |
| Afap1 | 0.31283217 | 0.32899819 | 0.31966265 | 0.00933554 | 0.44835005 | 0.90650187 |
| Adgrl3 | 0.11375197 | 0.13901066 | 0.12969132 | 0.00931934 | 0.37469197 | 0.90650187 |
| RP23-378D1( | 0.03493566 | 0.05117968 | 0.0418892 | 0.00929048 | 0.16207218 | 0.90650187 |
| Ptcd3 | 0.20654714 | 0.23270997 | 0.22342548 | 0.0092845 | 0.42671719 | 0.90650187 |
| Arhgap24 | 0.10493677 | 0.12953604 | 0.12025859 | 0.00927745 | 0.36581427 | 0.90650187 |
| Btd | 0.25367942 | 0.27673179 | 0.26745733 | 0.00927446 | 0.43874273 | 0.90650187 |
| Ptprg | 0.1805032 | 0.20734342 | 0.19809989 | 0.00924354 | 0.41783455 | 0.90650187 |
| Tal1 | 0.02084053 | 0.03445884 | 0.02522489 | 0.00923395 | 0.05182814 | 0.60884045 |
| Trim12a | 0.04745768 | 0.06563474 | 0.05643394 | 0.00920079 | 0.23428856 | 0.90650187 |
| Kcnmb1 | 0.05633031 | 0.0757868 | 0.06659531 | 0.00919149 | 0.26974558 | 0.90650187 |
| Gm15867 | 0.01969803 | 0.03305134 | 0.02386033 | 0.00919101 | 0.04340687 | 0.53126237 |
| Sc5d | 0.14404146 | 0.17056997 | 0.16137918 | 0.00919079 | 0.40006299 | 0.90650187 |
| Kif26a | 0.04628384 | 0.06426112 | 0.05508069 | 0.00918043 | 0.22935901 | 0.90650187 |
| Dhx16 | 0.1799173 | 0.2066857 | 0.19752169 | 0.00916401 | 0.4182964 | 0.90650187 |
| Ajuba | 0.09267035 | 0.11616119 | 0.10700934 | 0.00915185 | 0.35189645 | 0.90650187 |
| Ccl27a | 0.35534149 | 0.36437788 | 0.35523784 | 0.00914003 | 0.45446729 | 0.90650187 |
| Mmp15 | 0.07844234 | 0.10055084 | 0.09141226 | 0.00913859 | 0.32836504 | 0.90650187 |
| Gpr35 | 0.01106083 | 0.02259637 | 0.01347619 | 0.00912019 | 0.00131217 | 0.02972626 |
| 2310001H17 | 0.04127559 | 0.05840317 | 0.04928337 | 0.0091198 | 0.20535314 | 0.90650187 |
| Pdzrn3 | 0.13628295 | 0.16247986 | 0.15336725 | 0.00911261 | 0.39583433 | 0.90650187 |
| Rasl12 | 0.03670953 | 0.05307096 | 0.04396431 | 0.00910665 | 0.17856421 | 0.90650187 |
| Znrf1 | 0.28232255 | 0.30222889 | 0.29314398 | 0.00908491 | 0.44520907 | 0.90650187 |

| | | | | | | |
|---|---|---|---|---|---|---|
| Gm10116 | 0.41972252 | 0.41536256 | 0.40627975 | 0.00908282 | 0.4604158 | 0.90650187 |
| Pabpc4l | 0.09602766 | 0.11971378 | 0.11064837 | 0.00906541 | 0.35784385 | 0.90650187 |
| Sfrp4 | 0.02929728 | 0.04432468 | 0.0352607 | 0.00906398 | 0.12656692 | 0.90650187 |
| Gm10126 | 0.07970639 | 0.10186724 | 0.09280953 | 0.00905772 | 0.33219234 | 0.90650187 |
| Acad9 | 0.21128881 | 0.23700308 | 0.22795829 | 0.00904479 | 0.42999378 | 0.90650187 |
| Lnx2 | 0.05554896 | 0.07474721 | 0.06570521 | 0.009042 | 0.27034138 | 0.90650187 |
| Rftn1 | 0.09066694 | 0.11386135 | 0.10483036 | 0.00903099 | 0.35086622 | 0.90650187 |
| Gimap5 | 0.04524873 | 0.06289994 | 0.05388564 | 0.0090143 | 0.22853304 | 0.90650187 |
| Ggct | 0.21219352 | 0.23782711 | 0.2288205 | 0.00900661 | 0.43054622 | 0.90650187 |
| Il18r1 | 0.01137051 | 0.02285414 | 0.0138506 | 0.00900354 | 0.00192712 | 0.04141506 |
| Tdrd3 | 0.18282 | 0.20937615 | 0.20038254 | 0.00899361 | 0.42092461 | 0.90650187 |
| Bspry | 0.11612292 | 0.14120456 | 0.13221194 | 0.00899261 | 0.381183 | 0.90650187 |
| Laptm4b | 0.68717282 | 0.59671825 | 0.58772661 | 0.00899165 | 0.47288746 | 0.90650187 |
| Acad10 | 0.01969576 | 0.03284671 | 0.02385761 | 0.0089891 | 0.04696502 | 0.56346908 |
| Klf7 | 0.53635871 | 0.49993668 | 0.49095207 | 0.00898462 | 0.46757928 | 0.90650187 |
| Katna1 | 0.40816435 | 0.40633984 | 0.39735532 | 0.00898452 | 0.45996625 | 0.90650187 |
| Cwc25 | 0.28720951 | 0.30643091 | 0.29744908 | 0.00898182 | 0.44660628 | 0.90650187 |
| Lcp2 | 0.00936604 | 0.02038367 | 0.01142438 | 0.00895929 | 0.00024482 | 0.00670547 |
| H2-Ab1 | 0.02212617 | 0.03571559 | 0.02675792 | 0.00895767 | 0.06834349 | 0.73367892 |
| RP23-291B1. | 0.01965341 | 0.03276465 | 0.02380699 | 0.00895766 | 0.04719203 | 0.5657461 |
| Rita1 | 0.18433262 | 0.21082502 | 0.2018697 | 0.00895532 | 0.42182947 | 0.90650187 |
| Cldn34c1 | 0.08390549 | 0.10637153 | 0.09743488 | 0.00893665 | 0.34172952 | 0.90650187 |
| Cep68 | 0.19881568 | 0.22491543 | 0.21598402 | 0.00893141 | 0.42707124 | 0.90650187 |
| Gsdmd | 0.12056243 | 0.14583644 | 0.1369132 | 0.00892324 | 0.38600765 | 0.90650187 |
| Tex261 | 0.74730346 | 0.63180222 | 0.62287908 | 0.00892314 | 0.47461007 | 0.90650187 |
| Lmbr1l | 0.1025591 | 0.12662058 | 0.11769776 | 0.00892282 | 0.36804821 | 0.90650187 |
| Gnb1l | 0.06014651 | 0.07985003 | 0.07092956 | 0.00892047 | 0.28804581 | 0.90650187 |
| Foxn3 | 0.30112142 | 0.3184902 | 0.30958428 | 0.00890592 | 0.44911868 | 0.90650187 |
| Wnt6 | 0.02320822 | 0.03693425 | 0.02804615 | 0.0088881 | 0.07943884 | 0.81530674 |
| Rhod | 0.09674758 | 0.12029918 | 0.11142667 | 0.00887252 | 0.3616739 | 0.90650187 |

| | | | | | | |
|---|---|---|---|---|---|---|
| 2700062C07I | 0.25546469 | 0.27794382 | 0.26908129 | 0.00886253 | 0.44179429 | 0.90650187 |
| Syne1 | 0.2611893 | 0.2831297 | 0.274268 | 0.0088617 | 0.44289269 | 0.90650187 |
| Itgb3 | 0.06817652 | 0.08883138 | 0.07997939 | 0.00885199 | 0.31134824 | 0.90650187 |
| Pdlim3 | 0.04413154 | 0.06143019 | 0.05259399 | 0.00883619 | 0.22756322 | 0.90650187 |
| Gpr45 | 0.07992667 | 0.10188114 | 0.0930528 | 0.00882834 | 0.3365947 | 0.90650187 |
| Lmod1 | 0.01296956 | 0.02460293 | 0.01578139 | 0.00882154 | 0.00647719 | 0.11597151 |
| Hpca | 0.08059023 | 0.1026016 | 0.09378518 | 0.00881643 | 0.33800414 | 0.90650187 |
| Clptm1l | 0.31777528 | 0.33268999 | 0.32388052 | 0.00880946 | 0.45187747 | 0.90650187 |
| Col8a2 | 0.08606229 | 0.10860583 | 0.09980091 | 0.00880492 | 0.34745045 | 0.90650187 |
| 4930523C07I | 0.06961446 | 0.09039288 | 0.08158998 | 0.0088029 | 0.31573983 | 0.90650187 |
| Pdzd8 | 0.25709587 | 0.27934524 | 0.27056241 | 0.00878284 | 0.44262759 | 0.90650187 |
| Pus1 | 0.20762793 | 0.23322942 | 0.22446073 | 0.00876869 | 0.43106231 | 0.90650187 |
| Ccdc109b | 0.03256665 | 0.04787262 | 0.03911025 | 0.00876238 | 0.15962331 | 0.90650187 |
| Wbp1 | 0.61764537 | 0.55349343 | 0.54473273 | 0.00876071 | 0.47150118 | 0.90650187 |
| 9430038I01R | 0.2258897 | 0.25051405 | 0.24176966 | 0.00874439 | 0.43612946 | 0.90650187 |
| Spock1 | 0.11598766 | 0.14080735 | 0.13206833 | 0.00873903 | 0.38431478 | 0.90650187 |
| Fis1 | 1.70585651 | 0.92736305 | 0.9186305 | 0.00873254 | 0.48314566 | 0.90650187 |
| Tceanc | 0.10485153 | 0.12889081 | 0.12016691 | 0.0087239 | 0.37344491 | 0.90650187 |
| Tmem74b | 0.01222686 | 0.0236074 | 0.01488512 | 0.00872227 | 0.00459368 | 0.08761418 |
| Dcaf17 | 0.31151117 | 0.32725019 | 0.31853185 | 0.00871834 | 0.45157701 | 0.90650187 |
| Rps19bp1 | 0.71407019 | 0.61239073 | 0.60367415 | 0.00871658 | 0.47440906 | 0.90650187 |
| Polr2e | 1.11324164 | 0.80260897 | 0.79391228 | 0.00869669 | 0.48057992 | 0.90650187 |
| Prpf6 | 0.45851161 | 0.44419311 | 0.43549668 | 0.00869643 | 0.46463066 | 0.90650187 |
| Ccdc124 | 0.7435574 | 0.62943782 | 0.62074145 | 0.00869637 | 0.47516938 | 0.90650187 |
| Mut | 0.19063546 | 0.2167334 | 0.20803978 | 0.00869362 | 0.42631093 | 0.90650187 |
| Zfr | 1.05600682 | 0.78167682 | 0.77300494 | 0.00867188 | 0.48011196 | 0.90650187 |
| Icosl | 0.04794148 | 0.06565617 | 0.05699107 | 0.0086651 | 0.24954456 | 0.90650187 |
| Pisd | 0.37225705 | 0.37763193 | 0.36897038 | 0.00866155 | 0.45844173 | 0.90650187 |
| Eno2 | 0.05667366 | 0.07564666 | 0.06698615 | 0.00866051 | 0.28272552 | 0.90650187 |
| Stk17b | 0.0465173 | 0.06400544 | 0.05535 | 0.00865544 | 0.24346855 | 0.90650187 |

| | | | | | | |
|---|---|---|---|---|---|---|
| Epb41l4a | 0.16153331 | 0.18783894 | 0.17918395 | 0.00865498 | 0.41498787 | 0.90650187 |
| Peak1 | 0.19183864 | 0.21786506 | 0.20921275 | 0.00865231 | 0.4270637 | 0.90650187 |
| Prrx2 | 0.05621944 | 0.07511551 | 0.06646907 | 0.00864644 | 0.28153286 | 0.90650187 |
| Rbm33 | 0.24506226 | 0.26821461 | 0.2595746 | 0.00864001 | 0.44118197 | 0.90650187 |
| Ids | 0.34005765 | 0.35125848 | 0.34262599 | 0.00863249 | 0.45540877 | 0.90650187 |
| Xrcc3 | 0.05136321 | 0.0695503 | 0.06092134 | 0.00862896 | 0.26445193 | 0.90650187 |
| Zbtb7c | 0.12994868 | 0.15539193 | 0.14677332 | 0.00861861 | 0.39702752 | 0.90650187 |
| Tmf1 | 0.39659713 | 0.39694036 | 0.38832289 | 0.00861746 | 0.46070624 | 0.90650187 |
| Zfp143 | 0.11366729 | 0.13821715 | 0.12960115 | 0.00861599 | 0.38378778 | 0.90650187 |
| Cacng7 | 0.05084356 | 0.06893816 | 0.0603256 | 0.00861255 | 0.2628155 | 0.90650187 |
| Wbscr17 | 0.07271593 | 0.09365275 | 0.08505362 | 0.00859913 | 0.32654937 | 0.90650187 |
| Ormdl2 | 0.45874195 | 0.4442636 | 0.43566692 | 0.00859668 | 0.46504891 | 0.90650187 |
| Gigyf1 | 0.22150787 | 0.24624069 | 0.2376482 | 0.00859249 | 0.43615031 | 0.90650187 |
| Jup | 0.43802901 | 0.42877808 | 0.42020622 | 0.00857187 | 0.46387073 | 0.90650187 |
| Grk5 | 0.09866319 | 0.12206144 | 0.11349414 | 0.0085673 | 0.36859219 | 0.90650187 |
| Sp140 | 0.03171417 | 0.04667497 | 0.0381081 | 0.00856686 | 0.15880261 | 0.90650187 |
| Mmp9 | 0.01397625 | 0.02555244 | 0.01699479 | 0.00855765 | 0.01259212 | 0.20276782 |
| Asb3 | 0.17936196 | 0.20553069 | 0.19697331 | 0.00855738 | 0.42342605 | 0.90650187 |
| Cacna1g | 0.07263794 | 0.09352241 | 0.08496669 | 0.00855572 | 0.32720282 | 0.90650187 |
| Grin2a | 0.0142733 | 0.02588022 | 0.01735252 | 0.0085277 | 0.01445384 | 0.22511565 |
| Prdm10 | 0.02866603 | 0.04304283 | 0.03451549 | 0.00852733 | 0.13603165 | 0.90650187 |
| Ablim1 | 0.15519982 | 0.18130363 | 0.17277805 | 0.00852558 | 0.41318303 | 0.90650187 |
| Golm1 | 0.19983921 | 0.22549437 | 0.21697278 | 0.00852158 | 0.4306967 | 0.90650187 |
| Dbnl | 0.78838587 | 0.6544079 | 0.64589165 | 0.00851624 | 0.47662881 | 0.90650187 |
| Ctdsp2 | 0.54797878 | 0.50739865 | 0.49888752 | 0.00851113 | 0.469772 | 0.90650187 |
| Mmp19 | 0.03709837 | 0.05291638 | 0.04441853 | 0.00849785 | 0.19752281 | 0.90650187 |
| Adk | 0.57338701 | 0.52444034 | 0.51594513 | 0.00849521 | 0.4708241 | 0.90650187 |
| Lacc1 | 0.04890593 | 0.06657213 | 0.05810066 | 0.00847147 | 0.25842801 | 0.90650187 |
| Gm20324 | 0.00833573 | 0.01864197 | 0.01017473 | 0.00846724 | 0.00010799 | 0.00316896 |
| Echdc3 | 0.03556275 | 0.05107711 | 0.04262334 | 0.00845377 | 0.18896229 | 0.90650187 |

| | | | | | | |
|---|---|---|---|---|---|---|
| Mospd2 | 0.16520991 | 0.19132153 | 0.18288174 | 0.00843979 | 0.41872361 | 0.90650187 |
| Rep15 | 0.04909215 | 0.0667509 | 0.05831473 | 0.00843616 | 0.26007045 | 0.90650187 |
| Scrn1 | 0.19549545 | 0.22116813 | 0.21276823 | 0.0083999 | 0.43034003 | 0.90650187 |
| Pard3 | 0.46307662 | 0.44725038 | 0.43886366 | 0.00838672 | 0.46614824 | 0.90650187 |
| Nell1 | 0.03768791 | 0.05348889 | 0.04510675 | 0.00838214 | 0.20436742 | 0.90650187 |
| Usp19 | 0.48023246 | 0.45976001 | 0.45138464 | 0.00837536 | 0.46712956 | 0.90650187 |
| Dcun1d4 | 0.1085569 | 0.1325156 | 0.12414401 | 0.00837159 | 0.38216967 | 0.90650187 |
| Gprin1 | 0.09007427 | 0.11255224 | 0.10418468 | 0.00836756 | 0.36052807 | 0.90650187 |
| Slc25a14 | 0.22964854 | 0.25364213 | 0.24528932 | 0.00835281 | 0.43983569 | 0.90650187 |
| Vrk1 | 0.20654315 | 0.23177281 | 0.22342165 | 0.00835116 | 0.43401156 | 0.90650187 |
| Pum3 | 0.44625114 | 0.43472036 | 0.42638134 | 0.00833902 | 0.46535736 | 0.90650187 |
| Rbfa | 0.38929962 | 0.39090771 | 0.38256964 | 0.00833807 | 0.46140625 | 0.90650187 |
| Cfap126 | 0.1527016 | 0.17856684 | 0.17023866 | 0.00832818 | 0.41391649 | 0.90650187 |
| Creb3l2 | 0.44729307 | 0.43547843 | 0.42716013 | 0.0083183 | 0.46550607 | 0.90650187 |
| Sypl | 0.88234589 | 0.70396429 | 0.69564864 | 0.00831565 | 0.47880941 | 0.90650187 |
| Itgb8 | 0.03801822 | 0.0538072 | 0.04549211 | 0.00831509 | 0.20823177 | 0.90650187 |
| C1qtnf1 | 0.0384995 | 0.0543647 | 0.04605331 | 0.00831139 | 0.21118609 | 0.90650187 |
| 2210019I11R | 0.0670365 | 0.08700849 | 0.07870034 | 0.00830815 | 0.31942441 | 0.90650187 |
| Gtdc1 | 0.15017446 | 0.17595339 | 0.16766259 | 0.0082908 | 0.41300089 | 0.90650187 |
| Gcnt2 | 0.20058512 | 0.22597846 | 0.21769265 | 0.00828581 | 0.43281632 | 0.90650187 |
| Frmd4b | 0.15999422 | 0.18591372 | 0.17763146 | 0.00828227 | 0.41789506 | 0.90650187 |
| Ckmt1 | 0.15389835 | 0.17973069 | 0.17145602 | 0.00827467 | 0.41505902 | 0.90650187 |
| Nes | 0.08812572 | 0.1103183 | 0.10205838 | 0.00825992 | 0.35949926 | 0.90650187 |
| Khsrp | 0.27957582 | 0.29897372 | 0.29071463 | 0.00825909 | 0.44974803 | 0.90650187 |
| Slitrk2 | 0.01447833 | 0.02585755 | 0.01759936 | 0.00825819 | 0.01848828 | 0.27348002 |
| Lama5 | 0.1261779 | 0.15107714 | 0.14282519 | 0.00825195 | 0.3986465 | 0.90650187 |
| Dmtn | 0.02663923 | 0.04036705 | 0.03211856 | 0.00824849 | 0.12679067 | 0.90650187 |
| Cdh3 | 0.03702788 | 0.05257237 | 0.0443362 | 0.00823616 | 0.20444659 | 0.90650187 |
| Eef2k | 0.16808958 | 0.19399731 | 0.1857674 | 0.00822991 | 0.42193318 | 0.90650187 |
| Slc45a4 | 0.05212578 | 0.07001676 | 0.06179485 | 0.00822191 | 0.27709417 | 0.90650187 |

| | | | | | | |
|---|---|---|---|---|---|---|
| Ptges3l | 0.14240419 | 0.16791133 | 0.1596943 | 0.00821703 | 0.40953261 | 0.90650187 |
| Iars | 0.26083398 | 0.28216386 | 0.27394697 | 0.00821688 | 0.4469609 | 0.90650187 |
| Cnnm3 | 0.11640178 | 0.14071175 | 0.13250795 | 0.0082038 | 0.39156778 | 0.90650187 |
| Neurl1b | 0.11050241 | 0.13441242 | 0.12622535 | 0.00818707 | 0.38654124 | 0.90650187 |
| Rffl | 0.10304634 | 0.1264001 | 0.11822312 | 0.00817698 | 0.37923776 | 0.90650187 |
| Sema3b | 0.07082601 | 0.09110445 | 0.08294467 | 0.00815978 | 0.33093176 | 0.90650187 |
| Ppp1r16b | 0.04678595 | 0.06381425 | 0.0556598 | 0.00815444 | 0.25742521 | 0.90650187 |
| Cilp | 0.01275815 | 0.02368073 | 0.01552636 | 0.00815437 | 0.0097766 | 0.16457811 |
| Zfp385a | 0.198612 | 0.22392945 | 0.21578713 | 0.00814232 | 0.43339136 | 0.90650187 |
| Cpne5 | 0.05953731 | 0.07836677 | 0.07023911 | 0.00812766 | 0.30347922 | 0.90650187 |
| 6230400D17 | 0.01759503 | 0.02946094 | 0.02134309 | 0.00811785 | 0.04542906 | 0.54937718 |
| Rnf112 | 0.02872826 | 0.04270352 | 0.03458899 | 0.00811453 | 0.14848944 | 0.90650187 |
| Slc35a5 | 0.16580579 | 0.19157527 | 0.18347962 | 0.00809565 | 0.42224523 | 0.90650187 |
| Cbln4 | 0.01378152 | 0.02484477 | 0.0167602 | 0.00808456 | 0.01599997 | 0.24313303 |
| Glb1 | 0.31116703 | 0.32631819 | 0.31823701 | 0.00808117 | 0.45505896 | 0.90650187 |
| Brd1 | 0.15635998 | 0.18202858 | 0.17395488 | 0.0080737 | 0.41826574 | 0.90650187 |
| Ick | 0.1554547 | 0.18109512 | 0.17303672 | 0.0080584 | 0.41799169 | 0.90650187 |
| 2610044O15 | 0.12746926 | 0.15223156 | 0.14417921 | 0.00805235 | 0.40195682 | 0.90650187 |
| Zcchc3 | 0.44957092 | 0.43690644 | 0.42886001 | 0.00804642 | 0.46676276 | 0.90650187 |
| Pacs2 | 0.1932968 | 0.21867406 | 0.21063221 | 0.00804185 | 0.43261071 | 0.90650187 |
| Tfb1m | 0.0753206 | 0.09597419 | 0.08795171 | 0.00802248 | 0.34255344 | 0.90650187 |
| Tbc1d31 | 0.06790566 | 0.08768732 | 0.07967567 | 0.00801165 | 0.32743015 | 0.90650187 |
| Zcchc16 | 0.03797199 | 0.0534465 | 0.04543818 | 0.00800832 | 0.21666028 | 0.90650187 |
| Gm10250 | 0.35692479 | 0.36454061 | 0.35653312 | 0.00800749 | 0.46023357 | 0.90650187 |
| Zfp644 | 0.5490927 | 0.50762455 | 0.49964362 | 0.00798093 | 0.4716946 | 0.90650187 |
| Ebi3 | 0.03137256 | 0.04567561 | 0.0377062 | 0.00796941 | 0.17371064 | 0.90650187 |
| Slc44a2 | 0.19593767 | 0.22116402 | 0.21319722 | 0.0079668 | 0.43402955 | 0.90650187 |
| Bhlha15 | 0.02958175 | 0.04354899 | 0.03559632 | 0.00795267 | 0.160305 | 0.90650187 |
| Pir | 0.11233202 | 0.13612078 | 0.12817841 | 0.00794237 | 0.39147952 | 0.90650187 |
| Exo5 | 0.10254874 | 0.1256211 | 0.11768658 | 0.00793452 | 0.38219298 | 0.90650187 |

| | | | | | | |
|---|---|---|---|---|---|---|
| Dtna | 0.03587502 | 0.05092219 | 0.0429887 | 0.00793349 | 0.20598585 | 0.90650187 |
| Rnf123 | 0.11044921 | 0.13409808 | 0.12616849 | 0.00792959 | 0.38996783 | 0.90650187 |
| E430018J23F | 0.08394474 | 0.10538288 | 0.09747799 | 0.00790489 | 0.359232 | 0.90650187 |
| Tnfsf12 | 0.18554054 | 0.21095863 | 0.20305551 | 0.00790313 | 0.43131491 | 0.90650187 |
| Kcna6 | 0.05965079 | 0.07826677 | 0.07036776 | 0.00789901 | 0.30887783 | 0.90650187 |
| Adora2b | 0.07641847 | 0.09706596 | 0.08917033 | 0.00789563 | 0.34692395 | 0.90650187 |
| Fam219b | 0.18916705 | 0.21450133 | 0.20660613 | 0.0078952 | 0.43255094 | 0.90650187 |
| Gpsm2 | 0.18757816 | 0.21294104 | 0.20505224 | 0.0078888 | 0.43209926 | 0.90650187 |
| Pkhd1l1 | 0.01511561 | 0.02625438 | 0.01836613 | 0.00788826 | 0.02810595 | 0.37875759 |
| Flywch2 | 0.26131487 | 0.28226604 | 0.27438141 | 0.00788463 | 0.4491738 | 0.90650187 |
| Hddc3 | 0.25472727 | 0.27629466 | 0.26841088 | 0.00788378 | 0.44805509 | 0.90650187 |
| Fam184b | 0.01557205 | 0.02678722 | 0.0189149 | 0.00787232 | 0.03213907 | 0.42043551 |
| Arpc5l | 1.19219001 | 0.82795793 | 0.82008615 | 0.00787178 | 0.48298144 | 0.90650187 |
| Fam206a | 0.08565644 | 0.10722289 | 0.09935619 | 0.0078667 | 0.36242332 | 0.90650187 |
| Cdh26 | 0.00968002 | 0.01967046 | 0.01180486 | 0.00786559 | 0.00152768 | 0.0338024 |
| Zfp280d | 0.41315087 | 0.40906622 | 0.40121783 | 0.00784839 | 0.46535073 | 0.90650187 |
| Bcl6b | 0.05989846 | 0.07849351 | 0.07064849 | 0.00784502 | 0.31077549 | 0.90650187 |
| Syt5 | 0.18608316 | 0.21142892 | 0.20358768 | 0.00784124 | 0.43202387 | 0.90650187 |
| Oxsm | 0.04281401 | 0.05890339 | 0.05106826 | 0.00783513 | 0.24759862 | 0.90650187 |
| Mpzl2 | 0.05739803 | 0.0756353 | 0.06781015 | 0.00782515 | 0.30397098 | 0.90650187 |
| Frem2 | 0.08151155 | 0.10262515 | 0.094801 | 0.00782415 | 0.35684447 | 0.90650187 |
| Casq2 | 0.01886037 | 0.030678 | 0.02285852 | 0.00781948 | 0.06416004 | 0.70418539 |
| Spdya | 0.01448923 | 0.02542827 | 0.01761248 | 0.0078158 | 0.02425899 | 0.33584788 |
| Fas | 0.0990094 | 0.12168301 | 0.11386725 | 0.00781576 | 0.38012856 | 0.90650187 |
| Ggt1 | 0.04369748 | 0.05989591 | 0.05209163 | 0.00780427 | 0.25269513 | 0.90650187 |
| Crispld2 | 0.08454232 | 0.1059373 | 0.09813417 | 0.00780312 | 0.36185996 | 0.90650187 |
| Mall | 0.04049121 | 0.0561698 | 0.04837193 | 0.00779787 | 0.23679165 | 0.90650187 |
| Sh3bp1 | 0.18486515 | 0.21018374 | 0.20239268 | 0.00779106 | 0.43205974 | 0.90650187 |
| Il1rl2 | 0.05916867 | 0.07760658 | 0.06982103 | 0.00778554 | 0.31004711 | 0.90650187 |
| Xab2 | 0.33761411 | 0.3483587 | 0.34059107 | 0.00776763 | 0.45962124 | 0.90650187 |

| | | | | | | |
|---|---|---|---|---|---|---|
| Abca8a | 0.04753286 | 0.06428614 | 0.05652053 | 0.00776561 | 0.27066548 | 0.90650187 |
| Htatip2 | 0.17220293 | 0.19763839 | 0.18987324 | 0.00776515 | 0.42786599 | 0.90650187 |
| Tmem200a | 0.06501898 | 0.08418968 | 0.07643213 | 0.00775755 | 0.32591884 | 0.90650187 |
| Ctla2b | 0.02270506 | 0.0351919 | 0.02744734 | 0.00774456 | 0.10485283 | 0.90650187 |
| Slc35e1 | 0.12713239 | 0.15156346 | 0.14382619 | 0.00773727 | 0.40549242 | 0.90650187 |
| Pcp4l1 | 0.02395863 | 0.03666535 | 0.02893844 | 0.00772691 | 0.11760785 | 0.90650187 |
| A930019D19 | 0.02332457 | 0.03590431 | 0.02818456 | 0.00771975 | 0.11167811 | 0.90650187 |
| She | 0.0537989 | 0.07141855 | 0.06370828 | 0.00771027 | 0.295279 | 0.90650187 |
| Tek | 0.04381427 | 0.05993385 | 0.05222682 | 0.00770703 | 0.2559021 | 0.90650187 |
| Mettl8 | 0.07814059 | 0.09877596 | 0.09107837 | 0.00769759 | 0.35356044 | 0.90650187 |
| Heatr5a | 0.08523366 | 0.10659023 | 0.09889267 | 0.00769756 | 0.36465821 | 0.90650187 |
| Zfp110 | 0.18249978 | 0.20776045 | 0.20006739 | 0.00769306 | 0.43213392 | 0.90650187 |
| Oraov1 | 0.17594252 | 0.20127913 | 0.19358925 | 0.00768987 | 0.42991476 | 0.90650187 |
| Actn3 | 0.10601584 | 0.12910791 | 0.12141846 | 0.00768945 | 0.38914771 | 0.90650187 |
| Col7a1 | 0.02148006 | 0.03367294 | 0.02598782 | 0.00768512 | 0.09431282 | 0.90650187 |
| Gm26883 | 0.0190789 | 0.03080141 | 0.02311999 | 0.00768142 | 0.06983583 | 0.74653619 |
| Zkscan6 | 0.12793365 | 0.15233554 | 0.14466564 | 0.0076699 | 0.40683366 | 0.90650187 |
| Mr1 | 0.08424235 | 0.10546353 | 0.09780485 | 0.00765867 | 0.36387651 | 0.90650187 |
| Nrbf2 | 0.3629588 | 0.36910931 | 0.36145069 | 0.00765862 | 0.46247672 | 0.90650187 |
| Klhl6 | 0.03233871 | 0.04649762 | 0.0388424 | 0.00765522 | 0.19047198 | 0.90650187 |
| Cep89 | 0.23337761 | 0.25642061 | 0.24876705 | 0.00765355 | 0.445605 | 0.90650187 |
| Mill2 | 0.03234121 | 0.04648861 | 0.03884534 | 0.00764327 | 0.19086186 | 0.90650187 |
| Rnf126 | 0.33927489 | 0.34961659 | 0.34197468 | 0.00764191 | 0.46043281 | 0.90650187 |
| Epha1 | 0.06892588 | 0.08845889 | 0.0808191 | 0.00763979 | 0.33715468 | 0.90650187 |
| Uchl3 | 0.51058039 | 0.48067076 | 0.47303867 | 0.00763209 | 0.47140992 | 0.90650187 |
| Arid3a | 0.16528885 | 0.19059203 | 0.18296096 | 0.00763107 | 0.4264496 | 0.90650187 |
| Cks1brt | 0.06425015 | 0.08319379 | 0.0755662 | 0.00762759 | 0.32681119 | 0.90650187 |
| Relb | 0.09915737 | 0.12165009 | 0.11402668 | 0.00762342 | 0.3831503 | 0.90650187 |
| Npat | 0.13470477 | 0.15934588 | 0.15172884 | 0.00761704 | 0.41169894 | 0.90650187 |
| P2ry14 | 0.01268189 | 0.02305122 | 0.01543434 | 0.00761688 | 0.01412069 | 0.22103441 |

| | | | | | | |
|---|---|---|---|---|---|---|
| Tmem205 | 0.37267088 | 0.37691602 | 0.36930332 | 0.0076127 | 0.46349192 | 0.90650187 |
| Fgd5 | 0.03862938 | 0.05381283 | 0.04620469 | 0.00760815 | 0.23207775 | 0.90650187 |
| Slc16a7 | 0.02663047 | 0.03970256 | 0.03210818 | 0.00759438 | 0.14651539 | 0.90650187 |
| Gss | 0.2148924 | 0.2389729 | 0.23138755 | 0.00758535 | 0.4420649 | 0.90650187 |
| Zfp800 | 0.20533257 | 0.2298326 | 0.22226063 | 0.00757197 | 0.43980926 | 0.90650187 |
| 1500015A07 | 0.0678324 | 0.08716442 | 0.0795935 | 0.00757092 | 0.33619687 | 0.90650187 |
| Kcnk3 | 0.16588093 | 0.19112568 | 0.18355498 | 0.0075707 | 0.42725839 | 0.90650187 |
| G630016G05 | 0.10054331 | 0.12308464 | 0.1155211 | 0.00756353 | 0.3854997 | 0.90650187 |
| Gimap8 | 0.01826039 | 0.02969867 | 0.02214028 | 0.0075584 | 0.06454869 | 0.70630256 |
| Lsm14a | 0.69366699 | 0.59915852 | 0.59161095 | 0.00754758 | 0.47738589 | 0.90650187 |
| Hltf | 0.13316035 | 0.1576444 | 0.15012262 | 0.00752178 | 0.41186749 | 0.90650187 |
| Parp12 | 0.09028313 | 0.11193401 | 0.10441228 | 0.00752173 | 0.37438716 | 0.90650187 |
| Pdzd2 | 0.1201598 | 0.14400209 | 0.13648782 | 0.00751426 | 0.40332481 | 0.90650187 |
| Pgm2l1 | 0.11356523 | 0.13698931 | 0.12949249 | 0.00749682 | 0.3984453 | 0.90650187 |
| Diaph2 | 0.10283574 | 0.12548565 | 0.11799607 | 0.00748958 | 0.38890413 | 0.90650187 |
| Fbxo32 | 0.10017415 | 0.1226104 | 0.11512193 | 0.00748847 | 0.3862224 | 0.90650187 |
| Prl | 0.03144642 | 0.04528125 | 0.03779311 | 0.00748814 | 0.18920469 | 0.90650187 |
| Plekhb1 | 0.11473575 | 0.13820623 | 0.13073804 | 0.00746819 | 0.39976794 | 0.90650187 |
| Gatsl3 | 0.07348229 | 0.09337272 | 0.08590732 | 0.00746539 | 0.34962776 | 0.90650187 |
| Stk10 | 0.03164856 | 0.04548699 | 0.03803093 | 0.00745606 | 0.19172072 | 0.90650187 |
| Etf1 | 0.6664725 | 0.58265353 | 0.57519928 | 0.00745425 | 0.47702865 | 0.90650187 |
| Vopp1 | 0.14253199 | 0.16727989 | 0.15982594 | 0.00745395 | 0.41787454 | 0.90650187 |
| Adap2 | 0.0133321 | 0.02366849 | 0.01621856 | 0.00744993 | 0.02057228 | 0.29474013 |
| Tram2 | 0.09634436 | 0.11843469 | 0.11099084 | 0.00744385 | 0.38279254 | 0.90650187 |
| Ntf5 | 0.03062449 | 0.04426773 | 0.03682545 | 0.00744227 | 0.18447588 | 0.90650187 |
| Gm14418 | 0.06288607 | 0.08146464 | 0.0740277 | 0.00743694 | 0.32757735 | 0.90650187 |
| Klf15 | 0.0479896 | 0.06447589 | 0.05704647 | 0.00742942 | 0.28130412 | 0.90650187 |
| Kcna1 | 0.01074892 | 0.02052703 | 0.01309893 | 0.0074281 | 0.00585106 | 0.1065174 |
| Tmem209 | 0.1412654 | 0.16594633 | 0.15852056 | 0.00742577 | 0.41751653 | 0.90650187 |
| Usp46 | 0.23046808 | 0.253461 | 0.24605482 | 0.00740618 | 0.44677582 | 0.90650187 |

| | | | | | | |
|---|---|---|---|---|---|---|
| Dis3l2 | 0.15770092 | 0.182719 | 0.17531319 | 0.00740581 | 0.42551786 | 0.90650187 |
| Hspb8 | 0.31552533 | 0.32936382 | 0.32196333 | 0.00740049 | 0.45930511 | 0.90650187 |
| Myof | 0.10374145 | 0.12636746 | 0.11897208 | 0.00739538 | 0.39114282 | 0.90650187 |
| Prdm1 | 0.00938835 | 0.01884559 | 0.01145142 | 0.00739417 | 0.00204898 | 0.04365509 |
| Fitm2 | 0.25172443 | 0.27304836 | 0.26567545 | 0.0073729 | 0.45090609 | 0.90650187 |
| Dnmt1 | 0.28096903 | 0.29931698 | 0.29194772 | 0.00736926 | 0.4553266 | 0.90650187 |
| Pcdhb7 | 0.04432309 | 0.06018406 | 0.05281558 | 0.00736848 | 0.26755601 | 0.90650187 |
| Serf1 | 1.09372164 | 0.79437753 | 0.7870126 | 0.00736492 | 0.48340786 | 0.90650187 |
| Ddx28 | 0.18240942 | 0.20732912 | 0.19997844 | 0.00735068 | 0.43509831 | 0.90650187 |
| Ftsj2 | 0.1208859 | 0.14459978 | 0.1372548 | 0.00734497 | 0.40597949 | 0.90650187 |
| Rgs16 | 0.09069313 | 0.11220231 | 0.10485888 | 0.00734343 | 0.37777378 | 0.90650187 |
| Rell1 | 0.21601338 | 0.23979095 | 0.23245156 | 0.00733939 | 0.44418586 | 0.90650187 |
| Cd300a | 0.0087308 | 0.01799111 | 0.01065411 | 0.007337 | 0.00110121 | 0.02579286 |
| Fam60a | 0.42425534 | 0.41708651 | 0.40975099 | 0.00733552 | 0.46828288 | 0.90650187 |
| Chek1 | 0.06984889 | 0.08917822 | 0.08185227 | 0.00732594 | 0.34535536 | 0.90650187 |
| Gm26549 | 0.02666902 | 0.03947956 | 0.03215384 | 0.00732573 | 0.15556378 | 0.90650187 |
| Hoxb3 | 0.09394973 | 0.11572128 | 0.10839792 | 0.00732336 | 0.38195706 | 0.90650187 |
| Hdac11 | 0.18964727 | 0.21439858 | 0.20707524 | 0.00732334 | 0.43753513 | 0.90650187 |
| Fbxl22 | 0.02758509 | 0.04055817 | 0.03323796 | 0.00732021 | 0.1637691 | 0.90650187 |
| Jmjd7 | 0.16382705 | 0.18880639 | 0.1814927 | 0.00731369 | 0.42891134 | 0.90650187 |
| Tmeff2 | 0.04124486 | 0.05653771 | 0.04924768 | 0.00729003 | 0.25524549 | 0.90650187 |
| 1700019D03I | 0.04861891 | 0.0650553 | 0.05777059 | 0.00728471 | 0.28754253 | 0.90650187 |
| Stambp | 0.30980228 | 0.32434993 | 0.31706673 | 0.00728321 | 0.45933145 | 0.90650187 |
| Map9 | 0.45252303 | 0.43832695 | 0.43105759 | 0.00726935 | 0.47011902 | 0.90650187 |
| Morc4 | 0.11854665 | 0.14203932 | 0.13478156 | 0.00725776 | 0.40540218 | 0.90650187 |
| Fam171b | 0.11532687 | 0.13861462 | 0.1313664 | 0.00724821 | 0.40311641 | 0.90650187 |
| Nudt16 | 0.17903544 | 0.20389828 | 0.19665073 | 0.00724755 | 0.43492755 | 0.90650187 |
| Ccdc77 | 0.16363109 | 0.1885377 | 0.18129569 | 0.00724201 | 0.42952506 | 0.90650187 |
| Arhgef15 | 0.03910666 | 0.05398811 | 0.04676078 | 0.00722733 | 0.24600627 | 0.90650187 |
| Socs5 | 0.12444728 | 0.14822521 | 0.14100741 | 0.0072178 | 0.4099946 | 0.90650187 |

| | | | | | | |
|---|---|---|---|---|---|---|
| Prima1 | 0.01825455 | 0.02935061 | 0.02213329 | 0.00721732 | 0.07357891 | 0.77295992 |
| Fuom | 0.37499881 | 0.37838512 | 0.37117368 | 0.00721143 | 0.46558515 | 0.90650187 |
| Gm13212 | 0.05030097 | 0.06691114 | 0.05970313 | 0.007208 | 0.29573006 | 0.90650187 |
| Rbfox1 | 0.02726901 | 0.04007143 | 0.03286405 | 0.00720738 | 0.16478963 | 0.90650187 |
| Ptger4 | 0.0344508 | 0.04851211 | 0.04132115 | 0.00719096 | 0.21956842 | 0.90650187 |
| Kcnk2 | 0.05544443 | 0.07277524 | 0.06558607 | 0.00718917 | 0.31302219 | 0.90650187 |
| Nr3c1 | 0.49901612 | 0.47204757 | 0.46486091 | 0.00718666 | 0.47260299 | 0.90650187 |
| Lhfpl4 | 0.09106369 | 0.11243298 | 0.10526233 | 0.00717065 | 0.38100502 | 0.90650187 |
| Gsc | 0.04760969 | 0.06376523 | 0.05660903 | 0.00715621 | 0.28706159 | 0.90650187 |
| Tirap | 0.06604965 | 0.08474652 | 0.07759162 | 0.0071549 | 0.34092519 | 0.90650187 |
| C330027C09I | 0.0709797 | 0.09026664 | 0.08311637 | 0.00715028 | 0.35106624 | 0.90650187 |
| Neurl2 | 0.03529746 | 0.04944326 | 0.04231284 | 0.00713041 | 0.22688099 | 0.90650187 |
| Pcdh20 | 0.04962691 | 0.06605142 | 0.05892921 | 0.00712221 | 0.29553098 | 0.90650187 |
| Pah | 0.0456709 | 0.06149521 | 0.05437324 | 0.00712197 | 0.28018266 | 0.90650187 |
| Sim1 | 0.10052789 | 0.12262515 | 0.11550444 | 0.00712071 | 0.39201677 | 0.90650187 |
| Dnm1 | 0.10230734 | 0.12454675 | 0.11742619 | 0.00712057 | 0.39374339 | 0.90650187 |
| Mocs2 | 0.78340593 | 0.65025288 | 0.6431434 | 0.00710948 | 0.48040269 | 0.90650187 |
| Trps1 | 0.19132596 | 0.21581682 | 0.20871313 | 0.00710369 | 0.43986595 | 0.90650187 |
| Rras | 0.27756685 | 0.29603554 | 0.28893337 | 0.00710216 | 0.45649187 | 0.90650187 |
| Chrna3 | 0.01210005 | 0.02182551 | 0.01473201 | 0.0070935 | 0.01615428 | 0.24507108 |
| Luzp2 | 0.01568785 | 0.0261343 | 0.01905408 | 0.00708022 | 0.04927512 | 0.58609633 |
| Flnb | 0.18864856 | 0.21317916 | 0.20609936 | 0.00707979 | 0.43931246 | 0.90650187 |
| Ccdc79 | 0.02138504 | 0.03293662 | 0.02587451 | 0.00706212 | 0.11249393 | 0.90650187 |
| Pdia5 | 0.34045762 | 0.35001955 | 0.34295861 | 0.00706095 | 0.46353668 | 0.90650187 |
| Serpinb10 | 0.0086069 | 0.01755789 | 0.01050381 | 0.00705408 | 0.00141533 | 0.03163897 |
| Akap2 | 0.06395134 | 0.08228002 | 0.07522942 | 0.0070506 | 0.33846785 | 0.90650187 |
| Luzp1 | 0.52035165 | 0.48691871 | 0.47988022 | 0.00703849 | 0.47400558 | 0.90650187 |
| Brat1 | 0.06953861 | 0.08854272 | 0.0815051 | 0.00703762 | 0.35054144 | 0.90650187 |
| Ptafr | 0.01193422 | 0.02156358 | 0.01453173 | 0.00703185 | 0.01572976 | 0.2400684 |
| Boc | 0.19828869 | 0.2225046 | 0.21547446 | 0.00703014 | 0.44233814 | 0.90650187 |

| | | | | | | |
|---|---|---|---|---|---|---|
| Mturn | 0.0676031 | 0.08636297 | 0.07933627 | 0.00702669 | 0.34688631 | 0.90650187 |
| Klhdc4 | 0.24234497 | 0.26409292 | 0.25707331 | 0.00701961 | 0.45169059 | 0.90650187 |
| Pts | 0.48068094 | 0.45870432 | 0.45170921 | 0.00699511 | 0.47255686 | 0.90650187 |
| Hmg20b | 0.79160064 | 0.65464558 | 0.64765933 | 0.00698625 | 0.48087629 | 0.90650187 |
| Fcgr2b | 0.00548498 | 0.01368866 | 0.00670799 | 0.00698066 | 1.86E-06 | 7.61E-05 |
| Dtl | 0.16546672 | 0.19010734 | 0.18313946 | 0.00696787 | 0.43284279 | 0.90650187 |
| Arhgdig | 0.11227504 | 0.13508296 | 0.12811764 | 0.00696533 | 0.40450915 | 0.90650187 |
| Ciapin1 | 0.70711273 | 0.6065469 | 0.59958374 | 0.00696316 | 0.47941246 | 0.90650187 |
| Tmem132c | 0.06614646 | 0.08465369 | 0.07770044 | 0.00695325 | 0.34537853 | 0.90650187 |
| Hexa | 0.91864496 | 0.72073234 | 0.71378422 | 0.00694812 | 0.48274139 | 0.90650187 |
| Erdr1 | 0.31726305 | 0.33038287 | 0.32344443 | 0.00693844 | 0.46201208 | 0.90650187 |
| Sox7 | 0.03253746 | 0.04601227 | 0.03907595 | 0.00693632 | 0.21501679 | 0.90650187 |
| Zfp30 | 0.26920213 | 0.28838467 | 0.28147549 | 0.00690918 | 0.45655239 | 0.90650187 |
| Ddah1 | 0.08552177 | 0.10611296 | 0.09920857 | 0.00690439 | 0.37851224 | 0.90650187 |
| Cers4 | 0.28870148 | 0.30565876 | 0.29875905 | 0.00689971 | 0.45911271 | 0.90650187 |
| Sema6c | 0.09065754 | 0.1117087 | 0.10482012 | 0.00688858 | 0.38508343 | 0.90650187 |
| Rab15 | 0.06141286 | 0.07924909 | 0.07236304 | 0.00688605 | 0.33613202 | 0.90650187 |
| Anxa11 | 0.34824047 | 0.35628871 | 0.34940327 | 0.00688543 | 0.4650948 | 0.90650187 |
| Gbas | 0.55041843 | 0.50742795 | 0.50054251 | 0.00688543 | 0.47561848 | 0.90650187 |
| Clec4a1 | 0.00717391 | 0.01564864 | 0.00876348 | 0.00688516 | 0.00023903 | 0.00658246 |
| Chst1 | 0.07082622 | 0.08981278 | 0.0829449 | 0.00686788 | 0.3564 | 0.90650187 |
| Antxr1 | 0.13357433 | 0.15740753 | 0.15055344 | 0.00685409 | 0.41980576 | 0.90650187 |
| 1110002L01F | 0.07578829 | 0.09532491 | 0.08847104 | 0.00685387 | 0.36527203 | 0.90650187 |
| Nsun4 | 0.10378783 | 0.1258615 | 0.11902204 | 0.00683946 | 0.39918329 | 0.90650187 |
| Skil | 0.2573396 | 0.27762119 | 0.27078349 | 0.0068377 | 0.45530926 | 0.90650187 |
| Prkrip1 | 0.50660756 | 0.47707451 | 0.47023911 | 0.0068354 | 0.47423772 | 0.90650187 |
| Rabepk | 0.20685612 | 0.2305543 | 0.22372156 | 0.00683274 | 0.44599979 | 0.90650187 |
| Kcna2 | 0.01552832 | 0.02569368 | 0.01886235 | 0.00683133 | 0.05369289 | 0.62276574 |
| Slc35b1 | 0.79841286 | 0.65821884 | 0.65138902 | 0.00682982 | 0.48141114 | 0.90650187 |
| Dach1 | 0.22399319 | 0.24681423 | 0.23998839 | 0.00682583 | 0.44969046 | 0.90650187 |

| | | | | | | |
|---|---|---|---|---|---|---|
| Dpep1 | 0.02891302 | 0.04162774 | 0.03480714 | 0.00682059 | 0.19184058 | 0.90650187 |
| Mmd2 | 0.13472796 | 0.15857305 | 0.15175294 | 0.00682011 | 0.42082034 | 0.90650187 |
| Eif2ak1 | 0.21549164 | 0.2387704 | 0.23195649 | 0.00681391 | 0.44804853 | 0.90650187 |
| Fkbp5 | 0.28038184 | 0.29823784 | 0.29142823 | 0.0068096 | 0.4586333 | 0.90650187 |
| Osgin2 | 0.12060614 | 0.14374854 | 0.13695936 | 0.00678917 | 0.41279077 | 0.90650187 |
| 2310039H08I | 0.72598263 | 0.61740666 | 0.61061998 | 0.00678668 | 0.48029618 | 0.90650187 |
| Akr1e1 | 0.23063946 | 0.25299867 | 0.24621481 | 0.00678386 | 0.45125625 | 0.90650187 |
| Egln3 | 0.11060958 | 0.13312343 | 0.12633987 | 0.00678357 | 0.40566992 | 0.90650187 |
| Twf2 | 0.44558113 | 0.4326476 | 0.42588013 | 0.00676747 | 0.47184089 | 0.90650187 |
| Syt4 | 0.03240359 | 0.04568216 | 0.03891865 | 0.00676351 | 0.21988444 | 0.90650187 |
| Tfap2a | 0.01055654 | 0.01962183 | 0.01286617 | 0.00675567 | 0.00979091 | 0.16463683 |
| Arhgef39 | 0.04665636 | 0.06223373 | 0.05551037 | 0.00672336 | 0.29513421 | 0.90650187 |
| Fam21 | 0.38036173 | 0.38218596 | 0.37546611 | 0.00671985 | 0.46829179 | 0.90650187 |
| Kat5 | 0.28286451 | 0.30033826 | 0.29362249 | 0.00671577 | 0.45950513 | 0.90650187 |
| Gm525 | 0.01757384 | 0.02803216 | 0.02131769 | 0.00671447 | 0.08072068 | 0.82406653 |
| Dab2ip | 0.19205164 | 0.2161222 | 0.20942024 | 0.00670196 | 0.44343327 | 0.90650187 |
| Tnfrsf25 | 0.01114327 | 0.0202745 | 0.01357588 | 0.00669863 | 0.01413364 | 0.22103441 |
| C1qtnf6 | 0.59261027 | 0.53527686 | 0.52859069 | 0.00668617 | 0.47757821 | 0.90650187 |
| Klhl42 | 0.15676498 | 0.18104633 | 0.17436534 | 0.00668099 | 0.4323722 | 0.90650187 |
| Rapgef3 | 0.05409194 | 0.07072265 | 0.06404298 | 0.00667967 | 0.32143996 | 0.90650187 |
| Gm15417 | 0.07899167 | 0.09867915 | 0.09201977 | 0.00665938 | 0.37383087 | 0.90650187 |
| Ppl | 0.07232003 | 0.09126955 | 0.08461226 | 0.00665728 | 0.36325199 | 0.90650187 |
| Zfp106 | 0.59582254 | 0.5373344 | 0.53068185 | 0.00665255 | 0.47777866 | 0.90650187 |
| Ybey | 0.05612115 | 0.07299471 | 0.06635713 | 0.00663758 | 0.32827342 | 0.90650187 |
| Larp6 | 0.03581013 | 0.04953934 | 0.04291279 | 0.00662656 | 0.24620313 | 0.90650187 |
| C230035I16R | 0.03328075 | 0.04657214 | 0.03994884 | 0.0066233 | 0.23054375 | 0.90650187 |
| Uap1 | 0.48424173 | 0.46089869 | 0.45428131 | 0.00661738 | 0.47418341 | 0.90650187 |
| Mief1 | 0.08385174 | 0.10398734 | 0.09737583 | 0.00661152 | 0.38138591 | 0.90650187 |
| Insc | 0.03718992 | 0.0511198 | 0.04452544 | 0.00659435 | 0.25513806 | 0.90650187 |
| Abcf3 | 0.40333899 | 0.40018096 | 0.39359976 | 0.00658119 | 0.47037276 | 0.90650187 |

| | | | | | | |
|---|---|---|---|---|---|---|
| Fam168a | 0.27713284 | 0.29512287 | 0.28854807 | 0.0065748 | 0.45965747 | 0.90650187 |
| Rbm25 | 1.67630506 | 0.92295413 | 0.91639143 | 0.0065627 | 0.48730101 | 0.90650187 |
| Tsc2 | 0.16598476 | 0.19022179 | 0.18365911 | 0.00656268 | 0.43689169 | 0.90650187 |
| Uxt | 0.28734166 | 0.30412682 | 0.29756519 | 0.00656163 | 0.4609541 | 0.90650187 |
| Sh3bp5 | 0.15063689 | 0.17468734 | 0.16813452 | 0.00655281 | 0.43122296 | 0.90650187 |
| Pcdhb14 | 0.04536048 | 0.06055992 | 0.05401474 | 0.00654518 | 0.29504963 | 0.90650187 |
| Rdh10 | 0.12281885 | 0.14583782 | 0.13929345 | 0.00654437 | 0.41727635 | 0.90650187 |
| Gnai2 | 1.90244277 | 0.933556 | 0.92701626 | 0.00653974 | 0.48749042 | 0.90650187 |
| Cpm | 0.04909465 | 0.06484021 | 0.0583176 | 0.00652261 | 0.30951537 | 0.90650187 |
| Mvk | 0.22212723 | 0.2447336 | 0.23823204 | 0.00650156 | 0.451717 | 0.90650187 |
| 0610038B21 | 0.02878432 | 0.04115635 | 0.03465519 | 0.00650116 | 0.20214708 | 0.90650187 |
| DHRSX | 0.37580423 | 0.3782995 | 0.37181979 | 0.00647971 | 0.46912336 | 0.90650187 |
| Ankrd11 | 0.7000056 | 0.60183882 | 0.5953807 | 0.00645812 | 0.48076978 | 0.90650187 |
| Gm10687 | 0.0127848 | 0.02199276 | 0.01555851 | 0.00643425 | 0.03299522 | 0.42758767 |
| Plac9a | 0.01088204 | 0.01969371 | 0.01325996 | 0.00643375 | 0.01550229 | 0.23755076 |
| Sytl5 | 0.11326319 | 0.13558495 | 0.12917082 | 0.00641413 | 0.41264279 | 0.90650187 |
| Pgm2 | 0.38833883 | 0.38821696 | 0.38180903 | 0.00640793 | 0.47026215 | 0.90650187 |
| Entpd7 | 0.26472603 | 0.28384991 | 0.27745676 | 0.00639315 | 0.45920549 | 0.90650187 |
| BC031181 | 1.12389551 | 0.80403372 | 0.79764611 | 0.00638761 | 0.48580036 | 0.90650187 |
| Casp9 | 0.10214027 | 0.12362959 | 0.11724593 | 0.00638366 | 0.40437115 | 0.90650187 |
| Papd4 | 0.17946132 | 0.20345142 | 0.19707146 | 0.00637996 | 0.44278127 | 0.90650187 |
| A230050P20 | 0.14993858 | 0.17380072 | 0.16742176 | 0.00637896 | 0.43274785 | 0.90650187 |
| Phf10 | 0.19222047 | 0.21594749 | 0.20958467 | 0.00636282 | 0.44631975 | 0.90650187 |
| Mtdh | 1.37134655 | 0.8744056 | 0.86805584 | 0.00634976 | 0.48702898 | 0.90650187 |
| Haus5 | 0.09129692 | 0.1118538 | 0.10551615 | 0.00633765 | 0.39472771 | 0.90650187 |
| Noxo1 | 0.04639528 | 0.0615468 | 0.05520925 | 0.00633755 | 0.30491436 | 0.90650187 |
| Ccdc6 | 0.2355721 | 0.257132 | 0.25080714 | 0.00632486 | 0.45536835 | 0.90650187 |
| Mipep | 0.1435015 | 0.16714841 | 0.16082386 | 0.00632455 | 0.43060753 | 0.90650187 |
| Gsk3b | 1.31409803 | 0.86065364 | 0.85433011 | 0.00632353 | 0.48687508 | 0.90650187 |
| Scd1 | 0.04161778 | 0.05599916 | 0.0496807 | 0.00631846 | 0.28590138 | 0.90650187 |

| | | | | | | |
|---|---|---|---|---|---|---|
| Nudt4 | 0.22233577 | 0.24473654 | 0.23842853 | 0.006308 | 0.45318615 | 0.90650187 |
| Dhfr | 0.20408771 | 0.22736024 | 0.22106512 | 0.00629511 | 0.44963072 | 0.90650187 |
| Ndufb2 | 1.90847138 | 0.93335804 | 0.9270695 | 0.00628854 | 0.48797146 | 0.90650187 |
| H2afy2 | 0.71166186 | 0.60854354 | 0.60226094 | 0.00628261 | 0.48150559 | 0.90650187 |
| Kdf1 | 0.21210139 | 0.2349903 | 0.22873273 | 0.00625756 | 0.45159966 | 0.90650187 |
| Leprotl1 | 0.52047294 | 0.48621887 | 0.47996473 | 0.00625414 | 0.47690294 | 0.90650187 |
| Ushbp1 | 0.03285479 | 0.04570043 | 0.03944872 | 0.00625171 | 0.24055254 | 0.90650187 |
| Lrfn2 | 0.02347549 | 0.03461443 | 0.02836405 | 0.00625039 | 0.16363009 | 0.90650187 |
| Irx2 | 0.06675286 | 0.08462583 | 0.07838182 | 0.00624401 | 0.36161559 | 0.90650187 |
| Cirh1a | 0.20762568 | 0.23070066 | 0.22445857 | 0.0062421 | 0.45080396 | 0.90650187 |
| Prdm5 | 0.11948223 | 0.14200516 | 0.13577152 | 0.00623364 | 0.41913382 | 0.90650187 |
| Ndufa11 | 1.96772221 | 0.93359238 | 0.92736122 | 0.00623116 | 0.48808494 | 0.90650187 |
| Phlda2 | 0.02282757 | 0.03382355 | 0.02759318 | 0.00623037 | 0.15773971 | 0.90650187 |
| Cd79b | 0.01094516 | 0.01956512 | 0.0133363 | 0.00622882 | 0.01893057 | 0.27776409 |
| Dtx3l | 0.05305882 | 0.06908035 | 0.06286242 | 0.00621793 | 0.33006658 | 0.90650187 |
| Faf2 | 0.65608415 | 0.5750296 | 0.56882571 | 0.00620389 | 0.48066448 | 0.90650187 |
| Foxf1 | 0.03587151 | 0.04918458 | 0.04298459 | 0.00619999 | 0.26068878 | 0.90650187 |
| Vdr | 0.19618296 | 0.21963285 | 0.21343509 | 0.00619776 | 0.44864237 | 0.90650187 |
| Gpr85 | 0.10798756 | 0.12973091 | 0.12353403 | 0.00619688 | 0.41176529 | 0.90650187 |
| Lzts2 | 0.67754606 | 0.58812283 | 0.58192808 | 0.00619475 | 0.48112732 | 0.90650187 |
| Izumo4 | 0.10546894 | 0.12702251 | 0.12083079 | 0.00619172 | 0.40989857 | 0.90650187 |
| Snx24 | 0.24062251 | 0.26167199 | 0.25548402 | 0.00618797 | 0.45712666 | 0.90650187 |
| Angel2 | 0.4164404 | 0.4099423 | 0.40375567 | 0.00618663 | 0.47284555 | 0.90650187 |
| D630024D03 | 0.01046092 | 0.01893462 | 0.01275045 | 0.00618418 | 0.01553435 | 0.23780228 |
| Tnfrsf1b | 0.00658131 | 0.01422349 | 0.0080428 | 0.00618068 | 0.00031735 | 0.00853823 |
| Enc1 | 0.14639397 | 0.16997222 | 0.16379461 | 0.00617761 | 0.43342203 | 0.90650187 |
| Tmem86b | 0.02164162 | 0.03235608 | 0.02618045 | 0.00617563 | 0.14716702 | 0.90650187 |
| Eif3a | 1.66187246 | 0.92104295 | 0.91487405 | 0.00616891 | 0.48804298 | 0.90650187 |
| Rabac1 | 1.5736436 | 0.91103866 | 0.90487037 | 0.00616828 | 0.48791205 | 0.90650187 |
| Hoxb8 | 0.02617939 | 0.03773738 | 0.03157384 | 0.00616355 | 0.1927439 | 0.90650187 |

| | | | | | | |
|---|---|---|---|---|---|---|
| Ntrk3 | 0.06257691 | 0.07984137 | 0.07367863 | 0.00616274 | 0.35500377 | 0.90650187 |
| Col25a1 | 0.03233831 | 0.04500061 | 0.03884193 | 0.00615869 | 0.24044091 | 0.90650187 |
| Kif4 | 0.03747739 | 0.05101496 | 0.04486106 | 0.00615391 | 0.27098552 | 0.90650187 |
| Pla2g15 | 0.07577524 | 0.09460541 | 0.08845656 | 0.00614885 | 0.37865096 | 0.90650187 |
| Dock6 | 0.15611038 | 0.17984358 | 0.17370183 | 0.00614175 | 0.43754843 | 0.90650187 |
| Calm2 | 3.57169527 | 0.76495151 | 0.75882247 | 0.00612905 | 0.48567811 | 0.90650187 |
| Olfml2a | 0.02505902 | 0.03637084 | 0.03024526 | 0.00612558 | 0.18396167 | 0.90650187 |
| Sharpin | 0.26438109 | 0.28325348 | 0.27714629 | 0.00610719 | 0.46098071 | 0.90650187 |
| Tcirg1 | 0.08274711 | 0.10226652 | 0.09616141 | 0.00610511 | 0.38887844 | 0.90650187 |
| Aldh1a1 | 0.04645112 | 0.06137437 | 0.05527367 | 0.0061007 | 0.31182883 | 0.90650187 |
| Hmgcl | 0.50507649 | 0.475258 | 0.46915745 | 0.00610055 | 0.47695112 | 0.90650187 |
| Hpse2 | 0.01096458 | 0.01945533 | 0.01335979 | 0.00609554 | 0.02126204 | 0.30347813 |
| Hic2 | 0.07784844 | 0.09684465 | 0.09075498 | 0.00608967 | 0.38273678 | 0.90650187 |
| Hars | 0.49560276 | 0.46851358 | 0.46243034 | 0.00608324 | 0.4766825 | 0.90650187 |
| Pygl | 0.17872413 | 0.20241035 | 0.19634306 | 0.00606729 | 0.44536686 | 0.90650187 |
| Hnmt | 0.02508839 | 0.03634669 | 0.03028012 | 0.00606657 | 0.18655322 | 0.90650187 |
| Erbb3 | 0.05086806 | 0.06641784 | 0.06035371 | 0.00606413 | 0.3275534 | 0.90650187 |
| Myl1 | 0.0082993 | 0.01618588 | 0.01013052 | 0.00605537 | 0.00393852 | 0.07666679 |
| Gm3411 | 0.01296936 | 0.02183622 | 0.01578114 | 0.00605507 | 0.04402861 | 0.53714196 |
| Igf2bp1 | 0.2987832 | 0.31360114 | 0.30755738 | 0.00604376 | 0.46519274 | 0.90650187 |
| Pramef8 | 0.14386319 | 0.16723487 | 0.16119588 | 0.006039 | 0.4338625 | 0.90650187 |
| Adra1a | 0.03237464 | 0.04491919 | 0.03888463 | 0.00603456 | 0.24512216 | 0.90650187 |
| Itga5 | 0.14489607 | 0.16827146 | 0.16225739 | 0.00601406 | 0.43456008 | 0.90650187 |
| Itgb3bp | 0.42321514 | 0.41496236 | 0.40895576 | 0.0060066 | 0.47396936 | 0.90650187 |
| Mras | 0.13942986 | 0.16262746 | 0.15662548 | 0.00600198 | 0.43236424 | 0.90650187 |
| Trub1 | 0.28297102 | 0.29970815 | 0.2937165 | 0.00599165 | 0.46387031 | 0.90650187 |
| Them4 | 0.13508379 | 0.15809643 | 0.15212259 | 0.00597384 | 0.43070544 | 0.90650187 |
| Ak7 | 0.01053688 | 0.01880974 | 0.01284237 | 0.00596736 | 0.01942816 | 0.28258006 |
| Alx4 | 0.0052065 | 0.01233399 | 0.00636863 | 0.00596536 | 1.56E-05 | 0.00054364 |
| Hpse | 0.03919416 | 0.05281307 | 0.04686268 | 0.00595038 | 0.2862137 | 0.90650187 |

| | | | | | | |
|---|---|---|---|---|---|---|
| Cldn23 | 0.0586805 | 0.07521534 | 0.0692671 | 0.00594825 | 0.35131888 | 0.90650187 |
| Rab10os | 0.16473925 | 0.18835671 | 0.18240922 | 0.0059475 | 0.44237515 | 0.90650187 |
| Tnpo2 | 0.2152198 | 0.23761913 | 0.23169844 | 0.00592069 | 0.45477724 | 0.90650187 |
| Ednra | 0.12263469 | 0.14501905 | 0.13909941 | 0.00591964 | 0.42497149 | 0.90650187 |
| 5730508B09 | 0.04358051 | 0.05787113 | 0.05195621 | 0.00591493 | 0.30639056 | 0.90650187 |
| Zfp367 | 0.07402438 | 0.09240689 | 0.08651069 | 0.00589621 | 0.38094727 | 0.90650187 |
| Brinp1 | 0.03163082 | 0.0439011 | 0.03801006 | 0.00589104 | 0.2454089 | 0.90650187 |
| Saysd1 | 0.43908874 | 0.42688866 | 0.42100491 | 0.00588375 | 0.47522985 | 0.90650187 |
| Cenpi | 0.04061323 | 0.05439557 | 0.04851378 | 0.00588179 | 0.29494912 | 0.90650187 |
| Zkscan5 | 0.06152263 | 0.07836834 | 0.07248719 | 0.00588115 | 0.35916625 | 0.90650187 |
| Myo5a | 0.09275051 | 0.11297619 | 0.10709641 | 0.00587978 | 0.40358776 | 0.90650187 |
| Zfat | 0.03705019 | 0.05023656 | 0.04436226 | 0.0058743 | 0.27804081 | 0.90650187 |
| Gm26699 | 0.68605661 | 0.59292506 | 0.58705669 | 0.00586837 | 0.48227707 | 0.90650187 |
| Cln8 | 0.22275972 | 0.24469261 | 0.23882786 | 0.00586475 | 0.4565346 | 0.90650187 |
| Sgsh | 0.03784938 | 0.0511583 | 0.04529515 | 0.00586315 | 0.28249229 | 0.90650187 |
| C78197 | 0.0213102 | 0.0316327 | 0.02578525 | 0.00584745 | 0.15669043 | 0.90650187 |
| Plekha1 | 0.35513233 | 0.36090877 | 0.35506658 | 0.00584219 | 0.47084455 | 0.90650187 |
| Fbp1 | 0.03966527 | 0.05324436 | 0.04741116 | 0.0058332 | 0.29220241 | 0.90650187 |
| Mkrn2os | 0.05268913 | 0.06827248 | 0.06243958 | 0.00583289 | 0.33896353 | 0.90650187 |
| Kif15 | 0.07060032 | 0.08852267 | 0.08269247 | 0.0058302 | 0.37697479 | 0.90650187 |
| Cradd | 0.20707413 | 0.22974793 | 0.22393041 | 0.00581752 | 0.45402716 | 0.90650187 |
| Sag | 0.19272108 | 0.21588854 | 0.21007204 | 0.0058165 | 0.45101772 | 0.90650187 |
| Rhox5 | 0.01315821 | 0.02181831 | 0.01600891 | 0.0058094 | 0.05334541 | 0.62090631 |
| Arrb1 | 0.0418357 | 0.05574176 | 0.04993364 | 0.00580812 | 0.30254374 | 0.90650187 |
| Serpinb9 | 0.08287982 | 0.10210732 | 0.0963074 | 0.00579991 | 0.39445456 | 0.90650187 |
| Hdac6 | 0.33839821 | 0.34704373 | 0.3412446 | 0.00579913 | 0.46988901 | 0.90650187 |
| Tbc1d30 | 0.05771752 | 0.07395957 | 0.06817334 | 0.00578623 | 0.35296708 | 0.90650187 |
| Rasd2 | 0.02955388 | 0.04134208 | 0.03556344 | 0.00577864 | 0.23503686 | 0.90650187 |
| Ngf | 0.07340387 | 0.09158521 | 0.08582001 | 0.0057652 | 0.38260505 | 0.90650187 |
| Nin | 0.1542136 | 0.17753676 | 0.17177643 | 0.00576034 | 0.44074582 | 0.90650187 |

| | | | | | | |
|---|---|---|---|---|---|---|
| H2-Q4 | 0.14945222 | 0.17268352 | 0.166925 | 0.00575851 | 0.43905608 | 0.90650187 |
| Tfpi2 | 0.06108447 | 0.07774197 | 0.07199153 | 0.00575044 | 0.36125746 | 0.90650187 |
| Wdhd1 | 0.11153635 | 0.13307794 | 0.12732956 | 0.00574838 | 0.42046654 | 0.90650187 |
| Hoxa2 | 0.09560652 | 0.11593938 | 0.11019274 | 0.00574664 | 0.40833036 | 0.90650187 |
| Slc18a3 | 0.00832903 | 0.01590444 | 0.0101666 | 0.00573784 | 0.00605336 | 0.10975223 |
| Unc45b | 0.02180128 | 0.03210628 | 0.02637076 | 0.00573552 | 0.16679745 | 0.90650187 |
| Drc1 | 0.10713172 | 0.1283515 | 0.12261635 | 0.00573516 | 0.41763979 | 0.90650187 |
| Myl7 | 0.02103694 | 0.03117947 | 0.02545927 | 0.0057202 | 0.15893591 | 0.90650187 |
| Zfp13 | 0.08088938 | 0.0998344 | 0.09411515 | 0.00571925 | 0.39352111 | 0.90650187 |
| Strbp | 0.56412309 | 0.51548838 | 0.50977203 | 0.00571635 | 0.48012055 | 0.90650187 |
| Bicd2 | 0.19787466 | 0.22078979 | 0.21507385 | 0.00571593 | 0.45297474 | 0.90650187 |
| Gm15050 | 0.07582405 | 0.09422645 | 0.08851074 | 0.0057157 | 0.38702509 | 0.90650187 |
| Mat2b | 0.56788029 | 0.51799634 | 0.51228257 | 0.00571377 | 0.48022684 | 0.90650187 |
| Trp53i13 | 0.55870876 | 0.51185009 | 0.50613929 | 0.0057108 | 0.47999742 | 0.90650187 |
| Trappc6a | 0.6304978 | 0.55858743 | 0.55287939 | 0.00570804 | 0.48169604 | 0.90650187 |
| Dennd4b | 0.03800826 | 0.05118559 | 0.0454805 | 0.00570509 | 0.28853878 | 0.90650187 |
| Plec | 0.30143361 | 0.31555224 | 0.30985453 | 0.0056977 | 0.46742387 | 0.90650187 |
| Sypl2 | 0.01061975 | 0.01862294 | 0.01294264 | 0.0056803 | 0.02552274 | 0.35078764 |
| E530001K10I | 0.03555627 | 0.04829508 | 0.04261577 | 0.00567931 | 0.27677256 | 0.90650187 |
| Il4ra | 0.07878418 | 0.09746945 | 0.09179035 | 0.00567909 | 0.39163858 | 0.90650187 |
| Arhgap25 | 0.01890926 | 0.02858201 | 0.02291703 | 0.00566498 | 0.1358987 | 0.90650187 |
| Ptpn6 | 0.15455494 | 0.17777998 | 0.17212323 | 0.00565675 | 0.4419201 | 0.90650187 |
| Trim26 | 0.35775862 | 0.36285933 | 0.35721443 | 0.0056449 | 0.47199655 | 0.90650187 |
| Syk | 0.02874092 | 0.04024577 | 0.03460393 | 0.00564183 | 0.23428342 | 0.90650187 |
| Jrkl | 0.17705694 | 0.20030609 | 0.19469355 | 0.00561254 | 0.44901279 | 0.90650187 |
| Stx7 | 0.84663414 | 0.68281123 | 0.67720045 | 0.00561078 | 0.48530909 | 0.90650187 |
| Ptprd | 0.17088981 | 0.1941746 | 0.18856456 | 0.00561005 | 0.44738824 | 0.90650187 |
| Ccdc42 | 0.00984682 | 0.01761596 | 0.01200693 | 0.00560903 | 0.01891163 | 0.27775432 |
| D630003M21 | 0.04336792 | 0.05727988 | 0.05171002 | 0.00556986 | 0.31602226 | 0.90650187 |
| Fads2 | 0.04266382 | 0.05645789 | 0.05089416 | 0.00556373 | 0.31348646 | 0.90650187 |

| | | | | | | |
|---|---|---|---|---|---|---|
| Pde4a | 0.06689302 | 0.08410137 | 0.07853923 | 0.00556215 | 0.37644243 | 0.90650187 |
| 1700021F05I | 0.32551052 | 0.33599245 | 0.33043687 | 0.00555558 | 0.4702095 | 0.90650187 |
| Cbr3 | 0.03589073 | 0.04855734 | 0.04300707 | 0.00555027 | 0.28307632 | 0.90650187 |
| Gm10392 | 0.01873724 | 0.0282472 | 0.02271117 | 0.00553603 | 0.13925932 | 0.90650187 |
| Kcnma1 | 0.11422006 | 0.1357226 | 0.1301895 | 0.00553309 | 0.42507006 | 0.90650187 |
| Dhx37 | 0.05862302 | 0.074731 | 0.06920184 | 0.00552915 | 0.36122023 | 0.90650187 |
| Arhgap15 | 0.01498062 | 0.02373129 | 0.01820376 | 0.00552752 | 0.08852458 | 0.87809516 |
| Mas1 | 0.01279898 | 0.02110253 | 0.01557562 | 0.00552692 | 0.05734058 | 0.64902021 |
| Itga6 | 0.26472046 | 0.28297716 | 0.27745175 | 0.00552541 | 0.4647263 | 0.90650187 |
| Nfkb2 | 0.05694148 | 0.07280937 | 0.0672909 | 0.00551847 | 0.3577122 | 0.90650187 |
| Ajap1 | 0.00746502 | 0.01463132 | 0.0091173 | 0.00551403 | 0.0035871 | 0.07083898 |
| Impdh1 | 0.32877108 | 0.33869666 | 0.33318458 | 0.00551208 | 0.47068564 | 0.90650187 |
| Pcdhb19 | 0.0258839 | 0.03670995 | 0.03122362 | 0.00548633 | 0.21735976 | 0.90650187 |
| Ccl7 | 0.01384165 | 0.02230675 | 0.01683265 | 0.0054741 | 0.07412442 | 0.77654394 |
| Ddx19a | 0.35156376 | 0.35760189 | 0.35213904 | 0.00546285 | 0.47250826 | 0.90650187 |
| Cd300lg | 0.01385341 | 0.02230251 | 0.01684681 | 0.0054557 | 0.07498012 | 0.78120142 |
| Neurl3 | 0.03049107 | 0.04210689 | 0.03666828 | 0.00543861 | 0.25482955 | 0.90650187 |
| Taok3 | 0.44881927 | 0.43373458 | 0.42829949 | 0.00543509 | 0.47750584 | 0.90650187 |
| E130114P18I | 0.01577249 | 0.02457984 | 0.01915579 | 0.00542406 | 0.10405213 | 0.90650187 |
| Pdia2 | 0.02044579 | 0.03017011 | 0.02475367 | 0.00541644 | 0.16533654 | 0.90650187 |
| Slc26a11 | 0.04933837 | 0.06400217 | 0.05859771 | 0.00540447 | 0.34089629 | 0.90650187 |
| Ap3m1 | 0.39269737 | 0.39064928 | 0.38525372 | 0.00539555 | 0.47517727 | 0.90650187 |
| Srgap1 | 0.12196544 | 0.14378263 | 0.13839393 | 0.0053887 | 0.4312863 | 0.90650187 |
| Gm19590 | 0.06148632 | 0.07782361 | 0.07244612 | 0.00537749 | 0.37070532 | 0.90650187 |
| Gab2 | 0.05104737 | 0.06593298 | 0.06055931 | 0.00537368 | 0.34661447 | 0.90650187 |
| Miip | 0.06992505 | 0.0873043 | 0.08193747 | 0.00536682 | 0.38545573 | 0.90650187 |
| Plxnb3 | 0.00725983 | 0.01423092 | 0.00886793 | 0.00536299 | 0.00358832 | 0.07083898 |
| Ephx2 | 0.05751707 | 0.07329809 | 0.0679455 | 0.0053526 | 0.36308862 | 0.90650187 |
| Fam102b | 0.14082705 | 0.16342069 | 0.15806835 | 0.00535233 | 0.4401724 | 0.90650187 |
| Zfp174 | 0.09175981 | 0.11136848 | 0.1060197 | 0.00534878 | 0.41126801 | 0.90650187 |

| | | | | | | |
|---|---|---|---|---|---|---|
| Gm9885 | 0.0201122 | 0.02970282 | 0.02435525 | 0.00534757 | 0.1645071 | 0.90650187 |
| Fastkd1 | 0.05679067 | 0.07245574 | 0.06711931 | 0.00533644 | 0.36187347 | 0.90650187 |
| Chrna4 | 0.03185753 | 0.04361201 | 0.03827671 | 0.0053353 | 0.26774009 | 0.90650187 |
| Mlip | 0.00998538 | 0.01748783 | 0.01217474 | 0.0053131 | 0.0261851 | 0.35922146 |
| Cct3 | 1.658161 | 0.91978905 | 0.91447628 | 0.00531277 | 0.48969753 | 0.90650187 |
| Cep63 | 0.25575894 | 0.27465709 | 0.26934866 | 0.00530843 | 0.46509104 | 0.90650187 |
| Thap11 | 0.52099506 | 0.48563651 | 0.4803284 | 0.00530811 | 0.48040847 | 0.90650187 |
| Efr3b | 0.01881391 | 0.02810079 | 0.02280293 | 0.00529786 | 0.15083508 | 0.90650187 |
| Ppp2r4 | 0.59191306 | 0.53342769 | 0.52813586 | 0.00529183 | 0.48223531 | 0.90650187 |
| Galnt18 | 0.26982712 | 0.28732251 | 0.2820351 | 0.0052874 | 0.46678933 | 0.90650187 |
| Rars | 0.60174941 | 0.5398114 | 0.53452468 | 0.00528671 | 0.48246446 | 0.90650187 |
| 0610010K14I | 0.74024772 | 0.62413291 | 0.61884729 | 0.00528562 | 0.48485571 | 0.90650187 |
| Pycard | 0.33711306 | 0.34545843 | 0.34017318 | 0.00528525 | 0.4724665 | 0.90650187 |
| Kcns3 | 0.02888802 | 0.04006137 | 0.03477763 | 0.00528374 | 0.24970507 | 0.90650187 |
| Jph2 | 0.00922012 | 0.01653027 | 0.01124751 | 0.00528276 | 0.01839777 | 0.27310979 |
| Grik5 | 0.18424767 | 0.20706235 | 0.20178624 | 0.0052761 | 0.45373147 | 0.90650187 |
| Cln3 | 0.18762042 | 0.21036426 | 0.2050936 | 0.00527066 | 0.45452109 | 0.90650187 |
| Adipor1 | 0.95601668 | 0.73695055 | 0.73168986 | 0.00526069 | 0.4872508 | 0.90650187 |
| Hist1h4i | 0.13608254 | 0.15841895 | 0.15315935 | 0.0052596 | 0.43933113 | 0.90650187 |
| Zfp760 | 0.08933369 | 0.10862398 | 0.10337717 | 0.00524681 | 0.41074358 | 0.90650187 |
| Hist1h2ak | 0.01594026 | 0.02460382 | 0.01935736 | 0.00524646 | 0.11412086 | 0.90650187 |
| Cmklr1 | 0.04702116 | 0.06117543 | 0.05593094 | 0.00524448 | 0.33839344 | 0.90650187 |
| Gab1 | 0.14639127 | 0.16902534 | 0.16379185 | 0.00523349 | 0.44352164 | 0.90650187 |
| Ina | 0.01716528 | 0.0260478 | 0.02082782 | 0.00521998 | 0.13260068 | 0.90650187 |
| Art4 | 0.02168658 | 0.03145317 | 0.02623404 | 0.00521913 | 0.18823188 | 0.90650187 |
| Dcaf12l1 | 0.11558961 | 0.13685917 | 0.13164556 | 0.00521361 | 0.43012295 | 0.90650187 |
| Cmc2 | 0.37710124 | 0.37806741 | 0.37285916 | 0.00520825 | 0.47524237 | 0.90650187 |
| Rnf165 | 0.11159021 | 0.13259011 | 0.12738704 | 0.00520307 | 0.4279566 | 0.90650187 |
| B3galnt1 | 0.25393925 | 0.27289476 | 0.26769388 | 0.00520089 | 0.46558557 | 0.90650187 |
| Ccm2l | 0.01087018 | 0.01844517 | 0.01324562 | 0.00519955 | 0.04048226 | 0.50180941 |

| | | | | | | |
|---|---|---|---|---|---|---|
| Gm15952 | 0.02808034 | 0.03901713 | 0.03382351 | 0.00519362 | 0.24742181 | 0.90650187 |
| Gpr160 | 0.02209245 | 0.0319051 | 0.02671776 | 0.00518735 | 0.19403293 | 0.90650187 |
| Acad8 | 0.29460867 | 0.30908764 | 0.3039258 | 0.00516184 | 0.46990701 | 0.90650187 |
| Zfp442 | 0.09949944 | 0.11955491 | 0.11439509 | 0.00515982 | 0.42053688 | 0.90650187 |
| BC051077 | 0.03675503 | 0.04916275 | 0.04401748 | 0.00514527 | 0.30165251 | 0.90650187 |
| Nkrf | 0.04769551 | 0.06184682 | 0.05670787 | 0.00513896 | 0.34352269 | 0.90650187 |
| Fras1 | 0.11856573 | 0.13993818 | 0.13480176 | 0.00513642 | 0.43274368 | 0.90650187 |
| 1810043H04 | 0.66263997 | 0.57798982 | 0.57285453 | 0.00513529 | 0.48410551 | 0.90650187 |
| Tbc1d8 | 0.12574181 | 0.14749613 | 0.14236749 | 0.00512864 | 0.4363821 | 0.90650187 |
| Mcl1 | 0.36730561 | 0.37009975 | 0.36497495 | 0.0051248 | 0.47511298 | 0.90650187 |
| B630019K06 | 0.25522388 | 0.27398448 | 0.26886243 | 0.00512205 | 0.46625295 | 0.90650187 |
| Vwc2 | 0.01880281 | 0.02790998 | 0.02278965 | 0.00512034 | 0.15893836 | 0.90650187 |
| Zfp286 | 0.08777398 | 0.10678341 | 0.10167398 | 0.00510943 | 0.41161007 | 0.90650187 |
| Nrip3 | 0.01964148 | 0.02889231 | 0.02379273 | 0.00509958 | 0.17033494 | 0.90650187 |
| Rnpepl1 | 0.31423574 | 0.32595803 | 0.32086246 | 0.00509557 | 0.47185807 | 0.90650187 |
| Frmd4a | 0.2704244 | 0.28766232 | 0.28256958 | 0.00509275 | 0.46806968 | 0.90650187 |
| Nexn | 0.08289398 | 0.10140569 | 0.09632297 | 0.00508271 | 0.40726619 | 0.90650187 |
| Akna | 0.02402661 | 0.03410038 | 0.02901922 | 0.00508115 | 0.21816417 | 0.90650187 |
| C6 | 0.01299572 | 0.02089365 | 0.01581294 | 0.00508071 | 0.07659178 | 0.79364116 |
| Coa5 | 0.21094097 | 0.23270569 | 0.22762657 | 0.00507912 | 0.46049114 | 0.90650187 |
| Scnn1g | 0.00992965 | 0.01718622 | 0.01210724 | 0.00507898 | 0.0310957 | 0.40827162 |
| Il6 | 0.00782647 | 0.01462815 | 0.00955641 | 0.00507174 | 0.00915341 | 0.15563866 |
| Tcf3 | 0.79258326 | 0.6532691 | 0.64819858 | 0.00507052 | 0.48612932 | 0.90650187 |
| Pdgfc | 0.04114834 | 0.05420326 | 0.04913556 | 0.0050677 | 0.32329418 | 0.90650187 |
| Adgrb3 | 0.09305051 | 0.112485 | 0.10742218 | 0.00506282 | 0.41702044 | 0.90650187 |
| Gfra4 | 0.04921236 | 0.06350832 | 0.0584529 | 0.00505542 | 0.35030859 | 0.90650187 |
| Ovol2 | 0.17752769 | 0.20021121 | 0.19515962 | 0.00505159 | 0.45419411 | 0.90650187 |
| Sh3rf3 | 0.01266463 | 0.02046476 | 0.01541352 | 0.00505125 | 0.0725729 | 0.76707662 |
| Pabpn1l | 0.10252918 | 0.12271295 | 0.11766549 | 0.00504746 | 0.42437955 | 0.90650187 |
| Zfp46 | 0.11915072 | 0.14045992 | 0.13542087 | 0.00503905 | 0.4343058 | 0.90650187 |

| | | | | | | |
|---|---|---|---|---|---|---|
| Cldn1 | 0.01939932 | 0.02854033 | 0.02350321 | 0.00503712 | 0.17035419 | 0.90650187 |
| Kat2a | 0.26573114 | 0.28339587 | 0.27836081 | 0.00503505 | 0.46795435 | 0.90650187 |
| Zfp383 | 0.1848294 | 0.20739018 | 0.20235758 | 0.0050326 | 0.45598206 | 0.90650187 |
| Rtp4 | 0.04382221 | 0.05726663 | 0.05223602 | 0.00503061 | 0.33427611 | 0.90650187 |
| Plcl1 | 0.12966756 | 0.15150335 | 0.14647957 | 0.00502378 | 0.43940807 | 0.90650187 |
| Smim12 | 0.2792852 | 0.29546997 | 0.29045718 | 0.00501279 | 0.46942175 | 0.90650187 |
| Irf9 | 0.10112748 | 0.12116404 | 0.11615242 | 0.00501162 | 0.42394386 | 0.90650187 |
| Epha5 | 0.01925047 | 0.02833632 | 0.0233252 | 0.00501112 | 0.16976822 | 0.90650187 |
| Phactr1 | 0.16335057 | 0.1860181 | 0.18101359 | 0.00500452 | 0.45108977 | 0.90650187 |
| Mfap1b | 0.38173585 | 0.38156121 | 0.37656229 | 0.00499892 | 0.47646967 | 0.90650187 |
| Hps3 | 0.07368678 | 0.09113137 | 0.08613498 | 0.00499639 | 0.39825201 | 0.90650187 |
| Dus2 | 0.12056009 | 0.14190515 | 0.13691073 | 0.00499442 | 0.4355848 | 0.90650187 |
| Oma1 | 0.0897529 | 0.10881978 | 0.10383436 | 0.00498542 | 0.41548883 | 0.90650187 |
| Ept1 | 0.24158034 | 0.26134859 | 0.25636816 | 0.00498043 | 0.46558842 | 0.90650187 |
| Ccnyl1 | 0.05271782 | 0.0674517 | 0.0624724 | 0.0049793 | 0.36154469 | 0.90650187 |
| Lgals2 | 0.02314231 | 0.03294584 | 0.02796772 | 0.00497811 | 0.21438575 | 0.90650187 |
| Cym | 0.00965696 | 0.01675014 | 0.01177692 | 0.00497323 | 0.03023735 | 0.40178149 |
| Xpr1 | 0.32310236 | 0.3333733 | 0.32840162 | 0.00497167 | 0.47317063 | 0.90650187 |
| Serpina1c | 0.03397225 | 0.0457142 | 0.04076013 | 0.00495407 | 0.29448538 | 0.90650187 |
| Spdef | 0.01531844 | 0.02356268 | 0.01861003 | 0.00495265 | 0.11838481 | 0.90650187 |
| B3galt2 | 0.02186269 | 0.03139076 | 0.02644397 | 0.00494679 | 0.20281005 | 0.90650187 |
| Gm19426 | 0.07231693 | 0.089554 | 0.0846088 | 0.0049452 | 0.39749522 | 0.90650187 |
| Rab22a | 0.65263048 | 0.5716358 | 0.56669418 | 0.00494162 | 0.48453846 | 0.90650187 |
| Nr4a2 | 0.09810236 | 0.1178205 | 0.11288937 | 0.00493113 | 0.42301399 | 0.90650187 |
| Mospd3 | 0.39686755 | 0.3934659 | 0.38853522 | 0.00493068 | 0.47750494 | 0.90650187 |
| Helz2 | 0.04246719 | 0.05559121 | 0.05066619 | 0.00492502 | 0.33282132 | 0.90650187 |
| Fbxo2 | 0.06025088 | 0.07596939 | 0.07104779 | 0.0049216 | 0.37905843 | 0.90650187 |
| Spryd4 | 0.12138648 | 0.14269854 | 0.13778318 | 0.00491535 | 0.43699397 | 0.90650187 |
| Atp2b4 | 0.26995266 | 0.28706055 | 0.28214747 | 0.00491308 | 0.46914788 | 0.90650187 |
| Chrm3 | 0.05744173 | 0.07277284 | 0.06785983 | 0.004913 | 0.37377983 | 0.90650187 |

| | | | | | | |
|---|---|---|---|---|---|---|
| Pi16 | 0.06257391 | 0.07858796 | 0.07367524 | 0.00491271 | 0.3834486 | 0.90650187 |
| Tmem18 | 0.08882332 | 0.10773204 | 0.10282022 | 0.00491182 | 0.41590893 | 0.90650187 |
| RP24-308M1 | 0.01551078 | 0.02374402 | 0.01884126 | 0.00490276 | 0.12367546 | 0.90650187 |
| Frmd6 | 0.1419492 | 0.16411707 | 0.15922554 | 0.00489154 | 0.4456843 | 0.90650187 |
| P2ry12 | 0.01252258 | 0.02013309 | 0.0152421 | 0.00489099 | 0.07685668 | 0.79500368 |
| Zfp738 | 0.06086752 | 0.07663593 | 0.07174602 | 0.00488992 | 0.38094705 | 0.90650187 |
| Ccbe1 | 0.02492074 | 0.03497007 | 0.03008115 | 0.00488892 | 0.2349876 | 0.90650187 |
| Tmem221 | 0.03990416 | 0.05257697 | 0.04768915 | 0.00488782 | 0.32432271 | 0.90650187 |
| Garnl3 | 0.07466431 | 0.09210869 | 0.0872224 | 0.00488629 | 0.40166203 | 0.90650187 |
| Mt3 | 0.05185597 | 0.06637204 | 0.06148589 | 0.00488615 | 0.36193832 | 0.90650187 |
| A730017C20 | 0.01388729 | 0.02176953 | 0.01688763 | 0.00488189 | 0.09937157 | 0.90650187 |
| Rasa4 | 0.03238244 | 0.04377512 | 0.03889379 | 0.00488133 | 0.28844199 | 0.90650187 |
| Casp7 | 0.15929671 | 0.18180244 | 0.17692698 | 0.00487546 | 0.45124969 | 0.90650187 |
| Gm28707 | 0.01192058 | 0.01938477 | 0.01451526 | 0.00486951 | 0.06792961 | 0.73233904 |
| Dcun1d2 | 0.12024869 | 0.14145092 | 0.13658175 | 0.00486917 | 0.4370366 | 0.90650187 |
| Ano10 | 0.20986216 | 0.23146396 | 0.22659694 | 0.00486701 | 0.46196444 | 0.90650187 |
| Prosc | 0.41414339 | 0.40684603 | 0.4019844 | 0.00486163 | 0.47856101 | 0.90650187 |
| Guf1 | 0.43556524 | 0.42320189 | 0.41834616 | 0.00485572 | 0.47942374 | 0.90650187 |
| Alkbh1 | 0.56584822 | 0.51578098 | 0.5109258 | 0.00485518 | 0.48315157 | 0.90650187 |
| 5730455P16I | 0.23220216 | 0.2525227 | 0.24767233 | 0.00485036 | 0.46531119 | 0.90650187 |
| Cyb561 | 0.36327664 | 0.36654412 | 0.36170889 | 0.00483522 | 0.47630577 | 0.90650187 |
| Fmnl2 | 0.18625272 | 0.20856822 | 0.20375391 | 0.0048143 | 0.45817168 | 0.90650187 |
| Ccz1 | 0.66014241 | 0.57613659 | 0.57132234 | 0.00481424 | 0.48505877 | 0.90650187 |
| Lrrn4cl | 0.04873365 | 0.06270962 | 0.05790256 | 0.00480707 | 0.35603631 | 0.90650187 |
| Ccdc97 | 0.15198213 | 0.17431062 | 0.16950601 | 0.0048046 | 0.44986222 | 0.90650187 |
| Tctex1d2 | 0.51624658 | 0.48181541 | 0.47701384 | 0.00480158 | 0.48215367 | 0.90650187 |
| 4833420G17 | 0.3449349 | 0.35145714 | 0.34667271 | 0.00478443 | 0.47553871 | 0.90650187 |
| Nadk2 | 0.09756424 | 0.11708858 | 0.11230869 | 0.00477989 | 0.42496552 | 0.90650187 |
| Nek6 | 0.25323222 | 0.27180823 | 0.26705005 | 0.00475818 | 0.46843288 | 0.90650187 |
| Pold1 | 0.11509266 | 0.13587509 | 0.13111749 | 0.0047576 | 0.435925 | 0.90650187 |

| | | | | | | |
|---|---|---|---|---|---|---|
| Plce1 | 0.06281778 | 0.07870582 | 0.07395061 | 0.00475521 | 0.38749122 | 0.90650187 |
| Mag | 0.01280399 | 0.0203182 | 0.01558166 | 0.00473654 | 0.08828653 | 0.8765797 |
| Cog4 | 0.38845678 | 0.38663396 | 0.38190244 | 0.00473151 | 0.47803813 | 0.90650187 |
| Csmd1 | 0.01716744 | 0.02556034 | 0.02083041 | 0.00472993 | 0.15637745 | 0.90650187 |
| Parp16 | 0.07678427 | 0.09430515 | 0.08957598 | 0.00472918 | 0.40721832 | 0.90650187 |
| Gm17586 | 0.06220555 | 0.07797839 | 0.07325914 | 0.00471925 | 0.38729377 | 0.90650187 |
| Kcnk6 | 0.03223225 | 0.04343084 | 0.03871728 | 0.00471357 | 0.29417718 | 0.90650187 |
| Grin3a | 0.04674191 | 0.06032159 | 0.05560903 | 0.00471256 | 0.35318302 | 0.90650187 |
| Zfp418 | 0.01940832 | 0.02822175 | 0.02351398 | 0.00470777 | 0.18671625 | 0.90650187 |
| Zdhhc15 | 0.06751867 | 0.08394755 | 0.07924154 | 0.00470601 | 0.39588369 | 0.90650187 |
| Ap3s2 | 0.42305706 | 0.41353961 | 0.40883484 | 0.00470477 | 0.47959943 | 0.90650187 |
| Dapl1 | 0.01204937 | 0.01937544 | 0.01467081 | 0.00470464 | 0.07698926 | 0.79547148 |
| 2410021H03I | 0.05454528 | 0.06926184 | 0.06456051 | 0.00470133 | 0.3730706 | 0.90650187 |
| Klhl5 | 0.04581561 | 0.05924096 | 0.05454031 | 0.00470065 | 0.35080343 | 0.90650187 |
| Zfp180 | 0.09185415 | 0.11081632 | 0.10612229 | 0.00469403 | 0.42205486 | 0.90650187 |
| Ptk7 | 0.15909118 | 0.18140417 | 0.1767193 | 0.00468487 | 0.45309162 | 0.90650187 |
| Pus7l | 0.07637736 | 0.09380231 | 0.08912472 | 0.00467759 | 0.40775589 | 0.90650187 |
| Srcap | 0.1187737 | 0.13969877 | 0.1350219 | 0.00467687 | 0.43881026 | 0.90650187 |
| Kntc1 | 0.02289317 | 0.0323455 | 0.02767126 | 0.00467424 | 0.22634019 | 0.90650187 |
| Proser2 | 0.11829932 | 0.13919109 | 0.13451967 | 0.00467142 | 0.43865462 | 0.90650187 |
| Espl1 | 0.0185184 | 0.02711549 | 0.02244922 | 0.00466627 | 0.17772808 | 0.90650187 |
| 1700012C14I | 0.02528205 | 0.03517482 | 0.03050989 | 0.00466493 | 0.24833844 | 0.90650187 |
| Mob1a | 0.27211647 | 0.28874626 | 0.28408187 | 0.00466439 | 0.47090577 | 0.90650187 |
| Dapk2 | 0.02014646 | 0.02905319 | 0.02439617 | 0.00465702 | 0.19804409 | 0.90650187 |
| Atg5 | 0.39007718 | 0.3878329 | 0.38318467 | 0.00464823 | 0.47849646 | 0.90650187 |
| Gda | 0.02144302 | 0.03058342 | 0.02594365 | 0.00463977 | 0.21329118 | 0.90650187 |
| Amph | 0.037776 | 0.0498382 | 0.04520954 | 0.00462866 | 0.3245 | 0.90650187 |
| Gm26542 | 0.01850304 | 0.0270563 | 0.02243083 | 0.00462547 | 0.17964201 | 0.90650187 |
| Ache | 0.01220279 | 0.019472 | 0.01485606 | 0.00461594 | 0.08359379 | 0.84264534 |
| Sult4a1 | 0.06004274 | 0.07541104 | 0.07081198 | 0.00459905 | 0.3863938 | 0.90650187 |

| | | | | | | |
|---|---|---|---|---|---|---|
| Gstt2 | 0.17272506 | 0.19498706 | 0.19039307 | 0.004594 | 0.45728815 | 0.90650187 |
| Ier5l | 0.03421342 | 0.04563654 | 0.04104291 | 0.00459364 | 0.30939546 | 0.90650187 |
| Bmp5 | 0.01339933 | 0.02088139 | 0.01629961 | 0.00458177 | 0.10571437 | 0.90650187 |
| Ska1 | 0.05631871 | 0.07116123 | 0.0665821 | 0.00457913 | 0.37990019 | 0.90650187 |
| Trmt61a | 0.1617677 | 0.18399835 | 0.17942015 | 0.00457821 | 0.45484201 | 0.90650187 |
| Gyg | 0.44534486 | 0.43027141 | 0.42570332 | 0.00456809 | 0.4809759 | 0.90650187 |
| Adam19 | 0.1159558 | 0.13658903 | 0.13203449 | 0.00455454 | 0.43906053 | 0.90650187 |
| Hspb7 | 0.04004514 | 0.0523971 | 0.04785317 | 0.00454393 | 0.33646299 | 0.90650187 |
| Apaf1 | 0.12988707 | 0.15124397 | 0.14670895 | 0.00453502 | 0.44534891 | 0.90650187 |
| Dpp6 | 0.07314819 | 0.09006985 | 0.08553525 | 0.0045346 | 0.40684005 | 0.90650187 |
| Sec61a1 | 0.8551155 | 0.68616221 | 0.68163351 | 0.0045287 | 0.4882185 | 0.90650187 |
| Cerkl | 0.10826724 | 0.12836231 | 0.12383372 | 0.00452859 | 0.43542638 | 0.90650187 |
| Slc19a1 | 0.14137993 | 0.16315887 | 0.15863868 | 0.00452019 | 0.4496002 | 0.90650187 |
| Ccbl2 | 0.09332939 | 0.11224145 | 0.10772491 | 0.00451654 | 0.42606987 | 0.90650187 |
| Epsti1 | 0.02295041 | 0.03225479 | 0.02773938 | 0.00451541 | 0.23463867 | 0.90650187 |
| Ubxn4 | 1.35845181 | 0.86961216 | 0.86509906 | 0.00451311 | 0.49074851 | 0.90650187 |
| Pik3ap1 | 0.01763998 | 0.02590131 | 0.02139696 | 0.00450435 | 0.17467267 | 0.90650187 |
| Atp6v0a1 | 0.23868538 | 0.25819722 | 0.25369314 | 0.00450408 | 0.46854512 | 0.90650187 |
| Ppp1r37 | 0.49838227 | 0.46891063 | 0.46441015 | 0.00450048 | 0.48281838 | 0.90650187 |
| Ganc | 0.11542035 | 0.13595886 | 0.13146573 | 0.00449313 | 0.43961784 | 0.90650187 |
| Ccdc137 | 0.2428398 | 0.26202161 | 0.25752935 | 0.00449226 | 0.46909389 | 0.90650187 |
| Arrdc3 | 0.26065049 | 0.27827171 | 0.27378115 | 0.00449056 | 0.47093617 | 0.90650187 |
| Dtwd2 | 0.04312853 | 0.05592137 | 0.05143272 | 0.00448865 | 0.34901611 | 0.90650187 |
| Akr1b3 | 0.5522266 | 0.50625418 | 0.50176678 | 0.0044874 | 0.48414305 | 0.90650187 |
| Prr11 | 0.06874708 | 0.08510505 | 0.08061882 | 0.00448624 | 0.40230437 | 0.90650187 |
| Tlr13 | 0.00784564 | 0.01406198 | 0.0095797 | 0.00448228 | 0.0187594 | 0.27607614 |
| Cbr2 | 0.00784564 | 0.01406198 | 0.0095797 | 0.00448228 | 0.0187594 | 0.27607614 |
| 5031425E22I | 0.39394594 | 0.39071882 | 0.38623779 | 0.00448103 | 0.47943299 | 0.90650187 |
| C920006O11 | 0.05688708 | 0.07170579 | 0.06722901 | 0.00447678 | 0.38360245 | 0.90650187 |
| Ppp1r21 | 0.20234086 | 0.22385444 | 0.21938479 | 0.00446965 | 0.46391598 | 0.90650187 |

| | | | | | | |
|---|---|---|---|---|---|---|
| Nwd2 | 0.00710427 | 0.01313286 | 0.00867883 | 0.00445403 | 0.01125898 | 0.18462538 |
| Grem1 | 0.01669179 | 0.02471249 | 0.02025975 | 0.00445274 | 0.16426806 | 0.90650187 |
| Gabra1 | 0.05096942 | 0.06491296 | 0.06046994 | 0.00444302 | 0.37196994 | 0.90650187 |
| Adora3 | 0.01314549 | 0.02042816 | 0.01599357 | 0.00443459 | 0.10885329 | 0.90650187 |
| Atxn1 | 0.18473975 | 0.20670097 | 0.20226955 | 0.00443142 | 0.46120578 | 0.90650187 |
| Sh2b1 | 0.2426715 | 0.26180473 | 0.25737427 | 0.00443046 | 0.46949988 | 0.90650187 |
| Myrf | 0.17405817 | 0.19613124 | 0.19171888 | 0.00441236 | 0.45925353 | 0.90650187 |
| Bcl9l | 0.14863805 | 0.17048972 | 0.16609272 | 0.004397 | 0.45315693 | 0.90650187 |
| Slc35d2 | 0.04494123 | 0.05792332 | 0.05353031 | 0.00439301 | 0.35761825 | 0.90650187 |
| Itga1 | 0.0617326 | 0.07711497 | 0.0727246 | 0.00439037 | 0.39420326 | 0.90650187 |
| Nmrk1 | 0.17037642 | 0.19243197 | 0.18805239 | 0.00437958 | 0.4587694 | 0.90650187 |
| Trim59 | 0.10397641 | 0.12360026 | 0.11922511 | 0.00437516 | 0.43520462 | 0.90650187 |
| Acadsb | 0.34530105 | 0.35134179 | 0.34697571 | 0.00436609 | 0.47769471 | 0.90650187 |
| Il17re | 0.11276385 | 0.1330014 | 0.12863877 | 0.00436263 | 0.44007959 | 0.90650187 |
| Cib2 | 0.2471109 | 0.26581627 | 0.26145561 | 0.00436067 | 0.47044724 | 0.90650187 |
| Sgip1 | 0.02951884 | 0.03987844 | 0.03552211 | 0.00435633 | 0.29280938 | 0.90650187 |
| Mansc4 | 0.01045435 | 0.01709177 | 0.01274249 | 0.00434928 | 0.06458526 | 0.70630256 |
| Syt1 | 0.03096715 | 0.04157442 | 0.037229 | 0.00434542 | 0.3019162 | 0.90650187 |
| B230118H07 | 0.5094492 | 0.47658657 | 0.4722426 | 0.00434398 | 0.48369044 | 0.90650187 |
| Slc25a26 | 0.06138349 | 0.07667188 | 0.07232982 | 0.00434206 | 0.39478242 | 0.90650187 |
| Shisa8 | 0.01921077 | 0.0276125 | 0.02327772 | 0.00433478 | 0.20387468 | 0.90650187 |
| Stox1 | 0.06027583 | 0.07541004 | 0.07107605 | 0.00433398 | 0.39316534 | 0.90650187 |
| Armt1 | 0.16355855 | 0.18555018 | 0.18122274 | 0.00432744 | 0.45772883 | 0.90650187 |
| RP23-49I18.5 | 0.01484275 | 0.02236228 | 0.01803791 | 0.00432438 | 0.14326117 | 0.90650187 |
| Mblac1 | 0.07674207 | 0.09385068 | 0.08952918 | 0.0043215 | 0.4150452 | 0.90650187 |
| 1700112E06I | 0.02659084 | 0.03638131 | 0.03206126 | 0.00432005 | 0.27458072 | 0.90650187 |
| Kdm6a | 0.2722599 | 0.28852884 | 0.28420989 | 0.00431895 | 0.47306919 | 0.90650187 |
| Rhot1 | 0.42153302 | 0.41198133 | 0.40766809 | 0.00431324 | 0.48124236 | 0.90650187 |
| Hnrnpm | 1.66398425 | 0.9193835 | 0.91509899 | 0.00428451 | 0.49169685 | 0.90650187 |
| Dpy19l4 | 0.06960614 | 0.08586392 | 0.08158067 | 0.00428325 | 0.40772194 | 0.90650187 |

| | | | | | | |
|---|---|---|---|---|---|---|
| Appl1 | 0.43296683 | 0.42065776 | 0.41637972 | 0.00427804 | 0.48178432 | 0.90650187 |
| Cab39l | 0.19237582 | 0.21401234 | 0.20973594 | 0.0042764 | 0.46388797 | 0.90650187 |
| Atp6v0e2 | 0.31689708 | 0.32740715 | 0.32313273 | 0.00427443 | 0.47655297 | 0.90650187 |
| Arhgap11a | 0.05212436 | 0.06606565 | 0.06179322 | 0.00427244 | 0.37927974 | 0.90650187 |
| Ephb1 | 0.02464511 | 0.03402455 | 0.02975392 | 0.00427063 | 0.26170893 | 0.90650187 |
| Crybg3 | 0.0887489 | 0.10700428 | 0.10273898 | 0.0042653 | 0.42678575 | 0.90650187 |
| 5330434G04 | 0.02576818 | 0.03534778 | 0.03108644 | 0.00426133 | 0.27112822 | 0.90650187 |
| Evc | 0.09548874 | 0.11432325 | 0.11006527 | 0.00425798 | 0.43172568 | 0.90650187 |
| Btnl9 | 0.01178055 | 0.01860334 | 0.01434611 | 0.00425723 | 0.09354418 | 0.90571023 |
| Fzd4 | 0.12344414 | 0.14420886 | 0.13995196 | 0.00425689 | 0.44621853 | 0.90650187 |
| Adamts14 | 0.03548011 | 0.04678228 | 0.04252663 | 0.00425566 | 0.32820558 | 0.90650187 |
| Arhgap44 | 0.12526499 | 0.14611952 | 0.14186677 | 0.00425275 | 0.44699143 | 0.90650187 |
| Gspt2 | 0.14519081 | 0.16681022 | 0.16256007 | 0.00425015 | 0.45373478 | 0.90650187 |
| Rnf135 | 0.1596763 | 0.18155307 | 0.17731043 | 0.00424263 | 0.45764312 | 0.90650187 |
| Ublcp1 | 0.2118386 | 0.23272328 | 0.22848235 | 0.00424093 | 0.46711812 | 0.90650187 |
| Pirt | 0.00938397 | 0.01568541 | 0.01144611 | 0.00423929 | 0.04982861 | 0.58991024 |
| Trib3 | 0.02620661 | 0.03584247 | 0.03160609 | 0.00423638 | 0.2756295 | 0.90650187 |
| Arhgap28 | 0.12458791 | 0.14539032 | 0.14115528 | 0.00423504 | 0.44694635 | 0.90650187 |
| Rps24 | 4.76076867 | 0.57632317 | 0.57210016 | 0.00422301 | 0.4869108 | 0.90650187 |
| Gid8 | 0.41517114 | 0.40699571 | 0.4027774 | 0.0042183 | 0.48143235 | 0.90650187 |
| Gm1976 | 0.02911129 | 0.03924396 | 0.0350412 | 0.00420276 | 0.29694906 | 0.90650187 |
| Mirt1 | 0.01996928 | 0.02838694 | 0.02418449 | 0.00420245 | 0.21991033 | 0.90650187 |
| Mkx | 0.02926415 | 0.03941782 | 0.0352216 | 0.00419622 | 0.2981811 | 0.90650187 |
| A730046J19F | 0.02218761 | 0.03101573 | 0.02683112 | 0.00418461 | 0.24404743 | 0.90650187 |
| Jag2 | 0.02042305 | 0.02891029 | 0.02472651 | 0.00418378 | 0.22596324 | 0.90650187 |
| 1810012K16I | 0.01191323 | 0.01868494 | 0.01450638 | 0.00417856 | 0.10017483 | 0.90650187 |
| 4930403P22I | 0.02183974 | 0.03059176 | 0.02641661 | 0.00417515 | 0.2411435 | 0.90650187 |
| Map3k8 | 0.07448555 | 0.09119437 | 0.08702365 | 0.00417072 | 0.4156395 | 0.90650187 |
| Impact | 0.26165016 | 0.27883684 | 0.27468419 | 0.00415265 | 0.47320797 | 0.90650187 |
| Disp1 | 0.09260514 | 0.1110813 | 0.10693851 | 0.00414279 | 0.43163125 | 0.90650187 |

| | | | | | | |
|---|---|---|---|---|---|---|
| Txlna | 0.66855429 | 0.58060734 | 0.57646911 | 0.00413823 | 0.48727067 | 0.90650187 |
| Tmem17 | 0.14713813 | 0.16869273 | 0.16455733 | 0.00413539 | 0.45552273 | 0.90650187 |
| Tgfbr3 | 0.13975879 | 0.16109657 | 0.15696537 | 0.0041312 | 0.45342817 | 0.90650187 |
| 6430573F11I | 0.06270153 | 0.077947 | 0.07381936 | 0.00412764 | 0.40184358 | 0.90650187 |
| Aak1 | 0.31142091 | 0.32258027 | 0.31845453 | 0.00412574 | 0.47703557 | 0.90650187 |
| Igsf5 | 0.14785567 | 0.16941507 | 0.16529217 | 0.00412291 | 0.45585279 | 0.90650187 |
| Dpy19l1 | 0.09734761 | 0.11619497 | 0.11207481 | 0.00412016 | 0.43508909 | 0.90650187 |
| Kantr | 0.04674827 | 0.05972996 | 0.05561635 | 0.00411361 | 0.37114777 | 0.90650187 |
| Ptx3 | 0.0313438 | 0.04178497 | 0.03767235 | 0.00411262 | 0.31372526 | 0.90650187 |
| 1110032F04I | 0.0069658 | 0.01262194 | 0.00851046 | 0.00411149 | 0.01586811 | 0.24169445 |
| Nxph4 | 0.02146138 | 0.03007645 | 0.02596554 | 0.0041109 | 0.24076782 | 0.90650187 |
| Nradd | 0.11843728 | 0.13877223 | 0.13466575 | 0.00410648 | 0.44608306 | 0.90650187 |
| Casp12 | 0.09926142 | 0.11823776 | 0.11413876 | 0.00409901 | 0.4365773 | 0.90650187 |
| Efcc1 | 0.07108845 | 0.08733276 | 0.08323784 | 0.00409493 | 0.41344038 | 0.90650187 |
| A230065H16 | 0.00987381 | 0.01613212 | 0.01203961 | 0.00409251 | 0.06537484 | 0.71135498 |
| Akr1b10 | 0.07987342 | 0.09707571 | 0.092994 | 0.00408171 | 0.42264668 | 0.90650187 |
| Sox5 | 0.09367774 | 0.11218338 | 0.10810291 | 0.00408047 | 0.43336854 | 0.90650187 |
| Tacc2 | 0.16579777 | 0.187551 | 0.18347157 | 0.00407943 | 0.46063004 | 0.90650187 |
| Gm26532 | 0.02357442 | 0.03255138 | 0.0284817 | 0.00406968 | 0.26264166 | 0.90650187 |
| Socs4 | 0.23798268 | 0.25710997 | 0.25304259 | 0.00406738 | 0.47151653 | 0.90650187 |
| Cdr2 | 0.10057983 | 0.11959815 | 0.11556059 | 0.00403756 | 0.43828264 | 0.90650187 |
| Nxf1 | 1.16722074 | 0.81612698 | 0.81209207 | 0.00403491 | 0.49118882 | 0.90650187 |
| Mrap2 | 0.03049401 | 0.04069568 | 0.03667174 | 0.00402394 | 0.31284225 | 0.90650187 |
| Mrgbp | 0.21563467 | 0.23611616 | 0.23209224 | 0.00402393 | 0.46928146 | 0.90650187 |
| Irx1 | 0.04929154 | 0.06256668 | 0.0585439 | 0.00402278 | 0.3800021 | 0.90650187 |
| Gpatch2l | 0.05260594 | 0.06636613 | 0.0623444 | 0.00402173 | 0.38714112 | 0.90650187 |
| Prkx | 0.10068892 | 0.11969901 | 0.1156785 | 0.00402051 | 0.43860342 | 0.90650187 |
| Osbpl10 | 0.0198445 | 0.02805494 | 0.02403539 | 0.00401956 | 0.22860146 | 0.90650187 |
| Rasgef1a | 0.01321954 | 0.02009959 | 0.01608286 | 0.00401673 | 0.13343478 | 0.90650187 |
| Epm2aip1 | 0.38200592 | 0.38078712 | 0.37677756 | 0.00400956 | 0.48113355 | 0.90650187 |

| | | | | | | |
|---|---|---|---|---|---|---|
| Oit3 | 0.01018046 | 0.01641626 | 0.01241096 | 0.0040053 | 0.07568628 | 0.78632364 |
| Depdc7 | 0.03912794 | 0.05078213 | 0.04678555 | 0.00399657 | 0.35206246 | 0.90650187 |
| Hgsnat | 0.20038843 | 0.22149706 | 0.21750289 | 0.00399417 | 0.46746729 | 0.90650187 |
| Ccdc88b | 0.01256031 | 0.01928083 | 0.01528763 | 0.0039932 | 0.12277789 | 0.90650187 |
| Casp8ap2 | 0.37484993 | 0.37504672 | 0.3710542 | 0.00399252 | 0.48092407 | 0.90650187 |
| Gm9242 | 0.02299477 | 0.03178281 | 0.02779217 | 0.00399064 | 0.26162679 | 0.90650187 |
| Dnah1 | 0.02518646 | 0.03438617 | 0.03039648 | 0.00398969 | 0.27977583 | 0.90650187 |
| Inpp4b | 0.01982314 | 0.02798209 | 0.02400985 | 0.00397224 | 0.23102039 | 0.90650187 |
| BC030867 | 0.02435646 | 0.0333822 | 0.02941112 | 0.00397108 | 0.27417188 | 0.90650187 |
| Arhgef19 | 0.09252112 | 0.11081213 | 0.10684723 | 0.0039649 | 0.43448455 | 0.90650187 |
| Dnajc14 | 0.46014991 | 0.4406586 | 0.4367067 | 0.0039519 | 0.48395499 | 0.90650187 |
| Ficd | 0.1368869 | 0.15793724 | 0.15399347 | 0.00394377 | 0.45467761 | 0.90650187 |
| Tead4 | 0.01737924 | 0.0250213 | 0.02108439 | 0.00393691 | 0.20324403 | 0.90650187 |
| Il2 | 0.14826019 | 0.16963759 | 0.16570617 | 0.00393142 | 0.45800023 | 0.90650187 |
| Nup214 | 0.21110561 | 0.23168893 | 0.22778359 | 0.00390534 | 0.46962215 | 0.90650187 |
| Arsa | 0.1628795 | 0.18444378 | 0.18053966 | 0.00390412 | 0.4617067 | 0.90650187 |
| E2f3 | 0.07669089 | 0.09337062 | 0.08947244 | 0.00389819 | 0.42320981 | 0.90650187 |
| Gtf2e2 | 1.03522903 | 0.76883662 | 0.76494291 | 0.00389372 | 0.49097308 | 0.90650187 |
| Cnr1 | 0.02232908 | 0.03087975 | 0.02699964 | 0.00388011 | 0.26145173 | 0.90650187 |
| Patl1 | 0.14841159 | 0.16973644 | 0.16586108 | 0.00387536 | 0.45863542 | 0.90650187 |
| Zfp456 | 0.02438285 | 0.03329863 | 0.02944247 | 0.00385616 | 0.28019801 | 0.90650187 |
| Lrrc4b | 0.0106102 | 0.01678549 | 0.0129311 | 0.0038544 | 0.09256732 | 0.90084241 |
| Sumo3 | 1.29433873 | 0.85319663 | 0.84935008 | 0.00384655 | 0.4919685 | 0.90650187 |
| Msantd2 | 0.07972136 | 0.09667214 | 0.09282607 | 0.00384608 | 0.42693007 | 0.90650187 |
| Slc5a6 | 0.03099081 | 0.04108198 | 0.03725686 | 0.00382512 | 0.3240419 | 0.90650187 |
| Wnt5b | 0.05118257 | 0.06453428 | 0.0607143 | 0.00381998 | 0.38985164 | 0.90650187 |
| Mfsd5 | 0.14095609 | 0.1620185 | 0.15820149 | 0.00381701 | 0.45729081 | 0.90650187 |
| Rpl7a-ps5 | 0.28423024 | 0.29864266 | 0.29482714 | 0.00381552 | 0.47706028 | 0.90650187 |
| Slc2a10 | 0.04610167 | 0.05868396 | 0.0548705 | 0.00381347 | 0.37867332 | 0.90650187 |
| Poln | 0.01253456 | 0.01906216 | 0.01525656 | 0.00380561 | 0.1337334 | 0.90650187 |

| | | | | | | |
|---|---|---|---|---|---|---|
| Ccdc167 | 0.38368682 | 0.38191494 | 0.37811609 | 0.00379885 | 0.48218753 | 0.90650187 |
| Fam220a | 0.62914489 | 0.55582009 | 0.55202612 | 0.00379397 | 0.48781269 | 0.90650187 |
| Rbm17 | 0.8382239 | 0.67656263 | 0.67277455 | 0.00378807 | 0.49001508 | 0.90650187 |
| Rnf213 | 0.07540285 | 0.09183036 | 0.08804306 | 0.0037873 | 0.42417104 | 0.90650187 |
| Parp2 | 0.31921863 | 0.32888714 | 0.32510809 | 0.00377905 | 0.47939359 | 0.90650187 |
| Gm4737 | 0.0122999 | 0.01874746 | 0.0149733 | 0.00377416 | 0.13123846 | 0.90650187 |
| Pak1 | 0.15585703 | 0.17721077 | 0.17344489 | 0.00376588 | 0.46155218 | 0.90650187 |
| Pcsk9 | 0.01861437 | 0.02632292 | 0.0225641 | 0.00375882 | 0.22947786 | 0.90650187 |
| Susd3 | 0.00963364 | 0.01550723 | 0.01174867 | 0.00375856 | 0.07748261 | 0.79797634 |
| Gpr157 | 0.0480065 | 0.06082161 | 0.05706592 | 0.00375569 | 0.3849218 | 0.90650187 |
| Pdcd2 | 0.36740942 | 0.36881458 | 0.36505893 | 0.00375565 | 0.48176055 | 0.90650187 |
| Fam120aos | 0.05472824 | 0.068513 | 0.06476929 | 0.00374371 | 0.39860505 | 0.90650187 |
| Syde1 | 0.14043102 | 0.16140211 | 0.15765961 | 0.0037425 | 0.45797799 | 0.90650187 |
| Frem1 | 0.04700362 | 0.05965242 | 0.05591073 | 0.00374169 | 0.38303801 | 0.90650187 |
| Rasgef1b | 0.02451771 | 0.0333438 | 0.02960265 | 0.00374115 | 0.28711562 | 0.90650187 |
| Gm6169 | 0.03312393 | 0.04350537 | 0.03976475 | 0.00374063 | 0.33790287 | 0.90650187 |
| Il27ra | 0.01855422 | 0.02623125 | 0.0224921 | 0.00373915 | 0.22993798 | 0.90650187 |
| Col17a1 | 0.01960022 | 0.02748104 | 0.02374341 | 0.00373763 | 0.24202089 | 0.90650187 |
| Ltb | 0.01115879 | 0.01732751 | 0.01359464 | 0.00373287 | 0.11110129 | 0.90650187 |
| Gbf1 | 0.42319519 | 0.41267005 | 0.4089405 | 0.00372955 | 0.48382968 | 0.90650187 |
| Myh11 | 0.01882357 | 0.02654343 | 0.02281449 | 0.00372894 | 0.23373042 | 0.90650187 |
| Fundc1 | 0.42538042 | 0.41433742 | 0.41061022 | 0.00372721 | 0.48390553 | 0.90650187 |
| Mrps12 | 1.17724712 | 0.81905155 | 0.81532792 | 0.00372363 | 0.49190074 | 0.90650187 |
| Dab1 | 0.04786701 | 0.0606289 | 0.05690534 | 0.00372355 | 0.38556603 | 0.90650187 |
| Rnf125 | 0.03173771 | 0.04184657 | 0.03813579 | 0.00371078 | 0.33266092 | 0.90650187 |
| Raet1e | 0.03191633 | 0.04205074 | 0.03834586 | 0.00370488 | 0.33377124 | 0.90650187 |
| Cd1d1 | 0.16507361 | 0.18644903 | 0.18274493 | 0.0037041 | 0.46410024 | 0.90650187 |
| Gdap1 | 0.10771659 | 0.12694686 | 0.12324358 | 0.00370329 | 0.44686584 | 0.90650187 |
| Abcb9 | 0.29000834 | 0.30360025 | 0.29990483 | 0.00369542 | 0.47815743 | 0.90650187 |
| Fam214a | 0.07779887 | 0.09439382 | 0.09070009 | 0.00369373 | 0.42816585 | 0.90650187 |

| | | | | | | |
|---|---|---|---|---|---|---|
| Akr1cl | 0.01636658 | 0.02355544 | 0.01986936 | 0.00368608 | 0.20476234 | 0.90650187 |
| Mbd1 | 0.20348125 | 0.22416786 | 0.22048212 | 0.00368575 | 0.47037941 | 0.90650187 |
| Fam188b | 0.07187443 | 0.08778804 | 0.08411523 | 0.0036728 | 0.42304379 | 0.90650187 |
| Impad1 | 0.53828086 | 0.49593693 | 0.49227076 | 0.00366617 | 0.48679404 | 0.90650187 |
| Scml4 | 0.0317958 | 0.04186978 | 0.03820412 | 0.00366566 | 0.33485241 | 0.90650187 |
| Gm15915 | 0.06317772 | 0.07802089 | 0.07435688 | 0.00366401 | 0.41330097 | 0.90650187 |
| Abhd8 | 0.4926442 | 0.46397986 | 0.46031696 | 0.0036629 | 0.48589028 | 0.90650187 |
| Ampd3 | 0.01448774 | 0.02126666 | 0.01761068 | 0.00365598 | 0.17802732 | 0.90650187 |
| Tbx18 | 0.01915046 | 0.02686108 | 0.02320559 | 0.0036555 | 0.24186973 | 0.90650187 |
| Atg9a | 0.17726418 | 0.19855252 | 0.19489876 | 0.00365376 | 0.46678999 | 0.90650187 |
| Smg7 | 0.28465545 | 0.29885026 | 0.29520185 | 0.00364842 | 0.47809174 | 0.90650187 |
| Cdc14a | 0.05810505 | 0.07225461 | 0.06861366 | 0.00364095 | 0.4067535 | 0.90650187 |
| Cxcl5 | 0.01004403 | 0.01588585 | 0.01224575 | 0.00364009 | 0.09317203 | 0.90318385 |
| Unkl | 0.06234586 | 0.07705621 | 0.07341766 | 0.00363854 | 0.41281019 | 0.90650187 |
| Wdr38 | 0.01667611 | 0.02386452 | 0.02024093 | 0.00362359 | 0.21305495 | 0.90650187 |
| Tmem178 | 0.05305935 | 0.06647936 | 0.06286303 | 0.00361633 | 0.39907459 | 0.90650187 |
| Mgat1 | 0.37603447 | 0.37562009 | 0.37200439 | 0.0036157 | 0.48276744 | 0.90650187 |
| Wdr60 | 0.19808046 | 0.21888214 | 0.215273 | 0.00360914 | 0.47029339 | 0.90650187 |
| Rpain | 0.26734063 | 0.28341198 | 0.27980649 | 0.00360549 | 0.47715925 | 0.90650187 |
| Erp29 | 1.99627369 | 0.93074154 | 0.92713711 | 0.00360443 | 0.49310535 | 0.90650187 |
| Plbd1 | 0.00983763 | 0.01559259 | 0.01199579 | 0.0035968 | 0.09127201 | 0.89511343 |
| Agtr1a | 0.05753541 | 0.07155528 | 0.06796634 | 0.00358894 | 0.40720197 | 0.90650187 |
| Kcnn4 | 0.03545857 | 0.04609021 | 0.04250143 | 0.00358878 | 0.35368787 | 0.90650187 |
| Prelid2 | 0.1136023 | 0.13311876 | 0.12953197 | 0.00358679 | 0.45101374 | 0.90650187 |
| Chl1 | 0.01840057 | 0.02588807 | 0.02230814 | 0.00357993 | 0.23779459 | 0.90650187 |
| Wfdc16 | 0.04723832 | 0.05976047 | 0.05618122 | 0.00357925 | 0.38850201 | 0.90650187 |
| Hbegf | 0.07775455 | 0.09421642 | 0.09065102 | 0.00356541 | 0.43059848 | 0.90650187 |
| Cxcr4 | 0.02601762 | 0.03494229 | 0.03138213 | 0.00356017 | 0.30701312 | 0.90650187 |
| Ogdh | 0.51316707 | 0.47841164 | 0.47485595 | 0.00355569 | 0.48672229 | 0.90650187 |
| Tpm3 | 1.44614557 | 0.88719826 | 0.88365269 | 0.00354558 | 0.49288421 | 0.90650187 |

| | | | | | | |
|---|---|---|---|---|---|---|
| Insig2 | 0.46489606 | 0.4437377 | 0.44020158 | 0.00353612 | 0.48575624 | 0.90650187 |
| Mar-11 | 0.05008179 | 0.06298215 | 0.05945155 | 0.00353059 | 0.39588729 | 0.90650187 |
| Tram1l1 | 0.10225037 | 0.12088863 | 0.11736472 | 0.00352391 | 0.4469067 | 0.90650187 |
| Fam19a1 | 0.03129211 | 0.04113378 | 0.03761153 | 0.00352225 | 0.33858697 | 0.90650187 |
| Plxdc1 | 0.01479121 | 0.02149628 | 0.01797589 | 0.0035204 | 0.19197894 | 0.90650187 |
| Hal | 0.02755354 | 0.03672012 | 0.03320064 | 0.00351948 | 0.31872675 | 0.90650187 |
| Kcna4 | 0.00908311 | 0.01458965 | 0.01108139 | 0.00350825 | 0.07964953 | 0.81595645 |
| Fbxo44 | 0.19048085 | 0.2113929 | 0.20788893 | 0.00350396 | 0.47013497 | 0.90650187 |
| Dennd4c | 0.25030246 | 0.26788004 | 0.26437707 | 0.00350297 | 0.47651434 | 0.90650187 |
| Nkx2-5 | 0.01013733 | 0.01585979 | 0.01235873 | 0.00350107 | 0.10394615 | 0.90650187 |
| Lpcat3 | 0.57103904 | 0.51788543 | 0.51438568 | 0.00349975 | 0.48793513 | 0.90650187 |
| Mtap7d3 | 0.03083154 | 0.04056346 | 0.03706931 | 0.00349415 | 0.33759259 | 0.90650187 |
| Ngly1 | 0.27455589 | 0.28975067 | 0.28625668 | 0.00349399 | 0.47836314 | 0.90650187 |
| Nr4a1 | 0.09582972 | 0.11391834 | 0.11043425 | 0.0034841 | 0.44422935 | 0.90650187 |
| Tcte2 | 0.036917 | 0.04769072 | 0.0442067 | 0.00348402 | 0.36303152 | 0.90650187 |
| Fam89a | 0.06717893 | 0.08234234 | 0.07886025 | 0.00348209 | 0.422189 | 0.90650187 |
| Taf1a | 0.10528058 | 0.12410957 | 0.1206283 | 0.00348126 | 0.4489567 | 0.90650187 |
| Cr1l | 0.54867014 | 0.50283793 | 0.49935689 | 0.00348104 | 0.48763854 | 0.90650187 |
| Mier3 | 0.23657506 | 0.25521863 | 0.25173793 | 0.0034807 | 0.47549329 | 0.90650187 |
| Btg3 | 0.08709119 | 0.10440293 | 0.10092731 | 0.00347562 | 0.43916252 | 0.90650187 |
| Cypt3 | 0.01675945 | 0.02381221 | 0.02034094 | 0.00347127 | 0.22402749 | 0.90650187 |
| Adamts16 | 0.05762341 | 0.07152617 | 0.06806637 | 0.00345981 | 0.4106125 | 0.90650187 |
| Sphk2 | 0.30230042 | 0.31405679 | 0.31060445 | 0.00345234 | 0.48029527 | 0.90650187 |
| 2210404O09 | 0.06353868 | 0.07820722 | 0.07476411 | 0.00344311 | 0.41889017 | 0.90650187 |
| Dio1 | 0.05270122 | 0.06589471 | 0.06245341 | 0.0034413 | 0.40324322 | 0.90650187 |
| D15Ertd621e | 0.23988308 | 0.25824103 | 0.25480085 | 0.00344018 | 0.476069 | 0.90650187 |
| Magi1 | 0.11098532 | 0.13016643 | 0.12674124 | 0.0034252 | 0.45218494 | 0.90650187 |
| Syt6 | 0.02548008 | 0.03416025 | 0.0307448 | 0.00341545 | 0.31070109 | 0.90650187 |
| Akap17b | 0.18402731 | 0.20498093 | 0.20156973 | 0.0034112 | 0.47001434 | 0.90650187 |
| Rab6a | 0.92662328 | 0.7210789 | 0.71767324 | 0.00340566 | 0.49158444 | 0.90650187 |

| | | | | | | |
|---|---|---|---|---|---|---|
| Scn3b | 0.04668516 | 0.05894637 | 0.05554359 | 0.00340278 | 0.39267556 | 0.90650187 |
| Isg20l2 | 0.22215022 | 0.24165513 | 0.23825371 | 0.00340142 | 0.47469709 | 0.90650187 |
| Hoxa1 | 0.02177173 | 0.02972506 | 0.02633555 | 0.00338952 | 0.28360282 | 0.90650187 |
| Shc1 | 0.4803125 | 0.45482904 | 0.45144258 | 0.00338646 | 0.48669841 | 0.90650187 |
| Nsun7 | 0.02468286 | 0.03318322 | 0.02979875 | 0.00338447 | 0.30680694 | 0.90650187 |
| Cd177 | 0.01098419 | 0.01676699 | 0.01338351 | 0.00338348 | 0.13052973 | 0.90650187 |
| Fst | 0.01502424 | 0.02163308 | 0.01825623 | 0.00337685 | 0.20545204 | 0.90650187 |
| Nfatc1 | 0.06284905 | 0.07735817 | 0.07398591 | 0.00337226 | 0.41971235 | 0.90650187 |
| Zcchc6 | 0.38343162 | 0.38128525 | 0.37791301 | 0.00337224 | 0.48417827 | 0.90650187 |
| Tmem252 | 0.01171724 | 0.01763953 | 0.01426962 | 0.00336991 | 0.14688802 | 0.90650187 |
| Dnase2a | 0.32323385 | 0.33188063 | 0.32851288 | 0.00336775 | 0.48182485 | 0.90650187 |
| Gm17501 | 0.01465524 | 0.02117797 | 0.01781228 | 0.0033657 | 0.20045026 | 0.90650187 |
| Ubxn2a | 0.36348375 | 0.36524205 | 0.36187711 | 0.00336494 | 0.48351333 | 0.90650187 |
| Mtch1 | 0.69106296 | 0.59341989 | 0.59005642 | 0.00336347 | 0.48989145 | 0.90650187 |
| Ank3 | 0.459177 | 0.4393514 | 0.43598835 | 0.00336305 | 0.48632227 | 0.90650187 |
| Asphd1 | 0.05656816 | 0.0702253 | 0.06686608 | 0.00335922 | 0.41163594 | 0.90650187 |
| Ms4a8a | 0.00983896 | 0.01535109 | 0.0119974 | 0.00335369 | 0.10698889 | 0.90650187 |
| Hist1h2bk | 0.00956394 | 0.01501667 | 0.01166422 | 0.00335245 | 0.1006734 | 0.90650187 |
| Lrrc3b | 0.0426339 | 0.05420943 | 0.05085948 | 0.00334994 | 0.38483093 | 0.90650187 |
| Asah1 | 0.81335783 | 0.6628473 | 0.65950049 | 0.00334681 | 0.49100045 | 0.90650187 |
| Cep41 | 0.14047777 | 0.16105168 | 0.15770787 | 0.00334381 | 0.46245197 | 0.90650187 |
| Tex15 | 0.03058533 | 0.04012003 | 0.03677933 | 0.00334069 | 0.34318072 | 0.90650187 |
| Tbc1d24 | 0.15832312 | 0.17928307 | 0.17594276 | 0.00334032 | 0.46636883 | 0.90650187 |
| Riiad1 | 0.22570542 | 0.24492665 | 0.24159673 | 0.00332992 | 0.47557064 | 0.90650187 |
| Gm4924 | 0.04017856 | 0.05131535 | 0.04800836 | 0.00330698 | 0.37971516 | 0.90650187 |
| Rerg | 0.07310605 | 0.08879357 | 0.08548831 | 0.00330527 | 0.43176506 | 0.90650187 |
| Gpr107 | 0.3787279 | 0.37745675 | 0.37416082 | 0.00329592 | 0.48438116 | 0.90650187 |
| Tspan2 | 0.13490812 | 0.15523247 | 0.15194012 | 0.00329235 | 0.46162892 | 0.90650187 |
| Tmem238 | 0.38261347 | 0.38054702 | 0.37726162 | 0.0032854 | 0.4845589 | 0.90650187 |
| Ccdc64 | 0.09311954 | 0.11078122 | 0.10749713 | 0.00328409 | 0.44598354 | 0.90650187 |

| | | | | | | |
|---|---|---|---|---|---|---|
| Iqgap2 | 0.16276399 | 0.18370405 | 0.18042342 | 0.00328063 | 0.46778687 | 0.90650187 |
| Slc41a2 | 0.01100497 | 0.0166878 | 0.01340864 | 0.00327917 | 0.1384738 | 0.90650187 |
| Cdk14 | 0.29589993 | 0.30832612 | 0.305051 | 0.00327511 | 0.48096599 | 0.90650187 |
| Zfp300 | 0.01965528 | 0.02706915 | 0.02380923 | 0.00325992 | 0.27136689 | 0.90650187 |
| Nhlh1 | 0.00949851 | 0.01484431 | 0.01158493 | 0.00325938 | 0.10551202 | 0.90650187 |
| Rasgrf1 | 0.11214488 | 0.13123498 | 0.12797883 | 0.00325615 | 0.45497201 | 0.90650187 |
| Gjb3 | 0.01870181 | 0.02591084 | 0.02266877 | 0.00324208 | 0.26245038 | 0.90650187 |
| Vwa9 | 0.21675525 | 0.23639393 | 0.23315502 | 0.00323891 | 0.47537824 | 0.90650187 |
| Psmd4 | 1.5897215 | 0.91001562 | 0.90677839 | 0.00323723 | 0.49366866 | 0.90650187 |
| Sytl1 | 0.13692789 | 0.15726963 | 0.15403596 | 0.00323367 | 0.46282213 | 0.90650187 |
| Aff1 | 0.08003763 | 0.09640044 | 0.0931753 | 0.00322514 | 0.43885275 | 0.90650187 |
| Gm14391 | 0.02173317 | 0.02950512 | 0.02628958 | 0.00321554 | 0.2933061 | 0.90650187 |
| Eif2ak3 | 0.06236751 | 0.07665497 | 0.07344212 | 0.00321285 | 0.4228998 | 0.90650187 |
| Wfdc18 | 0.01811369 | 0.02517668 | 0.02196457 | 0.00321211 | 0.2578047 | 0.90650187 |
| AA474408 | 0.03281793 | 0.04260898 | 0.03940542 | 0.00320355 | 0.3588943 | 0.90650187 |
| Meis3 | 0.26586801 | 0.28167809 | 0.27848385 | 0.00319424 | 0.47966609 | 0.90650187 |
| Pus7 | 0.20189928 | 0.22215162 | 0.21895951 | 0.00319211 | 0.47416247 | 0.90650187 |
| AU021092 | 0.04748764 | 0.0596565 | 0.05646845 | 0.00318805 | 0.40091229 | 0.90650187 |
| Gm10600 | 0.01906131 | 0.0262863 | 0.02309894 | 0.00318737 | 0.26979309 | 0.90650187 |
| Blm | 0.07214561 | 0.08759485 | 0.08441775 | 0.00317709 | 0.43356228 | 0.90650187 |
| Serpina1d | 0.01716186 | 0.02399832 | 0.02082371 | 0.00317461 | 0.24896807 | 0.90650187 |
| Nlrx1 | 0.0322559 | 0.04191114 | 0.03874507 | 0.00316607 | 0.3582 | 0.90650187 |
| Pkhd1 | 0.0521023 | 0.06493168 | 0.06176796 | 0.00316372 | 0.40993928 | 0.90650187 |
| Armc8 | 0.33239574 | 0.33938952 | 0.33622845 | 0.00316108 | 0.48333078 | 0.90650187 |
| Srsf12 | 0.03835275 | 0.04904232 | 0.04588223 | 0.00316009 | 0.37973134 | 0.90650187 |
| Pdgfd | 0.05453536 | 0.06770817 | 0.06454918 | 0.00315899 | 0.41388424 | 0.90650187 |
| Car7 | 0.01093764 | 0.01648512 | 0.0133272 | 0.00315792 | 0.1460793 | 0.90650187 |
| Crygn | 0.01967875 | 0.02699428 | 0.02383728 | 0.00315699 | 0.27800687 | 0.90650187 |
| Ap1s3 | 0.13674092 | 0.15699745 | 0.15384215 | 0.0031553 | 0.46367507 | 0.90650187 |
| B3galt1 | 0.03097666 | 0.04039136 | 0.03724019 | 0.00315117 | 0.35339312 | 0.90650187 |

| | | | | | | |
|---|---|---|---|---|---|---|
| Gabpb1 | 0.27398317 | 0.28889407 | 0.28574658 | 0.00314749 | 0.48047231 | 0.90650187 |
| Lzts1 | 0.04346408 | 0.05496604 | 0.05182139 | 0.00314465 | 0.39366852 | 0.90650187 |
| Pla2r1 | 0.06121973 | 0.07528602 | 0.07214457 | 0.00314145 | 0.423253 | 0.90650187 |
| Tmem86a | 0.24843918 | 0.26581323 | 0.26267275 | 0.00314048 | 0.47880579 | 0.90650187 |
| Zfp710 | 0.1064695 | 0.12504366 | 0.12190564 | 0.00313803 | 0.45444599 | 0.90650187 |
| Plac1 | 0.03656477 | 0.04693282 | 0.04379515 | 0.00313767 | 0.37505211 | 0.90650187 |
| Zyg11b | 0.14138307 | 0.16176457 | 0.15864191 | 0.00312265 | 0.46513472 | 0.90650187 |
| Rbpms2 | 0.06919362 | 0.08424074 | 0.08111892 | 0.00312182 | 0.43207769 | 0.90650187 |
| 2900079G21 | 0.00701665 | 0.01168669 | 0.00857229 | 0.0031144 | 0.053141 | 0.61947578 |
| 2700069I18R | 0.01250225 | 0.0183294 | 0.01521756 | 0.00311184 | 0.18165349 | 0.90650187 |
| Smim3 | 0.05097789 | 0.06358685 | 0.06047964 | 0.00310721 | 0.40966841 | 0.90650187 |
| Fank1 | 0.03214722 | 0.04171763 | 0.03861732 | 0.00310031 | 0.3605811 | 0.90650187 |
| Bst1 | 0.0094527 | 0.0146289 | 0.01152941 | 0.00309949 | 0.11601893 | 0.90650187 |
| Cfap61 | 0.02569506 | 0.03408638 | 0.03099975 | 0.00308663 | 0.32901078 | 0.90650187 |
| Mrvi1 | 0.04423514 | 0.05578883 | 0.05271384 | 0.00307499 | 0.39769101 | 0.90650187 |
| Pparg | 0.01093843 | 0.01640061 | 0.01332816 | 0.00307245 | 0.15272538 | 0.90650187 |
| Negr1 | 0.03555969 | 0.0456917 | 0.04261976 | 0.00307194 | 0.37432145 | 0.90650187 |
| Hist1h3c | 0.02601254 | 0.03444548 | 0.0313761 | 0.00306938 | 0.33182008 | 0.90650187 |
| Gls2 | 0.00957134 | 0.01473162 | 0.01167318 | 0.00305844 | 0.1220553 | 0.90650187 |
| Suv39h2 | 0.06602767 | 0.08061784 | 0.07756691 | 0.00305093 | 0.43059538 | 0.90650187 |
| Gm14327 | 0.01533224 | 0.02167375 | 0.01862662 | 0.00304713 | 0.23353409 | 0.90650187 |
| Gm10644 | 0.00689125 | 0.01146299 | 0.00841981 | 0.00304318 | 0.05405119 | 0.62576701 |
| Gm37350 | 0.03189609 | 0.04136494 | 0.03832205 | 0.00304288 | 0.36204642 | 0.90650187 |
| Vipr1 | 0.01155811 | 0.01712014 | 0.01407734 | 0.0030428 | 0.16829772 | 0.90650187 |
| Rin1 | 0.01688884 | 0.02353589 | 0.0204962 | 0.00303969 | 0.25484967 | 0.90650187 |
| Fgfr4 | 0.08878544 | 0.10581174 | 0.10277887 | 0.00303287 | 0.44781491 | 0.90650187 |
| Lrig3 | 0.10541077 | 0.12379142 | 0.12076826 | 0.00302316 | 0.45569497 | 0.90650187 |
| Arl5c | 0.01009869 | 0.01532799 | 0.01231195 | 0.00301604 | 0.13806934 | 0.90650187 |
| Nudt15 | 0.01215714 | 0.01781637 | 0.01480094 | 0.00301543 | 0.18254363 | 0.90650187 |
| Gm16152 | 0.01629143 | 0.0227916 | 0.01977914 | 0.00301246 | 0.24917511 | 0.90650187 |

| | | | | | | |
|---|---|---|---|---|---|---|
| Chrd | 0.0520986 | 0.06477595 | 0.06176372 | 0.00301223 | 0.41417691 | 0.90650187 |
| Stk32b | 0.05291649 | 0.06571146 | 0.06269965 | 0.0030118 | 0.41545035 | 0.90650187 |
| Pla2g6 | 0.33316659 | 0.33988595 | 0.33687434 | 0.00301162 | 0.48414893 | 0.90650187 |
| Lrrc34 | 0.00918564 | 0.01420896 | 0.0112057 | 0.00300326 | 0.11673371 | 0.90650187 |
| Zc3h6 | 0.11008701 | 0.12878446 | 0.12578134 | 0.00300312 | 0.45773483 | 0.90650187 |
| Gm609 | 0.14495235 | 0.16531692 | 0.1623152 | 0.00300172 | 0.4672386 | 0.90650187 |
| Nod1 | 0.18055112 | 0.20114834 | 0.19814717 | 0.00300118 | 0.47315789 | 0.90650187 |
| Supt3 | 0.14022746 | 0.16045061 | 0.15744944 | 0.00300117 | 0.46623466 | 0.90650187 |
| Inhbb | 0.02109561 | 0.02852482 | 0.02552927 | 0.00299554 | 0.30096155 | 0.90650187 |
| Wwp1 | 0.21156844 | 0.23120886 | 0.22822487 | 0.00298399 | 0.47682441 | 0.90650187 |
| Frrs1 | 0.20384038 | 0.22379774 | 0.22082741 | 0.00297033 | 0.47615845 | 0.90650187 |
| Eftud2 | 0.45070039 | 0.43266981 | 0.42970153 | 0.00296828 | 0.4877507 | 0.90650187 |
| Necab2 | 0.22421451 | 0.24316084 | 0.24019646 | 0.00296438 | 0.47812284 | 0.90650187 |
| Cc2d1a | 0.17419215 | 0.19481244 | 0.19185199 | 0.00296045 | 0.47265415 | 0.90650187 |
| Csf2rb | 0.01212939 | 0.0177255 | 0.01476743 | 0.00295807 | 0.18659852 | 0.90650187 |
| Tubgcp2 | 0.23485815 | 0.25309997 | 0.25014395 | 0.00295602 | 0.47905122 | 0.90650187 |
| Sp6 | 0.00999101 | 0.01513248 | 0.01218156 | 0.00295092 | 0.14075644 | 0.90650187 |
| Catip | 0.0126762 | 0.0183738 | 0.01542748 | 0.00294631 | 0.1979367 | 0.90650187 |
| Aplf | 0.05600608 | 0.06917189 | 0.06622606 | 0.00294583 | 0.42162158 | 0.90650187 |
| Crebrf | 0.15013146 | 0.17056145 | 0.16761869 | 0.00294275 | 0.46889489 | 0.90650187 |
| Diras2 | 0.02464684 | 0.03269868 | 0.02975598 | 0.0029427 | 0.330097 | 0.90650187 |
| Trim23 | 0.16034532 | 0.18091374 | 0.17798585 | 0.00292789 | 0.47085104 | 0.90650187 |
| Wdr81 | 0.03373984 | 0.04341474 | 0.04048755 | 0.0029272 | 0.37394927 | 0.90650187 |
| Mrpl35 | 0.56521723 | 0.51342737 | 0.510504 | 0.00292337 | 0.48984502 | 0.90650187 |
| Abhd11os | 0.06310922 | 0.07719824 | 0.07427957 | 0.00291867 | 0.43066509 | 0.90650187 |
| Usp8 | 0.45243056 | 0.43390611 | 0.43098886 | 0.00291725 | 0.48799718 | 0.90650187 |
| Cd46 | 0.03120298 | 0.04041787 | 0.03750662 | 0.00291125 | 0.36502222 | 0.90650187 |
| Acat2 | 0.35749083 | 0.3599068 | 0.35699568 | 0.00291112 | 0.48554083 | 0.90650187 |
| 2810032G03 | 0.01901672 | 0.0259555 | 0.0230456 | 0.00290991 | 0.2872847 | 0.90650187 |
| Asb13 | 0.1774884 | 0.19802338 | 0.19512073 | 0.00290265 | 0.47363569 | 0.90650187 |

| | | | | | | |
|---|---|---|---|---|---|---|
| Tmem135 | 0.16008706 | 0.18062531 | 0.17772518 | 0.00290013 | 0.47108464 | 0.90650187 |
| Cpped1 | 0.15551896 | 0.17599467 | 0.17310192 | 0.00289275 | 0.47038914 | 0.90650187 |
| Mcoln3 | 0.05346819 | 0.06621612 | 0.0633304 | 0.00288572 | 0.41973584 | 0.90650187 |
| Sema4g | 0.05455011 | 0.06745026 | 0.06456602 | 0.00288424 | 0.42129152 | 0.90650187 |
| Syt2 | 0.01542834 | 0.02161986 | 0.01874215 | 0.00287771 | 0.24743377 | 0.90650187 |
| Tob1 | 0.61364109 | 0.5450443 | 0.5421748 | 0.0028695 | 0.49061427 | 0.90650187 |
| Rad54l | 0.07295215 | 0.08817973 | 0.08531686 | 0.00286287 | 0.44070747 | 0.90650187 |
| Abcb1a | 0.00834524 | 0.01304458 | 0.01018628 | 0.00285831 | 0.10611557 | 0.90650187 |
| Osgin1 | 0.04844255 | 0.06042531 | 0.05756772 | 0.00285759 | 0.41267302 | 0.90650187 |
| Cttnbp2nl | 0.17876466 | 0.1992388 | 0.19638312 | 0.00285568 | 0.47422823 | 0.90650187 |
| Arhgap22 | 0.01913922 | 0.02604671 | 0.02319214 | 0.00285457 | 0.29212718 | 0.90650187 |
| Gm16380 | 0.00932444 | 0.01422818 | 0.01137396 | 0.00285422 | 0.13229793 | 0.90650187 |
| Sema6d | 0.2255921 | 0.24434382 | 0.24149039 | 0.00285344 | 0.47905357 | 0.90650187 |
| Toporsos | 0.58087667 | 0.52373866 | 0.52089774 | 0.00284092 | 0.49032827 | 0.90650187 |
| Slc25a37 | 0.0558708 | 0.06891205 | 0.06607196 | 0.00284009 | 0.42422612 | 0.90650187 |
| Zfp775 | 0.07111041 | 0.0860964 | 0.08326236 | 0.00283404 | 0.43986276 | 0.90650187 |
| Sspn | 0.17036548 | 0.1908641 | 0.18804146 | 0.00282264 | 0.47339761 | 0.90650187 |
| Papola | 1.23743722 | 0.83635218 | 0.83353526 | 0.00281692 | 0.49400657 | 0.90650187 |
| Zfp142 | 0.11326209 | 0.1319831 | 0.12916964 | 0.00281346 | 0.46143064 | 0.90650187 |
| Napepld | 0.0677482 | 0.08229633 | 0.07949905 | 0.00279728 | 0.43784924 | 0.90650187 |
| Hlcs | 0.05202151 | 0.06447094 | 0.06167545 | 0.00279549 | 0.42015362 | 0.90650187 |
| Irak3 | 0.02035367 | 0.027432 | 0.02464366 | 0.00278833 | 0.30748111 | 0.90650187 |
| Prkcsh | 1.02901242 | 0.76526286 | 0.76247454 | 0.00278832 | 0.49351456 | 0.90650187 |
| D16Ertd472e | 0.03949418 | 0.04999514 | 0.04721201 | 0.00278312 | 0.39663528 | 0.90650187 |
| Arrb2 | 0.16701938 | 0.18746089 | 0.18469605 | 0.00276484 | 0.47347026 | 0.90650187 |
| Btbd7 | 0.1339428 | 0.15369633 | 0.15093673 | 0.0027596 | 0.46760974 | 0.90650187 |
| Sv2c | 0.04896303 | 0.06091336 | 0.0581663 | 0.00274707 | 0.41685095 | 0.90650187 |
| Plekha2 | 0.03957981 | 0.05005866 | 0.04731169 | 0.00274697 | 0.39815847 | 0.90650187 |
| Gm14703 | 0.03258228 | 0.04186753 | 0.0391286 | 0.00273893 | 0.3778305 | 0.90650187 |
| Zkscan14 | 0.11738176 | 0.13628362 | 0.13354745 | 0.00273617 | 0.46371322 | 0.90650187 |

| | | | | | | |
|---|---|---|---|---|---|---|
| Tnni2 | 0.02012592 | 0.02710494 | 0.02437163 | 0.0027333 | 0.3090379 | 0.90650187 |
| Naip2 | 0.00706219 | 0.01135898 | 0.00862766 | 0.00273132 | 0.07965788 | 0.81595645 |
| Tbc1d10c | 0.01550017 | 0.0215534 | 0.0188285 | 0.00272489 | 0.25999014 | 0.90650187 |
| St8sia4 | 0.02770215 | 0.03609874 | 0.0333764 | 0.00272233 | 0.35845036 | 0.90650187 |
| Gtf2h2 | 0.24239663 | 0.25984285 | 0.25712093 | 0.00272192 | 0.48123195 | 0.90650187 |
| N4bp2l1 | 0.15384854 | 0.17411974 | 0.17140538 | 0.00271436 | 0.47193755 | 0.90650187 |
| a | 0.0195089 | 0.02634714 | 0.02363423 | 0.00271291 | 0.30492077 | 0.90650187 |
| Adamts6 | 0.01147676 | 0.01669081 | 0.01397902 | 0.00271179 | 0.19423315 | 0.90650187 |
| Tdrkh | 0.22046003 | 0.23936899 | 0.23665953 | 0.00270945 | 0.47970396 | 0.90650187 |
| Gm37233 | 0.00914629 | 0.01386356 | 0.011158 | 0.00270556 | 0.14052617 | 0.90650187 |
| Ndufv1 | 0.88681281 | 0.70062151 | 0.69791778 | 0.00270373 | 0.49312964 | 0.90650187 |
| Mgat4a | 0.10730232 | 0.12550099 | 0.12279935 | 0.00270164 | 0.46104338 | 0.90650187 |
| Xrcc6bp1 | 0.08397446 | 0.10021121 | 0.09751065 | 0.00270056 | 0.45100557 | 0.90650187 |
| Cep72 | 0.0300128 | 0.03880397 | 0.03610463 | 0.00269933 | 0.36980418 | 0.90650187 |
| Rin2 | 0.06956009 | 0.0842269 | 0.08152914 | 0.00269776 | 0.44152539 | 0.90650187 |
| Cacnb4 | 0.02053951 | 0.02755567 | 0.02486556 | 0.00269011 | 0.31527518 | 0.90650187 |
| Gm20517 | 0.00705275 | 0.01129757 | 0.00861618 | 0.00268139 | 0.08325697 | 0.83980701 |
| Slc2a3 | 0.04606332 | 0.05750083 | 0.05482623 | 0.0026746 | 0.41415436 | 0.90650187 |
| Ccdc181 | 0.21988906 | 0.23878161 | 0.23612034 | 0.00266127 | 0.48001911 | 0.90650187 |
| Slc10a6 | 0.0231364 | 0.03062095 | 0.0279607 | 0.00266025 | 0.33615999 | 0.90650187 |
| Slc35e4 | 0.05405666 | 0.06665515 | 0.06400268 | 0.00265246 | 0.42691293 | 0.90650187 |
| Noc4l | 0.19751778 | 0.21738064 | 0.21472839 | 0.00265225 | 0.47810482 | 0.90650187 |
| Bms1 | 0.26333713 | 0.27885045 | 0.27620593 | 0.00264452 | 0.48302443 | 0.90650187 |
| Mtmr7 | 0.23660842 | 0.2544112 | 0.25176887 | 0.00264233 | 0.48139336 | 0.90650187 |
| 1700123O20 | 0.62029527 | 0.54904901 | 0.54641987 | 0.00262914 | 0.49146714 | 0.90650187 |
| Fam84a | 0.02651098 | 0.03459568 | 0.03196667 | 0.00262901 | 0.35732552 | 0.90650187 |
| Fam198a | 0.05564817 | 0.06844525 | 0.06581828 | 0.00262697 | 0.42958337 | 0.90650187 |
| Dpp7 | 0.10420705 | 0.12209576 | 0.11947341 | 0.00262235 | 0.46113382 | 0.90650187 |
| Gm26771 | 0.03281976 | 0.0420284 | 0.03940758 | 0.00262083 | 0.38374562 | 0.90650187 |
| Pdzd3 | 0.00913664 | 0.0137606 | 0.0111463 | 0.00261431 | 0.14854486 | 0.90650187 |

| | | | | | | |
|---|---|---|---|---|---|---|
| Cend1 | 0.01364681 | 0.01920101 | 0.01659789 | 0.00260312 | 0.24283133 | 0.90650187 |
| Rwdd2a | 0.04123972 | 0.05184066 | 0.04924171 | 0.00259895 | 0.40724554 | 0.90650187 |
| Wdr62 | 0.00775508 | 0.01206855 | 0.00946971 | 0.00259884 | 0.11122464 | 0.90650187 |
| Chst7 | 0.04249943 | 0.05329439 | 0.05070358 | 0.00259081 | 0.41015031 | 0.90650187 |
| Dscc1 | 0.04100029 | 0.05155097 | 0.04896358 | 0.0025874 | 0.40713525 | 0.90650187 |
| Gm4070 | 0.00904953 | 0.01362586 | 0.01104068 | 0.00258518 | 0.14895188 | 0.90650187 |
| Hoxb5os | 0.0632963 | 0.07707061 | 0.07449068 | 0.00257993 | 0.43881657 | 0.90650187 |
| Gm10827 | 0.03462413 | 0.04410016 | 0.04152427 | 0.00257589 | 0.39135994 | 0.90650187 |
| Blnk | 0.05820929 | 0.07129904 | 0.06873206 | 0.00256697 | 0.43406568 | 0.90650187 |
| Hmx3 | 0.00694435 | 0.01104384 | 0.00848438 | 0.00255946 | 0.08994493 | 0.88667504 |
| Ilf2 | 1.12339358 | 0.80002898 | 0.79747018 | 0.0025588 | 0.49430954 | 0.90650187 |
| Oxld1 | 0.1329908 | 0.15250182 | 0.1499461 | 0.00255571 | 0.46980031 | 0.90650187 |
| Gnb5 | 0.01178287 | 0.01690377 | 0.0143489 | 0.00255487 | 0.2143103 | 0.90650187 |
| Gm26545 | 0.01316605 | 0.01856796 | 0.01601835 | 0.00254961 | 0.23959716 | 0.90650187 |
| Mylpf | 0.05487188 | 0.06747756 | 0.06493316 | 0.00254439 | 0.43085403 | 0.90650187 |
| Slc22a5 | 0.02066534 | 0.02755991 | 0.02501579 | 0.00254411 | 0.32559165 | 0.90650187 |
| Lrrn3 | 0.03096848 | 0.0397695 | 0.03723056 | 0.00253893 | 0.38088107 | 0.90650187 |
| Gm9833 | 0.01064615 | 0.01551167 | 0.01297459 | 0.00253708 | 0.19234068 | 0.90650187 |
| Igf1r | 0.47272928 | 0.44846371 | 0.44593434 | 0.00252937 | 0.48994142 | 0.90650187 |
| Prrg3 | 0.08435088 | 0.10044765 | 0.09792401 | 0.00252364 | 0.45439319 | 0.90650187 |
| Prmt3 | 0.23859098 | 0.25612187 | 0.25360577 | 0.00251609 | 0.48240991 | 0.90650187 |
| Tmem237 | 0.29683281 | 0.3083702 | 0.30586291 | 0.00250728 | 0.48546477 | 0.90650187 |
| Fgf7 | 0.04239036 | 0.05308282 | 0.05057709 | 0.00250573 | 0.41283916 | 0.90650187 |
| 4933406P04I | 0.03078308 | 0.03951622 | 0.03701224 | 0.00250397 | 0.38180009 | 0.90650187 |
| Mum1l1 | 0.10579347 | 0.12367755 | 0.12117955 | 0.002498 | 0.46349129 | 0.90650187 |
| Fam212b | 0.01133044 | 0.01629441 | 0.01380216 | 0.00249225 | 0.21106215 | 0.90650187 |
| Actn1 | 0.62622674 | 0.5526633 | 0.55018223 | 0.00248107 | 0.4920027 | 0.90650187 |
| Wdr13 | 0.31000015 | 0.31970917 | 0.31723651 | 0.00247266 | 0.48617913 | 0.90650187 |
| Pcbp4 | 0.61649712 | 0.54646651 | 0.54400029 | 0.00246622 | 0.49196022 | 0.90650187 |
| Slc44a5 | 0.0143654 | 0.01992233 | 0.01746341 | 0.00245893 | 0.2656702 | 0.90650187 |

| | | | | | | |
|---|---|---|---|---|---|---|
| Edf1 | 2.25267328 | 0.91977829 | 0.9173303 | 0.00244799 | 0.49526724 | 0.90650187 |
| Mapk10 | 0.03603648 | 0.04561831 | 0.04317754 | 0.00244076 | 0.40078968 | 0.90650187 |
| Bnc1 | 0.01892991 | 0.0253817 | 0.02294173 | 0.00243998 | 0.31817297 | 0.90650187 |
| Khdrbs2 | 0.02456314 | 0.03209308 | 0.02965659 | 0.00243648 | 0.3574677 | 0.90650187 |
| Ccdc163 | 0.11697392 | 0.13554706 | 0.13311498 | 0.00243208 | 0.4676321 | 0.90650187 |
| Sun2 | 0.07274046 | 0.08751177 | 0.08508096 | 0.00243081 | 0.44946491 | 0.90650187 |
| Pstk | 0.2753792 | 0.28941928 | 0.28698943 | 0.00242985 | 0.48498752 | 0.90650187 |
| Reps2 | 0.03632663 | 0.04594523 | 0.0435168 | 0.00242842 | 0.40203488 | 0.90650187 |
| Zfp422 | 0.65578521 | 0.57106966 | 0.56864147 | 0.00242819 | 0.49242714 | 0.90650187 |
| Adgra1 | 0.01125148 | 0.01613378 | 0.01370671 | 0.00242707 | 0.21558512 | 0.90650187 |
| Tfap4 | 0.16539361 | 0.18548225 | 0.1830661 | 0.00241615 | 0.47660582 | 0.90650187 |
| Usp29 | 0.44330319 | 0.42658852 | 0.4241733 | 0.00241522 | 0.48990264 | 0.90650187 |
| Lap3 | 0.53783324 | 0.49437846 | 0.49196387 | 0.00241459 | 0.49129604 | 0.90650187 |
| Arhgef16 | 0.23423971 | 0.25198251 | 0.24956907 | 0.00241344 | 0.48285436 | 0.90650187 |
| Arhgef6 | 0.03256711 | 0.04151801 | 0.03911079 | 0.00240722 | 0.39218789 | 0.90650187 |
| Lrp4 | 0.06299099 | 0.07655158 | 0.07414613 | 0.00240545 | 0.44266164 | 0.90650187 |
| Hpcal4 | 0.00822479 | 0.01243771 | 0.01004007 | 0.00239763 | 0.14419972 | 0.90650187 |
| Pcdhb11 | 0.0180277 | 0.0242591 | 0.02186156 | 0.00239753 | 0.31293622 | 0.90650187 |
| Npm3 | 0.82874865 | 0.67014076 | 0.66774742 | 0.00239334 | 0.49364354 | 0.90650187 |
| Prdm8 | 0.00659851 | 0.01045366 | 0.00806373 | 0.00238993 | 0.09382011 | 0.90650187 |
| Gm10244 | 0.02834733 | 0.03652186 | 0.03413903 | 0.00238283 | 0.37816889 | 0.90650187 |
| Stambpl1 | 0.06106727 | 0.07435181 | 0.07197208 | 0.00237973 | 0.44156909 | 0.90650187 |
| Ywhag | 0.91201574 | 0.71289788 | 0.71052668 | 0.0023712 | 0.49408148 | 0.90650187 |
| Snx18 | 0.20823758 | 0.2274114 | 0.22504414 | 0.00236726 | 0.48135074 | 0.90650187 |
| Fam214b | 0.15294498 | 0.172853 | 0.17048637 | 0.00236663 | 0.47539595 | 0.90650187 |
| Gltscr1l | 0.0955769 | 0.11252492 | 0.11016069 | 0.00236423 | 0.46199453 | 0.90650187 |
| Ldhc | 0.00928685 | 0.01368998 | 0.0113284 | 0.00236158 | 0.17702642 | 0.90650187 |
| Trim5 | 0.00676208 | 0.01062197 | 0.0082627 | 0.00235927 | 0.10215572 | 0.90650187 |
| BC034090 | 0.0329147 | 0.04186341 | 0.03951908 | 0.00234433 | 0.39599807 | 0.90650187 |
| Plekho2 | 0.08906079 | 0.10542107 | 0.10307941 | 0.00234166 | 0.45977883 | 0.90650187 |

| | | | | | | |
|---|---|---|---|---|---|---|
| Rnasel | 0.08476742 | 0.10072218 | 0.09838122 | 0.00234097 | 0.45787746 | 0.90650187 |
| Lrg1 | 0.01556023 | 0.02123744 | 0.0189007 | 0.00233674 | 0.29129001 | 0.90650187 |
| Polk | 0.10243568 | 0.11989655 | 0.11756464 | 0.00233191 | 0.46486715 | 0.90650187 |
| Nxnl2 | 0.02746765 | 0.03542662 | 0.03309905 | 0.00232757 | 0.37728431 | 0.90650187 |
| Slc6a13 | 0.01084702 | 0.01553663 | 0.0132176 | 0.00231903 | 0.21770092 | 0.90650187 |
| Pde7b | 0.01876479 | 0.02506277 | 0.02274414 | 0.00231863 | 0.32520134 | 0.90650187 |
| Pard3b | 0.16090091 | 0.18086275 | 0.17854636 | 0.00231638 | 0.47700361 | 0.90650187 |
| Slc35f1 | 0.02288954 | 0.02997754 | 0.02766693 | 0.00231061 | 0.35521629 | 0.90650187 |
| Slc16a11 | 0.0325773 | 0.04143205 | 0.03912276 | 0.00230929 | 0.39650269 | 0.90650187 |
| Camsap3 | 0.07579831 | 0.09078705 | 0.08848217 | 0.00230488 | 0.45390393 | 0.90650187 |
| Afmid | 0.0108912 | 0.01557453 | 0.01327103 | 0.00230349 | 0.22016451 | 0.90650187 |
| Disc1 | 0.02186826 | 0.0287539 | 0.0264506 | 0.00230329 | 0.34933395 | 0.90650187 |
| Megf11 | 0.0176527 | 0.02371151 | 0.02141221 | 0.00229931 | 0.31654489 | 0.90650187 |
| AI427809 | 0.02887246 | 0.037058 | 0.03475926 | 0.00229875 | 0.38437787 | 0.90650187 |
| Fads1 | 0.13930041 | 0.15878699 | 0.15649168 | 0.00229532 | 0.4740053 | 0.90650187 |
| Mrpl24 | 0.99684015 | 0.75170226 | 0.7494187 | 0.00228356 | 0.494596 | 0.90650187 |
| Fmnl1 | 0.04928202 | 0.06081545 | 0.05853295 | 0.00228249 | 0.43118559 | 0.90650187 |
| Psmc3ip | 0.11733513 | 0.13577903 | 0.13349801 | 0.00228102 | 0.46972549 | 0.90650187 |
| Mapk1ip1l | 0.42926343 | 0.41584948 | 0.41356852 | 0.00228096 | 0.49021935 | 0.90650187 |
| Cyp27a1 | 0.01471613 | 0.02013821 | 0.01788556 | 0.00225266 | 0.28776804 | 0.90650187 |
| Trmt1l | 0.39680952 | 0.39073846 | 0.38848966 | 0.00224879 | 0.48973491 | 0.90650187 |
| Srr | 0.14642302 | 0.16606955 | 0.16382439 | 0.00224515 | 0.47570933 | 0.90650187 |
| Dusp26 | 0.03512298 | 0.04434758 | 0.04210856 | 0.00223902 | 0.40656735 | 0.90650187 |
| Nrxn1 | 0.03574582 | 0.04506951 | 0.04283755 | 0.00223195 | 0.40841339 | 0.90650187 |
| Hecw2 | 0.03681578 | 0.04631736 | 0.04408845 | 0.00222891 | 0.41108722 | 0.90650187 |
| Tle4 | 0.06661334 | 0.0804537 | 0.0782251 | 0.0022286 | 0.44960734 | 0.90650187 |
| Xlr3b | 0.0948598 | 0.11160845 | 0.10938428 | 0.00222417 | 0.46398666 | 0.90650187 |
| Dnajb1 | 0.69208963 | 0.59287361 | 0.59066972 | 0.00220389 | 0.49338291 | 0.90650187 |
| Tmigd3 | 0.03923011 | 0.04910685 | 0.04690454 | 0.00220231 | 0.41732779 | 0.90650187 |
| Atrx | 1.34122061 | 0.86323945 | 0.86104008 | 0.00219937 | 0.49546991 | 0.90650187 |

| | | | | | | |
|---|---|---|---|---|---|---|
| Rb1cc1 | 0.52038734 | 0.48209777 | 0.4799051 | 0.00219267 | 0.49189731 | 0.90650187 |
| Trpm7 | 0.38436129 | 0.38082409 | 0.37865256 | 0.00217153 | 0.48983004 | 0.90650187 |
| Slc10a4 | 0.00957054 | 0.01384297 | 0.01167222 | 0.00217075 | 0.20418185 | 0.90650187 |
| Grid1 | 0.0083872 | 0.01240453 | 0.0102372 | 0.00216733 | 0.17330526 | 0.90650187 |
| Gm14403 | 0.04121266 | 0.05137365 | 0.04921027 | 0.00216338 | 0.42252528 | 0.90650187 |
| Nxph3 | 0.01382485 | 0.01897213 | 0.01681241 | 0.00215973 | 0.28397135 | 0.90650187 |
| 9930012K11I | 0.04217871 | 0.05248922 | 0.05033162 | 0.0021576 | 0.42443008 | 0.90650187 |
| Mcf2l | 0.32187762 | 0.32952183 | 0.32736463 | 0.0021572 | 0.48831475 | 0.90650187 |
| Cers5 | 0.55084679 | 0.50298659 | 0.50083272 | 0.00215387 | 0.49237322 | 0.90650187 |
| Ube2u | 0.02142663 | 0.0280667 | 0.0259241 | 0.0021426 | 0.35665134 | 0.90650187 |
| 2700029M09 | 0.62716462 | 0.55291651 | 0.55077536 | 0.00214115 | 0.49310567 | 0.90650187 |
| Dlgap1 | 0.02921797 | 0.03730045 | 0.0351671 | 0.00213335 | 0.3937015 | 0.90650187 |
| Gm9958 | 0.02599191 | 0.03348354 | 0.03135166 | 0.00213188 | 0.38121275 | 0.90650187 |
| Stradb | 0.13600662 | 0.15520926 | 0.15308059 | 0.00212868 | 0.47535358 | 0.90650187 |
| Wipf1 | 0.16448622 | 0.18428227 | 0.18215508 | 0.00212719 | 0.47929806 | 0.90650187 |
| Hand1 | 0.01428135 | 0.01948865 | 0.01736221 | 0.00212643 | 0.29305755 | 0.90650187 |
| Gm17529 | 0.01441971 | 0.0196545 | 0.01752879 | 0.00212571 | 0.29490315 | 0.90650187 |
| Hspa14 | 0.48281198 | 0.45536767 | 0.45324957 | 0.0021181 | 0.49171258 | 0.90650187 |
| Rbm12b2 | 0.10600984 | 0.12353002 | 0.12141201 | 0.00211801 | 0.46909186 | 0.90650187 |
| Dct | 0.06633436 | 0.08002466 | 0.07791163 | 0.00211303 | 0.45201632 | 0.90650187 |
| Gm10101 | 0.02203395 | 0.02875972 | 0.02664805 | 0.00211167 | 0.36231295 | 0.90650187 |
| Cdh12 | 0.0183652 | 0.0243702 | 0.02226579 | 0.00210441 | 0.33718156 | 0.90650187 |
| Syce1 | 0.01134822 | 0.01592599 | 0.01382366 | 0.00210233 | 0.24948938 | 0.90650187 |
| Hsd11b1 | 0.0139488 | 0.019061 | 0.01696172 | 0.00209927 | 0.29108711 | 0.90650187 |
| Ablim2 | 0.02380315 | 0.03084517 | 0.02875363 | 0.00209154 | 0.37320632 | 0.90650187 |
| Ski | 0.3483755 | 0.35160504 | 0.34951461 | 0.00209044 | 0.48939376 | 0.90650187 |
| Fat3 | 0.00823346 | 0.01213533 | 0.01005059 | 0.00208474 | 0.17823123 | 0.90650187 |
| Taco1os | 0.03687756 | 0.04624296 | 0.04416063 | 0.00208233 | 0.41697966 | 0.90650187 |
| Aqp7 | 0.01188598 | 0.01655196 | 0.01447346 | 0.0020785 | 0.26159904 | 0.90650187 |
| Ccdc122 | 0.04645397 | 0.05735449 | 0.05527696 | 0.00207753 | 0.43365206 | 0.90650187 |

| | | | | | | |
|---|---|---|---|---|---|---|
| Myl4 | 0.04340927 | 0.0538248 | 0.05175791 | 0.00206689 | 0.42954609 | 0.90650187 |
| Slco4a1 | 0.02365451 | 0.03064349 | 0.02857692 | 0.00206657 | 0.37392025 | 0.90650187 |
| Inafm1 | 0.25113629 | 0.26720459 | 0.26513866 | 0.00206592 | 0.48618359 | 0.90650187 |
| Arl2bp | 0.6308815 | 0.55518373 | 0.55312121 | 0.00206252 | 0.49338699 | 0.90650187 |
| Pcbd2 | 0.70371214 | 0.59963569 | 0.59757581 | 0.00205988 | 0.49388674 | 0.90650187 |
| Cyp26b1 | 0.01462117 | 0.01981743 | 0.01777128 | 0.00204615 | 0.3043747 | 0.90650187 |
| Slc38a6 | 0.09096588 | 0.10719944 | 0.10515585 | 0.00204359 | 0.46557603 | 0.90650187 |
| Acsl3 | 0.17512606 | 0.19482215 | 0.19277933 | 0.00204282 | 0.48121328 | 0.90650187 |
| Lrrc8d | 0.18892871 | 0.20841095 | 0.20637321 | 0.00203774 | 0.48249361 | 0.90650187 |
| Lsm14b | 0.15895641 | 0.17861785 | 0.17658309 | 0.00203476 | 0.4795725 | 0.90650187 |
| Atp6v0a4 | 0.0079301 | 0.01171698 | 0.00968227 | 0.00203471 | 0.17509134 | 0.90650187 |
| Gm26645 | 0.00644563 | 0.00990846 | 0.00787772 | 0.00203074 | 0.12589796 | 0.90650187 |
| Rasal3 | 0.0210598 | 0.0275124 | 0.02548655 | 0.00202585 | 0.36190598 | 0.90650187 |
| Kcp | 0.03043437 | 0.03862332 | 0.03660148 | 0.00202184 | 0.40300682 | 0.90650187 |
| Arxes2 | 0.30480189 | 0.31478136 | 0.31276483 | 0.00201654 | 0.48856675 | 0.90650187 |
| Xlr3c | 0.01176684 | 0.01634167 | 0.01432954 | 0.00201213 | 0.26623392 | 0.90650187 |
| Col4a6 | 0.01996866 | 0.02619188 | 0.02418374 | 0.00200814 | 0.35600868 | 0.90650187 |
| Stk4 | 0.20074073 | 0.21984819 | 0.21784276 | 0.00200543 | 0.48367758 | 0.90650187 |
| Mybpc1 | 0.01109047 | 0.0155088 | 0.01351203 | 0.00199676 | 0.25560433 | 0.90650187 |
| Bfsp1 | 0.00746964 | 0.01111853 | 0.00912291 | 0.00199561 | 0.16540961 | 0.90650187 |
| Aim2 | 0.0063844 | 0.00979669 | 0.00780321 | 0.00199348 | 0.12803808 | 0.90650187 |
| Alpl | 0.00551341 | 0.00873027 | 0.00674263 | 0.00198764 | 0.09501213 | 0.90650187 |
| Sec22c | 0.07703032 | 0.09182358 | 0.08984871 | 0.00197487 | 0.46107961 | 0.90650187 |
| RP24-143K11 | 0.03386888 | 0.04260557 | 0.0406389 | 0.00196667 | 0.41482932 | 0.90650187 |
| Syap1 | 0.4199531 | 0.40841854 | 0.40645675 | 0.00196179 | 0.49144055 | 0.90650187 |
| Ccdc58 | 0.44474198 | 0.42721324 | 0.42525195 | 0.00196129 | 0.49182087 | 0.90650187 |
| Pcdhga3 | 0.01066023 | 0.01494595 | 0.01299163 | 0.00195432 | 0.25182765 | 0.90650187 |
| Gm26691 | 0.03463652 | 0.04349077 | 0.04153878 | 0.00195199 | 0.41726015 | 0.90650187 |
| Tnfaip1 | 0.62727715 | 0.55279704 | 0.55084649 | 0.00195054 | 0.49372016 | 0.90650187 |
| Rxrg | 0.01136673 | 0.01579499 | 0.01384602 | 0.00194897 | 0.26573489 | 0.90650187 |

| | | | | | | |
|---|---|---|---|---|---|---|
| Kcnk5 | 0.03658717 | 0.04576734 | 0.04382133 | 0.00194601 | 0.42174973 | 0.90650187 |
| Rho | 0.00658224 | 0.00998971 | 0.00804394 | 0.00194577 | 0.14110693 | 0.90650187 |
| Xlr4b | 0.02394768 | 0.03087012 | 0.02892543 | 0.0019447 | 0.38251424 | 0.90650187 |
| Dennd2c | 0.03052473 | 0.0386508 | 0.03670793 | 0.00194287 | 0.40698965 | 0.90650187 |
| Elovl2 | 0.03971256 | 0.04940424 | 0.0474662 | 0.00193804 | 0.42798223 | 0.90650187 |
| Eepd1 | 0.09767496 | 0.11434735 | 0.1124282 | 0.00191915 | 0.46975464 | 0.90650187 |
| Cyp1b1 | 0.01126861 | 0.01563595 | 0.01372742 | 0.00190853 | 0.26826269 | 0.90650187 |
| Slc50a1 | 0.61962844 | 0.54790419 | 0.54599568 | 0.00190851 | 0.49380091 | 0.90650187 |
| Htr1b | 0.00653151 | 0.00988381 | 0.00798221 | 0.0019016 | 0.14478252 | 0.90650187 |
| Ttc9b | 0.01831626 | 0.02410605 | 0.02220719 | 0.00189886 | 0.35192487 | 0.90650187 |
| Gm16104 | 0.02217535 | 0.02871176 | 0.02681651 | 0.00189525 | 0.37668775 | 0.90650187 |
| Yif1a | 0.89530573 | 0.70409063 | 0.70219753 | 0.0018931 | 0.49521871 | 0.90650187 |
| Mertk | 0.05730915 | 0.06959917 | 0.06770909 | 0.00189008 | 0.45061891 | 0.90650187 |
| Gpr162 | 0.02209673 | 0.02861098 | 0.02672285 | 0.00188813 | 0.37671927 | 0.90650187 |
| Gosr2 | 0.41733559 | 0.40633302 | 0.4044449 | 0.00188812 | 0.49172095 | 0.90650187 |
| Zfp768 | 0.25079769 | 0.26671681 | 0.26482948 | 0.00188734 | 0.48736279 | 0.90650187 |
| 4930579G24 | 0.09414957 | 0.11049788 | 0.1086146 | 0.00188327 | 0.46927895 | 0.90650187 |
| Gm26906 | 0.01794196 | 0.02363661 | 0.02175884 | 0.00187777 | 0.35061823 | 0.90650187 |
| Ift46 | 0.36059543 | 0.36140575 | 0.35952812 | 0.00187763 | 0.49073856 | 0.90650187 |
| Plpp5 | 0.34474674 | 0.34839294 | 0.34651696 | 0.00187598 | 0.4903993 | 0.90650187 |
| Cttn | 0.7304197 | 0.61505045 | 0.61318943 | 0.00186102 | 0.4946175 | 0.90650187 |
| Atp6v1g2 | 0.02169002 | 0.02809648 | 0.02623814 | 0.00185834 | 0.37643237 | 0.90650187 |
| Eif3g | 1.4315977 | 0.88266896 | 0.88081996 | 0.001849 | 0.49627707 | 0.90650187 |
| Gusb | 0.51976859 | 0.48132137 | 0.47947386 | 0.00184751 | 0.49316653 | 0.90650187 |
| Tmem54 | 0.01183079 | 0.01625202 | 0.01440679 | 0.00184523 | 0.28454357 | 0.90650187 |
| Mtmr2 | 0.4154716 | 0.40483738 | 0.40300909 | 0.00182829 | 0.49195469 | 0.90650187 |
| Igsf21 | 0.09391001 | 0.11018215 | 0.10835484 | 0.00182731 | 0.47011876 | 0.90650187 |
| Pdhb | 0.79047208 | 0.64886262 | 0.64703948 | 0.00182315 | 0.49500287 | 0.90650187 |
| Dnm3 | 0.06360667 | 0.07666125 | 0.07484078 | 0.00182047 | 0.45694337 | 0.90650187 |
| Ccrl2 | 0.0348972 | 0.04365447 | 0.04184415 | 0.00181031 | 0.42374132 | 0.90650187 |

| | | | | | | |
|---|---|---|---|---|---|---|
| Tsr2 | 0.29205411 | 0.30350441 | 0.30169532 | 0.00180909 | 0.48936639 | 0.90650187 |
| Il11 | 0.01769406 | 0.02326743 | 0.02146179 | 0.00180564 | 0.35419417 | 0.90650187 |
| Vps41 | 0.40242201 | 0.39467715 | 0.39288408 | 0.00179307 | 0.49190636 | 0.90650187 |
| Sfrp5 | 0.03914887 | 0.04860006 | 0.04680993 | 0.00179013 | 0.43250054 | 0.90650187 |
| Pcyox1l | 0.32747973 | 0.33388568 | 0.33209743 | 0.00178824 | 0.49045094 | 0.90650187 |
| Gm20467 | 0.02429329 | 0.03111893 | 0.02933608 | 0.00178285 | 0.39351305 | 0.90650187 |
| Hsf2bp | 0.00932947 | 0.01315914 | 0.01138006 | 0.00177908 | 0.24352875 | 0.90650187 |
| Plppr5 | 0.00860635 | 0.01227753 | 0.01050313 | 0.00177439 | 0.22631507 | 0.90650187 |
| Tmem51 | 0.25401474 | 0.26953578 | 0.26776259 | 0.0017732 | 0.48825682 | 0.90650187 |
| Gk5 | 0.08486564 | 0.10026104 | 0.09848899 | 0.00177206 | 0.46812366 | 0.90650187 |
| Egr4 | 0.01469188 | 0.01962826 | 0.01785636 | 0.00177189 | 0.32955639 | 0.90650187 |
| C2cd4d | 0.01351408 | 0.0182088 | 0.01643793 | 0.00177087 | 0.31599501 | 0.90650187 |
| Irf7 | 0.04559676 | 0.05605809 | 0.05428762 | 0.00177047 | 0.44236186 | 0.90650187 |
| Onecut1 | 0.04569657 | 0.05617168 | 0.05440287 | 0.00176881 | 0.44253682 | 0.90650187 |
| Xrcc2 | 0.03795451 | 0.04718395 | 0.0454178 | 0.00176615 | 0.43137492 | 0.90650187 |
| Wnt11 | 0.01584568 | 0.02100336 | 0.01924373 | 0.00175962 | 0.34218722 | 0.90650187 |
| Agrp | 0.00655026 | 0.00976117 | 0.00800503 | 0.00175615 | 0.16471053 | 0.90650187 |
| Rab9b | 0.02250069 | 0.02895838 | 0.02720401 | 0.00175437 | 0.38717346 | 0.90650187 |
| Nr2f1 | 0.05634048 | 0.06835035 | 0.06660689 | 0.00174346 | 0.45368153 | 0.90650187 |
| Mpg | 0.41019357 | 0.40066946 | 0.39892939 | 0.00174007 | 0.49226457 | 0.90650187 |
| Acot4 | 0.01275807 | 0.01726529 | 0.01552626 | 0.00173902 | 0.30926625 | 0.90650187 |
| Dhx34 | 0.08230788 | 0.09741421 | 0.09567804 | 0.00173617 | 0.46785228 | 0.90650187 |
| Fam71a | 0.01151106 | 0.01575451 | 0.01402047 | 0.00173404 | 0.2912185 | 0.90650187 |
| Oprk1 | 0.00887985 | 0.01256675 | 0.01083491 | 0.00173184 | 0.23867191 | 0.90650187 |
| Ptges | 0.06690816 | 0.08027722 | 0.07855623 | 0.00172099 | 0.46120699 | 0.90650187 |
| Edn3 | 0.08346786 | 0.09866541 | 0.09695399 | 0.00171141 | 0.46872588 | 0.90650187 |
| 9230110C19I | 0.06398513 | 0.07697866 | 0.07526751 | 0.00171115 | 0.45974843 | 0.90650187 |
| Dnah11 | 0.01921809 | 0.02499273 | 0.02328648 | 0.00170625 | 0.3723111 | 0.90650187 |
| Prtn3 | 0.01909696 | 0.02484778 | 0.02314159 | 0.00170619 | 0.37154371 | 0.90650187 |
| Prpf18 | 0.33244754 | 0.33797798 | 0.33627187 | 0.00170611 | 0.49100252 | 0.90650187 |

| | | | | | | |
|---|---|---|---|---|---|---|
| Fmo5 | 0.02363572 | 0.03025891 | 0.02855458 | 0.00170433 | 0.39537257 | 0.90650187 |
| Cep19 | 0.28662344 | 0.29863787 | 0.29693394 | 0.00170393 | 0.48982378 | 0.90650187 |
| Rel | 0.02847843 | 0.03599402 | 0.03429391 | 0.00170012 | 0.41278361 | 0.90650187 |
| Mybpc2 | 0.0053983 | 0.00829949 | 0.00660238 | 0.00169711 | 0.12657782 | 0.90650187 |
| Gm11423 | 0.067728 | 0.08116908 | 0.0794764 | 0.00169269 | 0.46228357 | 0.90650187 |
| Trib2 | 0.05201051 | 0.06335285 | 0.06166286 | 0.00168998 | 0.45151287 | 0.90650187 |
| Gm29609 | 0.00615829 | 0.00921633 | 0.00752801 | 0.00168832 | 0.15937654 | 0.90650187 |
| Ctnna2 | 0.02681699 | 0.03401724 | 0.03232904 | 0.0016882 | 0.40821193 | 0.90650187 |
| Fam134a | 0.56633045 | 0.51293542 | 0.511248 | 0.00168742 | 0.49414648 | 0.90650187 |
| Rhno1 | 0.31737882 | 0.3252278 | 0.32354301 | 0.00168478 | 0.49076548 | 0.90650187 |
| Gbp8 | 0.01422457 | 0.01897273 | 0.01729384 | 0.00167888 | 0.33302379 | 0.90650187 |
| Zfp101 | 0.14048575 | 0.15939371 | 0.15771611 | 0.0016776 | 0.48114216 | 0.90650187 |
| Sh3tc1 | 0.03464555 | 0.04322542 | 0.04154937 | 0.00167605 | 0.4288391 | 0.90650187 |
| Dbh | 0.00639794 | 0.00949355 | 0.00781968 | 0.00167387 | 0.17064454 | 0.90650187 |
| Dock2 | 0.00597911 | 0.00898218 | 0.00730986 | 0.00167232 | 0.15456478 | 0.90650187 |
| Rhbdl3 | 0.05385012 | 0.06543555 | 0.06376679 | 0.00166876 | 0.45369141 | 0.90650187 |
| Gm44148 | 0.01592358 | 0.02100343 | 0.01933732 | 0.00166611 | 0.35084719 | 0.90650187 |
| Gm28523 | 0.01352588 | 0.01811499 | 0.01645215 | 0.00166284 | 0.3265991 | 0.90650187 |
| Panx1 | 0.13968224 | 0.15854634 | 0.15688628 | 0.00166006 | 0.48124056 | 0.90650187 |
| Mgmt | 0.16235753 | 0.1816691 | 0.18001425 | 0.00165485 | 0.48370058 | 0.90650187 |
| Bend6 | 0.01901921 | 0.02470043 | 0.02304858 | 0.00165186 | 0.37501131 | 0.90650187 |
| Cdkn2b | 0.06026268 | 0.07270929 | 0.07106115 | 0.00164814 | 0.45893867 | 0.90650187 |
| 4930507D05I | 0.00593706 | 0.00890624 | 0.00725866 | 0.00164758 | 0.15647093 | 0.90650187 |
| Plxna3 | 0.07282087 | 0.08680889 | 0.08517057 | 0.00163832 | 0.46592623 | 0.90650187 |
| Hist1h2ac | 0.00366922 | 0.00612573 | 0.00449289 | 0.00163284 | 0.05308458 | 0.61929297 |
| Pxylp1 | 0.10609346 | 0.12312595 | 0.12150183 | 0.00162412 | 0.4763069 | 0.90650187 |
| Thap1 | 0.10584384 | 0.12285737 | 0.12123367 | 0.0016237 | 0.47626082 | 0.90650187 |
| Hoxc5 | 0.03313922 | 0.04139982 | 0.0397827 | 0.00161712 | 0.42829805 | 0.90650187 |
| Tmem9b | 0.53534891 | 0.49187353 | 0.49025838 | 0.00161516 | 0.49415728 | 0.90650187 |
| Glt1d1 | 0.01687935 | 0.02209883 | 0.02048481 | 0.00161402 | 0.36306644 | 0.90650187 |

| | | | | | | |
|---|---|---|---|---|---|---|
| Osmr | 0.01974566 | 0.02552664 | 0.02391726 | 0.00160938 | 0.38241546 | 0.90650187 |
| Gpd1 | 0.12517483 | 0.14337703 | 0.14177206 | 0.00160497 | 0.47993062 | 0.90650187 |
| Tmem44 | 0.05218884 | 0.06346737 | 0.06186704 | 0.00160033 | 0.4542244 | 0.90650187 |
| Ccdc66 | 0.15908729 | 0.17831208 | 0.17671537 | 0.00159672 | 0.48397944 | 0.90650187 |
| Ecm2 | 0.01501267 | 0.01983599 | 0.01824231 | 0.00159368 | 0.34886938 | 0.90650187 |
| Slc24a2 | 0.01882295 | 0.02440734 | 0.02281375 | 0.0015936 | 0.3780764 | 0.90650187 |
| Padi4 | 0.01051289 | 0.01440547 | 0.01281334 | 0.00159213 | 0.29033987 | 0.90650187 |
| Larp7 | 0.69604567 | 0.5946199 | 0.59302797 | 0.00159193 | 0.49523919 | 0.90650187 |
| Atp8a1 | 0.04067531 | 0.05017522 | 0.04858594 | 0.00158928 | 0.44219004 | 0.90650187 |
| Trrap | 0.32003847 | 0.32738985 | 0.32580456 | 0.00158528 | 0.49137108 | 0.90650187 |
| Tubd1 | 0.11286677 | 0.13032748 | 0.12874845 | 0.00157902 | 0.47825936 | 0.90650187 |
| Tmem255b | 0.01908796 | 0.02470605 | 0.02313082 | 0.00157522 | 0.38104091 | 0.90650187 |
| Cdh10 | 0.00363832 | 0.00602302 | 0.00445516 | 0.00156786 | 0.05885043 | 0.66167564 |
| Cep112os1 | 0.00608825 | 0.00900909 | 0.00744274 | 0.00156635 | 0.17474151 | 0.90650187 |
| Six4 | 0.0282287 | 0.03556501 | 0.03399885 | 0.00156616 | 0.41886905 | 0.90650187 |
| Rassf5 | 0.04207171 | 0.05176299 | 0.05020749 | 0.0015555 | 0.44522649 | 0.90650187 |
| Aqp5 | 0.00644491 | 0.00942946 | 0.00787684 | 0.00155262 | 0.19043907 | 0.90650187 |
| Dok5 | 0.02958968 | 0.03715508 | 0.03560567 | 0.00154941 | 0.42330162 | 0.90650187 |
| Kif27 | 0.02030254 | 0.02613052 | 0.0245826 | 0.00154792 | 0.38976473 | 0.90650187 |
| Rnf144b | 0.00362116 | 0.00596632 | 0.00443419 | 0.00153213 | 0.06226003 | 0.68780144 |
| Imp3 | 0.84415685 | 0.67743078 | 0.67590003 | 0.00153075 | 0.49598343 | 0.90650187 |
| A630033H20 | 0.01496583 | 0.01971287 | 0.01818596 | 0.0015269 | 0.35447934 | 0.90650187 |
| 2510017J16F | 0.01052435 | 0.0143511 | 0.0128272 | 0.0015239 | 0.29869856 | 0.90650187 |
| Vstm2l | 0.01421606 | 0.01880672 | 0.0172836 | 0.00152312 | 0.34761438 | 0.90650187 |
| Vmo1 | 0.00909595 | 0.01261838 | 0.01109696 | 0.00152142 | 0.27109609 | 0.90650187 |
| Zscan12 | 0.05671018 | 0.06854832 | 0.06702772 | 0.0015206 | 0.4598333 | 0.90650187 |
| Fancm | 0.0999725 | 0.11642051 | 0.11490433 | 0.00151618 | 0.47661131 | 0.90650187 |
| Nudt1 | 0.15734949 | 0.1764678 | 0.17495741 | 0.00151039 | 0.48469292 | 0.90650187 |
| L3mbtl3 | 0.13902827 | 0.15771963 | 0.15621034 | 0.00150929 | 0.48286944 | 0.90650187 |
| Il21r | 0.00360994 | 0.00592942 | 0.00442049 | 0.00150893 | 0.06457019 | 0.70630256 |

| | | | | | | |
|---|---|---|---|---|---|---|
| Paqr5 | 0.00520702 | 0.00787709 | 0.00636925 | 0.00150784 | 0.14629927 | 0.90650187 |
| 1110046J04F | 0.0471125 | 0.05753408 | 0.05603622 | 0.00149786 | 0.45270416 | 0.90650187 |
| Rapgefl1 | 0.01427972 | 0.01885712 | 0.01736026 | 0.00149686 | 0.3507426 | 0.90650187 |
| Cspp1 | 0.88125577 | 0.69658852 | 0.69509367 | 0.00149485 | 0.49618593 | 0.90650187 |
| Gm20661 | 0.0077023 | 0.0108942 | 0.00940559 | 0.00148861 | 0.24084156 | 0.90650187 |
| Mapk13 | 0.19187564 | 0.21072779 | 0.2092488 | 0.00147899 | 0.48746653 | 0.90650187 |
| Adrb2 | 0.01859222 | 0.0240072 | 0.02253758 | 0.00146962 | 0.38595149 | 0.90650187 |
| Psme2b | 0.15528841 | 0.17433685 | 0.17286796 | 0.00146889 | 0.48493353 | 0.90650187 |
| Xlr | 0.00873141 | 0.01212254 | 0.01065485 | 0.00146769 | 0.270145 | 0.90650187 |
| 9930111J21F | 0.01655378 | 0.02156149 | 0.0200941 | 0.00146739 | 0.37272529 | 0.90650187 |
| Thbs4 | 0.01604664 | 0.02095 | 0.01948515 | 0.00146485 | 0.36911017 | 0.90650187 |
| Hvcn1 | 0.03165143 | 0.03949531 | 0.0380343 | 0.00146101 | 0.43220288 | 0.90650187 |
| Mthfd1 | 0.26892116 | 0.28267307 | 0.28122377 | 0.0014493 | 0.4908608 | 0.90650187 |
| Cacng4 | 0.06212769 | 0.07461635 | 0.07317117 | 0.00144518 | 0.4650163 | 0.90650187 |
| Rbbp9 | 0.15602849 | 0.1750506 | 0.17361879 | 0.00143182 | 0.48537707 | 0.90650187 |
| Fam122b | 0.09027736 | 0.10582617 | 0.10440599 | 0.00142018 | 0.47589008 | 0.90650187 |
| Ccdc158 | 0.01625739 | 0.02115138 | 0.01973826 | 0.00141312 | 0.37513832 | 0.90650187 |
| Fam46b | 0.02116675 | 0.02702545 | 0.02561414 | 0.00141131 | 0.40324835 | 0.90650187 |
| Asl | 0.27612188 | 0.28906028 | 0.28764987 | 0.00141041 | 0.49130462 | 0.90650187 |
| Cdc6 | 0.02963124 | 0.03705814 | 0.03565469 | 0.00140345 | 0.43054423 | 0.90650187 |
| Smurf1 | 0.0858299 | 0.10094836 | 0.09954629 | 0.00140207 | 0.47503678 | 0.90650187 |
| Gm16499 | 0.0077656 | 0.01088094 | 0.00948248 | 0.00139846 | 0.25603126 | 0.90650187 |
| Fam26e | 0.03203954 | 0.03988308 | 0.03849073 | 0.00139235 | 0.43611998 | 0.90650187 |
| Pard6a | 0.21524604 | 0.23311117 | 0.23172336 | 0.00138781 | 0.48937936 | 0.90650187 |
| Lrr1 | 0.0523251 | 0.06340942 | 0.06202302 | 0.00138641 | 0.460421 | 0.90650187 |
| Slc9a1 | 0.08605019 | 0.10117282 | 0.09978765 | 0.00138517 | 0.4753967 | 0.90650187 |
| Phf24 | 0.00735191 | 0.01035731 | 0.00897985 | 0.00137746 | 0.24764119 | 0.90650187 |
| Gm7120 | 0.00829338 | 0.01150043 | 0.01012334 | 0.0013771 | 0.27267439 | 0.90650187 |
| Adamts20 | 0.02736988 | 0.03435811 | 0.03298339 | 0.00137473 | 0.42650077 | 0.90650187 |
| Gli1 | 0.03765242 | 0.04643969 | 0.04506533 | 0.00137435 | 0.44607775 | 0.90650187 |

| | | | | | | |
|---|---|---|---|---|---|---|
| Cer1 | 0.04292247 | 0.0525679 | 0.05119396 | 0.00137394 | 0.45251437 | 0.90650187 |
| Dkk3 | 0.11465224 | 0.13200785 | 0.13064923 | 0.00135862 | 0.48156356 | 0.90650187 |
| 4921504A21 | 0.01289793 | 0.01705242 | 0.01569499 | 0.00135743 | 0.35030678 | 0.90650187 |
| Supt4a | 0.70991126 | 0.60258925 | 0.6012319 | 0.00135734 | 0.49599611 | 0.90650187 |
| Tmem158 | 0.04604444 | 0.05615994 | 0.05480444 | 0.0013555 | 0.45622253 | 0.90650187 |
| Il7 | 0.0131409 | 0.01733893 | 0.01598803 | 0.0013509 | 0.35359658 | 0.90650187 |
| Gm17146 | 0.01096526 | 0.01470948 | 0.01336061 | 0.00134886 | 0.32678064 | 0.90650187 |
| Fam131c | 0.01452953 | 0.01900906 | 0.01766098 | 0.00134807 | 0.36717836 | 0.90650187 |
| Pcdh19 | 0.06525176 | 0.07803896 | 0.07669414 | 0.00134482 | 0.46893266 | 0.90650187 |
| Ndnf | 0.01166501 | 0.01554521 | 0.01420651 | 0.00133869 | 0.33763967 | 0.90650187 |
| Tnfrsf11a | 0.00971593 | 0.0131751 | 0.01184837 | 0.00132673 | 0.30931238 | 0.90650187 |
| Slfn8 | 0.00989727 | 0.01339176 | 0.01206802 | 0.00132373 | 0.31290426 | 0.90650187 |
| Gm26764 | 0.03729437 | 0.04596616 | 0.0446474 | 0.00131876 | 0.4477647 | 0.90650187 |
| Ak4 | 0.01943097 | 0.02485912 | 0.02354105 | 0.00131806 | 0.40171559 | 0.90650187 |
| Evc2 | 0.13329964 | 0.1515831 | 0.15026759 | 0.00131551 | 0.48447756 | 0.90650187 |
| Naaladl2 | 0.04461763 | 0.05446199 | 0.05315622 | 0.00130576 | 0.45652001 | 0.90650187 |
| Nppc | 0.01116124 | 0.01490334 | 0.0135976 | 0.00130574 | 0.33472671 | 0.90650187 |
| Ube2a | 0.6210279 | 0.54818685 | 0.54688563 | 0.00130121 | 0.49578027 | 0.90650187 |
| Fgf18 | 0.02508087 | 0.03156971 | 0.03027119 | 0.00129852 | 0.42438023 | 0.90650187 |
| Sema6b | 0.02082754 | 0.02649618 | 0.02520939 | 0.00128678 | 0.41024261 | 0.90650187 |
| Calb1 | 0.01573337 | 0.02039537 | 0.01910878 | 0.00128659 | 0.38234745 | 0.90650187 |
| Nlgn3 | 0.03947468 | 0.04847284 | 0.04718931 | 0.00128353 | 0.45187787 | 0.90650187 |
| Eno4 | 0.01857466 | 0.02379922 | 0.02251657 | 0.00128265 | 0.40004064 | 0.90650187 |
| 1700008J07R | 0.10413163 | 0.12066924 | 0.11939222 | 0.00127702 | 0.48103739 | 0.90650187 |
| Hrsp12 | 0.38350977 | 0.37924475 | 0.37797521 | 0.00126954 | 0.49404327 | 0.90650187 |
| Dsn1 | 0.0510799 | 0.06186388 | 0.0605966 | 0.00126728 | 0.4629627 | 0.90650187 |
| Tshr | 0.00789397 | 0.01089984 | 0.0096384 | 0.00126144 | 0.28034245 | 0.90650187 |
| Tmem81 | 0.01045222 | 0.0139978 | 0.01273992 | 0.00125789 | 0.3303524 | 0.90650187 |
| Gm42595 | 0.01181678 | 0.01564173 | 0.01438987 | 0.00125186 | 0.34947076 | 0.90650187 |
| Sorcs3 | 0.00812888 | 0.01115488 | 0.00992363 | 0.00123124 | 0.29062012 | 0.90650187 |

| | | | | | | |
|---|---|---|---|---|---|---|
| Scrg1 | 0.00598389 | 0.00854647 | 0.00731568 | 0.00123079 | 0.2272516 | 0.90650187 |
| Tnni1 | 0.01404415 | 0.01830315 | 0.01707657 | 0.00122657 | 0.37474245 | 0.90650187 |
| Sec62 | 1.94586369 | 0.92853892 | 0.92731524 | 0.00122368 | 0.49765967 | 0.90650187 |
| 4930426I24R | 0.0059386 | 0.0084828 | 0.00726053 | 0.00122227 | 0.22711204 | 0.90650187 |
| Gm2990 | 0.00582685 | 0.00834495 | 0.00712444 | 0.00122051 | 0.22315192 | 0.90650187 |
| Zfp804a | 0.01437571 | 0.01869507 | 0.01747582 | 0.00121925 | 0.37821925 | 0.90650187 |
| Ddo | 0.02275066 | 0.02871818 | 0.02750163 | 0.00121655 | 0.42204918 | 0.90650187 |
| Glt8d1 | 0.46781656 | 0.44355961 | 0.44234434 | 0.00121526 | 0.49512763 | 0.90650187 |
| Heatr5b | 0.08462721 | 0.09943516 | 0.09822734 | 0.00120782 | 0.47820315 | 0.90650187 |
| Rnf152 | 0.07260531 | 0.08613676 | 0.08493032 | 0.00120643 | 0.47482366 | 0.90650187 |
| Uqcc1 | 0.21474515 | 0.23245204 | 0.23124768 | 0.00120436 | 0.49076407 | 0.90650187 |
| Ankrd46 | 0.33246572 | 0.3374876 | 0.3362871 | 0.0012005 | 0.49366897 | 0.90650187 |
| Pcdhga2 | 0.01294632 | 0.01695237 | 0.01575336 | 0.00119901 | 0.36754592 | 0.90650187 |
| Zfp521 | 0.06866179 | 0.08171684 | 0.08052326 | 0.00119358 | 0.47373015 | 0.90650187 |
| Grik2 | 0.00695118 | 0.00968161 | 0.00849268 | 0.00118893 | 0.26685275 | 0.90650187 |
| Fam207a | 0.3573761 | 0.35808576 | 0.35690194 | 0.00118381 | 0.49411754 | 0.90650187 |
| Asb2 | 0.00580662 | 0.00828291 | 0.0070998 | 0.00118311 | 0.22940357 | 0.90650187 |
| Atp11c | 0.07470559 | 0.08844468 | 0.0872683 | 0.00117638 | 0.47610689 | 0.90650187 |
| Rnf24 | 0.04592341 | 0.05583784 | 0.05466476 | 0.00117308 | 0.46199832 | 0.90650187 |
| Hoxc6 | 0.0073424 | 0.01014025 | 0.00896829 | 0.00117196 | 0.28063993 | 0.90650187 |
| Gjc3 | 0.0059121 | 0.00839813 | 0.00722826 | 0.00116987 | 0.23591528 | 0.90650187 |
| Sema3e | 0.02022094 | 0.02565115 | 0.02448513 | 0.00116602 | 0.41616781 | 0.90650187 |
| Gm27200 | 0.00594105 | 0.00842756 | 0.00726352 | 0.00116404 | 0.23809366 | 0.90650187 |
| Ankrd13b | 0.04590061 | 0.05580043 | 0.05463843 | 0.00116199 | 0.46233831 | 0.90650187 |
| Hoxd8 | 0.01838183 | 0.02344745 | 0.0222857 | 0.00116175 | 0.40836694 | 0.90650187 |
| Chst10 | 0.03533743 | 0.0435186 | 0.04235963 | 0.00115897 | 0.45159472 | 0.90650187 |
| G6pc3 | 0.35153168 | 0.35326753 | 0.35211267 | 0.00115485 | 0.49418337 | 0.90650187 |
| Angptl1 | 0.02058481 | 0.02607369 | 0.02491965 | 0.00115405 | 0.41844272 | 0.90650187 |
| Mpped2 | 0.23505134 | 0.25147216 | 0.25032346 | 0.00114871 | 0.49186198 | 0.90650187 |
| Crlf1 | 0.02108081 | 0.02665878 | 0.02551162 | 0.00114717 | 0.42077815 | 0.90650187 |

| | | | | | | |
|---|---|---|---|---|---|---|
| Sgcd | 0.03983043 | 0.04874799 | 0.04760337 | 0.00114462 | 0.45743632 | 0.90650187 |
| Spr | 0.50145266 | 0.46772927 | 0.46659121 | 0.00113806 | 0.49567427 | 0.90650187 |
| Gm5468 | 0.01368837 | 0.01778065 | 0.01664797 | 0.00113269 | 0.38114762 | 0.90650187 |
| Pcdhb18 | 0.01090818 | 0.01442285 | 0.01329157 | 0.00113128 | 0.35257496 | 0.90650187 |
| A330048O09 | 0.00860009 | 0.01161464 | 0.01049554 | 0.0011191 | 0.31774369 | 0.90650187 |
| Slc35a1 | 0.58091726 | 0.52204206 | 0.52092449 | 0.00111757 | 0.49619519 | 0.90650187 |
| Lhx8 | 0.00888163 | 0.01195251 | 0.01083707 | 0.00111544 | 0.32362724 | 0.90650187 |
| AA386476 | 0.02373482 | 0.02978634 | 0.0286724 | 0.00111394 | 0.43143785 | 0.90650187 |
| Gm4221 | 0.00759995 | 0.01039344 | 0.00928125 | 0.00111219 | 0.29711237 | 0.90650187 |
| Cdh7 | 0.01318079 | 0.0171479 | 0.01603613 | 0.00111177 | 0.37896143 | 0.90650187 |
| RP23-449M8 | 0.00613245 | 0.00860255 | 0.00749654 | 0.00110601 | 0.25594879 | 0.90650187 |
| Zdhhc2 | 0.24058793 | 0.25655436 | 0.25545208 | 0.00110228 | 0.49234762 | 0.90650187 |
| Pcdhb16 | 0.03355299 | 0.04136801 | 0.04026833 | 0.00109968 | 0.45168524 | 0.90650187 |
| Nsmce4a | 0.58610036 | 0.52543174 | 0.52433248 | 0.00109925 | 0.49628186 | 0.90650187 |
| 4731419I09R | 0.00797927 | 0.01083591 | 0.00974199 | 0.00109392 | 0.308821 | 0.90650187 |
| RP23-4H17.3 | 0.08497401 | 0.09969843 | 0.09860787 | 0.00109056 | 0.48039327 | 0.90650187 |
| Eme1 | 0.0331411 | 0.0408709 | 0.03978491 | 0.00108599 | 0.4517069 | 0.90650187 |
| Wrb | 0.18619518 | 0.20478295 | 0.20369751 | 0.00108545 | 0.49055018 | 0.90650187 |
| Slc48a1 | 0.42809033 | 0.41375243 | 0.41267595 | 0.00107648 | 0.49537377 | 0.90650187 |
| Flt4 | 0.02984178 | 0.03697386 | 0.03590299 | 0.00107086 | 0.44725595 | 0.90650187 |
| Kcnq4 | 0.07595215 | 0.0897187 | 0.08865293 | 0.00106578 | 0.47868879 | 0.90650187 |
| Doc2g | 0.00585957 | 0.00822044 | 0.0071643 | 0.00105614 | 0.25611854 | 0.90650187 |
| Lims2 | 0.04669917 | 0.05661247 | 0.05555975 | 0.00105272 | 0.4664354 | 0.90650187 |
| Gm5141 | 0.06894979 | 0.08189292 | 0.08084588 | 0.00104704 | 0.47704354 | 0.90650187 |
| Cd274 | 0.01070013 | 0.0140844 | 0.0130399 | 0.00104449 | 0.36088565 | 0.90650187 |
| Ccdc78 | 0.02116726 | 0.02665895 | 0.02561474 | 0.00104421 | 0.42809391 | 0.90650187 |
| Inpp5b | 0.07946351 | 0.0935853 | 0.09254123 | 0.00104407 | 0.47999897 | 0.90650187 |
| Drp2 | 0.00853698 | 0.01146248 | 0.01041896 | 0.00104352 | 0.32806892 | 0.90650187 |
| Esyt2 | 0.19457097 | 0.21291267 | 0.21187071 | 0.00104195 | 0.49127865 | 0.90650187 |
| Begain | 0.050625 | 0.06111673 | 0.06007492 | 0.00104182 | 0.46927372 | 0.90650187 |

| | | | | | | |
|---|---|---|---|---|---|---|
| Stx11 | 0.02788463 | 0.03463314 | 0.03359216 | 0.00104098 | 0.44521343 | 0.90650187 |
| Fam180a | 0.01398573 | 0.01804589 | 0.01700621 | 0.00103968 | 0.39289457 | 0.90650187 |
| Arhgap4 | 0.00795425 | 0.01075 | 0.00971159 | 0.00103841 | 0.31727063 | 0.90650187 |
| Gm26786 | 0.00849389 | 0.01140214 | 0.01036667 | 0.00103547 | 0.32850445 | 0.90650187 |
| Trim12c | 0.1411117 | 0.15939549 | 0.15836203 | 0.00103346 | 0.48842767 | 0.90650187 |
| Pld5 | 0.01548149 | 0.01983107 | 0.01880605 | 0.00102502 | 0.40427088 | 0.90650187 |
| Zfp951 | 0.01393932 | 0.0179716 | 0.01695031 | 0.00102129 | 0.39440441 | 0.90650187 |
| Krtcap3 | 0.28220552 | 0.29405828 | 0.29304061 | 0.00101767 | 0.49384111 | 0.90650187 |
| 4930556M19 | 0.01152562 | 0.01504824 | 0.01403807 | 0.00101017 | 0.37452088 | 0.90650187 |
| Nuggc | 0.0059824 | 0.0083221 | 0.00731386 | 0.00100824 | 0.26999044 | 0.90650187 |
| Kdelr1 | 1.39454575 | 0.87419472 | 0.87319238 | 0.00100235 | 0.49796415 | 0.90650187 |
| Gm16685 | 0.00577646 | 0.00806429 | 0.00706307 | 0.00100122 | 0.26428671 | 0.90650187 |
| Phldb3 | 0.03774963 | 0.04617782 | 0.04517877 | 0.00099905 | 0.46084431 | 0.90650187 |
| Rasa3 | 0.07038458 | 0.08344973 | 0.08245134 | 0.00099839 | 0.47853486 | 0.90650187 |
| Syn3 | 0.01670677 | 0.02127467 | 0.02027772 | 0.00099695 | 0.41349331 | 0.90650187 |
| Abhd14a | 0.4786696 | 0.45124935 | 0.45025254 | 0.0009968 | 0.49607366 | 0.90650187 |
| L3hypdh | 0.13247155 | 0.15040196 | 0.14940533 | 0.00099663 | 0.48817109 | 0.90650187 |
| Cp | 0.0133417 | 0.01722502 | 0.01623014 | 0.00099488 | 0.39261564 | 0.90650187 |
| Unc80 | 0.22551452 | 0.24240566 | 0.24141757 | 0.00098809 | 0.49274152 | 0.90650187 |
| Traf6 | 0.05521912 | 0.06631624 | 0.0653292 | 0.00098704 | 0.47322424 | 0.90650187 |
| Gm26740 | 0.00480983 | 0.00686237 | 0.005885 | 0.00097737 | 0.23015966 | 0.90650187 |
| Coch | 0.01800673 | 0.02280507 | 0.02183644 | 0.00096863 | 0.42183558 | 0.90650187 |
| Adcyap1r1 | 0.07391086 | 0.08735213 | 0.08638438 | 0.00096776 | 0.48013936 | 0.90650187 |
| Elovl4 | 0.03448884 | 0.04233269 | 0.04136573 | 0.00096695 | 0.45861669 | 0.90650187 |
| C130026L21F | 0.00825836 | 0.01104735 | 0.01008082 | 0.00096654 | 0.33496682 | 0.90650187 |
| Ap3s1 | 1.22311743 | 0.83027918 | 0.8293132 | 0.00096598 | 0.49793419 | 0.90650187 |
| Gm15879 | 0.00987533 | 0.01300692 | 0.01204146 | 0.00096546 | 0.36075517 | 0.90650187 |
| Olfml1 | 0.05268652 | 0.06340005 | 0.06243659 | 0.00096346 | 0.47265406 | 0.90650187 |
| Troap | 0.03070679 | 0.03787537 | 0.0369224 | 0.00095297 | 0.45432508 | 0.90650187 |
| Atxn7l3 | 0.10846639 | 0.12499695 | 0.12404707 | 0.00094988 | 0.48642186 | 0.90650187 |

| | | | | | | |
|---|---|---|---|---|---|---|
| Dmrt2 | 0.02159957 | 0.02707071 | 0.02613031 | 0.0009404 | 0.43644375 | 0.90650187 |
| Wnt9a | 0.04363838 | 0.05296307 | 0.0520232 | 0.00093986 | 0.46799332 | 0.90650187 |
| Gm22 | 0.05388099 | 0.06473905 | 0.06380206 | 0.00093699 | 0.47397236 | 0.90650187 |
| Mgat3 | 0.06975856 | 0.08268749 | 0.08175122 | 0.00093627 | 0.47969697 | 0.90650187 |
| Man1a2 | 0.48209113 | 0.45366461 | 0.45272886 | 0.00093575 | 0.49633431 | 0.90650187 |
| Cd83 | 0.02180958 | 0.02731612 | 0.02638066 | 0.00093546 | 0.43736936 | 0.90650187 |
| Tmprss5 | 0.00554811 | 0.00771816 | 0.00678491 | 0.00093326 | 0.27043806 | 0.90650187 |
| Zfp438 | 0.05042622 | 0.06077487 | 0.05984686 | 0.000928 | 0.47252071 | 0.90650187 |
| Erbb2 | 0.10836774 | 0.12486815 | 0.1239414 | 0.00092675 | 0.48674104 | 0.90650187 |
| BC089491 | 0.00576582 | 0.00797575 | 0.00705012 | 0.00092564 | 0.2797176 | 0.90650187 |
| Gdap10 | 0.01747014 | 0.02211019 | 0.02119338 | 0.00091681 | 0.42374823 | 0.90650187 |
| Galnt5 | 0.01626361 | 0.02066109 | 0.01974573 | 0.00091536 | 0.41836199 | 0.90650187 |
| Tlr6 | 0.01198731 | 0.01550797 | 0.01459586 | 0.00091212 | 0.39057814 | 0.90650187 |
| Fam173b | 0.27898378 | 0.29110022 | 0.29019009 | 0.00091013 | 0.49443777 | 0.90650187 |
| Dnajc28 | 0.04404462 | 0.05340171 | 0.05249342 | 0.00090829 | 0.46934266 | 0.90650187 |
| Ston1 | 0.02626548 | 0.03257629 | 0.03167585 | 0.00090045 | 0.44971754 | 0.90650187 |
| 2410018L13I | 0.0605682 | 0.07230704 | 0.07140717 | 0.00089988 | 0.47766138 | 0.90650187 |
| Csrnp3 | 0.02268733 | 0.02832437 | 0.02742623 | 0.00089814 | 0.44212567 | 0.90650187 |
| 5033403F01I | 0.00859606 | 0.01138597 | 0.01049065 | 0.00089532 | 0.35219281 | 0.90650187 |
| Map1a | 0.27849004 | 0.2906444 | 0.28975239 | 0.00089201 | 0.49454029 | 0.90650187 |
| Cnksr2 | 0.00835957 | 0.01109406 | 0.01020367 | 0.00089039 | 0.3490327 | 0.90650187 |
| S100z | 0.0106578 | 0.01387433 | 0.01298868 | 0.00088565 | 0.38089598 | 0.90650187 |
| Slc12a2 | 0.15472532 | 0.17317423 | 0.17229628 | 0.00087795 | 0.49096357 | 0.90650187 |
| Tenm2 | 0.02457936 | 0.03055015 | 0.02967585 | 0.0008743 | 0.44789774 | 0.90650187 |
| Sec23ip | 0.39155758 | 0.38522159 | 0.38435433 | 0.00086726 | 0.49599824 | 0.90650187 |
| Mirg | 0.26483225 | 0.27841277 | 0.27755234 | 0.00086042 | 0.49450214 | 0.90650187 |
| Lpxn | 0.00722141 | 0.00967886 | 0.00882122 | 0.00085764 | 0.33279053 | 0.90650187 |
| Col4a4 | 0.00967448 | 0.01265474 | 0.01179815 | 0.00085659 | 0.37343603 | 0.90650187 |
| Ciz1 | 0.2847012 | 0.29609855 | 0.29524216 | 0.00085639 | 0.49485572 | 0.90650187 |
| Usp32 | 0.11210712 | 0.12879261 | 0.12793856 | 0.00085405 | 0.48816252 | 0.90650187 |

| | | | | | | |
|---|---|---|---|---|---|---|
| Rabggta | 0.16218069 | 0.18068553 | 0.17983617 | 0.00084936 | 0.4916242 | 0.90650187 |
| Ankrd61 | 0.04184296 | 0.05078084 | 0.04994206 | 0.00083879 | 0.47024074 | 0.90650187 |
| Ccdc189 | 0.17641179 | 0.19488946 | 0.19405443 | 0.00083504 | 0.49236872 | 0.90650187 |
| Abcb11 | 0.00758764 | 0.01009594 | 0.00926629 | 0.00082966 | 0.34530107 | 0.90650187 |
| Hspa4l | 0.38714037 | 0.38168752 | 0.38085926 | 0.00082826 | 0.49614314 | 0.90650187 |
| Igdcc3 | 0.00487386 | 0.00679028 | 0.00596308 | 0.00082719 | 0.26871918 | 0.90650187 |
| Hid1 | 0.30633595 | 0.31491409 | 0.31408695 | 0.00082713 | 0.49532956 | 0.90650187 |
| Fam65c | 0.02000521 | 0.0250537 | 0.02422742 | 0.00082628 | 0.43974412 | 0.90650187 |
| Foxc1 | 0.01467902 | 0.01866051 | 0.01784089 | 0.00081962 | 0.41908552 | 0.90650187 |
| Dolpp1 | 0.09612971 | 0.11157696 | 0.11075874 | 0.00081822 | 0.48690049 | 0.90650187 |
| Map4k1 | 0.01973508 | 0.02471598 | 0.02390462 | 0.00081136 | 0.44003069 | 0.90650187 |
| Sec14l2 | 0.0249165 | 0.0308868 | 0.03007611 | 0.00081068 | 0.45230938 | 0.90650187 |
| Gpr20 | 0.00818977 | 0.01080727 | 0.00999755 | 0.00080972 | 0.35940272 | 0.90650187 |
| Pou6f2 | 0.0359748 | 0.04391199 | 0.0431054 | 0.0008066 | 0.4668514 | 0.90650187 |
| Agpat9 | 0.01610794 | 0.0203591 | 0.01955877 | 0.00080032 | 0.42782685 | 0.90650187 |
| Chd5 | 0.00756089 | 0.01003177 | 0.00923379 | 0.00079798 | 0.35041892 | 0.90650187 |
| Pfkfb4 | 0.04605289 | 0.05560917 | 0.0548142 | 0.00079497 | 0.47429617 | 0.90650187 |
| Rpl28 | 4.31646768 | 0.63224134 | 0.63145 | 0.00079134 | 0.4977774 | 0.90650187 |
| Tppp | 0.01011631 | 0.0131211 | 0.01233328 | 0.00078782 | 0.38821424 | 0.90650187 |
| Ppp2r3c | 0.29182618 | 0.30228122 | 0.30149603 | 0.0007852 | 0.49538121 | 0.90650187 |
| RP24-274B1[illegible] | 0.00474174 | 0.00658537 | 0.00580196 | 0.00078341 | 0.2741638 | 0.90650187 |
| Adh1 | 0.04431049 | 0.05358306 | 0.05280101 | 0.00078205 | 0.47375048 | 0.90650187 |
| Scn1b | 0.01453331 | 0.01844584 | 0.01766553 | 0.00078031 | 0.42216102 | 0.90650187 |
| Aff3 | 0.03614646 | 0.04408621 | 0.04330615 | 0.00078006 | 0.46808773 | 0.90650187 |
| Polr3d | 0.43131969 | 0.41590533 | 0.41513063 | 0.00077469 | 0.49669035 | 0.90650187 |
| Trhde | 0.01387868 | 0.0176451 | 0.01687726 | 0.00076784 | 0.41985825 | 0.90650187 |
| Ccdc30 | 0.09233157 | 0.1073995 | 0.10664126 | 0.00075824 | 0.48739186 | 0.90650187 |
| Asap2 | 0.14513974 | 0.16324823 | 0.16250763 | 0.0007406 | 0.49191798 | 0.90650187 |
| Lynx1 | 0.09310946 | 0.10822645 | 0.10748618 | 0.00074027 | 0.4877874 | 0.90650187 |
| Ctnna3 | 0.01663156 | 0.02092594 | 0.02018746 | 0.00073848 | 0.43540656 | 0.90650187 |

| | | | | | | |
|---|---|---|---|---|---|---|
| Gm15747 | 0.01106832 | 0.01421893 | 0.01348525 | 0.00073368 | 0.40444109 | 0.90650187 |
| Pgpep1 | 0.29742694 | 0.30711274 | 0.30637945 | 0.00073328 | 0.49575532 | 0.90650187 |
| Tpbgl | 0.02305153 | 0.02858887 | 0.02785971 | 0.00072916 | 0.45368648 | 0.90650187 |
| Vps53 | 0.39610655 | 0.38866427 | 0.38793756 | 0.00072671 | 0.49667773 | 0.90650187 |
| Rgs7bp | 0.08092004 | 0.09487313 | 0.09414895 | 0.00072418 | 0.48636087 | 0.90650187 |
| Rab43 | 0.03578317 | 0.04359905 | 0.04288125 | 0.00071781 | 0.47033939 | 0.90650187 |
| Zdhhc14 | 0.05602005 | 0.06695962 | 0.06624198 | 0.00071764 | 0.48079346 | 0.90650187 |
| Oxtr | 0.01954172 | 0.02438752 | 0.02367348 | 0.00071404 | 0.44666614 | 0.90650187 |
| Aqp8 | 0.00764577 | 0.01004176 | 0.00933692 | 0.00070484 | 0.36858681 | 0.90650187 |
| Usp25 | 0.09527922 | 0.11054052 | 0.10983848 | 0.00070205 | 0.48866571 | 0.90650187 |
| Dnah8 | 0.00588263 | 0.00789077 | 0.00719237 | 0.0006984 | 0.33298627 | 0.90650187 |
| Nudt3 | 0.30779655 | 0.31604207 | 0.31534383 | 0.00069824 | 0.49607306 | 0.90650187 |
| Golph3 | 0.47649192 | 0.44936479 | 0.44867231 | 0.00069247 | 0.49726277 | 0.90650187 |
| Dchs1 | 0.10029078 | 0.11593432 | 0.11524806 | 0.00068626 | 0.48944042 | 0.90650187 |
| Apobec1 | 0.00826625 | 0.01077606 | 0.0100904 | 0.00068566 | 0.38129321 | 0.90650187 |
| Kbtbd13 | 0.0111235 | 0.01423722 | 0.01355197 | 0.00068525 | 0.41107144 | 0.90650187 |
| Pced1a | 0.0659522 | 0.07816647 | 0.07748205 | 0.00068441 | 0.484338 | 0.90650187 |
| Cyp2s1 | 0.00817654 | 0.0106593 | 0.0099815 | 0.0006778 | 0.38137086 | 0.90650187 |
| Gm12474 | 0.01177695 | 0.01501895 | 0.01434176 | 0.00067719 | 0.41686756 | 0.90650187 |
| Cxcl1 | 0.01669395 | 0.0209325 | 0.02026234 | 0.00067016 | 0.44155225 | 0.90650187 |
| D130043K22 | 0.01200642 | 0.015289 | 0.01461893 | 0.00067006 | 0.41926826 | 0.90650187 |
| Sdcbp2 | 0.03899336 | 0.04729553 | 0.0466288 | 0.00066674 | 0.47465755 | 0.90650187 |
| 4933415A04 | 0.02187382 | 0.02712247 | 0.02645723 | 0.00066525 | 0.45549849 | 0.90650187 |
| Has1 | 0.0101974 | 0.01309437 | 0.01243146 | 0.00066292 | 0.40630301 | 0.90650187 |
| Chordc1 | 0.5280779 | 0.48590596 | 0.48524457 | 0.00066139 | 0.49758266 | 0.90650187 |
| Zfp711 | 0.04664776 | 0.05615963 | 0.05550045 | 0.00065917 | 0.47894565 | 0.90650187 |
| Zfp607 | 0.04340187 | 0.05240528 | 0.05174935 | 0.00065594 | 0.47753179 | 0.90650187 |
| Mthfsl | 0.4559347 | 0.43424054 | 0.43358961 | 0.00065093 | 0.49733748 | 0.90650187 |
| Tnks1bp1 | 0.12835476 | 0.14575613 | 0.14510652 | 0.00064962 | 0.49206069 | 0.90650187 |
| Arrdc4 | 0.26340753 | 0.27691893 | 0.27626937 | 0.00064956 | 0.49583016 | 0.90650187 |

| | | | | | | |
|---|---|---|---|---|---|---|
| Hmox1 | 0.08566958 | 0.10001784 | 0.09937059 | 0.00064725 | 0.48844975 | 0.90650187 |
| Dgat2 | 0.09032871 | 0.10510858 | 0.10446193 | 0.00064665 | 0.48902264 | 0.90650187 |
| Ltbr | 0.28663641 | 0.29759163 | 0.29694535 | 0.00064628 | 0.49614008 | 0.90650187 |
| Adam11 | 0.014101 | 0.01778753 | 0.01714504 | 0.00064249 | 0.4338451 | 0.90650187 |
| Ky | 0.01152182 | 0.01467403 | 0.01403348 | 0.00064054 | 0.41960085 | 0.90650187 |
| Vill | 0.05723611 | 0.06826284 | 0.06762603 | 0.00063681 | 0.48330397 | 0.90650187 |
| Zfp526 | 0.02711981 | 0.03331644 | 0.03268749 | 0.00062895 | 0.46591621 | 0.90650187 |
| Dlx2 | 0.00572107 | 0.00762361 | 0.00699561 | 0.00062799 | 0.34491719 | 0.90650187 |
| Josd1 | 0.23610899 | 0.25192026 | 0.25130552 | 0.00061474 | 0.49566169 | 0.90650187 |
| Akt2 | 0.44502921 | 0.42607607 | 0.42546702 | 0.00060905 | 0.49746122 | 0.90650187 |
| Gm15594 | 0.02086877 | 0.02586569 | 0.0252586 | 0.00060709 | 0.45745399 | 0.90650187 |
| A830005F24 | 0.01308913 | 0.01653116 | 0.0159256 | 0.00060557 | 0.43288118 | 0.90650187 |
| Smim11 | 0.88770476 | 0.69896636 | 0.69836994 | 0.00059643 | 0.49848536 | 0.90650187 |
| Pih1d1 | 0.40927823 | 0.39881592 | 0.39821974 | 0.00059618 | 0.49734484 | 0.90650187 |
| Lipc | 0.00509535 | 0.00682783 | 0.00623313 | 0.0005947 | 0.3357273 | 0.90650187 |
| Lmntd2 | 0.01488179 | 0.01867784 | 0.01808487 | 0.00059297 | 0.44205406 | 0.90650187 |
| Cpa4 | 0.02135943 | 0.02643585 | 0.02584397 | 0.00059188 | 0.45945201 | 0.90650187 |
| Adcy4 | 0.06339183 | 0.07518892 | 0.07459845 | 0.00059047 | 0.4859648 | 0.90650187 |
| Lin7a | 0.05073846 | 0.0607932 | 0.06020507 | 0.00058813 | 0.48268003 | 0.90650187 |
| Krt17 | 0.02319313 | 0.02861472 | 0.02802819 | 0.00058653 | 0.46293962 | 0.90650187 |
| Phlpp1 | 0.04228403 | 0.05103713 | 0.05045378 | 0.00058335 | 0.47950335 | 0.90650187 |
| Trmt10a | 0.22484244 | 0.24136568 | 0.24078649 | 0.0005792 | 0.49573394 | 0.90650187 |
| Agbl3 | 0.05349426 | 0.06393875 | 0.06336019 | 0.00057856 | 0.48380973 | 0.90650187 |
| Best1 | 0.0079298 | 0.01025907 | 0.0096819 | 0.00057717 | 0.39549782 | 0.90650187 |
| Nckap5l | 0.06487389 | 0.07684291 | 0.07626878 | 0.00057412 | 0.48665198 | 0.90650187 |
| E2f4 | 0.44430053 | 0.42549171 | 0.42492123 | 0.00057049 | 0.49761892 | 0.90650187 |
| Taco1 | 0.04014435 | 0.04853604 | 0.04796857 | 0.00056747 | 0.47902861 | 0.90650187 |
| Sympk | 0.37396256 | 0.3709043 | 0.37034164 | 0.00056266 | 0.49730548 | 0.90650187 |
| Rbm42 | 0.88225729 | 0.69616491 | 0.69560355 | 0.00056136 | 0.49856875 | 0.90650187 |
| Mafk | 0.09667081 | 0.11190226 | 0.11134371 | 0.00055855 | 0.49110389 | 0.90650187 |

| | | | | | | |
|---|---|---|---|---|---|---|
| Las1l | 0.3361323 | 0.33991115 | 0.33935459 | 0.00055656 | 0.49709132 | 0.90650187 |
| Gm8773 | 0.02189352 | 0.02703624 | 0.0264807 | 0.00055554 | 0.46284671 | 0.90650187 |
| 2700016F22I | 0.00730109 | 0.00947135 | 0.00891808 | 0.00055328 | 0.39134875 | 0.90650187 |
| A330017A19 | 0.11816529 | 0.13492756 | 0.13437772 | 0.00054984 | 0.49274348 | 0.90650187 |
| Trabd2b | 0.03196757 | 0.03894707 | 0.03840611 | 0.00054097 | 0.47503519 | 0.90650187 |
| Abcb1b | 0.01374796 | 0.01725631 | 0.01671977 | 0.00053654 | 0.44327929 | 0.90650187 |
| Sfmbt2 | 0.01257227 | 0.0158377 | 0.01530207 | 0.00053563 | 0.43816862 | 0.90650187 |
| Bend4 | 0.0267301 | 0.03275786 | 0.03222617 | 0.00053169 | 0.47076501 | 0.90650187 |
| Chac2 | 0.20996104 | 0.22722302 | 0.22669136 | 0.00053165 | 0.49584064 | 0.90650187 |
| Zfp516 | 0.48888507 | 0.45815217 | 0.45762254 | 0.00052963 | 0.49794742 | 0.90650187 |
| Prrg1 | 0.08458312 | 0.09870738 | 0.09817895 | 0.00052843 | 0.49045521 | 0.90650187 |
| Slc23a3 | 0.00812725 | 0.01044994 | 0.00992166 | 0.00052828 | 0.40644206 | 0.90650187 |
| H2-DMb1 | 0.01703889 | 0.02120336 | 0.02067621 | 0.00052715 | 0.45487978 | 0.90650187 |
| Fam83b | 0.01067194 | 0.01352724 | 0.01300579 | 0.00052144 | 0.4292677 | 0.90650187 |
| Tekt4 | 0.0264398 | 0.03240347 | 0.03188236 | 0.00052111 | 0.47103728 | 0.90650187 |
| Fam78b | 0.00969004 | 0.01233724 | 0.01181699 | 0.00052024 | 0.4224158 | 0.90650187 |
| Cdh16 | 0.00859955 | 0.01101407 | 0.01049488 | 0.00051919 | 0.41296364 | 0.90650187 |
| Abcg4 | 0.01133528 | 0.01432571 | 0.01380801 | 0.0005177 | 0.43381203 | 0.90650187 |
| Utp15 | 0.07807385 | 0.09152071 | 0.0910045 | 0.00051621 | 0.48994099 | 0.90650187 |
| Fam84b | 0.07174202 | 0.08448166 | 0.08396748 | 0.00051418 | 0.48914103 | 0.90650187 |
| Hdac7 | 0.2068635 | 0.22423955 | 0.22372863 | 0.00051092 | 0.49594993 | 0.90650187 |
| Ryk | 0.39904325 | 0.39075083 | 0.3902415 | 0.00050934 | 0.49768522 | 0.90650187 |
| Neu3 | 0.02764399 | 0.03381148 | 0.03330762 | 0.00050385 | 0.47319153 | 0.90650187 |
| Gm15492 | 0.0119736 | 0.01508237 | 0.01457929 | 0.00050308 | 0.43904104 | 0.90650187 |
| 6330562C20I | 0.01488669 | 0.01859201 | 0.01809077 | 0.00050124 | 0.45098416 | 0.90650187 |
| Zfp385b | 0.0199409 | 0.02464877 | 0.02415058 | 0.00049819 | 0.46346587 | 0.90650187 |
| 4833403J16F | 0.00540305 | 0.00710628 | 0.00660816 | 0.00049811 | 0.36877351 | 0.90650187 |
| Coro2a | 0.00730478 | 0.00941269 | 0.00892257 | 0.00049012 | 0.40353764 | 0.90650187 |
| Ppp1r3d | 0.01637556 | 0.02036374 | 0.01988015 | 0.0004836 | 0.45694162 | 0.90650187 |
| A730098A19 | 0.02010199 | 0.02482653 | 0.02434305 | 0.00048348 | 0.46482136 | 0.90650187 |

| | | | | | | |
|---|---|---|---|---|---|---|
| Rab42 | 0.01253411 | 0.01573912 | 0.01525601 | 0.00048311 | 0.4440227 | 0.90650187 |
| Padi3 | 0.01440835 | 0.01799635 | 0.01751511 | 0.00048124 | 0.45139208 | 0.90650187 |
| Catsperd | 0.00303767 | 0.00419849 | 0.00372117 | 0.00047732 | 0.28425746 | 0.90650187 |
| Tbx1 | 0.00782904 | 0.01003648 | 0.00955953 | 0.00047695 | 0.41223448 | 0.90650187 |
| Pilra | 0.00737186 | 0.00947707 | 0.00900408 | 0.00047299 | 0.40767504 | 0.90650187 |
| F420014N23 | 0.00806796 | 0.01032052 | 0.00984967 | 0.00047085 | 0.41585307 | 0.90650187 |
| Zbtb40 | 0.06811456 | 0.08037047 | 0.07990992 | 0.00046054 | 0.48977973 | 0.90650187 |
| Amd2 | 0.02449652 | 0.03003789 | 0.02957748 | 0.00046041 | 0.47241446 | 0.90650187 |
| Itpripl1 | 0.09147275 | 0.10616698 | 0.10570746 | 0.00045952 | 0.49229068 | 0.90650187 |
| Sprtn | 0.00816429 | 0.01041539 | 0.00996662 | 0.00044876 | 0.42067367 | 0.90650187 |
| Isoc2a | 0.05014202 | 0.05996601 | 0.05952069 | 0.00044531 | 0.48673341 | 0.90650187 |
| Tmem28 | 0.01015755 | 0.01282673 | 0.01238321 | 0.00044352 | 0.436746 | 0.90650187 |
| Gm15834 | 0.01453762 | 0.01811361 | 0.01767072 | 0.00044289 | 0.4556404 | 0.90650187 |
| RP23-457J22 | 0.00301338 | 0.00413161 | 0.00369148 | 0.00044013 | 0.29803885 | 0.90650187 |
| Pknox1 | 0.27729284 | 0.28912493 | 0.28869014 | 0.0004348 | 0.49732889 | 0.90650187 |
| Gm16740 | 0.05172143 | 0.06176614 | 0.06133178 | 0.00043436 | 0.48744157 | 0.90650187 |
| Halr1 | 0.01752194 | 0.02168419 | 0.02125547 | 0.00042872 | 0.46427585 | 0.90650187 |
| Srcin1 | 0.01282404 | 0.0160325 | 0.01560585 | 0.00042664 | 0.45163289 | 0.90650187 |
| Gm17660 | 0.00529427 | 0.00690188 | 0.0064756 | 0.00042628 | 0.38489428 | 0.90650187 |
| Hist1h2ag | 0.05033838 | 0.06017075 | 0.05974607 | 0.00042469 | 0.48739544 | 0.90650187 |
| Zc3h12a | 0.0069542 | 0.0089144 | 0.00849636 | 0.00041804 | 0.41342784 | 0.90650187 |
| Fbxo31 | 0.22961785 | 0.24567599 | 0.24526063 | 0.00041535 | 0.4969965 | 0.90650187 |
| Pcdhb9 | 0.01959747 | 0.02414925 | 0.02374012 | 0.00040913 | 0.46946509 | 0.90650187 |
| Jakmip2 | 0.00968213 | 0.01220977 | 0.01180742 | 0.00040235 | 0.43979515 | 0.90650187 |
| Angel1 | 0.03790945 | 0.04576748 | 0.04536523 | 0.00040225 | 0.48427827 | 0.90650187 |
| Apcs | 0.01397046 | 0.01738976 | 0.01698782 | 0.00040195 | 0.45811364 | 0.90650187 |
| Acss1 | 0.0098262 | 0.0123805 | 0.01198195 | 0.00039855 | 0.44122119 | 0.90650187 |
| Gm29340 | 0.00565652 | 0.00731336 | 0.00691698 | 0.00039637 | 0.39945671 | 0.90650187 |
| Ccdc141 | 0.06213388 | 0.07357282 | 0.07317816 | 0.00039466 | 0.49043591 | 0.90650187 |
| Dhx58os | 0.00695713 | 0.00889233 | 0.00849992 | 0.00039241 | 0.41869344 | 0.90650187 |

| | | | | | | |
|---|---|---|---|---|---|---|
| Naa38 | 1.14185459 | 0.80427125 | 0.80388164 | 0.00038962 | 0.49914042 | 0.90650187 |
| St7 | 0.09090623 | 0.10547392 | 0.10509091 | 0.00038301 | 0.49353656 | 0.90650187 |
| Ifitm5 | 0.01999563 | 0.02459865 | 0.02421597 | 0.00038267 | 0.47199652 | 0.90650187 |
| Slitrk5 | 0.0176636 | 0.02180035 | 0.02142528 | 0.00037508 | 0.46898328 | 0.90650187 |
| Insl6 | 0.02089064 | 0.02565829 | 0.0252847 | 0.00037359 | 0.47381417 | 0.90650187 |
| Clspn | 0.0646528 | 0.07638944 | 0.07601982 | 0.00036962 | 0.49137753 | 0.90650187 |
| Esr1 | 0.01380922 | 0.01716206 | 0.01679358 | 0.00036848 | 0.46114732 | 0.90650187 |
| Cap1 | 0.75775081 | 0.62917194 | 0.62880615 | 0.00036579 | 0.4989683 | 0.90650187 |
| Gm26541 | 0.01410285 | 0.01751132 | 0.01714728 | 0.00036405 | 0.46240263 | 0.90650187 |
| D830044I16F | 0.01020855 | 0.01280566 | 0.01244496 | 0.00036069 | 0.44873933 | 0.90650187 |
| Thop1 | 0.31192907 | 0.31924623 | 0.31888975 | 0.00035648 | 0.49801741 | 0.90650187 |
| Gm2093 | 0.00942502 | 0.01185008 | 0.01149587 | 0.00035421 | 0.4455247 | 0.90650187 |
| Pcca | 0.17106408 | 0.18909077 | 0.18873835 | 0.00035242 | 0.49668842 | 0.90650187 |
| Exoc3l2 | 0.00295396 | 0.00397028 | 0.00361883 | 0.00035145 | 0.33296472 | 0.90650187 |
| Nsun3 | 0.05592452 | 0.06647984 | 0.06613316 | 0.00034669 | 0.49070352 | 0.90650187 |
| Adamts7 | 0.07302266 | 0.08573998 | 0.08539541 | 0.00034457 | 0.49284424 | 0.90650187 |
| Tiam2 | 0.00801719 | 0.01013127 | 0.00978802 | 0.00034325 | 0.43805614 | 0.90650187 |
| Slc25a31 | 0.00458845 | 0.00595813 | 0.00561497 | 0.00034316 | 0.39292893 | 0.90650187 |
| Chrnb1 | 0.0244402 | 0.02985176 | 0.02951059 | 0.00034117 | 0.4795054 | 0.90650187 |
| Gm3839 | 0.00706846 | 0.00897417 | 0.00863529 | 0.00033888 | 0.43075117 | 0.90650187 |
| Adrb3 | 0.00475289 | 0.00615372 | 0.00581555 | 0.00033817 | 0.39800826 | 0.90650187 |
| Eva1a | 0.04943215 | 0.05904239 | 0.05870547 | 0.00033692 | 0.48982236 | 0.90650187 |
| Arhgef10 | 0.11200734 | 0.12816842 | 0.12783212 | 0.0003363 | 0.49533426 | 0.90650187 |
| Lancl2 | 0.11681044 | 0.13327747 | 0.13294158 | 0.0003359 | 0.49551898 | 0.90650187 |
| Oas1g | 0.00753286 | 0.00953366 | 0.00919973 | 0.00033392 | 0.43590439 | 0.90650187 |
| Sh2b2 | 0.00507286 | 0.0065389 | 0.00620571 | 0.00033319 | 0.40567344 | 0.90650187 |
| Ccar2 | 0.25962972 | 0.27319077 | 0.27285807 | 0.0003327 | 0.49783752 | 0.90650187 |
| BC029214 | 0.41536797 | 0.40325416 | 0.40292919 | 0.00032497 | 0.49856961 | 0.90650187 |
| Otud7a | 0.00530766 | 0.00680966 | 0.00649192 | 0.00031774 | 0.41387569 | 0.90650187 |
| Car8 | 0.1249888 | 0.14189396 | 0.14157661 | 0.00031734 | 0.49602469 | 0.90650187 |

| | | | | | | |
|---|---|---|---|---|---|---|
| Akap6 | 0.01510893 | 0.01867476 | 0.01835809 | 0.00031668 | 0.4694363 | 0.90650187 |
| 1700029I15R | 0.01740934 | 0.0214357 | 0.02112048 | 0.00031522 | 0.4735493 | 0.90650187 |
| Gm26627 | 0.01037287 | 0.01295576 | 0.01264388 | 0.00031188 | 0.45634058 | 0.90650187 |
| Gm7676 | 0.00734879 | 0.00928571 | 0.00897605 | 0.00030967 | 0.43905341 | 0.90650187 |
| Pla2g7 | 0.00654594 | 0.008308 | 0.00799978 | 0.00030822 | 0.43200058 | 0.90650187 |
| Tnfaip3 | 0.02693421 | 0.03276694 | 0.03246781 | 0.00029912 | 0.483665 | 0.90650187 |
| Csf3 | 0.02168536 | 0.02652072 | 0.02623259 | 0.00028813 | 0.48052757 | 0.90650187 |
| Dync1i1 | 0.01875858 | 0.02302146 | 0.02273671 | 0.00028476 | 0.47779949 | 0.90650187 |
| Mak16 | 0.38514914 | 0.37956051 | 0.37927876 | 0.00028175 | 0.49868252 | 0.90650187 |
| Srrm4 | 0.01719919 | 0.02114655 | 0.02086848 | 0.00027806 | 0.4763821 | 0.90650187 |
| Pnpla3 | 0.00725339 | 0.00913516 | 0.0088601 | 0.00027507 | 0.44511384 | 0.90650187 |
| Msantd3 | 0.11028681 | 0.12626648 | 0.12599492 | 0.00027156 | 0.49617756 | 0.90650187 |
| RP23-474J16 | 0.01512429 | 0.01864513 | 0.01837657 | 0.00026857 | 0.47409858 | 0.90650187 |
| Gpatch4 | 0.25747222 | 0.27117224 | 0.27090377 | 0.00026847 | 0.49824241 | 0.90650187 |
| Bglap | 0.02001998 | 0.02450978 | 0.02424506 | 0.00026471 | 0.4806436 | 0.90650187 |
| Ghitm | 0.58602555 | 0.52454759 | 0.5242834 | 0.00026418 | 0.49910632 | 0.90650187 |
| Gm20512 | 0.00710367 | 0.00894111 | 0.00867809 | 0.00026302 | 0.4464096 | 0.90650187 |
| Foxc2 | 0.00980609 | 0.0122199 | 0.01195759 | 0.00026231 | 0.46115556 | 0.90650187 |
| Gm11149 | 0.00792234 | 0.0099332 | 0.00967285 | 0.00026035 | 0.45237724 | 0.90650187 |
| Rnf166 | 0.23343302 | 0.24907859 | 0.24881862 | 0.00025997 | 0.49814701 | 0.90650187 |
| Tpk1 | 0.13086263 | 0.1479861 | 0.14772769 | 0.00025842 | 0.49689761 | 0.90650187 |
| 2410015M20 | 2.04439458 | 0.92670753 | 0.92645429 | 0.00025324 | 0.49951522 | 0.90650187 |
| Lgals4 | 0.00956168 | 0.01191344 | 0.01166147 | 0.00025197 | 0.46173756 | 0.90650187 |
| 5730507C01I | 0.00832724 | 0.01041615 | 0.01016442 | 0.00025172 | 0.45616677 | 0.90650187 |
| Hcar1 | 0.02290162 | 0.0279263 | 0.02768131 | 0.00024499 | 0.48430772 | 0.90650187 |
| Cldn8 | 0.03358349 | 0.04054706 | 0.04030412 | 0.00024295 | 0.48931057 | 0.90650187 |
| Gm43101 | 0.03572448 | 0.04305455 | 0.04281258 | 0.00024197 | 0.48997718 | 0.90650187 |
| Pm20d1 | 0.00287628 | 0.0037642 | 0.00352385 | 0.00024035 | 0.38085973 | 0.90650187 |
| Siglec1 | 0.00732832 | 0.00919084 | 0.00895117 | 0.00023967 | 0.45262447 | 0.90650187 |
| Wt1os | 0.01643418 | 0.02018941 | 0.01995053 | 0.00023888 | 0.47877446 | 0.90650187 |

| | | | | | | |
|---|---|---|---|---|---|---|
| Klhl10 | 0.00823855 | 0.01029144 | 0.01005677 | 0.00023467 | 0.45869017 | 0.90650187 |
| Scube1 | 0.01737802 | 0.02131616 | 0.02108293 | 0.00023323 | 0.48038805 | 0.90650187 |
| Cd1d2 | 0.01386216 | 0.01708791 | 0.01685736 | 0.00023055 | 0.47575918 | 0.90650187 |
| Dusp2 | 0.01260424 | 0.01556845 | 0.01534064 | 0.0002278 | 0.47368249 | 0.90650187 |
| Itga10 | 0.00556545 | 0.00703132 | 0.00680604 | 0.00022528 | 0.44150701 | 0.90650187 |
| RP23-299D2. | 0.05796477 | 0.06867731 | 0.0684543 | 0.00022301 | 0.49422225 | 0.90650187 |
| 1700001O22 | 0.02342938 | 0.02852509 | 0.02830921 | 0.00021588 | 0.48647797 | 0.90650187 |
| Gpr182 | 0.02214037 | 0.0269887 | 0.02677484 | 0.00021386 | 0.48583716 | 0.90650187 |
| Fbxl17 | 0.14064917 | 0.15809658 | 0.15788478 | 0.00021179 | 0.4976209 | 0.90650187 |
| Nqo1 | 0.08553364 | 0.09943304 | 0.09922158 | 0.00021146 | 0.49622029 | 0.90650187 |
| Sh3rf1 | 0.08823871 | 0.10239037 | 0.10218182 | 0.00020855 | 0.49638034 | 0.90650187 |
| AW112010 | 0.01871561 | 0.02289067 | 0.02268528 | 0.00020539 | 0.48394651 | 0.90650187 |
| Parp14 | 0.00585087 | 0.0073589 | 0.0071537 | 0.0002052 | 0.44926461 | 0.90650187 |
| Padi2 | 0.03229292 | 0.0389906 | 0.03878859 | 0.00020201 | 0.4907643 | 0.90650187 |
| Myh8 | 0.0055423 | 0.00697614 | 0.00677783 | 0.00019831 | 0.44825517 | 0.90650187 |
| D630045J12F | 0.0851999 | 0.09905222 | 0.09885564 | 0.00019658 | 0.49647329 | 0.90650187 |
| Reln | 0.01288018 | 0.01586756 | 0.01567358 | 0.00019398 | 0.47806157 | 0.90650187 |
| Gm26910 | 0.00942762 | 0.01169286 | 0.01149902 | 0.00019385 | 0.47013014 | 0.90650187 |
| Tstd2 | 0.19885104 | 0.21621164 | 0.2160182 | 0.00019345 | 0.49841177 | 0.90650187 |
| Ankrd34b | 0.00949592 | 0.01177285 | 0.01158179 | 0.00019105 | 0.47076974 | 0.90650187 |
| A930029G22 | 0.02684047 | 0.03254702 | 0.03235683 | 0.00019019 | 0.48957651 | 0.90650187 |
| A4galt | 0.04457206 | 0.05329157 | 0.05310353 | 0.00018804 | 0.49372023 | 0.90650187 |
| St8sia2 | 0.02995529 | 0.03622122 | 0.03603683 | 0.00018439 | 0.49092622 | 0.90650187 |
| Cideb | 0.00283221 | 0.00364975 | 0.00346997 | 0.00017978 | 0.40891727 | 0.90650187 |
| Thada | 0.04838933 | 0.05768243 | 0.05750649 | 0.00017594 | 0.49457402 | 0.90650187 |
| Ica1l | 0.01247334 | 0.01535602 | 0.01518267 | 0.00017335 | 0.47975861 | 0.90650187 |
| Gm973 | 0.00681263 | 0.00849619 | 0.00832419 | 0.000172 | 0.46340511 | 0.90650187 |
| Zfp729b | 0.07607914 | 0.08895768 | 0.08879385 | 0.00016383 | 0.49672777 | 0.90650187 |
| B230208H11 | 0.01407487 | 0.01727563 | 0.01711357 | 0.00016206 | 0.48321034 | 0.90650187 |
| Plekho1 | 0.06388671 | 0.07531805 | 0.07515656 | 0.0001615 | 0.49618903 | 0.90650187 |

| | | | | | | |
|---|---|---|---|---|---|---|
| Otud3 | 0.05282876 | 0.06275999 | 0.06259931 | 0.00016068 | 0.49544784 | 0.90650187 |
| Mtus2 | 0.08954001 | 0.10375896 | 0.10360221 | 0.00015676 | 0.49731649 | 0.90650187 |
| Jade1 | 0.32769499 | 0.3324354 | 0.33227876 | 0.00015664 | 0.49916392 | 0.90650187 |
| Plet1 | 0.01076893 | 0.01327829 | 0.01312314 | 0.00015515 | 0.47904188 | 0.90650187 |
| Fignl1 | 0.11748745 | 0.13381189 | 0.13365949 | 0.0001524 | 0.49797782 | 0.90650187 |
| Spats2 | 0.15739261 | 0.17515249 | 0.17500107 | 0.00015142 | 0.49846547 | 0.90650187 |
| Npy1r | 0.03205106 | 0.03864577 | 0.03850427 | 0.00014149 | 0.49348298 | 0.90650187 |
| Slc6a20b | 0.01581408 | 0.01934576 | 0.01920576 | 0.00014001 | 0.4870736 | 0.90650187 |
| Psd2 | 0.00279828 | 0.00356283 | 0.00342848 | 0.00013435 | 0.43085051 | 0.90650187 |
| Slc22a17 | 0.35817134 | 0.35768578 | 0.35755145 | 0.00013433 | 0.4993337 | 0.90650187 |
| Pcdh12 | 0.02505982 | 0.03037565 | 0.03024621 | 0.00012944 | 0.49241028 | 0.90650187 |
| Zkscan2 | 0.0306408 | 0.03697316 | 0.03684467 | 0.0001285 | 0.49381495 | 0.90650187 |
| Agap3 | 0.12551779 | 0.14226038 | 0.14213227 | 0.00012811 | 0.49840149 | 0.90650187 |
| Gm26759 | 0.01295274 | 0.01588741 | 0.0157611 | 0.00012631 | 0.48579016 | 0.90650187 |
| Zfp52 | 0.02399491 | 0.02910664 | 0.02898156 | 0.00012509 | 0.49234574 | 0.90650187 |
| Zfyve28 | 0.00530519 | 0.00661137 | 0.00648891 | 0.00012246 | 0.46656976 | 0.90650187 |
| 1700026L06F | 0.09507782 | 0.1097385 | 0.10962041 | 0.00011809 | 0.49808939 | 0.90650187 |
| Astn1 | 0.00534596 | 0.00665618 | 0.0065386 | 0.00011758 | 0.46814163 | 0.90650187 |
| Phyhd1 | 0.01613612 | 0.01970969 | 0.01959263 | 0.00011706 | 0.48940504 | 0.90650187 |
| Dhrs9 | 0.00798492 | 0.00985796 | 0.00974884 | 0.00010912 | 0.48015663 | 0.90650187 |
| Il1r2 | 0.01354088 | 0.01657753 | 0.01647022 | 0.0001073 | 0.488447 | 0.90650187 |
| Tfdp1 | 0.01721819 | 0.02099793 | 0.02089127 | 0.00010666 | 0.49094575 | 0.90650187 |
| Tm6sf2 | 0.00789394 | 0.00974284 | 0.00963836 | 0.00010448 | 0.48078147 | 0.90650187 |
| Prrg4 | 0.007108 | 0.00878776 | 0.00868336 | 0.0001044 | 0.47868637 | 0.90650187 |
| Dusp8 | 0.07212047 | 0.08449368 | 0.08438971 | 0.00010397 | 0.49781491 | 0.90650187 |
| Pou3f1 | 0.01293808 | 0.015843 | 0.01574342 | 9.96E-05 | 0.48878338 | 0.90650187 |
| Plppr2 | 0.156087 | 0.17377717 | 0.17367812 | 9.90E-05 | 0.49898856 | 0.90650187 |
| Plekhh2 | 0.22374693 | 0.23985338 | 0.23975684 | 9.65E-05 | 0.49928584 | 0.90650187 |
| 2610021A01 | 0.04259668 | 0.05091181 | 0.05081633 | 9.55E-05 | 0.49666782 | 0.90650187 |
| 5730596B20 | 0.01043961 | 0.01281613 | 0.01272465 | 9.15E-05 | 0.4872531 | 0.90650187 |

| | | | | | | |
|---|---|---|---|---|---|---|
| Cd37 | 0.0084253 | 0.01037479 | 0.01028344 | 9.13E-05 | 0.4842499 | 0.90650187 |
| Megf8 | 0.08158238 | 0.09496945 | 0.09487905 | 9.04E-05 | 0.49831011 | 0.90650187 |
| 2310040G24 | 0.1007758 | 0.11586006 | 0.11577241 | 8.76E-05 | 0.49865737 | 0.90650187 |
| Gna15 | 0.00657294 | 0.00811987 | 0.00803262 | 8.72E-05 | 0.48074375 | 0.90650187 |
| Myo1h | 0.018385 | 0.02237411 | 0.0222895 | 8.46E-05 | 0.49326848 | 0.90650187 |
| Sptbn4 | 0.01002439 | 0.01230656 | 0.01222197 | 8.46E-05 | 0.48772672 | 0.90650187 |
| Zbtb48 | 0.09408404 | 0.10862633 | 0.10854356 | 8.28E-05 | 0.49864756 | 0.90650187 |
| Mical1 | 0.09629898 | 0.11102323 | 0.11094177 | 8.15E-05 | 0.49869783 | 0.90650187 |
| Kifc1 | 0.08699789 | 0.10090634 | 0.10082523 | 8.11E-05 | 0.49857328 | 0.90650187 |
| Ptgir | 0.00543065 | 0.00671524 | 0.0066418 | 7.34E-05 | 0.48039627 | 0.90650187 |
| Inpp5j | 0.02655108 | 0.03208706 | 0.03201417 | 7.29E-05 | 0.49596179 | 0.90650187 |
| Timm23 | 0.09380471 | 0.10831013 | 0.10824063 | 6.95E-05 | 0.4988612 | 0.90650187 |
| Olfr78 | 0.0052231 | 0.0064532 | 0.00638886 | 6.43E-05 | 0.48214712 | 0.90650187 |
| Gm37229 | 0.0102365 | 0.01253669 | 0.0124788 | 5.79E-05 | 0.49177299 | 0.90650187 |
| Ctr9 | 0.60184239 | 0.53464166 | 0.53458481 | 5.68E-05 | 0.49981141 | 0.90650187 |
| Igtp | 0.02744658 | 0.03313025 | 0.03307412 | 5.61E-05 | 0.49699034 | 0.90650187 |
| Nckap5 | 0.01971591 | 0.02393148 | 0.0238817 | 4.98E-05 | 0.49630264 | 0.90650187 |
| Cntrl | 0.16853096 | 0.18625677 | 0.18620888 | 4.79E-05 | 0.49954381 | 0.90650187 |
| A530016L24I | 0.01304626 | 0.01592133 | 0.0158739 | 4.74E-05 | 0.49470113 | 0.90650187 |
| A930041C12 | 0.00537565 | 0.00662024 | 0.00657478 | 4.55E-05 | 0.48773932 | 0.90650187 |
| Maml1 | 0.12063567 | 0.13703553 | 0.13699055 | 4.50E-05 | 0.49941763 | 0.90650187 |
| Seh1l | 0.47761309 | 0.44952695 | 0.44948629 | 4.07E-05 | 0.49983956 | 0.90650187 |
| 9330182L06I | 0.08825632 | 0.10224004 | 0.10220106 | 3.90E-05 | 0.4993235 | 0.90650187 |
| Cebpa | 0.02297963 | 0.02781281 | 0.02777415 | 3.87E-05 | 0.49753164 | 0.90650187 |
| Serpina1e | 0.0154163 | 0.01875973 | 0.01872768 | 3.20E-05 | 0.49696519 | 0.90650187 |
| Nkain3 | 0.01001237 | 0.01223437 | 0.01220742 | 2.69E-05 | 0.49608494 | 0.90650187 |
| Git1 | 0.14847277 | 0.16594616 | 0.16592366 | 2.25E-05 | 0.49975955 | 0.90650187 |
| Anxa8 | 0.00271009 | 0.00334178 | 0.00332061 | 2.12E-05 | 0.48869925 | 0.90650187 |
| Ppp1r36 | 0.00842421 | 0.01030153 | 0.01028212 | 1.94E-05 | 0.49665152 | 0.90650187 |
| Adap1 | 0.00684466 | 0.0083815 | 0.00836314 | 1.84E-05 | 0.49610776 | 0.90650187 |

| | | | | | | |
|---|---|---|---|---|---|---|
| B3gnt7 | 0.01739054 | 0.02111508 | 0.02109794 | 1.71E-05 | 0.49855918 | 0.90650187 |
| Tnfaip8l3 | 0.00869147 | 0.01062195 | 0.0106064 | 1.56E-05 | 0.49739902 | 0.90650187 |
| Arid4a | 0.4256268 | 0.41081194 | 0.41079825 | 1.37E-05 | 0.49994089 | 0.90650187 |
| Zfp618 | 0.25222254 | 0.26614264 | 0.26612981 | 1.28E-05 | 0.49991448 | 0.90650187 |
| Apoa2 | 0.03680891 | 0.04408917 | 0.04408042 | 8.75E-06 | 0.4996481 | 0.90650187 |
| Iqsec2 | 0.04517543 | 0.05380622 | 0.05380095 | 5.27E-06 | 0.49982633 | 0.90650187 |
| Cldn2 | 0.02264824 | 0.02738094 | 0.0273797 | 1.24E-06 | 0.49991974 | 0.90650187 |
| Hmcn2 | 0 | 0 | 0 | 0 | NA | NA |
| Gm26725 | 0 | 0 | 0 | 0 | NA | NA |
| Npm1 | 3.44064803 | 0.77974619 | 0.77974619 | 0 | 0.5 | 0.90650187 |
| Mcmdc2 | 0.00791535 | 0.00966354 | 0.00966435 | -8.11E-07 | 0.50014885 | 0.90650187 |
| Wbscr25 | 0.00982466 | 0.01197888 | 0.01198008 | -1.20E-06 | 0.50017792 | 0.90650187 |
| Pdp2 | 0.03149836 | 0.0378524 | 0.03785422 | -1.82E-06 | 0.50008537 | 0.90650187 |
| Gm14295 | 0.07925208 | 0.09230385 | 0.0923076 | -3.75E-06 | 0.50007206 | 0.90650187 |
| Kcnj12 | 0.0051802 | 0.00633138 | 0.00633657 | -5.18E-06 | 0.50145069 | 0.90650187 |
| Car12 | 0.01479928 | 0.01797991 | 0.0179856 | -5.69E-06 | 0.50056093 | 0.90650187 |
| Asic2 | 0.00512257 | 0.00626031 | 0.00626632 | -6.00E-06 | 0.50169924 | 0.90650187 |
| Cfap36 | 0.71031413 | 0.60146214 | 0.60146885 | -6.71E-06 | 0.50001978 | 0.90650187 |
| Siah2 | 0.07759273 | 0.09046047 | 0.09047182 | -1.13E-05 | 0.50022232 | 0.90650187 |
| Amy2a5 | 0.00994194 | 0.01211056 | 0.01212213 | -1.16E-05 | 0.50169311 | 0.90650187 |
| Dnajc27 | 0.0386724 | 0.04624248 | 0.04625482 | -1.23E-05 | 0.5004734 | 0.90650187 |
| Gpc2 | 0.00746626 | 0.00910557 | 0.00911881 | -1.32E-05 | 0.50257609 | 0.90650187 |
| Herpud2 | 0.31914559 | 0.32503137 | 0.32504601 | -1.46E-05 | 0.50007992 | 0.90650187 |
| Eea1 | 0.54260127 | 0.49520765 | 0.49522639 | -1.87E-05 | 0.5000671 | 0.90650187 |
| Aatk | 0.06487326 | 0.07624684 | 0.07626807 | -2.12E-05 | 0.50049374 | 0.90650187 |
| Hey2 | 0.0051715 | 0.00630313 | 0.00632597 | -2.28E-05 | 0.50640052 | 0.90650187 |
| Gm20939 | 0.01401737 | 0.01701924 | 0.01704432 | -2.51E-05 | 0.50261009 | 0.90650187 |
| Cplx1 | 0.01050317 | 0.01277046 | 0.01280158 | -3.11E-05 | 0.50431058 | 0.90650187 |
| Dact3 | 0.03745171 | 0.04479896 | 0.04483108 | -3.21E-05 | 0.50127074 | 0.90650187 |
| Bcl2l12 | 0.22410734 | 0.24006301 | 0.24009572 | -3.27E-05 | 0.50024156 | 0.90650187 |

| | | | | | | |
|---|---|---|---|---|---|---|
| Gm28557 | 0.01759162 | 0.02130381 | 0.021339 | -3.52E-05 | 0.50292474 | 0.90650187 |
| Siah1b | 0.12061306 | 0.13693001 | 0.13696668 | -3.67E-05 | 0.50047476 | 0.90650187 |
| Twist1 | 0.00951186 | 0.01156302 | 0.01160111 | -3.81E-05 | 0.50582217 | 0.90650187 |
| Ints5 | 0.17182141 | 0.18945342 | 0.1894932 | -3.98E-05 | 0.50037228 | 0.90650187 |
| Ubtd1 | 0.14311951 | 0.16039036 | 0.16043081 | -4.05E-05 | 0.50044718 | 0.90650187 |
| Ift88 | 0.2775209 | 0.28885102 | 0.28889259 | -4.16E-05 | 0.50025522 | 0.90650187 |
| Jdp2 | 0.0188106 | 0.02275459 | 0.02279897 | -4.44E-05 | 0.50345208 | 0.90650187 |
| Apba2 | 0.01510897 | 0.01830857 | 0.01835814 | -4.96E-05 | 0.50478899 | 0.90650187 |
| Tceal6 | 0.00783755 | 0.00951679 | 0.00956988 | -5.31E-05 | 0.50983736 | 0.90650187 |
| Opcml | 0.03375985 | 0.04045791 | 0.04051102 | -5.31E-05 | 0.50232527 | 0.90650187 |
| BC065403 | 0.01504773 | 0.01822525 | 0.01828448 | -5.92E-05 | 0.50574518 | 0.90650187 |
| Lamp3 | 0.00713526 | 0.00865724 | 0.00871649 | -5.93E-05 | 0.51205415 | 0.90650187 |
| Zfp335os | 0.03242804 | 0.03888533 | 0.03894737 | -6.20E-05 | 0.5028251 | 0.90650187 |
| Sema4a | 0.07656499 | 0.08926246 | 0.08933283 | -7.04E-05 | 0.50139695 | 0.90650187 |
| Gm26870 | 0.00645507 | 0.00781036 | 0.00788921 | -7.88E-05 | 0.51771994 | 0.90650187 |
| Pcdhb8 | 0.01797196 | 0.02171557 | 0.02179478 | -7.92E-05 | 0.50644557 | 0.90650187 |
| Mslnl | 0.0052107 | 0.00628953 | 0.00637374 | -8.42E-05 | 0.52341975 | 0.90650187 |
| Idh3a | 0.56184706 | 0.50816244 | 0.50824708 | -8.46E-05 | 0.50029535 | 0.90650187 |
| Pik3cb | 0.04085398 | 0.04870666 | 0.04879358 | -8.69E-05 | 0.50315937 | 0.90650187 |
| Zfp128 | 0.0126058 | 0.0152506 | 0.01534253 | -9.19E-05 | 0.51062608 | 0.90650187 |
| Cftr | 0.00447612 | 0.00538517 | 0.00547793 | -9.28E-05 | 0.53000436 | 0.90650187 |
| Eno1b | 0.07149666 | 0.08359791 | 0.08369363 | -9.57E-05 | 0.50202841 | 0.90650187 |
| Rsrc1 | 0.45162947 | 0.43029538 | 0.43039307 | -9.77E-05 | 0.50040257 | 0.90650187 |
| Gpr22 | 0.00682807 | 0.00824237 | 0.00834297 | -0.0001006 | 0.5213746 | 0.90650187 |
| 6030408B16 | 0.03317702 | 0.03972283 | 0.03982708 | -0.0001042 | 0.50464223 | 0.90650187 |
| Gm2885 | 0.01058229 | 0.01278698 | 0.01289732 | -0.0001103 | 0.51517052 | 0.90650187 |
| Cpxm2 | 0.02816479 | 0.03381284 | 0.03392332 | -0.0001105 | 0.50577602 | 0.90650187 |
| Dlk2 | 0.00259954 | 0.0030747 | 0.0031854 | -0.0001107 | 0.5613891 | 0.90650187 |
| Upk3bl | 0.0159327 | 0.01923746 | 0.01934828 | -0.0001108 | 0.51015668 | 0.90650187 |
| Prkch | 0.06084269 | 0.07160682 | 0.07171791 | -0.0001111 | 0.50274704 | 0.90650187 |

| | | | | | | |
|---|---|---|---|---|---|---|
| Gjc2 | 0.02093561 | 0.02522487 | 0.02533836 | -0.0001135 | 0.50794309 | 0.90650187 |
| Gm3716 | 0.00989762 | 0.01195223 | 0.01206845 | -0.0001162 | 0.51707398 | 0.90650187 |
| 9530046B11 | 0.01058886 | 0.01278285 | 0.01290527 | -0.0001224 | 0.51681836 | 0.90650187 |
| Dyx1c1 | 0.01552605 | 0.01873681 | 0.01885961 | -0.0001228 | 0.51154627 | 0.90650187 |
| Inmt | 0.00794052 | 0.00956988 | 0.00969493 | -0.000125 | 0.52286279 | 0.90650187 |
| Xlr4c | 0.01534236 | 0.01851344 | 0.01863879 | -0.0001253 | 0.51192532 | 0.90650187 |
| Gm9828 | 0.06358806 | 0.07469079 | 0.07481979 | -0.000129 | 0.50305785 | 0.90650187 |
| Chrdl1 | 0.00257561 | 0.00301836 | 0.00315613 | -0.0001378 | 0.57693527 | 0.90650187 |
| Apela | 0.00681663 | 0.0081898 | 0.00832906 | -0.0001393 | 0.52962583 | 0.90650187 |
| Zfp945 | 0.31652699 | 0.32267566 | 0.32281739 | -0.0001417 | 0.50077867 | 0.90650187 |
| Smad3 | 0.07938687 | 0.09231092 | 0.09245655 | -0.0001456 | 0.5027936 | 0.90650187 |
| Tmem8 | 0.02560146 | 0.03074263 | 0.03088876 | -0.0001461 | 0.50838958 | 0.90650187 |
| Grik4 | 0.00630983 | 0.00756564 | 0.00771245 | -0.0001468 | 0.53371992 | 0.90650187 |
| Rnf145 | 0.49519603 | 0.4619933 | 0.4621402 | -0.0001469 | 0.50056376 | 0.90650187 |
| Anxa6 | 1.45906209 | 0.88594546 | 0.88609288 | -0.0001474 | 0.50029508 | 0.90650187 |
| Wnt2b | 0.00975796 | 0.01174875 | 0.01189929 | -0.0001505 | 0.52242432 | 0.90650187 |
| Mapkapk2 | 0.15200474 | 0.16937498 | 0.16952904 | -0.0001541 | 0.50161174 | 0.90650187 |
| Col4a5 | 0.06501216 | 0.07626589 | 0.07642446 | -0.0001586 | 0.50367982 | 0.90650187 |
| Mettl7a1 | 0.12048361 | 0.13667034 | 0.13682994 | -0.0001596 | 0.50206869 | 0.90650187 |
| Tagap | 0.00780788 | 0.00936835 | 0.00953384 | -0.0001655 | 0.53075548 | 0.90650187 |
| A230005M16 | 0.00675744 | 0.00809056 | 0.00825706 | -0.0001665 | 0.53571501 | 0.90650187 |
| Gm15608 | 0.00519148 | 0.00618352 | 0.00635031 | -0.0001668 | 0.54647699 | 0.90650187 |
| Gm20721 | 0.18173951 | 0.19915187 | 0.19931871 | -0.0001668 | 0.5014845 | 0.90650187 |
| A830080D01 | 0.10580654 | 0.12102556 | 0.12119359 | -0.000168 | 0.50245906 | 0.90650187 |
| Palb2 | 0.03086761 | 0.03694258 | 0.03711179 | -0.0001692 | 0.50808556 | 0.90650187 |
| Sbk2 | 0.00254716 | 0.00295204 | 0.00312133 | -0.0001693 | 0.5952629 | 0.90650187 |
| Hexim2 | 0.06020606 | 0.07082701 | 0.07099702 | -0.00017 | 0.50424698 | 0.90650187 |
| Iqcb1 | 0.29106716 | 0.30065941 | 0.30083201 | -0.0001726 | 0.50101756 | 0.90650187 |
| Ccdc15 | 0.04741198 | 0.0562078 | 0.05638129 | -0.0001735 | 0.50545734 | 0.90650187 |
| Lypd6b | 0.02160726 | 0.02596376 | 0.02613949 | -0.0001757 | 0.51192128 | 0.90650187 |

| | | | | | | |
|---|---|---|---|---|---|---|
| Mir17hg | 0.03863222 | 0.04602674 | 0.046208 | -0.0001813 | 0.50695685 | 0.90650187 |
| Arntl2 | 0.01495952 | 0.01799375 | 0.01817838 | -0.0001846 | 0.51800703 | 0.90650187 |
| Psen2 | 0.26300165 | 0.27571708 | 0.27590352 | -0.0001864 | 0.50119849 | 0.90650187 |
| Gm26720 | 0.02266793 | 0.02721627 | 0.02740314 | -0.0001869 | 0.51209223 | 0.90650187 |
| Ankrd35 | 0.02265093 | 0.02719348 | 0.0273829 | -0.0001894 | 0.5122666 | 0.90650187 |
| Mapkapk3 | 0.0281475 | 0.03370776 | 0.03390288 | -0.0001951 | 0.51020599 | 0.90650187 |
| Rspo2 | 0.00706099 | 0.00842294 | 0.0086262 | -0.0002033 | 0.54171395 | 0.90650187 |
| Nup37 | 0.16521701 | 0.18268259 | 0.18288886 | -0.0002063 | 0.50200031 | 0.90650187 |
| Skida1 | 0.04731828 | 0.05606463 | 0.05627335 | -0.0002087 | 0.5065779 | 0.90650187 |
| Plcb1 | 0.10896017 | 0.1243653 | 0.12457583 | -0.0002105 | 0.50299711 | 0.90650187 |
| Rspo4 | 0.03746229 | 0.04463146 | 0.04484343 | -0.000212 | 0.50838266 | 0.90650187 |
| Casc1 | 0.02761343 | 0.03305177 | 0.03327148 | -0.0002197 | 0.51171002 | 0.90650187 |
| Egr2 | 0.04219367 | 0.05012324 | 0.05034897 | -0.0002257 | 0.50795083 | 0.90650187 |
| Adprh | 0.82000749 | 0.66284977 | 0.66307625 | -0.0002265 | 0.50060578 | 0.90650187 |
| Bcor | 0.25050113 | 0.26433089 | 0.26455859 | -0.0002277 | 0.50152645 | 0.90650187 |
| Adamtsl2 | 0.02383434 | 0.02855924 | 0.02879071 | -0.0002315 | 0.51425537 | 0.90650187 |
| Arhgef7 | 0.26177522 | 0.27456447 | 0.27479709 | -0.0002326 | 0.5015013 | 0.90650187 |
| Fgfr3 | 0.09930985 | 0.11395656 | 0.11419092 | -0.0002344 | 0.50363977 | 0.90650187 |
| Pnma2 | 0.03119978 | 0.03725975 | 0.03750285 | -0.0002431 | 0.51149471 | 0.90650187 |
| Zfp454 | 0.03956216 | 0.04704661 | 0.04729115 | -0.0002445 | 0.50916995 | 0.90650187 |
| Mppe1 | 0.01385217 | 0.01659432 | 0.01684533 | -0.000251 | 0.52640735 | 0.90650187 |
| 2010320M18 | 0.33977669 | 0.34214094 | 0.34239227 | -0.0002513 | 0.50130185 | 0.90650187 |
| Tep1 | 0.0834628 | 0.096696 | 0.09694843 | -0.0002524 | 0.50461775 | 0.90650187 |
| Paqr6 | 0.03783892 | 0.04502911 | 0.04528295 | -0.0002538 | 0.50994086 | 0.90650187 |
| Ppm1j | 0.00246394 | 0.00276231 | 0.00301953 | -0.0002572 | 0.64754157 | 0.90650187 |
| Zmat4 | 0.0074393 | 0.00882547 | 0.00908605 | -0.0002606 | 0.5507251 | 0.90650187 |
| Gm26810 | 0.04764379 | 0.05638753 | 0.0566483 | -0.0002608 | 0.50816368 | 0.90650187 |
| Rapsn | 0.01702507 | 0.02039391 | 0.02065963 | -0.0002657 | 0.5227987 | 0.90650187 |
| Sfxn4 | 0.10021196 | 0.11489583 | 0.11516282 | -0.000267 | 0.50411169 | 0.90650187 |
| Sorbs1 | 0.03280669 | 0.03912374 | 0.03939223 | -0.0002685 | 0.51208624 | 0.90650187 |

| | | | | | | |
|---|---|---|---|---|---|---|
| Gbp4 | 0.01149058 | 0.01372684 | 0.01399573 | -0.0002689 | 0.53403232 | 0.90650187 |
| Gm5148 | 0.01042616 | 0.01243597 | 0.01270837 | -0.0002724 | 0.53795774 | 0.90650187 |
| Tnfrsf21 | 0.1290054 | 0.14551156 | 0.14578728 | -0.0002757 | 0.50335415 | 0.90650187 |
| Arc | 0.08590505 | 0.09935249 | 0.09962863 | -0.0002761 | 0.50491569 | 0.90650187 |
| Dysf | 0.02042666 | 0.02445213 | 0.02473082 | -0.0002787 | 0.51997797 | 0.90650187 |
| Gm42567 | 0.00439845 | 0.00509874 | 0.00538317 | -0.0002844 | 0.59285307 | 0.90650187 |
| Erg | 0.02060632 | 0.02466091 | 0.02494534 | -0.0002844 | 0.52021334 | 0.90650187 |
| Eda | 0.02847357 | 0.03400318 | 0.03428817 | -0.000285 | 0.51473766 | 0.90650187 |
| Atcay | 0.00666113 | 0.00785263 | 0.00813991 | -0.0002873 | 0.56233699 | 0.90650187 |
| Aloxe3 | 0.00881975 | 0.01047107 | 0.01076201 | -0.0002909 | 0.54782969 | 0.90650187 |
| Gabrp | 0.00397833 | 0.00457917 | 0.00487038 | -0.0002912 | 0.60480895 | 0.90650187 |
| Ddias | 0.01772571 | 0.02120686 | 0.02149971 | -0.0002929 | 0.52414295 | 0.90650187 |
| Col9a1 | 0.0423189 | 0.05019989 | 0.05049422 | -0.0002943 | 0.51033662 | 0.90650187 |
| Efcab11 | 0.04248838 | 0.0503929 | 0.05069076 | -0.0002979 | 0.51041999 | 0.90650187 |
| Gm21887 | 0.00422686 | 0.00485947 | 0.00517376 | -0.0003143 | 0.60644308 | 0.90650187 |
| Rhobtb1 | 0.11235523 | 0.12788876 | 0.12820315 | -0.0003144 | 0.50434912 | 0.90650187 |
| Slfn9 | 0.03222534 | 0.03838678 | 0.03870915 | -0.0003224 | 0.51476668 | 0.90650187 |
| Gria1 | 0.00902089 | 0.01068346 | 0.01100595 | -0.0003225 | 0.55181966 | 0.90650187 |
| Gm33466 | 0.00700316 | 0.00823282 | 0.00855589 | -0.0003231 | 0.56665414 | 0.90650187 |
| Fabp3 | 0.23003335 | 0.24532569 | 0.24564883 | -0.0003231 | 0.50233302 | 0.90650187 |
| Fndc3c1 | 0.00401252 | 0.00458489 | 0.00491211 | -0.0003272 | 0.61644182 | 0.90650187 |
| Tor3a | 0.05940279 | 0.06975864 | 0.07008657 | -0.0003279 | 0.50829787 | 0.90650187 |
| Trappc12 | 0.20744452 | 0.22395559 | 0.22428513 | -0.0003295 | 0.50260584 | 0.90650187 |
| Hecw1 | 0.00757979 | 0.00892637 | 0.00925675 | -0.0003304 | 0.56303514 | 0.90650187 |
| AI846148 | 0.18968392 | 0.20677876 | 0.20711103 | -0.0003323 | 0.50284533 | 0.90650187 |
| Ssr1 | 1.44234823 | 0.88258318 | 0.88292002 | -0.0003368 | 0.50067663 | 0.90650187 |
| Angptl7 | 0.00937657 | 0.01109981 | 0.01143714 | -0.0003373 | 0.5521609 | 0.90650187 |
| 1700020I14R | 0.46827887 | 0.44234279 | 0.442683 | -0.0003402 | 0.501363 | 0.90650187 |
| Sirt7 | 0.33664228 | 0.33943987 | 0.33978035 | -0.0003405 | 0.5017772 | 0.90650187 |
| Pik3c2b | 0.02981406 | 0.03552859 | 0.0358703 | -0.0003417 | 0.51689018 | 0.90650187 |

| | | | | | | |
|---|---|---|---|---|---|---|
| Prrt2 | 0.00772966 | 0.00909514 | 0.00943882 | -0.0003437 | 0.5642965 | 0.90650187 |
| Popdc2 | 0.01221858 | 0.01453121 | 0.01487513 | -0.0003439 | 0.54093371 | 0.90650187 |
| Ifi44 | 0.00467079 | 0.00537149 | 0.00571542 | -0.0003439 | 0.6054645 | 0.90650187 |
| Liph | 0.00508937 | 0.00588036 | 0.00622585 | -0.0003455 | 0.59743043 | 0.90650187 |
| Gm43543 | 0.00687421 | 0.00805121 | 0.00839908 | -0.0003479 | 0.57304363 | 0.90650187 |
| Gm42670 | 0.01116076 | 0.01324787 | 0.01359703 | -0.0003492 | 0.54544347 | 0.90650187 |
| Zfp933 | 0.0275144 | 0.03279996 | 0.03315435 | -0.0003544 | 0.5189502 | 0.90650187 |
| Tmem110 | 0.15497601 | 0.17219445 | 0.17255084 | -0.0003564 | 0.50366298 | 0.90650187 |
| Enpp3 | 0.03373829 | 0.04012793 | 0.04048572 | -0.0003578 | 0.51566968 | 0.90650187 |
| Phka1 | 0.03779059 | 0.04486773 | 0.04522657 | -0.0003588 | 0.51406867 | 0.90650187 |
| Spata13 | 0.12487626 | 0.14109816 | 0.14145835 | -0.0003602 | 0.50451583 | 0.90650187 |
| D130017N08 | 0.05020961 | 0.05923235 | 0.05959827 | -0.0003659 | 0.51088777 | 0.90650187 |
| Cxcr2 | 0.02259853 | 0.02694945 | 0.02732051 | -0.0003711 | 0.5240733 | 0.90650187 |
| Adra2b | 0.00934606 | 0.01102863 | 0.01140016 | -0.0003715 | 0.55759814 | 0.90650187 |
| Gm38186 | 0.00770954 | 0.00904159 | 0.00941439 | -0.0003728 | 0.56986808 | 0.90650187 |
| Xylt2 | 0.17117935 | 0.1884779 | 0.18885329 | -0.0003754 | 0.50352522 | 0.90650187 |
| Lca5l | 0.02268275 | 0.02704466 | 0.02742079 | -0.0003761 | 0.52431243 | 0.90650187 |
| Spns2 | 0.0137912 | 0.01639377 | 0.01677187 | -0.0003781 | 0.53991472 | 0.90650187 |
| RP23-53O7.2 | 0.0192468 | 0.02293947 | 0.02332082 | -0.0003813 | 0.52897568 | 0.90650187 |
| Gm30074 | 0.03642835 | 0.04324937 | 0.04363571 | -0.0003863 | 0.51569851 | 0.90650187 |
| Rgs22 | 0.0070167 | 0.00818509 | 0.00857235 | -0.0003873 | 0.57958675 | 0.90650187 |
| Rilp | 0.01433911 | 0.01704229 | 0.01743176 | -0.0003895 | 0.53956027 | 0.90650187 |
| Lancl3 | 0.01722352 | 0.02050797 | 0.02089766 | -0.0003897 | 0.53303463 | 0.90650187 |
| Per3 | 0.08800255 | 0.10153177 | 0.10192379 | -0.000392 | 0.506821 | 0.90650187 |
| Gm26559 | 0.01761042 | 0.02096527 | 0.02136153 | -0.0003963 | 0.53286279 | 0.90650187 |
| Fryl | 0.37192155 | 0.36830224 | 0.36870035 | -0.0003981 | 0.501915 | 0.90650187 |
| Slc25a43 | 0.00826513 | 0.00968969 | 0.01008904 | -0.0003994 | 0.5698407 | 0.90650187 |
| Gm5426 | 0.00892922 | 0.01049394 | 0.01089478 | -0.0004008 | 0.56496259 | 0.90650187 |
| Adam9 | 0.14674788 | 0.16375515 | 0.16415743 | -0.0004023 | 0.50434613 | 0.90650187 |
| Gm15675 | 0.00230848 | 0.00242473 | 0.00282931 | -0.0004046 | 0.73751397 | 0.90650187 |

| | | | | | | |
|---|---|---|---|---|---|---|
| Ccdc28a | 0.06305634 | 0.07381279 | 0.07421989 | -0.0004071 | 0.50972698 | 0.90650187 |
| Edar | 0.00700614 | 0.00814919 | 0.00855952 | -0.0004103 | 0.58438136 | 0.90650187 |
| Fam57a | 0.10609411 | 0.12109027 | 0.12150253 | -0.0004123 | 0.50601738 | 0.90650187 |
| Itga7 | 0.03906319 | 0.04628998 | 0.04671013 | -0.0004202 | 0.51594869 | 0.90650187 |
| Sh3kbp1 | 0.18628648 | 0.20336659 | 0.20378701 | -0.0004204 | 0.5036588 | 0.90650187 |
| Bean1 | 0.00638668 | 0.00738336 | 0.00780599 | -0.0004226 | 0.59510272 | 0.90650187 |
| Rsl24d1 | 0.52831118 | 0.48498298 | 0.48540595 | -0.000423 | 0.5015454 | 0.90650187 |
| Tifa | 0.01891828 | 0.02250077 | 0.02292782 | -0.000427 | 0.53299562 | 0.90650187 |
| Cep85l | 0.0566841 | 0.06656997 | 0.06699803 | -0.0004281 | 0.51133001 | 0.90650187 |
| Bcas3os2 | 0.0073794 | 0.00858346 | 0.00901326 | -0.0004298 | 0.58394149 | 0.90650187 |
| Gm5868 | 0.00688748 | 0.00798148 | 0.00841522 | -0.0004337 | 0.59061937 | 0.90650187 |
| Igsf1 | 0.06297009 | 0.07368596 | 0.07412255 | -0.0004366 | 0.51044512 | 0.90650187 |
| Slc27a6 | 0.01531409 | 0.01816669 | 0.0186048 | -0.0004381 | 0.54168766 | 0.90650187 |
| Sdccag3 | 0.403335 | 0.39315553 | 0.39359665 | -0.0004411 | 0.50198769 | 0.90650187 |
| Fv1 | 0.00533414 | 0.00608181 | 0.0065242 | -0.0004424 | 0.61846111 | 0.90650187 |
| Mfsd7a | 0.0126624 | 0.01496691 | 0.01541082 | -0.0004439 | 0.55094776 | 0.90650187 |
| C2cd4a | 0.01545241 | 0.01832655 | 0.01877109 | -0.0004445 | 0.54192389 | 0.90650187 |
| 1700058P15I | 0.01141419 | 0.01345409 | 0.0139034 | -0.0004493 | 0.55711872 | 0.90650187 |
| Gm5424 | 0.0356257 | 0.04224728 | 0.04269701 | -0.0004497 | 0.51867378 | 0.90650187 |
| Rab17 | 0.01206126 | 0.01423377 | 0.01468516 | -0.0004514 | 0.55434622 | 0.90650187 |
| Clvs2 | 0.00487469 | 0.00551243 | 0.0059641 | -0.0004517 | 0.63181678 | 0.90650187 |
| Zfp185 | 0.03260401 | 0.03870235 | 0.03915414 | -0.0004518 | 0.52045535 | 0.90650187 |
| Dnaic2 | 0.00654588 | 0.00754702 | 0.0079997 | -0.0004527 | 0.59930988 | 0.90650187 |
| Wdr25 | 0.02708592 | 0.0321938 | 0.03264738 | -0.0004536 | 0.52462424 | 0.90650187 |
| Zfp941 | 0.01140849 | 0.01344273 | 0.01389651 | -0.0004538 | 0.55771109 | 0.90650187 |
| Fam217b | 0.0327411 | 0.03885899 | 0.03931518 | -0.0004562 | 0.52057005 | 0.90650187 |
| Maml2 | 0.07287011 | 0.08476903 | 0.08522544 | -0.0004564 | 0.50949714 | 0.90650187 |
| Ccdc171 | 0.02957508 | 0.03513089 | 0.03558844 | -0.0004576 | 0.52278986 | 0.90650187 |
| Cmtm1 | 0.00224161 | 0.00228629 | 0.00274748 | -0.0004612 | 0.77223477 | 0.90650187 |
| Cds1 | 0.00224027 | 0.00228356 | 0.00274584 | -0.0004623 | 0.77290344 | 0.90650187 |

| | | | | | | |
|---|---|---|---|---|---|---|
| Artn | 0.01278672 | 0.01509443 | 0.01556083 | -0.0004664 | 0.55300087 | 0.90650187 |
| Banf2os | 0.00441817 | 0.00493709 | 0.00540722 | -0.0004701 | 0.6504472 | 0.90650187 |
| Gm11770 | 0.00223037 | 0.00226342 | 0.00273372 | -0.0004703 | 0.77780726 | 0.90650187 |
| 4933405D12I | 0.01668883 | 0.01978518 | 0.0202562 | -0.000471 | 0.54116642 | 0.90650187 |
| Gbp3 | 0.00884523 | 0.0103199 | 0.01079292 | -0.000473 | 0.57723874 | 0.90650187 |
| Gulo | 0.01225377 | 0.014444 | 0.01491762 | -0.0004736 | 0.55612202 | 0.90650187 |
| Gm26648 | 0.01270335 | 0.01498176 | 0.01546024 | -0.0004785 | 0.55471615 | 0.90650187 |
| Dhh | 0.01155779 | 0.01359364 | 0.01407695 | -0.0004833 | 0.5606557 | 0.90650187 |
| 9530059O14 | 0.00749044 | 0.00865934 | 0.0091482 | -0.0004889 | 0.5938902 | 0.90650187 |
| Camk2a | 0.00756414 | 0.00874886 | 0.00923774 | -0.0004889 | 0.59300193 | 0.90650187 |
| Fgf16 | 0.00487205 | 0.00547007 | 0.00596088 | -0.0004908 | 0.64283187 | 0.90650187 |
| Cntnap1 | 0.01028065 | 0.01204134 | 0.01253224 | -0.0004909 | 0.56912182 | 0.90650187 |
| Extl3 | 0.12204305 | 0.13798432 | 0.13847577 | -0.0004914 | 0.50629401 | 0.90650187 |
| Dtx3 | 0.69521444 | 0.59203989 | 0.59253311 | -0.0004932 | 0.5014763 | 0.90650187 |
| Aldh1l1 | 0.01561498 | 0.01847098 | 0.0189665 | -0.0004955 | 0.54623181 | 0.90650187 |
| Best3 | 0.01615617 | 0.01911803 | 0.01961671 | -0.0004987 | 0.54498985 | 0.90650187 |
| Mettl24 | 0.01086653 | 0.01273865 | 0.0132412 | -0.0005026 | 0.56699409 | 0.90650187 |
| Pdyn | 0.00218936 | 0.00218095 | 0.00268353 | -0.0005026 | 0.79746044 | 0.9075779 |
| Ccdc178 | 0.0048467 | 0.00542378 | 0.00592996 | -0.0005062 | 0.64783151 | 0.90650187 |
| Fam20a | 0.09860723 | 0.11292691 | 0.11343381 | -0.0005069 | 0.5079248 | 0.90650187 |
| Traf1 | 0.00740805 | 0.0085398 | 0.00904807 | -0.0005083 | 0.59860257 | 0.90650187 |
| Hist1h2an | 0.01816436 | 0.02151586 | 0.02202527 | -0.0005094 | 0.54094698 | 0.90650187 |
| Slc28a2 | 0.00217854 | 0.00215944 | 0.00267028 | -0.0005108 | 0.80246837 | 0.90870058 |
| Gm5449 | 0.00476707 | 0.00531975 | 0.00583285 | -0.0005131 | 0.65212439 | 0.90650187 |
| Wif1 | 0.04822507 | 0.05680022 | 0.05731749 | -0.0005173 | 0.51600103 | 0.90650187 |
| Zfp808 | 0.00940653 | 0.01095569 | 0.01147346 | -0.0005178 | 0.57950202 | 0.90650187 |
| Gm43948 | 0.01186215 | 0.01392685 | 0.01444467 | -0.0005178 | 0.56331165 | 0.90650187 |
| RP24-259P13 | 0.00935206 | 0.01088848 | 0.01140744 | -0.000519 | 0.58013775 | 0.90650187 |
| Gm26514 | 0.00828754 | 0.0095963 | 0.01011624 | -0.0005199 | 0.59036591 | 0.90650187 |
| Ift140 | 0.04515176 | 0.05324898 | 0.0537736 | -0.0005246 | 0.51729728 | 0.90650187 |

| | | | | | | |
|---|---|---|---|---|---|---|
| Ppp1r42 | 0.01079656 | 0.01263151 | 0.01315656 | -0.0005251 | 0.57040899 | 0.90650187 |
| Dnaaf3 | 0.0021587 | 0.00212029 | 0.002646 | -0.0005257 | 0.81145299 | 0.9092537 |
| Chst3 | 0.00729007 | 0.00837884 | 0.00890469 | -0.0005259 | 0.60354255 | 0.90650187 |
| Frmd8os | 0.00759103 | 0.00874405 | 0.00927042 | -0.0005264 | 0.59964081 | 0.90650187 |
| Ttc39b | 0.09529437 | 0.10932652 | 0.10985488 | -0.0005284 | 0.50852947 | 0.90650187 |
| Gm17455 | 0.00214743 | 0.00209822 | 0.00263221 | -0.000534 | 0.81643934 | 0.9092537 |
| Xkr8 | 0.05763997 | 0.0675503 | 0.0680852 | -0.0005349 | 0.51393071 | 0.90650187 |
| Tiparp | 0.22599109 | 0.241329 | 0.24186479 | -0.0005358 | 0.50392878 | 0.90650187 |
| Nek2 | 0.02520854 | 0.02988343 | 0.03042268 | -0.0005392 | 0.53140387 | 0.90650187 |
| Rims1 | 0.00478219 | 0.00531181 | 0.00585129 | -0.0005395 | 0.65905186 | 0.90650187 |
| Epn2 | 0.26428374 | 0.27651516 | 0.27705864 | -0.0005435 | 0.50347895 | 0.90650187 |
| Nat1 | 0.00212393 | 0.00205254 | 0.00260345 | -0.0005509 | 0.82657452 | 0.90947782 |
| Kif14 | 0.02785684 | 0.03300642 | 0.03355931 | -0.0005529 | 0.52919299 | 0.90650187 |
| Gm11940 | 0.01884262 | 0.0222842 | 0.02283728 | -0.0005531 | 0.54286941 | 0.90650187 |
| Slc17a9 | 0.01424366 | 0.01676298 | 0.01731684 | -0.0005539 | 0.55653433 | 0.90650187 |
| Gm20692 | 0.00485651 | 0.00538636 | 0.00594192 | -0.0005556 | 0.66117015 | 0.90650187 |
| Jph1 | 0.00938869 | 0.01089589 | 0.01145183 | -0.0005559 | 0.58543514 | 0.90650187 |
| Gm43598 | 0.00621375 | 0.00703365 | 0.00759551 | -0.0005619 | 0.62886661 | 0.90650187 |
| Tbx15 | 0.00697762 | 0.00795972 | 0.00852483 | -0.0005651 | 0.61588786 | 0.90650187 |
| Pde4dip | 0.18810652 | 0.20500353 | 0.20556926 | -0.0005657 | 0.50488074 | 0.90650187 |
| Pitpnc1 | 0.36286323 | 0.36080616 | 0.36137302 | -0.0005669 | 0.50278205 | 0.90650187 |
| Pradc1 | 0.21177198 | 0.22785158 | 0.22841886 | -0.0005673 | 0.50440453 | 0.90650187 |
| 2900089D17 | 0.01450107 | 0.01705728 | 0.01762672 | -0.0005694 | 0.55709902 | 0.90650187 |
| Amhr2 | 0.00484115 | 0.00534931 | 0.00592319 | -0.0005739 | 0.66666324 | 0.90650187 |
| Tulp2 | 0.00577017 | 0.00648006 | 0.00705541 | -0.0005754 | 0.6415215 | 0.90650187 |
| Micalcl | 0.06309818 | 0.07369061 | 0.07426711 | -0.0005765 | 0.51376448 | 0.90650187 |
| Gm9938 | 0.0020868 | 0.0019814 | 0.002558 | -0.0005766 | 0.84184691 | 0.91251845 |
| 2810429I04R | 0.00433584 | 0.00472883 | 0.00530676 | -0.0005779 | 0.68586028 | 0.90650187 |
| Caskin1 | 0.00207712 | 0.00196307 | 0.00254615 | -0.0005831 | 0.84567664 | 0.9136218 |
| Svopl | 0.02145233 | 0.025371 | 0.02595475 | -0.0005838 | 0.53982266 | 0.90650187 |

| | | | | | | |
|---|---|---|---|---|---|---|
| Slc1a2 | 0.00682958 | 0.00775891 | 0.0083448 | -0.0005859 | 0.62252792 | 0.90650187 |
| Shmt1 | 0.0661391 | 0.07710561 | 0.07769218 | -0.0005866 | 0.51338747 | 0.90650187 |
| Pcbp3 | 0.16969623 | 0.18678371 | 0.18737335 | -0.0005896 | 0.50558096 | 0.90650187 |
| Cenpu | 0.04802633 | 0.05649582 | 0.05708875 | -0.0005929 | 0.5184136 | 0.90650187 |
| Ube2d2a | 1.58019728 | 0.90505639 | 0.9056559 | -0.0005995 | 0.50117402 | 0.90650187 |
| Clca3a1 | 0.03688362 | 0.0435679 | 0.0441677 | -0.0005998 | 0.52407013 | 0.90650187 |
| Mmp24 | 0.01148607 | 0.01339025 | 0.01399027 | -0.0006 | 0.575607 | 0.90650187 |
| Rslcan18 | 0.00481722 | 0.00529308 | 0.00589401 | -0.0006009 | 0.67481801 | 0.90650187 |
| Mmp28 | 0.00613041 | 0.00689001 | 0.00749407 | -0.0006041 | 0.63995418 | 0.90650187 |
| 1700007G11 | 0.00711503 | 0.00808424 | 0.0086919 | -0.0006077 | 0.62202298 | 0.90650187 |
| Rgag4 | 0.13555317 | 0.15199685 | 0.15260999 | -0.0006131 | 0.50712514 | 0.90650187 |
| Prex1 | 0.09690487 | 0.11098307 | 0.11159662 | -0.0006135 | 0.50974982 | 0.90650187 |
| Arf4os | 0.00667099 | 0.00753771 | 0.0081519 | -0.0006142 | 0.63116758 | 0.90650187 |
| Strn | 0.08503752 | 0.09806172 | 0.09867755 | -0.0006158 | 0.51106692 | 0.90650187 |
| Etv6 | 0.08814614 | 0.10146405 | 0.10208068 | -0.0006166 | 0.51071207 | 0.90650187 |
| Hcn1 | 0.00478725 | 0.00523762 | 0.00585746 | -0.0006198 | 0.68097893 | 0.90650187 |
| Popdc3 | 0.00201342 | 0.00184451 | 0.00246818 | -0.0006237 | 0.86935016 | 0.92141903 |
| Med9 | 0.41952244 | 0.40550082 | 0.40612611 | -0.0006253 | 0.50273061 | 0.90650187 |
| Dlg2 | 0.01432579 | 0.0167871 | 0.01741573 | -0.0006286 | 0.56374319 | 0.90650187 |
| Dkkl1 | 0.00473382 | 0.00516342 | 0.0057923 | -0.0006289 | 0.68533369 | 0.90650187 |
| Napsa | 0.00710887 | 0.00805305 | 0.00868441 | -0.0006314 | 0.62672767 | 0.90650187 |
| Hk1os | 0.01503696 | 0.01763986 | 0.01827153 | -0.0006317 | 0.56107221 | 0.90650187 |
| Prkg2 | 0.01383246 | 0.01618928 | 0.01682158 | -0.0006323 | 0.56635535 | 0.90650187 |
| Gm28151 | 0.00433481 | 0.00466977 | 0.0053055 | -0.0006357 | 0.70287853 | 0.90650187 |
| Ninl | 0.05129549 | 0.06020738 | 0.06084374 | -0.0006364 | 0.5185424 | 0.90650187 |
| Nox4 | 0.02157292 | 0.02546034 | 0.02609854 | -0.0006382 | 0.54328442 | 0.90650187 |
| Myo7a | 0.01606987 | 0.01887265 | 0.01951305 | -0.0006404 | 0.55799991 | 0.90650187 |
| Col6a5 | 0.02942636 | 0.03477127 | 0.03541301 | -0.0006417 | 0.53210439 | 0.90650187 |
| Snip1 | 0.22300961 | 0.23842019 | 0.23906316 | -0.000643 | 0.50476986 | 0.90650187 |
| Slc1a6 | 0.01067812 | 0.01236773 | 0.01301327 | -0.0006455 | 0.58727063 | 0.90650187 |

| | | | | | | |
|---|---|---|---|---|---|---|
| Galm | 0.05814842 | 0.06801705 | 0.06866292 | -0.0006459 | 0.51667776 | 0.90650187 |
| Bdp1 | 0.42385874 | 0.40879945 | 0.40944788 | -0.0006484 | 0.50280871 | 0.90650187 |
| 1700041G16 | 0.00750185 | 0.00851278 | 0.00916206 | -0.0006493 | 0.6236346 | 0.90650187 |
| Gm12031 | 0.00196626 | 0.00175912 | 0.00241044 | -0.0006513 | 0.88517221 | 0.92807528 |
| Phactr3 | 0.01328585 | 0.01551115 | 0.0161628 | -0.0006517 | 0.57112445 | 0.90650187 |
| Gm17430 | 0.00196309 | 0.00175344 | 0.00240656 | -0.0006531 | 0.88618576 | 0.92870801 |
| Arhgef33 | 0.00478239 | 0.005198 | 0.00585154 | -0.0006535 | 0.69023498 | 0.90650187 |
| Gpr88 | 0.00195723 | 0.001743 | 0.00239939 | -0.0006564 | 0.88803855 | 0.92986219 |
| Lgr5 | 0.00364959 | 0.00381148 | 0.00446892 | -0.0006574 | 0.74344742 | 0.90650187 |
| 5430402O13 | 0.00878892 | 0.01006661 | 0.01072462 | -0.000658 | 0.60748204 | 0.90650187 |
| Sorcs1 | 0.02707766 | 0.03197916 | 0.03263761 | -0.0006584 | 0.53573241 | 0.90650187 |
| Gm38399 | 0.03296898 | 0.03892143 | 0.03958281 | -0.0006614 | 0.5296067 | 0.90650187 |
| Cfap70 | 0.004099 | 0.00435152 | 0.00501769 | -0.0006662 | 0.72247806 | 0.90650187 |
| Ace | 0.01538273 | 0.01802027 | 0.01868733 | -0.0006671 | 0.56304334 | 0.90650187 |
| Gm15559 | 0.01445623 | 0.01690473 | 0.01757276 | -0.000668 | 0.56710096 | 0.90650187 |
| Mid2 | 0.03865333 | 0.04555992 | 0.0462326 | -0.0006727 | 0.52578692 | 0.90650187 |
| M5C1000I18 | 0.00476119 | 0.00515251 | 0.00582567 | -0.0006732 | 0.69626766 | 0.90650187 |
| Chdh | 0.01178979 | 0.01368305 | 0.01435727 | -0.0006742 | 0.58268406 | 0.90650187 |
| Tulp1 | 0.03155203 | 0.03723755 | 0.03791737 | -0.0006798 | 0.53176442 | 0.90650187 |
| Mrpl20 | 1.17835173 | 0.81500216 | 0.81568229 | -0.0006801 | 0.50147879 | 0.90650187 |
| Spg20 | 0.20848906 | 0.22460456 | 0.2252847 | -0.0006801 | 0.50535422 | 0.90650187 |
| Alox12 | 0.01730944 | 0.0203205 | 0.02100069 | -0.0006802 | 0.55724557 | 0.90650187 |
| Mrap | 0.00711388 | 0.00800637 | 0.00869051 | -0.0006841 | 0.63681984 | 0.90650187 |
| Fgd3 | 0.00451672 | 0.00484245 | 0.00552746 | -0.000685 | 0.70916319 | 0.90650187 |
| Hrc | 0.023277 | 0.02744143 | 0.02812797 | -0.0006865 | 0.54320316 | 0.90650187 |
| Fam109a | 0.12281001 | 0.13859733 | 0.13928413 | -0.0006868 | 0.50874458 | 0.90650187 |
| Tmem114 | 0.0047306 | 0.00510095 | 0.00578836 | -0.0006874 | 0.70123363 | 0.90650187 |
| Bmp2 | 0.03394876 | 0.04004073 | 0.04073259 | -0.0006919 | 0.53009572 | 0.90650187 |
| 2310040G07 | 0.01006831 | 0.01158226 | 0.01227516 | -0.0006929 | 0.59907178 | 0.90650187 |
| Gm10083 | 0.01086746 | 0.01254222 | 0.01324233 | -0.0007001 | 0.59290933 | 0.90650187 |

| | | | | | | |
|---|---|---|---|---|---|---|
| Gm11696 | 0.01612013 | 0.01887202 | 0.01957342 | -0.0007014 | 0.56328557 | 0.90650187 |
| 2610008E11I | 0.13265892 | 0.14889751 | 0.14960051 | -0.000703 | 0.50833353 | 0.90650187 |
| Fen1 | 0.20163851 | 0.21800438 | 0.21870827 | -0.0007039 | 0.50570776 | 0.90650187 |
| Aknad1 | 0.00654673 | 0.00728979 | 0.00800073 | -0.0007109 | 0.6535903 | 0.90650187 |
| Nckipsd | 0.05355939 | 0.06272282 | 0.06343462 | -0.0007118 | 0.51989279 | 0.90650187 |
| Gm9945 | 0.00655704 | 0.00729965 | 0.00801328 | -0.0007136 | 0.65391247 | 0.90650187 |
| 0610025J13F | 0.00471974 | 0.00506082 | 0.00577512 | -0.0007143 | 0.7087905 | 0.90650187 |
| Gm20707 | 0.01098767 | 0.01267142 | 0.01338772 | -0.0007163 | 0.59400519 | 0.90650187 |
| Cytl1 | 0.00714544 | 0.00801053 | 0.00872888 | -0.0007183 | 0.64276217 | 0.90650187 |
| Scube2 | 0.01697552 | 0.01988004 | 0.0206002 | -0.0007202 | 0.5617523 | 0.90650187 |
| Ttc21a | 0.00456014 | 0.00485956 | 0.00558043 | -0.0007209 | 0.7171113 | 0.90650187 |
| Hao2 | 0.00440193 | 0.00466347 | 0.00538741 | -0.0007239 | 0.72487264 | 0.90650187 |
| Tmtc1 | 0.04091327 | 0.04813456 | 0.04886247 | -0.0007279 | 0.52640145 | 0.90650187 |
| Gm26826 | 0.0080916 | 0.00915041 | 0.00987838 | -0.000728 | 0.62839758 | 0.90650187 |
| C530044C16I | 0.02524313 | 0.02973508 | 0.03046372 | -0.0007286 | 0.54234011 | 0.90650187 |
| Gm28154 | 0.00973065 | 0.01113371 | 0.0118662 | -0.0007325 | 0.60811974 | 0.90650187 |
| Gm9970 | 0.00383294 | 0.00395991 | 0.00469285 | -0.0007329 | 0.7562618 | 0.90650187 |
| Mb | 0.00089528 | 0.0003647 | 0.00109833 | -0.0007336 | 0.99850835 | 0.99857405 |
| Fan1 | 0.03832841 | 0.04511741 | 0.04585385 | -0.0007364 | 0.52846008 | 0.90650187 |
| Emx1 | 0.00469191 | 0.00500464 | 0.00574117 | -0.0007365 | 0.71577019 | 0.90650187 |
| Clic5 | 0.00393545 | 0.00407997 | 0.00481802 | -0.000738 | 0.7520636 | 0.90650187 |
| Pcdhga9 | 0.01946319 | 0.02284034 | 0.02357959 | -0.0007392 | 0.55542319 | 0.90650187 |
| Tmc3 | 0.00987702 | 0.01130273 | 0.0120435 | -0.0007408 | 0.60774238 | 0.90650187 |
| Ccl17 | 0.00688067 | 0.00766449 | 0.00840693 | -0.0007424 | 0.65269545 | 0.90650187 |
| Gpha2 | 0.00739785 | 0.00829287 | 0.00903567 | -0.0007428 | 0.64261622 | 0.90650187 |
| Prokr1 | 0.01749253 | 0.02047672 | 0.02122021 | -0.0007435 | 0.56188939 | 0.90650187 |
| Ttc28 | 0.24861535 | 0.26208999 | 0.26283403 | -0.000744 | 0.50502045 | 0.90650187 |
| Serp2 | 0.0850035 | 0.09789575 | 0.09864022 | -0.0007445 | 0.51338302 | 0.90650187 |
| Necab1 | 0.00677319 | 0.00753022 | 0.00827622 | -0.000746 | 0.65568441 | 0.90650187 |
| Numb | 0.13344929 | 0.14967497 | 0.15042333 | -0.0007484 | 0.50882272 | 0.90650187 |

| | | | | | | |
|---|---|---|---|---|---|---|
| Gm11508 | 0.00380401 | 0.0039072 | 0.00465752 | -0.0007503 | 0.7630608 | 0.90650187 |
| Kcnv2 | 0.00724314 | 0.00809655 | 0.00884764 | -0.0007511 | 0.64705865 | 0.90650187 |
| Tm4sf20 | 0.01121794 | 0.01291285 | 0.01366616 | -0.0007533 | 0.59679272 | 0.90650187 |
| Cyp4x1 | 0.00459868 | 0.00486788 | 0.00562745 | -0.0007596 | 0.72576384 | 0.90650187 |
| Lrrc4c | 0.00451648 | 0.0047622 | 0.00552717 | -0.000765 | 0.73081641 | 0.90650187 |
| Tmod4 | 0.00926043 | 0.01052849 | 0.01129637 | -0.0007679 | 0.61874765 | 0.90650187 |
| Acap1 | 0.0147435 | 0.01714901 | 0.01791849 | -0.0007695 | 0.57570112 | 0.90650187 |
| Fam229a | 0.00463322 | 0.00489738 | 0.00566959 | -0.0007722 | 0.72757778 | 0.90650187 |
| Rhof | 0.01664503 | 0.01942686 | 0.02020363 | -0.0007768 | 0.56785657 | 0.90650187 |
| Gm26669 | 0.00380642 | 0.00388271 | 0.00466046 | -0.0007778 | 0.77092558 | 0.90650187 |
| 6330403A02I | 0.00445652 | 0.00467625 | 0.00545402 | -0.0007778 | 0.73694786 | 0.90650187 |
| Nt5e | 0.02242993 | 0.02634141 | 0.02711975 | -0.0007783 | 0.55076333 | 0.90650187 |
| Mylk3 | 0.00428642 | 0.00446726 | 0.00524645 | -0.0007792 | 0.74545166 | 0.90650187 |
| Ccdc162 | 0.01675282 | 0.01955224 | 0.02033298 | -0.0007807 | 0.56777097 | 0.90650187 |
| Dnajc6 | 0.03076832 | 0.0362137 | 0.03699487 | -0.0007812 | 0.53739427 | 0.90650187 |
| Zfp563 | 0.03218973 | 0.03788558 | 0.03866729 | -0.0007817 | 0.53580652 | 0.90650187 |
| Tspan3 | 1.89763213 | 0.92618292 | 0.92696924 | -0.0007863 | 0.50150442 | 0.90650187 |
| Gm10605 | 0.00923601 | 0.01047972 | 0.01126676 | -0.000787 | 0.62192878 | 0.90650187 |
| Gm38250 | 0.00462855 | 0.00487537 | 0.00566389 | -0.0007885 | 0.73201352 | 0.90650187 |
| Sprn | 0.01440489 | 0.016722 | 0.01751095 | -0.000789 | 0.5793753 | 0.90650187 |
| B3gat1 | 0.00162396 | 0.00119994 | 0.00199127 | -0.0007913 | 0.96135893 | 0.97093994 |
| Camkv | 0.00674914 | 0.0074543 | 0.00824697 | -0.0007927 | 0.66541923 | 0.90650187 |
| Gm15893 | 0.00161797 | 0.00119111 | 0.00198394 | -0.0007928 | 0.96217996 | 0.97133123 |
| Mgat5b | 0.00431062 | 0.0044813 | 0.00527599 | -0.0007947 | 0.74844781 | 0.90650187 |
| D930048N14 | 0.01119333 | 0.01284068 | 0.01363641 | -0.0007957 | 0.6023424 | 0.90650187 |
| Sel1l | 0.54912951 | 0.49887192 | 0.4996686 | -0.0007967 | 0.50282773 | 0.90650187 |
| Kcnq5 | 0.00109633 | 0.00054689 | 0.00134479 | -0.0007979 | 0.9958266 | 0.99633807 |
| Myom1 | 0.00109633 | 0.00054689 | 0.00134479 | -0.0007979 | 0.9958266 | 0.99633807 |
| Foxn2 | 0.15259416 | 0.16933073 | 0.1701293 | -0.0007986 | 0.50832421 | 0.90650187 |
| Gabre | 0.01923346 | 0.02250168 | 0.02330487 | -0.0008032 | 0.56088569 | 0.90650187 |

| | | | | | | |
|---|---|---|---|---|---|---|
| Defb1 | 0.00426371 | 0.00441348 | 0.00521873 | -0.0008053 | 0.75363086 | 0.90650187 |
| Itln1 | 0.00402293 | 0.00411887 | 0.00492483 | -0.000806 | 0.76655107 | 0.90650187 |
| Slc38a3 | 0.01323036 | 0.01528989 | 0.0160959 | -0.000806 | 0.5880827 | 0.90650187 |
| Kcnn2 | 0.0105181 | 0.01201314 | 0.01281965 | -0.0008065 | 0.61013938 | 0.90650187 |
| Fosl2 | 0.10942571 | 0.12426689 | 0.12507406 | -0.0008072 | 0.51144403 | 0.90650187 |
| Fcor | 0.09829783 | 0.11228894 | 0.11310021 | -0.0008113 | 0.51271941 | 0.90650187 |
| AK157302 | 0.01162935 | 0.01335043 | 0.01416343 | -0.000813 | 0.6007092 | 0.90650187 |
| Sptssb | 0.00118754 | 0.00064166 | 0.00145658 | -0.0008149 | 0.99356211 | 0.99493242 |
| Deaf1 | 0.39912736 | 0.38949103 | 0.39030739 | -0.0008164 | 0.50370944 | 0.90650187 |
| Agmo | 0.00119882 | 0.00065391 | 0.0014704 | -0.0008165 | 0.99321731 | 0.99474162 |
| A230060F14 | 0.00120837 | 0.00066437 | 0.0014821 | -0.0008177 | 0.99291267 | 0.99467941 |
| 2210407C18I | 0.00120837 | 0.00066437 | 0.0014821 | -0.0008177 | 0.99291267 | 0.99467941 |
| Abcg8 | 0.00120837 | 0.00066437 | 0.0014821 | -0.0008177 | 0.99291267 | 0.99467941 |
| Stxbp2 | 0.33576002 | 0.3382252 | 0.33904366 | -0.0008185 | 0.50428129 | 0.90650187 |
| Colec11 | 0.03779764 | 0.04441456 | 0.04523479 | -0.0008202 | 0.5321242 | 0.90650187 |
| Glyctk | 0.00460617 | 0.00481622 | 0.00563659 | -0.0008204 | 0.74119428 | 0.90650187 |
| Gucy2c | 0.01330631 | 0.01536556 | 0.01618748 | -0.0008219 | 0.58929243 | 0.90650187 |
| Gal3st3 | 0.0012506 | 0.00071162 | 0.00153385 | -0.0008222 | 0.99141643 | 0.9936389 |
| Gm9530 | 0.00713143 | 0.0078891 | 0.00871185 | -0.0008227 | 0.66269881 | 0.90650187 |
| D330045A20 | 0.00143354 | 0.00093504 | 0.00175801 | -0.000823 | 0.98128803 | 0.98582773 |
| Gm12932 | 0.00143354 | 0.00093504 | 0.00175801 | -0.000823 | 0.98128803 | 0.98582773 |
| Lamc3 | 0.0014265 | 0.00092588 | 0.00174939 | -0.0008235 | 0.98181319 | 0.98622496 |
| Cacng2 | 0.00131285 | 0.00078422 | 0.00161013 | -0.0008259 | 0.98870662 | 0.99124979 |
| Wdr63 | 0.00137741 | 0.00086325 | 0.00168924 | -0.000826 | 0.98513879 | 0.98832462 |
| 1700026J14F | 0.00715245 | 0.00790948 | 0.00873739 | -0.0008279 | 0.66321074 | 0.90650187 |
| Gm12207 | 0.01244614 | 0.01432144 | 0.01514984 | -0.0008284 | 0.59603181 | 0.90650187 |
| D7Ertd443e | 0.0061888 | 0.00673118 | 0.00756515 | -0.000834 | 0.68795981 | 0.90650187 |
| Gm26767 | 0.0146998 | 0.01703006 | 0.0178659 | -0.0008358 | 0.58237864 | 0.90650187 |
| C1galt1 | 0.19301503 | 0.20951894 | 0.2103581 | -0.0008392 | 0.5070747 | 0.90650187 |
| Dnah7a | 0.00614282 | 0.00666976 | 0.00750917 | -0.0008394 | 0.69038722 | 0.90650187 |

| | | | | | | |
|---|---|---|---|---|---|---|
| C030034I22R | 0.04331181 | 0.05080534 | 0.05164503 | -0.0008397 | 0.52881072 | 0.90650187 |
| Xpa | 0.39405142 | 0.38548075 | 0.38632088 | -0.0008401 | 0.50385685 | 0.90650187 |
| Tatdn1 | 0.12152942 | 0.13708376 | 0.13793401 | -0.0008503 | 0.5109311 | 0.90650187 |
| Rd3 | 0.00675359 | 0.00740152 | 0.00825238 | -0.0008509 | 0.67665423 | 0.90650187 |
| Gm16867 | 0.0044071 | 0.00454148 | 0.00539372 | -0.0008522 | 0.75879621 | 0.90650187 |
| Noxred1 | 0.00443834 | 0.00457723 | 0.00543184 | -0.0008546 | 0.75786181 | 0.90650187 |
| Gm20635 | 0.00722394 | 0.00796612 | 0.0088243 | -0.0008582 | 0.66725371 | 0.90650187 |
| Tac2 | 0.00715608 | 0.00788287 | 0.00874181 | -0.0008589 | 0.66887618 | 0.90650187 |
| B230312C02 | 0.00717314 | 0.00790209 | 0.00876255 | -0.0008605 | 0.66878072 | 0.90650187 |
| Gm11744 | 0.01247937 | 0.01432774 | 0.01518995 | -0.0008622 | 0.59961045 | 0.90650187 |
| Krt222 | 0.00386239 | 0.00386642 | 0.00472881 | -0.0008624 | 0.79124369 | 0.90660917 |
| S100a7a | 0.00948591 | 0.01070585 | 0.01156966 | -0.0008638 | 0.63002329 | 0.90650187 |
| Gm17039 | 0.00534067 | 0.00566339 | 0.00653215 | -0.0008688 | 0.72282554 | 0.90650187 |
| Setd7 | 0.14464421 | 0.16112752 | 0.16199866 | -0.0008711 | 0.50953624 | 0.90650187 |
| Prr16 | 0.00998847 | 0.01130687 | 0.01217848 | -0.0008716 | 0.62482229 | 0.90650187 |
| Gm16033 | 0.00371908 | 0.00368172 | 0.0045538 | -0.0008721 | 0.80271611 | 0.90870476 |
| Eif1ax | 0.9359179 | 0.72129257 | 0.72217325 | -0.0008807 | 0.50216278 | 0.90650187 |
| Dchs2 | 0.01469982 | 0.01698218 | 0.01786592 | -0.0008837 | 0.58702625 | 0.90650187 |
| Bloc1s6 | 0.20213555 | 0.21830155 | 0.21918708 | -0.0008855 | 0.50716484 | 0.90650187 |
| Sycp3 | 0.00435426 | 0.00444103 | 0.00532924 | -0.0008882 | 0.77063279 | 0.90650187 |
| Nrbp2 | 0.08709704 | 0.10004541 | 0.10093371 | -0.0008883 | 0.51560465 | 0.90650187 |
| Adgrb2 | 0.04499242 | 0.05270114 | 0.05358948 | -0.0008883 | 0.52937267 | 0.90650187 |
| Fhod1 | 0.08492002 | 0.09765236 | 0.09854865 | -0.0008963 | 0.51612564 | 0.90650187 |
| Rnf167 | 0.54204502 | 0.49394988 | 0.4948465 | -0.0008966 | 0.50321347 | 0.90650187 |
| Gm26812 | 0.00445049 | 0.00454911 | 0.00544666 | -0.0008975 | 0.76809379 | 0.90650187 |
| Gabrb1 | 0.01675066 | 0.01942908 | 0.0203304 | -0.0009013 | 0.57812093 | 0.90650187 |
| Vdac3-ps1 | 0.03693041 | 0.04332005 | 0.04422237 | -0.0009023 | 0.53613789 | 0.90650187 |
| Gm37986 | 0.00428502 | 0.00433727 | 0.00524474 | -0.0009075 | 0.77911381 | 0.90650187 |
| Gm37584 | 0.0071828 | 0.00786329 | 0.00877429 | -0.000911 | 0.67780453 | 0.90650187 |
| Serpinf2 | 0.01719663 | 0.01995179 | 0.02086541 | -0.0009136 | 0.57716964 | 0.90650187 |

| | | | | | | |
|---|---|---|---|---|---|---|
| Gm26889 | 0.0067371 | 0.00731493 | 0.00823232 | -0.0009174 | 0.68984367 | 0.90650187 |
| Glp2r | 0.01224528 | 0.01398934 | 0.01490736 | -0.000918 | 0.60786876 | 0.90650187 |
| Igsf10 | 0.07858415 | 0.09064645 | 0.09156913 | -0.0009227 | 0.5178648 | 0.90650187 |
| Pip5k1b | 0.10544099 | 0.11987618 | 0.12080075 | -0.0009246 | 0.51357147 | 0.90650187 |
| Adgrg4 | 0.01084617 | 0.01229154 | 0.01321657 | -0.000925 | 0.62215658 | 0.90650187 |
| Gm15698 | 0.00378803 | 0.00371243 | 0.004638 | -0.0009256 | 0.81250853 | 0.9092537 |
| Ror1 | 0.10931103 | 0.12402366 | 0.12495135 | -0.0009277 | 0.51316498 | 0.90650187 |
| Capn10 | 0.35140622 | 0.351081 | 0.35200955 | -0.0009286 | 0.50467824 | 0.90650187 |
| Mgst2 | 0.00643242 | 0.00693273 | 0.00786164 | -0.0009289 | 0.70030633 | 0.90650187 |
| Mex3a | 0.23647362 | 0.25071412 | 0.25164383 | -0.0009297 | 0.50655216 | 0.90650187 |
| Gnmt | 0.01072486 | 0.01213853 | 0.01306982 | -0.0009313 | 0.6242911 | 0.90650187 |
| Zbtb39 | 0.04472734 | 0.05235072 | 0.05328307 | -0.0009324 | 0.53100231 | 0.90650187 |
| Plekhg1 | 0.10781691 | 0.12241793 | 0.12335112 | -0.0009332 | 0.51341496 | 0.90650187 |
| Nlgn1 | 0.01791046 | 0.02078708 | 0.02172111 | -0.000934 | 0.575802 | 0.90650187 |
| Rab9 | 0.4756413 | 0.44711788 | 0.44805419 | -0.0009363 | 0.50370615 | 0.90650187 |
| Adcy1 | 0.02530541 | 0.02960069 | 0.03053761 | -0.0009369 | 0.55424563 | 0.90650187 |
| Kcnab3 | 0.02568004 | 0.03004458 | 0.03098194 | -0.0009374 | 0.55349756 | 0.90650187 |
| Prr22 | 0.00445558 | 0.00451332 | 0.00545288 | -0.0009396 | 0.77816172 | 0.90650187 |
| Prtg | 0.02955416 | 0.03462339 | 0.03556377 | -0.0009404 | 0.5467884 | 0.90650187 |
| Nxf2 | 0.00620683 | 0.00664261 | 0.00758709 | -0.0009445 | 0.710009 | 0.90650187 |
| Ugt2b36 | 0.00445098 | 0.00450194 | 0.00544726 | -0.0009453 | 0.77979334 | 0.90650187 |
| Gm26760 | 0.00375488 | 0.00365098 | 0.00459751 | -0.0009465 | 0.81997296 | 0.9092537 |
| Tec | 0.0549471 | 0.06407149 | 0.06501896 | -0.0009475 | 0.52582653 | 0.90650187 |
| Ccdc63 | 0.00496041 | 0.00512089 | 0.00606861 | -0.0009477 | 0.75624153 | 0.90650187 |
| Thoc2 | 0.90150711 | 0.70435247 | 0.70530127 | -0.0009488 | 0.50238581 | 0.90650187 |
| Il15ra | 0.00713867 | 0.00776939 | 0.00872064 | -0.0009513 | 0.68613685 | 0.90650187 |
| Tmod1 | 0.06570671 | 0.0762521 | 0.07720599 | -0.0009539 | 0.52190136 | 0.90650187 |
| Gabra4 | 0.00922161 | 0.01029523 | 0.01124931 | -0.0009541 | 0.64692911 | 0.90650187 |
| Fgl1 | 0.02001666 | 0.02328654 | 0.0242411 | -0.0009546 | 0.56948306 | 0.90650187 |
| Ppargc1b | 0.01747018 | 0.02023866 | 0.02119343 | -0.0009548 | 0.57936712 | 0.90650187 |

| | | | | | | |
|---|---|---|---|---|---|---|
| Gm13075 | 0.0745389 | 0.08612624 | 0.08708297 | -0.0009567 | 0.51947707 | 0.90650187 |
| Pcdhga10 | 0.01476191 | 0.01698218 | 0.01794064 | -0.0009585 | 0.59386619 | 0.90650187 |
| Cntnap5a | 0.01066924 | 0.01204284 | 0.01300252 | -0.0009597 | 0.62858901 | 0.90650187 |
| Lrrk2 | 0.03195741 | 0.03743103 | 0.03839416 | -0.0009631 | 0.54439806 | 0.90650187 |
| Grik1 | 0.01732812 | 0.02005416 | 0.02102309 | -0.0009689 | 0.58117239 | 0.90650187 |
| Creg2 | 0.00399543 | 0.00392127 | 0.00489126 | -0.00097 | 0.81100592 | 0.9092537 |
| Lypd6 | 0.01762728 | 0.02041016 | 0.02138174 | -0.0009716 | 0.58004448 | 0.90650187 |
| Gm17296 | 0.01110503 | 0.01255712 | 0.01352964 | -0.0009725 | 0.62534643 | 0.90650187 |
| Shisa9 | 0.00717587 | 0.00779313 | 0.00876587 | -0.0009727 | 0.68910838 | 0.90650187 |
| Mmp17 | 0.02557499 | 0.02988306 | 0.03085737 | -0.0009743 | 0.55581539 | 0.90650187 |
| Gm17249 | 0.01417359 | 0.01625358 | 0.01723246 | -0.0009789 | 0.5996844 | 0.90650187 |
| Col22a1 | 0.00706265 | 0.00764927 | 0.00862822 | -0.0009789 | 0.69300749 | 0.90650187 |
| Vstm2b | 0.0093974 | 0.01048339 | 0.01146239 | -0.000979 | 0.64791498 | 0.90650187 |
| Gabbr1 | 0.24816271 | 0.26143768 | 0.26241957 | -0.0009819 | 0.50663574 | 0.90650187 |
| 1700022I11R | 0.00430338 | 0.00428465 | 0.00526714 | -0.0009825 | 0.79651842 | 0.9075779 |
| Gm12905 | 0.00626474 | 0.00667452 | 0.00765758 | -0.0009831 | 0.71590421 | 0.90650187 |
| Arhgef11 | 0.08051493 | 0.09271822 | 0.09370209 | -0.0009839 | 0.51861545 | 0.90650187 |
| Fndc5 | 0.04236644 | 0.04956357 | 0.05054936 | -0.0009858 | 0.53454358 | 0.90650187 |
| Chaf1b | 0.1077061 | 0.12224579 | 0.12323234 | -0.0009866 | 0.51419528 | 0.90650187 |
| Cntnap2 | 0.02647266 | 0.03093428 | 0.03192128 | -0.000987 | 0.55466536 | 0.90650187 |
| Kcnk10 | 0.00617599 | 0.00656171 | 0.00754955 | -0.0009878 | 0.71961549 | 0.90650187 |
| Slc44a4 | 0.00404509 | 0.00396185 | 0.00495187 | -0.00099 | 0.81294649 | 0.9092537 |
| Cabp7 | 0.02379189 | 0.02774923 | 0.02874024 | -0.000991 | 0.56091588 | 0.90650187 |
| Syn2 | 0.0639041 | 0.07418259 | 0.07517617 | -0.0009936 | 0.52342671 | 0.90650187 |
| Gm26981 | 0.01879003 | 0.02177888 | 0.02277435 | -0.0009955 | 0.57703682 | 0.90650187 |
| Apc2 | 0.00419194 | 0.00413534 | 0.00513114 | -0.0009958 | 0.80586645 | 0.9092537 |
| Lect1 | 0.01727301 | 0.01995586 | 0.02095702 | -0.0010012 | 0.58409271 | 0.90650187 |
| Agbl2 | 0.00434854 | 0.0043183 | 0.00532225 | -0.001004 | 0.7991507 | 0.90777025 |
| Vash1 | 0.01602051 | 0.01844876 | 0.01945376 | -0.001005 | 0.59082348 | 0.90650187 |
| Nrn1l | 0.0069103 | 0.00743737 | 0.00844297 | -0.0010056 | 0.70176767 | 0.90650187 |

| | | | | | | |
|---|---|---|---|---|---|---|
| Wdr55 | 0.10221581 | 0.11632161 | 0.11732743 | -0.0010058 | 0.51520057 | 0.90650187 |
| A230087F16 | 0.00710751 | 0.00767466 | 0.00868276 | -0.0010081 | 0.69712502 | 0.90650187 |
| Sdc3 | 0.18344633 | 0.19998828 | 0.20099863 | -0.0010103 | 0.50891422 | 0.90650187 |
| Kctd21 | 0.03244296 | 0.03795396 | 0.03896491 | -0.001011 | 0.54591296 | 0.90650187 |
| 2810030D12 | 0.00436415 | 0.00432956 | 0.00534131 | -0.0010117 | 0.8001293 | 0.90805731 |
| Sil1 | 0.33707814 | 0.33912775 | 0.34014405 | -0.0010163 | 0.50529893 | 0.90650187 |
| Tet1 | 0.13970666 | 0.15589418 | 0.15691151 | -0.0010173 | 0.51149708 | 0.90650187 |
| Rab11fip5 | 0.07680996 | 0.0885838 | 0.08960445 | -0.0010206 | 0.520193 | 0.90650187 |
| Stag3 | 0.00432491 | 0.00427085 | 0.00529342 | -0.0010226 | 0.8047728 | 0.90882608 |
| Stab2 | 0.00413775 | 0.00404085 | 0.005065 | -0.0010241 | 0.8156478 | 0.9092537 |
| Tinag | 0.00334279 | 0.0030697 | 0.00409409 | -0.0010244 | 0.86700555 | 0.92056229 |
| Tomm6os | 0.00435482 | 0.00430526 | 0.00532993 | -0.0010247 | 0.80362911 | 0.90882608 |
| Fam49a | 0.03962878 | 0.04634232 | 0.0473687 | -0.0010264 | 0.53836941 | 0.90650187 |
| Glra1 | 0.00658198 | 0.00701586 | 0.00804362 | -0.0010278 | 0.71499351 | 0.90650187 |
| 2210010C04I | 0.01002872 | 0.01119827 | 0.01222723 | -0.001029 | 0.64583876 | 0.90650187 |
| Hist1h1a | 0.00843515 | 0.00926526 | 0.0102954 | -0.0010301 | 0.67177499 | 0.90650187 |
| Fgf9 | 0.09169901 | 0.10492303 | 0.10595358 | -0.0010306 | 0.51724491 | 0.90650187 |
| Dpyd | 0.02011401 | 0.02332294 | 0.02435741 | -0.0010345 | 0.57487757 | 0.90650187 |
| 8030474K03I | 0.00380199 | 0.00361805 | 0.00465506 | -0.001037 | 0.83899725 | 0.91188312 |
| Lbx2 | 0.003569 | 0.00333305 | 0.00437048 | -0.0010374 | 0.85434788 | 0.91621818 |
| 3930402G23 | 0.01127471 | 0.01269706 | 0.0137348 | -0.0010377 | 0.63152218 | 0.90650187 |
| Gm26916 | 0.0043297 | 0.00426095 | 0.00529927 | -0.0010383 | 0.80813837 | 0.9092537 |
| Cdh6 | 0.00887304 | 0.00978619 | 0.01082664 | -0.0010405 | 0.66539315 | 0.90650187 |
| Cadps | 0.27016713 | 0.28129749 | 0.2823394 | -0.0010419 | 0.50654458 | 0.90650187 |
| Gm14399 | 0.010508 | 0.01176384 | 0.01280742 | -0.0010436 | 0.64141321 | 0.90650187 |
| Hpx | 0.00605904 | 0.00636263 | 0.00740718 | -0.0010445 | 0.73464118 | 0.90650187 |
| Rnf39 | 0.00930541 | 0.01030441 | 0.01135089 | -0.0010465 | 0.65904487 | 0.90650187 |
| Gm10638 | 0.0034408 | 0.00316555 | 0.00421385 | -0.0010483 | 0.86562638 | 0.92040222 |
| Gm26558 | 0.02388063 | 0.02779688 | 0.02884572 | -0.0010488 | 0.56420711 | 0.90650187 |
| Lnpep | 0.05942888 | 0.06906591 | 0.07011616 | -0.0010502 | 0.52654568 | 0.90650187 |

| | | | | | | |
|---|---|---|---|---|---|---|
| Kcnd3 | 0.01199492 | 0.01355141 | 0.01460504 | -0.0010536 | 0.62578629 | 0.90650187 |
| B230217O12 | 0.00662959 | 0.00704695 | 0.00810154 | -0.0010546 | 0.718601 | 0.90650187 |
| Anks4b | 0.02125708 | 0.0246673 | 0.02572189 | -0.0010546 | 0.57231437 | 0.90650187 |
| Tprg | 0.00686759 | 0.0073353 | 0.00839103 | -0.0010557 | 0.71203281 | 0.90650187 |
| Syt15 | 0.00582038 | 0.00605515 | 0.00711657 | -0.0010614 | 0.74635145 | 0.90650187 |
| 4930578C19l | 0.00328786 | 0.00296502 | 0.00402697 | -0.001062 | 0.87947185 | 0.92556435 |
| Pml | 0.1160344 | 0.13105543 | 0.13211796 | -0.0010625 | 0.51426029 | 0.90650187 |
| Samd13 | 0.01970834 | 0.02280955 | 0.02387265 | -0.0010631 | 0.57846678 | 0.90650187 |
| Lca5 | 0.11998148 | 0.13523566 | 0.13629937 | -0.0010637 | 0.51383823 | 0.90650187 |
| Tcp11 | 0.24120765 | 0.25495987 | 0.25602425 | -0.0010644 | 0.50737277 | 0.90650187 |
| Ncan | 0.00562745 | 0.00581499 | 0.00688157 | -0.0010666 | 0.75459834 | 0.90650187 |
| Gm27042 | 0.00652907 | 0.00690654 | 0.00797925 | -0.0010727 | 0.7249656 | 0.90650187 |
| Espn | 0.04352394 | 0.05081576 | 0.05189071 | -0.0010749 | 0.53668808 | 0.90650187 |
| H2-Q6 | 0.02995505 | 0.0349591 | 0.03603654 | -0.0010774 | 0.55287075 | 0.90650187 |
| Add2 | 0.00881929 | 0.00968323 | 0.01076145 | -0.0010782 | 0.67199157 | 0.90650187 |
| Gm26652 | 0.00796905 | 0.00864984 | 0.00972957 | -0.0010797 | 0.6891164 | 0.90650187 |
| Pola2 | 0.12636231 | 0.14193733 | 0.14301867 | -0.0010813 | 0.51340689 | 0.90650187 |
| Gm10330 | 0.00352728 | 0.0032373 | 0.00431951 | -0.0010822 | 0.86731871 | 0.92056229 |
| Zkscan4 | 0.02208436 | 0.02562587 | 0.02670811 | -0.0010822 | 0.57147881 | 0.90650187 |
| Col20a1 | 0.00755185 | 0.0081383 | 0.00922281 | -0.0010845 | 0.69942892 | 0.90650187 |
| Zfp935 | 0.00637817 | 0.00670998 | 0.00779563 | -0.0010856 | 0.73207871 | 0.90650187 |
| Cav3 | 0.00424945 | 0.0041125 | 0.00520133 | -0.0010888 | 0.82397681 | 0.90929265 |
| Dusp9 | 0.01838599 | 0.02120102 | 0.02229068 | -0.0010897 | 0.58602038 | 0.90650187 |
| Gm17638 | 0.0094042 | 0.01037236 | 0.01147064 | -0.0010983 | 0.66481988 | 0.90650187 |
| Brinp2 | 0.01082824 | 0.01209215 | 0.01319489 | -0.0011027 | 0.64487968 | 0.90650187 |
| Pianp | 0.15762394 | 0.17412947 | 0.17523527 | -0.0011058 | 0.51119024 | 0.90650187 |
| Calhm2 | 0.12349737 | 0.13890215 | 0.140008 | -0.0011059 | 0.51400539 | 0.90650187 |
| Slc35b3 | 0.16106862 | 0.17760709 | 0.17871549 | -0.0011084 | 0.51099821 | 0.90650187 |
| Pias1 | 0.3031905 | 0.31026232 | 0.3113738 | -0.0011115 | 0.50633057 | 0.90650187 |
| Drd4 | 0.00363331 | 0.00333702 | 0.00444903 | -0.001112 | 0.86674899 | 0.92056229 |

| | | | | | | |
|---|---|---|---|---|---|---|
| Erbb4 | 0.00324938 | 0.00286678 | 0.00397995 | -0.0011132 | 0.89314105 | 0.93181736 |
| Gm44257 | 0.00614727 | 0.00639956 | 0.00751459 | -0.001115 | 0.74525933 | 0.90650187 |
| Casp4 | 0.01358676 | 0.01541014 | 0.01652552 | -0.0011154 | 0.61793193 | 0.90650187 |
| Crispld1 | 0.02344915 | 0.02721696 | 0.02833272 | -0.0011158 | 0.56948778 | 0.90650187 |
| E230001N04 | 0.08220127 | 0.09444488 | 0.09556068 | -0.0011158 | 0.52069924 | 0.90650187 |
| Rhbg | 0.0107668 | 0.01200304 | 0.01312056 | -0.0011175 | 0.64752495 | 0.90650187 |
| A130051J06I | 0.00395803 | 0.00372396 | 0.00484559 | -0.0011216 | 0.84827028 | 0.91440801 |
| Mctp1 | 0.00410783 | 0.00390643 | 0.00502847 | -0.001122 | 0.83939557 | 0.91213511 |
| 4933406C10I | 0.00677121 | 0.00715154 | 0.00827381 | -0.0011223 | 0.7267488 | 0.90650187 |
| Kif5a | 0.01863706 | 0.02146777 | 0.02259126 | -0.0011235 | 0.58748694 | 0.90650187 |
| Lck | 0.01741263 | 0.02000051 | 0.02112442 | -0.0011239 | 0.59348777 | 0.90650187 |
| Gm9830 | 0.00418985 | 0.00400419 | 0.00512858 | -0.0011244 | 0.8351347 | 0.91066681 |
| Hist1h4j | 0.00411165 | 0.00390858 | 0.00503314 | -0.0011246 | 0.83971358 | 0.91213511 |
| Gm43597 | 0.0042161 | 0.00403575 | 0.00516062 | -0.0011249 | 0.8337322 | 0.91062493 |
| Padi1 | 0.00421059 | 0.00402655 | 0.00515391 | -0.0011274 | 0.83457986 | 0.91062493 |
| Il15 | 0.00509352 | 0.00510342 | 0.00623091 | -0.0011275 | 0.78942706 | 0.90650187 |
| Gm26753 | 0.00689089 | 0.00729056 | 0.00841937 | -0.0011288 | 0.72442358 | 0.90650187 |
| P2rx3 | 0.0054152 | 0.00549242 | 0.00662298 | -0.0011306 | 0.77603564 | 0.90650187 |
| Baiap2 | 0.19006362 | 0.20634845 | 0.20748175 | -0.0011333 | 0.50968643 | 0.90650187 |
| Lif | 0.00651373 | 0.00682669 | 0.00796059 | -0.0011339 | 0.73670712 | 0.90650187 |
| Doc2b | 0.02261807 | 0.02620962 | 0.02734378 | -0.0011342 | 0.57314782 | 0.90650187 |
| Myrip | 0.00519977 | 0.00522612 | 0.00636042 | -0.0011343 | 0.78605765 | 0.90650187 |
| Ube2v1 | 0.20874794 | 0.22439682 | 0.22553225 | -0.0011354 | 0.50892803 | 0.90650187 |
| Gm42504 | 0.00593087 | 0.00611493 | 0.00725112 | -0.0011362 | 0.75697083 | 0.90650187 |
| Defb42 | 0.00360528 | 0.00327553 | 0.00441479 | -0.0011393 | 0.87435403 | 0.9235672 |
| Car11 | 0.0867378 | 0.09939985 | 0.1005406 | -0.0011408 | 0.52011436 | 0.90650187 |
| Tigd5 | 0.01159503 | 0.01297985 | 0.01412195 | -0.0011421 | 0.64040199 | 0.90650187 |
| Gm10501 | 0.00625945 | 0.00650847 | 0.00765113 | -0.0011427 | 0.74663343 | 0.90650187 |
| Gm29394 | 0.02171504 | 0.02512483 | 0.02626798 | -0.0011432 | 0.57670351 | 0.90650187 |
| Trpc7 | 0.00404088 | 0.00380124 | 0.00494674 | -0.0011455 | 0.84836611 | 0.91440801 |

| | | | | | | |
|---|---|---|---|---|---|---|
| Zfp72 | 0.00912386 | 0.00998498 | 0.0111308 | -0.0011458 | 0.67639374 | 0.90650187 |
| Bcap31 | 1.16204643 | 0.80928985 | 0.81043625 | -0.0011464 | 0.50250875 | 0.90650187 |
| Cep128 | 0.01303179 | 0.01470658 | 0.01585645 | -0.0011499 | 0.62641928 | 0.90650187 |
| AI838599 | 0.01844198 | 0.0212068 | 0.02235772 | -0.0011509 | 0.59050685 | 0.90650187 |
| RP23-106P7. | 0.00577277 | 0.00590765 | 0.00705859 | -0.0011509 | 0.76573597 | 0.90650187 |
| Gal3st4 | 0.0058189 | 0.00596119 | 0.00711477 | -0.0011536 | 0.76448447 | 0.90650187 |
| C1qtnf3 | 0.01437972 | 0.01632519 | 0.01748065 | -0.0011555 | 0.61556427 | 0.90650187 |
| 8030443G20 | 0.00609311 | 0.00629304 | 0.00744865 | -0.0011556 | 0.75481104 | 0.90650187 |
| Gm29502 | 0.00946832 | 0.01039271 | 0.01154834 | -0.0011556 | 0.67179234 | 0.90650187 |
| Wdr66 | 0.01562253 | 0.01781891 | 0.01897557 | -0.0011567 | 0.60679835 | 0.90650187 |
| Gm26751 | 0.00332553 | 0.00291351 | 0.00407301 | -0.0011595 | 0.89716555 | 0.93404279 |
| Gm5741 | 0.00590349 | 0.00605772 | 0.00721777 | -0.0011601 | 0.76254296 | 0.90650187 |
| Gm28041 | 0.0064904 | 0.00676741 | 0.0079322 | -0.0011648 | 0.74306004 | 0.90650187 |
| Arhgap27 | 0.04358921 | 0.05080133 | 0.05196628 | -0.0011649 | 0.53969242 | 0.90650187 |
| Ecm1 | 0.05755037 | 0.06681733 | 0.06798334 | -0.001166 | 0.53038929 | 0.90650187 |
| Gart | 0.5278668 | 0.48392992 | 0.48509851 | -0.0011686 | 0.50427234 | 0.90650187 |
| 1700109H08 | 0.01210739 | 0.01357109 | 0.01474087 | -0.0011698 | 0.63787601 | 0.90650187 |
| Apobec2 | 0.00675136 | 0.00707801 | 0.00824966 | -0.0011716 | 0.73610472 | 0.90650187 |
| Mmel1 | 0.00413193 | 0.00388349 | 0.00505788 | -0.0011744 | 0.84901721 | 0.91444933 |
| Milr1 | 0.00685842 | 0.00720476 | 0.00837987 | -0.0011751 | 0.73349194 | 0.90650187 |
| Gm13306 | 0.00390788 | 0.00360786 | 0.00478435 | -0.0011765 | 0.86284508 | 0.91950555 |
| Rtp3 | 0.00693666 | 0.00729543 | 0.00847502 | -0.0011796 | 0.73196325 | 0.90650187 |
| Tmem213 | 0.00667656 | 0.00697817 | 0.00815868 | -0.0011805 | 0.73997061 | 0.90650187 |
| Pitpnm3 | 0.00658005 | 0.00686073 | 0.00804127 | -0.0011805 | 0.74301378 | 0.90650187 |
| Gm2237 | 0.01307531 | 0.01472794 | 0.01590894 | -0.001181 | 0.62930663 | 0.90650187 |
| Zfp868 | 0.11180033 | 0.12643024 | 0.12761127 | -0.001181 | 0.51640936 | 0.90650187 |
| Lrsam1 | 0.04960072 | 0.05771622 | 0.05889912 | -0.0011829 | 0.53557163 | 0.90650187 |
| 6820408C15I | 0.00412302 | 0.00386395 | 0.00504701 | -0.0011831 | 0.85131523 | 0.9150732 |
| Fcgbp | 0.00411597 | 0.00385385 | 0.00503841 | -0.0011846 | 0.85203148 | 0.91551074 |
| Strip2 | 0.00352863 | 0.00313657 | 0.00432116 | -0.0011846 | 0.8885209 | 0.93012921 |

| | | | | | | |
|---|---|---|---|---|---|---|
| Kcne2 | 0.00342101 | 0.00300392 | 0.00418967 | -0.0011858 | 0.89583775 | 0.93322571 |
| Lad1 | 0.01388035 | 0.01569306 | 0.01687927 | -0.0011862 | 0.62263982 | 0.90650187 |
| BC048403 | 0.01147357 | 0.01278615 | 0.01397516 | -0.001189 | 0.64737169 | 0.90650187 |
| Gm26614 | 0.00328824 | 0.00283709 | 0.00402743 | -0.0011903 | 0.90556652 | 0.93732239 |
| Cd2 | 0.00939363 | 0.01026626 | 0.01145782 | -0.0011916 | 0.67807557 | 0.90650187 |
| Gm29107 | 0.00349213 | 0.00308442 | 0.00427656 | -0.0011921 | 0.89237648 | 0.93159913 |
| Grik3 | 0.02082765 | 0.02401735 | 0.02520953 | -0.0011922 | 0.58325844 | 0.90650187 |
| Orc1 | 0.0244975 | 0.02838297 | 0.02957864 | -0.0011957 | 0.57130857 | 0.90650187 |
| Gm26527 | 0.00576821 | 0.00585531 | 0.00705302 | -0.0011977 | 0.77485571 | 0.90650187 |
| AI839979 | 0.00409964 | 0.0038198 | 0.00501847 | -0.0011987 | 0.85584573 | 0.9165432 |
| Zfhx2os | 0.00409726 | 0.00381604 | 0.00501556 | -0.0011995 | 0.85615641 | 0.9165432 |
| Capn12 | 0.00682913 | 0.00714273 | 0.00834426 | -0.0012015 | 0.73896139 | 0.90650187 |
| 8030462N17 | 0.01193464 | 0.01332761 | 0.01453224 | -0.0012046 | 0.64375473 | 0.90650187 |
| Pde6h | 0.00925171 | 0.01007743 | 0.0112858 | -0.0012084 | 0.68295985 | 0.90650187 |
| D930032P07 | 0.00338877 | 0.0029392 | 0.00415028 | -0.0012111 | 0.90272905 | 0.93635703 |
| Hist1h3f | 0.00528114 | 0.00524603 | 0.0064596 | -0.0012136 | 0.79819808 | 0.9075779 |
| Slmo1 | 0.00873753 | 0.00944745 | 0.01066228 | -0.0012148 | 0.69375474 | 0.90650187 |
| H2-DMa | 0.05318457 | 0.06178855 | 0.0630062 | -0.0012177 | 0.5342332 | 0.90650187 |
| Prr33 | 0.01129935 | 0.01254632 | 0.01376458 | -0.0012183 | 0.65301211 | 0.90650187 |
| Grifin | 0.00385189 | 0.00349684 | 0.00471599 | -0.0012192 | 0.87477396 | 0.92388237 |
| Bcas3os1 | 0.00610312 | 0.00623921 | 0.00746085 | -0.0012216 | 0.76666869 | 0.90650187 |
| Cbln3 | 0.00675902 | 0.00703701 | 0.00825899 | -0.001222 | 0.74465375 | 0.90650187 |
| Lcat | 0.01626561 | 0.01852412 | 0.01974813 | -0.001224 | 0.60855016 | 0.90650187 |
| BC024139 | 0.00404976 | 0.00373203 | 0.00495759 | -0.0012256 | 0.86411423 | 0.92013888 |
| Dock4 | 0.00323437 | 0.00273172 | 0.0039616 | -0.0012299 | 0.91622798 | 0.94231902 |
| Gm38393 | 0.00618103 | 0.00632153 | 0.00755568 | -0.0012342 | 0.7661274 | 0.90650187 |
| Myo7b | 0.01678367 | 0.01913451 | 0.02037001 | -0.0012355 | 0.60628037 | 0.90650187 |
| Icam5 | 0.00308464 | 0.00254172 | 0.00377859 | -0.0012369 | 0.92719458 | 0.94840756 |
| Itga11 | 0.00905916 | 0.00981121 | 0.01105236 | -0.0012411 | 0.69119053 | 0.90650187 |
| Lzts3 | 0.0114057 | 0.01265086 | 0.01389314 | -0.0012423 | 0.65450484 | 0.90650187 |

| | | | | | | |
|---|---|---|---|---|---|---|
| Ptprj | 0.12027979 | 0.13537219 | 0.13661461 | -0.0012424 | 0.51612477 | 0.90650187 |
| Far2 | 0.01304759 | 0.01463228 | 0.0158755 | -0.0012432 | 0.63613228 | 0.90650187 |
| Zfp759 | 0.01932305 | 0.02216862 | 0.02341201 | -0.0012434 | 0.59332333 | 0.90650187 |
| Gm43714 | 0.00683449 | 0.00710667 | 0.00835077 | -0.0012441 | 0.74611324 | 0.90650187 |
| Slc39a4 | 0.01170411 | 0.0130075 | 0.01425375 | -0.0012463 | 0.65124894 | 0.90650187 |
| Tekt5 | 0.00873177 | 0.00940793 | 0.0106553 | -0.0012474 | 0.69861613 | 0.90650187 |
| Slc38a7 | 0.05289038 | 0.06142037 | 0.06266978 | -0.0012494 | 0.53531179 | 0.90650187 |
| Kctd7 | 0.04133663 | 0.04810376 | 0.04935425 | -0.0012505 | 0.54484137 | 0.90650187 |
| Adam4 | 0.00401801 | 0.00366749 | 0.00491882 | -0.0012513 | 0.87096189 | 0.92222706 |
| Gm42556 | 0.01154463 | 0.01280499 | 0.01406105 | -0.0012561 | 0.65436043 | 0.90650187 |
| Gm16701.1 | 0.00395025 | 0.00357978 | 0.00483609 | -0.0012563 | 0.87592867 | 0.92433126 |
| Srpk3 | 0.00662607 | 0.00684087 | 0.00809726 | -0.0012564 | 0.75483824 | 0.90650187 |
| 5530401A14 | 0.00607336 | 0.00616595 | 0.00742461 | -0.0012587 | 0.77446758 | 0.90650187 |
| Lor | 0.00394337 | 0.00356873 | 0.00482769 | -0.001259 | 0.87683628 | 0.92473192 |
| Sv2b | 0.01712457 | 0.01951992 | 0.02077898 | -0.0012591 | 0.60617876 | 0.90650187 |
| Nr1i2 | 0.00524074 | 0.00514937 | 0.00641036 | -0.001261 | 0.80907665 | 0.9092537 |
| Apoa1 | 0.00395408 | 0.00357727 | 0.00484076 | -0.0012635 | 0.87704823 | 0.9247416 |
| Fut4 | 0.02068612 | 0.0237761 | 0.0250406 | -0.0012645 | 0.58881372 | 0.90650187 |
| Spag9 | 0.47365671 | 0.44534541 | 0.44661011 | -0.0012647 | 0.5050221 | 0.90650187 |
| Krt81 | 0.00682354 | 0.00707111 | 0.00833746 | -0.0012663 | 0.75023464 | 0.90650187 |
| Adamts17 | 0.05433605 | 0.06305406 | 0.06432169 | -0.0012676 | 0.53490778 | 0.90650187 |
| Sstr3 | 0.09364915 | 0.10680385 | 0.10807189 | -0.001268 | 0.52080007 | 0.90650187 |
| Xk | 0.0394501 | 0.04589096 | 0.0471607 | -0.0012697 | 0.54763661 | 0.90650187 |
| Akr1c14 | 0.01008504 | 0.01102517 | 0.01229542 | -0.0012702 | 0.67698266 | 0.90650187 |
| Kbtbd8 | 0.02364687 | 0.02729696 | 0.02856784 | -0.0012709 | 0.57838744 | 0.90650187 |
| Ypel2 | 0.27511906 | 0.28548607 | 0.28675797 | -0.0012719 | 0.50786598 | 0.90650187 |
| Cpa6 | 0.00397535 | 0.00359469 | 0.00486673 | -0.001272 | 0.87737536 | 0.92482217 |
| Synpo2l | 0.00580089 | 0.00582066 | 0.00709282 | -0.0012722 | 0.78737958 | 0.90650187 |
| Gm4319 | 0.00913168 | 0.00986808 | 0.01114028 | -0.0012722 | 0.69416082 | 0.90650187 |
| Pcdh8 | 0.03361472 | 0.03906705 | 0.04034076 | -0.0012737 | 0.55581395 | 0.90650187 |

| | | | | | | |
|---|---|---|---|---|---|---|
| Gm13594 | 0.01043664 | 0.01144644 | 0.01272106 | -0.0012746 | 0.67199911 | 0.90650187 |
| 3110021N24 | 0.02373832 | 0.02740131 | 0.02867656 | -0.0012753 | 0.57835932 | 0.90650187 |
| Shroom2 | 0.08473658 | 0.09707137 | 0.09834737 | -0.001276 | 0.52299782 | 0.90650187 |
| Gm14434 | 0.00573545 | 0.00573676 | 0.00701312 | -0.0012764 | 0.79076617 | 0.90650187 |
| Dydc2 | 0.00357117 | 0.00309618 | 0.00437312 | -0.0012769 | 0.90287503 | 0.93635703 |
| Slc34a2 | 0.00344188 | 0.00293778 | 0.00421517 | -0.0012774 | 0.91104712 | 0.94011454 |
| Gm15917 | 0.00954313 | 0.01035946 | 0.011639 | -0.0012795 | 0.68748333 | 0.90650187 |
| Fzd7 | 0.05325878 | 0.06180973 | 0.06309105 | -0.0012813 | 0.53597006 | 0.90650187 |
| Zkscan16 | 0.00659466 | 0.00677745 | 0.00805904 | -0.0012816 | 0.76020561 | 0.90650187 |
| Steap3 | 0.02169741 | 0.02496403 | 0.02624696 | -0.0012829 | 0.58601148 | 0.90650187 |
| Gm15892 | 0.00395808 | 0.00356099 | 0.00484565 | -0.0012847 | 0.88072114 | 0.92622922 |
| Zfp82 | 0.00671338 | 0.00691668 | 0.00820347 | -0.0012868 | 0.75720312 | 0.90650187 |
| Krtap16-1 | 0.00617088 | 0.00625561 | 0.00754333 | -0.0012877 | 0.77604623 | 0.90650187 |
| Pms1 | 0.04473931 | 0.05200792 | 0.0532969 | -0.001289 | 0.54281036 | 0.90650187 |
| 1700055D18 | 0.02102827 | 0.02415716 | 0.02544893 | -0.0012918 | 0.58926491 | 0.90650187 |
| Zfp54 | 0.05301269 | 0.06151378 | 0.06280966 | -0.0012959 | 0.53654042 | 0.90650187 |
| Ncapg2 | 0.07327735 | 0.08438211 | 0.08567911 | -0.001297 | 0.52682732 | 0.90650187 |
| Polr1b | 0.07818266 | 0.08982604 | 0.09112493 | -0.0012989 | 0.52526295 | 0.90650187 |
| Mfsd7c | 0.01414499 | 0.01589907 | 0.01719802 | -0.0012989 | 0.63147872 | 0.90650187 |
| Tub | 0.00315166 | 0.00256101 | 0.00386051 | -0.0012995 | 0.9327326 | 0.95188026 |
| Gm12519 | 0.00586257 | 0.00586837 | 0.00716795 | -0.0012996 | 0.78988086 | 0.90650187 |
| Rbm20 | 0.00339105 | 0.00285163 | 0.00415307 | -0.0013014 | 0.9182078 | 0.94321373 |
| Prkg1 | 0.01844237 | 0.02105565 | 0.02235819 | -0.0013025 | 0.60217903 | 0.90650187 |
| Bcat2 | 0.20340394 | 0.21910272 | 0.22040777 | -0.001305 | 0.51050002 | 0.90650187 |
| Lamb3 | 0.01159333 | 0.01281252 | 0.0141199 | -0.0013074 | 0.65969311 | 0.90650187 |
| Slc39a2 | 0.00845982 | 0.0090164 | 0.01032533 | -0.0013089 | 0.71347496 | 0.90650187 |
| Gm29019 | 0.00390613 | 0.0034728 | 0.00478222 | -0.0013094 | 0.88824533 | 0.92996882 |
| Npepps | 0.78122807 | 0.64062857 | 0.64193803 | -0.0013095 | 0.50361774 | 0.90650187 |
| Slc2a4 | 0.01864322 | 0.0212884 | 0.02259864 | -0.0013102 | 0.60169984 | 0.90650187 |
| Mab21l1 | 0.01075729 | 0.01179603 | 0.01310906 | -0.001313 | 0.67194092 | 0.90650187 |

| | | | | | | |
|---|---|---|---|---|---|---|
| Camkmt | 0.0182273 | 0.02078755 | 0.02210065 | -0.0013131 | 0.60416133 | 0.90650187 |
| Fbxo25 | 0.16978787 | 0.18615055 | 0.18746487 | -0.0013143 | 0.5124323 | 0.90650187 |
| Agpat1 | 0.31604817 | 0.32109473 | 0.32240924 | -0.0013145 | 0.50723054 | 0.90650187 |
| Cdh22 | 0.02255385 | 0.02595158 | 0.02726732 | -0.0013157 | 0.58492748 | 0.90650187 |
| Pnpla7 | 0.05727512 | 0.06635427 | 0.0676704 | -0.0013161 | 0.53445086 | 0.90650187 |
| Sec14l4 | 0.01044331 | 0.01141111 | 0.01272913 | -0.001318 | 0.67735532 | 0.90650187 |
| Rbm11 | 0.00939246 | 0.01013824 | 0.0114564 | -0.0013182 | 0.6955018 | 0.90650187 |
| Pld2 | 0.07128704 | 0.08214005 | 0.0834596 | -0.0013195 | 0.52801771 | 0.90650187 |
| Gsg2 | 0.04146477 | 0.04818202 | 0.04950306 | -0.001321 | 0.54721779 | 0.90650187 |
| Slc6a9 | 0.02637822 | 0.03048638 | 0.03180941 | -0.001323 | 0.57334793 | 0.90650187 |
| Nos1ap | 0.05344394 | 0.0619783 | 0.06330269 | -0.0013244 | 0.53705161 | 0.90650187 |
| Npw | 0.00649361 | 0.00661077 | 0.0079361 | -0.0013253 | 0.77108339 | 0.90650187 |
| Plk5 | 0.00387975 | 0.00342391 | 0.00475001 | -0.0013261 | 0.89271869 | 0.93179124 |
| Nmnat2 | 0.00555791 | 0.00546991 | 0.00679685 | -0.0013269 | 0.80727885 | 0.9092537 |
| Slc13a4 | 0.00784194 | 0.00824825 | 0.00957521 | -0.001327 | 0.73108018 | 0.90650187 |
| Ttc34 | 0.00672085 | 0.00688441 | 0.00821256 | -0.0013281 | 0.76391373 | 0.90650187 |
| Greb1l | 0.02647912 | 0.03060076 | 0.03192893 | -0.0013282 | 0.57335709 | 0.90650187 |
| Trp53cor1 | 0.0391872 | 0.04552495 | 0.04685458 | -0.0013296 | 0.55019597 | 0.90650187 |
| Lrrc36 | 0.0037347 | 0.0032432 | 0.00457287 | -0.0013297 | 0.90193684 | 0.93591959 |
| Gpr21 | 0.01083857 | 0.01187694 | 0.01320738 | -0.0013304 | 0.6728606 | 0.90650187 |
| Stk39 | 0.1243168 | 0.13953933 | 0.1408702 | -0.0013309 | 0.51675053 | 0.90650187 |
| Slc35f4 | 0.00670942 | 0.00686727 | 0.00819866 | -0.0013314 | 0.76482962 | 0.90650187 |
| Gm26789 | 0.00388021 | 0.0034186 | 0.00475057 | -0.001332 | 0.89370306 | 0.93189602 |
| Fgf2 | 0.03298773 | 0.03827099 | 0.03960483 | -0.0013338 | 0.55950816 | 0.90650187 |
| Lrrc2 | 0.00375446 | 0.00326106 | 0.004597 | -0.0013359 | 0.90181146 | 0.93591959 |
| Gm26734 | 0.01499896 | 0.01688885 | 0.01822582 | -0.001337 | 0.62782947 | 0.90650187 |
| Fam134b | 0.12844416 | 0.14386249 | 0.14520009 | -0.0013376 | 0.51633347 | 0.90650187 |
| Gm16279 | 0.00594306 | 0.00592787 | 0.00726597 | -0.0013381 | 0.79352133 | 0.90731056 |
| Nrsn2 | 0.00894768 | 0.00957847 | 0.01091717 | -0.0013387 | 0.70716968 | 0.90650187 |
| Nuak1 | 0.08081986 | 0.0926992 | 0.09403847 | -0.0013393 | 0.52524138 | 0.90650187 |

| | | | | | | |
|---|---|---|---|---|---|---|
| Lama1 | 0.00888582 | 0.00950245 | 0.01084215 | -0.0013397 | 0.70860687 | 0.90650187 |
| Gm17102 | 0.00669079 | 0.00683417 | 0.00817599 | -0.0013418 | 0.76718253 | 0.90650187 |
| Gli3 | 0.06073321 | 0.07024837 | 0.07159398 | -0.0013456 | 0.53329498 | 0.90650187 |
| Prr29 | 0.00676455 | 0.00691825 | 0.00826571 | -0.0013475 | 0.76568562 | 0.90650187 |
| Ooep | 0.01130391 | 0.01242213 | 0.01377009 | -0.001348 | 0.6682856 | 0.90650187 |
| Ccer2 | 0.00837306 | 0.00886936 | 0.01022004 | -0.0013507 | 0.72157627 | 0.90650187 |
| RP23-231J2.1 | 0.00357953 | 0.00303219 | 0.00438334 | -0.0013512 | 0.91471107 | 0.94165806 |
| 4933424G06 | 0.00357476 | 0.00302605 | 0.00437752 | -0.0013515 | 0.91504456 | 0.94181003 |
| Gm7932 | 0.00564645 | 0.00554931 | 0.00690471 | -0.0013554 | 0.80858114 | 0.9092537 |
| Ttll11 | 0.01252958 | 0.01389309 | 0.01525055 | -0.0013575 | 0.65383868 | 0.90650187 |
| 2810425M01 | 0.00721024 | 0.00744919 | 0.00880764 | -0.0013585 | 0.75354017 | 0.90650187 |
| B3galt5 | 0.00342547 | 0.00283434 | 0.00419512 | -0.0013608 | 0.92535301 | 0.94735258 |
| 9430037G07 | 0.00653928 | 0.00663 | 0.00799167 | -0.0013617 | 0.77561451 | 0.90650187 |
| Oacyl | 0.00650944 | 0.00659291 | 0.00795536 | -0.0013624 | 0.77677982 | 0.90650187 |
| Asprv1 | 0.00364747 | 0.00310359 | 0.00446633 | -0.0013627 | 0.91251636 | 0.9407366 |
| Prps1l3 | 0.0521736 | 0.06048483 | 0.06184959 | -0.0013648 | 0.53907181 | 0.90650187 |
| Taf4b | 0.00379361 | 0.00327741 | 0.00464482 | -0.0013674 | 0.90469244 | 0.93703811 |
| Wisp3 | 0.00380725 | 0.00329362 | 0.00466148 | -0.0013679 | 0.90397162 | 0.93662822 |
| A430106G13 | 0.00633668 | 0.00637584 | 0.00774514 | -0.0013693 | 0.78405406 | 0.90650187 |
| RP24-143K11 | 0.00378963 | 0.00326958 | 0.00463995 | -0.0013704 | 0.90540483 | 0.93721885 |
| C030034L19R | 0.00580832 | 0.00573026 | 0.00710187 | -0.0013716 | 0.80471898 | 0.90882608 |
| Bzrap1 | 0.00380309 | 0.0032837 | 0.0046564 | -0.0013727 | 0.90499645 | 0.93712658 |
| Grin1 | 0.00350804 | 0.00292288 | 0.004296 | -0.0013731 | 0.92233329 | 0.94557609 |
| Gm28800 | 0.00352833 | 0.00294669 | 0.0043208 | -0.0013741 | 0.92128886 | 0.94542037 |
| Adam8 | 0.00934469 | 0.01002414 | 0.0113985 | -0.0013744 | 0.7040294 | 0.90650187 |
| Fbxo41 | 0.00344673 | 0.00284608 | 0.00422109 | -0.001375 | 0.92621176 | 0.94759147 |
| Ugt2b35 | 0.00887583 | 0.00945364 | 0.01083003 | -0.0013764 | 0.71396075 | 0.90650187 |
| Gm16712 | 0.00588967 | 0.00582262 | 0.00720095 | -0.0013783 | 0.80259627 | 0.90870445 |
| Rasl10a | 0.01709912 | 0.0193691 | 0.02074846 | -0.0013794 | 0.61621096 | 0.90650187 |
| Bbs4 | 0.16784489 | 0.18414275 | 0.18552255 | -0.0013798 | 0.51318812 | 0.90650187 |

| | | | | | | |
|---|---|---|---|---|---|---|
| Eda2r | 0.01705435 | 0.01931292 | 0.02069476 | -0.0013818 | 0.61670728 | 0.90650187 |
| 9030612E09I | 0.01232173 | 0.01361623 | 0.01499966 | -0.0013834 | 0.65910563 | 0.90650187 |
| Gm42903 | 0.00333738 | 0.00270327 | 0.00408749 | -0.0013842 | 0.93390155 | 0.95257582 |
| Coprs | 0.30773619 | 0.31390726 | 0.31529192 | -0.0013847 | 0.50778834 | 0.90650187 |
| Nell2 | 0.01261343 | 0.01396624 | 0.01535174 | -0.0013855 | 0.6558697 | 0.90650187 |
| Gm572 | 0.01466275 | 0.01643555 | 0.01782131 | -0.0013858 | 0.63521055 | 0.90650187 |
| Lrrc7 | 0.00656868 | 0.00664051 | 0.00802744 | -0.0013869 | 0.77878116 | 0.90650187 |
| Ackr1 | 0.00852495 | 0.00901734 | 0.01040437 | -0.001387 | 0.72329264 | 0.90650187 |
| Dnah2os | 0.00350928 | 0.00291002 | 0.00429752 | -0.0013875 | 0.92440088 | 0.94682419 |
| D5Ertd605e | 0.01051688 | 0.01143052 | 0.01281817 | -0.0013876 | 0.68483569 | 0.90650187 |
| Cd109 | 0.01211961 | 0.01336598 | 0.01475562 | -0.0013896 | 0.66227335 | 0.90650187 |
| Gm12689 | 0.00371581 | 0.0031566 | 0.0045498 | -0.0013932 | 0.91328945 | 0.94082155 |
| Gm26660 | 0.00323468 | 0.00256817 | 0.00396198 | -0.0013938 | 0.94108485 | 0.95726919 |
| Plek2 | 0.04619099 | 0.0535775 | 0.05497356 | -0.0013961 | 0.54494394 | 0.90650187 |
| Vmn1r90 | 0.00608269 | 0.00603752 | 0.00743597 | -0.0013985 | 0.7984418 | 0.90770671 |
| Gm29417 | 0.00338338 | 0.00274323 | 0.0041437 | -0.0014005 | 0.93351622 | 0.95231059 |
| Ccdc65 | 0.00941725 | 0.01008449 | 0.01148644 | -0.001402 | 0.70629742 | 0.90650187 |
| RP23-36D15. | 0.00362633 | 0.00303769 | 0.0044405 | -0.0014028 | 0.91990333 | 0.94435798 |
| Tlx1 | 0.00859214 | 0.00908233 | 0.0104859 | -0.0014036 | 0.72409684 | 0.90650187 |
| Pxn | 0.23990809 | 0.25341957 | 0.25482397 | -0.0014044 | 0.50977344 | 0.90650187 |
| Cct6b | 0.00606064 | 0.00600344 | 0.00740912 | -0.0014057 | 0.80050718 | 0.90806412 |
| Gm3636 | 0.0037356 | 0.0031678 | 0.00457397 | -0.0014062 | 0.91414255 | 0.9413278 |
| Cfap53 | 0.0060812 | 0.00602706 | 0.00743416 | -0.0014071 | 0.79994987 | 0.90805731 |
| 1700034H15I | 0.0028202 | 0.00204716 | 0.00345528 | -0.0014081 | 0.96498418 | 0.97343716 |
| Gm27000 | 0.00662624 | 0.0066893 | 0.00809746 | -0.0014082 | 0.78026746 | 0.90650187 |
| Naip5 | 0.00620269 | 0.00616841 | 0.00758205 | -0.0014136 | 0.79640964 | 0.9075779 |
| Ccdc82 | 0.10655813 | 0.12058598 | 0.12200079 | -0.0014148 | 0.5205582 | 0.90650187 |
| Atp8b3 | 0.0048637 | 0.00453461 | 0.0059507 | -0.0014161 | 0.85495346 | 0.91627438 |
| Kif6 | 0.00338536 | 0.00272912 | 0.00414611 | -0.001417 | 0.93566257 | 0.9537321 |
| Il17d | 0.01988471 | 0.0226664 | 0.02408344 | -0.001417 | 0.60317517 | 0.90650187 |

| | | | | | | |
|---|---|---|---|---|---|---|
| Mep1a | 0.00620132 | 0.00616228 | 0.00758038 | -0.0014181 | 0.79719913 | 0.9075779 |
| Crb2 | 0.01359297 | 0.0151132 | 0.01653301 | -0.0014198 | 0.64868575 | 0.90650187 |
| Ephx1 | 0.08601877 | 0.09833057 | 0.09975323 | -0.0014227 | 0.52527694 | 0.90650187 |
| Pygm | 0.04444096 | 0.05152868 | 0.05295192 | -0.0014232 | 0.54755598 | 0.90650187 |
| Gm16222 | 0.00348093 | 0.00283854 | 0.00426289 | -0.0014243 | 0.93128098 | 0.95113723 |
| Pcdhb4 | 0.00863769 | 0.00911505 | 0.01054116 | -0.0014261 | 0.72622745 | 0.90650187 |
| Gucy1a2 | 0.01348222 | 0.01497245 | 0.01639953 | -0.0014271 | 0.65056759 | 0.90650187 |
| Gm26690 | 0.00620328 | 0.00615467 | 0.00758276 | -0.0014281 | 0.79877712 | 0.90777025 |
| Gm13420 | 0.01610852 | 0.01812859 | 0.01955948 | -0.0014309 | 0.62749295 | 0.90650187 |
| RP23-58K20. | 0.00652846 | 0.00654752 | 0.0079785 | -0.001431 | 0.78737387 | 0.90650187 |
| Cisd3 | 0.41493894 | 0.40116448 | 0.40259831 | -0.0014338 | 0.50631607 | 0.90650187 |
| Tmem88b | 0.01453167 | 0.01622758 | 0.01766356 | -0.001436 | 0.64110325 | 0.90650187 |
| 1700018L02F | 0.01601536 | 0.01801023 | 0.01944757 | -0.0014373 | 0.62875862 | 0.90650187 |
| Slc6a2 | 0.00307489 | 0.0023283 | 0.00376667 | -0.0014384 | 0.955211 | 0.96646449 |
| Cdkl3 | 0.05376412 | 0.0622298 | 0.06366855 | -0.0014387 | 0.54001005 | 0.90650187 |
| Pcolce2 | 0.01048664 | 0.01134137 | 0.01278157 | -0.0014402 | 0.69178697 | 0.90650187 |
| Acaa1b | 0.00661686 | 0.00664309 | 0.00808605 | -0.001443 | 0.78620406 | 0.90650187 |
| Asic1 | 0.00330038 | 0.00259756 | 0.00404226 | -0.0014447 | 0.94395439 | 0.95903292 |
| Chad | 0.00635026 | 0.00630926 | 0.00776167 | -0.0014524 | 0.79726318 | 0.9075779 |
| Pla2g3 | 0.00600538 | 0.00588923 | 0.00734184 | -0.0014526 | 0.81045673 | 0.9092537 |
| Abcc1 | 0.07628639 | 0.0875691 | 0.08902381 | -0.0014547 | 0.52895542 | 0.90650187 |
| Ermap | 0.00663302 | 0.00665084 | 0.00810571 | -0.0014549 | 0.78754476 | 0.90650187 |
| Gnb3 | 0.02609247 | 0.03001424 | 0.03147084 | -0.0014566 | 0.58151139 | 0.90650187 |
| Ddx60 | 0.00360525 | 0.0029557 | 0.00441476 | -0.0014591 | 0.92911785 | 0.94996709 |
| Tmc7 | 0.00289587 | 0.0020883 | 0.00354781 | -0.0014595 | 0.96628842 | 0.97444265 |
| Cdh24 | 0.01176478 | 0.01286326 | 0.01432705 | -0.0014638 | 0.67515929 | 0.90650187 |
| Steap1 | 0.09946481 | 0.11289149 | 0.1143578 | -0.0014663 | 0.52272828 | 0.90650187 |
| Gm17087 | 0.00579215 | 0.00561482 | 0.00708218 | -0.0014674 | 0.8214969 | 0.9092537 |
| Fbxo30 | 0.13615198 | 0.15176154 | 0.15323139 | -0.0014699 | 0.51700731 | 0.90650187 |
| Gm26513 | 0.00642335 | 0.00637903 | 0.00785061 | -0.0014716 | 0.79766801 | 0.9075779 |

| | | | | | | |
|---|---|---|---|---|---|---|
| Gm9913 | 0.00557594 | 0.00534721 | 0.00681882 | -0.0014716 | 0.83132961 | 0.9098597 |
| RP23-378L12 | 0.00316443 | 0.00240438 | 0.00387611 | -0.0014717 | 0.95429063 | 0.96585471 |
| Cfap74 | 0.00812476 | 0.00844478 | 0.00991864 | -0.0014739 | 0.7455635 | 0.90650187 |
| Fkbp6 | 0.0032895 | 0.00255495 | 0.00402897 | -0.001474 | 0.94807317 | 0.96173909 |
| Il34 | 0.0162583 | 0.01826432 | 0.01973936 | -0.001475 | 0.63013153 | 0.90650187 |
| Pmel | 0.00328739 | 0.00255118 | 0.0040264 | -0.0014752 | 0.94832404 | 0.96189773 |
| Zfp820 | 0.08902565 | 0.10156468 | 0.10304106 | -0.0014764 | 0.52539427 | 0.90650187 |
| Muc20 | 0.00643613 | 0.00638901 | 0.00786615 | -0.0014771 | 0.79809025 | 0.9075779 |
| Psg16 | 0.00324487 | 0.00249594 | 0.00397443 | -0.0014785 | 0.95091231 | 0.96348599 |
| Rnf180 | 0.00606643 | 0.00593619 | 0.00741617 | -0.00148 | 0.81250581 | 0.9092537 |
| Fstl4 | 0.00653489 | 0.00650599 | 0.00798632 | -0.0014803 | 0.79503827 | 0.90731056 |
| Gpatch3 | 0.0468648 | 0.05426997 | 0.0557507 | -0.0014807 | 0.54699569 | 0.90650187 |
| Acan | 0.00636376 | 0.00629715 | 0.00777809 | -0.0014809 | 0.80134537 | 0.90823524 |
| Sspo | 0.00279386 | 0.001942 | 0.00342307 | -0.0014811 | 0.97279127 | 0.97935527 |
| 4930515G01 | 0.01034211 | 0.01112507 | 0.01260664 | -0.0014816 | 0.6993246 | 0.90650187 |
| Shoc2 | 0.74658501 | 0.62098743 | 0.62246962 | -0.0014822 | 0.50422299 | 0.90650187 |
| Dlx1 | 0.00919093 | 0.00972894 | 0.01121212 | -0.0014832 | 0.72176061 | 0.90650187 |
| Dusp27 | 0.00272809 | 0.0018567 | 0.00334264 | -0.0014859 | 0.97593662 | 0.98135957 |
| Rbm41 | 0.03995414 | 0.04625979 | 0.0477473 | -0.0014875 | 0.55507633 | 0.90650187 |
| Areg | 0.00549592 | 0.00523253 | 0.00672132 | -0.0014888 | 0.83761703 | 0.91121886 |
| Sntb1 | 0.0138903 | 0.01539792 | 0.01689126 | -0.0014933 | 0.65285208 | 0.90650187 |
| Ccsap | 0.03623723 | 0.04191856 | 0.04341229 | -0.0014937 | 0.56078667 | 0.90650187 |
| Moxd1 | 0.00485945 | 0.00445132 | 0.00594551 | -0.0014942 | 0.86805537 | 0.9208814 |
| Pgc | 0.00640367 | 0.00632965 | 0.00782666 | -0.001497 | 0.80242379 | 0.90870058 |
| Slc6a8 | 0.0754764 | 0.08662711 | 0.08812474 | -0.0014976 | 0.53011168 | 0.90650187 |
| Nek11 | 0.00558948 | 0.00533355 | 0.00683532 | -0.0015018 | 0.83564832 | 0.91078297 |
| Gm9916 | 0.00333847 | 0.00258592 | 0.00408881 | -0.0015029 | 0.94887397 | 0.96195723 |
| B230334C09 | 0.00333847 | 0.00258592 | 0.00408881 | -0.0015029 | 0.94887397 | 0.96195723 |
| Lax1 | 0.00294651 | 0.00210569 | 0.00360972 | -0.001504 | 0.96800996 | 0.97564776 |
| Sox6os | 0.00482282 | 0.00439289 | 0.00590084 | -0.0015079 | 0.87203555 | 0.92254675 |

| | | | | | | |
|---|---|---|---|---|---|---|
| Caly | 0.01116796 | 0.01209761 | 0.01360573 | -0.0015081 | 0.68891355 | 0.90650187 |
| Rgs8 | 0.00568974 | 0.00544675 | 0.00695745 | -0.0015107 | 0.83280057 | 0.91035535 |
| Pfkfb1 | 0.0146734 | 0.01632284 | 0.01783413 | -0.0015113 | 0.64681156 | 0.90650187 |
| Gm5577 | 0.02559708 | 0.02937215 | 0.03088357 | -0.0015114 | 0.58611576 | 0.90650187 |
| Slc30a4 | 0.11256301 | 0.12691284 | 0.12842468 | -0.0015118 | 0.52086895 | 0.90650187 |
| Cyp4f17 | 0.0033195 | 0.00255305 | 0.00406563 | -0.0015126 | 0.95093073 | 0.96348599 |
| Ttc30a2 | 0.00615432 | 0.00600993 | 0.00752317 | -0.0015132 | 0.81439442 | 0.9092537 |
| Plcg1 | 0.12253792 | 0.13748265 | 0.13899745 | -0.0015148 | 0.51932043 | 0.90650187 |
| Gm10767 | 0.01604514 | 0.01796677 | 0.01948335 | -0.0015166 | 0.63534537 | 0.90650187 |
| 5031425F14I | 0.00655693 | 0.00649434 | 0.00801314 | -0.0015188 | 0.80027753 | 0.90806412 |
| Rab38 | 0.01258511 | 0.0137985 | 0.01531757 | -0.0015191 | 0.67034878 | 0.90650187 |
| Gm5087 | 0.00929929 | 0.00982288 | 0.01134348 | -0.0015206 | 0.72439055 | 0.90650187 |
| Wdr49 | 0.0090375 | 0.00950537 | 0.01102609 | -0.0015207 | 0.73010792 | 0.90650187 |
| Mei4 | 0.00291861 | 0.0020538 | 0.00357561 | -0.0015218 | 0.97076053 | 0.97777172 |
| A830082K12 | 0.01730369 | 0.01946933 | 0.02099379 | -0.0015245 | 0.62658363 | 0.90650187 |
| Slc22a15 | 0.08749629 | 0.09984562 | 0.10137038 | -0.0015248 | 0.52665684 | 0.90650187 |
| Gab3 | 0.03713853 | 0.0429406 | 0.04446543 | -0.0015248 | 0.56058429 | 0.90650187 |
| Aass | 0.00497291 | 0.00455872 | 0.00608386 | -0.0015251 | 0.86745681 | 0.92056229 |
| Xkr4 | 0.00890799 | 0.00934048 | 0.01086903 | -0.0015285 | 0.73408034 | 0.90650187 |
| Nlrc3 | 0.01358733 | 0.01499704 | 0.01652621 | -0.0015292 | 0.65959236 | 0.90650187 |
| Tekt1 | 0.00932523 | 0.0098447 | 0.01137492 | -0.0015302 | 0.72509599 | 0.90650187 |
| Natd1 | 0.08094721 | 0.09264869 | 0.09417891 | -0.0015302 | 0.52879147 | 0.90650187 |
| Kcnc1 | 0.0063321 | 0.0062087 | 0.00773956 | -0.0015309 | 0.810389 | 0.9092537 |
| Cpne4 | 0.0019887 | 0.00090672 | 0.00243791 | -0.0015312 | 0.99738229 | 0.99757917 |
| Nr1h3 | 0.03057653 | 0.03523689 | 0.03676896 | -0.0015321 | 0.57347852 | 0.90650187 |
| Elf5 | 0.01317399 | 0.01449358 | 0.01602793 | -0.0015343 | 0.66479167 | 0.90650187 |
| Gm16118 | 0.00527517 | 0.00491582 | 0.00645232 | -0.0015365 | 0.85511775 | 0.91636552 |
| Zar1l | 0.00198784 | 0.000897 | 0.00243686 | -0.0015399 | 0.99751676 | 0.99764802 |
| Mettl5os | 0.01599461 | 0.01788201 | 0.01942266 | -0.0015406 | 0.63781911 | 0.90650187 |
| Gm15821 | 0.00319204 | 0.00236907 | 0.00390986 | -0.0015408 | 0.96010568 | 0.9701425 |

| | | | | | | |
|---|---|---|---|---|---|---|
| Hapln3 | 0.00645489 | 0.00634513 | 0.00788898 | -0.0015439 | 0.80784863 | 0.9092537 |
| Apol7d | 0.00907088 | 0.0095225 | 0.01106657 | -0.0015441 | 0.73246198 | 0.90650187 |
| Necab3 | 0.00327895 | 0.00247106 | 0.00401608 | -0.001545 | 0.95639169 | 0.96707976 |
| Gckr | 0.00310883 | 0.00226258 | 0.00380816 | -0.0015456 | 0.9644072 | 0.97298426 |
| Hrasls | 0.00586955 | 0.00562989 | 0.00717645 | -0.0015466 | 0.83098086 | 0.9098597 |
| Klf1 | 0.00314751 | 0.00230817 | 0.00385544 | -0.0015473 | 0.96279718 | 0.97181141 |
| S100a3 | 0.00650991 | 0.00640844 | 0.00795594 | -0.0015475 | 0.80640031 | 0.9092537 |
| Car6 | 0.00992205 | 0.01055052 | 0.01209803 | -0.0015475 | 0.71520614 | 0.90650187 |
| Gm53 | 0.00203212 | 0.00094327 | 0.00249107 | -0.0015478 | 0.99712952 | 0.99739197 |
| Clvs1 | 0.00622993 | 0.00606716 | 0.00761521 | -0.0015481 | 0.81692869 | 0.9092537 |
| Gm13056 | 0.00203783 | 0.00094711 | 0.00249805 | -0.0015509 | 0.9971109 | 0.99739197 |
| Styx | 0.01338967 | 0.01473689 | 0.01628797 | -0.0015511 | 0.6639792 | 0.90650187 |
| Prlr | 0.01395684 | 0.01541893 | 0.01697141 | -0.0015525 | 0.65787404 | 0.90650187 |
| Gm10874 | 0.00806512 | 0.00829322 | 0.00984623 | -0.001553 | 0.75840695 | 0.90650187 |
| Cadm2 | 0.00265423 | 0.00169801 | 0.0032523 | -0.0015543 | 0.9831889 | 0.98715023 |
| Rab39 | 0.01717309 | 0.01928191 | 0.02083718 | -0.0015553 | 0.6299865 | 0.90650187 |
| Neb | 0.00848876 | 0.00880505 | 0.01036045 | -0.0015554 | 0.74774494 | 0.90650187 |
| Unc5d | 0.01081188 | 0.01161933 | 0.01317509 | -0.0015558 | 0.70019227 | 0.90650187 |
| Ak8 | 0.04468625 | 0.05167723 | 0.05323556 | -0.0015583 | 0.55176977 | 0.90650187 |
| Chrm1 | 0.00214056 | 0.00106448 | 0.0026238 | -0.0015593 | 0.99587926 | 0.99633807 |
| Gm43614 | 0.00625977 | 0.00608911 | 0.00765152 | -0.0015624 | 0.81800136 | 0.9092537 |
| Saraf | 1.36850982 | 0.86585208 | 0.86741506 | -0.001563 | 0.50319568 | 0.90650187 |
| Gm5218 | 0.00478722 | 0.00429442 | 0.00585742 | -0.001563 | 0.88224201 | 0.92700639 |
| Hnf1aos1 | 0.12449672 | 0.13949529 | 0.14105941 | -0.0015641 | 0.51965766 | 0.90650187 |
| Ctdp1 | 0.03738571 | 0.04318916 | 0.04475404 | -0.0015649 | 0.56176488 | 0.90650187 |
| Omd | 0.00243254 | 0.00141495 | 0.0029811 | -0.0015661 | 0.99024238 | 0.99265856 |
| Snap47 | 0.72812444 | 0.61029486 | 0.61186146 | -0.0015666 | 0.50454085 | 0.90650187 |
| Hoxaas3 | 0.02263156 | 0.02579205 | 0.02735985 | -0.0015678 | 0.6005404 | 0.90650187 |
| Aox1 | 0.00779264 | 0.00794748 | 0.00951532 | -0.0015678 | 0.76806953 | 0.90650187 |
| Slc1a4 | 0.03870951 | 0.04472967 | 0.04629808 | -0.0015684 | 0.55985467 | 0.90650187 |

| | | | | | | |
|---|---|---|---|---|---|---|
| P2ry2 | 0.01007867 | 0.01071902 | 0.01228771 | -0.0015687 | 0.71482629 | 0.90650187 |
| Gnal | 0.00611713 | 0.00590898 | 0.0074779 | -0.0015689 | 0.82451876 | 0.90934433 |
| Lhx9 | 0.00817046 | 0.00840505 | 0.00997411 | -0.0015691 | 0.75783471 | 0.90650187 |
| Fam19a2 | 0.0062824 | 0.0061092 | 0.00767907 | -0.0015699 | 0.81828233 | 0.9092537 |
| Lgi1 | 0.00296038 | 0.00205677 | 0.00362668 | -0.0015699 | 0.97284814 | 0.97935527 |
| Tmem35 | 0.01967835 | 0.02226584 | 0.02383681 | -0.001571 | 0.61523436 | 0.90650187 |
| Tcaf2 | 0.01526776 | 0.01697585 | 0.01854909 | -0.0015732 | 0.64693387 | 0.90650187 |
| Ptch2 | 0.03071353 | 0.03535454 | 0.03693034 | -0.0015758 | 0.57522447 | 0.90650187 |
| B430219N15 | 0.00555185 | 0.0052133 | 0.00678947 | -0.0015762 | 0.84897366 | 0.91444933 |
| Synpo2 | 0.0021678 | 0.00107964 | 0.00265715 | -0.0015775 | 0.99584601 | 0.99633807 |
| Angpt4 | 0.01091361 | 0.01172012 | 0.01329815 | -0.001578 | 0.70109035 | 0.90650187 |
| Cbfa2t2 | 0.24585155 | 0.2587216 | 0.26029979 | -0.0015782 | 0.51075162 | 0.90650187 |
| Dhx58 | 0.01363603 | 0.01500579 | 0.01658489 | -0.0015791 | 0.66395466 | 0.90650187 |
| Cep55 | 0.04897742 | 0.05660292 | 0.05818284 | -0.0015799 | 0.54804286 | 0.90650187 |
| Wfikkn1 | 0.00557926 | 0.0052429 | 0.00682286 | -0.00158 | 0.84836941 | 0.91440801 |
| Gm3550 | 0.11511021 | 0.12955565 | 0.13113615 | -0.0015805 | 0.52136505 | 0.90650187 |
| Prdm9 | 0.00295781 | 0.00204211 | 0.00362354 | -0.0015814 | 0.97382299 | 0.97994725 |
| Tmem8c | 0.0073949 | 0.00744792 | 0.00903209 | -0.0015842 | 0.78222524 | 0.90650187 |
| Hdac9 | 0.00952297 | 0.01002977 | 0.01161457 | -0.0015848 | 0.72794129 | 0.90650187 |
| Sgcz | 0.00217751 | 0.00108411 | 0.00266902 | -0.0015849 | 0.99585326 | 0.99633807 |
| Gm43062 | 0.02252369 | 0.02564592 | 0.02723141 | -0.0015855 | 0.60211903 | 0.90650187 |
| Snx20 | 0.00551814 | 0.00516205 | 0.0067484 | -0.0015864 | 0.85199622 | 0.91551074 |
| Lrriq1 | 0.00227609 | 0.00120329 | 0.00278968 | -0.0015864 | 0.99426511 | 0.99531274 |
| Rimkla | 0.00544814 | 0.00507347 | 0.00666311 | -0.0015896 | 0.85556467 | 0.9165432 |
| Fam151b | 0.00550895 | 0.00514756 | 0.0067372 | -0.0015896 | 0.85289784 | 0.91562204 |
| Ctrl | 0.00791401 | 0.00807272 | 0.00966273 | -0.00159 | 0.76777205 | 0.90650187 |
| Optc | 0.003064 | 0.002162 | 0.00375335 | -0.0015914 | 0.97027511 | 0.97741232 |
| Slc30a6 | 0.19037498 | 0.20619423 | 0.20778564 | -0.0015914 | 0.51358074 | 0.90650187 |
| Gm17705 | 0.00816247 | 0.00837272 | 0.00996442 | -0.0015917 | 0.76118945 | 0.90650187 |
| Rassf4 | 0.0701178 | 0.08056107 | 0.08215305 | -0.001592 | 0.53432556 | 0.90650187 |

| | | | | | | |
|---|---|---|---|---|---|---|
| Gm26562 | 0.00824239 | 0.00846864 | 0.01006144 | -0.0015928 | 0.75921313 | 0.90650187 |
| Chrna7 | 0.00599795 | 0.0057398 | 0.0073328 | -0.001593 | 0.83292209 | 0.91042272 |
| C330011M18 | 0.01795069 | 0.02017527 | 0.02176931 | -0.001594 | 0.62760833 | 0.90650187 |
| Lrrc55 | 0.0024333 | 0.00138767 | 0.00298204 | -0.0015944 | 0.99126985 | 0.9935575 |
| Trio | 0.29129848 | 0.29943919 | 0.30103443 | -0.0015952 | 0.50939749 | 0.90650187 |
| Gm26718 | 0.00629928 | 0.00610361 | 0.00769961 | -0.001596 | 0.82160605 | 0.9092537 |
| D430041D05 | 0.00229129 | 0.00121164 | 0.00280827 | -0.0015966 | 0.99425655 | 0.99531274 |
| Paqr9 | 0.00545813 | 0.00507811 | 0.00667528 | -0.0015972 | 0.85626448 | 0.9165432 |
| Gm7535 | 0.0126435 | 0.01378416 | 0.01538802 | -0.0016039 | 0.67844609 | 0.90650187 |
| Col8a1 | 0.0113617 | 0.01223545 | 0.01383995 | -0.0016045 | 0.6968598 | 0.90650187 |
| Rpusd2 | 0.02488855 | 0.02843577 | 0.03004294 | -0.0016072 | 0.59399018 | 0.90650187 |
| Hkdc1 | 0.00226343 | 0.00116579 | 0.00277418 | -0.0016084 | 0.99502327 | 0.99594052 |
| Gm15247 | 0.00805307 | 0.00822172 | 0.00983159 | -0.0016099 | 0.76667635 | 0.90650187 |
| Cbln2 | 0.0028199 | 0.00184452 | 0.00345491 | -0.0016104 | 0.980875 | 0.98560821 |
| Atp1a3 | 0.00601581 | 0.00574181 | 0.00735454 | -0.0016127 | 0.83518229 | 0.91066681 |
| Dnajb3 | 0.08762635 | 0.09989978 | 0.1015126 | -0.0016128 | 0.52815425 | 0.90650187 |
| Slc22a2 | 0.00838933 | 0.00862573 | 0.01023978 | -0.0016141 | 0.75826784 | 0.90650187 |
| Klhl35 | 0.00305376 | 0.00212547 | 0.00374084 | -0.0016154 | 0.97255263 | 0.97931721 |
| Gli2 | 0.06365238 | 0.07327604 | 0.07489233 | -0.0016163 | 0.53821714 | 0.90650187 |
| Gm16191 | 0.0023214 | 0.00122775 | 0.00284512 | -0.0016174 | 0.99425124 | 0.99531274 |
| Nutf2-ps1 | 0.01562609 | 0.01736207 | 0.01897986 | -0.0016178 | 0.64763029 | 0.90650187 |
| Gm16253 | 0.00276743 | 0.00177249 | 0.00339075 | -0.0016183 | 0.98306826 | 0.98709923 |
| Igsf23 | 0.00243993 | 0.00137031 | 0.00299015 | -0.0016198 | 0.99198633 | 0.99407899 |
| Il13ra1 | 0.12724455 | 0.14232367 | 0.14394374 | -0.0016201 | 0.51995273 | 0.90650187 |
| Kctd14 | 0.01877904 | 0.02113864 | 0.0227612 | -0.0016226 | 0.62434438 | 0.90650187 |
| Ikzf3 | 0.00238677 | 0.00130225 | 0.0029251 | -0.0016229 | 0.99317668 | 0.99474162 |
| Mlkl | 0.00272479 | 0.00171531 | 0.0033386 | -0.0016233 | 0.98467292 | 0.98818334 |
| Klhl40 | 0.01168052 | 0.01260131 | 0.01422525 | -0.0016239 | 0.69410014 | 0.90650187 |
| Gdpd2 | 0.012322 | 0.01337209 | 0.01499999 | -0.0016279 | 0.6852635 | 0.90650187 |
| Adra1b | 0.00551897 | 0.00512052 | 0.00674941 | -0.0016289 | 0.85834137 | 0.91744797 |

| | | | | | | |
|---|---|---|---|---|---|---|
| Lrrc43 | 0.00566111 | 0.00529359 | 0.00692257 | -0.001629 | 0.8522467 | 0.91557373 |
| Galnt13 | 0.00634088 | 0.00611999 | 0.00775025 | -0.0016303 | 0.8251393 | 0.90934433 |
| Slc6a15 | 0.0023803 | 0.00128636 | 0.00291719 | -0.0016308 | 0.9935275 | 0.99493242 |
| Lrp2bp | 0.00582814 | 0.00549362 | 0.00712602 | -0.0016324 | 0.84575224 | 0.9136218 |
| Hcn4 | 0.00529598 | 0.00484439 | 0.00647768 | -0.0016333 | 0.86883839 | 0.92132629 |
| Ppargc1a | 0.00240719 | 0.00131538 | 0.00295009 | -0.0016347 | 0.99311885 | 0.99474162 |
| Fam132b | 0.00243904 | 0.00135393 | 0.00298906 | -0.0016351 | 0.99249036 | 0.99445296 |
| Dclre1a | 0.11926949 | 0.13390999 | 0.13554651 | -0.0016365 | 0.52140255 | 0.90650187 |
| Glrb | 0.00871599 | 0.00899924 | 0.01063615 | -0.0016369 | 0.75306978 | 0.90650187 |
| Gpr161 | 0.02334134 | 0.02656601 | 0.02820449 | -0.0016385 | 0.60189629 | 0.90650187 |
| Wdr41 | 0.16814653 | 0.18418528 | 0.18582437 | -0.0016391 | 0.51563976 | 0.90650187 |
| Slc5a7 | 0.00569485 | 0.00532311 | 0.00696367 | -0.0016406 | 0.85252716 | 0.91557373 |
| Gm9949 | 0.00633619 | 0.00610273 | 0.00774453 | -0.0016418 | 0.82701813 | 0.90947782 |
| Nrg3 | 0.00262726 | 0.00157749 | 0.00321932 | -0.0016418 | 0.9883118 | 0.99098467 |
| Esyt3 | 0.00570119 | 0.00532546 | 0.00697139 | -0.0016459 | 0.85304827 | 0.91566617 |
| Setbp1 | 0.14002168 | 0.1555909 | 0.15723693 | -0.001646 | 0.51855955 | 0.90650187 |
| Rasl2-9 | 0.00635707 | 0.00612341 | 0.00776995 | -0.0016465 | 0.82692286 | 0.90947782 |
| Gm11733 | 0.00594619 | 0.00562313 | 0.00726977 | -0.0016466 | 0.84302248 | 0.91273395 |
| Tceal7 | 0.00597144 | 0.00565239 | 0.00730052 | -0.0016481 | 0.84221852 | 0.91251845 |
| Adgrg2 | 0.02461168 | 0.02806336 | 0.02971424 | -0.0016509 | 0.5975426 | 0.90650187 |
| Plppr4 | 0.00276172 | 0.00173286 | 0.00338377 | -0.0016509 | 0.98495749 | 0.98821581 |
| Colgalt2 | 0.00596907 | 0.00564578 | 0.00729764 | -0.0016519 | 0.84286086 | 0.91273395 |
| Atp1b2 | 0.07504273 | 0.08598974 | 0.087643 | -0.0016533 | 0.53341624 | 0.90650187 |
| Vpreb1 | 0.00860102 | 0.00884125 | 0.01049667 | -0.0016554 | 0.75838377 | 0.90650187 |
| Gm3294 | 0.00530713 | 0.00483498 | 0.00649127 | -0.0016563 | 0.8716716 | 0.92233607 |
| Trdmt1 | 0.06714247 | 0.07716233 | 0.07881932 | -0.001657 | 0.53723019 | 0.90650187 |
| Ggn | 0.00568489 | 0.00528755 | 0.00695154 | -0.001664 | 0.85636886 | 0.9165432 |
| Kcnmb4os2 | 0.00897663 | 0.00928732 | 0.01095228 | -0.001665 | 0.75042275 | 0.90650187 |
| 9430065F17I | 0.02218098 | 0.02515823 | 0.02682322 | -0.001665 | 0.60870675 | 0.90650187 |
| Bcl2l15 | 0.01319662 | 0.01438955 | 0.01605522 | -0.0016657 | 0.67767739 | 0.90650187 |

| | | | | | | |
|---|---|---|---|---|---|---|
| RP23-316J6.4 | 0.00625018 | 0.00597397 | 0.00763985 | -0.0016659 | 0.83381988 | 0.91062493 |
| Gm28874 | 0.01564584 | 0.01733432 | 0.0190036 | -0.0016693 | 0.65191703 | 0.90650187 |
| Prr15 | 0.02756522 | 0.03154508 | 0.03321446 | -0.0016694 | 0.5884031 | 0.90650187 |
| Mstn | 0.00543688 | 0.00497536 | 0.00664939 | -0.001674 | 0.86847478 | 0.92100496 |
| Gltscr1 | 0.00582781 | 0.00544539 | 0.00712562 | -0.0016802 | 0.85274462 | 0.91562204 |
| Gm10561 | 0.02614222 | 0.02984827 | 0.0315298 | -0.0016815 | 0.59370689 | 0.90650187 |
| Efhc2 | 0.00630166 | 0.00601626 | 0.00770252 | -0.0016863 | 0.83478444 | 0.91062493 |
| Tollip | 0.24579102 | 0.25855201 | 0.2602442 | -0.0016922 | 0.51153051 | 0.90650187 |
| Hyls1 | 0.0483027 | 0.05571325 | 0.05740681 | -0.0016936 | 0.55217179 | 0.90650187 |
| 2810433D01 | 0.03073598 | 0.03526294 | 0.03695678 | -0.0016938 | 0.58072741 | 0.90650187 |
| Tmem72 | 0.00499484 | 0.00441621 | 0.0061106 | -0.0016944 | 0.8911569 | 0.93109327 |
| Zfp641 | 0.00628182 | 0.00598191 | 0.00767836 | -0.0016965 | 0.83700256 | 0.91092327 |
| Gm10642 | 0.00628909 | 0.00599045 | 0.00768722 | -0.0016968 | 0.8367679 | 0.91092327 |
| Rps6ka4 | 0.20103216 | 0.21642667 | 0.21812381 | -0.0016971 | 0.51379649 | 0.90650187 |
| Kcnj5 | 0.00557766 | 0.00512349 | 0.00682092 | -0.0016974 | 0.8657055 | 0.92040222 |
| Fitm1 | 0.01385201 | 0.0151465 | 0.01684513 | -0.0016986 | 0.67302662 | 0.90650187 |
| Vmn1r4 | 0.00505825 | 0.00448895 | 0.0061879 | -0.001699 | 0.88887886 | 0.9302877 |
| Abhd3 | 0.02286698 | 0.02593877 | 0.02764009 | -0.0017013 | 0.60781898 | 0.90650187 |
| Heca | 0.02855416 | 0.03268204 | 0.03438337 | -0.0017013 | 0.58705427 | 0.90650187 |
| Wfdc3 | 0.0280247 | 0.03205589 | 0.03375775 | -0.0017019 | 0.58866839 | 0.90650187 |
| Plbd2 | 0.32384082 | 0.32731935 | 0.32902626 | -0.0017069 | 0.50919991 | 0.90650187 |
| Tmem82 | 0.00547421 | 0.00498708 | 0.00669488 | -0.0017078 | 0.87160868 | 0.9223337 |
| Fam150b | 0.00892518 | 0.00917972 | 0.01088988 | -0.0017102 | 0.75745662 | 0.90650187 |
| Gm32950 | 0.00955232 | 0.00993533 | 0.01165014 | -0.0017148 | 0.74355883 | 0.90650187 |
| Try4 | 0.00809676 | 0.00816882 | 0.00988464 | -0.0017158 | 0.77985051 | 0.90650187 |
| Gm19705 | 0.01309923 | 0.01422119 | 0.01593778 | -0.0017166 | 0.68396503 | 0.90650187 |
| Nrg2 | 0.00836866 | 0.00849775 | 0.0102147 | -0.001717 | 0.77254311 | 0.90650187 |
| Jph3 | 0.03320252 | 0.03813976 | 0.03985701 | -0.0017173 | 0.57594911 | 0.90650187 |
| Bcl6 | 0.00566412 | 0.00520709 | 0.00692624 | -0.0017191 | 0.8650811 | 0.92040222 |
| Syndig1 | 0.00740966 | 0.0073302 | 0.00905003 | -0.0017198 | 0.80089512 | 0.90813119 |

| | | | | | | |
|---|---|---|---|---|---|---|
| Acnat1 | 0.0062542 | 0.00592485 | 0.00764475 | -0.0017199 | 0.84138437 | 0.91251845 |
| Smim10l2a | 0.03452281 | 0.03968401 | 0.04140554 | -0.0017215 | 0.57332141 | 0.90650187 |
| Ppfibp2 | 0.04246299 | 0.04893971 | 0.05066132 | -0.0017216 | 0.56004149 | 0.90650187 |
| Tmprss6 | 0.01368114 | 0.01491641 | 0.01663926 | -0.0017228 | 0.67735078 | 0.90650187 |
| Mitf | 0.03171602 | 0.03638722 | 0.03811028 | -0.0017231 | 0.5796495 | 0.90650187 |
| Crym | 0.00620627 | 0.00586167 | 0.00758641 | -0.0017247 | 0.84391823 | 0.91300169 |
| Gm12981 | 0.0056331 | 0.00516189 | 0.00688846 | -0.0017266 | 0.86741919 | 0.92056229 |
| Gm15965 | 0.00621994 | 0.00587508 | 0.00760305 | -0.001728 | 0.84383923 | 0.91300169 |
| Stac3 | 0.0216645 | 0.02447892 | 0.02620772 | -0.0017288 | 0.61533652 | 0.90650187 |
| Sstr2 | 0.04472873 | 0.05155535 | 0.05328468 | -0.0017293 | 0.55736029 | 0.90650187 |
| 3110083C13I | 0.00617005 | 0.00581029 | 0.00754232 | -0.001732 | 0.84634798 | 0.91405582 |
| Gm28285 | 0.00580857 | 0.00537 | 0.00710218 | -0.0017322 | 0.86087409 | 0.91848494 |
| B4galnt4 | 0.00871472 | 0.00890164 | 0.01063461 | -0.001733 | 0.76560088 | 0.90650187 |
| Pcdhga5 | 0.00893016 | 0.00916231 | 0.01089592 | -0.0017336 | 0.76031819 | 0.90650187 |
| Sugct | 0.00541673 | 0.00489 | 0.00662484 | -0.0017348 | 0.8778219 | 0.92484883 |
| Gm14393 | 0.00835865 | 0.00846749 | 0.01020255 | -0.0017351 | 0.77518259 | 0.90650187 |
| Dusp14 | 0.03800626 | 0.043743 | 0.04547817 | -0.0017352 | 0.56734431 | 0.90650187 |
| Ifi47 | 0.01053939 | 0.01110876 | 0.01284541 | -0.0017366 | 0.72608902 | 0.90650187 |
| Raver2 | 0.01325327 | 0.01438227 | 0.01612353 | -0.0017413 | 0.68442267 | 0.90650187 |
| Cilp2 | 0.03183188 | 0.03650432 | 0.03824655 | -0.0017422 | 0.5802404 | 0.90650187 |
| Gm8898 | 0.00798242 | 0.00800219 | 0.0097458 | -0.0017436 | 0.78679765 | 0.90650187 |
| Ccdc107 | 0.46180014 | 0.43617917 | 0.43792364 | -0.0017445 | 0.50706452 | 0.90650187 |
| Kcnj11 | 0.00881854 | 0.00901465 | 0.01076055 | -0.0017459 | 0.76463677 | 0.90650187 |
| A830009L08I | 0.00508786 | 0.00447785 | 0.006224 | -0.0017462 | 0.89384162 | 0.93197657 |
| Slc16a4 | 0.01494626 | 0.01641421 | 0.01816243 | -0.0017482 | 0.665642 | 0.90650187 |
| Wdr93 | 0.00817426 | 0.0082302 | 0.00997873 | -0.0017485 | 0.78200551 | 0.90650187 |
| S100a14 | 0.00584672 | 0.00539882 | 0.00714864 | -0.0017498 | 0.86174314 | 0.9188337 |
| Gm17509 | 0.00538551 | 0.0048358 | 0.00658679 | -0.001751 | 0.8813565 | 0.92652145 |
| Zdhhc24 | 0.06914445 | 0.07931123 | 0.08106387 | -0.0017526 | 0.53828589 | 0.90650187 |
| Fanci | 0.00525811 | 0.00467872 | 0.00643153 | -0.0017528 | 0.88716298 | 0.92936447 |

| | | | | | | |
|---|---|---|---|---|---|---|
| Ar | 0.02172104 | 0.02452229 | 0.02627512 | -0.0017528 | 0.61660253 | 0.90650187 |
| Gm42912 | 0.00953675 | 0.00987718 | 0.01163126 | -0.0017541 | 0.74870884 | 0.90650187 |
| Gm26808 | 0.00780514 | 0.00777362 | 0.00953051 | -0.0017569 | 0.79375478 | 0.90731056 |
| Gm12996 | 0.01778782 | 0.01981553 | 0.02157414 | -0.0017586 | 0.64146717 | 0.90650187 |
| Reep6 | 0.17095212 | 0.18686703 | 0.1886267 | -0.0017597 | 0.51654035 | 0.90650187 |
| Gm340 | 0.03065324 | 0.03509666 | 0.03685933 | -0.0017627 | 0.58417856 | 0.90650187 |
| Rsad1 | 0.0238092 | 0.02699809 | 0.02876083 | -0.0017627 | 0.60736955 | 0.90650187 |
| D930028M14 | 0.01809635 | 0.02018086 | 0.02194381 | -0.001763 | 0.63951291 | 0.90650187 |
| Cables1 | 0.02974878 | 0.03402928 | 0.03579332 | -0.001764 | 0.58671282 | 0.90650187 |
| Smim18 | 0.00618604 | 0.00579568 | 0.00756178 | -0.0017661 | 0.85043392 | 0.91483663 |
| Eml6 | 0.01372083 | 0.01491993 | 0.01668708 | -0.0017671 | 0.68110304 | 0.90650187 |
| Cap2 | 0.01387695 | 0.01510624 | 0.01687518 | -0.0017689 | 0.67939616 | 0.90650187 |
| Rgs14 | 0.01806171 | 0.02013302 | 0.0219023 | -0.0017693 | 0.64024685 | 0.90650187 |
| Ak9 | 0.00987329 | 0.01026741 | 0.01203899 | -0.0017716 | 0.74350426 | 0.90650187 |
| Pbx4 | 0.03208098 | 0.03676553 | 0.03853945 | -0.0017739 | 0.58106751 | 0.90650187 |
| Abcb4 | 0.00612334 | 0.00571084 | 0.00748545 | -0.0017746 | 0.85404486 | 0.91608365 |
| Acsbg1 | 0.01506524 | 0.01652877 | 0.01830554 | -0.0017768 | 0.66694736 | 0.90650187 |
| Cebpz | 0.59730051 | 0.52986506 | 0.53164201 | -0.001777 | 0.50592764 | 0.90650187 |
| Nptx1 | 0.03786817 | 0.04353756 | 0.04531708 | -0.0017795 | 0.5692917 | 0.90650187 |
| Amz1 | 0.00543602 | 0.00486685 | 0.00664834 | -0.0017815 | 0.88322191 | 0.92745622 |
| Kcnt2 | 0.00616514 | 0.00575422 | 0.00753634 | -0.0017821 | 0.85342949 | 0.91590713 |
| 1700108N11 | 0.00606134 | 0.00562618 | 0.00740998 | -0.0017838 | 0.85773284 | 0.91722806 |
| Ddx23 | 0.17655257 | 0.19240944 | 0.19419393 | -0.0017845 | 0.51629289 | 0.90650187 |
| Ifnar1 | 0.31204476 | 0.31720426 | 0.3189888 | -0.0017845 | 0.5099208 | 0.90650187 |
| Bmper | 0.00565884 | 0.00513478 | 0.00691981 | -0.001785 | 0.87426413 | 0.9235672 |
| A530083I20R | 0.00604123 | 0.00560019 | 0.0073855 | -0.0017853 | 0.85873258 | 0.9176516 |
| Gm26511 | 0.00575593 | 0.00524927 | 0.00703807 | -0.0017888 | 0.87073968 | 0.92218448 |
| Sbno2 | 0.00572167 | 0.00520744 | 0.00699634 | -0.0017889 | 0.8721684 | 0.92254675 |
| 4930524O07 | 0.00531361 | 0.00470985 | 0.00649917 | -0.0017893 | 0.88951357 | 0.9302877 |
| Serpina7 | 0.00485812 | 0.00415353 | 0.00594389 | -0.0017904 | 0.9097244 | 0.9393235 |

| | | | | | | |
|---|---|---|---|---|---|---|
| 4930500F10l | 0.00522544 | 0.00459573 | 0.00639171 | -0.001796 | 0.89419577 | 0.93228189 |
| Efcab12 | 0.00566318 | 0.00512812 | 0.0069251 | -0.001797 | 0.87566388 | 0.92418014 |
| Iqcd | 0.00518888 | 0.00454968 | 0.00634715 | -0.0017975 | 0.89597841 | 0.93330829 |
| Slc25a38 | 0.18723382 | 0.20291621 | 0.20471513 | -0.0017989 | 0.51558085 | 0.90650187 |
| Abcc5 | 0.30328442 | 0.30965393 | 0.31145494 | -0.001801 | 0.51025448 | 0.90650187 |
| Spag8 | 0.01162347 | 0.01235367 | 0.01415632 | -0.0018027 | 0.71433684 | 0.90650187 |
| Actn2 | 0.01096068 | 0.01155146 | 0.01335507 | -0.0018036 | 0.72587436 | 0.90650187 |
| AA415398 | 0.01614365 | 0.01779777 | 0.01960167 | -0.0018039 | 0.65877424 | 0.90650187 |
| Mbd5 | 0.1236843 | 0.138399 | 0.14020476 | -0.0018058 | 0.52282976 | 0.90650187 |
| Dna2 | 0.04835903 | 0.05566335 | 0.05747163 | -0.0018083 | 0.55562105 | 0.90650187 |
| Gm26520 | 0.00864422 | 0.0087384 | 0.01054908 | -0.0018107 | 0.77728544 | 0.90650187 |
| Sptb | 0.00519765 | 0.00454578 | 0.00635784 | -0.0018121 | 0.89743107 | 0.93411734 |
| Gm3604 | 0.02796587 | 0.03187534 | 0.03368821 | -0.0018129 | 0.59453695 | 0.90650187 |
| Pdgfb | 0.02824727 | 0.03220593 | 0.03402079 | -0.0018149 | 0.59373142 | 0.90650187 |
| Hnf1a | 0.06294498 | 0.07227827 | 0.0740942 | -0.0018159 | 0.54338075 | 0.90650187 |
| Spp2 | 0.00580647 | 0.00528176 | 0.00709962 | -0.0018179 | 0.87250231 | 0.92276543 |
| Pou6f1 | 0.02693945 | 0.03065604 | 0.03247402 | -0.001818 | 0.59827175 | 0.90650187 |
| Hmgcll1 | 0.01856754 | 0.02068772 | 0.02250804 | -0.0018203 | 0.6404027 | 0.90650187 |
| Lingo1 | 0.24534824 | 0.25801693 | 0.25983742 | -0.0018205 | 0.51242389 | 0.90650187 |
| Rtn3 | 1.26936867 | 0.840831 | 0.84265183 | -0.0018208 | 0.50383226 | 0.90650187 |
| Phyhip | 0.03253044 | 0.0372464 | 0.03906771 | -0.0018213 | 0.58209302 | 0.90650187 |
| Kcnab3os | 0.00542809 | 0.00481567 | 0.00663868 | -0.001823 | 0.88891524 | 0.9302877 |
| Gm43689 | 0.00598707 | 0.00549646 | 0.00731956 | -0.0018231 | 0.86591311 | 0.9204717 |
| Gm16984 | 0.0083428 | 0.00835864 | 0.01018332 | -0.0018247 | 0.78715224 | 0.90650187 |
| Fam169a | 0.00590644 | 0.00539538 | 0.00722136 | -0.001826 | 0.86951683 | 0.92144928 |
| Gm16035 | 0.01060681 | 0.01110045 | 0.01292699 | -0.0018265 | 0.73504668 | 0.90650187 |
| Gm43140 | 0.00607737 | 0.0056014 | 0.00742949 | -0.0018281 | 0.86299622 | 0.91950555 |
| Polq | 0.00834079 | 0.00835133 | 0.01018087 | -0.0018295 | 0.78782443 | 0.90650187 |
| Hcn2 | 0.006028 | 0.00553855 | 0.00736939 | -0.0018308 | 0.86530263 | 0.92040222 |
| Gm43703 | 0.00609593 | 0.00561927 | 0.00745208 | -0.0018328 | 0.86288645 | 0.91950555 |

| | | | | | | |
|---|---|---|---|---|---|---|
| Gpr3 | 0.00525208 | 0.00458847 | 0.00642418 | -0.0018357 | 0.89801665 | 0.9343964 |
| Zfp459 | 0.0082892 | 0.0082809 | 0.01011826 | -0.0018374 | 0.79024459 | 0.90650187 |
| Cdhr4 | 0.02606549 | 0.02959884 | 0.03143886 | -0.00184 | 0.60264028 | 0.90650187 |
| Dym | 0.09554183 | 0.10828205 | 0.11012274 | -0.0018407 | 0.52961737 | 0.90650187 |
| Ifnlr1 | 0.0051254 | 0.00442757 | 0.00626976 | -0.0018422 | 0.90426078 | 0.93677582 |
| Tubb4a | 0.10208015 | 0.11533652 | 0.11718105 | -0.0018445 | 0.52789439 | 0.90650187 |
| Murc | 0.01155997 | 0.0122295 | 0.01407959 | -0.0018501 | 0.72044476 | 0.90650187 |
| Gm42851 | 0.00727455 | 0.00703427 | 0.00888582 | -0.0018516 | 0.82286464 | 0.9092537 |
| Mrps27 | 0.1440369 | 0.15951956 | 0.16137449 | -0.0018549 | 0.52037721 | 0.90650187 |
| Glt28d2 | 0.01725915 | 0.01908292 | 0.02094039 | -0.0018575 | 0.65333358 | 0.90650187 |
| Fhdc1 | 0.05128844 | 0.05897505 | 0.06083566 | -0.0018606 | 0.55407558 | 0.90650187 |
| Arhgef12 | 0.2717783 | 0.2819189 | 0.28377986 | -0.001861 | 0.51162881 | 0.90650187 |
| Fnip1 | 0.16491741 | 0.1807258 | 0.18258811 | -0.0018623 | 0.51808302 | 0.90650187 |
| Tspyl2 | 0.25719296 | 0.26878574 | 0.27065048 | -0.0018647 | 0.51221755 | 0.90650187 |
| Gm26722 | 0.03338739 | 0.03820921 | 0.04007401 | -0.0018648 | 0.58194478 | 0.90650187 |
| Gm26596 | 0.01047498 | 0.01089826 | 0.01276746 | -0.0018692 | 0.74242846 | 0.90650187 |
| Vstm5 | 0.00433443 | 0.00343256 | 0.00530504 | -0.0018725 | 0.9416914 | 0.95769392 |
| Cngb1 | 0.01016035 | 0.01051209 | 0.01238661 | -0.0018745 | 0.74945359 | 0.90650187 |
| Hoxb3os | 0.01339768 | 0.01442195 | 0.01629762 | -0.0018757 | 0.69554797 | 0.90650187 |
| Ccdc151 | 0.01479737 | 0.0161069 | 0.0179833 | -0.0018764 | 0.67862573 | 0.90650187 |
| Nfatc3 | 0.12124352 | 0.13575446 | 0.13763232 | -0.0018779 | 0.5241834 | 0.90650187 |
| Gm13563 | 0.00449119 | 0.00361601 | 0.00549631 | -0.0018803 | 0.9358515 | 0.95373281 |
| Mc4r | 0.00509549 | 0.00435143 | 0.00623331 | -0.0018819 | 0.91022795 | 0.93965196 |
| A630023P12 | 0.00590078 | 0.00533246 | 0.00721447 | -0.001882 | 0.87691669 | 0.92473192 |
| Engase | 0.04346208 | 0.04993682 | 0.05181907 | -0.0018822 | 0.56414206 | 0.90650187 |
| Rnf19b | 0.06463755 | 0.07411957 | 0.07600264 | -0.0018831 | 0.54385315 | 0.90650187 |
| Cfap45 | 0.01332944 | 0.01432723 | 0.01621536 | -0.0018881 | 0.69764851 | 0.90650187 |
| Gm15351 | 0.00586503 | 0.00528111 | 0.00717094 | -0.0018898 | 0.87932093 | 0.92549228 |
| AI317395 | 0.00869589 | 0.00871787 | 0.01061177 | -0.0018939 | 0.78623153 | 0.90650187 |
| Iglon5 | 0.01618479 | 0.01775418 | 0.01965107 | -0.0018969 | 0.66608473 | 0.90650187 |

| | | | | | | |
|---|---|---|---|---|---|---|
| 2310074N15 | 0.00521124 | 0.00447624 | 0.00637441 | -0.0018982 | 0.90721702 | 0.93808668 |
| Hist1h4c | 0.01226772 | 0.01303412 | 0.01493446 | -0.0019003 | 0.71419498 | 0.90650187 |
| Atp7b | 0.00596428 | 0.00538704 | 0.0072918 | -0.0019048 | 0.87723557 | 0.92481156 |
| Ssh3 | 0.13550225 | 0.15065229 | 0.15255712 | -0.0019048 | 0.52213316 | 0.90650187 |
| 1700022N22 | 0.00580764 | 0.00519534 | 0.00710105 | -0.0019057 | 0.88357925 | 0.92767795 |
| Gm16702 | 0.00457709 | 0.00369353 | 0.00560112 | -0.0019076 | 0.93499379 | 0.95330609 |
| Zhx2 | 0.03753862 | 0.0430233 | 0.04493252 | -0.0019092 | 0.57491353 | 0.90650187 |
| Gm26830 | 0.02272545 | 0.02556189 | 0.02747162 | -0.0019097 | 0.62135556 | 0.90650187 |
| Gm7694 | 0.00884061 | 0.00887679 | 0.01078731 | -0.0019105 | 0.78446322 | 0.90650187 |
| 5730522E02l | 0.00783097 | 0.00764945 | 0.00956188 | -0.0019124 | 0.81303739 | 0.9092537 |
| Ppp1r3e | 0.00594231 | 0.00535165 | 0.00726505 | -0.0019134 | 0.87916895 | 0.92549228 |
| Trim62 | 0.07180277 | 0.08211883 | 0.08403528 | -0.0019165 | 0.54037696 | 0.90650187 |
| Smad2 | 0.25920308 | 0.27055539 | 0.27247197 | -0.0019166 | 0.51247314 | 0.90650187 |
| Zfp772 | 0.04516778 | 0.05187409 | 0.05379211 | -0.001918 | 0.56297384 | 0.90650187 |
| Slc25a17 | 0.72770515 | 0.60970048 | 0.61161859 | -0.0019181 | 0.50556186 | 0.90650187 |
| Clec2l | 0.00506363 | 0.00427457 | 0.00619446 | -0.0019199 | 0.91587615 | 0.94228314 |
| Scml2 | 0.01954035 | 0.02175055 | 0.02367184 | -0.0019213 | 0.64088285 | 0.90650187 |
| Arhgef4 | 0.02965038 | 0.0337559 | 0.03567727 | -0.0019214 | 0.59460786 | 0.90650187 |
| Serpinb6b | 0.01992422 | 0.02220924 | 0.02413065 | -0.0019214 | 0.6383243 | 0.90650187 |
| Tlr5 | 0.00758651 | 0.00734294 | 0.00926492 | -0.001922 | 0.82179442 | 0.9092537 |
| Lhx6 | 0.01031496 | 0.01064504 | 0.01257377 | -0.0019287 | 0.7523576 | 0.90650187 |
| Spata9 | 0.01140719 | 0.01196565 | 0.01389493 | -0.0019293 | 0.73146971 | 0.90650187 |
| C130080G10 | 0.01070616 | 0.01111648 | 0.0130472 | -0.0019307 | 0.74468552 | 0.90650187 |
| Dennd3 | 0.01869557 | 0.0207302 | 0.02266129 | -0.0019311 | 0.64759417 | 0.90650187 |
| A930012L18l | 0.00538802 | 0.00465869 | 0.00658985 | -0.0019312 | 0.90367618 | 0.93651361 |
| Rtn4r | 0.00564946 | 0.00497698 | 0.00690839 | -0.0019314 | 0.89304338 | 0.93179124 |
| Piwil4 | 0.00562033 | 0.00493895 | 0.00687289 | -0.0019339 | 0.89452086 | 0.93247252 |
| Gm12108 | 0.00486521 | 0.00401849 | 0.00595253 | -0.001934 | 0.92569031 | 0.94746893 |
| Parpbp | 0.02381579 | 0.02683461 | 0.02876866 | -0.001934 | 0.61747946 | 0.90650187 |
| Pde1b | 0.0049688 | 0.00414055 | 0.00607885 | -0.0019383 | 0.9218355 | 0.94549985 |

| | | | | | | |
|---|---|---|---|---|---|---|
| Spice1 | 0.15583597 | 0.17148284 | 0.17342353 | -0.0019407 | 0.5198385 | 0.90650187 |
| Slc43a1 | 0.07417965 | 0.08474169 | 0.08668343 | -0.0019417 | 0.53966225 | 0.90650187 |
| Rab33a | 0.05406821 | 0.06207384 | 0.06401587 | -0.001942 | 0.55364078 | 0.90650187 |
| Trpm3 | 0.00554069 | 0.00483324 | 0.00677587 | -0.0019426 | 0.89876463 | 0.93440169 |
| 9430041J12F | 0.00553637 | 0.00482763 | 0.00677061 | -0.001943 | 0.89898185 | 0.93440169 |
| Gm6288 | 0.00636576 | 0.00583719 | 0.00778052 | -0.0019433 | 0.86658154 | 0.92056229 |
| Gm13572 | 0.00522943 | 0.00445237 | 0.00639657 | -0.0019442 | 0.91168737 | 0.94039219 |
| Edil3 | 0.00565089 | 0.00496482 | 0.00691012 | -0.0019453 | 0.89462399 | 0.93247252 |
| Gm15627 | 0.00813516 | 0.00798521 | 0.00993126 | -0.0019461 | 0.80815731 | 0.9092537 |
| Il17rb | 0.00573285 | 0.00506054 | 0.00700996 | -0.0019494 | 0.89182497 | 0.93134318 |
| 1700013F07I | 0.0054537 | 0.00471949 | 0.00666989 | -0.0019504 | 0.90319606 | 0.93635703 |
| Zfp558 | 0.00700248 | 0.00660236 | 0.00855507 | -0.0019527 | 0.84487903 | 0.91325304 |
| Cpsf7 | 0.3181851 | 0.32227517 | 0.32422926 | -0.0019541 | 0.51068761 | 0.90650187 |
| Gm12843 | 0.00507548 | 0.00425365 | 0.00620891 | -0.0019553 | 0.91923879 | 0.94389009 |
| Gm36266 | 0.01029232 | 0.01059059 | 0.01254637 | -0.0019558 | 0.75584431 | 0.90650187 |
| Trpc3 | 0.00489973 | 0.00403672 | 0.00599463 | -0.0019579 | 0.92674705 | 0.94801359 |
| Smco3 | 0.01353521 | 0.01450508 | 0.01646339 | -0.0019583 | 0.70152886 | 0.90650187 |
| Adcy8 | 0.00522463 | 0.00443171 | 0.00639072 | -0.001959 | 0.91351988 | 0.94094157 |
| Abcb10 | 0.18850055 | 0.20399474 | 0.20595464 | -0.0019599 | 0.51687232 | 0.90650187 |
| Rsph10b | 0.00498815 | 0.00414134 | 0.00610244 | -0.0019611 | 0.92344778 | 0.94620379 |
| Trpc6 | 0.00685748 | 0.0064166 | 0.00837874 | -0.0019621 | 0.85107968 | 0.91505486 |
| Ppm1n | 0.00583685 | 0.00517331 | 0.00713662 | -0.0019633 | 0.88933506 | 0.9302877 |
| Gm16230 | 0.00446387 | 0.00349827 | 0.00546299 | -0.0019647 | 0.94507019 | 0.95971754 |
| 1700042O10 | 0.00549574 | 0.00475636 | 0.0067211 | -0.0019647 | 0.90312365 | 0.93635703 |
| Ankrd45 | 0.00553038 | 0.00479675 | 0.00676331 | -0.0019666 | 0.90193291 | 0.93591959 |
| Cyp2c68 | 0.00525618 | 0.00446016 | 0.00642917 | -0.001969 | 0.91332562 | 0.94082155 |
| 2310022A10 | 0.13555762 | 0.15064538 | 0.15261461 | -0.0019692 | 0.52287195 | 0.90650187 |
| Dph1 | 0.01096405 | 0.01138753 | 0.01335915 | -0.0019716 | 0.74412257 | 0.90650187 |
| Gabrg3 | 0.00560769 | 0.00488563 | 0.00685749 | -0.0019719 | 0.8994336 | 0.93453795 |
| Gm14410 | 0.00586514 | 0.00519499 | 0.00717108 | -0.0019761 | 0.88972106 | 0.93029646 |

| | | | | | | |
|---|---|---|---|---|---|---|
| Slc18a2 | 0.01637755 | 0.01790634 | 0.01988254 | -0.0019762 | 0.67070695 | 0.90650187 |
| Parvaos | 0.00831293 | 0.00816937 | 0.01014706 | -0.0019777 | 0.80688081 | 0.9092537 |
| Rmst | 0.00538167 | 0.00460344 | 0.00658211 | -0.0019787 | 0.90929215 | 0.93919616 |
| Zkscan7 | 0.01116152 | 0.01161893 | 0.01359795 | -0.001979 | 0.74118526 | 0.90650187 |
| Gm26850 | 0.0072147 | 0.00683391 | 0.00881307 | -0.0019792 | 0.84094643 | 0.91251845 |
| Stra6 | 0.00452321 | 0.0035519 | 0.00553538 | -0.0019835 | 0.94441804 | 0.95931164 |
| Neu2 | 0.00729418 | 0.00692432 | 0.00890967 | -0.0019854 | 0.83906485 | 0.91189139 |
| Spaca1 | 0.01551821 | 0.01686445 | 0.01885019 | -0.0019857 | 0.68022021 | 0.90650187 |
| Nrg1 | 0.06299798 | 0.0721672 | 0.07415402 | -0.0019868 | 0.54740672 | 0.90650187 |
| Gm26584 | 0.00485606 | 0.00395428 | 0.00594137 | -0.0019871 | 0.93147067 | 0.95113723 |
| Gm3512 | 0.00547538 | 0.00470916 | 0.0066963 | -0.0019871 | 0.90645529 | 0.93785917 |
| 1700001L05I | 0.0055596 | 0.00481051 | 0.00679891 | -0.0019884 | 0.90322567 | 0.93635703 |
| Ddx3x | 0.85281378 | 0.67844454 | 0.68043336 | -0.0019888 | 0.50518367 | 0.90650187 |
| Asb5 | 0.00582437 | 0.00513214 | 0.00712142 | -0.0019893 | 0.89284899 | 0.93179124 |
| Plekhd1 | 0.00423187 | 0.00319014 | 0.00517988 | -0.0019897 | 0.95615229 | 0.96703067 |
| Kdm6bos | 0.00487064 | 0.00396553 | 0.00595915 | -0.0019936 | 0.93152776 | 0.95113723 |
| Gm26911 | 0.00500394 | 0.00412705 | 0.00612169 | -0.0019946 | 0.92626392 | 0.94759147 |
| Islr2 | 0.01105566 | 0.0114718 | 0.01346994 | -0.0019981 | 0.74520094 | 0.90650187 |
| Sp2 | 0.09853722 | 0.11135732 | 0.11335833 | -0.002001 | 0.5312747 | 0.90650187 |
| Gm9797 | 0.00541377 | 0.00461833 | 0.00662123 | -0.0020029 | 0.91065256 | 0.93983499 |
| Pmaip1 | 0.0055341 | 0.00476402 | 0.00676784 | -0.0020038 | 0.90595544 | 0.93759726 |
| Gm21992 | 0.00540029 | 0.00459827 | 0.0066048 | -0.0020065 | 0.9115838 | 0.94039219 |
| Etv2 | 0.00818268 | 0.00798228 | 0.00998895 | -0.0020067 | 0.81409084 | 0.9092537 |
| Ascl2 | 0.00432944 | 0.00329137 | 0.00529895 | -0.0020076 | 0.9539364 | 0.96556047 |
| Dennd2d | 0.01962774 | 0.0217683 | 0.02377631 | -0.002008 | 0.64633667 | 0.90650187 |
| Gm16263 | 0.00525089 | 0.00441459 | 0.00642273 | -0.0020081 | 0.91773072 | 0.94297078 |
| Pcx | 0.12853788 | 0.14328979 | 0.14529817 | -0.0020084 | 0.52449931 | 0.90650187 |
| Ccdc24 | 0.00558035 | 0.00481478 | 0.00682419 | -0.0020094 | 0.90473708 | 0.93703811 |
| Scn2a1 | 0.01109924 | 0.01150509 | 0.01352263 | -0.0020175 | 0.74642243 | 0.90650187 |
| Gm9917 | 0.01624979 | 0.01771056 | 0.01972913 | -0.0020186 | 0.67539046 | 0.90650187 |

| | | | | | | |
|---|---|---|---|---|---|---|
| Igf2os | 0.01260973 | 0.01332816 | 0.01534727 | -0.0020191 | 0.72068274 | 0.90650187 |
| Gm15423 | 0.00572215 | 0.00497622 | 0.00699693 | -0.0020207 | 0.90040991 | 0.93504361 |
| AU022754 | 0.00452503 | 0.00351326 | 0.0055376 | -0.0020243 | 0.9479348 | 0.9616629 |
| Gm6525 | 0.00770806 | 0.00738717 | 0.00941258 | -0.0020254 | 0.83061814 | 0.9098597 |
| Slc25a13 | 0.07016161 | 0.08017553 | 0.08220204 | -0.0020265 | 0.54363532 | 0.90650187 |
| Abca2 | 0.07310268 | 0.08345767 | 0.08548455 | -0.0020269 | 0.54197374 | 0.90650187 |
| Usp30 | 0.11167967 | 0.12545462 | 0.12748251 | -0.0020279 | 0.5281886 | 0.90650187 |
| Acot3 | 0.00508873 | 0.00419666 | 0.00622506 | -0.0020284 | 0.92627207 | 0.94759147 |
| Stx19 | 0.00458993 | 0.00358683 | 0.00561678 | -0.0020299 | 0.94593736 | 0.96040568 |
| Gm10848 | 0.03131755 | 0.0356111 | 0.03764146 | -0.0020304 | 0.59475465 | 0.90650187 |
| Mdga2 | 0.00470151 | 0.00372003 | 0.00575289 | -0.0020329 | 0.94189834 | 0.95777624 |
| Ercc6l | 0.02156203 | 0.02405235 | 0.02608555 | -0.0020332 | 0.63551872 | 0.90650187 |
| 1500002C15I | 0.03042319 | 0.03455496 | 0.03658831 | -0.0020334 | 0.59756883 | 0.90650187 |
| A330041J22I | 0.00494668 | 0.00401598 | 0.00605188 | -0.0020359 | 0.93261268 | 0.95188026 |
| Lrtm2 | 0.0079874 | 0.0077149 | 0.00975185 | -0.002037 | 0.82344999 | 0.9092537 |
| Kcnh1 | 0.00444559 | 0.00340319 | 0.00544068 | -0.0020375 | 0.95202816 | 0.96414257 |
| 4933407L21I | 0.00836812 | 0.00817213 | 0.01021404 | -0.0020419 | 0.81292627 | 0.9092537 |
| Ppara | 0.00970721 | 0.00979355 | 0.0118378 | -0.0020443 | 0.77866901 | 0.90650187 |
| Ly75 | 0.00453599 | 0.00350454 | 0.00555098 | -0.0020464 | 0.94938557 | 0.96236396 |
| A930031H19 | 0.00568087 | 0.00489952 | 0.00694665 | -0.0020471 | 0.90491723 | 0.93712658 |
| 4921507P07I | 0.00525188 | 0.00437647 | 0.00642394 | -0.0020475 | 0.9217479 | 0.94549985 |
| Rab10 | 1.17627463 | 0.81296647 | 0.81501559 | -0.0020491 | 0.50445897 | 0.90650187 |
| Gm34342 | 0.01368361 | 0.0145928 | 0.01664223 | -0.0020494 | 0.70796919 | 0.90650187 |
| Slc3a1 | 0.00548159 | 0.00465355 | 0.00670387 | -0.0020503 | 0.91302892 | 0.9407546 |
| Gm26624 | 0.00854194 | 0.00837231 | 0.01042498 | -0.0020527 | 0.80930487 | 0.9092537 |
| Slc9b2 | 0.01113708 | 0.01151507 | 0.0135684 | -0.0020533 | 0.74945065 | 0.90650187 |
| Tmem132e | 0.00808463 | 0.00781478 | 0.00986991 | -0.0020551 | 0.8226925 | 0.9092537 |
| Pik3cg | 0.00866566 | 0.0085197 | 0.01057509 | -0.0020554 | 0.80622107 | 0.9092537 |
| Zar1 | 0.00482871 | 0.00385142 | 0.00590803 | -0.0020566 | 0.93913256 | 0.95593613 |
| Gm15816 | 0.0048593 | 0.00388733 | 0.00594533 | -0.002058 | 0.93808107 | 0.9554285 |

| | | | | | | |
|---|---|---|---|---|---|---|
| Fignl2 | 0.02851787 | 0.03228052 | 0.0343405 | -0.00206 | 0.60514154 | 0.90650187 |
| Kcnab1 | 0.02416051 | 0.02711756 | 0.02917834 | -0.0020608 | 0.62323177 | 0.90650187 |
| Gm14226 | 0.01553456 | 0.01680845 | 0.01886984 | -0.0020614 | 0.68639197 | 0.90650187 |
| 2900055J20F | 0.01047303 | 0.01070314 | 0.0127651 | -0.002062 | 0.76365365 | 0.90650187 |
| Gabrd | 0.0056773 | 0.00487937 | 0.0069423 | -0.0020629 | 0.9067535 | 0.9379403 |
| Tceb3 | 0.43068816 | 0.4125852 | 0.41465119 | -0.002066 | 0.50883591 | 0.90650187 |
| Gp9 | 0.00500651 | 0.00405579 | 0.00612483 | -0.002069 | 0.93342248 | 0.95231059 |
| Sult1b1 | 0.00555951 | 0.00472643 | 0.0067988 | -0.0020724 | 0.91230428 | 0.94062208 |
| Kl | 0.0127843 | 0.0134834 | 0.0155579 | -0.0020745 | 0.72333573 | 0.90650187 |
| Myb | 0.00531217 | 0.00442116 | 0.00649741 | -0.0020763 | 0.92228328 | 0.94557609 |
| Gm26778 | 0.00701359 | 0.00649203 | 0.00856857 | -0.0020765 | 0.85934153 | 0.91797966 |
| Mfsd9 | 0.02146367 | 0.02388963 | 0.02596827 | -0.0020786 | 0.63902355 | 0.90650187 |
| Gm960 | 0.00780754 | 0.0074545 | 0.00953343 | -0.0020789 | 0.83383715 | 0.91062493 |
| Gm16052 | 0.00541454 | 0.00454216 | 0.00662217 | -0.00208 | 0.91869651 | 0.94359967 |
| 4930471D02I | 0.00531773 | 0.00442414 | 0.00650419 | -0.0020801 | 0.92244531 | 0.94558576 |
| Txlng | 0.17318479 | 0.18876957 | 0.19085052 | -0.0020809 | 0.51933029 | 0.90650187 |
| Pou5f1 | 0.00978547 | 0.00985123 | 0.01193261 | -0.0020814 | 0.78095886 | 0.90650187 |
| Cntnap4 | 0.00859509 | 0.00840546 | 0.01048948 | -0.002084 | 0.81144609 | 0.9092537 |
| Tmc8 | 0.02009539 | 0.02224931 | 0.02433516 | -0.0020859 | 0.64841663 | 0.90650187 |
| Scnn1a | 0.00516663 | 0.00423377 | 0.00632002 | -0.0020863 | 0.92888032 | 0.9498762 |
| Spata45 | 0.00485329 | 0.00385145 | 0.005938 | -0.0020865 | 0.94087235 | 0.95711708 |
| Trappc8 | 0.14385909 | 0.15910199 | 0.16119166 | -0.0020897 | 0.52297919 | 0.90650187 |
| Dpcd | 0.61791443 | 0.54281387 | 0.54490421 | -0.0020903 | 0.50680327 | 0.90650187 |
| Gm16794 | 0.01612817 | 0.01749142 | 0.01958308 | -0.0020917 | 0.68254672 | 0.90650187 |
| Ikbkb | 0.34531832 | 0.34489524 | 0.34698999 | -0.0020948 | 0.51070545 | 0.90650187 |
| Ccdc96 | 0.02818948 | 0.03185329 | 0.0339525 | -0.0020992 | 0.60828913 | 0.90650187 |
| Plcb4 | 0.19797793 | 0.21307326 | 0.21517379 | -0.0021005 | 0.5173079 | 0.90650187 |
| Tas1r2 | 0.00485628 | 0.00384102 | 0.00594165 | -0.0021006 | 0.94199142 | 0.95777624 |
| Car10 | 0.01493845 | 0.01605165 | 0.01815304 | -0.0021014 | 0.69659166 | 0.90650187 |
| Xkr5 | 0.00451869 | 0.00342794 | 0.00552987 | -0.0021019 | 0.9544665 | 0.96590409 |

| | | | | | | |
|---|---|---|---|---|---|---|
| Ppm1e | 0.00810347 | 0.0077906 | 0.00989278 | -0.0021022 | 0.82758759 | 0.90947782 |
| Slco1a6 | 0.01902283 | 0.02094985 | 0.0230529 | -0.0021031 | 0.65746746 | 0.90650187 |
| Ogfrl1 | 0.05912523 | 0.06766805 | 0.06977176 | -0.0021037 | 0.55331475 | 0.90650187 |
| Gprc5a | 0.03910661 | 0.04465696 | 0.04676071 | -0.0021037 | 0.57926208 | 0.90650187 |
| Epn3 | 0.03019615 | 0.03421651 | 0.03632075 | -0.0021042 | 0.60162538 | 0.90650187 |
| Aqp3 | 0.01935497 | 0.02134535 | 0.02345018 | -0.0021048 | 0.65506317 | 0.90650187 |
| Cdk15 | 0.0110744 | 0.01138759 | 0.0134926 | -0.002105 | 0.75602379 | 0.90650187 |
| Gm15406 | 0.00537162 | 0.00446265 | 0.00656987 | -0.0021072 | 0.92304864 | 0.945972 |
| 4933430I17R | 0.00554222 | 0.00467036 | 0.00677774 | -0.0021074 | 0.91655308 | 0.9423899 |
| E030044B06 | 0.00559731 | 0.00473708 | 0.00684485 | -0.0021078 | 0.91449404 | 0.94149841 |
| Jsrp1 | 0.01675485 | 0.01822758 | 0.02033542 | -0.0021078 | 0.67753019 | 0.90650187 |
| Ppm1h | 0.19024753 | 0.20555214 | 0.20766126 | -0.0021091 | 0.51800695 | 0.90650187 |
| Zpbp | 0.00542131 | 0.00451848 | 0.00663041 | -0.0021119 | 0.92161533 | 0.94549985 |
| Hist2h2ac | 0.06005511 | 0.06871235 | 0.07082601 | -0.0021137 | 0.55277274 | 0.90650187 |
| Polr3e | 0.1562713 | 0.17174956 | 0.17386498 | -0.0021154 | 0.52156818 | 0.90650187 |
| Gm26590 | 0.00452393 | 0.00341905 | 0.00553626 | -0.0021172 | 0.95544561 | 0.96651455 |
| Actrt3 | 0.00492813 | 0.00391197 | 0.00602926 | -0.0021173 | 0.94075686 | 0.95711708 |
| Msrb3 | 0.03492153 | 0.0397532 | 0.04187266 | -0.0021195 | 0.58901911 | 0.90650187 |
| Cd5 | 0.00762409 | 0.00718977 | 0.00931057 | -0.0021208 | 0.84438435 | 0.91316779 |
| Aldoart1 | 0.00532309 | 0.00438964 | 0.00651073 | -0.0021211 | 0.92623452 | 0.94759147 |
| Doc2a | 0.00702735 | 0.00646377 | 0.00858531 | -0.0021215 | 0.864021 | 0.92013334 |
| Cstad | 0.00833381 | 0.00805032 | 0.0101724 | -0.0021221 | 0.82314206 | 0.9092537 |
| Gm12666 | 0.00537557 | 0.0044522 | 0.00657467 | -0.0021225 | 0.9243789 | 0.94682419 |
| Crtac1 | 0.00795114 | 0.00758327 | 0.00970782 | -0.0021245 | 0.8347026 | 0.91062493 |
| Clstn2 | 0.00753911 | 0.00708213 | 0.00920732 | -0.0021252 | 0.84758184 | 0.9143112 |
| Krt23 | 0.01420652 | 0.01514558 | 0.01727212 | -0.0021265 | 0.70792835 | 0.90650187 |
| Anxa9 | 0.01021453 | 0.01032525 | 0.0124522 | -0.002127 | 0.77617855 | 0.90650187 |
| Klhl31 | 0.01446829 | 0.0154591 | 0.01758727 | -0.0021282 | 0.70469251 | 0.90650187 |
| Vwa3a | 0.00432676 | 0.00316696 | 0.00529568 | -0.0021287 | 0.96303221 | 0.9719225 |
| Ism2 | 0.01409175 | 0.01500463 | 0.0171339 | -0.0021293 | 0.70968568 | 0.90650187 |

| | | | | | | |
|---|---|---|---|---|---|---|
| Gm26564 | 0.00541654 | 0.00449376 | 0.00662461 | -0.0021308 | 0.9236351 | 0.94629143 |
| Vps37b | 0.33428903 | 0.33568295 | 0.33781392 | -0.002131 | 0.51118625 | 0.90650187 |
| Tcf24 | 0.00486634 | 0.00382233 | 0.00595392 | -0.0021316 | 0.9442619 | 0.95921713 |
| Gm6277 | 0.01278188 | 0.01342249 | 0.01555499 | -0.0021325 | 0.72889255 | 0.90650187 |
| RP23-235M8 | 0.00444617 | 0.00330581 | 0.00544139 | -0.0021356 | 0.95948708 | 0.96982267 |
| Gm14308 | 0.0044089 | 0.00326012 | 0.00539591 | -0.0021358 | 0.9607661 | 0.97053465 |
| Kif7 | 0.02435782 | 0.02727318 | 0.02941273 | -0.0021396 | 0.62679785 | 0.90650187 |
| Ubtf | 0.59934914 | 0.53083112 | 0.53297083 | -0.0021397 | 0.50711983 | 0.90650187 |
| Gm11491 | 0.01360594 | 0.01440789 | 0.01654863 | -0.0021407 | 0.71738365 | 0.90650187 |
| 6030498E09I | 0.00411025 | 0.00289006 | 0.00503143 | -0.0021414 | 0.97075724 | 0.97777172 |
| Cdh15 | 0.00849237 | 0.0082222 | 0.01036483 | -0.0021426 | 0.82095247 | 0.9092537 |
| Tmem117 | 0.00414507 | 0.00292887 | 0.00507393 | -0.0021451 | 0.96990722 | 0.97717124 |
| Platr26 | 0.00794283 | 0.00755181 | 0.00969774 | -0.0021459 | 0.83737586 | 0.91109492 |
| Opa1 | 0.21441776 | 0.22878915 | 0.23093664 | -0.0021475 | 0.51648754 | 0.90650187 |
| Arhgap8 | 0.00960052 | 0.00956049 | 0.01170854 | -0.0021481 | 0.79263343 | 0.90681274 |
| Ablim3 | 0.00528995 | 0.00432228 | 0.00647034 | -0.0021481 | 0.9300114 | 0.95045739 |
| Adamtsl4 | 0.03480254 | 0.0395835 | 0.04173328 | -0.0021498 | 0.59056694 | 0.90650187 |
| Smc1b | 0.00529831 | 0.00433066 | 0.00648052 | -0.0021499 | 0.92986647 | 0.95037317 |
| Gm12089 | 0.00483285 | 0.00376275 | 0.00591307 | -0.0021503 | 0.9470252 | 0.96112498 |
| Slc25a47 | 0.02031409 | 0.02244502 | 0.0245964 | -0.0021514 | 0.65130454 | 0.90650187 |
| Otoa | 0.00479132 | 0.00370766 | 0.00586243 | -0.0021548 | 0.94887118 | 0.96195723 |
| BC037039 | 0.01603275 | 0.01731278 | 0.01946846 | -0.0021557 | 0.68872794 | 0.90650187 |
| Gnl3l | 0.30065191 | 0.30701754 | 0.30917767 | -0.0021601 | 0.51238922 | 0.90650187 |
| Gm6712 | 0.01229728 | 0.01280915 | 0.01497014 | -0.002161 | 0.73947894 | 0.90650187 |
| Pcdhb5 | 0.01257613 | 0.01314515 | 0.01530672 | -0.0021616 | 0.73493115 | 0.90650187 |
| Slc2a9 | 0.00491352 | 0.00384549 | 0.00601144 | -0.002166 | 0.9453973 | 0.95992147 |
| Creb3l4 | 0.01413421 | 0.01501708 | 0.01718504 | -0.002168 | 0.71254345 | 0.90650187 |
| Mrs2 | 0.04362924 | 0.0498442 | 0.05201263 | -0.0021684 | 0.57351859 | 0.90650187 |
| Msantd1 | 0.00720614 | 0.00663356 | 0.00880266 | -0.0021691 | 0.86334221 | 0.91966818 |
| Oas1c | 0.0122182 | 0.01270555 | 0.01487467 | -0.0021691 | 0.74160052 | 0.90650187 |

| | | | | | | |
|---|---|---|---|---|---|---|
| 1700049E15I | 0.01495281 | 0.0160008 | 0.01817031 | -0.0021695 | 0.70222102 | 0.90650187 |
| Uckl1os | 0.01479954 | 0.01581588 | 0.01798592 | -0.00217 | 0.70414888 | 0.90650187 |
| Zfp599 | 0.03785821 | 0.04313522 | 0.04530546 | -0.0021702 | 0.58431882 | 0.90650187 |
| Scx | 0.02218178 | 0.02465324 | 0.02682418 | -0.0021709 | 0.64049883 | 0.90650187 |
| Efhd1 | 0.01045476 | 0.0105716 | 0.01274299 | -0.0021714 | 0.77563268 | 0.90650187 |
| Slc27a2 | 0.00472476 | 0.00360925 | 0.00578125 | -0.002172 | 0.95256069 | 0.96448911 |
| Cort | 0.01125931 | 0.01154143 | 0.01371617 | -0.0021747 | 0.75955268 | 0.90650187 |
| Pdk2 | 0.18112543 | 0.19653703 | 0.19871352 | -0.0021765 | 0.51941773 | 0.90650187 |
| Matk | 0.00743867 | 0.00690539 | 0.00908528 | -0.0021799 | 0.85693824 | 0.91689436 |
| Gm44195 | 0.00838793 | 0.00805513 | 0.01023809 | -0.002183 | 0.82840719 | 0.9096066 |
| Tssk6 | 0.0084238 | 0.00809745 | 0.01028161 | -0.0021842 | 0.82751681 | 0.90947782 |
| Zfp786 | 0.01805597 | 0.01971022 | 0.02189543 | -0.0021852 | 0.67136351 | 0.90650187 |
| Slc19a3 | 0.00768006 | 0.00719233 | 0.00937857 | -0.0021862 | 0.84997271 | 0.91472107 |
| Ankrd31 | 0.00843537 | 0.00810825 | 0.01029566 | -0.0021874 | 0.82754613 | 0.90947782 |
| Prr18 | 0.00446836 | 0.00328034 | 0.00546846 | -0.0021881 | 0.96236729 | 0.971442 |
| Dnd1 | 0.00478156 | 0.00365957 | 0.00585052 | -0.002191 | 0.9520266 | 0.96414257 |
| Gm11100 | 0.0120012 | 0.01242088 | 0.01461263 | -0.0021917 | 0.74754881 | 0.90650187 |
| 1700064M15 | 0.00407019 | 0.00279073 | 0.00498253 | -0.0021918 | 0.97474422 | 0.98054972 |
| Mgat4c | 0.01470374 | 0.01567786 | 0.01787065 | -0.0021928 | 0.7072923 | 0.90650187 |
| Tnfrsf19 | 0.03090267 | 0.03495831 | 0.03715307 | -0.0021948 | 0.60357697 | 0.90650187 |
| Gm11665 | 0.00514494 | 0.00409864 | 0.00629358 | -0.0021949 | 0.9394826 | 0.95608724 |
| Col19a1 | 0.00453919 | 0.00335953 | 0.00555489 | -0.0021954 | 0.96053716 | 0.97043237 |
| Fmo4 | 0.00455185 | 0.0033747 | 0.00557032 | -0.0021956 | 0.96013869 | 0.9701425 |
| Xrra1 | 0.00448934 | 0.0032983 | 0.00549406 | -0.0021958 | 0.96219365 | 0.97133123 |
| Apobec4 | 0.00436233 | 0.00314145 | 0.00533909 | -0.0021976 | 0.96636556 | 0.97444265 |
| Rbfox3 | 0.07745097 | 0.08811697 | 0.0903148 | -0.0021978 | 0.54307537 | 0.90650187 |
| Gm20109 | 0.00451832 | 0.00332974 | 0.00552942 | -0.0021997 | 0.96151301 | 0.97096654 |
| Rbm14 | 0.32988199 | 0.33191884 | 0.33411868 | -0.0021998 | 0.5116753 | 0.90650187 |
| Pgpep1l | 0.00514835 | 0.00409785 | 0.00629774 | -0.0021999 | 0.93977826 | 0.95632409 |
| Rgs6 | 0.00526953 | 0.0042432 | 0.00644545 | -0.0022023 | 0.93561319 | 0.9537321 |

| | | | | | | |
|---|---|---|---|---|---|---|
| Mak | 0.01042457 | 0.01050302 | 0.01270645 | -0.0022034 | 0.77962055 | 0.90650187 |
| Tspan2os | 0.01047716 | 0.0105653 | 0.0127701 | -0.0022048 | 0.77862167 | 0.90650187 |
| Rbm3os | 0.01621516 | 0.01748246 | 0.01968754 | -0.0022051 | 0.69073199 | 0.90650187 |
| A430105J06I | 0.03693713 | 0.04202494 | 0.04423021 | -0.0022053 | 0.58770806 | 0.90650187 |
| Zfp341 | 0.03141702 | 0.03555319 | 0.03775852 | -0.0022053 | 0.60243323 | 0.90650187 |
| Vwa3b | 0.00751758 | 0.00697495 | 0.00918117 | -0.0022062 | 0.85730089 | 0.9170242 |
| Gm27204 | 0.00527869 | 0.00424836 | 0.00645661 | -0.0022082 | 0.93580212 | 0.95373281 |
| Gm4724 | 0.00977108 | 0.00970689 | 0.01191518 | -0.0022083 | 0.79500821 | 0.90731056 |
| Gm17767 | 0.00651306 | 0.00575106 | 0.00795977 | -0.0022087 | 0.89132133 | 0.93118847 |
| 9030407P20I | 0.0136154 | 0.01435058 | 0.01656003 | -0.0022095 | 0.72345494 | 0.90650187 |
| Fam107a | 0.00458061 | 0.00339526 | 0.0056054 | -0.0022101 | 0.96018643 | 0.9701425 |
| Ptgfr | 0.004931 | 0.00382201 | 0.00603275 | -0.0022107 | 0.94835612 | 0.96189773 |
| Pcdhgb5 | 0.01001671 | 0.0100008 | 0.01221267 | -0.0022119 | 0.78963496 | 0.90650187 |
| Sftpd | 0.01582142 | 0.01699982 | 0.01921458 | -0.0022148 | 0.69582176 | 0.90650187 |
| Hist1h2bj | 0.00901776 | 0.00878693 | 0.01100215 | -0.0022152 | 0.81463216 | 0.9092537 |
| Pqlc3 | 0.03088529 | 0.03491671 | 0.03713261 | -0.0022159 | 0.60460788 | 0.90650187 |
| D630011A20 | 0.01143257 | 0.0117076 | 0.01392561 | -0.002218 | 0.76055157 | 0.90650187 |
| Lrrc71 | 0.00789167 | 0.00741264 | 0.0096356 | -0.002223 | 0.84746318 | 0.9143112 |
| Atcayos | 0.0046726 | 0.00349439 | 0.00571763 | -0.0022232 | 0.95806216 | 0.9685112 |
| Hdx | 0.00752044 | 0.00695743 | 0.00918465 | -0.0022272 | 0.85949029 | 0.91800029 |
| Amigo3 | 0.00952126 | 0.00938367 | 0.0116125 | -0.0022288 | 0.80324355 | 0.90882608 |
| Syngr3 | 0.00745464 | 0.00687579 | 0.00910469 | -0.0022289 | 0.86177305 | 0.9188337 |
| Abcc10 | 0.01084758 | 0.01098817 | 0.01321828 | -0.0022301 | 0.7733831 | 0.90650187 |
| 9330136K24I | 0.00810516 | 0.00766053 | 0.00989484 | -0.0022343 | 0.84227303 | 0.91251845 |
| Rqcd1 | 0.12538418 | 0.13975745 | 0.14199196 | -0.0022345 | 0.5278872 | 0.90650187 |
| Stk32c | 0.01115732 | 0.01135824 | 0.01359286 | -0.0022346 | 0.7675628 | 0.90650187 |
| Pddc1 | 0.67725236 | 0.57951525 | 0.58175043 | -0.0022352 | 0.50681391 | 0.90650187 |
| Slc39a5 | 0.00684986 | 0.00613366 | 0.00836947 | -0.0022358 | 0.88250308 | 0.92703446 |
| Gm17619 | 0.00520454 | 0.00412989 | 0.00636624 | -0.0022363 | 0.94081628 | 0.95711708 |
| Dlgap2 | 0.00845497 | 0.00808036 | 0.01031945 | -0.0022391 | 0.83262604 | 0.91029548 |

| | | | | | | |
|---|---|---|---|---|---|---|
| Ano9 | 0.00760709 | 0.00705079 | 0.00928992 | -0.0022391 | 0.85803156 | 0.91737198 |
| Rasl10b | 0.00921751 | 0.00900338 | 0.01124434 | -0.002241 | 0.81218988 | 0.9092537 |
| 4931429I11R | 0.00813121 | 0.00768548 | 0.00992646 | -0.002241 | 0.8422225 | 0.91251845 |
| Gm27010 | 0.00415713 | 0.00284547 | 0.00508865 | -0.0022432 | 0.97498538 | 0.98066253 |
| Gnat2 | 0.01194608 | 0.01230261 | 0.01454606 | -0.0022434 | 0.75353264 | 0.90650187 |
| Cbln1 | 0.00445499 | 0.00320824 | 0.00545215 | -0.0022439 | 0.96634892 | 0.97444265 |
| Rpl21-ps4 | 0.00835268 | 0.00794805 | 0.0101953 | -0.0022473 | 0.83643283 | 0.91085431 |
| Asgr2 | 0.00521412 | 0.00413023 | 0.00637791 | -0.0022477 | 0.94140812 | 0.95746988 |
| Hist1h3i | 0.00751257 | 0.0069258 | 0.00917508 | -0.0022493 | 0.86210941 | 0.91899895 |
| Gpbar1 | 0.00657896 | 0.00578719 | 0.00803994 | -0.0022528 | 0.89355102 | 0.93186533 |
| Cyp2j9 | 0.00819022 | 0.00774521 | 0.0099981 | -0.0022529 | 0.84176488 | 0.91251845 |
| Gm4349 | 0.01222974 | 0.01263436 | 0.0148886 | -0.0022542 | 0.7495571 | 0.90650187 |
| Kif26b | 0.02426165 | 0.02704155 | 0.0292985 | -0.0022569 | 0.63399683 | 0.90650187 |
| Gpld1 | 0.02217806 | 0.02456243 | 0.02681974 | -0.0022573 | 0.64586095 | 0.90650187 |
| Prss23os | 0.01025009 | 0.01023709 | 0.01249525 | -0.0022582 | 0.78913287 | 0.90650187 |
| Adgrb1 | 0.01065931 | 0.0107321 | 0.01299052 | -0.0022584 | 0.78020149 | 0.90650187 |
| Zfp825 | 0.10412346 | 0.11712462 | 0.11938343 | -0.0022588 | 0.53351706 | 0.90650187 |
| Sctr | 0.00436916 | 0.00308786 | 0.00534742 | -0.0022596 | 0.9698436 | 0.97717124 |
| Zfp846 | 0.04200744 | 0.04787226 | 0.05013292 | -0.0022607 | 0.57944245 | 0.90650187 |
| Fam131b | 0.01052563 | 0.01056518 | 0.01282875 | -0.0022636 | 0.78359902 | 0.90650187 |
| Krt20 | 0.00484189 | 0.0036597 | 0.00592409 | -0.0022644 | 0.95536518 | 0.96651455 |
| C77370 | 0.02117903 | 0.0233634 | 0.02562879 | -0.0022654 | 0.65282437 | 0.90650187 |
| Gal | 0.00506924 | 0.00393526 | 0.00620131 | -0.0022661 | 0.94786453 | 0.9616629 |
| Trpm2 | 0.02506712 | 0.027986 | 0.03025487 | -0.0022689 | 0.63057801 | 0.90650187 |
| Prkcg | 0.00734393 | 0.0066994 | 0.00897014 | -0.0022707 | 0.86978623 | 0.92149177 |
| Gm17399 | 0.01245788 | 0.01289085 | 0.01516401 | -0.0022732 | 0.74742843 | 0.90650187 |
| Gm43573 | 0.00679388 | 0.00602721 | 0.00830139 | -0.0022742 | 0.88836578 | 0.93003087 |
| Vgll4 | 0.52519038 | 0.48096845 | 0.4832442 | -0.0022758 | 0.50835156 | 0.90650187 |
| 4930480K23I | 0.01282202 | 0.01332399 | 0.01560341 | -0.0022794 | 0.74197169 | 0.90650187 |
| Trim6 | 0.01193162 | 0.01224884 | 0.01452859 | -0.0022797 | 0.75728019 | 0.90650187 |

| | | | | | | |
|---|---|---|---|---|---|---|
| R3hcc1 | 0.31022787 | 0.31515205 | 0.31743185 | -0.0022798 | 0.51273547 | 0.90650187 |
| Ambra1 | 0.12899966 | 0.14350046 | 0.14578127 | -0.0022808 | 0.52772543 | 0.90650187 |
| Cxx1c | 0.00803449 | 0.00752786 | 0.00980903 | -0.0022812 | 0.8493999 | 0.91455786 |
| Ankrd1 | 0.00427493 | 0.00295083 | 0.00523243 | -0.0022816 | 0.97371919 | 0.97994563 |
| Fmn1 | 0.00870366 | 0.00833851 | 0.01062119 | -0.0022827 | 0.8303223 | 0.90980532 |
| Mterf1b | 0.01264769 | 0.01310721 | 0.01539308 | -0.0022859 | 0.74542785 | 0.90650187 |
| 4631405K08I | 0.00491882 | 0.00373166 | 0.0060179 | -0.0022862 | 0.95438232 | 0.9658832 |
| Sidt1 | 0.00822352 | 0.00774545 | 0.01003853 | -0.0022931 | 0.84506645 | 0.91332066 |
| H2afj | 1.23701426 | 0.83111485 | 0.83341152 | -0.0022967 | 0.5048873 | 0.90650187 |
| A830039N20 | 0.01701616 | 0.01835213 | 0.02064895 | -0.0022968 | 0.68952059 | 0.90650187 |
| Cyp46a1 | 0.01795155 | 0.01947297 | 0.02177033 | -0.0022974 | 0.68051289 | 0.90650187 |
| Wfs1 | 0.37376369 | 0.36788374 | 0.37018187 | -0.0022981 | 0.51100894 | 0.90650187 |
| Gm38394 | 0.00487738 | 0.00366924 | 0.00596737 | -0.0022981 | 0.95655887 | 0.96718447 |
| Zfp384 | 0.06084022 | 0.0694142 | 0.07171511 | -0.0023009 | 0.55671011 | 0.90650187 |
| Slc12a5 | 0.02614787 | 0.02923532 | 0.03153649 | -0.0023012 | 0.62717822 | 0.90650187 |
| Tpcn2 | 0.04923801 | 0.05618112 | 0.05848238 | -0.0023013 | 0.56943387 | 0.90650187 |
| Ankrd29 | 0.01854469 | 0.02017904 | 0.02248069 | -0.0023016 | 0.67550161 | 0.90650187 |
| Gm29096 | 0.00969852 | 0.00952473 | 0.01182727 | -0.0023025 | 0.80661051 | 0.9092537 |
| Gm26682 | 0.01359709 | 0.01423222 | 0.01653797 | -0.0023058 | 0.73230989 | 0.90650187 |
| Abcg5 | 0.00439391 | 0.00307141 | 0.00537762 | -0.0023062 | 0.97170806 | 0.97859641 |
| Ankef1 | 0.00745499 | 0.00679786 | 0.00910511 | -0.0023072 | 0.87002797 | 0.92162336 |
| Bhlhe41 | 0.03016712 | 0.03397731 | 0.03628654 | -0.0023092 | 0.61137833 | 0.90650187 |
| Gpr135 | 0.00429997 | 0.00295197 | 0.00526298 | -0.002311 | 0.97453672 | 0.98047076 |
| Cyp4f14 | 0.00306434 | 0.00144221 | 0.00375377 | -0.0023116 | 0.99690545 | 0.9972991 |
| Ints4 | 0.16060184 | 0.17593276 | 0.17824468 | -0.0023119 | 0.52299096 | 0.90650187 |
| Spa17 | 0.24426095 | 0.25652342 | 0.25883774 | -0.0023143 | 0.51585343 | 0.90650187 |
| A930024N18 | 0.00481618 | 0.00357738 | 0.00589274 | -0.0023154 | 0.95966019 | 0.96993315 |
| 6430531B16 | 0.00994876 | 0.00981497 | 0.01213039 | -0.0023154 | 0.80193848 | 0.90850103 |
| Cacnb1 | 0.04835517 | 0.05515098 | 0.05746719 | -0.0023162 | 0.5711018 | 0.90650187 |
| Zfp493 | 0.01708697 | 0.01841714 | 0.02073388 | -0.0023167 | 0.69031306 | 0.90650187 |

| | | | | | | |
|---|---|---|---|---|---|---|
| Pxk | 0.24799391 | 0.25994717 | 0.26226497 | -0.0023178 | 0.51566983 | 0.90650187 |
| 1700028J19F | 0.00652601 | 0.0056558 | 0.00797553 | -0.0023197 | 0.90199984 | 0.93591959 |
| C330013E15I | 0.02322901 | 0.02575077 | 0.02807087 | -0.0023201 | 0.6433534 | 0.90650187 |
| Gm26876 | 0.00977453 | 0.00959886 | 0.01191935 | -0.0023205 | 0.8066125 | 0.9092537 |
| Hspbap1 | 0.04790512 | 0.05462851 | 0.05694921 | -0.0023207 | 0.57187892 | 0.90650187 |
| Gm17518 | 0.01331007 | 0.01387073 | 0.016192 | -0.0023213 | 0.73804154 | 0.90650187 |
| Gm26606 | 0.00430537 | 0.00294718 | 0.00526957 | -0.0023224 | 0.97495849 | 0.98066253 |
| Nagpa | 0.1187841 | 0.13270998 | 0.13503291 | -0.0023229 | 0.53047999 | 0.90650187 |
| Rcan3 | 0.30459025 | 0.31025782 | 0.31258226 | -0.0023244 | 0.51318607 | 0.90650187 |
| Glmn | 0.21680616 | 0.23087856 | 0.23320327 | -0.0023247 | 0.51767396 | 0.90650187 |
| Gm6878 | 0.00480152 | 0.00354741 | 0.00587486 | -0.0023274 | 0.96089941 | 0.97060482 |
| Kcnt1 | 0.0073997 | 0.00671034 | 0.00903792 | -0.0023276 | 0.87387421 | 0.9235672 |
| Tnfrsf10b | 0.03084169 | 0.03475354 | 0.03708126 | -0.0023277 | 0.60990331 | 0.90650187 |
| Adgrg3 | 0.00988159 | 0.00971982 | 0.01204904 | -0.0023292 | 0.80493681 | 0.90882608 |
| Dedd | 0.31406166 | 0.31838426 | 0.32071375 | -0.0023295 | 0.51287981 | 0.90650187 |
| Gm11716 | 0.01279683 | 0.0132431 | 0.01557302 | -0.0023299 | 0.7470129 | 0.90650187 |
| Cby3 | 0.00445229 | 0.00311588 | 0.00544886 | -0.002333 | 0.97150759 | 0.97845935 |
| Gm11266 | 0.02441174 | 0.02714202 | 0.02947678 | -0.0023348 | 0.63762637 | 0.90650187 |
| Myo5b | 0.04763068 | 0.05429691 | 0.05663321 | -0.0023363 | 0.57275604 | 0.90650187 |
| Hist4h4 | 0.00967201 | 0.00945726 | 0.01179516 | -0.0023379 | 0.81088408 | 0.9092537 |
| Ogdhl | 0.00426267 | 0.00287943 | 0.00521746 | -0.002338 | 0.97682254 | 0.98199057 |
| Cog3 | 0.09581548 | 0.10808076 | 0.11041884 | -0.0023381 | 0.53749871 | 0.90650187 |
| Fgf11 | 0.02325387 | 0.02576236 | 0.02810045 | -0.0023381 | 0.64426979 | 0.90650187 |
| Nog | 0.01299865 | 0.01347832 | 0.01581648 | -0.0023382 | 0.74447251 | 0.90650187 |
| 9130011E15I | 0.01913301 | 0.0208463 | 0.02318472 | -0.0023384 | 0.67306216 | 0.90650187 |
| P2rx6 | 0.00462526 | 0.00332035 | 0.00565988 | -0.0023395 | 0.96693967 | 0.97480297 |
| Kcnh6 | 0.05793784 | 0.06608406 | 0.0684237 | -0.0023396 | 0.56041055 | 0.90650187 |
| Ptchd1 | 0.00711504 | 0.0063519 | 0.00869192 | -0.00234 | 0.88431505 | 0.9278081 |
| Nccrp1 | 0.00441975 | 0.00306898 | 0.00540915 | -0.0023402 | 0.97277951 | 0.97935527 |
| RP23-2D23.2 | 0.00451108 | 0.0031802 | 0.00552059 | -0.0023404 | 0.97026207 | 0.97741232 |

| | | | | | | |
|---|---|---|---|---|---|---|
| Obfc1 | 0.20101502 | 0.21576352 | 0.21810728 | -0.0023438 | 0.51905104 | 0.90650187 |
| Tchh | 0.03627695 | 0.04111 | 0.04345872 | -0.0023487 | 0.59493654 | 0.90650187 |
| Depdc1b | 0.0056997 | 0.00461872 | 0.00696958 | -0.0023509 | 0.93313007 | 0.95210835 |
| Mrps22 | 0.26222609 | 0.27285296 | 0.27520401 | -0.0023511 | 0.51514763 | 0.90650187 |
| Lrig2 | 0.07938976 | 0.0901079 | 0.09245975 | -0.0023518 | 0.54501661 | 0.90650187 |
| Serpina3g | 0.00942072 | 0.00913736 | 0.01149066 | -0.0023533 | 0.81871105 | 0.9092537 |
| Golph3l | 0.23286242 | 0.2459336 | 0.24828742 | -0.0023538 | 0.51680853 | 0.90650187 |
| Sec16b | 0.06474356 | 0.07376746 | 0.07612203 | -0.0023546 | 0.554686 | 0.90650187 |
| Sstr1 | 0.01325814 | 0.01377475 | 0.01612939 | -0.0023546 | 0.74182748 | 0.90650187 |
| Dgkb | 0.01584924 | 0.01689109 | 0.01924801 | -0.0023569 | 0.70690506 | 0.90650187 |
| Mbnl3 | 0.01424241 | 0.01495766 | 0.01731534 | -0.0023577 | 0.72751656 | 0.90650187 |
| Hist1h1d | 0.00779365 | 0.00715758 | 0.00951655 | -0.002359 | 0.86476475 | 0.92040222 |
| Tmprss7 | 0.00925322 | 0.0089284 | 0.01128763 | -0.0023592 | 0.82360133 | 0.9092537 |
| Gm12088 | 0.00441574 | 0.00304411 | 0.00540426 | -0.0023601 | 0.97390002 | 0.9799599 |
| Hapln1 | 0.02378206 | 0.02636743 | 0.02872856 | -0.0023611 | 0.64258416 | 0.90650187 |
| Cc2d2b | 0.01292269 | 0.01336215 | 0.01572485 | -0.0023627 | 0.74792168 | 0.90650187 |
| Gm14305 | 0.24846484 | 0.26033039 | 0.26269624 | -0.0023659 | 0.51596833 | 0.90650187 |
| Cacna1e | 0.0106641 | 0.01062976 | 0.01299631 | -0.0023666 | 0.79089173 | 0.90650187 |
| Zpbp2 | 0.0032721 | 0.00163988 | 0.00400772 | -0.0023678 | 0.99568719 | 0.99633807 |
| Parp11 | 0.08405072 | 0.09522623 | 0.09759441 | -0.0023682 | 0.54295247 | 0.90650187 |
| Fbxl19 | 0.11326062 | 0.12679727 | 0.12916808 | -0.0023708 | 0.53251616 | 0.90650187 |
| Dll1 | 0.04077175 | 0.04632683 | 0.04869802 | -0.0023712 | 0.58568707 | 0.90650187 |
| Fadd | 0.11239394 | 0.12587262 | 0.12824443 | -0.0023718 | 0.53276381 | 0.90650187 |
| Gm28727 | 0.0126786 | 0.01305852 | 0.01543038 | -0.0023719 | 0.75280687 | 0.90650187 |
| Ak6 | 0.42157496 | 0.40532635 | 0.40770022 | -0.0023739 | 0.51032543 | 0.90650187 |
| A730056A06 | 0.00625042 | 0.00526626 | 0.00764014 | -0.0023739 | 0.91640696 | 0.94231902 |
| Macrod1 | 0.24365438 | 0.25590309 | 0.25827954 | -0.0023765 | 0.51631393 | 0.90650187 |
| Srsf4 | 0.47765353 | 0.44713624 | 0.44951564 | -0.0023794 | 0.50938692 | 0.90650187 |
| Map3k6 | 0.04117667 | 0.04678767 | 0.04916847 | -0.0023808 | 0.58521849 | 0.90650187 |
| Cebpb | 0.00702283 | 0.00619854 | 0.00857981 | -0.0023813 | 0.89137053 | 0.93118847 |

| | | | | | | |
|---|---|---|---|---|---|---|
| Magee2 | 0.0100272 | 0.0098435 | 0.01222538 | -0.0023819 | 0.80679339 | 0.9092537 |
| 4732491K20I | 0.03097487 | 0.03485344 | 0.03723809 | -0.0023847 | 0.61205825 | 0.90650187 |
| Lins1 | 0.08404247 | 0.09520048 | 0.09758534 | -0.0023849 | 0.54325798 | 0.90650187 |
| Fam124a | 0.02459891 | 0.02731409 | 0.02969907 | -0.002385 | 0.63945531 | 0.90650187 |
| Spaca7 | 0.00389515 | 0.00238293 | 0.00476882 | -0.0023859 | 0.98693194 | 0.98979696 |
| Rab26 | 0.01437256 | 0.01508571 | 0.01747203 | -0.0023863 | 0.72813386 | 0.90650187 |
| 2610020C07I | 0.02859464 | 0.03204382 | 0.03443118 | -0.0023874 | 0.62105241 | 0.90650187 |
| Ap5b1 | 0.00774975 | 0.00707486 | 0.00946323 | -0.0023884 | 0.8690688 | 0.92141903 |
| Armc4 | 0.00441621 | 0.00301506 | 0.00540483 | -0.0023898 | 0.97533043 | 0.98094468 |
| Tfcp2l1 | 0.01852032 | 0.02006141 | 0.02245151 | -0.0023901 | 0.68198682 | 0.90650187 |
| Rgma | 0.12403742 | 0.13818609 | 0.14057633 | -0.0023902 | 0.53012707 | 0.90650187 |
| Slc26a8 | 0.00424775 | 0.00280839 | 0.00519926 | -0.0023909 | 0.97953814 | 0.98452521 |
| Zfp365 | 0.00748419 | 0.00674876 | 0.00914059 | -0.0023918 | 0.87764329 | 0.92482217 |
| Prox2 | 0.02260571 | 0.02493373 | 0.02732906 | -0.0023953 | 0.65160166 | 0.90650187 |
| Rsph4a | 0.01088455 | 0.01086627 | 0.01326299 | -0.0023967 | 0.78911522 | 0.90650187 |
| Pcdh10 | 0.00994396 | 0.00972712 | 0.01212457 | -0.0023975 | 0.81031366 | 0.9092537 |
| RP23-455C1: | 0.00941188 | 0.00908152 | 0.01147994 | -0.0023984 | 0.82349996 | 0.9092537 |
| Prx | 0.01515077 | 0.01600907 | 0.01840841 | -0.0023993 | 0.71885339 | 0.90650187 |
| Ltv1 | 0.42435056 | 0.40742377 | 0.40982374 | -0.0024 | 0.51038487 | 0.90650187 |
| Gm43159 | 0.01082247 | 0.01078786 | 0.0131879 | -0.0024 | 0.79075588 | 0.90650187 |
| Gm26583 | 0.00361411 | 0.00202412 | 0.00442558 | -0.0024015 | 0.99207455 | 0.99410187 |
| Pura | 0.59361086 | 0.52683985 | 0.5292427 | -0.0024029 | 0.50805164 | 0.90650187 |
| Calr3 | 0.01290202 | 0.01329649 | 0.01569992 | -0.0024034 | 0.75192482 | 0.90650187 |
| Gm44210 | 0.0109788 | 0.01097088 | 0.01337699 | -0.0024061 | 0.78803733 | 0.90650187 |
| Apod | 0.01548857 | 0.01640755 | 0.01881456 | -0.002407 | 0.71523407 | 0.90650187 |
| B3gnt2 | 0.00974715 | 0.00947703 | 0.01188619 | -0.0024092 | 0.81622228 | 0.9092537 |
| Ppp2r2c | 0.00855562 | 0.00803094 | 0.01044158 | -0.0024106 | 0.84763711 | 0.9143112 |
| Eci3 | 0.01981197 | 0.02158362 | 0.02399651 | -0.0024129 | 0.67256716 | 0.90650187 |
| Reep1 | 0.05056941 | 0.05759749 | 0.06001115 | -0.0024137 | 0.57095366 | 0.90650187 |
| Gm20633 | 0.00427117 | 0.00281273 | 0.00522784 | -0.0024151 | 0.97999949 | 0.98492378 |

| | | | | | | |
|---|---|---|---|---|---|---|
| Tmem169 | 0.01994694 | 0.02174266 | 0.0241578 | -0.0024151 | 0.67163877 | 0.90650187 |
| Tnfrsf9 | 0.0036543 | 0.00205928 | 0.00447468 | -0.0024154 | 0.99179576 | 0.99395355 |
| Cndp1 | 0.01043387 | 0.01030183 | 0.0127177 | -0.0024159 | 0.80080301 | 0.90809448 |
| Tmem179 | 0.00759022 | 0.00685182 | 0.00926942 | -0.0024176 | 0.87687024 | 0.92473192 |
| Stac2 | 0.00386244 | 0.00231094 | 0.00472887 | -0.0024179 | 0.98848984 | 0.99109782 |
| Gm15537 | 0.0152606 | 0.01612148 | 0.01854048 | -0.002419 | 0.71905104 | 0.90650187 |
| Cyp2d26 | 0.00446509 | 0.00304515 | 0.00546448 | -0.0024193 | 0.97547954 | 0.98096484 |
| Slc25a48 | 0.00409004 | 0.00258719 | 0.00500675 | -0.0024196 | 0.98415845 | 0.98786269 |
| Gm5784 | 0.02863266 | 0.03205575 | 0.03447609 | -0.0024203 | 0.62251629 | 0.90650187 |
| Ccnf | 0.05431664 | 0.06187909 | 0.06429953 | -0.0024204 | 0.56645161 | 0.90650187 |
| Slc17a4 | 0.01112672 | 0.01113369 | 0.01355586 | -0.0024222 | 0.78650237 | 0.90650187 |
| Cpne7 | 0.00682394 | 0.00591422 | 0.00833795 | -0.0024237 | 0.90186945 | 0.93591959 |
| Ttc27 | 0.10796323 | 0.12108053 | 0.12350795 | -0.0024274 | 0.53481287 | 0.90650187 |
| Grem2 | 0.01093576 | 0.01089745 | 0.01332493 | -0.0024275 | 0.79099623 | 0.90650887 |
| Kiss1r | 0.00794508 | 0.00727242 | 0.00970046 | -0.002428 | 0.86709069 | 0.92056229 |
| Gm4262 | 0.00407032 | 0.00255257 | 0.00498268 | -0.0024301 | 0.98492714 | 0.98821581 |
| Npas2 | 0.00373892 | 0.00214681 | 0.00457803 | -0.0024312 | 0.99088453 | 0.99323677 |
| Gm29361 | 0.00425309 | 0.00277065 | 0.00520578 | -0.0024351 | 0.98121599 | 0.98582773 |
| Gm12002 | 0.00352807 | 0.00188497 | 0.00432048 | -0.0024355 | 0.99389553 | 0.99513937 |
| Slc26a6 | 0.0268628 | 0.0299474 | 0.03238328 | -0.0024359 | 0.63096102 | 0.90650187 |
| Zfp3 | 0.05062883 | 0.05764334 | 0.06007931 | -0.002436 | 0.57152227 | 0.90650187 |
| Pld6 | 0.01299546 | 0.01337386 | 0.01581262 | -0.0024388 | 0.75352977 | 0.90650187 |
| Tnnt3 | 0.00914257 | 0.00871461 | 0.01115349 | -0.0024389 | 0.83449943 | 0.91062493 |
| Yme1l1 | 0.76622963 | 0.63114016 | 0.63357907 | -0.0024389 | 0.50682678 | 0.90650187 |
| Adnp | 0.00423059 | 0.00273882 | 0.00517831 | -0.0024395 | 0.98188418 | 0.9862311 |
| Tmem183a | 0.53342899 | 0.48649445 | 0.4889377 | -0.0024432 | 0.50886174 | 0.90650187 |
| Magix | 0.00416603 | 0.00265538 | 0.00509951 | -0.0024441 | 0.98344411 | 0.98734125 |
| Dpysl5 | 0.02100567 | 0.02297642 | 0.02542196 | -0.0024455 | 0.66555033 | 0.90650187 |
| Fbxl21 | 0.01220624 | 0.01241362 | 0.01486023 | -0.0024466 | 0.76789443 | 0.90650187 |
| Timm8a1 | 0.04029562 | 0.04569715 | 0.0481445 | -0.0024473 | 0.58939351 | 0.90650187 |

| | | | | | | |
|---|---|---|---|---|---|---|
| Glod5 | 0.00945491 | 0.00908359 | 0.0115321 | -0.0024485 | 0.8273884 | 0.90947782 |
| Mtfr2 | 0.00941145 | 0.00903063 | 0.01147942 | -0.0024488 | 0.82852145 | 0.9096066 |
| Gm527 | 0.00410331 | 0.00257207 | 0.00502296 | -0.0024509 | 0.98496532 | 0.98821581 |
| Adamts19 | 0.01367393 | 0.01417945 | 0.01663057 | -0.0024511 | 0.74383735 | 0.90650187 |
| Gm26908 | 0.00642988 | 0.00540632 | 0.00785856 | -0.0024522 | 0.91731495 | 0.94283706 |
| Gm26852 | 0.03213549 | 0.03614995 | 0.03860354 | -0.0024536 | 0.61124146 | 0.90650187 |
| Alg9 | 0.1466025 | 0.16155339 | 0.16400841 | -0.002455 | 0.52652826 | 0.90650187 |
| Ccdc28b | 0.63607093 | 0.55392665 | 0.55638381 | -0.0024572 | 0.50783198 | 0.90650187 |
| Clcn4 | 0.55438653 | 0.50076765 | 0.50322659 | -0.0024589 | 0.50866545 | 0.90650187 |
| Nol11 | 0.3487545 | 0.34736423 | 0.34982702 | -0.0024628 | 0.51248374 | 0.90650187 |
| RP23-361O1 | 0.00423552 | 0.00272126 | 0.00518433 | -0.0024631 | 0.98266109 | 0.98681583 |
| Gm14443 | 0.01094859 | 0.01087736 | 0.01334045 | -0.0024631 | 0.79412195 | 0.90731056 |
| Amer3 | 0.00390727 | 0.00231623 | 0.00478361 | -0.0024674 | 0.98907719 | 0.99155592 |
| Ppil3 | 0.30008044 | 0.30621419 | 0.30868251 | -0.0024683 | 0.51417881 | 0.90650187 |
| Rasgrp1 | 0.00641484 | 0.00537017 | 0.00784025 | -0.0024701 | 0.9193323 | 0.94392235 |
| Corin | 0.00405706 | 0.0024942 | 0.00496649 | -0.0024723 | 0.98655111 | 0.98954559 |
| Gm12359 | 0.00416501 | 0.00262339 | 0.00509826 | -0.0024749 | 0.98453806 | 0.98811323 |
| Glb1l | 0.04993887 | 0.05681214 | 0.05928747 | -0.0024753 | 0.5736247 | 0.90650187 |
| Fam221b | 0.02690544 | 0.02995802 | 0.03243376 | -0.0024757 | 0.63282428 | 0.90650187 |
| Spred3 | 0.07637495 | 0.08664445 | 0.08912205 | -0.0024776 | 0.54917924 | 0.90650187 |
| Cabin1 | 0.32353662 | 0.32628992 | 0.32876901 | -0.0024791 | 0.51337094 | 0.90650187 |
| Gad1os | 0.00681494 | 0.00584645 | 0.008327 | -0.0024806 | 0.90730088 | 0.93809554 |
| Prodh2 | 0.02838822 | 0.03170363 | 0.03418733 | -0.0024837 | 0.62664215 | 0.90650187 |
| 2010308F09I | 0.02464848 | 0.02727367 | 0.02975793 | -0.0024843 | 0.64472938 | 0.90650187 |
| Col2a1 | 0.00637028 | 0.00530005 | 0.00778603 | -0.002486 | 0.9221127 | 0.94549985 |
| Gm16853 | 0.00958798 | 0.00920727 | 0.01169335 | -0.0024861 | 0.82771287 | 0.90947782 |
| Xpnpep2 | 0.02089646 | 0.02280453 | 0.02529165 | -0.0024871 | 0.66900589 | 0.90650187 |
| 3830408C21I | 0.0081079 | 0.0074082 | 0.00989817 | -0.00249 | 0.86828754 | 0.92093489 |
| Fam217a | 0.0139072 | 0.01442142 | 0.01691161 | -0.0024902 | 0.74363906 | 0.90650187 |
| 9230105E05I | 0.00713951 | 0.00622966 | 0.00872167 | -0.002492 | 0.89799815 | 0.9343964 |

| | | | | | | |
|---|---|---|---|---|---|---|
| Gm26802 | 0.00793054 | 0.00719078 | 0.00968281 | -0.002492 | 0.8737195 | 0.9235672 |
| Slc35f2 | 0.00691667 | 0.00595864 | 0.00845072 | -0.0024921 | 0.9050691 | 0.93712658 |
| Pcdhga11 | 0.0102574 | 0.01000998 | 0.0125041 | -0.0024941 | 0.81239055 | 0.9092537 |
| Lrrtm4 | 0.01266244 | 0.01291543 | 0.01541088 | -0.0024954 | 0.7641966 | 0.90650187 |
| Creb3 | 0.72601876 | 0.60814313 | 0.61064095 | -0.0024978 | 0.50725422 | 0.90650187 |
| Phldb1 | 0.16460637 | 0.1797778 | 0.18227576 | -0.002498 | 0.52429009 | 0.90650187 |
| Gm11772 | 0.01787089 | 0.01917194 | 0.0216737 | -0.0025018 | 0.69607652 | 0.90650187 |
| 4430402I18R | 0.00737222 | 0.00650081 | 0.00900453 | -0.0025037 | 0.89179008 | 0.93134318 |
| BC051628 | 0.00799857 | 0.00726088 | 0.00976542 | -0.0025045 | 0.8728911 | 0.92289911 |
| A430046D13 | 0.01912615 | 0.02066844 | 0.02317651 | -0.0025081 | 0.68477205 | 0.90650187 |
| Ttc39c | 0.0245327 | 0.02710919 | 0.02962044 | -0.0025112 | 0.64687693 | 0.90650187 |
| Gm15445 | 0.00766526 | 0.00684927 | 0.00936059 | -0.0025113 | 0.88350575 | 0.92767795 |
| Erich2os | 0.01136006 | 0.01132032 | 0.01383797 | -0.0025177 | 0.79069227 | 0.90650187 |
| Tex40 | 0.00693699 | 0.00595567 | 0.00847543 | -0.0025198 | 0.90686381 | 0.9379403 |
| Npm2 | 0.00715556 | 0.00621981 | 0.00874118 | -0.0025214 | 0.90013553 | 0.93499868 |
| RP24-274B19 | 0.01008756 | 0.0097767 | 0.01229847 | -0.0025218 | 0.81899978 | 0.9092537 |
| Cfap73 | 0.01251711 | 0.01271309 | 0.0152355 | -0.0025224 | 0.76914085 | 0.90650187 |
| Rrn3 | 0.21472745 | 0.22870652 | 0.23123087 | -0.0025244 | 0.51935417 | 0.90650187 |
| Acat3 | 0.00674944 | 0.00572264 | 0.00824733 | -0.0025247 | 0.91322717 | 0.94082155 |
| Ccdc116 | 0.01078966 | 0.01062102 | 0.01314821 | -0.0025272 | 0.80358004 | 0.90882608 |
| Slc25a36 | 0.43430376 | 0.41486401 | 0.41739211 | -0.0025281 | 0.51074083 | 0.90650187 |
| Acta1 | 0.00716839 | 0.00622797 | 0.00875678 | -0.0025288 | 0.90039672 | 0.93504361 |
| Pcdhb21 | 0.01818701 | 0.01952124 | 0.02205239 | -0.0025311 | 0.69506597 | 0.90650187 |
| 3300002I08R | 0.02438312 | 0.0269116 | 0.0294428 | -0.0025312 | 0.64883928 | 0.90650187 |
| Cntn1 | 0.02969463 | 0.0331936 | 0.03572946 | -0.0025359 | 0.62381739 | 0.90650187 |
| Gm9776 | 0.09134219 | 0.10302909 | 0.10556541 | -0.0025363 | 0.54253023 | 0.90650187 |
| Tmem198b | 0.15109867 | 0.16605809 | 0.16860555 | -0.0025475 | 0.5267763 | 0.90650187 |
| Sema3a | 0.0501417 | 0.05697259 | 0.05952033 | -0.0025477 | 0.57546 | 0.90650187 |
| Cyp2r1 | 0.00726667 | 0.00632746 | 0.00887624 | -0.0025488 | 0.89911805 | 0.93440169 |
| Zfp85os | 0.00738017 | 0.00646462 | 0.00901419 | -0.0025496 | 0.8956944 | 0.93320429 |

| | | | | | | |
|---|---|---|---|---|---|---|
| Lrif1 | 0.30518386 | 0.31054408 | 0.31309422 | -0.0025501 | 0.51444226 | 0.90650187 |
| Lrrc74b | 0.01256667 | 0.01274497 | 0.0152953 | -0.0025503 | 0.77073207 | 0.90650187 |
| Ifitm10 | 0.01001594 | 0.00965765 | 0.01221174 | -0.0025541 | 0.82376318 | 0.9092537 |
| Sdhaf1 | 0.46403096 | 0.43701061 | 0.43956572 | -0.0025551 | 0.51030807 | 0.90650187 |
| Exoc3l4 | 0.05211251 | 0.05922236 | 0.06177965 | -0.0025573 | 0.5730013 | 0.90650187 |
| Gata4 | 0.01017714 | 0.00984774 | 0.01240693 | -0.0025592 | 0.82042807 | 0.9092537 |
| P2rx4 | 0.24540644 | 0.2573295 | 0.25989091 | -0.0025614 | 0.51747389 | 0.90650187 |
| Arpp21 | 0.00993878 | 0.00955533 | 0.0121183 | -0.002563 | 0.826449 | 0.90947782 |
| Kcnh7 | 0.0275476 | 0.03062927 | 0.03319362 | -0.0025643 | 0.63436702 | 0.90650187 |
| Ifnk | 0.00602547 | 0.00480052 | 0.0073663 | -0.0025658 | 0.93924461 | 0.95593613 |
| Lef1 | 0.01236225 | 0.01248239 | 0.01504858 | -0.0025662 | 0.77580319 | 0.90650187 |
| Slc46a2 | 0.0105305 | 0.01026527 | 0.01283465 | -0.0025694 | 0.81325877 | 0.9092537 |
| Chn1os3 | 0.01283484 | 0.01304927 | 0.01561889 | -0.0025696 | 0.7677296 | 0.90650187 |
| Gsto2 | 0.02752771 | 0.03059922 | 0.0331701 | -0.0025709 | 0.63478749 | 0.90650187 |
| Parp3 | 0.02521375 | 0.02785639 | 0.03042886 | -0.0025725 | 0.64647977 | 0.90650187 |
| Gm16008 | 0.00753492 | 0.00662579 | 0.00920224 | -0.0025764 | 0.89337589 | 0.93181736 |
| Dnmt3bos | 0.00736707 | 0.00641704 | 0.00899827 | -0.0025812 | 0.89889105 | 0.93440169 |
| Sh3tc2 | 0.01417508 | 0.01465051 | 0.01723426 | -0.0025838 | 0.74744887 | 0.90650187 |
| Zcchc17 | 0.71405032 | 0.60107776 | 0.60366251 | -0.0025847 | 0.50759343 | 0.90650187 |
| Hnf4g | 0.00677184 | 0.00568677 | 0.00827457 | -0.0025878 | 0.9177852 | 0.94297078 |
| Cmya5 | 0.01240449 | 0.01250656 | 0.01509956 | -0.002593 | 0.77739636 | 0.90650187 |
| 4930590J08F | 0.00717869 | 0.00617157 | 0.0087693 | -0.0025977 | 0.90606767 | 0.93764958 |
| Dgkg | 0.01869505 | 0.02006132 | 0.02266067 | -0.0025994 | 0.69495494 | 0.90650187 |
| Lrp8os2 | 0.00701415 | 0.00596814 | 0.00856925 | -0.0026011 | 0.91139899 | 0.94034997 |
| Gm26901 | 0.00734521 | 0.00636522 | 0.0089717 | -0.0026065 | 0.90174314 | 0.93591959 |
| H2-Eb1 | 0.0147904 | 0.01536832 | 0.01797492 | -0.0026066 | 0.74043011 | 0.90650187 |
| Prr5l | 0.04214535 | 0.04768217 | 0.05029291 | -0.0026107 | 0.59125503 | 0.90650187 |
| Gramd1a | 0.45317148 | 0.4289282 | 0.43153949 | -0.0026113 | 0.51073057 | 0.90650187 |
| Ccdc40 | 0.02858506 | 0.031805 | 0.03441986 | -0.0026149 | 0.63221717 | 0.90650187 |
| Copg1 | 1.01101603 | 0.75261657 | 0.75523317 | -0.0026166 | 0.50614442 | 0.90650187 |

| | | | | | | |
|---|---|---|---|---|---|---|
| Hhat | 0.04057988 | 0.04585659 | 0.048475 | -0.0026184 | 0.59488629 | 0.90650187 |
| Tdrd5 | 0.0071392 | 0.00610248 | 0.00872129 | -0.0026188 | 0.90904839 | 0.93919616 |
| Fstl5 | 0.00711481 | 0.00607005 | 0.00869164 | -0.0026216 | 0.91002264 | 0.93950381 |
| Lrp1b | 0.00661529 | 0.00546255 | 0.00808414 | -0.0026216 | 0.92530096 | 0.94735258 |
| Ift80 | 0.12379811 | 0.13769739 | 0.14032453 | -0.0026271 | 0.53316574 | 0.90650187 |
| Snx8 | 0.17555007 | 0.19057134 | 0.19320004 | -0.0026287 | 0.5241163 | 0.90650187 |
| Efhc1 | 0.03911988 | 0.04414179 | 0.04677617 | -0.0026344 | 0.59884999 | 0.90650187 |
| Fancb | 0.02365618 | 0.02594176 | 0.02857891 | -0.0026372 | 0.65917891 | 0.90650187 |
| Stxbp3 | 0.30823398 | 0.31308123 | 0.31571988 | -0.0026386 | 0.51481907 | 0.90650187 |
| Rorb | 0.01070502 | 0.01040254 | 0.01304581 | -0.0026433 | 0.81613837 | 0.9092537 |
| Gm5616 | 0.00752481 | 0.00654413 | 0.00918996 | -0.0026458 | 0.89971118 | 0.93469858 |
| Actr2 | 0.79592623 | 0.64738339 | 0.65002995 | -0.0026466 | 0.50722047 | 0.90650187 |
| Pde7a | 0.11007906 | 0.12312581 | 0.12577284 | -0.002647 | 0.53727167 | 0.90650187 |
| C2cd4b | 0.04483416 | 0.05075905 | 0.05340655 | -0.0026475 | 0.5872124 | 0.90650187 |
| Nacad | 0.00688621 | 0.00576504 | 0.00841368 | -0.0026486 | 0.91916581 | 0.94387891 |
| Tmco4 | 0.01375168 | 0.01407499 | 0.01672425 | -0.0026493 | 0.75935229 | 0.90650187 |
| B230303O12 | 0.01007424 | 0.0096319 | 0.01228234 | -0.0026504 | 0.83130345 | 0.9098597 |
| Tmem91 | 0.02041098 | 0.02206026 | 0.0247121 | -0.0026518 | 0.68333851 | 0.90650187 |
| Gm2788 | 0.01306078 | 0.0132395 | 0.0158914 | -0.0026519 | 0.77091588 | 0.90650187 |
| Syngap1 | 0.02589033 | 0.02857908 | 0.03123126 | -0.0026522 | 0.64710774 | 0.90650187 |
| Tdo2 | 0.01018582 | 0.0097591 | 0.01241744 | -0.0026583 | 0.82938095 | 0.90974008 |
| C230037L18I | 0.01209423 | 0.01206535 | 0.01472498 | -0.0026596 | 0.78900336 | 0.90650187 |
| Mamstr | 0.06888608 | 0.07811196 | 0.08077452 | -0.0026626 | 0.55825258 | 0.90650187 |
| Ikbke | 0.05137135 | 0.05826805 | 0.06093067 | -0.0026626 | 0.57701753 | 0.90650187 |
| Gm9755 | 0.00606828 | 0.00475183 | 0.00741843 | -0.0026666 | 0.94497832 | 0.95968836 |
| Ppfia4 | 0.01124755 | 0.01103374 | 0.01370195 | -0.0026682 | 0.8066733 | 0.9092537 |
| Dnase1l2 | 0.00740248 | 0.006372 | 0.0090413 | -0.0026693 | 0.9053235 | 0.93719848 |
| Gm16062 | 0.0168846 | 0.01782175 | 0.02049111 | -0.0026694 | 0.71874764 | 0.90650187 |
| Scube3 | 0.01954612 | 0.02100881 | 0.02367873 | -0.0026699 | 0.6919092 | 0.90650187 |
| Gm26566 | 0.00712858 | 0.00603767 | 0.00870838 | -0.0026707 | 0.91362078 | 0.94098174 |

| | | | | | | |
|---|---|---|---|---|---|---|
| Brsk2 | 0.01186941 | 0.01178165 | 0.01445345 | -0.0026718 | 0.79440306 | 0.90731056 |
| Hykk | 0.02523576 | 0.02778316 | 0.03045498 | -0.0026718 | 0.65173708 | 0.90650187 |
| Gm15541 | 0.00953577 | 0.00895768 | 0.01163008 | -0.0026724 | 0.84649788 | 0.91408934 |
| Masp2 | 0.01416213 | 0.01454434 | 0.01721866 | -0.0026743 | 0.7550507 | 0.90650187 |
| Zscan25 | 0.01294527 | 0.01307744 | 0.01575209 | -0.0026747 | 0.77483028 | 0.90650187 |
| Greb1 | 0.0065443 | 0.00532266 | 0.00799777 | -0.0026751 | 0.93148927 | 0.95113723 |
| Mar-04 | 0.0533435 | 0.06051072 | 0.06318788 | -0.0026772 | 0.57469987 | 0.90650187 |
| Cacna2d4 | 0.0122031 | 0.01217596 | 0.01485644 | -0.0026805 | 0.78875299 | 0.90650187 |
| Wnt2 | 0.01016608 | 0.00971299 | 0.01239355 | -0.0026806 | 0.83185612 | 0.91013901 |
| Ccdc142os | 0.02760329 | 0.0305757 | 0.03325948 | -0.0026838 | 0.64009992 | 0.90650187 |
| Abcf1 | 1.04390872 | 0.76566071 | 0.76834848 | -0.0026878 | 0.5062038 | 0.90650187 |
| Slc25a32 | 0.06040074 | 0.06852932 | 0.07121753 | -0.0026882 | 0.56663215 | 0.90650187 |
| Gm17477 | 0.01219156 | 0.01215221 | 0.01484251 | -0.0026903 | 0.7898193 | 0.90650187 |
| Appbp2os | 0.0067969 | 0.00561471 | 0.00830505 | -0.0026903 | 0.92508194 | 0.94726646 |
| Sybu | 0.0240221 | 0.02632255 | 0.02901386 | -0.0026913 | 0.65996658 | 0.90650187 |
| Adcy5 | 0.02212677 | 0.02406352 | 0.02675864 | -0.0026951 | 0.67283716 | 0.90650187 |
| C1rl | 0.01900148 | 0.020329 | 0.02302737 | -0.0026984 | 0.69879546 | 0.90650187 |
| Gm12204 | 0.0073061 | 0.00622466 | 0.00892417 | -0.0026995 | 0.91065031 | 0.93983499 |
| Selenbp2 | 0.00833593 | 0.00747413 | 0.01017498 | -0.0027008 | 0.88100828 | 0.92635898 |
| Cntd1 | 0.00648868 | 0.00522841 | 0.0079301 | -0.0027017 | 0.93505851 | 0.95330814 |
| Gm7598 | 0.01296523 | 0.01307242 | 0.01577617 | -0.0027038 | 0.77693943 | 0.90650187 |
| Gm26982 | 0.01408942 | 0.01442454 | 0.0171311 | -0.0027066 | 0.75877476 | 0.90650187 |
| A930024E05 | 0.00702569 | 0.0058704 | 0.00858329 | -0.0027129 | 0.92000509 | 0.94435798 |
| 2410004I01R | 0.01743513 | 0.01843838 | 0.0211514 | -0.002713 | 0.71573679 | 0.90650187 |
| Hist1h2af | 0.02000101 | 0.02150928 | 0.0242224 | -0.0027131 | 0.69073939 | 0.90650187 |
| Ell3 | 0.00851203 | 0.00766994 | 0.01038868 | -0.0027187 | 0.87767151 | 0.92482217 |
| Gm12758 | 0.00670049 | 0.0054662 | 0.00818779 | -0.0027216 | 0.93025621 | 0.95057977 |
| Zfp592 | 0.11110833 | 0.12414961 | 0.12687261 | -0.002723 | 0.53800682 | 0.90650187 |
| Pan2 | 0.08172443 | 0.09230473 | 0.09503554 | -0.0027308 | 0.55082376 | 0.90650187 |
| Fsbp | 0.00666011 | 0.00540764 | 0.00813867 | -0.002731 | 0.93212152 | 0.95146248 |

| | | | | | | |
|---|---|---|---|---|---|---|
| Gm996 | 0.00674663 | 0.00551214 | 0.00824392 | -0.0027318 | 0.92964258 | 0.95027213 |
| Npas3 | 0.04218868 | 0.04761025 | 0.05034318 | -0.0027329 | 0.59535198 | 0.90650187 |
| 2210411M09 | 0.0060449 | 0.00465697 | 0.00738997 | -0.002733 | 0.94992177 | 0.96276378 |
| Erbb2ip | 0.22437541 | 0.2376131 | 0.24034768 | -0.0027346 | 0.52017005 | 0.90650187 |
| 1700003F12I | 0.00643379 | 0.00512755 | 0.00786332 | -0.0027358 | 0.93903115 | 0.95593613 |
| Lrrc23 | 0.00643809 | 0.00513076 | 0.00786854 | -0.0027378 | 0.93904489 | 0.95593613 |
| Micall2 | 0.01815245 | 0.01927127 | 0.022011 | -0.0027397 | 0.7099874 | 0.90650187 |
| Slc12a8 | 0.01275759 | 0.01278587 | 0.01552569 | -0.0027398 | 0.78363165 | 0.90650187 |
| Gpr4 | 0.00615852 | 0.00478763 | 0.00752828 | -0.0027406 | 0.94721284 | 0.96125124 |
| Tomm40l | 0.15820295 | 0.17307873 | 0.1758212 | -0.0027425 | 0.52764167 | 0.90650187 |
| Zfp947 | 0.01631383 | 0.01706333 | 0.01980603 | -0.0027427 | 0.73092675 | 0.90650187 |
| Lrrc20 | 0.01453682 | 0.01492627 | 0.01766975 | -0.0027435 | 0.75497995 | 0.90650187 |
| Cntn3 | 0.00704374 | 0.00586144 | 0.00860524 | -0.0027438 | 0.92183002 | 0.94549985 |
| Mlph | 0.00624916 | 0.00489356 | 0.00763862 | -0.0027451 | 0.94493405 | 0.95968836 |
| Gm42477 | 0.00983593 | 0.009247 | 0.01199372 | -0.0027467 | 0.84568591 | 0.9136218 |
| Hist1h3e | 0.01246591 | 0.01242285 | 0.0151737 | -0.0027508 | 0.78986302 | 0.90650187 |
| Tbc1d12 | 0.04251789 | 0.04797234 | 0.05072498 | -0.0027526 | 0.59531727 | 0.90650187 |
| Mast3 | 0.22769255 | 0.24070621 | 0.24345957 | -0.0027534 | 0.52004901 | 0.90650187 |
| Zfp131 | 0.30720804 | 0.31208417 | 0.31483763 | -0.0027535 | 0.51550686 | 0.90650187 |
| Nhs | 0.07866212 | 0.08890115 | 0.09165537 | -0.0027542 | 0.55313622 | 0.90650187 |
| Ppp1r14d | 0.01048007 | 0.01001723 | 0.01277362 | -0.0027564 | 0.83129854 | 0.9098597 |
| Ticrr | 0.03320571 | 0.03710381 | 0.03986076 | -0.0027569 | 0.62076024 | 0.90650187 |
| Cdk6 | 0.1395072 | 0.15394695 | 0.15670541 | -0.0027585 | 0.53118752 | 0.90650187 |
| Gm3336 | 0.00629178 | 0.0049318 | 0.00769048 | -0.0027587 | 0.94461023 | 0.95944274 |
| Gm15265 | 0.00713918 | 0.00596176 | 0.00872126 | -0.0027595 | 0.92023252 | 0.94452765 |
| Pcdhgc5 | 0.01773823 | 0.0187536 | 0.02151473 | -0.0027611 | 0.7158424 | 0.90650187 |
| Fam46d | 0.00680296 | 0.00554977 | 0.00831243 | -0.0027627 | 0.93023067 | 0.95057977 |
| Foxo6 | 0.01338021 | 0.01350983 | 0.01627656 | -0.0027667 | 0.77507819 | 0.90650187 |
| Sp7 | 0.01258359 | 0.01254373 | 0.01531573 | -0.002772 | 0.78947805 | 0.90650187 |
| Gm42469 | 0.01536851 | 0.01589676 | 0.01867023 | -0.0027735 | 0.74550036 | 0.90650187 |

| | | | | | | |
|---|---|---|---|---|---|---|
| Klf8 | 0.01470256 | 0.01509403 | 0.01786922 | -0.0027752 | 0.75503769 | 0.90650187 |
| Tmem231 | 0.10592499 | 0.11854476 | 0.12132086 | -0.0027761 | 0.54051276 | 0.90650187 |
| Gad1 | 0.0618674 | 0.07009979 | 0.07287699 | -0.0027772 | 0.56726421 | 0.90650187 |
| D130051D11 | 0.02175063 | 0.02353296 | 0.02631039 | -0.0027774 | 0.68057145 | 0.90650187 |
| Zfp870 | 0.08158963 | 0.09210877 | 0.09488704 | -0.0027783 | 0.55178254 | 0.90650187 |
| Spag4 | 0.00589908 | 0.00442928 | 0.00721241 | -0.0027831 | 0.95687061 | 0.96737098 |
| Has3 | 0.00562332 | 0.00409134 | 0.00687653 | -0.0027852 | 0.96411682 | 0.97282043 |
| RP24-267C3. | 0.01376447 | 0.0139544 | 0.01673966 | -0.0027853 | 0.77025589 | 0.90650187 |
| Tmem59l | 0.00691296 | 0.00565999 | 0.00844621 | -0.0027862 | 0.92874491 | 0.94980162 |
| Nat8l | 0.00499951 | 0.00332892 | 0.00611629 | -0.0027874 | 0.97861725 | 0.98366467 |
| Klhdc2 | 0.98810256 | 0.74300546 | 0.74579454 | -0.0027891 | 0.50663228 | 0.90650187 |
| Proca1 | 0.06764699 | 0.07659581 | 0.07938552 | -0.0027897 | 0.56207192 | 0.90650187 |
| Ccdc180 | 0.00970652 | 0.0090471 | 0.01183696 | -0.0027899 | 0.85263217 | 0.91557373 |
| Usp3 | 0.36854881 | 0.36319015 | 0.3659801 | -0.00279 | 0.51351755 | 0.90650187 |
| Gm16140 | 0.00666741 | 0.00535743 | 0.00814755 | -0.0027901 | 0.93604488 | 0.953802 |
| Chrm2 | 0.02097128 | 0.02258184 | 0.02538092 | -0.0027991 | 0.68802997 | 0.90650187 |
| B930036N10 | 0.05358991 | 0.06067025 | 0.0634695 | -0.0027992 | 0.57772151 | 0.90650187 |
| Spata2 | 0.07812264 | 0.08825671 | 0.0910585 | -0.0028018 | 0.55440053 | 0.90650187 |
| 1190005I06R | 0.03462484 | 0.03871808 | 0.0415251 | -0.002807 | 0.61810776 | 0.90650187 |
| Gm37459 | 0.00712843 | 0.00589761 | 0.00870819 | -0.0028106 | 0.9243313 | 0.94682419 |
| Ccdc130 | 0.12013125 | 0.13364562 | 0.13645765 | -0.002812 | 0.53649682 | 0.90650187 |
| Vav3 | 0.01873019 | 0.01988811 | 0.02270273 | -0.0028146 | 0.70923579 | 0.90650187 |
| Mrpl53 | 0.00626384 | 0.00484148 | 0.00765648 | -0.002815 | 0.94892104 | 0.96195723 |
| Ptpro | 0.01353596 | 0.01364605 | 0.0164643 | -0.0028183 | 0.77666243 | 0.90650187 |
| Pif1 | 0.00600515 | 0.00452315 | 0.00734157 | -0.0028184 | 0.95605753 | 0.96699916 |
| Sntg1 | 0.00945891 | 0.00871805 | 0.01153694 | -0.0028189 | 0.86130978 | 0.91859749 |
| 4933438K21I | 0.00917065 | 0.00836513 | 0.01118754 | -0.0028224 | 0.86897195 | 0.92140364 |
| Mettl21a | 0.19487634 | 0.20934389 | 0.21216728 | -0.0028234 | 0.52358741 | 0.90650187 |
| Stard6 | 0.01692723 | 0.01771343 | 0.02054226 | -0.0028288 | 0.72979566 | 0.90650187 |
| Rtca | 0.37589832 | 0.36906338 | 0.37189523 | -0.0028318 | 0.5135023 | 0.90650187 |

| | | | | | | |
|---|---|---|---|---|---|---|
| Snap91 | 0.02290526 | 0.02484979 | 0.02768564 | -0.0028358 | 0.67557664 | 0.90650187 |
| Magi2 | 0.06970219 | 0.07884873 | 0.08168815 | -0.0028394 | 0.5614024 | 0.90650187 |
| Sez6 | 0.00652776 | 0.00513776 | 0.00797765 | -0.0028399 | 0.94323804 | 0.95871682 |
| 1700010I14R | 0.00952396 | 0.00877586 | 0.01161577 | -0.0028399 | 0.86145839 | 0.91867924 |
| Rims3 | 0.07246696 | 0.08193593 | 0.08477609 | -0.0028402 | 0.55919795 | 0.90650187 |
| Tmem190 | 0.00657551 | 0.00519437 | 0.00803575 | -0.0028414 | 0.94201726 | 0.95777624 |
| Tslp | 0.02059805 | 0.02209217 | 0.02493546 | -0.0028433 | 0.69389294 | 0.90650187 |
| Spata2l | 0.05097859 | 0.05763583 | 0.06048045 | -0.0028446 | 0.5828122 | 0.90650187 |
| Aoc2 | 0.00805346 | 0.00698456 | 0.00983206 | -0.0028475 | 0.90104164 | 0.93544147 |
| Gm17546 | 0.01435423 | 0.01460119 | 0.01744996 | -0.0028488 | 0.76600779 | 0.90650187 |
| Ust | 0.01312751 | 0.01312162 | 0.01597188 | -0.0028503 | 0.78621022 | 0.90650187 |
| Sdr42e1 | 0.0143515 | 0.01459406 | 0.01744668 | -0.0028526 | 0.76635004 | 0.90650187 |
| Arg2 | 0.01008737 | 0.00944297 | 0.01229824 | -0.0028553 | 0.84899462 | 0.91444933 |
| F2 | 0.01970626 | 0.02101446 | 0.02387017 | -0.0028557 | 0.70258518 | 0.90650187 |
| Gm1818 | 0.00995365 | 0.00927901 | 0.01213631 | -0.0028573 | 0.85236924 | 0.91557373 |
| Hif3a | 0.17517274 | 0.18996728 | 0.19282566 | -0.0028584 | 0.52627128 | 0.90650187 |
| AA986860 | 0.01371412 | 0.01381946 | 0.016679 | -0.0028595 | 0.77702331 | 0.90650187 |
| Vma21-ps | 0.01493986 | 0.01529247 | 0.01815473 | -0.0028623 | 0.75831469 | 0.90650187 |
| Cyb5d1 | 0.01650234 | 0.01716942 | 0.02003236 | -0.0028629 | 0.73739908 | 0.90650187 |
| Kcnip4 | 0.0159963 | 0.01656127 | 0.01942468 | -0.0028634 | 0.74387233 | 0.90650187 |
| Gm43796 | 0.0931206 | 0.1046322 | 0.10749828 | -0.0028661 | 0.54717497 | 0.90650187 |
| Cdh23 | 0.0193393 | 0.02056314 | 0.02343145 | -0.0028683 | 0.70684773 | 0.90650187 |
| Zfp276 | 0.08546521 | 0.09627811 | 0.09914656 | -0.0028684 | 0.55116988 | 0.90650187 |
| Itih1 | 0.00593505 | 0.00438457 | 0.00725621 | -0.0028716 | 0.96074042 | 0.97053465 |
| Ltbp2 | 0.0098757 | 0.00917005 | 0.0120419 | -0.0028718 | 0.85547845 | 0.9165432 |
| Tnfrsf22 | 0.0162986 | 0.01691232 | 0.01978774 | -0.0028754 | 0.7408622 | 0.90650187 |
| Gm38037 | 0.00597247 | 0.00442592 | 0.00730177 | -0.0028759 | 0.96002128 | 0.9701425 |
| Gm6576 | 0.10204998 | 0.11427162 | 0.11714849 | -0.0028769 | 0.54346723 | 0.90650187 |
| Atp10a | 0.03727568 | 0.0417477 | 0.04462558 | -0.0028779 | 0.612827 | 0.90650187 |
| Ebf4 | 0.01180091 | 0.01149251 | 0.0143707 | -0.0028782 | 0.81336742 | 0.9092537 |

| | | | | | | |
|---|---|---|---|---|---|---|
| Spag5 | 0.04604203 | 0.05192312 | 0.05480166 | -0.0028785 | 0.5923183 | 0.90650187 |
| Alb | 0.0061274 | 0.00460968 | 0.0074904 | -0.0028807 | 0.95634186 | 0.96707976 |
| Kcnq2 | 0.00584753 | 0.00426878 | 0.00714963 | -0.0028809 | 0.96337757 | 0.97220358 |
| RP23-143D1: | 0.01904131 | 0.02019251 | 0.02307501 | -0.0028825 | 0.71066894 | 0.90650187 |
| Bub1 | 0.05990526 | 0.06776499 | 0.0706562 | -0.0028912 | 0.57217384 | 0.90650187 |
| 4930481A15I | 0.02789936 | 0.03071807 | 0.03360958 | -0.0028915 | 0.6489419 | 0.90650187 |
| Endov | 0.10899574 | 0.12172197 | 0.1246139 | -0.0028919 | 0.54108586 | 0.90650187 |
| Tmem144 | 0.00897408 | 0.00805529 | 0.01094919 | -0.0028939 | 0.87999959 | 0.92579928 |
| Aire | 0.0162068 | 0.01678241 | 0.0196775 | -0.0028951 | 0.74346607 | 0.90650187 |
| Unc13c | 0.00543146 | 0.00374528 | 0.00664279 | -0.0028975 | 0.97375691 | 0.97994563 |
| Hist1h4b | 0.00633494 | 0.00484011 | 0.00774302 | -0.0029029 | 0.95221155 | 0.96420588 |
| Stk26 | 0.03526343 | 0.03936367 | 0.042273 | -0.0029093 | 0.62018144 | 0.90650187 |
| Itga3 | 0.06621954 | 0.07487185 | 0.07778259 | -0.0029107 | 0.56606364 | 0.90650187 |
| Usp27x | 0.06421473 | 0.07261337 | 0.07552629 | -0.0029129 | 0.56806872 | 0.90650187 |
| Stmnd1 | 0.00580189 | 0.00417969 | 0.00709404 | -0.0029144 | 0.96609985 | 0.97436862 |
| Gm28172 | 0.04139908 | 0.04651122 | 0.04942678 | -0.0029156 | 0.60342986 | 0.90650187 |
| Acsl6 | 0.03234912 | 0.03593788 | 0.03885464 | -0.0029168 | 0.63070649 | 0.90650187 |
| Lpcat2 | 0.02011959 | 0.02144614 | 0.02436407 | -0.0029179 | 0.70278313 | 0.90650187 |
| RP24-159C1: | 0.00669874 | 0.00526747 | 0.00818566 | -0.0029182 | 0.94350122 | 0.95876476 |
| Osbp2 | 0.00645388 | 0.00496632 | 0.00788776 | -0.0029214 | 0.95017478 | 0.96290712 |
| Pgk1-rs7 | 0.0090978 | 0.00816957 | 0.0110992 | -0.0029296 | 0.87968545 | 0.92562058 |
| Cth | 0.05347133 | 0.06040138 | 0.06333399 | -0.0029326 | 0.58154565 | 0.90650187 |
| Gm15728 | 0.01026601 | 0.0095815 | 0.01251452 | -0.002933 | 0.8512746 | 0.9150732 |
| Caprin2 | 0.07878508 | 0.08885766 | 0.09179135 | -0.0029337 | 0.55649296 | 0.90650187 |
| Gria4 | 0.00951468 | 0.00866863 | 0.01160452 | -0.0029359 | 0.86964563 | 0.92144928 |
| Hadha | 0.84091451 | 0.67125733 | 0.67419417 | -0.0029368 | 0.50772519 | 0.90650187 |
| Rdm1 | 0.22064733 | 0.23389794 | 0.23683635 | -0.0029384 | 0.52199303 | 0.90650187 |
| Hscb | 0.17274909 | 0.18747641 | 0.19041698 | -0.0029406 | 0.52736697 | 0.90650187 |
| Crkl | 0.01903549 | 0.02012577 | 0.02306805 | -0.0029423 | 0.7146524 | 0.90650187 |
| Zfp7 | 0.05250932 | 0.0592901 | 0.06223385 | -0.0029437 | 0.58327668 | 0.90650187 |

| | | | | | | |
|---|---|---|---|---|---|---|
| Il1rapl1 | 0.01215247 | 0.01185152 | 0.01479531 | -0.0029438 | 0.81179546 | 0.9092537 |
| Gm26620 | 0.00548642 | 0.00376425 | 0.00670975 | -0.0029455 | 0.974505 | 0.98047076 |
| Capn9 | 0.00675015 | 0.00530069 | 0.0082482 | -0.0029475 | 0.94392974 | 0.95903292 |
| Bcas1 | 0.00774341 | 0.00650518 | 0.00945553 | -0.0029504 | 0.91729988 | 0.94283706 |
| Car5b | 0.00594045 | 0.00431057 | 0.00726279 | -0.0029522 | 0.96462486 | 0.97313927 |
| Tmc6 | 0.13343477 | 0.14745461 | 0.15040822 | -0.0029536 | 0.53478336 | 0.90650187 |
| Ank1 | 0.0099293 | 0.00915278 | 0.01210681 | -0.002954 | 0.86097637 | 0.91848494 |
| 4833422C13I | 0.00805661 | 0.00688066 | 0.0098359 | -0.0029552 | 0.90917611 | 0.93919616 |
| Slc38a11 | 0.01834363 | 0.01927988 | 0.02223997 | -0.0029601 | 0.72297427 | 0.90650187 |
| Zfp94 | 0.04007194 | 0.04492394 | 0.04788435 | -0.0029604 | 0.60828261 | 0.90650187 |
| Ccdc36 | 0.02744782 | 0.03011341 | 0.03307559 | -0.0029622 | 0.65473609 | 0.90650187 |
| Bbof1 | 0.04347901 | 0.04887468 | 0.05183867 | -0.002964 | 0.60032486 | 0.90650187 |
| Gm10371 | 0.00635509 | 0.00480323 | 0.00776754 | -0.0029643 | 0.95511058 | 0.96642721 |
| Iqsec3 | 0.00660971 | 0.00511303 | 0.00807736 | -0.0029643 | 0.94860766 | 0.96195723 |
| Acy3 | 0.09775298 | 0.1095459 | 0.1125124 | -0.0029665 | 0.54665428 | 0.90650187 |
| Gm20649 | 0.0088329 | 0.00781106 | 0.01077796 | -0.0029669 | 0.88947875 | 0.9302877 |
| Xpnpep3 | 0.06604654 | 0.07462094 | 0.07758811 | -0.0029672 | 0.56749964 | 0.90650187 |
| Plekhf1 | 0.05565069 | 0.06285347 | 0.06582116 | -0.0029677 | 0.57943164 | 0.90650187 |
| Klhdc7a | 0.00503505 | 0.00318955 | 0.00615962 | -0.0029701 | 0.98396737 | 0.98773611 |
| Rec8 | 0.01265804 | 0.01243295 | 0.01540557 | -0.0029726 | 0.80450226 | 0.90882608 |
| A730071L15I | 0.00632523 | 0.00475753 | 0.00773119 | -0.0029737 | 0.95635983 | 0.96707976 |
| Teddm2 | 0.0138752 | 0.01389709 | 0.01687307 | -0.002976 | 0.78350638 | 0.90650187 |
| Syne3 | 0.00601343 | 0.00437542 | 0.00735165 | -0.0029762 | 0.96404998 | 0.97281756 |
| Cntn4 | 0.01002387 | 0.00924342 | 0.01222135 | -0.0029779 | 0.86064979 | 0.91847356 |
| Gm8973 | 0.00596074 | 0.00430734 | 0.00728749 | -0.0029802 | 0.96546718 | 0.97379514 |
| Shisa7 | 0.00923842 | 0.00828469 | 0.01126969 | -0.002985 | 0.88050306 | 0.92607258 |
| 2600006K01I | 0.02564758 | 0.02795291 | 0.03094344 | -0.0029905 | 0.66627352 | 0.90650187 |
| B230369F24 | 0.01963528 | 0.02079441 | 0.02378533 | -0.0029909 | 0.71192522 | 0.90650187 |
| Trim34a | 0.01761352 | 0.01837278 | 0.02136525 | -0.0029925 | 0.73324665 | 0.90650187 |
| Slc52a3 | 0.01301273 | 0.01283766 | 0.01583346 | -0.0029958 | 0.79986514 | 0.90805731 |

| | | | | | | |
|---|---|---|---|---|---|---|
| Gtf2a1l | 0.00693 | 0.0054677 | 0.00846693 | -0.0029992 | 0.94234399 | 0.95797305 |
| Sema3f | 0.12542719 | 0.13903588 | 0.14203713 | -0.0030013 | 0.53741996 | 0.90650187 |
| Gm42893 | 0.00658392 | 0.00504315 | 0.00804598 | -0.0030028 | 0.95145608 | 0.96369163 |
| 1700112J16R | 0.00795494 | 0.00670757 | 0.00971244 | -0.0030049 | 0.91549548 | 0.94214655 |
| 1700040D17I | 0.00650718 | 0.00494255 | 0.00795261 | -0.0030101 | 0.95377972 | 0.96556047 |
| Hist2h2aa2 | 0.00981454 | 0.00895722 | 0.01196782 | -0.0030106 | 0.86828621 | 0.92093489 |
| E2f8 | 0.0263657 | 0.02878035 | 0.03179458 | -0.0030142 | 0.66328951 | 0.90650187 |
| Vwa2 | 0.01521139 | 0.01546644 | 0.01848131 | -0.0030149 | 0.76583949 | 0.90650187 |
| Kri1 | 0.1378879 | 0.15201189 | 0.15503043 | -0.0030185 | 0.53448889 | 0.90650187 |
| Cd59b | 0.03362221 | 0.03733065 | 0.04034954 | -0.0030189 | 0.63028716 | 0.90650187 |
| Ube2ql1 | 0.00551277 | 0.00372247 | 0.00674186 | -0.0030194 | 0.97675909 | 0.98199057 |
| Snx30 | 0.13795959 | 0.15208224 | 0.15510465 | -0.0030224 | 0.53451649 | 0.90650187 |
| B230354K17 | 0.20450872 | 0.21844465 | 0.22146962 | -0.003025 | 0.5242092 | 0.90650187 |
| Arl15 | 0.07212317 | 0.08136639 | 0.08439272 | -0.0030263 | 0.56333089 | 0.90650187 |
| Haus6 | 0.04989186 | 0.05620634 | 0.0592335 | -0.0030272 | 0.58986376 | 0.90650187 |
| Gm26721 | 0.01739108 | 0.01806974 | 0.02109858 | -0.0030288 | 0.73832739 | 0.90650187 |
| Pde4c | 0.00556684 | 0.00377833 | 0.00680774 | -0.0030294 | 0.97605076 | 0.98140942 |
| Tesk2 | 0.02024319 | 0.0214822 | 0.02451171 | -0.0030295 | 0.70865251 | 0.90650187 |
| Gm15420 | 0.00902725 | 0.00797958 | 0.01101366 | -0.0030341 | 0.88965448 | 0.93029083 |
| Slc12a4 | 0.14729365 | 0.16168194 | 0.16471666 | -0.0030347 | 0.53263897 | 0.90650187 |
| Abcd2 | 0.0119827 | 0.01155524 | 0.01459028 | -0.003035 | 0.82245592 | 0.9092537 |
| Zfp773 | 0.01000778 | 0.00916508 | 0.01220186 | -0.0030368 | 0.86572666 | 0.92040222 |
| Polr1c | 0.6050478 | 0.53361694 | 0.5366546 | -0.0030377 | 0.51003782 | 0.90650187 |
| H2-Q7 | 0.00961352 | 0.00868559 | 0.01172429 | -0.0030387 | 0.87538261 | 0.92418014 |
| Loxl4 | 0.00618579 | 0.00452191 | 0.00756147 | -0.0030396 | 0.96303511 | 0.9719225 |
| Slc6a19 | 0.01175279 | 0.0112729 | 0.01431257 | -0.0030397 | 0.82745256 | 0.90947782 |
| Tmem240 | 0.02948168 | 0.03243603 | 0.03547827 | -0.0030422 | 0.64847586 | 0.90650187 |
| Hist1h2bm | 0.00809616 | 0.00684119 | 0.00988391 | -0.0030427 | 0.91443155 | 0.94149783 |
| Chac1 | 0.01012435 | 0.00929777 | 0.01234301 | -0.0030452 | 0.86363801 | 0.91985436 |
| 1700001L19R | 0.00901611 | 0.00795456 | 0.01100015 | -0.0030456 | 0.89080973 | 0.9309865 |

| | | | | | | |
|---|---|---|---|---|---|---|
| Gm11110 | 0.03786192 | 0.04226333 | 0.04530978 | -0.0030464 | 0.6174936 | 0.90650187 |
| Rnasek | 0.00941195 | 0.00843236 | 0.01148003 | -0.0030477 | 0.88104016 | 0.92635898 |
| Bdh2 | 0.02039255 | 0.021642 | 0.02469009 | -0.0030481 | 0.70843767 | 0.90650187 |
| B230317F23 | 0.01240551 | 0.01205264 | 0.0151008 | -0.0030482 | 0.81523727 | 0.9092537 |
| Fam120c | 0.02030584 | 0.02153701 | 0.02458654 | -0.0030495 | 0.70931978 | 0.90650187 |
| Recql5 | 0.049144 | 0.05532164 | 0.05837433 | -0.0030527 | 0.59191863 | 0.90650187 |
| Tmem52 | 0.00593494 | 0.00420325 | 0.00725607 | -0.0030528 | 0.96928487 | 0.97673844 |
| Synm | 0.01860883 | 0.01950047 | 0.02255747 | -0.003057 | 0.72656962 | 0.90650187 |
| Acacb | 0.00976654 | 0.00885183 | 0.01190967 | -0.0030578 | 0.87315305 | 0.92306833 |
| 2810001G20 | 0.18478346 | 0.19925362 | 0.20231248 | -0.0030589 | 0.52679483 | 0.90650187 |
| Gm7221 | 0.00519376 | 0.00329119 | 0.0063531 | -0.0030619 | 0.98392666 | 0.98773611 |
| Ms4a4d | 0.01890631 | 0.01985075 | 0.02291349 | -0.0030627 | 0.72381898 | 0.90650187 |
| Gm20342 | 0.18416786 | 0.19864361 | 0.20170784 | -0.0030642 | 0.52692217 | 0.90650187 |
| Ccdc25 | 0.41167037 | 0.3970064 | 0.400073 | -0.0030666 | 0.51359172 | 0.90650187 |
| Trpc5 | 0.01634796 | 0.016779 | 0.01984701 | -0.003068 | 0.75402646 | 0.90650187 |
| Gm2861 | 0.02694511 | 0.02941214 | 0.03248071 | -0.0030686 | 0.66275487 | 0.90650187 |
| Rnft2 | 0.00989055 | 0.00899025 | 0.01205989 | -0.0030696 | 0.87108857 | 0.92225579 |
| Gm16845 | 0.04158842 | 0.04657249 | 0.04964661 | -0.0030741 | 0.60844653 | 0.90650187 |
| Zfp955a | 0.01897789 | 0.01992437 | 0.02299914 | -0.0030748 | 0.72385668 | 0.90650187 |
| 4930547M16 | 0.005993 | 0.00425153 | 0.00732677 | -0.0030752 | 0.96897511 | 0.97653493 |
| Il1rap | 0.02341714 | 0.02521698 | 0.02829466 | -0.0030777 | 0.68565083 | 0.90650187 |
| Cited1 | 0.12087447 | 0.13416271 | 0.13724273 | -0.00308 | 0.53973616 | 0.90650187 |
| Gm4876 | 0.0124062 | 0.0120208 | 0.01510163 | -0.0030808 | 0.81777901 | 0.9092537 |
| Pnrc2 | 0.63496885 | 0.55260812 | 0.55569215 | -0.003084 | 0.50984197 | 0.90650187 |
| Taf5 | 0.05627322 | 0.06344355 | 0.0665303 | -0.0030868 | 0.58170574 | 0.90650187 |
| Mrpl15 | 0.74963543 | 0.62111692 | 0.62420647 | -0.0030895 | 0.50877754 | 0.90650187 |
| Dusp19 | 0.25204082 | 0.26287387 | 0.26596408 | -0.0030902 | 0.52059742 | 0.90650187 |
| Adssl1 | 0.03197522 | 0.0353224 | 0.0384151 | -0.0030927 | 0.63979256 | 0.90650187 |
| Slc31a1 | 0.43787811 | 0.4169971 | 0.42009241 | -0.0030953 | 0.5130654 | 0.90650187 |
| A330074K22 | 0.00589611 | 0.00411312 | 0.0072088 | -0.0030957 | 0.97187381 | 0.9786985 |

| | | | | | | |
|---|---|---|---|---|---|---|
| Zfp869 | 0.00572273 | 0.00389987 | 0.00699763 | -0.0030978 | 0.97546657 | 0.98096484 |
| Tgm5 | 0.00615562 | 0.00442544 | 0.00752475 | -0.0030993 | 0.96645466 | 0.97446784 |
| 2610528J11F | 0.1934409 | 0.20767028 | 0.21077237 | -0.0031021 | 0.52608383 | 0.90650187 |
| Zfp407 | 0.08850328 | 0.09936717 | 0.1024708 | -0.0031036 | 0.55355505 | 0.90650187 |
| 2610020H08I | 0.03080745 | 0.033937 | 0.03704095 | -0.003104 | 0.64525163 | 0.90650187 |
| Slc6a20a | 0.01282375 | 0.01250121 | 0.0156055 | -0.0031043 | 0.81174197 | 0.9092537 |
| Bad | 0.82836565 | 0.664438 | 0.66754343 | -0.0031054 | 0.50824995 | 0.90650187 |
| Gm42809 | 0.01196406 | 0.0114615 | 0.01456777 | -0.0031063 | 0.828418 | 0.9096066 |
| Gm26533 | 0.00926968 | 0.00820002 | 0.01130759 | -0.0031076 | 0.88909826 | 0.9302877 |
| Rrp12 | 0.0595744 | 0.06717178 | 0.07028116 | -0.0031094 | 0.57796201 | 0.90650187 |
| Gm26664 | 0.00623499 | 0.00451125 | 0.00762137 | -0.0031101 | 0.96517321 | 0.97356323 |
| Sytl3 | 0.00895951 | 0.00781795 | 0.01093151 | -0.0031136 | 0.89728225 | 0.93409036 |
| Tspear | 0.00975524 | 0.00878145 | 0.01189599 | -0.0031145 | 0.87777273 | 0.92484883 |
| Kif16b | 0.12162036 | 0.13491372 | 0.13802995 | -0.0031162 | 0.53997325 | 0.90650187 |
| Trpv4 | 0.02623481 | 0.02852326 | 0.0316395 | -0.0031162 | 0.66925488 | 0.90650187 |
| Nol3 | 0.01471394 | 0.01476237 | 0.01788292 | -0.0031205 | 0.78105271 | 0.90650187 |
| Gm16090 | 0.00614102 | 0.00437977 | 0.00750698 | -0.0031272 | 0.96798156 | 0.97564776 |
| Zfp605 | 0.05563305 | 0.06267278 | 0.06580105 | -0.0031283 | 0.58369282 | 0.90650187 |
| Shank3 | 0.00515738 | 0.00317143 | 0.00630875 | -0.0031373 | 0.98647474 | 0.98953428 |
| Pik3cd | 0.00878824 | 0.00758592 | 0.0107238 | -0.0031379 | 0.9033403 | 0.93635703 |
| Gm37035 | 0.0099826 | 0.00903221 | 0.01217136 | -0.0031392 | 0.87422264 | 0.9235672 |
| Lgals12 | 0.02207103 | 0.02355068 | 0.02669224 | -0.0031416 | 0.69959218 | 0.90650187 |
| Dpp8 | 0.17951819 | 0.19398513 | 0.19712763 | -0.0031425 | 0.52824909 | 0.90650187 |
| Fam181b | 0.07691363 | 0.08657623 | 0.08971937 | -0.0031431 | 0.56188214 | 0.90650187 |
| Gm4577 | 0.02389273 | 0.02571555 | 0.02886012 | -0.0031446 | 0.68594486 | 0.90650187 |
| Slc5a9 | 0.01363752 | 0.01344093 | 0.01658669 | -0.0031458 | 0.80042358 | 0.90806412 |
| Ptger1 | 0.00908896 | 0.00793862 | 0.01108849 | -0.0031499 | 0.89667811 | 0.93371722 |
| 8030423F21I | 0.01666863 | 0.01708152 | 0.02023195 | -0.0031504 | 0.75561026 | 0.90650187 |
| Dnaja4 | 0.26203273 | 0.27187757 | 0.27502953 | -0.003152 | 0.52031674 | 0.90650187 |
| 2900026A02I | 0.24803156 | 0.25914533 | 0.26229946 | -0.0031541 | 0.52131652 | 0.90650187 |

| | | | | | | |
|---|---|---|---|---|---|---|
| Tmem40 | 0.0062302 | 0.00446031 | 0.00761554 | -0.0031552 | 0.9672537 | 0.97501482 |
| Adam5 | 0.01068726 | 0.00986834 | 0.01302433 | -0.003156 | 0.85931271 | 0.91797966 |
| Rnf122 | 0.09469365 | 0.10604543 | 0.10920429 | -0.0031589 | 0.55116068 | 0.90650187 |
| Anks1b | 0.0061335 | 0.00433489 | 0.00749782 | -0.0031629 | 0.96962886 | 0.9770203 |
| Vash2 | 0.153371 | 0.16775426 | 0.17091979 | -0.0031655 | 0.53280987 | 0.90650187 |
| Irgm2 | 0.00952539 | 0.0084512 | 0.01161751 | -0.0031663 | 0.88717363 | 0.92936447 |
| Hist1h3a | 0.00931362 | 0.00819436 | 0.01136085 | -0.0031665 | 0.89234174 | 0.93159913 |
| Frrs1l | 0.00928 | 0.00814905 | 0.0113201 | -0.003171 | 0.89349424 | 0.93186533 |
| Pcsk4 | 0.03753224 | 0.04175332 | 0.04492507 | -0.0031718 | 0.62318788 | 0.90650187 |
| Stap1 | 0.00865368 | 0.00738678 | 0.01056055 | -0.0031738 | 0.90923267 | 0.93919616 |
| 4732456N10 | 0.00955885 | 0.00848392 | 0.01165805 | -0.0031741 | 0.88693786 | 0.92936447 |
| Hspa13 | 0.47073779 | 0.44130651 | 0.44448126 | -0.0031748 | 0.51266557 | 0.90650187 |
| Cd22 | 0.00853971 | 0.00724484 | 0.01042227 | -0.0031774 | 0.9123434 | 0.94062208 |
| Hist2h2be | 0.03352716 | 0.03706036 | 0.04023802 | -0.0031777 | 0.6372346 | 0.90650187 |
| 5330439K02I | 0.01253289 | 0.0120731 | 0.01525454 | -0.0031814 | 0.82307837 | 0.9092537 |
| St6galnac5 | 0.13631931 | 0.15022305 | 0.15340496 | -0.0031819 | 0.53673451 | 0.90650187 |
| Barx2 | 0.01955905 | 0.02051192 | 0.02369419 | -0.0031823 | 0.72476995 | 0.90650187 |
| RP23-16L19.4 | 0.00862234 | 0.00734023 | 0.01052253 | -0.0031823 | 0.91060217 | 0.93983499 |
| Fam160a1 | 0.00877086 | 0.00752009 | 0.01070271 | -0.0031826 | 0.90691133 | 0.9379403 |
| Slc22a18 | 0.02039513 | 0.02150825 | 0.02469317 | -0.0031849 | 0.71681069 | 0.90650187 |
| 2900011O08 | 0.00869919 | 0.00742989 | 0.01061577 | -0.0031859 | 0.90892512 | 0.93913613 |
| Phf19 | 0.02096367 | 0.02218352 | 0.02537185 | -0.0031883 | 0.71180096 | 0.90650187 |
| A730011C13 | 0.00888964 | 0.00765777 | 0.01084678 | -0.003189 | 0.90439882 | 0.93681543 |
| Mapkap1 | 0.29583076 | 0.30179704 | 0.30499076 | -0.0031937 | 0.51856498 | 0.90650187 |
| D230017M19 | 0.01860724 | 0.01936089 | 0.02255557 | -0.0031947 | 0.73554079 | 0.90650187 |
| Sertad4 | 0.03973603 | 0.0442964 | 0.04749351 | -0.0031971 | 0.61763071 | 0.90650187 |
| Pdp1 | 0.07335285 | 0.08256319 | 0.0857632 | -0.0032 | 0.56587224 | 0.90650187 |
| Rinl | 0.02986723 | 0.03273284 | 0.035933 | -0.0032002 | 0.65391825 | 0.90650187 |
| Lrrc3 | 0.03832105 | 0.04264431 | 0.04584527 | -0.003201 | 0.62187012 | 0.90650187 |
| Cib3 | 0.00885816 | 0.00760548 | 0.0108086 | -0.0032031 | 0.90615663 | 0.9376778 |

| | | | | | | |
|---|---|---|---|---|---|---|
| Fzd9 | 0.00471257 | 0.00256309 | 0.00576637 | -0.0032033 | 0.99323652 | 0.99474162 |
| Fuk | 0.04234947 | 0.04732462 | 0.05052968 | -0.0032051 | 0.61102116 | 0.90650187 |
| Pabpc5 | 0.00855238 | 0.00723118 | 0.01043765 | -0.0032065 | 0.91398333 | 0.9412276 |
| Pcdhga7 | 0.01369179 | 0.01344497 | 0.01665209 | -0.0032071 | 0.80405735 | 0.90882608 |
| Med15 | 0.17827456 | 0.19268861 | 0.19589856 | -0.0032099 | 0.52903513 | 0.90650187 |
| Fam110c | 0.03058209 | 0.03356491 | 0.03677551 | -0.0032106 | 0.65103379 | 0.90650187 |
| Zfp652os | 0.00826295 | 0.00687314 | 0.01008639 | -0.0032132 | 0.92164853 | 0.94549985 |
| Mybl1 | 0.01143413 | 0.01071365 | 0.01392749 | -0.0032138 | 0.84751847 | 0.9143112 |
| Pigb | 0.08359437 | 0.093879 | 0.09709303 | -0.003214 | 0.55849754 | 0.90650187 |
| 6430550D23I | 0.01198761 | 0.01138073 | 0.01459622 | -0.0032155 | 0.8362959 | 0.91079271 |
| 6430573P05I | 0.01055132 | 0.00964318 | 0.01285984 | -0.0032167 | 0.86692862 | 0.92056229 |
| Calca | 0.01622546 | 0.01648294 | 0.01969991 | -0.003217 | 0.76606916 | 0.90650187 |
| Tarsl2 | 0.0737927 | 0.08303564 | 0.08625287 | -0.0032172 | 0.56585118 | 0.90650187 |
| Prob1 | 0.01417585 | 0.01401318 | 0.01723519 | -0.003222 | 0.79703501 | 0.9075779 |
| Adamtsl3 | 0.01628205 | 0.01654438 | 0.01976787 | -0.0032235 | 0.76575344 | 0.90650187 |
| Gm13544 | 0.01209841 | 0.01150633 | 0.01473003 | -0.0032237 | 0.8347062 | 0.91062493 |
| Gm43949 | 0.00825499 | 0.0068521 | 0.01007674 | -0.0032246 | 0.9225784 | 0.94559465 |
| Gm16365 | 0.01090278 | 0.01005873 | 0.01328505 | -0.0032263 | 0.85984657 | 0.91802142 |
| Gm14085 | 0.00905632 | 0.0078209 | 0.01104891 | -0.003228 | 0.90299616 | 0.93635703 |
| 2210406O10 | 0.00865098 | 0.00732909 | 0.01055728 | -0.0032282 | 0.91298532 | 0.9407546 |
| Phtf1os | 0.02712672 | 0.02946595 | 0.03269566 | -0.0032297 | 0.66972045 | 0.90650187 |
| Rnd1 | 0.01186095 | 0.01120756 | 0.01444323 | -0.0032357 | 0.84035927 | 0.91226452 |
| Frs2 | 0.09275811 | 0.10386683 | 0.10710466 | -0.0032378 | 0.55345424 | 0.90650187 |
| Cox7a1 | 0.06921331 | 0.07790093 | 0.08114096 | -0.00324 | 0.57044887 | 0.90650187 |
| Ces2e | 0.0052677 | 0.00320229 | 0.00644321 | -0.0032409 | 0.98732841 | 0.99012925 |
| Nyap2 | 0.01139263 | 0.01063449 | 0.01387733 | -0.0032428 | 0.85056155 | 0.91483663 |
| Nim1k | 0.02787618 | 0.03033924 | 0.03358217 | -0.0032429 | 0.66614738 | 0.90650187 |
| Gm17227 | 0.02191119 | 0.02325569 | 0.02650176 | -0.0032461 | 0.70695832 | 0.90650187 |
| Gm26899 | 0.01255233 | 0.0120318 | 0.01527801 | -0.0032462 | 0.82756452 | 0.90947782 |
| Gipc2 | 0.03987538 | 0.04440901 | 0.04765567 | -0.0032467 | 0.61900472 | 0.90650187 |

| | | | | | | |
|---|---|---|---|---|---|---|
| Cyp2j5 | 0.00473657 | 0.00254872 | 0.00579565 | -0.0032469 | 0.99362338 | 0.99493242 |
| Ppih | 0.33082104 | 0.33166024 | 0.33490744 | -0.0032472 | 0.51719058 | 0.90650187 |
| Fam234a | 0.3481792 | 0.34610252 | 0.34935275 | -0.0032502 | 0.51649554 | 0.90650187 |
| 4932438A13I | 0.37105408 | 0.36474888 | 0.36800177 | -0.0032529 | 0.51567284 | 0.90650187 |
| Etv4 | 0.00941384 | 0.00822675 | 0.01148232 | -0.0032556 | 0.89624782 | 0.93346099 |
| Zfp874b | 0.02232291 | 0.0237323 | 0.02699229 | -0.00326 | 0.70433746 | 0.90650187 |
| Dennd6b | 0.06997272 | 0.07872864 | 0.08199079 | -0.0032621 | 0.57019715 | 0.90650187 |
| Ybx2 | 0.0087506 | 0.00741356 | 0.01067813 | -0.0032646 | 0.91294782 | 0.9407546 |
| 2200002D01I | 0.02882365 | 0.03143683 | 0.03470162 | -0.0032648 | 0.66211795 | 0.90650187 |
| Gm10010 | 0.00935856 | 0.00814944 | 0.01141532 | -0.0032659 | 0.89829054 | 0.93440169 |
| Srsf2 | 0.8907114 | 0.69662045 | 0.69988838 | -0.0032679 | 0.50828046 | 0.90650187 |
| Tat | 0.00899543 | 0.00770672 | 0.01097508 | -0.0032684 | 0.90723059 | 0.93808668 |
| Crp | 0.01264576 | 0.0121209 | 0.01539075 | -0.0032698 | 0.82754131 | 0.90947782 |
| Gm16537 | 0.00912269 | 0.00785876 | 0.01112939 | -0.0032706 | 0.90429895 | 0.93677582 |
| Ccdc173 | 0.11856979 | 0.13153444 | 0.13480605 | -0.0032716 | 0.54295869 | 0.90650187 |
| Gm13373 | 0.00564982 | 0.00363548 | 0.00690882 | -0.0032733 | 0.98241036 | 0.98669441 |
| Sec61a2 | 0.37942609 | 0.37144482 | 0.37471888 | -0.0032741 | 0.51549219 | 0.90650187 |
| Atrip | 0.00964903 | 0.00849168 | 0.01176731 | -0.0032756 | 0.89205073 | 0.93145097 |
| Alpk2 | 0.01158961 | 0.01083852 | 0.0141154 | -0.0032769 | 0.8489744 | 0.91444933 |
| Paqr8 | 0.01075967 | 0.00983387 | 0.01311194 | -0.0032781 | 0.86680852 | 0.92056229 |
| Slc20a1 | 0.07965932 | 0.08947787 | 0.09275754 | -0.0032797 | 0.56245058 | 0.90650187 |
| Gm4978 | 0.04160137 | 0.0463792 | 0.04966164 | -0.0032824 | 0.61555871 | 0.90650187 |
| Pex11g | 0.15667461 | 0.17098976 | 0.17427378 | -0.003284 | 0.53338156 | 0.90650187 |
| Gm26609 | 0.02315291 | 0.02469402 | 0.02798034 | -0.0032863 | 0.69921396 | 0.90650187 |
| Cnot10 | 0.25968355 | 0.26962019 | 0.27290678 | -0.0032866 | 0.52134836 | 0.90650187 |
| Plxnc1 | 0.02472868 | 0.02656609 | 0.02985316 | -0.0032871 | 0.68775474 | 0.90650187 |
| Lrrc75a | 0.04488997 | 0.05018378 | 0.05347106 | -0.0032873 | 0.60769103 | 0.90650187 |
| Dmc1 | 0.00867906 | 0.00730319 | 0.01059135 | -0.0032882 | 0.91623274 | 0.94231902 |
| Slc25a35 | 0.05024209 | 0.05634362 | 0.05963555 | -0.0032919 | 0.59692709 | 0.90650187 |
| C130021I20R | 0.02122179 | 0.02238747 | 0.02567979 | -0.0032923 | 0.71564653 | 0.90650187 |

| | | | | | | |
|---|---|---|---|---|---|---|
| Atp8b5 | 0.00872497 | 0.00735359 | 0.01064704 | -0.0032934 | 0.91545887 | 0.94214655 |
| 1700008O03 | 0.00901828 | 0.00770447 | 0.01100279 | -0.0032983 | 0.90867896 | 0.93907328 |
| Galr2 | 0.01956113 | 0.02039813 | 0.02369667 | -0.0032985 | 0.73198333 | 0.90650187 |
| Gabra3 | 0.00864746 | 0.00725427 | 0.01055302 | -0.0032987 | 0.91768128 | 0.94297078 |
| Gm13248 | 0.01201407 | 0.01132912 | 0.01462817 | -0.0032991 | 0.84197459 | 0.91251845 |
| Ddhd2 | 0.26768093 | 0.27681273 | 0.28011185 | -0.0032991 | 0.52087893 | 0.90650187 |
| Pcdhgb4 | 0.02103902 | 0.02215997 | 0.02546175 | -0.0033018 | 0.7178577 | 0.90650187 |
| Dscam | 0.00888053 | 0.00752407 | 0.01083573 | -0.0033117 | 0.91287671 | 0.9407546 |
| Sh3bp5l | 0.24080427 | 0.25233941 | 0.25565186 | -0.0033125 | 0.52296686 | 0.90650187 |
| Mapk8 | 0.28203423 | 0.2895767 | 0.29288929 | -0.0033126 | 0.52005039 | 0.90650187 |
| Klhl3 | 0.00744302 | 0.00577638 | 0.00909057 | -0.0033142 | 0.94746502 | 0.96144298 |
| Tmem145 | 0.01166112 | 0.01088398 | 0.01420181 | -0.0033178 | 0.85050051 | 0.91483663 |
| Colq | 0.02052555 | 0.02152963 | 0.0248489 | -0.0033193 | 0.72368943 | 0.90650187 |
| Clybl | 0.08780338 | 0.09838581 | 0.10170612 | -0.0033203 | 0.55769665 | 0.90650187 |
| Tctn1 | 0.11191623 | 0.12441223 | 0.12773493 | -0.0033227 | 0.54603153 | 0.90650187 |
| Mycn | 0.04618685 | 0.05164391 | 0.05496879 | -0.0033249 | 0.60599657 | 0.90650187 |
| Hdhd2 | 0.41790632 | 0.40155633 | 0.40488401 | -0.0033277 | 0.51457322 | 0.90650187 |
| Bcat1 | 0.05666959 | 0.06365354 | 0.06698152 | -0.003328 | 0.58740712 | 0.90650187 |
| Acot11 | 0.01058404 | 0.00956979 | 0.01289944 | -0.0033296 | 0.8744158 | 0.92356828 |
| Gm29650 | 0.00822663 | 0.00671146 | 0.0100423 | -0.0033308 | 0.92982935 | 0.95037317 |
| Fibp | 0.46263448 | 0.43520674 | 0.43853819 | -0.0033315 | 0.51347052 | 0.90650187 |
| Cage1 | 0.01257478 | 0.01197221 | 0.01530509 | -0.0033329 | 0.83349997 | 0.91060043 |
| Gm8225 | 0.01146683 | 0.01063185 | 0.01396701 | -0.0033352 | 0.85578226 | 0.9165432 |
| Atp2c2 | 0.01358753 | 0.01319083 | 0.01652645 | -0.0033356 | 0.81521537 | 0.9092537 |
| Tns3 | 0.28040745 | 0.28811442 | 0.2914509 | -0.0033365 | 0.52029442 | 0.90650187 |
| Unc13d | 0.01640053 | 0.01657313 | 0.01991012 | -0.003337 | 0.77189347 | 0.90650187 |
| Styk1 | 0.01017336 | 0.00906382 | 0.01240235 | -0.0033385 | 0.88428659 | 0.9278081 |
| Ppp2r5e | 0.36937837 | 0.36331032 | 0.36665013 | -0.0033398 | 0.51615073 | 0.90650187 |
| Pla2g2c | 0.02345066 | 0.02499288 | 0.02833452 | -0.0033416 | 0.69996319 | 0.90650187 |
| 1700001C19I | 0.00871542 | 0.00729313 | 0.01063546 | -0.0033423 | 0.91880643 | 0.94363736 |

| | | | | | | |
|---|---|---|---|---|---|---|
| Grhl2 | 0.02103921 | 0.02211815 | 0.02546198 | -0.0033438 | 0.72033088 | 0.90650187 |
| Clp1 | 0.10723618 | 0.11938323 | 0.1227284 | -0.0033452 | 0.54822285 | 0.90650187 |
| Veph1 | 0.01402716 | 0.01370628 | 0.01705611 | -0.0033498 | 0.80870089 | 0.9092537 |
| Npb | 0.038814 | 0.04306658 | 0.04641983 | -0.0033532 | 0.62594756 | 0.90650187 |
| Nfkbie | 0.02712333 | 0.02933776 | 0.03269166 | -0.0033539 | 0.67583514 | 0.90650187 |
| A530095I07F | 0.01176954 | 0.01097596 | 0.0143328 | -0.0033568 | 0.85110647 | 0.91505486 |
| Slc41a1 | 0.07162145 | 0.08047249 | 0.08383292 | -0.0033604 | 0.57071767 | 0.90650187 |
| Slc9a5 | 0.02628581 | 0.02833715 | 0.03169994 | -0.0033628 | 0.68139306 | 0.90650187 |
| Bcl2l14 | 0.00892178 | 0.00751659 | 0.01088576 | -0.0033692 | 0.9155777 | 0.94216738 |
| Plekha6 | 0.09881996 | 0.11029363 | 0.11366311 | -0.0033695 | 0.5524238 | 0.90650187 |
| Gm15859 | 0.00909532 | 0.00772471 | 0.0110962 | -0.0033715 | 0.91161396 | 0.94039219 |
| Gm1661 | 0.01138876 | 0.01049493 | 0.01387266 | -0.0033777 | 0.86046753 | 0.9184725 |
| Atoh8 | 0.01591522 | 0.01594335 | 0.01932728 | -0.0033839 | 0.78182218 | 0.90650187 |
| Mmaa | 0.140998 | 0.15486004 | 0.15824473 | -0.0033847 | 0.53787715 | 0.90650187 |
| Gm17541 | 0.0332084 | 0.03647737 | 0.03986392 | -0.0033865 | 0.64716083 | 0.90650187 |
| Pecr | 0.16885599 | 0.18314305 | 0.18653383 | -0.0033908 | 0.53220402 | 0.90650187 |
| Arfgap2 | 0.66552358 | 0.57122858 | 0.57461949 | -0.0033909 | 0.51046472 | 0.90650187 |
| St6galnac6 | 0.1781044 | 0.19233839 | 0.19573026 | -0.0033919 | 0.53070378 | 0.90650187 |
| Kctd17 | 0.01142761 | 0.01052459 | 0.01391962 | -0.003395 | 0.86088284 | 0.91848494 |
| St8sia5 | 0.01428418 | 0.01396786 | 0.01736563 | -0.0033978 | 0.80780334 | 0.9092537 |
| Ces2g | 0.01702826 | 0.0172651 | 0.02066346 | -0.0033984 | 0.76765178 | 0.90650187 |
| Phactr4 | 0.24867015 | 0.25948356 | 0.2628842 | -0.0034006 | 0.52292966 | 0.90650187 |
| Pfn4 | 0.01952399 | 0.02024973 | 0.02365228 | -0.0034025 | 0.73876119 | 0.90650187 |
| Fam228b | 0.01220834 | 0.01145786 | 0.01486277 | -0.0034049 | 0.8457661 | 0.9136218 |
| Nostrin | 0.05499633 | 0.06166987 | 0.06507512 | -0.0034052 | 0.59197547 | 0.90650187 |
| 9030624G23 | 0.00973293 | 0.00846265 | 0.01186896 | -0.0034063 | 0.89899723 | 0.93440169 |
| Urb2 | 0.01782829 | 0.01821598 | 0.02162264 | -0.0034067 | 0.75816396 | 0.90650187 |
| Bid | 0.09114355 | 0.10193852 | 0.10534924 | -0.0034107 | 0.55722139 | 0.90650187 |
| Ccdc89 | 0.02055683 | 0.0214744 | 0.02488625 | -0.0034119 | 0.72889804 | 0.90650187 |
| Rsph14 | 0.01181618 | 0.01097662 | 0.01438915 | -0.0034125 | 0.85413154 | 0.91608365 |

| | | | | | | |
|---|---|---|---|---|---|---|
| Hist1h2ab | 0.01952436 | 0.02023884 | 0.02365272 | -0.0034139 | 0.73944963 | 0.90650187 |
| Gm43579 | 0.02501026 | 0.0267735 | 0.0301874 | -0.0034139 | 0.69243175 | 0.90650187 |
| A930007I19F | 0.01195715 | 0.01114359 | 0.01455942 | -0.0034158 | 0.85152511 | 0.91522542 |
| Lrrc6 | 0.00870644 | 0.00720613 | 0.01062456 | -0.0034184 | 0.92369398 | 0.94629143 |
| Sync | 0.01663015 | 0.01676503 | 0.02018577 | -0.0034207 | 0.7743849 | 0.90650187 |
| Gc | 0.00773699 | 0.00602429 | 0.00944772 | -0.0034234 | 0.94639814 | 0.96068101 |
| Zfp93 | 0.07645885 | 0.08578873 | 0.08921511 | -0.0034264 | 0.56778483 | 0.90650187 |
| B230307C23 | 0.0469591 | 0.05243155 | 0.05585942 | -0.0034279 | 0.6075001 | 0.90650187 |
| Slc2a13 | 0.01816049 | 0.01859268 | 0.02202063 | -0.003428 | 0.75554625 | 0.90650187 |
| 6030458C11I | 0.13184424 | 0.14532248 | 0.1487516 | -0.0034291 | 0.54081337 | 0.90650187 |
| Zfp382 | 0.01935129 | 0.02000851 | 0.02344577 | -0.0034373 | 0.74271786 | 0.90650187 |
| Hist1h2ai | 0.00754904 | 0.00578029 | 0.0092194 | -0.0034391 | 0.95137568 | 0.9636744 |
| Kbtbd11 | 0.01797024 | 0.01835325 | 0.02179273 | -0.0034395 | 0.75854737 | 0.90650187 |
| Zfp790 | 0.05456016 | 0.061136 | 0.06457749 | -0.0034415 | 0.59363953 | 0.90650187 |
| Gm13713 | 0.01224953 | 0.01146408 | 0.01491249 | -0.0034484 | 0.84802919 | 0.91440801 |
| Dancr.1 | 0.01020577 | 0.00898426 | 0.01244159 | -0.0034573 | 0.89165278 | 0.93134318 |
| Gfpt2 | 0.02794849 | 0.03021 | 0.03366766 | -0.0034577 | 0.67600711 | 0.90650187 |
| Myo15b | 0.01994871 | 0.02069926 | 0.02415991 | -0.0034606 | 0.73786768 | 0.90650187 |
| Naprt | 0.12449578 | 0.13759616 | 0.14105841 | -0.0034622 | 0.54344492 | 0.90650187 |
| Rltpr | 0.09652533 | 0.10772389 | 0.11118647 | -0.0034626 | 0.55505589 | 0.90650187 |
| Peo1 | 0.1037897 | 0.11555964 | 0.11902405 | -0.0034644 | 0.55147826 | 0.90650187 |
| WDR97 | 0.01139321 | 0.01041343 | 0.01387803 | -0.0034646 | 0.86646425 | 0.92056229 |
| Gm14419 | 0.05035371 | 0.05629432 | 0.05976365 | -0.0034693 | 0.60182425 | 0.90650187 |
| Sipa1l2 | 0.0248387 | 0.02651323 | 0.02998376 | -0.0034705 | 0.69657157 | 0.90650187 |
| Zfp960 | 0.04635998 | 0.0516978 | 0.05516854 | -0.0034707 | 0.61013827 | 0.90650187 |
| Peg12 | 0.04457789 | 0.04963885 | 0.05311027 | -0.0034714 | 0.61431214 | 0.90650187 |
| Gm4779 | 0.01174686 | 0.01083195 | 0.0143054 | -0.0034734 | 0.85980034 | 0.91802142 |
| Fsip1 | 0.00892177 | 0.00741152 | 0.01088574 | -0.0034742 | 0.92202588 | 0.94549985 |
| Glce | 0.09737879 | 0.10863149 | 0.11210847 | -0.003477 | 0.55483163 | 0.90650187 |
| Vangl2 | 0.29186074 | 0.29804729 | 0.30152625 | -0.003479 | 0.52045388 | 0.90650187 |

| | | | | | | |
|---|---|---|---|---|---|---|
| K230010J24F | 0.01802809 | 0.01838111 | 0.02186203 | -0.0034809 | 0.76047753 | 0.90650187 |
| Spata24 | 0.2079708 | 0.22130732 | 0.22478889 | -0.0034816 | 0.52744724 | 0.90650187 |
| Nipa1 | 0.03656391 | 0.04031111 | 0.04379414 | -0.003483 | 0.63816869 | 0.90650187 |
| Gcnt4 | 0.01471563 | 0.01440123 | 0.01788495 | -0.0034837 | 0.80673732 | 0.9092537 |
| Focad | 0.05011734 | 0.05600329 | 0.05949236 | -0.0034891 | 0.60284683 | 0.90650187 |
| Serpini1 | 0.09685376 | 0.10805124 | 0.11154139 | -0.0034902 | 0.55531595 | 0.90650187 |
| C730034F03I | 0.05897421 | 0.06610752 | 0.06960042 | -0.0034929 | 0.58827237 | 0.90650187 |
| E2f1 | 0.1137362 | 0.12618073 | 0.12967452 | -0.0034938 | 0.54767018 | 0.90650187 |
| Gm15222 | 0.01186238 | 0.0109367 | 0.01444495 | -0.0035083 | 0.85986376 | 0.91802142 |
| Fzd6 | 0.07456364 | 0.08360186 | 0.08711048 | -0.0035086 | 0.57105462 | 0.90650187 |
| Wdr78 | 0.03211147 | 0.03506641 | 0.0385753 | -0.0035089 | 0.65703391 | 0.90650187 |
| Dram1 | 0.01459058 | 0.01422311 | 0.01773446 | -0.0035113 | 0.81062729 | 0.9092537 |
| Impg2 | 0.01030679 | 0.00905218 | 0.01256389 | -0.0035117 | 0.89298957 | 0.93179124 |
| Il2rg | 0.02186944 | 0.02293809 | 0.026452 | -0.0035139 | 0.72259186 | 0.90650187 |
| Adnp2 | 0.06897788 | 0.07736243 | 0.08087734 | -0.0035149 | 0.5766008 | 0.90650187 |
| Nthl1 | 0.09421613 | 0.10517012 | 0.10868677 | -0.0035167 | 0.55718717 | 0.90650187 |
| Plcd3 | 0.01631085 | 0.01628554 | 0.01980246 | -0.0035169 | 0.78510158 | 0.90650187 |
| Shc4 | 0.01184916 | 0.01091202 | 0.01442899 | -0.003517 | 0.86072603 | 0.91848494 |
| Vipr2 | 0.02304224 | 0.02433059 | 0.02784866 | -0.0035181 | 0.71280727 | 0.90650187 |
| Pdk4 | 0.02278233 | 0.02401913 | 0.02753933 | -0.0035202 | 0.71506998 | 0.90650187 |
| Mamdc4 | 0.00655431 | 0.00448976 | 0.00800996 | -0.0035202 | 0.97463441 | 0.98050415 |
| Thsd4 | 0.073344 | 0.08223209 | 0.08575334 | -0.0035212 | 0.57242353 | 0.90650187 |
| Sncaip | 0.06326494 | 0.07093245 | 0.07445529 | -0.0035228 | 0.58330017 | 0.90650187 |
| Gm16432 | 0.01211018 | 0.01122092 | 0.01474424 | -0.0035233 | 0.85595785 | 0.9165432 |
| Slc36a1 | 0.13805817 | 0.15167989 | 0.15520671 | -0.0035268 | 0.54023238 | 0.90650187 |
| Cabp1 | 0.0204126 | 0.02118543 | 0.02471404 | -0.0035286 | 0.73719854 | 0.90650187 |
| Glis1 | 0.01597268 | 0.01586675 | 0.01939631 | -0.0035296 | 0.79073369 | 0.90650187 |
| Radil | 0.02095077 | 0.02182592 | 0.02535645 | -0.0035305 | 0.73203813 | 0.90650187 |
| Ak2 | 0.83612364 | 0.66813142 | 0.67166349 | -0.0035321 | 0.50932564 | 0.90650187 |
| Zfp146 | 0.11878606 | 0.13150183 | 0.13503498 | -0.0035332 | 0.54629976 | 0.90650187 |

| | | | | | | |
|---|---|---|---|---|---|---|
| Maats1 | 0.01097201 | 0.0098307 | 0.01336878 | -0.0035381 | 0.88031043 | 0.92599812 |
| Tmem87a | 0.17730267 | 0.19139497 | 0.19493687 | -0.0035419 | 0.53218923 | 0.90650187 |
| Polr3g | 0.04255569 | 0.04722361 | 0.05076881 | -0.0035452 | 0.62188589 | 0.90650187 |
| Adam28 | 0.01017738 | 0.00886106 | 0.01240722 | -0.0035462 | 0.89806798 | 0.9343964 |
| Gm2109 | 0.00746751 | 0.00557087 | 0.00912033 | -0.0035495 | 0.95819766 | 0.96858377 |
| Gm14322 | 0.0313674 | 0.03414941 | 0.03770012 | -0.0035507 | 0.66228239 | 0.90650187 |
| Lpar2 | 0.03190214 | 0.03477723 | 0.03832917 | -0.0035519 | 0.65981973 | 0.90650187 |
| Gm43254 | 0.00830415 | 0.00658203 | 0.0101364 | -0.0035544 | 0.94048642 | 0.95691661 |
| Tbc1d2 | 0.01200551 | 0.01105661 | 0.01461784 | -0.0035612 | 0.86060626 | 0.91847356 |
| Rb1 | 0.06649932 | 0.07453556 | 0.07809699 | -0.0035614 | 0.58032771 | 0.90650187 |
| Pvt1 | 0.02251879 | 0.02365368 | 0.02722557 | -0.0035719 | 0.72013626 | 0.90650187 |
| Palm3 | 0.01178824 | 0.01078165 | 0.01435539 | -0.0035737 | 0.86579304 | 0.92040843 |
| Limk1 | 0.12811011 | 0.14127247 | 0.14485041 | -0.0035779 | 0.54372025 | 0.90650187 |
| Ccdc159 | 0.00874984 | 0.00709741 | 0.01067721 | -0.0035798 | 0.93195377 | 0.95146248 |
| Il18bp | 0.01524856 | 0.01494588 | 0.01852601 | -0.0035801 | 0.80485979 | 0.90882608 |
| Cdkl5 | 0.06055903 | 0.06781435 | 0.07139678 | -0.0035824 | 0.5882573 | 0.90650187 |
| Eogt | 0.06488784 | 0.07270094 | 0.07628449 | -0.0035836 | 0.58271251 | 0.90650187 |
| Ada | 0.02383256 | 0.02520442 | 0.02878859 | -0.0035842 | 0.71003184 | 0.90650187 |
| Tmem79 | 0.02395204 | 0.02534468 | 0.0289306 | -0.0035859 | 0.70919305 | 0.90650187 |
| 9530053A07I | 0.01347254 | 0.01280095 | 0.01638786 | -0.0035869 | 0.8347325 | 0.91062493 |
| Prkaa2 | 0.04749634 | 0.05289106 | 0.05647847 | -0.0035874 | 0.61117258 | 0.90650187 |
| Rit2 | 0.01139378 | 0.01028957 | 0.01387873 | -0.0035892 | 0.87486027 | 0.92390933 |
| Gpr62 | 0.01167382 | 0.01062478 | 0.01421715 | -0.0035924 | 0.86934676 | 0.92141903 |
| Adgrv1 | 0.023901 | 0.02527588 | 0.02886995 | -0.0035941 | 0.7100191 | 0.90650187 |
| Bloc1s6os | 0.00815082 | 0.00635224 | 0.00995027 | -0.003598 | 0.94603179 | 0.9604315 |
| Chst15 | 0.06341726 | 0.07102516 | 0.07462714 | -0.003602 | 0.58494987 | 0.90650187 |
| Mia2 | 0.01457061 | 0.01410676 | 0.01771043 | -0.0036037 | 0.81715596 | 0.9092537 |
| Fn3k | 0.01034761 | 0.00900839 | 0.01261329 | -0.0036049 | 0.89805793 | 0.9343964 |
| Jak2 | 0.06129679 | 0.06862068 | 0.07223175 | -0.0036111 | 0.58793985 | 0.90650187 |
| Slc12a1 | 0.00875245 | 0.00706735 | 0.01068038 | -0.003613 | 0.93369495 | 0.95242899 |

| | | | | | | |
|---|---|---|---|---|---|---|
| Plppr3 | 0.04432573 | 0.04920456 | 0.05281864 | -0.0036141 | 0.6195076 | 0.90650187 |
| Panx2 | 0.00930287 | 0.00773077 | 0.01134781 | -0.003617 | 0.92175988 | 0.94549985 |
| Ano8 | 0.01447912 | 0.01398147 | 0.01760031 | -0.0036188 | 0.81966019 | 0.9092537 |
| Pcdhb20 | 0.03660434 | 0.04022132 | 0.0438414 | -0.0036201 | 0.64322138 | 0.90650187 |
| Nans | 0.74152163 | 0.6159564 | 0.61957701 | -0.0036206 | 0.51036285 | 0.90650187 |
| Pcmtd1 | 0.40487077 | 0.39117226 | 0.39479385 | -0.0036216 | 0.51626485 | 0.90650187 |
| Vamp1 | 0.03387624 | 0.03702455 | 0.04064753 | -0.003623 | 0.65403763 | 0.90650187 |
| Gm29440 | 0.00796066 | 0.00609576 | 0.00971938 | -0.0036236 | 0.95128353 | 0.9636744 |
| Mtif3 | 0.22005469 | 0.23265214 | 0.23627679 | -0.0036247 | 0.5271863 | 0.90650187 |
| Rdh12 | 0.00833524 | 0.00654749 | 0.01017414 | -0.0036266 | 0.94347915 | 0.95876476 |
| Armc7 | 0.04396283 | 0.04877082 | 0.05239877 | -0.0036279 | 0.62088331 | 0.90650187 |
| 4930447C04I | 0.0132124 | 0.01244354 | 0.01607425 | -0.0036307 | 0.84234319 | 0.91251845 |
| Stil | 0.02646463 | 0.02827823 | 0.03191177 | -0.0036335 | 0.69363678 | 0.90650187 |
| Zfp518b | 0.05662378 | 0.06329269 | 0.06692939 | -0.0036367 | 0.59543877 | 0.90650187 |
| P3h2 | 0.00996409 | 0.00850876 | 0.01214895 | -0.0036402 | 0.90857591 | 0.93903063 |
| Slc23a1 | 0.01833461 | 0.01858808 | 0.02222915 | -0.0036411 | 0.7667474 | 0.90650187 |
| Nipal1 | 0.10566842 | 0.11740405 | 0.12104518 | -0.0036411 | 0.55319085 | 0.90650187 |
| Zmym1 | 0.17967805 | 0.19364216 | 0.19728549 | -0.0036433 | 0.53271569 | 0.90650187 |
| H3f3a | 4.21081904 | 0.64585351 | 0.64949927 | -0.0036458 | 0.50995417 | 0.90650187 |
| Rsl1 | 0.01799754 | 0.01817699 | 0.02182543 | -0.0036484 | 0.77130445 | 0.90650187 |
| Stk11 | 0.77796921 | 0.63647683 | 0.64013042 | -0.0036536 | 0.51012154 | 0.90650187 |
| Chil1 | 0.0112643 | 0.01006823 | 0.01372221 | -0.003654 | 0.88175116 | 0.92669144 |
| Nudt17 | 0.04370176 | 0.04844144 | 0.05209659 | -0.0036551 | 0.62244471 | 0.90650187 |
| Hoxa4 | 0.02366598 | 0.0249287 | 0.02859055 | -0.0036619 | 0.71545415 | 0.90650187 |
| Ntf3 | 0.01736593 | 0.0174064 | 0.02106843 | -0.003662 | 0.7801553 | 0.90650187 |
| Gm17315 | 0.01132665 | 0.01013308 | 0.01379758 | -0.0036645 | 0.88114101 | 0.92635898 |
| Zfp84 | 0.13315053 | 0.14644659 | 0.1501124 | -0.0036658 | 0.54322566 | 0.90650187 |
| Pom121 | 0.2357246 | 0.24728191 | 0.25094873 | -0.0036668 | 0.52589646 | 0.90650187 |
| Gm26631 | 0.01030448 | 0.00889193 | 0.0125611 | -0.0036692 | 0.90295709 | 0.93635703 |
| Bora | 0.03278877 | 0.03570068 | 0.03937118 | -0.0036705 | 0.66072962 | 0.90650187 |

| | | | | | | |
|---|---|---|---|---|---|---|
| Dnajc17 | 0.23938189 | 0.25066698 | 0.2543375 | -0.0036705 | 0.5255776 | 0.90650187 |
| Gm20696 | 0.02663568 | 0.02844125 | 0.03211436 | -0.0036731 | 0.69443809 | 0.90650187 |
| Gpam | 0.0147754 | 0.01428275 | 0.01795687 | -0.0036741 | 0.81848477 | 0.9092537 |
| Zbtb42 | 0.03967536 | 0.04374516 | 0.04742291 | -0.0036778 | 0.63486505 | 0.90650187 |
| 1810062G17 | 0.02272819 | 0.02379712 | 0.02747488 | -0.0036778 | 0.72410633 | 0.90650187 |
| Recql4 | 0.01357556 | 0.01283399 | 0.01651202 | -0.003678 | 0.83897485 | 0.91188312 |
| Gm6297 | 0.01120214 | 0.00996722 | 0.01364706 | -0.0036798 | 0.88468336 | 0.92796521 |
| Nab1 | 0.10900866 | 0.12094552 | 0.12462773 | -0.0036822 | 0.55225009 | 0.90650187 |
| Mef2a | 0.49177783 | 0.45601322 | 0.45969681 | -0.0036836 | 0.51420851 | 0.90650187 |
| Dner | 0.06443431 | 0.072089 | 0.0757737 | -0.0036847 | 0.58557607 | 0.90650187 |
| Hist1h2bp | 0.00642357 | 0.00416077 | 0.00785088 | -0.0036901 | 0.98167066 | 0.98614695 |
| Cyp2u1 | 0.01336318 | 0.01256205 | 0.01625603 | -0.003694 | 0.8438023 | 0.91300169 |
| Pcdhb22 | 0.02071387 | 0.0213762 | 0.02507371 | -0.0036975 | 0.74395117 | 0.90650187 |
| Reep4 | 0.23134205 | 0.24317129 | 0.24687041 | -0.0036991 | 0.52655524 | 0.90650187 |
| Tbkbp1 | 0.0398304 | 0.04390318 | 0.04760333 | -0.0037001 | 0.63516011 | 0.90650187 |
| Kirrel3 | 0.01214806 | 0.01108698 | 0.01478999 | -0.003703 | 0.86715826 | 0.92056229 |
| 4930432K21I | 0.01374754 | 0.01301515 | 0.01671927 | -0.0037041 | 0.83766676 | 0.91121886 |
| Tnfrsf18 | 0.00827325 | 0.00639384 | 0.01009889 | -0.0037051 | 0.94855384 | 0.96195723 |
| Gm12454 | 0.02734527 | 0.02924893 | 0.03295427 | -0.0037053 | 0.69141314 | 0.90650187 |
| 6030419C18I | 0.01620292 | 0.01596597 | 0.01967284 | -0.0037069 | 0.79889114 | 0.90777025 |
| Plk3 | 0.01535488 | 0.01494531 | 0.01865385 | -0.0037085 | 0.81160417 | 0.9092537 |
| Inadl | 0.15291041 | 0.16674133 | 0.17045118 | -0.0037098 | 0.53854074 | 0.90650187 |
| Stag2 | 0.49324931 | 0.45703895 | 0.46074981 | -0.0037109 | 0.51428096 | 0.90650187 |
| Mgat4b | 0.11888781 | 0.13142781 | 0.13514268 | -0.0037149 | 0.54863092 | 0.90650187 |
| Timp4 | 0.01190305 | 0.01077687 | 0.01449409 | -0.0037172 | 0.87288615 | 0.92289911 |
| E130102H24I | 0.0200238 | 0.02052792 | 0.02424964 | -0.0037217 | 0.75247207 | 0.90650187 |
| D930030I03F | 0.00986203 | 0.00830086 | 0.01202534 | -0.0037245 | 0.91572699 | 0.94222861 |
| Mrps35 | 0.42340905 | 0.40537437 | 0.40910406 | -0.0037297 | 0.51616448 | 0.90650187 |
| Pidd1 | 0.0203916 | 0.0209584 | 0.02468896 | -0.0037306 | 0.74912576 | 0.90650187 |
| Gm10699 | 0.01616707 | 0.01589506 | 0.0196298 | -0.0037347 | 0.80117224 | 0.90817444 |

| | | | | | | |
|---|---|---|---|---|---|---|
| Trim71 | 0.00866113 | 0.00683211 | 0.0105696 | -0.0037375 | 0.94202438 | 0.95777624 |
| Prkab2 | 0.0774217 | 0.08654358 | 0.09028238 | -0.0037388 | 0.5730336 | 0.90650187 |
| Hist1h3h | 0.00633085 | 0.00399435 | 0.00773804 | -0.0037437 | 0.9842539 | 0.98789327 |
| Astn2 | 0.01709003 | 0.01699382 | 0.02073755 | -0.0037437 | 0.78888565 | 0.90650187 |
| Fam175b | 0.34935607 | 0.34657789 | 0.35032266 | -0.0037448 | 0.51895112 | 0.90650187 |
| Speg | 0.07775134 | 0.08690259 | 0.09064747 | -0.0037449 | 0.57285969 | 0.90650187 |
| Fam189a2 | 0.02157 | 0.02234812 | 0.02609506 | -0.0037469 | 0.73837284 | 0.90650187 |
| Ncoa4 | 0.03068441 | 0.033149 | 0.03689603 | -0.003747 | 0.67417846 | 0.90650187 |
| Naa50 | 0.60759046 | 0.53454266 | 0.5382923 | -0.0037496 | 0.51235215 | 0.90650187 |
| Zfp964 | 0.01053044 | 0.00908358 | 0.01283457 | -0.003751 | 0.90307249 | 0.93635703 |
| E130311K13I | 0.07540617 | 0.08429517 | 0.08804675 | -0.0037516 | 0.57511927 | 0.90650187 |
| Ttc26 | 0.06127447 | 0.06845001 | 0.0722065 | -0.0037565 | 0.59145106 | 0.90650187 |
| Trim16 | 0.03436162 | 0.03745933 | 0.04121664 | -0.0037573 | 0.65735724 | 0.90650187 |
| 2600014E21I | 0.0327282 | 0.03554129 | 0.03930003 | -0.0037587 | 0.66464988 | 0.90650187 |
| Rsph9 | 0.28687716 | 0.29339749 | 0.29715699 | -0.0037595 | 0.52242622 | 0.90650187 |
| Dclk3 | 0.01502346 | 0.01448928 | 0.01825529 | -0.003766 | 0.82045883 | 0.9092537 |
| Fbxw24 | 0.01439238 | 0.0137291 | 0.01749589 | -0.0037668 | 0.8307469 | 0.9098597 |
| Cdca4 | 0.24025625 | 0.25137848 | 0.25514569 | -0.0037672 | 0.52616744 | 0.90650187 |
| Dnaaf2 | 0.17813277 | 0.19198866 | 0.19575832 | -0.0037697 | 0.5341109 | 0.90650187 |
| Dhtkd1 | 0.01099464 | 0.00962574 | 0.01339615 | -0.0037704 | 0.89457604 | 0.93247252 |
| Arhgap27os2 | 0.01972415 | 0.02011547 | 0.02389154 | -0.0037761 | 0.75885707 | 0.90650187 |
| Plin5 | 0.0144775 | 0.01382177 | 0.01759836 | -0.0037766 | 0.82996351 | 0.90974008 |
| Hs1bp3 | 0.05428501 | 0.06048484 | 0.06426342 | -0.0037786 | 0.60310587 | 0.90650187 |
| Shox2 | 0.03663087 | 0.0400882 | 0.04387239 | -0.0037842 | 0.64930826 | 0.90650187 |
| Nphp3 | 0.03933882 | 0.04324599 | 0.04703113 | -0.0037851 | 0.63974968 | 0.90650187 |
| Ntn1 | 0.02402184 | 0.02522686 | 0.02901356 | -0.0037867 | 0.71911589 | 0.90650187 |
| Lrfn5 | 0.011694 | 0.01045251 | 0.01424154 | -0.003789 | 0.8815511 | 0.92661786 |
| Dsc2 | 0.04077394 | 0.04491028 | 0.04870057 | -0.0037903 | 0.63532661 | 0.90650187 |
| Mfsd3 | 0.0663814 | 0.07417317 | 0.0779645 | -0.0037913 | 0.58557825 | 0.90650187 |
| Gm21814 | 0.02523278 | 0.02665817 | 0.03045145 | -0.0037933 | 0.71013536 | 0.90650187 |

| | | | | | | |
|---|---|---|---|---|---|---|
| Sapcd2 | 0.0282047 | 0.03017721 | 0.03397049 | -0.0037933 | 0.6901988 | 0.90650187 |
| Gm9484 | 0.01035087 | 0.00882288 | 0.01261724 | -0.0037944 | 0.90937573 | 0.93921867 |
| Mmp11 | 0.09757921 | 0.10852933 | 0.11232485 | -0.0037955 | 0.5597043 | 0.90650187 |
| Rpl10-ps3 | 1.33769575 | 0.85638284 | 0.86017871 | -0.0037959 | 0.50782594 | 0.90650187 |
| Uprt | 0.00631309 | 0.00391942 | 0.00771643 | -0.003797 | 0.98564851 | 0.98877074 |
| Gm3739 | 0.01106714 | 0.00968492 | 0.01348383 | -0.0037989 | 0.89480523 | 0.93253978 |
| Foxred2 | 0.01825784 | 0.01833453 | 0.02213723 | -0.0038027 | 0.77746489 | 0.90650187 |
| 3110052M02 | 0.01816068 | 0.01821298 | 0.02202086 | -0.0038079 | 0.77897628 | 0.90650187 |
| 4732414G09 | 0.01872823 | 0.01888892 | 0.02270038 | -0.0038115 | 0.77229662 | 0.90650187 |
| Vtcn1 | 0.02536213 | 0.02679185 | 0.0306049 | -0.003813 | 0.710168 | 0.90650187 |
| B430203G13 | 0.01561061 | 0.01514807 | 0.01896125 | -0.0038132 | 0.8143469 | 0.9092537 |
| Pik3c2g | 0.01673743 | 0.01650097 | 0.02031451 | -0.0038135 | 0.79801464 | 0.9075779 |
| Ctf1 | 0.12773956 | 0.14064722 | 0.14446237 | -0.0038152 | 0.54673059 | 0.90650187 |
| Kcnq1 | 0.05141098 | 0.05715641 | 0.06097609 | -0.0038197 | 0.60967926 | 0.90650187 |
| Gm14325 | 0.02916403 | 0.0312824 | 0.03510345 | -0.003821 | 0.6857747 | 0.90650187 |
| Thrsp | 0.00717215 | 0.00494011 | 0.00876134 | -0.0038212 | 0.97374544 | 0.97994563 |
| Cpt1a | 0.10994004 | 0.12179982 | 0.12562419 | -0.0038244 | 0.5538275 | 0.90650187 |
| Spata33 | 0.05598132 | 0.06237316 | 0.06619786 | -0.0038247 | 0.60135388 | 0.90650187 |
| Nalcn | 0.03324635 | 0.03607768 | 0.03990846 | -0.0038308 | 0.66521206 | 0.90650187 |
| Ufsp1 | 0.08331429 | 0.0929543 | 0.09678518 | -0.0038309 | 0.56983822 | 0.90650187 |
| Pax6os1 | 0.06620715 | 0.07393662 | 0.07776866 | -0.003832 | 0.58669717 | 0.90650187 |
| Prkar2a | 0.32990966 | 0.33030275 | 0.33414192 | -0.0038392 | 0.52036849 | 0.90650187 |
| Ube2cbp | 0.0307946 | 0.03318439 | 0.03702581 | -0.0038414 | 0.67768343 | 0.90650187 |
| Pde6c | 0.02268483 | 0.02357537 | 0.02742327 | -0.0038479 | 0.73361583 | 0.90650187 |
| Ptpn13 | 0.12461227 | 0.13733147 | 0.1411809 | -0.0038494 | 0.54823902 | 0.90650187 |
| Cspg4 | 0.07498819 | 0.08373285 | 0.08758239 | -0.0038495 | 0.57745984 | 0.90650187 |
| Ccdc74a | 0.03476419 | 0.03783553 | 0.04168835 | -0.0038528 | 0.6594128 | 0.90650187 |
| Kif9 | 0.04788776 | 0.05307481 | 0.05692923 | -0.0038544 | 0.61828971 | 0.90650187 |
| Adcy7 | 0.0167743 | 0.01650328 | 0.02035876 | -0.0038555 | 0.80007683 | 0.90805731 |
| 5730409E04I | 0.07861979 | 0.0877511 | 0.09160855 | -0.0038575 | 0.57424642 | 0.90650187 |

| | | | | | | |
|---|---|---|---|---|---|---|
| Fgf1 | 0.04608977 | 0.05098896 | 0.05485676 | -0.0038678 | 0.62302972 | 0.90650187 |
| F2rl3 | 0.01200372 | 0.01074568 | 0.01461567 | -0.00387 | 0.88042677 | 0.92605642 |
| 6030443J06F | 0.0167904 | 0.01650759 | 0.02037808 | -0.0038705 | 0.80076888 | 0.90809448 |
| Prrc1 | 0.36695459 | 0.36081756 | 0.36469092 | -0.0038734 | 0.51882965 | 0.90650187 |
| Gm14286 | 0.01342997 | 0.01246272 | 0.01633654 | -0.0038738 | 0.85409704 | 0.91608365 |
| Defb25 | 0.01596638 | 0.01551469 | 0.01938874 | -0.003874 | 0.81280303 | 0.9092537 |
| Sh2d5 | 0.05529587 | 0.06154045 | 0.0654167 | -0.0038763 | 0.6038879 | 0.90650187 |
| Gdpgp1 | 0.05496594 | 0.06116218 | 0.06504045 | -0.0038783 | 0.60452827 | 0.90650187 |
| Gm10614 | 0.01067052 | 0.00912306 | 0.01300408 | -0.003881 | 0.90770844 | 0.93845311 |
| Ncam2 | 0.01541575 | 0.01484594 | 0.01872702 | -0.0038811 | 0.82156074 | 0.9092537 |
| Diras1 | 0.00703943 | 0.0047166 | 0.00859999 | -0.0038834 | 0.97764912 | 0.98275653 |
| Fgfr2 | 0.08148792 | 0.0908903 | 0.09477496 | -0.0038847 | 0.57229399 | 0.90650187 |
| Lrrc46 | 0.04306786 | 0.04747292 | 0.05136243 | -0.0038895 | 0.63181015 | 0.90650187 |
| Pcdhb17 | 0.04753878 | 0.05263473 | 0.05652736 | -0.0038926 | 0.62024975 | 0.90650187 |
| Nif3l1 | 0.2028539 | 0.21598564 | 0.21987863 | -0.003893 | 0.53136842 | 0.90650187 |
| Bcas3 | 0.08998744 | 0.10019191 | 0.10409005 | -0.0038981 | 0.56611294 | 0.90650187 |
| Gm10369 | 0.01129987 | 0.00986491 | 0.01376521 | -0.0039003 | 0.89610038 | 0.93337138 |
| 4930532G15 | 0.01368069 | 0.01273832 | 0.01663872 | -0.0039004 | 0.85132328 | 0.9150732 |
| 2200002J24F | 0.01293191 | 0.01183316 | 0.01573598 | -0.0039028 | 0.86489817 | 0.92040222 |
| Tsacc | 0.08304889 | 0.09259032 | 0.09649336 | -0.003903 | 0.57135277 | 0.90650187 |
| Lpcat4 | 0.07551511 | 0.08426439 | 0.08816773 | -0.0039033 | 0.57801383 | 0.90650187 |
| Rnf43 | 0.01008013 | 0.00838569 | 0.01228948 | -0.0039038 | 0.92104852 | 0.94523755 |
| Kif18b | 0.01525081 | 0.01462268 | 0.01852871 | -0.003906 | 0.82566621 | 0.90947782 |
| Sik1 | 0.05483963 | 0.0609883 | 0.06489637 | -0.0039081 | 0.60554059 | 0.90650187 |
| Rilpl1 | 0.22952653 | 0.2412656 | 0.24517529 | -0.0039097 | 0.52825807 | 0.90650187 |
| Stard8 | 0.07122066 | 0.07947094 | 0.08338549 | -0.0039145 | 0.58265864 | 0.90650187 |
| 9330159F19I | 0.00656968 | 0.00411157 | 0.00802865 | -0.0039171 | 0.9849571 | 0.98821581 |
| Zc3h12b | 0.00898489 | 0.00704027 | 0.0109623 | -0.003922 | 0.94414157 | 0.95915899 |
| Dennd2a | 0.27443533 | 0.28222507 | 0.28614933 | -0.0039243 | 0.52430733 | 0.90650187 |
| Gcfc2 | 0.11807791 | 0.13036028 | 0.13428516 | -0.0039249 | 0.55169155 | 0.90650187 |

| | | | | | | |
|---|---|---|---|---|---|---|
| Bard1 | 0.01772545 | 0.01757218 | 0.02149941 | -0.0039272 | 0.79162295 | 0.90674858 |
| Ogfod3 | 0.26544826 | 0.27417385 | 0.27810647 | -0.0039326 | 0.52506261 | 0.90650187 |
| Pprc1 | 0.10296075 | 0.11419793 | 0.11813084 | -0.0039329 | 0.55883133 | 0.90650187 |
| Tmem30a | 0.91094493 | 0.7060562 | 0.70999616 | -0.00394 | 0.50984086 | 0.90650187 |
| Gm26881 | 0.04784327 | 0.05293551 | 0.056878 | -0.0039425 | 0.62101475 | 0.90650187 |
| Lrrn2 | 0.0441526 | 0.04867523 | 0.05261836 | -0.0039431 | 0.63048801 | 0.90650187 |
| Eaf2 | 0.01919771 | 0.01931738 | 0.0232621 | -0.0039447 | 0.77453733 | 0.90650187 |
| 2500002B13I | 0.03279385 | 0.03543155 | 0.03937714 | -0.0039456 | 0.67200257 | 0.90650187 |
| Ube2r2 | 0.66492676 | 0.57030863 | 0.57425455 | -0.0039459 | 0.51218479 | 0.90650187 |
| Wtip | 0.03847513 | 0.04207859 | 0.0460249 | -0.0039463 | 0.6484651 | 0.90650187 |
| Rhbdf2 | 0.04335854 | 0.04774671 | 0.05169915 | -0.0039524 | 0.63302285 | 0.90650187 |
| Fam184a | 0.04964085 | 0.05499162 | 0.05894522 | -0.0039536 | 0.61721615 | 0.90650187 |
| Fam71e1 | 0.10958313 | 0.1212873 | 0.12524248 | -0.0039552 | 0.55582525 | 0.90650187 |
| Smarcal1 | 0.09914816 | 0.11005772 | 0.11401675 | -0.003959 | 0.56133936 | 0.90650187 |
| Itgb6 | 0.01464176 | 0.01383695 | 0.01779605 | -0.0039591 | 0.83867258 | 0.91185623 |
| Mat1a | 0.01421261 | 0.01331992 | 0.01727944 | -0.0039595 | 0.84582591 | 0.9136218 |
| Gm684 | 0.01557213 | 0.01495458 | 0.018915 | -0.0039604 | 0.82402653 | 0.90929265 |
| Secisbp2l | 0.20041389 | 0.21356127 | 0.21752746 | -0.0039662 | 0.53230168 | 0.90650187 |
| 2810459M11 | 0.00636788 | 0.0038159 | 0.00778311 | -0.0039672 | 0.98827468 | 0.99098467 |
| Ddx58 | 0.06243986 | 0.0695553 | 0.07352385 | -0.0039686 | 0.59481847 | 0.90650187 |
| Krba1 | 0.13474576 | 0.14779906 | 0.15177144 | -0.0039724 | 0.54631505 | 0.90650187 |
| Zfp879 | 0.01499355 | 0.01424243 | 0.01821931 | -0.0039769 | 0.83407151 | 0.91062493 |
| Map3k9 | 0.03033708 | 0.03250991 | 0.03648684 | -0.0039769 | 0.68600376 | 0.90650187 |
| Peli2 | 0.11874176 | 0.13100793 | 0.1349881 | -0.0039802 | 0.55214398 | 0.90650187 |
| Ctf2 | 0.01025534 | 0.00851567 | 0.0125016 | -0.0039859 | 0.92181803 | 0.94549985 |
| Dexi | 0.06319671 | 0.07039215 | 0.0743783 | -0.0039861 | 0.59415753 | 0.90650187 |
| Fam83e | 0.00561392 | 0.00287748 | 0.00686508 | -0.0039876 | 0.99509195 | 0.99594369 |
| Gm26945 | 0.01022352 | 0.00847507 | 0.01246308 | -0.003988 | 0.92256435 | 0.94559465 |
| Draxin | 0.01091967 | 0.00931536 | 0.01330547 | -0.0039901 | 0.908762 | 0.93909526 |
| Mfsd7b | 0.04614753 | 0.05093255 | 0.05492342 | -0.0039909 | 0.62666322 | 0.90650187 |

| | | | | | | |
|---|---|---|---|---|---|---|
| Dusp5 | 0.15997334 | 0.17360448 | 0.17761038 | -0.0040059 | 0.53993423 | 0.90650187 |
| Pip4k2c | 0.23513307 | 0.24638931 | 0.25039938 | -0.0040101 | 0.52837877 | 0.90650187 |
| Gm20632 | 0.01586149 | 0.01524798 | 0.01926273 | -0.0040147 | 0.82292132 | 0.9092537 |
| Ccdc69 | 0.02635121 | 0.02775956 | 0.03177742 | -0.0040179 | 0.71297298 | 0.90650187 |
| Fam166a | 0.03019244 | 0.03229769 | 0.03631638 | -0.0040187 | 0.68861917 | 0.90650187 |
| Gemin2 | 0.16810505 | 0.18176163 | 0.18578287 | -0.0040212 | 0.53832884 | 0.90650187 |
| Kcnj2 | 0.01880183 | 0.01876658 | 0.02278846 | -0.0040219 | 0.78365431 | 0.90650187 |
| Smim4 | 0.48588743 | 0.45144394 | 0.45546717 | -0.0040232 | 0.51566197 | 0.90650187 |
| Zfp59 | 0.02150511 | 0.0219896 | 0.02601769 | -0.0040281 | 0.7543598 | 0.90650187 |
| Arsg | 0.0363732 | 0.03954255 | 0.04357124 | -0.0040287 | 0.65948209 | 0.90650187 |
| Trpc4 | 0.02055951 | 0.02085828 | 0.02488945 | -0.0040312 | 0.76424521 | 0.90650187 |
| Pask | 0.03764959 | 0.04102778 | 0.04506203 | -0.0040342 | 0.65468485 | 0.90650187 |
| Trnp1 | 0.008883 | 0.00680341 | 0.01083872 | -0.0040353 | 0.95105038 | 0.96353751 |
| Chn2 | 0.02804821 | 0.02974754 | 0.03378553 | -0.004038 | 0.7024053 | 0.90650187 |
| Slc4a1ap | 0.19641962 | 0.20962613 | 0.21366452 | -0.0040384 | 0.53348164 | 0.90650187 |
| Rmnd1 | 0.19313307 | 0.20643289 | 0.21047295 | -0.0040401 | 0.53400203 | 0.90650187 |
| Gprasp1 | 0.48480084 | 0.45064058 | 0.45468439 | -0.0040438 | 0.51576916 | 0.90650187 |
| Cidea | 0.01404032 | 0.01302637 | 0.01707197 | -0.0040456 | 0.85394178 | 0.91608365 |
| Tmem170 | 0.01325921 | 0.01208499 | 0.01613068 | -0.0040457 | 0.86757288 | 0.92056229 |
| Prss16 | 0.01073085 | 0.0090262 | 0.01307707 | -0.0040509 | 0.91576117 | 0.94222861 |
| Gm10069 | 0.0282424 | 0.0299614 | 0.03401503 | -0.0040536 | 0.70187146 | 0.90650187 |
| Txnrd3 | 0.03931626 | 0.04294779 | 0.04700487 | -0.0040571 | 0.64940297 | 0.90650187 |
| Pyroxd2 | 0.04446834 | 0.04892477 | 0.05298358 | -0.0040588 | 0.63328087 | 0.90650187 |
| Ciart | 0.01661473 | 0.01610833 | 0.02016726 | -0.0040589 | 0.81453645 | 0.9092537 |
| Gm15912 | 0.01495487 | 0.01411051 | 0.01817279 | -0.0040623 | 0.83982845 | 0.91213511 |
| Gm13889 | 0.02335945 | 0.02415426 | 0.02822604 | -0.0040718 | 0.73933825 | 0.90650187 |
| Gm42726 | 0.0202734 | 0.02047522 | 0.02454779 | -0.0040726 | 0.76960366 | 0.90650187 |
| Sh3bp2 | 0.02433105 | 0.025307 | 0.02938094 | -0.0040739 | 0.73119268 | 0.90650187 |
| Snx22 | 0.01925309 | 0.01925359 | 0.02332833 | -0.0040747 | 0.7812762 | 0.90650187 |
| Pter | 0.06250585 | 0.06951881 | 0.07359837 | -0.0040796 | 0.59732149 | 0.90650187 |

| | | | | | | |
|---|---|---|---|---|---|---|
| 2010007H06I | 0.01416962 | 0.01314763 | 0.01722769 | -0.0040801 | 0.8537972 | 0.91608365 |
| Chrnb4 | 0.01567365 | 0.0149565 | 0.01903701 | -0.0040805 | 0.82967962 | 0.90974008 |
| Zcchc10 | 0.25240028 | 0.26221043 | 0.26629187 | -0.0040814 | 0.52716192 | 0.90650187 |
| 4933431E20I | 0.03573835 | 0.03874619 | 0.04282882 | -0.0040826 | 0.66413468 | 0.90650187 |
| Adgrl2 | 0.09130997 | 0.10144406 | 0.10553035 | -0.0040863 | 0.56833645 | 0.90650187 |
| Nr1d1 | 0.01005932 | 0.00817607 | 0.01226428 | -0.0040882 | 0.9308179 | 0.95091965 |
| Gm43270 | 0.02162682 | 0.02207239 | 0.02616281 | -0.0040904 | 0.75648682 | 0.90650187 |
| Caps2 | 0.0109364 | 0.00923517 | 0.01332571 | -0.0040905 | 0.9138182 | 0.94112131 |
| Mreg | 0.02763338 | 0.02919809 | 0.03329508 | -0.004097 | 0.70782268 | 0.90650187 |
| Pde8a | 0.05969138 | 0.06631485 | 0.07041378 | -0.0040989 | 0.60210056 | 0.90650187 |
| Tigd3 | 0.01050057 | 0.00869302 | 0.01279843 | -0.0041054 | 0.92307103 | 0.945972 |
| Sntg2 | 0.01359744 | 0.01243247 | 0.01653839 | -0.0041059 | 0.86513648 | 0.92040222 |
| Ebf3 | 0.04670146 | 0.0514503 | 0.05556238 | -0.0041121 | 0.62892682 | 0.90650187 |
| Unc5cl | 0.01208479 | 0.01059991 | 0.01471358 | -0.0041137 | 0.89305089 | 0.93179124 |
| Gm12999 | 0.00995407 | 0.00802005 | 0.01213682 | -0.0041168 | 0.93421429 | 0.95276695 |
| Kank4 | 0.01258539 | 0.01119342 | 0.0153179 | -0.0041245 | 0.88435048 | 0.9278081 |
| Cdan1 | 0.04026667 | 0.04397687 | 0.04811083 | -0.004134 | 0.64876632 | 0.90650187 |
| Pggt1b | 0.15194286 | 0.16532922 | 0.169466 | -0.0041368 | 0.5432085 | 0.90650187 |
| Top3a | 0.02911733 | 0.03090888 | 0.03504833 | -0.0041394 | 0.70022814 | 0.90650187 |
| Syt3 | 0.01047185 | 0.00861609 | 0.01276367 | -0.0041476 | 0.92571598 | 0.94746893 |
| Bche | 0.01068015 | 0.00886421 | 0.01301573 | -0.0041515 | 0.92190129 | 0.94549985 |
| AW011738 | 0.01789478 | 0.01754901 | 0.02170232 | -0.0041533 | 0.80255493 | 0.90870445 |
| Zfp9 | 0.23460635 | 0.24575595 | 0.24990994 | -0.004154 | 0.52945286 | 0.90650187 |
| Gm26601 | 0.06769602 | 0.07528624 | 0.07944053 | -0.0041543 | 0.59191735 | 0.90650187 |
| Zfp319 | 0.07101731 | 0.07900407 | 0.08315837 | -0.0041543 | 0.58787706 | 0.90650187 |
| Pnma1 | 0.01315051 | 0.01184477 | 0.01599962 | -0.0041549 | 0.87584302 | 0.92430504 |
| Yap1 | 0.1329422 | 0.14573748 | 0.14989551 | -0.004158 | 0.54907272 | 0.90650187 |
| Pced1b | 0.07855139 | 0.08737393 | 0.09153289 | -0.004159 | 0.58003928 | 0.90650187 |
| Ggact | 0.15677141 | 0.17020809 | 0.17437187 | -0.0041638 | 0.54227044 | 0.90650187 |
| Kcnk12 | 0.0094891 | 0.00740966 | 0.01157352 | -0.0041639 | 0.94513623 | 0.9597205 |

| | | | | | | |
|---|---|---|---|---|---|---|
| Gm12216 | 0.03001264 | 0.03193788 | 0.03610444 | -0.0041666 | 0.69603688 | 0.90650187 |
| Lats1 | 0.2611314 | 0.27004798 | 0.27421569 | -0.0041677 | 0.52693493 | 0.90650187 |
| Snn | 0.18823625 | 0.20152778 | 0.20569617 | -0.0041684 | 0.53589186 | 0.90650187 |
| Gm20878 | 0.01273112 | 0.01132376 | 0.01549375 | -0.00417 | 0.88424761 | 0.9278081 |
| Nanp | 0.01027361 | 0.0083489 | 0.01252372 | -0.0041748 | 0.93082418 | 0.95091965 |
| A330102I10F | 0.01572581 | 0.01492376 | 0.01909969 | -0.0041759 | 0.83447056 | 0.91062493 |
| 1700066B19 | 0.01028058 | 0.0083559 | 0.01253216 | -0.0041763 | 0.93075948 | 0.95091965 |
| Ccdc160 | 0.01945953 | 0.01939877 | 0.02357521 | -0.0041764 | 0.7845219 | 0.90650187 |
| Zfp429 | 0.04432871 | 0.04864359 | 0.05282209 | -0.0041785 | 0.63745725 | 0.90650187 |
| Usp43 | 0.01272171 | 0.0113003 | 0.01548239 | -0.0041821 | 0.88509411 | 0.92807528 |
| Npas4 | 0.01027962 | 0.00834837 | 0.012531 | -0.0041826 | 0.93107769 | 0.95109952 |
| 1700084E18I | 0.02814781 | 0.0297186 | 0.03390326 | -0.0041847 | 0.70840066 | 0.90650187 |
| Syt14 | 0.11481 | 0.12662929 | 0.13081699 | -0.0041877 | 0.55658329 | 0.90650187 |
| Mustn1 | 0.01939051 | 0.01930397 | 0.02349267 | -0.0041887 | 0.7860076 | 0.90650187 |
| Tnnc1 | 0.03863732 | 0.04202471 | 0.04621395 | -0.0041892 | 0.65652157 | 0.90650187 |
| Rpl6l | 2.11868647 | 0.91986258 | 0.92405459 | -0.004192 | 0.5080452 | 0.90650187 |
| Msh5 | 0.04851987 | 0.05346163 | 0.05765666 | -0.004195 | 0.62682545 | 0.90650187 |
| Fam129c | 0.01570022 | 0.0148735 | 0.01906894 | -0.0041954 | 0.835987 | 0.91079271 |
| 1700047I17R | 0.03669923 | 0.0397554 | 0.04395227 | -0.0041969 | 0.66439844 | 0.90650187 |
| Cpeb3 | 0.05503798 | 0.06092454 | 0.06512263 | -0.0041981 | 0.61278412 | 0.90650187 |
| Pde8b | 0.01103963 | 0.00925185 | 0.01345055 | -0.0041987 | 0.91739077 | 0.94283706 |
| Efhb | 0.01874113 | 0.01850599 | 0.02271583 | -0.0042098 | 0.79499796 | 0.90731056 |
| Nfkbid | 0.02586622 | 0.02699122 | 0.03120267 | -0.0042114 | 0.72575588 | 0.90650187 |
| 2810455O05 | 0.0120652 | 0.01047614 | 0.01468992 | -0.0042138 | 0.89888347 | 0.93440169 |
| 3830406C13I | 0.38283224 | 0.37322146 | 0.37743585 | -0.0042144 | 0.51979494 | 0.90650187 |
| Ispd | 0.07374704 | 0.08198115 | 0.08620205 | -0.0042209 | 0.58616086 | 0.90650187 |
| Gm9801 | 0.0148559 | 0.01383157 | 0.01805372 | -0.0042222 | 0.85075472 | 0.91491493 |
| Edem1 | 0.22132574 | 0.23324535 | 0.23747643 | -0.0042311 | 0.53156589 | 0.90650187 |
| Mtx1 | 0.41858676 | 0.40117333 | 0.40540721 | -0.0042339 | 0.51851537 | 0.90650187 |
| Tcea2 | 0.07312943 | 0.08127898 | 0.08551436 | -0.0042354 | 0.58713575 | 0.90650187 |

| | | | | | | |
|---|---|---|---|---|---|---|
| Plekhn1 | 0.01593115 | 0.01510786 | 0.01934641 | -0.0042386 | 0.83496574 | 0.91066681 |
| Tbk1 | 0.18072408 | 0.19407889 | 0.19831776 | -0.0042389 | 0.53785097 | 0.90650187 |
| Itpr2 | 0.12142967 | 0.1335885 | 0.13782876 | -0.0042403 | 0.55439282 | 0.90650187 |
| Clk2 | 0.10025127 | 0.11095693 | 0.11520533 | -0.0042484 | 0.56511061 | 0.90650187 |
| Apbb2 | 0.30414132 | 0.30794117 | 0.31219487 | -0.0042537 | 0.52414997 | 0.90650187 |
| Synj2 | 0.00945488 | 0.00727626 | 0.01153205 | -0.0042558 | 0.94956069 | 0.9624773 |
| Rasa1 | 0.34197066 | 0.33995898 | 0.34421561 | -0.0042566 | 0.52192087 | 0.90650187 |
| Fut1 | 0.0238404 | 0.02454059 | 0.02879791 | -0.0042573 | 0.74447827 | 0.90650187 |
| Smpd5 | 0.01002562 | 0.00796526 | 0.01222347 | -0.0042582 | 0.93927123 | 0.95593613 |
| Man2c1os | 0.02096199 | 0.02111153 | 0.02536984 | -0.0042583 | 0.77222396 | 0.90650187 |
| Gm42997 | 0.02515738 | 0.0261007 | 0.03036198 | -0.0042613 | 0.73366588 | 0.90650187 |
| 4933417E11I | 0.01598652 | 0.01514886 | 0.01941294 | -0.0042641 | 0.83558764 | 0.91078214 |
| Arhgap6 | 0.03534456 | 0.0381039 | 0.04236798 | -0.0042641 | 0.67271534 | 0.90650187 |
| Smpdl3b | 0.02022427 | 0.02021591 | 0.02448911 | -0.0042732 | 0.78104614 | 0.90650187 |
| Shpk | 0.02982995 | 0.03161356 | 0.03588904 | -0.0042755 | 0.7018076 | 0.90650187 |
| Tmem150c | 0.05709647 | 0.06319117 | 0.06746721 | -0.004276 | 0.61093654 | 0.90650187 |
| Pacsin3 | 0.2012262 | 0.21403374 | 0.21831088 | -0.0042771 | 0.53470329 | 0.90650187 |
| Kcnj13 | 0.03281305 | 0.03511794 | 0.0393997 | -0.0042818 | 0.68549682 | 0.90650187 |
| Gpr173 | 0.05849379 | 0.06477078 | 0.06905514 | -0.0042844 | 0.6086556 | 0.90650187 |
| B4galnt1 | 0.01623408 | 0.01542206 | 0.01971027 | -0.0042882 | 0.83327685 | 0.91056108 |
| Ttc9 | 0.03534062 | 0.03807334 | 0.04236336 | -0.00429 | 0.67371475 | 0.90650187 |
| Maneal | 0.01350758 | 0.01213423 | 0.0164301 | -0.0042959 | 0.87745596 | 0.92482217 |
| Tmem191c | 0.06464106 | 0.07170997 | 0.07600659 | -0.0042966 | 0.59921213 | 0.90650187 |
| Scyl1 | 0.48452171 | 0.45018313 | 0.45448318 | -0.0043 | 0.51677522 | 0.90650187 |
| Srprb | 0.57954266 | 0.51571674 | 0.52001803 | -0.0043013 | 0.51466642 | 0.90650187 |
| 1110035H17I | 0.0324484 | 0.03466998 | 0.03897131 | -0.0043013 | 0.68816963 | 0.90650187 |
| Adipor2 | 0.21679267 | 0.22888747 | 0.23319049 | -0.004303 | 0.53269014 | 0.90650187 |
| Pdpr | 0.02391493 | 0.02458263 | 0.0288865 | -0.0043039 | 0.74613029 | 0.90650187 |
| Ntm | 0.04826368 | 0.0530554 | 0.05736192 | -0.0043065 | 0.63071948 | 0.90650187 |
| Rnf19a | 0.24541464 | 0.2555903 | 0.25989844 | -0.0043081 | 0.52937208 | 0.90650187 |

| | | | | | | |
|---|---|---|---|---|---|---|
| Itpkc | 0.03888996 | 0.04219748 | 0.04650834 | -0.0043109 | 0.65985338 | 0.90650187 |
| Ttc36 | 0.02202529 | 0.02232618 | 0.02663773 | -0.0043116 | 0.7641024 | 0.90650187 |
| Gm43137 | 0.01638471 | 0.0155773 | 0.01989113 | -0.0043138 | 0.83250891 | 0.91029548 |
| Ago4 | 0.06246495 | 0.06923545 | 0.07355218 | -0.0043167 | 0.6029191 | 0.90650187 |
| Cmpk2 | 0.03826404 | 0.04146055 | 0.04577879 | -0.0043182 | 0.66251887 | 0.90650187 |
| Traf4 | 0.62221149 | 0.54331799 | 0.54763746 | -0.0043195 | 0.51398586 | 0.90650187 |
| Nek7 | 0.3725683 | 0.3648985 | 0.3692208 | -0.0043223 | 0.52075267 | 0.90650187 |
| Daam2 | 0.01772847 | 0.0171758 | 0.02150302 | -0.0043272 | 0.81450622 | 0.9092537 |
| Ascl1 | 0.0097916 | 0.00760062 | 0.01194003 | -0.0043394 | 0.9469185 | 0.96108087 |
| Stx1b | 0.02308857 | 0.0235637 | 0.02790379 | -0.0043401 | 0.75536147 | 0.90650187 |
| Dbndd2 | 0.05287767 | 0.0583116 | 0.06265525 | -0.0043436 | 0.62103375 | 0.90650187 |
| Elac2 | 0.18578576 | 0.19895164 | 0.20329605 | -0.0043444 | 0.53784338 | 0.90650187 |
| Mdga1 | 0.04261913 | 0.04649743 | 0.05084236 | -0.0043449 | 0.64799636 | 0.90650187 |
| Pigm | 0.1149965 | 0.12666764 | 0.13101527 | -0.0043476 | 0.55864056 | 0.90650187 |
| Gm14508 | 0.01210701 | 0.01038923 | 0.01474041 | -0.0043512 | 0.90528829 | 0.93719848 |
| Zfp74 | 0.13815826 | 0.15095746 | 0.15531031 | -0.0043528 | 0.54957831 | 0.90650187 |
| Nos3 | 0.02060238 | 0.02058185 | 0.02494062 | -0.0043588 | 0.78140335 | 0.90650187 |
| D730045B01 | 0.01768857 | 0.01708873 | 0.0214552 | -0.0043665 | 0.81720131 | 0.9092537 |
| Siah1a | 0.23590375 | 0.24674052 | 0.25111503 | -0.0043745 | 0.53086482 | 0.90650187 |
| Ap5z1 | 0.07763904 | 0.08614457 | 0.09052311 | -0.0043785 | 0.58512839 | 0.90650187 |
| Ism1 | 0.02407802 | 0.02470065 | 0.02908032 | -0.0043797 | 0.7484229 | 0.90650187 |
| Mpc2 | 1.25413902 | 0.83397248 | 0.83835239 | -0.0043799 | 0.50926491 | 0.90650187 |
| 9030622O22 | 0.02471627 | 0.02545642 | 0.02983842 | -0.004382 | 0.74308011 | 0.90650187 |
| Tmprss4 | 0.02932008 | 0.03090556 | 0.0352876 | -0.004382 | 0.70954604 | 0.90650187 |
| Guca2a | 0.0125843 | 0.01093338 | 0.01531659 | -0.0043832 | 0.89835175 | 0.93440169 |
| Gm14306 | 0.02035422 | 0.02025968 | 0.02464432 | -0.0043846 | 0.78551324 | 0.90650187 |
| Rnf150 | 0.02679932 | 0.02792056 | 0.03230812 | -0.0043876 | 0.726988 | 0.90650187 |
| Ddx59 | 0.05270028 | 0.05806056 | 0.06245234 | -0.0043918 | 0.62271693 | 0.90650187 |
| Usp51 | 0.17687596 | 0.19012178 | 0.19451431 | -0.0043925 | 0.53998297 | 0.90650187 |
| Ypel4 | 0.04075001 | 0.04428008 | 0.04867275 | -0.0043927 | 0.65586769 | 0.90650187 |

| | | | | | | |
|---|---|---|---|---|---|---|
| Fastkd5 | 0.07107798 | 0.0788322 | 0.08322614 | -0.0043939 | 0.59278177 | 0.90650187 |
| Ccdc57 | 0.02627477 | 0.02729152 | 0.03168685 | -0.0043953 | 0.73126941 | 0.90650187 |
| Aip | 0.5999239 | 0.52894703 | 0.53334321 | -0.0043962 | 0.51461544 | 0.90650187 |
| Socs1 | 0.10420604 | 0.11507569 | 0.11947233 | -0.0043966 | 0.56497708 | 0.90650187 |
| Wdr76 | 0.08459322 | 0.09379125 | 0.09819004 | -0.0043988 | 0.57893028 | 0.90650187 |
| Agpat4 | 0.41644147 | 0.39935764 | 0.40375649 | -0.0043988 | 0.51931483 | 0.90650187 |
| Ppp1r3b | 0.01574621 | 0.01472343 | 0.0191242 | -0.0044008 | 0.84684677 | 0.91420247 |
| Dnhd1 | 0.01421814 | 0.01288409 | 0.01728611 | -0.004402 | 0.87120465 | 0.92225579 |
| Mbl2 | 0.01344911 | 0.01195649 | 0.01635961 | -0.0044031 | 0.88425264 | 0.9278081 |
| Rapgef5 | 0.01097735 | 0.00896987 | 0.01337524 | -0.0044054 | 0.92843535 | 0.94954893 |
| Telo2 | 0.22304558 | 0.23469114 | 0.23909701 | -0.0044059 | 0.53264471 | 0.90650187 |
| D130020L05I | 0.091042 | 0.10083275 | 0.10523872 | -0.004406 | 0.57382536 | 0.90650187 |
| Hdac8 | 0.13081225 | 0.14326546 | 0.14767511 | -0.0044096 | 0.55280361 | 0.90650187 |
| Slc19a2 | 0.04369425 | 0.04767714 | 0.0520879 | -0.0044108 | 0.6467089 | 0.90650187 |
| Smyd5 | 0.12873428 | 0.14108977 | 0.14550367 | -0.0044139 | 0.55363827 | 0.90650187 |
| Ttc16 | 0.01644274 | 0.01554684 | 0.0199608 | -0.004414 | 0.83721222 | 0.91105039 |
| Gm4673 | 0.02494052 | 0.02569006 | 0.03010462 | -0.0044146 | 0.7427691 | 0.90650187 |
| 1010001B22I | 0.01239435 | 0.01066948 | 0.01508732 | -0.0044178 | 0.90349958 | 0.93645828 |
| Sft2d1 | 0.39589062 | 0.38334921 | 0.38776789 | -0.0044187 | 0.52020113 | 0.90650187 |
| Phkg1 | 0.01224163 | 0.01047446 | 0.01490296 | -0.0044285 | 0.90675575 | 0.9379403 |
| Gm9930 | 0.0543408 | 0.05989225 | 0.06432711 | -0.0044349 | 0.62038429 | 0.90650187 |
| Csrnp1 | 0.08747014 | 0.09690586 | 0.10134179 | -0.0044359 | 0.57714415 | 0.90650187 |
| Klhl29 | 0.01762426 | 0.01694187 | 0.02137812 | -0.0044363 | 0.82187221 | 0.9092537 |
| Npdc1 | 1.03841043 | 0.76176303 | 0.76620032 | -0.0044373 | 0.51026995 | 0.90650187 |
| Kcnh3 | 0.01647197 | 0.01555849 | 0.0199959 | -0.0044374 | 0.83806846 | 0.91139496 |
| Gm12728 | 0.01335973 | 0.01180836 | 0.01625187 | -0.0044435 | 0.88791191 | 0.92986219 |
| 4732463B04I | 0.00729597 | 0.00446603 | 0.00891185 | -0.0044458 | 0.98671449 | 0.98964416 |
| Mast1 | 0.01187663 | 0.01001266 | 0.01446216 | -0.0044495 | 0.91430629 | 0.94143263 |
| Rbm24 | 0.00946411 | 0.00708944 | 0.01154325 | -0.0044538 | 0.95685294 | 0.96737098 |
| BC025920 | 0.01961979 | 0.01931254 | 0.02376681 | -0.0044543 | 0.79762686 | 0.9075779 |

| | | | | | | |
|---|---|---|---|---|---|---|
| Hist2h4 | 0.02115307 | 0.02114277 | 0.02559782 | -0.004455 | 0.7804497 | 0.90650187 |
| Camk1d | 0.03369829 | 0.03597649 | 0.0404388 | -0.0044623 | 0.68813136 | 0.90650187 |
| 4933406I18R | 0.00817697 | 0.00551733 | 0.00998202 | -0.0044647 | 0.97661637 | 0.98191318 |
| Aunip | 0.02644753 | 0.02742647 | 0.03189151 | -0.004465 | 0.73316643 | 0.90650187 |
| Rell2 | 0.00916885 | 0.00671593 | 0.01118535 | -0.0044694 | 0.96216393 | 0.97133123 |
| Atp6v1b1 | 0.00886599 | 0.00634509 | 0.0108181 | -0.004473 | 0.96697941 | 0.97480297 |
| Rap2c | 0.09920539 | 0.10960266 | 0.1140784 | -0.0044757 | 0.56923171 | 0.90650187 |
| Gm17435 | 0.01248251 | 0.01071325 | 0.01519374 | -0.0044805 | 0.90506557 | 0.93712658 |
| 2310011J03F | 0.48880593 | 0.45308279 | 0.45756572 | -0.0044829 | 0.5173705 | 0.90650187 |
| Tmprss2 | 0.01399182 | 0.01252937 | 0.01701355 | -0.0044842 | 0.87934248 | 0.92549228 |
| Capza1 | 0.47741232 | 0.44485372 | 0.44934059 | -0.0044869 | 0.51770382 | 0.90650187 |
| Gm37065 | 0.0128415 | 0.01113977 | 0.01562692 | -0.0044871 | 0.89911442 | 0.93440169 |
| RP23-335G1 | 0.02239457 | 0.02259001 | 0.02707764 | -0.0044876 | 0.7693715 | 0.90650187 |
| Klhdc10 | 0.3133908 | 0.31564566 | 0.3201404 | -0.0044947 | 0.52488415 | 0.90650187 |
| Pex19 | 0.73776369 | 0.61292519 | 0.6174216 | -0.0044964 | 0.51291366 | 0.90650187 |
| Gm13154 | 0.03161122 | 0.03348569 | 0.037987 | -0.0045013 | 0.70082889 | 0.90650187 |
| Zfpm2 | 0.01927545 | 0.01885377 | 0.02335509 | -0.0045013 | 0.80422739 | 0.90882608 |
| B3gntl1 | 0.0289042 | 0.03029288 | 0.03479673 | -0.0045039 | 0.7174926 | 0.90650187 |
| Tstd1 | 0.0269887 | 0.02802617 | 0.03253231 | -0.0045061 | 0.73097773 | 0.90650187 |
| Zbtb32 | 0.00937627 | 0.0069292 | 0.01143678 | -0.0045076 | 0.96012568 | 0.9701425 |
| Sytl2 | 0.01291496 | 0.01120696 | 0.01571553 | -0.0045086 | 0.89891321 | 0.93440169 |
| Thoc3 | 0.74520315 | 0.61716542 | 0.62168138 | -0.004516 | 0.51288096 | 0.90650187 |
| Tcf25 | 1.4977223 | 0.88842568 | 0.89294235 | -0.0045167 | 0.50897014 | 0.90650187 |
| Igsf11 | 0.01810189 | 0.01743357 | 0.02195044 | -0.0045169 | 0.81985127 | 0.9092537 |
| Ldhd | 0.03891463 | 0.04201969 | 0.04653707 | -0.0045174 | 0.66696148 | 0.90650187 |
| Zbtb3 | 0.02343745 | 0.023801 | 0.02831881 | -0.0045178 | 0.76090729 | 0.90650187 |
| Sfxn5 | 0.03264732 | 0.03468671 | 0.03920502 | -0.0045183 | 0.69579619 | 0.90650187 |
| Prss27 | 0.00969015 | 0.00729779 | 0.01181713 | -0.0045193 | 0.95545123 | 0.96651455 |
| Zkscan17 | 0.1155094 | 0.12703901 | 0.13156036 | -0.0045213 | 0.56071514 | 0.90650187 |
| Gpbp1l1 | 0.42344362 | 0.40460864 | 0.4091305 | -0.0045219 | 0.51959392 | 0.90650187 |

| | | | | | | |
|---|---|---|---|---|---|---|
| Gm10311 | 0.01904379 | 0.018553 | 0.02307799 | -0.004525 | 0.80830501 | 0.9092537 |
| Ggt7 | 0.01272709 | 0.01096314 | 0.01548889 | -0.0045257 | 0.9030237 | 0.93635703 |
| Cbarp | 0.30118213 | 0.30510986 | 0.30963684 | -0.004527 | 0.52591142 | 0.90650187 |
| 4930522L14F | 0.03625008 | 0.03889804 | 0.04342731 | -0.0045293 | 0.67855293 | 0.90650187 |
| Cyp2e1 | 0.06874736 | 0.07608876 | 0.08061912 | -0.0045304 | 0.59863654 | 0.90650187 |
| Cecr2 | 0.05202425 | 0.05714407 | 0.0616786 | -0.0045345 | 0.6281037 | 0.90650187 |
| Sh3d21 | 0.05079018 | 0.05572739 | 0.06026439 | -0.004537 | 0.6310694 | 0.90650187 |
| Slc22a23 | 0.01887134 | 0.01833317 | 0.02287165 | -0.0045385 | 0.81115261 | 0.9092537 |
| Lrrtm2 | 0.01332794 | 0.01167394 | 0.01621355 | -0.0045396 | 0.89338243 | 0.93181736 |
| E2f7 | 0.03844935 | 0.04145458 | 0.04599485 | -0.0045403 | 0.66961044 | 0.90650187 |
| Ankrd17 | 0.71796469 | 0.60141068 | 0.60595286 | -0.0045422 | 0.51329187 | 0.90650187 |
| Abca8b | 0.01224907 | 0.01036723 | 0.01491195 | -0.0045447 | 0.91227326 | 0.94062208 |
| 4833412C05I | 0.01239383 | 0.01053631 | 0.0150867 | -0.0045504 | 0.91001833 | 0.93950381 |
| Ssc4d | 0.02400329 | 0.02444107 | 0.02899151 | -0.0045504 | 0.75734015 | 0.90650187 |
| 5033417F24I | 0.03576104 | 0.03830451 | 0.04285535 | -0.0045508 | 0.68156611 | 0.90650187 |
| Gm2447 | 0.01317465 | 0.01147788 | 0.01602872 | -0.0045508 | 0.89655955 | 0.93371722 |
| Rasgef1c | 0.01849155 | 0.01786519 | 0.02241707 | -0.0045519 | 0.81665698 | 0.9092537 |
| Ucn3 | 0.02368283 | 0.02405798 | 0.02861059 | -0.0045526 | 0.76034062 | 0.90650187 |
| Nanos3 | 0.04799127 | 0.05249363 | 0.05704839 | -0.0045548 | 0.63868214 | 0.90650187 |
| A130014A01 | 0.04763576 | 0.05207455 | 0.05663906 | -0.0045645 | 0.63992902 | 0.90650187 |
| Rhbdd1 | 0.08218375 | 0.09097369 | 0.0955414 | -0.0045677 | 0.5841566 | 0.90650187 |
| Slc7a11 | 0.01877447 | 0.01818795 | 0.02275572 | -0.0045678 | 0.81390237 | 0.9092537 |
| 4931428F04I | 0.07534733 | 0.08341273 | 0.08798139 | -0.0045687 | 0.5912843 | 0.90650187 |
| Usp13 | 0.01306365 | 0.0113248 | 0.01589487 | -0.0045701 | 0.89940931 | 0.93453795 |
| Slc25a45 | 0.02003989 | 0.01969762 | 0.02426886 | -0.0045712 | 0.79880679 | 0.90777025 |
| Mapk8ip3 | 0.28146326 | 0.28781259 | 0.29238473 | -0.0045721 | 0.52771125 | 0.90650187 |
| Cetn4 | 0.11321987 | 0.12454708 | 0.12912467 | -0.0045776 | 0.56261429 | 0.90650187 |
| Tusc2 | 0.28383905 | 0.2899042 | 0.29448226 | -0.0045781 | 0.52754974 | 0.90650187 |
| Cecr5 | 0.09987145 | 0.11021728 | 0.11479558 | -0.0045783 | 0.57036386 | 0.90650187 |
| Fxyd4 | 0.0289569 | 0.03027713 | 0.03485895 | -0.0045818 | 0.72049957 | 0.90650187 |

| | | | | | | |
|---|---|---|---|---|---|---|
| Brap | 0.25143545 | 0.26082672 | 0.26541175 | -0.004585 | 0.5306081 | 0.90650187 |
| Dok7 | 0.02545167 | 0.02612287 | 0.03071111 | -0.0045882 | 0.74671031 | 0.90650187 |
| A830011K09 | 0.03189468 | 0.03372732 | 0.0383204 | -0.0045931 | 0.70293132 | 0.90650187 |
| Abhd14b | 0.24358407 | 0.25362002 | 0.25821481 | -0.0045948 | 0.53152631 | 0.90650187 |
| Tldc1 | 0.01490704 | 0.01352007 | 0.01811525 | -0.0045952 | 0.87027405 | 0.92175556 |
| B3gnt4 | 0.0091291 | 0.00654018 | 0.01113716 | -0.004597 | 0.96674507 | 0.97469601 |
| Dll3 | 0.00865059 | 0.00595891 | 0.0105568 | -0.0045979 | 0.97358049 | 0.97994563 |
| Kcnn3 | 0.05133779 | 0.0562918 | 0.06089222 | -0.0046004 | 0.6315141 | 0.90650187 |
| Lmo1 | 0.07383226 | 0.08169628 | 0.08629691 | -0.0046006 | 0.59367253 | 0.90650187 |
| Ndst4 | 0.05112229 | 0.05603961 | 0.0606452 | -0.0046056 | 0.63217294 | 0.90650187 |
| Fut10 | 0.03857868 | 0.0415391 | 0.0461456 | -0.0046065 | 0.6714006 | 0.90650187 |
| Gm26733 | 0.01021492 | 0.00784606 | 0.01245267 | -0.0046066 | 0.94997 | 0.96276378 |
| A930038B10 | 0.02929729 | 0.03065064 | 0.03526072 | -0.0046101 | 0.71945709 | 0.90650187 |
| Fam76a | 0.48626154 | 0.45111619 | 0.45573649 | -0.0046203 | 0.51797425 | 0.90650187 |
| Dynlt1a | 0.26090833 | 0.26939268 | 0.27401416 | -0.0046215 | 0.5298843 | 0.90650187 |
| Faxc | 0.03408918 | 0.03626433 | 0.04089724 | -0.0046329 | 0.69273094 | 0.90650187 |
| 9530082P21I | 0.01993659 | 0.01950224 | 0.02414543 | -0.0046432 | 0.80369522 | 0.90882608 |
| Tap1 | 0.06495628 | 0.07171816 | 0.07636155 | -0.0046434 | 0.60654599 | 0.90650187 |
| Irs1 | 0.053983 | 0.05927089 | 0.06391857 | -0.0046477 | 0.62674735 | 0.90650187 |
| Myo5c | 0.01176056 | 0.00967135 | 0.01432195 | -0.0046506 | 0.92557003 | 0.94746893 |
| Alg13 | 0.07070799 | 0.07816137 | 0.0828128 | -0.0046514 | 0.59859067 | 0.90650187 |
| B9d1 | 0.70189055 | 0.59183367 | 0.59649787 | -0.0046642 | 0.51386505 | 0.90650187 |
| Zfp97 | 0.01323334 | 0.01143198 | 0.01609949 | -0.0046675 | 0.90127581 | 0.93562068 |
| Lrp3 | 0.05488476 | 0.06027856 | 0.06494785 | -0.0046693 | 0.62536729 | 0.90650187 |
| Ankrd40 | 0.20622659 | 0.2184439 | 0.2231182 | -0.0046743 | 0.53710181 | 0.90650187 |
| Fzd8 | 0.01108454 | 0.00882424 | 0.01350486 | -0.0046806 | 0.93831647 | 0.9555402 |
| Lrguk | 0.0120344 | 0.00996838 | 0.01465272 | -0.0046843 | 0.92237367 | 0.94557609 |
| Npc1 | 0.11430758 | 0.12559752 | 0.13028262 | -0.0046851 | 0.56350773 | 0.90650187 |
| Kit | 0.18379567 | 0.19665508 | 0.20134207 | -0.004687 | 0.54121225 | 0.90650187 |
| Pknox2 | 0.03397772 | 0.03607866 | 0.04076655 | -0.0046879 | 0.69540009 | 0.90650187 |

| | | | | | | |
|---|---|---|---|---|---|---|
| Aldh18a1 | 0.25089798 | 0.26022738 | 0.26492107 | -0.0046937 | 0.53138991 | 0.90650187 |
| Nfxl1 | 0.24857773 | 0.25810366 | 0.26279959 | -0.0046959 | 0.53165788 | 0.90650187 |
| Mkln1os | 0.01427144 | 0.01265034 | 0.01735028 | -0.0046999 | 0.8857539 | 0.92839449 |
| Dph2 | 0.10518624 | 0.11582616 | 0.12052687 | -0.0047007 | 0.56882534 | 0.90650187 |
| Malt1 | 0.01890934 | 0.01821449 | 0.02291713 | -0.0047026 | 0.81918053 | 0.9092537 |
| Pck2 | 0.41004623 | 0.3941114 | 0.3988152 | -0.0047038 | 0.52090837 | 0.90650187 |
| Apool | 0.31359725 | 0.31561296 | 0.32031688 | -0.0047039 | 0.52602629 | 0.90650187 |
| U2surp | 0.8619694 | 0.68048895 | 0.68519447 | -0.0047055 | 0.51217776 | 0.90650187 |
| Slc27a1 | 0.10440677 | 0.11498191 | 0.11968838 | -0.0047065 | 0.5693867 | 0.90650187 |
| 6430571L13F | 0.01674494 | 0.01561619 | 0.02032353 | -0.0047073 | 0.84842263 | 0.91440801 |
| Xlr4a | 0.02778612 | 0.02876588 | 0.0334757 | -0.0047098 | 0.73416823 | 0.90650187 |
| Ankrd33b | 0.01725594 | 0.01622632 | 0.02093654 | -0.0047102 | 0.84138258 | 0.91251845 |
| Aim1 | 0.01589334 | 0.01458832 | 0.01930099 | -0.0047127 | 0.86114383 | 0.91848494 |
| Ephb2 | 0.18037671 | 0.19325689 | 0.1979751 | -0.0047182 | 0.54218871 | 0.90650187 |
| 9030617O03 | 0.02777734 | 0.02874274 | 0.03346532 | -0.0047226 | 0.73478759 | 0.90650187 |
| Fancc | 0.05470298 | 0.06001437 | 0.06474046 | -0.0047261 | 0.62723221 | 0.90650187 |
| Aldh1l2 | 0.00834344 | 0.00545623 | 0.01018409 | -0.0047279 | 0.98048292 | 0.98527936 |
| Dubr | 0.13233617 | 0.14453639 | 0.14926429 | -0.0047279 | 0.55599131 | 0.90650187 |
| Bmp1 | 0.35366215 | 0.34913364 | 0.35386176 | -0.0047281 | 0.52368322 | 0.90650187 |
| Rdh13 | 0.12701442 | 0.13896983 | 0.14370253 | -0.0047327 | 0.55820183 | 0.90650187 |
| Kif13b | 0.02220855 | 0.02212133 | 0.02685607 | -0.0047347 | 0.78341055 | 0.90650187 |
| Stk38l | 0.05266187 | 0.05766076 | 0.06240839 | -0.0047476 | 0.63239109 | 0.90650187 |
| Mapk11 | 0.02618883 | 0.02683569 | 0.03158502 | -0.0047493 | 0.74808361 | 0.90650187 |
| Serpina9 | 0.02117637 | 0.02087027 | 0.02562562 | -0.0047553 | 0.79530603 | 0.907548 |
| Acot6 | 0.02526821 | 0.0257358 | 0.03049347 | -0.0047577 | 0.75603848 | 0.90650187 |
| 6330403L08F | 0.01839793 | 0.01754548 | 0.02230498 | -0.0047595 | 0.82859214 | 0.9096066 |
| Mterf3 | 0.33259808 | 0.33163597 | 0.33639804 | -0.0047621 | 0.52508975 | 0.90650187 |
| Gm26797 | 0.01533291 | 0.01386445 | 0.01862743 | -0.004763 | 0.87217416 | 0.92254675 |
| Maoa | 0.13588693 | 0.14819185 | 0.1529564 | -0.0047645 | 0.55506922 | 0.90650187 |
| Slc24a3 | 0.03786082 | 0.04054047 | 0.0453085 | -0.004768 | 0.68004767 | 0.90650187 |

| | | | | | | |
|---|---|---|---|---|---|---|
| Fah | 0.05104776 | 0.05579 | 0.06055976 | -0.0047698 | 0.63688349 | 0.90650187 |
| Arhgef9 | 0.12216629 | 0.13383221 | 0.13860571 | -0.0047735 | 0.56084192 | 0.90650187 |
| A630072M18 | 0.08467819 | 0.09350972 | 0.09828329 | -0.0047736 | 0.58547529 | 0.90650187 |
| Cdk16 | 0.54863658 | 0.49456025 | 0.49933411 | -0.0047739 | 0.51695077 | 0.90650187 |
| Ube3c | 0.39667406 | 0.38360429 | 0.38838331 | -0.004779 | 0.52181237 | 0.90650187 |
| Kcnc2 | 0.01166195 | 0.0094233 | 0.01420281 | -0.0047795 | 0.93267711 | 0.95188026 |
| Dpp10 | 0.01458885 | 0.01295183 | 0.01773238 | -0.0047806 | 0.8846409 | 0.92796521 |
| Mtm1 | 0.0966996 | 0.10659378 | 0.11137482 | -0.004781 | 0.57567402 | 0.90650187 |
| Iffo2 | 0.04909927 | 0.0535406 | 0.05832292 | -0.0047823 | 0.64226815 | 0.90650187 |
| Ap1m2 | 0.33204599 | 0.33114962 | 0.33593523 | -0.0047856 | 0.52524833 | 0.90650187 |
| Whrn | 0.05917814 | 0.06504536 | 0.06983178 | -0.0047864 | 0.61970711 | 0.90650187 |
| Sphk1 | 0.05822894 | 0.06396539 | 0.06875438 | -0.004789 | 0.62158747 | 0.90650187 |
| Ehd3 | 0.03276675 | 0.03455317 | 0.03934531 | -0.0047921 | 0.705905 | 0.90650187 |
| Upb1 | 0.15918636 | 0.17202295 | 0.17681548 | -0.0047925 | 0.54795523 | 0.90650187 |
| Tmem147os | 0.02195717 | 0.02176304 | 0.02655656 | -0.0047935 | 0.78885134 | 0.90650187 |
| Mettl22 | 0.08309252 | 0.09174073 | 0.09654134 | -0.0048006 | 0.58747798 | 0.90650187 |
| Cfap43 | 0.07853512 | 0.08671363 | 0.0915149 | -0.0048013 | 0.59221092 | 0.90650187 |
| Dagla | 0.01523108 | 0.01370361 | 0.01850499 | -0.0048014 | 0.87564235 | 0.92418014 |
| Gm17669 | 0.02007879 | 0.0195129 | 0.02431532 | -0.0048024 | 0.8100376 | 0.9092537 |
| Gm28050 | 0.01677569 | 0.01555597 | 0.02036043 | -0.0048045 | 0.85291811 | 0.91562204 |
| Eif5a2 | 0.0507876 | 0.0554558 | 0.06026143 | -0.0048056 | 0.63852571 | 0.90650187 |
| Cdip1 | 0.25416954 | 0.26309453 | 0.26790347 | -0.0048089 | 0.53180183 | 0.90650187 |
| Derl3 | 0.03472696 | 0.03683484 | 0.04164474 | -0.0048099 | 0.69618541 | 0.90650187 |
| 2810408A11 | 0.05045871 | 0.05507259 | 0.05988414 | -0.0048116 | 0.63952696 | 0.90650187 |
| Cutal | 0.01523461 | 0.01369529 | 0.01850923 | -0.0048139 | 0.87620594 | 0.92449549 |
| Polg2 | 0.08748575 | 0.09654082 | 0.10135886 | -0.004818 | 0.58368101 | 0.90650187 |
| Vsig10l | 0.01823944 | 0.01729624 | 0.02211519 | -0.0048189 | 0.83365437 | 0.91062493 |
| Zfp458 | 0.03765194 | 0.04023204 | 0.04506478 | -0.0048327 | 0.68322865 | 0.90650187 |
| Drg2 | 0.42251846 | 0.40358853 | 0.4084227 | -0.0048342 | 0.52098229 | 0.90650187 |
| Crybb3 | 0.02763602 | 0.02846133 | 0.0332982 | -0.0048369 | 0.74078439 | 0.90650187 |

| | | | | | | |
|---|---|---|---|---|---|---|
| Papd7 | 0.10957403 | 0.1203924 | 0.12523273 | -0.0048403 | 0.56821276 | 0.90650187 |
| Rbm8a2 | 0.19694324 | 0.20933116 | 0.21417195 | -0.0048408 | 0.54001882 | 0.90650187 |
| Clcn6 | 0.06094679 | 0.0669947 | 0.07183573 | -0.004841 | 0.6177553 | 0.90650187 |
| 9130019O22 | 0.05260668 | 0.05749926 | 0.06234525 | -0.004846 | 0.63515913 | 0.90650187 |
| 1700030J22F | 0.02288907 | 0.02282013 | 0.02766637 | -0.0048462 | 0.78192891 | 0.90650187 |
| Aqp4 | 0.03364405 | 0.0355221 | 0.04037518 | -0.0048531 | 0.70345377 | 0.90650187 |
| Rnf5 | 0.74374379 | 0.61599354 | 0.62084797 | -0.0048544 | 0.51386456 | 0.90650187 |
| 4930503L19F | 0.04497107 | 0.04870987 | 0.0535648 | -0.0048549 | 0.65650166 | 0.90650187 |
| Itga8 | 0.02755644 | 0.0283489 | 0.03320407 | -0.0048552 | 0.74216877 | 0.90650187 |
| Lrrc27 | 0.02782318 | 0.02866361 | 0.03351951 | -0.0048559 | 0.74022037 | 0.90650187 |
| Vsig2 | 0.02701762 | 0.02770712 | 0.03256655 | -0.0048594 | 0.74644757 | 0.90650187 |
| Galnt14 | 0.02668285 | 0.02730899 | 0.03217021 | -0.0048612 | 0.74913597 | 0.90650187 |
| C920021L13F | 0.0521948 | 0.05700858 | 0.06187387 | -0.0048653 | 0.63666861 | 0.90650187 |
| Car13 | 0.01885551 | 0.01798558 | 0.02285271 | -0.0048671 | 0.82813399 | 0.90957116 |
| Mnx1 | 0.05790953 | 0.06352366 | 0.06839153 | -0.0048679 | 0.62415916 | 0.90650187 |
| Sirt6 | 0.23110015 | 0.24177016 | 0.24664475 | -0.0048746 | 0.53500653 | 0.90650187 |
| 1010001N08 | 0.03984157 | 0.04274168 | 0.04761633 | -0.0048746 | 0.67548553 | 0.90650187 |
| Strada | 0.09959264 | 0.10961889 | 0.11449545 | -0.0048766 | 0.57508917 | 0.90650187 |
| Abcc12 | 0.01231112 | 0.01011012 | 0.01498685 | -0.0048767 | 0.92599596 | 0.94759147 |
| Gm6793 | 0.04785576 | 0.05201039 | 0.05689238 | -0.004882 | 0.64857732 | 0.90650187 |
| Angptl8 | 0.01214189 | 0.00989658 | 0.01478253 | -0.004886 | 0.92913384 | 0.94996709 |
| Plekhf2 | 0.21900179 | 0.23039564 | 0.2352818 | -0.0048862 | 0.53677933 | 0.90650187 |
| Foxj2 | 0.10527093 | 0.11572989 | 0.12061793 | -0.004888 | 0.57148578 | 0.90650187 |
| Hspa1b | 0.01714214 | 0.01591159 | 0.02080006 | -0.0048885 | 0.85194589 | 0.91551074 |
| Echdc2 | 0.2606412 | 0.26888143 | 0.27377275 | -0.0048913 | 0.53165341 | 0.90650187 |
| Atp8a2 | 0.01522947 | 0.01360704 | 0.01850306 | -0.004896 | 0.88026917 | 0.92599812 |
| Erc2 | 0.01776192 | 0.01664512 | 0.02154311 | -0.004898 | 0.84393045 | 0.91300169 |
| Sla2 | 0.01158781 | 0.00920525 | 0.01411322 | -0.004908 | 0.93894652 | 0.95593613 |
| Ss18 | 0.46154556 | 0.43282613 | 0.43773603 | -0.0049099 | 0.51988481 | 0.90650187 |
| Epha7 | 0.16849311 | 0.18125958 | 0.18617102 | -0.0049114 | 0.54668131 | 0.90650187 |

| | | | | | | |
|---|---|---|---|---|---|---|
| Gm15638 | 0.01407707 | 0.01220453 | 0.01711622 | -0.0049117 | 0.89897189 | 0.93440169 |
| Lrrc8e | 0.02215162 | 0.0218748 | 0.02678825 | -0.0049134 | 0.7925795 | 0.90681274 |
| Zbed5 | 0.02964323 | 0.03075393 | 0.03566883 | -0.0049149 | 0.72992014 | 0.90650187 |
| Zfp953 | 0.02560926 | 0.02598218 | 0.03089801 | -0.0049158 | 0.76030693 | 0.90650187 |
| Grb7 | 0.30017598 | 0.30384427 | 0.30876531 | -0.004921 | 0.52824268 | 0.90650187 |
| Vti1a | 0.07577634 | 0.08353046 | 0.08845778 | -0.0049273 | 0.59779017 | 0.90650187 |
| Mtcp1 | 0.04207942 | 0.0452866 | 0.05021643 | -0.0049298 | 0.66873843 | 0.90650187 |
| Trim14 | 0.01163947 | 0.00924418 | 0.01417564 | -0.0049315 | 0.93901427 | 0.95593613 |
| Gm16287 | 0.01056196 | 0.00794072 | 0.01287272 | -0.004932 | 0.95574141 | 0.96674375 |
| 9130221H12I | 0.02193906 | 0.02160125 | 0.02653497 | -0.0049337 | 0.79576367 | 0.9075779 |
| Ccdc73 | 0.05478994 | 0.05990535 | 0.06483968 | -0.0049343 | 0.63243637 | 0.90650187 |
| Rnf157 | 0.06941901 | 0.07643634 | 0.08137124 | -0.0049349 | 0.60627041 | 0.90650187 |
| Ulbp1 | 0.01488042 | 0.01314742 | 0.01808322 | -0.0049358 | 0.88751599 | 0.92965343 |
| 1810010H24I | 0.03346616 | 0.03521913 | 0.04016644 | -0.0049473 | 0.70800521 | 0.90650187 |
| Cecr6 | 0.05100926 | 0.05556692 | 0.06051562 | -0.0049487 | 0.64190008 | 0.90650187 |
| Wdpcp | 0.05780188 | 0.06331718 | 0.06826921 | -0.004952 | 0.62645163 | 0.90650187 |
| B230398E01 | 0.01715227 | 0.0158601 | 0.02081221 | -0.0049521 | 0.85492646 | 0.91627438 |
| Ankrd55 | 0.0121603 | 0.00985124 | 0.01480476 | -0.0049535 | 0.9315526 | 0.95113723 |
| AA543186 | 0.01551047 | 0.01388675 | 0.01884088 | -0.0049541 | 0.87878874 | 0.92542207 |
| E430024P14I | 0.12092327 | 0.13233942 | 0.13729425 | -0.0049548 | 0.56373205 | 0.90650187 |
| 1700016L21I | 0.01193869 | 0.00958123 | 0.01453713 | -0.0049559 | 0.93518549 | 0.95337366 |
| Rad54b | 0.02496116 | 0.02517064 | 0.03012912 | -0.0049585 | 0.76780386 | 0.90650187 |
| Polm | 0.0433371 | 0.04671577 | 0.05167432 | -0.0049586 | 0.66516112 | 0.90650187 |
| Lrrtm3 | 0.04055651 | 0.04348843 | 0.04844783 | -0.0049594 | 0.6754737 | 0.90650187 |
| Stx2 | 0.14585113 | 0.15827818 | 0.16323782 | -0.0049596 | 0.55372189 | 0.90650187 |
| Als2 | 0.03457781 | 0.03650572 | 0.04146999 | -0.0049643 | 0.7026967 | 0.90650187 |
| Gm10451 | 0.04347745 | 0.04687201 | 0.05183687 | -0.0049649 | 0.66487101 | 0.90650187 |
| Mst1r | 0.02058787 | 0.01995625 | 0.02492331 | -0.0049671 | 0.81218627 | 0.9092537 |
| Sema4c | 0.20756063 | 0.21942853 | 0.2243963 | -0.0049678 | 0.53919996 | 0.90650187 |
| Gm14091 | 0.03391338 | 0.03572332 | 0.0406911 | -0.0049678 | 0.70634635 | 0.90650187 |

| | | | | | | |
|---|---|---|---|---|---|---|
| H6pd | 0.08871135 | 0.09772954 | 0.10269799 | -0.0049684 | 0.58514551 | 0.90650187 |
| Paqr3 | 0.01745175 | 0.01620146 | 0.02117132 | -0.0049699 | 0.85166105 | 0.91530682 |
| Cyp4f15 | 0.01527428 | 0.01358442 | 0.01855693 | -0.0049725 | 0.88322109 | 0.92745622 |
| Chd6 | 0.55378828 | 0.49784802 | 0.50282254 | -0.0049745 | 0.51754031 | 0.90650187 |
| Syvn1 | 0.18037051 | 0.19299122 | 0.19796898 | -0.0049778 | 0.54450145 | 0.90650187 |
| Arrdc2 | 0.03819403 | 0.04071927 | 0.04569716 | -0.0049779 | 0.68590254 | 0.90650187 |
| Cdc7 | 0.11577274 | 0.12686192 | 0.13184009 | -0.0049782 | 0.56665415 | 0.90650187 |
| Naif1 | 0.01074595 | 0.00811088 | 0.01309534 | -0.0049845 | 0.95468875 | 0.96606468 |
| Smim5 | 0.0160629 | 0.01451925 | 0.01950468 | -0.0049854 | 0.87208559 | 0.92254675 |
| Epg5 | 0.02671526 | 0.02721653 | 0.03220859 | -0.0049921 | 0.75459865 | 0.90650187 |
| Txndc17 | 1.3368016 | 0.85496506 | 0.85995961 | -0.0049945 | 0.51029937 | 0.90650187 |
| Gm17690 | 0.01642117 | 0.01493841 | 0.01993491 | -0.0049965 | 0.8674138 | 0.92056229 |
| Mss51 | 0.04756422 | 0.05155648 | 0.05655666 | -0.0050002 | 0.65285357 | 0.90650187 |
| Nphs1 | 0.01149057 | 0.00899383 | 0.01399571 | -0.0050019 | 0.94394736 | 0.95903292 |
| Sord | 0.11958843 | 0.13088112 | 0.13588384 | -0.0050027 | 0.56500454 | 0.90650187 |
| Dock8 | 0.05988146 | 0.06562549 | 0.07062922 | -0.0050037 | 0.62359994 | 0.90650187 |
| Nxph1 | 0.05432109 | 0.05929659 | 0.06430461 | -0.005008 | 0.63541215 | 0.90650187 |
| Snx9 | 0.17628339 | 0.18891283 | 0.19392717 | -0.0050143 | 0.54575731 | 0.90650187 |
| Mob3b | 0.07349741 | 0.08090582 | 0.08592416 | -0.0050183 | 0.60243019 | 0.90650187 |
| Paupar | 0.0110448 | 0.00843841 | 0.01345681 | -0.0050184 | 0.95132947 | 0.9636744 |
| Large | 0.0793016 | 0.0873439 | 0.09236233 | -0.0050184 | 0.59543487 | 0.90650187 |
| Gstt3 | 0.07547783 | 0.08309906 | 0.08812634 | -0.0050273 | 0.60009967 | 0.90650187 |
| Gm37305 | 0.02609596 | 0.02644716 | 0.03147497 | -0.0050278 | 0.76119187 | 0.90650187 |
| Wars2 | 0.12356153 | 0.13502801 | 0.14007553 | -0.0050475 | 0.56363595 | 0.90650187 |
| Nphp4 | 0.01102931 | 0.0083872 | 0.01343807 | -0.0050509 | 0.95263308 | 0.96449816 |
| Tmem136 | 0.03487613 | 0.03676749 | 0.04181948 | -0.005052 | 0.70438331 | 0.90650187 |
| Kremen1 | 0.11994382 | 0.13120734 | 0.13625956 | -0.0050522 | 0.56546259 | 0.90650187 |
| Il17rd | 0.03696102 | 0.03920055 | 0.04425811 | -0.0050576 | 0.6942799 | 0.90650187 |
| Zfp449 | 0.12961545 | 0.14136617 | 0.14642511 | -0.0050589 | 0.56103515 | 0.90650187 |
| Ntn4 | 0.01950236 | 0.01856592 | 0.02362641 | -0.0050605 | 0.82950079 | 0.90974008 |

| | | | | | | |
|---|---|---|---|---|---|---|
| Mettl14 | 0.06804624 | 0.07477209 | 0.07983331 | -0.0050612 | 0.6109669 | 0.90650187 |
| Ddx11 | 0.02544919 | 0.0256465 | 0.03070816 | -0.0050617 | 0.76815204 | 0.90650187 |
| Jade2 | 0.01913134 | 0.01811946 | 0.02318271 | -0.0050632 | 0.8342128 | 0.91062493 |
| RP23-124B2. | 0.03364398 | 0.03531171 | 0.04037509 | -0.0050634 | 0.7114126 | 0.90650187 |
| Tmem140 | 0.0259044 | 0.02617949 | 0.03124793 | -0.0050684 | 0.76457004 | 0.90650187 |
| Plekhg6 | 0.02024911 | 0.0194501 | 0.02451878 | -0.0050687 | 0.82095856 | 0.9092537 |
| Gm4707 | 0.08658096 | 0.09529929 | 0.10036892 | -0.0050696 | 0.58883425 | 0.90650187 |
| Rasal1 | 0.02684868 | 0.02729644 | 0.03236655 | -0.0050701 | 0.7569076 | 0.90650187 |
| Spata7 | 0.113902 | 0.12477688 | 0.12985102 | -0.0050741 | 0.56895695 | 0.90650187 |
| Ncmap | 0.01046143 | 0.00767206 | 0.01275106 | -0.005079 | 0.96170082 | 0.97109168 |
| Copg2 | 0.36426565 | 0.35743049 | 0.36251184 | -0.0050814 | 0.52484375 | 0.90650187 |
| Shroom4 | 0.01378259 | 0.01167976 | 0.01676149 | -0.0050817 | 0.91114107 | 0.94014766 |
| Pank4 | 0.11595852 | 0.12695343 | 0.13203738 | -0.0050839 | 0.56795612 | 0.90650187 |
| 4930524J08F | 0.0164481 | 0.01488191 | 0.01996724 | -0.0050853 | 0.8712298 | 0.92225579 |
| Vps13c | 0.09928349 | 0.1090751 | 0.11416252 | -0.0050874 | 0.57852039 | 0.90650187 |
| Syt16 | 0.05954102 | 0.06515077 | 0.07024331 | -0.0050925 | 0.62638738 | 0.90650187 |
| Hoga1 | 0.01733007 | 0.0159276 | 0.02102543 | -0.0050978 | 0.85945686 | 0.91800029 |
| Adamts4 | 0.04248351 | 0.04558589 | 0.05068511 | -0.0050992 | 0.67265426 | 0.90650187 |
| Zbtb25 | 0.09480392 | 0.10422384 | 0.10932375 | -0.0050999 | 0.5821457 | 0.90650187 |
| Arhgef10l | 0.0639831 | 0.0701649 | 0.07526522 | -0.0051003 | 0.61838985 | 0.90650187 |
| Lss | 0.08062319 | 0.08871942 | 0.09382153 | -0.0051021 | 0.59551574 | 0.90650187 |
| Sestd1 | 0.18365185 | 0.19609777 | 0.2012007 | -0.0051029 | 0.54488609 | 0.90650187 |
| Pdcd2l | 0.30219046 | 0.30540605 | 0.31050935 | -0.0051033 | 0.52912269 | 0.90650187 |
| Fanca | 0.03166452 | 0.0329452 | 0.0380497 | -0.0051045 | 0.72454409 | 0.90650187 |
| RP23-218K1! | 0.22229441 | 0.23328415 | 0.23838957 | -0.0051054 | 0.53792536 | 0.90650187 |
| Samd1 | 0.10961932 | 0.12017189 | 0.12528119 | -0.0051093 | 0.57193542 | 0.90650187 |
| 2810405F17I | 0.01762986 | 0.01627415 | 0.02138483 | -0.0051107 | 0.85598162 | 0.9165432 |
| Ints2 | 0.18839095 | 0.20073666 | 0.20584746 | -0.0051108 | 0.54394439 | 0.90650187 |
| Gm7628 | 0.0167984 | 0.01527679 | 0.02038768 | -0.0051109 | 0.86745659 | 0.92056229 |
| Porcn | 0.05429102 | 0.05915622 | 0.06427028 | -0.0051141 | 0.63823397 | 0.90650187 |

| | | | | | | |
|---|---|---|---|---|---|---|
| Pde9a | 0.03113926 | 0.03231658 | 0.03743161 | -0.005115 | 0.72823795 | 0.90650187 |
| Carns1 | 0.02014367 | 0.01927647 | 0.02439283 | -0.0051164 | 0.82445057 | 0.90933631 |
| Mknk2 | 0.17347003 | 0.18601629 | 0.19113424 | -0.005118 | 0.54737771 | 0.90650187 |
| Zfp639 | 0.26708731 | 0.27445949 | 0.27957912 | -0.0051196 | 0.53244105 | 0.90650187 |
| Lrrc10b | 0.03129614 | 0.03248769 | 0.03761626 | -0.0051286 | 0.72777927 | 0.90650187 |
| Tube1 | 0.02712567 | 0.02756333 | 0.03269443 | -0.0051311 | 0.75731695 | 0.90650187 |
| Zbtb12 | 0.14859958 | 0.160909 | 0.16605338 | -0.0051444 | 0.55477157 | 0.90650187 |
| Hsbp1l1 | 0.05385873 | 0.05863107 | 0.06377663 | -0.0051456 | 0.64008129 | 0.90650187 |
| Arfgap1 | 0.54459424 | 0.49143666 | 0.49658549 | -0.0051488 | 0.51838239 | 0.90650187 |
| Gm4353 | 0.031933 | 0.03321532 | 0.03836546 | -0.0051501 | 0.7246704 | 0.90650187 |
| Zfp575 | 0.07954923 | 0.08748546 | 0.09263593 | -0.0051505 | 0.59761213 | 0.90650187 |
| Nek3 | 0.08712204 | 0.09580972 | 0.10096106 | -0.0051513 | 0.58972131 | 0.90650187 |
| Pawr | 0.02046274 | 0.01961853 | 0.0247739 | -0.0051554 | 0.8225476 | 0.9092537 |
| Batf3 | 0.0500144 | 0.05420456 | 0.05937418 | -0.0051696 | 0.65064881 | 0.90650187 |
| Pqlc2 | 0.10553763 | 0.11573389 | 0.12090462 | -0.0051707 | 0.57539449 | 0.90650187 |
| Gm7160 | 0.02042568 | 0.01955712 | 0.02472966 | -0.0051725 | 0.82377751 | 0.9092537 |
| Nrip2 | 0.02290483 | 0.02251253 | 0.02768513 | -0.0051726 | 0.79690081 | 0.9075779 |
| Polr2l | 0.97770821 | 0.73616698 | 0.74134226 | -0.0051753 | 0.51237901 | 0.90650187 |
| Hlf | 0.01311929 | 0.0107848 | 0.01596197 | -0.0051772 | 0.92533558 | 0.94735258 |
| Gm21988 | 0.13374922 | 0.14555258 | 0.15073539 | -0.0051828 | 0.56074385 | 0.90650187 |
| Bbc3 | 0.11025413 | 0.12077285 | 0.12596 | -0.0051872 | 0.57263012 | 0.90650187 |
| 6720427I07R | 0.05317428 | 0.05780438 | 0.06299444 | -0.0051901 | 0.64291799 | 0.90650187 |
| Agpat5 | 0.27180169 | 0.27860991 | 0.28380076 | -0.0051909 | 0.53240318 | 0.90650187 |
| Gm15491 | 0.01511651 | 0.01317434 | 0.01836721 | -0.0051929 | 0.89560213 | 0.93317211 |
| Auh | 0.55873386 | 0.50096138 | 0.50615618 | -0.0051948 | 0.51819592 | 0.90650187 |
| Ankrd54 | 0.58695763 | 0.51969767 | 0.52489465 | -0.005197 | 0.51755413 | 0.90650187 |
| Camsap2 | 0.16998923 | 0.18246395 | 0.18766592 | -0.005202 | 0.54903718 | 0.90650187 |
| Fam53b | 0.10685267 | 0.11711311 | 0.12231694 | -0.0052038 | 0.57500569 | 0.90650187 |
| Otop3 | 0.03509041 | 0.03686157 | 0.04207043 | -0.0052089 | 0.70898547 | 0.90650187 |
| Sowahb | 0.01423094 | 0.01209178 | 0.01730153 | -0.0052097 | 0.90965573 | 0.9393235 |

| | | | | | | |
|---|---|---|---|---|---|---|
| Lamb2 | 0.13198528 | 0.14368731 | 0.14889862 | -0.0052113 | 0.56182263 | 0.90650187 |
| Nnt | 0.21623124 | 0.22743526 | 0.2326582 | -0.0052229 | 0.53974815 | 0.90650187 |
| 2310015A10 | 0.11827567 | 0.12926447 | 0.13449462 | -0.0052302 | 0.56862647 | 0.90650187 |
| Abl1 | 0.22076051 | 0.23169571 | 0.23694317 | -0.0052475 | 0.53921429 | 0.90650187 |
| Gm30173 | 0.01787374 | 0.01642689 | 0.02167711 | -0.0052502 | 0.85919953 | 0.91797966 |
| Gm26521 | 0.01728632 | 0.01572202 | 0.02097297 | -0.0052509 | 0.86715338 | 0.92056229 |
| Zfp513 | 0.23718093 | 0.24704017 | 0.25229972 | -0.0052596 | 0.53691915 | 0.90650187 |
| AI429214 | 0.03258055 | 0.0338655 | 0.03912658 | -0.0052611 | 0.72500317 | 0.90650187 |
| 2010109A12 | 0.02066924 | 0.01975701 | 0.02502045 | -0.0052634 | 0.82515795 | 0.90934433 |
| Map10 | 0.0495279 | 0.05354554 | 0.05881547 | -0.0052699 | 0.65480698 | 0.90650187 |
| Xxylt1 | 0.015775 | 0.01388842 | 0.0191588 | -0.0052704 | 0.88932366 | 0.9302877 |
| Fam83f | 0.02916856 | 0.02983767 | 0.03510879 | -0.0052711 | 0.74775712 | 0.90650187 |
| Nadsyn1 | 0.04340973 | 0.04648249 | 0.05175844 | -0.005276 | 0.67478303 | 0.90650187 |
| Dach2 | 0.01651769 | 0.01477247 | 0.02005078 | -0.0052783 | 0.87905868 | 0.92549228 |
| Mms19 | 0.17504656 | 0.18742206 | 0.19270042 | -0.0052784 | 0.5484602 | 0.90650187 |
| Tiam1 | 0.02975963 | 0.03052648 | 0.03580612 | -0.0052796 | 0.7439298 | 0.90650187 |
| Zfp811 | 0.01960165 | 0.01846475 | 0.02374512 | -0.0052804 | 0.8385708 | 0.91181078 |
| Fam126a | 0.1398428 | 0.1517714 | 0.15705215 | -0.0052808 | 0.55941254 | 0.90650187 |
| Tvp23a | 0.06452714 | 0.07059177 | 0.07587827 | -0.0052865 | 0.62161642 | 0.90650187 |
| Adsl | 0.30299225 | 0.30591342 | 0.3112025 | -0.0052891 | 0.53011375 | 0.90650187 |
| Chst11 | 0.02810899 | 0.02856054 | 0.03385737 | -0.0052968 | 0.75662721 | 0.90650187 |
| Tatdn2 | 0.05397173 | 0.05860651 | 0.06390569 | -0.0052992 | 0.64380115 | 0.90650187 |
| Fbxw8 | 0.13375162 | 0.1454382 | 0.15073788 | -0.0052997 | 0.56210164 | 0.90650187 |
| Acsf2 | 0.06861276 | 0.07516712 | 0.08046832 | -0.0053012 | 0.61519064 | 0.90650187 |
| Cdyl2 | 0.0823366 | 0.09040726 | 0.09570967 | -0.0053024 | 0.5972714 | 0.90650187 |
| Proc | 0.02488559 | 0.0247344 | 0.03003942 | -0.005305 | 0.78380341 | 0.90650187 |
| Bsn | 0.01728115 | 0.01565788 | 0.02096677 | -0.0053089 | 0.86984313 | 0.92149177 |
| Prdm6 | 0.03297142 | 0.03427328 | 0.03958568 | -0.0053124 | 0.72461315 | 0.90650187 |
| Ppp6r2 | 0.01369135 | 0.01133597 | 0.01665156 | -0.0053156 | 0.92207203 | 0.94549985 |
| Ppp1r13l | 0.03317997 | 0.03451475 | 0.03983054 | -0.0053158 | 0.72351325 | 0.90650187 |

| | | | | | | |
|---|---|---|---|---|---|---|
| Itga2b | 0.03437708 | 0.03591692 | 0.04123475 | -0.0053178 | 0.71678828 | 0.90650187 |
| Slc12a9 | 0.13626629 | 0.14803185 | 0.15334997 | -0.0053181 | 0.56126292 | 0.90650187 |
| Zfpl1 | 0.51268195 | 0.46919383 | 0.47451545 | -0.0053216 | 0.51988178 | 0.90650187 |
| Nhej1 | 0.03837765 | 0.0405888 | 0.04591125 | -0.0053225 | 0.69685471 | 0.90650187 |
| Schip1.1 | 0.04148535 | 0.04419837 | 0.04952695 | -0.0053286 | 0.68378186 | 0.90650187 |
| Rnd2 | 0.03588677 | 0.0376731 | 0.04300244 | -0.0053293 | 0.70916585 | 0.90650187 |
| Furin | 0.26929029 | 0.27622334 | 0.28155445 | -0.0053311 | 0.53354158 | 0.90650187 |
| Ddx19b | 0.06662591 | 0.07290505 | 0.07823922 | -0.0053342 | 0.6190899 | 0.90650187 |
| Ccdc88c | 0.03002974 | 0.03078116 | 0.0361246 | -0.0053434 | 0.74459592 | 0.90650187 |
| Adora1 | 0.03060219 | 0.03145397 | 0.03679919 | -0.0053452 | 0.74077646 | 0.90650187 |
| Zfp11 | 0.02171631 | 0.02092207 | 0.02626948 | -0.0053474 | 0.81725345 | 0.9092537 |
| Ctns | 0.06750791 | 0.07387819 | 0.07922947 | -0.0053513 | 0.61801184 | 0.90650187 |
| Zc3h8 | 0.10528313 | 0.11527573 | 0.12063105 | -0.0053553 | 0.57822672 | 0.90650187 |
| Tox4 | 0.46100905 | 0.43197584 | 0.4373405 | -0.0053647 | 0.52174447 | 0.90650187 |
| Mansc1 | 0.06754163 | 0.07390172 | 0.07926731 | -0.0053656 | 0.6182631 | 0.90650187 |
| Kifc5b | 0.02565536 | 0.0255863 | 0.03095268 | -0.0053664 | 0.77957261 | 0.90650187 |
| Gm27217 | 0.01850461 | 0.01706597 | 0.02243271 | -0.0053667 | 0.8562345 | 0.9165432 |
| Bola2 | 1.16692678 | 0.80662891 | 0.81199667 | -0.0053678 | 0.51172242 | 0.90650187 |
| Gen1 | 0.00922136 | 0.00588005 | 0.01124901 | -0.005369 | 0.98307317 | 0.98709923 |
| Cnbd2 | 0.02582108 | 0.02577926 | 0.03114916 | -0.0053699 | 0.77827818 | 0.90650187 |
| Samd15 | 0.04598958 | 0.04937037 | 0.05474113 | -0.0053708 | 0.66864251 | 0.90650187 |
| Gramd1b | 0.019051 | 0.01771322 | 0.0230866 | -0.0053734 | 0.84959786 | 0.91464143 |
| Rab3c | 0.05809318 | 0.06322672 | 0.06860018 | -0.0053735 | 0.63616429 | 0.90650187 |
| Gm15614 | 0.03160991 | 0.03261148 | 0.03798546 | -0.005374 | 0.73530613 | 0.90650187 |
| Gm10762 | 0.0299701 | 0.03067716 | 0.03605429 | -0.0053771 | 0.74633995 | 0.90650187 |
| Cx3cl1 | 0.05149094 | 0.05568901 | 0.06106772 | -0.0053787 | 0.65230795 | 0.90650187 |
| Paip2b | 0.23788986 | 0.24757606 | 0.25295662 | -0.0053806 | 0.53766825 | 0.90650187 |
| Dbil5 | 0.01338416 | 0.01089937 | 0.01628132 | -0.005382 | 0.92915668 | 0.94996709 |
| Trpv2 | 0.01302328 | 0.01046089 | 0.01584619 | -0.0053853 | 0.93458422 | 0.95301635 |
| Zfp108 | 0.01918934 | 0.01786246 | 0.02325209 | -0.0053896 | 0.84860102 | 0.91440801 |

| | | | | | | |
|---|---|---|---|---|---|---|
| 1700086P04I | 0.01429557 | 0.01198457 | 0.01737934 | -0.0053948 | 0.9162024 | 0.94231902 |
| Ryr3 | 0.01588283 | 0.0138914 | 0.01928837 | -0.005397 | 0.89323185 | 0.93181736 |
| Gm20554 | 0.02953956 | 0.03014584 | 0.03554655 | -0.0054007 | 0.75030051 | 0.90650187 |
| Tradd | 0.24765122 | 0.25654704 | 0.261951 | -0.005404 | 0.53653629 | 0.90650187 |
| Nudt12 | 0.03842606 | 0.04056341 | 0.0459677 | -0.0054043 | 0.69939357 | 0.90650187 |
| Gm5422 | 0.01306267 | 0.01048725 | 0.01589368 | -0.0054064 | 0.93476219 | 0.95313389 |
| Eps8l1 | 0.02847708 | 0.0288849 | 0.03429232 | -0.0054074 | 0.75835156 | 0.90650187 |
| Gm37276 | 0.01443828 | 0.01213898 | 0.01755115 | -0.0054122 | 0.91479238 | 0.94167513 |
| Yjefn3 | 0.04175194 | 0.0444187 | 0.04983643 | -0.0054177 | 0.68555241 | 0.90650187 |
| Rdh5 | 0.03456949 | 0.03603867 | 0.04146024 | -0.0054216 | 0.71949163 | 0.90650187 |
| Card6 | 0.01825308 | 0.01670743 | 0.02213152 | -0.0054241 | 0.86204426 | 0.91899895 |
| Fndc3b | 0.34092339 | 0.33791619 | 0.34334576 | -0.0054296 | 0.52802319 | 0.90650187 |
| Exosc10 | 0.28537237 | 0.29040354 | 0.29583324 | -0.0054297 | 0.53251536 | 0.90650187 |
| Gins3 | 0.06115124 | 0.06663594 | 0.07206709 | -0.0054311 | 0.63119938 | 0.90650187 |
| Junos | 0.0187354 | 0.01727671 | 0.02270897 | -0.0054323 | 0.85620917 | 0.9165432 |
| Gm26534 | 0.0170011 | 0.01519359 | 0.02063087 | -0.0054373 | 0.87932967 | 0.92549228 |
| Ttll5 | 0.05595035 | 0.06072428 | 0.06616258 | -0.0054383 | 0.64259781 | 0.90650187 |
| Kiz | 0.32283143 | 0.32273255 | 0.32817234 | -0.0054398 | 0.52937162 | 0.90650187 |
| Zswim5 | 0.01286316 | 0.0102092 | 0.01565304 | -0.0054438 | 0.93895989 | 0.95593613 |
| Prss36 | 0.11171324 | 0.12207101 | 0.12751834 | -0.0054473 | 0.57530911 | 0.90650187 |
| Atxn3 | 0.06910725 | 0.07557135 | 0.08102222 | -0.0054509 | 0.61756186 | 0.90650187 |
| Tmem51os1 | 0.0175916 | 0.01588694 | 0.02133898 | -0.005452 | 0.87198774 | 0.92254675 |
| Rgp1 | 0.09289251 | 0.10179211 | 0.10725063 | -0.0054585 | 0.58950002 | 0.90650187 |
| Slc37a1 | 0.21242628 | 0.22357822 | 0.22904219 | -0.005464 | 0.54222993 | 0.90650187 |
| Thtpa | 0.20059623 | 0.21223812 | 0.21770337 | -0.0054653 | 0.54443091 | 0.90650187 |
| Raet1d | 0.02132271 | 0.02033364 | 0.02580016 | -0.0054665 | 0.82688647 | 0.90947782 |
| Tlr3 | 0.03613583 | 0.03782595 | 0.04329373 | -0.0054678 | 0.71275711 | 0.90650187 |
| Dusp23 | 0.17742019 | 0.18958435 | 0.19505321 | -0.0054689 | 0.54959771 | 0.90650187 |
| Trim65 | 0.04174413 | 0.04435678 | 0.04982735 | -0.0054706 | 0.68725533 | 0.90650187 |
| Zfp248 | 0.06713455 | 0.07333683 | 0.07881043 | -0.0054736 | 0.62124773 | 0.90650187 |

| | | | | | | |
|---|---|---|---|---|---|---|
| Arid1b | 0.29114479 | 0.29542624 | 0.30089994 | -0.0054737 | 0.5322275 | 0.90650187 |
| Sel1l3 | 0.02558834 | 0.02539608 | 0.0308732 | -0.0054771 | 0.78485075 | 0.90650187 |
| F730016J06F | 0.01587817 | 0.01379651 | 0.01928277 | -0.0054863 | 0.89703676 | 0.9340267 |
| Slain1 | 0.04430609 | 0.04730891 | 0.05279592 | -0.005487 | 0.67796873 | 0.90650187 |
| Zfp28 | 0.05736088 | 0.06227987 | 0.06776791 | -0.005488 | 0.64058326 | 0.90650187 |
| Slc38a9 | 0.08148813 | 0.08928705 | 0.0947752 | -0.0054881 | 0.60157718 | 0.90650187 |
| Rundc3b | 0.02521806 | 0.02494072 | 0.03043398 | -0.0054933 | 0.78884535 | 0.90650187 |
| Golt1a | 0.02004592 | 0.01878124 | 0.02427606 | -0.0054948 | 0.84285278 | 0.91273395 |
| Aph1c | 0.0278168 | 0.02801603 | 0.03351197 | -0.0054959 | 0.76702283 | 0.90650187 |
| Gne | 0.13154601 | 0.14294323 | 0.14844063 | -0.0054974 | 0.56538643 | 0.90650187 |
| Gm20045 | 0.05705879 | 0.06191602 | 0.06742436 | -0.0055083 | 0.64176872 | 0.90650187 |
| Lrrc29 | 0.02354915 | 0.02294176 | 0.02845164 | -0.0055099 | 0.80536057 | 0.90899065 |
| Igfals | 0.09201071 | 0.10078161 | 0.10629251 | -0.0055109 | 0.59114427 | 0.90650187 |
| Dclre1c | 0.05564287 | 0.06029617 | 0.06581224 | -0.0055161 | 0.6452803 | 0.90650187 |
| Kif21b | 0.04703954 | 0.05042654 | 0.05595213 | -0.0055256 | 0.66967977 | 0.90650187 |
| Hspa12a | 0.0241379 | 0.02362391 | 0.02915147 | -0.0055276 | 0.80037277 | 0.90806412 |
| Ctso | 0.41775736 | 0.3992375 | 0.40476942 | -0.0055319 | 0.52422378 | 0.90650187 |
| 9430060I03R | 0.0156715 | 0.01349106 | 0.01903443 | -0.0055434 | 0.90228594 | 0.93609395 |
| Syt9 | 0.02319654 | 0.0224856 | 0.02803225 | -0.0055466 | 0.81047189 | 0.9092537 |
| Vwa8 | 0.06157796 | 0.06700143 | 0.07254975 | -0.0055483 | 0.63306518 | 0.90650187 |
| Gemin6 | 0.37584726 | 0.366299 | 0.37185429 | -0.0055553 | 0.52647622 | 0.90650187 |
| Ccm2 | 0.34405694 | 0.34038798 | 0.34594572 | -0.0055577 | 0.52846833 | 0.90650187 |
| Mapk12 | 0.03822375 | 0.04017382 | 0.04573181 | -0.005558 | 0.70550322 | 0.90650187 |
| Asphd2 | 0.02030018 | 0.01902162 | 0.02457977 | -0.0055582 | 0.84261709 | 0.91268507 |
| Cenpt | 0.19178759 | 0.2036016 | 0.20916301 | -0.0055614 | 0.5470469 | 0.90650187 |
| Rpl7a-ps3 | 0.02223467 | 0.02132526 | 0.02688718 | -0.0055619 | 0.82111576 | 0.9092537 |
| Fhl3 | 0.04178713 | 0.04431323 | 0.04987727 | -0.005564 | 0.69002738 | 0.90650187 |
| Tpp2 | 0.51984095 | 0.47395964 | 0.47952431 | -0.0055647 | 0.52057203 | 0.90650187 |
| Ppp4r2 | 0.32762823 | 0.32665102 | 0.33222252 | -0.0055715 | 0.52971545 | 0.90650187 |
| Ticam1 | 0.14607672 | 0.15789771 | 0.16346925 | -0.0055715 | 0.56021725 | 0.90650187 |

| | | | | | | |
|---|---|---|---|---|---|---|
| Dgka | 0.05696821 | 0.0617438 | 0.06732131 | -0.0055775 | 0.64368065 | 0.90650187 |
| Zfp78 | 0.02189246 | 0.02090138 | 0.02647944 | -0.0055781 | 0.82549086 | 0.90947782 |
| Fermt1 | 0.06138982 | 0.06675702 | 0.07233698 | -0.00558 | 0.63417327 | 0.90650187 |
| Fdxacb1 | 0.0986035 | 0.1078477 | 0.11342979 | -0.0055821 | 0.58658791 | 0.90650187 |
| Casr | 0.01831338 | 0.01661839 | 0.02220374 | -0.0055853 | 0.86827878 | 0.92093489 |
| Mnd1 | 0.05756831 | 0.06241791 | 0.06800374 | -0.0055858 | 0.64250482 | 0.90650187 |
| Eif2s3x | 0.37007531 | 0.36162556 | 0.36721261 | -0.005587 | 0.52696341 | 0.90650187 |
| Gm26890 | 0.14214052 | 0.15383348 | 0.15942267 | -0.0055892 | 0.56192764 | 0.90650187 |
| Gna14 | 0.03231019 | 0.03321618 | 0.03880888 | -0.0055927 | 0.73912533 | 0.90650187 |
| Shank1 | 0.06138107 | 0.06673143 | 0.07232708 | -0.0055956 | 0.63455364 | 0.90650187 |
| Setd4 | 0.03703878 | 0.03875327 | 0.04434893 | -0.0055957 | 0.71257341 | 0.90650187 |
| Itgb4 | 0.02459993 | 0.02409811 | 0.02970028 | -0.0056022 | 0.79913793 | 0.90777025 |
| Ephb3 | 0.08581592 | 0.09392786 | 0.09953096 | -0.0056031 | 0.59880906 | 0.90650187 |
| Tgfa | 0.03091253 | 0.03155937 | 0.03716468 | -0.0056053 | 0.74873134 | 0.90650187 |
| Mfsd6 | 0.12597869 | 0.13700503 | 0.14261614 | -0.0056111 | 0.56942419 | 0.90650187 |
| Tmem243 | 0.15973308 | 0.17175593 | 0.17736777 | -0.0056118 | 0.55592955 | 0.90650187 |
| Gemin8 | 0.09579095 | 0.10477931 | 0.1103923 | -0.005613 | 0.5894151 | 0.90650187 |
| Cd40 | 0.051112 | 0.05502024 | 0.06063341 | -0.0056132 | 0.65966706 | 0.90650187 |
| Neurl1a | 0.05576514 | 0.06033483 | 0.06595158 | -0.0056167 | 0.6475107 | 0.90650187 |
| Tnfrsf23 | 0.01818613 | 0.01643231 | 0.02205134 | -0.005619 | 0.87135304 | 0.92225579 |
| Tmem130 | 0.02093242 | 0.01971529 | 0.02533455 | -0.0056193 | 0.83794482 | 0.9113909 |
| Uox | 0.01681884 | 0.01479253 | 0.02041221 | -0.0056197 | 0.88950859 | 0.9302877 |
| Ppp1r26 | 0.03372067 | 0.03484435 | 0.04046505 | -0.0056207 | 0.73154971 | 0.90650187 |
| Celf5 | 0.02398779 | 0.02334409 | 0.02897309 | -0.005629 | 0.80612611 | 0.9092537 |
| Zfp273 | 0.0426944 | 0.04529667 | 0.05092962 | -0.0056329 | 0.68853277 | 0.90650187 |
| Man1c1 | 0.13602892 | 0.14746996 | 0.15310372 | -0.0056338 | 0.56497107 | 0.90650187 |
| Cybrd1 | 0.04577506 | 0.04885756 | 0.0544935 | -0.0056359 | 0.67716395 | 0.90650187 |
| Gabpa | 0.14255885 | 0.15421314 | 0.15985359 | -0.0056405 | 0.56232394 | 0.90650187 |
| Ctu1 | 0.09391307 | 0.10271743 | 0.10835816 | -0.0056407 | 0.59150651 | 0.90650187 |
| Mypopos | 0.01895776 | 0.01733354 | 0.02297505 | -0.0056415 | 0.86249921 | 0.91935001 |

| | | | | | | |
|---|---|---|---|---|---|---|
| Tusc1 | 0.03600463 | 0.03749601 | 0.04314029 | -0.0056443 | 0.71959647 | 0.90650187 |
| Cbll1 | 0.25267468 | 0.26089574 | 0.26654203 | -0.0056463 | 0.53751428 | 0.90650187 |
| Fam65b | 0.03367893 | 0.03476495 | 0.04041609 | -0.0056511 | 0.73289839 | 0.90650187 |
| Ndufaf1 | 0.16809053 | 0.18011716 | 0.18576834 | -0.0056512 | 0.55378811 | 0.90650187 |
| Cdh8 | 0.01225929 | 0.00927129 | 0.01492428 | -0.005653 | 0.95389942 | 0.96556047 |
| Wscd2 | 0.0191248 | 0.01752169 | 0.0231749 | -0.0056532 | 0.86091649 | 0.91848494 |
| Gm27030 | 0.0169222 | 0.01487939 | 0.02053623 | -0.0056568 | 0.88963169 | 0.93029083 |
| Kcnip3 | 0.06395213 | 0.06957086 | 0.07523031 | -0.0056595 | 0.63097454 | 0.90650187 |
| Inca1 | 0.02054774 | 0.01921523 | 0.0248754 | -0.0056602 | 0.84412482 | 0.913082 |
| Irf1 | 0.11384206 | 0.12412477 | 0.12978722 | -0.0056624 | 0.57689478 | 0.90650187 |
| Aim1l | 0.01306857 | 0.0102376 | 0.01590081 | -0.0056632 | 0.94332791 | 0.95871682 |
| Syndig1l | 0.01973329 | 0.01823569 | 0.02390247 | -0.0056668 | 0.85404975 | 0.91608365 |
| Pfas | 0.14666934 | 0.15840127 | 0.16407693 | -0.0056757 | 0.56110836 | 0.90650187 |
| Slc11a2 | 0.07933225 | 0.08672045 | 0.0923962 | -0.0056757 | 0.60760652 | 0.90650187 |
| Gm26735 | 0.05166947 | 0.0555934 | 0.06127227 | -0.0056789 | 0.65984131 | 0.90650187 |
| Aldh16a1 | 0.10015083 | 0.10941603 | 0.11509671 | -0.0056807 | 0.58683697 | 0.90650187 |
| Actr5 | 0.03065908 | 0.03118449 | 0.0368662 | -0.0056817 | 0.75337331 | 0.90650187 |
| Mpdu1 | 0.46364057 | 0.43358688 | 0.43927861 | -0.0056917 | 0.52296702 | 0.90650187 |
| Stk16 | 0.80889617 | 0.65138627 | 0.65708902 | -0.0057027 | 0.51538839 | 0.90650187 |
| Sp3 | 0.49765587 | 0.45819028 | 0.46389323 | -0.005703 | 0.52179248 | 0.90650187 |
| Clk1 | 1.13711207 | 0.79654201 | 0.80224611 | -0.0057041 | 0.51260807 | 0.90650187 |
| Slc29a2 | 0.02540676 | 0.02495117 | 0.03065784 | -0.0057067 | 0.79602519 | 0.9075779 |
| Cspg5 | 0.04803539 | 0.05138396 | 0.05709918 | -0.0057152 | 0.67183046 | 0.90650187 |
| Macc1 | 0.01280394 | 0.0098598 | 0.01558161 | -0.0057218 | 0.94871243 | 0.96195723 |
| Mthfr | 0.07121748 | 0.07765853 | 0.08338194 | -0.0057234 | 0.61987405 | 0.90650187 |
| Def6 | 0.02371693 | 0.02292197 | 0.02865114 | -0.0057292 | 0.81298665 | 0.9092537 |
| Ccdc113 | 0.04636985 | 0.04944989 | 0.05517992 | -0.00573 | 0.67783145 | 0.90650187 |
| Cdk5rap2 | 0.11422875 | 0.12445414 | 0.13019875 | -0.0057446 | 0.57775304 | 0.90650187 |
| Cklf | 0.13872465 | 0.15014682 | 0.15589634 | -0.0057495 | 0.56511697 | 0.90650187 |
| Pvrl4 | 0.02126877 | 0.01998497 | 0.02573583 | -0.0057509 | 0.83974339 | 0.91213511 |

| | | | | | | |
|---|---|---|---|---|---|---|
| 9330104G04 | 0.02240795 | 0.02134046 | 0.02709357 | -0.0057531 | 0.82741267 | 0.90947782 |
| Tmem178b | 0.0609353 | 0.06606722 | 0.07182272 | -0.0057555 | 0.63917265 | 0.90650187 |
| Bcl11b | 0.02928925 | 0.02949568 | 0.03525123 | -0.0057555 | 0.76603285 | 0.90650187 |
| Bach2 | 0.09011908 | 0.09847325 | 0.10423351 | -0.0057603 | 0.59703412 | 0.90650187 |
| 4632411P08I | 0.01913565 | 0.01742686 | 0.02318787 | -0.005761 | 0.8653138 | 0.92040222 |
| Angpt1 | 0.0287537 | 0.02885725 | 0.03461903 | -0.0057618 | 0.77031961 | 0.90650187 |
| Nek5 | 0.01586392 | 0.0135033 | 0.01926565 | -0.0057624 | 0.90818809 | 0.93875746 |
| G730003C15 | 0.08238294 | 0.08999358 | 0.09576066 | -0.0057671 | 0.60554706 | 0.90650187 |
| Iyd | 0.02221362 | 0.02109413 | 0.02686211 | -0.005768 | 0.83010596 | 0.9097662 |
| Fam167a | 0.02302981 | 0.02206283 | 0.02783387 | -0.005771 | 0.82166963 | 0.9092537 |
| Kcnn1 | 0.0182776 | 0.01638922 | 0.02216088 | -0.0057717 | 0.87653341 | 0.92471264 |
| Ctc1 | 0.15518005 | 0.16698426 | 0.17275798 | -0.0057737 | 0.55905584 | 0.90650187 |
| Pdk3 | 0.26830366 | 0.27489572 | 0.28067034 | -0.0057746 | 0.53643871 | 0.90650187 |
| Mark4 | 0.20874028 | 0.21973892 | 0.22552493 | -0.005786 | 0.54540303 | 0.90650187 |
| Ppp4r4 | 0.0242815 | 0.02353233 | 0.02932208 | -0.0057897 | 0.80997406 | 0.9092537 |
| 3110040N11 | 0.23483719 | 0.2443293 | 0.25012447 | -0.0057952 | 0.54101885 | 0.90650187 |
| Kcnip1 | 0.04085441 | 0.04299853 | 0.04879408 | -0.0057956 | 0.70126091 | 0.90650187 |
| Ccdc18 | 0.03100862 | 0.03147774 | 0.03727782 | -0.0058001 | 0.75543547 | 0.90650187 |
| Zfp455 | 0.0276519 | 0.02751683 | 0.03331697 | -0.0058001 | 0.78051515 | 0.90650187 |
| Zfp821 | 0.25099455 | 0.25920616 | 0.26500925 | -0.0058031 | 0.5387752 | 0.90650187 |
| Prkab1 | 0.25365161 | 0.26162879 | 0.26743201 | -0.0058032 | 0.53842585 | 0.90650187 |
| Mkrn3 | 0.05736284 | 0.06196574 | 0.06777014 | -0.0058044 | 0.6483089 | 0.90650187 |
| Tmem125 | 0.01305404 | 0.01007773 | 0.01588328 | -0.0058056 | 0.94791117 | 0.9616629 |
| 9130604C24I | 0.02782859 | 0.02771825 | 0.03352591 | -0.0058077 | 0.77938211 | 0.90650187 |
| Vav2 | 0.04315568 | 0.04565321 | 0.05146418 | -0.005811 | 0.69215432 | 0.90650187 |
| Afm | 0.03056679 | 0.03094589 | 0.03675749 | -0.0058116 | 0.75893437 | 0.90650187 |
| Myo1e | 0.1108125 | 0.12074416 | 0.12655665 | -0.0058125 | 0.58089261 | 0.90650187 |
| Esyt1 | 0.38231187 | 0.37120658 | 0.37702136 | -0.0058148 | 0.52733179 | 0.90650187 |
| Slc35e3 | 0.03928811 | 0.04115532 | 0.04697209 | -0.0058168 | 0.70901855 | 0.90650187 |
| Cdc42bpb | 0.20971764 | 0.22064155 | 0.22645892 | -0.0058174 | 0.5454606 | 0.90650187 |

| | | | | | | |
|---|---|---|---|---|---|---|
| Rccd1 | 0.04766539 | 0.05085523 | 0.05667317 | -0.0058179 | 0.67594025 | 0.90650187 |
| Bend3 | 0.02195125 | 0.02073098 | 0.0265495 | -0.0058185 | 0.83504293 | 0.91066681 |
| Ero1l | 0.22783197 | 0.237771 | 0.24359011 | -0.0058191 | 0.54228852 | 0.90650187 |
| Celsr1 | 0.04337279 | 0.04589376 | 0.05171566 | -0.0058219 | 0.69162628 | 0.90650187 |
| Zfp316 | 0.06165133 | 0.06680764 | 0.07263271 | -0.0058251 | 0.63927933 | 0.90650187 |
| Alyref2 | 0.26843241 | 0.27495939 | 0.28078576 | -0.0058264 | 0.53674925 | 0.90650187 |
| Ypel1 | 0.13734266 | 0.14863562 | 0.15446576 | -0.0058301 | 0.56662744 | 0.90650187 |
| Fus | 2.23666307 | 0.91249067 | 0.91832478 | -0.0058341 | 0.51126578 | 0.90650187 |
| Mllt1 | 0.03348547 | 0.03435411 | 0.04018911 | -0.005835 | 0.74068231 | 0.90650187 |
| Zfp619 | 0.02298983 | 0.02194986 | 0.02778629 | -0.0058364 | 0.82479306 | 0.90934433 |
| Efnb3 | 0.12432716 | 0.13504407 | 0.14088109 | -0.005837 | 0.57306843 | 0.90650187 |
| Gba | 0.16914834 | 0.18098575 | 0.18682602 | -0.0058403 | 0.5552638 | 0.90650187 |
| Plch2 | 0.01679748 | 0.01454597 | 0.02038657 | -0.0058406 | 0.89860319 | 0.93440169 |
| Gm2694 | 0.04400438 | 0.04660601 | 0.05244685 | -0.0058408 | 0.68973284 | 0.90650187 |
| Cnnm4 | 0.04594184 | 0.04884308 | 0.05468602 | -0.0058429 | 0.68260366 | 0.90650187 |
| Herc6 | 0.04809343 | 0.05131361 | 0.05716599 | -0.0058524 | 0.67548851 | 0.90650187 |
| Lingo4 | 0.02052338 | 0.01899049 | 0.02484631 | -0.0058558 | 0.85262247 | 0.91557373 |
| Xndc1 | 0.12687695 | 0.13770091 | 0.1435584 | -0.0058575 | 0.57196948 | 0.90650187 |
| Ncbp1 | 0.2514512 | 0.25956469 | 0.26542612 | -0.0058614 | 0.53910245 | 0.90650187 |
| Cfap54 | 0.02339344 | 0.02240173 | 0.02826647 | -0.0058647 | 0.82183373 | 0.9092537 |
| Als2cl | 0.02123993 | 0.01983582 | 0.02570143 | -0.0058656 | 0.84484722 | 0.91325304 |
| Gng10 | 0.08961654 | 0.09781671 | 0.10368567 | -0.005869 | 0.59933889 | 0.90650187 |
| Farsb | 0.5143534 | 0.46981667 | 0.47568796 | -0.0058713 | 0.52187937 | 0.90650187 |
| Cdk18 | 0.02032526 | 0.0187364 | 0.02460973 | -0.0058733 | 0.85565142 | 0.9165432 |
| Zfp354b | 0.024136 | 0.02327032 | 0.02914921 | -0.0058789 | 0.8150343 | 0.9092537 |
| Akap5 | 0.03369205 | 0.03455119 | 0.04043149 | -0.0058803 | 0.7410426 | 0.90650187 |
| Gm4285 | 0.04823113 | 0.05144292 | 0.05732446 | -0.0058815 | 0.67584936 | 0.90650187 |
| Nmb | 0.05950471 | 0.06431879 | 0.07020215 | -0.0058834 | 0.64526514 | 0.90650187 |
| Folr1 | 0.03277274 | 0.03346746 | 0.03935234 | -0.0058849 | 0.74691412 | 0.90650187 |
| Dzip1l | 0.11459238 | 0.12469794 | 0.13058557 | -0.0058876 | 0.57943031 | 0.90650187 |

| | | | | | | |
|---|---|---|---|---|---|---|
| U2af1l4 | 0.02988424 | 0.03006488 | 0.03595305 | -0.0058882 | 0.7667161 | 0.90650187 |
| Mcm10 | 0.0213813 | 0.01998143 | 0.02587004 | -0.0058886 | 0.84421222 | 0.91310457 |
| Spaca6 | 0.11295558 | 0.12294613 | 0.12884309 | -0.005897 | 0.58061574 | 0.90650187 |
| Pctp | 0.01758646 | 0.01543455 | 0.02133282 | -0.0058983 | 0.89049326 | 0.93072624 |
| Ern1 | 0.11379796 | 0.12384022 | 0.12974027 | -0.0059 | 0.58010719 | 0.90650187 |
| Tsga10ip | 0.03316904 | 0.03391534 | 0.03981771 | -0.0059024 | 0.74504974 | 0.90650187 |
| Mettl6 | 0.2322415 | 0.24180633 | 0.247709 | -0.0059027 | 0.54218282 | 0.90650187 |
| Ttc21b | 0.08048163 | 0.08776031 | 0.09366535 | -0.005905 | 0.61036439 | 0.90650187 |
| Lurap1 | 0.01649281 | 0.01411517 | 0.02002091 | -0.0059057 | 0.90513167 | 0.93712753 |
| Nek4 | 0.07475632 | 0.08141791 | 0.08732469 | -0.0059068 | 0.61818142 | 0.90650187 |
| Dnajc21 | 0.25629135 | 0.26392456 | 0.26983222 | -0.0059077 | 0.53876835 | 0.90650187 |
| Rassf10 | 0.10624263 | 0.1157534 | 0.12166204 | -0.0059086 | 0.58546927 | 0.90650187 |
| Cep97 | 0.09692591 | 0.10570877 | 0.11161935 | -0.0059106 | 0.59305406 | 0.90650187 |
| Phf20 | 0.40148566 | 0.38622912 | 0.39215262 | -0.0059235 | 0.52676936 | 0.90650187 |
| Fam227a | 0.04039858 | 0.04233621 | 0.04826422 | -0.005928 | 0.70747667 | 0.90650187 |
| Pold4 | 0.47123425 | 0.43891438 | 0.44484376 | -0.0059294 | 0.52362592 | 0.90650187 |
| Oprl1 | 0.02540639 | 0.02472684 | 0.0306574 | -0.0059306 | 0.80510156 | 0.90885402 |
| Commd2 | 0.51070939 | 0.46719769 | 0.47312941 | -0.0059317 | 0.52222372 | 0.90650187 |
| Nxpe4 | 0.01969374 | 0.01792263 | 0.0238552 | -0.0059326 | 0.86554619 | 0.92040222 |
| Plekha4 | 0.02277492 | 0.02158922 | 0.02753051 | -0.0059413 | 0.83132025 | 0.9098597 |
| Pde4b | 0.07430695 | 0.08088338 | 0.08682503 | -0.0059417 | 0.6195214 | 0.90650187 |
| Oit1 | 0.02176547 | 0.02038468 | 0.02632808 | -0.0059434 | 0.8422067 | 0.91251845 |
| Igbp1 | 0.57375818 | 0.51024472 | 0.51619134 | -0.0059466 | 0.52042262 | 0.90650187 |
| Atg12 | 0.40433732 | 0.38842733 | 0.39437821 | -0.0059509 | 0.52674138 | 0.90650187 |
| Mvb12b | 0.16927536 | 0.18100067 | 0.18695294 | -0.0059523 | 0.55627864 | 0.90650187 |
| Kif1b | 0.71243292 | 0.5967612 | 0.60271371 | -0.0059525 | 0.51751018 | 0.90650187 |
| Vamp7 | 0.69631837 | 0.58723612 | 0.59319024 | -0.0059541 | 0.51779592 | 0.90650187 |
| Prps2 | 0.17015471 | 0.18187488 | 0.18783111 | -0.0059562 | 0.55605421 | 0.90650187 |
| Ppp1r12c | 0.09732057 | 0.10608807 | 0.11204561 | -0.0059575 | 0.59342953 | 0.90650187 |
| Wdr45b | 0.73605775 | 0.61047807 | 0.61644079 | -0.0059627 | 0.51714984 | 0.90650187 |

| | | | | | | |
|---|---|---|---|---|---|---|
| Sergef | 0.08971556 | 0.0978276 | 0.10379364 | -0.005966 | 0.60084412 | 0.90650187 |
| Orc5 | 0.12627183 | 0.13695629 | 0.14292374 | -0.0059675 | 0.57362705 | 0.90650187 |
| Nicn1 | 0.25253313 | 0.26044431 | 0.266413 | -0.0059687 | 0.53966857 | 0.90650187 |
| Shroom1 | 0.02719168 | 0.02680385 | 0.03277254 | -0.0059687 | 0.79093049 | 0.90650187 |
| Gm29666 | 0.03100071 | 0.0312991 | 0.03726851 | -0.0059694 | 0.76178841 | 0.90650187 |
| Gm16136 | 0.01231376 | 0.00901012 | 0.01499004 | -0.0059799 | 0.96192464 | 0.97125317 |
| Atm | 0.17519466 | 0.18686345 | 0.19284741 | -0.005984 | 0.55485809 | 0.90650187 |
| Rtf1 | 1.07898861 | 0.77564502 | 0.78163306 | -0.005988 | 0.51358433 | 0.90650187 |
| B230322F03 | 0.02548759 | 0.02476307 | 0.03075372 | -0.0059906 | 0.80674894 | 0.9092537 |
| Zbtb7b | 0.11959588 | 0.12990088 | 0.13589172 | -0.0059908 | 0.57768956 | 0.90650187 |
| Cox19 | 0.5041252 | 0.46249172 | 0.46848462 | -0.0059929 | 0.52267507 | 0.90650187 |
| Rgn | 0.02546514 | 0.02472915 | 0.03072709 | -0.0059979 | 0.80724383 | 0.9092537 |
| Mpp5 | 0.26475606 | 0.27148561 | 0.27748378 | -0.0059982 | 0.53827843 | 0.90650187 |
| Zfp184 | 0.05231046 | 0.0560062 | 0.06200626 | -0.0060001 | 0.66646785 | 0.90650187 |
| Zik1 | 0.02760214 | 0.02725506 | 0.03325813 | -0.0060031 | 0.78884759 | 0.90650187 |
| Exo1 | 0.04033727 | 0.04218228 | 0.04819294 | -0.0060107 | 0.71036766 | 0.90650187 |
| Dact1 | 0.14877681 | 0.160216 | 0.16623463 | -0.0060186 | 0.56393602 | 0.90650187 |
| Wdr59 | 0.04272907 | 0.04495004 | 0.05096981 | -0.0060198 | 0.70022456 | 0.90650187 |
| Gcdh | 0.32252235 | 0.32188333 | 0.32791069 | -0.0060274 | 0.53256336 | 0.90650187 |
| Lbhd1 | 0.03651878 | 0.03771347 | 0.0437414 | -0.0060279 | 0.7299434 | 0.90650187 |
| Gng7 | 0.02860454 | 0.02841387 | 0.03444287 | -0.006029 | 0.78176698 | 0.90650187 |
| Pgbd1 | 0.03106706 | 0.03131191 | 0.03734663 | -0.0060347 | 0.76372954 | 0.90650187 |
| Ftx | 0.02117901 | 0.01959238 | 0.02562877 | -0.0060364 | 0.85246902 | 0.91557373 |
| Mtx3 | 0.04316392 | 0.04543477 | 0.05147372 | -0.006039 | 0.69901281 | 0.90650187 |
| Gpr19 | 0.0512335 | 0.05473078 | 0.06077269 | -0.0060419 | 0.67074579 | 0.90650187 |
| Ltk | 0.02108477 | 0.01947402 | 0.02551633 | -0.0060423 | 0.85376715 | 0.91608365 |
| Akt1 | 0.41154376 | 0.39392611 | 0.39997502 | -0.0060489 | 0.52680146 | 0.90650187 |
| Gm12942 | 0.10937279 | 0.11896712 | 0.12501744 | -0.0060503 | 0.58517431 | 0.90650187 |
| Rps6ka1 | 0.05706372 | 0.06136841 | 0.06742996 | -0.0060616 | 0.65528679 | 0.90650187 |
| Tgfbr3l | 0.02404505 | 0.02297634 | 0.02904114 | -0.0060648 | 0.82339929 | 0.9092537 |

| | | | | | | |
|---|---|---|---|---|---|---|
| Igdcc4 | 0.10641812 | 0.11578411 | 0.12185048 | -0.0060664 | 0.58758076 | 0.90650187 |
| Rab35 | 0.12115674 | 0.13147192 | 0.13754072 | -0.0060688 | 0.57775623 | 0.90650187 |
| Ccndbp1 | 0.35691597 | 0.35045572 | 0.3565259 | -0.0060702 | 0.53016736 | 0.90650187 |
| Usp1 | 0.44521231 | 0.41953065 | 0.4256041 | -0.0060734 | 0.52529176 | 0.90650187 |
| Gpatch2 | 0.32654052 | 0.32522746 | 0.33130585 | -0.0060784 | 0.53250268 | 0.90650187 |
| Tmem8b | 0.04227451 | 0.0443594 | 0.05044273 | -0.0060833 | 0.70406738 | 0.90650187 |
| Cacna1c | 0.03540453 | 0.03634833 | 0.04243818 | -0.0060898 | 0.73824404 | 0.90650187 |
| Bmp3 | 0.02305561 | 0.02177061 | 0.02786457 | -0.006094 | 0.8345371 | 0.91062493 |
| Snx16 | 0.1828898 | 0.19435628 | 0.20045122 | -0.0060949 | 0.55376268 | 0.90650187 |
| Mfsd11 | 0.27047177 | 0.27651593 | 0.28261195 | -0.006096 | 0.53819722 | 0.90650187 |
| Kcnd1 | 0.02853314 | 0.0282568 | 0.03435854 | -0.0061017 | 0.78508889 | 0.90650187 |
| Enkur | 0.0149701 | 0.012087 | 0.01819111 | -0.0061041 | 0.93211675 | 0.95146248 |
| Slamf1 | 0.04070421 | 0.04250806 | 0.04861952 | -0.0061115 | 0.71185471 | 0.90650187 |
| A930015D03 | 0.03921159 | 0.04076981 | 0.04688298 | -0.0061132 | 0.71893185 | 0.90650187 |
| Nr1h4 | 0.02755249 | 0.02708332 | 0.0331994 | -0.0061161 | 0.79360139 | 0.90731056 |
| 4833418N02 | 0.02632242 | 0.02562554 | 0.03174331 | -0.0061178 | 0.80421816 | 0.90882608 |
| Mcph1 | 0.06626127 | 0.0717105 | 0.07782949 | -0.006119 | 0.63664897 | 0.90650187 |
| Ext2 | 0.56725455 | 0.50574593 | 0.51186504 | -0.0061191 | 0.52119188 | 0.90650187 |
| Crlf2 | 0.20904451 | 0.21969451 | 0.22581576 | -0.0061213 | 0.54795974 | 0.90650187 |
| Dmtf1 | 0.35003216 | 0.34475399 | 0.35087934 | -0.0061254 | 0.5309299 | 0.90650187 |
| Susd1 | 0.01869443 | 0.01653302 | 0.02265994 | -0.0061269 | 0.88532338 | 0.92812419 |
| Samd4 | 0.09967062 | 0.10844545 | 0.1145794 | -0.006134 | 0.59405688 | 0.90650187 |
| Cyp7b1 | 0.10456542 | 0.11371875 | 0.11985911 | -0.0061404 | 0.5900788 | 0.90650187 |
| Sfn | 0.02487953 | 0.02389123 | 0.03003222 | -0.006141 | 0.81833511 | 0.9092537 |
| Gpank1 | 0.17677487 | 0.18826747 | 0.19441418 | -0.0061467 | 0.5558891 | 0.90650187 |
| Syngr2 | 0.59577487 | 0.52449983 | 0.53065087 | -0.006151 | 0.52054888 | 0.90650187 |
| Snapc3 | 0.28671617 | 0.290863 | 0.29701547 | -0.0061525 | 0.5366858 | 0.90650187 |
| Rps6kl1 | 0.01163467 | 0.00801521 | 0.01416985 | -0.0061546 | 0.97325473 | 0.9796997 |
| Btaf1 | 0.11956356 | 0.129701 | 0.13585754 | -0.0061565 | 0.57982958 | 0.90650187 |
| Snph | 0.01881714 | 0.01664925 | 0.0228068 | -0.0061575 | 0.88498018 | 0.92807528 |

| | | | | | | |
|---|---|---|---|---|---|---|
| Mpp7 | 0.12826072 | 0.13884621 | 0.14500809 | -0.0061619 | 0.57491774 | 0.90650187 |
| C1galt1c1 | 0.44704837 | 0.42080931 | 0.4269773 | -0.006168 | 0.52560243 | 0.90650187 |
| Ffar3 | 0.02497209 | 0.02396997 | 0.0301421 | -0.0061721 | 0.81867212 | 0.9092537 |
| Zbed3 | 0.2213856 | 0.23136011 | 0.23753288 | -0.0061728 | 0.5459867 | 0.90650187 |
| Mettl21b | 0.02879667 | 0.02849509 | 0.03466977 | -0.0061747 | 0.78575042 | 0.90650187 |
| Pds5a | 0.43904394 | 0.41479473 | 0.42097117 | -0.0061764 | 0.52600273 | 0.90650187 |
| Gm17275 | 0.09532225 | 0.10370768 | 0.10988506 | -0.0061774 | 0.59867442 | 0.90650187 |
| Gpr146 | 0.0423421 | 0.04434187 | 0.05052113 | -0.0061793 | 0.70669111 | 0.90650187 |
| Mbd4 | 0.05220791 | 0.05570822 | 0.06188887 | -0.0061806 | 0.6714674 | 0.90650187 |
| D630036H23 | 0.03330513 | 0.03379362 | 0.03997746 | -0.0061838 | 0.75416675 | 0.90650187 |
| Rassf9 | 0.04040981 | 0.04209269 | 0.04827728 | -0.0061846 | 0.71549394 | 0.90650187 |
| Chrna1os | 0.07428254 | 0.08061283 | 0.08679788 | -0.006185 | 0.62429669 | 0.90650187 |
| 3222401L13F | 0.02275462 | 0.02131356 | 0.02750634 | -0.0061928 | 0.84155874 | 0.91251845 |
| Zfp119a | 0.06449473 | 0.06964736 | 0.07584177 | -0.0061944 | 0.64173366 | 0.90650187 |
| Rfx5 | 0.01985174 | 0.01784795 | 0.02404404 | -0.0061961 | 0.87402417 | 0.9235672 |
| Tubg2 | 0.04712709 | 0.04985402 | 0.05605304 | -0.006199 | 0.68851623 | 0.90650187 |
| Wasf3 | 0.02398973 | 0.02277557 | 0.0289754 | -0.0061998 | 0.82925413 | 0.90974008 |
| Lsg1 | 0.35397223 | 0.34791331 | 0.35411602 | -0.0062027 | 0.53103402 | 0.90650187 |
| Rgs7 | 0.02047089 | 0.01857617 | 0.02478363 | -0.0062075 | 0.86724758 | 0.92056229 |
| Zfp758 | 0.11387659 | 0.1236122 | 0.12982398 | -0.0062118 | 0.58422421 | 0.90650187 |
| B4galt7 | 0.23223218 | 0.24148359 | 0.2477003 | -0.0062167 | 0.54441967 | 0.90650187 |
| Slco2a1 | 0.07530548 | 0.08171676 | 0.08793491 | -0.0062182 | 0.6233772 | 0.90650187 |
| Fgfrl1 | 0.06199564 | 0.06680318 | 0.07302193 | -0.0062188 | 0.64750776 | 0.90650187 |
| Rfesd | 0.17876725 | 0.1901658 | 0.19638569 | -0.0062199 | 0.55598611 | 0.90650187 |
| Serpina10 | 0.04672955 | 0.04937201 | 0.05559476 | -0.0062228 | 0.69061669 | 0.90650187 |
| Ptpn3 | 0.02819935 | 0.02773881 | 0.03396416 | -0.0062254 | 0.7924191 | 0.90681274 |
| Nsg2 | 0.01859776 | 0.01631557 | 0.02254422 | -0.0062286 | 0.89032481 | 0.93072624 |
| Gm16973 | 0.02967924 | 0.0294789 | 0.03571131 | -0.0062324 | 0.7810832 | 0.90650187 |
| Inf2 | 0.06083585 | 0.06547577 | 0.07171017 | -0.0062344 | 0.65043534 | 0.90650187 |
| Shf | 0.06078077 | 0.06541103 | 0.07164782 | -0.0062368 | 0.65061481 | 0.90650187 |

| | | | | | | |
|---|---|---|---|---|---|---|
| Dcaf13 | 0.29419328 | 0.29732266 | 0.30356351 | -0.0062408 | 0.53641084 | 0.90650187 |
| Nqo2 | 0.17701538 | 0.18840554 | 0.19465239 | -0.0062469 | 0.55672457 | 0.90650187 |
| Slc8a2 | 0.02067638 | 0.01877797 | 0.02502897 | -0.006251 | 0.86656347 | 0.92056229 |
| Ascc1 | 0.51288203 | 0.46839781 | 0.47465591 | -0.0062581 | 0.52336986 | 0.90650187 |
| 4930518I15R | 0.01651767 | 0.01379245 | 0.02005076 | -0.0062583 | 0.91736686 | 0.94283706 |
| Tfip11 | 0.18946261 | 0.2006332 | 0.20689488 | -0.0062617 | 0.55351479 | 0.90650187 |
| Zfp141 | 0.05486088 | 0.05865788 | 0.06492062 | -0.0062627 | 0.66598624 | 0.90650187 |
| Kctd20 | 0.5019116 | 0.46065393 | 0.46691669 | -0.0062628 | 0.52377442 | 0.90650187 |
| Cyb561d1 | 0.02422612 | 0.02299189 | 0.02925628 | -0.0062644 | 0.82942596 | 0.90974008 |
| Kansl1l | 0.15677163 | 0.16810361 | 0.17437209 | -0.0062685 | 0.56348637 | 0.90650187 |
| Dus4l | 0.0542744 | 0.05798087 | 0.06425132 | -0.0062705 | 0.66780426 | 0.90650187 |
| Gpa33 | 0.0179672 | 0.01551612 | 0.02178908 | -0.006273 | 0.89970389 | 0.93469858 |
| Lyrm7 | 0.08429155 | 0.09158179 | 0.09785887 | -0.0062771 | 0.61223922 | 0.90650187 |
| 4930539J05F | 0.02277724 | 0.02125103 | 0.02753327 | -0.0062822 | 0.84479217 | 0.91325304 |
| Zbtb46 | 0.05075825 | 0.0539373 | 0.06022777 | -0.0062905 | 0.6787906 | 0.90650187 |
| Gm14326 | 0.03378429 | 0.03424782 | 0.04053969 | -0.0062919 | 0.75489373 | 0.90650187 |
| C8g | 0.02891931 | 0.02852198 | 0.03481457 | -0.0062926 | 0.78916435 | 0.90650187 |
| 1810032O08 | 0.10305336 | 0.11193692 | 0.11823067 | -0.0062938 | 0.59353665 | 0.90650187 |
| Nt5m | 0.35026387 | 0.34477593 | 0.35107004 | -0.0062941 | 0.53176303 | 0.90650187 |
| Bbs12 | 0.02720808 | 0.02649743 | 0.03279195 | -0.0062945 | 0.80326704 | 0.90882608 |
| Hinfp | 0.02055885 | 0.01859099 | 0.02488866 | -0.0062977 | 0.86968178 | 0.92144928 |
| Irf3 | 0.49546935 | 0.45603454 | 0.46233519 | -0.0063007 | 0.52415483 | 0.90650187 |
| Crebzf | 0.13708385 | 0.14789473 | 0.15419759 | -0.0063029 | 0.57209688 | 0.90650187 |
| Mb21d1 | 0.03588556 | 0.03669299 | 0.04300102 | -0.006308 | 0.74284706 | 0.90650187 |
| Wdr82 | 0.31583561 | 0.31591792 | 0.32222797 | -0.0063101 | 0.53468665 | 0.90650187 |
| Heatr6 | 0.25002207 | 0.25780227 | 0.26412081 | -0.0063185 | 0.54234842 | 0.90650187 |
| Parp10 | 0.06548122 | 0.07063297 | 0.07695233 | -0.0063194 | 0.64247254 | 0.90650187 |
| Kif18a | 0.05104204 | 0.05423192 | 0.0605532 | -0.0063213 | 0.67870704 | 0.90650187 |
| Rtn4ip1 | 0.03224885 | 0.03241438 | 0.03873678 | -0.0063224 | 0.76595383 | 0.90650187 |
| Erich3 | 0.01477054 | 0.01162655 | 0.01795102 | -0.0063245 | 0.94135683 | 0.95746988 |

| | | | | | | |
|---|---|---|---|---|---|---|
| Snhg12 | 0.31526336 | 0.31541413 | 0.32173981 | -0.0063257 | 0.53482498 | 0.90650187 |
| Memo1 | 0.2841222 | 0.28840268 | 0.2947319 | -0.0063292 | 0.53802816 | 0.90650187 |
| Gm16638 | 0.06247609 | 0.06723049 | 0.07356477 | -0.0063343 | 0.64906143 | 0.90650187 |
| Gpr142 | 0.02466469 | 0.02343884 | 0.02977717 | -0.0063383 | 0.82799877 | 0.90955407 |
| Kirrel | 0.02663862 | 0.02576951 | 0.03211784 | -0.0063483 | 0.81021971 | 0.9092537 |
| Slc25a15 | 0.07634861 | 0.08274244 | 0.08909283 | -0.0063504 | 0.62433092 | 0.90650187 |
| Pitpnm2 | 0.06834715 | 0.07381695 | 0.08017067 | -0.0063537 | 0.63770277 | 0.90650187 |
| 1810041L15F | 0.14035288 | 0.15121479 | 0.15757893 | -0.0063641 | 0.57124497 | 0.90650187 |
| Ssc5d | 0.05006606 | 0.05306274 | 0.05943349 | -0.0063708 | 0.6831518 | 0.90650187 |
| Lag3 | 0.04667189 | 0.049154 | 0.05552828 | -0.0063743 | 0.69508869 | 0.90650187 |
| Ppm1m | 0.17103897 | 0.18233752 | 0.18871331 | -0.0063758 | 0.55969551 | 0.90650187 |
| Hyal3 | 0.01772018 | 0.01511501 | 0.02149308 | -0.0063781 | 0.9064546 | 0.93785917 |
| Nubp1 | 0.29979771 | 0.30205585 | 0.30843741 | -0.0063816 | 0.53664285 | 0.90650187 |
| Timm21 | 0.33747515 | 0.33408905 | 0.34047519 | -0.0063861 | 0.53322698 | 0.90650187 |
| Ap1g2 | 0.05531232 | 0.05904634 | 0.06543546 | -0.0063891 | 0.66788084 | 0.90650187 |
| Cntrob | 0.07156011 | 0.07737233 | 0.08376446 | -0.0063921 | 0.6327885 | 0.90650187 |
| Ccdc68 | 0.02946957 | 0.02906813 | 0.03546398 | -0.0063959 | 0.78865277 | 0.90650187 |
| Fbxl8 | 0.06030356 | 0.06471124 | 0.07110746 | -0.0063962 | 0.65538079 | 0.90650187 |
| Fam134c | 0.33043968 | 0.3281904 | 0.33458721 | -0.0063968 | 0.53386664 | 0.90650187 |
| Clcn2 | 0.06567238 | 0.07076436 | 0.07716738 | -0.006403 | 0.64389041 | 0.90650187 |
| Fam161a | 0.03827018 | 0.03938097 | 0.04578596 | -0.006405 | 0.73299441 | 0.90650187 |
| Ric1 | 0.12518361 | 0.1353749 | 0.14178128 | -0.0064064 | 0.57960164 | 0.90650187 |
| Rasgrf2 | 0.02856142 | 0.02798144 | 0.03439194 | -0.0064105 | 0.79634579 | 0.9075779 |
| Brwd3 | 0.14808764 | 0.15911682 | 0.1655296 | -0.0064128 | 0.56837057 | 0.90650187 |
| Gm29095 | 0.01138881 | 0.00745664 | 0.01387272 | -0.0064161 | 0.9801124 | 0.98497214 |
| Lsm6 | 1.3680064 | 0.86088305 | 0.86729986 | -0.0064168 | 0.51311935 | 0.90650187 |
| 2010015M23 | 0.03311824 | 0.03334012 | 0.03975806 | -0.0064179 | 0.76350787 | 0.90650187 |
| Hyal1 | 0.11344243 | 0.12294323 | 0.12936172 | -0.0064185 | 0.5872887 | 0.90650187 |
| Klhl1 | 0.0303168 | 0.03004342 | 0.03646294 | -0.0064195 | 0.78309108 | 0.90650187 |
| Fam98b | 0.26012849 | 0.26688468 | 0.27330924 | -0.0064246 | 0.54161404 | 0.90650187 |

| | | | | | | |
|---|---|---|---|---|---|---|
| Rmdn2 | 0.03648227 | 0.03727384 | 0.04369873 | -0.0064249 | 0.74332293 | 0.90650187 |
| Fam58b | 0.24781574 | 0.25567362 | 0.26210175 | -0.0064281 | 0.54341058 | 0.90650187 |
| Dzip1 | 0.0639487 | 0.06879821 | 0.07522644 | -0.0064282 | 0.64798463 | 0.90650187 |
| Aifm3 | 0.02699867 | 0.02611312 | 0.03254412 | -0.006431 | 0.81016107 | 0.9092537 |
| Iba57 | 0.02596347 | 0.02488157 | 0.03131795 | -0.0064364 | 0.81954919 | 0.9092537 |
| Sh3bgr | 0.09050798 | 0.09821878 | 0.10465723 | -0.0064384 | 0.60776207 | 0.90650187 |
| Hs6st1 | 0.15662065 | 0.16777463 | 0.17421909 | -0.0064445 | 0.56530971 | 0.90650187 |
| Egfr | 0.06299265 | 0.0677034 | 0.074148 | -0.0064446 | 0.6503966 | 0.90650187 |
| Rad17 | 0.37095197 | 0.36146898 | 0.36791949 | -0.0064505 | 0.53106296 | 0.90650187 |
| Wnk4 | 0.13779994 | 0.14848309 | 0.15493936 | -0.0064563 | 0.57348225 | 0.90650187 |
| Gm20628 | 0.02547697 | 0.02428482 | 0.03074112 | -0.0064563 | 0.82476379 | 0.90934433 |
| Spef1 | 0.04022546 | 0.04159461 | 0.04806291 | -0.0064683 | 0.72517641 | 0.90650187 |
| Gcgr | 0.0248769 | 0.02355589 | 0.0300291 | -0.0064732 | 0.83104802 | 0.9098597 |
| Kdm5b | 0.85243881 | 0.67376433 | 0.68023763 | -0.0064733 | 0.51687239 | 0.90650187 |
| Rnf144a | 0.15077508 | 0.16180198 | 0.1682755 | -0.0064735 | 0.56789663 | 0.90650187 |
| Gm12802 | 0.02560388 | 0.02441804 | 0.03089162 | -0.0064736 | 0.82423152 | 0.90929265 |
| Ythdc2 | 0.12474098 | 0.13484203 | 0.14131619 | -0.0064742 | 0.58069339 | 0.90650187 |
| Gm26882 | 0.05080978 | 0.05381226 | 0.06028687 | -0.0064746 | 0.68347675 | 0.90650187 |
| Acss2 | 0.06724607 | 0.07245164 | 0.07893562 | -0.006484 | 0.64250906 | 0.90650187 |
| Ankle1 | 0.03373885 | 0.0340015 | 0.04048638 | -0.0064849 | 0.76179013 | 0.90650187 |
| F8 | 0.03171227 | 0.03160637 | 0.03810587 | -0.0064995 | 0.77585275 | 0.90650187 |
| C87436 | 0.2252769 | 0.23468786 | 0.2411945 | -0.0065066 | 0.54772985 | 0.90650187 |
| 0610009L18I | 0.16158788 | 0.17273141 | 0.17923895 | -0.0065075 | 0.56411255 | 0.90650187 |
| Cfi | 0.03838188 | 0.03940729 | 0.04591619 | -0.0065089 | 0.73571554 | 0.90650187 |
| Scnn1b | 0.04539829 | 0.04754783 | 0.05405841 | -0.0065106 | 0.70381733 | 0.90650187 |
| Pappa2 | 0.04496049 | 0.04703948 | 0.05355257 | -0.0065131 | 0.70563462 | 0.90650187 |
| Idua | 0.02492097 | 0.02356767 | 0.03008141 | -0.0065137 | 0.83213561 | 0.91015207 |
| Ccdc106 | 0.03117529 | 0.03095672 | 0.03747403 | -0.0065173 | 0.78028585 | 0.90650187 |
| Calr4 | 0.01533502 | 0.01210988 | 0.01862996 | -0.0065201 | 0.94012997 | 0.95661796 |
| Mapkbp1 | 0.03903888 | 0.04015257 | 0.04668182 | -0.0065293 | 0.73296146 | 0.90650187 |

| | | | | | | |
|---|---|---|---|---|---|---|
| Mtrf1 | 0.06395485 | 0.06869725 | 0.07523337 | -0.0065361 | 0.6503351 | 0.90650187 |
| Slc9a6 | 0.07256574 | 0.07834759 | 0.08488622 | -0.0065386 | 0.63398888 | 0.90650187 |
| Zfp217 | 0.08830317 | 0.09571168 | 0.10225224 | -0.0065406 | 0.61193386 | 0.90650187 |
| Cops2 | 0.67101184 | 0.57142431 | 0.57796526 | -0.0065409 | 0.52006304 | 0.90650187 |
| 2010300C02I | 0.0377209 | 0.03860291 | 0.04514525 | -0.0065423 | 0.74029208 | 0.90650187 |
| Ager | 0.01512706 | 0.01183523 | 0.0183799 | -0.0065447 | 0.94328704 | 0.95871682 |
| Osbpl6 | 0.05038813 | 0.05325758 | 0.05980316 | -0.0065456 | 0.68672216 | 0.90650187 |
| Map4k3 | 0.10847211 | 0.11750647 | 0.1240532 | -0.0065467 | 0.59274453 | 0.90650187 |
| 3110045C21I | 0.01739837 | 0.01455659 | 0.02110732 | -0.0065507 | 0.91616193 | 0.94231902 |
| Zfp280c | 0.08869889 | 0.09613103 | 0.10268438 | -0.0065534 | 0.61168753 | 0.90650187 |
| Acvr2b | 0.05432783 | 0.05775718 | 0.0643123 | -0.0065551 | 0.6747705 | 0.90650187 |
| 4933408B17I | 0.02359552 | 0.02194536 | 0.02850678 | -0.0065614 | 0.84690475 | 0.91420247 |
| Dld | 0.67451616 | 0.57353162 | 0.58009327 | -0.0065616 | 0.5200527 | 0.90650187 |
| Slc16a6 | 0.05420874 | 0.05761198 | 0.06417634 | -0.0065644 | 0.67534626 | 0.90650187 |
| Fam101a | 0.04338991 | 0.04516544 | 0.05173549 | -0.00657 | 0.71381385 | 0.90650187 |
| Mme | 0.02012954 | 0.01780386 | 0.02437595 | -0.0065721 | 0.88465775 | 0.92796521 |
| Sfmbt1 | 0.15230786 | 0.16325697 | 0.16983778 | -0.0065808 | 0.56838214 | 0.90650187 |
| Prepl | 0.15182012 | 0.16275586 | 0.16934095 | -0.0065851 | 0.56862491 | 0.90650187 |
| Akap1 | 0.11530876 | 0.12476011 | 0.13134716 | -0.0065871 | 0.58821163 | 0.90650187 |
| Vps16 | 0.18971377 | 0.20055303 | 0.20714019 | -0.0065872 | 0.55621214 | 0.90650187 |
| Galnt3 | 0.02877696 | 0.02805625 | 0.03464649 | -0.0065902 | 0.80111663 | 0.90817444 |
| Fcho2 | 0.26138212 | 0.26784863 | 0.27444215 | -0.0065935 | 0.54252874 | 0.90650187 |
| Cnnm2 | 0.04847077 | 0.05099694 | 0.05760019 | -0.0066032 | 0.69484856 | 0.90650187 |
| Fam179b | 0.21916141 | 0.22882899 | 0.23543271 | -0.0066037 | 0.54961796 | 0.90650187 |
| Ccdc125 | 0.0545631 | 0.05797638 | 0.06458084 | -0.0066045 | 0.67531466 | 0.90650187 |
| Ttk | 0.05484329 | 0.05829507 | 0.06490055 | -0.0066055 | 0.67453355 | 0.90650187 |
| Zfp830 | 0.36953272 | 0.3601685 | 0.36677473 | -0.0066062 | 0.53191034 | 0.90650187 |
| Mmgt2 | 0.22908501 | 0.23815612 | 0.24476256 | -0.0066064 | 0.54775537 | 0.90650187 |
| Tbcc | 0.51426302 | 0.46901717 | 0.47562461 | -0.0066074 | 0.52462259 | 0.90650187 |
| Erich5 | 0.04615941 | 0.04832834 | 0.05493713 | -0.0066088 | 0.70360409 | 0.90650187 |

| | | | | | | |
|---|---|---|---|---|---|---|
| Cenpl | 0.07720376 | 0.08342152 | 0.09004092 | -0.0066194 | 0.62809841 | 0.90650187 |
| Mpzl3 | 0.02565985 | 0.02433617 | 0.03095799 | -0.0066218 | 0.82917282 | 0.90974008 |
| Suclg2 | 0.50974355 | 0.4658218 | 0.47244982 | -0.006628 | 0.52486499 | 0.90650187 |
| Pxmp2 | 0.03368946 | 0.03379959 | 0.04042845 | -0.0066289 | 0.76697618 | 0.90650187 |
| Cdhr2 | 0.03286293 | 0.03282892 | 0.03945827 | -0.0066294 | 0.77243892 | 0.90650187 |
| Gpt | 0.0312504 | 0.03093034 | 0.03756244 | -0.0066321 | 0.78375119 | 0.90650187 |
| AV099323 | 0.04684222 | 0.04909241 | 0.05572467 | -0.0066323 | 0.70163505 | 0.90650187 |
| Zfp595 | 0.032327 | 0.03219606 | 0.03882864 | -0.0066326 | 0.77618834 | 0.90650187 |
| Ppa1 | 0.59778607 | 0.52532357 | 0.53195718 | -0.0066336 | 0.52210508 | 0.90650187 |
| Rhd | 0.01888103 | 0.01624696 | 0.02288325 | -0.0066363 | 0.90134519 | 0.93562881 |
| Wdr19 | 0.03884567 | 0.03981585 | 0.04645674 | -0.0066409 | 0.73744541 | 0.90650187 |
| AA474331 | 0.01923958 | 0.01666344 | 0.02331218 | -0.0066487 | 0.89758439 | 0.934149 |
| Tbc1d13 | 0.03858332 | 0.03949587 | 0.046151 | -0.0066551 | 0.73926192 | 0.90650187 |
| Capn5 | 0.03818659 | 0.03903306 | 0.04568848 | -0.0066554 | 0.74137469 | 0.90650187 |
| Eno3 | 0.04711277 | 0.0493785 | 0.05603653 | -0.006658 | 0.7013227 | 0.90650187 |
| Arhgef26 | 0.04365303 | 0.04537581 | 0.05204017 | -0.0066644 | 0.71542805 | 0.90650187 |
| Kif24 | 0.0264675 | 0.02524665 | 0.03191517 | -0.0066685 | 0.82352646 | 0.9092537 |
| Gm43848 | 0.0274048 | 0.02635477 | 0.0330247 | -0.0066699 | 0.8153725 | 0.9092537 |
| Gm27003 | 0.02975514 | 0.02912915 | 0.03580082 | -0.0066717 | 0.79629549 | 0.9075779 |
| Mamld1 | 0.13159769 | 0.14182192 | 0.14849453 | -0.0066726 | 0.57916723 | 0.90650187 |
| Gm16183 | 0.03425666 | 0.03442013 | 0.0410936 | -0.0066735 | 0.76483789 | 0.90650187 |
| Phf7 | 0.05302661 | 0.05614815 | 0.06282559 | -0.0066774 | 0.68171602 | 0.90650187 |
| Amot | 0.05812853 | 0.06195779 | 0.06864033 | -0.0066825 | 0.66742217 | 0.90650187 |
| 2810029C07I | 0.02383162 | 0.02210354 | 0.02878747 | -0.0066839 | 0.84900844 | 0.91444933 |
| Arl4d | 0.039725 | 0.04079648 | 0.04748067 | -0.0066842 | 0.73429002 | 0.90650187 |
| Odf2l | 0.17748233 | 0.18842629 | 0.19511472 | -0.0066884 | 0.56056159 | 0.90650187 |
| Rpp40 | 0.08778389 | 0.09499596 | 0.10168482 | -0.0066889 | 0.6150223 | 0.90650187 |
| Jak3 | 0.02694436 | 0.0257879 | 0.03247982 | -0.0066919 | 0.82015172 | 0.9092537 |
| Zc3h13 | 0.6528891 | 0.56015973 | 0.56685402 | -0.0066943 | 0.52093509 | 0.90650187 |
| Pdzrn4 | 0.05179512 | 0.0547121 | 0.06141619 | -0.0067041 | 0.68625951 | 0.90650187 |

| | | | | | | |
|---|---|---|---|---|---|---|
| Casp6 | 0.43014645 | 0.40753535 | 0.41423971 | -0.0067044 | 0.52867955 | 0.90650187 |
| Gm10053 | 0.22670779 | 0.23582536 | 0.2425369 | -0.0067115 | 0.54895435 | 0.90650187 |
| Rtkn2 | 0.04080143 | 0.04201842 | 0.04873251 | -0.0067141 | 0.72989349 | 0.90650187 |
| 4732440D04I | 0.05017162 | 0.05283955 | 0.05955467 | -0.0067151 | 0.69190834 | 0.90650187 |
| Polr1d | 1.54887595 | 0.89459697 | 0.90131449 | -0.0067175 | 0.51321583 | 0.90650187 |
| Tada2b | 0.0655974 | 0.07035948 | 0.07708303 | -0.0067235 | 0.65090572 | 0.90650187 |
| Snx33 | 0.06469082 | 0.0693388 | 0.07606264 | -0.0067238 | 0.65283511 | 0.90650187 |
| Csnk1g3 | 0.30267379 | 0.30420142 | 0.31092726 | -0.0067258 | 0.53830536 | 0.90650187 |
| Dhx29 | 0.19582818 | 0.20636127 | 0.21309103 | -0.0067298 | 0.5558278 | 0.90650187 |
| Ccdc17 | 0.02523987 | 0.02372913 | 0.03045986 | -0.0067307 | 0.83703547 | 0.91092327 |
| Mkl2 | 0.25159355 | 0.25882423 | 0.26555603 | -0.0067318 | 0.54486395 | 0.90650187 |
| Spin1 | 0.45923564 | 0.42929707 | 0.43603167 | -0.0067346 | 0.52737118 | 0.90650187 |
| Clmn | 0.11056722 | 0.1195464 | 0.1262946 | -0.0067482 | 0.59388049 | 0.90650187 |
| Plekhh1 | 0.019721 | 0.01713609 | 0.02388778 | -0.0067517 | 0.89553536 | 0.93316651 |
| Wipf3 | 0.02460256 | 0.02295034 | 0.0297034 | -0.0067531 | 0.84392156 | 0.91300169 |
| Clcn7 | 0.06014731 | 0.06417544 | 0.07093047 | -0.006755 | 0.66398907 | 0.90650187 |
| Zmynd12 | 0.01639049 | 0.01314116 | 0.01989807 | -0.0067569 | 0.9344301 | 0.95292311 |
| Trim67 | 0.02047942 | 0.01803621 | 0.02479383 | -0.0067576 | 0.88717899 | 0.92936447 |
| Rnf182 | 0.03640497 | 0.03684968 | 0.04360838 | -0.0067587 | 0.75459063 | 0.90650187 |
| Ripk2 | 0.08772897 | 0.09486459 | 0.10162478 | -0.0067602 | 0.61628058 | 0.90650187 |
| Usf1 | 0.22031443 | 0.22975985 | 0.23652208 | -0.0067622 | 0.55056979 | 0.90650187 |
| Lmtk3 | 0.0168529 | 0.01368982 | 0.02045308 | -0.0067633 | 0.92922417 | 0.9499722 |
| Sall4 | 0.04281678 | 0.04430804 | 0.05107147 | -0.0067634 | 0.72198032 | 0.90650187 |
| Elmod1 | 0.06510137 | 0.06976137 | 0.07652488 | -0.0067635 | 0.65280951 | 0.90650187 |
| Mfsd4 | 0.04588084 | 0.04785153 | 0.05461561 | -0.0067641 | 0.70904036 | 0.90650187 |
| Pomgnt2 | 0.23983131 | 0.24798868 | 0.254753 | -0.0067643 | 0.54698274 | 0.90650187 |
| Cep152 | 0.02771106 | 0.02662072 | 0.03338693 | -0.0067662 | 0.8161922 | 0.9092537 |
| 2700012I20R | 0.03928773 | 0.04020422 | 0.04697164 | -0.0067674 | 0.73907585 | 0.90650187 |
| Arhgef38 | 0.02534002 | 0.02381058 | 0.03057867 | -0.0067681 | 0.83743295 | 0.91109492 |
| Atad3aos | 0.02778135 | 0.02669938 | 0.03347005 | -0.0067707 | 0.81575449 | 0.9092537 |

| | | | | | | |
|---|---|---|---|---|---|---|
| Atp9a | 0.29603002 | 0.29838857 | 0.30516428 | -0.0067757 | 0.53931493 | 0.90650187 |
| 4632415L05F | 0.14440023 | 0.15496185 | 0.16174796 | -0.0067861 | 0.57397946 | 0.90650187 |
| Papolg | 0.12077278 | 0.13034484 | 0.13713535 | -0.0067905 | 0.58711609 | 0.90650187 |
| Apobr | 0.02417967 | 0.02240736 | 0.0292011 | -0.0067937 | 0.84949858 | 0.91459933 |
| Cep76 | 0.05188588 | 0.0547168 | 0.06152015 | -0.0068033 | 0.68850875 | 0.90650187 |
| Fancf | 0.02054582 | 0.0180674 | 0.0248731 | -0.0068057 | 0.88808231 | 0.92986219 |
| Grhl1 | 0.02719199 | 0.02596534 | 0.0327729 | -0.0068076 | 0.82211042 | 0.9092537 |
| Ksr1 | 0.04172859 | 0.04300115 | 0.04980932 | -0.0068082 | 0.72828899 | 0.90650187 |
| 2410022M11 | 0.08605253 | 0.09297662 | 0.09979022 | -0.0068136 | 0.61926178 | 0.90650187 |
| Rpn1 | 1.58413139 | 0.89930417 | 0.90612229 | -0.0068181 | 0.51334252 | 0.90650187 |
| Nars2 | 0.0835146 | 0.09018644 | 0.09700536 | -0.0068189 | 0.62267004 | 0.90650187 |
| Znfx1 | 0.09334921 | 0.1009219 | 0.10774642 | -0.0068245 | 0.61086681 | 0.90650187 |
| Cenpn | 0.05116309 | 0.05386729 | 0.06069197 | -0.0068247 | 0.69142695 | 0.90650187 |
| Me3 | 0.0235845 | 0.02166762 | 0.02849368 | -0.0068261 | 0.85656344 | 0.91662238 |
| Rab20 | 0.09699515 | 0.10486701 | 0.11169415 | -0.0068271 | 0.60708559 | 0.90650187 |
| Topbp1 | 0.17799401 | 0.1887888 | 0.19562106 | -0.0068323 | 0.5616947 | 0.90650187 |
| Rtn4rl1 | 0.17177379 | 0.1826074 | 0.18944575 | -0.0068383 | 0.56374509 | 0.90650187 |
| Parp8 | 0.08317728 | 0.08979125 | 0.09663454 | -0.0068433 | 0.62355113 | 0.90650187 |
| Zranb3 | 0.08119649 | 0.08760607 | 0.09445376 | -0.0068477 | 0.62638607 | 0.90650187 |
| Mroh7 | 0.0200343 | 0.01741326 | 0.02426218 | -0.0068489 | 0.89524996 | 0.93293307 |
| Fbxo33 | 0.11223562 | 0.12122638 | 0.1280756 | -0.0068492 | 0.5939594 | 0.90650187 |
| Ube2t | 0.05977259 | 0.06364724 | 0.07050583 | -0.0068586 | 0.66729466 | 0.90650187 |
| Nynrin | 0.17203897 | 0.18284684 | 0.18970993 | -0.0068631 | 0.56388545 | 0.90650187 |
| Gm4799 | 0.02367382 | 0.02173297 | 0.02859989 | -0.0068669 | 0.85710491 | 0.91700814 |
| Lrriq3 | 0.01419971 | 0.01039638 | 0.01726391 | -0.0068675 | 0.96150708 | 0.97096654 |
| Ccdc112 | 0.23484898 | 0.2432674 | 0.25013543 | -0.006868 | 0.54857591 | 0.90650187 |
| Trim9 | 0.03589711 | 0.03614376 | 0.04301453 | -0.0068708 | 0.76117986 | 0.90650187 |
| Hist2h2bb | 0.02618712 | 0.02470949 | 0.03158299 | -0.0068735 | 0.8333556 | 0.91056919 |
| Wdr31 | 0.06581261 | 0.07044532 | 0.07732509 | -0.0068798 | 0.65377584 | 0.90650187 |
| Klhl21 | 0.07291328 | 0.07839108 | 0.08527354 | -0.0068825 | 0.6401301 | 0.90650187 |

| | | | | | | |
|---|---|---|---|---|---|---|
| Gm13166 | 0.03109332 | 0.03048853 | 0.03737754 | -0.006889 | 0.79371053 | 0.90731056 |
| 1700001C02I | 0.03173527 | 0.03123281 | 0.03813292 | -0.0069001 | 0.78942515 | 0.90650187 |
| Tnks | 0.07388922 | 0.07945247 | 0.08636029 | -0.0069078 | 0.63892873 | 0.90650187 |
| Tstd3 | 0.06017061 | 0.06404466 | 0.07095686 | -0.0069122 | 0.66751613 | 0.90650187 |
| Lmo7 | 0.13059549 | 0.14053595 | 0.14744884 | -0.0069129 | 0.58255131 | 0.90650187 |
| Ahcy | 0.07552716 | 0.08126751 | 0.08818112 | -0.0069136 | 0.63628497 | 0.90650187 |
| Pgbd5 | 0.02990716 | 0.02905756 | 0.03598008 | -0.0069225 | 0.80381549 | 0.90882608 |
| Lhfpl5 | 0.01694597 | 0.01364062 | 0.02056474 | -0.0069241 | 0.93278128 | 0.95188026 |
| Tgif2 | 0.13477278 | 0.14487466 | 0.15179951 | -0.0069249 | 0.58035488 | 0.90650187 |
| Srfbp1 | 0.09129746 | 0.0985869 | 0.10551674 | -0.0069298 | 0.61484379 | 0.90650187 |
| Rabgap1l | 0.26845959 | 0.27386327 | 0.28081012 | -0.0069468 | 0.54378662 | 0.90650187 |
| Erich2 | 0.0307166 | 0.02998649 | 0.03693395 | -0.0069475 | 0.7984905 | 0.90770671 |
| Zfp932 | 0.16498269 | 0.17570313 | 0.18265365 | -0.0069505 | 0.56716801 | 0.90650187 |
| Camkk2 | 0.28779697 | 0.29100964 | 0.29796512 | -0.0069555 | 0.54132602 | 0.90650187 |
| Ngef | 0.02015219 | 0.0174446 | 0.02440302 | -0.0069584 | 0.89753767 | 0.934149 |
| Rad51c | 0.04777579 | 0.04983075 | 0.05680031 | -0.0069696 | 0.70729137 | 0.90650187 |
| Ptcra | 0.04591371 | 0.04768321 | 0.05465356 | -0.0069704 | 0.71463603 | 0.90650187 |
| D2hgdh | 0.09418792 | 0.10168226 | 0.10865618 | -0.0069739 | 0.61230588 | 0.90650187 |
| Xrn2 | 0.6718416 | 0.57149414 | 0.5784697 | -0.0069756 | 0.52137625 | 0.90650187 |
| Fancd2 | 0.02544955 | 0.02373296 | 0.03070859 | -0.0069756 | 0.84371493 | 0.91300169 |
| Lrrc75b | 0.02210403 | 0.01975251 | 0.02673155 | -0.006979 | 0.87710837 | 0.9247416 |
| Bcar1 | 0.29005895 | 0.29296899 | 0.29994917 | -0.0069802 | 0.54119898 | 0.90650187 |
| Pfkfb3 | 0.04799753 | 0.05007532 | 0.0570556 | -0.0069803 | 0.70673897 | 0.90650187 |
| Nudt18 | 0.21216428 | 0.22180846 | 0.22879264 | -0.0069842 | 0.55397385 | 0.90650187 |
| Mapk4 | 0.01697426 | 0.01361337 | 0.02059868 | -0.0069853 | 0.93416764 | 0.95276695 |
| Lefty1 | 0.07264145 | 0.07798487 | 0.08497061 | -0.0069857 | 0.64262694 | 0.90650187 |
| Purg | 0.05285424 | 0.05563961 | 0.06262845 | -0.0069888 | 0.69008606 | 0.90650187 |
| Tprgl | 0.78948665 | 0.63950206 | 0.64649765 | -0.0069956 | 0.51918364 | 0.90650187 |
| Zfp551 | 0.03092523 | 0.03018223 | 0.03717964 | -0.0069974 | 0.79861581 | 0.90777025 |
| Enthd2 | 0.07198562 | 0.07723988 | 0.08423929 | -0.0069994 | 0.64407913 | 0.90650187 |

| | | | | | | |
|---|---|---|---|---|---|---|
| Actr8 | 0.31508173 | 0.31458333 | 0.32158481 | -0.0070015 | 0.53855295 | 0.90650187 |
| 1110034G24 | 0.13354851 | 0.14352462 | 0.15052657 | -0.007002 | 0.58191451 | 0.90650187 |
| Lgi3 | 0.02641264 | 0.02484791 | 0.03185019 | -0.0070023 | 0.83580655 | 0.91079271 |
| Stard3 | 0.28922579 | 0.29221322 | 0.29921892 | -0.0070057 | 0.54144959 | 0.90650187 |
| Klhl18 | 0.07620608 | 0.08192515 | 0.08893471 | -0.0070096 | 0.6369767 | 0.90650187 |
| Rab11fip4 | 0.0220355 | 0.01963869 | 0.0266499 | -0.0070112 | 0.87891402 | 0.92548989 |
| Kif17 | 0.04516006 | 0.04677092 | 0.05378319 | -0.0070123 | 0.71891429 | 0.90650187 |
| Tnip2 | 0.07428915 | 0.07978776 | 0.08680523 | -0.0070175 | 0.64034833 | 0.90650187 |
| Helb | 0.03682289 | 0.03707507 | 0.04409676 | -0.0070217 | 0.76049344 | 0.90650187 |
| Klhdc8a | 0.06456971 | 0.06889705 | 0.07592623 | -0.0070292 | 0.65967281 | 0.90650187 |
| Cox11 | 0.22451876 | 0.23345198 | 0.2404824 | -0.0070304 | 0.55170327 | 0.90650187 |
| 1700105P06I | 0.02295577 | 0.02071278 | 0.02774576 | -0.007033 | 0.87010107 | 0.92163656 |
| Ttll4 | 0.05299947 | 0.05576152 | 0.06279455 | -0.007033 | 0.6907264 | 0.90650187 |
| Ctnnd2 | 0.02103294 | 0.01841942 | 0.0254545 | -0.0070351 | 0.8904021 | 0.93072624 |
| Atmin | 0.10528039 | 0.11359079 | 0.1206281 | -0.0070373 | 0.60231813 | 0.90650187 |
| Dppa5a | 0.0275748 | 0.0261881 | 0.03322579 | -0.0070377 | 0.82681217 | 0.90947782 |
| Sufu | 0.06429613 | 0.06857915 | 0.07561801 | -0.0070389 | 0.66049571 | 0.90650187 |
| Gfod2 | 0.27163919 | 0.27660905 | 0.28365558 | -0.0070465 | 0.54396863 | 0.90650187 |
| Abhd10 | 0.22326324 | 0.23225328 | 0.23930184 | -0.0070486 | 0.55209015 | 0.90650187 |
| Rrnad1 | 0.26075387 | 0.2668223 | 0.27387459 | -0.0070523 | 0.54556919 | 0.90650187 |
| Alg10b | 0.03267861 | 0.03218654 | 0.03924177 | -0.0070552 | 0.78793336 | 0.90650187 |
| Mtg1 | 0.17859273 | 0.18915727 | 0.19621316 | -0.0070559 | 0.56350646 | 0.90650187 |
| Ap5m1 | 0.07778063 | 0.08362043 | 0.0906799 | -0.0070595 | 0.63536344 | 0.90650187 |
| Ipo9 | 0.34581796 | 0.34034073 | 0.34740326 | -0.0070625 | 0.53600617 | 0.90650187 |
| Rcc2 | 0.27197226 | 0.27688895 | 0.2839531 | -0.0070641 | 0.54403212 | 0.90650187 |
| Rpusd1 | 0.21001224 | 0.21967445 | 0.22674025 | -0.0070658 | 0.55509177 | 0.90650187 |
| Trappc9 | 0.20908235 | 0.21878472 | 0.22585193 | -0.0070672 | 0.55531799 | 0.90650187 |
| Dsp | 0.16123845 | 0.17181569 | 0.17888673 | -0.007071 | 0.56974541 | 0.90650187 |
| Zfp963 | 0.04545389 | 0.0470469 | 0.05412263 | -0.0070757 | 0.71944608 | 0.90650187 |
| Nudcd1 | 0.15715826 | 0.16768565 | 0.17476375 | -0.0070781 | 0.57144391 | 0.90650187 |

| | | | | | | |
|---|---|---|---|---|---|---|
| Gm15706 | 0.05915657 | 0.06272862 | 0.06980731 | -0.0070787 | 0.67393293 | 0.90650187 |
| Gm26615 | 0.02456415 | 0.02257824 | 0.02965779 | -0.0070795 | 0.85570068 | 0.9165432 |
| Rhbdl2 | 0.02446573 | 0.02246045 | 0.02954091 | -0.0070805 | 0.8566835 | 0.91668632 |
| Nbn | 0.0812453 | 0.08742678 | 0.09450757 | -0.0070808 | 0.6304623 | 0.90650187 |
| Tmco6 | 0.11903527 | 0.12821772 | 0.13529872 | -0.007081 | 0.59198933 | 0.90650187 |
| Ltbp1 | 0.05208351 | 0.05466058 | 0.06174645 | -0.0070859 | 0.69503262 | 0.90650187 |
| Mmgt1 | 0.05649305 | 0.0596923 | 0.06678058 | -0.0070883 | 0.68149003 | 0.90650187 |
| Hdac3 | 0.66664976 | 0.56820043 | 0.57530749 | -0.0071071 | 0.5218984 | 0.90650187 |
| Gm9920 | 0.02727199 | 0.02575792 | 0.03286757 | -0.0071096 | 0.83188418 | 0.91013901 |
| Vps39 | 0.12207658 | 0.13139335 | 0.13851113 | -0.0071178 | 0.59035176 | 0.90650187 |
| Wdr45 | 0.21752987 | 0.22676789 | 0.23388893 | -0.007121 | 0.55383322 | 0.90650187 |
| Cobl | 0.03136668 | 0.03057582 | 0.03769928 | -0.0071235 | 0.7995511 | 0.90799325 |
| Tmem194b | 0.03392632 | 0.0335802 | 0.04070628 | -0.0071261 | 0.78178958 | 0.90650187 |
| Zfp746 | 0.07763414 | 0.08338452 | 0.09051768 | -0.0071332 | 0.63695504 | 0.90650187 |
| F5 | 0.0255806 | 0.02372987 | 0.03086402 | -0.0071341 | 0.8479291 | 0.91440801 |
| Stxbp4 | 0.13839201 | 0.14841769 | 0.15555221 | -0.0071345 | 0.58078457 | 0.90650187 |
| Rft1 | 0.13207779 | 0.14185171 | 0.14899504 | -0.0071433 | 0.58439018 | 0.90650187 |
| Dazap2 | 0.68016811 | 0.57636712 | 0.58351211 | -0.007145 | 0.52170592 | 0.90650187 |
| Zfp574 | 0.12267341 | 0.13198358 | 0.13914021 | -0.0071566 | 0.59043272 | 0.90650187 |
| Zfp583 | 0.02613552 | 0.02436332 | 0.03152185 | -0.0071585 | 0.84365316 | 0.91300169 |
| Zfp873 | 0.03433389 | 0.03402482 | 0.04118413 | -0.0071593 | 0.78018256 | 0.90650187 |
| Dguok | 0.62337403 | 0.54121145 | 0.54837517 | -0.0071637 | 0.52315569 | 0.90650187 |
| Tsc22d4 | 0.91833085 | 0.70646619 | 0.71363061 | -0.0071644 | 0.51779937 | 0.90650187 |
| D8Ertd82e | 0.05686682 | 0.06003957 | 0.06720596 | -0.0071664 | 0.68226783 | 0.90650187 |
| Gm42715 | 0.06640039 | 0.07081861 | 0.07798584 | -0.0071672 | 0.65857228 | 0.90650187 |
| Kdm1b | 0.04355473 | 0.04475084 | 0.05192635 | -0.0071755 | 0.73049859 | 0.90650187 |
| Zfp871 | 0.26678902 | 0.27212969 | 0.2793113 | -0.0071816 | 0.5455019 | 0.90650187 |
| Smim24 | 0.12047451 | 0.12963714 | 0.13682033 | -0.0071832 | 0.5922739 | 0.90650187 |
| Zfp658 | 0.03206033 | 0.03132891 | 0.03851517 | -0.0071863 | 0.7965816 | 0.9075779 |
| Cc2d2a | 0.21875285 | 0.22785936 | 0.23504639 | -0.007187 | 0.55406311 | 0.90650187 |

| | | | | | | |
|---|---|---|---|---|---|---|
| 2700038G22 | 0.10452684 | 0.11263005 | 0.11981759 | -0.0071875 | 0.60514227 | 0.90650187 |
| Cst6 | 0.0330702 | 0.03251071 | 0.03970166 | -0.007191 | 0.78965061 | 0.90650187 |
| Zfp472 | 0.06140963 | 0.06516579 | 0.07235938 | -0.0071936 | 0.67074059 | 0.90650187 |
| Fam159a | 0.03924134 | 0.03972349 | 0.04691763 | -0.0071941 | 0.75227604 | 0.90650187 |
| Tmem220 | 0.03501767 | 0.03478714 | 0.04198525 | -0.0071981 | 0.77702106 | 0.90650187 |
| Col11a2 | 0.03414202 | 0.03375134 | 0.04095921 | -0.0072079 | 0.78298757 | 0.90650187 |
| Pak7 | 0.03500472 | 0.03475822 | 0.04197008 | -0.0072119 | 0.77753753 | 0.90650187 |
| Gm10184 | 0.02799281 | 0.02650549 | 0.03372005 | -0.0072146 | 0.82923802 | 0.90974008 |
| Cfap52 | 0.02430878 | 0.02213096 | 0.02935448 | -0.0072235 | 0.86301617 | 0.91950555 |
| Lpin3 | 0.03203834 | 0.03126574 | 0.03848932 | -0.0072236 | 0.79795542 | 0.9075779 |
| Ttc22 | 0.03961174 | 0.04012145 | 0.04734886 | -0.0072274 | 0.75129952 | 0.90650187 |
| Slc18b1 | 0.0450277 | 0.04639992 | 0.05363024 | -0.0072303 | 0.72552941 | 0.90650187 |
| Ly6g6d | 0.04033439 | 0.04095484 | 0.04818959 | -0.0072347 | 0.74774907 | 0.90650187 |
| Mier2 | 0.15892944 | 0.1693183 | 0.17655582 | -0.0072375 | 0.57230205 | 0.90650187 |
| Nf1 | 0.26482667 | 0.27030129 | 0.27754733 | -0.007246 | 0.54619875 | 0.90650187 |
| Kirrel2 | 0.02303118 | 0.02058907 | 0.0278355 | -0.0072464 | 0.87643061 | 0.92466836 |
| Zfp940 | 0.03372071 | 0.03321818 | 0.04046511 | -0.0072469 | 0.78703257 | 0.90650187 |
| Gpr155 | 0.03776452 | 0.03794365 | 0.04519615 | -0.0072525 | 0.76219281 | 0.90650187 |
| Rimklb | 0.05184439 | 0.05421862 | 0.06147262 | -0.007254 | 0.70006876 | 0.90650187 |
| Meig1 | 0.06700134 | 0.07139432 | 0.07866087 | -0.0072665 | 0.65934522 | 0.90650187 |
| Atp2b2 | 0.01963194 | 0.01650855 | 0.02378132 | -0.0072728 | 0.91301372 | 0.9407546 |
| Gpatch1 | 0.21452636 | 0.22376262 | 0.23103982 | -0.0072772 | 0.5556803 | 0.90650187 |
| Samd5 | 0.18897735 | 0.19914181 | 0.20642075 | -0.0072789 | 0.56228453 | 0.90650187 |
| Slc26a1 | 0.0430177 | 0.04402344 | 0.05130431 | -0.0072809 | 0.73594699 | 0.90650187 |
| Med21 | 0.8615108 | 0.67767154 | 0.6849568 | -0.0072853 | 0.5188565 | 0.90650187 |
| Hist1h3d | 0.016014 | 0.01215527 | 0.01944594 | -0.0072907 | 0.95221753 | 0.96420588 |
| Tmem266 | 0.02242086 | 0.01981747 | 0.02710895 | -0.0072915 | 0.8840995 | 0.9278081 |
| Tbc1d2b | 0.13138094 | 0.14096815 | 0.14826848 | -0.0073003 | 0.58663191 | 0.90650187 |
| St3gal3 | 0.08089074 | 0.08681581 | 0.09411664 | -0.0073008 | 0.63489835 | 0.90650187 |
| Slx4ip | 0.07221964 | 0.07719292 | 0.08450031 | -0.0073074 | 0.64967585 | 0.90650187 |

| | | | | | | |
|---|---|---|---|---|---|---|
| Polr3a | 0.04243976 | 0.04332637 | 0.05063438 | -0.007308 | 0.73944324 | 0.90650187 |
| Lhx1os | 0.1385803 | 0.14843281 | 0.15574702 | -0.0073142 | 0.58268806 | 0.90650187 |
| Ubox5 | 0.0540467 | 0.0566763 | 0.06399131 | -0.007315 | 0.69433957 | 0.90650187 |
| Btbd19 | 0.03387135 | 0.03332335 | 0.04064181 | -0.0073185 | 0.78829821 | 0.90650187 |
| Map7d2 | 0.03707225 | 0.03706929 | 0.04438802 | -0.0073187 | 0.76821947 | 0.90650187 |
| Clcf1 | 0.03569632 | 0.03545898 | 0.04277963 | -0.0073207 | 0.77659776 | 0.90650187 |
| Ulk4 | 0.07587382 | 0.08124489 | 0.08856599 | -0.0073211 | 0.64337153 | 0.90650187 |
| Mrpl11 | 0.73381085 | 0.60782543 | 0.61514686 | -0.0073214 | 0.52109869 | 0.90650187 |
| Plcxd2 | 0.03917039 | 0.03950937 | 0.046835 | -0.0073256 | 0.75658296 | 0.90650187 |
| Vps33a | 0.15858453 | 0.1688804 | 0.17620713 | -0.0073267 | 0.57332638 | 0.90650187 |
| Rad51ap2 | 0.02229093 | 0.01962614 | 0.0269542 | -0.0073281 | 0.88659753 | 0.92907549 |
| Fbxl12os | 0.03525266 | 0.03493083 | 0.04226039 | -0.0073296 | 0.77965774 | 0.90650187 |
| Sgk2 | 0.02517349 | 0.02304652 | 0.0303811 | -0.0073346 | 0.85842132 | 0.91744797 |
| Atp6v0c | 0.02664117 | 0.02478388 | 0.03212086 | -0.007337 | 0.84505567 | 0.91332066 |
| 1810041H14 | 0.01908251 | 0.01578666 | 0.0231243 | -0.0073376 | 0.92082594 | 0.94507293 |
| Ccdc93 | 0.0485819 | 0.050387 | 0.05772802 | -0.007341 | 0.71407465 | 0.90650187 |
| Arhgap42 | 0.07661667 | 0.08204512 | 0.08939014 | -0.007345 | 0.64255152 | 0.90650187 |
| Chst14 | 0.10600607 | 0.11405907 | 0.12140796 | -0.0073489 | 0.60607172 | 0.90650187 |
| Zfp157 | 0.30646654 | 0.30684873 | 0.3141994 | -0.0073507 | 0.54141716 | 0.90650187 |
| Dhdh | 0.01941239 | 0.016168 | 0.02351884 | -0.0073508 | 0.91765633 | 0.94297078 |
| Zfp507 | 0.13697637 | 0.1467353 | 0.1540862 | -0.0073509 | 0.58397928 | 0.90650187 |
| Slc25a10 | 0.11919222 | 0.12811379 | 0.13546477 | -0.007351 | 0.59531553 | 0.90650187 |
| Zbtb8b | 0.01400244 | 0.00966895 | 0.01702633 | -0.0073574 | 0.97263578 | 0.97933606 |
| Bdh1 | 0.04258224 | 0.04343858 | 0.05079958 | -0.007361 | 0.74027105 | 0.90650187 |
| Arhgap19 | 0.02308618 | 0.02053579 | 0.02790094 | -0.0073651 | 0.87970795 | 0.92562058 |
| Klb | 0.03471496 | 0.03426527 | 0.04163068 | -0.0073654 | 0.78422094 | 0.90650187 |
| Anks6 | 0.06188595 | 0.06553247 | 0.07289796 | -0.0073655 | 0.6733479 | 0.90650187 |
| Dock9 | 0.11618641 | 0.12490924 | 0.13227935 | -0.0073701 | 0.59781374 | 0.90650187 |
| Usp35 | 0.02769015 | 0.02599106 | 0.03336221 | -0.0073712 | 0.83700585 | 0.91092327 |
| Cfap77 | 0.18171866 | 0.19192178 | 0.19929816 | -0.0073764 | 0.56534687 | 0.90650187 |

| | | | | | | |
|---|---|---|---|---|---|---|
| Pip4k2a | 0.04866791 | 0.05044885 | 0.05782695 | -0.0073781 | 0.71471469 | 0.90650187 |
| Fbxo7 | 0.18097601 | 0.19118778 | 0.1985662 | -0.0073784 | 0.56560365 | 0.90650187 |
| Grin2c | 0.04604284 | 0.04741946 | 0.05480259 | -0.0073831 | 0.72538773 | 0.90650187 |
| Zfp689 | 0.06064148 | 0.06410323 | 0.07149013 | -0.0073869 | 0.67700994 | 0.90650187 |
| Ppip5k1 | 0.13899149 | 0.14878358 | 0.1561723 | -0.0073887 | 0.58329396 | 0.90650187 |
| Ccdc9 | 0.2070786 | 0.21654555 | 0.2239347 | -0.0073891 | 0.5583121 | 0.90650187 |
| Tmem55a | 0.48023596 | 0.44399608 | 0.45138718 | -0.0073911 | 0.52901465 | 0.90650187 |
| Trim21 | 0.02120825 | 0.0182705 | 0.02566365 | -0.0073931 | 0.89984771 | 0.93477653 |
| Fggy | 0.02400029 | 0.02159477 | 0.02898795 | -0.0073932 | 0.87156508 | 0.9223337 |
| Etaa1os | 0.05723922 | 0.06022985 | 0.06762957 | -0.0073997 | 0.68666453 | 0.90650187 |
| Gorasp2 | 0.87705917 | 0.6855509 | 0.69295285 | -0.0074019 | 0.51893739 | 0.90650187 |
| Fbxo9 | 0.44674847 | 0.41934771 | 0.42675317 | -0.0074055 | 0.53074578 | 0.90650187 |
| Slc25a40 | 0.057865 | 0.06093175 | 0.06834093 | -0.0074092 | 0.68508651 | 0.90650187 |
| Fzd5 | 0.01953704 | 0.01625761 | 0.02366788 | -0.0074103 | 0.91802249 | 0.9430871 |
| Fabp3-ps1 | 0.02581564 | 0.02373112 | 0.03114271 | -0.0074116 | 0.85497243 | 0.91627438 |
| Rprm | 0.04526312 | 0.04649008 | 0.05390226 | -0.0074122 | 0.72950682 | 0.90650187 |
| Zswim6 | 0.08878297 | 0.09535951 | 0.10277618 | -0.0074167 | 0.62582266 | 0.90650187 |
| Tef | 0.21410958 | 0.22322513 | 0.23064374 | -0.0074186 | 0.55685176 | 0.90650187 |
| Slc15a2 | 0.03132265 | 0.03022463 | 0.03764746 | -0.0074228 | 0.80962801 | 0.9092537 |
| F8a | 0.26246775 | 0.26799871 | 0.27542204 | -0.0074233 | 0.54768733 | 0.90650187 |
| Perp | 0.08122109 | 0.08705482 | 0.09448087 | -0.0074261 | 0.63661227 | 0.90650187 |
| Catsper2 | 0.05100551 | 0.05308328 | 0.06051132 | -0.007428 | 0.70737007 | 0.90650187 |
| Ppp4r1 | 0.16338039 | 0.17361287 | 0.18104358 | -0.0074307 | 0.57239083 | 0.90650187 |
| Mtr | 0.0391862 | 0.03942106 | 0.04685341 | -0.0074323 | 0.75965896 | 0.90650187 |
| 5830454E08I | 0.03580077 | 0.03546831 | 0.04290184 | -0.0074335 | 0.77943399 | 0.90650187 |
| Hs6st3 | 0.02226655 | 0.01948048 | 0.02692516 | -0.0074447 | 0.89049947 | 0.93072624 |
| Fbxw17 | 0.13396427 | 0.1435141 | 0.15095907 | -0.007445 | 0.58677139 | 0.90650187 |
| Wdtc1 | 0.1777065 | 0.18789015 | 0.19533658 | -0.0074464 | 0.56728699 | 0.90650187 |
| Zfp763 | 0.09314354 | 0.10007551 | 0.10752318 | -0.0074477 | 0.62093153 | 0.90650187 |
| Dnal4 | 0.41761466 | 0.39720679 | 0.40465964 | -0.0074528 | 0.53262784 | 0.90650187 |

| | | | | | | |
|---|---|---|---|---|---|---|
| Rin3 | 0.0591179 | 0.06230714 | 0.06976344 | -0.0074563 | 0.68265829 | 0.90650187 |
| Dnmt3b | 0.06164043 | 0.06516221 | 0.07262039 | -0.0074582 | 0.67600917 | 0.90650187 |
| Gm16223 | 0.02748206 | 0.02565625 | 0.0331161 | -0.0074599 | 0.84169139 | 0.91251845 |
| Gm26632 | 0.02584777 | 0.02371823 | 0.0311808 | -0.0074626 | 0.8563304 | 0.9165432 |
| Gal3st1 | 0.03705618 | 0.03690451 | 0.04436925 | -0.0074647 | 0.77275031 | 0.90650187 |
| Fchsd2 | 0.17773794 | 0.18790094 | 0.1953677 | -0.0074668 | 0.56745822 | 0.90650187 |
| Tyw3 | 0.03540647 | 0.03497113 | 0.04244044 | -0.0074693 | 0.78301186 | 0.90650187 |
| Gk | 0.08816559 | 0.09462547 | 0.10210194 | -0.0074765 | 0.6276107 | 0.90650187 |
| Gabbr2 | 0.01465258 | 0.01033164 | 0.01780908 | -0.0074774 | 0.96901868 | 0.97653493 |
| Tkfc | 0.18669201 | 0.1967048 | 0.20418443 | -0.0074796 | 0.5646818 | 0.90650187 |
| Golga3 | 0.29725707 | 0.29874929 | 0.3062318 | -0.0074825 | 0.5432499 | 0.90650187 |
| Eif4ebp2 | 0.17121647 | 0.18140651 | 0.18889029 | -0.0074838 | 0.56990555 | 0.90650187 |
| Ss18l1 | 0.05235032 | 0.05456611 | 0.06205188 | -0.0074858 | 0.70412634 | 0.90650187 |
| Hmga1-rs1 | 0.0301141 | 0.02873715 | 0.03622404 | -0.0074869 | 0.82090811 | 0.9092537 |
| Lrch3 | 0.16564878 | 0.17583445 | 0.18332212 | -0.0074877 | 0.572043 | 0.90650187 |
| Gpd1l | 0.10360133 | 0.11132476 | 0.11882115 | -0.0074964 | 0.61044231 | 0.90650187 |
| Tnk2 | 0.05983704 | 0.0630618 | 0.07057888 | -0.0075171 | 0.68206604 | 0.90650187 |
| Paox | 0.10841975 | 0.11647189 | 0.1239971 | -0.0075252 | 0.60634206 | 0.90650187 |
| Camkk1 | 0.02175558 | 0.01878044 | 0.02631629 | -0.0075358 | 0.89849624 | 0.93440169 |
| Tmcc2 | 0.04350243 | 0.04432581 | 0.05186579 | -0.00754 | 0.74095072 | 0.90650187 |
| Pkn1 | 0.19912789 | 0.20874571 | 0.21628574 | -0.00754 | 0.56158155 | 0.90650187 |
| Il1r1 | 0.32148322 | 0.31949005 | 0.32703041 | -0.0075404 | 0.54082114 | 0.90650187 |
| Ttll13 | 0.01846401 | 0.01484115 | 0.0223841 | -0.007543 | 0.93294275 | 0.95198112 |
| Xylb | 0.01866725 | 0.01508392 | 0.0226274 | -0.0075435 | 0.93083905 | 0.95091965 |
| Mboat7 | 0.38690419 | 0.37312619 | 0.38067196 | -0.0075458 | 0.53511012 | 0.90650187 |
| Grip1 | 0.08159928 | 0.08734028 | 0.09489767 | -0.0075574 | 0.63834399 | 0.90650187 |
| Chic2 | 0.76438518 | 0.62498357 | 0.63254384 | -0.0075603 | 0.52118768 | 0.90650187 |
| Atg7 | 0.14979706 | 0.15971508 | 0.16727725 | -0.0075622 | 0.57964061 | 0.90650187 |
| Rptoros | 0.02185397 | 0.01886637 | 0.02643357 | -0.0075672 | 0.89842957 | 0.93440169 |
| Slc5a1 | 0.03621645 | 0.03582065 | 0.04338799 | -0.0075673 | 0.78093731 | 0.90650187 |

| | | | | | | |
|---|---|---|---|---|---|---|
| Snx19 | 0.08932694 | 0.0957988 | 0.10336981 | -0.007571 | 0.62763829 | 0.90650187 |
| Gm7008 | 0.02642536 | 0.02429283 | 0.03186526 | -0.0075724 | 0.85461862 | 0.91621818 |
| Vgf | 0.04154207 | 0.04201966 | 0.0495928 | -0.0075731 | 0.75139079 | 0.90650187 |
| Ncbp2 | 0.72437687 | 0.6021145 | 0.60968778 | -0.0075733 | 0.52201897 | 0.90650187 |
| Myo1b | 0.47111575 | 0.43718245 | 0.44475724 | -0.0075748 | 0.53017688 | 0.90650187 |
| Hddc2 | 0.27055066 | 0.27510755 | 0.28268252 | -0.007575 | 0.5474129 | 0.90650187 |
| Zfp661 | 0.0311709 | 0.02989079 | 0.03746886 | -0.0075781 | 0.81570645 | 0.9092537 |
| Smoc1 | 0.11627392 | 0.12478992 | 0.13237224 | -0.0075823 | 0.60050163 | 0.90650187 |
| Klhl23 | 0.11812583 | 0.12675314 | 0.13433592 | -0.0075828 | 0.59906929 | 0.90650187 |
| Myt1l | 0.10106008 | 0.10849667 | 0.1160796 | -0.0075829 | 0.61424892 | 0.90650187 |
| Dyrk1a | 0.18879129 | 0.1986522 | 0.2062389 | -0.0075867 | 0.56495178 | 0.90650187 |
| Etnk2 | 0.03136179 | 0.03010485 | 0.03769352 | -0.0075887 | 0.81461043 | 0.9092537 |
| Klhl11 | 0.0215147 | 0.01843766 | 0.02602912 | -0.0075915 | 0.90261103 | 0.93630341 |
| Tbl3 | 0.25974398 | 0.26536531 | 0.27296145 | -0.0075961 | 0.54922957 | 0.90650187 |
| Pik3r2 | 0.38250609 | 0.36956843 | 0.37717609 | -0.0076077 | 0.53572452 | 0.90650187 |
| Pgap3 | 0.04633865 | 0.04753499 | 0.05514392 | -0.0076089 | 0.73020031 | 0.90650187 |
| Bclaf1 | 1.13355333 | 0.79339831 | 0.80101361 | -0.0076153 | 0.51685618 | 0.90650187 |
| Tspyl3 | 0.05102773 | 0.05291428 | 0.06053679 | -0.0076225 | 0.71218174 | 0.90650187 |
| Zfp27 | 0.04995609 | 0.05168223 | 0.05930724 | -0.007625 | 0.71619082 | 0.90650187 |
| Clip4 | 0.06089178 | 0.06414752 | 0.07177347 | -0.007626 | 0.68166045 | 0.90650187 |
| Agt | 0.07477645 | 0.07971756 | 0.08734707 | -0.0076295 | 0.6511068 | 0.90650187 |
| Nt5c3b | 0.45989915 | 0.42888708 | 0.43652162 | -0.0076345 | 0.53098707 | 0.90650187 |
| Scn2b | 0.04080535 | 0.04110235 | 0.04873707 | -0.0076347 | 0.75691371 | 0.90650187 |
| Slc25a27 | 0.03876036 | 0.03872204 | 0.04635733 | -0.0076353 | 0.76798116 | 0.90650187 |
| Gm26843 | 0.02290557 | 0.02005035 | 0.02768601 | -0.0076357 | 0.8899148 | 0.9303978 |
| Dennd4a | 0.05814134 | 0.06099841 | 0.06865488 | -0.0076565 | 0.68997493 | 0.90650187 |
| Samd9l | 0.05297819 | 0.05510767 | 0.06277021 | -0.0076625 | 0.70632764 | 0.90650187 |
| Palm | 0.26305484 | 0.26828755 | 0.27595148 | -0.0076639 | 0.54913123 | 0.90650187 |
| Phtf2 | 0.03188901 | 0.03064248 | 0.03831373 | -0.0076713 | 0.81329603 | 0.9092537 |
| 5430403G16 | 0.08154954 | 0.08716932 | 0.09484286 | -0.0076735 | 0.64045862 | 0.90650187 |

| | | | | | | |
|---|---|---|---|---|---|---|
| Dmap1 | 0.3085448 | 0.30831035 | 0.31598698 | -0.0076766 | 0.54300302 | 0.90650187 |
| Smurf2 | 0.15255093 | 0.16240388 | 0.17008528 | -0.0076814 | 0.57956183 | 0.90650187 |
| Btc | 0.02120483 | 0.01797464 | 0.02565957 | -0.0076849 | 0.90847773 | 0.93899299 |
| 3110001I22R | 0.03704044 | 0.03666471 | 0.04435087 | -0.0076862 | 0.77948015 | 0.90650187 |
| Dusp4 | 0.03437534 | 0.03354565 | 0.04123271 | -0.0076871 | 0.79639107 | 0.9075779 |
| Pkd2 | 0.51571515 | 0.46895137 | 0.47664188 | -0.0076905 | 0.52859115 | 0.90650187 |
| Nap1l2 | 0.0565443 | 0.05914661 | 0.06683892 | -0.0076923 | 0.69554615 | 0.90650187 |
| Suco | 0.26346421 | 0.26862624 | 0.27632045 | -0.0076942 | 0.54925879 | 0.90650187 |
| Rev1 | 0.17759441 | 0.18752974 | 0.19522566 | -0.0076959 | 0.56955788 | 0.90650187 |
| H2-M3 | 0.05653658 | 0.05912574 | 0.06683014 | -0.0077044 | 0.69585079 | 0.90650187 |
| Kat2b | 0.11704052 | 0.125479 | 0.13318561 | -0.0077066 | 0.60150297 | 0.90650187 |
| Aldh3a2 | 0.16745562 | 0.17742613 | 0.18513292 | -0.0077068 | 0.57341041 | 0.90650187 |
| Klhl25 | 0.07546843 | 0.08039636 | 0.08811589 | -0.0077195 | 0.65153373 | 0.90650187 |
| Taf1c | 0.06209074 | 0.06540915 | 0.07312941 | -0.0077203 | 0.68058021 | 0.90650187 |
| Egf | 0.02095884 | 0.01763914 | 0.02536608 | -0.0077269 | 0.91216483 | 0.94062208 |
| Eif1b | 1.29715097 | 0.842335 | 0.8500664 | -0.0077314 | 0.51612605 | 0.90650187 |
| Ppp1r3f | 0.0401181 | 0.04020161 | 0.04793804 | -0.0077364 | 0.76345163 | 0.90650187 |
| Acox3 | 0.08241231 | 0.08804849 | 0.095793 | -0.0077445 | 0.64035608 | 0.90650187 |
| Cand2 | 0.10929943 | 0.11719431 | 0.12493893 | -0.0077446 | 0.60856118 | 0.90650187 |
| Trpm5 | 0.06577832 | 0.0695417 | 0.07728653 | -0.0077448 | 0.67201747 | 0.90650187 |
| A230072E10 | 0.0345737 | 0.03371798 | 0.04146517 | -0.0077472 | 0.79690009 | 0.9075779 |
| Gm16861 | 0.03378131 | 0.0327881 | 0.04053619 | -0.0077481 | 0.80226334 | 0.9086659 |
| Exoc8 | 0.11211315 | 0.12019601 | 0.12794499 | -0.007749 | 0.60613049 | 0.90650187 |
| Gm10643 | 0.02598736 | 0.02359613 | 0.03134626 | -0.0077501 | 0.86414727 | 0.92013888 |
| Fbxo4 | 0.12873087 | 0.13774941 | 0.14550011 | -0.0077507 | 0.59359965 | 0.90650187 |
| Kank1 | 0.12196125 | 0.13063742 | 0.13838952 | -0.0077521 | 0.59833019 | 0.90650187 |
| Ghdc | 0.1560826 | 0.1659185 | 0.17367366 | -0.0077552 | 0.57867787 | 0.90650187 |
| Rpl13 | 5.37550097 | 0.58148411 | 0.5892413 | -0.0077572 | 0.52333481 | 0.90650187 |
| Mrpl36 | 0.64161927 | 0.55209522 | 0.55985589 | -0.0077607 | 0.52456912 | 0.90650187 |
| Cd3eap | 0.31315065 | 0.31217236 | 0.31993505 | -0.0077627 | 0.54294868 | 0.90650187 |

| | | | | | | |
|---|---|---|---|---|---|---|
| Gfod1 | 0.03174681 | 0.03038297 | 0.0381465 | -0.0077635 | 0.81720486 | 0.9092537 |
| Acrbp | 0.04450688 | 0.04526423 | 0.05302815 | -0.0077639 | 0.74244036 | 0.90650187 |
| Gdpd5 | 0.09108124 | 0.0975169 | 0.10528143 | -0.0077645 | 0.62849292 | 0.90650187 |
| Srgap2 | 0.19610792 | 0.20559016 | 0.21336233 | -0.0077722 | 0.56432365 | 0.90650187 |
| Foxp2 | 0.06806177 | 0.07207444 | 0.07985073 | -0.0077763 | 0.66746968 | 0.90650187 |
| Churc1 | 1.1645346 | 0.80345479 | 0.81123287 | -0.0077781 | 0.5169995 | 0.90650187 |
| Slc25a19 | 0.09563209 | 0.10243927 | 0.11022041 | -0.0077811 | 0.62317977 | 0.90650187 |
| Pfn2 | 0.50086259 | 0.45838817 | 0.46617254 | -0.0077844 | 0.52958823 | 0.90650187 |
| Foxj1 | 0.02156188 | 0.01829633 | 0.02608537 | -0.007789 | 0.90782012 | 0.93850473 |
| Gm43672 | 0.03654117 | 0.03597696 | 0.04376758 | -0.0077906 | 0.78562042 | 0.90650187 |
| Zscan22 | 0.07548996 | 0.08033886 | 0.08813981 | -0.0078009 | 0.653012 | 0.90650187 |
| Prr14 | 0.16693536 | 0.17680749 | 0.18461189 | -0.0078044 | 0.57453672 | 0.90650187 |
| Cadps2 | 0.01906151 | 0.01529422 | 0.02309918 | -0.007805 | 0.93346726 | 0.95231059 |
| Tlcd1 | 0.14706883 | 0.15666875 | 0.16448633 | -0.0078176 | 0.58366857 | 0.90650187 |
| C030037D09 | 0.02316905 | 0.02017444 | 0.02799954 | -0.0078251 | 0.89296021 | 0.93179124 |
| Kcnb1 | 0.07764674 | 0.08270559 | 0.09053164 | -0.007826 | 0.64962278 | 0.90650187 |
| Thrb | 0.02828751 | 0.02623896 | 0.03406834 | -0.0078294 | 0.84652965 | 0.91408934 |
| Parn | 0.19285696 | 0.20237475 | 0.21020429 | -0.0078295 | 0.56575889 | 0.90650187 |
| Rnpc3 | 0.26791068 | 0.27247444 | 0.28031794 | -0.0078435 | 0.54949723 | 0.90650187 |
| Gm10273 | 0.02396872 | 0.02110627 | 0.02895043 | -0.0078442 | 0.88581021 | 0.92839449 |
| A530017D24 | 0.03910003 | 0.0389084 | 0.04675305 | -0.0078447 | 0.77214338 | 0.90650187 |
| Elac1 | 0.06317716 | 0.06650995 | 0.07435625 | -0.0078463 | 0.6805058 | 0.90650187 |
| Rrp1b | 0.20668703 | 0.21570965 | 0.22355954 | -0.0078499 | 0.56202267 | 0.90650187 |
| Immp2l | 0.10541998 | 0.1129249 | 0.12077817 | -0.0078533 | 0.61373351 | 0.90650187 |
| Cox18 | 0.29526294 | 0.29663832 | 0.30449612 | -0.0078578 | 0.54566759 | 0.90650187 |
| 4933404O12 | 0.05622894 | 0.05861596 | 0.06647988 | -0.0078639 | 0.70051121 | 0.90650187 |
| Camk1g | 0.05451233 | 0.056654 | 0.06452291 | -0.0078689 | 0.70614744 | 0.90650187 |
| Gmps | 0.70517299 | 0.5905634 | 0.59843909 | -0.0078757 | 0.52332716 | 0.90650187 |
| Fbxl15 | 0.39996747 | 0.38308571 | 0.39096522 | -0.0078795 | 0.53569615 | 0.90650187 |
| Fam109b | 0.07337443 | 0.07790593 | 0.08578723 | -0.0078813 | 0.65851689 | 0.90650187 |

| | | | | | | |
|---|---|---|---|---|---|---|
| Spidr | 0.09539504 | 0.10207981 | 0.10996386 | -0.0078841 | 0.6250365 | 0.90650187 |
| Nr1d2 | 0.04137683 | 0.0415147 | 0.04940094 | -0.0078862 | 0.76105054 | 0.90650187 |
| Agtrap | 0.44511225 | 0.41764157 | 0.42552919 | -0.0078876 | 0.53283723 | 0.90650187 |
| Drap1 | 1.17097369 | 0.80541943 | 0.81330737 | -0.0078879 | 0.51719552 | 0.90650187 |
| Shd | 0.04511916 | 0.0458435 | 0.05373593 | -0.0078924 | 0.74310449 | 0.90650187 |
| Zfp784 | 0.05158613 | 0.0532813 | 0.06117678 | -0.0078955 | 0.7169316 | 0.90650187 |
| Brca1 | 0.03624368 | 0.03552184 | 0.04341982 | -0.007898 | 0.79064081 | 0.90650187 |
| Ppil4 | 0.58491214 | 0.51565438 | 0.52355259 | -0.0078982 | 0.52673523 | 0.90650187 |
| Elmsan1 | 0.05402737 | 0.05606953 | 0.06396924 | -0.0078997 | 0.70849671 | 0.90650187 |
| Llgl1 | 0.14369191 | 0.1531182 | 0.16101973 | -0.0079015 | 0.58634539 | 0.90650187 |
| Fbxo17 | 0.08381167 | 0.08943014 | 0.0973318 | -0.0079017 | 0.64091528 | 0.90650187 |
| Csgalnact2 | 0.12488883 | 0.13356872 | 0.14147157 | -0.0079028 | 0.59806404 | 0.90650187 |
| Rap1gapos | 0.09425401 | 0.10082258 | 0.10872784 | -0.0079053 | 0.62673793 | 0.90650187 |
| Actr3b | 0.0356854 | 0.0348601 | 0.04276686 | -0.0079068 | 0.79443485 | 0.90731056 |
| Neil2 | 0.03097205 | 0.029326 | 0.03723477 | -0.0079088 | 0.82748139 | 0.90947782 |
| Dock5 | 0.02006161 | 0.01638248 | 0.02429481 | -0.0079123 | 0.92616973 | 0.94759147 |
| C330018D20 | 0.11259384 | 0.12054252 | 0.12845755 | -0.007915 | 0.60792696 | 0.90650187 |
| Ttll10 | 0.02444606 | 0.02159464 | 0.02951754 | -0.0079229 | 0.88361614 | 0.92767795 |
| Ufc1 | 1.13405889 | 0.7932649 | 0.80118897 | -0.0079241 | 0.51753536 | 0.90650187 |
| Ypel5 | 0.35494259 | 0.34698306 | 0.35491119 | -0.0079281 | 0.5395529 | 0.90650187 |
| Birc3 | 0.05697659 | 0.05940165 | 0.06733084 | -0.0079292 | 0.699699 | 0.90650187 |
| Ube2g1 | 0.76408949 | 0.62444844 | 0.63237769 | -0.0079292 | 0.52222652 | 0.90650187 |
| Kctd6 | 0.03309512 | 0.03179685 | 0.03973092 | -0.0079341 | 0.81266683 | 0.9092537 |
| Alg11 | 0.24652995 | 0.25298726 | 0.26092261 | -0.0079354 | 0.55377421 | 0.90650187 |
| Gm15972 | 0.02435074 | 0.0214634 | 0.02940433 | -0.0079409 | 0.88504296 | 0.92807528 |
| Apoo-ps | 0.0222077 | 0.01891312 | 0.02685505 | -0.0079419 | 0.90569699 | 0.93739361 |
| Arsk | 0.15199766 | 0.16157988 | 0.16952183 | -0.007942 | 0.58249204 | 0.90650187 |
| Mbd6 | 0.19707885 | 0.20636132 | 0.21430332 | -0.007942 | 0.56543062 | 0.90650187 |
| D10Wsu102e | 0.13566184 | 0.14477318 | 0.15272279 | -0.0079496 | 0.5915 | 0.90650187 |
| Lonrf2 | 0.03455923 | 0.03349467 | 0.04144822 | -0.0079536 | 0.8031906 | 0.90882608 |

| | | | | | | |
|---|---|---|---|---|---|---|
| Uevld | 0.04815275 | 0.04927899 | 0.05723425 | -0.0079553 | 0.73168455 | 0.90650187 |
| Cstb | 1.49499192 | 0.88451897 | 0.89247696 | -0.007958 | 0.51581006 | 0.90650187 |
| Acsl4 | 0.06331007 | 0.0665417 | 0.07450621 | -0.0079645 | 0.68268601 | 0.90650187 |
| Tmem56 | 0.02134583 | 0.01786115 | 0.02582774 | -0.0079666 | 0.91485154 | 0.94167513 |
| Exosc7 | 0.55562563 | 0.49609459 | 0.50406276 | -0.0079682 | 0.52801287 | 0.90650187 |
| Prmt2 | 0.59287597 | 0.5207954 | 0.52876388 | -0.0079685 | 0.52670732 | 0.90650187 |
| Dusp12 | 0.25875269 | 0.26409422 | 0.27206418 | -0.00797 | 0.55180831 | 0.90650187 |
| Ints6 | 0.03308293 | 0.03174637 | 0.03971662 | -0.0079702 | 0.81383966 | 0.9092537 |
| Mbtps1 | 0.71953087 | 0.59889184 | 0.60686695 | -0.0079751 | 0.52329364 | 0.90650187 |
| Pigv | 0.12627661 | 0.13495205 | 0.14292876 | -0.0079767 | 0.59797328 | 0.90650187 |
| Ext1 | 0.16566312 | 0.17535924 | 0.1833365 | -0.0079773 | 0.57669099 | 0.90650187 |
| B3galnt2 | 0.13380667 | 0.14281398 | 0.15079515 | -0.0079812 | 0.59300959 | 0.90650187 |
| Sult1c2 | 0.02442417 | 0.02150287 | 0.02949155 | -0.0079887 | 0.88574911 | 0.92839449 |
| Lamc2 | 0.03970491 | 0.03945692 | 0.0474573 | -0.0080004 | 0.77320473 | 0.90650187 |
| Tonsl | 1.19407203 | 0.81265471 | 0.8206552 | -0.0080005 | 0.51728467 | 0.90650187 |
| Zfp287 | 0.03932065 | 0.03900924 | 0.04700998 | -0.0080007 | 0.77535773 | 0.90650187 |
| Asnsd1 | 0.42513786 | 0.40242416 | 0.41042505 | -0.0080009 | 0.53453042 | 0.90650187 |
| Epb41l3 | 0.6508964 | 0.55761988 | 0.56562156 | -0.0080017 | 0.5250732 | 0.90650187 |
| Psph | 0.20337095 | 0.2123717 | 0.22037605 | -0.0080043 | 0.56413846 | 0.90650187 |
| Tdp2 | 0.36894131 | 0.35828817 | 0.36629719 | -0.008009 | 0.53871713 | 0.90650187 |
| Gm13205 | 0.02042975 | 0.01672458 | 0.02473451 | -0.0080099 | 0.92501655 | 0.94726331 |
| Kifc2 | 0.03401716 | 0.03279492 | 0.0408128 | -0.0080179 | 0.80876702 | 0.9092537 |
| Cdh4 | 0.03224096 | 0.03070655 | 0.03872751 | -0.008021 | 0.82140732 | 0.9092537 |
| Pptc7 | 0.11000903 | 0.11767588 | 0.12569796 | -0.0080221 | 0.61168752 | 0.90650187 |
| Abtb2 | 0.02922927 | 0.02715741 | 0.03518044 | -0.008023 | 0.84466981 | 0.91325304 |
| E130012A19 | 0.0197495 | 0.01589568 | 0.02392185 | -0.0080262 | 0.93209455 | 0.95146248 |
| Snx25 | 0.04773234 | 0.0487215 | 0.05675027 | -0.0080288 | 0.73530726 | 0.90650187 |
| Zfp202 | 0.0365621 | 0.03575813 | 0.04379203 | -0.0080339 | 0.79262758 | 0.90681274 |
| Zfp444 | 0.41817721 | 0.39705253 | 0.40509234 | -0.0080398 | 0.53515358 | 0.90650187 |
| Ccdc126 | 0.25518874 | 0.2607885 | 0.26883048 | -0.008042 | 0.55289894 | 0.90650187 |

| | | | | | | |
|---|---|---|---|---|---|---|
| Wrn | 0.12666978 | 0.13529861 | 0.14334117 | -0.0080426 | 0.59848718 | 0.90650187 |
| Tyro3 | 0.04770289 | 0.04867232 | 0.05671637 | -0.008044 | 0.73582195 | 0.90650187 |
| Zfp112 | 0.07971065 | 0.08476872 | 0.09281424 | -0.0080455 | 0.65001587 | 0.90650187 |
| Ubr4 | 0.43437942 | 0.40940301 | 0.41744935 | -0.0080463 | 0.53414324 | 0.90650187 |
| St14 | 0.15220079 | 0.16168164 | 0.16972874 | -0.0080471 | 0.58346782 | 0.90650187 |
| Lcorl | 0.30955517 | 0.3087993 | 0.31685465 | -0.0080554 | 0.54499268 | 0.90650187 |
| Fam171a2 | 0.05760398 | 0.05998697 | 0.06804429 | -0.0080573 | 0.70070078 | 0.90650187 |
| Cdhr5 | 0.02204072 | 0.01859843 | 0.02665612 | -0.0080577 | 0.91049842 | 0.93983499 |
| Pon3 | 0.15690074 | 0.16644288 | 0.1745029 | -0.00806 | 0.58134501 | 0.90650187 |
| Lcmt2 | 0.12931505 | 0.13804888 | 0.14611108 | -0.0080622 | 0.59688879 | 0.90650187 |
| Slc16a13 | 0.03803523 | 0.0374484 | 0.04551195 | -0.0080636 | 0.78454856 | 0.90650187 |
| Osbpl1a | 0.26840588 | 0.27269503 | 0.28076198 | -0.008067 | 0.55081974 | 0.90650187 |
| Cfap69 | 0.03958719 | 0.03924981 | 0.04732028 | -0.0080705 | 0.77583375 | 0.90650187 |
| Mrgpre | 0.03005754 | 0.02808517 | 0.03615738 | -0.0080722 | 0.83952087 | 0.91213511 |
| Lin7b | 0.0235294 | 0.02035465 | 0.02842816 | -0.0080735 | 0.89662234 | 0.93371722 |
| Cacna1b | 0.02506341 | 0.02217622 | 0.03025047 | -0.0080743 | 0.88230567 | 0.92700639 |
| 2610305D13 | 0.08109752 | 0.08626746 | 0.09434465 | -0.0080772 | 0.64825162 | 0.90650187 |
| Shroom3 | 0.09156031 | 0.09772497 | 0.10580272 | -0.0080777 | 0.63284954 | 0.90650187 |
| Gmeb2 | 0.07886048 | 0.0837968 | 0.09187472 | -0.0080779 | 0.65205413 | 0.90650187 |
| Brpf3 | 0.07347404 | 0.0778175 | 0.08589814 | -0.0080806 | 0.66210283 | 0.90650187 |
| Ehhadh | 0.03412423 | 0.03285293 | 0.04093834 | -0.0080854 | 0.81003269 | 0.9092537 |
| Dmpk | 0.10482995 | 0.11204603 | 0.1201437 | -0.0080977 | 0.61777155 | 0.90650187 |
| Tfeb | 0.06038999 | 0.06310139 | 0.07120535 | -0.008104 | 0.6935575 | 0.90650187 |
| Cct2 | 1.35661979 | 0.8565652 | 0.86467397 | -0.0081088 | 0.51662715 | 0.90650187 |
| Pvrl1 | 0.09877825 | 0.10550793 | 0.11361815 | -0.0081102 | 0.62450473 | 0.90650187 |
| Comtd1 | 0.16618288 | 0.17574577 | 0.18385777 | -0.008112 | 0.57775159 | 0.90650187 |
| Gm15234 | 0.03956051 | 0.03917237 | 0.04728923 | -0.0081169 | 0.77728534 | 0.90650187 |
| Ybx3 | 0.28175685 | 0.28452589 | 0.29264421 | -0.0081183 | 0.54907583 | 0.90650187 |
| Fgd4 | 0.1946276 | 0.20380654 | 0.21192571 | -0.0081192 | 0.56761975 | 0.90650187 |
| Apitd1 | 0.17765388 | 0.18716385 | 0.19528451 | -0.0081207 | 0.57333257 | 0.90650187 |

| | | | | | | |
|---|---|---|---|---|---|---|
| AI987944 | 0.13415535 | 0.14303521 | 0.15115776 | -0.0081225 | 0.59440341 | 0.90650187 |
| Mboat2 | 0.16311269 | 0.17264996 | 0.1807743 | -0.0081243 | 0.57917913 | 0.90650187 |
| Iqcg | 0.19707377 | 0.20617194 | 0.2142984 | -0.0081265 | 0.56693759 | 0.90650187 |
| Atg101 | 0.42010129 | 0.3984422 | 0.40657044 | -0.0081282 | 0.5354104 | 0.90650187 |
| Ccdc157 | 0.05124703 | 0.05265951 | 0.06078819 | -0.0081287 | 0.72390192 | 0.90650187 |
| Fig4 | 0.06713055 | 0.07067605 | 0.07880593 | -0.0081299 | 0.6767478 | 0.90650187 |
| Gca | 0.07688267 | 0.08154982 | 0.08968506 | -0.0081352 | 0.65662023 | 0.90650187 |
| Spsb1 | 0.12803478 | 0.136636 | 0.14477155 | -0.0081355 | 0.5986383 | 0.90650187 |
| Mettl4 | 0.03049146 | 0.02853165 | 0.03666874 | -0.0081371 | 0.83806012 | 0.91139496 |
| Neto2 | 0.04180563 | 0.04176003 | 0.04989874 | -0.0081387 | 0.76580407 | 0.90650187 |
| Yrdc | 0.33725108 | 0.33214833 | 0.34028831 | -0.00814 | 0.54234472 | 0.90650187 |
| Wwc1 | 0.0942616 | 0.10059476 | 0.10873606 | -0.0081413 | 0.63037698 | 0.90650187 |
| Slc39a8 | 0.03669969 | 0.03580659 | 0.04395282 | -0.0081462 | 0.79501604 | 0.90731056 |
| Enpp1 | 0.0359566 | 0.03493279 | 0.04308412 | -0.0081513 | 0.79985098 | 0.90805731 |
| Gtf2b | 0.65635348 | 0.56083202 | 0.56899167 | -0.0081597 | 0.52541633 | 0.90650187 |
| Unc79 | 0.04787365 | 0.04875147 | 0.05691298 | -0.0081615 | 0.73810516 | 0.90650187 |
| Vil1 | 0.05214884 | 0.05365601 | 0.06182125 | -0.0081652 | 0.72145447 | 0.90650187 |
| Prkdc | 0.06516603 | 0.06842836 | 0.07659766 | -0.0081693 | 0.6822975 | 0.90650187 |
| Fchsd1 | 0.03637626 | 0.03540463 | 0.04357482 | -0.0081702 | 0.7977305 | 0.9075779 |
| Adarb1 | 0.10968165 | 0.11717656 | 0.12534786 | -0.0081713 | 0.61401711 | 0.90650187 |
| Lsm12 | 0.57642191 | 0.50975865 | 0.51795597 | -0.0081973 | 0.52804543 | 0.90650187 |
| Fasn | 0.3485247 | 0.34143228 | 0.34963761 | -0.0082053 | 0.54154621 | 0.90650187 |
| Mrpl55 | 0.59434011 | 0.52151219 | 0.52971754 | -0.0082054 | 0.52745054 | 0.90650187 |
| Triobp | 0.55685292 | 0.49667797 | 0.50489005 | -0.0082121 | 0.52882166 | 0.90650187 |
| Fam120a | 0.56257328 | 0.50052047 | 0.50873398 | -0.0082135 | 0.52860925 | 0.90650187 |
| Ttc38 | 0.07260182 | 0.0767104 | 0.08492643 | -0.008216 | 0.66643157 | 0.90650187 |
| Eml2 | 0.34364314 | 0.33738443 | 0.34560286 | -0.0082184 | 0.54209631 | 0.90650187 |
| Daam1 | 0.43625251 | 0.41064633 | 0.41886547 | -0.0082191 | 0.53475704 | 0.90650187 |
| Gm5093 | 0.10770745 | 0.11501296 | 0.12323378 | -0.0082208 | 0.61659972 | 0.90650187 |
| Zfp239 | 0.04426012 | 0.04451299 | 0.05274274 | -0.0082298 | 0.75605662 | 0.90650187 |

| | | | | | | |
|---|---|---|---|---|---|---|
| Pde1a | 0.09515967 | 0.10147822 | 0.10970904 | -0.0082308 | 0.63063202 | 0.90650187 |
| Cacna2d2 | 0.02479101 | 0.02169466 | 0.02992715 | -0.0082325 | 0.88931941 | 0.9302877 |
| Clk3 | 0.36392104 | 0.35399413 | 0.36223216 | -0.008238 | 0.54026596 | 0.90650187 |
| Wdr5b | 0.0619903 | 0.06476945 | 0.0730159 | -0.0082464 | 0.69219745 | 0.90650187 |
| 9130023H24 | 0.05575046 | 0.05768807 | 0.06593484 | -0.0082468 | 0.71090561 | 0.90650187 |
| Gm16536 | 0.02185748 | 0.01818436 | 0.02643775 | -0.0082534 | 0.91740695 | 0.94283706 |
| Ncapd3 | 0.18168753 | 0.19101367 | 0.1992675 | -0.0082538 | 0.57304853 | 0.90650187 |
| Metap1d | 0.14806921 | 0.15725214 | 0.16551074 | -0.0082586 | 0.58777509 | 0.90650187 |
| Map3k3 | 0.04618289 | 0.04670468 | 0.05496422 | -0.0082595 | 0.74794794 | 0.90650187 |
| Zfp114 | 0.04139557 | 0.0411566 | 0.04942271 | -0.0082661 | 0.77142316 | 0.90650187 |
| Ubiad1 | 0.04736837 | 0.04806209 | 0.05633105 | -0.008269 | 0.74298661 | 0.90650187 |
| Zgrf1 | 0.04667273 | 0.04725756 | 0.05552926 | -0.0082717 | 0.74608548 | 0.90650187 |
| Gm17491 | 0.05141943 | 0.05271166 | 0.06098577 | -0.0082741 | 0.72679496 | 0.90650187 |
| Dlgap3 | 0.02496108 | 0.02184286 | 0.03012902 | -0.0082862 | 0.88926752 | 0.9302877 |
| Nupr1l | 0.0556404 | 0.057522 | 0.06580943 | -0.0082874 | 0.71220523 | 0.90650187 |
| Filip1 | 0.17158971 | 0.18096679 | 0.18926233 | -0.0082955 | 0.57724678 | 0.90650187 |
| Adam23 | 0.04186719 | 0.04167325 | 0.04997018 | -0.0082969 | 0.76978473 | 0.90650187 |
| Cnot2 | 0.50922697 | 0.46378743 | 0.4720861 | -0.0082987 | 0.53114487 | 0.90650187 |
| B130006D01 | 0.05801791 | 0.06021376 | 0.06851467 | -0.0083009 | 0.70492143 | 0.90650187 |
| Gabrb3 | 0.06615761 | 0.06941016 | 0.07771298 | -0.0083028 | 0.68259447 | 0.90650187 |
| Tmem194 | 0.05197094 | 0.05331465 | 0.06161756 | -0.0083029 | 0.72542728 | 0.90650187 |
| Cdk5r2 | 0.04099341 | 0.04064645 | 0.04895559 | -0.0083091 | 0.77473883 | 0.90650187 |
| Zfp169 | 0.01980095 | 0.01567208 | 0.02398334 | -0.0083113 | 0.93829227 | 0.9555402 |
| Tbcd | 0.2146357 | 0.22283151 | 0.23114371 | -0.0083122 | 0.56350799 | 0.90650187 |
| Atl1 | 0.03972974 | 0.03916735 | 0.0474862 | -0.0083188 | 0.78195158 | 0.90650187 |
| Npr2 | 0.05874008 | 0.06101196 | 0.06933472 | -0.0083228 | 0.70320444 | 0.90650187 |
| Tex26 | 0.03485976 | 0.03347414 | 0.0418003 | -0.0083262 | 0.81206088 | 0.9092537 |
| Tmem25 | 0.03075248 | 0.02864121 | 0.0369762 | -0.008335 | 0.84185392 | 0.91251845 |
| Onecut2 | 0.07731992 | 0.08182601 | 0.09016963 | -0.0083436 | 0.65959672 | 0.90650187 |
| Zfp85 | 0.0416995 | 0.04143108 | 0.04977555 | -0.0083445 | 0.77194578 | 0.90650187 |

| | | | | | | |
|---|---|---|---|---|---|---|
| Cstf3 | 0.52249304 | 0.47302356 | 0.4813708 | -0.0083472 | 0.53072377 | 0.90650187 |
| Afg3l1 | 0.57737313 | 0.51023761 | 0.51858511 | -0.0083475 | 0.52852376 | 0.90650187 |
| Zfand2b | 0.35732905 | 0.34850474 | 0.3568635 | -0.0083588 | 0.54146642 | 0.90650187 |
| Hspb6 | 0.14702795 | 0.15608234 | 0.16444445 | -0.0083621 | 0.5894234 | 0.90650187 |
| Gm4723 | 0.03949727 | 0.03885263 | 0.04721561 | -0.008363 | 0.78448188 | 0.90650187 |
| Stard13 | 0.11448612 | 0.12210735 | 0.13047254 | -0.0083652 | 0.61218871 | 0.90650187 |
| Plekha3 | 0.06822292 | 0.07166512 | 0.0800314 | -0.0083663 | 0.67893852 | 0.90650187 |
| 2900009J06F | 0.03060416 | 0.02843221 | 0.03680151 | -0.0083693 | 0.84399366 | 0.9130051 |
| Rhou | 0.13201174 | 0.1405548 | 0.1489262 | -0.0083714 | 0.59866582 | 0.90650187 |
| Zfp712 | 0.02399462 | 0.02060906 | 0.02898121 | -0.0083722 | 0.90047388 | 0.93504361 |
| Rdh11 | 0.15200458 | 0.16115566 | 0.16952888 | -0.0083732 | 0.58689836 | 0.90650187 |
| Gm43740 | 0.04673731 | 0.04722496 | 0.05560371 | -0.0083788 | 0.74853763 | 0.90650187 |
| Cgref1 | 0.04013476 | 0.0395784 | 0.04795742 | -0.008379 | 0.78134013 | 0.90650187 |
| Zfp597 | 0.07630784 | 0.08066456 | 0.0890476 | -0.008383 | 0.66221445 | 0.90650187 |
| Tsen15 | 0.49966167 | 0.45693585 | 0.46531973 | -0.0083839 | 0.53192056 | 0.90650187 |
| Prep | 0.13341553 | 0.14200082 | 0.15038821 | -0.0083874 | 0.59790897 | 0.90650187 |
| Aaed1 | 0.13324311 | 0.14181938 | 0.15020876 | -0.0083894 | 0.59804634 | 0.90650187 |
| Ccl25 | 0.14015669 | 0.14898244 | 0.15737636 | -0.0083939 | 0.59371563 | 0.90650187 |
| Krt10 | 0.40458587 | 0.3861737 | 0.39457189 | -0.0083982 | 0.53769232 | 0.90650187 |
| Mms22l | 0.07113796 | 0.07489017 | 0.08329312 | -0.008403 | 0.67310061 | 0.90650187 |
| Bco1 | 0.05282253 | 0.05418255 | 0.06259218 | -0.0084096 | 0.72484429 | 0.90650187 |
| Pusl1 | 0.20429266 | 0.21285168 | 0.22126206 | -0.0084104 | 0.56709446 | 0.90650187 |
| Mink1 | 0.15032552 | 0.15939767 | 0.16781678 | -0.0084191 | 0.58824363 | 0.90650187 |
| Tbc1d32 | 0.05461268 | 0.05621647 | 0.06463742 | -0.008421 | 0.71876376 | 0.90650187 |
| Nmnat1 | 0.05454189 | 0.05613219 | 0.06455664 | -0.0084245 | 0.71908939 | 0.90650187 |
| Gm17354 | 0.05922307 | 0.06145685 | 0.06988274 | -0.0084259 | 0.70402593 | 0.90650187 |
| Gm15446 | 0.09857415 | 0.10496575 | 0.11339814 | -0.0084324 | 0.62951894 | 0.90650187 |
| Trmt11 | 0.12621588 | 0.13443117 | 0.14286505 | -0.0084339 | 0.6035094 | 0.90650187 |
| Spire2 | 0.04798322 | 0.04860477 | 0.05703912 | -0.0084343 | 0.74452837 | 0.90650187 |
| Heatr1 | 0.02241379 | 0.0186656 | 0.02710053 | -0.0084349 | 0.91677129 | 0.94250239 |

| | | | | | | |
|---|---|---|---|---|---|---|
| Tspan1 | 0.03099443 | 0.02882191 | 0.03726112 | -0.0084392 | 0.84300313 | 0.91273395 |
| Stc2 | 0.04655034 | 0.0469407 | 0.05538811 | -0.0084474 | 0.75111833 | 0.90650187 |
| Ccdc64b | 0.02688912 | 0.02396603 | 0.03241443 | -0.0084484 | 0.87670858 | 0.92473192 |
| Zdhhc18 | 0.04107203 | 0.04059456 | 0.04904692 | -0.0084524 | 0.77819771 | 0.90650187 |
| Pde4d | 0.16374258 | 0.172954 | 0.18140778 | -0.0084538 | 0.58206128 | 0.90650187 |
| Gm26909 | 0.06269055 | 0.0653498 | 0.07380696 | -0.0084572 | 0.69476379 | 0.90650187 |
| Nptx2 | 0.02551932 | 0.02233368 | 0.03079135 | -0.0084577 | 0.88897677 | 0.9302877 |
| Zfyve1 | 0.03483838 | 0.03331459 | 0.04177526 | -0.0084607 | 0.81603587 | 0.9092537 |
| Zfyve26 | 0.03381782 | 0.032117 | 0.04057902 | -0.008462 | 0.82304987 | 0.9092537 |
| Arfgef3 | 0.0936485 | 0.09960182 | 0.10807118 | -0.0084694 | 0.63622814 | 0.90650187 |
| Zfp14 | 0.03137209 | 0.02922899 | 0.03770565 | -0.0084767 | 0.84120492 | 0.91251845 |
| Gm12976 | 0.02727728 | 0.02439617 | 0.03287383 | -0.0084777 | 0.87419613 | 0.9235672 |
| Pcsk6 | 0.0863039 | 0.09158748 | 0.10006554 | -0.0084781 | 0.64678486 | 0.90650187 |
| 4930427A07I | 0.02829065 | 0.02559387 | 0.03407205 | -0.0084782 | 0.86568091 | 0.92040222 |
| 3110056K07I | 0.09286098 | 0.0987344 | 0.10721638 | -0.008482 | 0.63746678 | 0.90650187 |
| Bcar3 | 0.03857891 | 0.03766236 | 0.04614587 | -0.0084835 | 0.79311728 | 0.9070651 |
| Slc25a25 | 0.17632082 | 0.18547105 | 0.19396427 | -0.0084932 | 0.5771713 | 0.90650187 |
| Itpka | 0.03268673 | 0.03075695 | 0.03925132 | -0.0084944 | 0.8319935 | 0.91015207 |
| Psd3 | 0.09703298 | 0.10323408 | 0.11173501 | -0.0085009 | 0.63240395 | 0.90650187 |
| Jakmip1 | 0.09524307 | 0.1012942 | 0.10979934 | -0.0085051 | 0.63471231 | 0.90650187 |
| Cdc45 | 0.06166683 | 0.06413425 | 0.07265024 | -0.008516 | 0.69885474 | 0.90650187 |
| Pvr | 0.05535003 | 0.05696241 | 0.06547846 | -0.008516 | 0.718432 | 0.90650187 |
| Csnk2a2 | 0.05686173 | 0.05867827 | 0.06720016 | -0.0085219 | 0.71354236 | 0.90650187 |
| Rspry1 | 0.32597364 | 0.32230575 | 0.3308277 | -0.008522 | 0.54558584 | 0.90650187 |
| Tor1b | 0.45917902 | 0.42746534 | 0.43598984 | -0.0085245 | 0.53463276 | 0.90650187 |
| Tcf7l1 | 0.10970919 | 0.11683847 | 0.12537732 | -0.0085388 | 0.61896693 | 0.90650187 |
| Sgpp2 | 0.07930948 | 0.0838302 | 0.09237104 | -0.0085408 | 0.65948248 | 0.90650187 |
| Senp1 | 0.15049039 | 0.15944327 | 0.16798503 | -0.0085418 | 0.58941943 | 0.90650187 |
| Ttc25 | 0.04385327 | 0.04372741 | 0.05227197 | -0.0085446 | 0.76629324 | 0.90650187 |
| Pars2 | 0.02709764 | 0.02410619 | 0.03266125 | -0.0085551 | 0.87787933 | 0.92484883 |

| | | | | | | |
|---|---|---|---|---|---|---|
| Rab11fip1 | 0.04986135 | 0.0506422 | 0.05919846 | -0.0085563 | 0.73974089 | 0.90650187 |
| Mcm9 | 0.03567995 | 0.03420009 | 0.04276049 | -0.0085604 | 0.81326267 | 0.9092537 |
| Hunk | 0.05410173 | 0.05548019 | 0.06405415 | -0.008574 | 0.72410051 | 0.90650187 |
| Mrm1 | 0.09027802 | 0.09582818 | 0.10440671 | -0.0085785 | 0.64254554 | 0.90650187 |
| Sult2b1 | 0.07250401 | 0.07623865 | 0.08481739 | -0.0085787 | 0.67351671 | 0.90650187 |
| Zfp275 | 0.13082063 | 0.13910275 | 0.14768385 | -0.0085811 | 0.60191607 | 0.90650187 |
| Per2 | 0.0765916 | 0.08078073 | 0.08936233 | -0.0085816 | 0.66528057 | 0.90650187 |
| Stap2 | 0.06976222 | 0.07317144 | 0.08175531 | -0.0085839 | 0.67966595 | 0.90650187 |
| Rictor | 0.11440791 | 0.12178793 | 0.13038936 | -0.0086014 | 0.61534003 | 0.90650187 |
| Arl14ep | 0.23965143 | 0.24598418 | 0.25458671 | -0.0086025 | 0.55970367 | 0.90650187 |
| Agpat3 | 0.36212435 | 0.35216926 | 0.36077239 | -0.0086031 | 0.54221355 | 0.90650187 |
| Med13l | 0.23316293 | 0.23996381 | 0.24856722 | -0.0086034 | 0.56114445 | 0.90650187 |
| Akap7 | 0.06469887 | 0.06746649 | 0.0760717 | -0.0086052 | 0.69247766 | 0.90650187 |
| Sbno1 | 0.83895182 | 0.66455371 | 0.6731592 | -0.0086055 | 0.52266027 | 0.90650187 |
| Sacs | 0.02566486 | 0.02235743 | 0.03096394 | -0.0086065 | 0.89170893 | 0.93134318 |
| Unc13b | 0.14537874 | 0.15413861 | 0.16275301 | -0.0086144 | 0.59301316 | 0.90650187 |
| Tex30 | 0.2514778 | 0.25683337 | 0.2654504 | -0.008617 | 0.55737333 | 0.90650187 |
| Zscan20 | 0.03785001 | 0.0366732 | 0.04529589 | -0.0086227 | 0.80130218 | 0.90823524 |
| Usp49 | 0.03431525 | 0.032537 | 0.04116228 | -0.0086253 | 0.82421481 | 0.90929265 |
| Arhgap26 | 0.07779915 | 0.08207248 | 0.0907004 | -0.0086279 | 0.66381247 | 0.90650187 |
| Gm12258 | 0.07749922 | 0.08171997 | 0.09036825 | -0.0086483 | 0.66474441 | 0.90650187 |
| Ror2 | 0.23089711 | 0.23780305 | 0.24645529 | -0.0086522 | 0.56201141 | 0.90650187 |
| Acadl | 0.57836961 | 0.51058975 | 0.5192436 | -0.0086538 | 0.52953125 | 0.90650187 |
| Fmr1 | 0.42484447 | 0.40153329 | 0.41020102 | -0.0086677 | 0.53742066 | 0.90650187 |
| Tab2 | 0.45994802 | 0.4278878 | 0.43655769 | -0.0086699 | 0.53517623 | 0.90650187 |
| Gm4425 | 0.02310211 | 0.01924775 | 0.02791989 | -0.0086721 | 0.91633604 | 0.94231902 |
| Nfatc2 | 0.13992949 | 0.14846909 | 0.15714171 | -0.0086726 | 0.59690811 | 0.90650187 |
| Col4a3bp | 0.3165282 | 0.31414491 | 0.32281842 | -0.0086735 | 0.54753854 | 0.90650187 |
| Ptgr2 | 0.36791079 | 0.35678663 | 0.3654644 | -0.0086778 | 0.54203374 | 0.90650187 |
| Enkd1 | 0.09071009 | 0.0961946 | 0.10487736 | -0.0086828 | 0.64358201 | 0.90650187 |

| | | | | | | |
|---|---|---|---|---|---|---|
| Adat3 | 0.03229739 | 0.03010991 | 0.03879384 | -0.0086839 | 0.84016608 | 0.91226452 |
| Hgfac | 0.02219114 | 0.01814976 | 0.02683533 | -0.0086856 | 0.92490659 | 0.94721452 |
| Bphl | 0.46593025 | 0.43227341 | 0.44096104 | -0.0086876 | 0.53489693 | 0.90650187 |
| Adgb | 0.02883256 | 0.02602272 | 0.03471215 | -0.0086894 | 0.86711567 | 0.92056229 |
| Nol10 | 0.093064 | 0.09874348 | 0.10743683 | -0.0086934 | 0.64047192 | 0.90650187 |
| Bivm | 0.08975258 | 0.09513008 | 0.10383401 | -0.0087039 | 0.6452974 | 0.90650187 |
| Fgd6 | 0.12176152 | 0.12945228 | 0.13817887 | -0.0087266 | 0.61055236 | 0.90650187 |
| Ankrd6 | 0.04513627 | 0.04502506 | 0.0537557 | -0.0087306 | 0.76486054 | 0.90650187 |
| Mettl20 | 0.06326987 | 0.06572547 | 0.07446086 | -0.0087354 | 0.69900365 | 0.90650187 |
| Zbtb16 | 0.05706005 | 0.058688 | 0.06742579 | -0.0087378 | 0.71773006 | 0.90650187 |
| Rad9b | 0.06224131 | 0.06455611 | 0.07329954 | -0.0087434 | 0.70204383 | 0.90650187 |
| Slx4 | 0.02815065 | 0.02516263 | 0.0339066 | -0.008744 | 0.87419578 | 0.9235672 |
| Slc9a7 | 0.04380887 | 0.04347178 | 0.05222057 | -0.0087488 | 0.77180247 | 0.90650187 |
| Acbd3 | 0.50477791 | 0.46019717 | 0.46894633 | -0.0087492 | 0.53305115 | 0.90650187 |
| Mzf1 | 0.04204296 | 0.04141927 | 0.05017413 | -0.0087549 | 0.78104044 | 0.90650187 |
| Garem | 0.02996002 | 0.0272837 | 0.03604241 | -0.0087587 | 0.86000286 | 0.91810545 |
| Zfp747 | 0.05180274 | 0.0526607 | 0.06142492 | -0.0087642 | 0.73705976 | 0.90650187 |
| Pim3 | 0.35769459 | 0.34839275 | 0.35716213 | -0.0087694 | 0.54345934 | 0.90650187 |
| Skiv2l2 | 0.28865251 | 0.28994521 | 0.29871609 | -0.0087709 | 0.55192706 | 0.90650187 |
| Tmem2 | 0.10675385 | 0.11343927 | 0.12221088 | -0.0087716 | 0.62516728 | 0.90650187 |
| Tex9 | 0.18552498 | 0.19426568 | 0.20304024 | -0.0087746 | 0.57617622 | 0.90650187 |
| Helq | 0.05447315 | 0.05569826 | 0.06447819 | -0.0087799 | 0.72752807 | 0.90650187 |
| Mbd3 | 0.93405785 | 0.7124946 | 0.72127532 | -0.0087807 | 0.52158033 | 0.90650187 |
| Zbtb2 | 0.08121857 | 0.08569699 | 0.0944781 | -0.0087811 | 0.66026612 | 0.90650187 |
| Kif13a | 0.0857707 | 0.09068584 | 0.09948142 | -0.0087956 | 0.65286063 | 0.90650187 |
| Slx1b | 0.16562787 | 0.1744867 | 0.18330114 | -0.0088144 | 0.58463944 | 0.90650187 |
| Wnt7b | 0.0317086 | 0.02928366 | 0.03810155 | -0.0088179 | 0.84822544 | 0.91440801 |
| Gm11992 | 0.03270211 | 0.03045037 | 0.03926938 | -0.008819 | 0.84095298 | 0.91251845 |
| Cep290 | 0.24402043 | 0.24979281 | 0.25861645 | -0.0088236 | 0.56027954 | 0.90650187 |
| Kcnh2 | 0.21705461 | 0.22461272 | 0.23343873 | -0.008826 | 0.56674077 | 0.90650187 |

| | | | | | | |
|---|---|---|---|---|---|---|
| B930041F14 | 0.02712545 | 0.02386627 | 0.03269417 | -0.0088279 | 0.88500358 | 0.92807528 |
| 6430503K07I | 0.06440636 | 0.06691195 | 0.07574221 | -0.0088303 | 0.69786941 | 0.90650187 |
| Tmem154 | 0.01959748 | 0.01490756 | 0.02374014 | -0.0088326 | 0.95093614 | 0.96348599 |
| Ccdc60 | 0.02552163 | 0.02196103 | 0.03079408 | -0.008833 | 0.89887873 | 0.93440169 |
| Dnajc22 | 0.02249352 | 0.01836226 | 0.02719548 | -0.0088332 | 0.9256241 | 0.94746893 |
| Rfx1 | 0.05709177 | 0.05862803 | 0.06746186 | -0.0088338 | 0.7197614 | 0.90650187 |
| Gtf2ird2 | 0.09523735 | 0.10095748 | 0.10979315 | -0.0088357 | 0.63973948 | 0.90650187 |
| Rassf7 | 0.2519898 | 0.25707565 | 0.26591754 | -0.0088419 | 0.55875701 | 0.90650187 |
| Cys1 | 0.05207935 | 0.05289873 | 0.06174169 | -0.008843 | 0.73784701 | 0.90650187 |
| Atp8b2 | 0.04536037 | 0.045169 | 0.05401461 | -0.0088456 | 0.76670157 | 0.90650187 |
| Hipk2 | 0.34360156 | 0.33672188 | 0.34556839 | -0.0088465 | 0.54530462 | 0.90650187 |
| Zfp653 | 0.03740073 | 0.03592194 | 0.04477156 | -0.0088496 | 0.8102252 | 0.9092537 |
| Ccp110 | 0.12672935 | 0.13454928 | 0.14340365 | -0.0088544 | 0.60814641 | 0.90650187 |
| Zfp329 | 0.37378396 | 0.36134161 | 0.37019815 | -0.0088565 | 0.54234996 | 0.90650187 |
| 1700040L02I | 0.02140018 | 0.0170278 | 0.02589257 | -0.0088648 | 0.93599926 | 0.953802 |
| Parp9 | 0.0818616 | 0.08631458 | 0.09518665 | -0.0088721 | 0.66069535 | 0.90650187 |
| Zscan18 | 0.03367305 | 0.03153522 | 0.04040919 | -0.008874 | 0.8355341 | 0.91078214 |
| Dctd | 0.13169913 | 0.1397263 | 0.14860031 | -0.008874 | 0.60467998 | 0.90650187 |
| Cryzl1 | 0.69182207 | 0.58163186 | 0.59050994 | -0.0088781 | 0.52664461 | 0.90650187 |
| Zfp608 | 0.13683372 | 0.14505892 | 0.15393835 | -0.0088794 | 0.60119076 | 0.90650187 |
| Zfp697 | 0.03745769 | 0.03595377 | 0.04483806 | -0.0088843 | 0.8108032 | 0.9092537 |
| Dnaja1 | 2.67776846 | 0.87141703 | 0.88030301 | -0.008886 | 0.51789653 | 0.90650187 |
| Gemin5 | 0.07545445 | 0.07921419 | 0.08810037 | -0.0088862 | 0.6730688 | 0.90650187 |
| Spag6l | 0.02137678 | 0.01697708 | 0.02586465 | -0.0088876 | 0.93669323 | 0.9542332 |
| Zc2hc1c | 0.09199719 | 0.09738887 | 0.10627781 | -0.0088889 | 0.64498841 | 0.90650187 |
| Ankrd26 | 0.21442701 | 0.22205605 | 0.23094543 | -0.0088894 | 0.56793408 | 0.90650187 |
| Gm2415 | 0.04408533 | 0.04364624 | 0.05254052 | -0.0088943 | 0.77414603 | 0.90650187 |
| Synrg | 0.39966258 | 0.38182783 | 0.39072654 | -0.0088987 | 0.54032305 | 0.90650187 |
| Szt2 | 0.05013104 | 0.05060712 | 0.05950809 | -0.008901 | 0.74696023 | 0.90650187 |
| Slc8a1 | 0.03439461 | 0.03234845 | 0.0412553 | -0.0089069 | 0.83141916 | 0.9098922 |

| | | | | | | |
|---|---|---|---|---|---|---|
| Nap1l3 | 0.08617527 | 0.09101759 | 0.09992466 | -0.0089071 | 0.65404781 | 0.90650187 |
| Rhpn2 | 0.12935847 | 0.13724206 | 0.14615648 | -0.0089144 | 0.60686129 | 0.90650187 |
| D17Wsu92e | 0.18721533 | 0.19578222 | 0.20469702 | -0.0089148 | 0.57675996 | 0.90650187 |
| Btbd8 | 0.04061916 | 0.03960323 | 0.04852066 | -0.0089174 | 0.79304757 | 0.9070651 |
| Ropn1l | 0.12624769 | 0.13397921 | 0.14289842 | -0.0089192 | 0.60929379 | 0.90650187 |
| Ptgfrn | 0.13401653 | 0.14209367 | 0.15101341 | -0.0089197 | 0.60356414 | 0.90650187 |
| Ammecr1 | 0.16389328 | 0.17263944 | 0.18155928 | -0.0089198 | 0.58644455 | 0.90650187 |
| Adam22 | 0.09470196 | 0.10029017 | 0.10921329 | -0.0089231 | 0.64178058 | 0.90650187 |
| Pigw | 0.06006348 | 0.06191169 | 0.07083549 | -0.0089238 | 0.71227865 | 0.90650187 |
| Nt5dc3 | 0.04887817 | 0.04914127 | 0.05806874 | -0.0089275 | 0.75284378 | 0.90650187 |
| 1600014C10I | 0.10372637 | 0.11002674 | 0.11895583 | -0.0089291 | 0.63069604 | 0.90650187 |
| Zfp959 | 0.05784147 | 0.0593812 | 0.0683142 | -0.008933 | 0.71948862 | 0.90650187 |
| Nufip2 | 0.32755849 | 0.32322821 | 0.33216378 | -0.0089356 | 0.54759668 | 0.90650187 |
| Ptpn2 | 0.31998485 | 0.31682084 | 0.32575903 | -0.0089382 | 0.54854204 | 0.90650187 |
| Gm26610 | 0.02779207 | 0.02454252 | 0.03348272 | -0.0089402 | 0.88238998 | 0.9270309 |
| Nptxr | 0.08553042 | 0.09027453 | 0.09921805 | -0.0089435 | 0.65568947 | 0.90650187 |
| BC067074 | 0.03331482 | 0.03104356 | 0.03998883 | -0.0089453 | 0.83999945 | 0.91225489 |
| 2210016L21F | 0.65659641 | 0.56019505 | 0.56914134 | -0.0089463 | 0.52785546 | 0.90650187 |
| Ttf2 | 0.03092802 | 0.02823272 | 0.03718292 | -0.0089502 | 0.85771054 | 0.91722806 |
| Zfp68 | 0.38237564 | 0.36812162 | 0.37707216 | -0.0089505 | 0.54202046 | 0.90650187 |
| Fcho1 | 0.02368259 | 0.01965945 | 0.0286103 | -0.0089509 | 0.91786104 | 0.94298498 |
| Gmeb1 | 0.16776299 | 0.17648853 | 0.18544059 | -0.0089521 | 0.58496421 | 0.90650187 |
| Lpin1 | 0.06632339 | 0.06894687 | 0.07789931 | -0.0089524 | 0.69529053 | 0.90650187 |
| Arnt2 | 0.02346992 | 0.01939573 | 0.02835742 | -0.0089617 | 0.91997876 | 0.94435798 |
| Slc39a14 | 0.06044414 | 0.06230489 | 0.07126668 | -0.0089618 | 0.71193125 | 0.90650187 |
| Tha1 | 0.04057382 | 0.03950169 | 0.04846796 | -0.0089663 | 0.79457811 | 0.90731056 |
| Zfp654 | 0.07676085 | 0.08057954 | 0.08955 | -0.0089705 | 0.67195878 | 0.90650187 |
| Lrrc73 | 0.22399225 | 0.23101306 | 0.23998752 | -0.0089745 | 0.56601825 | 0.90650187 |
| Kdm8 | 0.03553068 | 0.03360996 | 0.04258581 | -0.0089759 | 0.82562198 | 0.90947782 |
| Gm5113 | 0.03158243 | 0.02897627 | 0.03795314 | -0.0089769 | 0.85348493 | 0.91590713 |

| | | | | | | |
|---|---|---|---|---|---|---|
| Kdm5a | 0.57377798 | 0.50720945 | 0.51620448 | -0.008995 | 0.53087363 | 0.90650187 |
| Zfp677 | 0.02940678 | 0.02638941 | 0.0353899 | -0.0090005 | 0.87089317 | 0.92221854 |
| Chst9 | 0.0327304 | 0.03029828 | 0.03930261 | -0.0090043 | 0.84577989 | 0.9136218 |
| Exosc9 | 0.28045678 | 0.28248829 | 0.29149455 | -0.0090063 | 0.55462496 | 0.90650187 |
| 2310068J16F | 0.0389916 | 0.03761646 | 0.04662675 | -0.0090103 | 0.80485234 | 0.90882608 |
| Accs | 0.05903267 | 0.06065414 | 0.06966675 | -0.0090126 | 0.71739379 | 0.90650187 |
| Gm10138 | 0.1017976 | 0.10786214 | 0.11687609 | -0.0090139 | 0.63414915 | 0.90650187 |
| Tpbg | 0.18278186 | 0.19132769 | 0.20034501 | -0.0090173 | 0.57929571 | 0.90650187 |
| Cep135 | 0.07619183 | 0.07989793 | 0.0889189 | -0.009021 | 0.67401024 | 0.90650187 |
| Rsph3b | 0.03480955 | 0.03271262 | 0.0417415 | -0.0090289 | 0.83187603 | 0.91013901 |
| Gdf11 | 0.05777546 | 0.05920692 | 0.06823919 | -0.0090323 | 0.7218777 | 0.90650187 |
| Ate1 | 0.40176796 | 0.38333413 | 0.39237322 | -0.0090391 | 0.5407857 | 0.90650187 |
| Ankrd28 | 0.07143914 | 0.07458936 | 0.08362942 | -0.0090401 | 0.68458365 | 0.90650187 |
| Kdm5d | 0.06398639 | 0.06622607 | 0.07526893 | -0.0090429 | 0.70336316 | 0.90650187 |
| Pole | 0.02784443 | 0.02449342 | 0.03354463 | -0.0090512 | 0.88484107 | 0.92806659 |
| Ppp3cc | 0.04804048 | 0.04805317 | 0.05710504 | -0.0090519 | 0.7594969 | 0.90650187 |
| Tmem161b | 0.10988901 | 0.11651536 | 0.12556963 | -0.0090543 | 0.62572533 | 0.90650187 |
| Thoc5 | 0.2288599 | 0.23549443 | 0.24455204 | -0.0090576 | 0.56539202 | 0.90650187 |
| Ahcyl1 | 0.49424366 | 0.45240023 | 0.46146041 | -0.0090602 | 0.53477701 | 0.90650187 |
| Plxna2 | 0.07975776 | 0.08379967 | 0.09286627 | -0.0090666 | 0.66786592 | 0.90650187 |
| Sh3gl3 | 0.03672088 | 0.03490932 | 0.04397758 | -0.0090683 | 0.82034802 | 0.9092537 |
| Vps26b | 0.30409512 | 0.30308089 | 0.31215499 | -0.0090741 | 0.5514123 | 0.90650187 |
| Lrrc48 | 0.03924682 | 0.03784603 | 0.04692401 | -0.009078 | 0.80511994 | 0.90885402 |
| Ccdc114 | 0.02112787 | 0.01648857 | 0.02556775 | -0.0090792 | 0.94279159 | 0.95836398 |
| Chek2 | 0.0622062 | 0.06417839 | 0.07325987 | -0.0090815 | 0.70921367 | 0.90650187 |
| Plcl2 | 0.05294864 | 0.05365039 | 0.06273642 | -0.009086 | 0.74016405 | 0.90650187 |
| Slc26a2 | 0.10910329 | 0.11564086 | 0.12472902 | -0.0090882 | 0.62700117 | 0.90650187 |
| Cars | 0.2361051 | 0.24221246 | 0.25130191 | -0.0090895 | 0.56387265 | 0.90650187 |
| Gm13225 | 0.06556079 | 0.06795169 | 0.07704185 | -0.0090902 | 0.70004737 | 0.90650187 |
| B830012L14I | 0.17878834 | 0.18731589 | 0.19640652 | -0.0090906 | 0.58151236 | 0.90650187 |

| | | | | | | |
|---|---|---|---|---|---|---|
| Ribc1 | 0.0343379 | 0.03209591 | 0.04118883 | -0.0090929 | 0.8368073 | 0.91092327 |
| Zfp626 | 0.03940273 | 0.03800892 | 0.04710554 | -0.0090966 | 0.80469117 | 0.90882608 |
| Pot1a | 0.11494185 | 0.12185666 | 0.13095718 | -0.0091005 | 0.62131477 | 0.90650187 |
| Rps6ka5 | 0.03235489 | 0.02975909 | 0.03886141 | -0.0091023 | 0.85112562 | 0.91505486 |
| Farp2 | 0.06817183 | 0.07086904 | 0.07997413 | -0.0091051 | 0.6936188 | 0.90650187 |
| Traf3 | 0.1262578 | 0.13380225 | 0.14290903 | -0.0091068 | 0.61152387 | 0.90650187 |
| Gm21781 | 0.03245825 | 0.02987519 | 0.03898287 | -0.0091077 | 0.850514 | 0.91483663 |
| Srl | 0.05144679 | 0.05190865 | 0.06101713 | -0.0091085 | 0.7465359 | 0.90650187 |
| Cdk5rap1 | 0.09889704 | 0.10463014 | 0.11374618 | -0.009116 | 0.63918729 | 0.90650187 |
| Cyp4b1 | 0.05763233 | 0.05895746 | 0.06807651 | -0.009119 | 0.72424597 | 0.90650187 |
| Scrn3 | 0.15304949 | 0.16146983 | 0.17059271 | -0.0091229 | 0.59395874 | 0.90650187 |
| Elmod2 | 0.10757906 | 0.11397283 | 0.12309613 | -0.0091233 | 0.62910628 | 0.90650187 |
| Ap3b2 | 0.25387933 | 0.25851528 | 0.26763933 | -0.009124 | 0.56023081 | 0.90650187 |
| BC021891 | 0.01857189 | 0.01338451 | 0.02251325 | -0.0091287 | 0.96427585 | 0.97291631 |
| Elmo2 | 0.16375148 | 0.1722865 | 0.18141673 | -0.0091302 | 0.5885185 | 0.90650187 |
| Cmtr2 | 0.03895445 | 0.03745209 | 0.04658347 | -0.0091314 | 0.80824201 | 0.9092537 |
| Ccdc191 | 0.20935033 | 0.21697531 | 0.22610803 | -0.0091327 | 0.57125185 | 0.90650187 |
| Ube2m | 0.19683122 | 0.20492869 | 0.21406342 | -0.0091347 | 0.57523084 | 0.90650187 |
| BC068281 | 0.04251763 | 0.04157906 | 0.05072468 | -0.0091456 | 0.7885904 | 0.90650187 |
| Coq10a | 0.03691719 | 0.03506102 | 0.04420691 | -0.0091459 | 0.82114663 | 0.9092537 |
| 1110017D15 | 0.10239162 | 0.10836442 | 0.1175171 | -0.0091527 | 0.63541916 | 0.90650187 |
| Dock7 | 0.40686032 | 0.38717754 | 0.39634214 | -0.0091646 | 0.54093735 | 0.90650187 |
| Setdb2 | 0.05195485 | 0.05243062 | 0.06159912 | -0.0091685 | 0.74591526 | 0.90650187 |
| Zfp420 | 0.05878671 | 0.06020452 | 0.06938764 | -0.0091831 | 0.72185283 | 0.90650187 |
| Tfpt | 0.27919464 | 0.28119346 | 0.29037694 | -0.0091835 | 0.55590585 | 0.90650187 |
| Hdac10 | 0.09149542 | 0.09654413 | 0.10573212 | -0.009188 | 0.65036928 | 0.90650187 |
| Ocln | 0.25273817 | 0.25741069 | 0.26659989 | -0.0091892 | 0.56089221 | 0.90650187 |
| Uhrf1bp1 | 0.03993241 | 0.03852995 | 0.04772202 | -0.0091921 | 0.80408373 | 0.90882608 |
| Ap5s1 | 0.1858233 | 0.19414071 | 0.20333287 | -0.0091922 | 0.57964074 | 0.90650187 |
| Setdb1 | 0.17272659 | 0.18120151 | 0.19039459 | -0.0091931 | 0.58498109 | 0.90650187 |

| | | | | | | |
|---|---|---|---|---|---|---|
| Ugcg | 0.16550591 | 0.17398557 | 0.18317879 | -0.0091932 | 0.58827588 | 0.90650187 |
| Smim6 | 0.06898854 | 0.07169325 | 0.08088927 | -0.009196 | 0.693363 | 0.90650187 |
| Zfp955b | 0.05102006 | 0.05131884 | 0.060528 | -0.0092092 | 0.75060208 | 0.90650187 |
| Cited2 | 0.24786819 | 0.25293995 | 0.2621498 | -0.0092098 | 0.56205562 | 0.90650187 |
| BC051142 | 0.03252095 | 0.02984311 | 0.03905655 | -0.0092134 | 0.85284595 | 0.91562204 |
| Abhd4 | 0.17567418 | 0.18410785 | 0.19332314 | -0.0092153 | 0.58391226 | 0.90650187 |
| Fastkd2 | 0.12570726 | 0.13310875 | 0.14233121 | -0.0092225 | 0.61334826 | 0.90650187 |
| Inafm2 | 0.08117235 | 0.08518132 | 0.09442715 | -0.0092458 | 0.6683246 | 0.90650187 |
| Dpm1 | 0.69861304 | 0.58530649 | 0.59455422 | -0.0092477 | 0.52756379 | 0.90650187 |
| Slc7a5 | 0.26538624 | 0.26880073 | 0.27805069 | -0.00925 | 0.55878633 | 0.90650187 |
| 1700016K19I | 0.04216094 | 0.04106038 | 0.050311 | -0.0092506 | 0.79315238 | 0.9070651 |
| Ankrd9 | 0.03764667 | 0.03580724 | 0.04505863 | -0.0092514 | 0.81931758 | 0.9092537 |
| Serinc2 | 0.12252297 | 0.1297295 | 0.13898168 | -0.0092522 | 0.61636569 | 0.90650187 |
| Xkr6 | 0.06295551 | 0.06484622 | 0.07410608 | -0.0092599 | 0.71072268 | 0.90650187 |
| Fam57b | 0.03130368 | 0.02836436 | 0.03762514 | -0.0092608 | 0.86306856 | 0.91950555 |
| Lncpint | 0.0294132 | 0.0261361 | 0.03539747 | -0.0092614 | 0.87761479 | 0.92482217 |
| Arel1 | 0.04674208 | 0.04634533 | 0.05560921 | -0.0092639 | 0.7705288 | 0.90650187 |
| Pnmal1 | 0.30300973 | 0.3019518 | 0.31121761 | -0.0092658 | 0.5526494 | 0.90650187 |
| Gm4419 | 0.08955231 | 0.09434735 | 0.10361563 | -0.0092683 | 0.65455709 | 0.90650187 |
| Tmem123 | 0.35516002 | 0.3458199 | 0.35508925 | -0.0092694 | 0.54619329 | 0.90650187 |
| Lonrf1 | 0.05343313 | 0.05401957 | 0.06329033 | -0.0092708 | 0.7425387 | 0.90650187 |
| Pank1 | 0.08318273 | 0.08736663 | 0.09664053 | -0.0092739 | 0.66516937 | 0.90650187 |
| Zfp324 | 0.08289872 | 0.08705279 | 0.09632819 | -0.0092754 | 0.66569833 | 0.90650187 |
| Dynlt1b | 0.08170548 | 0.08573468 | 0.09501467 | -0.00928 | 0.66792768 | 0.90650187 |
| Ptprr | 0.03252854 | 0.02978541 | 0.03906547 | -0.0092801 | 0.85452983 | 0.91621818 |
| Acbd4 | 0.10264727 | 0.10851056 | 0.11779285 | -0.0092823 | 0.63695131 | 0.90650187 |
| Pou5f2 | 0.05609008 | 0.05703661 | 0.06632174 | -0.0092851 | 0.73315752 | 0.90650187 |
| Rom1 | 0.09311985 | 0.09821085 | 0.10749747 | -0.0092866 | 0.64953018 | 0.90650187 |
| Gm5294 | 0.03520304 | 0.03291214 | 0.04220231 | -0.0092902 | 0.83611857 | 0.91079271 |
| Guk1 | 1.00423733 | 0.74316978 | 0.7524628 | -0.009293 | 0.52189247 | 0.90650187 |

| | | | | | | |
|---|---|---|---|---|---|---|
| Gm8994 | 0.03521238 | 0.03291306 | 0.04221324 | -0.0093002 | 0.83631634 | 0.91079271 |
| Pank2 | 0.2885497 | 0.28932429 | 0.29862588 | -0.0093016 | 0.55506616 | 0.90650187 |
| Ppcdc | 0.11730639 | 0.12416377 | 0.13346755 | -0.0093038 | 0.62167956 | 0.90650187 |
| Zfp593 | 0.18076296 | 0.18904708 | 0.19835611 | -0.009309 | 0.5826339 | 0.90650187 |
| Scly | 0.33850759 | 0.33202595 | 0.34133572 | -0.0093098 | 0.54825423 | 0.90650187 |
| Mrps30 | 0.37474795 | 0.36165978 | 0.37097233 | -0.0093125 | 0.54442909 | 0.90650187 |
| Igsf9 | 0.13537946 | 0.14311452 | 0.15242965 | -0.0093151 | 0.60706421 | 0.90650187 |
| Tmem132b | 0.03003955 | 0.02681612 | 0.03613617 | -0.00932 | 0.87422388 | 0.9235672 |
| Fbxo10 | 0.0453572 | 0.04468972 | 0.05401094 | -0.0093212 | 0.77852571 | 0.90650187 |
| Cbx4 | 0.24632004 | 0.25140326 | 0.26072995 | -0.0093267 | 0.56317573 | 0.90650187 |
| Ddi2 | 0.25477233 | 0.25912504 | 0.26845186 | -0.0093268 | 0.56137395 | 0.90650187 |
| 6720489N17 | 0.04167253 | 0.04041529 | 0.04974425 | -0.009329 | 0.79778124 | 0.9075779 |
| Tspyl5 | 0.08758596 | 0.09213858 | 0.10146843 | -0.0093299 | 0.65864437 | 0.90650187 |
| Nsun6 | 0.07856603 | 0.08221687 | 0.09154909 | -0.0093322 | 0.67478717 | 0.90650187 |
| Atp4a | 0.03688589 | 0.03483653 | 0.04417035 | -0.0093338 | 0.82624241 | 0.90947782 |
| Phtf1 | 0.32504592 | 0.32070916 | 0.33004461 | -0.0093354 | 0.55003342 | 0.90650187 |
| Sh3bgrl2 | 0.03451405 | 0.03205317 | 0.04139528 | -0.0093421 | 0.84213878 | 0.91251845 |
| Gramd1c | 0.05899347 | 0.06027624 | 0.06962227 | -0.009346 | 0.72467083 | 0.90650187 |
| Pkia | 0.11047455 | 0.11684827 | 0.12619557 | -0.0093473 | 0.62903023 | 0.90650187 |
| Cbx7 | 0.02558362 | 0.0215187 | 0.0308676 | -0.0093489 | 0.91092044 | 0.94004764 |
| Arhgap33 | 0.05775291 | 0.05886121 | 0.06821356 | -0.0093523 | 0.72890675 | 0.90650187 |
| Nln | 0.07239207 | 0.07533906 | 0.08469259 | -0.0093535 | 0.68827857 | 0.90650187 |
| Atp6ap2 | 0.78566295 | 0.63502423 | 0.64439034 | -0.0093661 | 0.52576017 | 0.90650187 |
| Zfp41 | 0.06479161 | 0.06680529 | 0.07617614 | -0.0093709 | 0.70776905 | 0.90650187 |
| Gm12524 | 0.02499145 | 0.02079373 | 0.03016507 | -0.0093713 | 0.91637754 | 0.94231902 |
| Ccdc84 | 0.08395258 | 0.08811394 | 0.09748661 | -0.0093727 | 0.66546135 | 0.90650187 |
| Gm17259 | 0.05833166 | 0.05948567 | 0.06887103 | -0.0093854 | 0.72768376 | 0.90650187 |
| Mtss1l | 0.0561668 | 0.05702326 | 0.06640912 | -0.0093859 | 0.73510113 | 0.90650187 |
| Gm10031 | 0.06953869 | 0.07211265 | 0.08150519 | -0.0093925 | 0.69578128 | 0.90650187 |
| Leng1 | 0.27123201 | 0.27389243 | 0.28329172 | -0.0093993 | 0.55863135 | 0.90650187 |

| | | | | | | |
|---|---|---|---|---|---|---|
| Tnr | 0.05587733 | 0.05667851 | 0.06607939 | -0.0094009 | 0.73645714 | 0.90650187 |
| Nat2 | 0.03430457 | 0.03174716 | 0.04114977 | -0.0094026 | 0.84513966 | 0.91333485 |
| Setd1b | 0.37511486 | 0.36186044 | 0.37126681 | -0.0094064 | 0.54483935 | 0.90650187 |
| 1810009A15 | 0.03370345 | 0.03102422 | 0.04044485 | -0.0094206 | 0.84978093 | 0.91470896 |
| Urb1 | 0.0395756 | 0.03787612 | 0.04730679 | -0.0094307 | 0.81225511 | 0.9092537 |
| Ovca2 | 0.04701987 | 0.04647347 | 0.05592946 | -0.009456 | 0.77385975 | 0.90650187 |
| Cpne8 | 0.05593051 | 0.05668271 | 0.06613998 | -0.0094573 | 0.73750478 | 0.90650187 |
| Nedd1 | 0.18552672 | 0.19357765 | 0.20304195 | -0.0094643 | 0.58208136 | 0.90650187 |
| Piga | 0.08636137 | 0.09065734 | 0.10012848 | -0.0094711 | 0.66294319 | 0.90650187 |
| Cnksr1 | 0.03279187 | 0.02990357 | 0.03937482 | -0.0094713 | 0.85754401 | 0.91715518 |
| AU022252 | 0.26802444 | 0.27094583 | 0.28041996 | -0.0094741 | 0.55969544 | 0.90650187 |
| Rusc2 | 0.03608133 | 0.03374096 | 0.04322999 | -0.009489 | 0.83542195 | 0.91078214 |
| Dars2 | 0.0567835 | 0.05762184 | 0.06711115 | -0.0094893 | 0.73519306 | 0.90650187 |
| Bbs10 | 0.05576933 | 0.05646481 | 0.06595634 | -0.0094915 | 0.73883273 | 0.90650187 |
| Degs2 | 0.04310668 | 0.04191228 | 0.05140741 | -0.0094951 | 0.79421063 | 0.90731056 |
| Mterf1a | 0.09332615 | 0.09822554 | 0.10772139 | -0.0094959 | 0.65242993 | 0.90650187 |
| Fjx1 | 0.06986894 | 0.07237566 | 0.0818747 | -0.009499 | 0.69699466 | 0.90650187 |
| Zfp566 | 0.06481899 | 0.0667071 | 0.07620697 | -0.0094999 | 0.71027356 | 0.90650187 |
| Proscos | 0.03088741 | 0.02763414 | 0.03713511 | -0.009501 | 0.87231489 | 0.92263142 |
| Sowaha | 0.05003624 | 0.04989814 | 0.05939926 | -0.0095011 | 0.76148584 | 0.90650187 |
| Atg2b | 0.10622485 | 0.11214171 | 0.12164295 | -0.0095012 | 0.63579279 | 0.90650187 |
| Chmp2b | 0.60161507 | 0.52493502 | 0.5344378 | -0.0095028 | 0.5315023 | 0.90650187 |
| Lrrc28 | 0.06982647 | 0.07232365 | 0.08182719 | -0.0095035 | 0.6971845 | 0.90650187 |
| Ephx4 | 0.0421145 | 0.04075219 | 0.05025714 | -0.0095049 | 0.7997646 | 0.90805731 |
| Adat1 | 0.07902827 | 0.08255453 | 0.09206023 | -0.0095057 | 0.67689437 | 0.90650187 |
| Dclk2 | 0.02617365 | 0.02206049 | 0.03156703 | -0.0095065 | 0.90968429 | 0.9393235 |
| Tjp3 | 0.05691434 | 0.05774657 | 0.06726002 | -0.0095135 | 0.73525984 | 0.90650187 |
| Marveld1 | 0.0919106 | 0.09666603 | 0.10618367 | -0.0095176 | 0.65486044 | 0.90650187 |
| Kdsr | 0.20927955 | 0.21652063 | 0.22604039 | -0.0095198 | 0.57425905 | 0.90650187 |
| Pde5a | 0.07156422 | 0.07424834 | 0.08376905 | -0.0095207 | 0.69331271 | 0.90650187 |

| | | | | | | |
|---|---|---|---|---|---|---|
| Met | 0.08368453 | 0.08767006 | 0.09719212 | -0.0095221 | 0.66841578 | 0.90650187 |
| Zfp612 | 0.17495836 | 0.18309008 | 0.19261288 | -0.0095228 | 0.58698306 | 0.90650187 |
| Nol6 | 0.11763304 | 0.12428988 | 0.1338138 | -0.0095239 | 0.6241532 | 0.90650187 |
| Cdc42bpg | 0.04323179 | 0.04202574 | 0.05155235 | -0.0095266 | 0.79432639 | 0.90731056 |
| Ppfia3 | 0.03015558 | 0.02674223 | 0.03627294 | -0.0095307 | 0.87861325 | 0.92534436 |
| Abhd17c | 0.13234814 | 0.13973372 | 0.14927676 | -0.009543 | 0.61187183 | 0.90650187 |
| Gm14167 | 0.03735602 | 0.03517494 | 0.04471938 | -0.0095444 | 0.82864497 | 0.9096066 |
| Cox10 | 0.14755319 | 0.15542799 | 0.16498247 | -0.0095545 | 0.60158599 | 0.90650187 |
| 2210408I21R | 0.05616972 | 0.05684736 | 0.06641244 | -0.0095651 | 0.73900483 | 0.90650187 |
| Fry | 0.10579016 | 0.11161014 | 0.121176 | -0.0095659 | 0.63718494 | 0.90650187 |
| Sars2 | 0.09126004 | 0.09590929 | 0.10547602 | -0.0095667 | 0.65660644 | 0.90650187 |
| Aldh6a1 | 0.26596155 | 0.2689983 | 0.27856793 | -0.0095696 | 0.56069033 | 0.90650187 |
| Cdhr1 | 0.09011618 | 0.09465182 | 0.10423036 | -0.0095785 | 0.65856179 | 0.90650187 |
| Nolc1 | 0.5862086 | 0.51482279 | 0.52440348 | -0.0095807 | 0.53236645 | 0.90650187 |
| Ndst2 | 0.07825539 | 0.0816234 | 0.09120541 | -0.009582 | 0.67976941 | 0.90650187 |
| Ndufs3 | 1.31377029 | 0.84466267 | 0.85424851 | -0.0095858 | 0.5198933 | 0.90650187 |
| Lrrc51 | 0.07482269 | 0.07781262 | 0.08739847 | -0.0095858 | 0.68708095 | 0.90650187 |
| Wfikkn2 | 0.08043088 | 0.08401865 | 0.09360935 | -0.0095907 | 0.67561531 | 0.90650187 |
| Gpr39 | 0.02603578 | 0.02179918 | 0.03140365 | -0.0096045 | 0.9130284 | 0.9407546 |
| Nprl3 | 0.09359645 | 0.09840852 | 0.10801471 | -0.0096062 | 0.65371369 | 0.90650187 |
| Usp18 | 0.14990686 | 0.15776898 | 0.16738938 | -0.0096204 | 0.60083269 | 0.90650187 |
| Stk36 | 0.04318958 | 0.04186993 | 0.05150345 | -0.0096335 | 0.79716366 | 0.9075779 |
| Atxn7l2 | 0.08606693 | 0.09016791 | 0.09980599 | -0.0096381 | 0.66614902 | 0.90650187 |
| Tmem219 | 0.42749108 | 0.40257782 | 0.41221961 | -0.0096418 | 0.54140835 | 0.90650187 |
| Mctp2 | 0.0386688 | 0.03659846 | 0.04625063 | -0.0096522 | 0.82323546 | 0.9092537 |
| 2810021J22F | 0.0869452 | 0.0911149 | 0.10076757 | -0.0096527 | 0.66489183 | 0.90650187 |
| Rsph1 | 0.18556852 | 0.19342913 | 0.20308296 | -0.0096538 | 0.58368437 | 0.90650187 |
| Ttc17 | 0.04949864 | 0.04912736 | 0.05878186 | -0.0096545 | 0.76735355 | 0.90650187 |
| Synpr | 0.12518931 | 0.13212589 | 0.14178727 | -0.0096614 | 0.61902598 | 0.90650187 |
| Pinlyp | 0.02781152 | 0.02383898 | 0.03350573 | -0.0096668 | 0.90018457 | 0.93499868 |

| | | | | | | |
|---|---|---|---|---|---|---|
| Rxra | 0.09686567 | 0.10188317 | 0.11155426 | -0.0096711 | 0.65003225 | 0.90650187 |
| A830010M2( | 0.03680004 | 0.03439512 | 0.04407006 | -0.0096749 | 0.83545989 | 0.91078214 |
| Rttn | 0.02511712 | 0.02063855 | 0.03031421 | -0.0096757 | 0.92204237 | 0.94549985 |
| Gm8394 | 0.02897832 | 0.02519966 | 0.03488423 | -0.0096846 | 0.89143419 | 0.93119098 |
| Spg11 | 0.0512291 | 0.05107951 | 0.06076763 | -0.0096881 | 0.76076279 | 0.90650187 |
| Bre | 0.58040177 | 0.51088909 | 0.52058469 | -0.0096956 | 0.53299351 | 0.90650187 |
| Rgmb | 0.09818012 | 0.10327745 | 0.11297325 | -0.0096958 | 0.6485983 | 0.90650187 |
| Eid2 | 0.03940236 | 0.03740774 | 0.04710511 | -0.0096974 | 0.81995734 | 0.9092537 |
| Frk | 0.10992884 | 0.1159146 | 0.12561223 | -0.0096976 | 0.63428108 | 0.90650187 |
| Osbpl3 | 0.04565193 | 0.04465184 | 0.05435134 | -0.0096995 | 0.78621568 | 0.90650187 |
| Zfp334 | 0.10503913 | 0.11066671 | 0.12036868 | -0.009702 | 0.63995008 | 0.90650187 |
| Wdr47 | 0.19038007 | 0.19806946 | 0.20779061 | -0.0097211 | 0.58237792 | 0.90650187 |
| Ranbp17 | 0.0300491 | 0.02642007 | 0.03614742 | -0.0097274 | 0.88421471 | 0.9278081 |
| Rnaseh2a | 0.26915063 | 0.27170114 | 0.28142935 | -0.0097282 | 0.56106581 | 0.90650187 |
| Fam83g | 0.02649219 | 0.02221585 | 0.03194442 | -0.0097286 | 0.91211499 | 0.94062208 |
| Lekr1 | 0.11840102 | 0.12489323 | 0.13462736 | -0.0097341 | 0.62606004 | 0.90650187 |
| Lrp8 | 0.1913983 | 0.19904239 | 0.20878364 | -0.0097413 | 0.58215882 | 0.90650187 |
| Ino80dos | 0.0202986 | 0.01482854 | 0.02457789 | -0.0097493 | 0.96108727 | 0.97073008 |
| Timm17a | 1.1406603 | 0.79371951 | 0.80347051 | -0.009751 | 0.52151345 | 0.90650187 |
| Micu2 | 0.41396279 | 0.39209047 | 0.40184497 | -0.0097545 | 0.54296802 | 0.90650187 |
| Camk2n2 | 0.04028728 | 0.03837799 | 0.04813481 | -0.0097568 | 0.81623622 | 0.9092537 |
| Ric8 | 0.41184065 | 0.39044296 | 0.40020476 | -0.0097618 | 0.54317559 | 0.90650187 |
| Rpl21 | 4.39915163 | 0.61061904 | 0.62038305 | -0.009764 | 0.52789043 | 0.90650187 |
| Fhod3 | 0.05129446 | 0.05107632 | 0.06084255 | -0.0097662 | 0.76226058 | 0.90650187 |
| Pkmyt1 | 0.11329737 | 0.11943886 | 0.12920722 | -0.0097684 | 0.63160152 | 0.90650187 |
| Kif19a | 0.06744723 | 0.06938624 | 0.07916138 | -0.0097751 | 0.70848357 | 0.90650187 |
| Zbtb43 | 0.09261666 | 0.09717431 | 0.10695101 | -0.0097767 | 0.65777046 | 0.90650187 |
| Gatb | 0.07608686 | 0.07902532 | 0.08880243 | -0.0097771 | 0.68774174 | 0.90650187 |
| Zswim3 | 0.05456885 | 0.05480975 | 0.06458741 | -0.0097777 | 0.74952772 | 0.90650187 |
| Mpv17l | 0.1143881 | 0.1205772 | 0.13036828 | -0.0097911 | 0.63076471 | 0.90650187 |

| | | | | | | |
|---|---|---|---|---|---|---|
| Abl2 | 0.09422954 | 0.098908 | 0.10870131 | -0.0097933 | 0.65561377 | 0.90650187 |
| Fam161b | 0.0796269 | 0.08292783 | 0.09272173 | -0.0097939 | 0.68067071 | 0.90650187 |
| Gtpbp4 | 0.61073297 | 0.53051576 | 0.54031131 | -0.0097955 | 0.53211854 | 0.90650187 |
| Nup43 | 0.21248127 | 0.21929255 | 0.22909455 | -0.009802 | 0.57542731 | 0.90650187 |
| Alyref | 0.21033866 | 0.2172485 | 0.22705187 | -0.0098034 | 0.57610813 | 0.90650187 |
| Zfp691 | 0.11015476 | 0.116049 | 0.12585377 | -0.0098048 | 0.6354562 | 0.90650187 |
| Zfp346 | 0.06226307 | 0.0635176 | 0.07332413 | -0.0098065 | 0.72393294 | 0.90650187 |
| Slc25a42 | 0.03876325 | 0.03655404 | 0.0463607 | -0.0098067 | 0.82648796 | 0.90947782 |
| Trim7 | 0.0738886 | 0.07654469 | 0.0863596 | -0.0098149 | 0.69330855 | 0.90650187 |
| Jrk | 0.05041051 | 0.05001282 | 0.05982883 | -0.009816 | 0.76711622 | 0.90650187 |
| Utp20 | 0.10705089 | 0.11271226 | 0.12252963 | -0.0098174 | 0.63915192 | 0.90650187 |
| Kcnk16 | 0.10911429 | 0.11492233 | 0.12474079 | -0.0098185 | 0.63679977 | 0.90650187 |
| Mark1 | 0.06436606 | 0.06587223 | 0.0756968 | -0.0098246 | 0.71802647 | 0.90650187 |
| Yod1 | 0.25799818 | 0.26155376 | 0.2713806 | -0.0098268 | 0.5639446 | 0.90650187 |
| Cmtr1 | 0.07996281 | 0.08325849 | 0.0930927 | -0.0098342 | 0.68068989 | 0.90650187 |
| Gzf1 | 0.1476504 | 0.1552478 | 0.16508202 | -0.0098342 | 0.60443071 | 0.90650187 |
| Gtpbp8 | 0.15041019 | 0.15805391 | 0.16790319 | -0.0098493 | 0.6028695 | 0.90650187 |
| Gm561 | 0.64985784 | 0.55512809 | 0.56497839 | -0.0098503 | 0.53089043 | 0.90650187 |
| Gm7467 | 0.04204667 | 0.04032385 | 0.05017844 | -0.0098546 | 0.80868941 | 0.9092537 |
| Zfp277 | 0.28054914 | 0.28171771 | 0.29157628 | -0.0098586 | 0.55974063 | 0.90650187 |
| Ythdf3 | 0.30744346 | 0.3051811 | 0.31504016 | -0.0098591 | 0.5553237 | 0.90650187 |
| Prpf19 | 1.13608693 | 0.79202939 | 0.80189153 | -0.0098621 | 0.52180121 | 0.90650187 |
| Gm7293 | 0.03307006 | 0.02983918 | 0.0397015 | -0.0098623 | 0.86527727 | 0.92040222 |
| Shisa2 | 0.06399638 | 0.06541741 | 0.07528019 | -0.0098628 | 0.719864 | 0.90650187 |
| Mtpap | 0.23561036 | 0.24097573 | 0.25084267 | -0.0098669 | 0.56940855 | 0.90650187 |
| Stat6 | 0.06353792 | 0.0648935 | 0.07476324 | -0.0098697 | 0.72135761 | 0.90650187 |
| Ccdc71l | 0.03728509 | 0.03476319 | 0.04463656 | -0.0098734 | 0.83728107 | 0.9110601 |
| Tsc22d2 | 0.25621352 | 0.25988187 | 0.26976155 | -0.0098797 | 0.56466784 | 0.90650187 |
| Zfp946 | 0.1184396 | 0.12478295 | 0.13466821 | -0.0098853 | 0.62791184 | 0.90650187 |
| Arv1 | 0.02549723 | 0.02087664 | 0.03076515 | -0.0098885 | 0.92348394 | 0.94620379 |

| | | | | | | |
|---|---|---|---|---|---|---|
| Tm7sf2 | 0.06918239 | 0.07121241 | 0.08110635 | -0.0098939 | 0.70619663 | 0.90650187 |
| Eme2 | 0.03999912 | 0.03790508 | 0.04779963 | -0.0098946 | 0.82127714 | 0.9092537 |
| 1110020A21 | 0.09037734 | 0.09461936 | 0.10451492 | -0.0098956 | 0.66309092 | 0.90650187 |
| 1110006O24 | 0.05717162 | 0.05764749 | 0.06755268 | -0.0099052 | 0.74275274 | 0.90650187 |
| Gna13 | 0.30968696 | 0.307061 | 0.31696776 | -0.0099068 | 0.55525382 | 0.90650187 |
| Fam210a | 0.14965463 | 0.15721217 | 0.16713178 | -0.0099196 | 0.60405445 | 0.90650187 |
| Ccpg1 | 0.37960994 | 0.36494404 | 0.37486576 | -0.0099217 | 0.54683293 | 0.90650187 |
| Ptpn23 | 0.12260784 | 0.12914435 | 0.13907113 | -0.0099268 | 0.62450088 | 0.90650187 |
| Gdnf | 0.02546571 | 0.02080069 | 0.03072776 | -0.0099271 | 0.92453034 | 0.94689298 |
| Rps6ka3 | 0.23127243 | 0.23687444 | 0.24680547 | -0.009931 | 0.57098546 | 0.90650187 |
| Dlg3 | 0.04776101 | 0.04684474 | 0.05678329 | -0.0099386 | 0.78174397 | 0.90650187 |
| Tdp1 | 0.16359405 | 0.1713161 | 0.18125844 | -0.0099423 | 0.5963262 | 0.90650187 |
| AK010878 | 0.38514601 | 0.36933146 | 0.37927627 | -0.0099448 | 0.54639801 | 0.90650187 |
| Aqp11 | 0.10523974 | 0.11063513 | 0.1205844 | -0.0099493 | 0.64311641 | 0.90650187 |
| Ago2 | 0.47029762 | 0.43420804 | 0.44415973 | -0.0099517 | 0.53967171 | 0.90650187 |
| Arap2 | 0.04509256 | 0.04375257 | 0.0537052 | -0.0099526 | 0.79498987 | 0.90731056 |
| Trmt6 | 0.36537291 | 0.35345598 | 0.36340986 | -0.0099539 | 0.5484579 | 0.90650187 |
| Ugt8a | 0.05245511 | 0.05221432 | 0.06217181 | -0.0099575 | 0.76177133 | 0.90650187 |
| Sep-10 | 0.26788316 | 0.27033455 | 0.28029325 | -0.0099587 | 0.56275209 | 0.90650187 |
| Amacr | 0.16381051 | 0.17151362 | 0.18147607 | -0.0099624 | 0.59640373 | 0.90650187 |
| Mapkapk5 | 0.03991344 | 0.03773686 | 0.04769995 | -0.0099631 | 0.82344 | 0.9092537 |
| Ttc32 | 0.21021846 | 0.2169738 | 0.22693714 | -0.0099633 | 0.57737312 | 0.90650187 |
| Retsat | 0.07053992 | 0.07265319 | 0.08262497 | -0.0099718 | 0.70420316 | 0.90650187 |
| Mtg2 | 0.21879502 | 0.2251053 | 0.23508627 | -0.009981 | 0.57485386 | 0.90650187 |
| Dbndd1 | 0.03506214 | 0.03205189 | 0.04203732 | -0.0099854 | 0.85451518 | 0.91621818 |
| Setmar | 0.02913519 | 0.02507808 | 0.03506941 | -0.0099913 | 0.89734378 | 0.93409045 |
| Nupl1 | 0.06727833 | 0.06897233 | 0.07897182 | -0.0099995 | 0.71325097 | 0.90650187 |
| Rtel1 | 0.0390043 | 0.03663653 | 0.04664154 | -0.010005 | 0.8298621 | 0.90974008 |
| Arih2 | 0.23476724 | 0.24005327 | 0.25005947 | -0.0100062 | 0.57059615 | 0.90650187 |
| Plekhm1 | 0.04714424 | 0.04606246 | 0.0560728 | -0.0100103 | 0.78629964 | 0.90650187 |

| | | | | | | |
|---|---|---|---|---|---|---|
| Shank2 | 0.02670087 | 0.02218016 | 0.03219155 | -0.0100114 | 0.91659987 | 0.9423899 |
| Sav1 | 0.1044354 | 0.10970594 | 0.11971919 | -0.0100132 | 0.6449895 | 0.90650187 |
| Rbm12b1 | 0.04936046 | 0.04860902 | 0.05862309 | -0.0100141 | 0.77619468 | 0.90650187 |
| Scd3 | 0.02269996 | 0.01742533 | 0.02744127 | -0.0100159 | 0.94766536 | 0.96158208 |
| Ppip5k2 | 0.15391781 | 0.16145357 | 0.1714758 | -0.0100222 | 0.60250333 | 0.90650187 |
| Crhr2 | 0.07340546 | 0.07579553 | 0.08582178 | -0.0100263 | 0.69824663 | 0.90650187 |
| Gtf2h4 | 0.26008373 | 0.26324147 | 0.27326876 | -0.0100273 | 0.56479055 | 0.90650187 |
| Trub2 | 0.17823292 | 0.18582621 | 0.19585738 | -0.0100312 | 0.59005613 | 0.90650187 |
| Tecpr2 | 0.05436989 | 0.05432627 | 0.06436033 | -0.0100341 | 0.75587327 | 0.90650187 |
| Nampt | 0.19754256 | 0.20471496 | 0.21475239 | -0.0100374 | 0.5823018 | 0.90650187 |
| Pagr1a | 0.03342951 | 0.03008568 | 0.04012344 | -0.0100378 | 0.86696663 | 0.92056229 |
| Lipo1 | 0.12003692 | 0.12631819 | 0.13635796 | -0.0100398 | 0.62828752 | 0.90650187 |
| Lrch1 | 0.10917067 | 0.11475768 | 0.12480114 | -0.0100435 | 0.63973982 | 0.90650187 |
| Zfp148 | 0.63574012 | 0.5461279 | 0.55617627 | -0.0100484 | 0.53200791 | 0.90650187 |
| Letm2 | 0.06386355 | 0.06507524 | 0.07513045 | -0.0100552 | 0.72407321 | 0.90650187 |
| Cars2 | 0.15933569 | 0.16690646 | 0.17696637 | -0.0100599 | 0.5997565 | 0.90650187 |
| Hexdc | 0.0647358 | 0.06605244 | 0.07611329 | -0.0100608 | 0.72161019 | 0.90650187 |
| Rassf6 | 0.04302483 | 0.04124252 | 0.05131257 | -0.0100701 | 0.80851772 | 0.9092537 |
| Dph7 | 0.08652034 | 0.09021768 | 0.10030255 | -0.0100849 | 0.67255562 | 0.90650187 |
| Cep164 | 0.38360699 | 0.36796614 | 0.37805257 | -0.0100864 | 0.54720726 | 0.90650187 |
| Zfp629 | 0.08864959 | 0.09254241 | 0.10263056 | -0.0100881 | 0.66893853 | 0.90650187 |
| Alms1 | 0.07824075 | 0.08109456 | 0.09118921 | -0.0100946 | 0.68868583 | 0.90650187 |
| Gper1 | 0.04818735 | 0.04717859 | 0.05727408 | -0.0100955 | 0.78336564 | 0.90650187 |
| A1cf | 0.05430971 | 0.054196 | 0.06429162 | -0.0100956 | 0.75743869 | 0.90650187 |
| Ptpdc1 | 0.04636674 | 0.04508037 | 0.05517633 | -0.010096 | 0.79201723 | 0.90681274 |
| Caskin2 | 0.10489986 | 0.11010609 | 0.12021889 | -0.0101128 | 0.6457844 | 0.90650187 |
| Tmed4 | 1.05638094 | 0.76303233 | 0.77314804 | -0.0101157 | 0.52319155 | 0.90650187 |
| Naa60 | 0.15468251 | 0.16213204 | 0.1722528 | -0.0101208 | 0.60303193 | 0.90650187 |
| Gga3 | 0.11520368 | 0.12111329 | 0.13123549 | -0.0101222 | 0.63415938 | 0.90650187 |
| Sco2 | 0.18196681 | 0.18941981 | 0.19954261 | -0.0101228 | 0.58921483 | 0.90650187 |

| | | | | | | |
|---|---|---|---|---|---|---|
| Rnf113a1 | 0.04945474 | 0.04860836 | 0.05873143 | -0.0101231 | 0.77823732 | 0.90650187 |
| Pdxk | 0.06167073 | 0.06253037 | 0.07265466 | -0.0101243 | 0.73220214 | 0.90650187 |
| Plekhg5 | 0.07814416 | 0.08095348 | 0.09108232 | -0.0101288 | 0.68947939 | 0.90650187 |
| Polr3gl | 0.41484985 | 0.39239964 | 0.40252958 | -0.0101299 | 0.54453947 | 0.90650187 |
| Mrpl13 | 0.61982746 | 0.53599174 | 0.54612231 | -0.0101306 | 0.53286193 | 0.90650187 |
| Gins1 | 0.09638445 | 0.10090336 | 0.11103418 | -0.0101308 | 0.65748895 | 0.90650187 |
| Gm10282 | 0.23791591 | 0.24284874 | 0.25298074 | -0.010132 | 0.57065765 | 0.90650187 |
| Zfp523 | 0.24809606 | 0.25221771 | 0.26235854 | -0.0101408 | 0.568216 | 0.90650187 |
| Map3k5 | 0.03972154 | 0.03733347 | 0.04747665 | -0.0101432 | 0.82888914 | 0.90967755 |
| Ppp2r5b | 0.41066372 | 0.38914907 | 0.39929365 | -0.0101446 | 0.54496348 | 0.90650187 |
| Muc4 | 0.07911136 | 0.08200288 | 0.09215207 | -0.0101492 | 0.68779788 | 0.90650187 |
| Fnbp1 | 0.20981996 | 0.21640563 | 0.22655665 | -0.010151 | 0.57894226 | 0.90650187 |
| Fut11 | 0.2615125 | 0.26440237 | 0.2745599 | -0.0101575 | 0.5653187 | 0.90650187 |
| Psmb7 | 1.64981389 | 0.90340592 | 0.91357027 | -0.0101644 | 0.51972431 | 0.90650187 |
| Thap3 | 0.35472702 | 0.34456475 | 0.3547346 | -0.0101699 | 0.55070816 | 0.90650187 |
| Slc30a9 | 0.53760725 | 0.48163757 | 0.49180889 | -0.0101713 | 0.53662776 | 0.90650187 |
| Tmem170b | 0.07350684 | 0.07575625 | 0.08593466 | -0.0101784 | 0.7007482 | 0.90650187 |
| Otulin | 0.06304902 | 0.06402173 | 0.07421163 | -0.0101899 | 0.72920819 | 0.90650187 |
| Tmem63a | 0.22047046 | 0.22647939 | 0.23666938 | -0.01019 | 0.57589746 | 0.90650187 |
| Ubn2 | 0.74037065 | 0.60871105 | 0.61891777 | -0.0102067 | 0.52922171 | 0.90650187 |
| Rab31 | 0.12691283 | 0.13338904 | 0.14359602 | -0.010207 | 0.62399851 | 0.90650187 |
| Dnajb5 | 0.0505848 | 0.04981215 | 0.06002881 | -0.0102167 | 0.77536249 | 0.90650187 |
| Map3k2 | 0.24400619 | 0.24838501 | 0.25860334 | -0.0102183 | 0.56972008 | 0.90650187 |
| 2810402E24I | 0.14098566 | 0.14800808 | 0.15823201 | -0.0102239 | 0.61303812 | 0.90650187 |
| Vps25 | 0.10463595 | 0.10969518 | 0.11993499 | -0.0102398 | 0.64786292 | 0.90650187 |
| Apoo | 0.37476759 | 0.36074819 | 0.3709881 | -0.0102399 | 0.54883019 | 0.90650187 |
| Hist1h4h | 0.0593046 | 0.05973129 | 0.06997522 | -0.0102439 | 0.74241494 | 0.90650187 |
| Tnfsf9 | 0.07233654 | 0.07438635 | 0.08463067 | -0.0102443 | 0.70475604 | 0.90650187 |
| Nefh | 0.04126699 | 0.03901972 | 0.04927338 | -0.0102537 | 0.82254793 | 0.9092537 |
| Ears2 | 0.14857785 | 0.15576115 | 0.16603116 | -0.01027 | 0.60833691 | 0.90650187 |

| | | | | | | |
|---|---|---|---|---|---|---|
| Mphosph9 | 0.10481843 | 0.10985735 | 0.12013131 | -0.010274 | 0.64810172 | 0.90650187 |
| Ccny | 0.33937072 | 0.3317603 | 0.34205445 | -0.0102941 | 0.55321596 | 0.90650187 |
| Arhgef5 | 0.07065113 | 0.0724481 | 0.08274926 | -0.0103012 | 0.71001122 | 0.90650187 |
| Aarsd1 | 0.06452952 | 0.06557873 | 0.07588095 | -0.0103022 | 0.726936 | 0.90650187 |
| Rfc5 | 0.25373804 | 0.25720287 | 0.2675107 | -0.0103078 | 0.56800602 | 0.90650187 |
| Csdc2 | 0.04353489 | 0.04159494 | 0.05190339 | -0.0103084 | 0.81136412 | 0.9092537 |
| Gnpda1 | 0.14078847 | 0.14770471 | 0.15802854 | -0.0103238 | 0.61425512 | 0.90650187 |
| Frs3 | 0.07202989 | 0.07395707 | 0.08428866 | -0.0103316 | 0.70709422 | 0.90650187 |
| Spink10 | 0.05469102 | 0.05439367 | 0.06472682 | -0.0103331 | 0.76105725 | 0.90650187 |
| Nhlrc2 | 0.16504125 | 0.17236722 | 0.18271243 | -0.0103452 | 0.59936785 | 0.90650187 |
| Fam73a | 0.13665045 | 0.14339727 | 0.15374835 | -0.0103511 | 0.61764428 | 0.90650187 |
| Rnmt | 0.25638821 | 0.25956681 | 0.26992016 | -0.0103534 | 0.5676996 | 0.90650187 |
| Ndc1 | 0.10280912 | 0.107614 | 0.11796737 | -0.0103534 | 0.65179365 | 0.90650187 |
| Pklr | 0.02995816 | 0.02568581 | 0.03604021 | -0.0103544 | 0.89923933 | 0.93446384 |
| 9130230L23F | 0.02498542 | 0.01979368 | 0.03015792 | -0.0103642 | 0.93671915 | 0.9542332 |
| Sep-05 | 0.03491978 | 0.03150613 | 0.0418706 | -0.0103645 | 0.86443117 | 0.92024779 |
| Pcdh15 | 0.04251037 | 0.04035052 | 0.05071626 | -0.0103657 | 0.81822578 | 0.9092537 |
| Tmem104 | 0.04935944 | 0.04825427 | 0.05862193 | -0.0103677 | 0.78413456 | 0.90650187 |
| Ube2e1 | 1.01938476 | 0.74825113 | 0.75862201 | -0.0103709 | 0.52423058 | 0.90650187 |
| Gtf2f1 | 0.45594532 | 0.42322475 | 0.43359748 | -0.0103727 | 0.54234765 | 0.90650187 |
| Xpo5 | 0.04155615 | 0.03923358 | 0.04960915 | -0.0103756 | 0.82375804 | 0.9092537 |
| Faap100 | 0.287173 | 0.28703422 | 0.297417 | -0.0103828 | 0.56166634 | 0.90650187 |
| Mastl | 0.05626982 | 0.05614349 | 0.06652643 | -0.0103829 | 0.75610749 | 0.90650187 |
| Zfp867 | 0.08056297 | 0.08336854 | 0.0937551 | -0.0103866 | 0.68881763 | 0.90650187 |
| Mtbp | 0.05060571 | 0.04966244 | 0.06005279 | -0.0103904 | 0.77910678 | 0.90650187 |
| Jpx | 0.06836216 | 0.06979636 | 0.08018748 | -0.0103911 | 0.71772129 | 0.90650187 |
| Gm10269 | 0.09747767 | 0.10181846 | 0.11221523 | -0.0103968 | 0.65978881 | 0.90650187 |
| 5730559C18I | 0.0386758 | 0.03585851 | 0.04625879 | -0.0104003 | 0.84122308 | 0.91251845 |
| Dzank1 | 0.04169601 | 0.03936989 | 0.0497715 | -0.0104016 | 0.82357493 | 0.9092537 |
| Gm26641 | 0.04142944 | 0.03905835 | 0.04946203 | -0.0104037 | 0.8251249 | 0.90934433 |

| | | | | | | |
|---|---|---|---|---|---|---|
| 0610010F05I | 0.22210964 | 0.22780912 | 0.23821546 | -0.0104063 | 0.57699206 | 0.90650187 |
| Fdx1 | 0.62699381 | 0.54025949 | 0.55066737 | -0.0104079 | 0.53348137 | 0.90650187 |
| Ptprt | 0.03293292 | 0.02912456 | 0.03954047 | -0.0104159 | 0.8792167 | 0.92549228 |
| Gpr137b-ps | 0.03969944 | 0.0370292 | 0.04745093 | -0.0104217 | 0.83556582 | 0.91078214 |
| Bckdhb | 0.19876609 | 0.20551321 | 0.21593609 | -0.0104229 | 0.58495343 | 0.90650187 |
| Lrrn1 | 0.17600248 | 0.18322575 | 0.1936487 | -0.0104229 | 0.59455584 | 0.90650187 |
| Rab19 | 0.04407657 | 0.04210156 | 0.05253038 | -0.0104288 | 0.81126921 | 0.9092537 |
| Ppdpf | 1.43747801 | 0.87154358 | 0.88197265 | -0.0104291 | 0.52096195 | 0.90650187 |
| Tma16 | 0.20097133 | 0.20763482 | 0.21806515 | -0.0104303 | 0.58419574 | 0.90650187 |
| Axin1 | 0.09780416 | 0.10212052 | 0.11256763 | -0.0104471 | 0.66004467 | 0.90650187 |
| Dhx35 | 0.02520832 | 0.01997492 | 0.03042243 | -0.0104475 | 0.9365804 | 0.95421975 |
| Fgf12 | 0.1220987 | 0.12806215 | 0.13853445 | -0.0104723 | 0.63158684 | 0.90650187 |
| Pias3 | 0.27165512 | 0.27319353 | 0.28366982 | -0.0104763 | 0.56520601 | 0.90650187 |
| Chst8 | 0.07847958 | 0.08097443 | 0.09145345 | -0.010479 | 0.6947611 | 0.90650187 |
| Gclc | 0.07971687 | 0.08233648 | 0.09282111 | -0.0104846 | 0.69222057 | 0.90650187 |
| Fam65a | 0.18050663 | 0.18760987 | 0.19810328 | -0.0104934 | 0.59308226 | 0.90650187 |
| Med9os | 0.04609017 | 0.04436249 | 0.05485723 | -0.0104947 | 0.8024733 | 0.90870058 |
| Trmo | 0.04861082 | 0.04726604 | 0.05776128 | -0.0104952 | 0.79038789 | 0.90650187 |
| Ydjc | 0.10827658 | 0.11334794 | 0.12384374 | -0.0104958 | 0.64682649 | 0.90650187 |
| 2210016F16I | 0.48593449 | 0.44500296 | 0.45550105 | -0.0104981 | 0.54080399 | 0.90650187 |
| Chd9 | 0.35050955 | 0.34077386 | 0.35127219 | -0.0104983 | 0.55284954 | 0.90650187 |
| Ube2o | 0.13621652 | 0.14279137 | 0.15329835 | -0.010507 | 0.61970238 | 0.90650187 |
| Nr6a1 | 0.05098323 | 0.049974 | 0.06048576 | -0.0105118 | 0.78011977 | 0.90650187 |
| Gm15860 | 0.04967929 | 0.04847716 | 0.05898938 | -0.0105122 | 0.78588686 | 0.90650187 |
| Coa4 | 0.15586229 | 0.16293679 | 0.17345023 | -0.0105134 | 0.60621368 | 0.90650187 |
| Kif1a | 0.2245073 | 0.22995618 | 0.24047163 | -0.0105155 | 0.57706845 | 0.90650187 |
| Nceh1 | 0.03134449 | 0.02715257 | 0.03767316 | -0.0105206 | 0.89278539 | 0.93179124 |
| Smim8 | 0.3378387 | 0.33025758 | 0.34077831 | -0.0105207 | 0.55458251 | 0.90650187 |
| Morc2a | 0.16368935 | 0.1708332 | 0.18135426 | -0.0105211 | 0.60176052 | 0.90650187 |
| Gm14966 | 0.05110319 | 0.05009176 | 0.06062331 | -0.0105315 | 0.78003034 | 0.90650187 |

| | | | | | | |
|---|---|---|---|---|---|---|
| BC024978 | 0.07110658 | 0.07271348 | 0.08325809 | -0.0105446 | 0.71329422 | 0.90650187 |
| Usp31 | 0.045456 | 0.04357133 | 0.05412507 | -0.0105537 | 0.80698512 | 0.9092537 |
| Myo1d | 0.03912743 | 0.036231 | 0.04678497 | -0.010554 | 0.84203587 | 0.91251845 |
| 1810014B01 | 0.07322494 | 0.07506134 | 0.08562074 | -0.0105594 | 0.70824537 | 0.90650187 |
| B3galt4 | 0.10693934 | 0.11184855 | 0.12240994 | -0.0105614 | 0.64934833 | 0.90650187 |
| Tmem50b | 0.15941339 | 0.16648327 | 0.17704487 | -0.0105616 | 0.60457313 | 0.90650187 |
| Ttc37 | 0.09649293 | 0.10058598 | 0.11115145 | -0.0105655 | 0.6636976 | 0.90650187 |
| Atad5 | 0.09595515 | 0.10000329 | 0.11056994 | -0.0105666 | 0.66452538 | 0.90650187 |
| Cadm4 | 0.05596453 | 0.05560223 | 0.06617874 | -0.0105765 | 0.76129825 | 0.90650187 |
| Zfp956 | 0.0544506 | 0.05386923 | 0.06445245 | -0.0105832 | 0.76729791 | 0.90650187 |
| Urah | 0.04755107 | 0.04595513 | 0.05654152 | -0.0105864 | 0.79739705 | 0.9075779 |
| Urgcp | 0.20800764 | 0.21423619 | 0.22482415 | -0.010588 | 0.58291764 | 0.90650187 |
| Prodh | 0.04082917 | 0.03817638 | 0.04876474 | -0.0105884 | 0.83279811 | 0.91035535 |
| Tomm34 | 0.42169579 | 0.39720177 | 0.40779278 | -0.010591 | 0.54595947 | 0.90650187 |
| Ints10 | 0.23418955 | 0.23892972 | 0.24952242 | -0.0105927 | 0.57484561 | 0.90650187 |
| Pomk | 0.0605484 | 0.06078916 | 0.07138475 | -0.0105956 | 0.7453277 | 0.90650187 |
| Sac3d1 | 0.41620583 | 0.39297706 | 0.40357497 | -0.0105979 | 0.54646773 | 0.90650187 |
| Ap4e1 | 0.05135121 | 0.05030694 | 0.06090759 | -0.0106006 | 0.7804561 | 0.90650187 |
| Cables2 | 0.0755766 | 0.07763184 | 0.08823602 | -0.0106042 | 0.70342322 | 0.90650187 |
| Uros | 0.17326942 | 0.18032557 | 0.19093471 | -0.0106091 | 0.59755237 | 0.90650187 |
| Cep57 | 0.31912894 | 0.31442029 | 0.32503186 | -0.0106116 | 0.55769955 | 0.90650187 |
| B4galt2 | 0.12059771 | 0.12633578 | 0.13695046 | -0.0106147 | 0.63479032 | 0.90650187 |
| Smg8 | 0.06235556 | 0.06281378 | 0.07342862 | -0.0106148 | 0.73977661 | 0.90650187 |
| BC052040 | 0.05656769 | 0.05624884 | 0.06686554 | -0.0106167 | 0.75986226 | 0.90650187 |
| Plekha7 | 0.09141643 | 0.09502825 | 0.10564619 | -0.0106179 | 0.67249179 | 0.90650187 |
| Lhpp | 0.0386685 | 0.03562002 | 0.04625029 | -0.0106303 | 0.84655942 | 0.91408934 |
| Endog | 0.28480197 | 0.28469729 | 0.29533093 | -0.0106336 | 0.56358616 | 0.90650187 |
| Ptpru | 0.05618935 | 0.05579493 | 0.0664348 | -0.0106399 | 0.76176338 | 0.90650187 |
| R3hdm4 | 0.9071369 | 0.69746608 | 0.70810598 | -0.0106399 | 0.52662917 | 0.90650187 |
| Cachd1 | 0.05460842 | 0.05398463 | 0.06463257 | -0.0106479 | 0.76803605 | 0.90650187 |

| | | | | | | |
|---|---|---|---|---|---|---|
| B3glct | 0.10099171 | 0.10535523 | 0.11600574 | -0.0106505 | 0.65841888 | 0.90650187 |
| Zfp369 | 0.04504509 | 0.04298013 | 0.05365034 | -0.0106702 | 0.81169545 | 0.9092537 |
| Zfp39 | 0.05878292 | 0.05870758 | 0.06938334 | -0.0106758 | 0.7530215 | 0.90650187 |
| Golim4 | 0.78379617 | 0.6326804 | 0.64335915 | -0.0106787 | 0.52941128 | 0.90650187 |
| Ankrd50 | 0.12206729 | 0.12781844 | 0.13850133 | -0.0106829 | 0.63416275 | 0.90650187 |
| Zfp709 | 0.05243025 | 0.05145356 | 0.06214336 | -0.0106898 | 0.77778265 | 0.90650187 |
| Scamp5 | 0.0915857 | 0.09513091 | 0.10583033 | -0.0106994 | 0.67344634 | 0.90650187 |
| Gtse1 | 0.02956269 | 0.02487217 | 0.03557383 | -0.0107017 | 0.90944918 | 0.93923072 |
| Zfyve9 | 0.15314193 | 0.15998479 | 0.17068677 | -0.010702 | 0.60977689 | 0.90650187 |
| Snapc4 | 0.08737299 | 0.09053056 | 0.10123556 | -0.010705 | 0.68085601 | 0.90650187 |
| C2cd3 | 0.17141002 | 0.17837822 | 0.18908324 | -0.010705 | 0.59935952 | 0.90650187 |
| Otub2 | 0.05770507 | 0.05744394 | 0.06815918 | -0.0107152 | 0.75768957 | 0.90650187 |
| Tmppe | 0.08532356 | 0.08827096 | 0.09899125 | -0.0107203 | 0.68489749 | 0.90650187 |
| Ccdc136 | 0.10457508 | 0.10914837 | 0.1198695 | -0.0107211 | 0.65454308 | 0.90650187 |
| Secisbp2 | 0.2541415 | 0.257153 | 0.26787796 | -0.010725 | 0.57063363 | 0.90650187 |
| A930004D18 | 0.05194456 | 0.0508616 | 0.06158734 | -0.0107257 | 0.78060103 | 0.90650187 |
| Mtmr14 | 0.16395435 | 0.17089159 | 0.18162066 | -0.0107291 | 0.60357828 | 0.90650187 |
| Git2 | 0.44688144 | 0.4161214 | 0.42685255 | -0.0107312 | 0.54449449 | 0.90650187 |
| Gstm4 | 0.12797324 | 0.13396907 | 0.14470711 | -0.010738 | 0.62925756 | 0.90650187 |
| Adgra3 | 0.14128191 | 0.14779667 | 0.15853759 | -0.0107409 | 0.61836509 | 0.90650187 |
| Psme1 | 1.11584752 | 0.78407886 | 0.79482271 | -0.0107439 | 0.52395904 | 0.90650187 |
| Srd5a1 | 0.03999779 | 0.03705287 | 0.04779808 | -0.0107452 | 0.84119745 | 0.91251845 |
| Intu | 0.04739262 | 0.04560824 | 0.05635899 | -0.0107507 | 0.80178684 | 0.90839694 |
| Gm27029 | 0.04442962 | 0.04218501 | 0.0529388 | -0.0107538 | 0.81675406 | 0.9092537 |
| Nop14 | 0.35044159 | 0.34045903 | 0.35121627 | -0.0107572 | 0.55415356 | 0.90650187 |
| Zc3h12c | 0.03253259 | 0.02831286 | 0.03907023 | -0.0107574 | 0.88952909 | 0.9302877 |
| Sep-03 | 0.04285478 | 0.04035097 | 0.05111551 | -0.0107645 | 0.8254177 | 0.90947782 |
| Cenpv | 0.06774526 | 0.06872976 | 0.07949576 | -0.010766 | 0.72643364 | 0.90650187 |
| Slc25a12 | 0.1631653 | 0.17004971 | 0.18082723 | -0.0107775 | 0.60448123 | 0.90650187 |
| Ecd | 0.3689931 | 0.35555386 | 0.36633901 | -0.0107852 | 0.552065 | 0.90650187 |

| | | | | | | |
|---|---|---|---|---|---|---|
| Nudt6 | 0.05098916 | 0.04970703 | 0.06049257 | -0.0107855 | 0.78600334 | 0.90650187 |
| Tspan17 | 0.20154379 | 0.20782931 | 0.218617 | -0.0107877 | 0.58681873 | 0.90650187 |
| Six5 | 0.07832133 | 0.08049043 | 0.09127837 | -0.0107879 | 0.70035231 | 0.90650187 |
| Wars | 0.36305993 | 0.35074487 | 0.36153285 | -0.010788 | 0.55276686 | 0.90650187 |
| Mad2l1bp | 0.14731083 | 0.15394267 | 0.16473425 | -0.0107916 | 0.61456125 | 0.90650187 |
| Lin9 | 0.09499998 | 0.09874016 | 0.10953611 | -0.0107959 | 0.6693655 | 0.90650187 |
| Tpm3-rs7 | 1.37173881 | 0.85734409 | 0.868144 | -0.0107999 | 0.52205201 | 0.90650187 |
| Esrra | 0.13569324 | 0.14195446 | 0.15275539 | -0.0108009 | 0.62336749 | 0.90650187 |
| Lrig1 | 0.06318261 | 0.0635543 | 0.07436239 | -0.0108081 | 0.74090572 | 0.90650187 |
| Relt | 0.09553699 | 0.0993073 | 0.1101175 | -0.0108102 | 0.66873528 | 0.90650187 |
| Pigl | 0.04390259 | 0.0415106 | 0.05232905 | -0.0108185 | 0.82097313 | 0.9092537 |
| Itga4 | 0.04317641 | 0.04066945 | 0.05148819 | -0.0108187 | 0.82487738 | 0.90934433 |
| Orc6 | 0.61844565 | 0.53442128 | 0.54524266 | -0.0108214 | 0.53515367 | 0.90650187 |
| Asb8 | 0.26571792 | 0.26751838 | 0.27834893 | -0.0108305 | 0.5686658 | 0.90650187 |
| Slc17a5 | 0.1513599 | 0.15803436 | 0.16887189 | -0.0108375 | 0.61229343 | 0.90650187 |
| 2410089E03I | 0.19751125 | 0.20387723 | 0.21472206 | -0.0108448 | 0.58882814 | 0.90650187 |
| Cdkl2 | 0.12537364 | 0.1311174 | 0.14198089 | -0.0108635 | 0.63312824 | 0.90650187 |
| Med14 | 0.13777303 | 0.14404423 | 0.15491149 | -0.0108673 | 0.6224287 | 0.90650187 |
| Zbtb34 | 0.07468228 | 0.0763736 | 0.08724238 | -0.0108688 | 0.71015588 | 0.90650187 |
| Hirip3 | 0.25914541 | 0.26153803 | 0.27241977 | -0.0108817 | 0.57047302 | 0.90650187 |
| Rusc1 | 0.14970724 | 0.1563031 | 0.16718551 | -0.0108824 | 0.61385187 | 0.90650187 |
| Tanc1 | 0.15776868 | 0.1644984 | 0.17538178 | -0.0108834 | 0.60867751 | 0.90650187 |
| Fam110a | 0.15940726 | 0.16615339 | 0.17703868 | -0.0108853 | 0.60770391 | 0.90650187 |
| Cdr2l | 0.10839121 | 0.11307768 | 0.12396654 | -0.0108889 | 0.65191255 | 0.90650187 |
| Mpdz | 0.20563782 | 0.21165969 | 0.22255353 | -0.0108938 | 0.58613323 | 0.90650187 |
| E030030I06R | 0.13550738 | 0.14166074 | 0.15256245 | -0.0109017 | 0.62463282 | 0.90650187 |
| Ighmbp2 | 0.06562409 | 0.06620839 | 0.07711306 | -0.0109047 | 0.73521408 | 0.90650187 |
| Fars2 | 0.32467579 | 0.31882435 | 0.32973197 | -0.0109076 | 0.55845839 | 0.90650187 |
| Tdrp | 0.03464711 | 0.03063521 | 0.04155119 | -0.010916 | 0.87857891 | 0.92534436 |
| Fsd1 | 0.05366343 | 0.05263685 | 0.06355351 | -0.0109167 | 0.77745518 | 0.90650187 |

| | | | | | | |
|---|---|---|---|---|---|---|
| Gm26724 | 0.28087327 | 0.28093412 | 0.29186303 | -0.0109289 | 0.56610534 | 0.90650187 |
| Gm17484 | 0.0565151 | 0.05587431 | 0.06680569 | -0.0109314 | 0.76651929 | 0.90650187 |
| Gm26935 | 0.02741721 | 0.02210591 | 0.03303938 | -0.0109335 | 0.92937479 | 0.95006228 |
| Grk4 | 0.04531637 | 0.04302808 | 0.05396378 | -0.0109357 | 0.81617839 | 0.9092537 |
| Qk | 0.36059048 | 0.34857887 | 0.35952409 | -0.0109452 | 0.55382866 | 0.90650187 |
| Srebf1 | 0.21996164 | 0.22524351 | 0.23618891 | -0.0109454 | 0.58161119 | 0.90650187 |
| Cndp2 | 0.26021294 | 0.26243763 | 0.2733856 | -0.010948 | 0.57064962 | 0.90650187 |
| Mtfmt | 0.13808404 | 0.14428457 | 0.15523349 | -0.0109489 | 0.62307135 | 0.90650187 |
| E2f6 | 0.20927547 | 0.21508508 | 0.22603649 | -0.0109514 | 0.58526781 | 0.90650187 |
| Tti2 | 0.09693665 | 0.10066525 | 0.11163095 | -0.0109657 | 0.66883533 | 0.90650187 |
| Nepro | 0.0352022 | 0.0312354 | 0.04220132 | -0.0109659 | 0.87599186 | 0.92433377 |
| Zfp958 | 0.07403178 | 0.07555022 | 0.08651892 | -0.0109687 | 0.71348867 | 0.90650187 |
| Stxbp5 | 0.07507639 | 0.07671125 | 0.0876804 | -0.0109691 | 0.71094987 | 0.90650187 |
| Sall2 | 0.07416755 | 0.07569716 | 0.08666998 | -0.0109728 | 0.71322604 | 0.90650187 |
| Rbl1 | 0.06488741 | 0.06530693 | 0.07628401 | -0.0109771 | 0.73881991 | 0.90650187 |
| Celsr3 | 0.0636649 | 0.06392905 | 0.07490645 | -0.0109774 | 0.74263662 | 0.90650187 |
| Dip2b | 0.34584774 | 0.33644685 | 0.34742788 | -0.010981 | 0.55587176 | 0.90650187 |
| Zfp362 | 0.12827428 | 0.13403087 | 0.14502228 | -0.0109914 | 0.63191803 | 0.90650187 |
| Pex3 | 0.3159509 | 0.31133284 | 0.3223263 | -0.0109935 | 0.56025843 | 0.90650187 |
| Zfp90 | 0.1737979 | 0.18046632 | 0.19146024 | -0.0109939 | 0.6007439 | 0.90650187 |
| Slit1 | 0.07764861 | 0.0795363 | 0.09053371 | -0.0109974 | 0.705409 | 0.90650187 |
| Etl4 | 0.25937703 | 0.26162884 | 0.27262941 | -0.0110006 | 0.57118027 | 0.90650187 |
| Pak6 | 0.06035733 | 0.06015629 | 0.07116836 | -0.0110121 | 0.75423594 | 0.90650187 |
| Ppp1r12b | 0.11995512 | 0.12525733 | 0.1362715 | -0.0110142 | 0.64032106 | 0.90650187 |
| Mdn1 | 0.13998311 | 0.14617948 | 0.15719709 | -0.0110176 | 0.62232127 | 0.90650187 |
| Nup98 | 0.04770579 | 0.04570038 | 0.0567197 | -0.0110193 | 0.80611767 | 0.9092537 |
| Zfand4 | 0.03713455 | 0.03344093 | 0.04446078 | -0.0110198 | 0.86474045 | 0.92040222 |
| Adck3 | 0.03772436 | 0.03412779 | 0.04514929 | -0.0110215 | 0.86108949 | 0.91848494 |
| Lyst | 0.05147772 | 0.05002728 | 0.06105257 | -0.0110253 | 0.78896002 | 0.90650187 |
| Mipol1 | 0.082785 | 0.08517733 | 0.09620309 | -0.0110258 | 0.69480179 | 0.90650187 |

| | | | | | | |
|---|---|---|---|---|---|---|
| Pwp2 | 0.04615754 | 0.04390159 | 0.05493496 | -0.0110334 | 0.81403784 | 0.9092537 |
| Acsl5 | 0.30691915 | 0.30355075 | 0.31458904 | -0.0110383 | 0.56197847 | 0.90650187 |
| Zfc3h1 | 0.24751388 | 0.25078494 | 0.26182513 | -0.0110402 | 0.57434792 | 0.90650187 |
| Fbxo11 | 0.29354815 | 0.29195542 | 0.30300054 | -0.0110451 | 0.5643681 | 0.90650187 |
| Sap30l | 0.07261294 | 0.07389243 | 0.08493882 | -0.0110464 | 0.71842064 | 0.90650187 |
| Pcdhga8.1 | 0.03713405 | 0.03340983 | 0.0444602 | -0.0110504 | 0.86540598 | 0.92040222 |
| Map6 | 0.08492439 | 0.08750229 | 0.09855344 | -0.0110511 | 0.69093471 | 0.90650187 |
| Fam187b | 0.11486395 | 0.11982204 | 0.13087435 | -0.0110523 | 0.64632993 | 0.90650187 |
| Ptrh1 | 0.08846692 | 0.0913772 | 0.10243108 | -0.0110539 | 0.68429706 | 0.90650187 |
| Lsm11 | 0.09055855 | 0.09365826 | 0.10471231 | -0.011054 | 0.68057399 | 0.90650187 |
| Hebp2 | 0.0729933 | 0.07430326 | 0.08536271 | -0.0110594 | 0.71768068 | 0.90650187 |
| Pms2 | 0.13092275 | 0.13673059 | 0.14779043 | -0.0110598 | 0.6303141 | 0.90650187 |
| Dclre1b | 0.05734531 | 0.0566892 | 0.0677502 | -0.011061 | 0.76601822 | 0.90650187 |
| Pxmp4 | 0.29962275 | 0.29722356 | 0.30828571 | -0.0110622 | 0.56337074 | 0.90650187 |
| Rngtt | 0.23186156 | 0.23628793 | 0.24735487 | -0.0110669 | 0.57883008 | 0.90650187 |
| Zfp850 | 0.03352119 | 0.02916378 | 0.04023102 | -0.0110672 | 0.88932586 | 0.9302877 |
| Kctd2 | 0.04652264 | 0.04428557 | 0.05535616 | -0.0110706 | 0.81301727 | 0.9092537 |
| Ndrg4 | 0.07394238 | 0.07534762 | 0.08641945 | -0.0110718 | 0.71551161 | 0.90650187 |
| Cish | 0.14591919 | 0.15223138 | 0.16330765 | -0.0110763 | 0.61849154 | 0.90650187 |
| Plscr3 | 0.64032756 | 0.54796982 | 0.55904904 | -0.0110792 | 0.53510258 | 0.90650187 |
| Leng8 | 0.31718743 | 0.31229861 | 0.32338003 | -0.0110814 | 0.56054053 | 0.90650187 |
| Pcna-ps2 | 0.03999894 | 0.03671728 | 0.04779943 | -0.0110821 | 0.84865918 | 0.91440801 |
| Lmln | 0.05189437 | 0.05044269 | 0.06152987 | -0.0110872 | 0.7884521 | 0.90650187 |
| Traf2 | 0.20804734 | 0.21377467 | 0.22486213 | -0.0110875 | 0.58675408 | 0.90650187 |
| Mtmr9 | 0.08656593 | 0.08925946 | 0.10035246 | -0.011093 | 0.6884351 | 0.90650187 |
| Top1mt | 0.06874335 | 0.06952099 | 0.08061464 | -0.0110937 | 0.72965667 | 0.90650187 |
| Hus1 | 0.08860211 | 0.09148346 | 0.10257872 | -0.0110953 | 0.68468901 | 0.90650187 |
| Zbtb5 | 0.14280976 | 0.14901402 | 0.16011196 | -0.0110979 | 0.62101251 | 0.90650187 |
| Rnf32 | 0.08005287 | 0.08208983 | 0.09319213 | -0.0111023 | 0.70181245 | 0.90650187 |
| Gnb2l1 | 3.03170184 | 0.82145965 | 0.83256697 | -0.0111073 | 0.52364704 | 0.90650187 |

| | | | | | | |
|---|---|---|---|---|---|---|
| AI661453 | 0.07536587 | 0.07689412 | 0.08800198 | -0.0111079 | 0.71264796 | 0.90650187 |
| Ercc6 | 0.06439535 | 0.06462071 | 0.07572981 | -0.0111091 | 0.74284567 | 0.90650187 |
| Pepd | 0.37220283 | 0.35781682 | 0.36892674 | -0.0111099 | 0.55324957 | 0.90650187 |
| Fads3 | 0.0828758 | 0.08518994 | 0.09630299 | -0.011113 | 0.69602714 | 0.90650187 |
| Gm20498 | 0.08720609 | 0.08993903 | 0.10105301 | -0.011114 | 0.68755623 | 0.90650187 |
| AW209491 | 0.19262171 | 0.19885423 | 0.20997532 | -0.0111211 | 0.59307268 | 0.90650187 |
| Jmjd4 | 0.0635015 | 0.06360068 | 0.07472217 | -0.0111215 | 0.74591003 | 0.90650187 |
| Galnt10 | 0.10037707 | 0.10421964 | 0.11534137 | -0.0111217 | 0.66591649 | 0.90650187 |
| Phka2 | 0.06429511 | 0.06448554 | 0.07561687 | -0.0111313 | 0.74358066 | 0.90650187 |
| Trappc10 | 0.07356348 | 0.0748651 | 0.08599771 | -0.0111326 | 0.71752165 | 0.90650187 |
| Gm26670 | 0.08013151 | 0.08214131 | 0.09327895 | -0.0111376 | 0.70222533 | 0.90650187 |
| Rab40b | 0.10907508 | 0.11355068 | 0.12469883 | -0.0111482 | 0.65447848 | 0.90650187 |
| Phxr4 | 0.07308763 | 0.07430622 | 0.08546779 | -0.0111616 | 0.71923461 | 0.90650187 |
| Cdc25a | 0.19474472 | 0.20087355 | 0.21203946 | -0.0111659 | 0.59254782 | 0.90650187 |
| Slc6a17 | 0.03048004 | 0.02548907 | 0.03665529 | -0.0111662 | 0.91217256 | 0.94062208 |
| Lipt1 | 0.06854231 | 0.06922304 | 0.08038939 | -0.0111663 | 0.73155046 | 0.90650187 |
| Gtpbp10 | 0.10317982 | 0.10718568 | 0.11836698 | -0.0111813 | 0.66273729 | 0.90650187 |
| 4921524J17F | 0.7707041 | 0.62489786 | 0.63608344 | -0.0111856 | 0.53115614 | 0.90650187 |
| 0610030E20I | 0.33248703 | 0.32511878 | 0.33630496 | -0.0111862 | 0.55877726 | 0.90650187 |
| Fam63a | 0.2023583 | 0.20820769 | 0.21940158 | -0.0111939 | 0.58971629 | 0.90650187 |
| Fyco1 | 0.13187411 | 0.13758621 | 0.14878274 | -0.0111965 | 0.63101792 | 0.90650187 |
| Med12l | 0.04203063 | 0.03895285 | 0.05015982 | -0.011207 | 0.83970869 | 0.91213511 |
| Zfp764 | 0.0552805 | 0.0541876 | 0.06539918 | -0.0112116 | 0.77700684 | 0.90650187 |
| Pyroxd1 | 0.25358456 | 0.25615583 | 0.26737096 | -0.0112151 | 0.57396394 | 0.90650187 |
| Txnl1 | 1.37830301 | 0.85839063 | 0.8696126 | -0.011222 | 0.5228742 | 0.90650187 |
| Rad52 | 0.10343626 | 0.10741912 | 0.11864332 | -0.0112242 | 0.66296712 | 0.90650187 |
| Maml3 | 0.27770426 | 0.27782738 | 0.28905532 | -0.0112279 | 0.56854981 | 0.90650187 |
| Camsap1 | 0.21947399 | 0.22449817 | 0.23572817 | -0.01123 | 0.58386341 | 0.90650187 |
| Gigyf2 | 0.58420317 | 0.51185629 | 0.52308686 | -0.0112306 | 0.53801982 | 0.90650187 |
| Adora2a | 0.05319192 | 0.05178297 | 0.0630146 | -0.0112316 | 0.78592995 | 0.90650187 |

| | | | | | | |
|---|---|---|---|---|---|---|
| Chmp4c | 0.07583597 | 0.07728887 | 0.08852397 | -0.0112351 | 0.71369715 | 0.90650187 |
| Dctn3 | 1.60494324 | 0.89732318 | 0.90855889 | -0.0112357 | 0.52192148 | 0.90650187 |
| Gm11713 | 0.08987073 | 0.09272673 | 0.10396282 | -0.0112361 | 0.68455406 | 0.90650187 |
| Mrpl28 | 0.99967606 | 0.73934956 | 0.75058834 | -0.0112388 | 0.52653615 | 0.90650187 |
| Tbc1d5 | 0.06216169 | 0.06196994 | 0.07320959 | -0.0112397 | 0.75254513 | 0.90650187 |
| Mpv17 | 0.5917028 | 0.51674966 | 0.52799862 | -0.011249 | 0.53772871 | 0.90650187 |
| Pphln1 | 0.21730411 | 0.22241608 | 0.2336751 | -0.011259 | 0.58480443 | 0.90650187 |
| Mfsd12 | 0.08259302 | 0.08472319 | 0.09599187 | -0.0112687 | 0.69912263 | 0.90650187 |
| Gm26619 | 0.03500104 | 0.03069697 | 0.04196577 | -0.0112688 | 0.88371222 | 0.92771475 |
| Wdr44 | 0.06928217 | 0.06994193 | 0.08121806 | -0.0112761 | 0.73145451 | 0.90650187 |
| Clip1 | 0.40835943 | 0.38623049 | 0.39750677 | -0.0112763 | 0.55017793 | 0.90650187 |
| 2410002F23I | 0.51847536 | 0.46729457 | 0.47857177 | -0.0112772 | 0.54171588 | 0.90650187 |
| Ptpn21 | 0.09416153 | 0.09734457 | 0.10862758 | -0.011283 | 0.67787313 | 0.90650187 |
| Lgalsl | 0.18691253 | 0.19311125 | 0.20440046 | -0.0112892 | 0.59697877 | 0.90650187 |
| Kcnb2 | 0.22445689 | 0.22913208 | 0.24042425 | -0.0112922 | 0.58269776 | 0.90650187 |
| Vamp8 | 1.42014281 | 0.86725028 | 0.87854654 | -0.0112963 | 0.52279158 | 0.90650187 |
| Nfatc2ip | 0.04330579 | 0.04034041 | 0.05163807 | -0.0112977 | 0.83463245 | 0.91062493 |
| Gm7334 | 0.08915708 | 0.09188336 | 0.10318449 | -0.0113011 | 0.68683478 | 0.90650187 |
| Vasn | 0.12462509 | 0.12989208 | 0.14119436 | -0.0113023 | 0.63902754 | 0.90650187 |
| Qtrtd1 | 0.10147929 | 0.10522988 | 0.11653241 | -0.0113025 | 0.66683183 | 0.90650187 |
| Tmem201 | 0.1479637 | 0.15409785 | 0.16540275 | -0.0113049 | 0.61937784 | 0.90650187 |
| Cep126 | 0.1174871 | 0.1223529 | 0.13365912 | -0.0113062 | 0.64656153 | 0.90650187 |
| Dusp22 | 0.15073977 | 0.15693272 | 0.16823948 | -0.0113068 | 0.61744336 | 0.90650187 |
| Tcaim | 0.05363669 | 0.05221609 | 0.06352295 | -0.0113069 | 0.78561613 | 0.90650187 |
| Pemt | 0.05832646 | 0.05755373 | 0.06886513 | -0.0113114 | 0.76736917 | 0.90650187 |
| Golga7b | 0.03867677 | 0.03494765 | 0.04625993 | -0.0113123 | 0.8615073 | 0.91867924 |
| Map1lc3b | 1.61199144 | 0.89806996 | 0.9093845 | -0.0113145 | 0.52205511 | 0.90650187 |
| Utp3 | 0.72923988 | 0.60119228 | 0.61250713 | -0.0113149 | 0.53272585 | 0.90650187 |
| Usp21 | 0.12965638 | 0.13514938 | 0.14646788 | -0.0113185 | 0.6344039 | 0.90650187 |
| Ubxn11 | 0.05628216 | 0.05522124 | 0.06654048 | -0.0113192 | 0.77524881 | 0.90650187 |

| | | | | | | |
|---|---|---|---|---|---|---|
| D3Ertd751e | 0.12837887 | 0.13381178 | 0.14513175 | -0.01132 | 0.63560966 | 0.90650187 |
| Dmxl1 | 0.20676601 | 0.21230958 | 0.22363523 | -0.0113256 | 0.58906503 | 0.90650187 |
| Uvssa | 0.27926115 | 0.27910413 | 0.29043588 | -0.0113317 | 0.56885164 | 0.90650187 |
| Lactb2 | 0.14933511 | 0.15547343 | 0.16680535 | -0.0113319 | 0.6186787 | 0.90650187 |
| Shb | 0.04811037 | 0.04584959 | 0.05718548 | -0.0113359 | 0.81090974 | 0.9092537 |
| Slc12a6 | 0.12302187 | 0.12817127 | 0.13950731 | -0.011336 | 0.64103981 | 0.90650187 |
| Hist2h3c1 | 0.06379206 | 0.0637065 | 0.07504984 | -0.0113433 | 0.74918486 | 0.90650187 |
| C130013H08 | 0.04387349 | 0.04095057 | 0.05229536 | -0.0113448 | 0.83258141 | 0.91029548 |
| Rai14 | 0.09809575 | 0.10153103 | 0.11288224 | -0.0113512 | 0.67257726 | 0.90650187 |
| Alg8 | 0.11180333 | 0.1162613 | 0.12761447 | -0.0113532 | 0.65376359 | 0.90650187 |
| Mrpl12 | 0.83930472 | 0.66198697 | 0.6733456 | -0.0113586 | 0.52988975 | 0.90650187 |
| Mterf2 | 0.0403657 | 0.03686404 | 0.048226 | -0.011362 | 0.85253705 | 0.91557373 |
| Erp44 | 0.98580751 | 0.73345186 | 0.74481543 | -0.0113636 | 0.52703798 | 0.90650187 |
| Ttc7b | 0.05712476 | 0.05612896 | 0.06749939 | -0.0113704 | 0.77303301 | 0.90650187 |
| Calm3 | 1.55671024 | 0.89108594 | 0.90245836 | -0.0113724 | 0.52233777 | 0.90650187 |
| Rhebl1 | 0.03927606 | 0.03557884 | 0.04695806 | -0.0113792 | 0.85932493 | 0.91797966 |
| Cdk2 | 0.15035752 | 0.15647004 | 0.16784943 | -0.0113794 | 0.61844179 | 0.90650187 |
| Ska2 | 0.23502316 | 0.23891602 | 0.25029727 | -0.0113812 | 0.58009864 | 0.90650187 |
| Frat2 | 0.0600413 | 0.05942213 | 0.07081035 | -0.0113882 | 0.76268738 | 0.90650187 |
| Ube2v2 | 1.01103075 | 0.74383999 | 0.75523917 | -0.0113992 | 0.5267488 | 0.90650187 |
| 6330416G13 | 0.03583418 | 0.0315384 | 0.04294092 | -0.0114025 | 0.8810963 | 0.92635898 |
| Dcp1a | 0.06140176 | 0.06094572 | 0.07235049 | -0.0114048 | 0.75827698 | 0.90650187 |
| Adcy9 | 0.04811488 | 0.04577652 | 0.05719068 | -0.0114142 | 0.8125297 | 0.9092537 |
| Rnaseh2b | 0.28040898 | 0.2800337 | 0.29145225 | -0.0114186 | 0.56913428 | 0.90650187 |
| Satb1 | 0.12266351 | 0.12770249 | 0.13912978 | -0.0114273 | 0.64249491 | 0.90650187 |
| Taf6 | 0.40601704 | 0.38425882 | 0.39568625 | -0.0114274 | 0.55107952 | 0.90650187 |
| Xpo4 | 0.12465028 | 0.12979167 | 0.14122085 | -0.0114292 | 0.64049737 | 0.90650187 |
| Tmem192 | 0.21356944 | 0.21869369 | 0.23013015 | -0.0114365 | 0.58742541 | 0.90650187 |
| Lats2 | 0.14478975 | 0.15071028 | 0.16214818 | -0.0114379 | 0.62308457 | 0.90650187 |
| Tapbpl | 0.2002528 | 0.20592621 | 0.21737201 | -0.0114458 | 0.59254048 | 0.90650187 |

| | | | | | | |
|---|---|---|---|---|---|---|
| E130309D14I | 0.0329928 | 0.02816292 | 0.03961078 | -0.0114479 | 0.90057225 | 0.93508188 |
| Smarca4 | 0.85042999 | 0.66773725 | 0.67918812 | -0.0114509 | 0.52987329 | 0.90650187 |
| Zkscan8 | 0.09183447 | 0.09464962 | 0.10610089 | -0.0114513 | 0.68431736 | 0.90650187 |
| Whamm | 0.05789557 | 0.05692078 | 0.06837567 | -0.0114549 | 0.77179368 | 0.90650187 |
| Zfp445 | 0.07100793 | 0.07169047 | 0.0831479 | -0.0114574 | 0.72992485 | 0.90650187 |
| Grtp1 | 0.10309722 | 0.10681998 | 0.11827796 | -0.011458 | 0.66664242 | 0.90650187 |
| Mok | 0.04488354 | 0.04200552 | 0.05346363 | -0.0114581 | 0.82964527 | 0.90974008 |
| Map2k6 | 0.07292331 | 0.07382339 | 0.08528472 | -0.0114613 | 0.72489333 | 0.90650187 |
| Acap2 | 0.21346725 | 0.21855953 | 0.23003295 | -0.0114734 | 0.58773987 | 0.90650187 |
| Sdk1 | 0.11950842 | 0.12431338 | 0.13579922 | -0.0114858 | 0.64654428 | 0.90650187 |
| Nrtn | 0.17558487 | 0.18174401 | 0.19323455 | -0.0114905 | 0.60424665 | 0.90650187 |
| Spdl1 | 0.03769453 | 0.03362227 | 0.04511448 | -0.0114922 | 0.87127711 | 0.92225579 |
| 2410131K14I | 0.11919879 | 0.12397132 | 0.13547173 | -0.0115004 | 0.64705974 | 0.90650187 |
| 1700066M21 | 0.02771548 | 0.0218911 | 0.03339217 | -0.0115011 | 0.93713675 | 0.95459464 |
| Mcm8 | 0.03131837 | 0.02614008 | 0.03764243 | -0.0115023 | 0.91283755 | 0.9407546 |
| Kctd11 | 0.13354994 | 0.13902167 | 0.15052806 | -0.0115064 | 0.63300508 | 0.90650187 |
| Slc4a4 | 0.08891062 | 0.0914071 | 0.10291551 | -0.0115084 | 0.69044958 | 0.90650187 |
| Psmd10 | 0.40578691 | 0.3839841 | 0.39550717 | -0.0115231 | 0.55152784 | 0.90650187 |
| Ldlrad4 | 0.04592165 | 0.04313376 | 0.05466272 | -0.011529 | 0.82578172 | 0.90947782 |
| Slc27a3 | 0.07649982 | 0.07772831 | 0.08926055 | -0.0115322 | 0.71713824 | 0.90650187 |
| Tctn3 | 0.05926856 | 0.05840118 | 0.06993434 | -0.0115332 | 0.76826454 | 0.90650187 |
| Fam103a1 | 0.98807203 | 0.73424852 | 0.74578183 | -0.0115333 | 0.5274057 | 0.90650187 |
| Mcc | 0.11909925 | 0.12382788 | 0.13536641 | -0.0115385 | 0.64763385 | 0.90650187 |
| Arid3b | 0.05745163 | 0.05632417 | 0.06787109 | -0.0115469 | 0.77527477 | 0.90650187 |
| Tprn | 0.04438516 | 0.04133722 | 0.05288738 | -0.0115502 | 0.83419675 | 0.91062493 |
| Paqr7 | 0.05162175 | 0.04966731 | 0.0612176 | -0.0115503 | 0.79920329 | 0.90777025 |
| Rap1gds1 | 0.17318189 | 0.17929175 | 0.19084763 | -0.0115559 | 0.60610549 | 0.90650187 |
| Rpp38 | 0.16713482 | 0.17325341 | 0.18481168 | -0.0115583 | 0.609506 | 0.90650187 |
| Rmi2 | 0.06622982 | 0.06623315 | 0.07779415 | -0.011561 | 0.74558546 | 0.90650187 |
| Nup160 | 0.04910715 | 0.04676336 | 0.05833198 | -0.0115686 | 0.81102219 | 0.9092537 |

| | | | | | | |
|---|---|---|---|---|---|---|
| Rab8b | 0.12596991 | 0.13103518 | 0.14260692 | -0.0115717 | 0.64085144 | 0.90650187 |
| Hace1 | 0.13785316 | 0.1434225 | 0.15499447 | -0.011572 | 0.63002186 | 0.90650187 |
| Ang | 0.19801335 | 0.20363117 | 0.21520806 | -0.0115769 | 0.59450406 | 0.90650187 |
| Adcy3 | 0.1083418 | 0.11233145 | 0.1239136 | -0.0115822 | 0.66112264 | 0.90650187 |
| Sephs1 | 0.33629257 | 0.32790422 | 0.33948842 | -0.0115842 | 0.56028639 | 0.90650187 |
| Krr1 | 0.26200967 | 0.26342441 | 0.27500871 | -0.0115843 | 0.57427322 | 0.90650187 |
| Zbtb38 | 0.26768647 | 0.2685271 | 0.28011681 | -0.0115897 | 0.5729678 | 0.90650187 |
| Zfp874a | 0.10928846 | 0.11333238 | 0.12492719 | -0.0115948 | 0.66005316 | 0.90650187 |
| Apba1 | 0.03701934 | 0.03273088 | 0.04432623 | -0.0115953 | 0.87757119 | 0.92482217 |
| Trim39 | 0.11631864 | 0.12082078 | 0.13241971 | -0.0115989 | 0.65151003 | 0.90650187 |
| Pafah2 | 0.05752639 | 0.05635393 | 0.06795609 | -0.0116022 | 0.77607351 | 0.90650187 |
| Crebl2 | 0.03297948 | 0.027983 | 0.03959514 | -0.0116121 | 0.90384414 | 0.93655998 |
| Pdzd4 | 0.0500583 | 0.04780807 | 0.05942459 | -0.0116165 | 0.80758791 | 0.9092537 |
| Ubxn10 | 0.11406315 | 0.1183995 | 0.13002253 | -0.011623 | 0.65446511 | 0.90650187 |
| Sipa1l3 | 0.22498267 | 0.22928895 | 0.24091821 | -0.0116293 | 0.5849572 | 0.90650187 |
| Aebp2 | 0.25939648 | 0.26101275 | 0.27264701 | -0.0116343 | 0.5752282 | 0.90650187 |
| Nlgn2 | 0.13330173 | 0.13863177 | 0.15026977 | -0.011638 | 0.63468989 | 0.90650187 |
| Sorbs3 | 0.10531389 | 0.10901911 | 0.12066411 | -0.011645 | 0.66605107 | 0.90650187 |
| Vars2 | 0.08314078 | 0.08494025 | 0.09659441 | -0.0116542 | 0.70414666 | 0.90650187 |
| Bmyc | 0.85621818 | 0.67055281 | 0.6822077 | -0.0116549 | 0.53027019 | 0.90650187 |
| Tmem67 | 0.10468255 | 0.10832457 | 0.11998513 | -0.0116606 | 0.66714348 | 0.90650187 |
| Lmx1b | 0.04584267 | 0.04289773 | 0.05457156 | -0.0116738 | 0.8291964 | 0.90974008 |
| Etv3 | 0.10687 | 0.11065176 | 0.12233553 | -0.0116838 | 0.66442921 | 0.90650187 |
| Myo19 | 0.04815363 | 0.04554607 | 0.05723527 | -0.0116892 | 0.81804237 | 0.9092537 |
| Cep95 | 0.11269094 | 0.11687063 | 0.12856106 | -0.0116904 | 0.65698626 | 0.90650187 |
| Zwilch | 0.05610498 | 0.05464337 | 0.06633871 | -0.0116953 | 0.78340621 | 0.90650187 |
| Stat5b | 0.10404641 | 0.10760178 | 0.11930048 | -0.0116987 | 0.66855971 | 0.90650187 |
| Mal2 | 0.07162289 | 0.07212884 | 0.08383453 | -0.0117057 | 0.73261237 | 0.90650187 |
| Rpap1 | 0.11447725 | 0.11875244 | 0.13046311 | -0.0117107 | 0.65507099 | 0.90650187 |
| Thoc1 | 0.29193761 | 0.28987862 | 0.30159347 | -0.0117148 | 0.5685491 | 0.90650187 |

| | | | | | | |
|---|---|---|---|---|---|---|
| Smcr8 | 0.04706228 | 0.04426137 | 0.05597835 | -0.011717 | 0.82394938 | 0.90929265 |
| Rhobtb2 | 0.05206981 | 0.05001158 | 0.06173076 | -0.0117192 | 0.80065676 | 0.90806412 |
| Rbks | 0.04950387 | 0.04706758 | 0.05878787 | -0.0117203 | 0.81227264 | 0.9092537 |
| Fbxl12 | 0.15146377 | 0.15724773 | 0.16897779 | -0.0117301 | 0.62118859 | 0.90650187 |
| Strn4 | 0.7025323 | 0.58514615 | 0.59687781 | -0.0117317 | 0.53481472 | 0.90650187 |
| Rab11fip3 | 0.04824325 | 0.04560615 | 0.05733841 | -0.0117323 | 0.81849235 | 0.9092537 |
| Kcnu1 | 0.03682124 | 0.03236222 | 0.04409483 | -0.0117326 | 0.88157013 | 0.92661786 |
| Tm7sf3 | 0.43289326 | 0.40458805 | 0.41632396 | -0.0117359 | 0.54986446 | 0.90650187 |
| Dpf3 | 0.07489741 | 0.07573716 | 0.0874815 | -0.0117443 | 0.72468621 | 0.90650187 |
| Ppp1r9b | 0.18578637 | 0.19154913 | 0.20329665 | -0.0117475 | 0.60136806 | 0.90650187 |
| Arl10 | 0.04915515 | 0.04663602 | 0.05838715 | -0.0117511 | 0.81453497 | 0.9092537 |
| Rps6ka6 | 0.05281785 | 0.05083053 | 0.06258683 | -0.0117563 | 0.79815863 | 0.9075779 |
| Pbx2 | 0.34199436 | 0.33247376 | 0.34423529 | -0.0117615 | 0.56036458 | 0.90650187 |
| Zfp92 | 0.03101145 | 0.0255107 | 0.03728115 | -0.0117705 | 0.91977691 | 0.94431506 |
| Smpd4 | 0.09586281 | 0.09869627 | 0.11047005 | -0.0117738 | 0.68218156 | 0.90650187 |
| Pomt2 | 0.14581479 | 0.15142155 | 0.16320053 | -0.011779 | 0.62584217 | 0.90650187 |
| Ldlrap1 | 0.07559371 | 0.07647338 | 0.08825502 | -0.0117816 | 0.72356603 | 0.90650187 |
| Tesk1 | 0.10960547 | 0.11347937 | 0.12526637 | -0.011787 | 0.66214024 | 0.90650187 |
| Atxn1l | 0.11130441 | 0.11529044 | 0.12708197 | -0.0117915 | 0.66001108 | 0.90650187 |
| Galc | 0.0368649 | 0.03235217 | 0.04414584 | -0.0117937 | 0.88251533 | 0.92703446 |
| Supv3l1 | 0.06038166 | 0.05940126 | 0.07119592 | -0.0117947 | 0.76928092 | 0.90650187 |
| Dusp16 | 0.12515031 | 0.12994792 | 0.1417463 | -0.0117984 | 0.64432145 | 0.90650187 |
| Fam173a | 0.79727556 | 0.63896429 | 0.65076776 | -0.0118035 | 0.53213324 | 0.90650187 |
| Zmynd19 | 0.06087712 | 0.05995198 | 0.07175688 | -0.0118049 | 0.76771957 | 0.90650187 |
| Ttll3 | 0.09481935 | 0.09753406 | 0.10934047 | -0.0118064 | 0.68439678 | 0.90650187 |
| Fbxo21 | 0.58393315 | 0.5110989 | 0.52290941 | -0.0118105 | 0.53999022 | 0.90650187 |
| Tapt1 | 0.07727738 | 0.07830956 | 0.09012249 | -0.0118129 | 0.71995709 | 0.90650187 |
| Nsmaf | 0.15623468 | 0.1620133 | 0.17382786 | -0.0118146 | 0.61873291 | 0.90650187 |
| Athl1 | 0.15141228 | 0.1571107 | 0.1689253 | -0.0118146 | 0.62207109 | 0.90650187 |
| Atp6v1d | 0.75114142 | 0.61324447 | 0.62506235 | -0.0118179 | 0.5334924 | 0.90650187 |

| | | | | | | |
|---|---|---|---|---|---|---|
| Rasal2 | 0.07599366 | 0.07687856 | 0.088699 | -0.0118204 | 0.72322277 | 0.90650187 |
| C030014I23R | 0.05247011 | 0.05036379 | 0.06218897 | -0.0118252 | 0.80103625 | 0.90815573 |
| Bloc1s5 | 0.14666376 | 0.15224576 | 0.16407121 | -0.0118255 | 0.62567413 | 0.90650187 |
| Nudt10 | 0.04211456 | 0.0384276 | 0.0502572 | -0.0118296 | 0.85231449 | 0.91557373 |
| Prdm11 | 0.12011293 | 0.12460636 | 0.13643829 | -0.0118319 | 0.650075 | 0.90650187 |
| Hook3 | 0.54263822 | 0.48341717 | 0.49525162 | -0.0118344 | 0.54230063 | 0.90650187 |
| Ptpn12 | 0.14590714 | 0.15145424 | 0.16329529 | -0.011841 | 0.62641176 | 0.90650187 |
| Edc4 | 0.19554492 | 0.20095962 | 0.21281622 | -0.0118566 | 0.59780807 | 0.90650187 |
| Spsb4 | 0.08873915 | 0.09087062 | 0.10272834 | -0.0118577 | 0.6960758 | 0.90650187 |
| Dnajc16 | 0.06053616 | 0.05949105 | 0.07137088 | -0.0118798 | 0.77034409 | 0.90650187 |
| Tbc1d9 | 0.04298272 | 0.03938315 | 0.05126378 | -0.0118806 | 0.84856326 | 0.91440801 |
| Fhit | 0.0578869 | 0.05648037 | 0.06836582 | -0.0118855 | 0.78020096 | 0.90650187 |
| Atp6v1h | 0.55295697 | 0.49037263 | 0.50226072 | -0.0118881 | 0.54190087 | 0.90650187 |
| Trim45 | 0.10515041 | 0.10859573 | 0.12048834 | -0.0118926 | 0.66959551 | 0.90650187 |
| Slc35b4 | 0.14484468 | 0.15031152 | 0.16220461 | -0.0118931 | 0.62777215 | 0.90650187 |
| Eps15l1 | 0.15672099 | 0.16242694 | 0.17432077 | -0.0118938 | 0.61917767 | 0.90650187 |
| Zfp715 | 0.09745568 | 0.10029367 | 0.11219149 | -0.0118978 | 0.68134136 | 0.90650187 |
| Pitrm1 | 0.15462392 | 0.16029148 | 0.17219329 | -0.0119018 | 0.62068315 | 0.90650187 |
| Cenpj | 0.04828822 | 0.04548701 | 0.05739015 | -0.0119031 | 0.82174912 | 0.9092537 |
| Trim68 | 0.04317313 | 0.03958042 | 0.05148439 | -0.011904 | 0.84799948 | 0.91440801 |
| Mcoln1 | 0.11806918 | 0.12237193 | 0.13427591 | -0.011904 | 0.65325272 | 0.90650187 |
| Flrt1 | 0.13930862 | 0.14459538 | 0.15650016 | -0.0119048 | 0.63238324 | 0.90650187 |
| Pex16 | 0.2131691 | 0.21784183 | 0.22974929 | -0.0119075 | 0.59111232 | 0.90650187 |
| Tmem177 | 0.06260284 | 0.0617937 | 0.07370791 | -0.0119142 | 0.76380373 | 0.90650187 |
| Zfp182 | 0.05460786 | 0.05271586 | 0.06463193 | -0.0119161 | 0.79378595 | 0.90731056 |
| Zswim4 | 0.06124599 | 0.06025772 | 0.07217428 | -0.0119166 | 0.76852753 | 0.90650187 |
| Acad12 | 0.04926871 | 0.04659758 | 0.05851767 | -0.0119201 | 0.81741892 | 0.9092537 |
| Slc14a2 | 0.03576688 | 0.03092381 | 0.04286219 | -0.0119384 | 0.89218495 | 0.93152712 |
| Gcn1l1 | 0.32966372 | 0.32199496 | 0.33393523 | -0.0119403 | 0.56314908 | 0.90650187 |
| Pacsin1 | 0.08480453 | 0.08648131 | 0.09842193 | -0.0119406 | 0.70517531 | 0.90650187 |

| | | | | | | |
|---|---|---|---|---|---|---|
| Gnptg | 0.49459352 | 0.44976847 | 0.46171021 | -0.0119417 | 0.54577023 | 0.90650187 |
| Lrrc8b | 0.08793979 | 0.08990752 | 0.10185521 | -0.0119477 | 0.69898133 | 0.90650187 |
| Map3k14 | 0.07361849 | 0.07411109 | 0.08605896 | -0.0119479 | 0.7314484 | 0.90650187 |
| Trpt1 | 0.08653295 | 0.08836445 | 0.10031636 | -0.0119519 | 0.70182584 | 0.90650187 |
| Snhg20 | 0.36362262 | 0.35003389 | 0.36198988 | -0.011956 | 0.55836761 | 0.90650187 |
| Tnrc6c | 0.32322092 | 0.31654531 | 0.32850194 | -0.0119566 | 0.56427178 | 0.90650187 |
| Pex13 | 0.48227915 | 0.44089298 | 0.45286471 | -0.0119717 | 0.54677685 | 0.90650187 |
| Ndufaf4 | 0.42030829 | 0.39475527 | 0.40672924 | -0.011974 | 0.55206375 | 0.90650187 |
| Chchd1 | 1.12899045 | 0.78744888 | 0.79942677 | -0.0119779 | 0.52655348 | 0.90650187 |
| Gna12 | 0.10838624 | 0.11197914 | 0.12396122 | -0.0119821 | 0.66629734 | 0.90650187 |
| Fam199x | 0.0816548 | 0.08297127 | 0.09495884 | -0.0119876 | 0.71267341 | 0.90650187 |
| Tmem167b | 0.38675429 | 0.36856005 | 0.38055306 | -0.011993 | 0.55571024 | 0.90650187 |
| Ccdc39 | 0.04566899 | 0.04237692 | 0.05437104 | -0.0119941 | 0.83662705 | 0.91092327 |
| Ssu72 | 1.1757561 | 0.80285249 | 0.81484893 | -0.0119964 | 0.52609193 | 0.90650187 |
| Rhov | 0.05615688 | 0.05440015 | 0.06639782 | -0.0119977 | 0.78909742 | 0.90650187 |
| Hist2h3c2 | 0.0416053 | 0.03766228 | 0.0496662 | -0.0120039 | 0.85869338 | 0.9176516 |
| Nbas | 0.10774359 | 0.11126738 | 0.12327253 | -0.0120051 | 0.66747245 | 0.90650187 |
| Bmpr1a | 0.33837687 | 0.3292177 | 0.34122682 | -0.0120091 | 0.56216417 | 0.90650187 |
| Gm19412 | 0.14514783 | 0.15050516 | 0.16251594 | -0.0120108 | 0.62875346 | 0.90650187 |
| Magee1 | 0.1262354 | 0.13087059 | 0.14288552 | -0.0120149 | 0.64573139 | 0.90650187 |
| Fam149a | 0.1052449 | 0.10856713 | 0.12058994 | -0.0120228 | 0.67119914 | 0.90650187 |
| Smad1 | 0.33724089 | 0.3282478 | 0.34027982 | -0.012032 | 0.5624536 | 0.90650187 |
| Ptp4a1 | 1.09470477 | 0.77532895 | 0.78736351 | -0.0120346 | 0.52708705 | 0.90650187 |
| Agfg2 | 0.15669077 | 0.16225422 | 0.17429015 | -0.0120359 | 0.62057817 | 0.90650187 |
| Wnk1 | 0.90752114 | 0.69625669 | 0.70829696 | -0.0120403 | 0.53011957 | 0.90650187 |
| Dmd | 0.17924886 | 0.1848081 | 0.1968616 | -0.0120535 | 0.60726467 | 0.90650187 |
| Pigo | 0.09309697 | 0.09541848 | 0.10747263 | -0.0120542 | 0.69097646 | 0.90650187 |
| Mrpl9 | 0.5349091 | 0.47789498 | 0.48995604 | -0.0120611 | 0.54357158 | 0.90650187 |
| Lysmd3 | 0.26581662 | 0.26637004 | 0.27843765 | -0.0120676 | 0.57639306 | 0.90650187 |
| Hist1h2be | 0.04778459 | 0.04473624 | 0.05681044 | -0.0120742 | 0.82763146 | 0.90947782 |

| | | | | | | |
|---|---|---|---|---|---|---|
| Dgkq | 0.05086462 | 0.04827535 | 0.06034975 | -0.0120744 | 0.81311926 | 0.9092537 |
| Sik2 | 0.12077346 | 0.12506044 | 0.13713607 | -0.0120756 | 0.65227187 | 0.90650187 |
| Pex5l | 0.0817747 | 0.08301322 | 0.09509092 | -0.0120777 | 0.71384261 | 0.90650187 |
| Vat1l | 0.23244029 | 0.23581571 | 0.24789422 | -0.0120785 | 0.58574378 | 0.90650187 |
| Polr3b | 0.08521966 | 0.08679834 | 0.09887731 | -0.012079 | 0.70646281 | 0.90650187 |
| Cblc | 0.04072631 | 0.03656449 | 0.04864521 | -0.0120807 | 0.86521151 | 0.92040222 |
| Rgs9 | 0.04717502 | 0.04402674 | 0.05610827 | -0.0120815 | 0.83077994 | 0.9098597 |
| Tmem116 | 0.0478949 | 0.04485376 | 0.05693745 | -0.0120837 | 0.82728208 | 0.90947782 |
| Prim2 | 0.12417886 | 0.12863914 | 0.14072512 | -0.012086 | 0.64869682 | 0.90650187 |
| Rab21 | 0.09556329 | 0.09804957 | 0.11014596 | -0.0120964 | 0.68730407 | 0.90650187 |
| Bbs1 | 0.03981765 | 0.03548961 | 0.0475885 | -0.0120989 | 0.87081458 | 0.92219956 |
| Mar-09 | 0.07757037 | 0.07834701 | 0.09044706 | -0.0121001 | 0.72398982 | 0.90650187 |
| Mina | 0.1612703 | 0.16681726 | 0.17891883 | -0.0121016 | 0.61817399 | 0.90650187 |
| Tm9sf1 | 0.11648369 | 0.1204933 | 0.13259488 | -0.0121016 | 0.6575327 | 0.90650187 |
| Galr1 | 0.05806418 | 0.05645681 | 0.06856723 | -0.0121104 | 0.78382826 | 0.90650187 |
| Fto | 0.33243145 | 0.32414509 | 0.33625838 | -0.0121133 | 0.56361757 | 0.90650187 |
| Syt17 | 0.07269984 | 0.07292058 | 0.08503569 | -0.0121151 | 0.73675381 | 0.90650187 |
| Pls1 | 0.0612903 | 0.06010761 | 0.0722244 | -0.0121168 | 0.77211236 | 0.90650187 |
| Cyp4f13 | 0.16472567 | 0.17027197 | 0.18239558 | -0.0121236 | 0.61619169 | 0.90650187 |
| Ctif | 0.0764181 | 0.07704332 | 0.08916991 | -0.0121266 | 0.72726977 | 0.90650187 |
| Tbc1d7 | 0.28626287 | 0.28448397 | 0.29661686 | -0.0121329 | 0.57214749 | 0.90650187 |
| Bcl2l11 | 0.17427974 | 0.17977195 | 0.19193901 | -0.0121671 | 0.61095469 | 0.90650187 |
| Fkbp15 | 0.18437691 | 0.18974512 | 0.20191321 | -0.0121681 | 0.60561565 | 0.90650187 |
| Sarm1 | 0.04976377 | 0.0469181 | 0.0590864 | -0.0121683 | 0.82004375 | 0.9092537 |
| C2cd2l | 0.13475351 | 0.13960983 | 0.15177948 | -0.0121696 | 0.63924739 | 0.90650187 |
| Copb2 | 1.09340108 | 0.77472796 | 0.78689811 | -0.0121701 | 0.52740793 | 0.90650187 |
| Gtpbp3 | 0.14891512 | 0.15418979 | 0.16637606 | -0.0121863 | 0.62764409 | 0.90650187 |
| Card10 | 0.03699018 | 0.03207527 | 0.04429217 | -0.0122169 | 0.8899404 | 0.9303978 |
| Simc1 | 0.08138095 | 0.08243328 | 0.09465708 | -0.0122238 | 0.71704838 | 0.90650187 |
| Kalrn | 0.1275709 | 0.13205814 | 0.1442857 | -0.0122276 | 0.64681851 | 0.90650187 |

| | | | | | | |
|---|---|---|---|---|---|---|
| Nxt2 | 0.11044879 | 0.11393651 | 0.12616805 | -0.0122315 | 0.66676054 | 0.90650187 |
| Taf8 | 0.13901646 | 0.14396651 | 0.15619812 | -0.0122316 | 0.63612797 | 0.90650187 |
| 4931406C07I | 0.2594413 | 0.26045314 | 0.27268758 | -0.0122344 | 0.5790474 | 0.90650187 |
| Brca2 | 0.0829503 | 0.08412977 | 0.09638493 | -0.0122552 | 0.71404834 | 0.90650187 |
| Fuz | 0.2303472 | 0.23368547 | 0.24594196 | -0.0122565 | 0.58766645 | 0.90650187 |
| Pard6b | 0.05247769 | 0.04993746 | 0.06219765 | -0.0122602 | 0.8095672 | 0.9092537 |
| Wdr91 | 0.10821167 | 0.11151341 | 0.12377418 | -0.0122608 | 0.6701658 | 0.90650187 |
| Cdk19 | 0.08968397 | 0.09149547 | 0.1037592 | -0.0122637 | 0.70036293 | 0.90650187 |
| Plod3 | 0.375279 | 0.35913166 | 0.37139851 | -0.0122668 | 0.55836808 | 0.90650187 |
| Tmem241 | 0.03643307 | 0.03136676 | 0.04364122 | -0.0122745 | 0.89441676 | 0.93244834 |
| Gm26782 | 0.08081002 | 0.08175272 | 0.09402762 | -0.0122749 | 0.71916225 | 0.90650187 |
| Ripk1 | 0.20401764 | 0.20872242 | 0.22099779 | -0.0122754 | 0.59751961 | 0.90650187 |
| Gnb2 | 2.90227272 | 0.83825201 | 0.85052804 | -0.012276 | 0.52558071 | 0.90650187 |
| Spata5 | 0.10782065 | 0.11107834 | 0.12335513 | -0.0122768 | 0.67091611 | 0.90650187 |
| Zfand2a | 0.07102456 | 0.07088786 | 0.08316647 | -0.0122786 | 0.74419881 | 0.90650187 |
| Aasdh | 0.06494655 | 0.06407182 | 0.07635059 | -0.0122788 | 0.76267896 | 0.90650187 |
| Zfr2 | 0.10147835 | 0.10424761 | 0.1165314 | -0.0122838 | 0.68032925 | 0.90650187 |
| Klf5 | 0.07773996 | 0.07835033 | 0.09063486 | -0.0122845 | 0.72659657 | 0.90650187 |
| Bptf | 0.8651246 | 0.67452476 | 0.68682615 | -0.0123014 | 0.53173143 | 0.90650187 |
| 6330418K02I | 0.10868912 | 0.11197709 | 0.12428561 | -0.0123085 | 0.67012806 | 0.90650187 |
| Micall1 | 0.0657899 | 0.06498714 | 0.07729955 | -0.0123124 | 0.76055977 | 0.90650187 |
| Svop | 0.04969872 | 0.04669875 | 0.05901169 | -0.0123129 | 0.82318896 | 0.9092537 |
| Nxpe3 | 0.04794455 | 0.04467944 | 0.0569946 | -0.0123152 | 0.83162038 | 0.9100469 |
| Nrde2 | 0.11617942 | 0.11995664 | 0.13227193 | -0.0123153 | 0.66053061 | 0.90650187 |
| Ankrd42 | 0.03849178 | 0.03372832 | 0.04604431 | -0.012316 | 0.88280314 | 0.9272727 |
| Gla | 0.0467535 | 0.04329386 | 0.05562239 | -0.0123285 | 0.83777511 | 0.91127152 |
| Lias | 0.36539599 | 0.35109792 | 0.36342856 | -0.0123306 | 0.55994647 | 0.90650187 |
| Armc6 | 0.1832449 | 0.18846885 | 0.20080054 | -0.0123317 | 0.60757969 | 0.90650187 |
| Alg12 | 0.09774226 | 0.10015402 | 0.11250083 | -0.0123468 | 0.68718946 | 0.90650187 |
| 2900060B14I | 0.07760301 | 0.07813144 | 0.09048321 | -0.0123518 | 0.72802968 | 0.90650187 |

| | | | | | | |
|---|---|---|---|---|---|---|
| Nop2 | 0.20042371 | 0.20518049 | 0.21753692 | -0.0123564 | 0.59967937 | 0.90650187 |
| Mad1l1 | 0.11495797 | 0.11861775 | 0.13097432 | -0.0123566 | 0.66254313 | 0.90650187 |
| Uty | 0.07159562 | 0.07143757 | 0.0838041 | -0.0123665 | 0.74409256 | 0.90650187 |
| 1600002H07 | 0.06482176 | 0.06384166 | 0.07621009 | -0.0123684 | 0.76469742 | 0.90650187 |
| Sass6 | 0.06795216 | 0.06735532 | 0.07972781 | -0.0123725 | 0.75486767 | 0.90650187 |
| Myl6b | 0.06473625 | 0.06374117 | 0.07611379 | -0.0123726 | 0.7650533 | 0.90650187 |
| Ssfa2 | 0.11043734 | 0.11377547 | 0.12615581 | -0.0123803 | 0.66867987 | 0.90650187 |
| Dbt | 0.14683709 | 0.15186626 | 0.16424886 | -0.0123826 | 0.63124437 | 0.90650187 |
| Csnk1g2 | 0.20911465 | 0.21349988 | 0.22588281 | -0.0123829 | 0.59627179 | 0.90650187 |
| Fbxo45 | 0.06206572 | 0.06070481 | 0.07310114 | -0.0123963 | 0.77453808 | 0.90650187 |
| Inha | 0.08818196 | 0.08971431 | 0.10211982 | -0.0124055 | 0.7054193 | 0.90650187 |
| Ago3 | 0.16546648 | 0.17073367 | 0.18313922 | -0.0124055 | 0.61834538 | 0.90650187 |
| Abhd6 | 0.0954082 | 0.09756776 | 0.10997811 | -0.0124103 | 0.69204631 | 0.90650187 |
| Babam1 | 0.62581691 | 0.53750383 | 0.5499229 | -0.0124191 | 0.53998519 | 0.90650187 |
| Cenpo | 0.0460539 | 0.04239533 | 0.05481536 | -0.01242 | 0.84310052 | 0.91275344 |
| Uvrag | 0.2212251 | 0.22496104 | 0.23738151 | -0.0124205 | 0.5919666 | 0.90650187 |
| Gpr179 | 0.04528592 | 0.04150235 | 0.0539286 | -0.0124263 | 0.84716703 | 0.9143112 |
| 3110070M22 | 0.03744712 | 0.03239681 | 0.04482572 | -0.0124289 | 0.89114515 | 0.93109327 |
| Eif2ak4 | 0.0907483 | 0.09248309 | 0.10491896 | -0.0124359 | 0.70087908 | 0.90650187 |
| Zfp628 | 0.08476096 | 0.08593617 | 0.09837412 | -0.0124379 | 0.71296957 | 0.90650187 |
| Casc4 | 0.4748538 | 0.4350435 | 0.44748149 | -0.012438 | 0.54917133 | 0.90650187 |
| Hcn3 | 0.04401785 | 0.04002176 | 0.05246244 | -0.0124407 | 0.85410632 | 0.91608365 |
| Dcp1b | 0.13906489 | 0.14380255 | 0.15624819 | -0.0124456 | 0.63837027 | 0.90650187 |
| Fdft1 | 0.49287954 | 0.44803964 | 0.46048533 | -0.0124457 | 0.54781899 | 0.90650187 |
| Insrr | 0.15798379 | 0.16314852 | 0.17559946 | -0.0124509 | 0.62370169 | 0.90650187 |
| Vps18 | 0.13219467 | 0.13666384 | 0.14911685 | -0.012453 | 0.64477818 | 0.90650187 |
| Cacna1d | 0.08588809 | 0.08714848 | 0.09961006 | -0.0124616 | 0.71094971 | 0.90650187 |
| Tnik | 0.24305092 | 0.24525575 | 0.25772384 | -0.0124681 | 0.58514294 | 0.90650187 |
| Sh2b3 | 0.16708747 | 0.17229604 | 0.18476425 | -0.0124682 | 0.61791036 | 0.90650187 |
| Trim11 | 0.19379374 | 0.19864673 | 0.21111545 | -0.0124687 | 0.6035561 | 0.90650187 |

| | | | | | | |
|---|---|---|---|---|---|---|
| Smim14 | 1.26346844 | 0.82852566 | 0.84099497 | -0.0124693 | 0.526277 | 0.90650187 |
| Eif3b | 0.90713303 | 0.69563399 | 0.70810406 | -0.0124701 | 0.53120105 | 0.90650187 |
| Slc30a1 | 0.06800854 | 0.06731913 | 0.07979104 | -0.0124719 | 0.75643482 | 0.90650187 |
| Ddx46 | 0.71154234 | 0.5897188 | 0.60219073 | -0.0124719 | 0.53667982 | 0.90650187 |
| Lmbrd2 | 0.0951797 | 0.09725697 | 0.10973073 | -0.0124738 | 0.69334661 | 0.90650187 |
| Unc5a | 0.04943476 | 0.04623288 | 0.05870846 | -0.0124756 | 0.82759242 | 0.90947782 |
| Nup210 | 0.09119799 | 0.09293266 | 0.10540849 | -0.0124758 | 0.70061445 | 0.90650187 |
| Zscan26 | 0.2500748 | 0.25168979 | 0.26416901 | -0.0124792 | 0.58316943 | 0.90650187 |
| Mllt6 | 0.12043014 | 0.12428962 | 0.13677346 | -0.0124838 | 0.65754329 | 0.90650187 |
| Kdm4c | 0.13425447 | 0.13876889 | 0.15126082 | -0.0124919 | 0.64324347 | 0.90650187 |
| Ap3m2 | 0.06175853 | 0.06025874 | 0.07275392 | -0.0124952 | 0.7774217 | 0.90650187 |
| Aftph | 0.21124856 | 0.21542275 | 0.22791991 | -0.0124972 | 0.59629115 | 0.90650187 |
| Fam204a | 0.75419828 | 0.61429408 | 0.62679638 | -0.0125023 | 0.53532937 | 0.90650187 |
| Slc7a6 | 0.05781741 | 0.05578396 | 0.06828685 | -0.0125029 | 0.79216678 | 0.90681274 |
| 9330020H09I | 0.11064996 | 0.11387707 | 0.12638301 | -0.0125059 | 0.66999686 | 0.90650187 |
| Tmem246 | 0.20814796 | 0.21245221 | 0.2249584 | -0.0125062 | 0.59760245 | 0.90650187 |
| Prpf40b | 0.15321735 | 0.15825671 | 0.1707635 | -0.0125068 | 0.62763571 | 0.90650187 |
| Zbtb11os1 | 0.11981991 | 0.12361522 | 0.13612857 | -0.0125134 | 0.65860309 | 0.90650187 |
| Hnrnpa0 | 0.54357213 | 0.48337421 | 0.4958889 | -0.0125147 | 0.54466489 | 0.90650187 |
| Dgke | 0.05937132 | 0.05753275 | 0.07005088 | -0.0125181 | 0.78652799 | 0.90650187 |
| Ralgps1 | 0.08612829 | 0.08735179 | 0.09987321 | -0.0125214 | 0.71135918 | 0.90650187 |
| Rgs12 | 0.13210209 | 0.13649572 | 0.14902037 | -0.0125247 | 0.64566315 | 0.90650187 |
| Kcnmb4 | 0.0433531 | 0.03915496 | 0.05169286 | -0.0125379 | 0.85954201 | 0.91800029 |
| Entpd5 | 0.12000299 | 0.12378403 | 0.1363221 | -0.0125381 | 0.65868593 | 0.90650187 |
| Uhmk1 | 0.16375839 | 0.16886817 | 0.18142367 | -0.0125555 | 0.62082955 | 0.90650187 |
| 1700029J07F | 0.0784116 | 0.07881852 | 0.09137825 | -0.0125597 | 0.72941286 | 0.90650187 |
| Stk11ip | 0.09050335 | 0.0920903 | 0.10465218 | -0.0125619 | 0.70320001 | 0.90650187 |
| Sirt5 | 0.10772503 | 0.11068276 | 0.12325263 | -0.0125699 | 0.67486362 | 0.90650187 |
| Clec16a | 0.06386933 | 0.06256408 | 0.07513697 | -0.0125729 | 0.77153039 | 0.90650187 |
| Amer2 | 0.05175264 | 0.04879451 | 0.06136753 | -0.012573 | 0.8188047 | 0.9092537 |

| | | | | | | |
|---|---|---|---|---|---|---|
| Crls1 | 0.07686733 | 0.07709343 | 0.08966805 | -0.0125746 | 0.73349919 | 0.90650187 |
| Map3k12 | 0.19601421 | 0.20069041 | 0.21327145 | -0.012581 | 0.60343555 | 0.90650187 |
| Sdccag8 | 0.26335854 | 0.26364315 | 0.27622522 | -0.0125821 | 0.58023615 | 0.90650187 |
| Rfwd2 | 0.27509972 | 0.27415511 | 0.28674076 | -0.0125856 | 0.5773532 | 0.90650187 |
| Hmga1 | 0.0699136 | 0.06933643 | 0.08192466 | -0.0125882 | 0.7527266 | 0.90650187 |
| Tk2 | 0.093078 | 0.09485596 | 0.10745203 | -0.0125961 | 0.69886413 | 0.90650187 |
| Zswim1 | 0.08094454 | 0.08157609 | 0.09417597 | -0.0125999 | 0.72400525 | 0.90650187 |
| Pfdn4 | 1.08695476 | 0.7719569 | 0.78455981 | -0.0126029 | 0.52846537 | 0.90650187 |
| Ccnj | 0.06102331 | 0.05931038 | 0.07192233 | -0.012612 | 0.78217506 | 0.90650187 |
| Eif5 | 1.69091898 | 0.90490925 | 0.91752339 | -0.0126141 | 0.52436751 | 0.90650187 |
| Cldn9 | 0.03069208 | 0.0242873 | 0.03690507 | -0.0126178 | 0.9357375 | 0.95373281 |
| Syne4 | 0.13268951 | 0.13700619 | 0.14963237 | -0.0126262 | 0.64621673 | 0.90650187 |
| Pnpla6 | 0.12301513 | 0.12685905 | 0.13950022 | -0.0126412 | 0.65647066 | 0.90650187 |
| Tigar | 0.06493649 | 0.0636916 | 0.07633927 | -0.0126477 | 0.7692975 | 0.90650187 |
| Chrnb2 | 0.04955075 | 0.04619131 | 0.05884172 | -0.0126504 | 0.83040404 | 0.90982931 |
| 4932422M17 | 0.06386319 | 0.06247766 | 0.07513004 | -0.0126524 | 0.77297136 | 0.90650187 |
| Dixdc1 | 0.12969785 | 0.13385794 | 0.14651123 | -0.0126533 | 0.64948863 | 0.90650187 |
| Stoml1 | 0.13832215 | 0.14282161 | 0.15547991 | -0.0126583 | 0.64130024 | 0.90650187 |
| Slc35d1 | 0.06834274 | 0.0674979 | 0.08016572 | -0.0126678 | 0.7588152 | 0.90650187 |
| Zfp111 | 0.08258895 | 0.0833102 | 0.09598739 | -0.0126772 | 0.72144433 | 0.90650187 |
| Tbc1d25 | 0.05282065 | 0.04991041 | 0.06259003 | -0.0126796 | 0.81609953 | 0.9092537 |
| Gpr27 | 0.06190282 | 0.06023302 | 0.07291703 | -0.012684 | 0.78033314 | 0.90650187 |
| Zfp799 | 0.0686542 | 0.06782237 | 0.08051476 | -0.0126924 | 0.75828854 | 0.90650187 |
| AW146154 | 0.07057232 | 0.06996528 | 0.08266119 | -0.0126959 | 0.75263324 | 0.90650187 |
| Ttl | 0.14692819 | 0.15163619 | 0.16434222 | -0.012706 | 0.63446766 | 0.90650187 |
| Fbf1 | 0.08537702 | 0.08633795 | 0.09904987 | -0.0127119 | 0.71584523 | 0.90650187 |
| Zfp40 | 0.08364193 | 0.08443157 | 0.0971453 | -0.0127137 | 0.71965304 | 0.90650187 |
| Tpp1 | 0.36932614 | 0.35388364 | 0.36660796 | -0.0127243 | 0.56131308 | 0.90650187 |
| Nfyb | 0.51567834 | 0.46388621 | 0.47661611 | -0.0127299 | 0.54725827 | 0.90650187 |
| Ppt2 | 0.19529822 | 0.1998464 | 0.21257682 | -0.0127304 | 0.60496913 | 0.90650187 |

| | | | | | | |
|---|---|---|---|---|---|---|
| Mon1b | 0.0419012 | 0.03726939 | 0.05000965 | -0.0127403 | 0.87129719 | 0.92225579 |
| Slc27a4 | 0.2004683 | 0.20483969 | 0.21757995 | -0.0127403 | 0.60268759 | 0.90650187 |
| Gm4950 | 0.18844619 | 0.19314466 | 0.20590149 | -0.0127568 | 0.6085078 | 0.90650187 |
| Ube3b | 0.11993515 | 0.12349348 | 0.1362504 | -0.0127569 | 0.66138114 | 0.90650187 |
| Ddb2 | 0.05293935 | 0.04996787 | 0.06272579 | -0.0127579 | 0.81705546 | 0.9092537 |
| Zmat3 | 0.19427252 | 0.19882185 | 0.21158077 | -0.0127589 | 0.60568213 | 0.90650187 |
| Usp39 | 0.19521466 | 0.19973182 | 0.21249572 | -0.0127639 | 0.60527797 | 0.90650187 |
| Prkar2b | 0.31032579 | 0.30474781 | 0.31751584 | -0.012768 | 0.57093998 | 0.90650187 |
| Tbc1d17 | 0.40876648 | 0.38505094 | 0.39782271 | -0.0127718 | 0.55674536 | 0.90650187 |
| 2510046G10 | 0.12654962 | 0.13044116 | 0.14321516 | -0.012774 | 0.65414053 | 0.90650187 |
| Emc1 | 0.18262793 | 0.18741142 | 0.20019352 | -0.0127821 | 0.61173591 | 0.90650187 |
| Fgfr1op | 0.16452909 | 0.16941506 | 0.18219814 | -0.0127831 | 0.62244377 | 0.90650187 |
| Orai2 | 0.10637072 | 0.10901526 | 0.12179958 | -0.0127843 | 0.67961398 | 0.90650187 |
| Ttpal | 0.10518374 | 0.10773887 | 0.12052418 | -0.0127853 | 0.68139155 | 0.90650187 |
| Eya1 | 0.04176009 | 0.03705969 | 0.04984588 | -0.0127862 | 0.87293227 | 0.92289911 |
| Spag1 | 0.13372193 | 0.13791464 | 0.150707 | -0.0127924 | 0.64704448 | 0.90650187 |
| Ulk3 | 0.09870709 | 0.10074727 | 0.11354146 | -0.0127942 | 0.6917948 | 0.90650187 |
| Dram2 | 0.4238421 | 0.39663787 | 0.40943516 | -0.0127973 | 0.55525585 | 0.90650187 |
| Eif2a | 0.57213681 | 0.50231583 | 0.5151152 | -0.0127994 | 0.54397872 | 0.90650187 |
| Slc25a22 | 0.16917849 | 0.17403807 | 0.18685615 | -0.0128181 | 0.61980264 | 0.90650187 |
| Ranbp6 | 0.10785207 | 0.11056952 | 0.12338881 | -0.0128193 | 0.67791257 | 0.90650187 |
| Ppm1l | 0.22057434 | 0.22392391 | 0.23676744 | -0.0128435 | 0.5952819 | 0.90650187 |
| Mroh1 | 0.10295185 | 0.10527192 | 0.11812125 | -0.0128493 | 0.68566399 | 0.90650187 |
| Tsta3 | 0.38685502 | 0.36777933 | 0.38063296 | -0.0128536 | 0.55966665 | 0.90650187 |
| Gm6594 | 0.05043143 | 0.04699888 | 0.05985284 | -0.012854 | 0.83014435 | 0.9097662 |
| Entpd6 | 0.10602583 | 0.10856517 | 0.12142919 | -0.012864 | 0.68116559 | 0.90650187 |
| Mycbpap | 0.0393964 | 0.03423387 | 0.04709817 | -0.0128643 | 0.88767648 | 0.92975747 |
| Gm15545 | 0.10512311 | 0.10758992 | 0.12045899 | -0.0128691 | 0.6825852 | 0.90650187 |
| Dtnbp1 | 0.20694358 | 0.21093217 | 0.22380536 | -0.0128732 | 0.60091236 | 0.90650187 |
| Vegfb | 0.16299123 | 0.16777474 | 0.18065209 | -0.0128773 | 0.62433811 | 0.90650187 |

| | | | | | | |
|---|---|---|---|---|---|---|
| Med6 | 0.40573996 | 0.38258441 | 0.39547063 | -0.0128862 | 0.55758848 | 0.90650187 |
| Akip1 | 0.14372628 | 0.14816381 | 0.16105508 | -0.0128913 | 0.639019 | 0.90650187 |
| Ints1 | 0.08510549 | 0.08585638 | 0.0987521 | -0.0128957 | 0.71922388 | 0.90650187 |
| Pign | 0.0911062 | 0.09240816 | 0.1053086 | -0.0129004 | 0.70698301 | 0.90650187 |
| Nsmf | 0.29228393 | 0.28898664 | 0.30189623 | -0.0129096 | 0.57538522 | 0.90650187 |
| Zfp579 | 0.30689662 | 0.30165912 | 0.31456964 | -0.0129105 | 0.57238755 | 0.90650187 |
| Myo9a | 0.16692198 | 0.1716794 | 0.18459849 | -0.0129191 | 0.62214713 | 0.90650187 |
| Fiz1 | 0.34355252 | 0.33260689 | 0.34552774 | -0.0129209 | 0.56601651 | 0.90650187 |
| Mdfic | 0.09733524 | 0.09913838 | 0.11206145 | -0.0129231 | 0.69591002 | 0.90650187 |
| Hif1an | 0.20223942 | 0.20636102 | 0.21928711 | -0.0129261 | 0.6033587 | 0.90650187 |
| Bbs5 | 0.17738115 | 0.18207928 | 0.19501456 | -0.0129353 | 0.61595579 | 0.90650187 |
| Gm13111 | 0.03737238 | 0.03179775 | 0.04473847 | -0.0129407 | 0.90076165 | 0.93521466 |
| Arhgap23 | 0.11835654 | 0.12163246 | 0.13458026 | -0.0129478 | 0.6655675 | 0.90650187 |
| Anks1 | 0.09129627 | 0.09256015 | 0.10551545 | -0.0129553 | 0.70741077 | 0.90650187 |
| Iqce | 0.13122846 | 0.13515209 | 0.14810942 | -0.0129573 | 0.65133352 | 0.90650187 |
| Sgtb | 0.05286601 | 0.04968069 | 0.06264191 | -0.0129612 | 0.82117157 | 0.9092537 |
| Hsd17b11 | 0.24848869 | 0.24975368 | 0.26271808 | -0.0129644 | 0.58682223 | 0.90650187 |
| Ankrd13d | 0.07247195 | 0.07181322 | 0.08478166 | -0.0129684 | 0.75175158 | 0.90650187 |
| Cwf19l1 | 0.04303855 | 0.03835894 | 0.05132847 | -0.0129695 | 0.8693456 | 0.92141903 |
| Rps2-ps6 | 0.03718013 | 0.03154401 | 0.04451401 | -0.01297 | 0.90239367 | 0.93614183 |
| Zfp692 | 0.05400791 | 0.05097635 | 0.06394702 | -0.0129707 | 0.81639799 | 0.9092537 |
| 2900076A07 | 0.24265284 | 0.24438216 | 0.25735707 | -0.0129749 | 0.58867181 | 0.90650187 |
| Traip | 0.04744635 | 0.04344234 | 0.05642089 | -0.0129785 | 0.84675785 | 0.91417373 |
| Ak5 | 0.05286908 | 0.04965815 | 0.06264543 | -0.0129873 | 0.82164096 | 0.9092537 |
| Chn1 | 0.06744639 | 0.06617032 | 0.07916044 | -0.0129901 | 0.76715756 | 0.90650187 |
| Rpia | 0.41510757 | 0.38973305 | 0.40272838 | -0.0129953 | 0.55703331 | 0.90650187 |
| Lysmd4 | 0.0709166 | 0.07004294 | 0.08304587 | -0.0130029 | 0.75680827 | 0.90650187 |
| Fam19a5 | 0.06336657 | 0.06155735 | 0.07456995 | -0.0130126 | 0.78105661 | 0.90650187 |
| Zbtb14 | 0.17550749 | 0.18013549 | 0.1931578 | -0.0130223 | 0.61780224 | 0.90650187 |
| BC049715 | 0.0607044 | 0.05853716 | 0.07156137 | -0.0130242 | 0.79077204 | 0.90650187 |

| | | | | | | |
|---|---|---|---|---|---|---|
| 2510039O18 | 0.18730552 | 0.19175761 | 0.20478533 | -0.0130277 | 0.61134022 | 0.90650187 |
| Rwdd2b | 0.05473467 | 0.05174866 | 0.06477662 | -0.013028 | 0.81436747 | 0.9092537 |
| Eri2 | 0.08547275 | 0.08612013 | 0.09915483 | -0.0130347 | 0.72052803 | 0.90650187 |
| Rtkn | 0.16810383 | 0.17273883 | 0.18578165 | -0.0130428 | 0.62251934 | 0.90650187 |
| Itga9 | 0.04224804 | 0.03736486 | 0.05041204 | -0.0130472 | 0.87504561 | 0.92397668 |
| Mdc1 | 0.07080966 | 0.06987682 | 0.0829264 | -0.0130496 | 0.75790387 | 0.90650187 |
| Hcfc2 | 0.20612438 | 0.20996745 | 0.2230202 | -0.0130528 | 0.60264098 | 0.90650187 |
| Srgap3 | 0.06634186 | 0.06486213 | 0.07792007 | -0.0130579 | 0.77186411 | 0.90650187 |
| Senp5 | 0.1981025 | 0.20223553 | 0.21529433 | -0.0130588 | 0.60628551 | 0.90650187 |
| Zfp646 | 0.15502672 | 0.15954281 | 0.17260232 | -0.0130595 | 0.63170231 | 0.90650187 |
| Nup155 | 0.09643603 | 0.09802849 | 0.11108994 | -0.0130615 | 0.69940646 | 0.90650187 |
| 3110009E18I | 0.12612895 | 0.12971051 | 0.14277383 | -0.0130633 | 0.65790702 | 0.90650187 |
| Gan | 0.09237606 | 0.09362338 | 0.10668961 | -0.0130662 | 0.70693454 | 0.90650187 |
| Tmem180 | 0.06172108 | 0.05964291 | 0.07271158 | -0.0130687 | 0.78786117 | 0.90650187 |
| Zscan29 | 0.11031727 | 0.11295564 | 0.12602748 | -0.0130718 | 0.67763941 | 0.90650187 |
| Ssbp3 | 0.5971414 | 0.51846571 | 0.53153871 | -0.013073 | 0.54353282 | 0.90650187 |
| Rhot2 | 0.20833746 | 0.21206525 | 0.2251397 | -0.0130744 | 0.60186132 | 0.90650187 |
| Hiatl1 | 0.36055896 | 0.34641872 | 0.35949842 | -0.0130797 | 0.5642468 | 0.90650187 |
| Slc35d3 | 0.13052932 | 0.13429959 | 0.14737975 | -0.0130802 | 0.65341388 | 0.90650187 |
| Dapk3 | 0.12535071 | 0.12887598 | 0.1419568 | -0.0130808 | 0.65896756 | 0.90650187 |
| Prr14l | 0.18064542 | 0.18515772 | 0.19824019 | -0.0130825 | 0.6153837 | 0.90650187 |
| Otud5 | 0.37814087 | 0.36060721 | 0.37369132 | -0.0130841 | 0.56184741 | 0.90650187 |
| Mid1 | 0.08500402 | 0.08555307 | 0.0986408 | -0.0130877 | 0.72235309 | 0.90650187 |
| Chtf18 | 0.05549799 | 0.05255412 | 0.06564712 | -0.013093 | 0.81236762 | 0.9092537 |
| Hsdl2 | 0.3164701 | 0.3096656 | 0.3227689 | -0.0131033 | 0.57161066 | 0.90650187 |
| Daglb | 0.12719053 | 0.13077914 | 0.14388712 | -0.013108 | 0.65725778 | 0.90650187 |
| Ikzf5 | 0.27770653 | 0.27594918 | 0.28905735 | -0.0131082 | 0.579885 | 0.90650187 |
| A430005L14I | 0.40245224 | 0.37979719 | 0.39290768 | -0.0131105 | 0.55896294 | 0.90650187 |
| Ercc3 | 0.14179686 | 0.14595734 | 0.15906852 | -0.0131112 | 0.64297714 | 0.90650187 |
| Cep112 | 0.10667963 | 0.10901402 | 0.12213122 | -0.0131172 | 0.68348597 | 0.90650187 |

| | | | | | | |
|---|---|---|---|---|---|---|
| Bckdha | 0.45250524 | 0.41792183 | 0.43104437 | -0.0131225 | 0.55382858 | 0.90650187 |
| Tyw5 | 0.13949447 | 0.14356773 | 0.15669225 | -0.0131245 | 0.6451888 | 0.90650187 |
| RP23-54G8.4 | 0.08019178 | 0.08021785 | 0.09334547 | -0.0131276 | 0.73408246 | 0.90650187 |
| Uggt2 | 0.16144318 | 0.16596353 | 0.17909311 | -0.0131296 | 0.62775523 | 0.90650187 |
| Elovl7 | 0.07278975 | 0.07200447 | 0.08513589 | -0.0131314 | 0.7535468 | 0.90650187 |
| Hist1h4d | 0.04394779 | 0.03923388 | 0.05238136 | -0.0131475 | 0.86775172 | 0.92062353 |
| Tpd52l1 | 0.08426105 | 0.08467593 | 0.09782539 | -0.0131495 | 0.72493584 | 0.90650187 |
| Zfp119b | 0.06112895 | 0.05889198 | 0.07204186 | -0.0131499 | 0.79144956 | 0.90661829 |
| Ralgapa2 | 0.14532607 | 0.14954675 | 0.16269894 | -0.0131522 | 0.64034167 | 0.90650187 |
| Gmds | 0.3309772 | 0.3218862 | 0.33503854 | -0.0131523 | 0.56927064 | 0.90650187 |
| Aqr | 0.31573356 | 0.30898769 | 0.32214094 | -0.0131533 | 0.57201926 | 0.90650187 |
| Pgap1 | 0.25164571 | 0.25244894 | 0.26560363 | -0.0131547 | 0.58713445 | 0.90650187 |
| Gm13251 | 0.14218119 | 0.14630516 | 0.15946458 | -0.0131594 | 0.64313926 | 0.90650187 |
| Tnrc6a | 0.92919632 | 0.70576121 | 0.71892227 | -0.0131611 | 0.53243178 | 0.90650187 |
| Abhd11 | 0.37471156 | 0.35777953 | 0.37094313 | -0.0131636 | 0.56267722 | 0.90650187 |
| Pag1 | 0.07898813 | 0.07885051 | 0.09201585 | -0.0131653 | 0.73763251 | 0.90650187 |
| 2610318N02 | 0.03097702 | 0.02406428 | 0.03724061 | -0.0131763 | 0.94213216 | 0.95782176 |
| Ankrd52 | 0.04259183 | 0.03763234 | 0.05081071 | -0.0131784 | 0.87554927 | 0.92418014 |
| Zfp651 | 0.14354764 | 0.14768181 | 0.16087133 | -0.0131895 | 0.64225266 | 0.90650187 |
| Mlf1 | 0.07382442 | 0.07309375 | 0.08628818 | -0.0131944 | 0.75167901 | 0.90650187 |
| Plch1 | 0.05184016 | 0.04826576 | 0.06146778 | -0.013202 | 0.83016689 | 0.9097662 |
| Swsap1 | 0.10229418 | 0.10420794 | 0.11741198 | -0.013204 | 0.69144449 | 0.90650187 |
| Wdr92 | 0.19497315 | 0.19904381 | 0.21226127 | -0.0132175 | 0.60904346 | 0.90650187 |
| Cobll1 | 0.10475352 | 0.10684314 | 0.12006147 | -0.0132183 | 0.68773672 | 0.90650187 |
| Clasp2 | 0.14743235 | 0.15163617 | 0.16485872 | -0.0132226 | 0.63928991 | 0.90650187 |
| Snx29 | 0.08450697 | 0.08486971 | 0.09809537 | -0.0132257 | 0.72553909 | 0.90650187 |
| Shmt2 | 0.26617555 | 0.26552025 | 0.27876023 | -0.01324 | 0.5836142 | 0.90650187 |
| Ddx20 | 0.13432202 | 0.13808891 | 0.15133104 | -0.0132421 | 0.65136567 | 0.90650187 |
| Esf1 | 0.57963678 | 0.50681514 | 0.52008013 | -0.013265 | 0.5451386 | 0.90650187 |
| AU041133 | 0.08459251 | 0.08492366 | 0.09818926 | -0.0132656 | 0.72595075 | 0.90650187 |

| | | | | | | |
|---|---|---|---|---|---|---|
| Tdg | 0.70214408 | 0.58338167 | 0.59664799 | -0.0132663 | 0.53937013 | 0.90650187 |
| Gramd4 | 0.05436782 | 0.0510905 | 0.06435795 | -0.0132675 | 0.82028872 | 0.9092537 |
| Cox6c | 3.18665985 | 0.80082868 | 0.81410622 | -0.0132775 | 0.52889995 | 0.90650187 |
| G6pdx | 0.13881497 | 0.14270962 | 0.15598975 | -0.0132801 | 0.64746117 | 0.90650187 |
| Tor4a | 0.04810996 | 0.04390191 | 0.057185 | -0.0132831 | 0.84911334 | 0.91444933 |
| Naf1 | 0.11890308 | 0.12186986 | 0.13515884 | -0.013289 | 0.66897954 | 0.90650187 |
| Brsk1 | 0.05548189 | 0.05232919 | 0.06562877 | -0.0132996 | 0.81617813 | 0.9092537 |
| Slmap | 0.32184081 | 0.31402748 | 0.32733344 | -0.013306 | 0.57170314 | 0.90650187 |
| Cdk5r1 | 0.079393 | 0.07915571 | 0.09246333 | -0.0133076 | 0.73885714 | 0.90650187 |
| Mrpl39 | 0.36645071 | 0.35096799 | 0.36428303 | -0.013315 | 0.56454116 | 0.90650187 |
| Lyrm1 | 0.08154603 | 0.08151957 | 0.094839 | -0.0133194 | 0.73380188 | 0.90650187 |
| Agbl5 | 0.09167201 | 0.0926025 | 0.10592421 | -0.0133217 | 0.71195594 | 0.90650187 |
| Celsr2 | 0.1662645 | 0.1706139 | 0.1839396 | -0.0133257 | 0.62629889 | 0.90650187 |
| Zdhhc1 | 0.20994697 | 0.21335121 | 0.22667793 | -0.0133267 | 0.60309396 | 0.90650187 |
| Rgs11 | 0.04646789 | 0.04196155 | 0.05529301 | -0.0133315 | 0.85810882 | 0.91737198 |
| Zfp335 | 0.06495111 | 0.06302015 | 0.07635572 | -0.0133356 | 0.78125181 | 0.90650187 |
| Nr2c1 | 0.10252226 | 0.10432028 | 0.11765802 | -0.0133377 | 0.69285357 | 0.90650187 |
| Tmem147 | 1.27719215 | 0.83143435 | 0.8447731 | -0.0133388 | 0.52798077 | 0.90650187 |
| Slc25a33 | 0.13491415 | 0.1385925 | 0.15194638 | -0.0133539 | 0.65199208 | 0.90650187 |
| Itch | 0.52599936 | 0.47044877 | 0.48380517 | -0.0133564 | 0.54883943 | 0.90650187 |
| Cog5 | 0.04753263 | 0.04316008 | 0.05652027 | -0.0133602 | 0.85333554 | 0.91587612 |
| Opn3 | 0.06076878 | 0.05826761 | 0.07163425 | -0.0133666 | 0.79659784 | 0.9075779 |
| Ccdc62 | 0.05323501 | 0.04969127 | 0.06306386 | -0.0133726 | 0.82707873 | 0.90947782 |
| Efna2 | 0.29261337 | 0.2888112 | 0.30218414 | -0.0133729 | 0.57798336 | 0.90650187 |
| Derl1 | 0.68050533 | 0.57033812 | 0.58371558 | -0.0133775 | 0.54057538 | 0.90650187 |
| Spin4 | 0.06547421 | 0.06356621 | 0.07694446 | -0.0133782 | 0.78022472 | 0.90650187 |
| 3010003L21F | 0.12310677 | 0.12621082 | 0.13959672 | -0.0133859 | 0.66505113 | 0.90650187 |
| Clasrp | 0.14604822 | 0.15004877 | 0.16344001 | -0.0133912 | 0.64216245 | 0.90650187 |
| Tmem259 | 0.15260232 | 0.1567391 | 0.1701376 | -0.0133985 | 0.63686666 | 0.90650187 |
| Snrpa | 1.08880742 | 0.77183767 | 0.78523706 | -0.0133994 | 0.53023494 | 0.90650187 |

| | | | | | | |
|---|---|---|---|---|---|---|
| D17H6S53E | 0.0791037 | 0.07873938 | 0.0921436 | -0.0134042 | 0.74108917 | 0.90650187 |
| Nfrkb | 0.18138547 | 0.1855583 | 0.19896985 | -0.0134115 | 0.61778011 | 0.90650187 |
| Sos1 | 0.15846493 | 0.16267153 | 0.17608618 | -0.0134147 | 0.63257332 | 0.90650187 |
| Me1 | 0.13275971 | 0.13628976 | 0.14970548 | -0.0134157 | 0.6548283 | 0.90650187 |
| Apex2 | 0.06398749 | 0.06185341 | 0.07527017 | -0.0134168 | 0.7859422 | 0.90650187 |
| Prkd2 | 0.09599729 | 0.09718012 | 0.11061552 | -0.0134354 | 0.70538937 | 0.90650187 |
| Alkbh2 | 0.18729765 | 0.19133185 | 0.20477762 | -0.0134458 | 0.61481849 | 0.90650187 |
| Bend7 | 0.22868146 | 0.23093924 | 0.24438513 | -0.0134459 | 0.59661466 | 0.90650187 |
| RP23-60P18. | 0.09413488 | 0.09515101 | 0.10859868 | -0.0134477 | 0.70901007 | 0.90650187 |
| Ezh1 | 0.19586863 | 0.19968215 | 0.21313026 | -0.0134481 | 0.61045656 | 0.90650187 |
| Scai | 0.19526963 | 0.19909659 | 0.21254907 | -0.0134525 | 0.61078578 | 0.90650187 |
| Kif5c | 0.13437427 | 0.13793112 | 0.15138535 | -0.0134542 | 0.65361511 | 0.90650187 |
| Id4 | 0.06091009 | 0.05833444 | 0.0717942 | -0.0134598 | 0.79770413 | 0.9075779 |
| Rragb | 0.08443767 | 0.08455877 | 0.0980193 | -0.0134605 | 0.72923264 | 0.90650187 |
| Park2 | 0.0608552 | 0.05826684 | 0.07173207 | -0.0134652 | 0.79800308 | 0.9075779 |
| Haus3 | 0.15221134 | 0.15627366 | 0.16973949 | -0.0134658 | 0.63783594 | 0.90650187 |
| Jtb | 0.90617754 | 0.69416225 | 0.7076289 | -0.0134667 | 0.53371145 | 0.90650187 |
| Mettl13 | 0.14210685 | 0.14591829 | 0.15938798 | -0.0134697 | 0.64642742 | 0.90650187 |
| Rwdd3 | 0.12714687 | 0.13037111 | 0.14384136 | -0.0134702 | 0.6614105 | 0.90650187 |
| Ccdc94 | 0.08256126 | 0.08248323 | 0.09595692 | -0.0134737 | 0.73376012 | 0.90650187 |
| Ap4m1 | 0.3524326 | 0.33937388 | 0.35285278 | -0.0134789 | 0.56742461 | 0.90650187 |
| Zfp81 | 0.16575642 | 0.16994334 | 0.1834301 | -0.0134868 | 0.62811484 | 0.90650187 |
| Tmem102 | 0.05215246 | 0.04833106 | 0.06182539 | -0.0134943 | 0.83405671 | 0.91062493 |
| Sgk3 | 0.09397398 | 0.0949298 | 0.10842422 | -0.0134944 | 0.70997019 | 0.90650187 |
| Bahcc1 | 0.09809812 | 0.09937684 | 0.1128848 | -0.013508 | 0.7026258 | 0.90650187 |
| D10Jhu81e | 0.46827355 | 0.42916124 | 0.44267911 | -0.0135179 | 0.55399181 | 0.90650187 |
| Slc10a7 | 0.14056414 | 0.14427431 | 0.15779702 | -0.0135227 | 0.64838892 | 0.90650187 |
| Carnmt1 | 0.20634418 | 0.20970548 | 0.22323093 | -0.0135254 | 0.60617252 | 0.90650187 |
| Qpctl | 0.2380701 | 0.23959639 | 0.25312354 | -0.0135272 | 0.59389553 | 0.90650187 |
| Ankle2 | 0.20167391 | 0.20520861 | 0.21874238 | -0.0135338 | 0.6083624 | 0.90650187 |

| | | | | | | |
|---|---|---|---|---|---|---|
| Cep57l1 | 0.1716387 | 0.17577694 | 0.18931115 | -0.0135342 | 0.62469108 | 0.90650187 |
| F2rl1 | 0.05504743 | 0.0515975 | 0.0651334 | -0.0135359 | 0.82222497 | 0.9092537 |
| Fhad1 | 0.1268442 | 0.12998583 | 0.14352408 | -0.0135382 | 0.66251702 | 0.90650187 |
| Cog2 | 0.19130145 | 0.19515037 | 0.20868924 | -0.0135389 | 0.61348428 | 0.90650187 |
| Anapc1 | 0.25939564 | 0.25910392 | 0.27264626 | -0.0135423 | 0.5873811 | 0.90650187 |
| Mib2 | 0.15641037 | 0.16046101 | 0.17400596 | -0.013545 | 0.63534911 | 0.90650187 |
| AI467606 | 0.13849066 | 0.14210849 | 0.15565428 | -0.0135458 | 0.65057687 | 0.90650187 |
| 9430015G10 | 0.0477023 | 0.04316344 | 0.05671569 | -0.0135522 | 0.85594577 | 0.9165432 |
| Zfp446 | 0.07231222 | 0.07104957 | 0.08460356 | -0.013554 | 0.76183335 | 0.90650187 |
| Nkapl | 0.03716259 | 0.03093877 | 0.04449353 | -0.0135548 | 0.91218512 | 0.94062208 |
| Acer3 | 0.1112895 | 0.11351045 | 0.12706605 | -0.0135556 | 0.68234419 | 0.90650187 |
| Zrsr1 | 0.06538455 | 0.06328765 | 0.07684357 | -0.0135559 | 0.78355222 | 0.90650187 |
| Osgepl1 | 0.11207316 | 0.11434608 | 0.12790233 | -0.0135562 | 0.68124588 | 0.90650187 |
| Abcd4 | 0.12033011 | 0.12310667 | 0.13666778 | -0.0135611 | 0.67043818 | 0.90650187 |
| Thumpd2 | 0.09950339 | 0.10083817 | 0.11439935 | -0.0135612 | 0.70090084 | 0.90650187 |
| Aste1 | 0.06342648 | 0.06107292 | 0.07463754 | -0.0135646 | 0.79043868 | 0.90650187 |
| Fam63b | 0.28711203 | 0.2837985 | 0.29736342 | -0.0135649 | 0.58035301 | 0.90650187 |
| Ddx52 | 0.29742019 | 0.29280852 | 0.30637359 | -0.0135651 | 0.57802153 | 0.90650187 |
| Vps33b | 0.08852025 | 0.08891152 | 0.10248932 | -0.0135778 | 0.72205416 | 0.90650187 |
| Trmt5 | 0.05991058 | 0.05708309 | 0.07066223 | -0.0135791 | 0.80353366 | 0.90882608 |
| Ptpn9 | 0.30897824 | 0.30277689 | 0.31635932 | -0.0135824 | 0.57568457 | 0.90650187 |
| Slc35g1 | 0.14209619 | 0.14579042 | 0.159377 | -0.0135866 | 0.6476484 | 0.90650187 |
| Tap2 | 0.10985779 | 0.11194188 | 0.12553624 | -0.0135944 | 0.68488965 | 0.90650187 |
| Plcb3 | 0.11550666 | 0.11796189 | 0.13155744 | -0.0135955 | 0.6770346 | 0.90650187 |
| Rptor | 0.12810891 | 0.13125122 | 0.14484915 | -0.0135979 | 0.66178438 | 0.90650187 |
| Soga3 | 0.05485529 | 0.05131337 | 0.06491423 | -0.0136009 | 0.82418846 | 0.90929265 |
| Mad2l2 | 0.41205196 | 0.38675971 | 0.40036823 | -0.0136085 | 0.56005405 | 0.90650187 |
| Wdfy1 | 0.33555807 | 0.32526544 | 0.33887494 | -0.0136095 | 0.57085029 | 0.90650187 |
| Naa16 | 0.22723361 | 0.22941649 | 0.24302969 | -0.0136132 | 0.59832682 | 0.90650187 |
| Zfp354a | 0.06726662 | 0.06534008 | 0.07895868 | -0.0136186 | 0.77839004 | 0.90650187 |

| | | | | | | |
|---|---|---|---|---|---|---|
| Sirt1 | 0.05607301 | 0.05267704 | 0.0663023 | -0.0136253 | 0.81953157 | 0.9092537 |
| Lrba | 0.17045181 | 0.1744971 | 0.18812761 | -0.0136305 | 0.62631146 | 0.90650187 |
| Lmo3 | 0.05856141 | 0.05549061 | 0.0691319 | -0.0136413 | 0.80981608 | 0.9092537 |
| AI606181 | 0.05873019 | 0.05567086 | 0.0693235 | -0.0136526 | 0.80935517 | 0.9092537 |
| Rnf185 | 0.08948253 | 0.08987404 | 0.10353952 | -0.0136655 | 0.72131265 | 0.90650187 |
| Lta4h | 0.29736104 | 0.29265099 | 0.30632218 | -0.0136712 | 0.578637 | 0.90650187 |
| Pomt1 | 0.07913352 | 0.07849518 | 0.09217657 | -0.0136814 | 0.74532273 | 0.90650187 |
| Bod1l | 0.49520586 | 0.44846546 | 0.46214721 | -0.0136818 | 0.55235406 | 0.90650187 |
| Taf4a | 0.08026433 | 0.07973698 | 0.09342555 | -0.0136886 | 0.74259419 | 0.90650187 |
| Ap4s1 | 0.30251275 | 0.29709558 | 0.31078804 | -0.0136925 | 0.57764058 | 0.90650187 |
| Wdr54 | 0.07338417 | 0.07210283 | 0.08579807 | -0.0136952 | 0.76102874 | 0.90650187 |
| Pdss1 | 0.05444723 | 0.05075282 | 0.0644486 | -0.0136958 | 0.82759974 | 0.90947782 |
| Mccc1 | 0.16019139 | 0.16412925 | 0.17783049 | -0.0137012 | 0.63401978 | 0.90650187 |
| Fam105a | 0.10217583 | 0.10357562 | 0.11728429 | -0.0137087 | 0.69833603 | 0.90650187 |
| Abtb1 | 0.14315821 | 0.14676001 | 0.16047064 | -0.0137106 | 0.64796529 | 0.90650187 |
| Pja1 | 0.92600835 | 0.70366024 | 0.71737436 | -0.0137141 | 0.53386419 | 0.90650187 |
| Plpp7 | 0.08441979 | 0.08427619 | 0.09799967 | -0.0137235 | 0.73320808 | 0.90650187 |
| Prickle3 | 0.08568427 | 0.0856563 | 0.09938669 | -0.0137304 | 0.73044715 | 0.90650187 |
| Klhdc9 | 0.04568712 | 0.04065827 | 0.05439197 | -0.0137337 | 0.86917361 | 0.92141903 |
| 4632404H12I | 0.04207879 | 0.03647798 | 0.0502157 | -0.0137377 | 0.88804635 | 0.92986219 |
| Gm5595 | 0.07573252 | 0.07467015 | 0.08840914 | -0.013739 | 0.75517256 | 0.90650187 |
| Tmie | 0.10710231 | 0.10884309 | 0.12258479 | -0.0137417 | 0.69088147 | 0.90650187 |
| Slc25a1 | 0.23268943 | 0.23438409 | 0.2481263 | -0.0137422 | 0.59724214 | 0.90650187 |
| Ercc4 | 0.07189862 | 0.07039576 | 0.08414222 | -0.0137465 | 0.76616836 | 0.90650187 |
| Caprin1 | 1.19758407 | 0.80796518 | 0.82171381 | -0.0137486 | 0.52964692 | 0.90650187 |
| Pi4k2a | 0.07855546 | 0.07778079 | 0.0915374 | -0.0137566 | 0.74796616 | 0.90650187 |
| Rfx2 | 0.07713346 | 0.07620093 | 0.08996303 | -0.0137621 | 0.75176882 | 0.90650187 |
| Acot8 | 0.23506215 | 0.23656944 | 0.2503335 | -0.0137641 | 0.59655197 | 0.90650187 |
| Jarid2 | 0.40498476 | 0.38111507 | 0.39488264 | -0.0137676 | 0.56158773 | 0.90650187 |
| Capn7 | 0.35791263 | 0.34356215 | 0.35734021 | -0.0137781 | 0.56804944 | 0.90650187 |

| | | | | | | |
|---|---|---|---|---|---|---|
| Acap3 | 0.12908134 | 0.13208749 | 0.1458667 | -0.0137792 | 0.6627395 | 0.90650187 |
| Rabep1 | 0.5458064 | 0.48361911 | 0.49741076 | -0.0137917 | 0.54905053 | 0.90650187 |
| Depdc5 | 0.0761555 | 0.07508512 | 0.08887859 | -0.0137935 | 0.7548821 | 0.90650187 |
| Alkbh6 | 0.48367064 | 0.44007387 | 0.45386936 | -0.0137955 | 0.55374367 | 0.90650187 |
| Bzw1 | 1.57359395 | 0.89105575 | 0.90486438 | -0.0138086 | 0.52704426 | 0.90650187 |
| D430019H16 | 0.09280363 | 0.093344 | 0.1071541 | -0.0138101 | 0.71666173 | 0.90650187 |
| Yae1d1 | 0.36503223 | 0.34932114 | 0.36313366 | -0.0138125 | 0.56714003 | 0.90650187 |
| Gm12355 | 0.09954106 | 0.10062466 | 0.11443991 | -0.0138152 | 0.70425424 | 0.90650187 |
| Rmnd5a | 0.19387261 | 0.19737665 | 0.21119212 | -0.0138155 | 0.61440412 | 0.90650187 |
| 2700060E02l | 1.76219257 | 0.90848849 | 0.92230396 | -0.0138155 | 0.52654678 | 0.90650187 |
| Gm20522 | 0.04790131 | 0.04312723 | 0.05694483 | -0.0138176 | 0.85964403 | 0.91802142 |
| Pole2 | 0.06640057 | 0.0641681 | 0.07798604 | -0.0138179 | 0.78456197 | 0.90650187 |
| 4930578M01 | 0.05949434 | 0.0563699 | 0.07019039 | -0.0138205 | 0.8093055 | 0.9092537 |
| Acad11 | 0.09111508 | 0.09149677 | 0.10531825 | -0.0138215 | 0.72019501 | 0.90650187 |
| Zfp809 | 0.08470067 | 0.08447914 | 0.09830796 | -0.0138288 | 0.73413134 | 0.90650187 |
| Map3k11 | 0.14693058 | 0.1505156 | 0.16434467 | -0.0138291 | 0.64582881 | 0.90650187 |
| Dyrk3 | 0.06483999 | 0.06238681 | 0.07623061 | -0.0138438 | 0.79026503 | 0.90650187 |
| Cbr4 | 0.17768478 | 0.18146721 | 0.19531509 | -0.0138479 | 0.62369299 | 0.90650187 |
| Slc35f6 | 0.1069203 | 0.10853814 | 0.1223895 | -0.0138514 | 0.69256316 | 0.90650187 |
| Fbxw4 | 0.07062573 | 0.06886734 | 0.08272087 | -0.0138535 | 0.77171892 | 0.90650187 |
| Pus10 | 0.08780897 | 0.08785488 | 0.10171223 | -0.0138574 | 0.727634 | 0.90650187 |
| Eps8l2 | 0.2507508 | 0.25092746 | 0.26478665 | -0.0138592 | 0.59199789 | 0.90650187 |
| Msrb2 | 0.13414401 | 0.13728171 | 0.15114597 | -0.0138643 | 0.65828417 | 0.90650187 |
| Gprc5c | 0.20652514 | 0.2095335 | 0.22340439 | -0.0138709 | 0.60873442 | 0.90650187 |
| Tbcel | 0.15180151 | 0.15544946 | 0.16932199 | -0.0138725 | 0.64215608 | 0.90650187 |
| Zc3hav1l | 0.10704314 | 0.10864629 | 0.12252131 | -0.013875 | 0.6926747 | 0.90650187 |
| Pdxdc1 | 0.4903692 | 0.44481232 | 0.45868745 | -0.0138751 | 0.55348767 | 0.90650187 |
| Aff4 | 0.48404169 | 0.44025693 | 0.45413703 | -0.0138801 | 0.5540396 | 0.90650187 |
| Noc3l | 0.14343745 | 0.14686134 | 0.16075798 | -0.0138966 | 0.64962118 | 0.90650187 |
| Slc38a1 | 0.34738429 | 0.33479871 | 0.34869697 | -0.0138983 | 0.57032109 | 0.90650187 |

| | | | | | | |
|---|---|---|---|---|---|---|
| Clint1 | 0.5314334 | 0.47366388 | 0.48756252 | -0.0138986 | 0.55042221 | 0.90650187 |
| Sbf1 | 0.12080903 | 0.12327103 | 0.13717364 | -0.0139026 | 0.67384809 | 0.90650187 |
| Bbs7 | 0.08103537 | 0.08036762 | 0.09427613 | -0.0139085 | 0.74404227 | 0.90650187 |
| Fam83h | 0.047381 | 0.04243603 | 0.0563456 | -0.0139096 | 0.86377755 | 0.91993853 |
| Arhgef3 | 0.08541443 | 0.08518064 | 0.09909089 | -0.0139103 | 0.73370874 | 0.90650187 |
| Brd2 | 0.77272299 | 0.62329998 | 0.63721044 | -0.0139105 | 0.53865614 | 0.90650187 |
| Grb14 | 0.20455148 | 0.20759628 | 0.22151069 | -0.0139144 | 0.60997551 | 0.90650187 |
| Tpcn1 | 0.31963183 | 0.31153447 | 0.32545919 | -0.0139247 | 0.57542557 | 0.90650187 |
| Cep85 | 0.08535429 | 0.08509749 | 0.09902495 | -0.0139275 | 0.73409864 | 0.90650187 |
| Dnttip2 | 0.55413996 | 0.48912983 | 0.50306008 | -0.0139303 | 0.5489874 | 0.90650187 |
| BC055324 | 0.05120614 | 0.04681094 | 0.06074132 | -0.0139304 | 0.84602972 | 0.91377702 |
| Pabpc1l | 0.06981419 | 0.06787354 | 0.08181346 | -0.0139399 | 0.77561638 | 0.90650187 |
| Flot2 | 0.30036929 | 0.29498704 | 0.30893283 | -0.0139458 | 0.5795267 | 0.90650187 |
| Frmd5 | 0.0571758 | 0.0536114 | 0.06755744 | -0.013946 | 0.82061635 | 0.9092537 |
| Proser1 | 0.41965951 | 0.39228202 | 0.40623136 | -0.0139493 | 0.56066486 | 0.90650187 |
| Fbxl18 | 0.14228218 | 0.14561807 | 0.15956862 | -0.0139506 | 0.65123782 | 0.90650187 |
| Ogfod1 | 0.05546062 | 0.05164648 | 0.06560452 | -0.013958 | 0.82788721 | 0.90949726 |
| Setd6 | 0.09116053 | 0.0914 | 0.10536772 | -0.0139677 | 0.72217577 | 0.90650187 |
| Nifk | 0.52658157 | 0.47024073 | 0.48420863 | -0.0139679 | 0.55102118 | 0.90650187 |
| Rbm38 | 0.08524085 | 0.08492618 | 0.09890055 | -0.0139744 | 0.73504774 | 0.90650187 |
| Cyth1 | 0.18317399 | 0.18673698 | 0.2007308 | -0.0139938 | 0.62169011 | 0.90650187 |
| Sema4d | 0.08666403 | 0.08646276 | 0.10045985 | -0.0139971 | 0.73217583 | 0.90650187 |
| Mylip | 0.068872 | 0.06676136 | 0.08075875 | -0.0139974 | 0.77950658 | 0.90650187 |
| Aif1l | 0.07875751 | 0.07776006 | 0.09176086 | -0.0140008 | 0.7512115 | 0.90650187 |
| Tubgcp6 | 0.04927965 | 0.04452786 | 0.05853023 | -0.0140024 | 0.85623049 | 0.9165432 |
| Tmem29 | 0.36466954 | 0.34882819 | 0.36283952 | -0.0140113 | 0.56815173 | 0.90650187 |
| Fbxl4 | 0.14537304 | 0.14872769 | 0.16274716 | -0.0140195 | 0.64912392 | 0.90650187 |
| Lrfn4 | 0.16301089 | 0.16664779 | 0.18067187 | -0.0140241 | 0.63498142 | 0.90650187 |
| Cenpp | 0.07420199 | 0.07268259 | 0.08670828 | -0.0140257 | 0.76396308 | 0.90650187 |
| Nol4 | 0.22309427 | 0.22511545 | 0.23914283 | -0.0140274 | 0.60286338 | 0.90650187 |

| | | | | | | |
|---|---|---|---|---|---|---|
| Nup205 | 0.10832886 | 0.10986577 | 0.12389974 | -0.014034 | 0.69271058 | 0.90650187 |
| Poll | 0.07899732 | 0.07799121 | 0.09202601 | -0.0140348 | 0.75111269 | 0.90650187 |
| Glis2 | 0.20597654 | 0.20883945 | 0.22287842 | -0.014039 | 0.61027126 | 0.90650187 |
| Msi1 | 0.20111788 | 0.20416509 | 0.21820645 | -0.0140414 | 0.61258786 | 0.90650187 |
| Dcbld1 | 0.19589168 | 0.19910817 | 0.21315262 | -0.0140444 | 0.61520732 | 0.90650187 |
| Klhdc1 | 0.10503107 | 0.10631539 | 0.12036001 | -0.0140446 | 0.69803278 | 0.90650187 |
| Med22 | 0.2068978 | 0.20969778 | 0.22376149 | -0.0140637 | 0.61003586 | 0.90650187 |
| Fpgs | 0.19680775 | 0.19995381 | 0.21404068 | -0.0140869 | 0.61507959 | 0.90650187 |
| Pcgf1 | 0.12550062 | 0.12800831 | 0.14211425 | -0.0141059 | 0.67048788 | 0.90650187 |
| Ptar1 | 0.06878051 | 0.0665477 | 0.08065626 | -0.0141086 | 0.78160777 | 0.90650187 |
| Gdap2 | 0.14707059 | 0.15037907 | 0.16448813 | -0.0141091 | 0.64851874 | 0.90650187 |
| Rhpn1 | 0.05850952 | 0.05495888 | 0.06907299 | -0.0141141 | 0.81816605 | 0.9092537 |
| Dda1 | 0.63659825 | 0.54259563 | 0.55671451 | -0.0141189 | 0.54488381 | 0.90650187 |
| Kcnc3 | 0.1532354 | 0.15665151 | 0.17078186 | -0.0141303 | 0.64349839 | 0.90650187 |
| Diexf | 0.08913241 | 0.08902704 | 0.10315756 | -0.0141305 | 0.72872419 | 0.90650187 |
| Rnf208 | 0.12830887 | 0.13092689 | 0.14505848 | -0.0141316 | 0.66752565 | 0.90650187 |
| Rnf186 | 0.04658357 | 0.04128825 | 0.05542644 | -0.0141382 | 0.87159975 | 0.9223337 |
| Ift122 | 0.27307952 | 0.2708024 | 0.28494112 | -0.0141387 | 0.58729416 | 0.90650187 |
| Rpgrip1l | 0.09447586 | 0.09482624 | 0.1089683 | -0.0141421 | 0.71801569 | 0.90650187 |
| Stim1 | 0.08026589 | 0.07926472 | 0.09342728 | -0.0141626 | 0.74981424 | 0.90650187 |
| Nek1 | 0.27591106 | 0.2732998 | 0.28746245 | -0.0141627 | 0.58668486 | 0.90650187 |
| Cyb5r1 | 0.09516882 | 0.09553986 | 0.10971895 | -0.0141791 | 0.71718873 | 0.90650187 |
| Mon2 | 0.14575194 | 0.14895506 | 0.16313605 | -0.014181 | 0.65041639 | 0.90650187 |
| Cyp4f16 | 0.10969955 | 0.1111845 | 0.12536701 | -0.0141825 | 0.69249043 | 0.90650187 |
| Slc39a9 | 0.10981447 | 0.11130707 | 0.12548991 | -0.0141828 | 0.69232144 | 0.90650187 |
| Utp14b | 0.03824199 | 0.03154499 | 0.04575308 | -0.0142081 | 0.91628847 | 0.94231902 |
| Cyp4v3 | 0.04813322 | 0.04299697 | 0.05721177 | -0.0142148 | 0.86532331 | 0.92040222 |
| Nup85 | 0.25657072 | 0.25586832 | 0.27008584 | -0.0142175 | 0.59251535 | 0.90650187 |
| Myl6 | 3.54782812 | 0.74854432 | 0.76277023 | -0.0142259 | 0.53303924 | 0.90650187 |
| 3110062M04 | 0.19173414 | 0.19487984 | 0.20911094 | -0.0142311 | 0.61888238 | 0.90650187 |

| | | | | | | |
|---|---|---|---|---|---|---|
| Klhl26 | 0.13414239 | 0.13691098 | 0.15114428 | -0.0142333 | 0.66226194 | 0.90650187 |
| Ccser1 | 0.04007547 | 0.03364454 | 0.04788845 | -0.0142439 | 0.90696563 | 0.9379403 |
| Tagap1 | 0.10858591 | 0.10992996 | 0.12417508 | -0.0142451 | 0.69497126 | 0.90650187 |
| Pop1 | 0.07013742 | 0.0679112 | 0.08217499 | -0.0142638 | 0.7798422 | 0.90650187 |
| Ivns1abp | 0.56125587 | 0.4935798 | 0.50785047 | -0.0142707 | 0.54970743 | 0.90650187 |
| 2010315B03I | 0.11144483 | 0.11295962 | 0.12723187 | -0.0142722 | 0.6909998 | 0.90650187 |
| Atp5s | 0.10937835 | 0.11074717 | 0.12502338 | -0.0142762 | 0.69414582 | 0.90650187 |
| Kdm4b | 0.13509641 | 0.13784647 | 0.1521357 | -0.0142892 | 0.66186195 | 0.90650187 |
| Dcaf4 | 0.13993822 | 0.14285346 | 0.15715073 | -0.0142973 | 0.65706006 | 0.90650187 |
| Lin37 | 0.31816384 | 0.30991307 | 0.32421117 | -0.0142981 | 0.57771703 | 0.90650187 |
| Setd1a | 0.09588341 | 0.09618127 | 0.11049233 | -0.0143111 | 0.71762443 | 0.90650187 |
| 6430548M08 | 0.05921909 | 0.05556678 | 0.06987823 | -0.0143115 | 0.81871661 | 0.9092537 |
| Arntl | 0.07612828 | 0.07453276 | 0.08884839 | -0.0143156 | 0.7630953 | 0.90650187 |
| Lars2 | 0.10288566 | 0.10372361 | 0.11804989 | -0.0143263 | 0.7052323 | 0.90650187 |
| Vac14 | 0.11029141 | 0.11166499 | 0.12599985 | -0.0143349 | 0.69349141 | 0.90650187 |
| Rab40c | 0.18411597 | 0.18732164 | 0.20165685 | -0.0143352 | 0.6240084 | 0.90650187 |
| Ikzf4 | 0.12905938 | 0.13150189 | 0.14584374 | -0.0143418 | 0.66900518 | 0.90650187 |
| Cpd | 0.26537391 | 0.26369665 | 0.27803961 | -0.014343 | 0.59069423 | 0.90650187 |
| Usp12 | 0.16489699 | 0.16822303 | 0.18256761 | -0.0143446 | 0.63656692 | 0.90650187 |
| Fam69b | 0.19967595 | 0.20246946 | 0.21681514 | -0.0143457 | 0.61567676 | 0.90650187 |
| Nipal2 | 0.10195594 | 0.10269475 | 0.117047 | -0.0143523 | 0.70716392 | 0.90650187 |
| Gm37494 | 0.1534291 | 0.15661625 | 0.17097889 | -0.0143626 | 0.64559014 | 0.90650187 |
| Impa1 | 0.66523758 | 0.56007531 | 0.57444463 | -0.0143693 | 0.54427258 | 0.90650187 |
| Napb | 0.05373084 | 0.0492592 | 0.06363052 | -0.0143713 | 0.84232666 | 0.91251845 |
| Nanos1 | 0.08762644 | 0.08714057 | 0.10151269 | -0.0143721 | 0.73545871 | 0.90650187 |
| Haghl | 0.44692486 | 0.412512 | 0.426885 | -0.014373 | 0.55949195 | 0.90650187 |
| 1810044D09I | 0.10385628 | 0.10471714 | 0.11909575 | -0.0143786 | 0.70427108 | 0.90650187 |
| Micu3 | 0.20198454 | 0.20465766 | 0.21904163 | -0.014384 | 0.61483108 | 0.90650187 |
| Rp9 | 0.79498869 | 0.63513101 | 0.64951684 | -0.0143858 | 0.53921788 | 0.90650187 |
| Nit2 | 0.50909436 | 0.45759712 | 0.4719927 | -0.0143956 | 0.55392695 | 0.90650187 |

| | | | | | | |
|---|---|---|---|---|---|---|
| Usp6nl | 0.04530161 | 0.03954877 | 0.05394673 | -0.014398 | 0.88228698 | 0.92700639 |
| Fam135a | 0.40496953 | 0.38046813 | 0.39487078 | -0.0144026 | 0.56440632 | 0.90650187 |
| Aldh3b1 | 0.05423366 | 0.0497972 | 0.0642048 | -0.0144076 | 0.84076262 | 0.91251845 |
| Reep2 | 0.05774019 | 0.05378941 | 0.06819911 | -0.0144097 | 0.82621324 | 0.90947782 |
| Kat6a | 0.30879097 | 0.301752 | 0.31619847 | -0.0144465 | 0.58047577 | 0.90650187 |
| Ing4 | 0.79415112 | 0.6346108 | 0.64905811 | -0.0144473 | 0.5394127 | 0.90650187 |
| Shtn1 | 0.17460228 | 0.17780358 | 0.19225936 | -0.0144558 | 0.63090847 | 0.90650187 |
| Chd3os | 0.21764923 | 0.21954121 | 0.23400197 | -0.0144608 | 0.60823726 | 0.90650187 |
| Med26 | 0.07676872 | 0.07509409 | 0.08955873 | -0.0144646 | 0.76362454 | 0.90650187 |
| Rbm45 | 0.25076186 | 0.25033032 | 0.26479675 | -0.0144664 | 0.59594824 | 0.90650187 |
| Fbxo46 | 0.1419623 | 0.1447678 | 0.15923903 | -0.0144712 | 0.65689561 | 0.90650187 |
| Tcf20 | 0.12989386 | 0.1322438 | 0.14671605 | -0.0144723 | 0.66949537 | 0.90650187 |
| Bcl7b | 0.47683925 | 0.43444613 | 0.44892458 | -0.0144784 | 0.55700377 | 0.90650187 |
| Sap130 | 0.08822235 | 0.08768348 | 0.10216395 | -0.0144805 | 0.73568851 | 0.90650187 |
| Nfasc | 0.12579805 | 0.12792846 | 0.14242653 | -0.0144981 | 0.67455745 | 0.90650187 |
| Nt5c2 | 0.43738097 | 0.40521861 | 0.4197174 | -0.0144988 | 0.56102552 | 0.90650187 |
| Hacd2 | 0.38086719 | 0.36136885 | 0.37586951 | -0.0145007 | 0.56808742 | 0.90650187 |
| B230219D22 | 1.05427415 | 0.75783633 | 0.77234151 | -0.0145052 | 0.53326982 | 0.90650187 |
| Cox20 | 0.92706192 | 0.70337319 | 0.71788635 | -0.0145132 | 0.53580658 | 0.90650187 |
| Ston2 | 0.05476603 | 0.05029354 | 0.06481241 | -0.0145189 | 0.84034492 | 0.91226452 |
| Sipa1l1 | 0.11765189 | 0.11929822 | 0.13383377 | -0.0145356 | 0.6853928 | 0.90650187 |
| Mrps18a | 0.92721542 | 0.70342504 | 0.7179609 | -0.0145359 | 0.53585875 | 0.90650187 |
| Fancg | 0.07681481 | 0.07507241 | 0.08960982 | -0.0145374 | 0.76461024 | 0.90650187 |
| Nbeal2 | 0.0620885 | 0.05858919 | 0.07312688 | -0.0145377 | 0.81159541 | 0.9092537 |
| Wdr90 | 0.07414288 | 0.07209002 | 0.08664252 | -0.0145525 | 0.77237315 | 0.90650187 |
| Aldh4a1 | 0.10320372 | 0.10384022 | 0.11839273 | -0.0145525 | 0.70761894 | 0.90650187 |
| Marveld3 | 0.25453349 | 0.25367883 | 0.26823462 | -0.0145558 | 0.59531601 | 0.90650187 |
| Grpel2 | 0.29352417 | 0.28842022 | 0.3029796 | -0.0145594 | 0.58458188 | 0.90650187 |
| Zfp777 | 0.06181371 | 0.05824479 | 0.0728163 | -0.0145715 | 0.81316626 | 0.9092537 |
| Pex26 | 0.12112878 | 0.12292678 | 0.13751121 | -0.0145844 | 0.68135834 | 0.90650187 |

| | | | | | | |
|---|---|---|---|---|---|---|
| Podxl2 | 0.40261369 | 0.37844795 | 0.39303373 | -0.0145858 | 0.56552009 | 0.90650187 |
| Osbpl7 | 0.05600001 | 0.05163311 | 0.06621915 | -0.014586 | 0.83626685 | 0.91079271 |
| Eif2b1 | 0.54689391 | 0.48356333 | 0.4981504 | -0.0145871 | 0.55178732 | 0.90650187 |
| Fbxo42 | 0.10172578 | 0.10219242 | 0.11679855 | -0.0146061 | 0.71087442 | 0.90650187 |
| Prorsd1 | 0.42355476 | 0.39460809 | 0.40921549 | -0.0146074 | 0.56304377 | 0.90650187 |
| Pibf1 | 0.13717511 | 0.1396844 | 0.15429216 | -0.0146078 | 0.66308296 | 0.90650187 |
| Zfp263 | 0.25539227 | 0.25439551 | 0.26901547 | -0.01462 | 0.59545563 | 0.90650187 |
| Phf21b | 0.07385945 | 0.07170627 | 0.08632716 | -0.0146209 | 0.77425547 | 0.90650187 |
| Nktr | 1.48412766 | 0.87598428 | 0.89060693 | -0.0146226 | 0.52909347 | 0.90650187 |
| Zfp408 | 0.07985433 | 0.07834525 | 0.09297292 | -0.0146277 | 0.75786112 | 0.90650187 |
| Map3k1 | 0.16851599 | 0.17156291 | 0.18619391 | -0.014631 | 0.63658028 | 0.90650187 |
| Fa2h | 0.06691861 | 0.06392942 | 0.07856797 | -0.0146386 | 0.7962475 | 0.9075779 |
| Fam118a | 0.11893659 | 0.12053852 | 0.13519429 | -0.0146558 | 0.68507107 | 0.90650187 |
| Rbl2 | 0.07063264 | 0.06807239 | 0.08272859 | -0.0146562 | 0.78452991 | 0.90650187 |
| Mkl1 | 0.07689748 | 0.07504371 | 0.08970147 | -0.0146578 | 0.76621516 | 0.90650187 |
| Gorasp1 | 0.14543124 | 0.148148 | 0.1628069 | -0.0146589 | 0.65552341 | 0.90650187 |
| Rsph3a | 0.09494425 | 0.09481584 | 0.10947576 | -0.0146599 | 0.72418255 | 0.90650187 |
| Psmc6 | 1.17589864 | 0.80023013 | 0.81489475 | -0.0146646 | 0.53188214 | 0.90650187 |
| Gde1 | 0.82733129 | 0.65232592 | 0.6669922 | -0.0146663 | 0.53893579 | 0.90650187 |
| Spty2d1 | 0.3838279 | 0.36356054 | 0.37822833 | -0.0146678 | 0.56843915 | 0.90650187 |
| Rps6kb2 | 0.27303701 | 0.27023209 | 0.28490321 | -0.0146711 | 0.59053718 | 0.90650187 |
| Gm13830 | 0.0803639 | 0.07886283 | 0.09353544 | -0.0146726 | 0.75721392 | 0.90650187 |
| Plag1 | 0.12520545 | 0.12712752 | 0.14180423 | -0.0146767 | 0.67728477 | 0.90650187 |
| Abca7 | 0.13878747 | 0.14128356 | 0.15596131 | -0.0146778 | 0.66216634 | 0.90650187 |
| Pgk1 | 0.18981932 | 0.19256172 | 0.20724324 | -0.0146815 | 0.62359476 | 0.90650187 |
| Diablo | 0.73754334 | 0.60261292 | 0.61729499 | -0.0146821 | 0.54210433 | 0.90650187 |
| Usp45 | 0.06471757 | 0.0614008 | 0.07609275 | -0.014692 | 0.80465188 | 0.90882608 |
| Cul4a | 0.65541934 | 0.55371571 | 0.5684159 | -0.0147002 | 0.54576589 | 0.90650187 |
| Robo1 | 0.064704 | 0.06137202 | 0.07607747 | -0.0147055 | 0.80491713 | 0.90882608 |
| Sclt1 | 0.14010614 | 0.14261647 | 0.15732415 | -0.0147077 | 0.66115009 | 0.90650187 |

| | | | | | | |
|---|---|---|---|---|---|---|
| Chadl | 0.10504757 | 0.10566704 | 0.12037776 | -0.0147107 | 0.70652923 | 0.90650187 |
| Gys1 | 0.10417518 | 0.1047283 | 0.1194391 | -0.0147108 | 0.70799833 | 0.90650187 |
| Scfd2 | 0.07823381 | 0.07646361 | 0.09118153 | -0.0147179 | 0.76349193 | 0.90650187 |
| Ankzf1 | 0.14471337 | 0.14734856 | 0.16206971 | -0.0147212 | 0.6568217 | 0.90650187 |
| Tigd2 | 0.28158154 | 0.27774764 | 0.29248927 | -0.0147416 | 0.58864518 | 0.90650187 |
| Thoc6 | 0.2458262 | 0.24551863 | 0.26027651 | -0.0147579 | 0.5995063 | 0.90650187 |
| Fbxl20 | 0.1336839 | 0.13590692 | 0.15066744 | -0.0147605 | 0.6684093 | 0.90650187 |
| Evpl | 0.07358744 | 0.0712575 | 0.0860244 | -0.0147669 | 0.77730711 | 0.90650187 |
| Tbl2 | 0.1448818 | 0.14747127 | 0.16224274 | -0.0147715 | 0.65717018 | 0.90650187 |
| Ube2h | 0.28975922 | 0.28491255 | 0.29968654 | -0.014774 | 0.5867373 | 0.90650187 |
| Kdm3a | 0.15826797 | 0.16110975 | 0.17588697 | -0.0147772 | 0.64561167 | 0.90650187 |
| Soga1 | 0.07101738 | 0.06837393 | 0.08315845 | -0.0147845 | 0.7853449 | 0.90650187 |
| Ankrd13c | 0.38694958 | 0.36591822 | 0.38070796 | -0.0147897 | 0.56855751 | 0.90650187 |
| Csrnp2 | 0.05256457 | 0.04749356 | 0.06229707 | -0.0148035 | 0.85460705 | 0.91621818 |
| 2610035D17 | 0.05875292 | 0.05453983 | 0.06934929 | -0.0148095 | 0.82877984 | 0.90967191 |
| Sirt4 | 0.12499922 | 0.12677637 | 0.14158756 | -0.0148112 | 0.67905073 | 0.90650187 |
| Pwwp2b | 0.06096289 | 0.05703762 | 0.07185395 | -0.0148163 | 0.82034669 | 0.9092537 |
| Tmem151b | 0.04774268 | 0.04194526 | 0.05676219 | -0.0148169 | 0.87706906 | 0.9247416 |
| Slc12a7 | 0.10049751 | 0.10065044 | 0.11547159 | -0.0148211 | 0.71586725 | 0.90650187 |
| Dnaaf5 | 0.0413614 | 0.03454284 | 0.04938302 | -0.0148402 | 0.90921829 | 0.93919616 |
| Mxi1 | 0.09939555 | 0.0994372 | 0.11428322 | -0.014846 | 0.71820212 | 0.90650187 |
| Itfg1 | 0.77953405 | 0.62615021 | 0.640999 | -0.0148488 | 0.54101168 | 0.90650187 |
| Rabl3 | 0.13730855 | 0.13956825 | 0.15443042 | -0.0148622 | 0.66561582 | 0.90650187 |
| Aida | 0.47003774 | 0.42910687 | 0.44396983 | -0.014863 | 0.55915505 | 0.90650187 |
| Gm15013 | 0.05534411 | 0.05059739 | 0.06547171 | -0.0148743 | 0.84374852 | 0.91300169 |
| Dph5 | 0.15841295 | 0.16115883 | 0.17603361 | -0.0148748 | 0.64641227 | 0.90650187 |
| 1190002N15 | 0.16504466 | 0.16784026 | 0.18271587 | -0.0148756 | 0.6412993 | 0.90650187 |
| Kmt2c | 0.42753647 | 0.39737572 | 0.41225419 | -0.0148785 | 0.56373441 | 0.90650187 |
| Atl3 | 0.10795592 | 0.10861829 | 0.12350013 | -0.0148818 | 0.70391638 | 0.90650187 |
| Nop58 | 0.85529913 | 0.66684671 | 0.68172917 | -0.0148825 | 0.53865651 | 0.90650187 |

| | | | | | | |
|---|---|---|---|---|---|---|
| Zfp51 | 0.10631143 | 0.10684964 | 0.12173592 | -0.0148863 | 0.70664983 | 0.90650187 |
| Coro7 | 0.08313156 | 0.08169191 | 0.09658426 | -0.0148924 | 0.75347613 | 0.90650187 |
| Pih1d2 | 0.07699486 | 0.07491208 | 0.08980941 | -0.0148973 | 0.76956796 | 0.90650187 |
| Lgals8 | 0.23998416 | 0.23999609 | 0.25489427 | -0.0148982 | 0.60250596 | 0.90650187 |
| Rnf168 | 0.17456547 | 0.17732397 | 0.1922228 | -0.0148988 | 0.63479145 | 0.90650187 |
| Ngrn | 0.54122103 | 0.47937641 | 0.49428341 | -0.014907 | 0.55332806 | 0.90650187 |
| Prox1os | 0.05479354 | 0.04993144 | 0.06484379 | -0.0149124 | 0.84669746 | 0.91417346 |
| Pex2 | 0.36242625 | 0.34610182 | 0.36101786 | -0.014916 | 0.57286679 | 0.90650187 |
| Mfsd6l | 0.06369208 | 0.06001321 | 0.0749371 | -0.0149239 | 0.81201669 | 0.9092537 |
| Mthfsd | 0.31771615 | 0.30889535 | 0.32383019 | -0.0149348 | 0.58122511 | 0.90650187 |
| Txnl4b | 0.05309493 | 0.04796695 | 0.06290371 | -0.0149368 | 0.854431 | 0.91621818 |
| Rapgef4 | 0.27844447 | 0.27476791 | 0.28971198 | -0.0149441 | 0.59068815 | 0.90650187 |
| Casp2 | 0.27592282 | 0.27252532 | 0.2874729 | -0.0149476 | 0.59140317 | 0.90650187 |
| Gm20186 | 0.06462102 | 0.06102296 | 0.07598402 | -0.0149611 | 0.80930236 | 0.9092537 |
| Abcb6 | 0.10526949 | 0.10565439 | 0.12061638 | -0.014962 | 0.7093408 | 0.90650187 |
| Fam219a | 0.2618814 | 0.25993034 | 0.27489294 | -0.0149626 | 0.5956011 | 0.90650187 |
| BC026585 | 0.11079103 | 0.11157021 | 0.12653372 | -0.0149635 | 0.70046012 | 0.90650187 |
| Rab4b | 0.1219462 | 0.12339895 | 0.13837364 | -0.0149747 | 0.68477733 | 0.90650187 |
| Rbm19 | 0.09421058 | 0.09370345 | 0.10868075 | -0.0149773 | 0.72994593 | 0.90650187 |
| Megf9 | 0.14140462 | 0.14368514 | 0.15866414 | -0.014979 | 0.66264631 | 0.90650187 |
| Zbtb10 | 0.08066002 | 0.07887768 | 0.09386216 | -0.0149845 | 0.76105857 | 0.90650187 |
| Ddx51 | 0.08233899 | 0.08072613 | 0.09571229 | -0.0149862 | 0.75680789 | 0.90650187 |
| Reep3 | 0.39230378 | 0.36995443 | 0.38494326 | -0.0149888 | 0.56871435 | 0.90650187 |
| Hectd3 | 0.07493904 | 0.07253753 | 0.08752777 | -0.0149902 | 0.77678088 | 0.90650187 |
| Actr6 | 0.15752143 | 0.16014078 | 0.1751315 | -0.0149907 | 0.64822411 | 0.90650187 |
| Gm10320 | 0.3011249 | 0.29459591 | 0.30958729 | -0.0149914 | 0.58522277 | 0.90650187 |
| Tbc1d22a | 0.0674932 | 0.06421002 | 0.07921296 | -0.0150029 | 0.80010558 | 0.90805731 |
| Unc119b | 0.1591405 | 0.16176192 | 0.17676915 | -0.0150072 | 0.64706925 | 0.90650187 |
| Ankra2 | 0.50851669 | 0.45657335 | 0.47158571 | -0.0150124 | 0.55627062 | 0.90650187 |
| Abat | 0.1499129 | 0.15238084 | 0.16739554 | -0.0150147 | 0.65496235 | 0.90650187 |

| | | | | | | |
|---|---|---|---|---|---|---|
| Sh3bp4 | 0.17518629 | 0.17781169 | 0.1928391 | -0.0150274 | 0.63549155 | 0.90650187 |
| Cml1 | 0.09237342 | 0.0916558 | 0.10668674 | -0.0150309 | 0.73445244 | 0.90650187 |
| Oxsr1 | 0.09293234 | 0.09226245 | 0.10729387 | -0.0150314 | 0.73329557 | 0.90650187 |
| Gm13157 | 0.05974501 | 0.05543657 | 0.07047457 | -0.015038 | 0.82859094 | 0.9096066 |
| Rapgef6 | 0.25930906 | 0.25752857 | 0.27256789 | -0.0150393 | 0.59688509 | 0.90650187 |
| Galt | 0.24130326 | 0.24106742 | 0.25611249 | -0.0150451 | 0.60301263 | 0.90650187 |
| Gm9726 | 0.07776875 | 0.07562085 | 0.09066674 | -0.0150459 | 0.76966251 | 0.90650187 |
| Gas2 | 0.07689855 | 0.07465072 | 0.08970266 | -0.0150519 | 0.77215549 | 0.90650187 |
| Mlh3 | 0.09350374 | 0.09286183 | 0.10791412 | -0.0150523 | 0.73240096 | 0.90650187 |
| Zscan2 | 0.10831809 | 0.10883388 | 0.1238882 | -0.0150543 | 0.70547369 | 0.90650187 |
| Papd5 | 0.06602702 | 0.06250674 | 0.07756618 | -0.0150594 | 0.80596223 | 0.9092537 |
| Atr | 0.15813744 | 0.16069462 | 0.17575493 | -0.0150603 | 0.64837637 | 0.90650187 |
| Poc1b | 0.13118655 | 0.13298626 | 0.1480657 | -0.0150794 | 0.6746368 | 0.90650187 |
| Hps5 | 0.07340774 | 0.07074093 | 0.08582431 | -0.0150834 | 0.78268789 | 0.90650187 |
| Ago1 | 0.16531903 | 0.16790542 | 0.18299125 | -0.0150858 | 0.64300266 | 0.90650187 |
| 2310033P09I | 0.38605451 | 0.36491056 | 0.37999777 | -0.0150872 | 0.57005166 | 0.90650187 |
| Itpr3 | 0.22864358 | 0.22925389 | 0.2443497 | -0.0150958 | 0.60820649 | 0.90650187 |
| Ddx24 | 0.84516156 | 0.66133016 | 0.67642775 | -0.0150976 | 0.53951983 | 0.90650187 |
| 6330408A02I | 0.0844753 | 0.08295665 | 0.0980606 | -0.015104 | 0.75324673 | 0.90650187 |
| 2700046G09 | 0.12969186 | 0.13140046 | 0.14650496 | -0.0151045 | 0.67664491 | 0.90650187 |
| Zdhhc21 | 0.2729051 | 0.26967993 | 0.28478556 | -0.0151056 | 0.59320778 | 0.90650187 |
| Dpy19l3 | 0.08462883 | 0.08312153 | 0.09822912 | -0.0151076 | 0.75292778 | 0.90650187 |
| Mogs | 0.10238592 | 0.1023913 | 0.11751096 | -0.0151197 | 0.71633845 | 0.90650187 |
| 4930451G09 | 0.05657897 | 0.05174593 | 0.06687838 | -0.0151324 | 0.84276797 | 0.91273395 |
| Ttc8 | 0.43727345 | 0.40450285 | 0.41963626 | -0.0151334 | 0.56368654 | 0.90650187 |
| Zbed4 | 0.12679299 | 0.12833192 | 0.14347038 | -0.0151385 | 0.68049509 | 0.90650187 |
| Tdrd7 | 0.14570754 | 0.14794934 | 0.16309048 | -0.0151411 | 0.66009599 | 0.90650187 |
| 2810454H06I | 0.06151511 | 0.05733362 | 0.07247868 | -0.0151451 | 0.82354217 | 0.9092537 |
| Diaph3 | 0.06332057 | 0.05936788 | 0.07451805 | -0.0151502 | 0.81695749 | 0.9092537 |
| Celf6 | 0.06657853 | 0.063025 | 0.07818599 | -0.015161 | 0.80566854 | 0.90920323 |

| | | | | | | |
|---|---|---|---|---|---|---|
| St6galnac2 | 0.18620292 | 0.18854218 | 0.20370509 | -0.0151629 | 0.62964425 | 0.90650187 |
| Isoc2b | 0.08895635 | 0.0877976 | 0.10296543 | -0.0151678 | 0.74372808 | 0.90650187 |
| Mrps18c | 0.80335125 | 0.63891215 | 0.65408008 | -0.0151679 | 0.54105514 | 0.90650187 |
| Arhgef18 | 0.10056253 | 0.10036998 | 0.11554189 | -0.0151719 | 0.72030796 | 0.90650187 |
| Vps13d | 0.11055465 | 0.11110537 | 0.12628116 | -0.0151758 | 0.70341544 | 0.90650187 |
| Usp16 | 0.52446507 | 0.46755736 | 0.48274091 | -0.0151835 | 0.55560154 | 0.90650187 |
| Ice1 | 0.11904254 | 0.12011879 | 0.13530641 | -0.0151876 | 0.69111103 | 0.90650187 |
| Tet3 | 0.50121225 | 0.45123038 | 0.46642066 | -0.0151903 | 0.55755906 | 0.90650187 |
| Nom1 | 0.30849667 | 0.30075483 | 0.31594563 | -0.0151908 | 0.5846277 | 0.90650187 |
| Pigx | 0.49749041 | 0.44857672 | 0.46377544 | -0.0151987 | 0.55791702 | 0.90650187 |
| Usp28 | 0.07990444 | 0.0778275 | 0.09302825 | -0.0152007 | 0.7662059 | 0.90650187 |
| Exph5 | 0.04644925 | 0.04006401 | 0.05527151 | -0.0152075 | 0.88936758 | 0.9302877 |
| 1700052K11I | 0.12563688 | 0.12702473 | 0.14225732 | -0.0152326 | 0.68297193 | 0.90650187 |
| Rcor2 | 0.15792754 | 0.16030715 | 0.17554255 | -0.0152354 | 0.65019085 | 0.90650187 |
| Wasl | 0.52003146 | 0.46442066 | 0.4796571 | -0.0152364 | 0.55615032 | 0.90650187 |
| Ltbp3 | 0.23512175 | 0.23515002 | 0.25038887 | -0.0152388 | 0.60663636 | 0.90650187 |
| Myh14 | 0.12810031 | 0.12960116 | 0.14484015 | -0.015239 | 0.68001228 | 0.90650187 |
| Slc39a3 | 0.19566315 | 0.19768721 | 0.21293094 | -0.0152437 | 0.62485649 | 0.90650187 |
| Abcc4 | 0.04573377 | 0.03920194 | 0.05444583 | -0.0152439 | 0.89337771 | 0.93181736 |
| Tars | 0.37541734 | 0.35626372 | 0.37150949 | -0.0152458 | 0.57237966 | 0.90650187 |
| Yeats2 | 0.18047849 | 0.18282644 | 0.19807551 | -0.0152491 | 0.63391865 | 0.90650187 |
| Inpp5f | 0.08347819 | 0.0817144 | 0.09696534 | -0.0152509 | 0.75779138 | 0.90650187 |
| Aven | 0.04179946 | 0.03463778 | 0.04989158 | -0.0152538 | 0.91295766 | 0.9407546 |
| Zfp937 | 0.07580422 | 0.07322608 | 0.08848873 | -0.0152626 | 0.77839497 | 0.90650187 |
| Pik3r3 | 0.0603655 | 0.05591282 | 0.07117762 | -0.0152648 | 0.82980815 | 0.90974008 |
| Phlpp2 | 0.07701108 | 0.07455947 | 0.08982739 | -0.0152679 | 0.77506076 | 0.90650187 |
| Mapk1ip1 | 0.15290164 | 0.15516861 | 0.17044226 | -0.0152737 | 0.65482467 | 0.90650187 |
| Pou2f1 | 0.15080909 | 0.15303426 | 0.16831019 | -0.0152759 | 0.65670456 | 0.90650187 |
| Pigh | 0.16032313 | 0.16268517 | 0.17796346 | -0.0152783 | 0.64864264 | 0.90650187 |
| Elf1 | 0.25582129 | 0.25411768 | 0.2694053 | -0.0152876 | 0.59958363 | 0.90650187 |

| | | | | | | |
|---|---|---|---|---|---|---|
| Trmt44 | 0.08266161 | 0.0807776 | 0.09606735 | -0.0152897 | 0.76038677 | 0.90650187 |
| Zbtb44 | 0.1954592 | 0.197443 | 0.21273305 | -0.01529 | 0.62533579 | 0.90650187 |
| Taz | 0.4647087 | 0.42476772 | 0.44006391 | -0.0152962 | 0.56140206 | 0.90650187 |
| Ermp1 | 0.07912577 | 0.07686059 | 0.092168 | -0.0153074 | 0.76984412 | 0.90650187 |
| Zfp931 | 0.15386843 | 0.15611781 | 0.17142561 | -0.0153078 | 0.65430873 | 0.90650187 |
| Asns | 0.23059229 | 0.23085931 | 0.24617078 | -0.0153115 | 0.60892191 | 0.90650187 |
| Phldb2 | 0.46477925 | 0.42480341 | 0.44011575 | -0.0153123 | 0.56145918 | 0.90650187 |
| Hectd2 | 0.05107048 | 0.04526673 | 0.06058581 | -0.0153191 | 0.869508 | 0.92144928 |
| Irak2 | 0.08299709 | 0.08111153 | 0.09643639 | -0.0153249 | 0.7600485 | 0.90650187 |
| Stk35 | 0.07221985 | 0.06917075 | 0.08450055 | -0.0153298 | 0.7900246 | 0.90650187 |
| Smug1 | 0.13220123 | 0.13378621 | 0.14912369 | -0.0153375 | 0.67624849 | 0.90650187 |
| Emc9 | 0.20248014 | 0.20417715 | 0.21951888 | -0.0153417 | 0.62198352 | 0.90650187 |
| Mettl16 | 0.44289606 | 0.40852534 | 0.42386778 | -0.0153424 | 0.56391962 | 0.90650187 |
| Usp14 | 0.65512664 | 0.55289174 | 0.56823539 | -0.0153437 | 0.54777488 | 0.90650187 |
| Epb41l4b | 0.08966774 | 0.0883976 | 0.10374151 | -0.0153439 | 0.7445793 | 0.90650187 |
| Krt1 | 0.07615725 | 0.07352613 | 0.08888053 | -0.0153544 | 0.7787544 | 0.90650187 |
| Pigg | 0.06093698 | 0.05646221 | 0.07182463 | -0.0153624 | 0.82916268 | 0.90974008 |
| Zfp667 | 0.2170615 | 0.21807832 | 0.23344525 | -0.0153669 | 0.61510112 | 0.90650187 |
| Zbtb6 | 0.10840139 | 0.10860758 | 0.12397745 | -0.0153699 | 0.70922959 | 0.90650187 |
| Eml3 | 0.11282127 | 0.1133266 | 0.12869997 | -0.0153734 | 0.70230204 | 0.90650187 |
| Lyrm2 | 0.49538803 | 0.44690086 | 0.46227718 | -0.0153763 | 0.55877743 | 0.90650187 |
| Xylt1 | 0.04994899 | 0.04391624 | 0.05929909 | -0.0153828 | 0.87559407 | 0.92418014 |
| Fktn | 0.17309331 | 0.17537468 | 0.19075951 | -0.0153848 | 0.64003034 | 0.90650187 |
| Psmc2 | 1.43366264 | 0.86583886 | 0.88122584 | -0.015387 | 0.5309365 | 0.90650187 |
| Gm12576 | 0.06687905 | 0.06312813 | 0.07852355 | -0.0153954 | 0.80828961 | 0.9092537 |
| Dcun1d3 | 0.12693441 | 0.12822262 | 0.14361865 | -0.015396 | 0.68316664 | 0.90650187 |
| Lrrc61 | 0.07637933 | 0.0737274 | 0.08912691 | -0.0153995 | 0.77879205 | 0.90650187 |
| Upf3a | 0.2612932 | 0.25896187 | 0.27436185 | -0.0154 | 0.59852562 | 0.90650187 |
| Dip2a | 0.10504257 | 0.10496805 | 0.12037238 | -0.0154043 | 0.71529291 | 0.90650187 |
| Dhx33 | 0.08441722 | 0.08258189 | 0.09799685 | -0.015415 | 0.75781656 | 0.90650187 |

| | | | | | | |
|---|---|---|---|---|---|---|
| Gfm2 | 0.13046403 | 0.13189507 | 0.14731158 | -0.0154165 | 0.67912139 | 0.90650187 |
| Icmt | 0.07086951 | 0.06757141 | 0.08299327 | -0.0154219 | 0.79562205 | 0.9075779 |
| Kdm2a | 0.55064503 | 0.48527242 | 0.50069604 | -0.0154236 | 0.55446252 | 0.90650187 |
| Tmem98 | 0.29619758 | 0.28988652 | 0.30531016 | -0.0154236 | 0.58884809 | 0.90650187 |
| Nat10 | 0.09110873 | 0.08988747 | 0.10531135 | -0.0154239 | 0.74250983 | 0.90650187 |
| Ttf1 | 0.31084293 | 0.30253248 | 0.31795924 | -0.0154268 | 0.58538609 | 0.90650187 |
| Pld1 | 0.11865279 | 0.11946716 | 0.13489392 | -0.0154268 | 0.69441677 | 0.90650187 |
| Atat1 | 0.37227086 | 0.3535536 | 0.36898149 | -0.0154279 | 0.57373061 | 0.90650187 |
| Lims1 | 0.59242009 | 0.51303236 | 0.5284667 | -0.0154343 | 0.55165266 | 0.90650187 |
| Cbx2 | 0.0876027 | 0.08604838 | 0.10148674 | -0.0154384 | 0.75056676 | 0.90650187 |
| Cep170 | 0.28858229 | 0.28320745 | 0.29865448 | -0.015447 | 0.59092926 | 0.90650187 |
| Bicd1 | 0.34528722 | 0.33151168 | 0.34696426 | -0.0154526 | 0.57847431 | 0.90650187 |
| Det1 | 0.08354576 | 0.08158072 | 0.09703961 | -0.0154589 | 0.76059163 | 0.90650187 |
| A930011G23 | 0.04954317 | 0.04337197 | 0.05883301 | -0.015461 | 0.87865407 | 0.92534436 |
| Slc4a8 | 0.10261521 | 0.10229603 | 0.11775828 | -0.0154622 | 0.72029903 | 0.90650187 |
| Snx10 | 0.16409258 | 0.16627869 | 0.18175958 | -0.0154809 | 0.64752342 | 0.90650187 |
| Tmem120b | 0.08633555 | 0.08461204 | 0.10010021 | -0.0154882 | 0.75422748 | 0.90650187 |
| Supt16 | 0.71033978 | 0.58598565 | 0.60148393 | -0.0154983 | 0.54559861 | 0.90650187 |
| Ddx49 | 0.27063535 | 0.26725375 | 0.28275826 | -0.0155045 | 0.5962941 | 0.90650187 |
| Tmem43 | 0.21203206 | 0.2131605 | 0.22866668 | -0.0155062 | 0.61846898 | 0.90650187 |
| Sox12 | 0.14924207 | 0.15120105 | 0.16671028 | -0.0155092 | 0.66040889 | 0.90650187 |
| 2810006K23I | 0.17880226 | 0.18090225 | 0.19642029 | -0.015518 | 0.63728898 | 0.90650187 |
| Ola1 | 0.64166286 | 0.54436487 | 0.5598831 | -0.0155182 | 0.54903293 | 0.90650187 |
| Phf21a | 0.42330461 | 0.3934969 | 0.40902419 | -0.0155273 | 0.56700852 | 0.90650187 |
| Gatad2a | 0.22427634 | 0.22472557 | 0.24025457 | -0.015529 | 0.61307574 | 0.90650187 |
| Rabl2 | 0.23545696 | 0.23514341 | 0.25070022 | -0.0155568 | 0.60867418 | 0.90650187 |
| Qrsl1 | 0.07612551 | 0.07327864 | 0.08884532 | -0.0155667 | 0.78198626 | 0.90650187 |
| Sep-02 | 1.30414181 | 0.83626737 | 0.8518362 | -0.0155688 | 0.53237899 | 0.90650187 |
| BC003965 | 0.39197638 | 0.3691143 | 0.38468492 | -0.0155706 | 0.57140083 | 0.90650187 |
| Morn4 | 0.06973886 | 0.06615782 | 0.08172918 | -0.0155714 | 0.8015009 | 0.90834378 |

| | | | | | | |
|---|---|---|---|---|---|---|
| Rbm43 | 0.08868424 | 0.08709696 | 0.10266839 | -0.0155714 | 0.74992465 | 0.90650187 |
| Prkaca | 0.29950898 | 0.29261508 | 0.30818705 | -0.015572 | 0.58886489 | 0.90650187 |
| Mettl25 | 0.0805378 | 0.07815191 | 0.09372733 | -0.0155754 | 0.7699741 | 0.90650187 |
| Dpp9 | 0.33461869 | 0.32251029 | 0.33808968 | -0.0155794 | 0.58115783 | 0.90650187 |
| Pycrl | 0.53996725 | 0.47783876 | 0.49342581 | -0.0155871 | 0.55584158 | 0.90650187 |
| Foxred1 | 0.1469397 | 0.14876302 | 0.16435402 | -0.015591 | 0.66338545 | 0.90650187 |
| Jazf1 | 0.04926078 | 0.04291673 | 0.05850855 | -0.0155918 | 0.88193197 | 0.92680593 |
| Dyrk2 | 0.04160862 | 0.03406796 | 0.04967007 | -0.0156021 | 0.91870766 | 0.94359967 |
| Ankrd24 | 0.23581862 | 0.23542052 | 0.25103601 | -0.0156155 | 0.60893158 | 0.90650187 |
| Zak | 0.07144103 | 0.06800814 | 0.08363154 | -0.0156234 | 0.79687009 | 0.9075779 |
| Cflar | 0.37828329 | 0.35817696 | 0.37380525 | -0.0156283 | 0.57372462 | 0.90650187 |
| Zfp37 | 0.10201008 | 0.10147338 | 0.11710543 | -0.0156321 | 0.72355355 | 0.90650187 |
| Taf2 | 0.17157675 | 0.17361707 | 0.18924941 | -0.0156323 | 0.64327069 | 0.90650187 |
| Gtf3c5 | 0.11911774 | 0.11974854 | 0.13538597 | -0.0156374 | 0.69619185 | 0.90650187 |
| Ercc8 | 0.08612442 | 0.08423127 | 0.09986896 | -0.0156377 | 0.75681766 | 0.90650187 |
| Ttc30a1 | 0.07270647 | 0.06939957 | 0.08504307 | -0.0156435 | 0.793253 | 0.90711193 |
| Fam3a | 0.37660539 | 0.3568168 | 0.37246197 | -0.0156452 | 0.57406639 | 0.90650187 |
| Sdsl | 0.12944519 | 0.13059697 | 0.14624714 | -0.0156502 | 0.68286744 | 0.90650187 |
| Znhit6 | 0.3302967 | 0.31881416 | 0.33446711 | -0.015653 | 0.58240617 | 0.90650187 |
| Dbr1 | 0.13346445 | 0.13478388 | 0.15043911 | -0.0156552 | 0.67818358 | 0.90650187 |
| Farp1 | 0.15358023 | 0.15546247 | 0.17113259 | -0.0156701 | 0.65802222 | 0.90650187 |
| Pcmtd2 | 0.35260757 | 0.33731999 | 0.35299643 | -0.0156764 | 0.5782536 | 0.90650187 |
| Ice2 | 0.09727298 | 0.09631705 | 0.11199422 | -0.0156772 | 0.73312922 | 0.90650187 |
| Dgat1 | 0.14703082 | 0.14876994 | 0.16444739 | -0.0156774 | 0.66415072 | 0.90650187 |
| Ap1ar | 0.31595351 | 0.30663751 | 0.32232852 | -0.015691 | 0.58566708 | 0.90650187 |
| Mzt2 | 0.36202216 | 0.34498886 | 0.36068929 | -0.0157004 | 0.57672123 | 0.90650187 |
| Mettl17 | 0.11406628 | 0.114323 | 0.13002587 | -0.0157029 | 0.70432595 | 0.90650187 |
| Trim46 | 0.07509292 | 0.07195962 | 0.08769876 | -0.0157391 | 0.78751978 | 0.90650187 |
| Dcbld2 | 0.15068807 | 0.15244494 | 0.16818674 | -0.0157418 | 0.66133013 | 0.90650187 |
| Ctbp2 | 0.29719519 | 0.29043531 | 0.306178 | -0.0157427 | 0.59040187 | 0.90650187 |

| | | | | | | |
|---|---|---|---|---|---|---|
| Pum1 | 0.98907566 | 0.73045321 | 0.7461997 | -0.0157465 | 0.53737085 | 0.90650187 |
| BC037034 | 0.11872675 | 0.11921955 | 0.1349722 | -0.0157527 | 0.69806673 | 0.90650187 |
| BC005561 | 0.72121517 | 0.59209562 | 0.60784866 | -0.015753 | 0.54586169 | 0.90650187 |
| Nadk | 0.40119401 | 0.37616818 | 0.39192465 | -0.0157565 | 0.57092353 | 0.90650187 |
| Uba1 | 0.96460788 | 0.71990886 | 0.73566862 | -0.0157598 | 0.53793606 | 0.90650187 |
| Exoc6 | 0.15140901 | 0.15315878 | 0.16892196 | -0.0157632 | 0.66087334 | 0.90650187 |
| Zfp12 | 0.12015562 | 0.12071988 | 0.13648341 | -0.0157635 | 0.69618432 | 0.90650187 |
| Cpsf1 | 0.13289557 | 0.13407657 | 0.14984695 | -0.0157704 | 0.68006127 | 0.90650187 |
| Mbtps2 | 0.1923734 | 0.193961 | 0.20973359 | -0.0157726 | 0.63093196 | 0.90650187 |
| Bzw2 | 0.7763446 | 0.6234438 | 0.63922751 | -0.0157837 | 0.54370419 | 0.90650187 |
| Spopl | 0.20782826 | 0.20886642 | 0.22465248 | -0.0157861 | 0.6226269 | 0.90650187 |
| Txndc11 | 0.3325026 | 0.32052936 | 0.33631801 | -0.0157887 | 0.58265942 | 0.90650187 |
| A530020G20 | 0.06586921 | 0.06158638 | 0.07738874 | -0.0158024 | 0.81799979 | 0.9092537 |
| Mettl3 | 0.18505832 | 0.18677708 | 0.20258231 | -0.0158052 | 0.63564461 | 0.90650187 |
| Plxnb1 | 0.13599097 | 0.13725877 | 0.15306435 | -0.0158056 | 0.67690366 | 0.90650187 |
| Tmem185b | 0.24606633 | 0.24469039 | 0.26049702 | -0.0158066 | 0.60632515 | 0.90650187 |
| Gm21967 | 0.09183502 | 0.09029121 | 0.10610148 | -0.0158103 | 0.74615647 | 0.90650187 |
| Hmgxb3 | 0.08066033 | 0.07805086 | 0.0938625 | -0.0158116 | 0.77303716 | 0.90650187 |
| Cul4b | 0.18995586 | 0.19155963 | 0.20737655 | -0.0158169 | 0.63272267 | 0.90650187 |
| Emc10 | 1.4219158 | 0.86308098 | 0.87890085 | -0.0158199 | 0.53188898 | 0.90650187 |
| Tefm | 0.14120952 | 0.14264278 | 0.15846292 | -0.0158201 | 0.67141617 | 0.90650187 |
| Gramd3 | 0.24738479 | 0.24588616 | 0.26170678 | -0.0158206 | 0.60593683 | 0.90650187 |
| Sidt2 | 0.14228121 | 0.14374608 | 0.15956762 | -0.0158215 | 0.67031872 | 0.90650187 |
| Lysmd1 | 0.06772427 | 0.06364302 | 0.07947222 | -0.0158292 | 0.81204882 | 0.9092537 |
| Asb1 | 0.10643981 | 0.1060374 | 0.12187377 | -0.0158364 | 0.71825494 | 0.90650187 |
| Klhl8 | 0.10203844 | 0.10129901 | 0.11713604 | -0.015837 | 0.72609838 | 0.90650187 |
| Dusp11 | 0.55443175 | 0.48741805 | 0.50325711 | -0.0158391 | 0.55563721 | 0.90650187 |
| Akap10 | 0.12320115 | 0.12385433 | 0.13969612 | -0.0158418 | 0.69291839 | 0.90650187 |
| Blvra | 0.2935982 | 0.28720136 | 0.30304422 | -0.0158429 | 0.59189101 | 0.90650187 |
| Ccdc166 | 0.14188502 | 0.14331302 | 0.15915939 | -0.0158464 | 0.67097881 | 0.90650187 |

| | | | | | | |
|---|---|---|---|---|---|---|
| B4galt4 | 0.15925945 | 0.16102242 | 0.17688934 | -0.0158669 | 0.65496846 | 0.90650187 |
| Sv2a | 0.10033627 | 0.09943018 | 0.11529726 | -0.0158671 | 0.7296647 | 0.90650187 |
| Bbs9 | 0.09055603 | 0.08884241 | 0.10470956 | -0.0158672 | 0.74973772 | 0.90650187 |
| Peli1 | 0.41894144 | 0.38981226 | 0.40567979 | -0.0158675 | 0.5690213 | 0.90650187 |
| Faap24 | 0.09972116 | 0.09875803 | 0.11463381 | -0.0158758 | 0.73094626 | 0.90650187 |
| Ltn1 | 0.1724159 | 0.17420686 | 0.1900853 | -0.0158784 | 0.6448138 | 0.90650187 |
| Slc23a2 | 0.06034636 | 0.05527311 | 0.07115594 | -0.0158828 | 0.83947708 | 0.91213511 |
| Rufy2 | 0.22106037 | 0.22134188 | 0.23722611 | -0.0158842 | 0.61702237 | 0.90650187 |
| Nfkbib | 0.26585734 | 0.2625863 | 0.27847426 | -0.015888 | 0.60011232 | 0.90650187 |
| Bloc1s4 | 0.20928601 | 0.21015217 | 0.22604656 | -0.0158944 | 0.62270449 | 0.90650187 |
| Hipk3 | 0.22966458 | 0.22940777 | 0.24530431 | -0.0158965 | 0.61336113 | 0.90650187 |
| Sobp | 0.16584676 | 0.16761961 | 0.18352071 | -0.0159011 | 0.64995105 | 0.90650187 |
| Lclat1 | 0.06213427 | 0.05725999 | 0.0731786 | -0.0159186 | 0.83324225 | 0.91056108 |
| Mob4 | 1.11333289 | 0.77802422 | 0.7939442 | -0.01592 | 0.53551557 | 0.90650187 |
| Erlin2 | 0.22268567 | 0.22283772 | 0.23875813 | -0.0159204 | 0.61655042 | 0.90650187 |
| Arfgef2 | 0.06750496 | 0.06330219 | 0.07922617 | -0.015924 | 0.81421596 | 0.9092537 |
| Enpp4 | 0.1241933 | 0.12481512 | 0.1407403 | -0.0159252 | 0.69252963 | 0.90650187 |
| Slc4a2 | 0.26038598 | 0.25761235 | 0.27354206 | -0.0159297 | 0.60213982 | 0.90650187 |
| Scrn2 | 0.12718402 | 0.12794831 | 0.1438803 | -0.015932 | 0.68873432 | 0.90650187 |
| Usp5 | 0.51544859 | 0.46052275 | 0.47645525 | -0.0159325 | 0.55908884 | 0.90650187 |
| Nob1 | 0.44503896 | 0.40954001 | 0.42547432 | -0.0159343 | 0.56611475 | 0.90650187 |
| Gpr137b | 0.1166378 | 0.11681976 | 0.13275841 | -0.0159387 | 0.70323541 | 0.90650187 |
| Gm26526 | 0.09710588 | 0.09587129 | 0.11181375 | -0.0159425 | 0.73691293 | 0.90650187 |
| Twsg1 | 0.53110335 | 0.47137891 | 0.48733484 | -0.0159559 | 0.55786347 | 0.90650187 |
| Epb41l1 | 0.09718781 | 0.09594431 | 0.11190224 | -0.0159579 | 0.73694983 | 0.90650187 |
| 4931414P19I | 0.09229948 | 0.09064718 | 0.10660639 | -0.0159592 | 0.74714004 | 0.90650187 |
| Dnah9 | 0.14165264 | 0.14296031 | 0.15891986 | -0.0159596 | 0.67236413 | 0.90650187 |
| Fnip2 | 0.09897872 | 0.09787167 | 0.11383419 | -0.0159625 | 0.73348588 | 0.90650187 |
| Fbxo28 | 0.19217316 | 0.19357553 | 0.20953859 | -0.0159631 | 0.63257273 | 0.90650187 |
| 2510009E07I | 0.13731874 | 0.13847726 | 0.15444097 | -0.0159637 | 0.67706877 | 0.90650187 |

| | | | | | | |
|---|---|---|---|---|---|---|
| Atp13a2 | 0.29446092 | 0.2878285 | 0.30379696 | -0.0159685 | 0.59238097 | 0.90650187 |
| Wdr43 | 0.55735952 | 0.48926168 | 0.50523127 | -0.0159696 | 0.55587497 | 0.90650187 |
| Flywch1 | 0.18012009 | 0.18175176 | 0.19772187 | -0.0159701 | 0.64023014 | 0.90650187 |
| Kat8 | 0.23196959 | 0.23148136 | 0.24745558 | -0.0159742 | 0.61293664 | 0.90650187 |
| Aars2 | 0.07985582 | 0.07699138 | 0.09297456 | -0.0159832 | 0.77763804 | 0.90650187 |
| Ahctf1 | 0.21059466 | 0.21130365 | 0.22729618 | -0.0159925 | 0.62278075 | 0.90650187 |
| Gpalpp1 | 0.1097916 | 0.10946907 | 0.12546546 | -0.0159964 | 0.71457454 | 0.90650187 |
| Sdhaf3 | 0.19223853 | 0.19360499 | 0.20960225 | -0.0159973 | 0.63280721 | 0.90650187 |
| Ufl1 | 0.40808649 | 0.38129218 | 0.39729486 | -0.0160027 | 0.57105666 | 0.90650187 |
| Lace1 | 0.06519322 | 0.06060818 | 0.07662826 | -0.0160201 | 0.82366143 | 0.9092537 |
| Atp6v0d1 | 1.11934682 | 0.7800279 | 0.79604849 | -0.0160206 | 0.5356452 | 0.90650187 |
| Vti1b | 0.73387614 | 0.59916196 | 0.61518449 | -0.0160225 | 0.54608898 | 0.90650187 |
| Lancl1 | 0.37178799 | 0.35256703 | 0.36859283 | -0.0160258 | 0.5766331 | 0.90650187 |
| Zfpm1 | 0.05148104 | 0.04502315 | 0.06105638 | -0.0160332 | 0.87847622 | 0.92534436 |
| Hps1 | 0.1287849 | 0.12952024 | 0.14555662 | -0.0160364 | 0.68785693 | 0.90650187 |
| Zxdc | 0.1146348 | 0.11459123 | 0.13063068 | -0.0160394 | 0.70741629 | 0.90650187 |
| Cdk12 | 0.64511526 | 0.54599378 | 0.56203404 | -0.0160403 | 0.55048077 | 0.90650187 |
| Bbx | 0.73649653 | 0.60064494 | 0.61669319 | -0.0160482 | 0.54605022 | 0.90650187 |
| Vps37d | 0.09054264 | 0.08864332 | 0.10469498 | -0.0160517 | 0.75225184 | 0.90650187 |
| Mgme1 | 0.10246512 | 0.10154265 | 0.11759639 | -0.0160537 | 0.72803962 | 0.90650187 |
| Nxn | 0.14094503 | 0.14212883 | 0.15819008 | -0.0160613 | 0.67413796 | 0.90650187 |
| Mtmr12 | 0.06891497 | 0.06473986 | 0.08080688 | -0.016067 | 0.81163302 | 0.9092537 |
| Srek1 | 0.7025819 | 0.58083465 | 0.59690717 | -0.0160725 | 0.54764114 | 0.90650187 |
| Zc3h7a | 0.39930516 | 0.37436328 | 0.39044666 | -0.0160834 | 0.57264983 | 0.90650187 |
| C2cd2 | 0.09602494 | 0.09456168 | 0.11064543 | -0.0160837 | 0.74093364 | 0.90650187 |
| Dffa | 0.16027332 | 0.16182602 | 0.17791318 | -0.0160872 | 0.6561499 | 0.90650187 |
| Zdhhc3 | 0.48715807 | 0.44028209 | 0.45638153 | -0.0160994 | 0.56230837 | 0.90650187 |
| Tmem42 | 0.23199973 | 0.23138194 | 0.24748367 | -0.0161017 | 0.61380076 | 0.90650187 |
| Arl5b | 0.18466274 | 0.18609208 | 0.20219394 | -0.0161019 | 0.63834142 | 0.90650187 |
| Hist2h2aa1 | 0.10125541 | 0.10018772 | 0.11629061 | -0.0161029 | 0.73091688 | 0.90650187 |

| | | | | | | |
|---|---|---|---|---|---|---|
| Trmt10c | 0.31518901 | 0.30556831 | 0.32167637 | -0.0161081 | 0.58808256 | 0.90650187 |
| Smarce1 | 1.00930377 | 0.73841516 | 0.75453512 | -0.01612 | 0.53783321 | 0.90650187 |
| Zbtb26 | 0.08612622 | 0.08374876 | 0.09987094 | -0.0161222 | 0.76351501 | 0.90650187 |
| Bcl11a | 0.15882061 | 0.16031094 | 0.17644582 | -0.0161349 | 0.65782088 | 0.90650187 |
| Wdr7 | 0.05310127 | 0.04677481 | 0.06291096 | -0.0161362 | 0.87291149 | 0.92289911 |
| Zcrb1 | 1.21842916 | 0.81176852 | 0.8279161 | -0.0161476 | 0.53454764 | 0.90650187 |
| Slc35f5 | 0.44855463 | 0.41195342 | 0.42810204 | -0.0161486 | 0.56658824 | 0.90650187 |
| Stx12 | 0.78334676 | 0.62696131 | 0.64311067 | -0.0161494 | 0.54444361 | 0.90650187 |
| Snx21 | 0.33743802 | 0.32428117 | 0.34044423 | -0.0161631 | 0.58358025 | 0.90650187 |
| Dvl1 | 0.07608572 | 0.07263401 | 0.08880116 | -0.0161672 | 0.79084822 | 0.90650187 |
| Fbxo16 | 0.13962337 | 0.14065693 | 0.15682545 | -0.0161685 | 0.67664479 | 0.90650187 |
| Tmem30b | 0.17264536 | 0.17414476 | 0.19031373 | -0.016169 | 0.64717204 | 0.90650187 |
| Tmem62 | 0.05400816 | 0.04777461 | 0.0639473 | -0.0161727 | 0.86956198 | 0.92144928 |
| Spryd7 | 0.19312566 | 0.1942927 | 0.21046574 | -0.016173 | 0.63368096 | 0.90650187 |
| Rad9a | 0.09023017 | 0.08817967 | 0.10435457 | -0.0161749 | 0.75460933 | 0.90650187 |
| 1110037F02I | 0.30021358 | 0.29261288 | 0.3087979 | -0.016185 | 0.59212221 | 0.90650187 |
| Mgat5 | 0.15019602 | 0.15149753 | 0.16768459 | -0.0161871 | 0.66609183 | 0.90650187 |
| Ankib1 | 0.11217735 | 0.11182419 | 0.12801346 | -0.0161893 | 0.71301534 | 0.90650187 |
| Dnph1 | 0.13618794 | 0.1370719 | 0.1532687 | -0.0161968 | 0.68074857 | 0.90650187 |
| Coq2 | 0.53762858 | 0.47562504 | 0.49182352 | -0.0161985 | 0.5582045 | 0.90650187 |
| Gpcpd1 | 0.27722567 | 0.27242917 | 0.2886305 | -0.0162013 | 0.59852829 | 0.90650187 |
| Zfp810 | 0.1076628 | 0.10697867 | 0.12318591 | -0.0162072 | 0.7206914 | 0.90650187 |
| 1190007I07R | 0.19216297 | 0.19331725 | 0.20952867 | -0.0162114 | 0.634562 | 0.90650187 |
| Ccdc149 | 0.05103197 | 0.04432111 | 0.06054165 | -0.0162205 | 0.88318956 | 0.92745622 |
| Nfx1 | 0.4823366 | 0.43667577 | 0.45290622 | -0.0162305 | 0.56328918 | 0.90650187 |
| Nme7 | 0.13313172 | 0.13386129 | 0.15009282 | -0.0162315 | 0.68465728 | 0.90650187 |
| 2610037D02I | 0.0734976 | 0.06968663 | 0.08592437 | -0.0162377 | 0.79957882 | 0.90799325 |
| Map1s | 0.15066963 | 0.15190185 | 0.16816793 | -0.0162661 | 0.66640297 | 0.90650187 |
| Orc3 | 0.24124064 | 0.23978104 | 0.2560547 | -0.0162737 | 0.61123614 | 0.90650187 |
| Bcl9 | 0.34243889 | 0.32831944 | 0.34460423 | -0.0162848 | 0.58319885 | 0.90650187 |

| | | | | | | |
|---|---|---|---|---|---|---|
| Chfr | 0.3435657 | 0.3292467 | 0.34553867 | -0.016292 | 0.58301319 | 0.90650187 |
| Per1 | 0.1536684 | 0.15492525 | 0.17122224 | -0.016297 | 0.66390136 | 0.90650187 |
| Slc16a9 | 0.10755643 | 0.10676595 | 0.12307186 | -0.0163059 | 0.72207027 | 0.90650187 |
| C2cd4c | 0.05975287 | 0.0541758 | 0.07048348 | -0.0163077 | 0.84816028 | 0.91440801 |
| Ddx39 | 0.60648441 | 0.52126932 | 0.53758035 | -0.016311 | 0.55364926 | 0.90650187 |
| Pdpk1 | 0.43785072 | 0.40375296 | 0.42007176 | -0.0163188 | 0.56855692 | 0.90650187 |
| Lmf1 | 0.21417797 | 0.21438448 | 0.23070875 | -0.0163243 | 0.62345137 | 0.90650187 |
| St5 | 0.14916406 | 0.15030136 | 0.16663056 | -0.0163292 | 0.66845592 | 0.90650187 |
| Jmjd1c | 0.31067806 | 0.30148772 | 0.3178179 | -0.0163302 | 0.59034206 | 0.90650187 |
| Hdac5 | 0.3242418 | 0.31302428 | 0.32936525 | -0.016341 | 0.58728331 | 0.90650187 |
| Grasp | 0.1827962 | 0.18401767 | 0.20035912 | -0.0163414 | 0.64154334 | 0.90650187 |
| Pygo1 | 0.07835342 | 0.0749656 | 0.09131388 | -0.0163483 | 0.78695912 | 0.90650187 |
| Dnaja3 | 0.4102377 | 0.3826128 | 0.39896359 | -0.0163508 | 0.57228514 | 0.90650187 |
| Eml1 | 0.21071767 | 0.21106225 | 0.22741355 | -0.0163513 | 0.62538099 | 0.90650187 |
| Ldlr | 0.27093505 | 0.26667457 | 0.28302626 | -0.0163517 | 0.60134982 | 0.90650187 |
| 1700086O06 | 0.07322879 | 0.06927325 | 0.08562503 | -0.0163518 | 0.80205259 | 0.90856259 |
| Cdk20 | 0.07475052 | 0.07096264 | 0.08731824 | -0.0163556 | 0.79749515 | 0.9075779 |
| Sars | 0.97448729 | 0.72359495 | 0.73995374 | -0.0163588 | 0.53914613 | 0.90650187 |
| Nup133 | 0.10414997 | 0.10304158 | 0.11941197 | -0.0163704 | 0.72888885 | 0.90650187 |
| Adam17 | 0.17590915 | 0.17717623 | 0.19355616 | -0.0163799 | 0.64662145 | 0.90650187 |
| Slc25a29 | 0.06872139 | 0.06420377 | 0.08059004 | -0.0163863 | 0.81698096 | 0.9092537 |
| Oard1 | 0.41136483 | 0.38344961 | 0.39983653 | -0.0163869 | 0.57228669 | 0.90650187 |
| Klf16 | 0.1107669 | 0.11011699 | 0.12650794 | -0.016391 | 0.71769011 | 0.90650187 |
| Thyn1 | 0.61129898 | 0.52427219 | 0.54067437 | -0.0164022 | 0.5536404 | 0.90650187 |
| Prpf4 | 0.21424192 | 0.21436467 | 0.23076952 | -0.0164049 | 0.62400918 | 0.90650187 |
| A230046K03 | 0.23702104 | 0.23574345 | 0.2521515 | -0.0164081 | 0.61381851 | 0.90650187 |
| Mfsd8 | 0.10842845 | 0.10757618 | 0.12400642 | -0.0164302 | 0.72207644 | 0.90650187 |
| Ostm1 | 0.09561639 | 0.09377043 | 0.11020343 | -0.016433 | 0.74630501 | 0.90650187 |
| Lpcat1 | 0.25654974 | 0.25363365 | 0.2700668 | -0.0164332 | 0.60661539 | 0.90650187 |
| Nudt22 | 0.14206004 | 0.14290417 | 0.15933975 | -0.0164356 | 0.67672336 | 0.90650187 |

| | | | | | | |
|---|---|---|---|---|---|---|
| Crnkl1 | 0.40198 | 0.37610291 | 0.39253887 | -0.016436 | 0.57383314 | 0.90650187 |
| Wdr24 | 0.07508055 | 0.07124682 | 0.08768502 | -0.0164382 | 0.79769389 | 0.9075779 |
| Ebag9 | 0.23152029 | 0.23059576 | 0.24703666 | -0.0164409 | 0.6163338 | 0.90650187 |
| Arrdc1 | 0.20154674 | 0.20217676 | 0.21861984 | -0.0164431 | 0.63094899 | 0.90650187 |
| Tmem128 | 0.59792672 | 0.51559779 | 0.53204845 | -0.0164507 | 0.55466473 | 0.90650187 |
| Mre11a | 0.13079057 | 0.1311938 | 0.14765248 | -0.0164587 | 0.68989309 | 0.90650187 |
| Zfp24 | 0.46096056 | 0.42083897 | 0.43730475 | -0.0164658 | 0.56646831 | 0.90650187 |
| Gpt2 | 0.11384298 | 0.11332162 | 0.12978819 | -0.0164666 | 0.71363173 | 0.90650187 |
| Slc5a3 | 0.08709419 | 0.08446241 | 0.10093059 | -0.0164682 | 0.76588458 | 0.90650187 |
| Slc25a28 | 0.06050502 | 0.05486441 | 0.07133562 | -0.0164712 | 0.84766711 | 0.9143112 |
| Mthfs | 0.32395773 | 0.31264927 | 0.32912511 | -0.0164758 | 0.58805512 | 0.90650187 |
| Arhgef2 | 0.48065931 | 0.43519625 | 0.45169356 | -0.0164973 | 0.56449192 | 0.90650187 |
| 9530077C05l | 0.0622285 | 0.0567846 | 0.07328507 | -0.0165005 | 0.84157415 | 0.91251845 |
| Ccdc90b | 0.49981303 | 0.44892276 | 0.46542727 | -0.0165045 | 0.56263197 | 0.90650187 |
| Erlin1 | 0.15374333 | 0.15479367 | 0.17129843 | -0.0165048 | 0.66579759 | 0.90650187 |
| Scn8a | 0.0473659 | 0.03982217 | 0.05632821 | -0.016506 | 0.90366318 | 0.93651361 |
| Ttll12 | 0.18217325 | 0.18323871 | 0.19974591 | -0.0165072 | 0.64333569 | 0.90650187 |
| Ppp2cb | 0.26824982 | 0.26410657 | 0.28062206 | -0.0165155 | 0.60319973 | 0.90650187 |
| Tead3 | 0.0892812 | 0.08679848 | 0.1033199 | -0.0165214 | 0.76141968 | 0.90650187 |
| Fpgt | 0.11917944 | 0.11892227 | 0.13545125 | -0.016529 | 0.70626057 | 0.90650187 |
| Sft2d2 | 0.1559991 | 0.15705489 | 0.17358898 | -0.0165341 | 0.66401193 | 0.90650187 |
| 1110019D14 | 0.08165097 | 0.07841817 | 0.09495461 | -0.0165364 | 0.78059572 | 0.90650187 |
| Vps35 | 1.02724096 | 0.74522274 | 0.76176841 | -0.0165457 | 0.53846166 | 0.90650187 |
| Usp2 | 0.06731564 | 0.06246584 | 0.07901371 | -0.0165479 | 0.82408487 | 0.90929265 |
| Pdss2 | 0.09132484 | 0.08899683 | 0.10554654 | -0.0165497 | 0.75712091 | 0.90650187 |
| Swt1 | 0.48189741 | 0.43603759 | 0.45258887 | -0.0165513 | 0.5645742 | 0.90650187 |
| Rbm4 | 0.05360949 | 0.04693675 | 0.06349188 | -0.0165551 | 0.87680558 | 0.92473192 |
| Fam221a | 0.21815262 | 0.21791164 | 0.2344785 | -0.0165669 | 0.62327758 | 0.90650187 |
| D6Wsu163e | 0.11557601 | 0.11506223 | 0.13163112 | -0.0165689 | 0.7121186 | 0.90650187 |
| AW554918 | 0.0855681 | 0.08266176 | 0.09925936 | -0.0165976 | 0.77137105 | 0.90650187 |

| | | | | | | |
|---|---|---|---|---|---|---|
| Zfp664 | 0.68285328 | 0.56852649 | 0.58513064 | -0.0166042 | 0.55019428 | 0.90650187 |
| Ak3 | 0.35796754 | 0.34078039 | 0.35738505 | -0.0166047 | 0.5818192 | 0.90650187 |
| Pcsk7 | 0.09007833 | 0.08757708 | 0.1041891 | -0.016612 | 0.76077947 | 0.90650187 |
| Coq9 | 0.49465822 | 0.44511051 | 0.46175639 | -0.0166459 | 0.56366184 | 0.90650187 |
| Hemk1 | 0.10340554 | 0.10196028 | 0.11861022 | -0.0166499 | 0.73370368 | 0.90650187 |
| Gprin3 | 0.13411952 | 0.13446977 | 0.15112051 | -0.0166507 | 0.68787116 | 0.90650187 |
| Bmpr2 | 0.26213399 | 0.25846955 | 0.27512091 | -0.0166514 | 0.60606012 | 0.90650187 |
| Cacna1h | 0.19145078 | 0.19218245 | 0.2088348 | -0.0166523 | 0.63851405 | 0.90650187 |
| Lamtor4 | 1.17257494 | 0.79717081 | 0.8138244 | -0.0166536 | 0.53624261 | 0.90650187 |
| Brdt | 0.06674613 | 0.0617192 | 0.07837426 | -0.0166551 | 0.82759942 | 0.90947782 |
| Ppif | 0.16145389 | 0.16244792 | 0.1791039 | -0.016656 | 0.66035217 | 0.90650187 |
| Gm10094 | 0.37243153 | 0.3524499 | 0.36911077 | -0.0166609 | 0.57951977 | 0.90650187 |
| Stard3nl | 0.63385014 | 0.53832748 | 0.55498939 | -0.0166619 | 0.5530876 | 0.90650187 |
| Dennd6a | 0.29770243 | 0.28995534 | 0.30661885 | -0.0166635 | 0.59545531 | 0.90650187 |
| Cep83os | 0.03796609 | 0.02876462 | 0.04543131 | -0.0166667 | 0.94854344 | 0.96195723 |
| Rcor3 | 0.38056547 | 0.35895675 | 0.37562874 | -0.016672 | 0.57820936 | 0.90650187 |
| Tbl1xr1 | 0.21960755 | 0.21918177 | 0.23585438 | -0.0166726 | 0.62333865 | 0.90650187 |
| Mtmr1 | 0.0590372 | 0.05299697 | 0.06967189 | -0.0166749 | 0.85633307 | 0.9165432 |
| Vps52 | 0.2632744 | 0.25947065 | 0.27614939 | -0.0166787 | 0.60584415 | 0.90650187 |
| Sf3a1 | 0.46887987 | 0.42644192 | 0.44312304 | -0.0166811 | 0.56645358 | 0.90650187 |
| Tmem248 | 0.43424128 | 0.40066325 | 0.41734482 | -0.0166816 | 0.57051836 | 0.90650187 |
| Tmem229a | 0.0630739 | 0.0575453 | 0.07423971 | -0.0166944 | 0.84126995 | 0.91251845 |
| Srrd | 0.10338825 | 0.10189431 | 0.11859159 | -0.0166973 | 0.73431817 | 0.90650187 |
| Hipk1 | 0.34953262 | 0.33376463 | 0.35046807 | -0.0167034 | 0.58389936 | 0.90650187 |
| Khnyn | 0.0688844 | 0.06406705 | 0.08077264 | -0.0167056 | 0.82107007 | 0.9092537 |
| Sin3a | 0.26049464 | 0.25693118 | 0.27364028 | -0.0167091 | 0.60698091 | 0.90650187 |
| Zfp189 | 0.0614013 | 0.05563969 | 0.07234996 | -0.0167103 | 0.84773734 | 0.91432207 |
| Rcl1 | 0.22885188 | 0.22782863 | 0.24454454 | -0.0167159 | 0.61939055 | 0.90650187 |
| Mavs | 0.1182667 | 0.11774552 | 0.13448512 | -0.0167396 | 0.70998904 | 0.90650187 |
| Glcci1 | 0.06575854 | 0.06051182 | 0.07726428 | -0.0167525 | 0.8324513 | 0.91029548 |

| | | | | | | |
|---|---|---|---|---|---|---|
| Sfxn1 | 0.55913131 | 0.48966889 | 0.50642344 | -0.0167545 | 0.55846507 | 0.90650187 |
| Prkci | 0.08885078 | 0.08609501 | 0.1028502 | -0.0167552 | 0.76553707 | 0.90650187 |
| Spire1 | 0.05712093 | 0.0507387 | 0.06749503 | -0.0167563 | 0.86513125 | 0.92040222 |
| Trappc13 | 0.47727766 | 0.43248624 | 0.44924286 | -0.0167566 | 0.5658506 | 0.90650187 |
| Neo1 | 0.29888464 | 0.2908793 | 0.30764542 | -0.0167661 | 0.59571744 | 0.90650187 |
| Wapl | 0.5328777 | 0.47178421 | 0.48855805 | -0.0167738 | 0.56065602 | 0.90650187 |
| Zc3h3 | 0.14973541 | 0.15043849 | 0.16721429 | -0.0167758 | 0.67220355 | 0.90650187 |
| Urod | 0.51836154 | 0.46171066 | 0.4784923 | -0.0167816 | 0.56195037 | 0.90650187 |
| Nvl | 0.33082109 | 0.31811768 | 0.33490748 | -0.0167898 | 0.5881817 | 0.90650187 |
| Stam | 0.15709452 | 0.15790615 | 0.17469919 | -0.016793 | 0.66543268 | 0.90650187 |
| Cnot8 | 0.39819429 | 0.37277748 | 0.38957614 | -0.0167987 | 0.57600986 | 0.90650187 |
| Noa1 | 0.21331418 | 0.21308353 | 0.22988733 | -0.0168038 | 0.62739252 | 0.90650187 |
| Mar-06 | 0.30687088 | 0.29774107 | 0.31454749 | -0.0168064 | 0.5938775 | 0.90650187 |
| Pno1 | 0.41210013 | 0.38359157 | 0.40040549 | -0.0168139 | 0.57404459 | 0.90650187 |
| Mettl5 | 0.27638533 | 0.27106317 | 0.28788403 | -0.0168209 | 0.60247312 | 0.90650187 |
| Wnk2 | 0.08789673 | 0.0849871 | 0.10180815 | -0.0168211 | 0.76868307 | 0.90650187 |
| Chm | 0.1347179 | 0.13491919 | 0.15174248 | -0.0168233 | 0.68894871 | 0.90650187 |
| Fam129b | 0.32909453 | 0.31662927 | 0.33345666 | -0.0168274 | 0.58875397 | 0.90650187 |
| Decr1 | 0.25487551 | 0.25171596 | 0.26854569 | -0.0168297 | 0.60972628 | 0.90650187 |
| Oxnad1 | 0.2394222 | 0.23754422 | 0.25437477 | -0.0168305 | 0.61567481 | 0.90650187 |
| Hmg20a | 0.17211143 | 0.17294817 | 0.18978211 | -0.0168339 | 0.6533318 | 0.90650187 |
| Ino80c | 0.0964287 | 0.09424068 | 0.11108202 | -0.0168413 | 0.74984633 | 0.90650187 |
| Wsb2 | 0.2581482 | 0.25466356 | 0.27151656 | -0.016853 | 0.60870227 | 0.90650187 |
| Rad18 | 0.05262009 | 0.045507 | 0.06236059 | -0.0168536 | 0.88521568 | 0.92807528 |
| Gm14296 | 0.09134929 | 0.0887194 | 0.10557314 | -0.0168537 | 0.76105439 | 0.90650187 |
| Spast | 0.16383171 | 0.16463909 | 0.18149738 | -0.0168583 | 0.66017021 | 0.90650187 |
| Yipf1 | 0.46117705 | 0.42059853 | 0.43746438 | -0.0168658 | 0.56804314 | 0.90650187 |
| Rab29 | 0.07239981 | 0.06783025 | 0.08470122 | -0.016871 | 0.81205253 | 0.9092537 |
| Aldh7a1 | 0.45520713 | 0.41616597 | 0.43305032 | -0.0168844 | 0.56880451 | 0.90650187 |
| Zfp839 | 0.05050186 | 0.04304764 | 0.05993365 | -0.016886 | 0.89481121 | 0.93253978 |

| | | | | | | |
|---|---|---|---|---|---|---|
| Zfp35 | 0.09291974 | 0.0903932 | 0.1072802 | -0.016887 | 0.75796883 | 0.90650187 |
| Fam193b | 0.13253476 | 0.13258007 | 0.14947118 | -0.0168911 | 0.69229965 | 0.90650187 |
| Coq10b | 0.29372004 | 0.28625662 | 0.30315057 | -0.016894 | 0.59783391 | 0.90650187 |
| Saal1 | 0.18594403 | 0.18655591 | 0.20345126 | -0.0168954 | 0.64400311 | 0.90650187 |
| Klhl20 | 0.11139131 | 0.11027173 | 0.12717473 | -0.016903 | 0.72269719 | 0.90650187 |
| Eefsec | 0.26428487 | 0.26015633 | 0.27705966 | -0.0169033 | 0.60689104 | 0.90650187 |
| Ccpg1os | 0.22490587 | 0.22394032 | 0.24084607 | -0.0169057 | 0.62249916 | 0.90650187 |
| Prkag2 | 0.10023313 | 0.0982765 | 0.11518572 | -0.0169092 | 0.74299718 | 0.90650187 |
| Dnajc11 | 0.33321029 | 0.31999507 | 0.33691094 | -0.0169159 | 0.58831327 | 0.90650187 |
| Gmip | 0.1058874 | 0.10434706 | 0.12128047 | -0.0169334 | 0.7326028 | 0.90650187 |
| Rad23a | 0.76078915 | 0.6135844 | 0.63052041 | -0.016936 | 0.54752498 | 0.90650187 |
| Faah | 0.21558234 | 0.21509024 | 0.23204258 | -0.0169523 | 0.62732724 | 0.90650187 |
| Ikzf2 | 0.1220017 | 0.12147316 | 0.13843217 | -0.016959 | 0.70699298 | 0.90650187 |
| Eml5 | 0.17148547 | 0.17217894 | 0.18915843 | -0.0169795 | 0.65507272 | 0.90650187 |
| Med24 | 0.18317791 | 0.18374813 | 0.20073465 | -0.0169865 | 0.64661405 | 0.90650187 |
| Zfp719 | 0.12983843 | 0.12966332 | 0.14665813 | -0.0169948 | 0.69677899 | 0.90650187 |
| Ndufaf7 | 0.2604118 | 0.25657035 | 0.27356539 | -0.016995 | 0.60879509 | 0.90650187 |
| Mns1 | 0.10801425 | 0.10654763 | 0.12356263 | -0.017015 | 0.72978911 | 0.90650187 |
| Abcd1 | 0.11215726 | 0.11095612 | 0.12799203 | -0.0170359 | 0.72297966 | 0.90650187 |
| Hmbox1 | 0.20080337 | 0.20084809 | 0.21790317 | -0.0170551 | 0.63606251 | 0.90650187 |
| AI597479 | 0.18195266 | 0.18247238 | 0.19952867 | -0.0170563 | 0.64803594 | 0.90650187 |
| Dusp28 | 0.18633309 | 0.18677479 | 0.20383269 | -0.0170579 | 0.64506629 | 0.90650187 |
| Zfp2 | 0.07335856 | 0.06870126 | 0.08576956 | -0.0170683 | 0.81183638 | 0.9092537 |
| Polr1a | 0.15999412 | 0.16056006 | 0.17763135 | -0.0170713 | 0.66539982 | 0.90650187 |
| 1700049G17 | 0.05905138 | 0.05260513 | 0.06968797 | -0.0170828 | 0.8620919 | 0.91899895 |
| Adam10 | 0.387053 | 0.36370211 | 0.38078998 | -0.0170879 | 0.57906247 | 0.90650187 |
| Zc3h10 | 0.08732741 | 0.08409694 | 0.10118571 | -0.0170888 | 0.77361256 | 0.90650187 |
| Aar2 | 0.20791529 | 0.20763208 | 0.22473577 | -0.0171037 | 0.63244507 | 0.90650187 |
| Wdr20 | 0.21638003 | 0.21569327 | 0.2327993 | -0.017106 | 0.62803818 | 0.90650187 |
| Marveld2 | 0.0899591 | 0.08694978 | 0.10405915 | -0.0171094 | 0.76759375 | 0.90650187 |

| | | | | | | |
|---|---|---|---|---|---|---|
| Scnm1 | 0.28576275 | 0.27906496 | 0.29617686 | -0.0171119 | 0.60135237 | 0.90650187 |
| Tsga10 | 0.10349041 | 0.10157877 | 0.11870166 | -0.0171229 | 0.73933134 | 0.90650187 |
| Gbe1 | 0.10942325 | 0.107948 | 0.12507142 | -0.0171234 | 0.72861979 | 0.90650187 |
| Pcgf5 | 0.08526853 | 0.08179005 | 0.09893091 | -0.0171409 | 0.77942413 | 0.90650187 |
| Sptbn2 | 0.07054963 | 0.06548494 | 0.08263582 | -0.0171509 | 0.82191195 | 0.9092537 |
| Setd8 | 0.22814125 | 0.2267279 | 0.24387965 | -0.0171518 | 0.62272821 | 0.90650187 |
| Madd | 0.23755396 | 0.235492 | 0.25264543 | -0.0171534 | 0.61861136 | 0.90650187 |
| Unc119 | 0.38066184 | 0.35855112 | 0.37570565 | -0.0171545 | 0.58042628 | 0.90650187 |
| Sigirr | 0.06617674 | 0.06056887 | 0.07773448 | -0.0171656 | 0.83687493 | 0.91092327 |
| Ttbk2 | 0.1026287 | 0.10060436 | 0.11777282 | -0.0171685 | 0.74152787 | 0.90650187 |
| B9d2 | 0.36821053 | 0.34853683 | 0.36570672 | -0.0171699 | 0.58266687 | 0.90650187 |
| 4933427D14I | 0.0656234 | 0.05993784 | 0.07711228 | -0.0171744 | 0.83894358 | 0.91188312 |
| Mrps31 | 0.30272295 | 0.29379061 | 0.31096975 | -0.0171791 | 0.59700097 | 0.90650187 |
| Prkca | 0.27263969 | 0.26736694 | 0.28454881 | -0.0171819 | 0.60581901 | 0.90650187 |
| Plxna1 | 0.1055752 | 0.10375954 | 0.120945 | -0.0171855 | 0.73620544 | 0.90650187 |
| Cldn12 | 0.10175116 | 0.09963445 | 0.11682596 | -0.0171915 | 0.74350611 | 0.90650187 |
| Gm4944 | 0.08593006 | 0.082462 | 0.09965604 | -0.017194 | 0.77846445 | 0.90650187 |
| Snap29 | 0.17802241 | 0.17845412 | 0.19564916 | -0.017195 | 0.65199477 | 0.90650187 |
| Sorcs2 | 0.07822795 | 0.07397263 | 0.09117505 | -0.0172024 | 0.79920173 | 0.90777025 |
| Mettl15 | 0.090896 | 0.0878766 | 0.10507978 | -0.0172032 | 0.76663637 | 0.90650187 |
| Cmc4 | 0.16464762 | 0.16511095 | 0.1823172 | -0.0172062 | 0.66259492 | 0.90650187 |
| Hgh1 | 0.14995839 | 0.15023221 | 0.16744199 | -0.0172098 | 0.67613636 | 0.90650187 |
| Ankrd16 | 0.1600594 | 0.16048094 | 0.17769726 | -0.0172163 | 0.66666242 | 0.90650187 |
| Pcnx | 0.16013262 | 0.16055145 | 0.17777117 | -0.0172197 | 0.66662823 | 0.90650187 |
| Mtor | 0.08297545 | 0.07919135 | 0.09641259 | -0.0172212 | 0.78642419 | 0.90650187 |
| Wdfy3 | 0.17487689 | 0.17530803 | 0.192532 | -0.017224 | 0.65457628 | 0.90650187 |
| Cdadc1 | 0.1988455 | 0.19878098 | 0.21601285 | -0.0172319 | 0.63856917 | 0.90650187 |
| Atf2 | 0.46289847 | 0.4214997 | 0.43873254 | -0.0172328 | 0.56930994 | 0.90650187 |
| BC005537 | 0.58897373 | 0.50898004 | 0.52621504 | -0.017235 | 0.55788372 | 0.90650187 |
| Klf11 | 0.1817444 | 0.18208295 | 0.19932353 | -0.0172406 | 0.64970518 | 0.90650187 |

| | | | | | | |
|---|---|---|---|---|---|---|
| B630005N14 | 0.18731742 | 0.18755176 | 0.20479699 | -0.0172452 | 0.64592759 | 0.90650187 |
| Slc35a3 | 0.12966559 | 0.12922919 | 0.14647751 | -0.0172483 | 0.69968271 | 0.90650187 |
| Mocs1 | 0.07429428 | 0.06956014 | 0.08681094 | -0.0172508 | 0.81149461 | 0.9092537 |
| Usf2 | 0.68261716 | 0.56773759 | 0.58498846 | -0.0172509 | 0.55215094 | 0.90650187 |
| Xrcc5 | 0.32948425 | 0.31653201 | 0.33378436 | -0.0172524 | 0.59086887 | 0.90650187 |
| Iws1 | 0.4227771 | 0.39136739 | 0.40862064 | -0.0172532 | 0.57444702 | 0.90650187 |
| Herc1 | 0.43560988 | 0.4011212 | 0.41837991 | -0.0172587 | 0.5727528 | 0.90650187 |
| Zfp668 | 0.12356214 | 0.12281525 | 0.14007618 | -0.0172609 | 0.70808929 | 0.90650187 |
| Usb1 | 0.16021574 | 0.16058195 | 0.17785507 | -0.0172731 | 0.66703961 | 0.90650187 |
| Kdm1a | 0.40531916 | 0.37786218 | 0.39514305 | -0.0172809 | 0.57707674 | 0.90650187 |
| Enox2 | 0.07613767 | 0.07157009 | 0.08885881 | -0.0172887 | 0.80646821 | 0.9092537 |
| Ncoa2 | 0.25765028 | 0.25376887 | 0.27106523 | -0.0172964 | 0.61166798 | 0.90650187 |
| Rnf169 | 0.05924888 | 0.0526154 | 0.06991202 | -0.0172966 | 0.86430505 | 0.92017797 |
| Suv39h1 | 0.0978111 | 0.09527769 | 0.11257512 | -0.0172974 | 0.75272124 | 0.90650187 |
| Gm15441 | 0.07871673 | 0.07441723 | 0.09171576 | -0.0172985 | 0.79912138 | 0.90777025 |
| Prkcz | 0.21906998 | 0.21804229 | 0.23534628 | -0.017304 | 0.62811539 | 0.90650187 |
| Cd27 | 0.16080933 | 0.16114739 | 0.178454 | -0.0173066 | 0.66681607 | 0.90650187 |
| Atp5sl | 0.12810397 | 0.12751383 | 0.14484398 | -0.0173301 | 0.70260357 | 0.90650187 |
| BC003331 | 0.24306344 | 0.24039246 | 0.25773538 | -0.0173429 | 0.61758459 | 0.90650187 |
| Dapp1 | 0.07976532 | 0.07553129 | 0.09287461 | -0.0173433 | 0.79677832 | 0.9075779 |
| Gtf3c4 | 0.20108182 | 0.20081872 | 0.21817169 | -0.017353 | 0.63817885 | 0.90650187 |
| Kdelc1 | 0.21456655 | 0.2137104 | 0.23107801 | -0.0173676 | 0.63085839 | 0.90650187 |
| Hn1l | 0.48403029 | 0.43675377 | 0.45412881 | -0.017375 | 0.56753018 | 0.90650187 |
| Cep250 | 0.1332647 | 0.13285302 | 0.15023123 | -0.0173782 | 0.69646195 | 0.90650187 |
| Mkks | 0.43611059 | 0.40138005 | 0.41875826 | -0.0173782 | 0.57318545 | 0.90650187 |
| Tmem38a | 0.2270322 | 0.22545636 | 0.24284097 | -0.0173846 | 0.62485385 | 0.90650187 |
| Sox13 | 0.06737599 | 0.0616934 | 0.07908143 | -0.017388 | 0.83583369 | 0.91079271 |
| Mob3c | 0.21512958 | 0.2142162 | 0.23161278 | -0.0173966 | 0.63077712 | 0.90650187 |
| Plekhm3 | 0.05708371 | 0.05004723 | 0.06745269 | -0.0174055 | 0.87433859 | 0.9235672 |
| Ccdc22 | 0.12279152 | 0.12185598 | 0.13926466 | -0.0174087 | 0.71079876 | 0.90650187 |

| | | | | | | |
|---|---|---|---|---|---|---|
| Il17rc | 0.12657856 | 0.1258197 | 0.1432455 | -0.0174258 | 0.70567989 | 0.90650187 |
| Lman2 | 1.24038509 | 0.81695919 | 0.83439603 | -0.0174368 | 0.53700941 | 0.90650187 |
| Ccdc134 | 0.11062039 | 0.10891272 | 0.12635141 | -0.0174387 | 0.73025061 | 0.90650187 |
| Pigc | 0.27325573 | 0.26765418 | 0.28509824 | -0.0174441 | 0.60719264 | 0.90650187 |
| 5033430I15R | 0.06603511 | 0.06012684 | 0.07757527 | -0.0174484 | 0.84132481 | 0.91251845 |
| Fam163a | 0.07754876 | 0.0729734 | 0.09042312 | -0.0174497 | 0.80452805 | 0.90882608 |
| Terf2 | 0.14664482 | 0.14659973 | 0.1640518 | -0.0174521 | 0.68186844 | 0.90650187 |
| Zbtb33 | 0.10936496 | 0.10755265 | 0.12500905 | -0.0174564 | 0.73263092 | 0.90650187 |
| Mrps10 | 0.53204011 | 0.4705154 | 0.48798087 | -0.0174655 | 0.5632105 | 0.90650187 |
| Parva | 0.48708762 | 0.43885621 | 0.45633086 | -0.0174746 | 0.56758898 | 0.90650187 |
| Tia1 | 1.10623767 | 0.77397668 | 0.79145289 | -0.0174762 | 0.53909904 | 0.90650187 |
| Mbip | 0.20285689 | 0.20240213 | 0.2198815 | -0.0174794 | 0.6381062 | 0.90650187 |
| Atxn7 | 0.17767792 | 0.17782429 | 0.1953083 | -0.017484 | 0.65467419 | 0.90650187 |
| Tmem159 | 0.25207927 | 0.24850855 | 0.26599915 | -0.0174906 | 0.61497823 | 0.90650187 |
| Ccnt2 | 0.77562193 | 0.62132369 | 0.63882549 | -0.0175018 | 0.54846955 | 0.90650187 |
| Ppp2r3a | 0.25343162 | 0.24972453 | 0.26723168 | -0.0175072 | 0.61456806 | 0.90650187 |
| Insr | 0.14775758 | 0.14767809 | 0.16519175 | -0.0175137 | 0.68129532 | 0.90650187 |
| Slc36a4 | 0.0884952 | 0.08494278 | 0.10246197 | -0.0175192 | 0.77640865 | 0.90650187 |
| Smyd3 | 0.16049425 | 0.16061692 | 0.17813614 | -0.0175192 | 0.66902173 | 0.90650187 |
| Trpc4ap | 0.64897108 | 0.54689861 | 0.56442877 | -0.0175302 | 0.55490853 | 0.90650187 |
| Ccdc148 | 0.07806344 | 0.07345955 | 0.09099298 | -0.0175334 | 0.80417347 | 0.90882608 |
| D430042O09 | 0.26642704 | 0.26145261 | 0.27898618 | -0.0175336 | 0.6100292 | 0.90650187 |
| Zfyve16 | 0.15468898 | 0.15472221 | 0.17225937 | -0.0175372 | 0.67457887 | 0.90650187 |
| Borcs6 | 0.21462655 | 0.2135799 | 0.23113501 | -0.0175551 | 0.63218713 | 0.90650187 |
| Hars2 | 0.15021883 | 0.1501496 | 0.16770788 | -0.0175583 | 0.67919014 | 0.90650187 |
| Cdc14b | 0.18044755 | 0.18048249 | 0.19804499 | -0.0175625 | 0.65329564 | 0.90650187 |
| Zdhhc4 | 0.23065688 | 0.2286616 | 0.24623108 | -0.0175695 | 0.62445788 | 0.90650187 |
| Galnt1 | 0.39326696 | 0.36813297 | 0.3857028 | -0.0175698 | 0.58024085 | 0.90650187 |
| Ipo4 | 0.31126812 | 0.30075365 | 0.31832363 | -0.01757 | 0.59691761 | 0.90650187 |
| Gstcd | 0.0848768 | 0.08092323 | 0.09850123 | -0.017578 | 0.78620976 | 0.90650187 |

| | | | | | | |
|---|---|---|---|---|---|---|
| BC017158 | 0.05636227 | 0.04905107 | 0.0666317 | -0.0175806 | 0.8795948 | 0.92562058 |
| Mrpl27 | 0.73813468 | 0.60005198 | 0.61763471 | -0.0175827 | 0.55035441 | 0.90650187 |
| Crocc | 0.06745287 | 0.06156918 | 0.07916771 | -0.0175985 | 0.83848076 | 0.9117781 |
| Commd9 | 0.53066588 | 0.46943207 | 0.48703296 | -0.0176009 | 0.5638193 | 0.90650187 |
| Exosc5 | 0.40003549 | 0.3734171 | 0.39101845 | -0.0176014 | 0.57930471 | 0.90650187 |
| Nsun5 | 0.14122938 | 0.14087942 | 0.15848341 | -0.017604 | 0.68927925 | 0.90650187 |
| Nedd4l | 0.24898571 | 0.2455629 | 0.263173 | -0.0176101 | 0.61694834 | 0.90650187 |
| Zfp866 | 0.11161561 | 0.10980125 | 0.12741415 | -0.0176129 | 0.73056791 | 0.90650187 |
| Lin52 | 0.07794624 | 0.07324992 | 0.09086325 | -0.0176133 | 0.80558907 | 0.90918104 |
| 2700033N17 | 0.12461664 | 0.12356631 | 0.14118549 | -0.0176192 | 0.71047998 | 0.90650187 |
| Tysnd1 | 0.25322798 | 0.24942014 | 0.2670462 | -0.0176261 | 0.61540202 | 0.90650187 |
| Cutc | 0.11176178 | 0.10993628 | 0.12757013 | -0.0176339 | 0.73056097 | 0.90650187 |
| Zfp524 | 0.15239086 | 0.15228594 | 0.1699223 | -0.0176364 | 0.6777488 | 0.90650187 |
| Fam35a | 0.07435955 | 0.06924477 | 0.08688353 | -0.0176388 | 0.81661219 | 0.9092537 |
| Pex5 | 0.22460325 | 0.22292153 | 0.24056178 | -0.0176403 | 0.62778544 | 0.90650187 |
| Zcchc2 | 0.07863799 | 0.07398512 | 0.09162868 | -0.0176436 | 0.80400767 | 0.90882608 |
| Dhx57 | 0.12553163 | 0.12449575 | 0.14214681 | -0.0176511 | 0.70953703 | 0.90650187 |
| Golga2 | 0.24685025 | 0.24355606 | 0.26121652 | -0.0176605 | 0.6181253 | 0.90650187 |
| Arl6ip6 | 0.68104064 | 0.56637265 | 0.58403845 | -0.0176658 | 0.55348422 | 0.90650187 |
| Pkdcc | 0.30711369 | 0.29708935 | 0.31475645 | -0.0176671 | 0.59852435 | 0.90650187 |
| Elk4 | 0.16455467 | 0.16455643 | 0.18222384 | -0.0176674 | 0.66677356 | 0.90650187 |
| Ldlrad3 | 0.18597602 | 0.18580969 | 0.20348263 | -0.0176729 | 0.65029286 | 0.90650187 |
| Rnf115 | 0.46041767 | 0.41922772 | 0.43690429 | -0.0176766 | 0.57137008 | 0.90650187 |
| N6amt1 | 0.45902439 | 0.41819398 | 0.43587561 | -0.0176816 | 0.57155697 | 0.90650187 |
| Baz1a | 0.14160799 | 0.14119217 | 0.15887383 | -0.0176817 | 0.68961804 | 0.90650187 |
| Dis3 | 0.12787123 | 0.12689803 | 0.14460028 | -0.0177023 | 0.70686182 | 0.90650187 |
| Serinc5 | 0.06432624 | 0.05793749 | 0.07565194 | -0.0177144 | 0.85105545 | 0.91505486 |
| Capn1 | 0.07448646 | 0.06930817 | 0.08702466 | -0.0177165 | 0.81727715 | 0.9092537 |
| Shc2 | 0.05945513 | 0.0524294 | 0.07014592 | -0.0177165 | 0.86924134 | 0.92141903 |
| Pdk1 | 0.2073716 | 0.20649664 | 0.22421531 | -0.0177187 | 0.63732402 | 0.90650187 |

| | | | | | | |
|---|---|---|---|---|---|---|
| Cyfip1 | 0.44229082 | 0.40569418 | 0.42341337 | -0.0177192 | 0.57379398 | 0.90650187 |
| Cdc42se2 | 0.45360215 | 0.41413577 | 0.43185937 | -0.0177236 | 0.57238466 | 0.90650187 |
| Syde2 | 0.07703211 | 0.07212389 | 0.0898507 | -0.0177268 | 0.80977895 | 0.9092537 |
| Clstn3 | 0.05287176 | 0.04492104 | 0.06264849 | -0.0177274 | 0.89579671 | 0.93322571 |
| Slc25a11 | 1.06614305 | 0.75913525 | 0.77686313 | -0.0177279 | 0.54040257 | 0.90650187 |
| Aasdhppt | 0.47373676 | 0.42892799 | 0.4466684 | -0.0177404 | 0.57007576 | 0.90650187 |
| Ppp2r1b | 0.21721637 | 0.2158473 | 0.23359198 | -0.0177447 | 0.63220834 | 0.90650187 |
| Kansl3 | 0.30957476 | 0.29911019 | 0.31687147 | -0.0177613 | 0.59839122 | 0.90650187 |
| Gpsm1 | 0.29072796 | 0.28277208 | 0.30053509 | -0.017763 | 0.60363064 | 0.90650187 |
| Unc13a | 0.10655692 | 0.10423593 | 0.12199949 | -0.0177636 | 0.74127988 | 0.90650187 |
| Nudt13 | 0.14606908 | 0.14569326 | 0.16346141 | -0.0177682 | 0.68553546 | 0.90650187 |
| Ehbp1 | 0.16998348 | 0.16988733 | 0.18766017 | -0.0177728 | 0.6631344 | 0.90650187 |
| Dcxr | 0.09418761 | 0.090883 | 0.10865585 | -0.0177729 | 0.76643862 | 0.90650187 |
| Pold3 | 0.2025004 | 0.20176305 | 0.21953839 | -0.0177753 | 0.64055656 | 0.90650187 |
| Ormdl1 | 0.25276494 | 0.24884669 | 0.26662428 | -0.0177776 | 0.6165445 | 0.90650187 |
| Slc15a4 | 0.15169235 | 0.15142677 | 0.16921075 | -0.017784 | 0.67983282 | 0.90650187 |
| Tfrc | 0.1155017 | 0.11375841 | 0.13155217 | -0.0177938 | 0.72618433 | 0.90650187 |
| Aaas | 0.11144784 | 0.10943165 | 0.12723508 | -0.0178034 | 0.7330467 | 0.90650187 |
| Gadd45gip1 | 0.95483212 | 0.71332889 | 0.73113424 | -0.0178054 | 0.5431069 | 0.90650187 |
| Arhgap1 | 0.37893829 | 0.35652366 | 0.37432903 | -0.0178054 | 0.58373589 | 0.90650187 |
| Mtmr3 | 0.2237595 | 0.22195918 | 0.23976866 | -0.0178095 | 0.6293792 | 0.90650187 |
| Cdipt | 0.51982812 | 0.46169557 | 0.47951536 | -0.0178198 | 0.5656099 | 0.90650187 |
| Pelo | 0.29319232 | 0.28486459 | 0.30268986 | -0.0178253 | 0.60326211 | 0.90650187 |
| A430105I19F | 0.06911286 | 0.06319441 | 0.0810285 | -0.0178341 | 0.83607708 | 0.91079271 |
| Cipc | 0.09731522 | 0.09419998 | 0.11203984 | -0.0178399 | 0.76048519 | 0.90650187 |
| Fastk | 0.4275127 | 0.39437282 | 0.41223608 | -0.0178633 | 0.57637941 | 0.90650187 |
| Dynlrb2 | 0.0848876 | 0.08064943 | 0.09851307 | -0.0178636 | 0.78991716 | 0.90650187 |
| Scyl2 | 0.18138525 | 0.18110276 | 0.19896963 | -0.0178669 | 0.65512778 | 0.90650187 |
| Chpt1 | 0.18496047 | 0.18461081 | 0.20248626 | -0.0178754 | 0.6526398 | 0.90650187 |
| Lrfn1 | 0.06088194 | 0.05388104 | 0.07176233 | -0.0178813 | 0.86601042 | 0.92051076 |

| | | | | | | |
|---|---|---|---|---|---|---|
| Bcl2l2 | 0.12357102 | 0.12219921 | 0.14008553 | -0.0178863 | 0.71485391 | 0.90650187 |
| Ift52 | 0.33816076 | 0.32315756 | 0.34104675 | -0.0178892 | 0.59219257 | 0.90650187 |
| Rnf138 | 0.53764645 | 0.47394437 | 0.49183577 | -0.0178914 | 0.56423558 | 0.90650187 |
| D930016D06 | 0.16904152 | 0.16881759 | 0.18671928 | -0.0179017 | 0.66502697 | 0.90650187 |
| Elmo3 | 0.12827367 | 0.12710425 | 0.14502164 | -0.0179174 | 0.70858428 | 0.90650187 |
| 9330151L19I | 0.13254481 | 0.13156339 | 0.14948166 | -0.0179183 | 0.70294583 | 0.90650187 |
| Spsb3 | 0.22151045 | 0.21972231 | 0.23765063 | -0.0179283 | 0.63132911 | 0.90650187 |
| Atrnl1 | 0.11778987 | 0.11604998 | 0.13398 | -0.01793 | 0.7240576 | 0.90650187 |
| Grhpr | 0.48218985 | 0.43486899 | 0.45280019 | -0.0179312 | 0.56987233 | 0.90650187 |
| Ccdc86 | 0.4288641 | 0.39532573 | 0.41326479 | -0.0179391 | 0.57651097 | 0.90650187 |
| Vps13b | 0.15879531 | 0.15848092 | 0.17642024 | -0.0179393 | 0.67438437 | 0.90650187 |
| Homez | 0.13367854 | 0.13271711 | 0.15066185 | -0.0179447 | 0.70177021 | 0.90650187 |
| Trmt2b | 0.11243785 | 0.1103462 | 0.12829125 | -0.017945 | 0.73297635 | 0.90650187 |
| Hnrnph2 | 0.97346998 | 0.72156851 | 0.73951432 | -0.0179458 | 0.54295519 | 0.90650187 |
| Cpt1c | 0.18229598 | 0.18191852 | 0.19986676 | -0.0179482 | 0.65513443 | 0.90650187 |
| Zbtb24 | 0.1455975 | 0.14502706 | 0.16297756 | -0.0179505 | 0.68780682 | 0.90650187 |
| Zfp882 | 0.07448833 | 0.06907371 | 0.08702674 | -0.017953 | 0.82045456 | 0.9092537 |
| Stk40 | 0.15426542 | 0.15387166 | 0.17182908 | -0.0179574 | 0.67889069 | 0.90650187 |
| Lrp12 | 0.11992541 | 0.11828044 | 0.1362401 | -0.0179597 | 0.72107428 | 0.90650187 |
| Cep192 | 0.07835876 | 0.07335736 | 0.09131979 | -0.0179624 | 0.80906123 | 0.9092537 |
| Dnm2 | 0.6550967 | 0.55024998 | 0.56821693 | -0.0179669 | 0.55589499 | 0.90650187 |
| Add1 | 0.56692764 | 0.49366927 | 0.51164681 | -0.0179775 | 0.56206371 | 0.90650187 |
| BC051226 | 0.07867335 | 0.0736774 | 0.09166779 | -0.0179904 | 0.80852631 | 0.9092537 |
| Ascc2 | 0.15863552 | 0.15826615 | 0.17625869 | -0.0179925 | 0.67501676 | 0.90650187 |
| Topors | 0.61320184 | 0.52389752 | 0.54189363 | -0.0179961 | 0.55868542 | 0.90650187 |
| Cs | 0.46790487 | 0.42441269 | 0.44240905 | -0.0179964 | 0.57175295 | 0.90650187 |
| Top2b | 0.64786365 | 0.54574221 | 0.5637418 | -0.0179996 | 0.5564374 | 0.90650187 |
| 5031439G07 | 0.13789299 | 0.13703417 | 0.1550357 | -0.0180015 | 0.69713873 | 0.90650187 |
| Arhgap21 | 0.15605148 | 0.15563332 | 0.1736421 | -0.0180088 | 0.67762322 | 0.90650187 |
| Kcnh8 | 0.08408283 | 0.07962005 | 0.09762966 | -0.0180096 | 0.79391334 | 0.90731056 |

| | | | | | | |
|---|---|---|---|---|---|---|
| Zcchc8 | 0.23685469 | 0.23398178 | 0.25199727 | -0.0180155 | 0.62468929 | 0.90650187 |
| Ncor1 | 0.80617736 | 0.63759646 | 0.65561533 | -0.0180189 | 0.54862281 | 0.90650187 |
| Tab1 | 0.14200848 | 0.14125931 | 0.15928663 | -0.0180273 | 0.6925646 | 0.90650187 |
| Lrch2 | 0.09914119 | 0.09597426 | 0.11400924 | -0.018035 | 0.75904852 | 0.90650187 |
| Wdyhv1 | 0.09015044 | 0.08622964 | 0.10426769 | -0.018038 | 0.77907666 | 0.90650187 |
| D3Ertd254e | 0.15835777 | 0.15793879 | 0.1759778 | -0.018039 | 0.67569973 | 0.90650187 |
| Zfp1 | 0.15835555 | 0.15792752 | 0.17597556 | -0.018048 | 0.67578381 | 0.90650187 |
| Rap1gap | 0.16458081 | 0.16419671 | 0.1822501 | -0.0180534 | 0.670167 | 0.90650187 |
| Ddhd1 | 0.12665344 | 0.1252688 | 0.14332403 | -0.0180552 | 0.71227219 | 0.90650187 |
| Inpp5e | 0.11146208 | 0.10918933 | 0.12725029 | -0.018061 | 0.735972 | 0.90650187 |
| Mks1 | 0.10698817 | 0.10440034 | 0.12246233 | -0.018062 | 0.74398599 | 0.90650187 |
| Trpm4 | 0.08736013 | 0.08315645 | 0.1012215 | -0.018065 | 0.78623026 | 0.90650187 |
| Gcc1 | 0.11532247 | 0.11329217 | 0.13136173 | -0.0180696 | 0.72957183 | 0.90650187 |
| Zbtb37 | 0.11574837 | 0.1137446 | 0.13181421 | -0.0180696 | 0.72887733 | 0.90650187 |
| Pik3c3 | 0.09948812 | 0.09631265 | 0.11438291 | -0.0180703 | 0.75875978 | 0.90650187 |
| Gm12184 | 0.05745014 | 0.04979614 | 0.0678694 | -0.0180733 | 0.88176205 | 0.92669144 |
| Alg6 | 0.14859088 | 0.1479642 | 0.16604447 | -0.0180803 | 0.6858333 | 0.90650187 |
| Grb2 | 0.45319912 | 0.41347685 | 0.43156001 | -0.0180832 | 0.57388706 | 0.90650187 |
| Asap1 | 0.4836877 | 0.43579028 | 0.45388167 | -0.0180914 | 0.5703238 | 0.90650187 |
| Cchcr1 | 0.07586243 | 0.07046163 | 0.08855334 | -0.0180917 | 0.81812711 | 0.9092537 |
| Sco1 | 0.07678925 | 0.07148233 | 0.08958149 | -0.0180992 | 0.81545953 | 0.9092537 |
| Wdr48 | 0.17343102 | 0.17298598 | 0.19109543 | -0.0181095 | 0.66323008 | 0.90650187 |
| Plagl2 | 0.11638653 | 0.11437836 | 0.13249177 | -0.0181134 | 0.72833141 | 0.90650187 |
| Ranbp10 | 0.13429867 | 0.13318988 | 0.15130676 | -0.0181169 | 0.70274076 | 0.90650187 |
| Fxn | 0.25559843 | 0.25108554 | 0.26920282 | -0.0181173 | 0.61760176 | 0.90650187 |
| Ncdn | 0.10268594 | 0.09971384 | 0.11783455 | -0.0181207 | 0.75290171 | 0.90650187 |
| Fbxo38 | 0.18817029 | 0.18749819 | 0.20563165 | -0.0181335 | 0.65248274 | 0.90650187 |
| Vps37c | 0.0872585 | 0.08297301 | 0.10111034 | -0.0181373 | 0.78740798 | 0.90650187 |
| Tet2 | 0.23903371 | 0.23587802 | 0.25401545 | -0.0181374 | 0.62454092 | 0.90650187 |
| Herc4 | 0.09812188 | 0.09476929 | 0.11291042 | -0.0181411 | 0.7624693 | 0.90650187 |

| | | | | | | |
|---|---|---|---|---|---|---|
| Hist3h2a | 0.44311494 | 0.40588689 | 0.42403205 | -0.0181452 | 0.57543797 | 0.90650187 |
| Dhx8 | 0.19507244 | 0.19421174 | 0.21235767 | -0.0181459 | 0.64798138 | 0.90650187 |
| Lmtk2 | 0.11996784 | 0.11813263 | 0.13628495 | -0.0181523 | 0.72311746 | 0.90650187 |
| Ifrd2 | 0.2025653 | 0.20143963 | 0.21960086 | -0.0181612 | 0.64343533 | 0.90650187 |
| Galnt7 | 0.12253214 | 0.1208288 | 0.13899135 | -0.0181625 | 0.71935442 | 0.90650187 |
| Tango6 | 0.09064773 | 0.08664318 | 0.10480944 | -0.0181663 | 0.77951075 | 0.90650187 |
| Fahd1 | 0.21561554 | 0.21390776 | 0.23207409 | -0.0181663 | 0.63607756 | 0.90650187 |
| Ift172 | 0.06453137 | 0.05771176 | 0.07588304 | -0.0181713 | 0.85646344 | 0.91657988 |
| Mark2 | 0.45061562 | 0.41146338 | 0.4296384 | -0.018175 | 0.57458627 | 0.90650187 |
| 2700049A03I | 0.20564719 | 0.20438299 | 0.22256251 | -0.0181795 | 0.6417459 | 0.90650187 |
| Bahd1 | 0.08136549 | 0.07645621 | 0.09464003 | -0.0181838 | 0.80349299 | 0.90882608 |
| Aptx | 0.23168126 | 0.22900085 | 0.24718677 | -0.0181859 | 0.62819263 | 0.90650187 |
| Tada1 | 0.319739 | 0.30736379 | 0.32555022 | -0.0181864 | 0.59806738 | 0.90650187 |
| Armcx5 | 0.15983797 | 0.15928173 | 0.17747369 | -0.018192 | 0.67569619 | 0.90650187 |
| Ftsj3 | 0.29138443 | 0.28291454 | 0.30110963 | -0.0181951 | 0.60589425 | 0.90650187 |
| Noct | 0.20181452 | 0.20067076 | 0.21887785 | -0.0182071 | 0.64423517 | 0.90650187 |
| Ippk | 0.12935403 | 0.12793893 | 0.14615184 | -0.0182129 | 0.71020896 | 0.90650187 |
| Gm16124 | 0.05232098 | 0.04380313 | 0.06201829 | -0.0182152 | 0.90417313 | 0.93677316 |
| 1700007L15F | 0.08842325 | 0.08416593 | 0.10238339 | -0.0182175 | 0.78553485 | 0.90650187 |
| Bach1 | 0.13808751 | 0.13700507 | 0.15523707 | -0.018232 | 0.69920648 | 0.90650187 |
| Znrf3 | 0.17068851 | 0.17010969 | 0.18836377 | -0.0182541 | 0.66669954 | 0.90650187 |
| Wdr11 | 0.15961085 | 0.15898882 | 0.17724433 | -0.0182555 | 0.67648105 | 0.90650187 |
| Usp24 | 0.10970254 | 0.10711085 | 0.1253702 | -0.0182594 | 0.74133726 | 0.90650187 |
| Rad54l2 | 0.12244885 | 0.1206401 | 0.13890357 | -0.0182635 | 0.7205654 | 0.90650187 |
| Vmac | 0.2243357 | 0.22204678 | 0.24031037 | -0.0182636 | 0.63226796 | 0.90650187 |
| Pgp | 0.7434455 | 0.60240591 | 0.6206775 | -0.0182716 | 0.55206106 | 0.90650187 |
| Tank | 0.14894318 | 0.14813249 | 0.16640476 | -0.0182723 | 0.68727995 | 0.90650187 |
| Fam172a | 0.31894642 | 0.30660273 | 0.32487672 | -0.018274 | 0.59872988 | 0.90650187 |
| Fam126b | 0.17848246 | 0.17782723 | 0.19610414 | -0.0182769 | 0.66068396 | 0.90650187 |
| Rreb1 | 0.21599598 | 0.21415484 | 0.23243505 | -0.0182802 | 0.63669238 | 0.90650187 |

| | | | | | | |
|---|---|---|---|---|---|---|
| Fam117a | 0.10388832 | 0.10084947 | 0.11913025 | -0.0182808 | 0.75243973 | 0.90650187 |
| Kctd9 | 0.10632086 | 0.1034616 | 0.12174604 | -0.0182844 | 0.74782599 | 0.90650187 |
| AU040320 | 0.11476667 | 0.11248485 | 0.13077091 | -0.0182861 | 0.73291127 | 0.90650187 |
| Mtif2 | 0.23397001 | 0.23103105 | 0.24931825 | -0.0182872 | 0.62781808 | 0.90650187 |
| Ptch1 | 0.17672476 | 0.17607401 | 0.19436454 | -0.0182905 | 0.66215452 | 0.90650187 |
| Eral1 | 0.14073987 | 0.13968558 | 0.15797839 | -0.0182928 | 0.69664335 | 0.90650187 |
| Fzd3 | 0.15498894 | 0.15426601 | 0.17256396 | -0.0182979 | 0.68132002 | 0.90650187 |
| Sorl1 | 0.09119815 | 0.08710957 | 0.10540867 | -0.0182991 | 0.77987278 | 0.90650187 |
| Slc25a51 | 0.71147052 | 0.58383955 | 0.60214854 | -0.018309 | 0.55376265 | 0.90650187 |
| Tbp | 0.12214173 | 0.12026506 | 0.13857982 | -0.0183148 | 0.7215771 | 0.90650187 |
| Pi4k2b | 0.35689689 | 0.33819456 | 0.3565103 | -0.0183157 | 0.5903298 | 0.90650187 |
| Gtpbp2 | 0.42849602 | 0.39465811 | 0.41298474 | -0.0183266 | 0.57819518 | 0.90650187 |
| Snhg9 | 0.43540936 | 0.3998992 | 0.41822833 | -0.0183291 | 0.57723763 | 0.90650187 |
| Ubap1 | 0.27056756 | 0.26436575 | 0.28269763 | -0.0183319 | 0.61343415 | 0.90650187 |
| Naa40 | 0.13168379 | 0.1302461 | 0.14858431 | -0.0183382 | 0.70838574 | 0.90650187 |
| Prdm15 | 0.07794706 | 0.07252343 | 0.09086415 | -0.0183407 | 0.8152301 | 0.9092537 |
| Qser1 | 0.18790634 | 0.18703266 | 0.20537342 | -0.0183408 | 0.65432082 | 0.90650187 |
| Imp4 | 0.43155291 | 0.39696072 | 0.41530761 | -0.0183469 | 0.57784827 | 0.90650187 |
| Rgl3 | 0.09228996 | 0.08824822 | 0.10659604 | -0.0183478 | 0.77792348 | 0.90650187 |
| Mtmr10 | 0.05907577 | 0.05136678 | 0.06971565 | -0.0183489 | 0.87901264 | 0.92549228 |
| Tinf2 | 0.05580724 | 0.04763455 | 0.06599953 | -0.018365 | 0.89196223 | 0.93142254 |
| Polg | 0.18345019 | 0.18263405 | 0.20100243 | -0.0183684 | 0.65772301 | 0.90650187 |
| Fbxl2 | 0.10472743 | 0.10164662 | 0.12003341 | -0.0183868 | 0.75205816 | 0.90650187 |
| Nova2 | 0.06796033 | 0.06134702 | 0.07973699 | -0.01839 | 0.84738981 | 0.9143112 |
| Asf1a | 0.50293124 | 0.44924799 | 0.46763932 | -0.0183913 | 0.5693961 | 0.90650187 |
| Wrap73 | 0.24857904 | 0.24439916 | 0.2628008 | -0.0184016 | 0.62220872 | 0.90650187 |
| Chml | 0.05820681 | 0.05032324 | 0.06872924 | -0.018406 | 0.88308548 | 0.92745622 |
| Wdr36 | 0.19783479 | 0.19662292 | 0.21503526 | -0.0184123 | 0.64826995 | 0.90650187 |
| Spen | 0.23370522 | 0.23065828 | 0.24907192 | -0.0184136 | 0.62879339 | 0.90650187 |
| Map2k5 | 0.22629812 | 0.22373604 | 0.24215278 | -0.0184167 | 0.6323588 | 0.90650187 |

| | | | | | | |
|---|---|---|---|---|---|---|
| Pacs1 | 0.06214154 | 0.0547683 | 0.07318681 | -0.0184185 | 0.86838745 | 0.92097661 |
| Axin2 | 0.11904858 | 0.11688178 | 0.1353128 | -0.018431 | 0.72758927 | 0.90650187 |
| Rbbp8 | 0.16935048 | 0.16859632 | 0.187028 | -0.0184317 | 0.66934893 | 0.90650187 |
| Stk25 | 0.64594009 | 0.54410837 | 0.56254699 | -0.0184386 | 0.55792631 | 0.90650187 |
| Dok1 | 0.11521495 | 0.11280762 | 0.13124747 | -0.0184398 | 0.73388099 | 0.90650187 |
| Ivd | 0.33771034 | 0.32222545 | 0.3406713 | -0.0184458 | 0.59511004 | 0.90650187 |
| Gm18025 | 0.07511451 | 0.06927683 | 0.08772275 | -0.0184459 | 0.82505454 | 0.90934433 |
| Spsb2 | 0.17939836 | 0.17855718 | 0.19700927 | -0.0184521 | 0.661434 | 0.90650187 |
| Atf7 | 0.23884254 | 0.23536993 | 0.25383858 | -0.0184686 | 0.62682313 | 0.90650187 |
| Dirc2 | 0.1238213 | 0.12187574 | 0.14034893 | -0.0184732 | 0.72077503 | 0.90650187 |
| 1700030K09I | 0.10596483 | 0.10287622 | 0.12136366 | -0.0184874 | 0.75086147 | 0.90650187 |
| Rev3l | 0.42388524 | 0.39097401 | 0.40946814 | -0.0184941 | 0.57956912 | 0.90650187 |
| Herc3 | 0.06837289 | 0.0617023 | 0.08019951 | -0.0184972 | 0.84739735 | 0.9143112 |
| Aco1 | 0.19903153 | 0.19769329 | 0.21619263 | -0.0184993 | 0.64817762 | 0.90650187 |
| Alkbh7 | 0.36829334 | 0.34727104 | 0.36577364 | -0.0185026 | 0.5889638 | 0.90650187 |
| Fnbp1l | 0.34295968 | 0.32653277 | 0.34503624 | -0.0185035 | 0.59421739 | 0.90650187 |
| Aagab | 0.19470548 | 0.19349016 | 0.21200136 | -0.0185112 | 0.65105648 | 0.90650187 |
| Polr1e | 0.10588316 | 0.10276012 | 0.12127592 | -0.0185158 | 0.75134642 | 0.90650187 |
| 1700017B05I | 0.12264341 | 0.12058617 | 0.13910861 | -0.0185224 | 0.72305343 | 0.90650187 |
| L2hgdh | 0.06274265 | 0.05534121 | 0.07386578 | -0.0185246 | 0.86755337 | 0.92056229 |
| Tmem143 | 0.07088753 | 0.0644743 | 0.0830134 | -0.0185391 | 0.83960159 | 0.91213511 |
| Zw10 | 0.10752441 | 0.10449652 | 0.12303753 | -0.018541 | 0.74854743 | 0.90650187 |
| Tango2 | 0.18026419 | 0.17930887 | 0.19786407 | -0.0185552 | 0.66162345 | 0.90650187 |
| Zfp948 | 0.22833847 | 0.22550288 | 0.24406423 | -0.0185614 | 0.63235352 | 0.90650187 |
| Thnsl1 | 0.10591755 | 0.10274611 | 0.12131286 | -0.0185668 | 0.75187228 | 0.90650187 |
| Smad4 | 0.29295221 | 0.28391275 | 0.30248016 | -0.0185674 | 0.60753055 | 0.90650187 |
| Ndufb8 | 2.17440155 | 0.9031382 | 0.92171263 | -0.0185744 | 0.53569279 | 0.90650187 |
| Frg1 | 0.79147615 | 0.6290059 | 0.64759099 | -0.0185851 | 0.55076073 | 0.90650187 |
| 9830147E19I | 0.06873561 | 0.06201378 | 0.08060597 | -0.0185922 | 0.84741323 | 0.9143112 |
| 9330160F10I | 0.09575203 | 0.09174719 | 0.1103502 | -0.018603 | 0.77320617 | 0.90650187 |

| | | | | | | |
|---|---|---|---|---|---|---|
| Map2k3 | 0.32839576 | 0.31426527 | 0.33286876 | -0.0186035 | 0.59810983 | 0.90650187 |
| Umad1 | 0.19112173 | 0.18990089 | 0.20851403 | -0.0186131 | 0.65425723 | 0.90650187 |
| Mrpl50 | 0.6431762 | 0.54221053 | 0.56082673 | -0.0186162 | 0.55865831 | 0.90650187 |
| Abcb7 | 0.06828319 | 0.06148146 | 0.08009897 | -0.0186175 | 0.84926786 | 0.91448046 |
| Ncor2 | 0.30816944 | 0.29704537 | 0.31566441 | -0.018619 | 0.60342335 | 0.90650187 |
| Ints9 | 0.12286859 | 0.1207096 | 0.13934585 | -0.0186363 | 0.72393111 | 0.90650187 |
| Lipe | 0.07978284 | 0.07425417 | 0.09289397 | -0.0186398 | 0.81381443 | 0.9092537 |
| Fbrsl1 | 0.13319266 | 0.13150893 | 0.15015625 | -0.0186473 | 0.70955407 | 0.90650187 |
| Trim13 | 0.10236444 | 0.09883586 | 0.11748779 | -0.0186519 | 0.75983384 | 0.90650187 |
| Nck2 | 0.10447607 | 0.10110208 | 0.11976296 | -0.0186609 | 0.7557488 | 0.90650187 |
| Tti1 | 0.07665139 | 0.07076612 | 0.08942864 | -0.0186625 | 0.82322789 | 0.9092537 |
| Rtn2 | 0.17609558 | 0.17507672 | 0.193741 | -0.0186643 | 0.66577381 | 0.90650187 |
| Ret | 0.12210895 | 0.11987491 | 0.13854526 | -0.0186704 | 0.72544349 | 0.90650187 |
| Gapvd1 | 0.54220135 | 0.4762785 | 0.49495329 | -0.0186748 | 0.56660392 | 0.90650187 |
| Kif1c | 0.09817423 | 0.09429037 | 0.11296689 | -0.0186765 | 0.76882439 | 0.90650187 |
| 0610040B10 | 0.07231675 | 0.06592566 | 0.08460861 | -0.018683 | 0.83686681 | 0.91092327 |
| Hexim1 | 0.21728135 | 0.21496561 | 0.23365354 | -0.0186879 | 0.63891712 | 0.90650187 |
| Flcn | 0.11514976 | 0.1124901 | 0.13117818 | -0.0186881 | 0.73674286 | 0.90650187 |
| Eif2ak2 | 0.19317761 | 0.19181072 | 0.21051627 | -0.0187056 | 0.65358489 | 0.90650187 |
| Ogg1 | 0.10828869 | 0.10514807 | 0.1238567 | -0.0187086 | 0.74905268 | 0.90650187 |
| Fgfbp3 | 0.08556208 | 0.08054214 | 0.09925276 | -0.0187106 | 0.79900236 | 0.90777025 |
| Nhsl1 | 0.07724455 | 0.07137325 | 0.09008612 | -0.0187129 | 0.82211375 | 0.9092537 |
| Knop1 | 0.97910049 | 0.72322764 | 0.74194117 | -0.0187135 | 0.54463936 | 0.90650187 |
| Smg5 | 0.25887456 | 0.25345632 | 0.27217454 | -0.0187182 | 0.62009756 | 0.90650187 |
| Rnf149 | 0.29148328 | 0.28247369 | 0.30119611 | -0.0187224 | 0.6088565 | 0.90650187 |
| Rbm48 | 0.13189193 | 0.13006918 | 0.14880131 | -0.0187321 | 0.71213864 | 0.90650187 |
| Mesdc2 | 0.83054254 | 0.64996781 | 0.66870207 | -0.0187343 | 0.54955907 | 0.90650187 |
| Sugp2 | 0.12899292 | 0.12703859 | 0.14577423 | -0.0187356 | 0.71612619 | 0.90650187 |
| Zfp954 | 0.11464227 | 0.11190087 | 0.13063863 | -0.0187378 | 0.7381467 | 0.90650187 |
| D630039A03 | 0.08730145 | 0.08241604 | 0.10115732 | -0.0187413 | 0.79492544 | 0.90731056 |

| | | | | | | |
|---|---|---|---|---|---|---|
| Zfp160 | 0.09810262 | 0.09414723 | 0.11288965 | -0.0187424 | 0.76976665 | 0.90650187 |
| Fam171a1 | 0.16613804 | 0.16506823 | 0.18381281 | -0.0187446 | 0.67485067 | 0.90650187 |
| Ptk2b | 0.0934629 | 0.0891244 | 0.1078698 | -0.0187454 | 0.78010525 | 0.90650187 |
| Prr7 | 0.15712448 | 0.15597243 | 0.17472954 | -0.0187571 | 0.68340196 | 0.90650187 |
| Tsn | 1.26636371 | 0.82304722 | 0.84180939 | -0.0187622 | 0.53946392 | 0.90650187 |
| Tuft1 | 0.09381538 | 0.08948938 | 0.10825221 | -0.0187628 | 0.7795088 | 0.90650187 |
| BC030336 | 0.09260526 | 0.08817119 | 0.10693863 | -0.0187674 | 0.78236104 | 0.90650187 |
| D930015E06 | 0.1328528 | 0.13103281 | 0.14980242 | -0.0187696 | 0.71124162 | 0.90650187 |
| Mto1 | 0.22095962 | 0.21835197 | 0.23713105 | -0.0187791 | 0.63760341 | 0.90650187 |
| Msl2 | 0.20979839 | 0.20775523 | 0.22653604 | -0.0187808 | 0.64377214 | 0.90650187 |
| Agtpbp1 | 0.13669639 | 0.13500712 | 0.15379599 | -0.0187889 | 0.70647244 | 0.90650187 |
| Cbwd1 | 0.16264843 | 0.16151463 | 0.1803071 | -0.0187925 | 0.67844051 | 0.90650187 |
| Lyplal1 | 0.41669758 | 0.38516057 | 0.40395374 | -0.0187932 | 0.58192605 | 0.90650187 |
| Homer1 | 0.079174 | 0.07342314 | 0.09222131 | -0.0187982 | 0.81758015 | 0.9092537 |
| Sptlc1 | 0.26401167 | 0.25801524 | 0.27681364 | -0.0187984 | 0.61863623 | 0.90650187 |
| Dgcr14 | 0.17518409 | 0.17402296 | 0.19283692 | -0.018814 | 0.66775833 | 0.90650187 |
| Supt5 | 0.72058452 | 0.58866566 | 0.60748124 | -0.0188156 | 0.55475897 | 0.90650187 |
| Ubr2 | 0.31054111 | 0.29888428 | 0.31770049 | -0.0188162 | 0.60383898 | 0.90650187 |
| Scaper | 0.34266988 | 0.32597629 | 0.34479587 | -0.0188196 | 0.59586141 | 0.90650187 |
| Bcdin3d | 0.14173672 | 0.14018582 | 0.15900653 | -0.0188207 | 0.70062587 | 0.90650187 |
| Cmtm6 | 0.29267981 | 0.28337974 | 0.30224219 | -0.0188624 | 0.60928006 | 0.90650187 |
| Zgpat | 0.21193435 | 0.20970306 | 0.22857359 | -0.0188705 | 0.64319778 | 0.90650187 |
| Adar | 0.09805836 | 0.09396884 | 0.11284191 | -0.0188731 | 0.7714219 | 0.90650187 |
| Zfp26 | 0.10683772 | 0.1034252 | 0.12230088 | -0.0188757 | 0.75368327 | 0.90650187 |
| Zfp950 | 0.40522354 | 0.3761857 | 0.3950686 | -0.0188829 | 0.58413544 | 0.90650187 |
| Timm10 | 0.38431271 | 0.35972965 | 0.37861393 | -0.0188843 | 0.58774003 | 0.90650187 |
| Pwwp2a | 0.19277943 | 0.19124265 | 0.21012884 | -0.0188862 | 0.65526228 | 0.90650187 |
| Pik3ca | 0.24719727 | 0.24264815 | 0.26153482 | -0.0188867 | 0.62590931 | 0.90650187 |
| Ercc2 | 0.09993063 | 0.09596877 | 0.11485928 | -0.0188905 | 0.76765731 | 0.90650187 |
| Abhd15 | 0.09947598 | 0.09547932 | 0.11436983 | -0.0188905 | 0.76861174 | 0.90650187 |

| | | | | | | |
|---|---|---|---|---|---|---|
| Enpep | 0.06901322 | 0.06202621 | 0.08091692 | -0.0188907 | 0.85033308 | 0.91483663 |
| Poldip3 | 0.53711662 | 0.47257809 | 0.4914723 | -0.0188942 | 0.56785171 | 0.90650187 |
| Phkb | 0.20299455 | 0.20110594 | 0.22001396 | -0.018908 | 0.64878997 | 0.90650187 |
| Cblb | 0.10219324 | 0.09839245 | 0.11730308 | -0.0189106 | 0.76321545 | 0.90650187 |
| Siae | 0.08689318 | 0.08179857 | 0.10071065 | -0.0189121 | 0.79809155 | 0.9075779 |
| Btbd2 | 0.11374563 | 0.11075861 | 0.12968456 | -0.018926 | 0.74176118 | 0.90650187 |
| Camta1 | 0.38503092 | 0.36025577 | 0.37918482 | -0.018929 | 0.58781438 | 0.90650187 |
| Zc3h11a | 0.21275687 | 0.21042076 | 0.22935695 | -0.0189362 | 0.64320488 | 0.90650187 |
| Eif4e3 | 0.27264544 | 0.2656057 | 0.28455393 | -0.0189482 | 0.61639627 | 0.90650187 |
| Dtnb | 0.25889872 | 0.25324704 | 0.27219642 | -0.0189494 | 0.62152478 | 0.90650187 |
| Acsf3 | 0.12231968 | 0.11981775 | 0.13876742 | -0.0189497 | 0.72810075 | 0.90650187 |
| Ksr2 | 0.09030555 | 0.08547968 | 0.1044367 | -0.018957 | 0.790153 | 0.90650187 |
| Pfkfb2 | 0.07651786 | 0.07031789 | 0.08928056 | -0.0189627 | 0.82747163 | 0.90947782 |
| Ptprk | 0.10122696 | 0.09728896 | 0.11625989 | -0.0189709 | 0.76590287 | 0.90650187 |
| Hint2 | 0.88226652 | 0.67663119 | 0.69560825 | -0.0189771 | 0.54826603 | 0.90650187 |
| Hnf1b | 0.1126968 | 0.10958094 | 0.1285673 | -0.0189864 | 0.74425241 | 0.90650187 |
| Sertad3 | 0.22595571 | 0.22283727 | 0.24183159 | -0.0189943 | 0.63652061 | 0.90650187 |
| Primpol | 0.10116096 | 0.09718678 | 0.11618859 | -0.0190018 | 0.76640141 | 0.90650187 |
| Nars | 0.75368604 | 0.60749539 | 0.62650611 | -0.0190107 | 0.5536536 | 0.90650187 |
| Wdr5 | 0.23090824 | 0.2274537 | 0.24646568 | -0.019012 | 0.63417335 | 0.90650187 |
| Mapk8ip1 | 0.4286246 | 0.39406787 | 0.41308258 | -0.0190147 | 0.58107208 | 0.90650187 |
| Cdk10 | 0.1953218 | 0.19358021 | 0.21259971 | -0.0190195 | 0.65457832 | 0.90650187 |
| Zbtb22 | 0.117103 | 0.11423216 | 0.13325187 | -0.0190197 | 0.73713708 | 0.90650187 |
| Zfp61 | 0.19199204 | 0.19033174 | 0.20936219 | -0.0190305 | 0.65692801 | 0.90650187 |
| Proser3 | 0.08968857 | 0.08472774 | 0.10376422 | -0.0190365 | 0.79263166 | 0.90681274 |
| Daxx | 0.26082873 | 0.25489994 | 0.27394223 | -0.0190423 | 0.62134807 | 0.90650187 |
| Dedd2 | 0.11761697 | 0.11473976 | 0.13379677 | -0.019057 | 0.73669804 | 0.90650187 |
| Tmem138 | 0.34977226 | 0.33160765 | 0.35066539 | -0.0190577 | 0.59545766 | 0.90650187 |
| Sgpp1 | 0.14289742 | 0.14114176 | 0.16020221 | -0.0190604 | 0.70157188 | 0.90650187 |
| Gnaq | 0.50008369 | 0.44655824 | 0.46561953 | -0.0190613 | 0.57220545 | 0.90650187 |

| | | | | | | |
|---|---|---|---|---|---|---|
| Fads6 | 0.06558257 | 0.05800221 | 0.07706636 | -0.0190641 | 0.86427368 | 0.92017797 |
| Sf3b2 | 1.75473919 | 0.90282118 | 0.92188898 | -0.0190678 | 0.53663121 | 0.90650187 |
| Mttp | 0.11579541 | 0.11278624 | 0.13186417 | -0.0190779 | 0.73994778 | 0.90650187 |
| Mta2 | 0.51021854 | 0.45370042 | 0.47278411 | -0.0190837 | 0.57120571 | 0.90650187 |
| Cluh | 0.13236212 | 0.13020593 | 0.14929133 | -0.0190854 | 0.71509323 | 0.90650187 |
| Comt | 0.8290533 | 0.64880793 | 0.66790964 | -0.0191017 | 0.5505856 | 0.90650187 |
| Leng9 | 0.09376774 | 0.08909189 | 0.10820053 | -0.0191086 | 0.78380596 | 0.90650187 |
| Pus3 | 0.17263172 | 0.17118955 | 0.19030016 | -0.0191106 | 0.6723615 | 0.90650187 |
| Rfwd3 | 0.17528195 | 0.17381992 | 0.19293403 | -0.0191141 | 0.67018688 | 0.90650187 |
| Fancl | 0.14616828 | 0.14444641 | 0.16356317 | -0.0191168 | 0.69832486 | 0.90650187 |
| Tprkb | 0.19335282 | 0.19156671 | 0.2106867 | -0.01912 | 0.65668866 | 0.90650187 |
| Elmod3 | 0.1004225 | 0.09626845 | 0.11539049 | -0.019122 | 0.7693499 | 0.90650187 |
| Echdc1 | 0.18911444 | 0.18743114 | 0.20655472 | -0.0191236 | 0.65968011 | 0.90650187 |
| Pnp | 0.2923529 | 0.28282799 | 0.30195651 | -0.0191285 | 0.61088368 | 0.90650187 |
| Wrap53 | 0.1572468 | 0.15571954 | 0.17485343 | -0.0191339 | 0.68668458 | 0.90650187 |
| Ap3d1 | 0.48283797 | 0.43412826 | 0.45326834 | -0.0191401 | 0.57445341 | 0.90650187 |
| Slc10a3 | 0.09270108 | 0.08789856 | 0.10704272 | -0.0191442 | 0.7867161 | 0.90650187 |
| Pip5k1c | 0.24708756 | 0.24227495 | 0.2614342 | -0.0191592 | 0.62771103 | 0.90650187 |
| Glod4 | 0.59656978 | 0.51200331 | 0.53116745 | -0.0191641 | 0.56371474 | 0.90650187 |
| Drosha | 0.23535248 | 0.2314213 | 0.25060319 | -0.0191819 | 0.63317694 | 0.90650187 |
| Cptp | 0.11918127 | 0.11627049 | 0.13545318 | -0.0191827 | 0.73551555 | 0.90650187 |
| Zdhhc5 | 0.12232094 | 0.11957181 | 0.13876874 | -0.0191969 | 0.73072086 | 0.90650187 |
| Acy1 | 0.11080609 | 0.10734835 | 0.1265498 | -0.0192014 | 0.75001389 | 0.90650187 |
| Pex7 | 0.45659566 | 0.41486659 | 0.43407921 | -0.0192126 | 0.57799442 | 0.90650187 |
| Sltm | 0.8101321 | 0.63854449 | 0.65775788 | -0.0192134 | 0.55166063 | 0.90650187 |
| Nup62 | 0.27692188 | 0.26913621 | 0.28836072 | -0.0192245 | 0.61653036 | 0.90650187 |
| Zfp788 | 0.12763799 | 0.12512988 | 0.14435598 | -0.0192261 | 0.72310464 | 0.90650187 |
| Polrmt | 0.09573788 | 0.09110746 | 0.11033489 | -0.0192274 | 0.78074457 | 0.90650187 |
| Sh3gl2 | 0.09869529 | 0.09429633 | 0.11352874 | -0.0192324 | 0.77430827 | 0.90650187 |
| Ubqln1 | 0.61927964 | 0.52651593 | 0.5457737 | -0.0192578 | 0.56232413 | 0.90650187 |

| | | | | | | |
|---|---|---|---|---|---|---|
| Zbtb4 | 0.15649537 | 0.15483345 | 0.17409213 | -0.0192587 | 0.68856551 | 0.90650187 |
| Dennd1a | 0.179949 | 0.17827792 | 0.19755299 | -0.0192751 | 0.66776637 | 0.90650187 |
| Tanc2 | 0.09688301 | 0.09228529 | 0.111573 | -0.0192877 | 0.77890945 | 0.90650187 |
| Adss | 0.61081352 | 0.52107379 | 0.54036299 | -0.0192892 | 0.56304489 | 0.90650187 |
| Tbrg4 | 0.10782485 | 0.10407012 | 0.12335963 | -0.0192895 | 0.75651963 | 0.90650187 |
| Bloc1s3 | 0.09358278 | 0.08870904 | 0.10799988 | -0.0192908 | 0.78642267 | 0.90650187 |
| Fbxo8 | 0.22443674 | 0.22110655 | 0.24040532 | -0.0192988 | 0.6394074 | 0.90650187 |
| Klhl15 | 0.07287622 | 0.06593241 | 0.08523225 | -0.0192998 | 0.84294981 | 0.91273395 |
| Prr36 | 0.06995355 | 0.06266585 | 0.08196935 | -0.0193035 | 0.85243296 | 0.91557373 |
| Cox15 | 0.06633973 | 0.05861257 | 0.07791767 | -0.0193051 | 0.86465083 | 0.92040222 |
| Slc25a44 | 0.24221727 | 0.23765039 | 0.25695559 | -0.0193052 | 0.63081045 | 0.90650187 |
| Rangrf | 0.23964055 | 0.23526711 | 0.25457666 | -0.0193096 | 0.63201605 | 0.90650187 |
| Slc30a7 | 0.15915784 | 0.15747394 | 0.17678667 | -0.0193127 | 0.68639302 | 0.90650187 |
| Usp15 | 0.33450952 | 0.31867551 | 0.33799837 | -0.0193229 | 0.60030966 | 0.90650187 |
| Mex3d | 0.11908157 | 0.11602226 | 0.13534771 | -0.0193254 | 0.73720755 | 0.90650187 |
| Ubr3 | 0.26953908 | 0.26243831 | 0.28177723 | -0.0193389 | 0.61985907 | 0.90650187 |
| Fastkd3 | 0.07320785 | 0.066258 | 0.0856017 | -0.0193437 | 0.84245251 | 0.91257184 |
| Gm10399 | 0.05879392 | 0.05001633 | 0.06939583 | -0.0193795 | 0.89278597 | 0.93179124 |
| Ino80 | 0.12228836 | 0.11935325 | 0.13873441 | -0.0193812 | 0.73271657 | 0.90650187 |
| Steap2 | 0.12945377 | 0.12687346 | 0.1462561 | -0.0193826 | 0.72212159 | 0.90650187 |
| R3hdm1 | 0.84262868 | 0.65571381 | 0.67509657 | -0.0193828 | 0.55078238 | 0.90650187 |
| Cactin | 0.29956701 | 0.2888375 | 0.30823738 | -0.0193999 | 0.61018379 | 0.90650187 |
| Sgpl1 | 0.19856909 | 0.19633987 | 0.21574564 | -0.0194058 | 0.65537421 | 0.90650187 |
| Mphosph6 | 0.46869139 | 0.423575 | 0.44298507 | -0.0194101 | 0.5772217 | 0.90650187 |
| Esrp1 | 0.17896601 | 0.17715827 | 0.19658212 | -0.0194239 | 0.66976366 | 0.90650187 |
| Tomm70a | 0.48285271 | 0.4338481 | 0.45327898 | -0.0194309 | 0.57556934 | 0.90650187 |
| Mknk1 | 0.16616513 | 0.16440834 | 0.18383998 | -0.0194316 | 0.68078486 | 0.90650187 |
| Vps54 | 0.08939099 | 0.08400636 | 0.10343968 | -0.0194333 | 0.79819766 | 0.9075779 |
| Ccng2 | 0.2167701 | 0.2137351 | 0.23316909 | -0.019434 | 0.64450581 | 0.90650187 |
| Tmcc1 | 0.33103844 | 0.31563893 | 0.33508995 | -0.019451 | 0.60181766 | 0.90650187 |

| | | | | | | |
|---|---|---|---|---|---|---|
| Erich1 | 0.26506183 | 0.25829194 | 0.27775891 | -0.019467 | 0.6223179 | 0.90650187 |
| Hsd17b12 | 0.64120384 | 0.54012949 | 0.55959651 | -0.019467 | 0.56145215 | 0.90650187 |
| Golga1 | 0.25702247 | 0.25102526 | 0.27049582 | -0.0194706 | 0.6255155 | 0.90650187 |
| Manea | 0.09325746 | 0.0881753 | 0.10764684 | -0.0194715 | 0.78934105 | 0.90650187 |
| Pabpc1 | 2.63159675 | 0.86563834 | 0.88512109 | -0.0194827 | 0.53897588 | 0.90650187 |
| Zfp58 | 0.07168271 | 0.06441204 | 0.0839013 | -0.0194893 | 0.84911863 | 0.91444933 |
| Csnk1g1 | 0.28340946 | 0.2746128 | 0.29410337 | -0.0194906 | 0.6158562 | 0.90650187 |
| Mmab | 0.08754926 | 0.08193315 | 0.1014283 | -0.0194952 | 0.8035786 | 0.90882608 |
| Atp9b | 0.11407466 | 0.11053541 | 0.13003479 | -0.0194994 | 0.74749934 | 0.90650187 |
| Pofut1 | 0.21547665 | 0.21243525 | 0.23194226 | -0.019507 | 0.64575629 | 0.90650187 |
| Pcyt2 | 0.33700968 | 0.32057245 | 0.34008693 | -0.0195145 | 0.60067424 | 0.90650187 |
| Stk38 | 0.49018183 | 0.43903729 | 0.45855309 | -0.0195158 | 0.57503308 | 0.90650187 |
| Ssh2 | 0.19778954 | 0.19547045 | 0.21499147 | -0.019521 | 0.65676745 | 0.90650187 |
| Trp53rkb | 0.12573441 | 0.12283687 | 0.14235971 | -0.0195228 | 0.72895613 | 0.90650187 |
| Tmem186 | 0.15840057 | 0.15648602 | 0.17602109 | -0.0195351 | 0.68912787 | 0.90650187 |
| Zfp53 | 0.13167134 | 0.12902287 | 0.14857133 | -0.0195485 | 0.7207047 | 0.90650187 |
| Wbp1l | 0.29227252 | 0.2823348 | 0.30188626 | -0.0195515 | 0.61329473 | 0.90650187 |
| Med4 | 0.32375218 | 0.30939503 | 0.32895131 | -0.0195563 | 0.60422289 | 0.90650187 |
| Ccnk | 0.14777728 | 0.14564877 | 0.16521192 | -0.0195632 | 0.70070014 | 0.90650187 |
| 1700048O20 | 0.07530661 | 0.06836012 | 0.08793617 | -0.019576 | 0.83883117 | 0.91188312 |
| Invs | 0.11309534 | 0.10941366 | 0.12899201 | -0.0195784 | 0.7500834 | 0.90650187 |
| Fbxw7 | 0.14270757 | 0.140426 | 0.16000674 | -0.0195807 | 0.70678914 | 0.90650187 |
| Cacnb2 | 0.09254247 | 0.08728857 | 0.10687042 | -0.0195819 | 0.79234016 | 0.90681274 |
| Errfi1 | 0.35652524 | 0.33662314 | 0.35620644 | -0.0195833 | 0.5965425 | 0.90650187 |
| Ckap5 | 0.24956005 | 0.24410912 | 0.2636984 | -0.0195893 | 0.62939424 | 0.90650187 |
| Arhgap32 | 0.12149672 | 0.11830582 | 0.13789952 | -0.0195937 | 0.73619606 | 0.90650187 |
| Dennd1b | 0.13359431 | 0.13097036 | 0.15057423 | -0.0196039 | 0.71863512 | 0.90650187 |
| Hdac4 | 0.18799507 | 0.18585493 | 0.20546023 | -0.0196053 | 0.66429218 | 0.90650187 |
| Rfng | 0.1876644 | 0.1855282 | 0.20513665 | -0.0196085 | 0.66456104 | 0.90650187 |
| Inpp5k | 0.19920265 | 0.19674315 | 0.21635798 | -0.0196148 | 0.65653875 | 0.90650187 |

| | | | | | | |
|---|---|---|---|---|---|---|
| Glis3 | 0.13538448 | 0.13281991 | 0.15243486 | -0.019615 | 0.71635594 | 0.90650187 |
| Zfp568 | 0.2119688 | 0.20898088 | 0.22860641 | -0.0196255 | 0.64863908 | 0.90650187 |
| Letmd1 | 0.16247902 | 0.16051068 | 0.18013657 | -0.0196259 | 0.68593096 | 0.90650187 |
| Grk6 | 0.12810766 | 0.12521925 | 0.14484785 | -0.0196286 | 0.72655681 | 0.90650187 |
| Ulk1 | 0.13753594 | 0.13503284 | 0.15466596 | -0.0196331 | 0.71373192 | 0.90650187 |
| Tjp2 | 0.10779138 | 0.10368744 | 0.12332376 | -0.0196363 | 0.76048239 | 0.90650187 |
| Pkp3 | 0.06094208 | 0.05217998 | 0.07183041 | -0.0196504 | 0.88804011 | 0.92986219 |
| Blzf1 | 0.23991641 | 0.23517965 | 0.25483166 | -0.019652 | 0.6341385 | 0.90650187 |
| Pnkp | 0.18198914 | 0.17990668 | 0.1995646 | -0.0196579 | 0.6692743 | 0.90650187 |
| Nus1 | 0.45767563 | 0.4152202 | 0.43487851 | -0.0196583 | 0.57963501 | 0.90650187 |
| Irs2 | 0.10613206 | 0.10188118 | 0.12154328 | -0.0196621 | 0.7639811 | 0.90650187 |
| Mib1 | 0.32315834 | 0.30877248 | 0.32844899 | -0.0196765 | 0.60500539 | 0.90650187 |
| Rnf214 | 0.35621421 | 0.33624858 | 0.35595206 | -0.0197035 | 0.59719119 | 0.90650187 |
| Sepn1 | 0.19546371 | 0.19302847 | 0.21273743 | -0.019709 | 0.6597794 | 0.90650187 |
| Neil1 | 0.09867277 | 0.09379288 | 0.11350446 | -0.0197116 | 0.77995453 | 0.90650187 |
| Ddx26b | 0.11726269 | 0.11370895 | 0.13342121 | -0.0197123 | 0.74434981 | 0.90650187 |
| Mlh1 | 0.17382186 | 0.1717706 | 0.19148405 | -0.0197135 | 0.67640848 | 0.90650187 |
| Rnf44 | 0.21012967 | 0.20713446 | 0.22685237 | -0.0197179 | 0.65040358 | 0.90650187 |
| Extl2 | 0.1796547 | 0.17753511 | 0.19726243 | -0.0197273 | 0.67169011 | 0.90650187 |
| Zfp952 | 0.10537859 | 0.10100634 | 0.12073367 | -0.0197273 | 0.76620062 | 0.90650187 |
| Carf | 0.16903562 | 0.16698497 | 0.18671338 | -0.0197284 | 0.68072575 | 0.90650187 |
| Banp | 0.46695902 | 0.42197579 | 0.44171578 | -0.01974 | 0.57874005 | 0.90650187 |
| Yipf2 | 0.26109481 | 0.25444095 | 0.27418264 | -0.0197417 | 0.62555086 | 0.90650187 |
| Rpap3 | 0.26960076 | 0.26208378 | 0.28183246 | -0.0197487 | 0.62229521 | 0.90650187 |
| Ppp1r13b | 0.14626389 | 0.14390918 | 0.16366122 | -0.019752 | 0.70420518 | 0.90650187 |
| Ikbkg | 0.09045089 | 0.08479925 | 0.10459504 | -0.0197958 | 0.79993279 | 0.90805731 |
| Skp1a | 1.42199664 | 0.85911636 | 0.87891698 | -0.0198006 | 0.53988839 | 0.90650187 |
| Etaa1 | 0.17021329 | 0.1680843 | 0.18788958 | -0.0198053 | 0.68032486 | 0.90650187 |
| Gm12166 | 0.07036536 | 0.06262315 | 0.08242984 | -0.0198067 | 0.85728814 | 0.9170242 |
| Slc25a46 | 0.41973301 | 0.38647108 | 0.4062878 | -0.0198167 | 0.58583146 | 0.90650187 |

| | | | | | | |
|---|---|---|---|---|---|---|
| Slco3a1 | 0.17765004 | 0.1754587 | 0.19528071 | -0.019822 | 0.67409613 | 0.90650187 |
| Map4k5 | 0.28621906 | 0.27675512 | 0.29657832 | -0.0198232 | 0.6168213 | 0.90650187 |
| Klhl28 | 0.09724643 | 0.09213813 | 0.11196556 | -0.0198274 | 0.78443238 | 0.90650187 |
| Blmh | 0.95049966 | 0.70926482 | 0.72909759 | -0.0198328 | 0.54812616 | 0.90650187 |
| Rabep2 | 0.16805189 | 0.16589317 | 0.18572968 | -0.0198365 | 0.68253687 | 0.90650187 |
| Adrm1 | 0.95468618 | 0.71122237 | 0.73106575 | -0.0198434 | 0.5480228 | 0.90650187 |
| 9530034E10I | 0.10646405 | 0.10205566 | 0.12189979 | -0.0198441 | 0.76537579 | 0.90650187 |
| Pkp2 | 0.08249183 | 0.07603519 | 0.09588051 | -0.0198453 | 0.82125285 | 0.9092537 |
| Ccnh | 0.45568462 | 0.41355774 | 0.43340429 | -0.0198466 | 0.58065691 | 0.90650187 |
| Dnal1 | 0.08999254 | 0.08424656 | 0.1040956 | -0.019849 | 0.80169521 | 0.90839694 |
| Lrrc24 | 0.14741055 | 0.14498646 | 0.16483639 | -0.0198499 | 0.70379572 | 0.90650187 |
| Cdcp1 | 0.08185561 | 0.07530728 | 0.09518005 | -0.0198728 | 0.82334916 | 0.9092537 |
| Ppm1a | 0.15907926 | 0.15683249 | 0.17670725 | -0.0198748 | 0.69146591 | 0.90650187 |
| Trak2 | 0.0761044 | 0.06893722 | 0.08882189 | -0.0198847 | 0.84019113 | 0.91226452 |
| Arhgef17 | 0.18503989 | 0.18267625 | 0.20256422 | -0.019888 | 0.66875372 | 0.90650187 |
| Psen1 | 0.26653567 | 0.25919546 | 0.27908376 | -0.0198883 | 0.62430478 | 0.90650187 |
| Mblac2 | 0.08912074 | 0.08325625 | 0.10314483 | -0.0198886 | 0.80433567 | 0.90882608 |
| Tspan33 | 0.10325656 | 0.09854664 | 0.11844968 | -0.019903 | 0.77246614 | 0.90650187 |
| 1810022K09I | 1.03203834 | 0.7437675 | 0.76367789 | -0.0199104 | 0.54613592 | 0.90650187 |
| Tecr | 1.90772306 | 0.90714886 | 0.92706323 | -0.0199144 | 0.53803983 | 0.90650187 |
| Phc3 | 0.1603972 | 0.15811821 | 0.1780382 | -0.01992 | 0.69054691 | 0.90650187 |
| Pikfyve | 0.11447774 | 0.11052957 | 0.13046364 | -0.0199341 | 0.75151415 | 0.90650187 |
| Epdr1 | 0.13127962 | 0.12821068 | 0.14816279 | -0.0199521 | 0.72530039 | 0.90650187 |
| Uba6 | 0.1946504 | 0.19197798 | 0.21194786 | -0.0199699 | 0.66234381 | 0.90650187 |
| Rnf220 | 0.17498794 | 0.17266642 | 0.19264224 | -0.0199758 | 0.6775943 | 0.90650187 |
| Snapc2 | 0.25495685 | 0.24862765 | 0.26861965 | -0.019992 | 0.62962643 | 0.90650187 |
| Pold2 | 0.28362233 | 0.27429602 | 0.29429114 | -0.0199951 | 0.61869278 | 0.90650187 |
| Nme5 | 0.12703236 | 0.12372601 | 0.14372133 | -0.0199953 | 0.73187642 | 0.90650187 |
| Bmi1 | 0.2033062 | 0.20031646 | 0.22031377 | -0.0199973 | 0.65671569 | 0.90650187 |
| Zcchc4 | 0.0998434 | 0.09476615 | 0.11476539 | -0.0199992 | 0.78074151 | 0.90650187 |

| | | | | | | |
|---|---|---|---|---|---|---|
| Tbc1d14 | 0.13710984 | 0.13421754 | 0.15422452 | -0.020007 | 0.71793308 | 0.90650187 |
| 3110043O21 | 0.08424573 | 0.07779494 | 0.09780856 | -0.0200136 | 0.81849942 | 0.9092537 |
| Pcdhga12 | 0.10183481 | 0.09690214 | 0.11691626 | -0.0200141 | 0.77667566 | 0.90650187 |
| Plekhm2 | 0.08622042 | 0.07994847 | 0.09997412 | -0.0200257 | 0.81339915 | 0.9092537 |
| Fam8a1 | 0.15449289 | 0.15203257 | 0.1720602 | -0.0200276 | 0.69758384 | 0.90650187 |
| Nprl2 | 0.23553746 | 0.23073719 | 0.25077498 | -0.0200378 | 0.63878737 | 0.90650187 |
| Pex11a | 0.08174328 | 0.07501812 | 0.09505631 | -0.0200382 | 0.82565898 | 0.90947782 |
| Mmachc | 0.0794509 | 0.07248651 | 0.0925273 | -0.0200408 | 0.83219822 | 0.91015505 |
| Snx32 | 0.08034682 | 0.07347402 | 0.0935166 | -0.0200426 | 0.829651 | 0.90974008 |
| Rubcn | 0.09617768 | 0.09076731 | 0.11081061 | -0.0200433 | 0.78933514 | 0.90650187 |
| Vps13a | 0.11043377 | 0.10610604 | 0.126152 | -0.020046 | 0.7600368 | 0.90650187 |
| Wwp2 | 0.15772253 | 0.15528727 | 0.17533507 | -0.0200478 | 0.69438209 | 0.90650187 |
| Ric8b | 0.11735808 | 0.11347373 | 0.13352235 | -0.0200486 | 0.74777866 | 0.90650187 |
| Prkcb | 0.13067923 | 0.1274874 | 0.14753626 | -0.0200489 | 0.72711745 | 0.90650187 |
| Samd8 | 0.15503169 | 0.1525561 | 0.17260737 | -0.0200513 | 0.69722382 | 0.90650187 |
| Pop7 | 0.53935251 | 0.47292682 | 0.49300497 | -0.0200781 | 0.57183663 | 0.90650187 |
| Tmem94 | 0.13622703 | 0.13322293 | 0.15330925 | -0.0200863 | 0.71987017 | 0.90650187 |
| Trim41 | 0.21809812 | 0.21431732 | 0.23442693 | -0.0201096 | 0.64852935 | 0.90650187 |
| Man2a2 | 0.17445409 | 0.17199534 | 0.19211219 | -0.0201168 | 0.67921951 | 0.90650187 |
| Pwp1 | 0.23707804 | 0.23207838 | 0.25220435 | -0.020126 | 0.63861548 | 0.90650187 |
| Tmem164 | 0.20306676 | 0.19993122 | 0.22008344 | -0.0201522 | 0.65802077 | 0.90650187 |
| Ptpn4 | 0.11437696 | 0.11020084 | 0.13035643 | -0.0201556 | 0.75407693 | 0.90650187 |
| Zfp944 | 0.1019592 | 0.09689414 | 0.11705052 | -0.0201564 | 0.77802616 | 0.90650187 |
| Exd2 | 0.10408849 | 0.0991838 | 0.11934578 | -0.020162 | 0.77368263 | 0.90650187 |
| Dnajb4 | 0.35016361 | 0.3308255 | 0.35098753 | -0.020162 | 0.60078219 | 0.90650187 |
| Zfp770 | 0.09809387 | 0.09271559 | 0.11288021 | -0.0201646 | 0.78644631 | 0.90650187 |
| Gm10263 | 0.37915743 | 0.35433376 | 0.37450419 | -0.0201704 | 0.59461622 | 0.90650187 |
| Ddx31 | 0.10006572 | 0.0948207 | 0.11500465 | -0.0201839 | 0.78237174 | 0.90650187 |
| Rps3 | 4.49509979 | 0.5868468 | 0.60703686 | -0.0201901 | 0.55877375 | 0.90650187 |
| Prmt10 | 0.12475827 | 0.12112735 | 0.14133436 | -0.020207 | 0.73748211 | 0.90650187 |

| | | | | | | |
|---|---|---|---|---|---|---|
| Wdr4 | 0.18335766 | 0.1807001 | 0.20091144 | -0.0202113 | 0.67264199 | 0.90650187 |
| Fam114a2 | 0.61986069 | 0.52591538 | 0.54614345 | -0.0202281 | 0.56539281 | 0.90650187 |
| Plekhg3 | 0.15009128 | 0.14734812 | 0.16757767 | -0.0202296 | 0.70424955 | 0.90650187 |
| Atp11b | 0.25796424 | 0.251112 | 0.27134984 | -0.0202378 | 0.62989023 | 0.90650187 |
| Gorab | 0.15716957 | 0.15453345 | 0.1747752 | -0.0202417 | 0.69667875 | 0.90650187 |
| Tmem168 | 0.29703632 | 0.28578792 | 0.30603988 | -0.020252 | 0.61569163 | 0.90650187 |
| Cpeb2 | 0.06568964 | 0.05693422 | 0.07718679 | -0.0202526 | 0.87828481 | 0.92521187 |
| Fam188a | 0.24203759 | 0.23652816 | 0.25678991 | -0.0202618 | 0.63712187 | 0.90650187 |
| Prkaa1 | 0.13412412 | 0.13085793 | 0.15112529 | -0.0202674 | 0.72447876 | 0.90650187 |
| Aph1a | 0.57602532 | 0.49742178 | 0.51769351 | -0.0202717 | 0.56909874 | 0.90650187 |
| Map4k2 | 0.11190943 | 0.10745285 | 0.12772767 | -0.0202748 | 0.75980401 | 0.90650187 |
| Zfp865 | 0.23468691 | 0.22968677 | 0.24998481 | -0.020298 | 0.64093919 | 0.90650187 |
| Efcab2 | 0.09787391 | 0.09233787 | 0.1126429 | -0.020305 | 0.78854002 | 0.90650187 |
| Mlxip | 0.11276604 | 0.1083241 | 0.12864111 | -0.020317 | 0.75869766 | 0.90650187 |
| Usp47 | 0.38758362 | 0.36089236 | 0.38121067 | -0.0203183 | 0.59365151 | 0.90650187 |
| Zc3h14 | 0.66945771 | 0.55669048 | 0.57701948 | -0.020329 | 0.56222913 | 0.90650187 |
| Rps6ka2 | 0.09775488 | 0.09218335 | 0.11251445 | -0.0203311 | 0.7891024 | 0.90650187 |
| Ppp1r2 | 1.19489189 | 0.80057106 | 0.82090271 | -0.0203316 | 0.54383744 | 0.90650187 |
| Zfp426 | 0.13054039 | 0.12705325 | 0.14739131 | -0.0203381 | 0.73020562 | 0.90650187 |
| Lin54 | 0.1208825 | 0.11690547 | 0.13725121 | -0.0203457 | 0.74505348 | 0.90650187 |
| Rmdn1 | 0.23013987 | 0.22540238 | 0.24574833 | -0.0203459 | 0.64358565 | 0.90650187 |
| Heatr3 | 0.15606923 | 0.15331275 | 0.1736601 | -0.0203474 | 0.69877534 | 0.90650187 |
| Osbpl11 | 0.13908204 | 0.13591222 | 0.15626592 | -0.0203537 | 0.71871961 | 0.90650187 |
| Wbscr27 | 0.23105534 | 0.22624741 | 0.24660294 | -0.0203555 | 0.64317443 | 0.90650187 |
| Jmjd8 | 0.39972359 | 0.37037759 | 0.3907743 | -0.0203967 | 0.59174702 | 0.90650187 |
| Gopc | 0.3450058 | 0.32632979 | 0.34673139 | -0.0204016 | 0.6031768 | 0.90650187 |
| Ubqln4 | 0.18592172 | 0.18302544 | 0.20342938 | -0.0204039 | 0.672163 | 0.90650187 |
| Mplkip | 0.47105951 | 0.4243115 | 0.44471619 | -0.0204047 | 0.58081368 | 0.90650187 |
| Kbtbd4 | 0.17124391 | 0.1685094 | 0.18891765 | -0.0204082 | 0.6844739 | 0.90650187 |
| Rbbp5 | 0.11381318 | 0.10934374 | 0.12975647 | -0.0204127 | 0.75783764 | 0.90650187 |

| | | | | | | |
|---|---|---|---|---|---|---|
| Fam117b | 0.13135943 | 0.12782699 | 0.14824604 | -0.020419 | 0.72983928 | 0.90650187 |
| Ccdc51 | 0.07826019 | 0.07079087 | 0.09121072 | -0.0204198 | 0.84019567 | 0.91226452 |
| Pde3b | 0.19720479 | 0.19400427 | 0.21442531 | -0.020421 | 0.66399186 | 0.90650187 |
| Rere | 0.50628284 | 0.44958181 | 0.47000983 | -0.020428 | 0.57660633 | 0.90650187 |
| Rpap2 | 0.14418177 | 0.14108975 | 0.16152342 | -0.0204337 | 0.71307668 | 0.90650187 |
| Whsc1 | 0.39435904 | 0.36612845 | 0.38656312 | -0.0204347 | 0.59289779 | 0.90650187 |
| Cnksr3 | 0.20643081 | 0.20287511 | 0.22331397 | -0.0204389 | 0.65795383 | 0.90650187 |
| Cd276 | 0.14272215 | 0.13957442 | 0.16002175 | -0.0204473 | 0.7150004 | 0.90650187 |
| Dtwd1 | 0.12660503 | 0.12282544 | 0.14327327 | -0.0204478 | 0.7371139 | 0.90650187 |
| Fbxl14 | 0.11927758 | 0.11509694 | 0.13555507 | -0.0204581 | 0.74886945 | 0.90650187 |
| Arglu1 | 0.72235541 | 0.5880392 | 0.60851248 | -0.0204733 | 0.55944867 | 0.90650187 |
| Zufsp | 0.11021297 | 0.1054332 | 0.125916 | -0.0204828 | 0.76521171 | 0.90650187 |
| Slc35a4 | 0.40632321 | 0.3754317 | 0.39592445 | -0.0204928 | 0.59099345 | 0.90650187 |
| Mepce | 0.09231086 | 0.08611433 | 0.10661876 | -0.0205044 | 0.80371295 | 0.90882608 |
| Haus2 | 0.15815485 | 0.15526102 | 0.17577254 | -0.0205115 | 0.69804062 | 0.90650187 |
| Wdr53 | 0.14653503 | 0.14339127 | 0.16393923 | -0.020548 | 0.71130711 | 0.90650187 |
| Lnp | 0.2267694 | 0.22203488 | 0.24259466 | -0.0205598 | 0.6468251 | 0.90650187 |
| Plppr1 | 0.15697225 | 0.15400114 | 0.17457535 | -0.0205742 | 0.69983549 | 0.90650187 |
| Slc25a20 | 0.20124179 | 0.19773915 | 0.21832591 | -0.0205868 | 0.66246238 | 0.90650187 |
| Rybp | 0.20906317 | 0.20523103 | 0.2258336 | -0.0206026 | 0.65747 | 0.90650187 |
| Hspa2 | 0.21941897 | 0.21506914 | 0.23567617 | -0.020607 | 0.65125634 | 0.90650187 |
| Gtf3c3 | 0.21248685 | 0.2084927 | 0.22909986 | -0.0206072 | 0.65537574 | 0.90650187 |
| R3hcc1l | 0.13651344 | 0.13299455 | 0.15360629 | -0.0206117 | 0.72459143 | 0.90650187 |
| Ric3 | 0.08233536 | 0.07509417 | 0.09570829 | -0.0206141 | 0.83084674 | 0.9098597 |
| Crtc3 | 0.37318159 | 0.34908598 | 0.36971401 | -0.020628 | 0.59794853 | 0.90650187 |
| Tmub1 | 0.20995824 | 0.2060549 | 0.22668869 | -0.0206338 | 0.65713284 | 0.90650187 |
| Gemin7 | 0.92378431 | 0.69565394 | 0.71629219 | -0.0206382 | 0.55096094 | 0.90650187 |
| Mfn1 | 0.16526575 | 0.1622948 | 0.18293779 | -0.020643 | 0.69204285 | 0.90650187 |
| Pgap2 | 0.36030464 | 0.33863994 | 0.35929126 | -0.0206513 | 0.60084088 | 0.90650187 |
| Erc1 | 0.23035565 | 0.2252954 | 0.24594984 | -0.0206544 | 0.64555018 | 0.90650187 |

| | | | | | | |
|---|---|---|---|---|---|---|
| Sat2 | 0.17807361 | 0.17503974 | 0.1956998 | -0.0206601 | 0.68058076 | 0.90650187 |
| Mgrn1 | 0.33396111 | 0.31687862 | 0.33753954 | -0.0206609 | 0.60723406 | 0.90650187 |
| Bcap29 | 0.15933937 | 0.15630298 | 0.17697008 | -0.0206671 | 0.69817943 | 0.90650187 |
| Nat14 | 0.16640429 | 0.16340329 | 0.18407973 | -0.0206764 | 0.69123185 | 0.90650187 |
| Sgms2 | 0.20144396 | 0.1978144 | 0.21852078 | -0.0207064 | 0.66321464 | 0.90650187 |
| Elf2 | 0.55342544 | 0.48187001 | 0.50257737 | -0.0207074 | 0.57266744 | 0.90650187 |
| Ralgps2 | 0.29940753 | 0.28738711 | 0.30809906 | -0.020712 | 0.61747497 | 0.90650187 |
| Exoc4 | 0.34095558 | 0.32265781 | 0.34337251 | -0.0207147 | 0.60572387 | 0.90650187 |
| Kpna1 | 0.32166418 | 0.30645826 | 0.32718377 | -0.0207255 | 0.61087781 | 0.90650187 |
| Mb21d2 | 0.09731703 | 0.09131371 | 0.11204179 | -0.0207281 | 0.79458983 | 0.90731056 |
| E4f1 | 0.21793001 | 0.21353489 | 0.2342678 | -0.0207329 | 0.65300281 | 0.90650187 |
| Cd99l2 | 0.23791918 | 0.23224923 | 0.25298377 | -0.0207345 | 0.64220606 | 0.90650187 |
| Gatad1 | 0.59994801 | 0.51261849 | 0.53335882 | -0.0207403 | 0.56862438 | 0.90650187 |
| Tex2 | 0.0846588 | 0.07751811 | 0.09826202 | -0.0207439 | 0.82600715 | 0.90947782 |
| Tmem41b | 0.36390107 | 0.34147092 | 0.36221595 | -0.020745 | 0.60048827 | 0.90650187 |
| 0610037L13F | 0.62687085 | 0.52983765 | 0.55058963 | -0.020752 | 0.56653407 | 0.90650187 |
| Fam13b | 0.13792332 | 0.13428693 | 0.1550671 | -0.0207802 | 0.72432723 | 0.90650187 |
| Mfhas1 | 0.14100257 | 0.13746721 | 0.15824945 | -0.0207822 | 0.72033022 | 0.90650187 |
| Mpnd | 0.56469684 | 0.48937292 | 0.51015595 | -0.020783 | 0.57185848 | 0.90650187 |
| Neurl4 | 0.11012588 | 0.10503315 | 0.1258229 | -0.0207897 | 0.76869279 | 0.90650187 |
| Ttc5 | 0.47890596 | 0.42962875 | 0.45042386 | -0.0207951 | 0.58130932 | 0.90650187 |
| Iffo1 | 0.18729818 | 0.1839825 | 0.20477814 | -0.0207956 | 0.67417118 | 0.90650187 |
| Cox6a1 | 2.92376278 | 0.82651849 | 0.84732494 | -0.0208065 | 0.54346375 | 0.90650187 |
| Eftud1 | 0.1189531 | 0.1144034 | 0.13521176 | -0.0208084 | 0.75306137 | 0.90650187 |
| Brf1 | 0.15502112 | 0.15178434 | 0.17259663 | -0.0208123 | 0.70404365 | 0.90650187 |
| Qsox2 | 0.35360558 | 0.33300213 | 0.35381536 | -0.0208132 | 0.60315165 | 0.90650187 |
| Ranbp9 | 0.14884219 | 0.14547452 | 0.16630149 | -0.020827 | 0.7111512 | 0.90650187 |
| Lrp6 | 0.39869352 | 0.3691366 | 0.38996747 | -0.0208309 | 0.59385418 | 0.90650187 |
| Tnrc6b | 0.70207804 | 0.57577569 | 0.59660889 | -0.0208332 | 0.56168304 | 0.90650187 |
| Dcaf5 | 0.12882532 | 0.12475472 | 0.14559891 | -0.0208442 | 0.73775541 | 0.90650187 |

| | | | | | | |
|---|---|---|---|---|---|---|
| Dmxl2 | 0.11104318 | 0.10595647 | 0.12680303 | -0.0208466 | 0.76756845 | 0.90650187 |
| Gas8 | 0.3559325 | 0.33486927 | 0.35572158 | -0.0208523 | 0.60279964 | 0.90650187 |
| Traf3ip2 | 0.11069003 | 0.10557142 | 0.12642582 | -0.0208544 | 0.76831805 | 0.90650187 |
| Rfx7 | 0.16469023 | 0.16150552 | 0.18235999 | -0.0208545 | 0.69441152 | 0.90650187 |
| Zbtb18 | 0.07406424 | 0.06569966 | 0.08655504 | -0.0208554 | 0.85795257 | 0.91737198 |
| Fam32a | 1.32434259 | 0.83601711 | 0.85688145 | -0.0208643 | 0.54310002 | 0.90650187 |
| Rpusd3 | 0.10226468 | 0.09651555 | 0.11738016 | -0.0208646 | 0.78529957 | 0.90650187 |
| Fem1c | 0.20357635 | 0.19970664 | 0.22057357 | -0.0208669 | 0.66296463 | 0.90650187 |
| Gtf2e1 | 0.16079774 | 0.15757524 | 0.17844231 | -0.0208671 | 0.69842284 | 0.90650187 |
| Recql | 0.11403153 | 0.10911408 | 0.12998888 | -0.0208748 | 0.76236045 | 0.90650187 |
| Slc25a39 | 0.60398398 | 0.51508051 | 0.53596832 | -0.0208878 | 0.56877435 | 0.90650187 |
| Ppp6r1 | 0.08690526 | 0.07983261 | 0.10072387 | -0.0208913 | 0.82175352 | 0.9092537 |
| Mtmr4 | 0.14998412 | 0.14657498 | 0.16746826 | -0.0208933 | 0.71042688 | 0.90650187 |
| Nek9 | 0.17727147 | 0.17399308 | 0.19490598 | -0.0209129 | 0.68331964 | 0.90650187 |
| Rnf217 | 0.12358608 | 0.11917391 | 0.14010138 | -0.0209275 | 0.74667301 | 0.90650187 |
| Dhx15 | 0.81707027 | 0.64055987 | 0.66149935 | -0.0209395 | 0.55595593 | 0.90650187 |
| Gpaa1 | 0.46582716 | 0.41994231 | 0.44088536 | -0.0209431 | 0.58362517 | 0.90650187 |
| Gstm7 | 0.11911882 | 0.11444206 | 0.13538712 | -0.0209451 | 0.75419725 | 0.90650187 |
| Otud7b | 0.23885945 | 0.23289202 | 0.25385423 | -0.0209622 | 0.64322753 | 0.90650187 |
| Scyl3 | 0.12360684 | 0.11915416 | 0.14012323 | -0.0209691 | 0.74706212 | 0.90650187 |
| Adrbk2 | 0.31871864 | 0.30371277 | 0.32468306 | -0.0209703 | 0.61299291 | 0.90650187 |
| Uhrf1bp1l | 0.16058672 | 0.15725458 | 0.17822943 | -0.0209749 | 0.69957523 | 0.90650187 |
| Zfp780b | 0.15315947 | 0.14972581 | 0.17070461 | -0.0209788 | 0.70758534 | 0.90650187 |
| Pi4kb | 0.20556245 | 0.20150206 | 0.22248122 | -0.0209792 | 0.66246658 | 0.90650187 |
| Fam73b | 0.07551099 | 0.06718182 | 0.08816316 | -0.0209813 | 0.85496752 | 0.91627438 |
| Aph1b | 0.08047669 | 0.07267677 | 0.0936599 | -0.0209831 | 0.8403684 | 0.91226452 |
| Flad1 | 0.1823619 | 0.17894538 | 0.19993166 | -0.0209863 | 0.67962235 | 0.90650187 |
| Ngdn | 0.60262282 | 0.51410015 | 0.53508928 | -0.0209891 | 0.56921698 | 0.90650187 |
| Stx17 | 0.21711784 | 0.21250269 | 0.23349864 | -0.0209959 | 0.65532752 | 0.90650187 |
| Dtnbos | 0.14078801 | 0.13703134 | 0.15802807 | -0.0209967 | 0.72263079 | 0.90650187 |

| | | | | | | |
|---|---|---|---|---|---|---|
| Rraga | 1.21022387 | 0.80448599 | 0.82548839 | -0.0210024 | 0.54502708 | 0.90650187 |
| Ipo7 | 0.35234577 | 0.33177023 | 0.35278147 | -0.0210112 | 0.60440862 | 0.90650187 |
| Zrsr2 | 0.73439705 | 0.59447206 | 0.61548466 | -0.0210126 | 0.5603169 | 0.90650187 |
| Nr2f6 | 0.38923432 | 0.36150076 | 0.38251797 | -0.0210172 | 0.59648568 | 0.90650187 |
| Trp53inp1 | 0.1135072 | 0.10840606 | 0.12943069 | -0.0210246 | 0.76489618 | 0.90650187 |
| Camta2 | 0.1029894 | 0.09713434 | 0.11816173 | -0.0210274 | 0.78556149 | 0.90650187 |
| Sh3d19 | 0.17047589 | 0.16711805 | 0.18815164 | -0.0210336 | 0.69039738 | 0.90650187 |
| Atg4c | 0.08331785 | 0.07575483 | 0.09678909 | -0.0210343 | 0.83300832 | 0.9104515 |
| Pygo2 | 0.22608125 | 0.22090902 | 0.24194936 | -0.0210403 | 0.65047295 | 0.90650187 |
| Rims2 | 0.10412621 | 0.0983417 | 0.11938639 | -0.0210447 | 0.78337549 | 0.90650187 |
| Slc7a1 | 0.1414707 | 0.13768575 | 0.15873228 | -0.0210465 | 0.72221991 | 0.90650187 |
| Tmem175 | 0.1706102 | 0.1672381 | 0.18828565 | -0.0210476 | 0.69038892 | 0.90650187 |
| Zfp213 | 0.13286372 | 0.12875638 | 0.14981379 | -0.0210574 | 0.73396921 | 0.90650187 |
| St8sia3 | 0.1189596 | 0.11415905 | 0.13521864 | -0.0210596 | 0.75565053 | 0.90650187 |
| Tpra1 | 0.13333412 | 0.12923728 | 0.15030348 | -0.0210662 | 0.73338612 | 0.90650187 |
| Dsg2 | 0.12380168 | 0.11925972 | 0.14032828 | -0.0210686 | 0.74775857 | 0.90650187 |
| Borcs5 | 0.20175337 | 0.19774304 | 0.21881894 | -0.0210759 | 0.66574206 | 0.90650187 |
| Vsig10 | 0.20936296 | 0.20501663 | 0.22612009 | -0.0211035 | 0.6608932 | 0.90650187 |
| Tmem14a | 0.08442655 | 0.07689162 | 0.09800708 | -0.0211155 | 0.83091784 | 0.9098597 |
| Zfp748 | 0.06696513 | 0.05750227 | 0.0786202 | -0.0211179 | 0.88378468 | 0.92772674 |
| Rrp36 | 0.17784714 | 0.17435527 | 0.19547575 | -0.0211205 | 0.6845045 | 0.90650187 |
| Gmpr | 0.25494028 | 0.24748355 | 0.26860458 | -0.021121 | 0.63666885 | 0.90650187 |
| Sms | 0.59166149 | 0.50684537 | 0.52797166 | -0.0211263 | 0.57059403 | 0.90650187 |
| 3010026O09 | 0.10213213 | 0.09610325 | 0.11723714 | -0.0211339 | 0.78854777 | 0.90650187 |
| Mpp2 | 0.10908001 | 0.10356772 | 0.1247041 | -0.0211364 | 0.77442408 | 0.90650187 |
| Adpgk | 0.20617595 | 0.20193317 | 0.22306964 | -0.0211365 | 0.66320774 | 0.90650187 |
| Syngr1 | 0.12155723 | 0.11682024 | 0.13796335 | -0.0211431 | 0.75215811 | 0.90650187 |
| Gnl2 | 0.4425099 | 0.40243027 | 0.42357789 | -0.0211476 | 0.58782438 | 0.90650187 |
| Edc3 | 0.17930566 | 0.17576509 | 0.1969177 | -0.0211526 | 0.6835113 | 0.90650187 |
| Fbxo36 | 0.21024947 | 0.20581367 | 0.22696674 | -0.0211531 | 0.66068226 | 0.90650187 |

| | | | | | | |
|---|---|---|---|---|---|---|
| Mapk9 | 0.21244045 | 0.20790158 | 0.22905568 | -0.0211541 | 0.65930514 | 0.90650187 |
| Vkorc1l1 | 0.20777769 | 0.20344654 | 0.22460408 | -0.0211575 | 0.66230894 | 0.90650187 |
| Ppfibp1 | 0.26843697 | 0.25961212 | 0.28078984 | -0.0211777 | 0.63129925 | 0.90650187 |
| Cpeb4 | 0.22978004 | 0.22422752 | 0.2454122 | -0.0211847 | 0.64942083 | 0.90650187 |
| Tbck | 0.2990755 | 0.28661731 | 0.30781103 | -0.0211937 | 0.62023325 | 0.90650187 |
| Ik | 0.86814941 | 0.66718554 | 0.68838437 | -0.0211988 | 0.55444614 | 0.90650187 |
| Rapgef1 | 0.13209099 | 0.12780762 | 0.1490088 | -0.0212012 | 0.73647871 | 0.90650187 |
| Hnrnpr | 1.45696878 | 0.86447457 | 0.88570053 | -0.021226 | 0.54242292 | 0.90650187 |
| Osbpl5 | 0.15292712 | 0.14923968 | 0.17046819 | -0.0212285 | 0.71007857 | 0.90650187 |
| Mob1b | 0.28879793 | 0.27761042 | 0.29884367 | -0.0212332 | 0.62394803 | 0.90650187 |
| Mcmbp | 0.37191366 | 0.34745749 | 0.368694 | -0.0212365 | 0.60104919 | 0.90650187 |
| Poglut1 | 0.27714822 | 0.26731497 | 0.28856172 | -0.0212468 | 0.62829057 | 0.90650187 |
| Nabp2 | 0.61740241 | 0.52332846 | 0.54457784 | -0.0212494 | 0.56885788 | 0.90650187 |
| Foxo1 | 0.26752522 | 0.25871203 | 0.27997214 | -0.0212601 | 0.63216156 | 0.90650187 |
| Epb41 | 0.20951039 | 0.20499815 | 0.22626095 | -0.0212628 | 0.66194399 | 0.90650187 |
| Cbr1 | 0.68709824 | 0.56641604 | 0.58768187 | -0.0212658 | 0.56390152 | 0.90650187 |
| Med25 | 0.94707472 | 0.7062084 | 0.72748252 | -0.0212741 | 0.55171876 | 0.90650187 |
| Cdpf1 | 0.25618157 | 0.24845379 | 0.26973254 | -0.0212788 | 0.63709572 | 0.90650187 |
| Isca1 | 0.31979254 | 0.30431421 | 0.3255957 | -0.0212815 | 0.61431083 | 0.90650187 |
| Pbx3 | 0.1649736 | 0.16135591 | 0.18264453 | -0.0212886 | 0.69782996 | 0.90650187 |
| Taf5l | 0.18408711 | 0.18033947 | 0.2016285 | -0.021289 | 0.68060497 | 0.90650187 |
| Srbd1 | 0.09034025 | 0.08318243 | 0.10447451 | -0.0212921 | 0.81753791 | 0.9092537 |
| Lasp1 | 0.79625026 | 0.62891266 | 0.6502072 | -0.0212945 | 0.5578796 | 0.90650187 |
| Edrf1 | 0.09401525 | 0.08716981 | 0.10846897 | -0.0212992 | 0.8086541 | 0.9092537 |
| Snx14 | 0.31920672 | 0.30379832 | 0.32509797 | -0.0212996 | 0.61457587 | 0.90650187 |
| BC029722 | 0.23779958 | 0.23156304 | 0.25287299 | -0.0213099 | 0.64603487 | 0.90650187 |
| Rnf103 | 0.12233732 | 0.11747055 | 0.13878601 | -0.0213155 | 0.75262665 | 0.90650187 |
| Mbd2 | 0.20151726 | 0.19727136 | 0.21859143 | -0.0213201 | 0.66770959 | 0.90650187 |
| 1500026H17 | 0.10418514 | 0.09812957 | 0.11944983 | -0.0213203 | 0.78625192 | 0.90650187 |
| Bud13 | 0.10141967 | 0.09513951 | 0.11646803 | -0.0213285 | 0.79221008 | 0.90681274 |

| | | | | | | |
|---|---|---|---|---|---|---|
| Trabd | 0.51612703 | 0.45559888 | 0.47693018 | -0.0213313 | 0.57880419 | 0.90650187 |
| Rfxank | 0.13326802 | 0.12889916 | 0.15023469 | -0.0213355 | 0.73609124 | 0.90650187 |
| Fam89b | 0.08581905 | 0.07817529 | 0.0995344 | -0.0213591 | 0.82995412 | 0.90974008 |
| Haus1 | 0.22985434 | 0.22411794 | 0.24548161 | -0.0213637 | 0.65058126 | 0.90650187 |
| Nsf | 0.35100517 | 0.33030867 | 0.35167984 | -0.0213712 | 0.60647919 | 0.90650187 |
| Eif2d | 0.25247659 | 0.2449864 | 0.26636145 | -0.021375 | 0.63936136 | 0.90650187 |
| Usp42 | 0.15032404 | 0.1464388 | 0.16781527 | -0.0213765 | 0.71440015 | 0.90650187 |
| Plekhh3 | 0.19360504 | 0.18955301 | 0.21093199 | -0.021379 | 0.67385517 | 0.90650187 |
| Idh3b | 0.87580028 | 0.67092781 | 0.69230928 | -0.0213815 | 0.55460289 | 0.90650187 |
| Foxk2 | 0.13195977 | 0.12749004 | 0.14887204 | -0.021382 | 0.73842835 | 0.90650187 |
| Mrpl45 | 0.28932049 | 0.27791852 | 0.29930196 | -0.0213834 | 0.62461094 | 0.90650187 |
| Ddx47 | 0.51205981 | 0.45268524 | 0.47407857 | -0.0213933 | 0.57949941 | 0.90650187 |
| Nipsnap1 | 0.3772264 | 0.3515631 | 0.37295939 | -0.0213963 | 0.60065378 | 0.90650187 |
| Bcr | 0.13164658 | 0.1271473 | 0.14854551 | -0.0213982 | 0.73904255 | 0.90650187 |
| Dhx38 | 0.13672907 | 0.13242991 | 0.15382987 | -0.0214 | 0.73186001 | 0.90650187 |
| Tm9sf4 | 0.38096834 | 0.354549 | 0.37595021 | -0.0214012 | 0.59989257 | 0.90650187 |
| Zmym3 | 0.17945084 | 0.17565542 | 0.1970611 | -0.0214057 | 0.68541737 | 0.90650187 |
| Cep83 | 0.29687489 | 0.28448441 | 0.30589952 | -0.0214151 | 0.62218467 | 0.90650187 |
| Nipa2 | 0.42279658 | 0.38722023 | 0.40863554 | -0.0214153 | 0.59211165 | 0.90650187 |
| Dcaf6 | 0.25830203 | 0.25022244 | 0.27165595 | -0.0214335 | 0.63711427 | 0.90650187 |
| Tacc1 | 0.24885447 | 0.24161663 | 0.26305291 | -0.0214363 | 0.64142635 | 0.90650187 |
| Tbc1d23 | 0.15662827 | 0.15278711 | 0.17422682 | -0.0214397 | 0.70783122 | 0.90650187 |
| Afg3l2 | 0.33610973 | 0.31789492 | 0.33933575 | -0.0214408 | 0.61060431 | 0.90650187 |
| Lig4 | 0.09955657 | 0.09301175 | 0.11445661 | -0.0214449 | 0.79756053 | 0.9075779 |
| Srrm3 | 0.08860813 | 0.08113961 | 0.10258529 | -0.0214457 | 0.82364971 | 0.9092537 |
| Polr2i | 1.03837656 | 0.74473813 | 0.76618695 | -0.0214488 | 0.54952104 | 0.90650187 |
| Utp6 | 0.19088294 | 0.18682954 | 0.20828117 | -0.0214516 | 0.67647681 | 0.90650187 |
| Tmed8 | 0.3176376 | 0.30230817 | 0.32376333 | -0.0214552 | 0.6158509 | 0.90650187 |
| BC004004 | 0.67830672 | 0.56092622 | 0.58238797 | -0.0214618 | 0.56506608 | 0.90650187 |
| Pink1 | 0.41561967 | 0.38165577 | 0.40312324 | -0.0214675 | 0.59357128 | 0.90650187 |

| | | | | | | |
|---|---|---|---|---|---|---|
| Tbccd1 | 0.12723787 | 0.1224536 | 0.14393674 | -0.0214831 | 0.74650275 | 0.90650187 |
| Lemd3 | 0.07126759 | 0.06195283 | 0.08343789 | -0.0214851 | 0.87384013 | 0.9235672 |
| Atg14 | 0.09108953 | 0.08380039 | 0.10529045 | -0.0214901 | 0.81789185 | 0.9092537 |
| Cad | 0.11933608 | 0.11412567 | 0.13561695 | -0.0214913 | 0.75943847 | 0.90650187 |
| Rnf40 | 0.23786085 | 0.231427 | 0.25292974 | -0.0215027 | 0.64726311 | 0.90650187 |
| Acvr1b | 0.0709984 | 0.06162111 | 0.08313725 | -0.0215161 | 0.87503814 | 0.92397668 |
| Fam53c | 0.1397723 | 0.13545696 | 0.15697933 | -0.0215224 | 0.7289057 | 0.90650187 |
| Pik3ip1 | 0.12515735 | 0.12021306 | 0.1417537 | -0.0215406 | 0.75033563 | 0.90650187 |
| Psme3 | 0.36492564 | 0.34148555 | 0.36304723 | -0.0215617 | 0.60412093 | 0.90650187 |
| Crat | 0.15606814 | 0.15208527 | 0.173659 | -0.0215737 | 0.70962151 | 0.90650187 |
| Me2 | 0.15486318 | 0.15085749 | 0.17243628 | -0.0215788 | 0.71100598 | 0.90650187 |
| Hspbp1 | 0.3543393 | 0.33283597 | 0.35441691 | -0.0215809 | 0.60668879 | 0.90650187 |
| Zfp64 | 0.21455083 | 0.20946739 | 0.23106308 | -0.0215957 | 0.66111081 | 0.90650187 |
| Ralgapa1 | 0.18729048 | 0.1831749 | 0.20477061 | -0.0215957 | 0.68041038 | 0.90650187 |
| Tmem198 | 0.0825691 | 0.0743549 | 0.09596555 | -0.0216106 | 0.84161406 | 0.91251845 |
| Zfp606 | 0.19101098 | 0.18679072 | 0.20840604 | -0.0216153 | 0.67763172 | 0.90650187 |
| Zdhhc13 | 0.13334616 | 0.12868978 | 0.15031601 | -0.0216262 | 0.73878276 | 0.90650187 |
| Naa35 | 0.58937607 | 0.50485059 | 0.52647826 | -0.0216277 | 0.57245388 | 0.90650187 |
| Rps6kc1 | 0.10504545 | 0.09874596 | 0.12037547 | -0.0216295 | 0.78779767 | 0.90650187 |
| Ctnnd1 | 0.35818233 | 0.33592989 | 0.35756042 | -0.0216305 | 0.60601048 | 0.90650187 |
| Cdv3 | 0.734403 | 0.59384065 | 0.61548809 | -0.0216474 | 0.56212428 | 0.90650187 |
| Atg4b | 0.404003 | 0.37247006 | 0.39411761 | -0.0216475 | 0.59645509 | 0.90650187 |
| Dlg4 | 0.16390639 | 0.15991879 | 0.18157245 | -0.0216537 | 0.7020018 | 0.90650187 |
| Mark3 | 0.18813364 | 0.18391169 | 0.20559579 | -0.0216841 | 0.68042097 | 0.90650187 |
| Diaph1 | 0.25283132 | 0.24499531 | 0.26668478 | -0.0216895 | 0.64116002 | 0.90650187 |
| Sall1 | 0.06878624 | 0.05896689 | 0.08066269 | -0.0216958 | 0.88409944 | 0.9278081 |
| Hnrnpa2b1 | 2.7422144 | 0.85119209 | 0.87288851 | -0.0216964 | 0.54399337 | 0.90650187 |
| Arhgap17 | 0.2187964 | 0.21338186 | 0.23508758 | -0.0217057 | 0.65926733 | 0.90650187 |
| Nacc1 | 0.19123002 | 0.1869116 | 0.2086196 | -0.021708 | 0.67817065 | 0.90650187 |
| Man2b2 | 0.18863353 | 0.18437356 | 0.20608467 | -0.0217111 | 0.68023159 | 0.90650187 |

| | | | | | | |
|---|---|---|---|---|---|---|
| Pitpnm1 | 0.0755904 | 0.06653915 | 0.08825134 | -0.0217122 | 0.86296503 | 0.91950555 |
| Snupn | 0.11564907 | 0.1099902 | 0.13170873 | -0.0217185 | 0.7682429 | 0.90650187 |
| Cep350 | 0.1900991 | 0.18579627 | 0.20751639 | -0.0217201 | 0.67914465 | 0.90650187 |
| Cep131 | 0.12418026 | 0.11900555 | 0.14072659 | -0.021721 | 0.75369898 | 0.90650187 |
| Atn1.1 | 0.23104351 | 0.22486326 | 0.2465919 | -0.0217286 | 0.65237045 | 0.90650187 |
| Tmbim4 | 0.92319842 | 0.69427793 | 0.71600679 | -0.0217289 | 0.55365931 | 0.90650187 |
| Gm26518 | 0.22903801 | 0.22298836 | 0.24471861 | -0.0217302 | 0.65348847 | 0.90650187 |
| Donson | 0.08732095 | 0.07944166 | 0.10117864 | -0.021737 | 0.83023228 | 0.90977227 |
| Zfp949 | 0.15115268 | 0.14692358 | 0.16866062 | -0.021737 | 0.71666114 | 0.90650187 |
| Pogk | 0.14213886 | 0.13768114 | 0.15942097 | -0.0217398 | 0.72782096 | 0.90650187 |
| Esrp2 | 0.10270381 | 0.09610731 | 0.11785382 | -0.0217465 | 0.79397969 | 0.90731056 |
| Atp6v0a2 | 0.24786781 | 0.24040019 | 0.26214945 | -0.0217493 | 0.64387286 | 0.90650187 |
| Alkbh3 | 0.20586252 | 0.20101281 | 0.22276907 | -0.0217563 | 0.66791771 | 0.90650187 |
| Edem3 | 0.11769459 | 0.11210973 | 0.13387903 | -0.0217693 | 0.76512148 | 0.90650187 |
| C77080 | 0.06691141 | 0.05678876 | 0.07855988 | -0.0217711 | 0.89102564 | 0.93109327 |
| Hnrnpdl | 1.37522689 | 0.84714448 | 0.86892596 | -0.0217815 | 0.54436572 | 0.90650187 |
| Nr5a2 | 0.07577251 | 0.06667191 | 0.08845353 | -0.0217816 | 0.86318195 | 0.9195619 |
| Rab11fip2 | 0.42854712 | 0.39124185 | 0.41302363 | -0.0217818 | 0.59268201 | 0.90650187 |
| Tmem53 | 0.23926423 | 0.23244235 | 0.25422868 | -0.0217863 | 0.64838715 | 0.90650187 |
| Sbk1 | 0.32237546 | 0.30598768 | 0.32778631 | -0.0217986 | 0.61624908 | 0.90650187 |
| Nploc4 | 0.10655252 | 0.10019303 | 0.12199477 | -0.0218017 | 0.78654091 | 0.90650187 |
| Lig3 | 0.16026974 | 0.15610096 | 0.17790957 | -0.0218086 | 0.70710733 | 0.90650187 |
| Zbtb17 | 0.08699624 | 0.07900052 | 0.10082342 | -0.0218229 | 0.83203472 | 0.91015207 |
| Bag4 | 0.30506509 | 0.29115862 | 0.31299182 | -0.0218332 | 0.62176125 | 0.90650187 |
| Mypop | 0.15837871 | 0.15416161 | 0.17599899 | -0.0218374 | 0.70938749 | 0.90650187 |
| Zfp280b | 0.10030253 | 0.09340717 | 0.11526076 | -0.0218536 | 0.80035577 | 0.90806412 |
| Dcaf15 | 0.08285671 | 0.07442438 | 0.09628198 | -0.0218576 | 0.84356687 | 0.91300169 |
| Anxa7 | 0.49030196 | 0.43675831 | 0.45863923 | -0.0218809 | 0.5839826 | 0.90650187 |
| Agk | 0.2328355 | 0.22638128 | 0.24826235 | -0.0218811 | 0.65240509 | 0.90650187 |
| Trnt1 | 0.45182907 | 0.40865521 | 0.43054156 | -0.0218863 | 0.58939538 | 0.90650187 |

| | | | | | | |
|---|---|---|---|---|---|---|
| Slc25a16 | 0.12241902 | 0.11698217 | 0.13887213 | -0.02189 | 0.75826938 | 0.90650187 |
| Plpp6 | 0.11793464 | 0.11222962 | 0.13413338 | -0.0219038 | 0.76606839 | 0.90650187 |
| Slc30a5 | 0.47490565 | 0.4256121 | 0.44751921 | -0.0219071 | 0.58613849 | 0.90650187 |
| Tspyl4 | 0.24169178 | 0.23455794 | 0.25647097 | -0.021913 | 0.64796613 | 0.90650187 |
| Ccdc127 | 0.619707 | 0.52412491 | 0.54604566 | -0.0219208 | 0.57082182 | 0.90650187 |
| Fam208a | 0.34296572 | 0.32312027 | 0.34504125 | -0.021921 | 0.61119538 | 0.90650187 |
| Atp7a | 0.14099782 | 0.13631307 | 0.15824455 | -0.0219315 | 0.73109465 | 0.90650187 |
| Smn1 | 0.38931022 | 0.36060232 | 0.38257803 | -0.0219757 | 0.60077768 | 0.90650187 |
| Zfp433 | 0.13782016 | 0.13297811 | 0.1549603 | -0.0219822 | 0.73586202 | 0.90650187 |
| Fnbp4 | 0.317426 | 0.30159877 | 0.32358318 | -0.0219844 | 0.61868801 | 0.90650187 |
| Noc2l | 0.22627691 | 0.22014557 | 0.24213289 | -0.0219873 | 0.65678016 | 0.90650187 |
| Mcm3ap | 0.13010126 | 0.12494102 | 0.14693272 | -0.0219917 | 0.74709705 | 0.90650187 |
| Tbc1d22b | 0.17494225 | 0.17060194 | 0.19259689 | -0.021995 | 0.69416716 | 0.90650187 |
| G2e3 | 0.20749206 | 0.20232774 | 0.22433064 | -0.0220029 | 0.66859473 | 0.90650187 |
| Arl16 | 0.30009119 | 0.28668749 | 0.30869182 | -0.0220043 | 0.62433797 | 0.90650187 |
| Ccdc61 | 0.12321351 | 0.11768849 | 0.13970913 | -0.0220206 | 0.75825692 | 0.90650187 |
| Trp53bp2 | 0.16395127 | 0.15958299 | 0.18161756 | -0.0220346 | 0.70518039 | 0.90650187 |
| Lsm7 | 0.99156737 | 0.72519577 | 0.74723538 | -0.0220396 | 0.55216082 | 0.90650187 |
| Ipp | 0.09081439 | 0.08294538 | 0.10499092 | -0.0220455 | 0.8247111 | 0.90934433 |
| Coq5 | 0.37255652 | 0.34716206 | 0.36921132 | -0.0220493 | 0.60468445 | 0.90650187 |
| Btrc | 0.43409672 | 0.39516839 | 0.41723541 | -0.022067 | 0.59294305 | 0.90650187 |
| Khdrbs1 | 1.09983551 | 0.76711821 | 0.78918891 | -0.0220707 | 0.5494719 | 0.90650187 |
| Wdr37 | 0.15481995 | 0.15030566 | 0.17239238 | -0.0220867 | 0.71551398 | 0.90650187 |
| Zfp532 | 0.13804352 | 0.13309824 | 0.15519154 | -0.0220933 | 0.73659504 | 0.90650187 |
| Tmem260 | 0.10551937 | 0.09878983 | 0.12088499 | -0.0220952 | 0.79176496 | 0.90681274 |
| Eif3k | 1.99038933 | 0.90508244 | 0.92717917 | -0.0220967 | 0.54218846 | 0.90650187 |
| Dpp3 | 0.51464095 | 0.45379223 | 0.47588949 | -0.0220973 | 0.58177072 | 0.90650187 |
| Tpgs2 | 0.23584554 | 0.2289556 | 0.251061 | -0.0221054 | 0.65225893 | 0.90650187 |
| Snx11 | 0.09486362 | 0.08728096 | 0.10938842 | -0.0221075 | 0.81552974 | 0.9092537 |
| Glb1l2 | 0.11041337 | 0.10400516 | 0.12613019 | -0.022125 | 0.78225232 | 0.90650187 |

| | | | | | | |
|---|---|---|---|---|---|---|
| Tcta | 0.36105881 | 0.33776665 | 0.35990543 | -0.0221388 | 0.60775078 | 0.90650187 |
| Ubxn2b | 0.20695918 | 0.20167263 | 0.2238203 | -0.0221477 | 0.66999728 | 0.90650187 |
| Gatc | 0.20698372 | 0.20169051 | 0.2238438 | -0.0221533 | 0.67002102 | 0.90650187 |
| Dnajc19 | 0.80485612 | 0.63274447 | 0.65489801 | -0.0221535 | 0.55976901 | 0.90650187 |
| Klc4 | 0.16488457 | 0.16039198 | 0.18255514 | -0.0221632 | 0.70530496 | 0.90650187 |
| Dhrs7b | 0.26945034 | 0.2595243 | 0.28169778 | -0.0221735 | 0.63680445 | 0.90650187 |
| Gphn | 0.22628127 | 0.21995428 | 0.24213698 | -0.0221827 | 0.65809576 | 0.90650187 |
| Slc35e2 | 0.09300256 | 0.08516965 | 0.10737013 | -0.0222005 | 0.82100439 | 0.9092537 |
| Dlg5 | 0.14791502 | 0.14314697 | 0.16535292 | -0.0222059 | 0.72475276 | 0.90650187 |
| Dvl3 | 0.13862989 | 0.1335822 | 0.15579831 | -0.0222161 | 0.73693442 | 0.90650187 |
| 1810030O07 | 0.20957955 | 0.20410666 | 0.22632701 | -0.0222204 | 0.66874889 | 0.90650187 |
| Nup153 | 0.17153183 | 0.16697401 | 0.18920465 | -0.0222306 | 0.6992823 | 0.90650187 |
| Ttc33 | 0.41850419 | 0.38309568 | 0.40534374 | -0.0222481 | 0.59638674 | 0.90650187 |
| Mov10 | 0.28964159 | 0.27733412 | 0.29958345 | -0.0222493 | 0.6293621 | 0.90650187 |
| Med23 | 0.12433079 | 0.11863062 | 0.14088491 | -0.0222543 | 0.75873258 | 0.90650187 |
| Stox2 | 0.15641987 | 0.15176091 | 0.17401559 | -0.0222547 | 0.71516728 | 0.90650187 |
| Pigyl | 0.75788694 | 0.60662698 | 0.62888304 | -0.0222561 | 0.5625073 | 0.90650187 |
| Adck5 | 0.23341158 | 0.22653532 | 0.24879867 | -0.0222633 | 0.65461409 | 0.90650187 |
| Sap30bp | 0.25802802 | 0.24913771 | 0.27140765 | -0.0222699 | 0.64236119 | 0.90650187 |
| Zfp60 | 0.17839652 | 0.17374628 | 0.19601917 | -0.0222729 | 0.69326865 | 0.90650187 |
| B230217C12 | 0.13404922 | 0.12877092 | 0.1510474 | -0.0222765 | 0.74397158 | 0.90650187 |
| Gpkow | 0.11073293 | 0.10419071 | 0.12647164 | -0.0222809 | 0.78324439 | 0.90650187 |
| C330007P06I | 0.22504137 | 0.2186912 | 0.24097333 | -0.0222821 | 0.65949042 | 0.90650187 |
| Rbm15b | 0.16728331 | 0.16265876 | 0.18496039 | -0.0223016 | 0.70403103 | 0.90650187 |
| Sik3 | 0.15115157 | 0.14635517 | 0.16865949 | -0.0223043 | 0.72170294 | 0.90650187 |
| Fzr1 | 0.45902429 | 0.41357039 | 0.43587554 | -0.0223051 | 0.58998091 | 0.90650187 |
| Bcorl1 | 0.11280388 | 0.10636578 | 0.12868143 | -0.0223156 | 0.77963084 | 0.90650187 |
| Lemd2 | 0.08948461 | 0.08121591 | 0.10354179 | -0.0223259 | 0.83111237 | 0.9098597 |
| Ppat | 0.1952379 | 0.19018519 | 0.21251828 | -0.0223331 | 0.67981416 | 0.90650187 |
| Cramp1l | 0.30997009 | 0.29487102 | 0.31721071 | -0.0223397 | 0.62289142 | 0.90650187 |

| | | | | | | |
|---|---|---|---|---|---|---|
| Ppid | 0.93855214 | 0.70109721 | 0.72344268 | -0.0223455 | 0.55460888 | 0.90650187 |
| Larp1b | 0.10508118 | 0.09805844 | 0.1204139 | -0.0223555 | 0.79541346 | 0.9075779 |
| Tmtc2 | 0.14972044 | 0.14483619 | 0.16719899 | -0.0223628 | 0.72394395 | 0.90650187 |
| Scrib | 0.13587079 | 0.13057307 | 0.15293965 | -0.0223666 | 0.74220083 | 0.90650187 |
| Klhl36 | 0.09379277 | 0.08584828 | 0.10822768 | -0.0223794 | 0.82102179 | 0.9092537 |
| Foxo4 | 0.1257313 | 0.11996834 | 0.14235645 | -0.0223881 | 0.75777136 | 0.90650187 |
| Ldoc1l | 0.11658352 | 0.11029973 | 0.13270082 | -0.0224011 | 0.77351057 | 0.90650187 |
| Ubr7 | 0.20970343 | 0.20403075 | 0.22644535 | -0.0224146 | 0.6700483 | 0.90650187 |
| Scaf1 | 0.36939839 | 0.34423747 | 0.36666629 | -0.0224288 | 0.60716433 | 0.90650187 |
| Cask | 0.21021648 | 0.20450047 | 0.22693525 | -0.0224348 | 0.6698474 | 0.90650187 |
| Hmces | 0.28831094 | 0.27597887 | 0.29841635 | -0.0224375 | 0.63090779 | 0.90650187 |
| Sp1 | 0.16797336 | 0.16321187 | 0.18565111 | -0.0224392 | 0.70448033 | 0.90650187 |
| Dhx40 | 0.31655015 | 0.30039739 | 0.32283713 | -0.0224397 | 0.62134112 | 0.90650187 |
| Ermard | 0.1058299 | 0.09877596 | 0.1212187 | -0.0224427 | 0.79476613 | 0.90731056 |
| Rint1 | 0.42714895 | 0.38950396 | 0.41195895 | -0.022455 | 0.59573403 | 0.90650187 |
| Mea1 | 0.70574541 | 0.5763163 | 0.59877707 | -0.0224608 | 0.56622033 | 0.90650187 |
| Sgsm2 | 0.15738209 | 0.15252205 | 0.17499042 | -0.0224684 | 0.71593433 | 0.90650187 |
| 1700113A16 | 0.06670179 | 0.05584486 | 0.07832445 | -0.0224796 | 0.89900743 | 0.93440169 |
| Kif3b | 0.18173305 | 0.17682884 | 0.19931234 | -0.0224835 | 0.69198627 | 0.90650187 |
| Adi1 | 0.30565087 | 0.29101004 | 0.31349678 | -0.0224867 | 0.62509045 | 0.90650187 |
| Adcy6 | 0.14076245 | 0.13550026 | 0.15800169 | -0.0225014 | 0.73666923 | 0.90650187 |
| Grwd1 | 0.1837546 | 0.17879193 | 0.2013017 | -0.0225098 | 0.6904451 | 0.90650187 |
| Mus81 | 0.15449256 | 0.14954956 | 0.17205986 | -0.0225103 | 0.71958569 | 0.90650187 |
| Smarcd1 | 0.28616733 | 0.2740188 | 0.29653282 | -0.022514 | 0.63214097 | 0.90650187 |
| Wdr73 | 0.17599639 | 0.17111783 | 0.19364266 | -0.0225248 | 0.6974639 | 0.90650187 |
| St7l | 0.36085853 | 0.33721744 | 0.35974237 | -0.0225249 | 0.60963104 | 0.90650187 |
| Coil | 0.12539079 | 0.1194556 | 0.1419989 | -0.0225433 | 0.75983462 | 0.90650187 |
| Hivep1 | 0.12479625 | 0.11883024 | 0.14137427 | -0.022544 | 0.76081039 | 0.90650187 |
| D11Wsu47e | 0.10408536 | 0.09679145 | 0.11934241 | -0.022551 | 0.79955774 | 0.90799325 |
| Rilpl2 | 0.33764642 | 0.31805361 | 0.34061801 | -0.0225644 | 0.61581258 | 0.90650187 |

| | | | | | | |
|---|---|---|---|---|---|---|
| Fam53a | 0.28146873 | 0.26981794 | 0.29238956 | -0.0225716 | 0.63427142 | 0.90650187 |
| Zfp410 | 0.30823053 | 0.29314498 | 0.31571692 | -0.0225719 | 0.62469501 | 0.90650187 |
| Dlg1 | 0.22640126 | 0.21967697 | 0.2422495 | -0.0225725 | 0.66064957 | 0.90650187 |
| Gtf3c6 | 0.52660007 | 0.46164614 | 0.48422145 | -0.0225753 | 0.58209754 | 0.90650187 |
| Msl1 | 0.73438658 | 0.5928947 | 0.61547863 | -0.0225839 | 0.56478956 | 0.90650187 |
| Fbxo34 | 0.1190483 | 0.1127234 | 0.1353125 | -0.0225891 | 0.77100194 | 0.90650187 |
| Zfp113 | 0.13620053 | 0.13065546 | 0.15328176 | -0.0226263 | 0.74416094 | 0.90650187 |
| Zfp938 | 0.27661625 | 0.26546153 | 0.28808922 | -0.0226277 | 0.63652135 | 0.90650187 |
| Tubgcp5 | 0.17910259 | 0.17408659 | 0.19671708 | -0.0226305 | 0.69547362 | 0.90650187 |
| Sort1 | 0.14431333 | 0.13901891 | 0.16165865 | -0.0226397 | 0.73322367 | 0.90650187 |
| Amigo1 | 0.1161581 | 0.10960874 | 0.13224929 | -0.0226406 | 0.77669245 | 0.90650187 |
| 1700037C18I | 0.08426559 | 0.07518301 | 0.09783037 | -0.0226474 | 0.84829323 | 0.91440801 |
| Arxes1 | 0.28548711 | 0.27327595 | 0.29593425 | -0.0226583 | 0.63321455 | 0.90650187 |
| Mllt4 | 0.25776719 | 0.24849422 | 0.27117122 | -0.022677 | 0.64496753 | 0.90650187 |
| Mical2 | 0.18775217 | 0.18253988 | 0.20522255 | -0.0226827 | 0.68841421 | 0.90650187 |
| Manba | 0.1125498 | 0.10572026 | 0.12841061 | -0.0226903 | 0.78393307 | 0.90650187 |
| Usp37 | 0.22089042 | 0.21437535 | 0.23706575 | -0.0226904 | 0.66476561 | 0.90650187 |
| Cnst | 0.15161279 | 0.14643414 | 0.16912967 | -0.0226955 | 0.72459825 | 0.90650187 |
| Ssrp1 | 1.11773616 | 0.77278394 | 0.795481 | -0.0226971 | 0.55046815 | 0.90650187 |
| Ict1 | 0.79354202 | 0.6260207 | 0.64872432 | -0.0227036 | 0.56181966 | 0.90650187 |
| Dyrk1b | 0.08036498 | 0.07082136 | 0.09353663 | -0.0227153 | 0.85984221 | 0.91802142 |
| Lmnb2 | 0.22435583 | 0.21761253 | 0.24032928 | -0.0227167 | 0.66283542 | 0.90650187 |
| Cul9 | 0.11720552 | 0.11063202 | 0.13336059 | -0.0227286 | 0.7756734 | 0.90650187 |
| Rora | 0.16826195 | 0.16319353 | 0.18593982 | -0.0227463 | 0.7067234 | 0.90650187 |
| Med13 | 0.31236817 | 0.29651097 | 0.31926563 | -0.0227547 | 0.62432007 | 0.90650187 |
| Zmym2 | 0.26073603 | 0.25108997 | 0.27385846 | -0.0227685 | 0.64416202 | 0.90650187 |
| 1700007K13I | 0.13153395 | 0.12565407 | 0.14842806 | -0.022774 | 0.75241511 | 0.90650187 |
| Gm26917 | 1.36422686 | 0.84365377 | 0.86643252 | -0.0227788 | 0.54652093 | 0.90650187 |
| Sec63 | 0.38774434 | 0.35855282 | 0.38133804 | -0.0227852 | 0.60473721 | 0.90650187 |
| Tfe3 | 0.21990313 | 0.21334487 | 0.23613364 | -0.0227888 | 0.66605156 | 0.90650187 |

| | | | | | | |
|---|---|---|---|---|---|---|
| Dcaf11 | 0.46933755 | 0.42066886 | 0.44345795 | -0.0227891 | 0.5903547 | 0.90650187 |
| Gucd1 | 0.19161158 | 0.186195 | 0.20899151 | -0.0227965 | 0.68613375 | 0.90650187 |
| Zdhhc12 | 0.34084591 | 0.3204662 | 0.34328137 | -0.0228152 | 0.61618026 | 0.90650187 |
| Kbtbd7 | 0.08323901 | 0.07386824 | 0.09670241 | -0.0228342 | 0.85307969 | 0.91566617 |
| Nrf1 | 0.26117012 | 0.2513983 | 0.27425067 | -0.0228524 | 0.64447164 | 0.90650187 |
| Vezf1 | 0.55981566 | 0.48402898 | 0.5068834 | -0.0228544 | 0.57943333 | 0.90650187 |
| 2310057M21 | 0.21850558 | 0.21194749 | 0.23481248 | -0.022865 | 0.66745389 | 0.90650187 |
| Mios | 0.127423 | 0.12125824 | 0.14413074 | -0.0228725 | 0.75974591 | 0.90650187 |
| Ankrd27 | 0.14992603 | 0.14453389 | 0.16740895 | -0.0228751 | 0.7282257 | 0.90650187 |
| Tor1a | 0.48574735 | 0.43248602 | 0.4553663 | -0.0228803 | 0.58837755 | 0.90650187 |
| Ercc5 | 0.12524177 | 0.1189488 | 0.14184238 | -0.0228936 | 0.76347578 | 0.90650187 |
| Kctd15 | 0.11052408 | 0.10334911 | 0.12624851 | -0.0228994 | 0.78998224 | 0.90650187 |
| Harbi1 | 0.16629684 | 0.16105762 | 0.18397202 | -0.0229144 | 0.71011366 | 0.90650187 |
| Nme3 | 0.1112191 | 0.10407532 | 0.12699088 | -0.0229156 | 0.78878469 | 0.90650187 |
| Mlst8 | 0.0757189 | 0.06547638 | 0.08839401 | -0.0229176 | 0.87546233 | 0.92418014 |
| Snrpf | 2.00143238 | 0.90417401 | 0.9270958 | -0.0229218 | 0.5437614 | 0.90650187 |
| Bbs2 | 0.14499491 | 0.13942691 | 0.16235891 | -0.022932 | 0.73496823 | 0.90650187 |
| Smndc1 | 0.41437095 | 0.37921713 | 0.40216005 | -0.0229429 | 0.60010482 | 0.90650187 |
| Mon1a | 0.08642634 | 0.07725449 | 0.10019963 | -0.0229451 | 0.84566537 | 0.9136218 |
| Max | 0.52557603 | 0.46055644 | 0.48351167 | -0.0229552 | 0.58357972 | 0.90650187 |
| Letm1 | 0.38345164 | 0.35494561 | 0.37792895 | -0.0229833 | 0.60655631 | 0.90650187 |
| Kazn | 0.14717089 | 0.14160704 | 0.16459089 | -0.0229839 | 0.73263265 | 0.90650187 |
| Phf1 | 0.1920019 | 0.18636653 | 0.20937179 | -0.0230053 | 0.6873923 | 0.90650187 |
| Rlim | 0.44761893 | 0.40439765 | 0.42740353 | -0.0230059 | 0.59456119 | 0.90650187 |
| Lrrc47 | 0.14887668 | 0.1433273 | 0.16633676 | -0.0230095 | 0.7307105 | 0.90650187 |
| Ube4a | 0.26013564 | 0.25030554 | 0.2733157 | -0.0230102 | 0.64589861 | 0.90650187 |
| Acox1 | 0.22760354 | 0.22036178 | 0.24337621 | -0.0230144 | 0.66289996 | 0.90650187 |
| Brcc3 | 0.34452664 | 0.32331994 | 0.34633473 | -0.0230148 | 0.61616404 | 0.90650187 |
| Rpp30 | 0.26967784 | 0.25887496 | 0.28190147 | -0.0230265 | 0.64174601 | 0.90650187 |
| Dstyk | 0.13942865 | 0.13358968 | 0.15662423 | -0.0230345 | 0.74338296 | 0.90650187 |

| | | | | | | |
|---|---|---|---|---|---|---|
| Tfcp2 | 0.13591266 | 0.12993657 | 0.15298309 | -0.0230465 | 0.74848169 | 0.90650187 |
| Ganab | 0.45271989 | 0.40815009 | 0.43120392 | -0.0230538 | 0.59393509 | 0.90650187 |
| Irf2 | 0.51623954 | 0.45395272 | 0.47700891 | -0.0230562 | 0.5850688 | 0.90650187 |
| Atp13a3 | 0.26506889 | 0.25470496 | 0.27776525 | -0.0230603 | 0.64396515 | 0.90650187 |
| Dync1h1 | 0.65277485 | 0.54372094 | 0.56678341 | -0.0230625 | 0.57177387 | 0.90650187 |
| Zfp317 | 0.10563521 | 0.09793733 | 0.1210095 | -0.0230722 | 0.80167451 | 0.90839694 |
| Cln5 | 0.19319311 | 0.1874531 | 0.21053135 | -0.0230783 | 0.68698541 | 0.90650187 |
| Sppl3 | 0.26334077 | 0.2531301 | 0.2762092 | -0.0230791 | 0.64485246 | 0.90650187 |
| Zfp707 | 0.13766783 | 0.13171924 | 0.15480256 | -0.0230833 | 0.74630427 | 0.90650187 |
| Tm9sf2 | 0.84067475 | 0.65098225 | 0.67406785 | -0.0230856 | 0.5605067 | 0.90650187 |
| Pak4 | 0.35220734 | 0.32957492 | 0.35266778 | -0.0230929 | 0.6145128 | 0.90650187 |
| Zfp609 | 0.13104702 | 0.12482254 | 0.14792012 | -0.0230976 | 0.75621434 | 0.90650187 |
| Naa30 | 0.16620168 | 0.16076782 | 0.18387662 | -0.0231088 | 0.71181824 | 0.90650187 |
| 1700096K18I | 0.16608737 | 0.16065121 | 0.18376201 | -0.0231108 | 0.71195358 | 0.90650187 |
| Tut1 | 0.1001835 | 0.09201594 | 0.11513205 | -0.0231161 | 0.81396017 | 0.9092537 |
| Pex6 | 0.19958503 | 0.1936081 | 0.21672734 | -0.0231192 | 0.68233278 | 0.90650187 |
| Mief2 | 0.10098956 | 0.09285366 | 0.11600341 | -0.0231497 | 0.81250613 | 0.9092537 |
| Mtx2 | 0.74845492 | 0.60037975 | 0.62353483 | -0.0231551 | 0.56556279 | 0.90650187 |
| Emc2 | 0.59901256 | 0.50959313 | 0.53275267 | -0.0231595 | 0.57662119 | 0.90650187 |
| Psme4 | 0.31583154 | 0.29906375 | 0.32222451 | -0.0231608 | 0.62534148 | 0.90650187 |
| Macf1 | 0.52364302 | 0.45898711 | 0.48217007 | -0.023183 | 0.58462775 | 0.90650187 |
| Pnisr | 1.47061115 | 0.86503432 | 0.88823609 | -0.0232018 | 0.54622305 | 0.90650187 |
| Pnck | 0.12669959 | 0.12015242 | 0.14337243 | -0.02322 | 0.76423513 | 0.90650187 |
| Larp4b | 0.57042511 | 0.49072146 | 0.51397739 | -0.0232559 | 0.57970947 | 0.90650187 |
| Polr3c | 0.2217871 | 0.21465444 | 0.23791146 | -0.023257 | 0.66806631 | 0.90650187 |
| Kptn | 0.18457581 | 0.1788433 | 0.20210857 | -0.0232653 | 0.69558559 | 0.90650187 |
| Ccdc53 | 0.58518342 | 0.50046039 | 0.52373072 | -0.0232703 | 0.57829225 | 0.90650187 |
| Zzef1 | 0.22057899 | 0.2134948 | 0.23677183 | -0.023277 | 0.66896155 | 0.90650187 |
| Trim56 | 0.11987629 | 0.11289012 | 0.13618817 | -0.0232981 | 0.77652842 | 0.90650187 |
| Arl8a | 0.21010542 | 0.20352211 | 0.22682922 | -0.0233071 | 0.67609045 | 0.90650187 |

| | | | | | | |
|---|---|---|---|---|---|---|
| Thap2 | 0.23749832 | 0.22927773 | 0.25259388 | -0.0233161 | 0.65922905 | 0.90650187 |
| Alas1 | 0.31760166 | 0.30041635 | 0.32373274 | -0.0233164 | 0.62558724 | 0.90650187 |
| Uba3 | 0.51137017 | 0.45026265 | 0.47359399 | -0.0233313 | 0.58667898 | 0.90650187 |
| Cog7 | 0.40318236 | 0.37014331 | 0.39347756 | -0.0233343 | 0.60396983 | 0.90650187 |
| Vezt | 0.19622022 | 0.19013327 | 0.21347122 | -0.0233379 | 0.68652359 | 0.90650187 |
| Scaf4 | 0.16688136 | 0.1612078 | 0.18455779 | -0.02335 | 0.71309587 | 0.90650187 |
| D5Ertd579e | 0.31497575 | 0.2981424 | 0.32149435 | -0.0233519 | 0.62661809 | 0.90650187 |
| Gtpbp1 | 0.09287284 | 0.08387252 | 0.10722926 | -0.0233567 | 0.83356391 | 0.91060043 |
| Pbrm1 | 1.09631534 | 0.76454795 | 0.78793757 | -0.0233896 | 0.55249445 | 0.90650187 |
| Wiz | 0.17049049 | 0.16476741 | 0.1881662 | -0.0233988 | 0.70980631 | 0.90650187 |
| Got1 | 0.1461901 | 0.14017412 | 0.16358555 | -0.0234114 | 0.73768664 | 0.90650187 |
| Bet1l | 0.58713501 | 0.5015935 | 0.52501091 | -0.0234174 | 0.57859103 | 0.90650187 |
| Pgls | 1.57546711 | 0.8816412 | 0.90509002 | -0.0234488 | 0.54584701 | 0.90650187 |
| Msh6 | 0.13492457 | 0.12850437 | 0.15195721 | -0.0234528 | 0.75368375 | 0.90650187 |
| D730003I15F | 0.08674075 | 0.07709029 | 0.10054382 | -0.0234535 | 0.8501349 | 0.91476609 |
| Prkce | 0.17598964 | 0.17018018 | 0.19363597 | -0.0234558 | 0.704888 | 0.90650187 |
| Lypla2 | 0.35942738 | 0.33511131 | 0.35857626 | -0.0234649 | 0.61444259 | 0.90650187 |
| Slc25a23 | 0.21132735 | 0.20453008 | 0.22799504 | -0.023465 | 0.67635713 | 0.90650187 |
| Tnk1 | 0.11629328 | 0.10892381 | 0.13239279 | -0.023469 | 0.78467002 | 0.90650187 |
| Naa15 | 0.71134104 | 0.57860151 | 0.60207246 | -0.023471 | 0.56879444 | 0.90650187 |
| Snrnp35 | 0.18054584 | 0.17466986 | 0.19814195 | -0.0234721 | 0.70077506 | 0.90650187 |
| Lrrcc1 | 0.26960442 | 0.25835727 | 0.28183574 | -0.0234785 | 0.64443672 | 0.90650187 |
| Psmd6 | 1.16665975 | 0.78842872 | 0.81191151 | -0.0234828 | 0.55115485 | 0.90650187 |
| Acp6 | 0.20371868 | 0.19722714 | 0.22071041 | -0.0234833 | 0.68189563 | 0.90650187 |
| 2900093K20I | 0.1761289 | 0.17028989 | 0.19377404 | -0.0234842 | 0.70498016 | 0.90650187 |
| Mau2 | 0.42019052 | 0.38315068 | 0.4066389 | -0.0234882 | 0.60132821 | 0.90650187 |
| Hdhd3 | 0.15346496 | 0.14752582 | 0.17101536 | -0.0234895 | 0.72927412 | 0.90650187 |
| Zfp961 | 0.10557001 | 0.09744204 | 0.12093942 | -0.0234974 | 0.8061344 | 0.9092537 |
| Rnf215 | 0.25496361 | 0.24509547 | 0.2686258 | -0.0235303 | 0.65151494 | 0.90650187 |
| Wbp11 | 1.05450396 | 0.74888988 | 0.77242957 | -0.0235397 | 0.55388352 | 0.90650187 |

| | | | | | | |
|---|---|---|---|---|---|---|
| Cdk8 | 0.22116496 | 0.21377586 | 0.23732477 | -0.0235489 | 0.6704377 | 0.90650187 |
| Cgrrf1 | 0.21968878 | 0.21237609 | 0.23593114 | -0.0235551 | 0.6714221 | 0.90650187 |
| Sgf29 | 0.25301489 | 0.24328309 | 0.26685206 | -0.023569 | 0.65270947 | 0.90650187 |
| Mettl9 | 1.09137488 | 0.76260367 | 0.78617352 | -0.0235698 | 0.55301455 | 0.90650187 |
| Zfp414 | 0.36077521 | 0.33610037 | 0.35967453 | -0.0235742 | 0.6146191 | 0.90650187 |
| Slc2a8 | 0.10693749 | 0.09882306 | 0.12240795 | -0.0235849 | 0.80415461 | 0.90882608 |
| Tfb2m | 0.15354205 | 0.14750619 | 0.17109376 | -0.0235876 | 0.73002429 | 0.90650187 |
| Fam76b | 0.25421866 | 0.2443563 | 0.26794818 | -0.0235919 | 0.65225646 | 0.90650187 |
| Traf3ip1 | 0.12714178 | 0.12024062 | 0.14383603 | -0.0235954 | 0.76708298 | 0.90650187 |
| Dck | 0.0943528 | 0.08523922 | 0.10883492 | -0.0235957 | 0.8324318 | 0.91029548 |
| Gm26853 | 0.16525022 | 0.15932548 | 0.18292219 | -0.0235967 | 0.71683917 | 0.90650187 |
| Med17 | 0.17172964 | 0.16579585 | 0.18940177 | -0.0236059 | 0.71023593 | 0.90650187 |
| Chchd6 | 0.45651669 | 0.41040793 | 0.43402073 | -0.0236128 | 0.59555651 | 0.90650187 |
| Ints8 | 0.21059387 | 0.20367744 | 0.22729543 | -0.023618 | 0.67793692 | 0.90650187 |
| Cdkn2aip | 0.24931724 | 0.23985171 | 0.26347632 | -0.0236246 | 0.65491174 | 0.90650187 |
| Zfp652 | 0.25145264 | 0.24179636 | 0.26542743 | -0.0236311 | 0.65387129 | 0.90650187 |
| Taf7 | 0.51771593 | 0.45440951 | 0.4780413 | -0.0236318 | 0.58697361 | 0.90650187 |
| Cyhr1 | 0.53627694 | 0.46725931 | 0.49089592 | -0.0236366 | 0.5847482 | 0.90650187 |
| Senp7 | 0.33055278 | 0.3110369 | 0.33468219 | -0.0236453 | 0.62327064 | 0.90650187 |
| Parp6 | 0.15184365 | 0.14571756 | 0.16936493 | -0.0236474 | 0.73260556 | 0.90650187 |
| Ccdc50 | 0.41408949 | 0.37828074 | 0.40194279 | -0.0236621 | 0.60322721 | 0.90650187 |
| Nubpl | 0.1132459 | 0.10548346 | 0.1291524 | -0.0236689 | 0.79238313 | 0.90681274 |
| Samd10 | 0.24298139 | 0.2339907 | 0.25765979 | -0.0236691 | 0.65850315 | 0.90650187 |
| E130308A19 | 0.09701172 | 0.08804153 | 0.11171205 | -0.0236705 | 0.82689698 | 0.90947782 |
| Bcl7a | 0.31118665 | 0.29457754 | 0.31825382 | -0.0236763 | 0.629575 | 0.90650187 |
| Akt3 | 0.11482049 | 0.10715049 | 0.13082814 | -0.0236776 | 0.7894685 | 0.90650187 |
| Fsd1l | 0.09321402 | 0.08391702 | 0.10759969 | -0.0236827 | 0.83608086 | 0.91079271 |
| 1810024B03 | 0.12530432 | 0.11822334 | 0.14190808 | -0.0236847 | 0.77095046 | 0.90650187 |
| Hivep3 | 0.11220186 | 0.10435364 | 0.1280396 | -0.023686 | 0.7945724 | 0.90731056 |
| Dnajc2 | 0.52983734 | 0.4627628 | 0.48646086 | -0.0236981 | 0.5857278 | 0.90650187 |

| | | | | | | |
|---|---|---|---|---|---|---|
| Ndufaf5 | 0.1335104 | 0.12678441 | 0.15048692 | -0.0237025 | 0.75810108 | 0.90650187 |
| Ankfy1 | 0.2317891 | 0.22356048 | 0.24728731 | -0.0237268 | 0.66514608 | 0.90650187 |
| Ide | 0.16564879 | 0.15958625 | 0.18332213 | -0.0237359 | 0.71755745 | 0.90650187 |
| Ppp1r16a | 0.29235707 | 0.27822238 | 0.30196016 | -0.0237378 | 0.63663524 | 0.90650187 |
| Adck1 | 0.13203774 | 0.12520546 | 0.1489533 | -0.0237478 | 0.76076626 | 0.90650187 |
| Bysl | 0.19298405 | 0.18657917 | 0.21032796 | -0.0237488 | 0.69215473 | 0.90650187 |
| Hibch | 0.23372826 | 0.22534175 | 0.24909336 | -0.0237516 | 0.66418009 | 0.90650187 |
| Cggbp1 | 0.67609546 | 0.55728674 | 0.58105023 | -0.0237635 | 0.5721357 | 0.90650187 |
| Mthfd1l | 0.13169099 | 0.12482592 | 0.14859182 | -0.0237659 | 0.76146841 | 0.90650187 |
| Ptpn14 | 0.1182015 | 0.11063385 | 0.13441607 | -0.0237822 | 0.7842304 | 0.90650187 |
| Anapc10 | 0.37547055 | 0.34775331 | 0.37155217 | -0.0237989 | 0.61208305 | 0.90650187 |
| Tead1 | 0.32252872 | 0.30411645 | 0.32791608 | -0.0237996 | 0.6265218 | 0.90650187 |
| Cdc42bpa | 0.37576741 | 0.34798919 | 0.37179026 | -0.0238011 | 0.61202333 | 0.90650187 |
| Smg9 | 0.16248168 | 0.15633442 | 0.18013924 | -0.0238048 | 0.72155678 | 0.90650187 |
| Unk | 0.12006548 | 0.11257202 | 0.13638814 | -0.0238161 | 0.78121187 | 0.90650187 |
| Zfx | 0.26164759 | 0.25086273 | 0.27468187 | -0.0238191 | 0.6500674 | 0.90650187 |
| Arnt | 0.21602677 | 0.20864222 | 0.23246426 | -0.023822 | 0.67564992 | 0.90650187 |
| Hira | 0.09707202 | 0.08793474 | 0.11177718 | -0.0238424 | 0.828502 | 0.9096066 |
| Dnajb14 | 0.45320121 | 0.4077156 | 0.43156157 | -0.023846 | 0.59702047 | 0.90650187 |
| Jph4 | 0.0923706 | 0.08283597 | 0.10668368 | -0.0238477 | 0.83982915 | 0.91213511 |
| Pcgf6 | 0.22749146 | 0.21942057 | 0.24327123 | -0.0238507 | 0.66852856 | 0.90650187 |
| Fam131a | 0.1449604 | 0.13846979 | 0.16232347 | -0.0238537 | 0.74321488 | 0.90650187 |
| Tada2a | 0.14210174 | 0.13552497 | 0.15938272 | -0.0238578 | 0.74712058 | 0.90650187 |
| Hacl1 | 0.1026176 | 0.09390173 | 0.11776085 | -0.0238591 | 0.81612842 | 0.9092537 |
| Yeats4 | 0.60475103 | 0.51259813 | 0.53646322 | -0.0238651 | 0.57838636 | 0.90650187 |
| Trim3 | 0.10324075 | 0.09453974 | 0.11843264 | -0.0238929 | 0.81510527 | 0.9092537 |
| E130309D02l | 0.20632622 | 0.1992979 | 0.22321372 | -0.0239158 | 0.68307548 | 0.90650187 |
| Stk3 | 0.26422984 | 0.25309379 | 0.27701011 | -0.0239163 | 0.64944475 | 0.90650187 |
| Fopnl | 0.55770608 | 0.48153928 | 0.50546461 | -0.0239253 | 0.58333217 | 0.90650187 |
| Aars | 0.5649677 | 0.4864066 | 0.51033713 | -0.0239305 | 0.58256572 | 0.90650187 |

| | | | | | | |
|---|---|---|---|---|---|---|
| 1600002K03Rik | 0.14899072 | 0.14251658 | 0.16645336 | -0.0239368 | 0.73868587 | 0.90650187 |
| AI480526 | 0.14822488 | 0.14172774 | 0.16567005 | -0.0239423 | 0.73971602 | 0.90650187 |
| Dopey2 | 0.14572694 | 0.13916428 | 0.16311039 | -0.0239461 | 0.74301098 | 0.90650187 |
| AW549877 | 0.11161664 | 0.10345209 | 0.12741525 | -0.0239632 | 0.79844827 | 0.90770671 |
| Pcgf2 | 0.33426113 | 0.31379708 | 0.33779058 | -0.0239935 | 0.62391318 | 0.90650187 |
| Kin | 0.36620527 | 0.34009066 | 0.36408427 | -0.0239936 | 0.61522786 | 0.90650187 |
| Ints7 | 0.15693881 | 0.15052542 | 0.17454146 | -0.024016 | 0.72962969 | 0.90650187 |
| Phykpl | 0.22059701 | 0.21276752 | 0.23678885 | -0.0240213 | 0.67400158 | 0.90650187 |
| Tgfbrap1 | 0.09119867 | 0.08138419 | 0.10540923 | -0.024025 | 0.84452959 | 0.91320576 |
| Asxl2 | 0.43213312 | 0.39172121 | 0.41574773 | -0.0240265 | 0.60137839 | 0.90650187 |
| AI837181 | 0.08585375 | 0.07554506 | 0.09957243 | -0.0240274 | 0.8583081 | 0.91744797 |
| Ctnnbl1 | 0.58569268 | 0.50003426 | 0.52406499 | -0.0240307 | 0.58076504 | 0.90650187 |
| Btbd6 | 0.13909695 | 0.13223479 | 0.15628134 | -0.0240466 | 0.75302267 | 0.90650187 |
| Epb41l5 | 0.18769981 | 0.18112144 | 0.2051713 | -0.0240499 | 0.69885393 | 0.90650187 |
| Zbtb1 | 0.33066868 | 0.31071487 | 0.33477951 | -0.0240646 | 0.6253488 | 0.90650187 |
| Mnt | 0.11392175 | 0.10580716 | 0.12987204 | -0.0240649 | 0.79496117 | 0.90731056 |
| Mtfr1l | 0.77204697 | 0.6127667 | 0.63683328 | -0.0240666 | 0.56670993 | 0.90650187 |
| Srsf6 | 1.66008749 | 0.89061217 | 0.91468314 | -0.024071 | 0.54656655 | 0.90650187 |
| Pdcd7 | 0.09747955 | 0.08814365 | 0.11221726 | -0.0240736 | 0.82988263 | 0.90974008 |
| Cops3 | 0.9079178 | 0.68441528 | 0.70849405 | -0.0240788 | 0.56004672 | 0.90650187 |
| Xrcc4 | 0.257332 | 0.24669362 | 0.2707766 | -0.024083 | 0.65372393 | 0.90650187 |
| Dvl2 | 0.26321939 | 0.25201433 | 0.27609981 | -0.0240855 | 0.65092208 | 0.90650187 |
| Tmem69 | 0.09282633 | 0.083091 | 0.10717876 | -0.0240878 | 0.84113225 | 0.91251845 |
| Rad51d | 0.11274982 | 0.10453436 | 0.12862381 | -0.0240895 | 0.79746493 | 0.9075779 |
| Sart1 | 0.46874387 | 0.41893352 | 0.44302348 | -0.02409 | 0.59550741 | 0.90650187 |
| Arvcf | 0.21332157 | 0.20580116 | 0.22989436 | -0.0240932 | 0.67935818 | 0.90650187 |
| Ascc3 | 0.38290887 | 0.35340279 | 0.37749687 | -0.0240941 | 0.61169703 | 0.90650187 |
| Actl6b | 0.1208852 | 0.11315871 | 0.13725406 | -0.0240953 | 0.78243378 | 0.90650187 |
| Dgkz | 0.17946548 | 0.17297629 | 0.19707557 | -0.0240993 | 0.70665143 | 0.90650187 |
| Arhgap39 | 0.12051274 | 0.11275631 | 0.13686071 | -0.0241044 | 0.78317946 | 0.90650187 |

| | | | | | | |
|---|---|---|---|---|---|---|
| Nol8 | 0.1392526 | 0.13233723 | 0.15644226 | -0.024105 | 0.75332504 | 0.90650187 |
| Mef2d | 0.19528599 | 0.18845644 | 0.21256495 | -0.0241085 | 0.69294243 | 0.90650187 |
| Birc2 | 0.39840941 | 0.36563591 | 0.38974479 | -0.0241089 | 0.60834066 | 0.90650187 |
| Apobec3 | 0.19746564 | 0.19055733 | 0.2146779 | -0.0241206 | 0.69128473 | 0.90650187 |
| Sos2 | 0.14513665 | 0.13836757 | 0.16250446 | -0.0241369 | 0.74547219 | 0.90650187 |
| Eif1ad | 0.30147184 | 0.28574406 | 0.30988762 | -0.0241436 | 0.63546406 | 0.90650187 |
| Wwox | 0.19504338 | 0.18818525 | 0.21232945 | -0.0241442 | 0.69340126 | 0.90650187 |
| Mkrn2 | 0.21600929 | 0.20828908 | 0.23244768 | -0.0241586 | 0.6779729 | 0.90650187 |
| Get4 | 0.55060178 | 0.47649279 | 0.50066674 | -0.0241739 | 0.5849797 | 0.90650187 |
| Zc3h4 | 0.11104708 | 0.10262373 | 0.1268072 | -0.0241835 | 0.80173196 | 0.90839694 |
| Wdr75 | 0.22852151 | 0.22004296 | 0.2442355 | -0.0241925 | 0.67016027 | 0.90650187 |
| Ccdc92 | 0.20344579 | 0.19623499 | 0.22044802 | -0.024213 | 0.68732625 | 0.90650187 |
| Tmem129 | 0.20262361 | 0.19542648 | 0.21965699 | -0.0242305 | 0.68807382 | 0.90650187 |
| Tnip1 | 0.18259104 | 0.17591636 | 0.20015722 | -0.0242409 | 0.70485067 | 0.90650187 |
| Tmem39b | 0.21924184 | 0.21126716 | 0.23550874 | -0.0242416 | 0.67637971 | 0.90650187 |
| Pigq | 0.19580399 | 0.18881497 | 0.21306756 | -0.0242526 | 0.69358037 | 0.90650187 |
| Lrrc40 | 0.15201827 | 0.1452888 | 0.16954283 | -0.024254 | 0.73760151 | 0.90650187 |
| Slc9a8 | 0.12484986 | 0.11717623 | 0.14143061 | -0.0242544 | 0.77708665 | 0.90650187 |
| Akap11 | 0.39809299 | 0.36522145 | 0.38949672 | -0.0242753 | 0.60913723 | 0.90650187 |
| Socs7 | 0.18501584 | 0.17826468 | 0.20254061 | -0.0242759 | 0.70292697 | 0.90650187 |
| Mxd1 | 0.14302553 | 0.13604794 | 0.16033408 | -0.0242861 | 0.74964888 | 0.90650187 |
| Nop16 | 0.4272216 | 0.38771584 | 0.41201431 | -0.0242985 | 0.60340823 | 0.90650187 |
| Toe1 | 0.21026679 | 0.20267243 | 0.22698327 | -0.0243108 | 0.68301361 | 0.90650187 |
| Dis3l | 0.13394468 | 0.12662603 | 0.15093868 | -0.0243127 | 0.76302965 | 0.90650187 |
| Abce1 | 0.49000797 | 0.4341107 | 0.45842839 | -0.0243177 | 0.59321407 | 0.90650187 |
| Ifi35 | 0.25637283 | 0.2455825 | 0.2699062 | -0.0243237 | 0.65565521 | 0.90650187 |
| Tmem64 | 0.15030073 | 0.14346296 | 0.16779147 | -0.0243285 | 0.7404002 | 0.90650187 |
| Klhl12 | 0.17045055 | 0.16377314 | 0.18812636 | -0.0243532 | 0.71751998 | 0.90650187 |
| Arhgap35 | 0.22224229 | 0.21397774 | 0.23834046 | -0.0243627 | 0.67523735 | 0.90650187 |
| Spg7 | 0.38760952 | 0.35685826 | 0.38123119 | -0.0243729 | 0.61187802 | 0.90650187 |

| | | | | | | |
|---|---|---|---|---|---|---|
| Snrnp40 | 0.7561669 | 0.60352818 | 0.62791081 | -0.0243826 | 0.56852845 | 0.90650187 |
| Gm20604 | 0.33065025 | 0.31038113 | 0.33476404 | -0.0243829 | 0.62695554 | 0.90650187 |
| Clpp | 0.78833347 | 0.62146563 | 0.64586279 | -0.0243972 | 0.5666811 | 0.90650187 |
| Gpn2 | 0.13576071 | 0.12839782 | 0.15282541 | -0.0244276 | 0.76132931 | 0.90650187 |
| H1fx | 0.21754117 | 0.20945085 | 0.23389964 | -0.0244488 | 0.67892156 | 0.90650187 |
| Mpp3 | 0.12321597 | 0.11526235 | 0.13971173 | -0.0244494 | 0.78170827 | 0.90650187 |
| Mbtd1 | 0.40785691 | 0.37266352 | 0.39711656 | -0.024453 | 0.60785961 | 0.90650187 |
| Cep44 | 0.15811216 | 0.15127572 | 0.17572935 | -0.0244536 | 0.73191949 | 0.90650187 |
| Csrp2bp | 0.13436735 | 0.12691744 | 0.15137816 | -0.0244607 | 0.76372969 | 0.90650187 |
| Fubp3 | 0.47693465 | 0.42453252 | 0.44899384 | -0.0244613 | 0.59568664 | 0.90650187 |
| Zfp688 | 0.17620479 | 0.16936701 | 0.19384927 | -0.0244823 | 0.71275774 | 0.90650187 |
| Abca3 | 0.18400474 | 0.17705516 | 0.20154755 | -0.0244924 | 0.70548337 | 0.90650187 |
| Med29 | 0.6494782 | 0.54024963 | 0.56474314 | -0.0244935 | 0.57644637 | 0.90650187 |
| Alg1 | 0.20295681 | 0.19548193 | 0.21997765 | -0.0244957 | 0.68971503 | 0.90650187 |
| Fam120b | 0.11675267 | 0.1083841 | 0.13288028 | -0.0244962 | 0.79376011 | 0.90731056 |
| 1110032A03 | 0.37772712 | 0.34885767 | 0.37336024 | -0.0245026 | 0.61476206 | 0.90650187 |
| Adck2 | 0.19430519 | 0.18710903 | 0.21161252 | -0.0245035 | 0.69664493 | 0.90650187 |
| Lpin2 | 0.1655234 | 0.15869221 | 0.18319633 | -0.0245041 | 0.7239576 | 0.90650187 |
| Ppp2r5d | 0.21083384 | 0.2030172 | 0.22752438 | -0.0245072 | 0.68397613 | 0.90650187 |
| Zswim8 | 0.24336217 | 0.23349795 | 0.2580105 | -0.0245126 | 0.66361865 | 0.90650187 |
| Phf5a | 0.97500563 | 0.71565823 | 0.74017748 | -0.0245193 | 0.55853894 | 0.90650187 |
| Dync2li1 | 0.54882691 | 0.47493919 | 0.49946328 | -0.0245241 | 0.58639578 | 0.90650187 |
| Ubr5 | 0.27785793 | 0.26466724 | 0.2891917 | -0.0245245 | 0.64691421 | 0.90650187 |
| Cox16 | 0.18035779 | 0.17342617 | 0.19795643 | -0.0245303 | 0.70914485 | 0.90650187 |
| Sart3 | 0.27972254 | 0.26630585 | 0.29084457 | -0.0245387 | 0.64619891 | 0.90650187 |
| Dhrs13 | 0.13707801 | 0.12964291 | 0.15419154 | -0.0245486 | 0.76045873 | 0.90650187 |
| Gng4 | 0.15996818 | 0.15303882 | 0.17760517 | -0.0245663 | 0.730695 | 0.90650187 |
| Cep104 | 0.22380366 | 0.21523709 | 0.23981018 | -0.0245731 | 0.67563835 | 0.90650187 |
| Ginm1 | 0.7601151 | 0.60556643 | 0.63014046 | -0.024574 | 0.5688191 | 0.90650187 |
| Gfm1 | 0.21074162 | 0.20286082 | 0.2274364 | -0.0245756 | 0.68451742 | 0.90650187 |

| | | | | | | |
|---|---|---|---|---|---|---|
| Lyrm4 | 0.17995135 | 0.17296052 | 0.19755531 | -0.0245948 | 0.71002543 | 0.90650187 |
| Pola1 | 0.15900376 | 0.15202589 | 0.17663095 | -0.0246051 | 0.73213481 | 0.90650187 |
| Taok1 | 0.5614255 | 0.48335253 | 0.50796429 | -0.0246118 | 0.58527126 | 0.90650187 |
| Foxk1 | 0.08785564 | 0.07715017 | 0.10176323 | -0.0246131 | 0.85886655 | 0.91773024 |
| Ggnbp1 | 0.10588749 | 0.09665826 | 0.12128057 | -0.0246223 | 0.81661606 | 0.9092537 |
| Evi5l | 0.11602202 | 0.10747417 | 0.13210482 | -0.0246306 | 0.7964119 | 0.9075779 |
| P4htm | 0.1352712 | 0.12765974 | 0.15231723 | -0.0246575 | 0.76413458 | 0.90650187 |
| Cab39 | 0.27969002 | 0.26614991 | 0.29081578 | -0.0246659 | 0.64693514 | 0.90650187 |
| Brms1 | 0.27211313 | 0.25940622 | 0.28407889 | -0.0246727 | 0.65029143 | 0.90650187 |
| Ccs | 0.43185408 | 0.39084841 | 0.4155361 | -0.0246877 | 0.60415657 | 0.90650187 |
| Vma21 | 0.81493606 | 0.63564413 | 0.66035125 | -0.0247071 | 0.56605249 | 0.90650187 |
| L3mbtl2 | 0.10943189 | 0.10035498 | 0.12508067 | -0.0247257 | 0.81024469 | 0.9092537 |
| Skiv2l | 0.23012385 | 0.22100161 | 0.24573337 | -0.0247318 | 0.67271649 | 0.90650187 |
| Orai3 | 0.29695062 | 0.28122022 | 0.30596537 | -0.0247451 | 0.64040492 | 0.90650187 |
| Cep78 | 0.12886986 | 0.12088594 | 0.14564551 | -0.0247596 | 0.77509978 | 0.90650187 |
| Vps11 | 0.17322237 | 0.16612555 | 0.19088791 | -0.0247624 | 0.71792651 | 0.90650187 |
| Dgcr2 | 0.41883099 | 0.38082798 | 0.40559492 | -0.0247669 | 0.60698256 | 0.90650187 |
| Thoc7 | 1.35784311 | 0.84018862 | 0.86495793 | -0.0247693 | 0.55065101 | 0.90650187 |
| Inip | 0.12675119 | 0.11864508 | 0.14342655 | -0.0247815 | 0.77879169 | 0.90650187 |
| Slc39a11 | 0.15417754 | 0.14694429 | 0.17173979 | -0.0247955 | 0.73951439 | 0.90650187 |
| Srp9 | 1.68054434 | 0.89192849 | 0.91672474 | -0.0247963 | 0.54785668 | 0.90650187 |
| Smek2 | 0.81493914 | 0.63555209 | 0.6603529 | -0.0248008 | 0.56630052 | 0.90650187 |
| Eya3 | 0.11790596 | 0.10929988 | 0.134103 | -0.0248031 | 0.79452994 | 0.90731056 |
| Elavl1 | 1.22289258 | 0.80442029 | 0.82924636 | -0.0248261 | 0.55294012 | 0.90650187 |
| Fbxl16 | 0.20021159 | 0.19250117 | 0.21733224 | -0.0248311 | 0.69424813 | 0.90650187 |
| Ano6 | 0.27237109 | 0.25946891 | 0.28430913 | -0.0248402 | 0.65114542 | 0.90650187 |
| Rp2 | 0.20355132 | 0.19569629 | 0.2205495 | -0.0248532 | 0.69180217 | 0.90650187 |
| Farsa | 0.62849372 | 0.52673431 | 0.55161507 | -0.0248808 | 0.57946319 | 0.90650187 |
| Tpgs1 | 0.76627842 | 0.60872189 | 0.63360643 | -0.0248845 | 0.56930253 | 0.90650187 |
| Cluap1 | 0.30540258 | 0.28839183 | 0.31328278 | -0.0248909 | 0.63803538 | 0.90650187 |

| | | | | | | |
|---|---|---|---|---|---|---|
| Atf4 | 0.81458451 | 0.63525463 | 0.66016194 | -0.0249073 | 0.56660164 | 0.90650187 |
| Pgm3 | 0.29224944 | 0.27694908 | 0.30186608 | -0.024917 | 0.6431736 | 0.90650187 |
| Gpr108 | 0.36230099 | 0.33598848 | 0.36091602 | -0.0249275 | 0.62059604 | 0.90650187 |
| Bcs1l | 0.13800262 | 0.13022052 | 0.1551492 | -0.0249287 | 0.7624792 | 0.90650187 |
| Phrf1 | 0.29167654 | 0.27643243 | 0.30136516 | -0.0249327 | 0.64348755 | 0.90650187 |
| Kpna6 | 0.15297157 | 0.14557563 | 0.17051342 | -0.0249378 | 0.74221052 | 0.90650187 |
| Khdrbs3 | 0.17250395 | 0.16522625 | 0.19017296 | -0.0249467 | 0.72011075 | 0.90650187 |
| Gmppa | 0.48549549 | 0.43023229 | 0.45518491 | -0.0249526 | 0.59626924 | 0.90650187 |
| Gipc1 | 0.86753658 | 0.66311047 | 0.68806897 | -0.0249585 | 0.56405437 | 0.90650187 |
| Tmem87b | 0.30464111 | 0.28766615 | 0.31262614 | -0.02496 | 0.63868126 | 0.90650187 |
| Rnf8 | 0.3649309 | 0.33808872 | 0.3630515 | -0.0249628 | 0.62007282 | 0.90650187 |
| Syf2 | 0.87619248 | 0.66753209 | 0.69250984 | -0.0249778 | 0.56369579 | 0.90650187 |
| Ipmk | 0.11806796 | 0.1092958 | 0.13427463 | -0.0249788 | 0.79588339 | 0.9075779 |
| Lrrc57 | 0.18962068 | 0.18206541 | 0.20704927 | -0.0249839 | 0.70417089 | 0.90650187 |
| Nras | 0.69650228 | 0.56830777 | 0.59329966 | -0.0249919 | 0.57427345 | 0.90650187 |
| Pskh1 | 0.25303034 | 0.24186357 | 0.26686614 | -0.0250026 | 0.66148041 | 0.90650187 |
| Fahd2a | 0.23325873 | 0.22364509 | 0.2486564 | -0.0250113 | 0.67262205 | 0.90650187 |
| Rnf141 | 0.22125704 | 0.21239375 | 0.23741164 | -0.0250179 | 0.68027489 | 0.90650187 |
| Pdlim5 | 0.10331275 | 0.09348848 | 0.11851023 | -0.0250218 | 0.82603878 | 0.90947782 |
| Kat6b | 0.22739773 | 0.21815413 | 0.24318344 | -0.0250293 | 0.67636491 | 0.90650187 |
| Abr | 0.24654791 | 0.23589378 | 0.26093909 | -0.0250453 | 0.66519954 | 0.90650187 |
| Mta1 | 0.27869706 | 0.26487432 | 0.28993594 | -0.0250616 | 0.64961134 | 0.90650187 |
| Ube4b | 0.41437504 | 0.37710033 | 0.40216321 | -0.0250629 | 0.6091295 | 0.90650187 |
| Hook2 | 0.09323307 | 0.08254421 | 0.10762037 | -0.0250762 | 0.8498643 | 0.91471921 |
| Atad2b | 0.22775895 | 0.21843564 | 0.24352175 | -0.0250861 | 0.67650907 | 0.90650187 |
| Irf6 | 0.18791994 | 0.18028286 | 0.20538672 | -0.0251039 | 0.70656396 | 0.90650187 |
| Lrrc56 | 0.08271851 | 0.07101092 | 0.09612995 | -0.025119 | 0.87731108 | 0.92482217 |
| Kctd13 | 0.41410356 | 0.37682442 | 0.40195365 | -0.0251292 | 0.60946669 | 0.90650187 |
| Rnf41 | 0.34778592 | 0.32389704 | 0.34902837 | -0.0251313 | 0.62555376 | 0.90650187 |
| Gprc5b | 0.20712736 | 0.19884806 | 0.2239814 | -0.0251333 | 0.69105721 | 0.90650187 |

| | | | | | | |
|---|---|---|---|---|---|---|
| Ift22 | 0.56947425 | 0.488206 | 0.51334458 | -0.0251386 | 0.5861694 | 0.90650187 |
| Ppp1r15b | 0.23107934 | 0.22148495 | 0.24662533 | -0.0251404 | 0.67478836 | 0.90650187 |
| Nol9 | 0.1548094 | 0.14723407 | 0.17238166 | -0.0251476 | 0.74168187 | 0.90650187 |
| Trmt13 | 0.25671804 | 0.24506826 | 0.27021956 | -0.0251513 | 0.66048417 | 0.90650187 |
| Vipas39 | 0.20755285 | 0.19923474 | 0.22438885 | -0.0251541 | 0.69088302 | 0.90650187 |
| Pomc | 0.11036757 | 0.10091579 | 0.12608125 | -0.0251655 | 0.81254949 | 0.9092537 |
| Gm16586 | 0.13402787 | 0.12584496 | 0.1510252 | -0.0251802 | 0.77071708 | 0.90650187 |
| Eid2b | 0.18299705 | 0.17537644 | 0.20055674 | -0.0251803 | 0.71163064 | 0.90650187 |
| Tceal1 | 0.39352713 | 0.36072664 | 0.38590784 | -0.0251812 | 0.61412405 | 0.90650187 |
| Arl5a | 0.20840969 | 0.20000073 | 0.22520878 | -0.025208 | 0.69061883 | 0.90650187 |
| Poc5 | 0.12247638 | 0.11371997 | 0.13893258 | -0.0252126 | 0.79009784 | 0.90650187 |
| Atpaf1 | 0.18633509 | 0.17862127 | 0.20383465 | -0.0252134 | 0.70880617 | 0.90650187 |
| Rasa2 | 0.24798784 | 0.23704205 | 0.26225941 | -0.0252174 | 0.66547925 | 0.90650187 |
| Rab3gap2 | 0.217428 | 0.20857483 | 0.23379245 | -0.0252176 | 0.6842148 | 0.90650187 |
| Acvr1 | 0.13706179 | 0.12895707 | 0.15417473 | -0.0252177 | 0.76643227 | 0.90650187 |
| Ctdspl | 0.19935345 | 0.19128309 | 0.21650366 | -0.0252206 | 0.69772545 | 0.90650187 |
| Rnaseh1 | 0.17249454 | 0.16492319 | 0.19016359 | -0.0252404 | 0.72242651 | 0.90650187 |
| Calcoco1 | 0.22810603 | 0.21858863 | 0.24384669 | -0.0252581 | 0.6774154 | 0.90650187 |
| Dlgap4 | 0.62113748 | 0.5216745 | 0.54695527 | -0.0252808 | 0.58140145 | 0.90650187 |
| Tbc1d1 | 0.1740556 | 0.16642692 | 0.19171632 | -0.0252894 | 0.72120417 | 0.90650187 |
| Ikbkap | 0.20372987 | 0.19542332 | 0.22072117 | -0.0252979 | 0.69481002 | 0.90650187 |
| Ncoa6 | 0.20261435 | 0.19434919 | 0.21964807 | -0.0252989 | 0.69568894 | 0.90650187 |
| Hs2st1 | 0.20766075 | 0.19918743 | 0.22449215 | -0.0253047 | 0.69185244 | 0.90650187 |
| Fdxr | 0.10485202 | 0.09485502 | 0.12016744 | -0.0253124 | 0.82547542 | 0.90947782 |
| Pnpt1 | 0.19212538 | 0.1841554 | 0.20949206 | -0.0253367 | 0.70459626 | 0.90650187 |
| Phf6 | 0.15159697 | 0.1437734 | 0.16911354 | -0.0253401 | 0.74733735 | 0.90650187 |
| Cxxc1 | 0.48729955 | 0.4311198 | 0.45648327 | -0.0253635 | 0.59754983 | 0.90650187 |
| Smap1 | 0.27043654 | 0.25721189 | 0.28258044 | -0.0253685 | 0.65509119 | 0.90650187 |
| Supt20 | 0.55165604 | 0.47600911 | 0.50138068 | -0.0253716 | 0.58899589 | 0.90650187 |
| Ndst1 | 0.09914077 | 0.0886359 | 0.11400879 | -0.0253729 | 0.8387611 | 0.91188312 |

| | | | | | | |
|---|---|---|---|---|---|---|
| Serhl | 0.20949565 | 0.20087184 | 0.22624686 | -0.025375 | 0.69097045 | 0.90650187 |
| Ulk2 | 0.10720541 | 0.09731724 | 0.1226954 | -0.0253782 | 0.82108911 | 0.9092537 |
| Tusc3 | 1.30816195 | 0.82746325 | 0.85284694 | -0.0253837 | 0.55263309 | 0.90650187 |
| Mrrf | 0.26000807 | 0.24781134 | 0.27320033 | -0.025389 | 0.66024726 | 0.90650187 |
| Map7d1 | 0.26380528 | 0.2512378 | 0.27662774 | -0.0253899 | 0.65837587 | 0.90650187 |
| Pcnt | 0.1925507 | 0.18450725 | 0.2099062 | -0.0253989 | 0.70468547 | 0.90650187 |
| Nmd3 | 0.38861692 | 0.35662492 | 0.38202925 | -0.0254043 | 0.61624201 | 0.90650187 |
| Kif3c | 0.07649058 | 0.06384167 | 0.08925031 | -0.0254086 | 0.89717511 | 0.93404279 |
| Pmpca | 0.4585074 | 0.41008344 | 0.43549357 | -0.0254101 | 0.60233348 | 0.90650187 |
| Wdr3 | 0.15942572 | 0.15164555 | 0.17705733 | -0.0254118 | 0.73827821 | 0.90650187 |
| Smek1 | 0.3921394 | 0.35939814 | 0.38481356 | -0.0254154 | 0.61547388 | 0.90650187 |
| Tnpo3 | 0.54045196 | 0.46833384 | 0.49375748 | -0.0254236 | 0.59052857 | 0.90650187 |
| Timm10b | 1.24735742 | 0.81098831 | 0.83641384 | -0.0254255 | 0.55374891 | 0.90650187 |
| Rrp8 | 0.25641132 | 0.24450702 | 0.26994114 | -0.0254341 | 0.66234421 | 0.90650187 |
| Prune | 0.16899892 | 0.16122396 | 0.1866767 | -0.0254527 | 0.72779024 | 0.90650187 |
| Tle3 | 0.35809105 | 0.33202525 | 0.3574859 | -0.0254606 | 0.62423488 | 0.90650187 |
| Vhl | 0.23205813 | 0.22207411 | 0.2475381 | -0.025464 | 0.67627766 | 0.90650187 |
| Pogz | 0.23014882 | 0.22028388 | 0.24575669 | -0.0254728 | 0.67752515 | 0.90650187 |
| Arhgap12 | 0.19087149 | 0.18279651 | 0.20827001 | -0.0254735 | 0.70669045 | 0.90650187 |
| Brf2 | 0.10163174 | 0.09122059 | 0.11669703 | -0.0254764 | 0.83410933 | 0.91062493 |
| Fam222b | 0.2238109 | 0.2143329 | 0.239817 | -0.0254841 | 0.68168339 | 0.90650187 |
| Uggt1 | 0.3131466 | 0.29443873 | 0.31993159 | -0.0254929 | 0.63841888 | 0.90650187 |
| 5930430L01F | 0.11207863 | 0.10241451 | 0.12790817 | -0.0254937 | 0.81220835 | 0.9092537 |
| Crtc2 | 0.10648821 | 0.09642623 | 0.12192573 | -0.0254995 | 0.82375038 | 0.9092537 |
| Azi2 | 0.47271689 | 0.42041076 | 0.44592531 | -0.0255145 | 0.60039306 | 0.90650187 |
| Mettl2 | 0.16715462 | 0.15931311 | 0.18483151 | -0.0255184 | 0.73031767 | 0.90650187 |
| Suox | 0.15262053 | 0.1446356 | 0.17015614 | -0.0255205 | 0.74753911 | 0.90650187 |
| Tom1 | 0.12443333 | 0.11545846 | 0.14099275 | -0.0255343 | 0.78962339 | 0.90650187 |
| Huwe1 | 0.69934052 | 0.56943814 | 0.5949861 | -0.025548 | 0.57569363 | 0.90650187 |
| Dus3l | 0.21588436 | 0.20677098 | 0.23232917 | -0.0255582 | 0.68759844 | 0.90650187 |

| | | | | | | |
|---|---|---|---|---|---|---|
| Tmeff1 | 0.12114903 | 0.11196732 | 0.13753259 | -0.0255653 | 0.79570353 | 0.9075779 |
| Usp36 | 0.20067521 | 0.19220566 | 0.21777956 | -0.0255739 | 0.69918172 | 0.90650187 |
| Men1 | 0.33820127 | 0.31550539 | 0.3410805 | -0.0255751 | 0.6305624 | 0.90650187 |
| Tada3 | 0.48543907 | 0.42955944 | 0.45514427 | -0.0255848 | 0.59866723 | 0.90650187 |
| Nfkbil1 | 0.30441652 | 0.28684727 | 0.31243237 | -0.0255851 | 0.64208996 | 0.90650187 |
| Arl6ip5 | 0.94560643 | 0.70120243 | 0.72678877 | -0.0255863 | 0.5621829 | 0.90650187 |
| Acd | 0.44975271 | 0.40340362 | 0.42899551 | -0.0255919 | 0.60457398 | 0.90650187 |
| Sesn1 | 0.21968216 | 0.21033162 | 0.23592488 | -0.0255933 | 0.68519005 | 0.90650187 |
| Ift57 | 0.4917013 | 0.43404692 | 0.45964201 | -0.0255951 | 0.5977598 | 0.90650187 |
| Lppos | 0.14464758 | 0.13640152 | 0.16200212 | -0.0256006 | 0.75882345 | 0.90650187 |
| Zfp87 | 0.18464455 | 0.17657427 | 0.20217607 | -0.0256018 | 0.71326637 | 0.90650187 |
| Pin4 | 0.85857415 | 0.65782595 | 0.68343284 | -0.0256069 | 0.56614508 | 0.90650187 |
| Gba2 | 0.12751866 | 0.11861557 | 0.14423097 | -0.0256154 | 0.78510155 | 0.90650187 |
| Fam234b | 0.13837537 | 0.12991082 | 0.15553499 | -0.0256242 | 0.76804005 | 0.90650187 |
| Zfp428 | 0.3562704 | 0.33036893 | 0.35599802 | -0.0256291 | 0.62553445 | 0.90650187 |
| Tnpo1 | 0.50453387 | 0.44312981 | 0.46877374 | -0.0256439 | 0.59607232 | 0.90650187 |
| Ripk4 | 0.11887144 | 0.1094768 | 0.13512535 | -0.0256485 | 0.80061886 | 0.90806412 |
| Ubl4a | 0.62381246 | 0.5229866 | 0.54865319 | -0.0256666 | 0.58237391 | 0.90650187 |
| Agps | 0.19672791 | 0.18828544 | 0.21396331 | -0.0256779 | 0.70316321 | 0.90650187 |
| Ppard | 0.10217093 | 0.09159646 | 0.11727901 | -0.0256825 | 0.83485471 | 0.91063622 |
| Tmem216 | 0.2973697 | 0.28063624 | 0.3063297 | -0.0256935 | 0.64537965 | 0.90650187 |
| Zfp703 | 0.27351281 | 0.25962555 | 0.28532743 | -0.0257019 | 0.65558884 | 0.90650187 |
| Hccs | 0.24816253 | 0.23670833 | 0.26241942 | -0.0257111 | 0.66842471 | 0.90650187 |
| Samm50 | 0.75305721 | 0.60043788 | 0.62614961 | -0.0257117 | 0.57242481 | 0.90650187 |
| Utp23 | 0.15612678 | 0.147991 | 0.17371846 | -0.0257275 | 0.7448556 | 0.90650187 |
| Rnmtl1 | 0.11162076 | 0.10168627 | 0.12741965 | -0.0257334 | 0.81536045 | 0.9092537 |
| Ahcyl2 | 0.25766371 | 0.24532771 | 0.27107741 | -0.0257497 | 0.66359298 | 0.90650187 |
| Mex3c | 0.15316213 | 0.14495349 | 0.17070732 | -0.0257538 | 0.74879044 | 0.90650187 |
| Csad | 0.27624868 | 0.26199861 | 0.28776258 | -0.025764 | 0.65469383 | 0.90650187 |
| Abca5 | 0.10911358 | 0.09897317 | 0.12474003 | -0.0257669 | 0.82077023 | 0.9092537 |

| | | | | | | |
|---|---|---|---|---|---|---|
| Arfrp1 | 0.32793638 | 0.30671476 | 0.33248203 | -0.0257673 | 0.63477737 | 0.90650187 |
| Ube2q2 | 0.14249654 | 0.13401173 | 0.15978943 | -0.0257777 | 0.76336793 | 0.90650187 |
| Tmem19 | 0.20643041 | 0.1975353 | 0.2233136 | -0.0257783 | 0.6960874 | 0.90650187 |
| Ndufaf6 | 0.15575245 | 0.14755779 | 0.17333881 | -0.025781 | 0.74575915 | 0.90650187 |
| Lonp1 | 0.41859948 | 0.37961965 | 0.40541699 | -0.0257973 | 0.61136601 | 0.90650187 |
| Slc52a2 | 0.16146552 | 0.15331682 | 0.17911563 | -0.0257988 | 0.73901777 | 0.90650187 |
| 2610507B11 | 0.35170405 | 0.32645399 | 0.35225433 | -0.0258003 | 0.62764091 | 0.90650187 |
| Plekha8 | 0.11779363 | 0.10818239 | 0.13398398 | -0.0258016 | 0.80402881 | 0.90882608 |
| Sbf2 | 0.27632342 | 0.26202548 | 0.28782901 | -0.0258035 | 0.65488514 | 0.90650187 |
| Fam189b | 0.21941492 | 0.20986589 | 0.23567234 | -0.0258064 | 0.68679998 | 0.90650187 |
| Ube2i | 1.59839554 | 0.88197325 | 0.90778118 | -0.0258079 | 0.55028726 | 0.90650187 |
| Sfi1 | 0.11313243 | 0.10322178 | 0.12903151 | -0.0258097 | 0.81306281 | 0.9092537 |
| Trove2 | 0.29752512 | 0.28065363 | 0.30646478 | -0.0258111 | 0.64595364 | 0.90650187 |
| Ibtk | 0.26077608 | 0.24808304 | 0.27389465 | -0.0258116 | 0.66237382 | 0.90650187 |
| Ier3ip1 | 1.54774395 | 0.87530253 | 0.90113902 | -0.0258365 | 0.55071169 | 0.90650187 |
| Ubxn8 | 0.2119175 | 0.20272089 | 0.22855753 | -0.0258366 | 0.69235651 | 0.90650187 |
| Srsf9 | 0.86367034 | 0.66023221 | 0.68607525 | -0.025843 | 0.56649468 | 0.90650187 |
| Map7 | 0.24501658 | 0.23368655 | 0.25953261 | -0.0258461 | 0.67101743 | 0.90650187 |
| Trit1 | 0.13500702 | 0.1261893 | 0.15204285 | -0.0258536 | 0.77515631 | 0.90650187 |
| Ecsit | 0.20559017 | 0.19665144 | 0.22250782 | -0.0258564 | 0.69728189 | 0.90650187 |
| Parg | 0.24685502 | 0.23535999 | 0.26122089 | -0.0258609 | 0.67007331 | 0.90650187 |
| Fam175a | 0.11634572 | 0.10658181 | 0.13244846 | -0.0258666 | 0.80735998 | 0.9092537 |
| Ppm1k | 0.18405757 | 0.17571893 | 0.20159947 | -0.0258805 | 0.71590162 | 0.90650187 |
| Mfap3 | 0.34408577 | 0.320083 | 0.3459696 | -0.0258866 | 0.63029512 | 0.90650187 |
| N4bp1 | 0.20752413 | 0.19847101 | 0.22436135 | -0.0258903 | 0.69602571 | 0.90650187 |
| Gosr1 | 0.15684504 | 0.14855476 | 0.17444647 | -0.0258917 | 0.74531732 | 0.90650187 |
| Ubr1 | 0.27152582 | 0.25765516 | 0.28355429 | -0.0258991 | 0.65764725 | 0.90650187 |
| Clic6 | 0.08624277 | 0.07408566 | 0.0999986 | -0.0259129 | 0.87534017 | 0.92418014 |
| Arf2 | 0.34475164 | 0.32056771 | 0.34652102 | -0.0259533 | 0.63041835 | 0.90650187 |
| D230025D16 | 0.14692025 | 0.13837892 | 0.16433409 | -0.0259552 | 0.7587058 | 0.90650187 |

| | | | | | | |
|---|---|---|---|---|---|---|
| Lrrc16b | 0.16443387 | 0.15612744 | 0.1821025 | -0.0259751 | 0.73699885 | 0.90650187 |
| Erap1 | 0.11562543 | 0.10569668 | 0.13168362 | -0.0259869 | 0.80984161 | 0.9092537 |
| Zfp236 | 0.11572336 | 0.10578773 | 0.13178764 | -0.0259999 | 0.80977235 | 0.9092537 |
| Anks3 | 0.38323273 | 0.35174689 | 0.37775471 | -0.0260078 | 0.62022544 | 0.90650187 |
| Bod1 | 0.19962092 | 0.19073595 | 0.216762 | -0.0260261 | 0.70325054 | 0.90650187 |
| Rnf14 | 0.46284251 | 0.41265516 | 0.43869135 | -0.0260362 | 0.60405042 | 0.90650187 |
| Acbd5 | 0.42658176 | 0.3854797 | 0.41152664 | -0.0260469 | 0.61078965 | 0.90650187 |
| Lactb | 0.15366489 | 0.14515162 | 0.17121868 | -0.0260671 | 0.75073929 | 0.90650187 |
| Ube2k | 1.17415769 | 0.78826087 | 0.81433458 | -0.0260737 | 0.5565948 | 0.90650187 |
| Gak | 0.197323 | 0.18845497 | 0.21453979 | -0.0260848 | 0.70558047 | 0.90650187 |
| Cops7b | 0.23173726 | 0.22113284 | 0.24723898 | -0.0261061 | 0.68061337 | 0.90650187 |
| Crtc1 | 0.200591 | 0.19158773 | 0.21769833 | -0.0261106 | 0.70305354 | 0.90650187 |
| Hnrnpul2 | 0.7424495 | 0.59397712 | 0.62010799 | -0.0261309 | 0.57430076 | 0.90650187 |
| Gxylt1 | 0.16580005 | 0.15733252 | 0.18347386 | -0.0261413 | 0.73676692 | 0.90650187 |
| Znrf2 | 0.20792271 | 0.19857796 | 0.22474287 | -0.0261649 | 0.69761888 | 0.90650187 |
| Cln6 | 0.2558151 | 0.24322793 | 0.26939968 | -0.0261717 | 0.66708697 | 0.90650187 |
| Tamm41 | 0.09810989 | 0.0867241 | 0.11289749 | -0.0261734 | 0.84864503 | 0.91440801 |
| Zfp622 | 0.55059569 | 0.47448724 | 0.50066262 | -0.0261754 | 0.59189519 | 0.90650187 |
| Sap30 | 0.33696016 | 0.31386744 | 0.34004562 | -0.0261782 | 0.63391561 | 0.90650187 |
| Zfp518a | 0.1462309 | 0.1374416 | 0.16362739 | -0.0261858 | 0.76159538 | 0.90650187 |
| Itsn2 | 0.23617348 | 0.22517888 | 0.25136537 | -0.0261865 | 0.67836438 | 0.90650187 |
| 4930453N24 | 0.28937914 | 0.2731631 | 0.29935338 | -0.0261903 | 0.65134119 | 0.90650187 |
| Fut8 | 0.21130578 | 0.20177938 | 0.22797447 | -0.0261951 | 0.69526013 | 0.90650187 |
| Tmem38b | 0.14923534 | 0.14049596 | 0.1667034 | -0.0262074 | 0.75769081 | 0.90650187 |
| Nipal3 | 0.11825915 | 0.10826475 | 0.13447712 | -0.0262124 | 0.8069031 | 0.9092537 |
| Nfat5 | 0.5690329 | 0.48681293 | 0.51305067 | -0.0262377 | 0.58992466 | 0.90650187 |
| Zbtb21 | 0.13058534 | 0.12119605 | 0.14743824 | -0.0262422 | 0.78560476 | 0.90650187 |
| 0610012G03 | 1.06313681 | 0.74947607 | 0.77572293 | -0.0262469 | 0.55978285 | 0.90650187 |
| Gna11 | 0.76142029 | 0.60462422 | 0.63087597 | -0.0262517 | 0.57338105 | 0.90650187 |
| Mettl18 | 0.11086418 | 0.10034989 | 0.12661186 | -0.026262 | 0.82176505 | 0.9092537 |

| | | | | | | |
|---|---|---|---|---|---|---|
| 9030624J02F | 0.36943899 | 0.34043474 | 0.36669907 | -0.0262643 | 0.62491325 | 0.90650187 |
| 5430416N02 | 0.15078968 | 0.14201978 | 0.16829039 | -0.0262706 | 0.75615009 | 0.90650187 |
| Tbce | 0.20098224 | 0.19180404 | 0.21807567 | -0.0262716 | 0.70386979 | 0.90650187 |
| Smyd2 | 0.44937661 | 0.40243804 | 0.42871515 | -0.0262771 | 0.60737475 | 0.90650187 |
| Uspl1 | 0.2193444 | 0.20932393 | 0.23560569 | -0.0262818 | 0.69001975 | 0.90650187 |
| Jmy | 0.19879632 | 0.18968334 | 0.21596531 | -0.026282 | 0.7057495 | 0.90650187 |
| Zbtb41 | 0.14086132 | 0.13181911 | 0.15810371 | -0.0262846 | 0.77007019 | 0.90650187 |
| Wdsub1 | 0.17074067 | 0.16212481 | 0.1884158 | -0.026291 | 0.73247872 | 0.90650187 |
| Pdzd11 | 0.60791172 | 0.51219218 | 0.53849896 | -0.0263068 | 0.58596479 | 0.90650187 |
| Kpna4 | 0.32599602 | 0.30453798 | 0.33084659 | -0.0263086 | 0.63814712 | 0.90650187 |
| Cdkn2aipnl | 0.18746387 | 0.17862909 | 0.20494037 | -0.0263113 | 0.71591518 | 0.90650187 |
| Psmf1 | 0.28004565 | 0.26480895 | 0.29113066 | -0.0263217 | 0.65613456 | 0.90650187 |
| Galnt4 | 0.10747334 | 0.09663997 | 0.12298276 | -0.0263428 | 0.82951356 | 0.90974008 |
| Lztfl1 | 0.67241133 | 0.55245921 | 0.57881586 | -0.0263567 | 0.5802108 | 0.90650187 |
| Reps1 | 0.10841625 | 0.0976275 | 0.12399336 | -0.0263659 | 0.82774986 | 0.90947782 |
| Polb | 0.46809639 | 0.41618014 | 0.44254935 | -0.0263692 | 0.60445309 | 0.90650187 |
| Smox | 0.18371494 | 0.17487756 | 0.20126271 | -0.0263852 | 0.71998955 | 0.90650187 |
| Map3k7 | 0.17821454 | 0.16945329 | 0.1958392 | -0.0263859 | 0.7254028 | 0.90650187 |
| Trip11 | 0.57108519 | 0.48802962 | 0.51441637 | -0.0263867 | 0.59019059 | 0.90650187 |
| Nup188 | 0.09179823 | 0.07967366 | 0.10606147 | -0.0263878 | 0.86564868 | 0.92040222 |
| Slc7a6os | 0.26870535 | 0.25462958 | 0.28103039 | -0.0264008 | 0.66189234 | 0.90650187 |
| Ubn1 | 0.24945836 | 0.23719404 | 0.2636054 | -0.0264114 | 0.67199102 | 0.90650187 |
| Vps45 | 0.23360779 | 0.22256714 | 0.24898126 | -0.0264141 | 0.68140418 | 0.90650187 |
| Mrps21 | 1.64227773 | 0.88633804 | 0.91276741 | -0.0264294 | 0.55121174 | 0.90650187 |
| Mtrr | 0.12276884 | 0.11281118 | 0.13924076 | -0.0264296 | 0.80061778 | 0.90806412 |
| Nae1 | 0.61884968 | 0.51905955 | 0.54549997 | -0.0264404 | 0.58530299 | 0.90650187 |
| Pex10 | 0.17082732 | 0.16206075 | 0.18850223 | -0.0264415 | 0.73355208 | 0.90650187 |
| 1810013L24F | 0.15602641 | 0.14716265 | 0.17361667 | -0.026454 | 0.75091704 | 0.90650187 |
| Zdhhc17 | 0.1556266 | 0.1467562 | 0.17321114 | -0.0264549 | 0.75142721 | 0.90650187 |
| Plekha5 | 0.23864536 | 0.22719789 | 0.25365611 | -0.0264582 | 0.67857216 | 0.90650187 |

| | | | | | | |
|---|---|---|---|---|---|---|
| Ppp2r3d | 0.34580135 | 0.32092766 | 0.34738954 | -0.0264619 | 0.63255857 | 0.90650187 |
| Acvr2a | 0.0914744 | 0.07924307 | 0.10570925 | -0.0264662 | 0.86715355 | 0.92056229 |
| Krcc1 | 0.96865682 | 0.71094718 | 0.73742961 | -0.0264824 | 0.5634215 | 0.90650187 |
| 2010111I01R | 0.20370093 | 0.1942081 | 0.22069334 | -0.0264852 | 0.70316103 | 0.90650187 |
| Usp4 | 0.44706722 | 0.40050556 | 0.42699138 | -0.0264858 | 0.60863235 | 0.90650187 |
| Akt1s1 | 0.8222331 | 0.63777821 | 0.66426867 | -0.0264905 | 0.57035849 | 0.90650187 |
| Rbm7 | 0.63590025 | 0.52977342 | 0.55627674 | -0.0265033 | 0.58387119 | 0.90650187 |
| Mcrs1 | 0.57912161 | 0.49322759 | 0.51974016 | -0.0265126 | 0.58970092 | 0.90650187 |
| Adprhl2 | 0.41541239 | 0.37644542 | 0.40296344 | -0.026518 | 0.61506856 | 0.90650187 |
| Ppil1 | 0.4638154 | 0.4128865 | 0.4394072 | -0.0265207 | 0.6057725 | 0.90650187 |
| Rnf216 | 0.30507379 | 0.28647368 | 0.31299931 | -0.0265256 | 0.64682036 | 0.90650187 |
| Fam208b | 0.14042178 | 0.13111781 | 0.15765007 | -0.0265323 | 0.77282673 | 0.90650187 |
| Sssca1 | 0.59935509 | 0.50644204 | 0.53297469 | -0.0265327 | 0.58757527 | 0.90650187 |
| Kmt2b | 0.17173095 | 0.1628638 | 0.18940307 | -0.0265393 | 0.73333177 | 0.90650187 |
| Nsun2 | 0.56011249 | 0.48054163 | 0.50708282 | -0.0265412 | 0.59199783 | 0.90650187 |
| Decr2 | 0.18288668 | 0.17390688 | 0.20044815 | -0.0265413 | 0.72194901 | 0.90650187 |
| Cyb561d2 | 0.21969992 | 0.20939512 | 0.23594166 | -0.0265465 | 0.69152902 | 0.90650187 |
| Dimt1 | 0.27063766 | 0.25621354 | 0.28276033 | -0.0265468 | 0.66179726 | 0.90650187 |
| Pafah1b3 | 0.94311755 | 0.69906248 | 0.72561093 | -0.0265485 | 0.56460481 | 0.90650187 |
| Snrpa1 | 0.66979039 | 0.55066877 | 0.57722203 | -0.0265533 | 0.58102093 | 0.90650187 |
| Phf13 | 0.1955647 | 0.18628108 | 0.21283541 | -0.0265543 | 0.71043452 | 0.90650187 |
| Zscan21 | 0.17932494 | 0.17038209 | 0.19693675 | -0.0265547 | 0.72555991 | 0.90650187 |
| Pgd | 0.48955931 | 0.4315461 | 0.45810652 | -0.0265604 | 0.60170025 | 0.90650187 |
| Mcu | 0.14863217 | 0.13952189 | 0.16608671 | -0.0265648 | 0.76147489 | 0.90650187 |
| Ubxn7 | 0.26752006 | 0.25335209 | 0.27996751 | -0.0266154 | 0.66371608 | 0.90650187 |
| Ptdss2 | 0.36987991 | 0.34043121 | 0.36705494 | -0.0266237 | 0.62644594 | 0.90650187 |
| Mtf1 | 0.22985528 | 0.21885004 | 0.24548249 | -0.0266325 | 0.6852053 | 0.90650187 |
| Zdhhc7 | 0.16389978 | 0.15493242 | 0.18156581 | -0.0266334 | 0.74283606 | 0.90650187 |
| Trim33 | 0.14868151 | 0.13949527 | 0.16613718 | -0.0266419 | 0.76204659 | 0.90650187 |
| Ubp1 | 0.17310403 | 0.16412435 | 0.19077018 | -0.0266458 | 0.73268053 | 0.90650187 |

| | | | | | | |
|---|---|---|---|---|---|---|
| Pdcl | 0.38952125 | 0.35609471 | 0.38274499 | -0.0266503 | 0.62154638 | 0.90650187 |
| Dgcr8 | 0.20405022 | 0.19437093 | 0.2210291 | -0.0266582 | 0.70408346 | 0.90650187 |
| Tbc1d10b | 0.1593004 | 0.15026384 | 0.17693071 | -0.0266669 | 0.74858342 | 0.90650187 |
| Ppm1g | 0.93145806 | 0.69333371 | 0.7200181 | -0.0266844 | 0.56543253 | 0.90650187 |
| Exoc2 | 0.12186306 | 0.11157697 | 0.13828596 | -0.026709 | 0.80472999 | 0.90882608 |
| Prdm4 | 0.11526888 | 0.10456684 | 0.13130478 | -0.0267379 | 0.81733895 | 0.9092537 |
| Med27 | 0.29465828 | 0.27722275 | 0.30396905 | -0.0267463 | 0.65216418 | 0.90650187 |
| Kdm2b | 0.24964613 | 0.23702421 | 0.26377711 | -0.0267529 | 0.67396341 | 0.90650187 |
| Sys1 | 1.09745774 | 0.76158551 | 0.78834418 | -0.0267587 | 0.55997139 | 0.90650187 |
| Cinp | 0.28553291 | 0.2691974 | 0.29597457 | -0.0267772 | 0.65623129 | 0.90650187 |
| Gtf3c2 | 0.34994617 | 0.32401238 | 0.35080856 | -0.0267962 | 0.63291112 | 0.90650187 |
| D1Ertd622e | 0.20317443 | 0.19339008 | 0.22018702 | -0.0267969 | 0.70575788 | 0.90650187 |
| Limk2 | 0.30807176 | 0.28878051 | 0.31558044 | -0.0267999 | 0.64711111 | 0.90650187 |
| Slc44a1 | 0.2220272 | 0.21133197 | 0.23813778 | -0.0268058 | 0.69160908 | 0.90650187 |
| Atp11a | 0.16033249 | 0.15113868 | 0.1779729 | -0.0268342 | 0.74866519 | 0.90650187 |
| Atf6 | 0.19660916 | 0.18700922 | 0.21384824 | -0.026839 | 0.71155927 | 0.90650187 |
| Zfp787 | 0.49322173 | 0.4338826 | 0.46073008 | -0.0268475 | 0.60220269 | 0.90650187 |
| Gtf2h5 | 1.45374263 | 0.85824225 | 0.88509349 | -0.0268512 | 0.55364155 | 0.90650187 |
| Zbtb11 | 0.24182606 | 0.22972568 | 0.25659483 | -0.0268691 | 0.6792199 | 0.90650187 |
| Mri1 | 0.30032867 | 0.28201735 | 0.30889763 | -0.0268803 | 0.65056888 | 0.90650187 |
| Timm44 | 0.63080472 | 0.52618642 | 0.55307282 | -0.0268864 | 0.58555025 | 0.90650187 |
| Isy1 | 0.24632232 | 0.23384448 | 0.26073204 | -0.0268876 | 0.67668391 | 0.90650187 |
| Psmd2 | 1.26480966 | 0.81446651 | 0.84137258 | -0.0269061 | 0.55652518 | 0.90650187 |
| Cep295 | 0.19422723 | 0.18461354 | 0.21153676 | -0.0269232 | 0.71423925 | 0.90650187 |
| Tubgcp3 | 0.17678371 | 0.16748648 | 0.19442293 | -0.0269365 | 0.73102614 | 0.90650187 |
| Ing5 | 0.13077306 | 0.12069245 | 0.1476342 | -0.0269417 | 0.79139704 | 0.90661829 |
| Cdc73 | 0.27126956 | 0.25638284 | 0.28332529 | -0.0269424 | 0.66376082 | 0.90650187 |
| Usp22 | 0.41604874 | 0.37649941 | 0.40345393 | -0.0269545 | 0.61676994 | 0.90650187 |
| Fxr2 | 0.25283486 | 0.23973119 | 0.26668801 | -0.0269568 | 0.67341468 | 0.90650187 |
| Gm15800 | 0.14879084 | 0.1392903 | 0.16624899 | -0.0269587 | 0.76451225 | 0.90650187 |

| | | | | | | |
|---|---|---|---|---|---|---|
| Acbd6 | 0.71119909 | 0.57502164 | 0.60198906 | -0.0269674 | 0.57892757 | 0.90650187 |
| Snapin | 0.36314135 | 0.33462574 | 0.361599 | -0.0269733 | 0.6299109 | 0.90650187 |
| Sgsm1 | 0.11943445 | 0.10874762 | 0.13572099 | -0.0269734 | 0.81152388 | 0.9092537 |
| Cc2d1b | 0.21020043 | 0.19993799 | 0.22691992 | -0.0269819 | 0.70146033 | 0.90650187 |
| Nhlrc3 | 0.07900346 | 0.06504812 | 0.09203279 | -0.0269847 | 0.90379515 | 0.93655998 |
| Pcyt1a | 0.16012402 | 0.15076749 | 0.17776249 | -0.026995 | 0.75019686 | 0.90650187 |
| Wdr89 | 2.41342014 | 0.8779199 | 0.90492396 | -0.0270041 | 0.55276982 | 0.90650187 |
| Rexo4 | 0.31677799 | 0.29601771 | 0.32303127 | -0.0270136 | 0.64496603 | 0.90650187 |
| Dennd5a | 0.21103303 | 0.20069343 | 0.22771438 | -0.0270209 | 0.70108465 | 0.90650187 |
| Pfdn6 | 1.04689497 | 0.74247787 | 0.76950312 | -0.0270252 | 0.56203529 | 0.90650187 |
| Gabpb2 | 0.16853938 | 0.1591903 | 0.18621729 | -0.027027 | 0.7406081 | 0.90650187 |
| Rundc1 | 0.17178462 | 0.16242727 | 0.18945655 | -0.0270293 | 0.73703901 | 0.90650187 |
| Lrp5 | 0.11365445 | 0.10255792 | 0.12958748 | -0.0270296 | 0.82310721 | 0.9092537 |
| Rrm2b | 0.16199867 | 0.15261898 | 0.17965283 | -0.0270339 | 0.74824183 | 0.90650187 |
| Ctdsp1 | 0.33836735 | 0.31418291 | 0.34121888 | -0.027036 | 0.63767123 | 0.90650187 |
| Srd5a3 | 0.29440084 | 0.27670391 | 0.30374456 | -0.0270406 | 0.65386154 | 0.90650187 |
| 2700097O09 | 0.1994546 | 0.18955667 | 0.21660137 | -0.0270447 | 0.71057844 | 0.90650187 |
| Ist1 | 0.46097409 | 0.41026667 | 0.43731472 | -0.0270481 | 0.60832734 | 0.90650187 |
| Arl6ip4 | 0.62127504 | 0.51998923 | 0.54704268 | -0.0270535 | 0.58700721 | 0.90650187 |
| Zfp120 | 0.16663245 | 0.15725082 | 0.1843084 | -0.0270576 | 0.74300637 | 0.90650187 |
| Exosc3 | 0.38675789 | 0.35349254 | 0.38055591 | -0.0270634 | 0.62405602 | 0.90650187 |
| Zfp212 | 0.16753921 | 0.15814687 | 0.1852166 | -0.0270697 | 0.74206768 | 0.90650187 |
| Fbxo18 | 0.44353707 | 0.39727652 | 0.42434876 | -0.0270722 | 0.61164804 | 0.90650187 |
| Pomgnt1 | 0.20690979 | 0.19669332 | 0.22377299 | -0.0270797 | 0.70470479 | 0.90650187 |
| Lypla1 | 0.48011766 | 0.42419063 | 0.45130154 | -0.0271109 | 0.60528839 | 0.90650187 |
| Zfp687 | 0.24765506 | 0.2348253 | 0.26195452 | -0.0271292 | 0.67738814 | 0.90650187 |
| Crk | 0.43597066 | 0.39152015 | 0.41865255 | -0.0271324 | 0.61337051 | 0.90650187 |
| Emd | 0.65968071 | 0.54390566 | 0.57103874 | -0.0271331 | 0.58364778 | 0.90650187 |
| Naglu | 0.2410392 | 0.22873501 | 0.25586876 | -0.0271337 | 0.6813356 | 0.90650187 |
| Pak1ip1 | 0.63838716 | 0.5306965 | 0.55783531 | -0.0271388 | 0.58561514 | 0.90650187 |

| | | | | | | |
|---|---|---|---|---|---|---|
| Egln1 | 0.15070753 | 0.14105776 | 0.1682066 | -0.0271488 | 0.763476 | 0.90650187 |
| Usp11 | 0.21480537 | 0.20415282 | 0.23130489 | -0.0271521 | 0.69911407 | 0.90650187 |
| Dip2c | 0.10151089 | 0.08940065 | 0.11656654 | -0.0271659 | 0.8499108 | 0.91471921 |
| Strip1 | 0.23512785 | 0.2232261 | 0.25039454 | -0.0271684 | 0.68522468 | 0.90650187 |
| Enpp5 | 0.23709885 | 0.2250464 | 0.25222364 | -0.0271772 | 0.68403656 | 0.90650187 |
| Xpc | 0.14053713 | 0.13056924 | 0.15776914 | -0.0271999 | 0.77829172 | 0.90650187 |
| Nfya | 0.14654741 | 0.13674285 | 0.16395193 | -0.0272091 | 0.76967668 | 0.90650187 |
| Zfp637 | 0.58232365 | 0.49463752 | 0.52185077 | -0.0272133 | 0.59166405 | 0.90650187 |
| Armc5 | 0.1788136 | 0.169195 | 0.19643149 | -0.0272365 | 0.73118865 | 0.90650187 |
| Repin1 | 0.16144936 | 0.15186209 | 0.17909934 | -0.0272372 | 0.75050615 | 0.90650187 |
| Klhl2 | 0.12385098 | 0.11313839 | 0.14038017 | -0.0272418 | 0.80584933 | 0.9092537 |
| Stx1a | 0.24549949 | 0.23272388 | 0.2599764 | -0.0272525 | 0.6793991 | 0.90650187 |
| Zfp512 | 0.40215873 | 0.3654235 | 0.39267848 | -0.027255 | 0.6211719 | 0.90650187 |
| Twf1 | 0.56038145 | 0.4799813 | 0.50726347 | -0.0272822 | 0.59448528 | 0.90650187 |
| Tbc1d10a | 0.25300415 | 0.23955926 | 0.26684227 | -0.027283 | 0.67527859 | 0.90650187 |
| Zfp827 | 0.14822147 | 0.13838044 | 0.16566656 | -0.0272861 | 0.76798093 | 0.90650187 |
| Eif1a | 0.85693053 | 0.65528626 | 0.68257837 | -0.0272921 | 0.57054132 | 0.90650187 |
| Mvp | 0.22571511 | 0.21430781 | 0.24160583 | -0.027298 | 0.69226834 | 0.90650187 |
| Uck1 | 0.45557472 | 0.40602398 | 0.43332284 | -0.0272989 | 0.61028745 | 0.90650187 |
| A130010J15I | 0.1254554 | 0.1147676 | 0.14206676 | -0.0272992 | 0.80351828 | 0.90882608 |
| Nbea | 0.37700364 | 0.34548016 | 0.372781 | -0.0273008 | 0.62762763 | 0.90650187 |
| Cacfd1 | 0.56219794 | 0.48117806 | 0.50848237 | -0.0273043 | 0.5943381 | 0.90650187 |
| Sccpdh | 0.4189862 | 0.37839452 | 0.40571418 | -0.0273197 | 0.61766532 | 0.90650187 |
| Tex10 | 0.26519728 | 0.25055505 | 0.27788075 | -0.0273257 | 0.66900363 | 0.90650187 |
| Pnpla8 | 0.48914901 | 0.43048549 | 0.45781204 | -0.0273265 | 0.60463231 | 0.90650187 |
| Inpp4a | 0.22939178 | 0.21770251 | 0.24504935 | -0.0273468 | 0.69009422 | 0.90650187 |
| Faap20 | 0.46393193 | 0.41214476 | 0.43949289 | -0.0273481 | 0.60896973 | 0.90650187 |
| Pnpla2 | 0.22665063 | 0.21509675 | 0.24248332 | -0.0273866 | 0.69219997 | 0.90650187 |
| Gtf2h1 | 0.33388745 | 0.31008732 | 0.33747789 | -0.0273906 | 0.64088182 | 0.90650187 |
| Mcts2 | 0.2072727 | 0.19671829 | 0.2241206 | -0.0274023 | 0.70662306 | 0.90650187 |

| | | | | | | |
|---|---|---|---|---|---|---|
| Dohh | 0.71487998 | 0.57671464 | 0.6041487 | -0.0274341 | 0.57999183 | 0.90650187 |
| Fbxl3 | 0.24918816 | 0.23592388 | 0.26335824 | -0.0274344 | 0.67835523 | 0.90650187 |
| Ino80d | 0.41498934 | 0.37518995 | 0.40263719 | -0.0274472 | 0.61907454 | 0.90650187 |
| Elk1 | 0.1050914 | 0.09297632 | 0.12042489 | -0.0274486 | 0.84453961 | 0.91320576 |
| Slc35c2 | 0.23737335 | 0.22501545 | 0.25247807 | -0.0274626 | 0.68565047 | 0.90650187 |
| Atp5e | 3.23250316 | 0.78105411 | 0.80854932 | -0.0274952 | 0.56008139 | 0.90650187 |
| Rab23 | 0.24433689 | 0.23140598 | 0.2589076 | -0.0275016 | 0.68161519 | 0.90650187 |
| Vcpkmt | 0.14430195 | 0.13414372 | 0.16164695 | -0.0275032 | 0.77529301 | 0.90650187 |
| Trmt1 | 0.34482208 | 0.31907602 | 0.34657933 | -0.0275033 | 0.63787615 | 0.90650187 |
| Trappc11 | 0.1669312 | 0.15709787 | 0.18460772 | -0.0275099 | 0.74616651 | 0.90650187 |
| Lzic | 0.20083035 | 0.19040148 | 0.21792919 | -0.0275277 | 0.71278749 | 0.90650187 |
| Med31 | 0.57149888 | 0.48714934 | 0.51469135 | -0.027542 | 0.59401756 | 0.90650187 |
| Mast2 | 0.11044395 | 0.09861087 | 0.12616287 | -0.027552 | 0.83418875 | 0.91062493 |
| Ptrh2 | 0.38260195 | 0.34966309 | 0.37725244 | -0.0275894 | 0.62745397 | 0.90650187 |
| Asxl1 | 0.09028618 | 0.07682463 | 0.1044156 | -0.027591 | 0.8799456 | 0.92579928 |
| Gin1 | 0.19939878 | 0.18892281 | 0.21654745 | -0.0276246 | 0.71468349 | 0.90650187 |
| Arl6 | 0.38491059 | 0.35145916 | 0.3790892 | -0.02763 | 0.62703835 | 0.90650187 |
| Rsl1d1 | 0.93909892 | 0.69607261 | 0.72370414 | -0.0276315 | 0.56739126 | 0.90650187 |
| Gga1 | 0.38770955 | 0.35366063 | 0.38131047 | -0.0276498 | 0.62641108 | 0.90650187 |
| Cops6 | 1.40113983 | 0.84693584 | 0.87459037 | -0.0276545 | 0.55589518 | 0.90650187 |
| Osbpl8 | 0.1461832 | 0.13591959 | 0.16357847 | -0.0276589 | 0.77388159 | 0.90650187 |
| Zfp398 | 0.28137307 | 0.26464286 | 0.292305 | -0.0276621 | 0.66301573 | 0.90650187 |
| Rhbdd3 | 0.11663608 | 0.10508852 | 0.13275659 | -0.0276681 | 0.82291151 | 0.9092537 |
| Rab3gap1 | 0.28640308 | 0.26906046 | 0.29674018 | -0.0276797 | 0.66081302 | 0.90650187 |
| Arpin | 0.22305438 | 0.21142303 | 0.23910529 | -0.0276823 | 0.6966142 | 0.90650187 |
| Terf1 | 0.24479867 | 0.23163225 | 0.25933228 | -0.0277 | 0.68255238 | 0.90650187 |
| Bop1 | 0.16954164 | 0.15951365 | 0.18721896 | -0.0277053 | 0.74469119 | 0.90650187 |
| Enoph1 | 0.14891537 | 0.13865336 | 0.16637632 | -0.027723 | 0.77058134 | 0.90650187 |
| Zcchc11 | 0.7071199 | 0.57185841 | 0.59958797 | -0.0277296 | 0.58144792 | 0.90650187 |
| Dhdds | 0.21850598 | 0.2070802 | 0.23481286 | -0.0277327 | 0.70022581 | 0.90650187 |

| | | | | | | |
|---|---|---|---|---|---|---|
| Lsm10 | 0.4353286 | 0.39041765 | 0.41816728 | -0.0277496 | 0.61600725 | 0.90650187 |
| Rbm26 | 0.32168405 | 0.29944331 | 0.3272006 | -0.0277573 | 0.64696245 | 0.90650187 |
| Snapc1 | 0.1825543 | 0.17235923 | 0.20012105 | -0.0277618 | 0.73128939 | 0.90650187 |
| Dcaf10 | 0.22623657 | 0.21430404 | 0.24209506 | -0.027791 | 0.69508919 | 0.90650187 |
| Atxn2 | 0.32047777 | 0.29837611 | 0.32617751 | -0.0278014 | 0.64762513 | 0.90650187 |
| Usp7 | 0.30619771 | 0.28611899 | 0.3139679 | -0.0278489 | 0.6533293 | 0.90650187 |
| Champ1 | 0.14985714 | 0.13948713 | 0.1673386 | -0.0278515 | 0.77032479 | 0.90650187 |
| Asb7 | 0.18922891 | 0.17880629 | 0.20666658 | -0.0278603 | 0.72551455 | 0.90650187 |
| Gps2 | 0.67281305 | 0.55119625 | 0.57905984 | -0.0278636 | 0.58469398 | 0.90650187 |
| Ofd1 | 0.1350549 | 0.12422676 | 0.15209258 | -0.0278658 | 0.79232274 | 0.90681274 |
| Med10 | 0.76628071 | 0.60573584 | 0.63360772 | -0.0278719 | 0.57752214 | 0.90650187 |
| Cdc34 | 0.19758305 | 0.18691501 | 0.21479158 | -0.0278766 | 0.71802008 | 0.90650187 |
| Phc2 | 0.23466799 | 0.22207584 | 0.24996723 | -0.0278914 | 0.69006743 | 0.90650187 |
| Rnf38 | 0.10547331 | 0.09294274 | 0.12083549 | -0.0278927 | 0.84759898 | 0.9143112 |
| Senp2 | 0.14500835 | 0.13447047 | 0.16237271 | -0.0279022 | 0.77754732 | 0.90650187 |
| Cic | 0.24582209 | 0.23236929 | 0.26027274 | -0.0279034 | 0.6831789 | 0.90650187 |
| Pick1 | 0.41934025 | 0.37805079 | 0.40598618 | -0.0279354 | 0.62015826 | 0.90650187 |
| Herc2 | 0.38671571 | 0.35258651 | 0.38052246 | -0.027936 | 0.62792891 | 0.90650187 |
| Slc29a4 | 0.21717583 | 0.20560694 | 0.23355357 | -0.0279466 | 0.70262104 | 0.90650187 |
| Ptdss1 | 0.36534911 | 0.33543489 | 0.36339056 | -0.0279557 | 0.63382491 | 0.90650187 |
| Ankrd49 | 0.10496431 | 0.09232967 | 0.12028822 | -0.0279585 | 0.84926608 | 0.91448046 |
| Lamtor1 | 0.93881367 | 0.69560116 | 0.72356856 | -0.0279674 | 0.56821509 | 0.90650187 |
| Cstf2 | 0.18809818 | 0.17759096 | 0.20556111 | -0.0279701 | 0.72737784 | 0.90650187 |
| Ythdc1 | 0.74195273 | 0.59183308 | 0.61982376 | -0.0279907 | 0.57955671 | 0.90650187 |
| Tmem184c | 0.37067619 | 0.33968842 | 0.36769725 | -0.0280088 | 0.63255876 | 0.90650187 |
| Dnajc8 | 1.26127115 | 0.81235567 | 0.8403751 | -0.0280194 | 0.55891681 | 0.90650187 |
| 1110038F14I | 0.57979962 | 0.49216442 | 0.52018756 | -0.0280231 | 0.59463727 | 0.90650187 |
| Cdc27 | 0.2248482 | 0.21275842 | 0.2407919 | -0.0280335 | 0.69761962 | 0.90650187 |
| Fam98a | 0.23358065 | 0.22090891 | 0.24895601 | -0.0280471 | 0.69175705 | 0.90650187 |
| Fgd1 | 0.09494944 | 0.08142691 | 0.10948138 | -0.0280545 | 0.87268624 | 0.92289575 |

| | | | | | | |
|---|---|---|---|---|---|---|
| Map3k10 | 0.12520323 | 0.11373706 | 0.14180189 | -0.0280648 | 0.81053125 | 0.9092537 |
| Cep162 | 0.2254772 | 0.21330601 | 0.24138254 | -0.0280765 | 0.69745431 | 0.90650187 |
| Cherp | 0.23085391 | 0.21832514 | 0.24641497 | -0.0280898 | 0.69384275 | 0.90650187 |
| Asun | 0.22314107 | 0.21109451 | 0.23918687 | -0.0280924 | 0.69921148 | 0.90650187 |
| Cuedc1 | 0.37621009 | 0.34404935 | 0.37214517 | -0.0280958 | 0.63142492 | 0.90650187 |
| Ap4b1 | 0.11952897 | 0.10770743 | 0.13582096 | -0.0281135 | 0.8212644 | 0.9092537 |
| Atg13 | 0.13651549 | 0.1254654 | 0.15360841 | -0.028143 | 0.79231833 | 0.90681274 |
| Mthfd2l | 0.29563071 | 0.27666553 | 0.30481653 | -0.028151 | 0.65930572 | 0.90650187 |
| Triap1 | 0.50938783 | 0.44404401 | 0.47219938 | -0.0281554 | 0.60452371 | 0.90650187 |
| Uchl5 | 0.27645268 | 0.25977531 | 0.28794388 | -0.0281686 | 0.66818199 | 0.90650187 |
| Rela | 0.2251937 | 0.21294432 | 0.24111637 | -0.0281721 | 0.69826778 | 0.90650187 |
| Chmp3 | 0.66444192 | 0.54576813 | 0.57395795 | -0.0281898 | 0.58641985 | 0.90650187 |
| Trim8 | 0.24088299 | 0.227517 | 0.25572455 | -0.0282076 | 0.68806399 | 0.90650187 |
| Pdik1l | 0.19528287 | 0.18435105 | 0.21256192 | -0.0282109 | 0.72240927 | 0.90650187 |
| Tbcb | 1.28813165 | 0.8194907 | 0.84770648 | -0.0282158 | 0.55881731 | 0.90650187 |
| Ring1 | 0.31189474 | 0.2906343 | 0.31886035 | -0.0282261 | 0.65303691 | 0.90650187 |
| Parp4 | 0.10104114 | 0.08781517 | 0.11605914 | -0.028244 | 0.86034611 | 0.91840739 |
| Opa3 | 0.27649027 | 0.25972974 | 0.28797728 | -0.0282475 | 0.66860602 | 0.90650187 |
| Tchp | 0.13934408 | 0.12828885 | 0.15653682 | -0.028248 | 0.78879157 | 0.90650187 |
| Amn1 | 0.42055522 | 0.37866132 | 0.40691862 | -0.0282573 | 0.62122983 | 0.90650187 |
| Braf | 0.25460973 | 0.24004448 | 0.26830397 | -0.0282595 | 0.68019336 | 0.90650187 |
| Raf1 | 0.60533348 | 0.50857403 | 0.53683878 | -0.0282647 | 0.59253186 | 0.90650187 |
| Tle2 | 0.31739972 | 0.29526979 | 0.32356081 | -0.028291 | 0.65125401 | 0.90650187 |
| Cmc1 | 0.46985525 | 0.41553859 | 0.44383645 | -0.0282979 | 0.61158035 | 0.90650187 |
| Narfl | 0.14411398 | 0.13314414 | 0.16145373 | -0.0283096 | 0.78215756 | 0.90650187 |
| Vprbp | 0.18984637 | 0.17895394 | 0.20726966 | -0.0283157 | 0.72818317 | 0.90650187 |
| Gm5914 | 0.21088939 | 0.19924986 | 0.22757737 | -0.0283275 | 0.70999268 | 0.90650187 |
| Taf3 | 0.33522665 | 0.31026254 | 0.33859798 | -0.0283354 | 0.64506433 | 0.90650187 |
| Dhx30 | 0.56880233 | 0.48456034 | 0.51289709 | -0.0283367 | 0.59700894 | 0.90650187 |
| Nr2c2ap | 0.28309119 | 0.26546528 | 0.29382255 | -0.0283573 | 0.66605746 | 0.90650187 |

| | | | | | | |
|---|---|---|---|---|---|---|
| Fbxw11 | 0.21261039 | 0.20085939 | 0.2292175 | -0.0283581 | 0.70883937 | 0.90650187 |
| Smyd4 | 0.10012611 | 0.08670089 | 0.11506997 | -0.0283691 | 0.86346272 | 0.9197321 |
| Kif1bp | 0.49256888 | 0.43187106 | 0.46026307 | -0.028392 | 0.60804752 | 0.90650187 |
| Zfand3 | 0.41241315 | 0.37225215 | 0.40064758 | -0.0283954 | 0.62364846 | 0.90650187 |
| Gmfb | 0.35948604 | 0.33022814 | 0.35862409 | -0.028396 | 0.63758248 | 0.90650187 |
| Abhd13 | 0.18476856 | 0.17390139 | 0.20229784 | -0.0283965 | 0.7336953 | 0.90650187 |
| Hsd17b4 | 0.40674138 | 0.36785205 | 0.39624966 | -0.0283976 | 0.62498452 | 0.90650187 |
| Csnk2b | 1.00768069 | 0.72547338 | 0.75387235 | -0.028399 | 0.56649968 | 0.90650187 |
| Dhps | 0.40229385 | 0.36438471 | 0.392784 | -0.0283993 | 0.62605726 | 0.90650187 |
| Lman2l | 0.66524789 | 0.54603603 | 0.57445094 | -0.0284149 | 0.5870253 | 0.90650187 |
| Phf3 | 0.54619758 | 0.46925355 | 0.49767689 | -0.0284233 | 0.60021268 | 0.90650187 |
| Pef1 | 0.34082665 | 0.31483312 | 0.34326537 | -0.0284322 | 0.64364692 | 0.90650187 |
| Cnep1r1 | 0.42320629 | 0.38050627 | 0.40894899 | -0.0284427 | 0.62141356 | 0.90650187 |
| Ndufc1 | 1.87186862 | 0.89804713 | 0.92649356 | -0.0284464 | 0.55428497 | 0.90650187 |
| Setx | 0.15215033 | 0.14122637 | 0.16967735 | -0.028451 | 0.77199278 | 0.90650187 |
| Rlf | 0.2040311 | 0.19255305 | 0.22101072 | -0.0284577 | 0.71648224 | 0.90650187 |
| Mtcl1 | 0.13504848 | 0.12362805 | 0.15208592 | -0.0284579 | 0.79725286 | 0.9075779 |
| B4galt6 | 0.41485506 | 0.3740339 | 0.4025336 | -0.0284997 | 0.62352522 | 0.90650187 |
| Gmpr2 | 0.31754926 | 0.29518687 | 0.32368813 | -0.0285013 | 0.65226498 | 0.90650187 |
| Htt | 0.17444201 | 0.16359235 | 0.19210019 | -0.0285078 | 0.74528703 | 0.90650187 |
| Cdc40 | 0.30910848 | 0.28795754 | 0.31647117 | -0.0285136 | 0.65562176 | 0.90650187 |
| Ptrhd1 | 0.41089005 | 0.37093894 | 0.39946894 | -0.02853 | 0.62456954 | 0.90650187 |
| Brd7 | 0.57035769 | 0.48538556 | 0.51393254 | -0.028547 | 0.59752128 | 0.90650187 |
| Clasp1 | 0.32995258 | 0.30559799 | 0.33417799 | -0.02858 | 0.64810258 | 0.90650187 |
| Akap9 | 0.97662216 | 0.71228781 | 0.74087453 | -0.0285867 | 0.56809814 | 0.90650187 |
| Gas2l1 | 0.1732675 | 0.16234333 | 0.1909328 | -0.0285895 | 0.74718776 | 0.90650187 |
| Pex12 | 0.14375925 | 0.13248973 | 0.16108899 | -0.0285993 | 0.78501988 | 0.90650187 |
| 3110082I17R | 0.19806336 | 0.18665682 | 0.21525645 | -0.0285996 | 0.72262482 | 0.90650187 |
| Wdr35 | 0.15245726 | 0.14138588 | 0.16998991 | -0.028604 | 0.77278732 | 0.90650187 |
| Emsy | 0.46099841 | 0.40871697 | 0.43733265 | -0.0286157 | 0.61443084 | 0.90650187 |

| | | | | | | |
|---|---|---|---|---|---|---|
| Araf | 0.7279869 | 0.58313756 | 0.6117818 | -0.0286442 | 0.5824432 | 0.90650187 |
| Abhd18 | 0.17104062 | 0.16005446 | 0.18871495 | -0.0286605 | 0.75021372 | 0.90650187 |
| Msh3 | 0.23453095 | 0.2211695 | 0.24983984 | -0.0286703 | 0.69502855 | 0.90650187 |
| Prr3 | 0.27135171 | 0.25471775 | 0.28339871 | -0.028681 | 0.67361272 | 0.90650187 |
| Maea | 0.53807449 | 0.4634413 | 0.49212929 | -0.028688 | 0.60224042 | 0.90650187 |
| Akirin2 | 0.18409613 | 0.17294646 | 0.20163736 | -0.0286909 | 0.73649062 | 0.90650187 |
| Zkscan3 | 0.56561127 | 0.48206989 | 0.51076743 | -0.0286975 | 0.59862011 | 0.90650187 |
| Sp4 | 0.10763902 | 0.09441658 | 0.12316042 | -0.0287438 | 0.85025776 | 0.91483353 |
| Scaf8 | 0.17094841 | 0.15987053 | 0.188623 | -0.0287525 | 0.75100615 | 0.90650187 |
| Mdm2 | 0.4697361 | 0.41499605 | 0.44374936 | -0.0287533 | 0.61334913 | 0.90650187 |
| Ampd2 | 0.24422237 | 0.23002396 | 0.25880225 | -0.0287783 | 0.68946787 | 0.90650187 |
| Suds3 | 0.32025949 | 0.29720881 | 0.32599222 | -0.0287834 | 0.65266439 | 0.90650187 |
| Osbpl2 | 0.25799967 | 0.24258799 | 0.27138195 | -0.028794 | 0.68142393 | 0.90650187 |
| Msl3l2 | 0.16451969 | 0.15338785 | 0.1821887 | -0.0288009 | 0.75890132 | 0.90650187 |
| Ubfd1 | 0.62264236 | 0.51909428 | 0.54791096 | -0.0288167 | 0.59243416 | 0.90650187 |
| Synj1 | 0.15043847 | 0.13911346 | 0.16793205 | -0.0288186 | 0.77723978 | 0.90650187 |
| Cry2 | 0.27255943 | 0.25562988 | 0.28447719 | -0.0288473 | 0.67393503 | 0.90650187 |
| Atrn | 0.15554618 | 0.14426464 | 0.17312954 | -0.0288649 | 0.77071108 | 0.90650187 |
| Erf | 0.26892818 | 0.25236225 | 0.28123007 | -0.0288678 | 0.67592522 | 0.90650187 |
| Ado | 0.33847315 | 0.31241966 | 0.34130703 | -0.0288874 | 0.64664007 | 0.90650187 |
| Anapc4 | 0.44447158 | 0.39615862 | 0.42504942 | -0.0288908 | 0.61873873 | 0.90650187 |
| Wdr70 | 0.35345435 | 0.32479052 | 0.35369132 | -0.0289008 | 0.64179434 | 0.90650187 |
| Mob2 | 0.32726719 | 0.30300233 | 0.33191837 | -0.028916 | 0.65072995 | 0.90650187 |
| Phb2 | 1.71229374 | 0.89016526 | 0.91908875 | -0.0289235 | 0.55563134 | 0.90650187 |
| Dnajc5 | 0.64197651 | 0.53113722 | 0.56007877 | -0.0289415 | 0.59084632 | 0.90650187 |
| Rab33b | 0.25363733 | 0.23847562 | 0.26741901 | -0.0289434 | 0.68479807 | 0.90650187 |
| Zfp330 | 0.55763249 | 0.47646469 | 0.50541507 | -0.0289504 | 0.60050179 | 0.90650187 |
| Prkrir | 0.16024304 | 0.14891635 | 0.17788263 | -0.0289663 | 0.76544233 | 0.90650187 |
| Armcx4 | 0.1548276 | 0.14342349 | 0.17240015 | -0.0289767 | 0.77253108 | 0.90650187 |
| Ncoa1 | 0.18837931 | 0.17684158 | 0.20583608 | -0.0289945 | 0.73441385 | 0.90650187 |

| | | | | | | |
|---|---|---|---|---|---|---|
| Eif2b3 | 0.33661423 | 0.31075292 | 0.33975694 | -0.029004 | 0.64784464 | 0.90650187 |
| Dph6 | 0.27763212 | 0.25998252 | 0.2889913 | -0.0290088 | 0.67229075 | 0.90650187 |
| Gsk3a | 0.16732482 | 0.15597395 | 0.18500195 | -0.029028 | 0.75726892 | 0.90650187 |
| Tmem131 | 0.20985006 | 0.19755586 | 0.22658539 | -0.0290295 | 0.71551174 | 0.90650187 |
| Cacul1 | 0.2520084 | 0.23689834 | 0.26593451 | -0.0290362 | 0.68630202 | 0.90650187 |
| Zfhx2 | 0.28948423 | 0.27039625 | 0.29944551 | -0.0290493 | 0.6668636 | 0.90650187 |
| Rai1 | 0.17501182 | 0.16360661 | 0.19266594 | -0.0290593 | 0.74873684 | 0.90650187 |
| Fam195a | 0.12612118 | 0.11369898 | 0.14276567 | -0.0290667 | 0.81729722 | 0.9092537 |
| Wasf1 | 0.15435611 | 0.14284891 | 0.17192123 | -0.0290723 | 0.77390337 | 0.90650187 |
| Cdk5 | 0.38766796 | 0.35220017 | 0.38127751 | -0.0290773 | 0.63270831 | 0.90650187 |
| Tyw1 | 0.21746676 | 0.20474783 | 0.23382916 | -0.0290813 | 0.70983408 | 0.90650187 |
| Spns1 | 0.37243028 | 0.34002659 | 0.36910976 | -0.0290832 | 0.63693629 | 0.90650187 |
| Nlk | 0.21122375 | 0.19881239 | 0.22789626 | -0.0290839 | 0.71475953 | 0.90650187 |
| Mtfp1 | 0.22695938 | 0.21366307 | 0.24277272 | -0.0291097 | 0.70300063 | 0.90650187 |
| Rbm34 | 0.31727829 | 0.29432225 | 0.32345741 | -0.0291352 | 0.65558167 | 0.90650187 |
| Med19 | 0.39626499 | 0.35891267 | 0.38806202 | -0.0291494 | 0.63078479 | 0.90650187 |
| Msantd4 | 0.48214538 | 0.42361854 | 0.45276806 | -0.0291495 | 0.61264183 | 0.90650187 |
| Rpf1 | 0.48845577 | 0.4281634 | 0.45731423 | -0.0291508 | 0.61155674 | 0.90650187 |
| Usp40 | 0.25417845 | 0.23874872 | 0.26791158 | -0.0291629 | 0.68577716 | 0.90650187 |
| Surf6 | 0.10134047 | 0.08719423 | 0.11638249 | -0.0291883 | 0.86756196 | 0.92056229 |
| Rfc4 | 0.20672703 | 0.1943984 | 0.22359787 | -0.0291995 | 0.71922752 | 0.90650187 |
| Dgkd | 0.19599926 | 0.18405347 | 0.21325696 | -0.0292035 | 0.72866588 | 0.90650187 |
| Trpc1 | 0.13577067 | 0.12362662 | 0.15283575 | -0.0292091 | 0.80223178 | 0.9086659 |
| Pde12 | 0.19123244 | 0.17940376 | 0.20862196 | -0.0292182 | 0.7332335 | 0.90650187 |
| Mapk14 | 0.27018112 | 0.25311834 | 0.28235192 | -0.0292336 | 0.67734242 | 0.90650187 |
| Ttc39a | 0.21927437 | 0.20630364 | 0.2355395 | -0.0292359 | 0.70945791 | 0.90650187 |
| Nbeal1 | 0.38203877 | 0.34754683 | 0.37680374 | -0.0292569 | 0.63502007 | 0.90650187 |
| Cdk11b | 0.6897652 | 0.56001655 | 0.58928004 | -0.0292635 | 0.58736332 | 0.90650187 |
| Pkd1 | 0.18815544 | 0.17634495 | 0.20561712 | -0.0292722 | 0.73659655 | 0.90650187 |
| Rpusd4 | 0.16704035 | 0.15543823 | 0.18471706 | -0.0292788 | 0.7594881 | 0.90650187 |

| | | | | | | |
|---|---|---|---|---|---|---|
| Smarca5 | 0.6687012 | 0.54725789 | 0.57655864 | -0.0293008 | 0.58937053 | 0.90650187 |
| Senp3 | 0.26989801 | 0.25279437 | 0.28209856 | -0.0293042 | 0.67788986 | 0.90650187 |
| Rab36 | 0.15053283 | 0.13869313 | 0.16802835 | -0.0293352 | 0.78116561 | 0.90650187 |
| Foxp4 | 0.31849569 | 0.29515556 | 0.32449345 | -0.0293379 | 0.65613344 | 0.90650187 |
| Napg | 0.3222068 | 0.2982828 | 0.32764347 | -0.0293607 | 0.65482426 | 0.90650187 |
| Rab5b | 0.25054994 | 0.23523598 | 0.26460318 | -0.0293672 | 0.68913478 | 0.90650187 |
| Fat1 | 0.30020227 | 0.27941463 | 0.3087881 | -0.0293735 | 0.66381163 | 0.90650187 |
| Luc7l3 | 1.23212199 | 0.80259249 | 0.83197602 | -0.0293835 | 0.56238108 | 0.90650187 |
| BC005624 | 0.63090765 | 0.52375353 | 0.55313768 | -0.0293842 | 0.59334671 | 0.90650187 |
| Adck4 | 0.21503915 | 0.20214055 | 0.23152692 | -0.0293864 | 0.71370967 | 0.90650187 |
| Dmrtc1a | 0.17503511 | 0.16329919 | 0.19268906 | -0.0293899 | 0.75113495 | 0.90650187 |
| Casd1 | 0.28058528 | 0.26221528 | 0.29160826 | -0.029393 | 0.67295944 | 0.90650187 |
| Sep-15 | 2.39053998 | 0.87746164 | 0.90686049 | -0.0293988 | 0.55729669 | 0.90650187 |
| Mrpl19 | 0.47565546 | 0.41866287 | 0.44806448 | -0.0294016 | 0.61474848 | 0.90650187 |
| Leo1 | 0.42284457 | 0.37926635 | 0.40867227 | -0.0294059 | 0.62547133 | 0.90650187 |
| Fer | 0.19893258 | 0.18666312 | 0.21609701 | -0.0294339 | 0.72758386 | 0.90650187 |
| Ncbp3 | 0.36592556 | 0.33441789 | 0.36385769 | -0.0294398 | 0.64046238 | 0.90650187 |
| Nudcd3 | 0.86031936 | 0.65487101 | 0.68433893 | -0.0294679 | 0.57590551 | 0.90650187 |
| Mrps11 | 0.66861752 | 0.54703408 | 0.57650765 | -0.0294736 | 0.58989647 | 0.90650187 |
| Gnl1 | 0.41575924 | 0.3736913 | 0.40323082 | -0.0295395 | 0.62766383 | 0.90650187 |
| Kifap3 | 0.44153429 | 0.39330451 | 0.42284501 | -0.0295405 | 0.62193845 | 0.90650187 |
| Rnf113a2 | 0.22288478 | 0.20940461 | 0.23894562 | -0.029541 | 0.70870774 | 0.90650187 |
| Rab1b | 0.57676089 | 0.48863771 | 0.51818023 | -0.0295425 | 0.60004099 | 0.90650187 |
| Gaa | 0.69041676 | 0.56011998 | 0.58966995 | -0.02955 | 0.5881472 | 0.90650187 |
| Crbn | 0.2504192 | 0.23492964 | 0.26448373 | -0.0295541 | 0.69032206 | 0.90650187 |
| Ino80e | 0.46300333 | 0.40924023 | 0.43880972 | -0.0295695 | 0.61774749 | 0.90650187 |
| Snrnp48 | 0.39560407 | 0.35796113 | 0.38754268 | -0.0295816 | 0.63282243 | 0.90650187 |
| Katnb1 | 0.20752984 | 0.19476955 | 0.22436682 | -0.0295973 | 0.72121024 | 0.90650187 |
| Usp20 | 0.14408706 | 0.13181694 | 0.16142606 | -0.0296091 | 0.79258478 | 0.90681274 |
| Mpc1 | 0.91460214 | 0.68218893 | 0.7118042 | -0.0296153 | 0.5733712 | 0.90650187 |

| | | | | | | |
|---|---|---|---|---|---|---|
| Coq6 | 0.2447435 | 0.22965308 | 0.25928155 | -0.0296285 | 0.69427536 | 0.90650187 |
| 4930402H24I | 0.33174434 | 0.30603852 | 0.33568225 | -0.0296437 | 0.65268755 | 0.90650187 |
| Nudt16l1 | 0.69999955 | 0.56570248 | 0.59537711 | -0.0296746 | 0.58767831 | 0.90650187 |
| Brpf1 | 0.14995799 | 0.13776533 | 0.16744158 | -0.0296763 | 0.78462538 | 0.90650187 |
| Rbm12 | 0.16462736 | 0.1525998 | 0.18229685 | -0.0296971 | 0.76553193 | 0.90650187 |
| Osbp | 0.18759093 | 0.17534549 | 0.20506474 | -0.0297192 | 0.74030459 | 0.90650187 |
| Wipf2 | 0.14781174 | 0.13552204 | 0.1652472 | -0.0297252 | 0.78805446 | 0.90650187 |
| 1300002E11I | 0.27189429 | 0.25415057 | 0.28388346 | -0.0297329 | 0.67925488 | 0.90650187 |
| Tcof1 | 0.26270573 | 0.24589538 | 0.27563669 | -0.0297413 | 0.6842735 | 0.90650187 |
| Ctu2 | 0.25052754 | 0.23484008 | 0.26458271 | -0.0297426 | 0.69137285 | 0.90650187 |
| Ccdc32 | 0.28311926 | 0.26409777 | 0.29384733 | -0.0297496 | 0.67367362 | 0.90650187 |
| Rbsn | 0.19313524 | 0.18072226 | 0.21047505 | -0.0297528 | 0.73513943 | 0.90650187 |
| Fam136a | 0.62905652 | 0.52215649 | 0.55197035 | -0.0298139 | 0.59488246 | 0.90650187 |
| Cyld | 0.18708363 | 0.17474933 | 0.20456805 | -0.0298187 | 0.74151098 | 0.90650187 |
| Cry1 | 0.32939259 | 0.30386974 | 0.3337073 | -0.0298376 | 0.65449774 | 0.90650187 |
| Pcm1 | 1.00834983 | 0.72429071 | 0.75414571 | -0.029855 | 0.56984989 | 0.90650187 |
| Amdhd2 | 0.1954858 | 0.18288944 | 0.21275886 | -0.0298694 | 0.73372657 | 0.90650187 |
| Clptm1 | 0.72814743 | 0.58199896 | 0.61187477 | -0.0298758 | 0.58592081 | 0.90650187 |
| Picalm | 0.5187032 | 0.44881785 | 0.47873078 | -0.0299129 | 0.60940881 | 0.90650187 |
| Nutf2 | 0.49669453 | 0.43329538 | 0.4632086 | -0.0299132 | 0.61297817 | 0.90650187 |
| Rbm8a | 1.51572694 | 0.86605349 | 0.89596874 | -0.0299153 | 0.55899952 | 0.90650187 |
| Armc10 | 0.39039393 | 0.35350171 | 0.38343508 | -0.0299334 | 0.63572394 | 0.90650187 |
| Tasp1 | 0.16476065 | 0.1524933 | 0.18243071 | -0.0299374 | 0.76716333 | 0.90650187 |
| Eef2kmt | 0.35621596 | 0.32601571 | 0.35595349 | -0.0299378 | 0.64576124 | 0.90650187 |
| Ube2f | 0.43719831 | 0.38963372 | 0.41957956 | -0.0299458 | 0.62448733 | 0.90650187 |
| Pa2g4 | 1.20456706 | 0.79385676 | 0.82380592 | -0.0299492 | 0.56419682 | 0.90650187 |
| Fam213b | 0.36349536 | 0.3319275 | 0.36188654 | -0.029959 | 0.64357555 | 0.90650187 |
| Wdr83 | 0.37204126 | 0.3388158 | 0.36879671 | -0.0299809 | 0.64109998 | 0.90650187 |
| Rchy1 | 0.82031668 | 0.63325679 | 0.66324204 | -0.0299852 | 0.57964537 | 0.90650187 |
| Fntb | 0.161054 | 0.14870468 | 0.17870075 | -0.0299961 | 0.77223384 | 0.90650187 |

| | | | | | | |
|---|---|---|---|---|---|---|
| Paxbp1 | 0.16556244 | 0.15323781 | 0.1832355 | -0.0299977 | 0.76663057 | 0.90650187 |
| Tmco1 | 1.41505448 | 0.8475016 | 0.87749935 | -0.0299977 | 0.56039677 | 0.90650187 |
| Cops8 | 0.86665168 | 0.6576014 | 0.68761329 | -0.0300119 | 0.57692576 | 0.90650187 |
| Igf2bp3 | 0.58048423 | 0.49061419 | 0.52063906 | -0.0300249 | 0.60116928 | 0.90650187 |
| Eif2b5 | 0.38115871 | 0.3460752 | 0.37610207 | -0.0300269 | 0.63867673 | 0.90650187 |
| Mapk6 | 0.21583305 | 0.20225045 | 0.23228048 | -0.03003 | 0.7172676 | 0.90650187 |
| Creb1 | 0.3254284 | 0.30033328 | 0.33036755 | -0.0300343 | 0.65695158 | 0.90650187 |
| Zfp598 | 0.11004705 | 0.09569979 | 0.12573862 | -0.0300388 | 0.85589374 | 0.9165432 |
| Katnal1 | 0.17285624 | 0.16048171 | 0.19052362 | -0.0300419 | 0.75834426 | 0.90650187 |
| Creld1 | 0.33068959 | 0.30475362 | 0.33479707 | -0.0300434 | 0.65502843 | 0.90650187 |
| Trim24 | 0.27312732 | 0.25493858 | 0.28498375 | -0.0300452 | 0.68035433 | 0.90650187 |
| Smad5 | 0.32606086 | 0.3008394 | 0.33090129 | -0.0300619 | 0.65684832 | 0.90650187 |
| Numa1 | 0.36222762 | 0.33078485 | 0.36085637 | -0.0300715 | 0.64448352 | 0.90650187 |
| Nr1h2 | 0.30756612 | 0.28505594 | 0.31514566 | -0.0300897 | 0.6643852 | 0.90650187 |
| Med18 | 0.13585424 | 0.12278626 | 0.15292247 | -0.0301362 | 0.80950922 | 0.9092537 |
| Yy1 | 0.60772053 | 0.50823209 | 0.53837598 | -0.0301439 | 0.59828532 | 0.90650187 |
| Cbfb | 0.64472007 | 0.53160587 | 0.56178815 | -0.0301823 | 0.5943859 | 0.90650187 |
| Prpf3 | 0.23716211 | 0.22208572 | 0.25228228 | -0.0301966 | 0.70267521 | 0.90650187 |
| Utp18 | 0.38071329 | 0.34554738 | 0.3757467 | -0.0301993 | 0.63956641 | 0.90650187 |
| Ppp1r35 | 0.50734203 | 0.44055788 | 0.47075745 | -0.0301996 | 0.61225076 | 0.90650187 |
| Uimc1 | 0.42387695 | 0.37924351 | 0.4094618 | -0.0302183 | 0.62857663 | 0.90650187 |
| Pyurf | 0.42498147 | 0.38006373 | 0.41030564 | -0.0302419 | 0.62841827 | 0.90650187 |
| Psma2 | 1.94236538 | 0.89705092 | 0.92729723 | -0.0302463 | 0.55764675 | 0.90650187 |
| Gspt1 | 0.91355815 | 0.68103651 | 0.71129005 | -0.0302535 | 0.57498749 | 0.90650187 |
| Rprd1a | 0.25762377 | 0.24078431 | 0.27104119 | -0.0302569 | 0.69014877 | 0.90650187 |
| Zfyve27 | 0.23163017 | 0.21687616 | 0.24713913 | -0.030263 | 0.70690946 | 0.90650187 |
| Ipo13 | 0.18543131 | 0.17266842 | 0.20294835 | -0.0302799 | 0.74642607 | 0.90650187 |
| Rpl7l1 | 0.67823719 | 0.55205704 | 0.58234595 | -0.0302889 | 0.59142974 | 0.90650187 |
| Irf2bp1 | 0.42983322 | 0.38369834 | 0.41400169 | -0.0303034 | 0.6275614 | 0.90650187 |
| Prpf39 | 0.27391395 | 0.25537052 | 0.28568491 | -0.0303144 | 0.68143989 | 0.90650187 |

| | | | | | | |
|---|---|---|---|---|---|---|
| Atxn7l1 | 0.16195378 | 0.149292 | 0.17960761 | -0.0303156 | 0.77348253 | 0.90650187 |
| B4galt5 | 0.22780465 | 0.21322245 | 0.24356454 | -0.0303421 | 0.71014629 | 0.90650187 |
| Limd1 | 0.51245625 | 0.44401234 | 0.47435699 | -0.0303447 | 0.61194263 | 0.90650187 |
| Pcnxl3 | 0.15819323 | 0.14546454 | 0.17581137 | -0.0303468 | 0.77856531 | 0.90650187 |
| Alg5 | 0.51251686 | 0.44404648 | 0.47439955 | -0.0303531 | 0.61196306 | 0.90650187 |
| Scarb1 | 0.29592837 | 0.27471262 | 0.30507576 | -0.0303631 | 0.67092036 | 0.90650187 |
| Sh3glb2 | 0.38205474 | 0.34645175 | 0.37681647 | -0.0303647 | 0.63991701 | 0.90650187 |
| Pdhx | 0.17940559 | 0.16661833 | 0.19701641 | -0.0303981 | 0.75361888 | 0.90650187 |
| Epc2 | 0.25311465 | 0.23652858 | 0.26694296 | -0.0304144 | 0.69375207 | 0.90650187 |
| Msh2 | 0.32278758 | 0.29770063 | 0.32813523 | -0.0304346 | 0.6599515 | 0.90650187 |
| Camk2g | 0.25979195 | 0.24254834 | 0.27300484 | -0.0304565 | 0.6900363 | 0.90650187 |
| Rad1 | 0.2309325 | 0.2160251 | 0.24648832 | -0.0304632 | 0.70864399 | 0.90650187 |
| Tmem65 | 0.21690143 | 0.20281965 | 0.23329356 | -0.0304739 | 0.71928078 | 0.90650187 |
| Zfp358 | 0.35749136 | 0.32650366 | 0.35699612 | -0.0304925 | 0.64792222 | 0.90650187 |
| Misp | 0.17568823 | 0.16283452 | 0.19333708 | -0.0305026 | 0.75846637 | 0.90650187 |
| Metap1 | 0.41555211 | 0.37254741 | 0.40307116 | -0.0305237 | 0.6318121 | 0.90650187 |
| Cep170b | 0.25695408 | 0.23990374 | 0.27043377 | -0.03053 | 0.69212294 | 0.90650187 |
| Fgfr1op2 | 0.77683965 | 0.60896674 | 0.63950277 | -0.030536 | 0.58405419 | 0.90650187 |
| Atg16l1 | 0.2197598 | 0.2054588 | 0.23599824 | -0.0305394 | 0.71745084 | 0.90650187 |
| Nomo1 | 0.24360351 | 0.22769109 | 0.25823271 | -0.0305416 | 0.70048273 | 0.90650187 |
| Hmgxb4 | 0.33380105 | 0.30681761 | 0.33740557 | -0.030588 | 0.65653419 | 0.90650187 |
| Zkscan1 | 0.39420662 | 0.35585347 | 0.3864431 | -0.0305896 | 0.63754351 | 0.90650187 |
| Gga2 | 0.43955455 | 0.39075413 | 0.42135573 | -0.0306016 | 0.62660273 | 0.90650187 |
| Ddx54 | 0.52298256 | 0.45109904 | 0.48171113 | -0.0306121 | 0.61122463 | 0.90650187 |
| Rpl15 | 3.96811536 | 0.65885283 | 0.6894878 | -0.030635 | 0.57829162 | 0.90650187 |
| Alg3 | 0.17102991 | 0.15806534 | 0.18870427 | -0.0306389 | 0.76479437 | 0.90650187 |
| Pkig | 0.80770504 | 0.62580146 | 0.65644377 | -0.0306423 | 0.58219719 | 0.90650187 |
| Cyfip2 | 0.16952308 | 0.15655656 | 0.18720042 | -0.0306439 | 0.76660919 | 0.90650187 |
| Tulp3 | 0.34610421 | 0.31699557 | 0.34763985 | -0.0306443 | 0.65242578 | 0.90650187 |
| 1700088E04I | 0.17130358 | 0.15832283 | 0.18897713 | -0.0306543 | 0.76458505 | 0.90650187 |

| | | | | | | |
|---|---|---|---|---|---|---|
| Mrps2 | 0.19156343 | 0.17827427 | 0.20894459 | -0.0306703 | 0.74297889 | 0.90650187 |
| Zfp638 | 0.64076434 | 0.52863924 | 0.55932198 | -0.0306827 | 0.59633531 | 0.90650187 |
| Setd2 | 0.49881402 | 0.43402936 | 0.46471721 | -0.0306879 | 0.61545601 | 0.90650187 |
| Pacrgl | 0.19417667 | 0.18079653 | 0.21148763 | -0.0306911 | 0.74058398 | 0.90650187 |
| Nubp2 | 0.26216746 | 0.24444401 | 0.27515111 | -0.0307071 | 0.69009991 | 0.90650187 |
| Vps9d1 | 0.24269953 | 0.22669124 | 0.2574001 | -0.0307089 | 0.70207525 | 0.90650187 |
| Sppl2b | 0.18268108 | 0.16953663 | 0.20024583 | -0.0307092 | 0.75230688 | 0.90650187 |
| Mydgf | 1.45785221 | 0.85515339 | 0.88586626 | -0.0307129 | 0.56124583 | 0.90650187 |
| Ilf3 | 0.51210424 | 0.44338909 | 0.47410978 | -0.0307207 | 0.61334964 | 0.90650187 |
| Ercc6l2 | 0.16922659 | 0.15617955 | 0.18690422 | -0.0307247 | 0.76754965 | 0.90650187 |
| Tmem245 | 0.34046792 | 0.31223421 | 0.34296717 | -0.030733 | 0.65482001 | 0.90650187 |
| 1600020E01I | 0.20177892 | 0.18810495 | 0.21884355 | -0.0307386 | 0.73382625 | 0.90650187 |
| Cdc26 | 1.19808875 | 0.79111562 | 0.82186557 | -0.03075 | 0.56605198 | 0.90650187 |
| Rnf25 | 0.16793656 | 0.15486429 | 0.18561429 | -0.03075 | 0.76928264 | 0.90650187 |
| Rrp9 | 0.20792679 | 0.19399083 | 0.22474677 | -0.0307559 | 0.7285284 | 0.90650187 |
| Ppfia1 | 0.17333183 | 0.16021753 | 0.19099678 | -0.0307793 | 0.7631316 | 0.90650187 |
| Trak1 | 0.21299582 | 0.19879635 | 0.2295844 | -0.030788 | 0.72446948 | 0.90650187 |
| N6amt2 | 0.52361416 | 0.45135169 | 0.48215002 | -0.0307983 | 0.61178421 | 0.90650187 |
| Ankrd39 | 0.33671418 | 0.30903766 | 0.33984036 | -0.0308027 | 0.65650531 | 0.90650187 |
| Dtd2 | 0.33647267 | 0.30881041 | 0.33963878 | -0.0308284 | 0.65671685 | 0.90650187 |
| Mtfr1 | 0.28078122 | 0.26094605 | 0.2917816 | -0.0308356 | 0.68075563 | 0.90650187 |
| Rif1 | 0.37279306 | 0.33856504 | 0.36940159 | -0.0308366 | 0.64472074 | 0.90650187 |
| Sec16a | 0.24193485 | 0.22585142 | 0.25669517 | -0.0308437 | 0.70338846 | 0.90650187 |
| Dus1l | 0.2629769 | 0.24503211 | 0.27588121 | -0.0308491 | 0.69044382 | 0.90650187 |
| Akap8 | 0.38904357 | 0.35151681 | 0.38236701 | -0.0308502 | 0.64008321 | 0.90650187 |
| Zranb2 | 0.9361213 | 0.69141035 | 0.72227136 | -0.030861 | 0.5753261 | 0.90650187 |
| Hps4 | 0.20590424 | 0.19194038 | 0.22280908 | -0.0308687 | 0.73102272 | 0.90650187 |
| Zfand1 | 0.18625681 | 0.17287275 | 0.20375792 | -0.0308852 | 0.74979823 | 0.90650187 |
| Fam222a | 0.20866301 | 0.19456322 | 0.22545105 | -0.0308878 | 0.72876045 | 0.90650187 |
| Sdr39u1 | 0.19660717 | 0.18293275 | 0.21384631 | -0.0309136 | 0.73977615 | 0.90650187 |

| | | | | | | |
|---|---|---|---|---|---|---|
| Anapc2 | 0.52635775 | 0.45311134 | 0.48405355 | -0.0309422 | 0.61186264 | 0.90650187 |
| Prdm2 | 0.25697773 | 0.23950146 | 0.27045522 | -0.0309538 | 0.69455454 | 0.90650187 |
| Foxj3 | 0.22670681 | 0.21153716 | 0.24253599 | -0.0309988 | 0.71504988 | 0.90650187 |
| Gtf2h3 | 0.35523174 | 0.32414284 | 0.35514798 | -0.0310051 | 0.65103287 | 0.90650187 |
| Mrpl51 | 1.03667097 | 0.73449995 | 0.7655133 | -0.0310134 | 0.57146512 | 0.90650187 |
| Dnpep | 0.97309259 | 0.70832544 | 0.7393512 | -0.0310258 | 0.57399451 | 0.90650187 |
| Ptbp2 | 0.29631945 | 0.27437826 | 0.30541625 | -0.031038 | 0.67428831 | 0.90650187 |
| Prmt6 | 0.23336542 | 0.21768109 | 0.24875571 | -0.0310746 | 0.71067107 | 0.90650187 |
| Exog | 0.11467027 | 0.09958894 | 0.1306684 | -0.0310795 | 0.85483244 | 0.91627438 |
| Psmg1 | 0.44747042 | 0.39620257 | 0.42729261 | -0.03109 | 0.62682836 | 0.90650187 |
| Usp38 | 0.08990798 | 0.07290276 | 0.10400343 | -0.0311007 | 0.90814065 | 0.93875746 |
| Zranb1 | 0.36030764 | 0.32817643 | 0.3592937 | -0.0311173 | 0.64988857 | 0.90650187 |
| Irak1bp1 | 0.30086536 | 0.27821402 | 0.30936255 | -0.0311485 | 0.67278306 | 0.90650187 |
| Atad3a | 0.33770388 | 0.30951398 | 0.34066591 | -0.0311519 | 0.65782158 | 0.90650187 |
| Gtf2ird1 | 0.23159545 | 0.21595074 | 0.24710674 | -0.031156 | 0.71243629 | 0.90650187 |
| Dhrs1 | 0.43453223 | 0.3863095 | 0.41756498 | -0.0312555 | 0.63034287 | 0.90650187 |
| Acyp1 | 0.56902666 | 0.48179093 | 0.51304652 | -0.0312556 | 0.60674034 | 0.90650187 |
| 9430016H08 | 0.34190868 | 0.31284958 | 0.34416416 | -0.0313146 | 0.65707519 | 0.90650187 |
| Dctn6 | 0.46222159 | 0.4069165 | 0.43823413 | -0.0313176 | 0.62464277 | 0.90650187 |
| Nacc2 | 0.22894925 | 0.21331736 | 0.24463561 | -0.0313182 | 0.71536485 | 0.90650187 |
| Vps51 | 0.17468788 | 0.16100843 | 0.19234435 | -0.0313359 | 0.76554667 | 0.90650187 |
| Sec24a | 0.27827136 | 0.25821951 | 0.28955846 | -0.031339 | 0.68479455 | 0.90650187 |
| Scfd1 | 0.66525752 | 0.54310191 | 0.57445683 | -0.0313549 | 0.59586152 | 0.90650187 |
| Eif4e | 1.18618019 | 0.78681438 | 0.81818165 | -0.0313673 | 0.56766584 | 0.90650187 |
| Ncstn | 0.50664838 | 0.43889772 | 0.47026792 | -0.0313702 | 0.61659628 | 0.90650187 |
| Rxrb | 0.20752081 | 0.19298729 | 0.22435818 | -0.0313709 | 0.73289986 | 0.90650187 |
| Stx8 | 0.45327246 | 0.40024319 | 0.4316145 | -0.0313713 | 0.62669889 | 0.90650187 |
| Bsdc1 | 0.30813619 | 0.28423852 | 0.31563583 | -0.0313973 | 0.67083453 | 0.90650187 |
| Zdhhc8 | 0.12759191 | 0.11291033 | 0.14430771 | -0.0313974 | 0.83328828 | 0.91056108 |
| C330006A16 | 0.33260156 | 0.30498704 | 0.33640095 | -0.0314139 | 0.66098072 | 0.90650187 |

| | | | | | | |
|---|---|---|---|---|---|---|
| Taok2 | 0.38312691 | 0.34624937 | 0.37767047 | -0.0314211 | 0.64425723 | 0.90650187 |
| Dctn5 | 0.28937454 | 0.26791148 | 0.29934935 | -0.0314379 | 0.67970734 | 0.90650187 |
| Sacm1l | 0.42860436 | 0.38160651 | 0.41306718 | -0.0314607 | 0.63254193 | 0.90650187 |
| Bscl2 | 0.35294372 | 0.32181017 | 0.35327235 | -0.0314622 | 0.65391892 | 0.90650187 |
| Zer1 | 0.21755667 | 0.20244736 | 0.23391432 | -0.031467 | 0.72509279 | 0.90650187 |
| Rfxap | 0.22560614 | 0.21001303 | 0.24150356 | -0.0314905 | 0.71893439 | 0.90650187 |
| Tulp4 | 0.49486842 | 0.43041258 | 0.46190643 | -0.0314938 | 0.61909763 | 0.90650187 |
| Rprd2 | 0.30902343 | 0.28489085 | 0.31639812 | -0.0315073 | 0.67100806 | 0.90650187 |
| Eri3 | 0.73323842 | 0.58330082 | 0.61481682 | -0.031516 | 0.5901323 | 0.90650187 |
| Gm9803 | 0.23525904 | 0.21897947 | 0.2505164 | -0.0315369 | 0.71213989 | 0.90650187 |
| Tom1l1 | 0.1892654 | 0.17515538 | 0.20670223 | -0.0315469 | 0.7512687 | 0.90650187 |
| Hacd3 | 0.34438362 | 0.31464214 | 0.3462163 | -0.0315742 | 0.65741908 | 0.90650187 |
| Cpsf4 | 0.16887361 | 0.15495943 | 0.18655145 | -0.031592 | 0.77423063 | 0.90650187 |
| Tmem127 | 0.22589639 | 0.21016706 | 0.24177593 | -0.0316089 | 0.71944775 | 0.90650187 |
| Gnai3 | 0.96678027 | 0.70498786 | 0.73661428 | -0.0316264 | 0.57568683 | 0.90650187 |
| Prpf38b | 0.83277756 | 0.63825704 | 0.66988957 | -0.0316325 | 0.58313643 | 0.90650187 |
| Dnajb12 | 0.30604779 | 0.28219907 | 0.31383876 | -0.0316397 | 0.6729899 | 0.90650187 |
| Rbm10 | 0.47868712 | 0.4185921 | 0.45026524 | -0.0316731 | 0.62275279 | 0.90650187 |
| 2310035C23I | 0.22819067 | 0.21222607 | 0.2439259 | -0.0316998 | 0.71827911 | 0.90650187 |
| Ormdl3 | 0.52913549 | 0.45425147 | 0.48597591 | -0.0317244 | 0.61417169 | 0.90650187 |
| Pym1 | 0.33314428 | 0.30512971 | 0.33685565 | -0.0317259 | 0.66228182 | 0.90650187 |
| Wash1 | 0.45227941 | 0.39914612 | 0.43087648 | -0.0317304 | 0.62831004 | 0.90650187 |
| Dopey1 | 0.24306988 | 0.22599598 | 0.25774131 | -0.0317453 | 0.70800084 | 0.90650187 |
| Dcaf8 | 0.87638503 | 0.66086 | 0.69260829 | -0.0317483 | 0.58073768 | 0.90650187 |
| Eif4enif1 | 0.22064197 | 0.20507659 | 0.23683128 | -0.0317547 | 0.72443694 | 0.90650187 |
| Grsf1 | 0.29121117 | 0.26920051 | 0.30095803 | -0.0317575 | 0.68050336 | 0.90650187 |
| Mpi | 0.11576222 | 0.10004718 | 0.13182892 | -0.0317817 | 0.85808634 | 0.91737198 |
| D030056L22I | 0.29880552 | 0.27579124 | 0.30757675 | -0.0317855 | 0.6770325 | 0.90650187 |
| Sgsm3 | 0.16136905 | 0.14722489 | 0.17901838 | -0.0317935 | 0.78510046 | 0.90650187 |
| Dlat | 0.5165247 | 0.44539915 | 0.47720842 | -0.0318093 | 0.6165112 | 0.90650187 |

| | | | | | | |
|---|---|---|---|---|---|---|
| 0610009O20 | 0.29505596 | 0.27248209 | 0.30431574 | -0.0318336 | 0.67904963 | 0.90650187 |
| Znrd1as | 0.12462781 | 0.10935724 | 0.14119723 | -0.03184 | 0.84194522 | 0.91251845 |
| Rps7 | 4.61002214 | 0.55994499 | 0.59179732 | -0.0318523 | 0.59455423 | 0.90650187 |
| Kidins220 | 0.26216942 | 0.24328336 | 0.27515288 | -0.0318695 | 0.69669249 | 0.90650187 |
| Tarbp2 | 0.30462626 | 0.28073054 | 0.31261332 | -0.0318828 | 0.67486897 | 0.90650187 |
| Dolk | 0.18644444 | 0.17205734 | 0.20394183 | -0.0318845 | 0.75648201 | 0.90650187 |
| Eaf1 | 0.15957536 | 0.14528483 | 0.17720849 | -0.0319237 | 0.78839502 | 0.90650187 |
| Ssbp4 | 0.86820777 | 0.65647692 | 0.6884144 | -0.0319375 | 0.58169991 | 0.90650187 |
| Zmiz2 | 0.22443184 | 0.20846095 | 0.24040072 | -0.0319398 | 0.72262106 | 0.90650187 |
| Mar-08 | 0.30904451 | 0.28443019 | 0.31641623 | -0.031986 | 0.67342829 | 0.90650187 |
| Mzt1 | 0.4935688 | 0.42899143 | 0.46097824 | -0.0319868 | 0.62113997 | 0.90650187 |
| Supt7l | 0.18368742 | 0.16924179 | 0.20123567 | -0.0319939 | 0.76015278 | 0.90650187 |
| Src | 0.13368399 | 0.11866671 | 0.15066753 | -0.0320008 | 0.82747097 | 0.90947782 |
| Slc7a4 | 0.21223131 | 0.19681157 | 0.2288565 | -0.0320449 | 0.73318762 | 0.90650187 |
| Tmem68 | 0.17762207 | 0.16319481 | 0.19525303 | -0.0320582 | 0.76727908 | 0.90650187 |
| Zfp386 | 0.2270914 | 0.210837 | 0.24289644 | -0.0320594 | 0.72131996 | 0.90650187 |
| Golga5 | 0.18419654 | 0.1696577 | 0.20173602 | -0.0320783 | 0.76018629 | 0.90650187 |
| Fcf1 | 0.58361198 | 0.49061539 | 0.52269828 | -0.0320829 | 0.60752285 | 0.90650187 |
| Timm9 | 0.36382477 | 0.33005725 | 0.362154 | -0.0320968 | 0.65321035 | 0.90650187 |
| Zzz3 | 0.40060643 | 0.35936023 | 0.39146517 | -0.0321049 | 0.64229283 | 0.90650187 |
| Usp50 | 0.92503725 | 0.68477552 | 0.71690207 | -0.0321266 | 0.57895511 | 0.90650187 |
| Stk19 | 0.24741059 | 0.22959346 | 0.26173043 | -0.032137 | 0.70741911 | 0.90650187 |
| Xrcc1 | 0.44769525 | 0.39530564 | 0.42746053 | -0.0321549 | 0.63096572 | 0.90650187 |
| 1110051M20 | 0.30448119 | 0.28031558 | 0.31248817 | -0.0321726 | 0.67641697 | 0.90650187 |
| Ldah | 0.26337695 | 0.24406789 | 0.27624181 | -0.0321739 | 0.69769422 | 0.90650187 |
| Cdc5l | 0.84784234 | 0.64564978 | 0.67783376 | -0.032184 | 0.58358765 | 0.90650187 |
| Ppwd1 | 0.2668733 | 0.24719039 | 0.27938698 | -0.0321966 | 0.69578374 | 0.90650187 |
| Rce1 | 0.40135197 | 0.35984797 | 0.39204813 | -0.0322002 | 0.64249349 | 0.90650187 |
| Nat6 | 0.14346447 | 0.1285751 | 0.16078577 | -0.0322107 | 0.81342835 | 0.9092537 |
| Rcbtb1 | 0.27292945 | 0.2525814 | 0.28480728 | -0.0322259 | 0.69252464 | 0.90650187 |

| | | | | | | |
|---|---|---|---|---|---|---|
| Slain2 | 0.37134468 | 0.33599888 | 0.36823585 | -0.032237 | 0.65143136 | 0.90650187 |
| Rmi1 | 0.16870054 | 0.15413778 | 0.18637844 | -0.0322407 | 0.77906009 | 0.90650187 |
| Ipo8 | 0.19103854 | 0.17618249 | 0.20843291 | -0.0322504 | 0.75422955 | 0.90650187 |
| Aimp1 | 1.17263696 | 0.78157405 | 0.81384441 | -0.0322704 | 0.56996148 | 0.90650187 |
| Zfp740 | 0.17589993 | 0.16126276 | 0.19354702 | -0.0322843 | 0.77081736 | 0.90650187 |
| Rexo1 | 0.43870402 | 0.38842461 | 0.42071505 | -0.0322904 | 0.63352582 | 0.90650187 |
| Scmh1 | 0.28222268 | 0.26075796 | 0.29305577 | -0.0322978 | 0.68791617 | 0.90650187 |
| Pvrl3 | 0.23186898 | 0.2150388 | 0.24736179 | -0.032323 | 0.71934992 | 0.90650187 |
| Zfp553 | 0.29949297 | 0.27584429 | 0.30817317 | -0.0323289 | 0.6795232 | 0.90650187 |
| Zmpste24 | 0.38799287 | 0.34918335 | 0.38153497 | -0.0323516 | 0.64689753 | 0.90650187 |
| Tfg | 1.0481446 | 0.73760211 | 0.76998526 | -0.0323831 | 0.5741572 | 0.90650187 |
| Tmem101 | 0.37981562 | 0.34263797 | 0.37503005 | -0.0323921 | 0.64950179 | 0.90650187 |
| Irf2bpl | 0.76471651 | 0.60032905 | 0.63272998 | -0.0324009 | 0.59004133 | 0.90650187 |
| Oscp1 | 0.11323548 | 0.09673412 | 0.12914129 | -0.0324072 | 0.86770247 | 0.92062353 |
| Zmym4 | 0.23847583 | 0.22108015 | 0.2534992 | -0.0324191 | 0.71516303 | 0.90650187 |
| Mterf4 | 0.23216529 | 0.21520575 | 0.24763797 | -0.0324322 | 0.71979225 | 0.90650187 |
| Map3k4 | 0.1790117 | 0.16417816 | 0.19662727 | -0.0324491 | 0.76841953 | 0.90650187 |
| Chuk | 0.38918338 | 0.35002219 | 0.38247766 | -0.0324555 | 0.64700078 | 0.90650187 |
| Ccnl1 | 0.95038037 | 0.69658053 | 0.72904141 | -0.0324609 | 0.57845496 | 0.90650187 |
| Lrrc45 | 0.20048086 | 0.18512761 | 0.21759207 | -0.0324645 | 0.746424 | 0.90650187 |
| Crcp | 0.22575062 | 0.2091694 | 0.24163915 | -0.0324697 | 0.72486945 | 0.90650187 |
| Tsen34 | 1.54897219 | 0.86885282 | 0.90132864 | -0.0324758 | 0.56363031 | 0.90650187 |
| Mcat | 0.22194331 | 0.20558194 | 0.23805871 | -0.0324768 | 0.72790349 | 0.90650187 |
| Luc7l | 0.78253039 | 0.61017957 | 0.64265907 | -0.0324795 | 0.58888494 | 0.90650187 |
| Dmwd | 0.13739185 | 0.12200933 | 0.15451671 | -0.0325074 | 0.82517569 | 0.90934433 |
| 1810026J23F | 0.28894181 | 0.26645734 | 0.29896988 | -0.0325125 | 0.68561299 | 0.90650187 |
| Tmem9 | 0.9041147 | 0.67408311 | 0.70660187 | -0.0325188 | 0.58105484 | 0.90650187 |
| Oxa1l | 0.54700888 | 0.46567362 | 0.49822856 | -0.0325549 | 0.61427624 | 0.90650187 |
| Ndufaf3 | 0.62283597 | 0.51543641 | 0.54803383 | -0.0325974 | 0.60427479 | 0.90650187 |
| Tlk2 | 0.20880579 | 0.19296607 | 0.22558756 | -0.0326215 | 0.73984412 | 0.90650187 |

| | | | | | | |
|---|---|---|---|---|---|---|
| Ankrd13a | 0.17700736 | 0.16198783 | 0.19464445 | -0.0326566 | 0.77212549 | 0.90650187 |
| Ccdc91 | 0.25106917 | 0.23240524 | 0.26507738 | -0.0326721 | 0.70813472 | 0.90650187 |
| Sf3b3 | 0.44411246 | 0.39210622 | 0.42478024 | -0.032674 | 0.63380772 | 0.90650187 |
| Prpsap2 | 0.20062876 | 0.18505745 | 0.21773475 | -0.0326773 | 0.74767459 | 0.90650187 |
| Fam20b | 0.28935309 | 0.26664856 | 0.29933055 | -0.032682 | 0.68629918 | 0.90650187 |
| Rbmx2 | 0.25049098 | 0.23186442 | 0.26454931 | -0.0326849 | 0.70858363 | 0.90650187 |
| Dhx32 | 0.25334689 | 0.23445147 | 0.26715451 | -0.032703 | 0.70684726 | 0.90650187 |
| Trmu | 0.13934558 | 0.12381828 | 0.15653837 | -0.0327201 | 0.82361587 | 0.9092537 |
| Mrps5 | 0.63624015 | 0.52376876 | 0.55648995 | -0.0327212 | 0.6031073 | 0.90650187 |
| Mettl10 | 0.44771822 | 0.39475289 | 0.42747768 | -0.0327248 | 0.63319431 | 0.90650187 |
| Klc2 | 0.2831693 | 0.26116666 | 0.29389148 | -0.0327248 | 0.68970639 | 0.90650187 |
| Taf15 | 0.37355609 | 0.33727698 | 0.37001504 | -0.0327381 | 0.65296462 | 0.90650187 |
| Ssh1 | 0.15007008 | 0.13481324 | 0.16755602 | -0.0327428 | 0.80750351 | 0.9092537 |
| Trim28 | 0.6636051 | 0.54070029 | 0.57344574 | -0.0327454 | 0.60019707 | 0.90650187 |
| Cdk9 | 0.36180382 | 0.3277631 | 0.36051169 | -0.0327486 | 0.65683307 | 0.90650187 |
| Isoc1 | 0.3557656 | 0.32281732 | 0.355585 | -0.0327677 | 0.65897566 | 0.90650187 |
| Mrps9 | 0.40068316 | 0.358754 | 0.39152519 | -0.0327712 | 0.64509202 | 0.90650187 |
| Top3b | 0.2233707 | 0.20662134 | 0.23940294 | -0.0327816 | 0.72865156 | 0.90650187 |
| Ankmy2 | 0.25489039 | 0.23577524 | 0.26855922 | -0.032784 | 0.7063287 | 0.90650187 |
| Ints3 | 0.26058144 | 0.24089701 | 0.27371875 | -0.0328217 | 0.70300904 | 0.90650187 |
| Stau2 | 0.24849709 | 0.22990271 | 0.26272577 | -0.0328231 | 0.71069015 | 0.90650187 |
| Ppp1r10 | 0.39275557 | 0.35247653 | 0.38529962 | -0.0328231 | 0.64755005 | 0.90650187 |
| 1110059G10 | 0.39184573 | 0.35175861 | 0.3845818 | -0.0328232 | 0.6478129 | 0.90650187 |
| Rnf181 | 0.48027593 | 0.41858057 | 0.45141611 | -0.0328355 | 0.626792 | 0.90650187 |
| Hsf1 | 0.25086521 | 0.23202411 | 0.26489114 | -0.032867 | 0.70938869 | 0.90650187 |
| AI413582 | 0.28816411 | 0.26541524 | 0.29828747 | -0.0328722 | 0.68790472 | 0.90650187 |
| Tceanc2 | 0.17404868 | 0.15879655 | 0.19170945 | -0.0329129 | 0.77733599 | 0.90650187 |
| Psmd1 | 0.90738189 | 0.67530337 | 0.70822775 | -0.0329244 | 0.58186588 | 0.90650187 |
| Ddx17 | 0.81740998 | 0.62874475 | 0.66168192 | -0.0329372 | 0.58756791 | 0.90650187 |
| Smim13 | 0.13744724 | 0.12163035 | 0.15457408 | -0.0329437 | 0.82830127 | 0.9096066 |

| | | | | | | |
|---|---|---|---|---|---|---|
| Zc2hc1a | 0.31314373 | 0.28698394 | 0.31992914 | -0.0329452 | 0.67644947 | 0.90650187 |
| Adprm | 0.20211531 | 0.18621281 | 0.21916759 | -0.0329548 | 0.74807884 | 0.90650187 |
| 2810403A07 | 0.41633847 | 0.37070823 | 0.40367715 | -0.0329689 | 0.64172719 | 0.90650187 |
| Fam50a | 0.45932508 | 0.4031223 | 0.43609773 | -0.0329754 | 0.63162243 | 0.90650187 |
| Taf1 | 0.29144875 | 0.26817423 | 0.3011659 | -0.0329917 | 0.68687107 | 0.90650187 |
| Mrps14 | 1.21547569 | 0.79399983 | 0.8270406 | -0.0330408 | 0.57048328 | 0.90650187 |
| Eef1d | 2.17006736 | 0.88887803 | 0.92192136 | -0.0330433 | 0.56329895 | 0.90650187 |
| Fam102a | 0.20539415 | 0.18925206 | 0.22231973 | -0.0330677 | 0.74576957 | 0.90650187 |
| Ufsp2 | 0.69148159 | 0.55723541 | 0.59030656 | -0.0330711 | 0.59834252 | 0.90650187 |
| Tax1bp1 | 1.1860325 | 0.78504847 | 0.81813469 | -0.0330862 | 0.5713393 | 0.90650187 |
| Med12 | 0.15970917 | 0.14423868 | 0.17734363 | -0.0331049 | 0.79669291 | 0.9075779 |
| Fam168b | 0.22789733 | 0.21054172 | 0.24365132 | -0.0331096 | 0.72711477 | 0.90650187 |
| Tor2a | 0.74673006 | 0.58943693 | 0.62255231 | -0.0331154 | 0.59346786 | 0.90650187 |
| 2410016O06 | 0.16395116 | 0.14850139 | 0.18161745 | -0.0331161 | 0.79120526 | 0.90660917 |
| Cdk7 | 0.3642124 | 0.32935075 | 0.36246863 | -0.0331179 | 0.65769639 | 0.90650187 |
| Tmem80 | 0.25744034 | 0.23774658 | 0.27087486 | -0.0331283 | 0.70667681 | 0.90650187 |
| S100pbp | 0.22234854 | 0.20528597 | 0.23844057 | -0.0331546 | 0.73176209 | 0.90650187 |
| Immt | 0.83810157 | 0.63955372 | 0.67270989 | -0.0331562 | 0.58671887 | 0.90650187 |
| Zfp617 | 0.20472825 | 0.18851362 | 0.22168047 | -0.0331669 | 0.74701696 | 0.90650187 |
| Sh3yl1 | 0.14892383 | 0.13320538 | 0.16638497 | -0.0331796 | 0.81233098 | 0.9092537 |
| Fbxl5 | 0.3007285 | 0.27605537 | 0.30924402 | -0.0331887 | 0.68335841 | 0.90650187 |
| Zfp322a | 0.47255789 | 0.41259324 | 0.4458094 | -0.0332162 | 0.62976505 | 0.90650187 |
| Eif4g3 | 0.57983554 | 0.48698624 | 0.52021126 | -0.033225 | 0.61176928 | 0.90650187 |
| Csnk1e | 0.40828007 | 0.36421683 | 0.39744516 | -0.0332283 | 0.64493226 | 0.90650187 |
| Dicer1 | 0.35294107 | 0.3200386 | 0.35327018 | -0.0332316 | 0.66209638 | 0.90650187 |
| Ttc19 | 0.19226528 | 0.17638322 | 0.2096283 | -0.0332451 | 0.75960468 | 0.90650187 |
| Map2k2 | 1.3545098 | 0.83092943 | 0.86418313 | -0.0332537 | 0.5679143 | 0.90650187 |
| Yars2 | 0.17145248 | 0.15585713 | 0.18912556 | -0.0332684 | 0.7828961 | 0.90650187 |
| Surf2 | 0.25452649 | 0.23495928 | 0.26822825 | -0.033269 | 0.70931987 | 0.90650187 |
| Uqcc3 | 0.88044412 | 0.66140169 | 0.69468017 | -0.0332785 | 0.58432313 | 0.90650187 |

| | | | | | | |
|---|---|---|---|---|---|---|
| Cul5 | 0.60600321 | 0.50397299 | 0.5372704 | -0.0332974 | 0.60854049 | 0.90650187 |
| Actr1a | 0.45379726 | 0.39870115 | 0.43200425 | -0.0333031 | 0.63409191 | 0.90650187 |
| Ppm1d | 0.22880649 | 0.21118326 | 0.24450209 | -0.0333188 | 0.72768033 | 0.90650187 |
| Fbxw2 | 0.6833511 | 0.55209669 | 0.5854303 | -0.0333336 | 0.5999149 | 0.90650187 |
| Stat3 | 0.65614896 | 0.53552847 | 0.56886565 | -0.0333372 | 0.60277077 | 0.90650187 |
| Dnajc18 | 0.29748834 | 0.27309186 | 0.30643281 | -0.033341 | 0.68569968 | 0.90650187 |
| 2310061I04R | 0.26292356 | 0.24248816 | 0.27583312 | -0.033345 | 0.70451164 | 0.90650187 |
| 2610301B20 | 0.21309398 | 0.19632477 | 0.22967781 | -0.033353 | 0.740723 | 0.90650187 |
| Sft2d3 | 0.12973076 | 0.11318109 | 0.14654561 | -0.0333645 | 0.84426582 | 0.91310457 |
| Aup1 | 0.36161435 | 0.32699148 | 0.36035755 | -0.0333661 | 0.65969307 | 0.90650187 |
| Rgl2 | 0.35202359 | 0.31913987 | 0.35251686 | -0.033377 | 0.66309277 | 0.90650187 |
| Tbc1d19 | 0.23088214 | 0.21305357 | 0.24644132 | -0.0333877 | 0.72650895 | 0.90650187 |
| Trim25 | 0.17513045 | 0.15938202 | 0.19278369 | -0.0334017 | 0.77942272 | 0.90650187 |
| Aes | 1.63256593 | 0.87825785 | 0.91171335 | -0.0334555 | 0.5647929 | 0.90650187 |
| Arfgap3 | 0.28762159 | 0.26434524 | 0.29781109 | -0.0334659 | 0.69130933 | 0.90650187 |
| Cmtm4 | 0.29647623 | 0.27207599 | 0.3055527 | -0.0334767 | 0.68689416 | 0.90650187 |
| Snx13 | 0.14395876 | 0.12781469 | 0.16129415 | -0.0334795 | 0.82193504 | 0.9092537 |
| Zfp281 | 0.21371835 | 0.19674694 | 0.23027177 | -0.0335248 | 0.74125762 | 0.90650187 |
| Arl2 | 0.72277517 | 0.57522875 | 0.6087567 | -0.0335279 | 0.59671242 | 0.90650187 |
| Sec24b | 0.2444478 | 0.22543357 | 0.25900962 | -0.0335761 | 0.71779205 | 0.90650187 |
| Gle1 | 0.35073964 | 0.31785854 | 0.35146146 | -0.0336029 | 0.66459644 | 0.90650187 |
| Pgs1 | 0.29727782 | 0.2726041 | 0.30624984 | -0.0336457 | 0.68737054 | 0.90650187 |
| Tnks2 | 0.65430268 | 0.53406425 | 0.56772701 | -0.0336628 | 0.60395496 | 0.90650187 |
| Txndc12 | 0.82628636 | 0.63277157 | 0.66643488 | -0.0336633 | 0.58883873 | 0.90650187 |
| Rbm3 | 2.67525579 | 0.84689621 | 0.88056861 | -0.0336724 | 0.56749384 | 0.90650187 |
| Syncrip | 0.89970487 | 0.67072751 | 0.70440081 | -0.0336733 | 0.58414874 | 0.90650187 |
| Kdm5c | 0.36430938 | 0.32883081 | 0.36254733 | -0.0337165 | 0.66035665 | 0.90650187 |
| Ptges2 | 0.17198195 | 0.15590989 | 0.18965313 | -0.0337432 | 0.78551809 | 0.90650187 |
| Nucb1 | 1.18276429 | 0.78333734 | 0.81709365 | -0.0337563 | 0.57285984 | 0.90650187 |
| Mrpl10 | 0.5264587 | 0.45035712 | 0.4841235 | -0.0337664 | 0.62174603 | 0.90650187 |

| | | | | | | |
|---|---|---|---|---|---|---|
| Kbtbd3 | 0.15172465 | 0.13543451 | 0.16924367 | -0.0338092 | 0.81275159 | 0.9092537 |
| Ccni | 0.81445922 | 0.62625897 | 0.66009446 | -0.0338355 | 0.59012842 | 0.90650187 |
| Use1 | 1.14008534 | 0.76943249 | 0.80327241 | -0.0338399 | 0.57428047 | 0.90650187 |
| Iqgap1 | 0.82903449 | 0.63405219 | 0.66789963 | -0.0338474 | 0.58912391 | 0.90650187 |
| Wdr6 | 0.69980515 | 0.56139767 | 0.59526179 | -0.0338641 | 0.59983041 | 0.90650187 |
| Osbpl9 | 0.2716409 | 0.24978607 | 0.28365711 | -0.033871 | 0.70223615 | 0.90650187 |
| Orc4 | 0.42044712 | 0.37295373 | 0.40683572 | -0.033882 | 0.64439738 | 0.90650187 |
| Irgq | 0.16806144 | 0.15184556 | 0.18573924 | -0.0338937 | 0.79138428 | 0.90661829 |
| Nap1l4 | 1.07123336 | 0.74481316 | 0.77876081 | -0.0339477 | 0.57683077 | 0.90650187 |
| Ddx56 | 0.33865986 | 0.30751118 | 0.34146256 | -0.0339514 | 0.67076391 | 0.90650187 |
| Anapc15 | 0.17748412 | 0.1611226 | 0.1951165 | -0.0339939 | 0.78069113 | 0.90650187 |
| Abcb8 | 0.31277378 | 0.28559886 | 0.31961271 | -0.0340138 | 0.68193288 | 0.90650187 |
| Usp34 | 0.63393049 | 0.52102004 | 0.55503988 | -0.0340198 | 0.60737461 | 0.90650187 |
| Eif3m | 1.4818243 | 0.85615961 | 0.8902064 | -0.0340468 | 0.56750532 | 0.90650187 |
| Gskip | 0.32560301 | 0.29643893 | 0.33051494 | -0.034076 | 0.67664624 | 0.90650187 |
| Arl8b | 0.44042467 | 0.38786203 | 0.42201064 | -0.0341486 | 0.64047668 | 0.90650187 |
| Pfkm | 0.20128741 | 0.1842188 | 0.21836989 | -0.0341511 | 0.75655165 | 0.90650187 |
| Eed | 0.33229571 | 0.30194924 | 0.33614459 | -0.0341954 | 0.67445308 | 0.90650187 |
| Paf1 | 0.24003198 | 0.22072554 | 0.25493846 | -0.0342129 | 0.72461393 | 0.90650187 |
| Dnajc1 | 0.66800359 | 0.54191348 | 0.57613342 | -0.0342199 | 0.60412955 | 0.90650187 |
| Mkln1 | 0.69991077 | 0.56109894 | 0.59532444 | -0.0342255 | 0.60086249 | 0.90650187 |
| Ing3 | 0.24868225 | 0.22866573 | 0.26289528 | -0.0342295 | 0.71864729 | 0.90650187 |
| Nkiras1 | 0.27385945 | 0.25139702 | 0.28563636 | -0.0342393 | 0.70294736 | 0.90650187 |
| Kansl1 | 0.29822131 | 0.27280984 | 0.30706957 | -0.0342597 | 0.6900514 | 0.90650187 |
| Timm22 | 0.59770207 | 0.49759405 | 0.53190267 | -0.0343086 | 0.61284779 | 0.90650187 |
| Rnf111 | 0.20686603 | 0.18942021 | 0.22373106 | -0.0343109 | 0.75230697 | 0.90650187 |
| Eri1 | 0.26181859 | 0.24051644 | 0.27483624 | -0.0343198 | 0.71060023 | 0.90650187 |
| Rock2 | 0.36336837 | 0.32746266 | 0.3617834 | -0.0343207 | 0.66339058 | 0.90650187 |
| Iqcc | 0.19722388 | 0.18012045 | 0.21444379 | -0.0343233 | 0.76162911 | 0.90650187 |
| Pitpnb | 0.63066504 | 0.51861349 | 0.55298479 | -0.0343713 | 0.60884922 | 0.90650187 |

| | | | | | | |
|---|---|---|---|---|---|---|
| Tmem126b | 0.32100858 | 0.29224815 | 0.32662794 | -0.0343798 | 0.68008264 | 0.90650187 |
| Tmx2 | 0.4995462 | 0.43085588 | 0.46523768 | -0.0343818 | 0.62874729 | 0.90650187 |
| Asb6 | 0.14666541 | 0.12966387 | 0.1640729 | -0.034409 | 0.82441527 | 0.90933631 |
| Apip | 0.50446426 | 0.43428488 | 0.4687245 | -0.0344396 | 0.62803056 | 0.90650187 |
| Cog1 | 0.29842399 | 0.27279837 | 0.30724556 | -0.0344472 | 0.69090751 | 0.90650187 |
| Cstf2t | 0.35472925 | 0.32024605 | 0.35473643 | -0.0344904 | 0.66721585 | 0.90650187 |
| Ap1s1 | 0.95100665 | 0.69482666 | 0.72933628 | -0.0345096 | 0.58330292 | 0.90650187 |
| Snrnp27 | 1.38007021 | 0.83549129 | 0.8700058 | -0.0345145 | 0.56999611 | 0.90650187 |
| Cse1l | 0.28653019 | 0.26232834 | 0.29685195 | -0.0345236 | 0.69742885 | 0.90650187 |
| Zfp219 | 0.17525946 | 0.1583633 | 0.19291171 | -0.0345484 | 0.78703056 | 0.90650187 |
| Cstf1 | 0.14822865 | 0.13109832 | 0.1656739 | -0.0345756 | 0.82323992 | 0.9092537 |
| Ccdc43 | 0.27071078 | 0.24824588 | 0.28282572 | -0.0345798 | 0.70662252 | 0.90650187 |
| Ythdf1 | 0.19176726 | 0.17456326 | 0.2091432 | -0.0345799 | 0.7688427 | 0.90650187 |
| Hectd1 | 0.69455959 | 0.55753366 | 0.59214302 | -0.0346094 | 0.60250455 | 0.90650187 |
| Nr2c2 | 0.22448129 | 0.20583064 | 0.24044719 | -0.0346165 | 0.7389209 | 0.90650187 |
| Mycbp2 | 0.73411056 | 0.58069824 | 0.61531959 | -0.0346213 | 0.59875848 | 0.90650187 |
| Dazap1 | 1.20612841 | 0.78963842 | 0.8242714 | -0.034633 | 0.57408686 | 0.90650187 |
| Usp48 | 0.44073001 | 0.3875988 | 0.42224033 | -0.0346415 | 0.64234252 | 0.90650187 |
| Rc3h1 | 0.33726105 | 0.30565467 | 0.34029663 | -0.034642 | 0.67456706 | 0.90650187 |
| Ddb1 | 1.25704372 | 0.80453581 | 0.83917816 | -0.0346423 | 0.5728051 | 0.90650187 |
| Fem1a | 0.24696784 | 0.22666491 | 0.26132439 | -0.0346595 | 0.7222777 | 0.90650187 |
| Fra10ac1 | 0.27979569 | 0.25624298 | 0.29090935 | -0.0346664 | 0.70186188 | 0.90650187 |
| Epn1 | 0.83281171 | 0.63522692 | 0.6699077 | -0.0346808 | 0.59101115 | 0.90650187 |
| Uckl1 | 0.29665569 | 0.27102768 | 0.30570887 | -0.0346812 | 0.69298609 | 0.90650187 |
| Naa10 | 0.61472288 | 0.50815328 | 0.54286678 | -0.0347135 | 0.61189956 | 0.90650187 |
| Nagk | 0.75033918 | 0.58989239 | 0.62460656 | -0.0347142 | 0.59757544 | 0.90650187 |
| Trnau1ap | 0.43684291 | 0.38459305 | 0.4193113 | -0.0347182 | 0.64359638 | 0.90650187 |
| Eef1g | 3.52755051 | 0.73128358 | 0.76600381 | -0.0347202 | 0.57984756 | 0.90650187 |
| Api5 | 0.40891472 | 0.36321585 | 0.39793774 | -0.0347219 | 0.65095482 | 0.90650187 |
| Man2c1 | 0.28454546 | 0.26037957 | 0.29510494 | -0.0347254 | 0.69955464 | 0.90650187 |

| | | | | | | |
|---|---|---|---|---|---|---|
| Stx3 | 0.23108705 | 0.21185148 | 0.24663252 | -0.034781 | 0.73464942 | 0.90650187 |
| Zfp580 | 0.24004403 | 0.22016022 | 0.2549496 | -0.0347894 | 0.72795154 | 0.90650187 |
| Cdc37l1 | 0.29239167 | 0.26717316 | 0.3019904 | -0.0348172 | 0.69586527 | 0.90650187 |
| Ppp6r3 | 0.23937759 | 0.219506 | 0.25433352 | -0.0348275 | 0.72866001 | 0.90650187 |
| Sepsecs | 0.19852363 | 0.18086667 | 0.21570169 | -0.034835 | 0.76360562 | 0.90650187 |
| Zfand6 | 1.19362112 | 0.78567907 | 0.82051897 | -0.0348399 | 0.57486113 | 0.90650187 |
| Ctps2 | 0.37025184 | 0.33250655 | 0.36735502 | -0.0348485 | 0.66338691 | 0.90650187 |
| Cuta | 1.21477145 | 0.7919776 | 0.826833 | -0.0348554 | 0.5743288 | 0.90650187 |
| Paxip1 | 0.155358 | 0.13806307 | 0.17293859 | -0.0348755 | 0.81501233 | 0.9092537 |
| Suz12 | 0.23805778 | 0.21823063 | 0.25311213 | -0.0348815 | 0.72994647 | 0.90650187 |
| Bnip3l | 1.36672296 | 0.83212269 | 0.8670058 | -0.0348831 | 0.57097796 | 0.90650187 |
| Med8 | 0.48960898 | 0.42325314 | 0.45814215 | -0.034889 | 0.63252464 | 0.90650187 |
| Mrto4 | 0.50915525 | 0.43713724 | 0.47203559 | -0.0348984 | 0.62879775 | 0.90650187 |
| Ppie | 0.44371449 | 0.3895815 | 0.42448183 | -0.0349003 | 0.64263528 | 0.90650187 |
| Lrfn3 | 0.12629451 | 0.10804532 | 0.14294754 | -0.0349022 | 0.86113747 | 0.91848494 |
| Tmem251 | 0.28128446 | 0.25732428 | 0.29222665 | -0.0349024 | 0.70227857 | 0.90650187 |
| Nisch | 1.91498682 | 0.89220403 | 0.92711996 | -0.0349159 | 0.56648193 | 0.90650187 |
| Mrps34 | 0.84686034 | 0.64239564 | 0.67731907 | -0.0349234 | 0.59065147 | 0.90650187 |
| Bccip | 0.80631017 | 0.62074844 | 0.65568739 | -0.0349389 | 0.59362835 | 0.90650187 |
| Itpa | 0.68782566 | 0.55315676 | 0.58811813 | -0.0349614 | 0.6042157 | 0.90650187 |
| Lcmt1 | 0.35969766 | 0.32382595 | 0.35879661 | -0.0349707 | 0.66760082 | 0.90650187 |
| Emc3 | 0.6476815 | 0.52865005 | 0.56362874 | -0.0349787 | 0.60868504 | 0.90650187 |
| Vps50 | 0.21549492 | 0.19697531 | 0.23195961 | -0.0349843 | 0.74872705 | 0.90650187 |
| 1600012H06 | 0.40339646 | 0.35865194 | 0.3936446 | -0.0349927 | 0.6536479 | 0.90650187 |
| Smg1 | 0.25065657 | 0.22970367 | 0.26470059 | -0.0349969 | 0.72165596 | 0.90650187 |
| Etfdh | 0.27276406 | 0.24959187 | 0.28465976 | -0.0350679 | 0.70803882 | 0.90650187 |
| Socs6 | 0.29205524 | 0.26661904 | 0.30169631 | -0.0350773 | 0.69737883 | 0.90650187 |
| Taf10 | 0.55022849 | 0.4653066 | 0.50041379 | -0.0351072 | 0.62243697 | 0.90650187 |
| Map2k4 | 0.27142898 | 0.24835854 | 0.28346776 | -0.0351092 | 0.70905139 | 0.90650187 |
| Pam16 | 0.89204707 | 0.66544571 | 0.70056055 | -0.0351148 | 0.58816658 | 0.90650187 |

| | | | | | | |
|---|---|---|---|---|---|---|
| Brd3 | 0.95269871 | 0.69497808 | 0.73013223 | -0.0351541 | 0.58474421 | 0.90650187 |
| Atf7ip | 0.37149281 | 0.33315623 | 0.36835516 | -0.0351989 | 0.66451201 | 0.90650187 |
| Uri1 | 0.43690874 | 0.3841244 | 0.419361 | -0.0352366 | 0.64562666 | 0.90650187 |
| Uck2 | 0.2454327 | 0.22467092 | 0.25991503 | -0.0352441 | 0.7266851 | 0.90650187 |
| Triqk | 0.2713812 | 0.24817819 | 0.28342506 | -0.0352469 | 0.70981956 | 0.90650187 |
| Taf6l | 0.81065337 | 0.62277722 | 0.65803978 | -0.0352626 | 0.59414785 | 0.90650187 |
| Ap3b1 | 0.40088619 | 0.35641652 | 0.39168397 | -0.0352675 | 0.65552668 | 0.90650187 |
| 2510002D24I | 0.48612018 | 0.42035383 | 0.45563473 | -0.0352809 | 0.63466489 | 0.90650187 |
| Nub1 | 0.31261983 | 0.28419591 | 0.31948099 | -0.0352851 | 0.68828501 | 0.90650187 |
| Cpsf3 | 0.41764847 | 0.36938633 | 0.40468565 | -0.0352993 | 0.65090866 | 0.90650187 |
| Atg10 | 0.24356196 | 0.22288914 | 0.25819446 | -0.0353053 | 0.72836944 | 0.90650187 |
| Mtch2 | 1.30500545 | 0.81671925 | 0.85205376 | -0.0353345 | 0.57313388 | 0.90650187 |
| Hmox2 | 0.59807972 | 0.49678573 | 0.53214772 | -0.035362 | 0.61616233 | 0.90650187 |
| Rnf219 | 0.21530489 | 0.19641618 | 0.23177922 | -0.035363 | 0.75120148 | 0.90650187 |
| Dscr3 | 0.34486651 | 0.31117996 | 0.34661611 | -0.0354361 | 0.67526357 | 0.90650187 |
| Smc5 | 0.274785 | 0.25100664 | 0.28646065 | -0.035454 | 0.70891359 | 0.90650187 |
| Trp53inp2 | 0.24855127 | 0.22731977 | 0.26277537 | -0.0354556 | 0.7256916 | 0.90650187 |
| Cbl | 0.28001288 | 0.25560412 | 0.29110165 | -0.0354975 | 0.70612738 | 0.90650187 |
| Fkbpl | 0.15308804 | 0.13513255 | 0.17063194 | -0.0354994 | 0.82248928 | 0.9092537 |
| Zfyve19 | 0.26527051 | 0.24242665 | 0.27794662 | -0.03552 | 0.7150242 | 0.90650187 |
| Nit1 | 0.45749466 | 0.3992186 | 0.43474463 | -0.035526 | 0.64180276 | 0.90650187 |
| Yipf4 | 1.27525021 | 0.80872149 | 0.84424839 | -0.0355269 | 0.57419949 | 0.90650187 |
| Ralgapb | 0.14108268 | 0.12280271 | 0.15833209 | -0.0355294 | 0.84075959 | 0.91251845 |
| Smchd1 | 0.4401596 | 0.38625416 | 0.42181119 | -0.035557 | 0.64607573 | 0.90650187 |
| Nfyc | 0.41921869 | 0.37032631 | 0.4058928 | -0.0355665 | 0.6515645 | 0.90650187 |
| Ralbp1 | 1.21297048 | 0.79073452 | 0.82630134 | -0.0355668 | 0.5758759 | 0.90650187 |
| Rer1 | 1.31568736 | 0.81914617 | 0.85472535 | -0.0355792 | 0.57340697 | 0.90650187 |
| Slc39a10 | 0.27380158 | 0.24998939 | 0.28558479 | -0.0355954 | 0.71024549 | 0.90650187 |
| Dzip3 | 0.29445348 | 0.26818527 | 0.30379047 | -0.0356052 | 0.69883023 | 0.90650187 |
| Nt5c | 0.573959 | 0.48070541 | 0.51632453 | -0.0356191 | 0.62045814 | 0.90650187 |

| | | | | | | |
|---|---|---|---|---|---|---|
| Armc9 | 0.16453238 | 0.14658023 | 0.18220144 | -0.0356212 | 0.80761369 | 0.9092537 |
| Sugp1 | 0.15828891 | 0.14027361 | 0.17590815 | -0.0356345 | 0.81609079 | 0.9092537 |
| Fam193a | 0.33923338 | 0.30629066 | 0.34194012 | -0.0356495 | 0.67849096 | 0.90650187 |
| Mtmr6 | 0.36701638 | 0.32907897 | 0.36474092 | -0.035662 | 0.66809608 | 0.90650187 |
| Dxo | 0.2140365 | 0.19490803 | 0.23057427 | -0.0356662 | 0.75416841 | 0.90650187 |
| Mrpl14 | 1.09609061 | 0.75216523 | 0.78785753 | -0.0356923 | 0.5798068 | 0.90650187 |
| Midn | 0.44261198 | 0.38793953 | 0.42365453 | -0.035715 | 0.64608584 | 0.90650187 |
| Atp2c1 | 0.47275555 | 0.41021263 | 0.44595349 | -0.0357409 | 0.63918919 | 0.90650187 |
| Arfgef1 | 0.40944056 | 0.36260387 | 0.39834564 | -0.0357418 | 0.65501081 | 0.90650187 |
| Rassf3 | 0.15661301 | 0.13845942 | 0.17421135 | -0.0357519 | 0.81920366 | 0.9092537 |
| Selo | 0.21879758 | 0.1993312 | 0.23508869 | -0.0357575 | 0.75053876 | 0.90650187 |
| Zmat5 | 0.3606542 | 0.32381826 | 0.35957599 | -0.0357577 | 0.67078779 | 0.90650187 |
| Ccdc59 | 0.86485229 | 0.65091939 | 0.68668569 | -0.0357663 | 0.59155668 | 0.90650187 |
| Tmem63b | 0.32238639 | 0.29200507 | 0.32779556 | -0.0357905 | 0.68630204 | 0.90650187 |
| Cda | 0.32329415 | 0.29275694 | 0.3285639 | -0.035807 | 0.68597846 | 0.90650187 |
| R3hdm2 | 0.22206914 | 0.20233114 | 0.2381773 | -0.0358462 | 0.74827541 | 0.90650187 |
| Sirt3 | 0.21979789 | 0.20017132 | 0.23603423 | -0.0358629 | 0.75030914 | 0.90650187 |
| Txn2 | 0.85060214 | 0.64341156 | 0.67927813 | -0.0358666 | 0.5927918 | 0.90650187 |
| Bnip3 | 0.32221777 | 0.29175371 | 0.32765276 | -0.035899 | 0.6868991 | 0.90650187 |
| Tmem57 | 0.66015001 | 0.53541495 | 0.57132701 | -0.0359121 | 0.61004592 | 0.90650187 |
| Rufy1 | 0.2432626 | 0.22200184 | 0.25791881 | -0.035917 | 0.73206863 | 0.90650187 |
| Ddx10 | 0.24996074 | 0.22813783 | 0.26406475 | -0.0359269 | 0.72735734 | 0.90650187 |
| Med16 | 0.27765221 | 0.25306124 | 0.28900913 | -0.0359479 | 0.70985421 | 0.90650187 |
| Cnppd1 | 0.27964687 | 0.25473467 | 0.29077755 | -0.0360429 | 0.7091997 | 0.90650187 |
| Kdm4a | 0.17764991 | 0.15922572 | 0.19528059 | -0.0360549 | 0.79411911 | 0.90731056 |
| Ptpn11 | 0.44752753 | 0.39127442 | 0.42733527 | -0.0360608 | 0.64622336 | 0.90650187 |
| Carkd | 0.6815068 | 0.54825353 | 0.58431949 | -0.036066 | 0.60810992 | 0.90650187 |
| Pnn | 1.24549445 | 0.79976627 | 0.83587869 | -0.0361124 | 0.57615369 | 0.90650187 |
| Brd8 | 0.719478 | 0.5706744 | 0.60683611 | -0.0361617 | 0.60446301 | 0.90650187 |
| Cd320 | 0.39286177 | 0.34915166 | 0.38538336 | -0.0362317 | 0.662009 | 0.90650187 |

| | | | | | | |
|---|---|---|---|---|---|---|
| Kdm3b | 0.15850623 | 0.13987646 | 0.17612795 | -0.0362515 | 0.8199083 | 0.9092537 |
| Ndufs1 | 0.53868131 | 0.45628921 | 0.4925452 | -0.036256 | 0.62825631 | 0.90650187 |
| Chd8 | 0.39032154 | 0.34710961 | 0.38337785 | -0.0362682 | 0.66296269 | 0.90650187 |
| Mrpl43 | 0.804675 | 0.61851223 | 0.65479962 | -0.0362874 | 0.5973 | 0.90650187 |
| C2cd5 | 0.21663053 | 0.19673767 | 0.2330368 | -0.0362991 | 0.75567977 | 0.90650187 |
| Med20 | 0.25782978 | 0.23490538 | 0.27122796 | -0.0363226 | 0.72419567 | 0.90650187 |
| Tsen2 | 0.19233534 | 0.1733674 | 0.20969652 | -0.0363291 | 0.77940472 | 0.90650187 |
| Commd5 | 0.28748155 | 0.26132599 | 0.29768808 | -0.0363621 | 0.7064431 | 0.90650187 |
| Casc3 | 0.32661891 | 0.29500385 | 0.33137195 | -0.0363681 | 0.68719166 | 0.90650187 |
| mt-Atp8 | 1.09160373 | 0.74988011 | 0.78625543 | -0.0363753 | 0.58147625 | 0.90650187 |
| Ddx27 | 0.32087497 | 0.29013371 | 0.32651458 | -0.0363809 | 0.68981916 | 0.90650187 |
| Mccc2 | 0.21057085 | 0.19087095 | 0.22727345 | -0.0364025 | 0.76178465 | 0.90650187 |
| Ogfod2 | 0.32882619 | 0.29682627 | 0.33323094 | -0.0364047 | 0.68640002 | 0.90650187 |
| Mecr | 0.20465253 | 0.18519559 | 0.22160775 | -0.0364122 | 0.76744346 | 0.90650187 |
| Lamtor3 | 0.44523473 | 0.38919786 | 0.42562088 | -0.036423 | 0.64818987 | 0.90650187 |
| Ppil6 | 0.12318116 | 0.103249 | 0.13967507 | -0.0364261 | 0.87684856 | 0.92473192 |
| Dnttip1 | 0.26915358 | 0.24494072 | 0.28143199 | -0.0364913 | 0.71783768 | 0.90650187 |
| Patz1 | 0.1956877 | 0.17645942 | 0.21295475 | -0.0364953 | 0.77693125 | 0.90650187 |
| Lamp1 | 1.29191123 | 0.81220252 | 0.84871116 | -0.0365086 | 0.57582914 | 0.90650187 |
| Tiprl | 0.82150245 | 0.62735215 | 0.66387744 | -0.0365253 | 0.59661261 | 0.90650187 |
| Atg2a | 0.12893389 | 0.10917467 | 0.14571248 | -0.0365378 | 0.86752085 | 0.92056229 |
| Cltc | 0.9022363 | 0.6691005 | 0.70566524 | -0.0365647 | 0.59109145 | 0.90650187 |
| Gpn3 | 0.24458485 | 0.22256045 | 0.25913566 | -0.0365752 | 0.7348233 | 0.90650187 |
| Sqstm1 | 1.04334139 | 0.7315504 | 0.76812873 | -0.0365783 | 0.58382946 | 0.90650187 |
| Armc1 | 0.63580459 | 0.51963372 | 0.55621671 | -0.036583 | 0.61500671 | 0.90650187 |
| Gpi1 | 1.11926211 | 0.75943565 | 0.79601867 | -0.036583 | 0.58094395 | 0.90650187 |
| Vbp1 | 1.12530691 | 0.76155305 | 0.79814032 | -0.0365873 | 0.58074098 | 0.90650187 |
| Tmtc4 | 0.18523096 | 0.16615265 | 0.20275174 | -0.0365991 | 0.78886369 | 0.90650187 |
| Tgds | 0.3141339 | 0.28417435 | 0.32077546 | -0.0366011 | 0.69401117 | 0.90650187 |
| Whsc1l1 | 0.69109029 | 0.55347157 | 0.59007275 | -0.0366012 | 0.6086312 | 0.90650187 |

| | | | | | | |
|---|---|---|---|---|---|---|
| Rcor1 | 0.3005373 | 0.27245591 | 0.30907839 | -0.0366225 | 0.70081875 | 0.90650187 |
| Thumpd3 | 0.29852848 | 0.27067134 | 0.30733628 | -0.0366649 | 0.70206772 | 0.90650187 |
| Ube2e2 | 0.22308138 | 0.20244605 | 0.23913069 | -0.0366846 | 0.75237836 | 0.90650187 |
| Zfp672 | 0.24795806 | 0.22553129 | 0.26223213 | -0.0367008 | 0.73309063 | 0.90650187 |
| Mgea5 | 0.36518573 | 0.32654958 | 0.36325812 | -0.0367085 | 0.67337233 | 0.90650187 |
| Fam160b2 | 0.18117297 | 0.16203998 | 0.19876039 | -0.0367204 | 0.79426601 | 0.90731056 |
| Trp53rka | 0.28917682 | 0.26243435 | 0.29917598 | -0.0367416 | 0.70745412 | 0.90650187 |
| Ppp3r1 | 0.60692823 | 0.50112409 | 0.53786611 | -0.036742 | 0.61931514 | 0.90650187 |
| Lonp2 | 0.31509664 | 0.28481555 | 0.32159754 | -0.036782 | 0.69443324 | 0.90650187 |
| Fem1b | 0.34072455 | 0.30639153 | 0.3431805 | -0.036789 | 0.68316558 | 0.90650187 |
| Zmym5 | 0.37875942 | 0.33737732 | 0.37418602 | -0.0368087 | 0.6690584 | 0.90650187 |
| Egln2 | 0.7767231 | 0.60262531 | 0.63943798 | -0.0368127 | 0.60099953 | 0.90650187 |
| Stard7 | 0.49068314 | 0.42209894 | 0.45891252 | -0.0368136 | 0.63931289 | 0.90650187 |
| Gfer | 0.43123778 | 0.37823824 | 0.41506847 | -0.0368302 | 0.65338351 | 0.90650187 |
| Agfg1 | 0.23544624 | 0.21384794 | 0.25069026 | -0.0368423 | 0.74323209 | 0.90650187 |
| Ncln | 0.43397327 | 0.38028626 | 0.41714196 | -0.0368557 | 0.65275981 | 0.90650187 |
| Zfp292 | 0.62475358 | 0.51238374 | 0.54924961 | -0.0368659 | 0.61729765 | 0.90650187 |
| Fam160b1 | 0.2031876 | 0.1833281 | 0.22019968 | -0.0368716 | 0.77168261 | 0.90650187 |
| Apeh | 0.22515726 | 0.20417577 | 0.24108216 | -0.0369064 | 0.75192555 | 0.90650187 |
| Uhrf2 | 0.71148073 | 0.5652373 | 0.60215454 | -0.0369172 | 0.60740188 | 0.90650187 |
| Zfp451 | 0.43196298 | 0.37865432 | 0.41561869 | -0.0369644 | 0.6537203 | 0.90650187 |
| Ppp2r5a | 0.20688834 | 0.18677983 | 0.22375243 | -0.0369726 | 0.7687045 | 0.90650187 |
| Akr7a5 | 0.78262861 | 0.60572067 | 0.64271342 | -0.0369927 | 0.60097686 | 0.90650187 |
| Ccser2 | 0.23233526 | 0.21079516 | 0.24779636 | -0.0370012 | 0.74659802 | 0.90650187 |
| Ccdc12 | 1.26366949 | 0.80404739 | 0.84105161 | -0.0370042 | 0.57753674 | 0.90650187 |
| Slc38a10 | 0.72788529 | 0.57471668 | 0.61172295 | -0.0370063 | 0.60601113 | 0.90650187 |
| Rpp14 | 0.29756881 | 0.26945711 | 0.30650275 | -0.0370456 | 0.70447701 | 0.90650187 |
| Ctcf | 0.61300873 | 0.50462955 | 0.54176999 | -0.0371404 | 0.61972691 | 0.90650187 |
| Exoc6b | 0.23550184 | 0.21359698 | 0.2507419 | -0.0371449 | 0.74491482 | 0.90650187 |
| Emc6 | 0.98116919 | 0.70568324 | 0.74282962 | -0.0371464 | 0.58796357 | 0.90650187 |

| | | | | | | |
|---|---|---|---|---|---|---|
| Xpot | 0.23217132 | 0.2104965 | 0.2476436 | -0.0371471 | 0.74756634 | 0.90650187 |
| Pik3c2a | 0.36201629 | 0.32349515 | 0.36068451 | -0.0371894 | 0.67665895 | 0.90650187 |
| Tcerg1 | 0.44723527 | 0.38989212 | 0.42711695 | -0.0372248 | 0.65078885 | 0.90650187 |
| Nufip1 | 0.2185115 | 0.19757194 | 0.23481809 | -0.0372461 | 0.75964118 | 0.90650187 |
| Plrg1 | 0.25488887 | 0.23127634 | 0.26855784 | -0.0372815 | 0.73143012 | 0.90650187 |
| Fam96b | 0.79020569 | 0.60960557 | 0.64689308 | -0.0372875 | 0.60112065 | 0.90650187 |
| Tmem161a | 0.29438692 | 0.26643686 | 0.30373241 | -0.0372956 | 0.70742569 | 0.90650187 |
| Pten | 0.3999633 | 0.35366502 | 0.39096195 | -0.0372969 | 0.66425404 | 0.90650187 |
| Nudt21 | 0.98178952 | 0.7057905 | 0.7430957 | -0.0373052 | 0.58830235 | 0.90650187 |
| Rragc | 0.34487827 | 0.30931011 | 0.34662584 | -0.0373157 | 0.6838842 | 0.90650187 |
| Atg4d | 0.24356577 | 0.22087697 | 0.25819797 | -0.037321 | 0.73975368 | 0.90650187 |
| Eif2s1 | 1.03438954 | 0.72726192 | 0.76461046 | -0.0373485 | 0.58595517 | 0.90650187 |
| Zpr1 | 0.33998245 | 0.30520229 | 0.34256345 | -0.0373612 | 0.68610976 | 0.90650187 |
| Rnf20 | 0.54516285 | 0.45951441 | 0.49697273 | -0.0374583 | 0.63121713 | 0.90650187 |
| Prdx1 | 3.17593114 | 0.77792519 | 0.81542789 | -0.0375027 | 0.58100292 | 0.90650187 |
| Uba52 | 2.5465781 | 0.85589797 | 0.89340132 | -0.0375033 | 0.57402007 | 0.90650187 |
| Trap1 | 0.34639749 | 0.31036847 | 0.34788216 | -0.0375137 | 0.68416925 | 0.90650187 |
| Tnrc18 | 0.36350308 | 0.32435122 | 0.3618928 | -0.0375416 | 0.67766265 | 0.90650187 |
| Urm1 | 0.42733809 | 0.37455823 | 0.41210306 | -0.0375448 | 0.65726823 | 0.90650187 |
| Xpo7 | 0.23379404 | 0.21160574 | 0.24915455 | -0.0375488 | 0.74856304 | 0.90650187 |
| Ttc9c | 0.29013801 | 0.26246492 | 0.30001844 | -0.0375535 | 0.71105166 | 0.90650187 |
| 2410004B18I | 0.47099177 | 0.40711259 | 0.44466673 | -0.0375541 | 0.64633775 | 0.90650187 |
| Caap1 | 0.27503592 | 0.24910787 | 0.28668398 | -0.0375761 | 0.71994985 | 0.90650187 |
| Tsr3 | 0.47844091 | 0.41247067 | 0.45008675 | -0.0376161 | 0.64488318 | 0.90650187 |
| Faf1 | 0.2305291 | 0.2084316 | 0.24611179 | -0.0376802 | 0.75194746 | 0.90650187 |
| M6pr | 0.69804168 | 0.55653005 | 0.59421485 | -0.0376848 | 0.61100449 | 0.90650187 |
| Ndfip1 | 0.31327425 | 0.28234747 | 0.32004075 | -0.0376933 | 0.6997174 | 0.90650187 |
| Morn2 | 0.3837604 | 0.34045844 | 0.37817463 | -0.0377162 | 0.67125102 | 0.90650187 |
| Apopt1 | 0.53183452 | 0.45010893 | 0.48783914 | -0.0377302 | 0.63451286 | 0.90650187 |
| Odf2 | 0.41283598 | 0.36324263 | 0.40097447 | -0.0377318 | 0.66214821 | 0.90650187 |

| | | | | | | |
|---|---|---|---|---|---|---|
| Tubgcp4 | 0.24415336 | 0.22098441 | 0.25873875 | -0.0377543 | 0.74173027 | 0.90650187 |
| Commd10 | 0.36989924 | 0.32931539 | 0.36707054 | -0.0377552 | 0.67625583 | 0.90650187 |
| Zdhhc6 | 0.32930463 | 0.29584064 | 0.33363334 | -0.0377927 | 0.69272211 | 0.90650187 |
| Gps1 | 0.97788507 | 0.70360513 | 0.74141838 | -0.0378133 | 0.58968369 | 0.90650187 |
| AI314180 | 0.27112993 | 0.24536822 | 0.28320048 | -0.0378323 | 0.7237041 | 0.90650187 |
| Mrpl22 | 0.51690215 | 0.4396388 | 0.47747242 | -0.0378336 | 0.63767753 | 0.90650187 |
| Rab39b | 0.1117432 | 0.08967713 | 0.12755031 | -0.0378732 | 0.90658622 | 0.93793079 |
| Mrpl40 | 0.62140253 | 0.50923799 | 0.54712369 | -0.0378857 | 0.62089679 | 0.90650187 |
| Nsmce1 | 0.8559644 | 0.64417873 | 0.68207559 | -0.0378969 | 0.59754693 | 0.90650187 |
| Mis12 | 0.24858568 | 0.2248769 | 0.26280687 | -0.03793 | 0.73943951 | 0.90650187 |
| Cyb5b | 0.40962831 | 0.36053995 | 0.39849122 | -0.0379513 | 0.66399392 | 0.90650187 |
| Zc3h7b | 0.23993804 | 0.21689714 | 0.25485165 | -0.0379545 | 0.74603833 | 0.90650187 |
| Btbd9 | 0.28280639 | 0.25561382 | 0.29357119 | -0.0379574 | 0.71728533 | 0.90650187 |
| Uso1 | 0.6595062 | 0.53296566 | 0.57093152 | -0.0379659 | 0.61624151 | 0.90650187 |
| Mrpl21 | 0.86237141 | 0.64743514 | 0.68540274 | -0.0379676 | 0.5972605 | 0.90650187 |
| Rab28 | 0.73429938 | 0.57745997 | 0.61542839 | -0.0379684 | 0.60806087 | 0.90650187 |
| Bak1 | 0.39734473 | 0.35090792 | 0.38890976 | -0.0380018 | 0.66799989 | 0.90650187 |
| Zmiz1 | 0.50501281 | 0.43109544 | 0.46911243 | -0.038017 | 0.64067854 | 0.90650187 |
| Zbtb45 | 0.19592464 | 0.17514959 | 0.21318459 | -0.038035 | 0.78615652 | 0.90650187 |
| Acaa1a | 0.99806086 | 0.71187037 | 0.74992257 | -0.0380522 | 0.58923467 | 0.90650187 |
| Ammecr1l | 0.30326823 | 0.27336952 | 0.31144095 | -0.0380714 | 0.70658781 | 0.90650187 |
| Rsbn1l | 0.49383568 | 0.42308147 | 0.46116901 | -0.0380875 | 0.64324969 | 0.90650187 |
| Micu1 | 0.42351344 | 0.37108722 | 0.40918389 | -0.0380967 | 0.66052778 | 0.90650187 |
| Yipf6 | 0.33110645 | 0.29702006 | 0.33514703 | -0.038127 | 0.69348067 | 0.90650187 |
| Hras | 0.26389483 | 0.23853582 | 0.27670841 | -0.0381726 | 0.73015504 | 0.90650187 |
| Polr3h | 0.22875001 | 0.20625729 | 0.24444926 | -0.038192 | 0.75633677 | 0.90650187 |
| Arih1 | 0.23937969 | 0.21611544 | 0.25433545 | -0.03822 | 0.74795213 | 0.90650187 |
| Phf2 | 0.18361855 | 0.16292025 | 0.20116796 | -0.0382477 | 0.80101029 | 0.90815573 |
| Glmp | 0.85902082 | 0.64541381 | 0.68366486 | -0.0382511 | 0.59821584 | 0.90650187 |
| Ccdc71 | 0.2268101 | 0.20431135 | 0.24263281 | -0.0383215 | 0.75870398 | 0.90650187 |

| | | | | | | |
|---|---|---|---|---|---|---|
| Mical3 | 0.17346022 | 0.15278236 | 0.19112448 | -0.0383421 | 0.81376278 | 0.9092537 |
| Wdr34 | 0.27608732 | 0.24925474 | 0.28761916 | -0.0383644 | 0.72340352 | 0.90650187 |
| Gpatch8 | 0.57083078 | 0.47586746 | 0.5142472 | -0.0383797 | 0.62997546 | 0.90650187 |
| Ppox | 0.14037029 | 0.11921025 | 0.15759691 | -0.0383867 | 0.86055988 | 0.91847356 |
| Cbx8 | 0.14902894 | 0.12808463 | 0.16649243 | -0.0384078 | 0.84744763 | 0.9143112 |
| Zmynd8 | 0.66134582 | 0.53360702 | 0.57206101 | -0.038454 | 0.61746665 | 0.90650187 |
| Nudt8 | 0.19672714 | 0.17547669 | 0.21396257 | -0.0384859 | 0.7880404 | 0.90650187 |
| Higd2a | 1.28626183 | 0.80864209 | 0.84720779 | -0.0385657 | 0.58018555 | 0.90650187 |
| Gns | 0.33323925 | 0.29834411 | 0.33693519 | -0.0385911 | 0.69468708 | 0.90650187 |
| Mff | 0.99572434 | 0.71033975 | 0.74895761 | -0.0386179 | 0.59065299 | 0.90650187 |
| Cul7 | 0.27989212 | 0.25233365 | 0.29099474 | -0.0386611 | 0.72261784 | 0.90650187 |
| Chkb | 0.43810985 | 0.38158024 | 0.42026717 | -0.0386869 | 0.65881537 | 0.90650187 |
| Rnf139 | 0.16552351 | 0.14450716 | 0.18319644 | -0.0386893 | 0.82610079 | 0.90947782 |
| Mdh2 | 2.12719893 | 0.88501804 | 0.92372038 | -0.0387023 | 0.57388093 | 0.90650187 |
| Elp3 | 0.31976904 | 0.2868428 | 0.32557574 | -0.0387329 | 0.7015566 | 0.90650187 |
| Ep300 | 0.28891546 | 0.2601729 | 0.29894676 | -0.0387739 | 0.71789586 | 0.90650187 |
| Zfp942 | 0.20515596 | 0.18328793 | 0.22209112 | -0.0388032 | 0.78133946 | 0.90650187 |
| Rmdn3 | 0.28387576 | 0.25570214 | 0.29451463 | -0.0388125 | 0.72101602 | 0.90650187 |
| Ufd1l | 0.399389 | 0.35168496 | 0.39051231 | -0.0388273 | 0.67075972 | 0.90650187 |
| Gfpt1 | 0.71495639 | 0.56536467 | 0.60419347 | -0.0388288 | 0.61244572 | 0.90650187 |
| Thap4 | 0.24189361 | 0.21782105 | 0.25665713 | -0.0388361 | 0.74942682 | 0.90650187 |
| Ccdc115 | 0.47885508 | 0.41154181 | 0.45038699 | -0.0388452 | 0.64929829 | 0.90650187 |
| Mtf2 | 0.47647118 | 0.40980209 | 0.44865725 | -0.0388552 | 0.6498827 | 0.90650187 |
| Cant1 | 0.33345417 | 0.29824313 | 0.33711517 | -0.038872 | 0.69588917 | 0.90650187 |
| Bhlhb9 | 0.53958551 | 0.45426715 | 0.4931645 | -0.0388974 | 0.63707012 | 0.90650187 |
| Dhx36 | 0.34145084 | 0.30488444 | 0.34378397 | -0.0388995 | 0.69252671 | 0.90650187 |
| Scaf11 | 1.13287886 | 0.7618759 | 0.80077951 | -0.0389036 | 0.58549727 | 0.90650187 |
| Commd1 | 1.36059377 | 0.82668296 | 0.86559479 | -0.0389118 | 0.57919995 | 0.90650187 |
| Prrc2a | 0.61021655 | 0.5010578 | 0.5399799 | -0.0389221 | 0.62568343 | 0.90650187 |
| Ctbp1 | 1.03910175 | 0.72752332 | 0.76647304 | -0.0389497 | 0.58936489 | 0.90650187 |

| | | | | | | |
|---|---|---|---|---|---|---|
| Arid2 | 0.28699447 | 0.2582946 | 0.2972601 | -0.0389655 | 0.71996574 | 0.90650187 |
| Pank3 | 0.23746787 | 0.21359808 | 0.25256566 | -0.0389676 | 0.75361155 | 0.90650187 |
| Timm50 | 0.57649934 | 0.47902906 | 0.5180072 | -0.0389781 | 0.63100472 | 0.90650187 |
| Tmem184b | 0.38214753 | 0.33786707 | 0.37689041 | -0.0390233 | 0.67734932 | 0.90650187 |
| Ttc13 | 0.25297146 | 0.22778775 | 0.26681249 | -0.0390247 | 0.74222714 | 0.90650187 |
| Phc1 | 0.16560939 | 0.1442572 | 0.18328261 | -0.0390254 | 0.82807353 | 0.90957047 |
| Snrpd1 | 1.5256749 | 0.85855863 | 0.89760435 | -0.0390457 | 0.57667068 | 0.90650187 |
| Stx5a | 0.88814612 | 0.65950198 | 0.69859356 | -0.0390916 | 0.59822881 | 0.90650187 |
| Rpgr | 0.20453626 | 0.1823576 | 0.22149608 | -0.0391385 | 0.78393183 | 0.90650187 |
| Elp6 | 0.17512469 | 0.153639 | 0.19277797 | -0.039139 | 0.81662408 | 0.9092537 |
| Acp2 | 0.20750229 | 0.18516823 | 0.22434044 | -0.0391722 | 0.78119929 | 0.90650187 |
| Ppcs | 0.25181507 | 0.22656259 | 0.26575815 | -0.0391956 | 0.74397982 | 0.90650187 |
| Nup35 | 0.23428792 | 0.21040093 | 0.2496139 | -0.039213 | 0.75753029 | 0.90650187 |
| Atpaf2 | 0.22702331 | 0.20353483 | 0.24283264 | -0.0392978 | 0.76406388 | 0.90650187 |
| Psmb5 | 1.98810026 | 0.88789271 | 0.92720125 | -0.0393085 | 0.57474617 | 0.90650187 |
| Wdr46 | 0.21909601 | 0.19602572 | 0.23537088 | -0.0393452 | 0.771301 | 0.90650187 |
| Txnrd1 | 0.60003777 | 0.49406854 | 0.53341696 | -0.0393484 | 0.62852066 | 0.90650187 |
| Thumpd1 | 0.44434629 | 0.38560592 | 0.42495553 | -0.0393496 | 0.65970192 | 0.90650187 |
| Eapp | 0.60779833 | 0.49907463 | 0.53842602 | -0.0393514 | 0.62737599 | 0.90650187 |
| Polr2h | 0.56040855 | 0.46790742 | 0.50728167 | -0.0393742 | 0.63497521 | 0.90650187 |
| Mrpl3 | 0.37999135 | 0.33576096 | 0.3751704 | -0.0394094 | 0.67974458 | 0.90650187 |
| Fbxw5 | 0.40642379 | 0.35658467 | 0.39600268 | -0.039418 | 0.67094157 | 0.90650187 |
| Dhodh | 0.16170705 | 0.13991404 | 0.17935903 | -0.039445 | 0.83588529 | 0.91079271 |
| Abhd17b | 0.16579035 | 0.14400168 | 0.18346413 | -0.0394625 | 0.83052282 | 0.9098597 |
| Samd14 | 0.36523047 | 0.32379081 | 0.36329439 | -0.0395036 | 0.68559493 | 0.90650187 |
| Trappc5 | 0.51135785 | 0.43407508 | 0.47358534 | -0.0395103 | 0.64463947 | 0.90650187 |
| Polr2b | 0.34127504 | 0.30410592 | 0.34363794 | -0.039532 | 0.69547204 | 0.90650187 |
| Kif2a | 0.20982874 | 0.186935 | 0.22656502 | -0.03963 | 0.78160086 | 0.90650187 |
| Rcan1 | 0.16804334 | 0.14605329 | 0.18572113 | -0.0396678 | 0.82882415 | 0.90967191 |
| Mfn2 | 0.17772503 | 0.15566863 | 0.19535493 | -0.0396863 | 0.81676985 | 0.9092537 |

| | | | | | | |
|---|---|---|---|---|---|---|
| Fh1 | 0.70419806 | 0.55817672 | 0.59786307 | -0.0396864 | 0.61604159 | 0.90650187 |
| Thap7 | 0.34331646 | 0.30563771 | 0.34533207 | -0.0396944 | 0.69532531 | 0.90650187 |
| Dcakd | 0.67905403 | 0.5431344 | 0.58283951 | -0.0397051 | 0.61899812 | 0.90650187 |
| Clpb | 0.26359268 | 0.23671205 | 0.27643621 | -0.0397242 | 0.73853585 | 0.90650187 |
| Pex11b | 0.35792993 | 0.3176261 | 0.35735433 | -0.0397282 | 0.68942896 | 0.90650187 |
| Ppme1 | 0.36575826 | 0.32398923 | 0.36372215 | -0.0397329 | 0.68638771 | 0.90650187 |
| Mrps16 | 0.99131378 | 0.7073875 | 0.74713009 | -0.0397426 | 0.59346914 | 0.90650187 |
| Pigt | 0.8117817 | 0.61888728 | 0.65864959 | -0.0397623 | 0.60579611 | 0.90650187 |
| Ttc14 | 0.94109133 | 0.68484463 | 0.72465028 | -0.0398057 | 0.5964624 | 0.90650187 |
| B3galt6 | 0.31121981 | 0.27847235 | 0.31828224 | -0.0398099 | 0.71090979 | 0.90650187 |
| Pes1 | 0.33558932 | 0.29909105 | 0.33890105 | -0.03981 | 0.69924007 | 0.90650187 |
| Xpo6 | 0.2981989 | 0.26722803 | 0.30705011 | -0.0398221 | 0.71788216 | 0.90650187 |
| Pi4ka | 0.19981889 | 0.17711185 | 0.21695317 | -0.0398413 | 0.792862 | 0.90693771 |
| Rsf1 | 0.721481 | 0.56814919 | 0.60800348 | -0.0398543 | 0.61462999 | 0.90650187 |
| Brox | 0.37149102 | 0.32849498 | 0.36835371 | -0.0398587 | 0.68475923 | 0.90650187 |
| Coa7 | 0.23742517 | 0.21265036 | 0.25252609 | -0.0398757 | 0.75865854 | 0.90650187 |
| D2Wsu81e | 0.37105624 | 0.32810849 | 0.3680035 | -0.039895 | 0.68507755 | 0.90650187 |
| Atp5g3 | 1.83392795 | 0.885534 | 0.92543106 | -0.0398971 | 0.57599509 | 0.90650187 |
| Srf | 0.21936279 | 0.19572197 | 0.23562308 | -0.0399011 | 0.77422456 | 0.90650187 |
| Tmem106c | 0.47347982 | 0.40656731 | 0.44648127 | -0.039914 | 0.65447206 | 0.90650187 |
| Ruvbl1 | 0.55033135 | 0.4605606 | 0.5004835 | -0.0399229 | 0.6385629 | 0.90650187 |
| Nup88 | 0.34990981 | 0.31084707 | 0.35077863 | -0.0399316 | 0.6935977 | 0.90650187 |
| Ap1g1 | 0.2845873 | 0.25517809 | 0.2951418 | -0.0399637 | 0.72639892 | 0.90650187 |
| Gtpbp6 | 0.24023037 | 0.21514336 | 0.25512179 | -0.0399784 | 0.75698631 | 0.90650187 |
| Gm10020 | 0.69863608 | 0.55458341 | 0.5945679 | -0.0399845 | 0.61751683 | 0.90650187 |
| Sdf4 | 1.4248695 | 0.83946752 | 0.87948913 | -0.0400216 | 0.58015876 | 0.90650187 |
| Nsfl1c | 0.80766552 | 0.61638157 | 0.65642235 | -0.0400408 | 0.60687191 | 0.90650187 |
| Trim37 | 0.28456725 | 0.25506249 | 0.29512414 | -0.0400616 | 0.72690169 | 0.90650187 |
| Pspc1 | 0.29629351 | 0.26532044 | 0.30539367 | -0.0400732 | 0.72016939 | 0.90650187 |
| Qtrt1 | 0.31513048 | 0.28154259 | 0.32162642 | -0.0400838 | 0.71022749 | 0.90650187 |

| | | | | | | |
|---|---|---|---|---|---|---|
| Psmg3 | 0.41955748 | 0.36605997 | 0.40615302 | -0.040093 | 0.66961279 | 0.90650187 |
| Surf1 | 0.43228783 | 0.37576305 | 0.41586504 | -0.040102 | 0.66592602 | 0.90650187 |
| Med1 | 0.3857482 | 0.3396171 | 0.37975457 | -0.0401375 | 0.68077616 | 0.90650187 |
| Abhd16a | 0.70888691 | 0.56047485 | 0.60062907 | -0.0401542 | 0.61684492 | 0.90650187 |
| Ppp2ca | 0.61730269 | 0.50435666 | 0.54451426 | -0.0401576 | 0.62849154 | 0.90650187 |
| Unc50 | 0.80821846 | 0.61655647 | 0.65672197 | -0.0401655 | 0.60714896 | 0.90650187 |
| Polr2d | 0.54092132 | 0.4539001 | 0.4940785 | -0.0401784 | 0.64114316 | 0.90650187 |
| Parl | 0.52034267 | 0.4396946 | 0.47987397 | -0.0401794 | 0.6451378 | 0.90650187 |
| Efcab14 | 0.44021001 | 0.38160942 | 0.42184912 | -0.0402397 | 0.66423942 | 0.90650187 |
| Atp13a1 | 0.3077275 | 0.27504351 | 0.31528446 | -0.0402409 | 0.71478202 | 0.90650187 |
| Scamp1 | 0.30519261 | 0.27285331 | 0.31310176 | -0.0402484 | 0.71615999 | 0.90650187 |
| Mrpl49 | 0.6228545 | 0.50777785 | 0.54804559 | -0.0402677 | 0.62803054 | 0.90650187 |
| Znhit2 | 0.3708734 | 0.32757944 | 0.36785618 | -0.0402767 | 0.68678275 | 0.90650187 |
| Prr12 | 0.25311246 | 0.22665503 | 0.26694095 | -0.0402859 | 0.74886345 | 0.90650187 |
| Ash2l | 0.661974 | 0.53214563 | 0.57244629 | -0.0403007 | 0.62284985 | 0.90650187 |
| Zfp318 | 0.21030407 | 0.18670119 | 0.22701886 | -0.0403177 | 0.78509664 | 0.90650187 |
| Lcor | 0.24088743 | 0.21540973 | 0.25572865 | -0.0403189 | 0.75831953 | 0.90650187 |
| Trip4 | 0.30502839 | 0.27263721 | 0.31296017 | -0.040323 | 0.71660603 | 0.90650187 |
| Baiap2l1 | 0.22320517 | 0.19891672 | 0.2392472 | -0.0403305 | 0.77319388 | 0.90650187 |
| Tbc1d9b | 0.32844048 | 0.29251441 | 0.3329064 | -0.040392 | 0.70519161 | 0.90650187 |
| Ntmt1 | 0.47165462 | 0.40475732 | 0.44515056 | -0.0403932 | 0.65667193 | 0.90650187 |
| Trp53bp1 | 0.33920704 | 0.30149777 | 0.3419182 | -0.0404204 | 0.70039771 | 0.90650187 |
| Atic | 0.44217952 | 0.38289659 | 0.42332978 | -0.0404332 | 0.66443947 | 0.90650187 |
| Exoc5 | 0.42327312 | 0.36856505 | 0.40900011 | -0.0404351 | 0.66985284 | 0.90650187 |
| Gtf3c1 | 0.30390727 | 0.27154231 | 0.31199283 | -0.0404505 | 0.71782156 | 0.90650187 |
| Arhgap5 | 0.821913 | 0.62360018 | 0.6640973 | -0.0404971 | 0.60684148 | 0.90650187 |
| Echs1 | 0.95943651 | 0.69272569 | 0.73329099 | -0.0405653 | 0.59713132 | 0.90650187 |
| Shkbp1 | 0.24074331 | 0.21500962 | 0.25559557 | -0.0405859 | 0.75987999 | 0.90650187 |
| Gm6563 | 0.40408076 | 0.35358268 | 0.39417822 | -0.0405955 | 0.67646742 | 0.90650187 |
| Fam160a2 | 0.1802657 | 0.1572657 | 0.19786555 | -0.0405999 | 0.81916566 | 0.9092537 |

| | | | | | | |
|---|---|---|---|---|---|---|
| Tmem218 | 0.216192 | 0.19202109 | 0.23262099 | -0.0405999 | 0.78109664 | 0.90650187 |
| Eprs | 1.18885279 | 0.77841072 | 0.81903011 | -0.0406194 | 0.5872504 | 0.90650187 |
| Rprd1b | 0.23211387 | 0.20694941 | 0.24759005 | -0.0406406 | 0.76722044 | 0.90650187 |
| Gpx4 | 2.44512054 | 0.86157097 | 0.90223012 | -0.0406592 | 0.57939344 | 0.90650187 |
| Phospho2 | 0.21573605 | 0.19151459 | 0.23218844 | -0.0406739 | 0.78194054 | 0.90650187 |
| Apoa1bp | 1.08926579 | 0.7447295 | 0.78540443 | -0.0406749 | 0.591044 | 0.90650187 |
| Mysm1 | 0.41109023 | 0.35894212 | 0.39962394 | -0.0406818 | 0.67456862 | 0.90650187 |
| Rps28 | 5.42377901 | 0.55804513 | 0.59874364 | -0.0406985 | 0.61874335 | 0.90650187 |
| Tomm5 | 1.07857005 | 0.74077126 | 0.78147862 | -0.0407074 | 0.59156495 | 0.90650187 |
| Wdr33 | 0.39387482 | 0.34543919 | 0.38618177 | -0.0407426 | 0.68047095 | 0.90650187 |
| Ift27 | 0.53011023 | 0.44589146 | 0.48664934 | -0.0407579 | 0.64517591 | 0.90650187 |
| Lrrc14 | 0.21665492 | 0.19224329 | 0.23305992 | -0.0408166 | 0.78188501 | 0.90650187 |
| Xrcc6 | 0.29149041 | 0.26033926 | 0.30120234 | -0.0408631 | 0.7267856 | 0.90650187 |
| Dnlz | 0.6309911 | 0.51231756 | 0.55319026 | -0.0408727 | 0.62872017 | 0.90650187 |
| Ranbp2 | 0.26728576 | 0.23888358 | 0.27975725 | -0.0408737 | 0.74199923 | 0.90650187 |
| Pmm2 | 0.31280869 | 0.27875711 | 0.31964257 | -0.0408855 | 0.71519882 | 0.90650187 |
| Cdk13 | 0.25900505 | 0.23140291 | 0.2722927 | -0.0408898 | 0.74780232 | 0.90650187 |
| Tmem55b | 0.50950207 | 0.43138879 | 0.47227982 | -0.040891 | 0.6498481 | 0.90650187 |
| Fam118b | 0.22894118 | 0.20370132 | 0.24462805 | -0.0409267 | 0.77148865 | 0.90650187 |
| 1110001J03F | 0.82382787 | 0.62417959 | 0.6651218 | -0.0409422 | 0.60782458 | 0.90650187 |
| Cox7b | 2.19009975 | 0.87996812 | 0.92093332 | -0.0409652 | 0.57837991 | 0.90650187 |
| Vrk3 | 0.3736546 | 0.32903196 | 0.37009421 | -0.0410622 | 0.68907973 | 0.90650187 |
| Arcn1 | 0.73868499 | 0.5768872 | 0.61795071 | -0.0410635 | 0.61616173 | 0.90650187 |
| Zhx1 | 0.27774763 | 0.24801731 | 0.28909381 | -0.0410765 | 0.73619658 | 0.90650187 |
| Kbtbd2 | 0.25538045 | 0.22789042 | 0.26900473 | -0.0411143 | 0.75157687 | 0.90650187 |
| Rbck1 | 0.4692982 | 0.40229335 | 0.44342917 | -0.0411358 | 0.65997975 | 0.90650187 |
| Cds2 | 0.27832505 | 0.24846828 | 0.28960607 | -0.0411378 | 0.7361389 | 0.90650187 |
| Ppp2r2d | 0.18718301 | 0.16352088 | 0.20466537 | -0.0411445 | 0.81426244 | 0.9092537 |
| Wipi2 | 0.50212561 | 0.4259134 | 0.46706842 | -0.041155 | 0.65236661 | 0.90650187 |
| Rps13 | 4.45035653 | 0.57264283 | 0.61382514 | -0.0411823 | 0.61724817 | 0.90650187 |

| | | | | | | |
|---|---|---|---|---|---|---|
| Golgb1 | 1.22227137 | 0.7878381 | 0.82906162 | -0.0412235 | 0.58747299 | 0.90650187 |
| Ykt6 | 0.43214005 | 0.37445093 | 0.41575298 | -0.0413021 | 0.67062472 | 0.90650187 |
| Pet100 | 0.99944903 | 0.70918219 | 0.75049482 | -0.0413126 | 0.59666266 | 0.90650187 |
| Ndufab1 | 1.73022525 | 0.87898425 | 0.920318 | -0.0413337 | 0.57912797 | 0.90650187 |
| Appl2 | 0.34108663 | 0.30214403 | 0.34348141 | -0.0413374 | 0.70368229 | 0.90650187 |
| Phf20l1 | 0.55391359 | 0.46156287 | 0.50290719 | -0.0413443 | 0.64262197 | 0.90650187 |
| Cdc123 | 0.73278293 | 0.57319411 | 0.6145541 | -0.04136 | 0.61760357 | 0.90650187 |
| Zfp397 | 0.31777143 | 0.28249534 | 0.32387724 | -0.0413819 | 0.71498815 | 0.90650187 |
| Rbmxl1 | 0.86472671 | 0.64522223 | 0.68662091 | -0.0413987 | 0.60566594 | 0.90650187 |
| Taf12 | 0.31906798 | 0.28357086 | 0.32498005 | -0.0414092 | 0.71446023 | 0.90650187 |
| Irak1 | 0.39753923 | 0.34762083 | 0.38906237 | -0.0414415 | 0.6820825 | 0.90650187 |
| Epc1 | 0.31454657 | 0.27965495 | 0.32112794 | -0.041473 | 0.71706366 | 0.90650187 |
| Ppp1r12a | 0.70627293 | 0.55758968 | 0.5990884 | -0.0414987 | 0.62093832 | 0.90650187 |
| Ap2a1 | 0.53216353 | 0.44656675 | 0.48806595 | -0.0414992 | 0.64728483 | 0.90650187 |
| Klhl9 | 0.74904304 | 0.58233726 | 0.62386951 | -0.0415323 | 0.61636701 | 0.90650187 |
| Fbxw9 | 0.28985966 | 0.25823719 | 0.29977456 | -0.0415374 | 0.73105016 | 0.90650187 |
| Wrnip1 | 0.32065389 | 0.28476565 | 0.32632699 | -0.0415613 | 0.71437005 | 0.90650187 |
| Gnpda2 | 0.36004787 | 0.31749982 | 0.35908205 | -0.0415822 | 0.69665629 | 0.90650187 |
| Ttc30b | 0.15555014 | 0.13155065 | 0.17313356 | -0.0415829 | 0.85718064 | 0.9170242 |
| Pcbp1 | 1.25131485 | 0.79593242 | 0.8375469 | -0.0416145 | 0.58740904 | 0.90650187 |
| Wdr83os | 1.00474273 | 0.71103902 | 0.75266999 | -0.041631 | 0.59711675 | 0.90650187 |
| Nudt14 | 0.4818855 | 0.4109191 | 0.45258026 | -0.0416612 | 0.65881433 | 0.90650187 |
| Wdr12 | 0.31972492 | 0.28386169 | 0.32553826 | -0.0416766 | 0.71537025 | 0.90650187 |
| Sirt2 | 0.56414844 | 0.46810646 | 0.509789 | -0.0416825 | 0.64188125 | 0.90650187 |
| Nol4l | 0.24911711 | 0.2215953 | 0.26329323 | -0.0416979 | 0.75930034 | 0.90650187 |
| Srp19 | 1.11423392 | 0.75256084 | 0.79425925 | -0.0416984 | 0.59227196 | 0.90650187 |
| Kpna3 | 0.32336096 | 0.28691686 | 0.32862043 | -0.0417036 | 0.71368063 | 0.90650187 |
| Tsc1 | 0.30044088 | 0.26726663 | 0.30899485 | -0.0417282 | 0.72586589 | 0.90650187 |
| Gpr89 | 0.23052734 | 0.20433246 | 0.24611015 | -0.0417777 | 0.77477065 | 0.90650187 |
| Smpd2 | 0.44897181 | 0.38662075 | 0.42841328 | -0.0417925 | 0.66773862 | 0.90650187 |

| | | | | | | |
|---|---|---|---|---|---|---|
| Zfp704 | 0.32589216 | 0.28896296 | 0.33075896 | -0.041796 | 0.71286169 | 0.90650187 |
| Gm9774 | 0.22545264 | 0.19953074 | 0.24135948 | -0.0418287 | 0.77948189 | 0.90650187 |
| Dpf2 | 0.7160224 | 0.56297458 | 0.6048176 | -0.041843 | 0.62079135 | 0.90650187 |
| Fbxl6 | 0.38571892 | 0.33788602 | 0.37973133 | -0.0418453 | 0.68789524 | 0.90650187 |
| Cog6 | 0.32342194 | 0.28680298 | 0.32867201 | -0.041869 | 0.71441154 | 0.90650187 |
| Dr1 | 0.35760383 | 0.31521505 | 0.357088 | -0.0418729 | 0.69892176 | 0.90650187 |
| Sec22b | 1.0325311 | 0.7219801 | 0.76387351 | -0.0418934 | 0.59631183 | 0.90650187 |
| Nipsnap3b | 0.59154203 | 0.48594632 | 0.52789367 | -0.0419473 | 0.63805159 | 0.90650187 |
| 1110065P20I | 0.70250316 | 0.55490924 | 0.59686057 | -0.0419513 | 0.62265708 | 0.90650187 |
| Xpo1 | 0.51186965 | 0.43195703 | 0.47394498 | -0.041988 | 0.65315309 | 0.90650187 |
| Ube2d1 | 0.35524914 | 0.31316825 | 0.35516223 | -0.041994 | 0.70043202 | 0.90650187 |
| Eef1e1 | 0.4958987 | 0.42063434 | 0.46264138 | -0.042007 | 0.65676618 | 0.90650187 |
| Smpd1 | 0.44161035 | 0.38089405 | 0.42290217 | -0.0420081 | 0.67060892 | 0.90650187 |
| Ireb2 | 0.31359766 | 0.27830087 | 0.32031723 | -0.0420164 | 0.7200988 | 0.90650187 |
| Elovl5 | 0.27500937 | 0.24463082 | 0.28666035 | -0.0420295 | 0.74273664 | 0.90650187 |
| Hif1a | 0.56266298 | 0.46669552 | 0.50879411 | -0.0420986 | 0.64350296 | 0.90650187 |
| Ncoa5 | 0.29827099 | 0.26500309 | 0.30711271 | -0.0421096 | 0.72892247 | 0.90650187 |
| Stx18 | 0.36062219 | 0.31743373 | 0.35954991 | -0.0421162 | 0.698726 | 0.90650187 |
| Minpp1 | 0.50162381 | 0.42458743 | 0.46671261 | -0.0421252 | 0.65588475 | 0.90650187 |
| Rae1 | 0.59822543 | 0.49010243 | 0.53224225 | -0.0421398 | 0.63757225 | 0.90650187 |
| Copa | 1.06759047 | 0.73526181 | 0.77740374 | -0.0421419 | 0.59521853 | 0.90650187 |
| Coq4 | 0.25954773 | 0.23062948 | 0.27278389 | -0.0421544 | 0.75395789 | 0.90650187 |
| Map4k4 | 0.37069415 | 0.32554329 | 0.36771173 | -0.0421684 | 0.69490882 | 0.90650187 |
| Psme2 | 1.40525934 | 0.83325233 | 0.87545737 | -0.042205 | 0.58485079 | 0.90650187 |
| Myg1 | 0.47884083 | 0.4081704 | 0.45037666 | -0.0422063 | 0.66151896 | 0.90650187 |
| Slk | 0.19809457 | 0.17307944 | 0.21528665 | -0.0422072 | 0.80827754 | 0.9092537 |
| Keap1 | 0.58212273 | 0.47950378 | 0.52171852 | -0.0422147 | 0.64046996 | 0.90650187 |
| Galnt11 | 0.45440738 | 0.3902381 | 0.43245711 | -0.042219 | 0.66785807 | 0.90650187 |
| Map2k7 | 0.27728777 | 0.24643288 | 0.28868563 | -0.0422527 | 0.74237024 | 0.90650187 |
| Mrpl37 | 0.25828513 | 0.22934004 | 0.27164064 | -0.0423006 | 0.75561977 | 0.90650187 |

| | | | | | | |
|---|---|---|---|---|---|---|
| Klc1 | 0.54100588 | 0.45181059 | 0.49413632 | -0.0423257 | 0.64832147 | 0.90650187 |
| Pias4 | 0.35508785 | 0.3126899 | 0.35503016 | -0.0423403 | 0.70200469 | 0.90650187 |
| 2300009A05l | 0.44844676 | 0.38566364 | 0.42802155 | -0.0423579 | 0.67001207 | 0.90650187 |
| H13 | 1.00755224 | 0.71144442 | 0.75381985 | -0.0423754 | 0.59867016 | 0.90650187 |
| Trmt2a | 0.28711861 | 0.25495406 | 0.2973692 | -0.0424151 | 0.73699135 | 0.90650187 |
| Mycbp | 0.69660448 | 0.55092336 | 0.59336045 | -0.0424371 | 0.62473835 | 0.90650187 |
| Ube2z | 0.22748852 | 0.20082461 | 0.24326848 | -0.0424439 | 0.78101918 | 0.90650187 |
| Fyttd1 | 0.83229974 | 0.62717866 | 0.66963587 | -0.0424572 | 0.61097713 | 0.90650187 |
| Kctd5 | 0.36454329 | 0.32027471 | 0.3627371 | -0.0424624 | 0.69861019 | 0.90650187 |
| Pin1 | 1.17888154 | 0.77338403 | 0.8158521 | -0.0424681 | 0.59150185 | 0.90650187 |
| Maz | 0.53697116 | 0.44886875 | 0.49137249 | -0.0425037 | 0.64971279 | 0.90650187 |
| Nfe2l1 | 0.67493093 | 0.53781995 | 0.58034472 | -0.0425248 | 0.62769396 | 0.90650187 |
| Ahdc1 | 0.33664582 | 0.29724936 | 0.33978331 | -0.0425339 | 0.71106523 | 0.90650187 |
| Akap8l | 0.50767158 | 0.42843409 | 0.4709899 | -0.0425558 | 0.65603956 | 0.90650187 |
| Usp33 | 0.32685241 | 0.28893439 | 0.3315688 | -0.0426344 | 0.71621483 | 0.90650187 |
| Lrrc16a | 0.23450721 | 0.2071831 | 0.24981778 | -0.0426347 | 0.77598428 | 0.90650187 |
| Naa25 | 0.22420818 | 0.19749912 | 0.2401905 | -0.0426914 | 0.78528301 | 0.90650187 |
| Fnta | 0.95282208 | 0.68745034 | 0.73019022 | -0.0427399 | 0.60265009 | 0.90650187 |
| Os9 | 1.04611416 | 0.72644435 | 0.76920155 | -0.0427572 | 0.5975906 | 0.90650187 |
| Bckdk | 0.47429578 | 0.40429234 | 0.44707542 | -0.0427831 | 0.66473629 | 0.90650187 |
| Ube2w | 0.83500757 | 0.62822619 | 0.67107233 | -0.0428461 | 0.61173344 | 0.90650187 |
| Mynn | 0.22110187 | 0.19439268 | 0.23726525 | -0.0428726 | 0.78909857 | 0.90650187 |
| Prdx6 | 1.1305123 | 0.75708139 | 0.79995685 | -0.0428755 | 0.59416498 | 0.90650187 |
| Mta3 | 0.32417168 | 0.28641461 | 0.32930597 | -0.0428914 | 0.7187156 | 0.90650187 |
| Ugp2 | 0.66933796 | 0.53404887 | 0.57694655 | -0.0428977 | 0.6295055 | 0.90650187 |
| Kat7 | 0.25303299 | 0.22393376 | 0.26686855 | -0.0429348 | 0.76276509 | 0.90650187 |
| Zwint | 0.94610205 | 0.68403246 | 0.72702303 | -0.0429906 | 0.60367813 | 0.90650187 |
| Fkrp | 0.23772447 | 0.20980393 | 0.25280341 | -0.0429995 | 0.77522209 | 0.90650187 |
| Sfpq | 1.16358886 | 0.76792254 | 0.81093034 | -0.0430078 | 0.59319571 | 0.90650187 |
| Zfp512b | 0.2052847 | 0.17919398 | 0.22221468 | -0.0430207 | 0.80528959 | 0.90897803 |

| | | | | | | |
|---|---|---|---|---|---|---|
| Atxn2l | 0.49108913 | 0.41615533 | 0.45920348 | -0.0430482 | 0.66157104 | 0.90650187 |
| Otud4 | 0.17944636 | 0.15398578 | 0.19705668 | -0.0430709 | 0.8343954 | 0.91062493 |
| Tmem261 | 0.66140642 | 0.52898734 | 0.57209819 | -0.0431108 | 0.63118808 | 0.90650187 |
| Vamp4 | 0.46751045 | 0.39900167 | 0.44212003 | -0.0431184 | 0.66769742 | 0.90650187 |
| Lage3 | 0.83146067 | 0.62606834 | 0.66919014 | -0.0431218 | 0.6127407 | 0.90650187 |
| Dnajc13 | 0.24495409 | 0.21633341 | 0.25947517 | -0.0431418 | 0.77009227 | 0.90650187 |
| Polr2a | 0.30352753 | 0.26852226 | 0.31166493 | -0.0431427 | 0.73085125 | 0.90650187 |
| Trmt112 | 1.20551558 | 0.78093014 | 0.8240888 | -0.0431587 | 0.59205037 | 0.90650187 |
| Pgam5 | 0.28123892 | 0.24902414 | 0.29218639 | -0.0431623 | 0.74431704 | 0.90650187 |
| Chtf8 | 0.61772138 | 0.50158963 | 0.54478117 | -0.0431915 | 0.63775269 | 0.90650187 |
| Larp1 | 0.38895106 | 0.33910218 | 0.38229379 | -0.0431916 | 0.69225952 | 0.90650187 |
| Specc1l | 0.24519219 | 0.21647973 | 0.25969402 | -0.0432143 | 0.77028006 | 0.90650187 |
| Chd3 | 1.00861616 | 0.7110255 | 0.75425447 | -0.043229 | 0.60055862 | 0.90650187 |
| Ccdc117 | 0.1719193 | 0.14631559 | 0.18959072 | -0.0432751 | 0.84488372 | 0.91325304 |
| Wbscr22 | 0.43555792 | 0.37505545 | 0.41834064 | -0.0432852 | 0.67723519 | 0.90650187 |
| Smim10l1 | 0.95976734 | 0.69015663 | 0.73344565 | -0.043289 | 0.6034881 | 0.90650187 |
| Tra2b | 1.7519202 | 0.87842458 | 0.92171471 | -0.0432901 | 0.58269647 | 0.90650187 |
| Ylpm1 | 0.40024087 | 0.34784404 | 0.39117917 | -0.0433351 | 0.6888127 | 0.90650187 |
| Ccar1 | 0.82717146 | 0.62354063 | 0.66690699 | -0.0433664 | 0.61374008 | 0.90650187 |
| Tgfbr1 | 0.35909454 | 0.31493196 | 0.35830482 | -0.0433729 | 0.7047603 | 0.90650187 |
| Chmp6 | 0.43016792 | 0.37080128 | 0.41425603 | -0.0434548 | 0.67951268 | 0.90650187 |
| Coa6 | 0.41934367 | 0.36251231 | 0.40598881 | -0.0434765 | 0.68298802 | 0.90650187 |
| Sumf1 | 0.36515581 | 0.31974427 | 0.36323386 | -0.0434896 | 0.70272968 | 0.90650187 |
| Gtf2a1 | 0.17997145 | 0.15407352 | 0.19757515 | -0.0435016 | 0.83616632 | 0.91079271 |
| Agap1 | 0.19864372 | 0.17231172 | 0.21581779 | -0.0435061 | 0.81492081 | 0.9092537 |
| Tmx3 | 0.35906511 | 0.3147587 | 0.35828081 | -0.0435221 | 0.70541165 | 0.90650187 |
| Tmem33 | 0.81199352 | 0.61523229 | 0.658764 | -0.0435317 | 0.61553304 | 0.90650187 |
| Usp53 | 0.22624841 | 0.19855154 | 0.24210615 | -0.0435546 | 0.78807487 | 0.90650187 |
| Pfdn1 | 1.54750971 | 0.85753366 | 0.90110215 | -0.0435685 | 0.58509531 | 0.90650187 |
| Tial1 | 0.5923249 | 0.48482028 | 0.5284046 | -0.0435843 | 0.64307347 | 0.90650187 |

| | | | | | | |
|---|---|---|---|---|---|---|
| Zcchc14 | 0.20688667 | 0.18015546 | 0.22375083 | -0.0435954 | 0.8068035 | 0.9092537 |
| Atp6v1a | 0.51072942 | 0.42954326 | 0.47314349 | -0.0436002 | 0.65897346 | 0.90650187 |
| Chchd5 | 0.33880272 | 0.29797363 | 0.34158154 | -0.0436079 | 0.71482872 | 0.90650187 |
| Srpr | 1.14233916 | 0.7604245 | 0.8040483 | -0.0436238 | 0.59529888 | 0.90650187 |
| Idnk | 0.33132792 | 0.29169862 | 0.33533289 | -0.0436343 | 0.71852735 | 0.90650187 |
| Snrnp200 | 0.28678253 | 0.25342298 | 0.29707381 | -0.0436508 | 0.74319259 | 0.90650187 |
| Ppp4c | 1.11194468 | 0.74974137 | 0.79345823 | -0.0437169 | 0.59674772 | 0.90650187 |
| Snx27 | 0.30288173 | 0.26733283 | 0.31110699 | -0.0437742 | 0.73418407 | 0.90650187 |
| Med30 | 0.49070942 | 0.41515199 | 0.45893136 | -0.0437794 | 0.66424802 | 0.90650187 |
| Hnrnpll | 0.67720402 | 0.53793684 | 0.58172119 | -0.0437843 | 0.63103921 | 0.90650187 |
| Aimp2 | 0.44013666 | 0.37793965 | 0.42179393 | -0.0438543 | 0.6780359 | 0.90650187 |
| Nap1l1 | 1.88985593 | 0.88302421 | 0.92688461 | -0.0438604 | 0.58330944 | 0.90650187 |
| Dido1 | 0.37815301 | 0.32979454 | 0.37370103 | -0.0439065 | 0.69927573 | 0.90650187 |
| Psmd14 | 0.68909777 | 0.54497201 | 0.58888042 | -0.0439084 | 0.62985738 | 0.90650187 |
| Asna1 | 0.71024357 | 0.55751705 | 0.60142735 | -0.0439103 | 0.6272484 | 0.90650187 |
| Plxnb2 | 0.51966816 | 0.43545319 | 0.47940385 | -0.0439507 | 0.65820277 | 0.90650187 |
| Cnot11 | 0.2857821 | 0.25223286 | 0.29619388 | -0.043961 | 0.74531396 | 0.90650187 |
| Nup93 | 0.20008525 | 0.17324631 | 0.2172103 | -0.043964 | 0.81588757 | 0.9092537 |
| Ubl7 | 0.51923807 | 0.43510956 | 0.47910393 | -0.0439944 | 0.65844533 | 0.90650187 |
| Rps3a1 | 4.7796605 | 0.52536258 | 0.56935975 | -0.0439972 | 0.63440138 | 0.90650187 |
| Polr3f | 0.2061535 | 0.17900051 | 0.22304812 | -0.0440476 | 0.81000638 | 0.9092537 |
| Abi2 | 0.27420275 | 0.24184823 | 0.28594219 | -0.044094 | 0.75349794 | 0.90650187 |
| Katnbl1 | 0.27933848 | 0.24640676 | 0.29050439 | -0.0440976 | 0.75010769 | 0.90650187 |
| Rnf11 | 0.38674845 | 0.33644206 | 0.38054843 | -0.0441064 | 0.69681267 | 0.90650187 |
| Ptpra | 0.32987081 | 0.28994727 | 0.33410928 | -0.044162 | 0.72160392 | 0.90650187 |
| Srp68 | 0.77605194 | 0.59486778 | 0.63906473 | -0.044197 | 0.62075064 | 0.90650187 |
| Nf2 | 0.19869947 | 0.17162866 | 0.21587169 | -0.044243 | 0.81888647 | 0.9092537 |
| Ube2q1 | 0.28208802 | 0.2486925 | 0.29293681 | -0.0442443 | 0.74903345 | 0.90650187 |
| Eif4e2 | 1.05414143 | 0.72795184 | 0.77229064 | -0.0443388 | 0.60072782 | 0.90650187 |
| Sra1 | 1.1002447 | 0.7449927 | 0.78933407 | -0.0443414 | 0.59860382 | 0.90650187 |

| | | | | | | |
|---|---|---|---|---|---|---|
| Gm14698 | 0.33243179 | 0.29189366 | 0.33625867 | -0.044365 | 0.72124383 | 0.90650187 |
| Sec22a | 0.32786049 | 0.28801889 | 0.33241813 | -0.0443992 | 0.72366867 | 0.90650187 |
| Tpr | 1.35145977 | 0.8190494 | 0.86347126 | -0.0444219 | 0.59045176 | 0.90650187 |
| N4bp2 | 0.36101377 | 0.31543473 | 0.35986877 | -0.044434 | 0.70846705 | 0.90650187 |
| Atp5f1 | 2.31233332 | 0.86868829 | 0.91314686 | -0.0444586 | 0.58567921 | 0.90650187 |
| Tmx4 | 0.34438455 | 0.30172903 | 0.34621707 | -0.044488 | 0.7160847 | 0.90650187 |
| Ppp1r7 | 0.38147958 | 0.33178728 | 0.37635797 | -0.0445707 | 0.70072149 | 0.90650187 |
| Sfswap | 0.445668 | 0.38135762 | 0.42594514 | -0.0445875 | 0.67916379 | 0.90650187 |
| Ccdc174 | 0.49059344 | 0.4142502 | 0.45884822 | -0.044598 | 0.6671632 | 0.90650187 |
| Ldb1 | 0.41523277 | 0.35813294 | 0.40282493 | -0.044692 | 0.68907291 | 0.90650187 |
| Idh3g | 1.10041224 | 0.74462001 | 0.78939348 | -0.0447735 | 0.59953727 | 0.90650187 |
| Tor1aip1 | 0.41672118 | 0.35919292 | 0.40397191 | -0.044779 | 0.68891642 | 0.90650187 |
| Ipo5 | 0.42094443 | 0.36243042 | 0.40721702 | -0.0447866 | 0.68755696 | 0.90650187 |
| Caml | 0.52983462 | 0.44165664 | 0.48645899 | -0.0448023 | 0.65888997 | 0.90650187 |
| Scap | 0.23001953 | 0.20069175 | 0.24563593 | -0.0449442 | 0.79201014 | 0.90681274 |
| Samd4b | 0.30099769 | 0.2645188 | 0.30947715 | -0.0449583 | 0.74080263 | 0.90650187 |
| Gmcl1 | 0.39357084 | 0.34090945 | 0.38594228 | -0.0450328 | 0.69802404 | 0.90650187 |
| Ubac1 | 0.42167878 | 0.36274165 | 0.40777975 | -0.0450381 | 0.68828835 | 0.90650187 |
| Gm26825 | 0.28338421 | 0.24903883 | 0.29408109 | -0.0450423 | 0.752033 | 0.90650187 |
| Hcfc1 | 0.33472622 | 0.29312631 | 0.33817961 | -0.0450533 | 0.72316074 | 0.90650187 |
| Jkamp | 0.64514437 | 0.51693423 | 0.56205215 | -0.0451179 | 0.6394034 | 0.90650187 |
| Pdha1 | 0.67515185 | 0.53530972 | 0.58047861 | -0.0451689 | 0.63530114 | 0.90650187 |
| Trim32 | 0.3095457 | 0.27164015 | 0.31684653 | -0.0452064 | 0.73705132 | 0.90650187 |
| Rmnd5b | 0.22811991 | 0.19855237 | 0.24385968 | -0.0453073 | 0.79558674 | 0.9075779 |
| Phkg2 | 0.29562226 | 0.25943772 | 0.30480916 | -0.0453714 | 0.74593016 | 0.90650187 |
| Anapc11 | 1.40926017 | 0.83090448 | 0.87629473 | -0.0453903 | 0.59106025 | 0.90650187 |
| Ehmt2 | 0.62197035 | 0.50208241 | 0.54748435 | -0.0454019 | 0.64381186 | 0.90650187 |
| Phf14 | 0.62510931 | 0.50402797 | 0.54947493 | -0.045447 | 0.64344973 | 0.90650187 |
| Chd1 | 0.53442973 | 0.44416026 | 0.48962638 | -0.0454661 | 0.66012841 | 0.90650187 |
| Emc8 | 0.67048172 | 0.53216049 | 0.57764279 | -0.0454823 | 0.63684457 | 0.90650187 |

| | | | | | | |
|---|---|---|---|---|---|---|
| Prcc | 0.28362971 | 0.24879772 | 0.29429765 | -0.0454999 | 0.75405744 | 0.90650187 |
| Emc4 | 0.6626063 | 0.52728757 | 0.5728339 | -0.0455463 | 0.6381329 | 0.90650187 |
| Rtfdc1 | 0.58200923 | 0.47609694 | 0.52164379 | -0.0455468 | 0.65105266 | 0.90650187 |
| Etfb | 1.5745207 | 0.85942681 | 0.90497613 | -0.0455493 | 0.58852663 | 0.90650187 |
| Ddx18 | 0.70114563 | 0.55050153 | 0.59605658 | -0.045555 | 0.63298365 | 0.90650187 |
| Tbc1d16 | 0.47114937 | 0.39917981 | 0.44478179 | -0.045602 | 0.67573062 | 0.90650187 |
| Psmd5 | 0.44129998 | 0.37697267 | 0.4226689 | -0.0456962 | 0.68461031 | 0.90650187 |
| Cnih1 | 1.05530739 | 0.72703686 | 0.77273727 | -0.0457004 | 0.60369285 | 0.90650187 |
| Atf1 | 0.44877286 | 0.3825392 | 0.42826487 | -0.0457257 | 0.68248377 | 0.90650187 |
| Fbxo3 | 0.52929028 | 0.44035706 | 0.48608289 | -0.0457258 | 0.66209893 | 0.90650187 |
| Lyrm5 | 0.33609695 | 0.29358903 | 0.33932508 | -0.045736 | 0.72548108 | 0.90650187 |
| Cux1 | 0.5005713 | 0.42012502 | 0.46596577 | -0.0458407 | 0.66907107 | 0.90650187 |
| Rnf146 | 0.50834107 | 0.42558521 | 0.47146193 | -0.0458767 | 0.66734379 | 0.90650187 |
| Arpc2 | 2.14295349 | 0.87717611 | 0.92309295 | -0.0459168 | 0.58750635 | 0.90650187 |
| Pcif1 | 0.62783833 | 0.50526425 | 0.55120112 | -0.0459369 | 0.64449314 | 0.90650187 |
| Stau1 | 0.41421087 | 0.35607001 | 0.40203649 | -0.0459665 | 0.69437303 | 0.90650187 |
| Hint3 | 0.26545905 | 0.23212689 | 0.27811618 | -0.0459893 | 0.76886842 | 0.90650187 |
| Rab24 | 0.80360042 | 0.60820777 | 0.65421557 | -0.0460078 | 0.62272212 | 0.90650187 |
| Mdp1 | 0.40622505 | 0.34982251 | 0.39584809 | -0.0460256 | 0.697385 | 0.90650187 |
| Plaa | 0.38676976 | 0.33452302 | 0.38056533 | -0.0460423 | 0.70465874 | 0.90650187 |
| Glrx2 | 0.70816177 | 0.55410587 | 0.60020201 | -0.0460961 | 0.633609 | 0.90650187 |
| Mecp2 | 0.32757743 | 0.28607946 | 0.33217973 | -0.0461003 | 0.7313715 | 0.90650187 |
| Rbm6 | 0.62724672 | 0.50468479 | 0.55082725 | -0.0461425 | 0.64520459 | 0.90650187 |
| Macrod2 | 0.35596781 | 0.30957686 | 0.35575047 | -0.0461736 | 0.71803185 | 0.90650187 |
| Ubxn6 | 0.75133509 | 0.57894714 | 0.62517234 | -0.0462252 | 0.62881175 | 0.90650187 |
| Bola1 | 0.47733985 | 0.40301084 | 0.449288 | -0.0462772 | 0.67648872 | 0.90650187 |
| Cnot1 | 0.42972496 | 0.3676212 | 0.41391941 | -0.0462982 | 0.69049549 | 0.90650187 |
| Gnpat | 0.29184537 | 0.25520875 | 0.30151281 | -0.0463041 | 0.75260894 | 0.90650187 |
| Cul2 | 0.2669047 | 0.2330982 | 0.27941517 | -0.046317 | 0.76941495 | 0.90650187 |
| Mfap1a | 0.3390117 | 0.29541809 | 0.34175557 | -0.0463375 | 0.72666738 | 0.90650187 |

| | | | | | | |
|---|---|---|---|---|---|---|
| Fkbp1a | 2.77431667 | 0.82258187 | 0.8689461 | -0.0463642 | 0.59375056 | 0.90650187 |
| Vps4a | 0.46850687 | 0.3964769 | 0.44284997 | -0.0463731 | 0.67922114 | 0.90650187 |
| Eif3l | 1.20718367 | 0.77820795 | 0.82458553 | -0.0463776 | 0.59872043 | 0.90650187 |
| Hnrnpd | 0.44444101 | 0.37860327 | 0.42502652 | -0.0464233 | 0.68636344 | 0.90650187 |
| Snx12 | 0.34204191 | 0.29780367 | 0.34427476 | -0.0464711 | 0.72577332 | 0.90650187 |
| Mdm4 | 0.39907375 | 0.34379005 | 0.39026539 | -0.0464753 | 0.70174006 | 0.90650187 |
| Habp4 | 0.27983319 | 0.24437891 | 0.29094256 | -0.0465636 | 0.76161066 | 0.90650187 |
| Atraid | 1.34246399 | 0.81476427 | 0.86134304 | -0.0465788 | 0.5949917 | 0.90650187 |
| Ankhd1 | 0.3252026 | 0.28354291 | 0.33017692 | -0.046634 | 0.73496362 | 0.90650187 |
| Pigu | 0.2451719 | 0.21302429 | 0.25967537 | -0.0466511 | 0.78775619 | 0.90650187 |
| Flii | 0.36272526 | 0.31455038 | 0.3612609 | -0.0467105 | 0.71729032 | 0.90650187 |
| H2afy | 1.50351113 | 0.84718154 | 0.89392465 | -0.0467431 | 0.59190952 | 0.90650187 |
| Psmc4 | 1.17726395 | 0.76858617 | 0.81533333 | -0.0467472 | 0.60059556 | 0.90650187 |
| Tmub2 | 0.31416677 | 0.2740456 | 0.32080355 | -0.0467579 | 0.74149412 | 0.90650187 |
| Tvp23b | 1.06633376 | 0.73013391 | 0.77693441 | -0.0468005 | 0.60557019 | 0.90650187 |
| Chp1 | 0.43006609 | 0.36731794 | 0.41417865 | -0.0468607 | 0.69251145 | 0.90650187 |
| mt-Nd4l | 1.07942702 | 0.73493181 | 0.78179475 | -0.0468629 | 0.60506597 | 0.90650187 |
| Ufm1 | 1.09658473 | 0.74116477 | 0.7880335 | -0.0468687 | 0.60426604 | 0.90650187 |
| Rbfox2 | 0.79608768 | 0.6032467 | 0.65011828 | -0.0468716 | 0.62571097 | 0.90650187 |
| Atp5o | 2.21390422 | 0.87268643 | 0.91967742 | -0.046991 | 0.58984591 | 0.90650187 |
| Prpf4b | 0.96748921 | 0.68992714 | 0.73692246 | -0.0469953 | 0.61160592 | 0.90650187 |
| Cenpb | 0.21552552 | 0.18496031 | 0.23198865 | -0.0470283 | 0.81626049 | 0.9092537 |
| Mrpl42 | 1.00158232 | 0.7042582 | 0.75137274 | -0.0471145 | 0.6097855 | 0.90650187 |
| Rps9 | 4.74875156 | 0.52673653 | 0.57386274 | -0.0471262 | 0.64247337 | 0.90650187 |
| Capza2 | 1.54054086 | 0.85286373 | 0.89999886 | -0.0471351 | 0.59205222 | 0.90650187 |
| Mlycd | 0.1846373 | 0.15502922 | 0.20216896 | -0.0471397 | 0.85003466 | 0.91472298 |
| Rnf4 | 0.56839463 | 0.46543796 | 0.51262543 | -0.0471875 | 0.65881114 | 0.90650187 |
| Sbds | 0.5876435 | 0.4781324 | 0.5253441 | -0.0472117 | 0.65524436 | 0.90650187 |
| Tmem230 | 0.42157328 | 0.36046837 | 0.40769893 | -0.0472306 | 0.69672653 | 0.90650187 |
| Arid1a | 0.71654254 | 0.55779434 | 0.60512179 | -0.0473275 | 0.63596624 | 0.90650187 |

| | | | | | | |
|---|---|---|---|---|---|---|
| Gon4l | 0.50407947 | 0.42112351 | 0.46845226 | -0.0473288 | 0.67333803 | 0.90650187 |
| Amfr | 0.3798142 | 0.32764606 | 0.37502892 | -0.0473829 | 0.71283234 | 0.90650187 |
| Ddx42 | 0.58472285 | 0.47600972 | 0.52342827 | -0.0474186 | 0.65642936 | 0.90650187 |
| Mrpl2 | 0.62470942 | 0.50169813 | 0.54922163 | -0.0475235 | 0.64976018 | 0.90650187 |
| Zfp91 | 0.36974037 | 0.31936891 | 0.36694234 | -0.0475734 | 0.71781576 | 0.90650187 |
| Ptcd2 | 0.2658981 | 0.23093695 | 0.2785109 | -0.047574 | 0.77618678 | 0.90650187 |
| 2810428I15R | 0.99513201 | 0.70111688 | 0.74871264 | -0.0475958 | 0.61126123 | 0.90650187 |
| Pias2 | 0.31327649 | 0.27236148 | 0.32004266 | -0.0476812 | 0.74611801 | 0.90650187 |
| Sf3a2 | 0.50106625 | 0.4186186 | 0.46631707 | -0.0476985 | 0.67534845 | 0.90650187 |
| Pelp1 | 0.3367523 | 0.29209787 | 0.33987217 | -0.0477743 | 0.73398129 | 0.90650187 |
| Hbs1l | 0.40030741 | 0.34339832 | 0.39123124 | -0.0478329 | 0.70661792 | 0.90650187 |
| Mrps26 | 0.84938407 | 0.63080145 | 0.67864102 | -0.0478396 | 0.6230059 | 0.90650187 |
| Zfp62 | 0.30473152 | 0.26483064 | 0.31270413 | -0.0478735 | 0.75193795 | 0.90650187 |
| Dcun1d1 | 0.47223879 | 0.39767972 | 0.44557671 | -0.047897 | 0.68363139 | 0.90650187 |
| Apc | 0.2812758 | 0.24428663 | 0.29221899 | -0.0479324 | 0.76706277 | 0.90650187 |
| Ccnt1 | 0.57700425 | 0.47040046 | 0.51834119 | -0.0479407 | 0.65952555 | 0.90650187 |
| Fam195b | 1.44545929 | 0.83556657 | 0.88352175 | -0.0479552 | 0.59533696 | 0.90650187 |
| Raver1 | 0.34693861 | 0.30036923 | 0.34832907 | -0.0479598 | 0.72976264 | 0.90650187 |
| Kpnb1 | 0.69232662 | 0.54284874 | 0.59081121 | -0.0479625 | 0.64091196 | 0.90650187 |
| Haus7 | 0.25761468 | 0.22301358 | 0.27103294 | -0.0480194 | 0.78454507 | 0.90650187 |
| Dot1l | 0.2245838 | 0.19252176 | 0.24054351 | -0.0480217 | 0.81259894 | 0.9092537 |
| Nat9 | 0.27972063 | 0.2428104 | 0.29084288 | -0.0480325 | 0.76858307 | 0.90650187 |
| Mia3 | 0.59253315 | 0.48045469 | 0.52854041 | -0.0480857 | 0.6570605 | 0.90650187 |
| Tmed7 | 0.42815785 | 0.36459468 | 0.41272735 | -0.0481327 | 0.6979282 | 0.90650187 |
| Dctn4 | 0.30016763 | 0.26062121 | 0.30875808 | -0.0481369 | 0.75587389 | 0.90650187 |
| 9530068E07I | 1.08271332 | 0.73483756 | 0.78300447 | -0.0481669 | 0.60775545 | 0.90650187 |
| Tmem11 | 0.6851562 | 0.5383481 | 0.58651583 | -0.0481677 | 0.64248024 | 0.90650187 |
| Msto1 | 0.24340506 | 0.20983701 | 0.25804999 | -0.048213 | 0.79689977 | 0.9075779 |
| Rabgap1 | 0.475968 | 0.40004327 | 0.44829165 | -0.0482484 | 0.68384152 | 0.90650187 |
| Golga7 | 0.82742366 | 0.61876911 | 0.66704145 | -0.0482723 | 0.62616715 | 0.90650187 |

| | | | | | | |
|---|---|---|---|---|---|---|
| Kctd3 | 0.2787552 | 0.2416837 | 0.28998748 | -0.0483038 | 0.77050569 | 0.90650187 |
| Coq7 | 0.43454893 | 0.36917831 | 0.41757762 | -0.0483993 | 0.69681703 | 0.90650187 |
| Copb1 | 1.17547207 | 0.76629608 | 0.81475759 | -0.0484615 | 0.60427308 | 0.90650187 |
| Sdhaf2 | 0.34608246 | 0.29900993 | 0.34762187 | -0.0486119 | 0.73292384 | 0.90650187 |
| Prpf40a | 1.13412411 | 0.75257587 | 0.80121159 | -0.0486357 | 0.60636616 | 0.90650187 |
| Clpx | 0.48589124 | 0.40676144 | 0.45546991 | -0.0487085 | 0.68275554 | 0.90650187 |
| Znhit3 | 0.33003729 | 0.28550941 | 0.33424917 | -0.0487398 | 0.74158911 | 0.90650187 |
| Tmem199 | 0.39064164 | 0.33488076 | 0.38363085 | -0.0487501 | 0.71393856 | 0.90650187 |
| Zfp326 | 0.65659893 | 0.52039095 | 0.56914289 | -0.0487519 | 0.64832568 | 0.90650187 |
| Ap2m1 | 1.87025385 | 0.87759027 | 0.92645457 | -0.0488643 | 0.5926923 | 0.90650187 |
| Cd2bp2 | 0.36849393 | 0.31695108 | 0.36593576 | -0.0489847 | 0.72411004 | 0.90650187 |
| Brms1l | 0.21775928 | 0.18501909 | 0.23410617 | -0.0490871 | 0.82437023 | 0.90933631 |
| Cacnb3 | 0.2285776 | 0.19510573 | 0.24428797 | -0.0491822 | 0.81461429 | 0.9092537 |
| Ccdc47 | 0.82000602 | 0.61387342 | 0.66307547 | -0.049202 | 0.62925331 | 0.90650187 |
| Rpf2 | 0.48999705 | 0.40921359 | 0.45842056 | -0.049207 | 0.68338745 | 0.90650187 |
| Lztr1 | 0.3757653 | 0.32257924 | 0.37178857 | -0.0492093 | 0.7218725 | 0.90650187 |
| Apmap | 0.33714769 | 0.29092394 | 0.34020207 | -0.0492781 | 0.74019421 | 0.90650187 |
| Suv420h1 | 0.33692344 | 0.29068531 | 0.34001498 | -0.0493297 | 0.7405275 | 0.90650187 |
| Cfap20 | 1.13341021 | 0.75158102 | 0.80096395 | -0.0493829 | 0.60799269 | 0.90650187 |
| Ndufa10 | 1.15073157 | 0.7573888 | 0.80678576 | -0.049397 | 0.60726223 | 0.90650187 |
| Vcpip1 | 0.22288241 | 0.18951932 | 0.2389434 | -0.0494241 | 0.82109556 | 0.9092537 |
| Atp5b | 3.33588673 | 0.74541836 | 0.79485889 | -0.0494405 | 0.60892506 | 0.90650187 |
| Wdr18 | 0.52118371 | 0.43101576 | 0.48045976 | -0.049444 | 0.6763428 | 0.90650187 |
| Nfu1 | 0.83234867 | 0.62018674 | 0.66966186 | -0.0494751 | 0.62871229 | 0.90650187 |
| Fam98c | 0.28995009 | 0.2502727 | 0.29985379 | -0.0495811 | 0.76885613 | 0.90650187 |
| Ndufs4 | 1.41963734 | 0.82884803 | 0.87844536 | -0.0495973 | 0.59909314 | 0.90650187 |
| Nbr1 | 0.46672002 | 0.39193491 | 0.4415405 | -0.0496056 | 0.69126844 | 0.90650187 |
| Rpl27-ps3 | 1.18630837 | 0.76857286 | 0.81822239 | -0.0496495 | 0.60632689 | 0.90650187 |
| Ddx50 | 0.76924281 | 0.58549882 | 0.63526656 | -0.0497677 | 0.63618336 | 0.90650187 |
| Tsfm | 0.45915362 | 0.38619941 | 0.43597109 | -0.0497717 | 0.69410599 | 0.90650187 |

| | | | | | | |
|---|---|---|---|---|---|---|
| Ifrd1 | 0.36290819 | 0.31163471 | 0.36140957 | -0.0497749 | 0.72981968 | 0.90650187 |
| Nsmce2 | 0.46829403 | 0.39283576 | 0.4426941 | -0.0498583 | 0.69170381 | 0.90650187 |
| Nelfa | 0.38762431 | 0.33136706 | 0.38124292 | -0.0498759 | 0.71957978 | 0.90650187 |
| Pak2 | 0.73679351 | 0.56697879 | 0.61686397 | -0.0498852 | 0.64039388 | 0.90650187 |
| Trim44 | 0.37987114 | 0.32514375 | 0.3750744 | -0.0499306 | 0.72301038 | 0.90650187 |
| Rpl34 | 4.55903515 | 0.54863738 | 0.59857148 | -0.0499341 | 0.64462979 | 0.90650187 |
| Btbd1 | 0.43840246 | 0.37048945 | 0.42048777 | -0.0499983 | 0.70146098 | 0.90650187 |
| Avl9 | 0.37422455 | 0.32055252 | 0.37055209 | -0.0499996 | 0.72569937 | 0.90650187 |
| Actr3 | 1.53396017 | 0.84881688 | 0.8989458 | -0.0501289 | 0.59789618 | 0.90650187 |
| Atl2 | 0.28276801 | 0.24336284 | 0.29353731 | -0.0501745 | 0.7763397 | 0.90650187 |
| Zfp282 | 0.23711439 | 0.20191245 | 0.25223804 | -0.0503256 | 0.81245269 | 0.9092537 |
| Uba5 | 0.6055642 | 0.48663747 | 0.5369875 | -0.05035 | 0.66160129 | 0.90650187 |
| Hiat1 | 0.66320419 | 0.52279936 | 0.57320021 | -0.0504009 | 0.6520634 | 0.90650187 |
| Cul3 | 0.38602604 | 0.32955935 | 0.37997517 | -0.0504158 | 0.72235485 | 0.90650187 |
| Safb2 | 0.66653587 | 0.52481377 | 0.57523796 | -0.0504242 | 0.65161815 | 0.90650187 |
| Rnf34 | 0.34186148 | 0.29366925 | 0.34412498 | -0.0504557 | 0.7427402 | 0.90650187 |
| Psmd13 | 1.07229009 | 0.7286964 | 0.77915353 | -0.0504571 | 0.61328552 | 0.90650187 |
| Rps6kb1 | 0.64788178 | 0.51324935 | 0.56375305 | -0.0505037 | 0.65478026 | 0.90650187 |
| Mars | 0.30981499 | 0.26650491 | 0.31707763 | -0.0505727 | 0.76085573 | 0.90650187 |
| Atad1 | 0.64141214 | 0.50913971 | 0.55972658 | -0.0505869 | 0.65607858 | 0.90650187 |
| Crebbp | 0.63875403 | 0.50747073 | 0.55806494 | -0.0505942 | 0.65654023 | 0.90650187 |
| Rcn2 | 1.41527471 | 0.82690071 | 0.87754495 | -0.0506442 | 0.60124138 | 0.90650187 |
| Med7 | 0.41047099 | 0.34849024 | 0.39914435 | -0.0506541 | 0.71368329 | 0.90650187 |
| Polr2j | 1.14557524 | 0.75444622 | 0.80510688 | -0.0506607 | 0.61015997 | 0.90650187 |
| Dalrd3 | 0.46890089 | 0.39237532 | 0.44313842 | -0.0507631 | 0.69471506 | 0.90650187 |
| D8Ertd738e | 1.78692437 | 0.87275591 | 0.92359657 | -0.0508407 | 0.59666127 | 0.90650187 |
| Tbl1x | 0.71367515 | 0.55245351 | 0.60344254 | -0.050989 | 0.64640771 | 0.90650187 |
| Kmt2a | 0.56324729 | 0.45817746 | 0.5091856 | -0.0510081 | 0.67196396 | 0.90650187 |
| Cnpy2 | 1.84588809 | 0.87477546 | 0.92579732 | -0.0510219 | 0.59677294 | 0.90650187 |
| Brd4 | 0.56166975 | 0.45710034 | 0.50812815 | -0.0510278 | 0.67236085 | 0.90650187 |

| | | | | | | |
|---|---|---|---|---|---|---|
| Nelfb | 0.61298861 | 0.49066495 | 0.54175711 | -0.0510922 | 0.66248595 | 0.90650187 |
| Abt1 | 0.29897339 | 0.25662721 | 0.30772243 | -0.0510952 | 0.76979225 | 0.90650187 |
| Smarcad1 | 0.42926711 | 0.36239925 | 0.41357132 | -0.0511721 | 0.7088624 | 0.90650187 |
| Trip12 | 0.59374974 | 0.47807906 | 0.52933316 | -0.0512541 | 0.66656863 | 0.90650187 |
| 2610002M06 | 0.27143265 | 0.23200725 | 0.28347103 | -0.0514638 | 0.79019379 | 0.90650187 |
| Got2 | 0.57444361 | 0.46515406 | 0.51664582 | -0.0514918 | 0.67114373 | 0.90650187 |
| Rsbn1 | 0.30595254 | 0.26209478 | 0.3137567 | -0.0516619 | 0.76791442 | 0.90650187 |
| Prpf31 | 0.4308143 | 0.36308004 | 0.41474698 | -0.0516669 | 0.71014544 | 0.90650187 |
| Tm2d3 | 0.56840704 | 0.46080344 | 0.5126337 | -0.0518303 | 0.6734561 | 0.90650187 |
| Pigf | 0.27721244 | 0.2367847 | 0.28861875 | -0.0518341 | 0.78768139 | 0.90650187 |
| Ap1m1 | 0.45039033 | 0.37761799 | 0.42947061 | -0.0518526 | 0.70427823 | 0.90650187 |
| Cfap97 | 0.35107531 | 0.29977934 | 0.35173752 | -0.0519582 | 0.7443127 | 0.90650187 |
| Aifm1 | 0.36350461 | 0.30987606 | 0.36189405 | -0.052018 | 0.73858961 | 0.90650187 |
| Setd5 | 0.98381937 | 0.69194268 | 0.74396529 | -0.0520226 | 0.62204801 | 0.90650187 |
| Taf11 | 0.54707061 | 0.44621948 | 0.49827051 | -0.052051 | 0.67882053 | 0.90650187 |
| Eif2b2 | 0.44279038 | 0.37165832 | 0.42378845 | -0.0521301 | 0.70776019 | 0.90650187 |
| Zfp207 | 1.33612206 | 0.80765878 | 0.85979294 | -0.0521342 | 0.60625158 | 0.90650187 |
| Mrps24 | 1.18651949 | 0.76614753 | 0.8182895 | -0.052142 | 0.61151777 | 0.90650187 |
| Brix1 | 0.56711634 | 0.45961828 | 0.51177278 | -0.0521545 | 0.67474345 | 0.90650187 |
| Tmem115 | 0.43656387 | 0.36693397 | 0.41910062 | -0.0521667 | 0.70999063 | 0.90650187 |
| Tomm7 | 2.35140878 | 0.85793063 | 0.91010333 | -0.0521727 | 0.6005803 | 0.90650187 |
| Mrpl33 | 1.65891568 | 0.86222532 | 0.91455741 | -0.0523321 | 0.60040016 | 0.90650187 |
| Usp10 | 0.31816442 | 0.2718754 | 0.32421167 | -0.0523363 | 0.76350956 | 0.90650187 |
| Mapk8ip2 | 0.22739106 | 0.1907871 | 0.2431772 | -0.0523901 | 0.83090848 | 0.9098597 |
| Pcgf3 | 0.37617205 | 0.31971664 | 0.37211468 | -0.052398 | 0.73434009 | 0.90650187 |
| Coro1b | 0.58435476 | 0.47073113 | 0.52318647 | -0.0524553 | 0.67210101 | 0.90650187 |
| Strn3 | 0.59474981 | 0.47752313 | 0.52998417 | -0.052461 | 0.67005065 | 0.90650187 |
| Pex14 | 0.45155578 | 0.37777135 | 0.43033824 | -0.0525669 | 0.70644953 | 0.90650187 |
| Vapb | 1.02408528 | 0.70793783 | 0.76050748 | -0.0525696 | 0.62069248 | 0.90650187 |
| Mllt10 | 0.47693975 | 0.39630208 | 0.44899754 | -0.0526955 | 0.69907689 | 0.90650187 |

| | | | | | | |
|---|---|---|---|---|---|---|
| Ythdf2 | 0.63867176 | 0.50530467 | 0.55801345 | -0.0527088 | 0.66272902 | 0.90650187 |
| Pigk | 0.58799198 | 0.47282036 | 0.52557236 | -0.052752 | 0.67227653 | 0.90650187 |
| Pdcd10 | 0.84556207 | 0.62385657 | 0.67663799 | -0.0527814 | 0.63562221 | 0.90650187 |
| Rab12 | 0.2572465 | 0.2179065 | 0.27069905 | -0.0527925 | 0.80702777 | 0.9092537 |
| Cuedc2 | 0.81131506 | 0.605559 | 0.65839746 | -0.0528385 | 0.63936944 | 0.90650187 |
| Gm5900 | 0.35117033 | 0.2989741 | 0.35181564 | -0.0528415 | 0.74784235 | 0.90650187 |
| Smarcb1 | 1.03264227 | 0.7109284 | 0.76391763 | -0.0529892 | 0.6210996 | 0.90650187 |
| Krit1 | 0.76077189 | 0.57750052 | 0.63051068 | -0.0530102 | 0.64571012 | 0.90650187 |
| Acadvl | 0.66288056 | 0.51998897 | 0.57300195 | -0.053013 | 0.65957318 | 0.90650187 |
| Helz | 0.46160212 | 0.38473724 | 0.43777772 | -0.0530405 | 0.70492643 | 0.90650187 |
| Elp2 | 0.34814183 | 0.29623108 | 0.34932193 | -0.0530908 | 0.75037006 | 0.90650187 |
| Set | 1.87930624 | 0.87350405 | 0.9266671 | -0.0531631 | 0.60065591 | 0.90650187 |
| Timm17b | 0.85728286 | 0.62959271 | 0.68276163 | -0.0531689 | 0.63540161 | 0.90650187 |
| 2310009B15I | 0.49088328 | 0.40586048 | 0.45905597 | -0.0531955 | 0.69677857 | 0.90650187 |
| Cldnd1 | 0.96109199 | 0.6808649 | 0.73406449 | -0.0531996 | 0.62634367 | 0.90650187 |
| Pkn2 | 0.90226562 | 0.65247731 | 0.70567987 | -0.0532026 | 0.63124885 | 0.90650187 |
| Sf3a3 | 0.62782012 | 0.49795946 | 0.55118961 | -0.0532302 | 0.66615719 | 0.90650187 |
| Upf2 | 0.78205979 | 0.58916346 | 0.64239861 | -0.0532351 | 0.64371416 | 0.90650187 |
| Vdac1 | 1.58682084 | 0.85319269 | 0.90643891 | -0.0532462 | 0.60300955 | 0.90650187 |
| Ctdnep1 | 0.39456756 | 0.33347502 | 0.38672729 | -0.0532523 | 0.72978374 | 0.90650187 |
| Fxr1 | 0.88092051 | 0.64161647 | 0.6949229 | -0.0533064 | 0.633454 | 0.90650187 |
| Larp4 | 0.28476862 | 0.24188459 | 0.29530155 | -0.053417 | 0.78934878 | 0.90650187 |
| Dnajc4 | 0.28685811 | 0.24366939 | 0.29714025 | -0.0534709 | 0.78814284 | 0.90650187 |
| 9930021J03F | 0.24224626 | 0.20346831 | 0.25698232 | -0.053514 | 0.82271287 | 0.9092537 |
| Rpe | 0.32073891 | 0.27287274 | 0.32639914 | -0.0535264 | 0.76701072 | 0.90650187 |
| Wdr74 | 0.36263943 | 0.3076297 | 0.36119115 | -0.0535614 | 0.74513047 | 0.90650187 |
| Traf7 | 0.49446057 | 0.40804064 | 0.46161529 | -0.0535747 | 0.6970564 | 0.90650187 |
| Nfs1 | 0.41149894 | 0.34635973 | 0.39994033 | -0.0535806 | 0.72427358 | 0.90650187 |
| Rbm27 | 0.53927047 | 0.43932607 | 0.49294878 | -0.0536227 | 0.68566184 | 0.90650187 |
| Oaz2 | 1.36723093 | 0.81349838 | 0.86712224 | -0.0536239 | 0.60831175 | 0.90650187 |

| | | | | | | |
|---|---|---|---|---|---|---|
| Rtcb | 1.00756745 | 0.70010496 | 0.75382607 | -0.0537211 | 0.62430776 | 0.90650187 |
| Eif2b4 | 0.41541739 | 0.34915762 | 0.40296729 | -0.0538097 | 0.72362282 | 0.90650187 |
| Rpl36al | 3.21230344 | 0.75712412 | 0.81093611 | -0.053812 | 0.61600343 | 0.90650187 |
| Gm10093 | 0.5741766 | 0.46264113 | 0.51646882 | -0.0538277 | 0.67843691 | 0.90650187 |
| Tlk1 | 0.28346267 | 0.24030941 | 0.29415031 | -0.0538409 | 0.79209676 | 0.90681274 |
| Ubald1 | 0.33126177 | 0.28143597 | 0.33527738 | -0.0538414 | 0.76235846 | 0.90650187 |
| Rdh14 | 0.39758285 | 0.33524931 | 0.3890966 | -0.0538473 | 0.73079853 | 0.90650187 |
| Txnl4a | 0.82739715 | 0.61309724 | 0.66702731 | -0.0539301 | 0.64036456 | 0.90650187 |
| Clock | 0.46392328 | 0.38539525 | 0.43948653 | -0.0540913 | 0.7078649 | 0.90650187 |
| Arl1 | 1.53566879 | 0.84508775 | 0.89922027 | -0.0541325 | 0.60550517 | 0.90650187 |
| Ddx1 | 0.91867015 | 0.65965454 | 0.71379654 | -0.054142 | 0.6320179 | 0.90650187 |
| Srrt | 0.67335636 | 0.52523627 | 0.57938967 | -0.0541534 | 0.66111728 | 0.90650187 |
| Ppp2r5c | 0.29098086 | 0.2465348 | 0.30075648 | -0.0542217 | 0.78857108 | 0.90650187 |
| Prelid1 | 1.35139564 | 0.80910443 | 0.86345627 | -0.0543518 | 0.61019866 | 0.90650187 |
| Cyc1 | 1.30602332 | 0.7979403 | 0.85230988 | -0.0543696 | 0.61163733 | 0.90650187 |
| Umps | 0.43957433 | 0.36697598 | 0.42137063 | -0.0543947 | 0.71697633 | 0.90650187 |
| Ash1l | 0.96391794 | 0.68092324 | 0.73536787 | -0.0544446 | 0.62897529 | 0.90650187 |
| Ehmt1 | 0.48610805 | 0.40109556 | 0.455626 | -0.0545304 | 0.70265815 | 0.90650187 |
| Swi5 | 2.36847766 | 0.85410472 | 0.90871303 | -0.0546083 | 0.605325 | 0.90650187 |
| Mrpl38 | 0.69376357 | 0.53701322 | 0.59166854 | -0.0546553 | 0.65933995 | 0.90650187 |
| N4bp2l2 | 0.86027576 | 0.62960392 | 0.68431631 | -0.0547124 | 0.63886801 | 0.90650187 |
| Rap1a | 1.34032769 | 0.80607214 | 0.86082223 | -0.0547501 | 0.61131569 | 0.90650187 |
| Pomp | 1.85233654 | 0.87120647 | 0.92598369 | -0.0547772 | 0.60371816 | 0.90650187 |
| Ptges3 | 1.6869088 | 0.86237389 | 0.91721754 | -0.0548437 | 0.60481046 | 0.90650187 |
| Jmjd6 | 0.66917743 | 0.52199581 | 0.57684877 | -0.054853 | 0.66375601 | 0.90650187 |
| Nckap1 | 0.52806196 | 0.43030531 | 0.48523354 | -0.0549282 | 0.6926025 | 0.90650187 |
| Btf3l4 | 1.27633783 | 0.78950616 | 0.84454242 | -0.0550363 | 0.61397966 | 0.90650187 |
| Scoc | 0.67680507 | 0.52644205 | 0.5814798 | -0.0550377 | 0.66304281 | 0.90650187 |
| Ttc1 | 0.63299363 | 0.49938181 | 0.55445087 | -0.0550691 | 0.670591 | 0.90650187 |
| Ctdspl2 | 0.46996944 | 0.38875418 | 0.44391991 | -0.0551657 | 0.70968117 | 0.90650187 |

| | | | | | | |
|---|---|---|---|---|---|---|
| Ccnc | 0.31031634 | 0.26219969 | 0.31750773 | -0.055308 | 0.78065265 | 0.90650187 |
| Cisd2 | 0.76843034 | 0.57945824 | 0.63481195 | -0.0553537 | 0.65086011 | 0.90650187 |
| Isca2 | 0.50653101 | 0.41477139 | 0.47018506 | -0.0554137 | 0.69983885 | 0.90650187 |
| Ildr1 | 0.2759737 | 0.23205612 | 0.28751815 | -0.055462 | 0.8044306 | 0.90882608 |
| Rab8a | 0.62223369 | 0.49214573 | 0.54765156 | -0.0555058 | 0.67385136 | 0.90650187 |
| Xrn1 | 0.35666759 | 0.3007956 | 0.35632284 | -0.0555272 | 0.75577522 | 0.90650187 |
| Synj2bp | 0.54923504 | 0.44420893 | 0.49974019 | -0.0555313 | 0.68934582 | 0.90650187 |
| Nck1 | 0.59515112 | 0.47471306 | 0.53024525 | -0.0555322 | 0.67924374 | 0.90650187 |
| Exosc4 | 0.76925665 | 0.5797203 | 0.6352743 | -0.055554 | 0.6512743 | 0.90650187 |
| Pdcd11 | 0.40060083 | 0.3357651 | 0.39146079 | -0.0556957 | 0.7364717 | 0.90650187 |
| Kansl2 | 0.5799673 | 0.4645661 | 0.52029817 | -0.0557321 | 0.6830319 | 0.90650187 |
| Mat2a | 1.10082876 | 0.73375357 | 0.78954115 | -0.0557876 | 0.62328455 | 0.90650187 |
| Prpsap1 | 0.63969356 | 0.50280039 | 0.5586527 | -0.0558523 | 0.67164407 | 0.90650187 |
| Preb | 0.50599065 | 0.41390616 | 0.46980346 | -0.0558973 | 0.70157614 | 0.90650187 |
| Necap1 | 0.34776343 | 0.29299683 | 0.34900981 | -0.056013 | 0.76222575 | 0.90650187 |
| Phpt1 | 1.35722387 | 0.8087857 | 0.86481424 | -0.0560285 | 0.61333298 | 0.90650187 |
| Gm16286 | 0.85689267 | 0.6265054 | 0.68255868 | -0.0560533 | 0.64247539 | 0.90650187 |
| Commd7 | 0.68363259 | 0.52947493 | 0.58559969 | -0.0561248 | 0.66497291 | 0.90650187 |
| Mrpl16 | 0.30909026 | 0.26025384 | 0.31645552 | -0.0562017 | 0.78509831 | 0.90650187 |
| Aspscr1 | 0.49541001 | 0.40605592 | 0.46229286 | -0.0562369 | 0.70567678 | 0.90650187 |
| Eif2s2 | 2.35998174 | 0.85311031 | 0.90941468 | -0.0563044 | 0.60843397 | 0.90650187 |
| Son | 2.31581939 | 0.85657291 | 0.91288095 | -0.056308 | 0.60803933 | 0.90650187 |
| Pabpn1 | 0.59032727 | 0.47077372 | 0.52710018 | -0.0563265 | 0.68262861 | 0.90650187 |
| Actr10 | 0.9208633 | 0.65853482 | 0.714868 | -0.0563332 | 0.63695208 | 0.90650187 |
| l7Rn6 | 0.63852049 | 0.50155985 | 0.55791877 | -0.0563589 | 0.67331264 | 0.90650187 |
| Sdf2 | 1.03028318 | 0.70653573 | 0.76298033 | -0.0564446 | 0.62887756 | 0.90650187 |
| Emc7 | 1.05751664 | 0.71698151 | 0.77358211 | -0.0566006 | 0.62751209 | 0.90650187 |
| Prmt7 | 0.37481771 | 0.31440326 | 0.37102833 | -0.0566251 | 0.75126415 | 0.90650187 |
| Rhobtb3 | 0.27854576 | 0.23308618 | 0.28980179 | -0.0567156 | 0.80785708 | 0.9092537 |
| Tpd52l2 | 0.82645003 | 0.6097862 | 0.6665222 | -0.056736 | 0.64744103 | 0.90650187 |

| | | | | | | |
|---|---|---|---|---|---|---|
| Lpp | 0.70996837 | 0.54447331 | 0.60126549 | -0.0567922 | 0.66272337 | 0.90650187 |
| Ankrd10 | 0.44976628 | 0.37219579 | 0.42900563 | -0.0568098 | 0.72196761 | 0.90650187 |
| Lamtor5 | 1.01111436 | 0.69837068 | 0.75527322 | -0.0569025 | 0.63116238 | 0.90650187 |
| Ppp3cb | 0.38768468 | 0.32438146 | 0.38129077 | -0.0569093 | 0.74650317 | 0.90650187 |
| Wdr26 | 0.58688993 | 0.4679373 | 0.52485026 | -0.056913 | 0.68512057 | 0.90650187 |
| App | 2.12397017 | 0.866919 | 0.92384827 | -0.0569293 | 0.60793718 | 0.90650187 |
| Mar-07 | 0.65129148 | 0.50893104 | 0.56586608 | -0.056935 | 0.67267006 | 0.90650187 |
| Cyb5r4 | 0.50811043 | 0.41432828 | 0.47129936 | -0.0569711 | 0.70450046 | 0.90650187 |
| Rpp25l | 0.55988139 | 0.44995049 | 0.50692757 | -0.0569771 | 0.69134707 | 0.90650187 |
| Ing2 | 0.53153272 | 0.43049392 | 0.48763102 | -0.0571371 | 0.69878325 | 0.90650187 |
| Rbm5 | 1.13412717 | 0.74406307 | 0.80121265 | -0.0571496 | 0.62441614 | 0.90650187 |
| Bap1 | 0.24835929 | 0.20541153 | 0.2625996 | -0.0571881 | 0.83351514 | 0.91060043 |
| Yipf3 | 0.94892852 | 0.6711288 | 0.72835725 | -0.0572284 | 0.63656793 | 0.90650187 |
| Fubp1 | 1.69665567 | 0.86071689 | 0.91795468 | -0.0572378 | 0.60918659 | 0.90650187 |
| Hnrnpab | 1.52828116 | 0.84073388 | 0.89802821 | -0.0572943 | 0.61165315 | 0.90650187 |
| Snw1 | 0.97611257 | 0.68324268 | 0.74065491 | -0.0574122 | 0.63480372 | 0.90650187 |
| Polr3k | 0.77027931 | 0.57843363 | 0.63584608 | -0.0574124 | 0.65593979 | 0.90650187 |
| Dnajc7 | 0.96468822 | 0.67827591 | 0.73570362 | -0.0574277 | 0.63571039 | 0.90650187 |
| Ndrg3 | 0.4844224 | 0.39689125 | 0.45441158 | -0.0575203 | 0.71319194 | 0.90650187 |
| Vdac2 | 1.71023764 | 0.86137972 | 0.91894337 | -0.0575636 | 0.6096772 | 0.90650187 |
| Prkag1 | 0.51864549 | 0.4210765 | 0.47869051 | -0.057614 | 0.70369657 | 0.90650187 |
| Commd6 | 0.73342155 | 0.5572118 | 0.61492242 | -0.0577106 | 0.66174171 | 0.90650187 |
| Zmynd11 | 0.67498552 | 0.52266595 | 0.5803778 | -0.0577119 | 0.67077859 | 0.90650187 |
| Llph | 1.07840525 | 0.72366495 | 0.78141779 | -0.0577528 | 0.62875768 | 0.90650187 |
| Zc3h18 | 0.40112636 | 0.33400833 | 0.39187176 | -0.0578634 | 0.74422762 | 0.90650187 |
| Pgrmc2 | 0.4446003 | 0.36720456 | 0.42514584 | -0.0579413 | 0.72770025 | 0.90650187 |
| Magohb | 0.46408104 | 0.38161783 | 0.43960254 | -0.0579847 | 0.72119223 | 0.90650187 |
| Ndufb3 | 1.46948912 | 0.82997816 | 0.88802946 | -0.0580513 | 0.61432647 | 0.90650187 |
| Pum2 | 0.69862738 | 0.53648452 | 0.59456274 | -0.0580782 | 0.667949 | 0.90650187 |
| Anp32a | 1.87197743 | 0.86841364 | 0.92649618 | -0.0580825 | 0.60976147 | 0.90650187 |

| | | | | | | |
|---|---|---|---|---|---|---|
| Rrp7a | 0.60712939 | 0.47990322 | 0.5379956 | -0.0580924 | 0.68439817 | 0.90650187 |
| Purb | 0.63476998 | 0.49747235 | 0.55556728 | -0.0580949 | 0.67898812 | 0.90650187 |
| Gm6483 | 0.88132476 | 0.63700556 | 0.69512881 | -0.0581232 | 0.64494922 | 0.90650187 |
| Phax | 0.82584665 | 0.60797109 | 0.66620022 | -0.0582291 | 0.65120195 | 0.90650187 |
| Pgam1 | 1.96749834 | 0.8690509 | 0.92736261 | -0.0583117 | 0.6100831 | 0.90650187 |
| Anapc7 | 0.35453488 | 0.29625763 | 0.35457718 | -0.0583195 | 0.76767056 | 0.90650187 |
| Tmem60 | 0.41119707 | 0.34137679 | 0.39970666 | -0.0583299 | 0.74175158 | 0.90650187 |
| 1110004E09I | 0.42513417 | 0.35206172 | 0.41042223 | -0.0583605 | 0.73635515 | 0.90650187 |
| Pcf11 | 0.36795327 | 0.30702576 | 0.36549875 | -0.058473 | 0.76152424 | 0.90650187 |
| Rabggtb | 0.5854727 | 0.46541097 | 0.52392061 | -0.0585096 | 0.69021855 | 0.90650187 |
| Ube2j2 | 0.8234215 | 0.60636894 | 0.66490451 | -0.0585356 | 0.65223947 | 0.90650187 |
| Mrpl44 | 0.39191016 | 0.32603228 | 0.38463266 | -0.0586004 | 0.75089424 | 0.90650187 |
| Serinc1 | 1.42214564 | 0.82025793 | 0.87894671 | -0.0586888 | 0.61670597 | 0.90650187 |
| Hmbs | 0.42702357 | 0.35311386 | 0.41186342 | -0.0587496 | 0.73700395 | 0.90650187 |
| Aggf1 | 0.44731606 | 0.36839837 | 0.4271773 | -0.0587789 | 0.72963469 | 0.90650187 |
| Ebna1bp2 | 0.86648823 | 0.62871538 | 0.68752908 | -0.0588137 | 0.64813608 | 0.90650187 |
| Rpl27a | 4.45947167 | 0.55349232 | 0.61245032 | -0.058958 | 0.66565979 | 0.90650187 |
| Tmem14c | 1.39849468 | 0.81505501 | 0.87403109 | -0.0589761 | 0.61790057 | 0.90650187 |
| Unc45a | 0.36315413 | 0.30262329 | 0.36160938 | -0.0589861 | 0.76582672 | 0.90650187 |
| Zcchc7 | 0.50344508 | 0.40897459 | 0.46800322 | -0.0590286 | 0.71250528 | 0.90650187 |
| Ndufb7 | 1.9264551 | 0.86816694 | 0.92720797 | -0.059041 | 0.61144189 | 0.90650187 |
| Tmem242 | 0.87748396 | 0.63402275 | 0.69316986 | -0.0591471 | 0.64778108 | 0.90650187 |
| Tspyl1 | 0.66270657 | 0.51370448 | 0.57289534 | -0.0591909 | 0.67699592 | 0.90650187 |
| Hnrnpul1 | 0.94236231 | 0.66595479 | 0.72525305 | -0.0592983 | 0.64187771 | 0.90650187 |
| Wdr61 | 0.87476122 | 0.63247764 | 0.69177761 | -0.0593 | 0.64842899 | 0.90650187 |
| Psmb4 | 1.81282424 | 0.86535032 | 0.92475301 | -0.0594027 | 0.61239596 | 0.90650187 |
| Ep400 | 0.35234106 | 0.29319495 | 0.3527776 | -0.0595827 | 0.77362678 | 0.90650187 |
| Prrc2c | 1.63190271 | 0.85199455 | 0.91164056 | -0.059646 | 0.61442203 | 0.90650187 |
| Phf12 | 0.33968713 | 0.28265714 | 0.34231775 | -0.0596606 | 0.78077089 | 0.90650187 |
| Cnot4 | 0.62275215 | 0.48818535 | 0.54798064 | -0.0597953 | 0.68619789 | 0.90650187 |

| | | | | | | |
|---|---|---|---|---|---|---|
| Maf1 | 0.78913244 | 0.58649187 | 0.64630272 | -0.0598108 | 0.65961407 | 0.90650187 |
| Amz2 | 0.49487199 | 0.40209377 | 0.46190898 | -0.0598152 | 0.71758654 | 0.90650187 |
| Ybx1 | 3.45525718 | 0.71761339 | 0.77743228 | -0.0598189 | 0.63384846 | 0.90650187 |
| Srp54b | 1.29435662 | 0.78934585 | 0.84935464 | -0.0600088 | 0.62327441 | 0.90650187 |
| Nrd1 | 0.79260208 | 0.58806549 | 0.6482089 | -0.0601434 | 0.6600066 | 0.90650187 |
| Dnm1l | 0.50732814 | 0.41054611 | 0.47074765 | -0.0602015 | 0.71516112 | 0.90650187 |
| Igf2bp2 | 0.53099466 | 0.42687726 | 0.48725985 | -0.0603826 | 0.70915295 | 0.90650187 |
| Gripap1 | 0.44371936 | 0.3640339 | 0.42448548 | -0.0604516 | 0.73666748 | 0.90650187 |
| Prpf38a | 0.60591421 | 0.47669222 | 0.53721306 | -0.0605208 | 0.69175422 | 0.90650187 |
| Ndufb11 | 2.43050841 | 0.84293806 | 0.90347478 | -0.0605367 | 0.61710066 | 0.90650187 |
| Rc3h2 | 0.44986452 | 0.36846867 | 0.42907884 | -0.0606102 | 0.73498848 | 0.90650187 |
| Cops5 | 1.0493105 | 0.70982433 | 0.77043456 | -0.0606102 | 0.63673184 | 0.90650187 |
| Fbxo22 | 0.47473536 | 0.38669753 | 0.44739532 | -0.0606978 | 0.72678959 | 0.90650187 |
| Hnrnpf | 2.27614952 | 0.85499883 | 0.91576662 | -0.0607678 | 0.61600245 | 0.90650187 |
| Ip6k1 | 0.41421683 | 0.34127273 | 0.40204109 | -0.0607684 | 0.74919279 | 0.90650187 |
| Nelfcd | 0.405187 | 0.33422378 | 0.39504015 | -0.0608164 | 0.7531388 | 0.90650187 |
| Ilkap | 0.83021705 | 0.60769112 | 0.66852895 | -0.0608378 | 0.65710007 | 0.90650187 |
| Pmpcb | 0.66188639 | 0.51148439 | 0.57239257 | -0.0609082 | 0.68191342 | 0.90650187 |
| Snrpc | 1.38439336 | 0.80992955 | 0.87096386 | -0.0610343 | 0.6223025 | 0.90650187 |
| Kmt2e | 1.09352963 | 0.72586474 | 0.78694403 | -0.0610793 | 0.63497133 | 0.90650187 |
| Ergic2 | 1.00215221 | 0.69052681 | 0.75160695 | -0.0610801 | 0.64105379 | 0.90650187 |
| Lsm8 | 0.72700584 | 0.54997843 | 0.61121335 | -0.0612349 | 0.67197986 | 0.90650187 |
| Cox5a | 1.59574963 | 0.84623514 | 0.90747725 | -0.0612421 | 0.61791778 | 0.90650187 |
| Ergic3 | 1.60125707 | 0.84682307 | 0.90812234 | -0.0612993 | 0.61794323 | 0.90650187 |
| Fip1l1 | 1.00581761 | 0.69171434 | 0.75311028 | -0.0613959 | 0.64148088 | 0.90650187 |
| Meaf6 | 0.52302428 | 0.42018729 | 0.48174013 | -0.0615528 | 0.71499048 | 0.90650187 |
| Ndel1 | 0.59325855 | 0.4674336 | 0.52901319 | -0.0615796 | 0.69759266 | 0.90650187 |
| U2af2 | 0.80491173 | 0.59313781 | 0.65492822 | -0.0617904 | 0.66254866 | 0.90650187 |
| Chmp7 | 0.46292678 | 0.37689306 | 0.43875338 | -0.0618603 | 0.73460184 | 0.90650187 |
| Sec11a | 1.678373 | 0.85467569 | 0.91655453 | -0.0618788 | 0.61796244 | 0.90650187 |

| | | | | | | |
|---|---|---|---|---|---|---|
| Prpf8 | 0.4205274 | 0.34498072 | 0.40689728 | -0.0619166 | 0.7506311 | 0.90650187 |
| Hdgf | 0.79029699 | 0.58495962 | 0.64694326 | -0.0619836 | 0.66492187 | 0.90650187 |
| Dnajc30 | 0.4855709 | 0.39314907 | 0.45523922 | -0.0620901 | 0.72785448 | 0.90650187 |
| Slmo2 | 0.65767845 | 0.50759387 | 0.56980755 | -0.0622137 | 0.68629931 | 0.90650187 |
| Dync1i2 | 1.79871506 | 0.86186564 | 0.92416611 | -0.0623005 | 0.61779319 | 0.90650187 |
| Brwd1 | 0.58646791 | 0.46223795 | 0.52457356 | -0.0623356 | 0.70134608 | 0.90650187 |
| Dag1 | 0.56203232 | 0.44598849 | 0.50837132 | -0.0623828 | 0.70730392 | 0.90650187 |
| Sdhc | 0.95915546 | 0.67069447 | 0.73315957 | -0.0624651 | 0.64756884 | 0.90650187 |
| Arl3 | 1.20829822 | 0.7624151 | 0.82491689 | -0.0625018 | 0.63187821 | 0.90650187 |
| Nelfe | 0.48298602 | 0.39080442 | 0.45337523 | -0.0625708 | 0.73024134 | 0.90650187 |
| Pafah1b1 | 1.41260599 | 0.81438503 | 0.87699148 | -0.0626065 | 0.62451548 | 0.90650187 |
| Lamtor2 | 1.43106358 | 0.81808214 | 0.88071478 | -0.0626326 | 0.62405657 | 0.90650187 |
| Ppp2r1a | 1.16581288 | 0.74870378 | 0.81164126 | -0.0629375 | 0.63485048 | 0.90650187 |
| Glg1 | 0.78917746 | 0.58330538 | 0.6463275 | -0.0630221 | 0.66766777 | 0.90650187 |
| Actl6a | 0.770818 | 0.5731022 | 0.63614708 | -0.0630449 | 0.67024089 | 0.90650187 |
| Hnrnpu | 1.86704188 | 0.86331524 | 0.9263756 | -0.0630604 | 0.61891098 | 0.90650187 |
| Rdx | 1.40905987 | 0.81317664 | 0.87625292 | -0.0630763 | 0.62552075 | 0.90650187 |
| Ndufa8 | 1.50110354 | 0.83043927 | 0.89351718 | -0.0630779 | 0.62317793 | 0.90650187 |
| Arfip2 | 0.62363472 | 0.485439 | 0.54854049 | -0.0631015 | 0.69546501 | 0.90650187 |
| Sar1b | 0.86209485 | 0.62214983 | 0.68525947 | -0.0631096 | 0.65888518 | 0.90650187 |
| Naa20 | 0.90162015 | 0.64217176 | 0.70535771 | -0.0631859 | 0.65477241 | 0.90650187 |
| Sri | 1.33330444 | 0.79578172 | 0.85910035 | -0.0633186 | 0.62841426 | 0.90650187 |
| Cwc15 | 1.08359676 | 0.71999764 | 0.78332899 | -0.0633314 | 0.64036057 | 0.90650187 |
| Tm2d1 | 1.04518506 | 0.7054155 | 0.7688424 | -0.0634269 | 0.64309632 | 0.90650187 |
| Cnot6 | 0.57773096 | 0.45538281 | 0.51882163 | -0.0634388 | 0.7066367 | 0.90650187 |
| Cnot3 | 0.68601693 | 0.52353606 | 0.58703286 | -0.0634968 | 0.68469272 | 0.90650187 |
| Twistnb | 0.39693151 | 0.32494853 | 0.38858544 | -0.0636369 | 0.76670458 | 0.90650187 |
| Stub1 | 1.70798967 | 0.85504506 | 0.91878336 | -0.0637383 | 0.62111208 | 0.90650187 |
| Sucla2 | 0.73944891 | 0.55461104 | 0.61838914 | -0.0637781 | 0.6767039 | 0.90650187 |
| Prrc2b | 0.60838652 | 0.47496947 | 0.53880428 | -0.0638348 | 0.7007966 | 0.90650187 |

| | | | | | | |
|---|---|---|---|---|---|---|
| Prkra | 0.55088596 | 0.4369094 | 0.50085925 | -0.0639499 | 0.71485224 | 0.90650187 |
| Dgcr6 | 0.59581158 | 0.46666492 | 0.53067473 | -0.0640098 | 0.70409935 | 0.90650187 |
| Ndufb10 | 1.7352246 | 0.85659874 | 0.92064864 | -0.0640499 | 0.62144673 | 0.90650187 |
| 1110059E24I | 0.59564235 | 0.46650548 | 0.53056471 | -0.0640592 | 0.70428083 | 0.90650187 |
| Tspan31 | 0.75965516 | 0.56581991 | 0.6298811 | -0.0640612 | 0.67441386 | 0.90650187 |
| Mrpl46 | 0.45657236 | 0.36995871 | 0.43406195 | -0.0641032 | 0.7442607 | 0.90650187 |
| Elp5 | 0.49290712 | 0.39638923 | 0.46050506 | -0.0641158 | 0.73202831 | 0.90650187 |
| Cers2 | 0.75414948 | 0.56260365 | 0.62676873 | -0.0641651 | 0.67548724 | 0.90650187 |
| Tra2a | 1.25749325 | 0.77504524 | 0.83930571 | -0.0642605 | 0.63321206 | 0.90650187 |
| G3bp1 | 1.56281686 | 0.83901714 | 0.9033393 | -0.0643222 | 0.62420701 | 0.90650187 |
| Gpbp1 | 0.90758067 | 0.64392737 | 0.70832654 | -0.0643992 | 0.65696083 | 0.90650187 |
| Wdr77 | 0.37367386 | 0.30570946 | 0.37010969 | -0.0644002 | 0.78040129 | 0.90650187 |
| Atp6ap1 | 0.85696683 | 0.61794713 | 0.68259725 | -0.0646501 | 0.66314147 | 0.90650187 |
| Gtf2a2 | 1.43844579 | 0.81734537 | 0.88216143 | -0.0648161 | 0.62802883 | 0.90650187 |
| Ggnbp2 | 0.73416955 | 0.5504271 | 0.61535358 | -0.0649265 | 0.68048654 | 0.90650187 |
| Iars2 | 0.34304872 | 0.28010273 | 0.34511008 | -0.0650074 | 0.79881767 | 0.90777025 |
| Vapa | 1.5778478 | 0.84024456 | 0.90537553 | -0.065131 | 0.6254432 | 0.90650187 |
| Phf23 | 0.58041719 | 0.45534877 | 0.52059485 | -0.0652461 | 0.71129343 | 0.90650187 |
| Suv420h2 | 0.30053604 | 0.243657 | 0.3090773 | -0.0654203 | 0.82664025 | 0.90947782 |
| Mpv17l2 | 0.26750213 | 0.21449293 | 0.27995142 | -0.0654585 | 0.85070819 | 0.91491493 |
| Eif4a3 | 1.01028882 | 0.68944659 | 0.75493685 | -0.0654903 | 0.65012328 | 0.90650187 |
| Etfa | 0.99482545 | 0.68297841 | 0.74858579 | -0.0656074 | 0.65159151 | 0.90650187 |
| Rab18 | 1.19295249 | 0.75456777 | 0.82031684 | -0.0657491 | 0.63919917 | 0.90650187 |
| Spg21 | 0.68822961 | 0.52240887 | 0.58836028 | -0.0659514 | 0.69087214 | 0.90650187 |
| Dhx9 | 0.5342047 | 0.42342994 | 0.48947157 | -0.0660416 | 0.72568707 | 0.90650187 |
| Pfdn2 | 1.5441166 | 0.83442116 | 0.90056648 | -0.0661453 | 0.62798582 | 0.90650187 |
| Spop | 1.07011384 | 0.71205982 | 0.7783443 | -0.0662845 | 0.64750448 | 0.90650187 |
| Rbbp6 | 0.82162223 | 0.5976006 | 0.66394159 | -0.066341 | 0.67155268 | 0.90650187 |
| Bag1 | 1.63433942 | 0.845288 | 0.91190747 | -0.0666195 | 0.62732397 | 0.90650187 |
| Hibadh | 0.61115324 | 0.47394779 | 0.54058091 | -0.0666331 | 0.70814534 | 0.90650187 |

| | | | | | | |
|---|---|---|---|---|---|---|
| Csnk1d | 0.79053891 | 0.58032764 | 0.6470762 | -0.0667486 | 0.676733 | 0.90650187 |
| Nudt5 | 0.56345232 | 0.44254076 | 0.50932293 | -0.0667822 | 0.72002161 | 0.90650187 |
| Rab11b | 1.22284597 | 0.76243925 | 0.8292325 | -0.0667933 | 0.63986051 | 0.90650187 |
| 2010107E04I | 2.31265995 | 0.84632838 | 0.913122 | -0.0667936 | 0.62748136 | 0.90650187 |
| Snrpd2 | 2.25449968 | 0.85038349 | 0.9172108 | -0.0668273 | 0.62699451 | 0.90650187 |
| Mmadhc | 0.75145278 | 0.55831667 | 0.62523917 | -0.0669225 | 0.68290428 | 0.90650187 |
| Thrap3 | 1.10954497 | 0.72567291 | 0.79261648 | -0.0669436 | 0.64634501 | 0.90650187 |
| Rsrc2 | 1.75644165 | 0.8550245 | 0.92198478 | -0.0669603 | 0.62660225 | 0.90650187 |
| Tomm40 | 0.88905556 | 0.63193719 | 0.69905411 | -0.0671169 | 0.66524745 | 0.90650187 |
| Ssb | 1.78746691 | 0.85632167 | 0.92362338 | -0.0673017 | 0.6270076 | 0.90650187 |
| Eif5b | 1.69611486 | 0.85018445 | 0.91791433 | -0.0677299 | 0.62855427 | 0.90650187 |
| Ndufa9 | 0.90375589 | 0.63865878 | 0.70642307 | -0.0677643 | 0.66510962 | 0.90650187 |
| Arpc1a | 1.67990051 | 0.84884076 | 0.91667438 | -0.0678336 | 0.62891214 | 0.90650187 |
| Hdac2 | 1.27405481 | 0.77603765 | 0.8439248 | -0.0678871 | 0.63968399 | 0.90650187 |
| Eif6 | 1.09566395 | 0.71977862 | 0.78770551 | -0.0679269 | 0.64927424 | 0.90650187 |
| Chtop | 0.82491511 | 0.59774231 | 0.66570281 | -0.0679605 | 0.67502966 | 0.90650187 |
| Atf6b | 0.66623366 | 0.50706866 | 0.57505345 | -0.0679848 | 0.70040816 | 0.90650187 |
| Senp6 | 0.85548826 | 0.61376751 | 0.68182767 | -0.0680602 | 0.67139239 | 0.90650187 |
| Smc3 | 1.27668537 | 0.7765377 | 0.84463629 | -0.0680986 | 0.6399877 | 0.90650187 |
| Srsf10 | 1.05615918 | 0.70490515 | 0.77306322 | -0.0681581 | 0.65245331 | 0.90650187 |
| Arhgdia | 1.61456757 | 0.84145916 | 0.90968329 | -0.0682241 | 0.63058764 | 0.90650187 |
| Dnaja2 | 1.15443751 | 0.73975603 | 0.80798646 | -0.0682304 | 0.64632195 | 0.90650187 |
| Hnrnpa3 | 2.39718953 | 0.83790084 | 0.90630248 | -0.0684016 | 0.63138529 | 0.90650187 |
| Wtap | 0.93175331 | 0.65161013 | 0.72016101 | -0.0685509 | 0.66391422 | 0.90650187 |
| Mphosph8 | 0.59651336 | 0.46256137 | 0.5311308 | -0.0685694 | 0.71699291 | 0.90650187 |
| Matr3 | 1.2637095 | 0.77220134 | 0.84106288 | -0.0688615 | 0.64206408 | 0.90650187 |
| Mrps23 | 0.46167085 | 0.3688808 | 0.43782837 | -0.0689476 | 0.75806053 | 0.90650187 |
| Sun1 | 0.41393772 | 0.33281877 | 0.40182561 | -0.0690068 | 0.77740507 | 0.90650187 |
| Psmc3 | 1.38223537 | 0.80110781 | 0.87048631 | -0.0693785 | 0.63844994 | 0.90650187 |
| Ctage5 | 0.61485269 | 0.47328461 | 0.54294978 | -0.0696652 | 0.71580012 | 0.90650187 |

| | | | | | | |
|---|---|---|---|---|---|---|
| Hypk | 1.35212264 | 0.79393001 | 0.86362621 | -0.0696962 | 0.64011585 | 0.90650187 |
| Psma5 | 1.73469456 | 0.85082729 | 0.92061384 | -0.0697865 | 0.63194017 | 0.90650187 |
| Cfl1 | 3.44731336 | 0.70886317 | 0.77869051 | -0.0698273 | 0.65492438 | 0.90650187 |
| Trappc4 | 1.03305975 | 0.69406642 | 0.76408328 | -0.0700169 | 0.6581332 | 0.90650187 |
| Tsnax | 0.6877249 | 0.51798933 | 0.58805772 | -0.0700684 | 0.70184138 | 0.90650187 |
| Dph3 | 0.77695803 | 0.5694791 | 0.63956858 | -0.0700895 | 0.68693749 | 0.90650187 |
| Gatad2b | 0.73755311 | 0.54717329 | 0.6173006 | -0.0701273 | 0.69323287 | 0.90650187 |
| Dars | 0.83848887 | 0.60278006 | 0.67291459 | -0.0701345 | 0.67844089 | 0.90650187 |
| Psma1 | 1.57906198 | 0.83537892 | 0.90552059 | -0.0701417 | 0.63471106 | 0.90650187 |
| Slc25a3 | 2.3758354 | 0.83775392 | 0.90809996 | -0.070346 | 0.63471946 | 0.90650187 |
| Cops7a | 0.80491705 | 0.58450372 | 0.6549311 | -0.0704274 | 0.68369411 | 0.90650187 |
| Psmd12 | 1.11865267 | 0.72515058 | 0.79580403 | -0.0706535 | 0.65346488 | 0.90650187 |
| Hax1 | 0.69546559 | 0.52190872 | 0.59268267 | -0.0707739 | 0.70224303 | 0.90650187 |
| Eif3d | 1.11881352 | 0.72501155 | 0.79586069 | -0.0708491 | 0.65385782 | 0.90650187 |
| Nedd8 | 2.02565545 | 0.85559146 | 0.92680244 | -0.071211 | 0.63366602 | 0.90650187 |
| Actr1b | 0.5646182 | 0.43846908 | 0.51010334 | -0.0716343 | 0.73378483 | 0.90650187 |
| Rnps1 | 0.81990268 | 0.59087098 | 0.66302005 | -0.0721491 | 0.68572393 | 0.90650187 |
| Zfp706 | 1.51083272 | 0.82300532 | 0.89515445 | -0.0721491 | 0.63994593 | 0.90650187 |
| Mlf2 | 1.57601525 | 0.83298775 | 0.90515589 | -0.0721681 | 0.63849859 | 0.90650187 |
| Ap2s1 | 1.64603428 | 0.84085103 | 0.91316925 | -0.0723182 | 0.63760663 | 0.90650187 |
| Puf60 | 1.3997501 | 0.80195032 | 0.87429679 | -0.0723465 | 0.64351335 | 0.90650187 |
| Anapc13 | 1.74999971 | 0.8492073 | 0.92159504 | -0.0723877 | 0.63652489 | 0.90650187 |
| Eny2 | 1.29391071 | 0.77681699 | 0.84924083 | -0.0724238 | 0.64770225 | 0.90650187 |
| Cbx6 | 1.1482597 | 0.73354673 | 0.80598213 | -0.0724354 | 0.65525104 | 0.90650187 |
| Wac | 0.64558139 | 0.48979326 | 0.56232396 | -0.0725307 | 0.71681692 | 0.90650187 |
| Ube2n | 1.64625905 | 0.84037135 | 0.91319319 | -0.0728218 | 0.6385221 | 0.90650187 |
| Ywhaz | 1.29104588 | 0.77535376 | 0.84848153 | -0.0731278 | 0.64919642 | 0.90650187 |
| Mrpl34 | 0.92742091 | 0.64478554 | 0.7180607 | -0.0732752 | 0.67496177 | 0.90650187 |
| Nmt1 | 1.08740931 | 0.71121958 | 0.78472609 | -0.0735065 | 0.66145116 | 0.90650187 |
| Atp5c1 | 2.30075415 | 0.84039695 | 0.91402129 | -0.0736243 | 0.63986322 | 0.90650187 |

| | | | | | | |
|---|---|---|---|---|---|---|
| Cand1 | 0.53969896 | 0.41940964 | 0.49324217 | -0.0738325 | 0.74712018 | 0.90650187 |
| Plgrkt | 0.49380267 | 0.38728322 | 0.46114542 | -0.0738622 | 0.76178556 | 0.90650187 |
| Pigp | 1.01895405 | 0.68454488 | 0.75844881 | -0.0739039 | 0.66755939 | 0.90650187 |
| mt-Nd5 | 1.66915977 | 0.84164991 | 0.91564603 | -0.0739961 | 0.64030078 | 0.90650187 |
| Hdac1 | 1.14872007 | 0.73208932 | 0.80613197 | -0.0740426 | 0.65848274 | 0.90650187 |
| Smdt1 | 2.18969893 | 0.84689701 | 0.92095389 | -0.0740569 | 0.63963584 | 0.90650187 |
| Selk | 2.39483033 | 0.83240852 | 0.90650091 | -0.0740924 | 0.64183177 | 0.90650187 |
| Sf1 | 0.70955351 | 0.5268279 | 0.60102142 | -0.0741935 | 0.70842556 | 0.90650187 |
| Smu1 | 0.7941576 | 0.57470809 | 0.64906166 | -0.0743536 | 0.69471801 | 0.90650187 |
| Lrp10 | 0.62168954 | 0.47293072 | 0.54730601 | -0.0743753 | 0.72712096 | 0.90650187 |
| Rpl3 | 4.6162265 | 0.51661635 | 0.59100805 | -0.0743917 | 0.71211697 | 0.90650187 |
| Srsf5 | 1.66347355 | 0.84061481 | 0.91504468 | -0.0744299 | 0.64117686 | 0.90650187 |
| Rbm22 | 0.66783877 | 0.50112032 | 0.57603292 | -0.0749126 | 0.71841768 | 0.90650187 |
| Polr2g | 1.20973767 | 0.75037684 | 0.82534422 | -0.0749674 | 0.65682075 | 0.90650187 |
| Cdc23 | 0.43495242 | 0.34275177 | 0.41788284 | -0.0751311 | 0.78793442 | 0.90650187 |
| Tbca | 2.27239187 | 0.84078688 | 0.91602093 | -0.075234 | 0.64249046 | 0.90650187 |
| Tmem208 | 1.12467794 | 0.7224732 | 0.79792017 | -0.075447 | 0.66288592 | 0.90650187 |
| Hdgfrp2 | 0.45769189 | 0.35929155 | 0.43489054 | -0.075599 | 0.7801796 | 0.90650187 |
| Psmd11 | 1.21494948 | 0.75116289 | 0.8268855 | -0.0757226 | 0.65803622 | 0.90650187 |
| Srsf7 | 1.75172024 | 0.84567168 | 0.92170229 | -0.0760306 | 0.64308424 | 0.90650187 |
| Pdcl3 | 0.83029162 | 0.5925306 | 0.66856862 | -0.076038 | 0.69343484 | 0.90650187 |
| Eif3c | 1.5868558 | 0.83031373 | 0.90644302 | -0.0761293 | 0.64556437 | 0.90650187 |
| Ywhae | 3.31710137 | 0.72127601 | 0.79742072 | -0.0761447 | 0.66440137 | 0.90650187 |
| Sf3b6 | 1.62608704 | 0.83484019 | 0.9109979 | -0.0761577 | 0.64492117 | 0.90650187 |
| Morf4l2 | 2.43630384 | 0.8268066 | 0.90298225 | -0.0761757 | 0.64618155 | 0.90650187 |
| Park7 | 2.01685125 | 0.85074292 | 0.92694794 | -0.076205 | 0.64262205 | 0.90650187 |
| Eif4h | 1.54654178 | 0.82469822 | 0.90094964 | -0.0762514 | 0.64663508 | 0.90650187 |
| Sgta | 0.89132696 | 0.62380197 | 0.70019825 | -0.0763963 | 0.68617785 | 0.90650187 |
| Pdrg1 | 0.55382045 | 0.42632546 | 0.50284427 | -0.0765188 | 0.75063732 | 0.90650187 |
| Sec31a | 0.72212817 | 0.53180805 | 0.60838024 | -0.0765722 | 0.71210154 | 0.90650187 |

| | | | | | | |
|---|---|---|---|---|---|---|
| Selt | 0.87114655 | 0.61330355 | 0.6899247 | -0.0766212 | 0.68924694 | 0.90650187 |
| Tor1aip2 | 0.66956104 | 0.5004284 | 0.5770824 | -0.076654 | 0.72257595 | 0.90650187 |
| Kars | 0.62393398 | 0.4720035 | 0.54873022 | -0.0767267 | 0.73290119 | 0.90650187 |
| Snx5 | 0.82606566 | 0.58946107 | 0.66631711 | -0.076856 | 0.69594613 | 0.90650187 |
| Ppp5c | 0.702151 | 0.51974572 | 0.59665209 | -0.0769064 | 0.71668549 | 0.90650187 |
| Ube2l3 | 1.39212276 | 0.79518588 | 0.87266322 | -0.0774773 | 0.65346515 | 0.90650187 |
| Aurkaip1 | 0.94865665 | 0.65066588 | 0.72822905 | -0.0775632 | 0.68207119 | 0.90650187 |
| Timm13 | 1.89960096 | 0.84872264 | 0.92698897 | -0.0782663 | 0.64629831 | 0.90650187 |
| Glo1 | 1.14720377 | 0.72736509 | 0.80563816 | -0.0782731 | 0.66710023 | 0.90650187 |
| Bag6 | 0.77044832 | 0.55738525 | 0.63594053 | -0.0785553 | 0.70854826 | 0.90650187 |
| Adgrl1 | 0.41148414 | 0.32118605 | 0.39992887 | -0.0787428 | 0.80929592 | 0.9092537 |
| Ndufs8 | 1.23582121 | 0.75413434 | 0.83306218 | -0.0789278 | 0.66319386 | 0.90650187 |
| Ddx39b | 1.89426711 | 0.84795672 | 0.92693393 | -0.0789772 | 0.6475734 | 0.90650187 |
| Tm9sf3 | 1.4415646 | 0.80376914 | 0.88276803 | -0.0789989 | 0.6546249 | 0.90650187 |
| Vcp | 2.27356453 | 0.83673905 | 0.91594174 | -0.0792027 | 0.64966546 | 0.90650187 |
| Eif4g1 | 1.18113382 | 0.73732078 | 0.81657292 | -0.0792521 | 0.66693539 | 0.90650187 |
| Cpsf6 | 0.82350944 | 0.58567195 | 0.66495154 | -0.0792796 | 0.70195526 | 0.90650187 |
| Setd3 | 0.6154426 | 0.46402016 | 0.5433268 | -0.0793066 | 0.7417993 | 0.90650187 |
| Ppp1r8 | 0.41999689 | 0.32715396 | 0.40649034 | -0.0793364 | 0.80721171 | 0.9092537 |
| Srsf11 | 1.77123054 | 0.84332974 | 0.92279187 | -0.0794621 | 0.64907103 | 0.90650187 |
| Dap3 | 0.62903007 | 0.47248442 | 0.55195365 | -0.0794692 | 0.73893641 | 0.90650187 |
| Pde6d | 0.61856979 | 0.4658299 | 0.54532173 | -0.0794918 | 0.74151992 | 0.90650187 |
| Ewsr1 | 1.62486624 | 0.83128781 | 0.91086199 | -0.0795742 | 0.65113051 | 0.90650187 |
| Nono | 1.9527959 | 0.84767494 | 0.92734495 | -0.07967 | 0.64874361 | 0.90650187 |
| Rhoa | 2.23271174 | 0.83861293 | 0.91856386 | -0.0799509 | 0.65060019 | 0.90650187 |
| Psma4 | 1.76910851 | 0.84262227 | 0.92267881 | -0.0800565 | 0.65014967 | 0.90650187 |
| Mrpl30 | 1.14934111 | 0.72623569 | 0.80633398 | -0.0800983 | 0.67061442 | 0.90650187 |
| Ogfr | 0.67579587 | 0.50069825 | 0.5808688 | -0.0801705 | 0.73025197 | 0.90650187 |
| Chmp1a | 0.73311901 | 0.53455176 | 0.61474796 | -0.0801962 | 0.71902454 | 0.90650187 |
| Tcf12 | 0.60489071 | 0.45595116 | 0.5365533 | -0.0806021 | 0.74787835 | 0.90650187 |

| | | | | | | |
|---|---|---|---|---|---|---|
| Rpl7a | 3.5658161 | 0.67914473 | 0.75980604 | -0.0806613 | 0.68151781 | 0.90650187 |
| Clns1a | 0.79084566 | 0.56649002 | 0.64724472 | -0.0807547 | 0.7104379 | 0.90650187 |
| Polr2f | 1.566841 | 0.82299951 | 0.90391475 | -0.0809152 | 0.65466828 | 0.90650187 |
| Hint1 | 2.86539408 | 0.77470965 | 0.8556796 | -0.0809699 | 0.66300272 | 0.90650187 |
| Rps17 | 4.45475252 | 0.53203465 | 0.61316971 | -0.0811351 | 0.72181636 | 0.90650187 |
| Pdcd6ip | 0.69214803 | 0.50906934 | 0.59070459 | -0.0816352 | 0.73051849 | 0.90650187 |
| Sdha | 1.15076889 | 0.72505087 | 0.80679788 | -0.081747 | 0.67380384 | 0.90650187 |
| Snrnp70 | 1.35784899 | 0.78320898 | 0.86495929 | -0.0817503 | 0.66281926 | 0.90650187 |
| Cript | 0.88373705 | 0.61423203 | 0.69635617 | -0.0821241 | 0.69996109 | 0.90650187 |
| Wbp4 | 0.94472692 | 0.64383953 | 0.72637282 | -0.0825333 | 0.69326517 | 0.90650187 |
| Dctn1 | 0.57954067 | 0.43722954 | 0.52001672 | -0.0827872 | 0.76044816 | 0.90650187 |
| Smim7 | 1.16685157 | 0.72909344 | 0.81197269 | -0.0828793 | 0.67500267 | 0.90650187 |
| Esd | 1.77947652 | 0.8402293 | 0.92322254 | -0.0829932 | 0.6552886 | 0.90650187 |
| Chmp4b | 1.21931611 | 0.74512868 | 0.82818097 | -0.0830523 | 0.67213557 | 0.90650187 |
| Atxn10 | 1.53062064 | 0.81522453 | 0.89840721 | -0.0831827 | 0.65968901 | 0.90650187 |
| Pnkd | 0.88936713 | 0.61594782 | 0.69921117 | -0.0832634 | 0.70173288 | 0.90650187 |
| Ube2d3 | 2.42433792 | 0.82026936 | 0.90399601 | -0.0837267 | 0.65973679 | 0.90650187 |
| Tomm22 | 1.31064366 | 0.76974117 | 0.85346835 | -0.0837272 | 0.66862665 | 0.90650187 |
| Cetn3 | 1.35118916 | 0.77959645 | 0.86340797 | -0.0838115 | 0.66696124 | 0.90650187 |
| Rnf10 | 1.31557411 | 0.77085156 | 0.85469721 | -0.0838457 | 0.66862271 | 0.90650187 |
| Rps21 | 3.80566758 | 0.63303175 | 0.71703676 | -0.084005 | 0.69875689 | 0.90650187 |
| Zmat2 | 0.92821493 | 0.63442287 | 0.71844617 | -0.0840233 | 0.69844033 | 0.90650187 |
| Mtpn | 0.92393106 | 0.631454 | 0.71636365 | -0.0849097 | 0.70087957 | 0.90650187 |
| Ndufs5 | 2.11890397 | 0.83905623 | 0.92404617 | -0.0849899 | 0.65868924 | 0.90650187 |
| Bloc1s2 | 0.5032067 | 0.38276726 | 0.46783442 | -0.0850672 | 0.79055617 | 0.90650187 |
| Aamp | 1.52983753 | 0.812987 | 0.8982805 | -0.0852935 | 0.66353094 | 0.90650187 |
| Qrich1 | 0.65513828 | 0.48275855 | 0.56824257 | -0.085484 | 0.74818154 | 0.90650187 |
| Uqcrc2 | 1.2981737 | 0.76472091 | 0.85032628 | -0.0856054 | 0.67276282 | 0.90650187 |
| Atp5a1 | 2.70528905 | 0.79162155 | 0.87725728 | -0.0856357 | 0.66784464 | 0.90650187 |
| Eif3i | 1.74693885 | 0.83569256 | 0.92140274 | -0.0857102 | 0.66039322 | 0.90650187 |

| | | | | | | |
|---|---|---|---|---|---|---|
| Hnrnph1 | 1.32820587 | 0.77195236 | 0.85784098 | -0.0858886 | 0.67187708 | 0.90650187 |
| Luc7l2 | 1.65447337 | 0.82805911 | 0.91407797 | -0.0860189 | 0.66215498 | 0.90650187 |
| Pfn1 | 2.96092266 | 0.75571554 | 0.84206815 | -0.0863526 | 0.67576514 | 0.90650187 |
| Cct5 | 2.17274706 | 0.83540372 | 0.92179259 | -0.0863889 | 0.66152724 | 0.90650187 |
| Magoh | 1.08011094 | 0.6955777 | 0.78204684 | -0.0864691 | 0.68847793 | 0.90650187 |
| Acp1 | 1.08007511 | 0.69540625 | 0.78203364 | -0.0866274 | 0.68879883 | 0.90650187 |
| Ptov1 | 1.18639511 | 0.73138472 | 0.81824997 | -0.0868652 | 0.68151685 | 0.90650187 |
| Srpk1 | 0.6313058 | 0.46644353 | 0.55338852 | -0.086945 | 0.75755716 | 0.90650187 |
| Ndufb6 | 1.39957155 | 0.78723492 | 0.87425903 | -0.0870241 | 0.67094261 | 0.90650187 |
| Acin1 | 1.28311205 | 0.7592738 | 0.84636522 | -0.0870914 | 0.67632805 | 0.90650187 |
| AY036118 | 1.80708031 | 0.83741891 | 0.92455357 | -0.0871347 | 0.66238343 | 0.90650187 |
| Clta | 2.29704613 | 0.82710908 | 0.91428835 | -0.0871793 | 0.66417978 | 0.90650187 |
| Vps72 | 0.77937582 | 0.55346885 | 0.64091122 | -0.0874424 | 0.72791984 | 0.90650187 |
| Commd3 | 0.94649095 | 0.63972869 | 0.72720679 | -0.0874781 | 0.70359784 | 0.90650187 |
| Hadhb | 0.95763398 | 0.64471677 | 0.73244761 | -0.0877308 | 0.70280467 | 0.90650187 |
| Rps27rt | 2.97025613 | 0.75305461 | 0.84079869 | -0.0877441 | 0.67865272 | 0.90650187 |
| Gtf2i | 0.91470338 | 0.62385627 | 0.71185386 | -0.0879976 | 0.70868759 | 0.90650187 |
| Snrpg | 2.73339011 | 0.78571812 | 0.8739909 | -0.0882728 | 0.67328587 | 0.90650187 |
| Gtf3a | 0.71820635 | 0.51780927 | 0.60609398 | -0.0882847 | 0.74136354 | 0.90650187 |
| Tmbim6 | 2.22446765 | 0.83052028 | 0.91905697 | -0.0885367 | 0.66577064 | 0.90650187 |
| Strap | 1.1504889 | 0.71805031 | 0.80670696 | -0.0886567 | 0.68742681 | 0.90650187 |
| Pcmt1 | 0.97720152 | 0.65238792 | 0.7411241 | -0.0887362 | 0.70273424 | 0.90650187 |
| Dcun1d5 | 0.98699963 | 0.65646946 | 0.74532442 | -0.088855 | 0.70194043 | 0.90650187 |
| Rnf187 | 1.16642584 | 0.72293145 | 0.8118369 | -0.0889054 | 0.68681587 | 0.90650187 |
| Glyr1 | 0.69826051 | 0.50529292 | 0.59434484 | -0.0890519 | 0.74732463 | 0.90650187 |
| Rps27l | 2.51699419 | 0.8066677 | 0.89622609 | -0.0895584 | 0.67156622 | 0.90650187 |
| Eif4g2 | 2.41918272 | 0.81466789 | 0.90442897 | -0.0897611 | 0.67046923 | 0.90650187 |
| Cdc42 | 2.4283471 | 0.81379861 | 0.90365772 | -0.0898591 | 0.67078002 | 0.90650187 |
| Erh | 2.74908658 | 0.782072 | 0.87202735 | -0.0899553 | 0.67673694 | 0.90650187 |
| Tardbp | 1.02028227 | 0.66900709 | 0.75898269 | -0.0899756 | 0.70090881 | 0.90650187 |

| | | | | | | |
|---|---|---|---|---|---|---|
| Gabarap | 2.89904558 | 0.760673 | 0.85100785 | -0.0903348 | 0.68150199 | 0.90650187 |
| Gm10076 | 3.05294036 | 0.73907781 | 0.82982465 | -0.0907468 | 0.68657193 | 0.90650187 |
| Srp72 | 1.0986356 | 0.69775957 | 0.78876289 | -0.0910033 | 0.69599311 | 0.90650187 |
| Ndfip2 | 0.8804537 | 0.6036681 | 0.69468505 | -0.0910169 | 0.71987121 | 0.90650187 |
| Mrpl54 | 1.34648442 | 0.77118542 | 0.86230402 | -0.0911186 | 0.68073737 | 0.90650187 |
| Txndc15 | 0.76515827 | 0.54168715 | 0.63297808 | -0.0912909 | 0.73929233 | 0.90650187 |
| C1d | 1.21081603 | 0.73423261 | 0.82566388 | -0.0914313 | 0.68874341 | 0.90650187 |
| 2310036O22 | 1.6263196 | 0.81958124 | 0.91102375 | -0.0914425 | 0.67228099 | 0.90650187 |
| Uba2 | 0.98234481 | 0.65174099 | 0.74333375 | -0.0915928 | 0.70807943 | 0.90650187 |
| Sf3b4 | 0.84157941 | 0.58281824 | 0.67454434 | -0.0917261 | 0.7272523 | 0.90650187 |
| Mum1 | 0.59936296 | 0.4411891 | 0.53297979 | -0.0917907 | 0.77805172 | 0.90650187 |
| Rps18 | 5.18468825 | 0.47001219 | 0.56181594 | -0.0918038 | 0.76621424 | 0.90650187 |
| Mapk1 | 0.89493933 | 0.60970047 | 0.70201367 | -0.0923132 | 0.72058732 | 0.90650187 |
| Ociad1 | 1.23703356 | 0.74105307 | 0.83341717 | -0.0923641 | 0.68888338 | 0.90650187 |
| Mrps17 | 1.07311366 | 0.68676418 | 0.7794593 | -0.0926951 | 0.70148692 | 0.90650187 |
| Ap2b1 | 0.84500233 | 0.58344032 | 0.67634414 | -0.0929038 | 0.72928643 | 0.90650187 |
| Rab1a | 1.5732451 | 0.81190326 | 0.90482226 | -0.092919 | 0.67599667 | 0.90650187 |
| Rpl5 | 3.27394189 | 0.71021672 | 0.8032201 | -0.0930034 | 0.69663564 | 0.90650187 |
| Anapc5 | 1.69150042 | 0.8235007 | 0.91756743 | -0.0940667 | 0.67571603 | 0.90650187 |
| Fkbp8 | 1.52365391 | 0.80315307 | 0.89727447 | -0.0941214 | 0.67951063 | 0.90650187 |
| Sap18 | 1.22976498 | 0.7371231 | 0.8312816 | -0.0941585 | 0.69271109 | 0.90650187 |
| Snrpe | 2.07093938 | 0.83150092 | 0.92570331 | -0.0942024 | 0.67450901 | 0.90650187 |
| Psenen | 1.53743896 | 0.80516551 | 0.89950385 | -0.0943383 | 0.67948071 | 0.90650187 |
| Rps15 | 4.40146469 | 0.52574127 | 0.62012022 | -0.0943789 | 0.75066925 | 0.90650187 |
| Ndufa2 | 2.65281448 | 0.78849394 | 0.88292397 | -0.09443 | 0.68277192 | 0.90650187 |
| Adh5 | 0.92352457 | 0.62140979 | 0.71616568 | -0.0947559 | 0.72180038 | 0.90650187 |
| Sumo2 | 2.98182142 | 0.74445108 | 0.83922199 | -0.0947709 | 0.6921766 | 0.90650187 |
| Fam96a | 0.89365879 | 0.60643677 | 0.70137071 | -0.0949339 | 0.72632456 | 0.90650187 |
| Nipbl | 0.74381277 | 0.52591438 | 0.62088739 | -0.094973 | 0.75175246 | 0.90650187 |
| Psmd8 | 1.56943155 | 0.8092926 | 0.90428307 | -0.0949905 | 0.67974613 | 0.90650187 |

| | | | | | | |
|---|---|---|---|---|---|---|
| Hp1bp3 | 1.0078667 | 0.65889722 | 0.75394836 | -0.0950511 | 0.71241807 | 0.90650187 |
| Nsd1 | 0.8282596 | 0.57212428 | 0.66748693 | -0.0953627 | 0.73733006 | 0.90650187 |
| 1110004F10I | 1.85222229 | 0.83031692 | 0.92598046 | -0.0956635 | 0.67698272 | 0.90650187 |
| Csnk2a1 | 1.15191459 | 0.71148684 | 0.80716959 | -0.0956827 | 0.70089867 | 0.90650187 |
| Sdhd | 1.09453391 | 0.69114593 | 0.78730255 | -0.0961566 | 0.70642162 | 0.90650187 |
| Eif3h | 2.35845853 | 0.81330292 | 0.90953976 | -0.0962368 | 0.68095894 | 0.90650187 |
| Ppig | 1.31361626 | 0.75788073 | 0.85421015 | -0.0963294 | 0.69193117 | 0.90650187 |
| Ppp1cc | 1.4729936 | 0.79225794 | 0.88865757 | -0.0963996 | 0.68518571 | 0.90650187 |
| Uqcrc1 | 1.60200853 | 0.81177156 | 0.9082116 | -0.09644 | 0.68155963 | 0.90650187 |
| Snrpb | 2.14885938 | 0.82618055 | 0.92285546 | -0.0966749 | 0.67928674 | 0.90650187 |
| Rpl31 | 4.46588378 | 0.51473196 | 0.61147373 | -0.0967418 | 0.75907904 | 0.90650187 |
| Hnrnpl | 1.19009371 | 0.72244742 | 0.81942323 | -0.0969758 | 0.70059815 | 0.90650187 |
| Trappc3 | 0.82492226 | 0.56851248 | 0.66570663 | -0.0971942 | 0.74185185 | 0.90650187 |
| Rab14 | 1.73771271 | 0.82299613 | 0.92081122 | -0.0978151 | 0.68162362 | 0.90650187 |
| Rpl38 | 4.63682686 | 0.49061269 | 0.58848699 | -0.0978743 | 0.77015933 | 0.90650187 |
| Tmem50a | 1.4279174 | 0.7821256 | 0.88009358 | -0.097968 | 0.68965213 | 0.90650187 |
| Tm2d2 | 1.09476167 | 0.68909426 | 0.7873838 | -0.0982895 | 0.71053468 | 0.90650187 |
| Nhp2l1 | 1.64081481 | 0.81336382 | 0.91261003 | -0.0992462 | 0.6856151 | 0.90650187 |
| Slirp | 1.36910522 | 0.76824537 | 0.86755122 | -0.0993059 | 0.69457993 | 0.90650187 |
| Ndufa13 | 2.53740814 | 0.79469982 | 0.89428273 | -0.0995829 | 0.68971433 | 0.90650187 |
| Spcs1 | 2.25661927 | 0.81714917 | 0.91707166 | -0.0999225 | 0.68594378 | 0.90650187 |
| Rad23b | 1.17891512 | 0.71571941 | 0.81586286 | -0.1001435 | 0.70735633 | 0.90650187 |
| Srrm2 | 2.13503473 | 0.82319566 | 0.92340434 | -0.1002087 | 0.68525374 | 0.90650187 |
| Ndufv2 | 1.61317272 | 0.80909495 | 0.9095217 | -0.1004268 | 0.68824115 | 0.90650187 |
| Rps29 | 5.55524235 | 0.5244172 | 0.62493847 | -0.1005213 | 0.7627187 | 0.90650187 |
| Ndufb5 | 1.90201149 | 0.82590718 | 0.92701221 | -0.101105 | 0.68611295 | 0.90650187 |
| Ubxn1 | 1.2864762 | 0.74605928 | 0.84726502 | -0.1012057 | 0.70230008 | 0.90650187 |
| Pcnp | 1.21558915 | 0.7257479 | 0.82707403 | -0.1013261 | 0.7069988 | 0.90650187 |
| Dynlrb1 | 1.92020682 | 0.82576799 | 0.9271628 | -0.1013948 | 0.6865777 | 0.90650187 |
| Psma7 | 2.36797392 | 0.80716818 | 0.90875482 | -0.1015866 | 0.69039113 | 0.90650187 |

| | | | | | | |
|---|---|---|---|---|---|---|
| Rnf7 | 1.49914995 | 0.79134651 | 0.89318543 | -0.1018389 | 0.69387967 | 0.90650187 |
| Glrx3 | 1.40827451 | 0.77374301 | 0.87608886 | -0.1023458 | 0.69823871 | 0.90650187 |
| Ppia | 4.83903665 | 0.45914161 | 0.56156186 | -0.1024202 | 0.79126267 | 0.90660917 |
| Mrpl52 | 1.75736203 | 0.81878318 | 0.92203632 | -0.1032531 | 0.69070017 | 0.90650187 |
| Rpl10 | 5.013311 | 0.43967599 | 0.54380901 | -0.104133 | 0.80269342 | 0.90870476 |
| Paip2 | 1.78248387 | 0.81922315 | 0.9233749 | -0.1041517 | 0.69196947 | 0.90650187 |
| Brk1 | 1.51930693 | 0.79230192 | 0.89656045 | -0.1042585 | 0.69741049 | 0.90650187 |
| Gm9843 | 2.24988629 | 0.81312598 | 0.91751194 | -0.104386 | 0.69349434 | 0.90650187 |
| Rbbp4 | 1.40644439 | 0.77111516 | 0.87570588 | -0.1045907 | 0.70227848 | 0.90650187 |
| Tmed2 | 2.48111824 | 0.7945892 | 0.89925373 | -0.1046645 | 0.69757061 | 0.90650187 |
| St13 | 1.8762188 | 0.82136426 | 0.92659627 | -0.105232 | 0.6931785 | 0.90650187 |
| Psmd7 | 1.42127488 | 0.77333699 | 0.87877287 | -0.1054359 | 0.7031173 | 0.90650187 |
| Ndufs2 | 1.52046997 | 0.79121381 | 0.89675196 | -0.1055382 | 0.69958264 | 0.90650187 |
| Txndc9 | 1.05759384 | 0.66804633 | 0.7736116 | -0.1055653 | 0.72795417 | 0.90650187 |
| Ndufc2 | 2.06112726 | 0.81990672 | 0.92599266 | -0.1060859 | 0.69473215 | 0.90650187 |
| Ywhab | 1.67997406 | 0.81040214 | 0.91668014 | -0.106278 | 0.69686826 | 0.90650187 |
| Tmem234 | 1.37898663 | 0.76238268 | 0.86976481 | -0.1073821 | 0.70844907 | 0.90650187 |
| Spag7 | 1.19786019 | 0.71435531 | 0.82179685 | -0.1074415 | 0.71945311 | 0.90650187 |
| Atp5j2 | 3.04573227 | 0.72265975 | 0.83074476 | -0.108085 | 0.71850321 | 0.90650187 |
| Rbm39 | 3.15213533 | 0.709894 | 0.81818381 | -0.1082898 | 0.72186584 | 0.90650187 |
| Ndufs7 | 1.48215751 | 0.78041971 | 0.89026443 | -0.1098447 | 0.70833197 | 0.90650187 |
| Ei24 | 0.92181585 | 0.60496807 | 0.71533279 | -0.1103647 | 0.7536081 | 0.90650187 |
| Denr | 1.26413744 | 0.73077951 | 0.84118339 | -0.1104039 | 0.72021429 | 0.90650187 |
| Pdcd6 | 1.11003942 | 0.68226726 | 0.79279009 | -0.1105228 | 0.73229268 | 0.90650187 |
| Myeov2 | 2.07383371 | 0.81457713 | 0.9256136 | -0.1110365 | 0.70308505 | 0.90650187 |
| Arpp19 | 1.75782224 | 0.81089363 | 0.92206203 | -0.1111684 | 0.70401535 | 0.90650187 |
| Ube3a | 1.24094768 | 0.72253539 | 0.83455998 | -0.1120246 | 0.72466027 | 0.90650187 |
| Appbp2 | 0.68156417 | 0.47152214 | 0.58435406 | -0.1128319 | 0.80466267 | 0.90882608 |
| Bcas2 | 1.34692841 | 0.74914277 | 0.86240848 | -0.1132657 | 0.72034625 | 0.90650187 |
| Rheb | 1.49443991 | 0.77869298 | 0.8923827 | -0.1136897 | 0.71442922 | 0.90650187 |

| | | | | | | |
|---|---|---|---|---|---|---|
| Sdhb | 1.58318556 | 0.79186293 | 0.90601052 | -0.1141476 | 0.71229391 | 0.90650187 |
| Rps10 | 4.0559603 | 0.56059508 | 0.67502962 | -0.1144345 | 0.77446799 | 0.90650187 |
| Akr1a1 | 2.42735602 | 0.78927314 | 0.90374148 | -0.1144683 | 0.71331071 | 0.90650187 |
| Capzb | 1.66432117 | 0.80024835 | 0.91513478 | -0.1148864 | 0.71161312 | 0.90650187 |
| Rrp1 | 1.53413527 | 0.78396475 | 0.89897396 | -0.1150092 | 0.71523492 | 0.90650187 |
| Dctn2 | 1.25020258 | 0.72181042 | 0.83722896 | -0.1154185 | 0.73001603 | 0.90650187 |
| Vps29 | 1.11172907 | 0.67753663 | 0.79338268 | -0.1158461 | 0.74187179 | 0.90650187 |
| Csnk1a1 | 2.12750062 | 0.80646115 | 0.92370836 | -0.1172472 | 0.71371997 | 0.90650187 |
| Atp5j | 2.82853673 | 0.74353506 | 0.86094156 | -0.1174065 | 0.72782457 | 0.90650187 |
| Rpl6 | 4.8245983 | 0.44586959 | 0.56332671 | -0.1174571 | 0.82302025 | 0.9092537 |
| Eci2 | 0.95043902 | 0.6115677 | 0.72906903 | -0.1175013 | 0.76315304 | 0.90650187 |
| Hnrnpc | 1.93376015 | 0.80923914 | 0.9272524 | -0.1180133 | 0.71423552 | 0.90650187 |
| Srsf1 | 1.05195801 | 0.65338318 | 0.77145286 | -0.1180697 | 0.75187226 | 0.90650187 |
| Psma6 | 1.48339871 | 0.77212839 | 0.89048032 | -0.1183519 | 0.72269173 | 0.90650187 |
| Psma3 | 2.15070554 | 0.80428743 | 0.92278031 | -0.1184929 | 0.71595064 | 0.90650187 |
| Rala | 1.1418866 | 0.68518063 | 0.80389264 | -0.118712 | 0.74424531 | 0.90650187 |
| Psmd3 | 1.00750312 | 0.63375817 | 0.75379977 | -0.1200416 | 0.76051397 | 0.90650187 |
| Eif5a | 2.89803777 | 0.73090676 | 0.85115762 | -0.1202509 | 0.73502151 | 0.90650187 |
| Eif3f | 2.69869454 | 0.75695597 | 0.87799099 | -0.121035 | 0.73001121 | 0.90650187 |
| Srrm1 | 1.49330984 | 0.77009934 | 0.89218948 | -0.1220901 | 0.72852278 | 0.90650187 |
| Drg1 | 0.86105472 | 0.56173668 | 0.68472035 | -0.1229837 | 0.78770462 | 0.90650187 |
| Cfdp1 | 1.48819238 | 0.76780117 | 0.89131026 | -0.1235091 | 0.73106225 | 0.90650187 |
| Naca | 3.58584946 | 0.63262729 | 0.7564368 | -0.1238095 | 0.76658084 | 0.90650187 |
| Atp5d | 2.34320779 | 0.78608102 | 0.91075296 | -0.1246719 | 0.72858995 | 0.90650187 |
| Pafah1b2 | 1.18463742 | 0.69236119 | 0.81769076 | -0.1253296 | 0.75218748 | 0.90650187 |
| Sf3b1 | 1.57277574 | 0.7786635 | 0.90475609 | -0.1260926 | 0.73222892 | 0.90650187 |
| Ccnl2 | 1.17692658 | 0.68888519 | 0.81522501 | -0.1263398 | 0.75457567 | 0.90650187 |
| Rpl35a | 4.8613317 | 0.43186657 | 0.55852759 | -0.126661 | 0.84331112 | 0.91291642 |
| Gm10709 | 2.85636989 | 0.72942671 | 0.85689006 | -0.1274634 | 0.74578647 | 0.90650187 |
| Snx17 | 1.09377908 | 0.65949629 | 0.78703311 | -0.1275368 | 0.76436058 | 0.90650187 |

| | | | | | | |
|---|---|---|---|---|---|---|
| Ubap2l | 1.27790209 | 0.71738471 | 0.84496463 | -0.1275799 | 0.74896467 | 0.90650187 |
| Ndufa4 | 2.83598612 | 0.73211086 | 0.85985362 | -0.1277428 | 0.74551882 | 0.90650187 |
| Rpl8 | 4.4295922 | 0.4871637 | 0.61692698 | -0.1297633 | 0.82512662 | 0.90934433 |
| Mrpl17 | 1.28395383 | 0.71669991 | 0.84659071 | -0.1298908 | 0.75240611 | 0.90650187 |
| Grpel1 | 1.10119835 | 0.65939304 | 0.78967213 | -0.1302791 | 0.76835201 | 0.90650187 |
| Snrpd3 | 1.524317 | 0.76696599 | 0.89738282 | -0.1304168 | 0.74088652 | 0.90650187 |
| Cdc37 | 1.62275383 | 0.7801234 | 0.91062227 | -0.1304989 | 0.73796625 | 0.90650187 |
| Fam104a | 1.33764211 | 0.72929915 | 0.86016557 | -0.1308664 | 0.75059358 | 0.90650187 |
| Polr2m | 1.56511198 | 0.76944855 | 0.90366799 | -0.1342194 | 0.7454675 | 0.90650187 |
| Rpl30 | 3.56005489 | 0.62608161 | 0.76077012 | -0.1346885 | 0.78437677 | 0.90650187 |
| Csde1 | 1.34299323 | 0.72637054 | 0.86147184 | -0.1351013 | 0.75715721 | 0.90650187 |
| Rps6 | 4.67861893 | 0.44741606 | 0.58327235 | -0.1358563 | 0.84977628 | 0.91470896 |
| Sarnp | 1.64853234 | 0.77727596 | 0.91343466 | -0.1361587 | 0.74622983 | 0.90650187 |
| Hnrnpk | 3.26555944 | 0.66796837 | 0.80432756 | -0.1363592 | 0.77447695 | 0.90650187 |
| Tcp1 | 1.6843623 | 0.78053479 | 0.91702146 | -0.1364867 | 0.74590878 | 0.90650187 |
| Gdi2 | 1.86096931 | 0.78939767 | 0.92622116 | -0.1368235 | 0.74431792 | 0.90650187 |
| Psmb3 | 1.69831052 | 0.78101937 | 0.91807774 | -0.1370584 | 0.74655164 | 0.90650187 |
| Psmb1 | 2.68874617 | 0.74150185 | 0.87909325 | -0.1375914 | 0.75672442 | 0.90650187 |
| U2af1 | 1.54862561 | 0.76324637 | 0.90127768 | -0.1380313 | 0.75201609 | 0.90650187 |
| Bud31 | 1.25990006 | 0.70140733 | 0.83998752 | -0.1385802 | 0.76835224 | 0.90650187 |
| Rpl26 | 4.69660216 | 0.44126353 | 0.58100791 | -0.1397444 | 0.85752373 | 0.91715518 |
| Cox5b | 2.76837638 | 0.72915509 | 0.86966129 | -0.1405062 | 0.76369878 | 0.90650187 |
| Rbx1 | 2.42439689 | 0.76268038 | 0.90399105 | -0.1413107 | 0.7564508 | 0.90650187 |
| Ndufb9 | 2.06828772 | 0.7829037 | 0.92578285 | -0.1428791 | 0.75367662 | 0.90650187 |
| Vdac3 | 1.46663778 | 0.7445896 | 0.88750282 | -0.1429132 | 0.76296393 | 0.90650187 |
| Serbp1 | 2.75153312 | 0.72818841 | 0.87172022 | -0.1435318 | 0.76791107 | 0.90650187 |
| Cops4 | 1.08480486 | 0.63985493 | 0.78377228 | -0.1439174 | 0.79283811 | 0.90693771 |
| Plekhj1 | 1.15926692 | 0.66411601 | 0.80954375 | -0.1454277 | 0.78774342 | 0.90650187 |
| Tceb1 | 1.60544637 | 0.76230014 | 0.90861823 | -0.1463181 | 0.76297101 | 0.90650187 |
| Psmb2 | 1.89495757 | 0.77902159 | 0.92694133 | -0.1479197 | 0.76096881 | 0.90650187 |

| | | | | | | |
|---|---|---|---|---|---|---|
| Uqcrh | 2.86669095 | 0.70708907 | 0.85550539 | -0.1484163 | 0.77971784 | 0.90650187 |
| Rpl4 | 4.51009595 | 0.45615545 | 0.60486525 | -0.1487098 | 0.86279857 | 0.91950555 |
| Rpl14 | 4.91184098 | 0.40203698 | 0.55251279 | -0.1504758 | 0.88700486 | 0.92936447 |
| Cnbp | 2.18192546 | 0.76970463 | 0.92134481 | -0.1516402 | 0.76781904 | 0.90650187 |
| Atxn7l3b | 1.99915703 | 0.77446337 | 0.92711453 | -0.1526512 | 0.76790884 | 0.90650187 |
| Cox8a | 3.38695228 | 0.63106669 | 0.78756598 | -0.1564993 | 0.81148937 | 0.9092537 |
| Rpl36 | 4.72657363 | 0.41712054 | 0.5770635 | -0.159943 | 0.89105821 | 0.93109327 |
| Atp5k | 2.52149063 | 0.73570181 | 0.8958007 | -0.1600989 | 0.78655526 | 0.90650187 |
| Ndufs6 | 1.62049687 | 0.74976094 | 0.91036498 | -0.160604 | 0.78356249 | 0.90650187 |
| Ppp1ca | 2.54608756 | 0.73275744 | 0.89344721 | -0.1606898 | 0.78801673 | 0.90650187 |
| Tpt1 | 4.50041076 | 0.44550024 | 0.60626721 | -0.160767 | 0.88077386 | 0.92622922 |
| Rps14 | 5.17637127 | 0.39258344 | 0.56082917 | -0.1682457 | 0.90884202 | 0.9391141 |
| Tma7 | 2.73193217 | 0.70515709 | 0.87417266 | -0.1690156 | 0.80497562 | 0.90882608 |
| Rsrp1 | 2.45276482 | 0.73253093 | 0.90160707 | -0.1690761 | 0.79776751 | 0.9075779 |
| Rps16 | 4.77316754 | 0.40078413 | 0.57028566 | -0.1695015 | 0.90680477 | 0.9379403 |
| Ndufa7 | 2.32269559 | 0.74184341 | 0.91235288 | -0.1705095 | 0.79696759 | 0.9075779 |
| Rps25 | 4.1155302 | 0.49377979 | 0.66534375 | -0.171564 | 0.8741714 | 0.9235672 |
| Ndufa3 | 2.35080435 | 0.73771941 | 0.91015143 | -0.172432 | 0.80017286 | 0.90805731 |
| Cox7a2 | 2.77992494 | 0.69545381 | 0.86826933 | -0.1728155 | 0.81187553 | 0.9092537 |
| Rpl7 | 4.24860679 | 0.46906932 | 0.64341504 | -0.1743457 | 0.88582552 | 0.92839449 |
| Ndufa5 | 1.68510255 | 0.74073956 | 0.91707861 | -0.176339 | 0.80367365 | 0.90882608 |
| Rpl29 | 3.26873808 | 0.62630464 | 0.80390775 | -0.1776031 | 0.83698565 | 0.91092327 |
| Rpl11 | 4.65598144 | 0.40807263 | 0.58622859 | -0.178156 | 0.91165677 | 0.94039219 |
| Tceb2 | 3.07667265 | 0.64838729 | 0.82678477 | -0.1783975 | 0.83128106 | 0.9098597 |
| Cox7c | 3.5627607 | 0.57679552 | 0.7603173 | -0.1835218 | 0.85837776 | 0.91744797 |
| Atp5g2 | 2.79326357 | 0.68252188 | 0.8665772 | -0.1840553 | 0.82747079 | 0.90947782 |
| Psmb6 | 2.2642569 | 0.72841125 | 0.91656622 | -0.188155 | 0.81927485 | 0.9092537 |
| Cox4i1 | 3.58135658 | 0.56780461 | 0.7572067 | -0.1894021 | 0.86692902 | 0.92056229 |
| Tomm20 | 2.07788521 | 0.73276111 | 0.92548605 | -0.1927249 | 0.82271499 | 0.9092537 |
| Uqcr10 | 2.53678218 | 0.7007253 | 0.89434278 | -0.1936175 | 0.83208452 | 0.91015207 |

| | | | | | | |
|---|---|---|---|---|---|---|
| Pcbp2 | 2.14030372 | 0.72922864 | 0.92319805 | -0.1939694 | 0.82486053 | 0.90934433 |
| Arf1 | 2.61382114 | 0.69244377 | 0.88691308 | -0.1944693 | 0.83516219 | 0.91066681 |
| Rpl13a | 5.2711417 | 0.37603855 | 0.57214277 | -0.1961042 | 0.93621433 | 0.95391072 |
| Rab2a | 2.05419806 | 0.72919172 | 0.92618873 | -0.196997 | 0.82781461 | 0.90948322 |
| Rpl27 | 3.97019112 | 0.48865202 | 0.6891444 | -0.2004924 | 0.9020563 | 0.93591959 |
| Rpl18 | 4.37969243 | 0.4197803 | 0.62259567 | -0.2028154 | 0.92621814 | 0.94759147 |
| Rps27 | 5.51674832 | 0.41422642 | 0.61719991 | -0.2029735 | 0.92812805 | 0.94929851 |
| Hsp90ab1 | 4.4625696 | 0.40743156 | 0.61197837 | -0.2045468 | 0.93134633 | 0.95113723 |
| Shfm1 | 3.13399636 | 0.61545562 | 0.82027407 | -0.2048184 | 0.86651046 | 0.92056229 |
| Ubb | 5.39739407 | 0.38689304 | 0.59356305 | -0.20667 | 0.93917626 | 0.95593613 |
| Eef1a1 | 5.42753591 | 0.39220727 | 0.59947993 | -0.2072727 | 0.93786397 | 0.95527138 |
| Rplp2 | 4.68759625 | 0.37105685 | 0.58212941 | -0.2110726 | 0.94651023 | 0.96073063 |
| Eif1 | 4.28521443 | 0.42416814 | 0.6369265 | -0.2127584 | 0.93122893 | 0.95113723 |
| Fau | 4.50713763 | 0.37597142 | 0.60529326 | -0.2293218 | 0.95393482 | 0.96556047 |
| Btf3 | 3.27997548 | 0.57283261 | 0.8024222 | -0.2295896 | 0.89830988 | 0.93440169 |
| Cox6b1 | 3.10832463 | 0.59265186 | 0.82349483 | -0.230843 | 0.893654 | 0.93189602 |
| Oaz1 | 3.61204972 | 0.51028167 | 0.75195106 | -0.2416694 | 0.92346517 | 0.94620379 |
| Morf4l1 | 3.27598148 | 0.55701879 | 0.80295045 | -0.2459317 | 0.91334146 | 0.94082155 |
| Atp5h | 2.93976255 | 0.59783081 | 0.84506672 | -0.2472359 | 0.9033063 | 0.93635703 |
| Sumo1 | 2.02283658 | 0.65910895 | 0.9268503 | -0.2677413 | 0.90046617 | 0.93504361 |
| Pfdn5 | 2.75808861 | 0.59680141 | 0.87089584 | -0.2740944 | 0.91911635 | 0.94387891 |
| Eif4a1 | 3.55366367 | 0.4862677 | 0.76183989 | -0.2755722 | 0.94608915 | 0.9604315 |
| Rpl37a | 5.59760749 | 0.33286817 | 0.63361865 | -0.3007505 | 0.98257645 | 0.98679603 |
| Rpl19 | 4.68282551 | 0.24933665 | 0.58272402 | -0.3333874 | 0.99451126 | 0.99549359 |
| Rpl41 | 6.41765137 | 0.33663335 | 1.25395647 | -0.9173231 | 0.99942733 | 0.99942733 |

[illegible]

[illegible]Ms for all cellular genes.
[illegible] of log-CPMs for all cellular genes.
[illegible]logical component of the variance for all genes.
[illegible]echnical component of the variance for all genes.

| | mean | total | tech | bio | p.value | FDR |
|---|---|---|---|---|---|---|
| pTimer | 6.34342159 | 15.177556 | 1.48134956 | 13.6962064 | 0 | 0 |
| tdTomato | 6.68715591 | 13.2137418 | 2.19999791 | 11.0137439 | 1.27E-120 | 1.67E-117 |
| eGFP | 3.44139964 | 10.5570509 | 1.04235111 | 9.51469982 | 0 | 0 |
| Iapp | 4.75706566 | 8.2767467 | 0.77215614 | 7.50459056 | 0 | 0 |
| Ins2 | 5.05069627 | 7.63301892 | 0.6901717 | 6.94284723 | 0 | 0 |
| Ins1 | 2.91965817 | 6.973443 | 1.10155648 | 5.87188652 | 1.89E-136 | 2.75E-133 |
| Rbp4 | 3.75358641 | 6.47109126 | 1.00453975 | 5.46655151 | 4.14E-142 | 6.67E-139 |
| Col1a1 | 1.6761 | 6.32605499 | 1.07691236 | 5.24914264 | 1.83E-114 | 2.21E-111 |
| Pyy | 4.37806505 | 5.94307395 | 0.8685308 | 5.07454315 | 1.85E-163 | 3.83E-160 |
| Itm2a | 1.95310755 | 6.14679116 | 1.11434915 | 5.03244201 | 1.41E-98 | 1.58E-95 |
| Sparc | 2.3695728 | 6.07343672 | 1.12954244 | 4.94389428 | 9.90E-93 | 8.97E-90 |
| Col3a1 | 1.75244348 | 5.94597427 | 1.09058232 | 4.85539195 | 7.22E-96 | 6.98E-93 |
| Meg3 | 5.21665646 | 5.27117394 | 0.67235955 | 4.59881439 | 3.91E-223 | 1.14E-219 |
| Cdkn1c | 2.71846087 | 5.68549269 | 1.11581279 | 4.5696799 | 1.85E-81 | 1.22E-78 |
| Nnat | 2.9514909 | **5.61063371** | 1.09904951 | 4.51158419 | 7.83E-82 | **5.68E-79** |
| Ptn | 1.73491787 | 5.59430675 | 1.08767316 | 4.50663359 | 2.50E-83 | 1.91E-80 |
| Pcsk2 | 2.66265645 | 5.59876708 | 1.11919088 | 4.4795762 | 6.62E-78 | 3.84E-75 |
| Dlk1 | 3.79799121 | 5.28524374 | 0.99897706 | 4.28626668 | 3.37E-89 | 2.87E-86 |
| Chgb | 2.69605984 | 5.35350824 | 1.11719597 | 4.23631227 | 3.81E-70 | 2.05E-67 |
| Lgals1 | 2.2180535 | 5.31825829 | 1.12771599 | 4.19054231 | 1.90E-67 | 8.59E-65 |
| H19 | 2.04219981 | 5.30254772 | 1.12034356 | 4.18220416 | 4.77E-68 | 2.23E-65 |
| Chga | 3.07656974 | 5.19139255 | 1.0878699 | 4.10352265 | 1.95E-69 | 1.01E-66 |
| Sst | 2.34207427 | 5.21143866 | 1.12947429 | 4.08196437 | 6.47E-64 | 2.76E-61 |
| Dcn | 1.47050327 | 5.10406449 | 1.02729253 | 4.07677196 | 1.27E-76 | 7.08E-74 |
| Gcg | 2.28725092 | 5.04821143 | 1.12893683 | 3.9192746 | 3.83E-59 | 1.54E-56 |
| Col1a2 | 1.50674805 | 4.93559573 | 1.03739743 | 3.8981983 | 6.36E-69 | 3.18E-66 |
| Ttr | 3.01439223 | 4.89105593 | 1.09376588 | 3.79729005 | 3.80E-59 | 1.54E-56 |
| Mest | 1.77151434 | 4.51133459 | 1.09357501 | 3.41775957 | 2.50E-48 | 8.63E-46 |

mean variance for all transcripts = 0.343

**mean**: A numeric vector of mean log-CP
**total**: A numeric vector of the variances
**bio**: A numeric vector containing the bic
**tech**: A numeric vector containing the te

| | | | | | | |
|---|---|---|---|---|---|---|
| Vim | 2.00558658 | 4.36212688 | 1.11815447 | 3.24397241 | 6.25E-42 | 1.89E-39 |
| Xist | 1.73024074 | 4.04837111 | 1.08687344 | 2.96149767 | 3.17E-37 | 8.66E-35 |
| Tmsb4x | 4.04640361 | 3.64172988 | 0.95281776 | 2.68891211 | 8.76E-40 | 2.49E-37 |
| Tmsb15b2 | 1.56523543 | 3.57263754 | 1.05253902 | 2.52009852 | 3.39E-29 | 6.46E-27 |
| Igf2 | 1.51325583 | 3.54889512 | 1.03916482 | 2.5097303 | 1.13E-29 | 2.28E-27 |
| Serpinh1 | 2.22574749 | 3.61528623 | 1.12790325 | 2.48738298 | 4.59E-25 | 7.40E-23 |
| Pcsk1n | 2.34345161 | 3.60796718 | 1.12948255 | 2.47848463 | 7.79E-25 | 1.23E-22 |
| S100a6 | 1.1217641 | 3.32313464 | 0.89550376 | 2.42763088 | 7.22E-37 | 1.94E-34 |
| Sparcl1 | 1.17697758 | 3.28373495 | 0.92048387 | 2.36325108 | 2.83E-33 | 6.62E-31 |
| Fos | 1.95124765 | 3.41166797 | 1.11420302 | 2.29746495 | 3.75E-22 | 5.22E-20 |
| Lum | 1.06389014 | 3.15432928 | 0.86732232 | 2.28700696 | 5.45E-35 | 1.36E-32 |
| Spp1 | 1.07471298 | 3.11970212 | 0.87275535 | 2.24694676 | 1.90E-33 | 4.50E-31 |
| Ppy | 1.01276673 | 3.08739684 | 0.84110455 | 2.24629229 | 7.69E-36 | 2.03E-33 |
| Mafb | 1.78371292 | 3.30274916 | 1.09540434 | 2.20734482 | 3.05E-21 | 4.02E-19 |
| Mt1 | 0.88860146 | 2.76763275 | 0.77339966 | 1.99423309 | 1.51E-33 | 3.66E-31 |
| Gsn | 1.13186966 | 2.85206333 | 0.90019392 | 1.95186941 | 2.71E-24 | 4.14E-22 |
| Cpe | 3.17037481 | 2.99874019 | 1.0785133 | 1.9202269 | 5.43E-17 | 5.36E-15 |
| Slc38a5 | 1.34780987 | 2.8939239 | 0.98793796 | 1.90598594 | 1.26E-19 | 1.50E-17 |
| Ghrl | 1.13897523 | 2.80761958 | 0.90345951 | 1.90416007 | 4.65E-23 | 6.74E-21 |
| Col6a1 | 0.98619028 | 2.6992663 | 0.82735534 | 1.87191095 | 2.80E-26 | 4.77E-24 |
| Isl1 | 1.80796832 | 2.9470935 | 1.09886452 | 1.84822898 | 2.32E-15 | 1.94E-13 |
| Stmn1 | 1.62986249 | 2.89178933 | 1.0674236 | 1.82436573 | 8.41E-16 | 7.30E-14 |
| Igfbp4 | 1.23580351 | 2.76683269 | 0.94541225 | 1.82142044 | 1.41E-19 | 1.67E-17 |
| Cd24a | 1.3393435 | 2.78336489 | 0.9849534 | 1.79841149 | 8.95E-18 | 9.68E-16 |
| Gpc3 | 1.18126272 | 2.71232726 | 0.9223582 | 1.78996906 | 7.74E-20 | 9.51E-18 |
| Anxa2 | 1.43395934 | 2.80322662 | 1.01637945 | 1.78684716 | 1.30E-16 | 1.24E-14 |
| 1700086L19F | 1.7986961 | 2.79598145 | 1.09757148 | 1.69840997 | 2.81E-13 | 1.99E-11 |
| Fam183b | 1.95799439 | 2.80997132 | 1.11472786 | 1.69524346 | 6.94E-13 | 4.68E-11 |
| Fstl1 | 1.12524505 | 2.57512825 | 0.89712653 | 1.67800172 | 1.46E-18 | 1.62E-16 |
| Spint2 | 2.40149127 | 2.79822863 | 1.12918503 | 1.66904359 | 2.86E-12 | 1.80E-10 |

| | | | | | | |
|---|---|---|---|---|---|---|
| Pax6 | 1.79325593 | 2.7657464 | 1.09679577 | 1.66895063 | 6.74E-13 | 4.57E-11 |
| Gng12 | 2.22979637 | 2.78957931 | 1.12799541 | 1.6615839 | 3.38E-12 | 2.09E-10 |
| Mdk | 1.83159077 | 2.74746291 | 1.10198309 | 1.64547982 | 1.74E-12 | 1.13E-10 |
| Scg2 | 1.29824983 | 2.5777282 | 0.96990762 | 1.60782058 | 5.67E-15 | 4.60E-13 |
| Hba-a1 | 1.11710647 | 2.48440341 | 0.89331981 | 1.5910836 | 5.29E-17 | 5.29E-15 |
| Tmem27 | 1.36586758 | 2.54509572 | 0.9941552 | 1.55094052 | 1.82E-13 | 1.31E-11 |
| Scg3 | 1.71286499 | 2.6316874 | 1.08380745 | 1.54787995 | 1.43E-11 | 8.31E-10 |
| Ifitm2 | 2.49009268 | 2.64263827 | 1.12720476 | 1.51543351 | 1.88E-10 | 9.67E-09 |
| Mt2 | 0.61907271 | 2.08799517 | 0.58916494 | 1.49883023 | 1.05E-32 | 2.33E-30 |
| S100a10 | 1.59679804 | 2.54055591 | 1.06004388 | 1.48051203 | 3.84E-11 | 2.12E-09 |
| Neurog3 | 0.51508447 | 1.98194283 | 0.50469576 | 1.47724707 | 1.21E-42 | 3.82E-40 |
| Col6a2 | 0.91169633 | 2.26151162 | 0.78675436 | 1.47475726 | 1.24E-18 | 1.38E-16 |
| Hba-a2 | 0.45834399 | 1.92232865 | 0.45572216 | 1.46660649 | 4.05E-51 | 1.43E-48 |
| Hbb-bs | 2.09140672 | 2.5883594 | 1.12296075 | 1.46539865 | 6.02E-10 | 2.91E-08 |
| Krt8 | 2.02744399 | 2.56901903 | 1.11951366 | 1.44950537 | 8.09E-10 | 3.88E-08 |
| Scgn | 1.70821552 | 2.52969113 | 1.08296351 | 1.44672763 | 2.43E-10 | 1.23E-08 |
| Cryba2 | 1.59328159 | 2.48921487 | 1.05922891 | 1.42998596 | 1.59E-10 | 8.26E-09 |
| Hbb-bt | 0.47209709 | 1.87035475 | 0.46777187 | 1.40258288 | 1.20E-44 | 3.85E-42 |
| Dbpht2 | 1.43067095 | 2.40039992 | 1.0153584 | 1.38504152 | 1.04E-10 | 5.53E-09 |
| Meis2 | 2.69909509 | 2.50140536 | 1.11699643 | 1.38440893 | 3.86E-09 | 1.67E-07 |
| Ctrb1 | 1.35180792 | 2.36545449 | 0.98933119 | 1.3761233 | 4.57E-11 | 2.50E-09 |
| Aplp1 | 1.89572618 | 2.47188201 | 1.10925376 | 1.36262825 | 5.23E-09 | 2.22E-07 |
| Mfap2 | 1.33971404 | 2.34236418 | 0.98508492 | 1.35727926 | 6.85E-11 | 3.68E-09 |
| Epcam | 1.93477213 | 2.44810257 | 1.11285272 | 1.33524985 | 1.14E-08 | 4.60E-07 |
| mt-Atp6 | 5.78373489 | 2.02295118 | 0.69711951 | 1.32583167 | 3.98E-19 | 4.57E-17 |
| Bex2 | 2.16087636 | 2.44238815 | 1.12601679 | 1.31637135 | 2.57E-08 | 9.95E-07 |
| Hmgb2 | 1.47271942 | 2.33594461 | 1.02792819 | 1.30801642 | 1.53E-09 | 7.00E-08 |
| Crip1 | 1.42915555 | 2.32012224 | 1.0148858 | 1.30523644 | 1.04E-09 | 4.85E-08 |
| Tuba1a | 3.63036546 | 2.31601155 | 1.01847435 | 1.2975372 | 1.46E-09 | 6.76E-08 |
| Ldha | 1.27959909 | 2.25955112 | 0.96283571 | 1.29671541 | 1.75E-10 | 9.05E-09 |

| | | | | | | |
|---|---|---|---|---|---|---|
| Hmgn3 | 2.29841572 | 2.42305519 | 1.12905956 | 1.29399562 | 4.66E-08 | 1.73E-06 |
| Clu | 0.85105239 | 2.031121 | 0.75086589 | 1.28025511 | 9.81E-16 | 8.47E-14 |
| Rcn3 | 1.15549739 | 2.14615843 | 0.9109236 | 1.23523482 | 1.33E-10 | 6.95E-09 |
| Neurod1 | 1.59314805 | 2.28349196 | 1.05919784 | 1.22429413 | 3.62E-08 | 1.37E-06 |
| Pdx1 | 1.47533132 | 2.25261449 | 1.02867389 | 1.22394061 | 1.48E-08 | 5.94E-07 |
| Gpx3 | 1.80911843 | 2.31830536 | 1.09902202 | 1.21928334 | 1.18E-07 | 4.17E-06 |
| Cald1 | 0.93723563 | 1.99385754 | 0.80108419 | 1.19277336 | 2.00E-12 | 1.27E-10 |
| Ifitm1 | 0.76023383 | 1.86792709 | 0.69212474 | 1.17580235 | 1.24E-15 | 1.06E-13 |
| Map1b | 2.18540685 | 2.2936482 | 1.12688614 | 1.16676206 | 7.04E-07 | 2.25E-05 |
| Ppic | 1.21588067 | 2.10106134 | 0.9372263 | 1.16383504 | 3.62E-09 | 1.57E-07 |
| Pcsk1 | 1.33426003 | 2.14664649 | 0.98314077 | 1.16350572 | 1.76E-08 | 6.96E-07 |
| Mfap4 | 0.68281218 | 1.79978834 | 0.63737539 | 1.16241295 | 9.75E-18 | 1.05E-15 |
| mt-Co1 | 5.46866269 | 1.82641718 | 0.67036753 | 1.15604965 | 4.70E-16 | 4.24E-14 |
| Tubb3 | 1.33561766 | 2.13691541 | 0.98362638 | 1.15328902 | 2.35E-08 | 9.17E-07 |
| Peg3 | 2.70509753 | 2.25696711 | 1.11659796 | 1.14036915 | 9.77E-07 | 3.04E-05 |
| Ccnd2 | 1.29681495 | 2.10075637 | 0.96936518 | 1.13139118 | 2.70E-08 | 1.04E-06 |
| B2m | 1.15159417 | 2.03670725 | 0.90917655 | 1.1275307 | 3.78E-09 | 1.64E-07 |
| Fabp5 | 1.42269196 | 2.14027872 | 1.01286619 | 1.12741253 | 1.08E-07 | 3.84E-06 |
| Pcolce | 0.93069975 | 1.91915836 | 0.79746136 | 1.121697 | 2.82E-11 | 1.58E-09 |
| Krt7 | 1.73713322 | 2.20798841 | 1.08804851 | 1.11993991 | 8.11E-07 | 2.58E-05 |
| Bgn | 0.77883686 | 1.81962938 | 0.70464369 | 1.11498569 | 8.40E-14 | 6.12E-12 |
| Hadh | 2.08136713 | 2.23239673 | 1.12244735 | 1.10994938 | 2.04E-06 | 6.08E-05 |
| Igfbp7 | 1.53011126 | 2.10326699 | 1.04368256 | 1.05958443 | 1.12E-06 | 3.46E-05 |
| Nkx6-1 | 1.31224747 | 2.02326836 | 0.97513338 | 1.04813498 | 2.75E-07 | 9.32E-06 |
| Krt18 | 1.77341414 | 2.13516706 | 1.09386401 | 1.04130305 | 4.60E-06 | 0.0001288 |
| Cxcl12 | 0.87150112 | 1.78673996 | 0.76326477 | 1.02347519 | 2.09E-10 | 1.07E-08 |
| Pmp22 | 0.80667372 | 1.7461736 | 0.72290947 | 1.02326413 | 2.13E-11 | 1.22E-09 |
| Nrp1 | 0.86594764 | 1.78072104 | 0.75992706 | 1.02079398 | 1.95E-10 | 9.97E-09 |
| Postn | 0.66306147 | 1.63992341 | 0.62273512 | 1.0171883 | 1.37E-14 | 1.08E-12 |
| Tspan7 | 1.61280988 | 2.07470334 | 1.06367609 | 1.01102726 | 4.75E-06 | 0.00013238 |

| | | | | | | |
|---|---|---|---|---|---|---|
| Akap12 | 0.8070103 | 1.7323008 | 0.72312688 | 1.00917392 | 3.97E-11 | 2.18E-09 |
| Ppp1r1a | 0.91732875 | 1.79135512 | 0.789952 | 1.00140312 | 1.75E-09 | 7.99E-08 |
| Ociad2 | 1.13082559 | 1.90055234 | 0.89971126 | 1.00084107 | 1.09E-07 | 3.90E-06 |
| Ifitm3 | 0.96152044 | 1.81282268 | 0.81430296 | 0.99851972 | 5.55E-09 | 2.33E-07 |
| Tmsb10 | 4.86830855 | 1.73739945 | 0.74012523 | 0.99727421 | 1.72E-10 | 8.90E-09 |
| Igf1 | 0.65806604 | 1.60664448 | 0.61898917 | 0.9876553 | 5.28E-14 | 3.88E-12 |
| Scg5 | 1.45623132 | 2.00475252 | 1.02313352 | 0.981619 | 3.91E-06 | 0.0001108 |
| Ndn | 1.80150424 | 2.06919514 | 1.09796694 | 0.9712282 | 1.88E-05 | 0.00047034 |
| Cd34 | 0.71228098 | 1.62795826 | 0.65871425 | 0.96924401 | 3.56E-12 | 2.20E-10 |
| Lrpprc | 1.20631723 | 1.90228226 | 0.9331935 | 0.96908876 | 6.55E-07 | 2.11E-05 |
| Nfib | 1.21387296 | 1.89806542 | 0.93638732 | 0.9616781 | 8.56E-07 | 2.71E-05 |
| Kcnq1ot1 | 2.23374227 | 2.07864607 | 1.12808101 | 0.95056506 | 4.32E-05 | 0.00098989 |
| Cystm1 | 1.85239363 | 2.0524726 | 1.1045131 | 0.94795951 | 3.19E-05 | 0.00075602 |
| Sdf2l1 | 1.43061201 | 1.9608062 | 1.01534004 | 0.94546616 | 7.18E-06 | 0.00019429 |
| Slc25a5 | 3.45153425 | 1.97564765 | 1.04094854 | 0.9346991 | 1.44E-05 | 0.00036586 |
| Junb | 1.2582039 | 1.88898732 | 0.95446234 | 0.93452499 | 2.54E-06 | 7.42E-05 |
| Cdkn1a | 0.63263424 | 1.53344396 | 0.59965466 | 0.9337893 | 2.01E-13 | 1.44E-11 |
| Dbi | 1.50945018 | 1.97113029 | 1.03812382 | 0.93300646 | 1.41E-05 | 0.00036108 |
| Itih5 | 0.61433935 | 1.51404775 | 0.58547459 | 0.92857316 | 7.39E-14 | 5.41E-12 |
| Csrp2 | 1.27671657 | 1.87426861 | 0.96172589 | 0.91254271 | 4.92E-06 | 0.0001369 |
| Syt13 | 1.64730665 | 1.98060389 | 1.07112034 | 0.90948355 | 3.81E-05 | 0.00088581 |
| Gng11 | 0.62267533 | 1.50142226 | 0.59196348 | 0.90945878 | 4.10E-13 | 2.82E-11 |
| Pnliprp1 | 0.76999776 | 1.60031088 | 0.69872668 | 0.90158421 | 9.19E-10 | 4.32E-08 |
| Ccdc80 | 0.680049 | 1.52807927 | 0.63534318 | 0.89273608 | 2.95E-11 | 1.65E-09 |
| Jun | 1.71031382 | 1.9713818 | 1.08334561 | 0.88803619 | 6.70E-05 | 0.0014646 |
| Hist3h2ba | 1.46432831 | 1.91318399 | 1.02550697 | 0.88767701 | 2.76E-05 | 0.00066337 |
| Hspa5 | 3.69534978 | 1.89432505 | 1.0113304 | 0.88299465 | 2.37E-05 | 0.00058063 |
| Maged2 | 1.89429007 | 1.98355756 | 1.10910708 | 0.87445048 | 0.00011975 | 0.00244179 |
| Cthrc1 | 0.64433042 | 1.48138461 | 0.60860177 | 0.87278284 | 1.18E-11 | 6.98E-10 |
| Sfrp1 | 0.65922496 | 1.4852197 | 0.61985976 | 0.86535993 | 3.91E-11 | 2.15E-09 |

| | | | | | | |
|---|---|---|---|---|---|---|
| Hes1 | 0.76320651 | 1.55895383 | 0.69414204 | 0.86481178 | 3.23E-09 | 1.41E-07 |
| Tm4sf4 | 1.36967198 | 1.84764851 | 0.9954404 | 0.85220811 | 3.32E-05 | 0.00078459 |
| Ebf1 | 0.68898676 | 1.49220154 | 0.64189714 | 0.8503044 | 3.38E-10 | 1.70E-08 |
| Mlxipl | 1.14149903 | 1.75425428 | 0.90461136 | 0.84964292 | 6.05E-06 | 0.00016639 |
| Gch1 | 1.4283867 | 1.86158385 | 1.01464553 | 0.84693831 | 5.04E-05 | 0.00114245 |
| Col4a1 | 0.96628478 | 1.6548134 | 0.81684792 | 0.83796548 | 8.79E-07 | 2.77E-05 |
| Btg2 | 1.55281108 | 1.88593915 | 1.04948138 | 0.83645776 | 0.0001023 | 0.00212788 |
| Emp3 | 0.7959985 | 1.55128763 | 0.71597115 | 0.83531648 | 2.73E-08 | 1.05E-06 |
| Apoe | 0.6151126 | 1.41622169 | 0.58607849 | 0.8301432 | 2.07E-11 | 1.19E-09 |
| Ubqln2 | 1.69872664 | 1.90734083 | 1.08121017 | 0.82613066 | 0.00018557 | 0.00367033 |
| Gm42418 | 1.77427739 | 1.91895378 | 1.09399481 | 0.82495897 | 0.00022136 | 0.00432522 |
| Peg10 | 0.64880316 | 1.43185359 | 0.61199828 | 0.81985531 | 2.17E-10 | 1.10E-08 |
| Id3 | 0.44494609 | 1.26098173 | 0.44387472 | 0.81710701 | 4.89E-18 | 5.38E-16 |
| Gas1 | 0.72554401 | 1.48393053 | 0.66811875 | 0.81581178 | 6.40E-09 | 2.66E-07 |
| Sepp1 | 1.69874505 | 1.89556045 | 1.08121361 | 0.81434684 | 0.00022489 | 0.00438266 |
| S100a16 | 0.77537638 | 1.51409898 | 0.70233396 | 0.81176502 | 3.63E-08 | 1.37E-06 |
| Mid1ip1 | 1.62063844 | 1.85608945 | 1.06540877 | 0.79068069 | 0.00027253 | 0.00524017 |
| Manf | 2.93095882 | 1.88821377 | 1.10066484 | 0.78754893 | 0.00042869 | 0.00787732 |
| Slc30a8 | 0.89484273 | 1.55085891 | 0.77704664 | 0.77381227 | 1.75E-06 | 5.24E-05 |
| Prkcdbp | 0.7024788 | 1.4230616 | 0.65168389 | 0.77137772 | 1.75E-08 | 6.94E-07 |
| Pam | 1.59331361 | 1.82801752 | 1.05923636 | 0.76878116 | 0.00036058 | 0.0067107 |
| Atp1b1 | 1.50685307 | 1.80586493 | 1.03742573 | 0.7684392 | 0.0002793 | 0.00532801 |
| Sec61b | 3.90291136 | 1.74961146 | 0.98269274 | 0.76691872 | 0.00013847 | 0.00279222 |
| mt-Nd2 | 3.51638837 | 1.7872787 | 1.03238347 | 0.75489523 | 0.00032878 | 0.00616637 |
| Fbln1 | 0.70738314 | 1.40969867 | 0.65520962 | 0.75448905 | 4.05E-08 | 1.52E-06 |
| Gpx8 | 0.75531318 | 1.43560912 | 0.68877198 | 0.74683714 | 2.19E-07 | 7.50E-06 |
| mt-Co3 | 5.92601206 | 1.47814533 | 0.73833669 | 0.73980864 | 1.52E-06 | 4.61E-05 |
| Hist1h2bc | 0.99546664 | 1.57086453 | 0.83216065 | 0.73870388 | 1.77E-05 | 0.00044302 |
| Pak3 | 1.15002937 | 1.64308013 | 0.90847333 | 0.7346068 | 8.25E-05 | 0.001767 |
| Smoc2 | 0.47033364 | 1.19971989 | 0.46623293 | 0.73348696 | 1.15E-13 | 8.37E-12 |

| | | | | | | |
|---|---|---|---|---|---|---|
| Tubb4b | 2.25849824 | 1.85684279 | 1.12854336 | 0.72829943 | 0.00132084 | 0.02106654 |
| Hhex | 0.52826541 | 1.24333837 | 0.51578815 | 0.72755023 | 2.49E-11 | 1.41E-09 |
| Emb | 1.27430041 | 1.68436043 | 0.96079167 | 0.72356876 | 0.00022521 | 0.00438266 |
| Abcc8 | 1.21346426 | 1.65895444 | 0.93621621 | 0.72273822 | 0.0001612 | 0.00321916 |
| Gadd45a | 0.79570199 | 1.42666481 | 0.71577725 | 0.71088756 | 1.85E-06 | 5.55E-05 |
| Ube2c | 0.38961647 | 1.09155451 | 0.39384977 | 0.69770474 | 7.70E-17 | 7.45E-15 |
| Plagl1 | 0.77873773 | 1.39417707 | 0.70457765 | 0.68959941 | 2.56E-06 | 7.45E-05 |
| 2810417H13 | 0.44387418 | 1.12556621 | 0.4429222 | 0.68264402 | 3.47E-13 | 2.41E-11 |
| Clps | 0.98166152 | 1.50276988 | 0.8249867 | 0.67778319 | 6.47E-05 | 0.00142532 |
| mt-Nd1 | 4.61368622 | 1.48688998 | 0.81061033 | 0.67627965 | 5.08E-05 | 0.0011501 |
| Fscn1 | 0.7886126 | 1.38262531 | 0.71112206 | 0.67150325 | 5.43E-06 | 0.00015047 |
| Ccl21a | 0.14896758 | 0.82710074 | 0.1579942 | 0.66910654 | 5.88E-87 | 4.74E-84 |
| Mgst1 | 0.96013327 | 1.48249575 | 0.81355886 | 0.6689369 | 6.39E-05 | 0.00140958 |
| Tuba4a | 1.16961251 | 1.58053768 | 0.91723447 | 0.66330322 | 0.00037702 | 0.00696304 |
| Penk | 0.37310985 | 1.03767896 | 0.3785936 | 0.65908537 | 2.52E-16 | 2.31E-14 |
| Id2 | 0.72809676 | 1.32548443 | 0.66991441 | 0.65557002 | 2.57E-06 | 7.46E-05 |
| mt-Nd4 | 4.32856342 | 1.53243294 | 0.88206707 | 0.65036587 | 0.00029613 | 0.00561221 |
| Zbtb20 | 1.67200218 | 1.72363405 | 1.07610816 | 0.64752589 | 0.00252817 | 0.03691186 |
| Cpa1 | 0.41354417 | 1.06212665 | 0.41569738 | 0.64642927 | 2.16E-13 | 1.55E-11 |
| Mmp14 | 0.91317614 | 1.43302178 | 0.78759674 | 0.64542504 | 6.73E-05 | 0.00146692 |
| Cks2 | 0.54273586 | 1.17251231 | 0.52783892 | 0.6446734 | 6.35E-09 | 2.64E-07 |
| Mgp | 0.32515553 | 0.97444246 | 0.33344046 | 0.64100201 | 1.68E-19 | 1.98E-17 |
| Cyba | 1.03718337 | 1.49465351 | 0.85367066 | 0.64098286 | 0.00023421 | 0.00453357 |
| Cplx2 | 1.26633191 | 1.59602291 | 0.95768256 | 0.63834035 | 0.0009501 | 0.01578698 |
| Cav1 | 0.30524002 | 0.93718823 | 0.3143375 | 0.62285072 | 1.33E-20 | 1.68E-18 |
| Sytl4 | 0.75775182 | 1.31210457 | 0.69043567 | 0.6216689 | 1.36E-05 | 0.00035023 |
| Rbp1 | 0.83971941 | 1.36346167 | 0.74386269 | 0.61959898 | 5.21E-05 | 0.00117444 |
| Igfbp5 | 0.57941767 | 1.17553761 | 0.55778506 | 0.61775255 | 1.24E-07 | 4.35E-06 |
| Acta2 | 0.33829756 | 0.96186624 | 0.34593453 | 0.61593171 | 5.42E-17 | 5.36E-15 |
| Rarres2 | 0.53824931 | 1.12576073 | 0.52411705 | 0.60164367 | 4.44E-08 | 1.66E-06 |

| | | | | | | |
|---|---|---|---|---|---|---|
| Pdia6 | 2.93633565 | 1.70106924 | 1.10023608 | 0.60083316 | 0.00547605 | 0.07069615 |
| Auts2 | 1.15955827 | 1.51220001 | 0.9127416 | 0.59945841 | 0.00110728 | 0.01805781 |
| Anxa4 | 1.60380497 | 1.65466805 | 1.0616492 | 0.59301885 | 0.00462864 | 0.0617919 |
| Islr | 0.532395 | 1.11038502 | 0.51924098 | 0.59114404 | 5.66E-08 | 2.08E-06 |
| Nrk | 0.51397234 | 1.0942431 | 0.50375496 | 0.59048814 | 2.37E-08 | 9.22E-07 |
| Tuba1b | 2.7890895 | 1.70056363 | 1.11124749 | 0.58931614 | 0.00674193 | 0.08397299 |
| Krt19 | 0.56291627 | 1.13309573 | 0.54442187 | 0.58867385 | 2.36E-07 | 8.06E-06 |
| Tubb6 | 0.50334453 | 1.08196837 | 0.4947249 | 0.58724347 | 1.60E-08 | 6.39E-07 |
| Cd63 | 3.47994785 | 1.61961767 | 1.03700039 | 0.58261728 | 0.00442844 | 0.05955793 |
| Ccnd3 | 1.38006662 | 1.58025402 | 0.99895953 | 0.58129449 | 0.00335365 | 0.0471593 |
| Sdpr | 0.51654604 | 1.08714878 | 0.50593097 | 0.5812178 | 4.34E-08 | 1.62E-06 |
| Gpx1 | 1.32670731 | 1.55528697 | 0.98041905 | 0.57486792 | 0.00314979 | 0.0449023 |
| mt-Co2 | 4.91634259 | 1.29645364 | 0.72611382 | 0.57033983 | 0.00012638 | 0.00256612 |
| Ambp | 0.6906996 | 1.2114935 | 0.64314672 | 0.56834678 | 1.92E-05 | 0.00047789 |
| Crip2 | 0.71227045 | 1.22587602 | 0.65870674 | 0.56716929 | 3.02E-05 | 0.00071932 |
| Alas2 | 0.09069145 | 0.65857072 | 0.09697654 | 0.56159418 | 1.27E-160 | 2.29E-157 |
| Ier2 | 1.42308128 | 1.5707566 | 1.01298824 | 0.55776836 | 0.00515446 | 0.06738445 |
| Vwa5b2 | 1.0776086 | 1.43075823 | 0.8741956 | 0.55656263 | 0.00150778 | 0.02354648 |
| Zeb2 | 0.56068207 | 1.08854611 | 0.54259896 | 0.54594715 | 1.38E-06 | 4.22E-05 |
| Fxyd3 | 1.03382698 | 1.39735527 | 0.85193127 | 0.54542399 | 0.00142835 | 0.0225335 |
| Calr | 3.45675347 | 1.5829688 | 1.04022505 | 0.54274376 | 0.00753212 | 0.09262144 |
| Tpm4 | 1.09263795 | 1.41925198 | 0.88159929 | 0.53765269 | 0.00224647 | 0.03330193 |
| Cd81 | 3.32275336 | 1.59580459 | 1.05903111 | 0.53677349 | 0.00910304 | 0.10874825 |
| Plvap | 0.22051358 | 0.76718268 | 0.23092028 | 0.53626239 | 1.39E-27 | 2.49E-25 |
| H2afz | 3.13308687 | 1.61654388 | 1.08221484 | 0.53432904 | 0.01071591 | 0.12408885 |
| Pcbd1 | 1.45010755 | 1.5513374 | 1.02130514 | 0.53003226 | 0.0078057 | 0.0953387 |
| Tubb2a | 1.5978421 | 1.58881747 | 1.06028464 | 0.52853282 | 0.01010575 | 0.1184423 |
| Ssr2 | 3.13755102 | 1.59852377 | 1.08177759 | 0.51674618 | 0.01302403 | 0.14469148 |
| Mfap5 | 0.34997061 | 0.87321522 | 0.3569577 | 0.51625752 | 8.02E-12 | 4.81E-10 |
| Eln | 0.52728712 | 1.0261844 | 0.51496857 | 0.51121583 | 1.87E-06 | 5.60E-05 |

| | | | | | | |
|---|---|---|---|---|---|---|
| Fbn2 | 0.49924344 | 0.99370112 | 0.49122132 | 0.50247981 | 9.41E-07 | 2.94E-05 |
| Tpm2 | 0.52414337 | 1.01389513 | 0.51233071 | 0.50156442 | 2.55E-06 | 7.42E-05 |
| Rrbp1 | 2.07010567 | 1.62294595 | 1.12183333 | 0.50111262 | 0.0187121 | 0.19226646 |
| Sox4 | 1.46953775 | 1.52693455 | 1.02701473 | 0.49991982 | 0.01166972 | 0.13260663 |
| Top2a | 0.30078545 | 0.80588891 | 0.31003769 | 0.49585122 | 4.63E-14 | 3.46E-12 |
| Zcchc18 | 0.92516006 | 1.28755857 | 0.79436613 | 0.49319244 | 0.00191056 | 0.02894551 |
| Tinagl1 | 0.4369307 | 0.92690223 | 0.43673686 | 0.49016537 | 8.53E-08 | 3.10E-06 |
| Cenpa | 0.34010824 | 0.83691425 | 0.34764916 | 0.48926509 | 2.75E-11 | 1.55E-09 |
| Cldn7 | 1.20004672 | 1.41788391 | 0.93051437 | 0.48736954 | 0.0073399 | 0.09071937 |
| Ccnb2 | 0.29477085 | 0.78970139 | 0.30421615 | 0.48548524 | 5.23E-14 | 3.87E-12 |
| mt-Cytb | 5.13897555 | 1.16436784 | 0.67926236 | 0.48510548 | 0.00043853 | 0.00801747 |
| Tyrobp | 0.14896082 | 0.64158614 | 0.1579872 | 0.48359894 | 1.91E-46 | 6.45E-44 |
| Mif | 2.41465136 | 1.61050784 | 1.12900156 | 0.48150628 | 0.02346202 | 0.22557166 |
| Snca | 0.10047078 | 0.58850336 | 0.10740572 | 0.48109763 | 5.17E-97 | 5.35E-94 |
| S100a11 | 3.32982319 | 1.53720248 | 1.0581423 | 0.47906019 | 0.01746042 | 0.18264153 |
| Snhg18 | 0.67389177 | 1.10962776 | 0.63079665 | 0.47883111 | 0.00020269 | 0.0039765 |
| Cela2a | 0.17787209 | 0.66601748 | 0.18771181 | 0.47830567 | 8.33E-33 | 1.89E-30 |
| Laptm4a | 2.93400602 | 1.57821806 | 1.1004222 | 0.47779586 | 0.02154273 | 0.21203425 |
| Immp1l | 1.76117308 | 1.56879317 | 1.09197412 | 0.47681905 | 0.0209575 | 0.20789599 |
| Arpc1b | 1.50999281 | 1.51121577 | 1.03827273 | 0.47294305 | 0.01691082 | 0.17869756 |
| Ap1s2 | 1.39017582 | 1.4716333 | 1.00235709 | 0.4692762 | 0.01458472 | 0.15815205 |
| Igfbp3 | 0.38484381 | 0.85778966 | 0.38945414 | 0.46833552 | 1.06E-08 | 4.30E-07 |
| Egfl7 | 0.48546308 | 0.94584267 | 0.47937404 | 0.46646863 | 2.90E-06 | 8.34E-05 |
| Ero1lb | 0.83733701 | 1.20622502 | 0.74237858 | 0.46384644 | 0.00180161 | 0.02755246 |
| Rpl39 | 4.5038592 | 1.29442925 | 0.83838867 | 0.45604058 | 0.0056348 | 0.07233093 |
| Col5a2 | 0.5371912 | 0.97777398 | 0.52323739 | 0.45453659 | 2.59E-05 | 0.00062722 |
| Gas6 | 0.54630839 | 0.9840006 | 0.5307932 | 0.4532074 | 3.48E-05 | 0.00081537 |
| Rpl36a | 4.08572723 | 1.39639081 | 0.94381343 | 0.45257738 | 0.01273972 | 0.14251576 |
| Birc5 | 0.31361731 | 0.77383335 | 0.32239721 | 0.45143614 | 3.43E-11 | 1.91E-09 |
| Txnip | 1.43931609 | 1.46827265 | 1.01802968 | 0.45024297 | 0.01967513 | 0.19891914 |

| | | | | | | |
|---|---|---|---|---|---|---|
| Capn6 | 0.4471952 | 0.89184746 | 0.44587108 | 0.44597638 | 1.58E-06 | 4.77E-05 |
| Gnas | 5.90352776 | 1.15669574 | 0.71119388 | 0.44550186 | 0.00175901 | 0.02695792 |
| Col14a1 | 0.4360034 | 0.88115242 | 0.43590872 | 0.4452437 | 9.74E-07 | 3.04E-05 |
| Egr1 | 1.15332475 | 1.35405516 | 0.90995239 | 0.44410278 | 0.01148904 | 0.13125935 |
| Anxa5 | 2.261894 | 1.57235387 | 1.1285949 | 0.44375897 | 0.03348488 | 0.2936188 |
| Phgdh | 0.57329394 | 0.99571414 | 0.55284599 | 0.44286815 | 9.49E-05 | 0.00198602 |
| Col5a1 | 0.49045559 | 0.92569262 | 0.48367939 | 0.44201324 | 1.03E-05 | 0.00027185 |
| Pecam1 | 0.24259649 | 0.69370852 | 0.25298446 | 0.44072406 | 2.40E-16 | 2.22E-14 |
| Tpm1 | 1.22992869 | 1.37970205 | 0.94302467 | 0.43667738 | 0.01548838 | 0.16599537 |
| Timp3 | 0.51163966 | 0.93795552 | 0.50177909 | 0.43617643 | 2.56E-05 | 0.00062131 |
| Olfml3 | 0.5061794 | 0.93116772 | 0.49714057 | 0.43402716 | 2.38E-05 | 0.00058063 |
| Akr1c19 | 1.13029136 | 1.33310076 | 0.89946472 | 0.43363604 | 0.0123489 | 0.13867887 |
| Gadd45g | 0.9861531 | 1.25837006 | 0.82733595 | 0.43103411 | 0.0076066 | 0.09337756 |
| Lmna | 1.39388245 | 1.43352449 | 1.00358801 | 0.42993648 | 0.02297308 | 0.22241326 |
| Agtr2 | 0.32930438 | 0.76715216 | 0.3373944 | 0.42975776 | 1.48E-09 | 6.79E-08 |
| Mxra8 | 0.54726002 | 0.96024732 | 0.53157883 | 0.42866848 | 8.60E-05 | 0.0018278 |
| Ramp2 | 0.84767459 | 1.17446827 | 0.74878846 | 0.42567981 | 0.00404168 | 0.05512347 |
| Nusap1 | 0.20428608 | 0.63943904 | 0.21456843 | 0.42487061 | 1.41E-20 | 1.77E-18 |
| Gast | 0.28347544 | 0.71749282 | 0.29323454 | 0.42425828 | 7.88E-12 | 4.74E-10 |
| Copz2 | 0.68771867 | 1.06517086 | 0.64097069 | 0.42420016 | 0.00102349 | 0.01684291 |
| Cldn4 | 0.91020391 | 1.21003922 | 0.78590316 | 0.42413606 | 0.00596283 | 0.07569976 |
| Eng | 0.36167057 | 0.79107303 | 0.36793266 | 0.42314036 | 4.21E-08 | 1.58E-06 |
| Hnrnpa1 | 2.71801017 | 1.53816156 | 1.11583941 | 0.42232215 | 0.03892386 | 0.33039706 |
| Matn2 | 0.50834918 | 0.91797698 | 0.49898606 | 0.41899092 | 4.58E-05 | 0.00104312 |
| Fkbp7 | 0.60873394 | 0.99608291 | 0.58108464 | 0.41499827 | 0.00043843 | 0.00801747 |
| Ckb | 0.74050877 | 1.08855672 | 0.67858007 | 0.40997666 | 0.00244049 | 0.03584829 |
| Klf2 | 0.49146715 | 0.89346426 | 0.48454982 | 0.40891444 | 4.22E-05 | 0.00096743 |
| Ahnak | 0.52811715 | 0.92396491 | 0.51566398 | 0.40830093 | 0.0001126 | 0.00231882 |
| Ndufa1 | 1.7849442 | 1.50322931 | 1.09558608 | 0.40764324 | 0.04150444 | 0.34488185 |
| Sfrp2 | 0.26118319 | 0.67889312 | 0.27138267 | 0.40751045 | 1.32E-12 | 8.72E-11 |

| | | | | | | |
|---|---|---|---|---|---|---|
| Htra3 | 0.39238009 | 0.80336976 | 0.39638979 | 0.40697997 | 8.62E-07 | 2.72E-05 |
| Atp2a2 | 1.25736537 | 1.36099242 | 0.95412778 | 0.40686463 | 0.02347825 | 0.22557166 |
| Arg1 | 0.30047194 | 0.71615003 | 0.3097347 | 0.40641534 | 4.88E-10 | 2.39E-08 |
| Tpd52 | 1.46550364 | 1.43093651 | 1.02584846 | 0.40508805 | 0.03290414 | 0.29017292 |
| Hist1h1c | 0.76441604 | 1.09991008 | 0.69496102 | 0.40494906 | 0.00331646 | 0.04672692 |
| 6330403K07I | 0.49587392 | 0.89158841 | 0.48833482 | 0.40325359 | 5.97E-05 | 0.00133639 |
| Nid1 | 0.49175487 | 0.88600068 | 0.48479729 | 0.40120339 | 5.77E-05 | 0.00129548 |
| Bambi | 1.18756465 | 1.32545386 | 0.92510627 | 0.40034759 | 0.02188972 | 0.21501166 |
| Acly | 1.58160165 | 1.45648244 | 1.05648815 | 0.39999429 | 0.03887358 | 0.33016351 |
| Oaf | 0.74430375 | 1.08020879 | 0.68120772 | 0.39900107 | 0.00317765 | 0.04510328 |
| Tm4sf1 | 0.25037956 | 0.65948493 | 0.26070808 | 0.39877685 | 5.16E-13 | 3.51E-11 |
| Spats2l | 0.48317576 | 0.87536572 | 0.47739638 | 0.39796934 | 5.14E-05 | 0.00116293 |
| Hist1h2ap | 0.26217863 | 0.66954074 | 0.27236357 | 0.39717718 | 5.46E-12 | 3.31E-10 |
| Zfos1 | 1.33364781 | 1.37792904 | 0.98292143 | 0.39500761 | 0.03058302 | 0.27268917 |
| Tppp3 | 0.40607937 | 0.80362942 | 0.40891668 | 0.39471273 | 3.44E-06 | 9.81E-05 |
| Hsp90aa1 | 3.51661977 | 1.42515555 | 1.03235413 | 0.39280142 | 0.03814046 | 0.32603795 |
| Ccnd1 | 0.50488916 | 0.88786802 | 0.49604175 | 0.39182627 | 0.00011656 | 0.00238684 |
| Myl12a | 2.49048463 | 1.51847891 | 1.12719197 | 0.39128694 | 0.05290914 | 0.40810662 |
| Pcp4 | 0.50778929 | 0.88952168 | 0.49851014 | 0.39101155 | 0.00012895 | 0.00261467 |
| Ffar2 | 0.85923284 | 1.14589259 | 0.75586231 | 0.39003027 | 0.00810736 | 0.09836021 |
| Gypa | 0.06217078 | 0.45595224 | 0.0665096 | 0.38944264 | 3.65E-164 | 8.82E-161 |
| Cldn3 | 0.65306521 | 1.00269265 | 0.6152219 | 0.38747075 | 0.00167187 | 0.02573124 |
| Fcer1g | 0.15858976 | 0.55489658 | 0.16792438 | 0.3869722 | 3.44E-27 | 6.01E-25 |
| Ltb | 0.10524061 | 0.49905736 | 0.11242007 | 0.38663729 | 4.41E-58 | 1.73E-55 |
| C1qa | 0.35117222 | 0.74456288 | 0.35808826 | 0.38647462 | 2.47E-07 | 8.40E-06 |
| Rnase1 | 0.18395594 | 0.57915013 | 0.1939238 | 0.38522633 | 1.07E-20 | 1.38E-18 |
| Mef2c | 0.39768906 | 0.78538522 | 0.40125709 | 0.38412812 | 4.10E-06 | 0.00011533 |
| Marcks | 1.72250331 | 1.46952106 | 1.08552535 | 0.38399571 | 0.04966994 | 0.39157955 |
| Ppib | 2.83531614 | 1.48993906 | 1.1079093 | 0.38202976 | 0.05408225 | 0.41528696 |
| Rps4x | 4.89160875 | 1.11308109 | 0.73334621 | 0.37973488 | 0.00792281 | 0.09660636 |

| | | | | | | |
|---|---|---|---|---|---|---|
| Cela3b | 0.24064356 | 0.63060482 | 0.25104202 | 0.3795628 | 9.33E-13 | 6.27E-11 |
| Hn1 | 1.68598288 | 1.4583049 | 1.07881916 | 0.37948574 | 0.05062428 | 0.39690185 |
| Lmo4 | 0.66475681 | 1.00192229 | 0.62400246 | 0.37791983 | 0.00238899 | 0.03523452 |
| Cd52 | 0.12300412 | 0.50893011 | 0.1310206 | 0.37790951 | 1.81E-41 | 5.35E-39 |
| Rps12 | 3.70863213 | 1.38765054 | 1.00979261 | 0.37785793 | 0.0406366 | 0.34035205 |
| Prrx1 | 0.46653253 | 0.84016104 | 0.46290971 | 0.37725133 | 7.33E-05 | 0.00158328 |
| Resp18 | 0.4652872 | 0.83888087 | 0.46181925 | 0.37706162 | 7.12E-05 | 0.00155041 |
| Marcksl1 | 2.7691087 | 1.4879374 | 1.11263 | 0.3753074 | 0.05802941 | 0.4361381 |
| Id1 | 0.29884949 | 0.68228767 | 0.30816586 | 0.37412181 | 7.74E-09 | 3.20E-07 |
| Cdk1 | 0.37105177 | 0.74844948 | 0.37668083 | 0.37176865 | 2.13E-06 | 6.33E-05 |
| Myl9 | 0.43393127 | 0.80562628 | 0.43405635 | 0.37156993 | 3.33E-05 | 0.00078459 |
| Nid2 | 0.48474709 | 0.85009288 | 0.47875534 | 0.37133754 | 0.00015098 | 0.00302745 |
| Rph3al | 1.08911515 | 1.25066234 | 0.87987747 | 0.37078487 | 0.02480557 | 0.23486923 |
| H2-D1 | 1.55036547 | 1.41963244 | 1.04887159 | 0.37076084 | 0.0497926 | 0.39214348 |
| H2afv | 1.92084441 | 1.48158104 | 1.11164158 | 0.36993946 | 0.06051877 | 0.44948829 |
| Glul | 1.49666985 | 1.40330924 | 1.03465331 | 0.36865593 | 0.04845511 | 0.38472188 |
| Blvrb | 0.45559581 | 0.81919012 | 0.45330082 | 0.3658893 | 8.48E-05 | 0.00181007 |
| Mkrn1 | 0.91577306 | 1.15438209 | 0.78907114 | 0.36531095 | 0.015506 | 0.16603106 |
| Lsp1 | 0.23972451 | 0.61502337 | 0.2501273 | 0.36489607 | 5.35E-12 | 3.26E-10 |
| Il11ra1 | 0.8566863 | 1.11682446 | 0.75431222 | 0.36251224 | 0.01257718 | 0.14091491 |
| Rps20 | 4.2968994 | 1.25204663 | 0.89038395 | 0.36166268 | 0.02921866 | 0.26426212 |
| Grb10 | 1.95774797 | 1.47594377 | 1.1147083 | 0.36123547 | 0.06554858 | 0.47418188 |
| Lmnb1 | 0.82109284 | 1.09287961 | 0.73215013 | 0.36072948 | 0.01085354 | 0.12527997 |
| Col6a3 | 0.43754019 | 0.79780157 | 0.43728089 | 0.36052068 | 6.12E-05 | 0.00135549 |
| Lrrc17 | 0.38324936 | 0.74742477 | 0.38798289 | 0.35944188 | 7.94E-06 | 0.00021268 |
| Cd9 | 1.05579116 | 1.22261656 | 0.86320511 | 0.35941145 | 0.02619978 | 0.24404059 |
| Aplnr | 0.12786866 | 0.49535978 | 0.13609389 | 0.35926589 | 4.59E-35 | 1.17E-32 |
| Col4a2 | 0.44353331 | 0.80176256 | 0.44261916 | 0.35914341 | 7.83E-05 | 0.00168477 |
| Pf4 | 0.08235475 | 0.44665651 | 0.08807561 | 0.3585809 | 1.57E-80 | 9.91E-78 |
| Htra1 | 0.37730958 | 0.74081032 | 0.38248953 | 0.35832079 | 6.37E-06 | 0.0001739 |

| | | | | | | |
|---|---|---|---|---|---|---|
| Lpl | 0.50345091 | 0.85308574 | 0.49481564 | 0.3582701 | 0.00037137 | 0.00686752 |
| Sox18 | 0.16682292 | 0.53245238 | 0.17639186 | 0.35606052 | 2.62E-21 | 3.48E-19 |
| Esam | 0.19003122 | 0.55559886 | 0.20011159 | 0.35548727 | 6.35E-17 | 6.22E-15 |
| Runx1t1 | 0.90989251 | 1.14080173 | 0.78572535 | 0.35507638 | 0.01762749 | 0.18359434 |
| Rpl23 | 4.85771175 | 1.09829141 | 0.74321605 | 0.35507535 | 0.01301267 | 0.14467612 |
| Hepacam2 | 0.99409276 | 1.18601689 | 0.83145289 | 0.354564 | 0.0234748 | 0.22557166 |
| Ly6e | 1.08767037 | 1.23361345 | 0.87916892 | 0.35444453 | 0.03017059 | 0.27028518 |
| Plac8 | 0.36467909 | 0.72515634 | 0.37074329 | 0.35441305 | 4.22E-06 | 0.00011856 |
| Cldn6 | 0.81968042 | 1.08554124 | 0.73125163 | 0.35428961 | 0.01199637 | 0.13582249 |
| Osr1 | 0.40087626 | 0.75700975 | 0.40417145 | 0.3528383 | 2.38E-05 | 0.00058063 |
| Fbn1 | 0.44420466 | 0.79544716 | 0.44321594 | 0.35223122 | 0.00010673 | 0.0022073 |
| Calu | 0.77906582 | 1.05525152 | 0.70479621 | 0.35045532 | 0.010262 | 0.11998263 |
| Actb | 4.60312256 | 1.16249119 | 0.81323425 | 0.34925693 | 0.02270376 | 0.22091219 |
| Clec14a | 0.19579724 | 0.55353831 | 0.20596987 | 0.34756844 | 1.89E-15 | 1.60E-13 |
| Cmtm3 | 0.60922814 | 0.92846334 | 0.58147253 | 0.34699081 | 0.00271627 | 0.03949903 |
| Fam159b | 0.43347895 | 0.78002451 | 0.43365167 | 0.34637284 | 9.91E-05 | 0.00206462 |
| Hsp90b1 | 3.35136941 | 1.40010583 | 1.05529493 | 0.3448109 | 0.06396955 | 0.46698415 |
| Dpysl3 | 0.43530368 | 0.77962466 | 0.4352835 | 0.34434116 | 0.0001141 | 0.00233974 |
| Ecscr | 0.22260632 | 0.5773076 | 0.23302055 | 0.34428705 | 2.92E-12 | 1.82E-10 |
| Oat | 0.91568979 | 1.13315029 | 0.78902394 | 0.34412635 | 0.0210789 | 0.20845964 |
| Selm | 1.80845639 | 1.4416985 | 1.09893157 | 0.34276692 | 0.07308872 | 0.51316074 |
| Slc2a2 | 0.41300644 | 0.75771016 | 0.41521 | 0.34250016 | 6.07E-05 | 0.00134874 |
| Tagln2 | 1.51564576 | 1.38218063 | 1.0398146 | 0.34236603 | 0.06251629 | 0.45938225 |
| Igfbp2 | 0.30069874 | 0.6511176 | 0.3099539 | 0.34116371 | 1.46E-07 | 5.10E-06 |
| Ngfrap1 | 2.81741748 | 1.45017889 | 1.10921974 | 0.34095915 | 0.07605825 | 0.52409337 |
| Vcan | 0.41205306 | 0.75396379 | 0.41434549 | 0.3396183 | 6.71E-05 | 0.0014646 |
| Nkain4 | 0.3745935 | 0.71856489 | 0.37997105 | 0.33859385 | 1.65E-05 | 0.00041622 |
| Ddah2 | 1.22032076 | 1.27503175 | 0.93907257 | 0.33595918 | 0.04778049 | 0.38082549 |
| Prdx4 | 1.15668767 | 1.24459332 | 0.91145436 | 0.33313896 | 0.04429758 | 0.36120719 |
| Sep-11 | 0.65270747 | 0.94792333 | 0.6149518 | 0.33297153 | 0.00582407 | 0.07413294 |

| | | | | | | |
|---|---|---|---|---|---|---|
| Cenpf | 0.23208486 | 0.57499602 | 0.24250859 | 0.33248744 | 8.43E-11 | 4.51E-09 |
| Ypel3 | 1.76651457 | 1.42504756 | 1.09280689 | 0.33224067 | 0.07832411 | 0.53437317 |
| Maf | 0.37597684 | 0.71205849 | 0.38125425 | 0.33080424 | 2.64E-05 | 0.00063909 |
| Dab2 | 0.33866841 | 0.67672466 | 0.34628585 | 0.33043881 | 4.38E-06 | 0.00012282 |
| Cd74 | 0.09338851 | 0.4289577 | 0.09985519 | 0.32910251 | 1.64E-53 | 6.10E-51 |
| Plxdc2 | 0.43836903 | 0.76708835 | 0.43802036 | 0.32906799 | 0.00023256 | 0.0045135 |
| Enpp2 | 0.74084174 | 1.007252 | 0.67881103 | 0.32844097 | 0.01209127 | 0.13652587 |
| Fabp4 | 0.18900447 | 0.52716239 | 0.19906693 | 0.32809546 | 8.03E-15 | 6.44E-13 |
| Rgcc | 0.20987953 | 0.54727333 | 0.2202178 | 0.32705552 | 2.27E-12 | 1.44E-10 |
| Serpine2 | 0.7945691 | 1.04177524 | 0.7150358 | 0.32673944 | 0.0166301 | 0.17624506 |
| Osr2 | 0.32454471 | 0.65906144 | 0.33285757 | 0.32620387 | 2.49E-06 | 7.32E-05 |
| Ank | 0.91602879 | 1.11415911 | 0.78921606 | 0.32494305 | 0.02754152 | 0.25315361 |
| Des | 0.3632517 | 0.6941637 | 0.36941031 | 0.32475339 | 2.10E-05 | 0.00052058 |
| Adamts5 | 0.30855263 | 0.64161374 | 0.3175285 | 0.32408524 | 9.92E-07 | 3.07E-05 |
| Ndufa4l2 | 0.32175899 | 0.65406101 | 0.33019679 | 0.32386422 | 2.44E-06 | 7.20E-05 |
| Rac2 | 0.12578482 | 0.45678401 | 0.13392165 | 0.32286236 | 1.42E-29 | 2.78E-27 |
| Kdelr3 | 0.45338666 | 0.77328298 | 0.4513511 | 0.32193188 | 0.0004452 | 0.00812905 |
| Cldn5 | 0.16038571 | 0.49146227 | 0.16977379 | 0.32168848 | 5.34E-19 | 6.09E-17 |
| Cpa2 | 0.23618388 | 0.56555416 | 0.2465997 | 0.31895446 | 8.41E-10 | 4.01E-08 |
| Cdh5 | 0.17235271 | 0.49975848 | 0.18206348 | 0.317695 | 2.15E-16 | 2.03E-14 |
| Pttg1 | 0.44585011 | 0.7610929 | 0.44467751 | 0.31641539 | 0.00045801 | 0.00835166 |
| Cnn2 | 0.47498763 | 0.78631616 | 0.47029029 | 0.31602587 | 0.0008719 | 0.01471572 |
| Tshz2 | 1.50046879 | 1.3515114 | 1.03569417 | 0.31581723 | 0.07770716 | 0.53174815 |
| Bsg | 3.74957097 | 1.32062586 | 1.00501545 | 0.31561042 | 0.07172378 | 0.50699727 |
| Smim22 | 1.1156083 | 1.20808446 | 0.89261425 | 0.31547021 | 0.04982262 | 0.39214348 |
| Tagln | 0.14673827 | 0.47078485 | 0.15568842 | 0.31509643 | 2.07E-21 | 2.77E-19 |
| Sez6l2 | 0.75874898 | 1.00573231 | 0.69111471 | 0.3146176 | 0.01696513 | 0.1790105 |
| Lox | 0.36176365 | 0.68245414 | 0.36801967 | 0.31443447 | 3.44E-05 | 0.0008077 |
| Pdia4 | 1.76082681 | 1.40454892 | 1.09191971 | 0.31262921 | 0.0911169 | 0.58666938 |
| Kdelr2 | 1.40686365 | 1.31971424 | 1.00783094 | 0.3118833 | 0.07468566 | 0.51900827 |

| | | | | | | |
|---|---|---|---|---|---|---|
| Aebp1 | 0.40862357 | 0.72234113 | 0.41123133 | 0.3111098 | 0.00021203 | 0.00415406 |
| Eno1 | 2.11383238 | 1.43513349 | 1.12406247 | 0.31107102 | 0.0986457 | 0.61645056 |
| Btbd17 | 0.35877662 | 0.67567728 | 0.36522495 | 0.31045233 | 3.74E-05 | 0.00087401 |
| Fkbp11 | 0.49357134 | 0.79679545 | 0.48635841 | 0.31043704 | 0.00147093 | 0.02312962 |
| Car2 | 0.13390977 | 0.4526785 | 0.14238208 | 0.31029642 | 1.60E-24 | 2.50E-22 |
| Bex4 | 0.87592236 | 1.07397685 | 0.76590576 | 0.30807109 | 0.03046722 | 0.27199122 |
| Arx | 0.31068483 | 0.62757986 | 0.3195796 | 0.30800026 | 3.56E-06 | 0.00010117 |
| Cck | 0.29173182 | 0.60884113 | 0.30126777 | 0.30757336 | 9.85E-07 | 3.06E-05 |
| Mien1 | 1.72432239 | 1.39245108 | 1.08584482 | 0.30660626 | 0.09416606 | 0.59930619 |
| Bpgm | 0.41121544 | 0.72006677 | 0.41358552 | 0.30648126 | 0.00027777 | 0.00531989 |
| Sh3bgrl3 | 1.63116329 | 1.37410182 | 1.06770437 | 0.30639746 | 0.09061636 | 0.58467113 |
| Nr2f2 | 0.44505727 | 0.74934476 | 0.44397348 | 0.30537128 | 0.00067652 | 0.01183133 |
| 1110008P14I | 1.12641575 | 1.20273275 | 0.89767047 | 0.30506227 | 0.05667839 | 0.4290983 |
| Cxcl13 | 0.17210262 | 0.48381772 | 0.18180725 | 0.30201047 | 5.00E-15 | 4.07E-13 |
| Mageh1 | 1.59443409 | 1.361353 | 1.0594967 | 0.3018563 | 0.09219525 | 0.59196045 |
| Camk2n1 | 0.89938479 | 1.08148247 | 0.77968347 | 0.301799 | 0.03566446 | 0.30938746 |
| Fosb | 0.77178114 | 1.00157043 | 0.69992506 | 0.30164536 | 0.02232935 | 0.21839217 |
| Nupr1 | 0.29947502 | 0.61018332 | 0.30877088 | 0.30141244 | 2.71E-06 | 7.83E-05 |
| Fech | 0.40361472 | 0.70734372 | 0.40667085 | 0.30067286 | 0.00028596 | 0.00543353 |
| Hist1h2al | 0.81496016 | 1.02855615 | 0.72823828 | 0.30031786 | 0.02734703 | 0.25189151 |
| Cyr61 | 0.31078467 | 0.61953474 | 0.3196756 | 0.29985913 | 6.21E-06 | 0.00017015 |
| Cks1b | 1.21844614 | 1.23640692 | 0.93829461 | 0.29811231 | 0.06940652 | 0.49496101 |
| Prss53 | 0.3533576 | 0.65716246 | 0.36014244 | 0.29702002 | 6.09E-05 | 0.00135064 |
| Gm13305 | 0.53954218 | 0.82192566 | 0.52519091 | 0.29673475 | 0.00424022 | 0.05723896 |
| Cela1 | 0.17392902 | 0.48033194 | 0.18367791 | 0.29665404 | 2.65E-14 | 2.05E-12 |
| Fcgrt | 0.45918676 | 0.75229468 | 0.45646379 | 0.29583089 | 0.00126627 | 0.02028551 |
| Rcn1 | 1.25515869 | 1.24827191 | 0.95324526 | 0.29502664 | 0.07466123 | 0.51900827 |
| Rgs2 | 0.33899229 | 0.64047868 | 0.34659261 | 0.29388607 | 3.90E-05 | 0.00090471 |
| Rpsa | 4.14234009 | 1.22366694 | 0.93024921 | 0.29341774 | 0.07084646 | 0.50201954 |
| Gabarapl2 | 2.54722261 | 1.41779468 | 1.12503757 | 0.29275711 | 0.11268941 | 0.66875608 |

| | | | | | | |
|---|---|---|---|---|---|---|
| Fth1 | 3.91636007 | 1.2724727 | 0.98012294 | 0.29234976 | 0.08231565 | 0.5519946 |
| Cotl1 | 1.25697309 | 1.24452169 | 0.95397112 | 0.29055057 | 0.07795363 | 0.53259744 |
| Gpx7 | 0.51691885 | 0.79593453 | 0.50624583 | 0.28968871 | 0.00383818 | 0.05269503 |
| Pdlim2 | 0.36059958 | 0.65649765 | 0.36693112 | 0.28956653 | 0.00011816 | 0.00241627 |
| Adamts1 | 0.41594365 | 0.70727564 | 0.41787037 | 0.28940527 | 0.00062624 | 0.01106931 |
| Slc35b1 | 1.11234801 | 1.18015628 | 0.89107364 | 0.28908264 | 0.0653361 | 0.47362135 |
| Emp1 | 0.50103133 | 0.78122417 | 0.49275002 | 0.28847415 | 0.00319065 | 0.045218 |
| Ctsz | 1.37167214 | 1.28419717 | 0.99611267 | 0.2880845 | 0.08892466 | 0.57774473 |
| Zfp36l1 | 0.61947095 | 0.87505556 | 0.58947474 | 0.28558082 | 0.01200087 | 0.13582249 |
| Angptl2 | 0.28422544 | 0.57865537 | 0.2939656 | 0.28468977 | 3.21E-06 | 9.20E-05 |
| H1f0 | 1.90951053 | 1.39506461 | 1.1106041 | 0.28446051 | 0.11637579 | 0.68087821 |
| Ube2l6 | 0.16227465 | 0.4561198 | 0.17171755 | 0.28440225 | 5.99E-15 | 4.83E-13 |
| C1qb | 0.07077515 | 0.35903875 | 0.07570562 | 0.28333314 | 2.18E-68 | 1.06E-65 |
| Lurap1l | 0.507159 | 0.78118085 | 0.49797413 | 0.28320672 | 0.00402909 | 0.05505533 |
| Actg1 | 5.09200404 | 0.96569861 | 0.68416078 | 0.28153783 | 0.02760672 | 0.25331782 |
| Arl6ip1 | 1.44287341 | 1.30008104 | 1.01911663 | 0.28096441 | 0.0994936 | 0.62041211 |
| Bst2 | 0.32360546 | 0.61289495 | 0.3319609 | 0.28093405 | 4.03E-05 | 0.00092815 |
| Spint1 | 0.97058937 | 1.09972391 | 0.81913302 | 0.28059089 | 0.05525528 | 0.42140509 |
| Gpx2 | 0.51697842 | 0.78682053 | 0.50629612 | 0.28052441 | 0.0049202 | 0.06520384 |
| Coro1a | 0.12146473 | 0.40912985 | 0.12941334 | 0.27971651 | 3.75E-24 | 5.67E-22 |
| Nbl1 | 0.42913545 | 0.70924784 | 0.42975943 | 0.27948841 | 0.00122325 | 0.0197271 |
| Rab3a | 0.98717834 | 1.10733381 | 0.82787014 | 0.27946367 | 0.05789227 | 0.43561417 |
| Nfkbia | 0.80300945 | 0.99897849 | 0.72053727 | 0.27844122 | 0.03590029 | 0.30996718 |
| Vcam1 | 0.29831319 | 0.58606622 | 0.307647 | 0.27841923 | 1.24E-05 | 0.00032202 |
| Fam174b | 0.9315376 | 1.07589282 | 0.79792754 | 0.27796528 | 0.05229637 | 0.40455457 |
| Aspn | 0.22667248 | 0.51371215 | 0.2370957 | 0.27661646 | 2.73E-08 | 1.05E-06 |
| S100a13 | 0.91313015 | 1.0633483 | 0.78757058 | 0.27577772 | 0.05140438 | 0.40024739 |
| Calm1 | 4.0039474 | 1.23788534 | 0.96235216 | 0.27553318 | 0.09111638 | 0.58666938 |
| BC028528 | 0.39787791 | 0.67691195 | 0.40142994 | 0.27548201 | 0.00069383 | 0.01209028 |
| Ogn | 0.29907755 | 0.58253743 | 0.30838646 | 0.27415097 | 1.72E-05 | 0.0004323 |

| | | | | | | |
|---|---|---|---|---|---|---|
| Tsc22d1 | 2.24459168 | 1.40244135 | 1.12830109 | 0.27414026 | 0.12882346 | 0.71857996 |
| Alcam | 0.86629359 | 1.03422269 | 0.76013565 | 0.27408705 | 0.04648793 | 0.37319049 |
| Mmp2 | 0.35207783 | 0.63253076 | 0.35893981 | 0.27359096 | 0.00019179 | 0.00378301 |
| Bicc1 | 0.63791529 | 0.87677568 | 0.60370612 | 0.27306955 | 0.01754399 | 0.1828909 |
| Hebp1 | 0.49771164 | 0.76232767 | 0.48990998 | 0.27241769 | 0.00478995 | 0.06359404 |
| Reep5 | 1.44553902 | 1.29204732 | 1.01992646 | 0.27212086 | 0.10692913 | 0.6470194 |
| Icam2 | 0.17246634 | 0.45392004 | 0.18217989 | 0.27174015 | 1.84E-12 | 1.18E-10 |
| Gsta3 | 0.22329433 | 0.50535909 | 0.23371061 | 0.27164848 | 3.06E-08 | 1.17E-06 |
| Ntrk2 | 0.3417115 | 0.62075606 | 0.34916593 | 0.27159014 | 0.00014513 | 0.00292233 |
| Tmem176b | 1.93145859 | 1.38340391 | 1.11256893 | 0.27083498 | 0.12836879 | 0.71695178 |
| Dek | 1.00211043 | 1.10580644 | 0.8356073 | 0.27019914 | 0.06596951 | 0.4755972 |
| Msn | 0.39392488 | 0.66775959 | 0.39780772 | 0.26995187 | 0.00078465 | 0.01338341 |
| Uqcc2 | 2.31102594 | 1.39894651 | 1.12922484 | 0.26972167 | 0.13289412 | 0.72977796 |
| Igf2r | 0.58590812 | 0.83203346 | 0.56299267 | 0.26904079 | 0.0129938 | 0.14467612 |
| Ptrf | 0.40157349 | 0.67340164 | 0.40480822 | 0.26859341 | 0.0009966 | 0.01643766 |
| Cdc20 | 0.25034455 | 0.52898185 | 0.2606734 | 0.26830845 | 8.12E-07 | 2.58E-05 |
| Rps15a | 4.98591288 | 0.97342324 | 0.70526083 | 0.26816241 | 0.03824076 | 0.32651032 |
| Plpp3 | 0.39760152 | 0.66856642 | 0.40117697 | 0.26738945 | 0.00095062 | 0.01578698 |
| Atp5l | 4.30261563 | 1.1557628 | 0.88887726 | 0.26688554 | 0.08092931 | 0.54572704 |
| Ccndbp1 | 0.53097689 | 0.78480076 | 0.5180565 | 0.26674426 | 0.00822307 | 0.0994312 |
| Axl | 0.33794022 | 0.61196926 | 0.34559594 | 0.26637332 | 0.00016474 | 0.00328524 |
| Tcf4 | 1.00540356 | 1.10250288 | 0.83731475 | 0.26518814 | 0.07003298 | 0.49795891 |
| Creld2 | 1.16365635 | 1.17935368 | 0.91458083 | 0.26477284 | 0.08870509 | 0.57718099 |
| Gstm1 | 0.4263427 | 0.69129338 | 0.42725096 | 0.26404241 | 0.00199317 | 0.0300385 |
| Lhfp | 0.3203947 | 0.5927721 | 0.32889221 | 0.26387988 | 9.27E-05 | 0.00194868 |
| Igfbpl1 | 0.32349121 | 0.59493048 | 0.33185179 | 0.26307868 | 0.00011063 | 0.00228151 |
| Ece1 | 0.96576038 | 1.07918435 | 0.81656861 | 0.26261573 | 0.06702027 | 0.48149645 |
| Timp2 | 0.57707054 | 0.81809436 | 0.55589494 | 0.26219942 | 0.01399286 | 0.15310824 |
| Phlda3 | 0.44897174 | 0.70958759 | 0.44744583 | 0.26214176 | 0.00317153 | 0.04507926 |
| Basp1 | 0.52692385 | 0.77649999 | 0.51466408 | 0.26183591 | 0.00888824 | 0.10658534 |

| | | | | | | |
|---|---|---|---|---|---|---|
| Gap43 | 0.28731553 | 0.55867164 | 0.29697501 | 0.26169664 | 2.02E-05 | 0.00049993 |
| Scd2 | 1.0248007 | 1.10807217 | 0.84721541 | 0.26085676 | 0.07571721 | 0.52298621 |
| Tmsb15l | 0.5575873 | 0.80045512 | 0.54006848 | 0.26038664 | 0.01234416 | 0.13867887 |
| Pfn1 | 2.82287953 | 1.369052 | 1.1088198 | 0.2602322 | 0.13710457 | 0.7420427 |
| Wipi1 | 0.92970169 | 1.05692404 | 0.79690538 | 0.26001866 | 0.06423728 | 0.46799603 |
| Mfge8 | 0.59516276 | 0.82957095 | 0.57036939 | 0.25920156 | 0.01712004 | 0.18038254 |
| Cdo1 | 0.45950989 | 0.71523278 | 0.45674803 | 0.25848475 | 0.00418698 | 0.05667869 |
| Nefm | 0.33496805 | 0.60085421 | 0.34277741 | 0.2580768 | 0.00022599 | 0.00439193 |
| Prnp | 1.48888215 | 1.28967171 | 1.03249503 | 0.25717668 | 0.12293042 | 0.70139524 |
| Akr1c12 | 0.72114943 | 0.92200692 | 0.66501657 | 0.25699035 | 0.03589791 | 0.30996718 |
| Dtymk | 0.98852471 | 1.08506647 | 0.82857049 | 0.25649598 | 0.07461697 | 0.51900827 |
| Galk1 | 0.51321677 | 0.75936339 | 0.50311534 | 0.25624804 | 0.00882442 | 0.10590762 |
| Arhgdib | 0.18784039 | 0.4537815 | 0.19788199 | 0.25589951 | 8.46E-10 | 4.02E-08 |
| Stmn2 | 0.13488321 | 0.39915729 | 0.14339401 | 0.25576327 | 4.79E-17 | 4.82E-15 |
| Nkx2-2 | 0.9406859 | 1.05869907 | 0.802984 | 0.25571507 | 0.06894854 | 0.49315042 |
| 1700011H14 | 0.20168732 | 0.46682912 | 0.2119394 | 0.25488971 | 1.05E-08 | 4.30E-07 |
| Npepl1 | 0.96415116 | 1.07018731 | 0.81571026 | 0.25447704 | 0.07304951 | 0.51316074 |
| Fxyd6 | 1.19069896 | 1.18078494 | 0.92647367 | 0.25431127 | 0.10047487 | 0.62288099 |
| Lyve1 | 0.18336767 | 0.44755667 | 0.19332381 | 0.25423286 | 4.48E-10 | 2.21E-08 |
| Lyz2 | 0.13260975 | 0.39483099 | 0.14103007 | 0.25380092 | 2.55E-17 | 2.66E-15 |
| Fxyd1 | 0.41094907 | 0.66693488 | 0.41334375 | 0.25359113 | 0.00212949 | 0.03179548 |
| Serpinf1 | 0.39416041 | 0.65145647 | 0.39802378 | 0.25343269 | 0.00150617 | 0.02354648 |
| Emcn | 0.13208925 | 0.39373897 | 0.14048857 | 0.2532504 | 2.26E-17 | 2.38E-15 |
| Ifi27 | 0.40929747 | 0.66441657 | 0.4118438 | 0.25257276 | 0.00213696 | 0.03184132 |
| B630019K06 | 0.72844048 | 0.92250765 | 0.67015584 | 0.25235182 | 0.03968472 | 0.33567622 |
| Fkbp2 | 1.58303552 | 1.3091021 | 1.05682759 | 0.25227451 | 0.13304064 | 0.73006176 |
| Sep-07 | 2.06172863 | 1.37355438 | 1.12141285 | 0.25214153 | 0.14742781 | 0.77404405 |
| Serf2 | 3.4261068 | 1.29657773 | 1.0445198 | 0.25205793 | 0.13044909 | 0.72323168 |
| Slit2 | 0.36753786 | 0.62422865 | 0.37340962 | 0.25081904 | 0.00087579 | 0.01476425 |
| G0s2 | 0.27813783 | 0.53870841 | 0.2880244 | 0.25068401 | 2.51E-05 | 0.00060992 |

| | | | | | | |
|---|---|---|---|---|---|---|
| P4ha2 | 0.43060597 | 0.6786028 | 0.43107842 | 0.24752439 | 0.00373474 | 0.05172638 |
| Fev | 0.59934885 | 0.82108304 | 0.57368716 | 0.24739588 | 0.02226253 | 0.21793524 |
| Rhoj | 0.39939589 | 0.65017752 | 0.40281854 | 0.24735898 | 0.00211208 | 0.03163318 |
| Emilin1 | 0.48578795 | 0.72667767 | 0.47965467 | 0.247023 | 0.00821181 | 0.09937802 |
| Cst3 | 2.94152862 | 1.34668779 | 1.09981924 | 0.24686855 | 0.14783611 | 0.77404405 |
| Pdzk1ip1 | 0.57559866 | 0.80104906 | 0.55470777 | 0.24634128 | 0.01927311 | 0.19636093 |
| Lockd | 0.22008028 | 0.4768233 | 0.23048518 | 0.24633813 | 3.19E-07 | 1.07E-05 |
| Tgfb1i1 | 0.37559273 | 0.62687251 | 0.38089806 | 0.24597445 | 0.00131215 | 0.02095108 |
| Crabp1 | 0.16701207 | 0.4220636 | 0.17658607 | 0.24547754 | 4.69E-11 | 2.55E-09 |
| BC048546 | 0.49637893 | 0.73410691 | 0.48876788 | 0.24533903 | 0.00967873 | 0.11417597 |
| Dnajb9 | 0.82346433 | 0.97845087 | 0.73365544 | 0.24479543 | 0.06002933 | 0.44651732 |
| Sgk1 | 0.26729906 | 0.52196396 | 0.27740167 | 0.24456229 | 2.00E-05 | 0.00049666 |
| Srm | 0.85858184 | 0.99930899 | 0.7554665 | 0.24384249 | 0.06632031 | 0.47741402 |
| Gjd2 | 0.61655496 | 0.83033416 | 0.58720388 | 0.24313028 | 0.02686402 | 0.24854787 |
| Celf2 | 0.94332307 | 1.04745062 | 0.8044302 | 0.24302042 | 0.07964284 | 0.54059058 |
| Ctsb | 1.69698096 | 1.32308 | 1.08088713 | 0.24219288 | 0.14826063 | 0.77505178 |
| Surf4 | 1.4417324 | 1.25988695 | 1.01876877 | 0.24111819 | 0.13508745 | 0.73635034 |
| Cirbp | 1.56169476 | 1.29231099 | 1.05167018 | 0.24064081 | 0.14320086 | 0.76468733 |
| Ccna2 | 0.19939864 | 0.44918812 | 0.20962163 | 0.23956649 | 5.06E-08 | 1.87E-06 |
| Tyms | 0.61879223 | 0.82796237 | 0.58894669 | 0.23901568 | 0.02932784 | 0.26491901 |
| Cyb5r3 | 1.90352437 | 1.34823463 | 1.11003625 | 0.23819838 | 0.15871718 | 0.80513702 |
| Ccnb1 | 0.15021831 | 0.39717234 | 0.15928696 | 0.23788539 | 1.73E-12 | 1.12E-10 |
| Lpar4 | 0.34217087 | 0.58724592 | 0.34960026 | 0.23764566 | 0.00077015 | 0.01318263 |
| C1qc | 0.05391244 | 0.29402144 | 0.05768044 | 0.236341 | 1.54E-81 | 1.06E-78 |
| Gamt | 0.31003002 | 0.55465015 | 0.31894983 | 0.23570032 | 0.00028775 | 0.00546049 |
| Epha3 | 0.29216068 | 0.53738278 | 0.30168412 | 0.23569867 | 0.00013636 | 0.00275727 |
| Efemp2 | 0.69268697 | 0.8801137 | 0.64459398 | 0.23551972 | 0.04435245 | 0.36145129 |
| Tfpi | 0.42396326 | 0.66040659 | 0.42511013 | 0.23529646 | 0.00495861 | 0.06553327 |
| Smc2 | 0.40728275 | 0.64526967 | 0.41001195 | 0.23525773 | 0.00375635 | 0.05186631 |
| Gars | 1.67849453 | 1.31242162 | 1.07737859 | 0.23504302 | 0.15471859 | 0.79345952 |

| | | | | | | |
|---|---|---|---|---|---|---|
| Elk3 | 0.28154593 | 0.52565118 | 0.29135273 | 0.23429845 | 8.96E-05 | 0.0018941 |
| Serinc3 | 1.10570084 | 1.1215467 | 0.88791063 | 0.23363607 | 0.11011666 | 0.65698409 |
| Rspo3 | 0.22705444 | 0.47043846 | 0.23747812 | 0.23296035 | 2.44E-06 | 7.19E-05 |
| Nucb2 | 0.92146947 | 1.02478501 | 0.79229155 | 0.23249347 | 0.08579073 | 0.56639069 |
| Gatsl2 | 0.82579971 | 0.96737884 | 0.73513381 | 0.23224503 | 0.070529 | 0.50026519 |
| Try5 | 0.07155591 | 0.30875577 | 0.0765399 | 0.23221587 | 1.16E-45 | 3.82E-43 |
| Cdc42ep2 | 0.37700939 | 0.6134713 | 0.38221138 | 0.23125992 | 0.00240922 | 0.03546072 |
| Sat1 | 0.54225567 | 0.75841333 | 0.52744119 | 0.23097213 | 0.0206652 | 0.20548976 |
| Nudt19 | 0.54074101 | 0.75709033 | 0.52618568 | 0.23090465 | 0.02045354 | 0.20404477 |
| Stard10 | 1.07297374 | 1.10271967 | 0.87188757 | 0.2308321 | 0.10870322 | 0.65261673 |
| Tmed3 | 1.94045358 | 1.34407112 | 1.11332975 | 0.23074138 | 0.16712684 | 0.82737245 |
| Cryab | 0.26821862 | 0.5078931 | 0.2783051 | 0.22958799 | 6.07E-05 | 0.00134874 |
| Cdh11 | 0.33561924 | 0.57293286 | 0.34339525 | 0.22953761 | 0.00092222 | 0.01540355 |
| Ssr4 | 2.47509076 | 1.35639477 | 1.12765943 | 0.22873534 | 0.17232347 | 0.83696674 |
| Rps2 | 4.83219835 | 0.97809269 | 0.75062515 | 0.22746754 | 0.07899947 | 0.53796819 |
| Rasip1 | 0.17823522 | 0.41543357 | 0.18808299 | 0.22735059 | 8.93E-09 | 3.67E-07 |
| Prss23 | 0.37723228 | 0.60957122 | 0.38241791 | 0.2271533 | 0.00282548 | 0.04076221 |
| Dynll2 | 1.45545904 | 1.24901927 | 1.02290524 | 0.22611404 | 0.15153722 | 0.78379829 |
| Cd47 | 1.40154543 | 1.2319547 | 1.00610689 | 0.22584781 | 0.14782094 | 0.77404405 |
| F2r | 0.44811955 | 0.67239848 | 0.44669067 | 0.22570781 | 0.00928353 | 0.11050302 |
| H2afx | 1.02917521 | 1.07495495 | 0.84950786 | 0.22544708 | 0.10815208 | 0.65088783 |
| Rap1b | 1.88055787 | 1.33262617 | 1.10766379 | 0.22496238 | 0.17201984 | 0.83696674 |
| Rpl35 | 4.50012854 | 1.0642111 | 0.83926003 | 0.22495107 | 0.10587656 | 0.64279663 |
| Erp29 | 2.16525237 | 1.3511023 | 1.12618679 | 0.22491551 | 0.17606731 | 0.84551968 |
| Tubb5 | 3.33943057 | 1.28148081 | 1.05693098 | 0.22454983 | 0.16113267 | 0.81133695 |
| Rpn1 | 1.68870574 | 1.30372496 | 1.07933642 | 0.22438854 | 0.16637985 | 0.82496531 |
| Hist1h2ae | 0.25360873 | 0.48741806 | 0.26390425 | 0.2235138 | 3.98E-05 | 0.00091771 |
| Ppp2r2b | 0.48518694 | 0.70099661 | 0.47913546 | 0.22186115 | 0.01549122 | 0.16599537 |
| Mpp1 | 0.24995658 | 0.48207604 | 0.26028906 | 0.22178698 | 3.60E-05 | 0.00084244 |
| Wfdc2 | 0.3204593 | 0.55025376 | 0.32895401 | 0.22129975 | 0.00086148 | 0.01457386 |

| | | | | | | |
|---|---|---|---|---|---|---|
| Cxxc4 | 0.79821998 | 0.93837472 | 0.71742185 | 0.22095287 | 0.07566221 | 0.52298621 |
| Pkm | 2.84589355 | 1.32803322 | 1.10714792 | 0.2208853 | 0.17631557 | 0.84615131 |
| Thra | 0.77857473 | 0.9252587 | 0.70446904 | 0.22078966 | 0.07212139 | 0.50853143 |
| Ctsh | 0.34224734 | 0.57024114 | 0.34967255 | 0.22056859 | 0.00164745 | 0.02543604 |
| Clec3b | 0.16191267 | 0.3918761 | 0.17134518 | 0.22053092 | 1.01E-09 | 4.74E-08 |
| Tmem119 | 0.26696674 | 0.49700281 | 0.27707507 | 0.21992774 | 0.00010864 | 0.00224372 |
| Trappc2l | 1.61893134 | 1.2846952 | 1.06503127 | 0.21966392 | 0.16829821 | 0.82993631 |
| Sepw1 | 2.49263161 | 1.34674388 | 1.12712137 | 0.21962251 | 0.18199017 | 0.86140825 |
| Papss2 | 0.7161273 | 0.88076612 | 0.66145451 | 0.21931161 | 0.06120739 | 0.45297841 |
| Ehd2 | 0.3570137 | 0.58274438 | 0.36357325 | 0.21917112 | 0.00248843 | 0.03644173 |
| Fbln2 | 0.38266983 | 0.60599386 | 0.38744778 | 0.21854608 | 0.00429506 | 0.05787153 |
| Kdr | 0.1593956 | 0.38702203 | 0.16875437 | 0.21826766 | 8.41E-10 | 4.01E-08 |
| Mmp23 | 0.31572274 | 0.54219616 | 0.32441745 | 0.21777871 | 0.00088147 | 0.01482555 |
| Loxl2 | 0.36337408 | 0.58716047 | 0.36952464 | 0.21763583 | 0.00303535 | 0.04344182 |
| Ffar4 | 0.85779371 | 0.97171941 | 0.7549869 | 0.21673251 | 0.09054075 | 0.58444336 |
| Cldn10 | 0.22590701 | 0.45296877 | 0.23632911 | 0.21663966 | 9.74E-06 | 0.00025861 |
| Arf4 | 2.26874435 | 1.34332235 | 1.12868987 | 0.21463248 | 0.18782288 | 0.87574983 |
| Rgs17 | 0.63215431 | 0.81132413 | 0.59928552 | 0.21203862 | 0.04963305 | 0.39150162 |
| Prss2 | 0.11885164 | 0.3380366 | 0.12668298 | 0.21135362 | 3.84E-15 | 3.18E-13 |
| Rsu1 | 0.88029162 | 0.97939463 | 0.76850156 | 0.21089308 | 0.10053448 | 0.62288099 |
| Coq10b | 0.45079742 | 0.6597092 | 0.44906218 | 0.21064702 | 0.01442882 | 0.15669589 |
| Btg1 | 1.6396085 | 1.28012523 | 1.06950698 | 0.21061825 | 0.17944393 | 0.85521962 |
| Taldo1 | 1.84151115 | 1.31382239 | 1.10321466 | 0.21060772 | 0.18689167 | 0.87419127 |
| Ctss | 0.07685309 | 0.29260488 | 0.08219932 | 0.21040556 | 4.36E-33 | 1.00E-30 |
| Akr1c13 | 0.66333474 | 0.83314587 | 0.62293953 | 0.21020634 | 0.05796075 | 0.43584798 |
| Eif4ebp1 | 0.69382332 | 0.85489932 | 0.64542025 | 0.20947907 | 0.06525211 | 0.47348601 |
| Fibin | 0.16136016 | 0.3801978 | 0.17077671 | 0.20942109 | 5.54E-09 | 2.33E-07 |
| Plk2 | 0.43634659 | 0.64555495 | 0.43621527 | 0.20933968 | 0.0126816 | 0.14197515 |
| Gja4 | 0.08685338 | 0.30134492 | 0.09287924 | 0.20846568 | 6.81E-26 | 1.12E-23 |
| Gm10073 | 1.27226362 | 1.16837049 | 0.96000131 | 0.20836918 | 0.15595377 | 0.79725592 |

| | | | | | | |
|---|---|---|---|---|---|---|
| Sec11c | 0.85364723 | 0.9607831 | 0.75245609 | 0.20832702 | 0.09854495 | 0.61608654 |
| Gimap4 | 0.13813757 | 0.35505733 | 0.14677443 | 0.2082829 | 1.90E-11 | 1.10E-09 |
| Cdk2ap2 | 1.03678217 | 1.06165199 | 0.85346314 | 0.20818884 | 0.1278789 | 0.71595682 |
| Smarca1 | 0.79032372 | 0.92018229 | 0.71224901 | 0.20793328 | 0.08689491 | 0.57045578 |
| Atp6v0b | 1.82703748 | 1.30917422 | 1.10140017 | 0.20777405 | 0.18973107 | 0.88079447 |
| Tubb2b | 0.49185166 | 0.69203541 | 0.48488052 | 0.20715489 | 0.0232716 | 0.22432955 |
| Bex1 | 0.89117528 | 0.98201127 | 0.77490647 | 0.20710481 | 0.10653491 | 0.64598207 |
| Abcg2 | 0.37305233 | 0.58514121 | 0.37854017 | 0.20660104 | 0.00549882 | 0.07092692 |
| Cldn11 | 0.49819152 | 0.69692114 | 0.49032095 | 0.20660019 | 0.02481857 | 0.23486923 |
| Cd7 | 0.06999026 | 0.28080355 | 0.0748669 | 0.20593666 | 6.73E-38 | 1.87E-35 |
| Col18a1 | 0.45288302 | 0.65666094 | 0.45090619 | 0.20575474 | 0.01675415 | 0.17730051 |
| Ddost | 2.44121767 | 1.33398964 | 1.12851285 | 0.20547679 | 0.19813871 | 0.89054584 |
| Maged1 | 2.90093273 | 1.30803414 | 1.10301071 | 0.20502343 | 0.19324885 | 0.88774457 |
| Gimap6 | 0.17339294 | 0.3880041 | 0.18312898 | 0.20487512 | 9.33E-08 | 3.37E-06 |
| Colec12 | 0.34366749 | 0.55578668 | 0.35101453 | 0.20477215 | 0.00328491 | 0.0463726 |
| Rps26 | 4.47037402 | 1.05070978 | 0.8461138 | 0.20459598 | 0.12996361 | 0.72219719 |
| Higd1a | 1.36691306 | 1.19897024 | 0.99450924 | 0.204461 | 0.16907399 | 0.8322688 |
| Smc4 | 0.62367897 | 0.79685521 | 0.59274154 | 0.20411367 | 0.05432056 | 0.4159266 |
| Madcam1 | 0.11488308 | 0.32651257 | 0.12253144 | 0.20398113 | 4.39E-15 | 3.61E-13 |
| P3h1 | 0.46315245 | 0.66377349 | 0.45994776 | 0.20382573 | 0.01948002 | 0.19777702 |
| Ly6h | 0.36447491 | 0.57380297 | 0.37055268 | 0.20325029 | 0.00530244 | 0.06894593 |
| Plat | 0.28577961 | 0.49850627 | 0.29547978 | 0.20302649 | 0.00068429 | 0.01193849 |
| Acat1 | 1.28244911 | 1.16602557 | 0.96392795 | 0.20209762 | 0.16433509 | 0.81873889 |
| Aplp2 | 1.1219919 | 1.09768816 | 0.8956102 | 0.20207796 | 0.14658237 | 0.77203158 |
| Cntfr | 0.32060236 | 0.53103524 | 0.32909085 | 0.20194438 | 0.00212538 | 0.03176672 |
| Vkorc1 | 1.14891991 | 1.10974786 | 0.90797376 | 0.20177409 | 0.15025712 | 0.77912292 |
| H2-Aa | 0.05788704 | 0.2631908 | 0.0619301 | 0.20126069 | 4.37E-52 | 1.58E-49 |
| Filip1l | 0.36494847 | 0.57203032 | 0.37099472 | 0.2010356 | 0.00579151 | 0.07384814 |
| Plpp1 | 0.38698614 | 0.59177769 | 0.3914287 | 0.20034899 | 0.00854806 | 0.10293174 |
| 1500009L16I | 0.72524317 | 0.86823516 | 0.66790682 | 0.20032834 | 0.08115004 | 0.54670692 |

| | | | | | | |
|---|---|---|---|---|---|---|
| Cadm1 | 0.36925018 | 0.57516548 | 0.37500452 | 0.20016095 | 0.00644601 | 0.08063351 |
| Tspo | 0.57115232 | 0.75095752 | 0.55111335 | 0.19984416 | 0.04556944 | 0.36805888 |
| Tk1 | 0.22219719 | 0.43221917 | 0.23261011 | 0.19960906 | 3.19E-05 | 0.00075676 |
| Pdia3 | 3.32105012 | 1.25875388 | 1.05924491 | 0.19950897 | 0.19010419 | 0.88196176 |
| Sulf2 | 0.42527796 | 0.62548687 | 0.42629341 | 0.19919347 | 0.01474117 | 0.15949063 |
| Meis1 | 0.44787436 | 0.64549189 | 0.44647332 | 0.19901857 | 0.01891207 | 0.19349842 |
| Mrfap1 | 3.69787805 | 1.20995151 | 1.01103793 | 0.19891358 | 0.17967372 | 0.85590699 |
| Gip | 0.14896958 | 0.35492198 | 0.15799626 | 0.19692572 | 3.18E-09 | 1.40E-07 |
| Prps1 | 1.08261598 | 1.07352295 | 0.87667911 | 0.19684383 | 0.14776011 | 0.77404405 |
| Mcm3 | 0.36840517 | 0.56981997 | 0.37421765 | 0.19560232 | 0.00744151 | 0.09174069 |
| Kcnmb2 | 0.54462373 | 0.72498604 | 0.52940111 | 0.19558493 | 0.04260496 | 0.35075907 |
| Fam210b | 0.24649368 | 0.45202194 | 0.25685542 | 0.19516652 | 0.00020006 | 0.00393018 |
| Trf | 0.17985228 | 0.3843009 | 0.18973534 | 0.19456556 | 8.87E-07 | 2.79E-05 |
| Hmgn2 | 3.39097508 | 1.24397938 | 1.04959065 | 0.19438872 | 0.19410831 | 0.889076 |
| Tspan4 | 0.46066927 | 0.65178998 | 0.45776733 | 0.19402264 | 0.02415272 | 0.23040214 |
| Ctsd | 1.47221552 | 1.22111767 | 1.02778389 | 0.19333378 | 0.19041136 | 0.88259328 |
| Serping1 | 0.6030444 | 0.76945966 | 0.57660701 | 0.19285265 | 0.05958879 | 0.4439457 |
| Pnpo | 0.21369212 | 0.41588169 | 0.2240607 | 0.19182099 | 3.32E-05 | 0.00078459 |
| Add3 | 0.50456313 | 0.6875552 | 0.49576392 | 0.19179128 | 0.03574452 | 0.30938746 |
| Col15a1 | 0.23515431 | 0.43630712 | 0.24557284 | 0.19073428 | 0.00014812 | 0.0029785 |
| Myct1 | 0.142206 | 0.3417125 | 0.15099468 | 0.19071782 | 2.00E-09 | 9.07E-08 |
| Tusc3 | 1.45644442 | 1.21382438 | 1.02319646 | 0.19062792 | 0.19269991 | 0.88662751 |
| Gstz1 | 0.96783334 | 1.00781477 | 0.81767154 | 0.19014323 | 0.13931524 | 0.74945915 |
| H2-Ab1 | 0.0438344 | 0.23627123 | 0.04690252 | 0.18936871 | 3.10E-79 | 1.87E-76 |
| Gpd2 | 0.73882526 | 0.86652573 | 0.67741109 | 0.18911464 | 0.09668933 | 0.61116934 |
| Hagh | 0.94055918 | 0.99199159 | 0.80291438 | 0.18907721 | 0.13629279 | 0.73978766 |
| Fam212a | 0.33466563 | 0.53108446 | 0.34249041 | 0.18859405 | 0.00515182 | 0.06738445 |
| Plod2 | 0.41273407 | 0.60305548 | 0.41496308 | 0.1880924 | 0.01735259 | 0.18217078 |
| Pcna | 0.50978546 | 0.68821308 | 0.50020605 | 0.18800703 | 0.03996385 | 0.33713503 |
| Sdc1 | 0.44522414 | 0.63211069 | 0.44412168 | 0.187989 | 0.02430201 | 0.231188 |

| | | | | | | |
|---|---|---|---|---|---|---|
| Pid1 | 0.23509033 | 0.4331922 | 0.24550901 | 0.18768319 | 0.0001843 | 0.00365026 |
| Fam151a | 0.3712109 | 0.5644747 | 0.37682881 | 0.18764589 | 0.01017128 | 0.11901788 |
| Fkbp10 | 0.45601658 | 0.64130221 | 0.45367184 | 0.18763037 | 0.02699899 | 0.24963736 |
| Hmgb3 | 0.77500793 | 0.88956854 | 0.70208752 | 0.18748103 | 0.10673243 | 0.64658138 |
| Nt5c3 | 0.80938532 | 0.91188204 | 0.72465861 | 0.18722343 | 0.11435507 | 0.67449953 |
| Etv1 | 0.35243356 | 0.54639973 | 0.35927419 | 0.18712555 | 0.00762098 | 0.09339729 |
| 0610011F06I | 1.34263466 | 1.17312866 | 0.98611868 | 0.18700998 | 0.18847415 | 0.87748819 |
| Gmfg | 0.14330349 | 0.33876804 | 0.15213201 | 0.18663604 | 5.47E-09 | 2.31E-07 |
| Slbp | 1.2160284 | 1.12356506 | 0.93728793 | 0.18627713 | 0.17724363 | 0.84891909 |
| Glud1 | 1.53631854 | 1.23143647 | 1.04530832 | 0.18612814 | 0.20338998 | 0.89054584 |
| Lsr | 0.96579622 | 1.00255318 | 0.81658771 | 0.18596547 | 0.14434423 | 0.76687762 |
| Cnn3 | 1.88738799 | 1.29413055 | 1.1083909 | 0.18573965 | 0.2174817 | 0.89054584 |
| Klhl7 | 1.20964313 | 1.12003025 | 0.93460431 | 0.18542594 | 0.17765566 | 0.85005011 |
| Flt1 | 0.1305149 | 0.32412413 | 0.13885008 | 0.18527404 | 2.54E-10 | 1.28E-08 |
| Slc9a3r2 | 0.3736821 | 0.56394642 | 0.37912503 | 0.1848214 | 0.01156718 | 0.13173679 |
| Tgfb1 | 0.18823347 | 0.3828824 | 0.19828219 | 0.18460021 | 7.21E-06 | 0.00019457 |
| Anxa3 | 0.21818367 | 0.41282142 | 0.22857969 | 0.18424173 | 8.66E-05 | 0.00183536 |
| Rrm2 | 0.22376441 | 0.4182353 | 0.23418196 | 0.18405333 | 0.0001253 | 0.00254773 |
| Hic1 | 0.29901603 | 0.49197655 | 0.30832695 | 0.1836496 | 0.00275994 | 0.04005362 |
| 2010107G23 | 0.55943073 | 0.72520622 | 0.54157654 | 0.18362967 | 0.05708962 | 0.43153558 |
| Usp9x | 1.12251178 | 1.07922904 | 0.895853 | 0.18337605 | 0.17012816 | 0.83373276 |
| Nsg1 | 0.34100431 | 0.53153391 | 0.34849705 | 0.18303686 | 0.00720336 | 0.08918391 |
| Ctsl | 2.039363 | 1.30252512 | 1.12018968 | 0.18233544 | 0.22412228 | 0.89054584 |
| Stt3a | 1.35251515 | 1.17142226 | 0.98957665 | 0.18184561 | 0.19596245 | 0.89054584 |
| Wt1 | 0.15804955 | 0.34916886 | 0.16736785 | 0.18180101 | 2.09E-07 | 7.18E-06 |
| Smim1 | 0.44917432 | 0.62896198 | 0.44762528 | 0.18133669 | 0.02955492 | 0.26663797 |
| Laptm5 | 0.09974135 | 0.28786675 | 0.10663377 | 0.18123298 | 1.20E-15 | 1.03E-13 |
| Trim35 | 1.48771925 | 1.21301295 | 1.0321699 | 0.18084304 | 0.20717071 | 0.89054584 |
| Fxyd5 | 0.17056541 | 0.36104623 | 0.18023172 | 0.18081451 | 1.48E-06 | 4.49E-05 |
| Hand2 | 0.20229566 | 0.39319221 | 0.21255509 | 0.18063712 | 3.76E-05 | 0.00087591 |

| | | | | | | |
|---|---|---|---|---|---|---|
| Rasd1 | 0.2876459 | 0.47736446 | 0.29729647 | 0.18006799 | 0.00238737 | 0.03523452 |
| Cd93 | 0.13603684 | 0.32460542 | 0.14459281 | 0.18001261 | 3.31E-09 | 1.45E-07 |
| Atp5g1 | 3.14426581 | 1.26095292 | 1.08111727 | 0.17983565 | 0.21917589 | 0.89054584 |
| Txndc5 | 1.30492394 | 1.15146922 | 0.97241416 | 0.17905506 | 0.19548154 | 0.89054584 |
| Anxa1 | 0.18567665 | 0.37461132 | 0.19567796 | 0.17893336 | 1.02E-05 | 0.00026912 |
| Adk | 0.68066393 | 0.81471168 | 0.6357959 | 0.17891578 | 0.094921 | 0.60278786 |
| Rnase4 | 0.96333717 | 0.99406333 | 0.81527536 | 0.17878797 | 0.15346267 | 0.78925214 |
| Tmem45a | 0.31526998 | 0.5024628 | 0.3239832 | 0.1784796 | 0.00513548 | 0.06725758 |
| Cdca8 | 0.2119149 | 0.40071576 | 0.22227015 | 0.17844561 | 9.19E-05 | 0.00193341 |
| Pim2 | 0.65686561 | 0.79599794 | 0.61808641 | 0.17791153 | 0.08995353 | 0.58142948 |
| Arhgap31 | 0.33387188 | 0.51956401 | 0.34173688 | 0.17782713 | 0.00766813 | 0.09389579 |
| Apoc1 | 0.21388872 | 0.40178778 | 0.22425868 | 0.17752909 | 0.00011295 | 0.00232275 |
| Lig1 | 0.40086641 | 0.58162452 | 0.40416246 | 0.17746206 | 0.02039415 | 0.20391333 |
| Chchd10 | 0.85320201 | 0.92953676 | 0.7521836 | 0.17735316 | 0.13599187 | 0.73850761 |
| Dusp10 | 0.48181424 | 0.65340071 | 0.47621765 | 0.17718306 | 0.04151047 | 0.34488185 |
| Gsta4 | 0.25202522 | 0.43944326 | 0.26233753 | 0.17710573 | 0.00082964 | 0.01408441 |
| Rbm39 | 3.34293809 | 1.23310037 | 1.05648777 | 0.17661261 | 0.21803899 | 0.89054584 |
| Mcm6 | 0.49532548 | 0.66441733 | 0.48786432 | 0.17655301 | 0.04589576 | 0.36925458 |
| Cd302 | 0.30031122 | 0.48601845 | 0.30957935 | 0.17643911 | 0.00396215 | 0.05424287 |
| Cdca3 | 0.20562471 | 0.39193249 | 0.21592161 | 0.17601088 | 7.30E-05 | 0.00158192 |
| Pycr2 | 1.31453417 | 1.15184698 | 0.97597575 | 0.17587123 | 0.2005823 | 0.89054584 |
| Srpx | 0.2748263 | 0.46001459 | 0.28478493 | 0.17522966 | 0.00207407 | 0.03112823 |
| Kif23 | 0.15765878 | 0.34218697 | 0.16696518 | 0.17522179 | 5.06E-07 | 1.66E-05 |
| Ostf1 | 0.94939793 | 0.98212408 | 0.80775261 | 0.17437146 | 0.15727114 | 0.80068898 |
| Ctsf | 0.96437016 | 0.9897948 | 0.81582719 | 0.17396762 | 0.16023897 | 0.8080503 |
| Foxa2 | 0.75013778 | 0.85913532 | 0.685227 | 0.17390832 | 0.11851881 | 0.68869169 |
| Glipr2 | 0.34743612 | 0.52825643 | 0.3545705 | 0.17368593 | 0.01123946 | 0.12891584 |
| Rest | 0.37253077 | 0.55155869 | 0.37805564 | 0.17350305 | 0.0162522 | 0.17287186 |
| Fam111a | 0.28162706 | 0.46479911 | 0.29143189 | 0.17336721 | 0.00278981 | 0.04036595 |
| Figf | 0.23508683 | 0.41868034 | 0.24550553 | 0.17317481 | 0.00050752 | 0.00914035 |

| | | | | | | |
|---|---|---|---|---|---|---|
| Frzb | 0.42535185 | 0.59940888 | 0.42635988 | 0.173049 | 0.02931552 | 0.26491901 |
| Eef1b2 | 2.92931518 | 1.27327869 | 1.10079533 | 0.17248336 | 0.23269253 | 0.89054584 |
| Gnao1 | 0.88416135 | 0.94322981 | 0.77078884 | 0.17244097 | 0.14863657 | 0.77655241 |
| Stxbp5l | 0.50036223 | 0.66425151 | 0.49217815 | 0.17207336 | 0.05167341 | 0.40185388 |
| Lmo2 | 0.12955542 | 0.3099014 | 0.13785105 | 0.17205035 | 3.04E-09 | 1.34E-07 |
| Siva1 | 1.00815382 | 1.01065383 | 0.83873481 | 0.17191902 | 0.16979823 | 0.8336386 |
| Actg2 | 0.06379106 | 0.24008751 | 0.06824155 | 0.17184596 | 4.36E-32 | 9.30E-30 |
| Bcl2l1 | 0.34820043 | 0.52711022 | 0.35529075 | 0.17181947 | 0.0121272 | 0.13671864 |
| Uqcr11 | 2.83853859 | 1.27939193 | 1.10767872 | 0.17171321 | 0.23507572 | 0.89054584 |
| Gatm | 0.14191956 | 0.32228409 | 0.15069776 | 0.17158633 | 5.64E-08 | 2.08E-06 |
| Sox9 | 0.58150818 | 0.73097535 | 0.55946545 | 0.1715099 | 0.07661047 | 0.52664703 |
| Pabpc1 | 2.91752405 | 1.2732168 | 1.1017234 | 0.1714934 | 0.23416248 | 0.89054584 |
| Gm9493 | 2.2004461 | 1.29874754 | 1.12728215 | 0.17146539 | 0.23927121 | 0.89054584 |
| Ffar1 | 0.36147334 | 0.53899018 | 0.36774827 | 0.17124192 | 0.01502483 | 0.16195539 |
| Rps26-ps1 | 1.64055595 | 1.24071178 | 1.06970704 | 0.17100475 | 0.22820105 | 0.89054584 |
| Spc25 | 0.46437343 | 0.63175689 | 0.4610185 | 0.17073839 | 0.04222404 | 0.3490103 |
| Cd68 | 0.08270243 | 0.25913579 | 0.0884469 | 0.17068889 | 1.22E-19 | 1.48E-17 |
| Asns | 0.33031143 | 0.5083499 | 0.33835279 | 0.16999711 | 0.00962197 | 0.1136914 |
| Dusp1 | 0.64837312 | 0.78159705 | 0.61167232 | 0.16992474 | 0.09778602 | 0.61452179 |
| Syt7 | 0.7884966 | 0.88042286 | 0.71104559 | 0.16937727 | 0.1335425 | 0.73110935 |
| Gimap1 | 0.15086058 | 0.32915816 | 0.15995056 | 0.1692076 | 4.14E-07 | 1.38E-05 |
| Slc6a6 | 0.91383879 | 0.95638219 | 0.78797343 | 0.16840876 | 0.15969138 | 0.80704115 |
| Vps35 | 1.40856322 | 1.17635884 | 1.00837848 | 0.16798036 | 0.21884261 | 0.89054584 |
| Tcf7l2 | 0.63657774 | 0.77065002 | 0.6026818 | 0.16796822 | 0.09706584 | 0.61211857 |
| Isyna1 | 1.05328118 | 1.02979716 | 0.86192016 | 0.167877 | 0.18209069 | 0.86160273 |
| Anp32e | 1.19464413 | 1.09590053 | 0.92818587 | 0.16771466 | 0.19994215 | 0.89054584 |
| Tead2 | 0.55370281 | 0.70434016 | 0.53688337 | 0.16745679 | 0.07309125 | 0.51316074 |
| Prc1 | 0.16854435 | 0.34525382 | 0.17815878 | 0.16709504 | 6.22E-06 | 0.00017023 |
| Hspb1 | 0.18168072 | 0.35859553 | 0.19160244 | 0.16699309 | 2.45E-05 | 0.00059646 |
| Tfdp2 | 0.33040249 | 0.50464 | 0.33843942 | 0.16620058 | 0.0110714 | 0.1271895 |

| | | | | | | |
|---|---|---|---|---|---|---|
| Tsc22d3 | 0.41162221 | 0.57955073 | 0.41395463 | 0.1655961 | 0.03117933 | 0.27732391 |
| Tmem176a | 1.26812311 | 1.12397148 | 0.95838669 | 0.16558479 | 0.21042533 | 0.89054584 |
| Dnajb11 | 1.38959438 | 1.16764926 | 1.00216329 | 0.16548597 | 0.22085016 | 0.89054584 |
| Selenbp1 | 0.38898129 | 0.5586981 | 0.39326537 | 0.16543273 | 0.02500663 | 0.23590763 |
| Nfia | 0.55263681 | 0.7014104 | 0.53600757 | 0.16540283 | 0.07526385 | 0.52184374 |
| Hspa8 | 5.07728311 | 0.85073852 | 0.685852 | 0.16488653 | 0.13134371 | 0.72635204 |
| Kcnj8 | 0.22376378 | 0.39872276 | 0.23418133 | 0.16454142 | 0.00053115 | 0.0095305 |
| Riok3 | 0.76129051 | 0.85627622 | 0.69284255 | 0.16343367 | 0.13588596 | 0.73849629 |
| Kctd12 | 0.58807185 | 0.72802977 | 0.56472249 | 0.16330729 | 0.08894511 | 0.57774473 |
| Uchl1 | 0.50422868 | 0.65864368 | 0.49547885 | 0.16316483 | 0.06248732 | 0.45938225 |
| Hap1 | 0.67588178 | 0.79480509 | 0.63226899 | 0.1625361 | 0.11552382 | 0.67726017 |
| Sphkap | 0.43449986 | 0.59679411 | 0.43456489 | 0.16222922 | 0.04099552 | 0.34177863 |
| Knstrn | 0.18754343 | 0.35969308 | 0.19757962 | 0.16211347 | 6.60E-05 | 0.0014479 |
| Creg1 | 0.69938542 | 0.81133543 | 0.64945142 | 0.16188401 | 0.12275992 | 0.70117835 |
| Abhd5 | 0.30252331 | 0.47292317 | 0.31171637 | 0.1612068 | 0.00798914 | 0.09720403 |
| Cenpe | 0.1466177 | 0.31661765 | 0.15556366 | 0.16105399 | 7.06E-07 | 2.25E-05 |
| C1s1 | 0.15807016 | 0.32814326 | 0.16738908 | 0.16075418 | 3.83E-06 | 0.00010877 |
| Rhoc | 1.20206125 | 1.09196128 | 0.93137784 | 0.16058343 | 0.21090907 | 0.89054584 |
| Tmem88 | 0.15381895 | 0.32337478 | 0.16300528 | 0.1603695 | 2.29E-06 | 6.77E-05 |
| Acyp2 | 0.4261 | 0.58667381 | 0.42703275 | 0.15964105 | 0.04078143 | 0.34061128 |
| Aldoa | 2.4069823 | 1.28851454 | 1.12911299 | 0.15940155 | 0.25535839 | 0.89054584 |
| Cenpw | 0.3064149 | 0.47465411 | 0.31546989 | 0.15918422 | 0.00936469 | 0.11110414 |
| Sod2 | 1.35069972 | 1.14795508 | 0.98894596 | 0.15900912 | 0.22690007 | 0.89054584 |
| Sulf1 | 0.20879162 | 0.37791137 | 0.21912007 | 0.15879129 | 0.00036744 | 0.00680342 |
| Tmc4 | 0.92959042 | 0.95542566 | 0.79684335 | 0.15858231 | 0.17691379 | 0.84817993 |
| PISD | 1.23634608 | 1.10421352 | 0.94563165 | 0.15858187 | 0.21731254 | 0.89054584 |
| Tmod1 | 0.16585861 | 0.33388787 | 0.17540153 | 0.15848634 | 1.28E-05 | 0.00033049 |
| Npm1 | 3.71522846 | 1.16728028 | 1.00902775 | 0.15825253 | 0.23248344 | 0.89054584 |
| Sh3bgrl | 0.76667377 | 0.854571 | 0.6964869 | 0.15808411 | 0.14515166 | 0.76747222 |
| Mgst3 | 0.84734948 | 0.90653091 | 0.74858806 | 0.15794285 | 0.16280886 | 0.8160116 |

| | | | | | | |
|---|---|---|---|---|---|---|
| Qpct | 0.22157567 | 0.38961863 | 0.23198645 | 0.15763219 | 0.00077356 | 0.01322525 |
| Nkd1 | 0.22823831 | 0.39613056 | 0.23866299 | 0.15746757 | 0.00105631 | 0.01734363 |
| S100b | 0.20165119 | 0.36919192 | 0.21190283 | 0.15728909 | 0.00027191 | 0.00524017 |
| Mki67 | 0.18357773 | 0.35052311 | 0.19353807 | 0.15698505 | 7.87E-05 | 0.00169087 |
| Spink1 | 0.08686907 | 0.24953692 | 0.09289599 | 0.15664093 | 1.98E-15 | 1.67E-13 |
| Hjurp | 0.40284202 | 0.56165004 | 0.40596605 | 0.155684 | 0.03699637 | 0.31794511 |
| Psmb8 | 0.14069067 | 0.30470877 | 0.14942355 | 0.15528522 | 6.44E-07 | 2.08E-05 |
| Sema5a | 0.30577626 | 0.47012235 | 0.31485443 | 0.15526791 | 0.01079459 | 0.12490018 |
| Ptgis | 0.27547717 | 0.44057539 | 0.28542207 | 0.15515332 | 0.00566155 | 0.0725099 |
| Ets1 | 0.21225027 | 0.37768014 | 0.22260814 | 0.155072 | 0.00058625 | 0.01044166 |
| Esrp1 | 0.38456595 | 0.54381225 | 0.38919785 | 0.1546144 | 0.03209694 | 0.28357189 |
| Tcf21 | 0.26647251 | 0.43084541 | 0.27658925 | 0.15425616 | 0.00468391 | 0.06242235 |
| Ckap4 | 0.50924412 | 0.65398107 | 0.49974638 | 0.15423469 | 0.07523546 | 0.52184374 |
| Svbp | 0.72836951 | 0.82426186 | 0.670106 | 0.15415586 | 0.14191047 | 0.76031707 |
| Cmbl | 0.17938068 | 0.34292887 | 0.18925354 | 0.15367533 | 7.74E-05 | 0.00166834 |
| Rpl22 | 3.14510716 | 1.23456164 | 1.08103432 | 0.15352732 | 0.25409529 | 0.89054584 |
| Muc1 | 0.3066884 | 0.46865343 | 0.3157334 | 0.15292003 | 0.01202036 | 0.13593695 |
| Upk3b | 0.06318201 | 0.21970185 | 0.06759053 | 0.15211131 | 5.07E-26 | 8.45E-24 |
| Arhgap29 | 0.44855765 | 0.5991174 | 0.44707895 | 0.15203845 | 0.05655484 | 0.42838665 |
| C2cd4b | 0.33152825 | 0.49130396 | 0.33951009 | 0.15179387 | 0.01862572 | 0.19165059 |
| Cel | 0.11520937 | 0.27465936 | 0.122873 | 0.15178637 | 4.33E-09 | 1.86E-07 |
| Prdx6 | 1.49717999 | 1.18657853 | 1.03479354 | 0.15178499 | 0.24718333 | 0.89054584 |
| Npy | 0.06441958 | 0.22056108 | 0.06891335 | 0.15164773 | 5.78E-25 | 9.21E-23 |
| Flrt2 | 0.33774916 | 0.49634289 | 0.34541488 | 0.15092801 | 0.02088971 | 0.20743768 |
| Tnfaip8 | 0.56995331 | 0.70095438 | 0.55014199 | 0.15081239 | 0.10077073 | 0.6232762 |
| Irx2 | 0.1541557 | 0.31403415 | 0.1633528 | 0.15068135 | 8.64E-06 | 0.00023098 |
| Acsl1 | 0.13974273 | 0.29902734 | 0.14844026 | 0.15058707 | 1.14E-06 | 3.51E-05 |
| Fn1 | 0.262139 | 0.42276006 | 0.27232452 | 0.15043554 | 0.00503148 | 0.06629241 |
| Zfp667 | 0.3498408 | 0.50715865 | 0.35683551 | 0.15032314 | 0.02484286 | 0.23494573 |
| Gclm | 0.85980075 | 0.90616257 | 0.75620735 | 0.14995521 | 0.17777864 | 0.85016198 |

| | | | | | | |
|---|---|---|---|---|---|---|
| Gpm6b | 0.28648746 | 0.44586681 | 0.29616902 | 0.1496978 | 0.00926512 | 0.1103745 |
| Ormdl2 | 0.81910621 | 0.88042624 | 0.73088594 | 0.1495403 | 0.17023779 | 0.83373276 |
| Mfsd10 | 0.56434752 | 0.69512404 | 0.54558793 | 0.14953612 | 0.10081271 | 0.6232762 |
| Rgs5 | 0.21871754 | 0.37845695 | 0.22911622 | 0.14934073 | 0.0011957 | 0.01933526 |
| Rpn2 | 2.13785736 | 1.27441746 | 1.12511676 | 0.1493007 | 0.26821121 | 0.89054584 |
| Rplp0 | 5.23802789 | 0.81860373 | 0.66960186 | 0.14900187 | 0.14993273 | 0.77909236 |
| Loxl1 | 0.40860553 | 0.56017416 | 0.41121493 | 0.14895923 | 0.04573795 | 0.36880358 |
| Tmcc2 | 0.10576789 | 0.26176825 | 0.11297388 | 0.14879437 | 4.23E-10 | 2.10E-08 |
| Trim47 | 0.34756883 | 0.50309637 | 0.35469559 | 0.14840078 | 0.02563276 | 0.24037761 |
| Dcdc2a | 0.21038472 | 0.36860923 | 0.22072738 | 0.14788186 | 0.00090004 | 0.01510282 |
| Pros1 | 0.30117626 | 0.45789727 | 0.31041533 | 0.14748194 | 0.01343101 | 0.14854065 |
| Slc4a1 | 0.03803104 | 0.18808233 | 0.04069478 | 0.14738754 | 3.52E-64 | 1.54E-61 |
| Mapt | 0.37605438 | 0.52868893 | 0.38132616 | 0.14736277 | 0.03589567 | 0.30996718 |
| Ptprn2 | 0.71523829 | 0.80804451 | 0.66082208 | 0.14722243 | 0.14964653 | 0.77846267 |
| Emilin2 | 0.20497276 | 0.36219454 | 0.21526262 | 0.14693192 | 0.00073623 | 0.01267685 |
| Prss35 | 0.16817797 | 0.32467678 | 0.17778282 | 0.14689395 | 5.92E-05 | 0.00132624 |
| Rnf130 | 0.77693512 | 0.85022396 | 0.70337543 | 0.14684853 | 0.16535639 | 0.82185015 |
| Capg | 0.38804868 | 0.53921303 | 0.39240706 | 0.14680597 | 0.04066795 | 0.34041799 |
| Ripply3 | 0.34331759 | 0.49724844 | 0.35068399 | 0.14656446 | 0.02575823 | 0.24093087 |
| Hoxb2 | 0.22424656 | 0.38094415 | 0.23466532 | 0.14627883 | 0.00184123 | 0.02806955 |
| Minos1 | 3.01587683 | 1.2398902 | 1.09362689 | 0.14626331 | 0.26661143 | 0.89054584 |
| Plin3 | 0.70332716 | 0.79844161 | 0.65229496 | 0.14614665 | 0.14828075 | 0.77505178 |
| Ralb | 0.7106083 | 0.80356658 | 0.65751929 | 0.14604729 | 0.15037237 | 0.77944175 |
| Notch2 | 0.19476463 | 0.35072211 | 0.20492178 | 0.14580032 | 0.00045854 | 0.00835166 |
| G6pc2 | 0.243683 | 0.39976907 | 0.25406437 | 0.1457047 | 0.00377138 | 0.05202418 |
| Dnajc3 | 1.11519536 | 1.03803233 | 0.89241951 | 0.14561282 | 0.2235728 | 0.89054584 |
| Hmgb1 | 3.87852525 | 1.13236611 | 0.98720827 | 0.14515784 | 0.24665784 | 0.89054584 |
| Grn | 0.64600886 | 0.75499207 | 0.60987797 | 0.14511411 | 0.13381293 | 0.73180684 |
| Ptp4a1 | 1.29921622 | 1.11531694 | 0.97027224 | 0.1450447 | 0.24307075 | 0.89054584 |
| Chchd7 | 1.01943543 | 0.9894272 | 0.84450363 | 0.14492357 | 0.21199694 | 0.89054584 |

| | | | | | | |
|---|---|---|---|---|---|---|
| Prr13 | 1.11114169 | 1.03539171 | 0.89050182 | 0.14488989 | 0.22421436 | 0.89054584 |
| Cdc42ep5 | 0.38097416 | 0.5307095 | 0.38588101 | 0.14482849 | 0.04018102 | 0.33777651 |
| Serpinb6a | 0.74694878 | 0.82770204 | 0.68303302 | 0.14466902 | 0.16187485 | 0.81375228 |
| Sycn | 0.08799261 | 0.23875255 | 0.09409552 | 0.14465704 | 3.96E-13 | 2.74E-11 |
| Cnot6l | 0.95169236 | 0.95333856 | 0.80900065 | 0.1443379 | 0.20292318 | 0.89054584 |
| Ghitm | 0.76388612 | 0.83893951 | 0.69460234 | 0.14433717 | 0.16649224 | 0.82523228 |
| Tuba1c | 0.31476355 | 0.4676715 | 0.32349736 | 0.14417414 | 0.01893007 | 0.19354591 |
| Wisp1 | 0.16831727 | 0.32172142 | 0.17792577 | 0.14379565 | 8.32E-05 | 0.0017789 |
| C1qtnf6 | 0.54693612 | 0.67496715 | 0.53131145 | 0.1436557 | 0.10389205 | 0.6342008 |
| Entpd3 | 0.57245518 | 0.69552396 | 0.55216777 | 0.1433562 | 0.11322109 | 0.67029056 |
| Gcat | 0.47140824 | 0.61004392 | 0.46717094 | 0.14287298 | 0.07710391 | 0.52951666 |
| Abi3bp | 0.23092565 | 0.38393778 | 0.24135026 | 0.14258753 | 0.00295747 | 0.04245287 |
| Tacc3 | 0.19489966 | 0.34671302 | 0.20505886 | 0.14165415 | 0.00064457 | 0.01134105 |
| Ltbp4 | 0.34736298 | 0.49608541 | 0.35450156 | 0.14158384 | 0.03139134 | 0.27869665 |
| Pdgfrb | 0.21226226 | 0.36414261 | 0.22262022 | 0.14152239 | 0.0015295 | 0.02381816 |
| Ctla2a | 0.13639892 | 0.28607175 | 0.14496895 | 0.1411028 | 2.88E-06 | 8.31E-05 |
| Epb41 | 0.33846752 | 0.48707158 | 0.34609555 | 0.14097603 | 0.02886524 | 0.26204647 |
| Clic1 | 2.42203011 | 1.26974905 | 1.12888255 | 0.14086649 | 0.28049846 | 0.89054584 |
| Ncam1 | 0.68026792 | 0.77621369 | 0.63550438 | 0.1407093 | 0.15114081 | 0.78258554 |
| Trip6 | 0.38483372 | 0.52981031 | 0.38944483 | 0.14036548 | 0.04655685 | 0.37333028 |
| Ppp1r1b | 0.15944395 | 0.30895427 | 0.16880417 | 0.1401501 | 5.48E-05 | 0.00123275 |
| Bnip3l | 1.51443305 | 1.17945039 | 1.03948527 | 0.13996512 | 0.26522494 | 0.89054584 |
| Naa38 | 1.29206242 | 1.10691154 | 0.96757513 | 0.13933642 | 0.2511375 | 0.89054584 |
| Mrpl57 | 1.35813599 | 1.13074647 | 0.99151693 | 0.13922954 | 0.25648763 | 0.89054584 |
| Prr5 | 0.30782141 | 0.45589146 | 0.3168246 | 0.13906686 | 0.02042774 | 0.2039679 |
| Pcsk2os1 | 0.47464012 | 0.60861238 | 0.46998778 | 0.13862459 | 0.08469422 | 0.56163058 |
| Pop5 | 1.23189759 | 1.08239167 | 0.9438273 | 0.13856437 | 0.24699248 | 0.89054584 |
| Ldhb | 0.44730134 | 0.58405491 | 0.44596521 | 0.1380897 | 0.07456584 | 0.51900827 |
| Impdh2 | 1.24991337 | 1.08914567 | 0.95113531 | 0.13801036 | 0.24951541 | 0.89054584 |
| Eif4g2 | 2.59223809 | 1.26060419 | 1.12291828 | 0.13768591 | 0.28391404 | 0.89054584 |

| | | | | | | |
|---|---|---|---|---|---|---|
| Lamb1 | 0.6697112 | 0.76520866 | 0.62769457 | 0.13751409 | 0.15370501 | 0.79021819 |
| Malat1 | 7.68065191 | 6.14539037 | 6.00792113 | 0.13746925 | 0.45755205 | 0.89054584 |
| Gngt2 | 0.20279074 | 0.35010566 | 0.21305604 | 0.13704962 | 0.00136373 | 0.02163172 |
| Try10 | 0.07419899 | 0.21608164 | 0.0793639 | 0.13671774 | 5.04E-16 | 4.51E-14 |
| Wdr70 | 0.30926104 | 0.45481338 | 0.31821017 | 0.13660321 | 0.02274912 | 0.22105677 |
| Lama4 | 0.30708064 | 0.4526681 | 0.31611124 | 0.13655686 | 0.02207755 | 0.21656314 |
| Chmp5 | 1.65125268 | 1.20847622 | 1.07193705 | 0.13653917 | 0.2764441 | 0.89054584 |
| Tmem39a | 0.59132195 | 0.70367864 | 0.56731492 | 0.13636373 | 0.13138773 | 0.72635204 |
| Prr15l | 0.40263901 | 0.54208741 | 0.40578081 | 0.1363066 | 0.05879218 | 0.4395921 |
| Cd248 | 0.32196052 | 0.4664924 | 0.33038941 | 0.13610298 | 0.02747713 | 0.25292919 |
| Rab3b | 0.52623655 | 0.65018875 | 0.51408777 | 0.13610098 | 0.10870944 | 0.65261673 |
| Lman1 | 1.74067849 | 1.2246966 | 1.08864458 | 0.13605201 | 0.28020069 | 0.89054584 |
| Rpl34 | 4.69599214 | 0.92503618 | 0.78902528 | 0.1360109 | 0.21095826 | 0.89054584 |
| Calml4 | 0.30918519 | 0.45403435 | 0.3181372 | 0.13589715 | 0.02328788 | 0.22433735 |
| St3gal4 | 0.4041888 | 0.54249143 | 0.40719428 | 0.13529715 | 0.06080975 | 0.4511872 |
| Pgrmc1 | 1.63564786 | 1.20386179 | 1.06866594 | 0.13519585 | 0.27779717 | 0.89054584 |
| Cd164l2 | 0.63690333 | 0.73800952 | 0.60293126 | 0.13507825 | 0.14829588 | 0.77505178 |
| Lgals3 | 0.07830047 | 0.21864387 | 0.08374542 | 0.13489846 | 3.08E-14 | 2.36E-12 |
| Ramp1 | 0.44187155 | 0.57596845 | 0.4411408 | 0.13482766 | 0.0772332 | 0.52992284 |
| Mia | 0.36136583 | 0.50227541 | 0.36764774 | 0.13462766 | 0.04400016 | 0.36020009 |
| Col6a6 | 0.23381694 | 0.37867527 | 0.24423826 | 0.13443701 | 0.00516714 | 0.06748932 |
| Raly | 1.54974958 | 1.18309323 | 1.04871753 | 0.1343757 | 0.27526511 | 0.89054584 |
| Upp1 | 0.32075217 | 0.46360409 | 0.32923413 | 0.13436996 | 0.02862165 | 0.25999793 |
| Il1r1 | 0.3195159 | 0.4623706 | 0.32805137 | 0.13431923 | 0.02822254 | 0.25750178 |
| Cycs | 2.09338912 | 1.25727107 | 1.12305837 | 0.1342127 | 0.28884112 | 0.89054584 |
| Gck | 0.77306844 | 0.83496262 | 0.70078866 | 0.13417396 | 0.18619452 | 0.87252679 |
| Pax4 | 0.2029572 | 0.34699703 | 0.21322443 | 0.1337726 | 0.00173381 | 0.02662793 |
| Cenpm | 0.20423005 | 0.34827057 | 0.21451178 | 0.13375879 | 0.00183578 | 0.028016 |
| Nkx2-3 | 0.23983636 | 0.38384461 | 0.25023865 | 0.13360596 | 0.00643268 | 0.08053629 |
| Fopnl | 0.84532911 | 0.88088063 | 0.74734099 | 0.13353965 | 0.20256642 | 0.89054584 |

| | | | | | | |
|---|---|---|---|---|---|---|
| Lhfpl2 | 0.25840803 | 0.4020936 | 0.26864557 | 0.13344803 | 0.01032563 | 0.12043522 |
| Atox1 | 1.79845575 | 1.23093442 | 1.09753748 | 0.13339695 | 0.28560946 | 0.89054584 |
| Cd164 | 1.34808306 | 1.12135021 | 0.98803346 | 0.13331674 | 0.26479321 | 0.89054584 |
| Rps12-ps3 | 1.60742078 | 1.19547143 | 1.06246795 | 0.13300348 | 0.27987171 | 0.89054584 |
| Igfbp6 | 0.09760975 | 0.23724021 | 0.10435964 | 0.13288057 | 1.49E-09 | 6.86E-08 |
| Unc93b1 | 0.18842411 | 0.33098533 | 0.19847624 | 0.13250909 | 0.00093399 | 0.0155652 |
| Phyh | 0.74142143 | 0.81167695 | 0.67921294 | 0.13246401 | 0.1817771 | 0.86128898 |
| Smarca2 | 0.98772351 | 0.9605225 | 0.82815388 | 0.13236862 | 0.22823698 | 0.89054584 |
| Txnrd2 | 0.28428033 | 0.42611472 | 0.2940191 | 0.13209562 | 0.01816794 | 0.18800767 |
| Gm1673 | 1.07308513 | 1.0040191 | 0.87194321 | 0.13207588 | 0.24018613 | 0.89054584 |
| Nfix | 0.40906769 | 0.54337994 | 0.411635 | 0.13174495 | 0.06796666 | 0.48757084 |
| Idh2 | 1.89710786 | 1.24103289 | 1.10939412 | 0.13163877 | 0.29019271 | 0.89054584 |
| Sap30 | 0.53068423 | 0.6493621 | 0.5178119 | 0.1315502 | 0.11828477 | 0.68802988 |
| Ass1 | 0.22363236 | 0.36532994 | 0.23404956 | 0.13128038 | 0.00448442 | 0.06019921 |
| Adamtsl1 | 0.19494624 | 0.3362148 | 0.20510614 | 0.13110866 | 0.00145019 | 0.02285303 |
| Guk1 | 1.3282369 | 1.11197951 | 0.98097302 | 0.13100648 | 0.26690851 | 0.89054584 |
| Trem2 | 0.04814344 | 0.182368 | 0.0515112 | 0.1308568 | 1.28E-32 | 2.81E-30 |
| Maob | 0.30063267 | 0.44071796 | 0.30989004 | 0.13082792 | 0.02459795 | 0.23354365 |
| Serp1 | 1.79998629 | 1.22828195 | 1.09775359 | 0.13052835 | 0.28979926 | 0.89054584 |
| Gskip | 0.38732668 | 0.52204288 | 0.39174233 | 0.13030054 | 0.06061306 | 0.44995811 |
| Ndufc1 | 2.17211197 | 1.25667535 | 1.12644217 | 0.13023318 | 0.29506705 | 0.89054584 |
| Slc46a3 | 0.08493456 | 0.22087544 | 0.09083048 | 0.13004496 | 1.28E-11 | 7.46E-10 |
| Lgmn | 0.48554243 | 0.60948596 | 0.4794426 | 0.13004336 | 0.10317157 | 0.63193134 |
| Myo1c | 0.22702534 | 0.36737029 | 0.23744899 | 0.12992131 | 0.00539912 | 0.06995212 |
| Vamp5 | 0.17728029 | 0.31680356 | 0.1871068 | 0.12969677 | 0.00062017 | 0.01099174 |
| Psap | 2.03718466 | 1.24959736 | 1.12006969 | 0.12952767 | 0.29502221 | 0.89054584 |
| Cox17 | 1.59139166 | 1.18831595 | 1.05878832 | 0.12952763 | 0.28435414 | 0.89054584 |
| Klf6 | 0.57910475 | 0.68704171 | 0.55753329 | 0.12950842 | 0.13957801 | 0.74975989 |
| Stab1 | 0.09354518 | 0.22934655 | 0.1000224 | 0.12932416 | 8.52E-10 | 4.04E-08 |
| Csrp1 | 0.67805063 | 0.76312698 | 0.63387047 | 0.1292565 | 0.17104751 | 0.83552788 |

| | | | | | | |
|---|---|---|---|---|---|---|
| Atp5e | 3.26251224 | 1.19641088 | 1.0672457 | 0.12916518 | 0.28642604 | 0.89054584 |
| Rgs4 | 0.183481 | 0.32232131 | 0.19343941 | 0.1288819 | 0.00095433 | 0.01583051 |
| Ccdc85b | 0.37109732 | 0.50560287 | 0.37672318 | 0.12887968 | 0.05548404 | 0.42238211 |
| Rgs10 | 0.23315657 | 0.37238311 | 0.24357897 | 0.12880413 | 0.00687671 | 0.08550476 |
| Pdgfra | 0.2148805 | 0.3538309 | 0.22525721 | 0.12857369 | 0.00391564 | 0.05365682 |
| Hoxb6 | 0.13439503 | 0.27127791 | 0.14288657 | 0.12839134 | 1.42E-05 | 0.00036184 |
| Ndufv3 | 2.05557192 | 1.249439 | 1.12108927 | 0.12834974 | 0.29688303 | 0.89054584 |
| Fkbp9 | 1.09867235 | 1.01268257 | 0.8845294 | 0.12815317 | 0.24983762 | 0.89054584 |
| Anxa7 | 0.57612382 | 0.68316308 | 0.55513151 | 0.12803157 | 0.14129821 | 0.75759671 |
| Sep-01 | 0.21131824 | 0.34954937 | 0.2216687 | 0.12788067 | 0.00359683 | 0.04999704 |
| Racgap1 | 0.1768912 | 0.31457379 | 0.18670893 | 0.12786486 | 0.00070987 | 0.01232539 |
| Nfic | 0.72013857 | 0.79207425 | 0.66430104 | 0.12777321 | 0.18509658 | 0.86930035 |
| Adipor1 | 1.15561577 | 1.03871814 | 0.91097643 | 0.12774171 | 0.25678138 | 0.89054584 |
| Vwa1 | 0.13950602 | 0.2759334 | 0.14819467 | 0.12773873 | 2.96E-05 | 0.00070737 |
| Ifnar2 | 0.36824058 | 0.50160994 | 0.37406434 | 0.12754559 | 0.05607874 | 0.42589295 |
| Fam92b | 0.38598853 | 0.51795584 | 0.39050954 | 0.1274463 | 0.06419427 | 0.46799603 |
| Gm10260 | 4.43798482 | 0.98121282 | 0.85418816 | 0.12702466 | 0.2442106 | 0.89054584 |
| Slc35g2 | 0.37040277 | 0.50302421 | 0.37607716 | 0.12694705 | 0.05789961 | 0.43561417 |
| Rasgrp2 | 0.21292541 | 0.35018 | 0.22328841 | 0.1268916 | 0.00405316 | 0.05522805 |
| Ptprcap | 0.05338299 | 0.18390736 | 0.05711431 | 0.12679305 | 2.26E-25 | 3.68E-23 |
| Trp53i11 | 0.56910982 | 0.67622477 | 0.54945808 | 0.12676669 | 0.14121543 | 0.75743297 |
| Psip1 | 1.05212017 | 0.98781303 | 0.86132436 | 0.12648866 | 0.2469299 | 0.89054584 |
| Acot1 | 0.49184057 | 0.61120099 | 0.48487099 | 0.12633 | 0.11240117 | 0.66731865 |
| Itgb1 | 1.74089646 | 1.21494595 | 1.08868105 | 0.1262649 | 0.29448039 | 0.89054584 |
| Cyb5a | 1.23345007 | 1.07071107 | 0.94445843 | 0.12625264 | 0.26670974 | 0.89054584 |
| Pim1 | 0.2016185 | 0.33794023 | 0.21186973 | 0.12607049 | 0.00278355 | 0.04031556 |
| Serpinb1a | 0.25654894 | 0.39286052 | 0.26681012 | 0.12605039 | 0.01386638 | 0.15195374 |
| Ly6d | 0.03451074 | 0.1629459 | 0.03692879 | 0.12601711 | 3.25E-57 | 1.24E-54 |
| Fkbp4 | 2.06306674 | 1.24732779 | 1.12148154 | 0.12584625 | 0.30055681 | 0.89054584 |
| Tpx2 | 0.15593116 | 0.29088186 | 0.16518427 | 0.12569759 | 0.00019613 | 0.00385818 |

| | | | | | | |
|---|---|---|---|---|---|---|
| Tnfrsf1a | 0.38913908 | 0.51909647 | 0.39341056 | 0.1256859 | 0.06831905 | 0.48937232 |
| Rnaseh2c | 0.85524929 | 0.87899322 | 0.75343541 | 0.12555782 | 0.21875672 | 0.89054584 |
| Fam220a | 0.50254546 | 0.6195413 | 0.49404308 | 0.12549821 | 0.11831001 | 0.68802988 |
| Itm2b | 3.67529804 | 1.13865935 | 1.01338089 | 0.12527846 | 0.28232173 | 0.89054584 |
| Fap | 0.25348123 | 0.38899298 | 0.26377814 | 0.12521484 | 0.01349757 | 0.14890122 |
| Mmrn1 | 0.06035994 | 0.18933512 | 0.06457382 | 0.1247613 | 1.11E-19 | 1.36E-17 |
| Larp7 | 0.7250723 | 0.79244534 | 0.66778642 | 0.12465892 | 0.19223024 | 0.88530942 |
| Gstm2 | 0.30882633 | 0.44244286 | 0.31779189 | 0.12465097 | 0.03381779 | 0.29600565 |
| Rangap1 | 1.01117467 | 0.96482557 | 0.84028838 | 0.12453719 | 0.24494278 | 0.89054584 |
| Lima1 | 0.45531752 | 0.57756222 | 0.45305538 | 0.12450685 | 0.1002108 | 0.62272587 |
| Ddx21 | 0.56313498 | 0.66903195 | 0.54460015 | 0.1244318 | 0.14355381 | 0.76537874 |
| Bok | 0.15300366 | 0.28653986 | 0.16216374 | 0.12437612 | 0.00017625 | 0.00350037 |
| Kpna2 | 0.43814534 | 0.56201288 | 0.43782083 | 0.12419205 | 0.09315851 | 0.59577065 |
| Cdh1 | 0.74577729 | 0.80613153 | 0.68222521 | 0.12390632 | 0.19873152 | 0.89054584 |
| Gm43861 | 0.29685269 | 0.43006341 | 0.30623322 | 0.12383019 | 0.02978819 | 0.26824174 |
| Sapcd1 | 0.17981722 | 0.31348634 | 0.18969952 | 0.12378682 | 0.00118229 | 0.01917321 |
| Ezh2 | 0.56195828 | 0.66741895 | 0.54364064 | 0.12377831 | 0.14439799 | 0.76687762 |
| Hes6 | 0.9431621 | 0.92811716 | 0.80434207 | 0.12377509 | 0.23670715 | 0.89054584 |
| Isg20 | 0.36274512 | 0.49268335 | 0.36893696 | 0.12374638 | 0.05906499 | 0.44094965 |
| Lrp1 | 0.30690097 | 0.43949064 | 0.31593818 | 0.12355246 | 0.03423127 | 0.29872948 |
| St3gal5 | 0.34418599 | 0.47489135 | 0.35150423 | 0.12338712 | 0.05098162 | 0.39853675 |
| Ppp1cb | 1.81110191 | 1.22262571 | 1.09929191 | 0.12333381 | 0.30058991 | 0.89054584 |
| Ak1 | 0.4919378 | 0.60824701 | 0.4849546 | 0.12329241 | 0.11811439 | 0.68771984 |
| 1110001J03F | 0.95910621 | 0.93614171 | 0.81300702 | 0.12313469 | 0.24021099 | 0.89054584 |
| F13a1 | 0.0435734 | 0.16974246 | 0.04662335 | 0.12311911 | 4.37E-35 | 1.13E-32 |
| Gja1 | 0.25210755 | 0.38546819 | 0.26241902 | 0.12304918 | 0.01445933 | 0.15690974 |
| P2ry1 | 0.2819917 | 0.41479583 | 0.29178765 | 0.12300818 | 0.02476168 | 0.23463713 |
| Klhdc8b | 0.85035892 | 0.87344718 | 0.75044007 | 0.1230071 | 0.22253423 | 0.89054584 |
| Rpgrip1 | 0.26663252 | 0.39937984 | 0.27674655 | 0.12263329 | 0.0194855 | 0.19777702 |
| Rgs3 | 0.45162682 | 0.57226973 | 0.44979582 | 0.12247391 | 0.10229673 | 0.62949832 |

| | | | | | | |
|---|---|---|---|---|---|---|
| Enho | 0.47256326 | 0.59063761 | 0.46817838 | 0.12245923 | 0.11149471 | 0.66356743 |
| Ogt | 1.12041868 | 1.01728347 | 0.89487438 | 0.12240909 | 0.26196716 | 0.89054584 |
| Fabp7 | 0.04721335 | 0.17241706 | 0.05051648 | 0.12190057 | 1.26E-29 | 2.51E-27 |
| Rps18 | 5.22160161 | 0.79355623 | 0.67172026 | 0.12183597 | 0.19904586 | 0.89054584 |
| Phactr2 | 0.34256356 | 0.47173082 | 0.34997146 | 0.12175936 | 0.05251925 | 0.40587639 |
| Tmem163 | 0.49378346 | 0.60815448 | 0.48654058 | 0.12161391 | 0.12210426 | 0.70054116 |
| Usmg5 | 2.89898442 | 1.22461319 | 1.10316011 | 0.12145307 | 0.30400052 | 0.89054584 |
| Tmem215 | 0.17851381 | 0.30958567 | 0.1883677 | 0.12121798 | 0.00135829 | 0.02159264 |
| Gm561 | 0.84201706 | 0.86646445 | 0.74529015 | 0.1211743 | 0.22438054 | 0.89054584 |
| Pla2g16 | 0.65894572 | 0.74049294 | 0.61965008 | 0.12084286 | 0.18178675 | 0.86128898 |
| Camk1 | 0.61315071 | 0.70533096 | 0.58454547 | 0.12078549 | 0.1678531 | 0.82942543 |
| Orc6 | 0.90850108 | 0.90565186 | 0.78492996 | 0.12072191 | 0.23682799 | 0.89054584 |
| Rab25 | 0.72297518 | 0.78693966 | 0.66630705 | 0.12063261 | 0.19947853 | 0.89054584 |
| Col12a1 | 0.24235571 | 0.37269694 | 0.25274507 | 0.11995187 | 0.0135139 | 0.14890122 |
| Ctse | 0.0532359 | 0.17673167 | 0.05695702 | 0.11977465 | 5.78E-23 | 8.22E-21 |
| Cda | 0.44491122 | 0.56351606 | 0.44384374 | 0.11967232 | 0.1045259 | 0.63753323 |
| Rogdi | 0.91429245 | 0.90775959 | 0.78823114 | 0.11952845 | 0.23994228 | 0.89054584 |
| F3 | 0.62921723 | 0.71648283 | 0.59702302 | 0.11945981 | 0.17561166 | 0.84465613 |
| Sptssa | 1.22867716 | 1.06162426 | 0.94251321 | 0.11911105 | 0.27800498 | 0.89054584 |
| Asb4 | 0.2273704 | 0.3568134 | 0.23779441 | 0.11901899 | 0.00985412 | 0.11605605 |
| Hotairm1 | 0.14439641 | 0.27219246 | 0.15326414 | 0.11892832 | 0.00015006 | 0.00301325 |
| Wnk3 | 0.30944799 | 0.43727761 | 0.31839002 | 0.11888759 | 0.04095939 | 0.34167394 |
| Irf8 | 0.05479324 | 0.17748906 | 0.05862224 | 0.11886682 | 1.75E-21 | 2.39E-19 |
| Rras2 | 0.27028377 | 0.39900487 | 0.28033257 | 0.1186723 | 0.02429026 | 0.231188 |
| Rbbp7 | 1.72951194 | 1.20531707 | 1.08674793 | 0.11856914 | 0.3056174 | 0.89054584 |
| Snx5 | 1.09514322 | 1.00136583 | 0.88281872 | 0.11854711 | 0.26578399 | 0.89054584 |
| Klf10 | 0.58139191 | 0.67774757 | 0.55937207 | 0.1183755 | 0.1620825 | 0.81451372 |
| Uba7 | 0.11970535 | 0.24594324 | 0.12757529 | 0.11836795 | 7.71E-06 | 0.00020729 |
| Serpina6 | 0.09602326 | 0.22102179 | 0.10266686 | 0.11835494 | 3.92E-08 | 1.48E-06 |
| Tspan13 | 0.97437969 | 0.93949609 | 0.82114693 | 0.11834915 | 0.25095817 | 0.89054584 |

| | | | | | | |
|---|---|---|---|---|---|---|
| Rnf26 | 0.41239706 | 0.53263006 | 0.41465749 | 0.11797257 | 0.09250318 | 0.5928873 |
| H2-Eb1 | 0.0378177 | 0.15842601 | 0.04046657 | 0.11795944 | 2.60E-42 | 8.03E-40 |
| Lrp11 | 0.74048246 | 0.79649973 | 0.67856182 | 0.11793791 | 0.20904178 | 0.89054584 |
| Lrrc42 | 0.71595059 | 0.77918312 | 0.66132884 | 0.11785429 | 0.20319534 | 0.89054584 |
| Rps5 | 5.18336309 | 0.79388894 | 0.67631664 | 0.1175723 | 0.20899306 | 0.89054584 |
| Fzd2 | 0.33703862 | 0.46225307 | 0.34474134 | 0.11751173 | 0.05613302 | 0.42608192 |
| Amotl2 | 0.38900942 | 0.51078532 | 0.39329126 | 0.11749406 | 0.08198564 | 0.55080068 |
| Rsad2 | 0.05337591 | 0.17448041 | 0.05710674 | 0.11737367 | 5.06E-22 | 6.98E-20 |
| Ndrg2 | 0.37149108 | 0.49444374 | 0.37708931 | 0.11735443 | 0.07354114 | 0.51488639 |
| Chac2 | 0.30602482 | 0.4323793 | 0.315094 | 0.1172853 | 0.04144431 | 0.34472728 |
| Tax1bp3 | 0.9666874 | 0.93434488 | 0.81706223 | 0.11728265 | 0.25182646 | 0.89054584 |
| Chodl | 0.08117771 | 0.20407524 | 0.08681857 | 0.11725668 | 1.56E-10 | 8.15E-09 |
| Fkbp14 | 0.42189056 | 0.54045989 | 0.42324257 | 0.11721731 | 0.0984738 | 0.61600365 |
| Nenf | 1.98687601 | 1.23367566 | 1.1168685 | 0.11680716 | 0.31303947 | 0.89054584 |
| Ccdc34 | 0.74650932 | 0.79939728 | 0.6827301 | 0.11666718 | 0.2129767 | 0.89054584 |
| Ucp2 | 0.7466676 | 0.79913291 | 0.68283922 | 0.11629369 | 0.213755 | 0.89054584 |
| Pdk3 | 0.32661775 | 0.45106019 | 0.33483501 | 0.11622518 | 0.05292043 | 0.40810662 |
| Sox17 | 0.08708882 | 0.20916245 | 0.09313062 | 0.11603183 | 3.23E-09 | 1.41E-07 |
| Exoc1 | 0.24933566 | 0.3754615 | 0.2596738 | 0.1157877 | 0.01888202 | 0.19341421 |
| Mthfd2l | 0.48219298 | 0.592104 | 0.47654566 | 0.11555834 | 0.12928988 | 0.72010939 |
| Egflam | 0.19318462 | 0.31876155 | 0.20331721 | 0.11544434 | 0.00408013 | 0.05549132 |
| Zfp36l2 | 0.48380577 | 0.59335853 | 0.47794143 | 0.1154171 | 0.13027795 | 0.72297247 |
| Thbs2 | 0.20067339 | 0.32628594 | 0.21091287 | 0.11537307 | 0.005409 | 0.069967 |
| Celf3 | 0.66084312 | 0.73641679 | 0.62107378 | 0.11534301 | 0.19345424 | 0.88812525 |
| Fam46c | 0.31229318 | 0.43631103 | 0.3211255 | 0.11518553 | 0.04734795 | 0.37820971 |
| Pdcd4 | 0.81964217 | 0.84635388 | 0.73122728 | 0.1151266 | 0.2316214 | 0.89054584 |
| Itga6 | 0.31241791 | 0.43629608 | 0.32124532 | 0.11505076 | 0.04760304 | 0.37982877 |
| Vamp7 | 0.75551473 | 0.80393513 | 0.68890965 | 0.11502548 | 0.21831649 | 0.89054584 |
| Stx8 | 0.59344826 | 0.68389308 | 0.56900713 | 0.11488595 | 0.17343747 | 0.83899716 |
| Ppie | 0.52679198 | 0.62922456 | 0.51455353 | 0.11467103 | 0.14957214 | 0.77846267 |

| | | | | | | |
|---|---|---|---|---|---|---|
| Cir1 | 0.54446063 | 0.64386515 | 0.52926624 | 0.11459891 | 0.15654172 | 0.79857208 |
| Spock2 | 0.59620556 | 0.6857464 | 0.57119699 | 0.11454942 | 0.17507022 | 0.84296516 |
| Gm2a | 0.32639352 | 0.44895693 | 0.33462123 | 0.1143357 | 0.05570367 | 0.423293 |
| Derl3 | 0.11685641 | 0.23883525 | 0.12459649 | 0.11423876 | 9.70E-06 | 0.00025809 |
| Sipa1 | 0.34803123 | 0.46900658 | 0.35513134 | 0.11387524 | 0.06759818 | 0.4851675 |
| Tspan8 | 0.12576796 | 0.24760942 | 0.13390406 | 0.11370536 | 3.81E-05 | 0.00088581 |
| Hspa12b | 0.14782266 | 0.27051212 | 0.15681025 | 0.11370187 | 0.00036486 | 0.00676645 |
| Ctsk | 0.19404237 | 0.31773349 | 0.20418842 | 0.11354506 | 0.00478845 | 0.06359404 |
| Arap1 | 0.45030544 | 0.56207731 | 0.44862681 | 0.11345051 | 0.11936415 | 0.69138694 |
| Aldh2 | 0.85280398 | 0.86523674 | 0.75193986 | 0.11329687 | 0.24134717 | 0.89054584 |
| Pxdc1 | 0.10016188 | 0.22036627 | 0.10708071 | 0.11328556 | 4.13E-07 | 1.38E-05 |
| Rdx | 1.52690977 | 1.15586704 | 1.04283609 | 0.11303095 | 0.30678873 | 0.89054584 |
| Tssc4 | 0.8668395 | 0.87326559 | 0.76046461 | 0.11280098 | 0.24476125 | 0.89054584 |
| Zdbf2 | 0.59869024 | 0.68591648 | 0.57316595 | 0.11275054 | 0.17970623 | 0.85590699 |
| Atg101 | 0.48083769 | 0.58812054 | 0.4753715 | 0.11274904 | 0.13457723 | 0.73460117 |
| Nrp2 | 0.15415831 | 0.27604303 | 0.16335549 | 0.11268754 | 0.00065478 | 0.01149267 |
| Nek6 | 0.25191559 | 0.37489846 | 0.26222902 | 0.11266943 | 0.02265513 | 0.22058704 |
| Eef2 | 3.85757467 | 1.10315715 | 0.99089392 | 0.11226323 | 0.29880718 | 0.89054584 |
| Msl3 | 0.37960975 | 0.4968463 | 0.38461916 | 0.11222714 | 0.08700649 | 0.57045578 |
| Fam110b | 0.41159616 | 0.5260299 | 0.41393099 | 0.11209891 | 0.103526 | 0.63276555 |
| Ubald2 | 1.1879782 | 1.03733659 | 0.92528705 | 0.11204953 | 0.28631444 | 0.89054584 |
| Nt5c2 | 0.57222073 | 0.66383607 | 0.55197811 | 0.11185796 | 0.17255134 | 0.83703076 |
| Nrtn | 0.30798788 | 0.42876115 | 0.31698487 | 0.11177628 | 0.05020681 | 0.39431114 |
| Vimp | 1.44027623 | 1.12998507 | 1.01832375 | 0.11166131 | 0.30472364 | 0.89054584 |
| Lcp1 | 0.10494614 | 0.22375693 | 0.11211075 | 0.11164618 | 1.75E-06 | 5.24E-05 |
| Nme1 | 2.38480411 | 1.2409115 | 1.12939806 | 0.11151344 | 0.32275447 | 0.89054584 |
| Bcl2 | 0.329374 | 0.44891055 | 0.33746068 | 0.11144987 | 0.06194324 | 0.45632783 |
| Adgrb3 | 0.10982848 | 0.22842772 | 0.11723531 | 0.1111924 | 4.96E-06 | 0.00013783 |
| Nrarp | 0.14765708 | 0.26782423 | 0.15663899 | 0.11118524 | 0.0004715 | 0.00855546 |
| Gm9844 | 0.86774351 | 0.87204838 | 0.76100889 | 0.11103949 | 0.24831863 | 0.89054584 |

| | | | | | | |
|---|---|---|---|---|---|---|
| Nynrin | 0.21315437 | 0.33452892 | 0.22351907 | 0.11100986 | 0.01033812 | 0.12048402 |
| Phip | 1.44474076 | 1.13047953 | 1.01968436 | 0.11079516 | 0.30634989 | 0.89054584 |
| Limd2 | 1.05749222 | 0.97467488 | 0.86407353 | 0.11060136 | 0.27547266 | 0.89054584 |
| Zdhhc16 | 0.39959291 | 0.51356264 | 0.40299866 | 0.11056398 | 0.10059025 | 0.62288099 |
| Ehd1 | 0.59701706 | 0.68234019 | 0.57184051 | 0.11049968 | 0.18398799 | 0.86722786 |
| Coq2 | 0.81158624 | 0.8365728 | 0.72607443 | 0.11049837 | 0.23915427 | 0.89054584 |
| Rpa2 | 0.38095386 | 0.49631281 | 0.38586224 | 0.11045056 | 0.09116886 | 0.58666938 |
| Socs2 | 0.36134827 | 0.47806655 | 0.36763133 | 0.11043522 | 0.08082701 | 0.5452908 |
| Adamts2 | 0.16190107 | 0.28175758 | 0.17133324 | 0.11042434 | 0.00133803 | 0.02131727 |
| Creb5 | 0.18085365 | 0.30108335 | 0.19075805 | 0.1103253 | 0.00352463 | 0.04918201 |
| Cdca7 | 0.24504065 | 0.36572282 | 0.255413 | 0.11030982 | 0.02210253 | 0.21666161 |
| Tspo2 | 0.03586861 | 0.14857149 | 0.03838146 | 0.11019003 | 4.21E-41 | 1.22E-38 |
| Slc25a19 | 0.18180327 | 0.30181422 | 0.19172753 | 0.11008669 | 0.00373552 | 0.05172638 |
| Ythdc1 | 1.10208341 | 0.99567441 | 0.88617493 | 0.10949948 | 0.28241634 | 0.89054584 |
| Sdc4 | 0.2444302 | 0.3638364 | 0.25480671 | 0.10902969 | 0.02310213 | 0.22314101 |
| Fndc3a | 1.18044368 | 1.030987 | 0.92200088 | 0.10898612 | 0.29091445 | 0.89054584 |
| Slc9a3r1 | 0.40093813 | 0.51300891 | 0.40422797 | 0.10878094 | 0.10496501 | 0.63913594 |
| Ly86 | 0.05825726 | 0.17099741 | 0.06232591 | 0.1086715 | 2.27E-16 | 2.11E-14 |
| Smim15 | 0.50272662 | 0.60270401 | 0.49419769 | 0.10850632 | 0.1531729 | 0.78856652 |
| Ldb2 | 0.24852734 | 0.36733348 | 0.25887258 | 0.10846091 | 0.02547054 | 0.2394741 |
| Adamts9 | 0.26327801 | 0.3817935 | 0.27344633 | 0.10834717 | 0.03244579 | 0.28647936 |
| Pon2 | 0.30107119 | 0.41857607 | 0.31031381 | 0.10826226 | 0.05203749 | 0.40344362 |
| Becn1 | 0.7424114 | 0.78815858 | 0.67989875 | 0.10825983 | 0.2290921 | 0.89054584 |
| Edem2 | 0.77202762 | 0.80822885 | 0.70009051 | 0.10813834 | 0.23587465 | 0.89054584 |
| Cd82 | 0.36600332 | 0.47997899 | 0.37197893 | 0.10800006 | 0.08808002 | 0.57428752 |
| Ptms | 1.95093368 | 1.22202716 | 1.11417823 | 0.10784893 | 0.32600531 | 0.89054584 |
| Ndufb8 | 2.34568536 | 1.23718464 | 1.12949501 | 0.10768963 | 0.32844965 | 0.89054584 |
| Slc29a1 | 0.64903506 | 0.71980959 | 0.61217401 | 0.10763558 | 0.20634785 | 0.89054584 |
| Nap1l1 | 2.08395606 | 1.23007953 | 1.12258279 | 0.10749674 | 0.32775066 | 0.89054584 |
| Atp6v0d1 | 1.26087898 | 1.06284801 | 0.95552671 | 0.1073213 | 0.30039131 | 0.89054584 |

| | | | | | | |
|---|---|---|---|---|---|---|
| Clec4a2 | 0.04141412 | 0.15158107 | 0.0443137 | 0.10726737 | 8.46E-30 | 1.73E-27 |
| Smtnl2 | 0.25211366 | 0.36953795 | 0.26242506 | 0.10711289 | 0.02861045 | 0.25999793 |
| Wfdc15b | 0.20708889 | 0.32445744 | 0.21740092 | 0.10705651 | 0.01088834 | 0.12538461 |
| Naalad2 | 0.1557576 | 0.27201855 | 0.16500529 | 0.10701326 | 0.0012575 | 0.02018966 |
| Phlda1 | 0.57361015 | 0.66004913 | 0.55310163 | 0.10694749 | 0.18383381 | 0.86722786 |
| Vps8 | 0.26456771 | 0.38163004 | 0.27471579 | 0.10691425 | 0.03490259 | 0.30373211 |
| Pitx2 | 0.07212469 | 0.18395466 | 0.07714764 | 0.10680702 | 5.59E-11 | 3.01E-09 |
| Fhl1 | 0.15775044 | 0.27379922 | 0.16705964 | 0.10673958 | 0.00145665 | 0.02292991 |
| Pgf | 0.36930894 | 0.48178671 | 0.37505922 | 0.10672749 | 0.09246023 | 0.59287414 |
| Cpb1 | 0.07907294 | 0.19121912 | 0.08457053 | 0.10664859 | 2.11E-09 | 9.57E-08 |
| Acvrl1 | 0.20835224 | 0.32530994 | 0.21867658 | 0.10663337 | 0.01154803 | 0.13162217 |
| Odc1 | 0.67010049 | 0.73456483 | 0.62798394 | 0.10658089 | 0.21455619 | 0.89054584 |
| Fam162a | 0.94333657 | 0.91084859 | 0.80443759 | 0.106411 | 0.26885331 | 0.89054584 |
| Rsrc1 | 0.606 | 0.68533981 | 0.57893578 | 0.10640403 | 0.19592195 | 0.89054584 |
| Banf1 | 0.96112225 | 0.92044919 | 0.81408951 | 0.10635968 | 0.27136686 | 0.89054584 |
| Spc24 | 0.3477952 | 0.46124538 | 0.35490892 | 0.10633646 | 0.08137318 | 0.54770118 |
| Arhgdig | 0.14664142 | 0.26181463 | 0.15558821 | 0.10622642 | 0.00073422 | 0.01267228 |
| Rrm1 | 0.38045534 | 0.49159647 | 0.38540131 | 0.10619516 | 0.0996156 | 0.62085259 |
| Slc2a5 | 0.39456629 | 0.50447545 | 0.39839605 | 0.1060794 | 0.10739183 | 0.64845225 |
| Decr1 | 0.30533406 | 0.42040472 | 0.31442817 | 0.10597654 | 0.05817565 | 0.4365583 |
| Hoxa5 | 0.23408579 | 0.3504314 | 0.24450662 | 0.10592478 | 0.02177797 | 0.21405898 |
| Hmmr | 0.10582255 | 0.21881252 | 0.11303128 | 0.10578124 | 6.50E-06 | 0.00017714 |
| Celf4 | 0.31051069 | 0.4251314 | 0.31941214 | 0.10571926 | 0.0615344 | 0.45470217 |
| Rpl22l1 | 4.1403229 | 1.03649204 | 0.93077725 | 0.10571479 | 0.29835137 | 0.89054584 |
| Neurod2 | 0.1145018 | 0.22758474 | 0.12213227 | 0.10545247 | 2.88E-05 | 0.00068948 |
| Atp6v0e2 | 0.44997844 | 0.55365277 | 0.44833735 | 0.10531542 | 0.1368908 | 0.74192255 |
| Zadh2 | 0.27346184 | 0.38872585 | 0.28344861 | 0.10527724 | 0.04177911 | 0.34614674 |
| Irs4 | 0.06114774 | 0.17067282 | 0.06541599 | 0.10525683 | 3.28E-14 | 2.49E-12 |
| Anapc13 | 1.95932258 | 1.22007675 | 1.11483291 | 0.10524384 | 0.33003405 | 0.89054584 |
| Anapc11 | 1.61151416 | 1.16858908 | 1.06338693 | 0.10520215 | 0.32242992 | 0.89054584 |

| | | | | | | |
|---|---|---|---|---|---|---|
| Gfra3 | 0.41735441 | 0.52431757 | 0.41914666 | 0.10517091 | 0.12120053 | 0.69728784 |
| Nrep | 0.84431557 | 0.85183582 | 0.74671426 | 0.10512156 | 0.25595057 | 0.89054584 |
| Prrg2 | 0.80921913 | 0.82966498 | 0.72455156 | 0.10511341 | 0.24955522 | 0.89054584 |
| Dok4 | 0.21311571 | 0.32855243 | 0.22348011 | 0.10507232 | 0.0142452 | 0.15505027 |
| Gnl3 | 0.63025338 | 0.70280907 | 0.59782186 | 0.10498721 | 0.20662562 | 0.89054584 |
| Ndufb4 | 2.26061005 | 1.23355331 | 1.12857576 | 0.10497755 | 0.33237514 | 0.89054584 |
| Sptbn1 | 1.19298448 | 1.03241491 | 0.9274668 | 0.1049481 | 0.29903239 | 0.89054584 |
| Chd7 | 0.85570871 | 0.85865852 | 0.7537159 | 0.10494262 | 0.25827216 | 0.89054584 |
| Chd2 | 0.68399802 | 0.74317484 | 0.63824589 | 0.10492894 | 0.22185614 | 0.89054584 |
| Csf1r | 0.03988289 | 0.14749881 | 0.04267576 | 0.10482305 | 1.27E-30 | 2.66E-28 |
| Mad2l1bp | 0.2210292 | 0.33623713 | 0.23143795 | 0.10479918 | 0.01744423 | 0.18264153 |
| Rhd | 0.04182167 | 0.14947063 | 0.04474964 | 0.10472099 | 5.60E-28 | 1.04E-25 |
| Rbms1 | 0.81697145 | 0.83396116 | 0.72952426 | 0.1044369 | 0.25239801 | 0.89054584 |
| Jund | 1.47808491 | 1.13388546 | 1.02945596 | 0.1044295 | 0.31824581 | 0.89054584 |
| Cmtm7 | 0.33069007 | 0.44291781 | 0.33871298 | 0.10420482 | 0.07588318 | 0.52338458 |
| Gm21988 | 0.16713834 | 0.28090206 | 0.1767157 | 0.10418636 | 0.00300923 | 0.04311047 |
| 1110059G10 | 0.42573122 | 0.53079553 | 0.42670112 | 0.10409442 | 0.12786194 | 0.71595682 |
| Dhrs3 | 0.11712226 | 0.22893594 | 0.12487459 | 0.10406135 | 5.17E-05 | 0.00116776 |
| Camk2d | 0.26295904 | 0.37680443 | 0.27313223 | 0.10367219 | 0.03849837 | 0.32813014 |
| Gdi1 | 1.29689095 | 1.07305552 | 0.96939394 | 0.10366158 | 0.30917113 | 0.89054584 |
| Baiap3 | 0.74785569 | 0.7871771 | 0.6836577 | 0.10351939 | 0.2402631 | 0.89054584 |
| Clec11a | 0.20933659 | 0.32301436 | 0.21967002 | 0.10334434 | 0.01419657 | 0.15463706 |
| Mcm4 | 0.34040466 | 0.45115605 | 0.34792969 | 0.10322636 | 0.0834476 | 0.55675256 |
| Lrrc59 | 0.71853642 | 0.76637058 | 0.66316547 | 0.10320511 | 0.23421141 | 0.89054584 |
| Xlr4b | 0.10649102 | 0.21685793 | 0.11373322 | 0.10312471 | 1.20E-05 | 0.00031183 |
| Spon1 | 0.22358697 | 0.33705818 | 0.23400406 | 0.10305412 | 0.02009589 | 0.20204592 |
| Anxa11 | 0.37102784 | 0.47966581 | 0.37665857 | 0.10300723 | 0.10131212 | 0.62529723 |
| Rnf181 | 0.69954628 | 0.75234522 | 0.64956768 | 0.10277754 | 0.23051291 | 0.89054584 |
| Tpst1 | 0.65094846 | 0.71630674 | 0.61362246 | 0.10268428 | 0.2178036 | 0.89054584 |
| Lsg1 | 0.44103382 | 0.54287286 | 0.44039505 | 0.10247781 | 0.1391573 | 0.74916543 |

| | | | | | | |
|---|---|---|---|---|---|---|
| Ubxn8 | 0.21559771 | 0.32841163 | 0.22597902 | 0.10243261 | 0.01735085 | 0.18217078 |
| Pls3 | 0.41382311 | 0.5183455 | 0.41595014 | 0.10239536 | 0.1257111 | 0.70971942 |
| Eef1d | 2.30258886 | 1.23137898 | 1.12911856 | 0.10226042 | 0.33653279 | 0.89054584 |
| Urod | 0.77154633 | 0.80202452 | 0.69976741 | 0.10225711 | 0.24799531 | 0.89054584 |
| Pdia5 | 0.39326182 | 0.49932726 | 0.39719927 | 0.10212799 | 0.11547637 | 0.67725583 |
| Wasf3 | 0.04065514 | 0.14558764 | 0.04350184 | 0.1020858 | 4.00E-28 | 7.53E-26 |
| Ptger3 | 0.20884564 | 0.32121519 | 0.21917459 | 0.1020406 | 0.01503936 | 0.1619599 |
| Anpep | 0.24623581 | 0.35863251 | 0.25659951 | 0.102033 | 0.0319728 | 0.28281976 |
| Ang | 0.21955004 | 0.33184154 | 0.22995262 | 0.10188893 | 0.01949396 | 0.19777702 |
| Isoc1 | 0.51568005 | 0.60684391 | 0.50519926 | 0.10164465 | 0.17428525 | 0.84142109 |
| Pea15a | 0.76681795 | 0.7981998 | 0.69658421 | 0.10161559 | 0.24836869 | 0.89054584 |
| Il7r | 0.04046848 | 0.14452021 | 0.04330217 | 0.10121804 | 6.42E-28 | 1.18E-25 |
| Epha4 | 0.19504438 | 0.30628756 | 0.20520577 | 0.10108179 | 0.01086839 | 0.12527997 |
| P3h3 | 0.27468283 | 0.38566629 | 0.28464446 | 0.10102183 | 0.04911631 | 0.38865305 |
| Usp1 | 0.48801086 | 0.58258266 | 0.48157308 | 0.10100958 | 0.16423278 | 0.81857464 |
| Pdia2 | 0.05261566 | 0.15726815 | 0.05629379 | 0.10097436 | 3.22E-17 | 3.31E-15 |
| Ghr | 0.91526757 | 0.88972292 | 0.78878454 | 0.10093838 | 0.27552392 | 0.89054584 |
| Asb2 | 0.04051653 | 0.14422967 | 0.04335357 | 0.1008761 | 1.11E-27 | 2.00E-25 |
| Dap | 1.17492313 | 1.02033406 | 0.91958101 | 0.10075305 | 0.30486721 | 0.89054584 |
| S100a4 | 0.06295341 | 0.16802328 | 0.06734618 | 0.1006771 | 1.65E-12 | 1.08E-10 |
| Zim1 | 0.45847607 | 0.55643642 | 0.45583842 | 0.100598 | 0.15193453 | 0.78458375 |
| Rpl10a | 4.55600783 | 0.92593377 | 0.82558659 | 0.10034719 | 0.28560236 | 0.89054584 |
| Wls | 0.9115452 | 0.88699681 | 0.78666824 | 0.10032857 | 0.27619486 | 0.89054584 |
| Col26a1 | 0.17459207 | 0.28466504 | 0.18435669 | 0.10030836 | 0.00562354 | 0.07227842 |
| Xlr3a | 0.31537677 | 0.42436528 | 0.32408564 | 0.10027964 | 0.07470977 | 0.51900827 |
| 1810058I24R | 1.69440672 | 1.18063811 | 1.08040817 | 0.10022995 | 0.33279365 | 0.89054584 |
| Slc25a25 | 0.20508558 | 0.31560529 | 0.21537667 | 0.10022862 | 0.01507534 | 0.16213823 |
| Scfd1 | 0.85887574 | 0.85574783 | 0.75564524 | 0.10010259 | 0.26855671 | 0.89054584 |
| Cope | 2.23896848 | 1.22823771 | 1.12818876 | 0.10004894 | 0.3397448 | 0.89054584 |
| AI662270 | 0.06367899 | 0.16813153 | 0.06812176 | 0.10000978 | 3.96E-12 | 2.44E-10 |

| | | | | | | |
|---|---|---|---|---|---|---|
| Tie1 | 0.1043664 | 0.21117838 | 0.11150167 | 0.09967671 | 1.56E-05 | 0.00039484 |
| Tgfb2 | 0.19990773 | 0.30980963 | 0.21013739 | 0.09967224 | 0.01355861 | 0.14891872 |
| Klhl4 | 0.07133104 | 0.17577212 | 0.07629962 | 0.0994725 | 6.24E-10 | 3.01E-08 |
| Mdfi | 0.33734506 | 0.44444926 | 0.34503185 | 0.09941741 | 0.08972889 | 0.58049509 |
| Cav2 | 0.14421456 | 0.25238656 | 0.1530758 | 0.09931076 | 0.00125317 | 0.02014245 |
| Crtap | 0.37087526 | 0.47579232 | 0.37651667 | 0.09927566 | 0.10964525 | 0.65546703 |
| Ahi1 | 0.47606821 | 0.57044839 | 0.47123044 | 0.09921795 | 0.16331021 | 0.81700185 |
| Tspan6 | 0.5503308 | 0.63323289 | 0.53411048 | 0.09912241 | 0.19362221 | 0.88861501 |
| Abhd2 | 0.19277015 | 0.30196606 | 0.20289612 | 0.09906994 | 0.01145705 | 0.13110054 |
| Pdlim4 | 0.31317503 | 0.42104194 | 0.32197254 | 0.09906941 | 0.07585266 | 0.52338458 |
| Pamr1 | 0.24053624 | 0.34999905 | 0.25093524 | 0.09906382 | 0.03293925 | 0.29030594 |
| Smpd3 | 0.4303553 | 0.52986853 | 0.43085366 | 0.09901487 | 0.14215769 | 0.76136024 |
| Ap3b2 | 0.42406202 | 0.52404628 | 0.42519905 | 0.09884723 | 0.13938707 | 0.74952855 |
| Fmo2 | 0.15853592 | 0.26666406 | 0.16786893 | 0.09879513 | 0.0030542 | 0.04358243 |
| Cox6b2 | 0.29556642 | 0.4037745 | 0.30498723 | 0.09878727 | 0.06564171 | 0.47441352 |
| Myadm | 0.54690605 | 0.62985236 | 0.53128662 | 0.09856574 | 0.19370072 | 0.888694 |
| Try4 | 0.04214731 | 0.14322931 | 0.04509797 | 0.09813134 | 1.88E-24 | 2.90E-22 |
| Crlf3 | 0.27628271 | 0.38434045 | 0.28621031 | 0.09813013 | 0.05509212 | 0.4206472 |
| Gm16141 | 0.2209465 | 0.32948129 | 0.23135492 | 0.09812636 | 0.02407552 | 0.23009023 |
| Armt1 | 0.2187347 | 0.32718511 | 0.22913347 | 0.09805165 | 0.02309398 | 0.22314101 |
| Ptgr1 | 0.23103995 | 0.3391995 | 0.24146447 | 0.09773503 | 0.0296643 | 0.26745229 |
| Sct | 0.18355435 | 0.29122485 | 0.19351422 | 0.09771062 | 0.00932568 | 0.1108227 |
| Rab3il1 | 0.10684248 | 0.21170261 | 0.1141022 | 0.09760041 | 3.37E-05 | 0.00079359 |
| Pmf1 | 0.36295509 | 0.46670954 | 0.36913318 | 0.09757636 | 0.10906037 | 0.65364084 |
| Xab2 | 0.38872891 | 0.49039598 | 0.39303311 | 0.09736287 | 0.12422522 | 0.70545441 |
| Hps5 | 0.15795291 | 0.26439828 | 0.16726827 | 0.09713001 | 0.00341139 | 0.04792476 |
| Nucks1 | 0.95988098 | 0.91035697 | 0.81342337 | 0.0969336 | 0.28938092 | 0.89054584 |
| Hacd4 | 0.14303674 | 0.24854935 | 0.15185561 | 0.09669374 | 0.0015057 | 0.02354648 |
| Dnm3os | 0.20746286 | 0.31432405 | 0.2177786 | 0.09654545 | 0.01944266 | 0.19767162 |
| Thy1 | 0.12333363 | 0.22754208 | 0.13136452 | 0.09617756 | 0.00032361 | 0.0060853 |

| | | | | | | |
|---|---|---|---|---|---|---|
| Fam46a | 0.46229297 | 0.55536851 | 0.4591935 | 0.09617501 | 0.16458572 | 0.81899774 |
| Gnb4 | 0.30363411 | 0.40881907 | 0.31278855 | 0.09603052 | 0.07630672 | 0.5255557 |
| Tbcb | 1.49165773 | 1.12905487 | 1.03326805 | 0.09578682 | 0.3329082 | 0.89054584 |
| Rgmb | 0.1881508 | 0.29378916 | 0.19819802 | 0.09559114 | 0.0123196 | 0.13867203 |
| Man2a1 | 0.37563312 | 0.47640652 | 0.38093552 | 0.09547101 | 0.12147693 | 0.69832374 |
| Rab34 | 0.81911131 | 0.82633522 | 0.73088918 | 0.09544603 | 0.27145908 | 0.89054584 |
| Gnb2l1 | 3.2039559 | 1.17036808 | 1.07492415 | 0.09544393 | 0.33955662 | 0.89054584 |
| Tmem218 | 0.36112879 | 0.46279024 | 0.36742609 | 0.09536415 | 0.11329075 | 0.6704038 |
| Dnajb2 | 0.61383294 | 0.68017108 | 0.58507886 | 0.09509222 | 0.22446126 | 0.89054584 |
| Pclo | 0.57493196 | 0.64924547 | 0.55416956 | 0.0950759 | 0.21205546 | 0.89054584 |
| Plk1 | 0.1057723 | 0.20803585 | 0.11297851 | 0.09505734 | 4.43E-05 | 0.00101119 |
| Reg1 | 0.04907359 | 0.14747972 | 0.05250596 | 0.09497376 | 1.77E-17 | 1.87E-15 |
| Pgam1 | 2.2171219 | 1.22260245 | 1.12769223 | 0.09491022 | 0.3474879 | 0.89054584 |
| Srp54b | 1.65957809 | 1.16852639 | 1.07363548 | 0.09489091 | 0.34025391 | 0.89054584 |
| Tgm2 | 0.21664065 | 0.32167973 | 0.22702825 | 0.09465147 | 0.02604542 | 0.24285937 |
| Lyl1 | 0.06726124 | 0.16659837 | 0.07195049 | 0.09464787 | 4.43E-10 | 2.19E-08 |
| Gpc1 | 0.22378153 | 0.32875548 | 0.23419913 | 0.09455635 | 0.02998482 | 0.2691764 |
| 3830406C13I | 0.41201787 | 0.50886523 | 0.41431358 | 0.09455166 | 0.14383883 | 0.76611882 |
| Sp3os | 0.53864994 | 0.61898051 | 0.52444993 | 0.09453057 | 0.20052132 | 0.89054584 |
| Pcgf6 | 0.33790225 | 0.43983185 | 0.34555996 | 0.0942719 | 0.10186406 | 0.62763501 |
| Sult5a1 | 0.13164266 | 0.23426893 | 0.14002388 | 0.09424505 | 0.00085703 | 0.01451544 |
| Sox11 | 0.24708316 | 0.35156094 | 0.25744031 | 0.09412063 | 0.04425437 | 0.36105784 |
| Thbd | 0.1650878 | 0.26861657 | 0.17460965 | 0.09400693 | 0.00606574 | 0.07673749 |
| Tbc1d1 | 0.32181042 | 0.42413236 | 0.33024595 | 0.09388641 | 0.09266827 | 0.59368299 |
| Cd200 | 0.54949681 | 0.62728688 | 0.53342351 | 0.09386337 | 0.20616329 | 0.89054584 |
| Zfhx4 | 0.18228045 | 0.28594531 | 0.19221454 | 0.09373077 | 0.01154723 | 0.13162217 |
| Gins2 | 0.28053653 | 0.38404479 | 0.29036755 | 0.09367723 | 0.06641318 | 0.47760826 |
| Gnaz | 0.46094968 | 0.55162837 | 0.45801374 | 0.09361462 | 0.17048431 | 0.83412025 |
| Mvd | 0.36834955 | 0.4676521 | 0.37416585 | 0.09348625 | 0.1222023 | 0.70076598 |
| Maff | 0.2472218 | 0.3509567 | 0.25757785 | 0.09337885 | 0.04561062 | 0.36818637 |

| | | | | | | |
|---|---|---|---|---|---|---|
| Eef2k | 0.20368636 | 0.30731228 | 0.21396198 | 0.0933503 | 0.02104307 | 0.20845964 |
| Dusp23 | 0.26814059 | 0.37150353 | 0.27822846 | 0.09327507 | 0.0591566 | 0.44117919 |
| Eif4a2 | 2.54813466 | 1.21826328 | 1.12499792 | 0.09326536 | 0.34965992 | 0.89054584 |
| Ace2 | 0.13338551 | 0.23508874 | 0.14183693 | 0.09325181 | 0.00109544 | 0.01788478 |
| H3f3b | 4.70004075 | 0.88107464 | 0.78791287 | 0.09316176 | 0.29086244 | 0.89054584 |
| Creb3l1 | 0.36135245 | 0.46069305 | 0.36763523 | 0.09305782 | 0.11914008 | 0.6908542 |
| Mllt3 | 0.19433297 | 0.29749448 | 0.20448352 | 0.09301097 | 0.01703861 | 0.17965514 |
| Gm10941 | 0.28135161 | 0.38403862 | 0.29116311 | 0.09287551 | 0.0686245 | 0.49131753 |
| 1810037I17R | 2.03046538 | 1.21253551 | 1.11968955 | 0.09284596 | 0.34962806 | 0.89054584 |
| Srpx2 | 0.13855142 | 0.24004798 | 0.14720402 | 0.09284396 | 0.00164918 | 0.02543604 |
| Serpina3g | 0.04318303 | 0.13888693 | 0.04620581 | 0.09268111 | 4.59E-21 | 5.94E-19 |
| Cmc2 | 0.47019265 | 0.55878665 | 0.46610981 | 0.09267684 | 0.17713478 | 0.84867812 |
| Epn1 | 1.01678324 | 0.93580142 | 0.84315557 | 0.09264585 | 0.30435211 | 0.89054584 |
| Gng2 | 0.21299199 | 0.31588263 | 0.22335549 | 0.09252715 | 0.02680274 | 0.2484566 |
| Kank3 | 0.14758645 | 0.24907668 | 0.15656593 | 0.09251075 | 0.00295397 | 0.04245287 |
| Pdpn | 0.1305535 | 0.23132018 | 0.13889027 | 0.09242991 | 0.00096606 | 0.01597028 |
| Slc38a4 | 0.48533746 | 0.57142288 | 0.47926551 | 0.09215737 | 0.18516324 | 0.86933183 |
| Rbm38 | 0.09841849 | 0.19736741 | 0.10522249 | 0.09214492 | 2.25E-05 | 0.00055544 |
| C2cd4a | 0.11754481 | 0.21733226 | 0.12531655 | 0.09201571 | 0.00031207 | 0.00589877 |
| Ptpn18 | 0.22361276 | 0.32602744 | 0.23402991 | 0.09199753 | 0.03351767 | 0.2936188 |
| Zyx | 0.45142872 | 0.54149648 | 0.44962062 | 0.09187586 | 0.17054545 | 0.83412025 |
| Hpgd | 0.17876398 | 0.28048594 | 0.18862336 | 0.09186258 | 0.01163481 | 0.13240308 |
| Gucy1b3 | 0.11868413 | 0.21831633 | 0.12650787 | 0.09180846 | 0.00036104 | 0.0067107 |
| Nisch | 1.93682015 | 1.20478374 | 1.11302606 | 0.09175767 | 0.35045873 | 0.89054584 |
| Lxn | 0.33928834 | 0.43859709 | 0.34687297 | 0.09172413 | 0.10898002 | 0.65342944 |
| Stbd1 | 0.36140935 | 0.45918562 | 0.36768844 | 0.09149718 | 0.12315656 | 0.70193617 |
| Hmox1 | 0.12128495 | 0.22067023 | 0.12922558 | 0.09144466 | 0.00048891 | 0.00883811 |
| Lgals3bp | 0.48452212 | 0.56993532 | 0.47856087 | 0.09137446 | 0.18685131 | 0.87419127 |
| S100a1 | 0.68641202 | 0.73123354 | 0.64001488 | 0.09121866 | 0.25333891 | 0.89054584 |
| Palm | 0.36696934 | 0.46407438 | 0.37287972 | 0.09119465 | 0.12726182 | 0.71430193 |

| | | | | | | |
|---|---|---|---|---|---|---|
| Kdm6b | 0.69032366 | 0.73381449 | 0.64287263 | 0.09094186 | 0.25492806 | 0.89054584 |
| Bbs7 | 0.17089189 | 0.27149816 | 0.18056642 | 0.09093174 | 0.00948286 | 0.11223066 |
| Gm8797 | 2.42213068 | 1.21974464 | 1.12888085 | 0.09086378 | 0.35383066 | 0.89054584 |
| Grina | 0.60484151 | 0.668813 | 0.57802369 | 0.09078931 | 0.23215361 | 0.89054584 |
| Igtp | 0.08371157 | 0.18025802 | 0.08952454 | 0.09073348 | 1.17E-06 | 3.58E-05 |
| Dis3 | 0.18129747 | 0.28194425 | 0.1912112 | 0.09073305 | 0.01352661 | 0.14890122 |
| Prox1 | 0.60206057 | 0.66655666 | 0.57583053 | 0.09072613 | 0.23145949 | 0.89054584 |
| Specc1 | 0.13651038 | 0.23575204 | 0.14508473 | 0.09066731 | 0.00179851 | 0.0275341 |
| 1110032A03 | 0.53106563 | 0.60875642 | 0.51813066 | 0.09062576 | 0.20756732 | 0.89054584 |
| Rcsd1 | 0.16078689 | 0.26073543 | 0.17018674 | 0.09054869 | 0.00659103 | 0.0823055 |
| Mmd | 0.18942895 | 0.28998398 | 0.19949887 | 0.09048512 | 0.01729529 | 0.1818326 |
| Aes | 2.12040903 | 1.21484161 | 1.1243636 | 0.09047801 | 0.35386483 | 0.89054584 |
| Hells | 0.26255596 | 0.36321154 | 0.27273525 | 0.09047629 | 0.06110763 | 0.45270234 |
| Gmnn | 0.30340525 | 0.4030425 | 0.31256769 | 0.09047481 | 0.08873889 | 0.57718099 |
| Ush1c | 0.42162972 | 0.51344448 | 0.42300738 | 0.0904371 | 0.15961027 | 0.80704115 |
| Fez2 | 0.24441612 | 0.34520841 | 0.25479272 | 0.09041569 | 0.04913788 | 0.38865305 |
| Med9os | 0.07668938 | 0.17241824 | 0.08202444 | 0.0903938 | 1.42E-07 | 4.95E-06 |
| Macrod2 | 0.34611768 | 0.4437182 | 0.35332733 | 0.09039087 | 0.11665305 | 0.68192309 |
| P4hb | 1.51726609 | 1.13060616 | 1.0402534 | 0.09035276 | 0.34286399 | 0.89054584 |
| Cpsf3l | 0.51126501 | 0.59180602 | 0.50146143 | 0.09034459 | 0.20063172 | 0.89054584 |
| Tle6 | 0.40301847 | 0.49646094 | 0.40612702 | 0.09033392 | 0.15003627 | 0.77909236 |
| Gprasp2 | 0.48914294 | 0.5728476 | 0.48254891 | 0.09029868 | 0.19165303 | 0.88461814 |
| Atf5 | 0.28154155 | 0.38157389 | 0.29134846 | 0.09022544 | 0.07453964 | 0.51900827 |
| Dynlt1a | 0.45211266 | 0.54020483 | 0.45022537 | 0.08997946 | 0.17589851 | 0.8453343 |
| Mapk8ip2 | 0.32137182 | 0.41958417 | 0.32982666 | 0.08975751 | 0.10242348 | 0.62974371 |
| Samd1 | 0.13012764 | 0.22820412 | 0.13844689 | 0.08975723 | 0.00126194 | 0.02023858 |
| Lgi2 | 0.22899752 | 0.32905566 | 0.23942251 | 0.08963315 | 0.04056399 | 0.33994031 |
| Ptp4a3 | 0.52877399 | 0.60580218 | 0.51621397 | 0.08958821 | 0.20938624 | 0.89054584 |
| Rnf34 | 0.41757975 | 0.50893199 | 0.41935041 | 0.08958158 | 0.15980918 | 0.80728693 |
| Ywhah | 2.0452023 | 1.20985632 | 1.12051614 | 0.08934018 | 0.35514624 | 0.89054584 |

| | | | | | | |
|---|---|---|---|---|---|---|
| Tcp11 | 0.35277755 | 0.44889503 | 0.35959746 | 0.08929756 | 0.12364877 | 0.7035557 |
| Eif5 | 1.95628496 | 1.20379298 | 1.11459585 | 0.08919712 | 0.35463436 | 0.89054584 |
| H2-K1 | 0.61938688 | 0.67855585 | 0.58940935 | 0.08914651 | 0.24051349 | 0.89054584 |
| Ostc | 2.57923289 | 1.21251828 | 1.12355904 | 0.08895924 | 0.35610944 | 0.89054584 |
| Syp | 0.41873939 | 0.50924171 | 0.4203985 | 0.08884321 | 0.16241559 | 0.81501438 |
| 2310030G06 | 0.38904999 | 0.48212192 | 0.39332859 | 0.08879333 | 0.1464574 | 0.77188653 |
| Slc7a2 | 0.47994159 | 0.56330875 | 0.47459454 | 0.08871421 | 0.19191018 | 0.88486593 |
| Ttll7 | 0.61196728 | 0.67230351 | 0.58361947 | 0.08868404 | 0.23948798 | 0.89054584 |
| Degs1 | 1.11622597 | 0.98155194 | 0.89290532 | 0.08864662 | 0.32184909 | 0.89054584 |
| Pnmal2 | 0.84428084 | 0.83532085 | 0.74669277 | 0.08862808 | 0.29013541 | 0.89054584 |
| Rnf2 | 0.67645809 | 0.72118177 | 0.63269485 | 0.08848691 | 0.25733403 | 0.89054584 |
| Tmed6 | 0.25186319 | 0.3505967 | 0.26217716 | 0.08841954 | 0.05806426 | 0.43617392 |
| Akr1b3 | 0.78309884 | 0.79588251 | 0.70747645 | 0.08840606 | 0.28022273 | 0.89054584 |
| Camk2b | 0.53984951 | 0.61381932 | 0.52544601 | 0.0883733 | 0.21664495 | 0.89054584 |
| Gnptg | 0.73987069 | 0.76640718 | 0.67813724 | 0.08826994 | 0.2721138 | 0.89054584 |
| Nop58 | 1.08151122 | 0.96434214 | 0.87613263 | 0.08820951 | 0.31951168 | 0.89054584 |
| Ptges3l | 0.16852289 | 0.26632125 | 0.17813677 | 0.08818448 | 0.01054545 | 0.12270301 |
| Opn3 | 0.13541899 | 0.23211089 | 0.14395084 | 0.08816005 | 0.0021635 | 0.03217072 |
| Cd53 | 0.05925275 | 0.15147133 | 0.06339017 | 0.08808116 | 4.78E-11 | 2.59E-09 |
| Ermard | 0.19883473 | 0.29700437 | 0.20905021 | 0.08795417 | 0.02498814 | 0.23590763 |
| B3gat3 | 0.39431406 | 0.48611546 | 0.39816472 | 0.08795074 | 0.15171279 | 0.78420187 |
| Sep-09 | 0.3462368 | 0.44138596 | 0.35343968 | 0.08794627 | 0.12317048 | 0.70193617 |
| Rassf1 | 0.50838007 | 0.5869538 | 0.49901231 | 0.08794148 | 0.20580881 | 0.89054584 |
| Ctsc | 0.17459121 | 0.27203942 | 0.1843558 | 0.08768361 | 0.01334941 | 0.14807932 |
| Rbpjl | 0.06523227 | 0.15731086 | 0.06978198 | 0.08752888 | 2.55E-09 | 1.13E-07 |
| Kif5b | 2.07376376 | 1.20941056 | 1.12203723 | 0.08737333 | 0.35837978 | 0.89054584 |
| Ung | 0.18483058 | 0.28216556 | 0.1948156 | 0.08734995 | 0.01835652 | 0.18968837 |
| Pik3r1 | 1.04111676 | 0.94301948 | 0.85569927 | 0.0873202 | 0.31724251 | 0.89054584 |
| Aaed1 | 0.17458506 | 0.27164045 | 0.18434951 | 0.08729094 | 0.01369042 | 0.15025257 |
| Syce2 | 0.82521812 | 0.82192875 | 0.73476602 | 0.08716273 | 0.29024335 | 0.89054584 |

| | | | | | | |
|---|---|---|---|---|---|---|
| Efna1 | 0.13938385 | 0.23522659 | 0.14806791 | 0.08715868 | 0.00304924 | 0.04358243 |
| Prex2 | 0.09420967 | 0.18786714 | 0.10073154 | 0.0871356 | 2.79E-05 | 0.00066909 |
| Nes | 0.10131673 | 0.19534496 | 0.10829564 | 0.08704933 | 9.02E-05 | 0.00190412 |
| Ptpn11 | 0.60490609 | 0.66508317 | 0.57807456 | 0.08700861 | 0.24157717 | 0.89054584 |
| Shcbp1 | 0.12937666 | 0.22458066 | 0.13766488 | 0.08691578 | 0.00163335 | 0.02527254 |
| Hbq1b | 0.03310399 | 0.12233904 | 0.03542379 | 0.08691525 | 1.46E-30 | 3.03E-28 |
| Rab8b | 0.1461046 | 0.24186703 | 0.15503267 | 0.08683437 | 0.00453386 | 0.06063832 |
| Skp2 | 0.25660861 | 0.35337723 | 0.26686906 | 0.08650817 | 0.06549102 | 0.47418188 |
| Idh1 | 0.42498792 | 0.51249869 | 0.42603244 | 0.08646625 | 0.17218536 | 0.83696674 |
| Dnmt1 | 0.35439318 | 0.447277 | 0.36111494 | 0.08616206 | 0.13314997 | 0.73038525 |
| Slc48a1 | 0.47056211 | 0.55257289 | 0.46643242 | 0.08614047 | 0.19478139 | 0.88999075 |
| Rybp | 0.29477234 | 0.39029409 | 0.3042176 | 0.08607649 | 0.09371523 | 0.5972952 |
| Tcof1 | 0.33921474 | 0.4328591 | 0.34680327 | 0.08605582 | 0.12382617 | 0.70422347 |
| Cnrip1 | 0.28199227 | 0.37780535 | 0.2917882 | 0.08601715 | 0.08481033 | 0.56182274 |
| Cnih4 | 0.73614828 | 0.76153525 | 0.67554808 | 0.08598718 | 0.27658603 | 0.89054584 |
| Rem1 | 0.15131114 | 0.24635405 | 0.160416 | 0.08593805 | 0.00628212 | 0.07906091 |
| Procr | 0.16987131 | 0.26535356 | 0.17951999 | 0.08583356 | 0.01295457 | 0.14447334 |
| Rec114 | 0.03562594 | 0.12395077 | 0.03812186 | 0.08582892 | 4.84E-26 | 8.15E-24 |
| Rnft1 | 0.3882319 | 0.47826537 | 0.39257571 | 0.08568967 | 0.15459269 | 0.79309443 |
| Napsa | 0.04349683 | 0.13202037 | 0.04654146 | 0.08547892 | 5.81E-18 | 6.33E-16 |
| Acaa2 | 0.49139851 | 0.56986637 | 0.48449077 | 0.0853756 | 0.20582727 | 0.89054584 |
| Arid5b | 0.48642109 | 0.56556136 | 0.48020139 | 0.08535997 | 0.20378789 | 0.89054584 |
| Hace1 | 0.18122957 | 0.27648922 | 0.19114187 | 0.08534735 | 0.01875001 | 0.1925196 |
| Car8 | 0.22166793 | 0.31739266 | 0.23207903 | 0.08531364 | 0.04338805 | 0.35599321 |
| Smad5 | 0.4323601 | 0.51789196 | 0.43265012 | 0.08524183 | 0.17933129 | 0.85496384 |
| Spop | 1.10095156 | 0.9708118 | 0.88562978 | 0.08518202 | 0.32703752 | 0.89054584 |
| Mrps6 | 0.46073071 | 0.54300265 | 0.45782133 | 0.08518132 | 0.1930167 | 0.88724036 |
| Arsi | 0.1477203 | 0.2418566 | 0.15670438 | 0.08515221 | 0.00567652 | 0.07263736 |
| Egfl6 | 0.10084307 | 0.19279202 | 0.1077974 | 0.08499463 | 0.00011966 | 0.00244179 |
| Mycbp | 0.88623042 | 0.85692648 | 0.77200728 | 0.0849192 | 0.30415969 | 0.89054584 |

| | | | | | | |
|---|---|---|---|---|---|---|
| Gpsm1 | 0.40314613 | 0.49111202 | 0.40624348 | 0.08486854 | 0.16520165 | 0.82136266 |
| Arhgef40 | 0.2456635 | 0.34080075 | 0.25603142 | 0.08476933 | 0.06147283 | 0.45447887 |
| Anp32b | 2.2322894 | 1.21279379 | 1.12804997 | 0.08474382 | 0.36316969 | 0.89054584 |
| Nt5dc2 | 1.09482644 | 0.96735376 | 0.88266476 | 0.084689 | 0.32743348 | 0.89054584 |
| Ankrd44 | 0.6968965 | 0.73227565 | 0.64765027 | 0.08462538 | 0.2713412 | 0.89054584 |
| Lipa | 0.24010557 | 0.33507053 | 0.25050662 | 0.08456391 | 0.05789113 | 0.43561417 |
| Fcgr3 | 0.03042551 | 0.11697841 | 0.03255812 | 0.08442029 | 6.72E-34 | 1.65E-31 |
| Gm8730 | 4.39470984 | 0.94894273 | 0.86452436 | 0.08441836 | 0.32457892 | 0.89054584 |
| Acot7 | 0.21464259 | 0.30925971 | 0.22501772 | 0.08424199 | 0.04056112 | 0.33994031 |
| Lgals7 | 0.05198051 | 0.13962524 | 0.05561461 | 0.08401064 | 9.77E-13 | 6.53E-11 |
| Cpxm1 | 0.2634112 | 0.35753071 | 0.27357746 | 0.08395325 | 0.0764018 | 0.52596072 |
| Zfp334 | 0.15903499 | 0.25228104 | 0.16838299 | 0.08389805 | 0.01013395 | 0.11867689 |
| Qdpr | 0.94363763 | 0.88848655 | 0.80460236 | 0.08388418 | 0.31358284 | 0.89054584 |
| Cpn1 | 0.37816999 | 0.46713218 | 0.38328649 | 0.08384569 | 0.15406234 | 0.79121355 |
| Pxdn | 0.26531103 | 0.35926557 | 0.27544709 | 0.08381848 | 0.07813706 | 0.53336934 |
| Rai2 | 0.39459869 | 0.48215171 | 0.39842576 | 0.08372596 | 0.16378039 | 0.81785018 |
| Snx8 | 0.21956448 | 0.31368713 | 0.22996713 | 0.08372001 | 0.04493374 | 0.36388968 |
| Mcm5 | 0.29153278 | 0.3847938 | 0.3010745 | 0.0837193 | 0.09757317 | 0.61424915 |
| Pbx1 | 1.15786136 | 0.9954654 | 0.91197681 | 0.08348859 | 0.33486765 | 0.89054584 |
| Rgs1 | 0.0370217 | 0.12306531 | 0.03961504 | 0.08345027 | 4.89E-23 | 7.02E-21 |
| Gspt1 | 1.10142222 | 0.96930465 | 0.88585657 | 0.08344808 | 0.33037631 | 0.89054584 |
| Ppa2 | 0.69075206 | 0.72659495 | 0.64318496 | 0.08340999 | 0.27286057 | 0.89054584 |
| Gm27033 | 0.41752905 | 0.50259209 | 0.41930457 | 0.08328752 | 0.17737446 | 0.84926515 |
| Tgfb3 | 0.12530342 | 0.2166663 | 0.13341962 | 0.08324668 | 0.00182498 | 0.02788044 |
| Cited4 | 0.19244558 | 0.28576538 | 0.20256632 | 0.08319906 | 0.02783923 | 0.25472969 |
| Zcchc24 | 0.15471912 | 0.24712071 | 0.16393413 | 0.08318659 | 0.00903626 | 0.10809216 |
| Smagp | 0.10028352 | 0.19022532 | 0.1072087 | 0.08301662 | 0.0001545 | 0.00309388 |
| Naga | 0.46911749 | 0.5481644 | 0.4651705 | 0.0829939 | 0.20292185 | 0.89054584 |
| Fbxo34 | 0.21381562 | 0.30715524 | 0.22418507 | 0.08297017 | 0.04233054 | 0.3494921 |
| Creld1 | 0.40474996 | 0.49066454 | 0.40770574 | 0.0829588 | 0.17156777 | 0.83588128 |

| | | | | | | |
|---|---|---|---|---|---|---|
| Smarcd2 | 0.84590199 | 0.83059451 | 0.7476949 | 0.08289961 | 0.30273396 | 0.89054584 |
| Armcx2 | 0.73932749 | 0.76060581 | 0.67776004 | 0.08284577 | 0.28451383 | 0.89054584 |
| Decr2 | 0.27186528 | 0.36463276 | 0.28188382 | 0.08274894 | 0.0857086 | 0.56636427 |
| Ptx3 | 0.06383674 | 0.15101506 | 0.06829037 | 0.0827247 | 8.32E-09 | 3.43E-07 |
| Stom | 0.15596072 | 0.24793267 | 0.16521475 | 0.08271792 | 0.0098349 | 0.11592383 |
| Rgs7bp | 0.1042963 | 0.19398624 | 0.11142801 | 0.08255823 | 0.00027837 | 0.00532351 |
| Rbmx | 0.64193057 | 0.6893164 | 0.60677367 | 0.08254272 | 0.26311184 | 0.89054584 |
| Lst1 | 0.12804643 | 0.21882071 | 0.13627913 | 0.08254158 | 0.00238753 | 0.03523452 |
| Dnmt3a | 1.2937377 | 1.05067223 | 0.96820488 | 0.08246736 | 0.34574724 | 0.89054584 |
| Gpihbp1 | 0.06576248 | 0.15278731 | 0.07034867 | 0.08243863 | 2.39E-08 | 9.27E-07 |
| Kif20a | 0.12458474 | 0.21510217 | 0.13266999 | 0.08243217 | 0.00189729 | 0.02887441 |
| Scpep1 | 0.89027816 | 0.85673509 | 0.77438179 | 0.0823533 | 0.31013591 | 0.89054584 |
| Meox1 | 0.14107204 | 0.2321596 | 0.14981905 | 0.08234055 | 0.00522527 | 0.06800354 |
| Nudt4 | 0.42480095 | 0.50815292 | 0.42586419 | 0.08228872 | 0.1839973 | 0.86722786 |
| Ulk1 | 0.18962333 | 0.28194606 | 0.19969664 | 0.08224942 | 0.02749935 | 0.25297304 |
| Dusp6 | 0.25479642 | 0.34724284 | 0.26507857 | 0.08216427 | 0.07435619 | 0.51877575 |
| St6galnac3 | 0.19244347 | 0.28472316 | 0.20256418 | 0.08215898 | 0.0294029 | 0.26543163 |
| Rps7 | 5.09761489 | 0.76561435 | 0.68351748 | 0.08209687 | 0.2878821 | 0.89054584 |
| Clec4n | 0.03818096 | 0.12293035 | 0.04085517 | 0.08207518 | 4.01E-21 | 5.23E-19 |
| Ptprn | 0.17202552 | 0.26375276 | 0.18172824 | 0.08202452 | 0.0177394 | 0.18449484 |
| Sgol2a | 0.0905174 | 0.17878286 | 0.09679075 | 0.08199211 | 3.96E-05 | 0.00091638 |
| Bmp1 | 0.45480093 | 0.53454896 | 0.45259963 | 0.08194933 | 0.19945632 | 0.89054584 |
| Gstt1 | 0.2529802 | 0.34522507 | 0.26328252 | 0.08194255 | 0.07352616 | 0.51488639 |
| Dhrs11 | 0.14093013 | 0.23150499 | 0.14967189 | 0.0818331 | 0.00542838 | 0.07014319 |
| Srpk2 | 0.84023692 | 0.826013 | 0.74418454 | 0.08182846 | 0.30422607 | 0.89054584 |
| Acsl5 | 0.32743844 | 0.41739258 | 0.33561724 | 0.08177534 | 0.12814962 | 0.71651107 |
| Acot13 | 0.77941009 | 0.78664323 | 0.70502546 | 0.08161777 | 0.2948242 | 0.89054584 |
| Adam12 | 0.18915361 | 0.28082286 | 0.1992187 | 0.08160415 | 0.02816977 | 0.2571826 |
| Adgrg1 | 0.51632091 | 0.5873232 | 0.50574079 | 0.08158241 | 0.22616196 | 0.89054584 |
| Nav2 | 0.91668878 | 0.87116734 | 0.78958985 | 0.08157748 | 0.31513585 | 0.89054584 |

| | | | | | | |
|---|---|---|---|---|---|---|
| Lag3 | 0.0839391 | 0.17132446 | 0.08976752 | 0.08155694 | 1.15E-05 | 0.00030198 |
| Nop56 | 1.08461965 | 0.95921593 | 0.87766817 | 0.08154775 | 0.33255048 | 0.89054584 |
| Zfp428 | 0.42210517 | 0.50497852 | 0.42343606 | 0.08154246 | 0.18480945 | 0.8690805 |
| Mar-02 | 0.36420768 | 0.45184511 | 0.37030318 | 0.08154193 | 0.15246615 | 0.7857996 |
| Ehd4 | 0.26336157 | 0.35506465 | 0.27352859 | 0.08153605 | 0.08244948 | 0.55212586 |
| Nrros | 0.09536795 | 0.18344362 | 0.10196758 | 0.08147604 | 9.86E-05 | 0.00205604 |
| Mmrn2 | 0.09657483 | 0.18459588 | 0.1032554 | 0.08134049 | 0.00012121 | 0.00246821 |
| Rps19 | 5.48281225 | 0.7524913 | 0.67135017 | 0.08114113 | 0.2866862 | 0.89054584 |
| Lmcd1 | 0.13200017 | 0.22152152 | 0.14039589 | 0.08112563 | 0.0035502 | 0.0494436 |
| Atad3a | 0.34164631 | 0.43022907 | 0.34910428 | 0.08112479 | 0.13948324 | 0.74952855 |
| Gpsm2 | 0.21777324 | 0.30923451 | 0.22816712 | 0.08106739 | 0.04893042 | 0.38764659 |
| Pnmal1 | 0.35811985 | 0.44564866 | 0.36460981 | 0.08103884 | 0.15021662 | 0.77912292 |
| Cited2 | 0.6904428 | 0.7239074 | 0.6429595 | 0.08094789 | 0.27875112 | 0.89054584 |
| 4921524J17F | 0.90384468 | 0.86310792 | 0.78225782 | 0.08085011 | 0.31507208 | 0.89054584 |
| Zfp219 | 0.26973564 | 0.36042631 | 0.27979464 | 0.08063167 | 0.08969732 | 0.58049509 |
| Pou3f4 | 0.21005699 | 0.30100881 | 0.22039682 | 0.08061199 | 0.04418616 | 0.36077541 |
| Ddr2 | 0.16575855 | 0.25577283 | 0.17529875 | 0.08047408 | 0.01622709 | 0.17285842 |
| Rhob | 0.86208243 | 0.83806104 | 0.75759122 | 0.08046983 | 0.31034826 | 0.89054584 |
| Ckap2l | 0.09989202 | 0.18722218 | 0.1067945 | 0.08042768 | 0.00022517 | 0.00438266 |
| Cd37 | 0.049 | 0.13283169 | 0.05242726 | 0.08040443 | 4.50E-13 | 3.07E-11 |
| Fam216a | 1.00679061 | 0.91842344 | 0.83803161 | 0.08039184 | 0.32746307 | 0.89054584 |
| Sema3c | 0.25060134 | 0.34130431 | 0.26092775 | 0.08037656 | 0.07562261 | 0.52298621 |
| Cd55 | 0.15098347 | 0.24043565 | 0.16007752 | 0.08035813 | 0.00967378 | 0.11417597 |
| Ssx2ip | 0.17052296 | 0.26054036 | 0.18018819 | 0.08035217 | 0.01887386 | 0.19341421 |
| Hspa1a | 0.17697711 | 0.26708599 | 0.18679679 | 0.0802892 | 0.02261489 | 0.22034316 |
| Armcx1 | 0.5777691 | 0.63669673 | 0.55645786 | 0.08023886 | 0.25085566 | 0.89054584 |
| Sil1 | 0.42065213 | 0.50234123 | 0.42212552 | 0.08021571 | 0.18798941 | 0.87607539 |
| Ppp1r14b | 1.01330948 | 0.92153639 | 0.84138237 | 0.08015402 | 0.32858184 | 0.89054584 |
| Lfng | 0.1976778 | 0.2879542 | 0.20787742 | 0.08007678 | 0.03635251 | 0.31315434 |
| Mzt2 | 0.45394761 | 0.53178565 | 0.45184646 | 0.0799392 | 0.20490028 | 0.89054584 |

| | | | | | | |
|---|---|---|---|---|---|---|
| Prdm4 | 0.19835729 | 0.28845472 | 0.20856629 | 0.07988843 | 0.03716775 | 0.3190397 |
| Ramp3 | 0.14069547 | 0.22930129 | 0.14942854 | 0.07987275 | 0.0063817 | 0.08003619 |
| Dlat | 0.60739659 | 0.65989395 | 0.58003412 | 0.07985983 | 0.26061553 | 0.89054584 |
| Fndc4 | 0.21715368 | 0.30730532 | 0.2275442 | 0.07976112 | 0.05122402 | 0.39948674 |
| Tbxa2r | 0.18875498 | 0.27853928 | 0.19881302 | 0.07972626 | 0.03086098 | 0.2748295 |
| Zfp422 | 0.69175922 | 0.72362603 | 0.64391872 | 0.07970732 | 0.28206908 | 0.89054584 |
| Hk2 | 0.14062569 | 0.22901937 | 0.14935616 | 0.0796632 | 0.00647805 | 0.08096439 |
| Snap25 | 0.55438579 | 0.61708813 | 0.5374441 | 0.07964403 | 0.24496816 | 0.89054584 |
| Fgl2 | 0.03673616 | 0.11894619 | 0.03930957 | 0.07963662 | 1.89E-21 | 2.56E-19 |
| Klf3 | 0.57910839 | 0.63706177 | 0.55753621 | 0.07952556 | 0.2531725 | 0.89054584 |
| Sox13 | 0.08945417 | 0.17516517 | 0.0956558 | 0.07950937 | 5.39E-05 | 0.00121245 |
| Trp53 | 0.93603607 | 0.87992113 | 0.80042162 | 0.07949951 | 0.32177697 | 0.89054584 |
| Gm12216 | 0.04815217 | 0.13099795 | 0.05152054 | 0.07947741 | 3.31E-13 | 2.32E-11 |
| Dapp1 | 0.15522063 | 0.2437063 | 0.16445148 | 0.07925482 | 0.01237431 | 0.13885664 |
| Atp8b1 | 0.67061301 | 0.70757882 | 0.62836476 | 0.07921406 | 0.2784931 | 0.89054584 |
| Plp1 | 0.1050511 | 0.1913938 | 0.112221 | 0.0791728 | 0.00050645 | 0.00913246 |
| Pcdh17 | 0.10103598 | 0.18710461 | 0.10800033 | 0.07910427 | 0.00032195 | 0.00606182 |
| Maz | 0.62086092 | 0.66952799 | 0.59055515 | 0.07897284 | 0.26663449 | 0.89054584 |
| Ocrl | 0.68662954 | 0.7190389 | 0.64017408 | 0.07886482 | 0.28300154 | 0.89054584 |
| Xpot | 0.27270402 | 0.3615377 | 0.28270602 | 0.07883167 | 0.09694951 | 0.61165099 |
| Slfn2 | 0.04229726 | 0.12394968 | 0.04525836 | 0.07869132 | 2.74E-16 | 2.49E-14 |
| Spon2 | 0.1286718 | 0.21553395 | 0.13693068 | 0.07860327 | 0.00374311 | 0.05178201 |
| Fam189b | 0.27600677 | 0.36439263 | 0.28594033 | 0.0784523 | 0.10057976 | 0.62288099 |
| Tmed10 | 2.67953141 | 1.19669437 | 1.1182593 | 0.07843507 | 0.37191728 | 0.89054584 |
| Col9a2 | 0.66185679 | 0.70012428 | 0.62183335 | 0.07829093 | 0.27874407 | 0.89054584 |
| Chic1 | 0.53816433 | 0.60232561 | 0.52404644 | 0.07827917 | 0.24323728 | 0.89054584 |
| Cat | 0.48266531 | 0.55523306 | 0.4769546 | 0.07827846 | 0.22224537 | 0.89054584 |
| Ezr | 0.61532079 | 0.66446662 | 0.58624102 | 0.0782256 | 0.26707943 | 0.89054584 |
| Dnajc10 | 0.96901098 | 0.89651911 | 0.81829671 | 0.07822241 | 0.32803115 | 0.89054584 |
| Pvrl2 | 0.75350802 | 0.76575677 | 0.6875377 | 0.07821907 | 0.29804434 | 0.89054584 |

| | | | | | | |
|---|---|---|---|---|---|---|
| Ptma | 5.01033117 | 0.77705788 | 0.69889784 | 0.07816005 | 0.30117477 | 0.89054584 |
| Gyltl1b | 0.14146779 | 0.22838875 | 0.1502294 | 0.07815935 | 0.00767759 | 0.09393219 |
| Slc1a5 | 0.14346407 | 0.2304431 | 0.15229837 | 0.07814472 | 0.00841236 | 0.10146624 |
| Cbfa2t3 | 0.48431956 | 0.55647722 | 0.47838574 | 0.07809147 | 0.22347087 | 0.89054584 |
| Pi15 | 0.15502579 | 0.24219152 | 0.1642505 | 0.07794103 | 0.01352534 | 0.14890122 |
| Mis12 | 0.42447332 | 0.50349007 | 0.42556932 | 0.07792075 | 0.19681821 | 0.89054584 |
| Rqcd1 | 0.2255044 | 0.31379568 | 0.2359258 | 0.07786988 | 0.0620568 | 0.45670026 |
| Zfp874a | 0.13583746 | 0.22224078 | 0.14438566 | 0.07785513 | 0.00599943 | 0.07600027 |
| Ddit4 | 0.44442719 | 0.52125742 | 0.4434137 | 0.07784373 | 0.20670632 | 0.89054584 |
| Rps16 | 5.02415348 | 0.77355502 | 0.69583782 | 0.0777172 | 0.30140949 | 0.89054584 |
| Galm | 0.12875646 | 0.21471285 | 0.13701888 | 0.07769397 | 0.00412377 | 0.05592741 |
| Ip6k2 | 0.9234963 | 0.87107501 | 0.79343213 | 0.07764288 | 0.32422737 | 0.89054584 |
| Pacsin3 | 0.19733656 | 0.28511268 | 0.20753139 | 0.07758129 | 0.04078502 | 0.34061128 |
| Cdc14b | 0.25537245 | 0.34319019 | 0.26564786 | 0.07754233 | 0.08692446 | 0.57045578 |
| Krtdap | 0.044948 | 0.12551954 | 0.04809361 | 0.07742593 | 3.18E-14 | 2.43E-12 |
| Arnt | 0.31540296 | 0.40153088 | 0.32411076 | 0.07742012 | 0.13288037 | 0.72977796 |
| Tff2 | 0.13430347 | 0.22013812 | 0.14279139 | 0.07734673 | 0.00580736 | 0.07398521 |
| Cdh2 | 0.27323592 | 0.36051784 | 0.28322726 | 0.07729058 | 0.101795 | 0.62747616 |
| Mboat1 | 0.09333606 | 0.17700415 | 0.09979921 | 0.07720494 | 0.00015658 | 0.0031311 |
| Gpsm3 | 0.10500938 | 0.18937903 | 0.11217718 | 0.07720185 | 0.00067216 | 0.01178354 |
| Tspan14 | 0.25055908 | 0.33804091 | 0.26088589 | 0.07715502 | 0.08412535 | 0.55921567 |
| Fgfr1 | 0.29676235 | 0.38320509 | 0.30614574 | 0.07705935 | 0.12045946 | 0.69516408 |
| Dpy30 | 1.64405536 | 1.14745655 | 1.07044215 | 0.0770144 | 0.36873945 | 0.89054584 |
| Psmd10 | 0.52475584 | 0.58984128 | 0.51284512 | 0.07699617 | 0.24212867 | 0.89054584 |
| Dpp8 | 0.19710511 | 0.28428212 | 0.20729667 | 0.07698545 | 0.04179469 | 0.34614674 |
| Dctn3 | 1.85920436 | 1.18222964 | 1.1053012 | 0.07692844 | 0.37287056 | 0.89054584 |
| Cygb | 0.18504574 | 0.27188836 | 0.19503494 | 0.07685342 | 0.03318966 | 0.29198044 |
| Syngr2 | 0.61802911 | 0.66515923 | 0.58835262 | 0.07680661 | 0.27152661 | 0.89054584 |
| Kitl | 0.15943331 | 0.24540548 | 0.16879321 | 0.07661228 | 0.01723153 | 0.18129371 |
| Canx | 2.10288354 | 1.20010126 | 1.12353173 | 0.07656953 | 0.3754282 | 0.89054584 |

| | | | | | | |
|---|---|---|---|---|---|---|
| Ppih | 0.40512448 | 0.48456818 | 0.40804699 | 0.07652119 | 0.19114278 | 0.88366966 |
| Guf1 | 0.53060701 | 0.59415687 | 0.51774735 | 0.07640952 | 0.24586187 | 0.89054584 |
| Fbxo21 | 0.59316105 | 0.64514526 | 0.56877873 | 0.07636653 | 0.2658122 | 0.89054584 |
| Ms4a6c | 0.04750518 | 0.12711965 | 0.0508286 | 0.07629105 | 1.35E-12 | 8.88E-11 |
| Tmem158 | 0.21038259 | 0.2969943 | 0.22072522 | 0.07626907 | 0.05371598 | 0.41314285 |
| 8430408G22 | 0.08958307 | 0.17202208 | 0.0957934 | 0.07622869 | 0.0001047 | 0.00217011 |
| Med30 | 0.79430935 | 0.79098264 | 0.71486568 | 0.07611696 | 0.30992201 | 0.89054584 |
| Sgce | 0.57604079 | 0.63116105 | 0.55506453 | 0.07609652 | 0.2615035 | 0.89054584 |
| Angptl1 | 0.05111995 | 0.13074336 | 0.05469436 | 0.076049 | 4.65E-11 | 2.53E-09 |
| Dqx1 | 0.03323214 | 0.11154037 | 0.03556089 | 0.07597948 | 1.21E-23 | 1.77E-21 |
| Erp27 | 0.06483139 | 0.14531096 | 0.06935351 | 0.07595745 | 1.68E-07 | 5.81E-06 |
| Rad23a | 0.85694877 | 0.83037489 | 0.75447221 | 0.07590268 | 0.31963967 | 0.89054584 |
| 2210016L21F | 0.85739019 | 0.83059356 | 0.75474116 | 0.0758524 | 0.31981034 | 0.89054584 |
| Ubxn2a | 0.48326938 | 0.55332691 | 0.4774774 | 0.07584952 | 0.2296213 | 0.89054584 |
| Atp1a1 | 1.52612307 | 1.11840947 | 1.04262725 | 0.07578222 | 0.36744422 | 0.89054584 |
| Vps26b | 0.30793912 | 0.39270701 | 0.31693793 | 0.07576908 | 0.1326837 | 0.72977796 |
| Ppp1r15a | 0.52333174 | 0.58738954 | 0.51164866 | 0.07574089 | 0.24519865 | 0.89054584 |
| Comt | 0.97115773 | 0.89509395 | 0.81943568 | 0.07565827 | 0.33354007 | 0.89054584 |
| Ly6a | 0.06781304 | 0.14817629 | 0.07254022 | 0.07563607 | 5.94E-07 | 1.93E-05 |
| Snhg6 | 0.80386767 | 0.79669872 | 0.72109375 | 0.07560497 | 0.31260532 | 0.89054584 |
| Slitrk4 | 0.12732993 | 0.21110461 | 0.13553246 | 0.07557216 | 0.00469115 | 0.06245396 |
| Nme4 | 0.34405415 | 0.42683353 | 0.35137972 | 0.0754538 | 0.15854846 | 0.80483939 |
| Entpd2 | 0.15082184 | 0.23526939 | 0.15991054 | 0.07535885 | 0.01406189 | 0.15374752 |
| Bmp2k | 0.11087703 | 0.19366517 | 0.11833474 | 0.07533043 | 0.00150936 | 0.02354648 |
| AY036118 | 0.89267037 | 0.85095937 | 0.77578013 | 0.07517924 | 0.32581883 | 0.89054584 |
| B3gnt9 | 0.19609861 | 0.28136524 | 0.20627567 | 0.07508957 | 0.04494552 | 0.36388968 |
| Fbln5 | 0.16150298 | 0.24600979 | 0.17092366 | 0.07508612 | 0.02034597 | 0.20371258 |
| 1110034G24 | 0.1893458 | 0.27448986 | 0.19941427 | 0.0750756 | 0.03971518 | 0.33573803 |
| Lyplal1 | 0.65567223 | 0.69219787 | 0.61718796 | 0.07500991 | 0.28562068 | 0.89054584 |
| Xlr4a | 0.06905969 | 0.14882784 | 0.07387248 | 0.07495536 | 1.14E-06 | 3.50E-05 |

| | | | | | | |
|---|---|---|---|---|---|---|
| Lpar6 | 0.48225314 | 0.55149029 | 0.47659775 | 0.07489254 | 0.23205203 | 0.89054584 |
| Gpr119 | 0.1139368 | 0.1964187 | 0.12154067 | 0.07487804 | 0.00205072 | 0.0308097 |
| Hspd1 | 2.15969875 | 1.20076716 | 1.1259701 | 0.07479706 | 0.37847488 | 0.89054584 |
| N4bp3 | 0.11330263 | 0.19560759 | 0.12087649 | 0.07473111 | 0.0019861 | 0.0299631 |
| Dnajc12 | 0.36458638 | 0.44526932 | 0.37065674 | 0.07461258 | 0.17416407 | 0.84132624 |
| Ncapd2 | 0.17040894 | 0.25464256 | 0.18007129 | 0.07457127 | 0.02684282 | 0.24851031 |
| Prss1 | 0.04803639 | 0.12595675 | 0.05139672 | 0.07456003 | 6.97E-12 | 4.21E-10 |
| Cacna1a | 0.55976835 | 0.61633321 | 0.5418525 | 0.07448072 | 0.26095692 | 0.89054584 |
| Prkab1 | 0.31571268 | 0.39880559 | 0.32440781 | 0.07439778 | 0.14265784 | 0.76319314 |
| Fdps | 0.81781165 | 0.80439211 | 0.73006058 | 0.07433152 | 0.31762421 | 0.89054584 |
| Grrp1 | 0.08467315 | 0.16483795 | 0.09055134 | 0.07428661 | 6.62E-05 | 0.00144888 |
| Lta | 0.04214986 | 0.11937095 | 0.04510069 | 0.07427026 | 8.45E-15 | 6.73E-13 |
| BC023829 | 0.60677747 | 0.65365204 | 0.57954738 | 0.07410465 | 0.27568042 | 0.89054584 |
| Lgr4 | 0.09358218 | 0.17414247 | 0.10006188 | 0.07408059 | 0.00028109 | 0.00535506 |
| Slc24a5 | 0.44046648 | 0.51391413 | 0.43988983 | 0.0740243 | 0.21651922 | 0.89054584 |
| Rps23 | 5.27497394 | 0.74214569 | 0.66812549 | 0.07402021 | 0.30287335 | 0.89054584 |
| Palld | 0.12301399 | 0.20503308 | 0.1310309 | 0.07400217 | 0.00425389 | 0.05737009 |
| Pole2 | 0.11860526 | 0.20042668 | 0.12642542 | 0.07400126 | 0.00319524 | 0.04523882 |
| Mpst | 0.46086324 | 0.53186904 | 0.45793779 | 0.07393125 | 0.22597818 | 0.89054584 |
| Cyp39a1 | 0.09984278 | 0.18064215 | 0.10674197 | 0.07390017 | 0.00062876 | 0.01107621 |
| Ndufs5 | 2.29826895 | 1.20285924 | 1.12905741 | 0.07380182 | 0.38035988 | 0.89054584 |
| Usp18 | 0.19214136 | 0.2760519 | 0.20225715 | 0.07379475 | 0.04458023 | 0.36228932 |
| Sbspon | 0.11133514 | 0.19258229 | 0.11881495 | 0.07376734 | 0.00191067 | 0.02894551 |
| Pja2 | 1.0548642 | 0.93648423 | 0.86273106 | 0.07375317 | 0.34521056 | 0.89054584 |
| Pgm1 | 0.30122134 | 0.38412993 | 0.31045889 | 0.07367104 | 0.13445981 | 0.73423666 |
| 2700094K13I | 1.15414288 | 0.98393279 | 0.91031847 | 0.07361431 | 0.35317877 | 0.89054584 |
| Tox | 0.52956284 | 0.59048701 | 0.51687414 | 0.07361288 | 0.25349771 | 0.89054584 |
| Cxadr | 0.29760335 | 0.38052929 | 0.30696 | 0.0735693 | 0.13208026 | 0.72837567 |
| Mndal | 0.0881129 | 0.16779207 | 0.09422394 | 0.07356813 | 0.00013759 | 0.00277828 |
| Krtap17-1 | 0.05216378 | 0.12935255 | 0.05581058 | 0.07354198 | 4.15E-10 | 2.07E-08 |

| | | | | | | |
|---|---|---|---|---|---|---|
| Tmsb15b1 | 0.16520034 | 0.24825933 | 0.17472529 | 0.07353405 | 0.0249547 | 0.23584958 |
| Mad2l1 | 0.18040314 | 0.26382652 | 0.19029799 | 0.07352854 | 0.03591839 | 0.30996718 |
| Tcea3 | 0.08931162 | 0.16899941 | 0.09550363 | 0.07349578 | 0.00016832 | 0.00334752 |
| Tdg | 0.79402501 | 0.78808154 | 0.71467938 | 0.07340215 | 0.31614608 | 0.89054584 |
| Dpt | 0.12836165 | 0.20997547 | 0.13660757 | 0.0733679 | 0.00617156 | 0.07787232 |
| Aurkb | 0.10511521 | 0.18560218 | 0.11228835 | 0.07331383 | 0.00117563 | 0.01908652 |
| Rpl13-ps3 | 1.04488622 | 0.93090551 | 0.85763343 | 0.07327207 | 0.34530127 | 0.89054584 |
| Mfap1a | 0.44486267 | 0.51696985 | 0.44380061 | 0.07316923 | 0.22120859 | 0.89054584 |
| Prkab2 | 0.12540047 | 0.20667915 | 0.13352083 | 0.07315832 | 0.00534438 | 0.06934802 |
| Bik | 0.13919763 | 0.22102524 | 0.14787468 | 0.07315057 | 0.01059218 | 0.12300859 |
| Gucy1a3 | 0.1159192 | 0.19664878 | 0.1236159 | 0.07303288 | 0.00295688 | 0.04245287 |
| Kctd12b | 0.33680341 | 0.41752713 | 0.34451832 | 0.0730088 | 0.16174724 | 0.81339284 |
| Tnk1 | 0.18157927 | 0.264457 | 0.19149887 | 0.07295813 | 0.03795019 | 0.32460284 |
| Phactr1 | 0.26696783 | 0.34998982 | 0.27707614 | 0.07291368 | 0.11009582 | 0.65698409 |
| Hcls1 | 0.05127636 | 0.12768603 | 0.05486161 | 0.07282442 | 3.12E-10 | 1.57E-08 |
| Klhl36 | 0.17779643 | 0.26002125 | 0.18763447 | 0.07238678 | 0.03613964 | 0.31169095 |
| Bloc1s1 | 0.29812725 | 0.37983332 | 0.30746706 | 0.07236626 | 0.13642112 | 0.73997914 |
| Sep-04 | 0.14219609 | 0.22331611 | 0.15098441 | 0.0723317 | 0.01280924 | 0.14318303 |
| Tmf1 | 0.39011217 | 0.46653363 | 0.39430568 | 0.07222795 | 0.19671504 | 0.89054584 |
| Piga | 0.15519949 | 0.23657871 | 0.16442967 | 0.07214904 | 0.02046359 | 0.20404477 |
| Pgm3 | 0.41755572 | 0.49125672 | 0.41932868 | 0.07192804 | 0.21210035 | 0.89054584 |
| Rps11 | 4.7822938 | 0.83692316 | 0.76500685 | 0.07191631 | 0.33070221 | 0.89054584 |
| Osbpl2 | 0.33020828 | 0.41010715 | 0.33825465 | 0.0718525 | 0.16117054 | 0.81133695 |
| Ccser2 | 0.32731936 | 0.40725093 | 0.33550376 | 0.07174717 | 0.15954969 | 0.80704115 |
| Enox1 | 0.06373074 | 0.13991373 | 0.06817707 | 0.07173666 | 4.74E-07 | 1.56E-05 |
| Fuca2 | 0.38871077 | 0.46471994 | 0.39301641 | 0.07170353 | 0.19766356 | 0.89054584 |
| Rarb | 0.0791323 | 0.15633017 | 0.08463393 | 0.07169623 | 3.96E-05 | 0.00091638 |
| Fbxo28 | 0.31033217 | 0.39085098 | 0.31924045 | 0.07161053 | 0.14799486 | 0.77431593 |
| Aif1 | 0.02637685 | 0.09977062 | 0.02822625 | 0.07154437 | 1.76E-32 | 3.80E-30 |
| Tmem108 | 0.57068572 | 0.62225733 | 0.55073546 | 0.07152187 | 0.27257669 | 0.89054584 |

| | | | | | | |
|---|---|---|---|---|---|---|
| Gpm6a | 0.05757711 | 0.13305565 | 0.06159874 | 0.07145691 | 3.25E-08 | 1.24E-06 |
| Cyth3 | 0.28203644 | 0.36309094 | 0.29183129 | 0.07125965 | 0.12763859 | 0.71558556 |
| Srgn | 0.08734492 | 0.16459993 | 0.09340404 | 0.0711959 | 0.00019173 | 0.00378301 |
| Eva1b | 1.44496824 | 1.09078695 | 1.01975339 | 0.07103356 | 0.37276818 | 0.89054584 |
| Sac3d1 | 0.51764843 | 0.57783035 | 0.50686173 | 0.07096862 | 0.25709575 | 0.89054584 |
| Ppp1r18 | 0.15168908 | 0.23175299 | 0.16080635 | 0.07094663 | 0.01991575 | 0.20079178 |
| Bag3 | 0.14039495 | 0.21985111 | 0.14911685 | 0.07073426 | 0.01355292 | 0.14891872 |
| Sox6 | 0.2841087 | 0.3644858 | 0.29385181 | 0.07063398 | 0.13138187 | 0.72635204 |
| Clpp | 0.82579832 | 0.80574764 | 0.73513294 | 0.0706147 | 0.32724833 | 0.89054584 |
| Efnb1 | 0.4044151 | 0.47800522 | 0.40740056 | 0.07060466 | 0.20971257 | 0.89054584 |
| H2-Ke6 | 1.05609141 | 0.9339088 | 0.86335853 | 0.07055026 | 0.35170909 | 0.89054584 |
| Npnt | 0.23863223 | 0.31952222 | 0.24903966 | 0.07048256 | 0.0936579 | 0.5972952 |
| Sertad3 | 0.27289116 | 0.35335695 | 0.28288943 | 0.07046752 | 0.12291485 | 0.70139524 |
| Ccdc166 | 0.18288568 | 0.26325893 | 0.19283211 | 0.07042683 | 0.04441941 | 0.36159044 |
| Snrpd1 | 1.81193499 | 1.16981138 | 1.09940476 | 0.07040661 | 0.38271356 | 0.89054584 |
| Amigo2 | 0.42030131 | 0.49217851 | 0.42180893 | 0.07036959 | 0.2185079 | 0.89054584 |
| Klf5 | 0.15592877 | 0.23551117 | 0.16518181 | 0.07032936 | 0.02364829 | 0.22675454 |
| Abt1 | 0.3559635 | 0.43290496 | 0.36258851 | 0.07031645 | 0.18312922 | 0.86510504 |
| Fat4 | 0.1741585 | 0.25415777 | 0.18391285 | 0.07024492 | 0.03758156 | 0.32198646 |
| Samd4b | 0.41640719 | 0.48852611 | 0.41828986 | 0.07023625 | 0.21702025 | 0.89054584 |
| Rab3d | 0.60621289 | 0.64927154 | 0.57910329 | 0.07016825 | 0.28620346 | 0.89054584 |
| Dpcd | 0.80437475 | 0.79155061 | 0.72142229 | 0.07012832 | 0.32531459 | 0.89054584 |
| Katnb1 | 0.23793525 | 0.31839519 | 0.24834535 | 0.07004984 | 0.09440122 | 0.60001264 |
| Fcrls | 0.02826312 | 0.10026449 | 0.0302445 | 0.07001999 | 2.00E-27 | 3.54E-25 |
| Mapk7 | 0.2264923 | 0.30678549 | 0.23691528 | 0.06987022 | 0.08472107 | 0.56163058 |
| Sord | 0.13319568 | 0.21134161 | 0.14163951 | 0.06970209 | 0.01093243 | 0.12579236 |
| Arhgap10 | 0.17851393 | 0.25801656 | 0.18836782 | 0.06964874 | 0.04247709 | 0.35050247 |
| Plekhf1 | 0.07202785 | 0.14666691 | 0.07704417 | 0.06962274 | 1.28E-05 | 0.00033033 |
| Emid1 | 0.22709276 | 0.30711237 | 0.23751648 | 0.06959588 | 0.08610461 | 0.56742938 |
| Cd36 | 0.07563595 | 0.15049034 | 0.08089908 | 0.06959126 | 3.07E-05 | 0.00072973 |

| | | | | | | |
|---|---|---|---|---|---|---|
| Deptor | 0.46376178 | 0.52998815 | 0.46048223 | 0.06950592 | 0.24095684 | 0.89054584 |
| Tmem41b | 0.49897155 | 0.56047218 | 0.49098866 | 0.06948351 | 0.25484452 | 0.89054584 |
| Pdgfrl | 0.23026565 | 0.31012125 | 0.24069058 | 0.06943066 | 0.08948411 | 0.57968753 |
| Morc4 | 0.15022801 | 0.2287149 | 0.15929698 | 0.06941793 | 0.02116514 | 0.20902737 |
| Psme2b | 0.12579129 | 0.20299238 | 0.13392839 | 0.06906399 | 0.00814136 | 0.09869022 |
| Tst | 0.19535725 | 0.27456791 | 0.20552334 | 0.06904457 | 0.05877402 | 0.4395921 |
| Osbpl6 | 0.10253405 | 0.17860229 | 0.10957578 | 0.06902651 | 0.00166845 | 0.0257058 |
| Gstm5 | 0.88184295 | 0.83839047 | 0.76941983 | 0.06897064 | 0.33811008 | 0.89054584 |
| Cast | 0.19270876 | 0.27169922 | 0.20283374 | 0.06886548 | 0.05684885 | 0.43016419 |
| Atf1 | 0.57156198 | 0.62021163 | 0.55144502 | 0.06876661 | 0.2806263 | 0.89054584 |
| Tpm3-rs7 | 1.23490393 | 1.01373639 | 0.94504808 | 0.06868831 | 0.36744696 | 0.89054584 |
| Klhl24 | 0.25602952 | 0.33486525 | 0.26629706 | 0.06856819 | 0.11514358 | 0.67639855 |
| Gfm1 | 0.26734241 | 0.34599544 | 0.27744426 | 0.06855118 | 0.12483828 | 0.70660334 |
| Gm4890 | 0.11009924 | 0.18606698 | 0.11751925 | 0.06854773 | 0.00328875 | 0.04638159 |
| Marf1 | 0.30670151 | 0.38423594 | 0.31574603 | 0.06848991 | 0.15610584 | 0.79775203 |
| Gimap5 | 0.10117138 | 0.17658073 | 0.10814276 | 0.06843797 | 0.0015971 | 0.02479162 |
| Drap1 | 1.2583236 | 1.02288618 | 0.95451006 | 0.06837612 | 0.36928703 | 0.89054584 |
| Khsrp | 0.2151066 | 0.2937561 | 0.22548478 | 0.06827132 | 0.07917919 | 0.538911 |
| Sirt3 | 0.26205406 | 0.3404984 | 0.27224084 | 0.06825756 | 0.12138067 | 0.69804721 |
| Fhl2 | 0.23650737 | 0.31517566 | 0.24692224 | 0.06825342 | 0.09890625 | 0.61754641 |
| Smarcc1 | 1.01570623 | 0.91080133 | 0.84260673 | 0.0681946 | 0.35306391 | 0.89054584 |
| Tipin | 0.52335417 | 0.57984853 | 0.51166751 | 0.06818102 | 0.26736036 | 0.89054584 |
| Kif4 | 0.06095647 | 0.13333417 | 0.06521152 | 0.06812264 | 5.67E-07 | 1.85E-05 |
| Mar-02 | 0.80866322 | 0.79231479 | 0.72419334 | 0.06812145 | 0.33060448 | 0.89054584 |
| Ubl3 | 1.27368223 | 1.02862136 | 0.96055206 | 0.0680693 | 0.37064151 | 0.89054584 |
| Dnase2a | 0.45507471 | 0.52078048 | 0.45284118 | 0.0679393 | 0.24228346 | 0.89054584 |
| Rab20 | 0.10573381 | 0.1808657 | 0.11293809 | 0.06792761 | 0.00253807 | 0.03701911 |
| Cox6a1 | 3.21249777 | 1.14185522 | 1.07393733 | 0.06791789 | 0.38413426 | 0.89054584 |
| BC029722 | 0.2527259 | 0.33089797 | 0.26303092 | 0.06786704 | 0.11466307 | 0.67576633 |
| Srsf6 | 1.81136194 | 1.16717398 | 1.09932716 | 0.06784682 | 0.3868516 | 0.89054584 |

| | | | | | | |
|---|---|---|---|---|---|---|
| App | 1.89697792 | 1.1772089 | 1.10938095 | 0.06782795 | 0.38787979 | 0.89054584 |
| Tcf7 | 0.03690817 | 0.10713247 | 0.03949358 | 0.0676389 | 7.37E-16 | 6.43E-14 |
| Adgre1 | 0.0352382 | 0.10533376 | 0.03770705 | 0.0676267 | 3.25E-17 | 3.32E-15 |
| Ndufa6 | 2.69238973 | 1.18497983 | 1.11743549 | 0.06754434 | 0.38911989 | 0.89054584 |
| Prcp | 0.18334832 | 0.26077598 | 0.19330407 | 0.06747191 | 0.05195543 | 0.4030229 |
| Cd151 | 0.52862332 | 0.5835572 | 0.51608784 | 0.06746936 | 0.27123763 | 0.89054584 |
| Tesc | 0.22754483 | 0.30541349 | 0.23796901 | 0.06744448 | 0.09334682 | 0.59664859 |
| Sesn3 | 0.69291483 | 0.71216159 | 0.64475974 | 0.06740185 | 0.31311639 | 0.89054584 |
| Tram1 | 1.67785709 | 1.14463365 | 1.07725474 | 0.06737891 | 0.38537187 | 0.89054584 |
| Thnsl1 | 0.17256226 | 0.24964097 | 0.18227815 | 0.06736282 | 0.04255642 | 0.35075781 |
| Napa | 1.58984782 | 1.12573912 | 1.05842707 | 0.06731205 | 0.38350376 | 0.89054584 |
| Cd44 | 0.17311157 | 0.25013585 | 0.18284082 | 0.06729504 | 0.04319688 | 0.35462533 |
| Kcne1l | 0.17610135 | 0.25318592 | 0.18590108 | 0.06728484 | 0.04587361 | 0.36925458 |
| Nuf2 | 0.09665797 | 0.17060167 | 0.10334411 | 0.06725756 | 0.00121432 | 0.01960492 |
| Alkbh7 | 0.4256037 | 0.49375259 | 0.42658643 | 0.06716617 | 0.23161099 | 0.89054584 |
| Casc5 | 0.10603813 | 0.18037097 | 0.11325767 | 0.06711331 | 0.00288337 | 0.04151256 |
| Epas1 | 0.06401285 | 0.13553017 | 0.06847861 | 0.06705156 | 2.54E-06 | 7.42E-05 |
| Crot | 0.46418999 | 0.52785535 | 0.4608577 | 0.06699765 | 0.24910838 | 0.89054584 |
| Hopx | 0.16017199 | 0.23646648 | 0.16955378 | 0.0669127 | 0.0329867 | 0.2905475 |
| Cmc4 | 0.22925869 | 0.30659546 | 0.23968373 | 0.06691173 | 0.09669401 | 0.61116934 |
| Asl | 0.38699481 | 0.4583187 | 0.39143668 | 0.06688202 | 0.21300429 | 0.89054584 |
| Ctps | 0.44823375 | 0.51366721 | 0.4467919 | 0.06687531 | 0.24279343 | 0.89054584 |
| Slc43a3 | 0.25388828 | 0.33104108 | 0.26418071 | 0.06686037 | 0.11917688 | 0.6908542 |
| Ltn1 | 0.23792883 | 0.31517299 | 0.24833895 | 0.06683404 | 0.10495115 | 0.63913594 |
| Pold2 | 0.38878377 | 0.45980541 | 0.39308359 | 0.06672182 | 0.21452848 | 0.89054584 |
| 1810008I18R | 0.25493973 | 0.33193825 | 0.26522021 | 0.06671803 | 0.12060056 | 0.69533658 |
| Polr3gl | 0.43160936 | 0.49868674 | 0.43197768 | 0.06670906 | 0.23592669 | 0.89054584 |
| Mal | 0.02893119 | 0.09766365 | 0.03095929 | 0.06670436 | 5.18E-24 | 7.75E-22 |
| Card10 | 0.04748522 | 0.11743775 | 0.05080725 | 0.06663049 | 4.98E-10 | 2.42E-08 |
| Cracr2b | 0.16731697 | 0.24343526 | 0.17689909 | 0.06653616 | 0.03985733 | 0.33654725 |

| | | | | | | |
|---|---|---|---|---|---|---|
| Tmem97 | 0.39600033 | 0.46621538 | 0.39971058 | 0.06650479 | 0.21912089 | 0.89054584 |
| Vldlr | 0.30658205 | 0.3821354 | 0.31563094 | 0.06650446 | 0.16313516 | 0.81647508 |
| Gjc1 | 0.1840934 | 0.26052881 | 0.19406398 | 0.06646483 | 0.05528488 | 0.42140917 |
| Slc25a37 | 0.0847435 | 0.1570441 | 0.09062647 | 0.06641763 | 0.0003196 | 0.00603333 |
| Nudt9 | 0.33091421 | 0.40532888 | 0.33892617 | 0.0664027 | 0.18067815 | 0.85856175 |
| Kif22 | 0.17420512 | 0.25036142 | 0.18396058 | 0.06640084 | 0.0463182 | 0.37224016 |
| Ttyh2 | 0.19596276 | 0.27251873 | 0.20613783 | 0.0663809 | 0.06677079 | 0.47994192 |
| Wasf1 | 0.24386991 | 0.3206222 | 0.25425009 | 0.06637212 | 0.11195075 | 0.66518936 |
| Atic | 0.54116993 | 0.59286898 | 0.52654137 | 0.06632761 | 0.27864227 | 0.89054584 |
| Cwc27 | 0.37410391 | 0.4458393 | 0.37951663 | 0.06632267 | 0.20777199 | 0.89054584 |
| Arxes1 | 0.31109933 | 0.38618549 | 0.31997813 | 0.06620736 | 0.16752509 | 0.82808684 |
| Trp53rkb | 0.18340945 | 0.25957023 | 0.19336642 | 0.06620381 | 0.05534444 | 0.42164146 |
| Irf9 | 0.11047923 | 0.18409139 | 0.11791769 | 0.0661737 | 0.00446751 | 0.0600278 |
| Cd59a | 0.20899982 | 0.28548375 | 0.21933019 | 0.06615356 | 0.07997903 | 0.54094194 |
| Snai2 | 0.14650098 | 0.22157637 | 0.15544289 | 0.06613349 | 0.0237309 | 0.22739625 |
| Slc7a6os | 0.29581918 | 0.37135528 | 0.30523214 | 0.06612315 | 0.15642084 | 0.79826004 |
| 1190007I07R | 0.29042021 | 0.36606768 | 0.29999385 | 0.06607382 | 0.15241409 | 0.7857996 |
| Scara5 | 0.05316314 | 0.12287325 | 0.05687922 | 0.06599404 | 3.23E-08 | 1.24E-06 |
| Rptor | 0.19840033 | 0.27451412 | 0.20860992 | 0.0659042 | 0.07052987 | 0.50026519 |
| Lamtor2 | 1.57141365 | 1.1198318 | 1.05404599 | 0.06578581 | 0.38561097 | 0.89054584 |
| Slc2a1 | 0.32598665 | 0.39994724 | 0.33423325 | 0.06571399 | 0.17983349 | 0.85601281 |
| Zc3h3 | 0.26067382 | 0.3365161 | 0.27088056 | 0.06563554 | 0.12947386 | 0.72054099 |
| Ost4 | 2.00890998 | 1.18399217 | 1.11837012 | 0.06562204 | 0.39228427 | 0.89054584 |
| Apln | 0.05394393 | 0.12333464 | 0.05771411 | 0.06562053 | 5.88E-08 | 2.15E-06 |
| Ift43 | 0.29234882 | 0.36746675 | 0.30186674 | 0.06560001 | 0.15566018 | 0.79659772 |
| Nostrin | 0.06029086 | 0.13004969 | 0.06449998 | 0.06554972 | 1.10E-06 | 3.39E-05 |
| Ergic1 | 0.45684022 | 0.51991408 | 0.4543978 | 0.06551628 | 0.2508751 | 0.89054584 |
| Sla | 0.04763921 | 0.11644421 | 0.05097194 | 0.06547227 | 1.09E-09 | 5.07E-08 |
| Slc33a1 | 0.38407224 | 0.45419964 | 0.38874237 | 0.06545727 | 0.21637852 | 0.89054584 |
| Chchd5 | 0.40055494 | 0.46931744 | 0.4038779 | 0.06543954 | 0.22516078 | 0.89054584 |

| | | | | | | |
|---|---|---|---|---|---|---|
| Rnd3 | 0.28661905 | 0.36170254 | 0.29629712 | 0.06540541 | 0.15187449 | 0.78458375 |
| Ccdc102a | 0.24309049 | 0.31885899 | 0.25347553 | 0.06538347 | 0.11472752 | 0.6757765 |
| Hid1 | 0.44219108 | 0.50675929 | 0.44142519 | 0.06533409 | 0.2452365 | 0.89054584 |
| Kifc3 | 0.33968052 | 0.41255837 | 0.34724429 | 0.06531408 | 0.19043013 | 0.88259328 |
| Robo4 | 0.05446877 | 0.12358005 | 0.05827531 | 0.06530475 | 8.90E-08 | 3.22E-06 |
| Cdc25b | 0.11957725 | 0.19274542 | 0.12744141 | 0.06530401 | 0.00848511 | 0.10225864 |
| Cnih2 | 0.36739354 | 0.43856039 | 0.37327512 | 0.06528527 | 0.20758137 | 0.89054584 |
| Cmtm6 | 0.35592226 | 0.42781468 | 0.36254982 | 0.06526486 | 0.20082302 | 0.89054584 |
| Hprt | 1.17068895 | 0.98296552 | 0.91771158 | 0.06525394 | 0.37021865 | 0.89054584 |
| Ikbip | 0.41521607 | 0.48245825 | 0.4172117 | 0.06524656 | 0.2331236 | 0.89054584 |
| Baiap2 | 0.22745225 | 0.30309138 | 0.23787634 | 0.06521504 | 0.10075294 | 0.6232762 |
| Gm5294 | 0.06561083 | 0.13539577 | 0.07018659 | 0.06520918 | 7.50E-06 | 0.00020224 |
| Ppp2r1b | 0.29139269 | 0.36613022 | 0.30093847 | 0.06519175 | 0.15642549 | 0.79826004 |
| Gsto1 | 0.55878172 | 0.60622176 | 0.54104585 | 0.06517591 | 0.28732066 | 0.89054584 |
| Golt1b | 0.46455522 | 0.52627702 | 0.46117785 | 0.06509917 | 0.25538183 | 0.89054584 |
| Tpi1 | 1.97534259 | 1.18109267 | 1.11604959 | 0.06504307 | 0.39299485 | 0.89054584 |
| Sema3d | 0.13818239 | 0.21183606 | 0.14682097 | 0.06501509 | 0.01955313 | 0.1979618 |
| Man2b1 | 0.44111161 | 0.50546181 | 0.4404643 | 0.06499751 | 0.24588359 | 0.89054584 |
| Cbarp | 0.33626212 | 0.40893372 | 0.34400499 | 0.06492874 | 0.18960705 | 0.88050062 |
| Skap1 | 0.06934889 | 0.13906188 | 0.07418152 | 0.06488035 | 2.30E-05 | 0.00056587 |
| Clmp | 0.21061703 | 0.28578817 | 0.22096166 | 0.06482651 | 0.08583515 | 0.56642604 |
| Ethe1 | 0.37047318 | 0.44093932 | 0.37614267 | 0.06479665 | 0.21110938 | 0.89054584 |
| Col16a1 | 0.19376434 | 0.26870073 | 0.20390607 | 0.06479466 | 0.06937545 | 0.49496101 |
| C330027C09l | 0.09422152 | 0.16552865 | 0.10074418 | 0.06478447 | 0.00136785 | 0.02167325 |
| Tpst2 | 1.0100097 | 0.90446203 | 0.83969003 | 0.064772 | 0.35965491 | 0.89054584 |
| Pear1 | 0.20183377 | 0.27684459 | 0.21208764 | 0.06475695 | 0.07743773 | 0.53076426 |
| Bahd1 | 0.10892547 | 0.18101072 | 0.11628816 | 0.06472257 | 0.00475673 | 0.06326885 |
| Tmem123 | 0.45086657 | 0.51383305 | 0.44912336 | 0.06470968 | 0.25102741 | 0.89054584 |
| Cep85 | 0.14845005 | 0.22215843 | 0.15745909 | 0.06469934 | 0.02778829 | 0.25466161 |
| Dhrs7 | 0.54641928 | 0.59556309 | 0.53088476 | 0.06467833 | 0.28515181 | 0.89054584 |

| | | | | | | |
|---|---|---|---|---|---|---|
| C2 | 0.08331366 | 0.15377138 | 0.08909963 | 0.06467175 | 0.00036029 | 0.0067107 |
| Ralgps2 | 0.30429725 | 0.37806102 | 0.31342833 | 0.06463269 | 0.16834404 | 0.82993631 |
| Vat1 | 0.49108575 | 0.5487774 | 0.48422171 | 0.06455569 | 0.26725939 | 0.89054584 |
| Mrps25 | 0.50804578 | 0.56322275 | 0.49872819 | 0.06449456 | 0.2734256 | 0.89054584 |
| Tspan18 | 0.19696842 | 0.27161376 | 0.20715804 | 0.06445572 | 0.07358564 | 0.51488639 |
| Pdzd2 | 0.10586255 | 0.17748435 | 0.11307329 | 0.06441106 | 0.00397814 | 0.05441051 |
| Mllt11 | 0.58501735 | 0.62668558 | 0.56227963 | 0.06440595 | 0.29678999 | 0.89054584 |
| Spin1 | 0.51764888 | 0.57125973 | 0.50686211 | 0.06439763 | 0.27695014 | 0.89054584 |
| Stat1 | 0.10320672 | 0.17466042 | 0.11028293 | 0.06437749 | 0.00326756 | 0.04617255 |
| Arhgef25 | 0.23283195 | 0.30762335 | 0.24325481 | 0.06436854 | 0.10882159 | 0.65274944 |
| Rfesd | 0.30805599 | 0.38139802 | 0.31705045 | 0.06434757 | 0.17218546 | 0.83696674 |
| Shmt1 | 0.1306266 | 0.20326897 | 0.13896637 | 0.06430261 | 0.0155501 | 0.16638031 |
| Cep290 | 0.36110937 | 0.43169098 | 0.36740793 | 0.06428305 | 0.20749449 | 0.89054584 |
| Rmi1 | 0.24957954 | 0.32418097 | 0.25991547 | 0.0642655 | 0.12467116 | 0.70642606 |
| H2afy | 1.39283767 | 1.06750449 | 1.00324186 | 0.06426263 | 0.38268813 | 0.89054584 |
| Epb41l2 | 0.28581949 | 0.35976447 | 0.29551862 | 0.06424585 | 0.15556461 | 0.79659772 |
| Cep57 | 0.4515042 | 0.51386916 | 0.44968738 | 0.06418179 | 0.25304156 | 0.89054584 |
| Mfng | 0.42478068 | 0.48999717 | 0.42584596 | 0.06415122 | 0.24138873 | 0.89054584 |
| Epb41l3 | 0.66791105 | 0.69045353 | 0.62635504 | 0.06409849 | 0.3167608 | 0.89054584 |
| Tmem255a | 0.09122311 | 0.16138268 | 0.09754404 | 0.06383864 | 0.00114768 | 0.01865367 |
| Itfg1 | 0.96335648 | 0.8791007 | 0.81528568 | 0.06381502 | 0.35767879 | 0.89054584 |
| Trappc6b | 1.06803761 | 0.93320777 | 0.86941365 | 0.06379412 | 0.36623017 | 0.89054584 |
| Vangl1 | 0.15629751 | 0.22932059 | 0.16556202 | 0.06375857 | 0.03639254 | 0.31331294 |
| Arpc5 | 2.07859553 | 1.18604536 | 1.12229999 | 0.06374537 | 0.39565011 | 0.89054584 |
| Dhdds | 0.26727567 | 0.34105119 | 0.27737868 | 0.06367251 | 0.14243034 | 0.76225733 |
| Snap23 | 0.56399922 | 0.60890356 | 0.54530429 | 0.06359928 | 0.29343546 | 0.89054584 |
| Adgra2 | 0.14364886 | 0.2160868 | 0.15248982 | 0.06359697 | 0.02600537 | 0.24277262 |
| Kif21a | 0.69434044 | 0.70932617 | 0.64579596 | 0.0635302 | 0.32336001 | 0.89054584 |
| Plekho1 | 0.10781265 | 0.17862772 | 0.11512053 | 0.06350719 | 0.00508282 | 0.06674888 |
| Myzap | 0.10777998 | 0.1785447 | 0.11508625 | 0.06345846 | 0.00510056 | 0.06692124 |

| | | | | | | |
|---|---|---|---|---|---|---|
| Hexa | 0.99529696 | 0.89543312 | 0.83207331 | 0.06335981 | 0.36138321 | 0.89054584 |
| Ccl11 | 0.065237 | 0.13313231 | 0.06978704 | 0.06334527 | 1.17E-05 | 0.00030674 |
| Tmem175 | 0.16614089 | 0.23902552 | 0.17569147 | 0.06333406 | 0.04652964 | 0.37331857 |
| Brca2 | 0.1420516 | 0.2141525 | 0.15083463 | 0.06331787 | 0.02524754 | 0.23753332 |
| Rbm47 | 0.45225554 | 0.51365104 | 0.45035167 | 0.06329937 | 0.25628557 | 0.89054584 |
| Pex5l | 0.22733239 | 0.30103959 | 0.23775636 | 0.06328323 | 0.10747608 | 0.64845225 |
| Lhx1 | 0.21774398 | 0.29131625 | 0.2281377 | 0.06317854 | 0.0984892 | 0.61600365 |
| Afap1l2 | 0.10536111 | 0.17566523 | 0.11254664 | 0.06311859 | 0.00448973 | 0.06021472 |
| Ccl5 | 0.03067463 | 0.09587843 | 0.03282465 | 0.06305378 | 1.79E-19 | 2.09E-17 |
| Ebpl | 0.44534517 | 0.50720319 | 0.44422917 | 0.06297402 | 0.25448175 | 0.89054584 |
| Pigf | 0.46839492 | 0.52751045 | 0.46453884 | 0.06297161 | 0.26383943 | 0.89054584 |
| Ncoa3 | 0.15846181 | 0.23076139 | 0.16779258 | 0.06296881 | 0.04019768 | 0.33777651 |
| 2410015M2( | 2.23377385 | 1.19100741 | 1.12808168 | 0.06292574 | 0.39747767 | 0.89054584 |
| Psmc1 | 1.3655047 | 1.05695459 | 0.99403216 | 0.06292242 | 0.38403061 | 0.89054584 |
| Galnt18 | 0.35491373 | 0.42449452 | 0.36160357 | 0.06289095 | 0.20888557 | 0.89054584 |
| Ephb4 | 0.27080105 | 0.34364939 | 0.28084009 | 0.0628093 | 0.14871453 | 0.77667985 |
| Rassf5 | 0.0599362 | 0.12691287 | 0.06412083 | 0.06279203 | 2.53E-06 | 7.42E-05 |
| Traip | 0.06297996 | 0.13015819 | 0.06737456 | 0.06278363 | 7.08E-06 | 0.00019212 |
| Selplg | 0.04439493 | 0.11027673 | 0.04750206 | 0.06277467 | 3.71E-10 | 1.85E-08 |
| Wdr89 | 1.30314055 | 1.03452044 | 0.97174705 | 0.06277339 | 0.38172229 | 0.89054584 |
| Neurl1b | 0.08423235 | 0.15284446 | 0.09008065 | 0.06276381 | 0.00058494 | 0.01043109 |
| Smdt1 | 2.2449799 | 1.19102089 | 1.12830854 | 0.06271235 | 0.39783771 | 0.89054584 |
| Timm10b | 1.24807328 | 1.0131024 | 0.95039105 | 0.06271135 | 0.37926214 | 0.89054584 |
| Pole3 | 0.39284641 | 0.45952487 | 0.39681795 | 0.06270692 | 0.23079647 | 0.89054584 |
| Reep4 | 0.28792313 | 0.36027107 | 0.29756618 | 0.06270489 | 0.16310921 | 0.81647508 |
| Tbc1d10a | 0.31762532 | 0.38889786 | 0.32624109 | 0.06265677 | 0.1854522 | 0.8704066 |
| Rgs18 | 0.03765491 | 0.10289327 | 0.04029242 | 0.06260085 | 2.27E-13 | 1.61E-11 |
| Vash2 | 0.14805446 | 0.21962661 | 0.15704999 | 0.06257662 | 0.03170136 | 0.28093297 |
| Eif4e3 | 0.36290583 | 0.43165148 | 0.36908714 | 0.06256433 | 0.21483929 | 0.89054584 |
| Kif15 | 0.05363617 | 0.11985091 | 0.05738502 | 0.06246589 | 1.97E-07 | 6.80E-06 |

| | | | | | | |
|---|---|---|---|---|---|---|
| Tysnd1 | 0.34042321 | 0.41035131 | 0.34794725 | 0.06240406 | 0.20169636 | 0.89054584 |
| Neil1 | 0.10028647 | 0.16960118 | 0.1072118 | 0.06238938 | 0.0033524 | 0.0471593 |
| Miat | 0.18893153 | 0.26131097 | 0.1989927 | 0.06231827 | 0.07227785 | 0.50878997 |
| Zfp639 | 0.29615582 | 0.36781344 | 0.30555828 | 0.06225516 | 0.17125343 | 0.83568909 |
| Mettl25 | 0.09802655 | 0.16704962 | 0.10480433 | 0.06224529 | 0.00282844 | 0.04076221 |
| Afap1 | 0.30503342 | 0.37636863 | 0.31413831 | 0.06223032 | 0.17802205 | 0.85072631 |
| Dpp4 | 0.2153888 | 0.28794492 | 0.22576879 | 0.06217612 | 0.09973553 | 0.62085259 |
| Col27a1 | 0.30296261 | 0.37423366 | 0.31214047 | 0.06209319 | 0.17701793 | 0.84839864 |
| Rab30 | 0.10599351 | 0.17528918 | 0.11321081 | 0.06207836 | 0.005314 | 0.06903437 |
| Cnksr2 | 0.03224178 | 0.09656032 | 0.03450133 | 0.06205899 | 2.64E-17 | 2.74E-15 |
| Mgst2 | 0.03760843 | 0.10227535 | 0.0402427 | 0.06203265 | 3.45E-13 | 2.40E-11 |
| Cox4i2 | 0.05783127 | 0.12364934 | 0.06187047 | 0.06177887 | 1.64E-06 | 4.95E-05 |
| Coq9 | 0.62634555 | 0.65656946 | 0.59480541 | 0.06176405 | 0.31427095 | 0.89054584 |
| Ankrd37 | 0.34588293 | 0.41473042 | 0.35310588 | 0.06162455 | 0.20808463 | 0.89054584 |
| Tnfaip8l2 | 0.04225158 | 0.10683315 | 0.0452095 | 0.06162365 | 1.07E-10 | 5.68E-09 |
| Ppif | 0.19941637 | 0.2712623 | 0.20963959 | 0.06162271 | 0.08542511 | 0.56503846 |
| Rap1gapos | 0.12456185 | 0.19426519 | 0.1326461 | 0.06161909 | 0.01522247 | 0.1634781 |
| Zfp36 | 0.30166109 | 0.37246683 | 0.31088371 | 0.06158312 | 0.1780317 | 0.85072631 |
| Derl2 | 0.98791737 | 0.88982444 | 0.82825473 | 0.06156971 | 0.36454727 | 0.89054584 |
| Eif4e | 1.39995642 | 1.06715733 | 1.00558793 | 0.0615694 | 0.38772469 | 0.89054584 |
| Tmem80 | 0.32363311 | 0.3934553 | 0.3319873 | 0.061468 | 0.19417577 | 0.889076 |
| Zeb1 | 0.33124264 | 0.40065904 | 0.33923852 | 0.06142052 | 0.19946876 | 0.89054584 |
| Pld3 | 0.80463709 | 0.78296357 | 0.7215922 | 0.06137137 | 0.34596301 | 0.89054584 |
| Rpl12 | 3.27750818 | 1.1264626 | 1.06518868 | 0.06127392 | 0.39434928 | 0.89054584 |
| Rasl11a | 0.09888717 | 0.1669724 | 0.10572251 | 0.06124989 | 0.00347597 | 0.04859656 |
| Hspg2 | 0.26202702 | 0.33345902 | 0.2722142 | 0.06124482 | 0.1472723 | 0.77404405 |
| Dfna5 | 0.23678978 | 0.30842882 | 0.24720378 | 0.06122503 | 0.12427442 | 0.70545441 |
| Tmem186 | 0.23962805 | 0.31124188 | 0.25003128 | 0.0612106 | 0.12702576 | 0.7132531 |
| Amotl1 | 0.39481928 | 0.45983184 | 0.39862804 | 0.0612038 | 0.237207 | 0.89054584 |
| Csnk1d | 0.79626637 | 0.7773319 | 0.71614627 | 0.06118563 | 0.34529757 | 0.89054584 |

| | | | | | | |
|---|---|---|---|---|---|---|
| Atf3 | 0.09486054 | 0.16252359 | 0.10142612 | 0.06109747 | 0.00250437 | 0.03663805 |
| Zfp286 | 0.10399392 | 0.17218917 | 0.11111027 | 0.0610789 | 0.00521727 | 0.06800354 |
| Mcm7 | 0.42000437 | 0.48260558 | 0.4215409 | 0.06106468 | 0.24987055 | 0.89054584 |
| BC003331 | 0.33464338 | 0.40350289 | 0.34246929 | 0.06103361 | 0.20318482 | 0.89054584 |
| Erdr1 | 0.24510758 | 0.31649081 | 0.25547946 | 0.06101135 | 0.13293858 | 0.72977796 |
| Fam198b | 0.13690289 | 0.20644138 | 0.14549241 | 0.06094897 | 0.02548686 | 0.2394741 |
| Nat2 | 0.04971348 | 0.11412478 | 0.05319028 | 0.0609345 | 4.72E-08 | 1.75E-06 |
| Tc2n | 0.06201598 | 0.12713625 | 0.06634412 | 0.06079213 | 9.81E-06 | 0.00026014 |
| Spink4 | 0.15421016 | 0.2241929 | 0.16340899 | 0.06078391 | 0.04154736 | 0.34499059 |
| Pabpn1l | 0.15266924 | 0.22258104 | 0.16181847 | 0.06076257 | 0.0401088 | 0.33768723 |
| Gxylt2 | 0.12804815 | 0.19704277 | 0.13628093 | 0.06076184 | 0.0188905 | 0.19341421 |
| Capza1 | 0.5307321 | 0.57858887 | 0.51785192 | 0.06073695 | 0.29238515 | 0.89054584 |
| Rnf138rt1 | 0.3124402 | 0.38199817 | 0.32126674 | 0.06073142 | 0.18923631 | 0.87982135 |
| Zbed4 | 0.14231009 | 0.21176723 | 0.15110256 | 0.06066466 | 0.03070916 | 0.27364557 |
| Klhl8 | 0.13746689 | 0.2067081 | 0.14607812 | 0.06062998 | 0.02657491 | 0.24681805 |
| Fam69a | 0.21786473 | 0.28880724 | 0.2282591 | 0.06054815 | 0.10826021 | 0.6512683 |
| Cebpz | 0.65379755 | 0.67631777 | 0.61577458 | 0.06054319 | 0.32345148 | 0.89054584 |
| Galnt16 | 0.11364054 | 0.18176667 | 0.12123041 | 0.06053627 | 0.00999762 | 0.11755519 |
| Trip4 | 0.36406842 | 0.43066191 | 0.37017314 | 0.06048877 | 0.22323831 | 0.89054584 |
| Aacs | 0.59131993 | 0.62779386 | 0.56731331 | 0.06048056 | 0.30970601 | 0.89054584 |
| Dbr1 | 0.24603137 | 0.31687416 | 0.2563966 | 0.06047757 | 0.1358997 | 0.73849629 |
| Fen1 | 0.28045962 | 0.35075386 | 0.29029247 | 0.0604614 | 0.16593535 | 0.82390627 |
| Rtn1 | 0.06679534 | 0.13190551 | 0.07145257 | 0.06045295 | 4.04E-05 | 0.00093082 |
| Prickle1 | 0.12041291 | 0.18870414 | 0.12831463 | 0.06038951 | 0.01416571 | 0.15453307 |
| Spcs2 | 2.62164798 | 1.18175192 | 1.12138904 | 0.06036287 | 0.40099033 | 0.89054584 |
| Antxr1 | 0.10757045 | 0.17521695 | 0.11486634 | 0.06035061 | 0.00718655 | 0.08912799 |
| Rhog | 0.30948855 | 0.37872798 | 0.31842903 | 0.06029895 | 0.18882484 | 0.87855667 |
| Rpl6l | 2.08562835 | 1.18291123 | 1.12266915 | 0.06024208 | 0.4012947 | 0.89054584 |
| Sdc2 | 0.37312751 | 0.43882163 | 0.37861 | 0.06021163 | 0.22936879 | 0.89054584 |
| Kcnk1 | 0.34432426 | 0.41182432 | 0.35163478 | 0.06018953 | 0.21258799 | 0.89054584 |

| | | | | | | |
|---|---|---|---|---|---|---|
| Adm | 0.0730164 | 0.13828357 | 0.0781004 | 0.06018316 | 0.00016526 | 0.00329118 |
| Atp5b | 3.5157881 | 1.09261941 | 1.03245956 | 0.06015985 | 0.393015 | 0.89054584 |
| Eif2s3y | 0.41303038 | 0.4753607 | 0.41523171 | 0.06012899 | 0.24994803 | 0.89054584 |
| Prr7 | 0.32186199 | 0.3904068 | 0.33029524 | 0.06011156 | 0.19824778 | 0.89054584 |
| Cltb | 1.26228291 | 1.0160602 | 0.95608353 | 0.05997667 | 0.3850428 | 0.89054584 |
| Rrs1 | 0.48200401 | 0.53626199 | 0.47638202 | 0.05987997 | 0.27906656 | 0.89054584 |
| Aup1 | 0.39800544 | 0.46140552 | 0.40154665 | 0.05985887 | 0.24368061 | 0.89054584 |
| Wnt4 | 0.20761175 | 0.27778154 | 0.21792895 | 0.05985259 | 0.10035278 | 0.62288099 |
| Mcts2 | 0.33153833 | 0.39932862 | 0.33951967 | 0.05980895 | 0.2059064 | 0.89054584 |
| Vstm4 | 0.17233035 | 0.24181136 | 0.18204058 | 0.05977078 | 0.06304477 | 0.46198345 |
| Casp12 | 0.10072208 | 0.16729632 | 0.10767011 | 0.05962621 | 0.0049392 | 0.06533623 |
| Olfm1 | 0.28003899 | 0.34944821 | 0.28988177 | 0.05956644 | 0.16919684 | 0.8322688 |
| Lbh | 0.24749062 | 0.31739587 | 0.25784451 | 0.05955136 | 0.14095878 | 0.75633618 |
| Zfp931 | 0.17043869 | 0.23962371 | 0.18010179 | 0.05952192 | 0.06181241 | 0.45559551 |
| Gm10036 | 0.90837431 | 0.8443113 | 0.78485742 | 0.05945388 | 0.36207409 | 0.89054584 |
| Ptpre | 0.07020711 | 0.13452307 | 0.07509863 | 0.05942444 | 0.00011366 | 0.00233399 |
| Narf | 0.35493289 | 0.42103606 | 0.36162155 | 0.05941451 | 0.2219973 | 0.89054584 |
| Olfml2b | 0.1673441 | 0.23632796 | 0.17692695 | 0.05940101 | 0.05888749 | 0.43985104 |
| Oxct1 | 0.67990241 | 0.69455843 | 0.63523522 | 0.05932321 | 0.33174921 | 0.89054584 |
| Grap | 0.08534407 | 0.15056219 | 0.09126774 | 0.05929446 | 0.00123578 | 0.01990701 |
| Atg10 | 0.31529963 | 0.38329442 | 0.32401165 | 0.05928277 | 0.19698973 | 0.89054584 |
| Ccl9 | 0.03289397 | 0.09443941 | 0.0351991 | 0.05924032 | 2.23E-15 | 1.87E-13 |
| Dus4l | 0.0597256 | 0.12313262 | 0.06389568 | 0.05923694 | 7.83E-06 | 0.00021017 |
| Ptprc | 0.0421882 | 0.10435486 | 0.04514171 | 0.05921316 | 4.94E-10 | 2.41E-08 |
| Golim4 | 0.84785889 | 0.8080696 | 0.74890203 | 0.05916758 | 0.35640559 | 0.89054584 |
| Dll1 | 0.08106235 | 0.14586163 | 0.08669536 | 0.05916627 | 0.00073752 | 0.01268396 |
| Vnn1 | 0.10296007 | 0.16917493 | 0.11002366 | 0.05915127 | 0.00612681 | 0.07737503 |
| Ppp1r26 | 0.06694039 | 0.13071893 | 0.07160759 | 0.05911135 | 6.01E-05 | 0.00134153 |
| Hip1 | 0.20353876 | 0.27292383 | 0.21381271 | 0.05911112 | 0.09886934 | 0.61754641 |
| Dync1i2 | 1.77954973 | 1.1538695 | 1.09478662 | 0.05908288 | 0.40074032 | 0.89054584 |

| | | | | | | |
|---|---|---|---|---|---|---|
| Hdac10 | 0.06311642 | 0.12658591 | 0.06752042 | 0.05906549 | 2.30E-05 | 0.00056498 |
| Hey1 | 0.06107933 | 0.12431667 | 0.06534286 | 0.0589738 | 1.31E-05 | 0.00033582 |
| Sec22a | 0.45756713 | 0.51400629 | 0.45503816 | 0.05896812 | 0.27300514 | 0.89054584 |
| Amt | 0.12107004 | 0.18795377 | 0.1290011 | 0.05895267 | 0.01662275 | 0.17624506 |
| As3mt | 0.49410274 | 0.54575724 | 0.48681473 | 0.05894251 | 0.28634479 | 0.89054584 |
| Stx2 | 0.15471266 | 0.22282492 | 0.16392746 | 0.05889746 | 0.04707416 | 0.37685322 |
| Klhl32 | 0.48804461 | 0.5404719 | 0.48160218 | 0.05886972 | 0.28450977 | 0.89054584 |
| Samd4 | 0.09096242 | 0.15612057 | 0.09726578 | 0.05885479 | 0.002408 | 0.03546072 |
| Psme3 | 0.42775757 | 0.48735746 | 0.42852238 | 0.05883509 | 0.26119639 | 0.89054584 |
| Sh2d3c | 0.39733401 | 0.4597371 | 0.40093208 | 0.05880502 | 0.24719901 | 0.89054584 |
| Hacd1 | 0.58472289 | 0.62084717 | 0.5620438 | 0.05880337 | 0.31297403 | 0.89054584 |
| Xrcc2 | 0.06107199 | 0.12409747 | 0.06533502 | 0.05876245 | 1.39E-05 | 0.00035686 |
| Sssca1 | 0.71456334 | 0.71898444 | 0.66034155 | 0.05864289 | 0.33953002 | 0.89054584 |
| Kif20b | 0.14106575 | 0.20839307 | 0.14981253 | 0.05858054 | 0.0342453 | 0.29872948 |
| Rab13 | 0.31605185 | 0.38322494 | 0.32473304 | 0.0584919 | 0.20068342 | 0.89054584 |
| Aqp12 | 0.03157553 | 0.09227688 | 0.03378852 | 0.05848836 | 3.67E-16 | 3.33E-14 |
| Otud6b | 0.42719218 | 0.48648435 | 0.42801446 | 0.05846989 | 0.26224313 | 0.89054584 |
| Mtch1 | 1.30058872 | 1.02924575 | 0.97078911 | 0.05845663 | 0.38953129 | 0.89054584 |
| Srp9 | 2.15741894 | 1.18425818 | 1.12587856 | 0.05837962 | 0.40455369 | 0.89054584 |
| Fabp3 | 0.26009462 | 0.32865417 | 0.27030946 | 0.0583447 | 0.15730282 | 0.80068898 |
| Ttyh1 | 0.15293729 | 0.2204189 | 0.16209521 | 0.05832369 | 0.04683583 | 0.37515241 |
| Gss | 0.23968365 | 0.30836905 | 0.25008663 | 0.05828242 | 0.1387923 | 0.74747802 |
| Col5a3 | 0.14177943 | 0.2088187 | 0.15055249 | 0.0582662 | 0.0356875 | 0.30938746 |
| Il2rg | 0.05966654 | 0.12209166 | 0.06383255 | 0.05825911 | 1.06E-05 | 0.00027804 |
| Osbpl5 | 0.1899598 | 0.25829568 | 0.20003895 | 0.05825673 | 0.08742093 | 0.57117106 |
| Aldh1a2 | 0.05031573 | 0.1119923 | 0.05383433 | 0.05815797 | 2.41E-07 | 8.21E-06 |
| Nosip | 0.29074495 | 0.35844345 | 0.30030934 | 0.05813411 | 0.18355996 | 0.86657514 |
| Fam43a | 0.34634868 | 0.41160029 | 0.35354521 | 0.05805508 | 0.22212471 | 0.89054584 |
| Med1 | 0.36428301 | 0.42841742 | 0.37037352 | 0.0580439 | 0.23265345 | 0.89054584 |
| Tgfbi | 0.07584593 | 0.13909546 | 0.0811234 | 0.05797206 | 0.00043524 | 0.00798748 |

| | | | | | | |
|---|---|---|---|---|---|---|
| Col23a1 | 0.13307116 | 0.19947544 | 0.14151 | 0.05796544 | 0.02816981 | 0.2571826 |
| Foxs1 | 0.06467578 | 0.12707739 | 0.06918719 | 0.0578902 | 4.85E-05 | 0.001101 |
| Zfp770 | 0.14981877 | 0.21673891 | 0.15887405 | 0.05786486 | 0.0448604 | 0.36375062 |
| Cbr1 | 0.89020855 | 0.83218192 | 0.77434106 | 0.05784086 | 0.36391696 | 0.89054584 |
| Rgs16 | 0.09088289 | 0.15501678 | 0.09718088 | 0.05783589 | 0.00277962 | 0.04029886 |
| Pld4 | 0.02591525 | 0.08554297 | 0.02773235 | 0.05781063 | 1.34E-22 | 1.89E-20 |
| Fhl3 | 0.03653899 | 0.09686473 | 0.03909864 | 0.0577661 | 2.92E-12 | 1.82E-10 |
| Praf2 | 0.28427317 | 0.35174235 | 0.29401212 | 0.05773022 | 0.18014884 | 0.85660801 |
| Th | 0.09338326 | 0.15752652 | 0.09984958 | 0.05767694 | 0.00355996 | 0.04953196 |
| Fam64a | 0.09394521 | 0.15809866 | 0.10044931 | 0.05764935 | 0.00374946 | 0.05182043 |
| Dlc1 | 0.38063585 | 0.44317316 | 0.38556823 | 0.05760494 | 0.24319612 | 0.89054584 |
| Atpaf1 | 0.26949948 | 0.33715424 | 0.27956283 | 0.05759141 | 0.16858641 | 0.83078375 |
| Casp8 | 0.25966851 | 0.32746171 | 0.26988922 | 0.05757249 | 0.16015025 | 0.8080503 |
| Gtdc1 | 0.14286215 | 0.20916827 | 0.15167471 | 0.05749357 | 0.03869703 | 0.32924273 |
| Cib2 | 0.29194894 | 0.35891191 | 0.30147856 | 0.05743334 | 0.1873903 | 0.87468918 |
| Ifrd2 | 0.30063269 | 0.36719428 | 0.30989006 | 0.05730422 | 0.1944754 | 0.88943353 |
| Arhgap20 | 0.06653028 | 0.12846153 | 0.07116929 | 0.05729225 | 8.82E-05 | 0.00186754 |
| Slc39a1 | 0.35979079 | 0.42341881 | 0.36617437 | 0.05724443 | 0.23320357 | 0.89054584 |
| Ap5s1 | 0.14328915 | 0.20934138 | 0.15211715 | 0.05722423 | 0.03983321 | 0.33653955 |
| Scn9a | 0.17545464 | 0.24245254 | 0.18523944 | 0.0572131 | 0.07508068 | 0.52132173 |
| Mcm2 | 0.34615743 | 0.41056447 | 0.35336482 | 0.05719964 | 0.22537983 | 0.89054584 |
| Fam217b | 0.03101655 | 0.09035758 | 0.03319047 | 0.05716711 | 5.10E-16 | 4.53E-14 |
| Sec61a2 | 0.4866243 | 0.53750333 | 0.48037681 | 0.05712652 | 0.28977495 | 0.89054584 |
| Ggt5 | 0.09580135 | 0.15955527 | 0.10243007 | 0.0571252 | 0.00468448 | 0.06242235 |
| Klf14 | 0.12588319 | 0.19113944 | 0.13402422 | 0.05711522 | 0.02354801 | 0.22594241 |
| Bora | 0.04815994 | 0.10863515 | 0.05152885 | 0.0571063 | 1.21E-07 | 4.28E-06 |
| Ocln | 0.25643527 | 0.32379369 | 0.26669786 | 0.05709583 | 0.15928367 | 0.80659958 |
| Gse1 | 0.48917496 | 0.53965383 | 0.48257649 | 0.05707734 | 0.29080242 | 0.89054584 |
| Lpar1 | 0.17345984 | 0.24020337 | 0.18319749 | 0.05700588 | 0.07356679 | 0.51488639 |
| Zmym1 | 0.23936918 | 0.30674326 | 0.24977355 | 0.05696971 | 0.14397336 | 0.76637203 |

| | | | | | | |
|---|---|---|---|---|---|---|
| Ahr | 0.21146036 | 0.2787484 | 0.22181198 | 0.05693642 | 0.11586911 | 0.67896213 |
| Adgrf5 | 0.13033956 | 0.19555228 | 0.13866754 | 0.05688473 | 0.02798887 | 0.2558529 |
| Dlgap4 | 0.57478826 | 0.61081946 | 0.55405352 | 0.05676593 | 0.31656179 | 0.89054584 |
| Usp37 | 0.27117996 | 0.33796168 | 0.28121176 | 0.05674992 | 0.1735582 | 0.8390286 |
| Nsmce2 | 0.58899218 | 0.62219698 | 0.5654573 | 0.05673967 | 0.32007393 | 0.89054584 |
| Ddc | 0.36874465 | 0.4312153 | 0.37453382 | 0.05668148 | 0.24038075 | 0.89054584 |
| Chaf1a | 0.18017243 | 0.24672138 | 0.19006236 | 0.05665903 | 0.08243681 | 0.55212586 |
| Rfc4 | 0.32519981 | 0.39013971 | 0.33348271 | 0.056657 | 0.21431733 | 0.89054584 |
| Rbm15 | 0.18579341 | 0.25243586 | 0.19579695 | 0.0566389 | 0.08887599 | 0.57774473 |
| Wdr60 | 0.19754506 | 0.26437625 | 0.20774283 | 0.05663342 | 0.10202658 | 0.62810247 |
| Gmfb | 0.50775308 | 0.55510833 | 0.49847934 | 0.05662898 | 0.29830842 | 0.89054584 |
| Cox7a1 | 0.10273479 | 0.16633307 | 0.10978682 | 0.05654624 | 0.00820626 | 0.09937802 |
| Smim24 | 0.15238934 | 0.218023 | 0.16152947 | 0.05649353 | 0.05161161 | 0.40164524 |
| Gnb1l | 0.06086273 | 0.12155829 | 0.06511132 | 0.05644697 | 2.68E-05 | 0.00064752 |
| Nedd9 | 0.06628641 | 0.12735508 | 0.07090865 | 0.05644643 | 0.00010415 | 0.00216334 |
| Tmem47 | 0.09007696 | 0.15275368 | 0.09632061 | 0.05643307 | 0.0031704 | 0.04507926 |
| Zwint | 1.19984756 | 0.98686118 | 0.93042886 | 0.05643231 | 0.38875178 | 0.89054584 |
| Airn | 0.37304261 | 0.43494697 | 0.37853114 | 0.05641582 | 0.24372731 | 0.89054584 |
| Edem1 | 0.25340892 | 0.32011584 | 0.26370662 | 0.05640922 | 0.15948189 | 0.80704115 |
| Rdh10 | 0.12833734 | 0.19298037 | 0.13658224 | 0.05639813 | 0.02719045 | 0.25100767 |
| Nde1 | 0.19291746 | 0.25935601 | 0.20304579 | 0.05631022 | 0.09816934 | 0.61534939 |
| Vbp1 | 1.38644143 | 1.05738517 | 1.00110892 | 0.05627625 | 0.3967003 | 0.89054584 |
| Brd9 | 0.26245219 | 0.32889582 | 0.27263304 | 0.05626278 | 0.1681612 | 0.82993631 |
| Ttyh3 | 0.39499883 | 0.45502705 | 0.39879267 | 0.05623439 | 0.25560182 | 0.89054584 |
| Hpcal1 | 0.77699221 | 0.75955249 | 0.70341354 | 0.05613895 | 0.35501087 | 0.89054584 |
| Ocel1 | 0.26469829 | 0.33097976 | 0.27484428 | 0.05613549 | 0.17065906 | 0.83412025 |
| Arhgap36 | 0.05711321 | 0.11713418 | 0.06110276 | 0.05603141 | 9.67E-06 | 0.00025784 |
| Fbxo3 | 0.69442847 | 0.70181685 | 0.6458599 | 0.05595694 | 0.34323576 | 0.89054584 |
| Parl | 0.60387271 | 0.63314176 | 0.57726025 | 0.05588152 | 0.32599208 | 0.89054584 |
| Fam208a | 0.33699711 | 0.4005595 | 0.34470199 | 0.05585751 | 0.22513593 | 0.89054584 |

| | | | | | | |
|---|---|---|---|---|---|---|
| Klf13 | 0.45215434 | 0.50610601 | 0.45026222 | 0.0558438 | 0.28169099 | 0.89054584 |
| Smim14 | 1.5934678 | 1.11501934 | 1.05927222 | 0.05574712 | 0.40315481 | 0.89054584 |
| Srd5a3 | 0.34344712 | 0.40644243 | 0.35080636 | 0.05563607 | 0.22998874 | 0.89054584 |
| Ncapg | 0.11453216 | 0.17779898 | 0.12216405 | 0.05563493 | 0.01692997 | 0.17876959 |
| Rps6ka1 | 0.10326329 | 0.16596856 | 0.11034239 | 0.05562617 | 0.00942002 | 0.11157803 |
| Paics | 1.04394606 | 0.91277446 | 0.85715193 | 0.05562253 | 0.38120055 | 0.89054584 |
| Ppp1r14a | 0.19692261 | 0.26268356 | 0.20711157 | 0.05557198 | 0.10563468 | 0.64240417 |
| Fam213a | 0.73075767 | 0.72733955 | 0.6717812 | 0.05555835 | 0.3500044 | 0.89054584 |
| Col8a2 | 0.11504996 | 0.17821105 | 0.12270613 | 0.05550492 | 0.0175403 | 0.1828909 |
| Gm4319 | 0.03170127 | 0.0894121 | 0.03392305 | 0.05548905 | 1.26E-14 | 9.99E-13 |
| Dnajc25 | 0.36106092 | 0.42284943 | 0.36736262 | 0.05548681 | 0.2408126 | 0.89054584 |
| Asph | 0.63319306 | 0.65556269 | 0.60008427 | 0.05547842 | 0.33333453 | 0.89054584 |
| Rbp7 | 0.04200874 | 0.10042157 | 0.04494974 | 0.05547182 | 4.47E-09 | 1.91E-07 |
| Dtl | 0.19532288 | 0.26094433 | 0.20548845 | 0.05545588 | 0.10431807 | 0.63653341 |
| Ptbp3 | 1.15346416 | 0.96546896 | 0.9100148 | 0.05545417 | 0.38824161 | 0.89054584 |
| Mrpl45 | 0.46002242 | 0.51263612 | 0.45719873 | 0.05543739 | 0.28606497 | 0.89054584 |
| Sqstm1 | 1.21484988 | 0.99218014 | 0.93679587 | 0.05538426 | 0.39148871 | 0.89054584 |
| Mcur1 | 0.33497042 | 0.39815698 | 0.34277967 | 0.05537731 | 0.2258244 | 0.89054584 |
| Tmem14c | 1.3014331 | 1.02648046 | 0.97110653 | 0.05537393 | 0.39525065 | 0.89054584 |
| Rfc1 | 0.69831412 | 0.70401709 | 0.64867669 | 0.0553404 | 0.3455115 | 0.89054584 |
| Ebf2 | 0.10486387 | 0.16736232 | 0.11202432 | 0.055338 | 0.01068358 | 0.12381337 |
| Ndufaf4 | 0.53594657 | 0.57751919 | 0.52220174 | 0.05531745 | 0.31082006 | 0.89054584 |
| Stmn4 | 0.05493197 | 0.11404177 | 0.05877058 | 0.05527119 | 5.89E-06 | 0.00016241 |
| Atp6v1e1 | 1.7791784 | 1.15000158 | 1.09473125 | 0.05527034 | 0.4070195 | 0.89054584 |
| Sesn2 | 0.33966834 | 0.40250133 | 0.34723275 | 0.05526858 | 0.22917775 | 0.89054584 |
| Gpr83 | 0.03234081 | 0.08983398 | 0.03460728 | 0.0552267 | 5.24E-14 | 3.87E-12 |
| Dact2 | 0.31343459 | 0.37743961 | 0.32222178 | 0.05521783 | 0.21232394 | 0.89054584 |
| Me1 | 0.26461528 | 0.32997903 | 0.27476261 | 0.05521642 | 0.1745689 | 0.84202063 |
| Gypc | 0.12517828 | 0.1884878 | 0.1332891 | 0.05519869 | 0.02684086 | 0.24851031 |
| Arfip2 | 0.83934025 | 0.79873587 | 0.74362675 | 0.05510911 | 0.36494666 | 0.89054584 |

| | | | | | | |
|---|---|---|---|---|---|---|
| Cdhr1 | 0.13805686 | 0.20178301 | 0.14669065 | 0.05509237 | 0.04008065 | 0.3376463 |
| Josd1 | 0.2442641 | 0.30971917 | 0.25464171 | 0.05507746 | 0.15679909 | 0.79932251 |
| Mylk | 0.12291212 | 0.18599843 | 0.13092458 | 0.05507385 | 0.02500966 | 0.23590763 |
| Zfp114 | 0.07548683 | 0.13578945 | 0.08073977 | 0.05504968 | 0.00074521 | 0.01280106 |
| Ngfr | 0.11463325 | 0.17730295 | 0.12226989 | 0.05503306 | 0.01799829 | 0.18665181 |
| Antxr2 | 0.19422326 | 0.25938876 | 0.20437212 | 0.05501664 | 0.10488841 | 0.63913594 |
| Gadd45b | 0.37503628 | 0.4353899 | 0.38038189 | 0.05500801 | 0.25023783 | 0.89054584 |
| Prss3 | 0.03442281 | 0.09181777 | 0.03683472 | 0.05498305 | 1.77E-12 | 1.14E-10 |
| Sod1 | 0.9800512 | 0.87911288 | 0.82414088 | 0.054972 | 0.37799094 | 0.89054584 |
| Ctrl | 0.02588396 | 0.08265951 | 0.02769886 | 0.05496065 | 1.18E-20 | 1.50E-18 |
| Gnai1 | 0.54730286 | 0.58656394 | 0.5316142 | 0.05494975 | 0.31505693 | 0.89054584 |
| Vamp2 | 0.97876524 | 0.87834074 | 0.82346408 | 0.05487667 | 0.37809889 | 0.89054584 |
| H2-Q2 | 0.13599429 | 0.19938919 | 0.1445486 | 0.05484059 | 0.03856679 | 0.32845551 |
| Atp6v1f | 2.22047977 | 1.18258218 | 1.12777677 | 0.05480541 | 0.41044291 | 0.89054584 |
| Dph3 | 0.84416971 | 0.80142761 | 0.746624 | 0.05480361 | 0.36618483 | 0.89054584 |
| Usp15 | 0.45912107 | 0.51119086 | 0.456406 | 0.05478486 | 0.28799974 | 0.89054584 |
| Dctpp1 | 0.71330864 | 0.71416939 | 0.65944743 | 0.05472196 | 0.34952391 | 0.89054584 |
| Ccdc141 | 0.07618631 | 0.13620889 | 0.08148703 | 0.05472186 | 0.000878 | 0.01478428 |
| Col9a3 | 0.24599634 | 0.31106413 | 0.25636182 | 0.05470231 | 0.1600818 | 0.8080503 |
| Gipc2 | 0.06220825 | 0.12119687 | 0.06654965 | 0.05464722 | 6.52E-05 | 0.00143389 |
| Gipr | 0.30981092 | 0.37326082 | 0.31873911 | 0.05452171 | 0.21274524 | 0.89054584 |
| Sec23a | 0.4344044 | 0.48899094 | 0.43447953 | 0.05451141 | 0.27943257 | 0.89054584 |
| Klrb1b | 0.03602908 | 0.09299607 | 0.03855314 | 0.05444294 | 2.37E-11 | 1.35E-09 |
| Bmp4 | 0.073957 | 0.13353746 | 0.07910536 | 0.0544321 | 0.00067344 | 0.01179168 |
| Rock1 | 1.20961258 | 0.98902343 | 0.93459139 | 0.05443204 | 0.39306327 | 0.89054584 |
| Ctxn1 | 0.74404225 | 0.73535956 | 0.68102699 | 0.05433256 | 0.35506113 | 0.89054584 |
| B4gat1 | 0.3280301 | 0.39045302 | 0.33618095 | 0.05427207 | 0.22598767 | 0.89054584 |
| Ppp1r14c | 0.19883253 | 0.26328205 | 0.20904798 | 0.05423407 | 0.11339175 | 0.67072771 |
| Pak1 | 0.17323051 | 0.2371933 | 0.18296263 | 0.05423067 | 0.08365113 | 0.55785378 |
| Dusp5 | 0.28698682 | 0.35087845 | 0.2966551 | 0.05422335 | 0.19722562 | 0.89054584 |

| | | | | | | |
|---|---|---|---|---|---|---|
| Ednrb | 0.06961665 | 0.12866177 | 0.07446766 | 0.05419411 | 0.00034875 | 0.00651565 |
| Hist1h1b | 0.08197316 | 0.14183503 | 0.08766808 | 0.05416694 | 0.00199728 | 0.03006907 |
| Ap1m2 | 0.41460604 | 0.47081334 | 0.41665932 | 0.05415401 | 0.27241293 | 0.89054584 |
| 1810011O10 | 0.10241981 | 0.16359696 | 0.10945567 | 0.05414129 | 0.01059717 | 0.12300859 |
| Insm1 | 0.67019973 | 0.68216235 | 0.6280577 | 0.05410465 | 0.34408156 | 0.89054584 |
| Mtfr1 | 0.42327723 | 0.47854602 | 0.42449228 | 0.05405375 | 0.27650441 | 0.89054584 |
| Inafm2 | 0.10593778 | 0.16710815 | 0.11315229 | 0.05395585 | 0.01315577 | 0.14604317 |
| Inhba | 0.05510172 | 0.11290131 | 0.05895208 | 0.05394923 | 1.01E-05 | 0.00026565 |
| Cercam | 0.13069436 | 0.19292345 | 0.1390369 | 0.05388655 | 0.03548524 | 0.30806285 |
| Syne2 | 0.53897671 | 0.57859173 | 0.52472136 | 0.05387036 | 0.31621521 | 0.89054584 |
| Zscan21 | 0.24913635 | 0.31333952 | 0.25947627 | 0.05386325 | 0.16674102 | 0.82618294 |
| Tshz3 | 0.17521665 | 0.23883317 | 0.18499591 | 0.05383726 | 0.08757523 | 0.57192148 |
| Eftud1 | 0.20254351 | 0.26664025 | 0.21280589 | 0.05383436 | 0.11928055 | 0.69117879 |
| Glis1 | 0.04471668 | 0.10165008 | 0.04784619 | 0.05380389 | 8.07E-08 | 2.94E-06 |
| Adra2a | 0.30617155 | 0.36902897 | 0.3152354 | 0.05379357 | 0.21329849 | 0.89054584 |
| Fxyd2 | 0.1306896 | 0.19282031 | 0.13903194 | 0.05378837 | 0.03573803 | 0.30938746 |
| Smim4 | 0.55030089 | 0.58781806 | 0.53408585 | 0.05373221 | 0.31963589 | 0.89054584 |
| Peak1 | 0.19366645 | 0.25752402 | 0.20380665 | 0.05371737 | 0.10973129 | 0.65549414 |
| Aggf1 | 0.61107995 | 0.63663495 | 0.58292454 | 0.05371041 | 0.33386237 | 0.89054584 |
| Chst12 | 0.36132549 | 0.42131048 | 0.36761003 | 0.05370046 | 0.2480693 | 0.89054584 |
| Osbpl11 | 0.22660374 | 0.2906902 | 0.23702687 | 0.05366333 | 0.14575997 | 0.76928578 |
| Mrps27 | 0.21369288 | 0.27772181 | 0.22406146 | 0.05366035 | 0.1322609 | 0.72909452 |
| Mark4 | 0.24311762 | 0.30710331 | 0.25350249 | 0.05360082 | 0.16228967 | 0.81470761 |
| Armcx3 | 1.20140381 | 0.98468784 | 0.93109633 | 0.0535915 | 0.39428938 | 0.89054584 |
| Kctd13 | 0.60192324 | 0.62930368 | 0.5757221 | 0.05358158 | 0.33228925 | 0.89054584 |
| Ptk2 | 0.29676993 | 0.35970676 | 0.30615308 | 0.05355367 | 0.20754677 | 0.89054584 |
| Erlec1 | 0.85669716 | 0.80785533 | 0.75431885 | 0.05353648 | 0.37045055 | 0.89054584 |
| Gm266 | 0.14530808 | 0.20774193 | 0.15420815 | 0.05353378 | 0.05289983 | 0.40810662 |
| 2410089E03I | 0.31589145 | 0.37805627 | 0.32457924 | 0.05347704 | 0.22136303 | 0.89054584 |
| B230217C12 | 0.14586072 | 0.20815486 | 0.15478024 | 0.05337462 | 0.05407159 | 0.41528696 |

| | | | | | | |
|---|---|---|---|---|---|---|
| Scand1 | 1.79293814 | 1.15004508 | 1.09675007 | 0.05329501 | 0.41044725 | 0.89054584 |
| Tnfrsf11b | 0.03793787 | 0.0938764 | 0.04059512 | 0.05328128 | 4.83E-10 | 2.37E-08 |
| Prkx | 0.14431942 | 0.20631172 | 0.1531844 | 0.05312733 | 0.05306755 | 0.40902355 |
| Arhgap25 | 0.04426579 | 0.10044667 | 0.04736393 | 0.05308274 | 8.88E-08 | 3.22E-06 |
| Map3k15 | 0.20221601 | 0.26549007 | 0.21247449 | 0.05301558 | 0.1225211 | 0.70076598 |
| Spdya | 0.0308353 | 0.08599897 | 0.03299655 | 0.05300242 | 3.61E-14 | 2.71E-12 |
| Bbs2 | 0.1427945 | 0.2046017 | 0.1516046 | 0.05299711 | 0.05169385 | 0.40185388 |
| Ttll4 | 0.07139215 | 0.12934802 | 0.07636491 | 0.05298311 | 0.00061368 | 0.0109034 |
| Rnasel | 0.11608173 | 0.17670573 | 0.12378598 | 0.05291975 | 0.02319956 | 0.22378393 |
| Nfe2l2 | 0.7739428 | 0.7542748 | 0.70137455 | 0.05290025 | 0.36264621 | 0.89054584 |
| Pcdh18 | 0.14301498 | 0.2047182 | 0.15183307 | 0.05288513 | 0.05232031 | 0.40455457 |
| Sec24d | 0.4082579 | 0.46375934 | 0.41089888 | 0.05286046 | 0.27446852 | 0.89054584 |
| Serpinb9 | 0.11163101 | 0.17195975 | 0.11912506 | 0.05283469 | 0.01939786 | 0.19735881 |
| 2310035C23I | 0.25909216 | 0.32208919 | 0.26932066 | 0.05276854 | 0.18066447 | 0.85856175 |
| Slc1a3 | 0.0895989 | 0.14846919 | 0.0958103 | 0.05265889 | 0.00522413 | 0.06800354 |
| Mgll | 0.16161134 | 0.22368316 | 0.17103515 | 0.05264801 | 0.07576982 | 0.5231004 |
| Gdf10 | 0.06452406 | 0.12163111 | 0.06902503 | 0.05260608 | 0.00019209 | 0.00378383 |
| Rnf185 | 0.14486901 | 0.20628808 | 0.15375355 | 0.05253454 | 0.0557072 | 0.423293 |
| Mgat2 | 0.97248513 | 0.87264828 | 0.8201416 | 0.05250669 | 0.38274765 | 0.89054584 |
| Ifi203 | 0.09067345 | 0.14944543 | 0.09695733 | 0.05248809 | 0.00583234 | 0.07417302 |
| Slc40a1 | 0.28204244 | 0.344283 | 0.29183714 | 0.05244586 | 0.20122454 | 0.89054584 |
| Sh3tc1 | 0.06749827 | 0.12463828 | 0.07220382 | 0.05243446 | 0.00035807 | 0.00668116 |
| Ubr7 | 0.4172663 | 0.47149988 | 0.41906698 | 0.0524329 | 0.27997415 | 0.89054584 |
| Cited1 | 0.2106063 | 0.27333713 | 0.22095084 | 0.05238629 | 0.1346642 | 0.73479922 |
| Tmem245 | 0.29024969 | 0.35219221 | 0.29982818 | 0.05236404 | 0.20791723 | 0.89054584 |
| Pus3 | 0.24588973 | 0.30860303 | 0.256256 | 0.05234703 | 0.17062171 | 0.83412025 |
| Amdhd2 | 0.16747524 | 0.22939512 | 0.17706156 | 0.05233356 | 0.08425157 | 0.559798 |
| Mecom | 0.11067229 | 0.17044317 | 0.1181201 | 0.05232307 | 0.01952081 | 0.19784693 |
| Dcaf13 | 0.36232846 | 0.42085065 | 0.36854757 | 0.05230308 | 0.25424784 | 0.89054584 |
| Rora | 0.24996173 | 0.31257609 | 0.26029416 | 0.05228194 | 0.17469284 | 0.84226179 |

| | | | | | | |
|---|---|---|---|---|---|---|
| Bub1b | 0.1034147 | 0.16277616 | 0.11050154 | 0.05227462 | 0.01376249 | 0.15092931 |
| Plin2 | 0.45011524 | 0.50072193 | 0.44845845 | 0.05226348 | 0.29358061 | 0.89054584 |
| Tbc1d4 | 0.07593441 | 0.13347553 | 0.08121793 | 0.0522576 | 0.00136023 | 0.02159978 |
| Gm12942 | 0.19590927 | 0.25829238 | 0.20608355 | 0.05220883 | 0.11894289 | 0.69060234 |
| Pdlim1 | 0.23526951 | 0.29780174 | 0.24568776 | 0.05211397 | 0.1615201 | 0.81253242 |
| Ifngr1 | 0.37065108 | 0.42836272 | 0.37630815 | 0.05205457 | 0.25963548 | 0.89054584 |
| Ctla2b | 0.03395791 | 0.08837034 | 0.03633736 | 0.05203298 | 1.27E-11 | 7.44E-10 |
| Tspan12 | 0.20393708 | 0.26624139 | 0.21421554 | 0.05202585 | 0.12892184 | 0.71861164 |
| Arrdc3 | 0.2764917 | 0.33839902 | 0.28641477 | 0.05198426 | 0.19888866 | 0.89054584 |
| Dhx57 | 0.19167156 | 0.25373839 | 0.20177963 | 0.05195876 | 0.11513005 | 0.67639855 |
| 5730409E04l | 0.11838337 | 0.17809542 | 0.12619343 | 0.05190198 | 0.02767162 | 0.25375276 |
| Plxnd1 | 0.10735973 | 0.16653837 | 0.11464517 | 0.0518932 | 0.01747955 | 0.18270984 |
| Ly6g6d | 0.05343848 | 0.10906116 | 0.05717363 | 0.05188752 | 1.18E-05 | 0.00030714 |
| Mtmr12 | 0.11654835 | 0.17614719 | 0.12427421 | 0.05187298 | 0.02590696 | 0.24216579 |
| Caap1 | 0.31676546 | 0.37728726 | 0.32541714 | 0.05187012 | 0.22885722 | 0.89054584 |
| Prkar2a | 0.37916624 | 0.43598035 | 0.38420877 | 0.05177158 | 0.26507217 | 0.89054584 |
| Nfkbil1 | 0.31049784 | 0.37114099 | 0.31939978 | 0.05174121 | 0.22520675 | 0.89054584 |
| Rpp40 | 0.14252988 | 0.20307132 | 0.15133036 | 0.05174096 | 0.05558837 | 0.42283327 |
| Npr3 | 0.11895682 | 0.17840669 | 0.12679293 | 0.05161376 | 0.02894559 | 0.26244722 |
| Hsph1 | 0.76898804 | 0.74958216 | 0.69804715 | 0.05153502 | 0.36543909 | 0.89054584 |
| Rrp8 | 0.3052527 | 0.36588148 | 0.31434973 | 0.05153174 | 0.22250929 | 0.89054584 |
| Gem | 0.22058911 | 0.2824415 | 0.23099613 | 0.05144537 | 0.14972879 | 0.77861117 |
| Lrrcc1 | 0.32828045 | 0.38785408 | 0.33641941 | 0.05143467 | 0.23813993 | 0.89054584 |
| Mxi1 | 0.19906746 | 0.26068251 | 0.20928606 | 0.05139645 | 0.12628229 | 0.71147001 |
| Arhgef15 | 0.07301794 | 0.12938731 | 0.07810205 | 0.05128526 | 0.0011094 | 0.01807196 |
| Crb3 | 0.65656664 | 0.66914667 | 0.61786142 | 0.05128525 | 0.34948439 | 0.89054584 |
| Ggta1 | 0.19043209 | 0.25179138 | 0.20051933 | 0.05127204 | 0.11677306 | 0.68200227 |
| Pltp | 0.16129391 | 0.22195056 | 0.17070853 | 0.05124203 | 0.08098347 | 0.54583838 |
| Cntnap3 | 0.02740853 | 0.08050483 | 0.02933012 | 0.05117471 | 2.17E-16 | 2.03E-14 |
| St18 | 0.49308764 | 0.53707755 | 0.48594291 | 0.05113464 | 0.31197868 | 0.89054584 |

| | | | | | | |
|---|---|---|---|---|---|---|
| Fam101b | 0.32206072 | 0.38151796 | 0.33048518 | 0.05103278 | 0.23594009 | 0.89054584 |
| Tgfbr2 | 0.18599367 | 0.24693561 | 0.19600101 | 0.0509346 | 0.11300239 | 0.66962283 |
| Fam122a | 0.68689039 | 0.69126237 | 0.64036495 | 0.05089742 | 0.35557894 | 0.89054584 |
| Cdk7 | 0.46656989 | 0.51383816 | 0.46294241 | 0.05089575 | 0.30425462 | 0.89054584 |
| Gas7 | 0.068677 | 0.12433796 | 0.07346351 | 0.05087445 | 0.00062684 | 0.01106931 |
| Frmd4a | 0.27535339 | 0.33615877 | 0.28530091 | 0.05085786 | 0.20312672 | 0.89054584 |
| Rap1gap2 | 0.53096297 | 0.56888205 | 0.51804488 | 0.05083717 | 0.32376553 | 0.89054584 |
| Crybb1 | 0.07466537 | 0.13069773 | 0.07986218 | 0.05083555 | 0.00151043 | 0.02354648 |
| Fam126a | 0.18423535 | 0.24496474 | 0.19420873 | 0.05075601 | 0.11168731 | 0.66441926 |
| Lpxn | 0.03346068 | 0.08655459 | 0.03580539 | 0.0507492 | 2.01E-11 | 1.16E-09 |
| Rbmx2 | 0.32157644 | 0.38070345 | 0.33002229 | 0.05068116 | 0.23715979 | 0.89054584 |
| Zfp566 | 0.09069409 | 0.14762667 | 0.09697936 | 0.0506473 | 0.00748459 | 0.09211506 |
| Tmem184b | 0.43588996 | 0.48644171 | 0.43580738 | 0.05063433 | 0.29415055 | 0.89054584 |
| Zfp608 | 0.21067746 | 0.27160062 | 0.2210226 | 0.05057802 | 0.14318122 | 0.76468733 |
| Yif1b | 0.73782531 | 0.72726537 | 0.67671579 | 0.05054958 | 0.3639143 | 0.89054584 |
| Gstm4 | 0.184469 | 0.24499436 | 0.19444696 | 0.05054739 | 0.11292597 | 0.66962283 |
| Xkr6 | 0.11049287 | 0.1684504 | 0.11793198 | 0.05051842 | 0.02298076 | 0.22241326 |
| Kif2c | 0.08245387 | 0.13868408 | 0.08818146 | 0.05050262 | 0.00381256 | 0.0523929 |
| Psmc3ip | 0.17945171 | 0.23980677 | 0.18932611 | 0.05048066 | 0.107076 | 0.64763783 |
| Uhrf1 | 0.1642168 | 0.22419292 | 0.17371455 | 0.05047837 | 0.08789917 | 0.57377853 |
| Sirt6 | 0.19373684 | 0.25431216 | 0.20387814 | 0.05043402 | 0.12455827 | 0.70642606 |
| Elp6 | 0.24461011 | 0.30539816 | 0.25498541 | 0.05041275 | 0.17849607 | 0.85194992 |
| Arl13b | 0.32790205 | 0.38638191 | 0.33605896 | 0.05032295 | 0.24269792 | 0.89054584 |
| H2afj | 1.46627069 | 1.07636943 | 1.02607091 | 0.05029851 | 0.409674 | 0.89054584 |
| Krtcap2 | 2.36772628 | 1.17985076 | 1.12955641 | 0.05029434 | 0.41783201 | 0.89054584 |
| Hnf1b | 0.14474862 | 0.20390672 | 0.15362888 | 0.05027784 | 0.06366409 | 0.46498842 |
| Ifi30 | 0.56203114 | 0.59396775 | 0.54370008 | 0.05026767 | 0.33332821 | 0.89054584 |
| Gm10320 | 0.29301858 | 0.35276855 | 0.30251671 | 0.05025183 | 0.21949237 | 0.89054584 |
| Ahnak2 | 0.12777518 | 0.18622263 | 0.13599649 | 0.05022614 | 0.04265831 | 0.35099897 |
| Pfdn4 | 1.32138708 | 1.02869752 | 0.97848122 | 0.0502163 | 0.40551405 | 0.89054584 |

| | | | | | | |
|---|---|---|---|---|---|---|
| Skap2 | 0.33196701 | 0.39013333 | 0.3399272 | 0.05020614 | 0.24569164 | 0.89054584 |
| Cttnbp2nl | 0.19461315 | 0.2549485 | 0.20476799 | 0.05018051 | 0.12678485 | 0.71241644 |
| Gpatch1 | 0.26750504 | 0.3276923 | 0.27760407 | 0.05008823 | 0.20028183 | 0.89054584 |
| Lrch3 | 0.18049209 | 0.24039883 | 0.19038883 | 0.05001 | 0.11051802 | 0.65883644 |
| Cyp20a1 | 0.47922555 | 0.52397593 | 0.47397334 | 0.05000259 | 0.31153551 | 0.89054584 |
| Rbms3 | 0.68185801 | 0.68666334 | 0.63667425 | 0.04998909 | 0.35725669 | 0.89054584 |
| Trappc2 | 0.40870938 | 0.46128542 | 0.41130933 | 0.04997608 | 0.28566834 | 0.89054584 |
| Hivep2 | 0.0957618 | 0.15235001 | 0.10238786 | 0.04996215 | 0.01150041 | 0.13128581 |
| Tmem183a | 0.59620754 | 0.62109487 | 0.57119856 | 0.04989631 | 0.34201288 | 0.89054584 |
| Stx7 | 1.00135063 | 0.88507125 | 0.83521226 | 0.04985899 | 0.39045972 | 0.89054584 |
| Reln | 0.03648764 | 0.08888438 | 0.03904371 | 0.04984068 | 1.36E-09 | 6.33E-08 |
| Zcchc5 | 0.13593963 | 0.1943079 | 0.14449181 | 0.04981608 | 0.05410933 | 0.41528696 |
| Scarf2 | 0.1220157 | 0.17980447 | 0.1299887 | 0.04981577 | 0.03709392 | 0.31859456 |
| 9330020H09I | 0.13688415 | 0.19523586 | 0.14547295 | 0.04976291 | 0.05549993 | 0.42238211 |
| Dennd5b | 0.18625461 | 0.24599728 | 0.19626688 | 0.0497304 | 0.1189028 | 0.69060234 |
| Lama1 | 0.03110336 | 0.08299744 | 0.03328335 | 0.04971408 | 1.71E-12 | 1.12E-10 |
| Trappc9 | 0.28774847 | 0.34710864 | 0.29739627 | 0.04971238 | 0.21805389 | 0.89054584 |
| Sox7 | 0.04758812 | 0.10061795 | 0.05091729 | 0.04970066 | 2.71E-06 | 7.83E-05 |
| Mtrf1l | 0.23364171 | 0.29375649 | 0.24406334 | 0.04969316 | 0.17141307 | 0.83588128 |
| Ubox5 | 0.05255098 | 0.10589508 | 0.05622462 | 0.04967046 | 1.93E-05 | 0.00047966 |
| Casz1 | 0.3786948 | 0.43340668 | 0.3837724 | 0.04963428 | 0.27340308 | 0.89054584 |
| Tspan5 | 0.58854899 | 0.6147277 | 0.56510352 | 0.04962418 | 0.34122515 | 0.89054584 |
| 1810043G02 | 0.56552884 | 0.59617126 | 0.54654934 | 0.04962192 | 0.33615081 | 0.89054584 |
| Nudt13 | 0.18834886 | 0.24801806 | 0.19839965 | 0.04961841 | 0.12197517 | 0.70028846 |
| Actn3 | 0.13238578 | 0.19039516 | 0.14079708 | 0.04959808 | 0.05037914 | 0.39545033 |
| Ranbp1 | 2.46821103 | 1.177427 | 1.12785323 | 0.04957377 | 0.41887219 | 0.89054584 |
| C1qbp | 1.14268062 | 0.95472062 | 0.90514917 | 0.04957144 | 0.39930293 | 0.89054584 |
| RP23-4H17.3 | 0.03533079 | 0.08735242 | 0.0378061 | 0.04954632 | 5.12E-10 | 2.48E-08 |
| 1110002L01F | 0.15615131 | 0.21493072 | 0.16541128 | 0.04951944 | 0.08154492 | 0.54834798 |
| Ccdc114 | 0.03409433 | 0.08600261 | 0.03648331 | 0.0495193 | 1.28E-10 | 6.71E-09 |

| | | | | | | |
|---|---|---|---|---|---|---|
| Mbnl1 | 0.3143204 | 0.37255024 | 0.32307211 | 0.04947813 | 0.23776461 | 0.89054584 |
| Fam109a | 0.25142223 | 0.31121059 | 0.26174063 | 0.04946996 | 0.18927871 | 0.87982135 |
| Rgma | 0.11599191 | 0.17314472 | 0.12369199 | 0.04945272 | 0.03125438 | 0.27782097 |
| Cox7b | 2.35974927 | 1.17899535 | 1.129547 | 0.04944835 | 0.41919426 | 0.89054584 |
| Ogg1 | 0.14209408 | 0.20032231 | 0.15087867 | 0.04944364 | 0.06341076 | 0.46337157 |
| Timp1 | 0.17615085 | 0.23539295 | 0.18595171 | 0.04944123 | 0.10772292 | 0.64857426 |
| Polr3d | 0.4880762 | 0.53101207 | 0.48162943 | 0.04938264 | 0.3164345 | 0.89054584 |
| Prkar1a | 2.43453824 | 1.17801347 | 1.12865125 | 0.04936223 | 0.41926993 | 0.89054584 |
| Hs3st1 | 0.10120449 | 0.15753167 | 0.10817759 | 0.04935408 | 0.01677003 | 0.17733907 |
| Dse | 0.16829407 | 0.22724068 | 0.17790197 | 0.0493387 | 0.09816225 | 0.61534939 |
| Gramd3 | 0.34321157 | 0.39990047 | 0.35058383 | 0.04931664 | 0.25611367 | 0.89054584 |
| Irf2bpl | 0.77071406 | 0.74850635 | 0.69920829 | 0.04929805 | 0.37127395 | 0.89054584 |
| Gsk3b | 1.35889835 | 1.04107255 | 0.99177864 | 0.04929391 | 0.40843944 | 0.89054584 |
| Zdhhc6 | 0.49244571 | 0.53465257 | 0.48539126 | 0.04926131 | 0.31816872 | 0.89054584 |
| Alkbh4 | 0.10885115 | 0.165464 | 0.11621019 | 0.04925381 | 0.02415583 | 0.23040214 |
| Rnf38 | 0.14082029 | 0.19878145 | 0.14955798 | 0.04922347 | 0.0625911 | 0.45969899 |
| Clptm1l | 0.80787492 | 0.77289069 | 0.72368498 | 0.04920571 | 0.37570654 | 0.89054584 |
| Zfp959 | 0.10658655 | 0.16303337 | 0.11383351 | 0.04919986 | 0.02202389 | 0.21618304 |
| Gm26825 | 0.21043413 | 0.2699739 | 0.22077721 | 0.04919669 | 0.14959523 | 0.77846267 |
| Mbtps2 | 0.16567435 | 0.22437554 | 0.17521226 | 0.04916328 | 0.09556137 | 0.60579307 |
| Ptcd2 | 0.35873503 | 0.41434048 | 0.36518601 | 0.04915447 | 0.26529715 | 0.89054584 |
| Kif12 | 0.68893477 | 0.69096536 | 0.64185918 | 0.04910617 | 0.36075642 | 0.89054584 |
| Etfb | 1.8595491 | 1.15441706 | 1.10534059 | 0.04907647 | 0.41806186 | 0.89054584 |
| Rgl1 | 0.2150499 | 0.274486 | 0.22542772 | 0.04905828 | 0.15531683 | 0.79568319 |
| Fadd | 0.11408267 | 0.1706937 | 0.12169342 | 0.04900029 | 0.03033181 | 0.27111628 |
| Rars2 | 0.51748545 | 0.55568103 | 0.50672418 | 0.04895686 | 0.32631165 | 0.89054584 |
| Nubpl | 0.17407257 | 0.23277976 | 0.18382487 | 0.04895489 | 0.10735183 | 0.64845225 |
| Zscan26 | 0.27425046 | 0.33316983 | 0.28422108 | 0.04894875 | 0.21117064 | 0.89054584 |
| Morn4 | 0.08555455 | 0.14043658 | 0.09149248 | 0.0489441 | 0.00634575 | 0.07965421 |
| Grhpr | 0.68249141 | 0.68606768 | 0.63713976 | 0.04892792 | 0.36025608 | 0.89054584 |

| | | | | | | |
|---|---|---|---|---|---|---|
| Paip1 | 0.33897644 | 0.39544529 | 0.34657761 | 0.04886768 | 0.25561812 | 0.89054584 |
| Sh3pxd2a | 0.2084287 | 0.26760836 | 0.21875376 | 0.0488546 | 0.14905657 | 0.77749274 |
| Leprot | 0.78836163 | 0.75979873 | 0.7109566 | 0.04884213 | 0.37445937 | 0.89054584 |
| Rragd | 0.09073318 | 0.14583456 | 0.09702108 | 0.04881348 | 0.00953863 | 0.11279862 |
| Hmcn1 | 0.12740006 | 0.18441243 | 0.13560555 | 0.04880688 | 0.04678718 | 0.37496992 |
| Ccdc79 | 0.04329184 | 0.09510166 | 0.04632219 | 0.04877946 | 4.65E-07 | 1.53E-05 |
| Stat5a | 0.09815145 | 0.15371129 | 0.10493759 | 0.0487737 | 0.01517784 | 0.16311955 |
| Vcpip1 | 0.30414827 | 0.36202379 | 0.31328461 | 0.04873917 | 0.23428319 | 0.89054584 |
| Afap1l1 | 0.08680911 | 0.14156735 | 0.09283198 | 0.04873538 | 0.00722509 | 0.08937658 |
| Pkig | 1.06822163 | 0.91816025 | 0.86950617 | 0.04865407 | 0.39716361 | 0.89054584 |
| Cbs | 0.11299269 | 0.16919627 | 0.12055183 | 0.04864444 | 0.03005736 | 0.26949389 |
| Aldoc | 0.34718935 | 0.40296859 | 0.35433789 | 0.04863069 | 0.26127781 | 0.89054584 |
| Fbxo16 | 0.14408381 | 0.20157029 | 0.15294037 | 0.04862992 | 0.06925169 | 0.49496101 |
| Sirpa | 0.17285538 | 0.23120782 | 0.18257842 | 0.04862941 | 0.10732134 | 0.64845225 |
| Fam114a1 | 0.25250909 | 0.31143949 | 0.26281639 | 0.0486231 | 0.19436006 | 0.88918656 |
| Abcb10 | 0.24901808 | 0.30793906 | 0.25935904 | 0.04858002 | 0.19142508 | 0.88412897 |
| Rdm1 | 0.25490699 | 0.31366737 | 0.26518785 | 0.04847951 | 0.19718747 | 0.89054584 |
| Appl1 | 0.49443869 | 0.53556354 | 0.48710317 | 0.04846037 | 0.32150176 | 0.89054584 |
| Vopp1 | 0.1691729 | 0.22722883 | 0.17880365 | 0.04842518 | 0.10351479 | 0.63276555 |
| Nt5m | 0.51079402 | 0.54947974 | 0.50106195 | 0.04841779 | 0.32628488 | 0.89054584 |
| Nmt2 | 0.66306005 | 0.67113878 | 0.62273405 | 0.04840473 | 0.35862498 | 0.89054584 |
| Elp4 | 0.64640478 | 0.65853278 | 0.61017872 | 0.04835406 | 0.35598889 | 0.89054584 |
| Bbs10 | 0.05473911 | 0.10690254 | 0.05856437 | 0.04833817 | 6.02E-05 | 0.00134213 |
| Spa17 | 0.24691903 | 0.3055955 | 0.25727747 | 0.04831803 | 0.19079429 | 0.88318504 |
| Pla2g2f | 0.36468299 | 0.41906325 | 0.37074693 | 0.04831632 | 0.27187218 | 0.89054584 |
| Gpr155 | 0.07846635 | 0.13223758 | 0.08392261 | 0.04831497 | 0.00365707 | 0.05073697 |
| Traf1 | 0.03794086 | 0.08890518 | 0.04059832 | 0.04830685 | 1.48E-08 | 5.94E-07 |
| Zc3h14 | 0.82095432 | 0.78036306 | 0.73206208 | 0.04830099 | 0.37927176 | 0.89054584 |
| Gria3 | 0.18585205 | 0.24413608 | 0.19585671 | 0.04827937 | 0.12539237 | 0.7081958 |
| Map3k6 | 0.04053814 | 0.09152737 | 0.04337669 | 0.04815068 | 1.16E-07 | 4.12E-06 |

| | | | | | | |
|---|---|---|---|---|---|---|
| Wasf2 | 0.90329618 | 0.83008088 | 0.781942 | 0.04813888 | 0.38712527 | 0.89054584 |
| Tmem192 | 0.26941154 | 0.3275821 | 0.2794765 | 0.0481056 | 0.21129615 | 0.89054584 |
| Gulp1 | 0.13635237 | 0.19298599 | 0.14492059 | 0.0480654 | 0.06114681 | 0.45276125 |
| Cldn2 | 0.0522242 | 0.10391916 | 0.05587519 | 0.04804397 | 3.09E-05 | 0.00073399 |
| Lyn | 0.12380507 | 0.17988717 | 0.13185651 | 0.04803066 | 0.04484008 | 0.36375062 |
| Mlec | 0.95890673 | 0.86090016 | 0.81289975 | 0.04800042 | 0.39161857 | 0.89054584 |
| Smim3 | 0.07239086 | 0.12540984 | 0.07743204 | 0.0479778 | 0.00194631 | 0.02942399 |
| Mrpl49 | 0.74005783 | 0.72622387 | 0.67826715 | 0.04795672 | 0.37092316 | 0.89054584 |
| Flna | 0.4122235 | 0.46236194 | 0.41450008 | 0.04786185 | 0.29530141 | 0.89054584 |
| Fam178a | 0.31693873 | 0.37340798 | 0.3255832 | 0.04782478 | 0.24687712 | 0.89054584 |
| Pqlc1 | 0.30078655 | 0.35783145 | 0.31003875 | 0.0477927 | 0.23632343 | 0.89054584 |
| Tnfrsf21 | 0.16001968 | 0.21707622 | 0.16939698 | 0.04767924 | 0.0948737 | 0.60275147 |
| Vapb | 1.17338337 | 0.9665745 | 0.91890254 | 0.04767196 | 0.40450542 | 0.89054584 |
| Fam53c | 0.19482352 | 0.25261874 | 0.20498156 | 0.04763717 | 0.13946511 | 0.74952855 |
| Atad2 | 0.18148451 | 0.23903235 | 0.19140215 | 0.0476302 | 0.12315227 | 0.70193617 |
| Rplp1 | 4.68974448 | 0.83837147 | 0.790745 | 0.04762648 | 0.38950574 | 0.89054584 |
| Mapk6 | 0.29973002 | 0.3566366 | 0.30901746 | 0.04761914 | 0.23639834 | 0.89054584 |
| Stxbp6 | 0.17866292 | 0.23613295 | 0.18852008 | 0.04761287 | 0.11966399 | 0.69257027 |
| Zfand6 | 1.27996799 | 1.01058995 | 0.96297737 | 0.04761258 | 0.40890895 | 0.89054584 |
| Enthd2 | 0.12509259 | 0.18067974 | 0.13319973 | 0.04748 | 0.04838554 | 0.38438004 |
| Ptpn6 | 0.20236021 | 0.26007086 | 0.21262041 | 0.04745044 | 0.1492316 | 0.77797903 |
| Tmem230 | 0.56068405 | 0.58996944 | 0.54260058 | 0.04736887 | 0.3421053 | 0.89054584 |
| Ralgds | 0.3036138 | 0.36013342 | 0.31276895 | 0.04736447 | 0.24023982 | 0.89054584 |
| Isg20l2 | 0.27713647 | 0.33439745 | 0.2870454 | 0.04735205 | 0.22107851 | 0.89054584 |
| Ren1 | 0.01953079 | 0.06823598 | 0.02090075 | 0.04733524 | 2.50E-26 | 4.32E-24 |
| Clec12a | 0.01841902 | 0.06704077 | 0.01971107 | 0.04732971 | 2.36E-29 | 4.57E-27 |
| Usp35 | 0.09391188 | 0.14770637 | 0.10041373 | 0.04729264 | 0.01410844 | 0.15414028 |
| 4930515G01 | 0.02341748 | 0.07227544 | 0.02505971 | 0.04721573 | 8.32E-19 | 9.35E-17 |
| Efcab1 | 0.7335948 | 0.72097942 | 0.67376622 | 0.04721321 | 0.3720349 | 0.89054584 |
| Tlk2 | 0.35048439 | 0.40459472 | 0.35744119 | 0.04715353 | 0.26940747 | 0.89054584 |

| | | | | | | |
|---|---|---|---|---|---|---|
| Mras | 0.11969146 | 0.17470673 | 0.12756078 | 0.04714595 | 0.04254147 | 0.35075781 |
| Cacybp | 1.98086307 | 1.16359201 | 1.11644747 | 0.04714453 | 0.4220177 | 0.89054584 |
| Oser1 | 0.87262833 | 0.81108238 | 0.76393947 | 0.04714291 | 0.38686303 | 0.89054584 |
| Necab2 | 0.26227458 | 0.31959782 | 0.27245808 | 0.04713974 | 0.2101002 | 0.89054584 |
| Sult1d1 | 0.07599919 | 0.12842685 | 0.08128713 | 0.04713972 | 0.00344831 | 0.04830302 |
| Mirg | 0.2523174 | 0.30972701 | 0.2626267 | 0.04710031 | 0.20170425 | 0.89054584 |
| Gba2 | 0.15221653 | 0.20843764 | 0.16135104 | 0.0470866 | 0.08697744 | 0.57045578 |
| Ccdc167 | 0.36667816 | 0.41965863 | 0.37260825 | 0.04705037 | 0.27816593 | 0.89054584 |
| Fgfr3 | 0.11619616 | 0.17094173 | 0.12390571 | 0.04703602 | 0.03848141 | 0.32813014 |
| Gm5218 | 0.02328372 | 0.07195003 | 0.02491658 | 0.04703345 | 7.19E-19 | 8.15E-17 |
| Pcbp3 | 0.25219961 | 0.30954269 | 0.26251013 | 0.04703256 | 0.20193825 | 0.89054584 |
| Cfap126 | 0.16635134 | 0.22289353 | 0.17590761 | 0.04698592 | 0.10667075 | 0.64653533 |
| Adam10 | 0.49132582 | 0.53138398 | 0.48442825 | 0.04695573 | 0.32578142 | 0.89054584 |
| Ipo11 | 0.13665994 | 0.19216172 | 0.14524008 | 0.04692164 | 0.06614536 | 0.47662791 |
| Npc2 | 1.97377488 | 1.16284838 | 1.11593462 | 0.04691377 | 0.42235934 | 0.89054584 |
| Ncoa7 | 0.37919781 | 0.43113895 | 0.38423798 | 0.04690097 | 0.2847863 | 0.89054584 |
| Smco4 | 0.31889864 | 0.37435214 | 0.32746053 | 0.04689161 | 0.25233821 | 0.89054584 |
| Ccne2 | 0.13442036 | 0.1897958 | 0.14291291 | 0.04688289 | 0.06320963 | 0.46236792 |
| Rpp38 | 0.20288666 | 0.25997657 | 0.21315307 | 0.04682349 | 0.15305157 | 0.78825634 |
| Grpel1 | 1.41373136 | 1.05685241 | 1.01003328 | 0.04681914 | 0.41450915 | 0.89054584 |
| Eef1g | 3.76780625 | 1.04965977 | 1.00285333 | 0.04680643 | 0.41392963 | 0.89054584 |
| Zfp101 | 0.14970351 | 0.205551 | 0.15875493 | 0.04679608 | 0.08482756 | 0.56182274 |
| Gm7334 | 0.29395788 | 0.35022088 | 0.30342788 | 0.046793 | 0.23623175 | 0.89054584 |
| Rap2c | 0.11296468 | 0.16731363 | 0.12052248 | 0.04679114 | 0.03524355 | 0.30651529 |
| Klf4 | 0.25000115 | 0.30711269 | 0.26033322 | 0.04677947 | 0.20124891 | 0.89054584 |
| Ugcg | 0.19273498 | 0.249616 | 0.20286039 | 0.04675561 | 0.14145598 | 0.75816224 |
| Ccdc173 | 0.14595966 | 0.20159497 | 0.15488265 | 0.04671232 | 0.07999159 | 0.54094194 |
| Ssh1 | 0.1671883 | 0.223466 | 0.176767 | 0.046699 | 0.10919558 | 0.65377127 |
| Ipo4 | 0.36219765 | 0.41510444 | 0.36842532 | 0.04667912 | 0.27750042 | 0.89054584 |
| Aldh9a1 | 0.47473327 | 0.51665542 | 0.47006888 | 0.04658655 | 0.32213818 | 0.89054584 |

| | | | | | | |
|---|---|---|---|---|---|---|
| Agfg1 | 0.44439307 | 0.48995879 | 0.44338338 | 0.04657541 | 0.31227904 | 0.89054584 |
| Cntrl | 0.23040362 | 0.28738632 | 0.24082849 | 0.04655783 | 0.18387839 | 0.86722786 |
| Pycard | 0.3284169 | 0.38310024 | 0.33654937 | 0.04655087 | 0.25965289 | 0.89054584 |
| Rrp7a | 0.61767569 | 0.63461359 | 0.58807735 | 0.04653624 | 0.35618487 | 0.89054584 |
| Kiz | 0.44371384 | 0.48920705 | 0.44277966 | 0.04642739 | 0.31259416 | 0.89054584 |
| Inpp5a | 0.15484549 | 0.21048317 | 0.1640645 | 0.04641867 | 0.09372648 | 0.5972952 |
| Sun2 | 0.09580848 | 0.14884206 | 0.10243766 | 0.04640439 | 0.01740738 | 0.18261378 |
| Prss8 | 0.14402461 | 0.19927239 | 0.15287905 | 0.04639334 | 0.07870703 | 0.53648072 |
| Dio3 | 0.06124896 | 0.11172973 | 0.0655242 | 0.04620553 | 0.00050931 | 0.00916124 |
| Smo | 0.38102411 | 0.43206689 | 0.38592719 | 0.0461397 | 0.28876308 | 0.89054584 |
| Gbp7 | 0.05344599 | 0.10330382 | 0.05718167 | 0.04612216 | 8.57E-05 | 0.00182424 |
| Trappc6a | 0.77997109 | 0.75148855 | 0.70539886 | 0.04608969 | 0.38040847 | 0.89054584 |
| Gm43796 | 0.04900979 | 0.09842194 | 0.05243772 | 0.04598422 | 2.20E-05 | 0.00054299 |
| Nup62cl | 0.02903578 | 0.0770504 | 0.0310712 | 0.0459792 | 2.71E-12 | 1.71E-10 |
| Stra13 | 1.00108035 | 0.88102307 | 0.83507163 | 0.04595144 | 0.39883361 | 0.89054584 |
| Podxl | 0.07953062 | 0.1309974 | 0.08505939 | 0.04593801 | 0.00593217 | 0.07537653 |
| Ipo13 | 0.21158371 | 0.26786762 | 0.22193632 | 0.0459313 | 0.16747257 | 0.82808684 |
| Jade3 | 0.25580573 | 0.31198981 | 0.26607598 | 0.04591383 | 0.21071506 | 0.89054584 |
| Yipf2 | 0.40619019 | 0.45491649 | 0.40901757 | 0.04589891 | 0.30055166 | 0.89054584 |
| Desi1 | 0.76505389 | 0.74128001 | 0.69539248 | 0.04588753 | 0.37925652 | 0.89054584 |
| Gm27029 | 0.05256669 | 0.10207709 | 0.05624143 | 0.04583566 | 7.32E-05 | 0.00158328 |
| Tenm4 | 0.40331716 | 0.45216213 | 0.40639948 | 0.04576266 | 0.29992348 | 0.89054584 |
| Panx1 | 0.14724299 | 0.20195465 | 0.15621063 | 0.04574402 | 0.08623606 | 0.56777946 |
| Dgkq | 0.09319687 | 0.14531892 | 0.09965066 | 0.04566826 | 0.01637576 | 0.17380369 |
| Ublcp1 | 0.26413844 | 0.31982611 | 0.27429335 | 0.04553276 | 0.21964719 | 0.89054584 |
| Alox5ap | 0.08633868 | 0.13785149 | 0.09232971 | 0.04552178 | 0.01080878 | 0.12496469 |
| Inppl1 | 0.33167215 | 0.38507231 | 0.3396469 | 0.04542542 | 0.26660905 | 0.89054584 |
| Cd48 | 0.03608193 | 0.0840334 | 0.03860968 | 0.04542372 | 2.11E-08 | 8.30E-07 |
| Lgals9 | 0.17547286 | 0.23066685 | 0.18525808 | 0.04540876 | 0.12673595 | 0.71241644 |
| Cxx1b | 0.57993171 | 0.60358701 | 0.55819853 | 0.04538848 | 0.3524071 | 0.89054584 |

| | | | | | | |
|---|---|---|---|---|---|---|
| Kcnu1 | 0.06477874 | 0.11467351 | 0.06929724 | 0.04537627 | 0.00114153 | 0.01857448 |
| BC005537 | 0.65126915 | 0.65921146 | 0.61386498 | 0.04534648 | 0.36536371 | 0.89054584 |
| Bivm | 0.101321 | 0.15363019 | 0.10830013 | 0.04533006 | 0.02558541 | 0.24008888 |
| Tmem204 | 0.11678166 | 0.16984404 | 0.12451829 | 0.04532575 | 0.04495285 | 0.36388968 |
| Pnrc1 | 1.35430455 | 1.03550417 | 0.99019638 | 0.04530779 | 0.41559475 | 0.89054584 |
| Plppr1 | 0.24704482 | 0.30270662 | 0.25740228 | 0.04530434 | 0.20610764 | 0.89054584 |
| Gm26782 | 0.10134924 | 0.15362572 | 0.10832984 | 0.04529587 | 0.02570511 | 0.24074467 |
| Xlr | 0.02362203 | 0.07055133 | 0.02527858 | 0.04527275 | 3.59E-17 | 3.64E-15 |
| Btbd9 | 0.5379579 | 0.56913644 | 0.52387487 | 0.04526158 | 0.34365005 | 0.89054584 |
| Tac2 | 0.0262967 | 0.07339234 | 0.02814049 | 0.04525185 | 3.39E-14 | 2.56E-12 |
| Stc2 | 0.07732798 | 0.12794755 | 0.08270662 | 0.04524094 | 0.00540992 | 0.069967 |
| 2210010C04I | 0.02208675 | 0.06885056 | 0.02363578 | 0.04521478 | 2.50E-19 | 2.89E-17 |
| Hspbp1 | 0.37517262 | 0.42569122 | 0.38050837 | 0.04518285 | 0.29005694 | 0.89054584 |
| Dnaja4 | 0.3002024 | 0.35464708 | 0.30947415 | 0.04517292 | 0.2482363 | 0.89054584 |
| Birc6 | 0.78279828 | 0.7524481 | 0.70727711 | 0.04517099 | 0.38302404 | 0.89054584 |
| Ppox | 0.15740963 | 0.21186391 | 0.16670843 | 0.04515548 | 0.10348445 | 0.63276555 |
| Rufy3 | 0.49279663 | 0.53082949 | 0.48569286 | 0.04513663 | 0.33251942 | 0.89054584 |
| Snx15 | 0.66995703 | 0.67299624 | 0.62787732 | 0.04511892 | 0.36889213 | 0.89054584 |
| Acot11 | 0.05260378 | 0.10138839 | 0.05628108 | 0.04510731 | 9.42E-05 | 0.0019744 |
| Uba5 | 0.8509799 | 0.79592519 | 0.75082139 | 0.0451038 | 0.38978681 | 0.89054584 |
| Farsb | 0.66431648 | 0.66876826 | 0.62367348 | 0.04509478 | 0.36810925 | 0.89054584 |
| Igf2os | 0.03528525 | 0.08283516 | 0.03775738 | 0.04507779 | 1.33E-08 | 5.36E-07 |
| Incenp | 0.15946465 | 0.21389528 | 0.16882548 | 0.0450698 | 0.10679485 | 0.64658138 |
| Hcst | 0.03587779 | 0.08343507 | 0.03839128 | 0.04504379 | 2.30E-08 | 9.01E-07 |
| Gpi1 | 1.32152848 | 1.02354252 | 0.97853294 | 0.04500958 | 0.41515761 | 0.89054584 |
| Atp8b2 | 0.09432763 | 0.14586242 | 0.10085742 | 0.04500501 | 0.01881182 | 0.19301749 |
| Coa3 | 2.07701247 | 1.16721502 | 1.12221472 | 0.0450003 | 0.42590033 | 0.89054584 |
| Yae1d1 | 0.47572441 | 0.51589041 | 0.47093139 | 0.04495902 | 0.32823895 | 0.89054584 |
| Dkk2 | 0.08037392 | 0.13089704 | 0.08596009 | 0.04493695 | 0.00743495 | 0.09173777 |
| H2-M3 | 0.06145633 | 0.11064038 | 0.06574587 | 0.0448945 | 0.00073294 | 0.01266533 |

| | | | | | | |
|---|---|---|---|---|---|---|
| Upf3b | 0.75841533 | 0.73577948 | 0.69088758 | 0.0448919 | 0.38104985 | 0.89054584 |
| Ino80dos | 0.06377219 | 0.11310903 | 0.06822138 | 0.04488765 | 0.00108673 | 0.01778261 |
| Bhlhe40 | 0.06979538 | 0.11948364 | 0.07465866 | 0.04482499 | 0.00257719 | 0.03755186 |
| Sqrdl | 0.15555201 | 0.20961756 | 0.16479327 | 0.04482429 | 0.10253353 | 0.6298723 |
| Gabarap | 3.04644618 | 1.13553729 | 1.09072344 | 0.04481385 | 0.42409871 | 0.89054584 |
| Msi1 | 0.28221122 | 0.33679948 | 0.29200178 | 0.0447977 | 0.2373801 | 0.89054584 |
| Phf8 | 0.27937715 | 0.33403305 | 0.28923537 | 0.04479768 | 0.23527286 | 0.89054584 |
| 1600002H07 | 0.09887353 | 0.15049538 | 0.10570796 | 0.04478742 | 0.02419386 | 0.23046158 |
| Gm17501 | 0.03443919 | 0.08163759 | 0.03685225 | 0.04478534 | 7.49E-09 | 3.10E-07 |
| Med18 | 0.18042644 | 0.23505691 | 0.19032178 | 0.04473513 | 0.13673993 | 0.74138199 |
| Tln1 | 0.75397088 | 0.73257536 | 0.68785441 | 0.04472095 | 0.38098237 | 0.89054584 |
| 2300009A05 | 0.56157187 | 0.58803867 | 0.54332536 | 0.04471332 | 0.35070766 | 0.89054584 |
| Cela3a | 0.02295169 | 0.06924711 | 0.0245613 | 0.04468581 | 1.16E-17 | 1.24E-15 |
| Pfas | 0.16024477 | 0.21430769 | 0.16962871 | 0.04467898 | 0.10988656 | 0.65615128 |
| Zfp747 | 0.11595878 | 0.16828432 | 0.12365732 | 0.04462699 | 0.04634488 | 0.37224823 |
| Wwtr1 | 0.20598584 | 0.26091131 | 0.21628656 | 0.04462475 | 0.16821481 | 0.82993631 |
| Nudt18 | 0.22321359 | 0.27825242 | 0.23362964 | 0.04462278 | 0.18677369 | 0.87419127 |
| Nsun2 | 0.68462182 | 0.68329884 | 0.63870341 | 0.04459543 | 0.37247862 | 0.89054584 |
| Nsl1 | 0.04131831 | 0.08878628 | 0.04421122 | 0.04457506 | 1.32E-06 | 4.03E-05 |
| Rapgef4 | 0.33525964 | 0.38762661 | 0.3430541 | 0.04457251 | 0.27248008 | 0.89054584 |
| Jarid2 | 0.56211432 | 0.58832055 | 0.54376793 | 0.04455262 | 0.35133389 | 0.89054584 |
| Cdkn2d | 0.45457081 | 0.49694186 | 0.45239656 | 0.0445453 | 0.32320922 | 0.89054584 |
| Srl | 0.08057201 | 0.13069556 | 0.08617166 | 0.0445239 | 0.00803723 | 0.09759104 |
| Aasdhppt | 0.49547862 | 0.53251512 | 0.48799572 | 0.0445194 | 0.33540706 | 0.89054584 |
| Sec61a1 | 0.96885464 | 0.86269894 | 0.81821377 | 0.04448518 | 0.40001806 | 0.89054584 |
| Adamts15 | 0.110452 | 0.16236448 | 0.11788913 | 0.04447535 | 0.03940341 | 0.33407636 |
| Gabarapl1 | 0.45755712 | 0.49949696 | 0.45502935 | 0.04446761 | 0.32444898 | 0.89054584 |
| Foxp1 | 0.88402569 | 0.81514299 | 0.77070884 | 0.04443415 | 0.39411692 | 0.89054584 |
| Ppl | 0.09666492 | 0.14777668 | 0.10335152 | 0.04442516 | 0.02260885 | 0.22034316 |
| Acap2 | 0.24964494 | 0.30434295 | 0.25998028 | 0.04436267 | 0.21330817 | 0.89054584 |

| | | | | | | |
|---|---|---|---|---|---|---|
| Tril | 0.1235601 | 0.17593943 | 0.13160088 | 0.04433855 | 0.05824528 | 0.4366288 |
| Kcnh2 | 0.24377988 | 0.29849491 | 0.25416063 | 0.04433428 | 0.2082013 | 0.89054584 |
| Ccne1 | 0.17751802 | 0.23162854 | 0.18734985 | 0.04427869 | 0.13542553 | 0.73756551 |
| Plcb1 | 0.17078624 | 0.22467997 | 0.18045811 | 0.04422187 | 0.12679161 | 0.71241644 |
| Pdgfc | 0.0506792 | 0.09842423 | 0.05422302 | 0.04420121 | 7.30E-05 | 0.00158192 |
| Serpine1 | 0.04889718 | 0.09650798 | 0.0523173 | 0.04419068 | 4.16E-05 | 0.00095477 |
| Nkiras1 | 0.4216761 | 0.46723199 | 0.4230492 | 0.04418279 | 0.31327988 | 0.89054584 |
| Disp2 | 0.32853494 | 0.3808288 | 0.3366618 | 0.044167 | 0.27052987 | 0.89054584 |
| Arhgap30 | 0.03769238 | 0.0844942 | 0.04033251 | 0.04416169 | 1.69E-07 | 5.84E-06 |
| Pacsin1 | 0.15793144 | 0.21137251 | 0.16724615 | 0.04412636 | 0.10949489 | 0.65489146 |
| 2810408A11 | 0.0617705 | 0.11019973 | 0.06608171 | 0.04411802 | 0.00093404 | 0.0155652 |
| Cbx5 | 0.83699315 | 0.78625987 | 0.74216407 | 0.0440958 | 0.39096232 | 0.89054584 |
| Pir | 0.12034519 | 0.17231918 | 0.12824387 | 0.04407531 | 0.05466615 | 0.41779117 |
| Ttc39b | 0.1829541 | 0.23696928 | 0.19290191 | 0.04406737 | 0.1435943 | 0.76537874 |
| Tox3 | 0.61654355 | 0.63124987 | 0.58719499 | 0.04405488 | 0.36334137 | 0.89054584 |
| Dsc2 | 0.05946939 | 0.10767511 | 0.06362178 | 0.04405333 | 0.00062774 | 0.01107166 |
| Rffl | 0.09821982 | 0.14898807 | 0.10501053 | 0.04397754 | 0.02552104 | 0.23963991 |
| 5330417C22I | 0.32480089 | 0.37707165 | 0.33310206 | 0.04396959 | 0.26928333 | 0.89054584 |
| Kbtbd3 | 0.19861301 | 0.25278168 | 0.20882549 | 0.04395619 | 0.16337727 | 0.81705542 |
| Entpd5 | 0.14848355 | 0.20144061 | 0.15749373 | 0.04394688 | 0.09679603 | 0.61121462 |
| Rita1 | 0.21892241 | 0.27325624 | 0.22932208 | 0.04393416 | 0.18604298 | 0.87233221 |
| Taf6l | 1.09031568 | 0.92439157 | 0.88046518 | 0.04392639 | 0.40810029 | 0.89054584 |
| 1810055G02 | 0.17756118 | 0.23125498 | 0.18739397 | 0.04386101 | 0.13775503 | 0.7438695 |
| Bcam | 0.32319298 | 0.37537294 | 0.33156698 | 0.04380596 | 0.26910241 | 0.89054584 |
| Hist3h2a | 0.75895643 | 0.73505106 | 0.69125592 | 0.04379515 | 0.38393145 | 0.89054584 |
| Ift81 | 0.54369372 | 0.57238876 | 0.52863183 | 0.04375693 | 0.34988208 | 0.89054584 |
| Slc16a10 | 0.2529508 | 0.30695039 | 0.26325344 | 0.04369695 | 0.21966384 | 0.89054584 |
| Mcam | 0.10034496 | 0.15096263 | 0.10727335 | 0.04368928 | 0.02888429 | 0.26205532 |
| Zc3h15 | 1.37414929 | 1.04058202 | 0.99694325 | 0.04363877 | 0.41920259 | 0.89054584 |
| Mmp16 | 0.14794698 | 0.20055573 | 0.15693883 | 0.0436169 | 0.09769004 | 0.61445131 |

| | | | | | | |
|---|---|---|---|---|---|---|
| Rpa1 | 0.36940133 | 0.41875722 | 0.37514523 | 0.04361199 | 0.29404059 | 0.89054584 |
| Gmpr2 | 0.3267868 | 0.37858937 | 0.33499617 | 0.0435932 | 0.27216774 | 0.89054584 |
| Mylpf | 0.12325268 | 0.17487076 | 0.13128004 | 0.04359072 | 0.06093494 | 0.45165379 |
| Fcho1 | 0.06000367 | 0.10777609 | 0.06419296 | 0.04358313 | 0.00078039 | 0.01332637 |
| Ano1 | 0.14330719 | 0.19570887 | 0.15213584 | 0.04357303 | 0.09104274 | 0.58666938 |
| Arel1 | 0.11875444 | 0.17015253 | 0.12658137 | 0.04357116 | 0.05439279 | 0.41614069 |
| 3110002H16 | 0.18738525 | 0.24095079 | 0.19741854 | 0.04353225 | 0.15213231 | 0.78519553 |
| D1Ertd622e | 0.26193582 | 0.31564861 | 0.27212434 | 0.04352427 | 0.22808655 | 0.89054584 |
| Gcdh | 0.4532258 | 0.49471405 | 0.45120901 | 0.04350504 | 0.32664022 | 0.89054584 |
| Pbk | 0.11143 | 0.16241916 | 0.11891438 | 0.04350478 | 0.04414649 | 0.36077541 |
| Dnaaf5 | 0.1936021 | 0.24712079 | 0.20374128 | 0.04337951 | 0.16060927 | 0.80963603 |
| Xrcc1 | 0.58778251 | 0.60785565 | 0.56449135 | 0.04336429 | 0.36020892 | 0.89054584 |
| Atp6v1g1 | 2.30922234 | 1.17253237 | 1.12920357 | 0.04332879 | 0.42905964 | 0.89054584 |
| Papd4 | 0.22335696 | 0.27709826 | 0.23377341 | 0.04332485 | 0.19394942 | 0.88899106 |
| Rnase6 | 0.02811013 | 0.07339826 | 0.0300808 | 0.04331746 | 9.80E-12 | 5.82E-10 |
| Tex9 | 0.22384548 | 0.27752795 | 0.23426324 | 0.04326471 | 0.19477532 | 0.88999075 |
| Abca5 | 0.1804998 | 0.23361746 | 0.19039671 | 0.04322075 | 0.14511966 | 0.76747222 |
| Ccdc32 | 0.33649533 | 0.38742242 | 0.34422618 | 0.04319624 | 0.27939456 | 0.89054584 |
| Pfkfb2 | 0.09880708 | 0.14880534 | 0.10563707 | 0.04316828 | 0.02846404 | 0.25931659 |
| Zswim7 | 0.22618165 | 0.2797565 | 0.23660418 | 0.04315232 | 0.19774416 | 0.89054584 |
| Fam81a | 0.1444525 | 0.19645596 | 0.15332223 | 0.04313373 | 0.09498168 | 0.6029091 |
| Insig1 | 0.14794379 | 0.20005618 | 0.15693554 | 0.04312064 | 0.10025122 | 0.62272587 |
| Fermt3 | 0.0386946 | 0.08445786 | 0.04140462 | 0.04305323 | 6.35E-07 | 2.06E-05 |
| Cldn15 | 0.04487764 | 0.09105829 | 0.04801836 | 0.04303993 | 1.48E-05 | 0.00037679 |
| Atg4a | 0.14519267 | 0.19710307 | 0.15408867 | 0.0430144 | 0.09670454 | 0.61116934 |
| Mob4 | 1.39377041 | 1.04655462 | 1.00355092 | 0.0430037 | 0.42087998 | 0.89054584 |
| Asun | 0.25656107 | 0.3097985 | 0.26682211 | 0.04297639 | 0.22650558 | 0.89054584 |
| Cyp51 | 0.35249936 | 0.40230982 | 0.35933602 | 0.0429738 | 0.28870456 | 0.89054584 |
| Dus1l | 0.29808131 | 0.35036924 | 0.3074226 | 0.04294664 | 0.2575718 | 0.89054584 |
| Inadl | 0.20714696 | 0.26037835 | 0.21745957 | 0.04291878 | 0.17891462 | 0.85325796 |

| | | | | | | |
|---|---|---|---|---|---|---|
| Trpt1 | 0.09041883 | 0.1395676 | 0.09668554 | 0.04288206 | 0.01939828 | 0.19735881 |
| Aqr | 0.32167992 | 0.37299371 | 0.33012121 | 0.04287251 | 0.27257222 | 0.89054584 |
| Sncaip | 0.11236028 | 0.16272047 | 0.11988927 | 0.0428312 | 0.04801264 | 0.38170821 |
| Aoc3 | 0.0562901 | 0.10301853 | 0.06022271 | 0.04279582 | 0.00046521 | 0.00845186 |
| Pacsin2 | 0.33905869 | 0.38944705 | 0.34665549 | 0.04279156 | 0.28261009 | 0.89054584 |
| Kat8 | 0.28226792 | 0.33484165 | 0.29205709 | 0.04278455 | 0.24745953 | 0.89054584 |
| Ccdc109b | 0.03185262 | 0.07685471 | 0.03408498 | 0.04276973 | 2.52E-09 | 1.12E-07 |
| Eri1 | 0.33746578 | 0.38788536 | 0.34514628 | 0.04273907 | 0.28199965 | 0.89054584 |
| Gpr183 | 0.06498925 | 0.11223883 | 0.06952224 | 0.04271659 | 0.00210093 | 0.03149874 |
| Casp9 | 0.13236856 | 0.18348772 | 0.14077916 | 0.04270855 | 0.07876972 | 0.53665574 |
| Klhl21 | 0.07232893 | 0.12006684 | 0.07736586 | 0.04270098 | 0.00506389 | 0.06656052 |
| Wars2 | 0.1980923 | 0.25095814 | 0.20829767 | 0.04266046 | 0.16999734 | 0.83373276 |
| Umad1 | 0.16976139 | 0.222044 | 0.17940726 | 0.04263674 | 0.13410088 | 0.73282872 |
| Ptprz1 | 0.09139752 | 0.14036025 | 0.0977302 | 0.04263005 | 0.02106484 | 0.20845964 |
| Mob3c | 0.24961478 | 0.30255689 | 0.2599504 | 0.0426065 | 0.22254934 | 0.89054584 |
| Cfh | 0.03372488 | 0.0786887 | 0.03608805 | 0.04260065 | 1.90E-08 | 7.49E-07 |
| Crat | 0.17850576 | 0.23094886 | 0.18835948 | 0.04258938 | 0.14607453 | 0.77066541 |
| Ankrd46 | 0.40724336 | 0.45255884 | 0.40997611 | 0.04258273 | 0.31422585 | 0.89054584 |
| 4930503L19F | 0.07792447 | 0.12589946 | 0.08334379 | 0.04255567 | 0.00868281 | 0.10429449 |
| Fam167b | 0.03824251 | 0.08346816 | 0.04092101 | 0.04254715 | 6.36E-07 | 2.06E-05 |
| Clstn1 | 0.2125208 | 0.26541879 | 0.22288075 | 0.04253804 | 0.18695183 | 0.87419127 |
| Jag1 | 0.17349896 | 0.22573123 | 0.18323755 | 0.04249368 | 0.13997512 | 0.75161454 |
| Kcnk3 | 0.20308191 | 0.25581765 | 0.2133506 | 0.04246705 | 0.17687142 | 0.84817993 |
| Heg1 | 0.27508075 | 0.32747158 | 0.28503404 | 0.04243754 | 0.24395072 | 0.89054584 |
| Adam15 | 0.21662214 | 0.26943646 | 0.22700964 | 0.04242682 | 0.19195084 | 0.88486593 |
| Ropn1l | 0.18972596 | 0.24221911 | 0.19980105 | 0.04241806 | 0.16130709 | 0.81174254 |
| Chchd2 | 4.11663347 | 0.97862139 | 0.9362365 | 0.04238489 | 0.41647581 | 0.89054584 |
| Supt7l | 0.24574155 | 0.29844991 | 0.2561089 | 0.04234101 | 0.22057998 | 0.89054584 |
| Prss50 | 0.1765143 | 0.22865679 | 0.18632348 | 0.04233332 | 0.14490763 | 0.76730125 |
| Vat1l | 0.34874512 | 0.39807469 | 0.35580385 | 0.04227083 | 0.28996064 | 0.89054584 |

| | | | | | | |
|---|---|---|---|---|---|---|
| Ube2v2 | 1.1903709 | 0.96858411 | 0.92633085 | 0.04225326 | 0.41585422 | 0.89054584 |
| Tceal5 | 0.06737765 | 0.11430984 | 0.07207491 | 0.04223494 | 0.00316597 | 0.04507926 |
| Ntf3 | 0.05325953 | 0.09919719 | 0.05698229 | 0.0422149 | 0.0002787 | 0.00532351 |
| Flnc | 0.13396983 | 0.18463673 | 0.14244452 | 0.04219221 | 0.08379595 | 0.55825208 |
| C330013E15I | 0.04411323 | 0.0893895 | 0.04720076 | 0.04218875 | 1.56E-05 | 0.00039515 |
| Ckap2 | 0.11446117 | 0.16426075 | 0.12208973 | 0.04217102 | 0.05378138 | 0.41342654 |
| Upb1 | 0.19845189 | 0.25082573 | 0.20866218 | 0.04216355 | 0.17324636 | 0.83835974 |
| Vdac1 | 1.6689722 | 1.11765403 | 1.07551193 | 0.0421421 | 0.42757452 | 0.89054584 |
| Nop10 | 1.64926355 | 1.1136121 | 1.0715263 | 0.04208579 | 0.42740416 | 0.89054584 |
| 1700064H15I | 0.02721525 | 0.07117677 | 0.02912332 | 0.04205345 | 8.64E-12 | 5.15E-10 |
| Rpusd1 | 0.28928527 | 0.340925 | 0.29889085 | 0.04203415 | 0.25616805 | 0.89054584 |
| Plp2 | 0.75696231 | 0.73191776 | 0.68989751 | 0.04202025 | 0.38829463 | 0.89054584 |
| Gne | 0.18980129 | 0.24186401 | 0.19987769 | 0.04198632 | 0.16387466 | 0.81785018 |
| Chmp2a | 1.81470649 | 1.14173659 | 1.09977811 | 0.04195848 | 0.4294609 | 0.89054584 |
| H2-DMa | 0.07955882 | 0.12700211 | 0.08508951 | 0.04191261 | 0.01087062 | 0.12527997 |
| Ccl6 | 0.02210001 | 0.06555801 | 0.02364997 | 0.04190804 | 7.55E-17 | 7.35E-15 |
| Kazald1 | 0.05855312 | 0.10452891 | 0.06264221 | 0.0418867 | 0.00091887 | 0.01536534 |
| Rnf170 | 0.32894805 | 0.37894109 | 0.33705517 | 0.04188592 | 0.28130551 | 0.89054584 |
| Amer1 | 0.3833471 | 0.42993181 | 0.38807311 | 0.0418587 | 0.30764844 | 0.89054584 |
| Gm9830 | 0.02208889 | 0.06549137 | 0.02363806 | 0.0418533 | 7.98E-17 | 7.66E-15 |
| Mrpl12 | 1.09260414 | 0.92343052 | 0.8815828 | 0.04184772 | 0.41248604 | 0.89054584 |
| Bub1 | 0.08073976 | 0.12819143 | 0.08635082 | 0.0418406 | 0.01198989 | 0.13582249 |
| A330069E16 | 0.05553882 | 0.10124829 | 0.05941944 | 0.04182885 | 0.00051958 | 0.00933442 |
| Rtkn2 | 0.03999821 | 0.08462425 | 0.04279913 | 0.04182512 | 2.65E-06 | 7.67E-05 |
| Rpl5 | 3.40701719 | 1.0890346 | 1.04721674 | 0.04181786 | 0.42620567 | 0.89054584 |
| Ehf | 0.13098805 | 0.18112753 | 0.1393426 | 0.04178493 | 0.08119395 | 0.54674867 |
| Gm26523 | 0.09776588 | 0.14629711 | 0.10452622 | 0.04177089 | 0.03131485 | 0.27818792 |
| Nacc2 | 0.17205566 | 0.22350503 | 0.18175914 | 0.0417459 | 0.14229799 | 0.76183025 |
| Sgf29 | 0.28601222 | 0.33745098 | 0.2957063 | 0.04174468 | 0.25536549 | 0.89054584 |
| Elp5 | 0.52682905 | 0.55628078 | 0.51458461 | 0.04169617 | 0.35289791 | 0.89054584 |

| | | | | | | |
|---|---|---|---|---|---|---|
| Hspa4l | 0.57263203 | 0.59397608 | 0.55231081 | 0.04166527 | 0.36262102 | 0.89054584 |
| Llgl1 | 0.18308968 | 0.23469985 | 0.19304023 | 0.04165963 | 0.15734328 | 0.80068898 |
| Tnfaip8l1 | 0.10623858 | 0.15512144 | 0.11346816 | 0.04165329 | 0.04360878 | 0.3573997 |
| Sec23b | 0.74600392 | 0.72400824 | 0.68238156 | 0.04162668 | 0.38812665 | 0.89054584 |
| Hdlbp | 1.23410995 | 0.98635287 | 0.94472622 | 0.04162665 | 0.41867556 | 0.89054584 |
| Brap | 0.31925641 | 0.36938459 | 0.32780302 | 0.04158157 | 0.27726583 | 0.89054584 |
| Cbwd1 | 0.19147388 | 0.24314436 | 0.20157867 | 0.04156568 | 0.1683572 | 0.82993631 |
| Fam214b | 0.12983939 | 0.17969745 | 0.13814676 | 0.04155069 | 0.08056526 | 0.54416367 |
| Pomt1 | 0.09641739 | 0.14463194 | 0.1030874 | 0.04154453 | 0.03022019 | 0.27028518 |
| Tspan9 | 0.27736189 | 0.32877746 | 0.28726582 | 0.04151164 | 0.25039571 | 0.89054584 |
| Zbtb44 | 0.2790851 | 0.33045455 | 0.28895007 | 0.04150448 | 0.25168225 | 0.89054584 |
| Polrmt | 0.1277141 | 0.17742212 | 0.13593283 | 0.04148929 | 0.07751348 | 0.53084105 |
| Mre11a | 0.19249896 | 0.24410874 | 0.20262056 | 0.04148818 | 0.17005308 | 0.83373276 |
| Magel2 | 0.05818101 | 0.10369177 | 0.06224439 | 0.04144739 | 0.00096011 | 0.01590815 |
| Adam17 | 0.19392748 | 0.24548164 | 0.20407175 | 0.04140989 | 0.17223133 | 0.83696674 |
| Scnn1b | 0.07431104 | 0.12089285 | 0.07948361 | 0.04140924 | 0.00760778 | 0.09337756 |
| Rprm | 0.12414814 | 0.17361366 | 0.13221449 | 0.04139918 | 0.07230817 | 0.50878997 |
| Bola3 | 0.95134042 | 0.85020641 | 0.80880947 | 0.04139694 | 0.40576351 | 0.89054584 |
| Snrnp48 | 0.53173342 | 0.56006505 | 0.51868856 | 0.04137649 | 0.3550767 | 0.89054584 |
| Noct | 0.43709214 | 0.47823462 | 0.43688098 | 0.04135364 | 0.32960677 | 0.89054584 |
| Nefl | 0.07425241 | 0.12068704 | 0.07942097 | 0.04126607 | 0.00774483 | 0.09467498 |
| 1110017D15 | 0.10278535 | 0.15105022 | 0.10983997 | 0.04121025 | 0.04023529 | 0.33777651 |
| Tspan17 | 0.28227159 | 0.33324328 | 0.29206067 | 0.04118261 | 0.25560864 | 0.89054584 |
| Mxd3 | 0.08098609 | 0.1277905 | 0.08661391 | 0.04117659 | 0.01338418 | 0.14835161 |
| Col13a1 | 0.0899134 | 0.1373179 | 0.09614602 | 0.04117188 | 0.02301802 | 0.22262525 |
| Mettl7a1 | 0.15858136 | 0.20906999 | 0.16791574 | 0.04115425 | 0.1267574 | 0.71241644 |
| Flrt3 | 0.1631874 | 0.21379273 | 0.17265626 | 0.04113647 | 0.13349461 | 0.73110935 |
| Gm5422 | 0.04322793 | 0.08737753 | 0.04625383 | 0.0411237 | 1.72E-05 | 0.00043215 |
| Tro | 0.1910551 | 0.24224451 | 0.2011529 | 0.04109161 | 0.17061676 | 0.83412025 |
| Pycr1 | 0.17385234 | 0.22464824 | 0.1835994 | 0.04104885 | 0.1487897 | 0.7767926 |

| | | | | | | |
|---|---|---|---|---|---|---|
| Tamm41 | 0.13846964 | 0.18816306 | 0.14711914 | 0.04104391 | 0.09683976 | 0.61122455 |
| Foxn2 | 0.17091638 | 0.2216304 | 0.18059153 | 0.04103887 | 0.14486136 | 0.76730125 |
| Jakmip1 | 0.20649103 | 0.25783471 | 0.21679699 | 0.04103772 | 0.18891664 | 0.87870177 |
| Abcc9 | 0.10304292 | 0.15114668 | 0.11011075 | 0.04103594 | 0.04125642 | 0.3435586 |
| Slc30a9 | 0.64495146 | 0.6501082 | 0.6090742 | 0.04103399 | 0.37680687 | 0.89054584 |
| Irx1 | 0.09296013 | 0.14042915 | 0.09939799 | 0.04103115 | 0.02722778 | 0.25100767 |
| Polh | 0.03006502 | 0.07319241 | 0.03217242 | 0.04101999 | 1.42E-09 | 6.59E-08 |
| Twist2 | 0.13925244 | 0.18892641 | 0.14793155 | 0.04099486 | 0.09833657 | 0.61558016 |
| BC002059 | 0.10964619 | 0.15803327 | 0.11704414 | 0.04098913 | 0.05138481 | 0.40024739 |
| Slc25a24 | 0.19990716 | 0.2511011 | 0.21013681 | 0.0409643 | 0.18188125 | 0.86130002 |
| Bbx | 0.92786778 | 0.83682394 | 0.79588186 | 0.04094208 | 0.40529414 | 0.89054584 |
| Hist2h2be | 0.07324371 | 0.11926343 | 0.07834328 | 0.04092015 | 0.00747781 | 0.09210987 |
| Dnajb1 | 0.89913107 | 0.82044057 | 0.77953658 | 0.04090399 | 0.40343542 | 0.89054584 |
| Ube2g2 | 0.61377292 | 0.62592505 | 0.58503195 | 0.0408931 | 0.37234272 | 0.89054584 |
| Cxcl16 | 0.03950658 | 0.08316493 | 0.04227323 | 0.0408917 | 3.29E-06 | 9.41E-05 |
| Cdkn2aip | 0.34029937 | 0.38870745 | 0.34783005 | 0.0408774 | 0.29200931 | 0.89054584 |
| Rad51ap1 | 0.13488465 | 0.18427084 | 0.14339551 | 0.04087533 | 0.09208155 | 0.59149236 |
| Arf2 | 0.40715999 | 0.45075567 | 0.40990025 | 0.04085542 | 0.3211934 | 0.89054584 |
| Slc30a5 | 0.60516337 | 0.61911611 | 0.57827718 | 0.04083893 | 0.37106937 | 0.89054584 |
| Ugdh | 0.55234144 | 0.57659883 | 0.53576477 | 0.04083406 | 0.36126194 | 0.89054584 |
| Fgd2 | 0.52785425 | 0.55625407 | 0.51544377 | 0.0408103 | 0.35611154 | 0.89054584 |
| Zfp946 | 0.16953513 | 0.21993331 | 0.17917521 | 0.0407581 | 0.14461818 | 0.76727455 |
| Adam9 | 0.170063 | 0.22047119 | 0.17971657 | 0.04075462 | 0.14536632 | 0.76804699 |
| Hspb7 | 0.02722267 | 0.06984833 | 0.02913126 | 0.04071708 | 3.71E-11 | 2.06E-09 |
| Wdr48 | 0.20568861 | 0.25660457 | 0.21598619 | 0.04061838 | 0.19047129 | 0.88259328 |
| Samsn1 | 0.03290745 | 0.07581966 | 0.03521351 | 0.04060614 | 3.88E-08 | 1.47E-06 |
| Coro1c | 0.40084824 | 0.4446931 | 0.40414586 | 0.04054724 | 0.32009852 | 0.89054584 |
| Grasp | 0.26597192 | 0.31662488 | 0.27609707 | 0.04052781 | 0.24702616 | 0.89054584 |
| Rps6kc1 | 0.10682705 | 0.15459681 | 0.11408601 | 0.04051079 | 0.04902891 | 0.3882147 |
| Acd | 0.48098513 | 0.51600759 | 0.47549929 | 0.0405083 | 0.34572052 | 0.89054584 |

| | | | | | | |
|---|---|---|---|---|---|---|
| Aurka | 0.11238197 | 0.16039841 | 0.119912 | 0.04048641 | 0.05785726 | 0.43561417 |
| Bub3 | 1.10689715 | 0.92893921 | 0.88848207 | 0.04045714 | 0.41599677 | 0.89054584 |
| Myh10 | 0.46451017 | 0.50158926 | 0.46113836 | 0.04045089 | 0.34138701 | 0.89054584 |
| Mtus2 | 0.1287176 | 0.17739103 | 0.1369784 | 0.04041263 | 0.08463954 | 0.56160366 |
| Fam219a | 0.24589975 | 0.29664309 | 0.25626594 | 0.04037715 | 0.23145586 | 0.89054584 |
| Sumf1 | 0.38054891 | 0.42586206 | 0.38548784 | 0.04037422 | 0.31279049 | 0.89054584 |
| Pla2g1b | 0.13496656 | 0.18383475 | 0.14348065 | 0.04035411 | 0.09504163 | 0.6030256 |
| Nexn | 0.05436473 | 0.09851121 | 0.05816406 | 0.04034715 | 0.00061505 | 0.01091435 |
| Gm12355 | 0.08259921 | 0.12866699 | 0.08833667 | 0.04033032 | 0.01670793 | 0.17694052 |
| Hsf2 | 0.29979283 | 0.34940774 | 0.30907819 | 0.04032955 | 0.27162221 | 0.89054584 |
| Sfxn5 | 0.07940515 | 0.1252523 | 0.08492537 | 0.04032693 | 0.01347281 | 0.14876532 |
| Rab24 | 0.93689013 | 0.84121097 | 0.80089346 | 0.04031751 | 0.40728514 | 0.89054584 |
| Pomk | 0.08416952 | 0.13029507 | 0.09001357 | 0.0402815 | 0.01853926 | 0.19116794 |
| Svil | 0.56043604 | 0.58265591 | 0.54239802 | 0.04025788 | 0.36474798 | 0.89054584 |
| Hddc3 | 0.34898135 | 0.39628279 | 0.35602634 | 0.04025645 | 0.29916843 | 0.89054584 |
| Proser2 | 0.23779898 | 0.28843992 | 0.24820958 | 0.04023034 | 0.22508513 | 0.89054584 |
| Fam13a | 0.08303273 | 0.12900843 | 0.08879963 | 0.0402088 | 0.01744702 | 0.18264153 |
| Azi2 | 0.557086 | 0.57984971 | 0.53965799 | 0.04019173 | 0.36430292 | 0.89054584 |
| Dennd6b | 0.11874351 | 0.16675712 | 0.12656995 | 0.04018717 | 0.06953564 | 0.49563803 |
| Ackr3 | 0.09053113 | 0.13695022 | 0.09680541 | 0.04014481 | 0.02667703 | 0.2476079 |
| Acox1 | 0.32378949 | 0.37227832 | 0.33213662 | 0.04014169 | 0.28669194 | 0.89054584 |
| Cap2 | 0.02570341 | 0.06763678 | 0.02750567 | 0.04013111 | 5.32E-12 | 3.26E-10 |
| H2-Oa | 0.03365542 | 0.07614083 | 0.03601374 | 0.04012709 | 1.05E-07 | 3.74E-06 |
| Epm2a | 0.06887169 | 0.11378484 | 0.07367157 | 0.04011327 | 0.00559455 | 0.0719697 |
| Spata5 | 0.17583085 | 0.22571966 | 0.18562435 | 0.0400953 | 0.15712546 | 0.80068898 |
| Elof1 | 1.76257052 | 1.1322866 | 1.09219321 | 0.0400934 | 0.4321019 | 0.89054584 |
| Zfp707 | 0.13592027 | 0.18453511 | 0.14447169 | 0.04006342 | 0.09818374 | 0.61534939 |
| Klhl17 | 0.05549997 | 0.09943918 | 0.0593779 | 0.04006128 | 0.00083525 | 0.01416302 |
| Tns1 | 0.24415514 | 0.29458729 | 0.25453347 | 0.04005382 | 0.23173686 | 0.89054584 |
| Ccrl2 | 0.04216728 | 0.08516693 | 0.04511933 | 0.0400476 | 1.77E-05 | 0.00044316 |

| | | | | | | |
|---|---|---|---|---|---|---|
| Actr3 | 1.64544203 | 1.11077716 | 1.07073179 | 0.04004537 | 0.4308367 | 0.89054584 |
| Utp18 | 0.44684441 | 0.48559616 | 0.44555991 | 0.04003625 | 0.3377418 | 0.89054584 |
| Vegfa | 0.57761717 | 0.59636902 | 0.55633546 | 0.04003355 | 0.36871656 | 0.89054584 |
| Gm10282 | 0.19782258 | 0.24805246 | 0.20802421 | 0.04002824 | 0.18499863 | 0.86930035 |
| Mios | 0.1826004 | 0.23256907 | 0.19254103 | 0.04002803 | 0.16638147 | 0.82496531 |
| Cep104 | 0.29278435 | 0.34230822 | 0.30228943 | 0.04001879 | 0.26869089 | 0.89054584 |
| Gm16973 | 0.06338929 | 0.10782835 | 0.0678121 | 0.04001626 | 0.00298646 | 0.04282658 |
| Hnrnpa2b1 | 3.12134999 | 1.12346494 | 1.08347759 | 0.03998735 | 0.43174033 | 0.89054584 |
| Ppp1r16a | 0.32521903 | 0.37347895 | 0.33350105 | 0.0399779 | 0.28825729 | 0.89054584 |
| Oip5 | 0.07227933 | 0.11728093 | 0.07731287 | 0.03996806 | 0.00800889 | 0.09732843 |
| Asf1b | 0.12109447 | 0.16895355 | 0.12902662 | 0.03992693 | 0.07469401 | 0.51900827 |
| Tut1 | 0.13240552 | 0.18071872 | 0.14081762 | 0.03990111 | 0.09339658 | 0.59664859 |
| Gosr2 | 0.50986561 | 0.54016308 | 0.50027409 | 0.03988899 | 0.35514102 | 0.89054584 |
| 1700026L06I | 0.09843782 | 0.14509987 | 0.10524311 | 0.03985675 | 0.03883287 | 0.33001109 |
| Gipc1 | 0.96915486 | 0.85821704 | 0.81837301 | 0.03984402 | 0.41027825 | 0.89054584 |
| Scn8a | 0.12442351 | 0.17233513 | 0.13250178 | 0.03983335 | 0.08066774 | 0.54446969 |
| Prim1 | 0.27617316 | 0.32593139 | 0.28610313 | 0.03982826 | 0.25830861 | 0.89054584 |
| Itgb7 | 0.03632405 | 0.07868053 | 0.0388687 | 0.03981182 | 9.12E-07 | 2.86E-05 |
| Coa6 | 0.66655202 | 0.66508577 | 0.62534226 | 0.03974351 | 0.38357758 | 0.89054584 |
| Zfp316 | 0.06702481 | 0.11141526 | 0.07169782 | 0.03971745 | 0.00492788 | 0.06524606 |
| Myh14 | 0.12807842 | 0.17602121 | 0.13631247 | 0.03970874 | 0.08736184 | 0.57104236 |
| A130010J15I | 0.23751628 | 0.28761121 | 0.24792788 | 0.03968333 | 0.22792131 | 0.89054584 |
| Fam195a | 0.15984672 | 0.20889664 | 0.16921889 | 0.03967774 | 0.13732664 | 0.74234216 |
| Camk1d | 0.06299187 | 0.10704769 | 0.06738729 | 0.0396604 | 0.00305338 | 0.04358243 |
| Rnf123 | 0.12126843 | 0.16884391 | 0.12920832 | 0.03963559 | 0.07647851 | 0.52623897 |
| Pard6g | 0.30461503 | 0.35335983 | 0.31373484 | 0.03962499 | 0.27812236 | 0.89054584 |
| Socs1 | 0.15901166 | 0.20798288 | 0.16835896 | 0.03962392 | 0.13643014 | 0.73997914 |
| Inpp5e | 0.12047332 | 0.16797691 | 0.12837774 | 0.03959917 | 0.07534638 | 0.52216624 |
| H6pd | 0.11439363 | 0.16161043 | 0.12201902 | 0.03959141 | 0.06530728 | 0.47362135 |
| AU021092 | 0.05965998 | 0.10341028 | 0.06382554 | 0.03958475 | 0.00192936 | 0.02919817 |

| | | | | | | |
|---|---|---|---|---|---|---|
| Ufm1 | 1.39168612 | 1.04244165 | 1.0028596 | 0.03958205 | 0.42705199 | 0.89054584 |
| Mrpl37 | 0.36679278 | 0.41225125 | 0.37271512 | 0.03953613 | 0.31058183 | 0.89054584 |
| Cd3g | 0.01903054 | 0.05989232 | 0.02036544 | 0.03952688 | 7.66E-20 | 9.49E-18 |
| Zfp719 | 0.14799432 | 0.1964968 | 0.1569878 | 0.039509 | 0.12049575 | 0.69516408 |
| Esyt1 | 0.31032423 | 0.35873151 | 0.31923281 | 0.0394987 | 0.28215516 | 0.89054584 |
| Ecm1 | 0.05932124 | 0.10286684 | 0.0634634 | 0.03940344 | 0.00190994 | 0.02894551 |
| Syne4 | 0.15942465 | 0.20818166 | 0.16878429 | 0.03939736 | 0.13841178 | 0.74639314 |
| Zfp119b | 0.07400618 | 0.11848154 | 0.07915791 | 0.03932363 | 0.01032171 | 0.12043522 |
| Hoxc4 | 0.10042516 | 0.14662673 | 0.10735773 | 0.039269 | 0.04417801 | 0.36077541 |
| Srebf1 | 0.30660315 | 0.35489742 | 0.31565127 | 0.03924615 | 0.28120543 | 0.89054584 |
| Ppm1m | 0.19094982 | 0.24028914 | 0.20104585 | 0.03924329 | 0.18156809 | 0.86081561 |
| Pon3 | 0.16810366 | 0.21692018 | 0.17770657 | 0.03921361 | 0.15195966 | 0.78458375 |
| Ttpal | 0.12072771 | 0.16785419 | 0.1286435 | 0.03921069 | 0.07779643 | 0.53202481 |
| Wdr36 | 0.20151163 | 0.25096336 | 0.21176155 | 0.03920181 | 0.19421325 | 0.889076 |
| Prrc1 | 0.38131769 | 0.42539579 | 0.38619856 | 0.03919723 | 0.31815678 | 0.89054584 |
| Nob1 | 0.55054335 | 0.57347734 | 0.53428548 | 0.03919186 | 0.36626915 | 0.89054584 |
| Nkrf | 0.05564609 | 0.09872222 | 0.05953413 | 0.03918809 | 0.00108208 | 0.01772651 |
| Nrm | 0.21213228 | 0.26167715 | 0.22248923 | 0.03918792 | 0.20593766 | 0.89054584 |
| Mmgt2 | 0.31019148 | 0.35828023 | 0.31910514 | 0.0391751 | 0.2836755 | 0.89054584 |
| Chn2 | 0.03532032 | 0.07694094 | 0.03779491 | 0.03914604 | 6.98E-07 | 2.24E-05 |
| Agtpbp1 | 0.15666326 | 0.20506265 | 0.1659391 | 0.03912354 | 0.13600602 | 0.73850761 |
| Pde2a | 0.06044859 | 0.10372602 | 0.06466859 | 0.03905743 | 0.00244777 | 0.03588252 |
| Ugt2b34 | 0.10956263 | 0.15593914 | 0.1169565 | 0.03898264 | 0.06022749 | 0.44755418 |
| Wdr86 | 0.14415984 | 0.1919791 | 0.15301912 | 0.03895999 | 0.11777036 | 0.68682008 |
| Pkp4 | 0.40729272 | 0.44896418 | 0.41002102 | 0.03894316 | 0.32906408 | 0.89054584 |
| Maml3 | 0.4258345 | 0.4657049 | 0.42679401 | 0.0389109 | 0.33550706 | 0.89054584 |
| Ssr1 | 1.55178903 | 1.0880974 | 1.04922693 | 0.03887047 | 0.43148342 | 0.89054584 |
| Xlr4c | 0.03307302 | 0.07419107 | 0.03539065 | 0.03880041 | 1.63E-07 | 5.67E-06 |
| Ccdc77 | 0.20676086 | 0.25586654 | 0.21706958 | 0.03879696 | 0.2025084 | 0.89054584 |
| Cdk2 | 0.1436862 | 0.19130747 | 0.1525285 | 0.03877897 | 0.11810899 | 0.68771984 |

| | | | | | | |
|---|---|---|---|---|---|---|
| Spock3 | 0.1342084 | 0.18143276 | 0.14269256 | 0.0387402 | 0.10295801 | 0.63115652 |
| Gpr173 | 0.05296182 | 0.09536233 | 0.05666394 | 0.03869839 | 0.0007318 | 0.01266074 |
| Tmem165 | 0.58848228 | 0.60371853 | 0.56505025 | 0.03866828 | 0.37492888 | 0.89054584 |
| Thoc7 | 1.42905602 | 1.05344908 | 1.01485472 | 0.03859436 | 0.4296847 | 0.89054584 |
| Sympk | 0.45201226 | 0.4886629 | 0.45013661 | 0.03852629 | 0.34503892 | 0.89054584 |
| Dnajb5 | 0.09549868 | 0.14061861 | 0.10210708 | 0.03851152 | 0.03944157 | 0.33420447 |
| Hlx | 0.08988982 | 0.13459006 | 0.09612085 | 0.03846922 | 0.03111956 | 0.27696218 |
| Nudt11 | 0.12275133 | 0.16919127 | 0.13075673 | 0.03843453 | 0.08543001 | 0.56503846 |
| Fbxo2 | 0.06468017 | 0.10761756 | 0.06919188 | 0.03842568 | 0.00483599 | 0.06414659 |
| Ccbl2 | 0.12353455 | 0.16999723 | 0.13157422 | 0.038423 | 0.08683132 | 0.57045578 |
| Rara | 0.22845396 | 0.27723393 | 0.23887876 | 0.03835517 | 0.22721345 | 0.89054584 |
| S100pbp | 0.28035302 | 0.32835838 | 0.29018839 | 0.03816998 | 0.26999944 | 0.89054584 |
| Ddit4l | 0.0934218 | 0.13803886 | 0.09989072 | 0.03814814 | 0.03759988 | 0.32198646 |
| Rltpr | 0.08528314 | 0.12934663 | 0.09120268 | 0.03814394 | 0.0256763 | 0.24063027 |
| Lims2 | 0.02504865 | 0.0649346 | 0.02680508 | 0.03812952 | 1.71E-11 | 9.92E-10 |
| Spata24 | 0.20356641 | 0.2519495 | 0.21384068 | 0.03810882 | 0.20319139 | 0.89054584 |
| Thbs4 | 0.03968142 | 0.08052507 | 0.04246026 | 0.03806481 | 1.48E-05 | 0.00037622 |
| P3h4 | 0.59242898 | 0.60625202 | 0.56819632 | 0.0380557 | 0.37750555 | 0.89054584 |
| Alg8 | 0.14918733 | 0.19624858 | 0.15822138 | 0.0380272 | 0.13141271 | 0.72635204 |
| Gm11744 | 0.02136914 | 0.06089495 | 0.0228679 | 0.03802704 | 4.69E-15 | 3.84E-13 |
| Rfng | 0.17537792 | 0.22315243 | 0.18516094 | 0.03799148 | 0.1695549 | 0.83300813 |
| Mtap | 0.27184693 | 0.31985564 | 0.28186583 | 0.0379898 | 0.26502461 | 0.89054584 |
| Zfp810 | 0.11441041 | 0.15999242 | 0.12203658 | 0.03795584 | 0.07366534 | 0.51519545 |
| Rasa2 | 0.314249 | 0.36094618 | 0.32300359 | 0.03794259 | 0.29209437 | 0.89054584 |
| Tmprss4 | 0.06265893 | 0.10495966 | 0.0670314 | 0.03792826 | 0.0041925 | 0.05670042 |
| Mex3d | 0.15011625 | 0.19709324 | 0.15918148 | 0.03791176 | 0.13358454 | 0.73110935 |
| Rft1 | 0.16112397 | 0.20844217 | 0.17053365 | 0.03790852 | 0.15018217 | 0.77912292 |
| Top1mt | 0.08775769 | 0.13172147 | 0.09384471 | 0.03787675 | 0.03002682 | 0.26938665 |
| Tmem100 | 0.08357461 | 0.12725043 | 0.08937829 | 0.03787214 | 0.02418368 | 0.23046158 |
| Ror1 | 0.10212988 | 0.14697988 | 0.10915081 | 0.03782907 | 0.05319021 | 0.40975112 |

| | | | | | | |
|---|---|---|---|---|---|---|
| Eri3 | 0.76461257 | 0.73292107 | 0.69509399 | 0.03782708 | 0.39992548 | 0.89054584 |
| Fyb | 0.12971933 | 0.1758408 | 0.13802174 | 0.03781907 | 0.10087398 | 0.62338915 |
| Rftn1 | 0.09441439 | 0.13876593 | 0.10095001 | 0.03781592 | 0.0404714 | 0.33955693 |
| Gdpd5 | 0.11139378 | 0.15664739 | 0.11887642 | 0.03777097 | 0.06939664 | 0.49496101 |
| Armc2 | 0.07837852 | 0.12159374 | 0.08382879 | 0.03776495 | 0.01791474 | 0.18605151 |
| Ccnyl1 | 0.06634344 | 0.10872297 | 0.0709696 | 0.03775336 | 0.00659875 | 0.08233106 |
| Spsb4 | 0.15788274 | 0.2049389 | 0.16719596 | 0.03774293 | 0.14646571 | 0.77188653 |
| 4930523C07I | 0.14830371 | 0.19502377 | 0.15730776 | 0.03771602 | 0.1319918 | 0.72831857 |
| Cdkn2aipnl | 0.30839661 | 0.35508862 | 0.31737833 | 0.03771029 | 0.28993775 | 0.89054584 |
| Gria2 | 0.16779217 | 0.21508993 | 0.17738687 | 0.03770306 | 0.16102737 | 0.81128869 |
| Crk | 0.56892902 | 0.58700452 | 0.54931142 | 0.0376931 | 0.37460127 | 0.89054584 |
| Trim11 | 0.23274708 | 0.2808594 | 0.24317005 | 0.03768935 | 0.23511732 | 0.89054584 |
| Obsl1 | 0.33423754 | 0.37970755 | 0.34208405 | 0.0376235 | 0.3041837 | 0.89054584 |
| Pdap1 | 2.51865024 | 1.163762 | 1.12619645 | 0.03756555 | 0.43825132 | 0.89054584 |
| Fermt2 | 0.72322409 | 0.70404548 | 0.66648281 | 0.03756267 | 0.39643877 | 0.89054584 |
| Fbxl15 | 0.39711863 | 0.43825165 | 0.40073489 | 0.03751676 | 0.33135683 | 0.89054584 |
| Wbp2 | 0.88602405 | 0.80939426 | 0.77188589 | 0.03750837 | 0.41044803 | 0.89054584 |
| Ccl12 | 0.02038006 | 0.05928911 | 0.02180952 | 0.03747959 | 5.91E-16 | 5.22E-14 |
| Medag | 0.14482464 | 0.19116164 | 0.1537076 | 0.03745405 | 0.12813578 | 0.71651107 |
| 4930579G24 | 0.10766003 | 0.15240755 | 0.11496036 | 0.03744719 | 0.06455715 | 0.46985422 |
| Pcdh19 | 0.09684443 | 0.14098386 | 0.10354306 | 0.0374408 | 0.04602744 | 0.3701086 |
| Rhbdf1 | 0.17895712 | 0.22626096 | 0.18882074 | 0.03744022 | 0.17779633 | 0.85016198 |
| Bicd2 | 0.21987685 | 0.2677072 | 0.23028087 | 0.03742633 | 0.22446713 | 0.89054584 |
| Impact | 0.25310412 | 0.30079249 | 0.26340512 | 0.03738736 | 0.25421519 | 0.89054584 |
| Atp2a3 | 0.39842382 | 0.43930524 | 0.40192947 | 0.03737577 | 0.33242058 | 0.89054584 |
| Gpn1 | 0.24925093 | 0.29691115 | 0.25958982 | 0.03732133 | 0.251487 | 0.89054584 |
| Rpap2 | 0.1172036 | 0.16225711 | 0.12495967 | 0.03729744 | 0.0821762 | 0.5516435 |
| Abi3 | 0.07909667 | 0.12189159 | 0.08459588 | 0.03729571 | 0.01998865 | 0.20124687 |
| Ube2o | 0.19414503 | 0.24156999 | 0.20429268 | 0.03727731 | 0.19762986 | 0.89054584 |
| Wwp1 | 0.26186663 | 0.30932384 | 0.27205617 | 0.03726767 | 0.26166979 | 0.89054584 |

| | | | | | | |
|---|---|---|---|---|---|---|
| Scara3 | 0.19978176 | 0.2472061 | 0.21000978 | 0.03719632 | 0.20463527 | 0.89054584 |
| Rab26os | 0.18309595 | 0.23023358 | 0.19304662 | 0.03718696 | 0.18473642 | 0.8690805 |
| Dnajb4 | 0.38761064 | 0.42914828 | 0.3920038 | 0.03714448 | 0.32944007 | 0.89054584 |
| Abhd4 | 0.22877534 | 0.27631341 | 0.23920027 | 0.03711314 | 0.23488343 | 0.89054584 |
| Ckap5 | 0.41625062 | 0.45526093 | 0.41814818 | 0.03711275 | 0.33961854 | 0.89054584 |
| Tmbim6 | 2.41516973 | 1.16610558 | 1.12899357 | 0.03711201 | 0.43914094 | 0.89054584 |
| Nmi | 0.23743002 | 0.28491316 | 0.24784191 | 0.03707124 | 0.24294316 | 0.89054584 |
| Traf5 | 0.16555592 | 0.21211476 | 0.17509059 | 0.03702417 | 0.16227108 | 0.81470761 |
| Arpc3 | 1.93205328 | 1.14964379 | 1.11262016 | 0.03702363 | 0.43839834 | 0.89054584 |
| Mall | 0.06959223 | 0.11133782 | 0.07444156 | 0.03689625 | 0.01046736 | 0.12189214 |
| 3632451O06 | 0.10640884 | 0.15047739 | 0.11364693 | 0.03683046 | 0.0655396 | 0.47418188 |
| Apaf1 | 0.09814691 | 0.14175552 | 0.10493275 | 0.03682277 | 0.05103443 | 0.39853675 |
| Msantd3 | 0.12380671 | 0.1686789 | 0.13185823 | 0.03682067 | 0.09663213 | 0.61116934 |
| Tmem38a | 0.29267319 | 0.33900037 | 0.30218155 | 0.03681882 | 0.28513257 | 0.89054584 |
| Psrc1 | 0.09288151 | 0.13610946 | 0.09931409 | 0.03679537 | 0.04216341 | 0.34870801 |
| Sstr3 | 0.13773213 | 0.18311612 | 0.14635351 | 0.03676261 | 0.1209438 | 0.69664013 |
| Tma7 | 3.13199155 | 1.11904823 | 1.08232191 | 0.03672633 | 0.43719238 | 0.89054584 |
| Fzd1 | 0.10075568 | 0.14438646 | 0.10770546 | 0.03668099 | 0.05629206 | 0.42706556 |
| Vwa5a | 0.27841916 | 0.32487239 | 0.28829936 | 0.03657303 | 0.27725246 | 0.89054584 |
| B4galt1 | 0.27641279 | 0.32290867 | 0.28633757 | 0.0365711 | 0.27590803 | 0.89054584 |
| Hdhd3 | 0.18367716 | 0.2302046 | 0.19363949 | 0.03656511 | 0.18949685 | 0.88027083 |
| Uso1 | 0.82352314 | 0.77012879 | 0.73369272 | 0.03643607 | 0.40851421 | 0.89054584 |
| Gm5617 | 0.70660318 | 0.69107861 | 0.65464994 | 0.03642867 | 0.39772058 | 0.89054584 |
| Ndufb2 | 2.00567693 | 1.1545833 | 1.11816038 | 0.03642291 | 0.43968805 | 0.89054584 |
| Efcab2 | 0.20711325 | 0.25382758 | 0.21742552 | 0.03640206 | 0.2176912 | 0.89054584 |
| Pip5k1b | 0.16126609 | 0.20707834 | 0.1706799 | 0.03639843 | 0.16022221 | 0.8080503 |
| Ubl7 | 0.66915138 | 0.66363821 | 0.62727824 | 0.03635997 | 0.39355938 | 0.89054584 |
| Cep70 | 0.35790249 | 0.40076401 | 0.36440618 | 0.03635782 | 0.3210246 | 0.89054584 |
| Bbip1 | 1.02889973 | 0.88572102 | 0.84936388 | 0.03635714 | 0.42096452 | 0.89054584 |
| Slitrk6 | 0.28615906 | 0.33219343 | 0.29584929 | 0.03634414 | 0.28354717 | 0.89054584 |

| | | | | | | |
|---|---|---|---|---|---|---|
| Arhgap26 | 0.09670145 | 0.13970828 | 0.1033905 | 0.03631778 | 0.05086361 | 0.39839042 |
| Iscu | 1.12874529 | 0.93506682 | 0.89875016 | 0.03631666 | 0.42533708 | 0.89054584 |
| Svip | 0.41119368 | 0.4498731 | 0.41356577 | 0.03630733 | 0.34126511 | 0.89054584 |
| Gab3 | 0.03216466 | 0.07072485 | 0.03441882 | 0.03630602 | 4.45E-07 | 1.47E-05 |
| Fbl | 1.31825443 | 1.01355163 | 0.97733943 | 0.0362122 | 0.43147419 | 0.89054584 |
| Sgsm3 | 0.17703942 | 0.22305664 | 0.1868605 | 0.03619614 | 0.18340403 | 0.86612104 |
| Mrpl48 | 0.89929798 | 0.81582234 | 0.77963322 | 0.03618912 | 0.41439278 | 0.89054584 |
| Pik3cd | 0.04195342 | 0.08107358 | 0.04489057 | 0.03618301 | 8.66E-05 | 0.00183536 |
| Ewsr1 | 1.78969425 | 1.13245966 | 1.09628104 | 0.03617861 | 0.43890313 | 0.89054584 |
| Plod1 | 0.28794268 | 0.33376013 | 0.2975852 | 0.03617493 | 0.28557853 | 0.89054584 |
| Nt5dc1 | 0.15621509 | 0.20164357 | 0.16547704 | 0.03616652 | 0.15427856 | 0.79182956 |
| Rhobtb1 | 0.17079714 | 0.21663188 | 0.18046928 | 0.03616259 | 0.17526476 | 0.84334167 |
| Ric1 | 0.13627566 | 0.18099843 | 0.14484091 | 0.03615753 | 0.1224066 | 0.70076598 |
| Mon1b | 0.09446919 | 0.13713057 | 0.10100849 | 0.03612208 | 0.04784633 | 0.38114069 |
| Pdcd2l | 0.39547674 | 0.43534969 | 0.39923076 | 0.03611893 | 0.33669482 | 0.89054584 |
| Qprt | 0.0431347 | 0.08226614 | 0.04615412 | 0.03611202 | 0.00013356 | 0.00270442 |
| Rnf141 | 0.24447589 | 0.29094625 | 0.25485209 | 0.03609416 | 0.25467954 | 0.89054584 |
| Lsm8 | 1.07598886 | 0.90947461 | 0.87339066 | 0.03608395 | 0.42368187 | 0.89054584 |
| Dnajc9 | 0.55194429 | 0.571488 | 0.53543822 | 0.03604979 | 0.37688377 | 0.89054584 |
| Proscos | 0.04973335 | 0.08925881 | 0.05321153 | 0.03604728 | 0.00079941 | 0.01361901 |
| Pold4 | 0.55423774 | 0.57335528 | 0.53732258 | 0.0360327 | 0.37735792 | 0.89054584 |
| Spcs3 | 0.85675857 | 0.79038778 | 0.75435629 | 0.03603149 | 0.41194979 | 0.89054584 |
| Zfp273 | 0.06658464 | 0.10724987 | 0.07122739 | 0.03602248 | 0.00923127 | 0.11006164 |
| Pdcd6ip | 0.77001596 | 0.73476136 | 0.69873892 | 0.03602244 | 0.40509351 | 0.89054584 |
| Pdcd7 | 0.16622878 | 0.21179576 | 0.17578174 | 0.03601403 | 0.16991017 | 0.83373276 |
| Dock10 | 0.10685338 | 0.15012347 | 0.11411365 | 0.03600982 | 0.07075647 | 0.50162704 |
| Spin4 | 0.05928666 | 0.09943393 | 0.06342643 | 0.0360075 | 0.00408571 | 0.05551505 |
| Mapk10 | 0.04154349 | 0.08044286 | 0.04445209 | 0.03599077 | 8.09E-05 | 0.00173571 |
| Fam204a | 0.83983978 | 0.77987381 | 0.74393756 | 0.03593624 | 0.4109693 | 0.89054584 |
| Bhlhe41 | 0.05402734 | 0.09371689 | 0.05780329 | 0.0359136 | 0.00189801 | 0.02887441 |

| | | | | | | |
|---|---|---|---|---|---|---|
| Tifab | 0.01956731 | 0.05684555 | 0.02093983 | 0.03590572 | 6.82E-16 | 5.99E-14 |
| Tubg2 | 0.07407015 | 0.11506674 | 0.07922626 | 0.03584049 | 0.01753201 | 0.1828909 |
| Bud13 | 0.16759344 | 0.21301651 | 0.17718289 | 0.03583361 | 0.17303796 | 0.83791058 |
| Cdc45 | 0.09330833 | 0.13559987 | 0.09976961 | 0.03583026 | 0.04714736 | 0.3772309 |
| Erp44 | 1.26863108 | 0.99441007 | 0.95858534 | 0.03582473 | 0.43088704 | 0.89054584 |
| Med14 | 0.12538172 | 0.16930381 | 0.13350128 | 0.03580253 | 0.10575203 | 0.64250089 |
| Pou6f2 | 0.08500259 | 0.12669131 | 0.09090312 | 0.03578819 | 0.03331168 | 0.29269863 |
| Ccdc22 | 0.13855058 | 0.18294613 | 0.14720316 | 0.03574298 | 0.12897372 | 0.71862451 |
| Chmp2b | 0.6404138 | 0.64133585 | 0.60561621 | 0.03571964 | 0.39173979 | 0.89054584 |
| Dnmt3b | 0.12338026 | 0.1671313 | 0.13141319 | 0.03571811 | 0.10270279 | 0.63039166 |
| Plcxd2 | 0.13310399 | 0.17724176 | 0.14154415 | 0.03569761 | 0.1199992 | 0.69340312 |
| Tctex1d2 | 0.62968232 | 0.6330756 | 0.59738168 | 0.03569392 | 0.3903624 | 0.89054584 |
| Zfp58 | 0.11567738 | 0.15901816 | 0.12336284 | 0.03565532 | 0.08906022 | 0.57823337 |
| Thyn1 | 0.86463604 | 0.79478414 | 0.75913546 | 0.03564869 | 0.41340996 | 0.89054584 |
| Lrpap1 | 1.51187563 | 1.07443359 | 1.03878818 | 0.03564541 | 0.43649219 | 0.89054584 |
| Cmtm8 | 0.24541807 | 0.29140286 | 0.25578776 | 0.0356151 | 0.25826726 | 0.89054584 |
| Mblac1 | 0.08011049 | 0.12129309 | 0.08567873 | 0.03561436 | 0.02639727 | 0.24548275 |
| Dym | 0.09379837 | 0.1358745 | 0.10029261 | 0.0355819 | 0.04917528 | 0.38873676 |
| Man2c1os | 0.03471955 | 0.07272048 | 0.03715219 | 0.0355683 | 4.09E-06 | 0.00011533 |
| Pth1r | 0.12223966 | 0.16578715 | 0.13022255 | 0.0355646 | 0.10161799 | 0.62665146 |
| Gpr142 | 0.06718525 | 0.10742721 | 0.07186929 | 0.03555792 | 0.01058191 | 0.12300859 |
| Rhobtb3 | 0.41100256 | 0.44894601 | 0.4133923 | 0.0355537 | 0.34432379 | 0.89054584 |
| Rpl28 | 4.52600817 | 0.86876676 | 0.83323317 | 0.03553359 | 0.42125554 | 0.89054584 |
| Sema5b | 0.10346731 | 0.1460655 | 0.11055683 | 0.03550867 | 0.06727994 | 0.48312261 |
| Atp11b | 0.40726768 | 0.44549196 | 0.40999824 | 0.03549372 | 0.34335402 | 0.89054584 |
| Zfp346 | 0.05919211 | 0.09881826 | 0.06332535 | 0.03549291 | 0.00451038 | 0.06037993 |
| Park7 | 2.42778271 | 1.16424056 | 1.12878183 | 0.03545874 | 0.44182152 | 0.89054584 |
| Ttc7b | 0.0905815 | 0.13228431 | 0.09685918 | 0.03542512 | 0.04419486 | 0.36077541 |
| Acot9 | 0.20970046 | 0.25543132 | 0.22003715 | 0.03539417 | 0.22680327 | 0.89054584 |
| Rassf9 | 0.06740412 | 0.10748922 | 0.07210319 | 0.03538603 | 0.01111354 | 0.12757252 |

| | | | | | | |
|---|---|---|---|---|---|---|
| Snai1 | 0.13038242 | 0.17406675 | 0.13871216 | 0.03535458 | 0.11752341 | 0.68593175 |
| Slc28a3 | 0.13915923 | 0.18318745 | 0.14783483 | 0.03535263 | 0.13261376 | 0.72977796 |
| C1qtnf7 | 0.11918657 | 0.16233919 | 0.12703309 | 0.0353061 | 0.09768562 | 0.61445131 |
| Gchfr | 0.15001542 | 0.1943392 | 0.15907728 | 0.03526192 | 0.15086612 | 0.78172157 |
| Sike1 | 0.46392014 | 0.49588121 | 0.46062109 | 0.03526011 | 0.36068157 | 0.89054584 |
| Tiprl | 1.03469377 | 0.88759727 | 0.8523812 | 0.03521606 | 0.42368164 | 0.89054584 |
| Tmem37 | 0.13841214 | 0.18226952 | 0.14705946 | 0.03521006 | 0.13232281 | 0.72915851 |
| Tmem167b | 0.42057699 | 0.45725111 | 0.42205772 | 0.03519339 | 0.34882784 | 0.89054584 |
| Pofut2 | 1.1296907 | 0.93434409 | 0.8991873 | 0.03515679 | 0.42772969 | 0.89054584 |
| Kcnq1 | 0.04070786 | 0.07869456 | 0.04355824 | 0.03513633 | 8.56E-05 | 0.00182424 |
| Evi2a | 0.03127757 | 0.06859926 | 0.03346974 | 0.03512952 | 5.04E-07 | 1.65E-05 |
| Dph7 | 0.09194447 | 0.13344342 | 0.09831399 | 0.03512944 | 0.04798359 | 0.38170821 |
| Cse1l | 0.37673147 | 0.41708309 | 0.38195382 | 0.03512927 | 0.33414549 | 0.89054584 |
| Lrrc8a | 0.21740436 | 0.26291716 | 0.22779626 | 0.0351209 | 0.23628598 | 0.89054584 |
| Mknk1 | 0.26246889 | 0.30774235 | 0.27264949 | 0.03509286 | 0.27436815 | 0.89054584 |
| Klhl22 | 0.39227312 | 0.43137703 | 0.39629155 | 0.03508547 | 0.33999491 | 0.89054584 |
| Ntpcr | 0.32275661 | 0.36622061 | 0.33115013 | 0.03507048 | 0.31086303 | 0.89054584 |
| Zhx2 | 0.07638122 | 0.11674747 | 0.08169524 | 0.03505223 | 0.02280585 | 0.22145959 |
| Shisa5 | 0.8675291 | 0.79588437 | 0.76087985 | 0.03500452 | 0.4151425 | 0.89054584 |
| Paox | 0.12055698 | 0.16343774 | 0.12846514 | 0.0349726 | 0.10234133 | 0.62950557 |
| Ttc3 | 2.04718186 | 1.15559998 | 1.12062829 | 0.03497168 | 0.44220047 | 0.89054584 |
| Fam185a | 0.0884762 | 0.129568 | 0.09461179 | 0.03495621 | 0.04259469 | 0.35075907 |
| Gm12728 | 0.03495755 | 0.07230342 | 0.0374068 | 0.03489662 | 6.92E-06 | 0.0001883 |
| Rnf144a | 0.1563493 | 0.20050919 | 0.16561542 | 0.03489377 | 0.16314859 | 0.81647508 |
| Trim23 | 0.2061179 | 0.25130206 | 0.21642 | 0.03488206 | 0.22635129 | 0.89054584 |
| Tmem191c | 0.13239318 | 0.17562999 | 0.14080478 | 0.03482521 | 0.12459998 | 0.70642606 |
| Slc25a13 | 0.10958988 | 0.15180112 | 0.11698507 | 0.03481604 | 0.08278919 | 0.55312337 |
| Shc1 | 0.51225542 | 0.53707419 | 0.502301 | 0.03477319 | 0.37352627 | 0.89054584 |
| Tmem184a | 0.45756387 | 0.48978177 | 0.45503529 | 0.03474648 | 0.36101141 | 0.89054584 |
| Slc25a22 | 0.18791302 | 0.23268606 | 0.19795595 | 0.03473012 | 0.20685515 | 0.89054584 |

| | | | | | | |
|---|---|---|---|---|---|---|
| Mir143hg | 0.07688829 | 0.11696601 | 0.08223693 | 0.03472909 | 0.02456307 | 0.23336523 |
| Ccr2 | 0.02576841 | 0.06229606 | 0.02757522 | 0.03472084 | 2.23E-09 | 1.00E-07 |
| Golgb1 | 1.53688824 | 1.08013643 | 1.04545486 | 0.03468157 | 0.43858641 | 0.89054584 |
| Dcxr | 0.1581437 | 0.20213452 | 0.16746485 | 0.03466967 | 0.16739154 | 0.82799129 |
| Nudt16 | 0.20226933 | 0.24717881 | 0.21252845 | 0.03465036 | 0.22375055 | 0.89054584 |
| Syn1 | 0.08358587 | 0.12403968 | 0.08939031 | 0.03464937 | 0.03546751 | 0.30806285 |
| Gfra1 | 0.033753 | 0.07074053 | 0.03611813 | 0.03462239 | 3.98E-06 | 0.00011261 |
| Ncf2 | 0.0594692 | 0.09820787 | 0.06362158 | 0.03458628 | 0.00565911 | 0.0725099 |
| Dapk2 | 0.0447416 | 0.08240617 | 0.04787285 | 0.03453332 | 0.00038865 | 0.00716884 |
| Scarf1 | 0.05102301 | 0.08911071 | 0.05459069 | 0.03452003 | 0.00160931 | 0.02495384 |
| Ppard | 0.10189621 | 0.14342455 | 0.10890509 | 0.03451946 | 0.0698731 | 0.49755416 |
| Habp2 | 0.1050231 | 0.14669927 | 0.11219159 | 0.03450768 | 0.07593054 | 0.52346218 |
| Alkbh2 | 0.18998316 | 0.23455654 | 0.20006271 | 0.03449383 | 0.21090886 | 0.89054584 |
| Tnfrsf25 | 0.03330185 | 0.07009525 | 0.03563547 | 0.03445977 | 3.32E-06 | 9.46E-05 |
| Glipr1 | 0.05199997 | 0.09008387 | 0.05563542 | 0.03444845 | 0.00195873 | 0.02958091 |
| Nus1 | 0.4163404 | 0.45263878 | 0.41822942 | 0.03440936 | 0.35074526 | 0.89054584 |
| Uxs1 | 0.27674609 | 0.32105858 | 0.2866636 | 0.03439498 | 0.28808176 | 0.89054584 |
| Snx27 | 0.35232908 | 0.3935003 | 0.35917598 | 0.03432432 | 0.32807772 | 0.89054584 |
| Crhr2 | 0.12530216 | 0.16772421 | 0.1334183 | 0.0343059 | 0.11546718 | 0.67725583 |
| Muc4 | 0.11696377 | 0.15899289 | 0.1247088 | 0.0342841 | 0.10013116 | 0.62272587 |
| Abcd1 | 0.13042485 | 0.17302919 | 0.13875633 | 0.03427286 | 0.12491546 | 0.70660334 |
| Exoc2 | 0.15874131 | 0.20233009 | 0.1680805 | 0.03424959 | 0.17122189 | 0.83568909 |
| Taf13 | 0.39030805 | 0.42869116 | 0.39448579 | 0.03420537 | 0.34311704 | 0.89054584 |
| Brms1l | 0.3004755 | 0.34392766 | 0.30973814 | 0.03418952 | 0.30353395 | 0.89054584 |
| Xrn2 | 0.71107611 | 0.69203378 | 0.65785369 | 0.03418009 | 0.40436508 | 0.89054584 |
| Numbl | 0.17245904 | 0.21634713 | 0.18217241 | 0.03417472 | 0.19106014 | 0.88366966 |
| Gm26735 | 0.04665478 | 0.08408286 | 0.04991908 | 0.03416378 | 0.00071506 | 0.01240072 |
| 5430416N02 | 0.16767856 | 0.21142679 | 0.17727026 | 0.03415653 | 0.18467617 | 0.8690805 |
| Zfp386 | 0.34049487 | 0.38210133 | 0.34801506 | 0.03408628 | 0.32408023 | 0.89054584 |
| Synpo | 0.07621486 | 0.11559269 | 0.08151753 | 0.03407517 | 0.02573828 | 0.24089971 |

| | | | | | | |
|---|---|---|---|---|---|---|
| Wnt5a | 0.11250869 | 0.15410061 | 0.12004477 | 0.03405584 | 0.09313266 | 0.59577065 |
| Zbtb10 | 0.12368386 | 0.16578127 | 0.13173004 | 0.03405123 | 0.11423652 | 0.67417108 |
| Ndrg1 | 0.50624714 | 0.53123884 | 0.49719823 | 0.03404061 | 0.37487318 | 0.89054584 |
| Rbfa | 0.48804958 | 0.51559172 | 0.48160647 | 0.03398524 | 0.37116674 | 0.89054584 |
| Hspa4 | 1.03469687 | 0.88633428 | 0.85238281 | 0.03395147 | 0.42639042 | 0.89054584 |
| S1pr2 | 0.15430061 | 0.19741128 | 0.16350233 | 0.03390895 | 0.16696612 | 0.82701566 |
| Lrrc56 | 0.1012517 | 0.14211209 | 0.10822724 | 0.03388486 | 0.07232833 | 0.50878997 |
| Gps2 | 0.62757094 | 0.62962006 | 0.59575225 | 0.03386781 | 0.39556047 | 0.89054584 |
| Zfp593 | 0.23721622 | 0.28148273 | 0.24762884 | 0.0338539 | 0.2620838 | 0.89054584 |
| Folr2 | 0.01801926 | 0.0531122 | 0.01928328 | 0.03382892 | 1.50E-16 | 1.42E-14 |
| St8sia5 | 0.04736325 | 0.08447359 | 0.0506768 | 0.03379679 | 0.0009447 | 0.01572478 |
| Arhgef26 | 0.0511444 | 0.08851166 | 0.0547205 | 0.03379116 | 0.00200736 | 0.03018954 |
| Tacc2 | 0.21967516 | 0.26382262 | 0.2300783 | 0.03374432 | 0.24720823 | 0.89054584 |
| Hspb8 | 0.26091676 | 0.3048514 | 0.27112005 | 0.03373135 | 0.28107829 | 0.89054584 |
| Plbd1 | 0.02571135 | 0.06122993 | 0.02751417 | 0.03371576 | 5.68E-09 | 2.38E-07 |
| Mthfr | 0.0999138 | 0.14049786 | 0.10681774 | 0.03368012 | 0.07091836 | 0.50228355 |
| Dpysl2 | 0.75631502 | 0.72312069 | 0.68945598 | 0.03366472 | 0.41002273 | 0.89054584 |
| Mettl21a | 0.13972192 | 0.18206144 | 0.14841867 | 0.03364277 | 0.1454694 | 0.76831161 |
| Uap1 | 0.47019744 | 0.49973518 | 0.466114 | 0.03362118 | 0.36841537 | 0.89054584 |
| Rbpms | 0.88810037 | 0.80671914 | 0.77310574 | 0.0336134 | 0.41973818 | 0.89054584 |
| Sf3b6 | 1.89040239 | 1.14221016 | 1.10870597 | 0.03350418 | 0.44401817 | 0.89054584 |
| Stk32a | 0.07664962 | 0.11546522 | 0.08198196 | 0.03348325 | 0.02853234 | 0.25967473 |
| Limch1 | 0.17768621 | 0.22099773 | 0.1875218 | 0.03347593 | 0.20278839 | 0.89054584 |
| Tchp | 0.14025719 | 0.18244976 | 0.14897396 | 0.0334758 | 0.14757139 | 0.77404405 |
| Prpf3 | 0.29805188 | 0.34086551 | 0.30739412 | 0.03347138 | 0.30597199 | 0.89054584 |
| Prrg3 | 0.12644316 | 0.16807501 | 0.13460806 | 0.03346695 | 0.12336469 | 0.70249071 |
| Ecd | 0.44743993 | 0.47955077 | 0.44608812 | 0.03346265 | 0.36336311 | 0.89054584 |
| Tbx3 | 0.05399921 | 0.09120905 | 0.05777321 | 0.03343584 | 0.00350541 | 0.04896098 |
| Yap1 | 0.19679621 | 0.24041805 | 0.20698337 | 0.03343468 | 0.22585312 | 0.89054584 |
| Ttc39a | 0.32883114 | 0.37035117 | 0.33694385 | 0.03340732 | 0.32206761 | 0.89054584 |

| | | | | | | |
|---|---|---|---|---|---|---|
| Gm28557 | 0.04246586 | 0.07882439 | 0.04543871 | 0.03338569 | 0.00030954 | 0.0058586 |
| Il10rb | 0.35288677 | 0.39304833 | 0.35970009 | 0.03334823 | 0.33289393 | 0.89054584 |
| Tbc1d2b | 0.15116994 | 0.19361418 | 0.16027015 | 0.03334403 | 0.16620085 | 0.8246338 |
| Spi1 | 0.02883789 | 0.06419766 | 0.03085947 | 0.03333819 | 2.41E-07 | 8.21E-06 |
| Sri | 1.40502769 | 1.04050961 | 1.0072376 | 0.03327201 | 0.43884489 | 0.89054584 |
| Mzt1 | 0.546102 | 0.56388882 | 0.53062275 | 0.03326607 | 0.38511248 | 0.89054584 |
| Slc25a12 | 0.17280747 | 0.21575381 | 0.18252935 | 0.03322447 | 0.1982099 | 0.89054584 |
| Tmeff2 | 0.05511433 | 0.09218373 | 0.05896556 | 0.03321816 | 0.00433744 | 0.05838834 |
| Zbtb1 | 0.31249104 | 0.35450979 | 0.32131558 | 0.0331942 | 0.31515094 | 0.89054584 |
| Map3k12 | 0.2221271 | 0.26571878 | 0.23253978 | 0.03317899 | 0.25310775 | 0.89054584 |
| Kras | 0.89917679 | 0.81273232 | 0.77956306 | 0.03316927 | 0.42143211 | 0.89054584 |
| Rab27b | 0.09571183 | 0.13549247 | 0.10233453 | 0.03315793 | 0.06557695 | 0.47418188 |
| Zfp606 | 0.28972193 | 0.3324451 | 0.2993153 | 0.0331298 | 0.30304027 | 0.89054584 |
| Dusp2 | 0.03286892 | 0.06829727 | 0.03517229 | 0.03312497 | 5.73E-06 | 0.00015844 |
| Aatf | 0.28010282 | 0.32302433 | 0.28994409 | 0.03308024 | 0.2975199 | 0.89054584 |
| Rnmt | 0.35929374 | 0.39878285 | 0.36570913 | 0.03307372 | 0.33675276 | 0.89054584 |
| Ttll12 | 0.16786802 | 0.21053705 | 0.17746473 | 0.03307232 | 0.19263148 | 0.88659402 |
| Mboat4 | 0.08768822 | 0.12684188 | 0.09377055 | 0.03307132 | 0.05017787 | 0.39431114 |
| Rnf128 | 0.23085141 | 0.27433637 | 0.24127606 | 0.03306032 | 0.26161281 | 0.89054584 |
| Npl | 0.14765694 | 0.1896847 | 0.15663884 | 0.03304586 | 0.16283084 | 0.8160116 |
| Vhl | 0.27188843 | 0.31488906 | 0.28190652 | 0.03298253 | 0.29284673 | 0.89054584 |
| Plekha2 | 0.06696556 | 0.10460422 | 0.0716345 | 0.03296972 | 0.01600547 | 0.17074856 |
| Trio | 0.31860373 | 0.3600916 | 0.32717817 | 0.03291343 | 0.31964934 | 0.89054584 |
| Btd | 0.24469028 | 0.28796709 | 0.25506504 | 0.03290204 | 0.27392704 | 0.89054584 |
| E2f2 | 0.0455062 | 0.08158174 | 0.04869064 | 0.0328911 | 0.0008243 | 0.01401012 |
| Clec2d | 0.10032342 | 0.14013573 | 0.10725068 | 0.03288505 | 0.07657143 | 0.52662835 |
| C130013H08 | 0.05017988 | 0.0865131 | 0.05368905 | 0.03282404 | 0.00219732 | 0.03264011 |
| Cwf19l2 | 0.34933411 | 0.38918035 | 0.35635851 | 0.03282184 | 0.3339236 | 0.89054584 |
| Ado | 0.23946854 | 0.2826882 | 0.24987247 | 0.03281573 | 0.27031572 | 0.89054584 |
| Adal | 0.12451847 | 0.16537788 | 0.13260085 | 0.03277703 | 0.12473794 | 0.70642606 |

| | | | | | | |
|---|---|---|---|---|---|---|
| Anapc15 | 0.26789757 | 0.31076426 | 0.27798973 | 0.03277453 | 0.29140613 | 0.89054584 |
| Siglecg | 0.02054394 | 0.05475149 | 0.02198489 | 0.0327666 | 1.91E-12 | 1.22E-10 |
| Trub1 | 0.33027176 | 0.37106861 | 0.33831504 | 0.03275357 | 0.32597679 | 0.89054584 |
| Ntrk1 | 0.06762684 | 0.10509421 | 0.07234123 | 0.03275298 | 0.01745644 | 0.18264153 |
| Lcorl | 0.45493313 | 0.48539728 | 0.45271627 | 0.032681 | 0.36831368 | 0.89054584 |
| Gls | 0.36911711 | 0.40755542 | 0.37488063 | 0.03267479 | 0.34234346 | 0.89054584 |
| Lrrn4 | 0.02767897 | 0.06228785 | 0.02961948 | 0.03266837 | 1.38E-07 | 4.86E-06 |
| Eif1ax | 1.11013256 | 0.92268081 | 0.89002271 | 0.0326581 | 0.43213032 | 0.89054584 |
| Lix1l | 0.15982152 | 0.20184329 | 0.16919295 | 0.03265033 | 0.18430819 | 0.8681287 |
| Gm42637 | 0.0536823 | 0.09006463 | 0.05743435 | 0.03263028 | 0.00406176 | 0.05529331 |
| Fam126b | 0.30735617 | 0.34900612 | 0.3163766 | 0.03262951 | 0.31543592 | 0.89054584 |
| Atmin | 0.17563792 | 0.21805327 | 0.18542697 | 0.0326263 | 0.20617892 | 0.89054584 |
| Dcp1b | 0.14031502 | 0.18164163 | 0.14903393 | 0.0326077 | 0.15401906 | 0.79121355 |
| Jmjd6 | 0.84458897 | 0.77947985 | 0.74688339 | 0.03259646 | 0.41943793 | 0.89054584 |
| Rassf4 | 0.1098967 | 0.14988033 | 0.11730686 | 0.03257347 | 0.09788749 | 0.61489291 |
| Sh3yl1 | 0.19775807 | 0.24046136 | 0.20795881 | 0.03250255 | 0.23325753 | 0.89054584 |
| Hs3st6 | 0.08719333 | 0.1257424 | 0.09324219 | 0.03250021 | 0.05219791 | 0.40403915 |
| Cul2 | 0.40687495 | 0.44213456 | 0.40964087 | 0.03249369 | 0.35585581 | 0.89054584 |
| Nif3l1 | 0.23214483 | 0.27501571 | 0.2425685 | 0.03244721 | 0.26657526 | 0.89054584 |
| Rad54l2 | 0.13422916 | 0.17516094 | 0.14271414 | 0.0324468 | 0.14474648 | 0.76727455 |
| Sult1a1 | 0.09639706 | 0.13550961 | 0.10306572 | 0.0324439 | 0.07124505 | 0.50410481 |
| Chpf2 | 0.20058228 | 0.24326103 | 0.2108206 | 0.03244042 | 0.23671684 | 0.89054584 |
| Prkacb | 0.59098661 | 0.59942932 | 0.56704776 | 0.03238156 | 0.39509986 | 0.89054584 |
| Wrb | 0.2174824 | 0.26021203 | 0.22787472 | 0.03233731 | 0.25426055 | 0.89054584 |
| Acvr1c | 0.34858006 | 0.38798028 | 0.35564839 | 0.03233189 | 0.3359496 | 0.89054584 |
| Ercc4 | 0.09722718 | 0.13625534 | 0.10395145 | 0.03230389 | 0.07383516 | 0.51613412 |
| Ncoa4 | 0.03103468 | 0.06548516 | 0.03320987 | 0.03227529 | 2.98E-06 | 8.55E-05 |
| Capn15 | 0.17057017 | 0.21250221 | 0.18023659 | 0.03226561 | 0.20213069 | 0.89054584 |
| Slc39a3 | 0.19153572 | 0.23389057 | 0.20164154 | 0.03224904 | 0.22810076 | 0.89054584 |
| Ybx3 | 0.29779452 | 0.33935324 | 0.30714504 | 0.0322082 | 0.31257968 | 0.89054584 |

| | | | | | | |
|---|---|---|---|---|---|---|
| Msln | 0.02996467 | 0.06427032 | 0.03206505 | 0.03220527 | 1.44E-06 | 4.38E-05 |
| Serpini2 | 0.06027074 | 0.09665326 | 0.06447847 | 0.03217479 | 0.01004096 | 0.11777817 |
| Zfp277 | 0.31230081 | 0.35330428 | 0.32113283 | 0.03217146 | 0.32034364 | 0.89054584 |
| Dgkz | 0.23361467 | 0.27620501 | 0.24403635 | 0.03216866 | 0.26956219 | 0.89054584 |
| Surf6 | 0.15428214 | 0.19561625 | 0.16348326 | 0.03213299 | 0.17990558 | 0.85601281 |
| Rfk | 0.61818904 | 0.62060594 | 0.58847715 | 0.03212878 | 0.39960802 | 0.89054584 |
| Shmt2 | 0.31863508 | 0.35929179 | 0.32720819 | 0.0320836 | 0.32389984 | 0.89054584 |
| Napepld | 0.08158798 | 0.11930654 | 0.08725673 | 0.03204981 | 0.04351739 | 0.35685246 |
| Ifi27l2a | 0.05024918 | 0.08579832 | 0.05376316 | 0.03203516 | 0.00275107 | 0.03996501 |
| Eno3 | 0.05124564 | 0.08685313 | 0.05482876 | 0.03202436 | 0.00325234 | 0.04600238 |
| Haus4 | 0.11857297 | 0.15840329 | 0.12639166 | 0.03201162 | 0.11900307 | 0.69067516 |
| Fam109b | 0.07286581 | 0.10994974 | 0.07793951 | 0.03201023 | 0.0278449 | 0.25472969 |
| Tuft1 | 0.13678199 | 0.17736107 | 0.14536685 | 0.03199423 | 0.15258714 | 0.78614371 |
| Tmem263 | 0.59587489 | 0.60291928 | 0.57093464 | 0.03198464 | 0.39704584 | 0.89054584 |
| Zfp112 | 0.06754042 | 0.10422135 | 0.07224887 | 0.03197248 | 0.01961654 | 0.19846523 |
| Zfp503 | 0.21583276 | 0.25818463 | 0.22621553 | 0.0319691 | 0.25513795 | 0.89054584 |
| 2610001J05F | 1.06487978 | 0.89976862 | 0.86782239 | 0.03194623 | 0.43191348 | 0.89054584 |
| Slc37a4 | 0.3542436 | 0.39286308 | 0.36097451 | 0.03188857 | 0.34032659 | 0.89054584 |
| Pah | 0.06386961 | 0.10019155 | 0.0683255 | 0.03186604 | 0.01489528 | 0.16079806 |
| Plscr4 | 0.12357276 | 0.1634293 | 0.13161409 | 0.03181521 | 0.1300382 | 0.72233481 |
| 9330160F10I | 0.11400694 | 0.15342834 | 0.12161411 | 0.03181423 | 0.11146417 | 0.66356743 |
| Smarcad1 | 0.55921833 | 0.57321383 | 0.5414029 | 0.03181094 | 0.39214104 | 0.89054584 |
| Lcmt1 | 0.46648879 | 0.4946777 | 0.46287142 | 0.03180627 | 0.37443119 | 0.89054584 |
| Osgin2 | 0.19294898 | 0.23487864 | 0.20307782 | 0.03180083 | 0.23282849 | 0.89054584 |
| Thap1 | 0.13404659 | 0.17432022 | 0.14252433 | 0.0317959 | 0.14931683 | 0.77812345 |
| Dcun1d4 | 0.08786747 | 0.12573462 | 0.09396192 | 0.0317727 | 0.05758264 | 0.43458258 |
| Phox2b | 0.02066315 | 0.05387252 | 0.02211245 | 0.03176007 | 1.10E-11 | 6.53E-10 |
| Pde4d | 0.21672158 | 0.25883573 | 0.22710965 | 0.03172608 | 0.25757786 | 0.89054584 |
| Zfp580 | 0.22780305 | 0.26995022 | 0.23822744 | 0.03172278 | 0.26749963 | 0.89054584 |
| Supt4a | 0.85724575 | 0.78633999 | 0.75465317 | 0.03168682 | 0.42245288 | 0.89054584 |

| | | | | | | |
|---|---|---|---|---|---|---|
| Vps54 | 0.12000995 | 0.15956944 | 0.12789359 | 0.03167585 | 0.12427139 | 0.70545441 |
| Cdc42ep1 | 0.16732372 | 0.20855436 | 0.17690602 | 0.03164834 | 0.20228683 | 0.89054584 |
| Nudt17 | 0.05480479 | 0.09025719 | 0.05863459 | 0.0316226 | 0.00599152 | 0.07599742 |
| Onecut1 | 0.0847386 | 0.12223121 | 0.09062123 | 0.03160998 | 0.05207009 | 0.40348056 |
| Pin1 | 1.37988555 | 1.03049536 | 0.99889814 | 0.03159723 | 0.4414192 | 0.89054584 |
| Cdc34 | 0.29261814 | 0.33371136 | 0.30212812 | 0.03158323 | 0.31311974 | 0.89054584 |
| Pqbp1 | 0.68728189 | 0.67221854 | 0.64065133 | 0.03156722 | 0.4092157 | 0.89054584 |
| Eps8l2 | 0.36117025 | 0.39902406 | 0.36746487 | 0.03155919 | 0.34453169 | 0.89054584 |
| Gnai2 | 1.70244922 | 1.11343363 | 1.08190299 | 0.03153064 | 0.44599811 | 0.89054584 |
| Ppp2r3a | 0.24845296 | 0.29030793 | 0.25879883 | 0.0315091 | 0.28527818 | 0.89054584 |
| Apbb1 | 0.43841774 | 0.4695725 | 0.4380638 | 0.0315087 | 0.36877312 | 0.89054584 |
| Heyl | 0.06551999 | 0.10158434 | 0.0700895 | 0.03149484 | 0.01815212 | 0.18797814 |
| Fhod1 | 0.07870322 | 0.11565894 | 0.08417562 | 0.03148331 | 0.0407075 | 0.34053187 |
| Fam178b | 0.02160126 | 0.05459088 | 0.02311628 | 0.0314746 | 1.12E-10 | 5.92E-09 |
| Hnf1aos1 | 0.1895505 | 0.23109692 | 0.19962254 | 0.03147437 | 0.2312999 | 0.89054584 |
| Rbm12b2 | 0.14848211 | 0.18895834 | 0.15749224 | 0.0314661 | 0.1759704 | 0.8453343 |
| Atg4c | 0.08710241 | 0.12461037 | 0.09314512 | 0.03146525 | 0.05776316 | 0.43549156 |
| Slc1a4 | 0.08041404 | 0.11745947 | 0.08600294 | 0.03145653 | 0.044185 | 0.36077541 |
| Atp5k | 2.30934863 | 1.16066126 | 1.12920509 | 0.03145618 | 0.44836876 | 0.89054584 |
| Mrpl47 | 0.30858401 | 0.34900183 | 0.31755869 | 0.03144314 | 0.32228953 | 0.89054584 |
| Fra10ac1 | 0.40323656 | 0.43774515 | 0.40632596 | 0.03141919 | 0.35932895 | 0.89054584 |
| Gmcl1 | 0.41784174 | 0.45096685 | 0.41958726 | 0.03137958 | 0.36375967 | 0.89054584 |
| Slc25a51 | 0.84111927 | 0.77611153 | 0.74473284 | 0.03137869 | 0.42218746 | 0.89054584 |
| Plcl1 | 0.23292293 | 0.27471169 | 0.24334567 | 0.03136602 | 0.27408194 | 0.89054584 |
| Gm5595 | 0.14147663 | 0.18159223 | 0.15023856 | 0.03135367 | 0.16545406 | 0.82205381 |
| Tmem120a | 0.27718616 | 0.31839803 | 0.28709399 | 0.03130403 | 0.30572678 | 0.89054584 |
| Parp1 | 0.642963 | 0.63884776 | 0.60756062 | 0.03128714 | 0.40519677 | 0.89054584 |
| Prmt6 | 0.29760735 | 0.33825067 | 0.30696387 | 0.0312868 | 0.31744514 | 0.89054584 |
| Rbp2 | 0.17188682 | 0.21286823 | 0.18158613 | 0.03128211 | 0.21110224 | 0.89054584 |
| Pdlim3 | 0.06044462 | 0.09593772 | 0.06466435 | 0.03127337 | 0.01212372 | 0.13671864 |

| | | | | | | |
|---|---|---|---|---|---|---|
| Nnmt | 0.08009574 | 0.1169295 | 0.08566298 | 0.03126652 | 0.04451959 | 0.36199945 |
| Litaf | 0.5046619 | 0.52709048 | 0.49584809 | 0.03124239 | 0.38455072 | 0.89054584 |
| Lrig1 | 0.12546164 | 0.16479158 | 0.13358463 | 0.03120695 | 0.13821285 | 0.74574245 |
| Mrps21 | 1.80063062 | 1.1290405 | 1.09784427 | 0.03119623 | 0.44733871 | 0.89054584 |
| Plpp5 | 0.43655743 | 0.46759964 | 0.43640357 | 0.03119607 | 0.36955134 | 0.89054584 |
| Lss | 0.06631587 | 0.10213489 | 0.07094013 | 0.03119476 | 0.02024608 | 0.20313329 |
| Kif18a | 0.07955072 | 0.11626775 | 0.08508085 | 0.0311869 | 0.04384014 | 0.35909288 |
| Umps | 0.55327327 | 0.56768483 | 0.53653056 | 0.03115427 | 0.39337748 | 0.89054584 |
| Crem | 0.26202333 | 0.30336085 | 0.27221056 | 0.03115029 | 0.29696705 | 0.89054584 |
| Slit3 | 0.09272443 | 0.13028834 | 0.09914645 | 0.0311419 | 0.07168263 | 0.50695358 |
| Fkbp1b | 0.12827805 | 0.16765447 | 0.13652047 | 0.031134 | 0.14400691 | 0.76637203 |
| Capns1 | 1.91586129 | 1.14231886 | 1.11119148 | 0.03112738 | 0.44808182 | 0.89054584 |
| Sncg | 0.05727108 | 0.0923855 | 0.06127155 | 0.03111395 | 0.00899485 | 0.10777465 |
| Zbed3 | 0.28116372 | 0.32208321 | 0.29097975 | 0.03110346 | 0.30924046 | 0.89054584 |
| Zfp248 | 0.09643146 | 0.13420179 | 0.10310242 | 0.03109937 | 0.07996561 | 0.54094194 |
| Il16 | 0.04057536 | 0.07450949 | 0.04341651 | 0.03109298 | 0.00042416 | 0.00780391 |
| 2310022B05I | 0.87368918 | 0.79566522 | 0.76457359 | 0.03109163 | 0.4248672 | 0.89054584 |
| Tbcel | 0.15817987 | 0.19854523 | 0.16750212 | 0.03104311 | 0.19394783 | 0.88899106 |
| Tmem121 | 0.11417916 | 0.15282654 | 0.12179445 | 0.03103208 | 0.11760308 | 0.68612048 |
| 1700096K18I | 0.23540475 | 0.27684317 | 0.24582267 | 0.0310205 | 0.2782949 | 0.89054584 |
| Rhbdl3 | 0.04660582 | 0.08083055 | 0.04986673 | 0.03096382 | 0.00190862 | 0.02894551 |
| Ier3 | 0.15222717 | 0.19228259 | 0.16136202 | 0.03092056 | 0.18599364 | 0.87233221 |
| Set | 1.94518736 | 1.14462936 | 1.11371804 | 0.03091133 | 0.44855648 | 0.89054584 |
| Pole4 | 0.52802448 | 0.54649659 | 0.51558636 | 0.03091023 | 0.39000308 | 0.89054584 |
| Ect2 | 0.06467613 | 0.10009675 | 0.06918757 | 0.03090919 | 0.01870059 | 0.19226646 |
| Bcl9 | 0.3657179 | 0.40260702 | 0.37171268 | 0.03089434 | 0.34929635 | 0.89054584 |
| Xiap | 0.53850593 | 0.55522084 | 0.52433029 | 0.03089055 | 0.39185856 | 0.89054584 |
| Flt4 | 0.03891792 | 0.07253203 | 0.04164352 | 0.03088851 | 0.00027443 | 0.00526973 |
| Cnp | 0.27500616 | 0.31583585 | 0.28496102 | 0.03087484 | 0.30685491 | 0.89054584 |
| Cdkn2c | 0.06938245 | 0.10508237 | 0.07421739 | 0.03086498 | 0.02634055 | 0.24511247 |

| | | | | | | |
|---|---|---|---|---|---|---|
| Rab4b | 0.06285328 | 0.09809887 | 0.06723915 | 0.03085972 | 0.01624853 | 0.17287186 |
| Fam163a | 0.09755055 | 0.13514755 | 0.10429647 | 0.03085108 | 0.08408268 | 0.55918838 |
| Pdzrn3 | 0.07089571 | 0.10668172 | 0.07583444 | 0.03084727 | 0.02903802 | 0.26312076 |
| Sntg1 | 0.05005294 | 0.08439498 | 0.0535533 | 0.03084167 | 0.00364702 | 0.05064608 |
| 2410131K14I | 0.17774071 | 0.21841405 | 0.18757751 | 0.03083654 | 0.22186784 | 0.89054584 |
| Pdlim7 | 0.46963932 | 0.49644285 | 0.46562649 | 0.03081636 | 0.37891153 | 0.89054584 |
| Zfp397 | 0.44624005 | 0.47583067 | 0.44502363 | 0.03080704 | 0.3735301 | 0.89054584 |
| Tti2 | 0.1666712 | 0.20700229 | 0.17623607 | 0.03076622 | 0.20801457 | 0.89054584 |
| Ddah1 | 0.11251403 | 0.15081235 | 0.12005036 | 0.03076198 | 0.1162746 | 0.68083567 |
| Slc9a8 | 0.20218498 | 0.24316782 | 0.21244309 | 0.03072473 | 0.25021842 | 0.89054584 |
| Phox2a | 0.01336683 | 0.04502537 | 0.01430466 | 0.03072071 | 7.20E-24 | 1.07E-21 |
| Micalcl | 0.1030156 | 0.14079932 | 0.11008203 | 0.03071729 | 0.09679515 | 0.61121462 |
| Nedd1 | 0.27277742 | 0.31344802 | 0.28277797 | 0.03067005 | 0.30667125 | 0.89054584 |
| Tpd52l1 | 0.09817796 | 0.1356263 | 0.10496587 | 0.03066043 | 0.08677705 | 0.57045578 |
| Clasp1 | 0.22679329 | 0.26787236 | 0.23721666 | 0.0306557 | 0.27356084 | 0.89054584 |
| Brpf3 | 0.10608792 | 0.14396272 | 0.11330995 | 0.03065277 | 0.10377278 | 0.63373959 |
| Cdca4 | 0.31469238 | 0.35406987 | 0.32342907 | 0.0306408 | 0.32947059 | 0.89054584 |
| Slc7a8 | 0.19319209 | 0.23392171 | 0.2033248 | 0.03059691 | 0.2416231 | 0.89054584 |
| Zdhhc8 | 0.19171968 | 0.23240586 | 0.20182854 | 0.03057732 | 0.24014492 | 0.89054584 |
| Sep-06 | 0.1556709 | 0.19545301 | 0.16491588 | 0.03053714 | 0.19415488 | 0.889076 |
| Ndufaf2 | 0.49153282 | 0.51513795 | 0.48460631 | 0.03053164 | 0.38455962 | 0.89054584 |
| Pla2g15 | 0.06730683 | 0.10252688 | 0.07199922 | 0.03052766 | 0.02411193 | 0.23028641 |
| Ercc1 | 0.31704214 | 0.35620834 | 0.3256823 | 0.03052604 | 0.33117165 | 0.89054584 |
| Rnf8 | 0.53016363 | 0.54786019 | 0.51737665 | 0.03048354 | 0.39184935 | 0.89054584 |
| Anapc10 | 0.49068898 | 0.51436126 | 0.48388027 | 0.03048099 | 0.38457768 | 0.89054584 |
| Npr1 | 0.09780821 | 0.13504684 | 0.10457138 | 0.03047546 | 0.0872691 | 0.57076343 |
| Mical3 | 0.22663012 | 0.2675275 | 0.23705329 | 0.03047421 | 0.27461084 | 0.89054584 |
| 5830454E08I | 0.0582608 | 0.09279124 | 0.06232969 | 0.03046155 | 0.01139606 | 0.13050563 |
| Gm10053 | 0.22011941 | 0.26098059 | 0.23052448 | 0.03045611 | 0.26910451 | 0.89054584 |
| Spink2 | 0.04791353 | 0.08171837 | 0.05126532 | 0.03045306 | 0.00282402 | 0.04076221 |

| | | | | | | |
|---|---|---|---|---|---|---|
| Tnks1bp1 | 0.13669119 | 0.17572276 | 0.14527254 | 0.03045022 | 0.16439622 | 0.81876208 |
| Mta3 | 0.44921072 | 0.47809754 | 0.44765753 | 0.03044001 | 0.37569742 | 0.89054584 |
| Dock2 | 0.03557073 | 0.06849726 | 0.0380628 | 0.03043447 | 9.76E-05 | 0.00203832 |
| Oraov1 | 0.18840103 | 0.22887962 | 0.19845275 | 0.03042686 | 0.23751749 | 0.89054584 |
| Dolk | 0.24099047 | 0.28179854 | 0.2513872 | 0.03041134 | 0.28650937 | 0.89054584 |
| Fam45a | 0.29495106 | 0.33478347 | 0.30439084 | 0.03039263 | 0.32090015 | 0.89054584 |
| Mccc2 | 0.29985042 | 0.33951663 | 0.30913387 | 0.03038275 | 0.32351324 | 0.89054584 |
| Sh3bp2 | 0.04410602 | 0.07756462 | 0.04719304 | 0.03037157 | 0.00135741 | 0.02159264 |
| Mpv17 | 0.56198628 | 0.57401191 | 0.54366348 | 0.03034843 | 0.3974042 | 0.89054584 |
| 3110040N11 | 0.34597123 | 0.38350438 | 0.35318918 | 0.0303152 | 0.34461887 | 0.89054584 |
| 1810024B03 | 0.14625297 | 0.18548257 | 0.15518622 | 0.03029635 | 0.18153117 | 0.86081561 |
| Adgrl4 | 0.04766585 | 0.08129277 | 0.05100043 | 0.03029234 | 0.00282669 | 0.04076221 |
| Gstcd | 0.10453287 | 0.14195112 | 0.11167657 | 0.03027455 | 0.10329485 | 0.63241921 |
| Gmppa | 0.49930392 | 0.5215328 | 0.49127306 | 0.03025975 | 0.38706939 | 0.89054584 |
| Dnajb14 | 0.41938815 | 0.45123685 | 0.4209845 | 0.03025234 | 0.36888953 | 0.89054584 |
| Arv1 | 0.08723186 | 0.12352509 | 0.09328333 | 0.03024176 | 0.06547146 | 0.47418188 |
| Sfr1 | 2.3046516 | 1.15938198 | 1.12914614 | 0.03023584 | 0.45035863 | 0.89054584 |
| Hint3 | 0.35224828 | 0.38931769 | 0.35910004 | 0.03021765 | 0.34751364 | 0.89054584 |
| Ppil1 | 0.57999561 | 0.58843779 | 0.55824992 | 0.03018787 | 0.40054508 | 0.89054584 |
| Hars2 | 0.17891558 | 0.2189603 | 0.18877829 | 0.03018201 | 0.22817367 | 0.89054584 |
| Arhgap1 | 0.36704154 | 0.40312618 | 0.37294702 | 0.03017916 | 0.35308492 | 0.89054584 |
| 1600020E01 | 0.22399644 | 0.26456774 | 0.23441458 | 0.03015315 | 0.27449054 | 0.89054584 |
| Cenpk | 0.16589145 | 0.20557494 | 0.17543527 | 0.03013967 | 0.21173857 | 0.89054584 |
| Sh3bp5 | 0.16523181 | 0.20487234 | 0.17475762 | 0.03011472 | 0.21103311 | 0.89054584 |
| Stk19 | 0.32497021 | 0.36335013 | 0.33326364 | 0.03008649 | 0.33702304 | 0.89054584 |
| Chp1 | 0.6708544 | 0.65862513 | 0.62854406 | 0.03008106 | 0.41177977 | 0.89054584 |
| Orc2 | 0.22236767 | 0.26285885 | 0.23278114 | 0.03007771 | 0.27359272 | 0.89054584 |
| Top3b | 0.31507815 | 0.35387474 | 0.32379919 | 0.03007556 | 0.33260252 | 0.89054584 |
| Itsn1 | 0.17445923 | 0.2142738 | 0.18422071 | 0.03005309 | 0.22361458 | 0.89054584 |
| Dusp3 | 0.28891935 | 0.32858464 | 0.29853509 | 0.03004956 | 0.31955138 | 0.89054584 |

| | | | | | | |
|---|---|---|---|---|---|---|
| 0610040J01F | 0.64151171 | 0.63649061 | 0.6064542 | 0.03003641 | 0.40875543 | 0.89054584 |
| Adra1a | 0.04792371 | 0.08130797 | 0.0512762 | 0.03003177 | 0.00317944 | 0.04510328 |
| Aqp1 | 0.10490973 | 0.14202901 | 0.1120725 | 0.02995651 | 0.10650791 | 0.64598207 |
| Cfap20 | 1.34534059 | 1.0170052 | 0.98707191 | 0.02993329 | 0.44382281 | 0.89054584 |
| Tmem87b | 0.44037828 | 0.46974202 | 0.43981127 | 0.02993075 | 0.37559984 | 0.89054584 |
| Phf12 | 0.31597173 | 0.35458662 | 0.32465622 | 0.0299304 | 0.33377508 | 0.89054584 |
| Nmrk1 | 0.18044202 | 0.22026156 | 0.19033769 | 0.02992386 | 0.23194589 | 0.89054584 |
| Pxn | 0.26998919 | 0.30994201 | 0.28004349 | 0.02989852 | 0.30945125 | 0.89054584 |
| Orai3 | 0.33318938 | 0.3709678 | 0.34108871 | 0.02987909 | 0.34159298 | 0.89054584 |
| Mysm1 | 0.51522003 | 0.53467002 | 0.50481038 | 0.02985964 | 0.39143636 | 0.89054584 |
| Cacna2d1 | 0.63444824 | 0.63088772 | 0.60104847 | 0.02983925 | 0.40854293 | 0.89054584 |
| Ubl5 | 2.73485256 | 1.14466341 | 1.1148256 | 0.02983782 | 0.4503827 | 0.89054584 |
| Boc | 0.13012035 | 0.16826227 | 0.13843931 | 0.02982296 | 0.15777576 | 0.80204521 |
| Fzd3 | 0.15638814 | 0.19547236 | 0.16565547 | 0.02981689 | 0.20085289 | 0.89054584 |
| Ttc30a1 | 0.11745582 | 0.15503752 | 0.12522347 | 0.02981404 | 0.13366506 | 0.731274 |
| Notch3 | 0.10793701 | 0.14505784 | 0.11525104 | 0.0298068 | 0.11411733 | 0.67417108 |
| Pold3 | 0.24758898 | 0.28770438 | 0.25794207 | 0.02976231 | 0.29543841 | 0.89054584 |
| Lama2 | 0.07966176 | 0.11496126 | 0.08519946 | 0.0297618 | 0.05182001 | 0.40240304 |
| Eif2s3x | 0.37071626 | 0.40611404 | 0.37636879 | 0.02974525 | 0.35635969 | 0.89054584 |
| Xndc1 | 0.16545446 | 0.20472532 | 0.17498636 | 0.02973896 | 0.21424224 | 0.89054584 |
| Syngr1 | 0.23359341 | 0.27373479 | 0.24401513 | 0.02971966 | 0.28521015 | 0.89054584 |
| Kcnb2 | 0.25751728 | 0.29748554 | 0.26776627 | 0.02971927 | 0.30254526 | 0.89054584 |
| Msmo1 | 0.51973781 | 0.5383357 | 0.50862369 | 0.02971202 | 0.39274987 | 0.89054584 |
| Acsf3 | 0.14161496 | 0.18007329 | 0.15038199 | 0.0296913 | 0.17882356 | 0.85310432 |
| Rassf2 | 0.08436331 | 0.11990562 | 0.0902205 | 0.02968513 | 0.06264716 | 0.45987776 |
| Eno1b | 0.10335718 | 0.14005521 | 0.11044108 | 0.02961413 | 0.10578353 | 0.64250089 |
| Jam3 | 0.61769206 | 0.61769706 | 0.5880901 | 0.02960695 | 0.40727867 | 0.89054584 |
| Gm26682 | 0.02906096 | 0.06068405 | 0.03109814 | 0.02958591 | 4.66E-06 | 0.00013017 |
| Abcb9 | 0.34040439 | 0.3774924 | 0.34792943 | 0.02956297 | 0.34610331 | 0.89054584 |
| Trpm4 | 0.11736896 | 0.15466913 | 0.12513262 | 0.02953651 | 0.13573054 | 0.73849629 |

| | | | | | | |
|---|---|---|---|---|---|---|
| Eif5a2 | 0.05740741 | 0.09094821 | 0.0614173 | 0.02953091 | 0.01254122 | 0.14062071 |
| Pde4dip | 0.25022709 | 0.29004011 | 0.26055705 | 0.02948307 | 0.29903509 | 0.89054584 |
| Krcc1 | 1.0954834 | 0.91246534 | 0.88298398 | 0.02948136 | 0.43819219 | 0.89054584 |
| Zfp637 | 0.53572678 | 0.55149328 | 0.52201874 | 0.02947453 | 0.39625374 | 0.89054584 |
| Tmem106a | 0.05452552 | 0.08778665 | 0.05833598 | 0.02945067 | 0.00933523 | 0.11084537 |
| Unc5b | 0.08527632 | 0.12063261 | 0.0911954 | 0.02943721 | 0.06630713 | 0.47741402 |
| Npat | 0.16691951 | 0.20590264 | 0.17649103 | 0.02941161 | 0.21875753 | 0.89054584 |
| St6gal1 | 0.16093979 | 0.19971512 | 0.1703441 | 0.02937101 | 0.2109 | 0.89054584 |
| Tbxas1 | 0.03657375 | 0.0684819 | 0.03913582 | 0.02934608 | 0.00023835 | 0.00460748 |
| Erbb2 | 0.13625328 | 0.17414836 | 0.14481766 | 0.0293307 | 0.17268695 | 0.83703076 |
| Lta4h | 0.28655016 | 0.32554985 | 0.29623006 | 0.02931978 | 0.3223544 | 0.89054584 |
| Pknox1 | 0.36398228 | 0.39941224 | 0.3700927 | 0.02931953 | 0.35602987 | 0.89054584 |
| Il18r1 | 0.03919795 | 0.07125327 | 0.04194307 | 0.02931019 | 0.00056555 | 0.01012258 |
| Mum1l1 | 0.16422961 | 0.20303234 | 0.17372771 | 0.02930463 | 0.21596988 | 0.89054584 |
| Zfp281 | 0.32352014 | 0.36117492 | 0.33187942 | 0.0292955 | 0.34044396 | 0.89054584 |
| Gm11627 | 0.10608365 | 0.14259269 | 0.11330547 | 0.02928723 | 0.11424759 | 0.67417108 |
| Gm10146 | 0.0664224 | 0.10030186 | 0.07105399 | 0.02924787 | 0.02757108 | 0.25315361 |
| H2-DMb2 | 0.01814986 | 0.0486431 | 0.01942304 | 0.02922007 | 1.20E-12 | 7.99E-11 |
| Gtpbp3 | 0.18337057 | 0.22254411 | 0.19332677 | 0.02921734 | 0.24068444 | 0.89054584 |
| Yipf4 | 1.41814495 | 1.04064188 | 1.01143437 | 0.02920751 | 0.4464887 | 0.89054584 |
| Tcf19 | 0.17651618 | 0.21551046 | 0.1863254 | 0.02918506 | 0.23277064 | 0.89054584 |
| Krt17 | 0.02631091 | 0.05731708 | 0.02815569 | 0.02916139 | 6.99E-07 | 2.24E-05 |
| Wnt7b | 0.08151978 | 0.11633227 | 0.08718389 | 0.02914837 | 0.05966007 | 0.44424843 |
| Slc7a14 | 0.4040796 | 0.4362307 | 0.40709472 | 0.02913598 | 0.36940002 | 0.89054584 |
| Rasgrf1 | 0.19713272 | 0.23645993 | 0.20732468 | 0.02913525 | 0.25632433 | 0.89054584 |
| Sep-02 | 1.05118529 | 0.88996914 | 0.86084396 | 0.02912518 | 0.43737524 | 0.89054584 |
| Hmha1 | 0.05076235 | 0.08342033 | 0.05431194 | 0.0291084 | 0.00626329 | 0.07889247 |
| Efs | 0.1096951 | 0.14620148 | 0.11709544 | 0.02910605 | 0.12342007 | 0.70253011 |
| Slc6a13 | 0.031518 | 0.06281839 | 0.03372697 | 0.02909142 | 2.93E-05 | 0.00070032 |
| Leng9 | 0.12319709 | 0.16026582 | 0.13122202 | 0.0290438 | 0.15122847 | 0.78275985 |

| | | | | | | |
|---|---|---|---|---|---|---|
| Clybl | 0.13171565 | 0.16912609 | 0.14009984 | 0.02902626 | 0.16720936 | 0.82737245 |
| Col11a1 | 0.0730444 | 0.10715195 | 0.07813032 | 0.02902163 | 0.04176417 | 0.34614674 |
| Bcl7b | 0.67913295 | 0.66368857 | 0.63466843 | 0.02902014 | 0.41565196 | 0.89054584 |
| Stmn3 | 0.11399908 | 0.15060769 | 0.12160588 | 0.02900181 | 0.13326014 | 0.73071314 |
| Aldh1a1 | 0.07686319 | 0.11120054 | 0.08221012 | 0.02899043 | 0.05019967 | 0.39431114 |
| Hpgds | 0.08125102 | 0.11588698 | 0.08689686 | 0.02899012 | 0.06005717 | 0.44651732 |
| P4ha1 | 0.51626967 | 0.53467765 | 0.50569751 | 0.02898015 | 0.39473796 | 0.89054584 |
| Tmem79 | 0.03024803 | 0.06129694 | 0.03236823 | 0.02892872 | 1.56E-05 | 0.00039515 |
| Ptdss1 | 0.33002716 | 0.3669776 | 0.33808231 | 0.0288953 | 0.34524453 | 0.89054584 |
| Kdm3a | 0.17340249 | 0.21202857 | 0.18313876 | 0.02888982 | 0.23118809 | 0.89054584 |
| Mrgprf | 0.09516761 | 0.13062827 | 0.1017538 | 0.02887447 | 0.09307434 | 0.59575795 |
| Cry2 | 0.33984061 | 0.3762669 | 0.34739584 | 0.02887106 | 0.34930738 | 0.89054584 |
| Ctxn2 | 0.12604844 | 0.16304484 | 0.13419652 | 0.02884832 | 0.15828392 | 0.80406459 |
| Ccdc126 | 0.2344204 | 0.27367105 | 0.24484056 | 0.02883049 | 0.29163999 | 0.89054584 |
| Nudt10 | 0.06528404 | 0.09866646 | 0.06983732 | 0.02882914 | 0.02722556 | 0.25100767 |
| Parp9 | 0.11088855 | 0.14717104 | 0.11834682 | 0.02882422 | 0.1282466 | 0.71677685 |
| Adamts4 | 0.08747383 | 0.12235892 | 0.09354166 | 0.02881726 | 0.07560448 | 0.52298621 |
| Abca6 | 0.02269414 | 0.05310217 | 0.02428571 | 0.02881647 | 1.62E-08 | 6.45E-07 |
| Mypop | 0.25300928 | 0.29204847 | 0.26331129 | 0.02873718 | 0.30556332 | 0.89054584 |
| Cyth4 | 0.10976478 | 0.14589991 | 0.1171685 | 0.0287314 | 0.1266357 | 0.71241644 |
| Golm1 | 0.19821998 | 0.23715816 | 0.2084271 | 0.02873106 | 0.26036475 | 0.89054584 |
| Cacna2d2 | 0.04074119 | 0.07232128 | 0.04359389 | 0.02872739 | 0.0010698 | 0.01754524 |
| Lace1 | 0.08630796 | 0.12101463 | 0.09229691 | 0.02871772 | 0.07358422 | 0.51488639 |
| Isca1 | 0.3253233 | 0.36231504 | 0.33360052 | 0.02871452 | 0.34420407 | 0.89054584 |
| Otulin | 0.29096507 | 0.32922772 | 0.30052316 | 0.02870456 | 0.32815982 | 0.89054584 |
| Ltbp1 | 0.06089652 | 0.09385016 | 0.06514744 | 0.02870272 | 0.02005371 | 0.20176171 |
| Btg3 | 0.19361018 | 0.23244605 | 0.20374949 | 0.02869656 | 0.25585532 | 0.89054584 |
| Zbtb8a | 0.17868409 | 0.21721581 | 0.18854172 | 0.02867409 | 0.23930241 | 0.89054584 |
| Slc24a2 | 0.05732259 | 0.08999538 | 0.06132662 | 0.02866876 | 0.01470504 | 0.15926281 |
| Zbtb46 | 0.05063205 | 0.08281132 | 0.0541726 | 0.02863872 | 0.00688936 | 0.08558858 |

| | | | | | | |
|---|---|---|---|---|---|---|
| Gart | 0.56296702 | 0.57307265 | 0.54446325 | 0.0286094 | 0.40330216 | 0.89054584 |
| Usp28 | 0.08631184 | 0.12089094 | 0.09230105 | 0.02858989 | 0.0744977 | 0.51900827 |
| Sephs1 | 0.34503787 | 0.3808974 | 0.35230846 | 0.02858894 | 0.35269256 | 0.89054584 |
| Rpp25 | 0.06997244 | 0.10342647 | 0.07484786 | 0.02857861 | 0.03762895 | 0.32204514 |
| Gabpa | 0.20179933 | 0.24058928 | 0.21205278 | 0.0285365 | 0.26534079 | 0.89054584 |
| Sdccag8 | 0.38949691 | 0.42227227 | 0.39373978 | 0.02853249 | 0.36782787 | 0.89054584 |
| Rad51 | 0.1407293 | 0.17798033 | 0.14946362 | 0.02851671 | 0.18702942 | 0.87419127 |
| Batf3 | 0.06216534 | 0.09500643 | 0.06650378 | 0.02850265 | 0.02292558 | 0.22217582 |
| Endog | 0.33486422 | 0.37109593 | 0.34267888 | 0.02841705 | 0.3496194 | 0.89054584 |
| Scly | 0.3679606 | 0.40221101 | 0.37380352 | 0.02840749 | 0.36164701 | 0.89054584 |
| Trmt12 | 0.20933111 | 0.2480562 | 0.21966449 | 0.02839171 | 0.2735306 | 0.89054584 |
| Pgm5 | 0.08466603 | 0.11891709 | 0.09054374 | 0.02837335 | 0.07215127 | 0.50853143 |
| Mcph1 | 0.11631375 | 0.1523917 | 0.12402875 | 0.02836295 | 0.14334551 | 0.76511581 |
| Tnnt1 | 0.04939735 | 0.0812151 | 0.0528522 | 0.0283629 | 0.00620615 | 0.07824067 |
| 4930522L14I | 0.05720624 | 0.08954795 | 0.06120222 | 0.02834573 | 0.01547237 | 0.16599537 |
| Stambp | 0.25070153 | 0.28936173 | 0.26102698 | 0.02833474 | 0.30652213 | 0.89054584 |
| Pmepa1 | 0.22311415 | 0.26182096 | 0.23352991 | 0.02829104 | 0.28623802 | 0.89054584 |
| Cdc6 | 0.06009871 | 0.09258071 | 0.06429456 | 0.02828615 | 0.02019766 | 0.20278789 |
| Trp53i13 | 0.67274452 | 0.65818164 | 0.62994658 | 0.02823506 | 0.41729437 | 0.89054584 |
| Irak2 | 0.05932748 | 0.09169467 | 0.06347007 | 0.02822461 | 0.01914241 | 0.19547721 |
| Flt3 | 0.02452225 | 0.0544472 | 0.02624183 | 0.02820537 | 2.76E-07 | 9.32E-06 |
| Epha7 | 0.17011338 | 0.20794126 | 0.17976823 | 0.02817303 | 0.23265142 | 0.89054584 |
| Cdc14a | 0.08714487 | 0.12135411 | 0.09319045 | 0.02816366 | 0.07956447 | 0.54059058 |
| RP23-474J16 | 0.02194657 | 0.05163779 | 0.02348578 | 0.028152 | 1.17E-08 | 4.73E-07 |
| Pde10a | 0.13243303 | 0.16899283 | 0.14084623 | 0.0281466 | 0.17591665 | 0.8453343 |
| Ubxn7 | 0.27434235 | 0.31244193 | 0.28431106 | 0.02813087 | 0.32240854 | 0.89054584 |
| Rmnd1 | 0.27587173 | 0.3139208 | 0.28580819 | 0.02811261 | 0.32338218 | 0.89054584 |
| Fbxo6 | 0.25191616 | 0.2903271 | 0.26222958 | 0.02809751 | 0.30881971 | 0.89054584 |
| Zfp775 | 0.1284834 | 0.16482722 | 0.13673442 | 0.0280928 | 0.1692322 | 0.8322688 |
| Rab3gap2 | 0.26460228 | 0.30283748 | 0.27474981 | 0.02808766 | 0.31693625 | 0.89054584 |

| | | | | | | |
|---|---|---|---|---|---|---|
| Wiz | 0.24110217 | 0.27958302 | 0.25149833 | 0.02808469 | 0.30144063 | 0.89054584 |
| Gsr | 0.68898323 | 0.66997783 | 0.64189457 | 0.02808326 | 0.41924264 | 0.89054584 |
| Abcd3 | 0.88937985 | 0.80193637 | 0.77385586 | 0.02808051 | 0.43287607 | 0.89054584 |
| Bbs12 | 0.08046035 | 0.1141305 | 0.08605241 | 0.0280781 | 0.06423426 | 0.46799603 |
| Errfi1 | 0.47637441 | 0.49956414 | 0.47149671 | 0.02806743 | 0.39075984 | 0.89054584 |
| Gab1 | 0.19660478 | 0.23482817 | 0.20678919 | 0.02803898 | 0.26378639 | 0.89054584 |
| Sart3 | 0.31694757 | 0.35362577 | 0.32559168 | 0.02803409 | 0.34415702 | 0.89054584 |
| Khdrbs1 | 1.39354238 | 1.03150048 | 1.00347541 | 0.02802507 | 0.44823774 | 0.89054584 |
| Adam23 | 0.06813686 | 0.10090584 | 0.07288628 | 0.02801956 | 0.03664427 | 0.31529292 |
| Mapk1ip1l | 0.41590317 | 0.44584905 | 0.41783373 | 0.02801532 | 0.37737695 | 0.89054584 |
| Fam210a | 0.24889672 | 0.28724861 | 0.25923875 | 0.02800986 | 0.30734762 | 0.89054584 |
| Tbp | 0.14189182 | 0.17865664 | 0.150669 | 0.02798763 | 0.19340275 | 0.88812525 |
| Foxo3 | 0.28006685 | 0.31789356 | 0.28990897 | 0.02798459 | 0.32645532 | 0.89054584 |
| Scn3a | 0.15038659 | 0.18744469 | 0.15946084 | 0.02798385 | 0.20679293 | 0.89054584 |
| Cipc | 0.13930228 | 0.17593779 | 0.14798327 | 0.02795452 | 0.18940554 | 0.88012871 |
| Rxrb | 0.18302037 | 0.22092344 | 0.19296952 | 0.02795392 | 0.24986874 | 0.89054584 |
| Lysmd4 | 0.09609981 | 0.13069139 | 0.10274854 | 0.02794285 | 0.10257473 | 0.6298723 |
| Spry4 | 0.07899772 | 0.11242091 | 0.08449018 | 0.02793072 | 0.06176183 | 0.45559551 |
| Nab2 | 0.25091741 | 0.2891515 | 0.26124077 | 0.02791073 | 0.30932761 | 0.89054584 |
| Iqgap2 | 0.28347018 | 0.32113976 | 0.29322942 | 0.02791034 | 0.32872007 | 0.89054584 |
| Rabggta | 0.2236727 | 0.26197275 | 0.23409001 | 0.02788274 | 0.28947059 | 0.89054584 |
| Ank2 | 0.33120701 | 0.36708372 | 0.33920463 | 0.02787909 | 0.35089087 | 0.89054584 |
| Olfml1 | 0.09392656 | 0.12827032 | 0.10042941 | 0.02784091 | 0.09825727 | 0.61534939 |
| Phyhipl | 0.32091938 | 0.35722024 | 0.32939405 | 0.02782619 | 0.34694816 | 0.89054584 |
| Ces1d | 0.0644822 | 0.09678309 | 0.06898029 | 0.0278028 | 0.03020384 | 0.27028518 |
| Meis3 | 0.35876499 | 0.39300056 | 0.36521406 | 0.0277865 | 0.36149515 | 0.89054584 |
| Klhl13 | 0.26967862 | 0.3075239 | 0.27973867 | 0.02778523 | 0.32177092 | 0.89054584 |
| Pcdhb5 | 0.02379942 | 0.05319342 | 0.02546839 | 0.02772503 | 1.97E-07 | 6.80E-06 |
| Efemp1 | 0.05183494 | 0.08317243 | 0.05545893 | 0.02771349 | 0.00995263 | 0.11712117 |
| Sntb2 | 0.41200883 | 0.44199045 | 0.41430537 | 0.02768508 | 0.3777772 | 0.89054584 |

| | | | | | | |
|---|---|---|---|---|---|---|
| Csnk2a2 | 0.0577133 | 0.08942162 | 0.06174434 | 0.02767728 | 0.01838094 | 0.1898055 |
| Hscb | 0.1632165 | 0.20035137 | 0.17268619 | 0.02766517 | 0.22771738 | 0.89054584 |
| Snx4 | 0.93841507 | 0.82934508 | 0.80173461 | 0.02761048 | 0.43626464 | 0.89054584 |
| Mab21l2 | 0.03955358 | 0.06990913 | 0.0423235 | 0.02758563 | 0.00119628 | 0.01933526 |
| Klhl10 | 0.02084624 | 0.04987631 | 0.02230837 | 0.02756795 | 4.27E-09 | 1.84E-07 |
| Adck1 | 0.09611606 | 0.13032828 | 0.10276587 | 0.02756241 | 0.10573103 | 0.64250089 |
| Anln | 0.06400528 | 0.09602934 | 0.06847052 | 0.02755882 | 0.0303836 | 0.27141185 |
| Msh3 | 0.3186267 | 0.35471617 | 0.32720017 | 0.027516 | 0.34760479 | 0.89054584 |
| Cln8 | 0.26335881 | 0.30100039 | 0.27352588 | 0.0274745 | 0.31990279 | 0.89054584 |
| Fst | 0.01576909 | 0.04434206 | 0.01687538 | 0.02746668 | 1.69E-14 | 1.32E-12 |
| Ddx10 | 0.43299636 | 0.46068352 | 0.43321977 | 0.02746376 | 0.38386276 | 0.89054584 |
| Pla2r1 | 0.05499557 | 0.08622918 | 0.05883858 | 0.0273906 | 0.01504759 | 0.1619599 |
| Cyp2j6 | 0.13189496 | 0.16763096 | 0.14028641 | 0.02734455 | 0.18190789 | 0.86130002 |
| Smim20 | 0.53585532 | 0.54947001 | 0.52212577 | 0.02734424 | 0.40361798 | 0.89054584 |
| Ccdc124 | 0.81263681 | 0.75408845 | 0.7267491 | 0.02733935 | 0.4304364 | 0.89054584 |
| Rnaset2a | 0.83902817 | 0.77077054 | 0.74343249 | 0.02733806 | 0.43198526 | 0.89054584 |
| Lingo4 | 0.05297582 | 0.08401126 | 0.05667892 | 0.02733234 | 0.01233004 | 0.13867887 |
| Fam222b | 0.34471595 | 0.3793151 | 0.35200459 | 0.02731051 | 0.35887516 | 0.89054584 |
| Deaf1 | 0.47878273 | 0.5008944 | 0.47358901 | 0.02730539 | 0.3941121 | 0.89054584 |
| Stk39 | 0.17941816 | 0.21654323 | 0.18929183 | 0.02725139 | 0.25119808 | 0.89054584 |
| Ebi3 | 0.0502298 | 0.08098053 | 0.05374244 | 0.02723809 | 0.00910611 | 0.10874825 |
| Fbrs | 0.13189389 | 0.16752123 | 0.14028531 | 0.02723593 | 0.1828604 | 0.86439845 |
| Cdk6 | 0.15243041 | 0.18879062 | 0.16157187 | 0.02721875 | 0.2162692 | 0.89054584 |
| Ier3ip1 | 1.64342789 | 1.0975254 | 1.07031077 | 0.02721462 | 0.45285074 | 0.89054584 |
| B930041F14 | 0.06685319 | 0.09871567 | 0.0715144 | 0.02720127 | 0.0381908 | 0.32627589 |
| Snd1 | 1.16974461 | 0.94447848 | 0.91729306 | 0.02718543 | 0.44509055 | 0.89054584 |
| Capsl | 0.46757393 | 0.49098443 | 0.46382089 | 0.02716354 | 0.3924844 | 0.89054584 |
| Plek2 | 0.0534472 | 0.08434532 | 0.05718296 | 0.02716235 | 0.01344767 | 0.14860084 |
| Trip13 | 0.07267826 | 0.1048971 | 0.07773911 | 0.02715799 | 0.05180585 | 0.40240304 |
| Bdh1 | 0.06751976 | 0.09937216 | 0.07222679 | 0.02714537 | 0.03997345 | 0.33713503 |

| | | | | | | |
|---|---|---|---|---|---|---|
| Plcxd3 | 0.3810667 | 0.41309294 | 0.38596656 | 0.02712638 | 0.37166823 | 0.89054584 |
| St6galnac6 | 0.21078256 | 0.24823077 | 0.22112859 | 0.02710219 | 0.28399512 | 0.89054584 |
| Ankrd50 | 0.15648091 | 0.19281122 | 0.16575111 | 0.02706011 | 0.22344586 | 0.89054584 |
| Lrrfip1 | 0.56267871 | 0.5712764 | 0.5442282 | 0.0270482 | 0.40844383 | 0.89054584 |
| Clec10a | 0.01629773 | 0.04441331 | 0.01744108 | 0.02697223 | 2.90E-13 | 2.04E-11 |
| Mtss1 | 0.46915047 | 0.49212817 | 0.46519933 | 0.02692884 | 0.39369915 | 0.89054584 |
| Gpr153 | 0.08601077 | 0.11889588 | 0.0919796 | 0.02691628 | 0.08638441 | 0.56824007 |
| Fmo1 | 0.13698173 | 0.17248426 | 0.14557429 | 0.02690996 | 0.1945576 | 0.88952889 |
| Dnph1 | 0.09543596 | 0.12894072 | 0.10204016 | 0.02690056 | 0.10968154 | 0.65546703 |
| Mepce | 0.14366722 | 0.17940776 | 0.15250884 | 0.02689893 | 0.20561633 | 0.89054584 |
| Fam3a | 0.45340175 | 0.47821739 | 0.45136443 | 0.02685296 | 0.39082334 | 0.89054584 |
| Myl1 | 0.01544814 | 0.04338194 | 0.01653192 | 0.02685002 | 1.91E-14 | 1.49E-12 |
| Tiparp | 0.27556427 | 0.31232547 | 0.28550731 | 0.02681816 | 0.33083022 | 0.89054584 |
| Pkib | 0.18342422 | 0.22015469 | 0.19338149 | 0.0267732 | 0.25945737 | 0.89054584 |
| Bmp2 | 0.03370621 | 0.06280836 | 0.03606807 | 0.02674028 | 0.0002761 | 0.00529491 |
| H1fx | 0.43086646 | 0.45802022 | 0.43131193 | 0.02670829 | 0.38648302 | 0.89054584 |
| Cd83 | 0.02856953 | 0.05727604 | 0.03057234 | 0.02670369 | 2.36E-05 | 0.00057815 |
| Socs3 | 0.18570986 | 0.22239619 | 0.19571181 | 0.02668439 | 0.26264091 | 0.89054584 |
| Pygb | 0.23550048 | 0.27260197 | 0.24591816 | 0.02668381 | 0.30659434 | 0.89054584 |
| Galr1 | 0.05404656 | 0.08450141 | 0.05782385 | 0.02667756 | 0.01579957 | 0.16892491 |
| Eci1 | 0.96510407 | 0.84286624 | 0.81621877 | 0.02664747 | 0.439553 | 0.89054584 |
| Zfp952 | 0.15105728 | 0.18678188 | 0.16015377 | 0.02662811 | 0.2192812 | 0.89054584 |
| Barx1 | 0.09360335 | 0.12670787 | 0.10008447 | 0.0266234 | 0.1076132 | 0.64845225 |
| Cdt1 | 0.16018384 | 0.19618735 | 0.16956598 | 0.02662136 | 0.23225517 | 0.89054584 |
| Arsj | 0.08341143 | 0.11581311 | 0.08920404 | 0.02660907 | 0.08230451 | 0.5519946 |
| Tff3 | 0.04443134 | 0.07414487 | 0.047541 | 0.02660387 | 0.00456515 | 0.0610005 |
| Cdca2 | 0.08701821 | 0.11965856 | 0.09305522 | 0.02660334 | 0.09144098 | 0.5881594 |
| Itgav | 0.25538269 | 0.2922589 | 0.26565798 | 0.02660091 | 0.32042526 | 0.89054584 |
| Idh3g | 1.24072302 | 0.97399039 | 0.94739696 | 0.02659342 | 0.44797609 | 0.89054584 |
| Gm15829 | 0.01817134 | 0.04603805 | 0.01944602 | 0.02659203 | 9.39E-11 | 5.00E-09 |

| | | | | | | |
|---|---|---|---|---|---|---|
| Aspm | 0.05772792 | 0.08834536 | 0.06175998 | 0.02658539 | 0.02245469 | 0.2193554 |
| Rprd1a | 0.32218583 | 0.35718956 | 0.33060475 | 0.02658481 | 0.3539649 | 0.89054584 |
| Ppp1r17 | 0.0470245 | 0.07688955 | 0.05031451 | 0.02657505 | 0.00693228 | 0.08604817 |
| Senp5 | 0.24685722 | 0.28378986 | 0.25721615 | 0.02657371 | 0.31514169 | 0.89054584 |
| Hk1 | 0.13822616 | 0.17342703 | 0.14686641 | 0.02656063 | 0.19973608 | 0.89054584 |
| Psd | 0.10968986 | 0.14364735 | 0.11708994 | 0.02655742 | 0.14532329 | 0.76804699 |
| Cdc7 | 0.10621783 | 0.1399928 | 0.11344637 | 0.02654643 | 0.13781442 | 0.7438695 |
| Glrx | 0.66109723 | 0.64780052 | 0.62126426 | 0.02653626 | 0.42113181 | 0.89054584 |
| Cckar | 0.0354789 | 0.06448231 | 0.03796456 | 0.02651775 | 0.00056854 | 0.01015112 |
| Tdrkh | 0.3416433 | 0.37561152 | 0.34910143 | 0.02651008 | 0.36174778 | 0.89054584 |
| Nifk | 0.65654035 | 0.64433709 | 0.61784163 | 0.02649545 | 0.42082103 | 0.89054584 |
| Bicd1 | 0.3880912 | 0.41893266 | 0.39244621 | 0.02648645 | 0.37659536 | 0.89054584 |
| Xlr5a | 0.05598479 | 0.08638155 | 0.05989627 | 0.02648527 | 0.01969367 | 0.19896783 |
| Rbm34 | 0.3590943 | 0.39200387 | 0.36552241 | 0.02648146 | 0.36785789 | 0.89054584 |
| Gm9780 | 0.0852947 | 0.1176958 | 0.09121502 | 0.02648077 | 0.08810061 | 0.57428752 |
| Fbp2 | 0.01989246 | 0.04772972 | 0.02128776 | 0.02644196 | 3.58E-09 | 1.56E-07 |
| Gprc5c | 0.17882939 | 0.21512127 | 0.18869021 | 0.02643106 | 0.25700446 | 0.89054584 |
| Sh2d4a | 0.07510025 | 0.10674854 | 0.08032677 | 0.02642177 | 0.06270465 | 0.46006681 |
| Clec1a | 0.04023997 | 0.06946869 | 0.04305774 | 0.02641095 | 0.00213347 | 0.03182207 |
| Hoxd3os1 | 0.03365513 | 0.06240191 | 0.03601342 | 0.02638849 | 0.00032032 | 0.0060391 |
| Arhgef2 | 0.60085931 | 0.60125271 | 0.57488155 | 0.02637116 | 0.41538407 | 0.89054584 |
| A930004D18 | 0.08406937 | 0.11625713 | 0.08990662 | 0.02635051 | 0.08605147 | 0.5673371 |
| Dpy19l4 | 0.14167719 | 0.1767871 | 0.1504465 | 0.0263406 | 0.20733567 | 0.89054584 |
| Hoxa2 | 0.06491779 | 0.09576291 | 0.06944586 | 0.02631704 | 0.0387364 | 0.32938437 |
| Igsf3 | 0.29231088 | 0.32811384 | 0.30182992 | 0.02628393 | 0.34247803 | 0.89054584 |
| Aldh6a1 | 0.23286608 | 0.26953602 | 0.24328889 | 0.02624713 | 0.30761288 | 0.89054584 |
| Zfp110 | 0.15538405 | 0.19084998 | 0.16462003 | 0.02622995 | 0.22894112 | 0.89054584 |
| Bysl | 0.29018966 | 0.32598584 | 0.29976985 | 0.02621599 | 0.34184176 | 0.89054584 |
| Eva1a | 0.07817791 | 0.10982858 | 0.08361451 | 0.02621408 | 0.07205852 | 0.50853143 |
| Ccdc82 | 0.08343906 | 0.1154404 | 0.08923354 | 0.02620686 | 0.08561263 | 0.5659881 |

| | | | | | | |
|---|---|---|---|---|---|---|
| Fam221a | 0.33735366 | 0.37121811 | 0.34504001 | 0.0261781 | 0.36186706 | 0.89054584 |
| Heatr6 | 0.30168452 | 0.3370677 | 0.31090634 | 0.02616136 | 0.34751846 | 0.89054584 |
| Wrap53 | 0.19540206 | 0.23172555 | 0.20556882 | 0.02615673 | 0.27665511 | 0.89054584 |
| Adora2b | 0.08562662 | 0.11771965 | 0.09156943 | 0.02615022 | 0.09167822 | 0.58922032 |
| Tnxb | 0.07576775 | 0.10718542 | 0.08103989 | 0.02614554 | 0.06640748 | 0.47760826 |
| Mast2 | 0.15737668 | 0.19281595 | 0.16667446 | 0.02614149 | 0.23247641 | 0.89054584 |
| Gripap1 | 0.47618078 | 0.49745805 | 0.47132834 | 0.02612971 | 0.39809369 | 0.89054584 |
| Phldb2 | 0.48853046 | 0.50813059 | 0.48202106 | 0.02610953 | 0.40038147 | 0.89054584 |
| Ppfibp1 | 0.34604365 | 0.37935133 | 0.35325749 | 0.02609384 | 0.36537091 | 0.89054584 |
| Pcdh9 | 0.08478294 | 0.1167545 | 0.09066858 | 0.02608591 | 0.09005665 | 0.5818366 |
| Stard3 | 0.34864217 | 0.38178919 | 0.35570689 | 0.0260823 | 0.36631915 | 0.89054584 |
| Ptbp2 | 0.29002249 | 0.32568584 | 0.2996074 | 0.02607844 | 0.3425463 | 0.89054584 |
| Pomt2 | 0.15739578 | 0.19277159 | 0.16669415 | 0.02607744 | 0.23304994 | 0.89054584 |
| G6pc3 | 0.45385889 | 0.47783849 | 0.45176812 | 0.02607037 | 0.39402046 | 0.89054584 |
| Srsf3 | 2.61463133 | 1.14780745 | 1.12174926 | 0.02605819 | 0.45690781 | 0.89054584 |
| Chst2 | 0.06012427 | 0.09035314 | 0.06432189 | 0.02603125 | 0.02968208 | 0.26745229 |
| Megf6 | 0.08501139 | 0.1169372 | 0.09091251 | 0.02602469 | 0.09115564 | 0.58666938 |
| Lamb2 | 0.09750558 | 0.13025907 | 0.1042485 | 0.02601057 | 0.12253013 | 0.70076598 |
| Unc45b | 0.03497045 | 0.06342935 | 0.0374206 | 0.02600874 | 0.00060156 | 0.01070104 |
| Fras1 | 0.13624809 | 0.17078684 | 0.14481226 | 0.02597457 | 0.20167271 | 0.89054584 |
| Psat1 | 0.22369663 | 0.26007311 | 0.234114 | 0.02595911 | 0.3027197 | 0.89054584 |
| Calcrl | 0.07662343 | 0.10787777 | 0.08195398 | 0.02592378 | 0.07027825 | 0.49945789 |
| Map4 | 0.31573494 | 0.35032633 | 0.32442915 | 0.02589718 | 0.35498566 | 0.89054584 |
| Zbtb45 | 0.22436784 | 0.26068124 | 0.23478688 | 0.02589435 | 0.30368585 | 0.89054584 |
| Fars2 | 0.40785769 | 0.4364005 | 0.41053495 | 0.02586556 | 0.38455688 | 0.89054584 |
| Znrf3 | 0.10784063 | 0.14100989 | 0.11514989 | 0.02586 | 0.14771351 | 0.77404405 |
| Cib1 | 0.87119156 | 0.78893592 | 0.76307932 | 0.0258566 | 0.43728113 | 0.89054584 |
| Elmo1 | 0.09513167 | 0.12756258 | 0.10171545 | 0.02584712 | 0.11822769 | 0.68802988 |
| Ttc17 | 0.06980028 | 0.10050163 | 0.07466389 | 0.02583775 | 0.05345422 | 0.41156627 |
| Zfp800 | 0.33779031 | 0.37129012 | 0.34545387 | 0.02583625 | 0.36375529 | 0.89054584 |

| | | | | | | |
|---|---|---|---|---|---|---|
| Cacna1b | 0.07727759 | 0.10846246 | 0.08265279 | 0.02580967 | 0.07285819 | 0.5122687 |
| Tmem60 | 0.57228974 | 0.57784195 | 0.55203394 | 0.02580801 | 0.41378893 | 0.89054584 |
| Snx7 | 0.24724638 | 0.28340126 | 0.25760224 | 0.02579902 | 0.32039457 | 0.89054584 |
| Xylt2 | 0.16339248 | 0.19866512 | 0.17286714 | 0.02579799 | 0.24343949 | 0.89054584 |
| Mea1 | 0.87524269 | 0.79125993 | 0.76550071 | 0.02575922 | 0.43771144 | 0.89054584 |
| Rpl23a | 5.14675294 | 0.70439479 | 0.67865245 | 0.02574235 | 0.42986381 | 0.89054584 |
| Relt | 0.11120101 | 0.14441056 | 0.11867436 | 0.0257362 | 0.15616225 | 0.79775908 |
| Exosc8 | 0.39876018 | 0.42796899 | 0.40223717 | 0.02573182 | 0.38283607 | 0.89054584 |
| Upf1 | 0.22384355 | 0.25997131 | 0.23426131 | 0.02571 | 0.30456517 | 0.89054584 |
| Nrgn | 0.03769859 | 0.0660446 | 0.04033915 | 0.02570545 | 0.00149467 | 0.02347748 |
| Dtd1 | 0.51630829 | 0.53141975 | 0.50573014 | 0.02568962 | 0.40646003 | 0.89054584 |
| Nap1l5 | 0.10712051 | 0.14007707 | 0.11439407 | 0.025683 | 0.14778174 | 0.77404405 |
| Izumo4 | 0.12378446 | 0.15750466 | 0.13183501 | 0.02566965 | 0.18216522 | 0.86167416 |
| Cachd1 | 0.06481687 | 0.09500437 | 0.06933799 | 0.02566638 | 0.04230255 | 0.34946006 |
| Tapbpl | 0.27191633 | 0.30759558 | 0.28193388 | 0.0256617 | 0.33576171 | 0.89054584 |
| Gcsh | 1.00830625 | 0.86447173 | 0.83881336 | 0.02565837 | 0.44333786 | 0.89054584 |
| Cacna1g | 0.05224693 | 0.08153203 | 0.05589949 | 0.02563253 | 0.01632576 | 0.17339991 |
| Podnl1 | 0.02597523 | 0.05342649 | 0.02779652 | 0.02562997 | 8.70E-06 | 0.00023237 |
| Nfkb2 | 0.09761437 | 0.1299914 | 0.10436456 | 0.02562683 | 0.12631016 | 0.71147001 |
| Atp5g2 | 2.95152475 | 1.12465215 | 1.09904688 | 0.02560527 | 0.45678263 | 0.89054584 |
| Cyp4f16 | 0.15452001 | 0.18933279 | 0.1637287 | 0.02560408 | 0.23313218 | 0.89054584 |
| Rpap3 | 0.35610003 | 0.38828472 | 0.36271656 | 0.02556817 | 0.37129997 | 0.89054584 |
| Hdac6 | 0.40239509 | 0.43112047 | 0.40555823 | 0.02556224 | 0.38451193 | 0.89054584 |
| Hs6st2 | 0.11612432 | 0.1493902 | 0.12383055 | 0.02555965 | 0.16811418 | 0.82993631 |
| Haus8 | 0.16240195 | 0.19740215 | 0.17184849 | 0.02555366 | 0.24422419 | 0.89054584 |
| Farp2 | 0.04989483 | 0.07889527 | 0.05338421 | 0.02551105 | 0.01299377 | 0.14467612 |
| Hoxb8 | 0.03212054 | 0.0598821 | 0.03437162 | 0.02551048 | 0.00027226 | 0.00524017 |
| Cript | 1.01998478 | 0.87028754 | 0.84478223 | 0.02550531 | 0.44406895 | 0.89054584 |
| Dusp14 | 0.04985154 | 0.0788229 | 0.05333792 | 0.02548498 | 0.01300534 | 0.14467612 |
| Wdr33 | 0.40782019 | 0.4359848 | 0.41050083 | 0.02548396 | 0.3862035 | 0.89054584 |

| | | | | | | |
|---|---|---|---|---|---|---|
| Klf9 | 0.16646407 | 0.20149326 | 0.17602338 | 0.02546989 | 0.25011422 | 0.89054584 |
| Gm26532 | 0.02114071 | 0.04809315 | 0.02262346 | 0.02546968 | 7.81E-08 | 2.85E-06 |
| Upk1b | 0.02374625 | 0.05086645 | 0.0254115 | 0.02545496 | 1.53E-06 | 4.63E-05 |
| Unc5a | 0.08035643 | 0.1113937 | 0.08594142 | 0.02545229 | 0.08382637 | 0.55825208 |
| Hcn2 | 0.0225316 | 0.04955225 | 0.02411178 | 0.02544047 | 4.42E-07 | 1.47E-05 |
| Piezo1 | 0.12211951 | 0.15550657 | 0.1300971 | 0.02540947 | 0.18142572 | 0.86081561 |
| Stap2 | 0.10370973 | 0.13621577 | 0.11081161 | 0.02540415 | 0.14274062 | 0.76335429 |
| Tbc1d19 | 0.32532847 | 0.3589822 | 0.33360546 | 0.02537674 | 0.36152071 | 0.89054584 |
| Il17rc | 0.15819267 | 0.19288353 | 0.1675153 | 0.02536823 | 0.24023651 | 0.89054584 |
| B3galt4 | 0.10902675 | 0.14175794 | 0.11639441 | 0.02536353 | 0.15499789 | 0.79432997 |
| Nudt8 | 0.2793122 | 0.3144912 | 0.28917193 | 0.02531928 | 0.34166364 | 0.89054584 |
| Hoxb4 | 0.13956854 | 0.17357573 | 0.14825954 | 0.02531619 | 0.21314925 | 0.89054584 |
| Osgin1 | 0.07957214 | 0.11040688 | 0.08510373 | 0.02530315 | 0.0829953 | 0.55424501 |
| Erlin2 | 0.29016437 | 0.32504185 | 0.29974528 | 0.02529657 | 0.34709155 | 0.89054584 |
| Zfp84 | 0.14948915 | 0.18382984 | 0.15853336 | 0.02529648 | 0.22861765 | 0.89054584 |
| Dpep1 | 0.06091351 | 0.09045866 | 0.0651656 | 0.02529306 | 0.03528002 | 0.30664848 |
| Thoc2 | 1.0558108 | 0.88847852 | 0.86321515 | 0.02526338 | 0.44577175 | 0.89054584 |
| Mapk15 | 0.47889736 | 0.49893801 | 0.47368852 | 0.02524949 | 0.40193559 | 0.89054584 |
| Dnm3 | 0.09141681 | 0.12299519 | 0.09775078 | 0.02524441 | 0.11445244 | 0.67479929 |
| Hoxd4 | 0.05648426 | 0.08562402 | 0.06043031 | 0.02519371 | 0.02604817 | 0.24285937 |
| Micu1 | 0.42582117 | 0.45195334 | 0.42678201 | 0.02517133 | 0.3917421 | 0.89054584 |
| Kctd7 | 0.05372435 | 0.08262448 | 0.05747932 | 0.02514516 | 0.02076888 | 0.20637918 |
| Tmcc3 | 0.22750879 | 0.26307781 | 0.23793293 | 0.02514487 | 0.31123269 | 0.89054584 |
| Mphosph6 | 0.53427109 | 0.54592765 | 0.52080599 | 0.02512166 | 0.41109495 | 0.89054584 |
| 3010026O09 | 0.14883352 | 0.18295736 | 0.15785559 | 0.02510177 | 0.22939153 | 0.89054584 |
| 5730455P16I | 0.32653452 | 0.35985589 | 0.33475567 | 0.02510022 | 0.36342052 | 0.89054584 |
| Dcaf7 | 0.50894937 | 0.52458006 | 0.49949603 | 0.02508403 | 0.40750574 | 0.89054584 |
| Cers5 | 0.56677445 | 0.5726024 | 0.54756208 | 0.02504032 | 0.41564166 | 0.89054584 |
| Sned1 | 0.07082696 | 0.10080085 | 0.07576098 | 0.02503987 | 0.06180053 | 0.45559551 |
| Tmem55a | 0.56772267 | 0.57337037 | 0.54833234 | 0.02503803 | 0.41576596 | 0.89054584 |

| | | | | | | |
|---|---|---|---|---|---|---|
| 2510046G10 | 0.15909236 | 0.19343422 | 0.16844208 | 0.02499214 | 0.24470172 | 0.89054584 |
| Lepr | 0.04325515 | 0.07123784 | 0.04628295 | 0.02495489 | 0.00600223 | 0.07600027 |
| Pcyt1b | 0.21247217 | 0.24777932 | 0.22283174 | 0.02494758 | 0.300974 | 0.89054584 |
| Reck | 0.1605527 | 0.19487269 | 0.16994568 | 0.024927 | 0.24719087 | 0.89054584 |
| Mier1 | 0.74085133 | 0.70373834 | 0.67881768 | 0.02492066 | 0.43209678 | 0.89054584 |
| Slc45a3 | 0.01802188 | 0.0441891 | 0.01928609 | 0.02490301 | 8.95E-10 | 4.23E-08 |
| Tceal8 | 1.60048001 | 1.0857876 | 1.06089052 | 0.02489708 | 0.4564678 | 0.89054584 |
| Rela | 0.28362501 | 0.3182668 | 0.29338034 | 0.02488645 | 0.34634689 | 0.89054584 |
| Zswim4 | 0.04146482 | 0.06924346 | 0.04436794 | 0.02487553 | 0.00449936 | 0.06028811 |
| Adamts8 | 0.07351025 | 0.10349528 | 0.07862805 | 0.02486723 | 0.0703141 | 0.49946779 |
| Homez | 0.19435218 | 0.22934748 | 0.20450302 | 0.02484445 | 0.28569666 | 0.89054584 |
| Mpp3 | 0.22906387 | 0.26432648 | 0.23948888 | 0.0248376 | 0.31448297 | 0.89054584 |
| Gm10273 | 0.02825594 | 0.055064 | 0.03023681 | 0.02482719 | 6.53E-05 | 0.00143389 |
| Mrps33 | 2.30065361 | 1.15391666 | 1.12909173 | 0.02482493 | 0.45920593 | 0.89054584 |
| Apold1 | 0.06115894 | 0.09024829 | 0.06542797 | 0.02482032 | 0.03858185 | 0.32845551 |
| Gm15234 | 0.04874286 | 0.07693009 | 0.05215226 | 0.02477783 | 0.01343198 | 0.14854065 |
| Ppp1r21 | 0.14164023 | 0.17517335 | 0.15040818 | 0.02476518 | 0.22150828 | 0.89054584 |
| Ap2m1 | 2.11885683 | 1.14904391 | 1.12429372 | 0.02475019 | 0.45915535 | 0.89054584 |
| Arnt2 | 0.04000544 | 0.0675532 | 0.04280686 | 0.02474634 | 0.00353742 | 0.04931294 |
| Bnc2 | 0.08525921 | 0.11592277 | 0.09117712 | 0.02474564 | 0.10303513 | 0.63136232 |
| Cd4 | 0.03517237 | 0.0623819 | 0.03763662 | 0.02474528 | 0.00109508 | 0.01788478 |
| Klk8 | 0.09911277 | 0.13069441 | 0.10596319 | 0.02473122 | 0.13843642 | 0.74639314 |
| Vav3 | 0.03592902 | 0.06315318 | 0.03844609 | 0.02470709 | 0.00137658 | 0.02176563 |
| Edn3 | 0.07562901 | 0.10559827 | 0.08089166 | 0.02470661 | 0.07737203 | 0.53062428 |
| Spdl1 | 0.07493636 | 0.10484868 | 0.08015169 | 0.02469699 | 0.0755651 | 0.52298621 |
| Plk4 | 0.12928712 | 0.16225558 | 0.13757162 | 0.02468396 | 0.20159535 | 0.89054584 |
| Adgrd1 | 0.06695696 | 0.09630491 | 0.0716253 | 0.02467961 | 0.05421276 | 0.4156407 |
| Hdac7 | 0.1691337 | 0.20339476 | 0.17876344 | 0.02463132 | 0.26045521 | 0.89054584 |
| Tnni1 | 0.02073319 | 0.04680129 | 0.02218739 | 0.0246139 | 1.18E-07 | 4.17E-06 |
| Supv3l1 | 0.11389108 | 0.14610233 | 0.12149279 | 0.02460955 | 0.17265793 | 0.83703076 |

| | | | | | | |
|---|---|---|---|---|---|---|
| Tmem150a | 0.15362414 | 0.18740575 | 0.16280422 | 0.02460153 | 0.24071079 | 0.89054584 |
| Sharpin | 0.35072276 | 0.38225867 | 0.35766546 | 0.02459321 | 0.37435129 | 0.89054584 |
| Mrps11 | 0.63387573 | 0.62512992 | 0.60060882 | 0.02452111 | 0.4245719 | 0.89054584 |
| Gpbp1l1 | 0.53404577 | 0.54509141 | 0.52061816 | 0.02447325 | 0.41332197 | 0.89054584 |
| Zrsr1 | 0.08356735 | 0.11383482 | 0.08937054 | 0.02446428 | 0.1010948 | 0.62422163 |
| Robo2 | 0.15674681 | 0.19047683 | 0.16602523 | 0.0244516 | 0.2463086 | 0.89054584 |
| Ajuba | 0.11921954 | 0.15151114 | 0.12706755 | 0.02444359 | 0.18506608 | 0.86930035 |
| Gdap1l1 | 0.50976554 | 0.52462561 | 0.50018914 | 0.02443648 | 0.40997468 | 0.89054584 |
| Begain | 0.07839463 | 0.10828021 | 0.083846 | 0.02443421 | 0.0872798 | 0.57076343 |
| Thbs1 | 0.10253965 | 0.13397298 | 0.10958167 | 0.02439131 | 0.14986471 | 0.77903859 |
| Trib1 | 0.2089372 | 0.2436571 | 0.21926699 | 0.02439011 | 0.30214677 | 0.89054584 |
| Git1 | 0.17136223 | 0.20543201 | 0.18104852 | 0.02438349 | 0.26517793 | 0.89054584 |
| Ccdc8 | 0.84553833 | 0.77184352 | 0.74747027 | 0.02437325 | 0.43962614 | 0.89054584 |
| Gm2694 | 0.04497651 | 0.07248388 | 0.0481241 | 0.02435978 | 0.00917958 | 0.10953548 |
| Ina | 0.02484695 | 0.05094697 | 0.02658926 | 0.02435771 | 9.86E-06 | 0.00026098 |
| Cep170b | 0.21709543 | 0.25182514 | 0.22748562 | 0.02433952 | 0.30907389 | 0.89054584 |
| Lcmt2 | 0.22442579 | 0.25917951 | 0.23484496 | 0.02433455 | 0.31463387 | 0.89054584 |
| Gm26909 | 0.03508714 | 0.06187319 | 0.03754544 | 0.02432775 | 0.00126896 | 0.02030612 |
| Ttc33 | 0.41396356 | 0.44038852 | 0.41607738 | 0.02431114 | 0.39272683 | 0.89054584 |
| Adgrl2 | 0.17651853 | 0.21062367 | 0.1863278 | 0.02429587 | 0.27176156 | 0.89054584 |
| Cct5 | 2.46748624 | 1.15212899 | 1.1278731 | 0.02425588 | 0.46009494 | 0.89054584 |
| Gm5611 | 0.16094722 | 0.19460067 | 0.17035174 | 0.02424892 | 0.25360683 | 0.89054584 |
| Ncmap | 0.03193153 | 0.05840547 | 0.0341694 | 0.02423607 | 0.00047565 | 0.00860918 |
| Prpf6 | 0.48233969 | 0.50090853 | 0.4766727 | 0.02423583 | 0.4063757 | 0.89054584 |
| Dpm3 | 1.45655899 | 1.04744295 | 1.02323028 | 0.02421267 | 0.45610777 | 0.89054584 |
| Arntl | 0.09070619 | 0.12119412 | 0.09699227 | 0.02420185 | 0.12251313 | 0.70076598 |
| Sat2 | 0.1989105 | 0.23329167 | 0.20912699 | 0.02416467 | 0.29517058 | 0.89054584 |
| Lin37 | 0.36756744 | 0.39759669 | 0.37343718 | 0.02415951 | 0.38155089 | 0.89054584 |
| Lzic | 0.25212087 | 0.28657069 | 0.26243221 | 0.02413849 | 0.33413305 | 0.89054584 |
| Fry | 0.11047524 | 0.14204075 | 0.1179135 | 0.02412725 | 0.17021802 | 0.83373276 |

| | | | | | | |
|---|---|---|---|---|---|---|
| Sertad2 | 0.27853354 | 0.31252196 | 0.28841114 | 0.02411082 | 0.34845989 | 0.89054584 |
| Zfp599 | 0.0670894 | 0.09586238 | 0.07176685 | 0.02409553 | 0.05888209 | 0.43985104 |
| Abhd17a | 0.4102265 | 0.4367756 | 0.41268772 | 0.02408788 | 0.39283621 | 0.89054584 |
| Arhgap22 | 0.02417301 | 0.04995536 | 0.02586814 | 0.02408723 | 7.18E-06 | 0.00019429 |
| Tmem246 | 0.21715731 | 0.25159053 | 0.22754785 | 0.02404268 | 0.31126665 | 0.89054584 |
| Akr1cl | 0.05313335 | 0.08088828 | 0.05684737 | 0.02404091 | 0.02440318 | 0.2319982 |
| Kifc1 | 0.06552614 | 0.0941226 | 0.07009607 | 0.02402652 | 0.05514124 | 0.42075671 |
| Eya2 | 0.06907029 | 0.09790392 | 0.0738838 | 0.02402012 | 0.06493013 | 0.47185816 |
| Lrp3 | 0.06159303 | 0.08989932 | 0.065892 | 0.02400732 | 0.04480537 | 0.3637112 |
| Rps18-ps3 | 1.36203785 | 1.01683047 | 0.9928528 | 0.02397767 | 0.45520764 | 0.89054584 |
| Tgfbr3 | 0.07508684 | 0.10428253 | 0.08031245 | 0.02397008 | 0.0821872 | 0.5516435 |
| Gm14419 | 0.05091977 | 0.0784333 | 0.05448028 | 0.02395302 | 0.02026194 | 0.20315185 |
| Igsf8 | 0.62155552 | 0.61503848 | 0.59109456 | 0.02394392 | 0.42515483 | 0.89054584 |
| Slc35a5 | 0.16455362 | 0.19799469 | 0.17406073 | 0.02393396 | 0.26088456 | 0.89054584 |
| S1pr3 | 0.10676909 | 0.13795603 | 0.11402516 | 0.02393087 | 0.16408939 | 0.81835843 |
| Zfp319 | 0.08999066 | 0.12015545 | 0.09622848 | 0.02392696 | 0.12334354 | 0.70249071 |
| Gm3604 | 0.0572403 | 0.08515907 | 0.06123864 | 0.02392043 | 0.03439251 | 0.29983318 |
| Nat14 | 0.12640036 | 0.15846948 | 0.13456344 | 0.02390604 | 0.20392301 | 0.89054584 |
| Marveld1 | 0.056976 | 0.08486001 | 0.06095606 | 0.02390394 | 0.03384948 | 0.29600565 |
| Ddx58 | 0.04788716 | 0.07513414 | 0.05123711 | 0.02389703 | 0.01489264 | 0.16079806 |
| Mycn | 0.07064459 | 0.09946037 | 0.07556611 | 0.02389426 | 0.07035219 | 0.49949366 |
| C330006A16I | 0.22370251 | 0.25799792 | 0.23411991 | 0.02387802 | 0.31733363 | 0.89054584 |
| Ddo | 0.05082507 | 0.07824892 | 0.05437901 | 0.02386991 | 0.0204243 | 0.2039679 |
| Mtmr2 | 0.39102332 | 0.41897134 | 0.39514333 | 0.02382801 | 0.38937652 | 0.89054584 |
| Prkar1b | 0.3405156 | 0.37185191 | 0.34803467 | 0.02381724 | 0.37492856 | 0.89054584 |
| Gltp | 0.49937114 | 0.5151244 | 0.49133056 | 0.02379384 | 0.41074843 | 0.89054584 |
| Glb1l | 0.09011375 | 0.12013328 | 0.09635988 | 0.0237734 | 0.12518946 | 0.70787708 |
| Fam206a | 0.14426681 | 0.17690166 | 0.15312991 | 0.02377175 | 0.23476323 | 0.89054584 |
| Fastkd1 | 0.06946886 | 0.09807091 | 0.07430973 | 0.02376119 | 0.06814667 | 0.48857443 |
| Klhl6 | 0.04669497 | 0.07370202 | 0.04996207 | 0.02373996 | 0.01342331 | 0.14854065 |

| | | | | | | |
|---|---|---|---|---|---|---|
| Samd14 | 0.33637434 | 0.36784779 | 0.34411143 | 0.02373636 | 0.37396585 | 0.89054584 |
| Hlcs | 0.0946246 | 0.12490781 | 0.10117433 | 0.02373348 | 0.13722006 | 0.7420427 |
| Taf1 | 0.34916492 | 0.37992861 | 0.3561992 | 0.02372941 | 0.3781394 | 0.89054584 |
| Bccip | 1.12658324 | 0.92147704 | 0.89774823 | 0.02372881 | 0.45099705 | 0.89054584 |
| Xpr1 | 0.33250121 | 0.36415988 | 0.3404349 | 0.02372498 | 0.37271066 | 0.89054584 |
| D030056L22I | 0.38464944 | 0.41297863 | 0.38927487 | 0.02370376 | 0.38832295 | 0.89054584 |
| Meig1 | 0.11617385 | 0.14757517 | 0.12388238 | 0.0236928 | 0.1864556 | 0.8734195 |
| Tmem65 | 0.20705156 | 0.24105078 | 0.21736322 | 0.02368756 | 0.30582599 | 0.89054584 |
| Mn1 | 0.10958669 | 0.14066371 | 0.11698173 | 0.02368198 | 0.17279853 | 0.83703076 |
| Map10 | 0.06265938 | 0.09069768 | 0.06703189 | 0.0236658 | 0.05000014 | 0.3933272 |
| Pcdh20 | 0.03332707 | 0.05931831 | 0.03566245 | 0.02365586 | 0.00099943 | 0.01646567 |
| Arrb2 | 0.15606477 | 0.18897596 | 0.16532204 | 0.02365392 | 0.25251668 | 0.89054584 |
| Il17re | 0.18949181 | 0.22321446 | 0.19956283 | 0.02365163 | 0.29041775 | 0.89054584 |
| Mettl2 | 0.19571271 | 0.22953142 | 0.20588409 | 0.02364734 | 0.29628536 | 0.89054584 |
| Atg4d | 0.32912753 | 0.36087248 | 0.33722605 | 0.02364643 | 0.37195235 | 0.89054584 |
| Spata17 | 0.02442441 | 0.04973609 | 0.02613714 | 0.02359895 | 1.30E-05 | 0.00033401 |
| Cog1 | 0.29671282 | 0.32967253 | 0.30609777 | 0.02357475 | 0.35986563 | 0.89054584 |
| Slco2b1 | 0.03593498 | 0.06201077 | 0.03845248 | 0.02355829 | 0.00215627 | 0.03209611 |
| Nox4 | 0.03034977 | 0.05602537 | 0.03247708 | 0.02354829 | 0.00036497 | 0.00676645 |
| Ganc | 0.12555684 | 0.1571654 | 0.13368391 | 0.0234815 | 0.2065815 | 0.89054584 |
| Ncf4 | 0.03361672 | 0.05942462 | 0.03597233 | 0.02345229 | 0.00119309 | 0.01932674 |
| Tbpl1 | 0.61195844 | 0.60705666 | 0.58361255 | 0.02344411 | 0.4257704 | 0.89054584 |
| Gale | 0.27150406 | 0.3049468 | 0.28152962 | 0.02341718 | 0.34918447 | 0.89054584 |
| Adamtsl5 | 0.10828138 | 0.13902614 | 0.1156124 | 0.02341374 | 0.17270615 | 0.83703076 |
| Klc3 | 0.60608739 | 0.60241701 | 0.57900454 | 0.02341247 | 0.42528639 | 0.89054584 |
| Prx | 0.0274717 | 0.05279004 | 0.02939772 | 0.02339232 | 0.00010478 | 0.00217011 |
| Arl4a | 0.80079709 | 0.74247367 | 0.71910027 | 0.0233734 | 0.43981721 | 0.89054584 |
| Mrc1 | 0.02905821 | 0.05444911 | 0.03109521 | 0.0233539 | 0.00023344 | 0.00452457 |
| Ushbp1 | 0.05813733 | 0.08553319 | 0.06219768 | 0.02333551 | 0.0402359 | 0.33777651 |
| Lyrm4 | 0.21007694 | 0.2437421 | 0.22041694 | 0.02332517 | 0.31099821 | 0.89054584 |

| | | | | | | |
|---|---|---|---|---|---|---|
| Nxf1 | 1.27072052 | 0.98271679 | 0.95940081 | 0.02331598 | 0.45492625 | 0.89054584 |
| Tek | 0.06605347 | 0.09397225 | 0.07065969 | 0.02331256 | 0.06213295 | 0.45702869 |
| Saraf | 1.47245681 | 1.05115151 | 1.02785301 | 0.0232985 | 0.45794792 | 0.89054584 |
| Myrf | 0.16751065 | 0.20039166 | 0.17709791 | 0.02329375 | 0.27000692 | 0.89054584 |
| Trip10 | 0.18754484 | 0.22084802 | 0.19758106 | 0.02326696 | 0.29162881 | 0.89054584 |
| Cirh1a | 0.31063199 | 0.34279016 | 0.31952879 | 0.02326137 | 0.36724229 | 0.89054584 |
| Hoxb3 | 0.07543064 | 0.10393402 | 0.08067974 | 0.02325427 | 0.08966129 | 0.58049509 |
| Tceal1 | 0.498661 | 0.51396897 | 0.49072288 | 0.0232461 | 0.41266328 | 0.89054584 |
| Pcyox1 | 0.27280776 | 0.30604899 | 0.2828077 | 0.0232413 | 0.35090642 | 0.89054584 |
| Mis18bp1 | 0.09210652 | 0.12171733 | 0.09848695 | 0.02323039 | 0.13590206 | 0.73849629 |
| Fut10 | 0.06074706 | 0.08821258 | 0.06498766 | 0.02322492 | 0.04795782 | 0.38170821 |
| Stx18 | 0.40461262 | 0.43078672 | 0.40758058 | 0.02320614 | 0.39540345 | 0.89054584 |
| Smek1 | 0.48980615 | 0.50632629 | 0.48312021 | 0.02320608 | 0.41146172 | 0.89054584 |
| Fam102b | 0.08482906 | 0.11392173 | 0.09071782 | 0.02320391 | 0.11669549 | 0.68192309 |
| Trim30a | 0.0324206 | 0.05789494 | 0.03469265 | 0.02320229 | 0.00091699 | 0.01535164 |
| Trappc10 | 0.20983374 | 0.24336995 | 0.2201716 | 0.02319835 | 0.3117526 | 0.89054584 |
| Icam1 | 0.05883881 | 0.08612919 | 0.06294764 | 0.02318155 | 0.04310526 | 0.3540737 |
| Rgs12 | 0.1448471 | 0.17689573 | 0.15373086 | 0.02316487 | 0.24133007 | 0.89054584 |
| Uaca | 0.29512132 | 0.32771312 | 0.30455587 | 0.02315725 | 0.36157657 | 0.89054584 |
| Trim33 | 0.23243088 | 0.26600629 | 0.24285424 | 0.02315205 | 0.32846656 | 0.89054584 |
| Depdc1a | 0.06685182 | 0.09465355 | 0.07151294 | 0.02314061 | 0.06583235 | 0.47501237 |
| Chchd4 | 0.68042513 | 0.65875227 | 0.63562013 | 0.02313214 | 0.43268087 | 0.89054584 |
| Lsamp | 0.04416341 | 0.07037922 | 0.04725443 | 0.02312479 | 0.01130575 | 0.12957378 |
| Rabgef1 | 0.1699383 | 0.20270204 | 0.17958869 | 0.02311335 | 0.27438205 | 0.89054584 |
| Arc | 0.09847907 | 0.12839947 | 0.10528712 | 0.02311234 | 0.15322057 | 0.78856652 |
| Ogfod3 | 0.2716271 | 0.30475195 | 0.28165028 | 0.02310168 | 0.3511794 | 0.89054584 |
| Mnd1 | 0.08705994 | 0.1161932 | 0.09309977 | 0.02309342 | 0.12391186 | 0.70422347 |
| Stard5 | 0.29250708 | 0.32510554 | 0.30202034 | 0.0230852 | 0.36087894 | 0.89054584 |
| Med17 | 0.21081196 | 0.24424336 | 0.22115823 | 0.02308512 | 0.31337246 | 0.89054584 |
| Ccdc88a | 0.89154302 | 0.79819559 | 0.77512151 | 0.02307408 | 0.44484798 | 0.89054584 |

| | | | | | | |
|---|---|---|---|---|---|---|
| Unc5cl | 0.03767666 | 0.06337433 | 0.04031569 | 0.02305864 | 0.00385296 | 0.05284788 |
| Aco2 | 1.44356516 | 1.04238076 | 1.01932717 | 0.02305358 | 0.45804158 | 0.89054584 |
| AB124611 | 0.02288586 | 0.04754205 | 0.02449086 | 0.02305119 | 5.80E-06 | 0.0001601 |
| Mapkap1 | 0.43075293 | 0.45424771 | 0.43121016 | 0.02303755 | 0.40171694 | 0.89054584 |
| Upf3a | 0.43982365 | 0.4623537 | 0.43931714 | 0.02303655 | 0.40349857 | 0.89054584 |
| Tlk1 | 0.33855063 | 0.36920547 | 0.34617428 | 0.02303119 | 0.37829484 | 0.89054584 |
| Ltv1 | 0.47117741 | 0.48998481 | 0.46696951 | 0.02301529 | 0.4091924 | 0.89054584 |
| Epn3 | 0.05058963 | 0.0771118 | 0.05412724 | 0.02298456 | 0.02394348 | 0.22897929 |
| Pmm1 | 0.22411786 | 0.25752043 | 0.23453631 | 0.02298413 | 0.32399089 | 0.89054584 |
| Bbs4 | 0.25968788 | 0.2928917 | 0.26990832 | 0.02298338 | 0.34578671 | 0.89054584 |
| Ing1 | 0.55063302 | 0.55733311 | 0.5343593 | 0.02297381 | 0.42062191 | 0.89054584 |
| Gm4577 | 0.03249536 | 0.05774004 | 0.03477263 | 0.02296741 | 0.00104463 | 0.01717121 |
| Pabpc4 | 0.59388165 | 0.59231208 | 0.56935166 | 0.02296042 | 0.42548436 | 0.89054584 |
| Tvp23a | 0.05884922 | 0.08591197 | 0.06295877 | 0.02295319 | 0.04470357 | 0.36308815 |
| Nfs1 | 0.50565692 | 0.51958333 | 0.49669572 | 0.0228876 | 0.41500734 | 0.89054584 |
| Epb42 | 0.04739533 | 0.07359091 | 0.05071111 | 0.02287979 | 0.0177759 | 0.18474197 |
| Sla2 | 0.0181072 | 0.04225281 | 0.01937738 | 0.02287543 | 1.90E-08 | 7.49E-07 |
| Acat2 | 0.49867652 | 0.51359703 | 0.49073616 | 0.02286087 | 0.41409011 | 0.89054584 |
| Dok2 | 0.03899342 | 0.06458379 | 0.04172429 | 0.0228595 | 0.00534771 | 0.06934802 |
| Idi1 | 0.24926396 | 0.28244425 | 0.25960274 | 0.02284151 | 0.34093106 | 0.89054584 |
| Tti1 | 0.10478691 | 0.13476302 | 0.11194347 | 0.02281955 | 0.17112729 | 0.83563604 |
| Gm37350 | 0.04327673 | 0.06911855 | 0.04630603 | 0.02281251 | 0.01086049 | 0.12527997 |
| Taf1a | 0.11041874 | 0.14065742 | 0.11785427 | 0.02280315 | 0.18367722 | 0.86684645 |
| Zfp462 | 0.15333414 | 0.18530742 | 0.16250489 | 0.02280253 | 0.25664022 | 0.89054584 |
| Mospd2 | 0.25396279 | 0.28703943 | 0.26425439 | 0.02278504 | 0.34394797 | 0.89054584 |
| Elac1 | 0.07302032 | 0.10088315 | 0.07810459 | 0.02277856 | 0.0871142 | 0.57045578 |
| Sv2b | 0.07140773 | 0.09915624 | 0.07638156 | 0.02277468 | 0.08239266 | 0.55212586 |
| Acaca | 0.18201661 | 0.21470752 | 0.19194528 | 0.02276224 | 0.29030618 | 0.89054584 |
| Ifngr2 | 0.42087691 | 0.44508047 | 0.42232834 | 0.02275213 | 0.40091032 | 0.89054584 |
| Alg1 | 0.24135446 | 0.27450066 | 0.25174931 | 0.02275134 | 0.33686202 | 0.89054584 |

| | | | | | | |
|---|---|---|---|---|---|---|
| Aff3 | 0.04081229 | 0.06641873 | 0.04366994 | 0.02274879 | 0.00761293 | 0.09337756 |
| Adat1 | 0.08718763 | 0.11597734 | 0.09323611 | 0.02274123 | 0.12790234 | 0.71595682 |
| Zfp383 | 0.16680462 | 0.19909591 | 0.17637307 | 0.02272285 | 0.27417647 | 0.89054584 |
| C1ra | 0.05914459 | 0.08597468 | 0.06327455 | 0.02270013 | 0.04731902 | 0.37818696 |
| Tnc | 0.02519231 | 0.04964785 | 0.0269588 | 0.02268904 | 4.41E-05 | 0.00100803 |
| Nfatc4 | 0.11585926 | 0.14622316 | 0.12355318 | 0.02266998 | 0.19632005 | 0.89054584 |
| Tmem238 | 0.58960512 | 0.58861573 | 0.56594638 | 0.02266935 | 0.42598049 | 0.89054584 |
| Smarcd1 | 0.27581848 | 0.30841642 | 0.28575609 | 0.02266033 | 0.35589542 | 0.89054584 |
| Fibp | 0.467443 | 0.4863491 | 0.46370637 | 0.02264274 | 0.41001918 | 0.89054584 |
| Taf2 | 0.17273154 | 0.20508775 | 0.18245157 | 0.02263618 | 0.28162532 | 0.89054584 |
| Hdgfrp3 | 0.15358686 | 0.18539838 | 0.16276575 | 0.02263263 | 0.2585483 | 0.89054584 |
| Stip1 | 1.25787937 | 0.97691341 | 0.95433291 | 0.0225805 | 0.45611136 | 0.89054584 |
| Dand5 | 0.10188128 | 0.13145345 | 0.10888939 | 0.02256406 | 0.16716578 | 0.82737245 |
| Gfpt1 | 0.74340961 | 0.70314304 | 0.68058955 | 0.02255349 | 0.43865161 | 0.89054584 |
| Aamp | 1.69670256 | 1.10335898 | 1.08083548 | 0.02252351 | 0.46132843 | 0.89054584 |
| B3gnt8 | 0.15997526 | 0.19187142 | 0.16935124 | 0.02252018 | 0.26777968 | 0.89054584 |
| Ift80 | 0.13342548 | 0.16437861 | 0.14187849 | 0.02250012 | 0.22999941 | 0.89054584 |
| Jup | 0.56728387 | 0.5704732 | 0.54797596 | 0.02249723 | 0.42415566 | 0.89054584 |
| Ptprr | 0.02548122 | 0.04976146 | 0.02726794 | 0.02249353 | 6.07E-05 | 0.00134874 |
| Fam132a | 0.24001816 | 0.27286881 | 0.25041961 | 0.0224492 | 0.33809899 | 0.89054584 |
| Zfp748 | 0.1321342 | 0.16297076 | 0.14053534 | 0.02243542 | 0.22850925 | 0.89054584 |
| Dennd1c | 0.07791878 | 0.1057647 | 0.08333771 | 0.02242699 | 0.1049635 | 0.63913594 |
| Lrp8 | 0.34999662 | 0.37939702 | 0.35698217 | 0.02241485 | 0.38493916 | 0.89054584 |
| Pdha1 | 0.72364732 | 0.68919584 | 0.66678155 | 0.02241429 | 0.4377748 | 0.89054584 |
| Wdr20 | 0.31247876 | 0.34371552 | 0.32130378 | 0.02241174 | 0.37260083 | 0.89054584 |
| Jrkl | 0.20421127 | 0.2369022 | 0.21449279 | 0.02240941 | 0.31321814 | 0.89054584 |
| Uvssa | 0.35215041 | 0.38141211 | 0.35900804 | 0.02240406 | 0.3856238 | 0.89054584 |
| Triqk | 0.25145399 | 0.28415977 | 0.26177207 | 0.02238769 | 0.34514944 | 0.89054584 |
| Mycl | 0.21955761 | 0.25233905 | 0.22996022 | 0.02237883 | 0.325134 | 0.89054584 |
| Tatdn1 | 0.17938923 | 0.21161897 | 0.18926227 | 0.02235669 | 0.29104323 | 0.89054584 |

| | | | | | | |
|---|---|---|---|---|---|---|
| Setmar | 0.0476054 | 0.07327883 | 0.05093578 | 0.02234305 | 0.02049403 | 0.20420787 |
| Zfp950 | 0.55887898 | 0.56346266 | 0.5411254 | 0.02233725 | 0.42374671 | 0.89054584 |
| Rsph1 | 0.29598217 | 0.32772625 | 0.30539005 | 0.0223362 | 0.3666442 | 0.89054584 |
| Gm4924 | 0.05736519 | 0.0836674 | 0.06137217 | 0.02229523 | 0.0452757 | 0.36609429 |
| Fxyd7 | 0.04604696 | 0.07155364 | 0.049269 | 0.02228464 | 0.01754733 | 0.1828909 |
| Arl8a | 0.30318281 | 0.33463052 | 0.31235301 | 0.02227751 | 0.36983776 | 0.89054584 |
| Tagap | 0.03980263 | 0.06486393 | 0.04258991 | 0.02227401 | 0.00741364 | 0.09155279 |
| Adcy4 | 0.07366915 | 0.10104847 | 0.07879783 | 0.02225064 | 0.09415839 | 0.59930619 |
| Etv5 | 0.11985409 | 0.14996246 | 0.12773072 | 0.02223174 | 0.2087126 | 0.89054584 |
| Sik2 | 0.18016044 | 0.21227828 | 0.19005011 | 0.02222817 | 0.29290855 | 0.89054584 |
| Brdt | 0.07906967 | 0.10675408 | 0.08456704 | 0.02218704 | 0.11079308 | 0.66020472 |
| Ogfrl1 | 0.07535919 | 0.10276636 | 0.08060341 | 0.02216295 | 0.1000905 | 0.62272587 |
| Tmtc1 | 0.04440582 | 0.06965776 | 0.04751371 | 0.02214405 | 0.01495322 | 0.16130338 |
| Igdcc4 | 0.10460933 | 0.1338789 | 0.11175691 | 0.022122 | 0.17820547 | 0.85127607 |
| Dctd | 0.1269033 | 0.15718446 | 0.13508776 | 0.02209669 | 0.2230072 | 0.89054584 |
| Nup37 | 0.22856954 | 0.2610827 | 0.23899439 | 0.02208831 | 0.33338393 | 0.89054584 |
| Abhd11 | 0.35702657 | 0.38565965 | 0.36358532 | 0.02207434 | 0.38864281 | 0.89054584 |
| Adam33 | 0.05531411 | 0.08123758 | 0.05917918 | 0.0220584 | 0.04123156 | 0.34354894 |
| Slc4a7 | 0.45841844 | 0.47784534 | 0.4557877 | 0.02205764 | 0.41080788 | 0.89054584 |
| Lrrtm3 | 0.06885677 | 0.09571048 | 0.07365562 | 0.02205486 | 0.08150242 | 0.54831655 |
| Aph1b | 0.08510375 | 0.11305889 | 0.09101113 | 0.02204776 | 0.12952386 | 0.72054099 |
| Cldn8 | 0.06795403 | 0.09469529 | 0.0726909 | 0.02200439 | 0.07922313 | 0.538911 |
| Zfp945 | 0.31179451 | 0.34264424 | 0.32064633 | 0.02199791 | 0.37462583 | 0.89054584 |
| Lifr | 0.26060612 | 0.29280979 | 0.27081381 | 0.02199598 | 0.35256396 | 0.89054584 |
| Pcsk5 | 0.04667755 | 0.07192684 | 0.04994344 | 0.02198341 | 0.0201477 | 0.20242643 |
| Ercc3 | 0.15033333 | 0.18136547 | 0.15940581 | 0.02195966 | 0.26049725 | 0.89054584 |
| Uap1l1 | 0.12401335 | 0.15403269 | 0.13207385 | 0.02195884 | 0.21928684 | 0.89054584 |
| Ctnna1 | 1.05148692 | 0.88295231 | 0.86099902 | 0.02195329 | 0.4527204 | 0.89054584 |
| Mmadhc | 0.80055253 | 0.74089346 | 0.7189412 | 0.02195226 | 0.44343858 | 0.89054584 |
| Pde3a | 0.09144504 | 0.11972333 | 0.09778092 | 0.02194242 | 0.14789958 | 0.77409677 |

| | | | | | | |
|---|---|---|---|---|---|---|
| Cyp7b1 | 0.0629787 | 0.08931309 | 0.06737321 | 0.02193989 | 0.06461182 | 0.47001612 |
| Chd6 | 0.69755029 | 0.67004622 | 0.64812382 | 0.0219224 | 0.43739155 | 0.89054584 |
| Dysf | 0.05458636 | 0.08031859 | 0.05840103 | 0.02191755 | 0.04019114 | 0.33777651 |
| Ccdc106 | 0.03706479 | 0.06156876 | 0.03966112 | 0.02190763 | 0.00503435 | 0.06629241 |
| Rabep2 | 0.17394325 | 0.2055584 | 0.18369247 | 0.02186592 | 0.2895912 | 0.89054584 |
| Gm12840 | 0.03214796 | 0.05625021 | 0.03440096 | 0.02184926 | 0.00154291 | 0.02400117 |
| Ntrk3 | 0.0533754 | 0.07895159 | 0.05710619 | 0.0218454 | 0.03735577 | 0.32027435 |
| Eps8 | 0.48161132 | 0.49786779 | 0.47604188 | 0.02182591 | 0.41542696 | 0.89054584 |
| Tmem173 | 0.0836492 | 0.1112784 | 0.08945794 | 0.02182046 | 0.12789404 | 0.71595682 |
| Rfx5 | 0.06206217 | 0.08819106 | 0.0663935 | 0.02179756 | 0.06306147 | 0.46198345 |
| Tmem248 | 0.58370849 | 0.58302627 | 0.56123095 | 0.02179532 | 0.42821154 | 0.89054584 |
| Zg16 | 0.02054685 | 0.04377902 | 0.021988 | 0.02179102 | 1.94E-06 | 5.80E-05 |
| Ly6c1 | 0.04097018 | 0.06562147 | 0.04383884 | 0.02178264 | 0.01030827 | 0.12042647 |
| Traf3ip1 | 0.14869502 | 0.17946961 | 0.15771239 | 0.02175722 | 0.26020147 | 0.89054584 |
| Tfcp2 | 0.16486517 | 0.19611389 | 0.17438089 | 0.021733 | 0.28074104 | 0.89054584 |
| Ddx41 | 0.8017469 | 0.74143617 | 0.71971764 | 0.02171853 | 0.44409678 | 0.89054584 |
| Pus1 | 0.27549938 | 0.3071165 | 0.2854438 | 0.0216727 | 0.36176827 | 0.89054584 |
| Cpne2 | 0.09958478 | 0.12813444 | 0.10646674 | 0.02166771 | 0.17152144 | 0.83588128 |
| Mlycd | 0.33740705 | 0.36675414 | 0.34509062 | 0.02166352 | 0.38496317 | 0.89054584 |
| Xpo7 | 0.28666832 | 0.31799933 | 0.29634508 | 0.02165424 | 0.36676499 | 0.89054584 |
| Stk3 | 0.25456667 | 0.28649396 | 0.26485145 | 0.02164251 | 0.35170983 | 0.89054584 |
| Rab27a | 0.58452634 | 0.58348412 | 0.56188636 | 0.02159777 | 0.42893736 | 0.89054584 |
| Zfx | 0.27516845 | 0.30671582 | 0.28511988 | 0.02159594 | 0.36208774 | 0.89054584 |
| Emc3 | 0.6642826 | 0.64522451 | 0.62364817 | 0.02157635 | 0.43597372 | 0.89054584 |
| Srrt | 0.85701973 | 0.77603359 | 0.75451546 | 0.02151813 | 0.4471485 | 0.89054584 |
| Frem1 | 0.07405202 | 0.10071928 | 0.07920688 | 0.0215124 | 0.10287161 | 0.63089369 |
| Hsd11b2 | 0.07228566 | 0.09882758 | 0.07731963 | 0.02150795 | 0.09749216 | 0.61400582 |
| Rasgrp3 | 0.06187844 | 0.08770501 | 0.0661971 | 0.02150791 | 0.06504801 | 0.47228321 |
| Fmnl3 | 0.08501624 | 0.11242027 | 0.09091769 | 0.02150257 | 0.13525969 | 0.73693912 |
| Gm26917 | 0.71640365 | 0.6831502 | 0.66165098 | 0.02149922 | 0.43983627 | 0.89054584 |

| | | | | | | |
|---|---|---|---|---|---|---|
| Prr29 | 0.02927526 | 0.05279188 | 0.03132743 | 0.02146445 | 0.00070611 | 0.01228952 |
| Ska1 | 0.0704627 | 0.09683487 | 0.07537175 | 0.02146312 | 0.09230463 | 0.59240039 |
| Gng3 | 0.07945802 | 0.10642146 | 0.08498184 | 0.02143963 | 0.11992136 | 0.69330539 |
| Heatr5a | 0.1067136 | 0.1353984 | 0.1139669 | 0.0214315 | 0.19048382 | 0.88259328 |
| Samm50 | 0.92615592 | 0.81634975 | 0.79492422 | 0.02142553 | 0.45003533 | 0.89054584 |
| Snrnp35 | 0.23134158 | 0.26318957 | 0.24176592 | 0.02142366 | 0.33986053 | 0.89054584 |
| Dct | 0.13075539 | 0.16049297 | 0.13910043 | 0.02139254 | 0.23683884 | 0.89054584 |
| Gm4707 | 0.09411714 | 0.12202008 | 0.10063279 | 0.02138729 | 0.16104903 | 0.81128869 |
| Ccng2 | 0.24876244 | 0.28046952 | 0.25910564 | 0.02136388 | 0.35043702 | 0.89054584 |
| Cmip | 0.65870876 | 0.64082863 | 0.6194721 | 0.02135653 | 0.43619686 | 0.89054584 |
| Sclt1 | 0.24495759 | 0.27668344 | 0.25533051 | 0.02135292 | 0.34840859 | 0.89054584 |
| Cbx4 | 0.18735124 | 0.21871204 | 0.19738392 | 0.02132812 | 0.30733547 | 0.89054584 |
| Itga1 | 0.05009259 | 0.07492 | 0.05359571 | 0.02132429 | 0.03189335 | 0.28228924 |
| Gm26522 | 0.01944832 | 0.04210348 | 0.0208125 | 0.02129098 | 9.39E-07 | 2.94E-05 |
| Rarg | 0.12473044 | 0.15411027 | 0.13282198 | 0.02128829 | 0.22761616 | 0.89054584 |
| Ctrc | 0.04637451 | 0.07090357 | 0.04961933 | 0.02128424 | 0.02283366 | 0.22158129 |
| Ube2h | 0.3144899 | 0.34450206 | 0.32323477 | 0.02126729 | 0.37959836 | 0.89054584 |
| Lgi1 | 0.02477776 | 0.0477805 | 0.02651523 | 0.02126526 | 9.33E-05 | 0.0019577 |
| Mecr | 0.36236794 | 0.3898256 | 0.36858446 | 0.02124114 | 0.39416112 | 0.89054584 |
| Orc4 | 0.52213257 | 0.53188119 | 0.51064017 | 0.02124102 | 0.42316731 | 0.89054584 |
| Zkscan17 | 0.13596806 | 0.16576168 | 0.14452135 | 0.02124033 | 0.24675861 | 0.89054584 |
| Zfr2 | 0.1444294 | 0.17453714 | 0.1532983 | 0.02123884 | 0.25930909 | 0.89054584 |
| Cdca5 | 0.05711909 | 0.08234181 | 0.06110905 | 0.02123275 | 0.05274722 | 0.40742101 |
| Kcna5 | 0.05900931 | 0.08435339 | 0.06312992 | 0.02122348 | 0.0586419 | 0.43892115 |
| Spns2 | 0.04934728 | 0.07402001 | 0.05279865 | 0.02122136 | 0.03056402 | 0.27268752 |
| Mcoln3 | 0.09674557 | 0.12464848 | 0.10343758 | 0.0212109 | 0.1696973 | 0.83342527 |
| Carkd | 0.86062985 | 0.77791482 | 0.75671065 | 0.02120417 | 0.44806544 | 0.89054584 |
| Pdcl | 0.37053386 | 0.39737551 | 0.37619911 | 0.0211764 | 0.39656294 | 0.89054584 |
| Limk2 | 0.27681007 | 0.30790185 | 0.28672618 | 0.02117567 | 0.3653938 | 0.89054584 |
| Tubd1 | 0.08879579 | 0.11612705 | 0.09495297 | 0.02117408 | 0.14942031 | 0.77812345 |

| | | | | | | |
|---|---|---|---|---|---|---|
| Kctd20 | 0.62977016 | 0.61861836 | 0.5974494 | 0.02116896 | 0.43444184 | 0.89054584 |
| Arhgap42 | 0.0824989 | 0.10938095 | 0.08822955 | 0.0211514 | 0.13201967 | 0.72831857 |
| Rbpj | 0.37944998 | 0.40561268 | 0.38447133 | 0.02114135 | 0.39890347 | 0.89054584 |
| Pinx1 | 0.16821893 | 0.19894654 | 0.17782487 | 0.02112168 | 0.29000174 | 0.89054584 |
| Mrc2 | 0.08362879 | 0.11055245 | 0.08943615 | 0.0211163 | 0.13566645 | 0.73849629 |
| Nkd2 | 0.04851956 | 0.07302654 | 0.05191345 | 0.0211131 | 0.02906063 | 0.26316115 |
| Prkch | 0.08031138 | 0.106984 | 0.08589329 | 0.0210907 | 0.12631562 | 0.71147001 |
| Glp1r | 0.0183472 | 0.04071237 | 0.01963421 | 0.02107816 | 2.84E-07 | 9.59E-06 |
| Susd6 | 0.17303416 | 0.20383399 | 0.18276153 | 0.02107246 | 0.29557256 | 0.89054584 |
| Mns1 | 0.14141751 | 0.17120982 | 0.15017727 | 0.02103255 | 0.25704202 | 0.89054584 |
| Icosl | 0.04392423 | 0.06801989 | 0.04699861 | 0.02102128 | 0.01858799 | 0.19139826 |
| Aldh1b1 | 0.15162079 | 0.18173735 | 0.16073582 | 0.02100152 | 0.27135142 | 0.89054584 |
| Tnfrsf9 | 0.0286235 | 0.05161437 | 0.03063008 | 0.02098428 | 0.00070699 | 0.01229011 |
| Adgrl3 | 0.14125684 | 0.17098966 | 0.15001068 | 0.02097898 | 0.25734494 | 0.89054584 |
| Mrpl33 | 1.60359084 | 1.08257872 | 1.06160051 | 0.02097821 | 0.4633232 | 0.89054584 |
| Map1lc3b | 1.84441364 | 1.12453715 | 1.10356571 | 0.02097144 | 0.46472555 | 0.89054584 |
| Cdk5rap3 | 0.72964424 | 0.69196739 | 0.67100068 | 0.02096671 | 0.44212743 | 0.89054584 |
| Spred3 | 0.10780591 | 0.13605841 | 0.11511346 | 0.02094495 | 0.19830351 | 0.89054584 |
| Fbxl12os | 0.03205726 | 0.05524652 | 0.03430392 | 0.0209426 | 0.00222551 | 0.03302508 |
| Pard3b | 0.13664444 | 0.16614647 | 0.14522398 | 0.02092248 | 0.25104155 | 0.89054584 |
| Prpf31 | 0.54817971 | 0.55325056 | 0.5323377 | 0.02091285 | 0.42738872 | 0.89054584 |
| Lysmd3 | 0.3167348 | 0.34629135 | 0.32538776 | 0.02090359 | 0.3823555 | 0.89054584 |
| Mfsd6l | 0.06134702 | 0.08649313 | 0.06562902 | 0.02086411 | 0.06928691 | 0.49496101 |
| Syne1 | 0.29040866 | 0.3208254 | 0.29998264 | 0.02084276 | 0.37308156 | 0.89054584 |
| Hoxb7 | 0.10277796 | 0.13067174 | 0.10983221 | 0.02083953 | 0.18835193 | 0.87720084 |
| Chtop | 0.85253077 | 0.77259421 | 0.75177249 | 0.02082172 | 0.44866382 | 0.89054584 |
| Garnl3 | 0.06758786 | 0.0931169 | 0.07229957 | 0.02081733 | 0.08988615 | 0.58125307 |
| AI467606 | 0.23010505 | 0.26133651 | 0.24053002 | 0.02080649 | 0.34347012 | 0.89054584 |
| Mbnl2 | 1.5303951 | 1.06452911 | 1.04375735 | 0.02077176 | 0.46306405 | 0.89054584 |
| Magee1 | 0.15821371 | 0.18830308 | 0.16753698 | 0.0207661 | 0.28181003 | 0.89054584 |

| | | | | | | |
|---|---|---|---|---|---|---|
| Slc7a10 | 0.04442961 | 0.06827637 | 0.04753915 | 0.02073722 | 0.02106202 | 0.20845964 |
| Thsd1 | 0.03380597 | 0.05690419 | 0.03617481 | 0.02072938 | 0.00379561 | 0.05230825 |
| Actr2 | 0.76782179 | 0.71798625 | 0.69726136 | 0.02072489 | 0.44493093 | 0.89054584 |
| Lsm3 | 0.91941028 | 0.81185049 | 0.79112966 | 0.02072083 | 0.45143985 | 0.89054584 |
| Gm13373 | 0.03811065 | 0.06148136 | 0.04077995 | 0.02070141 | 0.00901382 | 0.10791272 |
| Sh3rf1 | 0.10024288 | 0.12785552 | 0.10716593 | 0.02068959 | 0.18420361 | 0.8679181 |
| Tprgl | 0.84419801 | 0.76732902 | 0.74664151 | 0.02068751 | 0.44864432 | 0.89054584 |
| Nacc1 | 0.18881201 | 0.21955141 | 0.19887106 | 0.02068035 | 0.31402359 | 0.89054584 |
| Zfp184 | 0.07558534 | 0.1015245 | 0.08084501 | 0.0206795 | 0.11668567 | 0.68192309 |
| Nfxl1 | 0.30816181 | 0.33783005 | 0.31715232 | 0.02067773 | 0.3806576 | 0.89054584 |
| Slc16a3 | 0.09006439 | 0.11698314 | 0.0963072 | 0.02067594 | 0.15860299 | 0.80483939 |
| Bckdha | 0.54497704 | 0.55033546 | 0.52969321 | 0.02064225 | 0.42796408 | 0.89054584 |
| Uri1 | 0.59712306 | 0.59256136 | 0.57192453 | 0.02063683 | 0.4332487 | 0.89054584 |
| Nova2 | 0.06390035 | 0.08898849 | 0.06835836 | 0.02063013 | 0.07985615 | 0.54094194 |
| Cypt3 | 0.03746164 | 0.06071093 | 0.04008567 | 0.02062526 | 0.00826129 | 0.09981018 |
| Icos | 0.02126265 | 0.04337643 | 0.02275395 | 0.02062248 | 1.21E-05 | 0.00031385 |
| Arhgef12 | 0.29070134 | 0.32088608 | 0.30026698 | 0.0206191 | 0.37451224 | 0.89054584 |
| Uros | 0.24333184 | 0.27431415 | 0.2537154 | 0.02059875 | 0.3526217 | 0.89054584 |
| Mogs | 0.24487112 | 0.27583924 | 0.25524464 | 0.02059459 | 0.35349191 | 0.89054584 |
| Pianp | 0.16332249 | 0.19337119 | 0.17279518 | 0.02057602 | 0.28952427 | 0.89054584 |
| Slc16a2 | 0.17230602 | 0.20257368 | 0.18201564 | 0.02055804 | 0.29937069 | 0.89054584 |
| Utp23 | 0.18504626 | 0.21556369 | 0.19503546 | 0.02052823 | 0.31193555 | 0.89054584 |
| Kat7 | 0.2691323 | 0.29972211 | 0.27920236 | 0.02051975 | 0.36602303 | 0.89054584 |
| Ascc1 | 0.71034755 | 0.67784916 | 0.65733283 | 0.02051633 | 0.44219263 | 0.89054584 |
| Fam184a | 0.05356068 | 0.07781087 | 0.05730431 | 0.02050656 | 0.04773541 | 0.38067544 |
| Slc30a4 | 0.10277329 | 0.13033001 | 0.1098273 | 0.02050271 | 0.19222193 | 0.88530942 |
| Tbcc | 0.71018362 | 0.67769507 | 0.65721558 | 0.02047949 | 0.44228547 | 0.89054584 |
| Sgip1 | 0.03685967 | 0.05990253 | 0.03944169 | 0.02046083 | 0.00782711 | 0.09551972 |
| Atl3 | 0.13851209 | 0.16761737 | 0.1471632 | 0.02045417 | 0.25863967 | 0.89054584 |
| BC068281 | 0.08564739 | 0.11204451 | 0.09159161 | 0.02045291 | 0.14908469 | 0.77749274 |

| | | | | | | |
|---|---|---|---|---|---|---|
| Slc35f6 | 0.1061433 | 0.13380305 | 0.1133681 | 0.02043495 | 0.20051347 | 0.89054584 |
| Hmbs | 0.5066785 | 0.51798253 | 0.49756534 | 0.02041719 | 0.42419389 | 0.89054584 |
| Haus1 | 0.29287579 | 0.3227928 | 0.30237816 | 0.02041464 | 0.37655475 | 0.89054584 |
| Zfp1 | 0.20705522 | 0.2377262 | 0.21736691 | 0.02035929 | 0.33128362 | 0.89054584 |
| Nr1h4 | 0.04382175 | 0.06724162 | 0.04688899 | 0.02035263 | 0.02157478 | 0.21220569 |
| Nipal2 | 0.21204665 | 0.2427484 | 0.22240294 | 0.02034546 | 0.33498075 | 0.89054584 |
| Tyw1 | 0.20907415 | 0.23972414 | 0.2194052 | 0.02031894 | 0.33306666 | 0.89054584 |
| Ino80c | 0.13406481 | 0.16285179 | 0.14254327 | 0.02030851 | 0.25341826 | 0.89054584 |
| N4bp2l1 | 0.21996765 | 0.25064569 | 0.23037207 | 0.02027362 | 0.34090127 | 0.89054584 |
| Snrpn | 0.42786366 | 0.44887356 | 0.42861767 | 0.02025589 | 0.41286733 | 0.89054584 |
| RP23-49I18.5 | 0.05766782 | 0.08194597 | 0.06169572 | 0.02025025 | 0.06310815 | 0.4620803 |
| Slco2a1 | 0.10965888 | 0.13730362 | 0.11705744 | 0.02024618 | 0.21017699 | 0.89054584 |
| Nme3 | 0.07491601 | 0.10036279 | 0.08012994 | 0.02023284 | 0.11972063 | 0.69262156 |
| Palmd | 0.08386031 | 0.10989715 | 0.08968337 | 0.02021378 | 0.14684044 | 0.77301841 |
| Cttnbp2 | 0.08256561 | 0.10849 | 0.08830079 | 0.02018922 | 0.14338665 | 0.76511581 |
| Bcat1 | 0.03639269 | 0.05909597 | 0.03894213 | 0.02015384 | 0.00795102 | 0.09686882 |
| Ms4a6b | 0.01912999 | 0.04059893 | 0.02047186 | 0.02012707 | 2.32E-06 | 6.86E-05 |
| Scin | 0.06416458 | 0.08876558 | 0.06864079 | 0.02012479 | 0.08597658 | 0.5671012 |
| Mrpl35 | 0.74492717 | 0.70173085 | 0.68163839 | 0.02009246 | 0.44538487 | 0.89054584 |
| Exoc3l2 | 0.0220455 | 0.04367744 | 0.02359163 | 0.0200858 | 3.65E-05 | 0.0008524 |
| Cox16 | 0.17041744 | 0.20016381 | 0.18008001 | 0.02008381 | 0.30167179 | 0.89054584 |
| Gm10020 | 0.93382209 | 0.81926795 | 0.79919599 | 0.02007196 | 0.45342598 | 0.89054584 |
| Ddx5 | 3.63820376 | 1.03764237 | 1.01757292 | 0.02006945 | 0.46339364 | 0.89054584 |
| Amica1 | 0.01754823 | 0.03884472 | 0.01877924 | 0.02006548 | 3.21E-07 | 1.08E-05 |
| Gm14403 | 0.0604826 | 0.08476695 | 0.06470495 | 0.020062 | 0.07429624 | 0.51860709 |
| Pced1a | 0.08621769 | 0.1122378 | 0.09220053 | 0.02003727 | 0.15565106 | 0.79659772 |
| Cpne8 | 0.08042596 | 0.1060215 | 0.08601568 | 0.02000582 | 0.13927276 | 0.74945915 |
| D5Ertd579e | 0.42446345 | 0.44555149 | 0.42556044 | 0.01999105 | 0.41338048 | 0.89054584 |
| Ncln | 0.38105864 | 0.40592134 | 0.38595911 | 0.01996223 | 0.4047909 | 0.89054584 |
| Sik1 | 0.07013662 | 0.09498328 | 0.0750233 | 0.01995998 | 0.1075777 | 0.64845225 |

| | | | | | | |
|---|---|---|---|---|---|---|
| Ggt1 | 0.06892118 | 0.09367482 | 0.07372446 | 0.01995036 | 0.10370084 | 0.63356715 |
| Ildr2 | 0.03438951 | 0.05674025 | 0.0367991 | 0.01994115 | 0.00579078 | 0.07384814 |
| Chst1 | 0.04774424 | 0.07096805 | 0.05108427 | 0.01988378 | 0.03488333 | 0.30373211 |
| Dkk3 | 0.06425758 | 0.08859513 | 0.06874019 | 0.01985494 | 0.08920129 | 0.57874406 |
| Nr4a1 | 0.09069623 | 0.11682096 | 0.09698164 | 0.01983932 | 0.17027747 | 0.83373276 |
| 9430020K01I | 0.05637909 | 0.08013183 | 0.06031786 | 0.01981397 | 0.0629554 | 0.46167296 |
| Bpnt1 | 0.65931451 | 0.63972637 | 0.619927 | 0.01979937 | 0.44085639 | 0.89054584 |
| Fbrsl1 | 0.1896918 | 0.21951079 | 0.19976631 | 0.01974448 | 0.3225859 | 0.89054584 |
| Rasal3 | 0.03798107 | 0.0603715 | 0.04064133 | 0.01973017 | 0.01185552 | 0.13459772 |
| Dbn1 | 1.09719295 | 0.90353162 | 0.8838133 | 0.01971832 | 0.45860717 | 0.89054584 |
| Cdc42ep4 | 0.29565104 | 0.32477816 | 0.30506923 | 0.01970893 | 0.38171109 | 0.89054584 |
| Cpox | 0.34093816 | 0.36813365 | 0.34843448 | 0.01969917 | 0.39612155 | 0.89054584 |
| Nagk | 0.82782355 | 0.75605576 | 0.73641173 | 0.01964403 | 0.45054725 | 0.89054584 |
| Apbb1ip | 0.03831916 | 0.06061049 | 0.041003 | 0.01960749 | 0.01294394 | 0.14446595 |
| Rundc3a | 0.34808739 | 0.37476906 | 0.35518426 | 0.01958481 | 0.39863056 | 0.89054584 |
| Arf6 | 0.56786971 | 0.56801449 | 0.54845173 | 0.01956277 | 0.43400759 | 0.89054584 |
| Chrm2 | 0.03233742 | 0.05416228 | 0.03460365 | 0.01955862 | 0.00422774 | 0.05712373 |
| Rp2 | 0.30746285 | 0.33603447 | 0.31647934 | 0.01955513 | 0.38672154 | 0.89054584 |
| Sumf2 | 0.33903778 | 0.36618454 | 0.3466357 | 0.01954884 | 0.39637347 | 0.89054584 |
| Zbtb33 | 0.08529359 | 0.11075556 | 0.09121384 | 0.01954172 | 0.15910699 | 0.8062012 |
| Pigm | 0.12821912 | 0.15599169 | 0.13645906 | 0.01953263 | 0.25242546 | 0.89054584 |
| Haus2 | 0.18463927 | 0.21414436 | 0.19462057 | 0.01952379 | 0.32011689 | 0.89054584 |
| Crim1 | 0.07462779 | 0.0993391 | 0.07982202 | 0.01951708 | 0.12732116 | 0.71435843 |
| Smim6 | 0.124385 | 0.1519664 | 0.13246161 | 0.01950479 | 0.24634944 | 0.89054584 |
| Mfsd12 | 0.10322581 | 0.12979334 | 0.11030299 | 0.01949036 | 0.20518941 | 0.89054584 |
| Arhgef10l | 0.06034603 | 0.08402943 | 0.06455896 | 0.01947047 | 0.07999583 | 0.54094194 |
| Fam58b | 0.27748755 | 0.30685621 | 0.2873887 | 0.01946752 | 0.37615552 | 0.89054584 |
| Arfgap3 | 0.41570167 | 0.4371146 | 0.41765134 | 0.01946326 | 0.41405953 | 0.89054584 |
| Guca2b | 0.02013883 | 0.04100967 | 0.02155139 | 0.01945828 | 1.30E-05 | 0.00033401 |
| Efhc1 | 0.08695972 | 0.11245013 | 0.09299277 | 0.01945735 | 0.16482625 | 0.81977734 |

| | | | | | | |
|---|---|---|---|---|---|---|
| Dmtn | 0.04378021 | 0.06625788 | 0.04684456 | 0.01941332 | 0.02675645 | 0.24818618 |
| Gm12166 | 0.34280986 | 0.36959506 | 0.35020423 | 0.01939083 | 0.39821664 | 0.89054584 |
| Ucn3 | 0.03932872 | 0.0614591 | 0.04208297 | 0.01937613 | 0.01597236 | 0.1705208 |
| Spag1 | 0.14774969 | 0.17609851 | 0.15673478 | 0.01936373 | 0.28244722 | 0.89054584 |
| Rpl23a-ps3 | 2.6178418 | 1.14094413 | 1.12158542 | 0.0193587 | 0.46795403 | 0.89054584 |
| Map2 | 0.39350168 | 0.41676778 | 0.39741941 | 0.01934838 | 0.41028159 | 0.89054584 |
| Paqr4 | 0.06511707 | 0.08899487 | 0.06965886 | 0.019336 | 0.09796482 | 0.61511216 |
| Armc6 | 0.14060973 | 0.16867493 | 0.14933961 | 0.01933532 | 0.27318683 | 0.89054584 |
| Acp5 | 0.06541186 | 0.0893088 | 0.06997393 | 0.01933487 | 0.09898847 | 0.61772836 |
| Dmrtc1a | 0.18445351 | 0.21374436 | 0.19443117 | 0.01931319 | 0.32176078 | 0.89054584 |
| Ccar2 | 0.27726963 | 0.30648723 | 0.28717561 | 0.01931162 | 0.37702693 | 0.89054584 |
| Gpn2 | 0.14948797 | 0.17784038 | 0.15853215 | 0.01930823 | 0.28521101 | 0.89054584 |
| Tmem30a | 0.91374887 | 0.80718934 | 0.78792233 | 0.01926701 | 0.45464873 | 0.89054584 |
| Rasl11b | 0.07674112 | 0.1013378 | 0.08207971 | 0.01925809 | 0.13717233 | 0.7420427 |
| Mvk | 0.20310604 | 0.23260549 | 0.213375 | 0.01923048 | 0.33728351 | 0.89054584 |
| Tmem141 | 0.16661907 | 0.19540835 | 0.17618254 | 0.01922582 | 0.30558493 | 0.89054584 |
| H2-T23 | 0.15780741 | 0.18633517 | 0.16711834 | 0.01921683 | 0.29607304 | 0.89054584 |
| Ppcs | 0.36669203 | 0.39182893 | 0.37262119 | 0.01920774 | 0.40510412 | 0.89054584 |
| Eepd1 | 0.10796581 | 0.13448719 | 0.11528126 | 0.01920592 | 0.21882068 | 0.89054584 |
| Pi4k2b | 0.48453359 | 0.49775044 | 0.47857078 | 0.01917965 | 0.42594161 | 0.89054584 |
| Efhd2 | 0.3309411 | 0.35810898 | 0.33895175 | 0.01915722 | 0.39615242 | 0.89054584 |
| Erg | 0.04627365 | 0.06866719 | 0.04951146 | 0.01915573 | 0.03573176 | 0.30938746 |
| Hnrnpu | 1.78568039 | 1.11484564 | 1.09569442 | 0.01915122 | 0.46754926 | 0.89054584 |
| Cd2ap | 0.61595842 | 0.60587595 | 0.58673861 | 0.01913734 | 0.43960995 | 0.89054584 |
| Dlx1 | 0.01108753 | 0.03097921 | 0.0118655 | 0.01911371 | 3.07E-14 | 2.36E-12 |
| Hmgcl | 0.45630663 | 0.47303898 | 0.45392755 | 0.01911143 | 0.42224539 | 0.89054584 |
| Igsf10 | 0.0899977 | 0.11533408 | 0.09623601 | 0.01909807 | 0.17759522 | 0.85004143 |
| Serpinb6b | 0.03494804 | 0.05649253 | 0.03739663 | 0.01909589 | 0.0086798 | 0.10429449 |
| Ier5 | 0.33552729 | 0.36235635 | 0.34330802 | 0.01904833 | 0.39801061 | 0.89054584 |
| Cybb | 0.01673426 | 0.03695107 | 0.0179082 | 0.01904286 | 3.63E-07 | 1.21E-05 |

| | | | | | | |
|---|---|---|---|---|---|---|
| Mpeg1 | 0.01673426 | 0.03695107 | 0.0179082 | 0.01904286 | 3.63E-07 | 1.21E-05 |
| Med9 | 0.5610975 | 0.56197115 | 0.54293816 | 0.01903299 | 0.43513259 | 0.89054584 |
| Cpz | 0.06904279 | 0.09288532 | 0.07385441 | 0.01903091 | 0.11496827 | 0.67619067 |
| Zfp763 | 0.09324252 | 0.11872434 | 0.09969938 | 0.01902495 | 0.18699243 | 0.87419127 |
| Kbtbd4 | 0.21573867 | 0.24513901 | 0.22612086 | 0.01901815 | 0.34758615 | 0.89054584 |
| Atrnl1 | 0.12007466 | 0.14694481 | 0.12796121 | 0.0189836 | 0.2447281 | 0.89054584 |
| Tmem199 | 0.54728392 | 0.55058051 | 0.53159856 | 0.01898195 | 0.43393753 | 0.89054584 |
| Kirrel2 | 0.05865297 | 0.08170309 | 0.06274896 | 0.01895413 | 0.07967029 | 0.54059058 |
| Rab5a | 0.94461069 | 0.82408908 | 0.80513622 | 0.01895285 | 0.45633509 | 0.89054584 |
| Efna5 | 0.13915903 | 0.16677597 | 0.14783461 | 0.01894136 | 0.27527759 | 0.89054584 |
| Fam181b | 0.09019378 | 0.11538498 | 0.09644531 | 0.01893967 | 0.18011934 | 0.85660801 |
| Gga3 | 0.15961794 | 0.1879221 | 0.16898333 | 0.01893877 | 0.30078298 | 0.89054584 |
| Sbsn | 0.1481038 | 0.17603265 | 0.15710102 | 0.01893163 | 0.28725197 | 0.89054584 |
| Obfc1 | 0.21693868 | 0.24625875 | 0.22732799 | 0.01893077 | 0.34901767 | 0.89054584 |
| Rin3 | 0.07493837 | 0.09906199 | 0.08015383 | 0.01890816 | 0.13587601 | 0.73849629 |
| Asxl3 | 0.1292775 | 0.15643546 | 0.1375616 | 0.01887386 | 0.26133972 | 0.89054584 |
| Wwc2 | 0.16165971 | 0.18994236 | 0.17108492 | 0.01885744 | 0.3037935 | 0.89054584 |
| Dhx34 | 0.09090127 | 0.11605215 | 0.09720051 | 0.01885164 | 0.18310843 | 0.86510504 |
| Impa2 | 0.10963357 | 0.13587754 | 0.1170309 | 0.01884664 | 0.22654446 | 0.89054584 |
| Dgat2 | 0.1104523 | 0.13673476 | 0.11788945 | 0.01884531 | 0.22820884 | 0.89054584 |
| Arhgef33 | 0.02576919 | 0.04641398 | 0.02757606 | 0.01883792 | 0.00072969 | 0.01263923 |
| Eif5a | 3.1629344 | 1.0980855 | 1.07926522 | 0.01882027 | 0.46762443 | 0.89054584 |
| Triobp | 0.67628692 | 0.65137563 | 0.6325684 | 0.01880723 | 0.44491573 | 0.89054584 |
| Usp11 | 0.21182096 | 0.24097134 | 0.22217546 | 0.01879588 | 0.34673777 | 0.89054584 |
| Camk4 | 0.0994493 | 0.12510607 | 0.10632221 | 0.01878386 | 0.20522802 | 0.89054584 |
| Lmf2 | 0.2290705 | 0.25825445 | 0.23949551 | 0.01875894 | 0.35758542 | 0.89054584 |
| Gm14966 | 0.06636 | 0.08974416 | 0.07098729 | 0.01875687 | 0.10915671 | 0.65377127 |
| Fam69b | 0.19819469 | 0.22713958 | 0.20840146 | 0.01873812 | 0.3376439 | 0.89054584 |
| Pcmtd1 | 0.49806431 | 0.50893999 | 0.49021202 | 0.01872797 | 0.42936567 | 0.89054584 |
| Lsm11 | 0.06028131 | 0.08320886 | 0.06448976 | 0.01871909 | 0.08813559 | 0.57428752 |

| | | | | | | |
|---|---|---|---|---|---|---|
| Capn3 | 0.0255977 | 0.04609235 | 0.02739257 | 0.01869978 | 0.00073519 | 0.01267387 |
| Bmf | 0.09553791 | 0.12084744 | 0.10214895 | 0.01869849 | 0.19687818 | 0.89054584 |
| Lrrc8c | 0.07923993 | 0.10341433 | 0.0847489 | 0.01866544 | 0.15242128 | 0.7857996 |
| Ncaph | 0.06313367 | 0.08619821 | 0.06753886 | 0.01865935 | 0.0990206 | 0.61772836 |
| Zbtb7b | 0.12641927 | 0.15323279 | 0.13458315 | 0.01864964 | 0.25926759 | 0.89054584 |
| Rpl13a-ps1 | 0.15604773 | 0.18395244 | 0.16530447 | 0.01864796 | 0.29959205 | 0.89054584 |
| Acsl4 | 0.06864497 | 0.09207566 | 0.07342929 | 0.01864637 | 0.11838969 | 0.68821721 |
| Glt8d2 | 0.07429702 | 0.09811176 | 0.07946864 | 0.01864312 | 0.13720251 | 0.7420427 |
| Zfp146 | 0.19804143 | 0.22686862 | 0.2082461 | 0.01862252 | 0.33847512 | 0.89054584 |
| Foxk2 | 0.18704665 | 0.2156938 | 0.19707371 | 0.0186201 | 0.32989965 | 0.89054584 |
| Efnb3 | 0.13503731 | 0.16215562 | 0.14355418 | 0.01860144 | 0.27302306 | 0.89054584 |
| Qtrtd1 | 0.09057973 | 0.11545138 | 0.09685728 | 0.01859409 | 0.18555498 | 0.87060715 |
| Cx3cr1 | 0.01575264 | 0.03544923 | 0.01685777 | 0.01859146 | 1.39E-07 | 4.86E-06 |
| Gpd1l | 0.14497253 | 0.17244866 | 0.15386073 | 0.01858792 | 0.28676955 | 0.89054584 |
| Clca3a1 | 0.0440479 | 0.06569551 | 0.04713088 | 0.01856463 | 0.03324286 | 0.29227103 |
| Ndfip1 | 0.43826028 | 0.45646399 | 0.43792336 | 0.01854063 | 0.42181654 | 0.89054584 |
| Haus7 | 0.40337704 | 0.42497743 | 0.40645409 | 0.01852334 | 0.41592791 | 0.89054584 |
| Rbfox3 | 0.10567906 | 0.13139196 | 0.11288059 | 0.01851138 | 0.22242664 | 0.89054584 |
| Dlg5 | 0.13527304 | 0.16230148 | 0.14379917 | 0.01850231 | 0.2744346 | 0.89054584 |
| Cfp | 0.03916442 | 0.06040403 | 0.04190721 | 0.01849682 | 0.01987472 | 0.20051753 |
| Cd46 | 0.02448274 | 0.04469312 | 0.02619956 | 0.01849357 | 0.0005034 | 0.00908871 |
| Rbm43 | 0.07441472 | 0.09804406 | 0.07959439 | 0.01844967 | 0.14008794 | 0.75194185 |
| Tsen2 | 0.24338868 | 0.27220487 | 0.2537719 | 0.01843297 | 0.36752834 | 0.89054584 |
| Nsmce1 | 0.95264483 | 0.82792007 | 0.80951761 | 0.01840246 | 0.45782691 | 0.89054584 |
| Nr4a2 | 0.08777176 | 0.1122246 | 0.09385973 | 0.01836486 | 0.1809945 | 0.85950156 |
| Fer | 0.31228793 | 0.33947547 | 0.32112045 | 0.01835502 | 0.3950034 | 0.89054584 |
| Selt | 1.02754364 | 0.86696531 | 0.84865437 | 0.01831094 | 0.45996442 | 0.89054584 |
| Them6 | 0.07167481 | 0.09497702 | 0.07666694 | 0.01831007 | 0.1329236 | 0.72977796 |
| Il6st | 0.39598098 | 0.41797286 | 0.39969286 | 0.01828 | 0.41563337 | 0.89054584 |
| Tnfrsf12a | 0.30705297 | 0.33434507 | 0.31608458 | 0.01826049 | 0.39390668 | 0.89054584 |

| | | | | | | |
|---|---|---|---|---|---|---|
| Magt1 | 0.95076514 | 0.82675119 | 0.80849676 | 0.01825443 | 0.45811226 | 0.89054584 |
| Melk | 0.06919429 | 0.09226267 | 0.07401631 | 0.01824636 | 0.12537835 | 0.7081958 |
| Lynx1 | 0.08183824 | 0.10575288 | 0.08752399 | 0.01822889 | 0.16594057 | 0.82390627 |
| Stard4 | 0.18737695 | 0.21563166 | 0.1974101 | 0.01822157 | 0.33358517 | 0.89054584 |
| Pbxip1 | 0.47509182 | 0.4885988 | 0.47038096 | 0.01821784 | 0.42840334 | 0.89054584 |
| Prpsap2 | 0.34043598 | 0.3661713 | 0.34795933 | 0.01821198 | 0.403675 | 0.89054584 |
| Smtn | 0.11539051 | 0.14126868 | 0.1230626 | 0.01820609 | 0.24533248 | 0.89054584 |
| Gm43194 | 0.12175526 | 0.14791521 | 0.12971674 | 0.01819847 | 0.25667797 | 0.89054584 |
| Usp51 | 0.22096409 | 0.24957011 | 0.23137258 | 0.01819753 | 0.35702317 | 0.89054584 |
| Cspg4 | 0.05179812 | 0.0736069 | 0.05541956 | 0.01818734 | 0.06313844 | 0.4620803 |
| Snx24 | 0.33220504 | 0.35832441 | 0.34015344 | 0.01817097 | 0.4017267 | 0.89054584 |
| Comtd1 | 0.18793311 | 0.21613755 | 0.19797639 | 0.01816116 | 0.33455014 | 0.89054584 |
| Magi1 | 0.11577488 | 0.14162536 | 0.12346488 | 0.01816049 | 0.24658203 | 0.89054584 |
| Lpin3 | 0.02639455 | 0.0463966 | 0.02824518 | 0.01815142 | 0.00137668 | 0.02176563 |
| Cd40 | 0.07675278 | 0.10024152 | 0.08209217 | 0.01814936 | 0.1514997 | 0.78379829 |
| Tec | 0.03110202 | 0.05142958 | 0.03328191 | 0.01814767 | 0.0055365 | 0.07134953 |
| Crispld2 | 0.06010541 | 0.08243915 | 0.06430172 | 0.01813743 | 0.09440019 | 0.60001264 |
| Ythdc2 | 0.18916253 | 0.21735886 | 0.19922778 | 0.01813108 | 0.33578465 | 0.89054584 |
| Myt1 | 0.35350531 | 0.37839939 | 0.36028118 | 0.01811821 | 0.40737843 | 0.89054584 |
| Clcn3 | 0.68139447 | 0.65444516 | 0.6363334 | 0.01811177 | 0.44725247 | 0.89054584 |
| Krr1 | 0.34789154 | 0.37309949 | 0.35499971 | 0.01809977 | 0.40611987 | 0.89054584 |
| Itgb1bp1 | 0.54697761 | 0.54943853 | 0.5313457 | 0.01809283 | 0.43697571 | 0.89054584 |
| Gm10334 | 0.02971709 | 0.04988248 | 0.03180016 | 0.01808232 | 0.00403432 | 0.05507498 |
| Prmt7 | 0.47551608 | 0.48880767 | 0.47075015 | 0.01805752 | 0.42908172 | 0.89054584 |
| Chsy1 | 0.10091066 | 0.12591076 | 0.1078685 | 0.01804227 | 0.21791259 | 0.89054584 |
| Arhgef16 | 0.19412785 | 0.22231743 | 0.20427523 | 0.01804219 | 0.34035609 | 0.89054584 |
| Itpa | 0.87980663 | 0.78625172 | 0.76821412 | 0.0180376 | 0.45644598 | 0.89054584 |
| Fam160a2 | 0.23110593 | 0.25955432 | 0.24153041 | 0.01802391 | 0.36404487 | 0.89054584 |
| Gm13524 | 0.02197076 | 0.04152639 | 0.02351166 | 0.01801473 | 0.0001787 | 0.00354409 |
| Ngf | 0.04943682 | 0.07088865 | 0.05289441 | 0.01799424 | 0.05649071 | 0.42812458 |

| | | | | | | |
|---|---|---|---|---|---|---|
| Tlx2 | 0.0137205 | 0.03266758 | 0.01468313 | 0.01798445 | 5.77E-09 | 2.41E-07 |
| Ttc23 | 0.12341667 | 0.14942813 | 0.13145119 | 0.01797694 | 0.2620151 | 0.89054584 |
| Commd9 | 0.66875869 | 0.64494123 | 0.62698607 | 0.01795516 | 0.44693089 | 0.89054584 |
| Nanos1 | 0.13295882 | 0.15933195 | 0.14139316 | 0.0179388 | 0.27723075 | 0.89054584 |
| C130074G19 | 0.04775919 | 0.06903535 | 0.05110026 | 0.01793509 | 0.05100481 | 0.39853675 |
| Dtnbos | 0.18047927 | 0.20830869 | 0.19037574 | 0.01793295 | 0.33038098 | 0.89054584 |
| Gpr22 | 0.01328976 | 0.03215233 | 0.01422219 | 0.01793015 | 2.13E-09 | 9.63E-08 |
| Kmt2d | 0.31669616 | 0.34326496 | 0.32535072 | 0.01791424 | 0.3987718 | 0.89054584 |
| Erich5 | 0.07965962 | 0.1031081 | 0.08519717 | 0.01791093 | 0.16367997 | 0.81785018 |
| Rubcn | 0.11396927 | 0.1394844 | 0.12157466 | 0.01790974 | 0.24625233 | 0.89054584 |
| Sh2b3 | 0.19197001 | 0.21995237 | 0.20208299 | 0.01786938 | 0.34018025 | 0.89054584 |
| Chrna4 | 0.02573745 | 0.04541118 | 0.0275421 | 0.01786908 | 0.00125273 | 0.02014245 |
| Sp2 | 0.11894818 | 0.14464965 | 0.1267839 | 0.01786575 | 0.25574659 | 0.89054584 |
| Gas2 | 0.13229663 | 0.15856784 | 0.14070433 | 0.01786351 | 0.27709645 | 0.89054584 |
| Thsd7a | 0.02449923 | 0.04405599 | 0.0262172 | 0.01783879 | 0.00076206 | 0.01307501 |
| P2rx1 | 0.03671213 | 0.05711782 | 0.03928386 | 0.01783396 | 0.01721281 | 0.18122826 |
| Rcan2 | 0.25023119 | 0.27839341 | 0.26056111 | 0.0178323 | 0.3749205 | 0.89054584 |
| 9330182L06F | 0.0828959 | 0.1064522 | 0.08865351 | 0.01779869 | 0.17480108 | 0.84226179 |
| Zfand5 | 1.02579463 | 0.86552739 | 0.84773742 | 0.01778996 | 0.46105788 | 0.89054584 |
| Men1 | 0.32365238 | 0.3497815 | 0.3320057 | 0.0177758 | 0.40150918 | 0.89054584 |
| Smc6 | 1.3882662 | 1.01949519 | 1.00171984 | 0.01777535 | 0.46705612 | 0.89054584 |
| Adgra3 | 0.1724739 | 0.1999373 | 0.18218764 | 0.01774966 | 0.32495109 | 0.89054584 |
| Ovol2 | 0.2501729 | 0.2782344 | 0.26050337 | 0.01773103 | 0.37558065 | 0.89054584 |
| Zfp9 | 0.25873588 | 0.28667983 | 0.26896912 | 0.01771072 | 0.37950705 | 0.89054584 |
| Itgb5 | 0.12908828 | 0.15505832 | 0.13736452 | 0.0176938 | 0.27421519 | 0.89054584 |
| Cdadc1 | 0.21594464 | 0.24401988 | 0.2263281 | 0.01769178 | 0.35786048 | 0.89054584 |
| Camsap2 | 0.21708983 | 0.24516231 | 0.22748 | 0.01768231 | 0.35862166 | 0.89054584 |
| Chka | 0.9323952 | 0.81607051 | 0.79840416 | 0.01766635 | 0.45894628 | 0.89054584 |
| Dmkn | 0.03029029 | 0.05007448 | 0.03241344 | 0.01766104 | 0.00556633 | 0.07167018 |
| Tnnt2 | 0.02379586 | 0.04310828 | 0.02546459 | 0.01764369 | 0.00062319 | 0.01103175 |

| | | | | | | |
|---|---|---|---|---|---|---|
| Atp7a | 0.2608317 | 0.28867103 | 0.27103621 | 0.01763482 | 0.38089484 | 0.89054584 |
| Ints3 | 0.2900349 | 0.31725223 | 0.29961946 | 0.01763277 | 0.39197265 | 0.89054584 |
| Kcnab1 | 0.04785928 | 0.06882821 | 0.0512073 | 0.01762091 | 0.0544469 | 0.41633498 |
| Arhgap23 | 0.08425207 | 0.10772131 | 0.09010172 | 0.01761959 | 0.18113024 | 0.8598645 |
| Celf5 | 0.04629258 | 0.06714586 | 0.04953171 | 0.01761415 | 0.04878171 | 0.38689129 |
| Runx2 | 0.01367609 | 0.03224387 | 0.01463561 | 0.01760826 | 1.04E-08 | 4.26E-07 |
| Pcdhga6 | 0.01560824 | 0.03430298 | 0.01670325 | 0.01759973 | 4.58E-07 | 1.51E-05 |
| Prkci | 0.1230965 | 0.14871531 | 0.13111703 | 0.01759828 | 0.26588241 | 0.89054584 |
| Nprl2 | 0.35952752 | 0.38351125 | 0.36592797 | 0.01758329 | 0.41143008 | 0.89054584 |
| Slc39a8 | 0.05154299 | 0.07272712 | 0.05514674 | 0.01758038 | 0.06874061 | 0.49190594 |
| Sh3gl1 | 0.65620861 | 0.63513252 | 0.6175919 | 0.01754062 | 0.44736498 | 0.89054584 |
| Alg3 | 0.2109205 | 0.23879228 | 0.22126768 | 0.0175246 | 0.35606675 | 0.89054584 |
| Gm11266 | 0.05511618 | 0.07648167 | 0.05896754 | 0.01751413 | 0.08321506 | 0.55545668 |
| Slc15a3 | 0.02333195 | 0.04247066 | 0.02496819 | 0.01750247 | 0.00054563 | 0.0097782 |
| Hivep1 | 0.14779699 | 0.17428243 | 0.1567837 | 0.01749873 | 0.30153613 | 0.89054584 |
| Zfp809 | 0.10117949 | 0.12564264 | 0.10815128 | 0.01749136 | 0.22557696 | 0.89054584 |
| Sar1a | 1.34678986 | 1.00505875 | 0.98758065 | 0.0174781 | 0.46714306 | 0.89054584 |
| Nrsn1 | 0.04018297 | 0.0604661 | 0.04299677 | 0.01746933 | 0.0291856 | 0.26412785 |
| A930033H14 | 0.06417177 | 0.08611164 | 0.06864848 | 0.01746316 | 0.11797589 | 0.68746563 |
| Igsf23 | 0.01424523 | 0.03266707 | 0.01524467 | 0.0174224 | 5.06E-08 | 1.87E-06 |
| Sec31a | 0.79717529 | 0.73413096 | 0.71674008 | 0.01739088 | 0.45499787 | 0.89054584 |
| Ccdc90b | 0.57030009 | 0.56780488 | 0.55042302 | 0.01738185 | 0.44151665 | 0.89054584 |
| Zgpat | 0.19842597 | 0.22601747 | 0.20863591 | 0.01738156 | 0.34895673 | 0.89054584 |
| Ebf3 | 0.06105959 | 0.08269389 | 0.06532176 | 0.01737213 | 0.10766693 | 0.64850649 |
| Dnaja1 | 3.18713705 | 1.09417193 | 1.0768061 | 0.01736583 | 0.47005344 | 0.89054584 |
| Ccdc138 | 0.02583285 | 0.04498205 | 0.02764418 | 0.01733787 | 0.00173899 | 0.02667926 |
| Kcne4 | 0.02981297 | 0.04922532 | 0.03190274 | 0.01732258 | 0.00570775 | 0.07297259 |
| Parm1 | 0.43687549 | 0.4540035 | 0.43668757 | 0.01731593 | 0.42671626 | 0.89054584 |
| Prkar2b | 0.41108363 | 0.43077212 | 0.41346588 | 0.01730624 | 0.42269374 | 0.89054584 |
| Neurl2 | 0.03714323 | 0.05704584 | 0.03974504 | 0.01730079 | 0.02127957 | 0.20972887 |

| | | | | | | |
|---|---|---|---|---|---|---|
| B3gnt5 | 0.03753673 | 0.05745381 | 0.040166 | 0.01728781 | 0.02246812 | 0.2193554 |
| Negr1 | 0.02138613 | 0.04016935 | 0.02288609 | 0.01728327 | 0.0002171 | 0.00424771 |
| Sema3a | 0.05146197 | 0.07231017 | 0.0550601 | 0.01725007 | 0.07219664 | 0.50860393 |
| Agfg2 | 0.19050549 | 0.21784081 | 0.20059399 | 0.01724683 | 0.34436832 | 0.89054584 |
| Gls2 | 0.0155199 | 0.0338522 | 0.01660871 | 0.01724348 | 6.59E-07 | 2.12E-05 |
| Pkp2 | 0.14273801 | 0.16877999 | 0.15154605 | 0.01723393 | 0.29811896 | 0.89054584 |
| 2410021H03I | 0.05752296 | 0.07876257 | 0.06154085 | 0.01722172 | 0.09615633 | 0.60929827 |
| Tns2 | 0.10553599 | 0.12995194 | 0.11273033 | 0.01722161 | 0.23831347 | 0.89054584 |
| Mccc1 | 0.19045525 | 0.21774725 | 0.20054289 | 0.01720436 | 0.34469407 | 0.89054584 |
| Ctnnb1 | 1.77211298 | 1.11080622 | 1.09366625 | 0.01713997 | 0.47089707 | 0.89054584 |
| Dlgap3 | 0.04530944 | 0.06561561 | 0.04848019 | 0.01713541 | 0.0498087 | 0.39214348 |
| Kif1a | 0.21315985 | 0.240633 | 0.22352458 | 0.01710842 | 0.36069858 | 0.89054584 |
| Tmprss2 | 0.03363666 | 0.0530711 | 0.03599367 | 0.01707743 | 0.01353647 | 0.14890122 |
| Naaa | 0.14464249 | 0.17059071 | 0.15351898 | 0.01707174 | 0.30219872 | 0.89054584 |
| Tnfrsf1b | 0.01428732 | 0.03235999 | 0.0152897 | 0.01707028 | 9.88E-08 | 3.55E-06 |
| Dll4 | 0.13703835 | 0.16269735 | 0.1456331 | 0.01706425 | 0.29256677 | 0.89054584 |
| Gm19412 | 0.13842783 | 0.16413891 | 0.14707574 | 0.01706317 | 0.29442177 | 0.89054584 |
| Tlr7 | 0.01426362 | 0.03231892 | 0.01526435 | 0.01705458 | 9.68E-08 | 3.48E-06 |
| Insrr | 0.18533781 | 0.21238475 | 0.19533264 | 0.01705211 | 0.34210851 | 0.89054584 |
| Arrb1 | 0.07124139 | 0.09323039 | 0.07620382 | 0.01702656 | 0.14894518 | 0.7773244 |
| Dgat1 | 0.20712545 | 0.23445778 | 0.21743785 | 0.01701994 | 0.35767573 | 0.89054584 |
| Myc | 0.07705347 | 0.09943269 | 0.08241337 | 0.01701931 | 0.1679934 | 0.82983589 |
| Smim10l1 | 0.97826699 | 0.84020978 | 0.82320153 | 0.01700826 | 0.46165748 | 0.89054584 |
| Syde2 | 0.07148615 | 0.09346469 | 0.07646535 | 0.01699934 | 0.15015966 | 0.77912292 |
| Cdca7l | 0.09993764 | 0.12384185 | 0.10684317 | 0.01699867 | 0.2292757 | 0.89054584 |
| Fbxo32 | 0.15002014 | 0.17607044 | 0.15908216 | 0.01698828 | 0.30940932 | 0.89054584 |
| Doc2b | 0.04794616 | 0.06826383 | 0.05130021 | 0.01696361 | 0.06170857 | 0.45559551 |
| Rnf144b | 0.02202043 | 0.04052719 | 0.02356482 | 0.01696237 | 0.00039886 | 0.00734765 |
| Alg6 | 0.16106964 | 0.18741777 | 0.17047774 | 0.01694003 | 0.32170023 | 0.89054584 |
| Blnk | 0.09004015 | 0.11322125 | 0.09628132 | 0.01693993 | 0.2061925 | 0.89054584 |

| | | | | | | |
|---|---|---|---|---|---|---|
| Tlx1 | 0.02791909 | 0.04680683 | 0.0298764 | 0.01693043 | 0.00414359 | 0.05614367 |
| Gm13199 | 0.01738692 | 0.03553193 | 0.01860663 | 0.01692531 | 1.13E-05 | 0.00029568 |
| Urah | 0.10431374 | 0.12836106 | 0.11144634 | 0.01691472 | 0.23974945 | 0.89054584 |
| Tjp3 | 0.06947922 | 0.09123422 | 0.0743208 | 0.01691342 | 0.14451538 | 0.76718564 |
| Rilpl1 | 0.23892379 | 0.26623384 | 0.24933004 | 0.0169038 | 0.37605372 | 0.89054584 |
| Fosl2 | 0.09109438 | 0.11429486 | 0.09740663 | 0.01688823 | 0.20961407 | 0.89054584 |
| Gpr160 | 0.03230101 | 0.05143513 | 0.0345647 | 0.01687043 | 0.01148456 | 0.13125935 |
| Pyurf | 0.46199359 | 0.47580097 | 0.45893066 | 0.0168703 | 0.43200857 | 0.89054584 |
| Sez6l | 0.09239084 | 0.11565842 | 0.0987904 | 0.01686802 | 0.21316332 | 0.89054584 |
| Smpdl3a | 0.32805894 | 0.35307353 | 0.33620842 | 0.01686511 | 0.40760705 | 0.89054584 |
| Tcf15 | 0.03586048 | 0.05522215 | 0.03837277 | 0.01684938 | 0.02039155 | 0.20391333 |
| Dscr3 | 0.32237035 | 0.34761884 | 0.33078108 | 0.01683776 | 0.40626856 | 0.89054584 |
| Plcb4 | 0.27193245 | 0.29874554 | 0.28194968 | 0.01679586 | 0.39068466 | 0.89054584 |
| Pusl1 | 0.21302928 | 0.24018123 | 0.22339305 | 0.01678817 | 0.36312309 | 0.89054584 |
| Kptn | 0.24452151 | 0.27168142 | 0.25489741 | 0.01678401 | 0.37950798 | 0.89054584 |
| Rnaset2b | 0.35631801 | 0.37968179 | 0.36292099 | 0.0167608 | 0.41481968 | 0.89054584 |
| Rraga | 1.44894581 | 1.03768779 | 1.02095566 | 0.01673213 | 0.46956874 | 0.89054584 |
| Cyb5r4 | 0.71763265 | 0.67925166 | 0.6625241 | 0.01672756 | 0.45318042 | 0.89054584 |
| Gtf2f2 | 0.52372057 | 0.52868499 | 0.51197546 | 0.01670953 | 0.43957157 | 0.89054584 |
| Hfe | 0.07137611 | 0.0930527 | 0.07634777 | 0.01670492 | 0.15401172 | 0.79121355 |
| Tcf24 | 0.01346604 | 0.03110178 | 0.01441083 | 0.01669094 | 3.41E-08 | 1.30E-06 |
| Sh3gl3 | 0.05662431 | 0.07726613 | 0.06058004 | 0.01668609 | 0.09970176 | 0.62085259 |
| Snhg12 | 0.3423174 | 0.36641203 | 0.34973878 | 0.01667325 | 0.412115 | 0.89054584 |
| Msr1 | 0.0126527 | 0.03019076 | 0.01354045 | 0.01665031 | 5.05E-09 | 2.15E-07 |
| Pou2f2 | 0.0226279 | 0.040862 | 0.02421483 | 0.01664717 | 0.00068021 | 0.01188147 |
| Fcgr1 | 0.01292358 | 0.03045714 | 0.01383033 | 0.01662681 | 1.07E-08 | 4.33E-07 |
| Kctd1 | 0.14732349 | 0.17291259 | 0.15629391 | 0.01661868 | 0.31016468 | 0.89054584 |
| Mcmdc2 | 0.02734542 | 0.04586782 | 0.0292626 | 0.01660522 | 0.00409977 | 0.05565396 |
| Fbxo10 | 0.05973009 | 0.08050498 | 0.06390049 | 0.01660449 | 0.11302021 | 0.66962283 |
| Gm29666 | 0.02673207 | 0.04518847 | 0.02860633 | 0.01658214 | 0.00346037 | 0.04842517 |

| | | | | | | |
|---|---|---|---|---|---|---|
| Rab7b | 0.07104511 | 0.09256327 | 0.07599409 | 0.01656919 | 0.15486211 | 0.79391473 |
| Klhl5 | 0.04412256 | 0.06377659 | 0.04721074 | 0.01656585 | 0.05104723 | 0.39853675 |
| Bnip2 | 0.595601 | 0.5872804 | 0.57071728 | 0.01656312 | 0.44622299 | 0.89054584 |
| Amot | 0.05673082 | 0.07724691 | 0.06069393 | 0.01655299 | 0.10193053 | 0.62777773 |
| Elavl3 | 0.0452072 | 0.06483886 | 0.04837084 | 0.01646802 | 0.05635238 | 0.42729956 |
| Hist2h2aa1 | 0.06425363 | 0.08518151 | 0.06873598 | 0.01644553 | 0.13249331 | 0.72954347 |
| Itga2 | 0.02877304 | 0.04722947 | 0.03079009 | 0.01643938 | 0.00643236 | 0.08053629 |
| Meox2 | 0.03650438 | 0.05548335 | 0.03906161 | 0.01642174 | 0.02507704 | 0.23638944 |
| Sugp1 | 0.18316036 | 0.2095241 | 0.19311233 | 0.01641177 | 0.34607354 | 0.89054584 |
| Zfp57 | 0.11350206 | 0.13749286 | 0.12108537 | 0.01640749 | 0.26392125 | 0.89054584 |
| Abca8a | 0.05174173 | 0.07176039 | 0.05535927 | 0.01640112 | 0.08374867 | 0.55824743 |
| Itga9 | 0.06536199 | 0.08630495 | 0.06992063 | 0.01638432 | 0.13747865 | 0.74288686 |
| Nrbf2 | 0.46757434 | 0.48019255 | 0.46382124 | 0.01637132 | 0.43469048 | 0.89054584 |
| Ptpn13 | 0.18472282 | 0.21100959 | 0.19470574 | 0.01630385 | 0.34822375 | 0.89054584 |
| Aifm3 | 0.05106142 | 0.07090763 | 0.05463176 | 0.01627587 | 0.0825701 | 0.55242329 |
| Itga4 | 0.07427068 | 0.0956964 | 0.0794405 | 0.01625591 | 0.17020413 | 0.83373276 |
| Krtcap3 | 0.36455144 | 0.38686133 | 0.37062412 | 0.0162372 | 0.41913368 | 0.89054584 |
| Ufd1l | 0.54457831 | 0.54559847 | 0.52936355 | 0.01623491 | 0.44319095 | 0.89054584 |
| Atad5 | 0.11084645 | 0.13450674 | 0.11830269 | 0.01620406 | 0.2616911 | 0.89054584 |
| Ube2v1 | 0.11220181 | 0.13592526 | 0.11972322 | 0.01620204 | 0.26418689 | 0.89054584 |
| Clec2i | 0.02068299 | 0.03833523 | 0.02213367 | 0.01620156 | 0.00032448 | 0.00609362 |
| 5033428I22R | 0.05879935 | 0.0791048 | 0.06290546 | 0.01619934 | 0.11511532 | 0.67639855 |
| Pomc | 0.16967668 | 0.19551829 | 0.17932039 | 0.0161979 | 0.33693622 | 0.89054584 |
| Sorbs3 | 0.10616166 | 0.12957731 | 0.11338739 | 0.01618992 | 0.25295395 | 0.89054584 |
| Dgkd | 0.27290857 | 0.29908935 | 0.28290649 | 0.01618287 | 0.39492647 | 0.89054584 |
| Gtf3c4 | 0.21960024 | 0.24617861 | 0.23000305 | 0.01617557 | 0.37158732 | 0.89054584 |
| Spns3 | 0.02044142 | 0.0380494 | 0.02187518 | 0.01617421 | 0.00028579 | 0.00543353 |
| Adprhl2 | 0.43134958 | 0.44791314 | 0.43174491 | 0.01616822 | 0.4307478 | 0.89054584 |
| Hpn | 0.19293699 | 0.21923322 | 0.20306563 | 0.01616758 | 0.35534357 | 0.89054584 |
| Gtf2f1 | 0.47883685 | 0.48979872 | 0.47363599 | 0.01616273 | 0.43684018 | 0.89054584 |

| | | | | | | |
|---|---|---|---|---|---|---|
| Plekhh1 | 0.07145851 | 0.09258589 | 0.07643581 | 0.01615007 | 0.16246321 | 0.81501438 |
| Ehbp1l1 | 0.10104435 | 0.12413229 | 0.10800914 | 0.01612315 | 0.24338129 | 0.89054584 |
| Ces2g | 0.02250584 | 0.04020097 | 0.02408422 | 0.01611675 | 0.00091139 | 0.01527557 |
| Snx13 | 0.09133512 | 0.11377751 | 0.09766359 | 0.01611392 | 0.22103635 | 0.89054584 |
| Zranb2 | 1.11301508 | 0.90750135 | 0.89138944 | 0.01611191 | 0.46644449 | 0.89054584 |
| Hgf | 0.0533111 | 0.07314626 | 0.05703743 | 0.01610883 | 0.09411981 | 0.59930619 |
| Pbdc1 | 0.91760019 | 0.80618936 | 0.79010575 | 0.0160836 | 0.46222149 | 0.89054584 |
| Telo2 | 0.22625072 | 0.25275638 | 0.23667336 | 0.01608303 | 0.37577502 | 0.89054584 |
| Eif2b4 | 0.37884973 | 0.39999796 | 0.38391582 | 0.01608214 | 0.42263308 | 0.89054584 |
| Shisa2 | 0.08500022 | 0.10697466 | 0.09090059 | 0.01607407 | 0.20501333 | 0.89054584 |
| Nol9 | 0.11966643 | 0.14360306 | 0.12753461 | 0.01606845 | 0.27860408 | 0.89054584 |
| Chmp4b | 1.09621954 | 0.89939024 | 0.88334133 | 0.01604891 | 0.46627159 | 0.89054584 |
| Hbegf | 0.0822953 | 0.10405333 | 0.08801212 | 0.01604121 | 0.19790032 | 0.89054584 |
| Slc52a3 | 0.02204261 | 0.03962915 | 0.02358855 | 0.0160406 | 0.00076705 | 0.01314494 |
| Osgep | 0.6867141 | 0.6562578 | 0.64023596 | 0.01602184 | 0.45359268 | 0.89054584 |
| Tlcd2 | 0.15055183 | 0.17563797 | 0.15963158 | 0.01600639 | 0.32019421 | 0.89054584 |
| Prelp | 0.08660744 | 0.10862286 | 0.09261666 | 0.0160062 | 0.21036143 | 0.89054584 |
| 2210019I11R | 0.09029899 | 0.11255784 | 0.09655762 | 0.01600021 | 0.22005238 | 0.89054584 |
| Arhgap9 | 0.03094577 | 0.04910179 | 0.03311475 | 0.01598705 | 0.01224891 | 0.1379834 |
| Spice1 | 0.18188079 | 0.20777357 | 0.19180665 | 0.01596692 | 0.34906981 | 0.89054584 |
| Piezo2 | 0.09297543 | 0.1153664 | 0.09941433 | 0.01595207 | 0.22735803 | 0.89054584 |
| Col2a1 | 0.0248713 | 0.0425619 | 0.02661531 | 0.01594659 | 0.00262396 | 0.03819499 |
| Car5b | 0.02568411 | 0.043428 | 0.02748502 | 0.01594298 | 0.00344134 | 0.0482684 |
| Ezh1 | 0.25244978 | 0.27869951 | 0.26275771 | 0.01594179 | 0.3887176 | 0.89054584 |
| Fitm2 | 0.28854939 | 0.31411538 | 0.29817533 | 0.01594006 | 0.40165687 | 0.89054584 |
| Mppe1 | 0.01854349 | 0.03577812 | 0.01984425 | 0.01593386 | 9.17E-05 | 0.00193214 |
| Gm14424 | 0.01582633 | 0.03283232 | 0.01693663 | 0.01589569 | 6.14E-06 | 0.00016856 |
| Tcirg1 | 0.10865481 | 0.1318876 | 0.11600421 | 0.01588339 | 0.26176751 | 0.89054584 |
| Sco2 | 0.16847367 | 0.193967 | 0.17808627 | 0.01588073 | 0.33890328 | 0.89054584 |
| Bcas3 | 0.08597593 | 0.10780267 | 0.0919424 | 0.01586028 | 0.21079143 | 0.89054584 |

| | | | | | | |
|---|---|---|---|---|---|---|
| Zeb2os | 0.06890036 | 0.08952793 | 0.07370221 | 0.01582571 | 0.15856051 | 0.80483939 |
| Asrgl1 | 0.20216139 | 0.22824053 | 0.21241921 | 0.01582132 | 0.36429366 | 0.89054584 |
| Clmn | 0.13970561 | 0.16421716 | 0.14840176 | 0.01581541 | 0.30976708 | 0.89054584 |
| Pde1a | 0.05569 | 0.07537954 | 0.05958109 | 0.01579845 | 0.10834849 | 0.65152897 |
| Ildr1 | 0.39039028 | 0.41035217 | 0.3945614 | 0.01579077 | 0.42604366 | 0.89054584 |
| Tnfaip6 | 0.04303996 | 0.06184194 | 0.04605278 | 0.01578916 | 0.05509788 | 0.4206472 |
| Tor3a | 0.09647072 | 0.11892875 | 0.10314431 | 0.01578444 | 0.23793217 | 0.89054584 |
| BC030336 | 0.10601133 | 0.1289961 | 0.11322953 | 0.01576657 | 0.25825612 | 0.89054584 |
| Ciapin1 | 0.98906788 | 0.84459874 | 0.82885266 | 0.01574608 | 0.46473648 | 0.89054584 |
| Mpzl2 | 0.0528852 | 0.07230222 | 0.05658201 | 0.01572021 | 0.09776427 | 0.61452179 |
| Pnpla2 | 0.28911605 | 0.31443792 | 0.29872633 | 0.01571158 | 0.40321346 | 0.89054584 |
| Kctd14 | 0.03281055 | 0.05081778 | 0.03510984 | 0.01570794 | 0.01856267 | 0.19127335 |
| Letm2 | 0.05026327 | 0.06946464 | 0.05377823 | 0.01568641 | 0.08708079 | 0.57045578 |
| Slc16a1 | 0.09560981 | 0.11789957 | 0.10222568 | 0.01567389 | 0.23750896 | 0.89054584 |
| Mmd2 | 0.18640045 | 0.21208176 | 0.19641547 | 0.01566629 | 0.35509468 | 0.89054584 |
| Atg16l2 | 0.10941054 | 0.13245854 | 0.11679698 | 0.01566157 | 0.26607487 | 0.89054584 |
| B4galt2 | 0.09897139 | 0.12147285 | 0.10581236 | 0.01566049 | 0.24524339 | 0.89054584 |
| Irf5 | 0.02382312 | 0.04114638 | 0.02549375 | 0.01565263 | 0.00211492 | 0.03164306 |
| Cdon | 0.17696229 | 0.20242725 | 0.18678162 | 0.01564563 | 0.34817476 | 0.89054584 |
| 0610009L18R | 0.20813481 | 0.23409427 | 0.21845708 | 0.0156372 | 0.36938304 | 0.89054584 |
| 0610030E20I | 0.31836782 | 0.34258647 | 0.32695227 | 0.0156342 | 0.4118529 | 0.89054584 |
| Tmem164 | 0.32796399 | 0.35173962 | 0.33611797 | 0.01562165 | 0.4142865 | 0.89054584 |
| Lix1 | 0.02317986 | 0.0404135 | 0.02480545 | 0.01560806 | 0.00168668 | 0.0259316 |
| Pcf11 | 0.30082688 | 0.32566994 | 0.31007773 | 0.01559221 | 0.40738611 | 0.89054584 |
| 1500011B03I | 0.89770723 | 0.79429671 | 0.77871136 | 0.01558535 | 0.4628543 | 0.89054584 |
| Arid3a | 0.25709916 | 0.28293688 | 0.26735347 | 0.01558341 | 0.39298062 | 0.89054584 |
| Prep | 0.22615086 | 0.25214242 | 0.23657335 | 0.01556907 | 0.37957071 | 0.89054584 |
| Mrps9 | 0.46714047 | 0.47900499 | 0.46344171 | 0.01556328 | 0.43783665 | 0.89054584 |
| Stxbp3 | 0.43524712 | 0.45078717 | 0.43523295 | 0.01555422 | 0.43388181 | 0.89054584 |
| Aldh5a1 | 0.2456561 | 0.27156253 | 0.25602408 | 0.01553845 | 0.38868139 | 0.89054584 |

| | | | | | | |
|---|---|---|---|---|---|---|
| Ubc | 3.19826585 | 1.09111757 | 1.07557912 | 0.01553844 | 0.47316914 | 0.89054584 |
| Fbxl20 | 0.18932917 | 0.21492772 | 0.19939735 | 0.01553037 | 0.35835146 | 0.89054584 |
| Cwc25 | 0.39463141 | 0.41398266 | 0.39845577 | 0.01552689 | 0.4279687 | 0.89054584 |
| Ptf1a | 0.01675715 | 0.03345425 | 0.0179327 | 0.01552155 | 2.76E-05 | 0.00066337 |
| Scrn1 | 0.20913374 | 0.23496763 | 0.21946534 | 0.01550229 | 0.37104313 | 0.89054584 |
| Top1 | 1.73777836 | 1.10365394 | 1.0881574 | 0.01549654 | 0.47355023 | 0.89054584 |
| Aplf | 0.114197 | 0.13729841 | 0.12181313 | 0.01548528 | 0.27683839 | 0.89054584 |
| Ripk3 | 0.08587562 | 0.10730236 | 0.0918353 | 0.01546707 | 0.21632434 | 0.89054584 |
| Ntm | 0.04730768 | 0.06607034 | 0.05061737 | 0.01545297 | 0.07746566 | 0.53076426 |
| Abcb4 | 0.0137865 | 0.03020429 | 0.01475376 | 0.01545053 | 5.33E-07 | 1.74E-05 |
| Mterf3 | 0.42886105 | 0.44495687 | 0.42951317 | 0.0154437 | 0.43348113 | 0.89054584 |
| St7 | 0.09221894 | 0.11405029 | 0.09860693 | 0.01544336 | 0.23279853 | 0.89054584 |
| Mex3b | 0.14223881 | 0.16645271 | 0.15102869 | 0.01542402 | 0.31710803 | 0.89054584 |
| Gcnt2 | 0.25893482 | 0.28458736 | 0.26916542 | 0.01542194 | 0.39475924 | 0.89054584 |
| Dusp22 | 0.17128697 | 0.19637751 | 0.18097138 | 0.01540613 | 0.34582512 | 0.89054584 |
| Rsad1 | 0.0470728 | 0.06576693 | 0.05036616 | 0.01540077 | 0.07713748 | 0.52951666 |
| Nsun4 | 0.10844234 | 0.13117259 | 0.11578129 | 0.0153913 | 0.26784852 | 0.89054584 |
| Slc2a8 | 0.14709714 | 0.1714499 | 0.15605974 | 0.01539016 | 0.32295503 | 0.89054584 |
| Rrm2b | 0.20370184 | 0.22936017 | 0.21397764 | 0.01538252 | 0.36884116 | 0.89054584 |
| 1110012L19F | 0.45758345 | 0.47043365 | 0.45505254 | 0.01538111 | 0.437435 | 0.89054584 |
| Jak3 | 0.04798753 | 0.06670842 | 0.05134446 | 0.01536396 | 0.08164231 | 0.54874839 |
| Mapk1ip1 | 0.29011292 | 0.31505725 | 0.29969527 | 0.01536197 | 0.40562607 | 0.89054584 |
| Cdk16 | 0.60759511 | 0.59550249 | 0.58019014 | 0.01531235 | 0.45106997 | 0.89054584 |
| Bzw1 | 1.62798943 | 1.08231617 | 1.06701786 | 0.01529831 | 0.47337152 | 0.89054584 |
| 2610020C07I | 0.06243916 | 0.08209074 | 0.06679648 | 0.01529426 | 0.14304271 | 0.76440591 |
| Mrm1 | 0.11102451 | 0.13378163 | 0.11848935 | 0.01529228 | 0.27382519 | 0.89054584 |
| Zmat5 | 0.33289621 | 0.35607651 | 0.3408102 | 0.01526631 | 0.41734381 | 0.89054584 |
| Gm20649 | 0.02237979 | 0.03920831 | 0.02394934 | 0.01525897 | 0.00149689 | 0.02348698 |
| Hecw2 | 0.03818632 | 0.05611864 | 0.0408609 | 0.01525774 | 0.04095756 | 0.34167394 |
| Acot2 | 0.12786149 | 0.15133926 | 0.13608643 | 0.01525284 | 0.3007713 | 0.89054584 |

| | | | | | | |
|---|---|---|---|---|---|---|
| Sash1 | 0.21151435 | 0.23711906 | 0.2218664 | 0.01525266 | 0.37437487 | 0.89054584 |
| Ankrd24 | 0.21887972 | 0.24449344 | 0.22927919 | 0.01521426 | 0.37860262 | 0.89054584 |
| Slc16a9 | 0.10848224 | 0.1310328 | 0.11582315 | 0.01520965 | 0.27033429 | 0.89054584 |
| Lipo1 | 0.10118679 | 0.12336147 | 0.10815897 | 0.01520251 | 0.25628309 | 0.89054584 |
| Adar | 0.12044342 | 0.14354769 | 0.12834651 | 0.01520118 | 0.29054319 | 0.89054584 |
| Lbp | 0.14100904 | 0.16495418 | 0.14975372 | 0.01520046 | 0.31814339 | 0.89054584 |
| Gpr182 | 0.05242808 | 0.07129099 | 0.05609321 | 0.01519778 | 0.10342327 | 0.63276555 |
| Ptprd | 0.18266331 | 0.20779529 | 0.19260522 | 0.01519007 | 0.3566478 | 0.89054584 |
| Dlgap1 | 0.01631086 | 0.03263625 | 0.01745513 | 0.01518111 | 2.54E-05 | 0.00061665 |
| Stat6 | 0.15314382 | 0.17748924 | 0.16230842 | 0.01518081 | 0.33150767 | 0.89054584 |
| Mettl10 | 0.57089999 | 0.56608848 | 0.55090901 | 0.01517947 | 0.44892804 | 0.89054584 |
| Sgol1 | 0.09228938 | 0.11385319 | 0.09868212 | 0.01517107 | 0.23691827 | 0.89054584 |
| Gimap9 | 0.11444334 | 0.13724132 | 0.12207107 | 0.01517026 | 0.28129961 | 0.89054584 |
| Rasgrf2 | 0.04357739 | 0.06179155 | 0.04662763 | 0.01516392 | 0.06486729 | 0.47163788 |
| Ppm1k | 0.13015758 | 0.15363366 | 0.13847807 | 0.0151556 | 0.30506313 | 0.89054584 |
| Itga7 | 0.0318093 | 0.04917478 | 0.03403863 | 0.01513615 | 0.01914593 | 0.19547721 |
| S100a14 | 0.01732972 | 0.03366891 | 0.01854541 | 0.0151235 | 7.26E-05 | 0.00157703 |
| Cenpi | 0.04345823 | 0.06160795 | 0.04650017 | 0.01510778 | 0.06505377 | 0.47228321 |
| Smpd5 | 0.03537102 | 0.05291598 | 0.03784915 | 0.01506683 | 0.03182584 | 0.28186377 |
| Mfsd5 | 0.09778628 | 0.11961343 | 0.10454798 | 0.01506545 | 0.25099648 | 0.89054584 |
| Guca2a | 0.02628368 | 0.04318709 | 0.02812656 | 0.01506053 | 0.00630396 | 0.07919825 |
| Fads1 | 0.15700999 | 0.18134916 | 0.16629652 | 0.01505263 | 0.33661742 | 0.89054584 |
| Dram1 | 0.01726371 | 0.03352736 | 0.01847478 | 0.01505258 | 7.35E-05 | 0.00158666 |
| Atp13a2 | 0.32619078 | 0.34947652 | 0.33442792 | 0.0150486 | 0.41697307 | 0.89054584 |
| 2310009B15 | 0.58414779 | 0.57662922 | 0.56158305 | 0.01504617 | 0.45033132 | 0.89054584 |
| Paxip1 | 0.20822878 | 0.23359127 | 0.21855194 | 0.01503933 | 0.37425746 | 0.89054584 |
| Csrnp1 | 0.09339669 | 0.11489796 | 0.09986392 | 0.01503404 | 0.24153246 | 0.89054584 |
| Sult1c2 | 0.05098807 | 0.06957765 | 0.05455332 | 0.01502433 | 0.09972818 | 0.62085259 |
| Lppos | 0.14038156 | 0.16411031 | 0.14910296 | 0.01500735 | 0.31956096 | 0.89054584 |
| Ptprb | 0.03375301 | 0.05111672 | 0.03611815 | 0.01499857 | 0.02651409 | 0.2464111 |

| | | | | | | |
|---|---|---|---|---|---|---|
| Rnf14 | 0.48880272 | 0.49723929 | 0.48225573 | 0.01498356 | 0.44245331 | 0.89054584 |
| Lrrn3 | 0.05454399 | 0.07333245 | 0.05835573 | 0.01497672 | 0.1159078 | 0.67896213 |
| Nfasc | 0.15395005 | 0.17811307 | 0.16314058 | 0.01497249 | 0.33447778 | 0.89054584 |
| Tmem258 | 2.5010487 | 1.14180716 | 1.12683611 | 0.01497105 | 0.4753219 | 0.89054584 |
| Adrb2 | 0.02217163 | 0.03869723 | 0.0237266 | 0.01497063 | 0.00164306 | 0.02539563 |
| Tbc1d31 | 0.08160421 | 0.10223239 | 0.08727406 | 0.01495833 | 0.21228475 | 0.89054584 |
| Mbd4 | 0.097199 | 0.11886674 | 0.10392138 | 0.01494535 | 0.25142217 | 0.89054584 |
| Kcnmb4 | 0.05550029 | 0.07431089 | 0.05937824 | 0.01493264 | 0.12066953 | 0.69533658 |
| Lrrc32 | 0.06916803 | 0.08887928 | 0.07398825 | 0.01489102 | 0.17420757 | 0.84132624 |
| Zc3h10 | 0.1492884 | 0.1732167 | 0.15832585 | 0.01489085 | 0.33062702 | 0.89054584 |
| Sestd1 | 0.11903615 | 0.14176179 | 0.12687586 | 0.01488594 | 0.29232041 | 0.89054584 |
| Ndst4 | 0.09041198 | 0.11155228 | 0.09667823 | 0.01487404 | 0.23675405 | 0.89054584 |
| Akap17b | 0.1907679 | 0.21573381 | 0.20086086 | 0.01487295 | 0.36505612 | 0.89054584 |
| Dock11 | 0.19986824 | 0.22495723 | 0.21009738 | 0.01485985 | 0.37088154 | 0.89054584 |
| Vrk2 | 0.12591453 | 0.14890172 | 0.1340569 | 0.01484482 | 0.30295899 | 0.89054584 |
| 1700113A16 | 0.09553453 | 0.11698271 | 0.10214534 | 0.01483736 | 0.24928375 | 0.89054584 |
| Dtx4 | 0.05723948 | 0.07605538 | 0.06123776 | 0.01481762 | 0.12980411 | 0.72158739 |
| Vasp | 1.27988742 | 0.97775022 | 0.96294644 | 0.01480378 | 0.47145062 | 0.89054584 |
| Vezt | 0.16582477 | 0.19014368 | 0.17536677 | 0.01477691 | 0.34731643 | 0.89054584 |
| Vsig10l | 0.02698741 | 0.04363392 | 0.02887953 | 0.01475439 | 0.008651 | 0.10408487 |
| Klf15 | 0.04440426 | 0.06226449 | 0.04751204 | 0.01475245 | 0.07400484 | 0.51682187 |
| Ubp1 | 0.26674782 | 0.29158542 | 0.27685989 | 0.01472553 | 0.4021453 | 0.89054584 |
| Ndufb10 | 1.78210144 | 1.10987925 | 1.0951655 | 0.01471376 | 0.475045 | 0.89054584 |
| Plxna2 | 0.09125404 | 0.11228905 | 0.09757705 | 0.014712 | 0.2412014 | 0.89054584 |
| Pde8a | 0.09848425 | 0.11999324 | 0.10529265 | 0.01470059 | 0.25769595 | 0.89054584 |
| Itga5 | 0.11464926 | 0.13696743 | 0.12228666 | 0.01468077 | 0.2879732 | 0.89054584 |
| Avp | 0.03348292 | 0.05048928 | 0.03582918 | 0.01466009 | 0.02830683 | 0.25810848 |
| Slc10a7 | 0.13981916 | 0.16315344 | 0.14851955 | 0.01463389 | 0.32309784 | 0.89054584 |
| Smad6 | 0.05699922 | 0.07560385 | 0.06098088 | 0.01462297 | 0.13195539 | 0.72831857 |
| Fes | 0.06751124 | 0.08683964 | 0.07221768 | 0.01462196 | 0.17276403 | 0.83703076 |

| | | | | | | |
|---|---|---|---|---|---|---|
| Rpusd3 | 0.16167026 | 0.18571311 | 0.17109577 | 0.01461734 | 0.34530392 | 0.89054584 |
| Upp2 | 0.10072112 | 0.12226404 | 0.10766911 | 0.01459493 | 0.26384498 | 0.89054584 |
| Cd1d1 | 0.18054344 | 0.20502782 | 0.19044128 | 0.01458654 | 0.36060428 | 0.89054584 |
| Svep1 | 0.11031074 | 0.13232515 | 0.11774102 | 0.01458413 | 0.28194092 | 0.89054584 |
| Sirt1 | 0.10071318 | 0.12223807 | 0.10766075 | 0.01457732 | 0.26407806 | 0.89054584 |
| Hoxb5 | 0.076109 | 0.09596997 | 0.08140444 | 0.01456553 | 0.20224909 | 0.89054584 |
| Atp1a2 | 0.06460649 | 0.08367605 | 0.06911313 | 0.01456292 | 0.16312582 | 0.81647508 |
| Traf3ip3 | 0.01971729 | 0.03565712 | 0.02110031 | 0.0145568 | 0.00065419 | 0.01149267 |
| AU019823 | 0.3899297 | 0.40868887 | 0.39413787 | 0.014551 | 0.43171824 | 0.89054584 |
| Mybpc2 | 0.01903271 | 0.03491658 | 0.02036776 | 0.01454882 | 0.0004375 | 0.00801747 |
| Apex2 | 0.13959999 | 0.16283902 | 0.14829217 | 0.01454685 | 0.32382728 | 0.89054584 |
| Efnb2 | 0.08422562 | 0.10461186 | 0.09007347 | 0.01453839 | 0.22603139 | 0.89054584 |
| Zfp330 | 0.69316743 | 0.65947005 | 0.64494345 | 0.0145266 | 0.4582128 | 0.89054584 |
| Ap4e1 | 0.08061017 | 0.10072292 | 0.08621242 | 0.0145105 | 0.21647576 | 0.89054584 |
| Tmem205 | 0.45338246 | 0.46584719 | 0.45134739 | 0.01449981 | 0.44051198 | 0.89054584 |
| Twsg1 | 0.65167625 | 0.6286449 | 0.61417274 | 0.01447216 | 0.45629124 | 0.89054584 |
| Nme5 | 0.17528842 | 0.19953337 | 0.18506936 | 0.01446402 | 0.35788509 | 0.89054584 |
| A1cf | 0.10744467 | 0.12918285 | 0.11473433 | 0.01444853 | 0.278702 | 0.89054584 |
| Acer3 | 0.10642349 | 0.12810668 | 0.11366231 | 0.01444437 | 0.27690355 | 0.89054584 |
| Sema6c | 0.08066954 | 0.10071686 | 0.08627583 | 0.01444103 | 0.2177474 | 0.89054584 |
| Snrpa | 1.29634276 | 0.98361309 | 0.9691864 | 0.01442669 | 0.47235551 | 0.89054584 |
| Pgm2l1 | 0.10876448 | 0.13054238 | 0.11611927 | 0.0144231 | 0.28140116 | 0.89054584 |
| Mlh1 | 0.22357651 | 0.24840474 | 0.23399356 | 0.01441117 | 0.38708126 | 0.89054584 |
| Csf2ra | 0.27015939 | 0.29461465 | 0.28021052 | 0.01440414 | 0.40536226 | 0.89054584 |
| Dhrs1 | 0.44842769 | 0.46135974 | 0.44696378 | 0.01439596 | 0.44035992 | 0.89054584 |
| Strn | 0.12098166 | 0.14330135 | 0.12890878 | 0.01439257 | 0.30147352 | 0.89054584 |
| Rab36 | 0.1967335 | 0.22129105 | 0.20691976 | 0.0143713 | 0.37312814 | 0.89054584 |
| 1700052K11I | 0.18759233 | 0.21199339 | 0.19762942 | 0.01436397 | 0.36744869 | 0.89054584 |
| Armcx6 | 0.3389223 | 0.36088777 | 0.34652633 | 0.01436144 | 0.42344611 | 0.89054584 |
| Nup62 | 0.3778969 | 0.39738183 | 0.38303359 | 0.01434824 | 0.4307278 | 0.89054584 |

| | | | | | | |
|---|---|---|---|---|---|---|
| Papd7 | 0.0917179 | 0.11240977 | 0.09807216 | 0.01433761 | 0.24790045 | 0.89054584 |
| Slc2a4 | 0.03127785 | 0.04778708 | 0.03347003 | 0.01431705 | 0.02313644 | 0.22332361 |
| Mrpl53 | 0.024823 | 0.04085845 | 0.02656363 | 0.01429482 | 0.00608564 | 0.07692212 |
| C8g | 0.04908447 | 0.06678254 | 0.0525176 | 0.01426494 | 0.10285117 | 0.63089369 |
| Zfp820 | 0.1201109 | 0.14225501 | 0.12799907 | 0.01425594 | 0.30191832 | 0.89054584 |
| Zfp811 | 0.04113151 | 0.05825218 | 0.0440114 | 0.01424078 | 0.06584195 | 0.47501237 |
| Cux2 | 0.04319044 | 0.06044368 | 0.04621374 | 0.01422995 | 0.07570588 | 0.52298621 |
| D8Ertd82e | 0.08322947 | 0.10322606 | 0.08900972 | 0.01421633 | 0.22840489 | 0.89054584 |
| Fnbp4 | 0.41789714 | 0.43383129 | 0.41963735 | 0.01419394 | 0.43739172 | 0.89054584 |
| Cog3 | 0.12138089 | 0.14351789 | 0.12932578 | 0.01419211 | 0.30458147 | 0.89054584 |
| P2ry14 | 0.03518906 | 0.05184224 | 0.03765448 | 0.01418776 | 0.03959262 | 0.33509273 |
| Lpcat3 | 0.68332779 | 0.65193561 | 0.63775401 | 0.0141816 | 0.45874345 | 0.89054584 |
| Gpr157 | 0.0333906 | 0.04988467 | 0.03573042 | 0.01415425 | 0.03247564 | 0.28656832 |
| Rbm45 | 0.04555065 | 0.06288936 | 0.04873818 | 0.01415117 | 0.08807132 | 0.57428752 |
| Mtap7d3 | 0.05188797 | 0.06964699 | 0.05551564 | 0.01413134 | 0.11782701 | 0.68687415 |
| Gbp2 | 0.04927823 | 0.06684642 | 0.05272481 | 0.01412161 | 0.10604612 | 0.64355657 |
| Abcc10 | 0.0587493 | 0.07696961 | 0.06285195 | 0.01411765 | 0.14766971 | 0.77404405 |
| Sall2 | 0.0859053 | 0.10597232 | 0.09186699 | 0.01410533 | 0.23720031 | 0.89054584 |
| Smco3 | 0.0376229 | 0.05436006 | 0.04025818 | 0.01410188 | 0.05134374 | 0.40020517 |
| Mettl14 | 0.09294584 | 0.11347499 | 0.09938274 | 0.01409225 | 0.25442564 | 0.89054584 |
| Calr3 | 0.01805738 | 0.0334119 | 0.01932408 | 0.01408782 | 0.00034126 | 0.00639227 |
| Thop1 | 0.48302496 | 0.49135053 | 0.47726589 | 0.01408463 | 0.44532159 | 0.89054584 |
| Plrg1 | 0.31757226 | 0.34025981 | 0.32619026 | 0.01406955 | 0.42036729 | 0.89054584 |
| Nckap5l | 0.05779119 | 0.07587124 | 0.06182762 | 0.01404362 | 0.14497324 | 0.76736841 |
| Epn2 | 0.31762572 | 0.34027758 | 0.32624147 | 0.01403611 | 0.42056637 | 0.89054584 |
| Kctd11 | 0.14921203 | 0.17227591 | 0.1582469 | 0.01402901 | 0.3397932 | 0.89054584 |
| Vegfb | 0.23798836 | 0.26242469 | 0.24839826 | 0.01402643 | 0.39624496 | 0.89054584 |
| Rspo1 | 0.03009687 | 0.04623268 | 0.0322065 | 0.01402619 | 0.02122883 | 0.20937119 |
| Auh | 0.67997627 | 0.649306 | 0.63528962 | 0.01401638 | 0.45906479 | 0.89054584 |
| Tubgcp3 | 0.26304239 | 0.28721759 | 0.27321432 | 0.01400328 | 0.40563477 | 0.89054584 |

| | | | | | | |
|---|---|---|---|---|---|---|
| Pkia | 0.09413038 | 0.11464254 | 0.10064692 | 0.01399562 | 0.25853887 | 0.89054584 |
| Mpp6 | 0.22106726 | 0.24544332 | 0.23147615 | 0.01396717 | 0.38930972 | 0.89054584 |
| Psd3 | 0.11136101 | 0.13278906 | 0.11884207 | 0.01394699 | 0.29227144 | 0.89054584 |
| Asb7 | 0.28466278 | 0.30833371 | 0.29439181 | 0.0139419 | 0.41268653 | 0.89054584 |
| Sgk3 | 0.13411623 | 0.15652793 | 0.14259674 | 0.01393119 | 0.32449636 | 0.89054584 |
| Pcdh12 | 0.03775953 | 0.05431973 | 0.04040434 | 0.01391539 | 0.05429585 | 0.4159266 |
| Hif3a | 0.11993845 | 0.14171166 | 0.12781888 | 0.01389278 | 0.30629292 | 0.89054584 |
| Plekhh2 | 0.23689343 | 0.26119985 | 0.24730711 | 0.01389274 | 0.3967683 | 0.89054584 |
| Ino80b | 0.4253348 | 0.44021944 | 0.42634454 | 0.0138749 | 0.43974335 | 0.89054584 |
| Ccdc15 | 0.04727056 | 0.064452 | 0.05057767 | 0.01387432 | 0.1006199 | 0.62288099 |
| Dsn1 | 0.04005837 | 0.05673185 | 0.04286348 | 0.01386837 | 0.06585563 | 0.47501237 |
| Prdm6 | 0.04159225 | 0.05836528 | 0.04450425 | 0.01386103 | 0.07338351 | 0.51488639 |
| Nup54 | 0.31881162 | 0.34123774 | 0.32737722 | 0.01386052 | 0.42181571 | 0.89054584 |
| Vtn | 0.05232893 | 0.06984272 | 0.05598718 | 0.01385554 | 0.12445906 | 0.70622602 |
| Pnpla7 | 0.04368481 | 0.060575 | 0.04674252 | 0.01383248 | 0.08399092 | 0.55909105 |
| Oxld1 | 0.18573311 | 0.20954372 | 0.1957355 | 0.01380821 | 0.37120397 | 0.89054584 |
| Lef1 | 0.03123325 | 0.04723041 | 0.03342232 | 0.01380809 | 0.02712755 | 0.25066618 |
| Edn1 | 0.01449038 | 0.02931365 | 0.015507 | 0.01380665 | 1.68E-05 | 0.00042193 |
| Csdc2 | 0.05740421 | 0.07521957 | 0.06141389 | 0.01380568 | 0.14747657 | 0.77404405 |
| Egln2 | 0.86272648 | 0.7717839 | 0.75798114 | 0.01380276 | 0.46619476 | 0.89054584 |
| Bco1 | 0.07797589 | 0.09720067 | 0.08339871 | 0.01380196 | 0.22034601 | 0.89054584 |
| B4galt7 | 0.28971613 | 0.31310441 | 0.29930966 | 0.01379475 | 0.4149913 | 0.89054584 |
| Akap2 | 0.05993855 | 0.07791139 | 0.06412334 | 0.01378805 | 0.15822408 | 0.80404231 |
| Naa40 | 0.20240522 | 0.2264447 | 0.21266596 | 0.01377874 | 0.38138121 | 0.89054584 |
| Rnf5 | 0.90360016 | 0.79588725 | 0.78211706 | 0.01377019 | 0.46731272 | 0.89054584 |
| Fam155a | 0.1254643 | 0.1473519 | 0.13358741 | 0.01376449 | 0.3155977 | 0.89054584 |
| Nell1 | 0.06018117 | 0.07814366 | 0.06438271 | 0.01376095 | 0.15967679 | 0.80704115 |
| Ppp1r3c | 0.09765537 | 0.11816332 | 0.10440831 | 0.01375501 | 0.26968022 | 0.89054584 |
| Fgfrl1 | 0.11231193 | 0.13359031 | 0.11983861 | 0.01375171 | 0.29645555 | 0.89054584 |
| Tgm7 | 0.02359399 | 0.03898156 | 0.02524857 | 0.01373299 | 0.0056376 | 0.07233093 |

| | | | | | | |
|---|---|---|---|---|---|---|
| Fam105a | 0.16572854 | 0.18899815 | 0.17526793 | 0.01373023 | 0.35756524 | 0.89054584 |
| Mnda | 0.02211555 | 0.03739408 | 0.0236666 | 0.01372748 | 0.00344251 | 0.0482684 |
| Rfx6 | 0.39961857 | 0.41674767 | 0.40302212 | 0.01372555 | 0.43696539 | 0.89054584 |
| Kcnn2 | 0.02728683 | 0.04291916 | 0.0291999 | 0.01371925 | 0.01430015 | 0.15553157 |
| Yy1 | 0.70102556 | 0.66435476 | 0.65063594 | 0.01371883 | 0.46087286 | 0.89054584 |
| Slc30a1 | 0.09279197 | 0.11292939 | 0.09921853 | 0.01371087 | 0.25984851 | 0.89054584 |
| Arhgap33 | 0.07607103 | 0.09506772 | 0.08136387 | 0.01370385 | 0.21631762 | 0.89054584 |
| Esco2 | 0.06585711 | 0.08413494 | 0.07044982 | 0.01368512 | 0.18272862 | 0.86405725 |
| Cmc1 | 0.4366124 | 0.45013465 | 0.43645266 | 0.013682 | 0.44194122 | 0.89054584 |
| Ndufa9 | 1.13675201 | 0.91612309 | 0.90244136 | 0.01368174 | 0.47184481 | 0.89054584 |
| Snx22 | 0.02966458 | 0.04542334 | 0.03174398 | 0.01367936 | 0.02233933 | 0.21839217 |
| Cryl1 | 0.24113472 | 0.2651969 | 0.25153071 | 0.01366619 | 0.4000839 | 0.89054584 |
| Ift74 | 0.62405271 | 0.60668782 | 0.5930311 | 0.01365672 | 0.45727975 | 0.89054584 |
| Tapbp | 0.59784039 | 0.58614942 | 0.57249295 | 0.01365647 | 0.45575418 | 0.89054584 |
| Creb1 | 0.36605658 | 0.38567161 | 0.37202861 | 0.013643 | 0.43216997 | 0.89054584 |
| Galnt11 | 0.63873666 | 0.61793638 | 0.60433453 | 0.01360185 | 0.45824366 | 0.89054584 |
| Tmem69 | 0.08050324 | 0.09969414 | 0.08609821 | 0.01359593 | 0.23095598 | 0.89054584 |
| Rtp4 | 0.05867032 | 0.0763528 | 0.06276751 | 0.01358529 | 0.15663771 | 0.79878068 |
| Clspn | 0.11630149 | 0.13759771 | 0.12401592 | 0.01358179 | 0.30494489 | 0.89054584 |
| Nek2 | 0.03421467 | 0.0501931 | 0.03661205 | 0.01358105 | 0.04197528 | 0.34735021 |
| Gclc | 0.11091863 | 0.13194615 | 0.11837835 | 0.01356779 | 0.29667803 | 0.89054584 |
| Peo1 | 0.11060568 | 0.13161096 | 0.11805027 | 0.01356069 | 0.29626172 | 0.89054584 |
| Plscr2 | 0.04325293 | 0.05982765 | 0.04628057 | 0.01354708 | 0.08632338 | 0.5680964 |
| Map3k2 | 0.28606811 | 0.30930751 | 0.29576073 | 0.01354678 | 0.41550969 | 0.89054584 |
| 4833412C05I | 0.02101871 | 0.03603201 | 0.02249292 | 0.0135391 | 0.00252096 | 0.03684364 |
| B430203G13 | 0.03486271 | 0.05083475 | 0.03730534 | 0.01352941 | 0.0455481 | 0.36805888 |
| Sema4c | 0.24679018 | 0.27067065 | 0.25714963 | 0.01352101 | 0.40324034 | 0.89054584 |
| Eif2b3 | 0.39423749 | 0.41161178 | 0.39809448 | 0.01351729 | 0.43715185 | 0.89054584 |
| Tacc1 | 0.29413912 | 0.31711692 | 0.30360364 | 0.01351328 | 0.41786139 | 0.89054584 |
| Gm10827 | 0.02687047 | 0.04224359 | 0.02875441 | 0.01348918 | 0.01442285 | 0.15669589 |

| | | | | | | |
|---|---|---|---|---|---|---|
| Pphln1 | 0.28098467 | 0.30429109 | 0.290805 | 0.01348609 | 0.41447107 | 0.89054584 |
| Gzf1 | 0.26933348 | 0.29288165 | 0.27939987 | 0.01348177 | 0.41106508 | 0.89054584 |
| Fgf7 | 0.02548477 | 0.04074494 | 0.02727173 | 0.01347321 | 0.01067653 | 0.12381337 |
| 3110062M04 | 0.19391209 | 0.21752813 | 0.20405612 | 0.013472 | 0.37919782 | 0.89054584 |
| Fam167a | 0.04176082 | 0.05813487 | 0.04468455 | 0.01345032 | 0.08040253 | 0.54343866 |
| Bvht | 0.08205697 | 0.10120399 | 0.0877576 | 0.0134464 | 0.2376592 | 0.89054584 |
| Dzip1 | 0.05975489 | 0.07735764 | 0.063927 | 0.01343064 | 0.16383627 | 0.81785018 |
| Khdrbs2 | 0.03596695 | 0.05190745 | 0.03848667 | 0.01342079 | 0.05212056 | 0.40365591 |
| Tox2 | 0.02188527 | 0.03683885 | 0.02342018 | 0.01341867 | 0.00379985 | 0.05230825 |
| Gdpd2 | 0.01866895 | 0.03337709 | 0.01997851 | 0.01339858 | 0.00089049 | 0.01495984 |
| Vti1b | 0.96962682 | 0.83201104 | 0.81862323 | 0.01338781 | 0.46963294 | 0.89054584 |
| Lrrk1 | 0.14297626 | 0.16516758 | 0.15179294 | 0.01337463 | 0.34071807 | 0.89054584 |
| Sp110 | 0.03361255 | 0.04933851 | 0.03596788 | 0.01337063 | 0.04164512 | 0.34560445 |
| Scd1 | 0.02060709 | 0.03541775 | 0.02205246 | 0.01336529 | 0.00237412 | 0.03512249 |
| Por | 0.87778977 | 0.78038028 | 0.76701692 | 0.01336336 | 0.46765319 | 0.89054584 |
| Sult2b1 | 0.09236536 | 0.11212578 | 0.09876321 | 0.01336257 | 0.26423264 | 0.89054584 |
| Pigh | 0.25254869 | 0.27621006 | 0.26285558 | 0.01335448 | 0.40644506 | 0.89054584 |
| Frmd4b | 0.15644048 | 0.17906018 | 0.16570943 | 0.01335075 | 0.35369709 | 0.89054584 |
| Brwd3 | 0.18436617 | 0.20768789 | 0.19434213 | 0.01334576 | 0.37450828 | 0.89054584 |
| Zfp385b | 0.03034462 | 0.04580746 | 0.03247157 | 0.01333589 | 0.02784843 | 0.25472969 |
| Elavl2 | 0.07206333 | 0.09041084 | 0.07708208 | 0.01332877 | 0.21023443 | 0.89054584 |
| Rabac1 | 1.47233944 | 1.04114232 | 1.02781939 | 0.01332293 | 0.47592223 | 0.89054584 |
| Tagln3 | 0.02837454 | 0.04366136 | 0.03036371 | 0.01329764 | 0.02065719 | 0.20548976 |
| Dmpk | 0.13308152 | 0.15481798 | 0.14152078 | 0.01329721 | 0.33078327 | 0.89054584 |
| Dbf4 | 0.18260658 | 0.20584162 | 0.19254734 | 0.01329428 | 0.37385027 | 0.89054584 |
| Ccdc189 | 0.22620935 | 0.24992056 | 0.23663192 | 0.01328863 | 0.39680181 | 0.89054584 |
| Smyd4 | 0.17901554 | 0.20216083 | 0.18888044 | 0.01328039 | 0.37161641 | 0.89054584 |
| Atp9b | 0.14946504 | 0.17176871 | 0.15850844 | 0.01326027 | 0.34836038 | 0.89054584 |
| Plbd2 | 0.27687863 | 0.30004985 | 0.28679323 | 0.01325662 | 0.41474604 | 0.89054584 |
| Copz1 | 1.31960335 | 0.99106477 | 0.97783182 | 0.01323295 | 0.47486365 | 0.89054584 |

| | | | | | | |
|---|---|---|---|---|---|---|
| Zfp651 | 0.10664973 | 0.12712857 | 0.11389985 | 0.01322872 | 0.29421739 | 0.89054584 |
| Hexb | 0.35646406 | 0.37627907 | 0.36305795 | 0.01322112 | 0.43263874 | 0.89054584 |
| Dsel | 0.09797467 | 0.11794406 | 0.10474898 | 0.01319508 | 0.27864213 | 0.89054584 |
| Sash3 | 0.02327402 | 0.03810004 | 0.0249062 | 0.01319384 | 0.00679288 | 0.0845349 |
| Pla2g4a | 0.11720296 | 0.1381504 | 0.12495901 | 0.01319139 | 0.31142168 | 0.89054584 |
| Ypel5 | 0.43107993 | 0.44467922 | 0.43150326 | 0.01317596 | 0.44343699 | 0.89054584 |
| Nipal3 | 0.14038003 | 0.16225817 | 0.14910137 | 0.01315679 | 0.34049677 | 0.89054584 |
| Calb1 | 0.05189707 | 0.06867389 | 0.05552538 | 0.01314851 | 0.13496016 | 0.73613709 |
| Peli2 | 0.13107692 | 0.15256145 | 0.13943511 | 0.01312634 | 0.33047928 | 0.89054584 |
| Cit | 0.16497624 | 0.18760515 | 0.17449502 | 0.01311012 | 0.36315636 | 0.89054584 |
| Gc | 0.03684035 | 0.05251513 | 0.03942103 | 0.0130941 | 0.0608696 | 0.45140021 |
| Ncapg2 | 0.08659683 | 0.10569762 | 0.09260533 | 0.01309229 | 0.2550543 | 0.89054584 |
| Col9a1 | 0.04754012 | 0.06395582 | 0.05086597 | 0.01308985 | 0.1152771 | 0.67690861 |
| Fli1 | 0.06786086 | 0.08568038 | 0.07259132 | 0.01308907 | 0.20043689 | 0.89054584 |
| Cfap61 | 0.02827057 | 0.04331414 | 0.03025247 | 0.01306168 | 0.02213502 | 0.21683347 |
| D930015E06 | 0.19846255 | 0.22173061 | 0.20867299 | 0.01305762 | 0.38532253 | 0.89054584 |
| Cd38 | 0.03681894 | 0.05244564 | 0.03939812 | 0.01304752 | 0.06142671 | 0.45436961 |
| Aff1 | 0.14026034 | 0.16201489 | 0.14897722 | 0.01303767 | 0.34173774 | 0.89054584 |
| Adipor2 | 0.34410929 | 0.364468 | 0.3514318 | 0.01303621 | 0.43139593 | 0.89054584 |
| Anapc2 | 0.37868739 | 0.39675509 | 0.38376554 | 0.01298955 | 0.43734893 | 0.89054584 |
| Itpkb | 0.30598002 | 0.32803537 | 0.31505082 | 0.01298455 | 0.42386551 | 0.89054584 |
| Zfp518b | 0.0668664 | 0.08451025 | 0.07152852 | 0.01298173 | 0.19890126 | 0.89054584 |
| Fam129a | 0.07574034 | 0.09399187 | 0.0810106 | 0.01298128 | 0.22766557 | 0.89054584 |
| Rps6kl1 | 0.03048643 | 0.04559097 | 0.03262329 | 0.01296768 | 0.0320188 | 0.28305401 |
| Adgrg3 | 0.0287127 | 0.04367971 | 0.03072553 | 0.01295418 | 0.02475029 | 0.23463713 |
| Gm4673 | 0.02743728 | 0.04230856 | 0.02936088 | 0.01294768 | 0.01996275 | 0.20112578 |
| Scn3b | 0.04281654 | 0.05874108 | 0.04581381 | 0.01292727 | 0.09432005 | 0.60001264 |
| Kansl1l | 0.19108975 | 0.21411264 | 0.20118813 | 0.01292451 | 0.38235758 | 0.89054584 |
| Dock1 | 0.15022248 | 0.17221554 | 0.15929127 | 0.01292427 | 0.35271221 | 0.89054584 |
| Pgs1 | 0.2897288 | 0.312241 | 0.29932197 | 0.01291903 | 0.42031624 | 0.89054584 |

| | | | | | | |
|---|---|---|---|---|---|---|
| Tmem59 | 2.93658438 | 1.11311797 | 1.10021617 | 0.0129018 | 0.47821521 | 0.89054584 |
| Tasp1 | 0.18491021 | 0.20779355 | 0.19489678 | 0.01289677 | 0.37892973 | 0.89054584 |
| Aqp8 | 0.01620004 | 0.03021957 | 0.01733654 | 0.01288303 | 0.00026796 | 0.00517302 |
| Crtc1 | 0.27250456 | 0.29539281 | 0.28251053 | 0.01288227 | 0.4158802 | 0.89054584 |
| Fam107b | 0.25881693 | 0.28192899 | 0.26904909 | 0.0128799 | 0.41175512 | 0.89054584 |
| Plaur | 0.04385429 | 0.059799 | 0.04692379 | 0.01287521 | 0.1005637 | 0.62288099 |
| G2e3 | 0.23844935 | 0.26172459 | 0.2488575 | 0.01286709 | 0.40482063 | 0.89054584 |
| Map3k8 | 0.06836705 | 0.08599796 | 0.07313228 | 0.01286568 | 0.20621725 | 0.89054584 |
| Ppp2r5b | 0.46371022 | 0.47329638 | 0.46043702 | 0.01285937 | 0.44823663 | 0.89054584 |
| Slitrk2 | 0.00957562 | 0.0231065 | 0.01024752 | 0.01285898 | 2.51E-09 | 1.12E-07 |
| Zfp536 | 0.00957562 | 0.0231065 | 0.01024752 | 0.01285898 | 2.51E-09 | 1.12E-07 |
| Sox10 | 0.00957562 | 0.0231065 | 0.01024752 | 0.01285898 | 2.51E-09 | 1.12E-07 |
| Hccs | 0.32213425 | 0.34338785 | 0.33055546 | 0.01283239 | 0.42823743 | 0.89054584 |
| Stard8 | 0.06116841 | 0.07825048 | 0.06543808 | 0.0128124 | 0.18083341 | 0.85901797 |
| Lrch1 | 0.14934717 | 0.17118087 | 0.1583866 | 0.01279427 | 0.35333083 | 0.89054584 |
| Tifa | 0.05245364 | 0.06890977 | 0.05612054 | 0.01278923 | 0.14418158 | 0.76653633 |
| Foxo4 | 0.14551266 | 0.16720358 | 0.15441994 | 0.01278363 | 0.34986321 | 0.89054584 |
| Trmt61a | 0.10764184 | 0.12771585 | 0.11494127 | 0.01277459 | 0.30230053 | 0.89054584 |
| Zfp445 | 0.09024057 | 0.10925231 | 0.09649525 | 0.01275705 | 0.26897023 | 0.89054584 |
| Pdcd2 | 0.42436109 | 0.4382152 | 0.42546829 | 0.01274691 | 0.44449571 | 0.89054584 |
| Smox | 0.25564613 | 0.27866359 | 0.26591829 | 0.01274529 | 0.411651 | 0.89054584 |
| Psd2 | 0.01420569 | 0.02794725 | 0.01520235 | 0.0127449 | 4.70E-05 | 0.00106862 |
| Ccdc60 | 0.06570289 | 0.08302657 | 0.07028499 | 0.01274158 | 0.1991688 | 0.89054584 |
| Cdk5r1 | 0.13166363 | 0.15277557 | 0.14004571 | 0.01272986 | 0.33596961 | 0.89054584 |
| Plaa | 0.45950457 | 0.46947185 | 0.45674335 | 0.0127285 | 0.44834842 | 0.89054584 |
| Wnt2b | 0.0167237 | 0.03059232 | 0.01789691 | 0.01269541 | 0.00047508 | 0.00860918 |
| Fam124b | 0.03378633 | 0.04882699 | 0.03615379 | 0.0126732 | 0.05122128 | 0.39948674 |
| Rims2 | 0.14971522 | 0.17143106 | 0.15876703 | 0.01266403 | 0.35508802 | 0.89054584 |
| Baz1a | 0.24411627 | 0.26712947 | 0.25449485 | 0.01263462 | 0.40854199 | 0.89054584 |
| Reep6 | 0.15952062 | 0.18151629 | 0.16888312 | 0.01263318 | 0.36372911 | 0.89054584 |

| | | | | | | |
|---|---|---|---|---|---|---|
| Slc35e4 | 0.06224194 | 0.07920073 | 0.06658566 | 0.01261507 | 0.18870856 | 0.8782975 |
| Arhgef28 | 0.12657732 | 0.14735511 | 0.13474794 | 0.01260717 | 0.33145617 | 0.89054584 |
| Fndc1 | 0.06100094 | 0.07783573 | 0.06525906 | 0.01257667 | 0.18462823 | 0.8690805 |
| Lrfn2 | 0.02495015 | 0.03926686 | 0.02669969 | 0.01256717 | 0.01415733 | 0.15453307 |
| Emc10 | 1.45209279 | 1.03446777 | 1.0219006 | 0.01256716 | 0.47715515 | 0.89054584 |
| Fam20c | 0.61516928 | 0.59868578 | 0.58612275 | 0.01256303 | 0.46022759 | 0.89054584 |
| Gimap8 | 0.0348207 | 0.04982011 | 0.0372604 | 0.01255971 | 0.05815717 | 0.4365583 |
| Rftn2 | 0.0802307 | 0.09836012 | 0.08580712 | 0.01255299 | 0.24775512 | 0.89054584 |
| Cmtm5 | 0.01732319 | 0.03108532 | 0.01853842 | 0.01254689 | 0.00080747 | 0.01374028 |
| Prdm5 | 0.09494559 | 0.11405704 | 0.10151688 | 0.01254017 | 0.28247346 | 0.89054584 |
| Ubxn2b | 0.29419686 | 0.31619679 | 0.30365963 | 0.01253716 | 0.42373279 | 0.89054584 |
| Tssc1 | 0.20125739 | 0.22402769 | 0.21150418 | 0.01252351 | 0.3913264 | 0.89054584 |
| Rasa3 | 0.08025688 | 0.0983474 | 0.08583509 | 0.01251231 | 0.24852432 | 0.89054584 |
| Dlx5 | 0.00948224 | 0.02265803 | 0.01014759 | 0.01251044 | 4.63E-09 | 1.97E-07 |
| 9230110C19I | 0.09757992 | 0.11682575 | 0.10432781 | 0.01249794 | 0.28838212 | 0.89054584 |
| Gm26526 | 0.07561562 | 0.09336865 | 0.08087736 | 0.01249128 | 0.23589783 | 0.89054584 |
| Camkk2 | 0.2937457 | 0.31571187 | 0.30322209 | 0.01248977 | 0.42390924 | 0.89054584 |
| Batf | 0.01831534 | 0.0320861 | 0.01960012 | 0.01248599 | 0.00149914 | 0.02349673 |
| She | 0.03954272 | 0.05479717 | 0.04231189 | 0.01248528 | 0.08460483 | 0.56160366 |
| Pus10 | 0.14550261 | 0.16688997 | 0.15440954 | 0.01248044 | 0.3532475 | 0.89054584 |
| Mus81 | 0.17838831 | 0.20070986 | 0.18823944 | 0.01247042 | 0.37879618 | 0.89054584 |
| Ei24 | 0.8408483 | 0.75702182 | 0.74456452 | 0.01245731 | 0.46893441 | 0.89054584 |
| Tenm3 | 0.2245251 | 0.24739167 | 0.2349445 | 0.01244717 | 0.4025211 | 0.89054584 |
| Gm29417 | 0.02877907 | 0.04323428 | 0.03079654 | 0.01243774 | 0.0299453 | 0.26898821 |
| Elf3 | 0.24201818 | 0.26483513 | 0.25240944 | 0.01242569 | 0.40929778 | 0.89054584 |
| Prpf40b | 0.16287804 | 0.18476358 | 0.17233814 | 0.01242544 | 0.36847089 | 0.89054584 |
| Arhgap18 | 0.18770892 | 0.21016808 | 0.19774813 | 0.01241995 | 0.38490868 | 0.89054584 |
| Lamc1 | 0.52365796 | 0.5243375 | 0.51192284 | 0.01241466 | 0.45502157 | 0.89054584 |
| Nudcd1 | 0.1829958 | 0.20534332 | 0.19294445 | 0.01239887 | 0.38232131 | 0.89054584 |
| Rnf31 | 0.10377402 | 0.12327514 | 0.11087918 | 0.01239596 | 0.3012337 | 0.89054584 |

| | | | | | | |
|---|---|---|---|---|---|---|
| Tmem198b | 0.14672056 | 0.1680311 | 0.1556701 | 0.012361 | 0.35571228 | 0.89054584 |
| Fgf18 | 0.06801313 | 0.08510438 | 0.07275405 | 0.01235033 | 0.21450803 | 0.89054584 |
| Efcab10 | 0.28987216 | 0.31180895 | 0.29946131 | 0.01234765 | 0.42383136 | 0.89054584 |
| Hps4 | 0.25201825 | 0.2746778 | 0.26233064 | 0.01234717 | 0.4132147 | 0.89054584 |
| Cln5 | 0.26195608 | 0.28448176 | 0.27214431 | 0.01233745 | 0.41636191 | 0.89054584 |
| Shf | 0.08592253 | 0.10420556 | 0.09188539 | 0.01232017 | 0.26609052 | 0.89054584 |
| Rmdn1 | 0.29255251 | 0.31437681 | 0.30206443 | 0.01231238 | 0.42469295 | 0.89054584 |
| Fbxo5 | 0.14702744 | 0.16829855 | 0.15598763 | 0.01231092 | 0.35655032 | 0.89054584 |
| Notch1 | 0.20523545 | 0.22783618 | 0.21552817 | 0.01230801 | 0.3950985 | 0.89054584 |
| Adam19 | 0.05364273 | 0.0696856 | 0.05739205 | 0.01229356 | 0.15914939 | 0.8062012 |
| Abcg1 | 0.16650984 | 0.1883573 | 0.17607038 | 0.01228692 | 0.37254505 | 0.89054584 |
| RP23-218K15 | 0.20790042 | 0.23050384 | 0.21822043 | 0.01228341 | 0.39656546 | 0.89054584 |
| Dhfr | 0.18707203 | 0.20937633 | 0.19709955 | 0.01227677 | 0.38583459 | 0.89054584 |
| Ap3m1 | 0.42311456 | 0.43661748 | 0.42434574 | 0.01227174 | 0.44641154 | 0.89054584 |
| Sepsecs | 0.20369703 | 0.22624382 | 0.21397277 | 0.01227104 | 0.39466381 | 0.89054584 |
| Dhx8 | 0.20131935 | 0.2238317 | 0.2115669 | 0.0122648 | 0.39354755 | 0.89054584 |
| B930036N10 | 0.02843742 | 0.04269565 | 0.03043099 | 0.01226466 | 0.03021095 | 0.27028518 |
| Sfswap | 0.43029631 | 0.44306363 | 0.43080078 | 0.01226285 | 0.44724778 | 0.89054584 |
| Nova1 | 0.02803823 | 0.04226378 | 0.03000388 | 0.0122599 | 0.0284751 | 0.25931659 |
| Tsga10ip | 0.04617398 | 0.06165987 | 0.04940485 | 0.01225501 | 0.12390943 | 0.70422347 |
| Irs2 | 0.10556022 | 0.12500001 | 0.11275577 | 0.01224424 | 0.30645701 | 0.89054584 |
| Mrpl30 | 1.36273187 | 1.00532239 | 0.99308946 | 0.01223293 | 0.47711763 | 0.89054584 |
| Khnyn | 0.07207135 | 0.08931168 | 0.07709064 | 0.01222104 | 0.23008295 | 0.89054584 |
| Gfra4 | 0.04729005 | 0.0628041 | 0.05059851 | 0.01220559 | 0.13053847 | 0.72345059 |
| Zbtb7c | 0.12526914 | 0.14558339 | 0.13338387 | 0.01219952 | 0.33501211 | 0.89054584 |
| Dhrs7b | 0.33695833 | 0.35683965 | 0.34466522 | 0.01217443 | 0.43464306 | 0.89054584 |
| Zfp11 | 0.03900115 | 0.05390675 | 0.04173255 | 0.01217419 | 0.08705689 | 0.57045578 |
| Fam20a | 0.11673126 | 0.13663742 | 0.12446557 | 0.01217186 | 0.32433461 | 0.89054584 |
| Nepro | 0.04848132 | 0.06404353 | 0.05187256 | 0.01217097 | 0.13716665 | 0.7420427 |
| Cfl2 | 1.09018055 | 0.89253191 | 0.88039908 | 0.01213283 | 0.47440344 | 0.89054584 |

| | | | | | | |
|---|---|---|---|---|---|---|
| Col17a1 | 0.03376021 | 0.04823754 | 0.03612585 | 0.01211169 | 0.05914703 | 0.44117919 |
| Nebl | 0.02633652 | 0.04028278 | 0.02818309 | 0.01209969 | 0.0227395 | 0.22105677 |
| Fanca | 0.02715539 | 0.04115773 | 0.02905927 | 0.01209847 | 0.02620852 | 0.24404059 |
| Pdhb | 0.92160008 | 0.80445518 | 0.79236514 | 0.01209004 | 0.4716643 | 0.89054584 |
| Steap3 | 0.02736246 | 0.04136725 | 0.02928083 | 0.01208643 | 0.02723376 | 0.25100767 |
| Ufc1 | 1.25521936 | 0.96535529 | 0.95326957 | 0.01208572 | 0.47644943 | 0.89054584 |
| Ppp1r16b | 0.04274901 | 0.05782461 | 0.04574157 | 0.01208304 | 0.10921741 | 0.65377127 |
| Jam2 | 0.04352891 | 0.05865367 | 0.04657577 | 0.0120779 | 0.113496 | 0.67107054 |
| Alad | 0.72249279 | 0.67803021 | 0.66596633 | 0.01206388 | 0.46637077 | 0.89054584 |
| Arsk | 0.18699889 | 0.20908612 | 0.19702506 | 0.01206106 | 0.38774506 | 0.89054584 |
| Zfp28 | 0.05850709 | 0.07464671 | 0.062593 | 0.01205371 | 0.18480999 | 0.8690805 |
| Osgepl1 | 0.20301265 | 0.22532779 | 0.21328053 | 0.01204726 | 0.39621263 | 0.89054584 |
| Tmem255b | 0.03075551 | 0.04493342 | 0.03291118 | 0.01202224 | 0.04438889 | 0.36154499 |
| Gm26830 | 0.03718554 | 0.05180058 | 0.03979031 | 0.01201027 | 0.07982488 | 0.54094194 |
| 4930589L23F | 0.0166797 | 0.02984756 | 0.01784982 | 0.01199774 | 0.00086958 | 0.01469359 |
| Gm26648 | 0.02222747 | 0.03577863 | 0.02378635 | 0.01199228 | 0.00941495 | 0.11157803 |
| Akna | 0.04640879 | 0.0616317 | 0.04965599 | 0.0119757 | 0.1305887 | 0.72345243 |
| Sec62 | 1.97068135 | 1.12767046 | 1.11570514 | 0.01196532 | 0.48007531 | 0.89054584 |
| Zfp672 | 0.26762425 | 0.28967637 | 0.2777212 | 0.01195517 | 0.42052318 | 0.89054584 |
| Bap1 | 0.30567897 | 0.32666724 | 0.31476066 | 0.01190658 | 0.4300544 | 0.89054584 |
| Ppp1r12a | 0.65214587 | 0.62643179 | 0.61452762 | 0.01190418 | 0.46404444 | 0.89054584 |
| Sel1l | 0.65661193 | 0.62979164 | 0.61789551 | 0.01189613 | 0.464264 | 0.89054584 |
| Galnt12 | 0.34096169 | 0.36034666 | 0.34845674 | 0.01188992 | 0.43684595 | 0.89054584 |
| Lefty1 | 0.10708592 | 0.12624443 | 0.11435776 | 0.01188667 | 0.31409938 | 0.89054584 |
| Gm10138 | 0.15832631 | 0.17953756 | 0.16765299 | 0.01188457 | 0.3706008 | 0.89054584 |
| Cfap69 | 0.04760991 | 0.06281421 | 0.0509406 | 0.01187361 | 0.13875281 | 0.74747802 |
| Uqcrq | 2.44866498 | 1.14020423 | 1.12834776 | 0.01185646 | 0.48047747 | 0.89054584 |
| Unk | 0.17465216 | 0.19627023 | 0.18441819 | 0.01185204 | 0.38231093 | 0.89054584 |
| Ints6 | 0.12301418 | 0.14286631 | 0.13103111 | 0.01183521 | 0.336946 | 0.89054584 |
| Fam65b | 0.0311736 | 0.04517591 | 0.0333585 | 0.01181741 | 0.04942528 | 0.39028741 |

| | | | | | | |
|---|---|---|---|---|---|---|
| Gm6576 | 0.14878756 | 0.1696252 | 0.15780807 | 0.01181714 | 0.36359146 | 0.89054584 |
| Cherp | 0.31035589 | 0.33107954 | 0.31926326 | 0.01181628 | 0.4315488 | 0.89054584 |
| Gapdh | 3.15570295 | 1.09180078 | 1.07998543 | 0.01181534 | 0.47967466 | 0.89054584 |
| Gal3st1 | 0.04842343 | 0.06362438 | 0.05181064 | 0.01181374 | 0.14404496 | 0.76637203 |
| Rab40c | 0.12973051 | 0.14984698 | 0.13803338 | 0.0118136 | 0.3450439 | 0.89054584 |
| Syt1 | 0.02243189 | 0.03581019 | 0.0240051 | 0.01180509 | 0.01097744 | 0.12621003 |
| Tmed5 | 0.45365354 | 0.46339172 | 0.45158679 | 0.01180493 | 0.45153291 | 0.89054584 |
| Syt2 | 0.02860579 | 0.04241449 | 0.03061114 | 0.01180336 | 0.03621206 | 0.31212984 |
| Crmp1 | 0.05417879 | 0.06975919 | 0.05796524 | 0.01179395 | 0.17158109 | 0.83588128 |
| Ikzf2 | 0.21125251 | 0.23339235 | 0.22160244 | 0.01178991 | 0.40211768 | 0.89054584 |
| Hoxa3 | 0.06827982 | 0.0848184 | 0.07303906 | 0.01177934 | 0.22621554 | 0.89054584 |
| Ikbke | 0.05106062 | 0.06640961 | 0.05463091 | 0.0117787 | 0.15757083 | 0.80128442 |
| Zfp131 | 0.2605713 | 0.28255412 | 0.27077949 | 0.01177462 | 0.41972766 | 0.89054584 |
| Dcbld2 | 0.09308085 | 0.11128484 | 0.09952684 | 0.011758 | 0.29102163 | 0.89054584 |
| Prickle2 | 0.06049924 | 0.0764805 | 0.06472274 | 0.01175776 | 0.19867599 | 0.89054584 |
| Epha5 | 0.0263182 | 0.03991644 | 0.02816349 | 0.01175295 | 0.0259339 | 0.24226138 |
| Senp2 | 0.2249591 | 0.24713026 | 0.23537944 | 0.01175082 | 0.40804077 | 0.89054584 |
| Tram2 | 0.04500551 | 0.05989792 | 0.04815513 | 0.01174279 | 0.12795764 | 0.71598989 |
| Smim5 | 0.04248436 | 0.05718109 | 0.04545849 | 0.01172259 | 0.11479359 | 0.6757765 |
| Tmem200a | 0.09374424 | 0.11195115 | 0.10023483 | 0.01171632 | 0.293022 | 0.89054584 |
| D930028M14 | 0.04247345 | 0.05716305 | 0.04544682 | 0.01171623 | 0.11486021 | 0.67582927 |
| Asah2 | 0.05861378 | 0.07440192 | 0.06270706 | 0.01169486 | 0.19245312 | 0.88605441 |
| 2500002B13 | 0.03447385 | 0.04857726 | 0.03688933 | 0.01168793 | 0.06995474 | 0.49764662 |
| Blvra | 0.45747967 | 0.46664638 | 0.45496113 | 0.01168524 | 0.45237606 | 0.89054584 |
| Rab11fip5 | 0.05442878 | 0.06991679 | 0.05823254 | 0.01168424 | 0.17494381 | 0.84263634 |
| 5033417F24 | 0.03694453 | 0.05121415 | 0.03953248 | 0.01168167 | 0.08430318 | 0.55988435 |
| Agpat2 | 0.32814728 | 0.3479664 | 0.33629257 | 0.01167382 | 0.43576035 | 0.89054584 |
| Zfp612 | 0.26065438 | 0.28253024 | 0.27086139 | 0.01166885 | 0.42046282 | 0.89054584 |
| Lmo3 | 0.06654957 | 0.08282576 | 0.0711899 | 0.01163586 | 0.22318065 | 0.89054584 |
| Rhoh | 0.02177054 | 0.03492424 | 0.02329741 | 0.01162682 | 0.01003346 | 0.11777817 |

| | | | | | | |
|---|---|---|---|---|---|---|
| Cxx1a | 0.83884302 | 0.75494373 | 0.74331719 | 0.01162654 | 0.4709538 | 0.89054584 |
| Lmntd1 | 0.03094802 | 0.04474263 | 0.03311715 | 0.01162548 | 0.05097421 | 0.39853675 |
| Cer1 | 0.09235814 | 0.11037624 | 0.0987555 | 0.01162073 | 0.29176846 | 0.89054584 |
| Ddi2 | 0.33110222 | 0.35072161 | 0.33910498 | 0.01161663 | 0.43659804 | 0.89054584 |
| Cpt1c | 0.17267529 | 0.19401046 | 0.18239394 | 0.01161652 | 0.38333861 | 0.89054584 |
| Phkg2 | 0.38322493 | 0.39953713 | 0.38796034 | 0.01157679 | 0.4447158 | 0.89054584 |
| Cdkn3 | 0.06003712 | 0.07579478 | 0.06422872 | 0.01156606 | 0.20074467 | 0.89054584 |
| Reps1 | 0.13366916 | 0.15369055 | 0.14213189 | 0.01155866 | 0.35238803 | 0.89054584 |
| Ubap1 | 0.40080719 | 0.41565989 | 0.40410836 | 0.01155153 | 0.44702676 | 0.89054584 |
| Pcdhb14 | 0.03074018 | 0.04444422 | 0.03289478 | 0.01154944 | 0.05094435 | 0.39853675 |
| Phtf1 | 0.29593741 | 0.31688806 | 0.30534668 | 0.01154138 | 0.43010889 | 0.89054584 |
| Bcap29 | 0.19019454 | 0.21180135 | 0.20027772 | 0.01152363 | 0.39432378 | 0.89054584 |
| Cd96 | 0.0213129 | 0.03432815 | 0.02280772 | 0.01152043 | 0.00930423 | 0.11065855 |
| Rnf152 | 0.05641372 | 0.07186302 | 0.06035489 | 0.01150814 | 0.1871786 | 0.87426397 |
| Gcnt1 | 0.06462816 | 0.08062163 | 0.06913629 | 0.01148533 | 0.21947391 | 0.89054584 |
| Trpm7 | 0.35551461 | 0.37363306 | 0.36216742 | 0.01146564 | 0.4413708 | 0.89054584 |
| Hk1os | 0.02325312 | 0.03634945 | 0.02488384 | 0.01146561 | 0.01590942 | 0.16997399 |
| Cd99l2 | 0.34326682 | 0.36209915 | 0.35063603 | 0.01146312 | 0.43947049 | 0.89054584 |
| Snapc1 | 0.22242037 | 0.2442898 | 0.23283401 | 0.01145579 | 0.40934624 | 0.89054584 |
| Ptprg | 0.12600026 | 0.14560158 | 0.13414628 | 0.0114553 | 0.34537181 | 0.89054584 |
| Smim13 | 0.18186414 | 0.2032103 | 0.19178965 | 0.01142064 | 0.3907252 | 0.89054584 |
| Slc16a7 | 0.03659404 | 0.05057748 | 0.03915753 | 0.01141994 | 0.08711471 | 0.57045578 |
| Rtkn | 0.2136528 | 0.23543399 | 0.2240211 | 0.0114129 | 0.4061916 | 0.89054584 |
| Plppr3 | 0.10142833 | 0.11981728 | 0.10841302 | 0.01140425 | 0.31203637 | 0.89054584 |
| Dner | 0.07509205 | 0.09169345 | 0.08031802 | 0.01137543 | 0.25467656 | 0.89054584 |
| Mesp2 | 0.04010228 | 0.05428226 | 0.04291045 | 0.01137181 | 0.10847509 | 0.65191723 |
| Rbfox1 | 0.01758432 | 0.03018403 | 0.01881786 | 0.01136617 | 0.00244612 | 0.03588252 |
| Tslp | 0.03829248 | 0.05233788 | 0.04097446 | 0.01136341 | 0.09816812 | 0.61534939 |
| Xkr8 | 0.05369793 | 0.06880836 | 0.05745106 | 0.01135729 | 0.17852282 | 0.85194992 |
| Gm20186 | 0.06064868 | 0.07623871 | 0.0648825 | 0.01135621 | 0.20740957 | 0.89054584 |

| | | | | | | |
|---|---|---|---|---|---|---|
| Bche | 0.02356336 | 0.03656553 | 0.0252158 | 0.01134973 | 0.01799647 | 0.18665181 |
| Tirap | 0.05539192 | 0.07060833 | 0.05926237 | 0.01134596 | 0.18620485 | 0.87252679 |
| Itgb3 | 0.05015744 | 0.06499591 | 0.05366505 | 0.01133086 | 0.16263387 | 0.81558832 |
| Sema6d | 0.2421872 | 0.26389718 | 0.25257752 | 0.01131967 | 0.417303 | 0.89054584 |
| Gm11837 | 0.13428375 | 0.1540878 | 0.14277089 | 0.01131691 | 0.35595331 | 0.89054584 |
| Kif5c | 0.20084387 | 0.22239525 | 0.2110855 | 0.01130975 | 0.40144006 | 0.89054584 |
| Nrxn3 | 0.05992773 | 0.07542032 | 0.06411177 | 0.01130855 | 0.20560069 | 0.89054584 |
| 6430503K07I | 0.1185071 | 0.13762368 | 0.12632279 | 0.01130089 | 0.33841567 | 0.89054584 |
| St6galnac5 | 0.14056968 | 0.16059519 | 0.14929807 | 0.01129712 | 0.36221898 | 0.89054584 |
| Pla2g2d | 0.17340571 | 0.19442623 | 0.18314205 | 0.01128418 | 0.38703406 | 0.89054584 |
| Prune2 | 0.11864843 | 0.13775334 | 0.12647055 | 0.01128279 | 0.33883773 | 0.89054584 |
| Gm26724 | 0.34302371 | 0.36167523 | 0.35040632 | 0.01126891 | 0.44044959 | 0.89054584 |
| C530044C16I | 0.04631955 | 0.06078391 | 0.04956055 | 0.01122336 | 0.14570068 | 0.76925292 |
| Sf3a2 | 0.52310797 | 0.52267593 | 0.51146054 | 0.01121539 | 0.45931405 | 0.89054584 |
| Vasn | 0.17008779 | 0.19094412 | 0.17974199 | 0.01120213 | 0.38577004 | 0.89054584 |
| Timm21 | 0.39058987 | 0.4059402 | 0.39474491 | 0.0111953 | 0.44744015 | 0.89054584 |
| Ppp1r35 | 0.54721766 | 0.54273657 | 0.53154387 | 0.0111927 | 0.46092518 | 0.89054584 |
| Gltscr2 | 1.74787221 | 1.10101839 | 1.08983676 | 0.01118163 | 0.48093768 | 0.89054584 |
| Gemin6 | 0.51950726 | 0.51960856 | 0.50842941 | 0.01117915 | 0.45920411 | 0.89054584 |
| Arhgap28 | 0.12015309 | 0.13922118 | 0.12804315 | 0.01117802 | 0.34210672 | 0.89054584 |
| Cdh13 | 0.03088103 | 0.04421304 | 0.03304547 | 0.01116757 | 0.05768968 | 0.4351639 |
| Fstl4 | 0.02017931 | 0.032762 | 0.02159471 | 0.01116729 | 0.00799194 | 0.09720403 |
| Mrps22 | 0.35770417 | 0.37538512 | 0.36422036 | 0.01116476 | 0.44321834 | 0.89054584 |
| Golt1a | 0.0237314 | 0.03655925 | 0.0253956 | 0.01116365 | 0.0202789 | 0.20318142 |
| Tex30 | 0.38310945 | 0.39899993 | 0.38785372 | 0.01114621 | 0.44674498 | 0.89054584 |
| Cox14 | 1.89965394 | 1.1207891 | 1.10965083 | 0.01113827 | 0.48135036 | 0.89054584 |
| Csgalnact1 | 0.03986895 | 0.05379196 | 0.04266086 | 0.0111311 | 0.11206602 | 0.66560148 |
| Tnfrsf13b | 0.0235473 | 0.03632828 | 0.02519862 | 0.01112966 | 0.01980714 | 0.19997491 |
| 1700025G04 | 0.28336405 | 0.30425458 | 0.29312596 | 0.01112862 | 0.42980206 | 0.89054584 |
| Lrrc27 | 0.03493029 | 0.0485062 | 0.03737764 | 0.01112856 | 0.08270242 | 0.55279838 |

| | | | | | | |
|---|---|---|---|---|---|---|
| Dll3 | 0.0560869 | 0.07113395 | 0.06000546 | 0.01112849 | 0.19378348 | 0.88879244 |
| Gm525 | 0.01915564 | 0.0316266 | 0.02049931 | 0.01112729 | 0.0057205 | 0.07307121 |
| Prdm1 | 0.01879312 | 0.03122691 | 0.02011138 | 0.01111553 | 0.00501214 | 0.06612012 |
| Ttk | 0.07244384 | 0.08858684 | 0.07748864 | 0.0110982 | 0.25229972 | 0.89054584 |
| Gm42726 | 0.02947194 | 0.04263021 | 0.03153787 | 0.01109235 | 0.05064619 | 0.39690185 |
| Ap5b1 | 0.02329406 | 0.03600904 | 0.02492764 | 0.0110814 | 0.0191748 | 0.19549667 |
| Dusp26 | 0.03637893 | 0.05000745 | 0.03892741 | 0.01108004 | 0.09240483 | 0.59278106 |
| Pcsk9 | 0.02849623 | 0.04157333 | 0.03049391 | 0.01107942 | 0.04525215 | 0.36609429 |
| Abhd11os | 0.06474268 | 0.08033768 | 0.0692587 | 0.01107898 | 0.22805475 | 0.89054584 |
| Hspb2 | 0.02307895 | 0.0357764 | 0.02469747 | 0.01107893 | 0.01831185 | 0.18936177 |
| Slc38a1 | 0.44720037 | 0.45695 | 0.44587566 | 0.01107433 | 0.45393906 | 0.89054584 |
| Cntln | 0.38858674 | 0.40397549 | 0.39290226 | 0.01107323 | 0.44776754 | 0.89054584 |
| Bcl9l | 0.13383603 | 0.1533775 | 0.1423054 | 0.0110721 | 0.35849302 | 0.89054584 |
| Nthl1 | 0.17022542 | 0.19095501 | 0.17988312 | 0.01107189 | 0.38714809 | 0.89054584 |
| Nhs | 0.06876397 | 0.08462384 | 0.07355645 | 0.01106739 | 0.24165475 | 0.89054584 |
| Tmem266 | 0.04181103 | 0.05579912 | 0.04473826 | 0.01106086 | 0.12469084 | 0.70642606 |
| Zfp825 | 0.13067503 | 0.15007325 | 0.13901677 | 0.01105648 | 0.35548953 | 0.89054584 |
| Smim10l2a | 0.06045516 | 0.07572998 | 0.06467562 | 0.01105436 | 0.21292702 | 0.89054584 |
| Parp8 | 0.1359976 | 0.15560578 | 0.14455203 | 0.01105375 | 0.3608212 | 0.89054584 |
| Tpcn2 | 0.02211007 | 0.03471255 | 0.02366074 | 0.01105181 | 0.01477136 | 0.15969813 |
| Ephx2 | 0.06914476 | 0.08499635 | 0.07396339 | 0.01103296 | 0.2435389 | 0.89054584 |
| Daam2 | 0.01591386 | 0.02806215 | 0.01703029 | 0.01103185 | 0.00127241 | 0.02033896 |
| Fam168b | 0.32335055 | 0.34274379 | 0.33171747 | 0.01102632 | 0.43846438 | 0.89054584 |
| Wdr53 | 0.18227808 | 0.20321971 | 0.19221212 | 0.01100758 | 0.39480858 | 0.89054584 |
| Pald1 | 0.08294209 | 0.09970966 | 0.08870283 | 0.01100683 | 0.28159427 | 0.89054584 |
| Bhlhe23 | 0.01626446 | 0.0284078 | 0.01740548 | 0.01100232 | 0.00161476 | 0.0250115 |
| Gm9801 | 0.04177251 | 0.05569455 | 0.04469706 | 0.01099749 | 0.12583349 | 0.71013386 |
| Ovca2 | 0.05736409 | 0.07235169 | 0.06137099 | 0.0109807 | 0.20225509 | 0.89054584 |
| Ablim3 | 0.02307265 | 0.03566817 | 0.02469073 | 0.01097744 | 0.01916282 | 0.19549667 |
| Tstd3 | 0.0962697 | 0.11390544 | 0.10292981 | 0.01097563 | 0.30966756 | 0.89054584 |

| | | | | | | |
|---|---|---|---|---|---|---|
| Prom1 | 0.18020555 | 0.20106913 | 0.19009619 | 0.01097295 | 0.3939923 | 0.89054584 |
| Bhlha15 | 0.09089406 | 0.1081628 | 0.09719281 | 0.01096999 | 0.29949727 | 0.89054584 |
| Plac9b | 0.05654939 | 0.07146733 | 0.06049994 | 0.01096739 | 0.19917542 | 0.89054584 |
| Gca | 0.14445482 | 0.16427507 | 0.15332463 | 0.01095044 | 0.3696647 | 0.89054584 |
| Scn2b | 0.04548749 | 0.05961756 | 0.04867063 | 0.01094693 | 0.14734633 | 0.77404405 |
| Usp12 | 0.20862299 | 0.22989315 | 0.21894987 | 0.01094329 | 0.407936 | 0.89054584 |
| Abat | 0.18866343 | 0.20965795 | 0.19871984 | 0.01093811 | 0.39880499 | 0.89054584 |
| Magix | 0.02702743 | 0.03985705 | 0.02892235 | 0.0109347 | 0.03908458 | 0.33156714 |
| Hoxb5os | 0.05357396 | 0.06824382 | 0.05731851 | 0.01092531 | 0.18726312 | 0.87437703 |
| Shb | 0.04968025 | 0.06407078 | 0.05315474 | 0.01091604 | 0.16934009 | 0.83251702 |
| Fzr1 | 0.45773151 | 0.46609795 | 0.45518292 | 0.01091503 | 0.45552321 | 0.89054584 |
| Mcc | 0.09860556 | 0.11632051 | 0.10542207 | 0.01089844 | 0.31503179 | 0.89054584 |
| Avpr1b | 0.03247109 | 0.04563691 | 0.03474667 | 0.01089023 | 0.07211807 | 0.50853143 |
| Neil3 | 0.0652572 | 0.08069516 | 0.06980864 | 0.01088652 | 0.23374994 | 0.89054584 |
| Ddx25 | 0.07142645 | 0.08728316 | 0.07640156 | 0.0108816 | 0.25348719 | 0.89054584 |
| Cgnl1 | 0.1331585 | 0.1524723 | 0.14160084 | 0.01087146 | 0.36028694 | 0.89054584 |
| Sccpdh | 0.45326764 | 0.46211701 | 0.45124597 | 0.01087105 | 0.45531681 | 0.89054584 |
| Polb | 0.52076669 | 0.52034467 | 0.50949035 | 0.01085432 | 0.46046778 | 0.89054584 |
| Arl5c | 0.02822262 | 0.04104439 | 0.03020117 | 0.01084322 | 0.04719175 | 0.37737782 |
| Zfp87 | 0.16304097 | 0.18333892 | 0.1725057 | 0.01083322 | 0.3849224 | 0.89054584 |
| Lipt2 | 0.13519318 | 0.1545424 | 0.14371617 | 0.01082623 | 0.36280886 | 0.89054584 |
| Ppargc1b | 0.03449322 | 0.04771951 | 0.03691005 | 0.01080946 | 0.08621838 | 0.56777946 |
| Slc20a1 | 0.09538221 | 0.11278934 | 0.1019828 | 0.01080654 | 0.31076555 | 0.89054584 |
| Ddx3y | 0.43730423 | 0.44787194 | 0.43707029 | 0.01080164 | 0.45416714 | 0.89054584 |
| Chad | 0.01655897 | 0.02851334 | 0.01772063 | 0.01079271 | 0.0022733 | 0.03366526 |
| Hpse | 0.03517603 | 0.04842005 | 0.03764054 | 0.01077951 | 0.09106434 | 0.58666938 |
| Inafm1 | 0.28290902 | 0.30344927 | 0.29268232 | 0.01076695 | 0.43195885 | 0.89054584 |
| Aifm1 | 0.40880262 | 0.42214907 | 0.41139409 | 0.01075499 | 0.45152962 | 0.89054584 |
| Srp72 | 1.30075675 | 0.98158913 | 0.97085231 | 0.01073682 | 0.47945398 | 0.89054584 |
| Usp49 | 0.03402955 | 0.04713262 | 0.036414 | 0.01071862 | 0.08512819 | 0.56355642 |

| | | | | | | |
|---|---|---|---|---|---|---|
| Snrpb | 2.22487333 | 1.13859455 | 1.12788278 | 0.01071177 | 0.48235373 | 0.89054584 |
| Chchd6 | 0.51374604 | 0.51424458 | 0.50356343 | 0.01068115 | 0.46064002 | 0.89054584 |
| Fam120b | 0.17166118 | 0.19202359 | 0.1813549 | 0.0106687 | 0.39201352 | 0.89054584 |
| Hnrnpf | 2.41276393 | 1.13969779 | 1.12903015 | 0.01066765 | 0.48244422 | 0.89054584 |
| Txndc16 | 0.08040323 | 0.09665655 | 0.08599139 | 0.01066515 | 0.28168978 | 0.89054584 |
| C030034I22R | 0.06856753 | 0.08400946 | 0.07334653 | 0.01066293 | 0.24910627 | 0.89054584 |
| Bin2 | 0.03163529 | 0.04451151 | 0.03385245 | 0.01065906 | 0.07119461 | 0.50399388 |
| Itpr1 | 0.58691079 | 0.57443384 | 0.56379466 | 0.01063919 | 0.46497117 | 0.89054584 |
| Xrcc3 | 0.05546502 | 0.06997347 | 0.05934053 | 0.01063294 | 0.20191151 | 0.89054584 |
| Fam53b | 0.10213805 | 0.11977069 | 0.10915941 | 0.01061128 | 0.32531353 | 0.89054584 |
| Zfp804a | 0.02195071 | 0.03409647 | 0.02349021 | 0.01060625 | 0.01770686 | 0.18428859 |
| Trp53cor1 | 0.05625205 | 0.07078826 | 0.06018203 | 0.01060624 | 0.20580236 | 0.89054584 |
| Nagpa | 0.13341511 | 0.1524675 | 0.14186771 | 0.01059979 | 0.36388349 | 0.89054584 |
| Mapk8ip1 | 0.61082064 | 0.59331942 | 0.58272135 | 0.01059806 | 0.46623681 | 0.89054584 |
| Borcs6 | 0.29848098 | 0.31839003 | 0.30780934 | 0.01058068 | 0.43638259 | 0.89054584 |
| Gm5771 | 0.01542878 | 0.02708497 | 0.0165112 | 0.01057377 | 0.00142447 | 0.02249665 |
| Mthfd2 | 0.32622399 | 0.34503017 | 0.33445959 | 0.01057058 | 0.44146909 | 0.89054584 |
| Itgb2 | 0.02568633 | 0.03805064 | 0.0274874 | 0.01056324 | 0.03669456 | 0.31553841 |
| Ctgf | 0.05936753 | 0.07406919 | 0.06351288 | 0.01055631 | 0.21936186 | 0.89054584 |
| Rerg | 0.05524711 | 0.06966319 | 0.05910754 | 0.01055565 | 0.20270118 | 0.89054584 |
| Taf1b | 0.16826347 | 0.18841954 | 0.17787057 | 0.01054897 | 0.39115583 | 0.89054584 |
| Stk17b | 0.05252238 | 0.06672413 | 0.05619405 | 0.01053008 | 0.19132364 | 0.88394202 |
| Rcan3 | 0.38529587 | 0.40039396 | 0.38987101 | 0.01052294 | 0.44996542 | 0.89054584 |
| Fzd4 | 0.07822483 | 0.09417704 | 0.08366463 | 0.01051241 | 0.27914169 | 0.89054584 |
| Pdgfb | 0.02605447 | 0.0383925 | 0.02788131 | 0.01051119 | 0.03950924 | 0.33458234 |
| Tmem132c | 0.02216796 | 0.03423021 | 0.02372268 | 0.01050753 | 0.01952814 | 0.19784693 |
| Psmb7 | 1.96458746 | 1.12570862 | 1.11524301 | 0.01046561 | 0.48256372 | 0.89054584 |
| Zfp52 | 0.03889019 | 0.05206256 | 0.04161386 | 0.0104487 | 0.12104087 | 0.69664572 |
| Lin7a | 0.0391054 | 0.05229002 | 0.04184408 | 0.01044594 | 0.12240332 | 0.70076598 |
| Arrdc1 | 0.33722261 | 0.35533829 | 0.34491578 | 0.01042251 | 0.44402117 | 0.89054584 |

| | | | | | | |
|---|---|---|---|---|---|---|
| Slc35d3 | 0.12309695 | 0.14151117 | 0.1311175 | 0.01039368 | 0.35594682 | 0.89054584 |
| Ankrd39 | 0.34084835 | 0.35874278 | 0.34834951 | 0.01039326 | 0.44472392 | 0.89054584 |
| Ssu72 | 1.5368601 | 1.05583932 | 1.04544763 | 0.0103917 | 0.48153173 | 0.89054584 |
| Kdelc1 | 0.17011062 | 0.19015152 | 0.1797654 | 0.01038612 | 0.39389753 | 0.89054584 |
| Polr3e | 0.20076694 | 0.22137945 | 0.2110076 | 0.01037185 | 0.40943241 | 0.89054584 |
| Zfp948 | 0.1676307 | 0.18758361 | 0.17722114 | 0.01036248 | 0.39265046 | 0.89054584 |
| Rbx1 | 2.72573863 | 1.12573799 | 1.11537906 | 0.01035893 | 0.48274344 | 0.89054584 |
| Agt | 0.09579353 | 0.11276991 | 0.10242172 | 0.01034819 | 0.31892147 | 0.89054584 |
| Snx10 | 0.36010275 | 0.37681162 | 0.36646631 | 0.01034531 | 0.44768141 | 0.89054584 |
| Ptgds | 0.09251967 | 0.10927307 | 0.09892791 | 0.01034516 | 0.31305898 | 0.89054584 |
| Adpgk | 0.29159771 | 0.31146767 | 0.30113755 | 0.01033013 | 0.4365119 | 0.89054584 |
| Prr5l | 0.05523894 | 0.06941651 | 0.0590988 | 0.0103177 | 0.20800117 | 0.89054584 |
| Gins1 | 0.08539164 | 0.10162645 | 0.09131853 | 0.01030792 | 0.29948047 | 0.89054584 |
| Rai1 | 0.16869034 | 0.18861515 | 0.17830858 | 0.01030657 | 0.39385115 | 0.89054584 |
| Tars | 0.40768005 | 0.42067836 | 0.41037337 | 0.01030499 | 0.45343325 | 0.89054584 |
| Zfp369 | 0.10535985 | 0.12281838 | 0.11254532 | 0.01027305 | 0.33532187 | 0.89054584 |
| 1110038F14I | 0.66980741 | 0.63802008 | 0.62776609 | 0.01025399 | 0.46966995 | 0.89054584 |
| Elmod1 | 0.06802989 | 0.08302535 | 0.07277196 | 0.0102534 | 0.25577252 | 0.89054584 |
| Tmem2 | 0.10723956 | 0.12476342 | 0.11451904 | 0.01024438 | 0.33842376 | 0.89054584 |
| Dnase1l1 | 0.05968735 | 0.07409783 | 0.06385479 | 0.01024304 | 0.22742691 | 0.89054584 |
| Cdx2 | 0.02690097 | 0.03902965 | 0.02878704 | 0.01024261 | 0.04869105 | 0.38638363 |
| Pld2 | 0.05755391 | 0.07180481 | 0.06157393 | 0.01023088 | 0.2194325 | 0.89054584 |
| Zfp459 | 0.01728923 | 0.02873188 | 0.01850208 | 0.0102298 | 0.00499856 | 0.06600098 |
| Tm7sf2 | 0.08416565 | 0.10021985 | 0.09000943 | 0.01021041 | 0.29857724 | 0.89054584 |
| Gm15050 | 0.05077979 | 0.06453948 | 0.0543306 | 0.01020888 | 0.19067028 | 0.88317499 |
| Man1a | 0.15867603 | 0.17821334 | 0.16801325 | 0.01020009 | 0.38864805 | 0.89054584 |
| Keap1 | 0.8150629 | 0.73849552 | 0.72830405 | 0.01019147 | 0.47400954 | 0.89054584 |
| Gria4 | 0.01658733 | 0.02792047 | 0.01775098 | 0.01016949 | 0.00380279 | 0.05230825 |
| Gm16124 | 0.10228081 | 0.1194766 | 0.10930951 | 0.01016709 | 0.3323849 | 0.89054584 |
| Cyp2d22 | 0.06162103 | 0.07607778 | 0.06592193 | 0.01015584 | 0.23645567 | 0.89054584 |

| | | | | | | |
|---|---|---|---|---|---|---|
| Sap30bp | 0.26981668 | 0.29002469 | 0.27987418 | 0.01015051 | 0.43290974 | 0.89054584 |
| Lmf1 | 0.23766239 | 0.25821628 | 0.24807348 | 0.0101428 | 0.42446385 | 0.89054584 |
| St8sia4 | 0.03114801 | 0.04347362 | 0.03333112 | 0.0101425 | 0.07814019 | 0.53336934 |
| Ttc38 | 0.09886557 | 0.11583725 | 0.10569947 | 0.01013777 | 0.32749344 | 0.89054584 |
| Scyl3 | 0.11075415 | 0.12833638 | 0.11820592 | 0.01013046 | 0.34484362 | 0.89054584 |
| Edc3 | 0.19596284 | 0.21625415 | 0.2061379 | 0.01011625 | 0.40957501 | 0.89054584 |
| Flad1 | 0.2535303 | 0.27393824 | 0.26382668 | 0.01011156 | 0.42914121 | 0.89054584 |
| Itpkc | 0.04503046 | 0.05828737 | 0.04818181 | 0.01010556 | 0.16424566 | 0.81857464 |
| Enpp5 | 0.26904858 | 0.2892051 | 0.27912017 | 0.01008493 | 0.43316071 | 0.89054584 |
| Gpt2 | 0.13737132 | 0.15605784 | 0.14597888 | 0.01007896 | 0.37385067 | 0.89054584 |
| Sep-05 | 0.07199768 | 0.08708895 | 0.07701193 | 0.01007701 | 0.2710559 | 0.89054584 |
| Mtg1 | 0.27174271 | 0.29183106 | 0.28176364 | 0.01006742 | 0.43389591 | 0.89054584 |
| Tmem151a | 0.03836067 | 0.05108153 | 0.04104741 | 0.01003411 | 0.12737526 | 0.7143855 |
| Cdk5rap1 | 0.13959088 | 0.15831621 | 0.14828272 | 0.01003349 | 0.37628771 | 0.89054584 |
| Cby1 | 0.60616237 | 0.58909679 | 0.57906354 | 0.01003325 | 0.46783073 | 0.89054584 |
| Dpp6 | 0.06678407 | 0.08147113 | 0.07144053 | 0.0100306 | 0.25651139 | 0.89054584 |
| Gm8113 | 0.02523866 | 0.03703798 | 0.02700839 | 0.01002959 | 0.04180597 | 0.34614674 |
| 1500026H17 | 0.12786231 | 0.1461019 | 0.13608727 | 0.01001462 | 0.36585564 | 0.89054584 |
| Gdpd3 | 0.04338396 | 0.05643047 | 0.04642072 | 0.01000975 | 0.15754173 | 0.80128442 |
| Pappa2 | 0.10021681 | 0.11714675 | 0.10713851 | 0.01000824 | 0.33170482 | 0.89054584 |
| Usp46 | 0.29105647 | 0.31061177 | 0.30061193 | 0.00999984 | 0.43841877 | 0.89054584 |
| Bcl10 | 0.66494317 | 0.63412805 | 0.62414165 | 0.00998641 | 0.47028872 | 0.89054584 |
| Trp53inp1 | 0.17859384 | 0.19842033 | 0.18844948 | 0.00997085 | 0.40264595 | 0.89054584 |
| Cfap43 | 0.04973156 | 0.06316884 | 0.05320961 | 0.00995923 | 0.19160061 | 0.88461814 |
| Dynlt1f | 0.79775239 | 0.72706002 | 0.7171168 | 0.00994322 | 0.47424672 | 0.89054584 |
| Ccdc122 | 0.03674338 | 0.04925237 | 0.03931729 | 0.00993508 | 0.1195445 | 0.69215504 |
| Gm13056 | 0.01272163 | 0.02353178 | 0.01361421 | 0.00991756 | 0.00034452 | 0.00644494 |
| Lhx1os | 0.14209284 | 0.16079483 | 0.15087738 | 0.00991745 | 0.37971059 | 0.89054584 |
| Gm10116 | 0.49399975 | 0.49663165 | 0.4867263 | 0.00990536 | 0.46223128 | 0.89054584 |
| Stk26 | 0.03839999 | 0.05099117 | 0.04108948 | 0.00990169 | 0.13078126 | 0.72424246 |

| | | | | | | |
|---|---|---|---|---|---|---|
| Fgl1 | 0.02415545 | 0.03573901 | 0.02584935 | 0.00988966 | 0.03733743 | 0.32027435 |
| Ccdc17 | 0.04318213 | 0.05607225 | 0.04620485 | 0.00986741 | 0.1598795 | 0.80736085 |
| Cerkl | 0.17858807 | 0.19830352 | 0.18844359 | 0.00985993 | 0.4037046 | 0.89054584 |
| Il18 | 0.04032203 | 0.05300254 | 0.04314552 | 0.00985702 | 0.14357806 | 0.76537874 |
| Il33 | 0.04294535 | 0.05580386 | 0.04595158 | 0.00985228 | 0.15892064 | 0.80588717 |
| Iqcb1 | 0.33792759 | 0.35543232 | 0.34558397 | 0.00984835 | 0.447188 | 0.89054584 |
| Rinl | 0.05497367 | 0.06865253 | 0.05881517 | 0.00983736 | 0.21791635 | 0.89054584 |
| Tmem208 | 1.17580818 | 0.92980519 | 0.91997029 | 0.0098349 | 0.48013842 | 0.89054584 |
| Qars | 0.51650945 | 0.51573263 | 0.50590006 | 0.00983257 | 0.46392499 | 0.89054584 |
| Gper1 | 0.03178203 | 0.04382642 | 0.03400945 | 0.00981696 | 0.08934139 | 0.57918151 |
| Cartpt | 0.02325395 | 0.0346997 | 0.02488473 | 0.00981498 | 0.033063 | 0.29104275 |
| Npdc1 | 1.11021999 | 0.89987167 | 0.89006425 | 0.00980742 | 0.47952896 | 0.89054584 |
| Kpna1 | 0.33805819 | 0.35550118 | 0.34570773 | 0.00979345 | 0.44749935 | 0.89054584 |
| Prps2 | 0.12431663 | 0.14218034 | 0.13239028 | 0.00979006 | 0.36522701 | 0.89054584 |
| Anks3 | 0.4073705 | 0.41987526 | 0.41009179 | 0.00978347 | 0.45574973 | 0.89054584 |
| Cdpf1 | 0.34185655 | 0.35908451 | 0.34930308 | 0.00978143 | 0.44810009 | 0.89054584 |
| Rab1b | 0.47909115 | 0.48363363 | 0.47385671 | 0.00977692 | 0.46171012 | 0.89054584 |
| Ndufa12 | 2.21250689 | 1.13734239 | 1.12757105 | 0.00977135 | 0.48389763 | 0.89054584 |
| Pdss1 | 0.09338467 | 0.10961381 | 0.09985109 | 0.00976272 | 0.32436838 | 0.89054584 |
| Fam219b | 0.21940759 | 0.239564 | 0.22980953 | 0.00975447 | 0.42161917 | 0.89054584 |
| Adcyap1r1 | 0.04009566 | 0.05265615 | 0.04290337 | 0.00975278 | 0.14478411 | 0.76727455 |
| Capn2 | 0.70539462 | 0.66352976 | 0.65378199 | 0.00974776 | 0.47231018 | 0.89054584 |
| Zfp641 | 0.02087683 | 0.03208031 | 0.0223411 | 0.00973921 | 0.02112831 | 0.20880589 |
| Slx1b | 0.13428438 | 0.15250925 | 0.14277155 | 0.0097377 | 0.37533269 | 0.89054584 |
| Cdkl1 | 0.0663434 | 0.08070661 | 0.07096956 | 0.00973705 | 0.2613437 | 0.89054584 |
| Rnls | 0.13365121 | 0.1518482 | 0.14211323 | 0.00973497 | 0.37480845 | 0.89054584 |
| Parp6 | 0.1799792 | 0.19959017 | 0.18986498 | 0.00972519 | 0.40569296 | 0.89054584 |
| Far1 | 0.39373762 | 0.40733566 | 0.3976359 | 0.00969976 | 0.45475833 | 0.89054584 |
| Dlg2 | 0.02239391 | 0.03364839 | 0.02396445 | 0.00968394 | 0.02987321 | 0.26867358 |
| Ltb4r1 | 0.0181266 | 0.02907807 | 0.01939815 | 0.00967992 | 0.01003947 | 0.11777817 |

| | | | | | | |
|---|---|---|---|---|---|---|
| Ptcd1 | 0.14760578 | 0.16624991 | 0.15658592 | 0.009664 | 0.38685122 | 0.89054584 |
| Coasy | 0.38131215 | 0.39585052 | 0.38619344 | 0.00965708 | 0.45362791 | 0.89054584 |
| Rgs14 | 0.02401144 | 0.03534907 | 0.02569526 | 0.00965381 | 0.04002618 | 0.33738342 |
| Dgcr8 | 0.22099058 | 0.24104195 | 0.23139918 | 0.00964277 | 0.42303101 | 0.89054584 |
| Frs2 | 0.13573989 | 0.15392502 | 0.14428428 | 0.00964074 | 0.37778642 | 0.89054584 |
| Cpeb3 | 0.04985091 | 0.06297262 | 0.05333724 | 0.00963537 | 0.19999544 | 0.89054584 |
| Man1b1 | 0.28425444 | 0.3036218 | 0.29399387 | 0.00962793 | 0.43936692 | 0.89054584 |
| Tax1bp1 | 1.42847748 | 1.0242945 | 1.01467392 | 0.00962058 | 0.48238305 | 0.89054584 |
| Poli | 0.05826817 | 0.07195781 | 0.06233757 | 0.00962024 | 0.23607337 | 0.89054584 |
| Mkl1 | 0.10888016 | 0.12585913 | 0.11624063 | 0.0096185 | 0.3499295 | 0.89054584 |
| Prdx2 | 3.31487672 | 1.06962864 | 1.06001874 | 0.0096099 | 0.4831549 | 0.89054584 |
| Gpr88 | 0.01844647 | 0.0293493 | 0.01974043 | 0.00960887 | 0.01167099 | 0.13260663 |
| Dedd | 0.34802544 | 0.36472821 | 0.35512588 | 0.00960233 | 0.4498762 | 0.89054584 |
| Trpm5 | 0.09006738 | 0.10590631 | 0.09631039 | 0.00959592 | 0.32125395 | 0.89054584 |
| Uggt2 | 0.12821156 | 0.14604289 | 0.1364512 | 0.0095917 | 0.3716466 | 0.89054584 |
| A730098A19 | 0.04792909 | 0.06087336 | 0.05128196 | 0.0095914 | 0.19177505 | 0.88486593 |
| Fbxo8 | 0.28802486 | 0.30723763 | 0.29766514 | 0.00957249 | 0.44045136 | 0.89054584 |
| Sgk2 | 0.03002083 | 0.04169374 | 0.03212514 | 0.0095686 | 0.08261654 | 0.55247906 |
| Hist1h4c | 0.01087183 | 0.0212028 | 0.01163466 | 0.00956814 | 6.37E-05 | 0.0014078 |
| Prkce | 0.26467345 | 0.28438696 | 0.27481984 | 0.00956711 | 0.4355787 | 0.89054584 |
| Gm21541 | 0.01845988 | 0.02931619 | 0.01975479 | 0.0095614 | 0.01206833 | 0.13637305 |
| 1110020A21 | 0.14869914 | 0.16727108 | 0.15771665 | 0.00955443 | 0.38888077 | 0.89054584 |
| Lrrc3b | 0.04677945 | 0.05959918 | 0.05005242 | 0.00954675 | 0.18710274 | 0.87419127 |
| Zbtb43 | 0.12765688 | 0.14541379 | 0.1358732 | 0.00954058 | 0.37178249 | 0.89054584 |
| Sarm1 | 0.05258477 | 0.06579848 | 0.05626075 | 0.00953773 | 0.2148172 | 0.89054584 |
| Nsun5 | 0.12675075 | 0.14446504 | 0.13492874 | 0.00953631 | 0.37097272 | 0.89054584 |
| Zfp764 | 0.06802436 | 0.08229509 | 0.07276605 | 0.00952904 | 0.27089475 | 0.89054584 |
| Tbc1d10c | 0.0300817 | 0.04171224 | 0.03219026 | 0.00952198 | 0.08408163 | 0.55918838 |
| Tmem54 | 0.0318267 | 0.04357845 | 0.03405725 | 0.00952121 | 0.09637693 | 0.61016274 |
| Mnt | 0.13897478 | 0.15715793 | 0.14764342 | 0.00951452 | 0.38199932 | 0.89054584 |

| | | | | | | |
|---|---|---|---|---|---|---|
| 4833418N02 | 0.05560477 | 0.06900397 | 0.05948995 | 0.00951402 | 0.22810917 | 0.89054584 |
| Apobec3 | 0.31722448 | 0.33536152 | 0.32585703 | 0.00950449 | 0.44595395 | 0.89054584 |
| Relb | 0.06593593 | 0.08003469 | 0.07053406 | 0.00950063 | 0.26515304 | 0.89054584 |
| Ccdc74a | 0.05044041 | 0.06346777 | 0.05396767 | 0.00950011 | 0.20607132 | 0.89054584 |
| Rbfox2 | 0.77079241 | 0.70875151 | 0.69926095 | 0.00949057 | 0.4747907 | 0.89054584 |
| Aldh4a1 | 0.10235249 | 0.11886478 | 0.10938488 | 0.0094799 | 0.34319104 | 0.89054584 |
| Derl1 | 0.80718438 | 0.73271468 | 0.72323929 | 0.00947539 | 0.47566435 | 0.89054584 |
| Cog2 | 0.21422561 | 0.23405686 | 0.22459792 | 0.00945895 | 0.42222234 | 0.89054584 |
| Adamtsl3 | 0.02323483 | 0.03431433 | 0.02486427 | 0.00945006 | 0.03830497 | 0.32686611 |
| Kcnh7 | 0.05682958 | 0.07024785 | 0.06079951 | 0.00944834 | 0.23453086 | 0.89054584 |
| Pzp | 0.01648354 | 0.02708757 | 0.01763992 | 0.00944765 | 0.00629338 | 0.07913392 |
| Fam187b | 0.16464029 | 0.18357936 | 0.1741498 | 0.00942956 | 0.40041837 | 0.89054584 |
| Rrp15 | 0.45752134 | 0.46442717 | 0.45499783 | 0.00942934 | 0.46154126 | 0.89054584 |
| Vps13c | 0.17133986 | 0.19045185 | 0.1810256 | 0.00942626 | 0.40415858 | 0.89054584 |
| H2-T22 | 0.12219274 | 0.13959032 | 0.13017356 | 0.00941676 | 0.36804798 | 0.89054584 |
| Arid3b | 0.09816827 | 0.11437195 | 0.10495554 | 0.00941642 | 0.33797662 | 0.89054584 |
| Gm26533 | 0.01953328 | 0.03030706 | 0.02090341 | 0.00940366 | 0.01804632 | 0.18701609 |
| Pde4b | 0.10440549 | 0.12094469 | 0.11154274 | 0.00940195 | 0.34726958 | 0.89054584 |
| Gna15 | 0.01322069 | 0.02354519 | 0.01414828 | 0.00939691 | 0.00098624 | 0.01628525 |
| Eda | 0.03896746 | 0.05108627 | 0.04169652 | 0.00938976 | 0.1470522 | 0.7738522 |
| Adgrg5 | 0.02231337 | 0.03326525 | 0.02387827 | 0.00938698 | 0.0335119 | 0.2936188 |
| Zfp759 | 0.0365259 | 0.04847132 | 0.03908463 | 0.00938669 | 0.13158999 | 0.72677778 |
| Klk1 | 0.03477005 | 0.04658238 | 0.03720621 | 0.00937617 | 0.12018165 | 0.69411919 |
| Lyrm7 | 0.04969016 | 0.06254144 | 0.05316534 | 0.0093761 | 0.20564132 | 0.89054584 |
| Wscd2 | 0.04364383 | 0.05607396 | 0.04669868 | 0.00937528 | 0.17480795 | 0.84226179 |
| Cacng8 | 0.01874017 | 0.02942936 | 0.02005472 | 0.00937464 | 0.01470913 | 0.15926281 |
| Cspp1 | 1.11652007 | 0.90241652 | 0.89304382 | 0.0093727 | 0.48050083 | 0.89054584 |
| Slc25a42 | 0.05890379 | 0.0723874 | 0.06301711 | 0.00937028 | 0.24423073 | 0.89054584 |
| Nxpe4 | 0.03465926 | 0.04644382 | 0.03708768 | 0.00935614 | 0.11993447 | 0.69330539 |
| Mdc1 | 0.10778471 | 0.12444412 | 0.11509121 | 0.00935291 | 0.35248883 | 0.89054584 |

| | | | | | | |
|---|---|---|---|---|---|---|
| Lect1 | 0.02819562 | 0.03950819 | 0.03017228 | 0.00933592 | 0.0747117 | 0.51900827 |
| Smek2 | 0.84781654 | 0.7582079 | 0.74887593 | 0.00933198 | 0.47685181 | 0.89054584 |
| Arhgap15 | 0.02650326 | 0.03766217 | 0.02836151 | 0.00930066 | 0.0632789 | 0.46264121 |
| Ptprt | 0.06623633 | 0.08014994 | 0.07085513 | 0.00929482 | 0.27054592 | 0.89054584 |
| 2610035D17I | 0.0620472 | 0.07566977 | 0.06637749 | 0.00929227 | 0.2571324 | 0.89054584 |
| Reps2 | 0.06537847 | 0.07921437 | 0.06993825 | 0.00927612 | 0.26831087 | 0.89054584 |
| AI464131 | 0.05057974 | 0.06338866 | 0.05411667 | 0.00927199 | 0.21236783 | 0.89054584 |
| E2f1 | 0.1436044 | 0.16171013 | 0.15244376 | 0.00926637 | 0.38851328 | 0.89054584 |
| Atg5 | 0.45923302 | 0.46575915 | 0.45650449 | 0.00925467 | 0.46237577 | 0.89054584 |
| Rpain | 0.27143366 | 0.29071174 | 0.28146058 | 0.00925116 | 0.43914711 | 0.89054584 |
| Ccdc130 | 0.12525437 | 0.14261287 | 0.13336846 | 0.00924441 | 0.37337276 | 0.89054584 |
| Gli2 | 0.05334277 | 0.06631477 | 0.05707129 | 0.00924348 | 0.22525076 | 0.89054584 |
| Tceanc | 0.09851556 | 0.11456435 | 0.10532605 | 0.0092383 | 0.34140101 | 0.89054584 |
| Arid2 | 0.36988926 | 0.38482154 | 0.37559936 | 0.00922217 | 0.45446352 | 0.89054584 |
| Troap | 0.04095291 | 0.05303413 | 0.04382036 | 0.00921376 | 0.16364202 | 0.81785018 |
| Apol7d | 0.02744371 | 0.03857724 | 0.02936777 | 0.00920947 | 0.07200734 | 0.50850579 |
| Clp1 | 0.138158 | 0.15600026 | 0.14679564 | 0.00920461 | 0.38509272 | 0.89054584 |
| Adat3 | 0.0535487 | 0.0664952 | 0.0572915 | 0.0092037 | 0.22709605 | 0.89054584 |
| Ttf2 | 0.09841657 | 0.11442347 | 0.10522045 | 0.00920303 | 0.34182379 | 0.89054584 |
| Ccdc36 | 0.02157276 | 0.0322868 | 0.02308578 | 0.00920102 | 0.03166576 | 0.2807891 |
| Lpp | 0.70961417 | 0.66600741 | 0.65680814 | 0.00919927 | 0.47398619 | 0.89054584 |
| 2010315B03I | 0.13172525 | 0.14930526 | 0.14010983 | 0.00919542 | 0.37989109 | 0.89054584 |
| Fam84b | 0.09484831 | 0.11059507 | 0.10141307 | 0.009182 | 0.336577 | 0.89054584 |
| Pknox2 | 0.03260223 | 0.04406229 | 0.03488697 | 0.00917532 | 0.11022963 | 0.6573876 |
| Lrrtm4 | 0.01820882 | 0.0286543 | 0.01948613 | 0.00916817 | 0.01418906 | 0.15463706 |
| Gfod2 | 0.34561383 | 0.36201999 | 0.35285199 | 0.009168 | 0.45182523 | 0.89054584 |
| Fam177a | 0.59075914 | 0.57603367 | 0.5668665 | 0.00916717 | 0.46997098 | 0.89054584 |
| E030030I06R | 0.2197662 | 0.23932559 | 0.23016974 | 0.00915585 | 0.42648646 | 0.89054584 |
| Mitf | 0.04720802 | 0.05965632 | 0.05051079 | 0.00914553 | 0.19945982 | 0.89054584 |
| 4933404O12 | 0.05858133 | 0.07181478 | 0.06267237 | 0.00914241 | 0.24836978 | 0.89054584 |

| | | | | | | |
|---|---|---|---|---|---|---|
| Dpp7 | 0.16292761 | 0.1815205 | 0.17238912 | 0.00913138 | 0.40253839 | 0.89054584 |
| Sv2a | 0.14301792 | 0.16096272 | 0.15183611 | 0.0091266 | 0.38972313 | 0.89054584 |
| Cdip1 | 0.374232 | 0.38876164 | 0.37963553 | 0.00912611 | 0.45541294 | 0.89054584 |
| Ace | 0.02155689 | 0.03214285 | 0.0230688 | 0.00907405 | 0.03343244 | 0.29340403 |
| Gm11713 | 0.12862771 | 0.14594463 | 0.13688475 | 0.00905987 | 0.37890521 | 0.89054584 |
| Dmd | 0.17086032 | 0.18957946 | 0.18053405 | 0.00904541 | 0.40771384 | 0.89054584 |
| Dctn4 | 0.27567553 | 0.2946226 | 0.28561619 | 0.00900641 | 0.44160089 | 0.89054584 |
| Rdh11 | 0.12461573 | 0.1416939 | 0.13270231 | 0.00899159 | 0.37612328 | 0.89054584 |
| Vps13d | 0.14793446 | 0.16591427 | 0.15692589 | 0.00898838 | 0.39479069 | 0.89054584 |
| Cyb5d1 | 0.02076544 | 0.03119708 | 0.02222191 | 0.00897517 | 0.02993891 | 0.26898821 |
| Zfp213 | 0.12839158 | 0.14560069 | 0.13663876 | 0.00896194 | 0.37996484 | 0.89054584 |
| A230107N01 | 0.0248922 | 0.03559943 | 0.02663768 | 0.00896175 | 0.05850892 | 0.43830347 |
| Mlxip | 0.13035013 | 0.14762152 | 0.13867854 | 0.00894298 | 0.38192017 | 0.89054584 |
| Tekt2 | 0.42747973 | 0.43720992 | 0.4282728 | 0.00893712 | 0.46127507 | 0.89054584 |
| Cyth2 | 1.10133501 | 0.89473858 | 0.88581456 | 0.00892402 | 0.48128217 | 0.89054584 |
| Dusp9 | 0.04938364 | 0.0617598 | 0.05283753 | 0.00892226 | 0.21572312 | 0.89054584 |
| Zfp654 | 0.11480324 | 0.13135799 | 0.12244786 | 0.00891013 | 0.36729882 | 0.89054584 |
| Zfp414 | 0.3534123 | 0.36909392 | 0.36019382 | 0.0089001 | 0.45417553 | 0.89054584 |
| Ascl1 | 0.01979135 | 0.03007599 | 0.02117956 | 0.00889642 | 0.02517511 | 0.2370057 |
| Hist1h3c | 0.02741619 | 0.03821825 | 0.02933832 | 0.00887993 | 0.07924943 | 0.538911 |
| Tnni2 | 0.01851925 | 0.028698 | 0.01981832 | 0.00887968 | 0.01842339 | 0.19010839 |
| Pdp2 | 0.04135371 | 0.05312078 | 0.04424909 | 0.00887169 | 0.17512851 | 0.84296586 |
| Fjx1 | 0.0826184 | 0.09722208 | 0.08835716 | 0.00886491 | 0.32009481 | 0.89054584 |
| Grtp1 | 0.11998091 | 0.13670607 | 0.12786325 | 0.00884283 | 0.37364862 | 0.89054584 |
| Sp100 | 0.01364863 | 0.02343934 | 0.01460622 | 0.00883312 | 0.00241999 | 0.03558314 |
| Ngef | 0.03359164 | 0.04477633 | 0.0359455 | 0.00883083 | 0.1261926 | 0.71147001 |
| Exoc3l | 0.03670553 | 0.04810354 | 0.0392768 | 0.00882674 | 0.1475472 | 0.77404405 |
| Cntnap1 | 0.02949629 | 0.04038978 | 0.03156391 | 0.00882587 | 0.09633378 | 0.61015603 |
| Slfn9 | 0.07047019 | 0.08419512 | 0.07537975 | 0.00881538 | 0.29292982 | 0.89054584 |
| Rab6b | 0.0334862 | 0.04464122 | 0.03583269 | 0.00880852 | 0.12604705 | 0.71106229 |

| | | | | | | |
|---|---|---|---|---|---|---|
| Gtf3c3 | 0.2242928 | 0.24350353 | 0.23471166 | 0.00879187 | 0.43073021 | 0.89054584 |
| Serpina1c | 0.05743236 | 0.07022989 | 0.06144398 | 0.00878591 | 0.25264641 | 0.89054584 |
| Azin2 | 0.09125435 | 0.10635334 | 0.09757738 | 0.00877596 | 0.33760143 | 0.89054584 |
| Cenpq | 0.17661195 | 0.19519819 | 0.18642335 | 0.00877484 | 0.41321066 | 0.89054584 |
| Miip | 0.11746098 | 0.13399798 | 0.12522887 | 0.00876911 | 0.37212109 | 0.89054584 |
| Fam83d | 0.04033677 | 0.0519162 | 0.04316128 | 0.00875492 | 0.17232221 | 0.83696674 |
| Plxna1 | 0.10775134 | 0.12380813 | 0.11505619 | 0.00875194 | 0.36152325 | 0.89054584 |
| Homer3 | 0.21132414 | 0.23040738 | 0.22167465 | 0.00873273 | 0.42718874 | 0.89054584 |
| Txnrd3 | 0.05656759 | 0.06924146 | 0.0605194 | 0.00872206 | 0.25096794 | 0.89054584 |
| Cpxm2 | 0.04451661 | 0.05634715 | 0.04763221 | 0.00871494 | 0.19699189 | 0.89054584 |
| D630003M21 | 0.04018455 | 0.05170447 | 0.04299846 | 0.00870601 | 0.17276156 | 0.83703076 |
| Afm | 0.05235256 | 0.06471477 | 0.05601245 | 0.00870233 | 0.23458431 | 0.89054584 |
| Tsr3 | 0.50071267 | 0.50116267 | 0.4924777 | 0.00868496 | 0.46725912 | 0.89054584 |
| Trps1 | 0.16732465 | 0.18558917 | 0.17690697 | 0.0086822 | 0.40957029 | 0.89054584 |
| Ubtd1 | 0.12630495 | 0.14312922 | 0.13446396 | 0.00866526 | 0.38199849 | 0.89054584 |
| Gm10126 | 0.03341802 | 0.04442247 | 0.03575975 | 0.00866272 | 0.12952944 | 0.72054099 |
| Gata2 | 0.03392119 | 0.04492953 | 0.03629807 | 0.00863147 | 0.13395927 | 0.73233089 |
| Tmtc2 | 0.10689716 | 0.12277848 | 0.11415961 | 0.00861887 | 0.36251559 | 0.89054584 |
| Ccnf | 0.06015363 | 0.07295652 | 0.06435327 | 0.00860325 | 0.2666935 | 0.89054584 |
| Lyrm5 | 0.38559812 | 0.39874469 | 0.39014968 | 0.00859502 | 0.45912566 | 0.89054584 |
| Tchh | 0.04359626 | 0.05523981 | 0.04664781 | 0.008592 | 0.19541179 | 0.89054584 |
| Ndc80 | 0.04309966 | 0.05470265 | 0.04611663 | 0.00858602 | 0.19286023 | 0.88708362 |
| Socs7 | 0.25365485 | 0.27252176 | 0.26394986 | 0.0085719 | 0.43986897 | 0.89054584 |
| Tor1aip1 | 0.50533862 | 0.50499557 | 0.49642464 | 0.00857092 | 0.46794443 | 0.89054584 |
| Pip5k1c | 0.18583563 | 0.20441078 | 0.19583998 | 0.0085708 | 0.41921762 | 0.89054584 |
| Camta2 | 0.09536883 | 0.110539 | 0.10196853 | 0.00857047 | 0.34768259 | 0.89054584 |
| Gm16638 | 0.10197617 | 0.11754464 | 0.10898918 | 0.00855546 | 0.35728741 | 0.89054584 |
| Psmb10 | 0.39977718 | 0.41170019 | 0.40316711 | 0.00853307 | 0.46072516 | 0.89054584 |
| Dusp28 | 0.19437164 | 0.2130515 | 0.20452278 | 0.00852872 | 0.42297814 | 0.89054584 |
| Gm28172 | 0.06909667 | 0.08243822 | 0.073912 | 0.00852623 | 0.2954822 | 0.89054584 |

| | | | | | | |
|---|---|---|---|---|---|---|
| Tescl | 0.01446149 | 0.02400214 | 0.01547609 | 0.00852605 | 0.00513365 | 0.06725758 |
| Hey2 | 0.02193962 | 0.0319801 | 0.02347834 | 0.00850176 | 0.04579665 | 0.36907161 |
| Slc39a2 | 0.02529886 | 0.0355664 | 0.02707281 | 0.00849359 | 0.07191786 | 0.50812144 |
| Nudt21 | 1.04103819 | 0.86414165 | 0.85565885 | 0.0084828 | 0.48158034 | 0.89054584 |
| Cnep1r1 | 0.49871088 | 0.49924489 | 0.49076557 | 0.00847931 | 0.46792142 | 0.89054584 |
| Frmd6 | 0.13622438 | 0.15325876 | 0.14478763 | 0.00847113 | 0.39258729 | 0.89054584 |
| Kremen1 | 0.10748928 | 0.12324387 | 0.11478115 | 0.00846272 | 0.36561117 | 0.89054584 |
| Shisa9 | 0.02793016 | 0.03834571 | 0.02988825 | 0.00845746 | 0.09369519 | 0.5972952 |
| D630033O11 | 0.04660229 | 0.05832039 | 0.04986295 | 0.00845744 | 0.21469948 | 0.89054584 |
| Slc37a1 | 0.24231787 | 0.26115961 | 0.25270745 | 0.00845217 | 0.43808547 | 0.89054584 |
| Nr1h2 | 0.36654786 | 0.38093854 | 0.37248677 | 0.00845178 | 0.45790542 | 0.89054584 |
| Ttc37 | 0.09922675 | 0.11453481 | 0.10608479 | 0.00845002 | 0.35528129 | 0.89054584 |
| Gm16152 | 0.02591016 | 0.03617211 | 0.0277269 | 0.00844521 | 0.07794282 | 0.53259744 |
| Slc12a4 | 0.09382484 | 0.10875855 | 0.10032085 | 0.00843769 | 0.3475846 | 0.89054584 |
| Pate2 | 0.01432821 | 0.02376974 | 0.01533346 | 0.00843627 | 0.00518429 | 0.06765245 |
| Ptpdc1 | 0.06876919 | 0.08199815 | 0.07356203 | 0.00843611 | 0.29657093 | 0.89054584 |
| Kcnq4 | 0.0826035 | 0.09676212 | 0.08834126 | 0.00842087 | 0.32848522 | 0.89054584 |
| Bsdc1 | 0.36341548 | 0.37798329 | 0.36956331 | 0.00841998 | 0.4577327 | 0.89054584 |
| Stambpl1 | 0.04742247 | 0.0591521 | 0.05074014 | 0.00841196 | 0.21994363 | 0.89054584 |
| Otud3 | 0.07906011 | 0.09295373 | 0.08455683 | 0.00839691 | 0.32180562 | 0.89054584 |
| Trim16 | 0.04366457 | 0.05510738 | 0.04672087 | 0.00838651 | 0.20149561 | 0.89054584 |
| Ptges | 0.03854986 | 0.04962993 | 0.0412498 | 0.00838013 | 0.17194926 | 0.83696674 |
| Camta1 | 0.53504259 | 0.529817 | 0.52144891 | 0.00836809 | 0.4702006 | 0.89054584 |
| Acsf2 | 0.0991785 | 0.11439707 | 0.10603332 | 0.00836374 | 0.35662675 | 0.89054584 |
| Hmgcs2 | 0.06426776 | 0.0770997 | 0.06875108 | 0.00834862 | 0.28578287 | 0.89054584 |
| Zfp932 | 0.18763011 | 0.20601306 | 0.19766789 | 0.00834518 | 0.42203431 | 0.89054584 |
| Rpl7l1 | 0.74281235 | 0.68851231 | 0.68017631 | 0.008336 | 0.47723342 | 0.89054584 |
| D16Ertd472e | 0.05116593 | 0.06304848 | 0.05474353 | 0.00830495 | 0.23984786 | 0.89054584 |
| Kif11 | 0.04856643 | 0.06025701 | 0.05196358 | 0.00829344 | 0.2285679 | 0.89054584 |
| Rbl1 | 0.08010092 | 0.09396134 | 0.08566851 | 0.00829283 | 0.32599762 | 0.89054584 |

| | | | | | | |
|---|---|---|---|---|---|---|
| Bag4 | 0.45786813 | 0.46359037 | 0.45530322 | 0.00828715 | 0.46621051 | 0.89054584 |
| Pcgf1 | 0.17582061 | 0.19389552 | 0.18561389 | 0.00828163 | 0.41766507 | 0.89054584 |
| Cpne7 | 0.01777381 | 0.02729459 | 0.01902063 | 0.00827396 | 0.02134977 | 0.21027779 |
| Slc35c2 | 0.33988678 | 0.35570942 | 0.34743954 | 0.00826988 | 0.45585028 | 0.89054584 |
| Pld1 | 0.13814003 | 0.15502626 | 0.14677699 | 0.00824926 | 0.39672046 | 0.89054584 |
| Tmub2 | 0.36106661 | 0.37560513 | 0.36736794 | 0.00823718 | 0.45840075 | 0.89054584 |
| Zfp354a | 0.04467449 | 0.05603776 | 0.04780107 | 0.00823669 | 0.21104798 | 0.89054584 |
| Rab29 | 0.08877963 | 0.10316961 | 0.09493571 | 0.0082339 | 0.34307833 | 0.89054584 |
| Dpp10 | 0.05611798 | 0.06825363 | 0.06003868 | 0.00821495 | 0.26190879 | 0.89054584 |
| Slc28a2 | 0.01950361 | 0.02908528 | 0.02087167 | 0.00821361 | 0.0333693 | 0.29302733 |
| Ankle1 | 0.02733725 | 0.03746398 | 0.02925386 | 0.00821013 | 0.09551519 | 0.60576518 |
| Tgfbr1 | 0.40454477 | 0.41572629 | 0.40751874 | 0.00820756 | 0.46262108 | 0.89054584 |
| Asb3 | 0.23712491 | 0.25573909 | 0.24753783 | 0.00820126 | 0.43866413 | 0.89054584 |
| Spred1 | 0.15509391 | 0.17250025 | 0.16432076 | 0.00817949 | 0.40830369 | 0.89054584 |
| Smap2 | 0.48734286 | 0.48916124 | 0.48099691 | 0.00816434 | 0.46848455 | 0.89054584 |
| Pnp | 0.3599912 | 0.37451949 | 0.36636192 | 0.00815756 | 0.45868869 | 0.89054584 |
| 6030408B16 | 0.03626522 | 0.04696176 | 0.03880576 | 0.008156 | 0.1637423 | 0.81785018 |
| Kdelc2 | 0.33944107 | 0.35517099 | 0.34701759 | 0.0081534 | 0.45641689 | 0.89054584 |
| Tm9sf4 | 0.52587295 | 0.52192963 | 0.51378275 | 0.00814688 | 0.47055484 | 0.89054584 |
| Hist2h2ac | 0.05161417 | 0.06335929 | 0.05522286 | 0.00813644 | 0.24621785 | 0.89054584 |
| Elavl4 | 0.22655596 | 0.24510954 | 0.23697903 | 0.00813052 | 0.43650211 | 0.89054584 |
| Lrig3 | 0.05623853 | 0.06828616 | 0.06016758 | 0.00811858 | 0.26479106 | 0.89054584 |
| Map2k4 | 0.26501041 | 0.28326305 | 0.27515137 | 0.00811169 | 0.44537732 | 0.89054584 |
| Ddx23 | 0.23034415 | 0.24886086 | 0.24076905 | 0.00809181 | 0.43778851 | 0.89054584 |
| Epha2 | 0.030006 | 0.0401997 | 0.03210927 | 0.00809043 | 0.12021888 | 0.69411919 |
| Myg1 | 0.6359081 | 0.61024922 | 0.60216852 | 0.0080807 | 0.47507443 | 0.89054584 |
| Cecr5 | 0.14352655 | 0.16044236 | 0.15236311 | 0.00807925 | 0.40243615 | 0.89054584 |
| Eid2b | 0.22064669 | 0.23912678 | 0.23105393 | 0.00807285 | 0.4353459 | 0.89054584 |
| Gm20498 | 0.07041802 | 0.08339465 | 0.07532401 | 0.00807064 | 0.30882438 | 0.89054584 |
| Xpa | 0.44810131 | 0.45474179 | 0.44667451 | 0.00806728 | 0.46647095 | 0.89054584 |

| | | | | | | |
|---|---|---|---|---|---|---|
| BC035044 | 0.01190624 | 0.02080514 | 0.01274164 | 0.0080635 | 0.00159714 | 0.02479162 |
| Ptch1 | 0.24319174 | 0.26163602 | 0.25357616 | 0.00805985 | 0.44113854 | 0.89054584 |
| Slitrk5 | 0.02699664 | 0.03693643 | 0.02888941 | 0.00804701 | 0.09719059 | 0.61237252 |
| 4931406C07I | 0.3865727 | 0.39908218 | 0.39104783 | 0.00803434 | 0.46187104 | 0.89054584 |
| Mctp2 | 0.0684999 | 0.08130102 | 0.07327425 | 0.00802677 | 0.30489967 | 0.89054584 |
| Tanc2 | 0.13096948 | 0.14734876 | 0.13932328 | 0.00802548 | 0.39420712 | 0.89054584 |
| Dctn5 | 0.36279019 | 0.376991 | 0.36897908 | 0.00801192 | 0.45971049 | 0.89054584 |
| Ppil4 | 0.73720213 | 0.68429245 | 0.6762821 | 0.00801035 | 0.47799607 | 0.89054584 |
| Fign | 0.07042419 | 0.08334053 | 0.07533059 | 0.00800993 | 0.31016344 | 0.89054584 |
| Lrfn5 | 0.01707719 | 0.02628491 | 0.01827518 | 0.00800973 | 0.02057871 | 0.20491081 |
| Atp10a | 0.02808798 | 0.03805372 | 0.03005711 | 0.00799662 | 0.10758029 | 0.64845225 |
| BC055324 | 0.06796012 | 0.08068542 | 0.0726974 | 0.00798802 | 0.30435136 | 0.89054584 |
| Ift88 | 0.29766983 | 0.31499457 | 0.30702435 | 0.00797021 | 0.45186769 | 0.89054584 |
| Spata2 | 0.07914427 | 0.09261499 | 0.08464673 | 0.00796826 | 0.330486 | 0.89054584 |
| Ptpn21 | 0.10937904 | 0.12471219 | 0.11676394 | 0.00794825 | 0.375569 | 0.89054584 |
| Sec16b | 0.06340576 | 0.07577113 | 0.0678297 | 0.00794143 | 0.29271724 | 0.89054584 |
| Myo10 | 0.38869034 | 0.40092659 | 0.39299761 | 0.00792899 | 0.46255567 | 0.89054584 |
| Faah | 0.36710576 | 0.38093313 | 0.37300689 | 0.00792624 | 0.4605689 | 0.89054584 |
| Espn | 0.05745174 | 0.06938823 | 0.0614647 | 0.00792353 | 0.27405567 | 0.89054584 |
| Fus | 2.35189298 | 1.13742166 | 1.12952357 | 0.0078981 | 0.48700585 | 0.89054584 |
| Fam134b | 0.2202652 | 0.23856008 | 0.23067088 | 0.0078892 | 0.43670014 | 0.89054584 |
| Ttll1 | 0.16131772 | 0.17861863 | 0.17073303 | 0.00788559 | 0.4148131 | 0.89054584 |
| Stap1 | 0.02407913 | 0.03365171 | 0.02576769 | 0.00788402 | 0.07701016 | 0.52914378 |
| St6galnac4 | 0.23359847 | 0.2518951 | 0.24402017 | 0.00787493 | 0.44024359 | 0.89054584 |
| Tank | 0.21059956 | 0.22881589 | 0.22094405 | 0.00787184 | 0.43408254 | 0.89054584 |
| Acot8 | 0.34484532 | 0.35998864 | 0.35212671 | 0.00786193 | 0.45857668 | 0.89054584 |
| Gna14 | 0.03472336 | 0.04501692 | 0.03715627 | 0.00786065 | 0.16215742 | 0.81460788 |
| Man1c1 | 0.09002972 | 0.10411558 | 0.09627018 | 0.0078454 | 0.35209374 | 0.89054584 |
| Spry2 | 0.08764463 | 0.10156722 | 0.09372401 | 0.00784321 | 0.34831282 | 0.89054584 |
| 1810044D09I | 0.11974824 | 0.13546315 | 0.12762011 | 0.00784304 | 0.38731625 | 0.89054584 |

| | | | | | | |
|---|---|---|---|---|---|---|
| Lyrm2 | 0.63221935 | 0.607173 | 0.59933555 | 0.00783745 | 0.47570968 | 0.89054584 |
| Aldh18a1 | 0.3800005 | 0.39281107 | 0.38498065 | 0.00783042 | 0.46225201 | 0.89054584 |
| Cpsf1 | 0.12302513 | 0.138869 | 0.13104254 | 0.00782646 | 0.39040895 | 0.89054584 |
| Nfkbie | 0.0384248 | 0.04892659 | 0.04111601 | 0.00781058 | 0.18806949 | 0.87616692 |
| Ppfia4 | 0.02837843 | 0.0381782 | 0.03036787 | 0.00781033 | 0.11541254 | 0.67725583 |
| Nckap5 | 0.04587648 | 0.05689081 | 0.04908667 | 0.00780414 | 0.22943446 | 0.89054584 |
| Capn7 | 0.4622834 | 0.46698763 | 0.45918509 | 0.00780254 | 0.46845053 | 0.89054584 |
| Enpep | 0.07447267 | 0.08745508 | 0.0796563 | 0.00779878 | 0.32414624 | 0.89054584 |
| Hoxa1 | 0.01708424 | 0.02607536 | 0.01828272 | 0.00779264 | 0.02352889 | 0.2259085 |
| Iyd | 0.02710134 | 0.03678944 | 0.02900144 | 0.007788 | 0.1054483 | 0.64180915 |
| Gdf11 | 0.09882631 | 0.11343665 | 0.10565758 | 0.00777906 | 0.36579336 | 0.89054584 |
| 2600006K01I | 0.04117169 | 0.05182884 | 0.04405438 | 0.00777446 | 0.20548637 | 0.89054584 |
| Trim25 | 0.17730532 | 0.19488712 | 0.18713238 | 0.00775474 | 0.42345368 | 0.89054584 |
| Chst15 | 0.07128916 | 0.08400369 | 0.07625486 | 0.00774883 | 0.31795324 | 0.89054584 |
| Ppan | 0.33442911 | 0.35001268 | 0.3422659 | 0.00774677 | 0.45800959 | 0.89054584 |
| Manea | 0.09827233 | 0.1128127 | 0.10506656 | 0.00774615 | 0.36561664 | 0.89054584 |
| Fbln7 | 0.02099256 | 0.03020804 | 0.02246494 | 0.0077431 | 0.05415677 | 0.41543112 |
| Nfe2l3 | 0.04884879 | 0.0600072 | 0.05226554 | 0.00774165 | 0.24506891 | 0.89054584 |
| Ssh2 | 0.20159566 | 0.21958804 | 0.21184662 | 0.00774142 | 0.4324069 | 0.89054584 |
| Mapkapk3 | 0.02639065 | 0.03597387 | 0.02824101 | 0.00773286 | 0.10103163 | 0.62409736 |
| Zfp141 | 0.05913285 | 0.07098115 | 0.063262 | 0.00771915 | 0.28485506 | 0.89054584 |
| Urb2 | 0.0610791 | 0.07305972 | 0.06534261 | 0.00771711 | 0.29108025 | 0.89054584 |
| Wdr83os | 1.22085804 | 0.94701114 | 0.93929513 | 0.00771601 | 0.48473552 | 0.89054584 |
| Inf2 | 0.04923288 | 0.0603766 | 0.05267631 | 0.00770029 | 0.24791988 | 0.89054584 |
| Nrbp2 | 0.07822649 | 0.09136389 | 0.0836664 | 0.00769749 | 0.33409524 | 0.89054584 |
| Ptar1 | 0.06723294 | 0.0796134 | 0.07192025 | 0.00769314 | 0.30911649 | 0.89054584 |
| S1pr1 | 0.03501765 | 0.04515989 | 0.0374711 | 0.00768878 | 0.1695414 | 0.83300813 |
| Ciao1 | 0.45570518 | 0.46107456 | 0.45339727 | 0.00767728 | 0.4685605 | 0.89054584 |
| Nespas | 0.25974642 | 0.27763859 | 0.26996606 | 0.00767253 | 0.44733053 | 0.89054584 |
| Myo1g | 0.01773135 | 0.02664527 | 0.01897519 | 0.00767008 | 0.0298353 | 0.26849919 |

| | | | | | | |
|---|---|---|---|---|---|---|
| 1810012K16I | 0.01682939 | 0.0256709 | 0.01801001 | 0.00766089 | 0.02375262 | 0.22745406 |
| Myl7 | 0.02076144 | 0.02986771 | 0.02221762 | 0.00765008 | 0.05433611 | 0.4159266 |
| Tm4sf5 | 0.01673583 | 0.0255519 | 0.01790989 | 0.00764201 | 0.02340943 | 0.22535852 |
| Cwf19l1 | 0.05003075 | 0.06116963 | 0.05352957 | 0.00764007 | 0.25304044 | 0.89054584 |
| Pias4 | 0.44073818 | 0.44776859 | 0.44013181 | 0.00763678 | 0.46778544 | 0.89054584 |
| Fam129c | 0.04808914 | 0.05908768 | 0.05145313 | 0.00763454 | 0.24469254 | 0.89054584 |
| Creb3l4 | 0.04109632 | 0.05160721 | 0.04397377 | 0.00763345 | 0.2093295 | 0.89054584 |
| Fam161b | 0.07845786 | 0.09154104 | 0.08391355 | 0.00762749 | 0.33597107 | 0.89054584 |
| Cep85l | 0.04656683 | 0.05745103 | 0.04982502 | 0.00762601 | 0.2378986 | 0.89054584 |
| Cep63 | 0.25695827 | 0.27482788 | 0.26721435 | 0.00761354 | 0.44719805 | 0.89054584 |
| Cep89 | 0.28247185 | 0.29985803 | 0.29225599 | 0.00760204 | 0.45177171 | 0.89054584 |
| Itga3 | 0.06753972 | 0.07984718 | 0.07224811 | 0.00759907 | 0.31205686 | 0.89054584 |
| Tab1 | 0.16273393 | 0.17977873 | 0.17218994 | 0.0075888 | 0.41865717 | 0.89054584 |
| Abcc3 | 0.07007344 | 0.08253967 | 0.07495579 | 0.00758387 | 0.31868395 | 0.89054584 |
| Il17d | 0.04265559 | 0.05321888 | 0.04564164 | 0.00757724 | 0.21962561 | 0.89054584 |
| Shprh | 0.19182441 | 0.20949462 | 0.201935 | 0.00755961 | 0.43077102 | 0.89054584 |
| Lmbr1l | 0.17089227 | 0.18812525 | 0.1805668 | 0.00755845 | 0.42268814 | 0.89054584 |
| Gm15675 | 0.01672328 | 0.02544835 | 0.01789646 | 0.00755189 | 0.02465136 | 0.23389751 |
| Sdk2 | 0.14600898 | 0.16248322 | 0.15493369 | 0.00754953 | 0.41020471 | 0.89054584 |
| Prickle3 | 0.1406855 | 0.15696756 | 0.1494182 | 0.00754936 | 0.40695182 | 0.89054584 |
| Polr1b | 0.09195198 | 0.10586464 | 0.098322 | 0.00754263 | 0.36039468 | 0.89054584 |
| Tnfrsf23 | 0.02014301 | 0.02908152 | 0.02155586 | 0.00752566 | 0.05191683 | 0.40293905 |
| Vsir | 0.04422862 | 0.05483766 | 0.04732417 | 0.00751349 | 0.229746 | 0.89054584 |
| Tfpi2 | 0.04835459 | 0.05924284 | 0.05173702 | 0.00750582 | 0.24955146 | 0.89054584 |
| Ttc8 | 0.48961464 | 0.49045497 | 0.48295527 | 0.0074997 | 0.47116263 | 0.89054584 |
| Msantd4 | 0.60561004 | 0.58612069 | 0.57862886 | 0.00749183 | 0.47594964 | 0.89054584 |
| Rspo4 | 0.02400812 | 0.03317957 | 0.0256917 | 0.00748786 | 0.08725539 | 0.57076343 |
| Zfp46 | 0.15969914 | 0.17655419 | 0.16906694 | 0.00748725 | 0.41826872 | 0.89054584 |
| Clint1 | 0.65228915 | 0.62212009 | 0.61463586 | 0.00748423 | 0.47737996 | 0.89054584 |
| Strip1 | 0.29090874 | 0.30795025 | 0.30046844 | 0.00748181 | 0.45382234 | 0.89054584 |

| | | | | | | |
|---|---|---|---|---|---|---|
| Gpcpd1 | 0.458002 | 0.46289838 | 0.4554211 | 0.00747728 | 0.46951373 | 0.89054584 |
| Dock8 | 0.06457271 | 0.07654425 | 0.06907702 | 0.00746723 | 0.30726013 | 0.89054584 |
| Abcb1b | 0.01659426 | 0.02521782 | 0.01775839 | 0.00745942 | 0.02517428 | 0.2370057 |
| Myof | 0.07595857 | 0.08870113 | 0.08124374 | 0.0074574 | 0.33445439 | 0.89054584 |
| Nsun6 | 0.07334893 | 0.08591181 | 0.07845569 | 0.00745612 | 0.32896697 | 0.89054584 |
| Crcp | 0.32442728 | 0.34019 | 0.33274549 | 0.00744451 | 0.45849165 | 0.89054584 |
| Cluap1 | 0.39276669 | 0.40418647 | 0.39674477 | 0.0074417 | 0.46518184 | 0.89054584 |
| Plxna3 | 0.11639736 | 0.13155163 | 0.12411624 | 0.00743539 | 0.39008341 | 0.89054584 |
| Anks6 | 0.06544553 | 0.07744295 | 0.07000992 | 0.00743302 | 0.31042556 | 0.89054584 |
| Trim59 | 0.20739356 | 0.22514019 | 0.21770862 | 0.00743157 | 0.43682075 | 0.89054584 |
| Ndufaf6 | 0.20180693 | 0.21947598 | 0.21206047 | 0.00741551 | 0.43529155 | 0.89054584 |
| Tmem240 | 0.03709513 | 0.04710847 | 0.03969358 | 0.00741489 | 0.19206691 | 0.88511955 |
| Pacs1 | 0.07360876 | 0.08614567 | 0.0787333 | 0.00741237 | 0.33046947 | 0.89054584 |
| Acsl3 | 0.19403131 | 0.21157915 | 0.20417719 | 0.00740196 | 0.43293813 | 0.89054584 |
| Jak1 | 0.82714159 | 0.74338239 | 0.73598145 | 0.00740094 | 0.4813165 | 0.89054584 |
| Csf1 | 0.10785106 | 0.12255139 | 0.11516084 | 0.00739055 | 0.38247319 | 0.89054584 |
| Zfp454 | 0.03289538 | 0.04259038 | 0.0352006 | 0.00738977 | 0.16402085 | 0.8182981 |
| Sspn | 0.10382116 | 0.11830996 | 0.11092872 | 0.00738124 | 0.37827759 | 0.89054584 |
| Plekho2 | 0.06615325 | 0.07814648 | 0.07076633 | 0.00738015 | 0.31352737 | 0.89054584 |
| Cdc73 | 0.38008915 | 0.39243829 | 0.38506265 | 0.00737565 | 0.46444587 | 0.89054584 |
| Gmip | 0.1010881 | 0.11542914 | 0.10805515 | 0.00737398 | 0.37526556 | 0.89054584 |
| Map4k1 | 0.03806643 | 0.04810033 | 0.04073265 | 0.00736768 | 0.19969689 | 0.89054584 |
| Pcdhb11 | 0.01598193 | 0.02446314 | 0.01710313 | 0.00736001 | 0.02248755 | 0.21939735 |
| Plcd3 | 0.03695962 | 0.04690438 | 0.03954862 | 0.00735576 | 0.19310004 | 0.88734213 |
| Ficd | 0.1528075 | 0.16931628 | 0.16196122 | 0.00735506 | 0.4162197 | 0.89054584 |
| Echdc1 | 0.25901347 | 0.27659041 | 0.26924302 | 0.00734739 | 0.44941552 | 0.89054584 |
| Cd79b | 0.02019452 | 0.028934 | 0.02161098 | 0.00732301 | 0.05720122 | 0.43192879 |
| Hmgcr | 0.74001225 | 0.68555524 | 0.67823551 | 0.00731972 | 0.47994935 | 0.89054584 |
| BC025920 | 0.03087946 | 0.04036218 | 0.0330438 | 0.00731839 | 0.15107342 | 0.78251605 |
| Fgf12 | 0.22434806 | 0.24208536 | 0.23476706 | 0.00731831 | 0.44226406 | 0.89054584 |

| | | | | | | |
|---|---|---|---|---|---|---|
| Depdc7 | 0.02290509 | 0.03181405 | 0.02451144 | 0.00730261 | 0.08256497 | 0.55242329 |
| Kcnh8 | 0.11186276 | 0.1266624 | 0.11936794 | 0.00729446 | 0.38793566 | 0.89054584 |
| Gm4285 | 0.06812186 | 0.08015441 | 0.07287025 | 0.00728416 | 0.32071127 | 0.89054584 |
| Syt16 | 0.05744083 | 0.06873334 | 0.06145303 | 0.0072803 | 0.29049414 | 0.89054584 |
| Specc1l | 0.24801127 | 0.26563536 | 0.25836087 | 0.00727449 | 0.44781701 | 0.89054584 |
| Zfyve21 | 0.78190753 | 0.71395361 | 0.70668596 | 0.00726765 | 0.48089273 | 0.89054584 |
| Pparg | 0.02446247 | 0.0334358 | 0.02617786 | 0.00725794 | 0.09822814 | 0.61534939 |
| Tmem231 | 0.10657851 | 0.1210665 | 0.11382507 | 0.00724142 | 0.3834636 | 0.89054584 |
| Zkscan2 | 0.01905805 | 0.02762227 | 0.02039488 | 0.00722739 | 0.04936904 | 0.3900558 |
| Trpc4 | 0.02817452 | 0.03736341 | 0.0301497 | 0.00721371 | 0.13248677 | 0.72954347 |
| Esm1 | 0.01269459 | 0.02079166 | 0.01358528 | 0.00720639 | 0.00672999 | 0.08389633 |
| Fzd6 | 0.07032252 | 0.0824222 | 0.07522196 | 0.00720024 | 0.32781547 | 0.89054584 |
| Cdc40 | 0.4288864 | 0.43672347 | 0.42953592 | 0.00718755 | 0.46893016 | 0.89054584 |
| Pacrg | 0.22471836 | 0.24229449 | 0.2351382 | 0.0071563 | 0.44362207 | 0.89054584 |
| Rnf165 | 0.07904871 | 0.09169938 | 0.08454466 | 0.00715473 | 0.34669076 | 0.89054584 |
| Coro2a | 0.04861103 | 0.05913905 | 0.05201127 | 0.00712778 | 0.26158219 | 0.89054584 |
| Zfp119a | 0.06521153 | 0.07688513 | 0.06975982 | 0.00712531 | 0.3170846 | 0.89054584 |
| Larp6 | 0.03265656 | 0.04203349 | 0.03494509 | 0.0070884 | 0.17231979 | 0.83696674 |
| Wash1 | 0.46875536 | 0.47193782 | 0.46485397 | 0.00708384 | 0.4717001 | 0.89054584 |
| Pcx | 0.17946081 | 0.19641744 | 0.1893354 | 0.00708204 | 0.43082803 | 0.89054584 |
| Syde1 | 0.14276971 | 0.15865954 | 0.15157891 | 0.00708063 | 0.4138584 | 0.89054584 |
| Lrfn4 | 0.17322303 | 0.19002587 | 0.18295497 | 0.0070709 | 0.42855267 | 0.89054584 |
| Slc39a11 | 0.22592644 | 0.2434161 | 0.23634857 | 0.00706753 | 0.44460011 | 0.89054584 |
| Acpt | 0.0242723 | 0.03303864 | 0.02597438 | 0.00706426 | 0.10256092 | 0.6298723 |
| 1700040D17 | 0.01195148 | 0.01985377 | 0.01279005 | 0.00706373 | 0.00504027 | 0.06631018 |
| Was | 0.02133299 | 0.02987656 | 0.02282922 | 0.00704735 | 0.07518748 | 0.52181337 |
| Klhl42 | 0.09895154 | 0.11283451 | 0.10579119 | 0.00704332 | 0.37821169 | 0.89054584 |
| Ccdc88b | 0.03231855 | 0.04162362 | 0.03458346 | 0.00704015 | 0.17145772 | 0.83588128 |
| Gm13139 | 0.03701361 | 0.04663676 | 0.03960638 | 0.00703038 | 0.20412089 | 0.89054584 |
| Foxc2 | 0.03023304 | 0.03937537 | 0.03235218 | 0.00702318 | 0.15591549 | 0.79725592 |

| | | | | | | |
|---|---|---|---|---|---|---|
| Vezf1 | 0.65812343 | 0.62604915 | 0.61903231 | 0.00701685 | 0.47894166 | 0.89054584 |
| Fmod | 0.05601946 | 0.06694726 | 0.05993334 | 0.00701392 | 0.2927978 | 0.89054584 |
| B3galt5 | 0.03585784 | 0.0453654 | 0.03836995 | 0.00699545 | 0.19782965 | 0.89054584 |
| Frs3 | 0.05285779 | 0.06354404 | 0.0565527 | 0.00699134 | 0.28232035 | 0.89054584 |
| Cdc42ep3 | 0.18812986 | 0.20516633 | 0.1981767 | 0.00698963 | 0.43473988 | 0.89054584 |
| Tmem151b | 0.04444629 | 0.05453508 | 0.04755699 | 0.00697809 | 0.24711031 | 0.89054584 |
| Fgf2 | 0.052253 | 0.0628824 | 0.05590599 | 0.00697641 | 0.28049193 | 0.89054584 |
| Ctnnal1 | 0.24698544 | 0.26430746 | 0.25734337 | 0.00696409 | 0.44983518 | 0.89054584 |
| Dchs1 | 0.06822776 | 0.07994656 | 0.07298342 | 0.00696314 | 0.32834194 | 0.89054584 |
| Fkbp3 | 1.9820567 | 1.12349298 | 1.11653207 | 0.00696091 | 0.48841407 | 0.89054584 |
| Entpd1 | 0.02434311 | 0.03298944 | 0.02605015 | 0.00693929 | 0.10729235 | 0.64845225 |
| Vsig10 | 0.30376079 | 0.31984907 | 0.31291079 | 0.00693829 | 0.45886067 | 0.89054584 |
| Spaca6 | 0.13503096 | 0.15047212 | 0.14354758 | 0.00692454 | 0.41109038 | 0.89054584 |
| Plcg2 | 0.01660646 | 0.02468831 | 0.01777145 | 0.00691685 | 0.03489179 | 0.30373211 |
| Parn | 0.21561281 | 0.23290873 | 0.22599422 | 0.00691451 | 0.44332486 | 0.89054584 |
| BC037039 | 0.01749289 | 0.02562215 | 0.01872002 | 0.00690213 | 0.04291965 | 0.35294902 |
| Trpc5 | 0.02481415 | 0.03345567 | 0.02655417 | 0.0069015 | 0.11297248 | 0.66962283 |
| Gstp1 | 0.82332601 | 0.74046872 | 0.73356776 | 0.00690096 | 0.48252056 | 0.89054584 |
| Megf11 | 0.03449198 | 0.04380856 | 0.03690873 | 0.00689984 | 0.19188873 | 0.88486593 |
| Gm9008 | 0.04384803 | 0.05381507 | 0.0469171 | 0.00689796 | 0.2466787 | 0.89054584 |
| Dubr | 0.0958753 | 0.10940066 | 0.10250897 | 0.0068917 | 0.37705623 | 0.89054584 |
| Soat2 | 0.01666421 | 0.02472441 | 0.01783325 | 0.00689116 | 0.03590526 | 0.30996718 |
| Acer2 | 0.53898296 | 0.5316156 | 0.52472656 | 0.00688905 | 0.47561332 | 0.89054584 |
| Fahd2a | 0.33515709 | 0.34984502 | 0.34295679 | 0.00688822 | 0.4627238 | 0.89054584 |
| Irak4 | 0.06842296 | 0.08006853 | 0.07319203 | 0.0068765 | 0.33079669 | 0.89054584 |
| Lrch4 | 0.02552582 | 0.03418631 | 0.02731565 | 0.00687065 | 0.1206277 | 0.69533658 |
| Il1rl2 | 0.05049835 | 0.06089216 | 0.05402963 | 0.00686253 | 0.27700831 | 0.89054584 |
| Otos | 0.012473 | 0.02020831 | 0.01334814 | 0.00686017 | 0.00832321 | 0.10047453 |
| Gpc6 | 0.11032843 | 0.12460964 | 0.11775957 | 0.00685007 | 0.39319151 | 0.89054584 |
| Lat | 0.02017163 | 0.02843568 | 0.0215865 | 0.00684919 | 0.06967214 | 0.49636689 |

| | | | | | | |
|---|---|---|---|---|---|---|
| AI413582 | 0.33846804 | 0.35294232 | 0.34609604 | 0.00684628 | 0.46328521 | 0.89054584 |
| Serinc2 | 0.15894093 | 0.1751273 | 0.16828611 | 0.00684119 | 0.4248913 | 0.89054584 |
| Gm10605 | 0.01935813 | 0.02755489 | 0.02071598 | 0.0068389 | 0.0620193 | 0.45665604 |
| Gm26691 | 0.03826627 | 0.04778282 | 0.04094643 | 0.0068364 | 0.21832734 | 0.89054584 |
| D030040B21 | 0.01626847 | 0.02423315 | 0.01740976 | 0.00682338 | 0.03392721 | 0.29648991 |
| Slc44a2 | 0.21847252 | 0.23569142 | 0.22886999 | 0.00682143 | 0.4447808 | 0.89054584 |
| Amy1 | 0.01926611 | 0.02743318 | 0.02061752 | 0.00681565 | 0.06176393 | 0.45559551 |
| Fgd5 | 0.02542311 | 0.03401621 | 0.02720575 | 0.00681046 | 0.12175051 | 0.69961907 |
| Ell | 0.14400912 | 0.1596672 | 0.152863 | 0.0068042 | 0.41785753 | 0.89054584 |
| BC049730 | 0.01999864 | 0.02820269 | 0.02140138 | 0.00680131 | 0.06935665 | 0.49496101 |
| Zbtb21 | 0.14985995 | 0.16571122 | 0.15891662 | 0.0067946 | 0.42105449 | 0.89054584 |
| 2210016F16I | 0.60055558 | 0.58142257 | 0.57464146 | 0.00678111 | 0.47807789 | 0.89054584 |
| Vrk3 | 0.41253893 | 0.42156538 | 0.41478614 | 0.00677923 | 0.46965171 | 0.89054584 |
| Erich1 | 0.31205199 | 0.3276729 | 0.32089376 | 0.00677915 | 0.46079781 | 0.89054584 |
| Wipf1 | 0.16359428 | 0.17985338 | 0.17307462 | 0.00677876 | 0.42760485 | 0.89054584 |
| Gltscr1l | 0.07053809 | 0.08222434 | 0.07545231 | 0.00677203 | 0.33791815 | 0.89054584 |
| Atr | 0.15379155 | 0.16973492 | 0.162977 | 0.00675792 | 0.42340695 | 0.89054584 |
| Kcnn3 | 0.0249432 | 0.03344925 | 0.02669225 | 0.006757 | 0.11912226 | 0.6908542 |
| Alas1 | 0.37517698 | 0.38726919 | 0.38051242 | 0.00675677 | 0.46703355 | 0.89054584 |
| Pde5a | 0.06767278 | 0.07914363 | 0.07239031 | 0.00675332 | 0.33191392 | 0.89054584 |
| Zbtb7a | 0.9144884 | 0.79508295 | 0.7883424 | 0.00674055 | 0.48411222 | 0.89054584 |
| Ccdc62 | 0.04789576 | 0.05798057 | 0.05124631 | 0.00673426 | 0.27019277 | 0.89054584 |
| D15Ertd621e | 0.29641608 | 0.31253921 | 0.30581038 | 0.00672883 | 0.45917516 | 0.89054584 |
| Cfi | 0.06990821 | 0.08149938 | 0.07477922 | 0.00672016 | 0.33772368 | 0.89054584 |
| Trmt44 | 0.07773302 | 0.08985533 | 0.08313928 | 0.00671605 | 0.35332749 | 0.89054584 |
| Tdh | 0.00941798 | 0.01678937 | 0.01007883 | 0.00671054 | 0.00096126 | 0.01590912 |
| Ints9 | 0.16099683 | 0.17710574 | 0.1704028 | 0.00670294 | 0.42729562 | 0.89054584 |
| Kank2 | 0.2605041 | 0.27741144 | 0.27071323 | 0.00669821 | 0.45411343 | 0.89054584 |
| Zcchc16 | 0.05165806 | 0.06195904 | 0.05526979 | 0.00668924 | 0.28642243 | 0.89054584 |
| Dgkg | 0.01513059 | 0.02285723 | 0.01619211 | 0.00666512 | 0.02757136 | 0.25315361 |

| | | | | | | |
|---|---|---|---|---|---|---|
| Slc2a10 | 0.02004469 | 0.02811419 | 0.02145066 | 0.00666353 | 0.07391008 | 0.5164088 |
| Aldh3b1 | 0.08296833 | 0.09538905 | 0.08873086 | 0.00665819 | 0.36332025 | 0.89054584 |
| Spata2l | 0.04126236 | 0.05080622 | 0.04415137 | 0.00665484 | 0.24126708 | 0.89054584 |
| Sfrp5 | 0.02655706 | 0.03506983 | 0.02841907 | 0.00665076 | 0.13778894 | 0.7438695 |
| Zdhhc1 | 0.25578498 | 0.27270541 | 0.26605548 | 0.00664993 | 0.45364873 | 0.89054584 |
| Rbm46 | 0.06737817 | 0.07871791 | 0.07207546 | 0.00664246 | 0.33382815 | 0.89054584 |
| Adamts17 | 0.08176151 | 0.09407202 | 0.08744206 | 0.00662996 | 0.36195176 | 0.89054584 |
| Shisa3 | 0.02507737 | 0.03345162 | 0.02683581 | 0.00661581 | 0.12536766 | 0.7081958 |
| Lrrc16b | 0.21764228 | 0.23464529 | 0.22803546 | 0.00660983 | 0.44628859 | 0.89054584 |
| Arap2 | 0.04513085 | 0.05489641 | 0.04828918 | 0.00660723 | 0.26191082 | 0.89054584 |
| Ap1m1 | 0.51774124 | 0.51354426 | 0.50694006 | 0.00660421 | 0.47580115 | 0.89054584 |
| Ttc21b | 0.08279073 | 0.09513777 | 0.08854119 | 0.00659657 | 0.3642564 | 0.89054584 |
| Vprbp | 0.22236871 | 0.23936825 | 0.23278219 | 0.00658605 | 0.44756555 | 0.89054584 |
| Cep135 | 0.12237108 | 0.13693744 | 0.13035976 | 0.00657768 | 0.40707317 | 0.89054584 |
| Bri3 | 1.16674854 | 0.92252348 | 0.91596138 | 0.0065621 | 0.4866868 | 0.89054584 |
| Rad21 | 0.96258812 | 0.82143412 | 0.81487473 | 0.00655939 | 0.48504218 | 0.89054584 |
| Pip4k2c | 0.27069879 | 0.28729552 | 0.28073976 | 0.00655576 | 0.45668281 | 0.89054584 |
| Mob2 | 0.37823133 | 0.38989771 | 0.38334329 | 0.00655442 | 0.46825432 | 0.89054584 |
| Cgn | 0.20620909 | 0.22306372 | 0.21651214 | 0.00655157 | 0.44394376 | 0.89054584 |
| Gm13251 | 0.12694439 | 0.14168112 | 0.13513059 | 0.00655052 | 0.41066118 | 0.89054584 |
| Slc38a3 | 0.02393628 | 0.03216361 | 0.02561484 | 0.00654878 | 0.11680312 | 0.68200227 |
| Rai14 | 0.058886 | 0.06954311 | 0.0629981 | 0.00654501 | 0.31418316 | 0.89054584 |
| RP24-308M1 | 0.02530427 | 0.03361571 | 0.0270786 | 0.00653711 | 0.13035246 | 0.72297247 |
| Bnip3 | 0.40983286 | 0.41885821 | 0.41233021 | 0.00652801 | 0.47060056 | 0.89054584 |
| Tmem56 | 0.01419908 | 0.02170911 | 0.01519528 | 0.00651383 | 0.02290337 | 0.22210909 |
| Efna4 | 0.14626475 | 0.16170807 | 0.15519841 | 0.00650966 | 0.42253393 | 0.89054584 |
| Dact1 | 0.10407866 | 0.11769034 | 0.11119932 | 0.00649102 | 0.39282802 | 0.89054584 |
| Efna3 | 0.06028349 | 0.07097719 | 0.0644921 | 0.00648509 | 0.31971846 | 0.89054584 |
| Zkscan5 | 0.07925679 | 0.09124315 | 0.0847669 | 0.00647625 | 0.36094045 | 0.89054584 |
| Rpl9-ps6 | 1.06694402 | 0.87533852 | 0.86886334 | 0.00647518 | 0.48615126 | 0.89054584 |

| | | | | | | |
|---|---|---|---|---|---|---|
| Agrn | 0.1972206 | 0.21388384 | 0.2074138 | 0.00647004 | 0.44222498 | 0.89054584 |
| Rab3c | 0.07082414 | 0.0822259 | 0.07575797 | 0.00646793 | 0.34540245 | 0.89054584 |
| Gm13111 | 0.06398755 | 0.07491889 | 0.06845157 | 0.00646732 | 0.3299042 | 0.89054584 |
| Elf5 | 0.03555943 | 0.04449886 | 0.0380507 | 0.00644816 | 0.2149057 | 0.89054584 |
| Wnt6 | 0.01990326 | 0.0277356 | 0.02129932 | 0.00643628 | 0.07958767 | 0.54059058 |
| Vegfc | 0.03266217 | 0.04138728 | 0.0349511 | 0.00643618 | 0.19546397 | 0.89054584 |
| Zc4h2 | 0.08768318 | 0.10018828 | 0.09376517 | 0.00642312 | 0.37480734 | 0.89054584 |
| E2f8 | 0.03195932 | 0.04062199 | 0.03419914 | 0.00642285 | 0.19079135 | 0.88318504 |
| Tarbp2 | 0.36844121 | 0.38065432 | 0.37425122 | 0.0064031 | 0.46823386 | 0.89054584 |
| Ankrd29 | 0.03911444 | 0.04824638 | 0.04185374 | 0.00639264 | 0.23835749 | 0.89054584 |
| Epsti1 | 0.02943609 | 0.03789092 | 0.03149951 | 0.00639141 | 0.17224703 | 0.83696674 |
| Susd2 | 0.04223298 | 0.05157673 | 0.04518961 | 0.00638712 | 0.25510944 | 0.89054584 |
| Alg2 | 0.88648336 | 0.77853361 | 0.77215601 | 0.0063776 | 0.48465234 | 0.89054584 |
| Hps6 | 0.10299597 | 0.11642419 | 0.11006139 | 0.00636279 | 0.39383425 | 0.89054584 |
| Tmem140 | 0.02446793 | 0.03254435 | 0.02618371 | 0.00636063 | 0.12886659 | 0.71857996 |
| Tanc1 | 0.16317725 | 0.17900579 | 0.17264584 | 0.00635996 | 0.43186545 | 0.89054584 |
| Parp16 | 0.082646 | 0.09474325 | 0.08838664 | 0.00635661 | 0.36878878 | 0.89054584 |
| Acy1 | 0.10590442 | 0.11946554 | 0.11311726 | 0.00634828 | 0.39686672 | 0.89054584 |
| Zw10 | 0.12932068 | 0.1439545 | 0.13760657 | 0.00634793 | 0.4149142 | 0.89054584 |
| Nudt1 | 0.1702337 | 0.18623108 | 0.17989161 | 0.00633947 | 0.43479343 | 0.89054584 |
| Arap3 | 0.03766565 | 0.04663878 | 0.04030391 | 0.00633487 | 0.23199848 | 0.89054584 |
| Arl4c | 0.14888702 | 0.1642426 | 0.15791091 | 0.0063317 | 0.42590563 | 0.89054584 |
| Fn3krp | 0.02918848 | 0.03756362 | 0.03123458 | 0.00632903 | 0.1725754 | 0.83703076 |
| Cog7 | 0.47653402 | 0.47795677 | 0.47163549 | 0.00632128 | 0.47510491 | 0.89054584 |
| Ubtd2 | 0.09306901 | 0.10582313 | 0.0995142 | 0.00630892 | 0.38385794 | 0.89054584 |
| Bcl11b | 0.0228019 | 0.03070806 | 0.02440102 | 0.00630704 | 0.11425289 | 0.67417108 |
| Nfx1 | 0.57978153 | 0.56438193 | 0.55807775 | 0.00630418 | 0.47901396 | 0.89054584 |
| Dgcr2 | 0.35430445 | 0.36731917 | 0.36103164 | 0.00628753 | 0.46766625 | 0.89054584 |
| Aldh16a1 | 0.12116639 | 0.13538457 | 0.12910174 | 0.00628283 | 0.41031658 | 0.89054584 |
| Esr1 | 0.01616928 | 0.0235735 | 0.01730362 | 0.00626988 | 0.04569178 | 0.36863632 |

| | | | | | | |
|---|---|---|---|---|---|---|
| Rbm24 | 0.02236113 | 0.0301977 | 0.02392938 | 0.00626833 | 0.11115385 | 0.66208238 |
| Eml5 | 0.21333318 | 0.22994981 | 0.22369917 | 0.00625064 | 0.44821167 | 0.89054584 |
| Ykt6 | 0.36645013 | 0.37864109 | 0.37239564 | 0.00624546 | 0.46886024 | 0.89054584 |
| Twf2 | 0.4875369 | 0.48740335 | 0.48116431 | 0.00623904 | 0.47591438 | 0.89054584 |
| Zfp738 | 0.08831973 | 0.100677 | 0.09444474 | 0.00623225 | 0.3792558 | 0.89054584 |
| Aox1 | 0.02325894 | 0.03111393 | 0.02489006 | 0.00622387 | 0.12201191 | 0.70028846 |
| Akr1b10 | 0.06384591 | 0.07450701 | 0.06830017 | 0.00620684 | 0.3360069 | 0.89054584 |
| Rcor2 | 0.22359453 | 0.24021733 | 0.23401164 | 0.0062057 | 0.45083604 | 0.89054584 |
| Tnik | 0.27320218 | 0.28939173 | 0.2831942 | 0.00619752 | 0.45939503 | 0.89054584 |
| Eme1 | 0.04606483 | 0.05548253 | 0.04928812 | 0.00619442 | 0.27909732 | 0.89054584 |
| Suclg2 | 0.66341848 | 0.62919375 | 0.62300217 | 0.00619158 | 0.4815348 | 0.89054584 |
| Fam13c | 0.02942948 | 0.03768288 | 0.03149243 | 0.00619045 | 0.17988456 | 0.85601281 |
| Pdss2 | 0.16840837 | 0.1842084 | 0.17801926 | 0.00618914 | 0.43566237 | 0.89054584 |
| Gm29394 | 0.02213407 | 0.02987369 | 0.02368641 | 0.00618728 | 0.11180368 | 0.66458783 |
| Tcf7l1 | 0.12406889 | 0.13830905 | 0.13213181 | 0.00617725 | 0.41378915 | 0.89054584 |
| Tradd | 0.28789442 | 0.30371386 | 0.29753826 | 0.0061756 | 0.46148257 | 0.89054584 |
| Cd200r1 | 0.01700064 | 0.02435828 | 0.01819326 | 0.00616502 | 0.05719767 | 0.43192879 |
| Primpol | 0.14373185 | 0.15873454 | 0.15257579 | 0.00615875 | 0.42541517 | 0.89054584 |
| Ednra | 0.10413345 | 0.11741314 | 0.1112569 | 0.00615624 | 0.39828221 | 0.89054584 |
| Lpar2 | 0.01617494 | 0.02346113 | 0.01730967 | 0.00615146 | 0.04889326 | 0.38756396 |
| Mospd1 | 0.12254341 | 0.1366857 | 0.13053967 | 0.00614603 | 0.41318839 | 0.89054584 |
| Tmem171 | 0.05285804 | 0.06269867 | 0.05655297 | 0.0061457 | 0.306325 | 0.89054584 |
| Nts | 0.01467955 | 0.02185361 | 0.01570944 | 0.00614418 | 0.03421411 | 0.29872948 |
| Slamf1 | 0.02754782 | 0.03561301 | 0.02947915 | 0.00613385 | 0.16617146 | 0.8246338 |
| Aard | 0.1091766 | 0.12268017 | 0.1165516 | 0.00612857 | 0.40323653 | 0.89054584 |
| Utrn | 0.31288296 | 0.32781685 | 0.32169204 | 0.00612481 | 0.46465888 | 0.89054584 |
| Homer2 | 0.1358481 | 0.15052089 | 0.14439671 | 0.00612418 | 0.42168061 | 0.89054584 |
| E130311K13I | 0.12656663 | 0.14085899 | 0.13473679 | 0.0061222 | 0.41617288 | 0.89054584 |
| Ism1 | 0.03672017 | 0.04541289 | 0.03929246 | 0.00612044 | 0.2340097 | 0.89054584 |
| C030014I23R | 0.08848434 | 0.10074064 | 0.09462047 | 0.00612017 | 0.38157531 | 0.89054584 |

| | | | | | | |
|---|---|---|---|---|---|---|
| Eid1 | 1.02808404 | 0.85502808 | 0.84893726 | 0.00609082 | 0.48666734 | 0.89054584 |
| Mettl8 | 0.10914737 | 0.12260065 | 0.11652093 | 0.00607971 | 0.40396802 | 0.89054584 |
| Dnajb12 | 0.25578082 | 0.27213005 | 0.26605137 | 0.00607868 | 0.45761406 | 0.89054584 |
| Pla2g12a | 0.43050596 | 0.43703053 | 0.43098875 | 0.00604178 | 0.47396319 | 0.89054584 |
| Usp6nl | 0.09332155 | 0.10582323 | 0.09978373 | 0.0060395 | 0.38897684 | 0.89054584 |
| Pcdh7 | 0.05002296 | 0.05955959 | 0.05352124 | 0.00603835 | 0.29957308 | 0.89054584 |
| Vgll3 | 0.08264562 | 0.0944237 | 0.08838623 | 0.00603747 | 0.37515059 | 0.89054584 |
| Zhx1 | 0.34452745 | 0.35784495 | 0.35182663 | 0.00601832 | 0.46823973 | 0.89054584 |
| Lrrtm1 | 0.01503706 | 0.02210992 | 0.01609202 | 0.0060179 | 0.04072853 | 0.34053187 |
| Zbtb16 | 0.04127206 | 0.05017092 | 0.04416175 | 0.00600917 | 0.26305638 | 0.89054584 |
| AI838599 | 0.01716533 | 0.02436381 | 0.01836949 | 0.00599431 | 0.06421773 | 0.46799603 |
| Ppil3 | 0.38084971 | 0.39174492 | 0.38576596 | 0.00597896 | 0.47121792 | 0.89054584 |
| Ext1 | 0.1118603 | 0.12534361 | 0.11936536 | 0.00597825 | 0.40774993 | 0.89054584 |
| Fndc3b | 0.33905176 | 0.3526164 | 0.34664894 | 0.00596746 | 0.46803816 | 0.89054584 |
| Sertad4 | 0.10496658 | 0.11809965 | 0.11213222 | 0.00596742 | 0.40209106 | 0.89054584 |
| Robo1 | 0.08550724 | 0.09740337 | 0.09144197 | 0.0059614 | 0.38066615 | 0.89054584 |
| Onecut2 | 0.1045885 | 0.11769572 | 0.11173502 | 0.0059607 | 0.40185859 | 0.89054584 |
| Wsb1 | 0.4983184 | 0.49638931 | 0.49042959 | 0.00595973 | 0.47742566 | 0.89054584 |
| Kdm1b | 0.05175025 | 0.06132732 | 0.05536838 | 0.00595894 | 0.30804063 | 0.89054584 |
| Hist1h2be | 0.06059446 | 0.07077655 | 0.06482453 | 0.00595202 | 0.33440859 | 0.89054584 |
| Rxrg | 0.02705322 | 0.03490084 | 0.02894995 | 0.00595089 | 0.1691115 | 0.8322688 |
| Bace2 | 0.046079 | 0.05525358 | 0.04930328 | 0.0059503 | 0.2869637 | 0.89054584 |
| Tgif2 | 0.11929528 | 0.13308816 | 0.12714671 | 0.00594145 | 0.41382826 | 0.89054584 |
| Slc17a9 | 0.03185931 | 0.04001686 | 0.03409214 | 0.00592472 | 0.20906878 | 0.89054584 |
| C1qtnf2 | 0.05401473 | 0.06371225 | 0.05778981 | 0.00592244 | 0.31651744 | 0.89054584 |
| Kcnip1 | 0.08899724 | 0.10108955 | 0.09516801 | 0.00592154 | 0.38595079 | 0.89054584 |
| Arid5a | 0.0597139 | 0.06980393 | 0.06388318 | 0.00592075 | 0.33294519 | 0.89054584 |
| Rhbdd1 | 0.09158222 | 0.10383995 | 0.09792734 | 0.00591261 | 0.38924191 | 0.89054584 |
| Trim6 | 0.01539255 | 0.02237901 | 0.01647244 | 0.00590658 | 0.04740309 | 0.37844164 |
| Gbp8 | 0.01477947 | 0.02170904 | 0.01581637 | 0.00589267 | 0.04130211 | 0.34374164 |

| | | | | | | |
|---|---|---|---|---|---|---|
| Nol4 | 0.29074731 | 0.30617318 | 0.30031163 | 0.00586155 | 0.4637724 | 0.89054584 |
| Urb1 | 0.03969662 | 0.04833674 | 0.04247652 | 0.00586022 | 0.26018822 | 0.89054584 |
| Nkain3 | 0.02412145 | 0.03167193 | 0.02581297 | 0.00585896 | 0.14514756 | 0.76747222 |
| Lrmp | 0.03233201 | 0.04045603 | 0.03459786 | 0.00585816 | 0.21509716 | 0.89054584 |
| Sdcbp | 1.2475618 | 0.95604002 | 0.95018379 | 0.00585623 | 0.48854624 | 0.89054584 |
| Slc2a12 | 0.01893633 | 0.02611344 | 0.02026463 | 0.0058488 | 0.08936605 | 0.57918151 |
| 2410002F23l | 0.47078874 | 0.47246649 | 0.46663027 | 0.00583621 | 0.47676676 | 0.89054584 |
| Etfa | 1.33052019 | 0.98762702 | 0.98179736 | 0.00582966 | 0.48896522 | 0.89054584 |
| Slc39a10 | 0.36040314 | 0.372576 | 0.36674736 | 0.00582865 | 0.47048782 | 0.89054584 |
| Nckap1l | 0.02022361 | 0.02745064 | 0.02164211 | 0.00580853 | 0.10557411 | 0.64230526 |
| Utp14a | 0.25027344 | 0.26640873 | 0.26060296 | 0.00580577 | 0.45866681 | 0.89054584 |
| Rrp9 | 0.25102162 | 0.26714669 | 0.26134397 | 0.00580272 | 0.45880518 | 0.89054584 |
| Asap3 | 0.03969639 | 0.04827275 | 0.04247627 | 0.00579648 | 0.26246071 | 0.89054584 |
| Slc35d2 | 0.03923459 | 0.04776997 | 0.04198227 | 0.0057877 | 0.26034446 | 0.89054584 |
| Lama5 | 0.15642911 | 0.17147378 | 0.16569771 | 0.00577607 | 0.43549287 | 0.89054584 |
| 1810032O08 | 0.09135566 | 0.10346156 | 0.09768551 | 0.00577604 | 0.39147439 | 0.89054584 |
| Syt4 | 0.02698316 | 0.03463612 | 0.02887499 | 0.00576113 | 0.17630188 | 0.84615131 |
| Prpsap1 | 0.65936909 | 0.62572838 | 0.61996797 | 0.00576041 | 0.48273583 | 0.89054584 |
| Tsku | 0.0308844 | 0.03880428 | 0.03304908 | 0.0057552 | 0.20859305 | 0.89054584 |
| Ppm1j | 0.01609941 | 0.02298084 | 0.01722886 | 0.00575198 | 0.05992282 | 0.44597591 |
| Tspan11 | 0.03572729 | 0.04397758 | 0.03823029 | 0.00574729 | 0.24184046 | 0.89054584 |
| Mtrr | 0.10121493 | 0.11392991 | 0.10818857 | 0.00574134 | 0.40236113 | 0.89054584 |
| Nhp2 | 0.59572303 | 0.57655211 | 0.57081413 | 0.00573798 | 0.48132317 | 0.89054584 |
| Ldlrad4 | 0.04546021 | 0.05437452 | 0.04864145 | 0.00573307 | 0.29146132 | 0.89054584 |
| Gdap10 | 0.02706309 | 0.03469115 | 0.02896051 | 0.00573063 | 0.17829063 | 0.85140236 |
| Sod3 | 0.08703495 | 0.09879098 | 0.0930731 | 0.00571788 | 0.38735522 | 0.89054584 |
| Ifi204 | 0.01220677 | 0.01875298 | 0.01306325 | 0.00568973 | 0.02121819 | 0.20937119 |
| Akap1 | 0.08871725 | 0.10055549 | 0.09486912 | 0.00568637 | 0.39002536 | 0.89054584 |
| Nptx1 | 0.04091364 | 0.04946448 | 0.04377835 | 0.00568613 | 0.27254819 | 0.89054584 |
| Gm12976 | 0.05274327 | 0.06211143 | 0.05643025 | 0.00568118 | 0.31951887 | 0.89054584 |

| | | | | | | |
|---|---|---|---|---|---|---|
| Ehmt2 | 0.62818907 | 0.60190711 | 0.59622961 | 0.0056775 | 0.48230712 | 0.89054584 |
| Ttll13 | 0.05649098 | 0.06610988 | 0.0604375 | 0.00567238 | 0.33095896 | 0.89054584 |
| Prl | 0.04318604 | 0.05186063 | 0.04620903 | 0.0056516 | 0.28440246 | 0.89054584 |
| Gm15860 | 0.05442222 | 0.0638731 | 0.05822552 | 0.00564758 | 0.32567288 | 0.89054584 |
| Gm16104 | 0.02336541 | 0.03065093 | 0.02500399 | 0.00564694 | 0.14635768 | 0.77187839 |
| Dcun1d1 | 0.57771513 | 0.56205368 | 0.55641439 | 0.00563929 | 0.48116948 | 0.89054584 |
| Dkc1 | 0.49836022 | 0.49610186 | 0.49046539 | 0.00563647 | 0.47865046 | 0.89054584 |
| Pot1a | 0.13811986 | 0.15238831 | 0.14675606 | 0.00563225 | 0.42904646 | 0.89054584 |
| Trim14 | 0.02422922 | 0.03156021 | 0.02592829 | 0.00563192 | 0.15577505 | 0.79690427 |
| Atp11c | 0.04802063 | 0.0570101 | 0.05137986 | 0.00563025 | 0.30483992 | 0.89054584 |
| AW551984 | 0.15782533 | 0.17275726 | 0.16713681 | 0.00562045 | 0.43775244 | 0.89054584 |
| Pglyrp1 | 0.05696473 | 0.06655641 | 0.06094401 | 0.0056124 | 0.33394462 | 0.89054584 |
| Dzip3 | 0.35011858 | 0.36269222 | 0.35709695 | 0.00559527 | 0.47090313 | 0.89054584 |
| Wwp2 | 0.16673357 | 0.18188662 | 0.17630011 | 0.0055865 | 0.44131739 | 0.89054584 |
| E2f5 | 0.23833212 | 0.25432042 | 0.24874073 | 0.00557969 | 0.45838299 | 0.89054584 |
| Fgf10 | 0.03376226 | 0.04169901 | 0.03612804 | 0.00557097 | 0.23625154 | 0.89054584 |
| Shank1 | 0.01436811 | 0.02094312 | 0.01537616 | 0.00556695 | 0.04582343 | 0.3690823 |
| Plekhh3 | 0.23623741 | 0.25221623 | 0.24665308 | 0.00556314 | 0.45815609 | 0.89054584 |
| Trim65 | 0.05513155 | 0.06454078 | 0.05898397 | 0.0055568 | 0.33036239 | 0.89054584 |
| Rapgef3 | 0.05672528 | 0.06623822 | 0.060688 | 0.00555022 | 0.33502345 | 0.89054584 |
| Celsr3 | 0.06930973 | 0.07966926 | 0.07413967 | 0.00552958 | 0.3641156 | 0.89054584 |
| Syt15 | 0.00726568 | 0.01330311 | 0.00777552 | 0.00552758 | 0.00046312 | 0.00842456 |
| 3110009E18I | 0.13503994 | 0.14908316 | 0.14355692 | 0.00552624 | 0.42883281 | 0.89054584 |
| Svopl | 0.02130153 | 0.02831688 | 0.02279556 | 0.00552132 | 0.12956617 | 0.72054099 |
| Gm15800 | 0.21739739 | 0.23329639 | 0.22778925 | 0.00550714 | 0.45515926 | 0.89054584 |
| Fgr | 0.0120919 | 0.01844372 | 0.01294032 | 0.00550341 | 0.02377276 | 0.22749665 |
| Frk | 0.1665195 | 0.18158138 | 0.17608029 | 0.00550109 | 0.44213659 | 0.89054584 |
| Ctbp2 | 0.26183609 | 0.2775198 | 0.27202608 | 0.00549371 | 0.46251889 | 0.89054584 |
| Hmgcs1 | 0.56307347 | 0.55004369 | 0.54455002 | 0.00549367 | 0.48125599 | 0.89054584 |
| Rif1 | 0.41043293 | 0.41836809 | 0.41287517 | 0.00549291 | 0.47528826 | 0.89054584 |

| | | | | | | |
|---|---|---|---|---|---|---|
| Ccdc157 | 0.03322432 | 0.04104332 | 0.03555252 | 0.00549079 | 0.23590564 | 0.89054584 |
| D10Wsu102e | 0.19294984 | 0.20855802 | 0.20307869 | 0.00547933 | 0.44998298 | 0.89054584 |
| Ppp1r3b | 0.02392822 | 0.03107857 | 0.02560621 | 0.00547235 | 0.15970486 | 0.80704115 |
| Cpne3 | 0.40918042 | 0.41720943 | 0.41173744 | 0.005472 | 0.4753143 | 0.89054584 |
| 4930447C04I | 0.01111319 | 0.01735466 | 0.01189295 | 0.00546171 | 0.01619487 | 0.17264208 |
| Acox3 | 0.09608567 | 0.10819185 | 0.10273345 | 0.0054584 | 0.40224622 | 0.89054584 |
| Gpr180 | 0.48720347 | 0.48632518 | 0.48087665 | 0.00544853 | 0.47895054 | 0.89054584 |
| Gm26734 | 0.01109243 | 0.01731697 | 0.01187074 | 0.00544624 | 0.01627864 | 0.17302624 |
| Hoxaas3 | 0.02938224 | 0.0368876 | 0.0314419 | 0.0054457 | 0.20985542 | 0.89054584 |
| Exoc3l4 | 0.037351 | 0.04539884 | 0.0399673 | 0.00543153 | 0.26331865 | 0.89054584 |
| Cped1 | 0.05154256 | 0.06057203 | 0.05514627 | 0.00542575 | 0.32333773 | 0.89054584 |
| Gm26513 | 0.01514526 | 0.02163243 | 0.0162078 | 0.00542463 | 0.05946112 | 0.44322231 |
| Nip7 | 0.58880862 | 0.57073249 | 0.56531079 | 0.0054217 | 0.48218027 | 0.89054584 |
| Borcs8 | 0.67790662 | 0.63918536 | 0.63376424 | 0.00542112 | 0.48410562 | 0.89054584 |
| Gm17586 | 0.0690015 | 0.07923034 | 0.07381029 | 0.00542005 | 0.36613182 | 0.89054584 |
| Fam129b | 0.33436241 | 0.34761549 | 0.34220259 | 0.0054129 | 0.47062683 | 0.89054584 |
| E130102H24I | 0.00721379 | 0.01311376 | 0.00771999 | 0.00539377 | 0.00056678 | 0.01013221 |
| Gm21781 | 0.04933585 | 0.0581762 | 0.05278643 | 0.00538977 | 0.31714326 | 0.89054584 |
| Mars | 0.3957428 | 0.40484201 | 0.39947461 | 0.0053674 | 0.47504322 | 0.89054584 |
| Fam217a | 0.03733921 | 0.04531024 | 0.0399547 | 0.00535554 | 0.26615418 | 0.89054584 |
| Inpp4b | 0.03323949 | 0.04091687 | 0.03556875 | 0.00534812 | 0.24180183 | 0.89054584 |
| Tstd2 | 0.22377706 | 0.23954264 | 0.23419465 | 0.00534799 | 0.45763642 | 0.89054584 |
| Cytl1 | 0.01140081 | 0.01752794 | 0.01220075 | 0.00532719 | 0.02096454 | 0.20789599 |
| Tm6sf1 | 0.04932392 | 0.0581006 | 0.05277367 | 0.00532693 | 0.31908104 | 0.89054584 |
| Ptpn14 | 0.16656114 | 0.18144958 | 0.17612305 | 0.00532653 | 0.4439739 | 0.89054584 |
| Wdr37 | 0.15415018 | 0.16867362 | 0.1633471 | 0.00532652 | 0.43962443 | 0.89054584 |
| Zmat1 | 0.10004643 | 0.11228449 | 0.10695922 | 0.00532527 | 0.40828501 | 0.89054584 |
| Ensa | 0.51383611 | 0.50894807 | 0.50363966 | 0.00530841 | 0.48041755 | 0.89054584 |
| Max | 0.65046982 | 0.61856612 | 0.61326037 | 0.00530576 | 0.48392384 | 0.89054584 |
| Dnajc21 | 0.30825954 | 0.32255028 | 0.31724639 | 0.00530389 | 0.46895755 | 0.89054584 |

| | | | | | | |
|---|---|---|---|---|---|---|
| Spire1 | 0.10044756 | 0.11267698 | 0.10738129 | 0.00529568 | 0.40913788 | 0.89054584 |
| Hdx | 0.02471507 | 0.03173929 | 0.02644815 | 0.00529114 | 0.17565445 | 0.84465613 |
| Cenph | 0.11956105 | 0.13270638 | 0.12742449 | 0.0052819 | 0.42343302 | 0.89054584 |
| Pitrm1 | 0.17680127 | 0.19189798 | 0.18661696 | 0.00528102 | 0.44755465 | 0.89054584 |
| Tk2 | 0.12834958 | 0.14187329 | 0.13659499 | 0.00527829 | 0.42856434 | 0.89054584 |
| Tbx1 | 0.0197822 | 0.02644624 | 0.02116977 | 0.00527647 | 0.12277623 | 0.70117835 |
| Zfp760 | 0.10710421 | 0.11965012 | 0.11437697 | 0.00527315 | 0.4149646 | 0.89054584 |
| Trim71 | 0.01024703 | 0.01621835 | 0.01096603 | 0.00525231 | 0.01282547 | 0.14325401 |
| Akap8 | 0.47790164 | 0.47807576 | 0.47282395 | 0.00525181 | 0.47936461 | 0.89054584 |
| Gm5426 | 0.01330487 | 0.01948638 | 0.01423836 | 0.00524802 | 0.04297063 | 0.35316795 |
| Abl1 | 0.23711236 | 0.25277322 | 0.24752533 | 0.0052479 | 0.46065794 | 0.89054584 |
| Hn1l | 0.63252581 | 0.60480598 | 0.59957127 | 0.00523471 | 0.48377706 | 0.89054584 |
| Nup107 | 0.1319088 | 0.14553158 | 0.14030082 | 0.00523076 | 0.43105186 | 0.89054584 |
| Rhof | 0.04194403 | 0.05010468 | 0.04488053 | 0.00522415 | 0.29380356 | 0.89054584 |
| Pms2 | 0.20316438 | 0.21865738 | 0.21343403 | 0.00522335 | 0.4546118 | 0.89054584 |
| Bmpr2 | 0.26226332 | 0.27764178 | 0.27244699 | 0.00519479 | 0.46460738 | 0.89054584 |
| Trim7 | 0.10129036 | 0.11346037 | 0.10826791 | 0.00519246 | 0.41159658 | 0.89054584 |
| Rnf180 | 0.02045948 | 0.02707612 | 0.02189451 | 0.00518162 | 0.13510084 | 0.73635034 |
| Hps1 | 0.14710613 | 0.16124505 | 0.15606904 | 0.00517601 | 0.43860253 | 0.89054584 |
| Dcaf17 | 0.4340087 | 0.43930161 | 0.43412562 | 0.00517599 | 0.4778511 | 0.89054584 |
| Mis18a | 0.21597483 | 0.2315326 | 0.22635847 | 0.00517413 | 0.45759496 | 0.89054584 |
| Tbc1d8b | 0.07553238 | 0.08595523 | 0.08078843 | 0.00516679 | 0.38286651 | 0.89054584 |
| Hcar1 | 0.02669619 | 0.03373367 | 0.02856793 | 0.00516574 | 0.19976993 | 0.89054584 |
| Fundc1 | 0.44636217 | 0.45029772 | 0.44513201 | 0.00516571 | 0.47844107 | 0.89054584 |
| Ptrh1 | 0.08717839 | 0.09839024 | 0.09322624 | 0.00516401 | 0.39817684 | 0.89054584 |
| Cnbd2 | 0.03695654 | 0.04470929 | 0.03954533 | 0.00516396 | 0.27146762 | 0.89054584 |
| Sec14l2 | 0.02027133 | 0.02684637 | 0.02169318 | 0.0051532 | 0.1342059 | 0.73312628 |
| Wfdc10 | 0.03048345 | 0.03776633 | 0.03262011 | 0.00514623 | 0.23116785 | 0.89054584 |
| Cand2 | 0.13721349 | 0.15095268 | 0.14581498 | 0.00513769 | 0.43480468 | 0.89054584 |
| E2f7 | 0.05351954 | 0.06239677 | 0.05726032 | 0.00513645 | 0.3380016 | 0.89054584 |

| | | | | | | |
|---|---|---|---|---|---|---|
| Ccdc91 | 0.3529022 | 0.36485071 | 0.35971459 | 0.00513612 | 0.47348117 | 0.89054584 |
| Tnfsf12 | 0.10683667 | 0.11923072 | 0.1140961 | 0.00513462 | 0.41696479 | 0.89054584 |
| Gsg2 | 0.049031 | 0.05759354 | 0.05246041 | 0.00513313 | 0.32424279 | 0.89054584 |
| Gm10184 | 0.06439594 | 0.07401454 | 0.06888808 | 0.00512646 | 0.36440637 | 0.89054584 |
| C8b | 0.02744341 | 0.0344884 | 0.02936744 | 0.00512096 | 0.20827964 | 0.89054584 |
| Gpam | 0.01310825 | 0.01914814 | 0.01402794 | 0.00512019 | 0.04451733 | 0.36199945 |
| Mob3b | 0.09007859 | 0.1014372 | 0.09632235 | 0.00511485 | 0.40230089 | 0.89054584 |
| Rad18 | 0.07726135 | 0.0877407 | 0.08263544 | 0.00510526 | 0.38673774 | 0.89054584 |
| Rpusd2 | 0.01952946 | 0.0259861 | 0.02089933 | 0.00508678 | 0.1284058 | 0.71695178 |
| Foxa3 | 0.34443933 | 0.35682595 | 0.35174344 | 0.00508251 | 0.47316374 | 0.89054584 |
| Cyyr1 | 0.02793345 | 0.03497204 | 0.02989177 | 0.00508028 | 0.21423474 | 0.89054584 |
| 4833439L19I | 0.78244785 | 0.71212334 | 0.70704461 | 0.00507873 | 0.48665175 | 0.89054584 |
| Ppm1f | 0.06908046 | 0.07896287 | 0.07389467 | 0.00506819 | 0.37465803 | 0.89054584 |
| Pvr | 0.05593092 | 0.064904 | 0.05983868 | 0.00506532 | 0.34665112 | 0.89054584 |
| Nov | 0.02554794 | 0.03239866 | 0.02733932 | 0.00505933 | 0.19429699 | 0.88917857 |
| Skil | 0.2865979 | 0.30133447 | 0.29627654 | 0.00505793 | 0.46830321 | 0.89054584 |
| Poc1a | 0.08250772 | 0.09329272 | 0.08823897 | 0.00505375 | 0.39479868 | 0.89054584 |
| Dvl2 | 0.18876286 | 0.2038633 | 0.19882104 | 0.00504226 | 0.45297268 | 0.89054584 |
| Gnpda1 | 0.14238249 | 0.15620576 | 0.1511776 | 0.00502815 | 0.43842805 | 0.89054584 |
| Gm26810 | 0.01931431 | 0.02569435 | 0.0206691 | 0.00502525 | 0.12866543 | 0.71801057 |
| Maoa | 0.18162903 | 0.19656829 | 0.19154967 | 0.00501862 | 0.45142385 | 0.89054584 |
| Sstr2 | 0.03630361 | 0.04386486 | 0.03884683 | 0.00501803 | 0.27364787 | 0.89054584 |
| Stab2 | 0.0150405 | 0.02110575 | 0.01609569 | 0.00501006 | 0.07350501 | 0.51488639 |
| H2-DMb1 | 0.01517854 | 0.02125074 | 0.01624342 | 0.00500732 | 0.07547276 | 0.52279223 |
| Rnd1 | 0.0199776 | 0.02636227 | 0.02137887 | 0.0049834 | 0.13874016 | 0.74747802 |
| Has2 | 0.01927694 | 0.02561021 | 0.02062911 | 0.0049811 | 0.13030586 | 0.72297247 |
| Syt17 | 0.07858507 | 0.08902888 | 0.08404942 | 0.00497946 | 0.39126772 | 0.89054584 |
| Poln | 0.02042267 | 0.02683105 | 0.02185512 | 0.00497593 | 0.14440446 | 0.76687762 |
| 1700016K19I | 0.09126349 | 0.102561 | 0.09758714 | 0.00497387 | 0.40615032 | 0.89054584 |
| Gm43672 | 0.04358634 | 0.05160658 | 0.04663719 | 0.00496939 | 0.30979543 | 0.89054584 |

| | | | | | | |
|---|---|---|---|---|---|---|
| Malt1 | 0.0355668 | 0.04302741 | 0.03805859 | 0.00496882 | 0.27150833 | 0.89054584 |
| D3Ertd254e | 0.18325223 | 0.19817362 | 0.19320606 | 0.00496757 | 0.45232587 | 0.89054584 |
| Gfra2 | 0.03925553 | 0.04697134 | 0.04200467 | 0.00496667 | 0.29085902 | 0.89054584 |
| Fcho2 | 0.37589892 | 0.38613939 | 0.381182 | 0.00495738 | 0.47584253 | 0.89054584 |
| Mrvi1 | 0.03130184 | 0.0384347 | 0.0334957 | 0.004939 | 0.24605107 | 0.89054584 |
| Pum3 | 0.45852926 | 0.46081259 | 0.45588524 | 0.00492735 | 0.47991963 | 0.89054584 |
| Wdr31 | 0.05835382 | 0.06734789 | 0.06242914 | 0.00491875 | 0.3567815 | 0.89054584 |
| Gli1 | 0.0381751 | 0.04576693 | 0.0408489 | 0.00491804 | 0.28742695 | 0.89054584 |
| 4833403J16F | 0.01839992 | 0.02460666 | 0.01969062 | 0.00491604 | 0.12238048 | 0.70076598 |
| Gm11696 | 0.01520073 | 0.02117754 | 0.01626717 | 0.00491038 | 0.07981171 | 0.54094194 |
| Osbpl7 | 0.05977149 | 0.06883073 | 0.06394475 | 0.00488598 | 0.3609251 | 0.89054584 |
| Ar | 0.01356403 | 0.01939869 | 0.0145157 | 0.004883 | 0.05852914 | 0.43830347 |
| Wdr38 | 0.01573668 | 0.02171881 | 0.01684069 | 0.00487813 | 0.08858449 | 0.57669418 |
| Wfikkn2 | 0.129238 | 0.14239764 | 0.13752046 | 0.00487718 | 0.43438155 | 0.89054584 |
| Gm15261 | 0.05502222 | 0.06374356 | 0.05886708 | 0.00487648 | 0.34977015 | 0.89054584 |
| Pcdhga10 | 0.01582803 | 0.02180754 | 0.01693845 | 0.00486909 | 0.09024458 | 0.58279103 |
| Nfkb1 | 0.35058471 | 0.36239271 | 0.35753558 | 0.00485713 | 0.47476706 | 0.89054584 |
| Enpp3 | 0.01995364 | 0.02620699 | 0.02135323 | 0.00485376 | 0.14479674 | 0.76727455 |
| A830010M2( | 0.06188307 | 0.07105575 | 0.06620205 | 0.0048537 | 0.36633456 | 0.89054584 |
| Nefh | 0.10204254 | 0.11390316 | 0.10905897 | 0.00484419 | 0.41802784 | 0.89054584 |
| Nin | 0.207088 | 0.2222431 | 0.21740002 | 0.00484308 | 0.45866858 | 0.89054584 |
| Tmem169 | 0.01843125 | 0.02456431 | 0.01972415 | 0.00484016 | 0.12646309 | 0.712024 |
| Ccdc68 | 0.02245519 | 0.02886346 | 0.02403002 | 0.00483343 | 0.17435196 | 0.84146294 |
| Rab33a | 0.06855114 | 0.07816053 | 0.07332901 | 0.00483152 | 0.37943334 | 0.89054584 |
| Angptl4 | 0.04246531 | 0.05025967 | 0.04543812 | 0.00482155 | 0.3105219 | 0.89054584 |
| Zfp768 | 0.20519722 | 0.22031094 | 0.21548952 | 0.00482142 | 0.45848926 | 0.89054584 |
| Rasgrp1 | 0.0264021 | 0.03306818 | 0.02825327 | 0.00481491 | 0.21360513 | 0.89054584 |
| Zbtb49 | 0.07224373 | 0.08208965 | 0.07727484 | 0.00481481 | 0.38579805 | 0.89054584 |
| Zfp423 | 0.05924652 | 0.06819615 | 0.06338352 | 0.00481262 | 0.36176423 | 0.89054584 |
| Cpne5 | 0.06598036 | 0.0753897 | 0.07058155 | 0.00480815 | 0.37547925 | 0.89054584 |

| | | | | | | |
|---|---|---|---|---|---|---|
| Dennd4a | 0.08137517 | 0.0918173 | 0.08702945 | 0.00478785 | 0.39885673 | 0.89054584 |
| Sgcd | 0.03149881 | 0.03849179 | 0.03370644 | 0.00478535 | 0.2541663 | 0.89054584 |
| Lat2 | 0.01094818 | 0.01650015 | 0.01171637 | 0.00478378 | 0.02856987 | 0.25985317 |
| Cyp27a1 | 0.01115331 | 0.01670047 | 0.01193589 | 0.00476458 | 0.03145963 | 0.27913202 |
| Mgmt | 0.18940973 | 0.20423828 | 0.19947932 | 0.00475896 | 0.45574948 | 0.89054584 |
| Wdr41 | 0.18351378 | 0.19822444 | 0.19347284 | 0.0047516 | 0.45445193 | 0.89054584 |
| Aoc2 | 0.01561125 | 0.02145452 | 0.01670646 | 0.00474805 | 0.0927362 | 0.59385576 |
| Map7d1 | 0.41195938 | 0.41899787 | 0.41426051 | 0.00473735 | 0.47875513 | 0.89054584 |
| Gpr85 | 0.0709868 | 0.08065355 | 0.07593178 | 0.00472177 | 0.38601769 | 0.89054584 |
| Polm | 0.06165454 | 0.07067271 | 0.06595776 | 0.00471495 | 0.36955138 | 0.89054584 |
| Tex15 | 0.01979443 | 0.02589463 | 0.02118286 | 0.00471176 | 0.15003087 | 0.77909236 |
| Amdhd1 | 0.00907659 | 0.01442487 | 0.00971349 | 0.00471139 | 0.0119183 | 0.13520465 |
| Slfn5 | 0.01808484 | 0.02406358 | 0.01935346 | 0.00471012 | 0.12842625 | 0.71695178 |
| Bloc1s2 | 0.71604037 | 0.666094 | 0.66139269 | 0.00470131 | 0.48679077 | 0.89054584 |
| Spry1 | 0.09920006 | 0.11074868 | 0.10605632 | 0.00469236 | 0.41834455 | 0.89054584 |
| Taok1 | 0.54731504 | 0.53631303 | 0.53162425 | 0.00468878 | 0.48361185 | 0.89054584 |
| Plekhg2 | 0.12240583 | 0.13508058 | 0.13039604 | 0.00468455 | 0.43353754 | 0.89054584 |
| Fgf13 | 0.01438423 | 0.02006882 | 0.01539341 | 0.0046754 | 0.07852703 | 0.5355056 |
| Akr1b8 | 0.07486405 | 0.08474657 | 0.08007443 | 0.00467214 | 0.39287362 | 0.89054584 |
| Gk | 0.1001586 | 0.11174833 | 0.10707725 | 0.00467108 | 0.419474 | 0.89054584 |
| Bcor | 0.25947615 | 0.27436525 | 0.26969947 | 0.00466578 | 0.4678803 | 0.89054584 |
| Pcp4l1 | 0.03610457 | 0.0432979 | 0.0386339 | 0.00466401 | 0.28690743 | 0.89054584 |
| Txk | 0.01748433 | 0.02337407 | 0.01871086 | 0.00466321 | 0.12279568 | 0.70117835 |
| Egr4 | 0.02230126 | 0.02852452 | 0.02386531 | 0.00465921 | 0.18152676 | 0.86081561 |
| Prss12 | 0.02572249 | 0.03217325 | 0.02752609 | 0.00464716 | 0.21577065 | 0.89054584 |
| Lncpint | 0.04201737 | 0.04960009 | 0.04495897 | 0.00464112 | 0.31527852 | 0.89054584 |
| Unc5c | 0.12051869 | 0.13306595 | 0.12842514 | 0.00464081 | 0.43315126 | 0.89054584 |
| Gm26901 | 0.01800656 | 0.02389984 | 0.0192697 | 0.00463014 | 0.13147319 | 0.72640942 |
| Fam83f | 0.04627682 | 0.05412824 | 0.04951485 | 0.00461339 | 0.33211452 | 0.89054584 |
| Gcnt4 | 0.02290315 | 0.0291036 | 0.02450935 | 0.00459424 | 0.1912466 | 0.88386776 |

| | | | | | | |
|---|---|---|---|---|---|---|
| Fam174a | 0.74892655 | 0.68898178 | 0.68439456 | 0.00458723 | 0.48754423 | 0.89054584 |
| Kif26a | 0.04853881 | 0.05651662 | 0.05193403 | 0.00458258 | 0.34050026 | 0.89054584 |
| 1110051M2C | 0.3510555 | 0.36255613 | 0.35797847 | 0.00457766 | 0.47624655 | 0.89054584 |
| Clic6 | 0.08071486 | 0.09089576 | 0.08632424 | 0.00457152 | 0.40256001 | 0.89054584 |
| Zfp839 | 0.05495791 | 0.06336372 | 0.05879832 | 0.0045654 | 0.35877182 | 0.89054584 |
| Rem2 | 0.29551467 | 0.30949531 | 0.30493708 | 0.00455823 | 0.47223917 | 0.89054584 |
| Nmnat3 | 0.09798267 | 0.10931049 | 0.10475752 | 0.00455298 | 0.41976796 | 0.89054584 |
| Zfp385a | 0.08436815 | 0.09476832 | 0.09022567 | 0.00454265 | 0.4072726 | 0.89054584 |
| Vav1 | 0.01879901 | 0.02465654 | 0.02011769 | 0.00453886 | 0.14659973 | 0.77203158 |
| Trim12a | 0.05182133 | 0.0599792 | 0.05544439 | 0.00453482 | 0.35158071 | 0.89054584 |
| Pigw | 0.06764622 | 0.07689469 | 0.07236193 | 0.00453275 | 0.38520591 | 0.89054584 |
| Akr1c14 | 0.02694569 | 0.03336066 | 0.02883489 | 0.00452577 | 0.23231565 | 0.89054584 |
| Hist1h4d | 0.04432466 | 0.05194775 | 0.0474269 | 0.00452085 | 0.32848464 | 0.89054584 |
| Wdr75 | 0.25144537 | 0.26628153 | 0.26176354 | 0.00451799 | 0.46795461 | 0.89054584 |
| Noxa1 | 0.0139684 | 0.01946351 | 0.01494842 | 0.00451509 | 0.07968286 | 0.54059058 |
| Itpr2 | 0.11706034 | 0.12932157 | 0.12480982 | 0.00451175 | 0.43312793 | 0.89054584 |
| Pygm | 0.05175951 | 0.05988274 | 0.05537827 | 0.00450447 | 0.35235944 | 0.89054584 |
| Nanp | 0.01575713 | 0.02136388 | 0.01686258 | 0.00450131 | 0.10681214 | 0.64658138 |
| Sft2d3 | 0.36687031 | 0.37728841 | 0.37278741 | 0.004501 | 0.47757066 | 0.89054584 |
| Gm10371 | 0.01290646 | 0.01831127 | 0.013812 | 0.00449926 | 0.06455119 | 0.46985422 |
| Pcdh8 | 0.02086485 | 0.02682455 | 0.02232828 | 0.00449627 | 0.17407703 | 0.84125625 |
| Scoc | 0.6644205 | 0.62824405 | 0.62375121 | 0.00449284 | 0.48661477 | 0.89054584 |
| Gm26767 | 0.02003075 | 0.02592481 | 0.02143574 | 0.00448907 | 0.16461301 | 0.81899774 |
| Dyrk3 | 0.09488971 | 0.10594306 | 0.10145725 | 0.00448581 | 0.41839952 | 0.89054584 |
| Cdk4 | 2.3019817 | 1.13359011 | 1.12911024 | 0.00447987 | 0.49262603 | 0.89054584 |
| Stard9 | 0.03595149 | 0.04294111 | 0.03847014 | 0.00447098 | 0.29409553 | 0.89054584 |
| Chrnb4 | 0.01854938 | 0.02431817 | 0.01985056 | 0.00446761 | 0.14719223 | 0.77404405 |
| Psen2 | 0.35117298 | 0.36254768 | 0.35808897 | 0.00445871 | 0.4768702 | 0.89054584 |
| Cep72 | 0.04606991 | 0.05374605 | 0.04929355 | 0.0044525 | 0.33694159 | 0.89054584 |
| Gal | 0.01363385 | 0.01903924 | 0.01459041 | 0.00444883 | 0.07771929 | 0.53174815 |

| | | | | | | |
|---|---|---|---|---|---|---|
| Arid4a | 0.57748843 | 0.56067904 | 0.55623173 | 0.00444732 | 0.48514272 | 0.89054584 |
| Dclk3 | 0.01511925 | 0.02062586 | 0.01617997 | 0.00444589 | 0.10024233 | 0.62272587 |
| Etl4 | 0.29350539 | 0.30743076 | 0.302989 | 0.00444176 | 0.47277371 | 0.89054584 |
| Rab37 | 0.21348823 | 0.22825498 | 0.22385535 | 0.00439963 | 0.46352133 | 0.89054584 |
| Glis3 | 0.19792306 | 0.21251362 | 0.20812609 | 0.00438753 | 0.46088052 | 0.89054584 |
| Chrdl1 | 0.01717267 | 0.02275781 | 0.01837735 | 0.00438046 | 0.13338885 | 0.73086605 |
| Fmo5 | 0.04150562 | 0.04878632 | 0.04441159 | 0.00437473 | 0.3231437 | 0.89054584 |
| Golga7 | 0.93682045 | 0.80522158 | 0.80085499 | 0.00436659 | 0.48986697 | 0.89054584 |
| Mfsd4 | 0.03942703 | 0.04655426 | 0.04218813 | 0.00436613 | 0.31484522 | 0.89054584 |
| Snx20 | 0.03573951 | 0.0426077 | 0.03824336 | 0.00436434 | 0.29747385 | 0.89054584 |
| Dscc1 | 0.07379577 | 0.08329095 | 0.07893311 | 0.00435784 | 0.39850563 | 0.89054584 |
| Pigt | 0.833469 | 0.7443171 | 0.73996057 | 0.00435653 | 0.48905856 | 0.89054584 |
| Pcdh1 | 0.21224437 | 0.22695542 | 0.22260219 | 0.00435323 | 0.46370235 | 0.89054584 |
| Churc1 | 1.15272557 | 0.9140307 | 0.90968399 | 0.00434671 | 0.49111961 | 0.89054584 |
| St8sia1 | 0.02475663 | 0.03083023 | 0.02649262 | 0.00433761 | 0.22279024 | 0.89054584 |
| Ecm2 | 0.01522959 | 0.020634 | 0.01629804 | 0.00433596 | 0.10758547 | 0.64845225 |
| Cpt2 | 0.57247085 | 0.55651542 | 0.55218044 | 0.00433498 | 0.48541163 | 0.89054584 |
| 1110008L16F | 0.11025246 | 0.12200229 | 0.11767992 | 0.00432238 | 0.43206367 | 0.89054584 |
| Evc2 | 0.0635629 | 0.07231572 | 0.06799766 | 0.00431806 | 0.38366972 | 0.89054584 |
| Myo9a | 0.19039991 | 0.20480322 | 0.20048661 | 0.00431661 | 0.46004907 | 0.89054584 |
| Ptger4 | 0.02377249 | 0.029756 | 0.02543957 | 0.00431643 | 0.21461801 | 0.89054584 |
| Clstn3 | 0.09112984 | 0.10175926 | 0.09744448 | 0.00431478 | 0.41828032 | 0.89054584 |
| Timm23 | 0.04493843 | 0.0523958 | 0.04808338 | 0.00431243 | 0.33803067 | 0.89054584 |
| Herpud1 | 0.87354142 | 0.76877779 | 0.76448532 | 0.00429247 | 0.48956515 | 0.89054584 |
| Myo1f | 0.011304 | 0.01638083 | 0.01209715 | 0.00428368 | 0.04949581 | 0.39063159 |
| Atp6v1a | 0.64988945 | 0.61709883 | 0.61282109 | 0.00427773 | 0.48702818 | 0.89054584 |
| Ube2e2 | 0.26594147 | 0.28034323 | 0.27606713 | 0.0042761 | 0.4712357 | 0.89054584 |
| Gm19590 | 0.11076501 | 0.12249197 | 0.11821731 | 0.00427466 | 0.43310902 | 0.89054584 |
| Zfp354c | 0.14606801 | 0.1592578 | 0.15499479 | 0.00426301 | 0.44901892 | 0.89054584 |
| Scube3 | 0.02064594 | 0.02634634 | 0.02209403 | 0.00425231 | 0.18494516 | 0.86930035 |

| | | | | | | |
|---|---|---|---|---|---|---|
| Kif9 | 0.08089524 | 0.09076699 | 0.08651688 | 0.00425011 | 0.40948553 | 0.89054584 |
| Rnf186 | 0.0874843 | 0.0978026 | 0.09355284 | 0.00424976 | 0.41619461 | 0.89054584 |
| Gm15879 | 0.01373826 | 0.01894837 | 0.01470214 | 0.00424623 | 0.08921872 | 0.57874406 |
| A830082K12 | 0.01851648 | 0.02406019 | 0.01981535 | 0.00424483 | 0.15913096 | 0.8062012 |
| Cxcr2 | 0.01318375 | 0.01835186 | 0.01410874 | 0.00424312 | 0.08058487 | 0.54416367 |
| Notum | 0.01125919 | 0.01629086 | 0.01204919 | 0.00424167 | 0.05049392 | 0.39613683 |
| Gpr19 | 0.08716574 | 0.09744922 | 0.09321274 | 0.00423649 | 0.41615214 | 0.89054584 |
| Nop14 | 0.38252169 | 0.39154639 | 0.38731097 | 0.00423542 | 0.47968361 | 0.89054584 |
| Btbd19 | 0.01886772 | 0.02442079 | 0.02019121 | 0.00422958 | 0.16454785 | 0.81899774 |
| Gm17546 | 0.01362584 | 0.01880758 | 0.01458184 | 0.00422575 | 0.08848578 | 0.57631032 |
| Osmr | 0.0242707 | 0.0301969 | 0.02597267 | 0.00422422 | 0.2243049 | 0.89054584 |
| Bckdk | 0.55927278 | 0.54565026 | 0.54144741 | 0.00420284 | 0.48557587 | 0.89054584 |
| 9130604C24I | 0.05075327 | 0.0584997 | 0.05430223 | 0.00419747 | 0.35937573 | 0.89054584 |
| Itga8 | 0.02139606 | 0.02709053 | 0.02289671 | 0.00419382 | 0.19673408 | 0.89054584 |
| Rims3 | 0.09732158 | 0.10824503 | 0.10405217 | 0.00419286 | 0.42554204 | 0.89054584 |
| B4galt4 | 0.1510273 | 0.16431512 | 0.1601228 | 0.00419233 | 0.45145739 | 0.89054584 |
| Crispld1 | 0.02901605 | 0.03524148 | 0.03105009 | 0.00419139 | 0.26470697 | 0.89054584 |
| Tead3 | 0.1042286 | 0.1155445 | 0.11135687 | 0.00418763 | 0.43046061 | 0.89054584 |
| Cdkl3 | 0.09369806 | 0.10437077 | 0.10018555 | 0.00418522 | 0.42284296 | 0.89054584 |
| Gm11232 | 0.02344403 | 0.02927263 | 0.02508811 | 0.00418452 | 0.2185565 | 0.89054584 |
| Wtip | 0.04103033 | 0.04808583 | 0.04390318 | 0.00418266 | 0.32857284 | 0.89054584 |
| Slc7a7 | 0.06082632 | 0.06925032 | 0.06507239 | 0.00417793 | 0.38242257 | 0.89054584 |
| Itga11 | 0.02935677 | 0.03559006 | 0.03141465 | 0.00417542 | 0.26788113 | 0.89054584 |
| Asb1 | 0.13691769 | 0.14968137 | 0.14550779 | 0.00417358 | 0.44684687 | 0.89054584 |
| Amer2 | 0.06741667 | 0.0762749 | 0.07211661 | 0.00415829 | 0.39410434 | 0.89054584 |
| Tmem136 | 0.02605598 | 0.0320389 | 0.02788292 | 0.00415598 | 0.24370941 | 0.89054584 |
| Nmnat2 | 0.01121525 | 0.01614987 | 0.01200218 | 0.0041477 | 0.0536956 | 0.41314285 |
| Zpbp | 0.01355437 | 0.01864759 | 0.01450535 | 0.00414224 | 0.0916872 | 0.58922032 |
| Mdga1 | 0.03607767 | 0.04274526 | 0.03860512 | 0.00414014 | 0.30866409 | 0.89054584 |
| Fam19a2 | 0.02069921 | 0.026274 | 0.02215104 | 0.00412296 | 0.1929255 | 0.88710241 |

| | | | | | | |
|---|---|---|---|---|---|---|
| Grb14 | 0.2593434 | 0.27368762 | 0.26956852 | 0.00411909 | 0.47162322 | 0.89054584 |
| Kynu | 0.01450096 | 0.01963038 | 0.01551832 | 0.00411205 | 0.10850298 | 0.65191723 |
| Parp3 | 0.03809956 | 0.04487842 | 0.04076808 | 0.00411034 | 0.31927552 | 0.89054584 |
| Ms4a4d | 0.02018988 | 0.02571562 | 0.02160602 | 0.0041096 | 0.187766 | 0.87574983 |
| Igf2bp1 | 0.42342322 | 0.42872573 | 0.42462378 | 0.00410195 | 0.48205113 | 0.89054584 |
| Dock6 | 0.17461452 | 0.1884757 | 0.18437967 | 0.00409603 | 0.45878338 | 0.89054584 |
| Bcorl1 | 0.11941121 | 0.13135147 | 0.12726788 | 0.00408359 | 0.44058369 | 0.89054584 |
| Armc3 | 0.01192555 | 0.01684259 | 0.0127623 | 0.00408029 | 0.06817396 | 0.48857443 |
| Fgfr4 | 0.07159039 | 0.08065663 | 0.07657674 | 0.00407988 | 0.40198178 | 0.89054584 |
| Ncf1 | 0.01884978 | 0.02425025 | 0.02017201 | 0.00407824 | 0.17311888 | 0.83802257 |
| Pygo1 | 0.09882235 | 0.10972053 | 0.10565336 | 0.00406717 | 0.42883221 | 0.89054584 |
| Calhm2 | 0.10358898 | 0.11474441 | 0.1106847 | 0.00405971 | 0.43215862 | 0.89054584 |
| Dok5 | 0.01337568 | 0.01836815 | 0.01431414 | 0.00405402 | 0.09350199 | 0.59691407 |
| Ttc32 | 0.26485767 | 0.27904617 | 0.27500109 | 0.00404507 | 0.47268197 | 0.89054584 |
| Pex12 | 0.15277515 | 0.16596391 | 0.16192782 | 0.00403609 | 0.45377659 | 0.89054584 |
| Npr2 | 0.06564839 | 0.07426001 | 0.07022674 | 0.00403327 | 0.3945142 | 0.89054584 |
| Fam101a | 0.0663396 | 0.074998 | 0.07096549 | 0.00403251 | 0.39560586 | 0.89054584 |
| Ttc28 | 0.41700563 | 0.42286265 | 0.41883123 | 0.00403142 | 0.48211576 | 0.89054584 |
| Epb41l5 | 0.26752813 | 0.28165689 | 0.27762675 | 0.00403013 | 0.47303973 | 0.89054584 |
| Ccnc | 0.29961361 | 0.31292507 | 0.3089049 | 0.00402018 | 0.47582583 | 0.89054584 |
| Zswim3 | 0.07184258 | 0.08085986 | 0.0768462 | 0.00401366 | 0.40387279 | 0.89054584 |
| Slc38a11 | 0.02092338 | 0.02639743 | 0.02239091 | 0.00400652 | 0.20223966 | 0.89054584 |
| Zfp869 | 0.02416818 | 0.02986506 | 0.02586297 | 0.00400209 | 0.23547493 | 0.89054584 |
| Ccm2l | 0.01428318 | 0.01927931 | 0.01528527 | 0.00399404 | 0.11172949 | 0.66441926 |
| Mir142hg | 0.01543463 | 0.02050905 | 0.01651746 | 0.00399159 | 0.13010953 | 0.72245421 |
| 3110045C21I | 0.02343096 | 0.02905019 | 0.02507413 | 0.00397606 | 0.23002057 | 0.89054584 |
| Lrp4 | 0.04283678 | 0.04980854 | 0.04583545 | 0.00397309 | 0.34316349 | 0.89054584 |
| Tmem53 | 0.21556805 | 0.22991999 | 0.22594918 | 0.00397081 | 0.46737277 | 0.89054584 |
| Lrrc28 | 0.08281034 | 0.0925277 | 0.08856214 | 0.00396556 | 0.41737465 | 0.89054584 |
| Gga2 | 0.586347 | 0.56730187 | 0.56334379 | 0.00395808 | 0.48694337 | 0.89054584 |

| | | | | | | |
|---|---|---|---|---|---|---|
| Vit | 0.00941898 | 0.01403267 | 0.01007989 | 0.00395278 | 0.03385138 | 0.29600565 |
| Abhd15 | 0.13661286 | 0.14914079 | 0.14519118 | 0.00394961 | 0.44957452 | 0.89054584 |
| Aff2 | 0.02745456 | 0.03331558 | 0.02937937 | 0.00393621 | 0.2662484 | 0.89054584 |
| Tagap1 | 0.16016576 | 0.1734787 | 0.16954737 | 0.00393133 | 0.45698676 | 0.89054584 |
| Gm17477 | 0.01467212 | 0.01963109 | 0.01570149 | 0.0039296 | 0.12180907 | 0.69967824 |
| Lsm14b | 0.26932004 | 0.28330812 | 0.27938668 | 0.00392144 | 0.47393081 | 0.89054584 |
| Rpl7a-ps3 | 0.0200574 | 0.02536393 | 0.02146426 | 0.00389967 | 0.19865215 | 0.89054584 |
| Ntn1 | 0.01916245 | 0.02440556 | 0.0205066 | 0.00389896 | 0.18785915 | 0.87574983 |
| Nmt1 | 1.00543472 | 0.84122909 | 0.83733086 | 0.00389823 | 0.49134766 | 0.89054584 |
| Synpr | 0.14776396 | 0.16064422 | 0.15674954 | 0.00389468 | 0.45392196 | 0.89054584 |
| Slc11a1 | 0.01470947 | 0.01963031 | 0.01574145 | 0.00388885 | 0.12487229 | 0.70660334 |
| Mtbp | 0.07370637 | 0.08271928 | 0.07883759 | 0.00388169 | 0.40928293 | 0.89054584 |
| Gm30074 | 0.03711116 | 0.04358711 | 0.03971074 | 0.00387637 | 0.32463238 | 0.89054584 |
| Irf2 | 0.53317456 | 0.52376198 | 0.51989156 | 0.00387042 | 0.48616576 | 0.89054584 |
| Pgk1 | 0.32374146 | 0.33594379 | 0.33209076 | 0.00385303 | 0.47844581 | 0.89054584 |
| Gstm3 | 0.01310349 | 0.01787192 | 0.01402285 | 0.00384907 | 0.10048149 | 0.62288099 |
| Wipi2 | 0.65226638 | 0.61846429 | 0.61461866 | 0.00384563 | 0.48837219 | 0.89054584 |
| Tbc1d5 | 0.0735464 | 0.08249629 | 0.07866668 | 0.00382961 | 0.41028796 | 0.89054584 |
| Dzip1l | 0.10267206 | 0.11354691 | 0.10972088 | 0.00382604 | 0.4354717 | 0.89054584 |
| Hist1h1e | 0.17052272 | 0.18401348 | 0.18018795 | 0.00382553 | 0.46060367 | 0.89054584 |
| Usp38 | 0.1246787 | 0.13658987 | 0.13276801 | 0.00382186 | 0.44665691 | 0.89054584 |
| Hhat | 0.05713579 | 0.06494362 | 0.0611269 | 0.00381672 | 0.3855636 | 0.89054584 |
| Ntng1 | 0.00997995 | 0.01449556 | 0.01068022 | 0.00381534 | 0.04802288 | 0.38170821 |
| Adamts19 | 0.01343695 | 0.01818503 | 0.0143797 | 0.00380534 | 0.10880499 | 0.65274944 |
| Galk2 | 0.52209785 | 0.51441282 | 0.51061095 | 0.00380186 | 0.48616382 | 0.89054584 |
| Eml2 | 0.46057818 | 0.46148524 | 0.45768728 | 0.00379795 | 0.48458051 | 0.89054584 |
| Tln2 | 0.18358476 | 0.19733647 | 0.19354524 | 0.00379123 | 0.4636426 | 0.89054584 |
| Wipf3 | 0.03995519 | 0.04653976 | 0.04275311 | 0.00378665 | 0.33993385 | 0.89054584 |
| Ipcef1 | 0.02140956 | 0.02669724 | 0.02291115 | 0.00378609 | 0.22068101 | 0.89054584 |
| Zscan29 | 0.13723426 | 0.14961692 | 0.14583654 | 0.00378038 | 0.45193693 | 0.89054584 |

| | | | | | | |
|---|---|---|---|---|---|---|
| Tsix | 0.16177764 | 0.17498053 | 0.17120625 | 0.00377428 | 0.4590976 | 0.89054584 |
| Irgm1 | 0.06622274 | 0.07461487 | 0.0708406 | 0.00377427 | 0.40198186 | 0.89054584 |
| Prpf38a | 0.68546192 | 0.6430888 | 0.63931914 | 0.00376966 | 0.4890421 | 0.89054584 |
| Flt3l | 0.03512982 | 0.04135629 | 0.03759111 | 0.00376519 | 0.32037613 | 0.89054584 |
| Spock1 | 0.13831214 | 0.1507075 | 0.14695565 | 0.00375185 | 0.45265943 | 0.89054584 |
| 2810433D01 | 0.0108113 | 0.01531671 | 0.01156989 | 0.00374682 | 0.06567964 | 0.4744511 |
| Nutf2 | 0.34052295 | 0.35177828 | 0.34804163 | 0.00373666 | 0.48005343 | 0.89054584 |
| Uchl4 | 0.04813676 | 0.05523552 | 0.05150406 | 0.00373146 | 0.36785515 | 0.89054584 |
| Rab38 | 0.01418317 | 0.01889717 | 0.01517825 | 0.00371892 | 0.12682762 | 0.71241644 |
| Ephb6 | 0.03669137 | 0.04297594 | 0.03926166 | 0.00371429 | 0.32969633 | 0.89054584 |
| Slc35f3 | 0.01604701 | 0.020881 | 0.01717278 | 0.00370822 | 0.15719989 | 0.80068898 |
| Fstl5 | 0.01330533 | 0.01793553 | 0.01423885 | 0.00369668 | 0.11322538 | 0.67029056 |
| Pnliprp2 | 0.01374163 | 0.01839856 | 0.01470575 | 0.00369281 | 0.12101587 | 0.69664572 |
| AW112010 | 0.01467125 | 0.01938692 | 0.01570056 | 0.00368636 | 0.13700357 | 0.7420427 |
| Lrrc49 | 0.14188589 | 0.1543445 | 0.15066286 | 0.00368165 | 0.45467936 | 0.89054584 |
| Cldn34c1 | 0.05668617 | 0.06432053 | 0.06064619 | 0.00367435 | 0.38886837 | 0.89054584 |
| Nell2 | 0.01914288 | 0.02415642 | 0.02048565 | 0.00367076 | 0.20190912 | 0.89054584 |
| Hip1r | 0.30068599 | 0.3136112 | 0.30994157 | 0.00366963 | 0.47800525 | 0.89054584 |
| Zc3hav1 | 0.10109668 | 0.11173257 | 0.10806418 | 0.00366839 | 0.4371676 | 0.89054584 |
| Cacna2d3 | 0.02338428 | 0.02868907 | 0.02502419 | 0.00366488 | 0.2475186 | 0.89054584 |
| Casp1 | 0.00647729 | 0.01057274 | 0.00693182 | 0.00364092 | 0.00720057 | 0.08918391 |
| Hvcn1 | 0.03324812 | 0.03921531 | 0.03557798 | 0.00363733 | 0.31692737 | 0.89054584 |
| Irs1 | 0.0259262 | 0.03135258 | 0.02774406 | 0.00360852 | 0.27226982 | 0.89054584 |
| Zfp808 | 0.02488608 | 0.030236 | 0.02663114 | 0.00360487 | 0.26413551 | 0.89054584 |
| Cacnb2 | 0.155737 | 0.1685842 | 0.16498405 | 0.00360015 | 0.45951184 | 0.89054584 |
| Ak5 | 0.05684007 | 0.0644095 | 0.06081073 | 0.00359877 | 0.39138336 | 0.89054584 |
| Evc | 0.050539 | 0.05766375 | 0.05407309 | 0.00359065 | 0.3785197 | 0.89054584 |
| Fhdc1 | 0.05162607 | 0.05882556 | 0.05523559 | 0.00358997 | 0.38102026 | 0.89054584 |
| Zfp652 | 0.24861161 | 0.26254036 | 0.25895612 | 0.00358424 | 0.47429204 | 0.89054584 |
| Serp2 | 0.10781586 | 0.11870698 | 0.1151239 | 0.00358308 | 0.44235413 | 0.89054584 |

| | | | | | | |
|---|---|---|---|---|---|---|
| Efr3b | 0.03964044 | 0.04598712 | 0.04241642 | 0.0035707 | 0.34745566 | 0.89054584 |
| Prss41 | 0.02536892 | 0.03070531 | 0.02714777 | 0.00355753 | 0.27075764 | 0.89054584 |
| Eme2 | 0.06070774 | 0.06849882 | 0.06494563 | 0.00355318 | 0.39940356 | 0.89054584 |
| Zfp446 | 0.10283841 | 0.1134411 | 0.10989576 | 0.00354535 | 0.44026314 | 0.89054584 |
| Wdr62 | 0.01610921 | 0.02078256 | 0.01723934 | 0.00354323 | 0.16914263 | 0.8322688 |
| 2510039O18 | 0.28692825 | 0.30013308 | 0.2965981 | 0.00353498 | 0.47785922 | 0.89054584 |
| Cyfip2 | 0.27508552 | 0.28857296 | 0.2850387 | 0.00353426 | 0.47696703 | 0.89054584 |
| 5930430L01I | 0.15765831 | 0.17049351 | 0.1669647 | 0.00352881 | 0.46078072 | 0.89054584 |
| Cdkal1 | 0.2727239 | 0.28625347 | 0.28272551 | 0.00352797 | 0.47682007 | 0.89054584 |
| Cd69 | 0.01477366 | 0.01933579 | 0.01581015 | 0.00352564 | 0.14941678 | 0.77812345 |
| 6030419C18I | 0.01907286 | 0.02393011 | 0.02041073 | 0.00351938 | 0.21089153 | 0.89054584 |
| Utp15 | 0.11178439 | 0.12279763 | 0.11928581 | 0.00351182 | 0.44545171 | 0.89054584 |
| Wwc1 | 0.16737355 | 0.18046196 | 0.17695718 | 0.00350478 | 0.46324014 | 0.89054584 |
| Tmod2 | 0.20168847 | 0.21544274 | 0.21194056 | 0.00350218 | 0.46931735 | 0.89054584 |
| Spred2 | 0.10112494 | 0.11156352 | 0.1080939 | 0.00346962 | 0.44056231 | 0.89054584 |
| Matn4 | 0.02802103 | 0.03344916 | 0.02998547 | 0.00346369 | 0.29523172 | 0.89054584 |
| Ankrd28 | 0.04693189 | 0.05367753 | 0.05021546 | 0.00346207 | 0.37402648 | 0.89054584 |
| Mto1 | 0.24686896 | 0.26068979 | 0.2572278 | 0.003462 | 0.47500106 | 0.89054584 |
| Haus3 | 0.16141619 | 0.17426743 | 0.17083436 | 0.00343307 | 0.46270333 | 0.89054584 |
| Ptk7 | 0.09053593 | 0.10024291 | 0.09681053 | 0.00343237 | 0.43440087 | 0.89054584 |
| Nxph1 | 0.05647593 | 0.06385212 | 0.0604214 | 0.00343072 | 0.39568412 | 0.89054584 |
| Nxnl2 | 0.03199659 | 0.03766696 | 0.03423901 | 0.00342795 | 0.32044836 | 0.89054584 |
| Cacng7 | 0.0242422 | 0.02936447 | 0.02594217 | 0.0034223 | 0.26940665 | 0.89054584 |
| Isoc2a | 0.02082848 | 0.02570685 | 0.02228936 | 0.00341749 | 0.23751248 | 0.89054584 |
| Ppil6 | 0.15205844 | 0.1646034 | 0.16118779 | 0.00341561 | 0.4606786 | 0.89054584 |
| Mcm10 | 0.02985285 | 0.03535832 | 0.03194541 | 0.0034129 | 0.30933354 | 0.89054584 |
| Gm26753 | 0.01390653 | 0.01829006 | 0.01488221 | 0.00340785 | 0.14302157 | 0.76440591 |
| 4833422C13I | 0.02609543 | 0.03132973 | 0.02792513 | 0.0034046 | 0.28501332 | 0.89054584 |
| Rimbp2 | 0.54593098 | 0.53388305 | 0.5304815 | 0.00340155 | 0.48808374 | 0.89054584 |
| Pkhd1 | 0.08901473 | 0.09858297 | 0.09518669 | 0.00339629 | 0.43398709 | 0.89054584 |

| | | | | | | |
|---|---|---|---|---|---|---|
| Wdhd1 | 0.09134246 | 0.10105479 | 0.09767142 | 0.00338337 | 0.43589428 | 0.89054584 |
| Inpp5d | 0.02355923 | 0.02859472 | 0.02521138 | 0.00338334 | 0.26591185 | 0.89054584 |
| Phf6 | 0.17220653 | 0.1852862 | 0.18191372 | 0.00337249 | 0.46558543 | 0.89054584 |
| D130017N08 | 0.03749473 | 0.04349318 | 0.04012106 | 0.00337212 | 0.34768514 | 0.89054584 |
| Snx25 | 0.03905625 | 0.04515523 | 0.0417915 | 0.00336373 | 0.35383377 | 0.89054584 |
| Serpina10 | 0.04417034 | 0.0506176 | 0.04726184 | 0.00335576 | 0.37039703 | 0.89054584 |
| Xpo6 | 0.3203617 | 0.33221553 | 0.32886065 | 0.00335488 | 0.48104609 | 0.89054584 |
| Podn | 0.01910252 | 0.0237933 | 0.02044246 | 0.00335084 | 0.22253103 | 0.89054584 |
| L3mbtl3 | 0.11978204 | 0.13100008 | 0.12765543 | 0.00334465 | 0.45142299 | 0.89054584 |
| Plekha4 | 0.03970124 | 0.0458219 | 0.04248146 | 0.00334044 | 0.35705347 | 0.89054584 |
| Prrg1 | 0.03828547 | 0.04429413 | 0.04096697 | 0.00332716 | 0.35257442 | 0.89054584 |
| Stx1b | 0.02530522 | 0.03038593 | 0.02707962 | 0.00330631 | 0.28473321 | 0.89054584 |
| Pkd1 | 0.16724009 | 0.1801231 | 0.17682017 | 0.00330293 | 0.46532491 | 0.89054584 |
| Cox20 | 1.18199011 | 0.92597527 | 0.92267519 | 0.00330008 | 0.49335259 | 0.89054584 |
| Wfdc1 | 0.03581984 | 0.04161863 | 0.0383293 | 0.00328933 | 0.34464483 | 0.89054584 |
| Gm15594 | 0.01609624 | 0.02051151 | 0.01722546 | 0.00328605 | 0.1870628 | 0.87419127 |
| Nalcn | 0.0233562 | 0.0282788 | 0.02499413 | 0.00328467 | 0.27018053 | 0.89054584 |
| Map2k3os | 0.01395636 | 0.01821161 | 0.01493554 | 0.00327607 | 0.15340802 | 0.78925105 |
| Cebpa | 0.02530459 | 0.03035448 | 0.02707894 | 0.00327554 | 0.28652777 | 0.89054584 |
| Cep55 | 0.04841558 | 0.05507681 | 0.05180224 | 0.00327457 | 0.38418599 | 0.89054584 |
| Timm10 | 0.4405277 | 0.44321732 | 0.43994435 | 0.00327297 | 0.48617533 | 0.89054584 |
| Phtf2 | 0.06745296 | 0.07542729 | 0.0721554 | 0.0032719 | 0.41634203 | 0.89054584 |
| Rmdn2 | 0.03440185 | 0.04007879 | 0.0368123 | 0.00326649 | 0.33965496 | 0.89054584 |
| Pcdhgb7 | 0.01220657 | 0.01631924 | 0.01306303 | 0.00325622 | 0.1227533 | 0.70117835 |
| Crtc2 | 0.13109071 | 0.14270239 | 0.13944946 | 0.00325293 | 0.45672857 | 0.89054584 |
| Aldh1l2 | 0.00894264 | 0.01281542 | 0.00957014 | 0.00324528 | 0.0570682 | 0.43153558 |
| Gm12027 | 0.01327259 | 0.01744679 | 0.01420381 | 0.00324298 | 0.14372794 | 0.76580953 |
| Tmem216 | 0.29015629 | 0.3029651 | 0.29973742 | 0.00322768 | 0.4799938 | 0.89054584 |
| Slc38a6 | 0.05741608 | 0.06464403 | 0.06142658 | 0.00321746 | 0.40360366 | 0.89054584 |
| Cc2d2b | 0.01245448 | 0.01653922 | 0.01332832 | 0.0032109 | 0.13085125 | 0.72435336 |

| | | | | | | |
|---|---|---|---|---|---|---|
| Psd4 | 0.0131826 | 0.01731493 | 0.01410751 | 0.00320743 | 0.14474575 | 0.76727455 |
| Gm6297 | 0.03211713 | 0.03757446 | 0.03436797 | 0.00320648 | 0.33190009 | 0.89054584 |
| Iffo1 | 0.17669363 | 0.18971084 | 0.18650688 | 0.00320396 | 0.46810474 | 0.89054584 |
| Cd5 | 0.01488744 | 0.01913339 | 0.01593191 | 0.00320149 | 0.17458367 | 0.84202063 |
| Tgif1 | 0.16019139 | 0.17277512 | 0.16957375 | 0.00320137 | 0.46495585 | 0.89054584 |
| Rnf24 | 0.03701112 | 0.04278897 | 0.03960372 | 0.00318526 | 0.35393799 | 0.89054584 |
| Cacnb4 | 0.03639109 | 0.04212555 | 0.03894042 | 0.00318514 | 0.35157245 | 0.89054584 |
| Rimklb | 0.02571323 | 0.03070096 | 0.02751619 | 0.00318478 | 0.29486228 | 0.89054584 |
| Fxr1 | 1.01049142 | 0.84310294 | 0.83993757 | 0.00316537 | 0.49299589 | 0.89054584 |
| Dnah1 | 0.03394544 | 0.03948785 | 0.03632402 | 0.00316383 | 0.34244658 | 0.89054584 |
| Rasa4 | 0.03001761 | 0.03528546 | 0.0321217 | 0.00316376 | 0.32316264 | 0.89054584 |
| Hlf | 0.0198793 | 0.02443139 | 0.02127368 | 0.0031577 | 0.24461373 | 0.89054584 |
| Sema6b | 0.00625072 | 0.00984601 | 0.00668934 | 0.00315666 | 0.01395577 | 0.15281779 |
| Proser3 | 0.10658616 | 0.11698871 | 0.1138331 | 0.00315561 | 0.44861865 | 0.89054584 |
| Otub2 | 0.0818803 | 0.09071462 | 0.08756892 | 0.0031457 | 0.43354293 | 0.89054584 |
| 2610028E06I | 0.01330976 | 0.01738353 | 0.01424359 | 0.00313995 | 0.15219943 | 0.78526239 |
| Degs2 | 0.05422293 | 0.06115049 | 0.05801243 | 0.00313806 | 0.40051452 | 0.89054584 |
| Slc19a2 | 0.018328 | 0.02274938 | 0.01961366 | 0.00313572 | 0.22818249 | 0.89054584 |
| Shox2 | 0.03861157 | 0.04444949 | 0.0413158 | 0.00313369 | 0.36190647 | 0.89054584 |
| Prodh | 0.06674767 | 0.07453409 | 0.07140162 | 0.00313246 | 0.41902333 | 0.89054584 |
| Phactr3 | 0.01640872 | 0.02069168 | 0.01755985 | 0.00313184 | 0.20300604 | 0.89054584 |
| Ankzf1 | 0.15473301 | 0.16707815 | 0.16394846 | 0.00312969 | 0.46456608 | 0.89054584 |
| 9030624J02F | 0.44347232 | 0.44569171 | 0.44256493 | 0.00312679 | 0.48687075 | 0.89054584 |
| Gm13470 | 0.02840517 | 0.03352149 | 0.03039648 | 0.003125 | 0.31597757 | 0.89054584 |
| Isg15 | 0.01419841 | 0.01831858 | 0.01519455 | 0.00312403 | 0.16906016 | 0.8322688 |
| Ftl1 | 5.27780555 | 0.67126411 | 0.66814205 | 0.00312205 | 0.4913157 | 0.89054584 |
| Crtac1 | 0.01065174 | 0.01451586 | 0.01139914 | 0.00311672 | 0.10136121 | 0.62533394 |
| Pdzd11 | 0.75422391 | 0.6911415 | 0.68802747 | 0.00311403 | 0.49158833 | 0.89054584 |
| Cry1 | 0.35169892 | 0.36169604 | 0.35858357 | 0.00311247 | 0.48387152 | 0.89054584 |
| Picalm | 0.50898686 | 0.50263757 | 0.49952787 | 0.0031097 | 0.48843101 | 0.89054584 |

| | | | | | | |
|---|---|---|---|---|---|---|
| Pappa | 0.0261046 | 0.03103428 | 0.02793494 | 0.00309934 | 0.30261231 | 0.89054584 |
| Ears2 | 0.18080991 | 0.19380878 | 0.19071339 | 0.00309539 | 0.46986178 | 0.89054584 |
| Pmaip1 | 0.01018293 | 0.01398494 | 0.01089744 | 0.00308751 | 0.09341925 | 0.59664859 |
| Rab1a | 1.72437838 | 1.08893644 | 1.08585462 | 0.00308182 | 0.49472503 | 0.89054584 |
| Cd2 | 0.01295679 | 0.01692857 | 0.01386587 | 0.0030627 | 0.15172343 | 0.78420187 |
| Mtss1l | 0.09047894 | 0.09981128 | 0.0967497 | 0.00306158 | 0.44139659 | 0.89054584 |
| Smurf1 | 0.10490289 | 0.115119 | 0.11206531 | 0.00305369 | 0.44948902 | 0.89054584 |
| Sav1 | 0.13403715 | 0.14555929 | 0.14251452 | 0.00304477 | 0.46035611 | 0.89054584 |
| Prtn3 | 0.01782559 | 0.02212039 | 0.01907603 | 0.00304435 | 0.22858261 | 0.89054584 |
| Cstf1 | 0.15283326 | 0.16502724 | 0.16198782 | 0.00303942 | 0.46517006 | 0.89054584 |
| Slc41a1 | 0.13360263 | 0.14509421 | 0.14206271 | 0.0030315 | 0.46040316 | 0.89054584 |
| Erc1 | 0.31906633 | 0.330651 | 0.32762107 | 0.00302993 | 0.48281603 | 0.89054584 |
| Slc2a3 | 0.03293656 | 0.03827134 | 0.03524466 | 0.00302668 | 0.34454411 | 0.89054584 |
| Iffo2 | 0.09089414 | 0.10021776 | 0.0971929 | 0.00302486 | 0.44235679 | 0.89054584 |
| Suv39h1 | 0.13948414 | 0.15119385 | 0.14817197 | 0.00302188 | 0.4621509 | 0.89054584 |
| Sema6a | 0.29148269 | 0.30404661 | 0.30102587 | 0.00302074 | 0.48135559 | 0.89054584 |
| Gm13889 | 0.01069193 | 0.01446113 | 0.01144215 | 0.00301899 | 0.10948878 | 0.65489146 |
| Asap1 | 0.53283404 | 0.52262625 | 0.51960742 | 0.00301883 | 0.48920288 | 0.89054584 |
| Ctnna3 | 0.02233221 | 0.02690742 | 0.02389843 | 0.00300899 | 0.27873764 | 0.89054584 |
| Pcdhga5 | 0.01830647 | 0.02259944 | 0.01959063 | 0.00300881 | 0.23713768 | 0.89054584 |
| Acap1 | 0.01913025 | 0.02347487 | 0.02047213 | 0.00300273 | 0.2471939 | 0.89054584 |
| Clec4a3 | 0.01006922 | 0.01377731 | 0.01077575 | 0.00300156 | 0.09718896 | 0.61237252 |
| Ndnf | 0.01867987 | 0.02298883 | 0.01999019 | 0.00299864 | 0.24231768 | 0.89054584 |
| Kif7 | 0.04195697 | 0.04789084 | 0.04489437 | 0.00299647 | 0.37791522 | 0.89054584 |
| Ptgs1 | 0.0665294 | 0.0741613 | 0.07116835 | 0.00299296 | 0.42233252 | 0.89054584 |
| Gli3 | 0.05451444 | 0.06131523 | 0.05832413 | 0.0029911 | 0.40558019 | 0.89054584 |
| Fblim1 | 0.13363736 | 0.14508054 | 0.14209883 | 0.00298171 | 0.46106132 | 0.89054584 |
| Rarres1 | 0.01823295 | 0.02248697 | 0.01951195 | 0.00297502 | 0.23874157 | 0.89054584 |
| Zfpm2 | 0.0186672 | 0.02294077 | 0.01997664 | 0.00296414 | 0.24469035 | 0.89054584 |
| Sqle | 0.54106041 | 0.52940909 | 0.52645056 | 0.00295853 | 0.48955602 | 0.89054584 |

| | | | | | | |
|---|---|---|---|---|---|---|
| 6720489N17 | 0.03855756 | 0.04421313 | 0.04125803 | 0.0029551 | 0.36930441 | 0.89054584 |
| Pthlh | 0.01066988 | 0.01437241 | 0.01141855 | 0.00295386 | 0.11405896 | 0.67412427 |
| Unkl | 0.04619345 | 0.05237022 | 0.04942568 | 0.00294454 | 0.39067612 | 0.89054584 |
| Kcna6 | 0.02934704 | 0.03434609 | 0.03140423 | 0.00294186 | 0.33126004 | 0.89054584 |
| Golga7b | 0.0298157 | 0.03483897 | 0.03190567 | 0.0029333 | 0.33420649 | 0.89054584 |
| Zfp61 | 0.17683212 | 0.18958136 | 0.18664852 | 0.00293284 | 0.4708208 | 0.89054584 |
| Hoxb3os | 0.03347048 | 0.03874682 | 0.03581588 | 0.00293094 | 0.35150611 | 0.89054584 |
| Lztr1 | 0.33548753 | 0.34619096 | 0.34327031 | 0.00292066 | 0.48419015 | 0.89054584 |
| Sox8 | 0.0234507 | 0.02800526 | 0.02509526 | 0.00291 | 0.29451492 | 0.89054584 |
| Rasl10a | 0.01739118 | 0.02152109 | 0.01861118 | 0.00290991 | 0.23317278 | 0.89054584 |
| Dcun1d2 | 0.18254789 | 0.19539491 | 0.19248745 | 0.00290746 | 0.47194887 | 0.89054584 |
| Cbr3 | 0.02213036 | 0.0265889 | 0.02368245 | 0.00290646 | 0.2837374 | 0.89054584 |
| Nudt22 | 0.15944959 | 0.17170638 | 0.16880997 | 0.0028964 | 0.46814367 | 0.89054584 |
| Ccdc151 | 0.02349909 | 0.02803831 | 0.02514703 | 0.00289128 | 0.29609598 | 0.89054584 |
| Slc39a13 | 0.16574689 | 0.17817103 | 0.17528678 | 0.00288425 | 0.46944682 | 0.89054584 |
| Gm15638 | 0.02521487 | 0.02986594 | 0.02698294 | 0.002883 | 0.30931744 | 0.89054584 |
| Eaf2 | 0.02715345 | 0.03193474 | 0.0290572 | 0.00287755 | 0.32226394 | 0.89054584 |
| Parpbp | 0.01990648 | 0.02417236 | 0.02130276 | 0.0028696 | 0.26513797 | 0.89054584 |
| Lcat | 0.01038458 | 0.01397891 | 0.01111323 | 0.00286568 | 0.11480463 | 0.6757765 |
| Zfp142 | 0.10184993 | 0.11172059 | 0.10885642 | 0.00286417 | 0.45121853 | 0.89054584 |
| Matk | 0.01425652 | 0.01811912 | 0.01525675 | 0.00286237 | 0.19103745 | 0.88366966 |
| Gata6 | 0.51358054 | 0.50628558 | 0.50342333 | 0.00286225 | 0.48943377 | 0.89054584 |
| Atoh8 | 0.03036777 | 0.03534831 | 0.03249633 | 0.00285198 | 0.34131281 | 0.89054584 |
| Sapcd2 | 0.02944903 | 0.03436133 | 0.03151335 | 0.00284797 | 0.33686113 | 0.89054584 |
| Ints5 | 0.17416099 | 0.18675966 | 0.1839154 | 0.00284426 | 0.47128072 | 0.89054584 |
| Apoo | 0.64017091 | 0.60827278 | 0.60543072 | 0.00284206 | 0.49127569 | 0.89054584 |
| Rasal2 | 0.1129707 | 0.12336304 | 0.1205288 | 0.00283424 | 0.45638101 | 0.89054584 |
| Eci2 | 1.0452511 | 0.86065392 | 0.85782015 | 0.00283377 | 0.49386028 | 0.89054584 |
| Gm4631 | 0.12873316 | 0.13982649 | 0.1369946 | 0.00283189 | 0.46163814 | 0.89054584 |
| Fndc3c1 | 0.01306742 | 0.0168071 | 0.01398426 | 0.00282285 | 0.17349383 | 0.83899716 |

| | | | | | | |
|---|---|---|---|---|---|---|
| Gm26609 | 0.0334994 | 0.03866873 | 0.03584682 | 0.00282191 | 0.35689992 | 0.89054584 |
| Ikzf1 | 0.02289992 | 0.0273223 | 0.0245059 | 0.0028164 | 0.29617293 | 0.89054584 |
| Mtr | 0.02223984 | 0.02661437 | 0.02379959 | 0.00281478 | 0.29081195 | 0.89054584 |
| Mar-05 | 1.01745463 | 0.8463084 | 0.8434973 | 0.0028111 | 0.49380599 | 0.89054584 |
| F11r | 0.60905803 | 0.58414698 | 0.58133903 | 0.00280795 | 0.49102321 | 0.89054584 |
| Ddit3 | 0.32299089 | 0.33418094 | 0.33137394 | 0.002807 | 0.48425986 | 0.89054584 |
| Lin52 | 0.09127947 | 0.10040652 | 0.0976042 | 0.00280232 | 0.44679495 | 0.89054584 |
| Serinc5 | 0.07193189 | 0.07972915 | 0.07694163 | 0.00278753 | 0.43298123 | 0.89054584 |
| Gsto2 | 0.03298993 | 0.03808818 | 0.03530176 | 0.00278643 | 0.35653424 | 0.89054584 |
| Micu3 | 0.17756453 | 0.19018372 | 0.1873974 | 0.00278632 | 0.47238672 | 0.89054584 |
| Gm10561 | 0.01470644 | 0.01851292 | 0.01573821 | 0.0027747 | 0.2057125 | 0.89054584 |
| Prpf18 | 0.36438493 | 0.37324226 | 0.37046867 | 0.00277359 | 0.48608769 | 0.89054584 |
| Fbxo27 | 0.05401648 | 0.06055593 | 0.05779169 | 0.00276424 | 0.41182923 | 0.89054584 |
| Gstt3 | 0.1054349 | 0.11538771 | 0.11262415 | 0.00276356 | 0.4544918 | 0.89054584 |
| Alox12 | 0.01862239 | 0.02269111 | 0.01992869 | 0.00276242 | 0.25920439 | 0.89054584 |
| Klhl3 | 0.03318407 | 0.03827023 | 0.03550946 | 0.00276077 | 0.35859371 | 0.89054584 |
| Ephb1 | 0.01439583 | 0.01815661 | 0.01540582 | 0.00275078 | 0.20273956 | 0.89054584 |
| Nrde2 | 0.11739333 | 0.12790297 | 0.12515812 | 0.00274485 | 0.45930866 | 0.89054584 |
| Tulp1 | 0.017017 | 0.02095499 | 0.01821077 | 0.00274422 | 0.24131878 | 0.89054584 |
| Dmrt2 | 0.01172129 | 0.01528511 | 0.01254371 | 0.0027414 | 0.15429152 | 0.79182956 |
| Ifih1 | 0.04013689 | 0.04568522 | 0.04294748 | 0.00273774 | 0.38323694 | 0.89054584 |
| Tldc1 | 0.02000726 | 0.02414218 | 0.02141061 | 0.00273157 | 0.27612597 | 0.89054584 |
| Vamp3 | 0.89446192 | 0.7795545 | 0.77682488 | 0.00272962 | 0.49346934 | 0.89054584 |
| Barx2 | 0.02116014 | 0.0253712 | 0.02264426 | 0.00272695 | 0.28737991 | 0.89054584 |
| Dpysl4 | 0.1064544 | 0.11642142 | 0.11369477 | 0.00272666 | 0.45551801 | 0.89054584 |
| Zcchc4 | 0.18433085 | 0.19702859 | 0.19430611 | 0.00272249 | 0.47397634 | 0.89054584 |
| Zscan2 | 0.16151803 | 0.17365301 | 0.17093914 | 0.00271387 | 0.47051856 | 0.89054584 |
| Pctp | 0.04214362 | 0.04780596 | 0.04509402 | 0.00271194 | 0.38966694 | 0.89054584 |
| Rad51ap2 | 0.01811388 | 0.02209586 | 0.01938454 | 0.00271132 | 0.25731437 | 0.89054584 |
| Fas | 0.0240933 | 0.02848163 | 0.02578285 | 0.00269878 | 0.31289255 | 0.89054584 |

| | | | | | | |
|---|---|---|---|---|---|---|
| Emp2 | 0.21039629 | 0.22343607 | 0.22073905 | 0.00269702 | 0.47730303 | 0.89054584 |
| Spata6 | 0.14061001 | 0.1520269 | 0.14933991 | 0.002687 | 0.46659738 | 0.89054584 |
| Mill2 | 0.02224022 | 0.02647968 | 0.0238 | 0.00267968 | 0.29994563 | 0.89054584 |
| Tbl3 | 0.24210711 | 0.25517511 | 0.25249788 | 0.00267723 | 0.4803008 | 0.89054584 |
| Ampd3 | 0.01390569 | 0.017558 | 0.01488131 | 0.00267669 | 0.20101544 | 0.89054584 |
| Glmp | 0.84002794 | 0.74672041 | 0.7440546 | 0.00266582 | 0.4933411 | 0.89054584 |
| Vsnl1 | 0.01353687 | 0.01715031 | 0.01448662 | 0.00266368 | 0.19581969 | 0.89054584 |
| Micall2 | 0.01907244 | 0.02307344 | 0.02041028 | 0.00266316 | 0.27162501 | 0.89054584 |
| Dcbld1 | 0.2277244 | 0.24079378 | 0.23814873 | 0.00264505 | 0.47936572 | 0.89054584 |
| Tmem189 | 0.12082674 | 0.13139061 | 0.12874696 | 0.00264365 | 0.46189317 | 0.89054584 |
| Ephb3 | 0.07906887 | 0.08720058 | 0.08456619 | 0.00263439 | 0.44230236 | 0.89054584 |
| Tor1b | 0.43070363 | 0.43379966 | 0.43116597 | 0.00263369 | 0.48864834 | 0.89054584 |
| Zscan18 | 0.03691665 | 0.04212819 | 0.03950266 | 0.00262554 | 0.37841138 | 0.89054584 |
| Foxk1 | 0.12298749 | 0.13362476 | 0.13100325 | 0.00262151 | 0.46286035 | 0.89054584 |
| Sec24a | 0.37506694 | 0.38302895 | 0.38041034 | 0.00261861 | 0.48720792 | 0.89054584 |
| Mkln1os | 0.01284649 | 0.01636434 | 0.01374782 | 0.00261652 | 0.1876202 | 0.87548043 |
| Gm14420 | 0.0083217 | 0.01151934 | 0.00890564 | 0.0026137 | 0.08575837 | 0.56639069 |
| Ncs1 | 0.18990966 | 0.20259035 | 0.19998794 | 0.00260241 | 0.47582856 | 0.89054584 |
| Lyst | 0.09769762 | 0.10705309 | 0.10445339 | 0.0025997 | 0.45384432 | 0.89054584 |
| Dffb | 0.08225187 | 0.09056392 | 0.08796574 | 0.00259819 | 0.44527507 | 0.89054584 |
| Naaladl2 | 0.07012685 | 0.07760743 | 0.07501286 | 0.00259456 | 0.43598978 | 0.89054584 |
| Slc25a20 | 0.24750757 | 0.26045155 | 0.25786133 | 0.00259022 | 0.48133666 | 0.89054584 |
| 8030423F21I | 0.02649679 | 0.03094419 | 0.02835459 | 0.0025896 | 0.33523703 | 0.89054584 |
| Gbp9 | 0.01360137 | 0.01714409 | 0.01455566 | 0.00258844 | 0.20369334 | 0.89054584 |
| Bcl2l12 | 0.16292002 | 0.17496481 | 0.17238131 | 0.0025835 | 0.47216684 | 0.89054584 |
| Slc5a3 | 0.10079691 | 0.11033021 | 0.10774883 | 0.00258138 | 0.45556394 | 0.89054584 |
| Dnaic1 | 0.04595247 | 0.05174378 | 0.04916795 | 0.00257583 | 0.40358654 | 0.89054584 |
| Snhg11 | 0.02705452 | 0.03152092 | 0.02895134 | 0.00256958 | 0.33961865 | 0.89054584 |
| Phtf1os | 0.03807867 | 0.04330344 | 0.04074574 | 0.0025577 | 0.38497055 | 0.89054584 |
| Ppp1r15b | 0.28902779 | 0.30119376 | 0.29864052 | 0.00255324 | 0.4841136 | 0.89054584 |

| | | | | | | |
|---|---|---|---|---|---|---|
| Ccdc3 | 0.02493792 | 0.02923788 | 0.02668661 | 0.00255128 | 0.32801487 | 0.89054584 |
| Maml2 | 0.05992835 | 0.06665426 | 0.06411244 | 0.00254182 | 0.42672826 | 0.89054584 |
| Parvg | 0.01468294 | 0.01825323 | 0.01571306 | 0.00254017 | 0.22567578 | 0.89054584 |
| Wdr3 | 0.17867648 | 0.19107174 | 0.18853393 | 0.00253781 | 0.4749976 | 0.89054584 |
| Ifit1 | 0.01842822 | 0.02225805 | 0.0197209 | 0.00253715 | 0.27445799 | 0.89054584 |
| Mfsd7c | 0.00924596 | 0.01242937 | 0.00989474 | 0.00253463 | 0.11635015 | 0.68087821 |
| Mrps16 | 1.11353968 | 0.89417098 | 0.89163757 | 0.0025334 | 0.49471919 | 0.89054584 |
| Gm21992 | 0.02891341 | 0.03347069 | 0.03094027 | 0.00253042 | 0.35159168 | 0.89054584 |
| Tbck | 0.35527901 | 0.36446481 | 0.36194635 | 0.00251846 | 0.48706959 | 0.89054584 |
| Gpr179 | 0.05569169 | 0.06207307 | 0.05958289 | 0.00249018 | 0.42280847 | 0.89054584 |
| Sorbs1 | 0.04201005 | 0.04743568 | 0.04495114 | 0.00248454 | 0.39839292 | 0.89054584 |
| Pcdh15 | 0.05929349 | 0.06591643 | 0.06343373 | 0.0024827 | 0.42765635 | 0.89054584 |
| Acvr2a | 0.12718375 | 0.1378604 | 0.13538009 | 0.00248031 | 0.46598885 | 0.89054584 |
| A930024E05 | 0.01389045 | 0.01734039 | 0.014865 | 0.00247539 | 0.21892492 | 0.89054584 |
| Mpp5 | 0.31710785 | 0.32821033 | 0.32574528 | 0.00246506 | 0.4859377 | 0.89054584 |
| Pcdhb19 | 0.02006112 | 0.02392518 | 0.02146823 | 0.00245695 | 0.29694891 | 0.89054584 |
| Herc6 | 0.04604778 | 0.05171111 | 0.04926989 | 0.00244122 | 0.40871934 | 0.89054584 |
| Plec | 0.22271281 | 0.23556683 | 0.23312737 | 0.00243945 | 0.48055874 | 0.89054584 |
| Acoxl | 0.01012666 | 0.0132709 | 0.01083722 | 0.00243368 | 0.14772436 | 0.77404405 |
| Cemip | 0.01826972 | 0.02198033 | 0.0195513 | 0.00242903 | 0.28135483 | 0.89054584 |
| Nr1d1 | 0.0136651 | 0.01705266 | 0.01462385 | 0.00242881 | 0.21952992 | 0.89054584 |
| Gfpt2 | 0.05282311 | 0.05894346 | 0.05651562 | 0.00242784 | 0.4206847 | 0.89054584 |
| Mrs2 | 0.09648494 | 0.10557721 | 0.10315949 | 0.00241772 | 0.4565257 | 0.89054584 |
| Sobp | 0.24066024 | 0.25347136 | 0.25105862 | 0.00241274 | 0.48214386 | 0.89054584 |
| 0610038B21 | 0.01455231 | 0.01798411 | 0.01557328 | 0.00241082 | 0.23538459 | 0.89054584 |
| Apobec1 | 0.01382089 | 0.01720128 | 0.01479056 | 0.00241072 | 0.22381783 | 0.89054584 |
| Metrn | 0.15026759 | 0.16174744 | 0.15933788 | 0.00240956 | 0.47191617 | 0.89054584 |
| Edem3 | 0.1612479 | 0.1730683 | 0.17066119 | 0.00240712 | 0.47380327 | 0.89054584 |
| Fignl1 | 0.11464299 | 0.12468709 | 0.12228009 | 0.002407 | 0.46346499 | 0.89054584 |
| Arhgap4 | 0.02983586 | 0.03432748 | 0.03192723 | 0.00240025 | 0.36307417 | 0.89054584 |

| | | | | | | |
|---|---|---|---|---|---|---|
| Ispd | 0.07818576 | 0.08601776 | 0.08362289 | 0.00239487 | 0.44692792 | 0.89054584 |
| Mnx1 | 0.17459767 | 0.1867548 | 0.18436242 | 0.00239238 | 0.47589595 | 0.89054584 |
| Cpq | 0.58588285 | 0.56536435 | 0.56297245 | 0.0023919 | 0.49210369 | 0.89054584 |
| Colec11 | 0.02326424 | 0.02728071 | 0.02489573 | 0.00238498 | 0.32768294 | 0.89054584 |
| Tmem19 | 0.28613574 | 0.29819411 | 0.29582658 | 0.00236753 | 0.48512848 | 0.89054584 |
| Ttll6 | 0.01751709 | 0.02110674 | 0.01874592 | 0.00236082 | 0.27869022 | 0.89054584 |
| Gm16183 | 0.0652321 | 0.07213662 | 0.06978181 | 0.00235481 | 0.43753682 | 0.89054584 |
| 2900089D17I | 0.01320549 | 0.01648644 | 0.01413201 | 0.00235443 | 0.21881757 | 0.89054584 |
| Hist1h2ab | 0.01377728 | 0.01709554 | 0.0147439 | 0.00235164 | 0.22871115 | 0.89054584 |
| Ppp4r4 | 0.02817257 | 0.03249078 | 0.03014762 | 0.00234316 | 0.35863611 | 0.89054584 |
| Plek | 0.01453819 | 0.01789958 | 0.01555816 | 0.00234142 | 0.24160586 | 0.89054584 |
| Zcchc12 | 0.02648789 | 0.03068617 | 0.02834506 | 0.00234111 | 0.35019403 | 0.89054584 |
| Gm8444 | 0.00915747 | 0.01214107 | 0.00980004 | 0.00234103 | 0.13287041 | 0.72977796 |
| Stag3 | 0.00915591 | 0.01213889 | 0.00979837 | 0.00234052 | 0.13288124 | 0.72977796 |
| Syt6 | 0.01740701 | 0.02096354 | 0.01862812 | 0.00233543 | 0.27957766 | 0.89054584 |
| Cyp2j9 | 0.00955457 | 0.01255524 | 0.010225 | 0.00233024 | 0.14417374 | 0.76653633 |
| Mettl5os | 0.01001597 | 0.01304252 | 0.01071877 | 0.00232375 | 0.15624209 | 0.79788579 |
| Stx11 | 0.01551921 | 0.01892819 | 0.01660797 | 0.00232022 | 0.25756274 | 0.89054584 |
| Pif1 | 0.02118116 | 0.02498155 | 0.02266675 | 0.00231479 | 0.31711397 | 0.89054584 |
| B3galt1 | 0.01364963 | 0.01692175 | 0.0146073 | 0.00231445 | 0.23020094 | 0.89054584 |
| Morn5 | 0.01289219 | 0.01609959 | 0.01379673 | 0.00230285 | 0.21839132 | 0.89054584 |
| Gm4262 | 0.01347912 | 0.01672663 | 0.01442483 | 0.0023018 | 0.2286084 | 0.89054584 |
| Slc22a21 | 0.00901608 | 0.01194901 | 0.00964873 | 0.00230028 | 0.13334812 | 0.73086605 |
| Tssk6 | 0.01248708 | 0.01564942 | 0.01336321 | 0.00228621 | 0.21270745 | 0.89054584 |
| Ankrd12 | 0.90224095 | 0.78361748 | 0.7813338 | 0.00228368 | 0.49456772 | 0.89054584 |
| Med4 | 0.34877133 | 0.35810739 | 0.35582854 | 0.00227885 | 0.48809831 | 0.89054584 |
| Nav3 | 0.04288576 | 0.04816242 | 0.04588784 | 0.00227458 | 0.40868241 | 0.89054584 |
| Tmem14a | 0.16938013 | 0.18129033 | 0.17901622 | 0.00227411 | 0.47640266 | 0.89054584 |
| Adarb1 | 0.12007896 | 0.13023054 | 0.1279657 | 0.00226484 | 0.46714141 | 0.89054584 |
| 1700112E06I | 0.0345294 | 0.03921249 | 0.03694876 | 0.00226373 | 0.38765408 | 0.89054584 |

| | | | | | | |
|---|---|---|---|---|---|---|
| Slc11a2 | 0.13223092 | 0.14289697 | 0.14063596 | 0.00226101 | 0.47014635 | 0.89054584 |
| 1110065P20I | 0.79643774 | 0.71851468 | 0.71625827 | 0.00225641 | 0.49414496 | 0.89054584 |
| Adam8 | 0.01332817 | 0.01651756 | 0.01426329 | 0.00225427 | 0.23076458 | 0.89054584 |
| Kbtbd11 | 0.02769181 | 0.03188331 | 0.02963323 | 0.00225009 | 0.36175981 | 0.89054584 |
| Ppara | 0.01997311 | 0.02362194 | 0.02137406 | 0.00224788 | 0.3120762 | 0.89054584 |
| Ntf5 | 0.01022008 | 0.01317067 | 0.0109372 | 0.00223347 | 0.17070181 | 0.83412025 |
| Gm16503 | 0.01389675 | 0.01709928 | 0.01487174 | 0.00222754 | 0.24264127 | 0.89054584 |
| Slc26a7 | 0.02113625 | 0.02484375 | 0.02261869 | 0.00222506 | 0.32336398 | 0.89054584 |
| Phxr4 | 0.07735361 | 0.08495346 | 0.082734 | 0.00221947 | 0.45026839 | 0.89054584 |
| Spink10 | 0.03767155 | 0.04252691 | 0.04031022 | 0.00221669 | 0.39889863 | 0.89054584 |
| Dck | 0.13309058 | 0.14374326 | 0.14153021 | 0.00221306 | 0.47096267 | 0.89054584 |
| Zdhhc21 | 0.25518499 | 0.26767529 | 0.26546261 | 0.00221267 | 0.48451175 | 0.89054584 |
| Ninl | 0.04698404 | 0.05247775 | 0.05027123 | 0.00220652 | 0.41898467 | 0.89054584 |
| Eogt | 0.07698204 | 0.08454254 | 0.08233708 | 0.00220546 | 0.45034361 | 0.89054584 |
| Gm26520 | 0.01270541 | 0.01579126 | 0.01359685 | 0.00219441 | 0.22605223 | 0.89054584 |
| Runx1 | 0.02702308 | 0.0311097 | 0.0289177 | 0.00219199 | 0.36198693 | 0.89054584 |
| Wdr73 | 0.23653811 | 0.24914303 | 0.2469529 | 0.00219013 | 0.48352102 | 0.89054584 |
| Gm11772 | 0.00819255 | 0.01095651 | 0.00876742 | 0.00218909 | 0.122361 | 0.70076598 |
| Zfp964 | 0.02662432 | 0.03067913 | 0.02849104 | 0.00218809 | 0.36024526 | 0.89054584 |
| 5430403G16 | 0.07780315 | 0.08538349 | 0.0832142 | 0.00216929 | 0.45166611 | 0.89054584 |
| Aim1l | 0.03691348 | 0.04165948 | 0.03949926 | 0.00216022 | 0.39943938 | 0.89054584 |
| Gm13620 | 0.01347895 | 0.01658182 | 0.01442464 | 0.00215718 | 0.24298406 | 0.89054584 |
| Tmem82 | 0.02375061 | 0.02756487 | 0.02541616 | 0.00214871 | 0.34683798 | 0.89054584 |
| Mgarp | 0.01417151 | 0.01731444 | 0.01516577 | 0.00214867 | 0.25460366 | 0.89054584 |
| Zfp865 | 0.27377151 | 0.28589889 | 0.28375198 | 0.00214691 | 0.48594007 | 0.89054584 |
| Eno2 | 0.05250603 | 0.05832025 | 0.05617656 | 0.00214369 | 0.42944603 | 0.89054584 |
| Rxra | 0.10424424 | 0.1135139 | 0.1113733 | 0.00214059 | 0.46432449 | 0.89054584 |
| Gm28727 | 0.01379801 | 0.01690507 | 0.01476608 | 0.00213899 | 0.24987376 | 0.89054584 |
| Mrpl3 | 0.52062722 | 0.51151026 | 0.50937288 | 0.00213739 | 0.49220139 | 0.89054584 |
| Pak7 | 0.02543618 | 0.02935121 | 0.02721974 | 0.00213146 | 0.3576221 | 0.89054584 |

| | | | | | | |
|---|---|---|---|---|---|---|
| Tpmt | 0.05125452 | 0.05696859 | 0.05483826 | 0.00213032 | 0.42818861 | 0.89054584 |
| Tnfrsf10b | 0.04165826 | 0.04668705 | 0.04457485 | 0.0021122 | 0.41263724 | 0.89054584 |
| Krt23 | 0.02216088 | 0.02581059 | 0.0237151 | 0.00209549 | 0.34029125 | 0.89054584 |
| Enox2 | 0.07886599 | 0.08644477 | 0.08434948 | 0.00209528 | 0.45393307 | 0.89054584 |
| Npm2 | 0.01318537 | 0.01620398 | 0.01411047 | 0.00209351 | 0.24471189 | 0.89054584 |
| Jph3 | 0.03067378 | 0.03491298 | 0.03282374 | 0.00208924 | 0.38340783 | 0.89054584 |
| Gna12 | 0.13451525 | 0.14509697 | 0.14301155 | 0.00208542 | 0.47291769 | 0.89054584 |
| Fermt1 | 0.08491878 | 0.09289411 | 0.09081362 | 0.00208048 | 0.45750022 | 0.89054584 |
| Samd12 | 0.01545749 | 0.01862103 | 0.01654192 | 0.00207911 | 0.27908256 | 0.89054584 |
| D130040H23 | 0.01337299 | 0.0163894 | 0.01431125 | 0.00207815 | 0.24935268 | 0.89054584 |
| Arhgef19 | 0.06149579 | 0.06785817 | 0.06578805 | 0.00207012 | 0.44172373 | 0.89054584 |
| Itga10 | 0.01289741 | 0.01586587 | 0.01380232 | 0.00206355 | 0.24304358 | 0.89054584 |
| Zfp958 | 0.09527573 | 0.1039254 | 0.10186918 | 0.00205621 | 0.46253867 | 0.89054584 |
| 4933415A04 | 0.01415918 | 0.01720289 | 0.01515257 | 0.00205032 | 0.2642137 | 0.89054584 |
| A930038B10 | 0.02016145 | 0.02362344 | 0.0215756 | 0.00204784 | 0.32917141 | 0.89054584 |
| Slc10a4 | 0.00567773 | 0.00812362 | 0.00607615 | 0.00204747 | 0.05821757 | 0.4366288 |
| Syngap1 | 0.01884554 | 0.02221404 | 0.02016748 | 0.00204656 | 0.31818474 | 0.89054584 |
| Kirrel | 0.03166996 | 0.03593482 | 0.03388955 | 0.00204527 | 0.38928915 | 0.89054584 |
| Gm12002 | 0.0101139 | 0.01286444 | 0.01082357 | 0.00204087 | 0.18984179 | 0.88102633 |
| Mypopos | 0.02008834 | 0.02351646 | 0.02149737 | 0.00201909 | 0.33084526 | 0.89054584 |
| Kntc1 | 0.02053914 | 0.0239968 | 0.02197975 | 0.00201705 | 0.3344912 | 0.89054584 |
| Tbc1d9 | 0.07214513 | 0.07918178 | 0.07716948 | 0.0020123 | 0.45165218 | 0.89054584 |
| Crbn | 0.38554801 | 0.3921157 | 0.39010348 | 0.00201222 | 0.49041369 | 0.89054584 |
| Lmbrd1 | 0.3808179 | 0.38774265 | 0.38573655 | 0.0020061 | 0.49033467 | 0.89054584 |
| Rrp1b | 0.23169188 | 0.24411903 | 0.24211596 | 0.00200307 | 0.48462686 | 0.89054584 |
| Trib2 | 0.0937872 | 0.1022753 | 0.10028068 | 0.00199461 | 0.46308373 | 0.89054584 |
| Ankrd33b | 0.05628339 | 0.06220887 | 0.06021554 | 0.00199333 | 0.43871558 | 0.89054584 |
| Rabif | 0.33262049 | 0.34253622 | 0.34054824 | 0.00198798 | 0.48915133 | 0.89054584 |
| Zfp420 | 0.04618217 | 0.05139847 | 0.04941361 | 0.00198486 | 0.42577489 | 0.89054584 |
| Cbx8 | 0.2104573 | 0.22278211 | 0.22080057 | 0.00198154 | 0.48332472 | 0.89054584 |

| | | | | | | |
|---|---|---|---|---|---|---|
| RP23-54G8.4 | 0.09374227 | 0.10221287 | 0.10023273 | 0.00198014 | 0.46333339 | 0.89054584 |
| C6 | 0.01086821 | 0.01360809 | 0.01163079 | 0.0019773 | 0.21416696 | 0.89054584 |
| Ssu2 | 0.02864453 | 0.0326168 | 0.03065259 | 0.00196422 | 0.38264378 | 0.89054584 |
| Prkcd | 0.22533823 | 0.23772296 | 0.23575932 | 0.00196364 | 0.48452319 | 0.89054584 |
| Ccdc190 | 0.01242324 | 0.015245 | 0.01329489 | 0.0019501 | 0.24718459 | 0.89054584 |
| Gm20324 | 0.01845073 | 0.02169376 | 0.019745 | 0.00194876 | 0.32282222 | 0.89054584 |
| Fpgs | 0.16217694 | 0.17355345 | 0.17161703 | 0.00193642 | 0.47903784 | 0.89054584 |
| Gm29440 | 0.01567455 | 0.01870836 | 0.01677421 | 0.00193415 | 0.29556452 | 0.89054584 |
| Parvb | 0.04246114 | 0.04736147 | 0.04543365 | 0.00192782 | 0.42164551 | 0.89054584 |
| Hdac8 | 0.17871628 | 0.19050226 | 0.18857461 | 0.00192765 | 0.48100765 | 0.89054584 |
| Synj1 | 0.14705029 | 0.15793763 | 0.15601126 | 0.00192637 | 0.47706278 | 0.89054584 |
| Rbpms2 | 0.04824187 | 0.05354044 | 0.05161647 | 0.00192397 | 0.43106679 | 0.89054584 |
| Slc18a2 | 0.01004724 | 0.01267082 | 0.01075223 | 0.0019186 | 0.2028941 | 0.89054584 |
| Rbms2 | 0.31301215 | 0.32373134 | 0.32181612 | 0.00191522 | 0.48894006 | 0.89054584 |
| Gm27204 | 0.01334916 | 0.01619693 | 0.01428576 | 0.00191118 | 0.26654955 | 0.89054584 |
| Srpk1 | 0.76990879 | 0.70057749 | 0.69866683 | 0.00191066 | 0.49491724 | 0.89054584 |
| Zfp729b | 0.0689304 | 0.07563677 | 0.07373432 | 0.00190246 | 0.45215922 | 0.89054584 |
| Vgf | 0.0256813 | 0.02938014 | 0.02748201 | 0.00189813 | 0.37380822 | 0.89054584 |
| Dpy19l1 | 0.06220464 | 0.0684425 | 0.06654579 | 0.00189671 | 0.4471795 | 0.89054584 |
| Tbx3os1 | 0.00559237 | 0.00788119 | 0.0059848 | 0.00189639 | 0.06993588 | 0.49764662 |
| Selo | 0.21506639 | 0.22732971 | 0.22544431 | 0.0018854 | 0.48445998 | 0.89054584 |
| Cd274 | 0.01292665 | 0.01571788 | 0.01383361 | 0.00188427 | 0.26284715 | 0.89054584 |
| Oit3 | 0.00986101 | 0.01243482 | 0.01055294 | 0.00188188 | 0.20303894 | 0.89054584 |
| Slc45a4 | 0.08448381 | 0.09223043 | 0.09034917 | 0.00188126 | 0.46135959 | 0.89054584 |
| BC026585 | 0.13374277 | 0.14408752 | 0.14220843 | 0.00187909 | 0.47545602 | 0.89054584 |
| Ctbs | 0.20201114 | 0.21414537 | 0.21226715 | 0.00187821 | 0.48355868 | 0.89054584 |
| Hist1h4i | 0.16940678 | 0.18091669 | 0.17904356 | 0.00187313 | 0.48056273 | 0.89054584 |
| Pdk4 | 0.02388876 | 0.02743024 | 0.02556398 | 0.00186626 | 0.36688361 | 0.89054584 |
| Kif16b | 0.16994709 | 0.18145835 | 0.17959771 | 0.00186064 | 0.48075176 | 0.89054584 |
| Gm10031 | 0.05869058 | 0.06464944 | 0.06278917 | 0.00186027 | 0.44510773 | 0.89054584 |

| | | | | | | |
|---|---|---|---|---|---|---|
| Gtf2a1 | 0.18882266 | 0.20073913 | 0.1988819 | 0.00185723 | 0.48264877 | 0.89054584 |
| Tmem40 | 0.01404701 | 0.01687984 | 0.01503254 | 0.00184729 | 0.28348493 | 0.89054584 |
| Fam229b | 0.19074573 | 0.20268001 | 0.20083831 | 0.00184171 | 0.48296121 | 0.89054584 |
| Gm13572 | 0.01241635 | 0.01512641 | 0.01328752 | 0.00183889 | 0.25954098 | 0.89054584 |
| Gm42715 | 0.02714347 | 0.03086831 | 0.02904651 | 0.00182179 | 0.38506365 | 0.89054584 |
| Steap1 | 0.1207618 | 0.13049792 | 0.12867912 | 0.0018188 | 0.47374808 | 0.89054584 |
| Zfp507 | 0.14804784 | 0.15885949 | 0.15704314 | 0.00181635 | 0.47851343 | 0.89054584 |
| Nr2f6 | 0.45532576 | 0.45485958 | 0.45306264 | 0.00179694 | 0.49262862 | 0.89054584 |
| Kcnc3 | 0.18614207 | 0.19794408 | 0.19615222 | 0.00179186 | 0.48302631 | 0.89054584 |
| Nudt15 | 0.01448951 | 0.01729595 | 0.01550608 | 0.00178987 | 0.2953634 | 0.89054584 |
| Drp2 | 0.01241783 | 0.01507519 | 0.0132891 | 0.00178609 | 0.26560081 | 0.89054584 |
| Sh3d21 | 0.07605991 | 0.08313531 | 0.08135199 | 0.00178332 | 0.45932716 | 0.89054584 |
| Gm15406 | 0.00989377 | 0.01236238 | 0.01058799 | 0.00177439 | 0.21746876 | 0.89054584 |
| Gm12524 | 0.01159732 | 0.01418187 | 0.01241105 | 0.00177082 | 0.25310723 | 0.89054584 |
| D630024D03 | 0.02050883 | 0.02371666 | 0.02194731 | 0.00176935 | 0.35360899 | 0.89054584 |
| Slc15a4 | 0.15639165 | 0.16742653 | 0.16565909 | 0.00176744 | 0.48017802 | 0.89054584 |
| Dkkl1 | 0.01386688 | 0.01660634 | 0.01483978 | 0.00176656 | 0.28958096 | 0.89054584 |
| Palb2 | 0.03108413 | 0.03502594 | 0.03326277 | 0.00176317 | 0.40247044 | 0.89054584 |
| Kazn | 0.16769149 | 0.17904628 | 0.17728354 | 0.00176274 | 0.48152593 | 0.89054584 |
| Spag4 | 0.00977291 | 0.01220989 | 0.01045866 | 0.00175123 | 0.21766378 | 0.89054584 |
| Spire2 | 0.05241208 | 0.05782375 | 0.0560761 | 0.00174765 | 0.44227674 | 0.89054584 |
| Lrrk2 | 0.01293447 | 0.0155812 | 0.01384198 | 0.00173922 | 0.27914323 | 0.89054584 |
| Gm43948 | 0.0350162 | 0.03920698 | 0.03746955 | 0.00173743 | 0.41448164 | 0.89054584 |
| Gas2l3 | 0.03639303 | 0.04067675 | 0.03894249 | 0.00173426 | 0.41781708 | 0.89054584 |
| Prrx2 | 0.02922435 | 0.03300546 | 0.03127296 | 0.0017325 | 0.3981643 | 0.89054584 |
| Chaf1b | 0.1280507 | 0.13801006 | 0.13628358 | 0.00172648 | 0.47646774 | 0.89054584 |
| Gm14325 | 0.03070467 | 0.03458166 | 0.03285679 | 0.00172487 | 0.40339133 | 0.89054584 |
| Dnaaf2 | 0.15678176 | 0.1677855 | 0.16606126 | 0.00172423 | 0.48070895 | 0.89054584 |
| Mcm9 | 0.05635082 | 0.06200966 | 0.06028764 | 0.00172202 | 0.44706702 | 0.89054584 |
| Tma16 | 0.24415719 | 0.25625597 | 0.2545355 | 0.00172046 | 0.48743904 | 0.89054584 |

| | | | | | | |
|---|---|---|---|---|---|---|
| 1810010H24[illegible] | 0.03818487 | 0.04257923 | 0.04085935 | 0.00171988 | 0.42226287 | 0.89054584 |
| Prkaa1 | 0.18108588 | 0.19271192 | 0.19099517 | 0.00171675 | 0.48329852 | 0.89054584 |
| Gm11963 | 0.01355583 | 0.01622343 | 0.01450692 | 0.00171652 | 0.29072632 | 0.89054584 |
| Trap1 | 0.37636793 | 0.38333325 | 0.38161685 | 0.0017164 | 0.49164094 | 0.89054584 |
| Adh1 | 0.05435064 | 0.05986437 | 0.05814899 | 0.00171538 | 0.44534248 | 0.89054584 |
| Zkscan6 | 0.10606479 | 0.1149964 | 0.11328567 | 0.00171073 | 0.47195558 | 0.89054584 |
| Grm4 | 0.00852033 | 0.01082853 | 0.0091182 | 0.00171034 | 0.1910874 | 0.88366966 |
| Mapkbp1 | 0.09834451 | 0.10684994 | 0.10514356 | 0.00170638 | 0.46986462 | 0.89054584 |
| Tarsl2 | 0.14054911 | 0.15097931 | 0.14927674 | 0.00170257 | 0.47881125 | 0.89054584 |
| Sag | 0.12270169 | 0.13240033 | 0.13070491 | 0.00169542 | 0.47590543 | 0.89054584 |
| P4htm | 0.21207784 | 0.22412755 | 0.22243437 | 0.00169317 | 0.48585491 | 0.89054584 |
| Med26 | 0.14555174 | 0.15614906 | 0.1544604 | 0.00168865 | 0.47968892 | 0.89054584 |
| Prkaa2 | 0.01195997 | 0.01448665 | 0.01279913 | 0.00168752 | 0.26952015 | 0.89054584 |
| Rin2 | 0.05036408 | 0.05556681 | 0.05388603 | 0.00168078 | 0.44222959 | 0.89054584 |
| Ankrd61 | 0.09420515 | 0.10240413 | 0.10072672 | 0.00167742 | 0.4690787 | 0.89054584 |
| Sptlc1 | 0.29831369 | 0.30932165 | 0.30764747 | 0.00167418 | 0.48988653 | 0.89054584 |
| Gm14418 | 0.03153662 | 0.03540734 | 0.0337469 | 0.00166045 | 0.40934366 | 0.89054584 |
| Dcdc2b | 0.01617103 | 0.0189586 | 0.0173055 | 0.00165311 | 0.32814353 | 0.89054584 |
| Gm15853 | 0.02633897 | 0.02983876 | 0.02818572 | 0.00165304 | 0.3923353 | 0.89054584 |
| Gm10311 | 0.02480093 | 0.02819217 | 0.02654002 | 0.00165215 | 0.38589839 | 0.89054584 |
| Pstk | 0.27701475 | 0.2885716 | 0.28692636 | 0.00164524 | 0.48934377 | 0.89054584 |
| Klb | 0.05218966 | 0.05748089 | 0.05583825 | 0.00164264 | 0.44549335 | 0.89054584 |
| Peg12 | 0.04388329 | 0.0485946 | 0.04695481 | 0.00163979 | 0.43537605 | 0.89054584 |
| Crkl | 0.02875627 | 0.03240869 | 0.03077215 | 0.00163654 | 0.40215446 | 0.89054584 |
| Lingo1 | 0.2455801 | 0.25758346 | 0.25594862 | 0.00163484 | 0.48812989 | 0.89054584 |
| Cerk | 0.10593448 | 0.11478151 | 0.11314883 | 0.00163269 | 0.47320074 | 0.89054584 |
| Nudt12 | 0.05764885 | 0.06330477 | 0.06167544 | 0.00162933 | 0.45102229 | 0.89054584 |
| Zfp335os | 0.03994489 | 0.04437004 | 0.04274209 | 0.00162794 | 0.42957825 | 0.89054584 |
| Eci3 | 0.00983285 | 0.01214632 | 0.0105228 | 0.00162352 | 0.23612913 | 0.89054584 |
| Rab43 | 0.03058475 | 0.03435058 | 0.03272849 | 0.00162209 | 0.40869372 | 0.89054584 |

| | | | | | | |
|---|---|---|---|---|---|---|
| Upk3bl | 0.03605131 | 0.04018617 | 0.03857692 | 0.00160925 | 0.42295098 | 0.89054584 |
| Mtg2 | 0.28503189 | 0.29634846 | 0.29475144 | 0.00159702 | 0.48993058 | 0.89054584 |
| Etv3 | 0.10169453 | 0.11028943 | 0.10869299 | 0.00159643 | 0.47272228 | 0.89054584 |
| Wfikkn1 | 0.00961794 | 0.01188793 | 0.01029282 | 0.00159512 | 0.23514321 | 0.89054584 |
| Gm17259 | 0.058789 | 0.06448708 | 0.0628944 | 0.00159268 | 0.45304222 | 0.89054584 |
| Ift140 | 0.07934473 | 0.08645299 | 0.08486083 | 0.00159215 | 0.4651725 | 0.89054584 |
| E2f6 | 0.11090953 | 0.11995639 | 0.11836881 | 0.00158757 | 0.47508789 | 0.89054584 |
| AI597479 | 0.20316941 | 0.21502111 | 0.21343912 | 0.001582 | 0.48622654 | 0.89054584 |
| Myl6b | 0.08560026 | 0.09311959 | 0.09154128 | 0.0015783 | 0.46798868 | 0.89054584 |
| Tmem35 | 0.02737657 | 0.03087014 | 0.02929593 | 0.00157422 | 0.40115898 | 0.89054584 |
| Dclk2 | 0.02029159 | 0.02328637 | 0.02171485 | 0.00157152 | 0.36799424 | 0.89054584 |
| Agps | 0.1753686 | 0.18671914 | 0.1851514 | 0.00156774 | 0.48426631 | 0.89054584 |
| Rbm3os | 0.02258459 | 0.02573383 | 0.02416849 | 0.00156534 | 0.38142167 | 0.89054584 |
| Hkdc1 | 0.01232146 | 0.01474723 | 0.01318598 | 0.00156125 | 0.29060076 | 0.89054584 |
| 4930590J08F | 0.02132291 | 0.02437691 | 0.02281843 | 0.00155848 | 0.37516583 | 0.89054584 |
| BC030867 | 0.03112205 | 0.03484216 | 0.03330335 | 0.00153881 | 0.41477823 | 0.89054584 |
| 1700084C06I | 0.01313398 | 0.0155902 | 0.01405548 | 0.00153472 | 0.30547747 | 0.89054584 |
| 2210013O21 | 1.22972904 | 0.94446746 | 0.94294315 | 0.00152431 | 0.49699544 | 0.89054584 |
| Gpr4 | 0.020706 | 0.02368073 | 0.0221583 | 0.00152243 | 0.37444558 | 0.89054584 |
| Gm3667 | 0.01373466 | 0.01621277 | 0.01469828 | 0.00151448 | 0.31559632 | 0.89054584 |
| Vsig2 | 0.0322512 | 0.03602505 | 0.03451141 | 0.00151364 | 0.41904519 | 0.89054584 |
| Obscn | 0.01363394 | 0.01610288 | 0.01459051 | 0.00151237 | 0.3145767 | 0.89054584 |
| Psen1 | 0.33095721 | 0.34047797 | 0.33896708 | 0.00151089 | 0.49171598 | 0.89054584 |
| Zcwpw1 | 0.11327482 | 0.12235819 | 0.12084736 | 0.00151083 | 0.47677644 | 0.89054584 |
| 2900026A02I | 0.28830066 | 0.29944292 | 0.29793342 | 0.0015095 | 0.4905839 | 0.89054584 |
| Simc1 | 0.11210105 | 0.12112423 | 0.11961765 | 0.00150658 | 0.47660386 | 0.89054584 |
| Mrpl1 | 0.63981117 | 0.60666236 | 0.6051559 | 0.00150645 | 0.49537325 | 0.89054584 |
| Sytl2 | 0.03610345 | 0.04013739 | 0.0386327 | 0.00150469 | 0.42800335 | 0.89054584 |
| Nppc | 0.01141372 | 0.01371475 | 0.01221457 | 0.00150018 | 0.28359098 | 0.89054584 |
| Gm9930 | 0.02637915 | 0.02972803 | 0.02822871 | 0.00149932 | 0.40227932 | 0.89054584 |

| | | | | | | |
|---|---|---|---|---|---|---|
| Slc1a6 | 0.0097392 | 0.01190532 | 0.01042258 | 0.00148273 | 0.25373334 | 0.89054584 |
| Lypd2 | 0.0132303 | 0.01563724 | 0.01415855 | 0.00147869 | 0.31328133 | 0.89054584 |
| Thumpd2 | 0.08925168 | 0.09691647 | 0.09543963 | 0.00147684 | 0.47126412 | 0.89054584 |
| Gsap | 0.03782067 | 0.04194516 | 0.04046974 | 0.00147541 | 0.43256322 | 0.89054584 |
| Neat1 | 0.13873564 | 0.14886453 | 0.14739522 | 0.00146931 | 0.48147872 | 0.89054584 |
| Ier5l | 0.03343009 | 0.03723555 | 0.03577266 | 0.00146289 | 0.4244497 | 0.89054584 |
| Wnk2 | 0.0851395 | 0.09250929 | 0.09104931 | 0.00145998 | 0.47022419 | 0.89054584 |
| Atp1b2 | 0.05735036 | 0.06281494 | 0.06135631 | 0.00145862 | 0.45590452 | 0.89054584 |
| Gxylt1 | 0.23022063 | 0.24210397 | 0.24064557 | 0.0014584 | 0.48873744 | 0.89054584 |
| Scai | 0.17148569 | 0.1826332 | 0.18117506 | 0.00145814 | 0.48504461 | 0.89054584 |
| Gm26847 | 0.01467988 | 0.01716022 | 0.01570979 | 0.00145043 | 0.33354576 | 0.89054584 |
| Dnal4 | 0.45153936 | 0.45116724 | 0.44971848 | 0.00144877 | 0.4940126 | 0.89054584 |
| Syt11 | 0.20576319 | 0.21750979 | 0.21606156 | 0.00144823 | 0.48754377 | 0.89054584 |
| Fam26f | 0.01663008 | 0.01924268 | 0.01779673 | 0.00144595 | 0.35251799 | 0.89054584 |
| Fam49b | 0.54914422 | 0.53457573 | 0.53313295 | 0.00144278 | 0.49497021 | 0.89054584 |
| Ice2 | 0.08275507 | 0.08994032 | 0.08850311 | 0.00143721 | 0.46984604 | 0.89054584 |
| Epg5 | 0.04768846 | 0.05246052 | 0.05102461 | 0.00143591 | 0.4478443 | 0.89054584 |
| A930029G22 | 0.07489469 | 0.08154307 | 0.08010717 | 0.0014359 | 0.46672283 | 0.89054584 |
| 2810006K23l | 0.25629586 | 0.26799491 | 0.26656016 | 0.00143476 | 0.4899969 | 0.89054584 |
| Usp32 | 0.1646674 | 0.17561018 | 0.17417766 | 0.00143253 | 0.48471727 | 0.89054584 |
| Prkrir | 0.19640288 | 0.20801542 | 0.20658437 | 0.00143105 | 0.48712701 | 0.89054584 |
| Cep120 | 0.12060239 | 0.1299349 | 0.12851259 | 0.00142232 | 0.47943845 | 0.89054584 |
| Unc13d | 0.03002832 | 0.03355428 | 0.03213315 | 0.00142112 | 0.41837732 | 0.89054584 |
| 2610305D13 | 0.03010825 | 0.03363477 | 0.03221867 | 0.0014161 | 0.41887441 | 0.89054584 |
| Zwilch | 0.05250883 | 0.05759462 | 0.05617955 | 0.00141507 | 0.45329097 | 0.89054584 |
| Atxn3 | 0.10534822 | 0.11394606 | 0.1125331 | 0.00141295 | 0.47667629 | 0.89054584 |
| Adamtsl4 | 0.03051027 | 0.03406126 | 0.03264879 | 0.00141247 | 0.42013134 | 0.89054584 |
| 3930402G23 | 0.02374288 | 0.02682004 | 0.02540789 | 0.00141214 | 0.39784126 | 0.89054584 |
| Zbtb18 | 0.10717773 | 0.11586489 | 0.11445414 | 0.00141075 | 0.47710309 | 0.89054584 |
| Mtor | 0.12046692 | 0.12978147 | 0.12837105 | 0.00141043 | 0.47958776 | 0.89054584 |

| | | | | | | |
|---|---|---|---|---|---|---|
| Elovl5 | 0.26962547 | 0.28109469 | 0.27968651 | 0.00140819 | 0.49064279 | 0.89054584 |
| Usp36 | 0.22585858 | 0.23767962 | 0.2362806 | 0.00139902 | 0.48899631 | 0.89054584 |
| Klhl20 | 0.15889365 | 0.16963463 | 0.16823741 | 0.00139721 | 0.48456777 | 0.89054584 |
| Sfn | 0.02945276 | 0.03290399 | 0.03151734 | 0.00138665 | 0.41879492 | 0.89054584 |
| Smoc1 | 0.14462789 | 0.15488362 | 0.15350386 | 0.00137976 | 0.48329852 | 0.89054584 |
| Susd4 | 0.04638012 | 0.05099742 | 0.04962533 | 0.00137209 | 0.44875211 | 0.89054584 |
| Tnfaip3 | 0.03279673 | 0.03646296 | 0.03509506 | 0.00136791 | 0.4279512 | 0.89054584 |
| Gigyf1 | 0.30583919 | 0.31627725 | 0.31491509 | 0.00136216 | 0.49196099 | 0.89054584 |
| Vstm2b | 0.02730925 | 0.03058225 | 0.02922389 | 0.00135836 | 0.41427837 | 0.89054584 |
| Scube1 | 0.01638966 | 0.01888591 | 0.01753945 | 0.00134646 | 0.36030051 | 0.89054584 |
| Rec8 | 0.02239471 | 0.0253019 | 0.02396531 | 0.00133659 | 0.39749425 | 0.89054584 |
| Kdelr1 | 1.56114663 | 1.05286632 | 1.05153632 | 0.00133001 | 0.49764915 | 0.89054584 |
| Gm38399 | 0.01213834 | 0.01431732 | 0.01299001 | 0.00132732 | 0.3170196 | 0.89054584 |
| Itgal | 0.01389503 | 0.01619443 | 0.0148699 | 0.00132453 | 0.33907363 | 0.89054584 |
| Fam192a | 0.63907719 | 0.60591816 | 0.60459492 | 0.00132324 | 0.49593216 | 0.89054584 |
| Gm37986 | 0.00976801 | 0.01177584 | 0.01045342 | 0.00132243 | 0.2778004 | 0.89054584 |
| Topbp1 | 0.22552314 | 0.23726294 | 0.23594458 | 0.00131836 | 0.48961582 | 0.89054584 |
| Ly96 | 0.13844731 | 0.14841427 | 0.14709596 | 0.00131831 | 0.48334722 | 0.89054584 |
| Adam11 | 0.02804495 | 0.03132225 | 0.03001106 | 0.00131119 | 0.41935235 | 0.89054584 |
| Amy2a5 | 0.00960498 | 0.01158819 | 0.01027895 | 0.00130924 | 0.27645236 | 0.89054584 |
| Ap2a2 | 0.85727764 | 0.75598173 | 0.7546726 | 0.00130913 | 0.49677584 | 0.89054584 |
| Dapl1 | 0.00920789 | 0.0111611 | 0.009854 | 0.00130711 | 0.26828894 | 0.89054584 |
| 2310010G23 | 0.04388961 | 0.04826494 | 0.04696157 | 0.00130336 | 0.44855871 | 0.89054584 |
| Nog | 0.01256698 | 0.01474907 | 0.01344872 | 0.00130036 | 0.32618446 | 0.89054584 |
| Dcakd | 0.7578177 | 0.69178009 | 0.69048055 | 0.00129953 | 0.49650194 | 0.89054584 |
| Map4k4 | 0.4997104 | 0.49291548 | 0.49162077 | 0.0012947 | 0.49510529 | 0.89054584 |
| Spag8 | 0.01250493 | 0.01467276 | 0.01338232 | 0.00129044 | 0.32662389 | 0.89054584 |
| Il15 | 0.0136721 | 0.01591169 | 0.01463134 | 0.00128035 | 0.34174998 | 0.89054584 |
| Car14 | 0.01331628 | 0.01552799 | 0.01425057 | 0.00127742 | 0.33811003 | 0.89054584 |
| Tmem194b | 0.01545166 | 0.01780708 | 0.01653569 | 0.0012714 | 0.36009051 | 0.89054584 |

| | | | | | | |
|---|---|---|---|---|---|---|
| Dhx16 | 0.14572272 | 0.15590081 | 0.1546374 | 0.00126341 | 0.48481822 | 0.89054584 |
| Colq | 0.00866705 | 0.01053565 | 0.00927522 | 0.00126044 | 0.26332859 | 0.89054584 |
| Asxl1 | 0.11226588 | 0.12104985 | 0.11979036 | 0.0012595 | 0.48046568 | 0.89054584 |
| 2010300C02I | 0.06738949 | 0.0733468 | 0.07208756 | 0.00125924 | 0.46756851 | 0.89054584 |
| Sh3pxd2b | 0.14277849 | 0.15284452 | 0.151588 | 0.00125652 | 0.48459743 | 0.89054584 |
| Akap11 | 0.4631599 | 0.46120638 | 0.4599543 | 0.00125208 | 0.49494053 | 0.89054584 |
| Gm13212 | 0.03290534 | 0.03646175 | 0.03521126 | 0.00125049 | 0.43429183 | 0.89054584 |
| Ncbp2 | 0.88112538 | 0.77024271 | 0.7689953 | 0.00124741 | 0.49698507 | 0.89054584 |
| Trank1 | 0.01343988 | 0.0156177 | 0.01438283 | 0.00123487 | 0.34457718 | 0.89054584 |
| Ccdc69 | 0.02124411 | 0.02396787 | 0.02273411 | 0.00123376 | 0.40019758 | 0.89054584 |
| Gm15496 | 0.01319084 | 0.0153471 | 0.01411633 | 0.00123077 | 0.34229696 | 0.89054584 |
| Hist1h4h | 0.04034997 | 0.04440441 | 0.04317541 | 0.00122901 | 0.4472475 | 0.89054584 |
| Serpina1e | 0.03901885 | 0.04297874 | 0.04175148 | 0.00122726 | 0.44553657 | 0.89054584 |
| Inpp5k | 0.15338541 | 0.16377198 | 0.16255782 | 0.00121416 | 0.48612036 | 0.89054584 |
| Rhpn1 | 0.06311092 | 0.06872786 | 0.06751454 | 0.00121332 | 0.46663672 | 0.89054584 |
| Bcs1l | 0.19301324 | 0.20435478 | 0.2031431 | 0.00121168 | 0.4889152 | 0.89054584 |
| Fam160b2 | 0.14956533 | 0.15982337 | 0.15861211 | 0.00121126 | 0.48580921 | 0.89054584 |
| Oasl2 | 0.02603449 | 0.0290704 | 0.02785993 | 0.00121047 | 0.41979262 | 0.89054584 |
| Tmem104 | 0.03874537 | 0.04266078 | 0.04145894 | 0.00120184 | 0.44628349 | 0.89054584 |
| Eda2r | 0.02122272 | 0.02390846 | 0.02271123 | 0.00119723 | 0.40299663 | 0.89054584 |
| B230322F03 | 0.03813667 | 0.04200183 | 0.04080779 | 0.00119404 | 0.44578372 | 0.89054584 |
| Zdhhc13 | 0.14348099 | 0.15350987 | 0.15231591 | 0.00119396 | 0.4854338 | 0.89054584 |
| Cobl | 0.05996547 | 0.06534493 | 0.06415212 | 0.00119281 | 0.46548444 | 0.89054584 |
| Card9 | 0.0175398 | 0.01996234 | 0.01877021 | 0.00119212 | 0.3836549 | 0.89054584 |
| Dmc1 | 0.01156618 | 0.01356553 | 0.01237772 | 0.00118781 | 0.32740614 | 0.89054584 |
| 2310001H17 | 0.05048978 | 0.05520622 | 0.05402046 | 0.00118576 | 0.45927336 | 0.89054584 |
| Rnasek | 0.00951701 | 0.01136834 | 0.0101848 | 0.00118354 | 0.29411619 | 0.89054584 |
| 6430573F11I | 0.08048894 | 0.0872646 | 0.08608294 | 0.00118166 | 0.47450383 | 0.89054584 |
| Tpgs2 | 0.2298855 | 0.2414904 | 0.24031053 | 0.00117988 | 0.49087521 | 0.89054584 |
| Syk | 0.02934874 | 0.0325838 | 0.03140605 | 0.00117775 | 0.43065251 | 0.89054584 |

| | | | | | | |
|---|---|---|---|---|---|---|
| 9330104G04 | 0.03845409 | 0.04232368 | 0.04114735 | 0.00117634 | 0.44702096 | 0.89054584 |
| Eya1 | 0.06247737 | 0.06801218 | 0.06683733 | 0.00117485 | 0.46736549 | 0.89054584 |
| Uty | 0.11025335 | 0.11884412 | 0.11768085 | 0.00116328 | 0.48163374 | 0.89054584 |
| Rasd2 | 0.01639815 | 0.01870492 | 0.01754854 | 0.00115638 | 0.37941908 | 0.89054584 |
| Fundc2 | 1.3104309 | 0.97561498 | 0.97446193 | 0.00115305 | 0.49780073 | 0.89054584 |
| Rnf122 | 0.06801449 | 0.07390585 | 0.0727555 | 0.00115035 | 0.47063913 | 0.89054584 |
| Col7a1 | 0.01878643 | 0.02125101 | 0.02010422 | 0.00114679 | 0.39521361 | 0.89054584 |
| AI504432 | 0.00948677 | 0.01129703 | 0.01015244 | 0.00114459 | 0.29970457 | 0.89054584 |
| Gm9484 | 0.01984733 | 0.02238347 | 0.02123947 | 0.001144 | 0.40093025 | 0.89054584 |
| Ehhadh | 0.01798884 | 0.02039139 | 0.01925073 | 0.00114066 | 0.39125253 | 0.89054584 |
| Plekhm3 | 0.05917875 | 0.06445126 | 0.06331107 | 0.00114019 | 0.46656622 | 0.89054584 |
| Syt3 | 0.03648983 | 0.04018561 | 0.03904605 | 0.00113956 | 0.44592212 | 0.89054584 |
| Saxo2 | 0.07408254 | 0.08037883 | 0.07923949 | 0.00113934 | 0.47329546 | 0.89054584 |
| Glyctk | 0.01420579 | 0.01632901 | 0.01520245 | 0.00112656 | 0.36495547 | 0.89054584 |
| Hck | 0.02509652 | 0.02798051 | 0.02685631 | 0.0011242 | 0.42268782 | 0.89054584 |
| Gm17249 | 0.02088549 | 0.02347029 | 0.02235037 | 0.00111993 | 0.40770636 | 0.89054584 |
| Tecr | 2.11796406 | 1.12537123 | 1.12425319 | 0.00111804 | 0.49815163 | 0.89054584 |
| BC022687 | 0.24961232 | 0.2610595 | 0.25994796 | 0.00111154 | 0.49205295 | 0.89054584 |
| Dtx1 | 0.03169003 | 0.03501869 | 0.03391102 | 0.00110766 | 0.43952301 | 0.89054584 |
| Invs | 0.21451103 | 0.22599104 | 0.22488528 | 0.00110576 | 0.49086184 | 0.89054584 |
| Cers4 | 0.22919301 | 0.24071405 | 0.23961804 | 0.001096 | 0.49149926 | 0.89054584 |
| Dpysl5 | 0.04520654 | 0.04946357 | 0.04837014 | 0.00109343 | 0.45806186 | 0.89054584 |
| Surf1 | 0.48719289 | 0.48196082 | 0.48086752 | 0.0010933 | 0.49577425 | 0.89054584 |
| Prodh2 | 0.06844242 | 0.07430297 | 0.07321282 | 0.00109014 | 0.47234683 | 0.89054584 |
| Slc6a9 | 0.0050924 | 0.00653501 | 0.00544976 | 0.00108525 | 0.17676261 | 0.848016 |
| Gm15915 | 0.0755785 | 0.08191838 | 0.0808377 | 0.00108067 | 0.47516881 | 0.89054584 |
| Hexim2 | 0.06489423 | 0.0705012 | 0.06942067 | 0.00108053 | 0.47109561 | 0.89054584 |
| Slc30a7 | 0.24076072 | 0.25223883 | 0.25115861 | 0.00108022 | 0.4920066 | 0.89054584 |
| Hist1h2ag | 0.02246158 | 0.02511158 | 0.02403686 | 0.00107473 | 0.41749404 | 0.89054584 |
| Fam57b | 0.05004933 | 0.0546213 | 0.05354944 | 0.00107187 | 0.4628504 | 0.89054584 |

| | | | | | | |
|---|---|---|---|---|---|---|
| Sema7a | 0.01345892 | 0.01546254 | 0.01440321 | 0.00105933 | 0.36592879 | 0.89054584 |
| Drc1 | 0.11896976 | 0.12785168 | 0.12680646 | 0.00104523 | 0.48468349 | 0.89054584 |
| Elovl4 | 0.03933312 | 0.04313087 | 0.04208767 | 0.0010432 | 0.45403303 | 0.89054584 |
| Gm21994 | 0.00905223 | 0.0107261 | 0.00968742 | 0.00103868 | 0.30870343 | 0.89054584 |
| Ccbe1 | 0.01946243 | 0.02185953 | 0.02082759 | 0.00103194 | 0.40872113 | 0.89054584 |
| Cox18 | 0.34599751 | 0.35423851 | 0.35321397 | 0.00102453 | 0.49460896 | 0.89054584 |
| Gm8225 | 0.01250181 | 0.01440248 | 0.01337897 | 0.00102351 | 0.36076506 | 0.89054584 |
| C1rb | 0.03281519 | 0.03613505 | 0.03511481 | 0.00102024 | 0.44616266 | 0.89054584 |
| Dnah11 | 0.01849095 | 0.02079908 | 0.01978804 | 0.00101104 | 0.40592408 | 0.89054584 |
| Cdc25c | 0.03604511 | 0.03957865 | 0.03857028 | 0.00100837 | 0.45152807 | 0.89054584 |
| Smad1 | 0.31804863 | 0.32765415 | 0.32664658 | 0.00100757 | 0.49426703 | 0.89054584 |
| Zfp182 | 0.0739461 | 0.08009752 | 0.07909371 | 0.00100381 | 0.47642486 | 0.89054584 |
| Pcna-ps2 | 0.03946103 | 0.04322272 | 0.0422245 | 0.00099823 | 0.4561485 | 0.89054584 |
| Mdga2 | 0.008484 | 0.01007617 | 0.00907932 | 0.00099685 | 0.30449396 | 0.89054584 |
| Jade2 | 0.02866922 | 0.03167565 | 0.030679 | 0.00099665 | 0.43984916 | 0.89054584 |
| Mitd1 | 0.36470787 | 0.37175757 | 0.37077015 | 0.00098742 | 0.49505023 | 0.89054584 |
| Slc41a3 | 0.05859621 | 0.06366909 | 0.06268828 | 0.00098081 | 0.47094556 | 0.89054584 |
| Gm10382 | 0.00501978 | 0.00634995 | 0.00537204 | 0.00097791 | 0.19819042 | 0.89054584 |
| Tmem203 | 0.32159288 | 0.33100609 | 0.330038 | 0.00096809 | 0.49454825 | 0.89054584 |
| Fam117a | 0.11958921 | 0.12841517 | 0.12745391 | 0.00096126 | 0.48598493 | 0.89054584 |
| Dcaf12 | 0.3592629 | 0.36664091 | 0.36568026 | 0.00096066 | 0.49511738 | 0.89054584 |
| Fancl | 0.16562674 | 0.17611833 | 0.17516335 | 0.00095498 | 0.48986781 | 0.89054584 |
| Abca1 | 0.25142102 | 0.26269171 | 0.26173944 | 0.00095227 | 0.49323812 | 0.89054584 |
| Gm17518 | 0.01494268 | 0.01694017 | 0.01599102 | 0.00094915 | 0.39106956 | 0.89054584 |
| A4galt | 0.02926676 | 0.03226509 | 0.03131834 | 0.00094675 | 0.44399859 | 0.89054584 |
| Oard1 | 0.52116338 | 0.51076672 | 0.50982442 | 0.0009423 | 0.49656472 | 0.89054584 |
| Rab31 | 0.07429032 | 0.08039786 | 0.07946147 | 0.00093638 | 0.47810852 | 0.89054584 |
| Ffar3 | 0.03331075 | 0.03657612 | 0.03564499 | 0.00093113 | 0.45156748 | 0.89054584 |
| Lca5l | 0.02457806 | 0.02723242 | 0.02630155 | 0.00093086 | 0.43451562 | 0.89054584 |
| Slmo1 | 0.01778012 | 0.01995764 | 0.01902738 | 0.00093026 | 0.4099093 | 0.89054584 |

| | | | | | | |
|---|---|---|---|---|---|---|
| Echs1 | 1.1873148 | 0.92592662 | 0.924997 | 0.00092963 | 0.49813206 | 0.89054584 |
| Slc43a1 | 0.04160238 | 0.04544137 | 0.04451508 | 0.00092629 | 0.46138476 | 0.89054584 |
| D930030I03F | 0.01342968 | 0.01529428 | 0.01437192 | 0.00092236 | 0.3824691 | 0.89054584 |
| Usp27x | 0.03488122 | 0.03824661 | 0.03732515 | 0.00092146 | 0.45421576 | 0.89054584 |
| Pcdhga12 | 0.06786604 | 0.07351734 | 0.07259686 | 0.00092048 | 0.47644727 | 0.89054584 |
| A230050P20 | 0.11810502 | 0.12682135 | 0.12590238 | 0.00091896 | 0.4864363 | 0.89054584 |
| Pcdhb18 | 0.00930141 | 0.01087212 | 0.00995408 | 0.00091804 | 0.33371311 | 0.89054584 |
| Ppp2r5c | 0.45019366 | 0.4494458 | 0.44852786 | 0.00091794 | 0.49619623 | 0.89054584 |
| AI427809 | 0.01724147 | 0.01936825 | 0.01845097 | 0.00091728 | 0.40841769 | 0.89054584 |
| Jmjd8 | 0.43766294 | 0.43830492 | 0.43739043 | 0.00091448 | 0.49611405 | 0.89054584 |
| Tmem116 | 0.07594475 | 0.08213559 | 0.08122897 | 0.00090662 | 0.47926442 | 0.89054584 |
| Zfp729a | 0.05725879 | 0.06216405 | 0.06125841 | 0.00090564 | 0.47254347 | 0.89054584 |
| Proca1 | 0.03817275 | 0.04175116 | 0.04084639 | 0.00090477 | 0.45890289 | 0.89054584 |
| Pcdh11x | 0.02925344 | 0.03220818 | 0.03130408 | 0.0009041 | 0.44648164 | 0.89054584 |
| Uvrag | 0.27498127 | 0.28583426 | 0.28493665 | 0.00089761 | 0.49414509 | 0.89054584 |
| Slc30a6 | 0.17624857 | 0.18694843 | 0.18605168 | 0.00089675 | 0.49104222 | 0.89054584 |
| Kcnh3 | 0.0124796 | 0.01425122 | 0.01335521 | 0.00089601 | 0.37730359 | 0.89054584 |
| Gdpgp1 | 0.09812259 | 0.10579363 | 0.10490679 | 0.00088684 | 0.4842918 | 0.89054584 |
| Adi1 | 0.22510125 | 0.2363989 | 0.23552189 | 0.00087701 | 0.4930793 | 0.89054584 |
| Tmem178 | 0.03478121 | 0.03809399 | 0.03721815 | 0.00087584 | 0.45634843 | 0.89054584 |
| Slc25a33 | 0.16612071 | 0.17654527 | 0.17567075 | 0.00087453 | 0.49074808 | 0.89054584 |
| Cd3e | 0.00494502 | 0.0061622 | 0.00529202 | 0.00087018 | 0.22181521 | 0.89054584 |
| Cldn18 | 0.00494502 | 0.0061622 | 0.00529202 | 0.00087018 | 0.22181521 | 0.89054584 |
| Acot6 | 0.0254994 | 0.02815658 | 0.02728739 | 0.00086919 | 0.44101265 | 0.89054584 |
| Plekhg3 | 0.18585183 | 0.19672478 | 0.19585648 | 0.00086831 | 0.49176047 | 0.89054584 |
| Tram1l1 | 0.09760744 | 0.10522528 | 0.10435717 | 0.00086811 | 0.48454252 | 0.89054584 |
| Kcnab2 | 0.01334988 | 0.01515089 | 0.01428652 | 0.00086436 | 0.38901946 | 0.89054584 |
| Gm42903 | 0.00919271 | 0.01069605 | 0.00983775 | 0.00085829 | 0.34219906 | 0.89054584 |
| Eml4 | 0.53687925 | 0.52383404 | 0.52297791 | 0.00085613 | 0.49695738 | 0.89054584 |
| Adgra1 | 0.01221927 | 0.01393202 | 0.01307662 | 0.0008554 | 0.38027341 | 0.89054584 |

| | | | | | | |
|---|---|---|---|---|---|---|
| Gm44036 | 0.00913886 | 0.01063455 | 0.00978012 | 0.00085443 | 0.34199536 | 0.89054584 |
| Abcc12 | 0.02066762 | 0.02297131 | 0.02211723 | 0.00085408 | 0.42861166 | 0.89054584 |
| Soga1 | 0.06747184 | 0.07302892 | 0.07217557 | 0.00085335 | 0.47803587 | 0.89054584 |
| Hspa9 | 1.24693941 | 0.9507846 | 0.94993138 | 0.00085322 | 0.49833059 | 0.89054584 |
| Adssl1 | 0.03665391 | 0.04005963 | 0.03922158 | 0.00083805 | 0.46035144 | 0.89054584 |
| Krt15 | 0.0131716 | 0.014925 | 0.01409574 | 0.00082926 | 0.39200896 | 0.89054584 |
| Bcl6b | 0.09355833 | 0.10086365 | 0.10003643 | 0.00082722 | 0.48463432 | 0.89054584 |
| Usp21 | 0.15649979 | 0.16659476 | 0.16577057 | 0.00082418 | 0.49075995 | 0.89054584 |
| Kcnk16 | 0.11276446 | 0.12113661 | 0.12031273 | 0.00082388 | 0.48727444 | 0.89054584 |
| Lrrc51 | 0.09152466 | 0.0986796 | 0.0978659 | 0.0008137 | 0.48455022 | 0.89054584 |
| Gmpr | 0.33060183 | 0.33944074 | 0.33862905 | 0.00081168 | 0.49554497 | 0.89054584 |
| Tmem86a | 0.27607552 | 0.2868106 | 0.28600759 | 0.00080301 | 0.49478171 | 0.89054584 |
| Srrm4 | 0.01275109 | 0.01444867 | 0.01364573 | 0.00080294 | 0.39198823 | 0.89054584 |
| Npas2 | 0.01722808 | 0.01923787 | 0.01843665 | 0.00080122 | 0.41977506 | 0.89054584 |
| Igsf21 | 0.08856142 | 0.09549329 | 0.09470276 | 0.00079052 | 0.48448901 | 0.89054584 |
| Meikin | 0.00855852 | 0.00994704 | 0.00915907 | 0.00078797 | 0.34427775 | 0.89054584 |
| Bcl11a | 0.21186063 | 0.22299942 | 0.22221545 | 0.00078397 | 0.49344303 | 0.89054584 |
| Gm15420 | 0.0131335 | 0.01483861 | 0.01405496 | 0.00078364 | 0.39752356 | 0.89054584 |
| Col25a1 | 0.02182736 | 0.02413837 | 0.02335822 | 0.00078015 | 0.43817205 | 0.89054584 |
| Adcy2 | 0.01321816 | 0.01489998 | 0.01414556 | 0.00075441 | 0.40188498 | 0.89054584 |
| Gm26797 | 0.01144761 | 0.01300243 | 0.01225083 | 0.0007516 | 0.3875044 | 0.89054584 |
| C1s2 | 0.03174795 | 0.03471842 | 0.03397299 | 0.00074543 | 0.45928865 | 0.89054584 |
| Col6a5 | 0.01281161 | 0.0144491 | 0.01371051 | 0.0007386 | 0.40091424 | 0.89054584 |
| Kif3c | 0.08245513 | 0.08892116 | 0.0881828 | 0.00073836 | 0.48444146 | 0.89054584 |
| Pnkp | 0.20334918 | 0.21435877 | 0.21362095 | 0.00073782 | 0.49358075 | 0.89054584 |
| Cdhr2 | 0.0469313 | 0.05094908 | 0.05021482 | 0.00073426 | 0.47284316 | 0.89054584 |
| Nap1l2 | 0.05159874 | 0.05592969 | 0.05520635 | 0.00072334 | 0.47566237 | 0.89054584 |
| Aim2 | 0.02174666 | 0.02399437 | 0.02327187 | 0.00072251 | 0.44249637 | 0.89054584 |
| Sema4g | 0.04904325 | 0.05318925 | 0.05247351 | 0.00071574 | 0.47466515 | 0.89054584 |
| Bloc1s5 | 0.21652096 | 0.22761131 | 0.22690786 | 0.00070345 | 0.49423814 | 0.89054584 |

| | | | | | | |
|---|---|---|---|---|---|---|
| Lrrc75b | 0.01344324 | 0.0150893 | 0.01438644 | 0.00070286 | 0.40997182 | 0.89054584 |
| Stk10 | 0.01561159 | 0.01740948 | 0.01670683 | 0.00070265 | 0.422327 | 0.89054584 |
| Jdp2 | 0.01725324 | 0.01916488 | 0.01846358 | 0.00070131 | 0.42976932 | 0.89054584 |
| Card6 | 0.02002557 | 0.02212941 | 0.02143019 | 0.00069922 | 0.43958932 | 0.89054584 |
| Atxn1 | 0.1848681 | 0.19554643 | 0.19485385 | 0.00069258 | 0.49339404 | 0.89054584 |
| Rbm12b1 | 0.05099158 | 0.05524913 | 0.05455707 | 0.00069206 | 0.47643659 | 0.89054584 |
| Morc3 | 0.36301399 | 0.36987812 | 0.36918821 | 0.00068991 | 0.49652674 | 0.89054584 |
| RP24-143K11 | 0.03042745 | 0.03324917 | 0.03256019 | 0.00068898 | 0.46073401 | 0.89054584 |
| Cfap97 | 0.48513875 | 0.47978269 | 0.47909382 | 0.00068887 | 0.49732753 | 0.89054584 |
| Atp8a1 | 0.03310726 | 0.03611208 | 0.03542728 | 0.0006848 | 0.4641215 | 0.89054584 |
| Fam19a1 | 0.03867718 | 0.04206904 | 0.041386 | 0.00068304 | 0.46935472 | 0.89054584 |
| Wdr6 | 0.80751757 | 0.72413514 | 0.72345438 | 0.00068076 | 0.49825104 | 0.89054584 |
| Paqr8 | 0.02418334 | 0.02655447 | 0.02587919 | 0.00067528 | 0.4516203 | 0.89054584 |
| Fbxl8 | 0.03846239 | 0.04182484 | 0.04115622 | 0.00066862 | 0.46983348 | 0.89054584 |
| Cdk20 | 0.07646923 | 0.08245366 | 0.08178926 | 0.0006644 | 0.48490529 | 0.89054584 |
| Necab3 | 0.0276496 | 0.03024845 | 0.02958806 | 0.00066039 | 0.45859062 | 0.89054584 |
| Rrp12 | 0.07258342 | 0.07829153 | 0.07763778 | 0.00065375 | 0.48435332 | 0.89054584 |
| RP24-267C3. | 0.01682989 | 0.01866226 | 0.01801054 | 0.00065172 | 0.43306147 | 0.89054584 |
| Rps6ka5 | 0.03380431 | 0.03681273 | 0.03617303 | 0.0006397 | 0.46716782 | 0.89054584 |
| Map1a | 0.26503726 | 0.27581606 | 0.27517778 | 0.00063828 | 0.4956889 | 0.89054584 |
| Lypd6b | 0.01568892 | 0.01742176 | 0.01678958 | 0.00063218 | 0.43037337 | 0.89054584 |
| Cxcr4 | 0.00869451 | 0.00993519 | 0.0093046 | 0.00063059 | 0.37609729 | 0.89054584 |
| Sdk1 | 0.1229666 | 0.1316116 | 0.13098144 | 0.00063015 | 0.49105877 | 0.89054584 |
| Dnd1 | 0.00476122 | 0.00571264 | 0.00509533 | 0.00061731 | 0.28622721 | 0.89054584 |
| Prr33 | 0.02327007 | 0.02551904 | 0.02490197 | 0.00061707 | 0.45404495 | 0.89054584 |
| Cd300a | 0.0047589 | 0.00570707 | 0.00509285 | 0.00061423 | 0.28709407 | 0.89054584 |
| Plk3 | 0.04057172 | 0.0440232 | 0.04341261 | 0.00061059 | 0.47387713 | 0.89054584 |
| Exd1 | 0.01107365 | 0.01245813 | 0.01185064 | 0.00060749 | 0.40561955 | 0.89054584 |
| Pfkp | 0.06568607 | 0.07087329 | 0.07026701 | 0.00060628 | 0.4839674 | 0.89054584 |
| Mmp19 | 0.02011902 | 0.02213396 | 0.0215302 | 0.00060376 | 0.44802655 | 0.89054584 |

| | | | | | | |
|---|---|---|---|---|---|---|
| Fbxl21 | 0.00900861 | 0.01023698 | 0.00964073 | 0.00059625 | 0.38661921 | 0.89054584 |
| Moxd1 | 0.00474475 | 0.0056732 | 0.00507771 | 0.00059549 | 0.29240341 | 0.89054584 |
| Epha1 | 0.05786345 | 0.06249721 | 0.06190488 | 0.00059232 | 0.48222177 | 0.89054584 |
| Adamts14 | 0.01455545 | 0.01616613 | 0.01557664 | 0.00058949 | 0.43002278 | 0.89054584 |
| Hhatl | 0.01340772 | 0.01493785 | 0.01434843 | 0.00058943 | 0.42411126 | 0.89054584 |
| Plxna4 | 0.03199707 | 0.03482875 | 0.03423952 | 0.00058923 | 0.46804869 | 0.89054584 |
| Hsdl2 | 0.38873153 | 0.39362344 | 0.39303552 | 0.00058793 | 0.49721974 | 0.89054584 |
| Gm14305 | 0.26921601 | 0.27987018 | 0.27928455 | 0.00058563 | 0.49610269 | 0.89054584 |
| Nrxn1 | 0.0483457 | 0.05231189 | 0.05172751 | 0.00058438 | 0.4790122 | 0.89054584 |
| Ube2u | 0.01479236 | 0.0164116 | 0.01583016 | 0.00058144 | 0.43206321 | 0.89054584 |
| Padi1 | 0.01319889 | 0.01470269 | 0.01412494 | 0.00057775 | 0.42443369 | 0.89054584 |
| Ccdc88c | 0.05232498 | 0.0565576 | 0.05598295 | 0.00057465 | 0.48092864 | 0.89054584 |
| Mfsd2a | 0.0258026 | 0.02817702 | 0.02761181 | 0.00056522 | 0.4620109 | 0.89054584 |
| Adcy7 | 0.01586384 | 0.01753627 | 0.01697677 | 0.0005595 | 0.43898504 | 0.89054584 |
| BC021891 | 0.0160795 | 0.01776564 | 0.01720755 | 0.00055809 | 0.43994692 | 0.89054584 |
| Zfp94 | 0.03920026 | 0.04250205 | 0.04194555 | 0.0005565 | 0.47535657 | 0.89054584 |
| Gm26624 | 0.0186415 | 0.0205054 | 0.01994914 | 0.00055626 | 0.44831864 | 0.89054584 |
| Nhlh1 | 0.01327279 | 0.0147602 | 0.01420403 | 0.00055617 | 0.42762468 | 0.89054584 |
| Fstl3 | 0.0695778 | 0.07497974 | 0.07442615 | 0.0005536 | 0.48617776 | 0.89054584 |
| Adora2a | 0.10476623 | 0.11247398 | 0.11192174 | 0.00055224 | 0.49082989 | 0.89054584 |
| Disp1 | 0.04215505 | 0.04565435 | 0.04510625 | 0.0005481 | 0.47742693 | 0.89054584 |
| Nxph4 | 0.01308846 | 0.01454797 | 0.01400677 | 0.00054119 | 0.4285718 | 0.89054584 |
| Ano9 | 0.02263006 | 0.02475719 | 0.02421714 | 0.00054005 | 0.45862621 | 0.89054584 |
| Csnk1g1 | 0.34217732 | 0.3501395 | 0.34960636 | 0.00053314 | 0.49716562 | 0.89054584 |
| Slc5a7 | 0.00828833 | 0.00940079 | 0.00886992 | 0.00053087 | 0.39018307 | 0.89054584 |
| Tnfrsf11a | 0.00893511 | 0.01008411 | 0.00956208 | 0.00052203 | 0.39961371 | 0.89054584 |
| Tle4 | 0.03833119 | 0.04153551 | 0.04101587 | 0.00051964 | 0.47646599 | 0.89054584 |
| Gak | 0.19882938 | 0.20956113 | 0.20904478 | 0.00051635 | 0.49540915 | 0.89054584 |
| Rnf125 | 0.01295603 | 0.01437861 | 0.01386505 | 0.00051356 | 0.4314958 | 0.89054584 |
| Tmem9 | 0.95822745 | 0.81304578 | 0.81253425 | 0.00051154 | 0.49882988 | 0.89054584 |

| | | | | | | |
|---|---|---|---|---|---|---|
| Brip1 | 0.06295155 | 0.06785443 | 0.0673442 | 0.00051024 | 0.48592079 | 0.89054584 |
| Ccdc180 | 0.01235927 | 0.01372946 | 0.01322644 | 0.00050302 | 0.42968067 | 0.89054584 |
| Pitpnm1 | 0.06888997 | 0.07419113 | 0.0736911 | 0.00050003 | 0.48739039 | 0.89054584 |
| St7l | 0.44943892 | 0.44835813 | 0.44785963 | 0.0004985 | 0.49793121 | 0.89054584 |
| Metap1d | 0.31023914 | 0.31964624 | 0.31915098 | 0.00049526 | 0.49711574 | 0.89054584 |
| Hrc | 0.01199939 | 0.01333597 | 0.01284131 | 0.00049466 | 0.42878617 | 0.89054584 |
| Rdh14 | 0.57919666 | 0.55809847 | 0.55760724 | 0.00049123 | 0.49836262 | 0.89054584 |
| AW549877 | 0.09900648 | 0.10634072 | 0.10584979 | 0.00049092 | 0.49138044 | 0.89054584 |
| Hnmt | 0.01808543 | 0.01984475 | 0.01935409 | 0.00049066 | 0.45298932 | 0.89054584 |
| Zfp467 | 0.35644994 | 0.36353341 | 0.36304471 | 0.0004887 | 0.49749808 | 0.89054584 |
| Mr1 | 0.05317971 | 0.05738159 | 0.05689693 | 0.00048466 | 0.48417189 | 0.89054584 |
| Slc6a20a | 0.02108822 | 0.02304812 | 0.0225673 | 0.00048082 | 0.46046441 | 0.89054584 |
| Zfp449 | 0.13279754 | 0.14169642 | 0.14122541 | 0.00047102 | 0.49380125 | 0.89054584 |
| Prkd2 | 0.08818651 | 0.09477239 | 0.09430252 | 0.00046987 | 0.49074004 | 0.89054584 |
| Mrpl44 | 0.48446753 | 0.47898006 | 0.47851367 | 0.00046638 | 0.49818848 | 0.89054584 |
| Tmem139 | 0.02368673 | 0.02581417 | 0.02534781 | 0.00046636 | 0.46584557 | 0.89054584 |
| Dnah2 | 0.02019324 | 0.02206932 | 0.02160962 | 0.0004597 | 0.46052556 | 0.89054584 |
| Alg10b | 0.03921304 | 0.04241392 | 0.04195922 | 0.0004547 | 0.47986705 | 0.89054584 |
| Zik1 | 0.03691093 | 0.03995107 | 0.03949653 | 0.00045454 | 0.47862024 | 0.89054584 |
| Arhgap11a | 0.06300966 | 0.06785843 | 0.0674063 | 0.00045213 | 0.48753519 | 0.89054584 |
| Oxtr | 0.03349791 | 0.03629692 | 0.03584522 | 0.0004517 | 0.4765917 | 0.89054584 |
| Angpt1 | 0.04550182 | 0.04913177 | 0.04868596 | 0.00044582 | 0.48298558 | 0.89054584 |
| Zswim1 | 0.0715357 | 0.07696329 | 0.07651829 | 0.000445 | 0.48919228 | 0.89054584 |
| Epb41l4aos | 0.38562358 | 0.39061426 | 0.39017314 | 0.00044112 | 0.49789867 | 0.89054584 |
| Igsf11 | 0.02308298 | 0.02514047 | 0.02470178 | 0.00043869 | 0.46702885 | 0.89054584 |
| Cfap70 | 0.02349876 | 0.02558199 | 0.02514668 | 0.00043532 | 0.46785966 | 0.89054584 |
| RP23-335G1 | 0.04356955 | 0.04705371 | 0.04661923 | 0.00043447 | 0.48268362 | 0.89054584 |
| Herc3 | 0.0490578 | 0.05292188 | 0.05248907 | 0.00043281 | 0.48467804 | 0.89054584 |
| 2310040G07 | 0.01211424 | 0.01338599 | 0.01296423 | 0.00042177 | 0.43976327 | 0.89054584 |
| Cdnf | 0.01084769 | 0.01202013 | 0.01160883 | 0.0004113 | 0.43444618 | 0.89054584 |

| | | | | | | |
|---|---|---|---|---|---|---|
| Olfml2a | 0.00858351 | 0.00959299 | 0.00918582 | 0.00040718 | 0.4181941 | 0.89054584 |
| E430024P14I | 0.05727447 | 0.06166981 | 0.06127517 | 0.00039464 | 0.48803137 | 0.89054584 |
| Pex26 | 0.12829857 | 0.13693012 | 0.13654185 | 0.00038827 | 0.49471489 | 0.89054584 |
| Rasal1 | 0.03148599 | 0.0340791 | 0.03369273 | 0.00038638 | 0.47869585 | 0.89054584 |
| Sdr39u1 | 0.25341546 | 0.26409921 | 0.26371309 | 0.00038612 | 0.49727862 | 0.89054584 |
| Prrc2c | 1.67981311 | 1.07801987 | 1.07763417 | 0.0003857 | 0.49933477 | 0.89054584 |
| Rasl2-9 | 0.01886709 | 0.02057566 | 0.02019054 | 0.00038512 | 0.46459415 | 0.89054584 |
| Cadm3 | 0.02871215 | 0.03110055 | 0.03072494 | 0.00037562 | 0.47729002 | 0.89054584 |
| Pik3cb | 0.03940412 | 0.04251804 | 0.04216362 | 0.00035441 | 0.48438079 | 0.89054584 |
| Zglp1 | 0.00812171 | 0.00904444 | 0.00869161 | 0.00035283 | 0.4249964 | 0.89054584 |
| Gm26610 | 0.01288436 | 0.01414068 | 0.01378836 | 0.00035232 | 0.45262043 | 0.89054584 |
| Rad1 | 0.27476311 | 0.28507538 | 0.28472307 | 0.00035231 | 0.49770014 | 0.89054584 |
| Pvt1 | 0.01703576 | 0.0185827 | 0.01823084 | 0.00035186 | 0.46417559 | 0.89054584 |
| Pcdhb22 | 0.04221966 | 0.04552703 | 0.04517536 | 0.00035167 | 0.48553446 | 0.89054584 |
| Sdhaf4 | 1.10022965 | 0.88563094 | 0.88528162 | 0.00034932 | 0.4992666 | 0.89054584 |
| Ppt2 | 0.21428204 | 0.22500199 | 0.22465473 | 0.00034725 | 0.49712706 | 0.89054584 |
| Slc2a9 | 0.01833182 | 0.01996272 | 0.01961775 | 0.00034497 | 0.46735275 | 0.89054584 |
| Nutf2-ps1 | 0.01483476 | 0.01621566 | 0.01587553 | 0.00034013 | 0.46024468 | 0.89054584 |
| Nop2 | 0.19916807 | 0.20972525 | 0.209388 | 0.00033725 | 0.4970064 | 0.89054584 |
| Arntl2 | 0.02030972 | 0.02206876 | 0.02173426 | 0.0003345 | 0.47141921 | 0.89054584 |
| Gm14308 | 0.00722704 | 0.00806133 | 0.00773417 | 0.00032715 | 0.42188554 | 0.89054584 |
| Swi5 | 2.777644 | 1.11237242 | 1.11204563 | 0.00032679 | 0.49945381 | 0.89054584 |
| Ralbp1 | 1.35922943 | 0.99221773 | 0.9918922 | 0.00032553 | 0.49939 | 0.89054584 |
| Pcgf5 | 0.11512012 | 0.12309953 | 0.12277957 | 0.00031996 | 0.49515652 | 0.89054584 |
| Zbtb8b | 0.0145918 | 0.01593124 | 0.01561553 | 0.00031571 | 0.46247798 | 0.89054584 |
| Amy2a2 | 0.00734229 | 0.0081731 | 0.0078575 | 0.0003156 | 0.42578054 | 0.89054584 |
| Sbf2 | 0.36409708 | 0.37051526 | 0.37019991 | 0.00031535 | 0.49841674 | 0.89054584 |
| Ccdc163 | 0.14826025 | 0.15757784 | 0.15726281 | 0.00031503 | 0.4962768 | 0.89054584 |
| Gm15706 | 0.09818816 | 0.10529024 | 0.10497675 | 0.00031348 | 0.49444987 | 0.89054584 |
| Tap1 | 0.05122179 | 0.05511013 | 0.05480326 | 0.00030687 | 0.48959377 | 0.89054584 |

| | | | | | | |
|---|---|---|---|---|---|---|
| Tbkbp1 | 0.04394582 | 0.04732767 | 0.0470217 | 0.00030597 | 0.48790759 | 0.89054584 |
| Aaas | 0.15000134 | 0.1593677 | 0.15906274 | 0.00030496 | 0.49643657 | 0.89054584 |
| Bhmt2 | 0.0066044 | 0.00737266 | 0.00706785 | 0.00030481 | 0.42037916 | 0.89054584 |
| Gm15965 | 0.00795991 | 0.00882196 | 0.00851846 | 0.0003035 | 0.43408124 | 0.89054584 |
| Grik2 | 0.00863091 | 0.00953685 | 0.00923654 | 0.00030031 | 0.43979963 | 0.89054584 |
| Eri2 | 0.09973595 | 0.10692378 | 0.10662801 | 0.00029578 | 0.49484443 | 0.89054584 |
| Gm29562 | 0.01173856 | 0.01285595 | 0.01256219 | 0.00029376 | 0.45662189 | 0.89054584 |
| E430018J23F | 0.07212599 | 0.07744 | 0.07714903 | 0.00029097 | 0.49299031 | 0.89054584 |
| A930007I19F | 0.0213967 | 0.02318433 | 0.02289739 | 0.00028694 | 0.47672161 | 0.89054584 |
| Asic2 | 0.01331371 | 0.01453213 | 0.01424782 | 0.00028432 | 0.46296405 | 0.89054584 |
| Pggt1b | 0.25105467 | 0.26166036 | 0.2613767 | 0.00028367 | 0.49798286 | 0.89054584 |
| Gm7808 | 0.01604182 | 0.01744669 | 0.01716722 | 0.00027947 | 0.4697719 | 0.89054584 |
| Scn7a | 0.01262875 | 0.01379354 | 0.01351482 | 0.00027872 | 0.46172788 | 0.89054584 |
| Nup160 | 0.09446754 | 0.10127759 | 0.10100673 | 0.00027086 | 0.49501594 | 0.89054584 |
| Irak3 | 0.02000732 | 0.02167418 | 0.02141067 | 0.00026351 | 0.47713737 | 0.89054584 |
| Zfp933 | 0.03379987 | 0.03642744 | 0.03616828 | 0.00025916 | 0.48668438 | 0.89054584 |
| Gm26632 | 0.06963648 | 0.07473814 | 0.07448885 | 0.0002493 | 0.49377977 | 0.89054584 |
| Hsdl1 | 0.2147722 | 0.22539475 | 0.2251482 | 0.00024655 | 0.49796466 | 0.89054584 |
| Ptchd1 | 0.01233822 | 0.01344668 | 0.01320391 | 0.00024277 | 0.46586812 | 0.89054584 |
| 4831440E17I | 0.01621991 | 0.01759858 | 0.0173578 | 0.00024077 | 0.47423627 | 0.89054584 |
| Dnah2os | 0.01280584 | 0.01394414 | 0.01370433 | 0.00023981 | 0.46751158 | 0.89054584 |
| BC089491 | 0.00749371 | 0.00825471 | 0.00801956 | 0.00023515 | 0.44566915 | 0.89054584 |
| Tm2d3 | 0.70290851 | 0.65222787 | 0.65199346 | 0.00023441 | 0.49933177 | 0.89054584 |
| Ifitm5 | 0.01286825 | 0.01400308 | 0.01377111 | 0.00023197 | 0.46872369 | 0.89054584 |
| BC034090 | 0.01974253 | 0.0213587 | 0.02112733 | 0.00023138 | 0.47965389 | 0.89054584 |
| Hapln1 | 0.03411876 | 0.03674053 | 0.03650944 | 0.00023109 | 0.48823723 | 0.89054584 |
| Gm20707 | 0.01891478 | 0.02046962 | 0.02024157 | 0.00022806 | 0.47906858 | 0.89054584 |
| Zp3 | 0.01249694 | 0.01359521 | 0.01337377 | 0.00022144 | 0.46925535 | 0.89054584 |
| Sap30l | 0.15109149 | 0.16040647 | 0.16018911 | 0.00021736 | 0.49747804 | 0.89054584 |
| Jsrp1 | 0.00851406 | 0.00932321 | 0.00911149 | 0.00021172 | 0.45689566 | 0.89054584 |

| | | | | | | |
|---|---|---|---|---|---|---|
| Ccdc171 | 0.00694239 | 0.00763043 | 0.00742955 | 0.00020088 | 0.44987731 | 0.89054584 |
| Tal1 | 0.02048856 | 0.02212435 | 0.02192562 | 0.00019872 | 0.4831592 | 0.89054584 |
| Mtus1 | 0.10696387 | 0.1144266 | 0.11422964 | 0.00019697 | 0.49679516 | 0.89054584 |
| Kif26b | 0.01989946 | 0.02149115 | 0.02129525 | 0.0001959 | 0.48290694 | 0.89054584 |
| Gm10501 | 0.00779289 | 0.00853382 | 0.00833973 | 0.00019409 | 0.45682797 | 0.89054584 |
| Chst13 | 0.01262343 | 0.01370213 | 0.01350912 | 0.00019301 | 0.47346381 | 0.89054584 |
| Ccdc65 | 0.04018039 | 0.04318582 | 0.042994 | 0.00019182 | 0.49170816 | 0.89054584 |
| Ick | 0.14134559 | 0.15029347 | 0.1501027 | 0.00019077 | 0.49763777 | 0.89054584 |
| Gm4349 | 0.01239153 | 0.01344015 | 0.01326096 | 0.00017919 | 0.47490088 | 0.89054584 |
| Palm3 | 0.02146853 | 0.02314727 | 0.02297425 | 0.00017302 | 0.48600561 | 0.89054584 |
| Loxl3 | 0.09675762 | 0.1036164 | 0.10345043 | 0.00016597 | 0.49701811 | 0.89054584 |
| Gm37459 | 0.00438924 | 0.00485489 | 0.00469725 | 0.00015764 | 0.43787809 | 0.89054584 |
| Ccdc85a | 0.02021628 | 0.02178678 | 0.02163427 | 0.00015252 | 0.4868993 | 0.89054584 |
| Wnt5b | 0.05830181 | 0.0625258 | 0.06237353 | 0.00015227 | 0.49546268 | 0.89054584 |
| Hspa12a | 0.04346402 | 0.04665037 | 0.04650636 | 0.00014402 | 0.49424444 | 0.89054584 |
| Gm13715 | 0.00837943 | 0.00911047 | 0.00896741 | 0.00014306 | 0.47037602 | 0.89054584 |
| Zfp384 | 0.04296662 | 0.04611622 | 0.04597433 | 0.00014189 | 0.49426384 | 0.89054584 |
| Mirt1 | 0.01210171 | 0.01308924 | 0.01295081 | 0.00013843 | 0.48014198 | 0.89054584 |
| Cops2 | 0.78875015 | 0.71134664 | 0.71121274 | 0.0001339 | 0.49965007 | 0.89054584 |
| Wbscr25 | 0.01217278 | 0.0131599 | 0.01302687 | 0.00013303 | 0.48102612 | 0.89054584 |
| B230307C23 | 0.06019637 | 0.06452853 | 0.06439896 | 0.00012958 | 0.49626028 | 0.89054584 |
| Lbr | 0.33955406 | 0.34724795 | 0.34712456 | 0.00012339 | 0.4993393 | 0.89054584 |
| Gm6277 | 0.00859886 | 0.00931908 | 0.00920224 | 0.00011684 | 0.47641567 | 0.89054584 |
| Fsd1 | 0.04912733 | 0.05267663 | 0.05256343 | 0.0001132 | 0.49599719 | 0.89054584 |
| P3h2 | 0.03247088 | 0.03485849 | 0.03474644 | 0.00011205 | 0.49400646 | 0.89054584 |
| Gm5093 | 0.03886502 | 0.04167612 | 0.04158693 | 8.92E-05 | 0.49601394 | 0.89054584 |
| Kcns3 | 0.03281232 | 0.03520091 | 0.03511174 | 8.92E-05 | 0.49528032 | 0.89054584 |
| 9430065F17l | 0.01614016 | 0.01735921 | 0.01727246 | 8.68E-05 | 0.49066577 | 0.89054584 |
| Gm26718 | 0.00773468 | 0.00836342 | 0.00827743 | 8.60E-05 | 0.48069847 | 0.89054584 |
| Zfp934 | 0.0210156 | 0.02257436 | 0.02248959 | 8.48E-05 | 0.49299447 | 0.89054584 |

| | | | | | | |
|---|---|---|---|---|---|---|
| Gnmt | 0.02069621 | 0.02223212 | 0.02214783 | 8.43E-05 | 0.49292688 | 0.89054584 |
| Gm43270 | 0.01787643 | 0.01921343 | 0.01913045 | 8.30E-05 | 0.49193835 | 0.89054584 |
| Zfp658 | 0.01870685 | 0.02010034 | 0.02001907 | 8.13E-05 | 0.49245454 | 0.89054584 |
| Dtx3 | 0.75531183 | 0.68885012 | 0.68877106 | 7.91E-05 | 0.49978665 | 0.89054584 |
| Xpnpep1 | 0.70800999 | 0.65573684 | 0.65565911 | 7.77E-05 | 0.49977964 | 0.89054584 |
| Sec14l1 | 0.30348754 | 0.31271967 | 0.31264711 | 7.26E-05 | 0.49956866 | 0.89054584 |
| Hsd11b1 | 0.00826254 | 0.0089141 | 0.00884232 | 7.18E-05 | 0.48491456 | 0.89054584 |
| Rap2b | 0.11519368 | 0.12292814 | 0.12285658 | 7.16E-05 | 0.49891726 | 0.89054584 |
| Il1r2 | 0.01721822 | 0.0184855 | 0.0184261 | 5.94E-05 | 0.49400804 | 0.89054584 |
| Kcnmb1 | 0.01992941 | 0.02138568 | 0.0213273 | 5.84E-05 | 0.49491223 | 0.89054584 |
| 1700109H08 | 0.01223118 | 0.01314111 | 0.01308936 | 5.17E-05 | 0.49265269 | 0.89054584 |
| Gm7694 | 0.01120877 | 0.01204107 | 0.01199524 | 4.58E-05 | 0.49289892 | 0.89054584 |
| Lck | 0.04545811 | 0.04867718 | 0.04863921 | 3.80E-05 | 0.49854882 | 0.89054584 |
| Gm12843 | 0.00855772 | 0.00919137 | 0.00915821 | 3.32E-05 | 0.49327173 | 0.89054584 |
| Milr1 | 0.0122391 | 0.01313054 | 0.01309784 | 3.27E-05 | 0.49536076 | 0.89054584 |
| Gm26737 | 0.01933331 | 0.02072205 | 0.02068943 | 3.26E-05 | 0.49706942 | 0.89054584 |
| Mettl24 | 0.00427403 | 0.00460336 | 0.00457395 | 2.94E-05 | 0.48805181 | 0.89054584 |
| Pcdhb7 | 0.01594605 | 0.01709408 | 0.01706474 | 2.93E-05 | 0.49680356 | 0.89054584 |
| Guca1b | 0.00693054 | 0.00743698 | 0.00741687 | 2.01E-05 | 0.49495918 | 0.89054584 |
| Apba2 | 0.00833553 | 0.00893491 | 0.00892043 | 1.45E-05 | 0.4969821 | 0.89054584 |
| Gata3 | 0.0165261 | 0.01769948 | 0.01768545 | 1.40E-05 | 0.49852612 | 0.89054584 |
| Krba1 | 0.17481 | 0.1845921 | 0.18457974 | 1.24E-05 | 0.4998756 | 0.89054584 |
| Rab42 | 0.01217574 | 0.01304084 | 0.01303003 | 1.08E-05 | 0.49845841 | 0.89054584 |
| Tmem51os1 | 0.02622561 | 0.02807516 | 0.02806442 | 1.07E-05 | 0.49928856 | 0.89054584 |
| Rad9a | 0.08728192 | 0.0933439 | 0.09333678 | 7.12E-06 | 0.49985819 | 0.89054584 |
| Upk3a | 0.00425257 | 0.00455725 | 0.00455099 | 6.26E-06 | 0.49744139 | 0.89054584 |
| Lzts1 | 0.03068441 | 0.03284138 | 0.03283512 | 6.26E-06 | 0.49964557 | 0.89054584 |
| B4galnt1 | 0.02693213 | 0.02882615 | 0.02882039 | 5.76E-06 | 0.49962845 | 0.89054584 |
| Itpripl2 | 0.09242888 | 0.09883446 | 0.098831 | 3.45E-06 | 0.49993503 | 0.89054584 |
| Ctnna2 | 0.02303491 | 0.02465074 | 0.02465035 | 3.99E-07 | 0.49996995 | 0.89054584 |

| | | | | | | |
|---|---|---|---|---|---|---|
| A830018L16I | 0 | 0 | 0 | 0 | NA | NA |
| Pm20d1 | 0 | 0 | 0 | 0 | NA | NA |
| Platr27 | 0 | 0 | 0 | 0 | NA | NA |
| A730011C13 | 0 | 0 | 0 | 0 | NA | NA |
| Gm28050 | 0 | 0 | 0 | 0 | NA | NA |
| Cplx1 | 0 | 0 | 0 | 0 | NA | NA |
| Sec61g | 3.88817147 | 0.98551264 | 0.98551264 | 0 | 0.5 | 0.89054584 |
| Grik4 | 0.00837584 | 0.00896356 | 0.00896357 | -5.73E-09 | 0.50000119 | 0.89054584 |
| Slc25a45 | 0.04726508 | 0.0505673 | 0.05057181 | -4.51E-06 | 0.50016582 | 0.89054584 |
| Pde1b | 0.0082923 | 0.00885921 | 0.00887417 | -1.50E-05 | 0.50313296 | 0.89054584 |
| Hoxc5 | 0.01600118 | 0.01710751 | 0.01712373 | -1.62E-05 | 0.5017609 | 0.89054584 |
| Klk1b3 | 0.01210764 | 0.01293875 | 0.01295716 | -1.84E-05 | 0.50264029 | 0.89054584 |
| Ccdc64b | 0.05064985 | 0.05416725 | 0.05419164 | -2.44E-05 | 0.50083673 | 0.89054584 |
| Fbxw15 | 0.00422171 | 0.00449136 | 0.00451797 | -2.66E-05 | 0.51094472 | 0.89054584 |
| Pi16 | 0.0483277 | 0.05167638 | 0.05170826 | -3.19E-05 | 0.50114602 | 0.89054584 |
| Zgrf1 | 0.03897551 | 0.04167301 | 0.04170513 | -3.21E-05 | 0.50143119 | 0.89054584 |
| Fam151b | 0.00843838 | 0.0089961 | 0.0090305 | -3.44E-05 | 0.50707997 | 0.89054584 |
| Angptl6 | 0.08676361 | 0.09274677 | 0.0927834 | -3.66E-05 | 0.50073388 | 0.89054584 |
| Gm20467 | 0.02459385 | 0.02627777 | 0.02631844 | -4.07E-05 | 0.5028724 | 0.89054584 |
| Arhgap27os2 | 0.00848067 | 0.00903468 | 0.00907576 | -4.11E-05 | 0.5084114 | 0.89054584 |
| Cyp4f17 | 0.01432795 | 0.015291 | 0.01533319 | -4.22E-05 | 0.50511387 | 0.89054584 |
| Ttl | 0.09494017 | 0.10146845 | 0.1015111 | -4.26E-05 | 0.50078082 | 0.89054584 |
| Gm26606 | 0.01134522 | 0.01209667 | 0.01214127 | -4.46E-05 | 0.50682663 | 0.89054584 |
| Cybrd1 | 0.0069443 | 0.00738032 | 0.0074316 | -5.13E-05 | 0.51282348 | 0.89054584 |
| Zfp953 | 0.01229406 | 0.01310114 | 0.01315665 | -5.55E-05 | 0.50784066 | 0.89054584 |
| Zfp563 | 0.03898694 | 0.04165492 | 0.04171735 | -6.24E-05 | 0.50278158 | 0.89054584 |
| Jph4 | 0.09839874 | 0.10513815 | 0.10520142 | -6.33E-05 | 0.50111791 | 0.89054584 |
| Elfn1 | 0.00741553 | 0.00787127 | 0.00793588 | -6.46E-05 | 0.51512853 | 0.89054584 |
| 4930518I15R | 0.03032404 | 0.03238257 | 0.03244955 | -6.70E-05 | 0.50383639 | 0.89054584 |
| Zfp799 | 0.11697637 | 0.12464753 | 0.12472198 | -7.45E-05 | 0.50110954 | 0.89054584 |

| | | | | | | |
|---|---|---|---|---|---|---|
| Tbc1d15 | 0.31103862 | 0.31983743 | 0.31991976 | -8.23E-05 | 0.50047835 | 0.89054584 |
| Slit1 | 0.11711965 | 0.12478829 | 0.12487185 | -8.36E-05 | 0.50124375 | 0.89054584 |
| Prelid2 | 0.15877098 | 0.16802338 | 0.16811106 | -8.77E-05 | 0.5009694 | 0.89054584 |
| Hes7 | 0.01530009 | 0.01628432 | 0.01637349 | -8.92E-05 | 0.51012086 | 0.89054584 |
| Prdm10 | 0.06033269 | 0.06445437 | 0.06454469 | -9.03E-05 | 0.50260077 | 0.89054584 |
| Ccl17 | 0.01258065 | 0.0133685 | 0.01346334 | -9.48E-05 | 0.51309055 | 0.89054584 |
| Pmm2 | 0.38250828 | 0.38720052 | 0.38729858 | -9.81E-05 | 0.50047057 | 0.89054584 |
| RP23-455C1: | 0.01118445 | 0.01186686 | 0.01196921 | -0.0001024 | 0.51588993 | 0.89054584 |
| Myt1l | 0.11956761 | 0.12732449 | 0.12743133 | -0.0001068 | 0.5015584 | 0.89054584 |
| Tbc1d32 | 0.06435986 | 0.06874263 | 0.06884952 | -0.0001069 | 0.50288554 | 0.89054584 |
| Lrp8os2 | 0.01931934 | 0.0205635 | 0.02067448 | -0.000111 | 0.50997673 | 0.89054584 |
| Trim12c | 0.11566505 | 0.12323893 | 0.12334993 | -0.000111 | 0.50167244 | 0.89054584 |
| C77370 | 0.04458014 | 0.04758273 | 0.04770015 | -0.0001174 | 0.50457556 | 0.89054584 |
| Kank1 | 0.11522219 | 0.12276884 | 0.12288641 | -0.0001176 | 0.50177829 | 0.89054584 |
| Stk32c | 0.02109168 | 0.02245278 | 0.022571 | -0.0001182 | 0.50973444 | 0.89054584 |
| Prr16 | 0.01624602 | 0.01726705 | 0.01738574 | -0.0001187 | 0.51268675 | 0.89054584 |
| Dynlt1b | 0.01681883 | 0.01787964 | 0.0179987 | -0.0001191 | 0.51229339 | 0.89054584 |
| L1cam | 0.02026035 | 0.02156159 | 0.02168143 | -0.0001198 | 0.51027216 | 0.89054584 |
| Asxl2 | 0.48764443 | 0.48112956 | 0.48125706 | -0.0001275 | 0.50049243 | 0.89054584 |
| D130051D11 | 0.02850816 | 0.03037633 | 0.03050668 | -0.0001304 | 0.50794123 | 0.89054584 |
| Gm28875 | 0.03647501 | 0.0388897 | 0.03903019 | -0.0001405 | 0.50669007 | 0.89054584 |
| Clic5 | 0.01976322 | 0.02100879 | 0.02114946 | -0.0001407 | 0.5123599 | 0.89054584 |
| Usp30 | 0.15502008 | 0.16410145 | 0.1642446 | -0.0001432 | 0.50161994 | 0.89054584 |
| Zbtb42 | 0.01143652 | 0.01209037 | 0.01223897 | -0.0001486 | 0.52255473 | 0.89054584 |
| 1500015O10 | 0.02405248 | 0.0255857 | 0.02573917 | -0.0001535 | 0.51108088 | 0.89054584 |
| Clasp2 | 0.05466577 | 0.05832777 | 0.05848595 | -0.0001582 | 0.50502679 | 0.89054584 |
| Gm16223 | 0.02353962 | 0.02502965 | 0.0251904 | -0.0001608 | 0.51185941 | 0.89054584 |
| Radil | 0.0351885 | 0.03748276 | 0.03765388 | -0.0001711 | 0.50844629 | 0.89054584 |
| Zfp956 | 0.04011106 | 0.04274778 | 0.04291985 | -0.0001721 | 0.5074508 | 0.89054584 |
| Ifitm10 | 0.0081051 | 0.00849675 | 0.00867384 | -0.0001771 | 0.5378882 | 0.89054584 |

| | | | | | | |
|---|---|---|---|---|---|---|
| RP23-299D2. | 0.03993735 | 0.04255523 | 0.04273402 | -0.0001788 | 0.50777575 | 0.89054584 |
| Sema4f | 0.00406655 | 0.00416729 | 0.00435192 | -0.0001846 | 0.57834313 | 0.89054584 |
| Rasgef1a | 0.00406655 | 0.00416729 | 0.00435192 | -0.0001846 | 0.57834313 | 0.89054584 |
| Fam228b | 0.0299307 | 0.03184371 | 0.0320287 | -0.000185 | 0.51073402 | 0.89054584 |
| 1700042O10 | 0.01132015 | 0.01192852 | 0.01211443 | -0.0001859 | 0.52849859 | 0.89054584 |
| Cyth1 | 0.23257785 | 0.24281101 | 0.24300103 | -0.00019 | 0.50145339 | 0.89054584 |
| Daglb | 0.16162388 | 0.17085518 | 0.17104806 | -0.0001929 | 0.50209589 | 0.89054584 |
| Zfp521 | 0.08376217 | 0.08937621 | 0.08957858 | -0.0002024 | 0.50419886 | 0.89054584 |
| Abca9 | 0.01155237 | 0.01215625 | 0.01236294 | -0.0002067 | 0.53104301 | 0.89054584 |
| Arhgef3 | 0.05770363 | 0.06152321 | 0.061734 | -0.0002108 | 0.50634611 | 0.89054584 |
| Ly9 | 0.01240602 | 0.01305857 | 0.01327646 | -0.0002179 | 0.53047405 | 0.89054584 |
| Gm21975 | 0.01240602 | 0.01305857 | 0.01327646 | -0.0002179 | 0.53047405 | 0.89054584 |
| Slc22a2 | 0.00831116 | 0.00867592 | 0.00889435 | -0.0002184 | 0.54554707 | 0.89054584 |
| Kctd17 | 0.01193607 | 0.01254345 | 0.01277356 | -0.0002301 | 0.5334434 | 0.89054584 |
| Hoxd9 | 0.0079015 | 0.00821754 | 0.00845595 | -0.0002384 | 0.55225321 | 0.89054584 |
| 1700047I17R | 0.04630278 | 0.04930039 | 0.04954262 | -0.0002422 | 0.50908656 | 0.89054584 |
| Nfkbid | 0.0146317 | 0.01541489 | 0.01565824 | -0.0002433 | 0.52885977 | 0.89054584 |
| P4ha3 | 0.08977453 | 0.09575199 | 0.09599777 | -0.0002458 | 0.50475868 | 0.89054584 |
| Rrad | 0.02053417 | 0.0217237 | 0.02197443 | -0.0002507 | 0.52119708 | 0.89054584 |
| Adck4 | 0.19289005 | 0.20276306 | 0.20301795 | -0.0002549 | 0.50233348 | 0.89054584 |
| Mipol1 | 0.11577976 | 0.1232139 | 0.12346998 | -0.0002561 | 0.50385477 | 0.89054584 |
| Vash1 | 0.0240257 | 0.025454 | 0.02571052 | -0.0002565 | 0.51853739 | 0.89054584 |
| Tinag | 0.00674158 | 0.00695773 | 0.00721465 | -0.0002569 | 0.56588621 | 0.89054584 |
| Iqcd | 0.00981726 | 0.01024591 | 0.01050612 | -0.0002602 | 0.54593184 | 0.89054584 |
| Fanci | 0.01202818 | 0.01261014 | 0.01287213 | -0.000262 | 0.53777266 | 0.89054584 |
| Mb | 0.01255924 | 0.01317754 | 0.01344044 | -0.0002629 | 0.53630578 | 0.89054584 |
| 4921507P07I | 0.00783056 | 0.00811558 | 0.00838004 | -0.0002645 | 0.5584439 | 0.89054584 |
| Wdr11 | 0.18026622 | 0.18989124 | 0.19015815 | -0.0002669 | 0.50260881 | 0.89054584 |
| Gm26947 | 0.01235357 | 0.0129515 | 0.01322033 | -0.0002688 | 0.53773879 | 0.89054584 |
| D2Wsu81e | 0.37272061 | 0.37795933 | 0.37823202 | -0.0002727 | 0.50134 | 0.89054584 |

| | | | | | | |
|---|---|---|---|---|---|---|
| Nrg1 | 0.03069795 | 0.03257422 | 0.0328496 | -0.0002754 | 0.51557711 | 0.89054584 |
| Sp140 | 0.01936882 | 0.02045158 | 0.02072742 | -0.0002758 | 0.52471954 | 0.89054584 |
| Zfp282 | 0.26353581 | 0.27341949 | 0.27370014 | -0.0002806 | 0.50190583 | 0.89054584 |
| Amy2a3 | 0.00694829 | 0.00715361 | 0.00743586 | -0.0002823 | 0.57018468 | 0.89054584 |
| Nr5a2 | 0.0969552 | 0.10337151 | 0.10366125 | -0.0002897 | 0.50519487 | 0.89054584 |
| Grin1 | 0.00821735 | 0.00848979 | 0.00879397 | -0.0003042 | 0.56401199 | 0.89054584 |
| Ppp1r3d | 0.03079237 | 0.03264398 | 0.03295062 | -0.0003066 | 0.51729166 | 0.89054584 |
| Rbm41 | 0.0234405 | 0.02477379 | 0.02508434 | -0.0003106 | 0.52299803 | 0.89054584 |
| Hddc2 | 0.31823153 | 0.32651107 | 0.32682176 | -0.0003107 | 0.50176687 | 0.89054584 |
| Vsig8 | 0.00393109 | 0.00389428 | 0.00420695 | -0.0003127 | 0.63542982 | 0.89054584 |
| Apod | 0.011472 | 0.01196315 | 0.01227693 | -0.0003138 | 0.5473922 | 0.89054584 |
| Ptpn7 | 0.01200642 | 0.01253333 | 0.01284884 | -0.0003155 | 0.54554115 | 0.89054584 |
| Klhdc1 | 0.10263472 | 0.10936518 | 0.10968162 | -0.0003164 | 0.50536205 | 0.89054584 |
| C1qtnf1 | 0.0154102 | 0.016169 | 0.01649132 | -0.0003223 | 0.53627682 | 0.89054584 |
| Gm26721 | 0.01236038 | 0.01290336 | 0.01322763 | -0.0003243 | 0.54546451 | 0.89054584 |
| Ryr3 | 0.01472353 | 0.01543044 | 0.0157565 | -0.0003261 | 0.53840216 | 0.89054584 |
| Plekhg6 | 0.02748915 | 0.0290903 | 0.02941638 | -0.0003261 | 0.52059374 | 0.89054584 |
| Srrm4os | 0.00818757 | 0.00842594 | 0.00876209 | -0.0003362 | 0.57092838 | 0.89054584 |
| Eya3 | 0.14681201 | 0.15542406 | 0.15576473 | -0.0003407 | 0.50406488 | 0.89054584 |
| Lrp2bp | 0.01545407 | 0.01619523 | 0.01653827 | -0.000343 | 0.53849177 | 0.89054584 |
| Mlkl | 0.03201549 | 0.03391041 | 0.03425923 | -0.0003488 | 0.51891702 | 0.89054584 |
| Diexf | 0.09256847 | 0.09862789 | 0.09897998 | -0.0003521 | 0.50661136 | 0.89054584 |
| Ksr1 | 0.01804977 | 0.01896297 | 0.01931593 | -0.000353 | 0.53392266 | 0.89054584 |
| Slc10a6 | 0.01663382 | 0.01744758 | 0.01780072 | -0.0003531 | 0.53682119 | 0.89054584 |
| Pabpc4l | 0.06896287 | 0.07341265 | 0.07376901 | -0.0003564 | 0.50897787 | 0.89054584 |
| Zfp941 | 0.02437239 | 0.02572457 | 0.02608148 | -0.0003569 | 0.52541784 | 0.89054584 |
| Tdo2 | 0.04147321 | 0.04401254 | 0.04437691 | -0.0003644 | 0.51525722 | 0.89054584 |
| Tpbgl | 0.01460757 | 0.01526581 | 0.01563241 | -0.0003666 | 0.54350146 | 0.89054584 |
| Gm4737 | 0.00813632 | 0.0083353 | 0.00870725 | -0.0003719 | 0.57887491 | 0.89054584 |
| Mapk3 | 0.89500257 | 0.77676388 | 0.77713969 | -0.0003758 | 0.5008988 | 0.89054584 |

| | | | | | | |
|---|---|---|---|---|---|---|
| Tmlhe | 0.01947364 | 0.02046366 | 0.02083959 | -0.0003759 | 0.5334889 | 0.89054584 |
| Gm7628 | 0.0118444 | 0.01229454 | 0.01267546 | -0.0003809 | 0.55567303 | 0.89054584 |
| Zfp874b | 0.03127912 | 0.03308892 | 0.0334714 | -0.0003825 | 0.52122882 | 0.89054584 |
| Serpina1a | 0.03378309 | 0.035765 | 0.03615033 | -0.0003853 | 0.51980321 | 0.89054584 |
| Islr2 | 0.0038493 | 0.00373391 | 0.00411942 | -0.0003855 | 0.66858288 | 0.89054584 |
| Sugp2 | 0.14059762 | 0.14893852 | 0.14932705 | -0.0003885 | 0.5048359 | 0.89054584 |
| Slc9a9 | 0.03016136 | 0.0318853 | 0.0322755 | -0.0003902 | 0.52245868 | 0.89054584 |
| Actn4 | 0.99421901 | 0.83112688 | 0.83151798 | -0.0003911 | 0.5008742 | 0.89054584 |
| 9930012K11I | 0.02795808 | 0.02952495 | 0.02991812 | -0.0003932 | 0.5244097 | 0.89054584 |
| Zfp160 | 0.1511617 | 0.15986669 | 0.16026163 | -0.0003949 | 0.50458032 | 0.89054584 |
| Mettl4 | 0.036722 | 0.03889825 | 0.03929442 | -0.0003962 | 0.51873186 | 0.89054584 |
| 1700001O22 | 0.02534045 | 0.02672086 | 0.02711731 | -0.0003965 | 0.52715268 | 0.89054584 |
| Pld5 | 0.00383558 | 0.00370735 | 0.00410474 | -0.0003974 | 0.67402228 | 0.89054584 |
| Cdh8 | 0.01058417 | 0.01092183 | 0.01132682 | -0.000405 | 0.56615032 | 0.89054584 |
| 5031425F14I | 0.00812898 | 0.00829399 | 0.0086994 | -0.0004054 | 0.58594102 | 0.89054584 |
| Etfdh | 0.3252487 | 0.33311848 | 0.33352935 | -0.0004109 | 0.50228959 | 0.89054584 |
| Gabra4 | 0.01167107 | 0.0120778 | 0.01248997 | -0.0004122 | 0.56109363 | 0.89054584 |
| Ctf2 | 0.00793279 | 0.00807615 | 0.00848944 | -0.0004133 | 0.58971435 | 0.89054584 |
| Gabrr2 | 0.01225364 | 0.01269765 | 0.0131134 | -0.0004157 | 0.55871322 | 0.89054584 |
| Lgals2 | 0.03986055 | 0.04222359 | 0.04265187 | -0.0004283 | 0.51865636 | 0.89054584 |
| Vac14 | 0.07344352 | 0.07812753 | 0.07855676 | -0.0004292 | 0.51015448 | 0.89054584 |
| Aagab | 0.27158905 | 0.28118256 | 0.28161297 | -0.0004304 | 0.5028407 | 0.89054584 |
| Kcnn1 | 0.0196855 | 0.02063449 | 0.0210663 | -0.0004318 | 0.53804039 | 0.89054584 |
| Oplah | 0.02700059 | 0.02846163 | 0.02889364 | -0.000432 | 0.52776754 | 0.89054584 |
| Aqp7 | 0.01018691 | 0.01046627 | 0.0109017 | -0.0004354 | 0.57381074 | 0.89054584 |
| Mxra7 | 0.26795489 | 0.27760835 | 0.27804604 | -0.0004377 | 0.50292574 | 0.89054584 |
| Rilp | 0.00661049 | 0.00663583 | 0.00707437 | -0.0004385 | 0.61363566 | 0.89054584 |
| Mme | 0.02823598 | 0.0297756 | 0.03021545 | -0.0004399 | 0.52703604 | 0.89054584 |
| Gm3550 | 0.07747401 | 0.08241389 | 0.0828626 | -0.0004487 | 0.51006386 | 0.89054584 |
| Trpv2 | 0.01567111 | 0.01631091 | 0.01677053 | -0.0004596 | 0.5508008 | 0.89054584 |

| | | | | | | |
|---|---|---|---|---|---|---|
| Elovl2 | 0.06174814 | 0.06559641 | 0.06605781 | -0.0004614 | 0.51298004 | 0.89054584 |
| Mid2 | 0.06628791 | 0.07044407 | 0.07091026 | -0.0004662 | 0.51221734 | 0.89054584 |
| Rel | 0.04110193 | 0.04351339 | 0.04397977 | -0.0004664 | 0.51970155 | 0.89054584 |
| AA474331 | 0.03856437 | 0.04079835 | 0.04126532 | -0.000467 | 0.52102333 | 0.89054584 |
| Utp14b | 0.08322266 | 0.08853162 | 0.08900245 | -0.0004708 | 0.50983141 | 0.89054584 |
| Cdr2 | 0.05986118 | 0.06355957 | 0.06404063 | -0.0004811 | 0.51395878 | 0.89054584 |
| Git2 | 0.50779823 | 0.49803556 | 0.49851773 | -0.0004822 | 0.50179772 | 0.89054584 |
| Pou6f1 | 0.05707281 | 0.06057288 | 0.06105957 | -0.0004867 | 0.51481128 | 0.89054584 |
| Elovl7 | 0.09749625 | 0.10375101 | 0.10423854 | -0.0004875 | 0.50869238 | 0.89054584 |
| Mamdc4 | 0.0075333 | 0.00756581 | 0.00806192 | -0.0004961 | 0.61282929 | 0.89054584 |
| Gm9917 | 0.01195869 | 0.0123013 | 0.01279776 | -0.0004965 | 0.57171191 | 0.89054584 |
| Plce1 | 0.03893008 | 0.04115966 | 0.04165653 | -0.0004969 | 0.52215823 | 0.89054584 |
| Ftx | 0.08035462 | 0.0854419 | 0.08593948 | -0.0004976 | 0.51076018 | 0.89054584 |
| Zfp526 | 0.02618722 | 0.02752314 | 0.02802335 | -0.0005002 | 0.5331383 | 0.89054584 |
| Alg9 | 0.13280702 | 0.14073019 | 0.14123527 | -0.0005051 | 0.50664656 | 0.89054584 |
| Gpd1 | 0.11194724 | 0.11894609 | 0.11945647 | -0.0005104 | 0.50794047 | 0.89054584 |
| Dclre1b | 0.07538516 | 0.08011864 | 0.08063116 | -0.0005125 | 0.5118125 | 0.89054584 |
| Spag5 | 0.03318633 | 0.03499665 | 0.03551188 | -0.0005152 | 0.52694587 | 0.89054584 |
| 2810414N06 | 0.01220206 | 0.01254094 | 0.01305821 | -0.0005173 | 0.57321025 | 0.89054584 |
| Ccdc57 | 0.04150867 | 0.04389316 | 0.04441485 | -0.0005217 | 0.52182055 | 0.89054584 |
| Tnnc1 | 0.02309552 | 0.02419274 | 0.0247152 | -0.0005225 | 0.53922752 | 0.89054584 |
| Cdhr4 | 0.00624717 | 0.00616022 | 0.00668555 | -0.0005253 | 0.64284986 | 0.89054584 |
| 1110006O24 | 0.07207517 | 0.07656505 | 0.07709472 | -0.0005297 | 0.51276749 | 0.89054584 |
| Dhx35 | 0.05500858 | 0.05832103 | 0.0588525 | -0.0005315 | 0.51677955 | 0.89054584 |
| Hyal3 | 0.02682965 | 0.02817872 | 0.02871074 | -0.000532 | 0.53439832 | 0.89054584 |
| Fam120c | 0.01672012 | 0.01736063 | 0.01789308 | -0.0005324 | 0.55513126 | 0.89054584 |
| BC030499 | 0.01076256 | 0.01098488 | 0.01151774 | -0.0005329 | 0.58532702 | 0.89054584 |
| Gtse1 | 0.02347827 | 0.02458506 | 0.02512475 | -0.0005397 | 0.53985817 | 0.89054584 |
| Ascl2 | 0.01115537 | 0.01139163 | 0.01193809 | -0.0005465 | 0.58443784 | 0.89054584 |
| Zdhhc15 | 0.05636901 | 0.05975668 | 0.06030708 | -0.0005504 | 0.51695816 | 0.89054584 |

| | | | | | | |
|---|---|---|---|---|---|---|
| Baiap2l2 | 0.00786526 | 0.00786527 | 0.00841717 | -0.0005519 | 0.61999902 | 0.89054584 |
| Ap1g2 | 0.05920289 | 0.06278494 | 0.06333687 | -0.0005519 | 0.51619214 | 0.89054584 |
| Usp16 | 0.55576651 | 0.53802435 | 0.53857674 | -0.0005524 | 0.50190632 | 0.89054584 |
| Erich6 | 0.00364178 | 0.00334217 | 0.00389734 | -0.0005552 | 0.74654712 | 0.89054584 |
| Nlrx1 | 0.04920715 | 0.05208698 | 0.0526488 | -0.0005618 | 0.51982565 | 0.89054584 |
| Nrip2 | 0.04074943 | 0.04303507 | 0.0436027 | -0.0005676 | 0.5241816 | 0.89054584 |
| Pik3ap1 | 0.00957912 | 0.00968305 | 0.01025127 | -0.0005682 | 0.60188936 | 0.89054584 |
| 4933427D14 | 0.08824513 | 0.09379665 | 0.09436511 | -0.0005685 | 0.511195 | 0.89054584 |
| Cacna2d4 | 0.01633907 | 0.01691669 | 0.01748531 | -0.0005686 | 0.56021205 | 0.89054584 |
| Npw | 0.00362102 | 0.00330417 | 0.00387512 | -0.000571 | 0.75378265 | 0.89054584 |
| Vtcn1 | 0.01747675 | 0.01812728 | 0.01870275 | -0.0005755 | 0.55699402 | 0.89054584 |
| Arhgef6 | 0.01864183 | 0.01937276 | 0.01994949 | -0.0005767 | 0.55357061 | 0.89054584 |
| Nqo1 | 0.12262924 | 0.13005067 | 0.13062927 | -0.0005786 | 0.50823197 | 0.89054584 |
| Gm12689 | 0.01176603 | 0.01200408 | 0.01259159 | -0.0005875 | 0.58604365 | 0.89054584 |
| Gm13205 | 0.01655489 | 0.01712855 | 0.01771627 | -0.0005877 | 0.56141417 | 0.89054584 |
| A730049H05 | 0.00793487 | 0.00790259 | 0.00849166 | -0.0005891 | 0.62672484 | 0.89054584 |
| Hsd17b11 | 0.19262452 | 0.20215717 | 0.20274815 | -0.000591 | 0.50541754 | 0.89054584 |
| Slc2a6 | 0.0077675 | 0.00771573 | 0.00831255 | -0.0005968 | 0.63099989 | 0.89054584 |
| Zfp85os | 0.01497317 | 0.01542364 | 0.01602364 | -0.0006 | 0.56924574 | 0.89054584 |
| Gm11508 | 0.01542048 | 0.01590092 | 0.01650232 | -0.0006014 | 0.56741135 | 0.89054584 |
| Lrrn4cl | 0.03779566 | 0.03984156 | 0.04044299 | -0.0006014 | 0.52761809 | 0.89054584 |
| Corin | 0.01012612 | 0.01023325 | 0.01083665 | -0.0006034 | 0.6023422 | 0.89054584 |
| Elk4 | 0.18813475 | 0.19757705 | 0.19818168 | -0.0006046 | 0.50567038 | 0.89054584 |
| Ces2e | 0.01191534 | 0.01214501 | 0.01275137 | -0.0006064 | 0.58766539 | 0.89054584 |
| Pi4kb | 0.10415982 | 0.11067779 | 0.1112846 | -0.0006068 | 0.51013371 | 0.89054584 |
| Hoxd8 | 0.01606366 | 0.01658329 | 0.01719059 | -0.0006073 | 0.56536684 | 0.89054584 |
| 1700105P06I | 0.01653355 | 0.01706911 | 0.01769343 | -0.0006243 | 0.56528897 | 0.89054584 |
| Tnfaip2 | 0.01194114 | 0.01215105 | 0.01277898 | -0.0006279 | 0.59053846 | 0.89054584 |
| P2ry12 | 0.00352631 | 0.00313358 | 0.00377376 | -0.0006402 | 0.78533652 | 0.89054584 |
| Parp14 | 0.01598016 | 0.01645869 | 0.01710124 | -0.0006426 | 0.569481 | 0.89054584 |

| | | | | | | |
|---|---|---|---|---|---|---|
| Gm15873 | 0.00592573 | 0.00569849 | 0.00634156 | -0.0006431 | 0.68169276 | 0.89054584 |
| Thumpd1 | 0.55189774 | 0.53475303 | 0.53539993 | -0.0006469 | 0.50224574 | 0.89054584 |
| Fancd2 | 0.02882403 | 0.03019714 | 0.03084464 | -0.0006475 | 0.53895524 | 0.89054584 |
| Gm43579 | 0.00777763 | 0.00767059 | 0.00832339 | -0.0006528 | 0.64259425 | 0.89054584 |
| Mterf1b | 0.00784012 | 0.00773535 | 0.00839026 | -0.0006549 | 0.64194357 | 0.89054584 |
| Timm8a1 | 0.04603271 | 0.0485986 | 0.04925376 | -0.0006552 | 0.52470741 | 0.89054584 |
| Impdh1 | 0.35062119 | 0.35691226 | 0.35756991 | -0.0006576 | 0.50341839 | 0.89054584 |
| 5730507C01I | 0.0117086 | 0.0118682 | 0.01253013 | -0.0006619 | 0.59720455 | 0.89054584 |
| Pdzrn4 | 0.05936527 | 0.06284659 | 0.06351047 | -0.0006639 | 0.51942097 | 0.89054584 |
| Slc26a9 | 0.00348436 | 0.00305948 | 0.00372888 | -0.0006694 | 0.79852389 | 0.89054584 |
| Rtn2 | 0.16143374 | 0.17018157 | 0.17085241 | -0.0006708 | 0.5072975 | 0.89054584 |
| Cdh16 | 0.01161838 | 0.01175736 | 0.01243359 | -0.0006762 | 0.60001503 | 0.89054584 |
| Hectd1 | 0.78085479 | 0.70530296 | 0.70598657 | -0.0006836 | 0.50179972 | 0.89054584 |
| Tbl2 | 0.17306137 | 0.18210201 | 0.18278941 | -0.0006874 | 0.5069893 | 0.89054584 |
| Gm17056 | 0.01057166 | 0.0106245 | 0.01131344 | -0.0006889 | 0.61168266 | 0.89054584 |
| Ugt8a | 0.04485433 | 0.04730378 | 0.04799342 | -0.0006896 | 0.52668775 | 0.89054584 |
| Usp39 | 0.27038882 | 0.27974102 | 0.28043565 | -0.0006946 | 0.50460374 | 0.89054584 |
| Colgalt2 | 0.00725161 | 0.00705968 | 0.00776047 | -0.0007008 | 0.66301765 | 0.89054584 |
| Pla2g6 | 0.3968286 | 0.39976775 | 0.40046931 | -0.0007016 | 0.50325601 | 0.89054584 |
| Sprn | 0.00343527 | 0.00297388 | 0.00367634 | -0.0007025 | 0.81332531 | 0.89054584 |
| Gp9 | 0.01144096 | 0.01153291 | 0.01224371 | -0.0007108 | 0.60660237 | 0.89054584 |
| Cfap54 | 0.02475713 | 0.02577435 | 0.02649316 | -0.0007188 | 0.55029469 | 0.89054584 |
| Fcgbp | 0.00340607 | 0.00292353 | 0.00364509 | -0.0007216 | 0.82180203 | 0.89054584 |
| Phlda2 | 0.00340607 | 0.00292353 | 0.00364509 | -0.0007216 | 0.82180203 | 0.89054584 |
| Clcn7 | 0.04409843 | 0.04646207 | 0.04718493 | -0.0007229 | 0.52844962 | 0.89054584 |
| Gm11131 | 0.00719609 | 0.00697005 | 0.00770105 | -0.000731 | 0.67084218 | 0.89054584 |
| Pde4c | 0.00338682 | 0.00289058 | 0.00362449 | -0.0007339 | 0.82725475 | 0.8919119 |
| Igsf1 | 0.05058876 | 0.05339046 | 0.0541263 | -0.0007358 | 0.52525135 | 0.89054584 |
| Slc12a8 | 0.00709468 | 0.00685599 | 0.00759253 | -0.0007365 | 0.67435006 | 0.89054584 |
| 6330408A02I | 0.10973758 | 0.11640291 | 0.11713998 | -0.0007371 | 0.5116933 | 0.89054584 |

| | | | | | | |
|---|---|---|---|---|---|---|
| Abca8b | 0.00763887 | 0.00743642 | 0.0081749 | -0.0007385 | 0.66307201 | 0.89054584 |
| Ccdc14 | 0.03869431 | 0.04066404 | 0.04140432 | -0.0007403 | 0.53319309 | 0.89054584 |
| Dgkb | 0.01414193 | 0.01439253 | 0.01513412 | -0.0007416 | 0.59029084 | 0.89054584 |
| Gpr21 | 0.00766162 | 0.0074575 | 0.00819925 | -0.0007417 | 0.66329439 | 0.89054584 |
| Borcs5 | 0.20475145 | 0.21429611 | 0.21503891 | -0.0007428 | 0.50642002 | 0.89054584 |
| Neurl3 | 0.01176854 | 0.01185122 | 0.01259428 | -0.0007431 | 0.60829356 | 0.89054584 |
| Vil1 | 0.06941418 | 0.07350369 | 0.07425129 | -0.0007476 | 0.5187071 | 0.89054584 |
| Gdf6 | 0.01205363 | 0.01215145 | 0.01289936 | -0.0007479 | 0.60646804 | 0.89054584 |
| Hectd2 | 0.05668784 | 0.05989872 | 0.06064797 | -0.0007492 | 0.5229491 | 0.89054584 |
| Sh3rf3 | 0.01090419 | 0.01091431 | 0.0116693 | -0.000755 | 0.61845618 | 0.89054584 |
| Hist1h2ad | 0.00707829 | 0.00681829 | 0.00757498 | -0.0007567 | 0.67917594 | 0.89054584 |
| Mylip | 0.08631094 | 0.09154333 | 0.09230009 | -0.0007568 | 0.51523525 | 0.89054584 |
| Csmd1 | 0.0033462 | 0.00282166 | 0.00358102 | -0.0007594 | 0.83840803 | 0.89497393 |
| Tshz1 | 0.73245616 | 0.67220312 | 0.67297015 | -0.000767 | 0.50211844 | 0.89054584 |
| Gm10101 | 0.00641503 | 0.00609446 | 0.00686519 | -0.0007707 | 0.6995275 | 0.89054584 |
| Pdk2 | 0.18974928 | 0.19905105 | 0.19982478 | -0.0007737 | 0.50719638 | 0.89054584 |
| Gm26541 | 0.01588979 | 0.01622602 | 0.01700454 | -0.0007785 | 0.58445321 | 0.89054584 |
| Nrg4 | 0.00767672 | 0.0074338 | 0.0082154 | -0.0007816 | 0.67120573 | 0.89054584 |
| Pkn3 | 0.01567759 | 0.01599481 | 0.01677746 | -0.0007826 | 0.58602574 | 0.89054584 |
| Pde7b | 0.02058383 | 0.02124091 | 0.02202757 | -0.0007867 | 0.56607199 | 0.89054584 |
| RP23-477O1: | 0.01157956 | 0.01160456 | 0.01239204 | -0.0007875 | 0.61640921 | 0.89054584 |
| Cytip | 0.13468867 | 0.14240268 | 0.14319181 | -0.0007891 | 0.51024194 | 0.89054584 |
| Als2cr12 | 0.00765744 | 0.00739868 | 0.00819477 | -0.0007961 | 0.67458048 | 0.89054584 |
| Nfatc2ip | 0.06406676 | 0.06773825 | 0.06853624 | -0.000798 | 0.52163028 | 0.89054584 |
| Tbc1d30 | 0.05688011 | 0.06005345 | 0.06085354 | -0.0008001 | 0.52442202 | 0.89054584 |
| Foxp2 | 0.06976286 | 0.07382063 | 0.0746239 | -0.0008033 | 0.51999858 | 0.89054584 |
| Rgag4 | 0.16689644 | 0.17566294 | 0.17646735 | -0.0008044 | 0.50847177 | 0.89054584 |
| Slc26a1 | 0.05250353 | 0.05536934 | 0.05617388 | -0.0008045 | 0.5266005 | 0.89054584 |
| 4930427A07 | 0.04120848 | 0.04328464 | 0.04409373 | -0.0008091 | 0.53406374 | 0.89054584 |
| Dixdc1 | 0.1697716 | 0.17860071 | 0.17941773 | -0.000817 | 0.50846316 | 0.89054584 |

| | | | | | | |
|---|---|---|---|---|---|---|
| Tmem154 | 0.01510664 | 0.01534638 | 0.01616648 | -0.0008201 | 0.59341597 | 0.89054584 |
| AW011738 | 0.0187832 | 0.01927971 | 0.02010077 | -0.0008211 | 0.57546451 | 0.89054584 |
| E130114P18I | 0.00765309 | 0.00736767 | 0.00819012 | -0.0008224 | 0.68005306 | 0.89054584 |
| Tyro3 | 0.01116073 | 0.0111191 | 0.01194383 | -0.0008247 | 0.62616172 | 0.89054584 |
| Magee2 | 0.0156733 | 0.01594506 | 0.01677286 | -0.0008278 | 0.59092897 | 0.89054584 |
| Bst1 | 0.00840097 | 0.0081622 | 0.00899046 | -0.0008283 | 0.66611618 | 0.89054584 |
| Ptcra | 0.03693517 | 0.03869395 | 0.03952246 | -0.0008285 | 0.5389012 | 0.89054584 |
| Tcte2 | 0.02552381 | 0.02648132 | 0.02731351 | -0.0008322 | 0.55643933 | 0.89054584 |
| Gm4353 | 0.01144818 | 0.01141895 | 0.01225145 | -0.0008325 | 0.62421845 | 0.89054584 |
| P2ry6 | 0.00760301 | 0.00730255 | 0.00813652 | -0.000834 | 0.68350631 | 0.89054584 |
| Ppp1r12b | 0.1373671 | 0.14513763 | 0.1459745 | -0.0008369 | 0.51065427 | 0.89054584 |
| Acsbg1 | 0.00321228 | 0.00260032 | 0.0034377 | -0.0008374 | 0.87178264 | 0.90667896 |
| 1700066B19I | 0.00321228 | 0.00260032 | 0.0034377 | -0.0008374 | 0.87178264 | 0.90667896 |
| Fam107a | 0.0032077 | 0.00259291 | 0.0034328 | -0.0008399 | 0.872832 | 0.90729956 |
| Dpf1 | 0.02143681 | 0.02209908 | 0.02294031 | -0.0008412 | 0.56782674 | 0.89054584 |
| Shisa4 | 0.22778231 | 0.23736513 | 0.23820668 | -0.0008415 | 0.50656603 | 0.89054584 |
| Nuak1 | 0.03074058 | 0.03205131 | 0.03289521 | -0.0008439 | 0.54756885 | 0.89054584 |
| Klf8 | 0.01104604 | 0.01097662 | 0.01182109 | -0.0008445 | 0.63036613 | 0.89054584 |
| Pfkfb3 | 0.02905017 | 0.0302396 | 0.0310866 | -0.000847 | 0.55050578 | 0.89054584 |
| Stox1 | 0.09965935 | 0.10569638 | 0.1065463 | -0.0008499 | 0.514823 | 0.89054584 |
| Pirb | 0.00757931 | 0.00726117 | 0.00811116 | -0.00085 | 0.68730438 | 0.89054584 |
| 9930111J21F | 0.00757931 | 0.00726117 | 0.00811116 | -0.00085 | 0.68730438 | 0.89054584 |
| 4930548H24I | 0.01193845 | 0.01192515 | 0.0127761 | -0.0008509 | 0.62183621 | 0.89054584 |
| Zfp296 | 0.00753357 | 0.00721032 | 0.00806221 | -0.0008519 | 0.68874103 | 0.89054584 |
| C130080G10 | 0.01197912 | 0.01196327 | 0.01281963 | -0.0008564 | 0.62218227 | 0.89054584 |
| Itpk1 | 0.06640454 | 0.07017441 | 0.0710349 | -0.0008605 | 0.52250303 | 0.89054584 |
| Syngr3 | 0.00758219 | 0.00725323 | 0.00811424 | -0.000861 | 0.68947655 | 0.89054584 |
| 6720427I07R | 0.04749649 | 0.04995828 | 0.0508193 | -0.000861 | 0.53145799 | 0.89054584 |
| Trim36 | 0.01129168 | 0.01122186 | 0.01208397 | -0.0008621 | 0.63019956 | 0.89054584 |
| Vps36 | 0.55774418 | 0.5393344 | 0.5401969 | -0.0008625 | 0.50296757 | 0.89054584 |

| | | | | | | |
|---|---|---|---|---|---|---|
| Sh3kbp1 | 0.19471371 | 0.20399927 | 0.20487009 | -0.0008708 | 0.50789989 | 0.89054584 |
| Ccdc71l | 0.03646218 | 0.03814493 | 0.03901647 | -0.0008715 | 0.54144306 | 0.89054584 |
| A830080D01 | 0.17065826 | 0.17945468 | 0.1803269 | -0.0008722 | 0.50898939 | 0.89054584 |
| Etohi1 | 0.03467091 | 0.03622752 | 0.03710015 | -0.0008726 | 0.54362996 | 0.89054584 |
| Ankrd31 | 0.00760313 | 0.00726323 | 0.00813665 | -0.0008734 | 0.69150105 | 0.89054584 |
| Il34 | 0.02040807 | 0.02096158 | 0.02183949 | -0.0008779 | 0.57428057 | 0.89054584 |
| Cdyl | 0.13005621 | 0.13749282 | 0.13837252 | -0.0008797 | 0.51181469 | 0.89054584 |
| Rad50 | 0.26973011 | 0.27890789 | 0.27978921 | -0.0008813 | 0.50585445 | 0.89054584 |
| Gm26760 | 0.01057804 | 0.01043454 | 0.01132026 | -0.0008857 | 0.64226764 | 0.89054584 |
| Hunk | 0.03553701 | 0.03713861 | 0.03802672 | -0.0008881 | 0.54332271 | 0.89054584 |
| Cwc22 | 0.25377193 | 0.26317453 | 0.26406565 | -0.0008911 | 0.50627194 | 0.89054584 |
| Lrrc34 | 0.00734202 | 0.00696566 | 0.00785722 | -0.0008916 | 0.70147603 | 0.89054584 |
| Cdkn2b | 0.06258777 | 0.06605721 | 0.06695534 | -0.0008981 | 0.5249155 | 0.89054584 |
| Gm2788 | 0.01351729 | 0.01356654 | 0.01446568 | -0.0008991 | 0.61393316 | 0.89054584 |
| Exosc9 | 0.30056023 | 0.3089193 | 0.30982003 | -0.0009007 | 0.50540342 | 0.89054584 |
| 1700003E16I | 0.01427581 | 0.01437346 | 0.01527739 | -0.0009039 | 0.60859548 | 0.89054584 |
| 2610318N02 | 0.04279359 | 0.04487817 | 0.04578926 | -0.0009111 | 0.53692943 | 0.89054584 |
| Gm17638 | 0.01034308 | 0.01015544 | 0.01106883 | -0.0009134 | 0.64967791 | 0.89054584 |
| Clec1b | 0.0117304 | 0.01163894 | 0.01255346 | -0.0009145 | 0.6328467 | 0.89054584 |
| Atp1a3 | 0.00305814 | 0.00235676 | 0.00327274 | -0.000916 | 0.90387615 | 0.92597487 |
| Slc25a32 | 0.08150395 | 0.08624958 | 0.08716699 | -0.0009174 | 0.51955394 | 0.89054584 |
| Pabpc5 | 0.00698417 | 0.00655456 | 0.00747426 | -0.0009197 | 0.71677212 | 0.89054584 |
| Piwil2 | 0.00303623 | 0.00232311 | 0.00324929 | -0.0009262 | 0.90791023 | 0.92866393 |
| Gm38037 | 0.00591512 | 0.00540195 | 0.0063302 | -0.0009283 | 0.7527536 | 0.89054584 |
| Hsbp1l1 | 0.07021814 | 0.07418201 | 0.07511042 | -0.0009284 | 0.52296123 | 0.89054584 |
| Chek1 | 0.03417358 | 0.03563961 | 0.03656809 | -0.0009285 | 0.54708212 | 0.89054584 |
| Tmem39b | 0.22653767 | 0.23603185 | 0.23696071 | -0.0009289 | 0.50728527 | 0.89054584 |
| 9530059O14 | 0.00745585 | 0.00705012 | 0.00797903 | -0.0009289 | 0.70622455 | 0.89054584 |
| Gm15834 | 0.00754354 | 0.0071431 | 0.00807288 | -0.0009298 | 0.70422187 | 0.89054584 |
| 6720483E21I | 0.01117884 | 0.01102991 | 0.01196321 | -0.0009333 | 0.6418707 | 0.89054584 |

| | | | | | | |
|---|---|---|---|---|---|---|
| Zar1l | 0.02521473 | 0.02604069 | 0.02698279 | -0.0009421 | 0.56460937 | 0.89054584 |
| Nyap2 | 0.00300029 | 0.00226843 | 0.00321083 | -0.0009424 | 0.91425431 | 0.93324361 |
| Mak | 0.01120941 | 0.01105199 | 0.01199592 | -0.0009439 | 0.64304145 | 0.89054584 |
| Lzts3 | 0.01545743 | 0.01559785 | 0.01654186 | -0.000944 | 0.60483333 | 0.89054584 |
| Aloxe3 | 0.00566586 | 0.00511891 | 0.00606345 | -0.0009445 | 0.76600609 | 0.89054584 |
| Cabp2 | 0.00299061 | 0.00225382 | 0.00320047 | -0.0009467 | 0.91590559 | 0.93425758 |
| Fam65c | 0.01984908 | 0.02029212 | 0.02124134 | -0.0009492 | 0.58246268 | 0.89054584 |
| Msh5 | 0.04375891 | 0.04587209 | 0.04682178 | -0.0009497 | 0.53764301 | 0.89054584 |
| Mmp15 | 0.10322496 | 0.109347 | 0.1103021 | -0.0009551 | 0.51608966 | 0.89054584 |
| Gm44257 | 0.01034851 | 0.01011862 | 0.01107464 | -0.000956 | 0.65622499 | 0.89054584 |
| Kif17 | 0.04321747 | 0.04528551 | 0.04624265 | -0.0009571 | 0.53841096 | 0.89054584 |
| P2rx6 | 0.00667334 | 0.0061795 | 0.00714162 | -0.0009621 | 0.73488383 | 0.89054584 |
| Dock9 | 0.12553151 | 0.13269218 | 0.1336575 | -0.0009653 | 0.51342115 | 0.89054584 |
| Trmt1 | 0.32509245 | 0.33241365 | 0.33338028 | -0.0009666 | 0.50538895 | 0.89054584 |
| Pot1b | 0.0214208 | 0.02195324 | 0.02292318 | -0.0009699 | 0.57813782 | 0.89054584 |
| Dnaaf3 | 0.01557259 | 0.01569197 | 0.0166651 | -0.0009731 | 0.60720788 | 0.89054584 |
| Gm13610 | 0.01140623 | 0.01123242 | 0.01220655 | -0.0009741 | 0.64498079 | 0.89054584 |
| Aatk | 0.06455262 | 0.06807842 | 0.06905555 | -0.0009771 | 0.52628059 | 0.89054584 |
| Gpr45 | 0.064976 | 0.06853079 | 0.06950807 | -0.0009773 | 0.52611392 | 0.89054584 |
| Fam171b | 0.08574407 | 0.09071714 | 0.09169483 | -0.0009777 | 0.51980958 | 0.89054584 |
| Il3ra | 0.25630909 | 0.26559512 | 0.26657323 | -0.0009781 | 0.50681942 | 0.89054584 |
| 5033403F01I | 0.00725328 | 0.0067825 | 0.00776225 | -0.0009798 | 0.72175149 | 0.89054584 |
| Gm14085 | 0.00725186 | 0.00677948 | 0.00776073 | -0.0009812 | 0.72209132 | 0.89054584 |
| Gramd4 | 0.01826546 | 0.0185626 | 0.01954674 | -0.0009841 | 0.59272788 | 0.89054584 |
| Slfn8 | 0.00747031 | 0.00701008 | 0.00799451 | -0.0009844 | 0.71691276 | 0.89054584 |
| Sgsm1 | 0.15046615 | 0.15855829 | 0.15954305 | -0.0009848 | 0.51147062 | 0.89054584 |
| RP23-53O7.2 | 0.02601899 | 0.02685731 | 0.02784335 | -0.000986 | 0.56552407 | 0.89054584 |
| Hs3st3b1 | 0.00747009 | 0.00700819 | 0.00799428 | -0.0009861 | 0.71724512 | 0.89054584 |
| Gm7102 | 0.01032137 | 0.01005414 | 0.01104559 | -0.0009915 | 0.66209465 | 0.89054584 |
| Msrb3 | 0.01577081 | 0.01588084 | 0.01687721 | -0.0009964 | 0.60836059 | 0.89054584 |

| | | | | | | |
|---|---|---|---|---|---|---|
| Hilpda | 0.26308796 | 0.27226195 | 0.27325918 | -0.0009972 | 0.50678264 | 0.89054584 |
| Gm28151 | 0.00743581 | 0.00696034 | 0.00795759 | -0.0009972 | 0.72034301 | 0.89054584 |
| Cage1 | 0.03228604 | 0.03354856 | 0.03454869 | -0.0010001 | 0.55364224 | 0.89054584 |
| Csrnp3 | 0.00139637 | 0.00049136 | 0.00149436 | -0.001003 | 0.99911709 | 0.99918601 |
| Kcnq2 | 0.00139637 | 0.00049136 | 0.00149436 | -0.001003 | 0.99911709 | 0.99918601 |
| Cyp2e1 | 0.03562434 | 0.03710983 | 0.03812015 | -0.0010103 | 0.54913562 | 0.89054584 |
| Chd5 | 0.00282726 | 0.00201433 | 0.00302566 | -0.0010113 | 0.94029453 | 0.95171671 |
| 1700001C02I | 0.03045805 | 0.03158076 | 0.03259292 | -0.0010122 | 0.55751877 | 0.89054584 |
| Adamts10 | 0.13300699 | 0.14042909 | 0.14144327 | -0.0010142 | 0.51332438 | 0.89054584 |
| Gm26981 | 0.02170701 | 0.02221392 | 0.02322944 | -0.0010155 | 0.5806957 | 0.89054584 |
| 2210406O10 | 0.00280627 | 0.00198454 | 0.0030032 | -0.0010187 | 0.94297757 | 0.95356691 |
| Fam122b | 0.04392951 | 0.04598533 | 0.04700425 | -0.0010189 | 0.54022206 | 0.89054584 |
| Gm11808 | 1.45926975 | 1.0230095 | 1.02402845 | -0.001019 | 0.50184943 | 0.89054584 |
| Gm15972 | 0.01798038 | 0.01822111 | 0.01924168 | -0.0010206 | 0.59758725 | 0.89054584 |
| Cryba4 | 0.00280056 | 0.00197647 | 0.00299709 | -0.0010206 | 0.94369174 | 0.95402293 |
| Slc2a13 | 0.0027896 | 0.00196104 | 0.00298536 | -0.0010243 | 0.94504062 | 0.9550536 |
| Ebag9 | 0.27313158 | 0.28210045 | 0.28312503 | -0.0010246 | 0.50672577 | 0.89054584 |
| Iqub | 0.00708485 | 0.00655605 | 0.00758201 | -0.001026 | 0.73579068 | 0.89054584 |
| Gm7879 | 0.06782944 | 0.07153071 | 0.07255775 | -0.001027 | 0.52628975 | 0.89054584 |
| Plch2 | 0.02388909 | 0.02453715 | 0.02556434 | -0.0010272 | 0.57424713 | 0.89054584 |
| Zxdb | 0.06442656 | 0.06788466 | 0.06892082 | -0.0010362 | 0.52792019 | 0.89054584 |
| Nkx6-3 | 0.01345751 | 0.01336535 | 0.0144017 | -0.0010363 | 0.63128418 | 0.89054584 |
| Nrn1 | 0.01467683 | 0.01466937 | 0.01570652 | -0.0010372 | 0.6208241 | 0.89054584 |
| Il17rd | 0.0300307 | 0.03109604 | 0.0321357 | -0.0010397 | 0.55990445 | 0.89054584 |
| Csad | 0.33400126 | 0.34081987 | 0.34185973 | -0.0010399 | 0.50565341 | 0.89054584 |
| Elmsan1 | 0.05248101 | 0.05510675 | 0.0561498 | -0.001043 | 0.53448344 | 0.89054584 |
| Amhr2 | 0.00735255 | 0.00681728 | 0.00786849 | -0.0010512 | 0.73316758 | 0.89054584 |
| Gm5580 | 0.0015441 | 0.00060083 | 0.00165245 | -0.0010516 | 0.99848642 | 0.99876198 |
| Mapk12 | 0.01964575 | 0.01997086 | 0.02102376 | -0.0010529 | 0.59224625 | 0.89054584 |
| G6b | 0.00269132 | 0.00182529 | 0.00288019 | -0.0010549 | 0.95603129 | 0.96387633 |

| | | | | | | |
|---|---|---|---|---|---|---|
| Gm15445 | 0.01406449 | 0.01399512 | 0.01505125 | -0.0010561 | 0.62813288 | 0.89054584 |
| Mfsd7a | 0.00267208 | 0.00179929 | 0.0028596 | -0.0010603 | 0.95795978 | 0.96541783 |
| 1010001B22 | 0.01526129 | 0.01527086 | 0.01633197 | -0.0010611 | 0.61893956 | 0.89054584 |
| Lrrc4 | 0.00266842 | 0.00179436 | 0.00285567 | -0.0010613 | 0.9583193 | 0.96571302 |
| Btbd16 | 0.00684629 | 0.00626489 | 0.00732671 | -0.0010618 | 0.75022464 | 0.89054584 |
| Pqlc3 | 0.02339176 | 0.02396923 | 0.02503219 | -0.001063 | 0.57841308 | 0.89054584 |
| Gfod1 | 0.04593582 | 0.0480849 | 0.04915014 | -0.0010652 | 0.54021441 | 0.89054584 |
| C920006O11 | 0.03599755 | 0.03745352 | 0.03851941 | -0.0010659 | 0.55128937 | 0.89054584 |
| Zfp947 | 0.01900417 | 0.01926947 | 0.02033723 | -0.0010678 | 0.59661933 | 0.89054584 |
| Cpa6 | 0.01169714 | 0.0114491 | 0.01251787 | -0.0010688 | 0.65460288 | 0.89054584 |
| Arpp21 | 0.00692305 | 0.00633323 | 0.00740885 | -0.0010756 | 0.75060239 | 0.89054584 |
| Tmem28 | 0.0025671 | 0.00166068 | 0.00274725 | -0.0010866 | 0.96731054 | 0.97301521 |
| Mrps36 | 1.33668257 | 0.98291801 | 0.98400651 | -0.0010885 | 0.50205602 | 0.89054584 |
| Gm807 | 0.00729606 | 0.00671907 | 0.00780803 | -0.001089 | 0.74207891 | 0.89054584 |
| Pde8b | 0.00255362 | 0.00164329 | 0.00273282 | -0.0010895 | 0.9683764 | 0.9736145 |
| Akap5 | 0.01417344 | 0.0140777 | 0.01516783 | -0.0010901 | 0.63112888 | 0.89054584 |
| Atp6v0c | 0.00170311 | 0.00073095 | 0.00182263 | -0.0010917 | 0.99736877 | 0.99778171 |
| Snph | 0.01430043 | 0.01421179 | 0.01530373 | -0.0010919 | 0.63021454 | 0.89054584 |
| Bdh2 | 0.03131438 | 0.03241609 | 0.03350912 | -0.001093 | 0.56039489 | 0.89054584 |
| Xpnpep2 | 0.01543461 | 0.01542397 | 0.01651744 | -0.0010935 | 0.62112157 | 0.89054584 |
| Dcaf12l2 | 0.01931859 | 0.01957929 | 0.02067368 | -0.0010944 | 0.59740119 | 0.89054584 |
| Edaradd | 0.01968411 | 0.01996909 | 0.02106482 | -0.0010957 | 0.59574313 | 0.89054584 |
| Me3 | 0.02698279 | 0.02777565 | 0.02887459 | -0.0010989 | 0.57037 | 0.89054584 |
| Efhd1 | 0.01241172 | 0.0121826 | 0.01328256 | -0.0011 | 0.6501848 | 0.89054584 |
| Rab17 | 0.00726975 | 0.00667465 | 0.00777988 | -0.0011052 | 0.74597024 | 0.89054584 |
| A830019P07 | 0.01053829 | 0.0101721 | 0.01127773 | -0.0011056 | 0.6760739 | 0.89054584 |
| Padi4 | 0.0024666 | 0.0015332 | 0.0026397 | -0.0011065 | 0.97458511 | 0.97870298 |
| Zmym6 | 0.05915974 | 0.06218225 | 0.06329074 | -0.0011085 | 0.5325166 | 0.89054584 |
| Dok3 | 0.0142393 | 0.01412735 | 0.01523832 | -0.001111 | 0.63294599 | 0.89054584 |
| Zfp455 | 0.01095347 | 0.01060754 | 0.01172203 | -0.0011145 | 0.67110095 | 0.89054584 |

| | | | | | | |
|---|---|---|---|---|---|---|
| Kif5a | 0.00729115 | 0.00668723 | 0.00780278 | -0.0011155 | 0.74731901 | 0.89054584 |
| Gm9885 | 0.01026424 | 0.00986838 | 0.01098446 | -0.0011161 | 0.68202645 | 0.89054584 |
| Gm2415 | 0.04272363 | 0.04459727 | 0.04571442 | -0.0011172 | 0.54532315 | 0.89054584 |
| Hrk | 0.0018519 | 0.00086424 | 0.00198186 | -0.0011176 | 0.99569651 | 0.99659013 |
| Cfap46 | 0.01059695 | 0.01022207 | 0.0113405 | -0.0011184 | 0.6770538 | 0.89054584 |
| Adam4 | 0.00185874 | 0.00087064 | 0.00198918 | -0.0011185 | 0.99560065 | 0.99656299 |
| Chst7 | 0.02300922 | 0.02350375 | 0.02462286 | -0.0011191 | 0.58384824 | 0.89054584 |
| Bend3 | 0.05264161 | 0.05519963 | 0.05632154 | -0.0011219 | 0.53697087 | 0.89054584 |
| Tbccd1 | 0.15425132 | 0.16232729 | 0.16345147 | -0.0011242 | 0.5127811 | 0.89054584 |
| Ostm1 | 0.13539102 | 0.1427969 | 0.14392177 | -0.0011249 | 0.5145237 | 0.89054584 |
| Slc35e2 | 0.09308147 | 0.09840124 | 0.0995275 | -0.0011263 | 0.52102273 | 0.89054584 |
| Lamp3 | 0.00231599 | 0.00135167 | 0.00247851 | -0.0011268 | 0.98291885 | 0.98584279 |
| Ccdc78 | 0.01111317 | 0.01076458 | 0.01189294 | -0.0011284 | 0.67076411 | 0.89054584 |
| Krt79 | 0.00227929 | 0.00130918 | 0.00243924 | -0.0011301 | 0.98455206 | 0.98700289 |
| Prox2 | 0.01563462 | 0.01560074 | 0.01673147 | -0.0011307 | 0.62356411 | 0.89054584 |
| Slc45a1 | 0.01164934 | 0.0113351 | 0.01246671 | -0.0011316 | 0.66381334 | 0.89054584 |
| Gm12474 | 0.00225627 | 0.00128287 | 0.0024146 | -0.0011317 | 0.98550653 | 0.98775483 |
| Pold1 | 0.11312313 | 0.11955666 | 0.12068847 | -0.0011318 | 0.51742465 | 0.89054584 |
| Gm10645 | 0.00200327 | 0.0010113 | 0.00214385 | -0.0011326 | 0.99307669 | 0.99479209 |
| Gm16432 | 0.01069788 | 0.01031587 | 0.01144851 | -0.0011326 | 0.67757329 | 0.89054584 |
| 1810041L15F | 0.17784612 | 0.18655127 | 0.18768526 | -0.001134 | 0.51122846 | 0.89054584 |
| Adgrg2 | 0.01455093 | 0.01443699 | 0.0155718 | -0.0011348 | 0.63289315 | 0.89054584 |
| Gabrg3 | 0.00212962 | 0.00114289 | 0.00227907 | -0.0011362 | 0.98989981 | 0.99188385 |
| Tssk4 | 0.00212962 | 0.00114289 | 0.00227907 | -0.0011362 | 0.98989981 | 0.99188385 |
| Gm5784 | 0.01546518 | 0.01541384 | 0.01655016 | -0.0011363 | 0.6254695 | 0.89054584 |
| F730016J06F | 0.0093941 | 0.00891647 | 0.01005327 | -0.0011368 | 0.70084157 | 0.89054584 |
| Ube2ql1 | 0.0116201 | 0.01129847 | 0.01243542 | -0.001137 | 0.66493135 | 0.89054584 |
| Tfdp1 | 0.01159862 | 0.01127311 | 0.01241244 | -0.0011393 | 0.66554357 | 0.89054584 |
| Phf11d | 0.01508622 | 0.01500367 | 0.01614462 | -0.001141 | 0.62901755 | 0.89054584 |
| Asb11 | 0.010938 | 0.01056276 | 0.01170548 | -0.0011427 | 0.67538003 | 0.89054584 |

| | | | | | | |
|---|---|---|---|---|---|---|
| Taf4b | 0.00653186 | 0.00584697 | 0.00699021 | -0.0011432 | 0.77695968 | 0.89054584 |
| Clcnka | 0.0060197 | 0.00529252 | 0.00644212 | -0.0011496 | 0.7971238 | 0.89054584 |
| A830052D11 | 0.01570147 | 0.01565064 | 0.01680301 | -0.0011524 | 0.62533208 | 0.89054584 |
| Gm12204 | 0.00698415 | 0.00631458 | 0.00747424 | -0.0011597 | 0.76511619 | 0.89054584 |
| Slc35e3 | 0.01962598 | 0.01984197 | 0.02100261 | -0.0011606 | 0.60158893 | 0.89054584 |
| P2ry2 | 0.01152939 | 0.0111722 | 0.01233835 | -0.0011662 | 0.67015326 | 0.89054584 |
| Zfp712 | 0.02386919 | 0.0243758 | 0.02554305 | -0.0011673 | 0.58429823 | 0.89054584 |
| Megf10 | 0.00720884 | 0.00654575 | 0.00771469 | -0.0011689 | 0.75988319 | 0.89054584 |
| Gaa | 0.81174886 | 0.72500935 | 0.72617892 | -0.0011696 | 0.50299347 | 0.89054584 |
| Magi3 | 0.10284443 | 0.10873085 | 0.10990209 | -0.0011712 | 0.51979969 | 0.89054584 |
| Agbl3 | 0.04299533 | 0.0448311 | 0.04600504 | -0.0011739 | 0.54731694 | 0.89054584 |
| Gm26945 | 0.01896376 | 0.01911401 | 0.02029398 | -0.00118 | 0.60676204 | 0.89054584 |
| 4921504A21 | 0.02729756 | 0.02802725 | 0.02921139 | -0.0011841 | 0.57489811 | 0.89054584 |
| Prox1os | 0.08508576 | 0.08980761 | 0.09099192 | -0.0011843 | 0.52417659 | 0.89054584 |
| B3gnt3 | 0.01124622 | 0.01084777 | 0.01203532 | -0.0011876 | 0.67713714 | 0.89054584 |
| Acrbp | 0.0546724 | 0.05730321 | 0.05849304 | -0.0011898 | 0.53775099 | 0.89054584 |
| Gm44148 | 0.00542736 | 0.00461693 | 0.00580822 | -0.0011913 | 0.83035461 | 0.89271879 |
| Slc8a2 | 0.03068109 | 0.0316402 | 0.03283156 | -0.0011914 | 0.56712511 | 0.89054584 |
| Pkd2l1 | 0.00720441 | 0.00651432 | 0.00770996 | -0.0011956 | 0.76500419 | 0.89054584 |
| Tmem132e | 0.01119784 | 0.01078775 | 0.01198355 | -0.0011958 | 0.67899859 | 0.89054584 |
| Gpr161 | 0.0224149 | 0.02279044 | 0.02398691 | -0.0011965 | 0.59188181 | 0.89054584 |
| Nicn1 | 0.29899966 | 0.30711353 | 0.30831112 | -0.0011976 | 0.50721928 | 0.89054584 |
| Pdgfd | 0.10408956 | 0.11001132 | 0.11121077 | -0.0011995 | 0.5200378 | 0.89054584 |
| Gtf2ird2 | 0.07611143 | 0.08020671 | 0.08140703 | -0.0012003 | 0.5273837 | 0.89054584 |
| RP23-124B2. | 0.01011488 | 0.00962392 | 0.01082462 | -0.0012007 | 0.69734603 | 0.89054584 |
| Gm9797 | 0.00694672 | 0.00623161 | 0.00743418 | -0.0012026 | 0.77446707 | 0.89054584 |
| Pik3r6 | 0.01548195 | 0.0153651 | 0.0165681 | -0.001203 | 0.63242525 | 0.89054584 |
| D930048G16 | 0.00681661 | 0.00609153 | 0.00729494 | -0.0012034 | 0.7789238 | 0.89054584 |
| Gatsl3 | 0.0485879 | 0.05078264 | 0.05198653 | -0.0012039 | 0.5429589 | 0.89054584 |
| Tlr4 | 0.01009413 | 0.00959667 | 0.01080241 | -0.0012057 | 0.69847558 | 0.89054584 |

| | | | | | | |
|---|---|---|---|---|---|---|
| Rgs22 | 0.00697523 | 0.00625875 | 0.00746469 | -0.0012059 | 0.7741735 | 0.89054584 |
| Gm12996 | 0.01156917 | 0.01117275 | 0.01238092 | -0.0012082 | 0.67531446 | 0.89054584 |
| Cilp2 | 0.01143718 | 0.01102743 | 0.01223967 | -0.0012122 | 0.67775523 | 0.89054584 |
| Wdr63 | 0.00717711 | 0.00646632 | 0.00768074 | -0.0012144 | 0.76932741 | 0.89054584 |
| Pcdhga9 | 0.00540226 | 0.00456576 | 0.00578135 | -0.0012156 | 0.83635681 | 0.89462889 |
| Gm16794 | 0.00860637 | 0.00799373 | 0.00921028 | -0.0012165 | 0.73084747 | 0.89054584 |
| Gm12500 | 0.01313229 | 0.01283609 | 0.01405367 | -0.0012176 | 0.65676207 | 0.89054584 |
| Cep112 | 0.1261416 | 0.13306986 | 0.13429366 | -0.0012238 | 0.51693243 | 0.89054584 |
| Fbxo47 | 0.01155199 | 0.0111378 | 0.01236254 | -0.0012247 | 0.67779928 | 0.89054584 |
| Pik3r5 | 0.00708752 | 0.00635974 | 0.00758487 | -0.0012251 | 0.7741315 | 0.89054584 |
| Cbln2 | 0.00618062 | 0.0053871 | 0.00661433 | -0.0012272 | 0.80632215 | 0.89054584 |
| Nlgn3 | 0.03324356 | 0.03434458 | 0.03557311 | -0.0012285 | 0.56391277 | 0.89054584 |
| Gm10600 | 0.00715742 | 0.0064311 | 0.00765966 | -0.0012286 | 0.77254818 | 0.89054584 |
| Tgfa | 0.01585986 | 0.01574362 | 0.0169725 | -0.0012289 | 0.63206514 | 0.89054584 |
| Fgf16 | 0.01146695 | 0.01104177 | 0.01227153 | -0.0012298 | 0.67970861 | 0.89054584 |
| Endod1 | 0.09861032 | 0.1041971 | 0.10542715 | -0.0012301 | 0.5216748 | 0.89054584 |
| Usp13 | 0.03104777 | 0.03199056 | 0.03322387 | -0.0012333 | 0.56865317 | 0.89054584 |
| Exoc5 | 0.46003469 | 0.4559723 | 0.45720952 | -0.0012372 | 0.50502941 | 0.89054584 |
| Gm11940 | 0.02741692 | 0.02810151 | 0.0293391 | -0.0012376 | 0.57790002 | 0.89054584 |
| Sftpd | 0.00696952 | 0.00621473 | 0.00745858 | -0.0012438 | 0.78140867 | 0.89054584 |
| Edil3 | 0.0108908 | 0.01041061 | 0.01165497 | -0.0012444 | 0.69055219 | 0.89054584 |
| Stxbp1 | 0.30234031 | 0.31028233 | 0.31153968 | -0.0012574 | 0.50750092 | 0.89054584 |
| Gm26569 | 0.01149936 | 0.01104762 | 0.01230622 | -0.0012586 | 0.68313528 | 0.89054584 |
| Insc | 0.01422454 | 0.01396374 | 0.01522252 | -0.0012588 | 0.64997704 | 0.89054584 |
| Slc27a4 | 0.26751023 | 0.27634673 | 0.27760917 | -0.0012624 | 0.50845163 | 0.89054584 |
| Stx3 | 0.32046817 | 0.32769707 | 0.32896249 | -0.0012654 | 0.50714932 | 0.89054584 |
| Slc39a4 | 0.00964546 | 0.00905504 | 0.01032226 | -0.0012672 | 0.71632591 | 0.89054584 |
| Kdm4b | 0.16001769 | 0.16812713 | 0.16939493 | -0.0012678 | 0.51390787 | 0.89054584 |
| A730071L15I | 0.0113779 | 0.01090712 | 0.01217623 | -0.0012691 | 0.68637238 | 0.89054584 |
| Whsc1 | 0.43348608 | 0.4323857 | 0.43365805 | -0.0012724 | 0.5054531 | 0.89054584 |

| | | | | | | |
|---|---|---|---|---|---|---|
| Zfand4 | 0.0486296 | 0.05075829 | 0.05203113 | -0.0012728 | 0.54536993 | 0.89054584 |
| Zfp951 | 0.02763998 | 0.02830449 | 0.02957776 | -0.0012733 | 0.57947805 | 0.89054584 |
| Gm11491 | 0.02298063 | 0.02331868 | 0.02459226 | -0.0012736 | 0.59532928 | 0.89054584 |
| Ltc4s | 0.00709343 | 0.00631744 | 0.00759119 | -0.0012737 | 0.78281479 | 0.89054584 |
| Prom2 | 0.00640457 | 0.00557883 | 0.006854 | -0.0012752 | 0.80696947 | 0.89054584 |
| Trim30d | 0.00709068 | 0.00631266 | 0.00758825 | -0.0012756 | 0.78323639 | 0.89054584 |
| Chek2 | 0.06423939 | 0.06744178 | 0.06872076 | -0.001279 | 0.53454827 | 0.89054584 |
| Fgd4 | 0.26609165 | 0.27493266 | 0.2762148 | -0.0012821 | 0.50862682 | 0.89054584 |
| Galnt4 | 0.0743842 | 0.07827383 | 0.07956178 | -0.0012879 | 0.53005926 | 0.89054584 |
| Gm11750 | 0.00707661 | 0.00628479 | 0.00757319 | -0.0012884 | 0.78599691 | 0.89054584 |
| Gpr34 | 0.00611292 | 0.00525085 | 0.00654188 | -0.001291 | 0.82106591 | 0.89054584 |
| Ikzf3 | 0.00820589 | 0.00749046 | 0.0087817 | -0.0012912 | 0.75333922 | 0.89054584 |
| Ceacam1 | 0.03268951 | 0.03368725 | 0.03498035 | -0.0012931 | 0.56836944 | 0.89054584 |
| Naif1 | 0.03911884 | 0.04056182 | 0.04185845 | -0.0012966 | 0.55737523 | 0.89054584 |
| Pgpep1l | 0.01107991 | 0.01055746 | 0.01185734 | -0.0012999 | 0.69523531 | 0.89054584 |
| Ccdc103 | 0.08305276 | 0.08751905 | 0.08882102 | -0.001302 | 0.52722339 | 0.89054584 |
| Ndst3 | 0.00705425 | 0.00624715 | 0.00754926 | -0.0013021 | 0.78918045 | 0.89054584 |
| Atl1 | 0.04288621 | 0.04458593 | 0.04588832 | -0.0013024 | 0.55259859 | 0.89054584 |
| Irf7 | 0.0475651 | 0.0495901 | 0.05089268 | -0.0013026 | 0.54745913 | 0.89054584 |
| Slc6a20b | 0.0114602 | 0.0109596 | 0.0122643 | -0.0013047 | 0.68992193 | 0.89054584 |
| Vps25 | 0.12755219 | 0.13445763 | 0.1357641 | -0.0013065 | 0.51787985 | 0.89054584 |
| Tcf3 | 0.95723185 | 0.81069067 | 0.81199793 | -0.0013073 | 0.50299227 | 0.89054584 |
| Pign | 0.12115342 | 0.12778019 | 0.1290882 | -0.001308 | 0.5188261 | 0.89054584 |
| Gm6410 | 0.01936231 | 0.01940944 | 0.02072046 | -0.001311 | 0.61591881 | 0.89054584 |
| Gm15396 | 0.00668776 | 0.0058439 | 0.00715705 | -0.0013132 | 0.80367105 | 0.89054584 |
| Znhit6 | 0.42148979 | 0.4215662 | 0.42288118 | -0.001315 | 0.50577943 | 0.89054584 |
| Gp1bb | 0.00991852 | 0.00929626 | 0.01061448 | -0.0013182 | 0.71856915 | 0.89054584 |
| RP23-32A8.8 | 0.01137026 | 0.010849 | 0.01216806 | -0.0013191 | 0.69323736 | 0.89054584 |
| 4930471D02 | 0.01131615 | 0.01078996 | 0.01211015 | -0.0013202 | 0.69423693 | 0.89054584 |
| B230303O12 | 0.01549491 | 0.01526085 | 0.01658197 | -0.0013211 | 0.64475194 | 0.89054584 |

| | | | | | | |
|---|---|---|---|---|---|---|
| Iglon5 | 0.0070278 | 0.00619963 | 0.00752095 | -0.0013213 | 0.7934654 | 0.89054584 |
| Tnk2 | 0.07144179 | 0.07508983 | 0.07641795 | -0.0013281 | 0.53226755 | 0.89054584 |
| Gemin8 | 0.13225306 | 0.13932952 | 0.140659 | -0.0013295 | 0.51756198 | 0.89054584 |
| Gm11716 | 0.00606983 | 0.00516223 | 0.00649576 | -0.0013335 | 0.83057602 | 0.89271879 |
| Lrrc20 | 0.05871599 | 0.06147963 | 0.06281634 | -0.0013367 | 0.53948671 | 0.89054584 |
| Crym | 0.0273476 | 0.02792818 | 0.02926493 | -0.0013367 | 0.58426183 | 0.89054584 |
| Sergef | 0.14338737 | 0.15087991 | 0.15221891 | -0.001339 | 0.51634514 | 0.89054584 |
| 9430041J12F | 0.00526525 | 0.00429208 | 0.00563473 | -0.0013426 | 0.86652908 | 0.90430169 |
| Pipox | 0.01304518 | 0.0126154 | 0.01396045 | -0.0013451 | 0.67323997 | 0.89054584 |
| Ctnnd2 | 0.03802288 | 0.03934043 | 0.04068606 | -0.0013456 | 0.56122975 | 0.89054584 |
| Bckdhb | 0.28067922 | 0.28915998 | 0.29050685 | -0.0013469 | 0.50861657 | 0.89054584 |
| Dcp1a | 0.09538167 | 0.10063404 | 0.10198222 | -0.0013482 | 0.52455544 | 0.89054584 |
| Gm14091 | 0.03284558 | 0.03379639 | 0.03514732 | -0.0013509 | 0.57105965 | 0.89054584 |
| Gm17767 | 0.0113976 | 0.01084579 | 0.01219732 | -0.0013515 | 0.69715504 | 0.89054584 |
| Cask | 0.20939552 | 0.2183769 | 0.21972948 | -0.0013526 | 0.51143963 | 0.89054584 |
| Gm43848 | 0.00695185 | 0.00608637 | 0.00743968 | -0.0013533 | 0.80163622 | 0.89054584 |
| Dbndd2 | 0.00901524 | 0.00829411 | 0.00964783 | -0.0013537 | 0.74335084 | 0.89054584 |
| Cyb5b | 0.51518389 | 0.50342532 | 0.50477982 | -0.0013545 | 0.50498728 | 0.89054584 |
| 9030624G23 | 0.01056711 | 0.00995279 | 0.01130857 | -0.0013558 | 0.71176921 | 0.89054584 |
| Zfp935 | 0.02406503 | 0.02439506 | 0.0257526 | -0.0013575 | 0.59700196 | 0.89054584 |
| Gk5 | 0.06210324 | 0.06507871 | 0.0664374 | -0.0013587 | 0.53795317 | 0.89054584 |
| Camkk1 | 0.01794218 | 0.01784113 | 0.01920081 | -0.0013597 | 0.62926865 | 0.89054584 |
| Gjb1 | 0.04766634 | 0.04964063 | 0.05100096 | -0.0013603 | 0.54944756 | 0.89054584 |
| Ube2b | 2.09362035 | 1.12170929 | 1.12306968 | -0.0013604 | 0.5022514 | 0.89054584 |
| Pramef8 | 0.12947931 | 0.1364099 | 0.13777178 | -0.0013619 | 0.51836637 | 0.89054584 |
| Dync2h1 | 0.49442869 | 0.48573251 | 0.48709459 | -0.0013621 | 0.50519726 | 0.89054584 |
| Fhl4 | 0.01132775 | 0.01076044 | 0.01212257 | -0.0013621 | 0.6996836 | 0.89054584 |
| Zfp418 | 0.00927177 | 0.00855593 | 0.00992236 | -0.0013664 | 0.73943144 | 0.89054584 |
| Gm20109 | 0.02815364 | 0.02875873 | 0.03012735 | -0.0013686 | 0.58380877 | 0.89054584 |
| Kcna2 | 0.00695132 | 0.00606816 | 0.0074391 | -0.0013709 | 0.80471679 | 0.89054584 |

| | | | | | | |
|---|---|---|---|---|---|---|
| Klhl23 | 0.15374449 | 0.16155743 | 0.16292844 | -0.001371 | 0.51563614 | 0.89054584 |
| Plpp2 | 0.16649435 | 0.17468334 | 0.17605447 | -0.0013711 | 0.51447215 | 0.89054584 |
| Chd3 | 1.14945876 | 0.90684434 | 0.9082165 | -0.0013722 | 0.50280808 | 0.89054584 |
| 4632415L05F | 0.17555804 | 0.18396923 | 0.18534524 | -0.001376 | 0.51379596 | 0.89054584 |
| Ulbp1 | 0.00692487 | 0.00603084 | 0.00741081 | -0.00138 | 0.80717674 | 0.89054584 |
| Asphd2 | 0.03207196 | 0.03293951 | 0.03431964 | -0.0013801 | 0.57430869 | 0.89054584 |
| Cpb2 | 0.02214082 | 0.02230486 | 0.02369363 | -0.0013888 | 0.60760296 | 0.89054584 |
| 9430060I03R | 0.00678075 | 0.00586745 | 0.00725657 | -0.0013891 | 0.81376616 | 0.89054584 |
| Mfsd2b | 0.00691259 | 0.00600292 | 0.00739766 | -0.0013947 | 0.81013328 | 0.89054584 |
| Neu2 | 0.00691832 | 0.00600816 | 0.00740379 | -0.0013956 | 0.81008818 | 0.89054584 |
| Cnr1 | 0.01573032 | 0.01543713 | 0.01683388 | -0.0013968 | 0.65046106 | 0.89054584 |
| Rnf182 | 0.02762793 | 0.02816713 | 0.02956488 | -0.0013977 | 0.58716631 | 0.89054584 |
| Gm5424 | 0.0231812 | 0.02340862 | 0.02480688 | -0.0013983 | 0.60357286 | 0.89054584 |
| 4930539J05F | 0.01307768 | 0.01259617 | 0.01399523 | -0.0013991 | 0.6792994 | 0.89054584 |
| 4732463B04 | 0.00947319 | 0.00873714 | 0.01013791 | -0.0014008 | 0.74012538 | 0.89054584 |
| Gpr162 | 0.02336778 | 0.02360176 | 0.02500652 | -0.0014048 | 0.60323146 | 0.89054584 |
| Glyr1 | 0.92406364 | 0.79234594 | 0.79375085 | -0.0014049 | 0.50328971 | 0.89054584 |
| Ptpro | 0.02036325 | 0.02038173 | 0.02179154 | -0.0014098 | 0.61844982 | 0.89054584 |
| Clcn5 | 0.30088227 | 0.30872089 | 0.31013125 | -0.0014104 | 0.50845179 | 0.89054584 |
| Cacng4 | 0.0433152 | 0.04493472 | 0.04634718 | -0.0014125 | 0.55645373 | 0.89054584 |
| Muc20 | 0.01130224 | 0.01067359 | 0.01209527 | -0.0014217 | 0.70802092 | 0.89054584 |
| 4930524O07 | 0.00870989 | 0.00789378 | 0.00932106 | -0.0014273 | 0.76219976 | 0.89054584 |
| Pkd1l1 | 0.00653811 | 0.00556939 | 0.00699691 | -0.0014275 | 0.82907642 | 0.89265604 |
| Tvp23bos | 0.00687141 | 0.00592595 | 0.00735359 | -0.0014276 | 0.81713295 | 0.89054584 |
| Ppm1e | 0.01125234 | 0.01061272 | 0.01204187 | -0.0014291 | 0.70984396 | 0.89054584 |
| Timd2 | 0.00647717 | 0.00550252 | 0.00693169 | -0.0014292 | 0.83161708 | 0.89335984 |
| Fmnl1 | 0.05548787 | 0.0579352 | 0.05936497 | -0.0014298 | 0.54467041 | 0.89054584 |
| Ugt2b35 | 0.01516741 | 0.01479917 | 0.0162315 | -0.0014323 | 0.65950884 | 0.89054584 |
| Slc36a1 | 0.15091058 | 0.15856541 | 0.16000222 | -0.0014368 | 0.51668571 | 0.89054584 |
| Akr1b7 | 0.00572012 | 0.00467857 | 0.00612152 | -0.0014429 | 0.86393991 | 0.90312213 |

| | | | | | | |
|---|---|---|---|---|---|---|
| Hist2h4 | 0.02430872 | 0.02457004 | 0.02601336 | -0.0014433 | 0.60198767 | 0.89054584 |
| Gm26982 | 0.00970651 | 0.00893781 | 0.0103876 | -0.0014498 | 0.7422325 | 0.89054584 |
| Cfap73 | 0.01044778 | 0.00972895 | 0.01118087 | -0.0014519 | 0.72740982 | 0.89054584 |
| Shroom4 | 0.02974809 | 0.03038064 | 0.03183333 | -0.0014527 | 0.58418341 | 0.89054584 |
| Myo7a | 0.01404675 | 0.01357749 | 0.01503226 | -0.0014548 | 0.67396077 | 0.89054584 |
| Tnfrsf22 | 0.00668802 | 0.00569686 | 0.00715734 | -0.0014605 | 0.82911479 | 0.89265604 |
| Sirt5 | 0.12134169 | 0.12782232 | 0.12928483 | -0.0014625 | 0.52101594 | 0.89054584 |
| Zfp930 | 0.0233141 | 0.02348258 | 0.02494909 | -0.0014665 | 0.60790125 | 0.89054584 |
| 4632428C04I | 0.00592683 | 0.00487487 | 0.00634273 | -0.0014679 | 0.85952464 | 0.90131997 |
| A730017C20 | 0.00652907 | 0.00551608 | 0.00698723 | -0.0014711 | 0.83668666 | 0.89462964 |
| Tsc22d2 | 0.27478441 | 0.28326551 | 0.28474392 | -0.0014784 | 0.50964934 | 0.89054584 |
| Gm22 | 0.01939974 | 0.01928192 | 0.02076051 | -0.0014786 | 0.62998579 | 0.89054584 |
| Tbx6 | 0.01460877 | 0.01415355 | 0.0156337 | -0.0014802 | 0.67042761 | 0.89054584 |
| Smg8 | 0.0492612 | 0.05122056 | 0.0527066 | -0.001486 | 0.55225365 | 0.89054584 |
| Cyb561d2 | 0.29439299 | 0.30235689 | 0.30384981 | -0.0014929 | 0.50913143 | 0.89054584 |
| Gm16751 | 0.00602974 | 0.00495937 | 0.00645286 | -0.0014935 | 0.85954947 | 0.90131997 |
| Hr | 0.00554086 | 0.00442935 | 0.00592968 | -0.0015003 | 0.8807617 | 0.91173336 |
| Gm26826 | 0.02860425 | 0.0291052 | 0.03060949 | -0.0015043 | 0.59055079 | 0.89054584 |
| 9030617O03 | 0.01105328 | 0.0103229 | 0.01182885 | -0.0015059 | 0.72345489 | 0.89054584 |
| Espnl | 0.00615976 | 0.00508604 | 0.006592 | -0.001506 | 0.85641405 | 0.9009252 |
| Gm17087 | 0.00553182 | 0.00441193 | 0.00592001 | -0.0015081 | 0.88235287 | 0.91235841 |
| Slc9a5 | 0.01103986 | 0.01030466 | 0.01181449 | -0.0015098 | 0.72420744 | 0.89054584 |
| Ankrd34b | 0.00613152 | 0.00504973 | 0.00656178 | -0.001512 | 0.85849265 | 0.90124516 |
| B230317F23 | 0.03130692 | 0.03198892 | 0.03350114 | -0.0015122 | 0.58328346 | 0.89054584 |
| BC049762 | 0.00673725 | 0.00569659 | 0.00721002 | -0.0015134 | 0.83594789 | 0.89462889 |
| Cd27 | 0.17939871 | 0.18775765 | 0.18927196 | -0.0015143 | 0.51486706 | 0.89054584 |
| Gm17619 | 0.02191664 | 0.02193813 | 0.02345375 | -0.0015156 | 0.61831918 | 0.89054584 |
| Zfp940 | 0.00939067 | 0.00852597 | 0.01004961 | -0.0015236 | 0.76001406 | 0.89054584 |
| D830044I16F | 0.02362817 | 0.02375751 | 0.02528515 | -0.0015276 | 0.61082737 | 0.89054584 |
| Adamtsl2 | 0.0230746 | 0.02316426 | 0.02469282 | -0.0015286 | 0.61348057 | 0.89054584 |

| | | | | | | |
|---|---|---|---|---|---|---|
| Zfp661 | 0.00644335 | 0.00536515 | 0.00689549 | -0.0015303 | 0.84942652 | 0.89883846 |
| Atp6v1b2 | 1.01590218 | 0.84117585 | 0.84270665 | -0.0015308 | 0.50337623 | 0.89054584 |
| 1700003F12I | 0.0104009 | 0.00959758 | 0.0111307 | -0.0015331 | 0.73946999 | 0.89054584 |
| Mapk11 | 0.01838664 | 0.01814279 | 0.01967641 | -0.0015336 | 0.64174492 | 0.89054584 |
| Shpk | 0.00585699 | 0.00473416 | 0.00626799 | -0.0015338 | 0.87287448 | 0.90729956 |
| Scd3 | 0.03856347 | 0.03972896 | 0.04126436 | -0.0015354 | 0.56881314 | 0.89054584 |
| 2200002D01I | 0.02868333 | 0.02915724 | 0.03069411 | -0.0015369 | 0.59222611 | 0.89054584 |
| Lrriq1 | 0.01090702 | 0.01013513 | 0.01167232 | -0.0015372 | 0.7302474 | 0.89054584 |
| Nhej1 | 0.04213665 | 0.04354824 | 0.04508657 | -0.0015383 | 0.56314991 | 0.89054584 |
| Adam28 | 0.01113705 | 0.01037565 | 0.01191849 | -0.0015428 | 0.72677866 | 0.89054584 |
| B930094E09 | 0.014784 | 0.01427558 | 0.01582121 | -0.0015456 | 0.67549921 | 0.89054584 |
| Cp | 0.00667697 | 0.00559726 | 0.00714551 | -0.0015483 | 0.84362654 | 0.89775379 |
| Gm29361 | 0.00662264 | 0.00553669 | 0.00708736 | -0.0015507 | 0.84598073 | 0.89825055 |
| Gm15265 | 0.006138 | 0.00501507 | 0.00656872 | -0.0015537 | 0.86475743 | 0.90358581 |
| Hist1h3e | 0.01013324 | 0.00928578 | 0.01084427 | -0.0015585 | 0.74843139 | 0.89054584 |
| Tcerg1 | 0.50796004 | 0.49709589 | 0.4986553 | -0.0015594 | 0.50581223 | 0.89054584 |
| Myd88 | 0.13938025 | 0.14650334 | 0.14806418 | -0.0015608 | 0.51958531 | 0.89054584 |
| Gm13166 | 0.01741469 | 0.01707332 | 0.01863634 | -0.001563 | 0.65200649 | 0.89054584 |
| Glt28d2 | 0.01041763 | 0.00958492 | 0.0111486 | -0.0015637 | 0.74326754 | 0.89054584 |
| Gm27217 | 0.01104651 | 0.01025723 | 0.01182159 | -0.0015644 | 0.73122568 | 0.89054584 |
| Scml4 | 0.03120976 | 0.03183196 | 0.03339718 | -0.0015652 | 0.58642213 | 0.89054584 |
| Klc2 | 0.34414019 | 0.34989539 | 0.35146097 | -0.0015656 | 0.50827873 | 0.89054584 |
| Sema3e | 0.01077586 | 0.0099634 | 0.01153196 | -0.0015686 | 0.73686182 | 0.89054584 |
| Fah | 0.02899168 | 0.02945472 | 0.03102401 | -0.0015693 | 0.59315346 | 0.89054584 |
| Gm14399 | 0.00935753 | 0.00844388 | 0.01001414 | -0.0015703 | 0.76747008 | 0.89054584 |
| Clvs1 | 0.00640615 | 0.00528281 | 0.00685569 | -0.0015729 | 0.85743983 | 0.9009252 |
| Egfr | 0.02249406 | 0.02249782 | 0.02407162 | -0.0015738 | 0.61966463 | 0.89054584 |
| Eefsec | 0.23197962 | 0.24082883 | 0.24240345 | -0.0015746 | 0.51207173 | 0.89054584 |
| Ccdc89 | 0.01678702 | 0.01638825 | 0.01796467 | -0.0015764 | 0.6586662 | 0.89054584 |
| Bzw2 | 0.96112255 | 0.81251195 | 0.81408967 | -0.0015777 | 0.50360205 | 0.89054584 |

| | | | | | | |
|---|---|---|---|---|---|---|
| Pfkfb1 | 0.01089944 | 0.0100849 | 0.01166421 | -0.0015793 | 0.73591795 | 0.89054584 |
| Wdr76 | 0.0987697 | 0.10401773 | 0.10559719 | -0.0015795 | 0.527778 | 0.89054584 |
| Wbscr17 | 0.04574725 | 0.04736396 | 0.04894846 | -0.0015845 | 0.55993851 | 0.89054584 |
| Glod5 | 0.00579058 | 0.00461197 | 0.00619693 | -0.001585 | 0.88328943 | 0.9130137 |
| Gm2885 | 0.00501506 | 0.00378113 | 0.00536698 | -0.0015859 | 0.91568796 | 0.93411512 |
| Mroh7 | 0.02019909 | 0.02002441 | 0.02161588 | -0.0015915 | 0.63420584 | 0.89054584 |
| Rsph4a | 0.01913006 | 0.01888024 | 0.02047194 | -0.0015917 | 0.64141077 | 0.89054584 |
| Nt5e | 0.00605731 | 0.0048888 | 0.00648237 | -0.0015936 | 0.87396046 | 0.90773126 |
| Pgp | 0.87523512 | 0.76390094 | 0.76549619 | -0.0015953 | 0.50387326 | 0.89054584 |
| Slc18b1 | 0.1235229 | 0.12996371 | 0.13156206 | -0.0015983 | 0.52256863 | 0.89054584 |
| Pagr1a | 0.01719865 | 0.01680679 | 0.01840515 | -0.0015984 | 0.65711304 | 0.89054584 |
| Necap2 | 0.37494623 | 0.37869539 | 0.38029834 | -0.0016029 | 0.50783364 | 0.89054584 |
| 1700034H15I | 0.0064764 | 0.00532717 | 0.00693086 | -0.0016037 | 0.85948448 | 0.90131997 |
| Gm10392 | 0.0091099 | 0.00814235 | 0.00974913 | -0.0016068 | 0.77871229 | 0.89054584 |
| Brf1 | 0.24110465 | 0.24988894 | 0.2515008 | -0.0016119 | 0.51191025 | 0.89054584 |
| Styk1 | 0.01484949 | 0.01427768 | 0.0158913 | -0.0016136 | 0.68191967 | 0.89054584 |
| Gm19705 | 0.01103747 | 0.01019285 | 0.01181193 | -0.0016191 | 0.73846043 | 0.89054584 |
| Lrrc4b | 0.01101314 | 0.01016602 | 0.01178589 | -0.0016199 | 0.73902225 | 0.89054584 |
| 5330434G04 | 0.01105157 | 0.01020321 | 0.01182701 | -0.0016238 | 0.73880067 | 0.89054584 |
| Dhtkd1 | 0.00889695 | 0.00789635 | 0.00952124 | -0.0016249 | 0.78672017 | 0.89054584 |
| Prex1 | 0.09351757 | 0.09836728 | 0.09999292 | -0.0016256 | 0.53018829 | 0.89054584 |
| Gm6976 | 0.00649365 | 0.00532269 | 0.00694932 | -0.0016266 | 0.86225846 | 0.90260095 |
| Cptp | 0.15576282 | 0.16337889 | 0.16501067 | -0.0016318 | 0.51837363 | 0.89054584 |
| 6330403A02I | 0.00948123 | 0.00851148 | 0.01014651 | -0.001635 | 0.77359919 | 0.89054584 |
| Gm11032 | 0.00516278 | 0.00388952 | 0.00552507 | -0.0016356 | 0.91607688 | 0.93431464 |
| Gm29650 | 0.01066792 | 0.0097804 | 0.01141645 | -0.0016361 | 0.74782393 | 0.89054584 |
| Mustn1 | 0.00650307 | 0.00532191 | 0.0069594 | -0.0016375 | 0.86350709 | 0.90290403 |
| Gltscr1 | 0.00489144 | 0.00359354 | 0.00523469 | -0.0016411 | 0.92794349 | 0.94303412 |
| Ncan | 0.00603795 | 0.00482028 | 0.00646165 | -0.0016414 | 0.88168501 | 0.91220076 |
| 2010308F09I | 0.00599412 | 0.00476977 | 0.00641474 | -0.001645 | 0.88390233 | 0.91351696 |

| | | | | | | |
|---|---|---|---|---|---|---|
| Helb | 0.04005647 | 0.04121274 | 0.04286145 | -0.0016487 | 0.57111406 | 0.89054584 |
| Art4 | 0.01103481 | 0.01015844 | 0.01180908 | -0.0016506 | 0.74254597 | 0.89054584 |
| Zfp954 | 0.13097747 | 0.13768083 | 0.13933159 | -0.0016508 | 0.52200951 | 0.89054584 |
| Nptx2 | 0.03792429 | 0.03892947 | 0.0405806 | -0.0016511 | 0.57517358 | 0.89054584 |
| Gm10033 | 0.00492519 | 0.00361952 | 0.00527081 | -0.0016513 | 0.92779954 | 0.94302003 |
| Cd22 | 0.01452266 | 0.01389016 | 0.01554155 | -0.0016514 | 0.68971439 | 0.89054584 |
| Dclk1 | 0.02559613 | 0.02573865 | 0.02739089 | -0.0016522 | 0.61065677 | 0.89054584 |
| Fbxo41 | 0.0183077 | 0.01793851 | 0.01959194 | -0.0016534 | 0.65290798 | 0.89054584 |
| Lrrc75aos2 | 0.01102453 | 0.01014401 | 0.01179808 | -0.0016541 | 0.74317806 | 0.89054584 |
| Idh3a | 0.88275921 | 0.76830614 | 0.76996135 | -0.0016552 | 0.50399553 | 0.89054584 |
| Spin2c | 0.11762623 | 0.12374541 | 0.1254017 | -0.0016563 | 0.52453326 | 0.89054584 |
| Zfp108 | 0.03892895 | 0.03999764 | 0.04165532 | -0.0016577 | 0.57354345 | 0.89054584 |
| Fmn2 | 0.21804017 | 0.22677699 | 0.22843545 | -0.0016585 | 0.51349133 | 0.89054584 |
| Mast1 | 0.04192303 | 0.04319715 | 0.04485806 | -0.0016609 | 0.56847836 | 0.89054584 |
| Rapgef2 | 0.09624381 | 0.10123628 | 0.10290219 | -0.0016659 | 0.53006167 | 0.89054584 |
| Gm9903 | 0.01382499 | 0.01312888 | 0.01479495 | -0.0016661 | 0.70008648 | 0.89054584 |
| Prph | 0.01140947 | 0.01054252 | 0.01221001 | -0.0016675 | 0.73769658 | 0.89054584 |
| Hrh3 | 0.00981962 | 0.00884078 | 0.01050865 | -0.0016679 | 0.77017973 | 0.89054584 |
| Gm37233 | 0.01357056 | 0.01285299 | 0.01452268 | -0.0016697 | 0.70389756 | 0.89054584 |
| Pcdhb13 | 0.00635608 | 0.00513235 | 0.0068021 | -0.0016697 | 0.8736166 | 0.90763953 |
| Upk1a | 0.00619394 | 0.00495679 | 0.00662859 | -0.0016718 | 0.88000917 | 0.91137745 |
| Frmd3 | 0.00636027 | 0.00513374 | 0.00680659 | -0.0016729 | 0.87390082 | 0.90773126 |
| Sorcs1 | 0.02814626 | 0.02844422 | 0.03011947 | -0.0016752 | 0.60223249 | 0.89054584 |
| Lamb3 | 0.00619048 | 0.00494798 | 0.00662488 | -0.0016769 | 0.88085592 | 0.91173336 |
| Gm16853 | 0.01517041 | 0.0145559 | 0.01623472 | -0.0016788 | 0.68501838 | 0.89054584 |
| A830039N20 | 0.01429871 | 0.0136214 | 0.01530189 | -0.0016805 | 0.69555418 | 0.89054584 |
| Tmprss11a | 0.01089183 | 0.00997386 | 0.01165606 | -0.0016822 | 0.74932885 | 0.89054584 |
| Pnma2 | 0.01806268 | 0.01764527 | 0.01932974 | -0.0016845 | 0.65762867 | 0.89054584 |
| Rnf32 | 0.09975998 | 0.10496878 | 0.10665364 | -0.0016849 | 0.52933549 | 0.89054584 |
| Cntd1 | 0.02168394 | 0.02151414 | 0.02320475 | -0.0016906 | 0.63285797 | 0.89054584 |

| | | | | | | |
|---|---|---|---|---|---|---|
| Socs5 | 0.12636949 | 0.13283573 | 0.13453125 | -0.0016955 | 0.5234115 | 0.89054584 |
| A930041C12 | 0.01077571 | 0.00983251 | 0.01153181 | -0.0016993 | 0.75381044 | 0.89054584 |
| Ttc22 | 0.02777953 | 0.02802586 | 0.02972708 | -0.0017012 | 0.6051199 | 0.89054584 |
| Tcaf2 | 0.01761687 | 0.01715022 | 0.01885269 | -0.0017025 | 0.66302022 | 0.89054584 |
| Rasl12 | 0.00584401 | 0.00454821 | 0.0062541 | -0.0017059 | 0.89809756 | 0.92211428 |
| Cep83 | 0.44496655 | 0.44218665 | 0.44389289 | -0.0017062 | 0.50714392 | 0.89054584 |
| a | 0.00550905 | 0.00418873 | 0.00589563 | -0.0017069 | 0.91130838 | 0.93102311 |
| 4933427E11I | 0.01062433 | 0.00966259 | 0.0113698 | -0.0017072 | 0.75789785 | 0.89054584 |
| Kcnk2 | 0.0225762 | 0.02244975 | 0.02415951 | -0.0017098 | 0.62919136 | 0.89054584 |
| Map9 | 0.5398125 | 0.52370266 | 0.5254153 | -0.0017126 | 0.50605818 | 0.89054584 |
| Pop4 | 0.20243792 | 0.21098566 | 0.21269904 | -0.0017134 | 0.51496871 | 0.89054584 |
| Dennd3 | 0.03382797 | 0.03448483 | 0.03619834 | -0.0017135 | 0.58727432 | 0.89054584 |
| Gpr135 | 0.00538752 | 0.00404994 | 0.00576558 | -0.0017156 | 0.91717865 | 0.93504367 |
| Hgfac | 0.01244358 | 0.01159912 | 0.01331666 | -0.0017175 | 0.72604485 | 0.89054584 |
| Nccrp1 | 0.01489991 | 0.01422398 | 0.01594525 | -0.0017213 | 0.69249136 | 0.89054584 |
| Lypd1 | 0.01262003 | 0.0117838 | 0.01350549 | -0.0017217 | 0.72371911 | 0.89054584 |
| Lgals4 | 0.01793288 | 0.01746571 | 0.01919085 | -0.0017251 | 0.66232287 | 0.89054584 |
| B230334C09 | 0.0063368 | 0.00505538 | 0.00678147 | -0.0017261 | 0.88215797 | 0.91223443 |
| RP24-274B19 | 0.00880694 | 0.0076985 | 0.00942492 | -0.0017264 | 0.80328296 | 0.89054584 |
| 1700112J16F | 0.0194436 | 0.0190799 | 0.02080745 | -0.0017275 | 0.65055141 | 0.89054584 |
| Samd15 | 0.07021807 | 0.07338205 | 0.07511034 | -0.0017283 | 0.54268569 | 0.89054584 |
| Fam72a | 0.0412708 | 0.04243071 | 0.0441604 | -0.0017297 | 0.57239827 | 0.89054584 |
| Kcnc1 | 0.01069457 | 0.00971487 | 0.01144497 | -0.0017301 | 0.75936995 | 0.89054584 |
| Kcp | 0.01921129 | 0.01882867 | 0.02055886 | -0.0017302 | 0.65250284 | 0.89054584 |
| A730090N16 | 0.01460577 | 0.01389929 | 0.01563049 | -0.0017312 | 0.69707772 | 0.89054584 |
| Lmln | 0.04149452 | 0.04266707 | 0.04439971 | -0.0017326 | 0.57213376 | 0.89054584 |
| Them7 | 0.01531032 | 0.01465051 | 0.01638443 | -0.0017339 | 0.68900945 | 0.89054584 |
| Col4a4 | 0.00536106 | 0.00400233 | 0.00573727 | -0.0017349 | 0.92056057 | 0.937526 |
| A530020G20 | 0.09344454 | 0.0981736 | 0.09991499 | -0.0017414 | 0.53235825 | 0.89054584 |
| Slc22a18 | 0.04232488 | 0.04354568 | 0.04528791 | -0.0017422 | 0.57112119 | 0.89054584 |

| | | | | | | |
|---|---|---|---|---|---|---|
| Metrnl | 0.08667322 | 0.09094444 | 0.09268689 | -0.0017425 | 0.53489675 | 0.89054584 |
| Slc7a3 | 0.00950503 | 0.00842682 | 0.01017198 | -0.0017452 | 0.7879455 | 0.89054584 |
| Fam131c | 0.01512392 | 0.01443971 | 0.01618497 | -0.0017453 | 0.69230138 | 0.89054584 |
| Adhfe1 | 0.01817932 | 0.01770793 | 0.01945457 | -0.0017466 | 0.66212939 | 0.89054584 |
| Vars2 | 0.03340416 | 0.03399473 | 0.03574493 | -0.0017502 | 0.59022288 | 0.89054584 |
| Capn12 | 0.00941532 | 0.00832424 | 0.01007598 | -0.0017517 | 0.79102073 | 0.89054584 |
| Gm13010 | 0.00521537 | 0.00382759 | 0.00558135 | -0.0017538 | 0.92839201 | 0.9434238 |
| Rcc1 | 0.13726896 | 0.144118 | 0.14587259 | -0.0017546 | 0.52234449 | 0.89054584 |
| Fam134c | 0.32523529 | 0.33176161 | 0.33351655 | -0.0017549 | 0.50977913 | 0.89054584 |
| Notch4 | 0.04388384 | 0.04519982 | 0.0469554 | -0.0017556 | 0.56914187 | 0.89054584 |
| Ankrd23 | 0.01090236 | 0.00991112 | 0.01166733 | -0.0017562 | 0.75843669 | 0.89054584 |
| Ngb | 0.01079801 | 0.00979891 | 0.01155566 | -0.0017568 | 0.76061334 | 0.89054584 |
| Vma21-ps | 0.00979993 | 0.00873064 | 0.01048757 | -0.0017569 | 0.78244869 | 0.89054584 |
| Fam162b | 0.01517274 | 0.01447918 | 0.01623721 | -0.001758 | 0.6930206 | 0.89054584 |
| Nsg2 | 0.01010719 | 0.00905666 | 0.01081639 | -0.0017597 | 0.77576413 | 0.89054584 |
| 4930426D05 | 0.01839423 | 0.01792464 | 0.01968453 | -0.0017599 | 0.6614893 | 0.89054584 |
| Pm20d2 | 0.01059438 | 0.00957681 | 0.01133775 | -0.0017609 | 0.76534815 | 0.89054584 |
| BC017643 | 0.3060222 | 0.31333048 | 0.31509148 | -0.001761 | 0.51038653 | 0.89054584 |
| Cmpk2 | 0.04202031 | 0.04319719 | 0.04496212 | -0.0017649 | 0.57255434 | 0.89054584 |
| Mslnl | 0.01032421 | 0.00928261 | 0.01104863 | -0.001766 | 0.77176953 | 0.89054584 |
| Fbxl2 | 0.13599981 | 0.14278803 | 0.14455434 | -0.0017663 | 0.52269859 | 0.89054584 |
| A930024N18 | 0.00924543 | 0.0081265 | 0.00989417 | -0.0017677 | 0.7973961 | 0.89054584 |
| Arg2 | 0.0053122 | 0.00391533 | 0.00568498 | -0.0017696 | 0.92650677 | 0.9423006 |
| Adamts7 | 0.0474275 | 0.04897544 | 0.05074551 | -0.0017701 | 0.56454802 | 0.89054584 |
| Ttc30a2 | 0.01067692 | 0.00965184 | 0.01142608 | -0.0017742 | 0.76529452 | 0.89054584 |
| Ska3 | 0.02772674 | 0.02789399 | 0.0296706 | -0.0017766 | 0.60986496 | 0.89054584 |
| Rims4 | 0.01060951 | 0.00957528 | 0.01135395 | -0.0017787 | 0.76725998 | 0.89054584 |
| Kcnj2 | 0.01637278 | 0.01574237 | 0.01752138 | -0.001779 | 0.68190788 | 0.89054584 |
| Poc1b | 0.13791708 | 0.14476171 | 0.14654554 | -0.0017838 | 0.52261229 | 0.89054584 |
| Mrps10 | 0.69692942 | 0.64588924 | 0.64767412 | -0.0017849 | 0.50512198 | 0.89054584 |

| | | | | | | |
|---|---|---|---|---|---|---|
| Lrrc7 | 0.00579492 | 0.00440773 | 0.00620157 | -0.0017938 | 0.91111041 | 0.93088645 |
| Gm15351 | 0.00609874 | 0.00473272 | 0.00652671 | -0.001794 | 0.89983186 | 0.92300013 |
| Glt1d1 | 0.00551749 | 0.00411038 | 0.00590467 | -0.0017943 | 0.92157439 | 0.93827146 |
| Dlec1 | 0.00986157 | 0.00875587 | 0.01055354 | -0.0017977 | 0.78628429 | 0.89054584 |
| 4931429I11R | 0.01073176 | 0.00968371 | 0.01148477 | -0.0018011 | 0.76749549 | 0.89054584 |
| Zfp382 | 0.00614849 | 0.004771 | 0.00657995 | -0.0018089 | 0.899873 | 0.92300013 |
| Styx | 0.00903896 | 0.00786274 | 0.00967322 | -0.0018105 | 0.80839232 | 0.89054584 |
| Hcar2 | 0.00489453 | 0.00341924 | 0.00523799 | -0.0018188 | 0.94713593 | 0.95662382 |
| Fam53a | 0.40644262 | 0.40742784 | 0.40924739 | -0.0018195 | 0.50826312 | 0.89054584 |
| Ppp4r1 | 0.18293023 | 0.19105672 | 0.19287756 | -0.0018208 | 0.51754073 | 0.89054584 |
| Gm12207 | 0.01517832 | 0.01442148 | 0.01624318 | -0.0018217 | 0.6993421 | 0.89054584 |
| 1700024P16I | 0.00547736 | 0.00403551 | 0.00586172 | -0.0018262 | 0.92667847 | 0.94234302 |
| Gm26916 | 0.00572593 | 0.00430087 | 0.00612774 | -0.0018269 | 0.91757973 | 0.93505805 |
| Tmc8 | 0.00572593 | 0.00430087 | 0.00612774 | -0.0018269 | 0.91757973 | 0.93505805 |
| Chdh | 0.00609503 | 0.00469542 | 0.00652273 | -0.0018273 | 0.90408402 | 0.92605695 |
| 2310047D07 | 0.01024934 | 0.00913665 | 0.01096851 | -0.0018319 | 0.78174224 | 0.89054584 |
| Reep1 | 0.03970498 | 0.0406529 | 0.04248546 | -0.0018326 | 0.57963421 | 0.89054584 |
| Pptc7 | 0.18301044 | 0.19112626 | 0.19295939 | -0.0018331 | 0.51765153 | 0.89054584 |
| Kl | 0.02010864 | 0.01968212 | 0.02151909 | -0.001837 | 0.65457739 | 0.89054584 |
| Cib3 | 0.00595392 | 0.0045338 | 0.00637173 | -0.0018379 | 0.91050466 | 0.93052985 |
| B3gnt4 | 0.01656916 | 0.01589067 | 0.01773153 | -0.0018409 | 0.68569559 | 0.89054584 |
| 1700028J19R | 0.00607146 | 0.00465417 | 0.00649751 | -0.0018433 | 0.9068723 | 0.9279951 |
| Atp8a2 | 0.0138607 | 0.01298846 | 0.01483316 | -0.0018447 | 0.71884144 | 0.89054584 |
| Atrip | 0.00598339 | 0.00455807 | 0.00640327 | -0.0018452 | 0.91028952 | 0.93044116 |
| Gm26881 | 0.06270476 | 0.06523438 | 0.06708039 | -0.001846 | 0.55100884 | 0.89054584 |
| St3gal3 | 0.08033788 | 0.0840712 | 0.0859216 | -0.0018504 | 0.53996063 | 0.89054584 |
| Hist1h2bg | 0.00526616 | 0.00378377 | 0.0056357 | -0.0018519 | 0.93711089 | 0.94935599 |
| Pdxk | 0.07745357 | 0.08098623 | 0.08284077 | -0.0018545 | 0.54153382 | 0.89054584 |
| Stard6 | 0.00970715 | 0.00853289 | 0.01038829 | -0.0018554 | 0.79732591 | 0.89054584 |
| Hist1h3d | 0.02568902 | 0.02563408 | 0.02749028 | -0.0018562 | 0.62346059 | 0.89054584 |

| | | | | | | |
|---|---|---|---|---|---|---|
| Acss3 | 0.01479528 | 0.01397522 | 0.01583328 | -0.0018581 | 0.7077193 | 0.89054584 |
| Engase | 0.01596648 | 0.0152237 | 0.0170866 | -0.0018629 | 0.69425597 | 0.89054584 |
| 1810064F22I | 0.02050609 | 0.02007959 | 0.02194438 | -0.0018648 | 0.6539136 | 0.89054584 |
| AY074887 | 0.00541377 | 0.00392557 | 0.00579367 | -0.0018681 | 0.9334795 | 0.94666941 |
| Cftr | 0.01015605 | 0.00899987 | 0.01086867 | -0.0018688 | 0.78845699 | 0.89054584 |
| Lpcat2 | 0.0134709 | 0.01254717 | 0.01441603 | -0.0018689 | 0.72706994 | 0.89054584 |
| Gm16712 | 0.01074758 | 0.00963133 | 0.0115017 | -0.0018704 | 0.77566108 | 0.89054584 |
| Rab11fip4 | 0.01723211 | 0.01656739 | 0.01844096 | -0.0018736 | 0.68201477 | 0.89054584 |
| Fcna | 0.01073407 | 0.00961333 | 0.01148724 | -0.0018739 | 0.7763758 | 0.89054584 |
| A930019D19 | 0.01830069 | 0.01770935 | 0.01958444 | -0.0018751 | 0.67222433 | 0.89054584 |
| Insr | 0.20320343 | 0.21159625 | 0.21347353 | -0.0018773 | 0.51634026 | 0.89054584 |
| Mex3c | 0.12768044 | 0.13401923 | 0.13589776 | -0.0018785 | 0.52567444 | 0.89054584 |
| 9530082P21I | 0.01756067 | 0.0169137 | 0.01879255 | -0.0018788 | 0.67931829 | 0.89054584 |
| Gm26882 | 0.02747675 | 0.02752412 | 0.02940312 | -0.001879 | 0.61704476 | 0.89054584 |
| Dlgap5 | 0.07000963 | 0.07300776 | 0.0748876 | -0.0018798 | 0.54654987 | 0.89054584 |
| Zfp51 | 0.13695984 | 0.14366887 | 0.14555156 | -0.0018827 | 0.52402682 | 0.89054584 |
| Ell3 | 0.00785817 | 0.00652489 | 0.00840958 | -0.0018847 | 0.85178718 | 0.89985795 |
| Prrg4 | 0.01328299 | 0.01232559 | 0.01421494 | -0.0018894 | 0.73211911 | 0.89054584 |
| Dnaic2 | 0.00603043 | 0.00456399 | 0.0064536 | -0.0018896 | 0.91373799 | 0.93295744 |
| 2310057J18F | 0.00932592 | 0.00808851 | 0.00998031 | -0.0018918 | 0.81141359 | 0.89054584 |
| Ptger1 | 0.01481573 | 0.01396306 | 0.01585516 | -0.0018921 | 0.71088905 | 0.89054584 |
| Cyb5r1 | 0.15972414 | 0.16719615 | 0.16909269 | -0.0018965 | 0.52083702 | 0.89054584 |
| Gm11789 | 0.01024353 | 0.00906412 | 0.0109623 | -0.0018982 | 0.79008616 | 0.89054584 |
| Car6 | 0.00566642 | 0.00416489 | 0.00606405 | -0.0018992 | 0.92773114 | 0.94301662 |
| Lad1 | 0.01454597 | 0.0136666 | 0.01556649 | -0.0018999 | 0.71519408 | 0.89054584 |
| E230013L22F | 0.00949726 | 0.00826173 | 0.01016367 | -0.0019019 | 0.80835013 | 0.89054584 |
| Diaph3 | 0.05074279 | 0.05238807 | 0.05429102 | -0.001903 | 0.56485906 | 0.89054584 |
| Rnf43 | 0.00535813 | 0.00383106 | 0.00573412 | -0.0019031 | 0.93897489 | 0.9506465 |
| Hist2h2bb | 0.06039856 | 0.06270581 | 0.06461511 | -0.0019093 | 0.55474795 | 0.89054584 |
| Mettl20 | 0.05159355 | 0.05328618 | 0.05520081 | -0.0019146 | 0.56418729 | 0.89054584 |

| | | | | | | |
|---|---|---|---|---|---|---|
| Crygn | 0.0051626 | 0.00360901 | 0.00552488 | -0.0019159 | 0.94690814 | 0.9564742 |
| Slc39a7 | 1.09970321 | 0.88311132 | 0.88502752 | -0.0019162 | 0.50402414 | 0.89054584 |
| A230065N10 | 0.00966604 | 0.00842335 | 0.01034429 | -0.0019209 | 0.80652602 | 0.89054584 |
| Dpyd | 0.01326279 | 0.01227175 | 0.01419332 | -0.0019216 | 0.73589911 | 0.89054584 |
| Pdf | 0.01019395 | 0.00898386 | 0.01090923 | -0.0019254 | 0.79453491 | 0.89054584 |
| Gm10643 | 0.02656874 | 0.02650467 | 0.02843157 | -0.0019269 | 0.62390478 | 0.89054584 |
| 2410004I01R | 0.01042045 | 0.00922385 | 0.01115162 | -0.0019278 | 0.78970128 | 0.89054584 |
| 4930451G09 | 0.0147913 | 0.01389941 | 0.01582903 | -0.0019296 | 0.71496294 | 0.89054584 |
| Per1 | 0.13861211 | 0.14533549 | 0.14726701 | -0.0019315 | 0.52436254 | 0.89054584 |
| Tulp2 | 0.00929232 | 0.00800958 | 0.00994435 | -0.0019348 | 0.81764932 | 0.89054584 |
| Kcnip2 | 0.00443765 | 0.00281368 | 0.00474906 | -0.0019354 | 0.97119488 | 0.97617744 |
| Dpep2 | 0.02108858 | 0.02063196 | 0.02256769 | -0.0019357 | 0.65528076 | 0.89054584 |
| Tnfaip8l3 | 0.01029011 | 0.00907444 | 0.01101214 | -0.0019377 | 0.7938325 | 0.89054584 |
| Kcnk6 | 0.02195656 | 0.02155719 | 0.02349647 | -0.0019393 | 0.64970542 | 0.89054584 |
| Gm43950 | 0.01279956 | 0.01175322 | 0.0136976 | -0.0019444 | 0.74580321 | 0.89054584 |
| Astn1 | 0.00994336 | 0.00869626 | 0.01064106 | -0.0019448 | 0.80275045 | 0.89054584 |
| Gm43573 | 0.02137141 | 0.02092181 | 0.02287032 | -0.0019485 | 0.65429177 | 0.89054584 |
| Gm9949 | 0.01047085 | 0.00925623 | 0.01120556 | -0.0019493 | 0.79116592 | 0.89054584 |
| 1700030K09I | 0.08276788 | 0.08656213 | 0.0885168 | -0.0019547 | 0.54097127 | 0.89054584 |
| Tmem88b | 0.00487384 | 0.00325836 | 0.00521586 | -0.0019575 | 0.9598101 | 0.96687929 |
| D630011A20 | 0.01363283 | 0.01262936 | 0.01458932 | -0.00196 | 0.73430683 | 0.89054584 |
| Heca | 0.02418575 | 0.02392099 | 0.02588178 | -0.0019608 | 0.63793984 | 0.89054584 |
| Ahcy | 0.05041578 | 0.05198034 | 0.05394133 | -0.001961 | 0.56724764 | 0.89054584 |
| A330102I10F | 0.01384519 | 0.01285514 | 0.01481657 | -0.0019614 | 0.73130127 | 0.89054584 |
| B430218F22 | 0.00964126 | 0.00835555 | 0.01031777 | -0.0019622 | 0.81219934 | 0.89054584 |
| Pcdhga11 | 0.0193734 | 0.01876205 | 0.02073232 | -0.0019703 | 0.67102953 | 0.89054584 |
| Irf1 | 0.09931947 | 0.10420189 | 0.1061837 | -0.0019818 | 0.53464612 | 0.89054584 |
| Xpnpep3 | 0.08730613 | 0.09138001 | 0.09336262 | -0.0019826 | 0.53940516 | 0.89054584 |
| Caly | 0.01433105 | 0.01335317 | 0.0153365 | -0.0019833 | 0.72657874 | 0.89054584 |
| Gm11110 | 0.00579116 | 0.00421347 | 0.00619755 | -0.0019841 | 0.9320862 | 0.94578568 |

| | | | | | | |
|---|---|---|---|---|---|---|
| Oprl1 | 0.01366759 | 0.01264128 | 0.01462652 | -0.0019852 | 0.73642162 | 0.89054584 |
| Mafk | 0.09259243 | 0.09701768 | 0.09900556 | -0.0019879 | 0.5372645 | 0.89054584 |
| Pcm1 | 1.14566569 | 0.90451515 | 0.90650376 | -0.0019886 | 0.50407727 | 0.89054584 |
| Med22 | 0.28569164 | 0.29340366 | 0.29539411 | -0.0019905 | 0.51252206 | 0.89054584 |
| Gm2237 | 0.0119229 | 0.01076673 | 0.01275946 | -0.0019927 | 0.76657644 | 0.89054584 |
| 2010109A12I | 0.01807184 | 0.01734508 | 0.01933955 | -0.0019945 | 0.68455402 | 0.89054584 |
| Ints7 | 0.18792496 | 0.19597212 | 0.1979681 | -0.001996 | 0.51873264 | 0.89054584 |
| Pde9a | 0.01503082 | 0.01408594 | 0.01608534 | -0.0019994 | 0.71874048 | 0.89054584 |
| Prox2os | 0.00454438 | 0.00285995 | 0.00486328 | -0.0020033 | 0.97251734 | 0.97730342 |
| Ptgfrn | 0.05736511 | 0.05936784 | 0.06137208 | -0.0020042 | 0.56046507 | 0.89054584 |
| Gm16253 | 0.01338521 | 0.01231983 | 0.01432433 | -0.0020045 | 0.74278547 | 0.89054584 |
| Gm7008 | 0.02936561 | 0.029418 | 0.0314241 | -0.0020061 | 0.61692882 | 0.89054584 |
| Kcnj13 | 0.0337384 | 0.03409062 | 0.03610251 | -0.0020119 | 0.60242526 | 0.89054584 |
| Adap2 | 0.00958259 | 0.00824198 | 0.01025498 | -0.002013 | 0.81977944 | 0.89054584 |
| D230017M19 | 0.01426146 | 0.01324219 | 0.01526203 | -0.0020198 | 0.73124548 | 0.89054584 |
| 4732414G09 | 0.02253561 | 0.02209347 | 0.02411608 | -0.0020226 | 0.6520065 | 0.89054584 |
| Adamts20 | 0.02413859 | 0.02380801 | 0.02583131 | -0.0020233 | 0.64241563 | 0.89054584 |
| Ctif | 0.05661857 | 0.05854933 | 0.06057391 | -0.0020246 | 0.56187217 | 0.89054584 |
| Chrd | 0.0473522 | 0.04864037 | 0.05066498 | -0.0020246 | 0.57384616 | 0.89054584 |
| Prkd1 | 0.02180428 | 0.02130673 | 0.02333353 | -0.0020268 | 0.6571454 | 0.89054584 |
| Serpina9 | 0.04593839 | 0.04712444 | 0.04915288 | -0.0020284 | 0.57623272 | 0.89054584 |
| Mmgt1 | 0.09113346 | 0.09541956 | 0.09744834 | -0.0020288 | 0.53863474 | 0.89054584 |
| Inmt | 0.01809978 | 0.01733607 | 0.01936945 | -0.0020334 | 0.68761067 | 0.89054584 |
| Akip1 | 0.21995376 | 0.22832365 | 0.23035812 | -0.0020345 | 0.51641048 | 0.89054584 |
| Lgi4 | 0.0102304 | 0.00891284 | 0.01094824 | -0.0020354 | 0.80679659 | 0.89054584 |
| Fam49a | 0.02651326 | 0.02633663 | 0.0283722 | -0.0020356 | 0.63090762 | 0.89054584 |
| Lrrc8e | 0.03473293 | 0.03512816 | 0.0371665 | -0.0020383 | 0.6008363 | 0.89054584 |
| Hnf4g | 0.00562413 | 0.00397952 | 0.00601879 | -0.0020393 | 0.94277809 | 0.95349821 |
| Gjc2 | 0.01026851 | 0.00894646 | 0.01098902 | -0.0020426 | 0.80674703 | 0.89054584 |
| Ankrd34c | 0.00444707 | 0.00271549 | 0.00475913 | -0.0020436 | 0.97728329 | 0.98100486 |

| | | | | | | |
|---|---|---|---|---|---|---|
| Slc1a1 | 0.00555748 | 0.00390172 | 0.00594746 | -0.0020457 | 0.94548111 | 0.9553025 |
| Gm21814 | 0.0055624 | 0.00390513 | 0.00595273 | -0.0020476 | 0.94548459 | 0.9553025 |
| C330021F23I | 0.01354068 | 0.01244161 | 0.0144907 | -0.0020491 | 0.74499057 | 0.89054584 |
| Mettl21b | 0.01380043 | 0.01271767 | 0.01476867 | -0.002051 | 0.74118759 | 0.89054584 |
| D2hgdh | 0.08263622 | 0.08632444 | 0.08837619 | -0.0020518 | 0.54306665 | 0.89054584 |
| Itpripl1 | 0.07468588 | 0.0778321 | 0.07988409 | -0.002052 | 0.54762959 | 0.89054584 |
| 1700088E04I | 0.25487783 | 0.26310214 | 0.26515903 | -0.0020569 | 0.51441475 | 0.89054584 |
| Hoxa7 | 0.01808497 | 0.01729534 | 0.0193536 | -0.0020583 | 0.68986885 | 0.89054584 |
| Heph | 0.03374789 | 0.03404962 | 0.03611266 | -0.002063 | 0.60494033 | 0.89054584 |
| Kctd6 | 0.06840074 | 0.07109885 | 0.07316828 | -0.0020694 | 0.55241677 | 0.89054584 |
| Dnajc30 | 0.44564187 | 0.44242273 | 0.44449263 | -0.0020699 | 0.50865463 | 0.89054584 |
| Mfap3l | 0.01025773 | 0.00890731 | 0.01097748 | -0.0020702 | 0.81019055 | 0.89054584 |
| 3300002I08R | 0.02958913 | 0.02959247 | 0.03166325 | -0.0020708 | 0.61970134 | 0.89054584 |
| Adcy5 | 0.00504392 | 0.00332664 | 0.00539787 | -0.0020712 | 0.96308746 | 0.96937253 |
| Maneal | 0.00406479 | 0.00227824 | 0.00435003 | -0.0020718 | 0.98675393 | 0.98893671 |
| Cbfb | 0.54801998 | 0.53013288 | 0.53220594 | -0.0020731 | 0.50723945 | 0.89054584 |
| Gm8773 | 0.0101584 | 0.00879712 | 0.01087119 | -0.0020741 | 0.81296028 | 0.89054584 |
| Gm15492 | 0.01045029 | 0.00910854 | 0.01118356 | -0.002075 | 0.8063231 | 0.89054584 |
| Gnat2 | 0.01792765 | 0.01710804 | 0.01918525 | -0.0020772 | 0.69302002 | 0.89054584 |
| 5730422E09I | 0.00955947 | 0.00815113 | 0.01023024 | -0.0020791 | 0.82814114 | 0.89229532 |
| Sacs | 0.04149125 | 0.0423168 | 0.0443962 | -0.0020794 | 0.58636849 | 0.89054584 |
| Grid1 | 0.00980605 | 0.00841429 | 0.01049412 | -0.0020798 | 0.82208889 | 0.89071405 |
| Lsm7 | 1.00298125 | 0.83397812 | 0.83605956 | -0.0020814 | 0.50462714 | 0.89054584 |
| Rapgef5 | 0.02374664 | 0.0233302 | 0.02541191 | -0.0020817 | 0.64864113 | 0.89054584 |
| Rhbg | 0.02286746 | 0.02238907 | 0.02447117 | -0.0020821 | 0.65409518 | 0.89054584 |
| Zfp790 | 0.06517013 | 0.06763222 | 0.06971557 | -0.0020834 | 0.55536418 | 0.89054584 |
| Gm609 | 0.15944517 | 0.16672173 | 0.16880542 | -0.0020837 | 0.52293002 | 0.89054584 |
| Gpr137b | 0.10431187 | 0.10936046 | 0.11144437 | -0.0020839 | 0.53471117 | 0.89054584 |
| 5330439K02I | 0.00460608 | 0.00284469 | 0.00492931 | -0.0020846 | 0.97559705 | 0.97954524 |
| Gm14327 | 0.004744 | 0.00299195 | 0.0050769 | -0.002085 | 0.97214569 | 0.97699766 |

| | | | | | | |
|---|---|---|---|---|---|---|
| Nr1h3 | 0.00812663 | 0.00661137 | 0.00869687 | -0.0020855 | 0.86804744 | 0.90466949 |
| Zfp458 | 0.04438285 | 0.04540351 | 0.04748914 | -0.0020856 | 0.58106246 | 0.89054584 |
| Esrrg | 0.00499388 | 0.00325309 | 0.00534432 | -0.0020912 | 0.96585133 | 0.97168223 |
| Gm26566 | 0.00876272 | 0.00728506 | 0.00937759 | -0.0020925 | 0.85073872 | 0.89942466 |
| Zkscan7 | 0.00876511 | 0.00728721 | 0.00938015 | -0.0020929 | 0.85071983 | 0.89942466 |
| U2af1l4 | 0.03518688 | 0.03555918 | 0.03765215 | -0.002093 | 0.60217332 | 0.89054584 |
| Bloc1s6os | 0.00978424 | 0.00837731 | 0.01047078 | -0.0020935 | 0.82419732 | 0.89120023 |
| Gm26669 | 0.00548093 | 0.00377125 | 0.00586554 | -0.0020943 | 0.95189101 | 0.96063732 |
| Emilin3 | 0.01484964 | 0.01379615 | 0.01589146 | -0.0020953 | 0.73048727 | 0.89054584 |
| Aqp3 | 0.01011605 | 0.00872765 | 0.01082587 | -0.0020982 | 0.81672994 | 0.89054584 |
| Tigd5 | 0.05421081 | 0.05589736 | 0.05799948 | -0.0021021 | 0.5670457 | 0.89054584 |
| 1190002N15 | 0.13248802 | 0.13880067 | 0.14090344 | -0.0021028 | 0.52771514 | 0.89054584 |
| C78197 | 0.00974504 | 0.00832533 | 0.01042883 | -0.0021035 | 0.82631699 | 0.89170439 |
| 2810405F17I | 0.01669483 | 0.0157622 | 0.01786601 | -0.0021038 | 0.70836462 | 0.89054584 |
| Tfrc | 0.15348566 | 0.16055569 | 0.16266129 | -0.0021056 | 0.52404508 | 0.89054584 |
| AA474408 | 0.0188559 | 0.01806821 | 0.02017856 | -0.0021104 | 0.68695975 | 0.89054584 |
| Chst10 | 0.03460228 | 0.03491583 | 0.03702672 | -0.0021109 | 0.60472893 | 0.89054584 |
| Nfyb | 0.73899271 | 0.67541435 | 0.67752745 | -0.0021131 | 0.50579663 | 0.89054584 |
| 1600002K03I | 0.15108327 | 0.15806609 | 0.16018062 | -0.0021145 | 0.52452033 | 0.89054584 |
| Selenbp2 | 0.00882177 | 0.00732472 | 0.00944079 | -0.0021161 | 0.85181752 | 0.89985795 |
| Acss1 | 0.00833388 | 0.00680232 | 0.00891867 | -0.0021163 | 0.86553715 | 0.90381429 |
| 2310068J16F | 0.02842029 | 0.02829532 | 0.03041267 | -0.0021173 | 0.62716667 | 0.89054584 |
| Slc14a2 | 0.05851731 | 0.06048448 | 0.06260393 | -0.0021194 | 0.56266434 | 0.89054584 |
| Gm26771 | 0.02152763 | 0.02091474 | 0.02303749 | -0.0021228 | 0.66614408 | 0.89054584 |
| Ccnk | 0.18613332 | 0.19402035 | 0.1961433 | -0.002123 | 0.52010856 | 0.89054584 |
| Gjb2 | 0.0033684 | 0.00147543 | 0.00360478 | -0.0021293 | 0.99703875 | 0.99752038 |
| Gm3336 | 0.00470532 | 0.00290542 | 0.00503551 | -0.0021301 | 0.97562652 | 0.97954524 |
| Cd109 | 0.01382863 | 0.0126686 | 0.01479885 | -0.0021302 | 0.7487746 | 0.89054584 |
| Prkcq | 0.00869595 | 0.00717538 | 0.00930614 | -0.0021308 | 0.85695213 | 0.9009252 |
| BC021785 | 0.00942026 | 0.0079502 | 0.01008126 | -0.0021311 | 0.83765078 | 0.89485554 |

| | | | | | | |
|---|---|---|---|---|---|---|
| RP23-235M8 | 0.01418519 | 0.01304666 | 0.01518041 | -0.0021338 | 0.74372063 | 0.89054584 |
| Ralyl | 0.00371236 | 0.00183654 | 0.00397287 | -0.0021363 | 0.99388169 | 0.99525464 |
| Otud1 | 0.03060261 | 0.03061077 | 0.0327476 | -0.0021368 | 0.61943709 | 0.89054584 |
| Pdzd4 | 0.03282387 | 0.03298662 | 0.0351241 | -0.0021375 | 0.61161056 | 0.89054584 |
| 4430402I18R | 0.00365264 | 0.0017679 | 0.00390896 | -0.0021411 | 0.99464226 | 0.99580993 |
| Gm27017 | 0.00945628 | 0.00797844 | 0.01011982 | -0.0021414 | 0.83789487 | 0.89493147 |
| Zfp97 | 0.00324082 | 0.00132104 | 0.00346824 | -0.0021472 | 0.998039 | 0.99838332 |
| Gm37584 | 0.00485233 | 0.00304348 | 0.00519284 | -0.0021494 | 0.97309581 | 0.97761368 |
| Ccdc84 | 0.08757047 | 0.09149488 | 0.09364484 | -0.00215 | 0.54259075 | 0.89054584 |
| Cog6 | 0.40802487 | 0.40853603 | 0.41068698 | -0.002151 | 0.50973364 | 0.89054584 |
| Hdc | 0.01179853 | 0.0104754 | 0.01262637 | -0.002151 | 0.78630711 | 0.89054584 |
| Dbil5 | 0.01415424 | 0.01299556 | 0.01514729 | -0.0021517 | 0.74595683 | 0.89054584 |
| Brinp3 | 0.00464307 | 0.0028171 | 0.00496889 | -0.0021518 | 0.9781817 | 0.98170278 |
| Gria1 | 0.01021409 | 0.0087775 | 0.01093079 | -0.0021533 | 0.82063254 | 0.89054584 |
| Sowahb | 0.02295573 | 0.02241207 | 0.02456562 | -0.0021535 | 0.65851954 | 0.89054584 |
| Kcnab3 | 0.02804736 | 0.02785803 | 0.03001364 | -0.0021556 | 0.63104049 | 0.89054584 |
| Gm19710 | 0.01245248 | 0.01116833 | 0.01332619 | -0.0021579 | 0.77469687 | 0.89054584 |
| D130043K22 | 0.01005529 | 0.00860085 | 0.01076085 | -0.00216 | 0.82515152 | 0.89156706 |
| Ncam2 | 0.00480812 | 0.00298313 | 0.00514553 | -0.0021624 | 0.97487978 | 0.97893108 |
| Klhdc9 | 0.00897574 | 0.00744244 | 0.00960556 | -0.0021631 | 0.85295017 | 0.9000052 |
| Fyn | 0.31538463 | 0.32192974 | 0.32409318 | -0.0021634 | 0.51240517 | 0.89054584 |
| 4930471M09 | 0.00918411 | 0.00766259 | 0.00982855 | -0.002166 | 0.84772155 | 0.89862623 |
| Hoxa4 | 0.0206408 | 0.01991831 | 0.02208854 | -0.0021702 | 0.67643242 | 0.89054584 |
| Sntg2 | 0.03406399 | 0.03428047 | 0.03645084 | -0.0021704 | 0.60926535 | 0.89054584 |
| Npy1r | 0.01021462 | 0.00875998 | 0.01093135 | -0.0021714 | 0.82263115 | 0.89073269 |
| A930031H19 | 0.00518648 | 0.0033787 | 0.00555043 | -0.0021717 | 0.96584185 | 0.97168223 |
| Wdr66 | 0.00491923 | 0.003092 | 0.00526442 | -0.0021724 | 0.97273265 | 0.97738429 |
| Aldh1a3 | 0.00491923 | 0.003092 | 0.00526442 | -0.0021724 | 0.97273265 | 0.97738429 |
| Gabre | 0.00741282 | 0.00576011 | 0.00793299 | -0.0021729 | 0.89904067 | 0.92252046 |
| Lor | 0.00962253 | 0.00812174 | 0.01029772 | -0.002176 | 0.83755693 | 0.89485554 |

| | | | | | | |
|---|---|---|---|---|---|---|
| Gm26549 | 0.02851936 | 0.02834212 | 0.03051866 | -0.0021765 | 0.63015662 | 0.89054584 |
| Acbd4 | 0.06019541 | 0.06222124 | 0.06439793 | -0.0021767 | 0.56256473 | 0.89054584 |
| Bsn | 0.04276918 | 0.04358074 | 0.04576314 | -0.0021824 | 0.5879135 | 0.89054584 |
| Crybg3 | 0.08251395 | 0.08606061 | 0.08824562 | -0.002185 | 0.54591929 | 0.89054584 |
| Thap8 | 0.00803032 | 0.00640878 | 0.00859381 | -0.002185 | 0.88190532 | 0.91223443 |
| Ssc5d | 0.02094462 | 0.0202266 | 0.02241364 | -0.002187 | 0.67530264 | 0.89054584 |
| A530095I07F | 0.00382338 | 0.00190294 | 0.00409169 | -0.0021887 | 0.99365207 | 0.99509343 |
| 4933431E20I | 0.03861763 | 0.03913177 | 0.04132229 | -0.0021905 | 0.59753532 | 0.89054584 |
| Gm43403 | 0.00824637 | 0.00663322 | 0.00882502 | -0.0021918 | 0.87638551 | 0.9091719 |
| Sult1b1 | 0.00896803 | 0.00740315 | 0.00959731 | -0.0021942 | 0.85659281 | 0.9009252 |
| Fam221b | 0.00421827 | 0.0023165 | 0.00451428 | -0.0021978 | 0.98834169 | 0.99045952 |
| Ltk | 0.01236253 | 0.01103164 | 0.01322993 | -0.0021983 | 0.7805734 | 0.89054584 |
| Arid1b | 0.32357066 | 0.32972923 | 0.33192766 | -0.0021984 | 0.51230829 | 0.89054584 |
| Gm43625 | 0.00494771 | 0.00309132 | 0.0052949 | -0.0022036 | 0.97374409 | 0.97799389 |
| Uox | 0.00467536 | 0.00279984 | 0.00500344 | -0.0022036 | 0.97990921 | 0.98311155 |
| Kalrn | 0.14239343 | 0.14898502 | 0.15118895 | -0.0022039 | 0.52707332 | 0.89054584 |
| Slc9b2 | 0.0129128 | 0.01161377 | 0.01381879 | -0.002205 | 0.77138393 | 0.89054584 |
| Cyp4x1 | 0.00349968 | 0.00153817 | 0.00374527 | -0.0022071 | 0.99697928 | 0.99752038 |
| Greb1 | 0.01368153 | 0.012431 | 0.01464143 | -0.0022104 | 0.75908604 | 0.89054584 |
| Def6 | 0.03973173 | 0.04030171 | 0.04251407 | -0.0022124 | 0.59578175 | 0.89054584 |
| Gm16062 | 0.02290641 | 0.02230027 | 0.02451284 | -0.0022126 | 0.66294859 | 0.89054584 |
| Nacad | 0.01291617 | 0.01160898 | 0.0138224 | -0.0022134 | 0.77218177 | 0.89054584 |
| Apoa2 | 0.02987375 | 0.02975306 | 0.03196777 | -0.0022147 | 0.62656508 | 0.89054584 |
| Fzd9 | 0.00854942 | 0.00693062 | 0.00914933 | -0.0022187 | 0.87071804 | 0.90602671 |
| Folr1 | 0.02351325 | 0.02294144 | 0.02516218 | -0.0022207 | 0.65953126 | 0.89054584 |
| Slc8b1 | 0.02328762 | 0.02269974 | 0.02492075 | -0.002221 | 0.66100941 | 0.89054584 |
| Hhipl2 | 0.01010142 | 0.00858906 | 0.01081021 | -0.0022211 | 0.83078252 | 0.89280428 |
| Ttc34 | 0.00806846 | 0.00641156 | 0.00863463 | -0.0022231 | 0.88483096 | 0.91404124 |
| A430078I02F | 0.00915114 | 0.00756822 | 0.00979326 | -0.002225 | 0.85509041 | 0.90088661 |
| Slc44a4 | 0.00457507 | 0.00266771 | 0.00489612 | -0.0022284 | 0.98301611 | 0.98586892 |

| | | | | | | |
|---|---|---|---|---|---|---|
| 5430402O13 | 0.00457037 | 0.00266166 | 0.0048911 | -0.0022294 | 0.98314891 | 0.98586892 |
| Lrguk | 0.01300931 | 0.01169117 | 0.01392207 | -0.0022309 | 0.77233473 | 0.89054584 |
| Ccdc158 | 0.00364847 | 0.00167093 | 0.00390449 | -0.0022336 | 0.99615226 | 0.99677103 |
| Synm | 0.00364847 | 0.00167093 | 0.00390449 | -0.0022336 | 0.99615226 | 0.99677103 |
| Dennd2c | 0.00383273 | 0.00186656 | 0.00410169 | -0.0022351 | 0.99443826 | 0.99567444 |
| Pogz | 0.28206159 | 0.28961833 | 0.29185582 | -0.0022375 | 0.51424617 | 0.89054584 |
| Phyhip | 0.01005429 | 0.0085205 | 0.01075977 | -0.0022393 | 0.83387566 | 0.89423396 |
| C1ql1 | 0.00917134 | 0.00757308 | 0.00981488 | -0.0022418 | 0.85636726 | 0.9009252 |
| Zfp942 | 0.27676534 | 0.28444001 | 0.28668243 | -0.0022424 | 0.51453508 | 0.89054584 |
| Gm26720 | 0.00892877 | 0.00731119 | 0.0095553 | -0.0022441 | 0.8630606 | 0.90285393 |
| Slc24a3 | 0.02796521 | 0.02767646 | 0.02992575 | -0.0022493 | 0.63689711 | 0.89054584 |
| Ddb2 | 0.02770639 | 0.02739953 | 0.02964882 | -0.0022493 | 0.63812399 | 0.89054584 |
| Lhx6 | 0.01007603 | 0.00853026 | 0.01078304 | -0.0022528 | 0.83480786 | 0.89453395 |
| 2610020H08I | 0.01239626 | 0.01101023 | 0.01326603 | -0.0022558 | 0.78588445 | 0.89054584 |
| A630033H20 | 0.01840281 | 0.01743757 | 0.01969372 | -0.0022562 | 0.70323851 | 0.89054584 |
| Spag6l | 0.03042152 | 0.03029707 | 0.03255385 | -0.0022568 | 0.6266443 | 0.89054584 |
| Clec12b | 0.01454415 | 0.01330439 | 0.01556454 | -0.0022602 | 0.75064886 | 0.89054584 |
| Fam199x | 0.11279266 | 0.11808097 | 0.12034228 | -0.0022613 | 0.5348806 | 0.89054584 |
| Xntrpc | 0.00389572 | 0.00190645 | 0.0041691 | -0.0022627 | 0.99427235 | 0.99557708 |
| Stac3 | 0.00398645 | 0.00200354 | 0.0042662 | -0.0022627 | 0.99326237 | 0.99484061 |
| Recql4 | 0.00398645 | 0.00200354 | 0.0042662 | -0.0022627 | 0.99326237 | 0.99484061 |
| Otop3 | 0.02705889 | 0.02669265 | 0.02895601 | -0.0022634 | 0.64213401 | 0.89054584 |
| Zfp324 | 0.08613505 | 0.08984204 | 0.09211229 | -0.0022703 | 0.54570882 | 0.89054584 |
| Erich2os | 0.00777815 | 0.0060531 | 0.00832395 | -0.0022708 | 0.898136 | 0.92211428 |
| Cep152 | 0.03511152 | 0.03530035 | 0.03757152 | -0.0022712 | 0.61088635 | 0.89054584 |
| Rwdd2b | 0.05673678 | 0.0584278 | 0.06070029 | -0.0022725 | 0.56923272 | 0.89054584 |
| Eif2b2 | 0.50371282 | 0.4927641 | 0.49503902 | -0.0022749 | 0.50854064 | 0.89054584 |
| Nceh1 | 0.05653711 | 0.05820783 | 0.06048681 | -0.002279 | 0.56967093 | 0.89054584 |
| Ppp1r27 | 0.01002391 | 0.00844782 | 0.01072726 | -0.0022794 | 0.83890824 | 0.89500145 |
| Nrxn2 | 0.05513502 | 0.0567065 | 0.05898769 | -0.0022812 | 0.57149107 | 0.89054584 |

| | | | | | | |
|---|---|---|---|---|---|---|
| Gm648 | 0.05092965 | 0.05220825 | 0.05449085 | -0.0022826 | 0.57736655 | 0.89054584 |
| Mapre1 | 1.44637165 | 1.01789579 | 1.0201786 | -0.0022828 | 0.50415895 | 0.89054584 |
| Plac1 | 0.04584314 | 0.04676647 | 0.04905101 | -0.0022845 | 0.58589145 | 0.89054584 |
| Tmem213 | 0.00974426 | 0.00814346 | 0.010428 | -0.0022845 | 0.84629484 | 0.89825055 |
| Gm5449 | 0.01453024 | 0.01326161 | 0.01554966 | -0.002288 | 0.75349772 | 0.89054584 |
| Gm14322 | 0.00812066 | 0.00639843 | 0.00869049 | -0.0022921 | 0.89042027 | 0.9171534 |
| Ttc21a | 0.0081127 | 0.00638755 | 0.00868197 | -0.0022944 | 0.89088273 | 0.91722772 |
| Rep15 | 0.02418574 | 0.0235859 | 0.02588176 | -0.0022959 | 0.66029721 | 0.89054584 |
| Ebf4 | 0.00978808 | 0.00817838 | 0.01047489 | -0.0022965 | 0.84647266 | 0.89825055 |
| Gm42912 | 0.00928307 | 0.00763681 | 0.00993445 | -0.0022976 | 0.85937613 | 0.90131997 |
| Adamts16 | 0.07438562 | 0.07726264 | 0.0795633 | -0.0023007 | 0.5535827 | 0.89054584 |
| Wscd1 | 0.07476536 | 0.07766394 | 0.079969 | -0.0023051 | 0.55341389 | 0.89054584 |
| Slc35f1 | 0.00867264 | 0.00697437 | 0.00928119 | -0.0023068 | 0.87656262 | 0.9091719 |
| Mc4r | 0.00959206 | 0.00795794 | 0.01026512 | -0.0023072 | 0.85248268 | 0.89988783 |
| 3110083C13I | 0.01288215 | 0.01147604 | 0.01378599 | -0.0023099 | 0.78249284 | 0.89054584 |
| Ticrr | 0.02034947 | 0.0194657 | 0.02177679 | -0.0023111 | 0.689501 | 0.89054584 |
| Smad7 | 0.18268234 | 0.19031287 | 0.19262464 | -0.0023118 | 0.52229477 | 0.89054584 |
| A930016O22 | 0.00954626 | 0.00790344 | 0.01021611 | -0.0023127 | 0.85421066 | 0.90035231 |
| Pwwp2b | 0.042029 | 0.04265239 | 0.04497142 | -0.002319 | 0.59493018 | 0.89054584 |
| Epor | 0.01401406 | 0.01267682 | 0.01499728 | -0.0023205 | 0.76450127 | 0.89054584 |
| Katnal2 | 0.06223788 | 0.06425999 | 0.06658133 | -0.0023213 | 0.56451739 | 0.89054584 |
| Arhgap24 | 0.11719484 | 0.12261964 | 0.12495051 | -0.0023309 | 0.53462833 | 0.89054584 |
| Dydc2 | 0.01442146 | 0.01309979 | 0.01543325 | -0.0023335 | 0.75941304 | 0.89054584 |
| Doc2g | 0.00886733 | 0.00715546 | 0.00948954 | -0.0023341 | 0.87408784 | 0.90775282 |
| Msl3l2 | 0.1953787 | 0.2032073 | 0.20554511 | -0.0023378 | 0.5211298 | 0.89054584 |
| Zmiz1os1 | 0.00843219 | 0.00668532 | 0.00902388 | -0.0023386 | 0.88635564 | 0.9147483 |
| Lcor | 0.36789753 | 0.3714059 | 0.37374475 | -0.0023389 | 0.51162957 | 0.89054584 |
| Gm26762 | 0.00900505 | 0.00729671 | 0.00963693 | -0.0023402 | 0.87105058 | 0.90617762 |
| Adap1 | 0.00984331 | 0.00819312 | 0.010534 | -0.0023409 | 0.8497395 | 0.89887961 |
| Gm17435 | 0.00865252 | 0.00691373 | 0.00925967 | -0.0023459 | 0.88106845 | 0.91182315 |

| | | | | | | |
|---|---|---|---|---|---|---|
| Pcdhb9 | 0.00934898 | 0.00765887 | 0.01000499 | -0.0023461 | 0.86269258 | 0.90267402 |
| Gm572 | 0.00969571 | 0.00802861 | 0.01037604 | -0.0023474 | 0.85406256 | 0.90035231 |
| Kif27 | 0.0086368 | 0.00689239 | 0.00924284 | -0.0023504 | 0.88194479 | 0.91223443 |
| Veph1 | 0.01002317 | 0.00837405 | 0.01072647 | -0.0023524 | 0.84655093 | 0.89825055 |
| Tdrd5 | 0.00812574 | 0.00634186 | 0.00869593 | -0.0023541 | 0.89638491 | 0.92122978 |
| D430019H16 | 0.08328371 | 0.0867128 | 0.08906765 | -0.0023548 | 0.54901658 | 0.89054584 |
| Tef | 0.1651228 | 0.17229069 | 0.17464561 | -0.0023549 | 0.52504553 | 0.89054584 |
| Rab19 | 0.03719057 | 0.03744044 | 0.03979568 | -0.0023552 | 0.60862324 | 0.89054584 |
| 4930426I24R | 0.00986225 | 0.00819895 | 0.01055426 | -0.0023553 | 0.85076028 | 0.89942466 |
| Gm10263 | 0.19680182 | 0.20463303 | 0.20698906 | -0.002356 | 0.52114597 | 0.89054584 |
| 5330416C01I | 0.01277915 | 0.01131557 | 0.01367577 | -0.0023602 | 0.7893171 | 0.89054584 |
| Arhgef17 | 0.25684978 | 0.26474542 | 0.26710722 | -0.0023618 | 0.5164298 | 0.89054584 |
| Gm12031 | 0.0083976 | 0.00662371 | 0.00898686 | -0.0023632 | 0.88973141 | 0.91679645 |
| H2-Q7 | 0.0218379 | 0.0210019 | 0.0233695 | -0.0023676 | 0.68153821 | 0.89054584 |
| Nufip2 | 0.40710411 | 0.40747604 | 0.40984941 | -0.0023734 | 0.51076182 | 0.89054584 |
| Pdzd3 | 0.01625961 | 0.0150233 | 0.01740028 | -0.002377 | 0.73775434 | 0.89054584 |
| Kcne2 | 0.0125312 | 0.01103337 | 0.01341043 | -0.0023771 | 0.79554659 | 0.89054584 |
| Ppp1r18os | 0.0097991 | 0.00810829 | 0.01048668 | -0.0023784 | 0.85466531 | 0.90070056 |
| Nubp1 | 0.27869046 | 0.28618134 | 0.28856448 | -0.0023831 | 0.51534607 | 0.89054584 |
| Cdkl4 | 0.00822429 | 0.00641717 | 0.00880139 | -0.0023842 | 0.89653746 | 0.92130916 |
| Fchsd1 | 0.04168872 | 0.04222306 | 0.04460744 | -0.0023844 | 0.59833219 | 0.89054584 |
| Zfp652os | 0.01417168 | 0.01277949 | 0.01516596 | -0.0023865 | 0.76825639 | 0.89054584 |
| Zfp335 | 0.03022899 | 0.02996062 | 0.03234785 | -0.0023872 | 0.63451064 | 0.89054584 |
| Kcnc2 | 0.02812233 | 0.02770574 | 0.03009385 | -0.0023881 | 0.64420188 | 0.89054584 |
| Slc7a6 | 0.08081227 | 0.08403731 | 0.08642826 | -0.002391 | 0.55127567 | 0.89054584 |
| P2rx3 | 0.0182017 | 0.0170875 | 0.01947851 | -0.002391 | 0.7163018 | 0.89054584 |
| Nupl2 | 0.07118556 | 0.07375299 | 0.07614417 | -0.0023912 | 0.55816004 | 0.89054584 |
| Catip | 0.01450826 | 0.0131339 | 0.01552614 | -0.0023922 | 0.76357275 | 0.89054584 |
| Gm26652 | 0.00943495 | 0.00770359 | 0.01009699 | -0.0023934 | 0.8652818 | 0.90377426 |
| 1700058P15I | 0.00946737 | 0.00773814 | 0.01013168 | -0.0023935 | 0.86447418 | 0.90335496 |

| | | | | | | |
|---|---|---|---|---|---|---|
| Ciz1 | 0.27921527 | 0.28668246 | 0.28907724 | -0.0023948 | 0.51539366 | 0.89054584 |
| Cd14 | 0.0272346 | 0.02674771 | 0.02914402 | -0.0023963 | 0.64916698 | 0.89054584 |
| Hs1bp3 | 0.01626096 | 0.01500392 | 0.01740173 | -0.0023978 | 0.73955125 | 0.89054584 |
| Mir17hg | 0.02267031 | 0.02186056 | 0.02426021 | -0.0023997 | 0.67753995 | 0.89054584 |
| H2-Q6 | 0.07985156 | 0.08299699 | 0.08540218 | -0.0024052 | 0.5521953 | 0.89054584 |
| Cntn3 | 0.01329375 | 0.01182035 | 0.01422645 | -0.0024061 | 0.78464013 | 0.89054584 |
| Adcy1 | 0.03410471 | 0.03408759 | 0.03649441 | -0.0024068 | 0.6206766 | 0.89054584 |
| M5C1000I18 | 0.00794826 | 0.00609879 | 0.008506 | -0.0024072 | 0.90632965 | 0.92763623 |
| Soat1 | 0.45989392 | 0.45467829 | 0.45708575 | -0.0024075 | 0.50978851 | 0.89054584 |
| Dhx38 | 0.14604807 | 0.15256529 | 0.15497415 | -0.0024089 | 0.52886492 | 0.89054584 |
| Cabp7 | 0.041949 | 0.04247693 | 0.04488584 | -0.0024089 | 0.59871942 | 0.89054584 |
| Ehd3 | 0.05551345 | 0.05698152 | 0.05939232 | -0.0024108 | 0.57499732 | 0.89054584 |
| Tmem26 | 0.02614307 | 0.02556404 | 0.0279761 | -0.0024121 | 0.65604332 | 0.89054584 |
| Eno4 | 0.01833869 | 0.01720796 | 0.0196251 | -0.0024171 | 0.71695623 | 0.89054584 |
| Gm17669 | 0.01643427 | 0.015166 | 0.01758719 | -0.0024212 | 0.73936369 | 0.89054584 |
| Adgre5 | 0.14865174 | 0.1552399 | 0.15766764 | -0.0024277 | 0.52859466 | 0.89054584 |
| Tjap1 | 0.27563528 | 0.2831455 | 0.2855768 | -0.0024313 | 0.51581974 | 0.89054584 |
| Srsf12 | 0.0468926 | 0.04774212 | 0.05017344 | -0.0024313 | 0.58930785 | 0.89054584 |
| Agbl2 | 0.0092326 | 0.00744859 | 0.00988044 | -0.0024318 | 0.87424582 | 0.90780805 |
| Zfp960 | 0.02235795 | 0.02149327 | 0.02392598 | -0.0024327 | 0.68214482 | 0.89054584 |
| Fam189a2 | 0.01097219 | 0.00930765 | 0.01174206 | -0.0024344 | 0.83295587 | 0.89393134 |
| Egfl8 | 0.03534793 | 0.03538927 | 0.03782444 | -0.0024352 | 0.61789114 | 0.89054584 |
| Tspan2os | 0.0082312 | 0.00637059 | 0.00880878 | -0.0024382 | 0.90139696 | 0.92418557 |
| Zfp385c | 0.00974733 | 0.00798806 | 0.01043128 | -0.0024432 | 0.86241213 | 0.90267402 |
| Tenm2 | 0.00970229 | 0.00793666 | 0.01038308 | -0.0024464 | 0.86383665 | 0.9030793 |
| Tm6sf2 | 0.0095898 | 0.0078131 | 0.0102627 | -0.0024496 | 0.86693952 | 0.90430169 |
| Rgs7 | 0.01819614 | 0.01701835 | 0.01947256 | -0.0024542 | 0.7214605 | 0.89054584 |
| Anxa6 | 1.6139659 | 1.06147757 | 1.06393335 | -0.0024558 | 0.50429007 | 0.89054584 |
| Serpina1d | 0.02145448 | 0.02050115 | 0.02295922 | -0.0024581 | 0.69103854 | 0.89054584 |
| Ankrd55 | 0.00940996 | 0.00760591 | 0.01007025 | -0.0024643 | 0.87288 | 0.90729956 |

| | | | | | | |
|---|---|---|---|---|---|---|
| Gm5577 | 0.04659328 | 0.04738799 | 0.04985331 | -0.0024653 | 0.59110621 | 0.89054584 |
| 2610206C17I | 0.00939946 | 0.00758854 | 0.01005901 | -0.0024705 | 0.87373488 | 0.90766755 |
| Gm17227 | 0.01426112 | 0.01279001 | 0.01526167 | -0.0024717 | 0.77473301 | 0.89054584 |
| Hist1h2bj | 0.00942356 | 0.00760893 | 0.0100848 | -0.0024759 | 0.87364531 | 0.90763953 |
| Bmp5 | 0.01439375 | 0.01292707 | 0.01540359 | -0.0024765 | 0.77308486 | 0.89054584 |
| 1810034E14I | 0.0230159 | 0.02215346 | 0.02463001 | -0.0024765 | 0.68027108 | 0.89054584 |
| Tbx15 | 0.00933646 | 0.0075141 | 0.00999159 | -0.0024775 | 0.87599952 | 0.90910816 |
| Fbxl22 | 0.02018438 | 0.01912224 | 0.02160013 | -0.0024779 | 0.70348736 | 0.89054584 |
| BC065403 | 0.00804617 | 0.00613197 | 0.00861077 | -0.0024788 | 0.91006894 | 0.93034688 |
| Nek5 | 0.02178927 | 0.02083667 | 0.02331746 | -0.0024808 | 0.68993785 | 0.89054584 |
| Zzef1 | 0.25757216 | 0.26533909 | 0.26782045 | -0.0024814 | 0.51721506 | 0.89054584 |
| Zfp879 | 0.00898421 | 0.00713181 | 0.00961463 | -0.0024828 | 0.88552966 | 0.91420299 |
| Cdc25a | 0.21302673 | 0.2209064 | 0.22339048 | -0.0024841 | 0.52065874 | 0.89054584 |
| Fitm1 | 0.01350849 | 0.0119709 | 0.01445626 | -0.0024854 | 0.78842828 | 0.89054584 |
| Fam160a1 | 0.01329521 | 0.01173594 | 0.01422802 | -0.0024921 | 0.79275761 | 0.89054584 |
| Pof1b | 0.01297083 | 0.01138711 | 0.01388089 | -0.0024938 | 0.79870512 | 0.89054584 |
| Muc16 | 0.00848829 | 0.00658577 | 0.00908391 | -0.0024981 | 0.89994553 | 0.92300013 |
| 6430550D23I | 0.00974233 | 0.00792715 | 0.01042593 | -0.0024988 | 0.86791836 | 0.90460678 |
| Wdr44 | 0.07992533 | 0.08298032 | 0.08548098 | -0.0025007 | 0.55420502 | 0.89054584 |
| Stxbp4 | 0.17883878 | 0.18619479 | 0.18869981 | -0.002505 | 0.52465809 | 0.89054584 |
| Arhgap27 | 0.06651035 | 0.06864054 | 0.07114798 | -0.0025074 | 0.56521062 | 0.89054584 |
| Gm17092 | 0.00927872 | 0.00742043 | 0.0099298 | -0.0025094 | 0.8804749 | 0.91159931 |
| Ercc6 | 0.07626321 | 0.07905879 | 0.08156918 | -0.0025104 | 0.5570068 | 0.89054584 |
| Tmem8 | 0.01968563 | 0.01855481 | 0.02106644 | -0.0025116 | 0.71070955 | 0.89054584 |
| Slc6a19 | 0.00789875 | 0.0059404 | 0.00845301 | -0.0025126 | 0.91694982 | 0.93494187 |
| Ophn1 | 0.09575139 | 0.09986324 | 0.10237675 | -0.0025135 | 0.54553352 | 0.89054584 |
| Gpr62 | 0.00897694 | 0.00709227 | 0.00960684 | -0.0025146 | 0.88866681 | 0.91609012 |
| Lims1 | 0.73690046 | 0.67355625 | 0.67607206 | -0.0025158 | 0.50691609 | 0.89054584 |
| B3galt2 | 0.00873342 | 0.00682664 | 0.00934624 | -0.0025196 | 0.89543728 | 0.92051689 |
| Iqgap3 | 0.02376352 | 0.02290939 | 0.02542998 | -0.0025206 | 0.67788379 | 0.89054584 |

| | | | | | | |
|---|---|---|---|---|---|---|
| Kif13b | 0.04579467 | 0.04647782 | 0.04899917 | -0.0025214 | 0.59473248 | 0.89054584 |
| Gm527 | 0.00899047 | 0.00709917 | 0.00962133 | -0.0025222 | 0.88901511 | 0.91631886 |
| Thada | 0.0432576 | 0.04376065 | 0.04628557 | -0.0025249 | 0.60031008 | 0.89054584 |
| Gm14443 | 0.00934386 | 0.00746964 | 0.0099995 | -0.0025299 | 0.88074214 | 0.91173336 |
| Etv4 | 0.03126253 | 0.0309217 | 0.03345365 | -0.0025319 | 0.63781071 | 0.89054584 |
| Mag | 0.02550456 | 0.0247577 | 0.02729291 | -0.0025352 | 0.66740677 | 0.89054584 |
| Dact3 | 0.02564283 | 0.02490365 | 0.02744085 | -0.0025372 | 0.666682 | 0.89054584 |
| Phyhd1 | 0.00955521 | 0.00768784 | 0.01022569 | -0.0025378 | 0.87621536 | 0.9091719 |
| Galnt6 | 0.00855489 | 0.00661619 | 0.00915519 | -0.002539 | 0.90183143 | 0.92431323 |
| 1700018L02F | 0.01386264 | 0.0122939 | 0.01483525 | -0.0025413 | 0.78759298 | 0.89054584 |
| Gm43570 | 0.00742664 | 0.0054039 | 0.00794778 | -0.0025439 | 0.93204661 | 0.94578568 |
| Cntn4 | 0.0170833 | 0.01573744 | 0.01828172 | -0.0025443 | 0.74163313 | 0.89054584 |
| Asic1 | 0.00956822 | 0.00768709 | 0.01023961 | -0.0025525 | 0.87725506 | 0.9096298 |
| Stx19 | 0.01422928 | 0.01267356 | 0.01522759 | -0.002554 | 0.78272104 | 0.89054584 |
| Lactb2 | 0.17440692 | 0.18161115 | 0.18416716 | -0.002556 | 0.52577773 | 0.89054584 |
| Smad3 | 0.06423642 | 0.06616043 | 0.06871758 | -0.0025572 | 0.56881997 | 0.89054584 |
| Srek1ip1 | 0.57058915 | 0.54809828 | 0.55065722 | -0.0025589 | 0.50863661 | 0.89054584 |
| Ager | 0.03265542 | 0.03238457 | 0.03494388 | -0.0025593 | 0.63353256 | 0.89054584 |
| Trim2 | 0.10909079 | 0.11390045 | 0.11646158 | -0.0025611 | 0.54080257 | 0.89054584 |
| Mob1b | 0.31607565 | 0.32219451 | 0.32475586 | -0.0025613 | 0.51465585 | 0.89054584 |
| Zfp54 | 0.05648528 | 0.05786977 | 0.06043139 | -0.0025616 | 0.57827689 | 0.89054584 |
| Ccdc142os | 0.03106547 | 0.03067913 | 0.03324282 | -0.0025637 | 0.64031352 | 0.89054584 |
| 4632411P08I | 0.00744341 | 0.00539968 | 0.00796573 | -0.002566 | 0.93329792 | 0.94660649 |
| Slc22a5 | 0.00887316 | 0.00691572 | 0.00949579 | -0.0025801 | 0.89721912 | 0.92169509 |
| Cdh6 | 0.00775629 | 0.00571572 | 0.00830055 | -0.0025848 | 0.9265847 | 0.94231376 |
| Gm10306 | 0.0089384 | 0.00697652 | 0.0095656 | -0.0025891 | 0.89634865 | 0.92122978 |
| Rimkla | 0.01410297 | 0.01250277 | 0.01509242 | -0.0025897 | 0.787974 | 0.89054584 |
| Cenpt | 0.24608281 | 0.25385532 | 0.25644764 | -0.0025923 | 0.51878135 | 0.89054584 |
| Intu | 0.05678195 | 0.05815349 | 0.06074859 | -0.0025951 | 0.57887768 | 0.89054584 |
| Jag2 | 0.01302925 | 0.011343 | 0.0139434 | -0.0026004 | 0.80754313 | 0.89054584 |

| | | | | | | |
|---|---|---|---|---|---|---|
| Dgke | 0.03981043 | 0.03999692 | 0.04259826 | -0.0026013 | 0.61198877 | 0.89054584 |
| Ankrd6 | 0.04993632 | 0.05082593 | 0.05342858 | -0.0026027 | 0.58976854 | 0.89054584 |
| Kif24 | 0.01532002 | 0.01378867 | 0.01639481 | -0.0026061 | 0.77052993 | 0.89054584 |
| Cadps | 0.31732924 | 0.32334937 | 0.32595742 | -0.002608 | 0.51486793 | 0.89054584 |
| Rmst | 0.02125611 | 0.02013874 | 0.02274695 | -0.0026082 | 0.70339981 | 0.89054584 |
| P2rx4 | 0.30439355 | 0.31091259 | 0.31352122 | -0.0026086 | 0.51546087 | 0.89054584 |
| Slc39a5 | 0.00845199 | 0.00643515 | 0.00904506 | -0.0026099 | 0.91057698 | 0.93053697 |
| Zfp937 | 0.1057404 | 0.11033251 | 0.112945 | -0.0026125 | 0.54290876 | 0.89054584 |
| 4930412M03 | 0.00838097 | 0.00635604 | 0.00896906 | -0.002613 | 0.91266039 | 0.9320103 |
| 6330416G13 | 0.05351318 | 0.05463927 | 0.05725351 | -0.0026142 | 0.58423169 | 0.89054584 |
| Ist1 | 0.45388731 | 0.44917834 | 0.45179322 | -0.0026149 | 0.51075614 | 0.89054584 |
| 1700040L02F | 0.04079388 | 0.04103522 | 0.04365024 | -0.002615 | 0.60991996 | 0.89054584 |
| Krt222 | 0.0122719 | 0.01051778 | 0.01313294 | -0.0026152 | 0.82322666 | 0.89080883 |
| Sap130 | 0.10954916 | 0.11432555 | 0.11694237 | -0.0026168 | 0.54151576 | 0.89054584 |
| Gm26660 | 0.00863552 | 0.0066245 | 0.00924147 | -0.002617 | 0.90646646 | 0.92771077 |
| Fam212b | 0.01251497 | 0.01077419 | 0.01339306 | -0.0026189 | 0.81885354 | 0.89054584 |
| Gm12359 | 0.01653935 | 0.01507999 | 0.01769963 | -0.0026196 | 0.7547612 | 0.89054584 |
| Als2 | 0.04309756 | 0.0434897 | 0.04611439 | -0.0026247 | 0.60456182 | 0.89054584 |
| Smc2os | 0.00792784 | 0.00585914 | 0.00848414 | -0.002625 | 0.92527557 | 0.94144468 |
| Gm10330 | 0.01691122 | 0.01547103 | 0.01809757 | -0.0026265 | 0.75053143 | 0.89054584 |
| Foxo6 | 0.00736743 | 0.00525741 | 0.00788441 | -0.002627 | 0.93970565 | 0.95118707 |
| Serpinf2 | 0.02570162 | 0.02487313 | 0.02750376 | -0.0026306 | 0.67206078 | 0.89054584 |
| Fbxo46 | 0.14230552 | 0.14846442 | 0.15109782 | -0.0026334 | 0.5323578 | 0.89054584 |
| Brpf1 | 0.17609301 | 0.1832589 | 0.18589255 | -0.0026336 | 0.52631345 | 0.89054584 |
| Hoga1 | 0.00936445 | 0.00738702 | 0.01002154 | -0.0026345 | 0.88966873 | 0.91679645 |
| Mat1a | 0.03223387 | 0.03185383 | 0.03449288 | -0.0026391 | 0.63925035 | 0.89054584 |
| C030010L15F | 0.00940519 | 0.00742534 | 0.01006514 | -0.0026398 | 0.88912945 | 0.91637156 |
| Gm29107 | 0.00903942 | 0.00703382 | 0.00967371 | -0.0026399 | 0.89820585 | 0.92212069 |
| Rapgefl1 | 0.03043208 | 0.02992233 | 0.03256514 | -0.0026428 | 0.64731977 | 0.89054584 |
| Gm13248 | 0.00927302 | 0.00727969 | 0.00992369 | -0.002644 | 0.8927524 | 0.91840803 |

| | | | | | | |
|---|---|---|---|---|---|---|
| Gm20554 | 0.02689982 | 0.02614089 | 0.02878581 | -0.0026449 | 0.66570272 | 0.89054584 |
| Sec14l4 | 0.01763342 | 0.01622527 | 0.01887039 | -0.0026451 | 0.7431407 | 0.89054584 |
| Prr14l | 0.29449411 | 0.30130004 | 0.30394785 | -0.0026478 | 0.516187 | 0.89054584 |
| Acaa1b | 0.01402892 | 0.01236481 | 0.01501318 | -0.0026484 | 0.79441948 | 0.89054584 |
| Zfp955b | 0.02604225 | 0.02521962 | 0.02786823 | -0.0026486 | 0.67104041 | 0.89054584 |
| Swap70 | 0.09119746 | 0.09486517 | 0.09751666 | -0.0026515 | 0.55040196 | 0.89054584 |
| Rasl10b | 0.01252251 | 0.01074903 | 0.01340113 | -0.0026521 | 0.82173856 | 0.89054584 |
| 1700001C19I | 0.0089949 | 0.00697373 | 0.00962607 | -0.0026523 | 0.90037825 | 0.92330484 |
| Hist2h3c1 | 0.05943909 | 0.06093575 | 0.06358939 | -0.0026536 | 0.57707698 | 0.89054584 |
| Cfap53 | 0.01695088 | 0.01548395 | 0.01814002 | -0.0026561 | 0.75243188 | 0.89054584 |
| Gm26802 | 0.00845772 | 0.00639226 | 0.00905119 | -0.0026589 | 0.91444404 | 0.93337156 |
| 1700016L21F | 0.01367377 | 0.01197396 | 0.01463313 | -0.0026592 | 0.80140061 | 0.89054584 |
| Mia2 | 0.01749971 | 0.01606699 | 0.01872732 | -0.0026603 | 0.74595958 | 0.89054584 |
| Lrtm2 | 0.01359656 | 0.01188969 | 0.0145505 | -0.0026608 | 0.80288429 | 0.89054584 |
| Pcdha11.1 | 0.00760141 | 0.00547084 | 0.0081348 | -0.002664 | 0.93645717 | 0.94880216 |
| Fam149a | 0.11042235 | 0.11519393 | 0.11785805 | -0.0026641 | 0.54193631 | 0.89054584 |
| Pinlyp | 0.04747499 | 0.04812908 | 0.0507963 | -0.0026672 | 0.5966296 | 0.89054584 |
| Acad10 | 0.02088888 | 0.01968259 | 0.02235399 | -0.0026714 | 0.71115553 | 0.89054584 |
| Trim67 | 0.0083828 | 0.0062939 | 0.00897102 | -0.0026771 | 0.91778344 | 0.93519991 |
| Stil | 0.0084732 | 0.0063904 | 0.00906777 | -0.0026774 | 0.91552829 | 0.93401795 |
| Gm15892 | 0.00775299 | 0.00561044 | 0.00829702 | -0.0026866 | 0.93429552 | 0.94723402 |
| Mfsd9 | 0.00779298 | 0.00565251 | 0.00833982 | -0.0026873 | 0.93335217 | 0.94660649 |
| Gm26876 | 0.00770906 | 0.00554773 | 0.00825001 | -0.0027023 | 0.93649876 | 0.94880216 |
| Usp2 | 0.0641816 | 0.06595455 | 0.06865899 | -0.0027044 | 0.57280218 | 0.89054584 |
| Gm10767 | 0.00702111 | 0.0048093 | 0.00751379 | -0.0027045 | 0.95322145 | 0.96191304 |
| Fam159a | 0.0622172 | 0.06384174 | 0.06655921 | -0.0027175 | 0.57542965 | 0.89054584 |
| Serpina1b | 0.02354878 | 0.02248102 | 0.02520021 | -0.0027192 | 0.69241772 | 0.89054584 |
| Trdmt1 | 0.06266862 | 0.06431602 | 0.06704176 | -0.0027257 | 0.57511823 | 0.89054584 |
| Timp4 | 0.0223748 | 0.02121447 | 0.02394401 | -0.0027295 | 0.70232667 | 0.89054584 |
| Dhdh | 0.02141071 | 0.02017818 | 0.02291238 | -0.0027342 | 0.71088272 | 0.89054584 |

| | | | | | | |
|---|---|---|---|---|---|---|
| Foxj1 | 0.01623255 | 0.01463626 | 0.01737132 | -0.0027351 | 0.7683846 | 0.89054584 |
| Dnmt3bos | 0.0122457 | 0.01036796 | 0.0131049 | -0.0027369 | 0.83472614 | 0.89453395 |
| Polq | 0.00912641 | 0.007028 | 0.0097668 | -0.0027388 | 0.90430173 | 0.92615843 |
| Ccdc13 | 0.01433984 | 0.01260707 | 0.01534591 | -0.0027388 | 0.79715363 | 0.89054584 |
| Cct4 | 1.83850525 | 1.10010855 | 1.10284747 | -0.0027389 | 0.50461584 | 0.89054584 |
| Dgka | 0.06331804 | 0.06499299 | 0.06773593 | -0.0027429 | 0.57482109 | 0.89054584 |
| Get4 | 0.64548167 | 0.60673422 | 0.60947733 | -0.0027431 | 0.50836472 | 0.89054584 |
| Ppp6r2 | 0.03657182 | 0.03639061 | 0.03913376 | -0.0027431 | 0.6280058 | 0.89054584 |
| Ankrd35 | 0.01387845 | 0.01210797 | 0.01485217 | -0.0027442 | 0.805332 | 0.89054584 |
| Gm7467 | 0.05604736 | 0.05721735 | 0.05996317 | -0.0027458 | 0.58446956 | 0.89054584 |
| Plk5 | 0.03033712 | 0.02971736 | 0.03246354 | -0.0027462 | 0.65325085 | 0.89054584 |
| Acot10 | 0.00678342 | 0.00451098 | 0.00725943 | -0.0027485 | 0.9611252 | 0.96773339 |
| Ephx1 | 0.04905887 | 0.04974012 | 0.05249021 | -0.0027501 | 0.59642087 | 0.89054584 |
| Ism2 | 0.00815346 | 0.00597219 | 0.00872559 | -0.0027534 | 0.92923969 | 0.94388825 |
| Slc46a1 | 0.08891113 | 0.09231794 | 0.0950761 | -0.0027582 | 0.55375567 | 0.89054584 |
| 4933429H19 | 0.00815739 | 0.00597022 | 0.0087298 | -0.0027596 | 0.9295893 | 0.94406031 |
| Atp8b5 | 0.01339819 | 0.01157338 | 0.01433822 | -0.0027648 | 0.8155093 | 0.89054584 |
| Pias2 | 0.35456302 | 0.35850913 | 0.36127438 | -0.0027653 | 0.51422338 | 0.89054584 |
| Aldh1l1 | 0.02541722 | 0.02443014 | 0.02719946 | -0.0027693 | 0.68237501 | 0.89054584 |
| Yod1 | 0.24903071 | 0.25660034 | 0.25937156 | -0.0027712 | 0.51985032 | 0.89054584 |
| Nhsl2 | 0.12712288 | 0.13254256 | 0.13531665 | -0.0027741 | 0.53804575 | 0.89054584 |
| Galr2 | 0.01706143 | 0.01548256 | 0.01825832 | -0.0027758 | 0.76061611 | 0.89054584 |
| Acsm3 | 0.00896383 | 0.00681272 | 0.00959282 | -0.0027801 | 0.91152606 | 0.93111426 |
| Zfp354b | 0.02595532 | 0.02499378 | 0.02777522 | -0.0027814 | 0.67958973 | 0.89054584 |
| Gab2 | 0.04741888 | 0.04795432 | 0.0507363 | -0.002782 | 0.60081593 | 0.89054584 |
| Anks4b | 0.01957116 | 0.01816191 | 0.02094394 | -0.002782 | 0.73199533 | 0.89054584 |
| Iqsec2 | 0.03269695 | 0.03220455 | 0.03498831 | -0.0027838 | 0.64456089 | 0.89054584 |
| Fam84a | 0.04100523 | 0.0410824 | 0.04387632 | -0.0027939 | 0.61664014 | 0.89054584 |
| Slc18a1 | 0.02537413 | 0.02435466 | 0.02715335 | -0.0027987 | 0.68445627 | 0.89054584 |
| Crlf1 | 0.037557 | 0.03738702 | 0.04018768 | -0.0028007 | 0.62728825 | 0.89054584 |

| | | | | | | |
|---|---|---|---|---|---|---|
| Cfap45 | 0.02058942 | 0.01923256 | 0.02203356 | -0.002801 | 0.72316315 | 0.89054584 |
| Fuk | 0.04031127 | 0.04033125 | 0.04313401 | -0.0028028 | 0.6189517 | 0.89054584 |
| Tmem98 | 0.2328446 | 0.24046404 | 0.24326745 | -0.0028034 | 0.52140874 | 0.89054584 |
| Nbn | 0.10297255 | 0.10723323 | 0.11003677 | -0.0028035 | 0.5472441 | 0.89054584 |
| Lin9 | 0.14100883 | 0.14694463 | 0.1497535 | -0.0028089 | 0.53481777 | 0.89054584 |
| Atg2a | 0.0990115 | 0.10304565 | 0.10585515 | -0.0028095 | 0.54920502 | 0.89054584 |
| Artn | 0.00699662 | 0.00467052 | 0.00748759 | -0.0028171 | 0.96018521 | 0.96718997 |
| Dtna | 0.01737222 | 0.01577272 | 0.0185909 | -0.0028182 | 0.75998163 | 0.89054584 |
| 1810041H14I | 0.01353799 | 0.01166876 | 0.01448783 | -0.0028191 | 0.81767627 | 0.89054584 |
| Atp6v1b1 | 0.01223346 | 0.01027108 | 0.01309181 | -0.0028207 | 0.84226326 | 0.89691028 |
| A330041J22F | 0.00913565 | 0.00695485 | 0.00977669 | -0.0028218 | 0.91063999 | 0.93053697 |
| Timeless | 0.11061455 | 0.1152375 | 0.11805956 | -0.0028221 | 0.54433698 | 0.89054584 |
| Bfsp1 | 0.01746755 | 0.0158706 | 0.0186929 | -0.0028223 | 0.75910314 | 0.89054584 |
| Star | 0.00798584 | 0.00572197 | 0.00854621 | -0.0028242 | 0.93817395 | 0.95016738 |
| Casr | 0.00828852 | 0.00604325 | 0.00887013 | -0.0028269 | 0.93120001 | 0.94534961 |
| Gm27042 | 0.01317264 | 0.01126971 | 0.01409685 | -0.0028271 | 0.82493968 | 0.89153749 |
| Sptbn4 | 0.01311307 | 0.01120341 | 0.01403311 | -0.0028297 | 0.82624948 | 0.89170439 |
| Acacb | 0.01694475 | 0.01530292 | 0.01813345 | -0.0028305 | 0.76645919 | 0.89054584 |
| Mlx | 0.42118275 | 0.41977315 | 0.42260425 | -0.0028311 | 0.51244938 | 0.89054584 |
| Wdr46 | 0.32383608 | 0.32934934 | 0.33218111 | -0.0028318 | 0.51584036 | 0.89054584 |
| Peli3 | 0.00776114 | 0.00547144 | 0.00830574 | -0.0028343 | 0.94406521 | 0.95433395 |
| Kif14 | 0.00818555 | 0.00592193 | 0.00875993 | -0.002838 | 0.934399 | 0.94723402 |
| Mtmr10 | 0.09359489 | 0.09723682 | 0.10007544 | -0.0028386 | 0.55256719 | 0.89054584 |
| Adam5 | 0.00830193 | 0.00604114 | 0.00888447 | -0.0028433 | 0.93202182 | 0.94578568 |
| 0610010F05I | 0.34510915 | 0.34952984 | 0.35237574 | -0.0028459 | 0.5150075 | 0.89054584 |
| Birc3 | 0.0439849 | 0.0442174 | 0.04706349 | -0.0028461 | 0.61092963 | 0.89054584 |
| Six4 | 0.03528841 | 0.03491383 | 0.03776076 | -0.0028469 | 0.6373017 | 0.89054584 |
| Myo7b | 0.02043738 | 0.01902375 | 0.02187086 | -0.0028471 | 0.72790583 | 0.89054584 |
| Cabp1 | 0.01354192 | 0.01164384 | 0.01449203 | -0.0028482 | 0.82007331 | 0.89054584 |
| Psmb9 | 0.06389415 | 0.06550316 | 0.06835174 | -0.0028486 | 0.57697556 | 0.89054584 |

| | | | | | | |
|---|---|---|---|---|---|---|
| Srcin1 | 0.00713896 | 0.00479119 | 0.00763991 | -0.0028487 | 0.95882377 | 0.96608709 |
| Exo1 | 0.07779629 | 0.08035639 | 0.08320687 | -0.0028505 | 0.56340382 | 0.89054584 |
| Gm5420 | 0.02111087 | 0.01973989 | 0.02259154 | -0.0028517 | 0.7217613 | 0.89054584 |
| Rmi2 | 0.05958528 | 0.06089292 | 0.06374568 | -0.0028528 | 0.58257975 | 0.89054584 |
| Slain1 | 0.05412861 | 0.05505672 | 0.05791158 | -0.0028549 | 0.59082631 | 0.89054584 |
| Cpsf7 | 0.34890635 | 0.35309759 | 0.35595571 | -0.0028581 | 0.5149204 | 0.89054584 |
| Il15ra | 0.01247507 | 0.01049182 | 0.01335036 | -0.0028585 | 0.84075319 | 0.89593824 |
| L3hypdh | 0.13171505 | 0.13723436 | 0.14009921 | -0.0028648 | 0.53794955 | 0.89054584 |
| Agpat9 | 0.00727629 | 0.00492038 | 0.00778688 | -0.0028665 | 0.95683165 | 0.964415 |
| Ak4 | 0.01706755 | 0.01539822 | 0.01826486 | -0.0028666 | 0.76767571 | 0.89054584 |
| Afp | 0.00822907 | 0.00593675 | 0.0088065 | -0.0028698 | 0.93551764 | 0.94807317 |
| Sowaha | 0.04491971 | 0.04518774 | 0.04806335 | -0.0028756 | 0.60977898 | 0.89054584 |
| Ankrd45 | 0.00743114 | 0.00507195 | 0.00795259 | -0.0028806 | 0.95425552 | 0.9627461 |
| Ttc16 | 0.00839961 | 0.00610635 | 0.00898901 | -0.0028827 | 0.93242053 | 0.9459925 |
| Ttc41 | 0.00758028 | 0.0052281 | 0.0081122 | -0.0028841 | 0.95117731 | 0.95999115 |
| Adcy8 | 0.01582059 | 0.01404582 | 0.01693048 | -0.0028847 | 0.78634477 | 0.89054584 |
| BC051019 | 0.00890746 | 0.0066477 | 0.00953249 | -0.0028848 | 0.92071881 | 0.9375979 |
| Gm16576 | 0.00743133 | 0.00506347 | 0.00795279 | -0.0028893 | 0.95473732 | 0.96297354 |
| Cyp2d26 | 0.01272889 | 0.01073058 | 0.01362198 | -0.0028914 | 0.83864554 | 0.89497393 |
| Ttll11 | 0.00780544 | 0.00546174 | 0.00835315 | -0.0028914 | 0.9465924 | 0.95624787 |
| Gstm6 | 0.00774093 | 0.00539223 | 0.00828411 | -0.0028919 | 0.9480654 | 0.95744303 |
| E530001K10I | 0.00787737 | 0.00553273 | 0.00843013 | -0.0028974 | 0.94534021 | 0.95528977 |
| Tm4sf20 | 0.01571146 | 0.0139126 | 0.0168137 | -0.0029011 | 0.7892645 | 0.89054584 |
| Vrk1 | 0.34068457 | 0.34529323 | 0.34819455 | -0.0029013 | 0.51548328 | 0.89054584 |
| Mtmr11 | 0.02019505 | 0.01870819 | 0.02161156 | -0.0029034 | 0.73430912 | 0.89054584 |
| Slc25a16 | 0.14241974 | 0.14830857 | 0.15121621 | -0.0029076 | 0.53569098 | 0.89054584 |
| Masp2 | 0.01371426 | 0.01176507 | 0.01467645 | -0.0029114 | 0.82230836 | 0.89073269 |
| Marveld2 | 0.15673828 | 0.16310353 | 0.16601644 | -0.0029129 | 0.53257541 | 0.89054584 |
| Gm13157 | 0.05456112 | 0.05545996 | 0.05837405 | -0.0029141 | 0.59195593 | 0.89054584 |
| Hist1h2bp | 0.00809385 | 0.00574761 | 0.0086618 | -0.0029142 | 0.94149694 | 0.95226891 |

| | | | | | | |
|---|---|---|---|---|---|---|
| Rbm8a2 | 0.13758873 | 0.14328474 | 0.14620463 | -0.0029199 | 0.53706596 | 0.89054584 |
| Wdr93 | 0.01522874 | 0.01337645 | 0.01629714 | -0.0029207 | 0.7981273 | 0.89054584 |
| Tmem179 | 0.02483356 | 0.0236542 | 0.02657493 | -0.0029207 | 0.69568998 | 0.89054584 |
| Mphosph9 | 0.10721785 | 0.11157506 | 0.11449625 | -0.0029212 | 0.54730893 | 0.89054584 |
| Pcdhgb4 | 0.02136011 | 0.01993532 | 0.02285823 | -0.0029229 | 0.72432696 | 0.89054584 |
| Tead4 | 0.00804274 | 0.00568294 | 0.0086071 | -0.0029242 | 0.94326882 | 0.95366188 |
| Fam124a | 0.01719942 | 0.01548021 | 0.01840598 | -0.0029258 | 0.77052414 | 0.89054584 |
| Spaca7 | 0.00731209 | 0.0048993 | 0.00782519 | -0.0029259 | 0.95924676 | 0.96637896 |
| Btc | 0.02029918 | 0.01879651 | 0.02172298 | -0.0029265 | 0.73488101 | 0.89054584 |
| Jph2 | 0.00806557 | 0.00570258 | 0.00863153 | -0.002929 | 0.94305267 | 0.95357634 |
| Crebl2 | 0.05195383 | 0.05265635 | 0.05558608 | -0.0029297 | 0.5969864 | 0.89054584 |
| Nr0b2 | 0.03653931 | 0.03616797 | 0.03909899 | -0.002931 | 0.63655023 | 0.89054584 |
| Atg9a | 0.17286267 | 0.17964789 | 0.18258588 | -0.002938 | 0.52987959 | 0.89054584 |
| Fbxw8 | 0.1802075 | 0.18715813 | 0.19009817 | -0.00294 | 0.5287208 | 0.89054584 |
| Prr15 | 0.01538961 | 0.01352704 | 0.01646929 | -0.0029422 | 0.79738649 | 0.89054584 |
| Orc1 | 0.01640466 | 0.01460978 | 0.0175555 | -0.0029457 | 0.78281832 | 0.89054584 |
| Diras1 | 0.00757464 | 0.00515942 | 0.00810616 | -0.0029467 | 0.95483018 | 0.96300021 |
| Ripk2 | 0.05881281 | 0.05996867 | 0.06291984 | -0.0029512 | 0.58648871 | 0.89054584 |
| Layn | 0.03731396 | 0.03697594 | 0.03992768 | -0.0029517 | 0.6347359 | 0.89054584 |
| Lars2 | 0.08248309 | 0.08525494 | 0.08821266 | -0.0029577 | 0.56206693 | 0.89054584 |
| Car13 | 0.02126341 | 0.01979505 | 0.02275476 | -0.0029597 | 0.72773859 | 0.89054584 |
| Dok7 | 0.04656499 | 0.04686173 | 0.04982305 | -0.0029613 | 0.60907626 | 0.89054584 |
| Kantr | 0.03861423 | 0.0383511 | 0.04131866 | -0.0029676 | 0.63104087 | 0.89054584 |
| Fez1 | 0.0636931 | 0.06516912 | 0.06813684 | -0.0029677 | 0.58040191 | 0.89054584 |
| Trim34a | 0.02899704 | 0.02805938 | 0.03102975 | -0.0029704 | 0.6721958 | 0.89054584 |
| Cenpl | 0.06587571 | 0.06749877 | 0.0704697 | -0.0029709 | 0.57785782 | 0.89054584 |
| Ackr4 | 0.0250813 | 0.02386717 | 0.02684002 | -0.0029728 | 0.6970839 | 0.89054584 |
| Alb | 0.00795302 | 0.00553762 | 0.00851109 | -0.0029735 | 0.94820219 | 0.95744779 |
| Rusc2 | 0.01709558 | 0.01532027 | 0.01829485 | -0.0029746 | 0.77562539 | 0.89054584 |
| Agtr1a | 0.01193974 | 0.00980057 | 0.01277749 | -0.0029769 | 0.86113777 | 0.90194881 |

| | | | | | | |
|---|---|---|---|---|---|---|
| Prss27 | 0.01213099 | 0.01000518 | 0.01298214 | -0.002977 | 0.85731902 | 0.9009252 |
| Gm4425 | 0.00717169 | 0.00468977 | 0.00767493 | -0.0029852 | 0.96501408 | 0.97104207 |
| Styxl1 | 0.00855942 | 0.00617323 | 0.00916004 | -0.0029868 | 0.93563605 | 0.94812689 |
| 6430531B16 | 0.02552994 | 0.02433311 | 0.02732006 | -0.0029869 | 0.69475339 | 0.89054584 |
| Kcnj5 | 0.00739203 | 0.00491601 | 0.00791074 | -0.0029947 | 0.96110955 | 0.96773339 |
| Prdm11 | 0.12295308 | 0.12797045 | 0.13096733 | -0.0029969 | 0.5424504 | 0.89054584 |
| Greb1l | 0.02952535 | 0.02859785 | 0.03159501 | -0.0029972 | 0.67073952 | 0.89054584 |
| Caps2 | 0.01023632 | 0.00795462 | 0.01095458 | -0.003 | 0.89899945 | 0.92252046 |
| 3110021N24 | 0.00772715 | 0.00526892 | 0.00826937 | -0.0030005 | 0.954529 | 0.9628974 |
| Dnajc6 | 0.02012556 | 0.01853661 | 0.02153719 | -0.0030006 | 0.74185963 | 0.89054584 |
| Arhgap19 | 0.03132543 | 0.03052029 | 0.03352095 | -0.0030007 | 0.66167855 | 0.89054584 |
| Ppp6r1 | 0.1385009 | 0.14414927 | 0.14715159 | -0.0030023 | 0.53786477 | 0.89054584 |
| Ablim2 | 0.01252198 | 0.01039617 | 0.01340056 | -0.0030044 | 0.85187978 | 0.89985815 |
| Gm26899 | 0.01226272 | 0.01011715 | 0.01312312 | -0.003006 | 0.85705323 | 0.9009252 |
| Fxyd4 | 0.01602289 | 0.01413619 | 0.01714697 | -0.0030108 | 0.79333531 | 0.89054584 |
| Mrpl16 | 0.35054613 | 0.35448843 | 0.35749929 | -0.0030109 | 0.5156495 | 0.89054584 |
| Ica1l | 0.01198806 | 0.00981357 | 0.01282919 | -0.0030156 | 0.86326979 | 0.90287732 |
| Gm42670 | 0.0254063 | 0.02416885 | 0.02718776 | -0.0030189 | 0.69753609 | 0.89054584 |
| Tgfbr3l | 0.01803377 | 0.01627708 | 0.01929881 | -0.0030217 | 0.76714573 | 0.89054584 |
| Serac1 | 0.00840063 | 0.00596708 | 0.0089901 | -0.003023 | 0.94139843 | 0.95223571 |
| 9530026P05 | 0.0062433 | 0.00365157 | 0.00668141 | -0.0030298 | 0.98268705 | 0.98567849 |
| Insl6 | 0.0240141 | 0.02266248 | 0.0256981 | -0.0030356 | 0.70895819 | 0.89054584 |
| Ndufaf5 | 0.12993932 | 0.13521463 | 0.13825081 | -0.0030362 | 0.54074734 | 0.89054584 |
| Fmn1 | 0.01597077 | 0.01405439 | 0.01709119 | -0.0030368 | 0.7961113 | 0.89054584 |
| Dab1 | 0.07727328 | 0.07961131 | 0.08264819 | -0.0030369 | 0.56796327 | 0.89054584 |
| Smpdl3b | 0.00763181 | 0.00512222 | 0.00816735 | -0.0030451 | 0.95881078 | 0.96608709 |
| Ttc36 | 0.0341289 | 0.03347151 | 0.03652029 | -0.0030488 | 0.65133867 | 0.89054584 |
| Mamstr | 0.06486803 | 0.06633926 | 0.06939267 | -0.0030534 | 0.58121493 | 0.89054584 |
| Psmf1 | 0.36848755 | 0.37123667 | 0.37429438 | -0.0030577 | 0.51518016 | 0.89054584 |
| Kifc5b | 0.02097042 | 0.01938003 | 0.02244125 | -0.0030612 | 0.73745816 | 0.89054584 |

| | | | | | | |
|---|---|---|---|---|---|---|
| Nrip3 | 0.01646172 | 0.01455304 | 0.01761657 | -0.0030635 | 0.79108483 | 0.89054584 |
| Gm26564 | 0.00760203 | 0.00507181 | 0.00813547 | -0.0030637 | 0.96032458 | 0.96726315 |
| Gm3294 | 0.00573944 | 0.00306824 | 0.0061422 | -0.003074 | 0.99014018 | 0.99205614 |
| Gm26631 | 0.05060341 | 0.05106788 | 0.05414197 | -0.0030741 | 0.604313 | 0.89054584 |
| Gm16861 | 0.00707652 | 0.00449808 | 0.00757309 | -0.003075 | 0.97073708 | 0.9758526 |
| Tmem74b | 0.01258459 | 0.01038823 | 0.01346756 | -0.0030793 | 0.85662047 | 0.9009252 |
| Ttf1 | 0.29627476 | 0.30259389 | 0.3056735 | -0.0030796 | 0.51871865 | 0.89054584 |
| Fam98c | 0.38529888 | 0.3867936 | 0.38987379 | -0.0030802 | 0.5146809 | 0.89054584 |
| Atp7b | 0.01451846 | 0.01245653 | 0.01553706 | -0.0030805 | 0.82218531 | 0.89073269 |
| Adgrb2 | 0.03283793 | 0.03205296 | 0.03513914 | -0.0030862 | 0.65879756 | 0.89054584 |
| Bbs9 | 0.09895318 | 0.1027051 | 0.10579294 | -0.0030878 | 0.55408262 | 0.89054584 |
| Mrps28 | 0.84161741 | 0.7419525 | 0.74504213 | -0.0030896 | 0.50770719 | 0.89054584 |
| Zfp655 | 0.30095824 | 0.3071111 | 0.31020466 | -0.0030936 | 0.51852897 | 0.89054584 |
| Gm26670 | 0.04004258 | 0.03975065 | 0.04284658 | -0.0030959 | 0.63180492 | 0.89054584 |
| Igf1r | 0.628562 | 0.59342077 | 0.59651747 | -0.0030967 | 0.50964787 | 0.89054584 |
| Stk36 | 0.06005739 | 0.0611523 | 0.06425039 | -0.0030981 | 0.58887397 | 0.89054584 |
| Erc2 | 0.02752567 | 0.02635087 | 0.02945546 | -0.0031046 | 0.68830446 | 0.89054584 |
| 2810021J22F | 0.08118069 | 0.08371588 | 0.08682174 | -0.0031059 | 0.56618264 | 0.89054584 |
| Slc5a6 | 0.01954667 | 0.01780443 | 0.02091774 | -0.0031133 | 0.75597624 | 0.89054584 |
| Kcnk10 | 0.01227474 | 0.01002027 | 0.01313598 | -0.0031157 | 0.86543119 | 0.90377426 |
| Tat | 0.01172118 | 0.00942772 | 0.01254359 | -0.0031159 | 0.87642478 | 0.9091719 |
| Gm19325 | 0.00700985 | 0.00438582 | 0.00750175 | -0.0031159 | 0.97351339 | 0.97796543 |
| Nradd | 0.12269099 | 0.12757546 | 0.13069374 | -0.0031183 | 0.54425519 | 0.89054584 |
| Zc3h12a | 0.0242758 | 0.02285562 | 0.02597813 | -0.0031225 | 0.71225783 | 0.89054584 |
| Kif18b | 0.01466826 | 0.01256719 | 0.01569736 | -0.0031302 | 0.82356224 | 0.8909122 |
| Cecr6 | 0.05614413 | 0.05693419 | 0.06006664 | -0.0031324 | 0.59598252 | 0.89054584 |
| Fnip1 | 0.15798669 | 0.16416968 | 0.16730308 | -0.0031334 | 0.53476631 | 0.89054584 |
| Zfp846 | 0.04131542 | 0.041074 | 0.04420813 | -0.0031341 | 0.62941169 | 0.89054584 |
| Ccdc160 | 0.01988087 | 0.01814104 | 0.02127536 | -0.0031343 | 0.75375783 | 0.89054584 |
| Atp6v0a4 | 0.01259746 | 0.01034677 | 0.01348134 | -0.0031346 | 0.86065324 | 0.90163673 |

| | | | | | | |
|---|---|---|---|---|---|---|
| Gm14226 | 0.01923303 | 0.01744477 | 0.02058212 | -0.0031373 | 0.76119945 | 0.89054584 |
| Alms1 | 0.09476611 | 0.09818661 | 0.10132535 | -0.0031387 | 0.55737591 | 0.89054584 |
| Zfp773 | 0.00661 | 0.00392876 | 0.00707384 | -0.0031451 | 0.98083924 | 0.9839612 |
| Sstr1 | 0.0070696 | 0.00441946 | 0.00756569 | -0.0031462 | 0.97365438 | 0.97799389 |
| Gm17315 | 0.00810362 | 0.00552163 | 0.00867225 | -0.0031506 | 0.95473337 | 0.96297354 |
| Neurl4 | 0.10232187 | 0.10620058 | 0.10935269 | -0.0031521 | 0.5534153 | 0.89054584 |
| Gm16586 | 0.13476762 | 0.14012039 | 0.14327388 | -0.0031535 | 0.54083769 | 0.89054584 |
| Fam171a2 | 0.10824339 | 0.11241883 | 0.11557254 | -0.0031537 | 0.55058195 | 0.89054584 |
| N4bp2l2 | 0.98219774 | 0.822113 | 0.82526793 | -0.0031549 | 0.50710509 | 0.89054584 |
| Gm29538 | 0.00575317 | 0.00299919 | 0.00615689 | -0.0031577 | 0.99156305 | 0.99341311 |
| Anks1b | 0.01336207 | 0.01113608 | 0.01429957 | -0.0031635 | 0.84865834 | 0.89883797 |
| Prdm9 | 0.01198768 | 0.00966214 | 0.01282879 | -0.0031666 | 0.87492991 | 0.90832322 |
| Gm26521 | 0.01231943 | 0.01001536 | 0.01318381 | -0.0031684 | 0.86857428 | 0.90498553 |
| 6230400D17 | 0.00642195 | 0.00370241 | 0.0068726 | -0.0031702 | 0.98418578 | 0.98670393 |
| Cndp1 | 0.01899316 | 0.01715414 | 0.02032544 | -0.0031713 | 0.76636135 | 0.89054584 |
| Kif21b | 0.03389497 | 0.03309663 | 0.03627001 | -0.0031734 | 0.65822593 | 0.89054584 |
| 4921507G05 | 0.01182679 | 0.00948147 | 0.01265661 | -0.0031751 | 0.87875402 | 0.91059794 |
| Thnsl2 | 0.01700247 | 0.01501839 | 0.01819522 | -0.0031768 | 0.79201577 | 0.89054584 |
| Champ1 | 0.13240036 | 0.13763044 | 0.14081225 | -0.0031818 | 0.54192077 | 0.89054584 |
| Garem | 0.02222732 | 0.02060307 | 0.02378619 | -0.0031831 | 0.73351207 | 0.89054584 |
| Rac3 | 0.28085542 | 0.28749318 | 0.29067884 | -0.0031857 | 0.52036077 | 0.89054584 |
| AV099323 | 0.06003087 | 0.06103577 | 0.06422203 | -0.0031863 | 0.59139906 | 0.89054584 |
| Tmem185b | 0.25641337 | 0.26348822 | 0.26667623 | -0.003188 | 0.52220792 | 0.89054584 |
| Fam166a | 0.02147235 | 0.01978725 | 0.02297834 | -0.0031911 | 0.74118543 | 0.89054584 |
| Xlr3c | 0.03255349 | 0.03163691 | 0.03483483 | -0.0031979 | 0.66556576 | 0.89054584 |
| Nolc1 | 0.60938919 | 0.57839967 | 0.58159891 | -0.0031992 | 0.51022286 | 0.89054584 |
| Mgat5b | 0.00775165 | 0.00509491 | 0.00829559 | -0.0032007 | 0.96387632 | 0.97003186 |
| Gm5475 | 0.01305991 | 0.01077146 | 0.01397622 | -0.0032048 | 0.85730709 | 0.9009252 |
| Zfp14 | 0.04013665 | 0.03973902 | 0.04294722 | -0.0032082 | 0.63609011 | 0.89054584 |
| Gm12927 | 0.01264909 | 0.01032805 | 0.01353659 | -0.0032085 | 0.86526707 | 0.90377426 |

| | | | | | | |
|---|---|---|---|---|---|---|
| Ypel2 | 0.30683909 | 0.3126671 | 0.31587857 | -0.0032115 | 0.51888938 | 0.89054584 |
| Spryd4 | 0.23860371 | 0.24579953 | 0.24901125 | -0.0032117 | 0.52395822 | 0.89054584 |
| Dsg2 | 0.13241278 | 0.13761291 | 0.14082517 | -0.0032123 | 0.54231655 | 0.89054584 |
| Slc38a7 | 0.05788033 | 0.05871004 | 0.06192293 | -0.0032129 | 0.59550557 | 0.89054584 |
| Bcl3 | 0.04049475 | 0.04011365 | 0.04333027 | -0.0032166 | 0.63527464 | 0.89054584 |
| Lrrc61 | 0.06428895 | 0.06555226 | 0.06877373 | -0.0032215 | 0.58637592 | 0.89054584 |
| Tmem110 | 0.18785321 | 0.19467316 | 0.19789505 | -0.0032219 | 0.5302313 | 0.89054584 |
| A830011K09 | 0.02028097 | 0.01847835 | 0.02170349 | -0.0032251 | 0.75563118 | 0.89054584 |
| Jak2 | 0.08649014 | 0.0892622 | 0.09249143 | -0.0032292 | 0.56460736 | 0.89054584 |
| Acpp | 0.00710562 | 0.00437419 | 0.00760423 | -0.00323 | 0.9760903 | 0.97994302 |
| Cdk17 | 0.10677857 | 0.11080349 | 0.11403511 | -0.0032316 | 0.55251929 | 0.89054584 |
| Lbhd1 | 0.03422772 | 0.03338924 | 0.03662601 | -0.0032368 | 0.65972988 | 0.89054584 |
| Gm29095 | 0.01712362 | 0.01508711 | 0.01832487 | -0.0032378 | 0.79479478 | 0.89054584 |
| Hypm | 0.01723982 | 0.01520824 | 0.01844921 | -0.003241 | 0.79344528 | 0.89054584 |
| Enkd1 | 0.05773611 | 0.05852756 | 0.06176873 | -0.0032412 | 0.59656545 | 0.89054584 |
| Pcsk4 | 0.01288854 | 0.01054614 | 0.01379283 | -0.0032467 | 0.86360672 | 0.90290403 |
| Gm42595 | 0.02381956 | 0.02224144 | 0.02548994 | -0.0032485 | 0.72365933 | 0.89054584 |
| Pradc1 | 0.18666262 | 0.1934326 | 0.19668254 | -0.0032499 | 0.53068164 | 0.89054584 |
| 4930403P22I | 0.0225202 | 0.02084568 | 0.02409959 | -0.0032539 | 0.73534078 | 0.89054584 |
| Parp12 | 0.14299395 | 0.14855638 | 0.15181128 | -0.0032549 | 0.53978402 | 0.89054584 |
| Gm14616 | 0.01276263 | 0.01040318 | 0.01365809 | -0.0032549 | 0.86656225 | 0.90430169 |
| Map3k10 | 0.10852929 | 0.11261495 | 0.11587252 | -0.0032576 | 0.5521039 | 0.89054584 |
| 1700012C14I | 0.01708013 | 0.01501926 | 0.01827833 | -0.0032591 | 0.79692866 | 0.89054584 |
| Erich3 | 0.01294252 | 0.01059044 | 0.01385059 | -0.0032602 | 0.86359652 | 0.90290403 |
| AW554918 | 0.12278871 | 0.12753214 | 0.13079576 | -0.0032636 | 0.54627279 | 0.89054584 |
| 1810062O18 | 0.00767884 | 0.00495158 | 0.00821767 | -0.0032661 | 0.96796431 | 0.97334573 |
| Vwa5b1 | 0.00733793 | 0.00458618 | 0.00785285 | -0.0032667 | 0.97369128 | 0.97799389 |
| Gm16174 | 0.00765359 | 0.00492307 | 0.00819065 | -0.0032676 | 0.96846095 | 0.97363199 |
| Angptl8 | 0.00556426 | 0.00268698 | 0.00595472 | -0.0032677 | 0.99471588 | 0.99581486 |
| Gm13544 | 0.01149695 | 0.00903262 | 0.01230363 | -0.003271 | 0.89225576 | 0.91809254 |

| | | | | | | |
|---|---|---|---|---|---|---|
| Tmco4 | 0.01252816 | 0.01013517 | 0.01340718 | -0.003272 | 0.87223307 | 0.90701739 |
| Mtmr4 | 0.19794316 | 0.20487227 | 0.20814647 | -0.0032742 | 0.52921085 | 0.89054584 |
| C3 | 0.01701052 | 0.01492527 | 0.01820384 | -0.0032786 | 0.79929062 | 0.89054584 |
| Zfp551 | 0.04387744 | 0.04366784 | 0.04694856 | -0.0032807 | 0.62762218 | 0.89054584 |
| Ppargc1a | 0.01640774 | 0.01427673 | 0.0175588 | -0.0032821 | 0.80808017 | 0.89054584 |
| Zfp185 | 0.03668418 | 0.03597041 | 0.03925396 | -0.0032836 | 0.65162648 | 0.89054584 |
| Ccdc39 | 0.03740995 | 0.03674261 | 0.04003037 | -0.0032878 | 0.649009 | 0.89054584 |
| Casc1 | 0.01163229 | 0.00916044 | 0.01244847 | -0.003288 | 0.89075888 | 0.91722772 |
| Vwc2 | 0.01195997 | 0.0095108 | 0.01279913 | -0.0032883 | 0.88434057 | 0.91373913 |
| Irak1bp1 | 0.31790351 | 0.32321582 | 0.32650758 | -0.0032918 | 0.51873149 | 0.89054584 |
| Ift122 | 0.32859925 | 0.33343114 | 0.33672303 | -0.0032919 | 0.51816435 | 0.89054584 |
| Gm15559 | 0.03874087 | 0.03815766 | 0.04145412 | -0.0032965 | 0.64448836 | 0.89054584 |
| Egln3 | 0.0944102 | 0.09764159 | 0.10094554 | -0.003304 | 0.56059873 | 0.89054584 |
| Gm5141 | 0.07998307 | 0.08223825 | 0.08554264 | -0.0033044 | 0.57141121 | 0.89054584 |
| Gm4724 | 0.01744165 | 0.01535718 | 0.01866519 | -0.003308 | 0.79551202 | 0.89054584 |
| Fam163b | 0.00755517 | 0.00477351 | 0.00808533 | -0.0033118 | 0.97182644 | 0.97674452 |
| Zcchc8 | 0.25998587 | 0.26688933 | 0.27020222 | -0.0033129 | 0.52277611 | 0.89054584 |
| Ppp1r3e | 0.01996068 | 0.01804392 | 0.02136076 | -0.0033168 | 0.76529154 | 0.89054584 |
| Wnt9a | 0.07802278 | 0.0801311 | 0.0834488 | -0.0033177 | 0.57347456 | 0.89054584 |
| Pcdhb20 | 0.04504746 | 0.04487874 | 0.04819999 | -0.0033212 | 0.6259051 | 0.89054584 |
| Gm12184 | 0.0313346 | 0.03020427 | 0.03353075 | -0.0033265 | 0.67803047 | 0.89054584 |
| Gm7676 | 0.01197639 | 0.00948944 | 0.01281671 | -0.0033273 | 0.88676078 | 0.91505129 |
| Pcnx | 0.17531675 | 0.18176815 | 0.18509834 | -0.0033302 | 0.53340064 | 0.89054584 |
| Uprt | 0.02618848 | 0.02469416 | 0.0280247 | -0.0033305 | 0.71010223 | 0.89054584 |
| Gm12981 | 0.02539544 | 0.02384517 | 0.02717615 | -0.003331 | 0.71601554 | 0.89054584 |
| Trnp1 | 0.02254678 | 0.02079684 | 0.02412803 | -0.0033312 | 0.73996182 | 0.89054584 |
| 2810429I04R | 0.02145553 | 0.01962302 | 0.02296034 | -0.0033373 | 0.75085583 | 0.89054584 |
| Carns1 | 0.01985774 | 0.01791048 | 0.0212506 | -0.0033401 | 0.76800195 | 0.89054584 |
| Tbx2 | 0.02297634 | 0.02124241 | 0.02458768 | -0.0033453 | 0.73691762 | 0.89054584 |
| Tnip2 | 0.06811337 | 0.06951565 | 0.07286118 | -0.0033455 | 0.58469607 | 0.89054584 |

| | | | | | | |
|---|---|---|---|---|---|---|
| 2010007H06I | 0.01138679 | 0.00883624 | 0.01218574 | -0.0033495 | 0.89983411 | 0.92300013 |
| Dtwd2 | 0.03639547 | 0.03559137 | 0.0389451 | -0.0033537 | 0.65586437 | 0.89054584 |
| Hist1h2an | 0.00587784 | 0.00293034 | 0.0062903 | -0.00336 | 0.9935871 | 0.99509343 |
| Platr22 | 0.00555189 | 0.00258043 | 0.00594149 | -0.0033611 | 0.99579966 | 0.99662457 |
| Xlr3b | 0.20647886 | 0.21342357 | 0.2167847 | -0.0033611 | 0.52879233 | 0.89054584 |
| Snta1 | 0.11879805 | 0.12326495 | 0.12662696 | -0.003362 | 0.54922256 | 0.89054584 |
| Xrcc6bp1 | 0.09050875 | 0.09341877 | 0.09678152 | -0.0033628 | 0.56429934 | 0.89054584 |
| Renbp | 0.12344589 | 0.12811775 | 0.13148169 | -0.0033639 | 0.54744081 | 0.89054584 |
| Gda | 0.01057312 | 0.00795053 | 0.011315 | -0.0033645 | 0.91702202 | 0.93494974 |
| Fbxw24 | 0.01753971 | 0.01540419 | 0.01877012 | -0.0033659 | 0.79826982 | 0.89054584 |
| Usp43 | 0.01989266 | 0.01792133 | 0.02128797 | -0.0033666 | 0.76937794 | 0.89054584 |
| Fank1 | 0.01118874 | 0.00860577 | 0.01197381 | -0.003368 | 0.90498443 | 0.92658646 |
| Sgsh | 0.03368929 | 0.03267507 | 0.03604997 | -0.0033749 | 0.66863876 | 0.89054584 |
| Gm9958 | 0.03120275 | 0.03000841 | 0.03338969 | -0.0033813 | 0.68146425 | 0.89054584 |
| Mif4gd | 0.15485037 | 0.16068663 | 0.16406954 | -0.0033829 | 0.53826419 | 0.89054584 |
| Gm5087 | 0.01500686 | 0.0126759 | 0.0160597 | -0.0033838 | 0.83686267 | 0.89468586 |
| 2810455O05 | 0.01160072 | 0.00903062 | 0.01241469 | -0.0033841 | 0.89795079 | 0.92211428 |
| Cenpn | 0.08121286 | 0.08347026 | 0.0868561 | -0.0033858 | 0.57205779 | 0.89054584 |
| Chst14 | 0.07704417 | 0.07901104 | 0.08240344 | -0.0033924 | 0.57605057 | 0.89054584 |
| Coch | 0.0206708 | 0.01872766 | 0.02212064 | -0.003393 | 0.7625754 | 0.89054584 |
| Raet1d | 0.01220741 | 0.00966778 | 0.01306393 | -0.0033961 | 0.88708207 | 0.91505129 |
| 9030612E09I | 0.02022526 | 0.01824482 | 0.02164388 | -0.0033991 | 0.76781298 | 0.89054584 |
| Mmp11 | 0.09681958 | 0.10011646 | 0.10351655 | -0.0034001 | 0.56081147 | 0.89054584 |
| Gm5160 | 0.01218946 | 0.00963517 | 0.01304472 | -0.0034096 | 0.88833619 | 0.91584392 |
| Brsk1 | 0.05055279 | 0.05067278 | 0.05408784 | -0.0034151 | 0.61568288 | 0.89054584 |
| Slc16a6 | 0.04648331 | 0.04632043 | 0.0497357 | -0.0034153 | 0.62548619 | 0.89054584 |
| Ptprf | 1.0020206 | 0.83214509 | 0.83556062 | -0.0034155 | 0.50759715 | 0.89054584 |
| Lmo1 | 0.07287062 | 0.0745283 | 0.07794464 | -0.0034163 | 0.58090259 | 0.89054584 |
| Gm9945 | 0.01714841 | 0.01493409 | 0.01835139 | -0.0034173 | 0.80718361 | 0.89054584 |
| Tmem145 | 0.02263377 | 0.02080043 | 0.02422111 | -0.0034207 | 0.74472129 | 0.89054584 |

| | | | | | | |
|---|---|---|---|---|---|---|
| Ikbkg | 0.13977419 | 0.14504989 | 0.1484729 | -0.003423 | 0.54276844 | 0.89054584 |
| Gm9755 | 0.01191523 | 0.00932798 | 0.01275125 | -0.0034233 | 0.89448992 | 0.91973864 |
| Tmem38b | 0.13308413 | 0.13809916 | 0.1415235 | -0.0034243 | 0.5448773 | 0.89054584 |
| Trmt5 | 0.0534315 | 0.05374176 | 0.05716618 | -0.0034244 | 0.60990978 | 0.89054584 |
| Tep1 | 0.07179561 | 0.07336524 | 0.07679601 | -0.0034308 | 0.58243736 | 0.89054584 |
| Cebpb | 0.01154227 | 0.00891854 | 0.01235213 | -0.0034336 | 0.90235192 | 0.9247401 |
| Tbc1d22b | 0.2232509 | 0.23022999 | 0.23366705 | -0.0034371 | 0.52731794 | 0.89054584 |
| Pitpnb | 0.73735218 | 0.67294825 | 0.67638655 | -0.0034383 | 0.50944724 | 0.89054584 |
| Tbc1d12 | 0.07248721 | 0.07409647 | 0.07753499 | -0.0034385 | 0.5818447 | 0.89054584 |
| Rgl3 | 0.09958939 | 0.10303238 | 0.10647166 | -0.0034393 | 0.55981285 | 0.89054584 |
| Tfe3 | 0.28407991 | 0.29038001 | 0.29382376 | -0.0034437 | 0.52177339 | 0.89054584 |
| Als2cl | 0.03072052 | 0.02942618 | 0.03287374 | -0.0034476 | 0.68743598 | 0.89054584 |
| Kank4 | 0.02087832 | 0.01888834 | 0.0223427 | -0.0034544 | 0.76433231 | 0.89054584 |
| Grin3a | 0.02317819 | 0.02134793 | 0.02480366 | -0.0034557 | 0.74186262 | 0.89054584 |
| Zbtb3 | 0.0334259 | 0.03230902 | 0.03576819 | -0.0034592 | 0.67385039 | 0.89054584 |
| Zfp687 | 0.23685483 | 0.24380653 | 0.24726863 | -0.0034621 | 0.52600516 | 0.89054584 |
| Col11a2 | 0.03375168 | 0.03264858 | 0.03611672 | -0.0034681 | 0.6726992 | 0.89054584 |
| Smarcd3 | 0.06444845 | 0.06546995 | 0.06894421 | -0.0034743 | 0.59280837 | 0.89054584 |
| Tfcp2l1 | 0.04689505 | 0.0467003 | 0.05017606 | -0.0034758 | 0.62655023 | 0.89054584 |
| Pan2 | 0.08774662 | 0.09034935 | 0.09383289 | -0.0034835 | 0.56865984 | 0.89054584 |
| Mybl1 | 0.01175343 | 0.00909439 | 0.0125781 | -0.0034837 | 0.90153895 | 0.92418557 |
| Slc26a11 | 0.06369348 | 0.06464936 | 0.06813724 | -0.0034879 | 0.59424838 | 0.89054584 |
| Taf1c | 0.07411716 | 0.07578788 | 0.07927648 | -0.0034886 | 0.58122136 | 0.89054584 |
| Pik3c2b | 0.02474217 | 0.02298133 | 0.02647715 | -0.0034958 | 0.73076461 | 0.89054584 |
| Tmem63a | 0.19107934 | 0.19767848 | 0.20117754 | -0.0034991 | 0.53229197 | 0.89054584 |
| Ly75 | 0.01737161 | 0.01509003 | 0.01859024 | -0.0035002 | 0.80980977 | 0.89054584 |
| Gm29019 | 0.01240815 | 0.00977604 | 0.01327874 | -0.0035027 | 0.89045444 | 0.9171534 |
| Brinp1 | 0.01500174 | 0.01255076 | 0.01605422 | -0.0035035 | 0.84535294 | 0.89825055 |
| Scml2 | 0.02086867 | 0.0188285 | 0.02233237 | -0.0035039 | 0.76760164 | 0.89054584 |
| Apobec4 | 0.01244033 | 0.00980135 | 0.01331319 | -0.0035118 | 0.89045739 | 0.9171534 |

| | | | | | | |
|---|---|---|---|---|---|---|
| Lrrc36 | 0.01035343 | 0.00756529 | 0.0110799 | -0.0035146 | 0.93027518 | 0.94460916 |
| Gm26739 | 0.01144935 | 0.00873516 | 0.01225269 | -0.0035175 | 0.90947005 | 0.92980021 |
| Ugt2b36 | 0.01446692 | 0.0119618 | 0.0154819 | -0.0035201 | 0.8552678 | 0.90089862 |
| A430105J06F | 0.0553047 | 0.05564655 | 0.05916911 | -0.0035226 | 0.60924984 | 0.89054584 |
| Gm9530 | 0.01209192 | 0.00941765 | 0.01294034 | -0.0035227 | 0.89765141 | 0.9219206 |
| Syn3 | 0.01430947 | 0.01177825 | 0.0153134 | -0.0035352 | 0.85893227 | 0.90131997 |
| Gm38393 | 0.01030546 | 0.00749317 | 0.01102857 | -0.0035354 | 0.93234711 | 0.94598421 |
| B230354K17 | 0.26248555 | 0.26912894 | 0.2726659 | -0.003537 | 0.52409527 | 0.89054584 |
| Gpr146 | 0.03389819 | 0.03273254 | 0.03627346 | -0.0035409 | 0.67537109 | 0.89054584 |
| Slc7a11 | 0.00931829 | 0.00642663 | 0.00997214 | -0.0035455 | 0.9511845 | 0.95999115 |
| Fn3k | 0.0113935 | 0.008646 | 0.01219293 | -0.0035469 | 0.91233723 | 0.93174592 |
| Gm27003 | 0.01234689 | 0.00966482 | 0.01321319 | -0.0035484 | 0.89455957 | 0.91974503 |
| Pde6c | 0.01091743 | 0.00813428 | 0.01168346 | -0.0035492 | 0.92150768 | 0.93827146 |
| Stc1 | 0.05859718 | 0.05913886 | 0.06268932 | -0.0035505 | 0.60405718 | 0.89054584 |
| Depdc5 | 0.10499129 | 0.10860761 | 0.11215817 | -0.0035506 | 0.5586262 | 0.89054584 |
| Apobr | 0.02315051 | 0.02122322 | 0.02477404 | -0.0035508 | 0.74785609 | 0.89054584 |
| Gm13420 | 0.01166886 | 0.00892412 | 0.0124876 | -0.0035635 | 0.90815549 | 0.92884926 |
| Pcdhga2 | 0.0565734 | 0.05696155 | 0.06052561 | -0.0035641 | 0.60808917 | 0.89054584 |
| Slc25a47 | 0.01631363 | 0.01389397 | 0.01745809 | -0.0035641 | 0.82923273 | 0.89271673 |
| Rab9b | 0.0112408 | 0.00846356 | 0.01202952 | -0.003566 | 0.91637127 | 0.93454915 |
| A630023P12 | 0.01477496 | 0.01224553 | 0.01581154 | -0.003566 | 0.85331145 | 0.90008483 |
| Zfp39 | 0.0302937 | 0.02884967 | 0.03241708 | -0.0035674 | 0.69592071 | 0.89054584 |
| Rassf10 | 0.10874504 | 0.11252985 | 0.11609887 | -0.003569 | 0.55694233 | 0.89054584 |
| Tmem130 | 0.01475361 | 0.01221548 | 0.01578869 | -0.0035732 | 0.85414696 | 0.90035231 |
| F8 | 0.01740084 | 0.01504685 | 0.01862152 | -0.0035747 | 0.81443309 | 0.89054584 |
| Proc | 0.03017138 | 0.02871071 | 0.03228622 | -0.0035755 | 0.69705526 | 0.89054584 |
| Nt5c3b | 0.56731104 | 0.54442251 | 0.54799804 | -0.0035755 | 0.51212526 | 0.89054584 |
| Pikfyve | 0.1306514 | 0.13541357 | 0.13899218 | -0.0035786 | 0.54773978 | 0.89054584 |
| Nr1d2 | 0.0613166 | 0.06201672 | 0.06559651 | -0.0035798 | 0.60034924 | 0.89054584 |
| 6430571L13F | 0.01667203 | 0.01426137 | 0.01784162 | -0.0035802 | 0.82508054 | 0.89156706 |

| | | | | | | |
|---|---|---|---|---|---|---|
| Ctdp1 | 0.16004009 | 0.16583708 | 0.16941799 | -0.0035809 | 0.53922194 | 0.89054584 |
| Tcp11l2 | 0.36167592 | 0.36435596 | 0.36793766 | -0.0035817 | 0.51808687 | 0.89054584 |
| C230037L18R | 0.02723614 | 0.02556235 | 0.02914566 | -0.0035833 | 0.71660765 | 0.89054584 |
| Nhlrc1 | 0.0321378 | 0.03080609 | 0.03439009 | -0.003584 | 0.68635177 | 0.89054584 |
| Cpeb1 | 0.01216464 | 0.00943406 | 0.01301815 | -0.0035841 | 0.90019691 | 0.92318418 |
| Fgd3 | 0.01885095 | 0.01658863 | 0.02017326 | -0.0035846 | 0.79612458 | 0.89054584 |
| Il4ra | 0.08761158 | 0.09010365 | 0.09368873 | -0.0035851 | 0.57074769 | 0.89054584 |
| Pip4k2a | 0.10469673 | 0.10826157 | 0.11184873 | -0.0035872 | 0.55938858 | 0.89054584 |
| Suv39h2 | 0.05790281 | 0.05835968 | 0.06194695 | -0.0035873 | 0.60634036 | 0.89054584 |
| Cfap57 | 0.01697797 | 0.01457998 | 0.01816901 | -0.003589 | 0.82129355 | 0.89054584 |
| Fancf | 0.04476179 | 0.04430496 | 0.04789445 | -0.0035895 | 0.63651852 | 0.89054584 |
| Mtx3 | 0.08996562 | 0.09261023 | 0.09620177 | -0.0035915 | 0.56904123 | 0.89054584 |
| Hist4h4 | 0.02140859 | 0.01931771 | 0.02291011 | -0.0035924 | 0.7674708 | 0.89054584 |
| Rundc3b | 0.02330849 | 0.02134479 | 0.02494309 | -0.0035983 | 0.74924042 | 0.89054584 |
| Elk1 | 0.09676461 | 0.09985917 | 0.1034579 | -0.0035987 | 0.56437024 | 0.89054584 |
| Dars | 0.98853926 | 0.82497287 | 0.82857805 | -0.0036052 | 0.50808651 | 0.89054584 |
| Mical1 | 0.06919912 | 0.07041415 | 0.07402148 | -0.0036073 | 0.58980627 | 0.89054584 |
| 3110001I22R | 0.0422015 | 0.04154512 | 0.04515594 | -0.0036108 | 0.64525625 | 0.89054584 |
| Wdr49 | 0.03225445 | 0.03090174 | 0.03451489 | -0.0036131 | 0.68712434 | 0.89054584 |
| Pigs | 0.47985417 | 0.47090168 | 0.47451872 | -0.003617 | 0.51416465 | 0.89054584 |
| Cdh4 | 0.02975805 | 0.02821992 | 0.03184398 | -0.0036241 | 0.70202034 | 0.89054584 |
| Atxn1l | 0.17515439 | 0.18130062 | 0.18493219 | -0.0036316 | 0.53644802 | 0.89054584 |
| Gm4792 | 0.01043594 | 0.00753515 | 0.01116819 | -0.003633 | 0.93518538 | 0.94786896 |
| Rpl27-ps3 | 0.80060821 | 0.71534135 | 0.71897742 | -0.0036361 | 0.50939885 | 0.89054584 |
| Prtg | 0.01180996 | 0.0089944 | 0.0126386 | -0.0036442 | 0.91042127 | 0.93051023 |
| Tmem221 | 0.02676154 | 0.02499297 | 0.02863786 | -0.0036449 | 0.72339866 | 0.89054584 |
| Krt1 | 0.09987565 | 0.10313023 | 0.10677704 | -0.0036468 | 0.56321253 | 0.89054584 |
| Ppp2r3c | 0.28291565 | 0.28904022 | 0.29268878 | -0.0036486 | 0.52315627 | 0.89054584 |
| Wdr19 | 0.06471888 | 0.06558371 | 0.06923326 | -0.0036495 | 0.59700032 | 0.89054584 |
| Ercc6l | 0.03076443 | 0.02926669 | 0.03292072 | -0.003654 | 0.69746314 | 0.89054584 |

| | | | | | | |
|---|---|---|---|---|---|---|
| F5 | 0.01479122 | 0.01217386 | 0.01582894 | -0.0036551 | 0.85899226 | 0.90131997 |
| Chuk | 0.497329 | 0.48592485 | 0.48958219 | -0.0036573 | 0.5138819 | 0.89054584 |
| Gm14286 | 0.02880785 | 0.02716948 | 0.03082733 | -0.0036579 | 0.70980427 | 0.89054584 |
| Fastkd5 | 0.0674654 | 0.0685104 | 0.07216869 | -0.0036583 | 0.59334792 | 0.89054584 |
| Sorbs2 | 0.51750279 | 0.50307796 | 0.50673881 | -0.0036609 | 0.51342497 | 0.89054584 |
| Zfp367 | 0.06635285 | 0.06731573 | 0.07097966 | -0.0036639 | 0.59502538 | 0.89054584 |
| Naa30 | 0.15142947 | 0.15687335 | 0.16053822 | -0.0036649 | 0.54235061 | 0.89054584 |
| Zfp93 | 0.09009856 | 0.09267292 | 0.09634367 | -0.0036708 | 0.57044554 | 0.89054584 |
| Draxin | 0.01577073 | 0.01320052 | 0.01687713 | -0.0036766 | 0.84493053 | 0.89801355 |
| Spats2 | 0.20282477 | 0.20941357 | 0.21309046 | -0.0036769 | 0.53203661 | 0.89054584 |
| Gm6594 | 0.02582514 | 0.02395771 | 0.02763592 | -0.0036782 | 0.73239845 | 0.89054584 |
| Tfeb | 0.02918698 | 0.02755326 | 0.03123298 | -0.0036797 | 0.70846065 | 0.89054584 |
| Gucy2c | 0.02176364 | 0.01960947 | 0.02329003 | -0.0036806 | 0.76921353 | 0.89054584 |
| Txndc11 | 0.4161673 | 0.41439075 | 0.41807278 | -0.003682 | 0.51636478 | 0.89054584 |
| Sema4a | 0.08981418 | 0.0923486 | 0.0960401 | -0.0036915 | 0.57106094 | 0.89054584 |
| Alpl | 0.01164086 | 0.00875872 | 0.01245764 | -0.0036989 | 0.91671852 | 0.93483379 |
| Ecsit | 0.21824419 | 0.22493881 | 0.22864051 | -0.0037017 | 0.53006313 | 0.89054584 |
| Fgfr2 | 0.11511754 | 0.1190738 | 0.12277687 | -0.0037031 | 0.5558747 | 0.89054584 |
| Tmem91 | 0.04263521 | 0.04191402 | 0.04561984 | -0.0037058 | 0.64745497 | 0.89054584 |
| Kcnip4 | 0.01529146 | 0.01265777 | 0.01636426 | -0.0037065 | 0.85434355 | 0.90042693 |
| Mlf1 | 0.09627875 | 0.09923038 | 0.10293947 | -0.0037091 | 0.56665715 | 0.89054584 |
| 4930480K23I | 0.019471 | 0.01712702 | 0.02083677 | -0.0037098 | 0.79658138 | 0.89054584 |
| Bmp7 | 0.11626419 | 0.12026531 | 0.1239769 | -0.0037116 | 0.55546382 | 0.89054584 |
| Gm9920 | 0.01978141 | 0.01745064 | 0.02116893 | -0.0037183 | 0.79341752 | 0.89054584 |
| Gm26759 | 0.04153389 | 0.04071664 | 0.04444182 | -0.0037252 | 0.65192369 | 0.89054584 |
| 2900009J06F | 0.01141156 | 0.00848288 | 0.01221225 | -0.0037294 | 0.92259474 | 0.93917838 |
| Cntnap2 | 0.01625292 | 0.01366319 | 0.01739312 | -0.0037299 | 0.84112789 | 0.89613993 |
| Sardh | 0.08005897 | 0.08189279 | 0.08562371 | -0.0037309 | 0.58043487 | 0.89054584 |
| Dnase1l2 | 0.01077676 | 0.00779823 | 0.01153292 | -0.0037347 | 0.93431241 | 0.94723402 |
| 1700064M15 | 0.01091813 | 0.00794866 | 0.01168421 | -0.0037356 | 0.93182308 | 0.94571726 |

| | | | | | | |
|---|---|---|---|---|---|---|
| Pde4a | 0.08253654 | 0.08453191 | 0.08826974 | -0.0037378 | 0.578198 | 0.89054584 |
| Rab39 | 0.01594316 | 0.01332221 | 0.01706165 | -0.0037394 | 0.84639913 | 0.89825055 |
| Zswim5 | 0.01143036 | 0.0084927 | 0.01223237 | -0.0037397 | 0.922824 | 0.93934581 |
| Tmtc3 | 0.14344792 | 0.14854016 | 0.15228165 | -0.0037415 | 0.54556663 | 0.89054584 |
| Ptpn1 | 0.58856073 | 0.56136747 | 0.56511289 | -0.0037454 | 0.51231667 | 0.89054584 |
| Sh3tc2 | 0.00959464 | 0.00652222 | 0.01026788 | -0.0037457 | 0.95539274 | 0.96336653 |
| Lgi3 | 0.01999296 | 0.01764951 | 0.0213953 | -0.0037458 | 0.79265444 | 0.89054584 |
| Lrriq3 | 0.01123244 | 0.0082706 | 0.01202057 | -0.00375 | 0.92694562 | 0.94249387 |
| Zfp365 | 0.02064792 | 0.01834581 | 0.02209615 | -0.0037503 | 0.78545632 | 0.89054584 |
| Gm9938 | 0.01099806 | 0.00801893 | 0.01176974 | -0.0037508 | 0.93119175 | 0.94534961 |
| Mbl2 | 0.0437122 | 0.04301991 | 0.04677181 | -0.0037519 | 0.64569616 | 0.89054584 |
| Angpt2 | 0.08211824 | 0.08406932 | 0.08782303 | -0.0037537 | 0.57891994 | 0.89054584 |
| Zfp677 | 0.05328355 | 0.05325239 | 0.05700797 | -0.0037556 | 0.62054881 | 0.89054584 |
| Camkmt | 0.01592225 | 0.01328247 | 0.01703927 | -0.0037568 | 0.84783672 | 0.89862623 |
| Nup155 | 0.11827794 | 0.12232494 | 0.1260832 | -0.0037583 | 0.55522458 | 0.89054584 |
| Tapt1 | 0.13101009 | 0.13560346 | 0.13936555 | -0.0037621 | 0.55004115 | 0.89054584 |
| Sphk1 | 0.09885148 | 0.10191509 | 0.10568444 | -0.0037694 | 0.56598698 | 0.89054584 |
| Wif1 | 0.02939441 | 0.02767892 | 0.03145491 | -0.003776 | 0.71201471 | 0.89054584 |
| Cetn4 | 0.14009503 | 0.14502919 | 0.14880574 | -0.0037766 | 0.54706111 | 0.89054584 |
| Creg2 | 0.01254248 | 0.00964474 | 0.0134225 | -0.0037778 | 0.90511451 | 0.9266542 |
| G430095P16 | 0.02261901 | 0.02042037 | 0.02420532 | -0.0037849 | 0.76684965 | 0.89054584 |
| Pcdhgc5 | 0.01609394 | 0.01343793 | 0.017223 | -0.0037851 | 0.84705612 | 0.89842854 |
| Lypd6 | 0.014329 | 0.0115467 | 0.01533431 | -0.0037876 | 0.87508653 | 0.90836342 |
| Rhbdl2 | 0.03448784 | 0.0331156 | 0.03690429 | -0.0037887 | 0.68378166 | 0.89054584 |
| Fut1 | 0.02428341 | 0.02219746 | 0.02598627 | -0.0037888 | 0.75151846 | 0.89054584 |
| Sigirr | 0.09664132 | 0.09953699 | 0.10332634 | -0.0037894 | 0.56783328 | 0.89054584 |
| Dph2 | 0.09784061 | 0.10081306 | 0.10460595 | -0.0037929 | 0.56707317 | 0.89054584 |
| Slc16a5 | 0.01544615 | 0.01273664 | 0.01652979 | -0.0037931 | 0.8574884 | 0.9009252 |
| Pecr | 0.30001914 | 0.30550213 | 0.30929699 | -0.0037949 | 0.52279193 | 0.89054584 |
| Wdr81 | 0.01691933 | 0.01431136 | 0.01810626 | -0.0037949 | 0.83558414 | 0.89459411 |

| | | | | | | |
|---|---|---|---|---|---|---|
| Zfp59 | 0.01927723 | 0.01683254 | 0.02062942 | -0.0037969 | 0.80441198 | 0.89054584 |
| Arhgef4 | 0.0297443 | 0.02803227 | 0.03182927 | -0.003797 | 0.71081873 | 0.89054584 |
| Mtfr2 | 0.01061426 | 0.00755934 | 0.01135902 | -0.0037997 | 0.9404371 | 0.95172812 |
| Mmachc | 0.08321429 | 0.08518475 | 0.08899352 | -0.0038088 | 0.57902285 | 0.89054584 |
| Pgam5 | 0.38170411 | 0.38274288 | 0.38655567 | -0.0038128 | 0.5183263 | 0.89054584 |
| Gm17634 | 0.01614038 | 0.01345464 | 0.0172727 | -0.0038181 | 0.84845709 | 0.89883797 |
| Cmklr1 | 0.01472578 | 0.01194061 | 0.01575891 | -0.0038183 | 0.87051645 | 0.90594699 |
| Slc36a4 | 0.04637793 | 0.04579989 | 0.04962299 | -0.0038231 | 0.64017963 | 0.89054584 |
| Syce1 | 0.02754875 | 0.02565496 | 0.02948015 | -0.0038252 | 0.7272509 | 0.89054584 |
| Ptpn4 | 0.14696183 | 0.15209212 | 0.15591974 | -0.0038276 | 0.54552812 | 0.89054584 |
| 9030407P20I | 0.01123389 | 0.0081886 | 0.01202213 | -0.0038335 | 0.93130885 | 0.94537357 |
| Gm10848 | 0.01590123 | 0.01317744 | 0.01701678 | -0.0038393 | 0.85340603 | 0.90009317 |
| Hs6st3 | 0.01086591 | 0.00778687 | 0.01162833 | -0.0038415 | 0.93810958 | 0.95016738 |
| Scx | 0.02511402 | 0.02303108 | 0.02687503 | -0.0038439 | 0.74741287 | 0.89054584 |
| Panx2 | 0.00927956 | 0.00608565 | 0.0099307 | -0.003845 | 0.96437542 | 0.97046678 |
| Eps8l1 | 0.03573765 | 0.03439397 | 0.03824136 | -0.0038474 | 0.68036802 | 0.89054584 |
| D930048N14 | 0.01040533 | 0.00728675 | 0.01113544 | -0.0038487 | 0.94632797 | 0.95608801 |
| Gm26778 | 0.01381711 | 0.01093726 | 0.01478652 | -0.0038493 | 0.88740136 | 0.9152899 |
| Apcs | 0.01666928 | 0.01398865 | 0.01783867 | -0.00385 | 0.8426751 | 0.89719494 |
| Hist1h2ak | 0.01085545 | 0.00776703 | 0.01161713 | -0.0038501 | 0.93871179 | 0.95044651 |
| Phf19 | 0.01973191 | 0.01726391 | 0.02111596 | -0.0038521 | 0.80230998 | 0.89054584 |
| Ldhd | 0.02726834 | 0.02532702 | 0.02918012 | -0.0038531 | 0.73078592 | 0.89054584 |
| Cmtr2 | 0.02544413 | 0.02337212 | 0.02722825 | -0.0038561 | 0.74531215 | 0.89054584 |
| Tatdn3 | 0.06982581 | 0.07083059 | 0.07469117 | -0.0038606 | 0.59514789 | 0.89054584 |
| Sostdc1 | 0.01983452 | 0.01736461 | 0.02122576 | -0.0038612 | 0.80164239 | 0.89054584 |
| Optn | 0.10097715 | 0.10407724 | 0.10793844 | -0.0038612 | 0.5661815 | 0.89054584 |
| Kcnn4 | 0.02714608 | 0.0251832 | 0.0290493 | -0.0038661 | 0.73238683 | 0.89054584 |
| 1700010I14R | 0.01561448 | 0.01283837 | 0.01670993 | -0.0038716 | 0.85980422 | 0.90132889 |
| Gm8898 | 0.01149261 | 0.00842053 | 0.01229899 | -0.0038785 | 0.92910879 | 0.94382141 |
| Gpt | 0.03888087 | 0.03772504 | 0.04160389 | -0.0038788 | 0.66798875 | 0.89054584 |

| | | | | | | |
|---|---|---|---|---|---|---|
| Cops7b | 0.34939741 | 0.35253775 | 0.35641811 | -0.0038804 | 0.52022664 | 0.89054584 |
| Calr4 | 0.01478442 | 0.01194063 | 0.01582166 | -0.003881 | 0.87344557 | 0.90763913 |
| Arl10 | 0.02130809 | 0.01892095 | 0.02280257 | -0.0038816 | 0.7861342 | 0.89054584 |
| Eml6 | 0.03078655 | 0.02905994 | 0.0329444 | -0.0038845 | 0.7086115 | 0.89054584 |
| 4930525G20 | 0.01083105 | 0.00770404 | 0.01159103 | -0.003887 | 0.94089742 | 0.95204654 |
| Gramd1c | 0.04026385 | 0.03919318 | 0.04308328 | -0.0038901 | 0.66300136 | 0.89054584 |
| Lmx1b | 0.03673527 | 0.03540717 | 0.03930862 | -0.0039014 | 0.6781055 | 0.89054584 |
| Gm10699 | 0.0201899 | 0.01770405 | 0.02160604 | -0.003902 | 0.79993578 | 0.89054584 |
| Zfp821 | 0.16445402 | 0.17005528 | 0.17395837 | -0.0039031 | 0.54162648 | 0.89054584 |
| Gdnf | 0.0309388 | 0.02920369 | 0.03310728 | -0.0039036 | 0.70860807 | 0.89054584 |
| 1500002C15I | 0.02981118 | 0.02799572 | 0.03190082 | -0.0039051 | 0.71576939 | 0.89054584 |
| Rassf8 | 0.05897243 | 0.05918287 | 0.0630905 | -0.0039076 | 0.61354095 | 0.89054584 |
| Inpp5j | 0.02437422 | 0.02217321 | 0.02608344 | -0.0039102 | 0.75754683 | 0.89054584 |
| Vav2 | 0.06811082 | 0.068948 | 0.07285845 | -0.0039105 | 0.5987273 | 0.89054584 |
| Nup98 | 0.07056834 | 0.07157404 | 0.07548463 | -0.0039106 | 0.59536314 | 0.89054584 |
| Slc38a9 | 0.10307262 | 0.1062256 | 0.11014197 | -0.0039164 | 0.56578777 | 0.89054584 |
| Sgsm2 | 0.1386188 | 0.14335067 | 0.14727396 | -0.0039233 | 0.54938652 | 0.89054584 |
| Ckmt1 | 0.13815937 | 0.14287179 | 0.14679707 | -0.0039253 | 0.54957106 | 0.89054584 |
| 9130008F23I | 0.02178051 | 0.0193769 | 0.02330808 | -0.0039312 | 0.78400313 | 0.89054584 |
| Ubash3b | 0.07565335 | 0.0769864 | 0.08091767 | -0.0039313 | 0.58953459 | 0.89054584 |
| 2600014E21I | 0.07381967 | 0.07502218 | 0.07895864 | -0.0039365 | 0.59183588 | 0.89054584 |
| Stim1 | 0.07361662 | 0.07480287 | 0.07874169 | -0.0039388 | 0.59213866 | 0.89054584 |
| B230369F24 | 0.01445457 | 0.0115261 | 0.01546868 | -0.0039426 | 0.88247415 | 0.91236613 |
| Paqr9 | 0.01107953 | 0.00791427 | 0.01185694 | -0.0039427 | 0.93933199 | 0.95094164 |
| Tet1 | 0.20093755 | 0.20723683 | 0.21118035 | -0.0039435 | 0.53466412 | 0.89054584 |
| Ipo9 | 0.30542256 | 0.31056357 | 0.31451349 | -0.0039499 | 0.5233291 | 0.89054584 |
| Rsph10b | 0.01647841 | 0.01367798 | 0.01763442 | -0.0039564 | 0.85205227 | 0.89988783 |
| Susd1 | 0.0162592 | 0.01344333 | 0.01739984 | -0.0039565 | 0.85528822 | 0.90089862 |
| Myo15b | 0.03343649 | 0.03182142 | 0.03577951 | -0.0039581 | 0.69686085 | 0.89054584 |
| Hdac9 | 0.01850046 | 0.0158346 | 0.01979821 | -0.0039636 | 0.82451812 | 0.89142971 |

| | | | | | | |
|---|---|---|---|---|---|---|
| Vamp1 | 0.02261819 | 0.02023902 | 0.02420444 | -0.0039654 | 0.77735067 | 0.89054584 |
| Vcpkmt | 0.15395097 | 0.15917317 | 0.16314153 | -0.0039684 | 0.54511439 | 0.89054584 |
| Ccdc162 | 0.02109442 | 0.01860376 | 0.02257393 | -0.0039702 | 0.7937172 | 0.89054584 |
| Slc15a2 | 0.073612 | 0.07476158 | 0.07873676 | -0.0039752 | 0.59297943 | 0.89054584 |
| Lancl3 | 0.02330899 | 0.02096687 | 0.02494362 | -0.0039767 | 0.77119024 | 0.89054584 |
| Slc22a15 | 0.09972254 | 0.10263103 | 0.10661371 | -0.0039827 | 0.56908306 | 0.89054584 |
| Ret | 0.13780153 | 0.14244209 | 0.14642557 | -0.0039835 | 0.5504291 | 0.89054584 |
| Cd59b | 0.02989484 | 0.02800685 | 0.03199033 | -0.0039835 | 0.71908923 | 0.89054584 |
| Kcnab3os | 0.014063 | 0.01106265 | 0.01504965 | -0.003987 | 0.89144753 | 0.91752139 |
| Prkag1 | 0.66394174 | 0.61940313 | 0.6233934 | -0.0039903 | 0.51189524 | 0.89054584 |
| Lrrc45 | 0.22819498 | 0.2346289 | 0.23861964 | -0.0039907 | 0.53105307 | 0.89054584 |
| 1700008O03 | 0.02824542 | 0.02623421 | 0.03022556 | -0.0039913 | 0.73079631 | 0.89054584 |
| Zfp715 | 0.10713315 | 0.11041236 | 0.11440734 | -0.003995 | 0.56461691 | 0.89054584 |
| Atp5sl | 0.21411385 | 0.22048638 | 0.22448539 | -0.003999 | 0.53307215 | 0.89054584 |
| Gm28874 | 0.01660649 | 0.01377213 | 0.01777148 | -0.0039993 | 0.85278719 | 0.89994913 |
| Tmem86b | 0.02098979 | 0.01846088 | 0.02246197 | -0.0040011 | 0.79669787 | 0.89054584 |
| Trpm3 | 0.00990276 | 0.00659209 | 0.01059761 | -0.0040055 | 0.96087368 | 0.96768176 |
| Dhx33 | 0.10426543 | 0.10738633 | 0.11139558 | -0.0040092 | 0.56658257 | 0.89054584 |
| Slc8a1 | 0.02427674 | 0.02196917 | 0.02597913 | -0.00401 | 0.7639671 | 0.89054584 |
| 2310033P09I | 0.44119355 | 0.43652421 | 0.44053725 | -0.004013 | 0.51692615 | 0.89054584 |
| 9530053A07I | 0.01934316 | 0.01668437 | 0.02069997 | -0.0040156 | 0.81694691 | 0.89054584 |
| Rap2a | 0.22564886 | 0.23205327 | 0.23607052 | -0.0040172 | 0.53159581 | 0.89054584 |
| Ccl27a | 0.35463461 | 0.35732417 | 0.36134158 | -0.0040174 | 0.5206553 | 0.89054584 |
| Gm37276 | 0.00977148 | 0.00643704 | 0.01045713 | -0.0040201 | 0.96335828 | 0.96957781 |
| Atad3aos | 0.03153083 | 0.02971471 | 0.0337407 | -0.004026 | 0.7108647 | 0.89054584 |
| Usp47 | 0.52230054 | 0.50675222 | 0.51078148 | -0.0040293 | 0.5146585 | 0.89054584 |
| Nhlrc3 | 0.09352478 | 0.09597027 | 0.10000062 | -0.0040304 | 0.57447183 | 0.89054584 |
| Prkg1 | 0.02335154 | 0.02095817 | 0.02498915 | -0.004031 | 0.77383331 | 0.89054584 |
| Gm1976 | 0.01786916 | 0.0150912 | 0.01912266 | -0.0040315 | 0.83700039 | 0.89476711 |
| Neu3 | 0.02757582 | 0.02547706 | 0.02950911 | -0.0040321 | 0.73780271 | 0.89054584 |

| | | | | | | |
|---|---|---|---|---|---|---|
| Gm20406 | 0.01124864 | 0.00800331 | 0.0120379 | -0.0040346 | 0.94079453 | 0.95202339 |
| Pcdhb16 | 0.01582501 | 0.01289631 | 0.01693522 | -0.0040389 | 0.86674148 | 0.90430169 |
| Tbc1d2 | 0.02270161 | 0.02025462 | 0.0242937 | -0.0040391 | 0.78071144 | 0.89054584 |
| Bcas1os1 | 0.01616655 | 0.01325987 | 0.0173007 | -0.0040408 | 0.86173844 | 0.90244755 |
| Calca | 0.04116923 | 0.04000074 | 0.04405176 | -0.004051 | 0.66583403 | 0.89054584 |
| Phf21a | 0.55031579 | 0.53004242 | 0.53409811 | -0.0040557 | 0.51411076 | 0.89054584 |
| Vwa2 | 0.01068201 | 0.00737293 | 0.01143153 | -0.0040586 | 0.95094522 | 0.9598833 |
| 4933407L21I | 0.00931875 | 0.0059106 | 0.00997263 | -0.004062 | 0.97113013 | 0.97617744 |
| Pla2g7 | 0.01371567 | 0.01060662 | 0.01467797 | -0.0040714 | 0.90187163 | 0.92431323 |
| Rgn | 0.02088232 | 0.01827461 | 0.02234697 | -0.0040724 | 0.80206351 | 0.89054584 |
| Tmem178b | 0.09288043 | 0.09523747 | 0.09931293 | -0.0040755 | 0.57581046 | 0.89054584 |
| 4732491K20I | 0.07393015 | 0.07500019 | 0.07907668 | -0.0040765 | 0.59490202 | 0.89054584 |
| Sirt4 | 0.15947893 | 0.16476283 | 0.16884018 | -0.0040774 | 0.54479032 | 0.89054584 |
| Tcn2 | 0.45530355 | 0.44896445 | 0.45304305 | -0.0040786 | 0.5167279 | 0.89054584 |
| AA386476 | 0.02834677 | 0.02625468 | 0.030334 | -0.0040793 | 0.73451802 | 0.89054584 |
| Gm5239 | 0.01937248 | 0.01664712 | 0.02073134 | -0.0040842 | 0.82064953 | 0.89054584 |
| Zfp236 | 0.05203836 | 0.05158539 | 0.05567646 | -0.0040911 | 0.63395106 | 0.89054584 |
| Coq3 | 0.08692432 | 0.08886307 | 0.09295498 | -0.0040919 | 0.58124834 | 0.89054584 |
| Rccd1 | 0.05502089 | 0.05477116 | 0.05886565 | -0.0040945 | 0.62705375 | 0.89054584 |
| Wfdc16 | 0.03161225 | 0.02973102 | 0.03382781 | -0.0040968 | 0.71370117 | 0.89054584 |
| Nphp1 | 0.27944973 | 0.28520916 | 0.28930627 | -0.0040971 | 0.52630279 | 0.89054584 |
| Zcchc11 | 0.87642622 | 0.76210506 | 0.76620583 | -0.0041008 | 0.50994654 | 0.89054584 |
| Ccdc67 | 0.01061007 | 0.00725276 | 0.01135455 | -0.0041018 | 0.95381492 | 0.96237794 |
| Efcab11 | 0.01442209 | 0.01132917 | 0.01543392 | -0.0041048 | 0.89234131 | 0.91811542 |
| Mstn | 0.01567933 | 0.01267361 | 0.01677932 | -0.0041057 | 0.87285498 | 0.90729956 |
| Reep3 | 0.31342567 | 0.31810264 | 0.32221321 | -0.0041106 | 0.52369742 | 0.89054584 |
| 1810062G17 | 0.01947505 | 0.01672921 | 0.0208411 | -0.0041119 | 0.82100338 | 0.89054584 |
| Mtcp1 | 0.04810643 | 0.04735843 | 0.05147162 | -0.0041132 | 0.64516703 | 0.89054584 |
| Actl6b | 0.17086826 | 0.17642727 | 0.18054219 | -0.0041149 | 0.54228286 | 0.89054584 |
| Fancg | 0.08229275 | 0.08389405 | 0.0880094 | -0.0041153 | 0.58622837 | 0.89054584 |

| | | | | | | |
|---|---|---|---|---|---|---|
| Csf3 | 0.01306987 | 0.00986366 | 0.01398687 | -0.0041232 | 0.91518866 | 0.93386861 |
| Gm10451 | 0.04249714 | 0.04134816 | 0.04547216 | -0.004124 | 0.66368168 | 0.89054584 |
| A430106G13 | 0.01523057 | 0.01217412 | 0.0162991 | -0.004125 | 0.88081766 | 0.91173336 |
| Rpia | 0.43913302 | 0.43457523 | 0.43870161 | -0.0041264 | 0.51747665 | 0.89054584 |
| Myo19 | 0.01976494 | 0.01702077 | 0.0211513 | -0.0041305 | 0.81854166 | 0.89054584 |
| Zufsp | 0.19540675 | 0.20143928 | 0.20557358 | -0.0041343 | 0.53732472 | 0.89054584 |
| Nxpe3 | 0.03306699 | 0.03122487 | 0.03538421 | -0.0041593 | 0.70803337 | 0.89054584 |
| Phf1 | 0.15763098 | 0.16277359 | 0.16693654 | -0.0041629 | 0.54624563 | 0.89054584 |
| Klhdc7a | 0.00929582 | 0.00577862 | 0.0099481 | -0.0041695 | 0.97457063 | 0.97870298 |
| Megf8 | 0.0953691 | 0.09779727 | 0.10196881 | -0.0041715 | 0.57557949 | 0.89054584 |
| Kbtbd8 | 0.01515311 | 0.0120439 | 0.01621621 | -0.0041723 | 0.8846799 | 0.91399482 |
| Gm15952 | 0.01471818 | 0.01157103 | 0.01575078 | -0.0041798 | 0.89183295 | 0.91778777 |
| Sall4 | 0.04811621 | 0.04730212 | 0.05148208 | -0.00418 | 0.64738544 | 0.89054584 |
| 4930532G15 | 0.01996472 | 0.01718356 | 0.02136508 | -0.0041815 | 0.81907226 | 0.89054584 |
| Wdr4 | 0.12856305 | 0.13263296 | 0.13681739 | -0.0041844 | 0.55665321 | 0.89054584 |
| Carnmt1 | 0.36055842 | 0.36270192 | 0.36689263 | -0.0041907 | 0.52121977 | 0.89054584 |
| Pld6 | 0.0141657 | 0.0109655 | 0.01515955 | -0.004194 | 0.90129127 | 0.92417574 |
| Gm26852 | 0.02305718 | 0.02047814 | 0.02467417 | -0.004196 | 0.78590411 | 0.89054584 |
| Arhgap39 | 0.12386819 | 0.12772104 | 0.13192238 | -0.0042013 | 0.55897607 | 0.89054584 |
| Paqr6 | 0.02689763 | 0.02457814 | 0.02878347 | -0.0042053 | 0.75196344 | 0.89054584 |
| Clcn4 | 0.58062119 | 0.55454123 | 0.55875283 | -0.0042116 | 0.51400668 | 0.89054584 |
| Tmem220 | 0.04637066 | 0.04540257 | 0.04961522 | -0.0042126 | 0.65379029 | 0.89054584 |
| Galc | 0.04746503 | 0.04657148 | 0.05078565 | -0.0042142 | 0.65047289 | 0.89054584 |
| Entpd7 | 0.2106644 | 0.21678968 | 0.22100944 | -0.0042198 | 0.53544061 | 0.89054584 |
| Rad54l | 0.03928476 | 0.03781157 | 0.04203594 | -0.0042244 | 0.68017815 | 0.89054584 |
| Hmgcll1 | 0.04254487 | 0.04129883 | 0.04552321 | -0.0042244 | 0.66725013 | 0.89054584 |
| Wfdc3 | 0.02082716 | 0.01806227 | 0.02228795 | -0.0042257 | 0.81146619 | 0.89054584 |
| Sel1l3 | 0.01566346 | 0.01253554 | 0.01676233 | -0.0042268 | 0.87996278 | 0.91137745 |
| Itpka | 0.01969968 | 0.01685452 | 0.02108147 | -0.004227 | 0.8248852 | 0.89153749 |
| Tmem150c | 0.09717528 | 0.09966587 | 0.10389608 | -0.0042302 | 0.57522485 | 0.89054584 |

| | | | | | | |
|---|---|---|---|---|---|---|
| Adat2 | 0.12923783 | 0.13328425 | 0.13752028 | -0.004236 | 0.55705596 | 0.89054584 |
| Lmnb2 | 0.17354148 | 0.17903992 | 0.18328109 | -0.0042412 | 0.54292641 | 0.89054584 |
| Gm28535 | 0.01042168 | 0.00691024 | 0.01115294 | -0.0042427 | 0.96182874 | 0.96837452 |
| Fbp1 | 0.05122324 | 0.05056208 | 0.05480481 | -0.0042427 | 0.64082818 | 0.89054584 |
| Acad9 | 0.18646997 | 0.19223441 | 0.19648628 | -0.0042519 | 0.5401523 | 0.89054584 |
| Paqr5 | 0.01618496 | 0.01306207 | 0.0173204 | -0.0042583 | 0.87398404 | 0.90773126 |
| 6430573P05I | 0.01437845 | 0.01112516 | 0.01538723 | -0.0042621 | 0.90155584 | 0.92418557 |
| Gng10 | 0.03666189 | 0.03496161 | 0.03923012 | -0.0042685 | 0.69389625 | 0.89054584 |
| Prob1 | 0.01476146 | 0.01152784 | 0.01579709 | -0.0042693 | 0.89600375 | 0.92096862 |
| Gm17146 | 0.01962567 | 0.01673027 | 0.02100227 | -0.004272 | 0.82834893 | 0.89229532 |
| Paqr3 | 0.02281339 | 0.02014078 | 0.02441331 | -0.0042725 | 0.79256493 | 0.89054584 |
| Gprc5a | 0.02091464 | 0.01810637 | 0.02238156 | -0.0042752 | 0.81324655 | 0.89054584 |
| Mamld1 | 0.14820718 | 0.15293181 | 0.15720793 | -0.0042761 | 0.55042104 | 0.89054584 |
| Clcnkb | 0.00856025 | 0.00488429 | 0.00916092 | -0.0042766 | 0.9851832 | 0.98749904 |
| Atg7 | 0.13015537 | 0.13419365 | 0.13847577 | -0.0042821 | 0.55727726 | 0.89054584 |
| Slc17a4 | 0.01940622 | 0.01648194 | 0.02076745 | -0.0042855 | 0.83182571 | 0.8933762 |
| Gm13075 | 0.03006715 | 0.02788058 | 0.03217469 | -0.0042941 | 0.73296144 | 0.89054584 |
| Cyp2j5 | 0.00911627 | 0.00546082 | 0.00975595 | -0.0042951 | 0.97987296 | 0.98311155 |
| Gulo | 0.02002341 | 0.01712764 | 0.02142789 | -0.0043002 | 0.82509986 | 0.89156706 |
| Nat9 | 0.35085054 | 0.35348162 | 0.35778568 | -0.0043041 | 0.5223473 | 0.89054584 |
| Rttn | 0.02020621 | 0.01731794 | 0.02162349 | -0.0043056 | 0.82320876 | 0.89080883 |
| Tmigd3 | 0.03476652 | 0.0328962 | 0.03720244 | -0.0043062 | 0.70515321 | 0.89054584 |
| Cidea | 0.01343698 | 0.01007319 | 0.01437973 | -0.0043065 | 0.91853615 | 0.93590112 |
| BC017158 | 0.09290623 | 0.09503311 | 0.09934048 | -0.0043074 | 0.58004548 | 0.89054584 |
| Tmem168 | 0.30754385 | 0.31224619 | 0.31655734 | -0.0043112 | 0.52529572 | 0.89054584 |
| Oit1 | 0.0398792 | 0.03835667 | 0.04267182 | -0.0043151 | 0.68122647 | 0.89054584 |
| Hpca | 0.05741189 | 0.05710407 | 0.06142209 | -0.004318 | 0.62836526 | 0.89054584 |
| Ttpa | 0.01305743 | 0.00965484 | 0.01397356 | -0.0043187 | 0.9250534 | 0.94128468 |
| 3222401L13I | 0.04936006 | 0.04848539 | 0.05281232 | -0.0043269 | 0.6486614 | 0.89054584 |
| Hspbap1 | 0.12219774 | 0.12584641 | 0.13017878 | -0.0043324 | 0.56160911 | 0.89054584 |

| | | | | | | |
|---|---|---|---|---|---|---|
| Lacc1 | 0.02507275 | 0.02249486 | 0.02683087 | -0.004336 | 0.77424795 | 0.89054584 |
| Pnma1 | 0.01359397 | 0.01021133 | 0.01454773 | -0.0043364 | 0.91754413 | 0.93505805 |
| Gen1 | 0.01378562 | 0.01041323 | 0.01475282 | -0.0043396 | 0.91472561 | 0.93359323 |
| Kctd2 | 0.08268531 | 0.0840848 | 0.08842862 | -0.0043438 | 0.59051038 | 0.89054584 |
| A330009N23 | 0.0321936 | 0.03010437 | 0.03444979 | -0.0043454 | 0.72162285 | 0.89054584 |
| Wdr78 | 0.0366885 | 0.03491243 | 0.03925858 | -0.0043462 | 0.69699329 | 0.89054584 |
| Slc6a17 | 0.02778596 | 0.02538549 | 0.02973396 | -0.0043485 | 0.75217494 | 0.89054584 |
| Tap2 | 0.14726033 | 0.15187386 | 0.15622856 | -0.0043547 | 0.55166255 | 0.89054584 |
| Aire | 0.02682724 | 0.0243533 | 0.02870815 | -0.0043549 | 0.76013375 | 0.89054584 |
| Pih1d1 | 0.43259653 | 0.42850475 | 0.43286183 | -0.0043571 | 0.51870183 | 0.89054584 |
| Tlr3 | 0.02480427 | 0.02218078 | 0.02654359 | -0.0043628 | 0.77809136 | 0.89054584 |
| Has3 | 0.00910474 | 0.00537951 | 0.00974361 | -0.0043641 | 0.9815427 | 0.98459877 |
| C030039L03I | 0.06466346 | 0.06480958 | 0.06917403 | -0.0043645 | 0.61560203 | 0.89054584 |
| Taco1os | 0.01779869 | 0.01468117 | 0.01904725 | -0.0043661 | 0.85722644 | 0.9009252 |
| A430046D13 | 0.02065875 | 0.01773907 | 0.02210774 | -0.0043687 | 0.82138188 | 0.89054584 |
| Map2k7 | 0.35926517 | 0.36131307 | 0.36568238 | -0.0043693 | 0.52219639 | 0.89054584 |
| Sipa1l2 | 0.03885633 | 0.03720517 | 0.04157764 | -0.0043725 | 0.68791622 | 0.89054584 |
| Add2 | 0.01532631 | 0.01202696 | 0.01640155 | -0.0043746 | 0.89299691 | 0.91859438 |
| Ntn4 | 0.02174969 | 0.01889914 | 0.02327511 | -0.004376 | 0.80946598 | 0.89054584 |
| Arhgap6 | 0.02864093 | 0.02627258 | 0.03064874 | -0.0043762 | 0.7470465 | 0.89054584 |
| Snip1 | 0.25183797 | 0.25777498 | 0.26215219 | -0.0043772 | 0.53100296 | 0.89054584 |
| Gm16845 | 0.03747104 | 0.03571851 | 0.04009573 | -0.0043772 | 0.6944883 | 0.89054584 |
| Ankle2 | 0.22386891 | 0.2299093 | 0.23428674 | -0.0043774 | 0.5346834 | 0.89054584 |
| Mmp24 | 0.01375679 | 0.01033858 | 0.01472197 | -0.0043834 | 0.91730565 | 0.93505805 |
| A430073D23 | 0.00783018 | 0.0039946 | 0.00837963 | -0.004385 | 0.99261572 | 0.99439903 |
| Scn1b | 0.0156815 | 0.01239481 | 0.01678164 | -0.0043868 | 0.88836481 | 0.91584392 |
| Zfp524 | 0.18958329 | 0.19526862 | 0.19965591 | -0.0043873 | 0.54077119 | 0.89054584 |
| 1810009A15I | 0.03262065 | 0.03051223 | 0.03490668 | -0.0043944 | 0.72123698 | 0.89054584 |
| Gnb3 | 0.0181822 | 0.01506175 | 0.01945765 | -0.0043959 | 0.85372734 | 0.90035231 |
| Cyp46a1 | 0.00742788 | 0.00354914 | 0.00794911 | -0.0044 | 0.99504287 | 0.99607343 |

| | | | | | | |
|---|---|---|---|---|---|---|
| Lrsam1 | 0.04827243 | 0.0472461 | 0.05164916 | -0.0044031 | 0.65437945 | 0.89054584 |
| Fam83g | 0.04605432 | 0.04486704 | 0.04927688 | -0.0044098 | 0.66163663 | 0.89054584 |
| Ccdc159 | 0.02892265 | 0.02653793 | 0.03095016 | -0.0044122 | 0.74671083 | 0.89054584 |
| Mybl2 | 0.06341751 | 0.06342799 | 0.06784226 | -0.0044143 | 0.61910903 | 0.89054584 |
| Taf9b | 0.19764866 | 0.20343359 | 0.20784787 | -0.0044143 | 0.5394097 | 0.89054584 |
| Ccdc28a | 0.09107288 | 0.09296353 | 0.09738368 | -0.0044202 | 0.58373779 | 0.89054584 |
| Dtx3l | 0.0406939 | 0.03911938 | 0.0435433 | -0.0044239 | 0.68201463 | 0.89054584 |
| Phkb | 0.27249493 | 0.27807207 | 0.28250109 | -0.004429 | 0.5291138 | 0.89054584 |
| Gsdmd | 0.10412341 | 0.10681535 | 0.11124634 | -0.004431 | 0.57360799 | 0.89054584 |
| Sufu | 0.09398842 | 0.09606071 | 0.10049543 | -0.0044347 | 0.58144524 | 0.89054584 |
| Gm14326 | 0.04611219 | 0.04490216 | 0.04933877 | -0.0044366 | 0.66236919 | 0.89054584 |
| Fam46d | 0.01406623 | 0.0106116 | 0.01505311 | -0.0044415 | 0.91538005 | 0.93399009 |
| Pars2 | 0.05442284 | 0.05378128 | 0.05822619 | -0.0044449 | 0.63895123 | 0.89054584 |
| Fzd8 | 0.01908183 | 0.01597009 | 0.02042032 | -0.0044502 | 0.84502567 | 0.89804883 |
| Ap3m2 | 0.09090013 | 0.09274513 | 0.09719929 | -0.0044542 | 0.58453003 | 0.89054584 |
| Noxo1 | 0.06421684 | 0.06423914 | 0.06869665 | -0.0044575 | 0.61879003 | 0.89054584 |
| Dcaf12l1 | 0.09250477 | 0.09445177 | 0.09891201 | -0.0044602 | 0.58319947 | 0.89054584 |
| Dagla | 0.01530834 | 0.01192181 | 0.01638231 | -0.0044605 | 0.897693 | 0.9219206 |
| Srr | 0.14308456 | 0.14743893 | 0.15190517 | -0.0044662 | 0.55447654 | 0.89054584 |
| Dock5 | 0.00966477 | 0.00587637 | 0.01034293 | -0.0044666 | 0.97788731 | 0.98147526 |
| Pus7l | 0.08849698 | 0.09016485 | 0.09463397 | -0.0044691 | 0.58707237 | 0.89054584 |
| Gm9242 | 0.04804405 | 0.04692989 | 0.05140491 | -0.004475 | 0.65747481 | 0.89054584 |
| Pxmp2 | 0.02363418 | 0.02081424 | 0.02529158 | -0.0044773 | 0.79524825 | 0.89054584 |
| Pms1 | 0.01979833 | 0.01670913 | 0.02118703 | -0.0044779 | 0.83760766 | 0.89485554 |
| Dnajc27 | 0.02233813 | 0.01940547 | 0.02390476 | -0.0044993 | 0.8097277 | 0.89054584 |
| Plin5 | 0.02251704 | 0.01959568 | 0.02409621 | -0.0045005 | 0.80789501 | 0.89054584 |
| Zpbp2 | 0.02808321 | 0.0255509 | 0.030052 | -0.0045011 | 0.75734949 | 0.89054584 |
| Myo5c | 0.02317324 | 0.0202941 | 0.02479836 | -0.0045043 | 0.80128756 | 0.89054584 |
| Cenpu | 0.04105989 | 0.03942986 | 0.04393479 | -0.0045049 | 0.68357285 | 0.89054584 |
| Kcnk12 | 0.02795611 | 0.025411 | 0.02991601 | -0.004505 | 0.75853035 | 0.89054584 |

| | | | | | | |
|---|---|---|---|---|---|---|
| Ccbl1 | 0.05033576 | 0.04934612 | 0.05385575 | -0.0045096 | 0.65177541 | 0.89054584 |
| Tmem120b | 0.04314693 | 0.04165672 | 0.04616719 | -0.0045105 | 0.67550745 | 0.89054584 |
| Fzd5 | 0.02752391 | 0.02494136 | 0.02945357 | -0.0045122 | 0.76230512 | 0.89054584 |
| Gad1 | 0.13362472 | 0.13757148 | 0.14208568 | -0.0045142 | 0.55883615 | 0.89054584 |
| Gm26619 | 0.03133557 | 0.02900827 | 0.03353179 | -0.0045235 | 0.73516267 | 0.89054584 |
| Erbb3 | 0.06772676 | 0.06791721 | 0.072448 | -0.0045308 | 0.61461343 | 0.89054584 |
| Fgf1 | 0.04147327 | 0.03984234 | 0.04437697 | -0.0045346 | 0.68298794 | 0.89054584 |
| Hinfp | 0.0315659 | 0.02923948 | 0.03377822 | -0.0045387 | 0.73434825 | 0.89054584 |
| Pou5f1 | 0.02268104 | 0.01973217 | 0.02427169 | -0.0045395 | 0.80822104 | 0.89054584 |
| Stk32b | 0.04563182 | 0.04427457 | 0.04882499 | -0.0045504 | 0.66793074 | 0.89054584 |
| Gm12454 | 0.02166662 | 0.01863576 | 0.02318622 | -0.0045505 | 0.81973372 | 0.89054584 |
| Ddx11 | 0.02608891 | 0.02336467 | 0.02791816 | -0.0045535 | 0.77633687 | 0.89054584 |
| Fzd7 | 0.07502594 | 0.0756886 | 0.08024739 | -0.0045588 | 0.60436853 | 0.89054584 |
| Pcdhb21 | 0.01951928 | 0.0163248 | 0.02088843 | -0.0045636 | 0.84562994 | 0.89825055 |
| Gm2115 | 0.09606916 | 0.09814937 | 0.10271583 | -0.0045665 | 0.58204338 | 0.89054584 |
| Ranbp6 | 0.11515076 | 0.11824371 | 0.12281165 | -0.0045679 | 0.56878735 | 0.89054584 |
| 4932422M17 | 0.07450613 | 0.07512283 | 0.07969205 | -0.0045692 | 0.60531349 | 0.89054584 |
| Rtel1 | 0.04673133 | 0.04542893 | 0.05000095 | -0.004572 | 0.66494871 | 0.89054584 |
| Helq | 0.07266728 | 0.07315496 | 0.07772738 | -0.0045724 | 0.60798423 | 0.89054584 |
| Zfp955a | 0.03806041 | 0.03614608 | 0.04072621 | -0.0045801 | 0.69984367 | 0.89054584 |
| Mkrn2os | 0.05686178 | 0.05624631 | 0.06083394 | -0.0045876 | 0.6373339 | 0.89054584 |
| Fam103a1 | 1.17750084 | 0.91612159 | 0.92071338 | -0.0045918 | 0.50926864 | 0.89054584 |
| Gm16536 | 0.02771803 | 0.02506489 | 0.02966128 | -0.0045964 | 0.76484105 | 0.89054584 |
| Fancb | 0.04086874 | 0.03913383 | 0.04373033 | -0.0045965 | 0.68782757 | 0.89054584 |
| Gm14703 | 0.0319985 | 0.02964234 | 0.03424105 | -0.0045987 | 0.73424889 | 0.89054584 |
| Zfp619 | 0.01875396 | 0.0154682 | 0.02006948 | -0.0046013 | 0.85727234 | 0.9009252 |
| Pkp3 | 0.09018367 | 0.09182141 | 0.09643452 | -0.0046131 | 0.58818112 | 0.89054584 |
| 1110035H17 | 0.0225759 | 0.01954288 | 0.02415919 | -0.0046163 | 0.81332752 | 0.89054584 |
| AA986860 | 0.02288221 | 0.01986818 | 0.02448696 | -0.0046188 | 0.81023912 | 0.89054584 |
| Gm2861 | 0.03629716 | 0.03421323 | 0.03883993 | -0.0046267 | 0.71054697 | 0.89054584 |

| | | | | | | |
|---|---|---|---|---|---|---|
| Klhdc8a | 0.04599524 | 0.04458401 | 0.04921369 | -0.0046297 | 0.66940842 | 0.89054584 |
| Atp6v1c1 | 0.76449457 | 0.69038057 | 0.69501415 | -0.0046336 | 0.51238943 | 0.89054584 |
| Bdp1 | 0.51235593 | 0.49775236 | 0.50238617 | -0.0046338 | 0.51713815 | 0.89054584 |
| Gm684 | 0.03575007 | 0.03361767 | 0.03825465 | -0.004637 | 0.71387035 | 0.89054584 |
| Mmaa | 0.12336665 | 0.12676087 | 0.13139899 | -0.0046381 | 0.56531205 | 0.89054584 |
| Ssh3 | 0.11468077 | 0.11767796 | 0.12231964 | -0.0046417 | 0.5701647 | 0.89054584 |
| Zfp280c | 0.03982166 | 0.03796672 | 0.04261027 | -0.0046436 | 0.69417483 | 0.89054584 |
| AI429214 | 0.02377955 | 0.02080207 | 0.02544713 | -0.0046451 | 0.80245746 | 0.89054584 |
| AU020206 | 0.04248704 | 0.04081624 | 0.04546136 | -0.0046451 | 0.68297634 | 0.89054584 |
| Kcnd1 | 0.01520838 | 0.01162916 | 0.01627535 | -0.0046462 | 0.90824167 | 0.92887188 |
| Slc29a2 | 0.01294085 | 0.00920052 | 0.01384881 | -0.0046483 | 0.94106217 | 0.95209067 |
| Gm26910 | 0.04213794 | 0.04043477 | 0.04508794 | -0.0046532 | 0.68467543 | 0.89054584 |
| Tesk2 | 0.02149176 | 0.01834412 | 0.02299911 | -0.004655 | 0.8271503 | 0.89186561 |
| Pla2g2c | 0.01908438 | 0.01576581 | 0.02042305 | -0.0046572 | 0.85597713 | 0.9009252 |
| Tspyl5 | 0.08835428 | 0.08982075 | 0.09448162 | -0.0046609 | 0.59088818 | 0.89054584 |
| Grhl2 | 0.04430587 | 0.04274549 | 0.04740681 | -0.0046613 | 0.67655771 | 0.89054584 |
| Ttc39c | 0.05471624 | 0.0538693 | 0.05853991 | -0.0046706 | 0.64494715 | 0.89054584 |
| Arhgef39 | 0.0449567 | 0.04342853 | 0.04810291 | -0.0046744 | 0.67462858 | 0.89054584 |
| Zbtb11os1 | 0.10349124 | 0.1059074 | 0.11058199 | -0.0046746 | 0.57806492 | 0.89054584 |
| Apba1 | 0.03575013 | 0.03357992 | 0.03825472 | -0.0046748 | 0.71543446 | 0.89054584 |
| Tmem147os | 0.03346953 | 0.03112982 | 0.03581486 | -0.004685 | 0.7288864 | 0.89054584 |
| Car11 | 0.08572267 | 0.08698289 | 0.09167199 | -0.0046891 | 0.59417934 | 0.89054584 |
| Fignl2 | 0.01317276 | 0.00940524 | 0.01409698 | -0.0046917 | 0.93949905 | 0.95104436 |
| Zkscan4 | 0.02217207 | 0.01903528 | 0.02372708 | -0.0046918 | 0.82154247 | 0.89054584 |
| Zfp68 | 0.40289023 | 0.40131246 | 0.40601003 | -0.0046976 | 0.52149432 | 0.89054584 |
| Amigo3 | 0.01793305 | 0.014491 | 0.01919103 | -0.0047 | 0.8730673 | 0.90740309 |
| Tmem125 | 0.02569831 | 0.02280002 | 0.02750022 | -0.0047002 | 0.78706625 | 0.89054584 |
| Spr | 0.53825998 | 0.5194177 | 0.52412593 | -0.0047082 | 0.51669138 | 0.89054584 |
| Zranb3 | 0.03565643 | 0.03344573 | 0.03815448 | -0.0047087 | 0.71734511 | 0.89054584 |
| Anxa9 | 0.0191906 | 0.01582762 | 0.02053672 | -0.0047091 | 0.85730806 | 0.9009252 |

| | | | | | | |
|---|---|---|---|---|---|---|
| 3110070M22 | 0.02374317 | 0.02069378 | 0.02540821 | -0.0047144 | 0.80633088 | 0.89054584 |
| Tstd1 | 0.0185318 | 0.01511586 | 0.01983175 | -0.0047159 | 0.86604163 | 0.90421083 |
| Abtb1 | 0.17682386 | 0.1819209 | 0.18664006 | -0.0047192 | 0.54688718 | 0.89054584 |
| Napb | 0.09695371 | 0.09893845 | 0.10365967 | -0.0047212 | 0.58402185 | 0.89054584 |
| Fcor | 0.05802208 | 0.05734882 | 0.06207447 | -0.0047256 | 0.6385849 | 0.89054584 |
| Mfsd7b | 0.0273645 | 0.02454657 | 0.02928301 | -0.0047364 | 0.77444655 | 0.89054584 |
| Exph5 | 0.11551799 | 0.11844994 | 0.12319602 | -0.0047461 | 0.5712212 | 0.89054584 |
| Lrrc46 | 0.03219028 | 0.02969954 | 0.03444623 | -0.0047467 | 0.73956435 | 0.89054584 |
| Thrb | 0.01428388 | 0.01053623 | 0.01528602 | -0.0047498 | 0.92614453 | 0.94206436 |
| Gm27030 | 0.02383205 | 0.02074725 | 0.02550331 | -0.0047561 | 0.80753146 | 0.89054584 |
| Stk24 | 0.33489898 | 0.33795398 | 0.34271187 | -0.0047579 | 0.52578576 | 0.89054584 |
| 2610002M06 | 0.30546657 | 0.30979247 | 0.31455591 | -0.0047634 | 0.52812291 | 0.89054584 |
| Snap91 | 0.0226131 | 0.01943286 | 0.02419899 | -0.0047661 | 0.82058742 | 0.89054584 |
| Tyk2 | 0.0364065 | 0.03418656 | 0.0389569 | -0.0047703 | 0.71582911 | 0.89054584 |
| Cdyl2 | 0.12022487 | 0.12334414 | 0.12811816 | -0.004774 | 0.5689118 | 0.89054584 |
| Lurap1 | 0.03195582 | 0.02941667 | 0.03419539 | -0.0047787 | 0.74250098 | 0.89054584 |
| Gm43949 | 0.01330246 | 0.00945578 | 0.01423578 | -0.00478 | 0.94113218 | 0.95209067 |
| Pou5f2 | 0.04806623 | 0.04664688 | 0.05142863 | -0.0047817 | 0.66755802 | 0.89054584 |
| Armc9 | 0.24462557 | 0.25020909 | 0.25500076 | -0.0047917 | 0.53488097 | 0.89054584 |
| S100z | 0.0287338 | 0.02595548 | 0.0307481 | -0.0047926 | 0.76613459 | 0.89054584 |
| 9230105E05I | 0.02486899 | 0.02181808 | 0.02661285 | -0.0047948 | 0.79937445 | 0.89054584 |
| C230035I16R | 0.03139128 | 0.02879056 | 0.0335914 | -0.0048008 | 0.7472464 | 0.89054584 |
| Mss51 | 0.03068064 | 0.02802553 | 0.03283108 | -0.0048055 | 0.75236084 | 0.89054584 |
| Shisa8 | 0.02162442 | 0.0183302 | 0.02314106 | -0.0048109 | 0.83361761 | 0.89411764 |
| Gm42997 | 0.01936558 | 0.01591276 | 0.02072396 | -0.0048112 | 0.86028556 | 0.90144139 |
| Mapk4 | 0.01452266 | 0.01072602 | 0.01554155 | -0.0048155 | 0.92556959 | 0.94167774 |
| 1700066M21 | 0.04917438 | 0.04779314 | 0.05261374 | -0.0048206 | 0.66526083 | 0.89054584 |
| Rab32 | 0.03628853 | 0.03400803 | 0.03883069 | -0.0048227 | 0.71857984 | 0.89054584 |
| Ada | 0.01852799 | 0.01500466 | 0.01982767 | -0.004823 | 0.87144939 | 0.90646242 |
| Synj2 | 0.01432248 | 0.01050309 | 0.01532733 | -0.0048242 | 0.92872905 | 0.94362101 |

| | | | | | | |
|---|---|---|---|---|---|---|
| Trim39 | 0.12621385 | 0.12954102 | 0.13436898 | -0.004828 | 0.56647192 | 0.89054584 |
| Zc3h12c | 0.05901572 | 0.05830059 | 0.06313678 | -0.0048362 | 0.63940484 | 0.89054584 |
| Rpgrip1l | 0.09498539 | 0.09672253 | 0.10155936 | -0.0048368 | 0.58779819 | 0.89054584 |
| Zbtb48 | 0.08932569 | 0.09068039 | 0.09551865 | -0.0048383 | 0.59327866 | 0.89054584 |
| Pfn4 | 0.02718488 | 0.02425069 | 0.02909082 | -0.0048401 | 0.78087586 | 0.89054584 |
| Zfp605 | 0.07048664 | 0.07055688 | 0.07539733 | -0.0048404 | 0.61756812 | 0.89054584 |
| Bmp3 | 0.012375 | 0.00840217 | 0.01324327 | -0.0048411 | 0.95572387 | 0.9636334 |
| Lemd3 | 0.04582298 | 0.04418508 | 0.04902945 | -0.0048444 | 0.67735984 | 0.89054584 |
| Gm14410 | 0.02282001 | 0.01957526 | 0.0244204 | -0.0048451 | 0.82235007 | 0.89073269 |
| Sim1 | 0.10817963 | 0.11066027 | 0.11550563 | -0.0048454 | 0.57747519 | 0.89054584 |
| Tspyl3 | 0.06946904 | 0.06946142 | 0.07430992 | -0.0048485 | 0.61942911 | 0.89054584 |
| Shroom1 | 0.01669479 | 0.01301745 | 0.01786597 | -0.0048485 | 0.8969491 | 0.9215483 |
| Rtl1 | 0.01324877 | 0.00932465 | 0.01417833 | -0.0048537 | 0.94463265 | 0.95470786 |
| Ccdc113 | 0.03288132 | 0.03033142 | 0.03518556 | -0.0048541 | 0.73980316 | 0.89054584 |
| Iqck | 0.01960564 | 0.0161253 | 0.02098084 | -0.0048555 | 0.85952927 | 0.90131997 |
| Rasgef1b | 0.0225792 | 0.01930697 | 0.02416272 | -0.0048557 | 0.82543098 | 0.89170439 |
| Jrk | 0.09149941 | 0.09297826 | 0.09783895 | -0.0048607 | 0.5915207 | 0.89054584 |
| Nr2f1 | 0.03427742 | 0.03181687 | 0.03667918 | -0.0048623 | 0.73158183 | 0.89054584 |
| Slc25a43 | 0.01759168 | 0.01396227 | 0.01882573 | -0.0048635 | 0.88562735 | 0.91420299 |
| Gm17491 | 0.0313502 | 0.02868086 | 0.03354744 | -0.0048666 | 0.75043273 | 0.89054584 |
| Guca1a | 0.06162266 | 0.06105585 | 0.06592367 | -0.0048678 | 0.63458346 | 0.89054584 |
| Il1rap | 0.02471949 | 0.02157747 | 0.02645288 | -0.0048754 | 0.8047381 | 0.89054584 |
| Fabp3-ps1 | 0.02899848 | 0.02615411 | 0.03103129 | -0.0048772 | 0.76799044 | 0.89054584 |
| Rnd2 | 0.03420045 | 0.03171765 | 0.03659683 | -0.0048792 | 0.73274674 | 0.89054584 |
| Mkrn3 | 0.05296136 | 0.05177805 | 0.05666345 | -0.0048854 | 0.65604178 | 0.89054584 |
| Erf | 0.19170619 | 0.19692878 | 0.20181483 | -0.004886 | 0.54490368 | 0.89054584 |
| Rad54b | 0.01887048 | 0.01530797 | 0.02019416 | -0.0048862 | 0.87018815 | 0.90589084 |
| Jpx | 0.06566435 | 0.06535437 | 0.0702438 | -0.0048894 | 0.62714192 | 0.89054584 |
| Mpz | 0.01062664 | 0.00648275 | 0.01137227 | -0.0048895 | 0.97741759 | 0.98107174 |
| Rab11fip1 | 0.02966371 | 0.02684861 | 0.03174304 | -0.0048944 | 0.76373167 | 0.89054584 |

| | | | | | | |
|---|---|---|---|---|---|---|
| Msx1 | 0.08073698 | 0.08145193 | 0.08634786 | -0.0048959 | 0.60417253 | 0.89054584 |
| Brat1 | 0.08000146 | 0.08066436 | 0.08556228 | -0.0048979 | 0.60514836 | 0.89054584 |
| Rragb | 0.11198575 | 0.11459454 | 0.11949682 | -0.0049023 | 0.57578823 | 0.89054584 |
| Actr5 | 0.18022772 | 0.18521551 | 0.19011883 | -0.0049033 | 0.54782095 | 0.89054584 |
| Tmem117 | 0.02203078 | 0.01867128 | 0.02357589 | -0.0049046 | 0.83378301 | 0.89422888 |
| Ppp1r14d | 0.02209017 | 0.01873226 | 0.02363944 | -0.0049072 | 0.83325894 | 0.89393134 |
| Tpra1 | 0.18051728 | 0.18550601 | 0.19041456 | -0.0049086 | 0.54779779 | 0.89054584 |
| Fam73b | 0.11602175 | 0.11881432 | 0.12372321 | -0.0049089 | 0.57332664 | 0.89054584 |
| Gramd1b | 0.03709104 | 0.03477412 | 0.03968921 | -0.0049151 | 0.71801687 | 0.89054584 |
| Smarce1 | 1.23817573 | 0.94145439 | 0.94637013 | -0.0049157 | 0.50965344 | 0.89054584 |
| Dio1 | 0.05529713 | 0.05424521 | 0.05916102 | -0.0049158 | 0.65066653 | 0.89054584 |
| Enkur | 0.02754889 | 0.02456264 | 0.0294803 | -0.0049177 | 0.78146994 | 0.89054584 |
| Mcf2l | 0.42317848 | 0.41948411 | 0.42440332 | -0.0049192 | 0.52153291 | 0.89054584 |
| Bloc1s6 | 0.22570781 | 0.23120903 | 0.23612958 | -0.0049205 | 0.53867038 | 0.89054584 |
| Fam13b | 0.14950211 | 0.15362329 | 0.15854676 | -0.0049235 | 0.55751729 | 0.89054584 |
| Cblc | 0.05380374 | 0.05263339 | 0.05756421 | -0.0049308 | 0.65508084 | 0.89054584 |
| Zfp229 | 0.05267523 | 0.05142587 | 0.05635749 | -0.0049316 | 0.6582476 | 0.89054584 |
| Ankrd13d | 0.04736534 | 0.04574491 | 0.05067904 | -0.0049341 | 0.67494005 | 0.89054584 |
| Spdef | 0.01621539 | 0.01241717 | 0.01735296 | -0.0049358 | 0.90744225 | 0.92844727 |
| Wdr25 | 0.01905963 | 0.01546035 | 0.02039657 | -0.0049362 | 0.87023894 | 0.90589084 |
| Smc5 | 0.46266687 | 0.45457862 | 0.45952168 | -0.0049431 | 0.51998503 | 0.89054584 |
| Uckl1os | 0.0217805 | 0.01836141 | 0.02330807 | -0.0049467 | 0.8386111 | 0.89497393 |
| Fam110c | 0.03395413 | 0.03138092 | 0.03633332 | -0.0049524 | 0.73729728 | 0.89054584 |
| Sgpp2 | 0.1312117 | 0.13462129 | 0.13957539 | -0.0049541 | 0.56567155 | 0.89054584 |
| 1700030J22F | 0.02684587 | 0.0237728 | 0.02872809 | -0.0049553 | 0.78919113 | 0.89054584 |
| Gm26843 | 0.01421731 | 0.01025819 | 0.01521478 | -0.0049566 | 0.93546312 | 0.94807317 |
| Tmco3 | 0.37233507 | 0.37291527 | 0.37787379 | -0.0049585 | 0.52437428 | 0.89054584 |
| Rwdd4a | 0.69529025 | 0.64151944 | 0.64648554 | -0.0049661 | 0.51427452 | 0.89054584 |
| Rapgef6 | 0.29569546 | 0.30014539 | 0.30511227 | -0.0049669 | 0.53022766 | 0.89054584 |
| Trim9 | 0.01420821 | 0.01023705 | 0.01520505 | -0.004968 | 0.93602449 | 0.94845422 |

| | | | | | | |
|---|---|---|---|---|---|---|
| Per3 | 0.13184624 | 0.135265 | 0.14023572 | -0.0049707 | 0.56558245 | 0.89054584 |
| Soga3 | 0.06806599 | 0.06783573 | 0.07281054 | -0.0049748 | 0.62488052 | 0.89054584 |
| Adgrv1 | 0.03190106 | 0.02915805 | 0.0341368 | -0.0049788 | 0.75158736 | 0.89054584 |
| Uqcc1 | 0.18701125 | 0.19205862 | 0.19703765 | -0.004979 | 0.54685877 | 0.89054584 |
| Arvcf | 0.32194494 | 0.3253926 | 0.33037453 | -0.0049819 | 0.52800469 | 0.89054584 |
| Gpa33 | 0.03117533 | 0.028372 | 0.03336035 | -0.0049883 | 0.75698838 | 0.89054584 |
| 5730559C18I | 0.03319711 | 0.03053337 | 0.03552341 | -0.00499 | 0.74358883 | 0.89054584 |
| Clpb | 0.26403431 | 0.26919905 | 0.27419086 | -0.0049918 | 0.53379726 | 0.89054584 |
| Inca1 | 0.02717344 | 0.02408638 | 0.02907858 | -0.0049922 | 0.78809976 | 0.89054584 |
| Igsf9 | 0.12052921 | 0.12344106 | 0.12843613 | -0.0049951 | 0.57189199 | 0.89054584 |
| Alyref2 | 0.2670601 | 0.2721691 | 0.27716683 | -0.0049977 | 0.5334748 | 0.89054584 |
| Slc26a6 | 0.04467208 | 0.04279828 | 0.0477985 | -0.0050002 | 0.68700315 | 0.89054584 |
| Cks1brt | 0.02131693 | 0.01781042 | 0.02281203 | -0.0050016 | 0.84648802 | 0.89825055 |
| Npas3 | 0.06504051 | 0.06456972 | 0.06957703 | -0.0050073 | 0.63129831 | 0.89054584 |
| Zfp623 | 0.1080999 | 0.11041184 | 0.11542198 | -0.0050101 | 0.58013195 | 0.89054584 |
| Dna2 | 0.06321983 | 0.06261576 | 0.06763096 | -0.0050152 | 0.63513535 | 0.89054584 |
| Csrnp2 | 0.04554861 | 0.04371769 | 0.048736 | -0.0050183 | 0.68428931 | 0.89054584 |
| Slco4a1 | 0.04000522 | 0.03778771 | 0.04280663 | -0.0050189 | 0.70755062 | 0.89054584 |
| Mertk | 0.05595962 | 0.05484734 | 0.05986937 | -0.005022 | 0.65202977 | 0.89054584 |
| Ush1g | 0.01328835 | 0.00919714 | 0.01422068 | -0.0050235 | 0.95009732 | 0.95922778 |
| Gm7535 | 0.01829245 | 0.01454566 | 0.01957563 | -0.00503 | 0.88436939 | 0.91373913 |
| Armc7 | 0.03310892 | 0.03039638 | 0.03542906 | -0.0050327 | 0.74595026 | 0.89054584 |
| Usp31 | 0.05389816 | 0.05263031 | 0.05766517 | -0.0050349 | 0.65791559 | 0.89054584 |
| Samhd1 | 0.22776623 | 0.23315324 | 0.23819059 | -0.0050373 | 0.5392439 | 0.89054584 |
| Gucd1 | 0.20433705 | 0.20958209 | 0.21461996 | -0.0050379 | 0.54354207 | 0.89054584 |
| Msrb2 | 0.19980517 | 0.20499386 | 0.2100335 | -0.0050396 | 0.54450452 | 0.89054584 |
| Ddias | 0.04105047 | 0.03888506 | 0.04392472 | -0.0050397 | 0.70351579 | 0.89054584 |
| Rras | 0.14827048 | 0.152225 | 0.1572734 | -0.0050484 | 0.55944001 | 0.89054584 |
| Acnat1 | 0.01037394 | 0.00604478 | 0.01110185 | -0.0050571 | 0.98309038 | 0.98586892 |
| Gm42893 | 0.01779307 | 0.01398264 | 0.01904124 | -0.0050586 | 0.89208979 | 0.91798693 |

| | | | | | | |
|---|---|---|---|---|---|---|
| Gstt2 | 0.20478188 | 0.21001028 | 0.21506967 | -0.0050594 | 0.54363621 | 0.89054584 |
| Shd | 0.02384741 | 0.02045205 | 0.02551974 | -0.0050677 | 0.82256107 | 0.89073269 |
| 2610021A01 | 0.05384293 | 0.05253623 | 0.05760611 | -0.0050699 | 0.65910836 | 0.89054584 |
| Chtf18 | 0.06454982 | 0.06398035 | 0.06905256 | -0.0050722 | 0.63390695 | 0.89054584 |
| Tmem190 | 0.02413544 | 0.02075356 | 0.02582794 | -0.0050744 | 0.81999213 | 0.89054584 |
| Itgb6 | 0.02162955 | 0.01807152 | 0.02314656 | -0.005075 | 0.84649221 | 0.89825055 |
| Serpinc1 | 0.02882275 | 0.02576754 | 0.03084327 | -0.0050757 | 0.77837012 | 0.89054584 |
| Aph1c | 0.02815404 | 0.02504969 | 0.03012778 | -0.0050781 | 0.78385344 | 0.89054584 |
| Sdr42e1 | 0.02518389 | 0.02187012 | 0.02694979 | -0.0050797 | 0.81006808 | 0.89054584 |
| Ttc12 | 0.02633651 | 0.02310098 | 0.02818308 | -0.0050821 | 0.79957967 | 0.89054584 |
| Nipal1 | 0.1674288 | 0.17193131 | 0.17701389 | -0.0050826 | 0.5532084 | 0.89054584 |
| Cnnm4 | 0.04332347 | 0.04127199 | 0.04635602 | -0.005084 | 0.69531169 | 0.89054584 |
| Dzank1 | 0.0320342 | 0.02919025 | 0.03427925 | -0.005089 | 0.75542234 | 0.89054584 |
| Ids | 0.3759389 | 0.37612533 | 0.38121908 | -0.0050937 | 0.52481875 | 0.89054584 |
| Gm16386 | 0.02360214 | 0.02016351 | 0.0252573 | -0.0050938 | 0.82628752 | 0.89170439 |
| Plscr1 | 0.20252753 | 0.20769355 | 0.21278971 | -0.0050962 | 0.54442115 | 0.89054584 |
| Xk | 0.0458478 | 0.04395982 | 0.049056 | -0.0050962 | 0.68580494 | 0.89054584 |
| 2210408I21R | 0.06074606 | 0.05989035 | 0.06498659 | -0.0050962 | 0.6425763 | 0.89054584 |
| Brsk2 | 0.02391777 | 0.02049428 | 0.02559503 | -0.0051007 | 0.82341661 | 0.89088762 |
| Pidd1 | 0.01528043 | 0.01125082 | 0.01635246 | -0.0051016 | 0.92695683 | 0.94249387 |
| Cntn1 | 0.05820635 | 0.05716706 | 0.06227147 | -0.0051044 | 0.64873033 | 0.89054584 |
| Arhgap8 | 0.02271452 | 0.01919429 | 0.02430752 | -0.0051132 | 0.83646593 | 0.89462889 |
| Tekt1 | 0.01202238 | 0.00775199 | 0.01286592 | -0.0051139 | 0.9679748 | 0.97334573 |
| Elmod3 | 0.12751507 | 0.13060945 | 0.13572541 | -0.005116 | 0.56970049 | 0.89054584 |
| Pigc | 0.42501675 | 0.42094241 | 0.42605838 | -0.005116 | 0.52230644 | 0.89054584 |
| Gm20045 | 0.0830403 | 0.08368981 | 0.08880771 | -0.0051179 | 0.6058389 | 0.89054584 |
| Rap1gds1 | 0.20174888 | 0.20688283 | 0.21200171 | -0.0051189 | 0.54478348 | 0.89054584 |
| Arhgef38 | 0.01205384 | 0.00778064 | 0.01289958 | -0.0051189 | 0.96775722 | 0.97334573 |
| Foxm1 | 0.0185323 | 0.01471123 | 0.01983228 | -0.0051211 | 0.88551631 | 0.91420299 |
| Rad51c | 0.05699344 | 0.0558501 | 0.06097471 | -0.0051246 | 0.65230826 | 0.89054584 |

| | | | | | | |
|---|---|---|---|---|---|---|
| Ddr1 | 0.75894661 | 0.68612439 | 0.69124923 | -0.0051248 | 0.51377708 | 0.89054584 |
| Amd2 | 0.02160341 | 0.01799295 | 0.02311858 | -0.0051256 | 0.8491829 | 0.89883846 |
| A630072M18 | 0.0619313 | 0.06112514 | 0.0662536 | -0.0051285 | 0.64081279 | 0.89054584 |
| Plet1 | 0.023216 | 0.01971493 | 0.02484412 | -0.0051292 | 0.83194041 | 0.8933762 |
| Esf1 | 0.88669988 | 0.76714364 | 0.77228329 | -0.0051397 | 0.51236758 | 0.89054584 |
| Gm43740 | 0.0182668 | 0.0144045 | 0.01954818 | -0.0051437 | 0.88988162 | 0.9168209 |
| Arsg | 0.02937911 | 0.02629202 | 0.03143855 | -0.0051465 | 0.77717123 | 0.89054584 |
| Rnft2 | 0.01275666 | 0.00850481 | 0.0136517 | -0.0051469 | 0.96049739 | 0.96737 |
| Rps8 | 5.22657273 | 0.66593124 | 0.67107833 | -0.0051471 | 0.51425259 | 0.89054584 |
| Bin1 | 0.49296026 | 0.48068188 | 0.48583346 | -0.0051516 | 0.51970035 | 0.89054584 |
| Prim2 | 0.11295714 | 0.11536147 | 0.12051459 | -0.0051531 | 0.57895201 | 0.89054584 |
| Tmem132b | 0.04376137 | 0.04165903 | 0.04682441 | -0.0051654 | 0.69635503 | 0.89054584 |
| Gata4 | 0.03227566 | 0.02935933 | 0.03453758 | -0.0051782 | 0.75757371 | 0.89054584 |
| Cntrob | 0.08770707 | 0.08861023 | 0.09379068 | -0.0051804 | 0.60153895 | 0.89054584 |
| Pih1d2 | 0.07441005 | 0.07440339 | 0.0795894 | -0.005186 | 0.61927386 | 0.89054584 |
| Zdhhc18 | 0.05279951 | 0.05130382 | 0.05649038 | -0.0051866 | 0.66558433 | 0.89054584 |
| 4930404N11 | 0.0233976 | 0.01985182 | 0.02503843 | -0.0051866 | 0.83274783 | 0.89393134 |
| Chrnb1 | 0.02388078 | 0.02036538 | 0.02555545 | -0.0051901 | 0.82797317 | 0.89229532 |
| 1700049E15I | 0.02496423 | 0.02152133 | 0.02671475 | -0.0051934 | 0.81745587 | 0.89054584 |
| Mtpn | 0.82936111 | 0.7321847 | 0.73738059 | -0.0051959 | 0.51309445 | 0.89054584 |
| Ranbp17 | 0.03412067 | 0.03131323 | 0.03651148 | -0.0051983 | 0.74643331 | 0.89054584 |
| Ttc25 | 0.03901201 | 0.03654538 | 0.04174417 | -0.0051988 | 0.71911738 | 0.89054584 |
| Plekhn1 | 0.02381718 | 0.02027303 | 0.0254874 | -0.0052144 | 0.82974427 | 0.89271879 |
| Col4a6 | 0.01779396 | 0.01382724 | 0.0190422 | -0.005215 | 0.89900661 | 0.92252046 |
| Prr11 | 0.04746024 | 0.04556252 | 0.05078053 | -0.005218 | 0.68393621 | 0.89054584 |
| 1700086P04I | 0.0310106 | 0.02796454 | 0.03318411 | -0.0052196 | 0.76816295 | 0.89054584 |
| Cldn9 | 0.03806096 | 0.03549619 | 0.04072679 | -0.0052306 | 0.72519823 | 0.89054584 |
| Cyp11a1 | 0.01060574 | 0.00611641 | 0.01134991 | -0.0052335 | 0.98415348 | 0.98670393 |
| Kdm8 | 0.02230811 | 0.0186352 | 0.02387264 | -0.0052374 | 0.84664049 | 0.89825055 |
| Cnnm2 | 0.08144906 | 0.08186955 | 0.08710836 | -0.0052388 | 0.6103361 | 0.89054584 |

| | | | | | | |
|---|---|---|---|---|---|---|
| Cdhr5 | 0.018249 | 0.01428863 | 0.01952912 | -0.0052405 | 0.89438546 | 0.91973864 |
| Strn4 | 0.55582932 | 0.533379 | 0.53862824 | -0.0052492 | 0.51810736 | 0.89054584 |
| Nadsyn1 | 0.0385272 | 0.0359752 | 0.04122555 | -0.0052504 | 0.72352585 | 0.89054584 |
| Wdr35 | 0.16553236 | 0.16981316 | 0.17506639 | -0.0052532 | 0.55559143 | 0.89054584 |
| Cacnb3 | 0.28283559 | 0.28734713 | 0.29261071 | -0.0052636 | 0.53339493 | 0.89054584 |
| Fa2h | 0.07573401 | 0.07571891 | 0.08100384 | -0.0052849 | 0.61942192 | 0.89054584 |
| Phka1 | 0.06170448 | 0.06072132 | 0.06601114 | -0.0052898 | 0.64555448 | 0.89054584 |
| Fam161a | 0.04906259 | 0.04720382 | 0.0524942 | -0.0052904 | 0.68065424 | 0.89054584 |
| Itgb8 | 0.03136086 | 0.02826721 | 0.03355885 | -0.0052916 | 0.76871858 | 0.89054584 |
| Mbd3 | 0.85991029 | 0.75097642 | 0.75627388 | -0.0052975 | 0.51301694 | 0.89054584 |
| Setd4 | 0.03731946 | 0.03463445 | 0.03993356 | -0.0052991 | 0.73178939 | 0.89054584 |
| Dbndd1 | 0.0350831 | 0.03223312 | 0.03754113 | -0.005308 | 0.74496742 | 0.89054584 |
| Nudt6 | 0.02241588 | 0.01867714 | 0.02398796 | -0.0053108 | 0.84883892 | 0.89883797 |
| Purg | 0.05999387 | 0.05887001 | 0.06418248 | -0.0053125 | 0.65011347 | 0.89054584 |
| BC051142 | 0.02803837 | 0.0246861 | 0.03000403 | -0.0053179 | 0.79552783 | 0.89054584 |
| Mvp | 0.2044457 | 0.2094114 | 0.21472981 | -0.0053184 | 0.5459329 | 0.89054584 |
| 6030443J06F | 0.01643321 | 0.01226749 | 0.01758606 | -0.0053186 | 0.92058354 | 0.937526 |
| C77080 | 0.08610612 | 0.08675991 | 0.0920814 | -0.0053215 | 0.60612963 | 0.89054584 |
| Nol4l | 0.19804615 | 0.20292537 | 0.20825089 | -0.0053255 | 0.54741829 | 0.89054584 |
| Mt3 | 0.06686938 | 0.06620598 | 0.0715317 | -0.0053257 | 0.6356556 | 0.89054584 |
| Brca1 | 0.04782936 | 0.04584717 | 0.0511753 | -0.0053281 | 0.68618531 | 0.89054584 |
| Bend5 | 0.16010703 | 0.16415651 | 0.16948691 | -0.0053304 | 0.55824639 | 0.89054584 |
| Stam2 | 0.33580664 | 0.33824253 | 0.34357301 | -0.0053305 | 0.52881155 | 0.89054584 |
| Tekt4 | 0.02541984 | 0.02186607 | 0.02720226 | -0.0053362 | 0.8196232 | 0.89054584 |
| Whamm | 0.09851338 | 0.09998685 | 0.10532373 | -0.0053369 | 0.59331244 | 0.89054584 |
| Dhrs13 | 0.12036205 | 0.12291358 | 0.12826148 | -0.0053479 | 0.5770123 | 0.89054584 |
| Ankrd13b | 0.03050809 | 0.02728473 | 0.03264647 | -0.0053617 | 0.77791427 | 0.89054584 |
| Carm1 | 0.16163012 | 0.16569134 | 0.17105448 | -0.0053631 | 0.55806842 | 0.89054584 |
| Taf1d | 0.80890471 | 0.71898415 | 0.72434898 | -0.0053648 | 0.51376319 | 0.89054584 |
| Kctd9 | 0.09157148 | 0.09254874 | 0.09791588 | -0.0053671 | 0.60078293 | 0.89054584 |

| | | | | | | |
|---|---|---|---|---|---|---|
| Gprasp1 | 0.56241136 | 0.53864081 | 0.5440102 | -0.0053694 | 0.51833846 | 0.89054584 |
| Zfp772 | 0.03904026 | 0.03639386 | 0.04177439 | -0.0053805 | 0.7257703 | 0.89054584 |
| Pask | 0.01753706 | 0.01338221 | 0.01876728 | -0.0053851 | 0.90936145 | 0.92975475 |
| Gm4723 | 0.06059424 | 0.05943782 | 0.06482429 | -0.0053865 | 0.65066912 | 0.89054584 |
| Mkrn2 | 0.21184635 | 0.21680186 | 0.22220106 | -0.0053992 | 0.54506645 | 0.89054584 |
| Slc4a4 | 0.09364947 | 0.09472638 | 0.10013369 | -0.0054073 | 0.59931987 | 0.89054584 |
| Etnk2 | 0.03202623 | 0.02886333 | 0.03427072 | -0.0054074 | 0.7688634 | 0.89054584 |
| Zfp583 | 0.03997043 | 0.03736031 | 0.04276941 | -0.0054091 | 0.72214399 | 0.89054584 |
| Aunip | 0.01712907 | 0.01292067 | 0.0183307 | -0.00541 | 0.91543655 | 0.93399009 |
| Zfp408 | 0.07998972 | 0.08013904 | 0.08554974 | -0.0054107 | 0.6158735 | 0.89054584 |
| Slc35f2 | 0.01763425 | 0.01345779 | 0.01887129 | -0.0054135 | 0.90930437 | 0.92975475 |
| Pcdhb17 | 0.03509491 | 0.03213894 | 0.03755376 | -0.0054148 | 0.74913356 | 0.89054584 |
| Cdh22 | 0.01606797 | 0.01176349 | 0.01719521 | -0.0054317 | 0.92944792 | 0.94403363 |
| Egr2 | 0.01292463 | 0.00839967 | 0.01383145 | -0.0054318 | 0.96634688 | 0.97211332 |
| Cep83os | 0.05991584 | 0.05866334 | 0.06409906 | -0.0054357 | 0.65361068 | 0.89054584 |
| 4931428F04I | 0.07171015 | 0.07126514 | 0.0767047 | -0.0054396 | 0.62944804 | 0.89054584 |
| St8sia2 | 0.04496959 | 0.04267378 | 0.04811671 | -0.0054429 | 0.70090866 | 0.89054584 |
| Tyw3 | 0.06028346 | 0.05904651 | 0.06449206 | -0.0054455 | 0.65298398 | 0.89054584 |
| Atrn | 0.19796087 | 0.20271707 | 0.20816443 | -0.0054474 | 0.54851792 | 0.89054584 |
| Six5 | 0.04235925 | 0.03987691 | 0.04532467 | -0.0054478 | 0.71225235 | 0.89054584 |
| Stx16 | 1.15386668 | 0.90474153 | 0.91019493 | -0.0054534 | 0.51113459 | 0.89054584 |
| Magi2 | 0.03339691 | 0.03028249 | 0.03573717 | -0.0054547 | 0.76149187 | 0.89054584 |
| Zfp202 | 0.0207213 | 0.01671955 | 0.02217468 | -0.0054551 | 0.87413002 | 0.90775282 |
| Arhgap21 | 0.16886514 | 0.17303158 | 0.17848792 | -0.0054563 | 0.55662705 | 0.89054584 |
| 2700012I20R | 0.02915327 | 0.02574049 | 0.03119691 | -0.0054564 | 0.7924241 | 0.89054584 |
| Bbs1 | 0.07537484 | 0.07516239 | 0.08062014 | -0.0054577 | 0.62376965 | 0.89054584 |
| Hnrnpc | 2.21096297 | 1.1220637 | 1.12752921 | -0.0054655 | 0.50900873 | 0.89054584 |
| Nos3 | 0.03123082 | 0.02795283 | 0.03341972 | -0.0054669 | 0.77700647 | 0.89054584 |
| Car9 | 0.04480249 | 0.04246649 | 0.04793797 | -0.0054715 | 0.70255251 | 0.89054584 |
| Fam46b | 0.02777018 | 0.02424536 | 0.02971707 | -0.0054717 | 0.80450857 | 0.89054584 |

| | | | | | | |
|---|---|---|---|---|---|---|
| Srd5a1 | 0.04469505 | 0.04235041 | 0.04782307 | -0.0054727 | 0.70303434 | 0.89054584 |
| Slc12a5 | 0.02108517 | 0.01708484 | 0.02256404 | -0.0054792 | 0.87104073 | 0.90617762 |
| Zfp85 | 0.01833709 | 0.01414244 | 0.01962339 | -0.005481 | 0.90341787 | 0.9256362 |
| Dnah7a | 0.02210296 | 0.01817156 | 0.02365312 | -0.0054816 | 0.85986204 | 0.90132889 |
| 9130019O22 | 0.08872327 | 0.08939393 | 0.09487555 | -0.0054816 | 0.60610418 | 0.89054584 |
| Rpl36-ps3 | 0.56881196 | 0.54372921 | 0.54921646 | -0.0054872 | 0.51856315 | 0.89054584 |
| Sema3b | 0.03102119 | 0.02770688 | 0.03319543 | -0.0054886 | 0.77944275 | 0.89054584 |
| Gm14964 | 0.05827926 | 0.05685666 | 0.06234943 | -0.0054928 | 0.65925718 | 0.89054584 |
| Gm10614 | 0.01656618 | 0.01223263 | 0.01772835 | -0.0054957 | 0.92566604 | 0.94170979 |
| Disc1 | 0.03093073 | 0.02759607 | 0.03309865 | -0.0055026 | 0.78069394 | 0.89054584 |
| Rab26 | 0.04261939 | 0.04009522 | 0.04560292 | -0.0055077 | 0.71317518 | 0.89054584 |
| Gm26641 | 0.0900904 | 0.09082414 | 0.09633496 | -0.0055108 | 0.60507814 | 0.89054584 |
| Ptprj | 0.10491179 | 0.10656047 | 0.11207466 | -0.0055142 | 0.59065284 | 0.89054584 |
| Mycbpap | 0.04992345 | 0.04790007 | 0.05341483 | -0.0055148 | 0.68474447 | 0.89054584 |
| Cep57l1 | 0.24950913 | 0.25432792 | 0.25984571 | -0.0055178 | 0.53940385 | 0.89054584 |
| Htatsf1 | 0.5379577 | 0.51835535 | 0.5238747 | -0.0055194 | 0.51957418 | 0.89054584 |
| Atg14 | 0.09876934 | 0.10007198 | 0.1055968 | -0.0055248 | 0.59629006 | 0.89054584 |
| Ppp1r10 | 0.61960575 | 0.58405211 | 0.58957957 | -0.0055275 | 0.51741977 | 0.89054584 |
| Pnpla6 | 0.09688533 | 0.09805455 | 0.1035867 | -0.0055322 | 0.59824823 | 0.89054584 |
| Hdac5 | 0.29692198 | 0.30076763 | 0.30630032 | -0.0055327 | 0.53353313 | 0.89054584 |
| Wdyhv1 | 0.15319119 | 0.15682157 | 0.16235733 | -0.0055358 | 0.56310723 | 0.89054584 |
| Ndrg4 | 0.06461575 | 0.0635862 | 0.06912303 | -0.0055368 | 0.64549507 | 0.89054584 |
| Dpy19l3 | 0.07442153 | 0.0740617 | 0.07960166 | -0.00554 | 0.62712332 | 0.89054584 |
| Dolpp1 | 0.10317527 | 0.10470797 | 0.11024987 | -0.0055419 | 0.59258116 | 0.89054584 |
| Cds1 | 0.02212689 | 0.01812079 | 0.02367872 | -0.0055579 | 0.86292578 | 0.90277803 |
| Aim1 | 0.015327 | 0.01084215 | 0.01640229 | -0.0055601 | 0.94286785 | 0.95352247 |
| Gpatch4 | 0.22852288 | 0.23338454 | 0.23894771 | -0.0055632 | 0.54318832 | 0.89054584 |
| Nim1k | 0.0318477 | 0.02851614 | 0.03407972 | -0.0055636 | 0.77654638 | 0.89054584 |
| Atp6v1g2 | 0.03052047 | 0.02709577 | 0.03265971 | -0.0055639 | 0.78631544 | 0.89054584 |
| 8030462N17 | 0.01652459 | 0.01211714 | 0.01768384 | -0.0055667 | 0.92875606 | 0.94362101 |

| | | | | | | |
|---|---|---|---|---|---|---|
| Gmppb | 0.46403133 | 0.45514987 | 0.46071859 | -0.0055687 | 0.52245369 | 0.89054584 |
| Trak1 | 0.20851485 | 0.21324925 | 0.21884072 | -0.0055915 | 0.54737731 | 0.89054584 |
| Lin54 | 0.13786962 | 0.14090468 | 0.14649627 | -0.0055916 | 0.57057026 | 0.89054584 |
| Tlr2 | 0.04314569 | 0.04057154 | 0.04616587 | -0.0055943 | 0.71381527 | 0.89054584 |
| Pdpr | 0.03470043 | 0.03152362 | 0.03713173 | -0.0056081 | 0.75917533 | 0.89054584 |
| Cutal | 0.0442868 | 0.04176806 | 0.04738641 | -0.0056183 | 0.70965774 | 0.89054584 |
| Acbd7 | 0.01787109 | 0.01349396 | 0.01912473 | -0.0056308 | 0.91492178 | 0.93370395 |
| Ccsap | 0.03493602 | 0.03175199 | 0.03738377 | -0.0056318 | 0.75861635 | 0.89054584 |
| Gng7 | 0.04176758 | 0.03905888 | 0.04469179 | -0.0056329 | 0.72146812 | 0.89054584 |
| Zfand2a | 0.08031438 | 0.08026358 | 0.0858965 | -0.0056329 | 0.62001643 | 0.89054584 |
| Hykk | 0.06810992 | 0.06722174 | 0.07285748 | -0.0056357 | 0.64071961 | 0.89054584 |
| 5033430I15R | 0.03851274 | 0.03557225 | 0.04121009 | -0.0056378 | 0.73806032 | 0.89054584 |
| Trpm2 | 0.01728802 | 0.01285934 | 0.01850078 | -0.0056414 | 0.9222908 | 0.9389349 |
| Igsf5 | 0.1701425 | 0.17414843 | 0.17979809 | -0.0056497 | 0.55819491 | 0.89054584 |
| Rnf150 | 0.02095826 | 0.01677846 | 0.02242823 | -0.0056498 | 0.87972405 | 0.91127746 |
| Mapkapk5 | 0.09568159 | 0.09665056 | 0.10230227 | -0.0056517 | 0.60155888 | 0.89054584 |
| Zfat | 0.05689452 | 0.05521644 | 0.06086894 | -0.0056525 | 0.66736387 | 0.89054584 |
| Gm17541 | 0.03639818 | 0.03329166 | 0.038948 | -0.0056563 | 0.75067286 | 0.89054584 |
| Tspan1 | 0.03716551 | 0.03411033 | 0.03976887 | -0.0056585 | 0.74630304 | 0.89054584 |
| Ypel4 | 0.03589948 | 0.03274562 | 0.03841449 | -0.0056689 | 0.754124 | 0.89054584 |
| Junos | 0.01456098 | 0.00991123 | 0.01558255 | -0.0056713 | 0.95502255 | 0.96306751 |
| Cmtr1 | 0.04194014 | 0.03919986 | 0.04487637 | -0.0056765 | 0.72217648 | 0.89054584 |
| Zfp558 | 0.02086524 | 0.01664536 | 0.0223287 | -0.0056833 | 0.882158 | 0.91223443 |
| Gm19426 | 0.04216067 | 0.03941116 | 0.04511226 | -0.0057011 | 0.72199479 | 0.89054584 |
| Mat2a | 1.44643459 | 1.01449075 | 1.02019765 | -0.0057069 | 0.51039593 | 0.89054584 |
| Gpld1 | 0.0169228 | 0.01240072 | 0.01810996 | -0.0057092 | 0.92904993 | 0.94382141 |
| AI606181 | 0.05635718 | 0.05458419 | 0.06029444 | -0.0057102 | 0.6704766 | 0.89054584 |
| Plekhm1 | 0.05634259 | 0.05456439 | 0.06027884 | -0.0057144 | 0.67063552 | 0.89054584 |
| Ggps1 | 0.21452887 | 0.21918766 | 0.22490324 | -0.0057156 | 0.54712466 | 0.89054584 |
| Vstm2l | 0.02906124 | 0.02538188 | 0.03109844 | -0.0057166 | 0.80411557 | 0.89054584 |

| | | | | | | |
|---|---|---|---|---|---|---|
| Klhl11 | 0.02347235 | 0.01939993 | 0.02511842 | -0.0057185 | 0.85557797 | 0.9009252 |
| Wdr54 | 0.04813731 | 0.04578075 | 0.05150465 | -0.0057239 | 0.69768899 | 0.89054584 |
| Cic | 0.21471461 | 0.21936074 | 0.22509022 | -0.0057295 | 0.54719962 | 0.89054584 |
| Hyou1 | 0.35547982 | 0.3564044 | 0.36213477 | -0.0057304 | 0.52938437 | 0.89054584 |
| Zfp688 | 0.20495151 | 0.20950826 | 0.21524114 | -0.0057329 | 0.54937774 | 0.89054584 |
| Acss2 | 0.09045799 | 0.09099362 | 0.09672733 | -0.0057337 | 0.60879068 | 0.89054584 |
| Slc16a13 | 0.04208228 | 0.03929148 | 0.04502841 | -0.0057369 | 0.72360389 | 0.89054584 |
| Pkmyt1 | 0.11925944 | 0.12136988 | 0.12710926 | -0.0057394 | 0.58330885 | 0.89054584 |
| Maats1 | 0.01568969 | 0.01104876 | 0.01679041 | -0.0057416 | 0.94443903 | 0.95457871 |
| Zfp35 | 0.11627663 | 0.11823833 | 0.12398991 | -0.0057516 | 0.58555157 | 0.89054584 |
| Dhx37 | 0.02763694 | 0.02382254 | 0.02957451 | -0.005752 | 0.8175644 | 0.89054584 |
| Zfp287 | 0.02538601 | 0.02140469 | 0.02716606 | -0.0057614 | 0.83844006 | 0.89497393 |
| Ppp1r3f | 0.0258233 | 0.02187198 | 0.02763395 | -0.005762 | 0.83433523 | 0.89435801 |
| Mia3 | 0.69544455 | 0.6408293 | 0.64659751 | -0.0057682 | 0.51657599 | 0.89054584 |
| Steap2 | 0.14627349 | 0.1494361 | 0.15520746 | -0.0057714 | 0.56876927 | 0.89054584 |
| AA415398 | 0.01627755 | 0.01164697 | 0.01741948 | -0.0057725 | 0.93869278 | 0.95044651 |
| Syt14 | 0.09498714 | 0.09578703 | 0.10156122 | -0.0057742 | 0.6044493 | 0.89054584 |
| Mettl13 | 0.18051785 | 0.18463842 | 0.19041514 | -0.0057767 | 0.55619949 | 0.89054584 |
| Ccdc28b | 0.84087306 | 0.73879518 | 0.7445799 | -0.0057847 | 0.51443689 | 0.89054584 |
| Tiam1 | 0.0258494 | 0.02187608 | 0.02766189 | -0.0057858 | 0.83508891 | 0.89459411 |
| Hsd17b7 | 0.08531082 | 0.08544419 | 0.09123224 | -0.0057881 | 0.61622357 | 0.89054584 |
| Crp | 0.01804145 | 0.01351742 | 0.01930703 | -0.0057896 | 0.91880497 | 0.93597769 |
| Mid1 | 0.03711533 | 0.03391856 | 0.0397152 | -0.0057966 | 0.75174667 | 0.89054584 |
| Tceanc2 | 0.1388525 | 0.14171906 | 0.14751652 | -0.0057975 | 0.57263947 | 0.89054584 |
| Tmod3 | 0.59017703 | 0.56060363 | 0.56640249 | -0.0057989 | 0.51902173 | 0.89054584 |
| Thoc1 | 0.39919363 | 0.39682954 | 0.40263359 | -0.005804 | 0.52677268 | 0.89054584 |
| Gprin3 | 0.16071369 | 0.16430195 | 0.17011139 | -0.0058094 | 0.56320759 | 0.89054584 |
| Fbxo45 | 0.10655294 | 0.10798323 | 0.11379823 | -0.005815 | 0.59408577 | 0.89054584 |
| 9130221H12 | 0.02723546 | 0.02332841 | 0.02914494 | -0.0058165 | 0.82376196 | 0.8909953 |
| 1700019G17 | 0.0256702 | 0.02164899 | 0.02747014 | -0.0058212 | 0.8382446 | 0.89497393 |

| | | | | | | |
|---|---|---|---|---|---|---|
| Lsm1 | 0.89173198 | 0.76940987 | 0.77523197 | -0.0058221 | 0.51395586 | 0.89054584 |
| Mbnl3 | 0.02939554 | 0.02561549 | 0.03145613 | -0.0058406 | 0.80649484 | 0.89054584 |
| Pola2 | 0.08500087 | 0.08506026 | 0.09090128 | -0.005841 | 0.61767035 | 0.89054584 |
| A130014A01 | 0.07686335 | 0.07636676 | 0.08221029 | -0.0058435 | 0.62973763 | 0.89054584 |
| Zfp472 | 0.07037075 | 0.06942788 | 0.07527349 | -0.0058456 | 0.64125124 | 0.89054584 |
| Gm26935 | 0.02879125 | 0.02496128 | 0.03080957 | -0.0058483 | 0.81174997 | 0.89054584 |
| Ribc1 | 0.02573061 | 0.02168553 | 0.02753478 | -0.0058492 | 0.83884079 | 0.89500145 |
| Reep2 | 0.03650617 | 0.03321395 | 0.03906353 | -0.0058496 | 0.75730327 | 0.89054584 |
| Gm6169 | 0.02500479 | 0.02090691 | 0.02675815 | -0.0058512 | 0.84584531 | 0.89825055 |
| Zfyve1 | 0.03831236 | 0.03513887 | 0.04099573 | -0.0058569 | 0.74716662 | 0.89054584 |
| Rnf13 | 0.35820277 | 0.35882979 | 0.36468749 | -0.0058577 | 0.52982625 | 0.89054584 |
| Gm5113 | 0.03900367 | 0.03587486 | 0.04173525 | -0.0058604 | 0.74350851 | 0.89054584 |
| Sv2c | 0.04023125 | 0.03718145 | 0.0430484 | -0.005867 | 0.73727127 | 0.89054584 |
| Dscam | 0.01636912 | 0.01164144 | 0.01751747 | -0.005876 | 0.94094874 | 0.95204654 |
| Polr3g | 0.06678111 | 0.0655602 | 0.07143736 | -0.0058772 | 0.64924821 | 0.89054584 |
| Adamts6 | 0.01574049 | 0.01096168 | 0.01684477 | -0.0058831 | 0.94814746 | 0.95744779 |
| Dennd4b | 0.03156663 | 0.02789413 | 0.033779 | -0.0058849 | 0.79150846 | 0.89054584 |
| Hcfc1r1 | 1.35808954 | 0.98561262 | 0.99150097 | -0.0058883 | 0.51103678 | 0.89054584 |
| Gm9828 | 0.07830022 | 0.07785638 | 0.08374516 | -0.0058888 | 0.62839535 | 0.89054584 |
| Dcaf15 | 0.08170892 | 0.08148666 | 0.08738589 | -0.0058992 | 0.62343496 | 0.89054584 |
| Sigmar1 | 0.62861382 | 0.59065487 | 0.59655746 | -0.0059026 | 0.51838375 | 0.89054584 |
| Zfp691 | 0.10886326 | 0.11031798 | 0.1162229 | -0.0059049 | 0.59355755 | 0.89054584 |
| Ppcdc | 0.14030249 | 0.14311268 | 0.14902094 | -0.0059083 | 0.57327331 | 0.89054584 |
| Cyp2u1 | 0.01555828 | 0.01074099 | 0.01664978 | -0.0059088 | 0.95087502 | 0.9598833 |
| Vps53 | 0.41266795 | 0.40899035 | 0.41490313 | -0.0059128 | 0.52646816 | 0.89054584 |
| Iba57 | 0.04310775 | 0.04020099 | 0.04612529 | -0.0059243 | 0.72521053 | 0.89054584 |
| Ccser1 | 0.03094863 | 0.0271909 | 0.0331178 | -0.0059269 | 0.79779851 | 0.89054584 |
| Gm340 | 0.01594868 | 0.01113951 | 0.01706755 | -0.005928 | 0.94718823 | 0.95662382 |
| Atp2b4 | 0.23300154 | 0.23749393 | 0.24342416 | -0.0059302 | 0.54518287 | 0.89054584 |
| Dnajb3 | 0.10910341 | 0.11054412 | 0.11647482 | -0.0059307 | 0.59375895 | 0.89054584 |

| | | | | | | |
|---|---|---|---|---|---|---|
| Fads3 | 0.07492419 | 0.07418738 | 0.08013868 | -0.0059513 | 0.63532298 | 0.89054584 |
| Usp29 | 0.52262156 | 0.50509727 | 0.51105151 | -0.0059542 | 0.5216444 | 0.89054584 |
| Helz2 | 0.02405605 | 0.01978349 | 0.025743 | -0.0059595 | 0.85960486 | 0.90131997 |
| 6330403L08F | 0.01397949 | 0.00899781 | 0.01496029 | -0.0059625 | 0.96833289 | 0.9736145 |
| Cep128 | 0.05313315 | 0.05087352 | 0.05684714 | -0.0059736 | 0.68778165 | 0.89054584 |
| Mpzl3 | 0.03139829 | 0.02762314 | 0.03359889 | -0.0059758 | 0.79633999 | 0.89054584 |
| Rfc5 | 0.41433354 | 0.41043531 | 0.41641253 | -0.0059772 | 0.5266593 | 0.89054584 |
| Gm20522 | 0.03944275 | 0.03622766 | 0.04220495 | -0.0059773 | 0.74531657 | 0.89054584 |
| Gemin5 | 0.07873023 | 0.07822685 | 0.08420448 | -0.0059776 | 0.6295779 | 0.89054584 |
| Zfp41 | 0.05084167 | 0.04841834 | 0.05439677 | -0.0059784 | 0.69568743 | 0.89054584 |
| Plcd1 | 0.12199733 | 0.12399073 | 0.12996953 | -0.0059788 | 0.58485056 | 0.89054584 |
| Tceal3 | 0.05546025 | 0.05335536 | 0.05933543 | -0.0059801 | 0.6806609 | 0.89054584 |
| Xaf1 | 0.0410642 | 0.03795849 | 0.0439394 | -0.0059809 | 0.73701266 | 0.89054584 |
| Paupar | 0.01646406 | 0.01163562 | 0.01761907 | -0.0059834 | 0.94319564 | 0.9536544 |
| Lmntd2 | 0.01876991 | 0.01409873 | 0.02008654 | -0.0059878 | 0.91755818 | 0.93505805 |
| Gm15614 | 0.03056112 | 0.02671413 | 0.0327032 | -0.0059891 | 0.80322883 | 0.89054584 |
| Xxylt1 | 0.02777696 | 0.0237351 | 0.02972433 | -0.0059892 | 0.82606816 | 0.89170439 |
| Tha1 | 0.04417718 | 0.04127569 | 0.04726916 | -0.0059935 | 0.72264871 | 0.89054584 |
| 4732440D04I | 0.06534531 | 0.06390905 | 0.0699028 | -0.0059938 | 0.65522906 | 0.89054584 |
| Il13ra1 | 0.13809407 | 0.1407353 | 0.14672928 | -0.005994 | 0.57547099 | 0.89054584 |
| Zfp595 | 0.02037129 | 0.01580473 | 0.02180014 | -0.0059954 | 0.89995378 | 0.92300013 |
| Shroom2 | 0.08794194 | 0.08804596 | 0.09404143 | -0.0059955 | 0.61677568 | 0.89054584 |
| Spaca1 | 0.02626944 | 0.02211488 | 0.02811132 | -0.0059964 | 0.83984021 | 0.89542723 |
| Lipt1 | 0.06687901 | 0.06554549 | 0.071542 | -0.0059965 | 0.65191776 | 0.89054584 |
| Slc25a40 | 0.07482408 | 0.07403394 | 0.08003173 | -0.0059978 | 0.63651299 | 0.89054584 |
| Tmem229a | 0.13614464 | 0.13870332 | 0.1447048 | -0.0060015 | 0.57660854 | 0.89054584 |
| Gm12802 | 0.03083865 | 0.02699301 | 0.03300013 | -0.0060071 | 0.80180407 | 0.89054584 |
| Tdp2 | 0.35564306 | 0.3562795 | 0.36228792 | -0.0060084 | 0.5307944 | 0.89054584 |
| Nfatc1 | 0.07044072 | 0.06933886 | 0.07534826 | -0.0060094 | 0.6448948 | 0.89054584 |
| 2200002J24F | 0.02053311 | 0.0159608 | 0.0219733 | -0.0060125 | 0.89881268 | 0.92252046 |

| | | | | | | |
|---|---|---|---|---|---|---|
| Gpr39 | 0.02472509 | 0.02044537 | 0.02645888 | -0.0060135 | 0.85517089 | 0.90089862 |
| Plcl2 | 0.04612281 | 0.04333613 | 0.04935013 | -0.006014 | 0.71489978 | 0.89054584 |
| Usp54 | 0.04090964 | 0.03775533 | 0.04377407 | -0.0060187 | 0.73910333 | 0.89054584 |
| Sncb | 0.03848979 | 0.03516603 | 0.04118554 | -0.0060195 | 0.75204265 | 0.89054584 |
| Inhbb | 0.01749815 | 0.0127009 | 0.01872565 | -0.0060247 | 0.93305772 | 0.9465065 |
| Shc4 | 0.0205723 | 0.01598889 | 0.02201523 | -0.0060263 | 0.89890135 | 0.92252046 |
| Mbd2 | 0.18242664 | 0.1863331 | 0.19236373 | -0.0060306 | 0.55806234 | 0.89054584 |
| St3gal1 | 0.0754532 | 0.07466899 | 0.08070384 | -0.0060349 | 0.63622459 | 0.89054584 |
| BC048403 | 0.01512881 | 0.01014898 | 0.0161902 | -0.0060412 | 0.95893336 | 0.96613036 |
| Zfp780b | 0.20824715 | 0.21252287 | 0.21857049 | -0.0060476 | 0.55128482 | 0.89054584 |
| Rapsn | 0.02057427 | 0.01596923 | 0.02201734 | -0.0060481 | 0.89969212 | 0.92300013 |
| Morn2 | 0.46622036 | 0.45658528 | 0.46263645 | -0.0060512 | 0.5242956 | 0.89054584 |
| Dlg3 | 0.06313124 | 0.06148029 | 0.06753627 | -0.006056 | 0.66194132 | 0.89054584 |
| Actr3b | 0.04020018 | 0.03695749 | 0.04301517 | -0.0060577 | 0.74412112 | 0.89054584 |
| Cd79a | 0.13357009 | 0.1359697 | 0.14202888 | -0.0060592 | 0.57877386 | 0.89054584 |
| Zc3h6 | 0.11170816 | 0.1131405 | 0.11920592 | -0.0060654 | 0.59369317 | 0.89054584 |
| Ifit2 | 0.03248804 | 0.02869216 | 0.03476481 | -0.0060726 | 0.79212434 | 0.89054584 |
| Esrp2 | 0.14058797 | 0.14324436 | 0.14931705 | -0.0060727 | 0.57514089 | 0.89054584 |
| Rgs9 | 0.0570561 | 0.05496721 | 0.0610417 | -0.0060745 | 0.67854305 | 0.89054584 |
| Usp45 | 0.12137941 | 0.12324936 | 0.12932423 | -0.0060749 | 0.58661606 | 0.89054584 |
| 4933417E11I | 0.0405712 | 0.03733555 | 0.04341206 | -0.0060765 | 0.74283967 | 0.89054584 |
| Adgrg6 | 0.04906961 | 0.04642205 | 0.0525017 | -0.0060797 | 0.70522948 | 0.89054584 |
| Rptoros | 0.02939459 | 0.02537451 | 0.03145511 | -0.0060806 | 0.81610536 | 0.89054584 |
| Slc17a5 | 0.2291701 | 0.23351058 | 0.23959513 | -0.0060845 | 0.54709072 | 0.89054584 |
| Gtpbp1 | 0.08514126 | 0.084965 | 0.09105119 | -0.0060862 | 0.6222594 | 0.89054584 |
| Gm17690 | 0.02303107 | 0.01855837 | 0.02464624 | -0.0060879 | 0.87509394 | 0.90836342 |
| Hcfc2 | 0.26620869 | 0.27023962 | 0.27632988 | -0.0060903 | 0.54089245 | 0.89054584 |
| Rsl1 | 0.028986 | 0.02490214 | 0.03101794 | -0.0061158 | 0.82084818 | 0.89054584 |
| Sesn1 | 0.29512401 | 0.29844203 | 0.30455848 | -0.0061165 | 0.53727287 | 0.89054584 |
| 2700038G22 | 0.13542632 | 0.13783941 | 0.14395845 | -0.006119 | 0.57848957 | 0.89054584 |

| | | | | | | |
|---|---|---|---|---|---|---|
| Atp4a | 0.02597059 | 0.0216675 | 0.02779155 | -0.0061241 | 0.84770245 | 0.89862623 |
| Zfp592 | 0.09675189 | 0.09731836 | 0.10344432 | -0.006126 | 0.60868843 | 0.89054584 |
| Ccnjl | 0.0515676 | 0.04904273 | 0.05517305 | -0.0061303 | 0.69765181 | 0.89054584 |
| Zfp850 | 0.03961683 | 0.03625549 | 0.04239116 | -0.0061357 | 0.7499497 | 0.89054584 |
| Wdr59 | 0.02941661 | 0.02533372 | 0.03147866 | -0.0061449 | 0.81844912 | 0.89054584 |
| Fnip2 | 0.11815399 | 0.11980817 | 0.1259536 | -0.0061454 | 0.5899107 | 0.89054584 |
| Pcsk6 | 0.06841966 | 0.06703972 | 0.0731885 | -0.0061488 | 0.65225337 | 0.89054584 |
| Top3a | 0.02049587 | 0.01578175 | 0.02193344 | -0.0061517 | 0.90434245 | 0.92615843 |
| Macc1 | 0.02052604 | 0.0158138 | 0.02196573 | -0.0061519 | 0.90402407 | 0.92605695 |
| Lats2 | 0.1471695 | 0.14998125 | 0.1561346 | -0.0061533 | 0.57284081 | 0.89054584 |
| Cox5a | 2.05139879 | 1.11470445 | 1.12086288 | -0.0061584 | 0.51021099 | 0.89054584 |
| Fam198a | 0.05564969 | 0.05337757 | 0.05953799 | -0.0061604 | 0.68511912 | 0.89054584 |
| A230072E10 | 0.03998473 | 0.03662153 | 0.04278471 | -0.0061632 | 0.74892964 | 0.89054584 |
| Clcn6 | 0.04005062 | 0.0366706 | 0.04285519 | -0.0061846 | 0.74931918 | 0.89054584 |
| Ing5 | 0.1167715 | 0.11832241 | 0.12450766 | -0.0061852 | 0.59151556 | 0.89054584 |
| Rnf157 | 0.06952071 | 0.06816588 | 0.07436514 | -0.0061993 | 0.65113291 | 0.89054584 |
| Dclre1c | 0.04428984 | 0.04118686 | 0.04738966 | -0.0062028 | 0.72900448 | 0.89054584 |
| Lpin2 | 0.2225832 | 0.22679065 | 0.23299736 | -0.0062067 | 0.54938486 | 0.89054584 |
| Mtf1 | 0.33315918 | 0.33485117 | 0.34106002 | -0.0062088 | 0.53379533 | 0.89054584 |
| Tmem180 | 0.04906599 | 0.0462849 | 0.05249783 | -0.0062129 | 0.70930959 | 0.89054584 |
| Slc5a9 | 0.04972631 | 0.0469865 | 0.05320399 | -0.0062175 | 0.70693428 | 0.89054584 |
| Panct2 | 0.01603404 | 0.01094129 | 0.0171589 | -0.0062176 | 0.95431261 | 0.9627461 |
| Fbxw5 | 0.43867825 | 0.43207248 | 0.43829613 | -0.0062237 | 0.52637293 | 0.89054584 |
| Zfp111 | 0.07991676 | 0.0792465 | 0.08547181 | -0.0062253 | 0.63282036 | 0.89054584 |
| Grik5 | 0.13253683 | 0.13472189 | 0.14095421 | -0.0062323 | 0.58160309 | 0.89054584 |
| Abca7 | 0.16451556 | 0.16778096 | 0.17402162 | -0.0062407 | 0.56634488 | 0.89054584 |
| Lmtk3 | 0.01256228 | 0.00719893 | 0.01344368 | -0.0062448 | 0.984773 | 0.98715612 |
| Ndst2 | 0.0683867 | 0.06690479 | 0.07315328 | -0.0062485 | 0.65466655 | 0.89054584 |
| Gm9833 | 0.01813095 | 0.01314123 | 0.0194028 | -0.0062616 | 0.93364606 | 0.94677209 |
| Mybbp1a | 0.39954343 | 0.39669137 | 0.40295343 | -0.0062621 | 0.52885894 | 0.89054584 |

| | | | | | | |
|---|---|---|---|---|---|---|
| Cdk19 | 0.11270092 | 0.11397831 | 0.12024616 | -0.0062679 | 0.59593848 | 0.89054584 |
| Vill | 0.06750016 | 0.06593603 | 0.07220584 | -0.0062698 | 0.65709496 | 0.89054584 |
| Osbpl10 | 0.01660004 | 0.01148957 | 0.01776458 | -0.006275 | 0.95008551 | 0.95922778 |
| Ttll10 | 0.04085023 | 0.03742973 | 0.04371053 | -0.0062808 | 0.74839461 | 0.89054584 |
| Gm10762 | 0.02907873 | 0.02483323 | 0.03111716 | -0.0062839 | 0.82660817 | 0.89181167 |
| Dst | 0.66454848 | 0.61755571 | 0.62384683 | -0.0062911 | 0.51873645 | 0.89054584 |
| Acadsb | 0.41724692 | 0.41275664 | 0.41904945 | -0.0062928 | 0.52788831 | 0.89054584 |
| Tnfsf9 | 0.07137069 | 0.07004751 | 0.07634198 | -0.0062945 | 0.64956061 | 0.89054584 |
| Fam19a5 | 0.05226407 | 0.04962262 | 0.05591783 | -0.0062952 | 0.70003505 | 0.89054584 |
| Samd9l | 0.08203966 | 0.08143585 | 0.08773911 | -0.0063033 | 0.63107529 | 0.89054584 |
| Gm10269 | 0.05216138 | 0.0495023 | 0.05580801 | -0.0063057 | 0.70069862 | 0.89054584 |
| Tmem219 | 0.46253293 | 0.4530963 | 0.45940413 | -0.0063078 | 0.52550271 | 0.89054584 |
| Tmem241 | 0.04445293 | 0.04125532 | 0.0475641 | -0.0063088 | 0.73169562 | 0.89054584 |
| Rabl6 | 0.58385661 | 0.55503888 | 0.56134968 | -0.0063108 | 0.52088575 | 0.89054584 |
| Sco1 | 0.08598405 | 0.08563753 | 0.09195107 | -0.0063135 | 0.62547481 | 0.89054584 |
| Bcl6 | 0.01608802 | 0.01089776 | 0.01721667 | -0.0063189 | 0.9563612 | 0.9641419 |
| Cmss1 | 0.1289707 | 0.13090872 | 0.13724205 | -0.0063333 | 0.58511516 | 0.89054584 |
| Slc25a27 | 0.02632086 | 0.02183301 | 0.02816634 | -0.0063333 | 0.852585 | 0.89988783 |
| Mcmbp | 0.366893 | 0.36647072 | 0.37280855 | -0.0063378 | 0.53156444 | 0.89054584 |
| Arrdc2 | 0.0372354 | 0.03349713 | 0.03984364 | -0.0063465 | 0.77098697 | 0.89054584 |
| Uhrf1bp1 | 0.03999482 | 0.03644549 | 0.0427955 | -0.00635 | 0.75530989 | 0.89054584 |
| 9130230L23I | 0.04687425 | 0.04380334 | 0.05015381 | -0.0063505 | 0.72237604 | 0.89054584 |
| Osbpl3 | 0.05019246 | 0.04734691 | 0.0537025 | -0.0063556 | 0.70931245 | 0.89054584 |
| Cacnb1 | 0.03048143 | 0.02626037 | 0.03261794 | -0.0063576 | 0.81808069 | 0.89054584 |
| Pfkfb4 | 0.04723036 | 0.04417299 | 0.05053468 | -0.0063617 | 0.72123121 | 0.89054584 |
| Dusp4 | 0.0306082 | 0.02638508 | 0.03275358 | -0.0063685 | 0.81749729 | 0.89054584 |
| Ankrd52 | 0.0761208 | 0.07504836 | 0.08141704 | -0.0063687 | 0.64223374 | 0.89054584 |
| Pgbd5 | 0.04292226 | 0.03955528 | 0.04592689 | -0.0063716 | 0.74097428 | 0.89054584 |
| Slc12a6 | 0.14443372 | 0.14692764 | 0.15330278 | -0.0063751 | 0.57681156 | 0.89054584 |
| Ggt7 | 0.02399735 | 0.01930065 | 0.02568018 | -0.0063795 | 0.87644245 | 0.9091719 |

| | | | | | | |
|---|---|---|---|---|---|---|
| Celf6 | 0.05256882 | 0.04986175 | 0.0562437 | -0.0063819 | 0.70147511 | 0.89054584 |
| Slc46a2 | 0.02042273 | 0.01547308 | 0.02185519 | -0.0063821 | 0.91316321 | 0.93245811 |
| Flii | 0.37018751 | 0.36948222 | 0.37587689 | -0.0063947 | 0.53158748 | 0.89054584 |
| Dhcr24 | 0.09437367 | 0.09450728 | 0.10090656 | -0.0063993 | 0.61617881 | 0.89054584 |
| Fads2 | 0.01505686 | 0.00971348 | 0.0161132 | -0.0063997 | 0.96787221 | 0.97334573 |
| Sox12 | 0.21910153 | 0.22309493 | 0.22950204 | -0.0064071 | 0.55174279 | 0.89054584 |
| Coro2b | 0.09695825 | 0.09725467 | 0.1036645 | -0.0064098 | 0.61335498 | 0.89054584 |
| Pter | 0.08301814 | 0.08236416 | 0.08878405 | -0.0064199 | 0.63189767 | 0.89054584 |
| Zfp341 | 0.02050094 | 0.01551115 | 0.02193887 | -0.0064277 | 0.91387229 | 0.93295744 |
| Mex3a | 0.29328423 | 0.29634628 | 0.30277445 | -0.0064282 | 0.53939647 | 0.89054584 |
| Atp2c2 | 0.0216979 | 0.01679011 | 0.02321968 | -0.0064296 | 0.90148692 | 0.92418557 |
| Arhgef18 | 0.11267623 | 0.11379061 | 0.1202203 | -0.0064297 | 0.59838628 | 0.89054584 |
| Atp2b2 | 0.0225336 | 0.01767272 | 0.02411392 | -0.0064412 | 0.89333761 | 0.9188173 |
| Chac1 | 0.02144364 | 0.01650189 | 0.02294761 | -0.0064457 | 0.90467244 | 0.92639787 |
| Cct7 | 2.39979787 | 1.1227563 | 1.12920594 | -0.0064496 | 0.51061471 | 0.89054584 |
| Kcnb1 | 0.08462818 | 0.08405026 | 0.09050333 | -0.0064531 | 0.63012778 | 0.89054584 |
| Jmjd4 | 0.03116859 | 0.02689835 | 0.03335314 | -0.0064548 | 0.81637559 | 0.89054584 |
| Mllt1 | 0.10191007 | 0.10246469 | 0.10891966 | -0.006455 | 0.60876651 | 0.89054584 |
| Flywch2 | 0.25483039 | 0.25865615 | 0.26511214 | -0.006456 | 0.54516477 | 0.89054584 |
| Frrs1l | 0.02694823 | 0.02235283 | 0.02883761 | -0.0064848 | 0.85260407 | 0.89988783 |
| Ppip5k1 | 0.09682605 | 0.09703635 | 0.10352345 | -0.0064871 | 0.61483512 | 0.89054584 |
| Gm6793 | 0.05140986 | 0.04851293 | 0.05500437 | -0.0064914 | 0.70878332 | 0.89054584 |
| Klhl25 | 0.06446143 | 0.06246525 | 0.06895808 | -0.0064928 | 0.6695491 | 0.89054584 |
| Ppp1r36 | 0.02650018 | 0.02186122 | 0.02835821 | -0.006497 | 0.85710063 | 0.9009252 |
| Bard1 | 0.01819027 | 0.01295625 | 0.01946628 | -0.00651 | 0.94039174 | 0.95172812 |
| Nanos3 | 0.03787859 | 0.03401829 | 0.0405317 | -0.0065134 | 0.7729773 | 0.89054584 |
| K230010J24F | 0.02804929 | 0.02350223 | 0.03001571 | -0.0065135 | 0.84399222 | 0.89775379 |
| 1700086O06 | 0.07826257 | 0.07719068 | 0.08370494 | -0.0065143 | 0.6415399 | 0.89054584 |
| Zfp169 | 0.03649712 | 0.03253614 | 0.03905385 | -0.0065177 | 0.78157731 | 0.89054584 |
| Pabpc1l | 0.05609076 | 0.05348926 | 0.06000958 | -0.0065203 | 0.69364742 | 0.89054584 |

| | | | | | | |
|---|---|---|---|---|---|---|
| Cth | 0.071638 | 0.0701001 | 0.0766276 | -0.0065275 | 0.65426853 | 0.89054584 |
| Nudt7 | 0.40041534 | 0.3972214 | 0.40375035 | -0.006529 | 0.53002735 | 0.89054584 |
| Gm15728 | 0.01929458 | 0.01411831 | 0.02064798 | -0.0065297 | 0.92966954 | 0.94406031 |
| 9430015G10 | 0.04423075 | 0.04079532 | 0.04732645 | -0.0065311 | 0.73986924 | 0.89054584 |
| Trmo | 0.06368444 | 0.06159518 | 0.06812758 | -0.0065324 | 0.67246212 | 0.89054584 |
| Eps15l1 | 0.19158807 | 0.19516187 | 0.20169476 | -0.0065329 | 0.55997383 | 0.89054584 |
| Ddx55 | 0.29950558 | 0.30226439 | 0.30880043 | -0.006536 | 0.53927633 | 0.89054584 |
| Cst6 | 0.04504636 | 0.04165374 | 0.04819882 | -0.0065451 | 0.73651975 | 0.89054584 |
| Polr3a | 0.0553771 | 0.05269731 | 0.05924652 | -0.0065492 | 0.69672616 | 0.89054584 |
| Nek3 | 0.06236908 | 0.06016685 | 0.06672157 | -0.0065547 | 0.67641373 | 0.89054584 |
| Zfp709 | 0.02608183 | 0.02134378 | 0.02791058 | -0.0065668 | 0.86349443 | 0.90290403 |
| Pklr | 0.0196565 | 0.01446483 | 0.02103526 | -0.0065704 | 0.92719798 | 0.94266789 |
| Traf6 | 0.12168892 | 0.12305456 | 0.12964747 | -0.0065929 | 0.59364028 | 0.89054584 |
| Abcd4 | 0.0966223 | 0.09670664 | 0.10330605 | -0.0065994 | 0.61700471 | 0.89054584 |
| Gm37305 | 0.03182043 | 0.02744494 | 0.03405054 | -0.0066056 | 0.81695086 | 0.89054584 |
| Spata33 | 0.040965 | 0.03722221 | 0.04383329 | -0.0066111 | 0.75887142 | 0.89054584 |
| Mocs1 | 0.06009791 | 0.05768053 | 0.0642937 | -0.0066132 | 0.6841064 | 0.89054584 |
| Capn5 | 0.07696942 | 0.07569439 | 0.08232359 | -0.0066292 | 0.64623167 | 0.89054584 |
| Gstk1 | 0.10775684 | 0.10842295 | 0.11506196 | -0.006639 | 0.60596529 | 0.89054584 |
| Lrrn1 | 0.19600017 | 0.19953381 | 0.20617578 | -0.006642 | 0.55965254 | 0.89054584 |
| Pcdhga7 | 0.02869215 | 0.02406109 | 0.03070354 | -0.0066425 | 0.84325415 | 0.8976137 |
| Erich2 | 0.02507287 | 0.02018506 | 0.026831 | -0.0066459 | 0.8757504 | 0.90897976 |
| Kdsr | 0.20155623 | 0.20515048 | 0.2118067 | -0.0066562 | 0.55820173 | 0.89054584 |
| Lyar | 0.73045775 | 0.6649072 | 0.67157104 | -0.0066638 | 0.51843634 | 0.89054584 |
| 9830147E19I | 0.08289789 | 0.08199125 | 0.08865564 | -0.0066644 | 0.63691326 | 0.89054584 |
| Zmym5 | 0.46114658 | 0.45152187 | 0.45818674 | -0.0066649 | 0.52701558 | 0.89054584 |
| Pitpnm2 | 0.05589076 | 0.05312622 | 0.05979574 | -0.0066695 | 0.69834645 | 0.89054584 |
| Sema3f | 0.11784445 | 0.11895857 | 0.12562991 | -0.0066713 | 0.59770242 | 0.89054584 |
| Fam89b | 0.02872985 | 0.02407153 | 0.03074387 | -0.0066723 | 0.84402282 | 0.89775379 |
| Gm15441 | 0.12838415 | 0.12995713 | 0.13663101 | -0.0066739 | 0.59001005 | 0.89054584 |

| | | | | | | |
|---|---|---|---|---|---|---|
| Gins3 | 0.03567788 | 0.03150136 | 0.03817742 | -0.0066761 | 0.79237964 | 0.89054584 |
| Ccdc40 | 0.03642859 | 0.03230098 | 0.03898054 | -0.0066796 | 0.78766304 | 0.89054584 |
| Gm20696 | 0.06111533 | 0.05869467 | 0.06538135 | -0.0066867 | 0.68313333 | 0.89054584 |
| Spg20 | 0.27542091 | 0.27867978 | 0.285367 | -0.0066872 | 0.54346872 | 0.89054584 |
| Ell2 | 0.2115669 | 0.21523199 | 0.22191938 | -0.0066874 | 0.55582568 | 0.89054584 |
| 4933408B17 | 0.03334426 | 0.02898728 | 0.03568084 | -0.0066936 | 0.80893966 | 0.89054584 |
| Gm17354 | 0.10476743 | 0.10522658 | 0.11192301 | -0.0066964 | 0.60978074 | 0.89054584 |
| Acvr1 | 0.09572075 | 0.09563609 | 0.10234406 | -0.006708 | 0.619955 | 0.89054584 |
| Hist2h3c2 | 0.03467687 | 0.03039712 | 0.03710653 | -0.0067094 | 0.80021948 | 0.89054584 |
| Plekhg5 | 0.04835902 | 0.04501952 | 0.05174176 | -0.0067222 | 0.7275051 | 0.89054584 |
| Dusp8 | 0.05264981 | 0.04960665 | 0.05633031 | -0.0067237 | 0.71092778 | 0.89054584 |
| Diras2 | 0.01744302 | 0.01194177 | 0.01866665 | -0.0067249 | 0.95336926 | 0.96199523 |
| Idua | 0.05616624 | 0.05336446 | 0.06009028 | -0.0067258 | 0.69897987 | 0.89054584 |
| Sep-10 | 0.22092259 | 0.22459809 | 0.23133093 | -0.0067328 | 0.55393023 | 0.89054584 |
| Arhgap35 | 0.18366031 | 0.18688917 | 0.1936223 | -0.0067331 | 0.56435203 | 0.89054584 |
| Arhgap12 | 0.24012843 | 0.24379461 | 0.25052937 | -0.0067348 | 0.54983411 | 0.89054584 |
| Tube1 | 0.041686 | 0.03786521 | 0.04460453 | -0.0067393 | 0.75925904 | 0.89054584 |
| Ap5m1 | 0.0866109 | 0.08588023 | 0.09262036 | -0.0067401 | 0.63270962 | 0.89054584 |
| Tada1 | 0.34307248 | 0.34370634 | 0.3504524 | -0.0067461 | 0.5357303 | 0.89054584 |
| Abcc4 | 0.05010267 | 0.04685353 | 0.05360648 | -0.0067529 | 0.72136424 | 0.89054584 |
| Map3k9 | 0.03000636 | 0.02535077 | 0.03210965 | -0.0067589 | 0.83662426 | 0.89462889 |
| Synj2bp | 0.66577662 | 0.61799549 | 0.62476384 | -0.0067683 | 0.52012702 | 0.89054584 |
| 4930452B06 | 0.04019612 | 0.03623237 | 0.04301083 | -0.0067785 | 0.76859981 | 0.89054584 |
| Tdrp | 0.04248006 | 0.03866954 | 0.04545389 | -0.0067843 | 0.75659244 | 0.89054584 |
| Il11 | 0.03826916 | 0.0341643 | 0.04094952 | -0.0067852 | 0.77993475 | 0.89054584 |
| Gm15417 | 0.08086056 | 0.07968731 | 0.08647985 | -0.0067925 | 0.64279334 | 0.89054584 |
| Zfp78 | 0.04888985 | 0.0455159 | 0.05230946 | -0.0067936 | 0.72743324 | 0.89054584 |
| Vps52 | 0.38397984 | 0.38186281 | 0.3886571 | -0.0067943 | 0.53245602 | 0.89054584 |
| Dnajc28 | 0.02337061 | 0.01821071 | 0.02500955 | -0.0067988 | 0.89733866 | 0.92175257 |
| Ube2cbp | 0.05429309 | 0.05128435 | 0.05808745 | -0.0068031 | 0.70734564 | 0.89054584 |

| | | | | | | |
|---|---|---|---|---|---|---|
| Lin7b | 0.02626547 | 0.021302 | 0.02810707 | -0.0068051 | 0.87033753 | 0.90589084 |
| Zfp784 | 0.03574821 | 0.03144364 | 0.03825266 | -0.006809 | 0.79653199 | 0.89054584 |
| Ggact | 0.16504869 | 0.16775632 | 0.17456946 | -0.0068131 | 0.57214212 | 0.89054584 |
| Zbed5 | 0.06348436 | 0.06109856 | 0.06791371 | -0.0068151 | 0.67993794 | 0.89054584 |
| Svop | 0.05120561 | 0.04796049 | 0.05478596 | -0.0068255 | 0.71918793 | 0.89054584 |
| Hps3 | 0.06674247 | 0.06456844 | 0.07139607 | -0.0068276 | 0.67203384 | 0.89054584 |
| Npb | 0.04048419 | 0.03648428 | 0.04331898 | -0.0068347 | 0.7688514 | 0.89054584 |
| Hyls1 | 0.03771478 | 0.03351728 | 0.04035647 | -0.0068392 | 0.78510434 | 0.89054584 |
| Gm30173 | 0.02569289 | 0.02065148 | 0.02749442 | -0.0068429 | 0.87688162 | 0.90943771 |
| Skiv2l2 | 0.26521007 | 0.26850308 | 0.27534778 | -0.0068447 | 0.5460999 | 0.89054584 |
| Iqsec1 | 0.12510481 | 0.12636173 | 0.13321247 | -0.0068507 | 0.59467844 | 0.89054584 |
| E130309D14 | 0.03431106 | 0.02985448 | 0.03671517 | -0.0068607 | 0.80800883 | 0.89054584 |
| Caskin2 | 0.10526905 | 0.10558168 | 0.11244995 | -0.0068683 | 0.61200964 | 0.89054584 |
| Cbx3 | 1.67608261 | 1.07001707 | 1.07690896 | -0.0068919 | 0.51189302 | 0.89054584 |
| Sppl3 | 0.23915698 | 0.24266576 | 0.24956225 | -0.0068965 | 0.55122104 | 0.89054584 |
| Nckap1 | 0.59851884 | 0.56612819 | 0.57303026 | -0.0069021 | 0.52237541 | 0.89054584 |
| Ddx59 | 0.05909087 | 0.05631496 | 0.06321712 | -0.0069022 | 0.69450885 | 0.89054584 |
| 9130023H24 | 0.08956998 | 0.08886641 | 0.09577942 | -0.006913 | 0.63166473 | 0.89054584 |
| Il23a | 0.02024177 | 0.0147373 | 0.02166154 | -0.0069242 | 0.93179041 | 0.94571726 |
| Dusp7 | 0.09645528 | 0.09620318 | 0.10312783 | -0.0069247 | 0.62279535 | 0.89054584 |
| Chst8 | 0.09729947 | 0.09710353 | 0.10402859 | -0.0069251 | 0.62177286 | 0.89054584 |
| Abtb2 | 0.03666707 | 0.03229404 | 0.03923566 | -0.0069416 | 0.79510575 | 0.89054584 |
| Cul4b | 0.2254403 | 0.22891373 | 0.23586159 | -0.0069479 | 0.55457949 | 0.89054584 |
| Ranbp9 | 0.20477353 | 0.20809989 | 0.21506122 | -0.0069613 | 0.55993551 | 0.89054584 |
| Vangl2 | 0.25487374 | 0.25819168 | 0.26515499 | -0.0069633 | 0.54868897 | 0.89054584 |
| Gcgr | 0.02906129 | 0.02413156 | 0.0310985 | -0.0069669 | 0.85169565 | 0.89985795 |
| Gprin1 | 0.07531213 | 0.07358292 | 0.08055313 | -0.0069702 | 0.65657571 | 0.89054584 |
| Abhd3 | 0.03022296 | 0.02537082 | 0.0323414 | -0.0069706 | 0.84234587 | 0.89691028 |
| Amph | 0.0294168 | 0.0245034 | 0.03147887 | -0.0069755 | 0.84905408 | 0.89883797 |
| Nipa1 | 0.04396928 | 0.04006782 | 0.04704679 | -0.006979 | 0.75525236 | 0.89054584 |

| | | | | | | |
|---|---|---|---|---|---|---|
| Ybey | 0.06950568 | 0.06736566 | 0.07434907 | -0.0069834 | 0.66916251 | 0.89054584 |
| Hmga2 | 0.17302212 | 0.17576564 | 0.1827492 | -0.0069836 | 0.57065284 | 0.89054584 |
| Neurl1a | 0.03607481 | 0.03161705 | 0.03860206 | -0.006985 | 0.80039477 | 0.89054584 |
| Tprkb | 0.29201239 | 0.29455445 | 0.30154016 | -0.0069857 | 0.54297547 | 0.89054584 |
| Gnptab | 0.07081045 | 0.0687465 | 0.07574334 | -0.0069968 | 0.66653788 | 0.89054584 |
| 1700019D03I | 0.10012445 | 0.10004308 | 0.10704132 | -0.0069982 | 0.61966312 | 0.89054584 |
| Met | 0.10538523 | 0.10557288 | 0.11257198 | -0.0069991 | 0.61396454 | 0.89054584 |
| Bcl2l15 | 0.0407255 | 0.03657319 | 0.0435771 | -0.0070039 | 0.77301321 | 0.89054584 |
| Elf4 | 0.07824256 | 0.07667697 | 0.08368357 | -0.0070066 | 0.65176147 | 0.89054584 |
| 3110052M02 | 0.04146139 | 0.03735676 | 0.04436427 | -0.0070075 | 0.76910357 | 0.89054584 |
| Wdr90 | 0.07941337 | 0.0779166 | 0.08493415 | -0.0070175 | 0.64985817 | 0.89054584 |
| Ssr3 | 1.69488746 | 1.07347985 | 1.08049785 | -0.007018 | 0.51207036 | 0.89054584 |
| Hmga1-rs1 | 0.02124252 | 0.01571051 | 0.02273241 | -0.0070219 | 0.92494127 | 0.94123665 |
| Pck2 | 0.44240606 | 0.4345914 | 0.4416165 | -0.0070251 | 0.52953977 | 0.89054584 |
| Pag1 | 0.09337752 | 0.09281576 | 0.09984346 | -0.0070277 | 0.62851763 | 0.89054584 |
| Gla | 0.06175782 | 0.05903359 | 0.06606816 | -0.0070346 | 0.69007328 | 0.89054584 |
| Zfp239 | 0.06857821 | 0.06632215 | 0.07335794 | -0.0070358 | 0.67250496 | 0.89054584 |
| Ptpn9 | 0.3717124 | 0.37025848 | 0.37729506 | -0.0070366 | 0.5346204 | 0.89054584 |
| Bend7 | 0.28038207 | 0.28317501 | 0.29021675 | -0.0070417 | 0.54500188 | 0.89054584 |
| Kifc2 | 0.04194064 | 0.03783094 | 0.04487691 | -0.007046 | 0.76775836 | 0.89054584 |
| Mrgpre | 0.04248741 | 0.03841455 | 0.04546175 | -0.0070472 | 0.7649145 | 0.89054584 |
| Lap3 | 0.69134294 | 0.63656637 | 0.64361553 | -0.0070492 | 0.5203479 | 0.89054584 |
| Dennd5a | 0.2593571 | 0.26252927 | 0.26958204 | -0.0070528 | 0.54850555 | 0.89054584 |
| Tm9sf1 | 0.13640856 | 0.13791933 | 0.14497897 | -0.0070596 | 0.58973504 | 0.89054584 |
| Atxn7l3 | 0.25586225 | 0.2590607 | 0.26613182 | -0.0070711 | 0.54925841 | 0.89054584 |
| Taz | 0.59410127 | 0.56244841 | 0.56952621 | -0.0070778 | 0.52308553 | 0.89054584 |
| Gmeb1 | 0.15930844 | 0.16157975 | 0.16866462 | -0.0070849 | 0.57757827 | 0.89054584 |
| Cenpo | 0.03764771 | 0.03319695 | 0.04028472 | -0.0070878 | 0.79380714 | 0.89054584 |
| Dyrk2 | 0.05313485 | 0.04975505 | 0.05684897 | -0.0070939 | 0.71950373 | 0.89054584 |
| Mllt6 | 0.12658129 | 0.12764312 | 0.13475208 | -0.007109 | 0.59707609 | 0.89054584 |

| | | | | | | |
|---|---|---|---|---|---|---|
| C730034F03I | 0.07011601 | 0.06789203 | 0.07500128 | -0.0071092 | 0.67061552 | 0.89054584 |
| Sytl3 | 0.03639038 | 0.03182926 | 0.03893966 | -0.0071104 | 0.80253886 | 0.89054584 |
| Ccdc112 | 0.18984675 | 0.19281194 | 0.19992394 | -0.007112 | 0.56581731 | 0.89054584 |
| Chrnb2 | 0.03030521 | 0.02531617 | 0.0324294 | -0.0071132 | 0.8465895 | 0.89825055 |
| Zfp617 | 0.20838814 | 0.21159923 | 0.21871281 | -0.0071136 | 0.56022156 | 0.89054584 |
| Snrnp200 | 0.34355268 | 0.34378974 | 0.35090608 | -0.0071163 | 0.53763719 | 0.89054584 |
| Bbof1 | 0.05300355 | 0.04959139 | 0.05670857 | -0.0071172 | 0.72063134 | 0.89054584 |
| Abhd18 | 0.22862741 | 0.23193481 | 0.23905229 | -0.0071175 | 0.55516193 | 0.89054584 |
| Dazap2 | 0.80896874 | 0.71727207 | 0.72439024 | -0.0071182 | 0.51825751 | 0.89054584 |
| Fut8 | 0.23749993 | 0.24078915 | 0.24791159 | -0.0071224 | 0.5532395 | 0.89054584 |
| Hist1h1a | 0.01742592 | 0.01152374 | 0.01864836 | -0.0071246 | 0.96245813 | 0.96887363 |
| Cfap52 | 0.03833122 | 0.03389079 | 0.04101591 | -0.0071251 | 0.79083811 | 0.89054584 |
| Snx32 | 0.07426141 | 0.07229583 | 0.07943059 | -0.0071348 | 0.66220342 | 0.89054584 |
| Ubxn10 | 0.14360267 | 0.1452991 | 0.15244197 | -0.0071429 | 0.58640254 | 0.89054584 |
| Nfya | 0.19881814 | 0.20188473 | 0.20903339 | -0.0071487 | 0.56329532 | 0.89054584 |
| Ankrd9 | 0.05928443 | 0.05626868 | 0.06342405 | -0.0071554 | 0.70042119 | 0.89054584 |
| Khk | 0.33650484 | 0.33707847 | 0.34423519 | -0.0071567 | 0.53858132 | 0.89054584 |
| Rbbp8 | 0.2199612 | 0.22319411 | 0.23036559 | -0.0071715 | 0.55765909 | 0.89054584 |
| Gm4978 | 0.03684332 | 0.03225103 | 0.0394242 | -0.0071732 | 0.80169388 | 0.89054584 |
| Mfn2 | 0.21295098 | 0.21614029 | 0.22331417 | -0.0071739 | 0.55948615 | 0.89054584 |
| Gm37035 | 0.03517307 | 0.03046296 | 0.03763738 | -0.0071744 | 0.81275257 | 0.89054584 |
| Orai2 | 0.11570863 | 0.11619805 | 0.12339554 | -0.0071975 | 0.60709267 | 0.89054584 |
| Klf11 | 0.17919147 | 0.18185841 | 0.18906021 | -0.0072018 | 0.57043087 | 0.89054584 |
| Sntb1 | 0.03407529 | 0.0292559 | 0.03646294 | -0.007207 | 0.82143748 | 0.89054584 |
| Ccdc181 | 0.27027156 | 0.27311277 | 0.28032059 | -0.0072078 | 0.54767698 | 0.89054584 |
| Paqr7 | 0.05845803 | 0.05532713 | 0.06254055 | -0.0072134 | 0.70449107 | 0.89054584 |
| Pik3c2g | 0.06311477 | 0.0603039 | 0.06751866 | -0.0072148 | 0.69069988 | 0.89054584 |
| 4933406I18R | 0.02858192 | 0.02336948 | 0.0305856 | -0.0072161 | 0.86415828 | 0.90322015 |
| Fgf11 | 0.02373915 | 0.01818525 | 0.02540391 | -0.0072187 | 0.9072258 | 0.92829131 |
| Dyx1c1 | 0.03702572 | 0.03239355 | 0.03961933 | -0.0072258 | 0.80225381 | 0.89054584 |

| | | | | | | |
|---|---|---|---|---|---|---|
| Ly6g6e | 0.04594434 | 0.04192479 | 0.04915925 | -0.0072345 | 0.75352603 | 0.89054584 |
| Tango6 | 0.06953724 | 0.06714643 | 0.0743828 | -0.0072364 | 0.67481488 | 0.89054584 |
| Capn9 | 0.02657455 | 0.02120034 | 0.02843778 | -0.0072374 | 0.88213061 | 0.91223443 |
| Fdx1l | 0.5018936 | 0.48624833 | 0.49348657 | -0.0072382 | 0.52724065 | 0.89054584 |
| Kin | 0.46250161 | 0.45212411 | 0.45937664 | -0.0072525 | 0.52931745 | 0.89054584 |
| Vti1a | 0.10194963 | 0.10170786 | 0.10896127 | -0.0072534 | 0.62177256 | 0.89054584 |
| Dppa5a | 0.03927777 | 0.03477355 | 0.04202846 | -0.0072549 | 0.78936558 | 0.89054584 |
| Gm15491 | 0.04795617 | 0.04405232 | 0.05131092 | -0.0072586 | 0.74507422 | 0.89054584 |
| Msrb1 | 0.46291202 | 0.45247526 | 0.45973681 | -0.0072616 | 0.52933092 | 0.89054584 |
| A330017A19 | 0.16298214 | 0.16518164 | 0.17244519 | -0.0072636 | 0.57778849 | 0.89054584 |
| Ccdc96 | 0.02862171 | 0.02336334 | 0.03062817 | -0.0072648 | 0.86543648 | 0.90377426 |
| 4930524J08F | 0.04278713 | 0.03851535 | 0.04578234 | -0.007267 | 0.77020106 | 0.89054584 |
| 2510017J16F | 0.03763365 | 0.03299786 | 0.04026968 | -0.0072718 | 0.79991071 | 0.89054584 |
| Kctd21 | 0.05779044 | 0.05454768 | 0.06182682 | -0.0072791 | 0.70833178 | 0.89054584 |
| Wdr82 | 0.44843236 | 0.43968444 | 0.44696792 | -0.0072835 | 0.53025816 | 0.89054584 |
| Tmem44 | 0.07802518 | 0.07616433 | 0.08345136 | -0.007287 | 0.65793059 | 0.89054584 |
| Polr1a | 0.1503226 | 0.15210538 | 0.15939472 | -0.0072893 | 0.58435989 | 0.89054584 |
| Kcnip3 | 0.05302886 | 0.04943278 | 0.05673563 | -0.0073029 | 0.72564215 | 0.89054584 |
| Fam149b | 0.08116491 | 0.0795019 | 0.08680489 | -0.007303 | 0.65245665 | 0.89054584 |
| Ppp1r2 | 1.34853911 | 0.98088695 | 0.98819279 | -0.0073058 | 0.51373852 | 0.89054584 |
| Rdh5 | 0.08385466 | 0.08236687 | 0.08967734 | -0.0073105 | 0.64795143 | 0.89054584 |
| Zfp758 | 0.13186587 | 0.13293951 | 0.14025615 | -0.0073166 | 0.59601273 | 0.89054584 |
| Mb21d1 | 0.04822429 | 0.04427923 | 0.05159767 | -0.0073184 | 0.74563261 | 0.89054584 |
| 4933406P04I | 0.02199128 | 0.0162117 | 0.02353362 | -0.0073219 | 0.92640334 | 0.94226152 |
| Naprt | 0.08090081 | 0.07919855 | 0.08652283 | -0.0073243 | 0.65335193 | 0.89054584 |
| 1010001N08 | 0.04394317 | 0.0396918 | 0.04701886 | -0.0073271 | 0.76608485 | 0.89054584 |
| Zfp629 | 0.1112263 | 0.11137166 | 0.11870087 | -0.0073292 | 0.6131992 | 0.89054584 |
| Tmem206 | 0.41885852 | 0.41317548 | 0.42050612 | -0.0073306 | 0.53236604 | 0.89054584 |
| Rassf6 | 0.05019049 | 0.04636483 | 0.0537004 | -0.0073356 | 0.73774865 | 0.89054584 |
| Faap24 | 0.17951508 | 0.18205377 | 0.18939085 | -0.0073371 | 0.57161584 | 0.89054584 |

| | | | | | | |
|---|---|---|---|---|---|---|
| Ncdn | 0.14121437 | 0.14261744 | 0.14996664 | -0.0073492 | 0.59029895 | 0.89054584 |
| Clcf1 | 0.0410317 | 0.03654663 | 0.04390464 | -0.007358 | 0.78253856 | 0.89054584 |
| A530017D24 | 0.04934579 | 0.04543405 | 0.05279706 | -0.007363 | 0.74206659 | 0.89054584 |
| Cdk18 | 0.0231991 | 0.01745257 | 0.02482603 | -0.0073735 | 0.91677934 | 0.93483379 |
| Efr3a | 0.42971184 | 0.42290227 | 0.43027657 | -0.0073743 | 0.53182066 | 0.89054584 |
| Rtn4ip1 | 0.03133373 | 0.02615402 | 0.03352982 | -0.0073758 | 0.84728579 | 0.89854206 |
| Zfp607 | 0.04805145 | 0.04403609 | 0.05141282 | -0.0073767 | 0.7480821 | 0.89054584 |
| Ss18l1 | 0.0492959 | 0.04536589 | 0.0527437 | -0.0073778 | 0.74270096 | 0.89054584 |
| Slc25a39 | 0.67550883 | 0.62460644 | 0.63199326 | -0.0073868 | 0.52171342 | 0.89054584 |
| Mcoln1 | 0.13623521 | 0.13740975 | 0.14479888 | -0.0073891 | 0.59396148 | 0.89054584 |
| Dhx29 | 0.22873631 | 0.23176505 | 0.23916123 | -0.0073962 | 0.55728134 | 0.89054584 |
| Kctd8 | 0.12339235 | 0.12402951 | 0.13142581 | -0.0073963 | 0.60341364 | 0.89054584 |
| Gm26534 | 0.02644382 | 0.02090139 | 0.02829791 | -0.0073965 | 0.88834157 | 0.91584392 |
| Mkx | 0.03165002 | 0.02647141 | 0.03386821 | -0.0073968 | 0.84554478 | 0.89825055 |
| Galnt3 | 0.03367453 | 0.02863624 | 0.03603418 | -0.0073979 | 0.8305893 | 0.89271879 |
| Fam227a | 0.03037231 | 0.02510132 | 0.03250119 | -0.0073999 | 0.85559763 | 0.9009252 |
| Wdtc1 | 0.15945216 | 0.16140674 | 0.16881262 | -0.0074059 | 0.58097541 | 0.89054584 |
| Ano8 | 0.03969436 | 0.03506348 | 0.0424741 | -0.0074106 | 0.79185188 | 0.89054584 |
| 9130011E15I | 0.02755894 | 0.02206928 | 0.02949105 | -0.0074218 | 0.87949727 | 0.91117275 |
| Tcea2 | 0.08950761 | 0.08828548 | 0.09571285 | -0.0074274 | 0.64115075 | 0.89054584 |
| Naa16 | 0.33941719 | 0.33956114 | 0.34699497 | -0.0074338 | 0.53975269 | 0.89054584 |
| Tex26 | 0.03639627 | 0.03150427 | 0.03894596 | -0.0074417 | 0.81332577 | 0.89054584 |
| D3Ertd751e | 0.27788457 | 0.28033389 | 0.28777685 | -0.007443 | 0.54795534 | 0.89054584 |
| Gldc | 0.06539511 | 0.06250954 | 0.06995603 | -0.0074465 | 0.69002544 | 0.89054584 |
| Slc25a29 | 0.12125455 | 0.12174007 | 0.12919382 | -0.0074537 | 0.60595604 | 0.89054584 |
| Leng1 | 0.35603063 | 0.35518538 | 0.36265147 | -0.0074661 | 0.53820635 | 0.89054584 |
| Oas1c | 0.02883156 | 0.02338525 | 0.0308527 | -0.0074675 | 0.87026182 | 0.90589084 |
| Fbxo42 | 0.14202264 | 0.14333333 | 0.15080461 | -0.0074713 | 0.59127149 | 0.89054584 |
| Map2k1 | 0.68003303 | 0.62785417 | 0.63533142 | -0.0074773 | 0.5218636 | 0.89054584 |
| Rcbtb2 | 0.48227484 | 0.46913452 | 0.47661655 | -0.007482 | 0.52915146 | 0.89054584 |

| | | | | | | |
|---|---|---|---|---|---|---|
| Slc22a17 | 0.32519779 | 0.3259953 | 0.33348078 | -0.0074855 | 0.54164426 | 0.89054584 |
| Rev3l | 0.43287865 | 0.42561293 | 0.43311439 | -0.0075015 | 0.53215654 | 0.89054584 |
| Mzf1 | 0.03733572 | 0.03244729 | 0.03995096 | -0.0075037 | 0.80922715 | 0.89054584 |
| Dnm1 | 0.05288021 | 0.04906949 | 0.05657668 | -0.0075072 | 0.7317774 | 0.89054584 |
| 3110056K07I | 0.16412511 | 0.16610871 | 0.17362031 | -0.0075116 | 0.57987234 | 0.89054584 |
| Mavs | 0.11391632 | 0.11400071 | 0.12151921 | -0.0075185 | 0.61342344 | 0.89054584 |
| Jazf1 | 0.06669535 | 0.06382601 | 0.0713457 | -0.0075197 | 0.68830188 | 0.89054584 |
| Homer1 | 0.04924089 | 0.04516275 | 0.05268487 | -0.0075221 | 0.74703372 | 0.89054584 |
| Setdb2 | 0.02284874 | 0.01692254 | 0.02445113 | -0.0075286 | 0.92428592 | 0.94063578 |
| Mterf1a | 0.08604773 | 0.08448732 | 0.09201906 | -0.0075317 | 0.64852195 | 0.89054584 |
| Gpr137b-ps | 0.07243711 | 0.06994847 | 0.07748146 | -0.007533 | 0.67471007 | 0.89054584 |
| Zfp13 | 0.07136286 | 0.06879247 | 0.07633361 | -0.0075411 | 0.67733718 | 0.89054584 |
| Cgref1 | 0.03424024 | 0.02909825 | 0.03663941 | -0.0075412 | 0.8311973 | 0.89310742 |
| Xpo1 | 0.57238486 | 0.54456588 | 0.55211088 | -0.007545 | 0.52538265 | 0.89054584 |
| 4930430F08I | 0.15034107 | 0.15186842 | 0.1594138 | -0.0075454 | 0.58726579 | 0.89054584 |
| Fbxl16 | 0.26262166 | 0.26525397 | 0.27279997 | -0.007546 | 0.55127067 | 0.89054584 |
| Gpkow | 0.10307714 | 0.10260037 | 0.11014672 | -0.0075463 | 0.62520965 | 0.89054584 |
| Gmeb2 | 0.08345387 | 0.08169881 | 0.08924935 | -0.0075505 | 0.65326454 | 0.89054584 |
| Fam173a | 1.03240872 | 0.84363808 | 0.85119397 | -0.0075559 | 0.51649419 | 0.89054584 |
| Cadps2 | 0.02717049 | 0.02151658 | 0.02907542 | -0.0075588 | 0.88709104 | 0.91505129 |
| Rbm14 | 0.39129604 | 0.38783276 | 0.39539396 | -0.0075612 | 0.53549636 | 0.89054584 |
| Sox5 | 0.0997128 | 0.09904209 | 0.10660332 | -0.0075612 | 0.62947084 | 0.89054584 |
| Tor4a | 0.0416622 | 0.03699358 | 0.04457907 | -0.0075855 | 0.78604005 | 0.89054584 |
| Tctn3 | 0.06280945 | 0.0596065 | 0.06719229 | -0.0075858 | 0.70054884 | 0.89054584 |
| Bcar3 | 0.05720633 | 0.05361062 | 0.06120232 | -0.0075917 | 0.7183368 | 0.89054584 |
| Med24 | 0.15539587 | 0.15702707 | 0.16463222 | -0.0076052 | 0.58520168 | 0.89054584 |
| Unc13a | 0.12168128 | 0.12203024 | 0.12963949 | -0.0076093 | 0.60774972 | 0.89054584 |
| Ppp1r7 | 0.49126609 | 0.47676353 | 0.48437686 | -0.0076133 | 0.52918774 | 0.89054584 |
| Map6 | 0.08188756 | 0.07995556 | 0.08757667 | -0.0076211 | 0.65741966 | 0.89054584 |
| Chst11 | 0.03282035 | 0.02749683 | 0.03512033 | -0.0076235 | 0.84406527 | 0.89775379 |

| | | | | | | |
|---|---|---|---|---|---|---|
| Strada | 0.13466316 | 0.1355368 | 0.14316529 | -0.0076285 | 0.59802934 | 0.89054584 |
| Trp53inp2 | 0.30153726 | 0.30313437 | 0.31076409 | -0.0076297 | 0.54553327 | 0.89054584 |
| Imp4 | 0.63275604 | 0.59211507 | 0.59974831 | -0.0076332 | 0.52364194 | 0.89054584 |
| Slc9a7 | 0.03618826 | 0.03108201 | 0.03872343 | -0.0076414 | 0.82104723 | 0.89054584 |
| Pacs2 | 0.12635235 | 0.12687125 | 0.13451339 | -0.0076421 | 0.60437557 | 0.89054584 |
| Gm7293 | 0.0301768 | 0.02464965 | 0.03229201 | -0.0076424 | 0.86490041 | 0.90367009 |
| Zfp786 | 0.04579254 | 0.04135453 | 0.0489969 | -0.0076424 | 0.76629066 | 0.89054584 |
| Zfp963 | 0.09451297 | 0.09340741 | 0.10105521 | -0.0076478 | 0.63780052 | 0.89054584 |
| Mtm1 | 0.09484569 | 0.09375928 | 0.10141027 | -0.007651 | 0.63739261 | 0.89054584 |
| Dguok | 0.87489177 | 0.75763944 | 0.76529143 | -0.007652 | 0.51857754 | 0.89054584 |
| Ccl25 | 0.16540257 | 0.16727882 | 0.17493306 | -0.0076542 | 0.58076582 | 0.89054584 |
| Gm13225 | 0.05301495 | 0.04906398 | 0.05672076 | -0.0076568 | 0.7352973 | 0.89054584 |
| RP23-378D1( | 0.01626083 | 0.00973646 | 0.01740159 | -0.0076651 | 0.97992447 | 0.98311155 |
| Kctd15 | 0.13400601 | 0.13481376 | 0.14248214 | -0.0076684 | 0.59899396 | 0.89054584 |
| Vdr | 0.28238994 | 0.28449919 | 0.29217611 | -0.0076769 | 0.5487142 | 0.89054584 |
| Dvl1 | 0.08067419 | 0.07860369 | 0.08628079 | -0.0076771 | 0.66076209 | 0.89054584 |
| Lactb | 0.14056932 | 0.14161913 | 0.1492977 | -0.0076786 | 0.59468581 | 0.89054584 |
| Bloc1s3 | 0.14019324 | 0.14121537 | 0.14890762 | -0.0076922 | 0.59509489 | 0.89054584 |
| Lrfn1 | 0.03113719 | 0.02562461 | 0.03331954 | -0.0076949 | 0.85902553 | 0.90131997 |
| Tex2 | 0.14009239 | 0.14110158 | 0.148803 | -0.0077014 | 0.59527182 | 0.89054584 |
| Cnnm3 | 0.1055579 | 0.10505085 | 0.11275334 | -0.0077025 | 0.62485806 | 0.89054584 |
| Grhl1 | 0.03612302 | 0.03094698 | 0.03865364 | -0.0077067 | 0.82352619 | 0.8909122 |
| Galnt14 | 0.03480656 | 0.02953577 | 0.03724527 | -0.0077095 | 0.83256884 | 0.89391861 |
| Ccdc94 | 0.05685075 | 0.05311157 | 0.06082214 | -0.0077106 | 0.72261441 | 0.89054584 |
| Icmt | 0.12549626 | 0.12591004 | 0.13362073 | -0.0077107 | 0.60597667 | 0.89054584 |
| Zfp235 | 0.08910221 | 0.08756685 | 0.09528008 | -0.0077132 | 0.64697131 | 0.89054584 |
| Phka2 | 0.09415564 | 0.09296031 | 0.10067388 | -0.0077136 | 0.63944097 | 0.89054584 |
| Apoo-ps | 0.02774595 | 0.02197705 | 0.02969114 | -0.0077141 | 0.88694598 | 0.91505129 |
| Snx33 | 0.03496422 | 0.02969038 | 0.03741394 | -0.0077236 | 0.83191735 | 0.8933762 |
| Nek4 | 0.06730135 | 0.06426847 | 0.07199336 | -0.0077249 | 0.69142894 | 0.89054584 |

| | | | | | | |
|---|---|---|---|---|---|---|
| Pbx4 | 0.03038863 | 0.02478797 | 0.03251866 | -0.0077307 | 0.86597707 | 0.90420854 |
| Cad | 0.09052758 | 0.08907088 | 0.09680162 | -0.0077307 | 0.64507999 | 0.89054584 |
| Gpalpp1 | 0.14446437 | 0.14559458 | 0.15333452 | -0.0077399 | 0.59296224 | 0.89054584 |
| Gm14167 | 0.06084087 | 0.05734125 | 0.06508795 | -0.0077467 | 0.71038234 | 0.89054584 |
| Cog5 | 0.03118068 | 0.02561809 | 0.03336607 | -0.007748 | 0.86034243 | 0.90144139 |
| Smim12 | 0.29878309 | 0.30035267 | 0.30810162 | -0.007749 | 0.54663944 | 0.89054584 |
| Klf7 | 0.54891599 | 0.52519178 | 0.53294483 | -0.0077531 | 0.52701813 | 0.89054584 |
| Traf4 | 0.80455985 | 0.71378859 | 0.72154218 | -0.0077536 | 0.51996443 | 0.89054584 |
| Arhgef5 | 0.12239283 | 0.12262303 | 0.13038247 | -0.0077594 | 0.60921246 | 0.89054584 |
| Arhgef1 | 0.58751486 | 0.55651522 | 0.56427749 | -0.0077623 | 0.52555032 | 0.89054584 |
| 2810029C07I | 0.02044164 | 0.01410931 | 0.02187542 | -0.0077661 | 0.95093587 | 0.9598833 |
| Ston2 | 0.03235354 | 0.02685467 | 0.03462091 | -0.0077662 | 0.85201245 | 0.89988783 |
| E130012A19I | 0.02363606 | 0.01752558 | 0.02529359 | -0.007768 | 0.92375992 | 0.94016646 |
| Polk | 0.13131794 | 0.13191687 | 0.13968595 | -0.0077691 | 0.60222941 | 0.89054584 |
| Myl4 | 0.04235822 | 0.03755277 | 0.04532357 | -0.0077708 | 0.7877918 | 0.89054584 |
| Asb8 | 0.32704694 | 0.3274586 | 0.33524413 | -0.0077855 | 0.54308017 | 0.89054584 |
| Lrr1 | 0.03504515 | 0.0297117 | 0.03750052 | -0.0077888 | 0.83339233 | 0.89400829 |
| Gid4 | 0.23918621 | 0.24180198 | 0.24959136 | -0.0077894 | 0.55780197 | 0.89054584 |
| Chrm3 | 0.0343064 | 0.02891814 | 0.03671018 | -0.007792 | 0.83864382 | 0.89497393 |
| Zfp217 | 0.10070527 | 0.09985702 | 0.10765242 | -0.0077954 | 0.63207976 | 0.89054584 |
| B3gntl1 | 0.04064342 | 0.03568931 | 0.04348931 | -0.0078 | 0.79831038 | 0.89054584 |
| Ccdc64 | 0.06459878 | 0.06130308 | 0.06910489 | -0.0078018 | 0.70055089 | 0.89054584 |
| Actr8 | 0.37899686 | 0.37624697 | 0.384052 | -0.007805 | 0.53771666 | 0.89054584 |
| F2 | 0.03449199 | 0.02910304 | 0.03690873 | -0.0078057 | 0.83776205 | 0.89485554 |
| Pitpnc1 | 0.38605216 | 0.38275853 | 0.39056818 | -0.0078097 | 0.53711113 | 0.89054584 |
| Ube2s | 1.0534008 | 0.85417105 | 0.8619815 | -0.0078104 | 0.51683628 | 0.89054584 |
| Ssc4d | 0.03381856 | 0.02837486 | 0.03618827 | -0.0078134 | 0.8427712 | 0.89720803 |
| Pink1 | 0.46925377 | 0.45747473 | 0.4652896 | -0.0078149 | 0.53118551 | 0.89054584 |
| Tada2a | 0.14815207 | 0.14932468 | 0.15715094 | -0.0078263 | 0.59173858 | 0.89054584 |
| mt-Nd3 | 2.09743067 | 1.1154211 | 1.12325358 | -0.0078325 | 0.51295813 | 0.89054584 |

| | | | | | | |
|---|---|---|---|---|---|---|
| Txnl4a | 0.97498496 | 0.81363139 | 0.82146756 | -0.0078362 | 0.51772424 | 0.89054584 |
| Zak | 0.0852292 | 0.08330335 | 0.09114509 | -0.0078417 | 0.65572933 | 0.89054584 |
| 1110046J04F | 0.05729333 | 0.05344345 | 0.06129534 | -0.0078519 | 0.72468187 | 0.89054584 |
| C920021L13F | 0.0361032 | 0.03077745 | 0.03863243 | -0.007855 | 0.82825322 | 0.89229532 |
| Lrrc48 | 0.0342125 | 0.02875264 | 0.03660973 | -0.0078571 | 0.84131826 | 0.89621103 |
| Rcbtb1 | 0.29290702 | 0.29454424 | 0.30240846 | -0.0078642 | 0.54821671 | 0.89054584 |
| Cdh3 | 0.03465333 | 0.02921379 | 0.03708134 | -0.0078676 | 0.83854327 | 0.89497393 |
| 2510009E07I | 0.14264905 | 0.14358616 | 0.15145386 | -0.0078677 | 0.59561849 | 0.89054584 |
| Perp | 0.07871279 | 0.07631591 | 0.08418584 | -0.0078699 | 0.6684115 | 0.89054584 |
| Pole | 0.05395589 | 0.04983926 | 0.0577269 | -0.0078876 | 0.73780214 | 0.89054584 |
| Mreg | 0.06141783 | 0.05781687 | 0.06570472 | -0.0078878 | 0.71202332 | 0.89054584 |
| Lpcat4 | 0.08154026 | 0.07931586 | 0.08720576 | -0.0078899 | 0.66331079 | 0.89054584 |
| B830012L14I | 0.17042614 | 0.17219893 | 0.18008893 | -0.00789 | 0.58086858 | 0.89054584 |
| Lrrc10b | 0.03928973 | 0.03414034 | 0.04204126 | -0.0079009 | 0.80936715 | 0.89054584 |
| Slc22a23 | 0.03356686 | 0.0280172 | 0.03591899 | -0.0079018 | 0.84729879 | 0.89854206 |
| F2rl1 | 0.10206231 | 0.10116537 | 0.10907976 | -0.0079144 | 0.63233121 | 0.89054584 |
| Kit | 0.23476752 | 0.23726522 | 0.24518694 | -0.0079217 | 0.55982494 | 0.89054584 |
| Col4a5 | 0.07761052 | 0.07507405 | 0.08300843 | -0.0079344 | 0.67195777 | 0.89054584 |
| Acy3 | 0.12105931 | 0.12104963 | 0.12898989 | -0.0079403 | 0.61286402 | 0.89054584 |
| Uimc1 | 0.44338802 | 0.43453899 | 0.44248997 | -0.007951 | 0.53335857 | 0.89054584 |
| 2900093K20I | 0.18640123 | 0.18845875 | 0.19641626 | -0.0079575 | 0.5748556 | 0.89054584 |
| Cables1 | 0.0272379 | 0.0211892 | 0.02914755 | -0.0079583 | 0.89832436 | 0.92217705 |
| Ltbp3 | 0.17969345 | 0.18160743 | 0.18957308 | -0.0079656 | 0.57760234 | 0.89054584 |
| Gm11992 | 0.03178701 | 0.02603521 | 0.03401478 | -0.0079796 | 0.86279152 | 0.90270271 |
| Gabbr2 | 0.03944343 | 0.03422394 | 0.04220567 | -0.0079817 | 0.81086188 | 0.89054584 |
| Nmb | 0.09211977 | 0.09051302 | 0.09850108 | -0.0079881 | 0.64721876 | 0.89054584 |
| Sf3b5 | 1.48159506 | 1.02245618 | 1.03044684 | -0.0079907 | 0.51440987 | 0.89054584 |
| Pik3ip1 | 0.15222156 | 0.15336226 | 0.16135623 | -0.007994 | 0.5912707 | 0.89054584 |
| Cyb561d1 | 0.046497 | 0.0417532 | 0.04975034 | -0.0079971 | 0.77304233 | 0.89054584 |
| Scrib | 0.1714536 | 0.17314136 | 0.18114216 | -0.0080008 | 0.58151827 | 0.89054584 |

| | | | | | | |
|---|---|---|---|---|---|---|
| Ccdc137 | 0.2507741 | 0.25309157 | 0.26109885 | -0.0080073 | 0.55680704 | 0.89054584 |
| Arhgap17 | 0.30789786 | 0.30888877 | 0.3168982 | -0.0080094 | 0.546868 | 0.89054584 |
| Morn1 | 0.05868226 | 0.05477061 | 0.06278028 | -0.0080097 | 0.72387748 | 0.89054584 |
| Map2k6 | 0.05818529 | 0.05423574 | 0.06224896 | -0.0080132 | 0.72565979 | 0.89054584 |
| Usp25 | 0.1127479 | 0.1122798 | 0.12029539 | -0.0080156 | 0.62188565 | 0.89054584 |
| Mks1 | 0.15372268 | 0.15488628 | 0.16290592 | -0.0080196 | 0.5907029 | 0.89054584 |
| 2010015M23 | 0.05504765 | 0.05087179 | 0.05889427 | -0.0080225 | 0.73716612 | 0.89054584 |
| Top2b | 0.82764639 | 0.72827482 | 0.73629998 | -0.0080252 | 0.52024928 | 0.89054584 |
| Tmed2 | 2.62124976 | 1.11338452 | 1.1214097 | -0.0080252 | 0.51329864 | 0.89054584 |
| Slc16a11 | 0.03835158 | 0.03299478 | 0.04103769 | -0.0080429 | 0.81940462 | 0.89054584 |
| Rngtt | 0.3185307 | 0.31905491 | 0.32710825 | -0.0080533 | 0.54565941 | 0.89054584 |
| Rwdd2a | 0.04451184 | 0.03957291 | 0.04762711 | -0.0080542 | 0.78461399 | 0.89054584 |
| Mfap3 | 0.44167087 | 0.43290357 | 0.44096216 | -0.0080586 | 0.53392585 | 0.89054584 |
| Cdan1 | 0.06670186 | 0.06328992 | 0.07135267 | -0.0080627 | 0.70071349 | 0.89054584 |
| Rhebl1 | 0.03132683 | 0.02545768 | 0.03352244 | -0.0080648 | 0.86882159 | 0.90515776 |
| Gm26722 | 0.05381005 | 0.04949846 | 0.05757096 | -0.0080725 | 0.74320836 | 0.89054584 |
| Ak8 | 0.08019326 | 0.07768994 | 0.08576714 | -0.0080772 | 0.6695824 | 0.89054584 |
| Skida1 | 0.03970788 | 0.0344083 | 0.04248856 | -0.0080803 | 0.81219503 | 0.89054584 |
| Adora1 | 0.03296092 | 0.02719015 | 0.03527072 | -0.0080806 | 0.85709755 | 0.9009252 |
| Frem2 | 0.04531258 | 0.04040164 | 0.04848355 | -0.0080819 | 0.78130757 | 0.89054584 |
| 1700049G17 | 0.04855193 | 0.04385757 | 0.05194807 | -0.0080905 | 0.76595599 | 0.89054584 |
| D17H6S53E | 0.06230899 | 0.05855201 | 0.06665734 | -0.0081053 | 0.71447748 | 0.89054584 |
| 2810454H06 | 0.10479237 | 0.10383053 | 0.11194921 | -0.0081187 | 0.63227001 | 0.89054584 |
| B630005N14 | 0.19752111 | 0.19959659 | 0.20771854 | -0.008122 | 0.5722746 | 0.89054584 |
| Hspb6 | 0.12935864 | 0.12950504 | 0.13764611 | -0.0081411 | 0.60855419 | 0.89054584 |
| A230046K03 | 0.28694454 | 0.2884641 | 0.29661395 | -0.0081499 | 0.55092967 | 0.89054584 |
| Itgb4 | 0.02576792 | 0.01941656 | 0.0275747 | -0.0081581 | 0.91595649 | 0.93425758 |
| Pard6b | 0.11230676 | 0.11167114 | 0.11983319 | -0.0081621 | 0.624503 | 0.89054584 |
| Ndufc2 | 2.41688517 | 1.12080135 | 1.12896675 | -0.0081654 | 0.51344037 | 0.89054584 |
| Otud7b | 0.21263901 | 0.21483218 | 0.22299985 | -0.0081677 | 0.56774661 | 0.89054584 |

| | | | | | | |
|---|---|---|---|---|---|---|
| Mfhas1 | 0.17639192 | 0.17801651 | 0.1861983 | -0.0081818 | 0.5811045 | 0.89054584 |
| Tprn | 0.0415584 | 0.03628601 | 0.04446804 | -0.008182 | 0.80434283 | 0.89054584 |
| Heatr1 | 0.04679199 | 0.04188326 | 0.05006583 | -0.0081826 | 0.7768029 | 0.89054584 |
| Lrrc73 | 0.23290358 | 0.23513797 | 0.24332635 | -0.0081884 | 0.56229163 | 0.89054584 |
| Mcm8 | 0.05847136 | 0.05436582 | 0.0625548 | -0.008189 | 0.72903479 | 0.89054584 |
| Zdhhc9 | 0.0916096 | 0.08975204 | 0.09795656 | -0.0082045 | 0.65181224 | 0.89054584 |
| Smcr8 | 0.05491529 | 0.05054371 | 0.05875274 | -0.008209 | 0.74246209 | 0.89054584 |
| Pqlc2 | 0.07363136 | 0.07054461 | 0.07875745 | -0.0082128 | 0.68645796 | 0.89054584 |
| Cpt1a | 0.09057223 | 0.08863631 | 0.09684929 | -0.008213 | 0.65361029 | 0.89054584 |
| Adcy3 | 0.08115004 | 0.07856883 | 0.08678901 | -0.0082202 | 0.67049101 | 0.89054584 |
| Sorcs2 | 0.07736846 | 0.07452732 | 0.08274986 | -0.0082225 | 0.67829663 | 0.89054584 |
| Ftsj3 | 0.35889879 | 0.35711365 | 0.36533936 | -0.0082257 | 0.54177132 | 0.89054584 |
| R3hcc1l | 0.17458153 | 0.17611426 | 0.1843459 | -0.0082316 | 0.58239987 | 0.89054584 |
| Ints2 | 0.19768938 | 0.19965657 | 0.20788917 | -0.0082326 | 0.57318871 | 0.89054584 |
| Taf5 | 0.05923372 | 0.05513559 | 0.06336984 | -0.0082342 | 0.72753686 | 0.89054584 |
| St8sia3 | 0.15760815 | 0.15865664 | 0.16691301 | -0.0082564 | 0.59113094 | 0.89054584 |
| Kif19a | 0.07557258 | 0.07257031 | 0.08083138 | -0.0082611 | 0.68301611 | 0.89054584 |
| Nrf1 | 0.31354778 | 0.31406331 | 0.32233045 | -0.0082671 | 0.54755749 | 0.89054584 |
| Adcy9 | 0.05451753 | 0.05005868 | 0.05832744 | -0.0082688 | 0.74552457 | 0.89054584 |
| Gm12576 | 0.08808734 | 0.08592763 | 0.09419665 | -0.008269 | 0.65872413 | 0.89054584 |
| Adam22 | 0.0839404 | 0.08149837 | 0.0897689 | -0.0082705 | 0.66612379 | 0.89054584 |
| Pxylp1 | 0.0936123 | 0.09181552 | 0.10009402 | -0.0082785 | 0.65000286 | 0.89054584 |
| Foxred2 | 0.02569313 | 0.01921587 | 0.02749467 | -0.0082788 | 0.91966686 | 0.93672405 |
| Rapgef1 | 0.15634213 | 0.15732521 | 0.16560803 | -0.0082828 | 0.59212514 | 0.89054584 |
| Srfbp1 | 0.08781784 | 0.08562069 | 0.09390893 | -0.0082882 | 0.65953344 | 0.89054584 |
| Cep76 | 0.0918726 | 0.0899452 | 0.09823727 | -0.0082921 | 0.65293438 | 0.89054584 |
| Spsb1 | 0.11240337 | 0.11163515 | 0.11993442 | -0.0082993 | 0.62642195 | 0.89054584 |
| Tmem62 | 0.06011393 | 0.05600582 | 0.06431082 | -0.008305 | 0.72629595 | 0.89054584 |
| Zfp74 | 0.12609473 | 0.12593205 | 0.13424478 | -0.0083127 | 0.61351494 | 0.89054584 |
| Rnf169 | 0.05862082 | 0.05439767 | 0.06271459 | -0.0083169 | 0.73166214 | 0.89054584 |

| | | | | | | |
|---|---|---|---|---|---|---|
| Cyb5d2 | 0.07736836 | 0.07442892 | 0.08274975 | -0.0083208 | 0.68027777 | 0.89054584 |
| Mafg | 0.66697113 | 0.61732979 | 0.62565473 | -0.0083249 | 0.52471529 | 0.89054584 |
| 4933406C10I | 0.02675759 | 0.02030631 | 0.02863363 | -0.0083273 | 0.91227965 | 0.93174592 |
| Caprin2 | 0.05718271 | 0.05284717 | 0.06117707 | -0.0083299 | 0.73707822 | 0.89054584 |
| Lgals8 | 0.22128538 | 0.2233578 | 0.23169509 | -0.0083373 | 0.56656943 | 0.89054584 |
| Tmem129 | 0.2399873 | 0.24204471 | 0.25038889 | -0.0083442 | 0.56169135 | 0.89054584 |
| Cdk8 | 0.47757603 | 0.4641912 | 0.47254111 | -0.0083499 | 0.53280567 | 0.89054584 |
| Blm | 0.0977979 | 0.09620929 | 0.10456038 | -0.0083511 | 0.64509191 | 0.89054584 |
| Zfp697 | 0.0354106 | 0.02953631 | 0.03789148 | -0.0083552 | 0.84786234 | 0.89862623 |
| Dync2li1 | 0.73046791 | 0.66321682 | 0.67157816 | -0.0083613 | 0.52312775 | 0.89054584 |
| Nr2c1 | 0.06871491 | 0.06514026 | 0.07350402 | -0.0083638 | 0.70198728 | 0.89054584 |
| Hic2 | 0.05167976 | 0.04692033 | 0.05529299 | -0.0083727 | 0.75974232 | 0.89054584 |
| Fam188b | 0.06314068 | 0.05916048 | 0.06754635 | -0.0083859 | 0.71850522 | 0.89054584 |
| Zfyve26 | 0.0312539 | 0.02505449 | 0.03344442 | -0.0083899 | 0.87874799 | 0.91059794 |
| Nop9 | 0.27469379 | 0.27626004 | 0.2846552 | -0.0083952 | 0.55464398 | 0.89054584 |
| Jmjd7 | 0.04781692 | 0.04276293 | 0.051162 | -0.0083991 | 0.77781678 | 0.89054584 |
| Fbxl19 | 0.06884888 | 0.0652475 | 0.0736472 | -0.0083997 | 0.70241771 | 0.89054584 |
| Traf3ip2 | 0.1101498 | 0.10917205 | 0.11757227 | -0.0084002 | 0.63038328 | 0.89054584 |
| Nmral1 | 0.49328436 | 0.47769089 | 0.48611191 | -0.008421 | 0.53216285 | 0.89054584 |
| Vdac3-ps1 | 0.0300591 | 0.023741 | 0.03216608 | -0.0084251 | 0.88882275 | 0.91618572 |
| Zfp81 | 0.11937474 | 0.118804 | 0.12722977 | -0.0084258 | 0.62116334 | 0.89054584 |
| Pdp1 | 0.06078506 | 0.05660017 | 0.06502828 | -0.0084281 | 0.72702205 | 0.89054584 |
| 0610009O20 | 0.33825713 | 0.3374576 | 0.34589622 | -0.0084386 | 0.54524692 | 0.89054584 |
| Accs | 0.06554657 | 0.06167879 | 0.07011792 | -0.0084391 | 0.71251008 | 0.89054584 |
| Btbd8 | 0.02621826 | 0.01961712 | 0.02805656 | -0.0084394 | 0.91945484 | 0.93657389 |
| Dmap1 | 0.32660319 | 0.32637685 | 0.33482113 | -0.0084443 | 0.54676798 | 0.89054584 |
| B3galnt2 | 0.14646934 | 0.1469658 | 0.15541014 | -0.0084443 | 0.59992297 | 0.89054584 |
| Actr6 | 0.16850572 | 0.16967168 | 0.17811915 | -0.0084475 | 0.58743617 | 0.89054584 |
| Fam213b | 0.41859827 | 0.41182229 | 0.42027099 | -0.0084487 | 0.53730983 | 0.89054584 |
| Cnksr1 | 0.04571906 | 0.04046324 | 0.0489183 | -0.0084551 | 0.78966325 | 0.89054584 |

| | | | | | | |
|---|---|---|---|---|---|---|
| Fgd6 | 0.15474153 | 0.15550183 | 0.16395724 | -0.0084554 | 0.59493783 | 0.89054584 |
| Zfp653 | 0.05243878 | 0.04764762 | 0.05610464 | -0.008457 | 0.75874556 | 0.89054584 |
| Npc1 | 0.10648526 | 0.10526011 | 0.11372717 | -0.0084671 | 0.63565219 | 0.89054584 |
| Nfatc2 | 0.21817111 | 0.22009673 | 0.22856706 | -0.0084703 | 0.56853788 | 0.89054584 |
| Lnx1 | 0.12040147 | 0.11983204 | 0.12830267 | -0.0084706 | 0.62080151 | 0.89054584 |
| Rnf113a1 | 0.10235872 | 0.10092079 | 0.10939144 | -0.0084706 | 0.64086117 | 0.89054584 |
| 2310022A10 | 0.18752782 | 0.18908947 | 0.19756373 | -0.0084743 | 0.57919699 | 0.89054584 |
| Med10 | 1.05146271 | 0.85250894 | 0.86098658 | -0.0084776 | 0.51829461 | 0.89054584 |
| Otud5 | 0.30605675 | 0.30664703 | 0.31512477 | -0.0084777 | 0.54987229 | 0.89054584 |
| Syt5 | 0.20845245 | 0.21029885 | 0.21877773 | -0.0084789 | 0.57164372 | 0.89054584 |
| Xpo5 | 0.02461027 | 0.01785405 | 0.02633602 | -0.008482 | 0.93325639 | 0.94660649 |
| Ggh | 0.48525545 | 0.47071214 | 0.47919466 | -0.0084825 | 0.53286381 | 0.89054584 |
| Kif1c | 0.11433769 | 0.1134728 | 0.12196044 | -0.0084876 | 0.62711862 | 0.89054584 |
| Adgb | 0.04736072 | 0.04218442 | 0.05067409 | -0.0084897 | 0.78246189 | 0.89054584 |
| Bcl2l14 | 0.03689972 | 0.03099117 | 0.03948454 | -0.0084934 | 0.84186855 | 0.89673133 |
| Pik3c3 | 0.1197845 | 0.1191632 | 0.127658 | -0.0084948 | 0.6217278 | 0.89054584 |
| Gpatch2l | 0.04317635 | 0.03769291 | 0.04619867 | -0.0085058 | 0.80449013 | 0.89054584 |
| Grk4 | 0.03832266 | 0.03250057 | 0.04100675 | -0.0085062 | 0.83308316 | 0.89393134 |
| Zscan20 | 0.02402333 | 0.01718861 | 0.02570798 | -0.0085194 | 0.93869697 | 0.95044651 |
| Tatdn2 | 0.07066642 | 0.06706857 | 0.07558944 | -0.0085209 | 0.70027168 | 0.89054584 |
| Plpp7 | 0.09803317 | 0.09627485 | 0.1048114 | -0.0085365 | 0.64782521 | 0.89054584 |
| Cyp4b1 | 0.03800493 | 0.03212502 | 0.04066686 | -0.0085418 | 0.83610729 | 0.89462889 |
| 9530077C05I | 0.04510161 | 0.03969986 | 0.0482579 | -0.008558 | 0.79565905 | 0.89054584 |
| Zfp2 | 0.08984688 | 0.0875082 | 0.09607501 | -0.0085668 | 0.66108571 | 0.89054584 |
| Zbtb39 | 0.04513487 | 0.03972148 | 0.04829348 | -0.008572 | 0.79586826 | 0.89054584 |
| Chml | 0.07538251 | 0.07205246 | 0.08062832 | -0.0085759 | 0.68988993 | 0.89054584 |
| Diaph2 | 0.08271727 | 0.0798817 | 0.08846275 | -0.008581 | 0.67433878 | 0.89054584 |
| Rnf112 | 0.04069479 | 0.03495564 | 0.04354425 | -0.0085886 | 0.82093218 | 0.89054584 |
| AI846148 | 0.10942032 | 0.10821431 | 0.11680724 | -0.0085929 | 0.63410115 | 0.89054584 |
| Heatr3 | 0.22760173 | 0.22942381 | 0.23802595 | -0.0086021 | 0.56685465 | 0.89054584 |

| | | | | | | |
|---|---|---|---|---|---|---|
| Pex3 | 0.36244317 | 0.36004135 | 0.36865476 | -0.0086134 | 0.54334065 | 0.89054584 |
| Taok3 | 0.59865585 | 0.56452416 | 0.57313872 | -0.0086146 | 0.52791361 | 0.89054584 |
| Raet1e | 0.03429129 | 0.02806511 | 0.03669402 | -0.0086289 | 0.86337065 | 0.90290403 |
| Tmem5 | 0.72391468 | 0.6583361 | 0.6669702 | -0.0086341 | 0.52404609 | 0.89054584 |
| Ctf1 | 0.08634901 | 0.08370033 | 0.09234073 | -0.0086404 | 0.66856024 | 0.89054584 |
| Srebf2 | 0.29421858 | 0.29503515 | 0.3036807 | -0.0086456 | 0.55275952 | 0.89054584 |
| Eif4enif1 | 0.24265909 | 0.2444 | 0.2530467 | -0.0086467 | 0.56324347 | 0.89054584 |
| Zfp128 | 0.0275813 | 0.02086608 | 0.02951498 | -0.0086489 | 0.91390962 | 0.93295744 |
| Pgbd1 | 0.03448823 | 0.02825042 | 0.03690471 | -0.0086543 | 0.86270184 | 0.90267402 |
| Ift20 | 1.58178466 | 1.04787471 | 1.05653153 | -0.0086568 | 0.51522536 | 0.89054584 |
| Mief2 | 0.11516235 | 0.11415478 | 0.12282378 | -0.008669 | 0.6288586 | 0.89054584 |
| Ipp | 0.20190234 | 0.20348186 | 0.21215703 | -0.0086752 | 0.57554357 | 0.89054584 |
| Mdm1 | 0.3525149 | 0.35066261 | 0.35935063 | -0.008688 | 0.54484175 | 0.89054584 |
| Hipk3 | 0.30352227 | 0.30398679 | 0.31268062 | -0.0086938 | 0.55153406 | 0.89054584 |
| Tubb4a | 0.11926255 | 0.11841004 | 0.12711251 | -0.0087025 | 0.62512296 | 0.89054584 |
| Snapc4 | 0.04403306 | 0.03841025 | 0.04711501 | -0.0087048 | 0.80531629 | 0.89054584 |
| Rpl10-ps3 | 1.38401963 | 0.9915881 | 1.00029515 | -0.008707 | 0.51617413 | 0.89054584 |
| Elovl6 | 0.2080529 | 0.20966423 | 0.21837438 | -0.0087102 | 0.5737102 | 0.89054584 |
| Necap1 | 0.58329375 | 0.55218609 | 0.56089841 | -0.0087123 | 0.52884482 | 0.89054584 |
| Myo5a | 0.09414071 | 0.09194464 | 0.10065794 | -0.0087133 | 0.65663414 | 0.89054584 |
| Egf | 0.04509804 | 0.0395355 | 0.04825409 | -0.0087186 | 0.8000447 | 0.89054584 |
| Nup133 | 0.12983925 | 0.12942361 | 0.13814661 | -0.008723 | 0.61568981 | 0.89054584 |
| Adnp2 | 0.08429606 | 0.08141728 | 0.09014868 | -0.0087314 | 0.67409335 | 0.89054584 |
| Retsat | 0.05149334 | 0.04635588 | 0.05509364 | -0.0087378 | 0.77001634 | 0.89054584 |
| Mturn | 0.07073437 | 0.06692014 | 0.07566204 | -0.0087419 | 0.70481098 | 0.89054584 |
| Sass6 | 0.09781703 | 0.09583721 | 0.10458079 | -0.0087436 | 0.6515526 | 0.89054584 |
| Gbe1 | 0.14549778 | 0.14566069 | 0.15440454 | -0.0087438 | 0.60404583 | 0.89054584 |
| Prkcb | 0.16178386 | 0.16246871 | 0.17121266 | -0.0087439 | 0.59403434 | 0.89054584 |
| Fdxacb1 | 0.05933917 | 0.05473566 | 0.06348257 | -0.0087469 | 0.73954058 | 0.89054584 |
| Zfp575 | 0.0567498 | 0.05196675 | 0.06071422 | -0.0087475 | 0.74896672 | 0.89054584 |

| | | | | | | |
|---|---|---|---|---|---|---|
| Fut4 | 0.06092743 | 0.05643193 | 0.06518048 | -0.0087486 | 0.73412116 | 0.89054584 |
| Rhbdd3 | 0.18103326 | 0.18219052 | 0.19094145 | -0.0087509 | 0.58453978 | 0.89054584 |
| Zfp873 | 0.04704963 | 0.04158892 | 0.05034138 | -0.0087525 | 0.79103388 | 0.89054584 |
| Lrch2 | 0.12639978 | 0.12580642 | 0.13456283 | -0.0087564 | 0.61912046 | 0.89054584 |
| Cadm4 | 0.05073234 | 0.04551614 | 0.05427985 | -0.0087637 | 0.77403617 | 0.89054584 |
| Sec11a | 1.9767887 | 1.10739067 | 1.11615487 | -0.0087642 | 0.51459111 | 0.89054584 |
| Samd10 | 0.29984877 | 0.3003614 | 0.30913227 | -0.0087709 | 0.55258142 | 0.89054584 |
| 1700048O20 | 0.0700479 | 0.06615123 | 0.0749285 | -0.0087773 | 0.70738351 | 0.89054584 |
| Dach2 | 0.03392408 | 0.02752075 | 0.03630117 | -0.0087804 | 0.8701057 | 0.90589084 |
| Dync1li2 | 0.32627488 | 0.3257211 | 0.33450811 | -0.008787 | 0.54870223 | 0.89054584 |
| Aptx | 0.34697779 | 0.34533654 | 0.35413844 | -0.0088019 | 0.54609259 | 0.89054584 |
| Map3k5 | 0.05092971 | 0.04568397 | 0.05449092 | -0.008807 | 0.77427182 | 0.89054584 |
| Lipe | 0.04671932 | 0.04116814 | 0.04998811 | -0.00882 | 0.79447006 | 0.89054584 |
| Mansc1 | 0.10193447 | 0.10010727 | 0.10894532 | -0.008838 | 0.64726636 | 0.89054584 |
| Cenpp | 0.06092395 | 0.05633834 | 0.06517676 | -0.0088384 | 0.73623625 | 0.89054584 |
| Tmem161b | 0.06966553 | 0.06567654 | 0.07451989 | -0.0088434 | 0.70982773 | 0.89054584 |
| Dnajc22 | 0.03413418 | 0.02767208 | 0.03652593 | -0.0088539 | 0.8706193 | 0.90598899 |
| Tsga10 | 0.10994425 | 0.10849564 | 0.11735672 | -0.0088611 | 0.63749686 | 0.89054584 |
| Setd6 | 0.16284088 | 0.16343875 | 0.17229993 | -0.0088612 | 0.59468129 | 0.89054584 |
| Eif2ak3 | 0.08971862 | 0.08707452 | 0.0959381 | -0.0088636 | 0.66655954 | 0.89054584 |
| Ppfibp2 | 0.04797551 | 0.04246503 | 0.0513316 | -0.0088666 | 0.78951653 | 0.89054584 |
| Zbtb2 | 0.1142944 | 0.11304783 | 0.12191512 | -0.0088673 | 0.6326426 | 0.89054584 |
| 2810001G20 | 0.23985311 | 0.2413824 | 0.25025532 | -0.0088729 | 0.56560076 | 0.89054584 |
| Lhpp | 0.06289289 | 0.05840329 | 0.06728148 | -0.0088782 | 0.73064845 | 0.89054584 |
| Gm14295 | 0.06990918 | 0.0659015 | 0.07478026 | -0.0088788 | 0.70992373 | 0.89054584 |
| Zfp7 | 0.06124877 | 0.05663906 | 0.065524 | -0.0088849 | 0.73622295 | 0.89054584 |
| Ltbr | 0.21362602 | 0.21510296 | 0.22399412 | -0.0088912 | 0.5733583 | 0.89054584 |
| Pbld2 | 0.06805159 | 0.06389739 | 0.07279516 | -0.0088978 | 0.715479 | 0.89054584 |
| C130021I20R | 0.03630828 | 0.02993838 | 0.03885183 | -0.0089134 | 0.85743396 | 0.9009252 |
| Galnt7 | 0.14480375 | 0.1447704 | 0.15368597 | -0.0089156 | 0.60652474 | 0.89054584 |

| | | | | | | |
|---|---|---|---|---|---|---|
| Hsbp1 | 2.28427539 | 1.11998082 | 1.12890293 | -0.0089221 | 0.51468624 | 0.89054584 |
| Rps2-ps6 | 0.05665575 | 0.05168385 | 0.06061366 | -0.0089298 | 0.75376041 | 0.89054584 |
| Yjefn3 | 0.04144058 | 0.03540966 | 0.04434201 | -0.0089324 | 0.8260081 | 0.89170439 |
| Osbpl8 | 0.09425087 | 0.09183154 | 0.1007755 | -0.008944 | 0.66037517 | 0.89054584 |
| Zhx3 | 0.12112113 | 0.12010669 | 0.12905446 | -0.0089478 | 0.62665993 | 0.89054584 |
| Scfd2 | 0.0855828 | 0.08256794 | 0.09152264 | -0.0089547 | 0.67574659 | 0.89054584 |
| Ndor1 | 0.035111 | 0.02860655 | 0.03757097 | -0.0089644 | 0.86684964 | 0.90430169 |
| Ap5z1 | 0.0953302 | 0.09295459 | 0.1019273 | -0.0089727 | 0.6591443 | 0.89054584 |
| Cdkl2 | 0.16552163 | 0.16607942 | 0.17505537 | -0.0089759 | 0.59440332 | 0.89054584 |
| Kdf1 | 0.31120483 | 0.31110235 | 0.32007955 | -0.0089772 | 0.55198111 | 0.89054584 |
| Nxn | 0.1772327 | 0.17806743 | 0.18705813 | -0.0089907 | 0.58859242 | 0.89054584 |
| C030037D09 | 0.03385157 | 0.02722385 | 0.03622359 | -0.0089997 | 0.87646847 | 0.9091719 |
| Kdm5d | 0.12750216 | 0.12669701 | 0.13571196 | -0.0090149 | 0.62152169 | 0.89054584 |
| Gm11423 | 0.04754272 | 0.04185112 | 0.05086874 | -0.0090176 | 0.79557013 | 0.89054584 |
| Vps33a | 0.16748349 | 0.16805206 | 0.17707003 | -0.009018 | 0.593778 | 0.89054584 |
| Zfp882 | 0.11301169 | 0.1115477 | 0.12057173 | -0.009024 | 0.63633991 | 0.89054584 |
| Cmas | 0.49342328 | 0.47720168 | 0.48623124 | -0.0090296 | 0.53447305 | 0.89054584 |
| Tpbg | 0.20988707 | 0.21118656 | 0.2202254 | -0.0090388 | 0.57582327 | 0.89054584 |
| Kif13a | 0.07706289 | 0.07338289 | 0.08242344 | -0.0090405 | 0.69532869 | 0.89054584 |
| Efcc1 | 0.06493294 | 0.06042098 | 0.06946206 | -0.0090411 | 0.72787564 | 0.89054584 |
| Amacr | 0.16873168 | 0.16930902 | 0.17835099 | -0.009042 | 0.59336016 | 0.89054584 |
| Rfwd3 | 0.22501157 | 0.22639001 | 0.23543202 | -0.009042 | 0.57100424 | 0.89054584 |
| Wdr55 | 0.14707817 | 0.14698624 | 0.15604011 | -0.0090539 | 0.60654466 | 0.89054584 |
| Unc79 | 0.0636269 | 0.05901123 | 0.06806607 | -0.0090548 | 0.73229857 | 0.89054584 |
| Cblb | 0.08834794 | 0.08541967 | 0.09447486 | -0.0090552 | 0.6723993 | 0.89054584 |
| Prkag2 | 0.10856716 | 0.1068458 | 0.11591225 | -0.0090665 | 0.64222563 | 0.89054584 |
| Slc25a26 | 0.06123585 | 0.05644362 | 0.06551018 | -0.0090666 | 0.74046966 | 0.89054584 |
| Eif2a | 0.7788773 | 0.69560376 | 0.70467064 | -0.0090669 | 0.5239006 | 0.89054584 |
| Srsf4 | 0.50231456 | 0.48477885 | 0.49384599 | -0.0090671 | 0.53408368 | 0.89054584 |
| Ilk | 0.96791214 | 0.80863084 | 0.8177134 | -0.0090826 | 0.5206353 | 0.89054584 |

| | | | | | | |
|---|---|---|---|---|---|---|
| Espl1 | 0.03659265 | 0.03004266 | 0.03915605 | -0.0091134 | 0.86089454 | 0.90182436 |
| Lnpep | 0.08058947 | 0.07707529 | 0.08619031 | -0.009115 | 0.68888935 | 0.89054584 |
| 2210404O09 | 0.08093349 | 0.0774352 | 0.08655774 | -0.0091225 | 0.6882929 | 0.89054584 |
| Ptpn3 | 0.05167719 | 0.04616161 | 0.05529024 | -0.0091286 | 0.77911554 | 0.89054584 |
| Trnt1 | 0.47772916 | 0.4635441 | 0.47267413 | -0.00913 | 0.53585271 | 0.89054584 |
| Rnf213 | 0.06873928 | 0.0643942 | 0.07353007 | -0.0091359 | 0.71865768 | 0.89054584 |
| Zfp867 | 0.03357659 | 0.02679335 | 0.0359294 | -0.0091361 | 0.88192523 | 0.91223443 |
| Ptprm | 0.04923834 | 0.04354515 | 0.05268215 | -0.009137 | 0.79046328 | 0.89054584 |
| Inpp4a | 0.26111942 | 0.26218133 | 0.27131981 | -0.0091385 | 0.56234623 | 0.89054584 |
| Catsper2 | 0.06257851 | 0.05780514 | 0.06694544 | -0.0091403 | 0.73764497 | 0.89054584 |
| Cnot7 | 1.06044223 | 0.85643372 | 0.86557493 | -0.0091412 | 0.51962101 | 0.89054584 |
| Casp7 | 0.14293182 | 0.1426025 | 0.15174689 | -0.0091444 | 0.61054951 | 0.89054584 |
| Clip2 | 0.08954345 | 0.0866063 | 0.09575111 | -0.0091448 | 0.67182456 | 0.89054584 |
| Arl4d | 0.04379246 | 0.03771227 | 0.04685766 | -0.0091454 | 0.81840509 | 0.89054584 |
| Syndig1l | 0.04857851 | 0.04282622 | 0.05197649 | -0.0091503 | 0.79394594 | 0.89054584 |
| B130006D01 | 0.03696026 | 0.03039604 | 0.0395493 | -0.0091533 | 0.85954156 | 0.90131997 |
| Ccdc93 | 0.04765508 | 0.04182884 | 0.05098892 | -0.0091601 | 0.79869562 | 0.89054584 |
| Ccnj | 0.10570207 | 0.10373337 | 0.11290475 | -0.0091714 | 0.64745217 | 0.89054584 |
| Gm10069 | 0.12552814 | 0.12447321 | 0.13365398 | -0.0091808 | 0.62552539 | 0.89054584 |
| Tctn1 | 0.06399226 | 0.05927528 | 0.0684566 | -0.0091813 | 0.73396591 | 0.89054584 |
| Uba52 | 2.08778577 | 1.113588 | 1.12277925 | -0.0091912 | 0.51521151 | 0.89054584 |
| Coro7 | 0.14307091 | 0.1426964 | 0.15189102 | -0.0091946 | 0.61103819 | 0.89054584 |
| Focad | 0.0436627 | 0.03751534 | 0.04671887 | -0.0092035 | 0.82063888 | 0.89054584 |
| Tubgcp6 | 0.03740583 | 0.03081626 | 0.04002596 | -0.0092097 | 0.85813802 | 0.90124396 |
| Tmem170 | 0.03999703 | 0.03358294 | 0.04279787 | -0.0092149 | 0.84210093 | 0.89691028 |
| Dusp16 | 0.07813153 | 0.07433303 | 0.08356497 | -0.0092319 | 0.6966194 | 0.89054584 |
| Sh3gl2 | 0.0795972 | 0.07589771 | 0.0851305 | -0.0092328 | 0.69332014 | 0.89054584 |
| Tnr | 0.0640354 | 0.05926983 | 0.06850271 | -0.0092329 | 0.73497774 | 0.89054584 |
| Enoph1 | 0.22866305 | 0.22985365 | 0.23908794 | -0.0092343 | 0.57140106 | 0.89054584 |
| Itga2b | 0.05130989 | 0.0456472 | 0.05489746 | -0.0092503 | 0.78378352 | 0.89054584 |

| | | | | | | |
|---|---|---|---|---|---|---|
| Gtpbp10 | 0.12482748 | 0.12366071 | 0.13292321 | -0.0092625 | 0.62727682 | 0.89054584 |
| B4galt3 | 0.56882531 | 0.53996305 | 0.54922729 | -0.0092642 | 0.53131904 | 0.89054584 |
| Dynlrb2 | 0.11612988 | 0.11456949 | 0.12383636 | -0.0092669 | 0.63631863 | 0.89054584 |
| Pwp2 | 0.07582035 | 0.07182332 | 0.08109607 | -0.0092728 | 0.70288501 | 0.89054584 |
| Btbd10 | 0.38188861 | 0.37745296 | 0.38672615 | -0.0092732 | 0.54447543 | 0.89054584 |
| Slc9a6 | 0.10257148 | 0.10034061 | 0.10961513 | -0.0092745 | 0.65328059 | 0.89054584 |
| Asphd1 | 0.05864347 | 0.05345919 | 0.0627388 | -0.0092796 | 0.75461923 | 0.89054584 |
| Sdcbp2 | 0.06605326 | 0.06136563 | 0.07065947 | -0.0092938 | 0.72999257 | 0.89054584 |
| Lgals12 | 0.04097267 | 0.03453218 | 0.0438415 | -0.0093093 | 0.83873712 | 0.89497393 |
| Klhl1 | 0.03101352 | 0.02387782 | 0.03318723 | -0.0093094 | 0.90437473 | 0.92615843 |
| Fbxo33 | 0.15343334 | 0.15329624 | 0.16260729 | -0.009311 | 0.60517894 | 0.89054584 |
| Fam208b | 0.15097516 | 0.15075311 | 0.16006893 | -0.0093158 | 0.60686017 | 0.89054584 |
| Eif4ebp2 | 0.12341597 | 0.12212956 | 0.13145047 | -0.0093209 | 0.62943483 | 0.89054584 |
| Qpctl | 0.28012325 | 0.28064202 | 0.28996404 | -0.009322 | 0.5595308 | 0.89054584 |
| Rbak | 0.14541998 | 0.14499569 | 0.154324 | -0.0093283 | 0.61088088 | 0.89054584 |
| 2610301B20 | 0.25540798 | 0.25634316 | 0.26568297 | -0.0093398 | 0.56504791 | 0.89054584 |
| Dennd1a | 0.15067021 | 0.15041239 | 0.15975389 | -0.0093415 | 0.6073537 | 0.89054584 |
| Vps13b | 0.15346084 | 0.15328494 | 0.16263567 | -0.0093507 | 0.60559865 | 0.89054584 |
| Lonrf2 | 0.03332894 | 0.02629275 | 0.03566445 | -0.0093717 | 0.88957098 | 0.91676145 |
| Smad4 | 0.33823496 | 0.33650317 | 0.34587521 | -0.009372 | 0.55022961 | 0.89054584 |
| Slc29a3 | 0.07312148 | 0.0688399 | 0.07821267 | -0.0093728 | 0.71168492 | 0.89054584 |
| Hmga1 | 0.14458973 | 0.14407768 | 0.15346435 | -0.0093867 | 0.61216448 | 0.89054584 |
| Gm4950 | 0.28000379 | 0.28045143 | 0.28984739 | -0.009396 | 0.56002337 | 0.89054584 |
| Zfp456 | 0.02652694 | 0.01899073 | 0.02838684 | -0.0093961 | 0.93847777 | 0.95040868 |
| Wdr45 | 0.3032727 | 0.30304233 | 0.31243977 | -0.0093974 | 0.55572129 | 0.89054584 |
| Commd10 | 0.47228514 | 0.4585345 | 0.46793587 | -0.0094014 | 0.53728794 | 0.89054584 |
| Tnfrsf19 | 0.07366924 | 0.06939614 | 0.07879792 | -0.0094018 | 0.71085416 | 0.89054584 |
| Gm20604 | 0.3884557 | 0.3833747 | 0.39278167 | -0.009407 | 0.54442173 | 0.89054584 |
| Ghdc | 0.18926905 | 0.18992763 | 0.19933617 | -0.0094085 | 0.58702512 | 0.89054584 |
| Uevld | 0.04859612 | 0.04258435 | 0.05199532 | -0.009411 | 0.80045635 | 0.89054584 |

| | | | | | | |
|---|---|---|---|---|---|---|
| Trim45 | 0.11051233 | 0.10854105 | 0.11795239 | -0.0094113 | 0.64495442 | 0.89054584 |
| Gm7160 | 0.04138361 | 0.03486509 | 0.04428107 | -0.009416 | 0.83907983 | 0.89500145 |
| Wbp4 | 1.05059715 | 0.85112347 | 0.86054534 | -0.0094219 | 0.52034099 | 0.89054584 |
| Ppp1r12c | 0.12185414 | 0.12039006 | 0.12982 | -0.0094299 | 0.6324763 | 0.89054584 |
| Slc25a35 | 0.04039606 | 0.03379129 | 0.0432247 | -0.0094334 | 0.84536919 | 0.89825055 |
| Pigg | 0.06307245 | 0.05803655 | 0.06747342 | -0.0094369 | 0.74267098 | 0.89054584 |
| Nmnat1 | 0.12839888 | 0.12720862 | 0.13664636 | -0.0094377 | 0.6261898 | 0.89054584 |
| Galnt10 | 0.13862206 | 0.13783472 | 0.14727735 | -0.0094426 | 0.61741784 | 0.89054584 |
| Ctns | 0.05013581 | 0.04419692 | 0.05364192 | -0.009445 | 0.79398402 | 0.89054584 |
| Dennd2d | 0.02906931 | 0.0216567 | 0.03110708 | -0.0094504 | 0.92152344 | 0.93827146 |
| Gm21967 | 0.06869221 | 0.06402834 | 0.07347976 | -0.0094514 | 0.72550089 | 0.89054584 |
| Tmem25 | 0.03347375 | 0.02635871 | 0.03581938 | -0.0094607 | 0.89074994 | 0.91722772 |
| Ubxn11 | 0.07889603 | 0.07490126 | 0.08438157 | -0.0094803 | 0.69966402 | 0.89054584 |
| Mbd1 | 0.21470194 | 0.21559519 | 0.22507747 | -0.0094823 | 0.57780305 | 0.89054584 |
| Gm8394 | 0.04672364 | 0.04051013 | 0.04999273 | -0.0094826 | 0.81157246 | 0.89054584 |
| Stat2 | 0.05410747 | 0.04838903 | 0.05788898 | -0.0095 | 0.77773382 | 0.89054584 |
| Hltf | 0.1718872 | 0.17208495 | 0.18158651 | -0.0095016 | 0.59629953 | 0.89054584 |
| Tmem237 | 0.3962345 | 0.39042253 | 0.39992513 | -0.0095026 | 0.54407324 | 0.89054584 |
| Gng13 | 0.0552742 | 0.04963186 | 0.05913651 | -0.0095046 | 0.77301194 | 0.89054584 |
| Igfals | 0.05488517 | 0.04921201 | 0.05872054 | -0.0095085 | 0.77470026 | 0.89054584 |
| Echdc3 | 0.04577885 | 0.03947026 | 0.04898225 | -0.009512 | 0.81719629 | 0.89054584 |
| Pax6os1 | 0.1121204 | 0.11012497 | 0.11963792 | -0.009513 | 0.64447771 | 0.89054584 |
| Mfsd3 | 0.07180805 | 0.06729167 | 0.07680931 | -0.0095176 | 0.71813208 | 0.89054584 |
| Ccp110 | 0.17503059 | 0.17527802 | 0.18480551 | -0.0095275 | 0.59490757 | 0.89054584 |
| Gm6710 | 0.12969091 | 0.12846096 | 0.13799214 | -0.0095312 | 0.62619598 | 0.89054584 |
| Hspa1b | 0.04176145 | 0.03515335 | 0.04468523 | -0.0095319 | 0.83984132 | 0.89542723 |
| Atxn7 | 0.20909582 | 0.2098934 | 0.21942706 | -0.0095337 | 0.58020638 | 0.89054584 |
| BC067074 | 0.02355906 | 0.01567517 | 0.0252112 | -0.009536 | 0.96098433 | 0.96772597 |
| Pla1a | 0.05605688 | 0.0504288 | 0.05997336 | -0.0095446 | 0.77079116 | 0.89054584 |
| Ccdc184 | 0.13485653 | 0.133821 | 0.14336629 | -0.0095453 | 0.62179217 | 0.89054584 |

| | | | | | | |
|---|---|---|---|---|---|---|
| Zbtb26 | 0.12149107 | 0.11989279 | 0.12944085 | -0.0095481 | 0.63444959 | 0.89054584 |
| Gm2000 | 0.97420618 | 0.81149375 | 0.82105497 | -0.0095612 | 0.52163342 | 0.89054584 |
| Mrps23 | 0.64093963 | 0.59644123 | 0.60601766 | -0.0095764 | 0.52934427 | 0.89054584 |
| Agpat4 | 0.44271849 | 0.43231777 | 0.44189447 | -0.0095767 | 0.54021215 | 0.89054584 |
| Mgea5 | 0.37488293 | 0.37066146 | 0.38023961 | -0.0095782 | 0.54671171 | 0.89054584 |
| Crybb3 | 0.04146657 | 0.03478834 | 0.04436981 | -0.0095815 | 0.84281115 | 0.89720803 |
| Aif1l | 0.03217321 | 0.02484223 | 0.03442797 | -0.0095857 | 0.90271606 | 0.92504788 |
| Mgrn1 | 0.33890273 | 0.33691449 | 0.34650779 | -0.0095933 | 0.55131561 | 0.89054584 |
| Athl1 | 0.13433311 | 0.13322744 | 0.1428222 | -0.0095948 | 0.62285464 | 0.89054584 |
| Msantd2 | 0.11277435 | 0.11071725 | 0.1203231 | -0.0096058 | 0.64503157 | 0.89054584 |
| Ndufs8 | 1.39288874 | 0.99364862 | 1.00325879 | -0.0096102 | 0.51779801 | 0.89054584 |
| Upf2 | 0.83214826 | 0.72952019 | 0.73913243 | -0.0096122 | 0.52415649 | 0.89054584 |
| Asb6 | 0.1764203 | 0.17661352 | 0.18622733 | -0.0096138 | 0.5950339 | 0.89054584 |
| 6430548M08 | 0.05681506 | 0.05116982 | 0.06078399 | -0.0096142 | 0.76940905 | 0.89054584 |
| Clcn2 | 0.06290896 | 0.05767344 | 0.06729867 | -0.0096252 | 0.7474011 | 0.89054584 |
| Tmppe | 0.10041819 | 0.09772279 | 0.10735039 | -0.0096276 | 0.66196507 | 0.89054584 |
| Bcdin3d | 0.19590344 | 0.19644219 | 0.20607763 | -0.0096354 | 0.58622114 | 0.89054584 |
| Pcdhga8.1 | 0.04295219 | 0.03632311 | 0.0459589 | -0.0096358 | 0.83566546 | 0.89459411 |
| Vps50 | 0.25070385 | 0.25138851 | 0.26102927 | -0.0096408 | 0.56830942 | 0.89054584 |
| Thg1l | 0.05565698 | 0.04989391 | 0.05954578 | -0.0096519 | 0.77492801 | 0.89054584 |
| E2f3 | 0.08222878 | 0.07828717 | 0.08794107 | -0.0096539 | 0.69548013 | 0.89054584 |
| Mdfic | 0.06361573 | 0.05840014 | 0.06805413 | -0.009654 | 0.74566374 | 0.89054584 |
| Plekha8 | 0.12226133 | 0.12058996 | 0.13024518 | -0.0096552 | 0.63509284 | 0.89054584 |
| Angel1 | 0.03771482 | 0.03069562 | 0.04035651 | -0.0096609 | 0.86763854 | 0.90444517 |
| Porcn | 0.04820738 | 0.0419141 | 0.05157959 | -0.0096655 | 0.80867895 | 0.89054584 |
| Nos1ap | 0.05304318 | 0.04707963 | 0.05675095 | -0.0096713 | 0.7863913 | 0.89054584 |
| Tns3 | 0.32851707 | 0.32696905 | 0.33664477 | -0.0096757 | 0.55326135 | 0.89054584 |
| Dffa | 0.18697717 | 0.18732128 | 0.19700294 | -0.0096817 | 0.59055098 | 0.89054584 |
| Arih2 | 0.23047211 | 0.23121439 | 0.24089696 | -0.0096826 | 0.5742718 | 0.89054584 |
| D730003I15F | 0.0732214 | 0.06863186 | 0.07831944 | -0.0096876 | 0.71778666 | 0.89054584 |

| | | | | | | |
|---|---|---|---|---|---|---|
| Tmem260 | 0.13522045 | 0.13405458 | 0.14374451 | -0.0096899 | 0.62326336 | 0.89054584 |
| Nphp4 | 0.03021898 | 0.02264295 | 0.03233715 | -0.0096942 | 0.91874525 | 0.93597769 |
| Rab3gap1 | 0.36460145 | 0.36094193 | 0.37067081 | -0.0097289 | 0.54866206 | 0.89054584 |
| Ikzf4 | 0.09972634 | 0.09688788 | 0.10661776 | -0.0097299 | 0.66464452 | 0.89054584 |
| Pdk1 | 0.20475711 | 0.20531294 | 0.21504463 | -0.0097317 | 0.58349283 | 0.89054584 |
| Arf3 | 0.61092131 | 0.57306777 | 0.58280024 | -0.0097325 | 0.5310072 | 0.89054584 |
| Yeats2 | 0.17245685 | 0.17243534 | 0.18217017 | -0.0097348 | 0.59830613 | 0.89054584 |
| Ablim1 | 0.18768226 | 0.18798338 | 0.19772099 | -0.0097376 | 0.5907402 | 0.89054584 |
| Gm10244 | 0.03547791 | 0.02822156 | 0.03796349 | -0.0097419 | 0.88406313 | 0.91361803 |
| Zfp3 | 0.03291135 | 0.02545008 | 0.03521769 | -0.0097676 | 0.90184814 | 0.92431323 |
| Ccdc149 | 0.05255473 | 0.04645706 | 0.05622864 | -0.0097716 | 0.79092779 | 0.89054584 |
| Snapc2 | 0.23277211 | 0.23340663 | 0.24319505 | -0.0097884 | 0.57437307 | 0.89054584 |
| Tle2 | 0.38068151 | 0.37581762 | 0.38561045 | -0.0097928 | 0.54709171 | 0.89054584 |
| Nek8 | 0.05843618 | 0.05272419 | 0.06251719 | -0.009793 | 0.76724364 | 0.89054584 |
| Zfp189 | 0.07180813 | 0.06699662 | 0.0768094 | -0.0098128 | 0.72414586 | 0.89054584 |
| Cspg5 | 0.03604749 | 0.02874455 | 0.03857283 | -0.0098283 | 0.88240375 | 0.91235841 |
| Zfp511 | 0.32150375 | 0.32012008 | 0.32995279 | -0.0098327 | 0.55521097 | 0.89054584 |
| Syvn1 | 0.2217842 | 0.22235027 | 0.23219572 | -0.0098454 | 0.57829985 | 0.89054584 |
| Numb | 0.23636832 | 0.23693358 | 0.24678361 | -0.00985 | 0.57376002 | 0.89054584 |
| Sos2 | 0.13065024 | 0.12912993 | 0.13899098 | -0.009861 | 0.62950384 | 0.89054584 |
| Slc6a8 | 0.07010971 | 0.06513209 | 0.07499455 | -0.0098625 | 0.72996029 | 0.89054584 |
| Zfp668 | 0.14145115 | 0.14033219 | 0.15021214 | -0.00988 | 0.62036298 | 0.89054584 |
| Vps9d1 | 0.25651883 | 0.25687518 | 0.26678039 | -0.0099052 | 0.56866657 | 0.89054584 |
| Nt5dc3 | 0.03958428 | 0.03244918 | 0.04235634 | -0.0099072 | 0.8620834 | 0.90260095 |
| Gm15013 | 0.07352406 | 0.06873471 | 0.0786428 | -0.0099081 | 0.72138915 | 0.89054584 |
| Pyroxd2 | 0.06149535 | 0.05587002 | 0.06578759 | -0.0099176 | 0.75876683 | 0.89054584 |
| Sowahc | 0.13630071 | 0.13494869 | 0.14486693 | -0.0099182 | 0.62512617 | 0.89054584 |
| Cobll1 | 0.10174617 | 0.09882902 | 0.1087473 | -0.0099183 | 0.66455182 | 0.89054584 |
| Mpzl1 | 1.22449838 | 0.93087837 | 0.94079822 | -0.0099198 | 0.51958985 | 0.89054584 |
| Gm9776 | 0.03114638 | 0.02339407 | 0.03332937 | -0.0099353 | 0.91755385 | 0.93505805 |

| | | | | | | |
|---|---|---|---|---|---|---|
| Spsb3 | 0.22754838 | 0.22803711 | 0.23797255 | -0.0099354 | 0.57711274 | 0.89054584 |
| Rexo2 | 0.84840573 | 0.73930268 | 0.74923884 | -0.0099362 | 0.52463312 | 0.89054584 |
| Zfp276 | 0.08836484 | 0.0845489 | 0.0944929 | -0.009944 | 0.68803404 | 0.89054584 |
| Ppp1r13l | 0.03199925 | 0.0242864 | 0.03424185 | -0.0099555 | 0.91221752 | 0.93174592 |
| Slc25a15 | 0.05748576 | 0.05153085 | 0.06150107 | -0.0099702 | 0.77496025 | 0.89054584 |
| Acadl | 0.67666331 | 0.62287332 | 0.63284645 | -0.0099731 | 0.52926443 | 0.89054584 |
| Wdr45b | 0.76327084 | 0.68420989 | 0.69418563 | -0.0099757 | 0.52668962 | 0.89054584 |
| Hus1 | 0.09540581 | 0.09202766 | 0.10200799 | -0.0099803 | 0.67574199 | 0.89054584 |
| 2700033N17 | 0.13061505 | 0.12895683 | 0.13895434 | -0.0099975 | 0.63126375 | 0.89054584 |
| Snx29 | 0.13606226 | 0.13462076 | 0.14461922 | -0.0099985 | 0.62631248 | 0.89054584 |
| D630045J12I | 0.06340671 | 0.05781534 | 0.06783071 | -0.0100154 | 0.75424339 | 0.89054584 |
| Arl15 | 0.10628108 | 0.10348424 | 0.11351278 | -0.0100285 | 0.6596854 | 0.89054584 |
| Ubiad1 | 0.05343286 | 0.04713674 | 0.05716763 | -0.0100309 | 0.79317259 | 0.89054584 |
| Mgat5 | 0.18100379 | 0.18087601 | 0.19091136 | -0.0100353 | 0.59673287 | 0.89054584 |
| Rps29 | 5.40513944 | 0.65916042 | 0.66920938 | -0.010049 | 0.52788703 | 0.89054584 |
| Rab40b | 0.12784336 | 0.12601042 | 0.13606753 | -0.0100571 | 0.63471021 | 0.89054584 |
| Eif4b | 1.0732028 | 0.86194305 | 0.87200198 | -0.0100589 | 0.52143001 | 0.89054584 |
| Gpr27 | 0.07247111 | 0.06745764 | 0.07751778 | -0.0100601 | 0.72728769 | 0.89054584 |
| Stxbp5 | 0.09438188 | 0.0908406 | 0.10091531 | -0.0100747 | 0.67907631 | 0.89054584 |
| Sacm1l | 0.41208069 | 0.40428991 | 0.41437055 | -0.0100806 | 0.54511984 | 0.89054584 |
| Xrn1 | 0.42022935 | 0.41165922 | 0.42174398 | -0.0100848 | 0.5443523 | 0.89054584 |
| Slc10a3 | 0.1011801 | 0.09805879 | 0.10815193 | -0.0100931 | 0.6681424 | 0.89054584 |
| Epb41l4a | 0.17379232 | 0.17344076 | 0.18353794 | -0.0100972 | 0.60114333 | 0.89054584 |
| Coa4 | 0.20587077 | 0.20607194 | 0.21617028 | -0.0100983 | 0.58614568 | 0.89054584 |
| Tmem87a | 0.23911564 | 0.23942 | 0.24952109 | -0.0101011 | 0.5747979 | 0.89054584 |
| Mms22l | 0.05907541 | 0.0530907 | 0.06320059 | -0.0101099 | 0.77194509 | 0.89054584 |
| Slmap | 0.41589906 | 0.40769947 | 0.41783001 | -0.0101305 | 0.54496841 | 0.89054584 |
| Sep-03 | 0.04599114 | 0.03907519 | 0.04920931 | -0.0101341 | 0.83133597 | 0.89319023 |
| Nck2 | 0.10559851 | 0.10265573 | 0.11279598 | -0.0101403 | 0.66233147 | 0.89054584 |
| 2310015A10I | 0.19991485 | 0.20000292 | 0.2101446 | -0.0101417 | 0.58894925 | 0.89054584 |

| | | | | | | |
|---|---|---|---|---|---|---|
| Lrrn2 | 0.04967188 | 0.04298349 | 0.05314579 | -0.0101623 | 0.81349847 | 0.89054584 |
| Chn1 | 0.14546162 | 0.14419523 | 0.15436711 | -0.0101719 | 0.62057699 | 0.89054584 |
| Clip4 | 0.03341521 | 0.02558103 | 0.03575675 | -0.0101757 | 0.90755523 | 0.92849733 |
| 0610040B10 | 0.08648442 | 0.08230445 | 0.09248531 | -0.0101809 | 0.6959754 | 0.89054584 |
| Ncapd3 | 0.23348595 | 0.2336906 | 0.24390784 | -0.0102172 | 0.5773669 | 0.89054584 |
| Mknk2 | 0.16151679 | 0.16071608 | 0.17093787 | -0.0102218 | 0.60972276 | 0.89054584 |
| Gm16740 | 0.04275087 | 0.03551333 | 0.04574356 | -0.0102302 | 0.85128022 | 0.89968367 |
| G730003C15 | 0.08056465 | 0.07592489 | 0.0861638 | -0.0102389 | 0.71008243 | 0.89054584 |
| L2hgdh | 0.10316422 | 0.09999627 | 0.11023825 | -0.010242 | 0.66743873 | 0.89054584 |
| Chst9 | 0.05918242 | 0.0530719 | 0.06331499 | -0.0102431 | 0.77449212 | 0.89054584 |
| Mok | 0.07150489 | 0.06623892 | 0.07648537 | -0.0102465 | 0.73373232 | 0.89054584 |
| Hnf4a | 0.13108364 | 0.12918976 | 0.1394421 | -0.0102523 | 0.63402914 | 0.89054584 |
| Eny2 | 1.61317738 | 1.05350377 | 1.06375794 | -0.0102542 | 0.51791055 | 0.89054584 |
| Ankrd42 | 0.05205334 | 0.04543597 | 0.05569249 | -0.0102565 | 0.80455545 | 0.89054584 |
| Iqcc | 0.22657226 | 0.22673708 | 0.23699535 | -0.0102583 | 0.57990907 | 0.89054584 |
| Tbc1d13 | 0.06337218 | 0.05750977 | 0.06779381 | -0.010284 | 0.76013655 | 0.89054584 |
| Grk5 | 0.13414738 | 0.13233816 | 0.14262912 | -0.010291 | 0.63162161 | 0.89054584 |
| Bnip1 | 0.35412995 | 0.35056726 | 0.3608678 | -0.0103005 | 0.55289697 | 0.89054584 |
| Mak16 | 0.47554866 | 0.46047553 | 0.4707785 | -0.010303 | 0.54060606 | 0.89054584 |
| Zfp27 | 0.07454056 | 0.06942308 | 0.07972883 | -0.0103057 | 0.72648434 | 0.89054584 |
| Nle1 | 0.1494036 | 0.14813489 | 0.15844493 | -0.01031 | 0.61911541 | 0.89054584 |
| Dis3l | 0.13421891 | 0.13239217 | 0.14270348 | -0.0103113 | 0.63180608 | 0.89054584 |
| Ap3b1 | 0.51048015 | 0.49048315 | 0.50079565 | -0.0103125 | 0.53821509 | 0.89054584 |
| Tlcd1 | 0.13065991 | 0.12868434 | 0.13900104 | -0.0103167 | 0.63524895 | 0.89054584 |
| Mkks | 0.48379792 | 0.46761539 | 0.47793464 | -0.0103192 | 0.54006314 | 0.89054584 |
| Rfx2 | 0.0564294 | 0.05004055 | 0.06037165 | -0.0103311 | 0.78735051 | 0.89054584 |
| Stradb | 0.19708806 | 0.19694802 | 0.20727938 | -0.0103314 | 0.59181415 | 0.89054584 |
| Cldn23 | 0.07273975 | 0.06747148 | 0.07780482 | -0.0103333 | 0.73196245 | 0.89054584 |
| Mterf2 | 0.08785007 | 0.08360164 | 0.09394335 | -0.0103417 | 0.69598133 | 0.89054584 |
| Dpf3 | 0.09381368 | 0.08996668 | 0.10030894 | -0.0103423 | 0.68451283 | 0.89054584 |

| | | | | | | |
|---|---|---|---|---|---|---|
| Pvrl4 | 0.0405326 | 0.0330192 | 0.04337076 | -0.0103516 | 0.86692597 | 0.90430169 |
| Me2 | 0.22815206 | 0.22822455 | 0.23857669 | -0.0103521 | 0.58010313 | 0.89054584 |
| Stard13 | 0.09046379 | 0.08637604 | 0.09673353 | -0.0103575 | 0.69105527 | 0.89054584 |
| Pml | 0.07087712 | 0.06545615 | 0.07581458 | -0.0103584 | 0.73778917 | 0.89054584 |
| Lrrc1 | 0.08951491 | 0.08535717 | 0.09572064 | -0.0103635 | 0.69301465 | 0.89054584 |
| Fam35a | 0.11199939 | 0.10914657 | 0.11951111 | -0.0103645 | 0.65691001 | 0.89054584 |
| Glcci1 | 0.04065741 | 0.03313727 | 0.04350427 | -0.010367 | 0.8665476 | 0.90430169 |
| Tom1 | 0.12225588 | 0.11985449 | 0.13023948 | -0.010385 | 0.64486487 | 0.89054584 |
| Pxk | 0.31834648 | 0.31654141 | 0.32693184 | -0.0103904 | 0.55885574 | 0.89054584 |
| Flnb | 0.19106568 | 0.1907602 | 0.20116365 | -0.0104034 | 0.59520052 | 0.89054584 |
| Cx3cl1 | 0.06626641 | 0.06048283 | 0.07088728 | -0.0104044 | 0.7529537 | 0.89054584 |
| Slc39a14 | 0.04749555 | 0.04040612 | 0.05081829 | -0.0104122 | 0.83010283 | 0.89271879 |
| Armcx5 | 0.15797979 | 0.1568726 | 0.16729597 | -0.0104234 | 0.61419709 | 0.89054584 |
| Camk1g | 0.05959082 | 0.05332357 | 0.0637516 | -0.010428 | 0.77699275 | 0.89054584 |
| Pced1b | 0.07690322 | 0.07182363 | 0.08225287 | -0.0104292 | 0.72264934 | 0.89054584 |
| Lrrc3 | 0.0475249 | 0.04041925 | 0.05084968 | -0.0104304 | 0.83037674 | 0.89271879 |
| Mars2 | 0.06092736 | 0.05474901 | 0.06518041 | -0.0104314 | 0.77204892 | 0.89054584 |
| Ddx56 | 0.35529939 | 0.3515323 | 0.36196548 | -0.0104332 | 0.55341256 | 0.89054584 |
| Phlpp1 | 0.05874537 | 0.05241405 | 0.06284775 | -0.0104337 | 0.78037394 | 0.89054584 |
| Aarsd1 | 0.06684503 | 0.06106913 | 0.07150568 | -0.0104365 | 0.75174508 | 0.89054584 |
| Enpp1 | 0.04092011 | 0.03334872 | 0.04378527 | -0.0104366 | 0.86660696 | 0.90430169 |
| Oscp1 | 0.19789992 | 0.19766485 | 0.20810263 | -0.0104378 | 0.59238257 | 0.89054584 |
| Lpin1 | 0.11495833 | 0.11216592 | 0.12261021 | -0.0104443 | 0.65426546 | 0.89054584 |
| Tusc1 | 0.03022032 | 0.02189153 | 0.03233858 | -0.0104471 | 0.93384844 | 0.94691109 |
| Spata1 | 0.05849176 | 0.05211661 | 0.06257661 | -0.01046 | 0.78194123 | 0.89054584 |
| Cebpg | 0.30247722 | 0.30119944 | 0.31167187 | -0.0104724 | 0.562198 | 0.89054584 |
| Zbtb22 | 0.04808739 | 0.04097403 | 0.05145126 | -0.0104772 | 0.8286184 | 0.89245298 |
| Mtmr9 | 0.07648979 | 0.07133231 | 0.08181122 | -0.0104789 | 0.72466192 | 0.89054584 |
| Rgl2 | 0.38678395 | 0.38075676 | 0.39124245 | -0.0104857 | 0.54968451 | 0.89054584 |
| Trim68 | 0.06604972 | 0.06016298 | 0.07065568 | -0.0104927 | 0.75549201 | 0.89054584 |

| | | | | | | |
|---|---|---|---|---|---|---|
| Zfp746 | 0.16680977 | 0.1658745 | 0.17637835 | -0.0105039 | 0.60928417 | 0.89054584 |
| Tmem17 | 0.14989168 | 0.14844185 | 0.15894941 | -0.0105076 | 0.62095334 | 0.89054584 |
| Tead1 | 0.34593327 | 0.34264085 | 0.35315337 | -0.0105125 | 0.55515061 | 0.89054584 |
| 1700037C18I | 0.11109545 | 0.10804977 | 0.11856371 | -0.0105139 | 0.66024923 | 0.89054584 |
| Fam57a | 0.16844985 | 0.16752557 | 0.17806182 | -0.0105363 | 0.60860243 | 0.89054584 |
| RP23-60P18. | 0.15692562 | 0.15567281 | 0.16620956 | -0.0105367 | 0.61613696 | 0.89054584 |
| BC052040 | 0.06055874 | 0.05424796 | 0.06478635 | -0.0105384 | 0.77572691 | 0.89054584 |
| Rab2b | 0.31039861 | 0.30876534 | 0.31930435 | -0.010539 | 0.56110598 | 0.89054584 |
| Rab11fip3 | 0.04670026 | 0.03942643 | 0.04996773 | -0.0105413 | 0.83716188 | 0.89483776 |
| Sult4a1 | 0.06915782 | 0.06343397 | 0.07397734 | -0.0105434 | 0.74665484 | 0.89054584 |
| Mrpl50 | 0.8508353 | 0.74018692 | 0.75073263 | -0.0105457 | 0.52609023 | 0.89054584 |
| Sema4b | 0.19174349 | 0.19130308 | 0.20185275 | -0.0105497 | 0.59618933 | 0.89054584 |
| Ssfa2 | 0.09294783 | 0.08882205 | 0.09938487 | -0.0105628 | 0.6897569 | 0.89054584 |
| G6pdx | 0.12791995 | 0.12557539 | 0.13614734 | -0.0105719 | 0.64123822 | 0.89054584 |
| Yars | 0.38331032 | 0.37746205 | 0.38803916 | -0.0105771 | 0.55052693 | 0.89054584 |
| Ank3 | 0.47475438 | 0.45950775 | 0.47008726 | -0.0105795 | 0.54175314 | 0.89054584 |
| BC049715 | 0.08419397 | 0.0794538 | 0.09003967 | -0.0105859 | 0.70806715 | 0.89054584 |
| Dbt | 0.17902208 | 0.17829467 | 0.18888712 | -0.0105924 | 0.60305593 | 0.89054584 |
| Ttll3 | 0.10181079 | 0.09821703 | 0.10881526 | -0.0105982 | 0.67500098 | 0.89054584 |
| Zfp251 | 0.08371829 | 0.0789299 | 0.08953172 | -0.0106018 | 0.70941783 | 0.89054584 |
| Fhod3 | 0.04532607 | 0.03789012 | 0.04849798 | -0.0106079 | 0.8459082 | 0.89825055 |
| Ncald | 0.23802929 | 0.23783084 | 0.24843905 | -0.0106082 | 0.57884284 | 0.89054584 |
| D630036H23 | 0.03521451 | 0.02706755 | 0.03768171 | -0.0106142 | 0.90529524 | 0.92670836 |
| 9030622O22 | 0.0422341 | 0.03456981 | 0.0451908 | -0.010621 | 0.86323499 | 0.90287732 |
| Rcl1 | 0.28546196 | 0.28453715 | 0.2951704 | -0.0106332 | 0.56664309 | 0.89054584 |
| Zfp523 | 0.23853099 | 0.23829471 | 0.24893882 | -0.0106441 | 0.57894945 | 0.89054584 |
| B4galt6 | 0.38889608 | 0.38252253 | 0.39318696 | -0.0106644 | 0.55027838 | 0.89054584 |
| Csgalnact2 | 0.14972491 | 0.14810038 | 0.15877705 | -0.0106767 | 0.62296671 | 0.89054584 |
| Lhfpl5 | 0.04940309 | 0.04216218 | 0.05285834 | -0.0106962 | 0.82709838 | 0.89186561 |
| Zfp710 | 0.15096931 | 0.14936563 | 0.16006289 | -0.0106973 | 0.62223827 | 0.89054584 |

| | | | | | | |
|---|---|---|---|---|---|---|
| Prdm16 | 0.19911641 | 0.19863581 | 0.20933566 | -0.0106999 | 0.59411149 | 0.89054584 |
| Ptges2 | 0.1951562 | 0.19461865 | 0.20531928 | -0.0107006 | 0.59592378 | 0.89054584 |
| Nkapl | 0.04194705 | 0.03417817 | 0.04488376 | -0.0107056 | 0.86676788 | 0.90430169 |
| Orai1 | 0.11193398 | 0.10871679 | 0.11944257 | -0.0107258 | 0.66216046 | 0.89054584 |
| Traf2 | 0.22502734 | 0.22471462 | 0.23544782 | -0.0107332 | 0.58409624 | 0.89054584 |
| Samd5 | 0.11153991 | 0.10826431 | 0.11902957 | -0.0107653 | 0.66325522 | 0.89054584 |
| Ptcd3 | 0.26536081 | 0.26473005 | 0.27549606 | -0.010766 | 0.57223418 | 0.89054584 |
| Whrn | 0.04029487 | 0.03234707 | 0.04311647 | -0.0107694 | 0.87772425 | 0.91005122 |
| Nabp1 | 0.11869971 | 0.11574785 | 0.12652415 | -0.0107763 | 0.65424659 | 0.89054584 |
| Lrig2 | 0.0798696 | 0.07464286 | 0.08542145 | -0.0107786 | 0.72169067 | 0.89054584 |
| Nek9 | 0.22725067 | 0.22689271 | 0.23767456 | -0.0107819 | 0.58369214 | 0.89054584 |
| Mdm4 | 0.47935911 | 0.4633003 | 0.47408923 | -0.0107889 | 0.54221849 | 0.89054584 |
| Dexi | 0.13247097 | 0.13008167 | 0.1408857 | -0.010804 | 0.63955807 | 0.89054584 |
| Sep-08 | 0.19944262 | 0.19885819 | 0.20966619 | -0.010808 | 0.59489768 | 0.89054584 |
| Ppfia3 | 0.03760538 | 0.02942611 | 0.04023943 | -0.0108133 | 0.89470998 | 0.91983444 |
| Arsa | 0.13212585 | 0.12969426 | 0.14052665 | -0.0108324 | 0.64025142 | 0.89054584 |
| Ss18 | 0.62857563 | 0.5856713 | 0.59652799 | -0.0108567 | 0.53378652 | 0.89054584 |
| Erlin1 | 0.15842441 | 0.15689206 | 0.16775405 | -0.010862 | 0.61854596 | 0.89054584 |
| Fan1 | 0.05248723 | 0.04529396 | 0.05615646 | -0.0108625 | 0.8162568 | 0.89054584 |
| 5031439G07 | 0.15101206 | 0.14924303 | 0.16010705 | -0.010864 | 0.62404857 | 0.89054584 |
| Mospd3 | 0.43189431 | 0.42136782 | 0.43223295 | -0.0108651 | 0.54661467 | 0.89054584 |
| Thsd4 | 0.07490763 | 0.06924725 | 0.08012099 | -0.0108737 | 0.73640313 | 0.89054584 |
| Ano6 | 0.30156132 | 0.2999118 | 0.31078733 | -0.0108755 | 0.56475361 | 0.89054584 |
| Faxc | 0.04513159 | 0.03741093 | 0.04828997 | -0.010879 | 0.85304738 | 0.90004228 |
| Unc13b | 0.12321831 | 0.12035509 | 0.13124417 | -0.0108891 | 0.65045295 | 0.89054584 |
| Nup210 | 0.10892441 | 0.10538412 | 0.11628705 | -0.0109029 | 0.66887778 | 0.89054584 |
| Ccdc125 | 0.05349289 | 0.04632535 | 0.05723182 | -0.0109065 | 0.81268609 | 0.89054584 |
| Myo5b | 0.0352139 | 0.02676532 | 0.03768106 | -0.0109157 | 0.91143397 | 0.9310858 |
| Stat3 | 0.77286421 | 0.68973538 | 0.70065173 | -0.0109164 | 0.52893286 | 0.89054584 |
| Polr1e | 0.10814426 | 0.10454926 | 0.11546852 | -0.0109193 | 0.67023846 | 0.89054584 |

| | | | | | | |
|---|---|---|---|---|---|---|
| Esrra | 0.05274358 | 0.04550107 | 0.05643058 | -0.0109295 | 0.81656354 | 0.89054584 |
| Gpr137 | 0.18076916 | 0.17973889 | 0.19067177 | -0.0109329 | 0.60531852 | 0.89054584 |
| Zc2hc1c | 0.10970256 | 0.10616969 | 0.11710325 | -0.0109336 | 0.66821541 | 0.89054584 |
| 0610010K14I | 0.65568795 | 0.60626283 | 0.6171998 | -0.010937 | 0.53289844 | 0.89054584 |
| Map4k2 | 0.13523352 | 0.13281575 | 0.1437581 | -0.0109423 | 0.63856352 | 0.89054584 |
| Rnf19a | 0.276542 | 0.27551759 | 0.28646397 | -0.0109464 | 0.5706494 | 0.89054584 |
| H2-Q4 | 0.21180611 | 0.21121055 | 0.2221605 | -0.0109499 | 0.59081112 | 0.89054584 |
| 4930578M01 | 0.0444082 | 0.03656486 | 0.04751626 | -0.0109514 | 0.85853889 | 0.90124516 |
| Gabrb3 | 0.04587864 | 0.03813015 | 0.04908898 | -0.0109588 | 0.85084903 | 0.89942466 |
| Frmd5 | 0.073057 | 0.06717871 | 0.07814378 | -0.0109651 | 0.74336001 | 0.89054584 |
| Lypla2 | 0.43889136 | 0.42751372 | 0.43848615 | -0.0109724 | 0.54640466 | 0.89054584 |
| Tmem201 | 0.07987993 | 0.07444649 | 0.08543248 | -0.010986 | 0.72544889 | 0.89054584 |
| Alg13 | 0.07584482 | 0.07012292 | 0.08112221 | -0.0109993 | 0.73620916 | 0.89054584 |
| Abhd17b | 0.24549254 | 0.24486051 | 0.25586169 | -0.0110012 | 0.57938425 | 0.89054584 |
| Trim3 | 0.06913828 | 0.06294625 | 0.07395646 | -0.0110102 | 0.7560322 | 0.89054584 |
| Mterf4 | 0.27049888 | 0.26949692 | 0.28054364 | -0.0110467 | 0.57277791 | 0.89054584 |
| Lin7c | 0.70911851 | 0.6453982 | 0.6564533 | -0.0110551 | 0.53126877 | 0.89054584 |
| Celsr1 | 0.03669174 | 0.02820312 | 0.03926205 | -0.0110589 | 0.90528762 | 0.92670836 |
| Large | 0.06149546 | 0.05472805 | 0.0657877 | -0.0110597 | 0.78325114 | 0.89054584 |
| Schip1.1 | 0.07021644 | 0.06404379 | 0.07510859 | -0.0110648 | 0.75375169 | 0.89054584 |
| Tmem43 | 0.15033486 | 0.14833463 | 0.15940739 | -0.0110728 | 0.62688688 | 0.89054584 |
| Pdlim5 | 0.11031365 | 0.10667083 | 0.11774408 | -0.0110732 | 0.66936131 | 0.89054584 |
| Hexdc | 0.05187047 | 0.04441338 | 0.05549693 | -0.0110836 | 0.82393324 | 0.89111407 |
| Trim21 | 0.04476442 | 0.03681063 | 0.04789726 | -0.0110866 | 0.85957014 | 0.90131997 |
| 4930550C14I | 0.06998131 | 0.06376614 | 0.07485734 | -0.0110912 | 0.754994 | 0.89054584 |
| Ube2t | 0.09015392 | 0.08531124 | 0.09640276 | -0.0110915 | 0.70403101 | 0.89054584 |
| Padi2 | 0.0556227 | 0.04841669 | 0.05950913 | -0.0110924 | 0.80741815 | 0.89054584 |
| Ptpn12 | 0.21261399 | 0.21187966 | 0.22297465 | -0.011095 | 0.59166268 | 0.89054584 |
| Slx4ip | 0.15098578 | 0.14897808 | 0.16007991 | -0.0111018 | 0.62669244 | 0.89054584 |
| Gm26615 | 0.03345195 | 0.02469007 | 0.03579605 | -0.011106 | 0.92583698 | 0.9418176 |

| | | | | | | |
|---|---|---|---|---|---|---|
| Nhsl1 | 0.13780706 | 0.13532429 | 0.14643131 | -0.011107 | 0.63810086 | 0.89054584 |
| Myo18a | 0.33775885 | 0.33431558 | 0.34542406 | -0.0111085 | 0.5595493 | 0.89054584 |
| Cdkn1b | 0.31761765 | 0.31512339 | 0.32623375 | -0.0111104 | 0.56303433 | 0.89054584 |
| Hacd3 | 0.43582983 | 0.42463474 | 0.43575366 | -0.0111189 | 0.5473148 | 0.89054584 |
| Tmem18 | 0.13388372 | 0.13122725 | 0.14235499 | -0.0111277 | 0.64213979 | 0.89054584 |
| Kcnh6 | 0.05646945 | 0.049261 | 0.06041447 | -0.0111535 | 0.80513698 | 0.89054584 |
| Dis3l2 | 0.16393194 | 0.16225375 | 0.17342174 | -0.011168 | 0.61792124 | 0.89054584 |
| Zscan12 | 0.08210368 | 0.07663125 | 0.08780748 | -0.0111762 | 0.72340778 | 0.89054584 |
| Tab3 | 0.06661166 | 0.06007617 | 0.07125627 | -0.0111801 | 0.7676065 | 0.89054584 |
| Vps11 | 0.19364228 | 0.19260032 | 0.2037821 | -0.0111818 | 0.60088628 | 0.89054584 |
| Arhgef7 | 0.20586152 | 0.2049653 | 0.21616093 | -0.0111956 | 0.595339 | 0.89054584 |
| Arl6 | 0.5483801 | 0.52130522 | 0.53250297 | -0.0111977 | 0.53902219 | 0.89054584 |
| Haus6 | 0.0649595 | 0.05828951 | 0.06949044 | -0.0112009 | 0.77366137 | 0.89054584 |
| Zfp60 | 0.16355781 | 0.16181976 | 0.17303712 | -0.0112174 | 0.61868239 | 0.89054584 |
| Sgms2 | 0.22836924 | 0.22757325 | 0.238794 | -0.0112207 | 0.58664386 | 0.89054584 |
| Neto2 | 0.08722307 | 0.08204959 | 0.09327394 | -0.0112244 | 0.71248036 | 0.89054584 |
| Smg7 | 0.2029218 | 0.20195154 | 0.21318863 | -0.0112371 | 0.59699282 | 0.89054584 |
| Dok1 | 0.13257516 | 0.12975292 | 0.14099409 | -0.0112412 | 0.64484793 | 0.89054584 |
| Pno1 | 0.48097361 | 0.46424366 | 0.47548931 | -0.0112456 | 0.54386951 | 0.89054584 |
| Tdrd7 | 0.17124259 | 0.16965924 | 0.1809259 | -0.0112667 | 0.61413879 | 0.89054584 |
| Bcr | 0.16238061 | 0.16055631 | 0.17182654 | -0.0112702 | 0.62003907 | 0.89054584 |
| 2610044O15 | 0.11467094 | 0.11102911 | 0.12230936 | -0.0112802 | 0.66628609 | 0.89054584 |
| Prdm15 | 0.13791831 | 0.13525737 | 0.14654682 | -0.0112894 | 0.64016811 | 0.89054584 |
| Utp20 | 0.06905123 | 0.06255791 | 0.07386343 | -0.0113055 | 0.76210695 | 0.89054584 |
| Map3k3 | 0.06323864 | 0.05634225 | 0.06765106 | -0.0113088 | 0.78195322 | 0.89054584 |
| Slc9a1 | 0.08149437 | 0.07584672 | 0.08715675 | -0.01131 | 0.72726916 | 0.89054584 |
| Btrc | 0.55341377 | 0.52533179 | 0.53664598 | -0.0113142 | 0.53912327 | 0.89054584 |
| Ermp1 | 0.06865194 | 0.06211961 | 0.07343673 | -0.0113171 | 0.76361378 | 0.89054584 |
| Wfs1 | 0.39086566 | 0.3836649 | 0.39499842 | -0.0113335 | 0.55317101 | 0.89054584 |
| Mettl15 | 0.07841858 | 0.07253803 | 0.08387158 | -0.0113336 | 0.73550943 | 0.89054584 |

| | | | | | | |
|---|---|---|---|---|---|---|
| Ankmy2 | 0.23358724 | 0.23266071 | 0.24400897 | -0.0113483 | 0.58576953 | 0.89054584 |
| Gm26890 | 0.14496881 | 0.1425075 | 0.15385688 | -0.0113494 | 0.63445303 | 0.89054584 |
| Rps6ka6 | 0.04597635 | 0.03782514 | 0.04919349 | -0.0113683 | 0.85918299 | 0.90131997 |
| Plekha6 | 0.13342025 | 0.13049848 | 0.14187305 | -0.0113746 | 0.64562197 | 0.89054584 |
| Rsph3b | 0.06740853 | 0.06072674 | 0.07210791 | -0.0113812 | 0.76893525 | 0.89054584 |
| Mtfp1 | 0.21069444 | 0.20965726 | 0.22103972 | -0.0113825 | 0.59480097 | 0.89054584 |
| Farp1 | 0.15869556 | 0.156641 | 0.16803338 | -0.0113924 | 0.6239487 | 0.89054584 |
| 4632404H12l | 0.05080932 | 0.04296579 | 0.05436217 | -0.0113964 | 0.83563901 | 0.89459411 |
| Bid | 0.11304355 | 0.10920732 | 0.1206051 | -0.0113978 | 0.67013753 | 0.89054584 |
| Pcnxl4 | 0.16047507 | 0.15846191 | 0.16986578 | -0.0114039 | 0.62277455 | 0.89054584 |
| Klhl18 | 0.05612813 | 0.04864304 | 0.06004954 | -0.0114065 | 0.81191422 | 0.89054584 |
| Mar-11 | 0.05627822 | 0.04879814 | 0.06021001 | -0.0114119 | 0.81138989 | 0.89054584 |
| Mdh1 | 2.03949202 | 1.1087843 | 1.12019674 | -0.0114124 | 0.51892859 | 0.89054584 |
| Megf9 | 0.1831393 | 0.18167748 | 0.19309084 | -0.0114134 | 0.60848947 | 0.89054584 |
| Nyap1 | 0.11110097 | 0.10715254 | 0.1185695 | -0.011417 | 0.67314182 | 0.89054584 |
| Heatr5b | 0.1005406 | 0.09605711 | 0.10747918 | -0.0114221 | 0.68974111 | 0.89054584 |
| Lnx2 | 0.08336061 | 0.07772714 | 0.08914977 | -0.0114226 | 0.72472735 | 0.89054584 |
| Sybu | 0.04258807 | 0.0341441 | 0.04556942 | -0.0114253 | 0.87861806 | 0.91059794 |
| Slc25a46 | 0.5091267 | 0.48821547 | 0.49964666 | -0.0114312 | 0.5424428 | 0.89054584 |
| St5 | 0.14337562 | 0.14076271 | 0.15220674 | -0.011444 | 0.63694087 | 0.89054584 |
| Ephx4 | 0.05726896 | 0.04981735 | 0.06126929 | -0.0114519 | 0.80807101 | 0.89054584 |
| Atxn7l2 | 0.09224505 | 0.08718041 | 0.0986348 | -0.0114544 | 0.70575988 | 0.89054584 |
| Zfp433 | 0.18794172 | 0.18652745 | 0.19798517 | -0.0114577 | 0.60627387 | 0.89054584 |
| Lonrf1 | 0.04259688 | 0.034118 | 0.04557884 | -0.0114608 | 0.87930001 | 0.91103349 |
| Zfp711 | 0.06690637 | 0.06010658 | 0.07157124 | -0.0114647 | 0.77225506 | 0.89054584 |
| Syt9 | 0.05012219 | 0.04216147 | 0.05362735 | -0.0114659 | 0.84040224 | 0.89576177 |
| Ppp3cc | 0.07902964 | 0.07305425 | 0.08452428 | -0.01147 | 0.73637957 | 0.89054584 |
| Chrna1os | 0.05080574 | 0.04288086 | 0.05435834 | -0.0114775 | 0.83737112 | 0.89485554 |
| Lpgat1 | 0.33556165 | 0.33186033 | 0.34334061 | -0.0114803 | 0.5618972 | 0.89054584 |
| Rbm27 | 0.61756315 | 0.57650568 | 0.58798967 | -0.011484 | 0.53625112 | 0.89054584 |

| | | | | | | |
|---|---|---|---|---|---|---|
| Rac1 | 1.69772854 | 1.06953497 | 1.08102564 | -0.0114907 | 0.51974828 | 0.89054584 |
| Tmem251 | 0.34522085 | 0.34098856 | 0.35248115 | -0.0114926 | 0.56036871 | 0.89054584 |
| Etaa1os | 0.04715675 | 0.03896321 | 0.05045595 | -0.0114927 | 0.85570115 | 0.9009252 |
| BC024978 | 0.09329426 | 0.08826056 | 0.0997546 | -0.011494 | 0.70430326 | 0.89054584 |
| Nnt | 0.25345276 | 0.25225173 | 0.26374998 | -0.0114983 | 0.58047464 | 0.89054584 |
| Ift172 | 0.09477616 | 0.08982894 | 0.10133607 | -0.0115071 | 0.70161187 | 0.89054584 |
| Zfp105 | 0.12713223 | 0.12381519 | 0.1353264 | -0.0115112 | 0.6540586 | 0.89054584 |
| Banp | 0.56238944 | 0.53248046 | 0.54399232 | -0.0115119 | 0.53926874 | 0.89054584 |
| 1700056E22I | 0.20416286 | 0.20292201 | 0.21444384 | -0.0115218 | 0.59882995 | 0.89054584 |
| Trim46 | 0.07720599 | 0.07104895 | 0.08257631 | -0.0115274 | 0.74227379 | 0.89054584 |
| Frg1 | 0.9141747 | 0.77662471 | 0.78816426 | -0.0115396 | 0.52719148 | 0.89054584 |
| Gm16136 | 0.03216049 | 0.02285533 | 0.03441437 | -0.011559 | 0.94118937 | 0.95209067 |
| Cacna1c | 0.06674501 | 0.05983859 | 0.07139878 | -0.0115602 | 0.77467448 | 0.89054584 |
| Coil | 0.17230332 | 0.17045107 | 0.18201288 | -0.0115618 | 0.61636397 | 0.89054584 |
| Prkd3 | 0.20131046 | 0.19998962 | 0.21155791 | -0.0115683 | 0.60054479 | 0.89054584 |
| Senp8 | 0.13078594 | 0.12753683 | 0.13913223 | -0.0115954 | 0.65109571 | 0.89054584 |
| Zfp180 | 0.06155603 | 0.05423354 | 0.06585245 | -0.0116189 | 0.79446629 | 0.89054584 |
| Dcaf4 | 0.13501575 | 0.13189952 | 0.14353177 | -0.0116323 | 0.64712678 | 0.89054584 |
| Appbp2 | 0.76841252 | 0.68602203 | 0.69765951 | -0.0116375 | 0.53097244 | 0.89054584 |
| Gm4799 | 0.04185772 | 0.03314847 | 0.04478821 | -0.0116397 | 0.88701107 | 0.91505129 |
| Rnf19b | 0.06662543 | 0.05961911 | 0.07127098 | -0.0116519 | 0.77687315 | 0.89054584 |
| Rsf1 | 0.8911337 | 0.7632127 | 0.77488214 | -0.0116694 | 0.52796762 | 0.89054584 |
| Ccdc73 | 0.05669815 | 0.04896918 | 0.06065899 | -0.0116898 | 0.81536489 | 0.89054584 |
| Ambra1 | 0.16299558 | 0.1607684 | 0.17245902 | -0.0116906 | 0.6239302 | 0.89054584 |
| Wbscr16 | 0.22351052 | 0.22222971 | 0.2339274 | -0.0116977 | 0.59210892 | 0.89054584 |
| Zfp12 | 0.09529801 | 0.09019324 | 0.10189295 | -0.0116997 | 0.70365997 | 0.89054584 |
| Klf16 | 0.04867497 | 0.04035717 | 0.05207965 | -0.0117225 | 0.85283495 | 0.89994913 |
| Sltm | 1.09309777 | 0.87008218 | 0.88182342 | -0.0117412 | 0.52473154 | 0.89054584 |
| Pls1 | 0.06911873 | 0.06219323 | 0.07393557 | -0.0117423 | 0.77032759 | 0.89054584 |
| Plag1 | 0.15376227 | 0.1511953 | 0.16294679 | -0.0117515 | 0.63156307 | 0.89054584 |

| | | | | | | |
|---|---|---|---|---|---|---|
| Gm42469 | 0.03812779 | 0.0290456 | 0.04079828 | -0.0117527 | 0.91021697 | 0.93043261 |
| Slc27a3 | 0.05075671 | 0.04254839 | 0.05430591 | -0.0117575 | 0.84343738 | 0.89774283 |
| Nlgn2 | 0.12706719 | 0.12349318 | 0.1352586 | -0.0117654 | 0.65735547 | 0.89054584 |
| Rhod | 0.11054672 | 0.10622107 | 0.11798844 | -0.0117674 | 0.67890946 | 0.89054584 |
| Myh9 | 0.49228169 | 0.47348235 | 0.48525027 | -0.0117679 | 0.5449788 | 0.89054584 |
| Dhcr7 | 0.31149682 | 0.30858417 | 0.32036022 | -0.011776 | 0.56798914 | 0.89054584 |
| Ccdc59 | 1.06983498 | 0.85853388 | 0.87031637 | -0.0117825 | 0.52514602 | 0.89054584 |
| Vars | 0.3868426 | 0.37950993 | 0.39129648 | -0.0117865 | 0.55580257 | 0.89054584 |
| Rab39b | 0.1592089 | 0.15677411 | 0.1685621 | -0.011788 | 0.62771663 | 0.89054584 |
| Oxsm | 0.06563629 | 0.05841337 | 0.07021381 | -0.0118004 | 0.78318689 | 0.89054584 |
| Nab1 | 0.15611762 | 0.15356754 | 0.16537654 | -0.011809 | 0.63031195 | 0.89054584 |
| Per2 | 0.09728582 | 0.09220126 | 0.10401401 | -0.0118128 | 0.70163565 | 0.89054584 |
| Ccdc186 | 0.37689807 | 0.37029278 | 0.38210823 | -0.0118154 | 0.55727443 | 0.89054584 |
| Eif2ak1 | 0.25228358 | 0.25076723 | 0.26259324 | -0.011826 | 0.58309495 | 0.89054584 |
| Pes1 | 0.36437524 | 0.35862829 | 0.37045963 | -0.0118313 | 0.55914123 | 0.89054584 |
| Fip1l1 | 1.15601638 | 0.89932298 | 0.91115514 | -0.0118322 | 0.52412148 | 0.89054584 |
| Srgap3 | 0.07038649 | 0.06345508 | 0.07529031 | -0.0118352 | 0.76802565 | 0.89054584 |
| Wdr77 | 0.45346121 | 0.43956527 | 0.45141694 | -0.0118517 | 0.54867623 | 0.89054584 |
| Zfp961 | 0.12761492 | 0.12396368 | 0.13582947 | -0.0118658 | 0.65799449 | 0.89054584 |
| Evpl | 0.06429097 | 0.05690633 | 0.06877589 | -0.0118696 | 0.78931734 | 0.89054584 |
| Fbxw4 | 0.04793992 | 0.03941177 | 0.05129355 | -0.0118818 | 0.85975281 | 0.90132889 |
| Rps6ka4 | 0.15548561 | 0.15283476 | 0.16472478 | -0.01189 | 0.63167306 | 0.89054584 |
| Nsmaf | 0.1534815 | 0.15076494 | 0.162657 | -0.0118921 | 0.63330558 | 0.89054584 |
| 1190005I06R | 0.03458254 | 0.02511224 | 0.03700561 | -0.0118934 | 0.93284946 | 0.94636145 |
| Usp33 | 0.40661723 | 0.39751257 | 0.40940632 | -0.0118938 | 0.5538316 | 0.89054584 |
| Axin2 | 0.0586352 | 0.05082109 | 0.06272997 | -0.0119089 | 0.81177863 | 0.89054584 |
| Bdnf | 0.08510449 | 0.0790994 | 0.09101192 | -0.0119125 | 0.72900519 | 0.89054584 |
| Zfp689 | 0.08351766 | 0.07740169 | 0.08931748 | -0.0119158 | 0.73287994 | 0.89054584 |
| Rps6ka3 | 0.28243643 | 0.28029991 | 0.29222145 | -0.0119215 | 0.57537179 | 0.89054584 |
| Ddt | 0.50835436 | 0.48706189 | 0.49899046 | -0.0119286 | 0.54434008 | 0.89054584 |

| | | | | | | |
|---|---|---|---|---|---|---|
| Tfap4 | 0.1818158 | 0.17978292 | 0.19174032 | -0.0119574 | 0.61429955 | 0.89054584 |
| Dnajc4 | 0.30915262 | 0.30614284 | 0.31810585 | -0.011963 | 0.56954226 | 0.89054584 |
| Dab2ip | 0.24111962 | 0.23955169 | 0.2515157 | -0.011964 | 0.58769308 | 0.89054584 |
| Ttc9 | 0.04108888 | 0.03199962 | 0.04396581 | -0.0119662 | 0.89760568 | 0.9219206 |
| A430105I19F | 0.11280711 | 0.10838703 | 0.12035741 | -0.0119704 | 0.67844834 | 0.89054584 |
| Ddx26b | 0.15245774 | 0.14962381 | 0.16160009 | -0.0119763 | 0.63505687 | 0.89054584 |
| Cox19 | 0.58889475 | 0.55339953 | 0.56537954 | -0.01198 | 0.53931963 | 0.89054584 |
| Ankrd49 | 0.0985582 | 0.09338733 | 0.10537155 | -0.0119842 | 0.70190068 | 0.89054584 |
| Klhl15 | 0.09533657 | 0.08993929 | 0.10193411 | -0.0119948 | 0.70823233 | 0.89054584 |
| Sgms1 | 0.33579552 | 0.33155914 | 0.34356246 | -0.0120033 | 0.56465148 | 0.89054584 |
| Tcaim | 0.05687262 | 0.04883992 | 0.06084554 | -0.0120056 | 0.82102272 | 0.89054584 |
| Zmiz1 | 0.56673341 | 0.53552145 | 0.54752872 | -0.0120073 | 0.54068925 | 0.89054584 |
| Ube3c | 0.46913064 | 0.45316379 | 0.465182 | -0.0120182 | 0.54790341 | 0.89054584 |
| Zc3h8 | 0.10955187 | 0.10491447 | 0.11694521 | -0.0120307 | 0.68413332 | 0.89054584 |
| Ldlrap1 | 0.10406193 | 0.09913912 | 0.11118174 | -0.0120426 | 0.69309125 | 0.89054584 |
| Pi4ka | 0.20194488 | 0.20015622 | 0.21220009 | -0.0120439 | 0.60427503 | 0.89054584 |
| Bcap31 | 1.18509102 | 0.9119783 | 0.92402264 | -0.0120443 | 0.524212 | 0.89054584 |
| Noc3l | 0.15900802 | 0.15629478 | 0.16835522 | -0.0120604 | 0.63071658 | 0.89054584 |
| 1600014C10I | 0.14178861 | 0.13849513 | 0.150562 | -0.0120669 | 0.64557212 | 0.89054584 |
| Tmx2 | 0.63264696 | 0.58757151 | 0.59966444 | -0.0120929 | 0.53742672 | 0.89054584 |
| Mapk8 | 0.32742142 | 0.3235071 | 0.33560102 | -0.0120939 | 0.56666606 | 0.89054584 |
| Exog | 0.16134108 | 0.15865477 | 0.17075706 | -0.0121023 | 0.6293753 | 0.89054584 |
| Slx4 | 0.05460461 | 0.04631248 | 0.05842056 | -0.0121081 | 0.83287746 | 0.89393134 |
| Zc3hc1 | 0.23297531 | 0.23128284 | 0.24339797 | -0.0121151 | 0.59169166 | 0.89054584 |
| Gm10399 | 0.08326255 | 0.07692877 | 0.08904505 | -0.0121163 | 0.73693934 | 0.89054584 |
| Lyrm1 | 0.07446569 | 0.0675225 | 0.07964884 | -0.0121263 | 0.76093463 | 0.89054584 |
| Zfp92 | 0.04494566 | 0.03595493 | 0.04809111 | -0.0121362 | 0.88014689 | 0.91145498 |
| Polg2 | 0.09910236 | 0.09381466 | 0.10595208 | -0.0121374 | 0.70322842 | 0.89054584 |
| Spata13 | 0.14392432 | 0.14063584 | 0.15277517 | -0.0121393 | 0.64438096 | 0.89054584 |
| Pygl | 0.12740298 | 0.12346508 | 0.13560859 | -0.0121435 | 0.66173395 | 0.89054584 |

| | | | | | | |
|---|---|---|---|---|---|---|
| Plch1 | 0.07751989 | 0.07076653 | 0.08291161 | -0.0121451 | 0.75252346 | 0.89054584 |
| Nr6a1 | 0.05626893 | 0.04803885 | 0.06020008 | -0.0121612 | 0.82669088 | 0.89183454 |
| Mast3 | 0.2221735 | 0.22040396 | 0.23258634 | -0.0121824 | 0.59639445 | 0.89054584 |
| Colgalt1 | 0.44268529 | 0.42968122 | 0.44186493 | -0.0121837 | 0.55110863 | 0.89054584 |
| Tmem194 | 0.04418801 | 0.03508536 | 0.04728074 | -0.0121954 | 0.88526134 | 0.91420299 |
| Fastkd2 | 0.11820159 | 0.11380241 | 0.12600336 | -0.0122009 | 0.67405096 | 0.89054584 |
| Lonp1 | 0.45489799 | 0.44048327 | 0.45268527 | -0.012202 | 0.54996795 | 0.89054584 |
| Fhit | 0.10308473 | 0.09795174 | 0.1101547 | -0.012203 | 0.69711412 | 0.89054584 |
| 2900060B14 | 0.03640697 | 0.02674762 | 0.0389574 | -0.0122098 | 0.92787908 | 0.94303412 |
| Vps33b | 0.09753832 | 0.09207333 | 0.10428342 | -0.0122101 | 0.70729351 | 0.89054584 |
| Cdk5r2 | 0.05643695 | 0.04816071 | 0.06037972 | -0.012219 | 0.82711527 | 0.89186561 |
| Taco1 | 0.05078836 | 0.04211778 | 0.05433976 | -0.012222 | 0.85265272 | 0.89988783 |
| Slc35a3 | 0.13918618 | 0.13562966 | 0.14786279 | -0.0122331 | 0.65004728 | 0.89054584 |
| Kri1 | 0.13966177 | 0.13611483 | 0.14835627 | -0.0122414 | 0.64966898 | 0.89054584 |
| Tfb1m | 0.12621757 | 0.12212074 | 0.13437286 | -0.0122521 | 0.66450966 | 0.89054584 |
| Sgtb | 0.08086084 | 0.07421813 | 0.08648014 | -0.012262 | 0.74556253 | 0.89054584 |
| Fam83h | 0.06741654 | 0.0598316 | 0.07211647 | -0.0122849 | 0.7862973 | 0.89054584 |
| Ric3 | 0.11572811 | 0.11111997 | 0.12341592 | -0.012296 | 0.67873743 | 0.89054584 |
| Snrk | 0.51032417 | 0.4883552 | 0.50066329 | -0.0123081 | 0.54559253 | 0.89054584 |
| Bcl2l2 | 0.15011494 | 0.14686742 | 0.15918013 | -0.0123127 | 0.64071577 | 0.89054584 |
| Zbtb34 | 0.079111 | 0.07229838 | 0.08461119 | -0.0123128 | 0.75110759 | 0.89054584 |
| Cyp4f13 | 0.15991379 | 0.15697152 | 0.16928795 | -0.0123164 | 0.63267936 | 0.89054584 |
| Zfp174 | 0.09468459 | 0.08891771 | 0.10123836 | -0.0123206 | 0.71463961 | 0.89054584 |
| Speg | 0.07001038 | 0.06256732 | 0.0748884 | -0.0123211 | 0.77831557 | 0.89054584 |
| Gm9774 | 0.14802019 | 0.14468922 | 0.15701455 | -0.0123253 | 0.64271207 | 0.89054584 |
| Pgap3 | 0.04894386 | 0.04000715 | 0.05236722 | -0.0123601 | 0.86425453 | 0.90325563 |
| Bscl2 | 0.36815696 | 0.36162636 | 0.37398645 | -0.0123601 | 0.56118564 | 0.89054584 |
| Nadk2 | 0.14377032 | 0.1402488 | 0.15261564 | -0.0123668 | 0.64710867 | 0.89054584 |
| Recql5 | 0.06454753 | 0.05667964 | 0.06905011 | -0.0123705 | 0.79804514 | 0.89054584 |
| Pskh1 | 0.22689116 | 0.22493811 | 0.23731466 | -0.0123765 | 0.59598775 | 0.89054584 |

| | | | | | | |
|---|---|---|---|---|---|---|
| Spef1 | 0.03696637 | 0.02717691 | 0.03955584 | -0.0123789 | 0.92758073 | 0.94292984 |
| Xylt1 | 0.06813 | 0.06049983 | 0.07287895 | -0.0123791 | 0.78563348 | 0.89054584 |
| Plcg1 | 0.11205758 | 0.10719191 | 0.11957209 | -0.0123802 | 0.68522998 | 0.89054584 |
| Rbm4 | 0.10151415 | 0.09611157 | 0.10850329 | -0.0123917 | 0.70266419 | 0.89054584 |
| Scrn2 | 0.14682656 | 0.14337609 | 0.15577978 | -0.0124037 | 0.64466634 | 0.89054584 |
| Psmd14 | 0.84832044 | 0.73677549 | 0.74918632 | -0.0124108 | 0.53075935 | 0.89054584 |
| Ruvbl1 | 0.55057906 | 0.52188915 | 0.53431488 | -0.0124257 | 0.5431393 | 0.89054584 |
| Slc25a28 | 0.14685474 | 0.14338255 | 0.15580894 | -0.0124264 | 0.64489323 | 0.89054584 |
| Zfand1 | 0.24377999 | 0.24173347 | 0.25416074 | -0.0124273 | 0.59009929 | 0.89054584 |
| Lpcat1 | 0.22416689 | 0.22215097 | 0.23458546 | -0.0124345 | 0.59752763 | 0.89054584 |
| Nupl1 | 0.07666491 | 0.06956019 | 0.08199829 | -0.0124381 | 0.7601243 | 0.89054584 |
| Mapk9 | 0.18976733 | 0.18738813 | 0.19984314 | -0.012455 | 0.61423094 | 0.89054584 |
| Vwa9 | 0.27437425 | 0.27188371 | 0.28434231 | -0.0124586 | 0.58087552 | 0.89054584 |
| Srsf7 | 2.17281081 | 1.11399496 | 1.12646743 | -0.0124725 | 0.52057019 | 0.89054584 |
| Vps37a | 0.78371338 | 0.69539769 | 0.70788381 | -0.0124861 | 0.5327472 | 0.89054584 |
| Txnl4b | 0.04097765 | 0.03135134 | 0.04384683 | -0.0124955 | 0.90786255 | 0.92866393 |
| Map3k14 | 0.04408383 | 0.0346663 | 0.04716931 | -0.012503 | 0.8915717 | 0.91758405 |
| Nphp3 | 0.05389148 | 0.04515246 | 0.05765803 | -0.0125056 | 0.8438691 | 0.89775379 |
| Maml1 | 0.08067286 | 0.07375906 | 0.08627937 | -0.0125203 | 0.75050383 | 0.89054584 |
| Cacna1h | 0.14378852 | 0.14010973 | 0.15263449 | -0.0125248 | 0.64888046 | 0.89054584 |
| Fndc5 | 0.06230022 | 0.05411535 | 0.06664796 | -0.0125326 | 0.8095057 | 0.89054584 |
| Zswim6 | 0.08204909 | 0.07521128 | 0.08774918 | -0.0125379 | 0.74719393 | 0.89054584 |
| Baiap2l1 | 0.30814049 | 0.30458938 | 0.31713179 | -0.0125424 | 0.57309481 | 0.89054584 |
| Cenpj | 0.07058712 | 0.06296195 | 0.0755047 | -0.0125427 | 0.78051609 | 0.89054584 |
| Pigl | 0.11272111 | 0.10771146 | 0.12026732 | -0.0125559 | 0.68665517 | 0.89054584 |
| Fam118b | 0.25556019 | 0.25325143 | 0.26583338 | -0.012582 | 0.5872625 | 0.89054584 |
| Mon1a | 0.07522951 | 0.06788102 | 0.08046487 | -0.0125838 | 0.76687943 | 0.89054584 |
| G630016G05 | 0.12220373 | 0.11758347 | 0.13018504 | -0.0126016 | 0.67399561 | 0.89054584 |
| Rb1 | 0.10015361 | 0.09446378 | 0.10707201 | -0.0126082 | 0.70836414 | 0.89054584 |
| Ifrd1 | 0.42459773 | 0.41306852 | 0.4256813 | -0.0126128 | 0.55489664 | 0.89054584 |

| | | | | | | |
|---|---|---|---|---|---|---|
| Cbx7 | 0.07222931 | 0.0646381 | 0.07725942 | -0.0126213 | 0.7767007 | 0.89054584 |
| Lrrfip2 | 0.45539055 | 0.44049627 | 0.45311979 | -0.0126235 | 0.5516353 | 0.89054584 |
| Rgs19 | 0.27777169 | 0.27504288 | 0.2876665 | -0.0126236 | 0.58099807 | 0.89054584 |
| Rpl7a-ps5 | 0.29314451 | 0.29001328 | 0.3026389 | -0.0126256 | 0.57705446 | 0.89054584 |
| Gm10110 | 0.07231791 | 0.06472227 | 0.07735409 | -0.0126318 | 0.7766114 | 0.89054584 |
| Gcfc2 | 0.11261996 | 0.1075232 | 0.12016134 | -0.0126381 | 0.68793693 | 0.89054584 |
| Brf2 | 0.11070828 | 0.10551029 | 0.11815783 | -0.0126475 | 0.69100111 | 0.89054584 |
| Zfp65 | 0.07612832 | 0.06877291 | 0.08142508 | -0.0126522 | 0.76544425 | 0.89054584 |
| Plekhg1 | 0.09217101 | 0.08590081 | 0.09855578 | -0.012655 | 0.72515595 | 0.89054584 |
| Slc23a2 | 0.07212469 | 0.06448861 | 0.07714764 | -0.012659 | 0.77770886 | 0.89054584 |
| Prkdc | 0.08180756 | 0.07480699 | 0.08749123 | -0.0126842 | 0.75030195 | 0.89054584 |
| Nrd1 | 1.03625403 | 0.8405007 | 0.85318981 | -0.0126891 | 0.52762074 | 0.89054584 |
| Cgrrf1 | 0.22204641 | 0.21976043 | 0.23245882 | -0.0126984 | 0.60044582 | 0.89054584 |
| Zfp597 | 0.08646916 | 0.07977054 | 0.09246902 | -0.0126985 | 0.73884792 | 0.89054584 |
| Mblac2 | 0.14170512 | 0.13777327 | 0.15047546 | -0.0127022 | 0.652943 | 0.89054584 |
| 9530034E10l | 0.08781172 | 0.0811945 | 0.0939024 | -0.0127079 | 0.73581596 | 0.89054584 |
| Zbtb40 | 0.10874412 | 0.10338867 | 0.11609791 | -0.0127092 | 0.69497937 | 0.89054584 |
| Dus3l | 0.25701003 | 0.25454971 | 0.26726546 | -0.0127157 | 0.58771052 | 0.89054584 |
| Kcnk5 | 0.05335608 | 0.0443642 | 0.05708553 | -0.0127213 | 0.85041861 | 0.89942466 |
| Rpl36al | 3.5834636 | 1.01109078 | 1.02382302 | -0.0127322 | 0.52310118 | 0.89054584 |
| Fbxo7 | 0.17962307 | 0.17676594 | 0.18950118 | -0.0127352 | 0.6228974 | 0.89054584 |
| Ptprk | 0.10894934 | 0.10356919 | 0.1163132 | -0.012744 | 0.69513648 | 0.89054584 |
| Ulk4 | 0.07419643 | 0.06661645 | 0.07936117 | -0.0127447 | 0.772826 | 0.89054584 |
| Zfp692 | 0.05868398 | 0.05003601 | 0.06278211 | -0.0127461 | 0.82789103 | 0.89229532 |
| Pnck | 0.12962586 | 0.12515196 | 0.1379244 | -0.0127724 | 0.66692569 | 0.89054584 |
| Agpat1 | 0.27922436 | 0.27629434 | 0.28908612 | -0.0127918 | 0.58166453 | 0.89054584 |
| 2610008E11l | 0.20610305 | 0.20360428 | 0.216405 | -0.0128007 | 0.6085663 | 0.89054584 |
| Zfp40 | 0.06761045 | 0.05951537 | 0.0723237 | -0.0128083 | 0.79533852 | 0.89054584 |
| Fbxo30 | 0.1645043 | 0.16119461 | 0.17401004 | -0.0128154 | 0.63424598 | 0.89054584 |
| A430033K04 | 0.10194848 | 0.09614205 | 0.10896005 | -0.012818 | 0.70817997 | 0.89054584 |

| | | | | | | |
|---|---|---|---|---|---|---|
| Tdrd3 | 0.26408955 | 0.26142352 | 0.27424522 | -0.0128217 | 0.58621459 | 0.89054584 |
| Cdk14 | 0.2490307 | 0.24654962 | 0.25937155 | -0.0128219 | 0.5910757 | 0.89054584 |
| Srf | 0.1548461 | 0.15124219 | 0.16406513 | -0.0128229 | 0.64212011 | 0.89054584 |
| Zfp280b | 0.11575239 | 0.11061807 | 0.12344133 | -0.0128233 | 0.68579905 | 0.89054584 |
| Nprl3 | 0.09506349 | 0.08880995 | 0.10164269 | -0.0128327 | 0.7218035 | 0.89054584 |
| Prps1l3 | 0.05390874 | 0.04483975 | 0.05767648 | -0.0128367 | 0.85011213 | 0.89917019 |
| Lhfpl4 | 0.08277119 | 0.07568294 | 0.08852033 | -0.0128374 | 0.75036815 | 0.89054584 |
| Mtmr1 | 0.1313674 | 0.12689492 | 0.13973743 | -0.0128425 | 0.66573927 | 0.89054584 |
| Akap10 | 0.13735801 | 0.13310872 | 0.14596506 | -0.0128563 | 0.65922573 | 0.89054584 |
| Abca2 | 0.08570136 | 0.07879206 | 0.09164923 | -0.0128572 | 0.74331124 | 0.89054584 |
| Phf21b | 0.06414225 | 0.05575757 | 0.06861692 | -0.0128594 | 0.80870186 | 0.89054584 |
| Sh3bgrl2 | 0.05318195 | 0.04403669 | 0.05689933 | -0.0128626 | 0.85387501 | 0.90035231 |
| Slc39a6 | 0.40885768 | 0.39857564 | 0.41144413 | -0.0128685 | 0.55792678 | 0.89054584 |
| Pip4k2b | 0.10201252 | 0.09615875 | 0.10902741 | -0.0128687 | 0.70880652 | 0.89054584 |
| Ndufb11 | 2.84070837 | 1.09465326 | 1.10752278 | -0.0128695 | 0.52158712 | 0.89054584 |
| Ttc27 | 0.08920531 | 0.08251008 | 0.09539014 | -0.0128801 | 0.73534973 | 0.89054584 |
| Mrpl21 | 1.07493204 | 0.85998045 | 0.87286451 | -0.0128841 | 0.52741328 | 0.89054584 |
| Wdpcp | 0.05000648 | 0.04061035 | 0.05350362 | -0.0128933 | 0.86921945 | 0.90542007 |
| Donson | 0.14174723 | 0.13761074 | 0.15051911 | -0.0129084 | 0.65525456 | 0.89054584 |
| Ctdsp1 | 0.31985208 | 0.31546035 | 0.32837308 | -0.0129127 | 0.57268149 | 0.89054584 |
| Spast | 0.1937225 | 0.19094766 | 0.20386358 | -0.0129159 | 0.61606809 | 0.89054584 |
| Stoml1 | 0.1347532 | 0.13032723 | 0.14325888 | -0.0129316 | 0.66295882 | 0.89054584 |
| Ccdc18 | 0.09503468 | 0.08867258 | 0.10161195 | -0.0129394 | 0.72350121 | 0.89054584 |
| Gusb | 0.60033874 | 0.56152918 | 0.57447001 | -0.0129408 | 0.54179222 | 0.89054584 |
| Dip2c | 0.10358368 | 0.09770142 | 0.11067913 | -0.0129777 | 0.70756512 | 0.89054584 |
| Pop1 | 0.06785624 | 0.05960862 | 0.07258639 | -0.0129778 | 0.7975707 | 0.89054584 |
| Ncoa2 | 0.38923222 | 0.38051794 | 0.39349627 | -0.0129783 | 0.5610617 | 0.89054584 |
| Mettl6 | 0.33830423 | 0.33294638 | 0.34594085 | -0.0129945 | 0.56946112 | 0.89054584 |
| Chd1l | 0.07266749 | 0.0647268 | 0.07772761 | -0.0130008 | 0.78208679 | 0.89054584 |
| Fbxo4 | 0.15352632 | 0.14970055 | 0.16270326 | -0.0130027 | 0.6451754 | 0.89054584 |

| | | | | | | |
|---|---|---|---|---|---|---|
| Ikzf5 | 0.23490017 | 0.23231473 | 0.24531929 | -0.0130046 | 0.59753585 | 0.89054584 |
| BC037034 | 0.06911335 | 0.06092116 | 0.07392982 | -0.0130087 | 0.79383079 | 0.89054584 |
| Mbip | 0.25773338 | 0.25496908 | 0.26797963 | -0.0130105 | 0.5894747 | 0.89054584 |
| Zfp429 | 0.07910962 | 0.0715988 | 0.08460972 | -0.0130109 | 0.76313767 | 0.89054584 |
| Yars2 | 0.26667372 | 0.26376936 | 0.27678705 | -0.0130177 | 0.58672044 | 0.89054584 |
| Ddx19b | 0.09898645 | 0.09278877 | 0.10582843 | -0.0130397 | 0.71703368 | 0.89054584 |
| Cish | 0.12149693 | 0.11640629 | 0.12944697 | -0.0130407 | 0.68058974 | 0.89054584 |
| Gm17484 | 0.06887234 | 0.06061957 | 0.07367227 | -0.0130527 | 0.79543804 | 0.89054584 |
| Capn1 | 0.04880277 | 0.03916259 | 0.05221633 | -0.0130537 | 0.87793083 | 0.91020031 |
| Macf1 | 0.62164614 | 0.5780956 | 0.5911649 | -0.0130693 | 0.54101794 | 0.89054584 |
| Dip2a | 0.1021791 | 0.09613243 | 0.10920257 | -0.0130701 | 0.71144635 | 0.89054584 |
| Alkbh3 | 0.23335117 | 0.23069771 | 0.24377327 | -0.0130756 | 0.59866663 | 0.89054584 |
| Aqp4 | 0.06448762 | 0.05590377 | 0.06898607 | -0.0130823 | 0.81151886 | 0.89054584 |
| Ptk2b | 0.11141682 | 0.10580893 | 0.11890057 | -0.0130916 | 0.696016 | 0.89054584 |
| Il2 | 0.08876932 | 0.08182226 | 0.09492471 | -0.0131024 | 0.73991166 | 0.89054584 |
| Ighmbp2 | 0.05884216 | 0.04984047 | 0.06295123 | -0.0131108 | 0.83405423 | 0.89427341 |
| Fgd1 | 0.1320888 | 0.12736941 | 0.1404881 | -0.0131187 | 0.66823658 | 0.89054584 |
| 4931414P19I | 0.13939819 | 0.13496315 | 0.14808279 | -0.0131196 | 0.66011111 | 0.89054584 |
| Bag5 | 0.54602723 | 0.51741883 | 0.530561 | -0.0131422 | 0.54593736 | 0.89054584 |
| Zfp827 | 0.12987413 | 0.12503981 | 0.13818293 | -0.0131431 | 0.67116428 | 0.89054584 |
| 5730508B09I | 0.09075924 | 0.08388869 | 0.0970489 | -0.0131602 | 0.73623164 | 0.89054584 |
| Tspan2 | 0.08258508 | 0.07515853 | 0.08832159 | -0.0131631 | 0.75626803 | 0.89054584 |
| Tada2b | 0.09230619 | 0.08553594 | 0.09870005 | -0.0131641 | 0.73282719 | 0.89054584 |
| Khdrbs3 | 0.18557197 | 0.18240654 | 0.19557128 | -0.0131647 | 0.62309264 | 0.89054584 |
| Patl1 | 0.16466429 | 0.161001 | 0.17417446 | -0.0131735 | 0.6377207 | 0.89054584 |
| Mettl22 | 0.10446257 | 0.09842362 | 0.11160272 | -0.0131791 | 0.70889911 | 0.89054584 |
| Pcbd2 | 0.75683814 | 0.67662551 | 0.68981284 | -0.0131873 | 0.53548534 | 0.89054584 |
| Klhl26 | 0.09226236 | 0.08544445 | 0.09865327 | -0.0132088 | 0.73361784 | 0.89054584 |
| Nckipsd | 0.05102645 | 0.04137762 | 0.05459437 | -0.0132167 | 0.87031534 | 0.90589084 |
| Tom1l2 | 0.06709299 | 0.05855364 | 0.07177068 | -0.013217 | 0.80454676 | 0.89054584 |

| | | | | | | |
|---|---|---|---|---|---|---|
| Pemt | 0.05807242 | 0.04890467 | 0.06212829 | -0.0132236 | 0.83931005 | 0.89504222 |
| Senp1 | 0.20943123 | 0.20654104 | 0.21976552 | -0.0132245 | 0.61039689 | 0.89054584 |
| Arhgef9 | 0.14702766 | 0.14276239 | 0.15598786 | -0.0132255 | 0.65358204 | 0.89054584 |
| Zfpm1 | 0.07072353 | 0.06242096 | 0.07565046 | -0.0132295 | 0.79238943 | 0.89054584 |
| Arpc4 | 1.36836096 | 0.9817642 | 0.99499848 | -0.0132343 | 0.5247057 | 0.89054584 |
| Ercc8 | 0.11795942 | 0.11250193 | 0.12575013 | -0.0132482 | 0.68822856 | 0.89054584 |
| Pex11g | 0.10837873 | 0.10246323 | 0.11571454 | -0.0132513 | 0.70316638 | 0.89054584 |
| Ap1s3 | 0.20423906 | 0.20126569 | 0.21452089 | -0.0132552 | 0.61327876 | 0.89054584 |
| A930011G23 | 0.07868081 | 0.0708862 | 0.08415169 | -0.0132655 | 0.7686551 | 0.89054584 |
| Lysmd1 | 0.07727707 | 0.0693839 | 0.08265223 | -0.0132683 | 0.77274256 | 0.89054584 |
| Tm2d1 | 1.11908527 | 0.88097901 | 0.89424943 | -0.0132704 | 0.52755988 | 0.89054584 |
| Gm15446 | 0.11757355 | 0.11207339 | 0.12534661 | -0.0132732 | 0.68911558 | 0.89054584 |
| Neil2 | 0.06278909 | 0.05389153 | 0.06717054 | -0.013279 | 0.82148271 | 0.89054584 |
| Dnal1 | 0.21045141 | 0.20751471 | 0.22079463 | -0.0132799 | 0.61034447 | 0.89054584 |
| Sbf1 | 0.12187616 | 0.11656268 | 0.12984299 | -0.0132803 | 0.68314643 | 0.89054584 |
| Ctsa | 0.6863819 | 0.62669693 | 0.63999284 | -0.0132959 | 0.53855338 | 0.89054584 |
| Usf1 | 0.20556943 | 0.20256577 | 0.21586574 | -0.0133 | 0.61296206 | 0.89054584 |
| Cdc42bpg | 0.0877911 | 0.08057916 | 0.09388039 | -0.0133012 | 0.74540241 | 0.89054584 |
| Pde6d | 0.85913151 | 0.74249657 | 0.75580073 | -0.0133042 | 0.53268066 | 0.89054584 |
| AI480526 | 0.1385819 | 0.13392422 | 0.14723566 | -0.0133114 | 0.66319936 | 0.89054584 |
| Lrrc8b | 0.05333528 | 0.0437509 | 0.05706329 | -0.0133124 | 0.86145765 | 0.90221867 |
| Prdx1 | 3.42938498 | 1.03071272 | 1.04405551 | -0.0133428 | 0.52373909 | 0.89054584 |
| Cox10 | 0.16742702 | 0.16366777 | 0.17701207 | -0.0133443 | 0.63728866 | 0.89054584 |
| Tmem143 | 0.05967807 | 0.05048943 | 0.06384488 | -0.0133555 | 0.83511695 | 0.89459411 |
| Gorasp1 | 0.18629361 | 0.18295017 | 0.19630662 | -0.0133565 | 0.62437381 | 0.89054584 |
| C2cd2l | 0.13063744 | 0.12561677 | 0.13897765 | -0.0133609 | 0.6728859 | 0.89054584 |
| Mar-09 | 0.07364093 | 0.06540027 | 0.07876767 | -0.0133674 | 0.78542686 | 0.89054584 |
| BC003965 | 0.47581943 | 0.45764593 | 0.47101405 | -0.0133681 | 0.55259805 | 0.89054584 |
| Ddx39 | 0.75182459 | 0.67301556 | 0.68638453 | -0.013369 | 0.53615198 | 0.89054584 |
| Limk1 | 0.11670543 | 0.11106735 | 0.12443855 | -0.0133712 | 0.69167839 | 0.89054584 |

| | | | | | | |
|---|---|---|---|---|---|---|
| Ube2a | 0.7838118 | 0.69457224 | 0.70794903 | -0.0133768 | 0.53507411 | 0.89054584 |
| 2310040G24 | 0.15706357 | 0.15295484 | 0.16635175 | -0.0133969 | 0.64624473 | 0.89054584 |
| Ift46 | 0.42099922 | 0.40904046 | 0.42243868 | -0.0133982 | 0.55873575 | 0.89054584 |
| Clec16a | 0.0697728 | 0.061236 | 0.07463452 | -0.0133985 | 0.79852975 | 0.89054584 |
| Slc35d1 | 0.06531896 | 0.05647191 | 0.06987464 | -0.0134027 | 0.81424175 | 0.89054584 |
| Wbscr27 | 0.24476789 | 0.24172637 | 0.25514212 | -0.0134158 | 0.59676174 | 0.89054584 |
| Nbeal2 | 0.11473028 | 0.10893828 | 0.12237147 | -0.0134332 | 0.695475 | 0.89054584 |
| Ppm1b | 0.70012637 | 0.63655222 | 0.64998677 | -0.0134346 | 0.53835705 | 0.89054584 |
| Plcb3 | 0.10315329 | 0.09678937 | 0.11022677 | -0.0134374 | 0.7149678 | 0.89054584 |
| Phc1 | 0.21603329 | 0.21297863 | 0.22641729 | -0.0134387 | 0.60892732 | 0.89054584 |
| Mmab | 0.08830908 | 0.08098818 | 0.09443338 | -0.0134452 | 0.74644007 | 0.89054584 |
| Fbxo44 | 0.14198699 | 0.13732138 | 0.15076766 | -0.0134463 | 0.66111551 | 0.89054584 |
| Sh2d5 | 0.08207382 | 0.07432628 | 0.08777559 | -0.0134493 | 0.76234329 | 0.89054584 |
| Shc2 | 0.12053867 | 0.11499599 | 0.12844601 | -0.01345 | 0.68717318 | 0.89054584 |
| Eml1 | 0.18559961 | 0.1821414 | 0.19559946 | -0.0134581 | 0.62572603 | 0.89054584 |
| Zfp703 | 0.26222009 | 0.25893865 | 0.2724044 | -0.0134658 | 0.59107272 | 0.89054584 |
| Lmo7 | 0.11026442 | 0.10422613 | 0.11769246 | -0.0134663 | 0.70300893 | 0.89054584 |
| 1810014B01 | 0.06068337 | 0.05145294 | 0.06491957 | -0.0134666 | 0.83308549 | 0.89393134 |
| Plekha3 | 0.10525948 | 0.09896855 | 0.1124399 | -0.0134713 | 0.7116408 | 0.89054584 |
| Ap1g1 | 0.28871539 | 0.28486493 | 0.29833676 | -0.0134718 | 0.58331492 | 0.89054584 |
| Mpp2 | 0.11992494 | 0.11432265 | 0.12780476 | -0.0134821 | 0.68845332 | 0.89054584 |
| Hnf1a | 0.08079359 | 0.07291651 | 0.08640832 | -0.0134918 | 0.7665242 | 0.89054584 |
| Rfxap | 0.209198 | 0.2060234 | 0.21953018 | -0.0135068 | 0.61280803 | 0.89054584 |
| Ttc9c | 0.35961743 | 0.35250243 | 0.36601212 | -0.0135097 | 0.56826706 | 0.89054584 |
| Huwe1 | 0.73234765 | 0.6593572 | 0.67289424 | -0.013537 | 0.53733693 | 0.89054584 |
| Tes | 0.62736186 | 0.58205152 | 0.59559073 | -0.0135392 | 0.54217271 | 0.89054584 |
| Ahctf1 | 0.29278145 | 0.28874253 | 0.30228662 | -0.0135441 | 0.58267668 | 0.89054584 |
| Nsun3 | 0.07155404 | 0.06299205 | 0.0765379 | -0.0135459 | 0.79518644 | 0.89054584 |
| Slc5a1 | 0.05302426 | 0.04317994 | 0.05673071 | -0.0135508 | 0.86711149 | 0.90441493 |
| Rpl17 | 5.12578561 | 0.66674081 | 0.68029854 | -0.0135577 | 0.53698801 | 0.89054584 |

| | | | | | | |
|---|---|---|---|---|---|---|
| Rhou | 0.17375825 | 0.16994524 | 0.18350306 | -0.0135578 | 0.63465887 | 0.89054584 |
| Secisbp2l | 0.21042201 | 0.20719119 | 0.22076498 | -0.0135738 | 0.61273561 | 0.89054584 |
| Myo1d | 0.06193487 | 0.05268179 | 0.06625742 | -0.0135756 | 0.83010522 | 0.89271879 |
| Flcn | 0.13489985 | 0.12983426 | 0.14341131 | -0.0135771 | 0.67041958 | 0.89054584 |
| Cep350 | 0.24928522 | 0.24604038 | 0.25962381 | -0.0135834 | 0.5962894 | 0.89054584 |
| Slc27a1 | 0.09808723 | 0.09128142 | 0.10486907 | -0.0135877 | 0.72696142 | 0.89054584 |
| Trpc1 | 0.23186707 | 0.2286953 | 0.242291 | -0.0135957 | 0.60311865 | 0.89054584 |
| Cmtm4 | 0.33358468 | 0.32785994 | 0.34146416 | -0.0136042 | 0.57362709 | 0.89054584 |
| Zfp626 | 0.04466367 | 0.03417172 | 0.04778949 | -0.0136178 | 0.90784163 | 0.92866393 |
| Aars2 | 0.08601316 | 0.07836296 | 0.09198215 | -0.0136192 | 0.75484587 | 0.89054584 |
| Zfp292 | 0.81574563 | 0.7151117 | 0.72874085 | -0.0136291 | 0.5347171 | 0.89054584 |
| Haus5 | 0.10467233 | 0.09817182 | 0.11182309 | -0.0136513 | 0.71524041 | 0.89054584 |
| Gm6483 | 0.86958807 | 0.74846542 | 0.76211757 | -0.0136522 | 0.53325619 | 0.89054584 |
| Mpv17l | 0.08685205 | 0.07922494 | 0.09287783 | -0.0136529 | 0.75328279 | 0.89054584 |
| Rnpc3 | 0.31632192 | 0.31133799 | 0.32499198 | -0.013654 | 0.57759218 | 0.89054584 |
| Nptxr | 0.12015364 | 0.11438318 | 0.12804373 | -0.0136606 | 0.69042195 | 0.89054584 |
| Arhgap44 | 0.05722095 | 0.04755447 | 0.06121796 | -0.0136635 | 0.85079485 | 0.89942466 |
| Micall1 | 0.06820561 | 0.05929478 | 0.07295976 | -0.013665 | 0.80855809 | 0.89054584 |
| Pigb | 0.08542734 | 0.07768458 | 0.09135665 | -0.0136721 | 0.75717324 | 0.89054584 |
| Brms1 | 0.29907112 | 0.29470086 | 0.30838025 | -0.0136794 | 0.5818643 | 0.89054584 |
| Tsacc | 0.10088646 | 0.09415339 | 0.10784304 | -0.0136897 | 0.72287666 | 0.89054584 |
| Lrp12 | 0.11217762 | 0.10600817 | 0.11969788 | -0.0136897 | 0.70292708 | 0.89054584 |
| Copb2 | 1.51043754 | 1.02470079 | 1.03839465 | -0.0136939 | 0.52449558 | 0.89054584 |
| Acp2 | 0.23220221 | 0.22893041 | 0.24262582 | -0.0136954 | 0.60371761 | 0.89054584 |
| Natd1 | 0.10627905 | 0.09981368 | 0.11351065 | -0.013697 | 0.71300316 | 0.89054584 |
| Inha | 0.05097094 | 0.04082764 | 0.054535 | -0.0137074 | 0.87920552 | 0.91100069 |
| Il6ra | 0.1566935 | 0.15226215 | 0.16597027 | -0.0137081 | 0.64980718 | 0.89054584 |
| Satb1 | 0.16144137 | 0.15713328 | 0.17086027 | -0.013727 | 0.64590994 | 0.89054584 |
| Rhov | 0.078521 | 0.07025165 | 0.08398099 | -0.0137293 | 0.77686573 | 0.89054584 |
| Tmed7 | 0.88432596 | 0.75713472 | 0.77088589 | -0.0137512 | 0.53311671 | 0.89054584 |

| | | | | | | |
|---|---|---|---|---|---|---|
| Cndp2 | 0.30897918 | 0.30418573 | 0.31793897 | -0.0137532 | 0.57985954 | 0.89054584 |
| Suox | 0.24284202 | 0.23946981 | 0.25322855 | -0.0137587 | 0.59991828 | 0.89054584 |
| Rhbdf2 | 0.05371817 | 0.04370884 | 0.05747271 | -0.0137639 | 0.86773496 | 0.90448066 |
| Chadl | 0.10378213 | 0.09711246 | 0.1108877 | -0.0137752 | 0.71862653 | 0.89054584 |
| Bspry | 0.07359299 | 0.06493892 | 0.07871645 | -0.0137775 | 0.79259031 | 0.89054584 |
| Atf7ip | 0.33044394 | 0.32468786 | 0.33847885 | -0.013791 | 0.5752765 | 0.89054584 |
| Fbxo31 | 0.24255961 | 0.23915629 | 0.2529478 | -0.0137915 | 0.60026 | 0.89054584 |
| Rnf121 | 0.41996459 | 0.40770202 | 0.42150498 | -0.013803 | 0.56062959 | 0.89054584 |
| Srgap1 | 0.2109966 | 0.20752614 | 0.22134442 | -0.0138183 | 0.61441788 | 0.89054584 |
| Hcfc1 | 0.37577823 | 0.36724462 | 0.38107008 | -0.0138255 | 0.56711322 | 0.89054584 |
| Dlst | 0.42958924 | 0.41634097 | 0.43016659 | -0.0138256 | 0.55951477 | 0.89054584 |
| Inpp5f | 0.12564158 | 0.11994182 | 0.13377227 | -0.0138304 | 0.68498289 | 0.89054584 |
| Arpin | 0.1937175 | 0.19001785 | 0.20385849 | -0.0138406 | 0.62411684 | 0.89054584 |
| Prr36 | 0.06332275 | 0.05388338 | 0.06774097 | -0.0138576 | 0.82972197 | 0.89271879 |
| Gpatch3 | 0.05284096 | 0.04266616 | 0.0565347 | -0.0138685 | 0.87345711 | 0.90763913 |
| Trak2 | 0.07297489 | 0.0641875 | 0.07805606 | -0.0138686 | 0.79610075 | 0.89054584 |
| Zfp777 | 0.06749977 | 0.0583274 | 0.07220542 | -0.013878 | 0.81472872 | 0.89054584 |
| Ormdl3 | 0.49391444 | 0.47275988 | 0.48665305 | -0.0138932 | 0.55290543 | 0.89054584 |
| Vwa8 | 0.05972978 | 0.05000193 | 0.06390015 | -0.0138982 | 0.84454521 | 0.89775379 |
| Cdkl5 | 0.07524693 | 0.0665791 | 0.08048348 | -0.0139044 | 0.78955606 | 0.89054584 |
| Ufsp2 | 0.81354759 | 0.71342894 | 0.72733335 | -0.0139044 | 0.53548482 | 0.89054584 |
| Cacul1 | 0.30815012 | 0.3032358 | 0.31714106 | -0.0139053 | 0.5809305 | 0.89054584 |
| Pik3r3 | 0.07507349 | 0.06637596 | 0.08029819 | -0.0139222 | 0.79038974 | 0.89054584 |
| Stk35 | 0.08344407 | 0.07529185 | 0.08923889 | -0.013947 | 0.76673619 | 0.89054584 |
| Atf7 | 0.25413333 | 0.25047178 | 0.26442302 | -0.0139512 | 0.59708562 | 0.89054584 |
| Armc10 | 0.40821723 | 0.39690612 | 0.4108619 | -0.0139558 | 0.56287035 | 0.89054584 |
| Zfp628 | 0.05670741 | 0.04670423 | 0.06066889 | -0.0139647 | 0.85822693 | 0.90124396 |
| Ccdc120 | 0.03824794 | 0.02694532 | 0.04092681 | -0.0139815 | 0.94426223 | 0.95446655 |
| Arl16 | 0.30296428 | 0.29814103 | 0.31214208 | -0.0140011 | 0.58276635 | 0.89054584 |
| Slc25a14 | 0.30414009 | 0.29927167 | 0.31327672 | -0.0140051 | 0.58249412 | 0.89054584 |

| | | | | | | |
|---|---|---|---|---|---|---|
| Ddx31 | 0.09437991 | 0.08690774 | 0.10091321 | -0.0140055 | 0.74105511 | 0.89054584 |
| Rdh13 | 0.11984535 | 0.11370267 | 0.12772159 | -0.0140189 | 0.69545483 | 0.89054584 |
| Eid2 | 0.04905627 | 0.03846706 | 0.05248743 | -0.0140204 | 0.89334037 | 0.9188173 |
| Tyw5 | 0.19011763 | 0.18616964 | 0.20019949 | -0.0140299 | 0.62797535 | 0.89054584 |
| Zc3h11a | 0.11446464 | 0.10805361 | 0.12209336 | -0.0140398 | 0.70393107 | 0.89054584 |
| Dohh | 0.7833742 | 0.69360913 | 0.70765901 | -0.0140499 | 0.5368491 | 0.89054584 |
| Taf4a | 0.08122379 | 0.07281109 | 0.08686777 | -0.0140567 | 0.77454412 | 0.89054584 |
| Hebp2 | 0.12440112 | 0.11841678 | 0.13247843 | -0.0140616 | 0.68952823 | 0.89054584 |
| Tmem160 | 0.86634094 | 0.74609621 | 0.76016419 | -0.014068 | 0.53435451 | 0.89054584 |
| Pam16 | 1.06925973 | 0.85595573 | 0.87002768 | -0.0140719 | 0.53003362 | 0.89054584 |
| Mdn1 | 0.128414 | 0.12257532 | 0.13666211 | -0.0140868 | 0.68446887 | 0.89054584 |
| Cep250 | 0.05782701 | 0.04777424 | 0.06186591 | -0.0140917 | 0.8557011 | 0.9009252 |
| Zbtb12 | 0.13073057 | 0.12496383 | 0.13907459 | -0.0141108 | 0.68178811 | 0.89054584 |
| Kpna4 | 0.2977414 | 0.29297611 | 0.30709363 | -0.0141175 | 0.58479585 | 0.89054584 |
| Ppm1h | 0.16905187 | 0.16455995 | 0.17867949 | -0.0141195 | 0.64362189 | 0.89054584 |
| 2410022M11 | 0.08436625 | 0.07610086 | 0.09022363 | -0.0141228 | 0.76708103 | 0.89054584 |
| Arih1 | 0.37567701 | 0.36685106 | 0.38097622 | -0.0141252 | 0.56857047 | 0.89054584 |
| Cc2d2a | 0.21138208 | 0.20760578 | 0.22173306 | -0.0141273 | 0.61670379 | 0.89054584 |
| Nup188 | 0.09957949 | 0.09232829 | 0.1064611 | -0.0141328 | 0.73187007 | 0.89054584 |
| Pak6 | 0.07176472 | 0.06262641 | 0.07676302 | -0.0141366 | 0.80454999 | 0.89054584 |
| Zbtb17 | 0.10334031 | 0.09628668 | 0.11042335 | -0.0141367 | 0.72456233 | 0.89054584 |
| Sp4 | 0.1107473 | 0.10405455 | 0.11819874 | -0.0141442 | 0.71141034 | 0.89054584 |
| Cinp | 0.27339888 | 0.26923141 | 0.28338692 | -0.0141555 | 0.59201047 | 0.89054584 |
| Wdr5b | 0.05934954 | 0.0493287 | 0.06349366 | -0.014165 | 0.8506847 | 0.89942466 |
| Szt2 | 0.06315542 | 0.05338353 | 0.06756211 | -0.0141786 | 0.83589486 | 0.89462889 |
| Slc31a2 | 0.31494491 | 0.3094925 | 0.32367137 | -0.0141789 | 0.58085891 | 0.89054584 |
| Rgs11 | 0.11862905 | 0.11225759 | 0.12645029 | -0.0141927 | 0.69948384 | 0.89054584 |
| Ago4 | 0.05025185 | 0.03956391 | 0.05376602 | -0.0142021 | 0.89077183 | 0.91722772 |
| Fbxo36 | 0.20042215 | 0.19645492 | 0.21065843 | -0.0142035 | 0.62328744 | 0.89054584 |
| Zfyve9 | 0.17858042 | 0.17423016 | 0.18843577 | -0.0142056 | 0.63728973 | 0.89054584 |

| | | | | | | |
|---|---|---|---|---|---|---|
| E330009J07F | 0.18006815 | 0.17574888 | 0.18995584 | -0.014207 | 0.6362474 | 0.89054584 |
| Ap1s1 | 1.21093607 | 0.92094044 | 0.93515086 | -0.0142104 | 0.52822018 | 0.89054584 |
| Lekr1 | 0.11622225 | 0.10971949 | 0.12393301 | -0.0142135 | 0.70344018 | 0.89054584 |
| Pex5 | 0.2478746 | 0.24399887 | 0.25822534 | -0.0142265 | 0.60128559 | 0.89054584 |
| 1110019D14 | 0.08733228 | 0.07913171 | 0.09339054 | -0.0142588 | 0.76155871 | 0.89054584 |
| Mthfs | 0.36228763 | 0.35424031 | 0.36850941 | -0.0142691 | 0.57158051 | 0.89054584 |
| Ttc26 | 0.05603578 | 0.04568088 | 0.0599508 | -0.0142699 | 0.86627513 | 0.90430169 |
| Dync1li1 | 0.62981645 | 0.58321239 | 0.59748509 | -0.0142727 | 0.54430784 | 0.89054584 |
| Med12l | 0.04670052 | 0.03569517 | 0.04996801 | -0.0142728 | 0.90836893 | 0.9289365 |
| Abracl | 1.14064239 | 0.88992061 | 0.90422086 | -0.0143003 | 0.52936795 | 0.89054584 |
| Fkbpl | 0.20265444 | 0.1986129 | 0.21291813 | -0.0143052 | 0.62286649 | 0.89054584 |
| Cradd | 0.16606411 | 0.16130734 | 0.17561261 | -0.0143053 | 0.64784678 | 0.89054584 |
| Slc26a2 | 0.08791532 | 0.07970483 | 0.094013 | -0.0143082 | 0.76085656 | 0.89054584 |
| Foxj2 | 0.07679351 | 0.06781581 | 0.08213568 | -0.0143199 | 0.79167848 | 0.89054584 |
| Lrrc57 | 0.21727289 | 0.21333674 | 0.22766407 | -0.0143273 | 0.61531379 | 0.89054584 |
| Ndst1 | 0.144358 | 0.13888849 | 0.15322436 | -0.0143359 | 0.66854418 | 0.89054584 |
| Vcl | 0.18916252 | 0.1848815 | 0.19922777 | -0.0143463 | 0.63137129 | 0.89054584 |
| Cys1 | 0.0520905 | 0.04137781 | 0.05573222 | -0.0143544 | 0.88492219 | 0.91404124 |
| Ogfod1 | 0.06376148 | 0.05385325 | 0.06820993 | -0.0143567 | 0.83660585 | 0.89462889 |
| Nol10 | 0.125747 | 0.11952376 | 0.13388221 | -0.0143584 | 0.69134164 | 0.89054584 |
| Gnb1 | 1.49169306 | 1.01891472 | 1.03327786 | -0.0143631 | 0.5258182 | 0.89054584 |
| Adck5 | 0.30848391 | 0.30309175 | 0.31746235 | -0.0143706 | 0.58351611 | 0.89054584 |
| Trim32 | 0.31339876 | 0.30781483 | 0.32218737 | -0.0143725 | 0.58231992 | 0.89054584 |
| Hemk1 | 0.10399586 | 0.09673287 | 0.11111231 | -0.0143794 | 0.72672247 | 0.89054584 |
| Nat10 | 0.08629391 | 0.07789727 | 0.09228191 | -0.0143846 | 0.7661489 | 0.89054584 |
| Gle1 | 0.32848404 | 0.32221187 | 0.33661332 | -0.0144014 | 0.57899578 | 0.89054584 |
| Slc16a12 | 0.38562389 | 0.37576249 | 0.39017343 | -0.0144109 | 0.56831135 | 0.89054584 |
| Shank2 | 0.08200307 | 0.07327683 | 0.08770003 | -0.0144232 | 0.77822501 | 0.89054584 |
| Manba | 0.10477991 | 0.09750398 | 0.11193612 | -0.0144321 | 0.72597541 | 0.89054584 |
| Phc3 | 0.17028634 | 0.16550383 | 0.17994558 | -0.0144418 | 0.64576403 | 0.89054584 |

| | | | | | | |
|---|---|---|---|---|---|---|
| Zfp866 | 0.2003772 | 0.19617114 | 0.2106129 | -0.0144418 | 0.62531284 | 0.89054584 |
| Epdr1 | 0.09614994 | 0.08835654 | 0.10280202 | -0.0144455 | 0.74365751 | 0.89054584 |
| Aamdc | 0.34769097 | 0.34036133 | 0.3548107 | -0.0144494 | 0.57524024 | 0.89054584 |
| Tmem94 | 0.1349871 | 0.12903355 | 0.143502 | -0.0144685 | 0.6807272 | 0.89054584 |
| Uggt1 | 0.39479635 | 0.38413047 | 0.39860702 | -0.0144766 | 0.56718142 | 0.89054584 |
| 1700023F06I | 0.25214231 | 0.24796693 | 0.26245342 | -0.0144865 | 0.60147116 | 0.89054584 |
| Mettl5 | 0.32476173 | 0.31856433 | 0.33306469 | -0.0145004 | 0.580367 | 0.89054584 |
| Nup85 | 0.24347225 | 0.23934959 | 0.25385495 | -0.0145054 | 0.60496259 | 0.89054584 |
| Srcap | 0.12423158 | 0.11779308 | 0.13230155 | -0.0145085 | 0.6952928 | 0.89054584 |
| Mad2l2 | 0.54029477 | 0.51130421 | 0.5258155 | -0.0145113 | 0.55115344 | 0.89054584 |
| Irgq | 0.17119539 | 0.16635369 | 0.18087751 | -0.0145238 | 0.64583377 | 0.89054584 |
| Celf1 | 0.87098042 | 0.74842677 | 0.76295279 | -0.014526 | 0.53534086 | 0.89054584 |
| Iqgap1 | 0.88761667 | 0.75829307 | 0.77282185 | -0.0145288 | 0.53489731 | 0.89054584 |
| Orc5 | 0.1457663 | 0.14012805 | 0.1546825 | -0.0145545 | 0.6694414 | 0.89054584 |
| Cep97 | 0.12777552 | 0.12139395 | 0.13599683 | -0.0146029 | 0.69155458 | 0.89054584 |
| Shroom3 | 0.11471112 | 0.10774552 | 0.12235142 | -0.0146059 | 0.71095226 | 0.89054584 |
| Grk6 | 0.15841785 | 0.1531378 | 0.16774729 | -0.0146095 | 0.65753967 | 0.89054584 |
| Fxn | 0.31353375 | 0.3077042 | 0.32231698 | -0.0146128 | 0.58364256 | 0.89054584 |
| Stk38 | 0.64937945 | 0.59781748 | 0.6124349 | -0.0146174 | 0.54427045 | 0.89054584 |
| Ak2 | 1.0670501 | 0.85429784 | 0.86891676 | -0.0146189 | 0.53123848 | 0.89054584 |
| 2700062C07I | 0.3157167 | 0.30979065 | 0.32441166 | -0.014621 | 0.58315653 | 0.89054584 |
| Aifm2 | 0.1336 | 0.12743746 | 0.14205998 | -0.0146225 | 0.68422777 | 0.89054584 |
| Tmem198 | 0.103492 | 0.09595499 | 0.11058279 | -0.0146278 | 0.73114537 | 0.89054584 |
| Eif2ak4 | 0.0821492 | 0.07322798 | 0.08785609 | -0.0146281 | 0.78104196 | 0.89054584 |
| Mier3 | 0.21470934 | 0.21045534 | 0.22508492 | -0.0146296 | 0.61898334 | 0.89054584 |
| Suv420h2 | 0.43879345 | 0.42375399 | 0.43839885 | -0.0146449 | 0.56183905 | 0.89054584 |
| Bach2 | 0.08280308 | 0.07390126 | 0.08855439 | -0.0146531 | 0.77962177 | 0.89054584 |
| Zbtb25 | 0.12995917 | 0.12360642 | 0.13827148 | -0.0146651 | 0.68939166 | 0.89054584 |
| Chpt1 | 0.16676558 | 0.16166012 | 0.17633298 | -0.0146729 | 0.650872 | 0.89054584 |
| B4galt5 | 0.20040396 | 0.19595925 | 0.21064001 | -0.0146808 | 0.62729944 | 0.89054584 |

| | | | | | | |
|---|---|---|---|---|---|---|
| Ilvbl | 0.29086371 | 0.28573269 | 0.30042471 | -0.014692 | 0.59011528 | 0.89054584 |
| Trmt11 | 0.11051438 | 0.10324825 | 0.11795454 | -0.0147063 | 0.71933456 | 0.89054584 |
| Traf3 | 0.14456085 | 0.13871179 | 0.15343443 | -0.0147226 | 0.6725783 | 0.89054584 |
| Zfp574 | 0.14528922 | 0.13946422 | 0.15418863 | -0.0147244 | 0.67180785 | 0.89054584 |
| Snn | 0.26240794 | 0.25785132 | 0.27258945 | -0.0147381 | 0.59943909 | 0.89054584 |
| Atl2 | 0.32922044 | 0.32256225 | 0.3373145 | -0.0147522 | 0.58072792 | 0.89054584 |
| Abhd6 | 0.10422624 | 0.09660086 | 0.11135439 | -0.0147535 | 0.73147202 | 0.89054584 |
| Xpo4 | 0.15495711 | 0.14942568 | 0.16417965 | -0.014754 | 0.66227306 | 0.89054584 |
| Tinf2 | 0.05293848 | 0.04187431 | 0.05663899 | -0.0147647 | 0.88772067 | 0.91537513 |
| Apba3 | 0.16036426 | 0.15498531 | 0.16975171 | -0.0147664 | 0.65736162 | 0.89054584 |
| Bcat2 | 0.21138911 | 0.20697214 | 0.22174014 | -0.014768 | 0.62182891 | 0.89054584 |
| Sfxn4 | 0.08746644 | 0.07876524 | 0.09353378 | -0.0147685 | 0.76902037 | 0.89054584 |
| Sh3bgr | 0.10950068 | 0.10210327 | 0.11689152 | -0.0147883 | 0.72220849 | 0.89054584 |
| Pgap2 | 0.35849314 | 0.35016572 | 0.36495947 | -0.0147938 | 0.57489551 | 0.89054584 |
| 1700020I14R | 0.5321921 | 0.5042409 | 0.51907159 | -0.0148307 | 0.5529481 | 0.89054584 |
| Frat2 | 0.05241189 | 0.04123814 | 0.05607589 | -0.0148377 | 0.89116765 | 0.91736357 |
| Btbd3 | 0.25675394 | 0.25216003 | 0.26701258 | -0.0148525 | 0.60224143 | 0.89054584 |
| Pafah2 | 0.06069568 | 0.05007275 | 0.06493274 | -0.01486 | 0.85683515 | 0.9009252 |
| Flrt1 | 0.1651075 | 0.1597691 | 0.17462989 | -0.0148608 | 0.65412085 | 0.89054584 |
| Nupr1l | 0.06032746 | 0.04967673 | 0.0645391 | -0.0148624 | 0.85833815 | 0.90124516 |
| Id4 | 0.10091628 | 0.09295994 | 0.10787441 | -0.0149145 | 0.74025588 | 0.89054584 |
| Crebrf | 0.1746222 | 0.1694694 | 0.18438752 | -0.0149181 | 0.64689015 | 0.89054584 |
| Ralgps1 | 0.13433135 | 0.12786398 | 0.14282037 | -0.0149564 | 0.68718694 | 0.89054584 |
| Arl8b | 0.46870587 | 0.44983358 | 0.46481071 | -0.0149771 | 0.5596652 | 0.89054584 |
| Ctu1 | 0.09700344 | 0.08872789 | 0.10371273 | -0.0149848 | 0.74957172 | 0.89054584 |
| 1500015A07 | 0.1475664 | 0.14154321 | 0.15654519 | -0.015002 | 0.67237249 | 0.89054584 |
| Zfp24 | 0.59569998 | 0.5557913 | 0.57079584 | -0.0150045 | 0.54873651 | 0.89054584 |
| Phf7 | 0.06239924 | 0.05174906 | 0.06675381 | -0.0150048 | 0.85250204 | 0.89988783 |
| Acad11 | 0.09309809 | 0.08452561 | 0.09954524 | -0.0150196 | 0.75895729 | 0.89054584 |
| Map3k11 | 0.19720724 | 0.19237137 | 0.20740025 | -0.0150289 | 0.63216831 | 0.89054584 |

| | | | | | | |
|---|---|---|---|---|---|---|
| Nxt2 | 0.10056954 | 0.09246874 | 0.10750963 | -0.0150409 | 0.74273439 | 0.89054584 |
| Raph1 | 0.43572053 | 0.4206001 | 0.43565601 | -0.0150559 | 0.56395687 | 0.89054584 |
| 6330418K02I | 0.13627125 | 0.12976554 | 0.14483633 | -0.0150708 | 0.68608417 | 0.89054584 |
| Gm26518 | 0.21253716 | 0.20780101 | 0.22289724 | -0.0150962 | 0.62382321 | 0.89054584 |
| Ric8b | 0.11642282 | 0.10904067 | 0.12414288 | -0.0151022 | 0.71456434 | 0.89054584 |
| AI314180 | 0.39639174 | 0.38496428 | 0.40006918 | -0.0151049 | 0.56981441 | 0.89054584 |
| Trmt2b | 0.14025697 | 0.13385985 | 0.14897372 | -0.0151139 | 0.68177381 | 0.89054584 |
| Casp8ap2 | 0.50688518 | 0.48262066 | 0.49774119 | -0.0151205 | 0.55627466 | 0.89054584 |
| Entpd6 | 0.11221704 | 0.1046138 | 0.11973919 | -0.0151254 | 0.72190653 | 0.89054584 |
| Ccdc92 | 0.23113756 | 0.22642604 | 0.24156202 | -0.015136 | 0.61482749 | 0.89054584 |
| Wdr24 | 0.08118826 | 0.07169106 | 0.08682983 | -0.0151388 | 0.79168613 | 0.89054584 |
| Tmem132a | 0.36451357 | 0.35544861 | 0.37058877 | -0.0151402 | 0.57547789 | 0.89054584 |
| Ccny | 0.45020702 | 0.43339844 | 0.44853969 | -0.0151413 | 0.56248429 | 0.89054584 |
| Edrf1 | 0.1114756 | 0.10379573 | 0.11896218 | -0.0151664 | 0.72373283 | 0.89054584 |
| Eral1 | 0.11170801 | 0.10403437 | 0.11920576 | -0.0151714 | 0.72339146 | 0.89054584 |
| Ammecr1 | 0.19114048 | 0.18606398 | 0.20123971 | -0.0151757 | 0.63733199 | 0.89054584 |
| Ankrd26 | 0.19695439 | 0.19196673 | 0.2071438 | -0.0151771 | 0.63358095 | 0.89054584 |
| Parp10 | 0.06169561 | 0.05082363 | 0.06600165 | -0.015178 | 0.85800296 | 0.90124396 |
| Lclat1 | 0.07881411 | 0.06911053 | 0.08429407 | -0.0151835 | 0.7993202 | 0.89054584 |
| Polr3b | 0.09422563 | 0.08555771 | 0.10074857 | -0.0151909 | 0.75880841 | 0.89054584 |
| Vps18 | 0.12658584 | 0.11956291 | 0.13475682 | -0.0151939 | 0.70031209 | 0.89054584 |
| Tbc1d25 | 0.07929983 | 0.06961279 | 0.08481288 | -0.0152001 | 0.79813314 | 0.89054584 |
| Sdc3 | 0.15225822 | 0.14618372 | 0.16139408 | -0.0152104 | 0.66969661 | 0.89054584 |
| Zcchc2 | 0.11524846 | 0.10768876 | 0.12291391 | -0.0152251 | 0.71806254 | 0.89054584 |
| Slc39a9 | 0.18652463 | 0.18130046 | 0.19654197 | -0.0152415 | 0.64105991 | 0.89054584 |
| Slc35b3 | 0.13846476 | 0.13183895 | 0.14711407 | -0.0152751 | 0.68571706 | 0.89054584 |
| Mastl | 0.07266807 | 0.06245066 | 0.07772823 | -0.0152776 | 0.82009314 | 0.89054584 |
| Acad12 | 0.07397275 | 0.06384443 | 0.07912219 | -0.0152778 | 0.81583298 | 0.89054584 |
| 2510002D24I | 0.73170884 | 0.65716434 | 0.67244726 | -0.0152829 | 0.54216331 | 0.89054584 |
| Znrf1 | 0.3058327 | 0.29959362 | 0.31490884 | -0.0153152 | 0.58962565 | 0.89054584 |

| | | | | | | |
|---|---|---|---|---|---|---|
| Calcoco1 | 0.20130551 | 0.19623343 | 0.21155289 | -0.0153195 | 0.6320826 | 0.89054584 |
| Pawr | 0.0822225 | 0.07260564 | 0.08793438 | -0.0153287 | 0.79164654 | 0.89054584 |
| Rhoq | 0.14435651 | 0.13789249 | 0.15322281 | -0.0153303 | 0.67944177 | 0.89054584 |
| Slc35e1 | 0.14012556 | 0.1334992 | 0.14883741 | -0.0153382 | 0.68442896 | 0.89054584 |
| Madd | 0.28844795 | 0.28273733 | 0.29807668 | -0.0153393 | 0.59473957 | 0.89054584 |
| C330018D20 | 0.14883643 | 0.14251051 | 0.1578586 | -0.0153481 | 0.67471627 | 0.89054584 |
| Abcb7 | 0.07356703 | 0.06333981 | 0.07868871 | -0.0153489 | 0.81826318 | 0.89054584 |
| Zscan22 | 0.05654763 | 0.04513918 | 0.06049806 | -0.0153589 | 0.88155259 | 0.91212885 |
| Bcl2l11 | 0.19024004 | 0.18495442 | 0.20032401 | -0.0153696 | 0.63962291 | 0.89054584 |
| Oxsr1 | 0.11527335 | 0.10756459 | 0.12293997 | -0.0153754 | 0.71994108 | 0.89054584 |
| Cdc16 | 0.73258314 | 0.65768128 | 0.67305897 | -0.0153777 | 0.54238536 | 0.89054584 |
| Slco3a1 | 0.20919996 | 0.20415398 | 0.21953215 | -0.0153782 | 0.62792312 | 0.89054584 |
| Eml3 | 0.10311844 | 0.09480406 | 0.11019013 | -0.0153861 | 0.74232732 | 0.89054584 |
| Cd320 | 0.3776907 | 0.3674551 | 0.3828426 | -0.0153875 | 0.57426991 | 0.89054584 |
| Vps13a | 0.13281369 | 0.12582616 | 0.14124221 | -0.0154161 | 0.69445069 | 0.89054584 |
| Fbf1 | 0.10128976 | 0.09283832 | 0.10826728 | -0.015429 | 0.74663455 | 0.89054584 |
| Zfp53 | 0.12338251 | 0.11598593 | 0.13141554 | -0.0154296 | 0.70781412 | 0.89054584 |
| Gfm2 | 0.20067912 | 0.19548129 | 0.21091867 | -0.0154374 | 0.63344569 | 0.89054584 |
| Slc31a1 | 0.61107931 | 0.56748618 | 0.58292404 | -0.0154379 | 0.54909887 | 0.89054584 |
| Ets2 | 0.44058727 | 0.42455681 | 0.43999741 | -0.0154406 | 0.56493515 | 0.89054584 |
| Kifap3 | 0.50558544 | 0.48119359 | 0.49663486 | -0.0154413 | 0.55758726 | 0.89054584 |
| Mfsd8 | 0.10150176 | 0.09303668 | 0.10849026 | -0.0154536 | 0.7465362 | 0.89054584 |
| Kidins220 | 0.29176948 | 0.2858505 | 0.30130433 | -0.0154538 | 0.59443016 | 0.89054584 |
| Ap4b1 | 0.16456191 | 0.15860588 | 0.17406924 | -0.0154634 | 0.66051718 | 0.89054584 |
| Csnk1e | 0.5336167 | 0.50479409 | 0.52026037 | -0.0154663 | 0.55507769 | 0.89054584 |
| Oxnad1 | 0.23997231 | 0.23490006 | 0.25037398 | -0.0154739 | 0.61330271 | 0.89054584 |
| Ttbk2 | 0.1381599 | 0.1313202 | 0.14679762 | -0.0154774 | 0.68836086 | 0.89054584 |
| Sidt2 | 0.12516026 | 0.11779116 | 0.13327031 | -0.0154791 | 0.70579068 | 0.89054584 |
| Ncbp1 | 0.34478428 | 0.33658802 | 0.35206909 | -0.0154811 | 0.58115985 | 0.89054584 |
| Extl3 | 0.08868594 | 0.07935433 | 0.0948357 | -0.0154814 | 0.7765354 | 0.89054584 |

| | | | | | | |
|---|---|---|---|---|---|---|
| Hat1 | 0.45238516 | 0.43498343 | 0.45046623 | -0.0154828 | 0.56361095 | 0.89054584 |
| Zdhhc20 | 0.77222869 | 0.6847365 | 0.70022545 | -0.0154889 | 0.54104053 | 0.89054584 |
| Nelfe | 0.53836302 | 0.5086923 | 0.52421155 | -0.0155192 | 0.5548512 | 0.89054584 |
| Elmo2 | 0.11857121 | 0.11086819 | 0.12638981 | -0.0155216 | 0.71639091 | 0.89054584 |
| Acvr1b | 0.11958684 | 0.11192813 | 0.12745143 | -0.0155233 | 0.71479529 | 0.89054584 |
| Mief1 | 0.10467674 | 0.09630222 | 0.11182773 | -0.0155255 | 0.74112589 | 0.89054584 |
| Setd1a | 0.09856836 | 0.08984657 | 0.10538239 | -0.0155358 | 0.75390711 | 0.89054584 |
| Rfx1 | 0.06071173 | 0.0494059 | 0.06494989 | -0.015544 | 0.8675729 | 0.90444517 |
| Sema4d | 0.11825157 | 0.11050385 | 0.12605562 | -0.0155518 | 0.71728141 | 0.89054584 |
| Lamc2 | 0.05810842 | 0.04660717 | 0.06216678 | -0.0155596 | 0.87820858 | 0.91042317 |
| Lrrc75a | 0.06108967 | 0.04978649 | 0.06535391 | -0.0155674 | 0.86645061 | 0.90430169 |
| Plekhf2 | 0.18710472 | 0.18156372 | 0.19713285 | -0.0155691 | 0.64354591 | 0.89054584 |
| Cables2 | 0.08479304 | 0.07510837 | 0.09067937 | -0.015571 | 0.78814738 | 0.89054584 |
| Map7d2 | 0.05655063 | 0.04490611 | 0.06050127 | -0.0155952 | 0.88510807 | 0.91411146 |
| AU040320 | 0.12558863 | 0.11810867 | 0.13371706 | -0.0156084 | 0.70671877 | 0.89054584 |
| Inpp1 | 0.13306485 | 0.1258928 | 0.14150344 | -0.0156106 | 0.69636481 | 0.89054584 |
| Cbfa2t2 | 0.30020353 | 0.29386303 | 0.30947525 | -0.0156122 | 0.59290785 | 0.89054584 |
| Coq10a | 0.07962401 | 0.06954213 | 0.08515914 | -0.015617 | 0.80355433 | 0.89054584 |
| Stk38l | 0.11521163 | 0.10724901 | 0.12287536 | -0.0156264 | 0.7232383 | 0.89054584 |
| Cul9 | 0.12121919 | 0.11353041 | 0.12915689 | -0.0156265 | 0.71351324 | 0.89054584 |
| Rpl32 | 5.40025773 | 0.65354466 | 0.66917572 | -0.0156311 | 0.54333004 | 0.89054584 |
| Rpusd4 | 0.10388083 | 0.09534469 | 0.11099143 | -0.0156467 | 0.74433959 | 0.89054584 |
| Rps6kb2 | 0.32875261 | 0.3212212 | 0.33686908 | -0.0156479 | 0.58566675 | 0.89054584 |
| Dstyk | 0.13104962 | 0.12375524 | 0.13940669 | -0.0156515 | 0.69953658 | 0.89054584 |
| Dnajc16 | 0.06350621 | 0.05228529 | 0.06793708 | -0.0156518 | 0.85844459 | 0.90124516 |
| Usp40 | 0.27759465 | 0.27183996 | 0.28749341 | -0.0156535 | 0.60012485 | 0.89054584 |
| Mrpl55 | 0.62968948 | 0.58173343 | 0.5973872 | -0.0156538 | 0.54858281 | 0.89054584 |
| Them4 | 0.1248719 | 0.11729214 | 0.13296954 | -0.0156774 | 0.70859993 | 0.89054584 |
| Man2b2 | 0.23580547 | 0.23052974 | 0.24622234 | -0.0156926 | 0.61674019 | 0.89054584 |
| Fbxl17 | 0.17926869 | 0.17343306 | 0.18913911 | -0.0157061 | 0.65057638 | 0.89054584 |

| | | | | | | |
|---|---|---|---|---|---|---|
| Lmtk2 | 0.10607754 | 0.0975904 | 0.11329906 | -0.0157087 | 0.74084461 | 0.89054584 |
| Pepd | 0.45295771 | 0.43525653 | 0.45097218 | -0.0157157 | 0.56448747 | 0.89054584 |
| Bbc3 | 0.17976069 | 0.1739224 | 0.18964178 | -0.0157194 | 0.65031795 | 0.89054584 |
| Smg5 | 0.33049451 | 0.32279985 | 0.33852696 | -0.0157271 | 0.58567865 | 0.89054584 |
| Mrrf | 0.32641881 | 0.31891397 | 0.33464535 | -0.0157314 | 0.58667988 | 0.89054584 |
| Stx1a | 0.17481093 | 0.16884661 | 0.1845807 | -0.0157341 | 0.65436746 | 0.89054584 |
| Zfp318 | 0.27132389 | 0.26559166 | 0.28135292 | -0.0157613 | 0.60295055 | 0.89054584 |
| Msi2 | 0.65530959 | 0.60115109 | 0.61691474 | -0.0157637 | 0.54738099 | 0.89054584 |
| Gatb | 0.07875391 | 0.06845868 | 0.08422976 | -0.0157711 | 0.80848716 | 0.89054584 |
| Ogdh | 0.44817568 | 0.43093164 | 0.44674043 | -0.0158088 | 0.56547518 | 0.89054584 |
| Wee1 | 0.0624407 | 0.05098527 | 0.06679812 | -0.0158129 | 0.86496311 | 0.90367048 |
| Wdr7 | 0.07153996 | 0.06069272 | 0.07652285 | -0.0158301 | 0.8324233 | 0.89382855 |
| 9130401M01 | 0.48606631 | 0.46404808 | 0.47989507 | -0.015847 | 0.5611346 | 0.89054584 |
| Srbd1 | 0.12070345 | 0.1127677 | 0.12861816 | -0.0158505 | 0.71706751 | 0.89054584 |
| Rfc3 | 0.42135022 | 0.40690289 | 0.42275531 | -0.0158524 | 0.56934242 | 0.89054584 |
| Ascc2 | 0.14252343 | 0.13545768 | 0.15132368 | -0.015866 | 0.68739546 | 0.89054584 |
| Adgrl1 | 0.38844538 | 0.37690584 | 0.39277218 | -0.0158663 | 0.57464068 | 0.89054584 |
| Ankrd54 | 0.65403334 | 0.60006446 | 0.61595249 | -0.015888 | 0.54782728 | 0.89054584 |
| Phldb3 | 0.0523987 | 0.04016324 | 0.05606178 | -0.0158985 | 0.90678733 | 0.92797365 |
| Rragc | 0.34746982 | 0.33868204 | 0.35460227 | -0.0159202 | 0.58284141 | 0.89054584 |
| Pdpk1 | 0.6033131 | 0.56089666 | 0.57681898 | -0.0159223 | 0.55116443 | 0.89054584 |
| Lemd2 | 0.16740434 | 0.16105503 | 0.17698879 | -0.0159338 | 0.66254908 | 0.89054584 |
| Scp2 | 1.54058326 | 1.03045606 | 1.04640113 | -0.0159451 | 0.52829834 | 0.89054584 |
| Plekha7 | 0.13938684 | 0.13212167 | 0.14807101 | -0.0159493 | 0.69210775 | 0.89054584 |
| Fam89a | 0.09222474 | 0.08266263 | 0.09861313 | -0.0159505 | 0.77444786 | 0.89054584 |
| Tfb2m | 0.21667407 | 0.21111053 | 0.22706186 | -0.0159513 | 0.62827783 | 0.89054584 |
| Tmeff1 | 0.1865772 | 0.18063728 | 0.19659552 | -0.0159582 | 0.6473516 | 0.89054584 |
| Smpd4 | 0.09690306 | 0.08763375 | 0.10360562 | -0.0159719 | 0.76369032 | 0.89054584 |
| Nub1 | 0.35356068 | 0.34435595 | 0.36033319 | -0.0159772 | 0.5818304 | 0.89054584 |
| Trappc4 | 1.14855362 | 0.89182418 | 0.90780865 | -0.0159845 | 0.53268996 | 0.89054584 |

| | | | | | | |
|---|---|---|---|---|---|---|
| Crocc | 0.06731609 | 0.05601604 | 0.07200912 | -0.0159931 | 0.84960515 | 0.89887961 |
| St3gal2 | 0.08044373 | 0.07003403 | 0.08603465 | -0.0160006 | 0.80688217 | 0.89054584 |
| Zfp438 | 0.06653227 | 0.05517023 | 0.07117142 | -0.0160012 | 0.85255421 | 0.89988783 |
| Dmwd | 0.14243638 | 0.13522048 | 0.15123345 | -0.016013 | 0.68909975 | 0.89054584 |
| Fbxl14 | 0.12693444 | 0.11908348 | 0.13512023 | -0.0160367 | 0.70984999 | 0.89054584 |
| Nubp2 | 0.33020066 | 0.32218071 | 0.3382474 | -0.0160667 | 0.58757014 | 0.89054584 |
| Serpini1 | 0.06371511 | 0.05208846 | 0.06816036 | -0.0160719 | 0.86401936 | 0.90314007 |
| 1700007K13I | 0.17575301 | 0.16946468 | 0.18554472 | -0.01608 | 0.65680619 | 0.89054584 |
| AU041133 | 0.09443981 | 0.08488726 | 0.10097714 | -0.0160899 | 0.77106667 | 0.89054584 |
| Evi5l | 0.08231107 | 0.07193752 | 0.08802896 | -0.0160914 | 0.80279327 | 0.89054584 |
| Cep126 | 0.10524898 | 0.09632798 | 0.11242887 | -0.0161009 | 0.7476794 | 0.89054584 |
| Gm26601 | 0.06823454 | 0.05688926 | 0.07299067 | -0.0161014 | 0.84796417 | 0.89866846 |
| Baz2a | 0.6537231 | 0.59960801 | 0.6157184 | -0.0161104 | 0.54851171 | 0.89054584 |
| Mul1 | 0.25890073 | 0.25301574 | 0.26913179 | -0.016116 | 0.60987178 | 0.89054584 |
| Zdhhc5 | 0.08710037 | 0.07702607 | 0.09314294 | -0.0161169 | 0.78992305 | 0.89054584 |
| Coa5 | 0.2158717 | 0.21013156 | 0.22625471 | -0.0161232 | 0.63005581 | 0.89054584 |
| Zfp395 | 0.43985502 | 0.42321087 | 0.4393451 | -0.0161342 | 0.56792428 | 0.89054584 |
| Mfsd6 | 0.18136181 | 0.17513874 | 0.19127688 | -0.0161381 | 0.65286893 | 0.89054584 |
| Rom1 | 0.11426332 | 0.10574182 | 0.12188258 | -0.0161408 | 0.73137525 | 0.89054584 |
| Nqo2 | 0.17946953 | 0.17320338 | 0.18934432 | -0.0161409 | 0.65437471 | 0.89054584 |
| Tsc22d4 | 1.0082058 | 0.82261756 | 0.8387616 | -0.016144 | 0.53572638 | 0.89054584 |
| Ino80 | 0.23280934 | 0.2270861 | 0.24323223 | -0.0161461 | 0.62144106 | 0.89054584 |
| Gng4 | 0.26500726 | 0.25899338 | 0.27514827 | -0.0161549 | 0.60778178 | 0.89054584 |
| C2cd2 | 0.13897673 | 0.13147868 | 0.14764544 | -0.0161668 | 0.69502365 | 0.89054584 |
| Rhobtb2 | 0.06904126 | 0.05768189 | 0.07385278 | -0.0161709 | 0.84616589 | 0.89825055 |
| C530008M17 | 0.10065553 | 0.09142541 | 0.1076001 | -0.0161747 | 0.7581434 | 0.89054584 |
| Qrsl1 | 0.10135904 | 0.09215824 | 0.10834014 | -0.0161819 | 0.75674458 | 0.89054584 |
| Gstm7 | 0.15399636 | 0.14699906 | 0.16318837 | -0.0161893 | 0.6780297 | 0.89054584 |
| mt-Nd6 | 0.12935855 | 0.12145062 | 0.13764602 | -0.0161954 | 0.70821234 | 0.89054584 |
| Gm18025 | 0.12620717 | 0.11814194 | 0.13436201 | -0.0162201 | 0.71308634 | 0.89054584 |

| | | | | | | |
|---|---|---|---|---|---|---|
| Mcu | 0.11769492 | 0.10925227 | 0.12547353 | -0.0162213 | 0.72651597 | 0.89054584 |
| 3110043O21 | 0.12540601 | 0.11730006 | 0.13352661 | -0.0162265 | 0.7143603 | 0.89054584 |
| Arsb | 0.19305635 | 0.18695908 | 0.2031869 | -0.0162278 | 0.6450883 | 0.89054584 |
| Sdad1 | 0.26879867 | 0.26264628 | 0.27887478 | -0.0162285 | 0.60684939 | 0.89054584 |
| Foxp4 | 0.36981914 | 0.35930267 | 0.37553411 | -0.0162314 | 0.57979537 | 0.89054584 |
| Mettl18 | 0.14117445 | 0.13368855 | 0.14992524 | -0.0162367 | 0.69306492 | 0.89054584 |
| Papd5 | 0.07312879 | 0.06198366 | 0.07822049 | -0.0162368 | 0.83325109 | 0.89393134 |
| Ephb2 | 0.15520908 | 0.14818917 | 0.16443957 | -0.0162504 | 0.67738905 | 0.89054584 |
| Rbks | 0.08754527 | 0.07736642 | 0.09361793 | -0.0162515 | 0.79067455 | 0.89054584 |
| Esyt2 | 0.16404839 | 0.15728405 | 0.17354144 | -0.0162574 | 0.66874497 | 0.89054584 |
| Abhd14b | 0.23876282 | 0.23290699 | 0.24916973 | -0.0162627 | 0.61946592 | 0.89054584 |
| Bhlhb9 | 0.74519306 | 0.66555502 | 0.68182198 | -0.016267 | 0.54425264 | 0.89054584 |
| Dnajc18 | 0.30384242 | 0.2967174 | 0.31298954 | -0.0162721 | 0.59569346 | 0.89054584 |
| Gm9726 | 0.08311312 | 0.07261197 | 0.08888547 | -0.0162735 | 0.80316401 | 0.89054584 |
| Mlst8 | 0.07826076 | 0.06741861 | 0.083703 | -0.0162844 | 0.81763674 | 0.89054584 |
| Itsn2 | 0.34148122 | 0.33266303 | 0.34894815 | -0.0162851 | 0.58606295 | 0.89054584 |
| Sytl1 | 0.18647119 | 0.18018237 | 0.19648753 | -0.0163052 | 0.65047894 | 0.89054584 |
| Gm11273 | 0.14872317 | 0.14143054 | 0.1577415 | -0.016311 | 0.68500782 | 0.89054584 |
| Lmbr1 | 0.13215042 | 0.12422928 | 0.14055221 | -0.0163229 | 0.70576786 | 0.89054584 |
| Pcca | 0.09453076 | 0.08474405 | 0.10107419 | -0.0163301 | 0.77419274 | 0.89054584 |
| Aig1 | 0.70972216 | 0.6405416 | 0.65688542 | -0.0163438 | 0.54614128 | 0.89054584 |
| Dock7 | 0.37701772 | 0.36586174 | 0.3822191 | -0.0163574 | 0.57901842 | 0.89054584 |
| Nol11 | 0.51804805 | 0.49082552 | 0.50719895 | -0.0163734 | 0.55977557 | 0.89054584 |
| Nln | 0.08232007 | 0.07165656 | 0.08803857 | -0.016382 | 0.80700835 | 0.89054584 |
| Prepl | 0.19557742 | 0.18936131 | 0.20574679 | -0.0163855 | 0.64469377 | 0.89054584 |
| Ppp6r3 | 0.2401454 | 0.23415825 | 0.25054626 | -0.016388 | 0.61971685 | 0.89054584 |
| Msh6 | 0.19715308 | 0.19095264 | 0.20734532 | -0.0163927 | 0.64368826 | 0.89054584 |
| 1110059E24I | 0.63266829 | 0.58327165 | 0.59968084 | -0.0164092 | 0.55072116 | 0.89054584 |
| Mar-04 | 0.1076407 | 0.09852782 | 0.11494007 | -0.0164122 | 0.7470542 | 0.89054584 |
| Mical2 | 0.24188417 | 0.23585312 | 0.25227618 | -0.0164231 | 0.61916739 | 0.89054584 |

| | | | | | | |
|---|---|---|---|---|---|---|
| Grin2c | 0.10635156 | 0.09714817 | 0.11358679 | -0.0164386 | 0.74992572 | 0.89054584 |
| Pigo | 0.10256201 | 0.09316294 | 0.10960517 | -0.0164422 | 0.75769363 | 0.89054584 |
| Mttp | 0.16471233 | 0.15776382 | 0.17422383 | -0.01646 | 0.67008907 | 0.89054584 |
| Magohb | 0.54712026 | 0.51500148 | 0.53146347 | -0.016462 | 0.55737212 | 0.89054584 |
| Aasdh | 0.0686729 | 0.05699614 | 0.07345913 | -0.016463 | 0.85178495 | 0.89985795 |
| Adcy6 | 0.1262455 | 0.11793372 | 0.13440198 | -0.0164683 | 0.7159525 | 0.89054584 |
| Ahsa1 | 1.62744941 | 1.0504299 | 1.06690056 | -0.0164707 | 0.52866884 | 0.89054584 |
| Zc3hav1l | 0.1244704 | 0.11607799 | 0.13255071 | -0.0164727 | 0.71870178 | 0.89054584 |
| Sfxn3 | 0.14776379 | 0.1402748 | 0.15674936 | -0.0164746 | 0.687813 | 0.89054584 |
| Fgf9 | 0.12568357 | 0.11734024 | 0.13381606 | -0.0164758 | 0.71688831 | 0.89054584 |
| Lsm2 | 0.86254636 | 0.74139614 | 0.75787213 | -0.016476 | 0.54033766 | 0.89054584 |
| Dhodh | 0.17876213 | 0.17214303 | 0.18862147 | -0.0164784 | 0.6580021 | 0.89054584 |
| Abcc1 | 0.05625895 | 0.04367144 | 0.06018941 | -0.016518 | 0.89947538 | 0.92290121 |
| Pik3r4 | 0.13465992 | 0.12663869 | 0.14316193 | -0.0165232 | 0.70461431 | 0.89054584 |
| Ipo7 | 0.44499027 | 0.42737923 | 0.44391396 | -0.0165347 | 0.5688843 | 0.89054584 |
| Prmt10 | 0.1329635 | 0.12486022 | 0.14139803 | -0.0165378 | 0.70709107 | 0.89054584 |
| Ppp2r5d | 0.11503208 | 0.10614809 | 0.12268742 | -0.0165393 | 0.73501954 | 0.89054584 |
| Mtrf1 | 0.08625311 | 0.07569771 | 0.09223835 | -0.0165406 | 0.79827189 | 0.89054584 |
| Kdm5c | 0.39911038 | 0.38599796 | 0.40255746 | -0.0165595 | 0.57599139 | 0.89054584 |
| Rab10os | 0.19284742 | 0.18641296 | 0.20297463 | -0.0165617 | 0.64808123 | 0.89054584 |
| Tmem8b | 0.05858296 | 0.04611057 | 0.06267411 | -0.0165635 | 0.89088856 | 0.91722772 |
| Ndc1 | 0.1284047 | 0.1200611 | 0.13665243 | -0.0165913 | 0.71418559 | 0.89054584 |
| Tbrg4 | 0.09913207 | 0.08936127 | 0.10598378 | -0.0166225 | 0.76752147 | 0.89054584 |
| Rad9b | 0.13808977 | 0.13009156 | 0.14672482 | -0.0166333 | 0.70130397 | 0.89054584 |
| Kdm6a | 0.31503659 | 0.30711442 | 0.32375932 | -0.0166449 | 0.59464969 | 0.89054584 |
| Agtrap | 0.37441661 | 0.36315604 | 0.37980687 | -0.0166508 | 0.5809206 | 0.89054584 |
| Iqcg | 0.14025195 | 0.1323118 | 0.14896851 | -0.0166567 | 0.69879333 | 0.89054584 |
| E230001N04 | 0.0745244 | 0.06302985 | 0.07971156 | -0.0166817 | 0.83522103 | 0.89459411 |
| Dennd2a | 0.19067345 | 0.18406641 | 0.2007648 | -0.0166984 | 0.65080782 | 0.89054584 |
| Snx9 | 0.2253918 | 0.21911066 | 0.23581299 | -0.0167023 | 0.62929504 | 0.89054584 |

| | | | | | | |
|---|---|---|---|---|---|---|
| Snx21 | 0.33429882 | 0.32542428 | 0.34214223 | -0.0167179 | 0.59003999 | 0.89054584 |
| Sort1 | 0.20733303 | 0.2009287 | 0.21764749 | -0.0167188 | 0.63978375 | 0.89054584 |
| Adck3 | 0.06058947 | 0.04809415 | 0.06481919 | -0.016725 | 0.88534314 | 0.91420299 |
| Unc80 | 0.31242864 | 0.30452505 | 0.32125563 | -0.0167306 | 0.59585462 | 0.89054584 |
| Gna11 | 0.64969982 | 0.59593795 | 0.61267752 | -0.0167396 | 0.5506452 | 0.89054584 |
| Gorab | 0.14309878 | 0.13517494 | 0.15191991 | -0.016745 | 0.69620592 | 0.89054584 |
| Raver1 | 0.37815004 | 0.36651227 | 0.38326802 | -0.0167558 | 0.58069824 | 0.89054584 |
| D130020L05I | 0.0651947 | 0.052986 | 0.06974183 | -0.0167558 | 0.86850127 | 0.90498553 |
| Sec16a | 0.26595409 | 0.25931956 | 0.27607954 | -0.01676 | 0.61134664 | 0.89054584 |
| Acsl6 | 0.0876189 | 0.07693439 | 0.09369655 | -0.0167622 | 0.79771207 | 0.89054584 |
| Pxmp4 | 0.32880268 | 0.3201516 | 0.33691676 | -0.0167652 | 0.59166549 | 0.89054584 |
| Furin | 0.24627492 | 0.23986183 | 0.25663832 | -0.0167765 | 0.61964773 | 0.89054584 |
| Galns | 0.19303003 | 0.18637375 | 0.20316016 | -0.0167864 | 0.64986346 | 0.89054584 |
| Gys1 | 0.07173083 | 0.05993725 | 0.0767268 | -0.0167896 | 0.84601277 | 0.89825055 |
| Plekhm2 | 0.09507881 | 0.08486931 | 0.10165904 | -0.0167897 | 0.77918929 | 0.89054584 |
| Inpp5b | 0.10049362 | 0.0906306 | 0.10742976 | -0.0167992 | 0.76685683 | 0.89054584 |
| Pa2g4 | 1.30188363 | 0.95447557 | 0.97127571 | -0.0168001 | 0.53211425 | 0.89054584 |
| Usb1 | 0.21952444 | 0.21311551 | 0.22992691 | -0.0168114 | 0.63331431 | 0.89054584 |
| Snupn | 0.12946546 | 0.12093807 | 0.13775736 | -0.0168193 | 0.7152631 | 0.89054584 |
| Slc4a1ap | 0.30736037 | 0.29955377 | 0.31638065 | -0.0168269 | 0.59785093 | 0.89054584 |
| Mier2 | 0.16377133 | 0.15642892 | 0.17325663 | -0.0168277 | 0.67454708 | 0.89054584 |
| Tmem259 | 0.14837214 | 0.14054334 | 0.15737853 | -0.0168352 | 0.69089156 | 0.89054584 |
| C87436 | 0.28166636 | 0.27463504 | 0.29147024 | -0.0168352 | 0.60607304 | 0.89054584 |
| Ccdc61 | 0.1439445 | 0.13592881 | 0.15279607 | -0.0168673 | 0.69648009 | 0.89054584 |
| Shtn1 | 0.20958787 | 0.20302533 | 0.21992356 | -0.0168982 | 0.63982016 | 0.89054584 |
| Ttll5 | 0.09250887 | 0.08201341 | 0.09891638 | -0.016903 | 0.78702076 | 0.89054584 |
| Poc5 | 0.14035395 | 0.13216192 | 0.14907431 | -0.0169124 | 0.70144262 | 0.89054584 |
| Bach1 | 0.12736678 | 0.11864588 | 0.13557087 | -0.016925 | 0.71959307 | 0.89054584 |
| Fancm | 0.10672418 | 0.09705072 | 0.11397801 | -0.0169273 | 0.75550515 | 0.89054584 |
| Tmed1 | 0.58383881 | 0.54440367 | 0.56133542 | -0.0169317 | 0.55587904 | 0.89054584 |

| | | | | | | |
|---|---|---|---|---|---|---|
| Sgta | 1.03808994 | 0.83719836 | 0.85413915 | -0.0169408 | 0.53681158 | 0.89054584 |
| Aldh3a2 | 0.16477816 | 0.15732806 | 0.17429148 | -0.0169634 | 0.6748858 | 0.89054584 |
| Mal2 | 0.09761211 | 0.08738731 | 0.10436215 | -0.0169748 | 0.77571225 | 0.89054584 |
| Ncoa6 | 0.21578144 | 0.20918807 | 0.22616389 | -0.0169758 | 0.63671783 | 0.89054584 |
| Btbd7 | 0.08569636 | 0.0746672 | 0.0916439 | -0.0169767 | 0.8059456 | 0.89054584 |
| Aftph | 0.25366658 | 0.24697977 | 0.26396146 | -0.0169817 | 0.61780776 | 0.89054584 |
| Tars2 | 0.15609511 | 0.14837068 | 0.16535333 | -0.0169826 | 0.68385191 | 0.89054584 |
| Immp2l | 0.08801053 | 0.07712468 | 0.09411465 | -0.01699 | 0.79984045 | 0.89054584 |
| Sorl1 | 0.12443226 | 0.11551877 | 0.13251092 | -0.0169922 | 0.72488845 | 0.89054584 |
| Hrsp12 | 0.51069794 | 0.48397795 | 0.50098044 | -0.0170025 | 0.56281775 | 0.89054584 |
| Sppl2b | 0.25835581 | 0.25158638 | 0.26859404 | -0.0170077 | 0.61600652 | 0.89054584 |
| Cep192 | 0.09554309 | 0.08514297 | 0.10215448 | -0.0170115 | 0.78107847 | 0.89054584 |
| Zfc3h1 | 0.26502861 | 0.25815382 | 0.27516927 | -0.0170154 | 0.61336172 | 0.89054584 |
| Mocs3 | 0.15513681 | 0.14734779 | 0.16436502 | -0.0170172 | 0.68522306 | 0.89054584 |
| 1700029J07R | 0.09106777 | 0.08034087 | 0.09737823 | -0.0170374 | 0.7925014 | 0.89054584 |
| Zfp317 | 0.0869051 | 0.075889 | 0.09293447 | -0.0170455 | 0.80358999 | 0.89054584 |
| Tnks | 0.12543201 | 0.11650573 | 0.13355372 | -0.017048 | 0.72398093 | 0.89054584 |
| Fam234b | 0.14278406 | 0.13454242 | 0.15159378 | -0.0170514 | 0.69987483 | 0.89054584 |
| Clasrp | 0.20753636 | 0.20079681 | 0.21785282 | -0.017056 | 0.64235293 | 0.89054584 |
| Myo9b | 0.13180263 | 0.12312167 | 0.14019035 | -0.0170687 | 0.71472545 | 0.89054584 |
| Ubald1 | 0.39002064 | 0.37715035 | 0.3942215 | -0.0170712 | 0.57994323 | 0.89054584 |
| Btbd2 | 0.24418504 | 0.23746764 | 0.25456317 | -0.0170955 | 0.62281312 | 0.89054584 |
| 3110082I17R | 0.22699023 | 0.22031609 | 0.23741383 | -0.0170977 | 0.63138407 | 0.89054584 |
| Setd7 | 0.10954365 | 0.0998328 | 0.11693659 | -0.0171038 | 0.75220425 | 0.89054584 |
| Ccpg1os | 0.32471477 | 0.31589387 | 0.33301988 | -0.017126 | 0.59467689 | 0.89054584 |
| Rab21 | 0.05908701 | 0.04608524 | 0.06321299 | -0.0171277 | 0.89658925 | 0.92130916 |
| Tmem138 | 0.38536425 | 0.37279734 | 0.38993406 | -0.0171367 | 0.58111624 | 0.89054584 |
| Rlim | 0.52905528 | 0.49930171 | 0.51644942 | -0.0171477 | 0.56146762 | 0.89054584 |
| Fgfr1op | 0.13874006 | 0.13025051 | 0.14739981 | -0.0171493 | 0.70610685 | 0.89054584 |
| Slc12a2 | 0.12060796 | 0.11136155 | 0.1285184 | -0.0171568 | 0.7330145 | 0.89054584 |

| | | | | | | |
|---|---|---|---|---|---|---|
| Kctd3 | 0.27557726 | 0.26835279 | 0.28552002 | -0.0171672 | 0.61030895 | 0.89054584 |
| Gtpbp8 | 0.16957894 | 0.16204312 | 0.17922014 | -0.017177 | 0.67239185 | 0.89054584 |
| Cog4 | 0.39581252 | 0.3823597 | 0.3995385 | -0.0171788 | 0.5793843 | 0.89054584 |
| Rabl3 | 0.23983776 | 0.23303999 | 0.25024004 | -0.0172001 | 0.62560251 | 0.89054584 |
| Dars2 | 0.09998105 | 0.08968749 | 0.10688947 | -0.017202 | 0.77330484 | 0.89054584 |
| Tbc1d10b | 0.11491128 | 0.1053389 | 0.12256096 | -0.0172221 | 0.7436588 | 0.89054584 |
| Lmbrd2 | 0.11251293 | 0.1028245 | 0.12004922 | -0.0172247 | 0.74808229 | 0.89054584 |
| Etaa1 | 0.26623453 | 0.25910631 | 0.27635528 | -0.017249 | 0.61439489 | 0.89054584 |
| Cpeb2 | 0.08505976 | 0.07370467 | 0.09096416 | -0.0172595 | 0.81164787 | 0.89054584 |
| Tmem30b | 0.13906276 | 0.13043044 | 0.14773471 | -0.0173043 | 0.70736569 | 0.89054584 |
| Zfp426 | 0.12544779 | 0.11626497 | 0.13357019 | -0.0173052 | 0.72694779 | 0.89054584 |
| Mettl17 | 0.11112282 | 0.10127903 | 0.1185924 | -0.0173134 | 0.75179888 | 0.89054584 |
| Cyp4v3 | 0.07028702 | 0.05784633 | 0.07518402 | -0.0173377 | 0.85867102 | 0.90127639 |
| Ccdc30 | 0.09386515 | 0.08301995 | 0.10036387 | -0.0173439 | 0.7896229 | 0.89054584 |
| Mrps12 | 1.42184579 | 0.99525218 | 1.01260063 | -0.0173484 | 0.53180964 | 0.89054584 |
| Zfp442 | 0.12952488 | 0.12045849 | 0.13781924 | -0.0173608 | 0.72135632 | 0.89054584 |
| Cecr2 | 0.06012463 | 0.04695652 | 0.06432227 | -0.0173658 | 0.8957721 | 0.92079579 |
| Slc4a3 | 0.0881217 | 0.07685884 | 0.09423333 | -0.0173745 | 0.80483064 | 0.89054584 |
| Nod1 | 0.21560018 | 0.20859976 | 0.22598151 | -0.0173818 | 0.63995945 | 0.89054584 |
| Snx12 | 0.52768006 | 0.49790862 | 0.51529784 | -0.0173892 | 0.56246443 | 0.89054584 |
| Fbxw17 | 0.12061215 | 0.11112924 | 0.12852278 | -0.0173935 | 0.73582102 | 0.89054584 |
| Med15 | 0.21045403 | 0.20339478 | 0.22079728 | -0.0174025 | 0.64326584 | 0.89054584 |
| Mpp7 | 0.12695322 | 0.1177158 | 0.1351398 | -0.017424 | 0.72597743 | 0.89054584 |
| Dbp | 0.13700294 | 0.1281642 | 0.14559632 | -0.0174321 | 0.71151335 | 0.89054584 |
| Supt6 | 0.46188228 | 0.44140041 | 0.45883292 | -0.0174325 | 0.57024889 | 0.89054584 |
| Nfrkb | 0.18772264 | 0.1803287 | 0.19776211 | -0.0174334 | 0.6593543 | 0.89054584 |
| Poll | 0.05850269 | 0.04514796 | 0.0625883 | -0.0174403 | 0.90289451 | 0.92516536 |
| Dcaf6 | 0.23695019 | 0.22991307 | 0.24736369 | -0.0174506 | 0.62879802 | 0.89054584 |
| Prss36 | 0.09545782 | 0.08459688 | 0.10206349 | -0.0174666 | 0.78736346 | 0.89054584 |
| Zfyve19 | 0.3991014 | 0.38507936 | 0.40254925 | -0.0174699 | 0.58011569 | 0.89054584 |

| | | | | | | |
|---|---|---|---|---|---|---|
| Arpc5l | 1.40332434 | 0.98920906 | 1.00668539 | -0.0174763 | 0.53223149 | 0.89054584 |
| Nfkbiz | 0.10041441 | 0.0898395 | 0.10734642 | -0.0175069 | 0.77631824 | 0.89054584 |
| Dcp2 | 0.2651433 | 0.25775904 | 0.2752821 | -0.0175231 | 0.61660049 | 0.89054584 |
| Mrps24 | 1.30430061 | 0.95464946 | 0.97218121 | -0.0175317 | 0.5334784 | 0.89054584 |
| Mipep | 0.25091232 | 0.24370314 | 0.26123574 | -0.0175326 | 0.6227383 | 0.89054584 |
| Ankrd10 | 0.54326391 | 0.51074291 | 0.5282761 | -0.0175332 | 0.56144259 | 0.89054584 |
| Fam8a1 | 0.23583134 | 0.22871434 | 0.24624815 | -0.0175338 | 0.62995509 | 0.89054584 |
| Mgat4a | 0.09722542 | 0.08641019 | 0.10394958 | -0.0175394 | 0.78409611 | 0.89054584 |
| Rbm5 | 1.3310878 | 0.96445258 | 0.98200179 | -0.0175492 | 0.53317731 | 0.89054584 |
| Pvrl1 | 0.13896419 | 0.13006758 | 0.14763243 | -0.0175648 | 0.71031539 | 0.89054584 |
| Sin3a | 0.29795764 | 0.28973643 | 0.30730291 | -0.0175665 | 0.60500398 | 0.89054584 |
| Mpdz | 0.24780975 | 0.24058567 | 0.25816103 | -0.0175754 | 0.62444508 | 0.89054584 |
| Zfp944 | 0.15977428 | 0.15156808 | 0.16914431 | -0.0175762 | 0.68585065 | 0.89054584 |
| AI837181 | 0.11171007 | 0.10161784 | 0.11920791 | -0.0175901 | 0.75410475 | 0.89054584 |
| Ftsj2 | 0.20687378 | 0.19958424 | 0.21718364 | -0.0175994 | 0.64711222 | 0.89054584 |
| Ska2 | 0.27967149 | 0.27190344 | 0.28952287 | -0.0176194 | 0.61161401 | 0.89054584 |
| Cfap77 | 0.19495836 | 0.18749352 | 0.20511845 | -0.0176249 | 0.65554049 | 0.89054584 |
| Aste1 | 0.09989913 | 0.08917117 | 0.10680209 | -0.0176309 | 0.77908286 | 0.89054584 |
| Zfp410 | 0.4521165 | 0.43256107 | 0.45022876 | -0.0176677 | 0.57253203 | 0.89054584 |
| Bcl2l13 | 0.17754865 | 0.16968991 | 0.18738116 | -0.0176913 | 0.66998278 | 0.89054584 |
| Rpap1 | 0.17382784 | 0.16587962 | 0.18357431 | -0.0176947 | 0.67331102 | 0.89054584 |
| Amigo1 | 0.11768058 | 0.10775329 | 0.12545854 | -0.0177052 | 0.74456693 | 0.89054584 |
| Fbxw11 | 0.21450319 | 0.20716171 | 0.22487739 | -0.0177157 | 0.64320095 | 0.89054584 |
| Nat6 | 0.15675611 | 0.1483185 | 0.16603482 | -0.0177163 | 0.69044788 | 0.89054584 |
| Fbxl4 | 0.12619244 | 0.11656869 | 0.13434666 | -0.017778 | 0.73122218 | 0.89054584 |
| Epb41l1 | 0.13019224 | 0.12071717 | 0.13851416 | -0.017797 | 0.72528152 | 0.89054584 |
| Tufm | 0.45094938 | 0.43139191 | 0.44919662 | -0.0178047 | 0.57325426 | 0.89054584 |
| Nuak2 | 0.06202139 | 0.04854192 | 0.06634991 | -0.017808 | 0.89443005 | 0.91973864 |
| Thap3 | 0.25789744 | 0.25032979 | 0.26814159 | -0.0178118 | 0.62152129 | 0.89054584 |
| Dach1 | 0.21483796 | 0.20738379 | 0.22521439 | -0.0178306 | 0.64388248 | 0.89054584 |

| | | | | | | |
|---|---|---|---|---|---|---|
| Asb13 | 0.26855297 | 0.26079079 | 0.27863349 | -0.0178427 | 0.61727895 | 0.89054584 |
| Cdr2l | 0.07248492 | 0.05968738 | 0.07753254 | -0.0178452 | 0.85821144 | 0.90124396 |
| Cdk2ap1 | 0.38832316 | 0.37481241 | 0.3926597 | -0.0178473 | 0.5838527 | 0.89054584 |
| Ccm2 | 0.33223284 | 0.32233157 | 0.34017985 | -0.0178483 | 0.59655559 | 0.89054584 |
| Mvb12b | 0.1045325 | 0.09379166 | 0.11167619 | -0.0178845 | 0.7722002 | 0.89054584 |
| Chmp4c | 0.10416642 | 0.09340125 | 0.11129153 | -0.0178903 | 0.77305088 | 0.89054584 |
| Lsm5 | 1.23068921 | 0.92543328 | 0.94333499 | -0.0179017 | 0.53522576 | 0.89054584 |
| Dapk3 | 0.66079619 | 0.60313152 | 0.62103859 | -0.0179071 | 0.55343165 | 0.89054584 |
| Isoc2b | 0.1173109 | 0.10716276 | 0.1250719 | -0.0179091 | 0.74765134 | 0.89054584 |
| Map3k7 | 0.17517913 | 0.16704347 | 0.18495751 | -0.017914 | 0.67409217 | 0.89054584 |
| Pde3b | 0.23784619 | 0.23033713 | 0.24825661 | -0.0179195 | 0.63167328 | 0.89054584 |
| Nrip1 | 0.45728993 | 0.43686658 | 0.45479401 | -0.0179274 | 0.57285592 | 0.89054584 |
| Cuedc1 | 0.49924863 | 0.47329712 | 0.49122576 | -0.0179286 | 0.56751102 | 0.89054584 |
| Fancc | 0.0517229 | 0.03740793 | 0.05533913 | -0.0179312 | 0.93442826 | 0.94723402 |
| Sall1 | 0.0654287 | 0.05205601 | 0.06999194 | -0.0179359 | 0.88373887 | 0.91341314 |
| Mina | 0.17876356 | 0.17068654 | 0.18862293 | -0.0179364 | 0.67112607 | 0.89054584 |
| Las1l | 0.46755488 | 0.4458449 | 0.46380422 | -0.0179593 | 0.57158167 | 0.89054584 |
| Mark1 | 0.07099907 | 0.05795549 | 0.07594489 | -0.0179894 | 0.86511316 | 0.90376211 |
| Rfwd2 | 0.31730912 | 0.30794739 | 0.32593814 | -0.0179907 | 0.60147187 | 0.89054584 |
| Sepn1 | 0.22164066 | 0.21405961 | 0.23205166 | -0.017992 | 0.64103604 | 0.89054584 |
| Opa3 | 0.30282764 | 0.29401386 | 0.31201018 | -0.0179963 | 0.60592795 | 0.89054584 |
| C2cd4c | 0.07360569 | 0.06073152 | 0.07873002 | -0.0179985 | 0.85658069 | 0.9009252 |
| Map3k1 | 0.2078251 | 0.20013806 | 0.21814438 | -0.0180063 | 0.6497196 | 0.89054584 |
| Pus7 | 0.22476292 | 0.2171764 | 0.23518285 | -0.0180065 | 0.63934385 | 0.89054584 |
| Sephs2 | 0.54617819 | 0.5126775 | 0.53068568 | -0.0180082 | 0.56280923 | 0.89054584 |
| Tefm | 0.15588647 | 0.14712558 | 0.16513819 | -0.0180126 | 0.69433599 | 0.89054584 |
| Lancl2 | 0.12596257 | 0.11609376 | 0.13410699 | -0.0180132 | 0.73427312 | 0.89054584 |
| Tm2d2 | 1.35849603 | 0.9736088 | 0.99164057 | -0.0180318 | 0.53375687 | 0.89054584 |
| Apex1 | 0.8817026 | 0.7512947 | 0.76933682 | -0.0180421 | 0.54350157 | 0.89054584 |
| Acadm | 0.63651564 | 0.58457341 | 0.60263422 | -0.0180608 | 0.55552281 | 0.89054584 |

| | | | | | | |
|---|---|---|---|---|---|---|
| Wdr18 | 0.63934042 | 0.5867332 | 0.60479615 | -0.018063 | 0.55533215 | 0.89054584 |
| Hmg20a | 0.1581567 | 0.14941526 | 0.16747825 | -0.018063 | 0.69233484 | 0.89054584 |
| Cpm | 0.07988576 | 0.06737163 | 0.08543871 | -0.0180671 | 0.83773487 | 0.89485554 |
| Kif3b | 0.19662145 | 0.18872684 | 0.2068061 | -0.0180793 | 0.65810277 | 0.89054584 |
| Trmu | 0.10489343 | 0.09397158 | 0.11205537 | -0.0180838 | 0.77393598 | 0.89054584 |
| Ago3 | 0.15448574 | 0.14560633 | 0.16369334 | -0.018087 | 0.6966472 | 0.89054584 |
| Scarb1 | 0.27168587 | 0.26361839 | 0.28170791 | -0.0180895 | 0.6175941 | 0.89054584 |
| Zfp870 | 0.08810337 | 0.07611559 | 0.09421376 | -0.0180982 | 0.81459791 | 0.89054584 |
| Ubac2 | 0.26626892 | 0.25828352 | 0.2763891 | -0.0181056 | 0.61989157 | 0.89054584 |
| Ncbp3 | 0.43951361 | 0.42092698 | 0.43904085 | -0.0181139 | 0.57621419 | 0.89054584 |
| Dph6 | 0.29424054 | 0.28558764 | 0.30370199 | -0.0181144 | 0.60944903 | 0.89054584 |
| Memo1 | 0.37807126 | 0.36507784 | 0.38319507 | -0.0181172 | 0.58717029 | 0.89054584 |
| Med16 | 0.33168653 | 0.32151867 | 0.33966057 | -0.0181419 | 0.59825875 | 0.89054584 |
| Ggcx | 0.26301756 | 0.25504793 | 0.27318986 | -0.0181419 | 0.62148751 | 0.89054584 |
| Zfp143 | 0.08960343 | 0.07763698 | 0.09581513 | -0.0181781 | 0.81162467 | 0.89054584 |
| Pacrgl | 0.22561574 | 0.21785427 | 0.23603735 | -0.0181831 | 0.64016522 | 0.89054584 |
| Slc20a2 | 0.12254993 | 0.11235618 | 0.13054648 | -0.0181903 | 0.74188745 | 0.89054584 |
| Stk11ip | 0.07449598 | 0.06148471 | 0.07968121 | -0.0181965 | 0.85632318 | 0.9009252 |
| Swsap1 | 0.10805188 | 0.09717221 | 0.11537158 | -0.0181994 | 0.7688083 | 0.89054584 |
| Ing3 | 0.30448682 | 0.29540444 | 0.31361118 | -0.0182067 | 0.60660277 | 0.89054584 |
| Adrbk2 | 0.34969873 | 0.33849221 | 0.35670178 | -0.0182096 | 0.59399673 | 0.89054584 |
| Habp4 | 0.28137214 | 0.27296975 | 0.29118314 | -0.0182134 | 0.61463304 | 0.89054584 |
| Egln1 | 0.17257287 | 0.16406969 | 0.18228903 | -0.0182193 | 0.67926683 | 0.89054584 |
| Rorc | 0.36778234 | 0.3554055 | 0.37363743 | -0.0182319 | 0.58991903 | 0.89054584 |
| Sik3 | 0.16247628 | 0.15369021 | 0.17192495 | -0.0182347 | 0.68939545 | 0.89054584 |
| Chd3os | 0.23959094 | 0.23174148 | 0.24999433 | -0.0182528 | 0.63313328 | 0.89054584 |
| Mib2 | 0.15661248 | 0.14762296 | 0.16588675 | -0.0182638 | 0.6960033 | 0.89054584 |
| Gdpd1 | 0.31402336 | 0.30451098 | 0.32278701 | -0.018276 | 0.60402793 | 0.89054584 |
| Exosc2 | 0.29460702 | 0.2857729 | 0.30405733 | -0.0182844 | 0.61032433 | 0.89054584 |
| Ppp1r8 | 0.45399381 | 0.43359651 | 0.45188724 | -0.0182907 | 0.57478772 | 0.89054584 |

| | | | | | | |
|---|---|---|---|---|---|---|
| Def8 | 0.19673168 | 0.18861196 | 0.20691792 | -0.018306 | 0.65989437 | 0.89054584 |
| Pdgfa | 0.40408798 | 0.38878538 | 0.40710236 | -0.018317 | 0.58301877 | 0.89054584 |
| Pcyt1a | 0.14376671 | 0.13427775 | 0.1526119 | -0.0183342 | 0.71215991 | 0.89054584 |
| Crlf2 | 0.18804632 | 0.17974396 | 0.19809166 | -0.0183477 | 0.66695546 | 0.89054584 |
| Vps16 | 0.13248895 | 0.12255324 | 0.14090441 | -0.0183512 | 0.72799921 | 0.89054584 |
| Bag2 | 0.30157668 | 0.29243464 | 0.31080217 | -0.0183675 | 0.60846873 | 0.89054584 |
| Tmbim1 | 0.40360182 | 0.38828692 | 0.40665909 | -0.0183722 | 0.58335466 | 0.89054584 |
| Gm5900 | 0.45855871 | 0.43753307 | 0.45591115 | -0.0183781 | 0.57448526 | 0.89054584 |
| Tmem177 | 0.13122259 | 0.12120864 | 0.13958672 | -0.0183781 | 0.73019386 | 0.89054584 |
| 1500011K16I | 0.77496802 | 0.68366453 | 0.70206082 | -0.0183963 | 0.54858196 | 0.89054584 |
| C2cd5 | 0.26454746 | 0.25628456 | 0.27469587 | -0.0184113 | 0.62257946 | 0.89054584 |
| Ercc5 | 0.14513532 | 0.13561401 | 0.15402929 | -0.0184153 | 0.71123943 | 0.89054584 |
| Ksr2 | 0.12351281 | 0.11313048 | 0.13155153 | -0.0184211 | 0.74292446 | 0.89054584 |
| Sertad1 | 0.21144026 | 0.20336555 | 0.22179171 | -0.0184262 | 0.65064365 | 0.89054584 |
| Ccdc58 | 0.56890556 | 0.53086072 | 0.54929239 | -0.0184317 | 0.56211438 | 0.89054584 |
| Btaf1 | 0.18189623 | 0.17337576 | 0.19182241 | -0.0184466 | 0.67293355 | 0.89054584 |
| Pfdn2 | 1.98703191 | 1.09843033 | 1.11687972 | -0.0184494 | 0.53067217 | 0.89054584 |
| Fam73a | 0.20478613 | 0.19662421 | 0.21507396 | -0.0184497 | 0.65529624 | 0.89054584 |
| Sfmbt1 | 0.12173954 | 0.11124932 | 0.12970033 | -0.018451 | 0.74626277 | 0.89054584 |
| Dirc2 | 0.15721711 | 0.14802296 | 0.16651 | -0.018487 | 0.69751494 | 0.89054584 |
| Zbtb4 | 0.18439207 | 0.17586212 | 0.19436853 | -0.0185064 | 0.67133116 | 0.89054584 |
| Fads6 | 0.08463017 | 0.07197494 | 0.09050545 | -0.0185305 | 0.82993116 | 0.89271879 |
| Foxred1 | 0.12642815 | 0.11604893 | 0.13459242 | -0.0185435 | 0.73952645 | 0.89054584 |
| Nbeal1 | 0.50785175 | 0.48001102 | 0.49856324 | -0.0185522 | 0.56881796 | 0.89054584 |
| Mapk14 | 0.27322942 | 0.26465259 | 0.28322089 | -0.0185683 | 0.61998664 | 0.89054584 |
| Phactr4 | 0.28118583 | 0.2724118 | 0.29100133 | -0.0185895 | 0.61700299 | 0.89054584 |
| Ptpru | 0.10825097 | 0.0969833 | 0.11558049 | -0.0185972 | 0.77326277 | 0.89054584 |
| Ddx20 | 0.17387899 | 0.1650271 | 0.18362668 | -0.0185996 | 0.68150274 | 0.89054584 |
| Carf | 0.17610916 | 0.16730622 | 0.18590907 | -0.0186029 | 0.6794615 | 0.89054584 |
| Tmem102 | 0.11995635 | 0.10923151 | 0.12783758 | -0.0186061 | 0.75114069 | 0.89054584 |

| | | | | | | |
|---|---|---|---|---|---|---|
| Cstf2 | 0.2343534 | 0.22615671 | 0.2447737 | -0.018617 | 0.63846158 | 0.89054584 |
| Papolg | 0.13083542 | 0.1205661 | 0.13918373 | -0.0186176 | 0.73342146 | 0.89054584 |
| Exoc6b | 0.26632614 | 0.25781401 | 0.27644535 | -0.0186313 | 0.62323739 | 0.89054584 |
| Inip | 0.18280254 | 0.17410855 | 0.19274728 | -0.0186387 | 0.67383262 | 0.89054584 |
| Park2 | 0.07031335 | 0.05656615 | 0.07521215 | -0.018646 | 0.87595643 | 0.90910816 |
| AW146154 | 0.09339684 | 0.08121321 | 0.09986408 | -0.0186509 | 0.80788141 | 0.89054584 |
| Zer1 | 0.24399618 | 0.23572462 | 0.25437554 | -0.0186509 | 0.63367236 | 0.89054584 |
| Cfap36 | 0.92959639 | 0.77819309 | 0.79684668 | -0.0186536 | 0.54342349 | 0.89054584 |
| Fpgt | 0.08957977 | 0.07713002 | 0.09578988 | -0.0186599 | 0.8179449 | 0.89054584 |
| Zbtb5 | 0.16210267 | 0.15287822 | 0.17154064 | -0.0186624 | 0.69387357 | 0.89054584 |
| Cchcr1 | 0.06877533 | 0.05490304 | 0.0735686 | -0.0186656 | 0.88140699 | 0.91204329 |
| Gm4944 | 0.09964553 | 0.08786399 | 0.10653156 | -0.0186676 | 0.79286083 | 0.89054584 |
| Fis1 | 1.79316054 | 1.07810956 | 1.09678206 | -0.0186725 | 0.53160998 | 0.89054584 |
| Mtfmt | 0.1477319 | 0.13804317 | 0.15671637 | -0.0186732 | 0.71059578 | 0.89054584 |
| Zdhhc24 | 0.09430243 | 0.08215579 | 0.10083053 | -0.0186747 | 0.80589785 | 0.89054584 |
| Ripk1 | 0.2475436 | 0.23919776 | 0.25789706 | -0.0186993 | 0.63224501 | 0.89054584 |
| Csrp2bp | 0.12733274 | 0.11680633 | 0.13553538 | -0.0187291 | 0.74014715 | 0.89054584 |
| Smurf2 | 0.21989044 | 0.21155871 | 0.23029453 | -0.0187358 | 0.64766807 | 0.89054584 |
| Slc35a4 | 0.5276766 | 0.49654571 | 0.51529494 | -0.0187492 | 0.56730521 | 0.89054584 |
| Gm4419 | 0.10836153 | 0.09694258 | 0.1156965 | -0.0187539 | 0.77493398 | 0.89054584 |
| Map2k5 | 0.28618735 | 0.27711146 | 0.29587683 | -0.0187654 | 0.61618778 | 0.89054584 |
| Bmpr1a | 0.37672855 | 0.36318393 | 0.38195112 | -0.0187672 | 0.59053339 | 0.89054584 |
| Prmt3 | 0.29714209 | 0.28774573 | 0.30651345 | -0.0187677 | 0.61227957 | 0.89054584 |
| Cnst | 0.16881031 | 0.15965976 | 0.17843167 | -0.0187719 | 0.68798408 | 0.89054584 |
| Zfp113 | 0.19296172 | 0.18431492 | 0.20309076 | -0.0187758 | 0.66666454 | 0.89054584 |
| Kbtbd7 | 0.13241264 | 0.12201814 | 0.14082502 | -0.0188069 | 0.73309202 | 0.89054584 |
| Llgl2 | 0.42044262 | 0.40312747 | 0.42193646 | -0.018809 | 0.58226261 | 0.89054584 |
| Cbx2 | 0.1350609 | 0.12475106 | 0.1435787 | -0.0188276 | 0.72937779 | 0.89054584 |
| Paxbp1 | 0.2581215 | 0.24952642 | 0.26836276 | -0.0188363 | 0.62817002 | 0.89054584 |
| Ngrn | 0.62861932 | 0.57771552 | 0.59656171 | -0.0188462 | 0.5585059 | 0.89054584 |

| | | | | | | |
|---|---|---|---|---|---|---|
| Rnf208 | 0.11469629 | 0.10348491 | 0.12233589 | -0.018851 | 0.76359221 | 0.89054584 |
| Gm12258 | 0.1267251 | 0.11603815 | 0.134902 | -0.0188639 | 0.74263075 | 0.89054584 |
| Asah1 | 0.75237502 | 0.66789099 | 0.6867618 | -0.0188708 | 0.55093271 | 0.89054584 |
| Sec22c | 0.11081391 | 0.09937937 | 0.11826857 | -0.0188892 | 0.771592 | 0.89054584 |
| Sytl5 | 0.12345328 | 0.11258305 | 0.1314894 | -0.0189063 | 0.74853637 | 0.89054584 |
| Eef1e1 | 0.64722988 | 0.59188977 | 0.61080514 | -0.0189154 | 0.55735943 | 0.89054584 |
| Zxdc | 0.09368905 | 0.08126008 | 0.10017593 | -0.0189159 | 0.81049744 | 0.89054584 |
| Zdhhc17 | 0.16487289 | 0.15545959 | 0.17438883 | -0.0189292 | 0.69346995 | 0.89054584 |
| Enc1 | 0.18015445 | 0.17106986 | 0.19004399 | -0.0189741 | 0.67908872 | 0.89054584 |
| Dennd6a | 0.30228774 | 0.29251269 | 0.31148892 | -0.0189762 | 0.61172874 | 0.89054584 |
| Zdhhc14 | 0.07587231 | 0.06216283 | 0.08115158 | -0.0189887 | 0.86217665 | 0.90260095 |
| Gtf2h2 | 0.2644243 | 0.25556792 | 0.27457468 | -0.0190068 | 0.62646445 | 0.89054584 |
| Tspan33 | 0.1242259 | 0.11328396 | 0.13229562 | -0.0190117 | 0.74841726 | 0.89054584 |
| Tigar | 0.11997171 | 0.10884112 | 0.12785364 | -0.0190125 | 0.75578541 | 0.89054584 |
| Ranbp2 | 0.40223076 | 0.38639412 | 0.40540825 | -0.0190141 | 0.58648407 | 0.89054584 |
| Arfgef3 | 0.11641475 | 0.10511788 | 0.12413443 | -0.0190165 | 0.76229922 | 0.89054584 |
| Ddx51 | 0.1079348 | 0.09623154 | 0.11524872 | -0.0190172 | 0.77898544 | 0.89054584 |
| Wdr91 | 0.09417586 | 0.08166942 | 0.10069546 | -0.019026 | 0.81064884 | 0.89054584 |
| Pik3ca | 0.25145863 | 0.24273519 | 0.26177667 | -0.0190415 | 0.63265316 | 0.89054584 |
| Pola1 | 0.18794982 | 0.17893524 | 0.19799341 | -0.0190582 | 0.67308708 | 0.89054584 |
| Acvr2b | 0.07175834 | 0.05769124 | 0.07675619 | -0.0190649 | 0.87640524 | 0.9091719 |
| Rbl2 | 0.11753374 | 0.10623502 | 0.12530497 | -0.0190699 | 0.76084899 | 0.89054584 |
| Tmem229b | 0.10362553 | 0.09165289 | 0.11072312 | -0.0190702 | 0.78884671 | 0.89054584 |
| Naglu | 0.18894125 | 0.1799234 | 0.19900259 | -0.0190792 | 0.67244393 | 0.89054584 |
| Tomm5 | 1.25482628 | 0.9340313 | 0.95311206 | -0.0190808 | 0.53715521 | 0.89054584 |
| Xrcc4 | 0.28742238 | 0.27799648 | 0.29707898 | -0.0190825 | 0.61762964 | 0.89054584 |
| Dph5 | 0.13500456 | 0.12441951 | 0.14352014 | -0.0191006 | 0.7323854 | 0.89054584 |
| Aida | 0.46981668 | 0.44667838 | 0.46578143 | -0.019103 | 0.57576725 | 0.89054584 |
| Cdk10 | 0.14976293 | 0.13970923 | 0.15881634 | -0.0191071 | 0.71243582 | 0.89054584 |
| Ralgapa2 | 0.1571891 | 0.14737195 | 0.16648114 | -0.0191092 | 0.70359443 | 0.89054584 |

| | | | | | | |
|---|---|---|---|---|---|---|
| Rbm12 | 0.18562373 | 0.17651475 | 0.19562403 | -0.0191093 | 0.67548237 | 0.89054584 |
| Hacl1 | 0.12223533 | 0.11110336 | 0.13021803 | -0.0191147 | 0.75297604 | 0.89054584 |
| Dcun1d5 | 1.05936884 | 0.84589877 | 0.86502931 | -0.0191305 | 0.54103231 | 0.89054584 |
| Stat5b | 0.14357461 | 0.13327562 | 0.1524129 | -0.0191373 | 0.72072157 | 0.89054584 |
| Spopl | 0.25230568 | 0.24347521 | 0.26261511 | -0.0191399 | 0.63290324 | 0.89054584 |
| Aldh7a1 | 0.5353437 | 0.50255515 | 0.52169973 | -0.0191446 | 0.56787517 | 0.89054584 |
| Mfsd1 | 0.27578211 | 0.26655671 | 0.28572049 | -0.0191638 | 0.62266345 | 0.89054584 |
| Rfc2 | 0.7960936 | 0.6968635 | 0.71603333 | -0.0191698 | 0.54963149 | 0.89054584 |
| Fam175a | 0.17637182 | 0.16698978 | 0.18617774 | -0.019188 | 0.68444414 | 0.89054584 |
| Abi2 | 0.25748498 | 0.24852937 | 0.26773437 | -0.019205 | 0.6308831 | 0.89054584 |
| Sypl | 0.87759473 | 0.74769139 | 0.76690099 | -0.0192096 | 0.5464506 | 0.89054584 |
| Med19 | 0.45082271 | 0.42986988 | 0.44908456 | -0.0192147 | 0.5790012 | 0.89054584 |
| Sars2 | 0.12175942 | 0.11049042 | 0.12972109 | -0.0192307 | 0.75511383 | 0.89054584 |
| Epb41l4b | 0.10207005 | 0.08983868 | 0.1090879 | -0.0192492 | 0.79448955 | 0.89054584 |
| 3010003L21F | 0.15947971 | 0.14957041 | 0.16884099 | -0.0192706 | 0.70254894 | 0.89054584 |
| Psmg3 | 0.50668877 | 0.47829818 | 0.49757407 | -0.0192759 | 0.57161454 | 0.89054584 |
| Spidr | 0.13702729 | 0.12634345 | 0.14562161 | -0.0192782 | 0.73130851 | 0.89054584 |
| Fam120aos | 0.09076386 | 0.0777731 | 0.09705383 | -0.0192807 | 0.82265904 | 0.89073269 |
| Ski | 0.39463428 | 0.37914264 | 0.3984584 | -0.0193158 | 0.58934015 | 0.89054584 |
| Tet2 | 0.25298595 | 0.24395464 | 0.26328821 | -0.0193336 | 0.63386659 | 0.89054584 |
| Ube2e1 | 1.28674567 | 0.94622364 | 0.96556507 | -0.0193414 | 0.537177 | 0.89054584 |
| Slc12a7 | 0.10054042 | 0.08811911 | 0.10747899 | -0.0193599 | 0.79932194 | 0.89054584 |
| Tbc1d24 | 0.19357343 | 0.18434854 | 0.20371216 | -0.0193636 | 0.67106294 | 0.89054584 |
| Stx12 | 0.79873364 | 0.69838162 | 0.71775677 | -0.0193752 | 0.55004049 | 0.89054584 |
| Sptbn2 | 0.09963538 | 0.08714319 | 0.10652072 | -0.0193775 | 0.80164816 | 0.89054584 |
| Dera | 0.48878662 | 0.46284671 | 0.48224185 | -0.0193951 | 0.57431729 | 0.89054584 |
| Snx19 | 0.09359523 | 0.08067688 | 0.10007581 | -0.0193989 | 0.81676356 | 0.89054584 |
| Zdhhc7 | 0.1761962 | 0.16659339 | 0.1859981 | -0.0194047 | 0.68653608 | 0.89054584 |
| Slc35c1 | 0.13402023 | 0.12308826 | 0.14249693 | -0.0194087 | 0.73714458 | 0.89054584 |
| C330007P06I | 0.2443135 | 0.23528016 | 0.25469079 | -0.0194106 | 0.63873128 | 0.89054584 |

| | | | | | | |
|---|---|---|---|---|---|---|
| Golga2 | 0.22273418 | 0.21372844 | 0.23314881 | -0.0194204 | 0.65101843 | 0.89054584 |
| Psma2 | 2.38284828 | 1.10999793 | 1.12941949 | -0.0194216 | 0.53192726 | 0.89054584 |
| Rbm15b | 0.15769655 | 0.14757878 | 0.16700411 | -0.0194253 | 0.70606032 | 0.89054584 |
| Samd8 | 0.19004488 | 0.1806947 | 0.20012549 | -0.0194308 | 0.67449183 | 0.89054584 |
| Pygo2 | 0.18645372 | 0.17703476 | 0.19646973 | -0.019435 | 0.67755296 | 0.89054584 |
| Slc7a5 | 0.24656936 | 0.23749013 | 0.25693052 | -0.0194404 | 0.63777362 | 0.89054584 |
| Ttc7 | 0.0688232 | 0.05415639 | 0.07361975 | -0.0194634 | 0.8909724 | 0.91722772 |
| Rwdd3 | 0.15205604 | 0.14172032 | 0.16118531 | -0.019465 | 0.71315349 | 0.89054584 |
| Efcab7 | 0.0814408 | 0.06763313 | 0.08709954 | -0.0194664 | 0.85112152 | 0.89958153 |
| Mapk13 | 0.26543727 | 0.25610413 | 0.27557126 | -0.0194671 | 0.62896778 | 0.89054584 |
| Dus2 | 0.09366574 | 0.08067728 | 0.10015105 | -0.0194738 | 0.81750645 | 0.89054584 |
| Setx | 0.20918238 | 0.20003828 | 0.21951442 | -0.0194761 | 0.66032805 | 0.89054584 |
| Impad1 | 0.70314346 | 0.63267606 | 0.65216268 | -0.0194866 | 0.55535756 | 0.89054584 |
| Pex13 | 0.5207578 | 0.48999502 | 0.50948287 | -0.0194879 | 0.57071928 | 0.89054584 |
| Trappc11 | 0.24401346 | 0.23488945 | 0.25439271 | -0.0195033 | 0.63952231 | 0.89054584 |
| Etv6 | 0.09109926 | 0.07789299 | 0.09741184 | -0.0195188 | 0.82472757 | 0.89148775 |
| Smndc1 | 0.44938876 | 0.42828816 | 0.44781521 | -0.0195271 | 0.58049261 | 0.89054584 |
| Tsc2 | 0.11921519 | 0.10753461 | 0.127063 | -0.0195284 | 0.76301525 | 0.89054584 |
| Ubr1 | 0.22597641 | 0.21686302 | 0.23639862 | -0.0195356 | 0.649884 | 0.89054584 |
| Cutc | 0.11865591 | 0.1069381 | 0.12647837 | -0.0195403 | 0.76417069 | 0.89054584 |
| Rnaseh2a | 0.30366224 | 0.29327475 | 0.31281569 | -0.0195409 | 0.6144875 | 0.89054584 |
| Nploc4 | 0.17425155 | 0.16446515 | 0.18400811 | -0.019543 | 0.68963368 | 0.89054584 |
| Sipa1l3 | 0.30791855 | 0.29737181 | 0.31691812 | -0.0195463 | 0.61307625 | 0.89054584 |
| Wdsub1 | 0.23250214 | 0.22336836 | 0.24292541 | -0.0195571 | 0.64619752 | 0.89054584 |
| Dyrk1b | 0.12605162 | 0.11462184 | 0.13419984 | -0.019578 | 0.751646 | 0.89054584 |
| Dgcr14 | 0.23754096 | 0.22836307 | 0.24795247 | -0.0195894 | 0.64359249 | 0.89054584 |
| Got1 | 0.25819651 | 0.24884434 | 0.26843681 | -0.0195925 | 0.63308807 | 0.89054584 |
| Thbs3 | 0.23795115 | 0.22876401 | 0.24836119 | -0.0195972 | 0.643421 | 0.89054584 |
| Pdzd8 | 0.29224519 | 0.28216695 | 0.30176615 | -0.0195992 | 0.61889893 | 0.89054584 |
| Cpped1 | 0.19212606 | 0.18262867 | 0.2022416 | -0.0196129 | 0.67429836 | 0.89054584 |

| | | | | | | |
|---|---|---|---|---|---|---|
| Ptprs | 0.36100641 | 0.34767909 | 0.36731164 | -0.0196326 | 0.59832627 | 0.89054584 |
| Cpeb4 | 0.27836404 | 0.26861217 | 0.28824549 | -0.0196333 | 0.62450546 | 0.89054584 |
| D230025D16 | 0.15607486 | 0.14569418 | 0.16533246 | -0.0196383 | 0.71000227 | 0.89054584 |
| Hira | 0.10113857 | 0.08846754 | 0.10810824 | -0.0196407 | 0.80134053 | 0.89054584 |
| Edc4 | 0.15632783 | 0.14595208 | 0.16559329 | -0.0196412 | 0.70973213 | 0.89054584 |
| Msra | 0.14044934 | 0.12952902 | 0.14917326 | -0.0196442 | 0.73023501 | 0.89054584 |
| Lins1 | 0.08461802 | 0.07084819 | 0.09049248 | -0.0196443 | 0.84408087 | 0.89775379 |
| Net1 | 0.07834701 | 0.0641502 | 0.08379514 | -0.0196449 | 0.86263678 | 0.90267402 |
| Brd2 | 0.93997158 | 0.7829461 | 0.8025914 | -0.0196453 | 0.54539635 | 0.89054584 |
| AI661453 | 0.09455687 | 0.08145413 | 0.10110206 | -0.0196479 | 0.81737519 | 0.89054584 |
| Apmap | 0.40693241 | 0.39003862 | 0.40969317 | -0.0196545 | 0.58842961 | 0.89054584 |
| Nfil3 | 0.18782486 | 0.17820998 | 0.19786619 | -0.0196562 | 0.67825454 | 0.89054584 |
| Rell1 | 0.22538453 | 0.21612288 | 0.23580571 | -0.0196828 | 0.6513191 | 0.89054584 |
| Adsl | 0.33038528 | 0.31873784 | 0.33842304 | -0.0196852 | 0.606804 | 0.89054584 |
| Golph3l | 0.27791576 | 0.26811965 | 0.28780733 | -0.0196877 | 0.62502241 | 0.89054584 |
| Fam193b | 0.17703239 | 0.16714181 | 0.18685332 | -0.0197115 | 0.68845664 | 0.89054584 |
| Smim8 | 0.49304037 | 0.46618442 | 0.48590229 | -0.0197179 | 0.57497689 | 0.89054584 |
| Ypel1 | 0.14216324 | 0.13121903 | 0.15095035 | -0.0197313 | 0.72873405 | 0.89054584 |
| Sptlc2 | 0.39742515 | 0.38126789 | 0.40101552 | -0.0197476 | 0.59073077 | 0.89054584 |
| Fbxl3 | 0.23177829 | 0.22242331 | 0.2422023 | -0.019779 | 0.64819932 | 0.89054584 |
| 1700007L15I | 0.11617142 | 0.10409831 | 0.12387983 | -0.0197815 | 0.77154794 | 0.89054584 |
| Srxn1 | 0.11397398 | 0.10179223 | 0.1215796 | -0.0197874 | 0.77585051 | 0.89054584 |
| Brd4 | 0.71271192 | 0.63922899 | 0.65902179 | -0.0197928 | 0.5556403 | 0.89054584 |
| Bms1 | 0.23332164 | 0.22394561 | 0.24374379 | -0.0197982 | 0.64744228 | 0.89054584 |
| Acbd5 | 0.50480225 | 0.47616584 | 0.49596769 | -0.0198019 | 0.57378212 | 0.89054584 |
| Exd2 | 0.21604428 | 0.20659772 | 0.22642834 | -0.0198306 | 0.65837441 | 0.89054584 |
| Fnbp1 | 0.18543142 | 0.17556578 | 0.19542804 | -0.0198623 | 0.68207531 | 0.89054584 |
| Zranb1 | 0.50741776 | 0.47833058 | 0.49819421 | -0.0198636 | 0.57368266 | 0.89054584 |
| Cyb5rl | 0.12128987 | 0.10936155 | 0.12923071 | -0.0198692 | 0.76310002 | 0.89054584 |
| Abcb8 | 0.34598433 | 0.33333206 | 0.35320154 | -0.0198695 | 0.60337411 | 0.89054584 |

| | | | | | | |
|---|---|---|---|---|---|---|
| Kti12 | 0.35008047 | 0.33718492 | 0.35706109 | -0.0198762 | 0.60231533 | 0.89054584 |
| Pex11a | 0.10397981 | 0.09120955 | 0.11109544 | -0.0198859 | 0.7978432 | 0.89054584 |
| Tspyl4 | 0.28822163 | 0.27794881 | 0.29785655 | -0.0199077 | 0.62224721 | 0.89054584 |
| Il17ra | 0.08755534 | 0.0737187 | 0.09362868 | -0.01991 | 0.83908782 | 0.89500145 |
| 2700081O15 | 0.20131675 | 0.19164538 | 0.21156427 | -0.0199189 | 0.66953948 | 0.89054584 |
| Mapre2 | 0.4310095 | 0.4115189 | 0.43144014 | -0.0199212 | 0.58516359 | 0.89054584 |
| Cml1 | 0.12051875 | 0.10848732 | 0.12842521 | -0.0199379 | 0.76525104 | 0.89054584 |
| Lig4 | 0.14809178 | 0.13714815 | 0.15708859 | -0.0199404 | 0.72287202 | 0.89054584 |
| Kdm4a | 0.23478377 | 0.22526042 | 0.24520316 | -0.0199427 | 0.64762584 | 0.89054584 |
| Tkt | 1.10884597 | 0.86946049 | 0.8894109 | -0.0199504 | 0.54161566 | 0.89054584 |
| Det1 | 0.12878696 | 0.11707925 | 0.13705066 | -0.0199714 | 0.75140384 | 0.89054584 |
| Zfp275 | 0.0841334 | 0.0699852 | 0.08997499 | -0.0199898 | 0.84968411 | 0.89887961 |
| Naf1 | 0.12532719 | 0.11343258 | 0.13344441 | -0.0200118 | 0.75762163 | 0.89054584 |
| Asap2 | 0.21573619 | 0.20610377 | 0.22611837 | -0.0200146 | 0.6599698 | 0.89054584 |
| 1810013L24F | 0.18352563 | 0.17345684 | 0.19348493 | -0.0200281 | 0.68518877 | 0.89054584 |
| Prr14 | 0.19113527 | 0.18119589 | 0.20123441 | -0.0200385 | 0.67865031 | 0.89054584 |
| Trim62 | 0.14854597 | 0.13751821 | 0.15755827 | -0.0200401 | 0.72326799 | 0.89054584 |
| Atp6v1h | 0.74648827 | 0.66267335 | 0.68271559 | -0.0200422 | 0.55439405 | 0.89054584 |
| Trmt10b | 0.15760682 | 0.14683876 | 0.16691164 | -0.0200729 | 0.71235807 | 0.89054584 |
| Abhd12 | 0.33494141 | 0.322661 | 0.34275214 | -0.0200911 | 0.60760938 | 0.89054584 |
| Akt2 | 0.47039635 | 0.4461936 | 0.46628769 | -0.0200941 | 0.57956129 | 0.89054584 |
| Fam193a | 0.38026718 | 0.36513054 | 0.3852273 | -0.0200968 | 0.59601679 | 0.89054584 |
| Nol8 | 0.18012571 | 0.16991532 | 0.19001463 | -0.0200993 | 0.68892736 | 0.89054584 |
| Tsfm | 0.69243581 | 0.62431097 | 0.64441124 | -0.0201003 | 0.55777096 | 0.89054584 |
| Rbm19 | 0.1045472 | 0.09157256 | 0.11169163 | -0.0201191 | 0.79932633 | 0.89054584 |
| Gm20721 | 0.109125 | 0.09636717 | 0.11649747 | -0.0201303 | 0.78960366 | 0.89054584 |
| Hcn3 | 0.08660033 | 0.07247191 | 0.09260907 | -0.0201372 | 0.84448243 | 0.89775379 |
| Ccdc97 | 0.15438662 | 0.14345061 | 0.16359107 | -0.0201405 | 0.71687533 | 0.89054584 |
| Gspt2 | 0.16697024 | 0.15638662 | 0.17654312 | -0.0201565 | 0.70261147 | 0.89054584 |
| Tmem106c | 0.55373198 | 0.51674782 | 0.53690733 | -0.0201595 | 0.56943324 | 0.89054584 |

| | | | | | | |
|---|---|---|---|---|---|---|
| Chordc1 | 0.64927741 | 0.59217646 | 0.6123576 | -0.0201811 | 0.56101468 | 0.89054584 |
| Ulk3 | 0.07907497 | 0.06439113 | 0.08457271 | -0.0201816 | 0.86687992 | 0.90430169 |
| Senp3 | 0.27419312 | 0.26398126 | 0.28416492 | -0.0201837 | 0.62964576 | 0.89054584 |
| Gm5786 | 0.0629646 | 0.04717175 | 0.06735814 | -0.0201864 | 0.91867681 | 0.93597769 |
| Phf10 | 0.18061319 | 0.17031438 | 0.19051251 | -0.0201981 | 0.68932595 | 0.89054584 |
| Tomm7 | 2.44047118 | 1.10832254 | 1.12852877 | -0.0202062 | 0.53324043 | 0.89054584 |
| Stk40 | 0.13782045 | 0.1262389 | 0.14644521 | -0.0202063 | 0.73983416 | 0.89054584 |
| Zfp90 | 0.16025948 | 0.14941954 | 0.16964385 | -0.0202243 | 0.71069681 | 0.89054584 |
| Nkiras2 | 0.68853484 | 0.62133294 | 0.64156711 | -0.0202342 | 0.55840903 | 0.89054584 |
| Lrrc41 | 0.52835023 | 0.49560846 | 0.51585918 | -0.0202507 | 0.57255892 | 0.89054584 |
| Rcan1 | 0.10658434 | 0.09356923 | 0.11383119 | -0.020262 | 0.79653042 | 0.89054584 |
| Cnih1 | 1.27102922 | 0.93921103 | 0.95952106 | -0.02031 | 0.53927805 | 0.89054584 |
| Tmem209 | 0.16793867 | 0.15722669 | 0.17753724 | -0.0203105 | 0.70297989 | 0.89054584 |
| 2410018L13F | 0.09019531 | 0.07613138 | 0.09644694 | -0.0203156 | 0.83679191 | 0.89467618 |
| Pfdn6 | 1.24044993 | 0.92694887 | 0.94728507 | -0.0203362 | 0.53983482 | 0.89054584 |
| Kat6b | 0.31316807 | 0.30161883 | 0.32196586 | -0.020347 | 0.61578438 | 0.89054584 |
| Ralgapa1 | 0.26078291 | 0.25063962 | 0.27098811 | -0.0203485 | 0.63677049 | 0.89054584 |
| Tor1aip2 | 0.6999069 | 0.62947537 | 0.64982825 | -0.0203529 | 0.55800768 | 0.89054584 |
| Rgp1 | 0.0840731 | 0.06955766 | 0.0899106 | -0.0203529 | 0.85420457 | 0.90035231 |
| Glce | 0.1294309 | 0.11736342 | 0.13772136 | -0.0203579 | 0.75448888 | 0.89054584 |
| C1qtnf4 | 0.47469746 | 0.44966869 | 0.47003771 | -0.020369 | 0.58000041 | 0.89054584 |
| Casp3 | 0.11666009 | 0.10401877 | 0.12439111 | -0.0203723 | 0.77727573 | 0.89054584 |
| Nol6 | 0.11392259 | 0.1011534 | 0.12152579 | -0.0203724 | 0.78260391 | 0.89054584 |
| Atad2b | 0.19346831 | 0.18322795 | 0.20360539 | -0.0203774 | 0.67949271 | 0.89054584 |
| Fam118a | 0.10849339 | 0.09541515 | 0.11583485 | -0.0204197 | 0.79425997 | 0.89054584 |
| Fbxo25 | 0.2329969 | 0.22299655 | 0.24341953 | -0.020423 | 0.65205956 | 0.89054584 |
| Nars2 | 0.10924107 | 0.09618953 | 0.11661922 | -0.0204297 | 0.79279768 | 0.89054584 |
| Mbtd1 | 0.3809605 | 0.36542539 | 0.38586839 | -0.020443 | 0.59747892 | 0.89054584 |
| Rps6ka2 | 0.08225597 | 0.06752388 | 0.08797011 | -0.0204462 | 0.8605609 | 0.90160514 |
| Riok1 | 0.34503558 | 0.33183653 | 0.3523063 | -0.0204698 | 0.60668712 | 0.89054584 |

| | | | | | | |
|---|---|---|---|---|---|---|
| Ano10 | 0.17941708 | 0.16881617 | 0.18929073 | -0.0204746 | 0.6928456 | 0.89054584 |
| Arfgef2 | 0.1071155 | 0.09388986 | 0.11438881 | -0.0204989 | 0.7981134 | 0.89054584 |
| Igf2bp2 | 0.72444101 | 0.64683855 | 0.66734142 | -0.0205029 | 0.55690925 | 0.89054584 |
| Znrd1as | 0.13837366 | 0.12651064 | 0.14701951 | -0.0205089 | 0.74212522 | 0.89054584 |
| Ulk2 | 0.13560619 | 0.12363119 | 0.14414535 | -0.0205142 | 0.7463479 | 0.89054584 |
| Gtpbp6 | 0.30632153 | 0.29485606 | 0.31537993 | -0.0205239 | 0.6191268 | 0.89054584 |
| Zfp512 | 0.49809932 | 0.46969995 | 0.490242 | -0.020542 | 0.57738896 | 0.89054584 |
| D630039A03 | 0.10184754 | 0.08831134 | 0.10885391 | -0.0205426 | 0.81035928 | 0.89054584 |
| Camsap1 | 0.25925101 | 0.24892763 | 0.26947738 | -0.0205497 | 0.63881029 | 0.89054584 |
| Lrfn3 | 0.14762884 | 0.13603581 | 0.15660977 | -0.020574 | 0.72974736 | 0.89054584 |
| Pank4 | 0.12769849 | 0.1153323 | 0.13591656 | -0.0205843 | 0.75977729 | 0.89054584 |
| Zfp512b | 0.24382331 | 0.23360482 | 0.25420379 | -0.020599 | 0.64710986 | 0.89054584 |
| Ubn2 | 0.81976232 | 0.71068927 | 0.73130377 | -0.0206145 | 0.55224265 | 0.89054584 |
| Mtmr14 | 0.19824467 | 0.18782352 | 0.20845214 | -0.0206286 | 0.67761971 | 0.89054584 |
| Abl2 | 0.13110585 | 0.11882853 | 0.13946522 | -0.0206367 | 0.75470923 | 0.89054584 |
| Mpi | 0.18284525 | 0.17214125 | 0.19279086 | -0.0206496 | 0.69111517 | 0.89054584 |
| Pcbp4 | 0.6632312 | 0.60219599 | 0.62286208 | -0.0206661 | 0.56142384 | 0.89054584 |
| Wdr34 | 0.29596251 | 0.2846974 | 0.305371 | -0.0206736 | 0.62377482 | 0.89054584 |
| Dnttip1 | 0.23780474 | 0.22750258 | 0.24821532 | -0.0207127 | 0.65127797 | 0.89054584 |
| Trp53bp2 | 0.23144984 | 0.22115463 | 0.2418741 | -0.0207195 | 0.6550885 | 0.89054584 |
| Tjp2 | 0.11268885 | 0.09950219 | 0.12023352 | -0.0207313 | 0.78910532 | 0.89054584 |
| Gas8 | 0.37370639 | 0.35839378 | 0.37914758 | -0.0207538 | 0.60064668 | 0.89054584 |
| Itpr3 | 0.29434833 | 0.28304049 | 0.30380651 | -0.020766 | 0.6249288 | 0.89054584 |
| Trim56 | 0.1678621 | 0.15669134 | 0.17745865 | -0.0207673 | 0.70719823 | 0.89054584 |
| Mgat3 | 0.10631121 | 0.09277133 | 0.11354442 | -0.0207731 | 0.80299261 | 0.89054584 |
| Dcaf11 | 0.54141495 | 0.50595393 | 0.5267445 | -0.0207906 | 0.57294937 | 0.89054584 |
| Ccdc134 | 0.10342198 | 0.08971672 | 0.11050919 | -0.0207925 | 0.80964442 | 0.89054584 |
| Creb3l2 | 0.59093406 | 0.54621307 | 0.56700589 | -0.0207928 | 0.56782885 | 0.89054584 |
| Rnmtl1 | 0.13207772 | 0.11967683 | 0.14047657 | -0.0207997 | 0.75484906 | 0.89054584 |
| Slc25a1 | 0.25044652 | 0.23997332 | 0.26077441 | -0.0208011 | 0.64491504 | 0.89054584 |

| | | | | | | |
|---|---|---|---|---|---|---|
| Dpagt1 | 0.21954426 | 0.20913635 | 0.22994682 | -0.0208105 | 0.66335603 | 0.89054584 |
| Zbtb14 | 0.19274169 | 0.18204631 | 0.2028672 | -0.0208209 | 0.68373238 | 0.89054584 |
| Rbm28 | 0.39642668 | 0.37927746 | 0.40010118 | -0.0208237 | 0.59579583 | 0.89054584 |
| C1galt1 | 0.23353578 | 0.22313181 | 0.24395759 | -0.0208258 | 0.65458059 | 0.89054584 |
| Mob3a | 0.08793815 | 0.07320992 | 0.09403738 | -0.0208275 | 0.84893237 | 0.89883797 |
| Lrif1 | 0.41521249 | 0.39636502 | 0.41720845 | -0.0208434 | 0.59202506 | 0.89054584 |
| Llph | 1.19972282 | 0.90951482 | 0.9303753 | -0.0208605 | 0.54159816 | 0.89054584 |
| Diablo | 0.99836614 | 0.81277554 | 0.83365655 | -0.020881 | 0.54644905 | 0.89054584 |
| Kcmf1 | 0.34544722 | 0.33181329 | 0.35269478 | -0.0208815 | 0.60866269 | 0.89054584 |
| Syn2 | 0.11481185 | 0.10156484 | 0.12245688 | -0.020892 | 0.78664941 | 0.89054584 |
| Frrs1 | 0.23119666 | 0.22071895 | 0.24162109 | -0.0209021 | 0.65653886 | 0.89054584 |
| Sh3d19 | 0.20357896 | 0.19294116 | 0.21385336 | -0.0209122 | 0.67565633 | 0.89054584 |
| St14 | 0.12575908 | 0.11297031 | 0.13389481 | -0.0209245 | 0.76671743 | 0.89054584 |
| Aven | 0.07664636 | 0.06105347 | 0.08197849 | -0.020925 | 0.88281857 | 0.91259206 |
| St3gal6 | 0.42027421 | 0.40085124 | 0.42178446 | -0.0209332 | 0.59142959 | 0.89054584 |
| Fam71e1 | 0.10077529 | 0.08678923 | 0.10772609 | -0.0209369 | 0.81739393 | 0.89054584 |
| Ibtk | 0.33275806 | 0.31973679 | 0.34067895 | -0.0209422 | 0.61271166 | 0.89054584 |
| Tango2 | 0.202764 | 0.19207715 | 0.21302898 | -0.0209518 | 0.67660146 | 0.89054584 |
| Trrap | 0.3322753 | 0.31925942 | 0.34022021 | -0.0209608 | 0.61295722 | 0.89054584 |
| Sdhaf3 | 0.22012335 | 0.20955389 | 0.23052844 | -0.0209745 | 0.66417775 | 0.89054584 |
| Kdm3b | 0.12097387 | 0.10792143 | 0.12890065 | -0.0209792 | 0.77585407 | 0.89054584 |
| Sh3bp5l | 0.20445702 | 0.19376058 | 0.21474126 | -0.0209807 | 0.67551314 | 0.89054584 |
| Cnot8 | 0.447194 | 0.42487174 | 0.44587001 | -0.0209983 | 0.58683585 | 0.89054584 |
| Frmd8 | 0.10307804 | 0.0891411 | 0.11014766 | -0.0210066 | 0.81286926 | 0.89054584 |
| Gas2l1 | 0.12670886 | 0.11386551 | 0.13488506 | -0.0210196 | 0.76608585 | 0.89054584 |
| Nap1l3 | 0.09725495 | 0.08294841 | 0.10398108 | -0.0210327 | 0.82700159 | 0.89186561 |
| Tmem170b | 0.09455352 | 0.08003985 | 0.10109848 | -0.0210586 | 0.83408866 | 0.89427341 |
| Znfx1 | 0.1039882 | 0.09004507 | 0.11110425 | -0.0210592 | 0.81140242 | 0.89054584 |
| Fem1a | 0.23671706 | 0.22607079 | 0.24713129 | -0.0210605 | 0.65432894 | 0.89054584 |
| Dmxl2 | 0.15126406 | 0.13929182 | 0.16036737 | -0.0210755 | 0.72982373 | 0.89054584 |

| | | | | | | |
|---|---|---|---|---|---|---|
| Siae | 0.10438247 | 0.09040237 | 0.11151855 | -0.0211162 | 0.81115931 | 0.89054584 |
| Ints12 | 0.18965944 | 0.17861409 | 0.19973338 | -0.0211193 | 0.68886102 | 0.89054584 |
| Eps15 | 0.23610632 | 0.22539667 | 0.24652237 | -0.0211257 | 0.65514449 | 0.89054584 |
| Zfp407 | 0.11995945 | 0.10671442 | 0.12784082 | -0.0211264 | 0.77932509 | 0.89054584 |
| Plpp6 | 0.2476128 | 0.23681901 | 0.25796569 | -0.0211467 | 0.64873806 | 0.89054584 |
| Fsd1l | 0.17069024 | 0.15921198 | 0.1803597 | -0.0211477 | 0.70756149 | 0.89054584 |
| Trmt6 | 0.45293021 | 0.42976301 | 0.45094788 | -0.0211849 | 0.58662444 | 0.89054584 |
| Dync1h1 | 0.62636307 | 0.57362875 | 0.59481895 | -0.0211902 | 0.56591093 | 0.89054584 |
| Glb1l2 | 0.11380619 | 0.10021072 | 0.12140389 | -0.0211932 | 0.79197686 | 0.89054584 |
| Rad52 | 0.11685792 | 0.10337554 | 0.12459807 | -0.0212225 | 0.78627069 | 0.89054584 |
| Plekhb1 | 0.11508052 | 0.10150806 | 0.12273812 | -0.0212301 | 0.78983788 | 0.89054584 |
| Rassf7 | 0.368454 | 0.35301605 | 0.37426313 | -0.0212471 | 0.60429886 | 0.89054584 |
| Mxd1 | 0.16769095 | 0.15602897 | 0.17728299 | -0.021254 | 0.71176516 | 0.89054584 |
| Rpl6 | 5.06561588 | 0.66648275 | 0.68774399 | -0.0212612 | 0.5572611 | 0.89054584 |
| Glis2 | 0.18370918 | 0.17232904 | 0.19367215 | -0.0213431 | 0.69617326 | 0.89054584 |
| Smyd5 | 0.12324007 | 0.10991832 | 0.13126688 | -0.0213486 | 0.77568646 | 0.89054584 |
| Tmtc4 | 0.22663713 | 0.21571149 | 0.23706031 | -0.0213488 | 0.66259937 | 0.89054584 |
| Ccdc66 | 0.18669763 | 0.1753679 | 0.1967182 | -0.0213503 | 0.69344745 | 0.89054584 |
| Fbxo38 | 0.17112983 | 0.15945453 | 0.18081032 | -0.0213558 | 0.70893458 | 0.89054584 |
| Rfxank | 0.1114982 | 0.0976268 | 0.11898586 | -0.0213591 | 0.79851338 | 0.89054584 |
| Rad51d | 0.13496174 | 0.12211052 | 0.14347564 | -0.0213651 | 0.75608624 | 0.89054584 |
| D6Wsu163e | 0.09546172 | 0.08069884 | 0.10206765 | -0.0213688 | 0.83531726 | 0.89459411 |
| Tpk1 | 0.14147508 | 0.12886512 | 0.15023696 | -0.0213718 | 0.74625595 | 0.89054584 |
| Pcmtd2 | 0.33453602 | 0.32097252 | 0.34236739 | -0.0213949 | 0.61452855 | 0.89054584 |
| Pias1 | 0.39618658 | 0.37847948 | 0.39988124 | -0.0214018 | 0.59845413 | 0.89054584 |
| Vps39 | 0.13086253 | 0.1178044 | 0.13921195 | -0.0214075 | 0.76313904 | 0.89054584 |
| Setbp1 | 0.21216365 | 0.20110097 | 0.22252085 | -0.0214199 | 0.67309304 | 0.89054584 |
| Gdap1 | 0.15090611 | 0.13857622 | 0.1599976 | -0.0214214 | 0.73360913 | 0.89054584 |
| Gm20342 | 0.20902734 | 0.19792982 | 0.21935796 | -0.0214281 | 0.67548564 | 0.89054584 |
| Usp24 | 0.09215016 | 0.07710155 | 0.09853353 | -0.021432 | 0.84455679 | 0.89775379 |

| | | | | | | |
|---|---|---|---|---|---|---|
| Rbm48 | 0.18117705 | 0.16961373 | 0.19108825 | -0.0214745 | 0.69971199 | 0.89054584 |
| Irf6 | 0.23220808 | 0.2211534 | 0.24263169 | -0.0214783 | 0.65998408 | 0.89054584 |
| Klhdc4 | 0.20970835 | 0.19856572 | 0.22004511 | -0.0214794 | 0.67536479 | 0.89054584 |
| Fam63a | 0.21322471 | 0.20207444 | 0.22358992 | -0.0215155 | 0.67303805 | 0.89054584 |
| Zbtb6 | 0.13242732 | 0.11931827 | 0.14084029 | -0.021522 | 0.76174898 | 0.89054584 |
| Txlng | 0.18555142 | 0.17401338 | 0.19555035 | -0.021537 | 0.69606392 | 0.89054584 |
| Klhl2 | 0.12317171 | 0.10965178 | 0.13119553 | -0.0215438 | 0.77787897 | 0.89054584 |
| Tgds | 0.35815038 | 0.34308447 | 0.36463842 | -0.021554 | 0.60849256 | 0.89054584 |
| B3glct | 0.12392359 | 0.11042118 | 0.13198019 | -0.021559 | 0.77668332 | 0.89054584 |
| Enpp4 | 0.19708517 | 0.18570533 | 0.20727645 | -0.0215711 | 0.68610964 | 0.89054584 |
| Erap1 | 0.1396625 | 0.1267812 | 0.14835702 | -0.0215758 | 0.75097397 | 0.89054584 |
| Pprc1 | 0.13092038 | 0.11767587 | 0.13927218 | -0.0215963 | 0.76498823 | 0.89054584 |
| Ubqln4 | 0.20639334 | 0.19508842 | 0.21669829 | -0.0216099 | 0.67889288 | 0.89054584 |
| Cep295 | 0.2606352 | 0.24923107 | 0.27084249 | -0.0216114 | 0.64496136 | 0.89054584 |
| Ddhd1 | 0.1103225 | 0.09613936 | 0.11775335 | -0.021614 | 0.80376951 | 0.89054584 |
| Fmnl2 | 0.15256868 | 0.14009747 | 0.16171464 | -0.0216172 | 0.73328644 | 0.89054584 |
| Lrwd1 | 0.20310843 | 0.19175591 | 0.21337743 | -0.0216215 | 0.68156875 | 0.89054584 |
| Ccng1 | 0.26307414 | 0.25161652 | 0.27324558 | -0.0216291 | 0.6438555 | 0.89054584 |
| Zc3h7a | 0.53407522 | 0.49899678 | 0.5206427 | -0.0216459 | 0.57679354 | 0.89054584 |
| Usf3 | 0.11363954 | 0.09958269 | 0.12122936 | -0.0216467 | 0.797267 | 0.89054584 |
| Slc4a8 | 0.11106227 | 0.09686874 | 0.11852892 | -0.0216602 | 0.80272168 | 0.89054584 |
| Spg11 | 0.07458557 | 0.05810986 | 0.07977692 | -0.0216671 | 0.89712668 | 0.92166544 |
| Hook2 | 0.10270025 | 0.08808206 | 0.10975051 | -0.0216684 | 0.82116967 | 0.89054584 |
| Rap1gap | 0.15993609 | 0.14761779 | 0.16931091 | -0.0216931 | 0.72472337 | 0.89054584 |
| Mbp | 0.29532407 | 0.2830478 | 0.30475238 | -0.0217046 | 0.6299842 | 0.89054584 |
| Fig4 | 0.08105206 | 0.06497675 | 0.08668437 | -0.0217076 | 0.87833396 | 0.91048804 |
| Mtfr1l | 0.96380679 | 0.79381172 | 0.81552633 | -0.0217146 | 0.54936267 | 0.89054584 |
| Adck2 | 0.21729089 | 0.20596214 | 0.22768217 | -0.02172 | 0.67163973 | 0.89054584 |
| Ccdc71 | 0.2764839 | 0.26468175 | 0.28640713 | -0.0217254 | 0.63810703 | 0.89054584 |
| Riiad1 | 0.25164478 | 0.24023043 | 0.26196096 | -0.0217305 | 0.65042736 | 0.89054584 |

| | | | | | | |
|---|---|---|---|---|---|---|
| Klhl28 | 0.12641426 | 0.11284341 | 0.13457794 | -0.0217345 | 0.77410241 | 0.89054584 |
| Cep68 | 0.2145122 | 0.20314651 | 0.22488645 | -0.0217399 | 0.67378298 | 0.89054584 |
| Ccdc148 | 0.12933678 | 0.11587907 | 0.13762334 | -0.0217443 | 0.7691667 | 0.89054584 |
| Tmem135 | 0.20634714 | 0.19488299 | 0.21665162 | -0.0217686 | 0.68015032 | 0.89054584 |
| Mad1l1 | 0.12483865 | 0.11116025 | 0.13293486 | -0.0217746 | 0.77730715 | 0.89054584 |
| Grwd1 | 0.22017612 | 0.20878928 | 0.23058143 | -0.0217921 | 0.67014532 | 0.89054584 |
| 4931406P16I | 0.33245377 | 0.31859169 | 0.34038982 | -0.0217981 | 0.61728296 | 0.89054584 |
| Apbb2 | 0.31408894 | 0.30104663 | 0.32284996 | -0.0218033 | 0.62348124 | 0.89054584 |
| Abcb6 | 0.12758007 | 0.11395527 | 0.13579315 | -0.0218379 | 0.77314313 | 0.89054584 |
| Nup205 | 0.12762687 | 0.11396671 | 0.13584192 | -0.0218752 | 0.77344773 | 0.89054584 |
| Trim24 | 0.27361363 | 0.26171734 | 0.28359732 | -0.02188 | 0.64036875 | 0.89054584 |
| Pank1 | 0.15021996 | 0.13740192 | 0.15928866 | -0.0218867 | 0.73896325 | 0.89054584 |
| Taf8 | 0.13887449 | 0.1256497 | 0.14753933 | -0.0218896 | 0.75528716 | 0.89054584 |
| Tmie | 0.08740828 | 0.07156366 | 0.09347169 | -0.021908 | 0.86257673 | 0.90267402 |
| Cdc42bpa | 0.32487484 | 0.31124549 | 0.33317263 | -0.0219271 | 0.62043354 | 0.89054584 |
| Atp2b1 | 1.53002544 | 1.02172561 | 1.04365994 | -0.0219343 | 0.5390004 | 0.89054584 |
| Yeats4 | 0.93180001 | 0.77613158 | 0.79807343 | -0.0219419 | 0.55096142 | 0.89054584 |
| Cnppd1 | 0.30830595 | 0.29533181 | 0.31729106 | -0.0219593 | 0.62643977 | 0.89054584 |
| Vgll4 | 0.53805525 | 0.50197843 | 0.52395578 | -0.0219774 | 0.57746774 | 0.89054584 |
| Saysd1 | 0.48223585 | 0.45459766 | 0.47658278 | -0.0219851 | 0.58508532 | 0.89054584 |
| N6amt1 | 0.56539306 | 0.52444471 | 0.54643888 | -0.0219942 | 0.57437448 | 0.89054584 |
| Ripk4 | 0.12606786 | 0.11222191 | 0.13421677 | -0.0219949 | 0.7774135 | 0.89054584 |
| Btbd6 | 0.14699276 | 0.13394674 | 0.15595174 | -0.022005 | 0.74453266 | 0.89054584 |
| Gcc1 | 0.15053693 | 0.13760384 | 0.15961618 | -0.0220123 | 0.73972731 | 0.89054584 |
| Eif2s2 | 2.42584982 | 1.10679687 | 1.12881645 | -0.0220196 | 0.53620641 | 0.89054584 |
| Lsm12 | 0.59191915 | 0.54576871 | 0.5677905 | -0.0220218 | 0.57169744 | 0.89054584 |
| Kctd18 | 0.10384613 | 0.0889313 | 0.11095496 | -0.0220237 | 0.82245513 | 0.89073269 |
| 2610528J11F | 0.29470387 | 0.28211449 | 0.30415122 | -0.0220367 | 0.6321508 | 0.89054584 |
| Mapk8ip3 | 0.33208184 | 0.31797156 | 0.34003634 | -0.0220648 | 0.61879447 | 0.89054584 |
| Ccdc9 | 0.2070871 | 0.19532742 | 0.21739911 | -0.0220717 | 0.68189492 | 0.89054584 |

| | | | | | | |
|---|---|---|---|---|---|---|
| Zfp740 | 0.16557272 | 0.15303064 | 0.17510785 | -0.0220772 | 0.72152879 | 0.89054584 |
| Filip1 | 0.28231867 | 0.27002708 | 0.29210659 | -0.0220795 | 0.63763939 | 0.89054584 |
| B3galnt1 | 0.28906103 | 0.27658429 | 0.29867284 | -0.0220886 | 0.6347856 | 0.89054584 |
| Cc2d1a | 0.15938771 | 0.14664025 | 0.16874625 | -0.022106 | 0.72917768 | 0.89054584 |
| Zfp37 | 0.10500564 | 0.09002107 | 0.11217325 | -0.0221522 | 0.82122828 | 0.89054584 |
| Fut11 | 0.22088453 | 0.20911 | 0.23129271 | -0.0221827 | 0.67250018 | 0.89054584 |
| Dtnb | 0.26335714 | 0.25129892 | 0.27352424 | -0.0222253 | 0.64749418 | 0.89054584 |
| Larp1b | 0.09794939 | 0.08248822 | 0.10472201 | -0.0222338 | 0.83870542 | 0.89497393 |
| Eif3j2 | 0.46140477 | 0.43617236 | 0.45841355 | -0.0222412 | 0.58941535 | 0.89054584 |
| Grip1 | 0.1235002 | 0.10925438 | 0.13153837 | -0.022284 | 0.78502398 | 0.89054584 |
| Ncaph2 | 0.53305472 | 0.49748812 | 0.51979157 | -0.0223035 | 0.57922373 | 0.89054584 |
| Ccdc191 | 0.25189949 | 0.23990846 | 0.26221308 | -0.0223046 | 0.65405939 | 0.89054584 |
| 9330151L19F | 0.11459284 | 0.09991477 | 0.12222759 | -0.0223128 | 0.80247547 | 0.89054584 |
| Anks1 | 0.10733574 | 0.09230251 | 0.11461999 | -0.0223175 | 0.81783226 | 0.89054584 |
| Gpr89 | 0.31855803 | 0.30481304 | 0.32713442 | -0.0223214 | 0.62471726 | 0.89054584 |
| Oma1 | 0.18475214 | 0.17239023 | 0.19473563 | -0.0223454 | 0.70353721 | 0.89054584 |
| Mtdh | 1.24116683 | 0.92521575 | 0.94757871 | -0.022363 | 0.54377608 | 0.89054584 |
| Phldb1 | 0.12586496 | 0.11162644 | 0.1340052 | -0.0223788 | 0.78172714 | 0.89054584 |
| Pafah1b1 | 1.62508925 | 1.04399629 | 1.06638621 | -0.0223899 | 0.53896219 | 0.89054584 |
| Zmat3 | 0.19033953 | 0.17801897 | 0.20042519 | -0.0224062 | 0.6987606 | 0.89054584 |
| Cnpy2 | 1.91941179 | 1.08908996 | 1.11151315 | -0.0224232 | 0.53744044 | 0.89054584 |
| Rictor | 0.10483191 | 0.08954664 | 0.11199074 | -0.0224441 | 0.82477067 | 0.89148775 |
| Zfp322a | 0.7056546 | 0.63152315 | 0.65396877 | -0.0224456 | 0.56352203 | 0.89054584 |
| Zfp362 | 0.14821889 | 0.13477367 | 0.15722005 | -0.0224464 | 0.7470256 | 0.89054584 |
| Trappc12 | 0.18892652 | 0.17653109 | 0.1989876 | -0.0224565 | 0.70047926 | 0.89054584 |
| Vcp | 2.53818314 | 1.10296121 | 1.12542015 | -0.0224589 | 0.53703792 | 0.89054584 |
| D8Ertd738e | 1.89715156 | 1.08692034 | 1.10939855 | -0.0224782 | 0.53760337 | 0.89054584 |
| Siah1b | 0.1264804 | 0.11216371 | 0.13464689 | -0.0224832 | 0.78169919 | 0.89054584 |
| Nampt | 0.20816169 | 0.1959947 | 0.21848421 | -0.0224895 | 0.68423148 | 0.89054584 |
| Ddhd2 | 0.28255558 | 0.26982401 | 0.29233766 | -0.0225136 | 0.64012635 | 0.89054584 |

| | | | | | | |
|---|---|---|---|---|---|---|
| 2810474O19 | 0.43757571 | 0.41477276 | 0.43731259 | -0.0225398 | 0.59488512 | 0.89054584 |
| Ldlrad3 | 0.20745259 | 0.19522157 | 0.21776823 | -0.0225467 | 0.68522634 | 0.89054584 |
| Map1s | 0.14468491 | 0.13101543 | 0.1535629 | -0.0225475 | 0.75303367 | 0.89054584 |
| Rab15 | 0.09533547 | 0.07937877 | 0.10193292 | -0.0225541 | 0.84869649 | 0.89883797 |
| Qsox1 | 0.4194885 | 0.39851722 | 0.42107513 | -0.0225579 | 0.59854765 | 0.89054584 |
| Med23 | 0.13995042 | 0.12609308 | 0.14865573 | -0.0225627 | 0.76025536 | 0.89054584 |
| Stk4 | 0.20543795 | 0.19316739 | 0.21573285 | -0.0225655 | 0.68698457 | 0.89054584 |
| Fam214a | 0.0953148 | 0.07933127 | 0.10191087 | -0.0225796 | 0.84902142 | 0.89883797 |
| AK010878 | 0.33895054 | 0.32396992 | 0.34655307 | -0.0225832 | 0.61928374 | 0.89054584 |
| Clcc1 | 0.40345371 | 0.38393182 | 0.40652402 | -0.0225922 | 0.60215016 | 0.89054584 |
| Man2a2 | 0.17874133 | 0.16600234 | 0.18860021 | -0.0225979 | 0.71165605 | 0.89054584 |
| Znrf2 | 0.20144417 | 0.18908074 | 0.21169327 | -0.0226125 | 0.69063677 | 0.89054584 |
| Pou2f1 | 0.20840879 | 0.19610811 | 0.21873366 | -0.0226255 | 0.68506695 | 0.89054584 |
| Cnpy4 | 0.34678086 | 0.3313242 | 0.35395277 | -0.0226286 | 0.61709152 | 0.89054584 |
| Mapre3 | 0.69006218 | 0.62004259 | 0.64268194 | -0.0226394 | 0.56518071 | 0.89054584 |
| Pdrg1 | 0.62000073 | 0.56724375 | 0.58988668 | -0.0226429 | 0.57096617 | 0.89054584 |
| AI987944 | 0.2342257 | 0.22198769 | 0.24464626 | -0.0226586 | 0.66694769 | 0.89054584 |
| Mast4 | 0.65045265 | 0.59057737 | 0.61324737 | -0.02267 | 0.56837076 | 0.89054584 |
| Endov | 0.1310956 | 0.11676997 | 0.13945455 | -0.0226846 | 0.77573059 | 0.89054584 |
| 2700046G09 | 0.19825036 | 0.18575886 | 0.2084579 | -0.022699 | 0.69403254 | 0.89054584 |
| Ccs | 0.52243655 | 0.48819387 | 0.5108959 | -0.022702 | 0.58200423 | 0.89054584 |
| Cdc27 | 0.23487877 | 0.22258777 | 0.24529794 | -0.0227102 | 0.66688728 | 0.89054584 |
| Rnpepl1 | 0.59428551 | 0.54694261 | 0.56967261 | -0.02273 | 0.57373539 | 0.89054584 |
| Herc4 | 0.12942841 | 0.11498324 | 0.13771878 | -0.0227355 | 0.77909164 | 0.89054584 |
| Fam21 | 0.3980646 | 0.37882663 | 0.40160079 | -0.0227742 | 0.6041876 | 0.89054584 |
| Dpm2 | 0.56209201 | 0.52096841 | 0.54374973 | -0.0227813 | 0.57737956 | 0.89054584 |
| Rpgr | 0.2394377 | 0.22705709 | 0.24984176 | -0.0227847 | 0.66453745 | 0.89054584 |
| Rsph3a | 0.11439909 | 0.09923868 | 0.12202474 | -0.0227861 | 0.8078439 | 0.89054584 |
| Tmem184c | 0.41689385 | 0.39592873 | 0.41873012 | -0.0228014 | 0.60013497 | 0.89054584 |
| Pbx3 | 0.20458967 | 0.19204913 | 0.21487536 | -0.0228262 | 0.68967161 | 0.89054584 |

| | | | | | | |
|---|---|---|---|---|---|---|
| Trappc8 | 0.12761333 | 0.11299972 | 0.13582781 | -0.0228281 | 0.78318988 | 0.89054584 |
| Hdac11 | 0.19296075 | 0.18024975 | 0.20308977 | -0.02284 | 0.69984585 | 0.89054584 |
| Plppr2 | 0.13374288 | 0.11936468 | 0.14220855 | -0.0228439 | 0.77288919 | 0.89054584 |
| Tecpr2 | 0.09145965 | 0.07493508 | 0.09779651 | -0.0228614 | 0.8619449 | 0.90253338 |
| Gosr1 | 0.17497775 | 0.16188738 | 0.18475142 | -0.022864 | 0.71788519 | 0.89054584 |
| Mtpap | 0.33942426 | 0.32411056 | 0.34700167 | -0.0228911 | 0.62070862 | 0.89054584 |
| Lgalsl | 0.23644007 | 0.22395673 | 0.24685514 | -0.0228984 | 0.66718958 | 0.89054584 |
| Pelp1 | 0.36576078 | 0.3488383 | 0.37175269 | -0.0229144 | 0.61300951 | 0.89054584 |
| Mphosph10 | 0.4379301 | 0.41470863 | 0.43762881 | -0.0229202 | 0.59638666 | 0.89054584 |
| Ticam1 | 0.13426947 | 0.11982475 | 0.14275605 | -0.0229313 | 0.77288409 | 0.89054584 |
| Zkscan8 | 0.10342733 | 0.08758278 | 0.11051481 | -0.022932 | 0.83316277 | 0.89393134 |
| Rsl24d1 | 0.56612612 | 0.52410015 | 0.54703508 | -0.0229349 | 0.57743279 | 0.89054584 |
| Spryd3 | 0.22589817 | 0.2133842 | 0.23632026 | -0.0229361 | 0.67442797 | 0.89054584 |
| Arfgef1 | 0.54535077 | 0.50705717 | 0.53000211 | -0.0229449 | 0.57992267 | 0.89054584 |
| Clk2 | 0.13164787 | 0.1170621 | 0.14002931 | -0.0229672 | 0.77760995 | 0.89054584 |
| Cars | 0.20343652 | 0.19074039 | 0.2137093 | -0.0229689 | 0.69171965 | 0.89054584 |
| Ppip5k2 | 0.16841538 | 0.15505167 | 0.17802645 | -0.0229748 | 0.72616276 | 0.89054584 |
| Kpna6 | 0.18986654 | 0.17694079 | 0.19994408 | -0.0230033 | 0.70402231 | 0.89054584 |
| Ttc30b | 0.13589303 | 0.12142645 | 0.14444339 | -0.0230169 | 0.77107704 | 0.89054584 |
| Ythdf3 | 0.47541779 | 0.44764209 | 0.47066463 | -0.0230225 | 0.59013486 | 0.89054584 |
| Gm13830 | 0.11018546 | 0.09457728 | 0.11760966 | -0.0230324 | 0.81921966 | 0.89054584 |
| Gyg | 0.70901445 | 0.63333171 | 0.65637879 | -0.0230471 | 0.56497182 | 0.89054584 |
| Kdm4c | 0.18806819 | 0.17505264 | 0.19811393 | -0.0230613 | 0.70620104 | 0.89054584 |
| Dtnbp1 | 0.22114808 | 0.2084921 | 0.23155727 | -0.0230652 | 0.678702 | 0.89054584 |
| Dmxl1 | 0.30925888 | 0.29512765 | 0.31820809 | -0.0230804 | 0.63229009 | 0.89054584 |
| Nbas | 0.13171889 | 0.11700694 | 0.1401032 | -0.0230963 | 0.77876672 | 0.89054584 |
| Fktn | 0.2212979 | 0.20860529 | 0.23170766 | -0.0231024 | 0.67886199 | 0.89054584 |
| Phf2 | 0.18875913 | 0.17569411 | 0.19881724 | -0.0231231 | 0.70603935 | 0.89054584 |
| Hgh1 | 0.11760805 | 0.10223922 | 0.12538269 | -0.0231435 | 0.80509441 | 0.89054584 |
| Cep131 | 0.21063665 | 0.19783213 | 0.22098145 | -0.0231493 | 0.68724487 | 0.89054584 |

| | | | | | | |
|---|---|---|---|---|---|---|
| Tollip | 0.34325474 | 0.32746337 | 0.35062461 | -0.0231612 | 0.62086605 | 0.89054584 |
| Kat2b | 0.11762302 | 0.10223257 | 0.12539834 | -0.0231658 | 0.80529314 | 0.89054584 |
| Rnf217 | 0.1595611 | 0.14575091 | 0.1689248 | -0.0231739 | 0.73863252 | 0.89054584 |
| Ggnbp1 | 0.13058062 | 0.11572574 | 0.1389185 | -0.0231928 | 0.78166259 | 0.89054584 |
| Arhgef11 | 0.12037818 | 0.10507007 | 0.12827834 | -0.0232083 | 0.80035813 | 0.89054584 |
| Smchd1 | 0.6235568 | 0.5694167 | 0.59264687 | -0.0232302 | 0.57245113 | 0.89054584 |
| Nup93 | 0.2901168 | 0.27646044 | 0.29969905 | -0.0232386 | 0.64104503 | 0.89054584 |
| Dedd2 | 0.18722898 | 0.17401336 | 0.1972594 | -0.023246 | 0.70850863 | 0.89054584 |
| Cmpk1 | 1.35386259 | 0.96678903 | 0.99004349 | -0.0232545 | 0.54356955 | 0.89054584 |
| Agbl5 | 0.10737829 | 0.09139788 | 0.11466466 | -0.0232668 | 0.8277606 | 0.89225823 |
| Kdm5b | 0.93320869 | 0.77558661 | 0.79885578 | -0.0232692 | 0.55397323 | 0.89054584 |
| Cab39l | 0.24709156 | 0.23413471 | 0.25744865 | -0.0233139 | 0.66345198 | 0.89054584 |
| Pan3 | 0.35928409 | 0.34238409 | 0.3657001 | -0.023316 | 0.61678239 | 0.89054584 |
| Zfp788 | 0.15197298 | 0.13777888 | 0.16109954 | -0.0233207 | 0.74997943 | 0.89054584 |
| Gm9803 | 0.16251778 | 0.14864613 | 0.17196763 | -0.0233215 | 0.7362495 | 0.89054584 |
| Nsd1 | 1.01612313 | 0.81946897 | 0.84281928 | -0.0233503 | 0.55135124 | 0.89054584 |
| Pde7a | 0.12364586 | 0.10832876 | 0.13169038 | -0.0233616 | 0.7957361 | 0.89054584 |
| St6galnac2 | 0.20514761 | 0.19206554 | 0.21543938 | -0.0233738 | 0.69338436 | 0.89054584 |
| Glb1 | 0.34974633 | 0.33336339 | 0.35674659 | -0.0233832 | 0.6199591 | 0.89054584 |
| Pex1 | 0.11207895 | 0.09620366 | 0.11959448 | -0.0233908 | 0.81890932 | 0.89054584 |
| Gm10094 | 0.37942764 | 0.36104982 | 0.38445066 | -0.0234008 | 0.61163452 | 0.89054584 |
| Mar-07 | 0.67051027 | 0.60488226 | 0.62828844 | -0.0234062 | 0.56889579 | 0.89054584 |
| Zbtb24 | 0.18716092 | 0.17377866 | 0.19719009 | -0.0234114 | 0.70991402 | 0.89054584 |
| Anapc16 | 1.100123 | 0.86180538 | 0.88523016 | -0.0234248 | 0.54905887 | 0.89054584 |
| Hmgxb3 | 0.09090264 | 0.07377445 | 0.09720197 | -0.0234275 | 0.86925859 | 0.90542007 |
| Noc2l | 0.17978927 | 0.16623598 | 0.18967097 | -0.023435 | 0.71757078 | 0.89054584 |
| Gm14296 | 0.08715522 | 0.06972834 | 0.0932015 | -0.0234732 | 0.87967677 | 0.91127746 |
| Aqp11 | 0.15493749 | 0.14067394 | 0.16415941 | -0.0234855 | 0.74746429 | 0.89054584 |
| Cldn12 | 0.14366613 | 0.12899096 | 0.15250771 | -0.0235167 | 0.76374791 | 0.89054584 |
| Trim13 | 0.13091186 | 0.11574008 | 0.13926331 | -0.0235232 | 0.7843446 | 0.89054584 |

| | | | | | | |
|---|---|---|---|---|---|---|
| Slc35b2 | 0.60128552 | 0.55168339 | 0.57521836 | -0.023535 | 0.57558828 | 0.89054584 |
| Scamp1 | 0.35733001 | 0.34032298 | 0.36386973 | -0.0235468 | 0.61847942 | 0.89054584 |
| 2310011J03F | 0.54592182 | 0.50692247 | 0.53047393 | -0.0235515 | 0.58193379 | 0.89054584 |
| Pin4 | 0.95261327 | 0.78594817 | 0.80950049 | -0.0235523 | 0.55391202 | 0.89054584 |
| Zfp943 | 0.19424583 | 0.18083898 | 0.20439503 | -0.023556 | 0.70434272 | 0.89054584 |
| Dcx | 0.19692909 | 0.18353852 | 0.20711814 | -0.0235796 | 0.70208371 | 0.89054584 |
| Spty2d1 | 0.41886772 | 0.39693084 | 0.42051443 | -0.0235836 | 0.60306406 | 0.89054584 |
| Iah1 | 0.40666942 | 0.3858659 | 0.40945382 | -0.0235879 | 0.60580177 | 0.89054584 |
| Mgat1 | 0.4231891 | 0.40081731 | 0.42441289 | -0.0235956 | 0.60218917 | 0.89054584 |
| Lrrc40 | 0.15158344 | 0.13709825 | 0.16069725 | -0.023599 | 0.75307019 | 0.89054584 |
| Tmem68 | 0.17891401 | 0.1651671 | 0.18877668 | -0.0236096 | 0.71994432 | 0.89054584 |
| Cep44 | 0.29364132 | 0.27949988 | 0.30312085 | -0.023621 | 0.64171709 | 0.89054584 |
| Pnrc2 | 0.75896807 | 0.66759442 | 0.69126384 | -0.0236694 | 0.5633727 | 0.89054584 |
| C2cd3 | 0.20524893 | 0.19185932 | 0.21554179 | -0.0236825 | 0.69563852 | 0.89054584 |
| Ifnar1 | 0.37460112 | 0.3562762 | 0.37997811 | -0.0237019 | 0.61432557 | 0.89054584 |
| Elmo3 | 0.08539069 | 0.06760603 | 0.09131752 | -0.0237115 | 0.88681058 | 0.91505129 |
| Slc25a10 | 0.13071403 | 0.11533093 | 0.13905737 | -0.0237264 | 0.78667127 | 0.89054584 |
| Stox2 | 0.18167777 | 0.16786521 | 0.19159942 | -0.0237342 | 0.71807156 | 0.89054584 |
| Clta | 2.60075155 | 1.09874784 | 1.12248354 | -0.0237357 | 0.53923894 | 0.89054584 |
| Tmem41a | 0.08589057 | 0.06811476 | 0.09185126 | -0.0237365 | 0.88570115 | 0.91420299 |
| Pard6a | 0.21771259 | 0.20436573 | 0.22810615 | -0.0237404 | 0.68612099 | 0.89054584 |
| Sos1 | 0.14882917 | 0.13401338 | 0.1578511 | -0.0238377 | 0.75914819 | 0.89054584 |
| M6pr | 0.72281745 | 0.64233987 | 0.66619566 | -0.0238558 | 0.56624887 | 0.89054584 |
| Blzf1 | 0.28380023 | 0.26969317 | 0.29355113 | -0.023858 | 0.64752571 | 0.89054584 |
| Misp | 0.20018462 | 0.18655943 | 0.21041786 | -0.0238584 | 0.70134002 | 0.89054584 |
| Mthfsd | 0.30448352 | 0.28973897 | 0.313608 | -0.023869 | 0.63855418 | 0.89054584 |
| Mdp1 | 0.52801388 | 0.49170433 | 0.51557748 | -0.0238731 | 0.58539943 | 0.89054584 |
| Exo5 | 0.10899217 | 0.09248394 | 0.11635813 | -0.0238742 | 0.83044233 | 0.89271879 |
| Tkfc | 0.22992101 | 0.21645622 | 0.24034602 | -0.0238898 | 0.67834911 | 0.89054584 |
| Smg6 | 0.45610178 | 0.42985389 | 0.45374696 | -0.0238931 | 0.5968984 | 0.89054584 |

| | | | | | | |
|---|---|---|---|---|---|---|
| Sh3glb1 | 1.4467416 | 0.99637724 | 1.02029051 | -0.0239133 | 0.54347603 | 0.89054584 |
| Morc2a | 0.13929771 | 0.12405874 | 0.14797852 | -0.0239198 | 0.77430153 | 0.89054584 |
| Sc5d | 0.16197995 | 0.14749227 | 0.17141439 | -0.0239221 | 0.74221489 | 0.89054584 |
| Phlpp2 | 0.13593693 | 0.12056657 | 0.14448901 | -0.0239224 | 0.77975357 | 0.89054584 |
| Rp9 | 0.63219785 | 0.57539109 | 0.59931902 | -0.0239279 | 0.57378119 | 0.89054584 |
| Rab12 | 0.31212481 | 0.29702834 | 0.32096372 | -0.0239354 | 0.63586715 | 0.89054584 |
| Ap3s2 | 0.54018648 | 0.50178846 | 0.52572566 | -0.0239372 | 0.58399707 | 0.89054584 |
| Paip2b | 0.29436816 | 0.27987562 | 0.30382574 | -0.0239501 | 0.64328638 | 0.89054584 |
| Etnk1 | 0.60819817 | 0.55670873 | 0.58066392 | -0.0239552 | 0.57620874 | 0.89054584 |
| Mb21d2 | 0.1228135 | 0.10686467 | 0.13082163 | -0.023957 | 0.80321956 | 0.89054584 |
| Tbc1d14 | 0.16548742 | 0.15103867 | 0.17502022 | -0.0239816 | 0.73838545 | 0.89054584 |
| Bahcc1 | 0.11017287 | 0.09360505 | 0.11759645 | -0.0239914 | 0.82906898 | 0.89265604 |
| Atg13 | 0.16212587 | 0.14756868 | 0.17156451 | -0.0239958 | 0.74267732 | 0.89054584 |
| Mpped2 | 0.33232184 | 0.31623855 | 0.34026444 | -0.0240259 | 0.62890902 | 0.89054584 |
| Fbxl6 | 0.37637499 | 0.35759629 | 0.38162339 | -0.0240271 | 0.61536432 | 0.89054584 |
| Rbm18 | 0.62170189 | 0.56718092 | 0.59120818 | -0.0240273 | 0.57508833 | 0.89054584 |
| Rusc1 | 0.16012352 | 0.14546866 | 0.16950388 | -0.0240352 | 0.74557387 | 0.89054584 |
| Snhg9 | 0.21588604 | 0.20221557 | 0.22626914 | -0.0240536 | 0.68979499 | 0.89054584 |
| Sh3bp4 | 0.17639493 | 0.1621315 | 0.18620138 | -0.0240699 | 0.72649679 | 0.89054584 |
| Vmac | 0.21435676 | 0.20064386 | 0.22472996 | -0.0240861 | 0.69122858 | 0.89054584 |
| Dvl3 | 0.10766333 | 0.09086032 | 0.11496382 | -0.0241035 | 0.83566616 | 0.89459411 |
| Akt3 | 0.09893194 | 0.08166086 | 0.10577028 | -0.0241094 | 0.85587433 | 0.9009252 |
| Camsap3 | 0.1048073 | 0.08784752 | 0.11196489 | -0.0241174 | 0.84220014 | 0.89691028 |
| Sfi1 | 0.10176285 | 0.08464282 | 0.10876485 | -0.024122 | 0.84926053 | 0.89883846 |
| Cant1 | 0.28327348 | 0.2688988 | 0.29303766 | -0.0241389 | 0.64942867 | 0.89054584 |
| Zbtb41 | 0.13391849 | 0.11825052 | 0.14239115 | -0.0241406 | 0.78519668 | 0.89054584 |
| Lrrc47 | 0.2842954 | 0.26987646 | 0.29403379 | -0.0241573 | 0.64905518 | 0.89054584 |
| Nav1 | 0.52123002 | 0.48572082 | 0.50988052 | -0.0241597 | 0.58735839 | 0.89054584 |
| Rpp14 | 0.46854688 | 0.44051166 | 0.46467171 | -0.0241601 | 0.59570139 | 0.89054584 |
| Smug1 | 0.11449884 | 0.09795238 | 0.12212917 | -0.0241768 | 0.82181078 | 0.89054584 |

| | | | | | | |
|---|---|---|---|---|---|---|
| Emc1 | 0.23550286 | 0.22173146 | 0.24592053 | -0.0241891 | 0.67661707 | 0.89054584 |
| Wrn | 0.14557157 | 0.13028973 | 0.15448093 | -0.0241912 | 0.76717491 | 0.89054584 |
| Rab33b | 0.20592539 | 0.19202388 | 0.21622547 | -0.0242016 | 0.69897817 | 0.89054584 |
| Slc7a1 | 0.15370008 | 0.13867998 | 0.1628826 | -0.0242026 | 0.75561579 | 0.89054584 |
| Bmi1 | 0.2548755 | 0.24092674 | 0.26515672 | -0.02423 | 0.66484892 | 0.89054584 |
| Arl14ep | 0.15197622 | 0.13687077 | 0.16110289 | -0.0242321 | 0.75827666 | 0.89054584 |
| Clpx | 0.57708827 | 0.53166763 | 0.55590923 | -0.0242416 | 0.58049619 | 0.89054584 |
| Slc52a2 | 0.10670107 | 0.08969473 | 0.11395374 | -0.024259 | 0.83935674 | 0.89504222 |
| Gm15545 | 0.09900989 | 0.08158718 | 0.10585344 | -0.0242663 | 0.85724744 | 0.9009252 |
| Ppm1a | 0.24334703 | 0.22943669 | 0.2537305 | -0.0242938 | 0.67222928 | 0.89054584 |
| Ankrd16 | 0.17977711 | 0.16535732 | 0.18965854 | -0.0243012 | 0.72473189 | 0.89054584 |
| Fgfbp3 | 0.08772128 | 0.06943468 | 0.09380585 | -0.0243712 | 0.88693986 | 0.91505129 |
| Gtf3c6 | 0.5725695 | 0.52788447 | 0.55226024 | -0.0243758 | 0.58146285 | 0.89054584 |
| Trmt13 | 0.29893787 | 0.28383509 | 0.30825135 | -0.0244163 | 0.64394719 | 0.89054584 |
| Supt3 | 0.20479756 | 0.19066788 | 0.21508551 | -0.0244176 | 0.70156513 | 0.89054584 |
| Slc7a4 | 0.22191068 | 0.20788636 | 0.23232263 | -0.0244363 | 0.68794681 | 0.89054584 |
| Setdb1 | 0.24103455 | 0.22690299 | 0.25143105 | -0.0245281 | 0.67526466 | 0.89054584 |
| Actl6a | 0.99542371 | 0.80760466 | 0.83213855 | -0.0245339 | 0.5546266 | 0.89054584 |
| Cdk5rap2 | 0.15354157 | 0.13817034 | 0.16271899 | -0.0245487 | 0.75893248 | 0.89054584 |
| Nsdhl | 0.23896744 | 0.22480442 | 0.2493735 | -0.0245691 | 0.67688792 | 0.89054584 |
| Dopey2 | 0.16585233 | 0.15082071 | 0.17539508 | -0.0245744 | 0.74304383 | 0.89054584 |
| Thap2 | 0.2838344 | 0.26900332 | 0.29358443 | -0.0245811 | 0.651762 | 0.89054584 |
| Ccdc43 | 0.30102113 | 0.28567502 | 0.31026544 | -0.0245904 | 0.64402905 | 0.89054584 |
| Rundc1 | 0.25780629 | 0.24342644 | 0.26805161 | -0.0246252 | 0.66567598 | 0.89054584 |
| Cep170 | 0.35537363 | 0.33740079 | 0.36203513 | -0.0246343 | 0.62438326 | 0.89054584 |
| Ercc2 | 0.13076104 | 0.11444948 | 0.13910631 | -0.0246568 | 0.79554275 | 0.89054584 |
| Zfp868 | 0.12196916 | 0.10527834 | 0.12994011 | -0.0246618 | 0.81171563 | 0.89054584 |
| Efcab14 | 0.60902815 | 0.55665157 | 0.58131558 | -0.024664 | 0.57834804 | 0.89054584 |
| Rbbp5 | 0.13980898 | 0.12384255 | 0.148509 | -0.0246664 | 0.78048194 | 0.89054584 |
| Alg12 | 0.14859483 | 0.13294012 | 0.15760879 | -0.0246687 | 0.76706331 | 0.89054584 |

| | | | | | | |
|---|---|---|---|---|---|---|
| Nfkbib | 0.30881065 | 0.29309979 | 0.31777681 | -0.024677 | 0.64124589 | 0.89054584 |
| Agap3 | 0.14073468 | 0.12478098 | 0.1494692 | -0.0246882 | 0.77921041 | 0.89054584 |
| Hsf1 | 0.25511839 | 0.24070063 | 0.26539679 | -0.0246962 | 0.66768581 | 0.89054584 |
| Slc12a9 | 0.16086658 | 0.14557071 | 0.17026876 | -0.024698 | 0.75041456 | 0.89054584 |
| Atf2 | 0.5178792 | 0.48235691 | 0.50705647 | -0.0246996 | 0.58976666 | 0.89054584 |
| Smarcal1 | 0.11137315 | 0.09415029 | 0.1188548 | -0.0247045 | 0.83357314 | 0.89411764 |
| Rnf103 | 0.18949032 | 0.17481971 | 0.19956131 | -0.0247416 | 0.71823807 | 0.89054584 |
| Gatc | 0.30382399 | 0.28822612 | 0.31297176 | -0.0247456 | 0.64370026 | 0.89054584 |
| Ap4m1 | 0.35942084 | 0.34107709 | 0.36582811 | -0.024751 | 0.6236998 | 0.89054584 |
| Rbm3 | 2.82641741 | 1.0837969 | 1.10855886 | -0.024762 | 0.54144196 | 0.89054584 |
| Daam1 | 0.58437543 | 0.53700301 | 0.56176545 | -0.0247624 | 0.58135637 | 0.89054584 |
| Lca5 | 0.1399222 | 0.12384676 | 0.14862645 | -0.0247797 | 0.78134993 | 0.89054584 |
| Zfp398 | 0.38028138 | 0.36042918 | 0.38524044 | -0.0248113 | 0.61793325 | 0.89054584 |
| Tbc1d20 | 0.26943538 | 0.25468468 | 0.27949991 | -0.0248152 | 0.6604316 | 0.89054584 |
| Zfp771 | 0.53739128 | 0.4985881 | 0.52340379 | -0.0248157 | 0.58741109 | 0.89054584 |
| Klc4 | 0.25692601 | 0.24235646 | 0.2671825 | -0.024826 | 0.66745596 | 0.89054584 |
| Tbc1d8 | 0.21162941 | 0.19715079 | 0.22198239 | -0.0248316 | 0.69887335 | 0.89054584 |
| Crls1 | 0.08916909 | 0.07051899 | 0.09535148 | -0.0248325 | 0.88749873 | 0.9152899 |
| Rangrf | 0.27034991 | 0.25555395 | 0.28039747 | -0.0248435 | 0.66011878 | 0.89054584 |
| Tmem67 | 0.13273022 | 0.11630693 | 0.14115538 | -0.0248485 | 0.79393247 | 0.89054584 |
| Tnrc6a | 0.8193782 | 0.70620513 | 0.73105919 | -0.0248541 | 0.56292587 | 0.89054584 |
| Letmd1 | 0.24252285 | 0.22805638 | 0.25291125 | -0.0248549 | 0.6764727 | 0.89054584 |
| Cept1 | 0.29816311 | 0.28264258 | 0.30750176 | -0.0248592 | 0.64677942 | 0.89054584 |
| Fryl | 0.52666633 | 0.48958659 | 0.51444818 | -0.0248616 | 0.58906922 | 0.89054584 |
| Tmem161a | 0.37040953 | 0.35122113 | 0.37608345 | -0.0248623 | 0.6209572 | 0.89054584 |
| Zfp704 | 0.30643538 | 0.29060566 | 0.31548962 | -0.024884 | 0.6433657 | 0.89054584 |
| Cacna1d | 0.13142045 | 0.11489547 | 0.13979264 | -0.0248972 | 0.79666294 | 0.89054584 |
| Eif1a | 0.99643406 | 0.80769939 | 0.83265819 | -0.0249588 | 0.55553215 | 0.89054584 |
| Ptpn2 | 0.37013586 | 0.35086965 | 0.37582883 | -0.0249592 | 0.6214933 | 0.89054584 |
| Med27 | 0.40182573 | 0.38006497 | 0.40503853 | -0.0249736 | 0.6130424 | 0.89054584 |

| | | | | | | |
|---|---|---|---|---|---|---|
| Toe1 | 0.20459 | 0.18989963 | 0.21487569 | -0.0249761 | 0.70592943 | 0.89054584 |
| D17Wsu92e | 0.3048712 | 0.28900364 | 0.31398188 | -0.0249782 | 0.64454467 | 0.89054584 |
| Rnf6 | 0.81966367 | 0.70625959 | 0.73124097 | -0.0249814 | 0.56322993 | 0.89054584 |
| Pi4k2a | 0.12088564 | 0.10381553 | 0.12880848 | -0.0249929 | 0.81699779 | 0.89054584 |
| Nup50 | 0.49590079 | 0.4633523 | 0.48835786 | -0.0250056 | 0.59427408 | 0.89054584 |
| Htatip2 | 0.17186298 | 0.15653798 | 0.1815617 | -0.0250237 | 0.73960175 | 0.89054584 |
| Ormdl1 | 0.28178036 | 0.2665535 | 0.29158146 | -0.025028 | 0.65538534 | 0.89054584 |
| Slc19a1 | 0.14214544 | 0.12586368 | 0.1509319 | -0.0250682 | 0.78047594 | 0.89054584 |
| Zfp26 | 0.14996527 | 0.13395463 | 0.15902546 | -0.0250708 | 0.76867668 | 0.89054584 |
| Gtf2h4 | 0.22765577 | 0.21299237 | 0.23808004 | -0.0250877 | 0.68826403 | 0.89054584 |
| Tmem115 | 0.4449628 | 0.41878652 | 0.44388956 | -0.025103 | 0.60390756 | 0.89054584 |
| Iqce | 0.18378337 | 0.16863837 | 0.19374781 | -0.0251094 | 0.72700904 | 0.89054584 |
| Hmgn5 | 0.50947653 | 0.4748338 | 0.49994375 | -0.02511 | 0.5925067 | 0.89054584 |
| Fastkd3 | 0.09324685 | 0.07457519 | 0.099704 | -0.0251288 | 0.87984495 | 0.91133758 |
| Ppp1r9a | 0.4209988 | 0.3973058 | 0.42243831 | -0.0251325 | 0.60917856 | 0.89054584 |
| Agk | 0.26808267 | 0.25303284 | 0.27817157 | -0.0251387 | 0.6631351 | 0.89054584 |
| Hectd3 | 0.12126433 | 0.10406323 | 0.12920404 | -0.0251408 | 0.8176768 | 0.89054584 |
| Ppp5c | 0.80362997 | 0.69578968 | 0.72093968 | -0.02515 | 0.56455468 | 0.89054584 |
| Ints10 | 0.3459003 | 0.3279628 | 0.35312227 | -0.0251595 | 0.63003387 | 0.89054584 |
| 1110004F10l | 2.05957936 | 1.09608769 | 1.12130129 | -0.0252136 | 0.54171729 | 0.89054584 |
| Cox15 | 0.14697375 | 0.13071565 | 0.15593208 | -0.0252164 | 0.77440026 | 0.89054584 |
| Atp11a | 0.13779311 | 0.12118856 | 0.14641683 | -0.0252283 | 0.78894219 | 0.89054584 |
| Tmc6 | 0.1348651 | 0.11814625 | 0.1433752 | -0.0252289 | 0.79383728 | 0.89054584 |
| Mbd6 | 0.24091294 | 0.22603327 | 0.25131006 | -0.0252768 | 0.68032103 | 0.89054584 |
| Atg16l1 | 0.19311901 | 0.17795509 | 0.20325055 | -0.0252955 | 0.71898425 | 0.89054584 |
| Mlh3 | 0.14685308 | 0.13046266 | 0.15580722 | -0.0253446 | 0.77573018 | 0.89054584 |
| Nfyc | 0.53142479 | 0.49306776 | 0.51843075 | -0.025363 | 0.59014875 | 0.89054584 |
| Dhx9 | 0.58269311 | 0.5350176 | 0.56041663 | -0.025399 | 0.5836153 | 0.89054584 |
| Mrpl15 | 0.83611529 | 0.71621524 | 0.74161604 | -0.0254008 | 0.5633907 | 0.89054584 |
| Slc44a1 | 0.25369088 | 0.23857429 | 0.26398549 | -0.0254112 | 0.67309262 | 0.89054584 |

| | | | | | | |
|---|---|---|---|---|---|---|
| Tusc2 | 0.36351199 | 0.34423952 | 0.36965346 | -0.0254139 | 0.62563165 | 0.89054584 |
| Rtn4rl1 | 0.16816843 | 0.15235265 | 0.17777303 | -0.0254204 | 0.74735781 | 0.89054584 |
| Lias | 0.4517819 | 0.42450946 | 0.44993294 | -0.0254235 | 0.60382246 | 0.89054584 |
| Cul7 | 0.33782574 | 0.3200613 | 0.34548745 | -0.0254262 | 0.63415368 | 0.89054584 |
| Fam110a | 0.27412866 | 0.25867152 | 0.2841018 | -0.0254303 | 0.66167075 | 0.89054584 |
| Casp2 | 0.27652228 | 0.2610053 | 0.28644468 | -0.0254394 | 0.66047664 | 0.89054584 |
| Mynn | 0.31181946 | 0.29522153 | 0.32067031 | -0.0254488 | 0.64421154 | 0.89054584 |
| Tbl1x | 0.76949662 | 0.6729382 | 0.69838951 | -0.0254513 | 0.56741044 | 0.89054584 |
| Rnf168 | 0.22258118 | 0.20753985 | 0.23299534 | -0.0254555 | 0.69462537 | 0.89054584 |
| Polr3c | 0.29612981 | 0.28006617 | 0.30553308 | -0.0254669 | 0.6511155 | 0.89054584 |
| D11Wsu47e | 0.09373715 | 0.07473512 | 0.10022727 | -0.0254921 | 0.88198554 | 0.91223443 |
| Abhd17c | 0.12379795 | 0.10635198 | 0.13184908 | -0.0254971 | 0.81619268 | 0.89054584 |
| Mar-06 | 0.33184106 | 0.31427616 | 0.33980747 | -0.0255313 | 0.63684865 | 0.89054584 |
| Fam171a1 | 0.18118309 | 0.16555408 | 0.19109442 | -0.0255403 | 0.73325303 | 0.89054584 |
| Zc3h4 | 0.17720812 | 0.1614826 | 0.187033 | -0.0255504 | 0.73775958 | 0.89054584 |
| Sp3 | 0.57138565 | 0.52573834 | 0.55130227 | -0.0255639 | 0.58551995 | 0.89054584 |
| Tbc1d22a | 0.1397404 | 0.12286924 | 0.14843784 | -0.0255686 | 0.78887049 | 0.89054584 |
| Zfp568 | 0.26360607 | 0.24819243 | 0.27376932 | -0.0255769 | 0.66831364 | 0.89054584 |
| Atg2b | 0.14235501 | 0.12553834 | 0.15114912 | -0.0256108 | 0.78506479 | 0.89054584 |
| Ext2 | 0.58448288 | 0.53623786 | 0.56185154 | -0.0256137 | 0.58409929 | 0.89054584 |
| Gsk3a | 0.19396134 | 0.17847418 | 0.20410614 | -0.025632 | 0.72075199 | 0.89054584 |
| Naa15 | 0.8021466 | 0.69433605 | 0.71997725 | -0.0256412 | 0.56589126 | 0.89054584 |
| Atm | 0.18129405 | 0.16556338 | 0.19120771 | -0.0256443 | 0.73396397 | 0.89054584 |
| Atp13a3 | 0.21213418 | 0.19684443 | 0.22249115 | -0.0256467 | 0.70437985 | 0.89054584 |
| Ftsj1 | 0.17921845 | 0.16338225 | 0.18908778 | -0.0257055 | 0.73675054 | 0.89054584 |
| Nt5c | 0.79987573 | 0.69278587 | 0.71850076 | -0.0257149 | 0.56621342 | 0.89054584 |
| Rhot2 | 0.10877541 | 0.09041342 | 0.11613073 | -0.0257173 | 0.84890049 | 0.89883797 |
| 2610037D02 | 0.15812104 | 0.14171853 | 0.1674415 | -0.025723 | 0.76291892 | 0.89054584 |
| Arl5b | 0.15389034 | 0.13735005 | 0.16307896 | -0.0257289 | 0.76884225 | 0.89054584 |
| Rae1 | 0.68619125 | 0.61412108 | 0.63985327 | -0.0257322 | 0.57431195 | 0.89054584 |

| | | | | | | |
|---|---|---|---|---|---|---|
| Gm37494 | 0.17012072 | 0.15404351 | 0.17977576 | -0.0257322 | 0.74756879 | 0.89054584 |
| Srrd | 0.07856264 | 0.05827986 | 0.08402546 | -0.0257456 | 0.92328366 | 0.93974772 |
| Zmynd19 | 0.14488096 | 0.12796524 | 0.15376592 | -0.0258007 | 0.78281416 | 0.89054584 |
| Ippk | 0.17501919 | 0.15898605 | 0.18479383 | -0.0258078 | 0.74236521 | 0.89054584 |
| Uqcc3 | 0.93780078 | 0.77558337 | 0.80139597 | -0.0258126 | 0.55964231 | 0.89054584 |
| Gm26853 | 0.18839666 | 0.17263379 | 0.19844831 | -0.0258145 | 0.72775705 | 0.89054584 |
| Dlg4 | 0.15613211 | 0.13955957 | 0.16539149 | -0.0258319 | 0.76659026 | 0.89054584 |
| Parp11 | 0.13101053 | 0.11353325 | 0.13936601 | -0.0258328 | 0.80608989 | 0.89054584 |
| Fance | 0.22530254 | 0.20988008 | 0.23572356 | -0.0258435 | 0.69524804 | 0.89054584 |
| E130308A19I | 0.13659276 | 0.11931565 | 0.1451703 | -0.0258547 | 0.79666043 | 0.89054584 |
| Zkscan14 | 0.15210032 | 0.13536385 | 0.16123103 | -0.0258672 | 0.7726069 | 0.89054584 |
| Tmem9b | 0.60232929 | 0.55016939 | 0.57604268 | -0.0258733 | 0.582877 | 0.89054584 |
| Cryz | 0.15754565 | 0.14097475 | 0.1668486 | -0.0258739 | 0.76500029 | 0.89054584 |
| Avl9 | 0.49646664 | 0.46296594 | 0.48884309 | -0.0258771 | 0.59740011 | 0.89054584 |
| Mon2 | 0.12690371 | 0.10917717 | 0.13508819 | -0.025911 | 0.8142382 | 0.89054584 |
| Rpl26 | 5.079846 | 0.65963639 | 0.6855572 | -0.0259208 | 0.56991347 | 0.89054584 |
| Eaf1 | 0.18893762 | 0.1730764 | 0.19899889 | -0.0259225 | 0.72803961 | 0.89054584 |
| Gdap2 | 0.21335739 | 0.19778256 | 0.22372356 | -0.025941 | 0.70547374 | 0.89054584 |
| Hyal1 | 0.13098659 | 0.11339233 | 0.13934109 | -0.0259488 | 0.80719604 | 0.89054584 |
| Phf13 | 0.23913884 | 0.22359 | 0.24954419 | -0.0259542 | 0.68600567 | 0.89054584 |
| Ube2q2 | 0.26300626 | 0.24722254 | 0.27317874 | -0.0259562 | 0.67099826 | 0.89054584 |
| Ppp1cc | 1.01176759 | 0.81461577 | 0.84059256 | -0.0259768 | 0.55723989 | 0.89054584 |
| Abcf3 | 0.52137348 | 0.48402411 | 0.5100013 | -0.0259772 | 0.59379017 | 0.89054584 |
| Srrm3 | 0.12005289 | 0.10192986 | 0.12793846 | -0.0260086 | 0.82821016 | 0.89229532 |
| Dennd1b | 0.1259991 | 0.10813161 | 0.13414508 | -0.0260135 | 0.81686007 | 0.89054584 |
| Prune | 0.1780359 | 0.16183669 | 0.18787925 | -0.0260426 | 0.74079273 | 0.89054584 |
| Gtf2ird1 | 0.21420333 | 0.19852401 | 0.22457548 | -0.0260515 | 0.70555735 | 0.89054584 |
| Brd1 | 0.19556515 | 0.17966936 | 0.20573433 | -0.026065 | 0.72248926 | 0.89054584 |
| 1700008J07F | 0.11435921 | 0.09589218 | 0.12198297 | -0.0260908 | 0.84049523 | 0.89579504 |
| Mcm3ap | 0.10354919 | 0.08451737 | 0.11064289 | -0.0261255 | 0.86435386 | 0.90329433 |

| | | | | | | |
|---|---|---|---|---|---|---|
| Ppp4c | 1.18851957 | 0.89939662 | 0.92552354 | -0.0261269 | 0.55231757 | 0.89054584 |
| Slc25a38 | 0.25538223 | 0.2395284 | 0.26565753 | -0.0261291 | 0.67660886 | 0.89054584 |
| Usp20 | 0.1956605 | 0.17967657 | 0.20583111 | -0.0261545 | 0.72307544 | 0.89054584 |
| Tox4 | 0.60607943 | 0.55280888 | 0.57899828 | -0.0261894 | 0.58345276 | 0.89054584 |
| Odf2l | 0.2083346 | 0.19246253 | 0.21865877 | -0.0261962 | 0.71163265 | 0.89054584 |
| Tnrc6b | 0.77799715 | 0.67787431 | 0.70408402 | -0.0262097 | 0.56884339 | 0.89054584 |
| Dld | 0.77291355 | 0.6744647 | 0.70068481 | -0.0262201 | 0.5692012 | 0.89054584 |
| Slc43a2 | 0.25234679 | 0.23640645 | 0.26265579 | -0.0262493 | 0.67925128 | 0.89054584 |
| Zbtb38 | 0.22957023 | 0.21373754 | 0.23999529 | -0.0262578 | 0.69488044 | 0.89054584 |
| Pura | 0.73595396 | 0.64914887 | 0.67541264 | -0.0262638 | 0.5718811 | 0.89054584 |
| Nufip1 | 0.25886839 | 0.24283424 | 0.26909987 | -0.0262656 | 0.67535127 | 0.89054584 |
| Nme7 | 0.14202553 | 0.12452783 | 0.15080761 | -0.0262798 | 0.79156646 | 0.89054584 |
| Casd1 | 0.25661806 | 0.24058405 | 0.26687839 | -0.0262943 | 0.67689184 | 0.89054584 |
| Jade1 | 0.45534151 | 0.42677886 | 0.45307654 | -0.0262977 | 0.60658003 | 0.89054584 |
| Scamp5 | 0.1085698 | 0.08960577 | 0.11591502 | -0.0263092 | 0.85484468 | 0.90074724 |
| Axin1 | 0.14243937 | 0.12492502 | 0.15123655 | -0.0263115 | 0.79118619 | 0.89054584 |
| Vkorc1l1 | 0.18561807 | 0.16928677 | 0.19561826 | -0.0263315 | 0.73471095 | 0.89054584 |
| Ankrd27 | 0.20537094 | 0.1893217 | 0.21566512 | -0.0263434 | 0.71535181 | 0.89054584 |
| Yaf2 | 0.76960206 | 0.67209734 | 0.69846046 | -0.0263631 | 0.56979409 | 0.89054584 |
| Naa60 | 0.17825052 | 0.16168776 | 0.18809861 | -0.0264109 | 0.74349572 | 0.89054584 |
| Kbtbd2 | 0.38199426 | 0.36040741 | 0.38682376 | -0.0264163 | 0.62481855 | 0.89054584 |
| Gpc4 | 0.46419893 | 0.43441595 | 0.46086553 | -0.0264496 | 0.60541239 | 0.89054584 |
| Fhad1 | 0.16153604 | 0.14449857 | 0.17095768 | -0.0264591 | 0.76456495 | 0.89054584 |
| 1110004E09I | 0.49996617 | 0.46534701 | 0.49183951 | -0.0264925 | 0.59907353 | 0.89054584 |
| Snx11 | 0.10857676 | 0.08942146 | 0.11592232 | -0.0265009 | 0.85657897 | 0.9009252 |
| Exoc7 | 0.72470661 | 0.64102588 | 0.66752868 | -0.0265028 | 0.573375 | 0.89054584 |
| Rab11a | 1.77279881 | 1.06724921 | 1.09377058 | -0.0265214 | 0.5449722 | 0.89054584 |
| Slc25a4 | 3.08133139 | 1.06091346 | 1.08743548 | -0.026522 | 0.54523417 | 0.89054584 |
| Dtwd1 | 0.1724057 | 0.15558251 | 0.18211777 | -0.0265353 | 0.75137581 | 0.89054584 |
| Stam | 0.19072427 | 0.17427027 | 0.20081648 | -0.0265462 | 0.73101047 | 0.89054584 |

| | | | | | | |
|---|---|---|---|---|---|---|
| Tspan15 | 0.43184214 | 0.40563567 | 0.43218622 | -0.0265505 | 0.61264261 | 0.89054584 |
| Srp19 | 1.38957447 | 0.9755285 | 1.00215665 | -0.0266281 | 0.54926001 | 0.89054584 |
| 2700049A03I | 0.3823701 | 0.36050587 | 0.38717095 | -0.0266651 | 0.62584506 | 0.89054584 |
| Gm5914 | 0.16722801 | 0.15008199 | 0.17680776 | -0.0267258 | 0.75935603 | 0.89054584 |
| Narfl | 0.13575397 | 0.11757192 | 0.14429891 | -0.026727 | 0.80591133 | 0.89054584 |
| Wdfy1 | 0.42174752 | 0.39638273 | 0.4231136 | -0.0267309 | 0.6157494 | 0.89054584 |
| Rnf41 | 0.28828326 | 0.27118281 | 0.2979165 | -0.0267337 | 0.66205292 | 0.89054584 |
| Tbcd | 0.15534464 | 0.13782853 | 0.16457939 | -0.0267509 | 0.7755553 | 0.89054584 |
| Prpf39 | 0.38870849 | 0.36625093 | 0.39301431 | -0.0267634 | 0.62447809 | 0.89054584 |
| Xpc | 0.18921339 | 0.17251205 | 0.19927954 | -0.0267675 | 0.73427532 | 0.89054584 |
| Ubac1 | 0.29180887 | 0.2745705 | 0.30134258 | -0.0267721 | 0.660531 | 0.89054584 |
| Mapk1 | 1.02104114 | 0.81854234 | 0.84531736 | -0.026775 | 0.55865882 | 0.89054584 |
| Camk2n2 | 0.13814902 | 0.11990542 | 0.14678633 | -0.0268809 | 0.80322292 | 0.89054584 |
| Fkbp5 | 0.33683854 | 0.31766541 | 0.34455164 | -0.0268862 | 0.64190254 | 0.89054584 |
| Acads | 0.60625294 | 0.55223675 | 0.57913481 | -0.0268981 | 0.58565641 | 0.89054584 |
| Arid4b | 1.35100472 | 0.96209345 | 0.98905206 | -0.0269586 | 0.55052538 | 0.89054584 |
| Tubgcp2 | 0.25811512 | 0.24139113 | 0.26835647 | -0.0269653 | 0.68015981 | 0.89054584 |
| Ap1ar | 0.34873513 | 0.32882386 | 0.35579445 | -0.0269706 | 0.63801787 | 0.89054584 |
| Ccdc107 | 0.50465118 | 0.46886428 | 0.49583896 | -0.0269747 | 0.60004215 | 0.89054584 |
| Cisd3 | 0.4530351 | 0.42405574 | 0.45104055 | -0.0269848 | 0.60977576 | 0.89054584 |
| Coa7 | 0.32619989 | 0.30745032 | 0.33443661 | -0.0269863 | 0.64651862 | 0.89054584 |
| Atxn7l1 | 0.23403499 | 0.21743026 | 0.24445591 | -0.0270257 | 0.69674656 | 0.89054584 |
| Ints8 | 0.20346069 | 0.18667654 | 0.21373374 | -0.0270572 | 0.72233413 | 0.89054584 |
| Clip3 | 0.30474199 | 0.28679617 | 0.31385727 | -0.0270611 | 0.65604717 | 0.89054584 |
| Timm8b | 1.74032815 | 1.06152309 | 1.08858593 | -0.0270628 | 0.54610391 | 0.89054584 |
| Tmem222 | 0.57035136 | 0.52339001 | 0.55046458 | -0.0270746 | 0.59062371 | 0.89054584 |
| Hmbox1 | 0.23466667 | 0.21798806 | 0.24508631 | -0.0270983 | 0.69676588 | 0.89054584 |
| Ube3b | 0.16738761 | 0.14987274 | 0.17697161 | -0.0270989 | 0.76220173 | 0.89054584 |
| Cep162 | 0.32507382 | 0.30624689 | 0.33336249 | -0.0271156 | 0.64764005 | 0.89054584 |
| Klc1 | 0.51626512 | 0.47855061 | 0.50569366 | -0.0271431 | 0.59873268 | 0.89054584 |

| | | | | | | |
|---|---|---|---|---|---|---|
| Polr2i | 1.15433644 | 0.88325905 | 0.91040502 | -0.027146 | 0.55524245 | 0.89054584 |
| Lsm6 | 1.5689612 | 1.02630355 | 1.05345015 | -0.0271466 | 0.54778113 | 0.89054584 |
| 2700097O09 | 0.25060575 | 0.23378215 | 0.26093212 | -0.02715 | 0.68607764 | 0.89054584 |
| Ern1 | 0.12830624 | 0.10938595 | 0.13654984 | -0.0271639 | 0.82298592 | 0.89075401 |
| Sf3a1 | 0.51818953 | 0.48009401 | 0.50731831 | -0.0272243 | 0.59871152 | 0.89054584 |
| Trit1 | 0.1923321 | 0.17521383 | 0.20245099 | -0.0272372 | 0.73460489 | 0.89054584 |
| 2900076A07 | 0.22525605 | 0.20843267 | 0.23567699 | -0.0272443 | 0.70490983 | 0.89054584 |
| Aak1 | 0.37298451 | 0.35122088 | 0.37847718 | -0.0272563 | 0.63138239 | 0.89054584 |
| Snapc3 | 0.36679763 | 0.34544229 | 0.37271964 | -0.0272774 | 0.63343427 | 0.89054584 |
| E2f4 | 0.3961174 | 0.37252976 | 0.39981785 | -0.0272881 | 0.62474951 | 0.89054584 |
| Herc2 | 0.52936023 | 0.48941503 | 0.51670462 | -0.0272896 | 0.59718228 | 0.89054584 |
| Lnp | 0.27588345 | 0.2584754 | 0.28581966 | -0.0273443 | 0.67209992 | 0.89054584 |
| Myef2 | 0.852017 | 0.72410584 | 0.75145761 | -0.0273518 | 0.5673288 | 0.89054584 |
| Hdgfrp2 | 0.43512282 | 0.40776609 | 0.43512185 | -0.0273558 | 0.61520218 | 0.89054584 |
| Eif2ak2 | 0.19563462 | 0.17844895 | 0.20580484 | -0.0273559 | 0.73213189 | 0.89054584 |
| Med11 | 0.13707222 | 0.11830357 | 0.14566828 | -0.0273647 | 0.80927062 | 0.89054584 |
| Gan | 0.12655812 | 0.10733867 | 0.13472792 | -0.0273893 | 0.82821383 | 0.89229532 |
| Wnk4 | 0.13210004 | 0.11307371 | 0.14049979 | -0.0274261 | 0.81844173 | 0.89054584 |
| Coq5 | 0.3955794 | 0.37189791 | 0.39932485 | -0.0274269 | 0.62551257 | 0.89054584 |
| Mms19 | 0.21074027 | 0.19364346 | 0.22108595 | -0.0274425 | 0.71846761 | 0.89054584 |
| Pttg1ip | 0.75549324 | 0.66144971 | 0.68889497 | -0.0274453 | 0.57362472 | 0.89054584 |
| Ccdc6 | 0.18682523 | 0.16934268 | 0.19684818 | -0.0275055 | 0.74247405 | 0.89054584 |
| Tomm34 | 0.38746667 | 0.3643626 | 0.39187124 | -0.0275086 | 0.62818463 | 0.89054584 |
| Stk11 | 0.9747716 | 0.79384558 | 0.82135456 | -0.027509 | 0.56199856 | 0.89054584 |
| Gpatch11 | 0.5191419 | 0.48059276 | 0.50812145 | -0.0275287 | 0.59963761 | 0.89054584 |
| Unc119b | 0.19111265 | 0.17367888 | 0.20121141 | -0.0275325 | 0.73810066 | 0.89054584 |
| Scaf1 | 0.35409227 | 0.33328649 | 0.36083242 | -0.0275459 | 0.63895366 | 0.89054584 |
| Cdk5 | 0.46431967 | 0.43340769 | 0.46097138 | -0.0275637 | 0.6097166 | 0.89054584 |
| Smarcc2 | 0.51869795 | 0.48013178 | 0.50774713 | -0.0276154 | 0.60001677 | 0.89054584 |
| Nme6 | 0.17791879 | 0.16009331 | 0.18775954 | -0.0276662 | 0.75379846 | 0.89054584 |

| | | | | | | |
|---|---|---|---|---|---|---|
| Evi5 | 0.4761511 | 0.44360567 | 0.47130252 | -0.0276969 | 0.60787707 | 0.89054584 |
| Tsen15 | 0.60402707 | 0.5496629 | 0.57738193 | -0.027719 | 0.58849182 | 0.89054584 |
| Fggy | 0.12463703 | 0.10499676 | 0.13272454 | -0.0277278 | 0.83480036 | 0.89453395 |
| Smap1 | 0.67598376 | 0.60460729 | 0.63234436 | -0.0277371 | 0.58096361 | 0.89054584 |
| Anapc1 | 0.27801804 | 0.26016801 | 0.28790731 | -0.0277393 | 0.67324084 | 0.89054584 |
| Ggct | 0.29314643 | 0.27489644 | 0.30264076 | -0.0277443 | 0.66534828 | 0.89054584 |
| Cpsf4 | 0.21089838 | 0.19348654 | 0.22124537 | -0.0277588 | 0.72057146 | 0.89054584 |
| Pex11b | 0.39416517 | 0.3702638 | 0.39802815 | -0.0277643 | 0.62740311 | 0.89054584 |
| Avpi1 | 0.22436604 | 0.20699127 | 0.23478508 | -0.0277938 | 0.70936293 | 0.89054584 |
| Leo1 | 0.55041655 | 0.50635916 | 0.53418108 | -0.0278219 | 0.59586294 | 0.89054584 |
| Ctnnbip1 | 0.94920588 | 0.77982503 | 0.80764798 | -0.027823 | 0.56375541 | 0.89054584 |
| Elac2 | 0.16560571 | 0.14731292 | 0.17514175 | -0.0278288 | 0.77043314 | 0.89054584 |
| Trpc4ap | 0.69770693 | 0.62039284 | 0.64823723 | -0.0278444 | 0.57930673 | 0.89054584 |
| Tspyl1 | 0.73227282 | 0.64494407 | 0.6728419 | -0.0278978 | 0.57658794 | 0.89054584 |
| Cacfd1 | 0.55077914 | 0.50657027 | 0.53447958 | -0.0279093 | 0.59610552 | 0.89054584 |
| Tsg101 | 1.51324971 | 1.01124803 | 1.03916315 | -0.0279151 | 0.54979899 | 0.89054584 |
| Gtf2e1 | 0.18690652 | 0.16900194 | 0.19693098 | -0.027929 | 0.74560975 | 0.89054584 |
| Recql | 0.1100375 | 0.0894964 | 0.11745451 | -0.0279581 | 0.86628148 | 0.90430169 |
| Ofd1 | 0.21956521 | 0.20199518 | 0.22996786 | -0.0279727 | 0.71454136 | 0.89054584 |
| Fam65a | 0.2017642 | 0.18402875 | 0.21201722 | -0.0279885 | 0.73073207 | 0.89054584 |
| Rnh1 | 1.1176199 | 0.865566 | 0.89356127 | -0.0279953 | 0.55802533 | 0.89054584 |
| Eef2kmt | 0.34758768 | 0.32671222 | 0.35471335 | -0.0280011 | 0.64348047 | 0.89054584 |
| Sfxn2 | 0.4887748 | 0.45422721 | 0.48223167 | -0.0280045 | 0.6066341 | 0.89054584 |
| Sdsl | 0.22974397 | 0.2121562 | 0.24016901 | -0.0280128 | 0.70657597 | 0.89054584 |
| Ercc6l2 | 0.20963752 | 0.19195876 | 0.21997365 | -0.0280149 | 0.72352407 | 0.89054584 |
| Katnal1 | 0.21000848 | 0.19231507 | 0.22034788 | -0.0280328 | 0.72331363 | 0.89054584 |
| Lancl1 | 0.39172531 | 0.36769873 | 0.39578838 | -0.0280897 | 0.62954614 | 0.89054584 |
| Papss1 | 1.05257818 | 0.83346083 | 0.86155951 | -0.0280987 | 0.56038504 | 0.89054584 |
| Zbtb37 | 0.14552639 | 0.12631074 | 0.15443416 | -0.0281234 | 0.80189851 | 0.89054584 |
| Hif1an | 0.20337406 | 0.18546963 | 0.21364612 | -0.0281765 | 0.7305375 | 0.89054584 |

| | | | | | | |
|---|---|---|---|---|---|---|
| BC051226 | 0.11324027 | 0.09256985 | 0.12081117 | -0.0282413 | 0.86194362 | 0.90253338 |
| Mgme1 | 0.1381996 | 0.11858453 | 0.14683884 | -0.0282543 | 0.81499645 | 0.89054584 |
| Rmnd5b | 0.31530087 | 0.29575734 | 0.32401284 | -0.0282555 | 0.65773245 | 0.89054584 |
| Pibf1 | 0.21110441 | 0.19319407 | 0.22145311 | -0.028259 | 0.72391618 | 0.89054584 |
| Smim7 | 1.15534824 | 0.88255106 | 0.91085703 | -0.028306 | 0.55755857 | 0.89054584 |
| Rpl7 | 4.55494325 | 0.79755356 | 0.82586993 | -0.0283164 | 0.56345698 | 0.89054584 |
| Ddx19a | 0.40962524 | 0.38382492 | 0.4121416 | -0.0283167 | 0.62555302 | 0.89054584 |
| Smc3 | 1.4123486 | 0.98127482 | 1.00959203 | -0.0283172 | 0.5519837 | 0.89054584 |
| Fchsd2 | 0.19249235 | 0.17426081 | 0.20261384 | -0.028353 | 0.74278487 | 0.89054584 |
| Bcl7c | 0.56139506 | 0.51482072 | 0.54318106 | -0.0283603 | 0.59609439 | 0.89054584 |
| Phf20 | 0.50298055 | 0.46603761 | 0.49441438 | -0.0283768 | 0.60541886 | 0.89054584 |
| Myo1e | 0.2058216 | 0.18774103 | 0.21612059 | -0.0283796 | 0.72965898 | 0.89054584 |
| Ammecr1l | 0.47079099 | 0.43823135 | 0.46663224 | -0.0284009 | 0.61162606 | 0.89054584 |
| Pex16 | 0.24821031 | 0.23013851 | 0.25855825 | -0.0284197 | 0.69570707 | 0.89054584 |
| Mcfd2 | 0.64182272 | 0.57824094 | 0.60669143 | -0.0284505 | 0.58647193 | 0.89054584 |
| Naa35 | 0.71089388 | 0.62926341 | 0.65772344 | -0.02846 | 0.57988323 | 0.89054584 |
| Tcp11l1 | 0.21758886 | 0.19951757 | 0.22798176 | -0.0284642 | 0.71960969 | 0.89054584 |
| Slf1 | 0.2549988 | 0.2368042 | 0.26527859 | -0.0284744 | 0.69149062 | 0.89054584 |
| Rhpn2 | 0.16707013 | 0.14815306 | 0.17664568 | -0.0284926 | 0.77381763 | 0.89054584 |
| Tbc1d23 | 0.14942206 | 0.12994039 | 0.15846401 | -0.0285236 | 0.79915653 | 0.89054584 |
| Ago1 | 0.21883114 | 0.20068483 | 0.22923037 | -0.0285455 | 0.71909909 | 0.89054584 |
| Rnf219 | 0.21560128 | 0.19743141 | 0.22598262 | -0.0285512 | 0.72194267 | 0.89054584 |
| Ccdc136 | 0.11654592 | 0.09571592 | 0.12427167 | -0.0285558 | 0.85781505 | 0.90120308 |
| Dpp9 | 0.37774898 | 0.35433267 | 0.38289658 | -0.0285639 | 0.63591274 | 0.89054584 |
| Apitd1 | 0.17947723 | 0.16078676 | 0.18935218 | -0.0285654 | 0.75892278 | 0.89054584 |
| Smim19 | 0.3493977 | 0.32784884 | 0.35641838 | -0.0285695 | 0.64559253 | 0.89054584 |
| Slc29a4 | 0.22533333 | 0.20717572 | 0.23575441 | -0.0285787 | 0.71388405 | 0.89054584 |
| Naa25 | 0.23179397 | 0.2136387 | 0.24221797 | -0.0285793 | 0.70874014 | 0.89054584 |
| Scyl2 | 0.20572561 | 0.18744406 | 0.21602358 | -0.0285795 | 0.73117458 | 0.89054584 |
| Plagl2 | 0.17341975 | 0.15456936 | 0.18315644 | -0.0285871 | 0.7664385 | 0.89054584 |

| | | | | | | |
|---|---|---|---|---|---|---|
| Zfyve27 | 0.27147322 | 0.25290549 | 0.28149938 | -0.0285939 | 0.68197963 | 0.89054584 |
| Zfyve16 | 0.17327329 | 0.15439597 | 0.18300645 | -0.0286105 | 0.76680329 | 0.89054584 |
| Chpf | 0.58352072 | 0.53246558 | 0.56108041 | -0.0286148 | 0.59390579 | 0.89054584 |
| Sgpp1 | 0.1857985 | 0.16715774 | 0.19580214 | -0.0286444 | 0.75224406 | 0.89054584 |
| Tgfbrap1 | 0.09581291 | 0.07379672 | 0.10244239 | -0.0286457 | 0.90367274 | 0.92583191 |
| Gadd45gip1 | 1.18213629 | 0.89400843 | 0.92273884 | -0.0287304 | 0.55766861 | 0.89054584 |
| Usp53 | 0.25600841 | 0.2375318 | 0.26627621 | -0.0287444 | 0.69249356 | 0.89054584 |
| Nsmf | 0.32882924 | 0.30819519 | 0.33694205 | -0.0287469 | 0.65449561 | 0.89054584 |
| Elp3 | 0.3776035 | 0.3540142 | 0.38276182 | -0.0287476 | 0.63679815 | 0.89054584 |
| Eif2b5 | 0.45214934 | 0.42150434 | 0.45025779 | -0.0287535 | 0.61696519 | 0.89054584 |
| Cep164 | 0.32086606 | 0.30055024 | 0.32934305 | -0.0287928 | 0.65810909 | 0.89054584 |
| Ankrd40 | 0.26748379 | 0.24878952 | 0.27758319 | -0.0287937 | 0.68554848 | 0.89054584 |
| Tbrg1 | 1.65299226 | 1.04349827 | 1.0722947 | -0.0287964 | 0.54978402 | 0.89054584 |
| Tsta3 | 0.45544046 | 0.42436079 | 0.45316381 | -0.028803 | 0.61643139 | 0.89054584 |
| 0610007P14I | 0.82226697 | 0.70407708 | 0.73289593 | -0.0288188 | 0.5726788 | 0.89054584 |
| Cd276 | 0.13819461 | 0.11798762 | 0.14683365 | -0.028846 | 0.81997418 | 0.89054584 |
| Prdx3 | 1.0272688 | 0.81963937 | 0.84851042 | -0.028871 | 0.5629777 | 0.89054584 |
| Snx2 | 1.13819575 | 0.87422273 | 0.90310291 | -0.0288802 | 0.55921826 | 0.89054584 |
| Tnpo2 | 0.26310442 | 0.24439115 | 0.27327539 | -0.0288842 | 0.68879357 | 0.89054584 |
| Parva | 0.47205648 | 0.43882709 | 0.46773645 | -0.0289094 | 0.61330935 | 0.89054584 |
| Tmem106b | 0.46130194 | 0.42938498 | 0.45832323 | -0.0289383 | 0.61568322 | 0.89054584 |
| Acap3 | 0.12158441 | 0.10059321 | 0.12953833 | -0.0289451 | 0.85106976 | 0.89958153 |
| Blmh | 1.07692845 | 0.84487746 | 0.87385781 | -0.0289803 | 0.56139527 | 0.89054584 |
| Kdm1a | 0.81165452 | 0.69711963 | 0.7261183 | -0.0289987 | 0.57380196 | 0.89054584 |
| Ppwd1 | 0.15471718 | 0.13490523 | 0.16393213 | -0.0290269 | 0.7952979 | 0.89054584 |
| Senp6 | 0.94807733 | 0.77800104 | 0.80703254 | -0.0290315 | 0.56654975 | 0.89054584 |
| Larp1 | 0.59047215 | 0.5376045 | 0.56663777 | -0.0290333 | 0.59433625 | 0.89054584 |
| Amn1 | 0.52122434 | 0.48081541 | 0.50987574 | -0.0290603 | 0.60470171 | 0.89054584 |
| Trmt10c | 0.4236578 | 0.39576668 | 0.42483506 | -0.0290684 | 0.62505239 | 0.89054584 |
| Fbxo17 | 0.10448876 | 0.08255662 | 0.11163023 | -0.0290736 | 0.88751176 | 0.9152899 |

| | | | | | | |
|---|---|---|---|---|---|---|
| Mesdc1 | 0.62597128 | 0.56543262 | 0.59451603 | -0.0290834 | 0.59014304 | 0.89054584 |
| Slc25a44 | 0.31909992 | 0.29850823 | 0.32765322 | -0.029145 | 0.66071532 | 0.89054584 |
| Glmn | 0.32182306 | 0.3011118 | 0.33025804 | -0.0291462 | 0.65952427 | 0.89054584 |
| E430025E21I | 0.17000642 | 0.15048738 | 0.17965855 | -0.0291712 | 0.77531705 | 0.89054584 |
| Abhd14a | 0.4487047 | 0.41801598 | 0.44720925 | -0.0291933 | 0.61948554 | 0.89054584 |
| Rab18 | 1.27301303 | 0.93107991 | 0.96029241 | -0.0292125 | 0.55635216 | 0.89054584 |
| Mcat | 0.18351232 | 0.16425733 | 0.19347136 | -0.029214 | 0.75912682 | 0.89054584 |
| Wars | 0.54109763 | 0.49726062 | 0.52648142 | -0.0292208 | 0.60202067 | 0.89054584 |
| Parp4 | 0.11368196 | 0.09204423 | 0.12127378 | -0.0292295 | 0.86926072 | 0.90542007 |
| Ints1 | 0.11321088 | 0.09152104 | 0.12078039 | -0.0292593 | 0.87047511 | 0.90594699 |
| Dpf2 | 0.67996941 | 0.6060196 | 0.63528457 | -0.029265 | 0.58496757 | 0.89054584 |
| Rc3h2 | 0.57215253 | 0.52265628 | 0.55192293 | -0.0292666 | 0.59756455 | 0.89054584 |
| Tmem63b | 0.43846129 | 0.40883057 | 0.43810264 | -0.0292721 | 0.62220982 | 0.89054584 |
| Rcn2 | 1.50038515 | 1.00639515 | 1.03567133 | -0.0292762 | 0.55238848 | 0.89054584 |
| Timm44 | 0.48138773 | 0.44656846 | 0.47584817 | -0.0292797 | 0.61281832 | 0.89054584 |
| Trove2 | 0.33107247 | 0.30979151 | 0.33907669 | -0.0292852 | 0.65629809 | 0.89054584 |
| Ubr4 | 0.47283616 | 0.43911398 | 0.46841628 | -0.0293023 | 0.61464459 | 0.89054584 |
| Dmtf1 | 0.39094556 | 0.36571585 | 0.39507186 | -0.029356 | 0.63539813 | 0.89054584 |
| Cnot3 | 0.68695564 | 0.61105362 | 0.64041268 | -0.0293591 | 0.58456442 | 0.89054584 |
| Pex6 | 0.35178216 | 0.32927737 | 0.35866184 | -0.0293845 | 0.64865729 | 0.89054584 |
| Gfer | 0.52902328 | 0.48701985 | 0.51642263 | -0.0294028 | 0.60459508 | 0.89054584 |
| Pigq | 0.24817183 | 0.22910784 | 0.25852009 | -0.0294122 | 0.70196319 | 0.89054584 |
| 1300002E11I | 0.41505226 | 0.3876197 | 0.41706336 | -0.0294437 | 0.62888808 | 0.89054584 |
| Kcnma1 | 0.17153771 | 0.15177806 | 0.18122836 | -0.0294503 | 0.77550377 | 0.89054584 |
| Nup153 | 0.18507477 | 0.1656129 | 0.19506453 | -0.0294516 | 0.75910474 | 0.89054584 |
| Atp6v0a2 | 0.31759129 | 0.29672637 | 0.32620849 | -0.0294821 | 0.66314675 | 0.89054584 |
| Zfp609 | 0.15656402 | 0.13630258 | 0.16583679 | -0.0295342 | 0.79665151 | 0.89054584 |
| Pwwp2a | 0.27671522 | 0.25709164 | 0.28663341 | -0.0295418 | 0.68444748 | 0.89054584 |
| Alg11 | 0.22407672 | 0.20494349 | 0.23449507 | -0.0295516 | 0.72144175 | 0.89054584 |
| Flywch1 | 0.24967997 | 0.23045741 | 0.26001499 | -0.0295576 | 0.70180952 | 0.89054584 |

| | | | | | | |
|---|---|---|---|---|---|---|
| Ankib1 | 0.16362177 | 0.14354105 | 0.17310287 | -0.0295618 | 0.78687807 | 0.89054584 |
| Lrrc14 | 0.2001833 | 0.18084972 | 0.21041652 | -0.0295668 | 0.7436547 | 0.89054584 |
| 6030458C11I | 0.18891786 | 0.16940744 | 0.19897879 | -0.0295714 | 0.75565421 | 0.89054584 |
| Cdcp1 | 0.11317585 | 0.09116795 | 0.12074369 | -0.0295757 | 0.87310478 | 0.90740309 |
| Mrpl22 | 0.63224362 | 0.56977348 | 0.59935422 | -0.0295807 | 0.59093043 | 0.89054584 |
| Fuz | 0.22461144 | 0.20544457 | 0.23503104 | -0.0295865 | 0.72122438 | 0.89054584 |
| Med6 | 0.5186635 | 0.47812812 | 0.50771808 | -0.02959 | 0.60700616 | 0.89054584 |
| Zfhx2 | 0.33592488 | 0.31405277 | 0.34368515 | -0.0296324 | 0.65604459 | 0.89054584 |
| Gde1 | 0.87120275 | 0.73344885 | 0.76308603 | -0.0296372 | 0.57179528 | 0.89054584 |
| Ech1 | 1.15683927 | 0.88188192 | 0.9115219 | -0.02964 | 0.56020733 | 0.89054584 |
| Rexo4 | 0.3455191 | 0.32309289 | 0.35276261 | -0.0296697 | 0.65241488 | 0.89054584 |
| Zkscan1 | 0.47648678 | 0.44191851 | 0.47159442 | -0.0296759 | 0.61530469 | 0.89054584 |
| Tsnax | 0.69471765 | 0.61638019 | 0.6460699 | -0.0296897 | 0.58476489 | 0.89054584 |
| Pofut1 | 0.22150044 | 0.20220997 | 0.23191094 | -0.029701 | 0.7246373 | 0.89054584 |
| Ntmt1 | 0.67114972 | 0.59905283 | 0.62876336 | -0.0297105 | 0.58712122 | 0.89054584 |
| Hspa13 | 0.38441129 | 0.35933605 | 0.38905518 | -0.0297191 | 0.63903738 | 0.89054584 |
| Ptpn23 | 0.12623974 | 0.10466776 | 0.13439597 | -0.0297282 | 0.84862439 | 0.89883797 |
| Dnajc1 | 0.76896422 | 0.66829992 | 0.69803111 | -0.0297312 | 0.57864878 | 0.89054584 |
| Plekhb2 | 0.21689072 | 0.19754128 | 0.22727976 | -0.0297385 | 0.72893671 | 0.89054584 |
| Ppfia1 | 0.29144431 | 0.27124954 | 0.30098859 | -0.0297391 | 0.67735845 | 0.89054584 |
| Rbm17 | 0.99004881 | 0.79960783 | 0.8293617 | -0.0297539 | 0.566371 | 0.89054584 |
| Polr3f | 0.29252064 | 0.2722626 | 0.3020335 | -0.0297709 | 0.67696318 | 0.89054584 |
| Prdx5 | 2.03879291 | 1.09038745 | 1.12015843 | -0.029771 | 0.54927224 | 0.89054584 |
| Txlna | 0.68607555 | 0.60996372 | 0.63976856 | -0.0298048 | 0.58591362 | 0.89054584 |
| Tmem181a | 0.2909821 | 0.27073253 | 0.3005397 | -0.0298072 | 0.67798382 | 0.89054584 |
| Ccdc51 | 0.08740861 | 0.06365949 | 0.09347203 | -0.0298125 | 0.93135403 | 0.94537357 |
| Dhx40 | 0.42574918 | 0.39689236 | 0.42671727 | -0.0298249 | 0.62764832 | 0.89054584 |
| Map4k3 | 0.15216054 | 0.13145425 | 0.16129322 | -0.029839 | 0.80562765 | 0.89054584 |
| Psmd13 | 1.27519759 | 0.93128483 | 0.96113899 | -0.0298542 | 0.55753106 | 0.89054584 |
| Lig3 | 0.13965346 | 0.11847124 | 0.14834764 | -0.0298764 | 0.82595111 | 0.89170439 |

| | | | | | | |
|---|---|---|---|---|---|---|
| 1700017B05 | 0.10634386 | 0.08370009 | 0.1135787 | -0.0298786 | 0.88982646 | 0.9168209 |
| Cers6 | 0.2871433 | 0.26692686 | 0.29680739 | -0.0298805 | 0.680476 | 0.89054584 |
| Rce1 | 0.40228898 | 0.37555807 | 0.40546138 | -0.0299033 | 0.6344276 | 0.89054584 |
| Elmod2 | 0.12456228 | 0.102734 | 0.13264656 | -0.0299126 | 0.85328294 | 0.90008483 |
| Snx17 | 1.21066633 | 0.90510963 | 0.93503692 | -0.0299273 | 0.55926917 | 0.89054584 |
| Rnaseh1 | 0.21476093 | 0.1951781 | 0.22513684 | -0.0299587 | 0.73235822 | 0.89054584 |
| Zfp157 | 0.34225085 | 0.31971122 | 0.34967587 | -0.0299646 | 0.65514067 | 0.89054584 |
| Fam188a | 0.34156066 | 0.31905516 | 0.34902328 | -0.0299681 | 0.65543271 | 0.89054584 |
| Slc35g1 | 0.12692223 | 0.10513311 | 0.1351075 | -0.0299744 | 0.84934151 | 0.89883846 |
| Ranbp10 | 0.11092666 | 0.08839963 | 0.11838677 | -0.0299871 | 0.88101994 | 0.91182315 |
| Poglut1 | 0.29399414 | 0.27346878 | 0.30346304 | -0.0299943 | 0.67741749 | 0.89054584 |
| N4bp2 | 0.4083588 | 0.38095295 | 0.41099062 | -0.0300377 | 0.63326132 | 0.89054584 |
| Bak1 | 0.42094711 | 0.39231539 | 0.42239167 | -0.0300763 | 0.62995668 | 0.89054584 |
| Cr1l | 0.71919941 | 0.6335575 | 0.66363561 | -0.0300781 | 0.5836181 | 0.89054584 |
| Exoc8 | 0.18074481 | 0.16056214 | 0.19064691 | -0.0300848 | 0.76889049 | 0.89054584 |
| Cuedc2 | 1.01405353 | 0.81165795 | 0.8417629 | -0.030105 | 0.56616672 | 0.89054584 |
| Pbx2 | 0.31340034 | 0.29207907 | 0.3221889 | -0.0301098 | 0.66836266 | 0.89054584 |
| Jmy | 0.23153195 | 0.21181695 | 0.24195615 | -0.0301392 | 0.7191572 | 0.89054584 |
| Fam50a | 0.61075777 | 0.55251773 | 0.58267209 | -0.0301544 | 0.5952646 | 0.89054584 |
| Psmd1 | 1.03048028 | 0.82003275 | 0.85018925 | -0.0301565 | 0.56562805 | 0.89054584 |
| Mroh1 | 0.13442456 | 0.11275435 | 0.14291728 | -0.0301629 | 0.83726449 | 0.8948515 |
| Yif1a | 0.94717004 | 0.7763627 | 0.80653709 | -0.0301744 | 0.56918578 | 0.89054584 |
| Cenpc1 | 0.31249438 | 0.29113269 | 0.32131879 | -0.0301861 | 0.66919137 | 0.89054584 |
| Wipf2 | 0.20022037 | 0.18026146 | 0.21045407 | -0.0301926 | 0.74805798 | 0.89054584 |
| Repin1 | 0.16814715 | 0.14755473 | 0.1777512 | -0.0301965 | 0.7856632 | 0.89054584 |
| Ydjc | 0.1683103 | 0.1476977 | 0.17791862 | -0.0302209 | 0.78563274 | 0.89054584 |
| Mov10 | 0.30378269 | 0.28269113 | 0.31293191 | -0.0302408 | 0.67372618 | 0.89054584 |
| Hibch | 0.24530781 | 0.22542069 | 0.25567828 | -0.0302576 | 0.70930329 | 0.89054584 |
| Timmdc1 | 0.34568244 | 0.32265334 | 0.35291673 | -0.0302634 | 0.65524293 | 0.89054584 |
| Rlf | 0.28754609 | 0.26691817 | 0.29719936 | -0.0302812 | 0.68249686 | 0.89054584 |

| | | | | | | |
|---|---|---|---|---|---|---|
| Scrn3 | 0.1725647 | 0.15195709 | 0.18228065 | -0.0303236 | 0.78084342 | 0.89054584 |
| Hbs1l | 0.49300698 | 0.45553019 | 0.48587361 | -0.0303434 | 0.61445781 | 0.89054584 |
| Ror2 | 0.16844799 | 0.14771095 | 0.17805991 | -0.030349 | 0.78642594 | 0.89054584 |
| Armcx4 | 0.2004911 | 0.18034249 | 0.21072826 | -0.0303858 | 0.74914186 | 0.89054584 |
| Rrn3 | 0.2251562 | 0.20518947 | 0.23557694 | -0.0303875 | 0.72606825 | 0.89054584 |
| Pdxdc1 | 0.45115636 | 0.41898804 | 0.44937972 | -0.0303917 | 0.62365165 | 0.89054584 |
| Ncoa5 | 0.28058066 | 0.26000554 | 0.29041063 | -0.0304051 | 0.68714595 | 0.89054584 |
| Rtca | 0.39478962 | 0.36818333 | 0.39860085 | -0.0304175 | 0.63890264 | 0.89054584 |
| Cdc42bpb | 0.24385166 | 0.2238072 | 0.25423196 | -0.0304248 | 0.71142428 | 0.89054584 |
| 2410004B18I | 0.6851032 | 0.60855619 | 0.63905628 | -0.0305001 | 0.58798172 | 0.89054584 |
| Dusp18 | 0.4191776 | 0.39026372 | 0.42079435 | -0.0305306 | 0.6323291 | 0.89054584 |
| Bcl7a | 0.34707938 | 0.32368813 | 0.35423422 | -0.0305461 | 0.6560648 | 0.89054584 |
| Pde1c | 0.3250032 | 0.30274855 | 0.33329511 | -0.0305466 | 0.66530775 | 0.89054584 |
| Cep95 | 0.15532518 | 0.13401042 | 0.16455932 | -0.0305489 | 0.80645005 | 0.89054584 |
| Tmem64 | 0.18258506 | 0.16197005 | 0.19252538 | -0.0305553 | 0.77017142 | 0.89054584 |
| Kansl2 | 0.6646515 | 0.59335521 | 0.62392379 | -0.0305686 | 0.59027816 | 0.89054584 |
| Trim41 | 0.21130531 | 0.19107669 | 0.22165567 | -0.030579 | 0.73980168 | 0.89054584 |
| Abhd13 | 0.27816329 | 0.25746579 | 0.28804929 | -0.0305835 | 0.68958024 | 0.89054584 |
| Rfx3 | 0.33858548 | 0.3156066 | 0.34620729 | -0.0306007 | 0.65975555 | 0.89054584 |
| Golga4 | 0.89760869 | 0.74804751 | 0.77865419 | -0.0306067 | 0.57265187 | 0.89054584 |
| Ice1 | 0.22118248 | 0.20097237 | 0.23159181 | -0.0306194 | 0.73104365 | 0.89054584 |
| Crebzf | 0.22831741 | 0.20808638 | 0.23874214 | -0.0306558 | 0.72515772 | 0.89054584 |
| Zfp949 | 0.20982986 | 0.18951105 | 0.22016769 | -0.0306566 | 0.74174077 | 0.89054584 |
| Map7 | 0.31630316 | 0.29428902 | 0.32497399 | -0.030685 | 0.66999899 | 0.89054584 |
| Zdhhc2 | 0.34519808 | 0.32176342 | 0.35245967 | -0.0306963 | 0.65753845 | 0.89054584 |
| Zfp518a | 0.14632965 | 0.12454714 | 0.15526558 | -0.0307184 | 0.82166944 | 0.89054584 |
| Pcsk7 | 0.11132548 | 0.08808011 | 0.11880483 | -0.0307247 | 0.88587373 | 0.91431604 |
| Vamp4 | 0.51051085 | 0.47009558 | 0.50082171 | -0.0307261 | 0.61249688 | 0.89054584 |
| Pigv | 0.18964091 | 0.16897884 | 0.19971452 | -0.0307357 | 0.76331363 | 0.89054584 |
| D930016D06 | 0.1739231 | 0.15292856 | 0.18367184 | -0.0307433 | 0.78225188 | 0.89054584 |

| | | | | | | |
|---|---|---|---|---|---|---|
| Mef2d | 0.21264234 | 0.19219593 | 0.22300322 | -0.0308073 | 0.74008905 | 0.89054584 |
| Srgap2 | 0.18090805 | 0.16000617 | 0.1908136 | -0.0308074 | 0.77403464 | 0.89054584 |
| Tmed8 | 0.31441973 | 0.2923421 | 0.32316744 | -0.0308253 | 0.67162086 | 0.89054584 |
| Ppp1r9b | 0.20051979 | 0.17991697 | 0.21075732 | -0.0308403 | 0.75230084 | 0.89054584 |
| Zcchc14 | 0.24985918 | 0.22935158 | 0.26019256 | -0.030841 | 0.70960482 | 0.89054584 |
| Cluh | 0.12636077 | 0.10367873 | 0.13452217 | -0.0308434 | 0.85728681 | 0.9009252 |
| Cars2 | 0.1585739 | 0.13705896 | 0.16790805 | -0.0308491 | 0.80399249 | 0.89054584 |
| Mbd5 | 0.15595194 | 0.13433317 | 0.1652057 | -0.0308725 | 0.80802231 | 0.89054584 |
| Hadhb | 1.03413144 | 0.82121223 | 0.85208937 | -0.0308771 | 0.56703321 | 0.89054584 |
| Nup214 | 0.19369551 | 0.17294939 | 0.20383615 | -0.0308868 | 0.75989263 | 0.89054584 |
| Ece2 | 0.29151517 | 0.2701575 | 0.3010574 | -0.0308999 | 0.68374039 | 0.89054584 |
| Ivns1abp | 0.68260582 | 0.60630605 | 0.63722381 | -0.0309178 | 0.58941849 | 0.89054584 |
| Pank2 | 0.24664335 | 0.22607067 | 0.25700394 | -0.0309333 | 0.71251773 | 0.89054584 |
| Numa1 | 0.42549432 | 0.39555351 | 0.42648804 | -0.0309345 | 0.63229116 | 0.89054584 |
| Gabbr1 | 0.191064 | 0.17017773 | 0.20116194 | -0.0309842 | 0.76349772 | 0.89054584 |
| Socs4 | 0.29827629 | 0.27662144 | 0.30761129 | -0.0309898 | 0.68059316 | 0.89054584 |
| Glt8d1 | 0.45748228 | 0.42396784 | 0.45496343 | -0.0309956 | 0.62453096 | 0.89054584 |
| Akap8l | 0.61831517 | 0.55757644 | 0.58857535 | -0.0309989 | 0.59691725 | 0.89054584 |
| Cpsf3 | 0.49422467 | 0.45591718 | 0.4869194 | -0.0310022 | 0.61662756 | 0.89054584 |
| Fam98b | 0.30398636 | 0.28211908 | 0.31312841 | -0.0310093 | 0.67773637 | 0.89054584 |
| Tsr1 | 0.22633774 | 0.20574542 | 0.23676051 | -0.0310151 | 0.72917199 | 0.89054584 |
| Mrps18b | 0.49598426 | 0.45740688 | 0.48842946 | -0.0310226 | 0.61635151 | 0.89054584 |
| Rhno1 | 0.40581468 | 0.37764252 | 0.40867565 | -0.0310331 | 0.63824835 | 0.89054584 |
| Tom1l1 | 0.25178503 | 0.23104827 | 0.26209979 | -0.0310515 | 0.70951032 | 0.89054584 |
| Grpel2 | 0.35969805 | 0.33500131 | 0.36608758 | -0.0310863 | 0.65380483 | 0.89054584 |
| Psme1 | 1.36707698 | 0.96347722 | 0.99456469 | -0.0310875 | 0.55789175 | 0.89054584 |
| C1d | 1.41806615 | 0.98031834 | 1.01140945 | -0.0310911 | 0.55694085 | 0.89054584 |
| Zfp938 | 0.27112542 | 0.2500472 | 0.28115827 | -0.0311111 | 0.69690758 | 0.89054584 |
| Helz | 0.56903122 | 0.51820452 | 0.54939432 | -0.0311898 | 0.60430032 | 0.89054584 |
| Zfp787 | 0.54085173 | 0.49508532 | 0.52627751 | -0.0311922 | 0.60877704 | 0.89054584 |

| | | | | | | |
|---|---|---|---|---|---|---|
| Serhl | 0.25563079 | 0.2347055 | 0.26590313 | -0.0311976 | 0.70768007 | 0.89054584 |
| Psmg2 | 0.53901526 | 0.49355522 | 0.52475338 | -0.0311982 | 0.60910537 | 0.89054584 |
| Arhgef10 | 0.14806974 | 0.12583731 | 0.1570658 | -0.0312285 | 0.82285767 | 0.89075401 |
| Extl2 | 0.17416443 | 0.15268225 | 0.18391892 | -0.0312367 | 0.78560722 | 0.89054584 |
| Metap2 | 2.00557349 | 1.08691363 | 1.11815361 | -0.03124 | 0.55178234 | 0.89054584 |
| Mrps7 | 0.8393628 | 0.71239133 | 0.74364079 | -0.0312495 | 0.5776085 | 0.89054584 |
| Camk2g | 0.25815013 | 0.23713804 | 0.26839102 | -0.031253 | 0.70626769 | 0.89054584 |
| Otub1 | 0.69788525 | 0.61710952 | 0.64836632 | -0.0312568 | 0.58885508 | 0.89054584 |
| Smyd3 | 0.23841718 | 0.21754095 | 0.24882546 | -0.0312845 | 0.72098251 | 0.89054584 |
| Cd3eap | 0.30235761 | 0.28027102 | 0.31155639 | -0.0312854 | 0.68004836 | 0.89054584 |
| Card19 | 0.64234991 | 0.57580324 | 0.60709339 | -0.0312902 | 0.59488376 | 0.89054584 |
| Coro1b | 0.51346055 | 0.47202882 | 0.50332175 | -0.0312929 | 0.61396162 | 0.89054584 |
| Hax1 | 0.86318482 | 0.7269615 | 0.75825845 | -0.0312969 | 0.57624512 | 0.89054584 |
| Tceb1 | 1.87720505 | 1.0760004 | 1.10730012 | -0.0312997 | 0.55238639 | 0.89054584 |
| Pdik1l | 0.23411908 | 0.21323306 | 0.24453984 | -0.0313068 | 0.72456358 | 0.89054584 |
| Sbk1 | 0.31748028 | 0.29479261 | 0.32610214 | -0.0313095 | 0.67267492 | 0.89054584 |
| Ccdc117 | 0.31411747 | 0.2915525 | 0.32287735 | -0.0313248 | 0.67436555 | 0.89054584 |
| Zfp516 | 0.59868901 | 0.54183843 | 0.57316497 | -0.0313265 | 0.6004979 | 0.89054584 |
| Ppp2r5a | 0.31196763 | 0.28945237 | 0.32081269 | -0.0313603 | 0.67559796 | 0.89054584 |
| Uba6 | 0.16577202 | 0.14394913 | 0.17531259 | -0.0313635 | 0.79771451 | 0.89054584 |
| Zmiz2 | 0.23802195 | 0.21702891 | 0.24843173 | -0.0314028 | 0.72203913 | 0.89054584 |
| Med12 | 0.13157114 | 0.10852752 | 0.13994946 | -0.0314219 | 0.85222909 | 0.89988783 |
| Arl6ip4 | 0.55345432 | 0.50521759 | 0.53667928 | -0.0314617 | 0.60761975 | 0.89054584 |
| Tmx4 | 0.36355867 | 0.33823085 | 0.36969707 | -0.0314662 | 0.65414623 | 0.89054584 |
| Msh2 | 0.42618823 | 0.39563095 | 0.42711208 | -0.0314811 | 0.63434988 | 0.89054584 |
| Ube2e3 | 1.70976172 | 1.05173849 | 1.08324527 | -0.0315068 | 0.55389482 | 0.89054584 |
| Foxn3 | 0.30973552 | 0.28711516 | 0.31866659 | -0.0315514 | 0.67770308 | 0.89054584 |
| Mboat2 | 0.30341824 | 0.28101209 | 0.31258024 | -0.0315682 | 0.68100697 | 0.89054584 |
| Mut | 0.30810181 | 0.28552175 | 0.31709455 | -0.0315728 | 0.67863542 | 0.89054584 |
| Rassf3 | 0.23180863 | 0.21061342 | 0.24223262 | -0.0316192 | 0.72845358 | 0.89054584 |

| | | | | | | |
|---|---|---|---|---|---|---|
| Akt1 | 0.49850876 | 0.45892205 | 0.49059255 | -0.0316705 | 0.61820155 | 0.89054584 |
| 4930453N24 | 0.4759893 | 0.43948477 | 0.47116181 | -0.031677 | 0.62294656 | 0.89054584 |
| Ctdspl2 | 0.53894383 | 0.49295416 | 0.52469405 | -0.0317399 | 0.6109629 | 0.89054584 |
| Iars | 0.23249699 | 0.21115675 | 0.24292027 | -0.0317635 | 0.72880042 | 0.89054584 |
| 1600012H06 | 0.47592079 | 0.43932909 | 0.47110222 | -0.0317731 | 0.62332256 | 0.89054584 |
| Ubxn6 | 0.81117522 | 0.69402522 | 0.72581026 | -0.031785 | 0.58083365 | 0.89054584 |
| Brip1os | 0.29692347 | 0.27451296 | 0.30630176 | -0.0317888 | 0.68563571 | 0.89054584 |
| Ptpra | 0.27680778 | 0.25493041 | 0.28672394 | -0.0317935 | 0.69728547 | 0.89054584 |
| Sft2d2 | 0.16218289 | 0.13982835 | 0.17162316 | -0.0317948 | 0.80596225 | 0.89054584 |
| Nek1 | 0.23956078 | 0.21816845 | 0.24996431 | -0.0317959 | 0.72328385 | 0.89054584 |
| Ak6 | 0.41790823 | 0.38785121 | 0.41964737 | -0.0317962 | 0.63795661 | 0.89054584 |
| Nelfcd | 0.53947878 | 0.49333315 | 0.52513827 | -0.0318051 | 0.61109338 | 0.89054584 |
| Cep78 | 0.13047223 | 0.10698815 | 0.13880566 | -0.0318175 | 0.85722584 | 0.9009252 |
| Rcc2 | 0.37049283 | 0.34429346 | 0.37616095 | -0.0318675 | 0.65346591 | 0.89054584 |
| Rhbdd2 | 0.31153006 | 0.28851797 | 0.32039217 | -0.0318742 | 0.67849502 | 0.89054584 |
| Epc2 | 0.22465518 | 0.20318552 | 0.23507488 | -0.0318894 | 0.73631113 | 0.89054584 |
| Chm | 0.18299055 | 0.16102464 | 0.1929391 | -0.0319145 | 0.77954137 | 0.89054584 |
| Nvl | 0.38849386 | 0.36089151 | 0.39281679 | -0.0319253 | 0.64752418 | 0.89054584 |
| Cln3 | 0.20510212 | 0.18346767 | 0.21539339 | -0.0319257 | 0.75507602 | 0.89054584 |
| Tsc1 | 0.30515246 | 0.28232691 | 0.31425309 | -0.0319262 | 0.68200745 | 0.89054584 |
| Josd2 | 0.8963802 | 0.74600912 | 0.7779409 | -0.0319318 | 0.57582862 | 0.89054584 |
| Cxxc5 | 0.94565966 | 0.77377775 | 0.80571096 | -0.0319332 | 0.57324841 | 0.89054584 |
| Glrx2 | 0.80880445 | 0.6923465 | 0.72428437 | -0.0319379 | 0.58138565 | 0.89054584 |
| Fastk | 0.47947901 | 0.44223885 | 0.47419327 | -0.0319544 | 0.62322111 | 0.89054584 |
| Ireb2 | 0.44188967 | 0.40920239 | 0.44115693 | -0.0319545 | 0.6321162 | 0.89054584 |
| Acad8 | 0.3265774 | 0.30283653 | 0.33479654 | -0.03196 | 0.67174879 | 0.89054584 |
| Cops8 | 0.95841028 | 0.78066759 | 0.81263265 | -0.0319651 | 0.57270312 | 0.89054584 |
| Pogk | 0.13771964 | 0.11437477 | 0.14634055 | -0.0319658 | 0.84558308 | 0.89825055 |
| Dhps | 0.4830543 | 0.44528221 | 0.47729128 | -0.0320091 | 0.62264969 | 0.89054584 |
| Cttn | 0.6723802 | 0.59763141 | 0.62967644 | -0.032045 | 0.59371017 | 0.89054584 |

| | | | | | | |
|---|---|---|---|---|---|---|
| 9430038I01R | 0.25297075 | 0.23121552 | 0.26327318 | -0.0320577 | 0.71474464 | 0.89054584 |
| Terf1 | 0.33472749 | 0.31048537 | 0.34254912 | -0.0320637 | 0.66861522 | 0.89054584 |
| Rad17 | 0.43420863 | 0.40222355 | 0.43430445 | -0.0320809 | 0.6346308 | 0.89054584 |
| Trafd1 | 0.46835193 | 0.43242003 | 0.46450125 | -0.0320812 | 0.6261881 | 0.89054584 |
| Dclre1a | 0.15163169 | 0.12861605 | 0.16074708 | -0.032131 | 0.82413928 | 0.89120023 |
| Ankfy1 | 0.21951313 | 0.19776301 | 0.22991554 | -0.0321525 | 0.74264766 | 0.89054584 |
| Sirt2 | 0.56731791 | 0.51583834 | 0.54800361 | -0.0321653 | 0.60774951 | 0.89054584 |
| 2310057M21 | 0.27107816 | 0.24892053 | 0.28111191 | -0.0321914 | 0.70316185 | 0.89054584 |
| Siah1a | 0.19603914 | 0.17399576 | 0.20621532 | -0.0322196 | 0.76666986 | 0.89054584 |
| Prkcz | 0.24465636 | 0.22279597 | 0.25503135 | -0.0322354 | 0.72202893 | 0.89054584 |
| Rit1 | 0.3805115 | 0.3532169 | 0.38545325 | -0.0322363 | 0.6515979 | 0.89054584 |
| Cpd | 0.21282768 | 0.19094622 | 0.22318995 | -0.0322437 | 0.74954744 | 0.89054584 |
| Polr2a | 0.41585803 | 0.38551798 | 0.41779288 | -0.0322749 | 0.64054189 | 0.89054584 |
| Fam102a | 0.21194673 | 0.1900219 | 0.22230223 | -0.0322803 | 0.75064507 | 0.89054584 |
| Atn1.1 | 0.20732664 | 0.18535011 | 0.21764104 | -0.0322909 | 0.75529165 | 0.89054584 |
| Fbxw7 | 0.18526389 | 0.16296274 | 0.1952573 | -0.0322946 | 0.77951759 | 0.89054584 |
| Cep41 | 0.14604932 | 0.12268019 | 0.15497545 | -0.0322953 | 0.83419462 | 0.89427341 |
| Ccdc86 | 0.53696199 | 0.49072556 | 0.52304674 | -0.0323212 | 0.61328638 | 0.89054584 |
| R3hdm2 | 0.30568571 | 0.28242871 | 0.31476716 | -0.0323384 | 0.68390575 | 0.89054584 |
| Zdhhc3 | 0.4825349 | 0.44449195 | 0.4768417 | -0.0323497 | 0.62402548 | 0.89054584 |
| Mtmr7 | 0.35816569 | 0.33225627 | 0.36465276 | -0.0323965 | 0.66052991 | 0.89054584 |
| Tob2 | 0.36067625 | 0.33459221 | 0.36700285 | -0.0324106 | 0.65962462 | 0.89054584 |
| Snx30 | 0.20455738 | 0.18239992 | 0.21484271 | -0.0324428 | 0.75913835 | 0.89054584 |
| Abca3 | 0.22058482 | 0.19854724 | 0.23099182 | -0.0324446 | 0.74356783 | 0.89054584 |
| E4f1 | 0.15944381 | 0.13634501 | 0.16880402 | -0.032459 | 0.81483601 | 0.89054584 |
| Ufsp1 | 0.12171193 | 0.09717763 | 0.1296715 | -0.0324939 | 0.8784888 | 0.91058344 |
| Mtmr3 | 0.25636343 | 0.23412354 | 0.26662691 | -0.0325034 | 0.71496622 | 0.89054584 |
| Ubxn4 | 1.58832629 | 1.02555956 | 1.05807103 | -0.0325115 | 0.55691645 | 0.89054584 |
| Tmem50b | 0.17012277 | 0.14726112 | 0.17977786 | -0.0325167 | 0.80029372 | 0.89054584 |
| Pex2 | 0.57957461 | 0.52538308 | 0.55791131 | -0.0325282 | 0.6070479 | 0.89054584 |

| | | | | | | |
|---|---|---|---|---|---|---|
| Pdhx | 0.19214138 | 0.1697256 | 0.20225716 | -0.0325316 | 0.77317873 | 0.89054584 |
| Fam135a | 0.59542934 | 0.53804679 | 0.57058102 | -0.0325342 | 0.60474578 | 0.89054584 |
| Armc5 | 0.17507815 | 0.15230421 | 0.18485417 | -0.03255 | 0.79399673 | 0.89054584 |
| Phospho2 | 0.27553865 | 0.25292765 | 0.28548224 | -0.0325546 | 0.70238628 | 0.89054584 |
| Dnajc19 | 1.02485532 | 0.8146865 | 0.84724412 | -0.0325576 | 0.57104381 | 0.89054584 |
| Snx16 | 0.23587203 | 0.21371137 | 0.24628873 | -0.0325774 | 0.73113595 | 0.89054584 |
| Rev1 | 0.17512339 | 0.15231894 | 0.18490047 | -0.0325815 | 0.79416478 | 0.89054584 |
| Fem1c | 0.20811782 | 0.18584748 | 0.21843993 | -0.0325924 | 0.75651616 | 0.89054584 |
| Stxbp2 | 0.38059633 | 0.35292632 | 0.38553169 | -0.0326054 | 0.65321617 | 0.89054584 |
| Atp6v0a1 | 0.23139393 | 0.20918288 | 0.24181823 | -0.0326354 | 0.73524765 | 0.89054584 |
| Map4k5 | 0.26319425 | 0.24071616 | 0.27336385 | -0.0326477 | 0.71103634 | 0.89054584 |
| Tmem256 | 1.99458587 | 1.08474786 | 1.11741325 | -0.0326654 | 0.55416645 | 0.89054584 |
| Fbxl18 | 0.15051024 | 0.12692307 | 0.15958861 | -0.0326655 | 0.82986188 | 0.89271879 |
| Rprd2 | 0.39606555 | 0.36709292 | 0.39977035 | -0.0326774 | 0.64833313 | 0.89054584 |
| Drosha | 0.24926921 | 0.22690334 | 0.25960794 | -0.0327046 | 0.72137091 | 0.89054584 |
| Fdxr | 0.12794638 | 0.10344547 | 0.13617488 | -0.0327294 | 0.86859385 | 0.90498553 |
| Cenpb | 0.21617045 | 0.19381642 | 0.22655527 | -0.0327389 | 0.74960597 | 0.89054584 |
| Uhrf1bp1l | 0.22518911 | 0.20286493 | 0.23560991 | -0.032745 | 0.74134515 | 0.89054584 |
| Qser1 | 0.25176017 | 0.22932805 | 0.26207518 | -0.0327471 | 0.71976706 | 0.89054584 |
| Plxnb1 | 0.13717388 | 0.11300379 | 0.14577384 | -0.0327701 | 0.85252702 | 0.89988783 |
| Ube4a | 0.31804147 | 0.29381423 | 0.32663973 | -0.0328255 | 0.68017849 | 0.89054584 |
| Unc119 | 0.17540809 | 0.15236559 | 0.18519181 | -0.0328262 | 0.79554739 | 0.89054584 |
| Osbp | 0.17600543 | 0.15297106 | 0.18580295 | -0.0328319 | 0.79481645 | 0.89054584 |
| Nae1 | 0.78859983 | 0.6782475 | 0.71111365 | -0.0328661 | 0.58524352 | 0.89054584 |
| BC023719 | 0.30292505 | 0.27923015 | 0.31210421 | -0.0328741 | 0.68819062 | 0.89054584 |
| Fam131a | 0.14239961 | 0.11831848 | 0.15119534 | -0.0328769 | 0.84448645 | 0.89775379 |
| Cnpy3 | 0.52521795 | 0.48034027 | 0.51323308 | -0.0328928 | 0.61737311 | 0.89054584 |
| Itfg2 | 0.38226335 | 0.3541773 | 0.38707235 | -0.0328951 | 0.6539241 | 0.89054584 |
| Scnm1 | 0.34767011 | 0.32189354 | 0.35479103 | -0.0328975 | 0.66712702 | 0.89054584 |
| Gabpb1 | 0.31573814 | 0.29153361 | 0.32443223 | -0.0328986 | 0.68169139 | 0.89054584 |

| | | | | | | |
|---|---|---|---|---|---|---|
| Prkca | 0.27451972 | 0.25155642 | 0.28448475 | -0.0329283 | 0.70514619 | 0.89054584 |
| Sh2b1 | 0.23941988 | 0.21687117 | 0.24982402 | -0.0329529 | 0.73056882 | 0.89054584 |
| Ldoc1l | 0.13616846 | 0.1117567 | 0.14472955 | -0.0329728 | 0.85574998 | 0.9009252 |
| Gphn | 0.24146939 | 0.21884406 | 0.25186364 | -0.0330196 | 0.72933132 | 0.89054584 |
| Gba | 0.17896679 | 0.15577538 | 0.18883062 | -0.0330552 | 0.79262368 | 0.89054584 |
| Adprm | 0.26582231 | 0.24288866 | 0.27594995 | -0.0330613 | 0.71164029 | 0.89054584 |
| Lrp5 | 0.19183937 | 0.16886508 | 0.20195021 | -0.0330851 | 0.77734754 | 0.89054584 |
| Phc2 | 0.23847692 | 0.21573257 | 0.24888496 | -0.0331524 | 0.73256462 | 0.89054584 |
| Mecp2 | 0.42483148 | 0.39272271 | 0.42589167 | -0.033169 | 0.64163927 | 0.89054584 |
| Ccdc25 | 0.5377958 | 0.49052885 | 0.52374012 | -0.0332113 | 0.61616788 | 0.89054584 |
| 2010111I01R | 0.22225806 | 0.19945918 | 0.23267118 | -0.033212 | 0.74698352 | 0.89054584 |
| Cenpv | 0.18579889 | 0.16258928 | 0.19580253 | -0.0332133 | 0.78531754 | 0.89054584 |
| Wdr5 | 0.26540888 | 0.24231587 | 0.27554334 | -0.0332275 | 0.71287952 | 0.89054584 |
| L3mbtl2 | 0.13499058 | 0.11025792 | 0.14350561 | -0.0332477 | 0.8597938 | 0.90132889 |
| Ralgapb | 0.13573346 | 0.11099507 | 0.1442776 | -0.0332825 | 0.85875515 | 0.90127639 |
| Aco1 | 0.30457141 | 0.28036835 | 0.31369277 | -0.0333244 | 0.68967582 | 0.89054584 |
| Ankrd13a | 0.18349164 | 0.16009009 | 0.19345026 | -0.0333602 | 0.78913587 | 0.89054584 |
| 1810022K09I | 1.60769115 | 1.02915623 | 1.06252891 | -0.0333727 | 0.55817012 | 0.89054584 |
| Elf1 | 0.47288951 | 0.43508874 | 0.46846278 | -0.033374 | 0.63002159 | 0.89054584 |
| 2810402E24I | 0.20492534 | 0.18182759 | 0.21521468 | -0.0333871 | 0.76508621 | 0.89054584 |
| Bola2 | 1.06194467 | 0.83292428 | 0.86633735 | -0.0334131 | 0.57130084 | 0.89054584 |
| Idnk | 0.4575792 | 0.42162301 | 0.45504879 | -0.0334258 | 0.63390892 | 0.89054584 |
| Tle1 | 0.32555471 | 0.30039088 | 0.33382128 | -0.0334304 | 0.67959546 | 0.89054584 |
| Gramd1a | 0.58748601 | 0.53076751 | 0.56425444 | -0.0334869 | 0.60891612 | 0.89054584 |
| Diaph1 | 0.33612661 | 0.31038788 | 0.34387647 | -0.0334886 | 0.67498249 | 0.89054584 |
| Nedd4 | 2.39366378 | 1.0957991 | 1.12929047 | -0.0334914 | 0.55494706 | 0.89054584 |
| Pfkm | 0.16371645 | 0.13970812 | 0.17320022 | -0.0334921 | 0.81618181 | 0.89054584 |
| Trp53rka | 0.29407234 | 0.26998906 | 0.30353888 | -0.0335498 | 0.69670521 | 0.89054584 |
| Cstb | 1.71465803 | 1.0505762 | 1.08413026 | -0.0335541 | 0.55732683 | 0.89054584 |
| Gm6133 | 0.98398541 | 0.79264256 | 0.82620401 | -0.0335614 | 0.57505235 | 0.89054584 |

| | | | | | | |
|---|---|---|---|---|---|---|
| Slc35b4 | 0.18098793 | 0.15727795 | 0.19089516 | -0.0336172 | 0.79402122 | 0.89054584 |
| Psmc2 | 1.71609791 | 1.05077033 | 1.08438842 | -0.0336181 | 0.55742189 | 0.89054584 |
| Tomm40l | 0.16046146 | 0.13623312 | 0.16985177 | -0.0336186 | 0.82177251 | 0.89054584 |
| Trmt10a | 0.25039685 | 0.22707338 | 0.26072521 | -0.0336518 | 0.72618988 | 0.89054584 |
| Ppp1r13b | 0.19192947 | 0.16837145 | 0.20204178 | -0.0336703 | 0.78124779 | 0.89054584 |
| Nek7 | 0.42574572 | 0.39303454 | 0.42671416 | -0.0336796 | 0.64345903 | 0.89054584 |
| Ctc1 | 0.11651202 | 0.09055304 | 0.1242362 | -0.0336832 | 0.89673082 | 0.92138933 |
| Lsm4 | 1.42723803 | 0.98059718 | 1.01428594 | -0.0336888 | 0.56148815 | 0.89054584 |
| Tbl1xr1 | 0.24592737 | 0.22258511 | 0.25629336 | -0.0337083 | 0.72998047 | 0.89054584 |
| Strap | 1.47389069 | 0.99454213 | 1.02826306 | -0.0337209 | 0.56071633 | 0.89054584 |
| Rfx7 | 0.20150772 | 0.17799852 | 0.2117576 | -0.0337591 | 0.77118148 | 0.89054584 |
| Mapkapk2 | 0.19954092 | 0.1759744 | 0.20976579 | -0.0337914 | 0.77352727 | 0.89054584 |
| Zfp513 | 0.24102036 | 0.21762288 | 0.25141694 | -0.0337941 | 0.73441757 | 0.89054584 |
| Nkap | 0.41680133 | 0.38483224 | 0.41864643 | -0.0338142 | 0.64665454 | 0.89054584 |
| Fntb | 0.14692481 | 0.12202095 | 0.15588143 | -0.0338605 | 0.84423403 | 0.89775379 |
| Gpr107 | 0.41328559 | 0.38156813 | 0.41546304 | -0.0338949 | 0.64806162 | 0.89054584 |
| Pgap1 | 0.28048462 | 0.25641039 | 0.29031688 | -0.0339065 | 0.70682188 | 0.89054584 |
| Babam1 | 0.65158922 | 0.58019889 | 0.61410696 | -0.0339081 | 0.60150502 | 0.89054584 |
| Rbsn | 0.20972118 | 0.18613788 | 0.22005805 | -0.0339202 | 0.76366395 | 0.89054584 |
| Slmo2 | 0.77818114 | 0.67028538 | 0.7042067 | -0.0339213 | 0.58878439 | 0.89054584 |
| Trabd | 0.67986755 | 0.60128724 | 0.63520955 | -0.0339223 | 0.5982434 | 0.89054584 |
| Ddx52 | 0.33624436 | 0.31005414 | 0.34398814 | -0.033934 | 0.67709788 | 0.89054584 |
| Anapc7 | 0.38878901 | 0.35911665 | 0.39308842 | -0.0339718 | 0.65639288 | 0.89054584 |
| Pard3 | 0.57342446 | 0.51897067 | 0.55295152 | -0.0339808 | 0.61267911 | 0.89054584 |
| Hook1 | 0.38285496 | 0.35361037 | 0.38761874 | -0.0340084 | 0.65864206 | 0.89054584 |
| Uhmk1 | 0.18298863 | 0.15891402 | 0.19293714 | -0.0340231 | 0.79433965 | 0.89054584 |
| Tnrc18 | 0.46802353 | 0.43018039 | 0.46421406 | -0.0340337 | 0.63366213 | 0.89054584 |
| Rab3ip | 0.47848904 | 0.43929924 | 0.47333405 | -0.0340348 | 0.63118688 | 0.89054584 |
| Zbtb11 | 0.24961668 | 0.22591593 | 0.25995228 | -0.0340364 | 0.72907226 | 0.89054584 |
| Tmem55b | 0.64207226 | 0.57284346 | 0.60688172 | -0.0340383 | 0.60307199 | 0.89054584 |

| | | | | | | |
|---|---|---|---|---|---|---|
| Rreb1 | 0.22788459 | 0.20425702 | 0.23830904 | -0.034052 | 0.74720398 | 0.89054584 |
| Alkbh8 | 0.15653055 | 0.13172524 | 0.16580229 | -0.034077 | 0.83085359 | 0.89280428 |
| Nemf | 0.62924789 | 0.56294726 | 0.59704666 | -0.0340994 | 0.60491439 | 0.89054584 |
| Atp5f1 | 2.80263642 | 1.07618238 | 1.11028404 | -0.0341017 | 0.55689296 | 0.89054584 |
| Tmem42 | 0.26623174 | 0.2422407 | 0.27635254 | -0.0341118 | 0.71738174 | 0.89054584 |
| Mrpl13 | 0.75566825 | 0.65489173 | 0.68901448 | -0.0341227 | 0.59123747 | 0.89054584 |
| Slc35a2 | 0.37143945 | 0.34286372 | 0.37704132 | -0.0341776 | 0.66360361 | 0.89054584 |
| Higd2a | 1.38314705 | 0.96578218 | 1.00000111 | -0.0342189 | 0.56333249 | 0.89054584 |
| Taf3 | 0.43244958 | 0.3984991 | 0.43273025 | -0.0342311 | 0.64376682 | 0.89054584 |
| Ints4 | 0.1649676 | 0.14025219 | 0.17448615 | -0.034234 | 0.81966214 | 0.89054584 |
| Smyd2 | 0.42249572 | 0.38955191 | 0.4237881 | -0.0342362 | 0.64668192 | 0.89054584 |
| Pkdcc | 0.33245184 | 0.30614243 | 0.34038798 | -0.0342456 | 0.68036656 | 0.89054584 |
| Zfp64 | 0.28411072 | 0.25960469 | 0.29385378 | -0.0342491 | 0.70643734 | 0.89054584 |
| Spata7 | 0.1777556 | 0.15333356 | 0.18759273 | -0.0342592 | 0.80257098 | 0.89054584 |
| Lats1 | 0.2677447 | 0.24355306 | 0.27783954 | -0.0342865 | 0.71733121 | 0.89054584 |
| Zfp598 | 0.14903164 | 0.12376738 | 0.15806042 | -0.034293 | 0.84394659 | 0.89775379 |
| Fuom | 0.35136164 | 0.32395845 | 0.35826641 | -0.034308 | 0.67225373 | 0.89054584 |
| Flot2 | 0.29987901 | 0.27482207 | 0.30916152 | -0.0343394 | 0.69759007 | 0.89054584 |
| Vps37c | 0.17198836 | 0.1473347 | 0.18169018 | -0.0343555 | 0.81082782 | 0.89054584 |
| Gmds | 0.40328137 | 0.37199837 | 0.40636683 | -0.0343685 | 0.65322048 | 0.89054584 |
| Fiz1 | 0.38770969 | 0.35772273 | 0.392095 | -0.0343723 | 0.65851632 | 0.89054584 |
| Dnah9 | 0.194798 | 0.17057364 | 0.20495566 | -0.034382 | 0.7827615 | 0.89054584 |
| 1700037H04 | 0.32770939 | 0.30148469 | 0.33587541 | -0.0343907 | 0.68333143 | 0.89054584 |
| Vps37d | 0.14198094 | 0.11636337 | 0.15076138 | -0.034398 | 0.85610746 | 0.9009252 |
| Pkd2 | 0.34189024 | 0.31493149 | 0.34933494 | -0.0344035 | 0.67682014 | 0.89054584 |
| Spsb2 | 0.1950791 | 0.17079168 | 0.20524102 | -0.0344493 | 0.78289125 | 0.89054584 |
| Nktr | 1.68973908 | 1.04507885 | 1.07953181 | -0.034453 | 0.5591004 | 0.89054584 |
| Lrba | 0.22701158 | 0.20293745 | 0.23743522 | -0.0344978 | 0.75076952 | 0.89054584 |
| Tcta | 0.40037767 | 0.36915081 | 0.40371593 | -0.0345651 | 0.65501162 | 0.89054584 |
| Tcp1 | 1.95170323 | 1.07965853 | 1.11423893 | -0.0345804 | 0.55748281 | 0.89054584 |

| | | | | | | |
|---|---|---|---|---|---|---|
| Polr3h | 0.24921091 | 0.22496319 | 0.25955016 | -0.034587 | 0.73264696 | 0.89054584 |
| Stx4a | 0.53089009 | 0.48339342 | 0.51798396 | -0.0345905 | 0.62214538 | 0.89054584 |
| Pcid2 | 0.16748713 | 0.14247934 | 0.17707377 | -0.0345944 | 0.81864279 | 0.89054584 |
| Rps9 | 5.01237819 | 0.66376405 | 0.69836711 | -0.0346031 | 0.59128188 | 0.89054584 |
| Ubqln1 | 0.70596736 | 0.61958418 | 0.65419341 | -0.0346092 | 0.59734275 | 0.89054584 |
| Ube2q1 | 0.45730755 | 0.42020014 | 0.45480952 | -0.0346094 | 0.63852869 | 0.89054584 |
| Mrto4 | 0.57610086 | 0.52048372 | 0.55511299 | -0.0346293 | 0.61433507 | 0.89054584 |
| Lrrc24 | 0.16256493 | 0.13736149 | 0.17201613 | -0.0346546 | 0.82603129 | 0.89170439 |
| Stim2 | 0.27991273 | 0.25508055 | 0.28975847 | -0.0346779 | 0.71143305 | 0.89054584 |
| Fam172a | 0.33679972 | 0.30983225 | 0.34451483 | -0.0346826 | 0.68047207 | 0.89054584 |
| Bbs5 | 0.16858942 | 0.14352119 | 0.17820503 | -0.0346838 | 0.81773434 | 0.89054584 |
| Gtf2h1 | 0.31234651 | 0.28648535 | 0.32117673 | -0.0346914 | 0.692598 | 0.89054584 |
| Zfp266 | 0.38613438 | 0.35593444 | 0.39064395 | -0.0347095 | 0.66054699 | 0.89054584 |
| Cnot10 | 0.23214897 | 0.2078441 | 0.24257263 | -0.0347285 | 0.74761687 | 0.89054584 |
| Fyco1 | 0.18845273 | 0.16376444 | 0.19850537 | -0.0347409 | 0.79257066 | 0.89054584 |
| Tesk1 | 0.15989286 | 0.13451597 | 0.1692664 | -0.0347504 | 0.83058553 | 0.89271879 |
| Sp1 | 0.2214599 | 0.19711928 | 0.23187025 | -0.034751 | 0.7574887 | 0.89054584 |
| Mrgbp | 0.23119873 | 0.20686394 | 0.24162316 | -0.0347592 | 0.74864184 | 0.89054584 |
| Gtf3c5 | 0.17462778 | 0.14963053 | 0.18439323 | -0.0347627 | 0.81011661 | 0.89054584 |
| Gnpda2 | 0.48381848 | 0.4431879 | 0.47795242 | -0.0347645 | 0.63264784 | 0.89054584 |
| Zfp120 | 0.23884435 | 0.21448546 | 0.24925092 | -0.0347655 | 0.74209836 | 0.89054584 |
| Ipmk | 0.17435384 | 0.14934099 | 0.18411283 | -0.0347718 | 0.81054193 | 0.89054584 |
| Ept1 | 0.25468449 | 0.2301882 | 0.26496792 | -0.0347797 | 0.72957671 | 0.89054584 |
| Ppt1 | 0.50389216 | 0.46039843 | 0.49519195 | -0.0347935 | 0.62829832 | 0.89054584 |
| Map2k3 | 0.33937549 | 0.31215683 | 0.34695549 | -0.0347987 | 0.67984973 | 0.89054584 |
| Snrnp25 | 0.27547815 | 0.25060413 | 0.28542302 | -0.0348189 | 0.71510014 | 0.89054584 |
| Dxo | 0.25656012 | 0.23200008 | 0.26682117 | -0.0348211 | 0.72840889 | 0.89054584 |
| Kmt2b | 0.17009672 | 0.14490154 | 0.17975115 | -0.0348496 | 0.81680643 | 0.89054584 |
| Paf1 | 0.30698254 | 0.28116443 | 0.31601675 | -0.0348523 | 0.69631016 | 0.89054584 |
| Fam134a | 0.66785851 | 0.59144402 | 0.62631591 | -0.0348719 | 0.60233653 | 0.89054584 |

| | | | | | | |
|---|---|---|---|---|---|---|
| Abcf2 | 0.47292482 | 0.4336145 | 0.46849356 | -0.0348791 | 0.63564999 | 0.89054584 |
| Sec24b | 0.24024103 | 0.21574313 | 0.25064144 | -0.0348983 | 0.7417314 | 0.89054584 |
| Kif2a | 0.22163302 | 0.19713482 | 0.232044 | -0.0349092 | 0.75831753 | 0.89054584 |
| Lonp2 | 0.41532947 | 0.38240233 | 0.41731437 | -0.034912 | 0.65164363 | 0.89054584 |
| Fyttd1 | 0.96182251 | 0.77950683 | 0.81446481 | -0.034958 | 0.57924732 | 0.89054584 |
| Zmym4 | 0.38883119 | 0.35809819 | 0.39312723 | -0.035029 | 0.66097602 | 0.89054584 |
| Cdk9 | 0.31103015 | 0.28487687 | 0.31991162 | -0.0350347 | 0.69505081 | 0.89054584 |
| Tubgcp4 | 0.3153455 | 0.28901193 | 0.32405565 | -0.0350437 | 0.69280704 | 0.89054584 |
| Ptp4a2 | 0.86899645 | 0.72669898 | 0.76176224 | -0.0350633 | 0.58490078 | 0.89054584 |
| Dnajc14 | 0.5231884 | 0.47644384 | 0.51152815 | -0.0350843 | 0.62534297 | 0.89054584 |
| Pdcd10 | 1.07284947 | 0.83672971 | 0.87182549 | -0.0350958 | 0.57438453 | 0.89054584 |
| Mrpl39 | 0.47374698 | 0.43409534 | 0.46920998 | -0.0351146 | 0.63632956 | 0.89054584 |
| Cisd1 | 1.27079005 | 0.92431068 | 0.9594279 | -0.0351172 | 0.56770245 | 0.89054584 |
| Zmym2 | 0.33953609 | 0.31197829 | 0.34710755 | -0.0351293 | 0.68136219 | 0.89054584 |
| A930015D03 | 0.15729817 | 0.13144455 | 0.16659355 | -0.035149 | 0.83718991 | 0.89483776 |
| Arhgap32 | 0.14273525 | 0.11639351 | 0.1515432 | -0.0351497 | 0.86006651 | 0.90134771 |
| Pank3 | 0.24344733 | 0.21865433 | 0.25383018 | -0.0351759 | 0.74074286 | 0.89054584 |
| Siah2 | 0.15282985 | 0.12680638 | 0.16198429 | -0.0351779 | 0.84417742 | 0.89775379 |
| Eif3a | 1.92242023 | 1.07660067 | 1.11178195 | -0.0351813 | 0.5586028 | 0.89054584 |
| Pomgnt1 | 0.19738812 | 0.17239243 | 0.20758368 | -0.0351913 | 0.78518389 | 0.89054584 |
| Anp32a | 2.17380461 | 1.09128269 | 1.12650311 | -0.0352204 | 0.5579063 | 0.89054584 |
| Nf1 | 0.24290957 | 0.21805646 | 0.2532957 | -0.0352392 | 0.74156105 | 0.89054584 |
| Casc3 | 0.36147842 | 0.33250388 | 0.36775302 | -0.0352491 | 0.67240349 | 0.89054584 |
| Ldah | 0.26135055 | 0.23628958 | 0.27154762 | -0.035258 | 0.72738459 | 0.89054584 |
| Tulp3 | 0.3600975 | 0.3311591 | 0.3664614 | -0.0353023 | 0.67321542 | 0.89054584 |
| Hs6st1 | 0.16613747 | 0.14036015 | 0.17568796 | -0.0353278 | 0.8255776 | 0.89170439 |
| Wdr12 | 0.34217377 | 0.31427465 | 0.349603 | -0.0353283 | 0.68110782 | 0.89054584 |
| Rab5c | 0.97275104 | 0.78495248 | 0.82028286 | -0.0353304 | 0.57951977 | 0.89054584 |
| 2410016O06 | 0.17206179 | 0.14642577 | 0.18176542 | -0.0353396 | 0.81748234 | 0.89054584 |
| Cbr4 | 0.22281384 | 0.1978706 | 0.23322872 | -0.0353581 | 0.76000182 | 0.89054584 |

| | | | | | | |
|---|---|---|---|---|---|---|
| Rps21 | 3.63987193 | 0.98202244 | 1.0173809 | -0.0353585 | 0.56431505 | 0.89054584 |
| B9d2 | 0.43187124 | 0.39685207 | 0.43221229 | -0.0353602 | 0.64845699 | 0.89054584 |
| Nudc | 1.82894975 | 1.06628507 | 1.10164603 | -0.035361 | 0.55943782 | 0.89054584 |
| Tbk1 | 0.28840061 | 0.26264298 | 0.29803064 | -0.0353877 | 0.70993498 | 0.89054584 |
| Ruvbl2 | 0.56751903 | 0.51277207 | 0.54816697 | -0.0353949 | 0.61822632 | 0.89054584 |
| Ube2j1 | 0.49479955 | 0.45196952 | 0.48741297 | -0.0354434 | 0.63261472 | 0.89054584 |
| Gopc | 0.36019961 | 0.33110745 | 0.36655694 | -0.0354495 | 0.67384782 | 0.89054584 |
| Dopey1 | 0.255048 | 0.22987094 | 0.26532722 | -0.0354563 | 0.73322134 | 0.89054584 |
| Rheb | 1.75198483 | 1.0550155 | 1.09050794 | -0.0354924 | 0.56026187 | 0.89054584 |
| Rab23 | 0.21674693 | 0.19164154 | 0.22713515 | -0.0354936 | 0.7667043 | 0.89054584 |
| Tigd2 | 0.33449823 | 0.30682638 | 0.34233152 | -0.0355051 | 0.68552497 | 0.89054584 |
| Vps37b | 0.40035971 | 0.36818883 | 0.40369952 | -0.0355107 | 0.65902926 | 0.89054584 |
| Prr12 | 0.31041678 | 0.28379613 | 0.31932182 | -0.0355257 | 0.6978838 | 0.89054584 |
| Flot1 | 0.48067146 | 0.4396992 | 0.47522741 | -0.0355282 | 0.63619423 | 0.89054584 |
| Fam60a | 0.58756033 | 0.52876514 | 0.56431383 | -0.0355487 | 0.61542536 | 0.89054584 |
| Dip2b | 0.29787527 | 0.27163973 | 0.30722319 | -0.0355835 | 0.70526761 | 0.89054584 |
| Rnf111 | 0.22127848 | 0.19608735 | 0.23168817 | -0.0356008 | 0.76296884 | 0.89054584 |
| Chkb | 0.46279966 | 0.42403285 | 0.45963821 | -0.0356054 | 0.6409128 | 0.89054584 |
| Mfn1 | 0.2137145 | 0.1884691 | 0.22408323 | -0.0356141 | 0.77048885 | 0.89054584 |
| Laptm4b | 0.8727559 | 0.72837846 | 0.76401577 | -0.0356373 | 0.58601858 | 0.89054584 |
| Dsp | 0.16727465 | 0.14118939 | 0.17685564 | -0.0356663 | 0.82627908 | 0.89170439 |
| Uspl1 | 0.27099069 | 0.24535641 | 0.28102612 | -0.0356697 | 0.72285495 | 0.89054584 |
| Tmem127 | 0.19125491 | 0.16567815 | 0.20135605 | -0.0356779 | 0.79545871 | 0.89054584 |
| Ptgr2 | 0.48691952 | 0.44492878 | 0.48063162 | -0.0357028 | 0.63535905 | 0.89054584 |
| Fkbp15 | 0.24017354 | 0.21486709 | 0.25057427 | -0.0357072 | 0.74662462 | 0.89054584 |
| Zmym3 | 0.24344648 | 0.2181003 | 0.25382934 | -0.035729 | 0.74402132 | 0.89054584 |
| Lamtor4 | 1.42728707 | 0.9785711 | 1.01430131 | -0.0357302 | 0.56518081 | 0.89054584 |
| Tubgcp5 | 0.28143891 | 0.25551211 | 0.2912483 | -0.0357362 | 0.7162213 | 0.89054584 |
| Tnrc6c | 0.3335591 | 0.30569531 | 0.34143986 | -0.0357446 | 0.6871308 | 0.89054584 |
| Pisd | 0.38218124 | 0.35125098 | 0.3869965 | -0.0357455 | 0.66652229 | 0.89054584 |

| | | | | | | |
|---|---|---|---|---|---|---|
| Cyld | 0.17349232 | 0.14743412 | 0.18323075 | -0.0357966 | 0.81863828 | 0.89054584 |
| Shq1 | 0.13952883 | 0.11241473 | 0.14821834 | -0.0358036 | 0.86979308 | 0.90589084 |
| Mob1a | 0.23695445 | 0.21155776 | 0.24736793 | -0.0358102 | 0.74998762 | 0.89054584 |
| Yrdc | 0.59020101 | 0.53060527 | 0.56642161 | -0.0358163 | 0.61584918 | 0.89054584 |
| Tfip11 | 0.17681949 | 0.15081614 | 0.1866356 | -0.0358195 | 0.81437991 | 0.89054584 |
| Pccb | 0.33708294 | 0.3089378 | 0.34478337 | -0.0358456 | 0.68593801 | 0.89054584 |
| Celsr2 | 0.24213912 | 0.21663608 | 0.25252971 | -0.0358936 | 0.74607991 | 0.89054584 |
| Aprt | 1.38686673 | 0.96533694 | 1.00125148 | -0.0359145 | 0.56636006 | 0.89054584 |
| Foxj3 | 0.2331214 | 0.20762536 | 0.24354385 | -0.0359185 | 0.75399422 | 0.89054584 |
| Mrpl46 | 0.58590065 | 0.52706423 | 0.56298669 | -0.0359225 | 0.61687089 | 0.89054584 |
| Mtcl1 | 0.20029616 | 0.17458332 | 0.21053083 | -0.0359475 | 0.78683872 | 0.89054584 |
| Ccpg1 | 0.37196973 | 0.34156839 | 0.37753426 | -0.0359659 | 0.67141901 | 0.89054584 |
| 2310039H08I | 0.92726269 | 0.75956016 | 0.79554361 | -0.0359834 | 0.58345096 | 0.89054584 |
| Rsl1d1 | 1.17168676 | 0.8821646 | 0.91815317 | -0.0359886 | 0.57244999 | 0.89054584 |
| Klhdc10 | 0.35079659 | 0.32174374 | 0.35773492 | -0.0359912 | 0.68036835 | 0.89054584 |
| Rtfdc1 | 0.67294407 | 0.59403039 | 0.63009451 | -0.0360641 | 0.60513428 | 0.89054584 |
| Hyal2 | 0.45083081 | 0.41302055 | 0.44909172 | -0.0360712 | 0.64587502 | 0.89054584 |
| Hspb11 | 0.69580229 | 0.61074977 | 0.64685704 | -0.0361073 | 0.60259124 | 0.89054584 |
| Qk | 0.38366931 | 0.35223814 | 0.38837053 | -0.0361324 | 0.66765555 | 0.89054584 |
| Ptges3 | 1.79974098 | 1.06155809 | 1.09771902 | -0.0361609 | 0.56098786 | 0.89054584 |
| Dido1 | 0.39314369 | 0.36090383 | 0.39709085 | -0.036187 | 0.66442523 | 0.89054584 |
| Wdr47 | 0.26463083 | 0.23856658 | 0.2747779 | -0.0362113 | 0.73038365 | 0.89054584 |
| Ndufb6 | 1.70618697 | 1.04635837 | 1.08259218 | -0.0362338 | 0.56195684 | 0.89054584 |
| Cbl | 0.30505694 | 0.27792149 | 0.31416099 | -0.0362395 | 0.70451262 | 0.89054584 |
| Ddx24 | 0.99767936 | 0.79702078 | 0.83329766 | -0.0362769 | 0.58036327 | 0.89054584 |
| Isy1 | 0.24193808 | 0.21604537 | 0.25232978 | -0.0362844 | 0.74855343 | 0.89054584 |
| Vps51 | 0.17995305 | 0.15352216 | 0.18983827 | -0.0363161 | 0.81360424 | 0.89054584 |
| Acp6 | 0.1952303 | 0.16904117 | 0.20539449 | -0.0363533 | 0.79520019 | 0.89054584 |
| Pcgf3 | 0.44368104 | 0.40639451 | 0.4427505 | -0.036356 | 0.64897847 | 0.89054584 |
| Oxr1 | 0.38371773 | 0.35205483 | 0.38841521 | -0.0363604 | 0.66863004 | 0.89054584 |

| | | | | | | |
|---|---|---|---|---|---|---|
| Mthfd1l | 0.18011053 | 0.15362521 | 0.18999913 | -0.0363739 | 0.81378198 | 0.89054584 |
| Ak3 | 0.32068386 | 0.29279123 | 0.3291688 | -0.0363776 | 0.69668002 | 0.89054584 |
| Ubr3 | 0.39363328 | 0.36115577 | 0.39754016 | -0.0363844 | 0.66509327 | 0.89054584 |
| Rint1 | 0.641717 | 0.5701674 | 0.60661079 | -0.0364434 | 0.61022115 | 0.89054584 |
| Ate1 | 0.46500132 | 0.42512519 | 0.46156878 | -0.0364436 | 0.64350785 | 0.89054584 |
| Zcchc10 | 0.24678977 | 0.22069446 | 0.25714923 | -0.0364548 | 0.74552547 | 0.89054584 |
| Dapk1 | 0.3807933 | 0.34923282 | 0.38571381 | -0.036481 | 0.67026492 | 0.89054584 |
| Clic4 | 0.45290816 | 0.41443724 | 0.45092841 | -0.0364912 | 0.6469216 | 0.89054584 |
| Enah | 0.705956 | 0.61768708 | 0.65418525 | -0.0364982 | 0.60254138 | 0.89054584 |
| Actr10 | 1.04974834 | 0.8236102 | 0.86011395 | -0.0365038 | 0.57837116 | 0.89054584 |
| Pik3r2 | 0.36966426 | 0.3388663 | 0.37538996 | -0.0365237 | 0.67483104 | 0.89054584 |
| Ppp2cb | 0.24845946 | 0.22222027 | 0.25880528 | -0.036585 | 0.74492149 | 0.89054584 |
| Deb1 | 0.91737373 | 0.75335585 | 0.78997748 | -0.0366216 | 0.58549768 | 0.89054584 |
| Hs2st1 | 0.23179669 | 0.20559876 | 0.24222068 | -0.0366219 | 0.75940704 | 0.89054584 |
| Fem1b | 0.38543568 | 0.3533659 | 0.38999992 | -0.036634 | 0.6691728 | 0.89054584 |
| Zmat2 | 0.92533806 | 0.75782948 | 0.79446594 | -0.0366365 | 0.58505597 | 0.89054584 |
| Phykpl | 0.22110188 | 0.19486923 | 0.2315109 | -0.0366417 | 0.76955418 | 0.89054584 |
| Cln6 | 0.30399971 | 0.27647929 | 0.3131413 | -0.036662 | 0.7072815 | 0.89054584 |
| Foxo1 | 0.49156211 | 0.44795491 | 0.4846315 | -0.0366766 | 0.63780044 | 0.89054584 |
| Nlk | 0.28880899 | 0.26173966 | 0.29842778 | -0.0366881 | 0.71659673 | 0.89054584 |
| Mrps34 | 0.95001155 | 0.77139761 | 0.80808677 | -0.0366892 | 0.58376219 | 0.89054584 |
| Dtx2 | 0.20710642 | 0.18071524 | 0.21741862 | -0.0367034 | 0.78421152 | 0.89054584 |
| Irak1 | 0.52935789 | 0.47997194 | 0.51670266 | -0.0367307 | 0.62974915 | 0.89054584 |
| Cep19 | 0.31169116 | 0.28379919 | 0.320547 | -0.0367478 | 0.7033657 | 0.89054584 |
| Rbm33 | 0.22076831 | 0.19442069 | 0.23117604 | -0.0367554 | 0.77057403 | 0.89054584 |
| Chchd1 | 1.31712969 | 0.94017041 | 0.97692804 | -0.0367576 | 0.56957649 | 0.89054584 |
| Smg1 | 0.29159614 | 0.26434052 | 0.30113602 | -0.0367955 | 0.71541398 | 0.89054584 |
| Nop16 | 0.42055829 | 0.38521006 | 0.42204084 | -0.0368308 | 0.65784093 | 0.89054584 |
| Eif3j1 | 0.8051313 | 0.68505032 | 0.72191213 | -0.0368618 | 0.5940175 | 0.89054584 |
| 0610009B22I | 1.1536009 | 0.87320585 | 0.910076 | -0.0368701 | 0.57485519 | 0.89054584 |

| | | | | | | |
|---|---|---|---|---|---|---|
| Sppl2a | 0.30611146 | 0.27829804 | 0.31517749 | -0.0368795 | 0.70717502 | 0.89054584 |
| Pwp1 | 0.21394251 | 0.18743013 | 0.22431285 | -0.0368827 | 0.77817612 | 0.89054584 |
| Actn1 | 0.65340807 | 0.57858884 | 0.61548067 | -0.0368918 | 0.60997594 | 0.89054584 |
| Tmem126b | 0.42196018 | 0.38640201 | 0.42330534 | -0.0369033 | 0.65768811 | 0.89054584 |
| Bcar1 | 0.3307504 | 0.30184446 | 0.33877037 | -0.0369259 | 0.69421156 | 0.89054584 |
| Nudt2 | 0.32316602 | 0.29457592 | 0.33154122 | -0.0369653 | 0.69827662 | 0.89054584 |
| Emc9 | 0.23591917 | 0.20932749 | 0.24633574 | -0.0370082 | 0.75801635 | 0.89054584 |
| Tmem51 | 0.33109245 | 0.30207637 | 0.33909569 | -0.0370193 | 0.6944907 | 0.89054584 |
| Gbf1 | 0.55336941 | 0.49956726 | 0.53660954 | -0.0370423 | 0.62612508 | 0.89054584 |
| Hivep3 | 0.16529492 | 0.1377749 | 0.17482246 | -0.0370476 | 0.83825248 | 0.89497393 |
| Elp2 | 0.45779638 | 0.41818471 | 0.45524004 | -0.0370553 | 0.64773978 | 0.89054584 |
| Gon4l | 0.58136026 | 0.52228738 | 0.55934664 | -0.0370593 | 0.62121603 | 0.89054584 |
| Znhit2 | 0.44526418 | 0.4070586 | 0.44415725 | -0.0370986 | 0.6514146 | 0.89054584 |
| Faim | 0.59284198 | 0.53141283 | 0.56852494 | -0.0371121 | 0.61948399 | 0.89054584 |
| Fcf1 | 0.67149688 | 0.59187762 | 0.62902108 | -0.0371435 | 0.60838364 | 0.89054584 |
| Zpr1 | 0.36696225 | 0.33572396 | 0.37287311 | -0.0371491 | 0.67873624 | 0.89054584 |
| Neo1 | 0.34030681 | 0.31067258 | 0.34783709 | -0.0371645 | 0.69068135 | 0.89054584 |
| Fkrp | 0.26651672 | 0.23946082 | 0.27663272 | -0.0371719 | 0.73435392 | 0.89054584 |
| Ogfr | 0.79818043 | 0.68021935 | 0.71739605 | -0.0371767 | 0.59539076 | 0.89054584 |
| Purb | 0.61372182 | 0.54780425 | 0.58499201 | -0.0371878 | 0.61644888 | 0.89054584 |
| Chmp1a | 0.61369143 | 0.54777303 | 0.58496825 | -0.0371952 | 0.61647619 | 0.89054584 |
| Thap7 | 0.38683348 | 0.35408368 | 0.39128808 | -0.0372044 | 0.6711102 | 0.89054584 |
| Mtf2 | 0.58275848 | 0.52319809 | 0.56046907 | -0.037271 | 0.62165025 | 0.89054584 |
| Zfp326 | 0.7757551 | 0.66530287 | 0.70258716 | -0.0372843 | 0.59763746 | 0.89054584 |
| Golga5 | 0.27824884 | 0.25081074 | 0.28813291 | -0.0373222 | 0.7269045 | 0.89054584 |
| Cetn2 | 1.24831706 | 0.9131622 | 0.95048977 | -0.0373276 | 0.57258751 | 0.89054584 |
| Agpat5 | 0.304706 | 0.27649195 | 0.31382257 | -0.0373306 | 0.71027975 | 0.89054584 |
| Akr1e1 | 0.53030701 | 0.48015791 | 0.51749654 | -0.0373386 | 0.63162258 | 0.89054584 |
| Fam117b | 0.15597504 | 0.12787204 | 0.16522952 | -0.0373575 | 0.85391215 | 0.90035231 |
| Wdr83 | 0.36497546 | 0.33365922 | 0.37101992 | -0.0373607 | 0.68051618 | 0.89054584 |

| | | | | | | |
|---|---|---|---|---|---|---|
| Ikbkap | 0.22880721 | 0.20186428 | 0.23923215 | -0.0373679 | 0.76660836 | 0.89054584 |
| Ero1l | 0.29178417 | 0.26394618 | 0.30131859 | -0.0373724 | 0.71831607 | 0.89054584 |
| Cdc42se2 | 0.6229462 | 0.55476012 | 0.59217354 | -0.0374134 | 0.61575517 | 0.89054584 |
| Ide | 0.20434507 | 0.17719147 | 0.21462807 | -0.0374366 | 0.79178714 | 0.89054584 |
| Utp6 | 0.21344724 | 0.18637571 | 0.22381407 | -0.0374384 | 0.78210484 | 0.89054584 |
| Prpf38b | 0.99602106 | 0.79500645 | 0.83244587 | -0.0374394 | 0.58298539 | 0.89054584 |
| Midn | 0.53736079 | 0.48592635 | 0.52337843 | -0.0374521 | 0.63057855 | 0.89054584 |
| Prmt5 | 0.45963577 | 0.41940577 | 0.45685874 | -0.037453 | 0.64874589 | 0.89054584 |
| Cox11 | 0.28566957 | 0.25788758 | 0.29537261 | -0.037485 | 0.72282523 | 0.89054584 |
| Pfn2 | 0.78945477 | 0.67417058 | 0.71167698 | -0.0375064 | 0.59697808 | 0.89054584 |
| Alg14 | 0.27201409 | 0.24450987 | 0.28202973 | -0.0375199 | 0.73230619 | 0.89054584 |
| Smg9 | 0.20476166 | 0.17749657 | 0.21504923 | -0.0375527 | 0.79205169 | 0.89054584 |
| Ppp3ca | 0.94774611 | 0.76927707 | 0.80685173 | -0.0375747 | 0.58588199 | 0.89054584 |
| Dnajc13 | 0.30425392 | 0.27579198 | 0.31338653 | -0.0375946 | 0.71188401 | 0.89054584 |
| Tmem126a | 0.79955065 | 0.68069412 | 0.71828909 | -0.037595 | 0.59632542 | 0.89054584 |
| Ppm1l | 0.25886058 | 0.23149508 | 0.26909217 | -0.0375971 | 0.74245715 | 0.89054584 |
| Dot1l | 0.34590246 | 0.31552403 | 0.35312431 | -0.0376003 | 0.69008058 | 0.89054584 |
| Dcaf10 | 0.20596253 | 0.17863097 | 0.216263 | -0.037632 | 0.79123234 | 0.89054584 |
| Tcf20 | 0.1371117 | 0.10805573 | 0.14570927 | -0.0376535 | 0.88569414 | 0.91420299 |
| Zfp329 | 0.46082834 | 0.42024803 | 0.45790713 | -0.0376591 | 0.64919929 | 0.89054584 |
| Tmpo | 0.92513525 | 0.75663251 | 0.79435222 | -0.0377197 | 0.58754302 | 0.89054584 |
| Usp42 | 0.18385498 | 0.15608794 | 0.19382084 | -0.0377329 | 0.81779645 | 0.89054584 |
| Rnf44 | 0.2402133 | 0.21287109 | 0.25061384 | -0.0377427 | 0.75854881 | 0.89054584 |
| Usp19 | 0.48560917 | 0.44172854 | 0.47950025 | -0.0377717 | 0.64319017 | 0.89054584 |
| Ppp2r3d | 0.29073107 | 0.26251983 | 0.30029586 | -0.037776 | 0.72108792 | 0.89054584 |
| Fto | 0.41972569 | 0.38347089 | 0.4212893 | -0.0378184 | 0.6621087 | 0.89054584 |
| Sec24c | 0.33918666 | 0.30895011 | 0.34677668 | -0.0378266 | 0.69434332 | 0.89054584 |
| Bmyc | 1.185616 | 0.88642613 | 0.92425295 | -0.0378268 | 0.57561032 | 0.89054584 |
| Pitpna | 0.45402822 | 0.41408366 | 0.45191762 | -0.037834 | 0.65174674 | 0.89054584 |
| Akap7 | 0.20520949 | 0.17766459 | 0.21550192 | -0.0378373 | 0.79332318 | 0.89054584 |

| | | | | | | |
|---|---|---|---|---|---|---|
| Akap13 | 0.54702568 | 0.49353272 | 0.53138539 | -0.0378527 | 0.63000803 | 0.89054584 |
| Gtpbp2 | 0.37976969 | 0.34691093 | 0.38476714 | -0.0378562 | 0.67666054 | 0.89054584 |
| Dhrs4 | 0.78717245 | 0.67229891 | 0.71017193 | -0.037873 | 0.59811013 | 0.89054584 |
| Tomm6 | 0.43343269 | 0.39570507 | 0.43361027 | -0.0379052 | 0.65809656 | 0.89054584 |
| Lysmd2 | 0.76451922 | 0.65710726 | 0.69503083 | -0.0379236 | 0.60033327 | 0.89054584 |
| Tbce | 0.35609245 | 0.32478461 | 0.36270945 | -0.0379248 | 0.68691978 | 0.89054584 |
| Add1 | 0.70531225 | 0.61577453 | 0.6537228 | -0.0379483 | 0.60659264 | 0.89054584 |
| Pias3 | 0.26399317 | 0.23618349 | 0.27415036 | -0.0379669 | 0.74060546 | 0.89054584 |
| Gabpb2 | 0.23044136 | 0.20289712 | 0.24086622 | -0.0379691 | 0.76865214 | 0.89054584 |
| Shkbp1 | 0.34275963 | 0.31213756 | 0.35015677 | -0.0380192 | 0.69352147 | 0.89054584 |
| Prosc | 0.41286328 | 0.3770448 | 0.41508022 | -0.0380354 | 0.66528009 | 0.89054584 |
| Med20 | 0.29012802 | 0.26167172 | 0.29970995 | -0.0380382 | 0.72283958 | 0.89054584 |
| Arl2bp | 0.58790772 | 0.52653717 | 0.56459138 | -0.0380542 | 0.62324663 | 0.89054584 |
| Tab2 | 0.58239043 | 0.52209031 | 0.56017374 | -0.0380834 | 0.6242789 | 0.89054584 |
| Cbll1 | 0.33338738 | 0.30318013 | 0.34127677 | -0.0380966 | 0.69849487 | 0.89054584 |
| Nomo1 | 0.29978941 | 0.27096905 | 0.30907489 | -0.0381058 | 0.71715166 | 0.89054584 |
| Akr7a5 | 0.88091536 | 0.73071323 | 0.76887098 | -0.0381577 | 0.59142622 | 0.89054584 |
| Dnaja3 | 0.52301057 | 0.47319398 | 0.51137864 | -0.0381847 | 0.63603609 | 0.89054584 |
| Usp10 | 0.36020558 | 0.32836836 | 0.36656252 | -0.0381942 | 0.6863181 | 0.89054584 |
| Katna1 | 0.55863126 | 0.50271439 | 0.54092278 | -0.0382084 | 0.62895505 | 0.89054584 |
| Ap1b1 | 0.36935843 | 0.33686766 | 0.37510529 | -0.0382376 | 0.68257975 | 0.89054584 |
| Mxd4 | 0.81432145 | 0.68957904 | 0.72782929 | -0.0382503 | 0.59671195 | 0.89054584 |
| Fahd1 | 0.21030051 | 0.18238812 | 0.22064245 | -0.0382543 | 0.79038366 | 0.89054584 |
| Wdr26 | 0.52495522 | 0.47475248 | 0.51301252 | -0.03826 | 0.63587707 | 0.89054584 |
| Slc25a23 | 0.26907841 | 0.2408497 | 0.27914946 | -0.0382998 | 0.73865823 | 0.89054584 |
| Ring1 | 0.40077082 | 0.36575431 | 0.40407514 | -0.0383208 | 0.67069566 | 0.89054584 |
| Rnf215 | 0.2357787 | 0.20786774 | 0.24619564 | -0.0383279 | 0.76586816 | 0.89054584 |
| Ube2g1 | 0.81049717 | 0.68703947 | 0.72537426 | -0.0383348 | 0.5972429 | 0.89054584 |
| Zfp646 | 0.20828287 | 0.18015906 | 0.21860654 | -0.0384475 | 0.79371875 | 0.89054584 |
| Gnpnat1 | 0.57710345 | 0.51745341 | 0.55592147 | -0.0384681 | 0.62641927 | 0.89054584 |

| | | | | | | |
|---|---|---|---|---|---|---|
| Arl5a | 0.20671471 | 0.17851145 | 0.21702297 | -0.0385115 | 0.79580951 | 0.89054584 |
| Klhl12 | 0.20448675 | 0.17625297 | 0.21477131 | -0.0385183 | 0.79829927 | 0.89054584 |
| Commd5 | 0.3817992 | 0.34809102 | 0.38664354 | -0.0385525 | 0.6788719 | 0.89054584 |
| Tsr2 | 0.24059907 | 0.21242437 | 0.25099775 | -0.0385734 | 0.76300031 | 0.89054584 |
| Fat1 | 0.3090542 | 0.27942751 | 0.31801116 | -0.0385837 | 0.7140515 | 0.89054584 |
| Nup43 | 0.24451999 | 0.2162886 | 0.25489589 | -0.0386073 | 0.75979933 | 0.89054584 |
| Nit1 | 0.43928825 | 0.40016636 | 0.43883999 | -0.0386736 | 0.6593091 | 0.89054584 |
| Sbds | 0.74202897 | 0.64093714 | 0.6796339 | -0.0386968 | 0.60459895 | 0.89054584 |
| Rnf145 | 0.57451432 | 0.51512915 | 0.55383227 | -0.0387031 | 0.62762814 | 0.89054584 |
| Atp5s | 0.21933085 | 0.19102531 | 0.22973243 | -0.0387071 | 0.78376586 | 0.89054584 |
| Osbpl1a | 0.23650577 | 0.208206 | 0.24692065 | -0.0387146 | 0.76744994 | 0.89054584 |
| Gprc5b | 0.2081553 | 0.17972333 | 0.21847776 | -0.0387544 | 0.79571733 | 0.89054584 |
| Ift27 | 0.57564239 | 0.51598663 | 0.55474307 | -0.0387564 | 0.6275953 | 0.89054584 |
| Cklf | 0.1636588 | 0.13436137 | 0.17314095 | -0.0387796 | 0.85164056 | 0.89985795 |
| Ptdss2 | 0.4203617 | 0.38308168 | 0.42186342 | -0.0387817 | 0.66578171 | 0.89054584 |
| Vps45 | 0.20827374 | 0.17979666 | 0.21859733 | -0.0388007 | 0.79586855 | 0.89054584 |
| Socs6 | 0.27029459 | 0.24154089 | 0.28034319 | -0.0388023 | 0.74048589 | 0.89054584 |
| Pick1 | 0.55662231 | 0.50047032 | 0.53927815 | -0.0388078 | 0.63128901 | 0.89054584 |
| Tmco6 | 0.17572415 | 0.14670099 | 0.1855152 | -0.0388142 | 0.83516077 | 0.89459411 |
| Nadk | 0.53817263 | 0.48523791 | 0.52405333 | -0.0388154 | 0.63498176 | 0.89054584 |
| Cdc42se1 | 0.64960764 | 0.57376317 | 0.61260772 | -0.0388446 | 0.61616217 | 0.89054584 |
| Dcaf5 | 0.17171885 | 0.1425253 | 0.181414 | -0.0388887 | 0.84103267 | 0.8961387 |
| Apeh | 0.28870905 | 0.25943244 | 0.29833059 | -0.0388982 | 0.72822749 | 0.89054584 |
| Fam222a | 0.21615167 | 0.18763037 | 0.22653639 | -0.038906 | 0.78818519 | 0.89054584 |
| Nr2c2 | 0.25942175 | 0.23072089 | 0.26964581 | -0.0389249 | 0.74938159 | 0.89054584 |
| Bloc1s4 | 0.3032099 | 0.27341984 | 0.31237916 | -0.0389593 | 0.719398 | 0.89054584 |
| Orc3 | 0.33201601 | 0.30099377 | 0.33997377 | -0.03898 | 0.70338983 | 0.89054584 |
| Apc | 0.2977292 | 0.26800312 | 0.30708182 | -0.0390787 | 0.7233722 | 0.89054584 |
| Zyg11b | 0.21330746 | 0.18458354 | 0.22367327 | -0.0390897 | 0.79223734 | 0.89054584 |
| Commd3 | 1.19838612 | 0.89069926 | 0.92980065 | -0.0391014 | 0.57766551 | 0.89054584 |

| | | | | | | |
|---|---|---|---|---|---|---|
| Ahsa2 | 0.6935015 | 0.60607258 | 0.64518634 | -0.0391138 | 0.61119829 | 0.89054584 |
| Msl2 | 0.19576574 | 0.1667852 | 0.2059379 | -0.0391527 | 0.81212468 | 0.89054584 |
| Dcun1d3 | 0.13773696 | 0.10718167 | 0.14635853 | -0.0391769 | 0.89381829 | 0.91924364 |
| Rnf146 | 0.52797095 | 0.47636097 | 0.51554153 | -0.0391806 | 0.63835832 | 0.89054584 |
| Trim37 | 0.38649301 | 0.35178857 | 0.39097442 | -0.0391858 | 0.6797313 | 0.89054584 |
| 4933434E20I | 0.8073316 | 0.68411027 | 0.72333434 | -0.0392241 | 0.59972649 | 0.89054584 |
| Gpatch2 | 0.4486636 | 0.40794768 | 0.44717282 | -0.0392251 | 0.65861031 | 0.89054584 |
| Crtc3 | 0.37948751 | 0.34525895 | 0.38450606 | -0.0392471 | 0.68280077 | 0.89054584 |
| Fam96a | 1.24663979 | 0.91051716 | 0.94980978 | -0.0392926 | 0.57641692 | 0.89054584 |
| Fnbp1l | 0.41798259 | 0.38036102 | 0.41971458 | -0.0393536 | 0.66888443 | 0.89054584 |
| Atp1b3 | 0.52687084 | 0.47526407 | 0.51461965 | -0.0393556 | 0.63918937 | 0.89054584 |
| Dnajc15 | 0.93286082 | 0.75930624 | 0.79866272 | -0.0393565 | 0.59079215 | 0.89054584 |
| Bop1 | 0.2825184 | 0.25290903 | 0.29230139 | -0.0393924 | 0.73495479 | 0.89054584 |
| N4bp1 | 0.3852716 | 0.35042208 | 0.38984864 | -0.0394266 | 0.68124128 | 0.89054584 |
| Toporsos | 0.46344316 | 0.42075832 | 0.46020278 | -0.0394445 | 0.65517228 | 0.89054584 |
| Prmt1 | 2.3161685 | 1.08982838 | 1.12928116 | -0.0394528 | 0.56464851 | 0.89054584 |
| Hsp90ab1 | 4.64015165 | 0.76446186 | 0.80394596 | -0.0394841 | 0.59049326 | 0.89054584 |
| Ddrgk1 | 0.87289474 | 0.72459511 | 0.76409879 | -0.0395037 | 0.59517047 | 0.89054584 |
| Sfxn1 | 0.64556388 | 0.57001427 | 0.60953982 | -0.0395256 | 0.61871536 | 0.89054584 |
| Tmub1 | 0.29238386 | 0.26237091 | 0.30190075 | -0.0395298 | 0.72907761 | 0.89054584 |
| Coq7 | 0.58493288 | 0.52265155 | 0.56221198 | -0.0395604 | 0.62847976 | 0.89054584 |
| Ndufaf1 | 0.21931705 | 0.19014251 | 0.22971856 | -0.0395761 | 0.7889101 | 0.89054584 |
| Rpp25l | 0.67183851 | 0.5896904 | 0.62927462 | -0.0395842 | 0.61526509 | 0.89054584 |
| Ash2l | 0.80870034 | 0.68461838 | 0.72421726 | -0.0395989 | 0.60053898 | 0.89054584 |
| Noa1 | 0.32777667 | 0.29633823 | 0.33593951 | -0.0396013 | 0.70856802 | 0.89054584 |
| Tm7sf3 | 0.57138494 | 0.51169803 | 0.5513017 | -0.0396037 | 0.63106779 | 0.89054584 |
| Dcaf8 | 0.86675005 | 0.72080331 | 0.76041072 | -0.0396074 | 0.59586935 | 0.89054584 |
| Dnajc11 | 0.38135902 | 0.34662858 | 0.38623676 | -0.0396082 | 0.68359288 | 0.89054584 |
| Ahdc1 | 0.33173813 | 0.30006481 | 0.33970962 | -0.0396448 | 0.70667878 | 0.89054584 |
| 1700021F05I | 0.38862901 | 0.35329204 | 0.39294115 | -0.0396491 | 0.68085936 | 0.89054584 |

| | | | | | | |
|---|---|---|---|---|---|---|
| Ube2i | 1.85107158 | 1.06461109 | 1.10435794 | -0.0397468 | 0.56658229 | 0.89054584 |
| Zfp532 | 0.16681537 | 0.13663526 | 0.17638411 | -0.0397488 | 0.85312039 | 0.9000538 |
| Terf2 | 0.15534254 | 0.12479786 | 0.16457722 | -0.0397794 | 0.86993752 | 0.90589084 |
| Rpl3 | 4.95793758 | 0.67375637 | 0.71354207 | -0.0397857 | 0.60248071 | 0.89054584 |
| Mink1 | 0.15848629 | 0.12802989 | 0.1678178 | -0.0397879 | 0.86533093 | 0.90377426 |
| Dyrk1a | 0.17639126 | 0.14639538 | 0.18619763 | -0.0398023 | 0.84035341 | 0.89576177 |
| Efna2 | 0.34623749 | 0.31360832 | 0.35344033 | -0.039832 | 0.70022692 | 0.89054584 |
| Dlg1 | 0.27387314 | 0.24398602 | 0.28385152 | -0.0398655 | 0.74354865 | 0.89054584 |
| Map3k4 | 0.21402225 | 0.18449465 | 0.22439315 | -0.0398985 | 0.79627485 | 0.89054584 |
| Sf3b3 | 0.53432312 | 0.48092539 | 0.52084936 | -0.039924 | 0.63949738 | 0.89054584 |
| Hdac4 | 0.26378808 | 0.23400798 | 0.27394849 | -0.0399405 | 0.75151142 | 0.89054584 |
| Snap29 | 0.19556648 | 0.16576629 | 0.20573568 | -0.0399694 | 0.81729854 | 0.89054584 |
| Supt20 | 0.65237563 | 0.57471721 | 0.61470119 | -0.039984 | 0.61907275 | 0.89054584 |
| Mark2 | 0.52270705 | 0.47108843 | 0.51112341 | -0.040035 | 0.64241552 | 0.89054584 |
| Cdk13 | 0.32996397 | 0.29795251 | 0.33802217 | -0.0400697 | 0.70962105 | 0.89054584 |
| Prcc | 0.3603571 | 0.32660971 | 0.36670429 | -0.0400946 | 0.6947632 | 0.89054584 |
| Eea1 | 0.57672717 | 0.5155211 | 0.55561812 | -0.040097 | 0.63164732 | 0.89054584 |
| Mthfd1 | 0.33201489 | 0.2998387 | 0.3399727 | -0.040134 | 0.70883726 | 0.89054584 |
| Capza2 | 1.57520367 | 1.01479169 | 1.05496071 | -0.040169 | 0.57040106 | 0.89054584 |
| Sec22b | 1.08003124 | 0.83522365 | 0.87539926 | -0.0401756 | 0.58465536 | 0.89054584 |
| Pnpt1 | 0.25029618 | 0.22043656 | 0.26062549 | -0.0401889 | 0.76374998 | 0.89054584 |
| Itgb3bp | 0.43222969 | 0.39228457 | 0.43253333 | -0.0402488 | 0.66768547 | 0.89054584 |
| Pom121 | 0.25036416 | 0.22038125 | 0.26069282 | -0.0403116 | 0.76436787 | 0.89054584 |
| Secisbp2 | 0.30708557 | 0.27578594 | 0.31611598 | -0.04033 | 0.72387343 | 0.89054584 |
| Zfp212 | 0.17838248 | 0.14787755 | 0.18823348 | -0.0403559 | 0.84106493 | 0.8961387 |
| Rnf135 | 0.17652668 | 0.14595866 | 0.18633614 | -0.0403775 | 0.84364536 | 0.89775379 |
| Thtpa | 0.20033237 | 0.17015626 | 0.2105675 | -0.0404112 | 0.81437243 | 0.89054584 |
| Mpv17l2 | 0.35702601 | 0.32316431 | 0.3635848 | -0.0404205 | 0.69775142 | 0.89054584 |
| Kdm7a | 0.53502983 | 0.48101521 | 0.52143828 | -0.0404231 | 0.64101441 | 0.89054584 |
| Pvrl3 | 0.25105264 | 0.22088184 | 0.26137469 | -0.0404928 | 0.76478369 | 0.89054584 |

| | | | | | | |
|---|---|---|---|---|---|---|
| Sdhaf2 | 0.48253588 | 0.43631434 | 0.47684256 | -0.0405282 | 0.65393891 | 0.89054584 |
| Cds2 | 0.25983695 | 0.22952452 | 0.27005535 | -0.0405308 | 0.75779576 | 0.89054584 |
| Ppa1 | 0.87111628 | 0.72246534 | 0.76303421 | -0.0405689 | 0.59781897 | 0.89054584 |
| Tdp1 | 0.19297472 | 0.16252508 | 0.20310397 | -0.0405789 | 0.82402848 | 0.8911506 |
| Gins4 | 0.39008001 | 0.35367395 | 0.3942761 | -0.0406021 | 0.68430558 | 0.89054584 |
| Ktn1 | 1.20521029 | 0.89209075 | 0.93272238 | -0.0406316 | 0.58041478 | 0.89054584 |
| Fam32a | 1.53505379 | 1.00433809 | 1.04497872 | -0.0406406 | 0.57189176 | 0.89054584 |
| Alkbh6 | 0.60111849 | 0.53438927 | 0.57508639 | -0.0406971 | 0.62918635 | 0.89054584 |
| Pnpla8 | 0.50396131 | 0.45454574 | 0.49525091 | -0.0407052 | 0.64911157 | 0.89054584 |
| Akt1s1 | 0.94246149 | 0.7632458 | 0.80395828 | -0.0407125 | 0.5932561 | 0.89054584 |
| Mfsd11 | 0.39500683 | 0.35807985 | 0.3988 | -0.0407202 | 0.6828592 | 0.89054584 |
| Emc2 | 0.78498176 | 0.66800278 | 0.70872374 | -0.040721 | 0.60553008 | 0.89054584 |
| Galt | 0.3023343 | 0.27078837 | 0.31153387 | -0.0407455 | 0.72885133 | 0.89054584 |
| Snx3 | 1.73564299 | 1.04704997 | 1.08779626 | -0.0407463 | 0.56926883 | 0.89054584 |
| Mettl1 | 0.59718847 | 0.53120402 | 0.57197638 | -0.0407724 | 0.63009489 | 0.89054584 |
| Ctdspl | 0.26688689 | 0.23621664 | 0.27699659 | -0.0407799 | 0.75361125 | 0.89054584 |
| Rbbp9 | 0.23901478 | 0.20863541 | 0.24942065 | -0.0407852 | 0.77691916 | 0.89054584 |
| Nrbp1 | 0.66284562 | 0.58176578 | 0.62257361 | -0.0408078 | 0.61996148 | 0.89054584 |
| Pcnxl3 | 0.16876569 | 0.13756525 | 0.17838589 | -0.0408206 | 0.85681554 | 0.9009252 |
| Ddx28 | 0.19009678 | 0.15934795 | 0.20017828 | -0.0408303 | 0.8290159 | 0.89265604 |
| Gemin2 | 0.23407368 | 0.2036062 | 0.24449453 | -0.0408883 | 0.7820524 | 0.89054584 |
| Hgsnat | 0.24095635 | 0.21042607 | 0.25135326 | -0.0409272 | 0.77595501 | 0.89054584 |
| Brcc3 | 0.44383969 | 0.40193959 | 0.44289154 | -0.040952 | 0.66668904 | 0.89054584 |
| Rpl30 | 3.78171588 | 0.96011203 | 1.00106825 | -0.0409562 | 0.57558404 | 0.89054584 |
| Carhsp1 | 0.7664013 | 0.65533163 | 0.69630294 | -0.0409713 | 0.60801063 | 0.89054584 |
| Exoc6 | 0.18152682 | 0.15046376 | 0.19144533 | -0.0409816 | 0.84069346 | 0.89593824 |
| Faap100 | 0.30546749 | 0.27356587 | 0.3145568 | -0.0409909 | 0.72811503 | 0.89054584 |
| Sipa1l1 | 0.14188908 | 0.10957541 | 0.15066617 | -0.0410908 | 0.89806806 | 0.92211428 |
| Nedd4l | 0.29184265 | 0.26027881 | 0.30137537 | -0.0410966 | 0.73738651 | 0.89054584 |
| Rnf166 | 0.25801446 | 0.22715502 | 0.26825711 | -0.0411021 | 0.76233644 | 0.89054584 |

| | | | | | | |
|---|---|---|---|---|---|---|
| Armc8 | 0.40003324 | 0.36229889 | 0.40340116 | -0.0411023 | 0.68249829 | 0.89054584 |
| Ncor2 | 0.30757942 | 0.27548832 | 0.31659159 | -0.0411033 | 0.72736833 | 0.89054584 |
| Nfatc3 | 0.1952703 | 0.16427057 | 0.20543509 | -0.0411645 | 0.82473091 | 0.89148775 |
| Braf | 0.39216106 | 0.35502246 | 0.39618864 | -0.0411662 | 0.68583886 | 0.89054584 |
| Zdhhc12 | 0.4595056 | 0.41557656 | 0.45674426 | -0.0411677 | 0.66272953 | 0.89054584 |
| Ip6k1 | 0.30532506 | 0.27320614 | 0.3144195 | -0.0412134 | 0.72929522 | 0.89054584 |
| Commd6 | 0.97935956 | 0.78256169 | 0.82377702 | -0.0412153 | 0.59215688 | 0.89054584 |
| Zfp263 | 0.37806947 | 0.34197645 | 0.3831934 | -0.041217 | 0.69185803 | 0.89054584 |
| Polr2b | 0.50596323 | 0.45573359 | 0.49695654 | -0.0412229 | 0.65042307 | 0.89054584 |
| Cggbp1 | 0.70400069 | 0.61154594 | 0.65277973 | -0.0412338 | 0.6157314 | 0.89054584 |
| Plscr3 | 0.7336059 | 0.63250698 | 0.67377398 | -0.041267 | 0.61231149 | 0.89054584 |
| Mrpl27 | 0.82237129 | 0.69167814 | 0.73296215 | -0.041284 | 0.60349898 | 0.89054584 |
| Prelid1 | 1.51888415 | 0.99935009 | 1.04069019 | -0.0413401 | 0.57341315 | 0.89054584 |
| Myo6 | 0.54109975 | 0.48511429 | 0.52648318 | -0.0413689 | 0.6428475 | 0.89054584 |
| Cnksr3 | 0.24265758 | 0.21164515 | 0.25304519 | -0.0414 | 0.77704066 | 0.89054584 |
| Hmgxb4 | 0.33778206 | 0.30402901 | 0.34544606 | -0.041417 | 0.71177612 | 0.89054584 |
| Rrnad1 | 0.33612526 | 0.30245437 | 0.34387519 | -0.0414208 | 0.71266386 | 0.89054584 |
| Emsy | 0.44798207 | 0.40514494 | 0.4465688 | -0.0414239 | 0.66718918 | 0.89054584 |
| Osbpl9 | 0.35297094 | 0.31835267 | 0.35977918 | -0.0414265 | 0.70417633 | 0.89054584 |
| Ola1 | 0.94510583 | 0.76390384 | 0.80540761 | -0.0415038 | 0.59486654 | 0.89054584 |
| Lamp2 | 1.14663737 | 0.86543888 | 0.90694342 | -0.0415045 | 0.58441747 | 0.89054584 |
| Snx18 | 0.34976106 | 0.3152405 | 0.35676046 | -0.04152 | 0.70616303 | 0.89054584 |
| Ccdc174 | 0.63853641 | 0.56265745 | 0.60418137 | -0.0415239 | 0.62559077 | 0.89054584 |
| Zfp30 | 0.36578489 | 0.33024755 | 0.37177517 | -0.0415276 | 0.69861037 | 0.89054584 |
| Zc3h18 | 0.49546748 | 0.44644166 | 0.48798616 | -0.0415445 | 0.65418242 | 0.89054584 |
| Tpp1 | 0.33404218 | 0.30034646 | 0.34189858 | -0.0415521 | 0.71437697 | 0.89054584 |
| Scyl1 | 0.50384209 | 0.45358746 | 0.49514926 | -0.0415618 | 0.65212422 | 0.89054584 |
| R3hcc1 | 0.29169674 | 0.25966979 | 0.30123371 | -0.0415639 | 0.73983468 | 0.89054584 |
| Minpp1 | 0.46976075 | 0.42411648 | 0.46573257 | -0.0416161 | 0.66140667 | 0.89054584 |
| Pigp | 1.292992 | 0.92624846 | 0.96792478 | -0.0416763 | 0.57949504 | 0.89054584 |

| | | | | | | |
|---|---|---|---|---|---|---|
| Rrp36 | 0.23176883 | 0.20044894 | 0.24219284 | -0.0417439 | 0.78901446 | 0.89054584 |
| Ldlr | 0.29153881 | 0.25928543 | 0.30108035 | -0.0417949 | 0.74109924 | 0.89054584 |
| Pcyt2 | 0.40196597 | 0.36328694 | 0.40516655 | -0.0418796 | 0.68494306 | 0.89054584 |
| Gpn3 | 0.38242259 | 0.34532791 | 0.38721944 | -0.0418915 | 0.69287983 | 0.89054584 |
| Utp11l | 0.82984686 | 0.69579105 | 0.73768632 | -0.0418953 | 0.60433898 | 0.89054584 |
| Fam234a | 0.37702575 | 0.34031727 | 0.38222654 | -0.0419093 | 0.69526498 | 0.89054584 |
| Bet1l | 0.75401055 | 0.64596945 | 0.68788154 | -0.0419121 | 0.61174333 | 0.89054584 |
| Fasn | 0.36790491 | 0.33177782 | 0.37375163 | -0.0419738 | 0.69958864 | 0.89054584 |
| Rc3h1 | 0.36082389 | 0.32514245 | 0.36714094 | -0.0419985 | 0.7029666 | 0.89054584 |
| Galnt1 | 0.48249215 | 0.43476811 | 0.47680469 | -0.0420366 | 0.65937087 | 0.89054584 |
| Slc37a3 | 0.32394939 | 0.29025155 | 0.33228929 | -0.0420377 | 0.72220357 | 0.89054584 |
| Txndc17 | 1.60424496 | 1.01968447 | 1.06174918 | -0.0420647 | 0.57322052 | 0.89054584 |
| Rnf40 | 0.25207726 | 0.22030159 | 0.26238904 | -0.0420874 | 0.77255832 | 0.89054584 |
| Myl6 | 3.66867377 | 0.97192537 | 1.01405633 | -0.042131 | 0.5767417 | 0.89054584 |
| Kdm5a | 0.74320025 | 0.63827006 | 0.68044472 | -0.0421747 | 0.61361975 | 0.89054584 |
| Bet1 | 0.91970756 | 0.74910571 | 0.79129759 | -0.0421919 | 0.59809299 | 0.89054584 |
| DHRSX | 0.30919317 | 0.27594229 | 0.31814487 | -0.0422026 | 0.73171836 | 0.89054584 |
| Ttc19 | 0.3218214 | 0.28804609 | 0.33025645 | -0.0422104 | 0.72423305 | 0.89054584 |
| Agpat3 | 0.45071225 | 0.40675509 | 0.44898682 | -0.0422317 | 0.66938669 | 0.89054584 |
| Ccar1 | 1.03674971 | 0.81121431 | 0.85344635 | -0.042232 | 0.59116531 | 0.89054584 |
| Cpsf2 | 0.45386583 | 0.40951968 | 0.45177425 | -0.0422546 | 0.66849169 | 0.89054584 |
| Rpl31 | 4.63447286 | 0.76321656 | 0.80547151 | -0.042255 | 0.59654241 | 0.89054584 |
| Taf5l | 0.18884157 | 0.15662746 | 0.19890115 | -0.0422737 | 0.83896023 | 0.89500145 |
| Rsph9 | 0.30818831 | 0.27489643 | 0.31717782 | -0.0422814 | 0.73271997 | 0.89054584 |
| Scaf4 | 0.22152941 | 0.18964824 | 0.23194002 | -0.0422918 | 0.80220064 | 0.89054584 |
| Hook3 | 0.52308304 | 0.46913982 | 0.51143957 | -0.0422998 | 0.65000285 | 0.89054584 |
| Gpbp1 | 1.07053027 | 0.82834945 | 0.87066499 | -0.0423155 | 0.58956691 | 0.89054584 |
| Rmdn3 | 0.37722339 | 0.34008416 | 0.38240967 | -0.0423255 | 0.69695285 | 0.89054584 |
| Abhd16a | 0.85691996 | 0.71210885 | 0.75445465 | -0.0423458 | 0.60314484 | 0.89054584 |
| Gpank1 | 0.22008852 | 0.18810216 | 0.23049346 | -0.0423913 | 0.80423642 | 0.89054584 |

| | | | | | | |
|---|---|---|---|---|---|---|
| Agap1 | 0.3009883 | 0.26783332 | 0.31023371 | -0.0424004 | 0.73785575 | 0.89054584 |
| Sphk2 | 0.32351234 | 0.28944153 | 0.33187198 | -0.0424304 | 0.72429687 | 0.89054584 |
| Jmjd1c | 0.38302621 | 0.34531691 | 0.38777686 | -0.04246 | 0.69502146 | 0.89054584 |
| Trappc1 | 1.11271582 | 0.84875912 | 0.8912478 | -0.0424887 | 0.58788466 | 0.89054584 |
| Aar2 | 0.26327421 | 0.23089968 | 0.27344258 | -0.0425429 | 0.76572797 | 0.89054584 |
| Rabl2 | 0.30013273 | 0.26685494 | 0.30940681 | -0.0425519 | 0.73915137 | 0.89054584 |
| Mpnd | 0.31155629 | 0.27778913 | 0.32041738 | -0.0426282 | 0.7323133 | 0.89054584 |
| Cc2d1b | 0.22871046 | 0.19649019 | 0.23913537 | -0.0426452 | 0.79696579 | 0.89054584 |
| Mtif3 | 0.26665846 | 0.23412402 | 0.27677206 | -0.042648 | 0.76359054 | 0.89054584 |
| Spryd7 | 0.23354961 | 0.20124858 | 0.24397139 | -0.0427228 | 0.79270587 | 0.89054584 |
| Tbc1d16 | 0.62114417 | 0.54803699 | 0.59077515 | -0.0427382 | 0.63195644 | 0.89054584 |
| Uba1 | 1.05051705 | 0.81772363 | 0.86050465 | -0.042781 | 0.5915853 | 0.89054584 |
| Szrd1 | 0.5556286 | 0.49559546 | 0.53846367 | -0.0428682 | 0.64464694 | 0.89054584 |
| Appl2 | 0.28388464 | 0.25074359 | 0.29363341 | -0.0428898 | 0.75190997 | 0.89054584 |
| Ifi35 | 0.24200572 | 0.20949425 | 0.25239705 | -0.0429028 | 0.78580022 | 0.89054584 |
| Azin1 | 0.72384185 | 0.62393476 | 0.66691881 | -0.042984 | 0.61801693 | 0.89054584 |
| Myo1b | 0.66274131 | 0.57949917 | 0.62249556 | -0.0429964 | 0.62619701 | 0.89054584 |
| Tmem128 | 0.6916731 | 0.60085853 | 0.643856 | -0.0429975 | 0.62214897 | 0.89054584 |
| Prorsd1 | 0.51938022 | 0.46530982 | 0.50832233 | -0.0430125 | 0.65329211 | 0.89054584 |
| Tmem57 | 0.51813539 | 0.46420188 | 0.50727263 | -0.0430707 | 0.65379049 | 0.89054584 |
| Podxl2 | 0.60939344 | 0.53851368 | 0.58160223 | -0.0430885 | 0.63501373 | 0.89054584 |
| Uchl5 | 0.45243041 | 0.40741507 | 0.45050623 | -0.0430912 | 0.6720678 | 0.89054584 |
| Rufy2 | 0.26584406 | 0.23281855 | 0.27597134 | -0.0431528 | 0.76684751 | 0.89054584 |
| Larp4b | 0.72474443 | 0.62437429 | 0.66755534 | -0.0431811 | 0.61843201 | 0.89054584 |
| Slc25a36 | 0.42233916 | 0.38046524 | 0.42364699 | -0.0431818 | 0.68256355 | 0.89054584 |
| Adrbk1 | 0.19596241 | 0.1629499 | 0.20613747 | -0.0431876 | 0.83548962 | 0.89459411 |
| Ddx18 | 0.72971177 | 0.62782423 | 0.67104804 | -0.0432238 | 0.61794711 | 0.89054584 |
| Ninj1 | 0.61977852 | 0.54646527 | 0.58971392 | -0.0432486 | 0.63370378 | 0.89054584 |
| Trub2 | 0.15922997 | 0.12529388 | 0.1685838 | -0.0432899 | 0.88421999 | 0.913715 |
| F8a | 0.35790642 | 0.32111379 | 0.36440986 | -0.0432961 | 0.71005166 | 0.89054584 |

| | | | | | | |
|---|---|---|---|---|---|---|
| Supt16 | 0.81195137 | 0.68298886 | 0.726309 | -0.0433201 | 0.60944736 | 0.89054584 |
| Trim27 | 0.89110909 | 0.73152283 | 0.77486775 | -0.0433449 | 0.60280491 | 0.89054584 |
| Tgs1 | 0.42958043 | 0.38681073 | 0.43015869 | -0.043348 | 0.6806406 | 0.89054584 |
| Ahcyl1 | 0.69083152 | 0.59987724 | 0.64324287 | -0.0433656 | 0.62327471 | 0.89054584 |
| Pcnt | 0.23847436 | 0.20549862 | 0.24888242 | -0.0433838 | 0.79163877 | 0.89054584 |
| Gna13 | 0.31022087 | 0.27572763 | 0.31913341 | -0.0434058 | 0.73685161 | 0.89054584 |
| Gnl3l | 0.27477297 | 0.2412644 | 0.28473272 | -0.0434683 | 0.76153552 | 0.89054584 |
| Fxr2 | 0.35715902 | 0.32023783 | 0.36370947 | -0.0434716 | 0.71118525 | 0.89054584 |
| Mrpl51 | 1.20156597 | 0.8876838 | 0.9311658 | -0.043482 | 0.58611219 | 0.89054584 |
| Wrnip1 | 0.32132442 | 0.2862956 | 0.32978135 | -0.0434857 | 0.73050433 | 0.89054584 |
| Golga3 | 0.36054351 | 0.32329436 | 0.36687867 | -0.0435843 | 0.71002959 | 0.89054584 |
| Pkn1 | 0.21053521 | 0.1772942 | 0.22087915 | -0.0435849 | 0.82103719 | 0.89054584 |
| Eif2d | 0.3265624 | 0.29117461 | 0.33478225 | -0.0436076 | 0.72802756 | 0.89054584 |
| Crnkl1 | 0.45205663 | 0.4065505 | 0.45017584 | -0.0436253 | 0.67418017 | 0.89054584 |
| Fam76a | 0.55237887 | 0.49212884 | 0.53579554 | -0.0436667 | 0.64791526 | 0.89054584 |
| Rsbn1 | 0.38521955 | 0.346056 | 0.38980064 | -0.0437446 | 0.69945753 | 0.89054584 |
| Pcgf2 | 0.29148651 | 0.25726355 | 0.30102958 | -0.043766 | 0.75090906 | 0.89054584 |
| Alg5 | 0.42648992 | 0.38360666 | 0.42738331 | -0.0437767 | 0.68339478 | 0.89054584 |
| Dnajc24 | 0.71761688 | 0.61871529 | 0.6625129 | -0.0437976 | 0.62095683 | 0.89054584 |
| Eif3m | 1.89759395 | 1.06558964 | 1.10944332 | -0.0438537 | 0.57305485 | 0.89054584 |
| Kat6a | 0.28592594 | 0.25173573 | 0.29562228 | -0.0438865 | 0.75541897 | 0.89054584 |
| Ppme1 | 0.44899938 | 0.40358312 | 0.44747032 | -0.0438872 | 0.67614353 | 0.89054584 |
| Zdhhc4 | 0.23214707 | 0.1985884 | 0.24257073 | -0.0439823 | 0.80087417 | 0.89054584 |
| Tnip1 | 0.1773977 | 0.14324255 | 0.18722684 | -0.0439843 | 0.86313263 | 0.90286413 |
| Mrps30 | 0.34652352 | 0.30971451 | 0.3537101 | -0.0439956 | 0.71887203 | 0.89054584 |
| Snap47 | 0.79914158 | 0.67401971 | 0.71802263 | -0.0440029 | 0.61237559 | 0.89054584 |
| Fnta | 1.03284414 | 0.80739988 | 0.85142048 | -0.0440206 | 0.59517557 | 0.89054584 |
| Tmx1 | 1.24454159 | 0.90490408 | 0.94895666 | -0.0440526 | 0.58561445 | 0.89054584 |
| Golph3 | 0.54124288 | 0.48252077 | 0.52660185 | -0.0440811 | 0.65172922 | 0.89054584 |
| Ccz1 | 0.63704264 | 0.55890105 | 0.60303798 | -0.0441369 | 0.63344533 | 0.89054584 |

| | | | | | | |
|---|---|---|---|---|---|---|
| Borcs7 | 0.56390956 | 0.50108207 | 0.54523126 | -0.0441492 | 0.64700607 | 0.89054584 |
| Tmem243 | 0.32882971 | 0.29279286 | 0.33694249 | -0.0441496 | 0.72922205 | 0.89054584 |
| Rps6 | 5.01888168 | 0.65278237 | 0.696969 | -0.0441866 | 0.61614391 | 0.89054584 |
| Tmem109 | 0.65211065 | 0.57030599 | 0.61450101 | -0.044195 | 0.63121456 | 0.89054584 |
| Fam160b1 | 0.40084617 | 0.35994161 | 0.40414396 | -0.0442024 | 0.69482063 | 0.89054584 |
| Uck2 | 0.23242155 | 0.19853781 | 0.24284492 | -0.0443071 | 0.80234435 | 0.89054584 |
| Tmed9 | 1.8925041 | 1.06456309 | 1.10892355 | -0.0443605 | 0.57392365 | 0.89054584 |
| Fam168a | 0.2858406 | 0.25117505 | 0.29553918 | -0.0443641 | 0.75783815 | 0.89054584 |
| Gnai3 | 1.13595587 | 0.85767517 | 0.90207596 | -0.0444008 | 0.59068844 | 0.89054584 |
| Kat5 | 0.37774661 | 0.33835413 | 0.38289438 | -0.0445403 | 0.70607426 | 0.89054584 |
| Esco1 | 0.39883522 | 0.3576928 | 0.40230581 | -0.044613 | 0.69729811 | 0.89054584 |
| Herpud2 | 0.24785703 | 0.21356539 | 0.25820792 | -0.0446425 | 0.78973488 | 0.89054584 |
| Selk | 2.82343135 | 1.06404697 | 1.1087792 | -0.0447322 | 0.57454552 | 0.89054584 |
| Ehbp1 | 0.24838344 | 0.2139802 | 0.25872991 | -0.0447497 | 0.78982279 | 0.89054584 |
| Rps27rt | 1.27125938 | 0.91484389 | 0.95961068 | -0.0447668 | 0.58602996 | 0.89054584 |
| Trappc13 | 0.55835857 | 0.49589424 | 0.5406997 | -0.0448055 | 0.65027626 | 0.89054584 |
| Qsox2 | 0.37784367 | 0.33816566 | 0.38298428 | -0.0448186 | 0.70719585 | 0.89054584 |
| Tmem70 | 0.79020635 | 0.66715341 | 0.71217178 | -0.0450184 | 0.61581386 | 0.89054584 |
| Ryk | 0.37083761 | 0.33141878 | 0.37648165 | -0.0450629 | 0.71145856 | 0.89054584 |
| Ssbp4 | 0.90764334 | 0.73932271 | 0.78443894 | -0.0451162 | 0.60563279 | 0.89054584 |
| Psph | 0.33364197 | 0.29635054 | 0.34151856 | -0.045168 | 0.73111044 | 0.89054584 |
| Pigu | 0.24120317 | 0.20641663 | 0.25159881 | -0.0451822 | 0.79860642 | 0.89054584 |
| Oxa1l | 0.61242713 | 0.53877825 | 0.5839794 | -0.0452011 | 0.64080493 | 0.89054584 |
| Spag9 | 0.49447755 | 0.44184752 | 0.48713653 | -0.045289 | 0.66754386 | 0.89054584 |
| Agl | 0.31408191 | 0.27745618 | 0.32284321 | -0.045387 | 0.74375933 | 0.89054584 |
| Slc38a2 | 0.70717018 | 0.60966208 | 0.65505684 | -0.0453948 | 0.62659929 | 0.89054584 |
| Tubg1 | 0.60996977 | 0.53663202 | 0.58205432 | -0.0454223 | 0.64191195 | 0.89054584 |
| Tmem11 | 0.78887595 | 0.66585094 | 0.71129565 | -0.0454447 | 0.61701863 | 0.89054584 |
| Smpd1 | 0.48741395 | 0.43552181 | 0.48105824 | -0.0455364 | 0.67039717 | 0.89054584 |
| Immt | 0.98326607 | 0.78027644 | 0.82582761 | -0.0455512 | 0.60140246 | 0.89054584 |

| | | | | | | |
|---|---|---|---|---|---|---|
| Afg3l1 | 0.58056674 | 0.51313625 | 0.55870906 | -0.0455728 | 0.648035 | 0.89054584 |
| Rock2 | 0.47429239 | 0.42411128 | 0.46968502 | -0.0455737 | 0.67438678 | 0.89054584 |
| Stx17 | 0.22374957 | 0.18857903 | 0.23416708 | -0.0455881 | 0.81779945 | 0.89054584 |
| Fam179b | 0.23657137 | 0.20131595 | 0.24698605 | -0.0456701 | 0.80551401 | 0.89054584 |
| Nipa2 | 0.42951554 | 0.38442053 | 0.43010047 | -0.0456799 | 0.68963463 | 0.89054584 |
| Lrrc16a | 0.23311636 | 0.19785861 | 0.24353882 | -0.0456802 | 0.80890563 | 0.89054584 |
| Gm6563 | 0.50847481 | 0.45340063 | 0.49909283 | -0.0456922 | 0.66513857 | 0.89054584 |
| Rab2a | 1.55114842 | 1.00335377 | 1.04906716 | -0.0457134 | 0.5804382 | 0.89054584 |
| Ppp1ca | 2.70593446 | 1.07076593 | 1.11654199 | -0.0457761 | 0.57574018 | 0.89054584 |
| Dcps | 0.82599238 | 0.68945572 | 0.7352556 | -0.0457999 | 0.61417241 | 0.89054584 |
| Copg2 | 0.40680023 | 0.36375996 | 0.40957288 | -0.0458129 | 0.69886103 | 0.89054584 |
| Hars | 0.51798162 | 0.4612962 | 0.50714289 | -0.0458467 | 0.66318717 | 0.89054584 |
| Med13l | 0.27383073 | 0.23790856 | 0.28380998 | -0.0459014 | 0.77442651 | 0.89054584 |
| Aga | 0.63579958 | 0.55611335 | 0.60208531 | -0.045972 | 0.63897914 | 0.89054584 |
| Plekha1 | 0.46372415 | 0.41445285 | 0.46044923 | -0.0459964 | 0.67917853 | 0.89054584 |
| Kdm2b | 0.28851837 | 0.25213701 | 0.29814516 | -0.0460082 | 0.76391189 | 0.89054584 |
| Scaf8 | 0.27801678 | 0.2418969 | 0.28790608 | -0.0460092 | 0.77172137 | 0.89054584 |
| Tmem159 | 0.32474036 | 0.28688695 | 0.3330443 | -0.0461573 | 0.74076089 | 0.89054584 |
| Abcc5 | 0.25603061 | 0.22013727 | 0.26629815 | -0.0461609 | 0.79033787 | 0.89054584 |
| Exoc4 | 0.42170087 | 0.37690312 | 0.42307154 | -0.0461684 | 0.69441886 | 0.89054584 |
| Cand1 | 0.55041367 | 0.48800943 | 0.53417871 | -0.0461693 | 0.65640601 | 0.89054584 |
| Baz2b | 0.86661894 | 0.71412623 | 0.76033173 | -0.0462055 | 0.61145937 | 0.89054584 |
| Tpd52l2 | 0.85599915 | 0.70764254 | 0.75389314 | -0.0462506 | 0.61249256 | 0.89054584 |
| Atp9a | 0.31878561 | 0.28109702 | 0.32735232 | -0.0462553 | 0.74483192 | 0.89054584 |
| Ube2d3 | 2.73604278 | 1.06849124 | 1.11475194 | -0.0462607 | 0.57665355 | 0.89054584 |
| Ncl | 2.55793051 | 1.07828234 | 1.12456282 | -0.0462805 | 0.5760251 | 0.89054584 |
| Gm10709 | 3.08464594 | 1.04083294 | 1.08713204 | -0.0462991 | 0.57864035 | 0.89054584 |
| 2610507B11 | 0.39962529 | 0.35671775 | 0.40302826 | -0.0463105 | 0.70379331 | 0.89054584 |
| E130309D02 | 0.32896247 | 0.2907362 | 0.33706889 | -0.0463327 | 0.73904601 | 0.89054584 |
| Arxes2 | 0.3443114 | 0.30524549 | 0.35162264 | -0.0463772 | 0.73055404 | 0.89054584 |

| | | | | | | |
|---|---|---|---|---|---|---|
| Cabin1 | 0.34737883 | 0.30804681 | 0.3545165 | -0.0464697 | 0.72929746 | 0.89054584 |
| Zrsr2 | 0.90251082 | 0.73501233 | 0.78148942 | -0.0464771 | 0.60914015 | 0.89054584 |
| Pex10 | 0.23278069 | 0.19671517 | 0.24320361 | -0.0464884 | 0.81341684 | 0.89054584 |
| Fubp3 | 0.50907703 | 0.45305919 | 0.49960447 | -0.0465453 | 0.66787276 | 0.89054584 |
| Rnf139 | 0.22518512 | 0.1890428 | 0.23560592 | -0.0465631 | 0.82141025 | 0.89054584 |
| Ep300 | 0.31492311 | 0.27707173 | 0.32365045 | -0.0465787 | 0.7487303 | 0.89054584 |
| Snf8 | 0.77126202 | 0.65294504 | 0.69957647 | -0.0466314 | 0.62192842 | 0.89054584 |
| Rprd1b | 0.37395512 | 0.33273206 | 0.37937851 | -0.0466465 | 0.71662339 | 0.89054584 |
| A430005L14I | 0.40077558 | 0.35743121 | 0.40407948 | -0.0466483 | 0.70465748 | 0.89054584 |
| Atf6 | 0.19970483 | 0.16327742 | 0.20993185 | -0.0466544 | 0.84975536 | 0.89887961 |
| Nsfl1c | 0.90708498 | 0.73742188 | 0.784119 | -0.0466971 | 0.60928531 | 0.89054584 |
| Trmt2a | 0.37704041 | 0.33553264 | 0.38224013 | -0.0467075 | 0.71542204 | 0.89054584 |
| Dtd2 | 0.39082915 | 0.34824818 | 0.39496487 | -0.0467167 | 0.70920455 | 0.89054584 |
| Nfe2l1 | 0.6802522 | 0.58877368 | 0.63549281 | -0.0467191 | 0.63401588 | 0.89054584 |
| Tfpt | 0.36752495 | 0.32665172 | 0.37339758 | -0.0467459 | 0.72013938 | 0.89054584 |
| Gid8 | 0.42886027 | 0.38275091 | 0.42951246 | -0.0467616 | 0.69400141 | 0.89054584 |
| Jagn1 | 1.09124999 | 0.83414003 | 0.8809219 | -0.0467819 | 0.59770676 | 0.89054584 |
| Chmp6 | 0.40299727 | 0.35929934 | 0.40610768 | -0.0468083 | 0.70436406 | 0.89054584 |
| Eif4a3 | 1.15589483 | 0.86428643 | 0.91110093 | -0.0468145 | 0.59459737 | 0.89054584 |
| Htt | 0.22685179 | 0.19045502 | 0.23727523 | -0.0468202 | 0.82103672 | 0.89054584 |
| Urgcp | 0.20220214 | 0.1656308 | 0.21246045 | -0.0468296 | 0.84776727 | 0.89862623 |
| Synrg | 0.43651912 | 0.38951223 | 0.43636936 | -0.0468571 | 0.6915592 | 0.89054584 |
| C1galt1c1 | 0.53944849 | 0.47820793 | 0.52511312 | -0.0469052 | 0.66135176 | 0.89054584 |
| Sgpl1 | 0.22763125 | 0.19114698 | 0.2380555 | -0.0469085 | 0.82070058 | 0.89054584 |
| Taf7 | 0.6659635 | 0.57799357 | 0.62490329 | -0.0469097 | 0.63673065 | 0.89054584 |
| Taf10 | 0.60395177 | 0.53039396 | 0.57732257 | -0.0469286 | 0.64754826 | 0.89054584 |
| Safb2 | 0.71649598 | 0.61478412 | 0.66171661 | -0.0469325 | 0.62946515 | 0.89054584 |
| 0610012G03 | 1.07867109 | 0.8277554 | 0.87472397 | -0.0469686 | 0.59876944 | 0.89054584 |
| Yipf6 | 0.41256128 | 0.36779746 | 0.41480641 | -0.0470089 | 0.70124542 | 0.89054584 |
| Rnf220 | 0.26726746 | 0.23035912 | 0.27737061 | -0.0470115 | 0.78513187 | 0.89054584 |

| | | | | | | |
|---|---|---|---|---|---|---|
| Acadvl | 0.74068679 | 0.63168796 | 0.67870356 | -0.0470156 | 0.62655287 | 0.89054584 |
| Dnajc17 | 0.24190156 | 0.20527342 | 0.25229347 | -0.04702 | 0.80738189 | 0.89054584 |
| Wdr13 | 0.23210732 | 0.19550127 | 0.24253102 | -0.0470298 | 0.81685009 | 0.89054584 |
| Aimp2 | 0.53973669 | 0.47827854 | 0.52535237 | -0.0470738 | 0.66182945 | 0.89054584 |
| Nhlrc2 | 0.19405651 | 0.15710737 | 0.20420279 | -0.0470954 | 0.85869946 | 0.90127639 |
| Ubr2 | 0.41622297 | 0.37100125 | 0.41812315 | -0.0471219 | 0.70022795 | 0.89054584 |
| Brd3 | 1.16923588 | 0.86989454 | 0.91706735 | -0.0471728 | 0.59469928 | 0.89054584 |
| Parp2 | 0.35129547 | 0.31099579 | 0.35820418 | -0.0472084 | 0.73039577 | 0.89054584 |
| Nbea | 0.4996912 | 0.44436864 | 0.49160435 | -0.0472357 | 0.67279835 | 0.89054584 |
| Kat2a | 0.35533803 | 0.31476382 | 0.36200174 | -0.0472379 | 0.72838943 | 0.89054584 |
| Cab39 | 0.31726823 | 0.27860079 | 0.32589896 | -0.0472982 | 0.75053003 | 0.89054584 |
| Nasp | 0.94126737 | 0.75599983 | 0.80330331 | -0.0473035 | 0.60809115 | 0.89054584 |
| Pcbp2 | 2.37092333 | 1.08218247 | 1.12953173 | -0.0473493 | 0.5774209 | 0.89054584 |
| Polr2e | 1.25369804 | 0.90531017 | 0.95265944 | -0.0473493 | 0.59155996 | 0.89054584 |
| Csk | 0.28806696 | 0.25031303 | 0.2977061 | -0.0473931 | 0.77085823 | 0.89054584 |
| Rbm6 | 0.7428827 | 0.63277314 | 0.680225 | -0.0474519 | 0.62741074 | 0.89054584 |
| Zkscan3 | 0.79953519 | 0.67079941 | 0.71827902 | -0.0474796 | 0.62094547 | 0.89054584 |
| Zfp579 | 0.31104702 | 0.2724233 | 0.31992784 | -0.0475045 | 0.75546342 | 0.89054584 |
| Polr2l | 0.74024768 | 0.63086463 | 0.67839891 | -0.0475343 | 0.62795587 | 0.89054584 |
| Cyb561 | 0.4971338 | 0.44186953 | 0.48941493 | -0.0475454 | 0.67458307 | 0.89054584 |
| Ndufaf3 | 0.77258173 | 0.65290909 | 0.70046229 | -0.0475532 | 0.62410791 | 0.89054584 |
| D10Jhu81e | 0.49414648 | 0.43927293 | 0.48685228 | -0.0475793 | 0.67555739 | 0.89054584 |
| Tcea1 | 1.02179568 | 0.7980839 | 0.8456991 | -0.0476152 | 0.60345926 | 0.89054584 |
| Ganab | 0.55525231 | 0.49049699 | 0.53815507 | -0.0476581 | 0.66004567 | 0.89054584 |
| Ubl4a | 0.62244499 | 0.54411601 | 0.59178482 | -0.0476688 | 0.64627462 | 0.89054584 |
| Hspa14 | 0.55274656 | 0.48842438 | 0.53609777 | -0.0476734 | 0.66067432 | 0.89054584 |
| 2610524H06 | 0.96928061 | 0.7707489 | 0.8184397 | -0.0476908 | 0.60698842 | 0.89054584 |
| Wrap73 | 0.27616939 | 0.23840584 | 0.28609944 | -0.0476936 | 0.78131998 | 0.89054584 |
| Mrpl23 | 1.39562767 | 0.95646845 | 1.00416482 | -0.0476964 | 0.58756782 | 0.89054584 |
| Macrod1 | 0.29529305 | 0.25702196 | 0.3047223 | -0.0477003 | 0.76709003 | 0.89054584 |

| | | | | | | |
|---|---|---|---|---|---|---|
| Fbxl5 | 0.39924823 | 0.35493873 | 0.40268352 | -0.0477448 | 0.70966093 | 0.89054584 |
| Tbc1d17 | 0.38744549 | 0.34401756 | 0.39185173 | -0.0478342 | 0.71522925 | 0.89054584 |
| Odf2 | 0.48132137 | 0.42794685 | 0.47579067 | -0.0478438 | 0.68028166 | 0.89054584 |
| Tbc1d9b | 0.30333142 | 0.26464958 | 0.31249644 | -0.0478469 | 0.76218182 | 0.89054584 |
| Pelo | 0.42582832 | 0.37892177 | 0.42678844 | -0.0478667 | 0.69934789 | 0.89054584 |
| Nudt3 | 0.33511242 | 0.29504073 | 0.34291441 | -0.0478737 | 0.74229164 | 0.89054584 |
| Plod3 | 0.27346409 | 0.23554938 | 0.28345082 | -0.0479014 | 0.78445628 | 0.89054584 |
| Atp6ap2 | 0.86825453 | 0.71337338 | 0.76131629 | -0.0479429 | 0.61538826 | 0.89054584 |
| Dnaja2 | 1.32099584 | 0.93035902 | 0.97833895 | -0.0479799 | 0.59036535 | 0.89054584 |
| Epc1 | 0.35246547 | 0.31132359 | 0.35930418 | -0.0479806 | 0.73307622 | 0.89054584 |
| Trp53bp1 | 0.47210509 | 0.41977253 | 0.46777885 | -0.0480063 | 0.68372077 | 0.89054584 |
| Rabgap1 | 0.55480359 | 0.48976818 | 0.53778696 | -0.0480188 | 0.66129317 | 0.89054584 |
| Clock | 0.53797602 | 0.4758635 | 0.52388992 | -0.0480264 | 0.66534598 | 0.89054584 |
| Rtn4 | 1.46787307 | 0.97850745 | 1.02653456 | -0.0480271 | 0.58627442 | 0.89054584 |
| Usp3 | 0.42554306 | 0.37849961 | 0.42653188 | -0.0480323 | 0.70008642 | 0.89054584 |
| Prr3 | 0.31257763 | 0.27334623 | 0.32139876 | -0.0480525 | 0.7569613 | 0.89054584 |
| Smad2 | 0.34657282 | 0.30569802 | 0.35375659 | -0.0480586 | 0.7366093 | 0.89054584 |
| Alyref | 0.47995082 | 0.42653393 | 0.47460254 | -0.0480686 | 0.68148882 | 0.89054584 |
| Mtx1 | 0.46857086 | 0.41660142 | 0.46469267 | -0.0480913 | 0.68515242 | 0.89054584 |
| Dhx36 | 0.57432497 | 0.50556173 | 0.55367931 | -0.0481176 | 0.65721931 | 0.89054584 |
| Mgat4b | 0.23371972 | 0.19598875 | 0.24414121 | -0.0481525 | 0.82092405 | 0.89054584 |
| Msto1 | 0.28711134 | 0.24855429 | 0.2967763 | -0.048222 | 0.77547894 | 0.89054584 |
| Ctps2 | 0.43641208 | 0.38803275 | 0.43627377 | -0.048241 | 0.69678039 | 0.89054584 |
| Dut | 1.05714128 | 0.81563983 | 0.86389452 | -0.0482547 | 0.60265883 | 0.89054584 |
| Srp14 | 2.51916771 | 1.07786364 | 1.12617678 | -0.0483131 | 0.57920858 | 0.89054584 |
| Phf20l1 | 0.59091115 | 0.51867321 | 0.56698764 | -0.0483144 | 0.65431628 | 0.89054584 |
| Srpr | 1.42320714 | 0.96470392 | 1.01302767 | -0.0483238 | 0.58793734 | 0.89054584 |
| Nf2 | 0.30475242 | 0.26546008 | 0.31386734 | -0.0484073 | 0.76378814 | 0.89054584 |
| Fbxl12 | 0.21972884 | 0.18171604 | 0.23013222 | -0.0484162 | 0.83649827 | 0.89462889 |
| Strbp | 0.73757284 | 0.62810774 | 0.67654012 | -0.0484324 | 0.6306313 | 0.89054584 |

| | | | | | | |
|---|---|---|---|---|---|---|
| Sec23ip | 0.39345751 | 0.34892799 | 0.39737887 | -0.0484509 | 0.7149984 | 0.89054584 |
| Col4a3bp | 0.37938744 | 0.33595628 | 0.38441346 | -0.0484572 | 0.72149288 | 0.89054584 |
| Saal1 | 0.25043145 | 0.21225712 | 0.26075948 | -0.0485024 | 0.80691464 | 0.89054584 |
| Cs | 0.45400844 | 0.4033845 | 0.45190015 | -0.0485157 | 0.69152595 | 0.89054584 |
| Ctr9 | 0.69298399 | 0.59619444 | 0.64481004 | -0.0486156 | 0.63730407 | 0.89054584 |
| Gatad1 | 0.6517073 | 0.56555949 | 0.61419621 | -0.0486367 | 0.64391018 | 0.89054584 |
| Ipo5 | 0.49366106 | 0.43779871 | 0.48643547 | -0.0486368 | 0.67933108 | 0.89054584 |
| Wdfy3 | 0.21227948 | 0.17398966 | 0.22263757 | -0.0486479 | 0.84566391 | 0.89825055 |
| Dalrd3 | 0.55449532 | 0.48886513 | 0.537534 | -0.0486689 | 0.66342372 | 0.89054584 |
| Skiv2l | 0.25394558 | 0.21552947 | 0.26423737 | -0.0487079 | 0.80477473 | 0.89054584 |
| Arrdc4 | 0.45190153 | 0.40130919 | 0.45003872 | -0.0487295 | 0.69303208 | 0.89054584 |
| Sgcb | 0.32460712 | 0.2841349 | 0.33291714 | -0.0487822 | 0.75259329 | 0.89054584 |
| Rmnd5a | 0.29074326 | 0.25147853 | 0.3003077 | -0.0488292 | 0.77563395 | 0.89054584 |
| Ndrg3 | 0.57370157 | 0.50433249 | 0.55317554 | -0.048843 | 0.65959743 | 0.89054584 |
| Zfp664 | 0.72095398 | 0.61596926 | 0.66487827 | -0.048909 | 0.63409382 | 0.89054584 |
| Mta2 | 0.69652698 | 0.59844756 | 0.64738248 | -0.0489349 | 0.63764245 | 0.89054584 |
| Ogfod2 | 0.39771607 | 0.35229578 | 0.40128182 | -0.048986 | 0.71523206 | 0.89054584 |
| Pex7 | 0.71226768 | 0.60971593 | 0.65870476 | -0.0489888 | 0.63551358 | 0.89054584 |
| Ssbp2 | 0.8635065 | 0.70944614 | 0.75845298 | -0.0490068 | 0.61830564 | 0.89054584 |
| Pithd1 | 0.44739301 | 0.39701913 | 0.44604651 | -0.0490274 | 0.69570578 | 0.89054584 |
| Chd8 | 0.53946889 | 0.47605262 | 0.52513007 | -0.0490774 | 0.66836898 | 0.89054584 |
| D430042O09 | 0.37343645 | 0.32981144 | 0.37889693 | -0.0490855 | 0.72693133 | 0.89054584 |
| Pigx | 0.73410835 | 0.62503906 | 0.67412496 | -0.0490859 | 0.63278452 | 0.89054584 |
| Tial1 | 0.71824446 | 0.61386852 | 0.66295834 | -0.0490898 | 0.63494486 | 0.89054584 |
| Wwox | 0.21265596 | 0.17389479 | 0.22301693 | -0.0491221 | 0.84759839 | 0.89862623 |
| Cdc37l1 | 0.37343933 | 0.32973602 | 0.37889959 | -0.0491636 | 0.7272491 | 0.89054584 |
| Stau1 | 0.59202491 | 0.51867756 | 0.5678747 | -0.0491971 | 0.65675456 | 0.89054584 |
| Rab4a | 0.40046247 | 0.35454024 | 0.40379342 | -0.0492532 | 0.7150775 | 0.89054584 |
| Rpe | 0.42905068 | 0.38042311 | 0.42968335 | -0.0492602 | 0.70336923 | 0.89054584 |
| Kctd10 | 0.50539953 | 0.44719641 | 0.49647652 | -0.0492801 | 0.67811893 | 0.89054584 |

| | | | | | | |
|---|---|---|---|---|---|---|
| Prdm2 | 0.26423894 | 0.22504236 | 0.27439225 | -0.0493499 | 0.79896168 | 0.89054584 |
| Hnrnph1 | 1.37270456 | 0.94709832 | 0.99645875 | -0.0493604 | 0.59125892 | 0.89054584 |
| Dusp19 | 0.23496203 | 0.19599511 | 0.24538101 | -0.0493859 | 0.82579256 | 0.89170439 |
| Scap | 0.23456054 | 0.19558897 | 0.24498041 | -0.0493914 | 0.82621345 | 0.89170439 |
| Tle3 | 0.38493816 | 0.34014006 | 0.38954116 | -0.0494011 | 0.72268646 | 0.89054584 |
| Cers2 | 0.78486631 | 0.65920698 | 0.70864733 | -0.0494404 | 0.62742511 | 0.89054584 |
| Glod4 | 0.69756108 | 0.59868546 | 0.64813163 | -0.0494462 | 0.63886707 | 0.89054584 |
| Cstf3 | 0.67854041 | 0.58477624 | 0.63423167 | -0.0494554 | 0.64180579 | 0.89054584 |
| Ncoa1 | 0.2536112 | 0.21443517 | 0.26390669 | -0.0494715 | 0.80876592 | 0.89054584 |
| Taf12 | 0.42224558 | 0.37408857 | 0.42356263 | -0.0494741 | 0.70684317 | 0.89054584 |
| Erbb2ip | 0.29484964 | 0.25481799 | 0.30429253 | -0.0494745 | 0.77562225 | 0.89054584 |
| Tcf25 | 1.46711485 | 0.97681107 | 1.02631535 | -0.0495043 | 0.58890295 | 0.89054584 |
| Ndel1 | 0.64776613 | 0.56168967 | 0.61121202 | -0.0495224 | 0.6470924 | 0.89054584 |
| Unc45a | 0.47653484 | 0.42206 | 0.4716362 | -0.0495762 | 0.68783618 | 0.89054584 |
| Kansl3 | 0.34631183 | 0.30387984 | 0.35351045 | -0.0496306 | 0.74347167 | 0.89054584 |
| Rpa3 | 0.83531985 | 0.69145062 | 0.74111897 | -0.0496683 | 0.62256833 | 0.89054584 |
| Rasa1 | 0.33203417 | 0.29024363 | 0.33999103 | -0.0497474 | 0.7522841 | 0.89054584 |
| Bin3 | 0.39301406 | 0.34718675 | 0.39697185 | -0.0497851 | 0.72048669 | 0.89054584 |
| Rabep1 | 0.60802335 | 0.53073659 | 0.5805266 | -0.04979 | 0.65526828 | 0.89054584 |
| Tmem33 | 0.93187723 | 0.74831257 | 0.79811636 | -0.0498038 | 0.61436942 | 0.89054584 |
| Zfp618 | 0.28054603 | 0.24054954 | 0.29037683 | -0.0498273 | 0.7879859 | 0.89054584 |
| Ehmt1 | 0.51057268 | 0.45104416 | 0.50087417 | -0.04983 | 0.67849692 | 0.89054584 |
| Tada3 | 0.47067014 | 0.41665439 | 0.46652674 | -0.0498723 | 0.69077462 | 0.89054584 |
| Blcap | 0.55227685 | 0.48580908 | 0.53571167 | -0.0499026 | 0.66785197 | 0.89054584 |
| Gnpat | 0.37445433 | 0.32992694 | 0.37984189 | -0.0499149 | 0.72980738 | 0.89054584 |
| Cactin | 0.32572906 | 0.28405191 | 0.33398758 | -0.0499357 | 0.75696554 | 0.89054584 |
| Tomm70a | 0.56169674 | 0.49346495 | 0.54342725 | -0.0499623 | 0.66579808 | 0.89054584 |
| Emc7 | 1.21463736 | 0.88673999 | 0.93670705 | -0.0499671 | 0.59813525 | 0.89054584 |
| Fam20b | 0.25406369 | 0.21436999 | 0.26435417 | -0.0499842 | 0.81081786 | 0.89054584 |
| Rab35 | 0.32942211 | 0.28749871 | 0.33750647 | -0.0500078 | 0.75499957 | 0.89054584 |

| | | | | | | |
|---|---|---|---|---|---|---|
| Tpcn1 | 0.45380267 | 0.40169834 | 0.45171848 | -0.0500201 | 0.69703738 | 0.89054584 |
| Tbc1d7 | 0.2589978 | 0.21920466 | 0.26922757 | -0.0500229 | 0.80665582 | 0.89054584 |
| Brox | 0.37325508 | 0.32864496 | 0.37872848 | -0.0500835 | 0.7310871 | 0.89054584 |
| Fbxw2 | 0.83260073 | 0.68929197 | 0.73941628 | -0.0501243 | 0.62393236 | 0.89054584 |
| Phf14 | 0.56704616 | 0.49765633 | 0.54778286 | -0.0501265 | 0.66506647 | 0.89054584 |
| Ascc3 | 0.43675806 | 0.38644375 | 0.43658272 | -0.050139 | 0.70369309 | 0.89054584 |
| Slc44a3 | 0.60715196 | 0.52970145 | 0.57984183 | -0.0501404 | 0.65647818 | 0.89054584 |
| Zfp644 | 0.70493839 | 0.60330542 | 0.65345411 | -0.0501487 | 0.63965839 | 0.89054584 |
| Gpaa1 | 0.48655917 | 0.43016878 | 0.4803206 | -0.0501518 | 0.6866777 | 0.89054584 |
| Slk | 0.22829479 | 0.18856545 | 0.2387195 | -0.0501541 | 0.83616694 | 0.89462889 |
| Noc4l | 0.25126408 | 0.21142319 | 0.26158405 | -0.0501609 | 0.81417574 | 0.89054584 |
| Fmr1 | 0.45405671 | 0.401734 | 0.45194278 | -0.0502088 | 0.69762679 | 0.89054584 |
| Baz1b | 0.96178523 | 0.76421316 | 0.81444484 | -0.0502317 | 0.61307583 | 0.89054584 |
| Src | 0.22270635 | 0.18288837 | 0.2331209 | -0.0502325 | 0.84228694 | 0.89691028 |
| AW209491 | 0.33168493 | 0.2893757 | 0.33965905 | -0.0502833 | 0.75481256 | 0.89054584 |
| Zbtb8os | 0.42263677 | 0.37359933 | 0.42391522 | -0.0503159 | 0.70986317 | 0.89054584 |
| Mrps2 | 0.19060565 | 0.15035288 | 0.20069586 | -0.050343 | 0.87872943 | 0.91059794 |
| Psmb5 | 2.20162956 | 1.07696675 | 1.12731064 | -0.0503439 | 0.58240927 | 0.89054584 |
| Pde12 | 0.32254228 | 0.28053653 | 0.33094536 | -0.0504088 | 0.76103601 | 0.89054584 |
| BC031181 | 1.41933327 | 0.9613855 | 1.0118097 | -0.0504242 | 0.59180144 | 0.89054584 |
| Klhdc3 | 0.32650434 | 0.28416928 | 0.33472689 | -0.0505576 | 0.75918799 | 0.89054584 |
| Mkln1 | 0.98672221 | 0.77707369 | 0.82763258 | -0.0505589 | 0.61202777 | 0.89054584 |
| Uckl1 | 0.31462358 | 0.27280126 | 0.32336305 | -0.0505618 | 0.76684093 | 0.89054584 |
| Rnf167 | 0.49329403 | 0.43555031 | 0.48612022 | -0.0505699 | 0.68604065 | 0.89054584 |
| 4932438A13I | 0.39416917 | 0.34745635 | 0.39803181 | -0.0505755 | 0.72306905 | 0.89054584 |
| Atp13a1 | 0.25704046 | 0.21669759 | 0.2672955 | -0.0505979 | 0.8110894 | 0.89054584 |
| Zmynd11 | 0.79977582 | 0.66783665 | 0.71843572 | -0.0505991 | 0.62859215 | 0.89054584 |
| Mrps18c | 0.97229803 | 0.76942993 | 0.82004217 | -0.0506122 | 0.61315269 | 0.89054584 |
| Cramp1l | 0.44153962 | 0.39019804 | 0.44084531 | -0.0506473 | 0.70376166 | 0.89054584 |
| Tfam | 0.41142998 | 0.36313043 | 0.41378021 | -0.0506498 | 0.71575916 | 0.89054584 |

| | | | | | | |
|---|---|---|---|---|---|---|
| Pfkl | 0.88986133 | 0.72343394 | 0.77413783 | -0.0507039 | 0.61987319 | 0.89054584 |
| Adrm1 | 1.08069539 | 0.824997 | 0.87572854 | -0.0507315 | 0.60637971 | 0.89054584 |
| Suv420h1 | 0.46607709 | 0.41176674 | 0.46251102 | -0.0507443 | 0.69537857 | 0.89054584 |
| 2700029M09 | 0.6477315 | 0.56043324 | 0.61118575 | -0.0507525 | 0.65057592 | 0.89054584 |
| Zfpl1 | 0.38852635 | 0.34208697 | 0.39284669 | -0.0507597 | 0.72640687 | 0.89054584 |
| Slc25a53 | 0.27049773 | 0.22975127 | 0.28054251 | -0.0507912 | 0.80052136 | 0.89054584 |
| Faf2 | 0.78731206 | 0.65944597 | 0.7102641 | -0.0508181 | 0.63056078 | 0.89054584 |
| Sars | 1.2000611 | 0.87968016 | 0.93052054 | -0.0508404 | 0.60046397 | 0.89054584 |
| Aimp1 | 1.23804881 | 0.89547094 | 0.94631897 | -0.050848 | 0.59883634 | 0.89054584 |
| Psmc5 | 1.67499821 | 1.02582449 | 1.07669691 | -0.0508724 | 0.58711444 | 0.89054584 |
| Clip1 | 0.43791896 | 0.38674158 | 0.43761887 | -0.0508773 | 0.70596864 | 0.89054584 |
| Htra2 | 0.31524876 | 0.27303994 | 0.32396285 | -0.0509229 | 0.76801559 | 0.89054584 |
| Ebp | 0.4430556 | 0.39125059 | 0.44219434 | -0.0509438 | 0.70427682 | 0.89054584 |
| Wdr92 | 0.23830013 | 0.19776197 | 0.24870886 | -0.0509469 | 0.83005007 | 0.89271879 |
| Pds5a | 0.45406445 | 0.40099929 | 0.45194961 | -0.0509503 | 0.70028609 | 0.89054584 |
| Nom1 | 0.33647078 | 0.29324619 | 0.34420289 | -0.0509567 | 0.75481544 | 0.89054584 |
| Exosc1 | 0.44484977 | 0.39283071 | 0.44378915 | -0.0509584 | 0.70366353 | 0.89054584 |
| Hipk2 | 0.40198423 | 0.3542211 | 0.40518322 | -0.0509621 | 0.72105575 | 0.89054584 |
| Ift57 | 0.64634848 | 0.55907631 | 0.61013596 | -0.0510597 | 0.65168986 | 0.89054584 |
| Katnbl1 | 0.31474579 | 0.27241129 | 0.32348032 | -0.051069 | 0.76898983 | 0.89054584 |
| Fam63b | 0.27892233 | 0.23770707 | 0.28879104 | -0.051084 | 0.79506306 | 0.89054584 |
| Ctnnbl1 | 0.67450345 | 0.58011544 | 0.6312495 | -0.0511341 | 0.64706008 | 0.89054584 |
| Ppat | 0.27959217 | 0.23826199 | 0.2894454 | -0.0511834 | 0.79498848 | 0.89054584 |
| Jkamp | 0.79313364 | 0.66286572 | 0.714095 | -0.0512293 | 0.63089818 | 0.89054584 |
| Angel2 | 0.46360325 | 0.40908793 | 0.4603432 | -0.0512553 | 0.69802656 | 0.89054584 |
| Zfp62 | 0.38272606 | 0.33622994 | 0.38749971 | -0.0512698 | 0.73119155 | 0.89054584 |
| Mtx2 | 1.03106529 | 0.79920766 | 0.85049431 | -0.0512866 | 0.61062342 | 0.89054584 |
| Setd1b | 0.41785259 | 0.3683039 | 0.41959707 | -0.0512932 | 0.71550074 | 0.89054584 |
| Srp68 | 0.78968671 | 0.6605313 | 0.71182973 | -0.0512984 | 0.63147006 | 0.89054584 |
| Elovl1 | 0.58203926 | 0.50857787 | 0.55989188 | -0.051314 | 0.66530715 | 0.89054584 |

| | | | | | | |
|---|---|---|---|---|---|---|
| Ddx49 | 0.32017413 | 0.27736202 | 0.32868121 | -0.0513192 | 0.76651918 | 0.89054584 |
| Zc2hc1a | 0.34573066 | 0.3016268 | 0.35296222 | -0.0513354 | 0.75098867 | 0.89054584 |
| Tmem261 | 0.79440339 | 0.66351331 | 0.71492727 | -0.051414 | 0.6312054 | 0.89054584 |
| Ttc4 | 0.40746013 | 0.35875348 | 0.41017332 | -0.0514198 | 0.72040689 | 0.89054584 |
| Akap9 | 1.14386198 | 0.85425332 | 0.90568596 | -0.0514326 | 0.60433147 | 0.89054584 |
| Elf2 | 0.77524523 | 0.65078512 | 0.70224625 | -0.0514611 | 0.63360277 | 0.89054584 |
| Twf1 | 0.59467436 | 0.51850037 | 0.56998153 | -0.0514812 | 0.66304911 | 0.89054584 |
| Rnf25 | 0.23573417 | 0.19460132 | 0.24615124 | -0.0515499 | 0.8353917 | 0.89459411 |
| Nup35 | 0.39488857 | 0.34712645 | 0.39869157 | -0.0515651 | 0.726602 | 0.89054584 |
| Rcor3 | 0.34241394 | 0.29826213 | 0.34983003 | -0.0515679 | 0.75388529 | 0.89054584 |
| Mtch2 | 1.51090155 | 0.98694439 | 1.03852175 | -0.0515774 | 0.59149121 | 0.89054584 |
| Rspry1 | 0.38072318 | 0.33402731 | 0.38564898 | -0.0516217 | 0.73356473 | 0.89054584 |
| Rpf1 | 0.6368141 | 0.55122661 | 0.60286291 | -0.0516363 | 0.65507062 | 0.89054584 |
| Eif1ad | 0.33393487 | 0.29011047 | 0.34179669 | -0.0516862 | 0.75944591 | 0.89054584 |
| Ube2m | 0.39076142 | 0.34318141 | 0.39490261 | -0.0517212 | 0.72913235 | 0.89054584 |
| Nit2 | 0.76755495 | 0.64532747 | 0.69708144 | -0.051754 | 0.63529011 | 0.89054584 |
| 4930402H24 | 0.40007487 | 0.35160441 | 0.40343922 | -0.0518348 | 0.72527735 | 0.89054584 |
| Gin1 | 0.25774866 | 0.21613704 | 0.26799471 | -0.0518577 | 0.8163433 | 0.89054584 |
| Gm14698 | 0.42204669 | 0.37152406 | 0.42338334 | -0.0518593 | 0.71588603 | 0.89054584 |
| Wdr43 | 0.65610276 | 0.56561647 | 0.6175122 | -0.0518957 | 0.65229999 | 0.89054584 |
| Man1a2 | 0.51810298 | 0.45525578 | 0.50724529 | -0.0519895 | 0.68350145 | 0.89054584 |
| Ube2f | 0.47225509 | 0.41588889 | 0.46790966 | -0.0520208 | 0.69775934 | 0.89054584 |
| Ppp4r2 | 0.57373101 | 0.50117838 | 0.55319933 | -0.052021 | 0.66934698 | 0.89054584 |
| Tra2b | 1.83563196 | 1.05046524 | 1.10249299 | -0.0520278 | 0.58700995 | 0.89054584 |
| Psma6 | 1.72146948 | 1.03330351 | 1.08534312 | -0.0520396 | 0.5883821 | 0.89054584 |
| Hspa2 | 0.28160364 | 0.23934552 | 0.29140904 | -0.0520635 | 0.79740065 | 0.89054584 |
| Ngly1 | 0.33594442 | 0.29159177 | 0.34370369 | -0.0521119 | 0.76002492 | 0.89054584 |
| Triap1 | 0.53535759 | 0.46958424 | 0.5217113 | -0.0521271 | 0.6792135 | 0.89054584 |
| Api5 | 0.51324855 | 0.45098113 | 0.50314225 | -0.0521611 | 0.68545062 | 0.89054584 |
| Meaf6 | 0.58517726 | 0.51014319 | 0.56240767 | -0.0522645 | 0.66747623 | 0.89054584 |

| | | | | | | |
|---|---|---|---|---|---|---|
| Slu7 | 0.48670167 | 0.4281227 | 0.4804436 | -0.0523209 | 0.69405043 | 0.89054584 |
| B3galt6 | 0.36233203 | 0.31620715 | 0.36855091 | -0.0523438 | 0.74591506 | 0.89054584 |
| Zfp638 | 0.7746836 | 0.64951068 | 0.70187051 | -0.0523598 | 0.63591427 | 0.89054584 |
| l7Rn6 | 0.85155391 | 0.69873519 | 0.75117362 | -0.0524384 | 0.62749821 | 0.89054584 |
| Cul3 | 0.36112629 | 0.31493369 | 0.36742375 | -0.0524901 | 0.74715866 | 0.89054584 |
| Nol12 | 0.51271656 | 0.45019123 | 0.50269169 | -0.0525005 | 0.68671979 | 0.89054584 |
| Ppil2 | 0.78301784 | 0.65491182 | 0.70742273 | -0.0525109 | 0.6352633 | 0.89054584 |
| Slc35f5 | 0.47753191 | 0.41998019 | 0.47250278 | -0.0525226 | 0.69772894 | 0.89054584 |
| Scmh1 | 0.37795999 | 0.33052299 | 0.38309202 | -0.052569 | 0.73869062 | 0.89054584 |
| Txndc15 | 0.95339829 | 0.75727987 | 0.80992608 | -0.0526462 | 0.61899298 | 0.89054584 |
| Eed | 0.45018048 | 0.395864 | 0.4485162 | -0.0526522 | 0.70778392 | 0.89054584 |
| Ttc13 | 0.34324379 | 0.29794919 | 0.35061426 | -0.0526651 | 0.7579771 | 0.89054584 |
| Psmd2 | 1.48626901 | 0.97909018 | 1.03176341 | -0.0526732 | 0.59399999 | 0.89054584 |
| Prrc2a | 0.56918564 | 0.49683715 | 0.54951957 | -0.0526824 | 0.67243701 | 0.89054584 |
| Wbp1l | 0.28771136 | 0.24466385 | 0.29736016 | -0.0526963 | 0.79549257 | 0.89054584 |
| Ubap2 | 0.4931984 | 0.43332819 | 0.48603807 | -0.0527099 | 0.69330944 | 0.89054584 |
| Ampd2 | 0.29585788 | 0.2525344 | 0.30526964 | -0.0527352 | 0.78954132 | 0.89054584 |
| Gm26699 | 0.67279545 | 0.57724155 | 0.62998434 | -0.0527428 | 0.65175029 | 0.89054584 |
| Cd2bp2 | 0.34883442 | 0.30310752 | 0.35588796 | -0.0527804 | 0.75520167 | 0.89054584 |
| Hdgf | 1.07182575 | 0.81848541 | 0.87131368 | -0.0528283 | 0.61121011 | 0.89054584 |
| Zswim8 | 0.25781606 | 0.2151439 | 0.26806125 | -0.0529174 | 0.82113805 | 0.89054584 |
| Daxx | 0.27533855 | 0.23235739 | 0.28528639 | -0.052929 | 0.80630776 | 0.89054584 |
| Rnf11 | 0.52721689 | 0.46189079 | 0.51490971 | -0.0530189 | 0.68428643 | 0.89054584 |
| Mrpl52 | 1.74920005 | 1.03703315 | 1.09005429 | -0.0530211 | 0.58963832 | 0.89054584 |
| Man2c1 | 0.24830953 | 0.20560318 | 0.25865662 | -0.0530534 | 0.8303634 | 0.89271879 |
| Ica1 | 0.49614062 | 0.43543685 | 0.48856354 | -0.0531267 | 0.69378787 | 0.89054584 |
| Zcchc3 | 0.56086229 | 0.48960194 | 0.54274613 | -0.0531442 | 0.67587353 | 0.89054584 |
| 2010320M18 | 0.37594268 | 0.32796354 | 0.38122259 | -0.053259 | 0.74243853 | 0.89054584 |
| Ttc14 | 0.94420622 | 0.75164203 | 0.8049144 | -0.0532724 | 0.6210908 | 0.89054584 |
| Prmt2 | 0.65071762 | 0.56015507 | 0.61344785 | -0.0532928 | 0.65716631 | 0.89054584 |

| | | | | | | |
|---|---|---|---|---|---|---|
| Hirip3 | 0.31874374 | 0.27401123 | 0.32731223 | -0.053301 | 0.77597888 | 0.89054584 |
| Tmcc1 | 0.40502714 | 0.35464432 | 0.4079583 | -0.053314 | 0.7286891 | 0.89054584 |
| Dnm1l | 0.5873335 | 0.51081705 | 0.56413256 | -0.0533155 | 0.6701441 | 0.89054584 |
| Mettl3 | 0.26959276 | 0.22624937 | 0.2796544 | -0.053405 | 0.81318875 | 0.89054584 |
| Med31 | 0.74940567 | 0.63128617 | 0.68472397 | -0.0534378 | 0.64192052 | 0.89054584 |
| Txn1 | 2.52395202 | 1.07251621 | 1.12599262 | -0.0534764 | 0.58755719 | 0.89054584 |
| Dhx32 | 0.29717877 | 0.25301811 | 0.30654897 | -0.0535309 | 0.79205254 | 0.89054584 |
| Mib1 | 0.31295405 | 0.26822927 | 0.32176032 | -0.053531 | 0.78086116 | 0.89054584 |
| Atp6v1d | 0.8629244 | 0.70445606 | 0.75810091 | -0.0536448 | 0.62917786 | 0.89054584 |
| Btbd1 | 0.74460124 | 0.62774395 | 0.68141327 | -0.0536693 | 0.64317054 | 0.89054584 |
| Ppp2r2d | 0.25948343 | 0.21598772 | 0.26970666 | -0.0537189 | 0.82328235 | 0.89080883 |
| Cdc123 | 0.87054446 | 0.70896484 | 0.76269143 | -0.0537266 | 0.62861665 | 0.89054584 |
| Ccdc53 | 0.5790691 | 0.50376821 | 0.5575046 | -0.0537364 | 0.67330687 | 0.89054584 |
| Amfr | 0.57668699 | 0.50180752 | 0.55558572 | -0.0537782 | 0.67399247 | 0.89054584 |
| Mvb12a | 0.60105743 | 0.52122524 | 0.57503813 | -0.0538129 | 0.66857815 | 0.89054584 |
| Tmem131 | 0.2595528 | 0.21595403 | 0.26977509 | -0.0538211 | 0.82367845 | 0.89097143 |
| Fbxw9 | 0.3724151 | 0.32411167 | 0.37794816 | -0.0538365 | 0.74653922 | 0.89054584 |
| Ndufs4 | 1.63285075 | 1.01421966 | 1.06806735 | -0.0538477 | 0.59285108 | 0.89054584 |
| Stk25 | 0.74451781 | 0.62750067 | 0.68135563 | -0.053855 | 0.64365543 | 0.89054584 |
| Ranbp3 | 0.48382248 | 0.42404718 | 0.47795588 | -0.0539087 | 0.7003761 | 0.89054584 |
| Csnk2a1 | 1.06219735 | 0.81255451 | 0.86646542 | -0.0539109 | 0.61404455 | 0.89054584 |
| Nbr1 | 0.62630235 | 0.54085322 | 0.59477201 | -0.0539188 | 0.66361678 | 0.89054584 |
| Rabepk | 0.30146965 | 0.25673639 | 0.31069877 | -0.0539624 | 0.79079082 | 0.89054584 |
| Plekha5 | 0.30174531 | 0.2569043 | 0.31096505 | -0.0540608 | 0.79101529 | 0.89054584 |
| Hexim1 | 0.32282325 | 0.27709886 | 0.33121379 | -0.0541149 | 0.7767297 | 0.89054584 |
| Peli1 | 0.52417761 | 0.45821375 | 0.51235947 | -0.0541457 | 0.68876511 | 0.89054584 |
| Kmt2a | 0.66784316 | 0.57213915 | 0.62630448 | -0.0541653 | 0.65649791 | 0.89054584 |
| Polr2f | 1.61175745 | 1.00924857 | 1.06344129 | -0.0541927 | 0.59383396 | 0.89054584 |
| Sms | 0.73713825 | 0.62198535 | 0.67623763 | -0.0542523 | 0.64571276 | 0.89054584 |
| Nme2 | 0.21306346 | 0.16912471 | 0.22342748 | -0.0543028 | 0.8712519 | 0.90632203 |

| | | | | | | |
|---|---|---|---|---|---|---|
| Pigyl | 0.82960266 | 0.68318793 | 0.73753264 | -0.0543447 | 0.63431047 | 0.89054584 |
| Ppp3r1 | 0.72963726 | 0.61654291 | 0.67099578 | -0.0544529 | 0.64731588 | 0.89054584 |
| Prpf8 | 0.46931265 | 0.41088807 | 0.46534106 | -0.054453 | 0.7071841 | 0.89054584 |
| Caprin1 | 1.3015014 | 0.91666148 | 0.97113218 | -0.0544707 | 0.60307694 | 0.89054584 |
| Gar1 | 0.61070597 | 0.5281284 | 0.58263149 | -0.0545031 | 0.66851897 | 0.89054584 |
| Terf2ip | 0.42429545 | 0.37086286 | 0.42540921 | -0.0545463 | 0.72487101 | 0.89054584 |
| Hdhd2 | 0.51018394 | 0.44589792 | 0.50054429 | -0.0546464 | 0.69449586 | 0.89054584 |
| Ppp2r4 | 0.67609489 | 0.57772641 | 0.63242649 | -0.0547001 | 0.65651226 | 0.89054584 |
| Tmem29 | 0.3183688 | 0.27218577 | 0.32695321 | -0.0547674 | 0.78242587 | 0.89054584 |
| 1810026J23F | 0.39733925 | 0.34616306 | 0.40093688 | -0.0547738 | 0.73776795 | 0.89054584 |
| Mrpl41 | 1.2063713 | 0.87840337 | 0.9332165 | -0.0548131 | 0.60782172 | 0.89054584 |
| Phf23 | 0.6535818 | 0.56073205 | 0.61561177 | -0.0548797 | 0.66104945 | 0.89054584 |
| Fam98a | 0.2395439 | 0.19505949 | 0.2499475 | -0.054888 | 0.84686848 | 0.89836096 |
| Larp4 | 0.37070441 | 0.32142408 | 0.37635776 | -0.0549337 | 0.75175575 | 0.89054584 |
| Atg12 | 0.45885005 | 0.40122769 | 0.45616754 | -0.0549399 | 0.71263976 | 0.89054584 |
| Ctu2 | 0.27075636 | 0.22584707 | 0.28079625 | -0.0549492 | 0.81903953 | 0.89054584 |
| Cdv3 | 0.83459527 | 0.68569939 | 0.74066577 | -0.0549664 | 0.63523468 | 0.89054584 |
| Sft2d1 | 0.39242664 | 0.34144574 | 0.39643254 | -0.0549868 | 0.74092962 | 0.89054584 |
| Iars2 | 0.37559481 | 0.32587064 | 0.38089998 | -0.0550293 | 0.74955373 | 0.89054584 |
| Ythdf1 | 0.33962104 | 0.29215698 | 0.34718798 | -0.055031 | 0.76988403 | 0.89054584 |
| Ube2d1 | 0.50900892 | 0.44449876 | 0.49954661 | -0.0550479 | 0.69616247 | 0.89054584 |
| Ccdc115 | 0.7555646 | 0.63389035 | 0.6889437 | -0.0550533 | 0.6451634 | 0.89054584 |
| Kpna3 | 0.37694365 | 0.3269878 | 0.38215046 | -0.0551627 | 0.74937009 | 0.89054584 |
| Vps4a | 0.417015 | 0.36362032 | 0.4188397 | -0.0552194 | 0.73046829 | 0.89054584 |
| Prkaca | 0.60422571 | 0.52231888 | 0.5775385 | -0.0552196 | 0.67200299 | 0.89054584 |
| Trim26 | 0.40554135 | 0.35320705 | 0.40842673 | -0.0552197 | 0.7356178 | 0.89054584 |
| Suds3 | 0.71492146 | 0.6053453 | 0.66059655 | -0.0552512 | 0.6516084 | 0.89054584 |
| Taf11 | 0.60540734 | 0.52319153 | 0.57846929 | -0.0552778 | 0.67191321 | 0.89054584 |
| Ctcf | 0.58122209 | 0.50393238 | 0.55923565 | -0.0553033 | 0.6775026 | 0.89054584 |
| Rnf114 | 0.45720087 | 0.39934535 | 0.45471555 | -0.0553702 | 0.7147496 | 0.89054584 |

| | | | | | | |
|---|---|---|---|---|---|---|
| Wapl | 0.64148772 | 0.55102611 | 0.6064359 | -0.0554098 | 0.66483148 | 0.89054584 |
| Harbi1 | 0.21435595 | 0.1692846 | 0.22472915 | -0.0554446 | 0.87481291 | 0.90828873 |
| Gtf2b | 0.86413017 | 0.70329791 | 0.75882992 | -0.055532 | 0.63342807 | 0.89054584 |
| Ppp2r5e | 0.43862398 | 0.3826982 | 0.43824774 | -0.0555495 | 0.72258506 | 0.89054584 |
| Metap1 | 0.48329085 | 0.42192073 | 0.47749597 | -0.0555752 | 0.70617656 | 0.89054584 |
| Rufy1 | 0.32021278 | 0.27313249 | 0.32871819 | -0.0555857 | 0.78459874 | 0.89054584 |
| Kif1bp | 0.58644106 | 0.5078303 | 0.56341902 | -0.0555887 | 0.67712204 | 0.89054584 |
| Rab11fip2 | 0.29989197 | 0.25356195 | 0.30917405 | -0.0556121 | 0.79898978 | 0.89054584 |
| Ppid | 1.25871949 | 0.89900241 | 0.95466783 | -0.0556654 | 0.60705699 | 0.89054584 |
| Gtf2h3 | 0.4532689 | 0.39555712 | 0.45124708 | -0.05569 | 0.71734623 | 0.89054584 |
| Cnot11 | 0.27798563 | 0.2321854 | 0.28787563 | -0.0556902 | 0.81628117 | 0.89054584 |
| Fbxo18 | 0.49355244 | 0.43061574 | 0.48634218 | -0.0557264 | 0.70327197 | 0.89054584 |
| Arfip1 | 0.41847834 | 0.3644228 | 0.42016263 | -0.0557398 | 0.73173434 | 0.89054584 |
| Atxn2l | 0.41389276 | 0.36020384 | 0.41601324 | -0.0558094 | 0.73401798 | 0.89054584 |
| Bace1 | 0.60054187 | 0.51881998 | 0.57463062 | -0.0558106 | 0.67454442 | 0.89054584 |
| Napg | 0.42593642 | 0.37106881 | 0.42688566 | -0.0558169 | 0.72879528 | 0.89054584 |
| Dr1 | 0.29970326 | 0.25310817 | 0.30899158 | -0.0558834 | 0.800275 | 0.89054584 |
| Srprb | 0.68183287 | 0.58074988 | 0.63665576 | -0.0559059 | 0.65877051 | 0.89054584 |
| Tecpr1 | 0.47091228 | 0.41082524 | 0.46673812 | -0.0559129 | 0.71161797 | 0.89054584 |
| Pik3c2a | 0.37722141 | 0.32649361 | 0.38240784 | -0.0559142 | 0.75213157 | 0.89054584 |
| Tpp2 | 0.60808296 | 0.52455104 | 0.58057343 | -0.0560224 | 0.67348766 | 0.89054584 |
| AU022252 | 0.28219339 | 0.23596108 | 0.29198439 | -0.0560233 | 0.81431639 | 0.89054584 |
| Ankrd13c | 0.37022617 | 0.31988952 | 0.37591286 | -0.0560233 | 0.75626393 | 0.89054584 |
| Coq4 | 0.29785929 | 0.25118247 | 0.30720773 | -0.0560253 | 0.80223954 | 0.89054584 |
| Alkbh1 | 0.53113691 | 0.46216112 | 0.51819023 | -0.0560291 | 0.6927801 | 0.89054584 |
| Mydgf | 1.45085535 | 0.96547901 | 1.0215297 | -0.0560507 | 0.60088295 | 0.89054584 |
| Phrf1 | 0.40303239 | 0.35006153 | 0.40613972 | -0.0560782 | 0.73998154 | 0.89054584 |
| Marveld3 | 0.35184756 | 0.3025841 | 0.35872333 | -0.0561392 | 0.76703332 | 0.89054584 |
| Grb7 | 0.38339953 | 0.33195903 | 0.38812151 | -0.0561625 | 0.74989661 | 0.89054584 |
| Creb3 | 0.67881327 | 0.57822041 | 0.63443282 | -0.0562124 | 0.66012137 | 0.89054584 |

| | | | | | | |
|---|---|---|---|---|---|---|
| Tnks2 | 0.67638707 | 0.57641708 | 0.63264239 | -0.0562253 | 0.66058403 | 0.89054584 |
| Ikbkb | 0.36084197 | 0.31089928 | 0.36715785 | -0.0562586 | 0.76234823 | 0.89054584 |
| Chchd3 | 0.94725919 | 0.75031954 | 0.80658579 | -0.0562662 | 0.62740982 | 0.89054584 |
| Polr2d | 0.79899277 | 0.66162644 | 0.71792566 | -0.0562992 | 0.64257525 | 0.89054584 |
| Ivd | 0.49250906 | 0.42914493 | 0.48544571 | -0.0563008 | 0.70551606 | 0.89054584 |
| Chmp3 | 0.73742353 | 0.62013272 | 0.67643621 | -0.0563035 | 0.65091399 | 0.89054584 |
| Pdcd11 | 0.42284322 | 0.36776952 | 0.42410126 | -0.0563317 | 0.73198556 | 0.89054584 |
| Tsen54 | 0.21056581 | 0.16457037 | 0.22091001 | -0.0563396 | 0.88262074 | 0.91245261 |
| Sun1 | 0.43194295 | 0.37592434 | 0.43227652 | -0.0563522 | 0.72818942 | 0.89054584 |
| 9930021J03F | 0.29644546 | 0.24943615 | 0.30583883 | -0.0564027 | 0.80488497 | 0.89054584 |
| Eif4e2 | 1.13500004 | 0.84523086 | 0.90163671 | -0.0564059 | 0.61465045 | 0.89054584 |
| Epm2aip1 | 0.47091308 | 0.41032042 | 0.46673881 | -0.0564184 | 0.71333803 | 0.89054584 |
| Klhdc2 | 1.31017545 | 0.917942 | 0.97436735 | -0.0564254 | 0.60634219 | 0.89054584 |
| Psma3 | 2.58968481 | 1.06658536 | 1.12304631 | -0.056461 | 0.5925958 | 0.89054584 |
| Chfr | 0.40269981 | 0.34927081 | 0.40583629 | -0.0565655 | 0.74194828 | 0.89054584 |
| Hmg20b | 0.74648539 | 0.62612142 | 0.68271361 | -0.0565922 | 0.65032339 | 0.89054584 |
| Rpl15 | 4.05013876 | 0.89524928 | 0.95195978 | -0.0567105 | 0.60931884 | 0.89054584 |
| Actr1b | 0.70642354 | 0.59776642 | 0.65452098 | -0.0567546 | 0.65688784 | 0.89054584 |
| Rab5b | 0.33049918 | 0.28175683 | 0.3385314 | -0.0567746 | 0.78269981 | 0.89054584 |
| Nfat5 | 0.72439498 | 0.61050414 | 0.66730897 | -0.0568048 | 0.65416634 | 0.89054584 |
| Gt(ROSA)26S | 0.49553239 | 0.43118638 | 0.48804185 | -0.0568555 | 0.70635013 | 0.89054584 |
| Atxn2 | 0.40418027 | 0.35031959 | 0.4071865 | -0.0568669 | 0.7423667 | 0.89054584 |
| Ube2z | 0.30926123 | 0.26129924 | 0.31821035 | -0.0569111 | 0.79764525 | 0.89054584 |
| Yme1l1 | 0.87734675 | 0.70982435 | 0.76675354 | -0.0569292 | 0.63529583 | 0.89054584 |
| Polg | 0.22756934 | 0.18103462 | 0.23799353 | -0.0569589 | 0.8675797 | 0.90444517 |
| Rtcb | 1.04556092 | 0.8009855 | 0.85797861 | -0.0569931 | 0.62152194 | 0.89054584 |
| Pgd | 0.45403525 | 0.39491597 | 0.45192383 | -0.0570079 | 0.72163364 | 0.89054584 |
| Irf2bp2 | 0.69168018 | 0.5868398 | 0.64386116 | -0.0570214 | 0.66005132 | 0.89054584 |
| Mrps31 | 0.42204183 | 0.36634864 | 0.42337896 | -0.0570303 | 0.73485793 | 0.89054584 |
| Zfp148 | 0.76281337 | 0.63681571 | 0.69387562 | -0.0570599 | 0.64918521 | 0.89054584 |

| | | | | | | |
|---|---|---|---|---|---|---|
| Ndufa11 | 2.2714458 | 1.07156772 | 1.12873131 | -0.0571636 | 0.59326347 | 0.89054584 |
| Casc4 | 0.57432973 | 0.49643242 | 0.55368315 | -0.0572507 | 0.68500242 | 0.89054584 |
| Rnf115 | 0.54946323 | 0.47606928 | 0.53339585 | -0.0573266 | 0.69171534 | 0.89054584 |
| Rab9 | 0.56634864 | 0.48986915 | 0.54721599 | -0.0573468 | 0.6873119 | 0.89054584 |
| Gorasp2 | 0.97923805 | 0.76633792 | 0.82371306 | -0.0573751 | 0.62722598 | 0.89054584 |
| Cflar | 0.44875343 | 0.38986717 | 0.44725242 | -0.0573852 | 0.72500361 | 0.89054584 |
| Usp4 | 0.44462141 | 0.38616619 | 0.44358627 | -0.0574201 | 0.72677135 | 0.89054584 |
| 9530068E07I | 1.2951969 | 0.91126218 | 0.96875199 | -0.0574898 | 0.60891072 | 0.89054584 |
| Otud4 | 0.20715508 | 0.15997682 | 0.21746778 | -0.057491 | 0.89096249 | 0.91722772 |
| 5031425E22I | 0.41783439 | 0.36206913 | 0.41958062 | -0.0575115 | 0.73845721 | 0.89054584 |
| Fam76b | 0.22938775 | 0.18229251 | 0.2398128 | -0.0575203 | 0.86810332 | 0.90466949 |
| Nol7 | 1.36640825 | 0.93679187 | 0.99433837 | -0.0575465 | 0.60627843 | 0.89054584 |
| Clk4 | 0.60380843 | 0.51966224 | 0.57720957 | -0.0575473 | 0.67885261 | 0.89054584 |
| Vta1 | 0.68166726 | 0.57897145 | 0.636534 | -0.0575626 | 0.66323702 | 0.89054584 |
| Zfp830 | 0.45127143 | 0.39188754 | 0.44948151 | -0.057594 | 0.72473626 | 0.89054584 |
| Sec13 | 1.09623115 | 0.82575175 | 0.88334696 | -0.0575952 | 0.61934775 | 0.89054584 |
| Fam207a | 0.37507436 | 0.32279931 | 0.38041722 | -0.0576179 | 0.75979469 | 0.89054584 |
| Xrcc5 | 0.4051262 | 0.35040311 | 0.40804855 | -0.0576454 | 0.74478664 | 0.89054584 |
| Phax | 0.85223422 | 0.69382391 | 0.75159077 | -0.0577669 | 0.63985965 | 0.89054584 |
| Pnkd | 0.98020074 | 0.76642238 | 0.82421951 | -0.0577971 | 0.62805302 | 0.89054584 |
| Thoc5 | 0.27211826 | 0.22423261 | 0.28213185 | -0.0578992 | 0.83049175 | 0.89271879 |
| Vps72 | 1.01273527 | 0.78317004 | 0.84108843 | -0.0579184 | 0.62582679 | 0.89054584 |
| Pcif1 | 0.70791244 | 0.59761627 | 0.65558918 | -0.0579729 | 0.65982448 | 0.89054584 |
| Eif3b | 1.04297506 | 0.79867516 | 0.856654 | -0.0579788 | 0.6237406 | 0.89054584 |
| Stau2 | 0.35683399 | 0.30542068 | 0.36340479 | -0.0579841 | 0.77137144 | 0.89054584 |
| Arl3 | 1.44982322 | 0.9631944 | 1.02121968 | -0.0580253 | 0.60438707 | 0.89054584 |
| Ccdc12 | 1.26552732 | 0.89931432 | 0.95736562 | -0.0580513 | 0.61122066 | 0.89054584 |
| Taok2 | 0.41843111 | 0.36202437 | 0.42011994 | -0.0580956 | 0.74029454 | 0.89054584 |
| Dad1 | 2.79323574 | 1.05285825 | 1.11095429 | -0.058096 | 0.59624279 | 0.89054584 |
| Spen | 0.33467339 | 0.28434327 | 0.34249777 | -0.0581545 | 0.78554732 | 0.89054584 |

| | | | | | | |
|---|---|---|---|---|---|---|
| Gnl1 | 0.44017525 | 0.38144324 | 0.43963041 | -0.0581872 | 0.73126159 | 0.89054584 |
| Rab28 | 0.84371012 | 0.68813601 | 0.74633951 | -0.0582035 | 0.64182037 | 0.89054584 |
| Scarb2 | 0.38176646 | 0.32834806 | 0.38661329 | -0.0582652 | 0.75870215 | 0.89054584 |
| Surf2 | 0.32137553 | 0.27143934 | 0.32983022 | -0.0583909 | 0.79525371 | 0.89054584 |
| Anapc5 | 1.85741012 | 1.04668023 | 1.1050954 | -0.0584152 | 0.59726373 | 0.89054584 |
| Amd1 | 0.92244213 | 0.73433489 | 0.79283928 | -0.0585044 | 0.63449708 | 0.89054584 |
| Suz12 | 0.33517462 | 0.28446111 | 0.34297343 | -0.0585123 | 0.78664368 | 0.89054584 |
| Rnf138 | 0.61706768 | 0.52906928 | 0.58760359 | -0.0585343 | 0.67871261 | 0.89054584 |
| Thap4 | 0.30814849 | 0.25858676 | 0.31713949 | -0.0585527 | 0.80515218 | 0.89054584 |
| Vps26a | 0.34911088 | 0.29754971 | 0.35614832 | -0.0585986 | 0.77832708 | 0.89054584 |
| Prpf4 | 0.37610536 | 0.32275422 | 0.38137342 | -0.0586192 | 0.76303673 | 0.89054584 |
| Abr | 0.32200797 | 0.2717565 | 0.33043476 | -0.0586783 | 0.79597497 | 0.89054584 |
| Rnaseh2b | 0.32208057 | 0.27178949 | 0.33050416 | -0.0587147 | 0.79607112 | 0.89054584 |
| Rbm22 | 0.78307713 | 0.6486937 | 0.70746205 | -0.0587683 | 0.65062737 | 0.89054584 |
| Ube2j2 | 0.9827255 | 0.76675208 | 0.82554452 | -0.0587924 | 0.6299775 | 0.89054584 |
| Dazap1 | 1.28845292 | 0.90739601 | 0.96621241 | -0.0588164 | 0.61164325 | 0.89054584 |
| Bag6 | 0.96712968 | 0.75842119 | 0.81729751 | -0.0588763 | 0.63142124 | 0.89054584 |
| Ctnnd1 | 0.43649795 | 0.3774713 | 0.43635045 | -0.0588791 | 0.73521297 | 0.89054584 |
| Pak4 | 0.36272441 | 0.31000698 | 0.36891761 | -0.0589106 | 0.77155065 | 0.89054584 |
| Pten | 0.38796182 | 0.33336437 | 0.39232712 | -0.0589627 | 0.75809601 | 0.89054584 |
| Ap3s1 | 1.17656194 | 0.8612821 | 0.92030143 | -0.0590193 | 0.61744602 | 0.89054584 |
| Parg | 0.30199535 | 0.25203992 | 0.31120655 | -0.0591666 | 0.81212616 | 0.89054584 |
| Adprh | 0.82847683 | 0.67765256 | 0.73682359 | -0.059171 | 0.6458495 | 0.89054584 |
| Gkap1 | 0.38341726 | 0.32896631 | 0.38813788 | -0.0591716 | 0.76122696 | 0.89054584 |
| Hmces | 0.38033598 | 0.32608585 | 0.38529093 | -0.0592051 | 0.76297604 | 0.89054584 |
| Med21 | 0.91768309 | 0.73090823 | 0.7901527 | -0.0592445 | 0.63657562 | 0.89054584 |
| Pnn | 1.4860522 | 0.97243472 | 1.03170253 | -0.0592678 | 0.60551192 | 0.89054584 |
| Bri3bp | 0.43140788 | 0.37247151 | 0.43179715 | -0.0593256 | 0.73894721 | 0.89054584 |
| Map1lc3a | 1.36065123 | 0.93295877 | 0.9923791 | -0.0594203 | 0.6098629 | 0.89054584 |
| Rrp1 | 1.59315938 | 0.99977749 | 1.05920047 | -0.059423 | 0.60309821 | 0.89054584 |

| | | | | | | |
|---|---|---|---|---|---|---|
| Med7 | 0.53279538 | 0.46014646 | 0.51957516 | -0.0594287 | 0.70294409 | 0.89054584 |
| Cog8 | 0.57661308 | 0.49608717 | 0.55552612 | -0.059439 | 0.6909296 | 0.89054584 |
| Dusp12 | 0.27059525 | 0.22119268 | 0.28063819 | -0.0594455 | 0.83814658 | 0.89497393 |
| Mfap1b | 0.42241356 | 0.36424054 | 0.42371405 | -0.0594735 | 0.74342478 | 0.89054584 |
| Herc1 | 0.51139004 | 0.44208936 | 0.50156744 | -0.0594781 | 0.70968948 | 0.89054584 |
| Nabp2 | 0.73631602 | 0.61615504 | 0.67566496 | -0.0595099 | 0.6592222 | 0.89054584 |
| Tmem107 | 0.60169045 | 0.51602729 | 0.57553824 | -0.059511 | 0.6850036 | 0.89054584 |
| Vipas39 | 0.26991193 | 0.2203819 | 0.27996767 | -0.0595858 | 0.83929548 | 0.89504222 |
| Zfp280d | 0.5183353 | 0.44784333 | 0.50744127 | -0.0595979 | 0.70787355 | 0.89054584 |
| Mpc1 | 1.18191631 | 0.86302462 | 0.92264304 | -0.0596184 | 0.61831059 | 0.89054584 |
| Snrpb2 | 1.2068426 | 0.87378614 | 0.93341683 | -0.0596307 | 0.61700845 | 0.89054584 |
| Dhx15 | 0.72889268 | 0.61083689 | 0.67047333 | -0.0596364 | 0.6607086 | 0.89054584 |
| Lsm14a | 0.61026329 | 0.52262807 | 0.58228448 | -0.0596564 | 0.68343268 | 0.89054584 |
| Ahcyl2 | 0.31022786 | 0.25942211 | 0.31914013 | -0.059718 | 0.80833814 | 0.89054584 |
| Ndufs1 | 0.77768424 | 0.64406032 | 0.70387533 | -0.059815 | 0.65391604 | 0.89054584 |
| Sdccag3 | 0.47711039 | 0.41229676 | 0.4721365 | -0.0598397 | 0.72256743 | 0.89054584 |
| Ppm1d | 0.31404021 | 0.26292527 | 0.32280319 | -0.0598779 | 0.80626198 | 0.89054584 |
| Fam173b | 0.31078444 | 0.25974713 | 0.31967538 | -0.0599282 | 0.8087755 | 0.89054584 |
| Hdac3 | 0.75542505 | 0.6287907 | 0.68884839 | -0.0600577 | 0.65769954 | 0.89054584 |
| Hipk1 | 0.33775508 | 0.28536087 | 0.34542049 | -0.0600596 | 0.79104959 | 0.89054584 |
| Rnpep | 0.60187039 | 0.51557921 | 0.57568036 | -0.0601012 | 0.68665624 | 0.89054584 |
| Leng8 | 0.29978992 | 0.24894031 | 0.30907537 | -0.0601351 | 0.81765484 | 0.89054584 |
| 2810004N23 | 0.65314954 | 0.55510853 | 0.61528555 | -0.060177 | 0.67568286 | 0.89054584 |
| Ing2 | 0.58599885 | 0.50284172 | 0.56306527 | -0.0602235 | 0.69086528 | 0.89054584 |
| Mrps18a | 1.14202973 | 0.84454136 | 0.90485302 | -0.0603117 | 0.62192291 | 0.89054584 |
| Tspyl2 | 0.34050191 | 0.28767966 | 0.34802172 | -0.0603421 | 0.79039563 | 0.89054584 |
| Ctdsp2 | 0.46037772 | 0.39715752 | 0.45751109 | -0.0603536 | 0.73058868 | 0.89054584 |
| Rbbp4 | 1.56281626 | 0.99151723 | 1.05194358 | -0.0604263 | 0.60550468 | 0.89054584 |
| Snhg20 | 0.37274097 | 0.31774476 | 0.37825093 | -0.0605062 | 0.77194225 | 0.89054584 |
| Apool | 0.34094576 | 0.2879062 | 0.34844166 | -0.0605355 | 0.7908595 | 0.89054584 |

| | | | | | | |
|---|---|---|---|---|---|---|
| Pspc1 | 0.35515778 | 0.30124696 | 0.3618326 | -0.0605856 | 0.78233344 | 0.89054584 |
| Ndufaf7 | 0.32350673 | 0.27125436 | 0.33186661 | -0.0606123 | 0.8025906 | 0.89054584 |
| Copa | 1.16252893 | 0.85337786 | 0.91407594 | -0.0606981 | 0.62148041 | 0.89054584 |
| Edf1 | 2.51087123 | 1.06577862 | 1.12648628 | -0.0607077 | 0.59912224 | 0.89054584 |
| Pgm2 | 0.46248362 | 0.39860619 | 0.45936084 | -0.0607547 | 0.73111457 | 0.89054584 |
| Usp48 | 0.46614592 | 0.4017812 | 0.46257128 | -0.0607901 | 0.72981963 | 0.89054584 |
| Snapin | 0.43685259 | 0.37587075 | 0.43666712 | -0.0607964 | 0.74171959 | 0.89054584 |
| Chid1 | 0.58958686 | 0.50513249 | 0.56593181 | -0.0607993 | 0.69164568 | 0.89054584 |
| Arf5 | 3.33188848 | 0.99697674 | 1.05788224 | -0.0609055 | 0.60573863 | 0.89054584 |
| Abhd8 | 0.41713186 | 0.35803579 | 0.41894539 | -0.0609096 | 0.75090927 | 0.89054584 |
| Bre | 0.56438198 | 0.48453582 | 0.54561598 | -0.0610802 | 0.69900999 | 0.89054584 |
| Mcee | 0.70515174 | 0.59248047 | 0.65360746 | -0.061127 | 0.66847851 | 0.89054584 |
| Mkl2 | 0.27627635 | 0.22503253 | 0.28620408 | -0.0611716 | 0.84032011 | 0.89576177 |
| Safb | 0.95538882 | 0.74981066 | 0.81100321 | -0.0611925 | 0.63740485 | 0.89054584 |
| Tm9sf3 | 1.41529957 | 0.94919258 | 1.01053238 | -0.0613398 | 0.61133492 | 0.89054584 |
| Nmd3 | 0.56026545 | 0.48090595 | 0.54225867 | -0.0613527 | 0.70094701 | 0.89054584 |
| Rilpl2 | 0.36142022 | 0.3063219 | 0.3676986 | -0.0613767 | 0.78162001 | 0.89054584 |
| Wdr74 | 0.43488467 | 0.37349186 | 0.43490896 | -0.0614171 | 0.7447077 | 0.89054584 |
| Farsa | 0.75061 | 0.62408448 | 0.68555125 | -0.0614668 | 0.66192493 | 0.89054584 |
| Capn10 | 0.46179668 | 0.39727987 | 0.45875775 | -0.0614779 | 0.73379856 | 0.89054584 |
| Hiatl1 | 0.31382841 | 0.26107615 | 0.32259988 | -0.0615237 | 0.81286863 | 0.89054584 |
| Hnrnpdl | 1.4757664 | 0.967255 | 1.02879774 | -0.0615427 | 0.60976175 | 0.89054584 |
| Gtpbp4 | 0.6611232 | 0.5597218 | 0.62128372 | -0.0615619 | 0.67783254 | 0.89054584 |
| Nars | 0.82258762 | 0.67149816 | 0.73309943 | -0.0616013 | 0.65228024 | 0.89054584 |
| Nudt14 | 0.62481879 | 0.53194668 | 0.59362433 | -0.0616777 | 0.68583001 | 0.89054584 |
| Zfp871 | 0.28172938 | 0.22979874 | 0.29153172 | -0.061733 | 0.83806812 | 0.89497393 |
| Dennd4c | 0.31556185 | 0.26244448 | 0.32426315 | -0.0618187 | 0.8127826 | 0.89054584 |
| Mphosph8 | 0.70694608 | 0.59306506 | 0.65489605 | -0.061831 | 0.66998317 | 0.89054584 |
| Ywhag | 0.97160984 | 0.75783271 | 0.81967625 | -0.0618435 | 0.63739759 | 0.89054584 |
| Lrrc8d | 0.22213963 | 0.17065821 | 0.23255235 | -0.0618941 | 0.89250931 | 0.91822311 |

| | | | | | | |
|---|---|---|---|---|---|---|
| Zmpste24 | 0.56481281 | 0.48405283 | 0.54596671 | -0.0619139 | 0.70136646 | 0.89054584 |
| Sdhc | 1.04998193 | 0.79820472 | 0.86023271 | -0.062028 | 0.63154116 | 0.89054584 |
| Zc3h13 | 0.69452652 | 0.58390046 | 0.64593111 | -0.0620307 | 0.6727111 | 0.89054584 |
| 1810030O07 | 0.32288151 | 0.26921523 | 0.33126946 | -0.0620542 | 0.80859338 | 0.89054584 |
| Pgpep1 | 0.29149627 | 0.23890826 | 0.30103905 | -0.0621308 | 0.83186181 | 0.8933762 |
| Slc50a1 | 0.66436641 | 0.56156256 | 0.62371079 | -0.0621482 | 0.67875835 | 0.89054584 |
| Nptn | 1.05031981 | 0.798248 | 0.86040444 | -0.0621564 | 0.63177809 | 0.89054584 |
| Lamtor3 | 0.6250178 | 0.53151315 | 0.59377837 | -0.0622652 | 0.68741961 | 0.89054584 |
| Pef1 | 0.45770211 | 0.39288461 | 0.45515703 | -0.0622724 | 0.73807245 | 0.89054584 |
| Krit1 | 0.9607277 | 0.75156997 | 0.81387789 | -0.0623079 | 0.63933201 | 0.89054584 |
| Nans | 0.85207043 | 0.68914422 | 0.75149037 | -0.0623461 | 0.65044468 | 0.89054584 |
| Impa1 | 0.7392148 | 0.61529801 | 0.67768175 | -0.0623837 | 0.66599367 | 0.89054584 |
| Dnajb6 | 1.80878698 | 1.03652873 | 1.09897676 | -0.062448 | 0.60439463 | 0.89054584 |
| Mta1 | 0.32375302 | 0.26957585 | 0.3321018 | -0.062526 | 0.80979875 | 0.89054584 |
| Drg2 | 0.5075901 | 0.43573601 | 0.49834076 | -0.0626048 | 0.7208223 | 0.89054584 |
| Fbxo22 | 0.61539936 | 0.52368645 | 0.58630235 | -0.0626159 | 0.69060483 | 0.89054584 |
| Tvp23b | 1.1585146 | 0.84961502 | 0.91227144 | -0.0626564 | 0.6255099 | 0.89054584 |
| Apoa1bp | 1.17405598 | 0.85653859 | 0.91919911 | -0.0626605 | 0.62460334 | 0.89054584 |
| Snapc5 | 0.53412222 | 0.4579345 | 0.52068189 | -0.0627474 | 0.71275431 | 0.89054584 |
| Sde2 | 0.68437706 | 0.57576616 | 0.63852392 | -0.0627578 | 0.67649045 | 0.89054584 |
| Commd8 | 0.73744187 | 0.6136813 | 0.67644898 | -0.0627677 | 0.6672394 | 0.89054584 |
| Mycbp2 | 1.03669824 | 0.79064977 | 0.85341972 | -0.06277 | 0.63407671 | 0.89054584 |
| Abhd10 | 0.35718765 | 0.30092159 | 0.36373631 | -0.0628147 | 0.78946509 | 0.89054584 |
| Ubtf | 0.66732868 | 0.56309679 | 0.6259212 | -0.0628244 | 0.67997295 | 0.89054584 |
| Letm1 | 0.47383565 | 0.40637182 | 0.46928723 | -0.0629154 | 0.73388486 | 0.89054584 |
| Rabgap1l | 0.29610254 | 0.24254099 | 0.30550667 | -0.0629657 | 0.83152756 | 0.89332987 |
| Xrcc6 | 0.36699395 | 0.30992699 | 0.37290266 | -0.0629757 | 0.78430063 | 0.89054584 |
| Atat1 | 0.48212787 | 0.41349929 | 0.47648928 | -0.06299 | 0.73101761 | 0.89054584 |
| Haghl | 0.41671614 | 0.35548455 | 0.41856937 | -0.0630848 | 0.75871463 | 0.89054584 |
| Clns1a | 0.81113228 | 0.66261737 | 0.72578265 | -0.0631653 | 0.65743409 | 0.89054584 |

| | | | | | | |
|---|---|---|---|---|---|---|
| Gns | 0.40848316 | 0.34790477 | 0.41110368 | -0.0631989 | 0.76307174 | 0.89054584 |
| Zzz3 | 0.49255781 | 0.42224592 | 0.48548761 | -0.0632417 | 0.72803921 | 0.89054584 |
| Zcchc17 | 0.76745307 | 0.63377082 | 0.69701272 | -0.0632419 | 0.66374974 | 0.89054584 |
| Atpaf2 | 0.27550535 | 0.22214351 | 0.28544964 | -0.0633061 | 0.84925545 | 0.89883846 |
| Ap4s1 | 0.29984508 | 0.24572491 | 0.3091287 | -0.0634038 | 0.8303556 | 0.89271879 |
| Pcmt1 | 1.21332804 | 0.87273556 | 0.93615916 | -0.0634236 | 0.62386111 | 0.89054584 |
| Ebna1bp2 | 1.00436214 | 0.77332204 | 0.83677562 | -0.0634536 | 0.63806538 | 0.89054584 |
| Acyp1 | 0.81105566 | 0.66211773 | 0.7257334 | -0.0636157 | 0.658506 | 0.89054584 |
| Csnk1g2 | 0.32019819 | 0.26507061 | 0.32870423 | -0.0636336 | 0.8164506 | 0.89054584 |
| Nxt1 | 0.39464714 | 0.33482582 | 0.39847019 | -0.0636444 | 0.77160244 | 0.89054584 |
| Exosc4 | 0.8850322 | 0.70765666 | 0.77130205 | -0.0636454 | 0.64967419 | 0.89054584 |
| Myl12b | 1.75579075 | 1.02747542 | 1.09112225 | -0.0636468 | 0.60709795 | 0.89054584 |
| Fbxo9 | 0.42708696 | 0.36422994 | 0.42791991 | -0.06369 | 0.75597669 | 0.89054584 |
| Atf6b | 0.78718276 | 0.6464794 | 0.71017874 | -0.0636993 | 0.66198359 | 0.89054584 |
| Leprotl1 | 0.59604042 | 0.50733704 | 0.57106598 | -0.0637289 | 0.69844078 | 0.89054584 |
| Mboat7 | 0.42272488 | 0.36020784 | 0.42399462 | -0.0637868 | 0.75831803 | 0.89054584 |
| 2010107E04I | 2.30921417 | 1.06520127 | 1.12920347 | -0.0640022 | 0.60413488 | 0.89054584 |
| Ttc5 | 0.62578184 | 0.53033833 | 0.59436952 | -0.0640312 | 0.69213313 | 0.89054584 |
| Tpm3 | 1.44077746 | 0.95443352 | 1.01847707 | -0.0640435 | 0.61522441 | 0.89054584 |
| Cdipt | 0.68137156 | 0.57227116 | 0.63631654 | -0.0640454 | 0.68043778 | 0.89054584 |
| Rpl37 | 5.08150079 | 0.62127459 | 0.68536694 | -0.0640923 | 0.6684665 | 0.89054584 |
| Ankrd11 | 0.89456798 | 0.7126698 | 0.77688665 | -0.0642168 | 0.64991998 | 0.89054584 |
| Calm2 | 3.88839998 | 0.92114843 | 0.98546889 | -0.0643205 | 0.61946803 | 0.89054584 |
| Gapvd1 | 0.61916696 | 0.52491176 | 0.58923827 | -0.0643265 | 0.69448776 | 0.89054584 |
| 2700060E02I | 2.18224201 | 1.06246409 | 1.12679472 | -0.0643306 | 0.60487535 | 0.89054584 |
| Aebp2 | 0.33509769 | 0.27856169 | 0.34290043 | -0.0643387 | 0.8089849 | 0.89054584 |
| Tonsl | 1.38786057 | 0.93720765 | 1.0015842 | -0.0643766 | 0.61770247 | 0.89054584 |
| Atpif1 | 3.46669868 | 0.97444451 | 1.03884424 | -0.0643997 | 0.6136388 | 0.89054584 |
| Pyroxd1 | 0.31907959 | 0.26312615 | 0.32763376 | -0.0645076 | 0.82050624 | 0.89054584 |
| Micu2 | 0.59213587 | 0.50343965 | 0.56796303 | -0.0645234 | 0.70169388 | 0.89054584 |

| | | | | | | |
|---|---|---|---|---|---|---|
| Gm16286 | 0.91291064 | 0.72287367 | 0.78744572 | -0.0645721 | 0.64878493 | 0.89054584 |
| Cstf2t | 0.37537835 | 0.31612214 | 0.38069922 | -0.0645771 | 0.78531927 | 0.89054584 |
| Setd2 | 0.60995932 | 0.51744894 | 0.58204613 | -0.0645972 | 0.69744392 | 0.89054584 |
| Bod1 | 0.3082671 | 0.25261312 | 0.31725367 | -0.0646405 | 0.82875603 | 0.89253491 |
| Rpl7a | 3.73966051 | 0.94151551 | 1.0061885 | -0.064673 | 0.61770336 | 0.89054584 |
| Mllt4 | 0.42761042 | 0.36363846 | 0.4283902 | -0.0647517 | 0.75934748 | 0.89054584 |
| Nsf | 0.4433268 | 0.37757983 | 0.44243553 | -0.0648557 | 0.75267902 | 0.89054584 |
| Pex14 | 0.42981303 | 0.36550227 | 0.43036735 | -0.0648651 | 0.75872179 | 0.89054584 |
| Ppp2r2a | 0.80306591 | 0.65563991 | 0.7205739 | -0.064934 | 0.66269776 | 0.89054584 |
| Nr2c2ap | 0.34822637 | 0.29034071 | 0.35531519 | -0.0649745 | 0.80288027 | 0.89054584 |
| Urm1 | 0.47512646 | 0.4054244 | 0.47041112 | -0.0649867 | 0.74009133 | 0.89054584 |
| Tob1 | 0.56846722 | 0.48389637 | 0.54893673 | -0.0650404 | 0.70952965 | 0.89054584 |
| Commd4 | 0.58762977 | 0.49931484 | 0.56436931 | -0.0650545 | 0.70437762 | 0.89054584 |
| Trim8 | 0.42161604 | 0.35791848 | 0.42299504 | -0.0650766 | 0.76324032 | 0.89054584 |
| Usp22 | 0.34204983 | 0.28438336 | 0.34948583 | -0.0651025 | 0.80726748 | 0.89054584 |
| Exosc10 | 0.33291388 | 0.27570566 | 0.340827 | -0.0651213 | 0.81331466 | 0.89054584 |
| Srek1 | 0.83798725 | 0.67765481 | 0.74278399 | -0.0651292 | 0.65854954 | 0.89054584 |
| Sdhaf1 | 0.53364192 | 0.45514595 | 0.5202814 | -0.0651355 | 0.72014273 | 0.89054584 |
| Pomgnt2 | 0.34455625 | 0.2866658 | 0.35185383 | -0.065188 | 0.80597605 | 0.89054584 |
| Nup88 | 0.49107104 | 0.41895836 | 0.48420905 | -0.0652507 | 0.73494109 | 0.89054584 |
| Ndufs2 | 1.58912165 | 0.99296595 | 1.05825673 | -0.0652908 | 0.61311249 | 0.89054584 |
| Coq6 | 0.2796549 | 0.22420689 | 0.28950667 | -0.0652998 | 0.85333605 | 0.90008483 |
| Rnf7 | 1.80959056 | 1.03377941 | 1.09908642 | -0.065307 | 0.60904518 | 0.89054584 |
| Pgrmc2 | 0.40330936 | 0.34104021 | 0.40639236 | -0.0653522 | 0.77313364 | 0.89054584 |
| Cnot1 | 0.42870465 | 0.36400682 | 0.42937279 | -0.065366 | 0.76091788 | 0.89054584 |
| Zcchc6 | 0.41626362 | 0.35267887 | 0.41815994 | -0.0654811 | 0.7671702 | 0.89054584 |
| Znhit3 | 0.36616547 | 0.30664121 | 0.37213016 | -0.065489 | 0.79386318 | 0.89054584 |
| Zcchc9 | 0.72509656 | 0.60225688 | 0.66780352 | -0.0655466 | 0.67626764 | 0.89054584 |
| Aars | 0.82267373 | 0.66755968 | 0.73315407 | -0.0655944 | 0.66159852 | 0.89054584 |
| Nipsnap1 | 0.51532627 | 0.43923286 | 0.50490021 | -0.0656673 | 0.7277234 | 0.89054584 |

| | | | | | | |
|---|---|---|---|---|---|---|
| Rps13 | 4.81406088 | 0.69026033 | 0.75593784 | -0.0656775 | 0.65717996 | 0.89054584 |
| Ndfip2 | 1.09309418 | 0.81610847 | 0.88182167 | -0.0657132 | 0.63577327 | 0.89054584 |
| Senp7 | 0.38080315 | 0.31995341 | 0.38572291 | -0.0657695 | 0.78651716 | 0.89054584 |
| Mrpl18 | 1.2580245 | 0.88862088 | 0.95439079 | -0.0657699 | 0.62591813 | 0.89054584 |
| Bud31 | 1.45317523 | 0.95638092 | 1.02222434 | -0.0658434 | 0.61794624 | 0.89054584 |
| Ankra2 | 0.4721207 | 0.40191581 | 0.46779246 | -0.0658767 | 0.74411794 | 0.89054584 |
| Tmem101 | 0.49315312 | 0.42002452 | 0.48599916 | -0.0659746 | 0.73645478 | 0.89054584 |
| Echdc2 | 0.38993278 | 0.32814554 | 0.3941407 | -0.0659952 | 0.78233282 | 0.89054584 |
| Os9 | 1.08044777 | 0.80960222 | 0.87560581 | -0.0660036 | 0.63727831 | 0.89054584 |
| Tor1a | 0.60142668 | 0.50931649 | 0.57532989 | -0.0660134 | 0.70352546 | 0.89054584 |
| Ctso | 0.52992284 | 0.45113941 | 0.51717527 | -0.0660359 | 0.72403815 | 0.89054584 |
| Tcaf1 | 0.67607315 | 0.56636951 | 0.63241043 | -0.0660409 | 0.68670097 | 0.89054584 |
| Uxt | 0.32469237 | 0.26692406 | 0.3329985 | -0.0660744 | 0.82237118 | 0.89073269 |
| Dynlt3 | 0.63576987 | 0.53586103 | 0.60206254 | -0.0662015 | 0.69577499 | 0.89054584 |
| Fam3c | 0.59853369 | 0.50675345 | 0.57304201 | -0.0662886 | 0.70503541 | 0.89054584 |
| Scaper | 0.39499358 | 0.3324973 | 0.39878786 | -0.0662906 | 0.78066952 | 0.89054584 |
| Dusp11 | 0.6176143 | 0.52173419 | 0.58802953 | -0.0662953 | 0.70029741 | 0.89054584 |
| Glrx3 | 1.72999262 | 1.02052678 | 1.08683074 | -0.066304 | 0.61188165 | 0.89054584 |
| Slain2 | 0.35128813 | 0.29175777 | 0.35819727 | -0.0664395 | 0.80624846 | 0.89054584 |
| Kmt2c | 0.57667559 | 0.48911086 | 0.55557653 | -0.0664657 | 0.71136142 | 0.89054584 |
| Gtf2e2 | 1.34714093 | 0.92122886 | 0.9877037 | -0.0664748 | 0.62307161 | 0.89054584 |
| Afg3l2 | 0.37019735 | 0.30933941 | 0.37588604 | -0.0665466 | 0.79526201 | 0.89054584 |
| Fam104a | 1.55306655 | 0.98296885 | 1.0495449 | -0.066576 | 0.61620617 | 0.89054584 |
| Akirin1 | 0.82752237 | 0.66960075 | 0.73622175 | -0.066621 | 0.66333765 | 0.89054584 |
| Arhgdia | 1.69594669 | 1.01402295 | 1.08069506 | -0.0666721 | 0.61310749 | 0.89054584 |
| Rpl18a | 5.34742386 | 0.60186353 | 0.6685615 | -0.066698 | 0.67895972 | 0.89054584 |
| Dicer1 | 0.45074809 | 0.38231651 | 0.44901853 | -0.066702 | 0.75555928 | 0.89054584 |
| Rnf126 | 0.35199225 | 0.29196342 | 0.35885936 | -0.0668959 | 0.80743577 | 0.89054584 |
| Qtrt1 | 0.40327159 | 0.3394182 | 0.40635792 | -0.0669397 | 0.77859945 | 0.89054584 |
| Ccdc127 | 0.76694221 | 0.62971598 | 0.69666807 | -0.0669521 | 0.67282942 | 0.89054584 |

| | | | | | | |
|---|---|---|---|---|---|---|
| Lyrm9 | 0.40373791 | 0.33976448 | 0.4067832 | -0.0670187 | 0.77862987 | 0.89054584 |
| Tcf12 | 0.70531551 | 0.58668228 | 0.65372515 | -0.0670429 | 0.68360313 | 0.89054584 |
| Naa10 | 0.61372881 | 0.51785482 | 0.58499747 | -0.0671427 | 0.70358051 | 0.89054584 |
| Setd8 | 0.32423465 | 0.26534397 | 0.33256162 | -0.0672176 | 0.826819 | 0.89184004 |
| Csnk1g3 | 0.41190766 | 0.34699256 | 0.4142136 | -0.067221 | 0.77519957 | 0.89054584 |
| 4833420G17 | 0.45709743 | 0.38731641 | 0.45462443 | -0.067308 | 0.75482907 | 0.89054584 |
| Pim3 | 0.42811237 | 0.36147913 | 0.42884102 | -0.0673619 | 0.76786125 | 0.89054584 |
| Gm8186 | 0.76779949 | 0.6298762 | 0.69724633 | -0.0673701 | 0.67370301 | 0.89054584 |
| Nipbl | 0.93212844 | 0.73085674 | 0.79825596 | -0.0673992 | 0.65297645 | 0.89054584 |
| Sirt7 | 0.33606614 | 0.27637816 | 0.34381913 | -0.067441 | 0.81960555 | 0.89054584 |
| Arfgap1 | 0.48120465 | 0.40824049 | 0.47568953 | -0.067449 | 0.74556596 | 0.89054584 |
| Med13 | 0.35693048 | 0.29604412 | 0.36349524 | -0.0674511 | 0.8063503 | 0.89054584 |
| Slc25a17 | 0.91075776 | 0.71875981 | 0.78621924 | -0.0674594 | 0.65532892 | 0.89054584 |
| Zfhx3 | 0.47156946 | 0.39984861 | 0.46731161 | -0.067463 | 0.74939391 | 0.89054584 |
| Scamp2 | 0.59346822 | 0.50154278 | 0.569023 | -0.0674802 | 0.70969779 | 0.89054584 |
| U2af1 | 1.7976784 | 1.0299059 | 1.09742733 | -0.0675214 | 0.61281012 | 0.89054584 |
| Tex10 | 0.32806388 | 0.26864431 | 0.33621313 | -0.0675688 | 0.82544276 | 0.89170439 |
| R3hdm4 | 0.89144851 | 0.70736166 | 0.77506625 | -0.0677046 | 0.65798641 | 0.89054584 |
| Ccdc47 | 0.9067775 | 0.71622115 | 0.78394272 | -0.0677216 | 0.6563297 | 0.89054584 |
| Mau2 | 0.42351145 | 0.35696371 | 0.42470325 | -0.0677395 | 0.77128767 | 0.89054584 |
| Rnf10 | 1.4871283 | 0.96422343 | 1.0320044 | -0.067781 | 0.62019554 | 0.89054584 |
| Shfm1 | 3.36866614 | 0.98503385 | 1.05283956 | -0.0678057 | 0.61792988 | 0.89054584 |
| Dimt1 | 0.35634731 | 0.29512447 | 0.36294846 | -0.067824 | 0.80801751 | 0.89054584 |
| Hnrnpa0 | 0.51998533 | 0.44096338 | 0.50883224 | -0.0678689 | 0.73283736 | 0.89054584 |
| Zc3h7b | 0.3137688 | 0.25467341 | 0.32254265 | -0.0678692 | 0.8365388 | 0.89462889 |
| Rab10 | 1.32473991 | 0.91177304 | 0.97970443 | -0.0679314 | 0.62666932 | 0.89054584 |
| Tnfaip1 | 0.70332485 | 0.58432576 | 0.65229329 | -0.0679675 | 0.68632192 | 0.89054584 |
| Hras | 0.45668609 | 0.38619597 | 0.45426198 | -0.068066 | 0.75743941 | 0.89054584 |
| Suco | 0.3015836 | 0.24272353 | 0.31080885 | -0.0680853 | 0.84627344 | 0.89825055 |
| Psmd4 | 1.81446917 | 1.03159742 | 1.09974627 | -0.0681489 | 0.61359617 | 0.89054584 |

| | | | | | | |
|---|---|---|---|---|---|---|
| Brd7 | 0.68514845 | 0.57090469 | 0.63908945 | -0.0681848 | 0.69042828 | 0.89054584 |
| Araf | 0.80554138 | 0.65392981 | 0.72217745 | -0.0682476 | 0.67013357 | 0.89054584 |
| Mar-08 | 0.30920409 | 0.24989929 | 0.31815538 | -0.0682561 | 0.84122737 | 0.89618006 |
| Usp7 | 0.40536304 | 0.34000202 | 0.40826431 | -0.0682623 | 0.78200445 | 0.89054584 |
| Atp5j2 | 3.20999936 | 1.00588426 | 1.07422641 | -0.0683421 | 0.61653816 | 0.89054584 |
| Mettl9 | 1.35594575 | 0.92238851 | 0.9907631 | -0.0683746 | 0.626093 | 0.89054584 |
| Snrpg | 2.24628274 | 1.05989022 | 1.12833325 | -0.068443 | 0.61125997 | 0.89054584 |
| Ylpm1 | 0.47780966 | 0.40422793 | 0.47274406 | -0.0685161 | 0.75023628 | 0.89054584 |
| Elavl1 | 1.31336445 | 0.90701619 | 0.97554525 | -0.0685291 | 0.62827055 | 0.89054584 |
| Eif4h | 1.60350129 | 0.99303477 | 1.06158014 | -0.0685454 | 0.61822567 | 0.89054584 |
| Isca2 | 0.5841107 | 0.49299717 | 0.56155333 | -0.0685562 | 0.71524656 | 0.89054584 |
| Preb | 0.60232053 | 0.5074758 | 0.57603577 | -0.06856 | 0.71038464 | 0.89054584 |
| Spns1 | 0.35119931 | 0.28949435 | 0.35811374 | -0.0686194 | 0.81399525 | 0.89054584 |
| Usp34 | 0.96697004 | 0.74856185 | 0.8172126 | -0.0686507 | 0.65224142 | 0.89054584 |
| Wbp1 | 0.79706303 | 0.64796884 | 0.71666677 | -0.0686979 | 0.67241615 | 0.89054584 |
| Gga1 | 0.3793327 | 0.31560672 | 0.38436281 | -0.0687561 | 0.79769232 | 0.89054584 |
| Ipo8 | 0.26251529 | 0.20380745 | 0.27269519 | -0.0688877 | 0.88038884 | 0.91159931 |
| Glrx5 | 1.43060909 | 0.9464274 | 1.01533913 | -0.0689117 | 0.6240767 | 0.89054584 |
| Ddx39b | 2.00971232 | 1.04949773 | 1.11842162 | -0.0689239 | 0.61298674 | 0.89054584 |
| Wdr1 | 0.82305279 | 0.66422195 | 0.73339451 | -0.0691726 | 0.66982271 | 0.89054584 |
| Cfdp1 | 1.53863783 | 0.97661279 | 1.04590382 | -0.069291 | 0.62120778 | 0.89054584 |
| Tmx3 | 0.36630867 | 0.30285735 | 0.37226372 | -0.0694064 | 0.80747577 | 0.89054584 |
| Snx14 | 0.35961781 | 0.29660311 | 0.36601248 | -0.0694094 | 0.81151848 | 0.89054584 |
| Smarca4 | 0.95171439 | 0.73959358 | 0.80901262 | -0.069419 | 0.65533713 | 0.89054584 |
| Ptrh2 | 0.43998766 | 0.36999791 | 0.43946328 | -0.0694654 | 0.76926627 | 0.89054584 |
| Thoc6 | 0.31905047 | 0.25810721 | 0.32760588 | -0.0694987 | 0.8385101 | 0.89497393 |
| H2afy2 | 0.80579402 | 0.65276364 | 0.72234086 | -0.0695772 | 0.67319685 | 0.89054584 |
| Ubr5 | 0.29046828 | 0.23045777 | 0.30004056 | -0.0695828 | 0.86003153 | 0.90134771 |
| Hsd17b4 | 0.46257022 | 0.38985183 | 0.45943686 | -0.069585 | 0.75979095 | 0.89054584 |
| Taf6 | 0.46616038 | 0.39294082 | 0.46258394 | -0.0696431 | 0.7584779 | 0.89054584 |

| | | | | | | |
|---|---|---|---|---|---|---|
| Mprip | 0.52072249 | 0.43965107 | 0.50945313 | -0.0698021 | 0.7383734 | 0.89054584 |
| Seh1l | 0.5354906 | 0.45198816 | 0.52182207 | -0.0698339 | 0.7335195 | 0.89054584 |
| Uhrf2 | 0.73330478 | 0.6037102 | 0.67356354 | -0.0698533 | 0.68551085 | 0.89054584 |
| Mef2a | 0.66803236 | 0.55658263 | 0.62644538 | -0.0698627 | 0.69832069 | 0.89054584 |
| Bphl | 0.61886506 | 0.51913822 | 0.58900337 | -0.0698651 | 0.70973967 | 0.89054584 |
| Ddx17 | 1.03949935 | 0.78498936 | 0.85486638 | -0.069877 | 0.64833277 | 0.89054584 |
| Syap1 | 0.58536027 | 0.49262257 | 0.56255419 | -0.0699316 | 0.71875846 | 0.89054584 |
| Rab11b | 1.30300035 | 0.90174045 | 0.97169452 | -0.0699541 | 0.63134048 | 0.89054584 |
| Hnrnpll | 0.64834074 | 0.54168223 | 0.61164777 | -0.0699655 | 0.70295893 | 0.89054584 |
| Dnajc2 | 0.64144615 | 0.5364007 | 0.60640418 | -0.0700035 | 0.70465266 | 0.89054584 |
| Psmb6 | 2.50088581 | 1.05683281 | 1.12684176 | -0.070009 | 0.61388258 | 0.89054584 |
| Ndufa5 | 1.82607777 | 1.03126475 | 1.1012762 | -0.0700114 | 0.61645481 | 0.89054584 |
| Zfp622 | 0.65891735 | 0.54960178 | 0.61962878 | -0.070027 | 0.70073915 | 0.89054584 |
| Asf1a | 0.66180122 | 0.5517273 | 0.62179173 | -0.0700644 | 0.70020025 | 0.89054584 |
| Mpg | 0.39383223 | 0.32765714 | 0.39772271 | -0.0700656 | 0.79410598 | 0.89054584 |
| Rbck1 | 0.49966647 | 0.42148461 | 0.4915832 | -0.0700986 | 0.7467681 | 0.89054584 |
| Arl2 | 0.70864154 | 0.58599038 | 0.65611169 | -0.0701213 | 0.69072974 | 0.89054584 |
| Fam96b | 0.97673264 | 0.75220568 | 0.82239187 | -0.0701862 | 0.65454208 | 0.89054584 |
| BC005624 | 0.78529576 | 0.63865455 | 0.70893153 | -0.070277 | 0.67790367 | 0.89054584 |
| Psmg1 | 0.55473667 | 0.4674445 | 0.53773205 | -0.0702876 | 0.72872962 | 0.89054584 |
| mt-Nd5 | 1.74540315 | 1.01903834 | 1.08943018 | -0.0703918 | 0.61830429 | 0.89054584 |
| Arfrp1 | 0.42602759 | 0.35642032 | 0.42696764 | -0.0705473 | 0.77928803 | 0.89054584 |
| Lsm10 | 0.45479969 | 0.38191491 | 0.45259853 | -0.0706836 | 0.76657086 | 0.89054584 |
| Cux1 | 0.552192 | 0.46491275 | 0.53564191 | -0.0707292 | 0.73078625 | 0.89054584 |
| Zfand2b | 0.48651046 | 0.40945991 | 0.48027855 | -0.0708186 | 0.75395109 | 0.89054584 |
| Gcc2 | 0.74256083 | 0.60910677 | 0.68000221 | -0.0708954 | 0.6864208 | 0.89054584 |
| Golga1 | 0.29644377 | 0.23491297 | 0.3058372 | -0.0709242 | 0.86002181 | 0.90134771 |
| Ythdf2 | 0.71745055 | 0.59146551 | 0.6623948 | -0.0709293 | 0.69106796 | 0.89054584 |
| Lzts2 | 0.74090243 | 0.60789297 | 0.67885312 | -0.0709602 | 0.68686963 | 0.89054584 |
| Xbp1 | 1.42453278 | 0.94243673 | 1.0134425 | -0.0710058 | 0.62794812 | 0.89054584 |

| | | | | | | |
|---|---|---|---|---|---|---|
| Ankhd1 | 0.40483352 | 0.33674971 | 0.40778188 | -0.0710322 | 0.79147457 | 0.89054584 |
| Wsb2 | 0.4102298 | 0.3416325 | 0.41269072 | -0.0710582 | 0.78877825 | 0.89054584 |
| Vps41 | 0.50286413 | 0.42322799 | 0.49431504 | -0.071087 | 0.74857066 | 0.89054584 |
| Rnf214 | 0.41591454 | 0.34666925 | 0.41784402 | -0.0711748 | 0.78628447 | 0.89054584 |
| Eif2b1 | 0.55320161 | 0.46527094 | 0.53647169 | -0.0712008 | 0.73182328 | 0.89054584 |
| Gbas | 0.58839677 | 0.49375657 | 0.56498197 | -0.0712254 | 0.72151139 | 0.89054584 |
| Pigk | 0.6333531 | 0.5289533 | 0.60020727 | -0.071254 | 0.70989879 | 0.89054584 |
| Rab22a | 0.51424958 | 0.43273182 | 0.50398956 | -0.0712577 | 0.74496069 | 0.89054584 |
| Uba3 | 0.56618611 | 0.47577328 | 0.54708386 | -0.0713106 | 0.72816668 | 0.89054584 |
| Ran | 2.83166646 | 1.03681629 | 1.10816899 | -0.0713527 | 0.61790361 | 0.89054584 |
| Rps6kb1 | 0.68043876 | 0.56426878 | 0.63563016 | -0.0713614 | 0.69953143 | 0.89054584 |
| Lrp6 | 0.46958519 | 0.39411197 | 0.46557919 | -0.0714672 | 0.76274355 | 0.89054584 |
| Rpf2 | 0.52996103 | 0.44571755 | 0.51720722 | -0.0714897 | 0.74020271 | 0.89054584 |
| 1110037F02I | 0.31739069 | 0.25445502 | 0.3260163 | -0.0715613 | 0.84676276 | 0.89831454 |
| Faap20 | 0.61066681 | 0.51101986 | 0.58260079 | -0.0715809 | 0.71648068 | 0.89054584 |
| Dctn1 | 0.68127891 | 0.56461894 | 0.6362484 | -0.0716295 | 0.70003741 | 0.89054584 |
| Srsf2 | 2.15303167 | 1.05406684 | 1.1257124 | -0.0716456 | 0.61658221 | 0.89054584 |
| Calm3 | 1.71895574 | 1.01323828 | 1.08489799 | -0.0716597 | 0.62085704 | 0.89054584 |
| Denr | 1.30254617 | 0.89983921 | 0.97152428 | -0.0716851 | 0.63448868 | 0.89054584 |
| Limd1 | 0.59102918 | 0.49529088 | 0.56708168 | -0.0717908 | 0.7223402 | 0.89054584 |
| Vamp8 | 1.63379191 | 0.9964443 | 1.06826919 | -0.0718249 | 0.62295201 | 0.89054584 |
| Tspan3 | 2.08687453 | 1.0508789 | 1.12273293 | -0.071854 | 0.61721275 | 0.89054584 |
| Itm2c | 1.65621956 | 1.00109711 | 1.07295435 | -0.0718572 | 0.62248572 | 0.89054584 |
| Atrx | 1.59731008 | 0.98829273 | 1.06016203 | -0.0718693 | 0.62393561 | 0.89054584 |
| Wasl | 0.69381943 | 0.57354776 | 0.64541743 | -0.0718697 | 0.69804665 | 0.89054584 |
| Ergic3 | 1.5733914 | 0.98265408 | 1.05452424 | -0.0718702 | 0.62457774 | 0.89054584 |
| Tceb3 | 0.50891411 | 0.4275661 | 0.49946607 | -0.0719 | 0.74878514 | 0.89054584 |
| Rpp30 | 0.33515437 | 0.2709993 | 0.34295421 | -0.0719549 | 0.83583627 | 0.89462889 |
| Tex264 | 0.55413664 | 0.4652797 | 0.53723958 | -0.0719599 | 0.73369803 | 0.89054584 |
| Opa1 | 0.29119224 | 0.22876626 | 0.3007438 | -0.0719775 | 0.86758175 | 0.90444517 |

| | | | | | | |
|---|---|---|---|---|---|---|
| Patz1 | 0.37755132 | 0.31063488 | 0.38271349 | -0.0720786 | 0.8098775 | 0.89054584 |
| Eftud2 | 0.45163765 | 0.37770179 | 0.4498054 | -0.0721036 | 0.77241575 | 0.89054584 |
| Dag1 | 0.47255837 | 0.39591055 | 0.46817411 | -0.0722636 | 0.76396538 | 0.89054584 |
| Sptan1 | 0.74581837 | 0.60993711 | 0.68225355 | -0.0723164 | 0.68928728 | 0.89054584 |
| Fam175b | 0.44711569 | 0.37347402 | 0.44580055 | -0.0723265 | 0.77513495 | 0.89054584 |
| Romo1 | 2.38153459 | 1.05707448 | 1.1294334 | -0.0723589 | 0.61733249 | 0.89054584 |
| Cct2 | 1.78891876 | 1.02380468 | 1.09616825 | -0.0723636 | 0.62079146 | 0.89054584 |
| Zfp553 | 0.36337032 | 0.29712221 | 0.36952112 | -0.0723989 | 0.81932853 | 0.89054584 |
| Polr2h | 0.69381836 | 0.57300761 | 0.64541665 | -0.072409 | 0.69940319 | 0.89054584 |
| Med25 | 0.96236237 | 0.74233268 | 0.8147539 | -0.0724212 | 0.66060705 | 0.89054584 |
| Ccdc50 | 0.49365767 | 0.41392303 | 0.48643255 | -0.0725095 | 0.75630934 | 0.89054584 |
| Psma4 | 2.12885691 | 1.05213692 | 1.12473114 | -0.0725942 | 0.61818026 | 0.89054584 |
| 9430016H08 | 0.38720632 | 0.31902503 | 0.39163149 | -0.0726065 | 0.80613565 | 0.89054584 |
| Aph1a | 0.61094084 | 0.51020884 | 0.58281554 | -0.0726067 | 0.71917998 | 0.89054584 |
| Adh5 | 1.23127459 | 0.87080272 | 0.9435736 | -0.0727709 | 0.64031832 | 0.89054584 |
| Cisd2 | 1.00519943 | 0.76442274 | 0.83720913 | -0.0727864 | 0.65727786 | 0.89054584 |
| Nudt16l1 | 0.83037342 | 0.66521452 | 0.73801752 | -0.072803 | 0.67709446 | 0.89054584 |
| 2310061I04R | 0.32465652 | 0.2601519 | 0.33296428 | -0.0728124 | 0.84585433 | 0.89825055 |
| Hif1a | 0.60022579 | 0.50152622 | 0.57438069 | -0.0728545 | 0.72271978 | 0.89054584 |
| Pabpn1 | 0.73955183 | 0.6049202 | 0.67791585 | -0.0729956 | 0.69204622 | 0.89054584 |
| Tnpo3 | 0.80294104 | 0.6474887 | 0.72049289 | -0.0730042 | 0.68156117 | 0.89054584 |
| Trmt1l | 0.50885874 | 0.42613294 | 0.49941904 | -0.0732861 | 0.75290682 | 0.89054584 |
| Eif3f | 3.05702735 | 1.01642659 | 1.08972939 | -0.0733028 | 0.62300891 | 0.89054584 |
| Rbm4b | 0.97623642 | 0.74876791 | 0.82212965 | -0.0733617 | 0.66119802 | 0.89054584 |
| Ccnh | 0.60625857 | 0.50571349 | 0.57913923 | -0.0734257 | 0.72263273 | 0.89054584 |
| Casp6 | 0.63809048 | 0.53040414 | 0.60384019 | -0.0734361 | 0.71450611 | 0.89054584 |
| Zfp444 | 0.41474502 | 0.34333179 | 0.41678518 | -0.0734534 | 0.79420036 | 0.89054584 |
| Gpat4 | 0.50529798 | 0.42288041 | 0.49639003 | -0.0735096 | 0.75488256 | 0.89054584 |
| Pex19 | 0.90071449 | 0.70685291 | 0.78045254 | -0.0735996 | 0.66979812 | 0.89054584 |
| Ctdnep1 | 0.38294669 | 0.314058 | 0.38770344 | -0.0736454 | 0.81191628 | 0.89054584 |

| | | | | | | |
|---|---|---|---|---|---|---|
| Gtf3a | 0.82969453 | 0.6639424 | 0.73759046 | -0.0736481 | 0.67910338 | 0.89054584 |
| Atraid | 1.62817302 | 0.99332014 | 1.06705771 | -0.0737376 | 0.62625462 | 0.89054584 |
| Ssb | 2.23458726 | 1.05434208 | 1.1280988 | -0.0737567 | 0.61966774 | 0.89054584 |
| Arl6ip5 | 1.17074334 | 0.84393649 | 0.91773567 | -0.0737992 | 0.64603813 | 0.89054584 |
| Dynlt1c | 0.61633974 | 0.51315126 | 0.58703605 | -0.0738848 | 0.72118918 | 0.89054584 |
| Maea | 0.59022176 | 0.49245144 | 0.56643815 | -0.0739867 | 0.728585 | 0.89054584 |
| Pts | 0.72466551 | 0.59350712 | 0.66749971 | -0.0739926 | 0.69722828 | 0.89054584 |
| Wbscr22 | 0.45031789 | 0.37463499 | 0.44863782 | -0.0740028 | 0.77890271 | 0.89054584 |
| Gm10250 | 0.31302771 | 0.24778675 | 0.32183106 | -0.0740443 | 0.85811569 | 0.90124396 |
| 1110008F13I | 1.14936748 | 0.83407152 | 0.90817539 | -0.0741039 | 0.64808408 | 0.89054584 |
| Rcor1 | 0.36445519 | 0.29634876 | 0.37053427 | -0.0741855 | 0.82453251 | 0.89142971 |
| Psme4 | 0.41244672 | 0.34048499 | 0.41470252 | -0.0742175 | 0.79780056 | 0.89054584 |
| Ppm1g | 1.060234 | 0.79118113 | 0.86546915 | -0.074288 | 0.65538593 | 0.89054584 |
| Npepps | 1.02642817 | 0.77377389 | 0.84806981 | -0.0742959 | 0.6584185 | 0.89054584 |
| Sep-15 | 2.74359897 | 1.03992832 | 1.11427986 | -0.0743515 | 0.62205139 | 0.89054584 |
| Gigyf2 | 0.69204694 | 0.56976861 | 0.6441282 | -0.0743596 | 0.70465572 | 0.89054584 |
| Irf3 | 0.56408448 | 0.47069245 | 0.54537373 | -0.0746813 | 0.73825554 | 0.89054584 |
| Lamtor5 | 1.28775392 | 0.89119901 | 0.96594759 | -0.0747486 | 0.64077311 | 0.89054584 |
| Acbd6 | 0.89293151 | 0.70113452 | 0.77593256 | -0.074798 | 0.67332414 | 0.89054584 |
| Rer1 | 1.41483217 | 0.93555255 | 1.01038377 | -0.0748312 | 0.63497211 | 0.89054584 |
| Birc2 | 0.50206175 | 0.41878052 | 0.49363015 | -0.0748496 | 0.7600427 | 0.89054584 |
| Txn2 | 0.9171262 | 0.71498041 | 0.78983741 | -0.074857 | 0.67059333 | 0.89054584 |
| Ift22 | 0.72176861 | 0.59058435 | 0.66545449 | -0.0748701 | 0.6999221 | 0.89054584 |
| Sae1 | 0.98098223 | 0.74971006 | 0.82463013 | -0.0749201 | 0.66395396 | 0.89054584 |
| Ddx27 | 0.42798782 | 0.35380503 | 0.42872917 | -0.0749241 | 0.79223206 | 0.89054584 |
| Slc4a2 | 0.30114841 | 0.23537938 | 0.31038842 | -0.075009 | 0.86989362 | 0.90589084 |
| Cops3 | 1.09969219 | 0.80999319 | 0.88502219 | -0.075029 | 0.65356685 | 0.89054584 |
| B9d1 | 0.87503003 | 0.69032435 | 0.76537389 | -0.0750495 | 0.67610604 | 0.89054584 |
| Ube2k | 1.37636898 | 0.9226518 | 0.99770198 | -0.0750502 | 0.63700314 | 0.89054584 |
| Hp1bp3 | 1.13353967 | 0.82582186 | 0.90096444 | -0.0751426 | 0.65120167 | 0.89054584 |

| | | | | | | |
|---|---|---|---|---|---|---|
| Hgs | 0.45418975 | 0.37688924 | 0.45206022 | -0.075171 | 0.78074555 | 0.89054584 |
| Atg4b | 0.40636862 | 0.33398896 | 0.40918002 | -0.0751911 | 0.80403668 | 0.89054584 |
| Hnrnpab | 1.48080311 | 0.95498541 | 1.03022387 | -0.0752385 | 0.6331651 | 0.89054584 |
| Pds5b | 0.41599582 | 0.34254309 | 0.41791759 | -0.0753745 | 0.79962267 | 0.89054584 |
| Abi1 | 0.42079957 | 0.34686105 | 0.42225856 | -0.0753975 | 0.7972644 | 0.89054584 |
| Rpl9 | 4.82033618 | 0.67864111 | 0.75409543 | -0.0754543 | 0.67945336 | 0.89054584 |
| Ppp2ca | 0.87280774 | 0.68852588 | 0.76404677 | -0.0755209 | 0.67742312 | 0.89054584 |
| Aen | 0.33701821 | 0.26919885 | 0.344722 | -0.0755231 | 0.84630182 | 0.89825055 |
| Insig2 | 0.57595276 | 0.4793986 | 0.55499351 | -0.0755949 | 0.73715142 | 0.89054584 |
| Sar1b | 1.14028849 | 0.82841356 | 0.9040594 | -0.0756458 | 0.65166889 | 0.89054584 |
| Asnsd1 | 0.59495629 | 0.49435675 | 0.57020545 | -0.0758487 | 0.73228292 | 0.89054584 |
| Gtf3c2 | 0.43171713 | 0.35602895 | 0.43207423 | -0.0760453 | 0.79388509 | 0.89054584 |
| Sec63 | 0.50096698 | 0.41660887 | 0.49269503 | -0.0760862 | 0.76407535 | 0.89054584 |
| Cnot2 | 0.6186573 | 0.51273017 | 0.58884168 | -0.0761115 | 0.72647871 | 0.89054584 |
| Lztfl1 | 0.80099761 | 0.64303169 | 0.71923066 | -0.076199 | 0.68920284 | 0.89054584 |
| Kctd5 | 0.50171392 | 0.41711273 | 0.49333313 | -0.0762204 | 0.76417914 | 0.89054584 |
| Ppp1r37 | 0.47009092 | 0.38974693 | 0.46602097 | -0.076274 | 0.77712895 | 0.89054584 |
| Med28 | 1.67840214 | 1.00101776 | 1.07736065 | -0.0763429 | 0.62935194 | 0.89054584 |
| Sart1 | 0.41550183 | 0.3411218 | 0.41747042 | -0.0763486 | 0.80290557 | 0.89054584 |
| Tmem147 | 1.43208806 | 0.9393434 | 1.01579916 | -0.0764558 | 0.63707931 | 0.89054584 |
| Ppia | 5.15447174 | 0.60149925 | 0.6780479 | -0.0765487 | 0.70054692 | 0.89054584 |
| Psmd5 | 0.51372562 | 0.42697745 | 0.50354614 | -0.0765687 | 0.760662 | 0.89054584 |
| Timm22 | 0.54588377 | 0.4538499 | 0.53044249 | -0.0765926 | 0.74943794 | 0.89054584 |
| Ufl1 | 0.67176415 | 0.55258902 | 0.62921944 | -0.0766304 | 0.71477742 | 0.89054584 |
| Csnk1a1 | 2.29633104 | 1.05232497 | 1.12902849 | -0.0767035 | 0.6241951 | 0.89054584 |
| Mdm2 | 0.66136517 | 0.54463601 | 0.62146505 | -0.076829 | 0.71768051 | 0.89054584 |
| Zcchc7 | 0.71842206 | 0.58621239 | 0.66308435 | -0.076872 | 0.70544121 | 0.89054584 |
| Dnm2 | 0.85138251 | 0.67419058 | 0.75106848 | -0.0768779 | 0.68327625 | 0.89054584 |
| Commd2 | 0.61718524 | 0.51078264 | 0.58769521 | -0.0769126 | 0.72897727 | 0.89054584 |
| Rb1cc1 | 0.74389362 | 0.6039704 | 0.68092425 | -0.0769538 | 0.70073815 | 0.89054584 |

| | | | | | | |
|---|---|---|---|---|---|---|
| Pgls | 1.86038105 | 1.02843017 | 1.10543546 | -0.0770053 | 0.62723713 | 0.89054584 |
| Vma21 | 0.88285708 | 0.69298625 | 0.77001916 | -0.0770329 | 0.67942148 | 0.89054584 |
| Mrps15 | 0.83020943 | 0.66085936 | 0.7379144 | -0.077055 | 0.68669341 | 0.89054584 |
| Acbd3 | 0.50746518 | 0.42116448 | 0.49823454 | -0.0770701 | 0.76444505 | 0.89054584 |
| Lamtor1 | 1.05604249 | 0.78615422 | 0.86333354 | -0.0771793 | 0.661476 | 0.89054584 |
| Sh3bp1 | 0.45052031 | 0.37162961 | 0.44881697 | -0.0771874 | 0.78850447 | 0.89054584 |
| Sbno1 | 1.02486983 | 0.77001787 | 0.84725174 | -0.0772339 | 0.66447242 | 0.89054584 |
| Ift52 | 0.40456246 | 0.33024539 | 0.40753486 | -0.0772895 | 0.81153683 | 0.89054584 |
| Mcts1 | 0.86947263 | 0.68468936 | 0.76204826 | -0.0773589 | 0.6818755 | 0.89054584 |
| Atxn7l3b | 2.15603575 | 1.04836373 | 1.12582229 | -0.0774586 | 0.62572177 | 0.89054584 |
| Rhot1 | 0.40465107 | 0.33012336 | 0.40761562 | -0.0774923 | 0.81211486 | 0.89054584 |
| Klhl9 | 0.75166712 | 0.60877392 | 0.68627656 | -0.0775026 | 0.70060589 | 0.89054584 |
| Ndufb3 | 1.68104886 | 1.00036131 | 1.07787295 | -0.0775116 | 0.63119962 | 0.89054584 |
| Yes1 | 0.40836644 | 0.33341937 | 0.41099757 | -0.0775782 | 0.81040845 | 0.89054584 |
| Fbxo11 | 0.3021585 | 0.23375805 | 0.31136412 | -0.0776061 | 0.87722369 | 0.9096298 |
| Stag2 | 0.57432821 | 0.47599714 | 0.55368192 | -0.0776848 | 0.74333987 | 0.89054584 |
| Ssrp1 | 1.16044757 | 0.83543688 | 0.91314165 | -0.0777048 | 0.65411627 | 0.89054584 |
| Rnf216 | 0.39837238 | 0.3241434 | 0.40188241 | -0.077739 | 0.81626234 | 0.89054584 |
| Smim11 | 0.90852728 | 0.70719859 | 0.78494495 | -0.0777464 | 0.67776343 | 0.89054584 |
| Hnrnpul2 | 0.56511501 | 0.46846486 | 0.54621265 | -0.0777478 | 0.74638363 | 0.89054584 |
| Nr3c1 | 0.62241105 | 0.51386449 | 0.59175849 | -0.077894 | 0.73014877 | 0.89054584 |
| Slc38a10 | 0.74503959 | 0.6036719 | 0.68171602 | -0.0780441 | 0.70310926 | 0.89054584 |
| Kdm2a | 0.57770949 | 0.47835865 | 0.55640985 | -0.0780512 | 0.7432958 | 0.89054584 |
| Got2 | 0.59069495 | 0.48876392 | 0.56681535 | -0.0780514 | 0.73941536 | 0.89054584 |
| Psmd9 | 0.82664589 | 0.65760124 | 0.73566849 | -0.0780672 | 0.68948632 | 0.89054584 |
| Wbp11 | 1.11234916 | 0.81295965 | 0.89107419 | -0.0781145 | 0.65851672 | 0.89054584 |
| Fdft1 | 0.47231923 | 0.38981919 | 0.4679656 | -0.0781464 | 0.78171692 | 0.89054584 |
| Ppp6c | 0.75909685 | 0.61316266 | 0.69135148 | -0.0781888 | 0.70087051 | 0.89054584 |
| Mrpl34 | 1.13802671 | 0.82483169 | 0.90302552 | -0.0781938 | 0.65668115 | 0.89054584 |
| Maf1 | 0.73773319 | 0.59839108 | 0.6766517 | -0.0782606 | 0.70500373 | 0.89054584 |

| | | | | | | |
|---|---|---|---|---|---|---|
| Tmed4 | 1.29949307 | 0.89205321 | 0.97037659 | -0.0783234 | 0.64655781 | 0.89054584 |
| Lrp10 | 0.57342136 | 0.47460519 | 0.55294901 | -0.0783438 | 0.74540378 | 0.89054584 |
| Cops6 | 1.71700947 | 1.00616066 | 1.08455136 | -0.0783907 | 0.63184558 | 0.89054584 |
| Bax | 1.71590905 | 1.00592794 | 1.08435461 | -0.0784267 | 0.63192686 | 0.89054584 |
| Exosc7 | 0.68629289 | 0.56149979 | 0.63992768 | -0.0784279 | 0.71599607 | 0.89054584 |
| Tmbim4 | 1.01097001 | 0.76132715 | 0.84018334 | -0.0788562 | 0.66904195 | 0.89054584 |
| Kxd1 | 0.47949648 | 0.39529969 | 0.47420842 | -0.0789087 | 0.78090468 | 0.89054584 |
| Tet3 | 0.58251467 | 0.48123383 | 0.56027344 | -0.0790396 | 0.74449058 | 0.89054584 |
| Ap3d1 | 0.49805998 | 0.41116186 | 0.49020831 | -0.0790464 | 0.7737512 | 0.89054584 |
| Tsn | 1.3703093 | 0.91657828 | 0.99565487 | -0.0790766 | 0.64431656 | 0.89054584 |
| Cap1 | 0.79900306 | 0.63883048 | 0.71793237 | -0.0791019 | 0.69613723 | 0.89054584 |
| Chd4 | 1.52256796 | 0.96235762 | 1.04167942 | -0.0793218 | 0.63861869 | 0.89054584 |
| Ccnt1 | 0.63188898 | 0.51970833 | 0.59908137 | -0.079373 | 0.73147133 | 0.89054584 |
| Eapp | 0.77026005 | 0.61949671 | 0.69890308 | -0.0794064 | 0.70171135 | 0.89054584 |
| Grsf1 | 0.48319604 | 0.39798815 | 0.47741393 | -0.0794258 | 0.78085754 | 0.89054584 |
| Rnf149 | 0.42599005 | 0.34748934 | 0.42693388 | -0.0794445 | 0.80701347 | 0.89054584 |
| Traf7 | 0.51172564 | 0.42235779 | 0.50185198 | -0.0794942 | 0.76973793 | 0.89054584 |
| Tjp1 | 0.38683577 | 0.31175395 | 0.39129019 | -0.0795362 | 0.82818246 | 0.89229532 |
| Cebpzos | 0.88239152 | 0.69017255 | 0.76974412 | -0.0795716 | 0.68495978 | 0.89054584 |
| Hsd17b10 | 1.21550724 | 0.857426 | 0.93707044 | -0.0796444 | 0.65393918 | 0.89054584 |
| Gnl2 | 0.59723802 | 0.4923526 | 0.57201565 | -0.0796631 | 0.74177845 | 0.89054584 |
| Matr3 | 1.35741551 | 0.91141272 | 0.99126927 | -0.0798566 | 0.64629018 | 0.89054584 |
| G3bp1 | 2.01347791 | 1.03879058 | 1.11866032 | -0.0798697 | 0.63029604 | 0.89054584 |
| Itch | 0.58505078 | 0.48240733 | 0.5623064 | -0.0798991 | 0.74601337 | 0.89054584 |
| Ddx1 | 0.98070676 | 0.74457613 | 0.82448544 | -0.0799093 | 0.67420181 | 0.89054584 |
| Ndufs3 | 1.48230572 | 0.95060003 | 1.03064663 | -0.0800466 | 0.64126537 | 0.89054584 |
| Kpnb1 | 0.79461833 | 0.63495129 | 0.71506805 | -0.0801168 | 0.69916178 | 0.89054584 |
| Eif4g3 | 0.78389665 | 0.62780671 | 0.70800525 | -0.0801985 | 0.70115891 | 0.89054584 |
| Mark3 | 0.44725175 | 0.36564816 | 0.44592123 | -0.0802731 | 0.799177 | 0.89054584 |
| Suclg1 | 1.44179244 | 0.93842514 | 1.01878709 | -0.080362 | 0.64337598 | 0.89054584 |

| | | | | | | |
|---|---|---|---|---|---|---|
| Drg1 | 0.96284838 | 0.73459756 | 0.81501397 | -0.0804164 | 0.67713132 | 0.89054584 |
| Gcn1l1 | 0.36037584 | 0.28615617 | 0.36672182 | -0.0805657 | 0.84697124 | 0.89840423 |
| Ino80d | 0.44180754 | 0.36038902 | 0.44108382 | -0.0806948 | 0.80298663 | 0.89054584 |
| Polr1d | 1.87429865 | 1.02619574 | 1.10698127 | -0.0807855 | 0.63307222 | 0.89054584 |
| Caml | 0.73091866 | 0.59109419 | 0.67189398 | -0.0807998 | 0.71235206 | 0.89054584 |
| Mpc2 | 1.38945042 | 0.92125055 | 1.00211527 | -0.0808647 | 0.6465225 | 0.89054584 |
| Pkn2 | 0.95387195 | 0.72930246 | 0.81018265 | -0.0808802 | 0.67907013 | 0.89054584 |
| Gatad2a | 0.36564668 | 0.29076498 | 0.37164624 | -0.0808813 | 0.84469049 | 0.89782425 |
| Aspscr1 | 0.64559495 | 0.52861681 | 0.60956344 | -0.0809466 | 0.73193686 | 0.89054584 |
| Rnf113a2 | 0.3595059 | 0.28490138 | 0.36590773 | -0.0810064 | 0.84882759 | 0.89883797 |
| Hnrnpm | 1.7749838 | 1.01308458 | 1.0941016 | -0.081017 | 0.63494887 | 0.89054584 |
| Gpatch8 | 0.68451693 | 0.55760411 | 0.63862649 | -0.0810224 | 0.72276616 | 0.89054584 |
| Mpdu1 | 0.5423289 | 0.4463814 | 0.52750186 | -0.0811205 | 0.76314717 | 0.89054584 |
| Nudcd2 | 0.98037703 | 0.74316068 | 0.82431217 | -0.0811515 | 0.67676126 | 0.89054584 |
| Mcl1 | 0.59656026 | 0.49029985 | 0.57147832 | -0.0811785 | 0.74595106 | 0.89054584 |
| Hnrnpul1 | 0.98121386 | 0.74353475 | 0.82475176 | -0.081217 | 0.67680637 | 0.89054584 |
| Aktip | 0.46709686 | 0.38213474 | 0.46340355 | -0.0812688 | 0.79305176 | 0.89054584 |
| Morf4l2 | 2.59464731 | 1.04149219 | 1.12279647 | -0.0813043 | 0.63207899 | 0.89054584 |
| Hspe1 | 2.71709008 | 1.03453932 | 1.11589366 | -0.0813543 | 0.63294393 | 0.89054584 |
| Thumpd3 | 0.38678098 | 0.30980502 | 0.39123972 | -0.0814347 | 0.8339111 | 0.89423396 |
| Cops5 | 1.13114763 | 0.81832698 | 0.89986021 | -0.0815332 | 0.66353516 | 0.89054584 |
| Armc1 | 0.80989126 | 0.64342021 | 0.72498437 | -0.0815642 | 0.6999139 | 0.89054584 |
| Ik | 1.08297965 | 0.79518709 | 0.87685883 | -0.0816717 | 0.66783387 | 0.89054584 |
| Usf2 | 0.84488583 | 0.66538463 | 0.74706698 | -0.0816823 | 0.6947628 | 0.89054584 |
| Ddx50 | 0.96684291 | 0.73546241 | 0.81714497 | -0.0816826 | 0.67928918 | 0.89054584 |
| Son | 2.42001721 | 1.04723225 | 1.12891616 | -0.0816839 | 0.63198021 | 0.89054584 |
| Fdx1 | 0.80905947 | 0.64271047 | 0.7244487 | -0.0817382 | 0.70043771 | 0.89054584 |
| Emc6 | 1.24069112 | 0.86564404 | 0.9473839 | -0.0817399 | 0.65614744 | 0.89054584 |
| Crebbp | 0.78612808 | 0.6276606 | 0.70948197 | -0.0818214 | 0.70446799 | 0.89054584 |
| BC004004 | 0.76286515 | 0.61205328 | 0.69391071 | -0.0818574 | 0.70870097 | 0.89054584 |

| | | | | | | |
|---|---|---|---|---|---|---|
| Serf1 | 1.27783622 | 0.88027966 | 0.96215759 | -0.0818779 | 0.65412001 | 0.89054584 |
| Setd3 | 0.82427346 | 0.6522792 | 0.7341681 | -0.0818889 | 0.69834867 | 0.89054584 |
| Trip12 | 0.65784922 | 0.53684531 | 0.61882619 | -0.0819809 | 0.73145104 | 0.89054584 |
| Yipf5 | 0.91151524 | 0.70466417 | 0.78665116 | -0.081987 | 0.68636322 | 0.89054584 |
| Mrpl10 | 0.57119664 | 0.46895791 | 0.55114924 | -0.0821913 | 0.75640178 | 0.89054584 |
| Ddx6 | 1.11298926 | 0.80912185 | 0.89137722 | -0.0822554 | 0.66637391 | 0.89054584 |
| Dnlz | 0.59428165 | 0.48723678 | 0.56966955 | -0.0824328 | 0.74989577 | 0.89054584 |
| Mrpl38 | 0.88745404 | 0.69028292 | 0.77272636 | -0.0824434 | 0.69043017 | 0.89054584 |
| Polr2g | 1.48104187 | 0.94770792 | 1.03029113 | -0.0825832 | 0.64558894 | 0.89054584 |
| Dram2 | 0.51306329 | 0.42036035 | 0.50298537 | -0.082625 | 0.7779597 | 0.89054584 |
| Pddc1 | 0.75220612 | 0.60388131 | 0.68664606 | -0.0827648 | 0.71279365 | 0.89054584 |
| Cul5 | 0.81639533 | 0.6463156 | 0.7291562 | -0.0828406 | 0.7017047 | 0.89054584 |
| Srsf9 | 0.99029854 | 0.74654974 | 0.82949118 | -0.0829414 | 0.67933893 | 0.89054584 |
| Pdcd5 | 1.72471575 | 1.00285086 | 1.0859137 | -0.0830628 | 0.63921751 | 0.89054584 |
| Pmpca | 0.57985618 | 0.47502892 | 0.55813779 | -0.0831089 | 0.75607557 | 0.89054584 |
| Clk3 | 0.46226074 | 0.37599881 | 0.4591652 | -0.0831664 | 0.80062385 | 0.89054584 |
| Chmp7 | 0.4883937 | 0.39865437 | 0.48190317 | -0.0832488 | 0.78954179 | 0.89054584 |
| Neu1 | 0.36178234 | 0.28477216 | 0.36803714 | -0.083265 | 0.85406817 | 0.90035231 |
| Ric8 | 0.40904276 | 0.32833592 | 0.41161234 | -0.0832764 | 0.82705359 | 0.89186561 |
| Idh3b | 1.07189322 | 0.78799616 | 0.87134743 | -0.0833513 | 0.67208022 | 0.89054584 |
| Znrd1 | 0.69878398 | 0.56551465 | 0.64901658 | -0.0835019 | 0.72555209 | 0.89054584 |
| Dhx30 | 0.64990557 | 0.52926271 | 0.6128333 | -0.0835706 | 0.73739292 | 0.89054584 |
| Hmgn1 | 3.21520461 | 0.98997044 | 1.07362373 | -0.0836533 | 0.64170116 | 0.89054584 |
| Shoc2 | 0.65840412 | 0.53554794 | 0.61924324 | -0.0836953 | 0.7355514 | 0.89054584 |
| Gnaq | 0.58498972 | 0.47855255 | 0.56225751 | -0.083705 | 0.75603048 | 0.89054584 |
| B230118H07 | 0.70135657 | 0.56712872 | 0.65087477 | -0.083746 | 0.72556411 | 0.89054584 |
| Proser1 | 0.44202466 | 0.35750138 | 0.44127708 | -0.0837757 | 0.81178467 | 0.89054584 |
| Brwd1 | 0.68260561 | 0.55335633 | 0.63722366 | -0.0838673 | 0.73012142 | 0.89054584 |
| Psma7 | 2.66910688 | 1.0349708 | 1.11883981 | -0.083869 | 0.63654448 | 0.89054584 |
| Dstn | 2.56805059 | 1.04021436 | 1.1240957 | -0.0838813 | 0.63595059 | 0.89054584 |

| | | | | | | |
|---|---|---|---|---|---|---|
| Pmvk | 0.49652066 | 0.40497789 | 0.4888894 | -0.0839115 | 0.78804223 | 0.89054584 |
| Nelfb | 0.73150463 | 0.58834596 | 0.67230432 | -0.0839584 | 0.71965452 | 0.89054584 |
| Rnf20 | 0.59479596 | 0.48586845 | 0.57007811 | -0.0842097 | 0.75433665 | 0.89054584 |
| Fam133b | 0.72815212 | 0.58562278 | 0.66995331 | -0.0843305 | 0.72121184 | 0.89054584 |
| Kansl1 | 0.40780433 | 0.32613824 | 0.41048641 | -0.0843482 | 0.83080102 | 0.89280428 |
| Arpc1a | 1.92643682 | 1.02777013 | 1.11213549 | -0.0843654 | 0.63811397 | 0.89054584 |
| Anapc4 | 0.54319247 | 0.44367085 | 0.52821697 | -0.0845461 | 0.77207796 | 0.89054584 |
| Zfp560 | 0.46602462 | 0.37765617 | 0.46246508 | -0.0848089 | 0.80355206 | 0.89054584 |
| Fam92a | 0.81515813 | 0.64351381 | 0.72836499 | -0.0848512 | 0.7063474 | 0.89054584 |
| Rps19bp1 | 0.70349777 | 0.56754637 | 0.65241778 | -0.0848714 | 0.7277659 | 0.89054584 |
| Emc4 | 0.82236196 | 0.64807807 | 0.73295622 | -0.0848782 | 0.70523491 | 0.89054584 |
| Syf2 | 0.91614144 | 0.70432028 | 0.78927989 | -0.0849596 | 0.69198944 | 0.89054584 |
| Mtmr6 | 0.40046288 | 0.31881694 | 0.40379379 | -0.0849769 | 0.83656954 | 0.89462889 |
| Zfand3 | 0.55600664 | 0.45375954 | 0.5387736 | -0.0850141 | 0.7688737 | 0.89054584 |
| Ssna1 | 1.00352026 | 0.75125969 | 0.83633922 | -0.0850795 | 0.68223117 | 0.89054584 |
| Npm3 | 0.99881393 | 0.7487692 | 0.83389037 | -0.0851212 | 0.68281017 | 0.89054584 |
| Mnat1 | 0.58502558 | 0.47702327 | 0.56228622 | -0.0852629 | 0.76005023 | 0.89054584 |
| Spg21 | 0.74449518 | 0.59604963 | 0.68134 | -0.0852904 | 0.72012329 | 0.89054584 |
| Zfp91 | 0.7285808 | 0.58492948 | 0.67025437 | -0.0853249 | 0.72344079 | 0.89054584 |
| Cdk11b | 0.83228161 | 0.65381754 | 0.7392161 | -0.0853986 | 0.70479021 | 0.89054584 |
| Rabggtb | 0.79774959 | 0.63170448 | 0.71711497 | -0.0854105 | 0.71051626 | 0.89054584 |
| Gtf2a2 | 1.75761019 | 1.00594502 | 1.09141165 | -0.0854666 | 0.64238232 | 0.89054584 |
| Snx1 | 0.74110151 | 0.59351097 | 0.67899116 | -0.0854802 | 0.72123956 | 0.89054584 |
| Sra1 | 1.22748198 | 0.85646339 | 0.94202383 | -0.0855604 | 0.66390852 | 0.89054584 |
| Atp2c1 | 0.59136384 | 0.48176699 | 0.56734828 | -0.0855813 | 0.75890202 | 0.89054584 |
| Ccnt2 | 0.8419109 | 0.65951282 | 0.74522428 | -0.0857115 | 0.70396713 | 0.89054584 |
| Desi2 | 0.71489554 | 0.57483214 | 0.6605781 | -0.085746 | 0.72732784 | 0.89054584 |
| Cdc23 | 0.60701559 | 0.49393496 | 0.57973462 | -0.0857997 | 0.75475028 | 0.89054584 |
| Arpp19 | 1.77353557 | 1.00808246 | 1.09388243 | -0.0858 | 0.64260463 | 0.89054584 |
| Fam136a | 0.79352309 | 0.62852507 | 0.71435039 | -0.0858253 | 0.71217383 | 0.89054584 |

| | | | | | | |
|---|---|---|---|---|---|---|
| R3hdm1 | 1.07798608 | 0.78847553 | 0.8743834 | -0.0859079 | 0.67643016 | 0.89054584 |
| Srrm2 | 2.22893744 | 1.04189358 | 1.12797623 | -0.0860826 | 0.63891188 | 0.89054584 |
| Pet100 | 0.95761553 | 0.72597653 | 0.8122047 | -0.0862282 | 0.68956551 | 0.89054584 |
| Ing4 | 1.01030144 | 0.75354324 | 0.83983997 | -0.0862967 | 0.68393243 | 0.89054584 |
| Ube4b | 0.45573372 | 0.36710096 | 0.45342244 | -0.0863215 | 0.81244927 | 0.89054584 |
| Cltc | 0.91236375 | 0.70077595 | 0.78713449 | -0.0863585 | 0.69537502 | 0.89054584 |
| Cox7a2l | 2.74693527 | 1.02769934 | 1.11406897 | -0.0863696 | 0.64102145 | 0.89054584 |
| Akirin2 | 0.37988186 | 0.29839189 | 0.3848709 | -0.086479 | 0.85241472 | 0.89988783 |
| Malsu1 | 0.57502227 | 0.4676858 | 0.55424248 | -0.0865567 | 0.76656837 | 0.89054584 |
| Pym1 | 0.34356745 | 0.2643343 | 0.35092003 | -0.0865857 | 0.87483403 | 0.90828873 |
| Mlf2 | 1.80224167 | 1.01145191 | 1.09807024 | -0.0866183 | 0.64337998 | 0.89054584 |
| Slc35a1 | 0.63056313 | 0.51125578 | 0.59806053 | -0.0868047 | 0.75054831 | 0.89054584 |
| Smpd2 | 0.49801343 | 0.40334974 | 0.49016845 | -0.0868187 | 0.7953688 | 0.89054584 |
| Spg7 | 0.47382979 | 0.38226679 | 0.46928212 | -0.0870153 | 0.80617067 | 0.89054584 |
| Dctn2 | 1.37252527 | 0.90936969 | 0.99639869 | -0.087029 | 0.65796974 | 0.89054584 |
| Eif3l | 1.48246288 | 0.94355873 | 1.03069108 | -0.0871324 | 0.65315655 | 0.89054584 |
| Lars | 0.62267387 | 0.50482525 | 0.59196235 | -0.0871371 | 0.75357978 | 0.89054584 |
| Manbal | 0.61229869 | 0.49672847 | 0.58387889 | -0.0871504 | 0.75659729 | 0.89054584 |
| Tfg | 1.52985484 | 0.95642792 | 1.04361496 | -0.087187 | 0.65144453 | 0.89054584 |
| Usp8 | 0.56119149 | 0.45582691 | 0.54301489 | -0.087188 | 0.77278577 | 0.89054584 |
| Ino80e | 0.61522782 | 0.49898027 | 0.58616844 | -0.0871882 | 0.75583977 | 0.89054584 |
| Stub1 | 1.90355905 | 1.02279277 | 1.11003958 | -0.0872468 | 0.64288579 | 0.89054584 |
| Mri1 | 0.3753962 | 0.29337675 | 0.38071578 | -0.087339 | 0.85741914 | 0.9009252 |
| Stt3b | 0.90359709 | 0.69474413 | 0.78211529 | -0.0873712 | 0.69862732 | 0.89054584 |
| Arcn1 | 0.88473727 | 0.68365879 | 0.77112831 | -0.0874695 | 0.70141213 | 0.89054584 |
| Uqcr10 | 2.65293173 | 1.03216112 | 1.11970826 | -0.0875471 | 0.64217209 | 0.89054584 |
| Mrpl40 | 0.7743427 | 0.61409497 | 0.70164233 | -0.0875474 | 0.71948726 | 0.89054584 |
| Bclaf1 | 1.29906965 | 0.88261399 | 0.97021698 | -0.087603 | 0.66300045 | 0.89054584 |
| 2810403A07 | 0.52878555 | 0.42854767 | 0.51622365 | -0.087676 | 0.78560958 | 0.89054584 |
| Coprs | 0.46983824 | 0.37807939 | 0.46580026 | -0.0877209 | 0.80986136 | 0.89054584 |

| | | | | | | |
|---|---|---|---|---|---|---|
| Cox6c | 3.36117416 | 0.96618177 | 1.05390444 | -0.0877227 | 0.65091468 | 0.89054584 |
| Pip5k1a | 0.49022578 | 0.39573339 | 0.48348155 | -0.0877482 | 0.80110138 | 0.89054584 |
| Arhgap5 | 1.06084696 | 0.77798286 | 0.86578046 | -0.0877976 | 0.68169963 | 0.89054584 |
| Sdhd | 1.33488947 | 0.89554537 | 0.98336605 | -0.0878207 | 0.66132146 | 0.89054584 |
| Arfgap2 | 0.71042471 | 0.56951574 | 0.65738801 | -0.0878723 | 0.73327728 | 0.89054584 |
| Ctage5 | 0.79791206 | 0.62934541 | 0.71722098 | -0.0878756 | 0.7159406 | 0.89054584 |
| Tardbp | 1.14158933 | 0.81672278 | 0.90465249 | -0.0879297 | 0.67466663 | 0.89054584 |
| Actr1a | 0.47873545 | 0.38555457 | 0.47354797 | -0.0879934 | 0.80667589 | 0.89054584 |
| Pycrl | 0.56917776 | 0.46144547 | 0.54951318 | -0.0880677 | 0.77236713 | 0.89054584 |
| Dnajc5 | 0.8076468 | 0.63527679 | 0.72353778 | -0.088261 | 0.71509198 | 0.89054584 |
| Tbca | 2.58513079 | 1.03500966 | 1.12327197 | -0.0882623 | 0.64284781 | 0.89054584 |
| Eif1b | 1.63894426 | 0.9810918 | 1.06936647 | -0.0882747 | 0.64972899 | 0.89054584 |
| Aff4 | 0.486381 | 0.39188099 | 0.48016678 | -0.0882858 | 0.8041713 | 0.89054584 |
| Pum2 | 0.87072764 | 0.67447105 | 0.76280126 | -0.0883302 | 0.70522633 | 0.89054584 |
| Wac | 0.86335792 | 0.66990991 | 0.75836314 | -0.0884532 | 0.70657455 | 0.89054584 |
| Ldb1 | 0.43295966 | 0.34467633 | 0.43318691 | -0.0885106 | 0.8294349 | 0.89271879 |
| Fubp1 | 1.89143374 | 1.02027802 | 1.10881296 | -0.0885349 | 0.64505379 | 0.89054584 |
| Gpr108 | 0.43194862 | 0.34372631 | 0.4322816 | -0.0885553 | 0.83006192 | 0.89271879 |
| Pafah1b3 | 1.07958186 | 0.78651241 | 0.87517628 | -0.0886639 | 0.68153521 | 0.89054584 |
| Mrpl4 | 0.73181028 | 0.58381498 | 0.67251826 | -0.0887033 | 0.73055797 | 0.89054584 |
| Tmem167 | 1.63587672 | 0.97998674 | 1.06871475 | -0.088728 | 0.65054777 | 0.89054584 |
| Pdcl3 | 1.08688806 | 0.78961497 | 0.87878467 | -0.0891697 | 0.68180055 | 0.89054584 |
| Arl6ip6 | 0.66530133 | 0.53512004 | 0.62440908 | -0.089289 | 0.74736402 | 0.89054584 |
| Glo1 | 1.79184158 | 1.00727137 | 1.09659202 | -0.0893207 | 0.64783594 | 0.89054584 |
| Slc25a3 | 2.69233688 | 1.02786714 | 1.11743893 | -0.0895718 | 0.64559393 | 0.89054584 |
| Polr1c | 0.68407765 | 0.54857791 | 0.63830431 | -0.0897264 | 0.74373621 | 0.89054584 |
| Timm9 | 0.46392564 | 0.37084955 | 0.46062592 | -0.0897764 | 0.81806922 | 0.89054584 |
| Ndufab1 | 2.15834387 | 1.03613936 | 1.12591588 | -0.0897765 | 0.64486272 | 0.89054584 |
| Luzp1 | 0.66204881 | 0.53219016 | 0.62197715 | -0.089787 | 0.74938399 | 0.89054584 |
| Hacd2 | 0.48197484 | 0.38654297 | 0.47635675 | -0.0898138 | 0.81013971 | 0.89054584 |

| | | | | | | |
|---|---|---|---|---|---|---|
| Rab6a | 1.09740791 | 0.79405694 | 0.88391744 | -0.0898605 | 0.68212012 | 0.89054584 |
| Atg3 | 0.8699258 | 0.67244723 | 0.76232029 | -0.0898731 | 0.70858705 | 0.89054584 |
| Taf15 | 0.46955144 | 0.37557909 | 0.46554971 | -0.0899706 | 0.81603879 | 0.89054584 |
| Rnps1 | 0.89644037 | 0.68798149 | 0.77797586 | -0.0899944 | 0.70503417 | 0.89054584 |
| Krt10 | 0.48824897 | 0.39172591 | 0.4817784 | -0.0900525 | 0.80807772 | 0.89054584 |
| Nck1 | 0.67669437 | 0.5427609 | 0.63286939 | -0.0901085 | 0.7464449 | 0.89054584 |
| Vmp1 | 0.84643215 | 0.65788397 | 0.74802221 | -0.0901382 | 0.71274171 | 0.89054584 |
| Mrpl17 | 1.41211342 | 0.91917423 | 1.00951687 | -0.0903426 | 0.66163635 | 0.89054584 |
| Pop7 | 0.649009 | 0.52179017 | 0.61215426 | -0.0903641 | 0.75419071 | 0.89054584 |
| Ptrhd1 | 0.37205391 | 0.28722332 | 0.3776125 | -0.0903892 | 0.86762013 | 0.90444517 |
| 1810043H04I | 0.76319881 | 0.60365265 | 0.69413683 | -0.0904842 | 0.72817942 | 0.89054584 |
| Uck1 | 0.48929504 | 0.39219243 | 0.48267996 | -0.0904875 | 0.80877938 | 0.89054584 |
| Ggnbp2 | 0.86292755 | 0.66749009 | 0.75810282 | -0.0906127 | 0.71118947 | 0.89054584 |
| 2810428I15R | 1.02506982 | 0.75664025 | 0.84735682 | -0.0907166 | 0.69103221 | 0.89054584 |
| Nelfa | 0.52525903 | 0.42245817 | 0.51326756 | -0.0908094 | 0.79510945 | 0.89054584 |
| Unc50 | 0.82928369 | 0.64639785 | 0.73733185 | -0.090934 | 0.71721232 | 0.89054584 |
| Sugt1 | 1.01170414 | 0.7494563 | 0.84056001 | -0.0911037 | 0.69320556 | 0.89054584 |
| Eif5b | 1.7762136 | 1.00316564 | 1.094287 | -0.0911214 | 0.65097362 | 0.89054584 |
| Rwdd1 | 1.4751535 | 0.93702498 | 1.02862324 | -0.0915983 | 0.66088359 | 0.89054584 |
| Mesdc2 | 0.9092977 | 0.69363255 | 0.78538551 | -0.091753 | 0.70687648 | 0.89054584 |
| Plgrkt | 0.73569703 | 0.58340726 | 0.67523353 | -0.0918263 | 0.73682192 | 0.89054584 |
| Arglu1 | 1.13321084 | 0.80889505 | 0.90081286 | -0.0919178 | 0.68274639 | 0.89054584 |
| Chic2 | 0.64499742 | 0.51713586 | 0.60910916 | -0.0919733 | 0.75912272 | 0.89054584 |
| Exosc3 | 0.54331938 | 0.43620954 | 0.52832202 | -0.0921125 | 0.79168517 | 0.89054584 |
| Exoc3 | 0.58160841 | 0.46731934 | 0.55954595 | -0.0922266 | 0.7787283 | 0.89054584 |
| Abce1 | 0.60122017 | 0.48293846 | 0.57516673 | -0.0922283 | 0.77248739 | 0.89054584 |
| Rtn3 | 1.50444948 | 0.94453669 | 1.03677641 | -0.0922397 | 0.66074447 | 0.89054584 |
| Vps4b | 0.72212992 | 0.57345597 | 0.6657099 | -0.0922539 | 0.74074212 | 0.89054584 |
| Ddx54 | 0.50038347 | 0.39991861 | 0.49219631 | -0.0922777 | 0.80879523 | 0.89054584 |
| Dctn6 | 0.54880307 | 0.44055934 | 0.53285174 | -0.0922924 | 0.79015243 | 0.89054584 |

| | | | | | | |
|---|---|---|---|---|---|---|
| Bfar | 0.60637774 | 0.48687328 | 0.57923298 | -0.0923597 | 0.7712217 | 0.89054584 |
| Psmg4 | 0.64366814 | 0.51568933 | 0.60809768 | -0.0924084 | 0.76052324 | 0.89054584 |
| Nudcd3 | 0.86368492 | 0.66605207 | 0.75856085 | -0.0925088 | 0.71504087 | 0.89054584 |
| Adss | 0.71301245 | 0.56668302 | 0.65923619 | -0.0925532 | 0.74347307 | 0.89054584 |
| Mrps35 | 0.46235537 | 0.36669055 | 0.45924827 | -0.0925577 | 0.82612727 | 0.89170439 |
| Tnpo1 | 0.45443429 | 0.35957028 | 0.45227607 | -0.0927058 | 0.83020397 | 0.89271879 |
| Ubfd1 | 0.72841074 | 0.57736011 | 0.67013495 | -0.0927748 | 0.74053422 | 0.89054584 |
| Use1 | 1.32417259 | 0.88663638 | 0.97949793 | -0.0928616 | 0.670644 | 0.89054584 |
| Ddx42 | 0.68176631 | 0.5437157 | 0.63660683 | -0.0928911 | 0.75168892 | 0.89054584 |
| Hnrnpr | 1.64101799 | 0.97689432 | 1.06980444 | -0.0929101 | 0.65712105 | 0.89054584 |
| Rbbp6 | 0.7857439 | 0.61625532 | 0.70922796 | -0.0929726 | 0.72931421 | 0.89054584 |
| Ap2a1 | 0.58185227 | 0.46674471 | 0.55974176 | -0.0929971 | 0.78054922 | 0.89054584 |
| Trappc5 | 0.54731078 | 0.43856094 | 0.53162073 | -0.0930598 | 0.79261951 | 0.89054584 |
| Rsrc2 | 1.8050835 | 1.00537766 | 1.09846614 | -0.0930885 | 0.65351124 | 0.89054584 |
| Snrpe | 2.46264959 | 1.03451371 | 1.12800306 | -0.0934893 | 0.65030163 | 0.89054584 |
| Sin3b | 1.49975967 | 0.94196315 | 1.03550047 | -0.0935373 | 0.66306575 | 0.89054584 |
| Ep400 | 0.46359373 | 0.3667035 | 0.46033485 | -0.0936314 | 0.82833879 | 0.89229532 |
| Qrich1 | 0.70447263 | 0.55930052 | 0.65311922 | -0.0938187 | 0.74832992 | 0.89054584 |
| Ten1 | 0.76485861 | 0.60138076 | 0.69526042 | -0.0938797 | 0.7353542 | 0.89054584 |
| Gtf2h5 | 1.66473488 | 0.98068147 | 1.07467077 | -0.0939893 | 0.65816672 | 0.89054584 |
| Cyfip1 | 0.54281389 | 0.43386875 | 0.52790354 | -0.0940348 | 0.79669987 | 0.89054584 |
| Snrnp70 | 1.49054796 | 0.93889293 | 1.03295947 | -0.0940665 | 0.6643144 | 0.89054584 |
| Alkbh5 | 0.70023952 | 0.55595151 | 0.65006849 | -0.094117 | 0.75001024 | 0.89054584 |
| Fam120a | 0.61903809 | 0.49500916 | 0.58913801 | -0.0941289 | 0.77167578 | 0.89054584 |
| Syncrip | 1.00999246 | 0.74541482 | 0.83968117 | -0.0942663 | 0.69952452 | 0.89054584 |
| Asna1 | 0.95854261 | 0.71828574 | 0.81270387 | -0.0944181 | 0.70583767 | 0.89054584 |
| Skp1a | 1.38578569 | 0.90636731 | 1.00088891 | -0.0945216 | 0.67002402 | 0.89054584 |
| Snrnp40 | 0.96386334 | 0.72096143 | 0.81555654 | -0.0945951 | 0.70553362 | 0.89054584 |
| Evl | 0.62159718 | 0.49651141 | 0.5911269 | -0.0946155 | 0.77207806 | 0.89054584 |
| Ddb1 | 1.44991031 | 0.92650586 | 1.02124586 | -0.09474 | 0.66720398 | 0.89054584 |

| | | | | | | |
|---|---|---|---|---|---|---|
| Sap18 | 1.65588659 | 0.97811903 | 1.07288652 | -0.0947675 | 0.65965501 | 0.89054584 |
| mt-Atp8 | 0.9283349 | 0.70130387 | 0.7961428 | -0.0948389 | 0.71054825 | 0.89054584 |
| Ndufs6 | 1.95771436 | 1.01977438 | 1.11470563 | -0.0949313 | 0.65423059 | 0.89054584 |
| Gtf3c1 | 0.40197244 | 0.31019217 | 0.40517245 | -0.0949803 | 0.86261557 | 0.90267402 |
| Apip | 0.55216522 | 0.44055698 | 0.53561989 | -0.0950629 | 0.79584699 | 0.89054584 |
| Slc3a2 | 1.01556393 | 0.74728533 | 0.84253415 | -0.0952488 | 0.70079734 | 0.89054584 |
| Timm50 | 0.57520733 | 0.45908938 | 0.5543919 | -0.0953025 | 0.78840333 | 0.89054584 |
| Txndc12 | 0.97973846 | 0.72845588 | 0.8239764 | -0.0955205 | 0.70543378 | 0.89054584 |
| Hiat1 | 0.78317177 | 0.61193185 | 0.70752481 | -0.095593 | 0.7354776 | 0.89054584 |
| Fuca1 | 1.05083124 | 0.76495711 | 0.86066422 | -0.0957071 | 0.69779914 | 0.89054584 |
| Bptf | 0.94088679 | 0.70730905 | 0.80309434 | -0.0957853 | 0.71078289 | 0.89054584 |
| Pak1ip1 | 0.81488 | 0.63231273 | 0.72818697 | -0.0958742 | 0.73019523 | 0.89054584 |
| Mrps26 | 0.8035451 | 0.62490941 | 0.72088466 | -0.0959752 | 0.73245993 | 0.89054584 |
| Dnajc8 | 1.50785397 | 0.941679 | 1.0376952 | -0.0960162 | 0.66679637 | 0.89054584 |
| Dnpep | 1.04407377 | 0.76116796 | 0.85721738 | -0.0960494 | 0.69917341 | 0.89054584 |
| Plxnb2 | 0.58759614 | 0.46826724 | 0.56434244 | -0.0960752 | 0.78615505 | 0.89054584 |
| Srsf5 | 1.7155799 | 0.98819006 | 1.08429565 | -0.0961056 | 0.66017513 | 0.89054584 |
| Stag1 | 0.46159815 | 0.36247108 | 0.4585834 | -0.0961123 | 0.83557811 | 0.89459411 |
| Psmd3 | 1.13858476 | 0.80716842 | 0.90328093 | -0.0961125 | 0.68995718 | 0.89054584 |
| Fkbp1a | 2.9753663 | 1.00095464 | 1.09716562 | -0.096211 | 0.65856313 | 0.89054584 |
| Srsf1 | 0.96538832 | 0.72014613 | 0.81637033 | -0.0962242 | 0.70854584 | 0.89054584 |
| Arid1a | 0.80711737 | 0.62693756 | 0.72319602 | -0.0962585 | 0.73240796 | 0.89054584 |
| Ptbp1 | 0.82293131 | 0.63698902 | 0.73331747 | -0.0963285 | 0.72973035 | 0.89054584 |
| BC029214 | 0.43521229 | 0.33886725 | 0.43520181 | -0.0963346 | 0.8487964 | 0.89883797 |
| Knop1 | 1.20110634 | 0.83462888 | 0.93096887 | -0.09634 | 0.68514103 | 0.89054584 |
| Med29 | 0.86473556 | 0.66281575 | 0.75919557 | -0.0963798 | 0.72289127 | 0.89054584 |
| Mrps14 | 1.51684224 | 0.94375078 | 1.04013876 | -0.096388 | 0.66703355 | 0.89054584 |
| Zfp106 | 0.7582295 | 0.59422099 | 0.69076105 | -0.0965401 | 0.74251836 | 0.89054584 |
| Hmox2 | 0.85538586 | 0.65684317 | 0.7535188 | -0.0966756 | 0.72499229 | 0.89054584 |
| Zfp451 | 0.49364336 | 0.38970244 | 0.48642027 | -0.0967178 | 0.82287181 | 0.89075401 |

| | | | | | | |
|---|---|---|---|---|---|---|
| Utp3 | 0.83225883 | 0.64240269 | 0.73920181 | -0.0967991 | 0.7290997 | 0.89054584 |
| Nras | 0.73816327 | 0.58013312 | 0.67695086 | -0.0968177 | 0.74739778 | 0.89054584 |
| Cops7a | 0.91522464 | 0.69170019 | 0.78876019 | -0.09706 | 0.71677946 | 0.89054584 |
| Smc1a | 1.04284209 | 0.75950306 | 0.85658577 | -0.0970827 | 0.70126061 | 0.89054584 |
| Trim44 | 0.57714082 | 0.45882437 | 0.55595158 | -0.0971272 | 0.79215968 | 0.89054584 |
| Ncstn | 0.54716853 | 0.43436127 | 0.53150331 | -0.097142 | 0.80275626 | 0.89054584 |
| Twistnb | 0.52008669 | 0.41174034 | 0.50891763 | -0.0971773 | 0.813165 | 0.89054584 |
| Gatad2b | 0.77852339 | 0.60716814 | 0.70443482 | -0.0972667 | 0.73998382 | 0.89054584 |
| Rad23b | 1.4352504 | 0.91938803 | 1.01677867 | -0.0973906 | 0.67229158 | 0.89054584 |
| Gm10093 | 0.76193761 | 0.59584041 | 0.69328173 | -0.0974413 | 0.74370754 | 0.89054584 |
| Mthfsl | 0.50185051 | 0.39598466 | 0.49344978 | -0.0974651 | 0.8212721 | 0.89054584 |
| Brix1 | 0.71216922 | 0.5611464 | 0.65863447 | -0.0974881 | 0.75477563 | 0.89054584 |
| Emd | 0.70870021 | 0.55866305 | 0.65615372 | -0.0974907 | 0.75560067 | 0.89054584 |
| Serbp1 | 2.9465515 | 1.00185234 | 1.09943237 | -0.09758 | 0.66038129 | 0.89054584 |
| Ndufa7 | 2.47817689 | 1.02992522 | 1.12756947 | -0.0976443 | 0.6566917 | 0.89054584 |
| Naa50 | 0.64530188 | 0.51153738 | 0.60934066 | -0.0978033 | 0.77270702 | 0.89054584 |
| 0610037L13F | 0.67467866 | 0.53350298 | 0.63137917 | -0.0978762 | 0.76492271 | 0.89054584 |
| Polr2j | 1.23391507 | 0.84675289 | 0.94464717 | -0.0978943 | 0.68538413 | 0.89054584 |
| Rpp21 | 0.91417537 | 0.68995016 | 0.78816464 | -0.0982145 | 0.7192309 | 0.89054584 |
| Hsd17b12 | 0.71310229 | 0.56108124 | 0.65930027 | -0.098219 | 0.756179 | 0.89054584 |
| Yipf1 | 0.68954972 | 0.54404608 | 0.64230807 | -0.098262 | 0.76199579 | 0.89054584 |
| Ube2l3 | 1.55962518 | 0.95289047 | 1.05116394 | -0.0982735 | 0.66842384 | 0.89054584 |
| Tm9sf2 | 0.96446922 | 0.71756298 | 0.81588006 | -0.0983171 | 0.71274527 | 0.89054584 |
| Ndnl2 | 0.82060751 | 0.63349518 | 0.73184156 | -0.0983464 | 0.73436814 | 0.89054584 |
| Mrpl36 | 0.73993566 | 0.57978953 | 0.67818235 | -0.0983928 | 0.75045865 | 0.89054584 |
| Mum1 | 0.6042234 | 0.47904836 | 0.57753668 | -0.0984883 | 0.78654699 | 0.89054584 |
| Mga | 0.57983112 | 0.45940031 | 0.55811763 | -0.0987173 | 0.79504512 | 0.89054584 |
| Cdc26 | 1.3306154 | 0.88310105 | 0.98183166 | -0.0987306 | 0.68028354 | 0.89054584 |
| Ywhaz | 1.13782897 | 0.80412467 | 0.90293498 | -0.0988103 | 0.69491785 | 0.89054584 |
| Esd | 1.96825253 | 1.01648724 | 1.11552256 | -0.0990353 | 0.6604229 | 0.89054584 |

| | | | | | | |
|---|---|---|---|---|---|---|
| Chd9 | 0.45839096 | 0.35669659 | 0.45576351 | -0.0990669 | 0.84439577 | 0.89775379 |
| Rbm10 | 0.47571137 | 0.37179584 | 0.47092005 | -0.0991242 | 0.83662053 | 0.89462889 |
| Bag1 | 1.83964008 | 1.00362126 | 1.10298653 | -0.0993653 | 0.66265216 | 0.89054584 |
| Tgoln1 | 0.54243451 | 0.4281915 | 0.52758934 | -0.0993978 | 0.80995884 | 0.89054584 |
| Ndufa10 | 1.4378902 | 0.91818753 | 1.01759199 | -0.0994045 | 0.6754861 | 0.89054584 |
| Hdac1 | 1.51233244 | 0.93949229 | 1.03891295 | -0.0994207 | 0.67214569 | 0.89054584 |
| Tor2a | 0.92235212 | 0.69328612 | 0.79278862 | -0.0995025 | 0.72063918 | 0.89054584 |
| Slirp | 1.35442543 | 0.890691 | 0.99023817 | -0.0995472 | 0.68023514 | 0.89054584 |
| Ccni | 0.72263286 | 0.56650064 | 0.66606528 | -0.0995646 | 0.75691925 | 0.89054584 |
| Uchl3 | 0.81159806 | 0.62639486 | 0.72608202 | -0.0996872 | 0.73879914 | 0.89054584 |
| Swt1 | 0.60451513 | 0.47803662 | 0.57776657 | -0.0997299 | 0.78935731 | 0.89054584 |
| Tex261 | 0.75696542 | 0.58984285 | 0.68989964 | -0.1000568 | 0.75038145 | 0.89054584 |
| Rnf4 | 0.69366785 | 0.54525015 | 0.64530726 | -0.1000571 | 0.76497104 | 0.89054584 |
| Commd1 | 1.51255449 | 0.93885751 | 1.03897356 | -0.1001161 | 0.67326184 | 0.89054584 |
| Phpt1 | 0.97749663 | 0.72265544 | 0.82279523 | -0.1001398 | 0.71465136 | 0.89054584 |
| Gnb2 | 3.19121549 | 0.97622969 | 1.076388 | -0.1001583 | 0.66768012 | 0.89054584 |
| Zmynd8 | 0.72025873 | 0.56408963 | 0.66438613 | -0.1002965 | 0.75907157 | 0.89054584 |
| Chtf8 | 0.69876316 | 0.54866694 | 0.64900152 | -0.1003346 | 0.76431899 | 0.89054584 |
| Cyhr1 | 0.6624435 | 0.52178573 | 0.62227266 | -0.1004869 | 0.7740777 | 0.89054584 |
| Tulp4 | 0.6292079 | 0.49625968 | 0.59701582 | -0.1007561 | 0.78414596 | 0.89054584 |
| Ndufv1 | 1.00302726 | 0.73532277 | 0.83608344 | -0.1007607 | 0.71276234 | 0.89054584 |
| Arpc2 | 2.4559281 | 1.02740432 | 1.12817591 | -0.1007716 | 0.6613494 | 0.89054584 |
| Mrpl28 | 1.19379251 | 0.82692813 | 0.92781711 | -0.100889 | 0.69378364 | 0.89054584 |
| Rbm42 | 0.86345902 | 0.65746973 | 0.75842428 | -0.1009545 | 0.73242242 | 0.89054584 |
| Ndufa13 | 2.63056265 | 1.01977534 | 1.12092004 | -0.1011447 | 0.66290089 | 0.89054584 |
| Srsf11 | 1.88021802 | 1.00646914 | 1.10762713 | -0.101158 | 0.66476184 | 0.89054584 |
| Ago2 | 0.59793069 | 0.47140407 | 0.57256449 | -0.1011604 | 0.79478566 | 0.89054584 |
| Ict1 | 1.03125473 | 0.74942712 | 0.85059305 | -0.1011659 | 0.71024983 | 0.89054584 |
| Pfdn5 | 3.00177194 | 0.99371943 | 1.09493967 | -0.1012202 | 0.66665297 | 0.89054584 |
| Mrpl2 | 0.68789727 | 0.53986217 | 0.64110124 | -0.1012391 | 0.76904748 | 0.89054584 |

| | | | | | | |
|---|---|---|---|---|---|---|
| Nsa2 | 1.30252228 | 0.87027011 | 0.97151532 | -0.1012452 | 0.6863482 | 0.89054584 |
| Rere | 0.59071124 | 0.46533063 | 0.56682833 | -0.1014977 | 0.7979276 | 0.89054584 |
| Ube2d2a | 1.92723375 | 1.01062751 | 1.11220491 | -0.1015774 | 0.66476386 | 0.89054584 |
| Snrpd3 | 1.80782861 | 0.99714112 | 1.09884531 | -0.1017042 | 0.66684241 | 0.89054584 |
| Znhit1 | 1.2617278 | 0.8541549 | 0.95586351 | -0.1017086 | 0.6899591 | 0.89054584 |
| Atxn10 | 1.67446736 | 0.97482194 | 1.07659291 | -0.101771 | 0.67018081 | 0.89054584 |
| Sfpq | 1.23839617 | 0.84457916 | 0.94645896 | -0.1018798 | 0.6919914 | 0.89054584 |
| Prpf19 | 1.31108759 | 0.87276845 | 0.97470489 | -0.1019364 | 0.68695612 | 0.89054584 |
| Cxxc1 | 0.51756216 | 0.4045663 | 0.50678892 | -0.1022226 | 0.82632421 | 0.89170439 |
| Mllt10 | 0.695433 | 0.54426032 | 0.64658913 | -0.1023288 | 0.76953649 | 0.89054584 |
| Uqcrc1 | 1.81881303 | 0.9978775 | 1.10032537 | -0.1024479 | 0.66777571 | 0.89054584 |
| Rsbn1l | 0.47743381 | 0.36995404 | 0.47241754 | -0.1024635 | 0.8438688 | 0.89775379 |
| Hnrnpl | 0.93419582 | 0.69687053 | 0.79940313 | -0.1025326 | 0.72493407 | 0.89054584 |
| Gemin7 | 1.02216109 | 0.74329755 | 0.84588383 | -0.1025863 | 0.71397078 | 0.89054584 |
| Nono | 2.27615514 | 1.02612849 | 1.12879913 | -0.1026706 | 0.66412847 | 0.89054584 |
| Mrpl54 | 1.59737246 | 0.95746535 | 1.06017641 | -0.1027111 | 0.67413616 | 0.89054584 |
| Copg1 | 1.08256997 | 0.77394359 | 0.87665637 | -0.1027128 | 0.70741936 | 0.89054584 |
| Sdha | 1.38938196 | 0.89929967 | 1.00209243 | -0.1027928 | 0.68364124 | 0.89054584 |
| Tmem242 | 0.81350757 | 0.62440555 | 0.72730769 | -0.1029021 | 0.74510475 | 0.89054584 |
| Mbtps1 | 0.73704863 | 0.57307059 | 0.67617524 | -0.1031046 | 0.76127359 | 0.89054584 |
| Stard7 | 0.54065964 | 0.42291671 | 0.52611819 | -0.1032015 | 0.81961042 | 0.89054584 |
| Trnau1ap | 0.55468031 | 0.43441183 | 0.5376858 | -0.103274 | 0.81456598 | 0.89054584 |
| Kmt2e | 1.23428319 | 0.84143112 | 0.94479649 | -0.1033654 | 0.69487302 | 0.89054584 |
| Emg1 | 1.37734907 | 0.89464616 | 0.99803608 | -0.1033899 | 0.68532269 | 0.89054584 |
| Dbnl | 0.94551407 | 0.70219486 | 0.80563124 | -0.1034364 | 0.72513656 | 0.89054584 |
| Bod1l | 0.74087617 | 0.57527689 | 0.67883491 | -0.103558 | 0.76137532 | 0.89054584 |
| Snrpa1 | 0.75997732 | 0.58833113 | 0.69195037 | -0.1036192 | 0.75730954 | 0.89054584 |
| Nsrp1 | 0.65809611 | 0.51535394 | 0.61901178 | -0.1036578 | 0.78235523 | 0.89054584 |
| Pbrm1 | 1.20849691 | 0.83042854 | 0.9341189 | -0.1036904 | 0.69747714 | 0.89054584 |
| Pja1 | 1.01860526 | 0.74035587 | 0.84408221 | -0.1037263 | 0.71651552 | 0.89054584 |

| | | | | | | |
|---|---|---|---|---|---|---|
| Mrps5 | 0.69308756 | 0.54113333 | 0.64488537 | -0.103752 | 0.77323749 | 0.89054584 |
| Arl1 | 1.87591061 | 1.00316011 | 1.10715853 | -0.1039984 | 0.66917167 | 0.89054584 |
| Kif3a | 0.56623244 | 0.44308317 | 0.54712153 | -0.1040384 | 0.81217101 | 0.89054584 |
| Mrpl11 | 0.90919382 | 0.68115204 | 0.78532613 | -0.1041741 | 0.7317163 | 0.89054584 |
| Cul4a | 0.75198494 | 0.58198836 | 0.68649446 | -0.1045061 | 0.76091152 | 0.89054584 |
| Stx6 | 0.70248075 | 0.54712184 | 0.65168529 | -0.1045635 | 0.77262845 | 0.89054584 |
| Ube2r2 | 0.85584595 | 0.64916205 | 0.75379966 | -0.1046376 | 0.74109476 | 0.89054584 |
| Ncor1 | 1.15148978 | 0.8044055 | 0.90912969 | -0.1047242 | 0.70425267 | 0.89054584 |
| Slc25a11 | 1.19242605 | 0.82246995 | 0.92722446 | -0.1047545 | 0.70067767 | 0.89054584 |
| Thrap3 | 1.30057602 | 0.86591171 | 0.97078433 | -0.1048726 | 0.69262319 | 0.89054584 |
| Vdac3 | 1.6742015 | 0.97166219 | 1.07654077 | -0.1048786 | 0.67504334 | 0.89054584 |
| Igbp1 | 0.68494908 | 0.53377188 | 0.63894332 | -0.1051714 | 0.7784211 | 0.89054584 |
| Rexo1 | 0.53470173 | 0.41598164 | 0.5211649 | -0.1051833 | 0.82646385 | 0.89172233 |
| Mtif2 | 0.45866923 | 0.35054985 | 0.45600843 | -0.1054586 | 0.8593599 | 0.90131997 |
| Prkrip1 | 0.55331584 | 0.43103328 | 0.53656553 | -0.1055323 | 0.82025185 | 0.89054584 |
| Rbm26 | 0.49419422 | 0.38123448 | 0.48689326 | -0.1056588 | 0.84399619 | 0.89775379 |
| Zfp358 | 0.46187793 | 0.35315251 | 0.4588291 | -0.1056766 | 0.85837295 | 0.90124516 |
| Serinc1 | 1.46984836 | 0.92129266 | 1.02710415 | -0.1058115 | 0.6843723 | 0.89054584 |
| Rpl36 | 4.63575669 | 0.699275 | 0.80512637 | -0.1058514 | 0.72990283 | 0.89054584 |
| Cct3 | 2.11237395 | 1.01796782 | 1.1239939 | -0.1060261 | 0.66984198 | 0.89054584 |
| Wtap | 0.96597506 | 0.71047202 | 0.81668298 | -0.106211 | 0.7277104 | 0.89054584 |
| Faf1 | 0.51280007 | 0.39653183 | 0.50276244 | -0.1062306 | 0.83754192 | 0.89485554 |
| Smarcb1 | 1.12461909 | 0.79054139 | 0.89683531 | -0.1062939 | 0.70958847 | 0.89054584 |
| Ssbp3 | 0.75174656 | 0.5800338 | 0.68633103 | -0.1062972 | 0.76471906 | 0.89054584 |
| Ccnl1 | 1.0686051 | 0.76318317 | 0.8696989 | -0.1065157 | 0.71586473 | 0.89054584 |
| Tmco1 | 1.54438107 | 0.94083955 | 1.04736623 | -0.1065267 | 0.68219896 | 0.89054584 |
| Ash1l | 1.17534421 | 0.8131125 | 0.91976628 | -0.1066538 | 0.70548393 | 0.89054584 |
| H13 | 1.15208759 | 0.80239191 | 0.90939796 | -0.107006 | 0.70822381 | 0.89054584 |
| Sh3glb2 | 0.5404463 | 0.41891116 | 0.52594122 | -0.1070301 | 0.82846165 | 0.89235044 |
| Exosc5 | 0.52975015 | 0.40991995 | 0.51703083 | -0.1071109 | 0.83277048 | 0.89393134 |

| | | | | | | |
|---|---|---|---|---|---|---|
| Mettl23 | 0.55960413 | 0.43457487 | 0.54171828 | -0.1071434 | 0.82159636 | 0.89054584 |
| Fam114a2 | 0.75489065 | 0.58132395 | 0.68848329 | -0.1071593 | 0.76581794 | 0.89054584 |
| Poldip3 | 0.63437996 | 0.49382157 | 0.60099604 | -0.1071745 | 0.79696236 | 0.89054584 |
| Zfp260 | 0.72704145 | 0.56179623 | 0.66917265 | -0.1073764 | 0.77264338 | 0.89054584 |
| Psmd8 | 1.74155174 | 0.98140835 | 1.08879054 | -0.1073822 | 0.67705848 | 0.89054584 |
| Zcrb1 | 1.4617542 | 0.9171576 | 1.02475639 | -0.1075988 | 0.68764498 | 0.89054584 |
| Mcrs1 | 0.57759117 | 0.44859245 | 0.55631452 | -0.1077221 | 0.8165074 | 0.89054584 |
| Atp6v0e | 1.96868643 | 1.00775347 | 1.11555533 | -0.1078019 | 0.67372319 | 0.89054584 |
| Rars | 0.73064876 | 0.56345739 | 0.67170489 | -0.1082475 | 0.77361469 | 0.89054584 |
| Cryzl1 | 0.82659231 | 0.62730323 | 0.73563464 | -0.1083314 | 0.75367102 | 0.89054584 |
| Yipf3 | 0.94769108 | 0.69846327 | 0.80682169 | -0.1083584 | 0.73424705 | 0.89054584 |
| N6amt2 | 0.66407103 | 0.51481309 | 0.62349005 | -0.108677 | 0.79162517 | 0.89054584 |
| Irf2bp1 | 0.53570038 | 0.41322447 | 0.52199677 | -0.1087723 | 0.83418048 | 0.89427341 |
| Cnot4 | 0.8118518 | 0.61745048 | 0.72624505 | -0.1087946 | 0.75738922 | 0.89054584 |
| Ddx3x | 0.87032383 | 0.65371506 | 0.7625591 | -0.108844 | 0.74697316 | 0.89054584 |
| Dpm1 | 0.80986961 | 0.61612465 | 0.72497044 | -0.1088458 | 0.7578756 | 0.89054584 |
| Topors | 0.5888247 | 0.45644335 | 0.56532363 | -0.1088803 | 0.81522147 | 0.89054584 |
| Puf60 | 1.38198869 | 0.89053941 | 0.99961008 | -0.1090707 | 0.69439676 | 0.89054584 |
| Sdf4 | 1.48769468 | 0.92301012 | 1.03216303 | -0.1091529 | 0.68888437 | 0.89054584 |
| Glg1 | 0.72099688 | 0.55571474 | 0.66490863 | -0.1091939 | 0.77789682 | 0.89054584 |
| Gps1 | 1.03244699 | 0.74179906 | 0.85121388 | -0.1094148 | 0.7253666 | 0.89054584 |
| Gm10076 | 2.7519289 | 1.00398899 | 1.11375056 | -0.1097616 | 0.67693487 | 0.89054584 |
| Myeov2 | 2.13666841 | 1.01517846 | 1.12506721 | -0.1098887 | 0.67546444 | 0.89054584 |
| Tomm22 | 1.59348355 | 0.94931586 | 1.05927589 | -0.10996 | 0.68567566 | 0.89054584 |
| Eif3h | 2.63178329 | 1.01082976 | 1.12085484 | -0.1100251 | 0.67628301 | 0.89054584 |
| Atp5g3 | 2.14082983 | 1.01512919 | 1.1252388 | -0.1101096 | 0.67576836 | 0.89054584 |
| Hibadh | 0.79583899 | 0.60567279 | 0.71586685 | -0.1101941 | 0.76336083 | 0.89054584 |
| Atad1 | 0.60283493 | 0.46624439 | 0.57644175 | -0.1101974 | 0.81343928 | 0.89054584 |
| Hnrnph2 | 0.92765213 | 0.68546122 | 0.79576134 | -0.1103001 | 0.74078704 | 0.89054584 |
| Atp6ap1 | 0.84442356 | 0.63638337 | 0.74678107 | -0.1103977 | 0.75450586 | 0.89054584 |

| | | | | | | |
|---|---|---|---|---|---|---|
| Polr2k | 1.5129022 | 0.92861137 | 1.03906842 | -0.1104571 | 0.68979302 | 0.89054584 |
| Rsrp1 | 2.8862425 | 0.99359498 | 1.10412722 | -0.1105322 | 0.67953484 | 0.89054584 |
| Hint2 | 0.86406989 | 0.64825668 | 0.75879351 | -0.1105368 | 0.75133244 | 0.89054584 |
| Mrpl9 | 0.58574829 | 0.45227108 | 0.56286477 | -0.1105937 | 0.82001059 | 0.89054584 |
| Med8 | 0.59772803 | 0.46172712 | 0.57240395 | -0.1106768 | 0.81615979 | 0.89054584 |
| Dnajc7 | 1.24414371 | 0.83806278 | 0.94879457 | -0.1107318 | 0.70668811 | 0.89054584 |
| Poldip2 | 0.66094204 | 0.51038754 | 0.62114793 | -0.1107604 | 0.79694599 | 0.89054584 |
| Dap3 | 0.70876971 | 0.54529875 | 0.6562035 | -0.1109048 | 0.78447795 | 0.89054584 |
| Timm13 | 2.20475289 | 1.01642586 | 1.12738397 | -0.1109581 | 0.67671667 | 0.89054584 |
| B230219D22 | 1.06269755 | 0.75553089 | 0.86671882 | -0.1111879 | 0.72497232 | 0.89054584 |
| St13 | 2.05641796 | 1.00983539 | 1.12113447 | -0.1112991 | 0.67814217 | 0.89054584 |
| Cuta | 1.40072866 | 0.89430895 | 1.00584067 | -0.1115317 | 0.69728314 | 0.89054584 |
| Cbx6 | 1.28316321 | 0.85260167 | 0.96420084 | -0.1115992 | 0.70513876 | 0.89054584 |
| Mrpl19 | 0.51721604 | 0.39466535 | 0.50649675 | -0.1118314 | 0.84818165 | 0.89876755 |
| Ankrd17 | 0.77720346 | 0.59165526 | 0.70355455 | -0.1118993 | 0.77065254 | 0.89054584 |
| Psmd7 | 1.61019107 | 0.95113535 | 1.06309081 | -0.1119555 | 0.68815913 | 0.89054584 |
| Snw1 | 1.12313533 | 0.78407839 | 0.89614397 | -0.1120656 | 0.71992376 | 0.89054584 |
| Cnot6 | 0.64137326 | 0.4942512 | 0.60634857 | -0.1120974 | 0.80546678 | 0.89054584 |
| Txnl1 | 1.78158065 | 0.98285965 | 1.0950884 | -0.1122287 | 0.68348473 | 0.89054584 |
| Stx5a | 0.9586594 | 0.70052573 | 0.81276671 | -0.112241 | 0.74001371 | 0.89054584 |
| Prpf4b | 1.10639659 | 0.77579493 | 0.88824308 | -0.1124481 | 0.72233897 | 0.89054584 |
| Uba2 | 1.23450575 | 0.83229456 | 0.94488672 | -0.1125922 | 0.71060627 | 0.89054584 |
| Ddx46 | 0.83123381 | 0.6259121 | 0.73855829 | -0.1126462 | 0.76133075 | 0.89054584 |
| Fh1 | 0.8306336 | 0.62550666 | 0.7381811 | -0.1126744 | 0.76149853 | 0.89054584 |
| Pcyox1l | 0.45457402 | 0.33971089 | 0.45239939 | -0.1126885 | 0.87707718 | 0.90957546 |
| Nudt5 | 0.68664245 | 0.52748256 | 0.64018353 | -0.112701 | 0.7939439 | 0.89054584 |
| Naa20 | 1.20282013 | 0.81896972 | 0.93170244 | -0.1127327 | 0.71352612 | 0.89054584 |
| Psmb3 | 2.06051862 | 1.00828524 | 1.12135023 | -0.113065 | 0.68073601 | 0.89054584 |
| Mat2b | 0.62511888 | 0.48078561 | 0.5938566 | -0.113071 | 0.81247881 | 0.89054584 |
| Commd7 | 0.79082477 | 0.59943166 | 0.71257859 | -0.1131469 | 0.7702805 | 0.89054584 |

| | | | | | | |
|---|---|---|---|---|---|---|
| Ywhaq | 2.93710337 | 0.98700059 | 1.10017462 | -0.113174 | 0.68412364 | 0.89054584 |
| Mplkip | 0.6415149 | 0.49325991 | 0.60645663 | -0.1131967 | 0.80774146 | 0.89054584 |
| Lasp1 | 0.74121299 | 0.5657847 | 0.67906846 | -0.1132838 | 0.78148412 | 0.89054584 |
| Atp5o | 2.47940489 | 1.01419465 | 1.12753316 | -0.1133385 | 0.68021929 | 0.89054584 |
| Lman2l | 0.87386008 | 0.6512484 | 0.76467567 | -0.1134273 | 0.75524192 | 0.89054584 |
| Rbmxl1 | 1.05673831 | 0.7500576 | 0.86368888 | -0.1136313 | 0.73004659 | 0.89054584 |
| Ywhab | 1.83562877 | 0.98885132 | 1.1024926 | -0.1136413 | 0.68446742 | 0.89054584 |
| Rpl4 | 4.75398329 | 0.65915673 | 0.77303425 | -0.1138775 | 0.75374425 | 0.89054584 |
| Usp14 | 0.71574667 | 0.54726415 | 0.6611838 | -0.1139197 | 0.78893149 | 0.89054584 |
| Mrpl43 | 0.85573956 | 0.63972607 | 0.75373472 | -0.1140087 | 0.75950265 | 0.89054584 |
| Fam195b | 1.56863438 | 0.93932558 | 1.05337051 | -0.1140449 | 0.69301269 | 0.89054584 |
| Fgfr1op2 | 0.89993487 | 0.66563863 | 0.78000178 | -0.1143632 | 0.75272493 | 0.89054584 |
| Cox5b | 2.91492942 | 0.98717649 | 1.10192572 | -0.1147492 | 0.68621845 | 0.89054584 |
| Gtf2i | 0.97085626 | 0.70416937 | 0.81927517 | -0.1151058 | 0.74362697 | 0.89054584 |
| Apopt1 | 0.59560577 | 0.45556444 | 0.57072107 | -0.1151566 | 0.82640477 | 0.89172233 |
| Amz2 | 0.5336441 | 0.40510954 | 0.52028322 | -0.1151737 | 0.84880863 | 0.89883797 |
| Psmc3 | 1.51096074 | 0.92326591 | 1.03853795 | -0.115272 | 0.69746277 | 0.89054584 |
| Smn1 | 0.52150719 | 0.39481341 | 0.51011386 | -0.1153005 | 0.85384214 | 0.90035231 |
| Tmem214 | 0.63798703 | 0.48838694 | 0.60376103 | -0.1153741 | 0.81334431 | 0.89054584 |
| Hnrnph3 | 1.17150603 | 0.80262488 | 0.91807324 | -0.1154484 | 0.72101692 | 0.89054584 |
| Snrpd2 | 2.49559137 | 1.01155031 | 1.1270226 | -0.1154723 | 0.68344177 | 0.89054584 |
| Phf5a | 0.98726913 | 0.71242236 | 0.82791741 | -0.1154951 | 0.74212983 | 0.89054584 |
| Rbm7 | 0.80060165 | 0.60318037 | 0.71897316 | -0.1157928 | 0.77347396 | 0.89054584 |
| Rtf1 | 0.96259021 | 0.69902929 | 0.81487585 | -0.1158466 | 0.74612232 | 0.89054584 |
| Bola1 | 0.57119662 | 0.43521104 | 0.55114923 | -0.1159382 | 0.83646721 | 0.89462889 |
| Usp50 | 0.68083812 | 0.51981712 | 0.6359241 | -0.116107 | 0.80251231 | 0.89054584 |
| Txnrd1 | 0.79018876 | 0.59587938 | 0.7121602 | -0.1162808 | 0.77658407 | 0.89054584 |
| Iws1 | 0.54859547 | 0.41638944 | 0.53268057 | -0.1162911 | 0.84544892 | 0.89825055 |
| Usp5 | 0.54394499 | 0.41245465 | 0.52883973 | -0.1163851 | 0.84739447 | 0.89854206 |
| Spag7 | 1.35712365 | 0.87451126 | 0.99116886 | -0.1166576 | 0.70827194 | 0.89054584 |

| | | | | | | |
|---|---|---|---|---|---|---|
| Jtb | 1.16561116 | 0.79872038 | 0.91545431 | -0.1167339 | 0.72377193 | 0.89054584 |
| Ppp2r1a | 1.27129174 | 0.84276853 | 0.95962328 | -0.1168547 | 0.71475369 | 0.89054584 |
| Aip | 0.69116168 | 0.52652224 | 0.64348347 | -0.1169612 | 0.80145262 | 0.89054584 |
| Ttc1 | 0.70207674 | 0.53441995 | 0.65139411 | -0.1169742 | 0.79859965 | 0.89054584 |
| Kars | 0.67157854 | 0.51196158 | 0.62908169 | -0.1171201 | 0.80713432 | 0.89054584 |
| Nipsnap3b | 0.6498729 | 0.49566426 | 0.61280857 | -0.1171443 | 0.8134286 | 0.89054584 |
| Cdk12 | 0.79928118 | 0.60091538 | 0.71811358 | -0.1171982 | 0.77647808 | 0.89054584 |
| Dpp3 | 0.63896035 | 0.48704401 | 0.60450558 | -0.1174616 | 0.81734137 | 0.89054584 |
| Hdac2 | 1.34164591 | 0.8680861 | 0.98576928 | -0.1176832 | 0.7109609 | 0.89054584 |
| Ilf3 | 0.54283689 | 0.41020514 | 0.52792257 | -0.1177174 | 0.8505655 | 0.89942466 |
| Clptm1 | 0.83290567 | 0.62179879 | 0.73960751 | -0.1178087 | 0.77098709 | 0.89054584 |
| Csnk2b | 1.16044096 | 0.79523275 | 0.91313868 | -0.1179059 | 0.72626979 | 0.89054584 |
| Fau | 4.70268141 | 0.66921434 | 0.78718816 | -0.1179738 | 0.75748123 | 0.89054584 |
| Sucla2 | 0.8959892 | 0.65961695 | 0.77771364 | -0.1180967 | 0.76036135 | 0.89054584 |
| Stard3nl | 0.6443982 | 0.49032061 | 0.60865335 | -0.1183327 | 0.81747353 | 0.89054584 |
| Prkra | 0.65409541 | 0.4973987 | 0.61599931 | -0.1186006 | 0.81514136 | 0.89054584 |
| Uqcrfs1 | 1.82230392 | 0.98213237 | 1.10078502 | -0.1186527 | 0.69223066 | 0.89054584 |
| Mrps17 | 1.19281665 | 0.8084673 | 0.92739399 | -0.1189267 | 0.72489684 | 0.89054584 |
| Rpl18 | 4.59623288 | 0.69590618 | 0.8149501 | -0.1190439 | 0.75192354 | 0.89054584 |
| Mrpl32 | 1.10220356 | 0.76709699 | 0.88623275 | -0.1191358 | 0.73444052 | 0.89054584 |
| Ptov1 | 2.5697418 | 1.00483828 | 1.12401591 | -0.1191776 | 0.68933991 | 0.89054584 |
| Kif1b | 0.94771261 | 0.68736533 | 0.80683344 | -0.1194681 | 0.75485607 | 0.89054584 |
| Chd1 | 0.70609983 | 0.53465047 | 0.65428852 | -0.1196381 | 0.80286446 | 0.89054584 |
| Ilkap | 0.8535861 | 0.63275023 | 0.75241868 | -0.1196684 | 0.77064765 | 0.89054584 |
| Cdc42 | 2.77096065 | 0.99281436 | 1.112504 | -0.1196896 | 0.69189745 | 0.89054584 |
| Pmpcb | 0.81389436 | 0.60769452 | 0.72755564 | -0.1198611 | 0.77861885 | 0.89054584 |
| Ube2n | 2.14293344 | 1.00545381 | 1.12532358 | -0.1198698 | 0.69014863 | 0.89054584 |
| Prrc2b | 0.62633658 | 0.47460012 | 0.59479848 | -0.1201984 | 0.82677321 | 0.89184004 |
| Nucb1 | 1.14946549 | 0.78786319 | 0.90821953 | -0.1203563 | 0.73151395 | 0.89054584 |
| Lypla1 | 0.58743059 | 0.44381295 | 0.56421015 | -0.1203972 | 0.8399311 | 0.89545709 |

| | | | | | | |
|---|---|---|---|---|---|---|
| 1700123O20 | 0.79139534 | 0.592055 | 0.7129537 | -0.1208987 | 0.78524696 | 0.89054584 |
| Thap11 | 0.61175259 | 0.46254126 | 0.58345138 | -0.1209101 | 0.83284887 | 0.89393134 |
| Tpgs1 | 0.73081677 | 0.55075659 | 0.6718226 | -0.121066 | 0.79942412 | 0.89054584 |
| Ube2w | 0.86229779 | 0.63632059 | 0.75772163 | -0.121401 | 0.7723018 | 0.89054584 |
| Grcc10 | 1.19515963 | 0.80696363 | 0.92840885 | -0.1214452 | 0.72888239 | 0.89054584 |
| Msl1 | 0.87797484 | 0.64556551 | 0.7671269 | -0.1215614 | 0.76982504 | 0.89054584 |
| Emc8 | 0.69954772 | 0.52797176 | 0.64956872 | -0.121597 | 0.80843324 | 0.89054584 |
| Nsmce4a | 0.48826214 | 0.36005037 | 0.48178975 | -0.1217394 | 0.88044784 | 0.91159931 |
| Rpl37a | 5.58003435 | 0.55603404 | 0.67817233 | -0.1221383 | 0.79928544 | 0.89054584 |
| Stk16 | 0.85813537 | 0.63298097 | 0.75519486 | -0.1222139 | 0.77456362 | 0.89054584 |
| Thoc3 | 0.85362593 | 0.63013742 | 0.75244305 | -0.1223056 | 0.77556079 | 0.89054584 |
| Ubn1 | 0.55601041 | 0.41646447 | 0.53877669 | -0.1223122 | 0.85489599 | 0.90074724 |
| Hadha | 0.86408982 | 0.63643621 | 0.75880555 | -0.1223693 | 0.77377223 | 0.89054584 |
| Btf3l4 | 1.42201686 | 0.89023169 | 1.01265435 | -0.1224227 | 0.71336175 | 0.89054584 |
| Timm17a | 1.44142583 | 0.89620263 | 1.01867518 | -0.1224725 | 0.71230524 | 0.89054584 |
| Morf4l1 | 3.4258789 | 0.92091183 | 1.04455206 | -0.1236402 | 0.70934209 | 0.89054584 |
| Ddx47 | 0.67004655 | 0.50359629 | 0.62794385 | -0.1243476 | 0.82188675 | 0.89056159 |
| Dynll1 | 4.11345396 | 0.8126031 | 0.93697113 | -0.124368 | 0.73184429 | 0.89054584 |
| Sf3a3 | 0.75479119 | 0.56403323 | 0.68841531 | -0.1243821 | 0.80004224 | 0.89054584 |
| Ndufa2 | 2.81771181 | 0.98388706 | 1.10919828 | -0.1253112 | 0.70067469 | 0.89054584 |
| Psmb4 | 1.95800679 | 0.98939926 | 1.11472885 | -0.1253296 | 0.69979373 | 0.89054584 |
| Ubap2l | 1.49239908 | 0.90804725 | 1.03347381 | -0.1254266 | 0.71410863 | 0.89054584 |
| Zfp207 | 1.48972744 | 0.90723334 | 1.03273089 | -0.1254976 | 0.71435578 | 0.89054584 |
| Rps10 | 4.28300451 | 0.76841682 | 0.89405604 | -0.1256392 | 0.74367216 | 0.89054584 |
| Dda1 | 0.70412746 | 0.52716243 | 0.65287094 | -0.1257085 | 0.81515824 | 0.89054584 |
| Lman2 | 1.3726458 | 0.87072791 | 0.99643907 | -0.1257112 | 0.72165797 | 0.89054584 |
| Chrac1 | 0.7912675 | 0.58686432 | 0.71286967 | -0.1260054 | 0.79488947 | 0.89054584 |
| Erh | 3.22728193 | 0.94611866 | 1.07221942 | -0.1261008 | 0.70812836 | 0.89054584 |
| Ilf2 | 1.25262343 | 0.8259386 | 0.95222759 | -0.126289 | 0.73167644 | 0.89054584 |
| Sumo3 | 1.56800623 | 0.92683555 | 1.0532173 | -0.1263817 | 0.7119375 | 0.89054584 |

| | | | | | | |
|---|---|---|---|---|---|---|
| Rnf187 | 1.46144018 | 0.89807105 | 1.02466457 | -0.1265935 | 0.71755551 | 0.89054584 |
| Mettl16 | 0.49433036 | 0.36005692 | 0.48701016 | -0.1269532 | 0.88771915 | 0.91537513 |
| Raf1 | 0.68614015 | 0.51278877 | 0.63981586 | -0.1270271 | 0.82250977 | 0.89073269 |
| Phf3 | 0.78111893 | 0.57892513 | 0.70616213 | -0.127237 | 0.79939233 | 0.89054584 |
| Mocs2 | 0.8112138 | 0.5983857 | 0.72583506 | -0.1274494 | 0.79333949 | 0.89054584 |
| Ugp2 | 0.70801853 | 0.5281916 | 0.65566523 | -0.1274736 | 0.81747533 | 0.89054584 |
| Brk1 | 1.80760913 | 0.97108146 | 1.09881511 | -0.1277336 | 0.70594834 | 0.89054584 |
| Rab8a | 0.82178219 | 0.604841 | 0.73258813 | -0.1277471 | 0.79172353 | 0.89054584 |
| Pak2 | 0.80332957 | 0.59235496 | 0.7207449 | -0.1283899 | 0.79670822 | 0.89054584 |
| Phb | 0.90394571 | 0.65360861 | 0.78231597 | -0.1287074 | 0.7783086 | 0.89054584 |
| Ubxn1 | 1.41274194 | 0.88074539 | 1.00971766 | -0.1289723 | 0.72410878 | 0.89054584 |
| Hbp1 | 0.78507283 | 0.57970357 | 0.70878401 | -0.1290804 | 0.80191037 | 0.89054584 |
| Uqcrc2 | 1.54610171 | 0.91844074 | 1.04780097 | -0.1293602 | 0.71741766 | 0.89054584 |
| Gm21092 | 0.79421552 | 0.58531872 | 0.71480421 | -0.1294855 | 0.80065378 | 0.89054584 |
| Ctbp1 | 0.78606656 | 0.57991292 | 0.7094413 | -0.1295284 | 0.80250982 | 0.89054584 |
| Ginm1 | 0.98399301 | 0.69667631 | 0.82620798 | -0.1295317 | 0.76743385 | 0.89054584 |
| Ndufb7 | 2.20972017 | 0.99774384 | 1.12749506 | -0.1297512 | 0.70407159 | 0.89054584 |
| Trip11 | 0.72845552 | 0.54025368 | 0.6701664 | -0.1299127 | 0.81677485 | 0.89054584 |
| Ndufv2 | 1.76477659 | 0.96261618 | 1.09253731 | -0.1299211 | 0.71021973 | 0.89054584 |
| Vps29 | 1.47292719 | 0.8979705 | 1.02798765 | -0.1300171 | 0.72215338 | 0.89054584 |
| Eif3i | 1.9098522 | 0.98055729 | 1.11063609 | -0.1300788 | 0.70735011 | 0.89054584 |
| Eprs | 1.34706484 | 0.85749775 | 0.98767704 | -0.1301793 | 0.73041374 | 0.89054584 |
| Trim28 | 0.97060549 | 0.68889392 | 0.81914161 | -0.1302477 | 0.77059124 | 0.89054584 |
| Strn3 | 0.79802931 | 0.58662497 | 0.71729747 | -0.1306725 | 0.80198564 | 0.89054584 |
| Ntan1 | 0.91504684 | 0.65760642 | 0.78865933 | -0.1310529 | 0.78058914 | 0.89054584 |
| Csde1 | 1.49824119 | 0.90401012 | 1.03508479 | -0.1310747 | 0.72239467 | 0.89054584 |
| Ppdpf | 1.46041043 | 0.89307377 | 1.0243631 | -0.1312893 | 0.72478688 | 0.89054584 |
| Stoml2 | 0.84404311 | 0.6152121 | 0.74654565 | -0.1313335 | 0.79378038 | 0.89054584 |
| Tpr | 1.75569534 | 0.95957558 | 1.09110704 | -0.1315315 | 0.71281558 | 0.89054584 |
| Ndufs7 | 1.58834121 | 0.92646033 | 1.05807452 | -0.1316142 | 0.71888319 | 0.89054584 |

| | | | | | | |
|---|---|---|---|---|---|---|
| Sf1 | 0.79337256 | 0.58250293 | 0.71425169 | -0.1317488 | 0.80493305 | 0.89054584 |
| Rchy1 | 0.97495667 | 0.68904757 | 0.82145258 | -0.132405 | 0.77365765 | 0.89054584 |
| Cyc1 | 1.52919364 | 0.91102005 | 1.0434405 | -0.1324204 | 0.72282509 | 0.89054584 |
| Gm9843 | 1.61868196 | 0.93218874 | 1.064976 | -0.1327873 | 0.71934718 | 0.89054584 |
| Bad | 0.94145979 | 0.67049924 | 0.80340893 | -0.1329097 | 0.7795691 | 0.89054584 |
| Rbm25 | 1.79475078 | 0.96394664 | 1.09701018 | -0.1330635 | 0.7140017 | 0.89054584 |
| Grb2 | 0.56492654 | 0.41250441 | 0.54605927 | -0.1335549 | 0.87274874 | 0.90729956 |
| Supt5 | 0.80205979 | 0.58625328 | 0.71992087 | -0.1336676 | 0.80648743 | 0.89054584 |
| Rpl13 | 5.65311119 | 0.55102385 | 0.68478844 | -0.1337646 | 0.81860564 | 0.89054584 |
| Aurkaip1 | 1.10456669 | 0.75352173 | 0.88736801 | -0.1338463 | 0.75888862 | 0.89054584 |
| Ppp3cb | 0.49748713 | 0.35582026 | 0.48971766 | -0.1338974 | 0.89863927 | 0.92243501 |
| Abcf1 | 1.07687368 | 0.73966968 | 0.87383059 | -0.1341609 | 0.76278702 | 0.89054584 |
| Ndufb9 | 2.34691812 | 0.99521496 | 1.12950139 | -0.1342864 | 0.71017689 | 0.89054584 |
| Sf3b4 | 1.10937774 | 0.7552587 | 0.88966391 | -0.1344052 | 0.75923552 | 0.89054584 |
| Dnttip2 | 0.72966616 | 0.5360009 | 0.67101605 | -0.1350152 | 0.82573018 | 0.89170439 |
| Cox7c | 3.67326822 | 0.87771937 | 1.01358796 | -0.1358686 | 0.73385623 | 0.89054584 |
| Sys1 | 1.32825866 | 0.84499937 | 0.98098089 | -0.1359815 | 0.74079967 | 0.89054584 |
| Timm17b | 0.99580043 | 0.6958868 | 0.83233239 | -0.1364456 | 0.77749121 | 0.89054584 |
| U2surp | 0.94581509 | 0.66900665 | 0.80579605 | -0.1367894 | 0.78549509 | 0.89054584 |
| Tspan31 | 0.95266748 | 0.67263316 | 0.80952989 | -0.1368967 | 0.78460984 | 0.89054584 |
| Rps28 | 4.88640331 | 0.59784967 | 0.73487766 | -0.137028 | 0.80750076 | 0.89054584 |
| Sdf2 | 1.26066239 | 0.81834574 | 0.9554407 | -0.137095 | 0.74809452 | 0.89054584 |
| Cox7a2 | 3.14946259 | 0.94286326 | 1.0806041 | -0.1377408 | 0.72369678 | 0.89054584 |
| Ergic2 | 1.23137405 | 0.80584048 | 0.94361412 | -0.1377736 | 0.75182214 | 0.89054584 |
| Lrrc58 | 0.86731733 | 0.62282712 | 0.76075237 | -0.1379253 | 0.80085279 | 0.89054584 |
| Rps27l | 2.98017072 | 0.95830277 | 1.09676538 | -0.1384626 | 0.72179234 | 0.89054584 |
| Gpx4 | 2.6483255 | 0.98146429 | 1.11994828 | -0.138484 | 0.71772215 | 0.89054584 |
| Ssbp1 | 1.11236338 | 0.75213679 | 0.89108092 | -0.1389441 | 0.76622087 | 0.89054584 |
| 2310036O22 | 1.94681901 | 0.97474476 | 1.11384995 | -0.1391052 | 0.7196629 | 0.89054584 |
| Copb1 | 1.35234625 | 0.84970355 | 0.98951805 | -0.1398145 | 0.74482336 | 0.89054584 |

| | | | | | | |
|---|---|---|---|---|---|---|
| Ndufa4 | 3.44523373 | 0.90199767 | 1.04182086 | -0.1398232 | 0.7341058 | 0.89054584 |
| Zfr | 1.28059277 | 0.82279017 | 0.9632171 | -0.1404269 | 0.75150223 | 0.89054584 |
| Whsc1l1 | 0.86172151 | 0.61683374 | 0.75737257 | -0.1405388 | 0.80634828 | 0.89054584 |
| Lage3 | 1.06138659 | 0.72542251 | 0.86605433 | -0.1406318 | 0.775334 | 0.89054584 |
| Smu1 | 0.95781924 | 0.67162957 | 0.81231444 | -0.1406849 | 0.79013299 | 0.89054584 |
| Psmd11 | 1.40810546 | 0.86749347 | 1.00823117 | -0.1407377 | 0.74226228 | 0.89054584 |
| Rps3a1 | 5.06721519 | 0.54654106 | 0.6874843 | -0.1409432 | 0.83024748 | 0.89271879 |
| Ngdn | 0.68640451 | 0.4986567 | 0.64000939 | -0.1413527 | 0.84825437 | 0.89877891 |
| Pafah1b2 | 1.24369398 | 0.80696591 | 0.94861124 | -0.1416453 | 0.75668121 | 0.89054584 |
| Plekhj1 | 1.17304078 | 0.77695908 | 0.91875138 | -0.1417923 | 0.76393616 | 0.89054584 |
| Cwc15 | 1.28996105 | 0.82454357 | 0.96678276 | -0.1422392 | 0.7534709 | 0.89054584 |
| Polr3k | 1.014421 | 0.69968461 | 0.84195069 | -0.1422661 | 0.78442655 | 0.89054584 |
| Txndc9 | 1.36527142 | 0.85075622 | 0.99395303 | -0.1431968 | 0.74895479 | 0.89054584 |
| Wdr61 | 1.15592841 | 0.7678496 | 0.91111591 | -0.1432663 | 0.76809402 | 0.89054584 |
| Naca | 4.09184502 | 0.79827881 | 0.9423003 | -0.1440215 | 0.76179056 | 0.89054584 |
| Rab7 | 1.6851054 | 0.93398866 | 1.07865172 | -0.1446631 | 0.73395952 | 0.89054584 |
| Eif3d | 1.15869336 | 0.76765217 | 0.91235202 | -0.1446999 | 0.77001987 | 0.89054584 |
| Psenen | 1.83544481 | 0.95745842 | 1.10246979 | -0.1450114 | 0.72999748 | 0.89054584 |
| Cpsf6 | 0.89386487 | 0.63146272 | 0.77647697 | -0.1450142 | 0.80787721 | 0.89054584 |
| Eif3g | 1.70317465 | 0.93672758 | 1.08203725 | -0.1453097 | 0.73423156 | 0.89054584 |
| Acin1 | 1.42630692 | 0.86855604 | 1.01399611 | -0.1454401 | 0.74801104 | 0.89054584 |
| Psmc4 | 1.62296169 | 0.92042199 | 1.0659202 | -0.1454982 | 0.73759417 | 0.89054584 |
| Mrpl20 | 1.43524515 | 0.87053826 | 1.01677704 | -0.1462388 | 0.74859532 | 0.89054584 |
| Tomm40 | 1.07014031 | 0.72419512 | 0.8704695 | -0.1462744 | 0.78315478 | 0.89054584 |
| Cbx1 | 1.70189485 | 0.9353607 | 1.08180022 | -0.1464395 | 0.73587008 | 0.89054584 |
| Mff | 1.33761697 | 0.83771577 | 0.9843395 | -0.1466237 | 0.75615243 | 0.89054584 |
| Tsen34 | 1.84065465 | 0.95560699 | 1.10311041 | -0.1475034 | 0.73334939 | 0.89054584 |
| Scamp3 | 0.72172186 | 0.51775696 | 0.66542143 | -0.1476645 | 0.84940182 | 0.89883846 |
| Vapa | 2.02876676 | 0.97172835 | 1.11959104 | -0.1478627 | 0.73082146 | 0.89054584 |
| Cfl1 | 3.60367644 | 0.8736349 | 1.02152447 | -0.1478896 | 0.75000032 | 0.89054584 |

| | | | | | | |
|---|---|---|---|---|---|---|
| Mrpl42 | 1.26826287 | 0.81055167 | 0.95844136 | -0.1478897 | 0.76389419 | 0.89054584 |
| Ap2b1 | 0.87695112 | 0.61858581 | 0.76651822 | -0.1479324 | 0.81571136 | 0.89054584 |
| Snx6 | 1.41949096 | 0.86386124 | 1.01185945 | -0.1479982 | 0.75220139 | 0.89054584 |
| Wnk1 | 0.94632023 | 0.65794344 | 0.80607247 | -0.148129 | 0.80404447 | 0.89054584 |
| Eif3e | 2.43694701 | 0.98028237 | 1.1286024 | -0.14832 | 0.72982187 | 0.89054584 |
| Psma1 | 1.94701932 | 0.96477401 | 1.11386607 | -0.1490921 | 0.7335561 | 0.89054584 |
| Nfu1 | 0.91792335 | 0.64094283 | 0.79028873 | -0.1493459 | 0.81068675 | 0.89054584 |
| Ccnl2 | 1.46904532 | 0.87749618 | 1.02687285 | -0.1493767 | 0.75102688 | 0.89054584 |
| Psmd12 | 1.40073163 | 0.85629432 | 1.00584164 | -0.1495473 | 0.75574661 | 0.89054584 |
| Rap1a | 1.53868739 | 0.89613976 | 1.04591651 | -0.1497768 | 0.7476673 | 0.89054584 |
| Lamp1 | 2.55403592 | 0.97467841 | 1.12473783 | -0.1500594 | 0.73289197 | 0.89054584 |
| Nap1l4 | 1.07859817 | 0.72369259 | 0.87468773 | -0.1509951 | 0.78937695 | 0.89054584 |
| Mrpl14 | 1.32371407 | 0.82803927 | 0.97933089 | -0.1512916 | 0.76415593 | 0.89054584 |
| Cul1 | 0.8443344 | 0.59541585 | 0.74672591 | -0.1513101 | 0.82742728 | 0.89203158 |
| Snrpf | 2.40196558 | 0.97779676 | 1.12917907 | -0.1513823 | 0.73388203 | 0.89054584 |
| Bcas2 | 1.71803555 | 0.93305614 | 1.08473432 | -0.1516782 | 0.74262482 | 0.89054584 |
| Luc7l | 0.91558029 | 0.63622336 | 0.78896187 | -0.1527385 | 0.81645678 | 0.89054584 |
| Psmb1 | 2.99838305 | 0.9421545 | 1.0952289 | -0.1530744 | 0.74252703 | 0.89054584 |
| Oaz2 | 1.45544841 | 0.86974151 | 1.02290209 | -0.1531606 | 0.75728315 | 0.89054584 |
| Rpl27a | 4.78584569 | 0.61032529 | 0.76399044 | -0.1536652 | 0.8256402 | 0.89170439 |
| Eif2s1 | 1.21785457 | 0.78422317 | 0.93804864 | -0.1538255 | 0.77756469 | 0.89054584 |
| Phb2 | 1.95216535 | 0.96034502 | 1.11427528 | -0.1539303 | 0.74008388 | 0.89054584 |
| G3bp2 | 1.22103361 | 0.78532316 | 0.93936782 | -0.1540447 | 0.77756891 | 0.89054584 |
| Setd5 | 1.13743771 | 0.7485729 | 0.90275573 | -0.1541828 | 0.78689873 | 0.89054584 |
| Prkcsh | 1.22957889 | 0.78844833 | 0.94288182 | -0.1544335 | 0.77729288 | 0.89054584 |
| U2af2 | 0.86968605 | 0.6075643 | 0.76217639 | -0.1546121 | 0.82769465 | 0.89225348 |
| Ppig | 1.4221356 | 0.85807952 | 1.01269163 | -0.1546121 | 0.76155118 | 0.89054584 |
| Magoh | 1.48403922 | 0.87639728 | 1.03113615 | -0.1547389 | 0.757771 | 0.89054584 |
| Ppp1r11 | 1.13048413 | 0.74426697 | 0.8995537 | -0.1552867 | 0.78937552 | 0.89054584 |
| Psmd6 | 1.45076911 | 0.86590929 | 1.02150381 | -0.1555945 | 0.76103805 | 0.89054584 |

| | | | | | | |
|---|---|---|---|---|---|---|
| Snrpc | 1.53643703 | 0.88964504 | 1.04533881 | -0.1556938 | 0.7561298 | 0.89054584 |
| Prpf40a | 1.42251864 | 0.85671025 | 1.01281183 | -0.1561016 | 0.76364241 | 0.89054584 |
| Rpl29 | 3.63129586 | 0.86219957 | 1.01836742 | -0.1561679 | 0.76252681 | 0.89054584 |
| Rpl14 | 5.21008401 | 0.51644355 | 0.67321032 | -0.1567668 | 0.86101745 | 0.90188794 |
| Rpl21 | 4.53628176 | 0.67406616 | 0.83085221 | -0.156786 | 0.81034435 | 0.89054584 |
| Luc7l2 | 1.94773637 | 0.95581403 | 1.11392368 | -0.1581096 | 0.74578595 | 0.89054584 |
| BC005561 | 0.92967971 | 0.63826042 | 0.79689312 | -0.1586327 | 0.82314764 | 0.89080883 |
| Sarnp | 1.73293604 | 0.92856674 | 1.0873355 | -0.1587688 | 0.75183691 | 0.89054584 |
| Pcnp | 1.17527763 | 0.76050208 | 0.919737 | -0.1592349 | 0.79005348 | 0.89054584 |
| Cldnd1 | 1.25454578 | 0.79322171 | 0.95299961 | -0.1597779 | 0.78263057 | 0.89054584 |
| Gdi2 | 1.99631486 | 0.95761402 | 1.11753255 | -0.1599185 | 0.74751585 | 0.89054584 |
| Srrm1 | 1.5675483 | 0.89263504 | 1.05310547 | -0.1604704 | 0.76112362 | 0.89054584 |
| Akr1a1 | 2.53416447 | 0.96447898 | 1.12558553 | -0.1611065 | 0.74756297 | 0.89054584 |
| Pum1 | 1.07427929 | 0.71110615 | 0.87253916 | -0.161433 | 0.80564936 | 0.89054584 |
| Ndufb5 | 2.33935774 | 0.96766728 | 1.12945671 | -0.1617894 | 0.74773262 | 0.89054584 |
| Hnrnpa3 | 2.80031372 | 0.94859814 | 1.11044886 | -0.1618507 | 0.75144768 | 0.89054584 |
| Cetn3 | 1.56113837 | 0.889375 | 1.0515343 | -0.1621593 | 0.76376471 | 0.89054584 |
| Atf4 | 0.89609573 | 0.61548714 | 0.77777557 | -0.1622884 | 0.83450491 | 0.89447377 |
| Pnisr | 1.55706536 | 0.88807235 | 1.0505347 | -0.1624623 | 0.76438914 | 0.89054584 |
| Brd8 | 0.78256668 | 0.54453774 | 0.70712347 | -0.1625857 | 0.85796236 | 0.90124396 |
| Etf1 | 0.97376906 | 0.65760092 | 0.82082319 | -0.1632223 | 0.82289117 | 0.89075401 |
| Smarca5 | 1.4438306 | 0.85610008 | 1.01940789 | -0.1633078 | 0.7722739 | 0.89054584 |
| Uqcrh | 3.05686712 | 0.92467517 | 1.08974451 | -0.1650693 | 0.7598171 | 0.89054584 |
| Hnrnpd | 1.03494396 | 0.6873679 | 0.85251098 | -0.1651431 | 0.81660457 | 0.89054584 |
| mt-Nd4l | 1.11365388 | 0.72601586 | 0.89169157 | -0.1656757 | 0.80665296 | 0.89054584 |
| Igf2bp3 | 0.79933554 | 0.55152252 | 0.71814899 | -0.1666265 | 0.86014645 | 0.90136633 |
| Cops4 | 1.31357812 | 0.80894657 | 0.97562395 | -0.1666774 | 0.78696714 | 0.89054584 |
| Ube3a | 1.46548337 | 0.85803817 | 1.02584258 | -0.1678044 | 0.77699875 | 0.89054584 |
| Tmem50a | 1.62822584 | 0.8990266 | 1.06706917 | -0.1680426 | 0.76843185 | 0.89054584 |
| Map2k2 | 1.57180852 | 0.88551835 | 1.05414163 | -0.1686233 | 0.77194168 | 0.89054584 |

| | | | | | | |
|---|---|---|---|---|---|---|
| Rab14 | 1.8795456 | 0.93807154 | 1.10755446 | -0.1694829 | 0.76205606 | 0.89054584 |
| Rpl24 | 4.76395197 | 0.6001815 | 0.77019799 | -0.1700165 | 0.84812684 | 0.89876755 |
| Dgcr6 | 0.73116811 | 0.50161811 | 0.6720687 | -0.1704506 | 0.88131862 | 0.91201694 |
| Atp5j | 3.1758033 | 0.90723807 | 1.0779624 | -0.1707243 | 0.76970285 | 0.89054584 |
| Tomm20 | 2.44933249 | 0.957566 | 1.12833242 | -0.1707664 | 0.75962696 | 0.89054584 |
| Hypk | 1.49458565 | 0.863301 | 1.03407894 | -0.1707779 | 0.77917896 | 0.89054584 |
| Ndufa3 | 2.17108448 | 0.95558227 | 1.12640477 | -0.1708225 | 0.76007435 | 0.89054584 |
| Fkbp8 | 1.47272161 | 0.85634732 | 1.02792882 | -0.1715815 | 0.78161783 | 0.89054584 |
| Pcbp1 | 2.28986136 | 0.95714692 | 1.12896475 | -0.1718178 | 0.76085237 | 0.89054584 |
| Cdc5l | 1.10240259 | 0.71322285 | 0.88632851 | -0.1731057 | 0.81856797 | 0.89054584 |
| Sdhb | 1.76487394 | 0.91905668 | 1.09255244 | -0.1734958 | 0.77029939 | 0.89054584 |
| Zfp706 | 1.81912118 | 0.92656555 | 1.10036615 | -0.1738006 | 0.76909518 | 0.89054584 |
| Ap2s1 | 1.7633142 | 0.91818672 | 1.09230944 | -0.1741227 | 0.77115983 | 0.89054584 |
| Polr2c | 1.40670282 | 0.83252623 | 1.00777904 | -0.1752528 | 0.79108452 | 0.89054584 |
| Rplp2 | 4.94574775 | 0.54180567 | 0.71718154 | -0.1753759 | 0.87270543 | 0.90729956 |
| Mrpl24 | 1.18968465 | 0.74987338 | 0.92603186 | -0.1761585 | 0.81226331 | 0.89054584 |
| Sf3b2 | 1.99176079 | 0.94095324 | 1.11721607 | -0.1762628 | 0.76884234 | 0.89054584 |
| Rpl38 | 4.54128044 | 0.65308673 | 0.82951479 | -0.1764281 | 0.83913331 | 0.89500145 |
| Eif4g1 | 1.26479778 | 0.78061808 | 0.9570779 | -0.1764598 | 0.8048257 | 0.89054584 |
| Ubb | 5.69040312 | 0.5108806 | 0.6881667 | -0.1772861 | 0.88497702 | 0.91404124 |
| Nhp2l1 | 1.94102398 | 0.93582366 | 1.11337698 | -0.1775533 | 0.77125138 | 0.89054584 |
| Vdac2 | 2.07038482 | 0.94340642 | 1.12184904 | -0.1784426 | 0.77067045 | 0.89054584 |
| Tia1 | 1.44643328 | 0.84167875 | 1.02019725 | -0.1785185 | 0.79253297 | 0.89054584 |
| Arf1 | 3.10194241 | 0.90611809 | 1.0855349 | -0.1794168 | 0.77935838 | 0.89054584 |
| Papola | 1.60442626 | 0.88140433 | 1.06179035 | -0.180386 | 0.78567408 | 0.89054584 |
| Rps24 | 5.08329749 | 0.50462004 | 0.68516043 | -0.1805404 | 0.89020803 | 0.91709202 |
| Cnbp | 2.69967891 | 0.93577192 | 1.1169579 | -0.181186 | 0.77509881 | 0.89054584 |
| Nedd8 | 2.26361421 | 0.94665216 | 1.12861988 | -0.1819677 | 0.77372256 | 0.89054584 |
| Clk1 | 1.29685303 | 0.78612556 | 0.96937959 | -0.183254 | 0.81077027 | 0.89054584 |
| Psma5 | 2.01499921 | 0.93519121 | 1.1187565 | -0.1835653 | 0.77769684 | 0.89054584 |

| | | | | | | |
|---|---|---|---|---|---|---|
| Pfdn1 | 1.71349862 | 0.89854135 | 1.08392169 | -0.1853803 | 0.78721836 | 0.89054584 |
| Ndufa8 | 1.77454327 | 0.90714406 | 1.09403503 | -0.186891 | 0.78694737 | 0.89054584 |
| Sumo1 | 2.29604509 | 0.94207601 | 1.12902415 | -0.1869481 | 0.77977848 | 0.89054584 |
| Psmc6 | 1.49745528 | 0.84737481 | 1.03486915 | -0.1874943 | 0.80069109 | 0.89054584 |
| Mdh2 | 2.43108369 | 0.94100921 | 1.12871921 | -0.18771 | 0.78077061 | 0.89054584 |
| Spcs1 | 2.71077265 | 0.9281103 | 1.11626297 | -0.1881527 | 0.78385876 | 0.89054584 |
| Acp1 | 1.24858734 | 0.76149799 | 0.95059919 | -0.1891012 | 0.82298372 | 0.89075401 |
| Uqcrb | 3.25413968 | 0.87846601 | 1.06838955 | -0.1899235 | 0.79622201 | 0.89054584 |
| Sf3b1 | 1.91216852 | 0.91978063 | 1.11085182 | -0.1910712 | 0.78853764 | 0.89054584 |
| Rpl10 | 5.41718434 | 0.47752017 | 0.66929296 | -0.1917728 | 0.90905011 | 0.92956753 |
| Rps27 | 5.36339168 | 0.4755267 | 0.66874557 | -0.1932189 | 0.91086413 | 0.93070041 |
| Gng5 | 3.20179055 | 0.88076056 | 1.07517363 | -0.1944131 | 0.80022648 | 0.89054584 |
| Tra2a | 1.52653015 | 0.84773449 | 1.04273536 | -0.1950009 | 0.80819494 | 0.89054584 |
| Cox6b1 | 3.37684321 | 0.85392864 | 1.05167503 | -0.1977464 | 0.80949053 | 0.89054584 |
| Eif6 | 1.283001 | 0.76415321 | 0.96413888 | -0.1999857 | 0.83307217 | 0.89393134 |
| Atp5d | 2.54173357 | 0.92241081 | 1.12527163 | -0.2028608 | 0.79951802 | 0.89054584 |
| Gmps | 0.91102048 | 0.58320625 | 0.78636909 | -0.2031628 | 0.88564016 | 0.91420299 |
| Rala | 1.37399901 | 0.79294679 | 0.99689177 | -0.203945 | 0.82973795 | 0.89271879 |
| Oaz1 | 4.09566005 | 0.73659884 | 0.94135769 | -0.2047589 | 0.84456228 | 0.89775379 |
| Pebp1 | 3.34861717 | 0.84986377 | 1.05568461 | -0.2058208 | 0.81814734 | 0.89054584 |
| Acaa1a | 1.32598469 | 0.77417366 | 0.98015684 | -0.2059832 | 0.83623269 | 0.89462889 |
| Trappc3 | 0.94101795 | 0.59629299 | 0.80316638 | -0.2068734 | 0.88493306 | 0.91404124 |
| Pdcd6 | 1.36766554 | 0.7864007 | 0.99476365 | -0.208363 | 0.83543325 | 0.89459411 |
| Eif3c | 1.78008594 | 0.88450642 | 1.09486647 | -0.2103601 | 0.81464298 | 0.89054584 |
| Rbm8a | 1.85205777 | 0.89093957 | 1.10447375 | -0.2135342 | 0.81613659 | 0.89054584 |
| Psmb2 | 2.37086108 | 0.91214379 | 1.12953223 | -0.2173884 | 0.8150483 | 0.89054584 |
| Imp3 | 1.13062643 | 0.6820953 | 0.89961938 | -0.2175241 | 0.87002553 | 0.90589084 |
| Capzb | 1.99431475 | 0.89775363 | 1.11739445 | -0.2196408 | 0.8201103 | 0.89054584 |
| Luc7l3 | 1.32800252 | 0.76060195 | 0.98088823 | -0.2202863 | 0.85228813 | 0.89988783 |
| Srsf10 | 1.18256333 | 0.70223583 | 0.92292474 | -0.2206889 | 0.86736935 | 0.90444517 |

| | | | | | | |
|---|---|---|---|---|---|---|
| Polr2m | 1.74080329 | 0.86524298 | 1.08866546 | -0.2234225 | 0.83049823 | 0.89271879 |
| Rps3 | 4.78980241 | 0.53821278 | 0.7628598 | -0.224647 | 0.9149629 | 0.93370395 |
| Trmt112 | 1.37086883 | 0.76977401 | 0.99584295 | -0.2260689 | 0.85488987 | 0.90074724 |
| H3f3a | 4.63692517 | 0.57854436 | 0.80481237 | -0.226268 | 0.90487373 | 0.92653855 |
| Snrnp27 | 1.36763418 | 0.76736395 | 0.99475306 | -0.2273891 | 0.85655759 | 0.9009252 |
| Paip2 | 2.05620037 | 0.88985345 | 1.12112286 | -0.2312694 | 0.83173996 | 0.8933762 |
| Rpl11 | 4.97083976 | 0.47610625 | 0.70971048 | -0.2336042 | 0.93742492 | 0.94960778 |
| Psme2 | 1.52804669 | 0.80942674 | 1.04313731 | -0.2337106 | 0.85171482 | 0.89985795 |
| Cct8 | 2.23816059 | 0.89258633 | 1.12817194 | -0.2355856 | 0.83469431 | 0.89453395 |
| Rhoa | 2.40017885 | 0.89043972 | 1.12920129 | -0.2387616 | 0.83771251 | 0.89485554 |
| Scaf11 | 1.34567791 | 0.74626736 | 0.98719043 | -0.2409231 | 0.87223317 | 0.90701739 |
| Wbp5 | 3.58786963 | 0.77882796 | 1.02332285 | -0.2444949 | 0.86717285 | 0.90441493 |
| Rpl8 | 4.70659289 | 0.54049678 | 0.78611592 | -0.2456191 | 0.92725802 | 0.94266789 |
| Tmem234 | 1.39302502 | 0.75699768 | 1.00330398 | -0.2463063 | 0.87363584 | 0.90763953 |
| Eif3k | 2.58204468 | 0.87616721 | 1.12342291 | -0.2472557 | 0.84741106 | 0.89854206 |
| Hnrnpk | 3.59172195 | 0.77476151 | 1.02288514 | -0.2481236 | 0.87078932 | 0.90603586 |
| Dynlrb1 | 2.20824639 | 0.87687913 | 1.12746281 | -0.2505837 | 0.84977541 | 0.89887961 |
| Cdc37 | 1.8270129 | 0.85051109 | 1.10139699 | -0.2508859 | 0.8557129 | 0.9009252 |
| Rpl19 | 4.97117058 | 0.4513353 | 0.70961251 | -0.2582772 | 0.95502977 | 0.96306751 |
| Rps27a | 5.45724783 | 0.40823624 | 0.66957667 | -0.2613404 | 0.9654998 | 0.9714634 |
| Atp5c1 | 2.65675205 | 0.85749712 | 1.11950673 | -0.2620096 | 0.86222703 | 0.90260095 |
| Atp5a1 | 3.10409967 | 0.81558131 | 1.08533411 | -0.2697528 | 0.87655774 | 0.9091719 |
| Pomp | 2.28073673 | 0.85867969 | 1.12885974 | -0.27018 | 0.8675889 | 0.90444517 |
| Rpl35a | 5.02034984 | 0.41996803 | 0.69665378 | -0.2766857 | 0.9678694 | 0.97334573 |
| Sumo2 | 3.38036001 | 0.77290606 | 1.05115693 | -0.2782509 | 0.89126144 | 0.91739498 |
| Rps17 | 4.81161995 | 0.47839367 | 0.75665573 | -0.2782621 | 0.95667476 | 0.9643239 |
| Ociad1 | 1.377815 | 0.71605615 | 0.99819471 | -0.2821386 | 0.90605458 | 0.92742017 |
| Rpl13a | 5.59470221 | 0.38885389 | 0.67949252 | -0.2906386 | 0.97685636 | 0.9806442 |
| Eif1 | 4.74081423 | 0.4808724 | 0.77679722 | -0.2959248 | 0.96203844 | 0.96851839 |
| Rps15 | 4.6126677 | 0.49997811 | 0.81086296 | -0.3108849 | 0.96296964 | 0.96932124 |

| | | | | | | |
|---|---|---|---|---|---|---|
| Tceb2 | 3.27188388 | 0.75368936 | 1.06596141 | -0.312272 | 0.9138469 | 0.93295744 |
| Rps25 | 4.44994286 | 0.5382688 | 0.85119864 | -0.3129298 | 0.95662411 | 0.9643239 |
| Cox4i1 | 3.9334674 | 0.65934568 | 0.97665375 | -0.3173081 | 0.93494368 | 0.94769024 |
| Rpl27 | 3.79279396 | 0.68182875 | 0.99964513 | -0.3178164 | 0.93072515 | 0.94499987 |
| Eef1a1 | 5.8692656 | 0.38397927 | 0.70675074 | -0.3227715 | 0.98332018 | 0.98597247 |
| Sub1 | 3.52741553 | 0.70639922 | 1.03098398 | -0.3245848 | 0.92878133 | 0.94362101 |
| Cox8a | 3.72013164 | 0.67535404 | 1.00845874 | -0.3331047 | 0.93808541 | 0.95016738 |
| Atp5h | 3.32641229 | 0.72387249 | 1.05857138 | -0.3346989 | 0.92963299 | 0.94406031 |
| Btf3 | 3.86525831 | 0.64696541 | 0.98954176 | -0.3425764 | 0.94661811 | 0.95624787 |
| Ybx1 | 4.00065535 | 0.60756154 | 0.9630942 | -0.3555327 | 0.95727281 | 0.96479257 |
| Rps14 | 5.80665866 | 0.30039213 | 0.6996829 | -0.3992908 | 0.99607828 | 0.99677103 |
| Eif4a1 | 4.11475564 | 0.52752381 | 0.93667032 | -0.4091465 | 0.97907808 | 0.98253436 |
| Ywhae | 3.80603718 | 0.55901051 | 0.99788685 | -0.4388763 | 0.97977036 | 0.98311155 |
| Hint1 | 3.2386589 | 0.62845937 | 1.07060135 | -0.442142 | 0.97282718 | 0.97741154 |
| Tpt1 | 4.92840701 | 0.25583634 | 0.72243542 | -0.4665991 | 0.99868975 | 0.99889645 |
| Rpl41 | 6.35352843 | 0.33100554 | 1.49864807 | -1.1676425 | 0.99985825 | 0.99985825 |

3

PMs for all cellular genes.
of log-CPMs for all cellular genes.
ological component of the variance for all genes.
echnical component of the variance for all genes.

| | mean | total | tech | bio | p.value | FDR |
|---|---|---|---|---|---|---|
| Ins2 | 3.58481023 | 10.2465926 | 0.25609704 | 9.99049554 | 0 | 0 |
| Ins1 | 2.85056265 | 8.84865412 | 0.35268456 | 8.49596955 | 0 | 0 |
| Iapp | 2.53402035 | 6.22868181 | 0.36871287 | 5.85996893 | 0 | 0 |
| Pyy | 1.54284025 | 3.3958754 | 0.40458158 | 2.99129382 | 0 | 0 |
| Nnat | 1.9969299 | **3.35141306** | 0.37074664 | 2.98066642 | 0 | **0** |
| Neurog3 | 1.25322908 | 2.94422153 | 0.42719791 | 2.51702362 | 0 | 0 |
| Chgb | 2.50667326 | 2.78210595 | 0.36964727 | 2.41245868 | 0 | 0 |
| Mdk | 1.48923273 | 2.20430932 | 0.41015194 | 1.79415738 | 1.08E-186 | 1.72E-183 |
| Rbp4 | 1.79994606 | 2.12487203 | 0.38041478 | 1.74445725 | 6.28E-205 | 1.11E-201 |
| Chga | 2.35191635 | 2.11169917 | 0.37066694 | 1.74103223 | 6.96E-215 | 1.39E-211 |
| Xist | 1.37309881 | 2.00847398 | 0.42044607 | 1.58802791 | 8.84E-140 | 1.17E-136 |
| Slc38a5 | 1.83017412 | 1.93759411 | 0.37826999 | 1.55932412 | 4.68E-166 | 6.78E-163 |
| Npy | 0.63793171 | 1.7181145 | 0.36427422 | 1.35384029 | 2.05E-135 | 2.52E-132 |
| Ttr | 2.51381498 | 1.71939817 | 0.36940555 | 1.34999261 | 5.25E-131 | 5.97E-128 |
| Tmsb4x | 2.28459114 | 1.71659661 | 0.3701723 | 1.34642431 | 8.50E-130 | 9.03E-127 |
| Pcsk1n | 1.93369308 | 1.66455654 | 0.37231042 | 1.29224612 | 2.14E-118 | 2.13E-115 |
| Pcsk2 | 1.5817426 | 1.64721786 | 0.40041192 | 1.24680594 | 9.96E-96 | 9.34E-93 |
| Cck | 0.80610434 | 1.64867125 | 0.40459753 | 1.24407372 | 2.11E-93 | 1.87E-90 |
| Gadd45a | 1.09335661 | 1.52574009 | 0.42965135 | 1.09608874 | 5.67E-65 | 4.75E-62 |
| Dlk1 | 1.73108503 | 1.31602465 | 0.38627221 | 0.92975244 | 4.50E-58 | 3.59E-55 |
| Cdkn1a | 0.8186486 | 1.33644201 | 0.40673875 | 0.92970327 | 1.41E-52 | 1.07E-49 |
| 1700086L19F | 1.26601887 | 1.31428903 | 0.42664779 | 0.88764124 | 6.49E-44 | 3.98E-41 |
| Hmgn3 | 1.7765666 | 1.24282156 | 0.38215694 | 0.86066462 | 4.21E-51 | 3.05E-48 |
| Hbb-bs | 2.34734057 | 1.20022843 | 0.37069262 | 0.82953581 | 1.76E-50 | 1.22E-47 |
| Hba-a1 | 1.75963776 | 1.18747571 | 0.38354734 | 0.80392837 | 1.53E-44 | 9.76E-42 |
| Cpe | 2.11977362 | 1.15707972 | 0.36846109 | 0.78861863 | 2.35E-46 | 1.56E-43 |
| Cryba2 | 1.42035351 | 1.19289766 | 0.41670234 | 0.77619531 | 1.33E-35 | 7.29E-33 |
| Manf | 1.81185359 | 1.13061107 | 0.37946532 | 0.75114576 | 6.02E-40 | 3.55E-37 |

mean variance for all transcripts = 0.086

**mean**: A numeric vector of mean log-CP
**total**: A numeric vector of the variances
**bio**: A numeric vector containing the bic
**tech**: A numeric vector containing the te

| | | | | | | |
|---|---|---|---|---|---|---|
| Sdf2l1 | 1.03235374 | 1.17426792 | 0.42802372 | 0.7462442 | 1.94E-31 | 1.03E-28 |
| Btbd17 | 0.94162867 | 1.15771819 | 0.42220643 | 0.73551176 | 2.16E-31 | 1.11E-28 |
| Cd24a | 0.98606391 | 1.15892173 | 0.4255244 | 0.73339734 | 9.09E-31 | 4.39E-28 |
| Fos | 1.28316304 | 1.15330655 | 0.42584211 | 0.72746444 | 2.91E-30 | 1.37E-27 |
| Igfbpl1 | 0.87669259 | 1.13715321 | 0.41527961 | 0.7218736 | 2.90E-31 | 1.45E-28 |
| Ambp | 1.01398657 | 1.11478466 | 0.42713835 | 0.68764631 | 4.28E-27 | 1.80E-24 |
| Scg2 | 0.84966504 | 1.09844036 | 0.41152436 | 0.686916 | 5.56E-29 | 2.46E-26 |
| Ppp1r1a | 0.83850282 | 1.03149684 | 0.4098275 | 0.62166935 | 2.85E-24 | 1.11E-21 |
| Sox4 | 1.0356187 | 1.0406408 | 0.42817622 | 0.61246458 | 8.55E-22 | 3.03E-19 |
| Hspa5 | 2.24036833 | 0.98104201 | 0.36961565 | 0.61142637 | 1.69E-28 | 7.28E-26 |
| Jun | 1.1213102 | 1.02466644 | 0.42992738 | 0.59473906 | 1.66E-20 | 5.63E-18 |
| Pdia6 | 1.7397555 | 0.97605963 | 0.38543328 | 0.59062635 | 9.91E-25 | 3.95E-22 |
| Isl1 | 0.90960058 | 0.99098539 | 0.41908233 | 0.57190307 | 5.25E-20 | 1.74E-17 |
| Marcksl1 | 1.69782346 | 0.94087651 | 0.38921124 | 0.55166527 | 1.96E-21 | 6.80E-19 |
| Lrpprc | 0.67479063 | 0.91908636 | 0.37556635 | 0.54352001 | 2.90E-22 | 1.07E-19 |
| Gpx3 | 1.33618943 | 0.96089858 | 0.42292165 | 0.53797694 | 1.26E-17 | 3.78E-15 |
| Rap1b | 1.39280176 | 0.94473975 | 0.41897464 | 0.52576511 | 3.33E-17 | 9.83E-15 |
| Clps | 1.30091595 | 0.94535156 | 0.42493317 | 0.52041839 | 1.79E-16 | 5.09E-14 |
| Scg5 | 1.39869157 | 0.92461583 | 0.41849962 | 0.50611621 | 4.14E-16 | 1.12E-13 |
| Fev | 0.56468732 | 0.82854158 | 0.33906792 | 0.48947366 | 3.66E-22 | 1.33E-19 |
| Mafb | 1.07209002 | 0.91860743 | 0.42935725 | 0.48925018 | 1.66E-14 | 4.28E-12 |
| Gnas | 3.26897941 | 0.77471802 | 0.28739553 | 0.48732249 | 7.62E-30 | 3.47E-27 |
| Gm10320 | 0.84198467 | 0.89395611 | 0.4103658 | 0.48359031 | 2.18E-15 | 5.79E-13 |
| Ctrb1 | 1.28678209 | 0.88661322 | 0.4256612 | 0.46095202 | 2.83E-13 | 6.85E-11 |
| Tuba1a | 1.92269354 | 0.82364301 | 0.37272188 | 0.45092113 | 4.04E-16 | 1.11E-13 |
| Scg3 | 1.22903357 | 0.86254324 | 0.42809986 | 0.43444337 | 7.14E-12 | 1.56E-09 |
| S100a11 | 1.71985369 | 0.79545094 | 0.38725546 | 0.40819548 | 1.14E-12 | 2.59E-10 |
| Stmn1 | 0.79024528 | 0.80540895 | 0.40168544 | 0.40372352 | 1.11E-11 | 2.34E-09 |
| Tm4sf4 | 1.12762572 | 0.83259371 | 0.42993568 | 0.40265803 | 2.27E-10 | 4.28E-08 |
| Hsp90b1 | 1.90437423 | 0.77356651 | 0.37408198 | 0.39948454 | 5.87E-13 | 1.36E-10 |

| | | | | | | |
|---|---|---|---|---|---|---|
| Btg2 | 1.20600362 | 0.82791646 | 0.42884606 | 0.3990704 | 2.93E-10 | 5.25E-08 |
| Krt7 | 1.6841358 | 0.78935129 | 0.39053951 | 0.39881179 | 5.33E-12 | 1.18E-09 |
| Spp1 | 0.46621469 | 0.69038131 | 0.29995123 | 0.39043008 | 2.27E-18 | 7.11E-16 |
| Gng12 | 1.16134584 | 0.81762965 | 0.42967848 | 0.38795117 | 9.28E-10 | 1.59E-07 |
| Cdkn1c | 0.66906947 | 0.76077129 | 0.37391786 | 0.38685343 | 2.86E-12 | 6.41E-10 |
| Cldn6 | 0.94012935 | 0.80682433 | 0.4220704 | 0.38475393 | 6.48E-10 | 1.12E-07 |
| Cldn4 | 0.88720284 | 0.79187543 | 0.41656867 | 0.37530676 | 1.00E-09 | 1.70E-07 |
| Hes6 | 1.17991842 | 0.79613549 | 0.42943711 | 0.36669838 | 6.58E-09 | 1.08E-06 |
| Pax4 | 0.56695987 | 0.70503317 | 0.33989802 | 0.36513514 | 4.32E-13 | 1.01E-10 |
| Hnrnpa1 | 1.70152285 | 0.75270364 | 0.38888098 | 0.36382265 | 2.37E-10 | 4.39E-08 |
| H19 | 0.41039702 | 0.63781216 | 0.27482245 | 0.36298972 | 7.36E-19 | 2.39E-16 |
| Hadh | 1.13118572 | 0.78438784 | 0.42993553 | 0.35445231 | 2.04E-08 | 3.18E-06 |
| Slc30a8 | 0.66342334 | 0.7221152 | 0.37222427 | 0.34989093 | 1.96E-10 | 3.77E-08 |
| Rps12 | 1.85533954 | 0.72399349 | 0.37662181 | 0.34737169 | 4.14E-10 | 7.34E-08 |
| Calr | 2.11608085 | 0.71559335 | 0.36846419 | 0.34712915 | 1.79E-10 | 3.49E-08 |
| Gch1 | 1.16286019 | 0.77563858 | 0.42966527 | 0.3459733 | 4.17E-08 | 6.38E-06 |
| Pax6 | 1.22506648 | 0.76672177 | 0.4282404 | 0.33848137 | 7.16E-08 | 1.05E-05 |
| Fam183b | 1.14277956 | 0.76755623 | 0.42986983 | 0.33768641 | 8.52E-08 | 1.24E-05 |
| Meis2 | 1.17397458 | 0.75291149 | 0.42954627 | 0.32336522 | 2.71E-07 | 3.69E-05 |
| Neurod1 | 1.44155762 | 0.73733948 | 0.414835 | 0.32250448 | 1.14E-07 | 1.62E-05 |
| Runx1t1 | 0.56710639 | 0.66119451 | 0.33995144 | 0.32124307 | 1.59E-10 | 3.12E-08 |
| Krt8 | 1.72225112 | 0.70709172 | 0.38704634 | 0.32004538 | 1.86E-08 | 2.96E-06 |
| Ybx1 | 1.82525291 | 0.69802247 | 0.37863765 | 0.31938482 | 9.84E-09 | 1.59E-06 |
| Tubb3 | 1.0948651 | 0.74779928 | 0.42967556 | 0.31812371 | 4.15E-07 | 5.56E-05 |
| Ttll6 | 0.43932855 | 0.60570797 | 0.2881322 | 0.31757576 | 1.10E-13 | 2.73E-11 |
| Cd81 | 1.55876793 | 0.71828996 | 0.40288856 | 0.3154014 | 9.39E-08 | 1.35E-05 |
| Gip | 0.20622184 | 0.47601093 | 0.16082814 | 0.31518278 | 3.40E-39 | 1.94E-36 |
| Gadd45g | 1.08948559 | 0.74432908 | 0.42958617 | 0.31474291 | 5.39E-07 | 7.04E-05 |
| Scgn | 1.04871985 | 0.74281005 | 0.42866316 | 0.31414688 | 5.35E-07 | 7.04E-05 |
| Ier2 | 1.0523598 | 0.73623131 | 0.42878739 | 0.30744392 | 9.09E-07 | 0.00011233 |

| | | | | | | |
|---|---|---|---|---|---|---|
| Tspan7 | 1.30952075 | 0.73179738 | 0.4244737 | 0.30732368 | 7.20E-07 | 9.19E-05 |
| Creld2 | 0.93421215 | 0.71816724 | 0.42152507 | 0.29664218 | 1.40E-06 | 0.00017222 |
| Mid1ip1 | 0.99086952 | 0.72065908 | 0.42583966 | 0.29481942 | 2.03E-06 | 0.00024322 |
| Ifitm2 | 1.03831761 | 0.72015303 | 0.4282847 | 0.29186833 | 2.86E-06 | 0.0003357 |
| Jund | 0.95944199 | 0.70489541 | 0.42360995 | 0.28128546 | 4.94E-06 | 0.00055806 |
| Junb | 0.79034162 | 0.67905473 | 0.40170366 | 0.27735108 | 2.16E-06 | 0.00025642 |
| Smarcd2 | 0.63796887 | 0.63642522 | 0.36428618 | 0.27213904 | 3.30E-07 | 4.46E-05 |
| Dbpht2 | 0.61027799 | 0.62600386 | 0.35513914 | 0.27086472 | 1.92E-07 | 2.68E-05 |
| Mfap4 | 0.32824347 | 0.503728 | 0.23343374 | 0.27029426 | 6.42E-15 | 1.68E-12 |
| Mtch1 | 0.77679019 | 0.66514201 | 0.39909526 | 0.26604675 | 4.55E-06 | 0.00051864 |
| Cystm1 | 1.60226962 | 0.66177705 | 0.3983644 | 0.26341265 | 5.38E-06 | 0.00060365 |
| Ffar2 | 0.98768477 | 0.68888815 | 0.42563259 | 0.26325556 | 1.92E-05 | 0.00197391 |
| Ptov1 | 1.0894881 | 0.688723 | 0.42958621 | 0.25913679 | 2.97E-05 | 0.00295786 |
| Tmed3 | 1.06485783 | 0.68589103 | 0.42917058 | 0.25672046 | 3.42E-05 | 0.00336668 |
| Apoe | 0.49544524 | 0.56278608 | 0.31222642 | 0.25055966 | 4.60E-08 | 6.99E-06 |
| Slc25a5 | 2.33021091 | 0.61961323 | 0.37043964 | 0.24917359 | 3.78E-06 | 0.0004365 |
| Rps4x | 2.81956781 | 0.60357892 | 0.35493552 | 0.24864341 | 1.56E-06 | 0.00018818 |
| Bex2 | 1.76253416 | 0.63186491 | 0.38329836 | 0.24856655 | 7.92E-06 | 0.000865 |
| Sct | 0.37731684 | 0.50658393 | 0.25881747 | 0.24776646 | 9.32E-11 | 1.88E-08 |
| Ssr4 | 1.6555932 | 0.63863996 | 0.39331188 | 0.24532808 | 1.65E-05 | 0.0017177 |
| Ppy | 0.20797935 | 0.40555529 | 0.16197916 | 0.24357613 | 6.92E-24 | 2.63E-21 |
| Hbb-bt | 0.72706212 | 0.63114423 | 0.38854304 | 0.24260119 | 1.62E-05 | 0.00169722 |
| Meg3 | 3.50435306 | 0.50227922 | 0.25986665 | 0.24241258 | 2.66E-10 | 4.88E-08 |
| Actg1 | 3.27617105 | 0.52232124 | 0.28616851 | 0.23615272 | 1.98E-08 | 3.12E-06 |
| Zcchc18 | 0.66645684 | 0.60890281 | 0.37313623 | 0.23576657 | 1.30E-05 | 0.00137308 |
| Cirbp | 1.28023529 | 0.66113119 | 0.42598658 | 0.2351446 | 0.00011925 | 0.01080063 |
| Cdk4 | 1.50429459 | 0.64169907 | 0.4086087 | 0.23309037 | 7.32E-05 | 0.00698692 |
| Sparc | 0.40487153 | 0.50381828 | 0.27220863 | 0.23160965 | 7.41E-09 | 1.21E-06 |
| Pdia4 | 1.17194226 | 0.66043661 | 0.42957385 | 0.23086276 | 0.0001736 | 0.01512177 |
| Rpl18a | 3.20782153 | 0.52558083 | 0.29908749 | 0.22649334 | 2.32E-07 | 3.21E-05 |

| | | | | | | |
|---|---|---|---|---|---|---|
| Actb | 2.93021631 | 0.56930816 | 0.34544095 | 0.22386721 | 8.03E-06 | 0.00087036 |
| Cacna2d1 | 0.63133181 | 0.58586563 | 0.36213733 | 0.2237283 | 1.96E-05 | 0.00200154 |
| Hbb-y | 0.30459621 | 0.44370409 | 0.22046152 | 0.22324257 | 7.90E-12 | 1.70E-09 |
| Ostc | 1.4684407 | 0.63433835 | 0.41222153 | 0.22211682 | 0.00016748 | 0.01475071 |
| Atp2a2 | 0.84941278 | 0.63299925 | 0.4114871 | 0.22151215 | 0.00016967 | 0.01486087 |
| Glud1 | 0.88295423 | 0.63676874 | 0.4160562 | 0.22071255 | 0.00020691 | 0.01792556 |
| Pdx1 | 1.02527812 | 0.64485881 | 0.42768122 | 0.21717759 | 0.00036228 | 0.02902041 |
| H3f3b | 3.09662355 | 0.53679955 | 0.31981642 | 0.21698313 | 3.15E-06 | 0.00036649 |
| Hmgn2 | 1.58481298 | 0.61656713 | 0.40010808 | 0.21645905 | 0.00015844 | 0.01403139 |
| Map1b | 1.5626451 | 0.61546382 | 0.40247185 | 0.21299197 | 0.00021366 | 0.01831167 |
| Serpina6 | 0.31632533 | 0.43949381 | 0.22695806 | 0.21253575 | 2.28E-10 | 4.28E-08 |
| Serpinh1 | 0.86994272 | 0.62608883 | 0.41438836 | 0.21170047 | 0.00033623 | 0.02748678 |
| Rpl36a | 2.04154066 | 0.58009872 | 0.36931852 | 0.21078019 | 7.27E-05 | 0.00697664 |
| Tsc22d1 | 1.25825473 | 0.63680732 | 0.42698643 | 0.20982089 | 0.00053597 | 0.04107652 |
| Stard10 | 1.12104562 | 0.63909436 | 0.42992679 | 0.20916757 | 0.00060092 | 0.04561526 |
| Pou3f4 | 0.3080854 | 0.42959677 | 0.22240693 | 0.20718984 | 2.81E-10 | 5.08E-08 |
| Tubb5 | 1.67550927 | 0.59718477 | 0.39139371 | 0.20579105 | 0.00023276 | 0.01973639 |
| Rps20 | 2.33937502 | 0.57566798 | 0.3705765 | 0.20509148 | 0.00011485 | 0.010462 |
| Mpc1 | 0.80465649 | 0.60674588 | 0.40433991 | 0.20240597 | 0.00043102 | 0.03368096 |
| Eef2 | 1.94670341 | 0.57402842 | 0.37181383 | 0.20221459 | 0.00014723 | 0.0131117 |
| Tff2 | 0.16229627 | 0.33209724 | 0.13096696 | 0.20113029 | 7.87E-25 | 3.22E-22 |
| Peg3 | 1.60902552 | 0.59174463 | 0.39774543 | 0.19399919 | 0.00058392 | 0.04453733 |
| Rpl12 | 1.76522194 | 0.57641094 | 0.38306681 | 0.19334414 | 0.00039022 | 0.03110216 |
| Pdia3 | 1.8806851 | 0.56846798 | 0.37535506 | 0.19311291 | 0.00030789 | 0.02556286 |
| Tff3 | 0.23142506 | 0.36891649 | 0.17702359 | 0.1918929 | 2.69E-13 | 6.59E-11 |
| Sec61b | 2.48442164 | 0.55925109 | 0.36989552 | 0.18935557 | 0.00032781 | 0.02693588 |
| Ccnd2 | 0.37125361 | 0.44274044 | 0.25578919 | 0.18695124 | 5.72E-07 | 7.42E-05 |
| Txnip | 0.70321169 | 0.56838374 | 0.38297066 | 0.18541308 | 0.00063508 | 0.0473075 |
| Fkbp2 | 0.97547624 | 0.60486318 | 0.42479447 | 0.18006871 | 0.00238921 | 0.14935824 |
| 1810008I18R | 0.44398889 | 0.46918323 | 0.2902178 | 0.17896543 | 2.02E-05 | 0.00205524 |

| | | | | | | |
|---|---|---|---|---|---|---|
| Gfra3 | 0.65905095 | 0.54844351 | 0.37090137 | 0.17754214 | 0.00072065 | 0.05318485 |
| Foxa3 | 0.58422418 | 0.52281078 | 0.34610499 | 0.17670579 | 0.00033886 | 0.02755978 |
| Gm8797 | 1.49796035 | 0.58592745 | 0.40927269 | 0.17665477 | 0.00203239 | 0.13223808 |
| Pcsk1 | 0.80749141 | 0.58147612 | 0.40484341 | 0.17663271 | 0.00184119 | 0.1217861 |
| Tecr | 1.60695736 | 0.57346428 | 0.39793536 | 0.17552893 | 0.00166176 | 0.11114947 |
| Gspt1 | 0.79366039 | 0.57724676 | 0.4023286 | 0.17491816 | 0.00190216 | 0.12478316 |
| Bambi | 0.47729067 | 0.47927463 | 0.30467131 | 0.17460332 | 6.82E-05 | 0.00658711 |
| Apoc1 | 0.31854711 | 0.40076114 | 0.22817457 | 0.17258657 | 2.39E-07 | 3.29E-05 |
| Rpl10a | 2.46262054 | 0.54222505 | 0.36984586 | 0.17237918 | 0.00095972 | 0.06829893 |
| Eef1a1 | 3.32802794 | 0.44841882 | 0.27632705 | 0.17209177 | 1.70E-05 | 0.00175518 |
| Hspa8 | 3.00837455 | 0.5070881 | 0.33518874 | 0.17189937 | 0.00032047 | 0.02646956 |
| Ypel3 | 1.38578397 | 0.59128187 | 0.41950639 | 0.17177548 | 0.00320941 | 0.18809274 |
| Krt19 | 0.44883051 | 0.46195785 | 0.29236825 | 0.1695896 | 5.64E-05 | 0.005487 |
| Eef1b2 | 1.59235317 | 0.56604937 | 0.39935828 | 0.16669109 | 0.00273182 | 0.16371397 |
| Neurod2 | 0.25279783 | 0.35637566 | 0.1902489 | 0.16612676 | 3.08E-09 | 5.17E-07 |
| Hsp90ab1 | 2.8926435 | 0.51432492 | 0.34934835 | 0.16497657 | 0.00083497 | 0.060316 |
| Pim2 | 0.72570526 | 0.55253211 | 0.38824137 | 0.16429074 | 0.00242563 | 0.15045522 |
| Nkx2-2 | 1.0039427 | 0.58844479 | 0.42663282 | 0.16181197 | 0.00579146 | 0.28587416 |
| Casp6 | 0.55535415 | 0.49578229 | 0.33562472 | 0.16015757 | 0.00074567 | 0.05452625 |
| Ddost | 1.54720432 | 0.56207722 | 0.40412299 | 0.15795423 | 0.00463838 | 0.24201117 |
| Emb | 1.07101622 | 0.5868331 | 0.42934152 | 0.15749158 | 0.00730928 | 0.34692483 |
| Ociad2 | 0.71167858 | 0.54189778 | 0.38500644 | 0.15689135 | 0.00333902 | 0.19215635 |
| Clu | 0.4218581 | 0.43674854 | 0.28017027 | 0.15657827 | 9.96E-05 | 0.00923102 |
| Pam | 0.7310478 | 0.54558981 | 0.38942357 | 0.15616624 | 0.00380018 | 0.20889214 |
| H1f0 | 1.28250146 | 0.58089607 | 0.42587491 | 0.15502116 | 0.00769739 | 0.3546359 |
| Sec61g | 2.26312696 | 0.52437774 | 0.36992861 | 0.15444913 | 0.00272548 | 0.16371397 |
| Ly6e | 0.7607395 | 0.54836563 | 0.39586133 | 0.1525043 | 0.0051686 | 0.25991397 |
| Ppp1r14a | 0.26872419 | 0.35225222 | 0.19980996 | 0.15244226 | 1.90E-07 | 2.68E-05 |
| Pak3 | 0.70703774 | 0.53588549 | 0.38389594 | 0.15198955 | 0.00420264 | 0.22406089 |
| Hsp90aa1 | 2.0813728 | 0.51938816 | 0.36863432 | 0.15075383 | 0.00324311 | 0.18937125 |

| | | | | | | |
|---|---|---|---|---|---|---|
| Cbfa2t3 | 0.44420722 | 0.43834843 | 0.29031513 | 0.1480333 | 0.00034426 | 0.02771603 |
| Pgrmc1 | 0.80508786 | 0.55244528 | 0.40441676 | 0.14802852 | 0.00741664 | 0.34875697 |
| Id2 | 0.30232979 | 0.3660608 | 0.21919209 | 0.14686871 | 4.10E-06 | 0.00046983 |
| Stxbp5l | 0.3540908 | 0.39364667 | 0.24705483 | 0.14659184 | 3.91E-05 | 0.00382789 |
| Fxyd3 | 0.50893164 | 0.46367056 | 0.31769545 | 0.14597511 | 0.00111234 | 0.07743181 |
| Hmgb3 | 0.45220223 | 0.43924794 | 0.29385609 | 0.14539185 | 0.00049495 | 0.03830099 |
| Rpl21 | 2.64769358 | 0.50965804 | 0.36448275 | 0.14517528 | 0.0040096 | 0.21763658 |
| Rps5 | 3.08180384 | 0.46608826 | 0.32274913 | 0.14333913 | 0.00155719 | 0.10608179 |
| Mfng | 0.53214179 | 0.46720033 | 0.32683033 | 0.14037 | 0.00212602 | 0.13610804 |
| Lpar6 | 0.43629414 | 0.42593466 | 0.28676548 | 0.13916918 | 0.00061814 | 0.04648012 |
| Rpl26 | 2.94693024 | 0.48261662 | 0.34356747 | 0.13904915 | 0.00353031 | 0.20012082 |
| Ubqln2 | 0.78064792 | 0.5385745 | 0.39984743 | 0.13872708 | 0.01046004 | 0.45065822 |
| Marcks | 0.81709991 | 0.54306889 | 0.4064945 | 0.13657439 | 0.01266249 | 0.51649113 |
| Ssr2 | 1.99779002 | 0.5066268 | 0.37074946 | 0.13587734 | 0.00735353 | 0.3478417 |
| Tmem258 | 1.46239951 | 0.54713638 | 0.41282426 | 0.13431212 | 0.01516928 | 0.56630796 |
| Gcg | 0.27797908 | 0.33943457 | 0.20525314 | 0.13418143 | 6.76E-06 | 0.00074838 |
| Btg1 | 0.90638401 | 0.55256176 | 0.41873829 | 0.13382347 | 0.01669808 | 0.59956436 |
| Ctsf | 0.72246429 | 0.52069094 | 0.38751687 | 0.13317407 | 0.01108261 | 0.4723742 |
| Eif3f | 1.76304794 | 0.51528905 | 0.38325414 | 0.13203492 | 0.01091875 | 0.46714787 |
| Tubb4b | 1.00586134 | 0.55632499 | 0.42674036 | 0.12958463 | 0.02162541 | 0.71171853 |
| Dusp1 | 0.41503393 | 0.40604622 | 0.27699778 | 0.12904844 | 0.00096408 | 0.06830379 |
| Qdpr | 0.85835922 | 0.54182505 | 0.41279136 | 0.1290337 | 0.01873074 | 0.64463872 |
| Slc2a2 | 0.43665854 | 0.41407614 | 0.28693 | 0.12714613 | 0.0015909 | 0.10791733 |
| Cotl1 | 0.80644084 | 0.52957797 | 0.40465726 | 0.12492071 | 0.01994591 | 0.67383848 |
| Zbtb20 | 0.79595509 | 0.52735272 | 0.40275766 | 0.12459507 | 0.01973864 | 0.67090317 |
| Krt18 | 1.00396566 | 0.55111106 | 0.42663416 | 0.1244769 | 0.02606298 | 0.81698086 |
| Cldn11 | 0.36919846 | 0.3780104 | 0.254756 | 0.1232544 | 0.00063998 | 0.04745073 |
| Rgs17 | 0.50611801 | 0.43974809 | 0.31656445 | 0.12318364 | 0.00479649 | 0.24664764 |
| Tead2 | 0.32528488 | 0.35474682 | 0.23183761 | 0.12290921 | 0.00020866 | 0.01797966 |
| Zdbf2 | 0.62028233 | 0.47982754 | 0.35849817 | 0.12132937 | 0.01213702 | 0.50170905 |

| | | | | | | |
|---|---|---|---|---|---|---|
| Egr1 | 0.61243475 | 0.4771611 | 0.35586791 | 0.12129319 | 0.01164214 | 0.49097191 |
| Calm2 | 2.63062863 | 0.48598924 | 0.36517322 | 0.12081601 | 0.01382461 | 0.53882175 |
| Tmsb10 | 2.28355789 | 0.49038214 | 0.37016261 | 0.12021952 | 0.01531611 | 0.56883004 |
| Grb10 | 0.84912265 | 0.53140662 | 0.41144422 | 0.1199624 | 0.02614396 | 0.81698086 |
| Dynll1 | 2.3594487 | 0.48923675 | 0.37062276 | 0.11861398 | 0.01657349 | 0.5977329 |
| Gm8773 | 0.11346178 | 0.21326239 | 0.09516224 | 0.11810015 | 7.23E-17 | 2.10E-14 |
| Igsf8 | 0.60346439 | 0.47087597 | 0.35281429 | 0.11806168 | 0.01296007 | 0.52009877 |
| Dbn1 | 0.7260116 | 0.50594134 | 0.38830957 | 0.11763177 | 0.0218775 | 0.71611751 |
| Cpa1 | 0.35097238 | 0.36279504 | 0.24544158 | 0.11735346 | 0.00072975 | 0.05360815 |
| Vwa5b2 | 0.88366108 | 0.53313795 | 0.41614199 | 0.11699596 | 0.03064501 | 0.89307523 |
| Npepl1 | 0.84614375 | 0.5279634 | 0.41100175 | 0.11696164 | 0.02909616 | 0.86533938 |
| Cited2 | 0.55456078 | 0.45083318 | 0.33532963 | 0.11550355 | 0.01093069 | 0.46714787 |
| Cyb5r3 | 1.76322073 | 0.49760972 | 0.38323926 | 0.11437046 | 0.02349095 | 0.75803498 |
| Rps8 | 2.99553249 | 0.45175595 | 0.33758235 | 0.1141736 | 0.01218504 | 0.50191641 |
| Sh3bgrl3 | 1.06579839 | 0.54278316 | 0.42919744 | 0.11358572 | 0.03906982 | 0.99999994 |
| Tmsb15b2 | 0.42000193 | 0.39238698 | 0.27931076 | 0.11307622 | 0.00352195 | 0.20012082 |
| Rnase4 | 0.63544234 | 0.47616154 | 0.36347093 | 0.1126906 | 0.01952125 | 0.66635616 |
| Hpcal1 | 0.47785483 | 0.41591088 | 0.30490948 | 0.1110014 | 0.00769164 | 0.3546359 |
| Rcn1 | 0.63341776 | 0.47217691 | 0.36281521 | 0.10936169 | 0.02240707 | 0.72452551 |
| Pdk3 | 0.3529438 | 0.35495395 | 0.24646238 | 0.10849156 | 0.00169428 | 0.11253555 |
| Rasd1 | 0.43723432 | 0.39517875 | 0.28718978 | 0.10798897 | 0.00615865 | 0.30100158 |
| Arx | 0.19038069 | 0.25812308 | 0.15030402 | 0.10781905 | 8.99E-07 | 0.00011198 |
| G6pc2 | 0.35920717 | 0.35699251 | 0.24968446 | 0.10730805 | 0.00211331 | 0.13583965 |
| Immp1l | 1.11459803 | 0.53603365 | 0.42989667 | 0.10613699 | 0.05015164 | 0.99999994 |
| Dnaja1 | 1.97956112 | 0.47688711 | 0.3707831 | 0.10610401 | 0.02840328 | 0.85915875 |
| Vim | 0.25336176 | 0.29553054 | 0.19059169 | 0.10493886 | 0.00012369 | 0.01113973 |
| Rpn2 | 1.11358695 | 0.53422063 | 0.42988911 | 0.10433152 | 0.05310575 | 0.99999994 |
| Sst | 0.14737134 | 0.22456854 | 0.12032524 | 0.1042433 | 4.04E-09 | 6.71E-07 |
| Mia | 0.29309023 | 0.31758637 | 0.21396768 | 0.10361869 | 0.00063319 | 0.0473075 |
| Sytl4 | 0.48580109 | 0.41170483 | 0.30824139 | 0.10346344 | 0.01273317 | 0.51649113 |

| | | | | | | |
|---|---|---|---|---|---|---|
| Ddc | 0.34636576 | 0.34556702 | 0.2430438 | 0.10252323 | 0.00249365 | 0.15230361 |
| Nono | 1.24671383 | 0.52926434 | 0.42746416 | 0.10180018 | 0.05645986 | 0.99999994 |
| Ppib | 1.69441063 | 0.490618 | 0.38951499 | 0.10110301 | 0.04201807 | 0.99999994 |
| Ube2i | 1.08160345 | 0.53055169 | 0.42948128 | 0.10107041 | 0.05862168 | 0.99999994 |
| Mgst1 | 0.29185672 | 0.31429338 | 0.21326427 | 0.10102912 | 0.00080718 | 0.05875474 |
| Tle6 | 0.34860639 | 0.34512937 | 0.24421225 | 0.10091712 | 0.00297393 | 0.1762357 |
| Bex1 | 0.74766309 | 0.49347367 | 0.39308962 | 0.10038405 | 0.04457987 | 0.99999994 |
| Ppp3ca | 0.596764 | 0.45040038 | 0.35050303 | 0.09989735 | 0.02890488 | 0.86317853 |
| Rpl9 | 2.85318001 | 0.45172121 | 0.35247483 | 0.09924638 | 0.03044926 | 0.89062697 |
| Mageh1 | 0.81008324 | 0.50366835 | 0.40530047 | 0.09836788 | 0.05309885 | 0.99999994 |
| Rbp1 | 0.24473418 | 0.28314786 | 0.18531266 | 0.0978352 | 0.00022054 | 0.01880003 |
| Cnih2 | 0.41558225 | 0.37454808 | 0.27725396 | 0.09729412 | 0.00974945 | 0.42696718 |
| Igfbp2 | 0.19095319 | 0.24760961 | 0.15068916 | 0.09692046 | 9.29E-06 | 0.00100089 |
| Rpl7a | 1.98079839 | 0.46750851 | 0.37076948 | 0.09673902 | 0.04121669 | 0.99999994 |
| Hhex | 0.11333412 | 0.19117378 | 0.09506484 | 0.09610893 | 8.51E-12 | 1.81E-09 |
| Rpsa | 2.4103008 | 0.46646704 | 0.37046565 | 0.09600139 | 0.04227192 | 0.99999994 |
| Rpl22 | 1.65223012 | 0.48928215 | 0.3936327 | 0.09564945 | 0.05289175 | 0.99999994 |
| Krtap17-1 | 0.14649425 | 0.21515942 | 0.11969183 | 0.09546759 | 5.51E-08 | 8.13E-06 |
| Ndufa1 | 1.10224955 | 0.52497804 | 0.42978451 | 0.09519353 | 0.07019701 | 0.99999994 |
| Prdx1 | 2.29287514 | 0.46536233 | 0.37024006 | 0.09512227 | 0.04361861 | 0.99999994 |
| Rps19 | 3.31027721 | 0.37483127 | 0.28006276 | 0.09476851 | 0.01214853 | 0.50170905 |
| Higd1a | 0.53903608 | 0.42368087 | 0.32947715 | 0.09420372 | 0.02850865 | 0.860713 |
| Cpt2 | 0.64254191 | 0.45988434 | 0.3657506 | 0.09413374 | 0.04334191 | 0.99999994 |
| Ube2e3 | 0.96255207 | 0.51697997 | 0.42384718 | 0.09313279 | 0.0717864 | 0.99999994 |
| Atp5g1 | 2.13898819 | 0.46145707 | 0.36849509 | 0.09296197 | 0.04655357 | 0.99999994 |
| Stx16 | 0.63827655 | 0.45697807 | 0.36438523 | 0.09259284 | 0.04537762 | 0.99999994 |
| Nefm | 0.14047161 | 0.20746225 | 0.11531728 | 0.09214498 | 5.22E-08 | 7.78E-06 |
| Nkx6-1 | 1.06533436 | 0.52118679 | 0.42918422 | 0.09200257 | 0.07680467 | 0.99999994 |
| Prps1 | 0.64286032 | 0.45701877 | 0.36585144 | 0.09116733 | 0.04858686 | 0.99999994 |
| Cited1 | 0.20507271 | 0.2507884 | 0.16007369 | 0.09071471 | 8.09E-05 | 0.00763169 |

| | | | | | | |
|---|---|---|---|---|---|---|
| Tagln2 | 0.65227024 | 0.4587967 | 0.36880729 | 0.08998941 | 0.05217093 | 0.99999994 |
| Pcbp4 | 0.61143207 | 0.44493272 | 0.35552942 | 0.0894033 | 0.04708075 | 0.99999994 |
| Aldh1b1 | 0.221779 | 0.26023954 | 0.17090345 | 0.08933609 | 0.00025121 | 0.02107615 |
| Hacd1 | 0.59347826 | 0.43852845 | 0.34935989 | 0.08916856 | 0.04466623 | 0.99999994 |
| Peg10 | 0.14720858 | 0.20913406 | 0.12020777 | 0.08892629 | 4.24E-07 | 5.63E-05 |
| Gm2000 | 0.5436374 | 0.41898966 | 0.33122734 | 0.08776232 | 0.03889335 | 0.99999994 |
| Tubb2a | 0.48430719 | 0.39524777 | 0.30761821 | 0.08762956 | 0.02896936 | 0.86317853 |
| Aplp2 | 0.46457382 | 0.38660774 | 0.29924453 | 0.08736321 | 0.02599018 | 0.81698086 |
| Krtcap2 | 1.27709738 | 0.51341494 | 0.42613692 | 0.08727802 | 0.0863935 | 0.99999994 |
| Celf3 | 0.60756291 | 0.44116045 | 0.3542158 | 0.08694465 | 0.05114539 | 0.99999994 |
| Pnrc1 | 0.60117741 | 0.43870087 | 0.35202821 | 0.08667265 | 0.05062143 | 0.99999994 |
| Ddah2 | 0.40707594 | 0.35942248 | 0.27325455 | 0.08616793 | 0.01790792 | 0.62329713 |
| Id1 | 0.14531895 | 0.20497707 | 0.11884157 | 0.0861355 | 7.02E-07 | 9.02E-05 |
| Spint1 | 0.87749255 | 0.50143553 | 0.41538395 | 0.08605157 | 0.08395643 | 0.99999994 |
| Mfap2 | 0.38523553 | 0.34857064 | 0.26272703 | 0.08584362 | 0.01481869 | 0.5624398 |
| Gdpd1 | 0.46881993 | 0.38630138 | 0.30106951 | 0.08523187 | 0.02975579 | 0.87840191 |
| Gpx2 | 0.4424052 | 0.37374275 | 0.2895108 | 0.08423195 | 0.02639472 | 0.81859587 |
| Ckb | 0.57584624 | 0.42710443 | 0.34311512 | 0.08398931 | 0.05161693 | 0.99999994 |
| Eif3h | 1.63645558 | 0.4785374 | 0.39505988 | 0.08347752 | 0.07978616 | 0.99999994 |
| Rpl32 | 3.2517449 | 0.3730752 | 0.29022076 | 0.08285444 | 0.02869636 | 0.86317853 |
| Lamp1 | 0.89092301 | 0.49977445 | 0.41701118 | 0.08276327 | 0.09323861 | 0.99999994 |
| Tox3 | 0.73107258 | 0.47199075 | 0.38942902 | 0.08256173 | 0.0790935 | 0.99999994 |
| Ffar4 | 0.56071465 | 0.4199483 | 0.33760871 | 0.08233959 | 0.0522495 | 0.99999994 |
| Phlda1 | 0.31621921 | 0.3091223 | 0.22689985 | 0.08222245 | 0.00793076 | 0.363288 |
| Fkbp11 | 0.37087283 | 0.33777277 | 0.25559802 | 0.08217475 | 0.01617662 | 0.58740677 |
| Resp18 | 0.3207893 | 0.31147061 | 0.22939791 | 0.0820727 | 0.00862174 | 0.38498356 |
| Mt1 | 0.34659464 | 0.32450507 | 0.24316334 | 0.08134172 | 0.01298534 | 0.52009877 |
| Srsf2 | 0.62629593 | 0.44147486 | 0.36049121 | 0.08098365 | 0.06741315 | 0.99999994 |
| Amotl2 | 0.23835424 | 0.26225786 | 0.18136189 | 0.08089597 | 0.00149361 | 0.10262764 |
| Srsf3 | 1.35948101 | 0.50212532 | 0.42141647 | 0.08070885 | 0.10118781 | 0.99999994 |

| | | | | | | |
|---|---|---|---|---|---|---|
| Ppp1r14b | 0.75045903 | 0.47438348 | 0.39369679 | 0.08068669 | 0.08625299 | 0.99999994 |
| Pla2g12a | 0.42102947 | 0.36003716 | 0.27978688 | 0.08025028 | 0.02811689 | 0.85536517 |
| Gast | 0.10007886 | 0.16505012 | 0.08484109 | 0.08020903 | 1.56E-10 | 3.10E-08 |
| Prrg2 | 0.75740545 | 0.47520755 | 0.39516513 | 0.08004242 | 0.08878509 | 0.99999994 |
| St3gal6 | 0.34183746 | 0.32062913 | 0.24066868 | 0.07996045 | 0.01349967 | 0.53399085 |
| Dad1 | 1.77365352 | 0.46223563 | 0.38235905 | 0.07987658 | 0.08218187 | 0.99999994 |
| Rpl9-ps6 | 0.58982781 | 0.42756818 | 0.34808171 | 0.07948647 | 0.06425325 | 0.99999994 |
| Prnp | 1.17325075 | 0.50876512 | 0.42955896 | 0.07920616 | 0.10984321 | 0.99999994 |
| Anxa5 | 0.62159081 | 0.43798903 | 0.35893349 | 0.07905554 | 0.07131448 | 0.99999994 |
| Prss53 | 0.32524478 | 0.3095386 | 0.23181593 | 0.07772267 | 0.0128165 | 0.51649113 |
| Zwint | 0.83571966 | 0.48651631 | 0.40939331 | 0.077123 | 0.10493029 | 0.99999994 |
| Anxa4 | 0.98546429 | 0.50245507 | 0.42548413 | 0.07697094 | 0.11426607 | 0.99999994 |
| Rcor2 | 0.36696187 | 0.33050575 | 0.25362777 | 0.07687798 | 0.02181484 | 0.71553568 |
| Hspa1a | 0.18733459 | 0.22484465 | 0.14824892 | 0.07659573 | 0.00029178 | 0.02435252 |
| Fxyd6 | 0.50133678 | 0.39109383 | 0.31463086 | 0.07646297 | 0.05286823 | 0.99999994 |
| Kirrel2 | 0.15217325 | 0.19929463 | 0.12377713 | 0.0755175 | 2.44E-05 | 0.00246388 |
| Rpl17 | 2.93408989 | 0.42044838 | 0.34495613 | 0.07549225 | 0.07259726 | 0.99999994 |
| Olfm1 | 0.24318994 | 0.25971984 | 0.18436008 | 0.07535976 | 0.00325543 | 0.18939691 |
| Rps9 | 3.12188498 | 0.39094426 | 0.31637968 | 0.07456458 | 0.0583502 | 0.99999994 |
| Ranbp1 | 1.52263432 | 0.48088459 | 0.4066739 | 0.07421069 | 0.11224677 | 0.99999994 |
| Trp53i11 | 0.43624291 | 0.36092875 | 0.28674234 | 0.07418641 | 0.04252181 | 0.99999994 |
| Mpzl1 | 0.78214134 | 0.47358394 | 0.40013667 | 0.07344727 | 0.11088959 | 0.99999994 |
| Rab3a | 0.77650673 | 0.47235936 | 0.39903971 | 0.07331964 | 0.11065717 | 0.99999994 |
| Tpst2 | 0.65176546 | 0.44191914 | 0.36864991 | 0.07326923 | 0.09292565 | 0.99999994 |
| Vasp | 0.74026694 | 0.46459981 | 0.39146448 | 0.07313533 | 0.10682824 | 0.99999994 |
| Gars | 1.1859879 | 0.50197048 | 0.42931665 | 0.07265383 | 0.12998452 | 0.99999994 |
| Lhx1 | 0.28053219 | 0.27873555 | 0.20674067 | 0.07199488 | 0.01022482 | 0.44524014 |
| Cdk2ap1 | 0.42581029 | 0.35395697 | 0.28199105 | 0.07196592 | 0.04468361 | 0.99999994 |
| Prox1 | 0.45203843 | 0.36573642 | 0.29378399 | 0.07195243 | 0.05152441 | 0.99999994 |
| Tmem238 | 0.42305226 | 0.35241162 | 0.28072176 | 0.07168986 | 0.04457673 | 0.99999994 |

| | | | | | | |
|---|---|---|---|---|---|---|
| Eif3e | 1.48310744 | 0.48168303 | 0.41076387 | 0.07091916 | 0.12523001 | 0.99999994 |
| Nucb2 | 0.51430763 | 0.39038868 | 0.31984239 | 0.07054629 | 0.07102971 | 0.99999994 |
| Hepacam2 | 0.9018795 | 0.48861989 | 0.41824801 | 0.07037188 | 0.13136572 | 0.99999994 |
| Fscn1 | 0.24764752 | 0.25712602 | 0.18710341 | 0.07002262 | 0.00636719 | 0.30757397 |
| Upp1 | 0.21466735 | 0.23589616 | 0.16632892 | 0.06956725 | 0.00268441 | 0.16209177 |
| Rph3al | 0.96977857 | 0.49394695 | 0.42438453 | 0.06956242 | 0.13762148 | 0.99999994 |
| Serpini1 | 0.28502853 | 0.27826127 | 0.20934543 | 0.06891584 | 0.0142168 | 0.54741557 |
| Epcam | 1.57545134 | 0.46968459 | 0.40109077 | 0.06859382 | 0.12748542 | 0.99999994 |
| Prr13 | 0.87936251 | 0.48337468 | 0.415617 | 0.06775767 | 0.13892093 | 0.99999994 |
| Foxa2 | 0.66325685 | 0.4397821 | 0.37217408 | 0.06760802 | 0.11329801 | 0.99999994 |
| Cer1 | 0.15142796 | 0.19045836 | 0.12324313 | 0.06721523 | 0.00014155 | 0.01267668 |
| Maged1 | 1.75901853 | 0.45071469 | 0.3836005 | 0.06711419 | 0.12209314 | 0.99999994 |
| Serp1 | 0.90263435 | 0.48544152 | 0.41833075 | 0.06711076 | 0.14279475 | 0.99999994 |
| Rps26 | 2.51682382 | 0.43625851 | 0.36930322 | 0.06695529 | 0.11375289 | 0.99999994 |
| Gpc3 | 0.2269456 | 0.24108875 | 0.1741933 | 0.06689545 | 0.00529049 | 0.26520646 |
| Klf2 | 0.1587155 | 0.19395386 | 0.12843726 | 0.0655166 | 0.00034258 | 0.02771603 |
| Cyba | 0.31398466 | 0.29072821 | 0.22567166 | 0.06505655 | 0.02749896 | 0.83977187 |
| Sox11 | 0.14110006 | 0.18016581 | 0.11577575 | 0.06439006 | 0.00010695 | 0.00983279 |
| Necab2 | 0.32977199 | 0.29829195 | 0.23425544 | 0.06403651 | 0.03441037 | 0.97085972 |
| Rps11 | 2.86452786 | 0.41559064 | 0.35156462 | 0.06402602 | 0.11271057 | 0.99999994 |
| Mgst3 | 0.46168933 | 0.36190009 | 0.29799774 | 0.06390236 | 0.07673474 | 0.99999994 |
| Prdx5 | 1.24604272 | 0.49139211 | 0.42749107 | 0.06390103 | 0.15987124 | 0.99999994 |
| Acaa2 | 0.21609945 | 0.2300039 | 0.16725429 | 0.06274961 | 0.00625718 | 0.30317865 |
| Swt1 | 0.41562352 | 0.33974296 | 0.27727323 | 0.06246972 | 0.06684878 | 0.99999994 |
| Prodh2 | 0.23085451 | 0.23911371 | 0.17666422 | 0.0624495 | 0.00931198 | 0.41348828 |
| Tuba4a | 0.90582864 | 0.48106846 | 0.41867829 | 0.06239017 | 0.16062064 | 0.99999994 |
| Ankrd44 | 0.4783128 | 0.36747617 | 0.30510265 | 0.06237352 | 0.08679011 | 0.99999994 |
| Cd164l2 | 0.51972349 | 0.38410703 | 0.32198603 | 0.062121 | 0.09953318 | 0.99999994 |
| Grhpr | 0.57420726 | 0.40438816 | 0.34252521 | 0.06186295 | 0.11464407 | 0.99999994 |
| Limd2 | 0.64701557 | 0.42892543 | 0.36716243 | 0.061763 | 0.13141798 | 0.99999994 |

| | | | | | | |
|---|---|---|---|---|---|---|
| Tuba1b | 0.9317535 | 0.48296959 | 0.42129447 | 0.06167511 | 0.16491586 | 0.99999994 |
| Lmna | 0.6379378 | 0.42569518 | 0.36427618 | 0.061419 | 0.13086763 | 0.99999994 |
| Ctsz | 0.6205311 | 0.41993742 | 0.35858101 | 0.06135642 | 0.12735943 | 0.99999994 |
| Spcs2 | 1.75512684 | 0.44513177 | 0.38393397 | 0.0611978 | 0.14434464 | 0.99999994 |
| Tshz2 | 0.60395187 | 0.4138662 | 0.35298147 | 0.06088472 | 0.12545635 | 0.99999994 |
| Foxp1 | 0.50713102 | 0.37768048 | 0.31697223 | 0.06070824 | 0.10117908 | 0.99999994 |
| Tmem171 | 0.12867394 | 0.16708009 | 0.10662307 | 0.06045701 | 8.02E-05 | 0.00761379 |
| Pcp4 | 0.2238589 | 0.23243032 | 0.17223119 | 0.06019913 | 0.00999423 | 0.43648767 |
| Tpm4 | 0.41708122 | 0.33812183 | 0.27795315 | 0.06016867 | 0.07480569 | 0.99999994 |
| Mlxipl | 0.64065737 | 0.42525929 | 0.36514999 | 0.0601093 | 0.13659653 | 0.99999994 |
| Dnajc3 | 0.69590507 | 0.44117917 | 0.38112676 | 0.06005241 | 0.14713231 | 0.99999994 |
| Dstn | 1.52694963 | 0.46626799 | 0.40623239 | 0.06003561 | 0.16262686 | 0.99999994 |
| Steap1 | 0.18717572 | 0.20807678 | 0.14814146 | 0.05993532 | 0.00354018 | 0.20012082 |
| Wfdc2 | 0.19431077 | 0.21282608 | 0.1529402 | 0.05988587 | 0.00457511 | 0.24149618 |
| S100a10 | 0.44789516 | 0.35155611 | 0.29195409 | 0.05960202 | 0.08709134 | 0.99999994 |
| Cald1 | 0.19301036 | 0.21150919 | 0.15206987 | 0.05943932 | 0.00463733 | 0.24201117 |
| Cd9 | 0.48632095 | 0.36772541 | 0.30845777 | 0.05926765 | 0.1004547 | 0.99999994 |
| Polr2k | 1.07326719 | 0.48861081 | 0.42937409 | 0.05923672 | 0.1792269 | 0.99999994 |
| Pkib | 0.23843842 | 0.24046428 | 0.18141428 | 0.05904999 | 0.01513148 | 0.56630796 |
| Ccnd3 | 0.43905543 | 0.3468133 | 0.2880095 | 0.0588038 | 0.08706547 | 0.99999994 |
| Cbx1 | 1.02329689 | 0.48622347 | 0.42758234 | 0.05864113 | 0.18064976 | 0.99999994 |
| Smarce1 | 0.77760231 | 0.45745889 | 0.39925421 | 0.05820469 | 0.16592673 | 0.99999994 |
| Rpl28 | 2.67487768 | 0.42147831 | 0.36343588 | 0.05804243 | 0.14387922 | 0.99999994 |
| Hnrnpk | 2.07959868 | 0.42664624 | 0.36867196 | 0.05797428 | 0.14761359 | 0.99999994 |
| Tmc4 | 0.73842924 | 0.44891152 | 0.39105638 | 0.05785514 | 0.16236607 | 0.99999994 |
| Npc2 | 1.25729867 | 0.48479903 | 0.42702707 | 0.05777196 | 0.18391903 | 0.99999994 |
| C2cd4b | 0.1496547 | 0.17956136 | 0.12196999 | 0.05759136 | 0.0008362 | 0.060316 |
| Snrk | 0.2640054 | 0.25425669 | 0.19700243 | 0.05725426 | 0.0265247 | 0.81943852 |
| Cldn3 | 0.45056246 | 0.35029908 | 0.2931335 | 0.05716559 | 0.09712766 | 0.99999994 |
| Set | 1.09577037 | 0.4864942 | 0.42968977 | 0.05680443 | 0.18943999 | 0.99999994 |

| | | | | | | |
|---|---|---|---|---|---|---|
| Hmgb2 | 0.55493863 | 0.3920609 | 0.33547022 | 0.05659068 | 0.13074739 | 0.99999994 |
| Traf4 | 0.43435574 | 0.3424309 | 0.28588869 | 0.05654221 | 0.09400741 | 0.99999994 |
| Lhx1os | 0.22679578 | 0.23060128 | 0.17409829 | 0.05650299 | 0.01537482 | 0.56883004 |
| Cpa2 | 0.27620109 | 0.26070644 | 0.20421364 | 0.05649281 | 0.03278243 | 0.94159421 |
| Rps3 | 2.91605802 | 0.40316521 | 0.34721591 | 0.0559493 | 0.14172867 | 0.99999994 |
| Rpl39 | 2.3104377 | 0.42614653 | 0.37034452 | 0.05580202 | 0.15794173 | 0.99999994 |
| Tmem184a | 0.51197502 | 0.37470315 | 0.3189131 | 0.05579005 | 0.12212071 | 0.99999994 |
| Dnajb11 | 0.91433852 | 0.47526142 | 0.41957431 | 0.05568711 | 0.18849631 | 0.99999994 |
| Pcyt1b | 0.22476479 | 0.22792773 | 0.17280804 | 0.05511969 | 0.01687089 | 0.60435699 |
| Lrpap1 | 0.97614874 | 0.47994001 | 0.42484204 | 0.05509798 | 0.19399551 | 0.99999994 |
| Dnajb9 | 0.63971287 | 0.41992312 | 0.36484696 | 0.05507616 | 0.15749102 | 0.99999994 |
| Rpl23a | 2.70613681 | 0.41662881 | 0.36182762 | 0.0548012 | 0.15669131 | 0.99999994 |
| Hmgn1 | 2.06985201 | 0.42354786 | 0.36887568 | 0.05467218 | 0.16192945 | 0.99999994 |
| Hpca | 0.21218617 | 0.21934579 | 0.16472063 | 0.05462516 | 0.013643 | 0.53455393 |
| Dbi | 0.5618786 | 0.3924097 | 0.33803726 | 0.05437244 | 0.14216152 | 0.99999994 |
| Hes1 | 0.10536796 | 0.14304318 | 0.08894703 | 0.05409615 | 2.58E-05 | 0.00258632 |
| Tpbg | 0.21766819 | 0.22182879 | 0.16826553 | 0.05356325 | 0.01704988 | 0.60693149 |
| Gapdh | 1.71270912 | 0.44118431 | 0.38787615 | 0.05330816 | 0.18014214 | 0.99999994 |
| Serping1 | 0.34590458 | 0.29589168 | 0.24280278 | 0.05308891 | 0.07277723 | 0.99999994 |
| Fam151a | 0.43152135 | 0.33754104 | 0.28460176 | 0.05293929 | 0.10782815 | 0.99999994 |
| Pbx1 | 0.55283256 | 0.38697531 | 0.33468555 | 0.05228976 | 0.14917939 | 0.99999994 |
| Tmem9 | 0.65829058 | 0.42271524 | 0.3706703 | 0.05204495 | 0.17499632 | 0.99999994 |
| Enpp2 | 0.42878619 | 0.33533416 | 0.28335438 | 0.05197978 | 0.11102942 | 0.99999994 |
| Chic2 | 0.50038256 | 0.36613554 | 0.31424301 | 0.05189254 | 0.13584085 | 0.99999994 |
| Glrx | 0.59805908 | 0.40275352 | 0.35095171 | 0.05180182 | 0.16292583 | 0.99999994 |
| Zfand5 | 0.41718441 | 0.32965877 | 0.27800123 | 0.05165754 | 0.10806786 | 0.99999994 |
| Lbp | 0.28662966 | 0.26136378 | 0.21026818 | 0.0510956 | 0.05288425 | 0.99999994 |
| Anp32a | 1.28614745 | 0.47663475 | 0.42569311 | 0.05094164 | 0.21285627 | 0.99999994 |
| Serpina1b | 0.10008668 | 0.13570927 | 0.08484719 | 0.05086208 | 3.30E-05 | 0.00326845 |
| Hbp1 | 0.55835395 | 0.38740539 | 0.33673708 | 0.05066831 | 0.15827517 | 0.99999994 |

| | | | | | | |
|---|---|---|---|---|---|---|
| Elovl1 | 0.44532976 | 0.34127697 | 0.29081502 | 0.05046196 | 0.12404415 | 0.99999994 |
| Pnliprp1 | 0.81769921 | 0.45669007 | 0.40658915 | 0.05010092 | 0.20604737 | 0.99999994 |
| Alcam | 0.300059 | 0.26791735 | 0.21791561 | 0.05000174 | 0.06333926 | 0.99999994 |
| Prr15l | 0.31265143 | 0.2749327 | 0.2249365 | 0.0499962 | 0.06950391 | 0.99999994 |
| Smarcc1 | 0.58706273 | 0.39693765 | 0.34710853 | 0.04982911 | 0.1696492 | 0.99999994 |
| Fam210b | 0.21269253 | 0.21420506 | 0.16504937 | 0.04915569 | 0.02371588 | 0.76220736 |
| Elavl1 | 0.61563036 | 0.40608822 | 0.35694302 | 0.04914521 | 0.17970983 | 0.99999994 |
| Maged2 | 0.7169518 | 0.43530483 | 0.38625435 | 0.04905048 | 0.19897401 | 0.99999994 |
| App | 0.92571544 | 0.46969794 | 0.42071812 | 0.04897982 | 0.21919044 | 0.99999994 |
| Sfpq | 0.63766449 | 0.4130649 | 0.36418814 | 0.04887675 | 0.18583953 | 0.99999994 |
| Cope | 1.47503539 | 0.46023659 | 0.41156925 | 0.04866735 | 0.21561087 | 0.99999994 |
| Syt13 | 1.21473216 | 0.47692981 | 0.4285899 | 0.04833992 | 0.22639881 | 0.99999994 |
| Cd79a | 0.18083963 | 0.19213151 | 0.14383329 | 0.04829822 | 0.01270346 | 0.51649113 |
| Arf5 | 2.12221978 | 0.41666256 | 0.36845864 | 0.04820392 | 0.19192519 | 0.99999994 |
| Casz1 | 0.34708916 | 0.29111627 | 0.24342149 | 0.04769478 | 0.09608044 | 0.99999994 |
| Ubc | 2.34050574 | 0.4181999 | 0.37059314 | 0.04760676 | 0.1962528 | 0.99999994 |
| Rps10 | 2.43212662 | 0.41794924 | 0.37044872 | 0.04750052 | 0.1966892 | 0.99999994 |
| Etv1 | 0.16121103 | 0.17765078 | 0.13020195 | 0.04744883 | 0.00763856 | 0.3546359 |
| Trim35 | 0.61310146 | 0.40335196 | 0.35609267 | 0.04725929 | 0.1885082 | 0.99999994 |
| Usp9x | 0.62534373 | 0.40738139 | 0.36017692 | 0.04720446 | 0.19150158 | 0.99999994 |
| Bri3 | 0.56502927 | 0.38639571 | 0.33919302 | 0.04720269 | 0.17714236 | 0.99999994 |
| Hmgn5 | 0.3005431 | 0.26520159 | 0.21818813 | 0.04701346 | 0.07574909 | 0.99999994 |
| Sms | 0.3514064 | 0.29254036 | 0.24566659 | 0.04687377 | 0.10203362 | 0.99999994 |
| Ppdpf | 1.01006705 | 0.47381396 | 0.4269492 | 0.04686476 | 0.23249777 | 0.99999994 |
| Insm1 | 0.70053559 | 0.42915702 | 0.38229862 | 0.0468584 | 0.20728702 | 0.99999994 |
| Cxxc5 | 0.65198731 | 0.41528745 | 0.3687191 | 0.04656836 | 0.20026294 | 0.99999994 |
| Arpc5 | 1.35917382 | 0.46727844 | 0.42143707 | 0.04584136 | 0.23452115 | 0.99999994 |
| Plcxd3 | 0.44397045 | 0.33590137 | 0.29020958 | 0.04569179 | 0.1473183 | 0.99999994 |
| Cyth2 | 0.81192177 | 0.45129619 | 0.4056228 | 0.0456734 | 0.22677484 | 0.99999994 |
| Pfn1 | 2.38536924 | 0.4160394 | 0.370453 | 0.0455864 | 0.20636366 | 0.99999994 |

| | | | | | | |
|---|---|---|---|---|---|---|
| Srsf7 | 1.13852187 | 0.47546881 | 0.42990667 | 0.04556213 | 0.24026497 | 0.99999994 |
| Atxn7l3b | 1.41307834 | 0.46283043 | 0.41731796 | 0.04551247 | 0.23393814 | 0.99999994 |
| Hnrnpm | 0.8836256 | 0.46154362 | 0.41613768 | 0.04540594 | 0.23382938 | 0.99999994 |
| Serpine2 | 0.19997528 | 0.20184315 | 0.15671022 | 0.04513294 | 0.02761593 | 0.84173146 |
| Hap1 | 0.36409624 | 0.29700251 | 0.25217636 | 0.04482615 | 0.11836196 | 0.99999994 |
| Uchl1 | 0.3082092 | 0.26709241 | 0.22247576 | 0.04461665 | 0.09095343 | 0.99999994 |
| Ndrg1 | 0.40754567 | 0.31772523 | 0.27347681 | 0.04424842 | 0.14074109 | 0.99999994 |
| Lrrc42 | 0.50650389 | 0.36064427 | 0.31671986 | 0.04392441 | 0.1779673 | 0.99999994 |
| Ubald2 | 0.86269026 | 0.45723598 | 0.41340896 | 0.04382702 | 0.24019815 | 0.99999994 |
| Fam43a | 0.19726369 | 0.1987042 | 0.15490979 | 0.04379441 | 0.0299307 | 0.88030488 |
| Abcc8 | 0.53902865 | 0.37326746 | 0.32947431 | 0.04379315 | 0.18814537 | 0.99999994 |
| Ube2b | 1.43751135 | 0.45899142 | 0.41520196 | 0.04378946 | 0.24133467 | 0.99999994 |
| Papss2 | 0.4490756 | 0.33619676 | 0.29247666 | 0.0437201 | 0.15986618 | 0.99999994 |
| Amigo2 | 0.2616493 | 0.23916769 | 0.19559269 | 0.043575 | 0.06904614 | 0.99999994 |
| Sec11c | 0.57729552 | 0.38718799 | 0.34363547 | 0.04355252 | 0.1994381 | 0.99999994 |
| Mob4 | 0.88552073 | 0.45986209 | 0.41636668 | 0.04349541 | 0.24341766 | 0.99999994 |
| Akirin2 | 0.25797216 | 0.23686737 | 0.19338192 | 0.04348545 | 0.06722221 | 0.99999994 |
| Dll3 | 0.10196951 | 0.12970965 | 0.08631288 | 0.04339677 | 0.00040888 | 0.03210805 |
| Pbxip1 | 0.33375679 | 0.27964788 | 0.23638713 | 0.04326076 | 0.11158019 | 0.99999994 |
| Kdm1a | 0.42931945 | 0.32675202 | 0.283598 | 0.04315402 | 0.15556052 | 0.99999994 |
| Psip1 | 0.53908641 | 0.3723131 | 0.32949636 | 0.04281674 | 0.19352841 | 0.99999994 |
| Rnf187 | 0.58329114 | 0.38852384 | 0.34577412 | 0.04274971 | 0.2052661 | 0.99999994 |
| Rbfox3 | 0.20401043 | 0.20210786 | 0.15937502 | 0.04273284 | 0.03715069 | 0.99999994 |
| Smarca1 | 0.24768093 | 0.22983747 | 0.1871239 | 0.04271358 | 0.06433038 | 0.99999994 |
| Pcbp1 | 0.82671281 | 0.45069582 | 0.40799251 | 0.04270331 | 0.24299515 | 0.99999994 |
| Rhob | 0.45446571 | 0.33743695 | 0.29485021 | 0.04258674 | 0.16817249 | 0.99999994 |
| Camk2n1 | 0.70302527 | 0.42544985 | 0.38292397 | 0.04252588 | 0.22988477 | 0.99999994 |
| Pla2g2d | 0.23703521 | 0.22305931 | 0.18053985 | 0.04251946 | 0.05848048 | 0.99999994 |
| Dnajc12 | 0.36651393 | 0.29587374 | 0.25340133 | 0.04247241 | 0.1322823 | 0.99999994 |
| Akap17b | 0.15343949 | 0.16696246 | 0.12468293 | 0.04227953 | 0.01199945 | 0.50170905 |

| | | | | | | |
|---|---|---|---|---|---|---|
| Igf2 | 0.17248213 | 0.18035798 | 0.13808238 | 0.0422756 | 0.02077873 | 0.69426319 |
| Col9a2 | 0.35359457 | 0.28877446 | 0.24679865 | 0.04197581 | 0.1287917 | 0.99999994 |
| Rem2 | 0.37501547 | 0.29957508 | 0.2576717 | 0.04190338 | 0.13952013 | 0.99999994 |
| Psmd4 | 1.38016987 | 0.4617264 | 0.41992635 | 0.04180005 | 0.2537988 | 0.99999994 |
| Litaf | 0.2086901 | 0.20390376 | 0.16244363 | 0.04146013 | 0.04467019 | 0.99999994 |
| Aplp1 | 1.63197554 | 0.43694906 | 0.39551457 | 0.04143449 | 0.24279926 | 0.99999994 |
| Ldha | 0.20468549 | 0.20094408 | 0.15981915 | 0.04112493 | 0.04337237 | 0.99999994 |
| Lpl | 0.22652897 | 0.21502816 | 0.17392903 | 0.04109913 | 0.05787261 | 0.99999994 |
| Uqcr11 | 1.78335663 | 0.42268137 | 0.38168347 | 0.0409979 | 0.23731017 | 0.99999994 |
| Kcnq1ot1 | 0.64106329 | 0.40620913 | 0.36528008 | 0.04092905 | 0.22788377 | 0.99999994 |
| St18 | 0.31000774 | 0.2643445 | 0.22347414 | 0.04087037 | 0.11173422 | 0.99999994 |
| Jup | 0.42281318 | 0.32148062 | 0.28061148 | 0.04086914 | 0.16615959 | 0.99999994 |
| Rps23 | 3.08015247 | 0.36374668 | 0.32307734 | 0.04066934 | 0.20104065 | 0.99999994 |
| Tspan13 | 0.70070507 | 0.42290213 | 0.38234129 | 0.04056083 | 0.24004977 | 0.99999994 |
| Parm1 | 0.20905975 | 0.20321005 | 0.16268499 | 0.04052506 | 0.04864784 | 0.99999994 |
| Dek | 0.35062167 | 0.28575937 | 0.24525965 | 0.04049972 | 0.13584822 | 0.99999994 |
| Akr1c13 | 0.4537351 | 0.33485851 | 0.29452985 | 0.04032866 | 0.18103561 | 0.99999994 |
| Tob1 | 0.40498509 | 0.31235306 | 0.2722626 | 0.04009046 | 0.16350761 | 0.99999994 |
| Usp18 | 0.17065647 | 0.17689649 | 0.13681561 | 0.04008088 | 0.02558709 | 0.80769072 |
| Ezh2 | 0.28425094 | 0.24891373 | 0.20889645 | 0.04001727 | 0.10113232 | 0.99999994 |
| Rpl8 | 2.74647253 | 0.39945792 | 0.35960772 | 0.0398502 | 0.23036985 | 0.99999994 |
| Ndufv3 | 1.26078701 | 0.46658883 | 0.42687788 | 0.03971095 | 0.26788649 | 0.99999994 |
| Ddit4 | 0.27954139 | 0.24581721 | 0.20616411 | 0.03965309 | 0.10022557 | 0.99999994 |
| Ppp1r14c | 0.12155222 | 0.14072101 | 0.1012929 | 0.03942811 | 0.00478519 | 0.24664764 |
| Gm8730 | 2.35992684 | 0.40980158 | 0.37061988 | 0.0391817 | 0.24080796 | 0.99999994 |
| Txndc12 | 0.64196276 | 0.40455854 | 0.36556706 | 0.03899148 | 0.23886168 | 0.99999994 |
| Rasl11b | 0.11531655 | 0.13538734 | 0.09657504 | 0.0388123 | 0.00373527 | 0.20603432 |
| Eno1 | 0.77643791 | 0.4377855 | 0.39902622 | 0.03875928 | 0.25895735 | 0.99999994 |
| Rplp0 | 2.32472678 | 0.40912013 | 0.37041494 | 0.03870519 | 0.24336085 | 0.99999994 |
| Snrpa1 | 0.5842155 | 0.38467955 | 0.34610192 | 0.03857764 | 0.22906209 | 0.99999994 |

| | | | | | | |
|---|---|---|---|---|---|---|
| Slc35d3 | 0.16756072 | 0.17319526 | 0.13465905 | 0.03853621 | 0.02839712 | 0.85915875 |
| Rtn4 | 0.50728052 | 0.35555461 | 0.31703236 | 0.03852226 | 0.20931314 | 0.99999994 |
| Mmp14 | 0.24255664 | 0.2223648 | 0.18396874 | 0.03839606 | 0.08237891 | 0.99999994 |
| Tox | 0.31194807 | 0.26280618 | 0.22454803 | 0.03825815 | 0.12837628 | 0.99999994 |
| Edem2 | 0.55288257 | 0.37291176 | 0.33470421 | 0.03820755 | 0.22367376 | 0.99999994 |
| Plk3 | 0.14302506 | 0.15522689 | 0.11717722 | 0.03804967 | 0.01533045 | 0.56883004 |
| Rpl5 | 1.78384608 | 0.41953353 | 0.38164922 | 0.03788431 | 0.254389 | 0.99999994 |
| Fubp1 | 1.0202013 | 0.46518306 | 0.42742755 | 0.03775551 | 0.27827653 | 0.99999994 |
| Spc25 | 0.18949799 | 0.18744413 | 0.14970953 | 0.0377346 | 0.04669806 | 0.99999994 |
| Arl4a | 0.38707077 | 0.30116754 | 0.26362612 | 0.03754142 | 0.17159216 | 0.99999994 |
| Irf2bpl | 0.3333549 | 0.27345413 | 0.23617275 | 0.03728138 | 0.14668699 | 0.99999994 |
| Miat | 0.15574167 | 0.16355813 | 0.12632505 | 0.03723308 | 0.02488727 | 0.78872359 |
| Hmga2 | 0.15282999 | 0.16129244 | 0.12424716 | 0.03704528 | 0.02359191 | 0.75975493 |
| Efna4 | 0.11924733 | 0.13648458 | 0.0995546 | 0.03692997 | 0.00677098 | 0.32316245 |
| Pgf | 0.16838857 | 0.17202688 | 0.1352368 | 0.03679007 | 0.03508529 | 0.98641032 |
| Kdm6b | 0.34022634 | 0.27657161 | 0.2398194 | 0.03675221 | 0.1538444 | 0.99999994 |
| Il11 | 0.13598004 | 0.14873244 | 0.11202692 | 0.03670552 | 0.01459357 | 0.55521746 |
| Csnk1e | 0.32443622 | 0.26808328 | 0.23137839 | 0.03670489 | 0.14549863 | 0.99999994 |
| Rpl7 | 2.68029282 | 0.39938903 | 0.36315881 | 0.03623022 | 0.25332412 | 0.99999994 |
| Dpy30 | 1.16861872 | 0.46582998 | 0.42960973 | 0.03622026 | 0.28733218 | 0.99999994 |
| Serinc2 | 0.24105314 | 0.21910936 | 0.18303809 | 0.03607127 | 0.09479944 | 0.99999994 |
| Fabp5 | 0.44224043 | 0.32550082 | 0.28943714 | 0.03606368 | 0.20344528 | 0.99999994 |
| Mt2 | 0.16353134 | 0.16788761 | 0.13183596 | 0.03605165 | 0.03436146 | 0.97085972 |
| Rap1gap2 | 0.40740117 | 0.30938891 | 0.27340846 | 0.03598045 | 0.190522 | 0.99999994 |
| Dpysl4 | 0.1316581 | 0.14467351 | 0.10883816 | 0.03583535 | 0.01420314 | 0.54741557 |
| Klhdc8b | 0.35703399 | 0.28440525 | 0.24857012 | 0.03583512 | 0.16862472 | 0.99999994 |
| Sphk1 | 0.09765851 | 0.11860647 | 0.08291803 | 0.03568845 | 0.00208545 | 0.13459169 |
| Ifitm1 | 0.10845393 | 0.12701461 | 0.09132639 | 0.03568823 | 0.00464559 | 0.24201117 |
| Mrps33 | 1.45587961 | 0.44912457 | 0.41347441 | 0.03565016 | 0.28301178 | 0.99999994 |
| Tmem163 | 0.35803246 | 0.28472583 | 0.24908258 | 0.03564325 | 0.1704186 | 0.99999994 |

| | | | | | | |
|---|---|---|---|---|---|---|
| F3 | 0.30694307 | 0.2572581 | 0.22177121 | 0.0354869 | 0.14341026 | 0.99999994 |
| Elavl4 | 0.186001 | 0.18278399 | 0.14734602 | 0.03543796 | 0.0546984 | 0.99999994 |
| Hnrnpab | 0.9763054 | 0.46019138 | 0.42485309 | 0.03533828 | 0.2899057 | 0.99999994 |
| Tac1 | 0.03563284 | 0.06681033 | 0.03150103 | 0.0353093 | 4.29E-14 | 1.09E-11 |
| Cbx3 | 0.49773451 | 0.34846088 | 0.31316342 | 0.03529746 | 0.22655045 | 0.99999994 |
| Ttc5 | 0.58426964 | 0.38130173 | 0.3461211 | 0.03518064 | 0.24933963 | 0.99999994 |
| Rpl23a-ps3 | 0.93244035 | 0.45644259 | 0.42135913 | 0.03508346 | 0.28971221 | 0.99999994 |
| Acot1 | 0.16710733 | 0.16937747 | 0.13434231 | 0.03503516 | 0.04128962 | 0.99999994 |
| Sumo1 | 1.46660314 | 0.44743427 | 0.41240236 | 0.03503191 | 0.28589043 | 0.99999994 |
| Hnrnpl | 0.79229431 | 0.43706129 | 0.40207199 | 0.03498931 | 0.28120929 | 0.99999994 |
| Ogt | 0.34974185 | 0.27976338 | 0.24480279 | 0.03496059 | 0.17090307 | 0.99999994 |
| Id3 | 0.10749507 | 0.1255139 | 0.09058837 | 0.03492553 | 0.00513951 | 0.25926879 |
| Rps18-ps3 | 1.56416578 | 0.43703171 | 0.40230864 | 0.03472307 | 0.28281234 | 0.99999994 |
| Rasgrp3 | 0.08318776 | 0.10590618 | 0.07128965 | 0.03461653 | 0.00061428 | 0.04640853 |
| Tekt2 | 0.32700672 | 0.26728489 | 0.23276744 | 0.03451745 | 0.16180128 | 0.99999994 |
| Oser1 | 0.626365 | 0.39501886 | 0.36051388 | 0.03450499 | 0.26203514 | 0.99999994 |
| D930028M14 | 0.10942763 | 0.12649695 | 0.09207466 | 0.03442229 | 0.0064143 | 0.30798308 |
| Anapc13 | 0.94866533 | 0.45715327 | 0.42277395 | 0.03437932 | 0.29415569 | 0.99999994 |
| Trp53 | 0.44231891 | 0.32372574 | 0.28947223 | 0.03425351 | 0.2154499 | 0.99999994 |
| Atg3 | 0.56891204 | 0.37483977 | 0.34060869 | 0.03423107 | 0.25175987 | 0.99999994 |
| Ccdc34 | 0.34343566 | 0.27566594 | 0.24150901 | 0.03415693 | 0.17324555 | 0.99999994 |
| Auts2 | 0.258597 | 0.22778751 | 0.1937585 | 0.03402902 | 0.12119596 | 0.99999994 |
| Cym | 0.04007016 | 0.06934785 | 0.03531971 | 0.03402814 | 7.14E-11 | 1.48E-08 |
| Cdk2ap2 | 0.66257566 | 0.40596482 | 0.37196858 | 0.03399625 | 0.27147404 | 0.99999994 |
| Lurap1l | 0.25468916 | 0.22539079 | 0.19139713 | 0.03399366 | 0.11855876 | 0.99999994 |
| Slc24a5 | 0.20906072 | 0.19654464 | 0.16268562 | 0.03385902 | 0.08297161 | 0.99999994 |
| Supt4a | 0.52270233 | 0.35683866 | 0.32315682 | 0.03368184 | 0.24391142 | 0.99999994 |
| Mif | 1.27610801 | 0.45983222 | 0.42618355 | 0.03364867 | 0.29960303 | 0.99999994 |
| Snw1 | 0.74467933 | 0.42600974 | 0.39243733 | 0.0335724 | 0.28452842 | 0.99999994 |
| Amer1 | 0.15341947 | 0.15815788 | 0.12466863 | 0.03348925 | 0.03688222 | 0.99999994 |

| | | | | | | |
|---|---|---|---|---|---|---|
| Fam204a | 0.67167468 | 0.40793577 | 0.37469376 | 0.03324202 | 0.27741513 | 0.99999994 |
| Sh3bgrl | 0.30990497 | 0.2565414 | 0.22341716 | 0.03312424 | 0.1618498 | 0.99999994 |
| Smco4 | 0.19673153 | 0.18763586 | 0.15455553 | 0.03308033 | 0.07712264 | 0.99999994 |
| Atp1b1 | 0.66771094 | 0.4065314 | 0.37351187 | 0.03301953 | 0.27811804 | 0.99999994 |
| Ghrl | 0.1088808 | 0.12458292 | 0.09165457 | 0.03292834 | 0.00839211 | 0.37684119 |
| Cartpt | 0.04602627 | 0.07331956 | 0.04041028 | 0.03290928 | 2.97E-08 | 4.59E-06 |
| Herpud1 | 0.46105657 | 0.33046269 | 0.29772347 | 0.03273922 | 0.23209412 | 0.99999994 |
| Agpat4 | 0.32848154 | 0.26629564 | 0.23356186 | 0.03273378 | 0.17543813 | 0.99999994 |
| Fam110a | 0.23706183 | 0.21325325 | 0.18055646 | 0.03269679 | 0.11402573 | 0.99999994 |
| Acin1 | 0.75875128 | 0.42812756 | 0.39544682 | 0.03268073 | 0.29112595 | 0.99999994 |
| Kdm5b | 0.61140547 | 0.38819076 | 0.35552043 | 0.03267033 | 0.2703741 | 0.99999994 |
| Lsm6 | 0.91580599 | 0.45210911 | 0.41972485 | 0.03238426 | 0.30377372 | 0.99999994 |
| Nsmce4a | 0.36014338 | 0.28249576 | 0.25016333 | 0.03233242 | 0.1948111 | 0.99999994 |
| Josd2 | 0.55673777 | 0.36830736 | 0.33613844 | 0.03216892 | 0.26205502 | 0.99999994 |
| Ndufb4 | 1.37774472 | 0.45222976 | 0.42010624 | 0.03212351 | 0.30538318 | 0.99999994 |
| Cbfb | 0.32308416 | 0.26272013 | 0.23064548 | 0.03207466 | 0.17731012 | 0.99999994 |
| Tmsb15l | 0.26706142 | 0.23088441 | 0.19882311 | 0.0320613 | 0.14155173 | 0.99999994 |
| Fam136a | 0.46039171 | 0.32932072 | 0.29743499 | 0.03188573 | 0.23774431 | 0.99999994 |
| Apex1 | 0.52370161 | 0.35526171 | 0.32354833 | 0.03171338 | 0.25705893 | 0.99999994 |
| Sox9 | 0.1949707 | 0.18505171 | 0.15338118 | 0.03167053 | 0.08465546 | 0.99999994 |
| Ilf2 | 0.72245787 | 0.41913702 | 0.38751542 | 0.03162159 | 0.29350826 | 0.99999994 |
| Klf10 | 0.24397395 | 0.21638353 | 0.18484399 | 0.03153954 | 0.12802848 | 0.99999994 |
| Bag1 | 0.93823639 | 0.45333559 | 0.42189742 | 0.03143817 | 0.30994501 | 0.99999994 |
| Snx15 | 0.51358432 | 0.35056633 | 0.3195546 | 0.03101174 | 0.25914683 | 0.99999994 |
| Csrp2 | 0.25747637 | 0.22400605 | 0.19308285 | 0.03092321 | 0.14320057 | 0.99999994 |
| Cct2 | 0.95546695 | 0.45419961 | 0.4233045 | 0.03089511 | 0.31354816 | 0.99999994 |
| Lman1 | 0.85140131 | 0.44266647 | 0.41178003 | 0.03088644 | 0.30879305 | 0.99999994 |
| Ctbp1 | 0.5599457 | 0.36818777 | 0.33732516 | 0.03086262 | 0.27126065 | 0.99999994 |
| Uqcc2 | 1.61513025 | 0.42794121 | 0.39718257 | 0.03075864 | 0.30310809 | 0.99999994 |
| Rac1 | 0.45554599 | 0.32596847 | 0.29532322 | 0.03064525 | 0.24487065 | 0.99999994 |

| | | | | | | |
|---|---|---|---|---|---|---|
| Smim22 | 0.78956144 | 0.43214034 | 0.40155599 | 0.03058435 | 0.30608482 | 0.99999994 |
| Wfdc15b | 0.12445653 | 0.13400418 | 0.10347403 | 0.03053016 | 0.02476692 | 0.78647307 |
| Rnf114 | 0.3358144 | 0.26789871 | 0.23748257 | 0.03041614 | 0.19696037 | 0.99999994 |
| Gnao1 | 0.50397858 | 0.34606644 | 0.31570106 | 0.03036538 | 0.26100994 | 0.99999994 |
| Canx | 0.95668867 | 0.45372277 | 0.42339795 | 0.03032483 | 0.31677182 | 0.99999994 |
| Snrpb2 | 0.62551415 | 0.39037151 | 0.36023321 | 0.0301383 | 0.28880031 | 0.99999994 |
| Suds3 | 0.31098291 | 0.25403806 | 0.22401426 | 0.03002379 | 0.18616251 | 0.99999994 |
| Mycl | 0.17875321 | 0.17239988 | 0.14240503 | 0.02999485 | 0.08045278 | 0.99999994 |
| Dazap1 | 0.88438437 | 0.44616037 | 0.41622955 | 0.02993082 | 0.31609216 | 0.99999994 |
| Tax1bp3 | 0.34494044 | 0.27210488 | 0.24229828 | 0.0298066 | 0.20643899 | 0.99999994 |
| Ctxn1 | 0.35701353 | 0.27832893 | 0.24855962 | 0.02976932 | 0.21266335 | 0.99999994 |
| Atf3 | 0.09144312 | 0.10769389 | 0.07794989 | 0.02974401 | 0.00554383 | 0.27530911 |
| Atp6v0b | 1.59650976 | 0.42863399 | 0.39894275 | 0.02969124 | 0.31015952 | 0.99999994 |
| Tmem86a | 0.18900616 | 0.17899651 | 0.14937791 | 0.02961859 | 0.09344757 | 0.99999994 |
| Nxf1 | 0.67271051 | 0.40459636 | 0.37500128 | 0.02959507 | 0.29968064 | 0.99999994 |
| Fxyd2 | 0.07324412 | 0.092675 | 0.06317272 | 0.02950228 | 0.00093993 | 0.06719025 |
| Stip1 | 0.78829577 | 0.43078262 | 0.40131581 | 0.02946682 | 0.31251058 | 0.99999994 |
| Ap1s2 | 0.53156634 | 0.35602657 | 0.32660797 | 0.0294186 | 0.2744001 | 0.99999994 |
| Ppp2ca | 0.46095473 | 0.32698788 | 0.2976793 | 0.02930859 | 0.25611656 | 0.99999994 |
| Gprasp2 | 0.21081889 | 0.19311539 | 0.16383164 | 0.02928375 | 0.11706699 | 0.99999994 |
| Glul | 0.8086795 | 0.43427143 | 0.40505332 | 0.02921812 | 0.31556047 | 0.99999994 |
| Cib1 | 0.78315702 | 0.42950626 | 0.40033278 | 0.02917348 | 0.3138136 | 0.99999994 |
| Cacybp | 1.02568953 | 0.45685769 | 0.42770159 | 0.02915611 | 0.32500112 | 0.99999994 |
| Sri | 0.57989932 | 0.37368836 | 0.34456716 | 0.0291212 | 0.28686584 | 0.99999994 |
| Klhdc2 | 0.71978803 | 0.41602257 | 0.38691439 | 0.02910818 | 0.30826709 | 0.99999994 |
| Sf3b1 | 1.08399959 | 0.4585915 | 0.4295082 | 0.02908329 | 0.32609494 | 0.99999994 |
| Peli1 | 0.31082403 | 0.25298903 | 0.22392632 | 0.0290627 | 0.19381919 | 0.99999994 |
| Pdlim4 | 0.21063151 | 0.19256271 | 0.16370965 | 0.02885306 | 0.1203674 | 0.99999994 |
| Dynll2 | 0.8933496 | 0.44599078 | 0.4172967 | 0.02869408 | 0.32358284 | 0.99999994 |
| Gm9493 | 0.75337133 | 0.42288786 | 0.39431534 | 0.02857252 | 0.31478684 | 0.99999994 |

| | | | | | | |
|---|---|---|---|---|---|---|
| Aldoa | 1.38513146 | 0.44810845 | 0.41955545 | 0.028553 | 0.32527405 | 0.99999994 |
| Aard | 0.073264 | 0.09161743 | 0.06318906 | 0.02842838 | 0.00137378 | 0.09480243 |
| Agtrap | 0.30640095 | 0.2498784 | 0.22146911 | 0.0284093 | 0.19659274 | 0.99999994 |
| Zfp703 | 0.21464274 | 0.19446377 | 0.166313 | 0.02815078 | 0.12993874 | 0.99999994 |
| Taf10 | 0.31836012 | 0.25618069 | 0.22807235 | 0.02810834 | 0.20600829 | 0.99999994 |
| Mtfp1 | 0.30898771 | 0.25099309 | 0.22290826 | 0.02808483 | 0.20083207 | 0.99999994 |
| Tmbim1 | 0.27363698 | 0.23074514 | 0.20270936 | 0.02803579 | 0.17862708 | 0.99999994 |
| Isyna1 | 0.70584169 | 0.41153966 | 0.38360787 | 0.02793179 | 0.31395451 | 0.99999994 |
| Snx6 | 0.60004635 | 0.37956832 | 0.35163838 | 0.02792994 | 0.29850632 | 0.99999994 |
| St6galnac2 | 0.21550925 | 0.1947437 | 0.16687318 | 0.02787051 | 0.13312945 | 0.99999994 |
| Pik3r1 | 0.42474539 | 0.30935161 | 0.28150163 | 0.02784999 | 0.25509627 | 0.99999994 |
| Bcl2 | 0.08004204 | 0.09650041 | 0.06873308 | 0.02776733 | 0.00358238 | 0.20108012 |
| Atf4 | 0.62788191 | 0.38870863 | 0.3610111 | 0.02769753 | 0.30478413 | 0.99999994 |
| Snrpe | 1.43091128 | 0.44344895 | 0.41579587 | 0.02765308 | 0.32899489 | 0.99999994 |
| Ube2g1 | 0.47510512 | 0.3313961 | 0.30374666 | 0.02764944 | 0.27228571 | 0.99999994 |
| Gata6 | 0.22656526 | 0.20125272 | 0.17395206 | 0.02730066 | 0.14808714 | 0.99999994 |
| Carhsp1 | 0.65786209 | 0.39774891 | 0.37053994 | 0.02720896 | 0.31249864 | 0.99999994 |
| Tspan6 | 0.23013363 | 0.20337743 | 0.17620969 | 0.02716774 | 0.15238282 | 0.99999994 |
| Dnajc7 | 0.6820732 | 0.40463711 | 0.37752605 | 0.02711106 | 0.31632183 | 0.99999994 |
| Nudt21 | 0.57457452 | 0.36969561 | 0.34265753 | 0.02703808 | 0.29971052 | 0.99999994 |
| Haus7 | 0.34666345 | 0.27022643 | 0.24319927 | 0.02702716 | 0.2297311 | 0.99999994 |
| Sulf2 | 0.18149741 | 0.17100668 | 0.14428259 | 0.02672409 | 0.10880769 | 0.99999994 |
| Dazap2 | 0.49535584 | 0.3388672 | 0.31218976 | 0.02667744 | 0.28474478 | 0.99999994 |
| Col9a3 | 0.13652428 | 0.13910163 | 0.11242689 | 0.02667474 | 0.05713275 | 0.99999994 |
| Epb42 | 0.09030708 | 0.10366881 | 0.07703754 | 0.02663127 | 0.0106944 | 0.45951322 |
| Mfge8 | 0.14317003 | 0.14390139 | 0.11728259 | 0.0266188 | 0.0654265 | 0.99999994 |
| Rhobtb3 | 0.22578738 | 0.20007277 | 0.1734582 | 0.02661457 | 0.15355171 | 0.99999994 |
| Tcp11 | 0.20479622 | 0.18649665 | 0.15989196 | 0.02660469 | 0.13402538 | 0.99999994 |
| Ythdc1 | 0.36883459 | 0.28112424 | 0.25457273 | 0.02655151 | 0.24376364 | 0.99999994 |
| Rpl23 | 2.834164 | 0.38051513 | 0.35396461 | 0.02655052 | 0.30878875 | 0.99999994 |

| | | | | | | |
|---|---|---|---|---|---|---|
| Rps7 | 2.83021741 | 0.38076508 | 0.35422739 | 0.02653769 | 0.30900418 | 0.99999994 |
| Tubb2b | 0.17983056 | 0.16966747 | 0.14314313 | 0.02652434 | 0.10870969 | 0.99999994 |
| Rrbp1 | 0.83331053 | 0.43542645 | 0.40901467 | 0.02641178 | 0.33366009 | 0.99999994 |
| Cnih4 | 0.35085721 | 0.27178934 | 0.24538185 | 0.02640749 | 0.23688963 | 0.99999994 |
| Cadm1 | 0.22620491 | 0.20004782 | 0.17372335 | 0.02632447 | 0.1565726 | 0.99999994 |
| Klc3 | 0.57108811 | 0.36770049 | 0.34139826 | 0.02630224 | 0.30403734 | 0.99999994 |
| Rab15 | 0.12112078 | 0.12723769 | 0.10096801 | 0.02626967 | 0.04165111 | 0.99999994 |
| Pex19 | 0.47987126 | 0.33202677 | 0.30575897 | 0.0262678 | 0.28371225 | 0.99999994 |
| Pdap1 | 1.50035137 | 0.43528379 | 0.40903032 | 0.02625347 | 0.33460373 | 0.99999994 |
| Zfand6 | 1.04133374 | 0.45461247 | 0.42839804 | 0.02621444 | 0.34188856 | 0.99999994 |
| Xlr3a | 0.12353396 | 0.1289947 | 0.1027823 | 0.0262124 | 0.04479539 | 0.99999994 |
| Mest | 0.07070637 | 0.08728501 | 0.06108439 | 0.02620061 | 0.00215132 | 0.13717669 |
| Rhoc | 0.31792494 | 0.25403264 | 0.22783434 | 0.0261983 | 0.22201558 | 0.99999994 |
| Dapk3 | 0.16928214 | 0.16189708 | 0.13585956 | 0.02603752 | 0.10103208 | 0.99999994 |
| Clic1 | 1.08977671 | 0.45557735 | 0.42959122 | 0.02598612 | 0.34360454 | 0.99999994 |
| Gdap1l1 | 0.47254257 | 0.32861559 | 0.30265841 | 0.02595718 | 0.28404203 | 0.99999994 |
| Serpinf2 | 0.06768241 | 0.08447339 | 0.05858696 | 0.02588643 | 0.00163531 | 0.11031784 |
| Ptprn2 | 0.48929788 | 0.33541622 | 0.30969337 | 0.02572285 | 0.29017584 | 0.99999994 |
| Mxd4 | 0.36383368 | 0.27765471 | 0.25204305 | 0.02561166 | 0.24939529 | 0.99999994 |
| Srsf11 | 1.11780289 | 0.45549499 | 0.42991798 | 0.025577 | 0.34605064 | 0.99999994 |
| Cox20 | 0.64716757 | 0.39277662 | 0.36721021 | 0.0255664 | 0.32152525 | 0.99999994 |
| Slc29a1 | 0.35736456 | 0.27424945 | 0.24873988 | 0.02550957 | 0.24741584 | 0.99999994 |
| Raly | 0.98047346 | 0.45062347 | 0.42514401 | 0.02547946 | 0.34497438 | 0.99999994 |
| Nit2 | 0.35271648 | 0.27177763 | 0.24634484 | 0.02543279 | 0.24597632 | 0.99999994 |
| Idh2 | 0.97764324 | 0.45015739 | 0.42494716 | 0.02521023 | 0.34646123 | 0.99999994 |
| Gclm | 0.27959509 | 0.23119462 | 0.20619538 | 0.02499924 | 0.20982736 | 0.99999994 |
| Fxr1 | 0.48354418 | 0.33215736 | 0.30729934 | 0.02485802 | 0.29513483 | 0.99999994 |
| Zfp326 | 0.46328445 | 0.3234319 | 0.29868792 | 0.02474399 | 0.29067021 | 0.99999994 |
| Mmd2 | 0.18263535 | 0.1697868 | 0.14505873 | 0.02472807 | 0.12824947 | 0.99999994 |
| Bnip3l | 0.88943148 | 0.44150974 | 0.41683446 | 0.02467528 | 0.34677746 | 0.99999994 |

| | | | | | | |
|---|---|---|---|---|---|---|
| Top2b | 0.39260265 | 0.29098308 | 0.2663202 | 0.02466288 | 0.26880886 | 0.99999994 |
| Magoh | 0.9896862 | 0.45042741 | 0.42576491 | 0.0246625 | 0.34990734 | 0.99999994 |
| Pla2g2f | 0.32590739 | 0.25678815 | 0.23217407 | 0.02461408 | 0.24019421 | 0.99999994 |
| Anp32b | 1.06432444 | 0.45371433 | 0.42915522 | 0.02455911 | 0.35163094 | 0.99999994 |
| Ndufc1 | 1.35098132 | 0.44648839 | 0.42198082 | 0.02450757 | 0.34953198 | 0.99999994 |
| Nr2f6 | 0.23037545 | 0.20083041 | 0.17636222 | 0.0244682 | 0.17787548 | 0.99999994 |
| Ubb | 3.25258732 | 0.31446057 | 0.29008212 | 0.02437844 | 0.28794471 | 0.99999994 |
| Fhl2 | 0.12439423 | 0.12757237 | 0.10342735 | 0.02414502 | 0.0601017 | 0.99999994 |
| Ptms | 0.56120154 | 0.36191735 | 0.33778807 | 0.02412927 | 0.31722074 | 0.99999994 |
| Espn | 0.08533213 | 0.09714661 | 0.07302648 | 0.02412013 | 0.01395504 | 0.54149729 |
| Cldnd1 | 0.54776093 | 0.35686033 | 0.3327849 | 0.02407543 | 0.31506059 | 0.99999994 |
| Homer2 | 0.13065272 | 0.13213241 | 0.10809318 | 0.02403923 | 0.06939239 | 0.99999994 |
| Npm1 | 2.06903014 | 0.39281124 | 0.36889262 | 0.02391862 | 0.33301918 | 0.99999994 |
| Asf1a | 0.45030936 | 0.31692535 | 0.2930218 | 0.02390355 | 0.29356522 | 0.99999994 |
| Ccng2 | 0.20086448 | 0.18110862 | 0.15729892 | 0.0238097 | 0.15683625 | 0.99999994 |
| Eef1g | 2.09319689 | 0.39229991 | 0.3684984 | 0.02380151 | 0.33362045 | 0.99999994 |
| Hnrnpd | 0.30296688 | 0.2433249 | 0.21954939 | 0.02377551 | 0.23550637 | 0.99999994 |
| Efcab10 | 0.19768842 | 0.17894592 | 0.15519232 | 0.0237536 | 0.15414505 | 0.99999994 |
| Glo1 | 0.75518577 | 0.41843425 | 0.39469856 | 0.02373569 | 0.34447205 | 0.99999994 |
| Ptbp3 | 0.58430916 | 0.36986311 | 0.3461351 | 0.02372801 | 0.32408643 | 0.99999994 |
| Hmgb1 | 1.89530935 | 0.39823305 | 0.37458426 | 0.02364878 | 0.33715557 | 0.99999994 |
| Phip | 0.72383679 | 0.41137947 | 0.38782436 | 0.0235551 | 0.34300193 | 0.99999994 |
| Tcp11l2 | 0.18068122 | 0.16726801 | 0.14372502 | 0.02354299 | 0.13777954 | 0.99999994 |
| Myt1 | 0.27995017 | 0.22988692 | 0.2064021 | 0.02348482 | 0.22441318 | 0.99999994 |
| Smpdl3a | 0.20585676 | 0.1839094 | 0.1605886 | 0.0233208 | 0.16686167 | 0.99999994 |
| Paics | 0.39619701 | 0.29135585 | 0.26805719 | 0.02329866 | 0.28144679 | 0.99999994 |
| Smim14 | 1.00376065 | 0.44978125 | 0.42662218 | 0.02315907 | 0.35892319 | 0.99999994 |
| Sult2b1 | 0.14392099 | 0.14085154 | 0.117828 | 0.02302354 | 0.09668926 | 0.99999994 |
| Gm10709 | 1.79595128 | 0.40373375 | 0.38073125 | 0.02300251 | 0.34378503 | 0.99999994 |
| Cks1b | 0.53173216 | 0.34960205 | 0.32667206 | 0.02292999 | 0.32016853 | 0.99999994 |

| | | | | | | |
|---|---|---|---|---|---|---|
| Syce2 | 0.49117867 | 0.33327302 | 0.31047099 | 0.02280203 | 0.31246852 | 0.99999994 |
| Rpl13a | 3.36577964 | 0.2935555 | 0.2708549 | 0.0227006 | 0.28846469 | 0.99999994 |
| Eci2 | 0.50842634 | 0.34017563 | 0.3174927 | 0.02268293 | 0.31719524 | 0.99999994 |
| Ccnl1 | 0.58647809 | 0.36958283 | 0.34690221 | 0.02268062 | 0.3317102 | 0.99999994 |
| Mad2l2 | 0.43919989 | 0.31073733 | 0.28807441 | 0.02266292 | 0.30025815 | 0.99999994 |
| Smdt1 | 1.46138293 | 0.43557702 | 0.41292597 | 0.02265105 | 0.35750555 | 0.99999994 |
| Pard3 | 0.36271263 | 0.27407965 | 0.25147321 | 0.02260644 | 0.27479314 | 0.99999994 |
| Paip2 | 1.37882249 | 0.44252729 | 0.42002641 | 0.02250088 | 0.36070231 | 0.99999994 |
| Serpinb1a | 0.12212699 | 0.12408341 | 0.10172537 | 0.02235804 | 0.07173513 | 0.99999994 |
| Gtf3c6 | 0.37834525 | 0.28161579 | 0.25932799 | 0.0222878 | 0.28363491 | 0.99999994 |
| Prss50 | 0.09032249 | 0.0992996 | 0.07704992 | 0.02224968 | 0.02729314 | 0.83669235 |
| Eml4 | 0.2467823 | 0.20867522 | 0.18657245 | 0.02210277 | 0.21518402 | 0.99999994 |
| Cct4 | 0.90298191 | 0.44037678 | 0.41836877 | 0.02200801 | 0.36311182 | 0.99999994 |
| Hsph1 | 0.3389087 | 0.2610954 | 0.23912321 | 0.02197219 | 0.27039155 | 0.99999994 |
| Srek1ip1 | 0.35231409 | 0.26809742 | 0.24613666 | 0.02196076 | 0.27629205 | 0.99999994 |
| Mex3a | 0.16179328 | 0.15244074 | 0.13061256 | 0.02182818 | 0.13297854 | 0.99999994 |
| Rogdi | 0.818224 | 0.42847832 | 0.40667189 | 0.02180643 | 0.36057394 | 0.99999994 |
| Mrpl1 | 0.50971835 | 0.33979918 | 0.31801077 | 0.02178841 | 0.32417403 | 0.99999994 |
| Yrdc | 0.19518754 | 0.17523805 | 0.15352598 | 0.02171207 | 0.17326 | 0.99999994 |
| Smc6 | 0.56472951 | 0.36073415 | 0.33908336 | 0.0216508 | 0.33541294 | 0.99999994 |
| Lingo1 | 0.225905 | 0.19517381 | 0.17353291 | 0.02164089 | 0.20324135 | 0.99999994 |
| Pttg1 | 0.11574422 | 0.11850189 | 0.09690014 | 0.02160175 | 0.06891928 | 0.99999994 |
| Pisd | 0.22891107 | 0.19691419 | 0.17543765 | 0.02147654 | 0.20757884 | 0.99999994 |
| Myef2 | 0.41902943 | 0.30031117 | 0.27885942 | 0.02145175 | 0.3043073 | 0.99999994 |
| Slbp | 0.57633871 | 0.36474142 | 0.34329207 | 0.02144935 | 0.33874225 | 0.99999994 |
| Pde1c | 0.2812524 | 0.22859083 | 0.2071592 | 0.02143163 | 0.24552726 | 0.99999994 |
| Btf3l4 | 0.98668515 | 0.44697668 | 0.42556598 | 0.0214107 | 0.36885446 | 0.99999994 |
| Rnaseh2c | 0.3867457 | 0.28480372 | 0.26346706 | 0.02133666 | 0.29492288 | 0.99999994 |
| Rbfox2 | 0.43835152 | 0.30901668 | 0.287693 | 0.02132368 | 0.31087687 | 0.99999994 |
| Ncl | 1.44904356 | 0.43530037 | 0.41415047 | 0.0211499 | 0.36695542 | 0.99999994 |

| | | | | | | |
|---|---|---|---|---|---|---|
| Qsox1 | 0.20187518 | 0.17900338 | 0.15796705 | 0.02103632 | 0.18769538 | 0.99999994 |
| Arrdc4 | 0.251214 | 0.21019385 | 0.18928434 | 0.02090952 | 0.23107768 | 0.99999994 |
| Fjx1 | 0.08750723 | 0.09556503 | 0.07478331 | 0.02078172 | 0.03217463 | 0.92747887 |
| Plk2 | 0.12985826 | 0.12825666 | 0.10750351 | 0.02075315 | 0.09939879 | 0.99999994 |
| Lsm7 | 0.40146956 | 0.29133611 | 0.27058732 | 0.02074879 | 0.30488098 | 0.99999994 |
| Btbd1 | 0.29043072 | 0.23315577 | 0.21244937 | 0.0207064 | 0.25824589 | 0.99999994 |
| Ahsa2 | 0.3904768 | 0.28591842 | 0.26528784 | 0.02063058 | 0.30235351 | 0.99999994 |
| Wsb1 | 0.19437737 | 0.17361241 | 0.15298472 | 0.02062768 | 0.18472357 | 0.99999994 |
| Ing4 | 0.50497566 | 0.3367035 | 0.3161038 | 0.02059969 | 0.33222478 | 0.99999994 |
| Sin3b | 1.00436277 | 0.44716223 | 0.42665732 | 0.02050491 | 0.37452178 | 0.99999994 |
| Rsl1d1 | 0.58187873 | 0.36569741 | 0.34527232 | 0.02042509 | 0.34687671 | 0.99999994 |
| Fblim1 | 0.11882995 | 0.11961281 | 0.09923913 | 0.02037368 | 0.08588431 | 0.99999994 |
| Dmrtc1a | 0.1037567 | 0.10805513 | 0.08769995 | 0.02035518 | 0.06118007 | 0.99999994 |
| P4hb | 0.83931625 | 0.43025434 | 0.40995374 | 0.0203006 | 0.3708444 | 0.99999994 |
| Tmed1 | 0.27132589 | 0.2215602 | 0.20134824 | 0.02021196 | 0.2520075 | 0.99999994 |
| Rpn1 | 0.90003013 | 0.43825546 | 0.41804431 | 0.02021115 | 0.37379668 | 0.99999994 |
| Gng3 | 0.06192864 | 0.07400836 | 0.05380778 | 0.02020058 | 0.00622831 | 0.30269945 |
| Ahsa1 | 0.94820755 | 0.44281679 | 0.42273761 | 0.02007918 | 0.37593921 | 0.99999994 |
| Rpl10-ps3 | 0.4697067 | 0.3214859 | 0.30144891 | 0.02003699 | 0.32908501 | 0.99999994 |
| Trp53inp1 | 0.16177919 | 0.15063135 | 0.13060262 | 0.02002873 | 0.153676 | 0.99999994 |
| Bckdha | 0.40049098 | 0.29014682 | 0.27011933 | 0.02002749 | 0.31082147 | 0.99999994 |
| Prkaca | 0.28121356 | 0.22713568 | 0.20713664 | 0.01999903 | 0.26021877 | 0.99999994 |
| F11r | 0.4397281 | 0.30828979 | 0.28831162 | 0.01997817 | 0.32231199 | 0.99999994 |
| Dera | 0.39205514 | 0.28596885 | 0.26605474 | 0.0199141 | 0.30916236 | 0.99999994 |
| Tecpr1 | 0.25666542 | 0.21248135 | 0.19259315 | 0.0198882 | 0.24592446 | 0.99999994 |
| Nasp | 0.40669633 | 0.29294457 | 0.2730747 | 0.01986987 | 0.31407231 | 0.99999994 |
| Bnip2 | 0.2504797 | 0.20870011 | 0.18883632 | 0.01986379 | 0.24190562 | 0.99999994 |
| Alyref | 0.18104634 | 0.16383566 | 0.14397454 | 0.01986112 | 0.17924744 | 0.99999994 |
| Snrpn | 0.29439283 | 0.23449501 | 0.21470899 | 0.01978602 | 0.2698054 | 0.99999994 |
| Fkbp3 | 1.05097103 | 0.448501 | 0.42874049 | 0.01976051 | 0.37950187 | 0.99999994 |

| | | | | | | |
|---|---|---|---|---|---|---|
| Yaf2 | 0.27443618 | 0.22276825 | 0.20317888 | 0.01958937 | 0.26051163 | 0.99999994 |
| Rbm18 | 0.35890565 | 0.26907535 | 0.24953008 | 0.01954527 | 0.30105059 | 0.99999994 |
| Supt16 | 0.41646866 | 0.29715843 | 0.27766763 | 0.0194908 | 0.32016364 | 0.99999994 |
| Bcl7a | 0.26623387 | 0.21778167 | 0.19833095 | 0.01945072 | 0.25694178 | 0.99999994 |
| Mrpl33 | 0.72277147 | 0.40698114 | 0.38758577 | 0.01939536 | 0.36953167 | 0.99999994 |
| Ppm1b | 0.58093845 | 0.36426481 | 0.34493763 | 0.01932719 | 0.3545883 | 0.99999994 |
| Ttc1 | 0.46913075 | 0.32051556 | 0.30120259 | 0.01931297 | 0.33476156 | 0.99999994 |
| Trf | 0.09844344 | 0.10284355 | 0.08354265 | 0.0193009 | 0.06204609 | 0.99999994 |
| Psmc2 | 1.21696143 | 0.44770782 | 0.42851655 | 0.01919127 | 0.3828106 | 0.99999994 |
| Ube2v2 | 0.82486602 | 0.426881 | 0.40770805 | 0.01917295 | 0.37713157 | 0.99999994 |
| Cnot6l | 0.70407351 | 0.40235241 | 0.38318059 | 0.01917182 | 0.36955201 | 0.99999994 |
| Gm28172 | 0.08270521 | 0.09005209 | 0.07089815 | 0.01915394 | 0.0360655 | 0.99999994 |
| Kctd12b | 0.15366418 | 0.14397126 | 0.12484347 | 0.01912779 | 0.15390022 | 0.99999994 |
| Plscr3 | 0.33291645 | 0.25504365 | 0.23593871 | 0.01910494 | 0.29494579 | 0.99999994 |
| Stk11 | 0.43247993 | 0.3041365 | 0.28503764 | 0.01909886 | 0.32779597 | 0.99999994 |
| Gm1673 | 0.62143043 | 0.37794825 | 0.35888018 | 0.01906807 | 0.3617944 | 0.99999994 |
| Smarca5 | 0.57407221 | 0.36142575 | 0.34247653 | 0.01894922 | 0.3563262 | 0.99999994 |
| Tmem59 | 1.9351415 | 0.39113113 | 0.37225566 | 0.01887547 | 0.36786434 | 0.99999994 |
| Capza2 | 0.96266785 | 0.44270271 | 0.42385594 | 0.01884676 | 0.38362455 | 0.99999994 |
| Hmgcr | 0.43351328 | 0.30427361 | 0.28550678 | 0.01876683 | 0.33086233 | 0.99999994 |
| Rgs16 | 0.1365729 | 0.13113627 | 0.1124626 | 0.01867368 | 0.13452681 | 0.99999994 |
| Wasf2 | 0.39796052 | 0.28757242 | 0.2689058 | 0.01866661 | 0.32201776 | 0.99999994 |
| Numbl | 0.16263498 | 0.14983933 | 0.13120545 | 0.01863388 | 0.17224269 | 0.99999994 |
| Grb7 | 0.26850739 | 0.21825715 | 0.19968149 | 0.01857566 | 0.26788658 | 0.99999994 |
| Gpd2 | 0.44967145 | 0.31128512 | 0.29274007 | 0.01854504 | 0.33662954 | 0.99999994 |
| Chd7 | 0.64050955 | 0.38363298 | 0.36510259 | 0.01853039 | 0.36774326 | 0.99999994 |
| Nrtn | 0.196433 | 0.17285759 | 0.15435667 | 0.01850092 | 0.21248761 | 0.99999994 |
| Ash2l | 0.44705004 | 0.31007664 | 0.29157936 | 0.01849728 | 0.33641481 | 0.99999994 |
| Zfp422 | 0.30136784 | 0.23713995 | 0.21865191 | 0.01848804 | 0.28677667 | 0.99999994 |
| Muc1 | 0.13198333 | 0.12753929 | 0.1090789 | 0.01846039 | 0.12997391 | 0.99999994 |

| | | | | | | |
|---|---|---|---|---|---|---|
| Prom1 | 0.10764802 | 0.10909797 | 0.09070617 | 0.0183918 | 0.08856265 | 0.99999994 |
| Lipc | 0.04586102 | 0.05866139 | 0.04026959 | 0.0183918 | 0.00118261 | 0.08196535 |
| Lynx1 | 0.09268599 | 0.09731341 | 0.07894651 | 0.01836691 | 0.06073871 | 0.99999994 |
| Sde2 | 0.29561901 | 0.23376587 | 0.21540541 | 0.01836046 | 0.28523262 | 0.99999994 |
| Crip2 | 0.16402255 | 0.15051667 | 0.1321811 | 0.01833557 | 0.17791497 | 0.99999994 |
| Tex30 | 0.23407663 | 0.19701957 | 0.17868943 | 0.01833015 | 0.24736151 | 0.99999994 |
| Sfr1 | 1.17800265 | 0.44776344 | 0.42947325 | 0.01829019 | 0.38840537 | 0.99999994 |
| Cd40 | 0.08639547 | 0.09215191 | 0.07388595 | 0.01826596 | 0.04992581 | 0.99999994 |
| Cplx2 | 0.59964076 | 0.3697237 | 0.35149841 | 0.01822529 | 0.36499679 | 0.99999994 |
| Tomm7 | 1.25779679 | 0.44521987 | 0.42700592 | 0.01821394 | 0.38823314 | 0.99999994 |
| Fundc2 | 0.5682121 | 0.35855534 | 0.34035414 | 0.01820119 | 0.36093393 | 0.99999994 |
| Halr1 | 0.06874336 | 0.07765062 | 0.0594643 | 0.01818632 | 0.02088756 | 0.69426319 |
| Rps21 | 1.86478795 | 0.39442163 | 0.37624222 | 0.01817941 | 0.37386868 | 0.99999994 |
| Ube2j2 | 0.49012978 | 0.32818278 | 0.31003761 | 0.01814518 | 0.34842795 | 0.99999994 |
| Nfix | 0.13587285 | 0.13008634 | 0.1119481 | 0.01813823 | 0.14040822 | 0.99999994 |
| Slc38a1 | 0.22608739 | 0.19161638 | 0.17364874 | 0.01796763 | 0.24549314 | 0.99999994 |
| Srsf9 | 0.62213668 | 0.37703999 | 0.35911482 | 0.01792517 | 0.36984978 | 0.99999994 |
| Capsl | 0.36083319 | 0.26840058 | 0.25051561 | 0.01788497 | 0.31731686 | 0.99999994 |
| Ptn | 0.08516207 | 0.09077023 | 0.07288891 | 0.01788131 | 0.05123908 | 0.99999994 |
| Serpinf1 | 0.08716375 | 0.09230873 | 0.0745062 | 0.01780252 | 0.05586432 | 0.99999994 |
| Vps37a | 0.62077626 | 0.37645671 | 0.35866261 | 0.0177941 | 0.3706102 | 0.99999994 |
| Insig1 | 0.13664859 | 0.13027491 | 0.11251819 | 0.01775672 | 0.1467549 | 0.99999994 |
| Pcolce | 0.12595524 | 0.12228781 | 0.10459555 | 0.01769227 | 0.13009999 | 0.99999994 |
| Snrpd3 | 1.10319807 | 0.44743873 | 0.42979467 | 0.01764406 | 0.39232713 | 0.99999994 |
| Lxn | 0.14755872 | 0.13810274 | 0.12046044 | 0.0176423 | 0.16481201 | 0.99999994 |
| Rpl35 | 2.0019025 | 0.38835133 | 0.3707623 | 0.01758903 | 0.37608521 | 0.99999994 |
| Crybb1 | 0.11251656 | 0.1120043 | 0.09444062 | 0.01756368 | 0.10787251 | 0.99999994 |
| Egr4 | 0.02993565 | 0.04404923 | 0.02656499 | 0.01748424 | 5.91E-06 | 0.00065845 |
| Tpst1 | 0.24462338 | 0.20271544 | 0.18524439 | 0.01747105 | 0.2650725 | 0.99999994 |
| Kdm5a | 0.35483527 | 0.2648826 | 0.24743878 | 0.01744381 | 0.31944314 | 0.99999994 |

| | | | | | | |
|---|---|---|---|---|---|---|
| Atrx | 0.70774512 | 0.40141694 | 0.38406595 | 0.01735099 | 0.3818159 | 0.99999994 |
| Eps8l2 | 0.2259272 | 0.19087775 | 0.17354701 | 0.01733074 | 0.25311584 | 0.99999994 |
| Arglu1 | 0.34761256 | 0.26101356 | 0.2436945 | 0.01731906 | 0.31808552 | 0.99999994 |
| Tmem184c | 0.19267545 | 0.16900877 | 0.15184543 | 0.01716335 | 0.22591143 | 0.99999994 |
| Nt5dc2 | 0.89244342 | 0.43433574 | 0.41719036 | 0.01714538 | 0.39221213 | 0.99999994 |
| Ppp1r17 | 0.04792954 | 0.05915649 | 0.04202855 | 0.01712794 | 0.00333714 | 0.19215635 |
| Rbm22 | 0.38771623 | 0.28106702 | 0.26394172 | 0.0171253 | 0.33291297 | 0.99999994 |
| Eif2s3y | 0.21714246 | 0.1850257 | 0.16792692 | 0.01709878 | 0.24895922 | 0.99999994 |
| Yy1 | 0.41911588 | 0.29597983 | 0.27889957 | 0.01708026 | 0.34176674 | 0.99999994 |
| Guk1 | 0.73544934 | 0.40744873 | 0.39039096 | 0.01705777 | 0.38558557 | 0.99999994 |
| Nhlh1 | 0.05533688 | 0.06528102 | 0.04828823 | 0.01699279 | 0.00958021 | 0.42187319 |
| Psmd10 | 0.34741532 | 0.26049958 | 0.24359164 | 0.01690794 | 0.32203187 | 0.99999994 |
| Eif4a1 | 2.567437 | 0.38401264 | 0.36712424 | 0.0168884 | 0.37972432 | 0.99999994 |
| Rbms3 | 0.15978472 | 0.14602657 | 0.12919422 | 0.01683235 | 0.19290575 | 0.99999994 |
| Sesn2 | 0.18045389 | 0.16038972 | 0.1435696 | 0.01682012 | 0.21774366 | 0.99999994 |
| Rtf1 | 0.59814951 | 0.36770821 | 0.350983 | 0.01672521 | 0.37554897 | 0.99999994 |
| Fuca1 | 0.50097038 | 0.3311998 | 0.31448204 | 0.01671776 | 0.36172575 | 0.99999994 |
| Rai2 | 0.1780808 | 0.15859847 | 0.1419437 | 0.01665476 | 0.21739707 | 0.99999994 |
| N6amt1 | 0.36005958 | 0.26660889 | 0.25012049 | 0.0164884 | 0.33040353 | 0.99999994 |
| Fam32a | 1.033665 | 0.44447941 | 0.42808539 | 0.01639402 | 0.39939518 | 0.99999994 |
| Slc25a33 | 0.09211937 | 0.09483649 | 0.07849235 | 0.01634414 | 0.08286942 | 0.99999994 |
| Tmed10 | 1.64719264 | 0.41040067 | 0.39409074 | 0.01630993 | 0.39147312 | 0.99999994 |
| Tcf3 | 0.39622876 | 0.28432646 | 0.26807249 | 0.01625397 | 0.34325591 | 0.99999994 |
| Enc1 | 0.12545866 | 0.12043118 | 0.10422425 | 0.01620693 | 0.15031875 | 0.99999994 |
| Mat2a | 0.39597352 | 0.28412196 | 0.26794948 | 0.01617248 | 0.3439325 | 0.99999994 |
| Pola1 | 0.11275882 | 0.11076415 | 0.09462568 | 0.01613847 | 0.12813618 | 0.99999994 |
| Smc1a | 0.44832384 | 0.30823185 | 0.29214398 | 0.01608787 | 0.35697576 | 0.99999994 |
| Chd3 | 0.47700023 | 0.32062593 | 0.30454862 | 0.01607731 | 0.36264667 | 0.99999994 |
| Pfn2 | 0.33591621 | 0.2536098 | 0.23753667 | 0.01607312 | 0.32620806 | 0.99999994 |
| Acads | 0.34317166 | 0.25743993 | 0.24137035 | 0.01606957 | 0.32882626 | 0.99999994 |

| | | | | | | |
|---|---|---|---|---|---|---|
| Hnrnpdl | 0.66462193 | 0.38863718 | 0.37258517 | 0.01605201 | 0.38714253 | 0.99999994 |
| Tcea3 | 0.0426265 | 0.05354997 | 0.03750948 | 0.01604049 | 0.00221004 | 0.13925008 |
| Jam3 | 0.19860127 | 0.171821 | 0.15579889 | 0.01602212 | 0.24682042 | 0.99999994 |
| Megf11 | 0.08909289 | 0.09207159 | 0.07606096 | 0.01601063 | 0.08058526 | 0.99999994 |
| Ndn | 0.49508899 | 0.32806114 | 0.31208031 | 0.01598083 | 0.36660636 | 0.99999994 |
| Gatad1 | 0.27978146 | 0.22227799 | 0.2063039 | 0.01597409 | 0.30313639 | 0.99999994 |
| Chd1 | 0.30431612 | 0.2362658 | 0.22030489 | 0.0159609 | 0.31481505 | 0.99999994 |
| Acat1 | 0.46887841 | 0.31699448 | 0.30109455 | 0.01589992 | 0.36260557 | 0.99999994 |
| Urah | 0.09079196 | 0.09327637 | 0.0774271 | 0.01584927 | 0.08651191 | 0.99999994 |
| Fam89a | 0.11814891 | 0.11448096 | 0.09872391 | 0.01575705 | 0.14402637 | 0.99999994 |
| Gabpb1 | 0.20009323 | 0.17250949 | 0.15678835 | 0.01572113 | 0.25224828 | 0.99999994 |
| Smap1 | 0.21919905 | 0.1848597 | 0.16924991 | 0.01560979 | 0.26963715 | 0.99999994 |
| Chd4 | 0.74957397 | 0.40909911 | 0.39350501 | 0.0155941 | 0.3959736 | 0.99999994 |
| Ewsr1 | 0.97427749 | 0.44028512 | 0.42470927 | 0.01557584 | 0.40357044 | 0.99999994 |
| Rnf138rt1 | 0.18396182 | 0.16149783 | 0.14596168 | 0.01553616 | 0.23931658 | 0.99999994 |
| Zranb2 | 0.5536898 | 0.3505174 | 0.33500525 | 0.01551215 | 0.37895847 | 0.99999994 |
| Psmd14 | 0.5500887 | 0.34915959 | 0.33365927 | 0.01550033 | 0.37857527 | 0.99999994 |
| Brd3 | 0.62321772 | 0.37495515 | 0.35947345 | 0.0154817 | 0.38718092 | 0.99999994 |
| Eif2b2 | 0.43626204 | 0.30219418 | 0.28675098 | 0.0154432 | 0.35999076 | 0.99999994 |
| Synpr | 0.13413547 | 0.12611083 | 0.11066871 | 0.01544212 | 0.17649823 | 0.99999994 |
| Ssbp2 | 0.2999664 | 0.23329097 | 0.21786347 | 0.01542751 | 0.31869368 | 0.99999994 |
| Gadd45gip1 | 0.70713708 | 0.39934364 | 0.38391983 | 0.01542381 | 0.39457434 | 0.99999994 |
| Ssrp1 | 0.52145293 | 0.33803993 | 0.32266644 | 0.0153735 | 0.37556745 | 0.99999994 |
| Hba-a2 | 0.32631111 | 0.24776125 | 0.2323921 | 0.01536915 | 0.32989062 | 0.99999994 |
| Rps26-ps1 | 0.3782788 | 0.2746516 | 0.25929503 | 0.01535657 | 0.34671 | 0.99999994 |
| Klf3 | 0.22233821 | 0.18658414 | 0.17126096 | 0.01532318 | 0.27573388 | 0.99999994 |
| Ptprf | 0.67899322 | 0.39202241 | 0.37670077 | 0.01532164 | 0.39329718 | 0.99999994 |
| Dnajb1 | 0.38366351 | 0.27710455 | 0.2619548 | 0.01514975 | 0.35013344 | 0.99999994 |
| Bzw1 | 0.71678448 | 0.40136356 | 0.38621498 | 0.01514858 | 0.39701401 | 0.99999994 |
| Mospd1 | 0.11254819 | 0.10958328 | 0.09446479 | 0.01511849 | 0.14336825 | 0.99999994 |

| | | | | | | |
|---|---|---|---|---|---|---|
| Gsr | 0.35570177 | 0.26299141 | 0.24788511 | 0.0151063 | 0.34250231 | 0.99999994 |
| Klhl24 | 0.16995621 | 0.15142023 | 0.13632874 | 0.01509149 | 0.23060554 | 0.99999994 |
| Inppl1 | 0.17134099 | 0.15236383 | 0.13729102 | 0.01507281 | 0.23245568 | 0.99999994 |
| Gstz1 | 0.78770022 | 0.41623666 | 0.40120252 | 0.01503414 | 0.40151344 | 0.99999994 |
| Pabpc1 | 1.33887231 | 0.43778828 | 0.42275539 | 0.01503289 | 0.40644651 | 0.99999994 |
| Zfp644 | 0.36399797 | 0.26714762 | 0.25212647 | 0.01502115 | 0.34584174 | 0.99999994 |
| Dtd1 | 0.26698321 | 0.21379757 | 0.19877662 | 0.01502095 | 0.30748185 | 0.99999994 |
| Dusp10 | 0.20379436 | 0.17423483 | 0.15923276 | 0.01500207 | 0.26528809 | 0.99999994 |
| Rfx6 | 0.27559019 | 0.21885273 | 0.2038558 | 0.01499693 | 0.31217874 | 0.99999994 |
| Cfl2 | 0.45865232 | 0.31166758 | 0.29667883 | 0.01498875 | 0.36832566 | 0.99999994 |
| Plac8 | 0.06250656 | 0.06927492 | 0.05428943 | 0.01498549 | 0.03307894 | 0.94678472 |
| Nudt11 | 0.08127522 | 0.08464689 | 0.06973654 | 0.01491034 | 0.07734004 | 0.99999994 |
| Prpsap1 | 0.43925022 | 0.30294992 | 0.28809702 | 0.0148529 | 0.36573533 | 0.99999994 |
| Lpcat3 | 0.42644713 | 0.2971225 | 0.28228334 | 0.01483916 | 0.36320183 | 0.99999994 |
| Dpysl2 | 0.35085047 | 0.26019484 | 0.24537835 | 0.01481649 | 0.34386936 | 0.99999994 |
| Lmo4 | 0.10877501 | 0.1063391 | 0.09157326 | 0.01476584 | 0.14156554 | 0.99999994 |
| Ythdf2 | 0.39468961 | 0.28205761 | 0.26732994 | 0.01472767 | 0.35691757 | 0.99999994 |
| Mbd3 | 0.68751007 | 0.39368173 | 0.37897037 | 0.01471137 | 0.39805097 | 0.99999994 |
| Brd4 | 0.34972617 | 0.2594998 | 0.24479464 | 0.01470516 | 0.34463102 | 0.99999994 |
| Ly6h | 0.06983214 | 0.07506743 | 0.06036339 | 0.01470404 | 0.05246217 | 0.99999994 |
| Fbxo44 | 0.17852788 | 0.15694515 | 0.14225049 | 0.01469466 | 0.24584905 | 0.99999994 |
| Ttc28 | 0.30112921 | 0.23318979 | 0.21851779 | 0.014672 | 0.32746293 | 0.99999994 |
| B630019K06 | 0.23318991 | 0.1928001 | 0.17813313 | 0.01466697 | 0.29182347 | 0.99999994 |
| Ube2m | 0.20417798 | 0.1741475 | 0.1594853 | 0.0146622 | 0.27028575 | 0.99999994 |
| Dync1i2 | 0.8651257 | 0.42837756 | 0.41374038 | 0.01463718 | 0.40691557 | 0.99999994 |
| Fus | 1.12299143 | 0.44454095 | 0.42993066 | 0.01461029 | 0.41052204 | 0.99999994 |
| Dnajc8 | 0.9418062 | 0.43677207 | 0.42222098 | 0.01455108 | 0.40927927 | 0.99999994 |
| Eif1b | 0.87955212 | 0.43012643 | 0.41564033 | 0.0144861 | 0.40827265 | 0.99999994 |
| Sgce | 0.09882013 | 0.09832491 | 0.08384217 | 0.01448273 | 0.12511187 | 0.99999994 |
| Asph | 0.27215054 | 0.2162819 | 0.20183449 | 0.0144474 | 0.31687152 | 0.99999994 |

| | | | | | | |
|---|---|---|---|---|---|---|
| Cdc34 | 0.21354662 | 0.18004228 | 0.16560325 | 0.01443902 | 0.28083229 | 0.99999994 |
| Upk3bl | 0.05373608 | 0.06137433 | 0.0469406 | 0.01443373 | 0.02034092 | 0.68362409 |
| Cbfa2t2 | 0.20342007 | 0.17341228 | 0.15898622 | 0.01442606 | 0.27292738 | 0.99999994 |
| Fabp3 | 0.10875386 | 0.10595404 | 0.091557 | 0.01439704 | 0.1476212 | 0.99999994 |
| Sap30 | 0.27290642 | 0.21662839 | 0.20227963 | 0.01434875 | 0.31840261 | 0.99999994 |
| Rps3a1 | 2.94744222 | 0.35784383 | 0.34351214 | 0.01433169 | 0.39061739 | 0.99999994 |
| Spink4 | 0.07708946 | 0.08058248 | 0.06632403 | 0.01425845 | 0.07621587 | 0.99999994 |
| Wnk3 | 0.12600357 | 0.11887881 | 0.10463167 | 0.01424714 | 0.18237238 | 0.99999994 |
| Sh2d3c | 0.22789229 | 0.18903874 | 0.17479314 | 0.0142456 | 0.29373993 | 0.99999994 |
| Zyx | 0.12735988 | 0.11986699 | 0.1056442 | 0.01422279 | 0.18508901 | 0.99999994 |
| St13 | 0.9311035 | 0.43544931 | 0.42123312 | 0.01421619 | 0.4111276 | 0.99999994 |
| Commd3 | 0.712111 | 0.39929917 | 0.38510932 | 0.01418985 | 0.40312674 | 0.99999994 |
| Ddx46 | 0.40346423 | 0.28572726 | 0.271539 | 0.01418826 | 0.36399398 | 0.99999994 |
| Gprasp1 | 0.2305834 | 0.19065172 | 0.17649334 | 0.01415838 | 0.29667901 | 0.99999994 |
| Ids | 0.26798389 | 0.2134281 | 0.19937095 | 0.01405716 | 0.31941923 | 0.99999994 |
| Fdft1 | 0.37232281 | 0.27037453 | 0.25632539 | 0.01404914 | 0.3576178 | 0.99999994 |
| Cir1 | 0.37330785 | 0.27085072 | 0.25681857 | 0.01403215 | 0.35804382 | 0.99999994 |
| Mat2b | 0.32551971 | 0.24595019 | 0.23196457 | 0.01398562 | 0.34409029 | 0.99999994 |
| Glrx3 | 1.07164265 | 0.44331144 | 0.42935074 | 0.0139607 | 0.41432351 | 0.99999994 |
| Nisch | 1.12203919 | 0.44388386 | 0.4299289 | 0.01395496 | 0.4144716 | 0.99999994 |
| Erp29 | 1.29496246 | 0.43919343 | 0.42524289 | 0.01395055 | 0.41357084 | 0.99999994 |
| Galk1 | 0.0703668 | 0.07474657 | 0.06080444 | 0.01394213 | 0.06347166 | 0.99999994 |
| Hspd1 | 1.09196771 | 0.443531 | 0.42962847 | 0.01390252 | 0.41472936 | 0.99999994 |
| Hnrnpa2b1 | 1.90790687 | 0.38769213 | 0.37382113 | 0.013871 | 0.40245771 | 0.99999994 |
| Rcn2 | 0.5486675 | 0.34692527 | 0.33312593 | 0.01379933 | 0.39137716 | 0.99999994 |
| Ttyh2 | 0.1433787 | 0.13121616 | 0.11743421 | 0.01378195 | 0.21734741 | 0.99999994 |
| Coa3 | 1.43576314 | 0.42909529 | 0.41535984 | 0.01373545 | 0.41288975 | 0.99999994 |
| mt-Nd3 | 0.73472874 | 0.40394304 | 0.39022937 | 0.01371367 | 0.40752288 | 0.99999994 |
| Tprgl | 0.58489544 | 0.36005525 | 0.34634272 | 0.01371253 | 0.39606683 | 0.99999994 |
| Klf13 | 0.18777299 | 0.16225189 | 0.14854531 | 0.01370658 | 0.26954303 | 0.99999994 |

| | | | | | | |
|---|---|---|---|---|---|---|
| Lyar | 0.35399728 | 0.2607128 | 0.24700656 | 0.01370624 | 0.35593075 | 0.99999994 |
| R3hcc1 | 0.18751474 | 0.16207037 | 0.14837074 | 0.01369963 | 0.26940721 | 0.99999994 |
| Rtn1 | 0.03619092 | 0.0456801 | 0.03198254 | 0.01369756 | 0.00218046 | 0.13848085 |
| Brix1 | 0.4515397 | 0.30722284 | 0.29356436 | 0.01365849 | 0.378396 | 0.99999994 |
| Kit | 0.17071582 | 0.15051399 | 0.13685685 | 0.01365713 | 0.25326608 | 0.99999994 |
| Ifitm3 | 0.12709888 | 0.11909958 | 0.10544953 | 0.01365005 | 0.19444266 | 0.99999994 |
| Kdm7a | 0.36500826 | 0.26627701 | 0.25263901 | 0.013638 | 0.3596761 | 0.99999994 |
| Mapk13 | 0.18629739 | 0.16116093 | 0.14754686 | 0.01361407 | 0.26954872 | 0.99999994 |
| S100a1 | 0.44959677 | 0.3062807 | 0.29270707 | 0.01357363 | 0.37878491 | 0.99999994 |
| Acvr2b | 0.08145082 | 0.08344394 | 0.0698793 | 0.01356464 | 0.09816115 | 0.99999994 |
| Arf6 | 0.17150005 | 0.15094614 | 0.13740141 | 0.01354473 | 0.25585767 | 0.99999994 |
| Pdcd10 | 0.60587367 | 0.36718098 | 0.35363927 | 0.0135417 | 0.39940479 | 0.99999994 |
| Hspa4l | 0.19130987 | 0.16445646 | 0.15092892 | 0.01352755 | 0.27538726 | 0.99999994 |
| Pdlim7 | 0.25287839 | 0.20382133 | 0.19029789 | 0.01352344 | 0.31809542 | 0.99999994 |
| Creb3l1 | 0.12188453 | 0.11500401 | 0.10154298 | 0.01346103 | 0.18878053 | 0.99999994 |
| Tmem53 | 0.19329136 | 0.16569928 | 0.15225809 | 0.01344119 | 0.27839484 | 0.99999994 |
| 2310033P09I | 0.28089874 | 0.22034457 | 0.20695375 | 0.01339083 | 0.33334473 | 0.99999994 |
| Fam107b | 0.10032954 | 0.09840942 | 0.08503649 | 0.01337292 | 0.14759929 | 0.99999994 |
| Map1lc3b | 1.23561138 | 0.44123945 | 0.42787214 | 0.01336731 | 0.41763258 | 0.99999994 |
| Zfp428 | 0.28751683 | 0.22413798 | 0.21077845 | 0.01335953 | 0.33655181 | 0.99999994 |
| Eva1b | 0.86244973 | 0.42673193 | 0.41337609 | 0.01335584 | 0.41485969 | 0.99999994 |
| Bhlhe23 | 0.02866734 | 0.03880463 | 0.02546102 | 0.01334361 | 0.0002429 | 0.0204874 |
| Gc | 0.07164673 | 0.07517213 | 0.06185902 | 0.0133131 | 0.07599179 | 0.99999994 |
| Cenpt | 0.12633837 | 0.11818608 | 0.10488182 | 0.01330426 | 0.19923439 | 0.99999994 |
| Top1 | 1.11671171 | 0.44321137 | 0.42991153 | 0.01329984 | 0.41842568 | 0.99999994 |
| Ppp1r15a | 0.33749354 | 0.25167334 | 0.23837387 | 0.01329947 | 0.35517926 | 0.99999994 |
| Tspan12 | 0.08746872 | 0.08805006 | 0.07475225 | 0.01329781 | 0.11818446 | 0.99999994 |
| Tpm1 | 0.44067266 | 0.3020188 | 0.28873531 | 0.0132835 | 0.37971438 | 0.99999994 |
| Zfp451 | 0.28646534 | 0.223452 | 0.21017359 | 0.01327841 | 0.33704651 | 0.99999994 |
| Elof1 | 1.16031985 | 0.44286225 | 0.42968709 | 0.01317516 | 0.41913817 | 0.99999994 |

| | | | | | | |
|---|---|---|---|---|---|---|
| Exoc7 | 0.53935163 | 0.3427029 | 0.32959758 | 0.01310532 | 0.39563287 | 0.99999994 |
| Myl12a | 0.94883926 | 0.43587505 | 0.42278774 | 0.01308731 | 0.41837737 | 0.99999994 |
| Akr1c12 | 0.37744878 | 0.27191337 | 0.25888302 | 0.01303036 | 0.36879958 | 0.99999994 |
| Map1a | 0.12370749 | 0.11592402 | 0.10291249 | 0.01301153 | 0.20001063 | 0.99999994 |
| Basp1 | 0.09551499 | 0.09420383 | 0.08120911 | 0.01299472 | 0.14341055 | 0.99999994 |
| Rnf220 | 0.18866957 | 0.16213351 | 0.14915083 | 0.01298268 | 0.28116106 | 0.99999994 |
| Uqcrq | 1.68335668 | 0.40359857 | 0.39061688 | 0.01298169 | 0.41246204 | 0.99999994 |
| Crip1 | 0.27135543 | 0.21434719 | 0.20136567 | 0.01298153 | 0.33391859 | 0.99999994 |
| Ocrl | 0.25000183 | 0.2015221 | 0.18854447 | 0.01297762 | 0.32341841 | 0.99999994 |
| Igf2bp1 | 0.15687874 | 0.14004553 | 0.12713387 | 0.01291166 | 0.24951504 | 0.99999994 |
| Kras | 0.58100675 | 0.35786971 | 0.34496195 | 0.01290775 | 0.4016545 | 0.99999994 |
| Sdcbp | 0.55448915 | 0.34820869 | 0.33530297 | 0.01290572 | 0.39889767 | 0.99999994 |
| Dnase2a | 0.31292654 | 0.23799182 | 0.22508832 | 0.0129035 | 0.35138466 | 0.99999994 |
| Gtpbp4 | 0.37765538 | 0.27186622 | 0.25898562 | 0.0128806 | 0.37030234 | 0.99999994 |
| Ldb1 | 0.31396351 | 0.23853175 | 0.22566001 | 0.01287174 | 0.35209092 | 0.99999994 |
| Spsb4 | 0.1462451 | 0.13233906 | 0.11951175 | 0.01282731 | 0.23747971 | 0.99999994 |
| Slc39a2 | 0.0934148 | 0.09235516 | 0.07953019 | 0.01282497 | 0.1415461 | 0.99999994 |
| Dcakd | 0.45664436 | 0.30861718 | 0.29580331 | 0.01281387 | 0.38654094 | 0.99999994 |
| Syt5 | 0.20176706 | 0.17070327 | 0.15789563 | 0.01280765 | 0.29462332 | 0.99999994 |
| Adss | 0.25319203 | 0.2032951 | 0.19048857 | 0.01280653 | 0.32725475 | 0.99999994 |
| Bad | 0.56193348 | 0.35078684 | 0.33805745 | 0.0127294 | 0.40104541 | 0.99999994 |
| Timm22 | 0.33595264 | 0.25027391 | 0.23755604 | 0.01271787 | 0.36078672 | 0.99999994 |
| Hras | 0.24543117 | 0.19842992 | 0.18574184 | 0.01268808 | 0.32466334 | 0.99999994 |
| Prkra | 0.31827757 | 0.24069343 | 0.22802721 | 0.01266621 | 0.355788 | 0.99999994 |
| Polr3k | 0.67942925 | 0.38947028 | 0.37681792 | 0.01265236 | 0.41157337 | 0.99999994 |
| Nfe2l2 | 0.29667915 | 0.22864196 | 0.21600644 | 0.01263552 | 0.34850099 | 0.99999994 |
| Abcf1 | 0.6893739 | 0.3920725 | 0.37945434 | 0.01261815 | 0.41241126 | 0.99999994 |
| Nhp2 | 0.36022367 | 0.26280018 | 0.25020437 | 0.01259582 | 0.36877663 | 0.99999994 |
| Alas1 | 0.22070503 | 0.18280924 | 0.17021595 | 0.01259329 | 0.31119518 | 0.99999994 |
| Fyttd1 | 0.55240674 | 0.34708817 | 0.33452657 | 0.0125616 | 0.40131352 | 0.99999994 |

| | | | | | | |
|---|---|---|---|---|---|---|
| Tmod3 | 0.20619726 | 0.17337041 | 0.16081202 | 0.01255839 | 0.30159483 | 0.99999994 |
| Gm2694 | 0.05937477 | 0.06422965 | 0.05167497 | 0.01255468 | 0.05291817 | 0.99999994 |
| Gyg | 0.40762429 | 0.28606762 | 0.273514 | 0.01255362 | 0.3799887 | 0.99999994 |
| Ccdc174 | 0.34106594 | 0.25281441 | 0.24026226 | 0.01255215 | 0.3640133 | 0.99999994 |
| Gsta3 | 0.06037702 | 0.0650286 | 0.05251282 | 0.01251578 | 0.05631751 | 0.99999994 |
| Dmpk | 0.09538076 | 0.09358063 | 0.08110193 | 0.0124787 | 0.15287601 | 0.99999994 |
| mt-Atp6 | 3.97237306 | 0.24088259 | 0.22842917 | 0.01245342 | 0.35834393 | 0.99999994 |
| Ypel5 | 0.30903405 | 0.23538321 | 0.22293399 | 0.01244922 | 0.35505492 | 0.99999994 |
| Myl12b | 0.79535746 | 0.4150679 | 0.40264615 | 0.01242175 | 0.41864914 | 0.99999994 |
| Rnf6 | 0.36536178 | 0.26522207 | 0.25281815 | 0.01240391 | 0.37199348 | 0.99999994 |
| Rpf2 | 0.38530806 | 0.27515954 | 0.26276261 | 0.01239693 | 0.3767447 | 0.99999994 |
| Pamr1 | 0.12742568 | 0.11807229 | 0.10569326 | 0.01237903 | 0.21781104 | 0.99999994 |
| AW551984 | 0.06683944 | 0.07023545 | 0.05788901 | 0.01234645 | 0.07785415 | 0.99999994 |
| Dhrs4 | 0.76259409 | 0.40858143 | 0.39624622 | 0.01233522 | 0.41792151 | 0.99999994 |
| Ppic | 0.1643456 | 0.1447283 | 0.13240793 | 0.01232038 | 0.26783794 | 0.99999994 |
| Elp4 | 0.33643173 | 0.25004244 | 0.23781053 | 0.01223191 | 0.36603499 | 0.99999994 |
| Tpm3 | 0.88362701 | 0.42834659 | 0.41613786 | 0.01220873 | 0.42258473 | 0.99999994 |
| Usp3 | 0.23855129 | 0.19369072 | 0.18148452 | 0.0122062 | 0.32718842 | 0.99999994 |
| Adrm1 | 0.7327074 | 0.4019301 | 0.38978775 | 0.01214234 | 0.41786696 | 0.99999994 |
| Jarid2 | 0.26195877 | 0.20791917 | 0.19577816 | 0.01214101 | 0.33988086 | 0.99999994 |
| Hnrnpf | 1.13004251 | 0.4420365 | 0.42993596 | 0.01210054 | 0.42569584 | 0.99999994 |
| 1500011B03l | 0.67453454 | 0.38759415 | 0.37549691 | 0.01209724 | 0.41509992 | 0.99999994 |
| Actl6a | 0.51570652 | 0.33247999 | 0.32039804 | 0.01208196 | 0.40090472 | 0.99999994 |
| Cox7a2l | 1.83347321 | 0.39003587 | 0.37799105 | 0.01204482 | 0.4160117 | 0.99999994 |
| Bco1 | 0.10720327 | 0.10240726 | 0.09036354 | 0.01204372 | 0.18749526 | 0.99999994 |
| Zfp329 | 0.29366287 | 0.22631708 | 0.21429376 | 0.01202333 | 0.35440057 | 0.99999994 |
| Ddx3y | 0.16846623 | 0.14725572 | 0.13529096 | 0.01196476 | 0.27804132 | 0.99999994 |
| Ctnna1 | 0.42322561 | 0.29275916 | 0.28080171 | 0.01195746 | 0.38841613 | 0.99999994 |
| Ccdc115 | 0.46872128 | 0.31295499 | 0.30102724 | 0.01192774 | 0.39598705 | 0.99999994 |
| Stc2 | 0.06859716 | 0.07126117 | 0.05934346 | 0.01191771 | 0.09064897 | 0.99999994 |

| | | | | | | |
|---|---|---|---|---|---|---|
| Fars2 | 0.22481245 | 0.18474823 | 0.17283837 | 0.01190987 | 0.3232344 | 0.99999994 |
| Cox6b2 | 0.1245721 | 0.11542191 | 0.10356061 | 0.0118613 | 0.22291506 | 0.99999994 |
| Asb4 | 0.03968291 | 0.0468419 | 0.03498734 | 0.01185456 | 0.01205616 | 0.50170905 |
| Apbb1 | 0.31471148 | 0.23792368 | 0.22607172 | 0.01185196 | 0.36355823 | 0.99999994 |
| Bbx | 0.34595697 | 0.25466664 | 0.24283017 | 0.01183647 | 0.37279629 | 0.99999994 |
| Rbm4b | 0.28335856 | 0.22019817 | 0.2083804 | 0.01181777 | 0.35290157 | 0.99999994 |
| Vps37b | 0.29484857 | 0.226783 | 0.21496799 | 0.01181501 | 0.35724043 | 0.99999994 |
| Gnpda2 | 0.29792628 | 0.228524 | 0.21671218 | 0.01181182 | 0.35837623 | 0.99999994 |
| Prox1os | 0.08043497 | 0.08082793 | 0.06905299 | 0.01177494 | 0.12817834 | 0.99999994 |
| 1810037I17R | 1.21053318 | 0.44049891 | 0.42872496 | 0.01177395 | 0.42747681 | 0.99999994 |
| Cks2 | 0.16427758 | 0.14412902 | 0.13236018 | 0.01176884 | 0.27697488 | 0.99999994 |
| Ftsj1 | 0.18895506 | 0.16110288 | 0.14934345 | 0.01175943 | 0.30009506 | 0.99999994 |
| Nfib | 0.12904619 | 0.11864889 | 0.10689999 | 0.01174891 | 0.23221525 | 0.99999994 |
| Gm4950 | 0.13011706 | 0.1194303 | 0.10769569 | 0.01173461 | 0.23413947 | 0.99999994 |
| Lrrc58 | 0.41654725 | 0.28942118 | 0.27770428 | 0.0117169 | 0.3894146 | 0.99999994 |
| Naa35 | 0.42513629 | 0.29332595 | 0.28168138 | 0.01164457 | 0.39159259 | 0.99999994 |
| Map1lc3a | 0.8239731 | 0.41919655 | 0.40756995 | 0.0116266 | 0.42470018 | 0.99999994 |
| Ssbp4 | 0.46006691 | 0.30890801 | 0.29729395 | 0.01161406 | 0.39741774 | 0.99999994 |
| Pnn | 0.74662147 | 0.404453 | 0.39286242 | 0.01159058 | 0.42215568 | 0.99999994 |
| Hdac6 | 0.26504153 | 0.2091694 | 0.19762072 | 0.01154868 | 0.34864238 | 0.99999994 |
| Sema6a | 0.09965927 | 0.09600655 | 0.08450892 | 0.01149763 | 0.18257038 | 0.99999994 |
| Fosb | 0.16904145 | 0.14713713 | 0.1356919 | 0.01144523 | 0.28724701 | 0.99999994 |
| Ass1 | 0.1308782 | 0.11969894 | 0.10826037 | 0.01143857 | 0.24093534 | 0.99999994 |
| Tdp2 | 0.25186926 | 0.20111648 | 0.18968369 | 0.01143279 | 0.3441371 | 0.99999994 |
| Smarca4 | 0.59923643 | 0.36277608 | 0.35135879 | 0.0114173 | 0.41437839 | 0.99999994 |
| Ccdc28b | 0.49241898 | 0.32233642 | 0.31098253 | 0.01135389 | 0.40399427 | 0.99999994 |
| Rsad2 | 0.05078684 | 0.05579447 | 0.04445037 | 0.01134409 | 0.04468256 | 0.99999994 |
| Cxcl12 | 0.0392042 | 0.04584277 | 0.03457624 | 0.01126653 | 0.01504333 | 0.56557946 |
| Psmd5 | 0.32051196 | 0.24049356 | 0.22924683 | 0.01124673 | 0.37200131 | 0.99999994 |
| Rfc3 | 0.19144153 | 0.16224807 | 0.15101739 | 0.01123068 | 0.31029576 | 0.99999994 |

| | | | | | | |
|---|---|---|---|---|---|---|
| Eif5b | 1.18467558 | 0.44055273 | 0.42934346 | 0.01120927 | 0.43101772 | 0.99999994 |
| Ppp1cb | 0.94537335 | 0.43371561 | 0.42251082 | 0.01120479 | 0.42994133 | 0.99999994 |
| Gm15915 | 0.10140843 | 0.09704564 | 0.08587657 | 0.01116907 | 0.19332092 | 0.99999994 |
| Msmo1 | 0.25552308 | 0.20306133 | 0.19190228 | 0.01115905 | 0.34935409 | 0.99999994 |
| Stox1 | 0.09244246 | 0.0898988 | 0.07875136 | 0.01114744 | 0.17303918 | 0.99999994 |
| Abtb1 | 0.18004942 | 0.15443477 | 0.14329292 | 0.01114185 | 0.30237906 | 0.99999994 |
| Rsu1 | 0.23453336 | 0.19010962 | 0.17897565 | 0.01113396 | 0.33940487 | 0.99999994 |
| Gm6133 | 0.15680278 | 0.13813964 | 0.12707988 | 0.01105976 | 0.28119255 | 0.99999994 |
| Nkd1 | 0.0558419 | 0.05976206 | 0.04871279 | 0.01104927 | 0.06554374 | 0.99999994 |
| Ube2d1 | 0.28916026 | 0.22261611 | 0.21172179 | 0.01089432 | 0.36598419 | 0.99999994 |
| Acat2 | 0.18074263 | 0.15465489 | 0.143767 | 0.01088789 | 0.30709365 | 0.99999994 |
| Habp2 | 0.07148015 | 0.07259014 | 0.06172188 | 0.01086827 | 0.12058215 | 0.99999994 |
| Smarcad1 | 0.31069623 | 0.23468697 | 0.22385556 | 0.0108314 | 0.37369891 | 0.99999994 |
| Atxn10 | 1.21211032 | 0.43947906 | 0.4286747 | 0.01080436 | 0.4333831 | 0.99999994 |
| Tbpl1 | 0.35491287 | 0.25827293 | 0.24747878 | 0.01079415 | 0.38578308 | 0.99999994 |
| Tcf15 | 0.03843992 | 0.04470519 | 0.03391936 | 0.01078583 | 0.01714614 | 0.60730275 |
| Tmeff1 | 0.15627684 | 0.13743372 | 0.12670591 | 0.01072781 | 0.28652216 | 0.99999994 |
| Coq7 | 0.33096079 | 0.24561922 | 0.23489283 | 0.01072639 | 0.38057808 | 0.99999994 |
| Rorc | 0.23109703 | 0.18750113 | 0.17681701 | 0.01068412 | 0.34376531 | 0.99999994 |
| Ptger3 | 0.04185334 | 0.0475133 | 0.03684797 | 0.01066533 | 0.02701307 | 0.83130387 |
| Gm10269 | 0.13196457 | 0.11971425 | 0.10906501 | 0.01064924 | 0.25786721 | 0.99999994 |
| Mycbp2 | 0.4320489 | 0.29548423 | 0.28484172 | 0.0106425 | 0.4017962 | 0.99999994 |
| Ube2r2 | 0.46993479 | 0.31218881 | 0.30154639 | 0.01064242 | 0.40713483 | 0.99999994 |
| Rab3b | 0.23381771 | 0.18916559 | 0.17852707 | 0.01063852 | 0.34581067 | 0.99999994 |
| Rprm | 0.07916721 | 0.07862454 | 0.06802025 | 0.01060429 | 0.14970037 | 0.99999994 |
| Ywhah | 1.00533621 | 0.43731395 | 0.42671386 | 0.01060009 | 0.43433344 | 0.99999994 |
| Hypk | 0.73266085 | 0.40035737 | 0.38977756 | 0.01057981 | 0.42831137 | 0.99999994 |
| Etfb | 1.00712322 | 0.43734506 | 0.42680366 | 0.0105414 | 0.4347074 | 0.99999994 |
| Dusp23 | 0.18816295 | 0.15934921 | 0.14880878 | 0.01054043 | 0.31864782 | 0.99999994 |
| Tspan15 | 0.28803367 | 0.22156945 | 0.21107538 | 0.01049406 | 0.37034681 | 0.99999994 |

| | | | | | | |
|---|---|---|---|---|---|---|
| Ppp5c | 0.54319549 | 0.34154407 | 0.33105981 | 0.01048426 | 0.41652214 | 0.99999994 |
| Klf6 | 0.2010431 | 0.1678813 | 0.15741708 | 0.01046422 | 0.32907162 | 0.99999994 |
| Plin2 | 0.13061535 | 0.11852796 | 0.10806546 | 0.01046249 | 0.2596439 | 0.99999994 |
| Snapc3 | 0.2631439 | 0.20691904 | 0.19648758 | 0.01043146 | 0.36190035 | 0.99999994 |
| Hadha | 0.46096085 | 0.30810643 | 0.29768195 | 0.01042447 | 0.40784271 | 0.99999994 |
| Cacna1a | 0.29879475 | 0.22760787 | 0.21720282 | 0.01040505 | 0.37491202 | 0.99999994 |
| Mtf2 | 0.27080051 | 0.2114237 | 0.20103812 | 0.01038559 | 0.36547349 | 0.99999994 |
| Donson | 0.05253227 | 0.05630431 | 0.04592532 | 0.01037899 | 0.06624973 | 0.99999994 |
| Cfap77 | 0.09554456 | 0.09159169 | 0.08123271 | 0.01035898 | 0.19798685 | 0.99999994 |
| Chd6 | 0.30678837 | 0.23202626 | 0.22168503 | 0.01034123 | 0.37808746 | 0.99999994 |
| Mboat4 | 0.0266269 | 0.03399834 | 0.02368105 | 0.01031729 | 0.00186568 | 0.12289594 |
| Cdo1 | 0.10290912 | 0.09733429 | 0.08704263 | 0.01029167 | 0.21563266 | 0.99999994 |
| Fa2h | 0.07423994 | 0.07425577 | 0.06399032 | 0.01026545 | 0.14279946 | 0.99999994 |
| Dut | 0.47688398 | 0.31469935 | 0.3044995 | 0.01019985 | 0.41177996 | 0.99999994 |
| Tax1bp1 | 0.83305507 | 0.41913654 | 0.40897437 | 0.01016218 | 0.4343158 | 0.99999994 |
| Dynlt3 | 0.45517359 | 0.3052781 | 0.29516025 | 0.01011785 | 0.40975533 | 0.99999994 |
| Ypel1 | 0.10036244 | 0.09517987 | 0.08506213 | 0.01011774 | 0.21425553 | 0.99999994 |
| Echs1 | 0.43201827 | 0.29494023 | 0.2848278 | 0.01011243 | 0.40659023 | 0.99999994 |
| Mdfi | 0.08863474 | 0.08578764 | 0.07569207 | 0.01009558 | 0.1873229 | 0.99999994 |
| Ccs | 0.42770547 | 0.29295173 | 0.28286002 | 0.01009171 | 0.40614166 | 0.99999994 |
| Med19 | 0.28936429 | 0.22193017 | 0.21183874 | 0.01009143 | 0.37558725 | 0.99999994 |
| Lsm3 | 0.39735239 | 0.27869664 | 0.26861344 | 0.0100832 | 0.40134542 | 0.99999994 |
| Cops2 | 0.36651835 | 0.26347895 | 0.25340356 | 0.01007538 | 0.39563634 | 0.99999994 |
| Tnnc1 | 0.07767066 | 0.07686683 | 0.06679896 | 0.01006787 | 0.15787263 | 0.99999994 |
| Igsf21 | 0.06671135 | 0.06784179 | 0.05778289 | 0.0100589 | 0.12327994 | 0.99999994 |
| Itm2c | 0.74689998 | 0.40297023 | 0.39292322 | 0.01004701 | 0.43242546 | 0.99999994 |
| Lrrn2 | 0.05789525 | 0.06045951 | 0.05043611 | 0.0100234 | 0.09294286 | 0.99999994 |
| Hnrnpr | 0.84884372 | 0.42136512 | 0.41140295 | 0.00996217 | 0.43597406 | 0.99999994 |
| Oxa1l | 0.2940889 | 0.2244766 | 0.21453616 | 0.00994044 | 0.37888159 | 0.99999994 |
| Btf3 | 2.13483335 | 0.37836002 | 0.36844094 | 0.00991908 | 0.42888995 | 0.99999994 |

| | | | | | | |
|---|---|---|---|---|---|---|
| Kdelr2 | 0.62849804 | 0.37109768 | 0.3612127 | 0.00988498 | 0.42772916 | 0.99999994 |
| Tgm7 | 0.03881031 | 0.04410908 | 0.03423779 | 0.00987129 | 0.02748381 | 0.83977187 |
| Ccar1 | 0.45104297 | 0.30320374 | 0.29334544 | 0.0098583 | 0.41149667 | 0.99999994 |
| Taf13 | 0.2408507 | 0.19269998 | 0.18291261 | 0.00978737 | 0.36085606 | 0.99999994 |
| Rnd3 | 0.07651719 | 0.07562148 | 0.06585604 | 0.00976544 | 0.16181245 | 0.99999994 |
| Igfbp5 | 0.06403904 | 0.06527889 | 0.05556488 | 0.00971402 | 0.12227601 | 0.99999994 |
| Lsm14a | 0.40201776 | 0.2805566 | 0.27084918 | 0.00970742 | 0.40572009 | 0.99999994 |
| Glod5 | 0.03588382 | 0.04140811 | 0.03171762 | 0.00969049 | 0.02099209 | 0.69426319 |
| Pole3 | 0.16270024 | 0.14093311 | 0.13125138 | 0.00968173 | 0.31171154 | 0.99999994 |
| Dpysl5 | 0.0637265 | 0.06498547 | 0.05530496 | 0.00968051 | 0.12198511 | 0.99999994 |
| Cryl1 | 0.12619401 | 0.11444221 | 0.10477397 | 0.00966824 | 0.26953179 | 0.99999994 |
| Pet100 | 0.47127726 | 0.31174777 | 0.30211943 | 0.00962834 | 0.41600143 | 0.99999994 |
| Zrsr2 | 0.43464073 | 0.29564034 | 0.28601777 | 0.00962257 | 0.41140139 | 0.99999994 |
| Adam23 | 0.04756066 | 0.0513122 | 0.04171523 | 0.00959698 | 0.06284051 | 0.99999994 |
| Arl2 | 0.58159704 | 0.35475339 | 0.34517211 | 0.00958128 | 0.42670588 | 0.99999994 |
| Ctbp2 | 0.1479298 | 0.13024856 | 0.12072807 | 0.00952049 | 0.29982168 | 0.99999994 |
| Fkbp4 | 1.29360062 | 0.43480332 | 0.42531342 | 0.0094899 | 0.44096565 | 0.99999994 |
| Plk5 | 0.05169957 | 0.05468142 | 0.04522208 | 0.00945933 | 0.08190911 | 0.99999994 |
| Calcoco1 | 0.13615028 | 0.12160511 | 0.11215207 | 0.00945304 | 0.28738194 | 0.99999994 |
| Scin | 0.04733676 | 0.0509664 | 0.04152497 | 0.00944143 | 0.06508406 | 0.99999994 |
| Ddx42 | 0.37419907 | 0.26669409 | 0.25726411 | 0.00942998 | 0.40362014 | 0.99999994 |
| Tmie | 0.08131813 | 0.07919507 | 0.06977143 | 0.00942364 | 0.18431635 | 0.99999994 |
| Arc | 0.11774156 | 0.10783097 | 0.09841546 | 0.0094155 | 0.26212096 | 0.99999994 |
| Capza1 | 0.21223851 | 0.17413861 | 0.16475462 | 0.00938398 | 0.35229625 | 0.99999994 |
| Pdzk1ip1 | 0.36471686 | 0.26187396 | 0.25249126 | 0.0093827 | 0.40231729 | 0.99999994 |
| Ndufa12 | 1.37184942 | 0.42991914 | 0.42053674 | 0.0093824 | 0.44097134 | 0.99999994 |
| Rad23a | 0.29572503 | 0.22484518 | 0.21546556 | 0.00937961 | 0.38599856 | 0.99999994 |
| Odc1 | 0.28471975 | 0.21852491 | 0.20916721 | 0.0093577 | 0.38293114 | 0.99999994 |
| Gamt | 0.12271325 | 0.11147908 | 0.10216607 | 0.00931301 | 0.27200361 | 0.99999994 |
| Ces1d | 0.03972872 | 0.0443396 | 0.03502666 | 0.00931294 | 0.03838008 | 0.99999994 |

| | | | | | | |
|---|---|---|---|---|---|---|
| Rpa3 | 0.52687702 | 0.3340977 | 0.32478829 | 0.00930941 | 0.42434461 | 0.99999994 |
| Puf60 | 0.87052132 | 0.42368148 | 0.41446552 | 0.00921596 | 0.44116771 | 0.99999994 |
| Bbs4 | 0.15562894 | 0.1354409 | 0.12624478 | 0.00919612 | 0.31388335 | 0.99999994 |
| Erich1 | 0.20660179 | 0.170257 | 0.16107729 | 0.00917971 | 0.35221678 | 0.99999994 |
| Ctsa | 0.41490604 | 0.28611162 | 0.276938 | 0.00917363 | 0.41274353 | 0.99999994 |
| Cdc14b | 0.23169293 | 0.18634862 | 0.17719219 | 0.00915643 | 0.36543515 | 0.99999994 |
| Cat | 0.12982486 | 0.11663278 | 0.10747871 | 0.00915407 | 0.28538125 | 0.99999994 |
| Pnrc2 | 0.32505816 | 0.24083029 | 0.23171499 | 0.00911531 | 0.39671827 | 0.99999994 |
| Terf1 | 0.17102081 | 0.14617835 | 0.13706871 | 0.00910963 | 0.32910572 | 0.99999994 |
| Romo1 | 1.45589765 | 0.4225675 | 0.41347262 | 0.00909489 | 0.44179667 | 0.99999994 |
| Mbip | 0.15893551 | 0.13766698 | 0.12859312 | 0.00907386 | 0.31928846 | 0.99999994 |
| Zfp292 | 0.4766411 | 0.31346107 | 0.30439683 | 0.00906424 | 0.42143998 | 0.99999994 |
| Cnppd1 | 0.17558549 | 0.1492894 | 0.14022719 | 0.00906221 | 0.33353663 | 0.99999994 |
| Eif4e | 0.86440447 | 0.42270317 | 0.4136425 | 0.00906067 | 0.44203775 | 0.99999994 |
| Tspan1 | 0.05416297 | 0.05635897 | 0.04730025 | 0.00905872 | 0.10119108 | 0.99999994 |
| Armcx3 | 0.47984195 | 0.31479129 | 0.30574664 | 0.00904465 | 0.42194914 | 0.99999994 |
| Snn | 0.11087479 | 0.10222468 | 0.09318459 | 0.00904009 | 0.25922012 | 0.99999994 |
| Serpina1a | 0.03212907 | 0.03750944 | 0.02846981 | 0.00903964 | 0.01727883 | 0.60803923 |
| Sult1d1 | 0.05735488 | 0.05898287 | 0.04998304 | 0.00899984 | 0.11535555 | 0.99999994 |
| Ss18 | 0.22425394 | 0.18146384 | 0.17248285 | 0.008981 | 0.36444991 | 0.99999994 |
| Dcun1d5 | 0.59872628 | 0.36010126 | 0.35118249 | 0.00891876 | 0.4328797 | 0.99999994 |
| Samd14 | 0.19032097 | 0.15916877 | 0.15026383 | 0.00890494 | 0.34661721 | 0.99999994 |
| Mapre3 | 0.54296106 | 0.33985901 | 0.33097089 | 0.00888813 | 0.42906522 | 0.99999994 |
| Surf4 | 0.71112633 | 0.3937347 | 0.38487489 | 0.00885981 | 0.4391088 | 0.99999994 |
| Phf23 | 0.38945217 | 0.27364119 | 0.26478889 | 0.0088523 | 0.4119498 | 0.99999994 |
| Gt(ROSA)26S | 0.25081567 | 0.19788142 | 0.18904137 | 0.00884006 | 0.37779834 | 0.99999994 |
| Hmces | 0.28067228 | 0.21565868 | 0.20682212 | 0.00883657 | 0.38805354 | 0.99999994 |
| Serpina1c | 0.02506458 | 0.03113022 | 0.02231487 | 0.00881535 | 0.00427487 | 0.2271524 |
| Rrm1 | 0.10570451 | 0.09801175 | 0.0892071 | 0.00880465 | 0.2555982 | 0.99999994 |
| Malat1 | 5.36172884 | 0.40082057 | 0.39203013 | 0.00879044 | 0.44067633 | 0.99999994 |

| | | | | | | |
|---|---|---|---|---|---|---|
| Dctn6 | 0.37696916 | 0.26741847 | 0.25864469 | 0.00877378 | 0.41067929 | 0.99999994 |
| Bloc1s2 | 0.48608328 | 0.31711994 | 0.30835887 | 0.00876107 | 0.42499937 | 0.99999994 |
| Nanos1 | 0.09113951 | 0.08645814 | 0.07770619 | 0.00875195 | 0.22671849 | 0.99999994 |
| Fmo1 | 0.15997508 | 0.13806893 | 0.12932885 | 0.00874008 | 0.32641344 | 0.99999994 |
| 6330403K07l | 0.08311518 | 0.07995554 | 0.07123078 | 0.00872476 | 0.20744774 | 0.99999994 |
| Chchd5 | 0.29419609 | 0.22331114 | 0.21459712 | 0.00871402 | 0.39346822 | 0.99999994 |
| Ift20 | 0.83309599 | 0.41768508 | 0.40898082 | 0.00870426 | 0.44367193 | 0.99999994 |
| Fgfr4 | 0.0848867 | 0.08136183 | 0.07266609 | 0.00869574 | 0.21285706 | 0.99999994 |
| Cdc42se1 | 0.38916115 | 0.27332164 | 0.26464703 | 0.00867461 | 0.41364356 | 0.99999994 |
| Psme2b | 0.08260766 | 0.07948769 | 0.07081898 | 0.00866871 | 0.20759843 | 0.99999994 |
| Pbld2 | 0.08943625 | 0.08498223 | 0.07633728 | 0.00864494 | 0.22548149 | 0.99999994 |
| Dennd2a | 0.12322018 | 0.11117709 | 0.10254679 | 0.0086303 | 0.28767267 | 0.99999994 |
| Errfi1 | 0.2040623 | 0.16802676 | 0.15940916 | 0.0086176 | 0.35948327 | 0.99999994 |
| Exosc8 | 0.18530929 | 0.15545147 | 0.14687695 | 0.00857452 | 0.34878932 | 0.99999994 |
| Psmd9 | 0.41857606 | 0.28720388 | 0.27864877 | 0.00855511 | 0.4190344 | 0.99999994 |
| Ap1m2 | 0.3616458 | 0.25947153 | 0.25093 | 0.00854154 | 0.41037545 | 0.99999994 |
| Nop10 | 0.80638563 | 0.41318346 | 0.40464746 | 0.008536 | 0.44416596 | 0.99999994 |
| Ak6 | 0.30763896 | 0.23069366 | 0.22215862 | 0.00853504 | 0.39908024 | 0.99999994 |
| Adamts1 | 0.05421902 | 0.05587743 | 0.04734746 | 0.00852997 | 0.11522727 | 0.99999994 |
| Calm3 | 1.24281174 | 0.43613676 | 0.42761202 | 0.00852474 | 0.44721567 | 0.99999994 |
| Apool | 0.2468973 | 0.19516206 | 0.18664306 | 0.008519 | 0.38063324 | 0.99999994 |
| Srl | 0.05533327 | 0.05679528 | 0.04828519 | 0.00851009 | 0.1203661 | 0.99999994 |
| Ptpn1 | 0.33037414 | 0.24308043 | 0.23457846 | 0.00850198 | 0.40468045 | 0.99999994 |
| Bdnf | 0.04054682 | 0.04422272 | 0.03572858 | 0.00849414 | 0.05676904 | 0.99999994 |
| Rbmxl1 | 0.39648822 | 0.27668135 | 0.26819749 | 0.00848386 | 0.41661542 | 0.99999994 |
| Ccnjl | 0.06795955 | 0.06729491 | 0.05881625 | 0.00847866 | 0.16864145 | 0.99999994 |
| Acvr1c | 0.2056074 | 0.16886461 | 0.16042491 | 0.0084397 | 0.3631016 | 0.99999994 |
| Sim1 | 0.08387737 | 0.08025473 | 0.07184872 | 0.00840602 | 0.21805832 | 0.99999994 |
| Baz1a | 0.12844272 | 0.1148554 | 0.10645099 | 0.00840441 | 0.29960869 | 0.99999994 |
| Ctps2 | 0.27026354 | 0.20910838 | 0.20072088 | 0.0083875 | 0.39044921 | 0.99999994 |

| | | | | | | |
|---|---|---|---|---|---|---|
| Trib1 | 0.1714653 | 0.14574593 | 0.13737729 | 0.00836864 | 0.34255994 | 0.99999994 |
| Ccdc50 | 0.28676591 | 0.21870611 | 0.21034659 | 0.00835952 | 0.39568401 | 0.99999994 |
| Gpbp1 | 0.53115924 | 0.33478122 | 0.32645053 | 0.00833069 | 0.43255868 | 0.99999994 |
| Cdk9 | 0.22064372 | 0.17849534 | 0.17017667 | 0.00831867 | 0.37244686 | 0.99999994 |
| Rsl24d1 | 0.2381433 | 0.18951206 | 0.18123058 | 0.00828148 | 0.38049998 | 0.99999994 |
| Grina | 0.45615648 | 0.30387125 | 0.29559017 | 0.00828109 | 0.4260339 | 0.99999994 |
| Kin | 0.34471595 | 0.25045252 | 0.24218065 | 0.00827186 | 0.41007464 | 0.99999994 |
| Gm10250 | 0.14104469 | 0.1239555 | 0.11573538 | 0.00822011 | 0.31818923 | 0.99999994 |
| B2m | 0.19022452 | 0.15836176 | 0.15019891 | 0.00816285 | 0.35876844 | 0.99999994 |
| Megf6 | 0.04605391 | 0.04858066 | 0.04043382 | 0.00814685 | 0.08993384 | 0.99999994 |
| Pigx | 0.37287094 | 0.26473391 | 0.25659991 | 0.008134 | 0.41644331 | 0.99999994 |
| Zc3h13 | 0.3349788 | 0.24513969 | 0.23703814 | 0.00810155 | 0.4100164 | 0.99999994 |
| Tgfbr1 | 0.27299617 | 0.21042599 | 0.20233245 | 0.00809354 | 0.39501856 | 0.99999994 |
| Stt3a | 0.65742298 | 0.37847355 | 0.37040627 | 0.00806728 | 0.44236628 | 0.99999994 |
| Dock11 | 0.06266528 | 0.06247999 | 0.05442164 | 0.00805835 | 0.16215889 | 0.99999994 |
| Rrp1 | 0.9613308 | 0.43181028 | 0.42375446 | 0.00805582 | 0.44965175 | 0.99999994 |
| Vdr | 0.18721991 | 0.15621938 | 0.14817135 | 0.00804802 | 0.35884628 | 0.99999994 |
| Anxa2 | 0.0781263 | 0.07521735 | 0.06717105 | 0.0080463 | 0.21262291 | 0.99999994 |
| Sh3bp1 | 0.09849053 | 0.09162587 | 0.08358009 | 0.00804577 | 0.2608363 | 0.99999994 |
| Fmr1 | 0.22079372 | 0.1783141 | 0.17027277 | 0.00804133 | 0.37662605 | 0.99999994 |
| Serbp1 | 1.63004252 | 0.40374221 | 0.39571025 | 0.00803196 | 0.44626329 | 0.99999994 |
| Brd7 | 0.27302989 | 0.21036766 | 0.2023523 | 0.00801537 | 0.39601916 | 0.99999994 |
| Krtcap3 | 0.31329172 | 0.23330325 | 0.22528975 | 0.00801349 | 0.40641965 | 0.99999994 |
| Cdc42ep3 | 0.06501973 | 0.0643876 | 0.05637974 | 0.00800786 | 0.17221931 | 0.99999994 |
| Spp2 | 0.01936576 | 0.02526557 | 0.01730722 | 0.00795834 | 0.00110381 | 0.07717468 |
| Mcfd2 | 0.50156632 | 0.32267751 | 0.31472403 | 0.00795348 | 0.43320717 | 0.99999994 |
| Zcchc3 | 0.2158067 | 0.17500062 | 0.1670653 | 0.00793532 | 0.37593832 | 0.99999994 |
| Arl6ip4 | 0.37548392 | 0.26582874 | 0.25790534 | 0.0079234 | 0.41898241 | 0.99999994 |
| Kcnmb2 | 0.27707928 | 0.21264671 | 0.20472744 | 0.00791927 | 0.39840361 | 0.99999994 |
| St8sia2 | 0.02623324 | 0.03123464 | 0.02333708 | 0.00789756 | 0.01214234 | 0.50170905 |

| | | | | | | |
|---|---|---|---|---|---|---|
| Lypd1 | 0.02268472 | 0.0281249 | 0.0202283 | 0.0078966 | 0.00468201 | 0.24311392 |
| Fcf1 | 0.4055462 | 0.28040211 | 0.27252912 | 0.00787298 | 0.42375644 | 0.99999994 |
| Strn3 | 0.3747409 | 0.26538866 | 0.25753468 | 0.00785398 | 0.41956851 | 0.99999994 |
| Pou3f1 | 0.02833286 | 0.033023 | 0.02516957 | 0.00785343 | 0.01890432 | 0.64807274 |
| Rbm17 | 0.48639245 | 0.31633658 | 0.30848751 | 0.00784907 | 0.43275576 | 0.99999994 |
| Dync1li1 | 0.35088921 | 0.25323612 | 0.24539845 | 0.00783768 | 0.4158216 | 0.99999994 |
| Mcm6 | 0.0914301 | 0.08577527 | 0.07793944 | 0.00783584 | 0.25167965 | 0.99999994 |
| Gpatch11 | 0.28275583 | 0.21586601 | 0.20803134 | 0.00783467 | 0.40102874 | 0.99999994 |
| Tmem97 | 0.20359265 | 0.16693426 | 0.15909991 | 0.00783435 | 0.37154266 | 0.99999994 |
| Crlf1 | 0.04890622 | 0.0506832 | 0.0428574 | 0.00782581 | 0.11209573 | 0.99999994 |
| Msx1 | 0.03038469 | 0.03476047 | 0.0269554 | 0.00780507 | 0.02696614 | 0.83130387 |
| Tiparp | 0.10820623 | 0.09893596 | 0.09113585 | 0.00780012 | 0.28443958 | 0.99999994 |
| Hnrnpc | 1.34478493 | 0.43017432 | 0.4223846 | 0.00778971 | 0.45114935 | 0.99999994 |
| Med6 | 0.29345359 | 0.22196237 | 0.21417462 | 0.00778775 | 0.404376 | 0.99999994 |
| Rtkn | 0.10731495 | 0.09820993 | 0.0904496 | 0.00776033 | 0.28396675 | 0.99999994 |
| Slc25a11 | 0.80029203 | 0.41131514 | 0.40355739 | 0.00775775 | 0.44909112 | 0.99999994 |
| Nelfcd | 0.30250199 | 0.22703882 | 0.2192887 | 0.00775011 | 0.40700759 | 0.99999994 |
| Tlcd2 | 0.10774594 | 0.09849588 | 0.09078157 | 0.00771431 | 0.28582108 | 0.99999994 |
| Phyhipl | 0.19353886 | 0.16012131 | 0.1524238 | 0.00769751 | 0.36837858 | 0.99999994 |
| Ggh | 0.16200164 | 0.13845666 | 0.1307594 | 0.00769725 | 0.34759129 | 0.99999994 |
| Cd151 | 0.31623826 | 0.23460612 | 0.2269103 | 0.00769582 | 0.41069602 | 0.99999994 |
| Hdac1 | 0.59902437 | 0.35896967 | 0.35128552 | 0.00768414 | 0.44211719 | 0.99999994 |
| Cd1d1 | 0.07260998 | 0.07033544 | 0.06265153 | 0.00768391 | 0.20714424 | 0.99999994 |
| Rabl2 | 0.12183132 | 0.1091757 | 0.10150295 | 0.00767275 | 0.30742481 | 0.99999994 |
| Proser1 | 0.18790973 | 0.15629015 | 0.14863772 | 0.00765244 | 0.36591406 | 0.99999994 |
| Slc38a11 | 0.03940994 | 0.04240187 | 0.03475296 | 0.00764891 | 0.07145719 | 0.99999994 |
| Zfp664 | 0.38499878 | 0.27022624 | 0.26261085 | 0.00761539 | 0.4234692 | 0.99999994 |
| Elovl7 | 0.06913515 | 0.06739357 | 0.05978798 | 0.0076056 | 0.19856671 | 0.99999994 |
| Spc24 | 0.09864189 | 0.09130121 | 0.08370047 | 0.00760075 | 0.27277008 | 0.99999994 |
| Dact2 | 0.16829464 | 0.14277044 | 0.13517128 | 0.00759916 | 0.3541233 | 0.99999994 |

| | | | | | | |
|---|---|---|---|---|---|---|
| Blnk | 0.07188605 | 0.06962766 | 0.06205602 | 0.00757164 | 0.20834937 | 0.99999994 |
| Elf3 | 0.2109752 | 0.17149425 | 0.16393337 | 0.00756089 | 0.37942072 | 0.99999994 |
| Gse1 | 0.39741178 | 0.27620047 | 0.268642 | 0.00755847 | 0.42571965 | 0.99999994 |
| Pnck | 0.11090348 | 0.10074718 | 0.09320657 | 0.00754061 | 0.29511125 | 0.99999994 |
| Adgrb3 | 0.05300088 | 0.0538582 | 0.04632073 | 0.00753747 | 0.13937194 | 0.99999994 |
| Magohb | 0.26042261 | 0.20239272 | 0.19485662 | 0.0075361 | 0.39842133 | 0.99999994 |
| Bzw2 | 0.42544193 | 0.28934893 | 0.28182185 | 0.00752708 | 0.42944698 | 0.99999994 |
| Poc1a | 0.04982829 | 0.05115518 | 0.04363893 | 0.00751625 | 0.12579966 | 0.99999994 |
| Fstl1 | 0.07674671 | 0.07353338 | 0.06604378 | 0.0074896 | 0.22516811 | 0.99999994 |
| Dll4 | 0.08124749 | 0.07719502 | 0.06971399 | 0.00748102 | 0.23752229 | 0.99999994 |
| Ddx5 | 2.14350288 | 0.3760292 | 0.36855292 | 0.00747628 | 0.44629506 | 0.99999994 |
| Ttyh1 | 0.07521216 | 0.07221237 | 0.06478752 | 0.00742485 | 0.2227795 | 0.99999994 |
| Frat2 | 0.05365727 | 0.05428693 | 0.04687419 | 0.00741275 | 0.14625178 | 0.99999994 |
| Cdv3 | 0.25223855 | 0.19730108 | 0.18990857 | 0.00739251 | 0.39777517 | 0.99999994 |
| Parl | 0.40903995 | 0.28156659 | 0.27418277 | 0.00738383 | 0.42886766 | 0.99999994 |
| Phldb2 | 0.27882786 | 0.21308296 | 0.20574834 | 0.00733462 | 0.40621621 | 0.99999994 |
| Gm4319 | 0.03612575 | 0.03925733 | 0.03192632 | 0.00733101 | 0.06320035 | 0.99999994 |
| Gprc5b | 0.10983486 | 0.09971442 | 0.09238726 | 0.00732716 | 0.29878069 | 0.99999994 |
| Drg1 | 0.65634838 | 0.37739421 | 0.37007394 | 0.00732027 | 0.44762403 | 0.99999994 |
| Mlf1 | 0.03095434 | 0.03476621 | 0.02745033 | 0.00731587 | 0.03803109 | 0.99999994 |
| Lipg | 0.03349537 | 0.03695675 | 0.02965351 | 0.00730324 | 0.05056835 | 0.99999994 |
| Plekhf2 | 0.09703159 | 0.08970386 | 0.0824187 | 0.00728516 | 0.27814215 | 0.99999994 |
| Sel1l | 0.39566656 | 0.27508378 | 0.26780147 | 0.00728231 | 0.42818146 | 0.99999994 |
| Vps36 | 0.38150817 | 0.26816998 | 0.2608929 | 0.00727708 | 0.42635345 | 0.99999994 |
| Esf1 | 0.38444264 | 0.26961148 | 0.26233777 | 0.0072737 | 0.42678813 | 0.99999994 |
| Cd2 | 0.042749 | 0.04488786 | 0.03761422 | 0.00727364 | 0.09901723 | 0.99999994 |
| Chordc1 | 0.3626094 | 0.25869341 | 0.25142069 | 0.00727272 | 0.42365764 | 0.99999994 |
| Gm36266 | 0.01176994 | 0.01783148 | 0.01057275 | 0.00725873 | 2.44E-06 | 0.00028806 |
| Tmem119 | 0.02703527 | 0.03129082 | 0.02403767 | 0.00725315 | 0.02229537 | 0.72237903 |
| Dnajc9 | 0.29141701 | 0.22026494 | 0.21301319 | 0.00725175 | 0.41036473 | 0.99999994 |

| | | | | | | |
|---|---|---|---|---|---|---|
| Fam98c | 0.17779002 | 0.14896754 | 0.14174406 | 0.00722348 | 0.36722289 | 0.99999994 |
| Edn3 | 0.03045787 | 0.03421578 | 0.027019 | 0.00719678 | 0.03811541 | 0.99999994 |
| Lima1 | 0.14272988 | 0.12415316 | 0.1169626 | 0.00719056 | 0.34119068 | 0.99999994 |
| Tmem98 | 0.16891714 | 0.14278463 | 0.13560529 | 0.00717934 | 0.36226652 | 0.99999994 |
| Snrnp48 | 0.27671282 | 0.21169141 | 0.20451312 | 0.00717829 | 0.40763447 | 0.99999994 |
| Leprot | 0.3009352 | 0.22558033 | 0.2184087 | 0.00717163 | 0.41349368 | 0.99999994 |
| Abcg2 | 0.05115803 | 0.05191705 | 0.04476432 | 0.00715273 | 0.14375592 | 0.99999994 |
| Fermt2 | 0.23995222 | 0.18950215 | 0.18235521 | 0.00714693 | 0.39709311 | 0.99999994 |
| Elavl3 | 0.04302154 | 0.04498375 | 0.0378472 | 0.00713655 | 0.10471462 | 0.99999994 |
| Sf3b2 | 1.11628875 | 0.43704181 | 0.42990866 | 0.00713316 | 0.45602863 | 0.99999994 |
| Spop | 0.53979419 | 0.33689859 | 0.32976639 | 0.0071322 | 0.44276446 | 0.99999994 |
| Wbp4 | 0.58279117 | 0.35272322 | 0.34559662 | 0.00712659 | 0.445412 | 0.99999994 |
| Usp14 | 0.34799963 | 0.25095268 | 0.24389624 | 0.00705644 | 0.42364292 | 0.99999994 |
| Tpi1 | 1.14705382 | 0.4368797 | 0.42982798 | 0.00705171 | 0.45652055 | 0.99999994 |
| Chst1 | 0.02936908 | 0.03312272 | 0.02607207 | 0.00705065 | 0.03592446 | 0.99942728 |
| Cfi | 0.04472492 | 0.04634542 | 0.03930146 | 0.00704396 | 0.11643229 | 0.99999994 |
| Cldn12 | 0.09223354 | 0.08562131 | 0.07858389 | 0.00703742 | 0.27555541 | 0.99999994 |
| Kcnb2 | 0.14302003 | 0.12420562 | 0.11717356 | 0.00703206 | 0.34477161 | 0.99999994 |
| Drap1 | 0.93514962 | 0.42862813 | 0.42161237 | 0.00701576 | 0.45590177 | 0.99999994 |
| Commd10 | 0.25402618 | 0.19799969 | 0.19099506 | 0.00700463 | 0.40356974 | 0.99999994 |
| Rpl3 | 2.72244931 | 0.36804165 | 0.36103822 | 0.00700343 | 0.44863127 | 0.99999994 |
| Nsrp1 | 0.38435878 | 0.26929024 | 0.26229658 | 0.00699366 | 0.42956584 | 0.99999994 |
| Maff | 0.11819037 | 0.10574748 | 0.09875529 | 0.00699219 | 0.31871622 | 0.99999994 |
| Abhd10 | 0.18245426 | 0.15190826 | 0.14493531 | 0.00697294 | 0.37439234 | 0.99999994 |
| Prlr | 0.07602142 | 0.07242265 | 0.06545033 | 0.00697232 | 0.2391339 | 0.99999994 |
| Ghr | 0.33349216 | 0.24321364 | 0.23624598 | 0.00696767 | 0.42218073 | 0.99999994 |
| Rock1 | 0.62066644 | 0.36558963 | 0.35862606 | 0.00696357 | 0.44858035 | 0.99999994 |
| Iffo2 | 0.05537969 | 0.05526998 | 0.04832422 | 0.00694575 | 0.16935089 | 0.99999994 |
| Shf | 0.07683446 | 0.07305201 | 0.06611554 | 0.00693648 | 0.24247969 | 0.99999994 |
| Proser2 | 0.08285243 | 0.07793545 | 0.07101761 | 0.00691784 | 0.2583641 | 0.99999994 |

| | | | | | | |
|---|---|---|---|---|---|---|
| Arl2bp | 0.38319868 | 0.26862717 | 0.2617261 | 0.00690107 | 0.43033875 | 0.99999994 |
| Afm | 0.03693115 | 0.03950729 | 0.03262066 | 0.00688663 | 0.07997478 | 0.99999994 |
| Rbm7 | 0.37746767 | 0.26577659 | 0.2588924 | 0.00688419 | 0.42975469 | 0.99999994 |
| Ubald1 | 0.32302854 | 0.23748561 | 0.2306153 | 0.00687031 | 0.42140474 | 0.99999994 |
| Rnf149 | 0.18746774 | 0.15520888 | 0.14833896 | 0.00686992 | 0.37893793 | 0.99999994 |
| Sds | 0.01549674 | 0.02070901 | 0.01388555 | 0.00682347 | 0.00053587 | 0.04107652 |
| Flrt2 | 0.02978482 | 0.033253 | 0.02643381 | 0.00681919 | 0.04297528 | 0.99999994 |
| Prmt6 | 0.13428978 | 0.11758369 | 0.11078249 | 0.00680121 | 0.34139828 | 0.99999994 |
| Tro | 0.05035775 | 0.05088428 | 0.04408726 | 0.00679702 | 0.15239319 | 0.99999994 |
| Sh3bp5 | 0.05436304 | 0.05423756 | 0.04746875 | 0.00676882 | 0.1712684 | 0.99999994 |
| Cela3b | 0.28070435 | 0.21360441 | 0.20684076 | 0.00676364 | 0.41384652 | 0.99999994 |
| Atg101 | 0.36384299 | 0.25880984 | 0.25204777 | 0.00676207 | 0.42913369 | 0.99999994 |
| B9d2 | 0.30265778 | 0.22613461 | 0.21937608 | 0.00675853 | 0.41875916 | 0.99999994 |
| Lman2l | 0.28141898 | 0.21400555 | 0.20725594 | 0.00674961 | 0.41419206 | 0.99999994 |
| Xlr3b | 0.05207231 | 0.05228382 | 0.04553697 | 0.00674686 | 0.16201248 | 0.99999994 |
| Thoc3 | 0.54677168 | 0.3391467 | 0.33241217 | 0.00673453 | 0.44636329 | 0.99999994 |
| Smn1 | 0.32071947 | 0.23609431 | 0.22935987 | 0.00673444 | 0.42252295 | 0.99999994 |
| Myl7 | 0.02314252 | 0.02735803 | 0.0206302 | 0.00672783 | 0.01497539 | 0.56435639 |
| Pdcd7 | 0.0805858 | 0.07589229 | 0.06917575 | 0.00671654 | 0.25904528 | 0.99999994 |
| Ccnb2 | 0.02306935 | 0.02727711 | 0.02056598 | 0.00671113 | 0.01492352 | 0.56373427 |
| Usf2 | 0.50844847 | 0.32420537 | 0.31750159 | 0.00670378 | 0.44411529 | 0.99999994 |
| Luc7l | 0.46390262 | 0.30564868 | 0.29895492 | 0.00669376 | 0.44076128 | 0.99999994 |
| Fat1 | 0.18520979 | 0.15346662 | 0.14680943 | 0.00665719 | 0.38138786 | 0.99999994 |
| Ankrd2 | 0.03268123 | 0.03559406 | 0.02894843 | 0.00664563 | 0.06324553 | 0.99999994 |
| Tmem114 | 0.01680543 | 0.0216904 | 0.01504492 | 0.00664548 | 0.00164013 | 0.11031784 |
| Tmem200a | 0.03682128 | 0.03916704 | 0.03252598 | 0.00664106 | 0.08706124 | 0.99999994 |
| Lfng | 0.04887828 | 0.04947046 | 0.0428337 | 0.00663676 | 0.15118767 | 0.99999994 |
| Atp6v1d | 0.6819892 | 0.3841091 | 0.37750361 | 0.00660548 | 0.45363938 | 0.99999994 |
| Dner | 0.0675814 | 0.06509812 | 0.05850336 | 0.00659476 | 0.22652658 | 0.99999994 |
| Pmvk | 0.39552485 | 0.27431869 | 0.26773312 | 0.00658557 | 0.43497187 | 0.99999994 |

| | | | | | | |
|---|---|---|---|---|---|---|
| Pgm2l1 | 0.06837362 | 0.06572444 | 0.05915868 | 0.00656576 | 0.23002648 | 0.99999994 |
| Cryzl1 | 0.44838177 | 0.29873406 | 0.29216963 | 0.00656444 | 0.44055812 | 0.99999994 |
| Bhlhb9 | 0.3393238 | 0.2458955 | 0.23934269 | 0.00655282 | 0.42769708 | 0.99999994 |
| Spcs1 | 1.84771521 | 0.38346702 | 0.37692304 | 0.00654399 | 0.45399865 | 0.99999994 |
| Lbh | 0.13536111 | 0.11811075 | 0.11157164 | 0.00653911 | 0.34822279 | 0.99999994 |
| Nab2 | 0.02925748 | 0.03250928 | 0.02597493 | 0.00653435 | 0.04701634 | 0.99999994 |
| Hspb11 | 0.38069736 | 0.2669735 | 0.26049249 | 0.00648101 | 0.43423227 | 0.99999994 |
| Fbln1 | 0.05780423 | 0.05682549 | 0.05035982 | 0.00646567 | 0.19638469 | 0.99999994 |
| Chac2 | 0.13101207 | 0.11481954 | 0.1083596 | 0.00645994 | 0.34574857 | 0.99999994 |
| Onecut2 | 0.06413836 | 0.06210476 | 0.05564745 | 0.00645731 | 0.21993828 | 0.99999994 |
| Wrap73 | 0.21502116 | 0.1729845 | 0.16655774 | 0.00642677 | 0.39865077 | 0.99999994 |
| F2r | 0.0687167 | 0.06586772 | 0.05944226 | 0.00642546 | 0.23590899 | 0.99999994 |
| Wac | 0.39607586 | 0.27441527 | 0.26799881 | 0.00641646 | 0.43668972 | 0.99999994 |
| Mplkip | 0.33811177 | 0.24510823 | 0.23870144 | 0.00640679 | 0.42910314 | 0.99999994 |
| Gm14964 | 0.06362839 | 0.06162777 | 0.05522335 | 0.00640442 | 0.22006947 | 0.99999994 |
| Grin3a | 0.03119839 | 0.03406385 | 0.02766226 | 0.00640159 | 0.06173001 | 0.99999994 |
| Dlgap4 | 0.31468616 | 0.23243117 | 0.2260578 | 0.00637337 | 0.42556917 | 0.99999994 |
| Sugt1 | 0.60323388 | 0.35910485 | 0.35273519 | 0.00636965 | 0.45216245 | 0.99999994 |
| Mnx1 | 0.07771095 | 0.07319698 | 0.06683187 | 0.00636511 | 0.26305492 | 0.99999994 |
| Prdx4 | 0.38756521 | 0.27018039 | 0.2638679 | 0.00631248 | 0.43674016 | 0.99999994 |
| Emid1 | 0.14660544 | 0.12607831 | 0.11977219 | 0.00630612 | 0.3629953 | 0.99999994 |
| Hint2 | 0.57944979 | 0.35070581 | 0.3444067 | 0.00629912 | 0.45155121 | 0.99999994 |
| Ube2e2 | 0.15061538 | 0.12894877 | 0.12266018 | 0.0062886 | 0.36645353 | 0.99999994 |
| Atad2b | 0.12809964 | 0.11248232 | 0.10619552 | 0.0062868 | 0.34676902 | 0.99999994 |
| Mrpl37 | 0.23961938 | 0.18843203 | 0.18214853 | 0.0062835 | 0.40919289 | 0.99999994 |
| Arhgef2 | 0.17576811 | 0.14661468 | 0.14035307 | 0.00626161 | 0.38324775 | 0.99999994 |
| Rpl15 | 1.53433832 | 0.41172123 | 0.40546959 | 0.00625164 | 0.45912849 | 0.99999994 |
| Zfp36l1 | 0.05756799 | 0.0564093 | 0.05016176 | 0.00624754 | 0.20354215 | 0.99999994 |
| Naf1 | 0.08054512 | 0.07536658 | 0.06914264 | 0.00622393 | 0.27452716 | 0.99999994 |
| Mast4 | 0.31798038 | 0.23408787 | 0.22786467 | 0.0062232 | 0.42787323 | 0.99999994 |

| | | | | | | |
|---|---|---|---|---|---|---|
| Bod1 | 0.11299155 | 0.10102063 | 0.09480339 | 0.00621725 | 0.33122731 | 0.99999994 |
| Phf14 | 0.30796626 | 0.22855681 | 0.22234068 | 0.00621614 | 0.42618453 | 0.99999994 |
| Glmn | 0.13148238 | 0.11489716 | 0.10870804 | 0.00618912 | 0.3523553 | 0.99999994 |
| Zfp36 | 0.16012628 | 0.13562434 | 0.12943575 | 0.00618859 | 0.37514597 | 0.99999994 |
| Ada | 0.03072503 | 0.03343706 | 0.02725115 | 0.00618591 | 0.06539829 | 0.99999994 |
| Hspa2 | 0.15477585 | 0.13182091 | 0.12563689 | 0.00618403 | 0.37159367 | 0.99999994 |
| Idnk | 0.37051138 | 0.26159204 | 0.25541644 | 0.0061756 | 0.43606986 | 0.99999994 |
| Ripply3 | 0.1454261 | 0.1250878 | 0.11891916 | 0.00616864 | 0.36494114 | 0.99999994 |
| Homer3 | 0.07984312 | 0.07473652 | 0.06857107 | 0.00616545 | 0.27475488 | 0.99999994 |
| Rbbp7 | 0.74873914 | 0.3994792 | 0.39332376 | 0.00615544 | 0.45851694 | 0.99999994 |
| Srrm2 | 1.32983184 | 0.42945562 | 0.4233106 | 0.00614502 | 0.46151113 | 0.99999994 |
| Gm5900 | 0.46182436 | 0.30419966 | 0.29805623 | 0.00614343 | 0.44543691 | 0.99999994 |
| Efcc1 | 0.04827291 | 0.04844223 | 0.04232006 | 0.00612216 | 0.16779099 | 0.99999994 |
| Nme1 | 1.77307338 | 0.38849282 | 0.38239923 | 0.00609359 | 0.45776338 | 0.99999994 |
| Ubn1 | 0.20575748 | 0.16661515 | 0.16052344 | 0.00609171 | 0.40028769 | 0.99999994 |
| Khdrbs1 | 0.56491404 | 0.34523913 | 0.33915087 | 0.00608826 | 0.452443 | 0.99999994 |
| Desi2 | 0.29265363 | 0.21979681 | 0.21371886 | 0.00607795 | 0.42492882 | 0.99999994 |
| Stxbp2 | 0.37066893 | 0.26156207 | 0.2554956 | 0.00606648 | 0.43720939 | 0.99999994 |
| Ifi204 | 0.01870144 | 0.02277467 | 0.01672098 | 0.00605369 | 0.0079789 | 0.36342269 |
| Dyrk1b | 0.09137499 | 0.08394375 | 0.07789521 | 0.00604854 | 0.30262543 | 0.99999994 |
| Pdgfa | 0.17913268 | 0.14870745 | 0.14266515 | 0.0060423 | 0.38900363 | 0.99999994 |
| Sbds | 0.36681959 | 0.2595979 | 0.25355586 | 0.00604203 | 0.43698582 | 0.99999994 |
| Dll1 | 0.05478497 | 0.05386413 | 0.04782393 | 0.0060402 | 0.20025718 | 0.99999994 |
| Echdc2 | 0.24198228 | 0.18965303 | 0.18361348 | 0.00603955 | 0.41334607 | 0.99999994 |
| Dexi | 0.05845739 | 0.05691607 | 0.05090709 | 0.00600898 | 0.21602048 | 0.99999994 |
| Mapk15 | 0.36784569 | 0.26005943 | 0.25407409 | 0.00598534 | 0.43769857 | 0.99999994 |
| Ctsc | 0.05207548 | 0.05152114 | 0.04553964 | 0.0059815 | 0.19097946 | 0.99999994 |
| Caprin1 | 0.52907732 | 0.33162047 | 0.3256438 | 0.00597667 | 0.45138347 | 0.99999994 |
| Mrpl45 | 0.18708249 | 0.154028 | 0.14807839 | 0.00594961 | 0.39456344 | 0.99999994 |
| Timm50 | 0.31692734 | 0.23323583 | 0.22728811 | 0.00594772 | 0.43086006 | 0.99999994 |

| | | | | | | |
|---|---|---|---|---|---|---|
| Plppr3 | 0.05382417 | 0.05293574 | 0.04701483 | 0.00592091 | 0.2009361 | 0.99999994 |
| Clk1 | 0.52398722 | 0.32958086 | 0.32366011 | 0.00592074 | 0.45154244 | 0.99999994 |
| Xab2 | 0.17663133 | 0.1468659 | 0.14094757 | 0.00591833 | 0.38993136 | 0.99999994 |
| Safb | 0.4504383 | 0.29899311 | 0.29307871 | 0.00591441 | 0.44657198 | 0.99999994 |
| Egfl7 | 0.19296487 | 0.15795245 | 0.15203939 | 0.00591306 | 0.39786518 | 0.99999994 |
| Avp | 0.01722031 | 0.02132042 | 0.01541204 | 0.00590839 | 0.00535851 | 0.26777407 |
| Akr1c19 | 0.13846268 | 0.11975276 | 0.11384857 | 0.0059042 | 0.36497226 | 0.99999994 |
| 1600014C10I | 0.10224713 | 0.09242846 | 0.0865286 | 0.00589986 | 0.32496606 | 0.99999994 |
| Pcdh9 | 0.02140961 | 0.02499746 | 0.01910759 | 0.00588987 | 0.02009431 | 0.67721644 |
| Ndc1 | 0.05550923 | 0.05431404 | 0.04843315 | 0.00588089 | 0.20947808 | 0.99999994 |
| Spry2 | 0.03260174 | 0.03474846 | 0.02887955 | 0.00586891 | 0.08807448 | 0.99999994 |
| Msrb2 | 0.08930641 | 0.08209034 | 0.0762328 | 0.00585754 | 0.3045152 | 0.99999994 |
| Bex4 | 0.55429626 | 0.34105522 | 0.33523116 | 0.00582406 | 0.45396788 | 0.99999994 |
| D030056L22I | 0.20410854 | 0.16525873 | 0.1594396 | 0.00581914 | 0.40402592 | 0.99999994 |
| Zfp706 | 1.02163989 | 0.4333113 | 0.42749864 | 0.00581266 | 0.46394261 | 0.99999994 |
| Ift74 | 0.27399404 | 0.20871916 | 0.2029192 | 0.00579996 | 0.4245542 | 0.99999994 |
| Dusp16 | 0.05701821 | 0.05548423 | 0.0497006 | 0.00578363 | 0.21928894 | 0.99999994 |
| Necap2 | 0.1750292 | 0.14562684 | 0.13984351 | 0.00578333 | 0.39155114 | 0.99999994 |
| 1700049E15I | 0.03537229 | 0.03705717 | 0.0312761 | 0.00578107 | 0.10928111 | 0.99999994 |
| Lmnb1 | 0.15612227 | 0.13236378 | 0.12659595 | 0.00576783 | 0.38084204 | 0.99999994 |
| Arid5b | 0.11417664 | 0.10146452 | 0.09570726 | 0.00575726 | 0.34442644 | 0.99999994 |
| Paip1 | 0.21980439 | 0.1753923 | 0.1696385 | 0.0057538 | 0.41068994 | 0.99999994 |
| Scn3a | 0.03820248 | 0.03944687 | 0.03371515 | 0.00573172 | 0.12889943 | 0.99999994 |
| Dynlt1f | 0.52003464 | 0.32782701 | 0.32210858 | 0.00571843 | 0.45296604 | 0.99999994 |
| Tbc1d9 | 0.06100136 | 0.05873623 | 0.0530342 | 0.00570203 | 0.23709893 | 0.99999994 |
| Ush1c | 0.21816222 | 0.17428148 | 0.16858347 | 0.00569801 | 0.41099725 | 0.99999994 |
| Acadl | 0.22630776 | 0.17948174 | 0.17378864 | 0.0056931 | 0.41369355 | 0.99999994 |
| Kcne1l | 0.02927669 | 0.03165513 | 0.02599165 | 0.00566348 | 0.07347429 | 0.99999994 |
| Tmem192 | 0.18700868 | 0.15366613 | 0.14802844 | 0.00563768 | 0.3999377 | 0.99999994 |
| Zfp768 | 0.13170788 | 0.1145038 | 0.10887502 | 0.00562879 | 0.36537405 | 0.99999994 |

| | | | | | | |
|---|---|---|---|---|---|---|
| Mthfd2l | 0.19810311 | 0.16109534 | 0.15546798 | 0.00562736 | 0.40480262 | 0.99999994 |
| Spats2 | 0.08760411 | 0.08048171 | 0.07486144 | 0.00562027 | 0.30863242 | 0.99999994 |
| Ece2 | 0.13439176 | 0.11647721 | 0.11085767 | 0.00561954 | 0.36789992 | 0.99999994 |
| Nefl | 0.02027246 | 0.02371559 | 0.01810651 | 0.00560907 | 0.01960291 | 0.66771373 |
| Cenpa | 0.04567663 | 0.04571499 | 0.04011256 | 0.00560243 | 0.1762686 | 0.99999994 |
| Calml4 | 0.09467748 | 0.08612905 | 0.08054013 | 0.00558891 | 0.32207511 | 0.99999994 |
| Trp53cor1 | 0.03327715 | 0.03504573 | 0.0294646 | 0.00558113 | 0.10368405 | 0.99999994 |
| Vps72 | 0.56394439 | 0.34437483 | 0.33879589 | 0.00557894 | 0.45635931 | 0.99999994 |
| Maf1 | 0.46985046 | 0.30707464 | 0.30151035 | 0.00556428 | 0.45111647 | 0.99999994 |
| Clk3 | 0.29276243 | 0.21934489 | 0.21378088 | 0.00556401 | 0.43123017 | 0.99999994 |
| Gpx7 | 0.09031272 | 0.08259542 | 0.07704207 | 0.00555335 | 0.31568339 | 0.99999994 |
| Sac3d1 | 0.32618523 | 0.23787711 | 0.23232414 | 0.00555297 | 0.43679538 | 0.99999994 |
| Cnot7 | 0.77222091 | 0.40373974 | 0.398195 | 0.00554474 | 0.46307587 | 0.99999994 |
| Cnksr3 | 0.13919708 | 0.11992865 | 0.11438603 | 0.00554262 | 0.37352328 | 0.99999994 |
| Abca1 | 0.09869611 | 0.08928097 | 0.08374357 | 0.0055374 | 0.32991771 | 0.99999994 |
| Nt5c | 0.4128752 | 0.28152386 | 0.27598704 | 0.00553682 | 0.44688351 | 0.99999994 |
| Gemin6 | 0.23004009 | 0.18168281 | 0.17615068 | 0.00553214 | 0.41720554 | 0.99999994 |
| Zbed3 | 0.11640735 | 0.10292141 | 0.09740378 | 0.00551762 | 0.3530638 | 0.99999994 |
| Tgif1 | 0.05225198 | 0.05120422 | 0.04568869 | 0.00551553 | 0.21082716 | 0.99999994 |
| Efna3 | 0.04269035 | 0.04307912 | 0.03756407 | 0.00551505 | 0.1642184 | 0.99999994 |
| Tgif2 | 0.05128996 | 0.05037586 | 0.04487587 | 0.00549999 | 0.2073062 | 0.99999994 |
| Kcnh2 | 0.2125177 | 0.17040843 | 0.16493589 | 0.00547253 | 0.41260216 | 0.99999994 |
| Mllt11 | 0.33338465 | 0.24165787 | 0.23618862 | 0.00546925 | 0.43875108 | 0.99999994 |
| Stard5 | 0.17709989 | 0.14671954 | 0.14126983 | 0.0054497 | 0.39867456 | 0.99999994 |
| Lrrc8a | 0.11163034 | 0.09919694 | 0.09376304 | 0.0054339 | 0.34983736 | 0.99999994 |
| Anpep | 0.05968692 | 0.05736434 | 0.05193603 | 0.00542831 | 0.24330318 | 0.99999994 |
| Gm10263 | 0.09077038 | 0.0828352 | 0.07740977 | 0.00542543 | 0.3204199 | 0.99999994 |
| Rpgr | 0.10387304 | 0.09321375 | 0.0877901 | 0.00542365 | 0.34045273 | 0.99999994 |
| Pcgf6 | 0.16137296 | 0.13573728 | 0.13031619 | 0.00542109 | 0.39092787 | 0.99999994 |
| Naa20 | 0.69521553 | 0.38637096 | 0.38095128 | 0.00541968 | 0.46227733 | 0.99999994 |

| | | | | | | |
|---|---|---|---|---|---|---|
| Fgfrl1 | 0.0470504 | 0.04670097 | 0.04128156 | 0.00541942 | 0.19110193 | 0.99999994 |
| Afp | 0.02480717 | 0.02750616 | 0.0220895 | 0.00541666 | 0.05131524 | 0.99999994 |
| Ak8 | 0.08488794 | 0.07807377 | 0.0726671 | 0.00540667 | 0.31021008 | 0.99999994 |
| Pip5k1a | 0.24409844 | 0.19032298 | 0.18492078 | 0.0054022 | 0.42290937 | 0.99999994 |
| Dcx | 0.18050755 | 0.14900442 | 0.14360629 | 0.00539813 | 0.40121211 | 0.99999994 |
| Hba-x | 0.03127004 | 0.03312075 | 0.02772447 | 0.00539628 | 0.09755815 | 0.99999994 |
| 1110004F10I | 1.20132988 | 0.43435643 | 0.42896599 | 0.00539043 | 0.46666953 | 0.99999994 |
| Rbbp9 | 0.10160172 | 0.09140984 | 0.08602692 | 0.00538291 | 0.33852047 | 0.99999994 |
| Mettl6 | 0.19775431 | 0.1606056 | 0.15523613 | 0.00536947 | 0.40895371 | 0.99999994 |
| Lmo1 | 0.0297105 | 0.03173044 | 0.02636915 | 0.00536128 | 0.08797234 | 0.99999994 |
| Bloc1s5 | 0.12296318 | 0.1077146 | 0.10235381 | 0.00536079 | 0.36368461 | 0.99999994 |
| Cct3 | 1.14649504 | 0.43519273 | 0.42983373 | 0.005359 | 0.46693018 | 0.99999994 |
| Rab13 | 0.09681548 | 0.08759745 | 0.08224648 | 0.00535097 | 0.33248415 | 0.99999994 |
| Gm5611 | 0.03266153 | 0.03427745 | 0.02893136 | 0.00534609 | 0.10934914 | 0.99999994 |
| Rfc4 | 0.1123884 | 0.09967281 | 0.09434269 | 0.00533011 | 0.35343372 | 0.99999994 |
| Ccnl2 | 0.58083482 | 0.35020623 | 0.34490071 | 0.00530552 | 0.45922234 | 0.99999994 |
| Rnf7 | 0.75703524 | 0.4003785 | 0.39508748 | 0.00529101 | 0.46448455 | 0.99999994 |
| Ramp1 | 0.12313578 | 0.10776935 | 0.10248343 | 0.00528593 | 0.36567738 | 0.99999994 |
| Setbp1 | 0.10303308 | 0.09242347 | 0.08713882 | 0.00528465 | 0.3432226 | 0.99999994 |
| Rbpj | 0.16669451 | 0.13933703 | 0.13405371 | 0.00528333 | 0.39652948 | 0.99999994 |
| Lage3 | 0.46367346 | 0.30413446 | 0.29885597 | 0.00527849 | 0.45320548 | 0.99999994 |
| Tppp3 | 0.07725632 | 0.0717328 | 0.06646041 | 0.00527239 | 0.29872966 | 0.99999994 |
| Rit1 | 0.27792961 | 0.21049596 | 0.20522425 | 0.00527171 | 0.43211752 | 0.99999994 |
| Gpx8 | 0.0423771 | 0.04256634 | 0.03729616 | 0.00527017 | 0.17345912 | 0.99999994 |
| Atp6v1a | 0.27866019 | 0.2109102 | 0.20565057 | 0.00525963 | 0.43241058 | 0.99999994 |
| Thoc6 | 0.22960463 | 0.18111946 | 0.17587582 | 0.00524364 | 0.42134424 | 0.99999994 |
| Syf2 | 0.62098137 | 0.36397135 | 0.35873085 | 0.0052405 | 0.46126831 | 0.99999994 |
| Mxi1 | 0.10022621 | 0.09019389 | 0.08495596 | 0.00523793 | 0.34075799 | 0.99999994 |
| Nme5 | 0.13106596 | 0.11363075 | 0.10839953 | 0.00523121 | 0.37401928 | 0.99999994 |
| Dnajb6 | 1.20658116 | 0.43405811 | 0.42883088 | 0.00522723 | 0.46766628 | 0.99999994 |

| | | | | | | |
|---|---|---|---|---|---|---|
| Yes1 | 0.1593122 | 0.1340607 | 0.12885985 | 0.00520085 | 0.39409793 | 0.99999994 |
| Alad | 0.44145779 | 0.29428441 | 0.289087 | 0.00519741 | 0.45237124 | 0.99999994 |
| Ppil2 | 0.44398394 | 0.29540628 | 0.2902156 | 0.00519069 | 0.45261665 | 0.99999994 |
| Ankrd11 | 0.31390031 | 0.23080061 | 0.22562519 | 0.00517542 | 0.43932353 | 0.99999994 |
| Rpl27-ps3 | 0.5064173 | 0.32185527 | 0.316685 | 0.00517028 | 0.45673077 | 0.99999994 |
| Vil1 | 0.07146043 | 0.0668741 | 0.06170563 | 0.00516847 | 0.28858031 | 0.99999994 |
| Mcm7 | 0.14864808 | 0.12641157 | 0.12124565 | 0.00516593 | 0.38835533 | 0.99999994 |
| Polb | 0.38515583 | 0.26784978 | 0.26268792 | 0.00516185 | 0.44796741 | 0.99999994 |
| Ptgds | 0.05291353 | 0.05140162 | 0.04624705 | 0.00515458 | 0.22907375 | 0.99999994 |
| Gal | 0.04931805 | 0.04835914 | 0.04320658 | 0.00515256 | 0.21365658 | 0.99999994 |
| Tipin | 0.20409411 | 0.16454969 | 0.1594301 | 0.00511959 | 0.41537188 | 0.99999994 |
| Notum | 0.02886718 | 0.03075161 | 0.02563509 | 0.00511651 | 0.0919993 | 0.99999994 |
| Mkrn1 | 0.57243289 | 0.34700045 | 0.34188482 | 0.00511564 | 0.46033119 | 0.99999994 |
| Mcm5 | 0.05123888 | 0.0499455 | 0.04483269 | 0.00511281 | 0.22389488 | 0.99999994 |
| Capn2 | 0.20167718 | 0.16294879 | 0.15783625 | 0.00511254 | 0.41464632 | 0.99999994 |
| Rbm26 | 0.176279 | 0.14579903 | 0.14070504 | 0.005094 | 0.40478431 | 0.99999994 |
| Ctxn2 | 0.03428213 | 0.035428 | 0.03033416 | 0.00509385 | 0.1318337 | 0.99999994 |
| Prmt5 | 0.19611504 | 0.15923656 | 0.15414475 | 0.00509181 | 0.41298348 | 0.99999994 |
| Irf2bp1 | 0.28919703 | 0.21683413 | 0.21174288 | 0.00509125 | 0.43642131 | 0.99999994 |
| Ndufb8 | 1.64166515 | 0.39967721 | 0.39459088 | 0.00508633 | 0.46581218 | 0.99999994 |
| Ptprk | 0.06834409 | 0.06421193 | 0.05913427 | 0.00507767 | 0.28381007 | 0.99999994 |
| Ostf1 | 0.41188811 | 0.28059449 | 0.27552371 | 0.00507078 | 0.4512497 | 0.99999994 |
| Pmpca | 0.30163595 | 0.22385755 | 0.21880255 | 0.00505499 | 0.4388909 | 0.99999994 |
| 9130230L23F | 0.02529078 | 0.02756498 | 0.02251284 | 0.00505213 | 0.06761902 | 0.99999994 |
| Slc25a17 | 0.45425022 | 0.29979829 | 0.29475576 | 0.00504253 | 0.45466899 | 0.99999994 |
| Sf3a2 | 0.32807155 | 0.23837265 | 0.2333412 | 0.00503146 | 0.44293631 | 0.99999994 |
| Nptx2 | 0.03883599 | 0.03928215 | 0.03425986 | 0.00502229 | 0.16458596 | 0.99999994 |
| Smox | 0.1198693 | 0.10504272 | 0.10002432 | 0.0050184 | 0.36920432 | 0.99999994 |
| Hebp2 | 0.06944086 | 0.06505014 | 0.06004043 | 0.00500971 | 0.28931061 | 0.99999994 |
| Tmx1 | 0.76305684 | 0.40135031 | 0.39634198 | 0.00500832 | 0.46648362 | 0.99999994 |

| | | | | | | |
|---|---|---|---|---|---|---|
| Strbp | 0.41592223 | 0.28241753 | 0.27741269 | 0.00500484 | 0.45220665 | 0.99999994 |
| Stk3 | 0.11385313 | 0.10045622 | 0.0954607 | 0.00499552 | 0.36379555 | 0.99999994 |
| Wif1 | 0.02107719 | 0.02380528 | 0.0188151 | 0.00499017 | 0.03874742 | 0.99999994 |
| Slc39a1 | 0.10063233 | 0.09025334 | 0.08527241 | 0.00498093 | 0.34870808 | 0.99999994 |
| Asap2 | 0.0926981 | 0.08393038 | 0.07895621 | 0.00497417 | 0.33748221 | 0.99999994 |
| Triobp | 0.27666768 | 0.20945496 | 0.20448671 | 0.00496824 | 0.43576162 | 0.99999994 |
| Sccpdh | 0.28696985 | 0.21542773 | 0.21046393 | 0.0049638 | 0.43762618 | 0.99999994 |
| Rnf167 | 0.36253103 | 0.25633823 | 0.25138081 | 0.00495742 | 0.44778144 | 0.99999994 |
| Irf8 | 0.01699574 | 0.02016472 | 0.01521335 | 0.00495137 | 0.01514059 | 0.56630796 |
| Tmem159 | 0.18621879 | 0.1524433 | 0.14749361 | 0.00494969 | 0.41162048 | 0.99999994 |
| Ppp4r2 | 0.21535079 | 0.17169455 | 0.1667708 | 0.00492375 | 0.42210058 | 0.99999994 |
| Bag2 | 0.14001251 | 0.11989985 | 0.11498206 | 0.00491779 | 0.38793965 | 0.99999994 |
| Pinlyp | 0.05234692 | 0.05068651 | 0.04576885 | 0.00491766 | 0.237244 | 0.99999994 |
| Epc2 | 0.14443738 | 0.1231143 | 0.11820267 | 0.00491164 | 0.39104766 | 0.99999994 |
| Egr2 | 0.05952671 | 0.05671175 | 0.05180206 | 0.00490969 | 0.26406096 | 0.99999994 |
| Alg13 | 0.04057615 | 0.04065334 | 0.03575372 | 0.00489961 | 0.18083878 | 0.99999994 |
| Col13a1 | 0.02233197 | 0.02481477 | 0.01991845 | 0.00489632 | 0.05089309 | 0.99999994 |
| 3010003L21F | 0.09063953 | 0.08219709 | 0.07730466 | 0.00489243 | 0.33678086 | 0.99999994 |
| Map9 | 0.32074675 | 0.23426338 | 0.22937473 | 0.00488865 | 0.44359269 | 0.99999994 |
| Jrkl | 0.06947783 | 0.06495872 | 0.06007095 | 0.00488778 | 0.29404495 | 0.99999994 |
| Ccdc28a | 0.06282953 | 0.05943068 | 0.05455844 | 0.00487224 | 0.27611107 | 0.99999994 |
| Rsrc1 | 0.26092934 | 0.20002577 | 0.19516085 | 0.00486492 | 0.43410678 | 0.99999994 |
| Ctcf | 0.30891877 | 0.22772113 | 0.22286998 | 0.00485114 | 0.44240006 | 0.99999994 |
| Cacna1h | 0.11480256 | 0.10102391 | 0.09618395 | 0.00483995 | 0.36883278 | 0.99999994 |
| Ncs1 | 0.06712927 | 0.06294042 | 0.05812906 | 0.00481136 | 0.29083445 | 0.99999994 |
| A1cf | 0.03629224 | 0.03686531 | 0.03206992 | 0.0047954 | 0.15978991 | 0.99999994 |
| Aldoc | 0.14461555 | 0.1231259 | 0.11833187 | 0.00479403 | 0.39370629 | 0.99999994 |
| Fbxo32 | 0.08097605 | 0.0742799 | 0.06949325 | 0.00478665 | 0.32330191 | 0.99999994 |
| Ube2f | 0.27283892 | 0.20701873 | 0.2022399 | 0.00477883 | 0.43750957 | 0.99999994 |
| Pink1 | 0.3438458 | 0.24649519 | 0.24172432 | 0.00477087 | 0.44773916 | 0.99999994 |

| | | | | | | |
|---|---|---|---|---|---|---|
| Speg | 0.04174002 | 0.04152043 | 0.03675096 | 0.00476947 | 0.19383514 | 0.99999994 |
| Nxph1 | 0.0254051 | 0.02738181 | 0.02261288 | 0.00476892 | 0.08019123 | 0.99999994 |
| Ebpl | 0.24423808 | 0.18976824 | 0.18500689 | 0.00476135 | 0.43199055 | 0.99999994 |
| Pon3 | 0.0626536 | 0.05917256 | 0.05441192 | 0.00476064 | 0.28015453 | 0.99999994 |
| Ppp2r2a | 0.34766743 | 0.24848219 | 0.2437231 | 0.00475909 | 0.44829267 | 0.99999994 |
| Tmem108 | 0.21029178 | 0.16824648 | 0.1634884 | 0.00475808 | 0.4231963 | 0.99999994 |
| Pim3 | 0.22333403 | 0.17665046 | 0.17189659 | 0.00475387 | 0.42697362 | 0.99999994 |
| Rhbdd2 | 0.23127981 | 0.18168406 | 0.17693213 | 0.00475193 | 0.42905829 | 0.99999994 |
| Ppp1r7 | 0.25074042 | 0.19374484 | 0.18899545 | 0.00474939 | 0.43357807 | 0.99999994 |
| Srrm4 | 0.05703646 | 0.05445277 | 0.04971591 | 0.00473686 | 0.26297264 | 0.99999994 |
| Rsrc2 | 1.00709149 | 0.43152782 | 0.42680208 | 0.00472575 | 0.47062367 | 0.99999994 |
| Fgd2 | 0.45635497 | 0.30039987 | 0.2956769 | 0.00472296 | 0.45766241 | 0.99999994 |
| Mrps31 | 0.24339047 | 0.18920024 | 0.18448391 | 0.00471634 | 0.4324382 | 0.99999994 |
| Tubb6 | 0.05454414 | 0.05233157 | 0.04762121 | 0.00471036 | 0.25514056 | 0.99999994 |
| Snai1 | 0.01953475 | 0.02216524 | 0.01745627 | 0.00470897 | 0.03627865 | 0.99999994 |
| Abhd11os | 0.02689092 | 0.02862054 | 0.02391164 | 0.0047089 | 0.09495715 | 0.99999994 |
| Nrip3 | 0.03342972 | 0.03429786 | 0.02959668 | 0.00470118 | 0.14518608 | 0.99999994 |
| Pygl | 0.06140625 | 0.0580606 | 0.05337209 | 0.00468851 | 0.27936382 | 0.99999994 |
| Evi5 | 0.28557646 | 0.21434947 | 0.20966147 | 0.004688 | 0.44084187 | 0.99999994 |
| Med10 | 0.62049677 | 0.36325649 | 0.35856958 | 0.00468691 | 0.46533331 | 0.99999994 |
| Rab3c | 0.04488166 | 0.04411736 | 0.03943511 | 0.00468226 | 0.21466739 | 0.99999994 |
| Mphosph8 | 0.47890682 | 0.31003343 | 0.305353 | 0.00468043 | 0.45936715 | 0.99999994 |
| Pkn2 | 0.4349325 | 0.29082159 | 0.28614985 | 0.00467174 | 0.45673093 | 0.99999994 |
| Slc25a13 | 0.05333739 | 0.05127313 | 0.04660453 | 0.0046686 | 0.25245022 | 0.99999994 |
| Fxn | 0.22663959 | 0.1786626 | 0.17399922 | 0.00466338 | 0.42920511 | 0.99999994 |
| Svbp | 0.15805503 | 0.13263099 | 0.12796902 | 0.00466196 | 0.40419877 | 0.99999994 |
| Utp3 | 0.43785577 | 0.29212998 | 0.28746989 | 0.00466009 | 0.45703581 | 0.99999994 |
| Gsta4 | 0.05476261 | 0.05246422 | 0.04780511 | 0.00465911 | 0.25825579 | 0.99999994 |
| Gm27033 | 0.17531525 | 0.14469238 | 0.14004084 | 0.00465154 | 0.41250935 | 0.99999994 |
| Elmod1 | 0.04897039 | 0.04756258 | 0.04291182 | 0.00465076 | 0.23532772 | 0.99999994 |

| | | | | | | |
|---|---|---|---|---|---|---|
| Lrp1b | 0.01706491 | 0.01991158 | 0.01527454 | 0.00463703 | 0.02165382 | 0.71171853 |
| Cdc42ep4 | 0.09932438 | 0.08887883 | 0.08424291 | 0.00463592 | 0.35706994 | 0.99999994 |
| Cbx6 | 0.3116002 | 0.22898801 | 0.22435574 | 0.00463227 | 0.44534399 | 0.99999994 |
| Mcm3 | 0.0578866 | 0.0550538 | 0.05042886 | 0.00462494 | 0.27077557 | 0.99999994 |
| Gnai1 | 0.16355989 | 0.13645821 | 0.13185603 | 0.00460218 | 0.4081419 | 0.99999994 |
| Pea15a | 0.42071922 | 0.28424381 | 0.27964321 | 0.0046006 | 0.45639967 | 0.99999994 |
| Drc1 | 0.04184367 | 0.04144012 | 0.03683969 | 0.00460043 | 0.20292282 | 0.99999994 |
| Fbrsl1 | 0.12427253 | 0.1079362 | 0.10333616 | 0.00460005 | 0.38349653 | 0.99999994 |
| Dusp6 | 0.06038168 | 0.05711105 | 0.05251672 | 0.00459433 | 0.28017578 | 0.99999994 |
| 1700128E19I | 0.02671364 | 0.02834775 | 0.02375681 | 0.00459094 | 0.09916538 | 0.99999994 |
| Gng2 | 0.08095632 | 0.07406689 | 0.0694772 | 0.00458969 | 0.33006934 | 0.99999994 |
| Galr1 | 0.06700397 | 0.06261124 | 0.05802529 | 0.00458595 | 0.2994176 | 0.99999994 |
| Sftpc | 0.0271729 | 0.0287408 | 0.02415782 | 0.00458297 | 0.1033349 | 0.99999994 |
| Il17re | 0.06606295 | 0.06181393 | 0.05724543 | 0.0045685 | 0.29763527 | 0.99999994 |
| Fmo2 | 0.03858994 | 0.03859778 | 0.03404835 | 0.00454942 | 0.18689417 | 0.99999994 |
| Timp3 | 0.02127151 | 0.02353409 | 0.01898609 | 0.004548 | 0.05541329 | 0.99999994 |
| Zfp503 | 0.02576225 | 0.02746433 | 0.02292531 | 0.00453902 | 0.09376789 | 0.99999994 |
| Pxmp4 | 0.19337026 | 0.15684873 | 0.15231092 | 0.00453781 | 0.42139997 | 0.99999994 |
| Pclo | 0.61420436 | 0.36100059 | 0.35646395 | 0.00453664 | 0.46624439 | 0.99999994 |
| Baiap3 | 0.33962347 | 0.24403683 | 0.23950105 | 0.00453578 | 0.44984176 | 0.99999994 |
| Gm15441 | 0.03585048 | 0.03621807 | 0.03168885 | 0.00452922 | 0.17070619 | 0.99999994 |
| Mpdu1 | 0.30826965 | 0.22703349 | 0.22250936 | 0.00452413 | 0.44617182 | 0.99999994 |
| Rassf3 | 0.10026531 | 0.08950537 | 0.08498643 | 0.00451893 | 0.36169397 | 0.99999994 |
| Adam10 | 0.24534004 | 0.19019566 | 0.18568575 | 0.00450991 | 0.43578338 | 0.99999994 |
| Kcnip3 | 0.03005757 | 0.03116322 | 0.02667102 | 0.0044922 | 0.13111622 | 0.99999994 |
| Mlycd | 0.10882427 | 0.09609459 | 0.09161113 | 0.00448346 | 0.37230197 | 0.99999994 |
| Xrn2 | 0.38832357 | 0.26871895 | 0.26423838 | 0.00448057 | 0.45506727 | 0.99999994 |
| Kat5 | 0.19845711 | 0.16018163 | 0.15570316 | 0.00447847 | 0.42408418 | 0.99999994 |
| Zfpm1 | 0.05241463 | 0.05030356 | 0.04582601 | 0.00447755 | 0.25772434 | 0.99999994 |
| Trp53i13 | 0.29116365 | 0.21733317 | 0.21286844 | 0.00446473 | 0.44448357 | 0.99999994 |

| | | | | | | |
|---|---|---|---|---|---|---|
| Irx1 | 0.01429992 | 0.01728294 | 0.0128235 | 0.00445944 | 0.01031239 | 0.44671149 |
| Fam53b | 0.05682111 | 0.05398404 | 0.04953519 | 0.00444885 | 0.27497988 | 0.99999994 |
| Samd5 | 0.04188178 | 0.04131746 | 0.03687231 | 0.00444515 | 0.21114354 | 0.99999994 |
| Smc2 | 0.07116045 | 0.06589252 | 0.06145857 | 0.00443395 | 0.31553355 | 0.99999994 |
| Mmp11 | 0.03056001 | 0.03153776 | 0.02710776 | 0.00442999 | 0.13834335 | 0.99999994 |
| Habp4 | 0.12880465 | 0.11114932 | 0.10672033 | 0.00442899 | 0.39117994 | 0.99999994 |
| Actn1 | 0.27851728 | 0.20999295 | 0.20556722 | 0.00442573 | 0.44302379 | 0.99999994 |
| Tgs1 | 0.16366682 | 0.13635604 | 0.13193118 | 0.00442486 | 0.41167092 | 0.99999994 |
| Mcm2 | 0.10972995 | 0.09670499 | 0.09230675 | 0.00439825 | 0.37555991 | 0.99999994 |
| Bhmt2 | 0.02591044 | 0.02744843 | 0.02305489 | 0.00439354 | 0.10231114 | 0.99999994 |
| Dhfr | 0.08123526 | 0.07408911 | 0.06970405 | 0.00438505 | 0.3376999 | 0.99999994 |
| Ctr9 | 0.31660158 | 0.23149096 | 0.22710955 | 0.00438141 | 0.44891035 | 0.99999994 |
| Dab1 | 0.07663053 | 0.07032896 | 0.06594875 | 0.00438021 | 0.32920679 | 0.99999994 |
| Prss8 | 0.10318965 | 0.09163878 | 0.08726029 | 0.0043785 | 0.36919019 | 0.99999994 |
| Rnf121 | 0.22427481 | 0.17687275 | 0.17249614 | 0.00437661 | 0.43294296 | 0.99999994 |
| Tmem109 | 0.29294502 | 0.21826152 | 0.21388494 | 0.00437658 | 0.44582985 | 0.99999994 |
| Srpk1 | 0.39866175 | 0.2736145 | 0.26924258 | 0.00437193 | 0.4569641 | 0.99999994 |
| Atp2c2 | 0.04367079 | 0.04276481 | 0.03840189 | 0.00436292 | 0.22475192 | 0.99999994 |
| Cast | 0.03993333 | 0.03955709 | 0.03520229 | 0.00435481 | 0.2051275 | 0.99999994 |
| Pop4 | 0.07260413 | 0.06699613 | 0.06264672 | 0.00434941 | 0.3219925 | 0.99999994 |
| Gcc2 | 0.38814646 | 0.26849181 | 0.26415189 | 0.00433991 | 0.45645791 | 0.99999994 |
| Aars | 0.28837057 | 0.21560866 | 0.21126881 | 0.00433985 | 0.44562063 | 0.99999994 |
| Tfeb | 0.05361374 | 0.05116885 | 0.04683749 | 0.00433136 | 0.26909398 | 0.99999994 |
| Trim13 | 0.10704652 | 0.09457219 | 0.09024272 | 0.00432946 | 0.37473292 | 0.99999994 |
| Vtn | 0.02044336 | 0.02258265 | 0.01825707 | 0.00432558 | 0.05739006 | 0.99999994 |
| Mtpn | 0.5221834 | 0.32727593 | 0.32295326 | 0.00432267 | 0.46450354 | 0.99999994 |
| Soat1 | 0.21067694 | 0.16805816 | 0.16373923 | 0.00431893 | 0.4303143 | 0.99999994 |
| Ddx20 | 0.09824081 | 0.08769946 | 0.08338145 | 0.004318 | 0.36515727 | 0.99999994 |
| Nsa2 | 0.61060293 | 0.35956455 | 0.3552491 | 0.00431544 | 0.46777698 | 0.99999994 |
| Shisa4 | 0.08147712 | 0.07420657 | 0.06990067 | 0.0043059 | 0.34089074 | 0.99999994 |

| | | | | | | |
|---|---|---|---|---|---|---|
| Ctnnal1 | 0.14969296 | 0.12630013 | 0.1219975 | 0.00430263 | 0.40719839 | 0.99999994 |
| Thoc1 | 0.15925404 | 0.13311515 | 0.12881868 | 0.00429647 | 0.41215368 | 0.99999994 |
| Dpp6 | 0.03148109 | 0.03220203 | 0.02790767 | 0.00429436 | 0.15285545 | 0.99999994 |
| Lama5 | 0.07809844 | 0.07143742 | 0.06714831 | 0.00428912 | 0.33535385 | 0.99999994 |
| Prmt2 | 0.36034693 | 0.254546 | 0.25026735 | 0.00427865 | 0.45469848 | 0.99999994 |
| Cdc23 | 0.21577277 | 0.17131639 | 0.16704339 | 0.004273 | 0.4323985 | 0.99999994 |
| Txnrd1 | 0.34767659 | 0.2479815 | 0.24372788 | 0.00425362 | 0.45375931 | 0.99999994 |
| Pik3c2g | 0.02785006 | 0.02899684 | 0.02474865 | 0.00424819 | 0.1266049 | 0.99999994 |
| Rps15a | 2.86657823 | 0.35564415 | 0.35140008 | 0.00424408 | 0.46796237 | 0.99999994 |
| Slc25a35 | 0.04398995 | 0.04291717 | 0.03867438 | 0.00424279 | 0.23262222 | 0.99999994 |
| Fyn | 0.09438469 | 0.08454886 | 0.08030609 | 0.00424277 | 0.36254224 | 0.99999994 |
| Hmgcs2 | 0.01948661 | 0.02165556 | 0.01741381 | 0.00424175 | 0.05246699 | 0.99999994 |
| Rps12-ps3 | 0.46912975 | 0.30544098 | 0.30120217 | 0.00423882 | 0.4626837 | 0.99999994 |
| Cxadr | 0.16537396 | 0.13736725 | 0.13312922 | 0.00423803 | 0.41609323 | 0.99999994 |
| Riiad1 | 0.16341242 | 0.13597904 | 0.13175236 | 0.00422668 | 0.41545316 | 0.99999994 |
| Tarbp2 | 0.20350471 | 0.16325332 | 0.15904198 | 0.00421133 | 0.4300463 | 0.99999994 |
| Dcun1d1 | 0.32236964 | 0.23446383 | 0.23025752 | 0.00420631 | 0.45160899 | 0.99999994 |
| Gm6563 | 0.15072874 | 0.12694569 | 0.12274155 | 0.00420414 | 0.40982569 | 0.99999994 |
| 9930012K11I | 0.0158824 | 0.01842802 | 0.01422742 | 0.0042006 | 0.02469158 | 0.78564569 |
| Pcbd2 | 0.44107016 | 0.29311237 | 0.28891342 | 0.00419895 | 0.46146616 | 0.99999994 |
| Tspan8 | 0.02143085 | 0.02332432 | 0.01912627 | 0.00419805 | 0.07200594 | 0.99999994 |
| Ccnt2 | 0.4084968 | 0.27812252 | 0.27392635 | 0.00419617 | 0.45939183 | 0.99999994 |
| Cmtm8 | 0.06820212 | 0.06320057 | 0.05901687 | 0.0041837 | 0.31851021 | 0.99999994 |
| Gps2 | 0.36879085 | 0.25873206 | 0.25455069 | 0.00418137 | 0.45646618 | 0.99999994 |
| Zmym5 | 0.24948842 | 0.19240886 | 0.18823068 | 0.00417818 | 0.44126918 | 0.99999994 |
| Arid4b | 0.75101239 | 0.39799104 | 0.39381465 | 0.00417639 | 0.47186189 | 0.99999994 |
| Iqcb1 | 0.19881202 | 0.16011171 | 0.1559388 | 0.00417291 | 0.42931288 | 0.99999994 |
| 6430503K07I | 0.07800965 | 0.07124755 | 0.06707581 | 0.00417174 | 0.33944155 | 0.99999994 |
| Slco4a1 | 0.06326495 | 0.05908833 | 0.05492094 | 0.00416739 | 0.30675022 | 0.99999994 |
| Pkn1 | 0.10907118 | 0.09596705 | 0.09180087 | 0.00416618 | 0.38129388 | 0.99999994 |

| | | | | | | |
|---|---|---|---|---|---|---|
| Tsen15 | 0.28039189 | 0.21081088 | 0.20665909 | 0.00415179 | 0.44680937 | 0.99999994 |
| Igbp1 | 0.28755981 | 0.21495227 | 0.21080315 | 0.00414911 | 0.44788244 | 0.99999994 |
| Lrrc10b | 0.04381427 | 0.04266163 | 0.0385244 | 0.00413723 | 0.2373532 | 0.99999994 |
| Gpcpd1 | 0.16025058 | 0.13366052 | 0.12952362 | 0.0041369 | 0.41581959 | 0.99999994 |
| Igf2bp2 | 0.27865951 | 0.20978282 | 0.20565017 | 0.00413265 | 0.44679502 | 0.99999994 |
| Sod3 | 0.02409159 | 0.0255916 | 0.0214626 | 0.004129 | 0.10017437 | 0.99999994 |
| Sf3a3 | 0.37474469 | 0.2616599 | 0.25753657 | 0.00412332 | 0.45756406 | 0.99999994 |
| Dennd1c | 0.09631763 | 0.08596857 | 0.08184956 | 0.00411901 | 0.3688216 | 0.99999994 |
| Polr3gl | 0.18252784 | 0.14910275 | 0.14498547 | 0.00411729 | 0.42503594 | 0.99999994 |
| Gtf2f2 | 0.31880853 | 0.23242913 | 0.22831742 | 0.00411171 | 0.45229206 | 0.99999994 |
| Adamtsl2 | 0.03442249 | 0.0345657 | 0.03045551 | 0.00411019 | 0.18450641 | 0.99999994 |
| Afap1 | 0.122696 | 0.10625694 | 0.1021531 | 0.00410384 | 0.3945773 | 0.99999994 |
| Plbd1 | 0.0217578 | 0.02350148 | 0.01941381 | 0.00408767 | 0.08052839 | 0.99999994 |
| Siah2 | 0.11378223 | 0.09948808 | 0.09540664 | 0.00408144 | 0.38791597 | 0.99999994 |
| Sf3b4 | 0.58447257 | 0.35025371 | 0.34619299 | 0.00406072 | 0.46888354 | 0.99999994 |
| Tmem44 | 0.04094394 | 0.04012482 | 0.03606902 | 0.0040558 | 0.22708617 | 0.99999994 |
| Slc28a3 | 0.04826611 | 0.04636979 | 0.04231429 | 0.00405549 | 0.26174993 | 0.99999994 |
| Syt11 | 0.06591181 | 0.0611681 | 0.05712008 | 0.00404801 | 0.31856115 | 0.99999994 |
| Slc14a2 | 0.04095215 | 0.04011671 | 0.03607606 | 0.00404065 | 0.22797378 | 0.99999994 |
| Rgs19 | 0.12986855 | 0.11154312 | 0.10751116 | 0.00403196 | 0.40143615 | 0.99999994 |
| Kif21a | 0.51087344 | 0.3225017 | 0.31847304 | 0.00402866 | 0.46644775 | 0.99999994 |
| Ctsl | 0.7734891 | 0.40247154 | 0.39844589 | 0.00402565 | 0.47319064 | 0.99999994 |
| Usf1 | 0.24638032 | 0.19034986 | 0.18632552 | 0.00402434 | 0.44284227 | 0.99999994 |
| Capn15 | 0.10560737 | 0.09315357 | 0.08913205 | 0.00402152 | 0.38196433 | 0.99999994 |
| Hells | 0.04143352 | 0.04050714 | 0.03648849 | 0.00401865 | 0.23174994 | 0.99999994 |
| Ppp2r2d | 0.1377742 | 0.1173519 | 0.11334412 | 0.00400778 | 0.4069628 | 0.99999994 |
| Sertad3 | 0.14277323 | 0.12099251 | 0.11699413 | 0.00399839 | 0.41002239 | 0.99999994 |
| Xlr4b | 0.02038613 | 0.02220319 | 0.01820666 | 0.00399653 | 0.07198958 | 0.99999994 |
| Klc1 | 0.2830275 | 0.21218259 | 0.20818871 | 0.00399388 | 0.44919512 | 0.99999994 |
| Sytl5 | 0.07921878 | 0.07205616 | 0.06806229 | 0.00399387 | 0.34804932 | 0.99999994 |

| | | | | | | |
|---|---|---|---|---|---|---|
| Rab7 | 1.03390974 | 0.43208923 | 0.42809683 | 0.0039924 | 0.47525088 | 0.99999994 |
| Marf1 | 0.1472817 | 0.12423341 | 0.12026054 | 0.00397287 | 0.41297603 | 0.99999994 |
| Lactb | 0.12753347 | 0.10974188 | 0.10577363 | 0.00396825 | 0.40140082 | 0.99999994 |
| Kcnk1 | 0.18850646 | 0.15300414 | 0.14904074 | 0.00396341 | 0.42974983 | 0.99999994 |
| Pabpn1l | 0.06891684 | 0.06356952 | 0.05960764 | 0.00396188 | 0.329092 | 0.99999994 |
| Nupr1 | 0.07642918 | 0.06973332 | 0.06578403 | 0.00394929 | 0.34472228 | 0.99999994 |
| Tmem176a | 0.69141607 | 0.38392877 | 0.37997975 | 0.00394902 | 0.47242412 | 0.99999994 |
| Midn | 0.24778487 | 0.19113515 | 0.18718763 | 0.00394752 | 0.44418254 | 0.99999994 |
| Ago1 | 0.15501747 | 0.12974868 | 0.12580915 | 0.00393954 | 0.41744489 | 0.99999994 |
| Apom | 0.01590932 | 0.01817822 | 0.01425128 | 0.00392694 | 0.03331514 | 0.95175039 |
| Cit | 0.05414438 | 0.05120738 | 0.0472846 | 0.00392277 | 0.2903996 | 0.99999994 |
| Sertad1 | 0.13458709 | 0.11490821 | 0.11100163 | 0.00390658 | 0.40739034 | 0.99999994 |
| Plxdc2 | 0.05003882 | 0.04772368 | 0.04381724 | 0.00390645 | 0.27644475 | 0.99999994 |
| Trappc2 | 0.34575657 | 0.24662787 | 0.24272539 | 0.00390248 | 0.45738687 | 0.99999994 |
| Psmc6 | 0.84294998 | 0.41441485 | 0.41051409 | 0.00390076 | 0.47478385 | 0.99999994 |
| Grin2c | 0.07056311 | 0.0648665 | 0.0609663 | 0.0039002 | 0.33511657 | 0.99999994 |
| Zbtb7a | 0.47423101 | 0.30726957 | 0.30337595 | 0.00389362 | 0.46595986 | 0.99999994 |
| Cecr2 | 0.05296713 | 0.05018366 | 0.04629226 | 0.0038914 | 0.28789526 | 0.99999994 |
| Gsdmd | 0.07097961 | 0.06519193 | 0.06130958 | 0.00388235 | 0.33669293 | 0.99999994 |
| Snhg6 | 0.19927198 | 0.16012191 | 0.156244 | 0.00387791 | 0.43439009 | 0.99999994 |
| Bod1l | 0.28782907 | 0.21483449 | 0.21095787 | 0.00387662 | 0.45132313 | 0.99999994 |
| Arg1 | 0.026785 | 0.02769141 | 0.02381914 | 0.00387227 | 0.13959929 | 0.99999994 |
| Slu7 | 0.25072044 | 0.19285335 | 0.18898326 | 0.00387009 | 0.44578727 | 0.99999994 |
| Cnot2 | 0.25948625 | 0.198154 | 0.19429379 | 0.00386022 | 0.44739402 | 0.99999994 |
| Aqp7 | 0.06105341 | 0.05693598 | 0.05307765 | 0.00385833 | 0.31424078 | 0.99999994 |
| Bend5 | 0.08281523 | 0.07484188 | 0.07098743 | 0.00385445 | 0.35889103 | 0.99999994 |
| Smad1 | 0.18510678 | 0.15059369 | 0.14673952 | 0.00385417 | 0.43060588 | 0.99999994 |
| Ptpra | 0.1448791 | 0.12236797 | 0.11852291 | 0.00384506 | 0.41451641 | 0.99999994 |
| Cdx2 | 0.01399685 | 0.01639557 | 0.01255428 | 0.00384129 | 0.02084124 | 0.69426319 |
| Zfp286 | 0.03886987 | 0.03812485 | 0.03428898 | 0.00383588 | 0.22824442 | 0.99999994 |

| | | | | | | |
|---|---|---|---|---|---|---|
| Kmt2a | 0.26363578 | 0.20061201 | 0.19678162 | 0.00383039 | 0.44845447 | 0.99999994 |
| Ccdc58 | 0.31452354 | 0.22978521 | 0.22596836 | 0.00381685 | 0.45523998 | 0.99999994 |
| Ap1g2 | 0.05175225 | 0.04908144 | 0.04526659 | 0.00381485 | 0.28741068 | 0.99999994 |
| Haus3 | 0.05058243 | 0.04808958 | 0.04427743 | 0.00381215 | 0.28329116 | 0.99999994 |
| Cd200 | 0.2281815 | 0.17878579 | 0.17497621 | 0.00380958 | 0.4423862 | 0.99999994 |
| Snap23 | 0.22306757 | 0.17553208 | 0.17172661 | 0.00380547 | 0.44136666 | 0.99999994 |
| Slc25a53 | 0.19190148 | 0.15511332 | 0.15132629 | 0.00378703 | 0.43385056 | 0.99999994 |
| Cdca8 | 0.03760555 | 0.03698515 | 0.03320148 | 0.00378367 | 0.22405633 | 0.99999994 |
| Cep290 | 0.17984861 | 0.14693434 | 0.14315548 | 0.00377886 | 0.43026208 | 0.99999994 |
| Lyrm4 | 0.11797983 | 0.10237317 | 0.0985959 | 0.00377727 | 0.39935805 | 0.99999994 |
| Phlda2 | 0.01077277 | 0.01346064 | 0.00968353 | 0.00377711 | 0.00471105 | 0.24382735 |
| Ssbp3 | 0.34224616 | 0.24465757 | 0.24088378 | 0.00377379 | 0.45847299 | 0.99999994 |
| Tgm5 | 0.02913639 | 0.02963335 | 0.02586952 | 0.00376383 | 0.16640939 | 0.99999994 |
| 1110059E24I | 0.30609942 | 0.22505776 | 0.22130097 | 0.0037568 | 0.45501603 | 0.99999994 |
| Jmy | 0.1777937 | 0.14549155 | 0.14174658 | 0.00374497 | 0.43020116 | 0.99999994 |
| E2f1 | 0.06594263 | 0.06088524 | 0.05714564 | 0.0037396 | 0.33156695 | 0.99999994 |
| Tle1 | 0.15275462 | 0.12792137 | 0.12419324 | 0.00372813 | 0.42081229 | 0.99999994 |
| Zfp358 | 0.27944241 | 0.20983004 | 0.20610647 | 0.00372358 | 0.45214033 | 0.99999994 |
| Gm3764 | 0.01996477 | 0.02155697 | 0.01783538 | 0.00372159 | 0.08242594 | 0.99999994 |
| Serpina10 | 0.05059596 | 0.04800825 | 0.04428887 | 0.00371937 | 0.28808093 | 0.99999994 |
| Shd | 0.03768207 | 0.03696903 | 0.03326735 | 0.00370168 | 0.22945032 | 0.99999994 |
| Wasf1 | 0.07120428 | 0.06518487 | 0.06149467 | 0.0036902 | 0.34478548 | 0.99999994 |
| Nfyb | 0.32050693 | 0.23293349 | 0.22924408 | 0.00368941 | 0.45734446 | 0.99999994 |
| Mapk7 | 0.0960599 | 0.08532629 | 0.08164397 | 0.00368232 | 0.38200628 | 0.99999994 |
| Spag1 | 0.09222356 | 0.08225024 | 0.07857589 | 0.00367435 | 0.37780017 | 0.99999994 |
| Ago2 | 0.19896243 | 0.15971281 | 0.15603862 | 0.00367419 | 0.43772688 | 0.99999994 |
| Hoxb4 | 0.01208253 | 0.01452407 | 0.01085126 | 0.00367282 | 0.01213038 | 0.50170905 |
| Smoc2 | 0.01898398 | 0.02063445 | 0.01697037 | 0.00366408 | 0.07533385 | 0.99999994 |
| Wipi2 | 0.28084213 | 0.21058294 | 0.20692084 | 0.00366209 | 0.4531113 | 0.99999994 |
| Ppp1r12a | 0.26972277 | 0.20405746 | 0.20040113 | 0.00365633 | 0.45166899 | 0.99999994 |

| | | | | | | |
|---|---|---|---|---|---|---|
| Bcl11a | 0.0956256 | 0.08495265 | 0.0812974 | 0.00365524 | 0.38236366 | 0.99999994 |
| Imp4 | 0.29983394 | 0.22144313 | 0.21778885 | 0.00365428 | 0.45553581 | 0.99999994 |
| Hey1 | 0.01944905 | 0.02103476 | 0.01738069 | 0.00365408 | 0.08084389 | 0.99999994 |
| Nt5c2 | 0.42254372 | 0.28414037 | 0.28048712 | 0.00365325 | 0.46545623 | 0.99999994 |
| Igfbp4 | 0.05760327 | 0.05383672 | 0.05019134 | 0.00364538 | 0.31438842 | 0.99999994 |
| 4921524J17F | 0.35574586 | 0.25154923 | 0.2479078 | 0.00364142 | 0.46105639 | 0.99999994 |
| Efs | 0.0425241 | 0.04106318 | 0.0374219 | 0.00364128 | 0.25859379 | 0.99999994 |
| Pspc1 | 0.19397152 | 0.15634875 | 0.15271332 | 0.00363542 | 0.43704788 | 0.99999994 |
| Mef2a | 0.22574561 | 0.17705638 | 0.17343166 | 0.00362472 | 0.44467857 | 0.99999994 |
| Gpr85 | 0.04803118 | 0.04573447 | 0.04211485 | 0.00361962 | 0.2836305 | 0.99999994 |
| Syngr1 | 0.16082024 | 0.13354149 | 0.12992607 | 0.00361542 | 0.4265264 | 0.99999994 |
| Mcm4 | 0.06343613 | 0.05867705 | 0.05506339 | 0.00361365 | 0.33111451 | 0.99999994 |
| Ing3 | 0.11567779 | 0.10044649 | 0.09684966 | 0.00359683 | 0.40237455 | 0.99999994 |
| Gtf2e2 | 0.85536613 | 0.41595123 | 0.4123589 | 0.00359232 | 0.47687916 | 0.99999994 |
| Chd2 | 0.3232809 | 0.23433869 | 0.23075222 | 0.00358646 | 0.45880038 | 0.99999994 |
| Vps37d | 0.08892092 | 0.07950529 | 0.07592252 | 0.00358277 | 0.37671777 | 0.99999994 |
| Fbxo28 | 0.11216637 | 0.09774804 | 0.09417299 | 0.00357505 | 0.40025293 | 0.99999994 |
| Ptp4a1 | 0.40146242 | 0.2741541 | 0.27058391 | 0.00357018 | 0.46500731 | 0.99999994 |
| Hnrnpll | 0.28544993 | 0.21315464 | 0.20958852 | 0.00356612 | 0.45491341 | 0.99999994 |
| Plcl1 | 0.06489065 | 0.05982967 | 0.05627255 | 0.00355712 | 0.3369638 | 0.99999994 |
| Khdrbs2 | 0.01660682 | 0.01841421 | 0.0148691 | 0.00354511 | 0.05625453 | 0.99999994 |
| Derl2 | 0.63735865 | 0.36763366 | 0.36408959 | 0.00354408 | 0.47416921 | 0.99999994 |
| Luzp1 | 0.26643807 | 0.20198777 | 0.19845246 | 0.00353531 | 0.4528044 | 0.99999994 |
| Kif3a | 0.19228661 | 0.15511594 | 0.15158469 | 0.00353124 | 0.4383857 | 0.99999994 |
| Rpl7a-ps5 | 0.08384816 | 0.0753464 | 0.07182505 | 0.00352135 | 0.37208367 | 0.99999994 |
| Rragb | 0.04772448 | 0.04535338 | 0.04185439 | 0.003499 | 0.28894574 | 0.99999994 |
| Uty | 0.05850557 | 0.05444536 | 0.05094745 | 0.00349792 | 0.32383275 | 0.99999994 |
| Actr6 | 0.11785416 | 0.10199646 | 0.09850075 | 0.00349572 | 0.40662749 | 0.99999994 |
| Otop3 | 0.02766205 | 0.02807967 | 0.02458466 | 0.003495 | 0.17200271 | 0.99999994 |
| Zscan29 | 0.0728736 | 0.06636217 | 0.06286825 | 0.00349392 | 0.35571764 | 0.99999994 |

| | | | | | | |
|---|---|---|---|---|---|---|
| Hif1a | 0.24439659 | 0.18859286 | 0.18510461 | 0.00348824 | 0.45008872 | 0.99999994 |
| Bbc3 | 0.090857 | 0.08096583 | 0.07747934 | 0.0034865 | 0.38226797 | 0.99999994 |
| Pbrm1 | 0.70562651 | 0.38704139 | 0.38355596 | 0.00348543 | 0.47588377 | 0.99999994 |
| Tmem41a | 0.08981131 | 0.08012433 | 0.07663897 | 0.00348535 | 0.38105352 | 0.99999994 |
| Coq2 | 0.227394 | 0.17796274 | 0.17447753 | 0.00348521 | 0.44711187 | 0.99999994 |
| Klhl13 | 0.04727783 | 0.04495137 | 0.04147488 | 0.00347649 | 0.28844056 | 0.99999994 |
| Tfpi | 0.05630472 | 0.05257436 | 0.04910163 | 0.00347274 | 0.31890103 | 0.99999994 |
| Pak1 | 0.10125958 | 0.08923242 | 0.08576076 | 0.00347167 | 0.39378981 | 0.99999994 |
| Wwc2 | 0.04181747 | 0.04028724 | 0.03681726 | 0.00346998 | 0.26521362 | 0.99999994 |
| Msi2 | 0.36100863 | 0.25406543 | 0.25060512 | 0.00346031 | 0.46338488 | 0.99999994 |
| Tubb4a | 0.10860018 | 0.09489587 | 0.09143885 | 0.00345702 | 0.40065347 | 0.99999994 |
| Gprc5a | 0.02304791 | 0.02400162 | 0.02054716 | 0.00345446 | 0.1315509 | 0.99999994 |
| Tubgcp4 | 0.10150693 | 0.08940734 | 0.0859532 | 0.00345414 | 0.3945443 | 0.99999994 |
| Smarcd1 | 0.12061007 | 0.10403084 | 0.10058314 | 0.0034477 | 0.40976065 | 0.99999994 |
| Cdc20 | 0.05575603 | 0.0520856 | 0.04864061 | 0.00344498 | 0.31866402 | 0.99999994 |
| Insrr | 0.1349316 | 0.11469746 | 0.11125542 | 0.00344204 | 0.41842063 | 0.99999994 |
| Bcl7c | 0.29700031 | 0.21961346 | 0.21618831 | 0.00342515 | 0.45800579 | 0.99999994 |
| Birc5 | 0.03994108 | 0.03862375 | 0.03520894 | 0.00341482 | 0.25927523 | 0.99999994 |
| Sulf1 | 0.03651895 | 0.03565594 | 0.03226539 | 0.00339055 | 0.24212896 | 0.99999994 |
| Pdcl3 | 0.43504054 | 0.28958916 | 0.28619875 | 0.00339041 | 0.46857458 | 0.99999994 |
| Ppie | 0.26279385 | 0.19966086 | 0.19627817 | 0.00338269 | 0.45433487 | 0.99999994 |
| Fam181b | 0.03210727 | 0.03182446 | 0.02845089 | 0.00337357 | 0.2149748 | 0.99999994 |
| Cmtm3 | 0.05258948 | 0.04934705 | 0.0459736 | 0.00337345 | 0.31262271 | 0.99999994 |
| Cog4 | 0.1980093 | 0.15877702 | 0.15540564 | 0.00337138 | 0.4425909 | 0.99999994 |
| Mapt | 0.14049702 | 0.11870104 | 0.11533583 | 0.00336522 | 0.42300327 | 0.99999994 |
| Slc35d2 | 0.03121232 | 0.0310357 | 0.02767436 | 0.00336134 | 0.20940573 | 0.99999994 |
| Rbms1 | 0.17748894 | 0.14489532 | 0.14153723 | 0.00335809 | 0.43725692 | 0.99999994 |
| Wapl | 0.32896314 | 0.23717319 | 0.23382087 | 0.00335232 | 0.46198543 | 0.99999994 |
| Caap1 | 0.18898432 | 0.15270983 | 0.14936318 | 0.00334665 | 0.44072037 | 0.99999994 |
| Psat1 | 0.05042558 | 0.04747896 | 0.04414468 | 0.00333428 | 0.30756679 | 0.99999994 |

| | | | | | | |
|---|---|---|---|---|---|---|
| Rnf128 | 0.14009893 | 0.11837863 | 0.11504518 | 0.00333345 | 0.42353083 | 0.99999994 |
| Dapk1 | 0.15777615 | 0.13110286 | 0.12777117 | 0.0033317 | 0.43110253 | 0.99999994 |
| Rdm1 | 0.06493989 | 0.05963939 | 0.05631344 | 0.00332595 | 0.34711041 | 0.99999994 |
| Fam221a | 0.19618282 | 0.15750825 | 0.15418993 | 0.00331832 | 0.44304583 | 0.99999994 |
| Il10rb | 0.15066148 | 0.12600983 | 0.12269327 | 0.00331655 | 0.42860385 | 0.99999994 |
| Nipsnap1 | 0.27706262 | 0.20803257 | 0.2047177 | 0.00331487 | 0.45708414 | 0.99999994 |
| Tmem138 | 0.17129892 | 0.14056751 | 0.13726181 | 0.00330569 | 0.43632007 | 0.99999994 |
| Ndrg2 | 0.05343653 | 0.04998145 | 0.04668812 | 0.00329333 | 0.31934455 | 0.99999994 |
| Lrp8os2 | 0.02569424 | 0.02615425 | 0.02286582 | 0.00328843 | 0.16921338 | 0.99999994 |
| Cacng4 | 0.01760593 | 0.019015 | 0.01575308 | 0.00326193 | 0.08405437 | 0.99999994 |
| Efna2 | 0.16470891 | 0.13592465 | 0.13266289 | 0.00326176 | 0.43499993 | 0.99999994 |
| Gstm1 | 0.117167 | 0.10123694 | 0.09798007 | 0.00325687 | 0.41244576 | 0.99999994 |
| Prkrip1 | 0.35809675 | 0.25235757 | 0.24911555 | 0.00324202 | 0.46548414 | 0.99999994 |
| Tmem229a | 0.0539645 | 0.05037487 | 0.04713307 | 0.0032418 | 0.32354008 | 0.99999994 |
| Itpka | 0.02996115 | 0.02982837 | 0.02658717 | 0.0032412 | 0.20854932 | 0.99999994 |
| Fitm1 | 0.03598395 | 0.03504168 | 0.031804 | 0.00323767 | 0.24900538 | 0.99999994 |
| Akirin1 | 0.49344472 | 0.31463615 | 0.31140482 | 0.00323133 | 0.47246671 | 0.99999994 |
| Syncrip | 0.45670605 | 0.29905878 | 0.29583026 | 0.00322853 | 0.47104478 | 0.99999994 |
| Hnrnpu | 0.74203959 | 0.39507499 | 0.3918565 | 0.00321849 | 0.47820004 | 0.99999994 |
| Nop58 | 0.49958386 | 0.31713432 | 0.31391786 | 0.00321645 | 0.47281237 | 0.99999994 |
| Gm19590 | 0.03294913 | 0.03239388 | 0.02918053 | 0.00321335 | 0.23178025 | 0.99999994 |
| Carnmt1 | 0.1127915 | 0.09786143 | 0.09465064 | 0.00321079 | 0.41067825 | 0.99999994 |
| Rbbp4 | 0.86595031 | 0.41705923 | 0.41385202 | 0.00320722 | 0.47942985 | 0.99999994 |
| Snx18 | 0.06687101 | 0.06111983 | 0.05791516 | 0.00320467 | 0.35631653 | 0.99999994 |
| Flnc | 0.02540044 | 0.02580621 | 0.0226088 | 0.0031974 | 0.17325966 | 0.99999994 |
| Pi16 | 0.0378349 | 0.03659387 | 0.03339889 | 0.00319498 | 0.2621418 | 0.99999994 |
| Rrad | 0.01476629 | 0.01642499 | 0.01323755 | 0.00318744 | 0.05449287 | 0.99999994 |
| Rhou | 0.10106903 | 0.0887942 | 0.08561245 | 0.00318175 | 0.40230677 | 0.99999994 |
| Scn9a | 0.10338203 | 0.09058781 | 0.0874095 | 0.00317831 | 0.40437728 | 0.99999994 |
| Lekr1 | 0.07018357 | 0.06383035 | 0.06065332 | 0.00317703 | 0.36367183 | 0.99999994 |

| | | | | | | |
|---|---|---|---|---|---|---|
| Gng11 | 0.0259177 | 0.02623805 | 0.02306124 | 0.00317681 | 0.17958478 | 0.99999994 |
| Fbln2 | 0.05611594 | 0.05211812 | 0.04894305 | 0.00317507 | 0.33293755 | 0.99999994 |
| Fermt1 | 0.04977292 | 0.04676546 | 0.04359203 | 0.00317343 | 0.31398994 | 0.99999994 |
| Hsf2 | 0.13245665 | 0.1125938 | 0.10942902 | 0.00316478 | 0.42367228 | 0.99999994 |
| Kif22 | 0.07497659 | 0.06775156 | 0.06459445 | 0.00315712 | 0.37246337 | 0.99999994 |
| Eef1e1 | 0.31552467 | 0.22967309 | 0.22651862 | 0.00315447 | 0.46307267 | 0.99999994 |
| Rgs3 | 0.07899672 | 0.07102925 | 0.06788124 | 0.00314801 | 0.37877869 | 0.99999994 |
| Efhd2 | 0.13709454 | 0.11598757 | 0.11284559 | 0.00314197 | 0.42648362 | 0.99999994 |
| Lsm12 | 0.30420396 | 0.2233802 | 0.22024215 | 0.00313805 | 0.4622206 | 0.99999994 |
| Cypt3 | 0.00690155 | 0.00935232 | 0.00621999 | 0.00313233 | 0.00040104 | 0.0316483 |
| Evl | 0.1580455 | 0.13109051 | 0.12796226 | 0.00312825 | 0.43536725 | 0.99999994 |
| Gm27216 | 0.00929529 | 0.01149173 | 0.00836378 | 0.00312794 | 0.00639811 | 0.30798308 |
| Ece1 | 0.53959988 | 0.33281809 | 0.32969229 | 0.0031258 | 0.47483997 | 0.99999994 |
| B3gnt8 | 0.0629334 | 0.05776794 | 0.05464493 | 0.00312301 | 0.35181794 | 0.99999994 |
| Prr5 | 0.04959725 | 0.04654534 | 0.0434432 | 0.00310214 | 0.31728313 | 0.99999994 |
| Dand5 | 0.0545086 | 0.05068975 | 0.0475913 | 0.00309845 | 0.33237471 | 0.99999994 |
| Siah1a | 0.10932337 | 0.0950915 | 0.09199459 | 0.00309691 | 0.41134765 | 0.99999994 |
| Tlcd1 | 0.05196043 | 0.04852928 | 0.04544246 | 0.00308681 | 0.3255794 | 0.99999994 |
| Rdh5 | 0.02283513 | 0.0234301 | 0.02036037 | 0.00306973 | 0.15779105 | 0.99999994 |
| Kcnk12 | 0.02629459 | 0.02646016 | 0.0233907 | 0.00306947 | 0.19119885 | 0.99999994 |
| Trmt6 | 0.31693138 | 0.23035659 | 0.22729032 | 0.00306627 | 0.46422393 | 0.99999994 |
| Aif1l | 0.03625676 | 0.03509993 | 0.03203932 | 0.00306061 | 0.26243406 | 0.99999994 |
| Cd55 | 0.03560273 | 0.0345329 | 0.03147504 | 0.00305786 | 0.25892008 | 0.99999994 |
| Nqo1 | 0.06491334 | 0.05933627 | 0.05629139 | 0.00304487 | 0.35940457 | 0.99999994 |
| Col9a1 | 0.02729736 | 0.02730339 | 0.02426645 | 0.00303694 | 0.20241106 | 0.99999994 |
| Ccer2 | 0.02101444 | 0.02179603 | 0.01875988 | 0.00303615 | 0.14067699 | 0.99999994 |
| Rnf165 | 0.0388618 | 0.03731118 | 0.03428204 | 0.00302914 | 0.27821541 | 0.99999994 |
| Cnot4 | 0.40061703 | 0.27320862 | 0.27017965 | 0.00302897 | 0.4702569 | 0.99999994 |
| Nkpd1 | 0.02122362 | 0.02196117 | 0.01894396 | 0.00301721 | 0.14453553 | 0.99999994 |
| Bcam | 0.112268 | 0.09726713 | 0.09425068 | 0.00301646 | 0.41565014 | 0.99999994 |

| | | | | | | |
|---|---|---|---|---|---|---|
| Gnb3 | 0.02746576 | 0.02742403 | 0.02441342 | 0.00301061 | 0.20586612 | 0.99999994 |
| Neto2 | 0.03718216 | 0.03584354 | 0.0328369 | 0.00300664 | 0.27110502 | 0.99999994 |
| Ctf1 | 0.03009255 | 0.02970566 | 0.02670143 | 0.00300422 | 0.22695338 | 0.99999994 |
| Prpf38a | 0.36158575 | 0.25390068 | 0.25089939 | 0.00300129 | 0.46826812 | 0.99999994 |
| Pip4k2a | 0.02104949 | 0.0217897 | 0.01879073 | 0.00299897 | 0.14403963 | 0.99999994 |
| Klk8 | 0.0192483 | 0.02019905 | 0.0172036 | 0.00299545 | 0.12323057 | 0.99999994 |
| Nosip | 0.1692001 | 0.13879729 | 0.13580241 | 0.00299488 | 0.44164725 | 0.99999994 |
| Kctd10 | 0.14375249 | 0.1206947 | 0.11770568 | 0.00298902 | 0.43288586 | 0.99999994 |
| Gm10036 | 0.33723993 | 0.24122479 | 0.2382394 | 0.00298538 | 0.4667624 | 0.99999994 |
| Ehmt1 | 0.26593354 | 0.20113619 | 0.19815218 | 0.00298401 | 0.46007722 | 0.99999994 |
| Gm266 | 0.02124698 | 0.02194686 | 0.01896452 | 0.00298234 | 0.14760208 | 0.99999994 |
| Psmd12 | 0.87971159 | 0.41863463 | 0.41565993 | 0.0029747 | 0.48100288 | 0.99999994 |
| Cdkn2aip | 0.16762709 | 0.13767087 | 0.1347054 | 0.00296548 | 0.4417489 | 0.99999994 |
| Immp2l | 0.05889242 | 0.05422673 | 0.05127134 | 0.0029554 | 0.35060501 | 0.99999994 |
| Pcmtd1 | 0.20114292 | 0.16042969 | 0.15748309 | 0.0029466 | 0.45044218 | 0.99999994 |
| Draxin | 0.02801415 | 0.02783333 | 0.02489174 | 0.00294158 | 0.21575338 | 0.99999994 |
| Serpina1d | 0.0172988 | 0.01842294 | 0.01548146 | 0.00294148 | 0.10298817 | 0.99999994 |
| Bmp2 | 0.01845117 | 0.01943692 | 0.01649998 | 0.00293694 | 0.11804653 | 0.99999994 |
| Thumpd1 | 0.24653076 | 0.18935375 | 0.18641795 | 0.0029358 | 0.45825631 | 0.99999994 |
| Reep3 | 0.12160778 | 0.10427008 | 0.10133472 | 0.00293535 | 0.42355232 | 0.99999994 |
| 5430403G16 | 0.0518315 | 0.04826311 | 0.04533355 | 0.00292956 | 0.3335436 | 0.99999994 |
| Lmo3 | 0.02175686 | 0.02233961 | 0.01941298 | 0.00292663 | 0.15781307 | 0.99999994 |
| Asxl3 | 0.03256875 | 0.03177663 | 0.02885096 | 0.00292567 | 0.24983867 | 0.99999994 |
| Ntf3 | 0.0215582 | 0.0221562 | 0.01923828 | 0.00291792 | 0.15634679 | 0.99999994 |
| Ints10 | 0.2053959 | 0.16319871 | 0.16028602 | 0.0029127 | 0.45186198 | 0.99999994 |
| Nedd4l | 0.15311627 | 0.12735974 | 0.12445189 | 0.00290785 | 0.43820275 | 0.99999994 |
| Vash2 | 0.04216342 | 0.0400172 | 0.03711335 | 0.00290385 | 0.30124875 | 0.99999994 |
| Plbd2 | 0.09771839 | 0.0858677 | 0.0829657 | 0.002902 | 0.40794735 | 0.99999994 |
| Gm15567 | 0.01903962 | 0.01992084 | 0.01701948 | 0.00290136 | 0.12824472 | 0.99999994 |
| Kctd5 | 0.26423977 | 0.20004208 | 0.19714238 | 0.00289971 | 0.46100333 | 0.99999994 |

| | | | | | | |
|---|---|---|---|---|---|---|
| Pdcd4 | 0.46142759 | 0.30078255 | 0.29788432 | 0.00289823 | 0.47418165 | 0.99999994 |
| Hyls1 | 0.04580709 | 0.04312138 | 0.04022366 | 0.00289772 | 0.3157826 | 0.99999994 |
| Baz1b | 0.32571897 | 0.23496214 | 0.23207227 | 0.00288987 | 0.46697033 | 0.99999994 |
| Prdm16 | 0.04377645 | 0.04138025 | 0.03849211 | 0.00288814 | 0.30873476 | 0.99999994 |
| Mrpl47 | 0.19878697 | 0.15880785 | 0.15592217 | 0.00288567 | 0.4509783 | 0.99999994 |
| Haus1 | 0.12126024 | 0.10395364 | 0.10107305 | 0.00288059 | 0.42476944 | 0.99999994 |
| Bik | 0.04669051 | 0.04385522 | 0.04097551 | 0.0028797 | 0.31996393 | 0.99999994 |
| Clic6 | 0.05713278 | 0.05267625 | 0.04979673 | 0.00287952 | 0.3501531 | 0.99999994 |
| Rab35 | 0.08128407 | 0.07262269 | 0.06974374 | 0.00287895 | 0.39174746 | 0.99999994 |
| Rexo1 | 0.24745917 | 0.18986601 | 0.18698788 | 0.00287812 | 0.4591978 | 0.99999994 |
| Lzts2 | 0.24885635 | 0.19071741 | 0.187844 | 0.00287341 | 0.4594494 | 0.99999994 |
| Tbk1 | 0.08102129 | 0.07238277 | 0.06953004 | 0.00285273 | 0.39238741 | 0.99999994 |
| 2310040G24 | 0.13092104 | 0.11114052 | 0.10829212 | 0.0028484 | 0.43050771 | 0.99999994 |
| Pawr | 0.02164634 | 0.02216364 | 0.0193158 | 0.00284784 | 0.16320161 | 0.99999994 |
| Mal2 | 0.05312837 | 0.04926897 | 0.04642827 | 0.0028407 | 0.34190581 | 0.99999994 |
| Fras1 | 0.02441033 | 0.02458257 | 0.02174191 | 0.00284065 | 0.19223896 | 0.99999994 |
| Ccdc43 | 0.17681918 | 0.14391729 | 0.14107679 | 0.00284049 | 0.44669274 | 0.99999994 |
| Pik3ip1 | 0.09375246 | 0.08263322 | 0.07980042 | 0.00283281 | 0.40660321 | 0.99999994 |
| Renbp | 0.03523715 | 0.03398886 | 0.03115941 | 0.00282945 | 0.27277724 | 0.99999994 |
| Metrn | 0.05482104 | 0.05068182 | 0.04785428 | 0.00282753 | 0.34704902 | 0.99999994 |
| AI467606 | 0.16706405 | 0.13713801 | 0.13431206 | 0.00282595 | 0.44430985 | 0.99999994 |
| Pdlim1 | 0.05052471 | 0.04703922 | 0.04422858 | 0.00281064 | 0.33614797 | 0.99999994 |
| Ehbp1l1 | 0.02593557 | 0.02588214 | 0.02307686 | 0.00280528 | 0.20921046 | 0.99999994 |
| Gprc5c | 0.09422289 | 0.08297643 | 0.08017672 | 0.00279971 | 0.40809997 | 0.99999994 |
| Vdac3 | 0.83303305 | 0.41176402 | 0.40897089 | 0.00279313 | 0.48187008 | 0.99999994 |
| Akr1b8 | 0.02182182 | 0.02225366 | 0.01947009 | 0.00278356 | 0.17064129 | 0.99999994 |
| Zfand3 | 0.24154787 | 0.186124 | 0.18334457 | 0.00277943 | 0.45981185 | 0.99999994 |
| Nhsl1 | 0.08971859 | 0.07934191 | 0.0765644 | 0.00277751 | 0.40459491 | 0.99999994 |
| Msantd4 | 0.26265305 | 0.19897072 | 0.19619392 | 0.00277681 | 0.46247117 | 0.99999994 |
| Pacsin3 | 0.04171664 | 0.0395068 | 0.03673093 | 0.00277587 | 0.30746781 | 0.99999994 |

| | | | | | | |
|---|---|---|---|---|---|---|
| Pde6d | 0.42932859 | 0.28637687 | 0.28360217 | 0.0027747 | 0.47403751 | 0.99999994 |
| Sun2 | 0.04476335 | 0.04209922 | 0.03933423 | 0.00276498 | 0.31992604 | 0.99999994 |
| Nsmce2 | 0.34174057 | 0.24336874 | 0.24061767 | 0.00275106 | 0.46966801 | 0.99999994 |
| Dedd2 | 0.12824074 | 0.10904485 | 0.1063006 | 0.00274425 | 0.4317817 | 0.99999994 |
| Fam171a2 | 0.02830617 | 0.02788866 | 0.02514631 | 0.00274235 | 0.23394545 | 0.99999994 |
| Ell2 | 0.14611811 | 0.12215765 | 0.11941993 | 0.00273772 | 0.43935755 | 0.99999994 |
| Atp6v1b2 | 0.70334807 | 0.38574038 | 0.38300481 | 0.00273557 | 0.4810405 | 0.99999994 |
| Ppp1r9a | 0.14333646 | 0.12013439 | 0.11740352 | 0.00273087 | 0.43847739 | 0.99999994 |
| Ppp2r5b | 0.24776826 | 0.18990227 | 0.18717744 | 0.00272483 | 0.461403 | 0.99999994 |
| Srsf10 | 0.63839404 | 0.36714451 | 0.36442304 | 0.00272147 | 0.48017715 | 0.99999994 |
| Marveld2 | 0.09320197 | 0.0820789 | 0.07935979 | 0.00271911 | 0.40979692 | 0.99999994 |
| Hsf1 | 0.12986687 | 0.11022329 | 0.10750991 | 0.00271338 | 0.43329328 | 0.99999994 |
| Igll1 | 0.00311721 | 0.00552789 | 0.00281659 | 0.0027113 | 7.39E-11 | 1.51E-08 |
| Smoc1 | 0.05851282 | 0.05365617 | 0.05095351 | 0.00270266 | 0.36202021 | 0.99999994 |
| Crbn | 0.1897171 | 0.15255294 | 0.14985717 | 0.00269576 | 0.45234441 | 0.99999994 |
| Elfn1 | 0.01366334 | 0.01495315 | 0.0122579 | 0.00269525 | 0.07165271 | 0.99999994 |
| C1ql1 | 0.00860871 | 0.01044331 | 0.0077496 | 0.00269371 | 0.01034192 | 0.44677666 |
| Hprt | 0.53786472 | 0.33172183 | 0.32902954 | 0.00269229 | 0.47828203 | 0.99999994 |
| Flna | 0.05667129 | 0.05209904 | 0.04940944 | 0.00268961 | 0.35854912 | 0.99999994 |
| Ppp1r35 | 0.35291779 | 0.24912922 | 0.24644893 | 0.00268029 | 0.47114489 | 0.99999994 |
| Mtfr1 | 0.22421665 | 0.17513738 | 0.1724591 | 0.00267828 | 0.45883349 | 0.99999994 |
| Upf1 | 0.1366897 | 0.11522099 | 0.11254838 | 0.00267261 | 0.43720318 | 0.99999994 |
| Csnk1g3 | 0.1530789 | 0.12709654 | 0.12442517 | 0.00267136 | 0.44318082 | 0.99999994 |
| Rnf34 | 0.24980965 | 0.1910978 | 0.18842704 | 0.00267075 | 0.46241687 | 0.99999994 |
| Stap2 | 0.05651554 | 0.05194825 | 0.04927867 | 0.00266959 | 0.35920053 | 0.99999994 |
| Rfc1 | 0.19485091 | 0.15596772 | 0.15330117 | 0.00266655 | 0.45391261 | 0.99999994 |
| Abcf2 | 0.25038029 | 0.19144177 | 0.18877563 | 0.00266614 | 0.46255057 | 0.99999994 |
| Med27 | 0.30292848 | 0.22218895 | 0.21952787 | 0.00266108 | 0.4678453 | 0.99999994 |
| Col2a1 | 0.01207974 | 0.01350719 | 0.01084877 | 0.00265842 | 0.05143501 | 0.99999994 |
| Rhov | 0.07024472 | 0.06335937 | 0.06070376 | 0.00265561 | 0.38545078 | 0.99999994 |

| | | | | | | |
|---|---|---|---|---|---|---|
| Ltbp1 | 0.02120779 | 0.0215824 | 0.01893003 | 0.00265237 | 0.17549973 | 0.99999994 |
| H2-T23 | 0.05751847 | 0.05276326 | 0.05012024 | 0.00264302 | 0.36278888 | 0.99999994 |
| Ica1 | 0.34057626 | 0.24264632 | 0.24000404 | 0.00264228 | 0.47079093 | 0.99999994 |
| Nkx6-2 | 0.00871333 | 0.01047979 | 0.00784323 | 0.00263656 | 0.01262422 | 0.51649113 |
| Itfg2 | 0.17098585 | 0.13967215 | 0.13704443 | 0.00262772 | 0.44922079 | 0.99999994 |
| Fibp | 0.26685523 | 0.20132788 | 0.19870054 | 0.00262734 | 0.46493259 | 0.99999994 |
| Cox17 | 0.89540421 | 0.42015077 | 0.41752861 | 0.00262216 | 0.48332779 | 0.99999994 |
| Slc5a9 | 0.01182593 | 0.0132396 | 0.01062264 | 0.00261695 | 0.05051977 | 0.99999994 |
| Sat1 | 0.14253412 | 0.11943672 | 0.11682021 | 0.00261651 | 0.44074239 | 0.99999994 |
| Fgfbp3 | 0.03653843 | 0.03489565 | 0.03228219 | 0.00261346 | 0.29498562 | 0.99999994 |
| Kctd15 | 0.08488711 | 0.07527892 | 0.07266643 | 0.00261249 | 0.40543344 | 0.99999994 |
| Rbm38 | 0.06959838 | 0.06277817 | 0.06017047 | 0.00260771 | 0.38649054 | 0.99999994 |
| 3830403N18 | 0.00514968 | 0.00724943 | 0.00464664 | 0.00260279 | 9.63E-05 | 0.0089777 |
| Triqk | 0.125881 | 0.10714104 | 0.10454006 | 0.00260098 | 0.43423132 | 0.99999994 |
| Mex3b | 0.03221601 | 0.03113985 | 0.02854519 | 0.00259466 | 0.2725758 | 0.99999994 |
| Dgkb | 0.00885469 | 0.01054671 | 0.00796971 | 0.00257701 | 0.01568549 | 0.57349171 |
| Pcdhb7 | 0.01481918 | 0.01585541 | 0.0132845 | 0.00257092 | 0.09884011 | 0.99999994 |
| St6galnac4 | 0.05402315 | 0.04975128 | 0.04718248 | 0.0025688 | 0.35852706 | 0.99999994 |
| Tbcc | 0.37417381 | 0.25981978 | 0.25725149 | 0.00256828 | 0.47350817 | 0.99999994 |
| Exosc5 | 0.30832308 | 0.22510032 | 0.22253906 | 0.00256126 | 0.46946696 | 0.99999994 |
| Elf2 | 0.26536051 | 0.20037169 | 0.19781086 | 0.00256083 | 0.46566476 | 0.99999994 |
| Lin9 | 0.07164114 | 0.0644112 | 0.06185443 | 0.00255677 | 0.39160323 | 0.99999994 |
| 2410004B18 | 0.34745946 | 0.24617013 | 0.24361466 | 0.00255547 | 0.47216687 | 0.99999994 |
| Hectd2 | 0.03030177 | 0.02943652 | 0.02688333 | 0.00255319 | 0.26363642 | 0.99999994 |
| 2900060B14 | 0.05164579 | 0.04772764 | 0.04517664 | 0.00255101 | 0.35350784 | 0.99999994 |
| Tmem63b | 0.26343568 | 0.19921271 | 0.19666203 | 0.00255068 | 0.46560118 | 0.99999994 |
| Arhgap27 | 0.02757297 | 0.0270548 | 0.02450695 | 0.00254785 | 0.2444614 | 0.99999994 |
| Enho | 0.39125711 | 0.26819995 | 0.26566727 | 0.00253268 | 0.47470134 | 0.99999994 |
| Dcbld1 | 0.09349098 | 0.08211843 | 0.07959116 | 0.00252727 | 0.4163032 | 0.99999994 |
| Tsr3 | 0.29429964 | 0.21718244 | 0.214656 | 0.00252643 | 0.46877751 | 0.99999994 |

| | | | | | | |
|---|---|---|---|---|---|---|
| Pon2 | 0.09355091 | 0.08216226 | 0.07963913 | 0.00252314 | 0.41648751 | 0.99999994 |
| Mrpl35 | 0.38980527 | 0.26747841 | 0.26496093 | 0.00251748 | 0.474786 | 0.99999994 |
| Bptf | 0.48135875 | 0.30888647 | 0.30638388 | 0.00250259 | 0.47832017 | 0.99999994 |
| Macrod2 | 0.22919939 | 0.17810755 | 0.17561986 | 0.00248769 | 0.46243997 | 0.99999994 |
| AI987944 | 0.08677252 | 0.07666806 | 0.07419043 | 0.00247763 | 0.41204286 | 0.99999994 |
| Aaas | 0.05999175 | 0.05466814 | 0.05219088 | 0.00247726 | 0.37602153 | 0.99999994 |
| Fam83f | 0.0306693 | 0.02967388 | 0.02720273 | 0.00247115 | 0.27269716 | 0.99999994 |
| Frk | 0.03353598 | 0.03215957 | 0.02968867 | 0.0024709 | 0.28979304 | 0.99999994 |
| Klc4 | 0.15409524 | 0.12761739 | 0.12515129 | 0.0024661 | 0.44782314 | 0.99999994 |
| Gm6710 | 0.04330701 | 0.04054397 | 0.03809115 | 0.00245282 | 0.33409785 | 0.99999994 |
| Brd9 | 0.11891014 | 0.10174743 | 0.09929975 | 0.00244768 | 0.43483601 | 0.99999994 |
| Dctpp1 | 0.32176201 | 0.23237105 | 0.22992726 | 0.00244378 | 0.47179948 | 0.99999994 |
| Irx2 | 0.01860414 | 0.01907407 | 0.01663507 | 0.002439 | 0.16454657 | 0.99999994 |
| Fbxo36 | 0.11660274 | 0.09998461 | 0.09755208 | 0.00243253 | 0.43408589 | 0.99999994 |
| Cthrc1 | 0.03148088 | 0.03033917 | 0.02790748 | 0.00243169 | 0.28095911 | 0.99999994 |
| Lgi3 | 0.03075652 | 0.02970996 | 0.02727851 | 0.00243146 | 0.27648644 | 0.99999994 |
| Calcoco2 | 0.01230479 | 0.01347506 | 0.01104921 | 0.00242585 | 0.0719525 | 0.99999994 |
| Celf4 | 0.18509583 | 0.14915317 | 0.14673209 | 0.00242108 | 0.45627213 | 0.99999994 |
| Rnf111 | 0.13618888 | 0.11460094 | 0.11218044 | 0.0024205 | 0.44289895 | 0.99999994 |
| Dqx1 | 0.00666134 | 0.00842402 | 0.00600448 | 0.00241954 | 0.00365677 | 0.20382044 |
| Sdc1 | 0.04561374 | 0.04247719 | 0.04005899 | 0.0024182 | 0.34390892 | 0.99999994 |
| Tmem17 | 0.09576936 | 0.08382564 | 0.08141215 | 0.00241349 | 0.42178479 | 0.99999994 |
| Cul1 | 0.36449053 | 0.25478831 | 0.25237646 | 0.00241185 | 0.47463965 | 0.99999994 |
| Eif4a2 | 1.36076401 | 0.42374164 | 0.42133024 | 0.0024114 | 0.48480542 | 0.99999994 |
| Pde1a | 0.01461001 | 0.01550592 | 0.01309883 | 0.00240709 | 0.11062735 | 0.99999994 |
| Zfp638 | 0.3557242 | 0.2503033 | 0.24789665 | 0.00240665 | 0.47423762 | 0.99999994 |
| Klhdc8a | 0.03487308 | 0.03324932 | 0.03084493 | 0.00240439 | 0.30192627 | 0.99999994 |
| Taf5 | 0.03872933 | 0.03657077 | 0.03416818 | 0.00240259 | 0.31987377 | 0.99999994 |
| Prcc | 0.1611913 | 0.1325905 | 0.13018803 | 0.00240247 | 0.45111866 | 0.99999994 |
| Dusp9 | 0.02325829 | 0.02313361 | 0.02073179 | 0.00240182 | 0.22030807 | 0.99999994 |

| | | | | | | |
|---|---|---|---|---|---|---|
| Bcl2l12 | 0.06734676 | 0.0607044 | 0.05830914 | 0.00239526 | 0.39225981 | 0.99999994 |
| Kif26a | 0.02136054 | 0.02145335 | 0.01906442 | 0.00238893 | 0.20211296 | 0.99999994 |
| Camk2n2 | 0.08237103 | 0.07300314 | 0.07062688 | 0.00237626 | 0.41139612 | 0.99999994 |
| Gsx1 | 0.00204682 | 0.00422632 | 0.00185077 | 0.00237554 | 6.50E-18 | 1.99E-15 |
| Mbnl2 | 0.54119623 | 0.33266496 | 0.3303004 | 0.00236456 | 0.48099692 | 0.99999994 |
| Ndst4 | 0.01706846 | 0.01763773 | 0.01527769 | 0.00236004 | 0.15191643 | 0.99999994 |
| Meis1 | 0.09293468 | 0.08150323 | 0.07914574 | 0.00235749 | 0.42141642 | 0.99999994 |
| Klb | 0.02406047 | 0.02379195 | 0.02143532 | 0.00235663 | 0.23214235 | 0.99999994 |
| Bmp7 | 0.02713563 | 0.02648156 | 0.02412529 | 0.00235627 | 0.25780912 | 0.99999994 |
| Btbd2 | 0.09423625 | 0.08253877 | 0.0801874 | 0.00235136 | 0.42262339 | 0.99999994 |
| Exoc6b | 0.13731787 | 0.11535756 | 0.11300947 | 0.0023481 | 0.44499972 | 0.99999994 |
| Nrp1 | 0.01969922 | 0.01993276 | 0.0176013 | 0.00233146 | 0.18896977 | 0.99999994 |
| Adgre5 | 0.04278213 | 0.03997344 | 0.03764255 | 0.00233089 | 0.34010622 | 0.99999994 |
| Rhbg | 0.02705384 | 0.02638344 | 0.02405389 | 0.00232955 | 0.25957731 | 0.99999994 |
| Trim9 | 0.03313085 | 0.03166685 | 0.02933792 | 0.00232894 | 0.29861005 | 0.99999994 |
| Zc3hc1 | 0.09037901 | 0.07941617 | 0.07709534 | 0.00232083 | 0.4205922 | 0.99999994 |
| Tbc1d17 | 0.22872371 | 0.17763889 | 0.1753192 | 0.00231968 | 0.46490987 | 0.99999994 |
| Brcc3 | 0.29925675 | 0.21978048 | 0.21746355 | 0.00231693 | 0.47173107 | 0.99999994 |
| Msi1 | 0.12226381 | 0.10414042 | 0.10182825 | 0.00231216 | 0.43993162 | 0.99999994 |
| Ldlrad3 | 0.07966127 | 0.07073405 | 0.06842292 | 0.00231113 | 0.41105469 | 0.99999994 |
| Taf11 | 0.24705166 | 0.18903763 | 0.18673782 | 0.0022998 | 0.46733226 | 0.99999994 |
| Tram1l1 | 0.02932934 | 0.02833642 | 0.02603748 | 0.00229894 | 0.27836214 | 0.99999994 |
| Snapin | 0.23822336 | 0.18357747 | 0.18128042 | 0.00229705 | 0.4663913 | 0.99999994 |
| Gpr160 | 0.02107502 | 0.02110896 | 0.01881319 | 0.00229577 | 0.20831704 | 0.99999994 |
| Gfra1 | 0.01816694 | 0.01854035 | 0.01624891 | 0.00229143 | 0.17394636 | 0.99999994 |
| Gm7932 | 0.01339906 | 0.01431405 | 0.01202294 | 0.0022911 | 0.10231992 | 0.99999994 |
| Tcf7l2 | 0.12202501 | 0.10393207 | 0.10164866 | 0.00228341 | 0.44056902 | 0.99999994 |
| Mrs2 | 0.04333826 | 0.04039778 | 0.03811785 | 0.00227993 | 0.34526553 | 0.99999994 |
| Obscn | 0.01510207 | 0.01580212 | 0.01353551 | 0.00226661 | 0.13249985 | 0.99999994 |
| Zfp711 | 0.02421479 | 0.02383662 | 0.02157057 | 0.00226605 | 0.24219205 | 0.99999994 |

| | | | | | | |
|---|---|---|---|---|---|---|
| Dbp | 0.10197783 | 0.08858281 | 0.08631934 | 0.00226347 | 0.43071912 | 0.99999994 |
| Arpc4 | 0.59367186 | 0.3516878 | 0.34942747 | 0.00226034 | 0.4828277 | 0.99999994 |
| Psme1 | 0.94447028 | 0.42469734 | 0.42243791 | 0.00225944 | 0.48579988 | 0.99999994 |
| Arid3a | 0.14253602 | 0.11907602 | 0.1168216 | 0.00225442 | 0.44889467 | 0.99999994 |
| Dixdc1 | 0.05709975 | 0.0520227 | 0.04976902 | 0.00225369 | 0.38154742 | 0.99999994 |
| Hkdc1 | 0.01838189 | 0.01869221 | 0.0164388 | 0.00225342 | 0.1807666 | 0.99999994 |
| Tsc22d3 | 0.08408143 | 0.07426157 | 0.07201406 | 0.00224751 | 0.41771569 | 0.99999994 |
| N4bp2l2 | 0.54101537 | 0.33247871 | 0.33023158 | 0.00224713 | 0.48193625 | 0.99999994 |
| Stx4a | 0.25100739 | 0.19139917 | 0.18915833 | 0.00224084 | 0.46857448 | 0.99999994 |
| Prss23 | 0.06136435 | 0.05557417 | 0.05333713 | 0.00223704 | 0.39005444 | 0.99999994 |
| Etfa | 0.75358058 | 0.39659487 | 0.39435962 | 0.00223525 | 0.48495205 | 0.99999994 |
| Fzd2 | 0.04503505 | 0.04179831 | 0.03956588 | 0.00223243 | 0.35361754 | 0.99999994 |
| Sft2d2 | 0.11557041 | 0.09899974 | 0.09676804 | 0.0022317 | 0.43899752 | 0.99999994 |
| Zfp105 | 0.05210975 | 0.04780001 | 0.04556858 | 0.00223142 | 0.3722304 | 0.99999994 |
| Ccndbp1 | 0.36685838 | 0.25580259 | 0.25357547 | 0.00222712 | 0.47669038 | 0.99999994 |
| Rwdd3 | 0.06362606 | 0.05744832 | 0.05522142 | 0.00222691 | 0.39418402 | 0.99999994 |
| Phyhip | 0.03166745 | 0.03029489 | 0.02806939 | 0.00222551 | 0.29883448 | 0.99999994 |
| L3mbtl3 | 0.05309784 | 0.0486237 | 0.04640251 | 0.00222119 | 0.37500541 | 0.99999994 |
| Tgm2 | 0.03786397 | 0.03564008 | 0.03342391 | 0.00221617 | 0.32947974 | 0.99999994 |
| Lor | 0.00550393 | 0.00717329 | 0.0049651 | 0.00220819 | 0.00153638 | 0.10511378 |
| Alb | 0.01839106 | 0.01865292 | 0.0164469 | 0.00220602 | 0.18597778 | 0.99999994 |
| Efcab6 | 0.00913513 | 0.01042658 | 0.00822056 | 0.00220602 | 0.03702859 | 0.99999994 |
| Nmnat3 | 0.04497979 | 0.04171851 | 0.03951878 | 0.00219973 | 0.35550017 | 0.99999994 |
| Msantd3 | 0.05143992 | 0.04720076 | 0.04500265 | 0.00219812 | 0.37254353 | 0.99999994 |
| Car3 | 0.00380931 | 0.0056313 | 0.00344033 | 0.00219097 | 1.12E-05 | 0.0011925 |
| Eif2ak2 | 0.1004959 | 0.08735597 | 0.08516613 | 0.00218985 | 0.43205184 | 0.99999994 |
| Tssc4 | 0.65328295 | 0.37130816 | 0.36912264 | 0.00218551 | 0.48428129 | 0.99999994 |
| Slc17a7 | 0.0101598 | 0.01131941 | 0.00913632 | 0.00218309 | 0.05585946 | 0.99999994 |
| Tmem266 | 0.02622782 | 0.02551123 | 0.02333234 | 0.00217889 | 0.26710085 | 0.99999994 |
| Pdhx | 0.11404966 | 0.09778873 | 0.0956105 | 0.00217823 | 0.43973248 | 0.99999994 |

| | | | | | | |
|---|---|---|---|---|---|---|
| Hp1bp3 | 0.61493549 | 0.35888654 | 0.35670971 | 0.00217683 | 0.48379918 | 0.99999994 |
| Plod2 | 0.03233018 | 0.030815 | 0.02864417 | 0.00217083 | 0.30696927 | 0.99999994 |
| Glrx2 | 0.54674209 | 0.33457143 | 0.33240101 | 0.00217042 | 0.4826663 | 0.99999994 |
| Steap2 | 0.11416939 | 0.09787193 | 0.09570174 | 0.00217019 | 0.44001008 | 0.99999994 |
| Nrip1 | 0.09554868 | 0.0834043 | 0.081236 | 0.0021683 | 0.4294922 | 0.99999994 |
| Nt5c3b | 0.26326524 | 0.19872311 | 0.19656014 | 0.00216297 | 0.47080463 | 0.99999994 |
| Ncam1 | 0.17240164 | 0.1401871 | 0.13802661 | 0.00216049 | 0.45850921 | 0.99999994 |
| Cidea | 0.01557351 | 0.01611167 | 0.01395361 | 0.00215806 | 0.15162976 | 0.99999994 |
| Vps13a | 0.11738803 | 0.10030524 | 0.09814761 | 0.00215763 | 0.44183033 | 0.99999994 |
| Cggbp1 | 0.32200517 | 0.23221501 | 0.23005947 | 0.00215554 | 0.47513542 | 0.99999994 |
| Cntn5 | 0.00505565 | 0.00671079 | 0.00456209 | 0.0021487 | 0.00085902 | 0.06168322 |
| Ppp2r2b | 0.12858751 | 0.10869825 | 0.10655875 | 0.00213949 | 0.44684095 | 0.99999994 |
| Npy1r | 0.01354987 | 0.01429624 | 0.01215703 | 0.00213921 | 0.12074148 | 0.99999994 |
| Cox19 | 0.36260995 | 0.25355774 | 0.25142097 | 0.00213677 | 0.47744351 | 0.99999994 |
| Ndufaf1 | 0.13841507 | 0.11594213 | 0.1138137 | 0.00212843 | 0.45046749 | 0.99999994 |
| Anapc15 | 0.10688034 | 0.09224204 | 0.09011461 | 0.00212743 | 0.43756575 | 0.99999994 |
| Mok | 0.04878903 | 0.04487971 | 0.042758 | 0.00212171 | 0.3705874 | 0.99999994 |
| Pdrg1 | 0.55309294 | 0.33690208 | 0.3347827 | 0.00211938 | 0.48319401 | 0.99999994 |
| Zfp518a | 0.11027745 | 0.09484485 | 0.09272676 | 0.00211809 | 0.43957524 | 0.99999994 |
| Rnf146 | 0.26971684 | 0.20251291 | 0.20039762 | 0.00211529 | 0.47199298 | 0.99999994 |
| Ptpn12 | 0.04464398 | 0.04134503 | 0.03923243 | 0.00211259 | 0.36000975 | 0.99999994 |
| Cadm4 | 0.04474601 | 0.04143197 | 0.03931945 | 0.00211252 | 0.36031112 | 0.99999994 |
| Rnf44 | 0.12397546 | 0.10521066 | 0.10311345 | 0.0020972 | 0.44615489 | 0.99999994 |
| Timm44 | 0.25083459 | 0.19114666 | 0.18905292 | 0.00209374 | 0.47061721 | 0.99999994 |
| Skp2 | 0.02505331 | 0.02439851 | 0.022305 | 0.00209351 | 0.26606647 | 0.99999994 |
| Bend7 | 0.16475929 | 0.13478833 | 0.13269824 | 0.00209009 | 0.45825043 | 0.99999994 |
| Sfrp1 | 0.01756924 | 0.01780998 | 0.01572064 | 0.00208935 | 0.18816975 | 0.99999994 |
| Pmf1 | 0.12812418 | 0.10830191 | 0.1062138 | 0.00208811 | 0.44794282 | 0.99999994 |
| Galnt9 | 0.01243126 | 0.01324932 | 0.01116182 | 0.0020875 | 0.10658778 | 0.99999994 |
| Kif20b | 0.01764058 | 0.01787025 | 0.01578371 | 0.00208654 | 0.18944466 | 0.99999994 |

| | | | | | | |
|---|---|---|---|---|---|---|
| Repin1 | 0.07395831 | 0.06583799 | 0.0637592 | 0.00207879 | 0.41409544 | 0.99999994 |
| Adra2c | 0.00821379 | 0.00947124 | 0.00739607 | 0.00207516 | 0.03090768 | 0.89706031 |
| Gm26737 | 0.00951919 | 0.01063834 | 0.00856395 | 0.00207439 | 0.05344539 | 0.99999994 |
| Sema5a | 0.02304274 | 0.02261552 | 0.02054262 | 0.0020729 | 0.250895 | 0.99999994 |
| Plpp1 | 0.05044464 | 0.04623357 | 0.04416081 | 0.00207276 | 0.37735772 | 0.99999994 |
| Cbr4 | 0.11920884 | 0.10159185 | 0.09952551 | 0.00206634 | 0.4450418 | 0.99999994 |
| Atoh8 | 0.01554611 | 0.01599059 | 0.01392932 | 0.00206127 | 0.16230941 | 0.99999994 |
| Lhfpl2 | 0.02449995 | 0.02388154 | 0.02182043 | 0.00206111 | 0.26475798 | 0.99999994 |
| Gm5426 | 0.03865093 | 0.03615757 | 0.03410079 | 0.00205678 | 0.34403439 | 0.99999994 |
| Cox4i2 | 0.01398591 | 0.01459908 | 0.01254456 | 0.00205452 | 0.13782112 | 0.99999994 |
| Pqbp1 | 0.39774455 | 0.27085498 | 0.268802 | 0.00205298 | 0.47972733 | 0.99999994 |
| Olfml2b | 0.03151298 | 0.02997807 | 0.02793534 | 0.00204273 | 0.3132229 | 0.99999994 |
| Dbndd1 | 0.04978654 | 0.04564452 | 0.04360356 | 0.00204096 | 0.37768552 | 0.99999994 |
| Acot9 | 0.01478422 | 0.01528672 | 0.01325347 | 0.00203325 | 0.15358642 | 0.99999994 |
| Sox13 | 0.04723611 | 0.0434708 | 0.04143943 | 0.00203137 | 0.37210021 | 0.99999994 |
| Fbxo2 | 0.05869327 | 0.05313554 | 0.05110462 | 0.00203092 | 0.39568727 | 0.99999994 |
| Zswim4 | 0.03476867 | 0.03278341 | 0.03075471 | 0.0020287 | 0.33030219 | 0.99999994 |
| Fmnl1 | 0.02328923 | 0.02278373 | 0.02075894 | 0.0020248 | 0.25808798 | 0.99999994 |
| Hpgd | 0.02366058 | 0.0231088 | 0.0210847 | 0.00202409 | 0.26141206 | 0.99999994 |
| Tgfb1 | 0.02211896 | 0.02175396 | 0.01973128 | 0.00202268 | 0.24750822 | 0.99999994 |
| Lzts1 | 0.03633586 | 0.03413018 | 0.03210753 | 0.00202265 | 0.33748937 | 0.99999994 |
| Eed | 0.13406417 | 0.11263052 | 0.11061613 | 0.00201439 | 0.45175981 | 0.99999994 |
| Snx20 | 0.00676429 | 0.00811044 | 0.00609685 | 0.00201359 | 0.01396119 | 0.54149729 |
| Gulo | 0.05266989 | 0.04805379 | 0.04604147 | 0.00201232 | 0.38555332 | 0.99999994 |
| Aldh5a1 | 0.19552264 | 0.15576042 | 0.15374965 | 0.00201077 | 0.46531455 | 0.99999994 |
| Vsig8 | 0.01033888 | 0.01130517 | 0.00929623 | 0.00200894 | 0.07515268 | 0.99999994 |
| Hnrnph3 | 0.61599809 | 0.35907441 | 0.35706638 | 0.00200803 | 0.48506975 | 0.99999994 |
| Plpp6 | 0.09784522 | 0.08507422 | 0.08306666 | 0.00200756 | 0.43609727 | 0.99999994 |
| Ppm1d | 0.10378249 | 0.08972686 | 0.08771994 | 0.00200692 | 0.43947951 | 0.99999994 |
| Cdc40 | 0.19221945 | 0.15353876 | 0.15153965 | 0.00199911 | 0.46501351 | 0.99999994 |

| | | | | | | |
|---|---|---|---|---|---|---|
| Col18a1 | 0.07100872 | 0.06333034 | 0.06133356 | 0.00199678 | 0.41421913 | 0.99999994 |
| Dnaaf5 | 0.04268486 | 0.03954429 | 0.03755938 | 0.00198491 | 0.36250534 | 0.99999994 |
| Gm5141 | 0.01691006 | 0.01712041 | 0.01513752 | 0.00198289 | 0.19162437 | 0.99999994 |
| Rbm24 | 0.02206434 | 0.02166497 | 0.01968328 | 0.00198169 | 0.25137958 | 0.99999994 |
| Etnk2 | 0.0241849 | 0.02352401 | 0.02154437 | 0.00197964 | 0.2703913 | 0.99999994 |
| Ssna1 | 0.58245721 | 0.34745317 | 0.34547798 | 0.00197519 | 0.48482147 | 0.99999994 |
| Dusp4 | 0.04374871 | 0.04044152 | 0.03846842 | 0.0019731 | 0.36639606 | 0.99999994 |
| Vdac3-ps1 | 0.02774486 | 0.02662755 | 0.0246569 | 0.00197065 | 0.29736444 | 0.99999994 |
| Srgap2 | 0.04385916 | 0.04053236 | 0.03856273 | 0.00196963 | 0.36693562 | 0.99999994 |
| Sall4 | 0.0409973 | 0.0380837 | 0.03611475 | 0.00196895 | 0.35833999 | 0.99999994 |
| Prr33 | 0.01572164 | 0.0160518 | 0.01408493 | 0.00196687 | 0.17631055 | 0.99999994 |
| Sephs1 | 0.13796532 | 0.11545079 | 0.11348421 | 0.00196658 | 0.4540844 | 0.99999994 |
| Baz2a | 0.28693429 | 0.21240642 | 0.21044347 | 0.00196294 | 0.47524631 | 0.99999994 |
| Nod1 | 0.12139751 | 0.10313788 | 0.10117643 | 0.00196145 | 0.44866183 | 0.99999994 |
| Ubxn11 | 0.03194031 | 0.03026615 | 0.0283061 | 0.00196006 | 0.32242797 | 0.99999994 |
| Tspyl1 | 0.30418145 | 0.22218374 | 0.22022956 | 0.00195418 | 0.47645038 | 0.99999994 |
| Fnbp1l | 0.21107894 | 0.16595328 | 0.16400087 | 0.00195241 | 0.46841963 | 0.99999994 |
| Med26 | 0.05257342 | 0.04790661 | 0.04596005 | 0.00194657 | 0.38900261 | 0.99999994 |
| Phpt1 | 0.41101045 | 0.27705522 | 0.27511115 | 0.00194407 | 0.48124186 | 0.99999994 |
| Try5 | 0.03540101 | 0.0332441 | 0.0313009 | 0.0019432 | 0.3397173 | 0.99999994 |
| Rps27rt | 0.82070598 | 0.40900287 | 0.40706149 | 0.00194138 | 0.48733735 | 0.99999994 |
| Maoa | 0.03789933 | 0.03539333 | 0.03345433 | 0.00193899 | 0.34982293 | 0.99999994 |
| Slc25a34 | 0.02152764 | 0.02113829 | 0.01921141 | 0.00192688 | 0.25218668 | 0.99999994 |
| Ebf1 | 0.01119586 | 0.01198282 | 0.01006097 | 0.00192185 | 0.10177488 | 0.99999994 |
| Acta1 | 0.01069643 | 0.01153472 | 0.00961541 | 0.00191932 | 0.09197901 | 0.99999994 |
| D16Ertd472e | 0.01560081 | 0.01589287 | 0.01397782 | 0.00191505 | 0.18089406 | 0.99999994 |
| Pth1r | 0.01359094 | 0.01410745 | 0.01219354 | 0.00191392 | 0.14805967 | 0.99999994 |
| Pard6g | 0.09591844 | 0.08344337 | 0.08153111 | 0.00191226 | 0.43796903 | 0.99999994 |
| Nkain3 | 0.00761729 | 0.0087644 | 0.00686173 | 0.00190267 | 0.03246607 | 0.93419069 |
| Pot1a | 0.06840948 | 0.06108811 | 0.05918833 | 0.00189978 | 0.41540978 | 0.99999994 |

| | | | | | | |
|---|---|---|---|---|---|---|
| Pgc | 0.00750717 | 0.00866211 | 0.00676304 | 0.00189907 | 0.03080343 | 0.89605388 |
| Larp7 | 0.20551764 | 0.16226454 | 0.16036597 | 0.00189857 | 0.46859404 | 0.99999994 |
| Lig1 | 0.06611426 | 0.05918483 | 0.05728797 | 0.00189685 | 0.4127807 | 0.99999994 |
| Acer2 | 0.35204605 | 0.24788568 | 0.24599792 | 0.00188776 | 0.47963078 | 0.99999994 |
| Dmd | 0.05238295 | 0.04768091 | 0.04579927 | 0.00188164 | 0.39224468 | 0.99999994 |
| Gstt1 | 0.06353292 | 0.0570203 | 0.05514393 | 0.00187637 | 0.41040849 | 0.99999994 |
| Gria2 | 0.02445239 | 0.02364777 | 0.02177876 | 0.001869 | 0.28392087 | 0.99999994 |
| E130311K13I | 0.08585302 | 0.07531217 | 0.07344765 | 0.00186452 | 0.43290759 | 0.99999994 |
| Nr4a2 | 0.0493491 | 0.04509653 | 0.04323289 | 0.00186364 | 0.38708078 | 0.99999994 |
| Bcl11b | 0.03982883 | 0.03697492 | 0.0351126 | 0.00186232 | 0.36202826 | 0.99999994 |
| Cnn3 | 0.82583514 | 0.40970796 | 0.40785752 | 0.00185044 | 0.48795387 | 0.99999994 |
| Clcn3 | 0.43210754 | 0.28671818 | 0.28486839 | 0.00184979 | 0.48276188 | 0.99999994 |
| Gimap9 | 0.02881485 | 0.0274352 | 0.02558951 | 0.00184569 | 0.31557792 | 0.99999994 |
| Dnaja4 | 0.18470518 | 0.14831093 | 0.14646685 | 0.00184408 | 0.46660521 | 0.99999994 |
| Iqcc | 0.16315255 | 0.1334127 | 0.13156962 | 0.00184307 | 0.46285466 | 0.99999994 |
| Krt15 | 0.00989576 | 0.01074022 | 0.00890046 | 0.00183975 | 0.08442792 | 0.99999994 |
| Plpp2 | 0.08846394 | 0.07739112 | 0.07555449 | 0.00183663 | 0.43572885 | 0.99999994 |
| Wfikkn1 | 0.00554929 | 0.00683286 | 0.00500586 | 0.001827 | 0.00756222 | 0.35248365 |
| Nptx1 | 0.03937786 | 0.03653926 | 0.0347254 | 0.00181385 | 0.3640368 | 0.99999994 |
| Eya2 | 0.0582363 | 0.05253437 | 0.0507219 | 0.00181247 | 0.40599669 | 0.99999994 |
| Klk10 | 0.00886905 | 0.00979054 | 0.00798256 | 0.00180799 | 0.0658269 | 0.99999994 |
| Gap43 | 0.01989379 | 0.01957437 | 0.01777282 | 0.00180155 | 0.24992562 | 0.99999994 |
| Tst | 0.0553742 | 0.05011819 | 0.04831961 | 0.00179858 | 0.40215726 | 0.99999994 |
| Elavl2 | 0.02932111 | 0.02782381 | 0.02603032 | 0.0017935 | 0.32325181 | 0.99999994 |
| Orc4 | 0.2899614 | 0.21397173 | 0.21218077 | 0.00179097 | 0.47759732 | 0.99999994 |
| Rps17 | 2.65175913 | 0.36611787 | 0.36432741 | 0.00179046 | 0.48695209 | 0.99999994 |
| Appl1 | 0.17863036 | 0.14411115 | 0.14232078 | 0.00179036 | 0.46663337 | 0.99999994 |
| Sertad2 | 0.12144989 | 0.10300531 | 0.10121586 | 0.00178944 | 0.4531602 | 0.99999994 |
| Pde10a | 0.09152338 | 0.07980041 | 0.07801429 | 0.00178612 | 0.4394374 | 0.99999994 |
| Gas6 | 0.04229783 | 0.03901197 | 0.03722835 | 0.00178363 | 0.37489766 | 0.99999994 |

| | | | | | | |
|---|---|---|---|---|---|---|
| Lrrtm3 | 0.00547546 | 0.00672017 | 0.00493951 | 0.00178066 | 0.00820699 | 0.37272847 |
| Prpf40b | 0.08716988 | 0.07628552 | 0.07451116 | 0.00177437 | 0.43702712 | 0.99999994 |
| Polr2m | 0.86913722 | 0.41605503 | 0.4142807 | 0.00177433 | 0.48862825 | 0.99999994 |
| Rnf126 | 0.23249165 | 0.17946723 | 0.17769451 | 0.00177272 | 0.47352763 | 0.99999994 |
| Acsl3 | 0.05624614 | 0.05081777 | 0.04905243 | 0.00176534 | 0.40533765 | 0.99999994 |
| Tmem54 | 0.01817122 | 0.01801705 | 0.0162527 | 0.00176435 | 0.23496424 | 0.99999994 |
| Ednra | 0.03072777 | 0.02901576 | 0.02725353 | 0.00176223 | 0.33345076 | 0.99999994 |
| Col16a1 | 0.01220508 | 0.01272026 | 0.01096041 | 0.00175985 | 0.14258668 | 0.99999994 |
| Sec61a2 | 0.21273991 | 0.16683872 | 0.16508011 | 0.0017586 | 0.47173454 | 0.99999994 |
| Zc3h8 | 0.07507874 | 0.06643266 | 0.06467817 | 0.00175449 | 0.42835523 | 0.99999994 |
| Clcn2 | 0.05332248 | 0.0483462 | 0.04659196 | 0.00175425 | 0.40105378 | 0.99999994 |
| Ly6g6e | 0.03015916 | 0.02850914 | 0.02675935 | 0.00174979 | 0.33168778 | 0.99999994 |
| Trim47 | 0.03239157 | 0.03044484 | 0.02869739 | 0.00174745 | 0.34262046 | 0.99999994 |
| Ehbp1 | 0.09896246 | 0.08569554 | 0.08395532 | 0.00174022 | 0.445131 | 0.99999994 |
| Dhrs3 | 0.01078121 | 0.01143127 | 0.00969106 | 0.00174021 | 0.1159901 | 0.99999994 |
| Sema3g | 0.02807801 | 0.02668659 | 0.02494742 | 0.00173917 | 0.32131035 | 0.99999994 |
| Prcp | 0.06053812 | 0.05438567 | 0.05264739 | 0.00173828 | 0.41302294 | 0.99999994 |
| Tmem45a | 0.03647824 | 0.03396815 | 0.0322303 | 0.00173785 | 0.35983132 | 0.99999994 |
| Med4 | 0.20985656 | 0.16494091 | 0.16320477 | 0.00173615 | 0.47177478 | 0.99999994 |
| Tpm3-rs7 | 0.03226151 | 0.03032004 | 0.02858464 | 0.00173541 | 0.34306375 | 0.99999994 |
| Mmrn2 | 0.02387148 | 0.02300502 | 0.02126964 | 0.00173538 | 0.29353389 | 0.99999994 |
| Slc43a1 | 0.02784388 | 0.02647745 | 0.02474326 | 0.00173418 | 0.32042006 | 0.99999994 |
| Leng1 | 0.155908 | 0.12817634 | 0.12644346 | 0.00173288 | 0.46365724 | 0.99999994 |
| Sft2d3 | 0.095996 | 0.0833234 | 0.08159299 | 0.00173041 | 0.44386916 | 0.99999994 |
| Pim1 | 0.07394395 | 0.06547657 | 0.06374741 | 0.00172915 | 0.42835869 | 0.99999994 |
| Igdcc4 | 0.0165024 | 0.01650268 | 0.01477665 | 0.00172604 | 0.21842665 | 0.99999994 |
| Psrc1 | 0.01009116 | 0.01079928 | 0.00907501 | 0.00172427 | 0.1029853 | 0.99999994 |
| Irak2 | 0.01905481 | 0.01875215 | 0.01703288 | 0.00171927 | 0.25082992 | 0.99999994 |
| Gm13373 | 0.01708362 | 0.01700579 | 0.0152911 | 0.00171468 | 0.22770766 | 0.99999994 |
| Peak1 | 0.03176398 | 0.02986346 | 0.02815313 | 0.00171032 | 0.34296704 | 0.99999994 |

| | | | | | | |
|---|---|---|---|---|---|---|
| Lyrm1 | 0.05918033 | 0.05321903 | 0.05151229 | 0.00170674 | 0.41272403 | 0.99999994 |
| C130021I20R | 0.03918436 | 0.0362611 | 0.0345592 | 0.0017019 | 0.37153164 | 0.99999994 |
| S100a16 | 0.06760868 | 0.06022404 | 0.05852594 | 0.0016981 | 0.42342815 | 0.99999994 |
| Tpmt | 0.05119694 | 0.04649488 | 0.04479722 | 0.00169765 | 0.40042307 | 0.99999994 |
| Galr2 | 0.02384407 | 0.02294078 | 0.02124561 | 0.00169517 | 0.29767273 | 0.99999994 |
| Acy3 | 0.04698173 | 0.04291833 | 0.04122317 | 0.00169516 | 0.39215015 | 0.99999994 |
| Tinagl1 | 0.0325167 | 0.03050082 | 0.02880585 | 0.00169497 | 0.34765186 | 0.99999994 |
| Ocln | 0.13702651 | 0.11448937 | 0.11279566 | 0.00169371 | 0.46019203 | 0.99999994 |
| Asf1b | 0.01567356 | 0.01573602 | 0.01404232 | 0.0016937 | 0.21103047 | 0.99999994 |
| Rap2b | 0.06152497 | 0.05516227 | 0.05347114 | 0.00169113 | 0.41663053 | 0.99999994 |
| Cenpk | 0.01852936 | 0.01825905 | 0.01656903 | 0.00169001 | 0.24858896 | 0.99999994 |
| Dok4 | 0.07519552 | 0.06646341 | 0.06477388 | 0.00168953 | 0.4310816 | 0.99999994 |
| Snrnp25 | 0.1718925 | 0.13935232 | 0.13767366 | 0.00167866 | 0.46765681 | 0.99999994 |
| Galt | 0.18304189 | 0.14700816 | 0.14533567 | 0.00167248 | 0.46947104 | 0.99999994 |
| Frmd8 | 0.03399599 | 0.03175661 | 0.03008669 | 0.00166992 | 0.3558957 | 0.99999994 |
| Stau2 | 0.2008461 | 0.15895601 | 0.15728676 | 0.00166924 | 0.47184125 | 0.99999994 |
| Ypel2 | 0.17839329 | 0.14382494 | 0.14215816 | 0.00166678 | 0.46889621 | 0.99999994 |
| D130040H23 | 0.00487325 | 0.00606434 | 0.00439803 | 0.00166631 | 0.00583569 | 0.28711953 |
| Zdhhc4 | 0.11357798 | 0.09691664 | 0.09525088 | 0.00166576 | 0.45366488 | 0.99999994 |
| Pcdhb19 | 0.02193052 | 0.02123027 | 0.01956565 | 0.00166461 | 0.28559087 | 0.99999994 |
| Taf9b | 0.08360425 | 0.07329124 | 0.07162736 | 0.00166388 | 0.43855866 | 0.99999994 |
| Rai14 | 0.01986886 | 0.01941274 | 0.01775085 | 0.00166189 | 0.26658048 | 0.99999994 |
| Cdca5 | 0.01715241 | 0.01701332 | 0.01535196 | 0.00166136 | 0.23565853 | 0.99999994 |
| Rabgap1 | 0.24848933 | 0.18927986 | 0.1876193 | 0.00166056 | 0.4765105 | 0.99999994 |
| Zfp513 | 0.19882546 | 0.15760661 | 0.15594772 | 0.00165889 | 0.47177576 | 0.99999994 |
| Baiap2l2 | 0.01544045 | 0.01549285 | 0.01383564 | 0.00165721 | 0.21264136 | 0.99999994 |
| Klk11 | 0.01320145 | 0.01350384 | 0.01184721 | 0.00165664 | 0.17598308 | 0.99999994 |
| Tsga10ip | 0.03176581 | 0.02980955 | 0.02815472 | 0.00165483 | 0.34781174 | 0.99999994 |
| Faf1 | 0.170483 | 0.13834953 | 0.13669505 | 0.00165448 | 0.46789402 | 0.99999994 |
| Nmb | 0.02862209 | 0.02707569 | 0.0254216 | 0.00165409 | 0.33246955 | 0.99999994 |

| | | | | | | |
|---|---|---|---|---|---|---|
| Ptprz1 | 0.00965674 | 0.0103353 | 0.00868689 | 0.00164841 | 0.10327679 | 0.99999994 |
| Gjc1 | 0.02168751 | 0.0209983 | 0.019352 | 0.0016463 | 0.2856067 | 0.99999994 |
| Gm13305 | 0.07795682 | 0.06867644 | 0.06703267 | 0.00164377 | 0.43517007 | 0.99999994 |
| Myo1b | 0.24132552 | 0.18484562 | 0.18320686 | 0.00163876 | 0.47626087 | 0.99999994 |
| Eif2b3 | 0.26716363 | 0.20052197 | 0.19888385 | 0.00163812 | 0.47813874 | 0.99999994 |
| Nsg2 | 0.01693917 | 0.01680077 | 0.01516328 | 0.00163749 | 0.2361245 | 0.99999994 |
| Ecscr | 0.00760095 | 0.0084801 | 0.00684708 | 0.00163302 | 0.05619614 | 0.99999994 |
| Tmem170 | 0.02365073 | 0.02270171 | 0.02107606 | 0.00162565 | 0.30382829 | 0.99999994 |
| Aldh6a1 | 0.06597332 | 0.05879404 | 0.0571711 | 0.00162294 | 0.42506326 | 0.99999994 |
| Camk1g | 0.04373558 | 0.04007828 | 0.03845722 | 0.00162107 | 0.38951563 | 0.99999994 |
| Eif4b | 0.35994743 | 0.25168003 | 0.25006316 | 0.00161687 | 0.48283523 | 0.99999994 |
| Foxo3 | 0.15444067 | 0.1270129 | 0.12539781 | 0.00161509 | 0.46583972 | 0.99999994 |
| Trim28 | 0.40086307 | 0.27190577 | 0.27029736 | 0.00160841 | 0.4842025 | 0.99999994 |
| Zscan21 | 0.11914257 | 0.10108227 | 0.09947544 | 0.00160683 | 0.45718816 | 0.99999994 |
| Tekt1 | 0.02938972 | 0.02769443 | 0.02609003 | 0.0016044 | 0.34114859 | 0.99999994 |
| Smad7 | 0.06021723 | 0.05397693 | 0.05237931 | 0.00159762 | 0.41955756 | 0.99999994 |
| Nacc2 | 0.08713896 | 0.07608186 | 0.0744862 | 0.00159566 | 0.44330539 | 0.99999994 |
| Acer3 | 0.05121192 | 0.04640261 | 0.0448099 | 0.00159272 | 0.40648642 | 0.99999994 |
| Gmfg | 0.01474597 | 0.01480412 | 0.01321952 | 0.0015846 | 0.21246773 | 0.99999994 |
| P3h4 | 0.24561076 | 0.18743238 | 0.18585234 | 0.00158004 | 0.47743609 | 0.99999994 |
| Rbm8a2 | 0.0354615 | 0.0329327 | 0.03135312 | 0.00157958 | 0.36868057 | 0.99999994 |
| Zkscan14 | 0.08973722 | 0.07815635 | 0.07657939 | 0.00157696 | 0.44548738 | 0.99999994 |
| Me1 | 0.06045812 | 0.05415613 | 0.05258057 | 0.00157556 | 0.42095328 | 0.99999994 |
| Gpt2 | 0.0493425 | 0.04480242 | 0.0432273 | 0.00157512 | 0.40417909 | 0.99999994 |
| Kansl1l | 0.0581462 | 0.05221856 | 0.05064641 | 0.00157215 | 0.41815141 | 0.99999994 |
| Cyp7b1 | 0.02781844 | 0.02629005 | 0.02472107 | 0.00156898 | 0.336344 | 0.99999994 |
| Surf2 | 0.16798826 | 0.13652418 | 0.13495753 | 0.00156665 | 0.46920436 | 0.99999994 |
| Proca1 | 0.01321107 | 0.01342084 | 0.01185576 | 0.00156507 | 0.1897812 | 0.99999994 |
| Akap1 | 0.0319386 | 0.02986946 | 0.02830461 | 0.00156484 | 0.35643583 | 0.99999994 |
| Nrk | 0.00801273 | 0.0087743 | 0.00721601 | 0.00155829 | 0.07529777 | 0.99999994 |

| | | | | | | |
|---|---|---|---|---|---|---|
| Pwwp2a | 0.13068757 | 0.10967502 | 0.10811902 | 0.00155599 | 0.46184173 | 0.99999994 |
| Macc1 | 0.0083662 | 0.00908255 | 0.00753253 | 0.00155002 | 0.08538606 | 0.99999994 |
| Gm14226 | 0.01099541 | 0.0114242 | 0.00988217 | 0.00154202 | 0.14948051 | 0.99999994 |
| Phtf1 | 0.11810551 | 0.10023217 | 0.09869105 | 0.00154112 | 0.45860737 | 0.99999994 |
| Trim59 | 0.036379 | 0.03368384 | 0.03214473 | 0.00153911 | 0.37497291 | 0.99999994 |
| Kynu | 0.0132268 | 0.01340777 | 0.01186976 | 0.00153801 | 0.1942094 | 0.99999994 |
| Snapc5 | 0.31852964 | 0.2296954 | 0.22816502 | 0.00153038 | 0.48219455 | 0.99999994 |
| Pter | 0.02343843 | 0.02241832 | 0.02088984 | 0.00152848 | 0.31311584 | 0.99999994 |
| Ccdc90b | 0.25545574 | 0.1933864 | 0.19186151 | 0.00152488 | 0.47890435 | 0.99999994 |
| Zfp277 | 0.16698163 | 0.13577911 | 0.13425445 | 0.00152466 | 0.46987153 | 0.99999994 |
| Cmtm7 | 0.05071684 | 0.04591538 | 0.04439115 | 0.00152423 | 0.40960758 | 0.99999994 |
| Pnma1 | 0.01339014 | 0.01353722 | 0.01201501 | 0.00152221 | 0.19952831 | 0.99999994 |
| Ckm | 0.0059209 | 0.00686091 | 0.00533973 | 0.00152117 | 0.02896324 | 0.86317853 |
| Adat2 | 0.05062987 | 0.04583772 | 0.04431757 | 0.00152016 | 0.40969769 | 0.99999994 |
| Snai2 | 0.0146222 | 0.01462973 | 0.01310965 | 0.00152008 | 0.22011295 | 0.99999994 |
| Riok1 | 0.14289878 | 0.11860423 | 0.11708542 | 0.00151881 | 0.46559607 | 0.99999994 |
| Hic2 | 0.03436793 | 0.03191817 | 0.03040835 | 0.00150983 | 0.37051111 | 0.99999994 |
| Cdyl | 0.03248102 | 0.03028134 | 0.02877492 | 0.00150641 | 0.36374347 | 0.99999994 |
| Tcea2 | 0.04507333 | 0.04110321 | 0.03959851 | 0.0015047 | 0.40015947 | 0.99999994 |
| Rnf2 | 0.22274173 | 0.17302231 | 0.17151866 | 0.00150365 | 0.47673318 | 0.99999994 |
| Bcl10 | 0.26509352 | 0.19915333 | 0.19765171 | 0.00150162 | 0.47983392 | 0.99999994 |
| Vpreb3 | 0.00136791 | 0.00273837 | 0.00123746 | 0.0015009 | 3.42E-16 | 9.56E-14 |
| Pah | 0.02031358 | 0.01964268 | 0.01814274 | 0.00149994 | 0.29105355 | 0.99999994 |
| Cbx2 | 0.04426333 | 0.04040667 | 0.03890771 | 0.00149896 | 0.39880427 | 0.99999994 |
| Vnn1 | 0.02263033 | 0.02167935 | 0.02018054 | 0.00149881 | 0.31052299 | 0.99999994 |
| Cops6 | 0.96910246 | 0.42583196 | 0.42433507 | 0.00149689 | 0.49063331 | 0.99999994 |
| Flad1 | 0.13776866 | 0.11483656 | 0.11334006 | 0.0014965 | 0.46498278 | 0.99999994 |
| Syt17 | 0.06810544 | 0.06042926 | 0.05893692 | 0.00149234 | 0.4330769 | 0.99999994 |
| Cgnl1 | 0.02776819 | 0.02616949 | 0.02467725 | 0.00149225 | 0.34365315 | 0.99999994 |
| Cdyl2 | 0.03731571 | 0.03444082 | 0.03295192 | 0.0014889 | 0.38179854 | 0.99999994 |

| | | | | | | |
|---|---|---|---|---|---|---|
| Akna | 0.01067522 | 0.01108533 | 0.00959647 | 0.00148886 | 0.15086918 | 0.99999994 |
| Tmem222 | 0.21034927 | 0.16501368 | 0.16352584 | 0.00148783 | 0.47585377 | 0.99999994 |
| Smndc1 | 0.27364263 | 0.20419507 | 0.20271268 | 0.0014824 | 0.48058851 | 0.99999994 |
| Crb3 | 0.46722854 | 0.30186851 | 0.30038697 | 0.00148154 | 0.48690511 | 0.99999994 |
| Rsf1 | 0.43567627 | 0.28796499 | 0.28648629 | 0.0014787 | 0.48629632 | 0.99999994 |
| Adgrg1 | 0.3390354 | 0.24066724 | 0.23919022 | 0.00147702 | 0.48360665 | 0.99999994 |
| Lrch2 | 0.03656839 | 0.03378348 | 0.03230802 | 0.00147546 | 0.38056914 | 0.99999994 |
| Cenpq | 0.08157865 | 0.0714579 | 0.0699832 | 0.0014747 | 0.44422566 | 0.99999994 |
| Trap1 | 0.20714679 | 0.16290877 | 0.16143441 | 0.00147437 | 0.47576246 | 0.99999994 |
| Tpp1 | 0.18937344 | 0.15109558 | 0.14962558 | 0.00147 | 0.47392959 | 0.99999994 |
| Ccnb1 | 0.02079007 | 0.0200256 | 0.01856238 | 0.00146322 | 0.2998948 | 0.99999994 |
| Gk | 0.01347543 | 0.01355107 | 0.01209085 | 0.00146022 | 0.21072936 | 0.99999994 |
| Nsmf | 0.10991639 | 0.09390994 | 0.09244982 | 0.00146012 | 0.4581371 | 0.99999994 |
| Col26a1 | 0.02513777 | 0.02383422 | 0.02237894 | 0.00145529 | 0.33255889 | 0.99999994 |
| Ltk | 0.02185567 | 0.02095121 | 0.01949985 | 0.00145135 | 0.31014978 | 0.99999994 |
| Brms1l | 0.11858957 | 0.10050721 | 0.09905734 | 0.00144987 | 0.46119361 | 0.99999994 |
| Plk4 | 0.02312066 | 0.02205957 | 0.02061101 | 0.00144856 | 0.31995897 | 0.99999994 |
| Dnph1 | 0.05193448 | 0.04686525 | 0.04542055 | 0.0014447 | 0.41616253 | 0.99999994 |
| Ccdc180 | 0.0124505 | 0.01262001 | 0.01117895 | 0.00144106 | 0.19543028 | 0.99999994 |
| Znrf1 | 0.14747191 | 0.12183647 | 0.12039781 | 0.00143866 | 0.46830217 | 0.99999994 |
| Shox2 | 0.03291963 | 0.03059203 | 0.02915497 | 0.00143706 | 0.37142041 | 0.99999994 |
| Zfp296 | 0.01289977 | 0.01300132 | 0.01157883 | 0.00142249 | 0.20674941 | 0.99999994 |
| Sbsn | 0.03158676 | 0.02941875 | 0.02799936 | 0.00141939 | 0.36789466 | 0.99999994 |
| Lin7a | 0.00789091 | 0.00852619 | 0.00710689 | 0.0014193 | 0.0918702 | 0.99999994 |
| Rchy1 | 0.5111494 | 0.32000051 | 0.31858335 | 0.00141716 | 0.48818922 | 0.99999994 |
| Gm28050 | 0.01456249 | 0.01447369 | 0.01305665 | 0.00141704 | 0.23501992 | 0.99999994 |
| Actr3 | 0.7858102 | 0.40225591 | 0.4008419 | 0.00141401 | 0.49063332 | 0.99999994 |
| Kcnh7 | 0.02164766 | 0.02072552 | 0.01931696 | 0.00140856 | 0.31370703 | 0.99999994 |
| Nrsn1 | 0.0069639 | 0.00768128 | 0.00627592 | 0.00140536 | 0.06803999 | 0.99999994 |
| Slc17a5 | 0.1753401 | 0.14146144 | 0.14005799 | 0.00140345 | 0.47341023 | 0.99999994 |

| | | | | | | |
|---|---|---|---|---|---|---|
| 1700016K19I | 0.05138936 | 0.04636309 | 0.04495991 | 0.00140318 | 0.41771497 | 0.99999994 |
| Creb3l4 | 0.0166428 | 0.01630388 | 0.01490095 | 0.00140293 | 0.26542879 | 0.99999994 |
| Gm3739 | 0.00804077 | 0.00864402 | 0.00724112 | 0.0014029 | 0.09859341 | 0.99999994 |
| Nradd | 0.03615298 | 0.03335019 | 0.03194982 | 0.00140037 | 0.38523876 | 0.99999994 |
| Prtn3 | 0.03119578 | 0.02905977 | 0.02765999 | 0.00139978 | 0.36811261 | 0.99999994 |
| Tstd3 | 0.04920047 | 0.04450623 | 0.0431069 | 0.00139933 | 0.41446387 | 0.99999994 |
| Klf16 | 0.030774 | 0.02869102 | 0.02729369 | 0.00139733 | 0.36663454 | 0.99999994 |
| Zfp821 | 0.13635636 | 0.11369961 | 0.11230352 | 0.00139609 | 0.467026 | 0.99999994 |
| Nek6 | 0.08128565 | 0.07113905 | 0.06974502 | 0.00139403 | 0.4470791 | 0.99999994 |
| Sbk1 | 0.20295449 | 0.16007126 | 0.15867934 | 0.00139191 | 0.4767195 | 0.99999994 |
| Cept1 | 0.13664542 | 0.11390183 | 0.11251586 | 0.00138597 | 0.46732625 | 0.99999994 |
| Amotl1 | 0.13206353 | 0.11052292 | 0.10913824 | 0.00138468 | 0.46634855 | 0.99999994 |
| Sec14l2 | 0.01214269 | 0.01228331 | 0.01090485 | 0.00137846 | 0.20005667 | 0.99999994 |
| Ppp2r3c | 0.22520502 | 0.17446416 | 0.17308807 | 0.00137609 | 0.47889799 | 0.99999994 |
| Pdk2 | 0.13183543 | 0.11034138 | 0.10896944 | 0.00137194 | 0.46660607 | 0.99999994 |
| Cd248 | 0.00901945 | 0.00948231 | 0.0081171 | 0.00136521 | 0.13145651 | 0.99999994 |
| Ist1 | 0.21192609 | 0.16591299 | 0.16455168 | 0.00136131 | 0.47804249 | 0.99999994 |
| Akap5 | 0.01973991 | 0.01899773 | 0.01763717 | 0.00136056 | 0.30380701 | 0.99999994 |
| Zfp820 | 0.02222898 | 0.02118412 | 0.01982797 | 0.00135616 | 0.32445794 | 0.99999994 |
| Klk1b9 | 0.00646846 | 0.00718162 | 0.00583139 | 0.00135023 | 0.06162762 | 0.99999994 |
| Gdi1 | 0.93003644 | 0.42248149 | 0.42113203 | 0.00134945 | 0.4914915 | 0.99999994 |
| Setmar | 0.02476504 | 0.02340195 | 0.02205261 | 0.00134933 | 0.34190026 | 0.99999994 |
| 1110019D14I | 0.03483579 | 0.03216152 | 0.03081271 | 0.00134881 | 0.38538119 | 0.99999994 |
| Tada2a | 0.06058115 | 0.05403206 | 0.05268334 | 0.00134873 | 0.43234495 | 0.99999994 |
| Gbe1 | 0.0627197 | 0.05581466 | 0.05446697 | 0.00134768 | 0.43459038 | 0.99999994 |
| Gm43672 | 0.02531525 | 0.02387978 | 0.02253426 | 0.00134553 | 0.34551679 | 0.99999994 |
| Palm3 | 0.0102505 | 0.01056159 | 0.00921732 | 0.00134426 | 0.16583122 | 0.99999994 |
| Slc18a2 | 0.00649246 | 0.00719432 | 0.00585293 | 0.0013414 | 0.06356351 | 0.99999994 |
| Mrpl27 | 0.48115927 | 0.30764149 | 0.30630017 | 0.00134132 | 0.48837292 | 0.99999994 |
| Dhx8 | 0.08913491 | 0.07743501 | 0.07609478 | 0.00134023 | 0.45333646 | 0.99999994 |

| | | | | | | |
|---|---|---|---|---|---|---|
| Uba7 | 0.01051544 | 0.01079074 | 0.00945386 | 0.00133688 | 0.17328077 | 0.99999994 |
| Tinf2 | 0.0212275 | 0.02028137 | 0.01894737 | 0.001334 | 0.31966207 | 0.99999994 |
| Prmt3 | 0.08932443 | 0.07757668 | 0.0762473 | 0.00132938 | 0.45380473 | 0.99999994 |
| Smtn | 0.0150755 | 0.01483908 | 0.01351194 | 0.00132714 | 0.2566243 | 0.99999994 |
| D030040B21 | 0.00918727 | 0.00959288 | 0.00826719 | 0.00132569 | 0.14289946 | 0.99999994 |
| Tmem67 | 0.06328346 | 0.0562604 | 0.05493634 | 0.00132406 | 0.43627133 | 0.99999994 |
| Armcx6 | 0.08165714 | 0.07137011 | 0.07004699 | 0.00132312 | 0.44997189 | 0.99999994 |
| Ppp4c | 0.8019272 | 0.40517461 | 0.40385161 | 0.001323 | 0.49130138 | 0.99999994 |
| Rnf138 | 0.27135267 | 0.20268586 | 0.20136404 | 0.00132182 | 0.48257395 | 0.99999994 |
| Rbmx | 0.25847058 | 0.1950028 | 0.19368234 | 0.00132046 | 0.48190188 | 0.99999994 |
| Pxdn | 0.04718752 | 0.04271494 | 0.04139812 | 0.00131682 | 0.41615912 | 0.99999994 |
| Cep126 | 0.05957594 | 0.05315964 | 0.05184323 | 0.00131641 | 0.43289061 | 0.99999994 |
| Crabp1 | 0.00637138 | 0.00705978 | 0.00574425 | 0.00131553 | 0.06370158 | 0.99999994 |
| Gata4 | 0.01263044 | 0.01265444 | 0.01133914 | 0.0013153 | 0.22002356 | 0.99999994 |
| Fdxr | 0.08685612 | 0.07557064 | 0.07425792 | 0.00131272 | 0.45316447 | 0.99999994 |
| Sox10 | 0.00464477 | 0.00550255 | 0.00419248 | 0.00131007 | 0.01876371 | 0.64463872 |
| Lefty1 | 0.03241953 | 0.03002812 | 0.02872163 | 0.00130649 | 0.38102714 | 0.99999994 |
| Spred3 | 0.03109073 | 0.02887183 | 0.02756878 | 0.00130305 | 0.37652645 | 0.99999994 |
| Fam114a2 | 0.45757366 | 0.29751184 | 0.29620886 | 0.00130298 | 0.48832048 | 0.99999994 |
| Smtnl2 | 0.03620523 | 0.03329437 | 0.03199488 | 0.00129949 | 0.39344446 | 0.99999994 |
| Ap4m1 | 0.16841309 | 0.13655239 | 0.1352539 | 0.00129849 | 0.47452357 | 0.99999994 |
| Tet3 | 0.25899777 | 0.1952961 | 0.19399984 | 0.00129626 | 0.48226236 | 0.99999994 |
| Rps6ka1 | 0.06761856 | 0.05982686 | 0.05853412 | 0.00129274 | 0.44156312 | 0.99999994 |
| Tmprss6 | 0.01740167 | 0.01686297 | 0.01557245 | 0.00129052 | 0.29060196 | 0.99999994 |
| Otop1 | 0.00467675 | 0.00551142 | 0.00422126 | 0.00129016 | 0.02095527 | 0.69426319 |
| Padi3 | 0.00830174 | 0.0087641 | 0.00747482 | 0.00128927 | 0.12546343 | 0.99999994 |
| Celf6 | 0.04453966 | 0.04043228 | 0.03914345 | 0.00128883 | 0.41326034 | 0.99999994 |
| Cryz | 0.09285886 | 0.08036947 | 0.07908501 | 0.00128447 | 0.45695425 | 0.99999994 |
| B930036N10 | 0.03399756 | 0.03137225 | 0.03008805 | 0.0012842 | 0.38816544 | 0.99999994 |
| Ckap4 | 0.13241289 | 0.11067814 | 0.10939667 | 0.00128148 | 0.46892485 | 0.99999994 |

| | | | | | | |
|---|---|---|---|---|---|---|
| Chn2 | 0.01715882 | 0.01663671 | 0.01535764 | 0.00127908 | 0.28965861 | 0.99999994 |
| Dusp26 | 0.03605659 | 0.03313712 | 0.03186667 | 0.00127044 | 0.39536143 | 0.99999994 |
| Cnot6 | 0.29477057 | 0.21619395 | 0.21492368 | 0.00127027 | 0.48430914 | 0.99999994 |
| Picalm | 0.24949108 | 0.18950117 | 0.18823231 | 0.00126886 | 0.48210545 | 0.99999994 |
| Gm26910 | 0.0100197 | 0.01027823 | 0.00901119 | 0.00126704 | 0.17465327 | 0.99999994 |
| Fhl3 | 0.01490466 | 0.01462597 | 0.01336036 | 0.00126561 | 0.26416776 | 0.99999994 |
| Hoxa3 | 0.0043802 | 0.00521391 | 0.00395439 | 0.00125953 | 0.01699693 | 0.60693149 |
| Rfwd3 | 0.09973385 | 0.08582565 | 0.08456814 | 0.0012575 | 0.46057782 | 0.99999994 |
| Iws1 | 0.23793444 | 0.18235793 | 0.18110052 | 0.00125741 | 0.48156895 | 0.99999994 |
| Mctp2 | 0.00528304 | 0.00602373 | 0.00476654 | 0.00125719 | 0.03957582 | 0.99999994 |
| Chrna3 | 0.00770553 | 0.0081953 | 0.0069408 | 0.0012545 | 0.11447051 | 0.99999994 |
| Arap2 | 0.00983859 | 0.01010195 | 0.00884939 | 0.00125256 | 0.17305704 | 0.99999994 |
| Ctss | 0.01026097 | 0.01047405 | 0.00922667 | 0.00124738 | 0.18408823 | 0.99999994 |
| Derl3 | 0.17301415 | 0.13969734 | 0.13845083 | 0.00124651 | 0.47610608 | 0.99999994 |
| Eif2s3x | 0.19544149 | 0.15494063 | 0.15369549 | 0.00124514 | 0.47849734 | 0.99999994 |
| Kcnh8 | 0.0811274 | 0.07086102 | 0.06961634 | 0.00124468 | 0.45263387 | 0.99999994 |
| Plac1 | 0.01454532 | 0.01428559 | 0.0130414 | 0.00124419 | 0.26270175 | 0.99999994 |
| Gpr153 | 0.0131682 | 0.01305928 | 0.01181763 | 0.00124165 | 0.24216129 | 0.99999994 |
| Chtf18 | 0.01312133 | 0.01301663 | 0.01177594 | 0.00124069 | 0.24155744 | 0.99999994 |
| Sft2d1 | 0.20153866 | 0.15898226 | 0.15774471 | 0.00123755 | 0.47917631 | 0.99999994 |
| Tfap2b | 0.00294287 | 0.00389378 | 0.00265938 | 0.00123441 | 0.0010018 | 0.07066263 |
| Dynlrb2 | 0.05708965 | 0.05099438 | 0.04976055 | 0.00123384 | 0.43445325 | 0.99999994 |
| Cdk6 | 0.02946624 | 0.02738285 | 0.02615662 | 0.00122623 | 0.37750092 | 0.99999994 |
| Mms22l | 0.01524923 | 0.01489157 | 0.01366605 | 0.00122551 | 0.27528226 | 0.99999994 |
| Stk24 | 0.08709157 | 0.07566829 | 0.07444796 | 0.00122033 | 0.45655781 | 0.99999994 |
| Cenpf | 0.02277459 | 0.02152613 | 0.02030721 | 0.00121892 | 0.34474744 | 0.99999994 |
| Apcs | 0.01094183 | 0.01105307 | 0.00983437 | 0.0012187 | 0.20472192 | 0.99999994 |
| Rmdn1 | 0.19211378 | 0.1526866 | 0.15146875 | 0.00121786 | 0.47865913 | 0.99999994 |
| Mcoln3 | 0.04018516 | 0.03663245 | 0.03541838 | 0.00121407 | 0.40975837 | 0.99999994 |
| Brf2 | 0.0502724 | 0.04522871 | 0.04401502 | 0.00121369 | 0.4271848 | 0.99999994 |

| | | | | | | |
|---|---|---|---|---|---|---|
| Tnfrsf12a | 0.14878366 | 0.12255427 | 0.12134328 | 0.00121099 | 0.47351793 | 0.99999994 |
| Fst | 0.00102423 | 0.00213479 | 0.00092677 | 0.00120802 | 2.04E-18 | 6.51E-16 |
| Ubqln4 | 0.08694857 | 0.07553896 | 0.07433255 | 0.00120641 | 0.45698508 | 0.99999994 |
| Pcdhb5 | 0.01010851 | 0.01029607 | 0.00909051 | 0.00120556 | 0.18868543 | 0.99999994 |
| Lmx1b | 0.05718444 | 0.05104425 | 0.04984007 | 0.00120418 | 0.43611607 | 0.99999994 |
| Cdh3 | 0.01406303 | 0.01381411 | 0.01261307 | 0.00120104 | 0.26309564 | 0.99999994 |
| Klf7 | 0.32888303 | 0.23497635 | 0.2337778 | 0.00119855 | 0.48638816 | 0.99999994 |
| Gimap6 | 0.01829233 | 0.01755621 | 0.01635969 | 0.00119652 | 0.31318765 | 0.99999994 |
| Cdca7l | 0.02393602 | 0.02252206 | 0.02132622 | 0.00119583 | 0.35448308 | 0.99999994 |
| Chrna7 | 0.00261729 | 0.003559 | 0.00236568 | 0.00119331 | 0.00039305 | 0.0311723 |
| Kctd20 | 0.22973559 | 0.17715023 | 0.17595849 | 0.00119173 | 0.48202091 | 0.99999994 |
| Mars2 | 0.01376409 | 0.01353904 | 0.01234744 | 0.00119159 | 0.26031555 | 0.99999994 |
| 2010308F09I | 0.01783374 | 0.01714404 | 0.01595446 | 0.00118958 | 0.30984046 | 0.99999994 |
| Pls1 | 0.03276234 | 0.03020642 | 0.02901871 | 0.00118772 | 0.39264172 | 0.99999994 |
| Mmp2 | 0.00674466 | 0.00726642 | 0.00607924 | 0.00118717 | 0.09682198 | 0.99999994 |
| Dock10 | 0.01646873 | 0.01592118 | 0.01474683 | 0.00117435 | 0.29803101 | 0.99999994 |
| Gabrb3 | 0.02259652 | 0.02132505 | 0.02015084 | 0.0011742 | 0.34905598 | 0.99999994 |
| Galm | 0.02733566 | 0.02547346 | 0.02429988 | 0.00117358 | 0.37392617 | 0.99999994 |
| Nt5e | 0.016021 | 0.0155185 | 0.01435024 | 0.00116826 | 0.29394416 | 0.99999994 |
| Celf2 | 0.15597919 | 0.12766098 | 0.12649413 | 0.00116686 | 0.47551947 | 0.99999994 |
| Pcdh7 | 0.00624585 | 0.00679765 | 0.00563155 | 0.0011661 | 0.08405445 | 0.99999994 |
| Prc1 | 0.01197348 | 0.01192013 | 0.01075411 | 0.00116601 | 0.23523515 | 0.99999994 |
| Cdk5rap2 | 0.05351824 | 0.04791784 | 0.046757 | 0.00116084 | 0.43437026 | 0.99999994 |
| Ebf2 | 0.00522389 | 0.00587209 | 0.00471337 | 0.00115873 | 0.05087927 | 0.99999994 |
| Adrb2 | 0.01558153 | 0.01511877 | 0.01396072 | 0.00115804 | 0.29042442 | 0.99999994 |
| Pld1 | 0.02357777 | 0.0221657 | 0.02101207 | 0.00115363 | 0.35738581 | 0.99999994 |
| C130013H08I | 0.02737117 | 0.02548361 | 0.02433087 | 0.00115274 | 0.37624272 | 0.99999994 |
| Pank1 | 0.05303538 | 0.04750058 | 0.04634983 | 0.00115075 | 0.43436925 | 0.99999994 |
| Raph1 | 0.07015731 | 0.06178159 | 0.06063166 | 0.00114993 | 0.4497696 | 0.99999994 |
| Zbtb18 | 0.06946591 | 0.0612108 | 0.0600611 | 0.0011497 | 0.44930528 | 0.99999994 |

| | | | | | | |
|---|---|---|---|---|---|---|
| Ccdc125 | 0.0415692 | 0.03775364 | 0.03660469 | 0.00114895 | 0.41725166 | 0.99999994 |
| Tead3 | 0.02540455 | 0.02375798 | 0.0226124 | 0.00114559 | 0.36797429 | 0.99999994 |
| Adamts10 | 0.01995154 | 0.01896743 | 0.01782373 | 0.00114371 | 0.33464504 | 0.99999994 |
| Ccdc103 | 0.02187488 | 0.02065607 | 0.01951675 | 0.00113932 | 0.34879523 | 0.99999994 |
| Brwd3 | 0.08711487 | 0.07560509 | 0.07446676 | 0.00113832 | 0.45947703 | 0.99999994 |
| Sla2 | 0.01081341 | 0.0108575 | 0.0097198 | 0.00113771 | 0.21795195 | 0.99999994 |
| Zfp867 | 0.03134431 | 0.02892569 | 0.02778894 | 0.00113675 | 0.39270046 | 0.99999994 |
| Aggf1 | 0.17830599 | 0.14323339 | 0.14209826 | 0.00113513 | 0.47879689 | 0.99999994 |
| Bhlhe22 | 0.0056814 | 0.00625533 | 0.00512457 | 0.00113076 | 0.07094975 | 0.99999994 |
| Mthfs | 0.15392883 | 0.12616135 | 0.12503249 | 0.00112886 | 0.47603911 | 0.99999994 |
| Slc31a2 | 0.24621411 | 0.18735128 | 0.18622337 | 0.00112791 | 0.48392055 | 0.99999994 |
| Cdh16 | 0.00961118 | 0.00977266 | 0.00864617 | 0.00112648 | 0.19290655 | 0.99999994 |
| Nr2f2 | 0.01807922 | 0.01729525 | 0.01617141 | 0.00112385 | 0.32182927 | 0.99999994 |
| Gm15611 | 0.00114819 | 0.00216111 | 0.00103885 | 0.00112226 | 3.22E-13 | 7.66E-11 |
| Nfatc4 | 0.01529714 | 0.01482731 | 0.01370854 | 0.00111877 | 0.29348524 | 0.99999994 |
| Pxn | 0.04239396 | 0.03842806 | 0.03731058 | 0.00111748 | 0.42098938 | 0.99999994 |
| Clvs1 | 0.00839244 | 0.00867293 | 0.00755602 | 0.00111691 | 0.16257629 | 0.99999994 |
| Sybu | 0.01398747 | 0.01366222 | 0.01254594 | 0.00111628 | 0.27684155 | 0.99999994 |
| Fastkd2 | 0.06968319 | 0.0613566 | 0.06024046 | 0.00111614 | 0.45092329 | 0.99999994 |
| Gem | 0.08440966 | 0.07339498 | 0.07227991 | 0.00111507 | 0.45910518 | 0.99999994 |
| Fbxo30 | 0.05611434 | 0.05005255 | 0.04894171 | 0.00111084 | 0.4399558 | 0.99999994 |
| Lypd6b | 0.01103619 | 0.01102566 | 0.00991855 | 0.00110711 | 0.22874634 | 0.99999994 |
| Ddx27 | 0.15045554 | 0.12364968 | 0.12254542 | 0.00110427 | 0.4760854 | 0.99999994 |
| Otud5 | 0.2105835 | 0.16477669 | 0.16367839 | 0.0010983 | 0.48218721 | 0.99999994 |
| Myl9 | 0.01833303 | 0.01749225 | 0.01639564 | 0.00109661 | 0.32808554 | 0.99999994 |
| Siah1b | 0.07515966 | 0.06583402 | 0.0647445 | 0.00108952 | 0.45540632 | 0.99999994 |
| Pcdh15 | 0.01159186 | 0.01150324 | 0.01041404 | 0.0010892 | 0.24315572 | 0.99999994 |
| 1700124L16I | 0.00332612 | 0.00409283 | 0.00300493 | 0.0010879 | 0.0079793 | 0.36342269 |
| Lpar2 | 0.01586416 | 0.01529736 | 0.01421125 | 0.00108611 | 0.30547451 | 0.99999994 |
| Epha3 | 0.00992417 | 0.01001073 | 0.00892585 | 0.00108488 | 0.20924604 | 0.99999994 |

| | | | | | | |
|---|---|---|---|---|---|---|
| Rimklb | 0.0156481 | 0.01510427 | 0.01401974 | 0.00108453 | 0.30330502 | 0.99999994 |
| Slc13a4 | 0.00599857 | 0.00649306 | 0.0054095 | 0.00108356 | 0.09121418 | 0.99999994 |
| Timeless | 0.03178551 | 0.02925495 | 0.02817181 | 0.00108313 | 0.39900626 | 0.99999994 |
| Sass6 | 0.04587064 | 0.04135448 | 0.04027778 | 0.0010767 | 0.42938617 | 0.99999994 |
| Trip13 | 0.01555963 | 0.01501771 | 0.01394131 | 0.0010764 | 0.30364911 | 0.99999994 |
| Otop2 | 0.00337857 | 0.00412805 | 0.0030522 | 0.00107585 | 0.00948151 | 0.41984649 |
| Prpf4b | 0.42430706 | 0.28237493 | 0.28129993 | 0.00107499 | 0.48985306 | 0.99999994 |
| Col4a2 | 0.03061979 | 0.02823454 | 0.02715971 | 0.00107483 | 0.3961127 | 0.99999994 |
| Syce1 | 0.01302136 | 0.01276071 | 0.01168701 | 0.0010737 | 0.27042548 | 0.99999994 |
| Cntrl | 0.09038327 | 0.07817055 | 0.07709877 | 0.00107178 | 0.46313742 | 0.99999994 |
| Rcbtb2 | 0.17458174 | 0.14060529 | 0.13953465 | 0.00107064 | 0.47963329 | 0.99999994 |
| Coil | 0.10433109 | 0.08921528 | 0.08814489 | 0.00107039 | 0.46778797 | 0.99999994 |
| Irs4 | 0.00356684 | 0.00429025 | 0.00322188 | 0.00106838 | 0.01364804 | 0.53455393 |
| Slco2a1 | 0.03029375 | 0.0279424 | 0.02687636 | 0.00106604 | 0.39588129 | 0.99999994 |
| Tmem91 | 0.01286735 | 0.0126141 | 0.01154998 | 0.00106412 | 0.26985168 | 0.99999994 |
| Jkamp | 0.50942881 | 0.31895804 | 0.31789476 | 0.00106328 | 0.49111876 | 0.99999994 |
| Xlr4c | 0.00965223 | 0.00974101 | 0.00868286 | 0.00105815 | 0.20862864 | 0.99999994 |
| G430095P16 | 0.01168673 | 0.0115566 | 0.01049859 | 0.00105801 | 0.25117281 | 0.99999994 |
| Tbrg4 | 0.05768364 | 0.05131266 | 0.05025873 | 0.00105393 | 0.44449437 | 0.99999994 |
| Xpc | 0.10038007 | 0.08612966 | 0.08507587 | 0.00105379 | 0.46714479 | 0.99999994 |
| Bmf | 0.04697533 | 0.04227005 | 0.04121774 | 0.00105232 | 0.43252804 | 0.99999994 |
| Rgs12 | 0.04270022 | 0.03862474 | 0.03757251 | 0.00105222 | 0.42606082 | 0.99999994 |
| Akap7 | 0.04017434 | 0.03645662 | 0.03540909 | 0.00104752 | 0.42194581 | 0.99999994 |
| Galnt3 | 0.02397349 | 0.0224062 | 0.02135907 | 0.00104713 | 0.37208758 | 0.99999994 |
| Prss41 | 0.00860419 | 0.00879183 | 0.00774556 | 0.00104627 | 0.18428863 | 0.99999994 |
| Trp53rkb | 0.10577319 | 0.09030592 | 0.08926016 | 0.00104576 | 0.46891986 | 0.99999994 |
| Cenpj | 0.05415941 | 0.04834244 | 0.04729726 | 0.00104519 | 0.44152903 | 0.99999994 |
| Nudt4 | 0.12225661 | 0.10286753 | 0.10182284 | 0.00104469 | 0.472776 | 0.99999994 |
| Adora2a | 0.03192088 | 0.02932459 | 0.02828924 | 0.00103534 | 0.40376538 | 0.99999994 |
| Prtg | 0.00576009 | 0.00623016 | 0.00519527 | 0.00103488 | 0.09243189 | 0.99999994 |

| | | | | | | |
|---|---|---|---|---|---|---|
| Madcam1 | 0.00893025 | 0.00907113 | 0.00803731 | 0.00103382 | 0.19594662 | 0.99999994 |
| Lrrcc1 | 0.098954 | 0.08498232 | 0.08394859 | 0.00103373 | 0.46733705 | 0.99999994 |
| Ajap1 | 0.01035732 | 0.01034595 | 0.0093127 | 0.00103325 | 0.23009812 | 0.99999994 |
| Dnajb5 | 0.03092356 | 0.02845664 | 0.0274236 | 0.00103304 | 0.40100638 | 0.99999994 |
| Zfp382 | 0.00629942 | 0.0067121 | 0.00567964 | 0.00103245 | 0.11313942 | 0.99999994 |
| Hoxb2 | 0.00834182 | 0.00854146 | 0.00751071 | 0.00103075 | 0.18049006 | 0.99999994 |
| Sntb2 | 0.0757191 | 0.06623234 | 0.0652028 | 0.00102954 | 0.45814706 | 0.99999994 |
| Foxc1 | 0.00814634 | 0.0083626 | 0.00733567 | 0.00102693 | 0.17571122 | 0.99999994 |
| Prl | 0.01484294 | 0.01433105 | 0.01330558 | 0.00102547 | 0.30397193 | 0.99999994 |
| Fgf18 | 0.00382001 | 0.00447439 | 0.00344997 | 0.00102442 | 0.02404859 | 0.76825358 |
| Zbtb12 | 0.09978746 | 0.08563381 | 0.08461071 | 0.0010231 | 0.46792481 | 0.99999994 |
| Mad2l1 | 0.03523296 | 0.03217586 | 0.03115579 | 0.00102007 | 0.41373997 | 0.99999994 |
| Sfn | 0.00888615 | 0.00901691 | 0.00799786 | 0.00101906 | 0.19818287 | 0.99999994 |
| Rhod | 0.02795902 | 0.02586245 | 0.02484367 | 0.00101879 | 0.39244075 | 0.99999994 |
| Gldc | 0.02470386 | 0.02301749 | 0.02199903 | 0.00101846 | 0.37898003 | 0.99999994 |
| Krtap4-16 | 0.0059884 | 0.00641862 | 0.00540036 | 0.00101826 | 0.10472476 | 0.99999994 |
| Trpv2 | 0.01015665 | 0.01015067 | 0.00913351 | 0.00101716 | 0.22925775 | 0.99999994 |
| Pthlh | 0.00910751 | 0.00921296 | 0.00819586 | 0.0010171 | 0.20438824 | 0.99999994 |
| Rara | 0.0349615 | 0.03193508 | 0.03092132 | 0.00101376 | 0.41362515 | 0.99999994 |
| Sirt5 | 0.07803509 | 0.06810687 | 0.06709658 | 0.00101029 | 0.46008229 | 0.99999994 |
| Tut1 | 0.05846206 | 0.0519202 | 0.05091101 | 0.00100919 | 0.44751306 | 0.99999994 |
| Smpd3 | 0.17653549 | 0.14188982 | 0.14088162 | 0.00100821 | 0.48100332 | 0.99999994 |
| Ncam2 | 0.00736722 | 0.00764306 | 0.00663759 | 0.00100547 | 0.15665155 | 0.99999994 |
| Xlr4a | 0.00852039 | 0.00867453 | 0.00767056 | 0.00100397 | 0.19181637 | 0.99999994 |
| Nadk2 | 0.03532973 | 0.03223893 | 0.03123935 | 0.00099958 | 0.41566875 | 0.99999994 |
| Reep5 | 0.64409699 | 0.36724144 | 0.36624256 | 0.00099887 | 0.4927579 | 0.99999994 |
| Nup62cl | 0.00386905 | 0.00449082 | 0.00349414 | 0.00099668 | 0.02880271 | 0.86317853 |
| Trim71 | 0.00946427 | 0.00950975 | 0.00851486 | 0.00099489 | 0.21836013 | 0.99999994 |
| Bbof1 | 0.02440389 | 0.02272962 | 0.02173627 | 0.00099334 | 0.38049005 | 0.99999994 |
| Map4k1 | 0.00993016 | 0.00991866 | 0.0089312 | 0.00098746 | 0.23088213 | 0.99999994 |

| | | | | | | |
|---|---|---|---|---|---|---|
| Rpl9-ps1 | 0.00638347 | 0.00673812 | 0.00575509 | 0.00098303 | 0.12777455 | 0.99999994 |
| Tmem29 | 0.20571949 | 0.16148042 | 0.1604985 | 0.00098192 | 0.48375827 | 0.99999994 |
| Gm5422 | 0.00740104 | 0.00764859 | 0.0066679 | 0.00098069 | 0.16379266 | 0.99999994 |
| Sema3a | 0.0046471 | 0.00517207 | 0.00419458 | 0.00097749 | 0.06042973 | 0.99999994 |
| Il1rap | 0.023933 | 0.0222984 | 0.02132358 | 0.00097483 | 0.38044903 | 0.99999994 |
| Prss27 | 0.01130232 | 0.01113062 | 0.0101559 | 0.00097472 | 0.26146057 | 0.99999994 |
| Ing2 | 0.32359184 | 0.23189523 | 0.23092086 | 0.00097437 | 0.48879653 | 0.99999994 |
| Ston2 | 0.02564593 | 0.02379755 | 0.02282357 | 0.00097398 | 0.38818433 | 0.99999994 |
| Srfbp1 | 0.04668218 | 0.0419421 | 0.04096844 | 0.00097366 | 0.43715128 | 0.99999994 |
| Mgll | 0.01761798 | 0.01673641 | 0.01576373 | 0.00097268 | 0.34063858 | 0.99999994 |
| Prr29 | 0.01714784 | 0.01631954 | 0.01534792 | 0.00097163 | 0.33673451 | 0.99999994 |
| Rtn4rl1 | 0.02727219 | 0.02521488 | 0.02424449 | 0.00097039 | 0.39495682 | 0.99999994 |
| BC055324 | 0.01103483 | 0.01087958 | 0.00991734 | 0.00096225 | 0.2591901 | 0.99999994 |
| Ifngr1 | 0.14493595 | 0.1195258 | 0.11856411 | 0.00096169 | 0.47847128 | 0.99999994 |
| Zbtb16 | 0.01062956 | 0.01051683 | 0.00955573 | 0.0009611 | 0.25159216 | 0.99999994 |
| Ppp1r16b | 0.008225 | 0.00836688 | 0.00740611 | 0.00096077 | 0.19392998 | 0.99999994 |
| Tbc1d25 | 0.03713392 | 0.03375605 | 0.03279535 | 0.0009607 | 0.42270007 | 0.99999994 |
| Abhd2 | 0.12556719 | 0.10526586 | 0.10430542 | 0.00096044 | 0.47556366 | 0.99999994 |
| Acrbp | 0.02531355 | 0.02348994 | 0.02253277 | 0.00095716 | 0.38868281 | 0.99999994 |
| Nudt10 | 0.02595555 | 0.02405117 | 0.02309433 | 0.00095684 | 0.39135636 | 0.99999994 |
| Slc29a4 | 0.17969025 | 0.14400207 | 0.14304708 | 0.000955 | 0.48227744 | 0.99999994 |
| 2610037D02I | 0.04228445 | 0.03817072 | 0.03721691 | 0.00095381 | 0.43227235 | 0.99999994 |
| Actr5 | 0.0356066 | 0.03243098 | 0.03147838 | 0.0009526 | 0.42017887 | 0.99999994 |
| Mag | 0.01953529 | 0.01840884 | 0.01745674 | 0.0009521 | 0.35828711 | 0.99999994 |
| Exd2 | 0.04529582 | 0.04073716 | 0.03978814 | 0.00094903 | 0.43692601 | 0.99999994 |
| Pgp | 0.46884777 | 0.30202818 | 0.30108143 | 0.00094675 | 0.49165037 | 0.99999994 |
| Tmx2 | 0.34139125 | 0.24137721 | 0.24043368 | 0.00094353 | 0.48958024 | 0.99999994 |
| Col11a2 | 0.03786946 | 0.03437198 | 0.03342863 | 0.00094334 | 0.42550102 | 0.99999994 |
| Adamts5 | 0.00413296 | 0.00467222 | 0.00373181 | 0.00094041 | 0.04673301 | 0.99999994 |
| Tatdn3 | 0.05923211 | 0.05249464 | 0.05155561 | 0.00093903 | 0.45175132 | 0.99999994 |

| | | | | | | |
|---|---|---|---|---|---|---|
| Trpv6 | 0.00941706 | 0.00940979 | 0.00847265 | 0.00093714 | 0.23079114 | 0.99999994 |
| Fgfbp1 | 0.00305329 | 0.00369568 | 0.00275896 | 0.00093672 | 0.01191153 | 0.50100705 |
| Cdc7 | 0.01091302 | 0.01074451 | 0.00980866 | 0.00093584 | 0.26268705 | 0.99999994 |
| Nhej1 | 0.03550857 | 0.0323296 | 0.03139376 | 0.00093583 | 0.42135711 | 0.99999994 |
| Enpp3 | 0.01328634 | 0.01285491 | 0.0119227 | 0.00093221 | 0.30137602 | 0.99999994 |
| Zfp467 | 0.0778833 | 0.06790365 | 0.06697264 | 0.00093101 | 0.46313754 | 0.99999994 |
| Acan | 0.00254763 | 0.00323376 | 0.00230283 | 0.00093093 | 0.00356337 | 0.20071964 |
| Psg16 | 0.00471003 | 0.00517983 | 0.00425121 | 0.00092862 | 0.07297408 | 0.99999994 |
| Uba2 | 0.44023775 | 0.28946878 | 0.2885403 | 0.00092848 | 0.49145562 | 0.99999994 |
| Sstr2 | 0.01671861 | 0.01589423 | 0.01496807 | 0.00092616 | 0.34021794 | 0.99999994 |
| Tfap2c | 0.00572924 | 0.00609264 | 0.00516755 | 0.00092508 | 0.11670811 | 0.99999994 |
| Alpl | 0.00613428 | 0.00645495 | 0.00553137 | 0.00092358 | 0.1331923 | 0.99999994 |
| Vill | 0.01086809 | 0.01068969 | 0.00976859 | 0.0009211 | 0.26511898 | 0.99999994 |
| Zcwpw1 | 0.02015473 | 0.0189225 | 0.01800279 | 0.00091971 | 0.36690765 | 0.99999994 |
| Ggt5 | 0.00823032 | 0.00832803 | 0.00741087 | 0.00091715 | 0.20503343 | 0.99999994 |
| Atpaf2 | 0.14807216 | 0.1217468 | 0.1208307 | 0.0009161 | 0.47987533 | 0.99999994 |
| Aurka | 0.0274088 | 0.02527855 | 0.02436371 | 0.00091483 | 0.40131689 | 0.99999994 |
| Dbf4 | 0.02773014 | 0.02555631 | 0.02464406 | 0.00091225 | 0.40268653 | 0.99999994 |
| Vgll4 | 0.16984517 | 0.13716187 | 0.13625148 | 0.00091039 | 0.48226261 | 0.99999994 |
| Dcp1b | 0.07239202 | 0.0633824 | 0.0624723 | 0.0009101 | 0.46137496 | 0.99999994 |
| Spdef | 0.01458253 | 0.01398262 | 0.01307444 | 0.00090818 | 0.32190885 | 0.99999994 |
| Mob3b | 0.01477296 | 0.01415153 | 0.01324348 | 0.00090805 | 0.32405208 | 0.99999994 |
| Dpysl3 | 0.01814583 | 0.01713791 | 0.01623026 | 0.00090765 | 0.35485511 | 0.99999994 |
| Rbms2 | 0.05771989 | 0.05119307 | 0.05028911 | 0.00090396 | 0.4523803 | 0.99999994 |
| Ldhb | 0.05621817 | 0.04993174 | 0.04902893 | 0.00090281 | 0.45122405 | 0.99999994 |
| Als2cl | 0.00780279 | 0.00792937 | 0.00702794 | 0.00090143 | 0.19661554 | 0.99999994 |
| Ppp2r3d | 0.18099205 | 0.14483714 | 0.14393745 | 0.00089969 | 0.48340635 | 0.99999994 |
| Dnajc22 | 0.0260488 | 0.02407461 | 0.02317586 | 0.00089875 | 0.39815266 | 0.99999994 |
| Lyrm9 | 0.22721747 | 0.17526417 | 0.17436566 | 0.00089851 | 0.48631877 | 0.99999994 |
| Trit1 | 0.13012236 | 0.10859442 | 0.10769962 | 0.0008948 | 0.47794864 | 0.99999994 |

| | | | | | | |
|---|---|---|---|---|---|---|
| Gpr157 | 0.00695359 | 0.00716147 | 0.00626667 | 0.00089479 | 0.17094556 | 0.99999994 |
| Mycn | 0.01432833 | 0.013743 | 0.01284872 | 0.00089428 | 0.321579 | 0.99999994 |
| 4933429H19 | 0.00568967 | 0.00602613 | 0.005132 | 0.00089413 | 0.12308355 | 0.99999994 |
| Dars | 0.40553546 | 0.27341698 | 0.27252403 | 0.00089296 | 0.49129962 | 0.99999994 |
| Nrg1 | 0.01461866 | 0.01399929 | 0.01310651 | 0.00089278 | 0.32512555 | 0.99999994 |
| Msh2 | 0.16266196 | 0.1321153 | 0.13122444 | 0.00089086 | 0.48197833 | 0.99999994 |
| Rdh10 | 0.0143936 | 0.01379687 | 0.01290669 | 0.00089018 | 0.32308432 | 0.99999994 |
| Asap3 | 0.00933355 | 0.00928763 | 0.008398 | 0.00088963 | 0.24036293 | 0.99999994 |
| Esrp2 | 0.07359797 | 0.06434681 | 0.06346336 | 0.00088344 | 0.46308684 | 0.99999994 |
| Egfl6 | 0.01319316 | 0.01272182 | 0.01183983 | 0.00088199 | 0.30999933 | 0.99999994 |
| Trim7 | 0.02760344 | 0.02541485 | 0.02453354 | 0.00088131 | 0.40550855 | 0.99999994 |
| Mterf4 | 0.15357045 | 0.12565634 | 0.12477651 | 0.00087983 | 0.48128224 | 0.99999994 |
| Zfp442 | 0.04577431 | 0.04107484 | 0.04019575 | 0.00087909 | 0.44212825 | 0.99999994 |
| Kcnd2 | 0.00576732 | 0.00608077 | 0.00520177 | 0.000879 | 0.13033858 | 0.99999994 |
| Ptpn18 | 0.03998674 | 0.03612699 | 0.03524813 | 0.00087886 | 0.43409158 | 0.99999994 |
| Ascl1 | 0.00785271 | 0.00795152 | 0.00707267 | 0.00087885 | 0.20408397 | 0.99999994 |
| Art5 | 0.0034103 | 0.00395927 | 0.0030808 | 0.00087847 | 0.02884672 | 0.86317853 |
| Fen1 | 0.11081413 | 0.09401618 | 0.09313812 | 0.00087806 | 0.47498162 | 0.99999994 |
| Acot2 | 0.03612341 | 0.03280203 | 0.03192431 | 0.00087772 | 0.42739532 | 0.99999994 |
| Gimd1 | 0.00514806 | 0.00552216 | 0.00464519 | 0.00087697 | 0.10443688 | 0.99999994 |
| Gpr39 | 0.03475649 | 0.03161939 | 0.03074419 | 0.00087519 | 0.42485575 | 0.99999994 |
| Wnt7b | 0.00516407 | 0.00553322 | 0.00465958 | 0.00087363 | 0.10601327 | 0.99999994 |
| Morn4 | 0.04259194 | 0.0383516 | 0.03747992 | 0.00087168 | 0.43848594 | 0.99999994 |
| Dvl3 | 0.06113062 | 0.05401304 | 0.05314209 | 0.00087095 | 0.4565647 | 0.99999994 |
| Praf2 | 0.10659713 | 0.09076554 | 0.08989619 | 0.00086935 | 0.47433738 | 0.99999994 |
| Shc2 | 0.05843284 | 0.05174894 | 0.05088653 | 0.00086241 | 0.45509061 | 0.99999994 |
| Arpc1b | 0.36841607 | 0.25522414 | 0.25436179 | 0.00086235 | 0.49099797 | 0.99999994 |
| Zfp618 | 0.11370152 | 0.09620698 | 0.09534509 | 0.00086189 | 0.47600958 | 0.99999994 |
| Rab11fip1 | 0.01746036 | 0.01648486 | 0.01562435 | 0.0008605 | 0.35696023 | 0.99999994 |
| Taok3 | 0.1979518 | 0.15622474 | 0.15536742 | 0.00085732 | 0.48535023 | 0.99999994 |

| | | | | | | |
|---|---|---|---|---|---|---|
| Haus6 | 0.01400645 | 0.01341823 | 0.01256281 | 0.00085541 | 0.32518911 | 0.99999994 |
| BC051019 | 0.00232851 | 0.00295937 | 0.00210508 | 0.00085429 | 0.00345349 | 0.19802917 |
| Lrrk1 | 0.00358756 | 0.00409471 | 0.00324055 | 0.00085416 | 0.03967076 | 0.99999994 |
| Ehf | 0.01712029 | 0.01617699 | 0.01532355 | 0.00085344 | 0.35542143 | 0.99999994 |
| Eif5a2 | 0.01273417 | 0.01228425 | 0.01143146 | 0.00085279 | 0.30974741 | 0.99999994 |
| Cd79b | 0.00189212 | 0.0025624 | 0.00171107 | 0.00085134 | 0.0004634 | 0.03603423 |
| Mzt1 | 0.23832234 | 0.18218845 | 0.18134204 | 0.00084642 | 0.48760737 | 0.99999994 |
| Aldob | 0.00326736 | 0.00379743 | 0.00295196 | 0.00084547 | 0.02829553 | 0.85915875 |
| Gm10645 | 0.0104534 | 0.01024394 | 0.00939848 | 0.00084546 | 0.27465708 | 0.99999994 |
| Tm4sf5 | 0.00428484 | 0.00471133 | 0.00386855 | 0.00084277 | 0.07351454 | 0.99999994 |
| Tpgs1 | 0.54553519 | 0.33278803 | 0.33194546 | 0.00084257 | 0.49325988 | 0.99999994 |
| Efcab12 | 0.00434799 | 0.00476745 | 0.00392539 | 0.00084206 | 0.07666086 | 0.99999994 |
| Prr7 | 0.0961873 | 0.0825873 | 0.0817456 | 0.0008417 | 0.47267886 | 0.99999994 |
| Zfp184 | 0.03001559 | 0.02747524 | 0.02663451 | 0.00084073 | 0.41679104 | 0.99999994 |
| Kitl | 0.00324715 | 0.00377076 | 0.00293374 | 0.00083703 | 0.02877387 | 0.86317853 |
| Otud3 | 0.04959885 | 0.04428058 | 0.04344455 | 0.00083604 | 0.44903752 | 0.99999994 |
| Nos1ap | 0.00754521 | 0.00763304 | 0.00679713 | 0.00083592 | 0.20650606 | 0.99999994 |
| A930031H19 | 0.00412454 | 0.00455974 | 0.00372423 | 0.00083551 | 0.06767614 | 0.99999994 |
| Ccdc136 | 0.0820434 | 0.07119297 | 0.07036081 | 0.00083216 | 0.46862554 | 0.99999994 |
| Trpc5 | 0.0065331 | 0.00672026 | 0.0058894 | 0.00083086 | 0.17384978 | 0.99999994 |
| Fam118b | 0.18869505 | 0.14999582 | 0.14916802 | 0.0008278 | 0.48526678 | 0.99999994 |
| Epha5 | 0.00654737 | 0.00672815 | 0.00590221 | 0.00082595 | 0.17580224 | 0.99999994 |
| Sema4c | 0.08675564 | 0.07500055 | 0.0741768 | 0.00082375 | 0.47053702 | 0.99999994 |
| Epha7 | 0.02214022 | 0.02057112 | 0.01974996 | 0.00082116 | 0.39098385 | 0.99999994 |
| Zfp101 | 0.06830953 | 0.05992652 | 0.05910569 | 0.00082083 | 0.46317433 | 0.99999994 |
| Arl4c | 0.05279498 | 0.04696757 | 0.04614703 | 0.00082054 | 0.45289231 | 0.99999994 |
| Slc52a3 | 0.03081972 | 0.02815369 | 0.02733341 | 0.00082028 | 0.42083441 | 0.99999994 |
| Haus4 | 0.01760403 | 0.01657137 | 0.0157514 | 0.00081997 | 0.36447988 | 0.99999994 |
| Smpd4 | 0.03333994 | 0.03033834 | 0.02951897 | 0.00081938 | 0.42670684 | 0.99999994 |
| Lpcat4 | 0.01561322 | 0.01480765 | 0.01398882 | 0.00081883 | 0.34840561 | 0.99999994 |

| | | | | | | |
|---|---|---|---|---|---|---|
| Kcnj13 | 0.00410863 | 0.00452644 | 0.0037099 | 0.00081654 | 0.07145448 | 0.99999994 |
| Rbpjl | 0.01480111 | 0.01408493 | 0.01326846 | 0.00081647 | 0.34105134 | 0.99999994 |
| Med18 | 0.05339615 | 0.04747026 | 0.04665407 | 0.00081619 | 0.45364821 | 0.99999994 |
| Nudt17 | 0.01397261 | 0.01334791 | 0.01253275 | 0.00081516 | 0.33252715 | 0.99999994 |
| Spire2 | 0.03504974 | 0.03181178 | 0.03099754 | 0.00081424 | 0.43059933 | 0.99999994 |
| Rgma | 0.00794503 | 0.00796863 | 0.00715537 | 0.00081326 | 0.22466279 | 0.99999994 |
| Ptgfrn | 0.0186646 | 0.01750163 | 0.01668845 | 0.00081317 | 0.37283979 | 0.99999994 |
| Arhgef15 | 0.00800074 | 0.00801827 | 0.00720527 | 0.000813 | 0.22630637 | 0.99999994 |
| D2hgdh | 0.03871301 | 0.0349642 | 0.03415415 | 0.00081005 | 0.4372791 | 0.99999994 |
| Gm8973 | 0.0126595 | 0.01216858 | 0.01136501 | 0.00080357 | 0.31894717 | 0.99999994 |
| Dlat | 0.29994149 | 0.21865103 | 0.21784944 | 0.00080159 | 0.49022985 | 0.99999994 |
| Zmat2 | 0.63539243 | 0.36425481 | 0.3634548 | 0.00080001 | 0.49415509 | 0.99999994 |
| Tcf24 | 0.00661526 | 0.00676232 | 0.00596313 | 0.00079919 | 0.1861688 | 0.99999994 |
| Eif2ak4 | 0.05016822 | 0.04472451 | 0.04392681 | 0.0007977 | 0.45189415 | 0.99999994 |
| Ccsap | 0.0140583 | 0.01340624 | 0.01260888 | 0.00079736 | 0.33690175 | 0.99999994 |
| Loxl1 | 0.02493214 | 0.02299568 | 0.02219893 | 0.00079675 | 0.40558948 | 0.99999994 |
| Sh3bgrl2 | 0.01350665 | 0.01291253 | 0.01211861 | 0.00079392 | 0.3313914 | 0.99999994 |
| Gm20449 | 0.00220603 | 0.00278781 | 0.00199452 | 0.00079329 | 0.00405483 | 0.21763658 |
| Ninl | 0.01833542 | 0.01718939 | 0.01639775 | 0.00079164 | 0.37397267 | 0.99999994 |
| Wdr75 | 0.11888597 | 0.10007197 | 0.09928148 | 0.00079049 | 0.47886653 | 0.99999994 |
| Sp3os | 0.27535653 | 0.20450718 | 0.20371884 | 0.00078834 | 0.48972504 | 0.99999994 |
| Igfn1 | 0.0036628 | 0.00409552 | 0.00330834 | 0.00078718 | 0.05661988 | 0.99999994 |
| Ncan | 0.00486961 | 0.00518168 | 0.00439476 | 0.00078691 | 0.11665659 | 0.99999994 |
| Ptpn5 | 0.00759779 | 0.00763076 | 0.00684425 | 0.00078651 | 0.22216105 | 0.99999994 |
| Nr4a1 | 0.06932073 | 0.06072639 | 0.05994123 | 0.00078516 | 0.46526004 | 0.99999994 |
| Tagln | 0.00671168 | 0.0068332 | 0.00604965 | 0.00078355 | 0.19430898 | 0.99999994 |
| Fam111a | 0.02080215 | 0.01935575 | 0.01857301 | 0.00078274 | 0.38953734 | 0.99999994 |
| Zbtb2 | 0.03860125 | 0.03484016 | 0.03405807 | 0.00078209 | 0.43925736 | 0.99999994 |
| Gm29202 | 0.00331798 | 0.00377902 | 0.00299759 | 0.00078143 | 0.04135087 | 0.99999994 |
| Fgf9 | 0.04638562 | 0.04149724 | 0.04071613 | 0.00078111 | 0.44919397 | 0.99999994 |

| | | | | | | |
|---|---|---|---|---|---|---|
| Bok | 0.00937907 | 0.00921924 | 0.00843869 | 0.00078054 | 0.26905187 | 0.99999994 |
| Lef1 | 0.00132972 | 0.00198311 | 0.00120294 | 0.00078017 | 7.91E-06 | 0.000865 |
| Gm20045 | 0.01810563 | 0.01697391 | 0.01619474 | 0.00077917 | 0.37438768 | 0.99999994 |
| Gfra4 | 0.00553926 | 0.00577466 | 0.00499684 | 0.00077781 | 0.15006946 | 0.99999994 |
| 4930579G24 | 0.03460717 | 0.03139264 | 0.03061515 | 0.00077749 | 0.43288191 | 0.99999994 |
| Hpd | 0.00734478 | 0.0073924 | 0.00661747 | 0.00077494 | 0.21784569 | 0.99999994 |
| Tenm4 | 0.06804083 | 0.05965612 | 0.05888348 | 0.00077264 | 0.46520021 | 0.99999994 |
| Bcar3 | 0.01784604 | 0.01673753 | 0.01596534 | 0.0007722 | 0.37374608 | 0.99999994 |
| Cldn23 | 0.08004841 | 0.06950915 | 0.06873827 | 0.00077087 | 0.47024716 | 0.99999994 |
| Chst9 | 0.02139874 | 0.01986848 | 0.01909803 | 0.00077045 | 0.39414595 | 0.99999994 |
| Neat1 | 0.0694949 | 0.06085403 | 0.06008504 | 0.00076899 | 0.46605502 | 0.99999994 |
| Tbxas1 | 0.00479974 | 0.00510007 | 0.00433191 | 0.00076816 | 0.11893272 | 0.99999994 |
| Fam89b | 0.01170017 | 0.01127857 | 0.01051057 | 0.000768 | 0.31335101 | 0.99999994 |
| Fndc4 | 0.00903769 | 0.00889977 | 0.00813341 | 0.00076636 | 0.26526919 | 0.99999994 |
| Hdhd3 | 0.11887527 | 0.1000371 | 0.09927339 | 0.0007637 | 0.47958029 | 0.99999994 |
| Spag6l | 0.01191591 | 0.01146403 | 0.01070282 | 0.0007612 | 0.31795945 | 0.99999994 |
| Rasl11a | 0.006012 | 0.00618163 | 0.00542155 | 0.00076008 | 0.17536054 | 0.99999994 |
| Hdac2 | 0.78795177 | 0.40200954 | 0.40125039 | 0.00075915 | 0.49497604 | 0.99999994 |
| Rpusd2 | 0.01324404 | 0.01264399 | 0.01188509 | 0.0007589 | 0.3354086 | 0.99999994 |
| Gpr142 | 0.01134308 | 0.01095095 | 0.01019225 | 0.0007587 | 0.31012531 | 0.99999994 |
| Ccne2 | 0.02310704 | 0.02135734 | 0.02059905 | 0.00075828 | 0.40321629 | 0.99999994 |
| Tmem183a | 0.30503571 | 0.22146533 | 0.22070716 | 0.00075817 | 0.49087869 | 0.99999994 |
| Dsc2 | 0.02647413 | 0.02430559 | 0.02354758 | 0.00075801 | 0.4151674 | 0.99999994 |
| Kdm2b | 0.09197662 | 0.07913559 | 0.07837788 | 0.00075771 | 0.47434582 | 0.99999994 |
| Gsg1l2 | 0.00816945 | 0.00811321 | 0.00735636 | 0.00075685 | 0.24672562 | 0.99999994 |
| Trim68 | 0.01750357 | 0.01641848 | 0.01566257 | 0.00075591 | 0.37401016 | 0.99999994 |
| Slc18a3 | 0.00433329 | 0.00466795 | 0.00391216 | 0.00075578 | 0.0992336 | 0.99999994 |
| Shcbp1 | 0.00873353 | 0.00861709 | 0.00786131 | 0.00075578 | 0.26110557 | 0.99999994 |
| Dusp5 | 0.14305488 | 0.11795315 | 0.1171989 | 0.00075425 | 0.48291543 | 0.99999994 |
| Gsdma2 | 0.00322327 | 0.00366589 | 0.00291221 | 0.00075368 | 0.04247355 | 0.99999994 |

| | | | | | | |
|---|---|---|---|---|---|---|
| Dmgdh | 0.00898115 | 0.00883467 | 0.00808284 | 0.00075183 | 0.26790903 | 0.99999994 |
| Akr1cl | 0.00750679 | 0.00751288 | 0.00676269 | 0.00075019 | 0.23013643 | 0.99999994 |
| Spock3 | 0.01912185 | 0.01784092 | 0.01709204 | 0.00074888 | 0.38527926 | 0.99999994 |
| Spag8 | 0.01230032 | 0.01179361 | 0.01104523 | 0.00074838 | 0.32599253 | 0.99999994 |
| 3425401B19I | 0.00187287 | 0.00244055 | 0.00169369 | 0.00074686 | 0.00166644 | 0.11114947 |
| Tle4 | 0.02288886 | 0.02115386 | 0.02040754 | 0.00074632 | 0.40383655 | 0.99999994 |
| Pabpc4l | 0.01361748 | 0.01296196 | 0.01221713 | 0.00074482 | 0.34244208 | 0.99999994 |
| Pls3 | 0.04534781 | 0.04057439 | 0.03983243 | 0.00074196 | 0.45066207 | 0.99999994 |
| Ntn4 | 0.01177901 | 0.0113227 | 0.01058084 | 0.00074186 | 0.32035617 | 0.99999994 |
| Sapcd1 | 0.01107076 | 0.01069088 | 0.00994938 | 0.0007415 | 0.30991839 | 0.99999994 |
| Mocs3 | 0.03252169 | 0.02954968 | 0.02881017 | 0.00073951 | 0.43216822 | 0.99999994 |
| Rab28 | 0.39994099 | 0.27059524 | 0.26985598 | 0.00073926 | 0.49272576 | 0.99999994 |
| Oas3 | 0.00929164 | 0.00909946 | 0.00836052 | 0.00073894 | 0.27816008 | 0.99999994 |
| Gabre | 0.00692257 | 0.00697715 | 0.00623885 | 0.00073831 | 0.21543154 | 0.99999994 |
| Prim1 | 0.07005135 | 0.06128096 | 0.06054426 | 0.0007367 | 0.46772329 | 0.99999994 |
| Guca2b | 0.01113132 | 0.01073691 | 0.01000341 | 0.00073351 | 0.31274621 | 0.99999994 |
| Ttll8 | 0.00331763 | 0.00373062 | 0.00299727 | 0.00073335 | 0.05169579 | 0.99999994 |
| Rapsn | 0.00624153 | 0.0063597 | 0.00562767 | 0.00073203 | 0.19328794 | 0.99999994 |
| Gm27153 | 0.00376878 | 0.00413533 | 0.00340382 | 0.00073151 | 0.07628617 | 0.99999994 |
| Bcl2l15 | 0.01136111 | 0.01093931 | 0.01020832 | 0.00073099 | 0.31680858 | 0.99999994 |
| Zmat5 | 0.23430539 | 0.1795632 | 0.17883281 | 0.00073039 | 0.48915578 | 0.99999994 |
| Cfap45 | 0.00594635 | 0.00609162 | 0.00536259 | 0.00072903 | 0.18275516 | 0.99999994 |
| 4933407L21F | 0.01105689 | 0.01066399 | 0.00993701 | 0.00072697 | 0.31314125 | 0.99999994 |
| Iqsec1 | 0.03993265 | 0.0359274 | 0.0352017 | 0.0007257 | 0.44542703 | 0.99999994 |
| Slc6a19 | 0.00367052 | 0.00403939 | 0.00331529 | 0.00072409 | 0.07299843 | 0.99999994 |
| Emx1 | 0.00453369 | 0.00481597 | 0.00409254 | 0.00072344 | 0.1196692 | 0.99999994 |
| Zfp661 | 0.01013473 | 0.00983696 | 0.00911393 | 0.00072304 | 0.29872488 | 0.99999994 |
| Car14 | 0.00412648 | 0.00444787 | 0.00372598 | 0.00072189 | 0.0985882 | 0.99999994 |
| Tmsb15b1 | 0.09962747 | 0.08520324 | 0.08448366 | 0.00071958 | 0.47739427 | 0.99999994 |
| Tmem123 | 0.11191853 | 0.09469986 | 0.09398349 | 0.00071638 | 0.47976743 | 0.99999994 |

| | | | | | | |
|---|---|---|---|---|---|---|
| A630023P12 | 0.0095151 | 0.0092751 | 0.00856029 | 0.0007148 | 0.28916645 | 0.99999994 |
| Nyap1 | 0.05458095 | 0.0483666 | 0.04765221 | 0.00071439 | 0.46025534 | 0.99999994 |
| Gm17092 | 0.00565101 | 0.0058115 | 0.00509727 | 0.00071423 | 0.17548916 | 0.99999994 |
| Spats2l | 0.00563821 | 0.00579977 | 0.00508576 | 0.00071401 | 0.17502023 | 0.99999994 |
| Fndc5 | 0.02993724 | 0.0272795 | 0.02656638 | 0.00071312 | 0.42909567 | 0.99999994 |
| Tmem139 | 0.01004144 | 0.00974361 | 0.0090306 | 0.00071301 | 0.29959923 | 0.99999994 |
| Wdyhv1 | 0.06163013 | 0.05427176 | 0.05355885 | 0.00071291 | 0.46469941 | 0.99999994 |
| Apela | 0.00433117 | 0.0046231 | 0.00391026 | 0.00071284 | 0.11247886 | 0.99999994 |
| Cd22 | 0.01492538 | 0.01409154 | 0.01337874 | 0.0007128 | 0.36142897 | 0.99999994 |
| Ramp3 | 0.00420165 | 0.00450624 | 0.00379366 | 0.00071259 | 0.10559442 | 0.99999994 |
| Gm15893 | 0.00526844 | 0.0054658 | 0.00475342 | 0.00071238 | 0.15924337 | 0.99999994 |
| Icosl | 0.00996549 | 0.00967465 | 0.00896276 | 0.00071189 | 0.29850713 | 0.99999994 |
| Zc3hav1l | 0.03049071 | 0.02775795 | 0.02704755 | 0.0007104 | 0.43060757 | 0.99999994 |
| Atpaf1 | 0.1111879 | 0.09413366 | 0.09342439 | 0.00070927 | 0.47984814 | 0.99999994 |
| Ccp110 | 0.06524542 | 0.05727542 | 0.05656713 | 0.0007083 | 0.46678792 | 0.99999994 |
| Gm26801 | 0.00867247 | 0.00851444 | 0.00780666 | 0.00070777 | 0.27309341 | 0.99999994 |
| E2f7 | 0.00951539 | 0.00926732 | 0.00856055 | 0.00070677 | 0.29131249 | 0.99999994 |
| Gm10827 | 0.00931997 | 0.0090919 | 0.00838585 | 0.00070606 | 0.28758945 | 0.99999994 |
| Gm4425 | 0.01432774 | 0.01355378 | 0.0128482 | 0.00070558 | 0.35735335 | 0.99999994 |
| Whrn | 0.0546595 | 0.0484235 | 0.04771833 | 0.00070517 | 0.46082073 | 0.99999994 |
| Armcx5 | 0.0885338 | 0.07631507 | 0.07561077 | 0.00070431 | 0.47528004 | 0.99999994 |
| Gimap3 | 0.00085113 | 0.0014742 | 0.00077023 | 0.00070397 | 5.87E-10 | 1.03E-07 |
| Apobr | 0.00844019 | 0.00829789 | 0.00759877 | 0.00069912 | 0.27013145 | 0.99999994 |
| Gcnt3 | 0.00952759 | 0.00926722 | 0.00857145 | 0.00069577 | 0.29448936 | 0.99999994 |
| Rnf38 | 0.0589628 | 0.05202558 | 0.05133025 | 0.00069533 | 0.4640766 | 0.99999994 |
| Ahr | 0.020432 | 0.01894221 | 0.01824706 | 0.00069515 | 0.39990842 | 0.99999994 |
| Gm28901 | 0.00328138 | 0.00365803 | 0.0029646 | 0.00069343 | 0.05974155 | 0.99999994 |
| Fgf21 | 0.0051564 | 0.00534522 | 0.00465268 | 0.00069254 | 0.16089742 | 0.99999994 |
| Xpnpep2 | 0.0106552 | 0.01027087 | 0.00957861 | 0.00069227 | 0.31523375 | 0.99999994 |
| Gata1 | 0.00351137 | 0.00386394 | 0.00317189 | 0.00069205 | 0.07320794 | 0.99999994 |

| | | | | | | |
|---|---|---|---|---|---|---|
| Nppc | 0.00360549 | 0.0039444 | 0.0032567 | 0.00068769 | 0.07992535 | 0.99999994 |
| Cideb | 0.00726326 | 0.00723185 | 0.00654438 | 0.00068747 | 0.24220318 | 0.99999994 |
| Kcna5 | 0.00897652 | 0.00876585 | 0.0080787 | 0.00068715 | 0.28563933 | 0.99999994 |
| Gm4673 | 0.01221962 | 0.01165879 | 0.01097336 | 0.00068543 | 0.33878663 | 0.99999994 |
| Rab24 | 0.46491017 | 0.30007443 | 0.29938954 | 0.00068489 | 0.49392547 | 0.99999994 |
| Tigd5 | 0.01170645 | 0.0112001 | 0.01051617 | 0.00068393 | 0.33254253 | 0.99999994 |
| Fcor | 0.02355487 | 0.02167474 | 0.02099198 | 0.00068276 | 0.41430069 | 0.99999994 |
| Cntnap4 | 0.00234813 | 0.00280439 | 0.00212279 | 0.0006816 | 0.01628808 | 0.58877176 |
| Cntln | 0.08498491 | 0.07342502 | 0.07274557 | 0.00067944 | 0.47521364 | 0.99999994 |
| Sult5a1 | 0.01706009 | 0.01594925 | 0.01527028 | 0.00067896 | 0.38362936 | 0.99999994 |
| Wnt3 | 0.00172939 | 0.00224253 | 0.00156408 | 0.00067845 | 0.00194257 | 0.12691219 |
| Brap | 0.08692096 | 0.07498771 | 0.07431026 | 0.00067746 | 0.47580573 | 0.99999994 |
| Cenpe | 0.01065441 | 0.01025038 | 0.0095779 | 0.00067248 | 0.32012282 | 0.99999994 |
| Ogfrl1 | 0.01757619 | 0.01639922 | 0.01572678 | 0.00067244 | 0.38797119 | 0.99999994 |
| Rims4 | 0.01142722 | 0.01093891 | 0.01026727 | 0.00067165 | 0.33162364 | 0.99999994 |
| Fzd9 | 0.00868894 | 0.00849192 | 0.00782141 | 0.00067051 | 0.28412281 | 0.99999994 |
| Hoga1 | 0.00957762 | 0.00928582 | 0.00861617 | 0.00066964 | 0.30246393 | 0.99999994 |
| Mcts2 | 0.23053562 | 0.17713198 | 0.17646322 | 0.00066876 | 0.48993722 | 0.99999994 |
| Spon2 | 0.00781844 | 0.00770977 | 0.00704196 | 0.00066781 | 0.2639417 | 0.99999994 |
| Amot | 0.00726016 | 0.00720932 | 0.0065416 | 0.00066772 | 0.24843313 | 0.99999994 |
| Prex1 | 0.07213711 | 0.06292859 | 0.06226261 | 0.00066599 | 0.47161969 | 0.99999994 |
| Mbl2 | 0.02607183 | 0.0238611 | 0.02319599 | 0.00066512 | 0.42431653 | 0.99999994 |
| Ccdc85a | 0.00391361 | 0.00419862 | 0.00353428 | 0.00066434 | 0.10542941 | 0.99999994 |
| Arhgap4 | 0.00412492 | 0.00438813 | 0.00372457 | 0.00066356 | 0.1178351 | 0.99999994 |
| E130308A19 | 0.05138034 | 0.04561348 | 0.04495228 | 0.00066119 | 0.46100323 | 0.99999994 |
| Efna5 | 0.01743136 | 0.01625782 | 0.01559871 | 0.00065911 | 0.38925629 | 0.99999994 |
| Bin1 | 0.1726008 | 0.13882297 | 0.13816459 | 0.00065837 | 0.48734827 | 0.99999994 |
| Serpinb8 | 0.00228642 | 0.00272412 | 0.00206708 | 0.00065704 | 0.0171817 | 0.60730275 |
| Scube1 | 0.00899851 | 0.00875536 | 0.00809837 | 0.000657 | 0.29459505 | 0.99999994 |
| Sh3gl3 | 0.01414381 | 0.01334157 | 0.01268483 | 0.00065673 | 0.36518961 | 0.99999994 |

| | | | | | | |
|---|---|---|---|---|---|---|
| Zfp383 | 0.03172807 | 0.02877852 | 0.02812198 | 0.00065654 | 0.43825325 | 0.99999994 |
| Stk39 | 0.04765389 | 0.04244878 | 0.04179442 | 0.00065436 | 0.45849893 | 0.99999994 |
| Aldh8a1 | 0.00277991 | 0.00316647 | 0.0025124 | 0.00065408 | 0.04155532 | 0.99999994 |
| Gnpda1 | 0.06315084 | 0.05547819 | 0.05482595 | 0.00065224 | 0.46844168 | 0.99999994 |
| Il27ra | 0.00626111 | 0.00629738 | 0.00564525 | 0.00065213 | 0.22096611 | 0.99999994 |
| Trafd1 | 0.1758169 | 0.1410383 | 0.1403867 | 0.0006516 | 0.48767648 | 0.99999994 |
| Vps11 | 0.07145068 | 0.06234808 | 0.0616976 | 0.00065048 | 0.47202601 | 0.99999994 |
| Proser3 | 0.05174407 | 0.0459096 | 0.04525968 | 0.00064992 | 0.46192529 | 0.99999994 |
| Naaladl2 | 0.00769975 | 0.00758478 | 0.00693562 | 0.00064917 | 0.26663344 | 0.99999994 |
| Tmc3 | 0.00174211 | 0.00222435 | 0.00157558 | 0.00064877 | 0.0030638 | 0.18022151 |
| Lrrc47 | 0.12026411 | 0.10097097 | 0.10032224 | 0.00064873 | 0.48283354 | 0.99999994 |
| Tmem37 | 0.01684554 | 0.01572882 | 0.01508042 | 0.0006484 | 0.38736387 | 0.99999994 |
| Grsf1 | 0.22865689 | 0.17592296 | 0.17527695 | 0.00064601 | 0.49021376 | 0.99999994 |
| Hsbp1l1 | 0.07997893 | 0.06932719 | 0.06868169 | 0.0006455 | 0.4750588 | 0.99999994 |
| Greb1 | 0.00944165 | 0.00913961 | 0.00849464 | 0.00064497 | 0.30664004 | 0.99999994 |
| Cpb2 | 0.02057079 | 0.01901249 | 0.0183693 | 0.00064319 | 0.40785345 | 0.99999994 |
| Gm2990 | 0.00348193 | 0.0037878 | 0.00314536 | 0.00064244 | 0.08698573 | 0.99999994 |
| Rfx2 | 0.0314181 | 0.02849509 | 0.02785299 | 0.0006421 | 0.43902154 | 0.99999994 |
| Rhbdf1 | 0.01487916 | 0.013978 | 0.01333772 | 0.00064028 | 0.37465918 | 0.99999994 |
| Etv4 | 0.00396553 | 0.00422017 | 0.00358104 | 0.00063913 | 0.11741742 | 0.99999994 |
| Adamts9 | 0.02824864 | 0.02573411 | 0.02509617 | 0.00063795 | 0.43281766 | 0.99999994 |
| Gm13166 | 0.00587416 | 0.00593549 | 0.00529775 | 0.00063773 | 0.21148445 | 0.99999994 |
| Rad18 | 0.01278293 | 0.0121102 | 0.01147486 | 0.00063534 | 0.35623134 | 0.99999994 |
| Slc25a20 | 0.15445189 | 0.12604082 | 0.12540582 | 0.000635 | 0.48655614 | 0.99999994 |
| Colgalt2 | 0.0012613 | 0.00177559 | 0.0011411 | 0.00063448 | 0.00010733 | 0.00983279 |
| 5730507C01I | 0.00732624 | 0.00723424 | 0.00660084 | 0.00063339 | 0.26150344 | 0.99999994 |
| Zik1 | 0.01037132 | 0.00995849 | 0.0093252 | 0.0006333 | 0.32561633 | 0.99999994 |
| Katnb1 | 0.09817981 | 0.08396457 | 0.08333292 | 0.00063165 | 0.47988029 | 0.99999994 |
| Cadm3 | 0.00657244 | 0.00655592 | 0.00592471 | 0.00063122 | 0.23911039 | 0.99999994 |
| Pdzrn3 | 0.00935285 | 0.00904528 | 0.00841525 | 0.00063003 | 0.30911963 | 0.99999994 |

| | | | | | | |
|---|---|---|---|---|---|---|
| Pcdhb9 | 0.00632383 | 0.00633081 | 0.00570156 | 0.00062926 | 0.23128169 | 0.99999994 |
| Fbln5 | 0.00614704 | 0.0061707 | 0.00554282 | 0.00062787 | 0.22542111 | 0.99999994 |
| Eci3 | 0.00403495 | 0.00427105 | 0.00364356 | 0.00062749 | 0.12582454 | 0.99999994 |
| AU022754 | 0.00172323 | 0.00218587 | 0.00155852 | 0.00062735 | 0.00368798 | 0.2041321 |
| Clcn5 | 0.07006692 | 0.06118298 | 0.0605571 | 0.00062589 | 0.47257578 | 0.99999994 |
| Gpsm3 | 0.01246485 | 0.01181715 | 0.01119173 | 0.00062542 | 0.35495638 | 0.99999994 |
| Sh3bgr | 0.05036499 | 0.04471834 | 0.04409339 | 0.00062495 | 0.46241823 | 0.99999994 |
| H2-Oa | 0.00283045 | 0.00318208 | 0.00255799 | 0.00062409 | 0.05218772 | 0.99999994 |
| Gm17322 | 0.00700238 | 0.00693407 | 0.00631044 | 0.00062364 | 0.25532646 | 0.99999994 |
| Tspan11 | 0.01186708 | 0.01128278 | 0.01065931 | 0.00062347 | 0.34851436 | 0.99999994 |
| Gm3558 | 0.00366338 | 0.00393014 | 0.00330886 | 0.00062127 | 0.10568571 | 0.99999994 |
| Rgs22 | 0.00494528 | 0.00508396 | 0.00446282 | 0.00062113 | 0.17711156 | 0.99999994 |
| 1700112D23I | 0.00370454 | 0.00396688 | 0.00334595 | 0.00062093 | 0.10836495 | 0.99999994 |
| Flywch1 | 0.09146363 | 0.07858677 | 0.07796634 | 0.00062042 | 0.47887851 | 0.99999994 |
| Tshz3 | 0.03293834 | 0.02979137 | 0.02917118 | 0.00062019 | 0.44373105 | 0.99999994 |
| Acvrl1 | 0.00639734 | 0.00638729 | 0.00576755 | 0.00061974 | 0.23722974 | 0.99999994 |
| Bid | 0.07935748 | 0.06879479 | 0.06817536 | 0.00061944 | 0.47588705 | 0.99999994 |
| Cnksr2 | 0.00791766 | 0.00774954 | 0.00713085 | 0.00061868 | 0.28179546 | 0.99999994 |
| Lrrc43 | 0.00312023 | 0.00343784 | 0.00281931 | 0.00061853 | 0.07209845 | 0.99999994 |
| Ell3 | 0.00469961 | 0.00485965 | 0.00424183 | 0.00061781 | 0.16615241 | 0.99999994 |
| Itga4 | 0.02031905 | 0.01876494 | 0.01814756 | 0.00061738 | 0.41042642 | 0.99999994 |
| Gins2 | 0.04784454 | 0.04257326 | 0.04195636 | 0.0006169 | 0.46101714 | 0.99999994 |
| H2-Q2 | 0.08184757 | 0.07081813 | 0.07020173 | 0.0006164 | 0.47669668 | 0.99999994 |
| Cldn9 | 0.02073187 | 0.01912745 | 0.01851113 | 0.00061632 | 0.4123049 | 0.99999994 |
| Sh2b3 | 0.03335306 | 0.03014546 | 0.02953032 | 0.00061514 | 0.44486031 | 0.99999994 |
| Pak6 | 0.01935544 | 0.01791195 | 0.01729812 | 0.00061384 | 0.40663666 | 0.99999994 |
| Hmgcll1 | 0.0314381 | 0.02848396 | 0.02787035 | 0.00061361 | 0.44174318 | 0.99999994 |
| Serpinb12 | 0.00352549 | 0.00379799 | 0.00318461 | 0.00061338 | 0.0999105 | 0.99999994 |
| Krtdap | 0.00246655 | 0.00284231 | 0.00222966 | 0.00061265 | 0.0337019 | 0.95936079 |
| 5031425F14I | 0.01187783 | 0.01128094 | 0.01066889 | 0.00061206 | 0.3512811 | 0.99999994 |

| | | | | | | |
|---|---|---|---|---|---|---|
| Zdhhc20 | 0.35178028 | 0.24647022 | 0.2458603 | 0.00060992 | 0.49341265 | 0.99999994 |
| Card9 | 0.01831387 | 0.01698724 | 0.01637872 | 0.00060852 | 0.40233567 | 0.99999994 |
| Pde4d | 0.03191205 | 0.02888983 | 0.02828159 | 0.00060824 | 0.44308364 | 0.99999994 |
| Wdr3 | 0.0713686 | 0.06223674 | 0.06163001 | 0.00060673 | 0.47387599 | 0.99999994 |
| F5 | 0.015363 | 0.01437337 | 0.01376695 | 0.00060642 | 0.38468253 | 0.99999994 |
| Htr5b | 0.00438496 | 0.00456297 | 0.00395867 | 0.0006043 | 0.15479075 | 0.99999994 |
| Zfhx4 | 0.01016591 | 0.00974555 | 0.00914177 | 0.00060378 | 0.33010339 | 0.99999994 |
| Csmd3 | 0.00181863 | 0.00224598 | 0.00164469 | 0.00060129 | 0.0074762 | 0.34949606 |
| Bloc1s1 | 0.0283722 | 0.02580469 | 0.02520386 | 0.00060084 | 0.43695986 | 0.99999994 |
| Trps1 | 0.0410993 | 0.03680217 | 0.03620216 | 0.00060001 | 0.4560775 | 0.99999994 |
| Kcnmb1 | 0.00984961 | 0.00945849 | 0.00885924 | 0.00059925 | 0.32626725 | 0.99999994 |
| Kcnj1 | 0.00294024 | 0.00325579 | 0.00265701 | 0.00059878 | 0.06679997 | 0.99999994 |
| Rcsd1 | 0.00424551 | 0.00443083 | 0.00383314 | 0.0005977 | 0.14965473 | 0.99999994 |
| Snta1 | 0.02489397 | 0.02276305 | 0.0221655 | 0.00059755 | 0.42879397 | 0.99999994 |
| Ciart | 0.00925952 | 0.00892905 | 0.0083318 | 0.00059725 | 0.31662827 | 0.99999994 |
| Pknox2 | 0.0024018 | 0.0027675 | 0.00217122 | 0.00059627 | 0.03377442 | 0.95971117 |
| Oplah | 0.01033732 | 0.00989101 | 0.00929484 | 0.00059616 | 0.33471449 | 0.99999994 |
| Bicd2 | 0.05321642 | 0.04709824 | 0.04650252 | 0.00059572 | 0.46602276 | 0.99999994 |
| Trim26 | 0.22385204 | 0.17282184 | 0.17222682 | 0.00059502 | 0.49082647 | 0.99999994 |
| Upk3a | 0.01005575 | 0.00963783 | 0.00904338 | 0.00059445 | 0.33085935 | 0.99999994 |
| Gm21781 | 0.00932252 | 0.00898159 | 0.00838813 | 0.00059346 | 0.31884411 | 0.99999994 |
| Taf9 | 0.00401211 | 0.00421642 | 0.00362299 | 0.00059343 | 0.13779388 | 0.99999994 |
| Ssc5d | 0.00210196 | 0.00249267 | 0.00190056 | 0.00059211 | 0.01905163 | 0.65172122 |
| Mapk3 | 0.85280692 | 0.41257788 | 0.41198604 | 0.00059184 | 0.49618528 | 0.99999994 |
| Slc46a2 | 0.00752782 | 0.00737293 | 0.00678154 | 0.00059139 | 0.28079645 | 0.99999994 |
| Cep295 | 0.0815133 | 0.07052086 | 0.06993009 | 0.00059078 | 0.47757791 | 0.99999994 |
| Rad54l | 0.00591861 | 0.00592845 | 0.00533768 | 0.00059077 | 0.23064646 | 0.99999994 |
| Ptrh1 | 0.04621898 | 0.04116453 | 0.04057432 | 0.00059021 | 0.46143218 | 0.99999994 |
| Cyp4b1 | 0.02150821 | 0.01978407 | 0.01919432 | 0.00058975 | 0.41897374 | 0.99999994 |
| Alkbh4 | 0.06322388 | 0.05547383 | 0.05488675 | 0.00058707 | 0.47162047 | 0.99999994 |

| | | | | | | |
|---|---|---|---|---|---|---|
| Fam217a | 0.00676533 | 0.00668444 | 0.00609779 | 0.00058666 | 0.26095819 | 0.99999994 |
| Zfp456 | 0.0145189 | 0.01360368 | 0.01301794 | 0.00058573 | 0.38228051 | 0.99999994 |
| Neil1 | 0.07572706 | 0.06579496 | 0.06520932 | 0.00058564 | 0.47616544 | 0.99999994 |
| Aif1 | 0.00541761 | 0.00547307 | 0.00488751 | 0.00058556 | 0.21258784 | 0.99999994 |
| Barx1 | 0.00593581 | 0.0059379 | 0.00535313 | 0.00058477 | 0.23357257 | 0.99999994 |
| 1700109G15 | 0.00223544 | 0.00260489 | 0.00202107 | 0.00058383 | 0.02725063 | 0.83669235 |
| Xrcc4 | 0.18393693 | 0.14652808 | 0.14594475 | 0.00058332 | 0.48938753 | 0.99999994 |
| Mgat5b | 0.00863212 | 0.0083534 | 0.00777056 | 0.00058284 | 0.30879478 | 0.99999994 |
| Zfp60 | 0.08332139 | 0.0719807 | 0.07139802 | 0.00058268 | 0.4783391 | 0.99999994 |
| Sptbn2 | 0.04269174 | 0.03814743 | 0.03756527 | 0.00058217 | 0.45891924 | 0.99999994 |
| Nepn | 0.0008011 | 0.00130598 | 0.00072498 | 0.000581 | 4.79E-08 | 7.21E-06 |
| Ubxn7 | 0.12677001 | 0.10578401 | 0.10520411 | 0.0005799 | 0.48536578 | 0.99999994 |
| Gm5093 | 0.02102289 | 0.01934608 | 0.01876732 | 0.00057876 | 0.4186789 | 0.99999994 |
| Rccd1 | 0.04655504 | 0.041439 | 0.04086028 | 0.00057872 | 0.46244501 | 0.99999994 |
| Pdpn | 0.01349577 | 0.01268582 | 0.01210893 | 0.00057689 | 0.37557726 | 0.99999994 |
| Ephb3 | 0.03578553 | 0.03220824 | 0.0316328 | 0.00057544 | 0.45181128 | 0.99999994 |
| Heph | 0.00225915 | 0.00261672 | 0.00204247 | 0.00057426 | 0.03063876 | 0.89307523 |
| Prrg3 | 0.0146832 | 0.01373723 | 0.0131638 | 0.00057343 | 0.38592432 | 0.99999994 |
| Spata6 | 0.01910004 | 0.01764532 | 0.01707279 | 0.00057253 | 0.41168252 | 0.99999994 |
| Stard9 | 0.00893049 | 0.00861005 | 0.00803753 | 0.00057252 | 0.31769958 | 0.99999994 |
| Kdm8 | 0.01092669 | 0.01039228 | 0.00982087 | 0.00057141 | 0.3492701 | 0.99999994 |
| Nxn | 0.03436124 | 0.03097359 | 0.03040256 | 0.00057103 | 0.45025334 | 0.99999994 |
| Dsn1 | 0.01191239 | 0.01127063 | 0.01069969 | 0.00057095 | 0.36122297 | 0.99999994 |
| Them4 | 0.07440942 | 0.0647001 | 0.06412936 | 0.00057073 | 0.47638064 | 0.99999994 |
| Jag1 | 0.00845633 | 0.00817935 | 0.00761321 | 0.00056614 | 0.31030659 | 0.99999994 |
| Lins1 | 0.04785567 | 0.04253054 | 0.04196581 | 0.00056473 | 0.46431298 | 0.99999994 |
| Iqck | 0.00804181 | 0.00780399 | 0.00724206 | 0.00056193 | 0.30275689 | 0.99999994 |
| Cd83 | 0.00542227 | 0.00545318 | 0.0048917 | 0.00056149 | 0.22242191 | 0.99999994 |
| Ttc41 | 0.00482281 | 0.00491391 | 0.00435266 | 0.00056124 | 0.19536612 | 0.99999994 |
| Plekho1 | 0.01722889 | 0.0159777 | 0.01541962 | 0.00055808 | 0.40481223 | 0.99999994 |

| | | | | | | |
|---|---|---|---|---|---|---|
| Slc4a4 | 0.01935976 | 0.01785981 | 0.01730193 | 0.00055788 | 0.41502997 | 0.99999994 |
| Fam20c | 0.46009948 | 0.29786547 | 0.29730809 | 0.00055738 | 0.49502171 | 0.99999994 |
| Tmem158 | 0.01094674 | 0.01039593 | 0.00983875 | 0.00055717 | 0.35310399 | 0.99999994 |
| Grik2 | 0.00682954 | 0.00671243 | 0.0061554 | 0.00055703 | 0.27346496 | 0.99999994 |
| Ascl2 | 0.00821812 | 0.00795673 | 0.00739995 | 0.00055678 | 0.3082451 | 0.99999994 |
| Enpp1 | 0.00890632 | 0.00857194 | 0.0080159 | 0.00055604 | 0.32213814 | 0.99999994 |
| Krt80 | 0.00271913 | 0.00301346 | 0.00245756 | 0.0005559 | 0.06607716 | 0.99999994 |
| Cnn2 | 0.00951176 | 0.00911172 | 0.00855731 | 0.00055442 | 0.33314083 | 0.99999994 |
| 1700016L21F | 0.01009043 | 0.00962846 | 0.00907436 | 0.00055409 | 0.34220703 | 0.99999994 |
| Gpr4 | 0.00368417 | 0.00388084 | 0.00332759 | 0.00055325 | 0.13421189 | 0.99999994 |
| Htra2 | 0.15312743 | 0.12501274 | 0.12445988 | 0.00055286 | 0.48820579 | 0.99999994 |
| Rnf144a | 0.00973121 | 0.00930605 | 0.00875344 | 0.00055261 | 0.33716193 | 0.99999994 |
| Gm28707 | 0.00852361 | 0.00822449 | 0.00767344 | 0.00055105 | 0.31632163 | 0.99999994 |
| Igdcc3 | 0.00368087 | 0.00387333 | 0.00332462 | 0.00054872 | 0.13596967 | 0.99999994 |
| 1700064M15 | 0.00404054 | 0.00419729 | 0.00364859 | 0.0005487 | 0.15840826 | 0.99999994 |
| Vwde | 0.00673961 | 0.00662247 | 0.00607472 | 0.00054775 | 0.27418728 | 0.99999994 |
| Dnah1 | 0.00734134 | 0.00716171 | 0.00661438 | 0.00054732 | 0.29088654 | 0.99999994 |
| Gm28209 | 0.00311949 | 0.00336592 | 0.00281865 | 0.00054727 | 0.09810834 | 0.99999994 |
| Cd44 | 0.01545081 | 0.01439046 | 0.01384482 | 0.00054564 | 0.39653184 | 0.99999994 |
| N4bp3 | 0.01718232 | 0.01592399 | 0.01537842 | 0.00054557 | 0.40666067 | 0.99999994 |
| Tmem126b | 0.23961646 | 0.1826916 | 0.18214671 | 0.00054489 | 0.49205665 | 0.99999994 |
| Ndufa10 | 0.77461178 | 0.39921202 | 0.39866733 | 0.00054468 | 0.49637195 | 0.99999994 |
| Nrxn2 | 0.02995338 | 0.02712505 | 0.02658041 | 0.00054464 | 0.44575625 | 0.99999994 |
| Cpxm1 | 0.01004419 | 0.00957452 | 0.00903306 | 0.00054147 | 0.34494637 | 0.99999994 |
| Gm26744 | 0.00167513 | 0.0020554 | 0.00151506 | 0.00054034 | 0.00879956 | 0.39182637 |
| A330009N23 | 0.04213151 | 0.03762482 | 0.03708604 | 0.00053878 | 0.46148144 | 0.99999994 |
| Podn | 0.00500635 | 0.00505528 | 0.00451775 | 0.00053754 | 0.21418222 | 0.99999994 |
| Adgrg4 | 0.00402046 | 0.00416756 | 0.0036305 | 0.00053705 | 0.16239646 | 0.99999994 |
| Nr5a2 | 0.02310239 | 0.02113136 | 0.02059498 | 0.00053638 | 0.43118413 | 0.99999994 |
| Cntnap1 | 0.00524501 | 0.00526712 | 0.00473236 | 0.00053476 | 0.22597282 | 0.99999994 |

| | | | | | | |
|---|---|---|---|---|---|---|
| Klhl7 | 0.46581787 | 0.30031449 | 0.2997805 | 0.000534 | 0.49526986 | 0.99999994 |
| Rgmb | 0.04676003 | 0.04156788 | 0.04103465 | 0.00053323 | 0.46553569 | 0.99999994 |
| Arhgap22 | 0.00264526 | 0.00292402 | 0.00239092 | 0.0005331 | 0.06888483 | 0.99999994 |
| Chrac1 | 0.46220817 | 0.29875442 | 0.29822242 | 0.000532 | 0.49526293 | 0.99999994 |
| Defa24 | 0.00314437 | 0.003372 | 0.00284108 | 0.00053092 | 0.10677035 | 0.99999994 |
| Phospho2 | 0.16887233 | 0.13610451 | 0.13557406 | 0.00053046 | 0.48961102 | 0.99999994 |
| Nfe2l3 | 0.02826817 | 0.02564365 | 0.02511319 | 0.00053046 | 0.44409309 | 0.99999994 |
| Crim1 | 0.00521797 | 0.00523698 | 0.00470804 | 0.00052894 | 0.22728289 | 0.99999994 |
| Dpf1 | 0.01259866 | 0.01183796 | 0.01131085 | 0.00052711 | 0.37820363 | 0.99999994 |
| Zfp647 | 0.0017767 | 0.00213373 | 0.00160682 | 0.00052691 | 0.01452687 | 0.55516698 |
| Kif15 | 0.0050022 | 0.00504017 | 0.00451402 | 0.00052615 | 0.21891594 | 0.99999994 |
| Itga11 | 0.00297973 | 0.00321756 | 0.00269262 | 0.00052494 | 0.09719826 | 0.99999994 |
| Anapc10 | 0.27218218 | 0.20237798 | 0.20185313 | 0.00052485 | 0.49309561 | 0.99999994 |
| Ccdc61 | 0.06761757 | 0.05905742 | 0.0585333 | 0.00052412 | 0.47623597 | 0.99999994 |
| Srd5a1 | 0.01077396 | 0.01020846 | 0.00968459 | 0.00052387 | 0.35939867 | 0.99999994 |
| Mturn | 0.03626799 | 0.03257257 | 0.032049 | 0.00052356 | 0.45670405 | 0.99999994 |
| Tc2n | 0.00320832 | 0.00342186 | 0.00289874 | 0.00052312 | 0.11482797 | 0.99999994 |
| Gpr20 | 0.00530855 | 0.00531244 | 0.00478948 | 0.00052296 | 0.23367121 | 0.99999994 |
| Nckap5 | 0.00427905 | 0.00438481 | 0.00386334 | 0.00052146 | 0.18447028 | 0.99999994 |
| Pde4a | 0.0192754 | 0.01774863 | 0.01722751 | 0.00052113 | 0.42021146 | 0.99999994 |
| B3gnt4 | 0.00516265 | 0.00517859 | 0.0046583 | 0.00052029 | 0.22860141 | 0.99999994 |
| Sh2d4a | 0.0050409 | 0.00506846 | 0.00454883 | 0.00051963 | 0.22351261 | 0.99999994 |
| Gsg1l | 0.01205751 | 0.01134791 | 0.01082897 | 0.00051894 | 0.37487049 | 0.99999994 |
| Kif2c | 0.00266838 | 0.00293068 | 0.00241178 | 0.0005189 | 0.07605396 | 0.99999994 |
| Arap3 | 0.00307345 | 0.0032949 | 0.00277714 | 0.00051777 | 0.10730168 | 0.99999994 |
| Gm5577 | 0.01452356 | 0.0135374 | 0.01302208 | 0.00051532 | 0.39611769 | 0.99999994 |
| Cyp2e1 | 0.00680285 | 0.00664621 | 0.00613145 | 0.00051476 | 0.28814077 | 0.99999994 |
| Fbxo46 | 0.03846363 | 0.03445434 | 0.03393975 | 0.00051459 | 0.45980557 | 0.99999994 |
| Ccdc14 | 0.0089982 | 0.0086118 | 0.0080981 | 0.0005137 | 0.33642334 | 0.99999994 |
| Snx31 | 0.00704927 | 0.0068661 | 0.00635249 | 0.00051361 | 0.29522777 | 0.99999994 |

| | | | | | | |
|---|---|---|---|---|---|---|
| Oprl1 | 0.01513283 | 0.0140763 | 0.0135628 | 0.0005135 | 0.40051446 | 0.99999994 |
| Msantd2 | 0.0399893 | 0.03576322 | 0.03525032 | 0.0005129 | 0.46142246 | 0.99999994 |
| Dnah11 | 0.00867882 | 0.0083246 | 0.00781235 | 0.00051226 | 0.33125193 | 0.99999994 |
| Ccdc96 | 0.02029299 | 0.01863677 | 0.01812461 | 0.00051217 | 0.42540049 | 0.99999994 |
| St3gal1 | 0.03165425 | 0.02856939 | 0.02805793 | 0.00051146 | 0.45171259 | 0.99999994 |
| Ptf1a | 0.01439832 | 0.01342183 | 0.01291088 | 0.00051095 | 0.39611099 | 0.99999994 |
| Grk6 | 0.04386147 | 0.03907552 | 0.0385647 | 0.00051082 | 0.46487127 | 0.99999994 |
| Bcan | 0.00144466 | 0.0018175 | 0.00130683 | 0.00051068 | 0.00464559 | 0.24201117 |
| Antxr2 | 0.01915317 | 0.01762995 | 0.01711968 | 0.00051027 | 0.42136615 | 0.99999994 |
| Otulin | 0.0567682 | 0.05000055 | 0.04949078 | 0.00050977 | 0.47266886 | 0.99999994 |
| Ppp1r18 | 0.00800828 | 0.00772163 | 0.00721202 | 0.00050961 | 0.31905469 | 0.99999994 |
| Gm33466 | 0.0026554 | 0.00290872 | 0.00240007 | 0.00050865 | 0.07916947 | 0.99999994 |
| Fibcd1 | 0.00204975 | 0.00236111 | 0.00185342 | 0.00050769 | 0.03412784 | 0.96800495 |
| Zc4h2 | 0.03294433 | 0.02968364 | 0.02917637 | 0.00050727 | 0.45393359 | 0.99999994 |
| Mbnl3 | 0.00714724 | 0.00694753 | 0.00644035 | 0.00050719 | 0.30007022 | 0.99999994 |
| Nox4 | 0.00287365 | 0.00310291 | 0.00259695 | 0.00050597 | 0.09733868 | 0.99999994 |
| Ccdc74a | 0.01882451 | 0.01733526 | 0.01682963 | 0.00050563 | 0.42074633 | 0.99999994 |
| P3h3 | 0.01166482 | 0.01098222 | 0.01047907 | 0.00050315 | 0.37463442 | 0.99999994 |
| Grm8 | 0.00076613 | 0.00119447 | 0.00069336 | 0.00050111 | 7.52E-07 | 9.51E-05 |
| Fignl2 | 0.01266349 | 0.01186877 | 0.01136856 | 0.00050022 | 0.38480734 | 0.99999994 |
| 4732465J04F | 0.00076562 | 0.00119288 | 0.0006929 | 0.00049998 | 7.81E-07 | 9.80E-05 |
| Gemin2 | 0.12505449 | 0.1044215 | 0.10392182 | 0.00049968 | 0.48723395 | 0.99999994 |
| Gm9767 | 0.00906013 | 0.00865266 | 0.00815348 | 0.00049917 | 0.34181492 | 0.99999994 |
| Foxm1 | 0.00174596 | 0.00207798 | 0.00157905 | 0.00049893 | 0.01772376 | 0.61823741 |
| Sf3a1 | 0.23232208 | 0.17808603 | 0.17758792 | 0.0004981 | 0.49255218 | 0.99999994 |
| Zfp462 | 0.01599171 | 0.01482196 | 0.01432428 | 0.00049768 | 0.40855431 | 0.99999994 |
| Pipox | 0.0038503 | 0.00397248 | 0.00347725 | 0.00049523 | 0.17156542 | 0.99999994 |
| Arhgap19 | 0.00646909 | 0.00632655 | 0.00583195 | 0.0004946 | 0.28620129 | 0.99999994 |
| BC049762 | 0.00462019 | 0.00466456 | 0.00417037 | 0.00049419 | 0.21511882 | 0.99999994 |
| Sycp3 | 0.0055351 | 0.00548726 | 0.00499311 | 0.00049415 | 0.25502706 | 0.99999994 |

| | | | | | | |
|---|---|---|---|---|---|---|
| Arhgap9 | 0.00336667 | 0.00353513 | 0.00304148 | 0.00049365 | 0.13998791 | 0.99999994 |
| Tnks | 0.0441471 | 0.03930174 | 0.03880852 | 0.00049322 | 0.46629126 | 0.99999994 |
| 4732463B04 | 0.00202147 | 0.00232049 | 0.00182789 | 0.0004926 | 0.03641925 | 0.99999994 |
| Ifit1 | 0.00269175 | 0.00292477 | 0.00243287 | 0.00049191 | 0.08917189 | 0.99999994 |
| Gm8251 | 0.00117505 | 0.00155504 | 0.00106313 | 0.0004919 | 0.00103559 | 0.07272363 |
| Hebp1 | 0.09288586 | 0.07959853 | 0.07910663 | 0.00049189 | 0.48349251 | 0.99999994 |
| Zfp57 | 0.00669072 | 0.00652241 | 0.00603084 | 0.00049157 | 0.29371962 | 0.99999994 |
| Slc19a2 | 0.01057954 | 0.01000258 | 0.00951108 | 0.0004915 | 0.36543036 | 0.99999994 |
| Psmd7 | 1.07679495 | 0.4299127 | 0.42942202 | 0.00049068 | 0.49696573 | 0.99999994 |
| Cenph | 0.01591606 | 0.01474528 | 0.01425725 | 0.00048803 | 0.4098816 | 0.99999994 |
| Pcdhga3 | 0.0045267 | 0.00457343 | 0.00408625 | 0.00048719 | 0.21371063 | 0.99999994 |
| Timp2 | 0.05331419 | 0.0470716 | 0.04658497 | 0.00048663 | 0.47228257 | 0.99999994 |
| Thpo | 0.00266416 | 0.00289288 | 0.00240798 | 0.00048491 | 0.09005387 | 0.99999994 |
| Gm7694 | 0.00218142 | 0.00245716 | 0.0019723 | 0.00048486 | 0.0508824 | 0.99999994 |
| Frg1 | 0.52945669 | 0.32627505 | 0.32579101 | 0.00048405 | 0.49605463 | 0.99999994 |
| Plxna4 | 0.00254618 | 0.00278541 | 0.00230152 | 0.00048388 | 0.0808373 | 0.99999994 |
| Zfp418 | 0.0123679 | 0.01158897 | 0.01110541 | 0.00048357 | 0.38596963 | 0.99999994 |
| Ager | 0.00834651 | 0.00799796 | 0.00751491 | 0.00048305 | 0.3343749 | 0.99999994 |
| Vwa3b | 0.00744834 | 0.00719129 | 0.0067103 | 0.00048099 | 0.31663642 | 0.99999994 |
| Hoxa5 | 0.00075694 | 0.00116598 | 0.00068504 | 0.00048094 | 1.48E-06 | 0.00018049 |
| Tm6sf1 | 0.01548369 | 0.01435482 | 0.01387398 | 0.00048084 | 0.40877595 | 0.99999994 |
| Pcdhb21 | 0.00977062 | 0.00926931 | 0.00878865 | 0.00048065 | 0.35791462 | 0.99999994 |
| Zfp53 | 0.05471831 | 0.04824832 | 0.04776783 | 0.00048049 | 0.47330847 | 0.99999994 |
| Rad51ap1 | 0.00939771 | 0.00893538 | 0.00845535 | 0.00048003 | 0.35275446 | 0.99999994 |
| Sncg | 0.00122586 | 0.00158726 | 0.00110906 | 0.0004782 | 0.00205188 | 0.13296325 |
| Zfp128 | 0.01518173 | 0.01408243 | 0.01360618 | 0.00047625 | 0.4078848 | 0.99999994 |
| Mcmbp | 0.13893049 | 0.11466602 | 0.114191 | 0.00047502 | 0.48895487 | 0.99999994 |
| Gm10146 | 0.03846076 | 0.03441138 | 0.03393728 | 0.0004741 | 0.46295649 | 0.99999994 |
| Camk1d | 0.02184946 | 0.01996849 | 0.01949439 | 0.00047409 | 0.43570084 | 0.99999994 |
| Lrig3 | 0.00309927 | 0.00327394 | 0.00280042 | 0.00047353 | 0.13018099 | 0.99999994 |

| | | | | | | |
|---|---|---|---|---|---|---|
| Klhl36 | 0.06048054 | 0.05307241 | 0.0525993 | 0.00047311 | 0.47612922 | 0.99999994 |
| Mirt1 | 0.00854461 | 0.00816518 | 0.00769223 | 0.00047294 | 0.34117685 | 0.99999994 |
| Oaz3 | 0.00217092 | 0.00243559 | 0.00196282 | 0.00047277 | 0.05443711 | 0.99999994 |
| Capn6 | 0.02295719 | 0.02093874 | 0.02046753 | 0.00047121 | 0.43910248 | 0.99999994 |
| Zfp385b | 0.00257575 | 0.00279878 | 0.00232821 | 0.00047057 | 0.08925351 | 0.99999994 |
| Tor4a | 0.0116145 | 0.010904 | 0.01043422 | 0.00046978 | 0.38220789 | 0.99999994 |
| Ccdc106 | 0.00527512 | 0.00522857 | 0.00475942 | 0.00046915 | 0.25586848 | 0.99999994 |
| Slc6a11 | 0.00491173 | 0.00490169 | 0.00443265 | 0.00046904 | 0.24060808 | 0.99999994 |
| Rnf8 | 0.25666156 | 0.19305879 | 0.19259082 | 0.00046797 | 0.49354779 | 0.99999994 |
| Gdnf | 0.00640865 | 0.00624471 | 0.0057777 | 0.00046701 | 0.29527697 | 0.99999994 |
| A230052G05 | 0.0015551 | 0.00187345 | 0.00140662 | 0.00046682 | 0.01358447 | 0.53455393 |
| Gab1 | 0.02343931 | 0.02135631 | 0.02089062 | 0.0004657 | 0.4410197 | 0.99999994 |
| Pcm1 | 0.57866636 | 0.34459142 | 0.34412654 | 0.00046488 | 0.49641274 | 0.99999994 |
| Adcy10 | 0.00300656 | 0.00317944 | 0.00271682 | 0.00046262 | 0.12851125 | 0.99999994 |
| A230083N12 | 0.0063667 | 0.00620249 | 0.00574005 | 0.00046245 | 0.29588639 | 0.99999994 |
| Nsl1 | 0.00218547 | 0.00243822 | 0.00197596 | 0.00046226 | 0.05971108 | 0.99999994 |
| Spry1 | 0.01309555 | 0.01221454 | 0.01175301 | 0.00046153 | 0.39689651 | 0.99999994 |
| Isg15 | 0.00218958 | 0.00244072 | 0.00197966 | 0.00046105 | 0.06054354 | 0.99999994 |
| Scrib | 0.08007123 | 0.06921726 | 0.06875685 | 0.00046041 | 0.48222425 | 0.99999994 |
| Tgfbi | 0.00743569 | 0.0071586 | 0.00669896 | 0.00045964 | 0.3239364 | 0.99999994 |
| Hdac11 | 0.07922878 | 0.06852945 | 0.06807045 | 0.000459 | 0.48209989 | 0.99999994 |
| Gm10784 | 0.00277233 | 0.00296396 | 0.00250556 | 0.0004584 | 0.11165003 | 0.99999994 |
| Col22a1 | 0.00317385 | 0.00332563 | 0.00286766 | 0.00045797 | 0.14388485 | 0.99999994 |
| Rest | 0.01416711 | 0.01316277 | 0.01270553 | 0.00045724 | 0.40534226 | 0.99999994 |
| Stk17b | 0.00509349 | 0.00505195 | 0.00459612 | 0.00045583 | 0.2545745 | 0.99999994 |
| Cfp | 0.00802372 | 0.00768167 | 0.00722586 | 0.00045581 | 0.33728387 | 0.99999994 |
| Tpx2 | 0.00586685 | 0.00574659 | 0.00529118 | 0.00045541 | 0.28335301 | 0.99999994 |
| Zfp143 | 0.05408106 | 0.04768664 | 0.04723126 | 0.00045537 | 0.4744151 | 0.99999994 |
| Zfp608 | 0.03896813 | 0.03482854 | 0.03437341 | 0.00045513 | 0.46488474 | 0.99999994 |
| Cercam | 0.01034401 | 0.00975497 | 0.00930081 | 0.00045416 | 0.37257769 | 0.99999994 |

| | | | | | | |
|---|---|---|---|---|---|---|
| Stra6 | 0.00552951 | 0.00543995 | 0.00498809 | 0.00045186 | 0.27325807 | 0.99999994 |
| Bfsp1 | 0.00306181 | 0.00321835 | 0.00276664 | 0.00045171 | 0.13856371 | 0.99999994 |
| Kdm1b | 0.01383409 | 0.01286067 | 0.01240965 | 0.00045102 | 0.40442131 | 0.99999994 |
| Bglap2 | 0.00425713 | 0.00429418 | 0.0038436 | 0.00045058 | 0.21760496 | 0.99999994 |
| Itpr2 | 0.02594685 | 0.02353719 | 0.02308672 | 0.00045047 | 0.44833139 | 0.99999994 |
| Smo | 0.05057597 | 0.04472023 | 0.04427195 | 0.00044828 | 0.47313203 | 0.99999994 |
| Stk38 | 0.25178415 | 0.19007936 | 0.18963184 | 0.00044752 | 0.49373345 | 0.99999994 |
| Tox2 | 0.01532716 | 0.01418209 | 0.01373517 | 0.00044692 | 0.41426534 | 0.99999994 |
| Plch2 | 0.01712843 | 0.0157771 | 0.01533075 | 0.00044635 | 0.42316784 | 0.99999994 |
| Wdr86 | 0.00323853 | 0.00337197 | 0.00292597 | 0.000446 | 0.15514454 | 0.99999994 |
| Bmpr1b | 0.0030194 | 0.00317398 | 0.0027284 | 0.00044558 | 0.13850553 | 0.99999994 |
| Adgrb2 | 0.00683799 | 0.00660839 | 0.00616298 | 0.00044541 | 0.31523333 | 0.99999994 |
| Ercc4 | 0.02729903 | 0.02471313 | 0.02426791 | 0.00044522 | 0.45140314 | 0.99999994 |
| A4galt | 0.00212573 | 0.00236627 | 0.00192202 | 0.00044425 | 0.06195982 | 0.99999994 |
| Gm6793 | 0.0083684 | 0.00797779 | 0.0075345 | 0.00044329 | 0.34766872 | 0.99999994 |
| Tm4sf20 | 0.00923489 | 0.00875293 | 0.00830977 | 0.00044315 | 0.36130297 | 0.99999994 |
| Hk2 | 0.01157457 | 0.01084096 | 0.01039863 | 0.00044233 | 0.38853299 | 0.99999994 |
| Ccdc89 | 0.0116088 | 0.01087036 | 0.01042914 | 0.00044122 | 0.38912204 | 0.99999994 |
| Wfdc10 | 0.01064284 | 0.01000756 | 0.00956758 | 0.00043998 | 0.37976246 | 0.99999994 |
| Oaf | 0.06463924 | 0.05650296 | 0.05606371 | 0.00043924 | 0.47920439 | 0.99999994 |
| Phyhd1 | 0.005214 | 0.0051431 | 0.00470447 | 0.00043863 | 0.26742546 | 0.99999994 |
| Gm38037 | 0.00463735 | 0.00462402 | 0.00418581 | 0.00043821 | 0.24294833 | 0.99999994 |
| Klf11 | 0.09773486 | 0.08341593 | 0.08297881 | 0.00043712 | 0.48601407 | 0.99999994 |
| Grid2ip | 0.00296166 | 0.00311296 | 0.00267633 | 0.00043663 | 0.13874984 | 0.99999994 |
| Gins1 | 0.02745006 | 0.02483474 | 0.02439972 | 0.00043502 | 0.45276592 | 0.99999994 |
| Bpi | 0.00205483 | 0.00229111 | 0.00185801 | 0.0004331 | 0.0603795 | 0.99999994 |
| Gm3252 | 0.00169432 | 0.00196542 | 0.00153241 | 0.00043302 | 0.02999255 | 0.88049966 |
| 2810408A11 | 0.02990605 | 0.02697213 | 0.02653925 | 0.00043287 | 0.45677177 | 0.99999994 |
| Cnbp | 1.45871643 | 0.41362303 | 0.41319217 | 0.00043086 | 0.49723095 | 0.99999994 |
| Amz1 | 0.00346437 | 0.00356001 | 0.00312953 | 0.00043048 | 0.17993513 | 0.99999994 |

| | | | | | | |
|---|---|---|---|---|---|---|
| Cep85l | 0.01613927 | 0.0148854 | 0.01445502 | 0.00043038 | 0.42145075 | 0.99999994 |
| Ephb1 | 0.00326625 | 0.00338113 | 0.00295096 | 0.00043017 | 0.16594515 | 0.99999994 |
| Ncapg | 0.00544983 | 0.00534534 | 0.00491647 | 0.00042886 | 0.28074308 | 0.99999994 |
| Slit3 | 0.00213871 | 0.00236234 | 0.00193374 | 0.0004286 | 0.07006078 | 0.99999994 |
| Egflam | 0.0058736 | 0.0057246 | 0.00529724 | 0.00042736 | 0.29563128 | 0.99999994 |
| Thnsl2 | 0.00913689 | 0.00864914 | 0.00822214 | 0.000427 | 0.36479107 | 0.99999994 |
| Cdk2 | 0.023674 | 0.02152195 | 0.02109647 | 0.00042548 | 0.44660339 | 0.99999994 |
| Gm42893 | 0.00351004 | 0.00359596 | 0.00317069 | 0.00042527 | 0.18598605 | 0.99999994 |
| Ttc12 | 0.01586824 | 0.01463951 | 0.01421487 | 0.00042464 | 0.42119281 | 0.99999994 |
| Islr | 0.00307252 | 0.0032009 | 0.0027763 | 0.0004246 | 0.15433613 | 0.99999994 |
| Vwa2 | 0.00348954 | 0.00357673 | 0.00315222 | 0.00042451 | 0.18501463 | 0.99999994 |
| Arhgap27os2 | 0.02597725 | 0.0235371 | 0.0231133 | 0.0004238 | 0.45143052 | 0.99999994 |
| Ier5l | 0.00867737 | 0.00823462 | 0.00781105 | 0.00042357 | 0.35906689 | 0.99999994 |
| BC016579 | 0.0007299 | 0.00108415 | 0.00066058 | 0.00042357 | 9.86E-06 | 0.00105454 |
| Pcdhga10 | 0.00738533 | 0.00707718 | 0.00665382 | 0.00042336 | 0.33595906 | 0.99999994 |
| Ripk2 | 0.05004829 | 0.04424763 | 0.04382526 | 0.00042238 | 0.47442451 | 0.99999994 |
| B4galnt4 | 0.00537683 | 0.00527269 | 0.00485086 | 0.00042183 | 0.28134904 | 0.99999994 |
| Sstr5 | 0.0047516 | 0.00471022 | 0.0042886 | 0.00042162 | 0.25642964 | 0.99999994 |
| Paqr7 | 0.01336562 | 0.01241477 | 0.01199321 | 0.00042156 | 0.40750437 | 0.99999994 |
| Gm6525 | 0.00355326 | 0.00363084 | 0.00320964 | 0.0004212 | 0.19119114 | 0.99999994 |
| Dtl | 0.0140571 | 0.01302842 | 0.01260781 | 0.00042061 | 0.41213207 | 0.99999994 |
| Pik3c2b | 0.00928449 | 0.0087743 | 0.00835412 | 0.00042018 | 0.36889278 | 0.99999994 |
| Suv39h2 | 0.02135363 | 0.0194783 | 0.01905834 | 0.00041996 | 0.44169374 | 0.99999994 |
| BC001981 | 0.00287233 | 0.0030152 | 0.00259576 | 0.00041945 | 0.14105463 | 0.99999994 |
| Fxyd1 | 0.02018537 | 0.01844908 | 0.01802979 | 0.00041929 | 0.43849103 | 0.99999994 |
| Nkd2 | 0.00169296 | 0.00195041 | 0.00153117 | 0.00041924 | 0.03418774 | 0.96800495 |
| Arhgap8 | 0.01062995 | 0.00997529 | 0.00955607 | 0.00041922 | 0.38513879 | 0.99999994 |
| Gm44386 | 0.0037548 | 0.0038083 | 0.00339122 | 0.00041708 | 0.20649258 | 0.99999994 |
| E030030I06R | 0.1264143 | 0.10535519 | 0.10493853 | 0.00041666 | 0.4894574 | 0.99999994 |
| Taco1 | 0.03063204 | 0.02758675 | 0.02717036 | 0.00041639 | 0.4593746 | 0.99999994 |

| | | | | | | |
|---|---|---|---|---|---|---|
| Catsper3 | 0.00189093 | 0.00212602 | 0.00171 | 0.00041603 | 0.05267642 | 0.99999994 |
| Nap1l2 | 0.02642801 | 0.02392238 | 0.02350728 | 0.00041509 | 0.45321649 | 0.99999994 |
| Gja1 | 0.00764178 | 0.0072942 | 0.00688367 | 0.00041053 | 0.34569417 | 0.99999994 |
| Tmem82 | 0.02228021 | 0.02028322 | 0.01987298 | 0.00041024 | 0.4453542 | 0.99999994 |
| Rit2 | 0.01006537 | 0.00946198 | 0.00905198 | 0.00041 | 0.38152031 | 0.99999994 |
| Ppp1r36 | 0.00531416 | 0.00520427 | 0.00479452 | 0.00040976 | 0.28472002 | 0.99999994 |
| Depdc1b | 0.00166575 | 0.00191623 | 0.00150659 | 0.00040964 | 0.03515903 | 0.98674321 |
| Chaf1b | 0.02785353 | 0.02516123 | 0.02475168 | 0.00040955 | 0.45614959 | 0.99999994 |
| E2f6 | 0.07969933 | 0.06886332 | 0.06845393 | 0.00040938 | 0.48412311 | 0.99999994 |
| Spin2c | 0.01916215 | 0.01753698 | 0.0171276 | 0.00040938 | 0.43679588 | 0.99999994 |
| Gabrg3 | 0.00699796 | 0.00671535 | 0.00630647 | 0.00040888 | 0.33302875 | 0.99999994 |
| Noxa1 | 0.00537049 | 0.00525396 | 0.00484515 | 0.00040881 | 0.2871854 | 0.99999994 |
| Jmjd4 | 0.01406154 | 0.01302041 | 0.01261175 | 0.00040866 | 0.41461543 | 0.99999994 |
| Gm21814 | 0.00904995 | 0.00855258 | 0.00814438 | 0.0004082 | 0.36933221 | 0.99999994 |
| Ccdc8 | 0.28377633 | 0.20903004 | 0.20862214 | 0.0004079 | 0.49480808 | 0.99999994 |
| Pros1 | 0.006511 | 0.00627628 | 0.00586957 | 0.00040671 | 0.32231564 | 0.99999994 |
| Uevld | 0.01616106 | 0.01488086 | 0.01447433 | 0.00040653 | 0.42584848 | 0.99999994 |
| 2610020C07l | 0.00386479 | 0.00389623 | 0.00349031 | 0.00040592 | 0.21942546 | 0.99999994 |
| Cacna2d3 | 0.00202371 | 0.0022358 | 0.0018299 | 0.0004059 | 0.06990825 | 0.99999994 |
| Stk36 | 0.03739652 | 0.03342707 | 0.03302151 | 0.00040556 | 0.46742232 | 0.99999994 |
| Wars2 | 0.05541069 | 0.04875555 | 0.0483503 | 0.00040525 | 0.47775411 | 0.99999994 |
| Palld | 0.02995274 | 0.02698473 | 0.02657986 | 0.00040488 | 0.4596194 | 0.99999994 |
| Gm6712 | 0.00320804 | 0.00330123 | 0.00289848 | 0.00040275 | 0.17750744 | 0.99999994 |
| Serpina7 | 0.00200653 | 0.00221712 | 0.00181439 | 0.00040274 | 0.06977157 | 0.99999994 |
| Padi4 | 0.00425006 | 0.00423949 | 0.00383724 | 0.00040225 | 0.24265999 | 0.99999994 |
| Slc1a3 | 0.00439155 | 0.00436551 | 0.0039646 | 0.00040091 | 0.25043914 | 0.99999994 |
| Six1 | 0.00121054 | 0.00149589 | 0.00109522 | 0.00040068 | 0.00744226 | 0.34893274 |
| Gm28557 | 0.00344417 | 0.00351149 | 0.00311132 | 0.00040017 | 0.19596575 | 0.99999994 |
| Eva1a | 0.00638536 | 0.00615561 | 0.00575679 | 0.00039881 | 0.32235117 | 0.99999994 |
| Kcnj6 | 0.00211911 | 0.00231486 | 0.00191604 | 0.00039881 | 0.08295242 | 0.99999994 |

| | | | | | | |
|---|---|---|---|---|---|---|
| 1700064H15 | 0.00168377 | 0.00192164 | 0.00152287 | 0.00039878 | 0.04066478 | 0.99999994 |
| Aldh1a3 | 0.00251448 | 0.00267047 | 0.00227292 | 0.00039756 | 0.12215712 | 0.99999994 |
| Kcnip4 | 0.00389802 | 0.00391613 | 0.00352024 | 0.00039589 | 0.22705406 | 0.99999994 |
| Bub1 | 0.00307848 | 0.00317663 | 0.00278167 | 0.00039496 | 0.17230041 | 0.99999994 |
| Gulp1 | 0.00430076 | 0.00427648 | 0.00388289 | 0.0003936 | 0.2499204 | 0.99999994 |
| Gm11827 | 0.00519301 | 0.00507852 | 0.0046856 | 0.00039292 | 0.28836007 | 0.99999994 |
| Krt20 | 0.00120717 | 0.00148385 | 0.00109217 | 0.00039168 | 0.00848999 | 0.38016558 |
| A230087F16 | 0.00251937 | 0.00266877 | 0.00227733 | 0.00039144 | 0.12628403 | 0.99999994 |
| Msn | 0.00831683 | 0.00787918 | 0.00748833 | 0.00039085 | 0.3641368 | 0.99999994 |
| Edil3 | 0.00374985 | 0.00377718 | 0.00338676 | 0.00039041 | 0.2214453 | 0.99999994 |
| Gm9930 | 0.00700452 | 0.00670211 | 0.00631235 | 0.00038976 | 0.34053522 | 0.99999994 |
| Actc1 | 0.00209285 | 0.00228126 | 0.00189234 | 0.00038892 | 0.08564987 | 0.99999994 |
| Cdhr5 | 0.01200036 | 0.01116663 | 0.01077806 | 0.00038857 | 0.40517516 | 0.99999994 |
| Pask | 0.00612537 | 0.0059119 | 0.00552337 | 0.00038853 | 0.31980917 | 0.99999994 |
| Sh2b2 | 0.0040054 | 0.00400505 | 0.00361695 | 0.00038811 | 0.23753874 | 0.99999994 |
| Rasgef1a | 0.00374154 | 0.00376735 | 0.00337928 | 0.00038807 | 0.22231283 | 0.99999994 |
| Gm20707 | 0.00586847 | 0.00568012 | 0.00529264 | 0.00038748 | 0.31301708 | 0.99999994 |
| Gm16090 | 0.00447339 | 0.00442547 | 0.00403826 | 0.00038721 | 0.26165685 | 0.99999994 |
| Dhx36 | 0.16674349 | 0.13447482 | 0.13408796 | 0.00038686 | 0.4923391 | 0.99999994 |
| Slc22a3 | 0.00166358 | 0.00189083 | 0.00150463 | 0.0003862 | 0.04376977 | 0.99999994 |
| Adcyap1r1 | 0.00303676 | 0.00312979 | 0.00274405 | 0.00038574 | 0.17471537 | 0.99999994 |
| Grhl2 | 0.0085715 | 0.00810124 | 0.0077163 | 0.00038494 | 0.36992095 | 0.99999994 |
| Samd15 | 0.05386454 | 0.04743306 | 0.04704885 | 0.00038421 | 0.47832536 | 0.99999994 |
| Clmp | 0.00716827 | 0.00684321 | 0.00645921 | 0.000384 | 0.34615524 | 0.99999994 |
| Map3k6 | 0.0048179 | 0.0047318 | 0.00434825 | 0.00038355 | 0.27855563 | 0.99999994 |
| Dkkl1 | 0.00559726 | 0.00543143 | 0.00504897 | 0.00038247 | 0.30705038 | 0.99999994 |
| Rsph10b | 0.00697082 | 0.00666446 | 0.00628213 | 0.00038234 | 0.34269765 | 0.99999994 |
| Phgr1 | 0.01044796 | 0.00977554 | 0.00939362 | 0.00038192 | 0.39333854 | 0.99999994 |
| S1pr3 | 0.00706394 | 0.0067472 | 0.00636564 | 0.00038155 | 0.34495548 | 0.99999994 |
| B3gnt5 | 0.0011985 | 0.00146572 | 0.00108433 | 0.00038139 | 0.00961081 | 0.42205476 |

| | | | | | | |
|---|---|---|---|---|---|---|
| Gm10440 | 0.001669 | 0.0018901 | 0.00150953 | 0.00038057 | 0.04666143 | 0.99999994 |
| Gys2 | 0.00207561 | 0.00225693 | 0.00187677 | 0.00038016 | 0.08878022 | 0.99999994 |
| Rxfp2 | 0.00294901 | 0.00304497 | 0.00266492 | 0.00038005 | 0.17123883 | 0.99999994 |
| Cfap206 | 0.00737516 | 0.00702427 | 0.0066447 | 0.00037958 | 0.3518832 | 0.99999994 |
| Scn3b | 0.00707123 | 0.00675056 | 0.00637219 | 0.00037838 | 0.34632874 | 0.99999994 |
| Gm16793 | 0.00329804 | 0.00335719 | 0.00297962 | 0.00037757 | 0.19948359 | 0.99999994 |
| Atp6v1e2 | 0.00202091 | 0.00220491 | 0.00182738 | 0.00037753 | 0.08453769 | 0.99999994 |
| Ammecr1 | 0.04184535 | 0.03721833 | 0.03684113 | 0.0003772 | 0.47283264 | 0.99999994 |
| Rbm45 | 0.02227094 | 0.02024145 | 0.01986483 | 0.00037662 | 0.44978717 | 0.99999994 |
| Fiz1 | 0.2267959 | 0.17447418 | 0.17409837 | 0.00037581 | 0.49426803 | 0.99999994 |
| Grm2 | 0.00439015 | 0.00433865 | 0.00396334 | 0.00037531 | 0.26424019 | 0.99999994 |
| Rnf17 | 0.00101529 | 0.00129389 | 0.00091868 | 0.00037521 | 0.00327817 | 0.19002633 |
| Exosc9 | 0.18210105 | 0.14506942 | 0.14469448 | 0.00037494 | 0.49311919 | 0.99999994 |
| Fkbp10 | 0.01018967 | 0.00953767 | 0.00916299 | 0.00037467 | 0.39274345 | 0.99999994 |
| Nudt12 | 0.02873898 | 0.02589742 | 0.02552343 | 0.00037399 | 0.46115079 | 0.99999994 |
| Gm13524 | 0.00550591 | 0.00534083 | 0.00496687 | 0.00037396 | 0.30812791 | 0.99999994 |
| Pwwp2b | 0.01417261 | 0.01308409 | 0.01271042 | 0.00037368 | 0.42242523 | 0.99999994 |
| Trip6 | 0.0205634 | 0.01873601 | 0.01836279 | 0.00037322 | 0.44619153 | 0.99999994 |
| Rsad1 | 0.01530948 | 0.01409198 | 0.01371949 | 0.00037249 | 0.4282926 | 0.99999994 |
| Npr2 | 0.01680889 | 0.01541923 | 0.01504798 | 0.00037125 | 0.43477928 | 0.99999994 |
| Prpf6 | 0.22773929 | 0.17506673 | 0.17469626 | 0.00037047 | 0.49436877 | 0.99999994 |
| Reep4 | 0.1067204 | 0.09036155 | 0.08999126 | 0.00037029 | 0.48907475 | 0.99999994 |
| Tagln3 | 0.01962958 | 0.01790875 | 0.01753989 | 0.00036886 | 0.4443371 | 0.99999994 |
| Epn3 | 0.00898311 | 0.00845334 | 0.00808459 | 0.00036875 | 0.38071501 | 0.99999994 |
| Mier3 | 0.06849589 | 0.05962797 | 0.05925976 | 0.00036821 | 0.48350488 | 0.99999994 |
| Chrnb4 | 0.01582984 | 0.01454903 | 0.01418083 | 0.0003682 | 0.43139326 | 0.99999994 |
| Ndc80 | 0.00564683 | 0.00546083 | 0.00509351 | 0.00036732 | 0.31560504 | 0.99999994 |
| Pcdhga6 | 0.00339406 | 0.0034328 | 0.00306616 | 0.00036664 | 0.21303533 | 0.99999994 |
| Rpain | 0.10911649 | 0.09220201 | 0.09183568 | 0.00036633 | 0.48940854 | 0.99999994 |
| Plekhg6 | 0.01467024 | 0.0135186 | 0.0131523 | 0.0003663 | 0.42646408 | 0.99999994 |

| | | | | | | |
|---|---|---|---|---|---|---|
| Cyp2d26 | 0.00535888 | 0.00520087 | 0.00483472 | 0.00036615 | 0.30709098 | 0.99999994 |
| Pm20d2 | 0.00408597 | 0.00405501 | 0.0036895 | 0.00036551 | 0.25480978 | 0.99999994 |
| Chrng | 0.00209204 | 0.00225694 | 0.0018916 | 0.00036534 | 0.0992942 | 0.99999994 |
| Rapgef5 | 0.00297328 | 0.00305184 | 0.0026868 | 0.00036504 | 0.18290465 | 0.99999994 |
| Cntn2 | 0.00209034 | 0.00225403 | 0.00189007 | 0.00036396 | 0.09996192 | 0.99999994 |
| Ccm2l | 0.00210467 | 0.00226513 | 0.00190301 | 0.00036212 | 0.1026454 | 0.99999994 |
| Endou | 0.00233185 | 0.00246901 | 0.00210809 | 0.00036092 | 0.12722436 | 0.99999994 |
| Zfp773 | 0.00935809 | 0.00877948 | 0.00841993 | 0.00035954 | 0.388115 | 0.99999994 |
| Tnfsf9 | 0.03232504 | 0.0289975 | 0.02863971 | 0.00035779 | 0.46686386 | 0.99999994 |
| Dydc2 | 0.00666648 | 0.00636593 | 0.00600909 | 0.00035684 | 0.34631916 | 0.99999994 |
| Tcerg1l | 0.00203046 | 0.00219261 | 0.001836 | 0.00035661 | 0.09802545 | 0.99999994 |
| Abcc12 | 0.00412228 | 0.00407668 | 0.0037222 | 0.00035449 | 0.26306501 | 0.99999994 |
| Csmd2 | 0.00069524 | 0.00098365 | 0.00062923 | 0.00035442 | 8.87E-05 | 0.00831716 |
| Gm43403 | 0.00814323 | 0.00768727 | 0.00733289 | 0.00035438 | 0.37384521 | 0.99999994 |
| Gltpd2 | 0.00117984 | 0.00142149 | 0.00106747 | 0.00035402 | 0.01363795 | 0.53455393 |
| Acp5 | 0.01603815 | 0.01471912 | 0.01436543 | 0.00035369 | 0.43491151 | 0.99999994 |
| Gjb2 | 0.00159625 | 0.00179674 | 0.0014438 | 0.00035294 | 0.05185009 | 0.99999994 |
| B3gnt2 | 0.00640523 | 0.00612581 | 0.00577462 | 0.00035119 | 0.34280691 | 0.99999994 |
| Il11ra1 | 0.0281066 | 0.02532281 | 0.02497234 | 0.00035047 | 0.46278638 | 0.99999994 |
| Ccdc102a | 0.00709358 | 0.0067423 | 0.00639223 | 0.00035007 | 0.35772845 | 0.99999994 |
| H3f3aos | 0.00254564 | 0.00265106 | 0.00230103 | 0.00035002 | 0.15564074 | 0.99999994 |
| Coch | 0.00259201 | 0.00269194 | 0.00234288 | 0.00034906 | 0.1606669 | 0.99999994 |
| Gm3550 | 0.02915923 | 0.02623807 | 0.0258894 | 0.00034868 | 0.46428375 | 0.99999994 |
| Styx | 0.00349248 | 0.00350348 | 0.00315486 | 0.00034861 | 0.23100923 | 0.99999994 |
| Mcmdc2 | 0.00316985 | 0.00321201 | 0.00286406 | 0.00034795 | 0.20934968 | 0.99999994 |
| Nr2f1 | 0.0020952 | 0.00224075 | 0.00189446 | 0.00034629 | 0.11185301 | 0.99999994 |
| Gm37469 | 0.00201725 | 0.00216974 | 0.00182407 | 0.00034567 | 0.10358324 | 0.99999994 |
| Ntan1 | 0.32703497 | 0.23312822 | 0.23278268 | 0.00034554 | 0.49605824 | 0.99999994 |
| Fcgr1 | 0.00209427 | 0.00223831 | 0.00189361 | 0.0003447 | 0.11281839 | 0.99999994 |
| Plk1 | 0.00917396 | 0.00859992 | 0.00825529 | 0.00034463 | 0.39055249 | 0.99999994 |

| | | | | | | |
|---|---|---|---|---|---|---|
| Socs5 | 0.05689858 | 0.0499448 | 0.04960021 | 0.00034459 | 0.48155798 | 0.99999994 |
| Rtp4 | 0.00397667 | 0.00393538 | 0.00359107 | 0.0003443 | 0.26167371 | 0.99999994 |
| Dbil5 | 0.00677682 | 0.00645139 | 0.0061081 | 0.00034329 | 0.35416184 | 0.99999994 |
| Ptx3 | 0.00117441 | 0.00140576 | 0.00106256 | 0.0003432 | 0.01577826 | 0.5755633 |
| Fam72a | 0.01319757 | 0.01218684 | 0.01184376 | 0.00034308 | 0.42355152 | 0.99999994 |
| Cnr1 | 0.00568209 | 0.00546742 | 0.00512519 | 0.00034223 | 0.32835041 | 0.99999994 |
| Klhdc7a | 0.00232094 | 0.00244027 | 0.00209825 | 0.00034202 | 0.13895838 | 0.99999994 |
| Jam2 | 0.00117342 | 0.00140265 | 0.00106166 | 0.000341 | 0.01625973 | 0.58877176 |
| Gpank1 | 0.10810406 | 0.09139813 | 0.09105723 | 0.0003409 | 0.49005942 | 0.99999994 |
| Sel1l3 | 0.02198648 | 0.01995548 | 0.01961484 | 0.00034063 | 0.45398686 | 0.99999994 |
| Adgra1 | 0.00673658 | 0.00641254 | 0.00607199 | 0.00034055 | 0.35445205 | 0.99999994 |
| Lzts3 | 0.00343271 | 0.00344109 | 0.003101 | 0.00034009 | 0.23269485 | 0.99999994 |
| Gm11716 | 0.00955088 | 0.00893169 | 0.00859227 | 0.00033941 | 0.39629823 | 0.99999994 |
| Faap24 | 0.0616352 | 0.05390216 | 0.05356308 | 0.00033908 | 0.48319445 | 0.99999994 |
| Prrt1 | 0.00341776 | 0.00342473 | 0.00308752 | 0.00033721 | 0.23362015 | 0.99999994 |
| Aire | 0.00373887 | 0.00371301 | 0.00337688 | 0.00033613 | 0.25380406 | 0.99999994 |
| A330074K22 | 0.00347034 | 0.00347081 | 0.00313491 | 0.0003359 | 0.23785692 | 0.99999994 |
| Gm29417 | 0.00240911 | 0.00251372 | 0.00217783 | 0.00033589 | 0.15229627 | 0.99999994 |
| D430019H16 | 0.0505098 | 0.04455175 | 0.04421596 | 0.00033579 | 0.47984165 | 0.99999994 |
| Bcas1 | 0.00283585 | 0.00289824 | 0.00256286 | 0.00033538 | 0.19185696 | 0.99999994 |
| Rnf112 | 0.02783869 | 0.02507362 | 0.02473873 | 0.00033489 | 0.46410136 | 0.99999994 |
| Dscaml1 | 0.00116887 | 0.00139197 | 0.00105755 | 0.00033442 | 0.01765018 | 0.61810138 |
| Rab33a | 0.00645651 | 0.00615408 | 0.00582066 | 0.00033342 | 0.35149354 | 0.99999994 |
| Ica1l | 0.00797691 | 0.00751726 | 0.00718392 | 0.00033334 | 0.37871502 | 0.99999994 |
| Trpc4 | 0.00862438 | 0.00809665 | 0.00776363 | 0.00033302 | 0.38762037 | 0.99999994 |
| Prph | 0.00116365 | 0.00138564 | 0.00105283 | 0.00033282 | 0.01768109 | 0.61810138 |
| 1700029F12I | 0.00292705 | 0.00297755 | 0.00264511 | 0.00033244 | 0.20141399 | 0.99999994 |
| Camk2g | 0.16926406 | 0.13617799 | 0.13584696 | 0.00033103 | 0.49352949 | 0.99999994 |
| Ascl4 | 0.00132643 | 0.00153011 | 0.00119997 | 0.00033014 | 0.03352633 | 0.95607013 |
| Sap30l | 0.05067595 | 0.04468559 | 0.04435655 | 0.00032904 | 0.4803093 | 0.99999994 |

| | | | | | | |
|---|---|---|---|---|---|---|
| Cep152 | 0.01402639 | 0.01290871 | 0.01258053 | 0.00032818 | 0.43107414 | 0.99999994 |
| Taf7l | 0.00114312 | 0.00136224 | 0.00103426 | 0.00032798 | 0.0173936 | 0.61072978 |
| Socs3 | 0.05670856 | 0.04976851 | 0.04944072 | 0.00032779 | 0.48239972 | 0.99999994 |
| Gm527 | 0.00527655 | 0.00508798 | 0.00476071 | 0.00032728 | 0.32362116 | 0.99999994 |
| Selplg | 0.00413945 | 0.00406456 | 0.00373765 | 0.00032691 | 0.2802193 | 0.99999994 |
| Nmnat1 | 0.04607154 | 0.0407745 | 0.04044882 | 0.00032568 | 0.47862914 | 0.99999994 |
| Gm17634 | 0.00327967 | 0.00328797 | 0.00296305 | 0.00032492 | 0.23272277 | 0.99999994 |
| Tbxa2r | 0.00877551 | 0.00822349 | 0.00789887 | 0.00032462 | 0.39221188 | 0.99999994 |
| Crtac1 | 0.00402662 | 0.00396015 | 0.00363606 | 0.00032409 | 0.27649099 | 0.99999994 |
| Gm17056 | 0.00870073 | 0.00815599 | 0.00783195 | 0.00032403 | 0.39150667 | 0.99999994 |
| Gm17541 | 0.01694575 | 0.01549291 | 0.0151691 | 0.00032381 | 0.44350462 | 0.99999994 |
| Slamf6 | 0.00116054 | 0.00137279 | 0.00105002 | 0.00032278 | 0.02037021 | 0.68362409 |
| Amhr2 | 0.00207752 | 0.0022001 | 0.00187849 | 0.00032161 | 0.12722527 | 0.99999994 |
| Dhrs2 | 0.00327295 | 0.00327844 | 0.002957 | 0.00032144 | 0.23466526 | 0.99999994 |
| Fstl5 | 0.00475666 | 0.00461378 | 0.00429315 | 0.00032063 | 0.30954961 | 0.99999994 |
| Grid2 | 0.00151868 | 0.00169394 | 0.00137372 | 0.00032022 | 0.06037356 | 0.99999994 |
| Mcam | 0.00707655 | 0.00669583 | 0.00637696 | 0.00031887 | 0.36962657 | 0.99999994 |
| Nkain4 | 0.0058936 | 0.00563407 | 0.00531521 | 0.00031886 | 0.34483148 | 0.99999994 |
| Gm13179 | 0.00115479 | 0.00136325 | 0.00104482 | 0.00031843 | 0.02124512 | 0.70117707 |
| B230369F24 | 0.00930528 | 0.00869099 | 0.00837271 | 0.00031828 | 0.4001212 | 0.99999994 |
| Elmo1 | 0.00804668 | 0.00756415 | 0.00724642 | 0.00031774 | 0.3851952 | 0.99999994 |
| Mrpl53 | 0.00115827 | 0.00136518 | 0.00104796 | 0.00031721 | 0.0219599 | 0.71665561 |
| S1pr4 | 0.0013499 | 0.00153808 | 0.00122118 | 0.0003169 | 0.04205201 | 0.99999994 |
| Thsd7a | 0.00452763 | 0.00440364 | 0.00408708 | 0.00031656 | 0.30308284 | 0.99999994 |
| Zfyve9 | 0.07029992 | 0.06106539 | 0.06074928 | 0.00031611 | 0.48618496 | 0.99999994 |
| Glcci1 | 0.00832498 | 0.00781137 | 0.00749562 | 0.00031574 | 0.38958977 | 0.99999994 |
| Epb41l5 | 0.11902194 | 0.09969931 | 0.09938427 | 0.00031503 | 0.49158305 | 0.99999994 |
| Hapln1 | 0.00767621 | 0.00722937 | 0.00691453 | 0.00031484 | 0.38091091 | 0.99999994 |
| Gm16576 | 0.00317103 | 0.00317918 | 0.00286511 | 0.00031407 | 0.23280115 | 0.99999994 |
| Mfsd9 | 0.00690856 | 0.00654001 | 0.00622628 | 0.00031374 | 0.36865868 | 0.99999994 |

| | | | | | | |
|---|---|---|---|---|---|---|
| Rbpms2 | 0.00272694 | 0.00277742 | 0.00246461 | 0.00031281 | 0.19910133 | 0.99999994 |
| Gm42875 | 0.0020636 | 0.00217754 | 0.00186592 | 0.00031162 | 0.13314686 | 0.99999994 |
| Katnbl1 | 0.15025087 | 0.12270967 | 0.12239843 | 0.00031124 | 0.49324784 | 0.99999994 |
| Gm6356 | 0.0068393 | 0.00647525 | 0.00616415 | 0.0003111 | 0.36845635 | 0.99999994 |
| Mtnr1a | 0.00285868 | 0.00289385 | 0.00258345 | 0.0003104 | 0.21192252 | 0.99999994 |
| Onecut3 | 0.00115348 | 0.00135375 | 0.00104363 | 0.00031012 | 0.02396701 | 0.76718503 |
| Cav2 | 0.00705976 | 0.0066711 | 0.0063619 | 0.00030921 | 0.37315056 | 0.99999994 |
| Art1 | 0.00335948 | 0.00334388 | 0.003035 | 0.00030888 | 0.24906148 | 0.99999994 |
| Chaf1a | 0.01713245 | 0.01564183 | 0.01533431 | 0.00030752 | 0.44690315 | 0.99999994 |
| Man1a2 | 0.22802466 | 0.17518421 | 0.17487694 | 0.00030727 | 0.49533426 | 0.99999994 |
| Tmem179 | 0.02228516 | 0.02018448 | 0.01987733 | 0.00030715 | 0.45903836 | 0.99999994 |
| Zbtb43 | 0.08097918 | 0.06980248 | 0.0694958 | 0.00030668 | 0.48828307 | 0.99999994 |
| Dnmt3aos | 0.00459838 | 0.00445726 | 0.00415075 | 0.00030652 | 0.31152066 | 0.99999994 |
| Ngfr | 0.00636142 | 0.00604167 | 0.0057353 | 0.00030636 | 0.36108431 | 0.99999994 |
| Hsd17b11 | 0.09600979 | 0.08190999 | 0.081604 | 0.00030598 | 0.49004391 | 0.99999994 |
| Psmc3ip | 0.06893305 | 0.05992653 | 0.05962103 | 0.00030551 | 0.48639543 | 0.99999994 |
| Doc2a | 0.00648149 | 0.00614849 | 0.00584308 | 0.00030541 | 0.36395057 | 0.99999994 |
| Magi3 | 0.01233826 | 0.0113844 | 0.01107902 | 0.00030539 | 0.42721126 | 0.99999994 |
| 4732419C18I | 0.00114576 | 0.00134148 | 0.00103665 | 0.00030483 | 0.02515298 | 0.79556275 |
| Slc25a31 | 0.00293612 | 0.00295774 | 0.00265329 | 0.00030445 | 0.22249952 | 0.99999994 |
| Slc4a10 | 0.00189109 | 0.00201458 | 0.00171014 | 0.00030445 | 0.11800993 | 0.99999994 |
| Npas1 | 0.00332743 | 0.00330855 | 0.00300611 | 0.00030244 | 0.2515306 | 0.99999994 |
| Cenpl | 0.02625971 | 0.02366253 | 0.02336021 | 0.00030232 | 0.46567634 | 0.99999994 |
| Il9r | 0.00114538 | 0.00133834 | 0.00103631 | 0.00030204 | 0.0261889 | 0.81698086 |
| Rfx8 | 0.00114674 | 0.00133951 | 0.00103754 | 0.00030197 | 0.02635393 | 0.81859587 |
| 6330409D20I | 0.00114642 | 0.00133922 | 0.00103725 | 0.00030197 | 0.02632266 | 0.81859587 |
| Kctd17 | 0.00114682 | 0.00133923 | 0.00103761 | 0.00030162 | 0.0265 | 0.81943852 |
| Artn | 0.00588815 | 0.00561193 | 0.00531031 | 0.00030161 | 0.35269086 | 0.99999994 |
| 6820408C15I | 0.00117795 | 0.00136727 | 0.00106575 | 0.00030152 | 0.02983625 | 0.87914907 |
| Cxcr2 | 0.00158412 | 0.00173405 | 0.00143284 | 0.00030122 | 0.08085853 | 0.99999994 |

| | | | | | | |
|---|---|---|---|---|---|---|
| Dhx30 | 0.28128129 | 0.20747655 | 0.20717598 | 0.00030057 | 0.49614748 | 0.99999994 |
| Wee1 | 0.01157848 | 0.01070247 | 0.01040212 | 0.00030035 | 0.42379444 | 0.99999994 |
| Slc35f3 | 0.00622349 | 0.00591053 | 0.00561147 | 0.00029906 | 0.3613907 | 0.99999994 |
| Fgd6 | 0.11488906 | 0.09654885 | 0.09624979 | 0.00029905 | 0.49174979 | 0.99999994 |
| Gm19325 | 0.00525773 | 0.00504246 | 0.00474379 | 0.00029867 | 0.33757449 | 0.99999994 |
| Dcst2 | 0.00159339 | 0.00173978 | 0.00144122 | 0.00029856 | 0.08395856 | 0.99999994 |
| 4933411E06I | 0.00159693 | 0.00174282 | 0.00144442 | 0.0002984 | 0.08454507 | 0.99999994 |
| Amd2 | 0.00375318 | 0.00368816 | 0.00338976 | 0.0002984 | 0.2789539 | 0.99999994 |
| Gins4 | 0.19488561 | 0.15362239 | 0.15332435 | 0.00029804 | 0.49483818 | 0.99999994 |
| Gm10425 | 0.00146164 | 0.00161994 | 0.00132217 | 0.00029777 | 0.06692356 | 0.99999994 |
| Nkain1 | 0.00160588 | 0.00174984 | 0.0014525 | 0.00029734 | 0.08649943 | 0.99999994 |
| Pctp | 0.02045094 | 0.01856069 | 0.01826374 | 0.00029695 | 0.45690828 | 0.99999994 |
| Asic4 | 0.00250603 | 0.00256204 | 0.00226529 | 0.00029675 | 0.1916121 | 0.99999994 |
| Zfp804a | 0.00265601 | 0.00269709 | 0.00240062 | 0.00029648 | 0.2055198 | 0.99999994 |
| Far2os1 | 0.00224791 | 0.00232864 | 0.00203232 | 0.00029632 | 0.16589642 | 0.99999994 |
| Gm15318 | 0.00575674 | 0.00548839 | 0.00519227 | 0.00029613 | 0.35211172 | 0.99999994 |
| L2hgdh | 0.03380082 | 0.03021335 | 0.02991785 | 0.0002955 | 0.47379034 | 0.99999994 |
| Chodl | 0.00200895 | 0.00211198 | 0.00181657 | 0.00029541 | 0.13952762 | 0.99999994 |
| Itgam | 0.00416861 | 0.00405862 | 0.00376391 | 0.00029471 | 0.30111989 | 0.99999994 |
| Tspan31 | 0.55310296 | 0.33508076 | 0.33478644 | 0.00029432 | 0.49766546 | 0.99999994 |
| Scn11a | 0.00288054 | 0.00289684 | 0.00260317 | 0.00029368 | 0.22634257 | 0.99999994 |
| Gpm6b | 0.00940733 | 0.00875676 | 0.00846395 | 0.00029281 | 0.40893842 | 0.99999994 |
| Ptcra | 0.00421616 | 0.00409737 | 0.00380672 | 0.00029065 | 0.30564657 | 0.99999994 |
| Espnl | 0.00113873 | 0.00132078 | 0.0010303 | 0.00029048 | 0.03028044 | 0.88731712 |
| Gm17705 | 0.00354846 | 0.00349563 | 0.00320531 | 0.00029032 | 0.27329113 | 0.99999994 |
| A230083G16 | 0.00112919 | 0.00131164 | 0.00102167 | 0.00028997 | 0.02943393 | 0.87375476 |
| Nepro | 0.01242489 | 0.01144582 | 0.01115615 | 0.00028967 | 0.43139197 | 0.99999994 |
| Peli2 | 0.02035648 | 0.01846849 | 0.01818054 | 0.00028795 | 0.45801899 | 0.99999994 |
| Mdfic | 0.00238966 | 0.0024476 | 0.00216027 | 0.00028733 | 0.18798546 | 0.99999994 |
| Sfxn4 | 0.06771446 | 0.05889968 | 0.05861348 | 0.0002862 | 0.48703572 | 0.99999994 |

| | | | | | | |
|---|---|---|---|---|---|---|
| Fancg | 0.03211755 | 0.02874555 | 0.02845981 | 0.00028574 | 0.47335807 | 0.99999994 |
| Pdcd1 | 0.00526735 | 0.0050378 | 0.00475243 | 0.00028536 | 0.34469411 | 0.99999994 |
| Usp43 | 0.00598806 | 0.00568484 | 0.00540006 | 0.00028478 | 0.36278142 | 0.99999994 |
| Iqcd | 0.00720847 | 0.00678002 | 0.00649525 | 0.00028476 | 0.38521032 | 0.99999994 |
| Prkaa2 | 0.01685316 | 0.01537173 | 0.01508716 | 0.00028457 | 0.45004361 | 0.99999994 |
| Krtap16-1 | 0.00573367 | 0.00545579 | 0.00517154 | 0.00028426 | 0.35723064 | 0.99999994 |
| Topbp1 | 0.04575263 | 0.04046135 | 0.04017728 | 0.00028406 | 0.48123193 | 0.99999994 |
| Phxr4 | 0.00328524 | 0.00324853 | 0.00296808 | 0.00028045 | 0.26469137 | 0.99999994 |
| Gm17501 | 0.00198989 | 0.0020798 | 0.00179937 | 0.00028043 | 0.14977612 | 0.99999994 |
| Car13 | 0.00250322 | 0.002543 | 0.00226276 | 0.00028024 | 0.20486421 | 0.99999994 |
| Klk14 | 0.00159304 | 0.00172091 | 0.0014409 | 0.00028002 | 0.09790923 | 0.99999994 |
| Rpgrip1 | 0.00885901 | 0.00825344 | 0.00797358 | 0.00027986 | 0.40763731 | 0.99999994 |
| Sox8 | 0.00642287 | 0.00607029 | 0.00579047 | 0.00027982 | 0.37385222 | 0.99999994 |
| Tnfrsf10b | 0.00817852 | 0.00764423 | 0.00736449 | 0.00027974 | 0.4001959 | 0.99999994 |
| Gm3289 | 0.00157742 | 0.00170638 | 0.00142679 | 0.00027959 | 0.09605386 | 0.99999994 |
| Wscd1 | 0.04631353 | 0.0409338 | 0.04065478 | 0.00027902 | 0.48178136 | 0.99999994 |
| Crabp2 | 0.00659887 | 0.00622586 | 0.00594843 | 0.00027744 | 0.3781076 | 0.99999994 |
| Atp2b3 | 0.00112751 | 0.00129659 | 0.00102016 | 0.00027644 | 0.03563945 | 0.99497107 |
| Rasgrp4 | 0.0029359 | 0.00292918 | 0.00265309 | 0.00027609 | 0.24425099 | 0.99999994 |
| Gm14302 | 0.00204074 | 0.00212078 | 0.00184528 | 0.00027549 | 0.16016669 | 0.99999994 |
| Lrrc3b | 0.00274453 | 0.00275579 | 0.00248048 | 0.00027531 | 0.23001372 | 0.99999994 |
| Arl5c | 0.00175775 | 0.00186438 | 0.0015897 | 0.00027468 | 0.12504341 | 0.99999994 |
| Nlgn3 | 0.00280503 | 0.00280968 | 0.00253505 | 0.00027462 | 0.23542799 | 0.99999994 |
| Rasal1 | 0.02153794 | 0.01949507 | 0.01922047 | 0.0002746 | 0.46211848 | 0.99999994 |
| Osgin2 | 0.05224827 | 0.04595988 | 0.04568556 | 0.00027433 | 0.48405892 | 0.99999994 |
| Gm9242 | 0.00708797 | 0.00666149 | 0.0063872 | 0.00027429 | 0.38749691 | 0.99999994 |
| Rps6ka6 | 0.02532781 | 0.02281947 | 0.02254526 | 0.00027422 | 0.4677362 | 0.99999994 |
| Polq | 0.00173622 | 0.00184362 | 0.00157025 | 0.00027337 | 0.12326221 | 0.99999994 |
| Phox2b | 0.00112325 | 0.0012894 | 0.0010163 | 0.0002731 | 0.03683167 | 0.99999994 |
| B9d1os | 0.00112499 | 0.00128959 | 0.00101787 | 0.00027172 | 0.03779302 | 0.99999994 |

| | | | | | | |
|---|---|---|---|---|---|---|
| Ly6g5b | 0.00201493 | 0.00209354 | 0.00182198 | 0.00027156 | 0.16057177 | 0.99999994 |
| Cyyr1 | 0.00155108 | 0.00167443 | 0.00140299 | 0.00027144 | 0.09890307 | 0.99999994 |
| Nckap5l | 0.01247893 | 0.01147536 | 0.01120426 | 0.00027109 | 0.43602528 | 0.99999994 |
| Gm9857 | 0.0013963 | 0.00153415 | 0.00126312 | 0.00027103 | 0.07660518 | 0.99999994 |
| Phtf2 | 0.0202436 | 0.01835206 | 0.01808109 | 0.00027097 | 0.46026935 | 0.99999994 |
| Acot7 | 0.01652729 | 0.01506962 | 0.01479868 | 0.00027094 | 0.45150255 | 0.99999994 |
| Hck | 0.00402939 | 0.00390892 | 0.00363855 | 0.00027037 | 0.31043687 | 0.99999994 |
| Fbxl12os | 0.02027368 | 0.01837754 | 0.01810759 | 0.00026995 | 0.46047626 | 0.99999994 |
| 1700003F12I | 0.00200008 | 0.00207845 | 0.00180856 | 0.00026989 | 0.16027642 | 0.99999994 |
| Pygb | 0.01406642 | 0.01288584 | 0.01261609 | 0.00026976 | 0.44341119 | 0.99999994 |
| Pdlim5 | 0.0602016 | 0.05263531 | 0.05236626 | 0.00026906 | 0.48635864 | 0.99999994 |
| Grap2 | 0.00409421 | 0.00396585 | 0.00369692 | 0.00026893 | 0.31411677 | 0.99999994 |
| Apoc4 | 0.00132589 | 0.00146836 | 0.00119948 | 0.00026888 | 0.06783382 | 0.99999994 |
| Zfp91 | 0.25681396 | 0.19295169 | 0.1926829 | 0.00026879 | 0.49629564 | 0.99999994 |
| Cers4 | 0.04437592 | 0.03927199 | 0.03900377 | 0.00026821 | 0.48174563 | 0.99999994 |
| Rnf224 | 0.00112347 | 0.00128469 | 0.0010165 | 0.00026819 | 0.03952958 | 0.99999994 |
| 0610025J13F | 0.00199059 | 0.002068 | 0.0018 | 0.000268 | 0.16082844 | 0.99999994 |
| Mapk4 | 0.00547362 | 0.00520552 | 0.00493785 | 0.00026767 | 0.35911286 | 0.99999994 |
| A330041J22F | 0.00221324 | 0.00226734 | 0.00200103 | 0.00026632 | 0.18783557 | 0.99999994 |
| Fas | 0.00111253 | 0.00127244 | 0.00100661 | 0.00026583 | 0.03938863 | 0.99999994 |
| Acot3 | 0.00111754 | 0.00127691 | 0.00101114 | 0.00026577 | 0.04009437 | 0.99999994 |
| Iqsec3 | 0.00154381 | 0.0016619 | 0.00139642 | 0.00026549 | 0.10284564 | 0.99999994 |
| Gm15410 | 0.00157616 | 0.00169113 | 0.00142565 | 0.00026548 | 0.10757269 | 0.99999994 |
| Rad51 | 0.01305002 | 0.01197782 | 0.01171251 | 0.00026531 | 0.44007409 | 0.99999994 |
| Arl5b | 0.08520254 | 0.07318669 | 0.07292165 | 0.00026504 | 0.49034914 | 0.99999994 |
| Zfp772 | 0.02533184 | 0.02281354 | 0.02254878 | 0.00026476 | 0.46885167 | 0.99999994 |
| Kifc1 | 0.00407126 | 0.00394101 | 0.00367626 | 0.00026476 | 0.31583519 | 0.99999994 |
| Tmem88 | 0.00326973 | 0.00321873 | 0.0029541 | 0.00026464 | 0.27548899 | 0.99999994 |
| 4930448E22I | 0.00149219 | 0.00161438 | 0.00134977 | 0.00026461 | 0.09595877 | 0.99999994 |
| Ints7 | 0.03576653 | 0.03188039 | 0.03161641 | 0.00026398 | 0.47783953 | 0.99999994 |

| | | | | | | |
|---|---|---|---|---|---|---|
| Pex5 | 0.1041013 | 0.08823055 | 0.08796694 | 0.00026362 | 0.49204264 | 0.99999994 |
| Hoxa7 | 0.00111936 | 0.00127619 | 0.00101278 | 0.00026341 | 0.04170685 | 0.99999994 |
| Tbx18 | 0.0013413 | 0.0014768 | 0.00121341 | 0.0002634 | 0.07424101 | 0.99999994 |
| Wnt9a | 0.01084658 | 0.01001237 | 0.00974939 | 0.00026297 | 0.42875494 | 0.99999994 |
| Cd46 | 0.00519371 | 0.00494914 | 0.00468623 | 0.00026291 | 0.35440816 | 0.99999994 |
| Rab43 | 0.0253331 | 0.02281269 | 0.02254988 | 0.00026281 | 0.46908231 | 0.99999994 |
| 1700022N22 | 0.00592107 | 0.00560204 | 0.00533988 | 0.00026215 | 0.37191517 | 0.99999994 |
| 1700016H13 | 0.00175601 | 0.00184943 | 0.00158813 | 0.00026131 | 0.13671035 | 0.99999994 |
| Siglece | 0.00233574 | 0.0023727 | 0.00211161 | 0.00026109 | 0.20524509 | 0.99999994 |
| Gata5os | 0.0032289 | 0.00317741 | 0.00291728 | 0.00026013 | 0.27641197 | 0.99999994 |
| Krt14 | 0.00267395 | 0.00267679 | 0.00241681 | 0.00025998 | 0.23698419 | 0.99999994 |
| Rasl10b | 0.00939269 | 0.0087105 | 0.00845086 | 0.00025963 | 0.41898053 | 0.99999994 |
| Nanos3 | 0.02242216 | 0.02025711 | 0.01999769 | 0.00025943 | 0.46559334 | 0.99999994 |
| Gm29019 | 0.00600342 | 0.0056731 | 0.00541385 | 0.00025925 | 0.37495606 | 0.99999994 |
| Nlgn1 | 0.00111733 | 0.00127016 | 0.00101095 | 0.00025921 | 0.04394141 | 0.99999994 |
| Gfpt2 | 0.00275874 | 0.00275227 | 0.0024933 | 0.00025898 | 0.24466016 | 0.99999994 |
| Lrrc75a | 0.02760014 | 0.02478944 | 0.02453065 | 0.00025878 | 0.47200939 | 0.99999994 |
| 4930430F08I | 0.07954883 | 0.06858973 | 0.06833131 | 0.00025842 | 0.48995814 | 0.99999994 |
| Uts2b | 0.00111365 | 0.001266 | 0.00100762 | 0.00025838 | 0.0439233 | 0.99999994 |
| 2010007H06 | 0.00234009 | 0.00237388 | 0.00211554 | 0.00025834 | 0.20814939 | 0.99999994 |
| Tmub2 | 0.2099704 | 0.16353669 | 0.16327897 | 0.00025771 | 0.49580877 | 0.99999994 |
| 2810455O05 | 0.00559475 | 0.0053041 | 0.00504671 | 0.00025739 | 0.36712327 | 0.99999994 |
| Cmya5 | 0.00379333 | 0.00368324 | 0.00342593 | 0.00025731 | 0.30856218 | 0.99999994 |
| Slc34a3 | 0.00228465 | 0.00232266 | 0.00206549 | 0.00025717 | 0.20361762 | 0.99999994 |
| Fign | 0.00843226 | 0.00784883 | 0.00759167 | 0.00025716 | 0.41080457 | 0.99999994 |
| Vrk2 | 0.02287119 | 0.02064905 | 0.02039203 | 0.00025701 | 0.46657037 | 0.99999994 |
| Ly6a | 0.00126972 | 0.00140563 | 0.00114871 | 0.00025692 | 0.06827444 | 0.99999994 |
| Tpgs2 | 0.11619511 | 0.09749896 | 0.09724264 | 0.00025631 | 0.49300102 | 0.99999994 |
| Hyou1 | 0.32121964 | 0.2298855 | 0.2296322 | 0.0002533 | 0.49707081 | 0.99999994 |
| Piwil4 | 0.00434979 | 0.0041801 | 0.00392701 | 0.00025308 | 0.33396881 | 0.99999994 |

| | | | | | | |
|---|---|---|---|---|---|---|
| Gm38250 | 0.00244075 | 0.00245943 | 0.00220638 | 0.00025305 | 0.22261026 | 0.99999994 |
| Bax | 1.31030068 | 0.42468404 | 0.42443137 | 0.00025267 | 0.49841915 | 0.99999994 |
| Ralb | 0.22156913 | 0.17102174 | 0.1707692 | 0.00025255 | 0.49607287 | 0.99999994 |
| Map2k1 | 0.41118024 | 0.27544353 | 0.27519101 | 0.00025252 | 0.49756325 | 0.99999994 |
| C130074G19 | 0.00155874 | 0.00166177 | 0.00140991 | 0.00025186 | 0.11720569 | 0.99999994 |
| Rasip1 | 0.01803201 | 0.01638146 | 0.01612969 | 0.00025177 | 0.45862435 | 0.99999994 |
| Mapk11 | 0.00110266 | 0.00124864 | 0.00099768 | 0.00025095 | 0.04703516 | 0.99999994 |
| Rhox3c | 0.00110915 | 0.00125449 | 0.00100355 | 0.00025094 | 0.04801169 | 0.99999994 |
| Zfp65 | 0.0289481 | 0.0259561 | 0.02570556 | 0.00025053 | 0.47413708 | 0.99999994 |
| Aldh1a7 | 0.00227346 | 0.00230506 | 0.00205539 | 0.00024967 | 0.20938727 | 0.99999994 |
| Hsd17b7 | 0.04533657 | 0.04007216 | 0.03982286 | 0.00024931 | 0.48338011 | 0.99999994 |
| Gpr6 | 0.00296318 | 0.00292657 | 0.00267769 | 0.00024888 | 0.26806293 | 0.99999994 |
| Gm4799 | 0.00873817 | 0.00811415 | 0.00786545 | 0.00024869 | 0.41665239 | 0.99999994 |
| 6720483E21I | 0.00185582 | 0.00192679 | 0.00167828 | 0.00024851 | 0.16216013 | 0.99999994 |
| Lrrc4b | 0.00265307 | 0.00264569 | 0.00239797 | 0.00024772 | 0.24584184 | 0.99999994 |
| Gm37233 | 0.00258425 | 0.00258353 | 0.00233588 | 0.00024766 | 0.24017742 | 0.99999994 |
| Gm10647 | 0.00063496 | 0.00082047 | 0.0005747 | 0.00024577 | 0.00220927 | 0.13925008 |
| Cutal | 0.06258017 | 0.05459539 | 0.05435075 | 0.00024464 | 0.48804897 | 0.99999994 |
| Adamtsl5 | 0.00258234 | 0.00257867 | 0.00233415 | 0.00024452 | 0.24280197 | 0.99999994 |
| Phf24 | 0.00063393 | 0.0008178 | 0.00057376 | 0.00024404 | 0.00231903 | 0.14554169 |
| Chil5 | 0.00371975 | 0.00360263 | 0.00335965 | 0.00024298 | 0.3151137 | 0.99999994 |
| Gm13814 | 0.00110427 | 0.00124202 | 0.00099915 | 0.00024287 | 0.05282818 | 0.99999994 |
| Gm6741 | 0.00063318 | 0.00081587 | 0.00057309 | 0.00024278 | 0.00240166 | 0.14955019 |
| Akr1c14 | 0.00381843 | 0.00369132 | 0.00344854 | 0.00024278 | 0.31967236 | 0.99999994 |
| Trabd2b | 0.00063281 | 0.00081491 | 0.00057275 | 0.00024216 | 0.00244388 | 0.15099967 |
| Lsamp | 0.00223453 | 0.00226238 | 0.00202025 | 0.00024213 | 0.2124969 | 0.99999994 |
| 6030445D17I | 0.00063262 | 0.00081443 | 0.00057258 | 0.00024185 | 0.00246522 | 0.15173012 |
| Galnt7 | 0.04831196 | 0.04259504 | 0.04235322 | 0.00024182 | 0.48484168 | 0.99999994 |
| Selp | 0.00063253 | 0.00081419 | 0.00057249 | 0.00024169 | 0.00247595 | 0.15180428 |
| Ccl17 | 0.00321177 | 0.00314326 | 0.00290185 | 0.00024141 | 0.28987567 | 0.99999994 |

| | | | | | | |
|---|---|---|---|---|---|---|
| 5330439B14 | 0.00151429 | 0.00161023 | 0.00136975 | 0.00024048 | 0.12127662 | 0.99999994 |
| Gdf15 | 0.00221217 | 0.00224047 | 0.00200006 | 0.00024041 | 0.211827 | 0.99999994 |
| Pde9a | 0.01737716 | 0.01579094 | 0.01555077 | 0.00024017 | 0.45905927 | 0.99999994 |
| Gm15917 | 0.00274971 | 0.00272522 | 0.00248515 | 0.00024007 | 0.26010889 | 0.99999994 |
| Ppp1r42 | 0.00498251 | 0.00473573 | 0.00449631 | 0.00023943 | 0.3615019 | 0.99999994 |
| Slc28a2 | 0.00063113 | 0.0008106 | 0.00057123 | 0.00023937 | 0.00264156 | 0.16011051 |
| Gm15991 | 0.00063113 | 0.0008106 | 0.00057123 | 0.00023937 | 0.00264156 | 0.16011051 |
| Cyth4 | 0.02674054 | 0.0240189 | 0.0237803 | 0.00023859 | 0.47337641 | 0.99999994 |
| Gm37245 | 0.00379874 | 0.00366923 | 0.00343081 | 0.00023842 | 0.32183522 | 0.99999994 |
| Atp8a2 | 0.00695774 | 0.00650875 | 0.0062704 | 0.00023835 | 0.40012618 | 0.99999994 |
| Gm43660 | 0.00109868 | 0.00123236 | 0.00099409 | 0.00023827 | 0.05530391 | 0.99999994 |
| Gm36937 | 0.0006303 | 0.00080845 | 0.00057048 | 0.00023798 | 0.00274524 | 0.163902 |
| Ccdc189 | 0.10602457 | 0.08969225 | 0.0894543 | 0.00023796 | 0.49293644 | 0.99999994 |
| Chst5 | 0.00209145 | 0.00212891 | 0.00189107 | 0.00023784 | 0.20124955 | 0.99999994 |
| 2900089D17 | 0.00490016 | 0.00465983 | 0.00442225 | 0.00023758 | 0.36031695 | 0.99999994 |
| Cd3e | 0.00062974 | 0.00080703 | 0.00056997 | 0.00023706 | 0.0028162 | 0.16751162 |
| Adap2os | 0.00142757 | 0.00152835 | 0.00129138 | 0.00023696 | 0.110965 | 0.99999994 |
| 4930500M09 | 0.00155179 | 0.00164044 | 0.00140363 | 0.00023681 | 0.13071873 | 0.99999994 |
| Hddc2 | 0.24496251 | 0.18569002 | 0.18545331 | 0.00023671 | 0.49661055 | 0.99999994 |
| Usp27x | 0.01608939 | 0.01464746 | 0.01441083 | 0.00023663 | 0.45648186 | 0.99999994 |
| Ttbk1 | 0.00232429 | 0.00233763 | 0.00210127 | 0.00023636 | 0.22700558 | 0.99999994 |
| Ifi203 | 0.00169092 | 0.00176533 | 0.00152933 | 0.000236 | 0.15216382 | 0.99999994 |
| Mamld1 | 0.06247082 | 0.05449536 | 0.05425966 | 0.00023571 | 0.48846587 | 0.99999994 |
| Tiam1 | 0.01204512 | 0.01105266 | 0.01081793 | 0.00023473 | 0.44258085 | 0.99999994 |
| Gm15969 | 0.00062827 | 0.00080326 | 0.00056864 | 0.00023462 | 0.00301287 | 0.17788196 |
| Pcdhgb7 | 0.00197942 | 0.00202449 | 0.00178991 | 0.00023457 | 0.19151098 | 0.99999994 |
| Tmem88b | 0.00784968 | 0.00730448 | 0.00706995 | 0.00023453 | 0.41262021 | 0.99999994 |
| Tprkb | 0.13574302 | 0.11208678 | 0.11185262 | 0.00023416 | 0.4944409 | 0.99999994 |
| Gm3667 | 0.00211825 | 0.00214894 | 0.00191527 | 0.00023367 | 0.20836271 | 0.99999994 |
| Clpb | 0.1455443 | 0.11923652 | 0.11900474 | 0.00023177 | 0.49482828 | 0.99999994 |

| | | | | | | |
|---|---|---|---|---|---|---|
| Hamp | 0.00223962 | 0.00225637 | 0.00202484 | 0.00023153 | 0.22329312 | 0.99999994 |
| Cd36 | 0.0020534 | 0.00208823 | 0.00185671 | 0.00023152 | 0.20327075 | 0.99999994 |
| Cfap54 | 0.01131869 | 0.01040066 | 0.0101705 | 0.00023016 | 0.44013201 | 0.99999994 |
| Npr3 | 0.00108816 | 0.0012143 | 0.00098457 | 0.00022973 | 0.06019354 | 0.99999994 |
| Matn3 | 0.00062471 | 0.00079418 | 0.00056542 | 0.00022876 | 0.00353987 | 0.20012082 |
| Frmd3 | 0.00062435 | 0.00079326 | 0.00056509 | 0.00022817 | 0.00359794 | 0.20124465 |
| Gm26511 | 0.00171531 | 0.00177932 | 0.00155136 | 0.00022796 | 0.16401251 | 0.99999994 |
| 2310081O03 | 0.00149609 | 0.00158125 | 0.0013533 | 0.00022794 | 0.13110931 | 0.99999994 |
| Csf3 | 0.00533139 | 0.00503772 | 0.00481 | 0.00022771 | 0.37633437 | 0.99999994 |
| Gabra1 | 0.0006238 | 0.00079188 | 0.0005646 | 0.00022728 | 0.00368649 | 0.2041321 |
| Cp | 0.00062295 | 0.0007897 | 0.00056382 | 0.00022588 | 0.00383033 | 0.20952724 |
| Gm2155 | 0.0006229 | 0.00078959 | 0.00056378 | 0.00022581 | 0.00383802 | 0.20952724 |
| Chst8 | 0.06964398 | 0.06043381 | 0.0602081 | 0.00022571 | 0.49004601 | 0.99999994 |
| Unc13c | 0.00131175 | 0.00141221 | 0.0011867 | 0.0002255 | 0.10295738 | 0.99999994 |
| Maf | 0.00810972 | 0.00752836 | 0.00730288 | 0.00022549 | 0.41858061 | 0.99999994 |
| Brat1 | 0.02598254 | 0.0233434 | 0.02311793 | 0.00022547 | 0.47411889 | 0.99999994 |
| Agmat | 0.00154076 | 0.001619 | 0.00139366 | 0.00022535 | 0.14089925 | 0.99999994 |
| Stat4 | 0.00152015 | 0.0015998 | 0.00137504 | 0.00022476 | 0.13829076 | 0.99999994 |
| Il16 | 0.00108068 | 0.00120251 | 0.00097781 | 0.00022469 | 0.06306185 | 0.99999994 |
| Pdcd2 | 0.2147622 | 0.16661495 | 0.16639027 | 0.00022468 | 0.49641433 | 0.99999994 |
| Mreg | 0.01440018 | 0.01313674 | 0.01291253 | 0.00022421 | 0.45399241 | 0.99999994 |
| Xntrpc | 0.00109142 | 0.00121154 | 0.00098752 | 0.00022402 | 0.06552233 | 0.99999994 |
| Gm6938 | 0.00062173 | 0.00078663 | 0.00056273 | 0.0002239 | 0.00404257 | 0.21763658 |
| Sval2 | 0.00062173 | 0.00078663 | 0.00056273 | 0.0002239 | 0.00404257 | 0.21763658 |
| Gm16208 | 0.00062173 | 0.00078663 | 0.00056273 | 0.0002239 | 0.00404257 | 0.21763658 |
| P2ry14 | 0.0010849 | 0.00120544 | 0.00098163 | 0.00022381 | 0.06454999 | 0.99999994 |
| Synj2 | 0.00980172 | 0.00904009 | 0.00881645 | 0.00022365 | 0.43295709 | 0.99999994 |
| Kif7 | 0.00287964 | 0.0028259 | 0.00260235 | 0.00022355 | 0.28372916 | 0.99999994 |
| Gm26555 | 0.00108898 | 0.00120879 | 0.00098532 | 0.00022347 | 0.06556317 | 0.99999994 |
| Gm28592 | 0.00107312 | 0.00119372 | 0.00097097 | 0.00022274 | 0.06338228 | 0.99999994 |

| | | | | | | |
|---|---|---|---|---|---|---|
| Gm34342 | 0.00062101 | 0.00078482 | 0.00056208 | 0.00022274 | 0.00417279 | 0.22321601 |
| Hic1 | 0.00269447 | 0.00265737 | 0.00243532 | 0.00022205 | 0.27194916 | 0.99999994 |
| 4933408N05 | 0.01438008 | 0.01311668 | 0.01289468 | 0.000222 | 0.45438187 | 0.99999994 |
| Lhpp | 0.01990513 | 0.01800473 | 0.01778282 | 0.00022191 | 0.46689977 | 0.99999994 |
| Gm12280 | 0.00062021 | 0.00078278 | 0.00056135 | 0.00022143 | 0.00432333 | 0.22896418 |
| Asic2 | 0.004791 | 0.00454538 | 0.00432405 | 0.00022133 | 0.36665998 | 0.99999994 |
| Pdzd2 | 0.00316176 | 0.00307777 | 0.00285676 | 0.00022101 | 0.30328993 | 0.99999994 |
| Pcdhb12 | 0.00258236 | 0.00255502 | 0.00233417 | 0.00022085 | 0.2644149 | 0.99999994 |
| Ren1 | 0.0022925 | 0.00229291 | 0.00207257 | 0.00022034 | 0.23958461 | 0.99999994 |
| Mlkl | 0.00302758 | 0.00295563 | 0.00273577 | 0.00021986 | 0.29634884 | 0.99999994 |
| Clspn | 0.01105561 | 0.01015543 | 0.00993587 | 0.00021956 | 0.44153048 | 0.99999994 |
| Tgm1 | 0.00198658 | 0.00201555 | 0.00179637 | 0.00021917 | 0.20835727 | 0.99999994 |
| Hook3 | 0.20066765 | 0.15738751 | 0.15716868 | 0.00021883 | 0.49630277 | 0.99999994 |
| Pcdhb10 | 0.0051257 | 0.00484359 | 0.00462508 | 0.00021851 | 0.37657839 | 0.99999994 |
| Daw1 | 0.00141084 | 0.00149454 | 0.00127626 | 0.00021828 | 0.12747144 | 0.99999994 |
| Nhlh2 | 0.00239812 | 0.00238578 | 0.00216791 | 0.00021787 | 0.25176503 | 0.99999994 |
| Prss44 | 0.0010652 | 0.00118142 | 0.00096382 | 0.0002176 | 0.06644978 | 0.99999994 |
| Mid1 | 0.03251874 | 0.02902514 | 0.02880762 | 0.00021752 | 0.47995722 | 0.99999994 |
| Fgf7 | 0.00061755 | 0.00077607 | 0.00055894 | 0.00021713 | 0.00485726 | 0.24896956 |
| Ttpa | 0.00730999 | 0.00680301 | 0.00658628 | 0.00021673 | 0.41330904 | 0.99999994 |
| A930014E10 | 0.00145164 | 0.0015298 | 0.00131313 | 0.00021666 | 0.13603907 | 0.99999994 |
| Gm11232 | 0.0006171 | 0.00077496 | 0.00055854 | 0.00021642 | 0.00495121 | 0.25216358 |
| Adrb3 | 0.0006171 | 0.00077496 | 0.00055854 | 0.00021642 | 0.00495121 | 0.25216358 |
| Gja6 | 0.00192063 | 0.00195302 | 0.00173682 | 0.00021621 | 0.20365881 | 0.99999994 |
| Rptoros | 0.02079233 | 0.01878034 | 0.01856436 | 0.00021598 | 0.46913645 | 0.99999994 |
| Gm16499 | 0.00061675 | 0.00077408 | 0.00055822 | 0.00021586 | 0.00502742 | 0.2544193 |
| 4930433N12 | 0.00061675 | 0.00077408 | 0.00055822 | 0.00021586 | 0.00502742 | 0.2544193 |
| Vgll2 | 0.00107335 | 0.00118659 | 0.00097119 | 0.0002154 | 0.0699286 | 0.99999994 |
| Zfp831 | 0.00108467 | 0.00119673 | 0.00098142 | 0.00021531 | 0.07210481 | 0.99999994 |
| Gm16143 | 0.00234135 | 0.00233186 | 0.00211667 | 0.0002152 | 0.24928792 | 0.99999994 |

| | | | | | | |
|---|---|---|---|---|---|---|
| Zfp68 | 0.15405022 | 0.12533399 | 0.12511916 | 0.00021483 | 0.4954405 | 0.99999994 |
| Angel2 | 0.21948331 | 0.16964706 | 0.16943243 | 0.00021462 | 0.49663626 | 0.99999994 |
| Fam163b | 0.0070448 | 0.00656188 | 0.00634848 | 0.0002134 | 0.41147558 | 0.99999994 |
| Lama1 | 0.00236106 | 0.0023478 | 0.00213446 | 0.00021334 | 0.25292438 | 0.99999994 |
| Adamts15 | 0.00198637 | 0.00200949 | 0.00179619 | 0.00021331 | 0.21462181 | 0.99999994 |
| Cap2 | 0.00108305 | 0.00119297 | 0.00097995 | 0.00021302 | 0.07395587 | 0.99999994 |
| 3110082J24F | 0.00061473 | 0.00076902 | 0.00055639 | 0.00021262 | 0.00548418 | 0.27319783 |
| Gm10522 | 0.00104929 | 0.00116185 | 0.00094943 | 0.00021242 | 0.06821513 | 0.99999994 |
| Zfp11 | 0.00677436 | 0.00631712 | 0.00610589 | 0.00021123 | 0.40893979 | 0.99999994 |
| Rag1 | 0.00061345 | 0.00076582 | 0.00055524 | 0.00021058 | 0.00579244 | 0.28587416 |
| Pcdh19 | 0.00126156 | 0.00135177 | 0.00114134 | 0.00021044 | 0.10985713 | 0.99999994 |
| Dchs1 | 0.00701069 | 0.0065283 | 0.00631789 | 0.00021041 | 0.4122818 | 0.99999994 |
| Colq | 0.00108059 | 0.00118781 | 0.00097773 | 0.00021008 | 0.07632691 | 0.99999994 |
| Rundc1 | 0.09263408 | 0.07911476 | 0.07890492 | 0.00020985 | 0.49293808 | 0.99999994 |
| Pgm5 | 0.00303015 | 0.00294722 | 0.00273809 | 0.00020913 | 0.30558918 | 0.99999994 |
| Gm26596 | 0.00153533 | 0.00159785 | 0.00138876 | 0.0002091 | 0.15812363 | 0.99999994 |
| Gm26877 | 0.00153337 | 0.00159574 | 0.00138698 | 0.00020876 | 0.15820059 | 0.99999994 |
| Drd4 | 0.00107952 | 0.00118518 | 0.00097677 | 0.00020841 | 0.07776503 | 0.99999994 |
| Fam25c | 0.00061195 | 0.00076207 | 0.00055387 | 0.00020819 | 0.00617449 | 0.30100158 |
| Bcl2a1a | 0.00061195 | 0.00076207 | 0.00055387 | 0.00020819 | 0.00617449 | 0.30100158 |
| Gm36186 | 0.00607 | 0.00568153 | 0.00547364 | 0.00020789 | 0.40020974 | 0.99999994 |
| Itgb7 | 0.00149066 | 0.001556 | 0.00134839 | 0.00020761 | 0.15271822 | 0.99999994 |
| Gimap1 | 0.00564147 | 0.00529553 | 0.00508869 | 0.00020684 | 0.39336344 | 0.99999994 |
| Nog | 0.00102669 | 0.00113479 | 0.000929 | 0.0002058 | 0.0701632 | 0.99999994 |
| 5730596B20 | 0.00061014 | 0.00075756 | 0.00055223 | 0.00020532 | 0.00666403 | 0.31901281 |
| Camk2d | 0.03910265 | 0.03469321 | 0.034489 | 0.00020421 | 0.4842805 | 0.99999994 |
| Ccr7 | 0.0010653 | 0.00116781 | 0.0009639 | 0.00020391 | 0.07955074 | 0.99999994 |
| C330011M18 | 0.00701808 | 0.00652778 | 0.00632452 | 0.00020326 | 0.41530292 | 0.99999994 |
| Cdc6 | 0.00722825 | 0.00671583 | 0.00651299 | 0.00020284 | 0.4178858 | 0.99999994 |
| 4930507D05 | 0.00436558 | 0.00414355 | 0.00394123 | 0.00020232 | 0.36629028 | 0.99999994 |

| | | | | | | |
|---|---|---|---|---|---|---|
| 4930547M16 | 0.00545679 | 0.00512489 | 0.00492273 | 0.00020216 | 0.3922901 | 0.99999994 |
| Hnf1aos1 | 0.06839438 | 0.0593778 | 0.05917585 | 0.00020196 | 0.49093807 | 0.99999994 |
| Gm13874 | 0.00060791 | 0.00075204 | 0.00055022 | 0.00020182 | 0.00731239 | 0.34692483 |
| Ccr5 | 0.00107218 | 0.00117168 | 0.00097013 | 0.00020155 | 0.08334792 | 0.99999994 |
| Nenf | 0.95731815 | 0.42364733 | 0.42344587 | 0.00020146 | 0.49873661 | 0.99999994 |
| Cacng6 | 0.00060757 | 0.0007512 | 0.00054991 | 0.00020128 | 0.00741642 | 0.34875697 |
| Bgn | 0.02035952 | 0.01838443 | 0.01818322 | 0.00020121 | 0.47064092 | 0.99999994 |
| Mamdc4 | 0.00185105 | 0.00187446 | 0.00167397 | 0.00020049 | 0.21266124 | 0.99999994 |
| Ttll10 | 0.01394277 | 0.01270632 | 0.01250623 | 0.0002001 | 0.4575926 | 0.99999994 |
| Gm21693 | 0.00060672 | 0.00074909 | 0.00054914 | 0.00019995 | 0.00768164 | 0.3546359 |
| F2 | 0.01372516 | 0.01251266 | 0.01231284 | 0.00019981 | 0.45699005 | 0.99999994 |
| Mastl | 0.00787571 | 0.00729305 | 0.00709327 | 0.00019978 | 0.42564286 | 0.99999994 |
| Plscr4 | 0.00290897 | 0.0028282 | 0.0026288 | 0.0001994 | 0.30681857 | 0.99999994 |
| 5430427N15 | 0.00060604 | 0.00074742 | 0.00054853 | 0.00019889 | 0.00789917 | 0.36288384 |
| Gm12766 | 0.00142232 | 0.00148517 | 0.00128663 | 0.00019854 | 0.15217779 | 0.99999994 |
| Stap1 | 0.00106834 | 0.00116484 | 0.00096666 | 0.00019818 | 0.08618497 | 0.99999994 |
| Spag4 | 0.00847303 | 0.00782626 | 0.00762817 | 0.00019809 | 0.43138225 | 0.99999994 |
| D11Wsu47e | 0.06362636 | 0.05541899 | 0.05522166 | 0.00019733 | 0.49051189 | 0.99999994 |
| Arhgef33 | 0.00793098 | 0.0073401 | 0.00714279 | 0.00019731 | 0.42705534 | 0.99999994 |
| Fbxo48 | 0.00060502 | 0.00074492 | 0.00054761 | 0.00019731 | 0.00823455 | 0.37291738 |
| Hfm1 | 0.00104917 | 0.00114609 | 0.00094932 | 0.00019677 | 0.08384089 | 0.99999994 |
| Gm15475 | 0.0006046 | 0.00074388 | 0.00054723 | 0.00019665 | 0.00837755 | 0.37684119 |
| Muc2 | 0.0006046 | 0.00074388 | 0.00054723 | 0.00019665 | 0.00837755 | 0.37684119 |
| Cabp4 | 0.00148657 | 0.00154073 | 0.00134469 | 0.00019604 | 0.1659246 | 0.99999994 |
| Fut1 | 0.0247237 | 0.02221161 | 0.02201641 | 0.0001952 | 0.47646929 | 0.99999994 |
| Zeb2os | 0.0018407 | 0.00185982 | 0.00166462 | 0.00019519 | 0.21754168 | 0.99999994 |
| Nab1 | 0.02148464 | 0.01936877 | 0.01917358 | 0.00019519 | 0.47298778 | 0.99999994 |
| Clca3a1 | 0.00359622 | 0.00344301 | 0.00324835 | 0.00019466 | 0.34498647 | 0.99999994 |
| Bbs1 | 0.02129066 | 0.01919732 | 0.01900294 | 0.00019438 | 0.47285855 | 0.99999994 |
| Tmem213 | 0.00548191 | 0.00513965 | 0.00494531 | 0.00019434 | 0.39682111 | 0.99999994 |

| | | | | | | |
|---|---|---|---|---|---|---|
| Tescl | 0.00467155 | 0.00440993 | 0.00421658 | 0.00019335 | 0.38009806 | 0.99999994 |
| Kdm5d | 0.0461646 | 0.04072073 | 0.04052804 | 0.00019269 | 0.48737637 | 0.99999994 |
| Kcnj2 | 0.00297264 | 0.00287877 | 0.00268623 | 0.00019254 | 0.31664564 | 0.99999994 |
| Dtna | 0.00306979 | 0.00296619 | 0.00277384 | 0.00019235 | 0.32218718 | 0.99999994 |
| Lpar1 | 0.00918972 | 0.00846143 | 0.00826938 | 0.00019205 | 0.43857341 | 0.99999994 |
| Eras | 0.00060142 | 0.00073607 | 0.00054435 | 0.00019172 | 0.00952928 | 0.42079308 |
| Flywch2 | 0.13513611 | 0.11159769 | 0.11140602 | 0.00019167 | 0.49543138 | 0.99999994 |
| Acd | 0.32142157 | 0.22993358 | 0.22974208 | 0.0001915 | 0.49778654 | 0.99999994 |
| Pcp4l1 | 0.00299753 | 0.00289989 | 0.00270867 | 0.00019121 | 0.31921722 | 0.99999994 |
| Fam83d | 0.00404169 | 0.00384047 | 0.00364963 | 0.00019084 | 0.36389623 | 0.99999994 |
| Dnajc2 | 0.35445578 | 0.24743391 | 0.24724312 | 0.00019078 | 0.49795093 | 0.99999994 |
| Srgn | 0.0085311 | 0.00787081 | 0.00768014 | 0.00019067 | 0.43437179 | 0.99999994 |
| Syn3 | 0.00260915 | 0.00254867 | 0.00235834 | 0.00019033 | 0.29555945 | 0.99999994 |
| Gm5148 | 0.0040356 | 0.00383421 | 0.00364414 | 0.00019007 | 0.36422836 | 0.99999994 |
| Gm37637 | 0.00059959 | 0.00073161 | 0.0005427 | 0.00018891 | 0.01025049 | 0.44524014 |
| Fzd4 | 0.00366072 | 0.00349462 | 0.00330646 | 0.00018815 | 0.35242762 | 0.99999994 |
| Lrrc40 | 0.08090673 | 0.06962494 | 0.06943686 | 0.00018808 | 0.49280758 | 0.99999994 |
| Tceal7 | 0.00793097 | 0.00733074 | 0.00714277 | 0.00018797 | 0.43047464 | 0.99999994 |
| Lrrc14b | 0.00105014 | 0.00113768 | 0.0009502 | 0.00018748 | 0.09453368 | 0.99999994 |
| Gm5218 | 0.00332593 | 0.00319217 | 0.00300476 | 0.00018742 | 0.33900601 | 0.99999994 |
| Tlx2 | 0.00141967 | 0.0014716 | 0.00128424 | 0.00018736 | 0.16575088 | 0.99999994 |
| Cep72 | 0.01170626 | 0.01070312 | 0.010516 | 0.00018712 | 0.45285889 | 0.99999994 |
| Pcdhb2 | 0.00207864 | 0.00206652 | 0.00187951 | 0.00018701 | 0.2538911 | 0.99999994 |
| Gm44107 | 0.00106027 | 0.00114622 | 0.00095936 | 0.00018686 | 0.09740698 | 0.99999994 |
| Runx2 | 0.00059721 | 0.00072579 | 0.00054054 | 0.00018526 | 0.01126525 | 0.47887837 |
| Gm21917 | 0.00059712 | 0.00072559 | 0.00054046 | 0.00018513 | 0.01130158 | 0.47914493 |
| Gm15640 | 0.0010446 | 0.00113009 | 0.00094519 | 0.00018489 | 0.09644469 | 0.99999994 |
| Plcd3 | 0.00148804 | 0.00153067 | 0.00134602 | 0.00018464 | 0.18059331 | 0.99999994 |
| Gm3839 | 0.0005968 | 0.0007248 | 0.00054017 | 0.00018463 | 0.0114478 | 0.48405669 |
| Exo1 | 0.00441788 | 0.00417287 | 0.0039883 | 0.00018457 | 0.37902772 | 0.99999994 |

| | | | | | | |
|---|---|---|---|---|---|---|
| Prss3 | 0.00161187 | 0.00164238 | 0.00145791 | 0.00018447 | 0.19982968 | 0.99999994 |
| Ccdc88c | 0.01017544 | 0.00933396 | 0.00915029 | 0.00018368 | 0.44685342 | 0.99999994 |
| Tulp2 | 0.00228474 | 0.00224878 | 0.00206557 | 0.00018321 | 0.27746105 | 0.99999994 |
| Mmp19 | 0.00148927 | 0.00153008 | 0.00134713 | 0.00018294 | 0.18300992 | 0.99999994 |
| Tril | 0.00320153 | 0.00307554 | 0.00289261 | 0.00018293 | 0.33689872 | 0.99999994 |
| Six2 | 0.00059565 | 0.00072202 | 0.00053913 | 0.00018289 | 0.01197111 | 0.50170905 |
| Krt13 | 0.00213018 | 0.00210877 | 0.00192604 | 0.00018273 | 0.26385055 | 0.99999994 |
| Mir142hg | 0.00142229 | 0.0014691 | 0.0012866 | 0.0001825 | 0.17254424 | 0.99999994 |
| Armc3 | 0.0028042 | 0.00271657 | 0.0025343 | 0.00018227 | 0.31606747 | 0.99999994 |
| Stmn4 | 0.00104634 | 0.00112894 | 0.00094676 | 0.00018218 | 0.10012477 | 0.99999994 |
| Tssk4 | 0.00304059 | 0.00292959 | 0.00274751 | 0.00018209 | 0.32955393 | 0.99999994 |
| Gm4131 | 0.00059508 | 0.00072064 | 0.00053862 | 0.00018203 | 0.01223958 | 0.5028638 |
| D030025P21 | 0.00105812 | 0.00113901 | 0.00095741 | 0.0001816 | 0.1033721 | 0.99999994 |
| Barhl2 | 0.00228265 | 0.0022452 | 0.00206368 | 0.00018152 | 0.27910995 | 0.99999994 |
| Trp73 | 0.00360174 | 0.00343468 | 0.00325332 | 0.00018136 | 0.35529213 | 0.99999994 |
| Gm11205 | 0.00149654 | 0.00153507 | 0.0013537 | 0.00018136 | 0.18625404 | 0.99999994 |
| Slc9b1 | 0.00101439 | 0.00109923 | 0.00091787 | 0.00018136 | 0.09422379 | 0.99999994 |
| 2210417A02 | 0.00130898 | 0.00136523 | 0.00118419 | 0.00018104 | 0.1544321 | 0.99999994 |
| Slc25a43 | 0.0065355 | 0.00607259 | 0.00589156 | 0.00018103 | 0.41896756 | 0.99999994 |
| Gpc2 | 0.00569104 | 0.00531378 | 0.00513323 | 0.00018055 | 0.40744565 | 0.99999994 |
| Hs6st2 | 0.00278841 | 0.00270061 | 0.00252007 | 0.00018054 | 0.31672466 | 0.99999994 |
| Myb | 0.00290971 | 0.00280979 | 0.00262947 | 0.00018031 | 0.32403151 | 0.99999994 |
| Mis18a | 0.05519157 | 0.04834632 | 0.04816602 | 0.00018031 | 0.49006039 | 0.99999994 |
| Gria1 | 0.00059387 | 0.0007177 | 0.00053752 | 0.00018018 | 0.01283047 | 0.51649113 |
| Sfta2 | 0.00059387 | 0.0007177 | 0.00053752 | 0.00018018 | 0.01283047 | 0.51649113 |
| Kng1 | 0.00059387 | 0.0007177 | 0.00053752 | 0.00018018 | 0.01283047 | 0.51649113 |
| Creb3l3 | 0.00251187 | 0.00245059 | 0.00227057 | 0.00018002 | 0.29883666 | 0.99999994 |
| Ldoc1 | 0.00137594 | 0.00142459 | 0.00124472 | 0.00017987 | 0.1680479 | 0.99999994 |
| Cd300a | 0.00236364 | 0.00231664 | 0.00213679 | 0.00017985 | 0.28765051 | 0.99999994 |
| Hoxa2 | 0.00100644 | 0.00109023 | 0.00091068 | 0.00017955 | 0.09469463 | 0.99999994 |

| | | | | | | |
|---|---|---|---|---|---|---|
| Brs3 | 0.0005933 | 0.00071633 | 0.000537 | 0.00017933 | 0.01311359 | 0.52260953 |
| Gm15637 | 0.0005933 | 0.00071633 | 0.000537 | 0.00017933 | 0.01311359 | 0.52260953 |
| Abcg5 | 0.0023563 | 0.00230946 | 0.00213017 | 0.0001793 | 0.2876478 | 0.99999994 |
| 1500009L16I | 0.30698692 | 0.22197488 | 0.22179563 | 0.00017925 | 0.49785387 | 0.99999994 |
| Rftn1 | 0.00352195 | 0.00336065 | 0.00318143 | 0.00017923 | 0.35383429 | 0.99999994 |
| D17H6S53E | 0.06382328 | 0.05556467 | 0.05538545 | 0.00017921 | 0.49140822 | 0.99999994 |
| 4930444P10I | 0.00059298 | 0.00071555 | 0.00053671 | 0.00017884 | 0.01327752 | 0.52782269 |
| Wdr62 | 0.00347045 | 0.00331381 | 0.00313502 | 0.00017879 | 0.35211349 | 0.99999994 |
| Gm10373 | 0.00059282 | 0.00071516 | 0.00053657 | 0.0001786 | 0.01336006 | 0.52978304 |
| Tns1 | 0.00964013 | 0.00885034 | 0.00867204 | 0.0001783 | 0.44557146 | 0.99999994 |
| Lrrn3 | 0.0035511 | 0.00338595 | 0.00320769 | 0.00017825 | 0.35573124 | 0.99999994 |
| Mmp23 | 0.00773056 | 0.00714145 | 0.00696323 | 0.00017822 | 0.4323615 | 0.99999994 |
| Kcnj8 | 0.00611296 | 0.00569043 | 0.00551222 | 0.00017821 | 0.41480687 | 0.99999994 |
| Tprg | 0.00144741 | 0.00148747 | 0.00130931 | 0.00017816 | 0.18254019 | 0.99999994 |
| Pla2g4a | 0.00129957 | 0.00135359 | 0.0011757 | 0.00017789 | 0.15692863 | 0.99999994 |
| Smad6 | 0.00536212 | 0.00501552 | 0.00483763 | 0.00017788 | 0.40332126 | 0.99999994 |
| Col20a1 | 0.00440862 | 0.00415777 | 0.00397997 | 0.0001778 | 0.38309614 | 0.99999994 |
| Fgf2 | 0.00099406 | 0.00107717 | 0.00089949 | 0.00017768 | 0.09427863 | 0.99999994 |
| Gm29570 | 0.00059209 | 0.00071342 | 0.00053591 | 0.00017751 | 0.0137364 | 0.53669581 |
| Gm28196 | 0.00105362 | 0.00113075 | 0.00095335 | 0.0001774 | 0.10774315 | 0.99999994 |
| 4632415L05I | 0.0668448 | 0.05807079 | 0.05789345 | 0.00017734 | 0.4918663 | 0.99999994 |
| Dnajb13 | 0.00148524 | 0.00152042 | 0.0013435 | 0.00017693 | 0.19035664 | 0.99999994 |
| Cbln2 | 0.00187135 | 0.00186866 | 0.00169231 | 0.00017635 | 0.24395991 | 0.99999994 |
| Ephb4 | 0.01157532 | 0.01057564 | 0.0103993 | 0.00017634 | 0.45506511 | 0.99999994 |
| Gm29590 | 0.00059129 | 0.00071148 | 0.00053518 | 0.0001763 | 0.01416398 | 0.54741557 |
| Gm12784 | 0.00199646 | 0.00198099 | 0.0018053 | 0.00017569 | 0.25856315 | 0.99999994 |
| Gm20633 | 0.00104882 | 0.00112447 | 0.00094901 | 0.00017547 | 0.10921312 | 0.99999994 |
| Tmprss3 | 0.00059073 | 0.00071013 | 0.00053468 | 0.00017546 | 0.01446928 | 0.55516698 |
| BC028528 | 0.01283404 | 0.0116958 | 0.01152034 | 0.00017545 | 0.45962597 | 0.99999994 |
| Rbp7 | 0.00141911 | 0.00145915 | 0.00128374 | 0.00017541 | 0.1815355 | 0.99999994 |

| | | | | | | |
|---|---|---|---|---|---|---|
| Unc5d | 0.00059057 | 0.00070975 | 0.00053453 | 0.00017522 | 0.01455742 | 0.55516698 |
| Drc7 | 0.00059057 | 0.00070975 | 0.00053453 | 0.00017522 | 0.01455742 | 0.55516698 |
| Fgf14 | 0.00182962 | 0.00182971 | 0.00165462 | 0.00017509 | 0.24060529 | 0.99999994 |
| Sidt1 | 0.0026895 | 0.00260584 | 0.00243083 | 0.00017501 | 0.31589136 | 0.99999994 |
| Bmp4 | 0.00230574 | 0.00225913 | 0.00208453 | 0.0001746 | 0.288578 | 0.99999994 |
| Zar1l | 0.00288797 | 0.00278445 | 0.00260986 | 0.00017459 | 0.32805639 | 0.99999994 |
| Slc25a32 | 0.0230599 | 0.02073228 | 0.02055769 | 0.00017459 | 0.47745962 | 0.99999994 |
| A330033J07R | 0.00059001 | 0.00070841 | 0.00053403 | 0.00017438 | 0.01486913 | 0.56301374 |
| Tulp1 | 0.00186627 | 0.00186189 | 0.00168772 | 0.00017417 | 0.24606231 | 0.99999994 |
| Gm12781 | 0.0010485 | 0.00112259 | 0.00094872 | 0.00017387 | 0.11125224 | 0.99999994 |
| Rspo1 | 0.00360467 | 0.00342922 | 0.00325596 | 0.00017326 | 0.36159173 | 0.99999994 |
| Dnah7b | 0.00058905 | 0.00070611 | 0.00053316 | 0.00017295 | 0.01541525 | 0.56883004 |
| Gm28178 | 0.00058905 | 0.00070611 | 0.00053316 | 0.00017295 | 0.01541525 | 0.56883004 |
| Panct2 | 0.0065635 | 0.00608957 | 0.00591669 | 0.00017288 | 0.42289345 | 0.99999994 |
| Cpne6 | 0.0012505 | 0.00130415 | 0.00113133 | 0.00017282 | 0.15462622 | 0.99999994 |
| Gm42722 | 0.0005889 | 0.00070573 | 0.00053302 | 0.00017272 | 0.01550773 | 0.5696055 |
| Kcnf1 | 0.0005889 | 0.00070573 | 0.00053302 | 0.00017272 | 0.01550773 | 0.5696055 |
| Frmpd3 | 0.00058866 | 0.00070516 | 0.0005328 | 0.00017236 | 0.01564724 | 0.5734084 |
| Olfml1 | 0.00105034 | 0.00112238 | 0.00095038 | 0.000172 | 0.11416223 | 0.99999994 |
| Gm15853 | 0.00865856 | 0.00796621 | 0.00779422 | 0.000172 | 0.44161045 | 0.99999994 |
| Trem2 | 0.00490051 | 0.00459453 | 0.00442256 | 0.00017197 | 0.39788167 | 0.99999994 |
| Gm26852 | 0.00730379 | 0.00675251 | 0.00658071 | 0.0001718 | 0.43102106 | 0.99999994 |
| 4933405L10R | 0.00058826 | 0.00070421 | 0.00053244 | 0.00017177 | 0.01588186 | 0.57801989 |
| Hes5 | 0.0023433 | 0.00229016 | 0.00211843 | 0.00017173 | 0.29473552 | 0.99999994 |
| Egln3 | 0.02164545 | 0.01948595 | 0.01931502 | 0.00017093 | 0.47651365 | 0.99999994 |
| Paqr8 | 0.00467093 | 0.00438691 | 0.00421602 | 0.00017089 | 0.39365369 | 0.99999994 |
| Crmp1 | 0.02823605 | 0.02525603 | 0.02508519 | 0.00017084 | 0.48192072 | 0.99999994 |
| A830082K12 | 0.00253586 | 0.00246296 | 0.00229221 | 0.00017075 | 0.30999984 | 0.99999994 |
| Oprk1 | 0.00145791 | 0.00148917 | 0.00131879 | 0.00017037 | 0.19491299 | 0.99999994 |
| Zglp1 | 0.00103475 | 0.00110633 | 0.00093629 | 0.00017004 | 0.11335223 | 0.99999994 |

| | | | | | | |
|---|---|---|---|---|---|---|
| Gm2093 | 0.00214308 | 0.00210753 | 0.00193769 | 0.00016984 | 0.27979665 | 0.99999994 |
| Mycs | 0.00058692 | 0.000701 | 0.00053123 | 0.00016977 | 0.01669949 | 0.59956436 |
| Mmp17 | 0.00149826 | 0.00152498 | 0.00135526 | 0.00016972 | 0.20225474 | 0.99999994 |
| Gm26543 | 0.00103459 | 0.00110566 | 0.00093614 | 0.00016952 | 0.1140291 | 0.99999994 |
| Rpa2 | 0.11938416 | 0.09982748 | 0.09965798 | 0.0001695 | 0.49548354 | 0.99999994 |
| Prr19 | 0.00146323 | 0.00149304 | 0.00132361 | 0.00016943 | 0.19708518 | 0.99999994 |
| C230034O21 | 0.00102761 | 0.00109924 | 0.00092982 | 0.00016942 | 0.1125984 | 0.99999994 |
| Plcxd1 | 0.00278337 | 0.00268442 | 0.00251552 | 0.0001689 | 0.32746234 | 0.99999994 |
| Hamp2 | 0.00058629 | 0.0006995 | 0.00053066 | 0.00016884 | 0.01709505 | 0.60693149 |
| BC065403 | 0.00058629 | 0.0006995 | 0.00053066 | 0.00016884 | 0.01709505 | 0.60693149 |
| Plvap | 0.0027358 | 0.0026413 | 0.0024726 | 0.0001687 | 0.3248624 | 0.99999994 |
| Rnf144b | 0.00058605 | 0.00069893 | 0.00053044 | 0.00016849 | 0.01724519 | 0.60803923 |
| Slc35f2 | 0.00123231 | 0.00128245 | 0.0011149 | 0.00016756 | 0.15856093 | 0.99999994 |
| Zfp521 | 0.00830146 | 0.00764139 | 0.00747457 | 0.00016682 | 0.44094943 | 0.99999994 |
| E2f8 | 0.0014956 | 0.0015196 | 0.00135285 | 0.00016675 | 0.20598039 | 0.99999994 |
| Tmem95 | 0.00058456 | 0.00069539 | 0.0005291 | 0.00016629 | 0.01821918 | 0.63000423 |
| Coro6 | 0.00058456 | 0.00069539 | 0.0005291 | 0.00016629 | 0.01821918 | 0.63000423 |
| Ctsw | 0.00058456 | 0.00069539 | 0.0005291 | 0.00016629 | 0.01821918 | 0.63000423 |
| Hrc | 0.00189981 | 0.00188363 | 0.00171802 | 0.00016562 | 0.26054359 | 0.99999994 |
| Agmo | 0.00271652 | 0.00262053 | 0.00245521 | 0.00016532 | 0.32700567 | 0.99999994 |
| B3gat1 | 0.00147815 | 0.00150236 | 0.00133708 | 0.00016528 | 0.20531339 | 0.99999994 |
| Sln | 0.00058378 | 0.00069353 | 0.00052839 | 0.00016514 | 0.01874805 | 0.64463872 |
| Gm12666 | 0.00145607 | 0.00148221 | 0.00131714 | 0.00016507 | 0.20207767 | 0.99999994 |
| Gm2109 | 0.00101925 | 0.00108713 | 0.00092227 | 0.00016486 | 0.1170507 | 0.99999994 |
| Rac2 | 0.00587154 | 0.00545954 | 0.0052954 | 0.00016415 | 0.41826503 | 0.99999994 |
| Rnf150 | 0.00297154 | 0.00284928 | 0.00268523 | 0.00016404 | 0.34213378 | 0.99999994 |
| Gm26632 | 0.00567661 | 0.00528395 | 0.00512027 | 0.00016368 | 0.41574711 | 0.99999994 |
| Rab9b | 0.00857196 | 0.00788027 | 0.00771672 | 0.00016355 | 0.4439042 | 0.99999994 |
| Dcdc2b | 0.00301 | 0.00288343 | 0.00271991 | 0.00016351 | 0.34451894 | 0.99999994 |
| 3110083C13I | 0.01083054 | 0.00989841 | 0.00973508 | 0.00016333 | 0.45553962 | 0.99999994 |

| | | | | | | |
|---|---|---|---|---|---|---|
| Nanp | 0.00216003 | 0.00211627 | 0.00195299 | 0.00016328 | 0.28892908 | 0.99999994 |
| Rho | 0.00076629 | 0.00085656 | 0.0006935 | 0.00016306 | 0.05877859 | 0.99999994 |
| Lamc3 | 0.00058223 | 0.00068984 | 0.00052699 | 0.00016286 | 0.01984006 | 0.6729158 |
| Slmapos2 | 0.00058207 | 0.00068948 | 0.00052685 | 0.00016263 | 0.01995181 | 0.67383848 |
| Gli3 | 0.00200609 | 0.00197642 | 0.001814 | 0.00016242 | 0.27558844 | 0.99999994 |
| Rgs4 | 0.00770808 | 0.00710465 | 0.00694309 | 0.00016157 | 0.43845276 | 0.99999994 |
| Slc1a6 | 0.00058099 | 0.00068692 | 0.00052587 | 0.00016105 | 0.02074716 | 0.69426319 |
| Tsku | 0.00490139 | 0.00458426 | 0.00442335 | 0.00016091 | 0.40433404 | 0.99999994 |
| Usp25 | 0.02046126 | 0.01843347 | 0.01827283 | 0.00016064 | 0.47666848 | 0.99999994 |
| Klhl4 | 0.00058068 | 0.00068619 | 0.00052559 | 0.0001606 | 0.02097865 | 0.69426319 |
| Tomm6os | 0.00341172 | 0.0032425 | 0.00308208 | 0.00016042 | 0.36449875 | 0.99999994 |
| Gtse1 | 0.0013828 | 0.00141134 | 0.00125092 | 0.00016042 | 0.1966596 | 0.99999994 |
| Kcnj14 | 0.00121858 | 0.00126184 | 0.00110248 | 0.00015936 | 0.16798965 | 0.99999994 |
| Hsf2bp | 0.00402752 | 0.003796 | 0.00363686 | 0.00015914 | 0.3854245 | 0.99999994 |
| 4930533D04 | 0.00299026 | 0.00286113 | 0.00270212 | 0.00015901 | 0.34763838 | 0.99999994 |
| Akr1d1 | 0.00057938 | 0.00068311 | 0.00052441 | 0.0001587 | 0.02198385 | 0.71665561 |
| Cadm2 | 0.00272629 | 0.00262246 | 0.00246402 | 0.00015844 | 0.33432126 | 0.99999994 |
| Kcnj16 | 0.00094262 | 0.00101132 | 0.00085297 | 0.00015835 | 0.10828369 | 0.99999994 |
| Nxf2 | 0.00057907 | 0.00068239 | 0.00052413 | 0.00015826 | 0.02222544 | 0.72157992 |
| Prr16 | 0.00057907 | 0.00068239 | 0.00052413 | 0.00015826 | 0.02222544 | 0.72157992 |
| Rnf39 | 0.00577767 | 0.00536899 | 0.00521106 | 0.00015792 | 0.42006709 | 0.99999994 |
| Pcdhga8 | 0.00216508 | 0.00211531 | 0.00195755 | 0.00015777 | 0.29581807 | 0.99999994 |
| Cstad | 0.00582963 | 0.00541532 | 0.00525774 | 0.00015758 | 0.42093858 | 0.99999994 |
| Gjb1 | 0.01276361 | 0.01161521 | 0.01145767 | 0.00015754 | 0.46353809 | 0.99999994 |
| 2700049A03 | 0.14353046 | 0.11770192 | 0.11754445 | 0.00015747 | 0.49644247 | 0.99999994 |
| Ier3ip1 | 0.97364669 | 0.4248217 | 0.42466423 | 0.00015746 | 0.49901535 | 0.99999994 |
| Pln | 0.00103791 | 0.00109578 | 0.00093914 | 0.00015664 | 0.13345502 | 0.99999994 |
| Nsun7 | 0.0021503 | 0.00210035 | 0.00194421 | 0.00015615 | 0.29646515 | 0.99999994 |
| Pfkfb1 | 0.00748379 | 0.00689789 | 0.00674208 | 0.00015581 | 0.43887287 | 0.99999994 |
| Caln1 | 0.00142998 | 0.00144891 | 0.00129355 | 0.00015535 | 0.2120231 | 0.99999994 |

| | | | | | | |
|---|---|---|---|---|---|---|
| Syne3 | 0.00057701 | 0.00067755 | 0.00052227 | 0.00015528 | 0.02390752 | 0.76682042 |
| Ckap2l | 0.00211857 | 0.00207076 | 0.00191555 | 0.00015521 | 0.29482998 | 0.99999994 |
| Kcnk5 | 0.00257752 | 0.00248488 | 0.0023298 | 0.00015508 | 0.32885961 | 0.99999994 |
| Larp6 | 0.00189523 | 0.00186888 | 0.00171388 | 0.00015501 | 0.27358172 | 0.99999994 |
| Slit2 | 0.00816748 | 0.00750959 | 0.0073546 | 0.000155 | 0.44421918 | 0.99999994 |
| Zfp317 | 0.02681297 | 0.02399845 | 0.02384357 | 0.00015488 | 0.48275587 | 0.99999994 |
| Zcchc12 | 0.00500047 | 0.00466724 | 0.00451246 | 0.00015478 | 0.40969884 | 0.99999994 |
| Fem1c | 0.06908788 | 0.05990366 | 0.05974893 | 0.00015473 | 0.49312359 | 0.99999994 |
| Grm1 | 0.00294315 | 0.00281429 | 0.00265963 | 0.00015466 | 0.34935022 | 0.99999994 |
| Dnah12 | 0.00057656 | 0.00067648 | 0.00052186 | 0.00015462 | 0.02429362 | 0.77452909 |
| 4921514A10 | 0.00103437 | 0.00109048 | 0.00093594 | 0.00015455 | 0.13585681 | 0.99999994 |
| Dlg2 | 0.00118066 | 0.00122267 | 0.00106821 | 0.00015447 | 0.16788971 | 0.99999994 |
| Entpd1 | 0.00099629 | 0.00105562 | 0.00090151 | 0.00015411 | 0.1275856 | 0.99999994 |
| Qprt | 0.00225776 | 0.00219482 | 0.00204121 | 0.00015361 | 0.30821779 | 0.99999994 |
| Ifit2 | 0.00142619 | 0.00144362 | 0.00129013 | 0.00015349 | 0.21420965 | 0.99999994 |
| Tln2 | 0.03920626 | 0.03473148 | 0.03457801 | 0.00015347 | 0.4882159 | 0.99999994 |
| Col8a2 | 0.00126375 | 0.00129673 | 0.00114331 | 0.00015341 | 0.18588134 | 0.99999994 |
| Wt1 | 0.00384082 | 0.0036218 | 0.00346872 | 0.00015308 | 0.38447172 | 0.99999994 |
| Galnt13 | 0.00233105 | 0.00226011 | 0.00210737 | 0.00015273 | 0.31475242 | 0.99999994 |
| Izumo1 | 0.00143923 | 0.00145446 | 0.00130191 | 0.00015255 | 0.21770675 | 0.99999994 |
| Nkx2-3 | 0.00146813 | 0.00148011 | 0.00132803 | 0.00015208 | 0.22295647 | 0.99999994 |
| Zfp82 | 0.00585349 | 0.00543121 | 0.00527918 | 0.00015203 | 0.42399322 | 0.99999994 |
| Dgkk | 0.0005746 | 0.00067188 | 0.00052008 | 0.0001518 | 0.02601942 | 0.81698086 |
| Trap1a | 0.00057452 | 0.0006717 | 0.00052001 | 0.00015169 | 0.02608752 | 0.81698086 |
| Afmid | 0.00212569 | 0.00207344 | 0.00192199 | 0.00015146 | 0.29995158 | 0.99999994 |
| Kdm4c | 0.07881549 | 0.06788468 | 0.06773344 | 0.00015125 | 0.49407063 | 0.99999994 |
| Slc16a1 | 0.00306876 | 0.00292366 | 0.0027729 | 0.00015076 | 0.35871671 | 0.99999994 |
| Rin3 | 0.00311629 | 0.00296632 | 0.00281576 | 0.00015056 | 0.36095099 | 0.99999994 |
| Arhgap42 | 0.01040607 | 0.00950676 | 0.00935622 | 0.00015054 | 0.45735623 | 0.99999994 |
| Cyp2b13 | 0.00148372 | 0.00149252 | 0.00134212 | 0.0001504 | 0.22785113 | 0.99999994 |

| | | | | | | |
|---|---|---|---|---|---|---|
| Dpep2 | 0.00453994 | 0.0042478 | 0.00409816 | 0.00014964 | 0.40398077 | 0.99999994 |
| Tmem45a2 | 0.00146811 | 0.00147764 | 0.00132801 | 0.00014963 | 0.22663027 | 0.99999994 |
| Rapgef3 | 0.00341405 | 0.00323344 | 0.00308418 | 0.00014927 | 0.37367029 | 0.99999994 |
| Mzb1 | 0.00364276 | 0.0034395 | 0.00329028 | 0.00014921 | 0.38137891 | 0.99999994 |
| Kdm3a | 0.0730549 | 0.06316639 | 0.06301726 | 0.00014913 | 0.49371591 | 0.99999994 |
| Ankrd33 | 0.00103137 | 0.0010818 | 0.00093322 | 0.00014857 | 0.14463475 | 0.99999994 |
| Wt1os | 0.00142653 | 0.00143888 | 0.00129044 | 0.00014844 | 0.22192562 | 0.99999994 |
| Cxcl15 | 0.00229511 | 0.00222284 | 0.00207493 | 0.00014792 | 0.31756635 | 0.99999994 |
| Lysmd4 | 0.03691257 | 0.03275241 | 0.03260464 | 0.00014777 | 0.48796668 | 0.99999994 |
| Calb2 | 0.00103019 | 0.00107956 | 0.00093216 | 0.0001474 | 0.14627028 | 0.99999994 |
| Gm10101 | 0.00472089 | 0.00440777 | 0.00426098 | 0.00014679 | 0.4093135 | 0.99999994 |
| Gad2 | 0.00125838 | 0.00128487 | 0.00113846 | 0.00014641 | 0.19599794 | 0.99999994 |
| Inhba | 0.0020336 | 0.00198515 | 0.00183883 | 0.00014632 | 0.29817473 | 0.99999994 |
| 1110032F04I | 0.00057072 | 0.00066283 | 0.00051657 | 0.00014626 | 0.02973472 | 0.87840191 |
| Gm8113 | 0.00057072 | 0.00066283 | 0.00051657 | 0.00014626 | 0.02973472 | 0.87840191 |
| 0610038B21 | 0.00188788 | 0.00185285 | 0.00170724 | 0.00014561 | 0.28511463 | 0.99999994 |
| Jsrp1 | 0.00097918 | 0.0010316 | 0.00088603 | 0.00014557 | 0.13706581 | 0.99999994 |
| Srsf6 | 1.01401929 | 0.42728521 | 0.42713991 | 0.0001453 | 0.49909668 | 0.99999994 |
| Magee2 | 0.00202001 | 0.00197168 | 0.00182657 | 0.00014512 | 0.29846282 | 0.99999994 |
| Abcg4 | 0.00101174 | 0.00106022 | 0.00091548 | 0.00014474 | 0.14631661 | 0.99999994 |
| Aadac | 0.00056953 | 0.00066009 | 0.0005155 | 0.00014459 | 0.03095058 | 0.89706031 |
| Siae | 0.02797079 | 0.02499813 | 0.02485393 | 0.0001442 | 0.48459717 | 0.99999994 |
| Wdfy2 | 0.0013737 | 0.00138668 | 0.00124269 | 0.00014399 | 0.22028175 | 0.99999994 |
| Arrb1 | 0.0005688 | 0.00065838 | 0.00051483 | 0.00014355 | 0.03172824 | 0.91793073 |
| Htra3 | 0.00483331 | 0.00450559 | 0.00436211 | 0.00014348 | 0.41334555 | 0.99999994 |
| Gm44081 | 0.00056865 | 0.00065804 | 0.0005147 | 0.00014334 | 0.03188542 | 0.92080701 |
| Erich2 | 0.00736574 | 0.00677958 | 0.00663626 | 0.00014332 | 0.44284717 | 0.99999994 |
| 9330198N18 | 0.00161738 | 0.00160593 | 0.00146289 | 0.00014304 | 0.2575751 | 0.99999994 |
| Klk9 | 0.00141756 | 0.00142535 | 0.00128233 | 0.00014302 | 0.22891776 | 0.99999994 |
| Gpm6a | 0.00367839 | 0.00346483 | 0.00332239 | 0.00014245 | 0.38767376 | 0.99999994 |

| | | | | | | |
|---|---|---|---|---|---|---|
| Zfpm2 | 0.00056755 | 0.0006555 | 0.00051371 | 0.00014179 | 0.03308193 | 0.94678472 |
| Ska1 | 0.00224726 | 0.00217283 | 0.00203174 | 0.0001411 | 0.321949 | 0.99999994 |
| Prex2 | 0.01180718 | 0.01074669 | 0.01060594 | 0.00014074 | 0.46480655 | 0.99999994 |
| Fam83e | 0.00318304 | 0.00301661 | 0.00287595 | 0.00014067 | 0.37237201 | 0.99999994 |
| Slc38a9 | 0.04268272 | 0.0376982 | 0.03755755 | 0.00014065 | 0.49005628 | 0.99999994 |
| Mef2c | 0.00176404 | 0.0017358 | 0.00159538 | 0.00014041 | 0.2789923 | 0.99999994 |
| Gm15884 | 0.00088368 | 0.00094007 | 0.00079967 | 0.0001404 | 0.12127063 | 0.99999994 |
| Slc25a27 | 0.01992993 | 0.01794479 | 0.01780467 | 0.00014011 | 0.47911219 | 0.99999994 |
| Gad1os | 0.00227019 | 0.00219243 | 0.00205244 | 0.00013999 | 0.32491303 | 0.99999994 |
| Fam169b | 0.00142313 | 0.00142704 | 0.00128736 | 0.00013967 | 0.23508829 | 0.99999994 |
| Letmd1 | 0.10341355 | 0.0875735 | 0.08743394 | 0.00013956 | 0.49576131 | 0.99999994 |
| Gm11724 | 0.0014108 | 0.0014156 | 0.00127622 | 0.00013937 | 0.23363433 | 0.99999994 |
| Aldoart1 | 0.0005658 | 0.00065146 | 0.00051212 | 0.00013934 | 0.03506179 | 0.98641032 |
| Plekho2 | 0.00329624 | 0.00311732 | 0.002978 | 0.00013932 | 0.37774283 | 0.99999994 |
| Fcgr3 | 0.00700472 | 0.00645173 | 0.00631254 | 0.0001392 | 0.44165263 | 0.99999994 |
| Pcdhb6 | 0.00345608 | 0.00326121 | 0.00312207 | 0.00013914 | 0.3833663 | 0.99999994 |
| 2010109A12I | 0.02755109 | 0.02462667 | 0.02448787 | 0.00013881 | 0.48495103 | 0.99999994 |
| Nphs2 | 0.00056536 | 0.00065046 | 0.00051173 | 0.00013873 | 0.03556948 | 0.99475976 |
| A830019P07 | 0.00056536 | 0.00065046 | 0.00051173 | 0.00013873 | 0.03556948 | 0.99475976 |
| B230208H11 | 0.00314635 | 0.0029815 | 0.00284287 | 0.00013864 | 0.37273828 | 0.99999994 |
| Dkk3 | 0.00243573 | 0.00234023 | 0.00220185 | 0.00013838 | 0.33784873 | 0.99999994 |
| Gm11454 | 0.00056507 | 0.00064979 | 0.00051147 | 0.00013833 | 0.03591078 | 0.99942728 |
| Slc16a2 | 0.02784846 | 0.0248855 | 0.02474725 | 0.00013825 | 0.48516882 | 0.99999994 |
| Ppt2 | 0.03875632 | 0.03432958 | 0.03419138 | 0.0001382 | 0.48926779 | 0.99999994 |
| Gm6169 | 0.00099524 | 0.00103845 | 0.00090056 | 0.0001379 | 0.15404501 | 0.99999994 |
| Dnd1 | 0.00503349 | 0.00467984 | 0.00454216 | 0.00013768 | 0.42004756 | 0.99999994 |
| Igfbp1 | 0.00121294 | 0.00123505 | 0.00109738 | 0.00013766 | 0.20185142 | 0.99999994 |
| Aox4 | 0.00056457 | 0.00064863 | 0.00051101 | 0.00013762 | 0.03651357 | 0.99999994 |
| Serpina1f | 0.00056421 | 0.0006478 | 0.00051068 | 0.00013712 | 0.03694844 | 0.99999994 |
| Cntnap5a | 0.00056413 | 0.00064763 | 0.00051062 | 0.00013702 | 0.03703584 | 0.99999994 |

| | | | | | | |
|---|---|---|---|---|---|---|
| Plet1 | 0.01017342 | 0.00928537 | 0.00914848 | 0.00013689 | 0.46033056 | 0.99999994 |
| U2surp | 0.53902612 | 0.32960956 | 0.32947334 | 0.00013622 | 0.4989021 | 0.99999994 |
| B3glct | 0.03713201 | 0.03292912 | 0.0327937 | 0.00013542 | 0.48903562 | 0.99999994 |
| Bub1b | 0.00169324 | 0.00166676 | 0.00153143 | 0.00013533 | 0.27819556 | 0.99999994 |
| Cytip | 0.00289489 | 0.00275107 | 0.00261611 | 0.00013496 | 0.36565594 | 0.99999994 |
| Dscam | 0.00168605 | 0.00165989 | 0.00152493 | 0.00013496 | 0.27790001 | 0.99999994 |
| Heyl | 0.00297618 | 0.00282409 | 0.00268942 | 0.00013467 | 0.36944531 | 0.99999994 |
| Rad9b | 0.0414666 | 0.03665147 | 0.03651682 | 0.00013465 | 0.49020888 | 0.99999994 |
| Dao | 0.00056234 | 0.00064351 | 0.00050899 | 0.00013452 | 0.03926802 | 0.99999994 |
| Scn7a | 0.00056219 | 0.00064318 | 0.00050886 | 0.00013432 | 0.03945053 | 0.99999994 |
| Igsf23 | 0.00293851 | 0.00278961 | 0.00265545 | 0.00013416 | 0.3683223 | 0.99999994 |
| Mapk12 | 0.00145737 | 0.00145225 | 0.00131831 | 0.00013394 | 0.24943735 | 0.99999994 |
| Mlip | 0.00145376 | 0.00144894 | 0.00131504 | 0.0001339 | 0.24896751 | 0.99999994 |
| Iqca | 0.00184098 | 0.00179861 | 0.00166488 | 0.00013373 | 0.29643534 | 0.99999994 |
| Cnpy4 | 0.09231917 | 0.07878604 | 0.07865253 | 0.00013351 | 0.49549258 | 0.99999994 |
| Cd300lb | 0.00056148 | 0.00064155 | 0.00050821 | 0.00013334 | 0.04037186 | 0.99999994 |
| Gm20379 | 0.00056148 | 0.00064155 | 0.00050821 | 0.00013334 | 0.04037186 | 0.99999994 |
| Naip1 | 0.00056148 | 0.00064155 | 0.00050821 | 0.00013334 | 0.04037186 | 0.99999994 |
| Ankrd28 | 0.01721059 | 0.01553633 | 0.01540344 | 0.00013289 | 0.47710283 | 0.99999994 |
| Arhgap33os | 0.0009593 | 0.00100077 | 0.00086806 | 0.00013271 | 0.15441858 | 0.99999994 |
| Gata5 | 0.00316081 | 0.00298862 | 0.00285591 | 0.00013271 | 0.37853894 | 0.99999994 |
| Aloxe3 | 0.00647421 | 0.0059692 | 0.00583654 | 0.00013266 | 0.43987259 | 0.99999994 |
| 4930434B07 | 0.00056083 | 0.00064008 | 0.00050763 | 0.00013245 | 0.04121366 | 0.99999994 |
| Gm9484 | 0.01618078 | 0.01462415 | 0.0144918 | 0.00013235 | 0.47576273 | 0.99999994 |
| Ksr1 | 0.00219977 | 0.00212111 | 0.00198887 | 0.00013224 | 0.32903628 | 0.99999994 |
| Yap1 | 0.00857026 | 0.0078473 | 0.0077152 | 0.0001321 | 0.45462898 | 0.99999994 |
| 1700019N19 | 0.00175835 | 0.00172221 | 0.00159025 | 0.00013196 | 0.29035032 | 0.99999994 |
| Agfg1 | 0.15386677 | 0.12512011 | 0.12498817 | 0.00013194 | 0.49719681 | 0.99999994 |
| Arhgap20 | 0.00089332 | 0.00093976 | 0.00080839 | 0.00013137 | 0.13969739 | 0.99999994 |
| Cdkn2c | 0.00265408 | 0.00253003 | 0.00239888 | 0.00013115 | 0.35795856 | 0.99999994 |

| | | | | | | |
|---|---|---|---|---|---|---|
| Gm11431 | 0.00055977 | 0.00063765 | 0.00050666 | 0.00013098 | 0.04264333 | 0.99999994 |
| St6gal2 | 0.00055977 | 0.00063765 | 0.00050666 | 0.00013098 | 0.04264333 | 0.99999994 |
| Slc22a8 | 0.00100987 | 0.00104476 | 0.00091378 | 0.00013098 | 0.17002026 | 0.99999994 |
| Gm16401 | 0.00249851 | 0.00238941 | 0.00225851 | 0.0001309 | 0.349824 | 0.99999994 |
| Gpr161 | 0.00237738 | 0.00227999 | 0.00214919 | 0.00013079 | 0.34270645 | 0.99999994 |
| Mroh2a | 0.00298994 | 0.00283236 | 0.00270183 | 0.00013053 | 0.37388311 | 0.99999994 |
| Sned1 | 0.00560109 | 0.0051828 | 0.00505241 | 0.00013039 | 0.43180234 | 0.99999994 |
| Fanci | 0.00200957 | 0.0019474 | 0.00181713 | 0.00013027 | 0.31661545 | 0.99999994 |
| Prickle2 | 0.00436868 | 0.00407379 | 0.00394402 | 0.00012977 | 0.41332037 | 0.99999994 |
| Sorcs2 | 0.00673092 | 0.00619663 | 0.00606691 | 0.00012972 | 0.44341166 | 0.99999994 |
| Aspa | 0.00055885 | 0.00063555 | 0.00050583 | 0.00012972 | 0.04390952 | 0.99999994 |
| Pla2g3 | 0.00130699 | 0.00131184 | 0.0011824 | 0.00012944 | 0.2330929 | 0.99999994 |
| BC030343 | 0.00142389 | 0.00141747 | 0.00128805 | 0.00012942 | 0.25180579 | 0.99999994 |
| Ldhc | 0.00055835 | 0.00063443 | 0.00050538 | 0.00012904 | 0.04460183 | 0.99999994 |
| Msr1 | 0.00079697 | 0.00085021 | 0.00072125 | 0.00012896 | 0.1169822 | 0.99999994 |
| Sox5 | 0.01935998 | 0.01743071 | 0.01730212 | 0.00012859 | 0.48027264 | 0.99999994 |
| Plg | 0.00140868 | 0.00140239 | 0.00127431 | 0.00012809 | 0.25172778 | 0.99999994 |
| Acsl6 | 0.02236084 | 0.02007171 | 0.01994382 | 0.00012789 | 0.48297652 | 0.99999994 |
| H60b | 0.00055751 | 0.00063251 | 0.00050462 | 0.00012789 | 0.04580584 | 0.99999994 |
| Mettl24 | 0.00196881 | 0.00190798 | 0.00178033 | 0.00012765 | 0.31658367 | 0.99999994 |
| Ccr1 | 0.00117947 | 0.00119459 | 0.00106713 | 0.00012746 | 0.21328849 | 0.99999994 |
| Smc1b | 0.00176866 | 0.00172694 | 0.00159955 | 0.00012739 | 0.29801461 | 0.99999994 |
| Adap2 | 0.00307362 | 0.00290461 | 0.00277728 | 0.00012733 | 0.38011754 | 0.99999994 |
| Samsn1 | 0.00055702 | 0.00063139 | 0.00050417 | 0.00012722 | 0.04651825 | 0.99999994 |
| Tpsg1 | 0.00055702 | 0.00063139 | 0.00050417 | 0.00012722 | 0.04651825 | 0.99999994 |
| Cckar | 0.01763589 | 0.01590678 | 0.01577957 | 0.00012721 | 0.47860255 | 0.99999994 |
| Lims2 | 0.00055695 | 0.00063123 | 0.00050411 | 0.00012712 | 0.04662063 | 0.99999994 |
| 5730403I07R | 0.00055691 | 0.00063115 | 0.00050408 | 0.00012707 | 0.04667187 | 0.99999994 |
| Zfp811 | 0.00098683 | 0.0010199 | 0.00089296 | 0.00012694 | 0.17200709 | 0.99999994 |
| Cd302 | 0.00276951 | 0.0026299 | 0.00250302 | 0.00012688 | 0.36789792 | 0.99999994 |

| | | | | | | |
|---|---|---|---|---|---|---|
| 1700030O20 | 0.00309725 | 0.00292539 | 0.0027986 | 0.00012679 | 0.38148936 | 0.99999994 |
| Tnc | 0.00055666 | 0.0006306 | 0.00050386 | 0.00012674 | 0.04703167 | 0.99999994 |
| Nnmt | 0.00055666 | 0.0006306 | 0.00050386 | 0.00012674 | 0.04703167 | 0.99999994 |
| She | 0.00100982 | 0.00104043 | 0.00091374 | 0.00012669 | 0.17802743 | 0.99999994 |
| Klk1b26 | 0.0009876 | 0.00102031 | 0.00089365 | 0.00012666 | 0.1727282 | 0.99999994 |
| Gm6594 | 0.01457448 | 0.01319395 | 0.01306729 | 0.00012666 | 0.47427835 | 0.99999994 |
| Atp1a3 | 0.00055652 | 0.00063028 | 0.00050373 | 0.00012655 | 0.04723811 | 0.99999994 |
| Pcdh10 | 0.00181614 | 0.00176879 | 0.00164244 | 0.00012635 | 0.3043113 | 0.99999994 |
| Gucy1a2 | 0.00101221 | 0.00104216 | 0.0009159 | 0.00012625 | 0.17942712 | 0.99999994 |
| Tnfaip2 | 0.00142928 | 0.00141914 | 0.00129292 | 0.00012622 | 0.25789675 | 0.99999994 |
| Gm38336 | 0.00139606 | 0.00138862 | 0.0012629 | 0.00012572 | 0.25377539 | 0.99999994 |
| Best1 | 0.00179637 | 0.0017496 | 0.00162459 | 0.00012502 | 0.30424663 | 0.99999994 |
| C130026I21R | 0.00221544 | 0.00212725 | 0.00200301 | 0.00012424 | 0.3398463 | 0.99999994 |
| Kifc5b | 0.00251339 | 0.00239614 | 0.00227193 | 0.00012421 | 0.35796424 | 0.99999994 |
| Ralyl | 0.00055478 | 0.00062633 | 0.00050215 | 0.00012418 | 0.04987022 | 0.99999994 |
| Dcaf12l2 | 0.00178894 | 0.00174173 | 0.00161787 | 0.00012386 | 0.30517323 | 0.99999994 |
| Pakap | 0.00055436 | 0.00062539 | 0.00050177 | 0.00012362 | 0.05051622 | 0.99999994 |
| Ifitm7 | 0.00055436 | 0.00062539 | 0.00050177 | 0.00012362 | 0.05051622 | 0.99999994 |
| Scube3 | 0.00055436 | 0.00062539 | 0.00050177 | 0.00012362 | 0.05051622 | 0.99999994 |
| Akt1s1 | 0.53420775 | 0.3277502 | 0.32762672 | 0.00012348 | 0.49899916 | 0.99999994 |
| Krt5 | 0.00145759 | 0.00144195 | 0.0013185 | 0.00012345 | 0.266575 | 0.99999994 |
| Tk1 | 0.01345389 | 0.01219505 | 0.0120717 | 0.00012336 | 0.47288528 | 0.99999994 |
| Nrarp | 0.00797874 | 0.00730889 | 0.00718557 | 0.00012333 | 0.4545218 | 0.99999994 |
| Angptl2 | 0.00260861 | 0.0024809 | 0.00235785 | 0.00012305 | 0.36415747 | 0.99999994 |
| Piezo2 | 0.00055367 | 0.00062383 | 0.00050115 | 0.00012268 | 0.05160526 | 0.99999994 |
| Zfp341 | 0.01205704 | 0.01095114 | 0.01082855 | 0.00012259 | 0.46996505 | 0.99999994 |
| Thbs1 | 0.00224122 | 0.00214879 | 0.00202628 | 0.00012251 | 0.34368013 | 0.99999994 |
| Gpr137b-ps | 0.02457937 | 0.0220122 | 0.02189 | 0.0001222 | 0.48517884 | 0.99999994 |
| Gm44210 | 0.00459055 | 0.00426581 | 0.0041437 | 0.00012211 | 0.42224592 | 0.99999994 |
| Rnase6 | 0.00098471 | 0.00101312 | 0.00089103 | 0.00012209 | 0.18086799 | 0.99999994 |

| | | | | | | |
|---|---|---|---|---|---|---|
| Smim5 | 0.04187833 | 0.03699135 | 0.03686936 | 0.00012199 | 0.49121422 | 0.99999994 |
| Rhag | 0.00055312 | 0.00062258 | 0.00050065 | 0.00012194 | 0.05248765 | 0.99999994 |
| Gm26586 | 0.00055298 | 0.00062227 | 0.00050052 | 0.00012175 | 0.0527098 | 0.99999994 |
| Dio3 | 0.00394525 | 0.0036845 | 0.00356277 | 0.00012173 | 0.41004516 | 0.99999994 |
| Rps6kb2 | 0.17577491 | 0.14047931 | 0.14035776 | 0.00012156 | 0.49770017 | 0.99999994 |
| Zyg11a | 0.00079282 | 0.00083886 | 0.0007175 | 0.00012136 | 0.13011445 | 0.99999994 |
| H2-T24 | 0.00097842 | 0.00100668 | 0.00088535 | 0.00012134 | 0.18081819 | 0.99999994 |
| Gm13539 | 0.00055236 | 0.00062087 | 0.00049996 | 0.00012091 | 0.05371718 | 0.99999994 |
| Gm43101 | 0.00055236 | 0.00062087 | 0.00049996 | 0.00012091 | 0.05371718 | 0.99999994 |
| Zmynd15 | 0.00099998 | 0.00102569 | 0.00090485 | 0.00012085 | 0.18700144 | 0.99999994 |
| Gm14102 | 0.00055222 | 0.00062056 | 0.00049983 | 0.00012073 | 0.05394276 | 0.99999994 |
| Stac3 | 0.0052857 | 0.00488876 | 0.00476894 | 0.00011983 | 0.43358717 | 0.99999994 |
| Glyr1 | 0.40074418 | 0.27036017 | 0.27024049 | 0.00011968 | 0.49882395 | 0.99999994 |
| 4930524B15 | 0.00100621 | 0.00103016 | 0.00091048 | 0.00011968 | 0.19080429 | 0.99999994 |
| Glipr1 | 0.00284155 | 0.0026876 | 0.002568 | 0.0001196 | 0.37827508 | 0.99999994 |
| 4930455G09 | 0.00433853 | 0.00403614 | 0.00391688 | 0.00011926 | 0.41969887 | 0.99999994 |
| Eef1a2 | 0.00100481 | 0.00102846 | 0.00090922 | 0.00011924 | 0.19134303 | 0.99999994 |
| Gm14409 | 0.00099134 | 0.00101619 | 0.00089703 | 0.00011916 | 0.18829254 | 0.99999994 |
| Gfi1 | 0.00140372 | 0.00138836 | 0.00126983 | 0.00011853 | 0.2671911 | 0.99999994 |
| Cep128 | 0.00567498 | 0.00523729 | 0.0051188 | 0.00011849 | 0.43877423 | 0.99999994 |
| Gm37170 | 0.00258732 | 0.00245644 | 0.00233864 | 0.00011779 | 0.36871285 | 0.99999994 |
| Heatr6 | 0.11805844 | 0.0987728 | 0.09865542 | 0.00011739 | 0.49684038 | 0.99999994 |
| Gm12184 | 0.01009897 | 0.00919927 | 0.00908199 | 0.00011728 | 0.46575075 | 0.99999994 |
| BC051077 | 0.00204575 | 0.00196665 | 0.00184981 | 0.00011685 | 0.33707235 | 0.99999994 |
| Rasl12 | 0.00130211 | 0.00129425 | 0.00117798 | 0.00011627 | 0.25559066 | 0.99999994 |
| Jakmip2 | 0.00307006 | 0.00289013 | 0.00277407 | 0.00011606 | 0.39032373 | 0.99999994 |
| Gm15638 | 0.00733839 | 0.00672741 | 0.00661174 | 0.00011567 | 0.45364622 | 0.99999994 |
| Pcdhga1 | 0.00246245 | 0.00234163 | 0.00222596 | 0.00011566 | 0.36471862 | 0.99999994 |
| Ang4 | 0.00054812 | 0.00061139 | 0.00049613 | 0.00011526 | 0.06100217 | 0.99999994 |
| Cd3g | 0.00747122 | 0.00684601 | 0.00673081 | 0.00011521 | 0.45464576 | 0.99999994 |

| | | | | | | |
|---|---|---|---|---|---|---|
| Oas2 | 0.00054798 | 0.00061108 | 0.000496 | 0.00011508 | 0.06124727 | 0.99999994 |
| S1pr2 | 0.00815228 | 0.00745593 | 0.00734099 | 0.00011494 | 0.45849794 | 0.99999994 |
| Vsig2 | 0.0183284 | 0.0165064 | 0.01639155 | 0.00011485 | 0.48140067 | 0.99999994 |
| Ptgdr | 0.00054758 | 0.00061018 | 0.00049564 | 0.00011454 | 0.06198639 | 0.99999994 |
| Bend6 | 0.00196413 | 0.00189035 | 0.0017761 | 0.00011424 | 0.33427001 | 0.99999994 |
| Cyp27a1 | 0.00177629 | 0.00172057 | 0.00160645 | 0.00011413 | 0.31814909 | 0.99999994 |
| Gm12089 | 0.00180398 | 0.00174553 | 0.00163146 | 0.00011407 | 0.32082473 | 0.99999994 |
| Prima1 | 0.00139299 | 0.00137409 | 0.00126012 | 0.00011396 | 0.27358783 | 0.99999994 |
| Hbb-bh1 | 0.0018138 | 0.00175429 | 0.00164033 | 0.00011395 | 0.3218954 | 0.99999994 |
| Fabp2 | 0.00118649 | 0.00118709 | 0.00107347 | 0.00011362 | 0.24055903 | 0.99999994 |
| P2ry13 | 0.00054684 | 0.00060853 | 0.00049496 | 0.00011356 | 0.06335631 | 0.99999994 |
| Gm31121 | 0.00054684 | 0.00060853 | 0.00049496 | 0.00011356 | 0.06335631 | 0.99999994 |
| Oxt | 0.0005467 | 0.00060823 | 0.00049484 | 0.00011338 | 0.06360745 | 0.99999994 |
| Gm15408 | 0.002294 | 0.0021871 | 0.00207393 | 0.00011317 | 0.35821859 | 0.99999994 |
| Ncf2 | 0.00300279 | 0.00282651 | 0.00271342 | 0.00011309 | 0.39073143 | 0.99999994 |
| Ehd2 | 0.00134857 | 0.00133256 | 0.00121998 | 0.00011258 | 0.26952379 | 0.99999994 |
| Slc16a3 | 0.01848763 | 0.01664407 | 0.01653219 | 0.00011189 | 0.48203419 | 0.99999994 |
| Slc15a3 | 0.00199575 | 0.00191629 | 0.00180466 | 0.00011163 | 0.34026576 | 0.99999994 |
| Bcl6b | 0.02198231 | 0.01972224 | 0.01961118 | 0.00011106 | 0.48496456 | 0.99999994 |
| Gm15867 | 0.00054476 | 0.0006039 | 0.00049308 | 0.00011082 | 0.06732072 | 0.99999994 |
| Sdr16c6 | 0.00182184 | 0.00175815 | 0.0016476 | 0.00011056 | 0.32756377 | 0.99999994 |
| Ctsg | 0.00054442 | 0.00060316 | 0.00049278 | 0.00011038 | 0.06797452 | 0.99999994 |
| Begain | 0.00224248 | 0.00213772 | 0.00202742 | 0.0001103 | 0.35862617 | 0.99999994 |
| Gm17597 | 0.00099149 | 0.00100729 | 0.00089717 | 0.00011013 | 0.20694658 | 0.99999994 |
| 4930426I24R | 0.00214205 | 0.00204649 | 0.00193676 | 0.00010973 | 0.3530355 | 0.99999994 |
| Cdk10 | 0.10559528 | 0.08923226 | 0.08912271 | 0.00010955 | 0.49673586 | 0.99999994 |
| Nup210 | 0.0530055 | 0.04643377 | 0.04632463 | 0.00010914 | 0.49374408 | 0.99999994 |
| Mei4 | 0.0009605 | 0.0009782 | 0.00086914 | 0.00010905 | 0.2018088 | 0.99999994 |
| Phox2a | 0.00054336 | 0.0006008 | 0.00049181 | 0.00010899 | 0.07009356 | 0.99999994 |
| Hecw2 | 0.00206371 | 0.00197501 | 0.00186602 | 0.00010899 | 0.34872198 | 0.99999994 |

| | | | | | | |
|---|---|---|---|---|---|---|
| Cldn13 | 0.00117438 | 0.00117136 | 0.00106253 | 0.00010883 | 0.24768487 | 0.99999994 |
| Tekt4 | 0.00624831 | 0.00574257 | 0.00563376 | 0.00010881 | 0.44885258 | 0.99999994 |
| Gm26850 | 0.00157858 | 0.00153665 | 0.00142784 | 0.00010881 | 0.3059918 | 0.99999994 |
| Fmn1 | 0.00532289 | 0.0049111 | 0.00480237 | 0.00010873 | 0.44010362 | 0.99999994 |
| Mettl18 | 0.06212806 | 0.05408257 | 0.05397402 | 0.00010855 | 0.4946595 | 0.99999994 |
| 3300002P13I | 0.00054282 | 0.00059963 | 0.00049133 | 0.0001083 | 0.07116847 | 0.99999994 |
| Parpbp | 0.00054282 | 0.00059963 | 0.00049133 | 0.0001083 | 0.07116847 | 0.99999994 |
| Adgre1 | 0.00054263 | 0.00059919 | 0.00049115 | 0.00010804 | 0.0715742 | 0.99999994 |
| Serpinc1 | 0.00224785 | 0.00214004 | 0.00203227 | 0.00010777 | 0.36205403 | 0.99999994 |
| Pifo | 0.00095023 | 0.00096756 | 0.00085986 | 0.0001077 | 0.20221848 | 0.99999994 |
| Gpr63 | 0.00135179 | 0.00133052 | 0.0012229 | 0.00010763 | 0.27899351 | 0.99999994 |
| Gm11772 | 0.01174321 | 0.01065578 | 0.01054894 | 0.00010684 | 0.47312537 | 0.99999994 |
| Pkp3 | 0.04419172 | 0.0389533 | 0.03884659 | 0.00010671 | 0.49270564 | 0.99999994 |
| Dbx2 | 0.00073706 | 0.00077303 | 0.00066706 | 0.00010598 | 0.14514245 | 0.99999994 |
| Gm13589 | 0.00054091 | 0.00059541 | 0.0004896 | 0.0001058 | 0.07515105 | 0.99999994 |
| Slc26a10 | 0.00054091 | 0.00059541 | 0.0004896 | 0.0001058 | 0.07515105 | 0.99999994 |
| Cntnap5b | 0.00054078 | 0.00059512 | 0.00048948 | 0.00010563 | 0.07543069 | 0.99999994 |
| Ahrr | 0.00054078 | 0.00059512 | 0.00048948 | 0.00010563 | 0.07543069 | 0.99999994 |
| Tnfaip8l3 | 0.00526818 | 0.00485833 | 0.00475319 | 0.00010514 | 0.44147174 | 0.99999994 |
| Rbfox1 | 0.00099323 | 0.00100387 | 0.00089874 | 0.00010513 | 0.21810541 | 0.99999994 |
| Atg9b | 0.00129859 | 0.00127978 | 0.00117481 | 0.00010497 | 0.27600528 | 0.99999994 |
| Cfap57 | 0.00267856 | 0.00252589 | 0.00242097 | 0.00010492 | 0.38649293 | 0.99999994 |
| Klf15 | 0.00639721 | 0.00587232 | 0.00576743 | 0.00010489 | 0.45182103 | 0.99999994 |
| Gm43689 | 0.00572775 | 0.00527105 | 0.00516621 | 0.00010483 | 0.44627856 | 0.99999994 |
| Nfam1 | 0.00094776 | 0.00096242 | 0.00085762 | 0.0001048 | 0.20799328 | 0.99999994 |
| Nfia | 0.04610741 | 0.04058385 | 0.04047935 | 0.0001045 | 0.49314485 | 0.99999994 |
| Cyp3a13 | 0.00069624 | 0.00073377 | 0.00063013 | 0.00010364 | 0.13679545 | 0.99999994 |
| Cyp4f18 | 0.00091614 | 0.00093231 | 0.00082903 | 0.00010329 | 0.20346303 | 0.99999994 |
| Il20ra | 0.00096931 | 0.00098019 | 0.00087711 | 0.00010308 | 0.21701806 | 0.99999994 |
| Plin1 | 0.00227306 | 0.00215802 | 0.00205502 | 0.00010299 | 0.36933894 | 0.99999994 |

| | | | | | | |
|---|---|---|---|---|---|---|
| Crp | 0.00977757 | 0.00889764 | 0.00879487 | 0.00010277 | 0.46900084 | 0.99999994 |
| Tmem47 | 0.00267477 | 0.00252031 | 0.00241754 | 0.00010277 | 0.38860439 | 0.99999994 |
| Stil | 0.00120982 | 0.00119709 | 0.00109457 | 0.00010252 | 0.26648909 | 0.99999994 |
| Gm11131 | 0.00276554 | 0.00260177 | 0.00249943 | 0.00010234 | 0.39260509 | 0.99999994 |
| Ggn | 0.00053797 | 0.00058894 | 0.00048694 | 0.00010201 | 0.08159819 | 0.99999994 |
| Ranbp3l | 0.0005379 | 0.0005888 | 0.00048688 | 0.00010192 | 0.08174515 | 0.99999994 |
| Naalad2 | 0.00053777 | 0.00058852 | 0.00048676 | 0.00010176 | 0.08203956 | 0.99999994 |
| Prss12 | 0.00362724 | 0.00337796 | 0.0032763 | 0.00010167 | 0.41817987 | 0.99999994 |
| Fgd3 | 0.00175574 | 0.00168948 | 0.00158788 | 0.0001016 | 0.33508616 | 0.99999994 |
| Hspb2 | 0.00082055 | 0.0008438 | 0.00074257 | 0.00010122 | 0.182114 | 0.99999994 |
| Glt1d1 | 0.00348711 | 0.00325114 | 0.00315003 | 0.00010111 | 0.4154047 | 0.99999994 |
| Ttn | 0.00053725 | 0.00058738 | 0.00048629 | 0.00010109 | 0.08322358 | 0.99999994 |
| Rnf125 | 0.00053725 | 0.00058738 | 0.00048629 | 0.00010109 | 0.08322358 | 0.99999994 |
| Trpm6 | 0.00053725 | 0.00058738 | 0.00048629 | 0.00010109 | 0.08322358 | 0.99999994 |
| C030034L19I | 0.00053654 | 0.00058583 | 0.00048565 | 0.00010018 | 0.08486834 | 0.99999994 |
| Asgr2 | 0.00235642 | 0.00223043 | 0.00213027 | 0.00010017 | 0.37714475 | 0.99999994 |
| Capn11 | 0.00337256 | 0.00314683 | 0.00304679 | 0.00010004 | 0.41349359 | 0.99999994 |
| Lsm11 | 0.02124711 | 0.01906466 | 0.01896463 | 0.00010003 | 0.48599561 | 0.99999994 |
| Btnl10 | 0.00053628 | 0.00058526 | 0.00048541 | 9.98E-05 | 0.08547125 | 0.99999994 |
| Hecw1 | 0.00053628 | 0.00058526 | 0.00048541 | 9.98E-05 | 0.08547125 | 0.99999994 |
| 1700021N21 | 0.00053583 | 0.00058428 | 0.00048501 | 9.93E-05 | 0.08653248 | 0.99999994 |
| Acsm3 | 0.00053583 | 0.00058428 | 0.00048501 | 9.93E-05 | 0.08653248 | 0.99999994 |
| Slc30a3 | 0.00092856 | 0.00093915 | 0.00084026 | 9.89E-05 | 0.21669527 | 0.99999994 |
| Ovol1 | 0.00143225 | 0.00139371 | 0.00129561 | 9.81E-05 | 0.30712968 | 0.99999994 |
| Slc7a11 | 0.00187899 | 0.00179731 | 0.00169921 | 9.81E-05 | 0.35038331 | 0.99999994 |
| Wnt16 | 0.00053474 | 0.00058189 | 0.00048402 | 9.79E-05 | 0.08914236 | 0.99999994 |
| Pdp2 | 0.02611719 | 0.02333352 | 0.02323564 | 9.79E-05 | 0.48881549 | 0.99999994 |
| Gm16984 | 0.00095265 | 0.00095965 | 0.00086205 | 9.76E-05 | 0.22552193 | 0.99999994 |
| Pcdhga11 | 0.00257449 | 0.0024244 | 0.00232707 | 9.73E-05 | 0.39034952 | 0.99999994 |
| Stk33 | 0.00180753 | 0.00173181 | 0.00163467 | 9.71E-05 | 0.34621686 | 0.99999994 |

| | | | | | | |
|---|---|---|---|---|---|---|
| Gm28876 | 0.00053365 | 0.00057953 | 0.00048303 | 9.65E-05 | 0.09179833 | 0.99999994 |
| Pi15 | 0.00053346 | 0.00057911 | 0.00048286 | 9.63E-05 | 0.0922718 | 0.99999994 |
| Rrh | 0.00264272 | 0.0024848 | 0.00238863 | 9.62E-05 | 0.39434672 | 0.99999994 |
| Capn1 | 0.0166625 | 0.01501435 | 0.0149184 | 9.60E-05 | 0.48292567 | 0.99999994 |
| Hmx2 | 0.0005332 | 0.00057856 | 0.00048263 | 9.59E-05 | 0.09290533 | 0.99999994 |
| Capns2 | 0.00053307 | 0.00057828 | 0.00048251 | 9.58E-05 | 0.09322304 | 0.99999994 |
| 4833422C13I | 0.00151071 | 0.00146193 | 0.00136651 | 9.54E-05 | 0.32103059 | 0.99999994 |
| A530095I07F | 0.00053269 | 0.00057745 | 0.00048217 | 9.53E-05 | 0.09417999 | 0.99999994 |
| Fndc1 | 0.00290619 | 0.00272136 | 0.0026263 | 9.51E-05 | 0.40480473 | 0.99999994 |
| Gm12474 | 0.00471181 | 0.00434717 | 0.0042528 | 9.44E-05 | 0.4412889 | 0.99999994 |
| Gm17590 | 0.00053193 | 0.0005758 | 0.00048148 | 9.43E-05 | 0.09611101 | 0.99999994 |
| Dkk1 | 0.0005318 | 0.00057553 | 0.00048136 | 9.42E-05 | 0.09643507 | 0.99999994 |
| Prr18 | 0.00092426 | 0.00093013 | 0.00083637 | 9.38E-05 | 0.2277754 | 0.99999994 |
| Kremen2 | 0.0008848 | 0.00089429 | 0.00080069 | 9.36E-05 | 0.2182504 | 0.99999994 |
| Asb17 | 0.00089523 | 0.00090361 | 0.00081012 | 9.35E-05 | 0.2211788 | 0.99999994 |
| Zfp930 | 0.00817593 | 0.00745546 | 0.00736217 | 9.33E-05 | 0.46639022 | 0.99999994 |
| Gm16010 | 0.0074223 | 0.00678004 | 0.00668696 | 9.31E-05 | 0.46309001 | 0.99999994 |
| Gm17029 | 0.00185372 | 0.00176942 | 0.00167639 | 9.30E-05 | 0.35590916 | 0.99999994 |
| Shisa8 | 0.00312167 | 0.0029136 | 0.00282061 | 9.30E-05 | 0.4131513 | 0.99999994 |
| Gm17163 | 0.00053085 | 0.00057347 | 0.0004805 | 9.30E-05 | 0.09888559 | 0.99999994 |
| Rab32 | 0.00322292 | 0.00300471 | 0.0029119 | 9.28E-05 | 0.41598964 | 0.99999994 |
| Rnf151 | 0.00308622 | 0.00288135 | 0.00278865 | 9.27E-05 | 0.41244013 | 0.99999994 |
| Vat1 | 0.14760226 | 0.12058422 | 0.12049185 | 9.24E-05 | 0.49796419 | 0.99999994 |
| Cdkl4 | 0.00222523 | 0.00210413 | 0.00201185 | 9.23E-05 | 0.38006105 | 0.99999994 |
| Gm42556 | 0.00139362 | 0.00135294 | 0.0012607 | 9.22E-05 | 0.31311772 | 0.99999994 |
| BC002059 | 0.03941547 | 0.03484877 | 0.0347577 | 9.11E-05 | 0.49304295 | 0.99999994 |
| Lcn3 | 0.00131914 | 0.00128424 | 0.00119338 | 9.09E-05 | 0.3061521 | 0.99999994 |
| Tex29 | 0.00052915 | 0.00056981 | 0.00047897 | 9.08E-05 | 0.10338569 | 0.99999994 |
| Creg2 | 0.00772799 | 0.00705175 | 0.00696092 | 9.08E-05 | 0.46539473 | 0.99999994 |
| 4930505N22 | 0.00147134 | 0.00142159 | 0.00133094 | 9.07E-05 | 0.32513689 | 0.99999994 |

| | | | | | | |
|---|---|---|---|---|---|---|
| L3mbtl2 | 0.06036628 | 0.05259451 | 0.05250385 | 9.07E-05 | 0.49541513 | 0.99999994 |
| Kif18b | 0.00052872 | 0.00056887 | 0.00047857 | 9.03E-05 | 0.10457102 | 0.99999994 |
| Kif6 | 0.00283887 | 0.00265552 | 0.00256558 | 8.99E-05 | 0.40774211 | 0.99999994 |
| Amy1 | 0.00208147 | 0.00197179 | 0.00188206 | 8.97E-05 | 0.37548646 | 0.99999994 |
| Gm38287 | 0.00052822 | 0.00056779 | 0.00047812 | 8.97E-05 | 0.10593502 | 0.99999994 |
| 6030498E09I | 0.00141371 | 0.00136846 | 0.00127885 | 8.96E-05 | 0.32046318 | 0.99999994 |
| Styxl1 | 0.00157321 | 0.00151233 | 0.00142298 | 8.93E-05 | 0.33799259 | 0.99999994 |
| Aunip | 0.00073615 | 0.00075548 | 0.00066623 | 8.92E-05 | 0.18629735 | 0.99999994 |
| Asic1 | 0.00908758 | 0.0082672 | 0.00817804 | 8.92E-05 | 0.47107529 | 0.99999994 |
| Gabrg1 | 0.00052778 | 0.00056685 | 0.00047772 | 8.91E-05 | 0.10713669 | 0.99999994 |
| S100a7a | 0.00052765 | 0.00056658 | 0.00047761 | 8.90E-05 | 0.10748142 | 0.99999994 |
| Chrd | 0.01429717 | 0.01290993 | 0.01282105 | 8.89E-05 | 0.48159837 | 0.99999994 |
| Serpinb6b | 0.00257463 | 0.00241602 | 0.0023272 | 8.88E-05 | 0.39972312 | 0.99999994 |
| AW495222 | 0.00266167 | 0.00249447 | 0.00240573 | 8.87E-05 | 0.40302211 | 0.99999994 |
| Ildr2 | 0.00077429 | 0.00078939 | 0.00070073 | 8.87E-05 | 0.19982755 | 0.99999994 |
| Piezo1 | 0.00207339 | 0.00196341 | 0.00187477 | 8.86E-05 | 0.3764876 | 0.99999994 |
| Gm29395 | 0.00137722 | 0.00133441 | 0.00124587 | 8.85E-05 | 0.31809532 | 0.99999994 |
| Lcp1 | 0.00455175 | 0.00419722 | 0.00410879 | 8.84E-05 | 0.44304458 | 0.99999994 |
| Procr | 0.00232269 | 0.00218778 | 0.00209982 | 8.80E-05 | 0.39019711 | 0.99999994 |
| Pde6h | 0.00052678 | 0.00056471 | 0.00047682 | 8.79E-05 | 0.10991186 | 0.99999994 |
| Rln1 | 0.00052678 | 0.00056471 | 0.00047682 | 8.79E-05 | 0.10991186 | 0.99999994 |
| Kcnk6 | 0.00163323 | 0.00156509 | 0.00147721 | 8.79E-05 | 0.3460664 | 0.99999994 |
| Prss23os | 0.0015248 | 0.00146701 | 0.00137924 | 8.78E-05 | 0.33592816 | 0.99999994 |
| Gm42413 | 0.00091707 | 0.00091762 | 0.00082987 | 8.77E-05 | 0.24076964 | 0.99999994 |
| Fndc8 | 0.00133726 | 0.00129742 | 0.00120976 | 8.77E-05 | 0.31479366 | 0.99999994 |
| Hoxb7 | 0.00052641 | 0.00056392 | 0.00047648 | 8.74E-05 | 0.11096273 | 0.99999994 |
| Mir124-2hg | 0.00052623 | 0.00056352 | 0.00047631 | 8.72E-05 | 0.11149025 | 0.99999994 |
| Gdf10 | 0.00052616 | 0.00056339 | 0.00047626 | 8.71E-05 | 0.1116664 | 0.99999994 |
| Sap30bpos | 0.0005261 | 0.00056325 | 0.0004762 | 8.71E-05 | 0.11184269 | 0.99999994 |
| Sox30 | 0.00075727 | 0.00077234 | 0.00068534 | 8.70E-05 | 0.19906097 | 0.99999994 |

| | | | | | | |
|---|---|---|---|---|---|---|
| Adnp | 0.00124713 | 0.00121528 | 0.00112829 | 8.70E-05 | 0.30391007 | 0.99999994 |
| Cbln4 | 0.00052579 | 0.00056259 | 0.00047592 | 8.67E-05 | 0.11272649 | 0.99999994 |
| Vcam1 | 0.00052579 | 0.00056259 | 0.00047592 | 8.67E-05 | 0.11272649 | 0.99999994 |
| Plek | 0.00077705 | 0.00078987 | 0.00070323 | 8.66E-05 | 0.20608208 | 0.99999994 |
| Lrrn4cl | 0.00052567 | 0.00056233 | 0.00047581 | 8.65E-05 | 0.11308109 | 0.99999994 |
| Serpina3b | 0.00214558 | 0.00202645 | 0.00193994 | 8.65E-05 | 0.38329652 | 0.99999994 |
| Cd37 | 0.0021234 | 0.00200622 | 0.00191992 | 8.63E-05 | 0.38239419 | 0.99999994 |
| Gm17210 | 0.00052548 | 0.00056193 | 0.00047564 | 8.63E-05 | 0.11361412 | 0.99999994 |
| Ccdc18 | 0.01504338 | 0.01356972 | 0.01348344 | 8.63E-05 | 0.48301207 | 0.99999994 |
| Serpinb5 | 0.001224 | 0.00119342 | 0.00110738 | 8.60E-05 | 0.30252603 | 0.99999994 |
| Slc4a1 | 0.00438458 | 0.004044 | 0.00395833 | 8.57E-05 | 0.44272395 | 0.99999994 |
| Spag5 | 0.00517542 | 0.00475544 | 0.00466979 | 8.57E-05 | 0.45141404 | 0.99999994 |
| Vmn1r4 | 0.00339208 | 0.00314992 | 0.00306438 | 8.55E-05 | 0.42629588 | 0.99999994 |
| B230398E01 | 0.00480797 | 0.00442479 | 0.00433931 | 8.55E-05 | 0.44784302 | 0.99999994 |
| Rnf225 | 0.0023619 | 0.00222062 | 0.00213522 | 8.54E-05 | 0.39502798 | 0.99999994 |
| Mgst2 | 0.02096744 | 0.0188039 | 0.01871852 | 8.54E-05 | 0.4878884 | 0.99999994 |
| Frmpd1 | 0.00073661 | 0.00075197 | 0.00066665 | 8.53E-05 | 0.19714171 | 0.99999994 |
| Rgs8 | 0.00052468 | 0.00056022 | 0.00047492 | 8.53E-05 | 0.11593982 | 0.99999994 |
| Myog | 0.0008964 | 0.00089644 | 0.00081117 | 8.53E-05 | 0.24206581 | 0.99999994 |
| Rfx4 | 0.00070986 | 0.00072719 | 0.00064245 | 8.47E-05 | 0.18998455 | 0.99999994 |
| B3gnt7 | 0.00276291 | 0.00258158 | 0.00249706 | 8.45E-05 | 0.41087115 | 0.99999994 |
| Mpzl3 | 0.01460969 | 0.013183 | 0.01309855 | 8.45E-05 | 0.48288408 | 0.99999994 |
| 1700084C06I | 0.00467997 | 0.00430843 | 0.00422416 | 8.43E-05 | 0.4471763 | 0.99999994 |
| Gm6288 | 0.0033419 | 0.0031034 | 0.00301915 | 8.42E-05 | 0.42632367 | 0.99999994 |
| Zswim6 | 0.04483295 | 0.03947762 | 0.03939358 | 8.40E-05 | 0.49433523 | 0.99999994 |
| Ccl21a | 0.00083037 | 0.00083518 | 0.00075146 | 8.37E-05 | 0.22916353 | 0.99999994 |
| Aoc1 | 0.00052327 | 0.00055721 | 0.00047364 | 8.36E-05 | 0.12011735 | 0.99999994 |
| Slc26a1 | 0.00827597 | 0.00753521 | 0.00745175 | 8.35E-05 | 0.4702872 | 0.99999994 |
| Myl3 | 0.00175337 | 0.00166917 | 0.00158575 | 8.34E-05 | 0.3630954 | 0.99999994 |
| Skil | 0.12859 | 0.10664383 | 0.10656061 | 8.32E-05 | 0.49792608 | 0.99999994 |

| | | | | | | |
|---|---|---|---|---|---|---|
| Tmem190 | 0.0160085 | 0.01442233 | 0.01433916 | 8.32E-05 | 0.4846016 | 0.99999994 |
| Car12 | 0.00096665 | 0.00095785 | 0.0008747 | 8.31E-05 | 0.26344994 | 0.99999994 |
| Tmem26 | 0.00091456 | 0.00091066 | 0.0008276 | 8.31E-05 | 0.25204541 | 0.99999994 |
| Gm12925 | 0.00139537 | 0.00134533 | 0.00126228 | 8.31E-05 | 0.33070833 | 0.99999994 |
| Gm13362 | 0.00069913 | 0.00071518 | 0.00063275 | 8.24E-05 | 0.19290949 | 0.99999994 |
| Fabp12 | 0.00052223 | 0.000555 | 0.0004727 | 8.23E-05 | 0.12325627 | 0.99999994 |
| Inhbc | 0.00052223 | 0.000555 | 0.0004727 | 8.23E-05 | 0.12325627 | 0.99999994 |
| Neto1 | 0.00052223 | 0.000555 | 0.0004727 | 8.23E-05 | 0.12325627 | 0.99999994 |
| Hsh2d | 0.00052223 | 0.000555 | 0.0004727 | 8.23E-05 | 0.12325627 | 0.99999994 |
| Accs | 0.02030748 | 0.01821962 | 0.01813737 | 8.23E-05 | 0.4879588 | 0.99999994 |
| Camk2a | 0.00052193 | 0.00055435 | 0.00047243 | 8.19E-05 | 0.1241877 | 0.99999994 |
| Gm26911 | 0.00052168 | 0.00055384 | 0.00047221 | 8.16E-05 | 0.12493552 | 0.99999994 |
| Arhgef39 | 0.00089927 | 0.00089462 | 0.00081377 | 8.08E-05 | 0.25422527 | 0.99999994 |
| Elovl4 | 0.00454877 | 0.00418694 | 0.0041061 | 8.08E-05 | 0.44787286 | 0.99999994 |
| Gm973 | 0.00265742 | 0.00248244 | 0.00240189 | 8.06E-05 | 0.41167476 | 0.99999994 |
| Bcas1os1 | 0.00165956 | 0.00158153 | 0.001501 | 8.05E-05 | 0.36050993 | 0.99999994 |
| Ate1 | 0.20862793 | 0.16248351 | 0.16240303 | 8.05E-05 | 0.49868398 | 0.99999994 |
| Myl1 | 0.00094855 | 0.00093879 | 0.00085834 | 8.05E-05 | 0.26634584 | 0.99999994 |
| Alas2 | 0.03171693 | 0.02819262 | 0.02811231 | 8.03E-05 | 0.49241428 | 0.99999994 |
| Ssmem1 | 0.00198105 | 0.00187165 | 0.00179138 | 8.03E-05 | 0.38275031 | 0.99999994 |
| Dact3 | 0.0127197 | 0.01149839 | 0.01141859 | 7.98E-05 | 0.48144969 | 0.99999994 |
| Ska3 | 0.00281455 | 0.0026234 | 0.00254364 | 7.98E-05 | 0.41733246 | 0.99999994 |
| Bhmt | 0.00124844 | 0.00120908 | 0.00112947 | 7.96E-05 | 0.31948588 | 0.99999994 |
| Trim50 | 0.00177204 | 0.00168169 | 0.00160261 | 7.91E-05 | 0.37127916 | 0.99999994 |
| Spaca7 | 0.00177296 | 0.00168247 | 0.00160344 | 7.90E-05 | 0.37142926 | 0.99999994 |
| Klk13 | 0.00245493 | 0.00229723 | 0.00221918 | 7.81E-05 | 0.40744641 | 0.99999994 |
| Boc | 0.00461767 | 0.00424586 | 0.0041681 | 7.78E-05 | 0.450586 | 0.99999994 |
| Sh3tc2 | 0.00051848 | 0.00054706 | 0.00046931 | 7.78E-05 | 0.13506624 | 0.99999994 |
| Col4a5 | 0.01732387 | 0.01558129 | 0.01550364 | 7.76E-05 | 0.48670259 | 0.99999994 |
| Notch3 | 0.00952458 | 0.00864598 | 0.00856876 | 7.72E-05 | 0.47608538 | 0.99999994 |

| | | | | | | |
|---|---|---|---|---|---|---|
| Sowaha | 0.01700431 | 0.01529809 | 0.01522093 | 7.72E-05 | 0.48654174 | 0.99999994 |
| Gm14401 | 0.0009533 | 0.00093974 | 0.00086263 | 7.71E-05 | 0.27593764 | 0.99999994 |
| Wfdc1 | 0.00051777 | 0.00054555 | 0.00046866 | 7.69E-05 | 0.13741655 | 0.99999994 |
| Arl6ip6 | 0.34417525 | 0.24197394 | 0.24189717 | 7.68E-05 | 0.49915717 | 0.99999994 |
| Ankrd22 | 0.00096169 | 0.00094678 | 0.00087022 | 7.66E-05 | 0.2790502 | 0.99999994 |
| 2810429I04R | 0.00263334 | 0.00245639 | 0.00238017 | 7.62E-05 | 0.4156036 | 0.99999994 |
| Gm4951 | 0.00051681 | 0.00054354 | 0.0004678 | 7.57E-05 | 0.14058224 | 0.99999994 |
| Dmrta1 | 0.00118981 | 0.00115202 | 0.00107648 | 7.55E-05 | 0.32020473 | 0.99999994 |
| Kcnk13 | 0.00092957 | 0.00091669 | 0.00084117 | 7.55E-05 | 0.27505676 | 0.99999994 |
| Kif11 | 0.00361007 | 0.0033363 | 0.00326083 | 7.55E-05 | 0.43878668 | 0.99999994 |
| Slco5a1 | 0.00250733 | 0.0023418 | 0.00226647 | 7.53E-05 | 0.41245858 | 0.99999994 |
| Gm8444 | 0.00051639 | 0.00054266 | 0.00046742 | 7.52E-05 | 0.14197861 | 0.99999994 |
| Pex11a | 0.05721025 | 0.04993668 | 0.04986173 | 7.50E-05 | 0.49600815 | 0.99999994 |
| Zfp276 | 0.0198691 | 0.0178258 | 0.01775106 | 7.47E-05 | 0.48881971 | 0.99999994 |
| Hsd3b6 | 0.00051598 | 0.00054179 | 0.00046704 | 7.47E-05 | 0.14338187 | 0.99999994 |
| Ms4a4c | 0.00051598 | 0.00054179 | 0.00046704 | 7.47E-05 | 0.14338187 | 0.99999994 |
| Platr23 | 0.00091631 | 0.00090374 | 0.00082918 | 7.46E-05 | 0.27472928 | 0.99999994 |
| Igf2os | 0.00089472 | 0.00088421 | 0.00080966 | 7.45E-05 | 0.26998292 | 0.99999994 |
| Sox18 | 0.00051568 | 0.00054116 | 0.00046678 | 7.44E-05 | 0.14438839 | 0.99999994 |
| Xpa | 0.23512707 | 0.17942168 | 0.17934742 | 7.43E-05 | 0.49890042 | 0.99999994 |
| Gm525 | 0.00051539 | 0.00054054 | 0.00046651 | 7.40E-05 | 0.1453984 | 0.99999994 |
| Serpina3f | 0.00051539 | 0.00054054 | 0.00046651 | 7.40E-05 | 0.1453984 | 0.99999994 |
| Lrrtm1 | 0.00051515 | 0.00054005 | 0.00046629 | 7.38E-05 | 0.1462089 | 0.99999994 |
| Vps33a | 0.07571373 | 0.06527177 | 0.0651984 | 7.34E-05 | 0.49701176 | 0.99999994 |
| Kif12 | 0.4820473 | 0.30674585 | 0.30667265 | 7.32E-05 | 0.49936621 | 0.99999994 |
| Gm10874 | 0.00051444 | 0.00053856 | 0.00046565 | 7.29E-05 | 0.14865365 | 0.99999994 |
| Mrc2 | 0.00051426 | 0.00053819 | 0.00046549 | 7.27E-05 | 0.14926793 | 0.99999994 |
| BB019430 | 0.00051426 | 0.00053819 | 0.00046549 | 7.27E-05 | 0.14926793 | 0.99999994 |
| Fbxo25 | 0.10862911 | 0.09153376 | 0.0914611 | 7.27E-05 | 0.49789047 | 0.99999994 |
| Fgf5 | 0.00051421 | 0.00053807 | 0.00046544 | 7.26E-05 | 0.14947296 | 0.99999994 |

| | | | | | | |
|---|---|---|---|---|---|---|
| Fzd3 | 0.12202413 | 0.10172055 | 0.101648 | 7.25E-05 | 0.49810477 | 0.99999994 |
| Kcnn3 | 0.00204751 | 0.00192367 | 0.00185139 | 7.23E-05 | 0.3974872 | 0.99999994 |
| Evc | 0.00242627 | 0.00226553 | 0.00219331 | 7.22E-05 | 0.41326399 | 0.99999994 |
| Gm26671 | 0.00051385 | 0.00053733 | 0.00046512 | 7.22E-05 | 0.15070603 | 0.99999994 |
| Pcdh8 | 0.00818819 | 0.00744527 | 0.00737315 | 7.21E-05 | 0.47404389 | 0.99999994 |
| Ankrd23 | 0.00051373 | 0.00053708 | 0.00046501 | 7.21E-05 | 0.15111814 | 0.99999994 |
| Wdr66 | 0.00127071 | 0.00122082 | 0.00114961 | 7.12E-05 | 0.34005916 | 0.99999994 |
| Ctse | 0.00615909 | 0.00562474 | 0.00555364 | 7.11E-05 | 0.46604486 | 0.99999994 |
| BC107364 | 0.00093646 | 0.00091848 | 0.00084741 | 7.11E-05 | 0.28832069 | 0.99999994 |
| Gm12840 | 0.00075814 | 0.00075715 | 0.00068613 | 7.10E-05 | 0.24539746 | 0.99999994 |
| Klk1b8 | 0.00269994 | 0.00251104 | 0.00244026 | 7.08E-05 | 0.42344744 | 0.99999994 |
| 6030407O03 | 0.00051215 | 0.00053378 | 0.00046358 | 7.02E-05 | 0.15673443 | 0.99999994 |
| Gm42788 | 0.00051215 | 0.00053378 | 0.00046358 | 7.02E-05 | 0.15673443 | 0.99999994 |
| Psd2 | 0.00051215 | 0.00053378 | 0.00046358 | 7.02E-05 | 0.15673443 | 0.99999994 |
| Zfyve28 | 0.00088423 | 0.00086984 | 0.00080017 | 6.97E-05 | 0.28110018 | 0.99999994 |
| Dennd4b | 0.00879547 | 0.00798638 | 0.00791672 | 6.97E-05 | 0.47664927 | 0.99999994 |
| Gm28536 | 0.00051169 | 0.00053281 | 0.00046316 | 6.96E-05 | 0.1584172 | 0.99999994 |
| Gm12052 | 0.00094621 | 0.00092577 | 0.00085622 | 6.96E-05 | 0.2943501 | 0.99999994 |
| Casp1 | 0.00073792 | 0.00073712 | 0.00066784 | 6.93E-05 | 0.24492776 | 0.99999994 |
| Kif16bos | 0.00072557 | 0.00072589 | 0.00065666 | 6.92E-05 | 0.24141908 | 0.99999994 |
| Optc | 0.00302014 | 0.00279808 | 0.00272906 | 6.90E-05 | 0.43315428 | 0.99999994 |
| Slc24a4 | 0.0005111 | 0.0005316 | 0.00046263 | 6.90E-05 | 0.16053252 | 0.99999994 |
| Ets1 | 0.00544313 | 0.00497937 | 0.00491044 | 6.89E-05 | 0.46277947 | 0.99999994 |
| Slc35f6 | 0.05525447 | 0.04828775 | 0.04821892 | 6.88E-05 | 0.4962095 | 0.99999994 |
| Il1rapl1 | 0.00051099 | 0.00053136 | 0.00046253 | 6.88E-05 | 0.16095716 | 0.99999994 |
| Adcy4 | 0.00051099 | 0.00053136 | 0.00046253 | 6.88E-05 | 0.16095716 | 0.99999994 |
| Tmem200c | 0.00051099 | 0.00053136 | 0.00046253 | 6.88E-05 | 0.16095716 | 0.99999994 |
| A430110L20I | 0.00095277 | 0.00093095 | 0.00086216 | 6.88E-05 | 0.2976638 | 0.99999994 |
| Siglech | 0.00466265 | 0.00427708 | 0.00420857 | 6.85E-05 | 0.45685412 | 0.99999994 |
| Sez6 | 0.00345938 | 0.00319322 | 0.00312503 | 6.82E-05 | 0.44225828 | 0.99999994 |

| | | | | | | |
|---|---|---|---|---|---|---|
| Lsmem1 | 0.0007504 | 0.00074694 | 0.00067913 | 6.78E-05 | 0.25312172 | 0.99999994 |
| Stx19 | 0.00632326 | 0.00576882 | 0.00570104 | 6.78E-05 | 0.46846318 | 0.99999994 |
| Rtn4r | 0.00128365 | 0.00122883 | 0.0011613 | 6.75E-05 | 0.34934569 | 0.99999994 |
| Prr11 | 0.00740916 | 0.00674271 | 0.00667518 | 6.75E-05 | 0.47315593 | 0.99999994 |
| Sval1 | 0.00131843 | 0.00126022 | 0.00119274 | 6.75E-05 | 0.35323902 | 0.99999994 |
| Sox6 | 0.03912824 | 0.03457842 | 0.03451098 | 6.74E-05 | 0.49481114 | 0.99999994 |
| Itga5 | 0.00252791 | 0.00235219 | 0.00228504 | 6.72E-05 | 0.42245672 | 0.99999994 |
| Gm26581 | 0.00050948 | 0.00052823 | 0.00046117 | 6.71E-05 | 0.16652455 | 0.99999994 |
| Gm10825 | 0.00050948 | 0.00052823 | 0.00046117 | 6.71E-05 | 0.16652455 | 0.99999994 |
| Foxq1 | 0.00050948 | 0.00052823 | 0.00046117 | 6.71E-05 | 0.16652455 | 0.99999994 |
| D030018L15I | 0.00050948 | 0.00052823 | 0.00046117 | 6.71E-05 | 0.16652455 | 0.99999994 |
| Col23a1 | 0.00126026 | 0.0012071 | 0.00114016 | 6.69E-05 | 0.347989 | 0.99999994 |
| Npm3 | 0.48867808 | 0.3095033 | 0.3094366 | 6.67E-05 | 0.49942757 | 0.99999994 |
| Cyp4x1 | 0.00248232 | 0.00231055 | 0.0022439 | 6.67E-05 | 0.42163118 | 0.99999994 |
| Abi3 | 0.00094251 | 0.00091947 | 0.00085287 | 6.66E-05 | 0.30161698 | 0.99999994 |
| Gpr22 | 0.00050902 | 0.00052727 | 0.00046075 | 6.65E-05 | 0.168255 | 0.99999994 |
| Gm8225 | 0.00089718 | 0.00087811 | 0.00081189 | 6.62E-05 | 0.29357592 | 0.99999994 |
| Gm43703 | 0.00050868 | 0.00052656 | 0.00046043 | 6.61E-05 | 0.16955815 | 0.99999994 |
| Odf3l2 | 0.00050862 | 0.00052644 | 0.00046038 | 6.61E-05 | 0.16977579 | 0.99999994 |
| Lingo2 | 0.00092616 | 0.00090381 | 0.00083809 | 6.57E-05 | 0.30082748 | 0.99999994 |
| Slc4a11 | 0.00050799 | 0.00052513 | 0.00045981 | 6.53E-05 | 0.17217799 | 0.99999994 |
| Gm11627 | 0.00093729 | 0.0009133 | 0.00084815 | 6.52E-05 | 0.30456457 | 0.99999994 |
| 8030462N17 | 0.00423436 | 0.00388816 | 0.00382311 | 6.51E-05 | 0.45490687 | 0.99999994 |
| Platr3 | 0.00050741 | 0.00052395 | 0.00045929 | 6.47E-05 | 0.17437484 | 0.99999994 |
| Lpcat2 | 0.00844064 | 0.00766375 | 0.00759917 | 6.46E-05 | 0.47744565 | 0.99999994 |
| A130051J06I | 0.00133253 | 0.00126987 | 0.00120548 | 6.44E-05 | 0.3610975 | 0.99999994 |
| Syndig1 | 0.00267416 | 0.00248135 | 0.00241699 | 6.44E-05 | 0.42966127 | 0.99999994 |
| F7 | 0.00050678 | 0.00052265 | 0.00045872 | 6.39E-05 | 0.17680555 | 0.99999994 |
| Zfp775 | 0.02640612 | 0.02355184 | 0.02348816 | 6.37E-05 | 0.49280034 | 0.99999994 |
| Paqr5 | 0.00440851 | 0.00404344 | 0.00397987 | 6.36E-05 | 0.45766449 | 0.99999994 |

| | | | | | | |
|---|---|---|---|---|---|---|
| Dpf2 | 0.43816729 | 0.28767349 | 0.28761011 | 6.34E-05 | 0.49941476 | 0.99999994 |
| Dnaaf3 | 0.00251018 | 0.00233233 | 0.00226904 | 6.33E-05 | 0.42635533 | 0.99999994 |
| Trp63 | 0.00050621 | 0.00052148 | 0.00045821 | 6.33E-05 | 0.17902803 | 0.99999994 |
| Hoxc4 | 0.0009238 | 0.00089905 | 0.00083595 | 6.31E-05 | 0.30769811 | 0.99999994 |
| A530016L24I | 0.00153136 | 0.00144806 | 0.00138517 | 6.29E-05 | 0.38123414 | 0.99999994 |
| 2510017J16F | 0.00288093 | 0.0026663 | 0.00260352 | 6.28E-05 | 0.43623655 | 0.99999994 |
| Pmm1 | 0.05571275 | 0.04866691 | 0.04860424 | 6.27E-05 | 0.49657587 | 0.99999994 |
| Ccbe1 | 0.00197735 | 0.00185037 | 0.00178804 | 6.23E-05 | 0.40826344 | 0.99999994 |
| Pou5f1 | 0.00883426 | 0.00801375 | 0.00795143 | 6.23E-05 | 0.47919539 | 0.99999994 |
| Smim19 | 0.08340954 | 0.07153176 | 0.0714695 | 6.23E-05 | 0.4976869 | 0.99999994 |
| Adgrb1 | 0.0009526 | 0.00092395 | 0.000862 | 6.20E-05 | 0.31617448 | 0.99999994 |
| G0s2 | 0.00891123 | 0.00808195 | 0.00802029 | 6.17E-05 | 0.47959476 | 0.99999994 |
| Lgals2 | 0.00766641 | 0.00696733 | 0.00690574 | 6.16E-05 | 0.47633104 | 0.99999994 |
| Gm6658 | 0.00050474 | 0.00051844 | 0.00045687 | 6.16E-05 | 0.18486234 | 0.99999994 |
| Gm20755 | 0.00050451 | 0.00051798 | 0.00045667 | 6.13E-05 | 0.18576698 | 0.99999994 |
| Olfm2 | 0.00084145 | 0.00082257 | 0.00076148 | 6.11E-05 | 0.29665959 | 0.99999994 |
| Stra6l | 0.00281558 | 0.00260554 | 0.00254457 | 6.10E-05 | 0.43663893 | 0.99999994 |
| Carns1 | 0.01122795 | 0.01015038 | 0.01008958 | 6.08E-05 | 0.48400296 | 0.99999994 |
| Gm15918 | 0.00113707 | 0.00108946 | 0.0010288 | 6.07E-05 | 0.34733662 | 0.99999994 |
| Stx1b | 0.00083851 | 0.00081945 | 0.00075882 | 6.06E-05 | 0.29739642 | 0.99999994 |
| Pygo1 | 0.02525188 | 0.02253938 | 0.02247881 | 6.06E-05 | 0.49284523 | 0.99999994 |
| Lrrc23 | 0.00225327 | 0.00209763 | 0.00203717 | 6.05E-05 | 0.42169053 | 0.99999994 |
| Kcnj9 | 0.00084279 | 0.00082302 | 0.00076269 | 6.03E-05 | 0.2992592 | 0.99999994 |
| Ppfibp2 | 0.00075072 | 0.00073962 | 0.00067942 | 6.02E-05 | 0.27763594 | 0.99999994 |
| Mansc4 | 0.00095143 | 0.00092081 | 0.00086094 | 5.99E-05 | 0.32172701 | 0.99999994 |
| C1ra | 0.00076483 | 0.00075166 | 0.00069217 | 5.95E-05 | 0.28366078 | 0.99999994 |
| AB124611 | 0.00165331 | 0.00155481 | 0.00149535 | 5.95E-05 | 0.39563649 | 0.99999994 |
| 1700001O22 | 0.00422063 | 0.00387007 | 0.00381074 | 5.93E-05 | 0.45873252 | 0.99999994 |
| Klhl2 | 0.03992656 | 0.0352558 | 0.03519648 | 5.93E-05 | 0.49552439 | 0.99999994 |
| Ern2 | 0.00050277 | 0.0005144 | 0.00045509 | 5.93E-05 | 0.19284021 | 0.99999994 |

| | | | | | | |
|---|---|---|---|---|---|---|
| Fgf20 | 0.00050277 | 0.0005144 | 0.00045509 | 5.93E-05 | 0.19284021 | 0.99999994 |
| Ptpn7 | 0.00050254 | 0.00051394 | 0.00045489 | 5.91E-05 | 0.19376079 | 0.99999994 |
| 4930573C15I | 0.00050254 | 0.00051394 | 0.00045489 | 5.91E-05 | 0.19376079 | 0.99999994 |
| Scn4b | 0.00050254 | 0.00051394 | 0.00045489 | 5.91E-05 | 0.19376079 | 0.99999994 |
| Ntng1 | 0.0037793 | 0.00347213 | 0.0034133 | 5.88E-05 | 0.45432894 | 0.99999994 |
| Capn10 | 0.26528221 | 0.19782275 | 0.19776419 | 5.86E-05 | 0.49921367 | 0.99999994 |
| Cxcr6 | 0.00077051 | 0.00075542 | 0.00069731 | 5.81E-05 | 0.28956197 | 0.99999994 |
| Notumos | 0.00256394 | 0.00237564 | 0.00231755 | 5.81E-05 | 0.4337473 | 0.99999994 |
| Bfsp2 | 0.0006805 | 0.00067375 | 0.00061589 | 5.79E-05 | 0.2658965 | 0.99999994 |
| Itgb6 | 0.00379228 | 0.00348263 | 0.00342499 | 5.76E-05 | 0.45539659 | 0.99999994 |
| Veph1 | 0.00148237 | 0.00139828 | 0.0013409 | 5.74E-05 | 0.38787106 | 0.99999994 |
| Crispld1 | 0.00094764 | 0.00091467 | 0.00085752 | 5.72E-05 | 0.3286366 | 0.99999994 |
| Strip2 | 0.00205737 | 0.00191737 | 0.0018603 | 5.71E-05 | 0.41910024 | 0.99999994 |
| Adam12 | 0.00094521 | 0.00091206 | 0.00085532 | 5.67E-05 | 0.32939488 | 0.99999994 |
| Inmt | 0.00322607 | 0.00297147 | 0.00291474 | 5.67E-05 | 0.4484551 | 0.99999994 |
| Ranbp6 | 0.04501569 | 0.03960582 | 0.03954938 | 5.64E-05 | 0.49621042 | 0.99999994 |
| Atp8b3 | 0.00217795 | 0.00202546 | 0.00196917 | 5.63E-05 | 0.42454982 | 0.99999994 |
| Fam136b-ps | 0.00094118 | 0.00090781 | 0.00085167 | 5.61E-05 | 0.33040822 | 0.99999994 |
| A230005M16 | 0.00455336 | 0.00416635 | 0.00411024 | 5.61E-05 | 0.4637945 | 0.99999994 |
| Tbx3 | 0.00049965 | 0.00050803 | 0.00045227 | 5.58E-05 | 0.20588697 | 0.99999994 |
| Gm9821 | 0.00094445 | 0.00091035 | 0.00085463 | 5.57E-05 | 0.33215198 | 0.99999994 |
| Atp2b1 | 0.70975342 | 0.38460278 | 0.38454716 | 5.56E-05 | 0.49961594 | 0.99999994 |
| Gm37065 | 0.00119517 | 0.00113689 | 0.00108132 | 5.56E-05 | 0.36614588 | 0.99999994 |
| Chrna1 | 0.00137765 | 0.00130182 | 0.00124626 | 5.56E-05 | 0.38332686 | 0.99999994 |
| Thrb | 0.00925125 | 0.00837991 | 0.00832441 | 5.55E-05 | 0.48230219 | 0.99999994 |
| Gm28578 | 0.00088329 | 0.00085471 | 0.00079932 | 5.54E-05 | 0.32232007 | 0.99999994 |
| Gm26520 | 0.00132169 | 0.00125105 | 0.00119569 | 5.54E-05 | 0.37896056 | 0.99999994 |
| Fbxo15 | 0.00071341 | 0.00070053 | 0.00064567 | 5.49E-05 | 0.28584963 | 0.99999994 |
| Dennd2c | 0.00115002 | 0.00109527 | 0.00104051 | 5.48E-05 | 0.3630326 | 0.99999994 |
| AI854703 | 0.00091701 | 0.00088451 | 0.00082981 | 5.47E-05 | 0.3304046 | 0.99999994 |

| | | | | | | |
|---|---|---|---|---|---|---|
| Ticrr | 0.00230605 | 0.00213945 | 0.00208481 | 5.46E-05 | 0.43074965 | 0.99999994 |
| Timp4 | 0.00160392 | 0.00150525 | 0.00145073 | 5.45E-05 | 0.40123215 | 0.99999994 |
| Gm44170 | 0.00049838 | 0.00050546 | 0.00045112 | 5.43E-05 | 0.21134114 | 0.99999994 |
| Prr22 | 0.00239505 | 0.00221941 | 0.00216514 | 5.43E-05 | 0.43374196 | 0.99999994 |
| Atrip | 0.00147868 | 0.00139172 | 0.00133756 | 5.42E-05 | 0.39376299 | 0.99999994 |
| Cdh23 | 0.0021585 | 0.00200577 | 0.00195161 | 5.42E-05 | 0.42672714 | 0.99999994 |
| 9430002A10 | 0.00223541 | 0.00207514 | 0.00202104 | 5.41E-05 | 0.42928536 | 0.99999994 |
| Adam19 | 0.00149577 | 0.00140704 | 0.00135301 | 5.40E-05 | 0.39517993 | 0.99999994 |
| Xkr5 | 0.00049766 | 0.00050401 | 0.00045047 | 5.35E-05 | 0.21444752 | 0.99999994 |
| Gm20732 | 0.00049761 | 0.0005039 | 0.00045042 | 5.35E-05 | 0.21468716 | 0.99999994 |
| Lrrc69 | 0.00049739 | 0.00050345 | 0.00045022 | 5.32E-05 | 0.21564672 | 0.99999994 |
| Ccr4 | 0.00049739 | 0.00050345 | 0.00045022 | 5.32E-05 | 0.21564672 | 0.99999994 |
| Ercc6l | 0.00251895 | 0.00232964 | 0.00227695 | 5.27E-05 | 0.43879012 | 0.99999994 |
| Nol3 | 0.00503501 | 0.00459609 | 0.00454353 | 5.26E-05 | 0.46931292 | 0.99999994 |
| Pgpep1l | 0.00678144 | 0.00616457 | 0.00611224 | 5.23E-05 | 0.47727889 | 0.99999994 |
| Ackr4 | 0.00049646 | 0.00050157 | 0.00044938 | 5.22E-05 | 0.21974217 | 0.99999994 |
| Sec31b | 0.00149818 | 0.00140696 | 0.00135519 | 5.18E-05 | 0.39963985 | 0.99999994 |
| Zbtb46 | 0.00373253 | 0.00342264 | 0.00337116 | 5.15E-05 | 0.45951955 | 0.99999994 |
| Gm13572 | 0.00224034 | 0.00207686 | 0.00202549 | 5.14E-05 | 0.43297047 | 0.99999994 |
| Mad2l1bp | 0.11173119 | 0.09389154 | 0.0938402 | 5.13E-05 | 0.49854705 | 0.99999994 |
| Spdl1 | 0.00644849 | 0.0058647 | 0.00581346 | 5.12E-05 | 0.47660735 | 0.99999994 |
| Lrp2 | 0.00092998 | 0.00089274 | 0.00084155 | 5.12E-05 | 0.34275977 | 0.99999994 |
| Ak9 | 0.00162867 | 0.00152416 | 0.00147309 | 5.11E-05 | 0.40875266 | 0.99999994 |
| Rfc5 | 0.1253318 | 0.10417981 | 0.10412935 | 5.05E-05 | 0.49871324 | 0.99999994 |
| Sox6os | 0.00179459 | 0.00167326 | 0.00162298 | 5.03E-05 | 0.41831869 | 0.99999994 |
| Gm9801 | 0.00259872 | 0.00239907 | 0.00234893 | 5.01E-05 | 0.44350891 | 0.99999994 |
| Cyb5d1 | 0.00722973 | 0.0065643 | 0.00651431 | 5.00E-05 | 0.47963309 | 0.99999994 |
| Crocc2 | 0.0004944 | 0.00049741 | 0.00044751 | 4.99E-05 | 0.2289949 | 0.99999994 |
| Gm6569 | 0.0004944 | 0.00049741 | 0.00044751 | 4.99E-05 | 0.2289949 | 0.99999994 |
| Snhg18 | 0.09450469 | 0.08045189 | 0.08040202 | 4.99E-05 | 0.49835297 | 0.99999994 |

| | | | | | | |
|---|---|---|---|---|---|---|
| Lgr6 | 0.00049434 | 0.0004973 | 0.00044747 | 4.98E-05 | 0.22924017 | 0.99999994 |
| Gm17202 | 0.00049429 | 0.00049719 | 0.00044742 | 4.98E-05 | 0.22948553 | 0.99999994 |
| Gm29292 | 0.00049429 | 0.00049719 | 0.00044742 | 4.98E-05 | 0.22948553 | 0.99999994 |
| Ms4a4a | 0.00049429 | 0.00049719 | 0.00044742 | 4.98E-05 | 0.22948553 | 0.99999994 |
| Pla2g12b | 0.00131344 | 0.00123789 | 0.00118823 | 4.97E-05 | 0.3904414 | 0.99999994 |
| Reck | 0.00863444 | 0.00782207 | 0.00777264 | 4.94E-05 | 0.48311603 | 0.99999994 |
| Pcdhb17 | 0.02424981 | 0.02165065 | 0.02160126 | 4.94E-05 | 0.49392865 | 0.99999994 |
| Gm5420 | 0.00127468 | 0.00120243 | 0.00115319 | 4.92E-05 | 0.38811885 | 0.99999994 |
| Anln | 0.00329905 | 0.00302943 | 0.00298053 | 4.89E-05 | 0.4565165 | 0.99999994 |
| Fgfr3 | 0.02936447 | 0.0261163 | 0.02606805 | 4.82E-05 | 0.49508511 | 0.99999994 |
| Lrrc39 | 0.00093862 | 0.00089746 | 0.00084935 | 4.81E-05 | 0.35309333 | 0.99999994 |
| Gpr75 | 0.00329527 | 0.0030251 | 0.00297712 | 4.80E-05 | 0.45728004 | 0.99999994 |
| 6530409C15I | 0.00049257 | 0.00049373 | 0.00044586 | 4.79E-05 | 0.23738311 | 0.99999994 |
| Gm10447 | 0.00049257 | 0.00049373 | 0.00044586 | 4.79E-05 | 0.23738311 | 0.99999994 |
| Rab36 | 0.0788094 | 0.06777629 | 0.06772847 | 4.78E-05 | 0.49812518 | 0.99999994 |
| Platr5 | 0.00049235 | 0.0004933 | 0.00044566 | 4.76E-05 | 0.23837645 | 0.99999994 |
| Wnt7a | 0.00049235 | 0.0004933 | 0.00044566 | 4.76E-05 | 0.23837645 | 0.99999994 |
| Rgr | 0.00049235 | 0.0004933 | 0.00044566 | 4.76E-05 | 0.23837645 | 0.99999994 |
| Islr2 | 0.00049235 | 0.0004933 | 0.00044566 | 4.76E-05 | 0.23837645 | 0.99999994 |
| Prrt4 | 0.00087068 | 0.00083553 | 0.00078791 | 4.76E-05 | 0.34372404 | 0.99999994 |
| Kcne2 | 0.00092053 | 0.00088058 | 0.000833 | 4.76E-05 | 0.35188154 | 0.99999994 |
| Tmem147os | 0.001333 | 0.00125308 | 0.00120591 | 4.72E-05 | 0.39729162 | 0.99999994 |
| Ppbp | 0.00049176 | 0.00049212 | 0.00044513 | 4.70E-05 | 0.241115 | 0.99999994 |
| Gm9754 | 0.00049176 | 0.00049212 | 0.00044513 | 4.70E-05 | 0.241115 | 0.99999994 |
| Gm10840 | 0.00135646 | 0.00127402 | 0.00122711 | 4.69E-05 | 0.39957422 | 0.99999994 |
| Gm15856 | 0.0004916 | 0.0004918 | 0.00044499 | 4.68E-05 | 0.24186359 | 0.99999994 |
| Pltp | 0.00369446 | 0.00338365 | 0.00333687 | 4.68E-05 | 0.46282441 | 0.99999994 |
| Clec4a1 | 0.00114425 | 0.00108188 | 0.00103529 | 4.66E-05 | 0.38227225 | 0.99999994 |
| Pcdhb20 | 0.01129474 | 0.01019563 | 0.01014914 | 4.65E-05 | 0.48783877 | 0.99999994 |
| Pcsk9 | 0.00741945 | 0.00673083 | 0.0066844 | 4.64E-05 | 0.48156236 | 0.99999994 |

| | | | | | | |
|---|---|---|---|---|---|---|
| Gm13306 | 0.00094776 | 0.000904 | 0.00085762 | 4.64E-05 | 0.35944567 | 0.99999994 |
| Rnf207 | 0.00169078 | 0.00157536 | 0.0015292 | 4.62E-05 | 0.42038604 | 0.99999994 |
| Ptger4 | 0.00200667 | 0.00186057 | 0.00181451 | 4.61E-05 | 0.43292231 | 0.99999994 |
| Lmx1a | 0.00177819 | 0.00165418 | 0.00160817 | 4.60E-05 | 0.42448941 | 0.99999994 |
| Vwce | 0.00129651 | 0.00121867 | 0.00117293 | 4.57E-05 | 0.39759252 | 0.99999994 |
| Dgkq | 0.02303748 | 0.02058373 | 0.020538 | 4.57E-05 | 0.49408765 | 0.99999994 |
| Elf5 | 0.00665366 | 0.00604325 | 0.00599759 | 4.57E-05 | 0.47979117 | 0.99999994 |
| Tvp23bos | 0.00137533 | 0.00128982 | 0.00124417 | 4.57E-05 | 0.40351838 | 0.99999994 |
| Gm43301 | 0.00073607 | 0.00071097 | 0.00066616 | 4.48E-05 | 0.32715829 | 0.99999994 |
| Gm29361 | 0.00130734 | 0.00122747 | 0.00118271 | 4.48E-05 | 0.40055481 | 0.99999994 |
| Rbm46 | 0.00048926 | 0.00048714 | 0.00044287 | 4.43E-05 | 0.25292462 | 0.99999994 |
| Hoxb3os | 0.00113255 | 0.00106887 | 0.00102471 | 4.42E-05 | 0.38709539 | 0.99999994 |
| Gm13470 | 0.00048916 | 0.00048692 | 0.00044277 | 4.42E-05 | 0.2534309 | 0.99999994 |
| Pitx2 | 0.0004891 | 0.00048682 | 0.00044273 | 4.41E-05 | 0.25368416 | 0.99999994 |
| Ftcd | 0.0004891 | 0.00048682 | 0.00044273 | 4.41E-05 | 0.25368416 | 0.99999994 |
| Cabyr | 0.00122507 | 0.00115234 | 0.00110835 | 4.40E-05 | 0.3958109 | 0.99999994 |
| Mterf1b | 0.00942833 | 0.00852665 | 0.00848273 | 4.39E-05 | 0.48625303 | 0.99999994 |
| Stx11 | 0.00081228 | 0.00077875 | 0.0007351 | 4.37E-05 | 0.34631184 | 0.99999994 |
| Gxylt2 | 0.00166272 | 0.00154734 | 0.00150386 | 4.35E-05 | 0.42368979 | 0.99999994 |
| Pcdhb22 | 0.01382792 | 0.01244758 | 0.01240417 | 4.34E-05 | 0.49070864 | 0.99999994 |
| Il34 | 0.00126778 | 0.00119013 | 0.00114696 | 4.32E-05 | 0.40107184 | 0.99999994 |
| Gm2449 | 0.00070886 | 0.0006845 | 0.00064155 | 4.30E-05 | 0.32790815 | 0.99999994 |
| Gm36975 | 0.000488 | 0.00048462 | 0.00044172 | 4.29E-05 | 0.25901924 | 0.99999994 |
| Ndufs2 | 1.01303251 | 0.42713566 | 0.42709279 | 4.29E-05 | 0.49973344 | 0.99999994 |
| Trim12a | 0.00104153 | 0.00098516 | 0.00094241 | 4.27E-05 | 0.38135549 | 0.99999994 |
| Gm5475 | 0.00196214 | 0.00181697 | 0.0017743 | 4.27E-05 | 0.43641604 | 0.99999994 |
| Gm11491 | 0.0048359 | 0.00440674 | 0.00436444 | 4.23E-05 | 0.47428094 | 0.99999994 |
| Kcnv2 | 0.00080037 | 0.00076644 | 0.00072432 | 4.21E-05 | 0.34934667 | 0.99999994 |
| Gm26636 | 0.00048716 | 0.00048295 | 0.00044096 | 4.20E-05 | 0.26310494 | 0.99999994 |
| Gm3716 | 0.00658486 | 0.00597765 | 0.00593585 | 4.18E-05 | 0.48130808 | 0.99999994 |

| | | | | | | |
|---|---|---|---|---|---|---|
| Hgsnat | 0.08444293 | 0.07234863 | 0.07230686 | 4.18E-05 | 0.49846575 | 0.99999994 |
| Gm43573 | 0.00181133 | 0.00167971 | 0.0016381 | 4.16E-05 | 0.43286869 | 0.99999994 |
| Ncaph | 0.00967163 | 0.00874178 | 0.0087002 | 4.16E-05 | 0.48730887 | 0.99999994 |
| Zbtb10 | 0.0363642 | 0.03217248 | 0.03213197 | 4.05E-05 | 0.49665225 | 0.99999994 |
| Stmn2 | 0.00390726 | 0.00356896 | 0.00352855 | 4.04E-05 | 0.46961829 | 0.99999994 |
| Amer3 | 0.00048538 | 0.00047944 | 0.00043936 | 4.01E-05 | 0.27184378 | 0.99999994 |
| Col15a1 | 0.00222933 | 0.00205547 | 0.00201556 | 3.99E-05 | 0.44756723 | 0.99999994 |
| Ooep | 0.00171033 | 0.00158673 | 0.00154687 | 3.99E-05 | 0.43190539 | 0.99999994 |
| Hapln3 | 0.00048497 | 0.00047862 | 0.00043898 | 3.96E-05 | 0.27391066 | 0.99999994 |
| Lta | 0.00048497 | 0.00047862 | 0.00043898 | 3.96E-05 | 0.27391066 | 0.99999994 |
| 0610031O16 | 0.00048497 | 0.00047862 | 0.00043898 | 3.96E-05 | 0.27391066 | 0.99999994 |
| Trim29 | 0.00132597 | 0.00123918 | 0.00119955 | 3.96E-05 | 0.41298434 | 0.99999994 |
| Rhoh | 0.00090843 | 0.00086131 | 0.00082205 | 3.93E-05 | 0.37527476 | 0.99999994 |
| B230312C02 | 0.00048455 | 0.0004778 | 0.00043861 | 3.92E-05 | 0.27598145 | 0.99999994 |
| Robo4 | 0.00048455 | 0.0004778 | 0.00043861 | 3.92E-05 | 0.27598145 | 0.99999994 |
| Myo7a | 0.00088447 | 0.0008394 | 0.00080039 | 3.90E-05 | 0.37279959 | 0.99999994 |
| Clic3 | 0.0011875 | 0.00111318 | 0.00107439 | 3.88E-05 | 0.40502982 | 0.99999994 |
| P3h2 | 0.00048357 | 0.00047587 | 0.00043772 | 3.81E-05 | 0.28091471 | 0.99999994 |
| Slc35f4 | 0.00184238 | 0.00170425 | 0.00166615 | 3.81E-05 | 0.43950707 | 0.99999994 |
| Tgfbr2 | 0.00610357 | 0.00554182 | 0.00550379 | 3.80E-05 | 0.48165772 | 0.99999994 |
| 6720489N17 | 0.00972718 | 0.00878771 | 0.00874984 | 3.79E-05 | 0.48850991 | 0.99999994 |
| Igfbp3 | 0.00681542 | 0.00618056 | 0.00614273 | 3.78E-05 | 0.4836528 | 0.99999994 |
| Cyp2j9 | 0.00092941 | 0.00087851 | 0.00084102 | 3.75E-05 | 0.38334156 | 0.99999994 |
| Bmyc | 0.59452684 | 0.34976307 | 0.34972566 | 3.74E-05 | 0.49971594 | 0.99999994 |
| Usp44 | 0.00048285 | 0.00047445 | 0.00043707 | 3.74E-05 | 0.28456288 | 0.99999994 |
| Rbm11 | 0.00265718 | 0.00243896 | 0.00240167 | 3.73E-05 | 0.45884045 | 0.99999994 |
| 2310026L22F | 0.00075086 | 0.00071683 | 0.00067954 | 3.73E-05 | 0.35745312 | 0.99999994 |
| Atg4a-ps | 0.00085465 | 0.00081067 | 0.00077342 | 3.73E-05 | 0.37425544 | 0.99999994 |
| Cspg4 | 0.00048244 | 0.00047365 | 0.0004367 | 3.69E-05 | 0.28665236 | 0.99999994 |
| Cdh17 | 0.00048234 | 0.00047344 | 0.0004366 | 3.68E-05 | 0.28717526 | 0.99999994 |

| | | | | | | |
|---|---|---|---|---|---|---|
| Irs3 | 0.00228952 | 0.00210661 | 0.00206989 | 3.67E-05 | 0.45299067 | 0.99999994 |
| 4921504A21 | 0.00214743 | 0.00197826 | 0.00194161 | 3.66E-05 | 0.45000963 | 0.99999994 |
| Pgam2 | 0.00109414 | 0.00102611 | 0.00098998 | 3.61E-05 | 0.40403754 | 0.99999994 |
| Celf5 | 0.02259904 | 0.02018903 | 0.02015305 | 3.60E-05 | 0.49525914 | 0.99999994 |
| Ptf1aos | 0.00048111 | 0.00047104 | 0.00043549 | 3.55E-05 | 0.29346618 | 0.99999994 |
| Gm13074 | 0.00048101 | 0.00047084 | 0.0004354 | 3.54E-05 | 0.29399172 | 0.99999994 |
| Gm16008 | 0.00048101 | 0.00047084 | 0.0004354 | 3.54E-05 | 0.29399172 | 0.99999994 |
| Taf3 | 0.1688574 | 0.13559871 | 0.13556366 | 3.51E-05 | 0.49931336 | 0.99999994 |
| Prm1 | 0.0004806 | 0.00047004 | 0.00043503 | 3.50E-05 | 0.29609583 | 0.99999994 |
| Cenpp | 0.01866824 | 0.01672634 | 0.01669167 | 3.47E-05 | 0.49448371 | 0.99999994 |
| Zfp872 | 0.00048014 | 0.00046915 | 0.00043462 | 3.45E-05 | 0.29846656 | 0.99999994 |
| Cfap99 | 0.00048014 | 0.00046915 | 0.00043462 | 3.45E-05 | 0.29846656 | 0.99999994 |
| Arhgap25 | 0.00048009 | 0.00046905 | 0.00043457 | 3.45E-05 | 0.2987302 | 0.99999994 |
| Spib | 0.00048009 | 0.00046905 | 0.00043457 | 3.45E-05 | 0.2987302 | 0.99999994 |
| Irf4 | 0.00048009 | 0.00046905 | 0.00043457 | 3.45E-05 | 0.2987302 | 0.99999994 |
| Kctd6 | 0.01677645 | 0.01505371 | 0.01501927 | 3.44E-05 | 0.49391095 | 0.99999994 |
| Cplx1 | 0.00142589 | 0.0013243 | 0.00128986 | 3.44E-05 | 0.42947288 | 0.99999994 |
| Angpt4 | 0.00130538 | 0.00121538 | 0.00118095 | 3.44E-05 | 0.42304847 | 0.99999994 |
| Stambpl1 | 0.00727952 | 0.00659329 | 0.00655895 | 3.43E-05 | 0.48610015 | 0.99999994 |
| Fscn3 | 0.00214973 | 0.00197766 | 0.00194369 | 3.40E-05 | 0.45369916 | 0.99999994 |
| Gm16140 | 0.00091037 | 0.00085734 | 0.00082381 | 3.35E-05 | 0.39324117 | 0.99999994 |
| Pdf | 0.00161097 | 0.00149055 | 0.0014571 | 3.34E-05 | 0.43928005 | 0.99999994 |
| 1700011M02 | 0.00171785 | 0.00158691 | 0.00155366 | 3.32E-05 | 0.44336119 | 0.99999994 |
| Pkp1 | 0.00047888 | 0.00046668 | 0.00043347 | 3.32E-05 | 0.30507102 | 0.99999994 |
| Zmynd12 | 0.00767449 | 0.00694615 | 0.00691299 | 3.32E-05 | 0.48726244 | 0.99999994 |
| Hipk4 | 0.00047868 | 0.00046628 | 0.00043329 | 3.30E-05 | 0.30613023 | 0.99999994 |
| Sox21 | 0.00047863 | 0.00046619 | 0.00043325 | 3.29E-05 | 0.30639513 | 0.99999994 |
| Aspg | 0.00047863 | 0.00046619 | 0.00043325 | 3.29E-05 | 0.30639513 | 0.99999994 |
| Kcna4 | 0.00047858 | 0.00046609 | 0.0004332 | 3.29E-05 | 0.30666008 | 0.99999994 |
| Foxd2 | 0.00047858 | 0.00046609 | 0.0004332 | 3.29E-05 | 0.30666008 | 0.99999994 |

| | | | | | | |
|---|---|---|---|---|---|---|
| Smpx | 0.00130983 | 0.00121773 | 0.00118496 | 3.28E-05 | 0.4269837 | 0.99999994 |
| 1700113H08I | 0.00047837 | 0.00046569 | 0.00043302 | 3.27E-05 | 0.30772027 | 0.99999994 |
| Gm37240 | 0.00047812 | 0.0004652 | 0.00043279 | 3.24E-05 | 0.30904641 | 0.99999994 |
| Tacr1 | 0.00047812 | 0.0004652 | 0.00043279 | 3.24E-05 | 0.30904641 | 0.99999994 |
| Dlx5 | 0.00047802 | 0.00046501 | 0.0004327 | 3.23E-05 | 0.30957714 | 0.99999994 |
| Sfrp2 | 0.00317308 | 0.00289925 | 0.00286696 | 3.23E-05 | 0.47012737 | 0.99999994 |
| Rasl10a | 0.00168918 | 0.00155998 | 0.00152776 | 3.22E-05 | 0.44418823 | 0.99999994 |
| Ccdc146 | 0.00385364 | 0.00351247 | 0.00348026 | 3.22E-05 | 0.47543719 | 0.99999994 |
| 2310043L19I | 0.00047792 | 0.00046481 | 0.00043261 | 3.22E-05 | 0.31010803 | 0.99999994 |
| Slc24a3 | 0.00127114 | 0.00118192 | 0.00115 | 3.19E-05 | 0.42670844 | 0.99999994 |
| Gm43936 | 0.00072911 | 0.00069147 | 0.00065986 | 3.16E-05 | 0.37493337 | 0.99999994 |
| Ino80b | 0.25675823 | 0.19268078 | 0.19264923 | 3.15E-05 | 0.49956515 | 0.99999994 |
| Atp10a | 0.00163382 | 0.00150888 | 0.00147774 | 3.11E-05 | 0.44422518 | 0.99999994 |
| Emcn | 0.00071383 | 0.00067696 | 0.00064604 | 3.09E-05 | 0.3750212 | 0.99999994 |
| Pllp | 0.00069162 | 0.00065662 | 0.00062595 | 3.07E-05 | 0.37213292 | 0.99999994 |
| Ccl24 | 0.00047632 | 0.0004617 | 0.00043116 | 3.05E-05 | 0.31862251 | 0.99999994 |
| Ptger1 | 0.00172473 | 0.00159036 | 0.00155988 | 3.05E-05 | 0.44825586 | 0.99999994 |
| Jph1 | 0.0024591 | 0.00225334 | 0.00222294 | 3.04E-05 | 0.46372995 | 0.99999994 |
| Itgb4 | 0.00047617 | 0.00046141 | 0.00043102 | 3.04E-05 | 0.31942259 | 0.99999994 |
| Prdx6b | 0.00091352 | 0.00085692 | 0.00082665 | 3.03E-05 | 0.4037194 | 0.99999994 |
| Tmem132d | 0.00047587 | 0.00046083 | 0.00043075 | 3.01E-05 | 0.32102365 | 0.99999994 |
| Rxrg | 0.00170622 | 0.00157315 | 0.00154315 | 3.00E-05 | 0.44853214 | 0.99999994 |
| Slc45a3 | 0.00246554 | 0.00225827 | 0.00222876 | 2.95E-05 | 0.46488015 | 0.99999994 |
| Afap1l2 | 0.00412342 | 0.00375239 | 0.00372322 | 2.92E-05 | 0.47920573 | 0.99999994 |
| Cfap74 | 0.00047498 | 0.0004591 | 0.00042994 | 2.92E-05 | 0.32583368 | 0.99999994 |
| Ankle1 | 0.00122041 | 0.00113316 | 0.00110414 | 2.90E-05 | 0.4305569 | 0.99999994 |
| Tmem198b | 0.02803982 | 0.02494308 | 0.02491413 | 2.90E-05 | 0.49691432 | 0.99999994 |
| Igsf3 | 0.04385951 | 0.03859164 | 0.03856303 | 2.86E-05 | 0.49802988 | 0.99999994 |
| Recql4 | 0.0010451 | 0.00097416 | 0.00094564 | 2.85E-05 | 0.42045155 | 0.99999994 |
| Mdga1 | 0.0024894 | 0.00227856 | 0.00225028 | 2.83E-05 | 0.4666729 | 0.99999994 |

| | | | | | | |
|---|---|---|---|---|---|---|
| A730061H03 | 0.00047409 | 0.00045738 | 0.00042914 | 2.82E-05 | 0.33065326 | 0.99999994 |
| Pald1 | 0.0100318 | 0.00904997 | 0.009022 | 2.80E-05 | 0.49176552 | 0.99999994 |
| Clec3b | 0.00227503 | 0.00208465 | 0.00205681 | 2.78E-05 | 0.46409746 | 0.99999994 |
| Gm20324 | 0.00657677 | 0.00595628 | 0.00592859 | 2.77E-05 | 0.48759754 | 0.99999994 |
| Peg12 | 0.0006227 | 0.00059054 | 0.00056361 | 2.69E-05 | 0.37518115 | 0.99999994 |
| Hormad2 | 0.00047271 | 0.00045472 | 0.00042789 | 2.68E-05 | 0.33816724 | 0.99999994 |
| Pou4f1 | 0.00267258 | 0.00244231 | 0.00241557 | 2.67E-05 | 0.47062747 | 0.99999994 |
| Pnlip | 0.00102209 | 0.00095158 | 0.00092484 | 2.67E-05 | 0.4236964 | 0.99999994 |
| Capn3 | 0.00091673 | 0.00085624 | 0.00082956 | 2.67E-05 | 0.41525665 | 0.99999994 |
| Cldn5 | 0.00267587 | 0.00244493 | 0.00241854 | 2.64E-05 | 0.47104471 | 0.99999994 |
| 1700003E16I | 0.0022135 | 0.00202735 | 0.00200126 | 2.61E-05 | 0.46542458 | 0.99999994 |
| Zfp108 | 0.00739224 | 0.00668607 | 0.00666001 | 2.61E-05 | 0.48961072 | 0.99999994 |
| Cfap53 | 0.00125056 | 0.00115707 | 0.00113139 | 2.57E-05 | 0.43995605 | 0.99999994 |
| Ttll11 | 0.00324919 | 0.0029612 | 0.00293558 | 2.56E-05 | 0.47683918 | 0.99999994 |
| Prickle4 | 0.00134012 | 0.0012378 | 0.00121235 | 2.55E-05 | 0.44443519 | 0.99999994 |
| Etl4 | 0.12252555 | 0.10205045 | 0.10202501 | 2.54E-05 | 0.49933782 | 0.99999994 |
| Nat14 | 0.08364179 | 0.07168319 | 0.07165779 | 2.54E-05 | 0.49905876 | 0.99999994 |
| Gm15873 | 0.00047109 | 0.00045162 | 0.00042643 | 2.52E-05 | 0.34704526 | 0.99999994 |
| Ptpro | 0.00256095 | 0.00233994 | 0.00231485 | 2.51E-05 | 0.471248 | 0.99999994 |
| Bcas3os2 | 0.00521521 | 0.00473041 | 0.00470556 | 2.48E-05 | 0.48598033 | 0.99999994 |
| Endov | 0.06747803 | 0.05844254 | 0.0584178 | 2.47E-05 | 0.49887549 | 0.99999994 |
| Anxa9 | 0.00220828 | 0.00202126 | 0.00199655 | 2.47E-05 | 0.46717047 | 0.99999994 |
| Tnfaip8l2 | 0.00237155 | 0.00216842 | 0.00214393 | 2.45E-05 | 0.46969709 | 0.99999994 |
| Psma8 | 0.00064561 | 0.00060868 | 0.00058433 | 2.44E-05 | 0.39071961 | 0.99999994 |
| Rad54l2 | 0.06875875 | 0.05950134 | 0.05947701 | 2.43E-05 | 0.49891375 | 0.99999994 |
| Shank2 | 0.01202461 | 0.01082392 | 0.01079966 | 2.43E-05 | 0.49403542 | 0.99999994 |
| Glipr2 | 0.00461704 | 0.00419162 | 0.00416754 | 2.41E-05 | 0.48466225 | 0.99999994 |
| Zpld1 | 0.00046968 | 0.00044893 | 0.00042515 | 2.38E-05 | 0.35486292 | 0.99999994 |
| Ubqln1 | 0.32683286 | 0.23269711 | 0.23267367 | 2.34E-05 | 0.49973244 | 0.99999994 |
| Trim62 | 0.03851931 | 0.03401104 | 0.03398763 | 2.34E-05 | 0.49817126 | 0.99999994 |

| | | | | | | |
|---|---|---|---|---|---|---|
| Lect2 | 0.0004693 | 0.00044819 | 0.0004248 | 2.34E-05 | 0.35702159 | 0.99999994 |
| 6330562C20I | 0.00294059 | 0.00268063 | 0.00265733 | 2.33E-05 | 0.47672071 | 0.99999994 |
| Gm26608 | 0.00128537 | 0.00118555 | 0.00116285 | 2.27E-05 | 0.44832261 | 0.99999994 |
| Slc26a3 | 0.00123251 | 0.00113715 | 0.00111508 | 2.21E-05 | 0.44757695 | 0.99999994 |
| Troap | 0.00046776 | 0.00044525 | 0.00042341 | 2.18E-05 | 0.36566306 | 0.99999994 |
| Gm13596 | 0.00046766 | 0.00044507 | 0.00042332 | 2.17E-05 | 0.36620344 | 0.99999994 |
| Robo3 | 0.00046756 | 0.00044488 | 0.00042323 | 2.16E-05 | 0.36674385 | 0.99999994 |
| F12 | 0.00088513 | 0.00082261 | 0.00080098 | 2.16E-05 | 0.42866305 | 0.99999994 |
| Kcnq2 | 0.00115879 | 0.00107003 | 0.00104843 | 2.16E-05 | 0.44545678 | 0.99999994 |
| Ednrb | 0.00257105 | 0.00234549 | 0.00232397 | 2.15E-05 | 0.47542466 | 0.99999994 |
| Ifit1bl1 | 0.00112048 | 0.00103526 | 0.00101379 | 2.15E-05 | 0.44396205 | 0.99999994 |
| Gm28529 | 0.00084308 | 0.00078435 | 0.00076295 | 2.14E-05 | 0.42594345 | 0.99999994 |
| Bche | 0.008126 | 0.00733868 | 0.00731745 | 2.12E-05 | 0.49229905 | 0.99999994 |
| Anxa3 | 0.00587211 | 0.00531711 | 0.00529591 | 2.12E-05 | 0.4893699 | 0.99999994 |
| Bricd5 | 0.00065327 | 0.00061232 | 0.00059125 | 2.11E-05 | 0.40624794 | 0.99999994 |
| mt-Co3 | 4.10515902 | 0.22242529 | 0.22240427 | 2.10E-05 | 0.49974904 | 0.99999994 |
| 9330102E08I | 0.00088108 | 0.00081833 | 0.00079732 | 2.10E-05 | 0.43038556 | 0.99999994 |
| Itih5 | 0.00120774 | 0.00111368 | 0.00109269 | 2.10E-05 | 0.44911765 | 0.99999994 |
| Mmab | 0.04034884 | 0.03557971 | 0.03555879 | 2.09E-05 | 0.49843775 | 0.99999994 |
| Hmg20b | 0.46907599 | 0.30119982 | 0.30117916 | 2.07E-05 | 0.49981788 | 0.99999994 |
| Gpr146 | 0.00847237 | 0.00764805 | 0.00762758 | 2.05E-05 | 0.49287384 | 0.99999994 |
| Slc43a3 | 0.00206643 | 0.00188809 | 0.00186848 | 1.96E-05 | 0.47215091 | 0.99999994 |
| Angptl3 | 0.00046485 | 0.00043973 | 0.00042078 | 1.90E-05 | 0.38215195 | 0.99999994 |
| Mfap5 | 0.00374558 | 0.00340186 | 0.00338292 | 1.89E-05 | 0.48513854 | 0.99999994 |
| Rgs9bp | 0.0030951 | 0.00281547 | 0.00279665 | 1.88E-05 | 0.48213411 | 0.99999994 |
| Ybx2 | 0.00298373 | 0.00271496 | 0.00269622 | 1.87E-05 | 0.48155301 | 0.99999994 |
| Cygb | 0.00169824 | 0.00155463 | 0.00153594 | 1.87E-05 | 0.46773028 | 0.99999994 |
| Hoxaas3 | 0.00302974 | 0.00275606 | 0.00273772 | 1.83E-05 | 0.48221043 | 0.99999994 |
| Kif17 | 0.00927644 | 0.00836508 | 0.00834693 | 1.82E-05 | 0.49422383 | 0.99999994 |
| Iglon5 | 0.00304284 | 0.00276763 | 0.00274953 | 1.81E-05 | 0.48252799 | 0.99999994 |

| | | | | | | |
|---|---|---|---|---|---|---|
| AI593442 | 0.00046367 | 0.0004375 | 0.00041971 | 1.78E-05 | 0.38890971 | 0.99999994 |
| A930017K11 | 0.00046367 | 0.0004375 | 0.00041971 | 1.78E-05 | 0.38890971 | 0.99999994 |
| Gm17203 | 0.00046353 | 0.00043723 | 0.00041958 | 1.77E-05 | 0.38972047 | 0.99999994 |
| Slc15a1 | 0.00582281 | 0.00526927 | 0.00525162 | 1.76E-05 | 0.49107671 | 0.99999994 |
| 4933428P19I | 0.00046348 | 0.00043714 | 0.00041954 | 1.76E-05 | 0.38999071 | 0.99999994 |
| Gm26646 | 0.00077197 | 0.00071619 | 0.00069863 | 1.76E-05 | 0.43358476 | 0.99999994 |
| Hgf | 0.00113906 | 0.00104805 | 0.00103059 | 1.75E-05 | 0.45510581 | 0.99999994 |
| Hnf1b | 0.05915337 | 0.05150711 | 0.05148973 | 1.74E-05 | 0.49910356 | 0.99999994 |
| Adgra2 | 0.00134825 | 0.00123699 | 0.00121969 | 1.73E-05 | 0.46239376 | 0.99999994 |
| Nat8l | 0.0004631 | 0.00043644 | 0.0004192 | 1.72E-05 | 0.39215242 | 0.99999994 |
| Gm11520 | 0.00250383 | 0.00228053 | 0.00226331 | 1.72E-05 | 0.47979983 | 0.99999994 |
| 9230104M06 | 0.00046273 | 0.00043573 | 0.00041886 | 1.69E-05 | 0.3943137 | 0.99999994 |
| Zfp536 | 0.00046268 | 0.00043564 | 0.00041882 | 1.68E-05 | 0.39458383 | 0.99999994 |
| Agtr1b | 0.00046254 | 0.00043537 | 0.00041869 | 1.67E-05 | 0.39539415 | 0.99999994 |
| Gm15774 | 0.00090544 | 0.00083599 | 0.00081935 | 1.66E-05 | 0.44624626 | 0.99999994 |
| Nr4a3 | 0.00187042 | 0.00170778 | 0.00169147 | 1.63E-05 | 0.47440949 | 0.99999994 |
| 6230400D17 | 0.0073058 | 0.00659875 | 0.00658252 | 1.62E-05 | 0.4934518 | 0.99999994 |
| Gm15839 | 0.00156665 | 0.00143315 | 0.00141706 | 1.61E-05 | 0.46988336 | 0.99999994 |
| Cyp2c70 | 0.00046184 | 0.00043406 | 0.00041805 | 1.60E-05 | 0.39944457 | 0.99999994 |
| Gstp2 | 0.00116418 | 0.00106891 | 0.0010533 | 1.56E-05 | 0.46072209 | 0.99999994 |
| Dleu7 | 0.00119905 | 0.00110006 | 0.00108483 | 1.52E-05 | 0.46275437 | 0.99999994 |
| Htr3a | 0.00131573 | 0.00120546 | 0.0011903 | 1.52E-05 | 0.46620433 | 0.99999994 |
| Rasd2 | 0.00067937 | 0.00063001 | 0.00061487 | 1.51E-05 | 0.43492085 | 0.99999994 |
| Adamts14 | 0.00046072 | 0.00043196 | 0.00041704 | 1.49E-05 | 0.40592015 | 0.99999994 |
| Rasal3 | 0.00658928 | 0.00595467 | 0.00593982 | 1.49E-05 | 0.49335982 | 0.99999994 |
| Ppm1m | 0.02055377 | 0.01836903 | 0.01835431 | 1.47E-05 | 0.49786968 | 0.99999994 |
| Inf2 | 0.00138805 | 0.00127028 | 0.00125567 | 1.46E-05 | 0.46912319 | 0.99999994 |
| Gm12359 | 0.00110107 | 0.00101083 | 0.00099625 | 1.46E-05 | 0.46119266 | 0.99999994 |
| Gm15050 | 0.00185574 | 0.00169269 | 0.00167822 | 1.45E-05 | 0.4771116 | 0.99999994 |
| Rassf2 | 0.00147961 | 0.00135262 | 0.00133841 | 1.42E-05 | 0.47181543 | 0.99999994 |

| | | | | | | |
|---|---|---|---|---|---|---|
| Adad1 | 0.00045989 | 0.00043039 | 0.00041629 | 1.41E-05 | 0.41077176 | 0.99999994 |
| Map3k13 | 0.00085136 | 0.00078427 | 0.00077044 | 1.38E-05 | 0.4524611 | 0.99999994 |
| Slc9b2 | 0.0007656 | 0.00070613 | 0.00069288 | 1.33E-05 | 0.44933924 | 0.99999994 |
| Erich2os | 0.00129893 | 0.0011883 | 0.00117511 | 1.32E-05 | 0.47023254 | 0.99999994 |
| 1700102P08I | 0.00089542 | 0.00082346 | 0.00081029 | 1.32E-05 | 0.45692128 | 0.99999994 |
| Esco2 | 0.00093842 | 0.0008623 | 0.00084918 | 1.31E-05 | 0.4590183 | 0.99999994 |
| Klk1b3 | 0.00219491 | 0.0019975 | 0.00198448 | 1.30E-05 | 0.4825888 | 0.99999994 |
| Itpripl2 | 0.00090155 | 0.00082879 | 0.00081583 | 1.30E-05 | 0.45790268 | 0.99999994 |
| Foxf2 | 0.00045869 | 0.00042815 | 0.0004152 | 1.29E-05 | 0.41777028 | 0.99999994 |
| Dph1 | 0.0006654 | 0.00061513 | 0.00060223 | 1.29E-05 | 0.44331043 | 0.99999994 |
| Gm12207 | 0.00662505 | 0.00598457 | 0.00597192 | 1.27E-05 | 0.49437423 | 0.99999994 |
| Fgl1 | 0.00104853 | 0.00096129 | 0.00094875 | 1.25E-05 | 0.46494721 | 0.99999994 |
| Epor | 0.00045804 | 0.00042695 | 0.00041462 | 1.23E-05 | 0.42153342 | 0.99999994 |
| Rrs1 | 0.17265661 | 0.13821508 | 0.13820325 | 1.18E-05 | 0.49977274 | 0.99999994 |
| Exoc3l | 0.00591648 | 0.00534759 | 0.00533576 | 1.18E-05 | 0.49411576 | 0.99999994 |
| Plscr2 | 0.00128199 | 0.00117141 | 0.0011598 | 1.16E-05 | 0.47344027 | 0.99999994 |
| Fam149a | 0.01332157 | 0.01196551 | 0.01195404 | 1.15E-05 | 0.49745141 | 0.99999994 |
| Myct1 | 0.00107973 | 0.00098831 | 0.00097695 | 1.14E-05 | 0.46915382 | 0.99999994 |
| Nxpe3 | 0.01415052 | 0.01270167 | 0.0126908 | 1.09E-05 | 0.49772482 | 0.99999994 |
| Gm5724 | 0.00089828 | 0.00082353 | 0.00081287 | 1.07E-05 | 0.46523205 | 0.99999994 |
| Cdh6 | 0.00080378 | 0.0007377 | 0.00072741 | 1.03E-05 | 0.46248633 | 0.99999994 |
| Gm4221 | 0.00088443 | 0.00081033 | 0.00080035 | 9.98E-06 | 0.46691836 | 0.99999994 |
| Sntb1 | 0.0012022 | 0.00109754 | 0.00108768 | 9.86E-06 | 0.47593994 | 0.99999994 |
| Lrrc9 | 0.00504079 | 0.00455858 | 0.00454872 | 9.86E-06 | 0.49424584 | 0.99999994 |
| Twist1 | 0.00045513 | 0.00042153 | 0.00041198 | 9.55E-06 | 0.43867845 | 0.99999994 |
| Mb | 0.00169045 | 0.0015384 | 0.00152891 | 9.50E-06 | 0.48351317 | 0.99999994 |
| Gimap8 | 0.00308216 | 0.00279447 | 0.00278499 | 9.48E-06 | 0.4909607 | 0.99999994 |
| Gm26920 | 0.00045499 | 0.00042128 | 0.00041186 | 9.42E-06 | 0.43947946 | 0.99999994 |
| Prl8a8 | 0.0004549 | 0.00042111 | 0.00041178 | 9.34E-06 | 0.44001332 | 0.99999994 |
| Ifi44 | 0.00045477 | 0.00042086 | 0.00041165 | 9.21E-06 | 0.44081389 | 0.99999994 |

| | | | | | | |
|---|---|---|---|---|---|---|
| Ackr2 | 0.00089689 | 0.00082077 | 0.00081162 | 9.15E-06 | 0.47007508 | 0.99999994 |
| Lrrc8c | 0.0247043 | 0.02200845 | 0.02199942 | 9.03E-06 | 0.49891005 | 0.99999994 |
| Kdm4d | 0.00089937 | 0.00082264 | 0.00081386 | 8.77E-06 | 0.47140265 | 0.99999994 |
| D630023F18 | 0.00068808 | 0.00063148 | 0.00062275 | 8.74E-06 | 0.462801 | 0.99999994 |
| Dpp4 | 0.0313849 | 0.02783283 | 0.02782417 | 8.66E-06 | 0.49917357 | 0.99999994 |
| Sirt7 | 0.22592586 | 0.17355463 | 0.17354616 | 8.47E-06 | 0.49987043 | 0.99999994 |
| Aplnr | 0.00086151 | 0.0007879 | 0.00077962 | 8.28E-06 | 0.47181691 | 0.99999994 |
| Trmt10c | 0.18625653 | 0.14752741 | 0.14751918 | 8.23E-06 | 0.49985187 | 0.99999994 |
| Lcp2 | 0.00202535 | 0.00183944 | 0.00183138 | 8.06E-06 | 0.48831684 | 0.99999994 |
| Sgk1 | 0.01373538 | 0.01232989 | 0.01232193 | 7.95E-06 | 0.49828592 | 0.99999994 |
| Mtfr2 | 0.00124682 | 0.00113571 | 0.00112801 | 7.70E-06 | 0.48187086 | 0.99999994 |
| BC037039 | 0.00108859 | 0.0009926 | 0.00098497 | 7.64E-06 | 0.47941513 | 0.99999994 |
| Mamdc2 | 0.00089728 | 0.00081887 | 0.00081197 | 6.90E-06 | 0.47744669 | 0.99999994 |
| Platr4 | 0.00045198 | 0.00041573 | 0.00040913 | 6.59E-06 | 0.45729416 | 0.99999994 |
| Irx3 | 0.00045194 | 0.00041564 | 0.00040909 | 6.55E-06 | 0.45755888 | 0.99999994 |
| Calhm2 | 0.00381102 | 0.00344842 | 0.00344187 | 6.55E-06 | 0.49494961 | 0.99999994 |
| Tcaf2 | 0.00045176 | 0.00041532 | 0.00040893 | 6.38E-06 | 0.45861738 | 0.99999994 |
| Nipal4 | 0.00045176 | 0.00041532 | 0.00040893 | 6.38E-06 | 0.45861738 | 0.99999994 |
| Alk | 0.00045167 | 0.00041515 | 0.00040885 | 6.30E-06 | 0.45914641 | 0.99999994 |
| Stac | 0.00045154 | 0.00041491 | 0.00040873 | 6.18E-06 | 0.45993967 | 0.99999994 |
| Npffr2 | 0.00068602 | 0.00062701 | 0.00062089 | 6.12E-06 | 0.47383514 | 0.99999994 |
| Mgam | 0.00222436 | 0.00201671 | 0.00201106 | 5.64E-06 | 0.49254731 | 0.99999994 |
| Kcnj11 | 0.00291332 | 0.00263822 | 0.00263273 | 5.49E-06 | 0.49446088 | 0.99999994 |
| Gm10273 | 0.00581147 | 0.00524666 | 0.00524143 | 5.23E-06 | 0.49735046 | 0.99999994 |
| Ccnf | 0.00497861 | 0.00449802 | 0.0044928 | 5.22E-06 | 0.49691711 | 0.99999994 |
| Fam78b | 0.00087707 | 0.00079874 | 0.00079369 | 5.05E-06 | 0.48309468 | 0.99999994 |
| Bmp8b | 0.00045025 | 0.00041255 | 0.00040757 | 4.98E-06 | 0.46758976 | 0.99999994 |
| Gm17227 | 0.0048125 | 0.00434819 | 0.00434339 | 4.79E-06 | 0.49707042 | 0.99999994 |
| Fcrl6 | 0.00125784 | 0.00114262 | 0.00113797 | 4.65E-06 | 0.48915829 | 0.99999994 |
| Tspan14 | 0.10211483 | 0.08643029 | 0.08642581 | 4.49E-06 | 0.49986213 | 0.99999994 |

| | | | | | | |
|---|---|---|---|---|---|---|
| 8030443G20 | 0.00236348 | 0.00214092 | 0.00213664 | 4.27E-06 | 0.49468746 | 0.99999994 |
| C130050O18 | 0.00044932 | 0.00041085 | 0.00040673 | 4.12E-06 | 0.47310805 | 0.99999994 |
| Banf2os | 0.00116844 | 0.00106067 | 0.00105716 | 3.52E-06 | 0.49116215 | 0.99999994 |
| Pla2g2e | 0.00080527 | 0.00073194 | 0.00072876 | 3.18E-06 | 0.48841478 | 0.99999994 |
| Gm42477 | 0.00390075 | 0.00352569 | 0.0035227 | 2.99E-06 | 0.49774336 | 0.99999994 |
| S1pr5 | 0.00044775 | 0.00040797 | 0.0004053 | 2.67E-06 | 0.48252303 | 0.99999994 |
| Gm13565 | 0.00089041 | 0.00080834 | 0.00080575 | 2.59E-06 | 0.49146419 | 0.99999994 |
| Kcne1 | 0.0007223 | 0.00065603 | 0.00065371 | 2.33E-06 | 0.49054924 | 0.99999994 |
| Hsf4 | 0.004117 | 0.00371898 | 0.00371744 | 1.54E-06 | 0.49890283 | 0.99999994 |
| 4930599N23 | 0.00152541 | 0.00138118 | 0.00137979 | 1.39E-06 | 0.4973275 | 0.99999994 |
| Pstpip1 | 0.00289201 | 0.00261471 | 0.00261351 | 1.20E-06 | 0.49878564 | 0.99999994 |
| Rem1 | 0.00118969 | 0.00107742 | 0.00107637 | 1.05E-06 | 0.49741173 | 0.99999994 |
| Cacna1f | 0.00044587 | 0.00040457 | 0.00040361 | 9.59E-07 | 0.49368868 | 0.99999994 |
| Lrfn2 | 0.00044587 | 0.00040457 | 0.00040361 | 9.59E-07 | 0.49368868 | 0.99999994 |
| Osr2 | 0.00122091 | 0.00110547 | 0.00110459 | 8.83E-07 | 0.49787695 | 0.99999994 |
| Gm12843 | 0.00087623 | 0.00079378 | 0.00079294 | 8.47E-07 | 0.497165 | 0.99999994 |
| L1td1 | 0.00300533 | 0.00271632 | 0.00271571 | 6.11E-07 | 0.49940222 | 0.99999994 |
| Tenm1 | 0.0004451 | 0.00040316 | 0.0004029 | 2.53E-07 | 0.49833511 | 0.99999994 |
| Sox17 | 0 | 0 | 0 | 0 | NA | NA |
| Rgs20 | 0 | 0 | 0 | 0 | NA | NA |
| Npbwr1 | 0 | 0 | 0 | 0 | NA | NA |
| Xkr9 | 0 | 0 | 0 | 0 | NA | NA |
| Gm9947 | 0 | 0 | 0 | 0 | NA | NA |
| Msc | 0 | 0 | 0 | 0 | NA | NA |
| Gm16070 | 0 | 0 | 0 | 0 | NA | NA |
| Gm28287 | 0 | 0 | 0 | 0 | NA | NA |
| Rims1 | 0 | 0 | 0 | 0 | NA | NA |
| Gm37724 | 0 | 0 | 0 | 0 | NA | NA |
| Gm28306 | 0 | 0 | 0 | 0 | NA | NA |
| Cfc1 | 0 | 0 | 0 | 0 | NA | NA |

| | | | | | | |
|---|---|---|---|---|---|---|
| Gm33280 | 0 | 0 | 0 | 0 | NA | NA |
| Gm33533 | 0 | 0 | 0 | 0 | NA | NA |
| 4930556I23R | 0 | 0 | 0 | 0 | NA | NA |
| Npas2 | 0 | 0 | 0 | 0 | NA | NA |
| 1700066B17I | 0 | 0 | 0 | 0 | NA | NA |
| Il1rl1 | 0 | 0 | 0 | 0 | NA | NA |
| Il18rap | 0 | 0 | 0 | 0 | NA | NA |
| Slc9a2 | 0 | 0 | 0 | 0 | NA | NA |
| 4930448I06R | 0 | 0 | 0 | 0 | NA | NA |
| Pantr1 | 0 | 0 | 0 | 0 | NA | NA |
| Pou3f3 | 0 | 0 | 0 | 0 | NA | NA |
| Gm29157 | 0 | 0 | 0 | 0 | NA | NA |
| Mettl21c | 0 | 0 | 0 | 0 | NA | NA |
| Dnah7c | 0 | 0 | 0 | 0 | NA | NA |
| Gm28055 | 0 | 0 | 0 | 0 | NA | NA |
| Gm28551 | 0 | 0 | 0 | 0 | NA | NA |
| Gm10561 | 0 | 0 | 0 | 0 | NA | NA |
| Boll | 0 | 0 | 0 | 0 | NA | NA |
| 4930558J18F | 0 | 0 | 0 | 0 | NA | NA |
| Aox3 | 0 | 0 | 0 | 0 | NA | NA |
| Aox2 | 0 | 0 | 0 | 0 | NA | NA |
| 9530026F06I | 0 | 0 | 0 | 0 | NA | NA |
| Pard3bos1 | 0 | 0 | 0 | 0 | NA | NA |
| Cryge | 0 | 0 | 0 | 0 | NA | NA |
| Abca12 | 0 | 0 | 0 | 0 | NA | NA |
| Gm29185 | 0 | 0 | 0 | 0 | NA | NA |
| Pinc | 0 | 0 | 0 | 0 | NA | NA |
| D530049I02F | 0 | 0 | 0 | 0 | NA | NA |
| Gm29183 | 0 | 0 | 0 | 0 | NA | NA |
| Plcd4 | 0 | 0 | 0 | 0 | NA | NA |

| | | | | | | |
|---|---|---|---|---|---|---|
| Prkag3 | 0 | 0 | 0 | 0 | NA | NA |
| Wnt10a | 0 | 0 | 0 | 0 | NA | NA |
| Gm15179 | 0 | 0 | 0 | 0 | NA | NA |
| Gm28386 | 0 | 0 | 0 | 0 | NA | NA |
| BC035947 | 0 | 0 | 0 | 0 | NA | NA |
| 5730419F03I | 0 | 0 | 0 | 0 | NA | NA |
| Fam124b | 0 | 0 | 0 | 0 | NA | NA |
| Slc16a14 | 0 | 0 | 0 | 0 | NA | NA |
| Htr2b | 0 | 0 | 0 | 0 | NA | NA |
| Gm21972 | 0 | 0 | 0 | 0 | NA | NA |
| Gm16341 | 0 | 0 | 0 | 0 | NA | NA |
| Gm28626 | 0 | 0 | 0 | 0 | NA | NA |
| Gm29374 | 0 | 0 | 0 | 0 | NA | NA |
| Prss56 | 0 | 0 | 0 | 0 | NA | NA |
| Ugt1a7c | 0 | 0 | 0 | 0 | NA | NA |
| Ugt1a6a | 0 | 0 | 0 | 0 | NA | NA |
| Glrp1 | 0 | 0 | 0 | 0 | NA | NA |
| Gm19589 | 0 | 0 | 0 | 0 | NA | NA |
| Gm29336 | 0 | 0 | 0 | 0 | NA | NA |
| Gm28723 | 0 | 0 | 0 | 0 | NA | NA |
| Gm28722 | 0 | 0 | 0 | 0 | NA | NA |
| Rbm44 | 0 | 0 | 0 | 0 | NA | NA |
| Gm29100 | 0 | 0 | 0 | 0 | NA | NA |
| Otos | 0 | 0 | 0 | 0 | NA | NA |
| Gm29483 | 0 | 0 | 0 | 0 | NA | NA |
| Gm29480 | 0 | 0 | 0 | 0 | NA | NA |
| Ankmy1 | 0 | 0 | 0 | 0 | NA | NA |
| Gpr35 | 0 | 0 | 0 | 0 | NA | NA |
| Agxt | 0 | 0 | 0 | 0 | NA | NA |
| Gm10550 | 0 | 0 | 0 | 0 | NA | NA |

| | | | | | | |
|---|---|---|---|---|---|---|
| 1810006J02F | 0 | 0 | 0 | 0 | NA | NA |
| Cdh20 | 0 | 0 | 0 | 0 | NA | NA |
| D630008O14 | 0 | 0 | 0 | 0 | NA | NA |
| Serpinb3a | 0 | 0 | 0 | 0 | NA | NA |
| Serpinb2 | 0 | 0 | 0 | 0 | NA | NA |
| Gm26831 | 0 | 0 | 0 | 0 | NA | NA |
| 3830432H09 | 0 | 0 | 0 | 0 | NA | NA |
| Cfap221 | 0 | 0 | 0 | 0 | NA | NA |
| Marco | 0 | 0 | 0 | 0 | NA | NA |
| Celrr | 0 | 0 | 0 | 0 | NA | NA |
| Acmsd | 0 | 0 | 0 | 0 | NA | NA |
| Thsd7b | 0 | 0 | 0 | 0 | NA | NA |
| Il19 | 0 | 0 | 0 | 0 | NA | NA |
| Il10 | 0 | 0 | 0 | 0 | NA | NA |
| Gm28913 | 0 | 0 | 0 | 0 | NA | NA |
| Lemd1 | 0 | 0 | 0 | 0 | NA | NA |
| Lax1 | 0 | 0 | 0 | 0 | NA | NA |
| Gm1627 | 0 | 0 | 0 | 0 | NA | NA |
| Chit1 | 0 | 0 | 0 | 0 | NA | NA |
| Mybph | 0 | 0 | 0 | 0 | NA | NA |
| Platr1 | 0 | 0 | 0 | 0 | NA | NA |
| Syt2 | 0 | 0 | 0 | 0 | NA | NA |
| Lmod1 | 0 | 0 | 0 | 0 | NA | NA |
| Gm4793 | 0 | 0 | 0 | 0 | NA | NA |
| Mroh3 | 0 | 0 | 0 | 0 | NA | NA |
| Gm26568 | 0 | 0 | 0 | 0 | NA | NA |
| Platr22 | 0 | 0 | 0 | 0 | NA | NA |
| Lhx9 | 0 | 0 | 0 | 0 | NA | NA |
| Kcnt2 | 0 | 0 | 0 | 0 | NA | NA |
| Rgs18 | 0 | 0 | 0 | 0 | NA | NA |

| | | | | | | |
|---|---|---|---|---|---|---|
| Gm29514 | 0 | 0 | 0 | 0 | NA | NA |
| Brinp3 | 0 | 0 | 0 | 0 | NA | NA |
| Ptgs2 | 0 | 0 | 0 | 0 | NA | NA |
| Ptgs2os2 | 0 | 0 | 0 | 0 | NA | NA |
| Prg4 | 0 | 0 | 0 | 0 | NA | NA |
| Nmnat2 | 0 | 0 | 0 | 0 | NA | NA |
| Rgsl1 | 0 | 0 | 0 | 0 | NA | NA |
| Gm29291 | 0 | 0 | 0 | 0 | NA | NA |
| A830008E24 | 0 | 0 | 0 | 0 | NA | NA |
| Gm5532 | 0 | 0 | 0 | 0 | NA | NA |
| 4930439D14 | 0 | 0 | 0 | 0 | NA | NA |
| Gm28694 | 0 | 0 | 0 | 0 | NA | NA |
| 4930562F07I | 0 | 0 | 0 | 0 | NA | NA |
| Tnfsf18 | 0 | 0 | 0 | 0 | NA | NA |
| Fasl | 0 | 0 | 0 | 0 | NA | NA |
| Gm16548 | 0 | 0 | 0 | 0 | NA | NA |
| Dpt | 0 | 0 | 0 | 0 | NA | NA |
| Gm26665 | 0 | 0 | 0 | 0 | NA | NA |
| Mael | 0 | 0 | 0 | 0 | NA | NA |
| 5330438I03R | 0 | 0 | 0 | 0 | NA | NA |
| Gm16701 | 0 | 0 | 0 | 0 | NA | NA |
| Gm2453 | 0 | 0 | 0 | 0 | NA | NA |
| Sh2d1b1 | 0 | 0 | 0 | 0 | NA | NA |
| Fcgr4 | 0 | 0 | 0 | 0 | NA | NA |
| Gm17224 | 0 | 0 | 0 | 0 | NA | NA |
| Aim2 | 0 | 0 | 0 | 0 | NA | NA |
| Ifi205 | 0 | 0 | 0 | 0 | NA | NA |
| Grem2 | 0 | 0 | 0 | 0 | NA | NA |
| Kmo | 0 | 0 | 0 | 0 | NA | NA |
| Gm10518 | 0 | 0 | 0 | 0 | NA | NA |

| | | | | | | |
|---|---|---|---|---|---|---|
| Mixl1 | 0 | 0 | 0 | 0 | NA | NA |
| Lefty2 | 0 | 0 | 0 | 0 | NA | NA |
| Cnih3 | 0 | 0 | 0 | 0 | NA | NA |
| 1700047M11 | 0 | 0 | 0 | 0 | NA | NA |
| Gm37885 | 0 | 0 | 0 | 0 | NA | NA |
| Capn8 | 0 | 0 | 0 | 0 | NA | NA |
| Gm37800 | 0 | 0 | 0 | 0 | NA | NA |
| Gm37662 | 0 | 0 | 0 | 0 | NA | NA |
| Gm34882 | 0 | 0 | 0 | 0 | NA | NA |
| Gm2061 | 0 | 0 | 0 | 0 | NA | NA |
| A430027H14 | 0 | 0 | 0 | 0 | NA | NA |
| Gm29678 | 0 | 0 | 0 | 0 | NA | NA |
| Gm37168 | 0 | 0 | 0 | 0 | NA | NA |
| Cr2 | 0 | 0 | 0 | 0 | NA | NA |
| Gm13180 | 0 | 0 | 0 | 0 | NA | NA |
| Gm13267 | 0 | 0 | 0 | 0 | NA | NA |
| Gm13383 | 0 | 0 | 0 | 0 | NA | NA |
| Gm13391 | 0 | 0 | 0 | 0 | NA | NA |
| Gm13262 | 0 | 0 | 0 | 0 | NA | NA |
| Prkcq | 0 | 0 | 0 | 0 | NA | NA |
| Gm13293 | 0 | 0 | 0 | 0 | NA | NA |
| E030013I19R | 0 | 0 | 0 | 0 | NA | NA |
| C1ql3 | 0 | 0 | 0 | 0 | NA | NA |
| Cubn | 0 | 0 | 0 | 0 | NA | NA |
| St8sia6 | 0 | 0 | 0 | 0 | NA | NA |
| 4921504E06I | 0 | 0 | 0 | 0 | NA | NA |
| Gm13375 | 0 | 0 | 0 | 0 | NA | NA |
| Nxph2 | 0 | 0 | 0 | 0 | NA | NA |
| Il1rn | 0 | 0 | 0 | 0 | NA | NA |
| Pax8 | 0 | 0 | 0 | 0 | NA | NA |

| | | | | | | |
|---|---|---|---|---|---|---|
| Entpd8 | 0 | 0 | 0 | 0 | NA | NA |
| Cysrt1 | 0 | 0 | 0 | 0 | NA | NA |
| Fcnaos | 0 | 0 | 0 | 0 | NA | NA |
| Lcn8 | 0 | 0 | 0 | 0 | NA | NA |
| Obp2a | 0 | 0 | 0 | 0 | NA | NA |
| Sohlh1 | 0 | 0 | 0 | 0 | NA | NA |
| Snhg7os | 0 | 0 | 0 | 0 | NA | NA |
| Stkld1 | 0 | 0 | 0 | 0 | NA | NA |
| Dbh | 0 | 0 | 0 | 0 | NA | NA |
| Sardhos | 0 | 0 | 0 | 0 | NA | NA |
| Gbgt1 | 0 | 0 | 0 | 0 | NA | NA |
| Gfi1b | 0 | 0 | 0 | 0 | NA | NA |
| Ntng2 | 0 | 0 | 0 | 0 | NA | NA |
| Gm13402 | 0 | 0 | 0 | 0 | NA | NA |
| Gm14487 | 0 | 0 | 0 | 0 | NA | NA |
| Gm13523 | 0 | 0 | 0 | 0 | NA | NA |
| Gm13528 | 0 | 0 | 0 | 0 | NA | NA |
| Gm13403 | 0 | 0 | 0 | 0 | NA | NA |
| C230014O12 | 0 | 0 | 0 | 0 | NA | NA |
| Gm13431 | 0 | 0 | 0 | 0 | NA | NA |
| Nr5a1 | 0 | 0 | 0 | 0 | NA | NA |
| Gm13479 | 0 | 0 | 0 | 0 | NA | NA |
| Gm13481 | 0 | 0 | 0 | 0 | NA | NA |
| Gm13499 | 0 | 0 | 0 | 0 | NA | NA |
| Gm13490 | 0 | 0 | 0 | 0 | NA | NA |
| Galnt5 | 0 | 0 | 0 | 0 | NA | NA |
| Ermn | 0 | 0 | 0 | 0 | NA | NA |
| Gm13546 | 0 | 0 | 0 | 0 | NA | NA |
| Gm13571 | 0 | 0 | 0 | 0 | NA | NA |
| Tbr1 | 0 | 0 | 0 | 0 | NA | NA |

| | | | | | | |
|---|---|---|---|---|---|---|
| Csrnp3 | 0 | 0 | 0 | 0 | NA | NA |
| Scn1a | 0 | 0 | 0 | 0 | NA | NA |
| B3galt1 | 0 | 0 | 0 | 0 | NA | NA |
| Myo3b | 0 | 0 | 0 | 0 | NA | NA |
| Sp5 | 0 | 0 | 0 | 0 | NA | NA |
| Dlx1as | 0 | 0 | 0 | 0 | NA | NA |
| Dlx1 | 0 | 0 | 0 | 0 | NA | NA |
| Sp9 | 0 | 0 | 0 | 0 | NA | NA |
| Hoxd9 | 0 | 0 | 0 | 0 | NA | NA |
| Hoxd3os1 | 0 | 0 | 0 | 0 | NA | NA |
| Hoxd8 | 0 | 0 | 0 | 0 | NA | NA |
| Hoxd3 | 0 | 0 | 0 | 0 | NA | NA |
| Haglr | 0 | 0 | 0 | 0 | NA | NA |
| Gm14424 | 0 | 0 | 0 | 0 | NA | NA |
| Gm14426 | 0 | 0 | 0 | 0 | NA | NA |
| Hoxd1 | 0 | 0 | 0 | 0 | NA | NA |
| Gm14461 | 0 | 0 | 0 | 0 | NA | NA |
| Ppp1r1c | 0 | 0 | 0 | 0 | NA | NA |
| Gm28675 | 0 | 0 | 0 | 0 | NA | NA |
| Calcrl | 0 | 0 | 0 | 0 | NA | NA |
| Gm13710 | 0 | 0 | 0 | 0 | NA | NA |
| Smtnl1 | 0 | 0 | 0 | 0 | NA | NA |
| Prg2 | 0 | 0 | 0 | 0 | NA | NA |
| P2rx3 | 0 | 0 | 0 | 0 | NA | NA |
| Pramel7 | 0 | 0 | 0 | 0 | NA | NA |
| Alx4 | 0 | 0 | 0 | 0 | NA | NA |
| Lrrc4c | 0 | 0 | 0 | 0 | NA | NA |
| Gm13920 | 0 | 0 | 0 | 0 | NA | NA |
| 4931422A03 | 0 | 0 | 0 | 0 | NA | NA |
| A930018P22 | 0 | 0 | 0 | 0 | NA | NA |

| | | | | | | |
|---|---|---|---|---|---|---|
| Ano3 | 0 | 0 | 0 | 0 | NA | NA |
| Chrm5 | 0 | 0 | 0 | 0 | NA | NA |
| Tmco5b | 0 | 0 | 0 | 0 | NA | NA |
| Gm13974 | 0 | 0 | 0 | 0 | NA | NA |
| 3110099E03I | 0 | 0 | 0 | 0 | NA | NA |
| Gm13977 | 0 | 0 | 0 | 0 | NA | NA |
| Gpr176 | 0 | 0 | 0 | 0 | NA | NA |
| Ankrd63 | 0 | 0 | 0 | 0 | NA | NA |
| Plcb2 | 0 | 0 | 0 | 0 | NA | NA |
| Pla2g4e | 0 | 0 | 0 | 0 | NA | NA |
| Pla2g4d | 0 | 0 | 0 | 0 | NA | NA |
| Serinc4 | 0 | 0 | 0 | 0 | NA | NA |
| Mageb3 | 0 | 0 | 0 | 0 | NA | NA |
| Patl2 | 0 | 0 | 0 | 0 | NA | NA |
| Trim69 | 0 | 0 | 0 | 0 | NA | NA |
| Duoxa2 | 0 | 0 | 0 | 0 | NA | NA |
| Gm9913 | 0 | 0 | 0 | 0 | NA | NA |
| Atp8b4 | 0 | 0 | 0 | 0 | NA | NA |
| 4933427J07F | 0 | 0 | 0 | 0 | NA | NA |
| Gm14029 | 0 | 0 | 0 | 0 | NA | NA |
| A730036I17F | 0 | 0 | 0 | 0 | NA | NA |
| Il1a | 0 | 0 | 0 | 0 | NA | NA |
| Il1bos | 0 | 0 | 0 | 0 | NA | NA |
| Pdyn | 0 | 0 | 0 | 0 | NA | NA |
| Prokr2 | 0 | 0 | 0 | 0 | NA | NA |
| 4921508D12I | 0 | 0 | 0 | 0 | NA | NA |
| 1700026D11I | 0 | 0 | 0 | 0 | NA | NA |
| Gm14055 | 0 | 0 | 0 | 0 | NA | NA |
| Sptlc3 | 0 | 0 | 0 | 0 | NA | NA |
| Otor | 0 | 0 | 0 | 0 | NA | NA |

| | | | | | | |
|---|---|---|---|---|---|---|
| Nkx2-2os | 0 | 0 | 0 | 0 | NA | NA |
| Cst12 | 0 | 0 | 0 | 0 | NA | NA |
| Cst13 | 0 | 0 | 0 | 0 | NA | NA |
| Cst3 | 1.9281702 | 0.37251798 | 0.37251798 | 0 | 0.5 | 0.99999994 |
| Gm14139 | 0 | 0 | 0 | 0 | NA | NA |
| Gm14124 | 0 | 0 | 0 | 0 | NA | NA |
| Gm10130 | 0 | 0 | 0 | 0 | NA | NA |
| Zfp345 | 0 | 0 | 0 | 0 | NA | NA |
| Cst7 | 0 | 0 | 0 | 0 | NA | NA |
| Acss1 | 0 | 0 | 0 | 0 | NA | NA |
| Nrsn2 | 0 | 0 | 0 | 0 | NA | NA |
| Gm14161 | 0 | 0 | 0 | 0 | NA | NA |
| Defb36 | 0 | 0 | 0 | 0 | NA | NA |
| Defb25 | 0 | 0 | 0 | 0 | NA | NA |
| Mylk2 | 0 | 0 | 0 | 0 | NA | NA |
| Efcab8 | 0 | 0 | 0 | 0 | NA | NA |
| Bpifa6 | 0 | 0 | 0 | 0 | NA | NA |
| Bpifb1 | 0 | 0 | 0 | 0 | NA | NA |
| Bpifb5 | 0 | 0 | 0 | 0 | NA | NA |
| Gm14214 | 0 | 0 | 0 | 0 | NA | NA |
| Gdf5 | 0 | 0 | 0 | 0 | NA | NA |
| Mroh8 | 0 | 0 | 0 | 0 | NA | NA |
| Ghrh | 0 | 0 | 0 | 0 | NA | NA |
| Platr27 | 0 | 0 | 0 | 0 | NA | NA |
| 2010009K17I | 0 | 0 | 0 | 0 | NA | NA |
| 1700060C20I | 0 | 0 | 0 | 0 | NA | NA |
| Emilin3 | 0 | 0 | 0 | 0 | NA | NA |
| Gtsf1l | 0 | 0 | 0 | 0 | NA | NA |
| Jph2 | 0 | 0 | 0 | 0 | NA | NA |
| Hnf4aos | 0 | 0 | 0 | 0 | NA | NA |

| | | | | | | |
|---|---|---|---|---|---|---|
| Kcns1 | 0 | 0 | 0 | 0 | NA | NA |
| Svs4 | 0 | 0 | 0 | 0 | NA | NA |
| Gm14317 | 0 | 0 | 0 | 0 | NA | NA |
| Wfdc6a | 0 | 0 | 0 | 0 | NA | NA |
| Wfdc8 | 0 | 0 | 0 | 0 | NA | NA |
| Wfdc9 | 0 | 0 | 0 | 0 | NA | NA |
| Gm11457 | 0 | 0 | 0 | 0 | NA | NA |
| Mmp9 | 0 | 0 | 0 | 0 | NA | NA |
| Zfp663 | 0 | 0 | 0 | 0 | NA | NA |
| Ocstamp | 0 | 0 | 0 | 0 | NA | NA |
| Slc13a3 | 0 | 0 | 0 | 0 | NA | NA |
| Gm14235 | 0 | 0 | 0 | 0 | NA | NA |
| Gm9873 | 0 | 0 | 0 | 0 | NA | NA |
| Gm16796 | 0 | 0 | 0 | 0 | NA | NA |
| Gm14275 | 0 | 0 | 0 | 0 | NA | NA |
| Cass4 | 0 | 0 | 0 | 0 | NA | NA |
| Fam209 | 0 | 0 | 0 | 0 | NA | NA |
| Spo11 | 0 | 0 | 0 | 0 | NA | NA |
| Gm14453 | 0 | 0 | 0 | 0 | NA | NA |
| Pmepa1os | 0 | 0 | 0 | 0 | NA | NA |
| 1700010B08 | 0 | 0 | 0 | 0 | NA | NA |
| Tubb1 | 0 | 0 | 0 | 0 | NA | NA |
| Gm14393 | 0 | 0 | 0 | 0 | NA | NA |
| Gm14434 | 0 | 0 | 0 | 0 | NA | NA |
| Gm14308 | 0 | 0 | 0 | 0 | NA | NA |
| Sycp2 | 0 | 0 | 0 | 0 | NA | NA |
| Rbbp8nl | 0 | 0 | 0 | 0 | NA | NA |
| Ntsr1 | 0 | 0 | 0 | 0 | NA | NA |
| Gm14342 | 0 | 0 | 0 | 0 | NA | NA |
| Birc7 | 0 | 0 | 0 | 0 | NA | NA |

| | | | | | | |
|---|---|---|---|---|---|---|
| Ccnb3 | 0 | 0 | 0 | 0 | NA | NA |
| Foxp3 | 0 | 0 | 0 | 0 | NA | NA |
| Gm14820 | 0 | 0 | 0 | 0 | NA | NA |
| Gm14493 | 0 | 0 | 0 | 0 | NA | NA |
| Gm14635 | 0 | 0 | 0 | 0 | NA | NA |
| Gpr34 | 0 | 0 | 0 | 0 | NA | NA |
| Ndp | 0 | 0 | 0 | 0 | NA | NA |
| Zfp300 | 0 | 0 | 0 | 0 | NA | NA |
| Gm14549 | 0 | 0 | 0 | 0 | NA | NA |
| Akap14 | 0 | 0 | 0 | 0 | NA | NA |
| Rhox5 | 0 | 0 | 0 | 0 | NA | NA |
| Sash3 | 0 | 0 | 0 | 0 | NA | NA |
| Usp26 | 0 | 0 | 0 | 0 | NA | NA |
| Gm28730 | 0 | 0 | 0 | 0 | NA | NA |
| Gm773 | 0 | 0 | 0 | 0 | NA | NA |
| Mcf2 | 0 | 0 | 0 | 0 | NA | NA |
| Slitrk4 | 0 | 0 | 0 | 0 | NA | NA |
| Slitrk2 | 0 | 0 | 0 | 0 | NA | NA |
| Fmr1nb | 0 | 0 | 0 | 0 | NA | NA |
| Gm6812 | 0 | 0 | 0 | 0 | NA | NA |
| 1700020N15 | 0 | 0 | 0 | 0 | NA | NA |
| Gpr50 | 0 | 0 | 0 | 0 | NA | NA |
| Gm1141 | 0 | 0 | 0 | 0 | NA | NA |
| Gabra3 | 0 | 0 | 0 | 0 | NA | NA |
| Pnma5 | 0 | 0 | 0 | 0 | NA | NA |
| Tktl1 | 0 | 0 | 0 | 0 | NA | NA |
| Fthl17a | 0 | 0 | 0 | 0 | NA | NA |
| Gm14762 | 0 | 0 | 0 | 0 | NA | NA |
| Nr0b1 | 0 | 0 | 0 | 0 | NA | NA |
| Gm27000 | 0 | 0 | 0 | 0 | NA | NA |

| | | | | | | |
|---|---|---|---|---|---|---|
| Gm14798 | 0 | 0 | 0 | 0 | NA | NA |
| Asb12 | 0 | 0 | 0 | 0 | NA | NA |
| F630028O10 | 0 | 0 | 0 | 0 | NA | NA |
| Vsig4 | 0 | 0 | 0 | 0 | NA | NA |
| Hsf3 | 0 | 0 | 0 | 0 | NA | NA |
| Gpr165 | 0 | 0 | 0 | 0 | NA | NA |
| Ar | 0 | 0 | 0 | 0 | NA | NA |
| Gm14808 | 0 | 0 | 0 | 0 | NA | NA |
| Gdpd2 | 0 | 0 | 0 | 0 | NA | NA |
| Slc7a3 | 0 | 0 | 0 | 0 | NA | NA |
| Cxcr3 | 0 | 0 | 0 | 0 | NA | NA |
| Gm4779 | 0 | 0 | 0 | 0 | NA | NA |
| 8030474K03I | 0 | 0 | 0 | 0 | NA | NA |
| Fgf16 | 0 | 0 | 0 | 0 | NA | NA |
| Fndc3c1 | 0 | 0 | 0 | 0 | NA | NA |
| Gm5127 | 0 | 0 | 0 | 0 | NA | NA |
| P2ry10 | 0 | 0 | 0 | 0 | NA | NA |
| Gpr174 | 0 | 0 | 0 | 0 | NA | NA |
| Tex16 | 0 | 0 | 0 | 0 | NA | NA |
| Tnmd | 0 | 0 | 0 | 0 | NA | NA |
| Nox1 | 0 | 0 | 0 | 0 | NA | NA |
| Cenpi | 0 | 0 | 0 | 0 | NA | NA |
| Drp2 | 0 | 0 | 0 | 0 | NA | NA |
| Tceal6 | 0 | 0 | 0 | 0 | NA | NA |
| Nxf3 | 0 | 0 | 0 | 0 | NA | NA |
| Kir3dl1 | 0 | 0 | 0 | 0 | NA | NA |
| Il1rapl2 | 0 | 0 | 0 | 0 | NA | NA |
| 4930513O06 | 0 | 0 | 0 | 0 | NA | NA |
| Ripply1 | 0 | 0 | 0 | 0 | NA | NA |
| Eif2c5 | 0 | 0 | 0 | 0 | NA | NA |

| | | | | | | |
|---|---|---|---|---|---|---|
| Vsig1 | 0 | 0 | 0 | 0 | NA | NA |
| Gucy2f | 0 | 0 | 0 | 0 | NA | NA |
| Htr2c | 0 | 0 | 0 | 0 | NA | NA |
| Gm15104 | 0 | 0 | 0 | 0 | NA | NA |
| Gm15138 | 0 | 0 | 0 | 0 | NA | NA |
| Gpr143 | 0 | 0 | 0 | 0 | NA | NA |
| Ptchd1 | 0 | 0 | 0 | 0 | NA | NA |
| Gm15155 | 0 | 0 | 0 | 0 | NA | NA |
| Ppef1 | 0 | 0 | 0 | 0 | NA | NA |
| Rs1 | 0 | 0 | 0 | 0 | NA | NA |
| Bmx | 0 | 0 | 0 | 0 | NA | NA |
| Gm8817 | 0 | 0 | 0 | 0 | NA | NA |
| Gm15232 | 0 | 0 | 0 | 0 | NA | NA |
| Tlr8 | 0 | 0 | 0 | 0 | NA | NA |
| Tlr7 | 0 | 0 | 0 | 0 | NA | NA |
| Frmpd4 | 0 | 0 | 0 | 0 | NA | NA |
| Gm17308 | 0 | 0 | 0 | 0 | NA | NA |
| Il7 | 0 | 0 | 0 | 0 | NA | NA |
| Gm16685 | 0 | 0 | 0 | 0 | NA | NA |
| Slc10a5 | 0 | 0 | 0 | 0 | NA | NA |
| Gm2464 | 0 | 0 | 0 | 0 | NA | NA |
| Car1 | 0 | 0 | 0 | 0 | NA | NA |
| Gm5150 | 0 | 0 | 0 | 0 | NA | NA |
| Trim55 | 0 | 0 | 0 | 0 | NA | NA |
| Gm43674 | 0 | 0 | 0 | 0 | NA | NA |
| Gm33051 | 0 | 0 | 0 | 0 | NA | NA |
| Gm37867 | 0 | 0 | 0 | 0 | NA | NA |
| Egfem1 | 0 | 0 | 0 | 0 | NA | NA |
| Lrrc31 | 0 | 0 | 0 | 0 | NA | NA |
| Gm43666 | 0 | 0 | 0 | 0 | NA | NA |

| | | | | | | |
|---|---|---|---|---|---|---|
| Sox2 | 0 | 0 | 0 | 0 | NA | NA |
| Trpc3 | 0 | 0 | 0 | 0 | NA | NA |
| Gm29230 | 0 | 0 | 0 | 0 | NA | NA |
| Gm20402 | 0 | 0 | 0 | 0 | NA | NA |
| Stoml3 | 0 | 0 | 0 | 0 | NA | NA |
| Sertm1 | 0 | 0 | 0 | 0 | NA | NA |
| Ccdc169 | 0 | 0 | 0 | 0 | NA | NA |
| Gm43549 | 0 | 0 | 0 | 0 | NA | NA |
| Mab21l1 | 0 | 0 | 0 | 0 | NA | NA |
| Gm37696 | 0 | 0 | 0 | 0 | NA | NA |
| 9330121J05F | 0 | 0 | 0 | 0 | NA | NA |
| A330015K06 | 0 | 0 | 0 | 0 | NA | NA |
| Gm35584 | 0 | 0 | 0 | 0 | NA | NA |
| Gm17641 | 0 | 0 | 0 | 0 | NA | NA |
| Arl14 | 0 | 0 | 0 | 0 | NA | NA |
| Sptssb | 0 | 0 | 0 | 0 | NA | NA |
| Slitrk3 | 0 | 0 | 0 | 0 | NA | NA |
| Platr10 | 0 | 0 | 0 | 0 | NA | NA |
| Gm37256 | 0 | 0 | 0 | 0 | NA | NA |
| Rxfp1 | 0 | 0 | 0 | 0 | NA | NA |
| Gm36569 | 0 | 0 | 0 | 0 | NA | NA |
| A330069K06 | 0 | 0 | 0 | 0 | NA | NA |
| A830029E22 | 0 | 0 | 0 | 0 | NA | NA |
| Fga | 0 | 0 | 0 | 0 | NA | NA |
| Gm10710 | 0 | 0 | 0 | 0 | NA | NA |
| Dchs2 | 0 | 0 | 0 | 0 | NA | NA |
| Cd5l | 0 | 0 | 0 | 0 | NA | NA |
| Fcrl1 | 0 | 0 | 0 | 0 | NA | NA |
| AI849053 | 0 | 0 | 0 | 0 | NA | NA |
| Gm16069 | 0 | 0 | 0 | 0 | NA | NA |

| | | | | | | |
|---|---|---|---|---|---|---|
| 4731419I09R | 0 | 0 | 0 | 0 | NA | NA |
| Dcst1 | 0 | 0 | 0 | 0 | NA | NA |
| Gm16048 | 0 | 0 | 0 | 0 | NA | NA |
| S100a2 | 0 | 0 | 0 | 0 | NA | NA |
| S100a5 | 0 | 0 | 0 | 0 | NA | NA |
| Sprr2b | 0 | 0 | 0 | 0 | NA | NA |
| Sprr2i | 0 | 0 | 0 | 0 | NA | NA |
| Lce6a | 0 | 0 | 0 | 0 | NA | NA |
| Lce3a | 0 | 0 | 0 | 0 | NA | NA |
| Lce3c | 0 | 0 | 0 | 0 | NA | NA |
| Crct1 | 0 | 0 | 0 | 0 | NA | NA |
| Selenbp2 | 0 | 0 | 0 | 0 | NA | NA |
| Adam30 | 0 | 0 | 0 | 0 | NA | NA |
| Reg4 | 0 | 0 | 0 | 0 | NA | NA |
| Hsd3b1 | 0 | 0 | 0 | 0 | NA | NA |
| Cd101 | 0 | 0 | 0 | 0 | NA | NA |
| Gm16160 | 0 | 0 | 0 | 0 | NA | NA |
| Gm43064 | 0 | 0 | 0 | 0 | NA | NA |
| Gm5546 | 0 | 0 | 0 | 0 | NA | NA |
| Wnt2b | 0 | 0 | 0 | 0 | NA | NA |
| Kcnd3os | 0 | 0 | 0 | 0 | NA | NA |
| Kcnd3 | 0 | 0 | 0 | 0 | NA | NA |
| I830077J02R | 0 | 0 | 0 | 0 | NA | NA |
| Chia1 | 0 | 0 | 0 | 0 | NA | NA |
| Chil3 | 0 | 0 | 0 | 0 | NA | NA |
| AI504432 | 0 | 0 | 0 | 0 | NA | NA |
| Kcnc4 | 0 | 0 | 0 | 0 | NA | NA |
| Gstm6 | 0 | 0 | 0 | 0 | NA | NA |
| Gm12498 | 0 | 0 | 0 | 0 | NA | NA |
| Gm12525 | 0 | 0 | 0 | 0 | NA | NA |

| | | | | | | |
|---|---|---|---|---|---|---|
| Mybphl | 0 | 0 | 0 | 0 | NA | NA |
| Aknad1 | 0 | 0 | 0 | 0 | NA | NA |
| Fndc7 | 0 | 0 | 0 | 0 | NA | NA |
| Gm9889 | 0 | 0 | 0 | 0 | NA | NA |
| Gpr88 | 0 | 0 | 0 | 0 | NA | NA |
| Plppr5 | 0 | 0 | 0 | 0 | NA | NA |
| Gm43569 | 0 | 0 | 0 | 0 | NA | NA |
| Ndst3 | 0 | 0 | 0 | 0 | NA | NA |
| Arsj | 0 | 0 | 0 | 0 | NA | NA |
| Gm15551 | 0 | 0 | 0 | 0 | NA | NA |
| 6330410L21I | 0 | 0 | 0 | 0 | NA | NA |
| 4930534D22I | 0 | 0 | 0 | 0 | NA | NA |
| Gm29865 | 0 | 0 | 0 | 0 | NA | NA |
| Gm29811 | 0 | 0 | 0 | 0 | NA | NA |
| Tacr3 | 0 | 0 | 0 | 0 | NA | NA |
| Gm16559 | 0 | 0 | 0 | 0 | NA | NA |
| Stpg2 | 0 | 0 | 0 | 0 | NA | NA |
| Gbp5 | 0 | 0 | 0 | 0 | NA | NA |
| Gbp2b | 0 | 0 | 0 | 0 | NA | NA |
| 4930519L02I | 0 | 0 | 0 | 0 | NA | NA |
| Gm33651 | 0 | 0 | 0 | 0 | NA | NA |
| Gm16233 | 0 | 0 | 0 | 0 | NA | NA |
| Clca2 | 0 | 0 | 0 | 0 | NA | NA |
| Lpar3 | 0 | 0 | 0 | 0 | NA | NA |
| Gm10636 | 0 | 0 | 0 | 0 | NA | NA |
| Gm30382 | 0 | 0 | 0 | 0 | NA | NA |
| 4930555A03I | 0 | 0 | 0 | 0 | NA | NA |
| Ptgfr | 0 | 0 | 0 | 0 | NA | NA |
| AI115009 | 0 | 0 | 0 | 0 | NA | NA |
| 5730460C07I | 0 | 0 | 0 | 0 | NA | NA |

| | | | | | | |
|---|---|---|---|---|---|---|
| Asb17os | 0 | 0 | 0 | 0 | NA | NA |
| Msh4 | 0 | 0 | 0 | 0 | NA | NA |
| 1810013D15 | 0 | 0 | 0 | 0 | NA | NA |
| Gm11802 | 0 | 0 | 0 | 0 | NA | NA |
| Gm11795 | 0 | 0 | 0 | 0 | NA | NA |
| Gdf6 | 0 | 0 | 0 | 0 | NA | NA |
| Gm11840 | 0 | 0 | 0 | 0 | NA | NA |
| Slc26a7 | 0 | 0 | 0 | 0 | NA | NA |
| Necab1 | 0 | 0 | 0 | 0 | NA | NA |
| Slc7a13 | 0 | 0 | 0 | 0 | NA | NA |
| Atp6v0d2 | 0 | 0 | 0 | 0 | NA | NA |
| Fut9 | 0 | 0 | 0 | 0 | NA | NA |
| Gm11915 | 0 | 0 | 0 | 0 | NA | NA |
| Bach2os | 0 | 0 | 0 | 0 | NA | NA |
| Gabrr1 | 0 | 0 | 0 | 0 | NA | NA |
| Cga | 0 | 0 | 0 | 0 | NA | NA |
| Gm12371 | 0 | 0 | 0 | 0 | NA | NA |
| 1700009N14 | 0 | 0 | 0 | 0 | NA | NA |
| Arid3c | 0 | 0 | 0 | 0 | NA | NA |
| Gm10600 | 0 | 0 | 0 | 0 | NA | NA |
| Gm2564 | 0 | 0 | 0 | 0 | NA | NA |
| Ccl27b | 0 | 0 | 0 | 0 | NA | NA |
| Ccl19 | 0 | 0 | 0 | 0 | NA | NA |
| 1700022I11R | 0 | 0 | 0 | 0 | NA | NA |
| Hrct1 | 0 | 0 | 0 | 0 | NA | NA |
| Acnat2 | 0 | 0 | 0 | 0 | NA | NA |
| Acnat1 | 0 | 0 | 0 | 0 | NA | NA |
| Baat | 0 | 0 | 0 | 0 | NA | NA |
| 4930522O17 | 0 | 0 | 0 | 0 | NA | NA |
| Gm12511 | 0 | 0 | 0 | 0 | NA | NA |

| | | | | | | |
|---|---|---|---|---|---|---|
| Gm11214 | 0 | 0 | 0 | 0 | NA | NA |
| Tnfsf15 | 0 | 0 | 0 | 0 | NA | NA |
| Tnfsf8 | 0 | 0 | 0 | 0 | NA | NA |
| Astn2 | 0 | 0 | 0 | 0 | NA | NA |
| Tlr4 | 0 | 0 | 0 | 0 | NA | NA |
| Saxo1 | 0 | 0 | 0 | 0 | NA | NA |
| Tek | 0 | 0 | 0 | 0 | NA | NA |
| Eqtn | 0 | 0 | 0 | 0 | NA | NA |
| Kank4 | 0 | 0 | 0 | 0 | NA | NA |
| Kank4os | 0 | 0 | 0 | 0 | NA | NA |
| Gm12688 | 0 | 0 | 0 | 0 | NA | NA |
| Foxd3 | 0 | 0 | 0 | 0 | NA | NA |
| Ube2uos | 0 | 0 | 0 | 0 | NA | NA |
| Raver2 | 0 | 0 | 0 | 0 | NA | NA |
| 0610043K17I | 0 | 0 | 0 | 0 | NA | NA |
| Gm44037 | 0 | 0 | 0 | 0 | NA | NA |
| 2700068H02I | 0 | 0 | 0 | 0 | NA | NA |
| Skint8 | 0 | 0 | 0 | 0 | NA | NA |
| Tal1 | 0 | 0 | 0 | 0 | NA | NA |
| Cyp4x1os | 0 | 0 | 0 | 0 | NA | NA |
| Mpl | 0 | 0 | 0 | 0 | NA | NA |
| Tie1 | 0 | 0 | 0 | 0 | NA | NA |
| Gm12866 | 0 | 0 | 0 | 0 | NA | NA |
| Gm12867 | 0 | 0 | 0 | 0 | NA | NA |
| 9530002B09I | 0 | 0 | 0 | 0 | NA | NA |
| Oxct2b | 0 | 0 | 0 | 0 | NA | NA |
| Bmp8a | 0 | 0 | 0 | 0 | NA | NA |
| Epha10 | 0 | 0 | 0 | 0 | NA | NA |
| 2610028E06I | 0 | 0 | 0 | 0 | NA | NA |
| Tlr12 | 0 | 0 | 0 | 0 | NA | NA |

| | | | | | | |
|---|---|---|---|---|---|---|
| Fam167b | 0 | 0 | 0 | 0 | NA | NA |
| Spocd1 | 0 | 0 | 0 | 0 | NA | NA |
| Hcrtr1 | 0 | 0 | 0 | 0 | NA | NA |
| Gm10570 | 0 | 0 | 0 | 0 | NA | NA |
| Oprd1 | 0 | 0 | 0 | 0 | NA | NA |
| Smpdl3b | 0 | 0 | 0 | 0 | NA | NA |
| Themis2 | 0 | 0 | 0 | 0 | NA | NA |
| E130218I03R | 0 | 0 | 0 | 0 | NA | NA |
| Trim63 | 0 | 0 | 0 | 0 | NA | NA |
| Slc30a2 | 0 | 0 | 0 | 0 | NA | NA |
| Il22ra1 | 0 | 0 | 0 | 0 | NA | NA |
| Cnr2 | 0 | 0 | 0 | 0 | NA | NA |
| Gm13008 | 0 | 0 | 0 | 0 | NA | NA |
| Fam43b | 0 | 0 | 0 | 0 | NA | NA |
| AB041806 | 0 | 0 | 0 | 0 | NA | NA |
| Htr6 | 0 | 0 | 0 | 0 | NA | NA |
| Tas1r2 | 0 | 0 | 0 | 0 | NA | NA |
| Gm13016 | 0 | 0 | 0 | 0 | NA | NA |
| Padi6 | 0 | 0 | 0 | 0 | NA | NA |
| Spata21 | 0 | 0 | 0 | 0 | NA | NA |
| Gm13056 | 0 | 0 | 0 | 0 | NA | NA |
| Gm29367 | 0 | 0 | 0 | 0 | NA | NA |
| Fv1 | 0 | 0 | 0 | 0 | NA | NA |
| Nppb | 0 | 0 | 0 | 0 | NA | NA |
| Nppa | 0 | 0 | 0 | 0 | NA | NA |
| Gm13054 | 0 | 0 | 0 | 0 | NA | NA |
| Ube4bos1 | 0 | 0 | 0 | 0 | NA | NA |
| Slc45a1 | 0 | 0 | 0 | 0 | NA | NA |
| Gm13049 | 0 | 0 | 0 | 0 | NA | NA |
| Tas1r1 | 0 | 0 | 0 | 0 | NA | NA |

| | | | | | | |
|---|---|---|---|---|---|---|
| Gm13134 | 0 | 0 | 0 | 0 | NA | NA |
| Gm13112 | 0 | 0 | 0 | 0 | NA | NA |
| 1700109H08 | 0 | 0 | 0 | 0 | NA | NA |
| Gm43825 | 0 | 0 | 0 | 0 | NA | NA |
| Steap4 | 0 | 0 | 0 | 0 | NA | NA |
| Abcb1a | 0 | 0 | 0 | 0 | NA | NA |
| Grm3 | 0 | 0 | 0 | 0 | NA | NA |
| Gm8953 | 0 | 0 | 0 | 0 | NA | NA |
| Gm21149 | 0 | 0 | 0 | 0 | NA | NA |
| Gm21083 | 0 | 0 | 0 | 0 | NA | NA |
| A930003O13 | 0 | 0 | 0 | 0 | NA | NA |
| Lhfpl3 | 0 | 0 | 0 | 0 | NA | NA |
| Gbx1 | 0 | 0 | 0 | 0 | NA | NA |
| Asb10 | 0 | 0 | 0 | 0 | NA | NA |
| Gm10062 | 0 | 0 | 0 | 0 | NA | NA |
| 9530036O11 | 0 | 0 | 0 | 0 | NA | NA |
| Il6 | 0 | 0 | 0 | 0 | NA | NA |
| Otof | 0 | 0 | 0 | 0 | NA | NA |
| Cib4 | 0 | 0 | 0 | 0 | NA | NA |
| Tcf23 | 0 | 0 | 0 | 0 | NA | NA |
| AI839979 | 0 | 0 | 0 | 0 | NA | NA |
| Poln | 0 | 0 | 0 | 0 | NA | NA |
| Hmx1 | 0 | 0 | 0 | 0 | NA | NA |
| Clnk | 0 | 0 | 0 | 0 | NA | NA |
| 4930513D17 | 0 | 0 | 0 | 0 | NA | NA |
| Nkx3-2 | 0 | 0 | 0 | 0 | NA | NA |
| Gm15866 | 0 | 0 | 0 | 0 | NA | NA |
| Gm42555 | 0 | 0 | 0 | 0 | NA | NA |
| Gm42984 | 0 | 0 | 0 | 0 | NA | NA |
| 9630001P10I | 0 | 0 | 0 | 0 | NA | NA |

| | | | | | | |
|---|---|---|---|---|---|---|
| Slc34a2 | 0 | 0 | 0 | 0 | NA | NA |
| Gm17182 | 0 | 0 | 0 | 0 | NA | NA |
| Nwd2 | 0 | 0 | 0 | 0 | NA | NA |
| Gm26761 | 0 | 0 | 0 | 0 | NA | NA |
| Tmem156 | 0 | 0 | 0 | 0 | NA | NA |
| Gm26725 | 0 | 0 | 0 | 0 | NA | NA |
| Gm43698 | 0 | 0 | 0 | 0 | NA | NA |
| Gm26756 | 0 | 0 | 0 | 0 | NA | NA |
| Shisa3 | 0 | 0 | 0 | 0 | NA | NA |
| Gm15477 | 0 | 0 | 0 | 0 | NA | NA |
| Gabrb1 | 0 | 0 | 0 | 0 | NA | NA |
| Lrrc66 | 0 | 0 | 0 | 0 | NA | NA |
| Spink2 | 0 | 0 | 0 | 0 | NA | NA |
| Tecrl | 0 | 0 | 0 | 0 | NA | NA |
| Gm42794 | 0 | 0 | 0 | 0 | NA | NA |
| Odam | 0 | 0 | 0 | 0 | NA | NA |
| Amtn | 0 | 0 | 0 | 0 | NA | NA |
| Adamts3 | 0 | 0 | 0 | 0 | NA | NA |
| Gm42530 | 0 | 0 | 0 | 0 | NA | NA |
| Epgn | 0 | 0 | 0 | 0 | NA | NA |
| U90926 | 0 | 0 | 0 | 0 | NA | NA |
| Ppef2 | 0 | 0 | 0 | 0 | NA | NA |
| Art3 | 0 | 0 | 0 | 0 | NA | NA |
| Gm42604 | 0 | 0 | 0 | 0 | NA | NA |
| 1700016F12I | 0 | 0 | 0 | 0 | NA | NA |
| Gm33370 | 0 | 0 | 0 | 0 | NA | NA |
| Gm8013 | 0 | 0 | 0 | 0 | NA | NA |
| A730035I17F | 0 | 0 | 0 | 0 | NA | NA |
| Prkg2 | 0 | 0 | 0 | 0 | NA | NA |
| 2310034O05 | 0 | 0 | 0 | 0 | NA | NA |

| | | | | | | |
|---|---|---|---|---|---|---|
| 4930458D05I | 0 | 0 | 0 | 0 | NA | NA |
| Gm29707 | 0 | 0 | 0 | 0 | NA | NA |
| Slc10a6 | 0 | 0 | 0 | 0 | NA | NA |
| Gm26703 | 0 | 0 | 0 | 0 | NA | NA |
| Abcg3 | 0 | 0 | 0 | 0 | NA | NA |
| Gbp10 | 0 | 0 | 0 | 0 | NA | NA |
| 1700028K03I | 0 | 0 | 0 | 0 | NA | NA |
| Gm42517 | 0 | 0 | 0 | 0 | NA | NA |
| Gm10419 | 0 | 0 | 0 | 0 | NA | NA |
| Rnf212 | 0 | 0 | 0 | 0 | NA | NA |
| P2rx2 | 0 | 0 | 0 | 0 | NA | NA |
| C130026L21I | 0 | 0 | 0 | 0 | NA | NA |
| E130006D01I | 0 | 0 | 0 | 0 | NA | NA |
| Gm42488 | 0 | 0 | 0 | 0 | NA | NA |
| 1700028D13I | 0 | 0 | 0 | 0 | NA | NA |
| Gm42865 | 0 | 0 | 0 | 0 | NA | NA |
| Myo18b | 0 | 0 | 0 | 0 | NA | NA |
| Crybb2 | 0 | 0 | 0 | 0 | NA | NA |
| 1700069L16I | 0 | 0 | 0 | 0 | NA | NA |
| Myo1h | 0 | 0 | 0 | 0 | NA | NA |
| Hnf1aos2 | 0 | 0 | 0 | 0 | NA | NA |
| Gm13842 | 0 | 0 | 0 | 0 | NA | NA |
| C330018A13I | 0 | 0 | 0 | 0 | NA | NA |
| B230112J18I | 0 | 0 | 0 | 0 | NA | NA |
| 4930562A09I | 0 | 0 | 0 | 0 | NA | NA |
| Hrk | 0 | 0 | 0 | 0 | NA | NA |
| Gm7538 | 0 | 0 | 0 | 0 | NA | NA |
| Tbx3os1 | 0 | 0 | 0 | 0 | NA | NA |
| Tbx3os2 | 0 | 0 | 0 | 0 | NA | NA |
| Tbx5 | 0 | 0 | 0 | 0 | NA | NA |

| | | | | | | |
|---|---|---|---|---|---|---|
| Gm10390 | 0 | 0 | 0 | 0 | NA | NA |
| Oas1f | 0 | 0 | 0 | 0 | NA | NA |
| Gm42878 | 0 | 0 | 0 | 0 | NA | NA |
| Myl2 | 0 | 0 | 0 | 0 | NA | NA |
| Morn3 | 0 | 0 | 0 | 0 | NA | NA |
| Dnah10 | 0 | 0 | 0 | 0 | NA | NA |
| Tmem132cos | 0 | 0 | 0 | 0 | NA | NA |
| Fzd10 | 0 | 0 | 0 | 0 | NA | NA |
| Gm43604 | 0 | 0 | 0 | 0 | NA | NA |
| Gm8066 | 0 | 0 | 0 | 0 | NA | NA |
| Lrch4 | 0 | 0 | 0 | 0 | NA | NA |
| Gm15498 | 0 | 0 | 0 | 0 | NA | NA |
| Pilrb1 | 0 | 0 | 0 | 0 | NA | NA |
| Pilrb2 | 0 | 0 | 0 | 0 | NA | NA |
| Nxpe5 | 0 | 0 | 0 | 0 | NA | NA |
| Gal3st4 | 0 | 0 | 0 | 0 | NA | NA |
| Gm17135 | 0 | 0 | 0 | 0 | NA | NA |
| Ocm | 0 | 0 | 0 | 0 | NA | NA |
| Kpna7 | 0 | 0 | 0 | 0 | NA | NA |
| Urad | 0 | 0 | 0 | 0 | NA | NA |
| Flt3 | 0 | 0 | 0 | 0 | NA | NA |
| D5Ertd605e | 0 | 0 | 0 | 0 | NA | NA |
| Gm15409 | 0 | 0 | 0 | 0 | NA | NA |
| Gm43332 | 0 | 0 | 0 | 0 | NA | NA |
| Gm29264 | 0 | 0 | 0 | 0 | NA | NA |
| Gm19719 | 0 | 0 | 0 | 0 | NA | NA |
| BC028471 | 0 | 0 | 0 | 0 | NA | NA |
| Wdr95 | 0 | 0 | 0 | 0 | NA | NA |
| Vmn2r18 | 0 | 0 | 0 | 0 | NA | NA |
| Gm8579 | 0 | 0 | 0 | 0 | NA | NA |

| | | | | | | |
|---|---|---|---|---|---|---|
| Calcr | 0 | 0 | 0 | 0 | NA | NA |
| Gngt1 | 0 | 0 | 0 | 0 | NA | NA |
| 1700019G24 | 0 | 0 | 0 | 0 | NA | NA |
| Gm20619 | 0 | 0 | 0 | 0 | NA | NA |
| Gm20618 | 0 | 0 | 0 | 0 | NA | NA |
| Dlx6os1 | 0 | 0 | 0 | 0 | NA | NA |
| Dlx6 | 0 | 0 | 0 | 0 | NA | NA |
| Tfec | 0 | 0 | 0 | 0 | NA | NA |
| Gm4876 | 0 | 0 | 0 | 0 | NA | NA |
| D830026I12F | 0 | 0 | 0 | 0 | NA | NA |
| Gm15594 | 0 | 0 | 0 | 0 | NA | NA |
| Ankrd7 | 0 | 0 | 0 | 0 | NA | NA |
| Opn1sw | 0 | 0 | 0 | 0 | NA | NA |
| Gm26627 | 0 | 0 | 0 | 0 | NA | NA |
| Smkr-ps | 0 | 0 | 0 | 0 | NA | NA |
| Gm44296 | 0 | 0 | 0 | 0 | NA | NA |
| Tsga13 | 0 | 0 | 0 | 0 | NA | NA |
| Klf14 | 0 | 0 | 0 | 0 | NA | NA |
| Gm43154 | 0 | 0 | 0 | 0 | NA | NA |
| Plxna4os1 | 0 | 0 | 0 | 0 | NA | NA |
| 9330158H04 | 0 | 0 | 0 | 0 | NA | NA |
| Chrm2 | 0 | 0 | 0 | 0 | NA | NA |
| Dgki | 0 | 0 | 0 | 0 | NA | NA |
| Atp6v0a4 | 0 | 0 | 0 | 0 | NA | NA |
| Klrg2 | 0 | 0 | 0 | 0 | NA | NA |
| Gm42962 | 0 | 0 | 0 | 0 | NA | NA |
| Wee2 | 0 | 0 | 0 | 0 | NA | NA |
| Gm4744 | 0 | 0 | 0 | 0 | NA | NA |
| 1810009J06F | 0 | 0 | 0 | 0 | NA | NA |
| Clcn1 | 0 | 0 | 0 | 0 | NA | NA |

| | | | | | | |
|---|---|---|---|---|---|---|
| Fam131b | 0 | 0 | 0 | 0 | NA | NA |
| Nobox | 0 | 0 | 0 | 0 | NA | NA |
| Gimap7 | 0 | 0 | 0 | 0 | NA | NA |
| 4833403J16F | 0 | 0 | 0 | 0 | NA | NA |
| Gpnmb | 0 | 0 | 0 | 0 | NA | NA |
| Gm6559 | 0 | 0 | 0 | 0 | NA | NA |
| Hoxa1 | 0 | 0 | 0 | 0 | NA | NA |
| Gm29430 | 0 | 0 | 0 | 0 | NA | NA |
| Hoxa4 | 0 | 0 | 0 | 0 | NA | NA |
| Hoxa6 | 0 | 0 | 0 | 0 | NA | NA |
| Cpvl | 0 | 0 | 0 | 0 | NA | NA |
| 9530036M11 | 0 | 0 | 0 | 0 | NA | NA |
| Mmrn1 | 0 | 0 | 0 | 0 | NA | NA |
| Gm16838 | 0 | 0 | 0 | 0 | NA | NA |
| Fabp1 | 0 | 0 | 0 | 0 | NA | NA |
| Smyd1 | 0 | 0 | 0 | 0 | NA | NA |
| Cd8b1 | 0 | 0 | 0 | 0 | NA | NA |
| Sh2d6 | 0 | 0 | 0 | 0 | NA | NA |
| Gm15402 | 0 | 0 | 0 | 0 | NA | NA |
| Gm44166 | 0 | 0 | 0 | 0 | NA | NA |
| Gm44287 | 0 | 0 | 0 | 0 | NA | NA |
| Nat8 | 0 | 0 | 0 | 0 | NA | NA |
| Gm19265 | 0 | 0 | 0 | 0 | NA | NA |
| D6Ertd527e | 0 | 0 | 0 | 0 | NA | NA |
| Gata2 | 0 | 0 | 0 | 0 | NA | NA |
| Gm44264 | 0 | 0 | 0 | 0 | NA | NA |
| Gm26588 | 0 | 0 | 0 | 0 | NA | NA |
| Vmn1r53 | 0 | 0 | 0 | 0 | NA | NA |
| Cfap100 | 0 | 0 | 0 | 0 | NA | NA |
| Gm44421 | 0 | 0 | 0 | 0 | NA | NA |

| | | | | | | |
|---|---|---|---|---|---|---|
| 4930471M09 | 0 | 0 | 0 | 0 | NA | NA |
| A730049H05 | 0 | 0 | 0 | 0 | NA | NA |
| Gm15737 | 0 | 0 | 0 | 0 | NA | NA |
| Gm32592 | 0 | 0 | 0 | 0 | NA | NA |
| Gm44169 | 0 | 0 | 0 | 0 | NA | NA |
| Cntn6 | 0 | 0 | 0 | 0 | NA | NA |
| Gm15631 | 0 | 0 | 0 | 0 | NA | NA |
| Il5ra | 0 | 0 | 0 | 0 | NA | NA |
| Gm20387 | 0 | 0 | 0 | 0 | NA | NA |
| Grm7 | 0 | 0 | 0 | 0 | NA | NA |
| 5031434C07I | 0 | 0 | 0 | 0 | NA | NA |
| Cidec | 0 | 0 | 0 | 0 | NA | NA |
| Fancd2os | 0 | 0 | 0 | 0 | NA | NA |
| Gm44206 | 0 | 0 | 0 | 0 | NA | NA |
| Alox5 | 0 | 0 | 0 | 0 | NA | NA |
| D030044L04I | 0 | 0 | 0 | 0 | NA | NA |
| Gm4640 | 0 | 0 | 0 | 0 | NA | NA |
| Gm9946 | 0 | 0 | 0 | 0 | NA | NA |
| Ninj2 | 0 | 0 | 0 | 0 | NA | NA |
| A2m | 0 | 0 | 0 | 0 | NA | NA |
| Klrg1 | 0 | 0 | 0 | 0 | NA | NA |
| 1700063H04I | 0 | 0 | 0 | 0 | NA | NA |
| Aicda | 0 | 0 | 0 | 0 | NA | NA |
| Dppa3 | 0 | 0 | 0 | 0 | NA | NA |
| Nanog | 0 | 0 | 0 | 0 | NA | NA |
| Clec4a3 | 0 | 0 | 0 | 0 | NA | NA |
| Clec4b1 | 0 | 0 | 0 | 0 | NA | NA |
| Cd163 | 0 | 0 | 0 | 0 | NA | NA |
| 1700013D24I | 0 | 0 | 0 | 0 | NA | NA |
| C1s2 | 0 | 0 | 0 | 0 | NA | NA |

| | | | | | | |
|---|---|---|---|---|---|---|
| Cd4 | 0 | 0 | 0 | 0 | NA | NA |
| Tuba3a | 0 | 0 | 0 | 0 | NA | NA |
| Kcna1 | 0 | 0 | 0 | 0 | NA | NA |
| Gm26673 | 0 | 0 | 0 | 0 | NA | NA |
| Platr31 | 0 | 0 | 0 | 0 | NA | NA |
| Gm43126 | 0 | 0 | 0 | 0 | NA | NA |
| Cracr2a | 0 | 0 | 0 | 0 | NA | NA |
| Gm26770 | 0 | 0 | 0 | 0 | NA | NA |
| Clec2h | 0 | 0 | 0 | 0 | NA | NA |
| Klrb1 | 0 | 0 | 0 | 0 | NA | NA |
| Klrb1b | 0 | 0 | 0 | 0 | NA | NA |
| Gm44066 | 0 | 0 | 0 | 0 | NA | NA |
| Gm44067 | 0 | 0 | 0 | 0 | NA | NA |
| Clec2i | 0 | 0 | 0 | 0 | NA | NA |
| Gm15987 | 0 | 0 | 0 | 0 | NA | NA |
| Klrb1f | 0 | 0 | 0 | 0 | NA | NA |
| Gm44000 | 0 | 0 | 0 | 0 | NA | NA |
| Clec1a | 0 | 0 | 0 | 0 | NA | NA |
| Olr1 | 0 | 0 | 0 | 0 | NA | NA |
| Klre1 | 0 | 0 | 0 | 0 | NA | NA |
| Klrk1 | 0 | 0 | 0 | 0 | NA | NA |
| Klra17 | 0 | 0 | 0 | 0 | NA | NA |
| Klra8 | 0 | 0 | 0 | 0 | NA | NA |
| Klra7 | 0 | 0 | 0 | 0 | NA | NA |
| Klra2 | 0 | 0 | 0 | 0 | NA | NA |
| 5430401F13I | 0 | 0 | 0 | 0 | NA | NA |
| Gm6619 | 0 | 0 | 0 | 0 | NA | NA |
| Gsg1 | 0 | 0 | 0 | 0 | NA | NA |
| Smco3 | 0 | 0 | 0 | 0 | NA | NA |
| Art4 | 0 | 0 | 0 | 0 | NA | NA |

| | | | | | | |
|---|---|---|---|---|---|---|
| Rergl | 0 | 0 | 0 | 0 | NA | NA |
| Gm30524 | 0 | 0 | 0 | 0 | NA | NA |
| Spx | 0 | 0 | 0 | 0 | NA | NA |
| Gm15687 | 0 | 0 | 0 | 0 | NA | NA |
| 2010013B24 | 0 | 0 | 0 | 0 | NA | NA |
| Gm15704 | 0 | 0 | 0 | 0 | NA | NA |
| Far2os2 | 0 | 0 | 0 | 0 | NA | NA |
| Gm43913 | 0 | 0 | 0 | 0 | NA | NA |
| Prkcg | 0 | 0 | 0 | 0 | NA | NA |
| Cacng7 | 0 | 0 | 0 | 0 | NA | NA |
| Gm44257 | 0 | 0 | 0 | 0 | NA | NA |
| Cacng8 | 0 | 0 | 0 | 0 | NA | NA |
| Oscar | 0 | 0 | 0 | 0 | NA | NA |
| Gm15929 | 0 | 0 | 0 | 0 | NA | NA |
| Pirb | 0 | 0 | 0 | 0 | NA | NA |
| Lilra6 | 0 | 0 | 0 | 0 | NA | NA |
| Lilra5 | 0 | 0 | 0 | 0 | NA | NA |
| Ncr1 | 0 | 0 | 0 | 0 | NA | NA |
| Gp6 | 0 | 0 | 0 | 0 | NA | NA |
| Tnni3 | 0 | 0 | 0 | 0 | NA | NA |
| Tmem150b | 0 | 0 | 0 | 0 | NA | NA |
| Slc27a5 | 0 | 0 | 0 | 0 | NA | NA |
| Sult2a1 | 0 | 0 | 0 | 0 | NA | NA |
| Sult2a8 | 0 | 0 | 0 | 0 | NA | NA |
| Vmn1r90 | 0 | 0 | 0 | 0 | NA | NA |
| Crxos | 0 | 0 | 0 | 0 | NA | NA |
| C5ar2 | 0 | 0 | 0 | 0 | NA | NA |
| Gng8 | 0 | 0 | 0 | 0 | NA | NA |
| Ptgir | 0 | 0 | 0 | 0 | NA | NA |
| Ceacam3 | 0 | 0 | 0 | 0 | NA | NA |

| | | | | | | |
|---|---|---|---|---|---|---|
| Ceacam12 | 0 | 0 | 0 | 0 | NA | NA |
| Exoc3l2 | 0 | 0 | 0 | 0 | NA | NA |
| Gm26600 | 0 | 0 | 0 | 0 | NA | NA |
| Nlrp5 | 0 | 0 | 0 | 0 | NA | NA |
| Lypd5 | 0 | 0 | 0 | 0 | NA | NA |
| Tex101 | 0 | 0 | 0 | 0 | NA | NA |
| Dmrtc2 | 0 | 0 | 0 | 0 | NA | NA |
| Erich4 | 0 | 0 | 0 | 0 | NA | NA |
| Cyp2s1 | 0 | 0 | 0 | 0 | NA | NA |
| Gm26707 | 0 | 0 | 0 | 0 | NA | NA |
| Cyp2b10 | 0 | 0 | 0 | 0 | NA | NA |
| Cyp2b23 | 0 | 0 | 0 | 0 | NA | NA |
| Cyp2g1 | 0 | 0 | 0 | 0 | NA | NA |
| Cyp2a5 | 0 | 0 | 0 | 0 | NA | NA |
| Cyp2f2 | 0 | 0 | 0 | 0 | NA | NA |
| Gm21983 | 0 | 0 | 0 | 0 | NA | NA |
| Prx | 0 | 0 | 0 | 0 | NA | NA |
| Catsperg2 | 0 | 0 | 0 | 0 | NA | NA |
| Nphs1os | 0 | 0 | 0 | 0 | NA | NA |
| Etv2 | 0 | 0 | 0 | 0 | NA | NA |
| Gm17077 | 0 | 0 | 0 | 0 | NA | NA |
| Gm6096 | 0 | 0 | 0 | 0 | NA | NA |
| Slc7a10 | 0 | 0 | 0 | 0 | NA | NA |
| Tdrd12 | 0 | 0 | 0 | 0 | NA | NA |
| Vstm2b | 0 | 0 | 0 | 0 | NA | NA |
| Siglecf | 0 | 0 | 0 | 0 | NA | NA |
| Zfp819 | 0 | 0 | 0 | 0 | NA | NA |
| Klk7 | 0 | 0 | 0 | 0 | NA | NA |
| Klk5 | 0 | 0 | 0 | 0 | NA | NA |
| Klk4 | 0 | 0 | 0 | 0 | NA | NA |

| | | | | | | |
|---|---|---|---|---|---|---|
| Klk1b22 | 0 | 0 | 0 | 0 | NA | NA |
| Mybpc2 | 0 | 0 | 0 | 0 | NA | NA |
| Tsks | 0 | 0 | 0 | 0 | NA | NA |
| Lhb | 0 | 0 | 0 | 0 | NA | NA |
| Hsd17b14 | 0 | 0 | 0 | 0 | NA | NA |
| Fut2 | 0 | 0 | 0 | 0 | NA | NA |
| Sec1 | 0 | 0 | 0 | 0 | NA | NA |
| Abcc6 | 0 | 0 | 0 | 0 | NA | NA |
| Myod1 | 0 | 0 | 0 | 0 | NA | NA |
| Saa3 | 0 | 0 | 0 | 0 | NA | NA |
| Saa1 | 0 | 0 | 0 | 0 | NA | NA |
| Saa2 | 0 | 0 | 0 | 0 | NA | NA |
| Mrgpra2a | 0 | 0 | 0 | 0 | NA | NA |
| Mrgpra2b | 0 | 0 | 0 | 0 | NA | NA |
| Csrp3 | 0 | 0 | 0 | 0 | NA | NA |
| Ano5 | 0 | 0 | 0 | 0 | NA | NA |
| Oca2 | 0 | 0 | 0 | 0 | NA | NA |
| Gm9962 | 0 | 0 | 0 | 0 | NA | NA |
| Gabra5 | 0 | 0 | 0 | 0 | NA | NA |
| Trpm1 | 0 | 0 | 0 | 0 | NA | NA |
| Gm20457 | 0 | 0 | 0 | 0 | NA | NA |
| Apba2 | 0 | 0 | 0 | 0 | NA | NA |
| Cers3 | 0 | 0 | 0 | 0 | NA | NA |
| Gm16157 | 0 | 0 | 0 | 0 | NA | NA |
| Gm28258 | 0 | 0 | 0 | 0 | NA | NA |
| Gm9885 | 0 | 0 | 0 | 0 | NA | NA |
| Gm26633 | 0 | 0 | 0 | 0 | NA | NA |
| Rlbp1 | 0 | 0 | 0 | 0 | NA | NA |
| Gm15880 | 0 | 0 | 0 | 0 | NA | NA |
| Slc28a1 | 0 | 0 | 0 | 0 | NA | NA |

| | | | | | | |
|---|---|---|---|---|---|---|
| Adamtsl3 | 0 | 0 | 0 | 0 | NA | NA |
| Cfap161 | 0 | 0 | 0 | 0 | NA | NA |
| Grm5 | 0 | 0 | 0 | 0 | NA | NA |
| Ccdc81 | 0 | 0 | 0 | 0 | NA | NA |
| Sytl2 | 0 | 0 | 0 | 0 | NA | NA |
| Gm9934 | 0 | 0 | 0 | 0 | NA | NA |
| Lrrc32 | 0 | 0 | 0 | 0 | NA | NA |
| Wnt11 | 0 | 0 | 0 | 0 | NA | NA |
| Tpbgl | 0 | 0 | 0 | 0 | NA | NA |
| Gm10605 | 0 | 0 | 0 | 0 | NA | NA |
| Chrdl2 | 0 | 0 | 0 | 0 | NA | NA |
| Kcne3 | 0 | 0 | 0 | 0 | NA | NA |
| P2ry2 | 0 | 0 | 0 | 0 | NA | NA |
| Chrna10 | 0 | 0 | 0 | 0 | NA | NA |
| Hbb-bh2 | 0 | 0 | 0 | 0 | NA | NA |
| Trim5 | 0 | 0 | 0 | 0 | NA | NA |
| Trim30c | 0 | 0 | 0 | 0 | NA | NA |
| Trim30d | 0 | 0 | 0 | 0 | NA | NA |
| Cckbr | 0 | 0 | 0 | 0 | NA | NA |
| Gvin1 | 0 | 0 | 0 | 0 | NA | NA |
| Nlrp10 | 0 | 0 | 0 | 0 | NA | NA |
| Tub | 0 | 0 | 0 | 0 | NA | NA |
| Trim66 | 0 | 0 | 0 | 0 | NA | NA |
| Ascl3 | 0 | 0 | 0 | 0 | NA | NA |
| 1600010M07 | 0 | 0 | 0 | 0 | NA | NA |
| Hs3st2 | 0 | 0 | 0 | 0 | NA | NA |
| Slc5a11 | 0 | 0 | 0 | 0 | NA | NA |
| Gm27040 | 0 | 0 | 0 | 0 | NA | NA |
| Hs3st4 | 0 | 0 | 0 | 0 | NA | NA |
| 3100003L05I | 0 | 0 | 0 | 0 | NA | NA |

| | | | | | | |
|---|---|---|---|---|---|---|
| 4930451I11R | 0 | 0 | 0 | 0 | NA | NA |
| Prrt2 | 0 | 0 | 0 | 0 | NA | NA |
| Spn | 0 | 0 | 0 | 0 | NA | NA |
| 1700120K04I | 0 | 0 | 0 | 0 | NA | NA |
| Gm15533 | 0 | 0 | 0 | 0 | NA | NA |
| Itgax | 0 | 0 | 0 | 0 | NA | NA |
| Itgad | 0 | 0 | 0 | 0 | NA | NA |
| Slc5a2 | 0 | 0 | 0 | 0 | NA | NA |
| Plpp4 | 0 | 0 | 0 | 0 | NA | NA |
| Hmx3 | 0 | 0 | 0 | 0 | NA | NA |
| Gm15582 | 0 | 0 | 0 | 0 | NA | NA |
| Gm15718 | 0 | 0 | 0 | 0 | NA | NA |
| D7Ertd443e | 0 | 0 | 0 | 0 | NA | NA |
| 4930544L04F | 0 | 0 | 0 | 0 | NA | NA |
| 5830432E09I | 0 | 0 | 0 | 0 | NA | NA |
| Ebf3 | 0 | 0 | 0 | 0 | NA | NA |
| Utf1 | 0 | 0 | 0 | 0 | NA | NA |
| Sprn | 0 | 0 | 0 | 0 | NA | NA |
| Nlrp6 | 0 | 0 | 0 | 0 | NA | NA |
| Ifitm10 | 0 | 0 | 0 | 0 | NA | NA |
| Syt8 | 0 | 0 | 0 | 0 | NA | NA |
| Tnnt3 | 0 | 0 | 0 | 0 | NA | NA |
| Tspan32os | 0 | 0 | 0 | 0 | NA | NA |
| 4933417O13 | 0 | 0 | 0 | 0 | NA | NA |
| Fgf15 | 0 | 0 | 0 | 0 | NA | NA |
| H60c | 0 | 0 | 0 | 0 | NA | NA |
| Vip | 0 | 0 | 0 | 0 | NA | NA |
| Gm16234 | 0 | 0 | 0 | 0 | NA | NA |
| Nmbr | 0 | 0 | 0 | 0 | NA | NA |
| Gm20655 | 0 | 0 | 0 | 0 | NA | NA |

| | | | | | | |
|---|---|---|---|---|---|---|
| Txlnb | 0 | 0 | 0 | 0 | NA | NA |
| Gm26577 | 0 | 0 | 0 | 0 | NA | NA |
| 1700021A07I | 0 | 0 | 0 | 0 | NA | NA |
| Gm26740 | 0 | 0 | 0 | 0 | NA | NA |
| Eya4 | 0 | 0 | 0 | 0 | NA | NA |
| Gm9996 | 0 | 0 | 0 | 0 | NA | NA |
| Nkain2 | 0 | 0 | 0 | 0 | NA | NA |
| Trdn | 0 | 0 | 0 | 0 | NA | NA |
| Clvs2 | 0 | 0 | 0 | 0 | NA | NA |
| Cdk19os | 0 | 0 | 0 | 0 | NA | NA |
| F930017D23I | 0 | 0 | 0 | 0 | NA | NA |
| Gm15934 | 0 | 0 | 0 | 0 | NA | NA |
| Lin28b | 0 | 0 | 0 | 0 | NA | NA |
| Edar | 0 | 0 | 0 | 0 | NA | NA |
| Oit3 | 0 | 0 | 0 | 0 | NA | NA |
| Gm10322 | 0 | 0 | 0 | 0 | NA | NA |
| Gm17059 | 0 | 0 | 0 | 0 | NA | NA |
| Gm20625 | 0 | 0 | 0 | 0 | NA | NA |
| Hk1os | 0 | 0 | 0 | 0 | NA | NA |
| 1700023F02I | 0 | 0 | 0 | 0 | NA | NA |
| Gm26576 | 0 | 0 | 0 | 0 | NA | NA |
| Gm16222 | 0 | 0 | 0 | 0 | NA | NA |
| Dnmt3l | 0 | 0 | 0 | 0 | NA | NA |
| Gm9978 | 0 | 0 | 0 | 0 | NA | NA |
| Ccdc105 | 0 | 0 | 0 | 0 | NA | NA |
| Grin3b | 0 | 0 | 0 | 0 | NA | NA |
| Amh | 0 | 0 | 0 | 0 | NA | NA |
| Atcay | 0 | 0 | 0 | 0 | NA | NA |
| 4930463O16 | 0 | 0 | 0 | 0 | NA | NA |
| Bpifc | 0 | 0 | 0 | 0 | NA | NA |

| | | | | | | |
|---|---|---|---|---|---|---|
| Stab2 | 0 | 0 | 0 | 0 | NA | NA |
| Igf1os | 0 | 0 | 0 | 0 | NA | NA |
| Slc5a8 | 0 | 0 | 0 | 0 | NA | NA |
| Ano4 | 0 | 0 | 0 | 0 | NA | NA |
| Slc17a8 | 0 | 0 | 0 | 0 | NA | NA |
| Epyc | 0 | 0 | 0 | 0 | NA | NA |
| Ccer1 | 0 | 0 | 0 | 0 | NA | NA |
| Csl | 0 | 0 | 0 | 0 | NA | NA |
| Mgat4c | 0 | 0 | 0 | 0 | NA | NA |
| Alx1 | 0 | 0 | 0 | 0 | NA | NA |
| Otogl | 0 | 0 | 0 | 0 | NA | NA |
| Trhde | 0 | 0 | 0 | 0 | NA | NA |
| 4930473D10 | 0 | 0 | 0 | 0 | NA | NA |
| Myrfl | 0 | 0 | 0 | 0 | NA | NA |
| Lyz1 | 0 | 0 | 0 | 0 | NA | NA |
| Ifng | 0 | 0 | 0 | 0 | NA | NA |
| Msrb3 | 0 | 0 | 0 | 0 | NA | NA |
| Avpr1a | 0 | 0 | 0 | 0 | NA | NA |
| Agap2 | 0 | 0 | 0 | 0 | NA | NA |
| Nxph4 | 0 | 0 | 0 | 0 | NA | NA |
| Gm16229 | 0 | 0 | 0 | 0 | NA | NA |
| Rdh1 | 0 | 0 | 0 | 0 | NA | NA |
| Mip | 0 | 0 | 0 | 0 | NA | NA |
| Gm17201 | 0 | 0 | 0 | 0 | NA | NA |
| Mettl7b | 0 | 0 | 0 | 0 | NA | NA |
| Tespa1 | 0 | 0 | 0 | 0 | NA | NA |
| Vmn2r87 | 0 | 0 | 0 | 0 | NA | NA |
| Pcp2 | 0 | 0 | 0 | 0 | NA | NA |
| Fcer2a | 0 | 0 | 0 | 0 | NA | NA |
| Clec4g | 0 | 0 | 0 | 0 | NA | NA |

| | | | | | | |
|---|---|---|---|---|---|---|
| Cd209f | 0 | 0 | 0 | 0 | NA | NA |
| 4932443L11F | 0 | 0 | 0 | 0 | NA | NA |
| Slc10a2 | 0 | 0 | 0 | 0 | NA | NA |
| Gm7676 | 0 | 0 | 0 | 0 | NA | NA |
| Gm15056 | 0 | 0 | 0 | 0 | NA | NA |
| Defb50 | 0 | 0 | 0 | 0 | NA | NA |
| Defb2 | 0 | 0 | 0 | 0 | NA | NA |
| Thsd1 | 0 | 0 | 0 | 0 | NA | NA |
| Gm26714 | 0 | 0 | 0 | 0 | NA | NA |
| Adam3 | 0 | 0 | 0 | 0 | NA | NA |
| 2310008N11 | 0 | 0 | 0 | 0 | NA | NA |
| Gm26795 | 0 | 0 | 0 | 0 | NA | NA |
| Gm29243 | 0 | 0 | 0 | 0 | NA | NA |
| Zfp42 | 0 | 0 | 0 | 0 | NA | NA |
| Gm2366 | 0 | 0 | 0 | 0 | NA | NA |
| Enpp6 | 0 | 0 | 0 | 0 | NA | NA |
| Cldn22 | 0 | 0 | 0 | 0 | NA | NA |
| Galntl6 | 0 | 0 | 0 | 0 | NA | NA |
| BC030500 | 0 | 0 | 0 | 0 | NA | NA |
| 4930512H18 | 0 | 0 | 0 | 0 | NA | NA |
| Tll1 | 0 | 0 | 0 | 0 | NA | NA |
| Npy5r | 0 | 0 | 0 | 0 | NA | NA |
| Gm43263 | 0 | 0 | 0 | 0 | NA | NA |
| Isx | 0 | 0 | 0 | 0 | NA | NA |
| Nr3c2 | 0 | 0 | 0 | 0 | NA | NA |
| Ttc29 | 0 | 0 | 0 | 0 | NA | NA |
| Pou4f2 | 0 | 0 | 0 | 0 | NA | NA |
| Hhip | 0 | 0 | 0 | 0 | NA | NA |
| Frem3 | 0 | 0 | 0 | 0 | NA | NA |
| Gm27048 | 0 | 0 | 0 | 0 | NA | NA |

| | | | | | | |
|---|---|---|---|---|---|---|
| Il15 | 0 | 0 | 0 | 0 | NA | NA |
| Podnl1 | 0 | 0 | 0 | 0 | NA | NA |
| Gm26887 | 0 | 0 | 0 | 0 | NA | NA |
| Zfp791 | 0 | 0 | 0 | 0 | NA | NA |
| Nod2 | 0 | 0 | 0 | 0 | NA | NA |
| Gm6625 | 0 | 0 | 0 | 0 | NA | NA |
| Irx3os | 0 | 0 | 0 | 0 | NA | NA |
| Ces1b | 0 | 0 | 0 | 0 | NA | NA |
| Ces1e | 0 | 0 | 0 | 0 | NA | NA |
| Ces1f | 0 | 0 | 0 | 0 | NA | NA |
| Ces1h | 0 | 0 | 0 | 0 | NA | NA |
| Adgrg5 | 0 | 0 | 0 | 0 | NA | NA |
| Cmtm2b | 0 | 0 | 0 | 0 | NA | NA |
| Ces3a | 0 | 0 | 0 | 0 | NA | NA |
| Plekhg4 | 0 | 0 | 0 | 0 | NA | NA |
| Dpep3 | 0 | 0 | 0 | 0 | NA | NA |
| D030068K23 | 0 | 0 | 0 | 0 | NA | NA |
| Pmfbp1 | 0 | 0 | 0 | 0 | NA | NA |
| Gm17344 | 0 | 0 | 0 | 0 | NA | NA |
| Chst4 | 0 | 0 | 0 | 0 | NA | NA |
| Gm15895 | 0 | 0 | 0 | 0 | NA | NA |
| Clec18a | 0 | 0 | 0 | 0 | NA | NA |
| Gm16118 | 0 | 0 | 0 | 0 | NA | NA |
| Pkd1l2 | 0 | 0 | 0 | 0 | NA | NA |
| Adad2 | 0 | 0 | 0 | 0 | NA | NA |
| Foxc2 | 0 | 0 | 0 | 0 | NA | NA |
| Gm26815 | 0 | 0 | 0 | 0 | NA | NA |
| Gm26747 | 0 | 0 | 0 | 0 | NA | NA |
| Cpne7 | 0 | 0 | 0 | 0 | NA | NA |
| 1810008B01 | 0 | 0 | 0 | 0 | NA | NA |

| | | | | | | |
|---|---|---|---|---|---|---|
| Gm15775 | 0 | 0 | 0 | 0 | NA | NA |
| Gm2888 | 0 | 0 | 0 | 0 | NA | NA |
| Gm5796 | 0 | 0 | 0 | 0 | NA | NA |
| Gm2974 | 0 | 0 | 0 | 0 | NA | NA |
| 2610042L04F | 0 | 0 | 0 | 0 | NA | NA |
| Gm3269 | 0 | 0 | 0 | 0 | NA | NA |
| Gm3526 | 0 | 0 | 0 | 0 | NA | NA |
| Gm3248 | 0 | 0 | 0 | 0 | NA | NA |
| Gm3453 | 0 | 0 | 0 | 0 | NA | NA |
| Gm3636 | 0 | 0 | 0 | 0 | NA | NA |
| Gm10406 | 0 | 0 | 0 | 0 | NA | NA |
| Gm3696 | 0 | 0 | 0 | 0 | NA | NA |
| Fhitos | 0 | 0 | 0 | 0 | NA | NA |
| Gm2237 | 0 | 0 | 0 | 0 | NA | NA |
| Synpo2l | 0 | 0 | 0 | 0 | NA | NA |
| Plau | 0 | 0 | 0 | 0 | NA | NA |
| Gm10248 | 0 | 0 | 0 | 0 | NA | NA |
| Zcchc24 | 0 | 0 | 0 | 0 | NA | NA |
| 4933413J09F | 0 | 0 | 0 | 0 | NA | NA |
| Hesx1 | 0 | 0 | 0 | 0 | NA | NA |
| Gm2670 | 0 | 0 | 0 | 0 | NA | NA |
| Itih4 | 0 | 0 | 0 | 0 | NA | NA |
| Itih3 | 0 | 0 | 0 | 0 | NA | NA |
| Chat | 0 | 0 | 0 | 0 | NA | NA |
| Gm15512 | 0 | 0 | 0 | 0 | NA | NA |
| Zfp488 | 0 | 0 | 0 | 0 | NA | NA |
| Opn4 | 0 | 0 | 0 | 0 | NA | NA |
| Nrg3 | 0 | 0 | 0 | 0 | NA | NA |
| Dydc1 | 0 | 0 | 0 | 0 | NA | NA |
| Sftpa1 | 0 | 0 | 0 | 0 | NA | NA |

| | | | | | | |
|---|---|---|---|---|---|---|
| Mbl1 | 0 | 0 | 0 | 0 | NA | NA |
| Ear1 | 0 | 0 | 0 | 0 | NA | NA |
| Gm15601 | 0 | 0 | 0 | 0 | NA | NA |
| Gm15217 | 0 | 0 | 0 | 0 | NA | NA |
| Gm15221 | 0 | 0 | 0 | 0 | NA | NA |
| Gm10371 | 0 | 0 | 0 | 0 | NA | NA |
| Ccnb1ip1 | 0 | 0 | 0 | 0 | NA | NA |
| Rnase12 | 0 | 0 | 0 | 0 | NA | NA |
| Rnase2a | 0 | 0 | 0 | 0 | NA | NA |
| Gm16617 | 0 | 0 | 0 | 0 | NA | NA |
| Tmem253 | 0 | 0 | 0 | 0 | NA | NA |
| Gm43434 | 0 | 0 | 0 | 0 | NA | NA |
| Psmb11 | 0 | 0 | 0 | 0 | NA | NA |
| 4930579G18 | 0 | 0 | 0 | 0 | NA | NA |
| Cebpe | 0 | 0 | 0 | 0 | NA | NA |
| Cmtm5 | 0 | 0 | 0 | 0 | NA | NA |
| Zfhx2os | 0 | 0 | 0 | 0 | NA | NA |
| Ltb4r2 | 0 | 0 | 0 | 0 | NA | NA |
| Ltb4r1 | 0 | 0 | 0 | 0 | NA | NA |
| Cma2 | 0 | 0 | 0 | 0 | NA | NA |
| Mcpt1 | 0 | 0 | 0 | 0 | NA | NA |
| Mcpt2 | 0 | 0 | 0 | 0 | NA | NA |
| Mcpt4 | 0 | 0 | 0 | 0 | NA | NA |
| Gzmc | 0 | 0 | 0 | 0 | NA | NA |
| Gzmb | 0 | 0 | 0 | 0 | NA | NA |
| Gja3 | 0 | 0 | 0 | 0 | NA | NA |
| Gjb6 | 0 | 0 | 0 | 0 | NA | NA |
| Phf11a | 0 | 0 | 0 | 0 | NA | NA |
| Arl11 | 0 | 0 | 0 | 0 | NA | NA |
| Gm26916 | 0 | 0 | 0 | 0 | NA | NA |

| | | | | | | |
|---|---|---|---|---|---|---|
| Blk | 0 | 0 | 0 | 0 | NA | NA |
| Mir124a-1hg | 0 | 0 | 0 | 0 | NA | NA |
| Adam2 | 0 | 0 | 0 | 0 | NA | NA |
| Adamdec1 | 0 | 0 | 0 | 0 | NA | NA |
| Piwil2 | 0 | 0 | 0 | 0 | NA | NA |
| Fgf17 | 0 | 0 | 0 | 0 | NA | NA |
| Gfra2 | 0 | 0 | 0 | 0 | NA | NA |
| Lrrc63 | 0 | 0 | 0 | 0 | NA | NA |
| Siah3 | 0 | 0 | 0 | 0 | NA | NA |
| Slc25a30 | 0 | 0 | 0 | 0 | NA | NA |
| Tnfsf11 | 0 | 0 | 0 | 0 | NA | NA |
| Olfm4 | 0 | 0 | 0 | 0 | NA | NA |
| Pcdh20 | 0 | 0 | 0 | 0 | NA | NA |
| 1700128A07I | 0 | 0 | 0 | 0 | NA | NA |
| Slitrk5 | 0 | 0 | 0 | 0 | NA | NA |
| Gm27198 | 0 | 0 | 0 | 0 | NA | NA |
| 1700006F04I | 0 | 0 | 0 | 0 | NA | NA |
| Zic5 | 0 | 0 | 0 | 0 | NA | NA |
| Gria4 | 0 | 0 | 0 | 0 | NA | NA |
| Casp4 | 0 | 0 | 0 | 0 | NA | NA |
| Mmp13 | 0 | 0 | 0 | 0 | NA | NA |
| Mmp8 | 0 | 0 | 0 | 0 | NA | NA |
| Mmp20 | 0 | 0 | 0 | 0 | NA | NA |
| Pgr | 0 | 0 | 0 | 0 | NA | NA |
| Gm16485 | 0 | 0 | 0 | 0 | NA | NA |
| Gm16833 | 0 | 0 | 0 | 0 | NA | NA |
| Gpr83 | 0 | 0 | 0 | 0 | NA | NA |
| Vstm5 | 0 | 0 | 0 | 0 | NA | NA |
| Fat3 | 0 | 0 | 0 | 0 | NA | NA |
| Col5a3 | 0 | 0 | 0 | 0 | NA | NA |

| | | | | | | |
|---|---|---|---|---|---|---|
| Bmper | 0 | 0 | 0 | 0 | NA | NA |
| Dpy19l2 | 0 | 0 | 0 | 0 | NA | NA |
| Tbx20 | 0 | 0 | 0 | 0 | NA | NA |
| Gm15520 | 0 | 0 | 0 | 0 | NA | NA |
| Slc37a2 | 0 | 0 | 0 | 0 | NA | NA |
| Tecta | 0 | 0 | 0 | 0 | NA | NA |
| Thy1 | 0 | 0 | 0 | 0 | NA | NA |
| Cxcr5 | 0 | 0 | 0 | 0 | NA | NA |
| Apoa5 | 0 | 0 | 0 | 0 | NA | NA |
| Htr3b | 0 | 0 | 0 | 0 | NA | NA |
| Isl2 | 0 | 0 | 0 | 0 | NA | NA |
| Gm17226 | 0 | 0 | 0 | 0 | NA | NA |
| Peak1os | 0 | 0 | 0 | 0 | NA | NA |
| Odf3l1 | 0 | 0 | 0 | 0 | NA | NA |
| Sema7a | 0 | 0 | 0 | 0 | NA | NA |
| Tbc1d21 | 0 | 0 | 0 | 0 | NA | NA |
| 4930461G14 | 0 | 0 | 0 | 0 | NA | NA |
| Tmem202 | 0 | 0 | 0 | 0 | NA | NA |
| Nr2e3 | 0 | 0 | 0 | 0 | NA | NA |
| Iqch | 0 | 0 | 0 | 0 | NA | NA |
| Lctl | 0 | 0 | 0 | 0 | NA | NA |
| Kbtbd13 | 0 | 0 | 0 | 0 | NA | NA |
| Gm28379 | 0 | 0 | 0 | 0 | NA | NA |
| Gm27253 | 0 | 0 | 0 | 0 | NA | NA |
| Gm27241 | 0 | 0 | 0 | 0 | NA | NA |
| Gm27204 | 0 | 0 | 0 | 0 | NA | NA |
| Bmp5 | 0 | 0 | 0 | 0 | NA | NA |
| Gfral | 0 | 0 | 0 | 0 | NA | NA |
| Hcrtr2 | 0 | 0 | 0 | 0 | NA | NA |
| Gm19541 | 0 | 0 | 0 | 0 | NA | NA |

| | | | | | | |
|---|---|---|---|---|---|---|
| Cd109 | 0 | 0 | 0 | 0 | NA | NA |
| D430036J16F | 0 | 0 | 0 | 0 | NA | NA |
| Ripply2 | 0 | 0 | 0 | 0 | NA | NA |
| Bcl2a1d | 0 | 0 | 0 | 0 | NA | NA |
| Gm26611 | 0 | 0 | 0 | 0 | NA | NA |
| Trim43b | 0 | 0 | 0 | 0 | NA | NA |
| Prr23a3 | 0 | 0 | 0 | 0 | NA | NA |
| Cpne4 | 0 | 0 | 0 | 0 | NA | NA |
| Nek11 | 0 | 0 | 0 | 0 | NA | NA |
| Col6a4 | 0 | 0 | 0 | 0 | NA | NA |
| Gm29123 | 0 | 0 | 0 | 0 | NA | NA |
| Gm28959 | 0 | 0 | 0 | 0 | NA | NA |
| Iqcf1 | 0 | 0 | 0 | 0 | NA | NA |
| Gnat1 | 0 | 0 | 0 | 0 | NA | NA |
| Mst1r | 0 | 0 | 0 | 0 | NA | NA |
| Gm20661 | 0 | 0 | 0 | 0 | NA | NA |
| 4921523L03F | 0 | 0 | 0 | 0 | NA | NA |
| Tmem89 | 0 | 0 | 0 | 0 | NA | NA |
| Fbxw22 | 0 | 0 | 0 | 0 | NA | NA |
| Fbxw16 | 0 | 0 | 0 | 0 | NA | NA |
| Fbxw26 | 0 | 0 | 0 | 0 | NA | NA |
| Spink8 | 0 | 0 | 0 | 0 | NA | NA |
| Prss45 | 0 | 0 | 0 | 0 | NA | NA |
| Prss46 | 0 | 0 | 0 | 0 | NA | NA |
| 2900079G21 | 0 | 0 | 0 | 0 | NA | NA |
| Susd5 | 0 | 0 | 0 | 0 | NA | NA |
| Gadl1 | 0 | 0 | 0 | 0 | NA | NA |
| Slc22a13 | 0 | 0 | 0 | 0 | NA | NA |
| Scn5a | 0 | 0 | 0 | 0 | NA | NA |
| Cyp8b1 | 0 | 0 | 0 | 0 | NA | NA |

| | | | | | | |
|---|---|---|---|---|---|---|
| Topaz1 | 0 | 0 | 0 | 0 | NA | NA |
| Ccr3 | 0 | 0 | 0 | 0 | NA | NA |
| Gm12735 | 0 | 0 | 0 | 0 | NA | NA |
| Lif | 0 | 0 | 0 | 0 | NA | NA |
| Ankrd36 | 0 | 0 | 0 | 0 | NA | NA |
| Gm11973 | 0 | 0 | 0 | 0 | NA | NA |
| Gm11983 | 0 | 0 | 0 | 0 | NA | NA |
| Pkd1l1 | 0 | 0 | 0 | 0 | NA | NA |
| Gm11998 | 0 | 0 | 0 | 0 | NA | NA |
| Gm12002 | 0 | 0 | 0 | 0 | NA | NA |
| Egfros | 0 | 0 | 0 | 0 | NA | NA |
| 4933406G16 | 0 | 0 | 0 | 0 | NA | NA |
| Gm12031 | 0 | 0 | 0 | 0 | NA | NA |
| Gm12056 | 0 | 0 | 0 | 0 | NA | NA |
| 4933430M04 | 0 | 0 | 0 | 0 | NA | NA |
| 2810471M01 | 0 | 0 | 0 | 0 | NA | NA |
| Gm12088 | 0 | 0 | 0 | 0 | NA | NA |
| Gm12108 | 0 | 0 | 0 | 0 | NA | NA |
| Tlx3 | 0 | 0 | 0 | 0 | NA | NA |
| D130052B06 | 0 | 0 | 0 | 0 | NA | NA |
| Fbll1 | 0 | 0 | 0 | 0 | NA | NA |
| Atp10b | 0 | 0 | 0 | 0 | NA | NA |
| Gm12146 | 0 | 0 | 0 | 0 | NA | NA |
| Gm12144 | 0 | 0 | 0 | 0 | NA | NA |
| Adra1b | 0 | 0 | 0 | 0 | NA | NA |
| Havcr2 | 0 | 0 | 0 | 0 | NA | NA |
| Dppa1 | 0 | 0 | 0 | 0 | NA | NA |
| Timd2 | 0 | 0 | 0 | 0 | NA | NA |
| Havcr1 | 0 | 0 | 0 | 0 | NA | NA |
| Gm5431 | 0 | 0 | 0 | 0 | NA | NA |

| | | | | | | |
|---|---|---|---|---|---|---|
| 9930111J21F | 0 | 0 | 0 | 0 | NA | NA |
| 9930111J21F | 0 | 0 | 0 | 0 | NA | NA |
| Ifi47 | 0 | 0 | 0 | 0 | NA | NA |
| Btnl9 | 0 | 0 | 0 | 0 | NA | NA |
| Grm6 | 0 | 0 | 0 | 0 | NA | NA |
| 9230009I02R | 0 | 0 | 0 | 0 | NA | NA |
| A630014C17 | 0 | 0 | 0 | 0 | NA | NA |
| Il4 | 0 | 0 | 0 | 0 | NA | NA |
| Gm17334 | 0 | 0 | 0 | 0 | NA | NA |
| Slc36a3os | 0 | 0 | 0 | 0 | NA | NA |
| Glra1 | 0 | 0 | 0 | 0 | NA | NA |
| Hand1 | 0 | 0 | 0 | 0 | NA | NA |
| 4930438A08I | 0 | 0 | 0 | 0 | NA | NA |
| Trim58 | 0 | 0 | 0 | 0 | NA | NA |
| Trim17 | 0 | 0 | 0 | 0 | NA | NA |
| Cdrt4 | 0 | 0 | 0 | 0 | NA | NA |
| Tekt3 | 0 | 0 | 0 | 0 | NA | NA |
| Gm12290 | 0 | 0 | 0 | 0 | NA | NA |
| Hs3st3a1 | 0 | 0 | 0 | 0 | NA | NA |
| Gm12292 | 0 | 0 | 0 | 0 | NA | NA |
| Myocd | 0 | 0 | 0 | 0 | NA | NA |
| Myh3 | 0 | 0 | 0 | 0 | NA | NA |
| Myh2 | 0 | 0 | 0 | 0 | NA | NA |
| Myh4 | 0 | 0 | 0 | 0 | NA | NA |
| Glp2r | 0 | 0 | 0 | 0 | NA | NA |
| Pik3r6 | 0 | 0 | 0 | 0 | NA | NA |
| Ccdc42os | 0 | 0 | 0 | 0 | NA | NA |
| Alox8 | 0 | 0 | 0 | 0 | NA | NA |
| Gucy2e | 0 | 0 | 0 | 0 | NA | NA |
| Tnfsfm13 | 0 | 0 | 0 | 0 | NA | NA |

| | | | | | | |
|---|---|---|---|---|---|---|
| Slc35g3 | 0 | 0 | 0 | 0 | NA | NA |
| Spem1 | 0 | 0 | 0 | 0 | NA | NA |
| Asgr1 | 0 | 0 | 0 | 0 | NA | NA |
| Mgl2 | 0 | 0 | 0 | 0 | NA | NA |
| Alox15 | 0 | 0 | 0 | 0 | NA | NA |
| Vmo1 | 0 | 0 | 0 | 0 | NA | NA |
| Chrne | 0 | 0 | 0 | 0 | NA | NA |
| Gp1ba | 0 | 0 | 0 | 0 | NA | NA |
| Gm12319 | 0 | 0 | 0 | 0 | NA | NA |
| Nlrp1a | 0 | 0 | 0 | 0 | NA | NA |
| Slc13a5 | 0 | 0 | 0 | 0 | NA | NA |
| P2rx5 | 0 | 0 | 0 | 0 | NA | NA |
| Scarf1 | 0 | 0 | 0 | 0 | NA | NA |
| Gm16075 | 0 | 0 | 0 | 0 | NA | NA |
| Bhlha9 | 0 | 0 | 0 | 0 | NA | NA |
| Gm10277 | 0 | 0 | 0 | 0 | NA | NA |
| Slc13a2 | 0 | 0 | 0 | 0 | NA | NA |
| Sebox | 0 | 0 | 0 | 0 | NA | NA |
| Nos2 | 0 | 0 | 0 | 0 | NA | NA |
| Rab11fip4os | 0 | 0 | 0 | 0 | NA | NA |
| Spaca3 | 0 | 0 | 0 | 0 | NA | NA |
| Ccl1 | 0 | 0 | 0 | 0 | NA | NA |
| Unc45b | 0 | 0 | 0 | 0 | NA | NA |
| Slfn5 | 0 | 0 | 0 | 0 | NA | NA |
| Slfn8 | 0 | 0 | 0 | 0 | NA | NA |
| Slfn3 | 0 | 0 | 0 | 0 | NA | NA |
| 1700020L24R | 0 | 0 | 0 | 0 | NA | NA |
| Mmp28 | 0 | 0 | 0 | 0 | NA | NA |
| Bcas3os1 | 0 | 0 | 0 | 0 | NA | NA |
| Gm11444 | 0 | 0 | 0 | 0 | NA | NA |

| | | | | | | |
|---|---|---|---|---|---|---|
| Tbx4 | 0 | 0 | 0 | 0 | NA | NA |
| Lpo | 0 | 0 | 0 | 0 | NA | NA |
| Ccdc182 | 0 | 0 | 0 | 0 | NA | NA |
| Gm11496 | 0 | 0 | 0 | 0 | NA | NA |
| 4930405D11 | 0 | 0 | 0 | 0 | NA | NA |
| Cdc34b | 0 | 0 | 0 | 0 | NA | NA |
| Dlx3 | 0 | 0 | 0 | 0 | NA | NA |
| Dlx4 | 0 | 0 | 0 | 0 | NA | NA |
| Dlx4os | 0 | 0 | 0 | 0 | NA | NA |
| Gm9796 | 0 | 0 | 0 | 0 | NA | NA |
| Zfp652os | 0 | 0 | 0 | 0 | NA | NA |
| B4galnt2 | 0 | 0 | 0 | 0 | NA | NA |
| Hoxb9 | 0 | 0 | 0 | 0 | NA | NA |
| Hoxb6 | 0 | 0 | 0 | 0 | NA | NA |
| Gm11525 | 0 | 0 | 0 | 0 | NA | NA |
| Gm11592 | 0 | 0 | 0 | 0 | NA | NA |
| 1700001P01I | 0 | 0 | 0 | 0 | NA | NA |
| Stac2 | 0 | 0 | 0 | 0 | NA | NA |
| Pnmt | 0 | 0 | 0 | 0 | NA | NA |
| Krt24 | 0 | 0 | 0 | 0 | NA | NA |
| Krt28 | 0 | 0 | 0 | 0 | NA | NA |
| Krt36 | 0 | 0 | 0 | 0 | NA | NA |
| Krt9 | 0 | 0 | 0 | 0 | NA | NA |
| Hspb9 | 0 | 0 | 0 | 0 | NA | NA |
| Gm11615 | 0 | 0 | 0 | 0 | NA | NA |
| Aoc3 | 0 | 0 | 0 | 0 | NA | NA |
| G6pc | 0 | 0 | 0 | 0 | NA | NA |
| Sost | 0 | 0 | 0 | 0 | NA | NA |
| Cd300lg | 0 | 0 | 0 | 0 | NA | NA |
| Nags | 0 | 0 | 0 | 0 | NA | NA |

| | | | | | | |
|---|---|---|---|---|---|---|
| Gm11636 | 0 | 0 | 0 | 0 | NA | NA |
| Meioc | 0 | 0 | 0 | 0 | NA | NA |
| Fam187a | 0 | 0 | 0 | 0 | NA | NA |
| Gfap | 0 | 0 | 0 | 0 | NA | NA |
| Arhgap27os3 | 0 | 0 | 0 | 0 | NA | NA |
| Arhgap27os1 | 0 | 0 | 0 | 0 | NA | NA |
| Efcab3 | 0 | 0 | 0 | 0 | NA | NA |
| Ace | 0 | 0 | 0 | 0 | NA | NA |
| Tcam1 | 0 | 0 | 0 | 0 | NA | NA |
| Apoh | 0 | 0 | 0 | 0 | NA | NA |
| E030025P04I | 0 | 0 | 0 | 0 | NA | NA |
| Gm11697 | 0 | 0 | 0 | 0 | NA | NA |
| Abca6 | 0 | 0 | 0 | 0 | NA | NA |
| 4732490B19I | 0 | 0 | 0 | 0 | NA | NA |
| Cpsf4l | 0 | 0 | 0 | 0 | NA | NA |
| Cd300c | 0 | 0 | 0 | 0 | NA | NA |
| Ict1os | 0 | 0 | 0 | 0 | NA | NA |
| Qrich2 | 0 | 0 | 0 | 0 | NA | NA |
| Gm11730 | 0 | 0 | 0 | 0 | NA | NA |
| 6030468B19I | 0 | 0 | 0 | 0 | NA | NA |
| Gm11725 | 0 | 0 | 0 | 0 | NA | NA |
| Dnah17 | 0 | 0 | 0 | 0 | NA | NA |
| Gm11738 | 0 | 0 | 0 | 0 | NA | NA |
| Gm11766 | 0 | 0 | 0 | 0 | NA | NA |
| Gm11770 | 0 | 0 | 0 | 0 | NA | NA |
| Gm17178 | 0 | 0 | 0 | 0 | NA | NA |
| Cbr2 | 0 | 0 | 0 | 0 | NA | NA |
| Gm28192 | 0 | 0 | 0 | 0 | NA | NA |
| Sectm1b | 0 | 0 | 0 | 0 | NA | NA |
| Sectm1a | 0 | 0 | 0 | 0 | NA | NA |

| | | | | | | |
|---|---|---|---|---|---|---|
| Tex19.1 | 0 | 0 | 0 | 0 | NA | NA |
| Akr1c6 | 0 | 0 | 0 | 0 | NA | NA |
| Gm26645 | 0 | 0 | 0 | 0 | NA | NA |
| Sugct | 0 | 0 | 0 | 0 | NA | NA |
| Sfrp4 | 0 | 0 | 0 | 0 | NA | NA |
| Gpr141 | 0 | 0 | 0 | 0 | NA | NA |
| Aoah | 0 | 0 | 0 | 0 | NA | NA |
| Gpx6 | 0 | 0 | 0 | 0 | NA | NA |
| Btn2a2 | 0 | 0 | 0 | 0 | NA | NA |
| Slc17a2 | 0 | 0 | 0 | 0 | NA | NA |
| Cmah | 0 | 0 | 0 | 0 | NA | NA |
| 4932702P03I | 0 | 0 | 0 | 0 | NA | NA |
| Prl3b1 | 0 | 0 | 0 | 0 | NA | NA |
| Gm5447 | 0 | 0 | 0 | 0 | NA | NA |
| Mylk4 | 0 | 0 | 0 | 0 | NA | NA |
| Serpinb9b | 0 | 0 | 0 | 0 | NA | NA |
| Serpinb1b | 0 | 0 | 0 | 0 | NA | NA |
| Serpinb6d | 0 | 0 | 0 | 0 | NA | NA |
| Fam50b | 0 | 0 | 0 | 0 | NA | NA |
| Ppp1r3g | 0 | 0 | 0 | 0 | NA | NA |
| Nrn1 | 0 | 0 | 0 | 0 | NA | NA |
| Ly86 | 0 | 0 | 0 | 0 | NA | NA |
| Bmp6 | 0 | 0 | 0 | 0 | NA | NA |
| Gm26688 | 0 | 0 | 0 | 0 | NA | NA |
| Gm5083 | 0 | 0 | 0 | 0 | NA | NA |
| Gm9817 | 0 | 0 | 0 | 0 | NA | NA |
| Phf2os1 | 0 | 0 | 0 | 0 | NA | NA |
| Ecm2 | 0 | 0 | 0 | 0 | NA | NA |
| Msx2 | 0 | 0 | 0 | 0 | NA | NA |
| Hk3 | 0 | 0 | 0 | 0 | NA | NA |

| | | | | | | |
|---|---|---|---|---|---|---|
| Pitx1 | 0 | 0 | 0 | 0 | NA | NA |
| Cxcl14 | 0 | 0 | 0 | 0 | NA | NA |
| Slc25a48 | 0 | 0 | 0 | 0 | NA | NA |
| 1700014D04 | 0 | 0 | 0 | 0 | NA | NA |
| 4930486L24F | 0 | 0 | 0 | 0 | NA | NA |
| Gm16133 | 0 | 0 | 0 | 0 | NA | NA |
| Rslcan18 | 0 | 0 | 0 | 0 | NA | NA |
| 8030423J24F | 0 | 0 | 0 | 0 | NA | NA |
| Tert | 0 | 0 | 0 | 0 | NA | NA |
| Fam81b | 0 | 0 | 0 | 0 | NA | NA |
| Gm28526 | 0 | 0 | 0 | 0 | NA | NA |
| Thbs4 | 0 | 0 | 0 | 0 | NA | NA |
| Crhbp | 0 | 0 | 0 | 0 | NA | NA |
| F2rl2 | 0 | 0 | 0 | 0 | NA | NA |
| Ankdd1b | 0 | 0 | 0 | 0 | NA | NA |
| Lncenc1 | 0 | 0 | 0 | 0 | NA | NA |
| 5330416C01I | 0 | 0 | 0 | 0 | NA | NA |
| Zfp366 | 0 | 0 | 0 | 0 | NA | NA |
| Naip6 | 0 | 0 | 0 | 0 | NA | NA |
| 1700099I09R | 0 | 0 | 0 | 0 | NA | NA |
| 2610204G07 | 0 | 0 | 0 | 0 | NA | NA |
| Gapt | 0 | 0 | 0 | 0 | NA | NA |
| Il31ra | 0 | 0 | 0 | 0 | NA | NA |
| Ddx4 | 0 | 0 | 0 | 0 | NA | NA |
| Gm10735 | 0 | 0 | 0 | 0 | NA | NA |
| Esm1 | 0 | 0 | 0 | 0 | NA | NA |
| Hspb3 | 0 | 0 | 0 | 0 | NA | NA |
| Fgf10 | 0 | 0 | 0 | 0 | NA | NA |
| BC147527 | 0 | 0 | 0 | 0 | NA | NA |
| Fam228a | 0 | 0 | 0 | 0 | NA | NA |

| | | | | | | |
|---|---|---|---|---|---|---|
| Apob | 0 | 0 | 0 | 0 | NA | NA |
| Kcns3 | 0 | 0 | 0 | 0 | NA | NA |
| Slc26a4 | 0 | 0 | 0 | 0 | NA | NA |
| Cdhr3 | 0 | 0 | 0 | 0 | NA | NA |
| Meox2 | 0 | 0 | 0 | 0 | NA | NA |
| Insm2 | 0 | 0 | 0 | 0 | NA | NA |
| Foxa1 | 0 | 0 | 0 | 0 | NA | NA |
| Lrfn5 | 0 | 0 | 0 | 0 | NA | NA |
| Gm9887 | 0 | 0 | 0 | 0 | NA | NA |
| Tomm20l | 0 | 0 | 0 | 0 | NA | NA |
| Gpr135 | 0 | 0 | 0 | 0 | NA | NA |
| Kcnh5 | 0 | 0 | 0 | 0 | NA | NA |
| Esr2 | 0 | 0 | 0 | 0 | NA | NA |
| Rad51b | 0 | 0 | 0 | 0 | NA | NA |
| Gm26777 | 0 | 0 | 0 | 0 | NA | NA |
| Galnt16 | 0 | 0 | 0 | 0 | NA | NA |
| Plekhd1os | 0 | 0 | 0 | 0 | NA | NA |
| Plekhd1 | 0 | 0 | 0 | 0 | NA | NA |
| Ccdc177 | 0 | 0 | 0 | 0 | NA | NA |
| Gm28370 | 0 | 0 | 0 | 0 | NA | NA |
| Gm5435 | 0 | 0 | 0 | 0 | NA | NA |
| Rgs6 | 0 | 0 | 0 | 0 | NA | NA |
| Papln | 0 | 0 | 0 | 0 | NA | NA |
| Gm26571 | 0 | 0 | 0 | 0 | NA | NA |
| Prox2os | 0 | 0 | 0 | 0 | NA | NA |
| Jdp2 | 0 | 0 | 0 | 0 | NA | NA |
| Gm26531 | 0 | 0 | 0 | 0 | NA | NA |
| Gm805 | 0 | 0 | 0 | 0 | NA | NA |
| Lrrc74a | 0 | 0 | 0 | 0 | NA | NA |
| Dio2 | 0 | 0 | 0 | 0 | NA | NA |

| | | | | | | |
|---|---|---|---|---|---|---|
| Tshr | 0 | 0 | 0 | 0 | NA | NA |
| Kcnk10 | 0 | 0 | 0 | 0 | NA | NA |
| Gm26839 | 0 | 0 | 0 | 0 | NA | NA |
| Gm16339 | 0 | 0 | 0 | 0 | NA | NA |
| Gm20069 | 0 | 0 | 0 | 0 | NA | NA |
| Fam181a | 0 | 0 | 0 | 0 | NA | NA |
| 9330161L09F | 0 | 0 | 0 | 0 | NA | NA |
| B430119L08I | 0 | 0 | 0 | 0 | NA | NA |
| Serpina1e | 0 | 0 | 0 | 0 | NA | NA |
| Serpina11 | 0 | 0 | 0 | 0 | NA | NA |
| Serpina3i | 0 | 0 | 0 | 0 | NA | NA |
| Gsc | 0 | 0 | 0 | 0 | NA | NA |
| Bdkrb1 | 0 | 0 | 0 | 0 | NA | NA |
| Gm16087 | 0 | 0 | 0 | 0 | NA | NA |
| Ccdc85c | 0 | 0 | 0 | 0 | NA | NA |
| Hhipl1 | 0 | 0 | 0 | 0 | NA | NA |
| A730018C14 | 0 | 0 | 0 | 0 | NA | NA |
| Tex22 | 0 | 0 | 0 | 0 | NA | NA |
| Tmem196 | 0 | 0 | 0 | 0 | NA | NA |
| Osmr | 0 | 0 | 0 | 0 | NA | NA |
| 4930556M19 | 0 | 0 | 0 | 0 | NA | NA |
| C1qtnf3 | 0 | 0 | 0 | 0 | NA | NA |
| Slc45a2 | 0 | 0 | 0 | 0 | NA | NA |
| 1810049J17F | 0 | 0 | 0 | 0 | NA | NA |
| Cdh9 | 0 | 0 | 0 | 0 | NA | NA |
| Cdh10 | 0 | 0 | 0 | 0 | NA | NA |
| Acot10 | 0 | 0 | 0 | 0 | NA | NA |
| Cdh12 | 0 | 0 | 0 | 0 | NA | NA |
| Cdh18 | 0 | 0 | 0 | 0 | NA | NA |
| Gm5468 | 0 | 0 | 0 | 0 | NA | NA |

| | | | | | | |
|---|---|---|---|---|---|---|
| Fbxl7 | 0 | 0 | 0 | 0 | NA | NA |
| Dnah5 | 0 | 0 | 0 | 0 | NA | NA |
| Gm10384 | 0 | 0 | 0 | 0 | NA | NA |
| Gm28221 | 0 | 0 | 0 | 0 | NA | NA |
| Odf1 | 0 | 0 | 0 | 0 | NA | NA |
| Baalc | 0 | 0 | 0 | 0 | NA | NA |
| Abra | 0 | 0 | 0 | 0 | NA | NA |
| Angpt1 | 0 | 0 | 0 | 0 | NA | NA |
| Rspo2 | 0 | 0 | 0 | 0 | NA | NA |
| Tmem74 | 0 | 0 | 0 | 0 | NA | NA |
| Pkhd1l1 | 0 | 0 | 0 | 0 | NA | NA |
| Kcnv1 | 0 | 0 | 0 | 0 | NA | NA |
| Colec10 | 0 | 0 | 0 | 0 | NA | NA |
| Has2 | 0 | 0 | 0 | 0 | NA | NA |
| Fam83a | 0 | 0 | 0 | 0 | NA | NA |
| Gsdmc4 | 0 | 0 | 0 | 0 | NA | NA |
| Adcy8 | 0 | 0 | 0 | 0 | NA | NA |
| Oc90 | 0 | 0 | 0 | 0 | NA | NA |
| Tg | 0 | 0 | 0 | 0 | NA | NA |
| Gm17140 | 0 | 0 | 0 | 0 | NA | NA |
| Gm17035 | 0 | 0 | 0 | 0 | NA | NA |
| Mroh5 | 0 | 0 | 0 | 0 | NA | NA |
| Gm28068 | 0 | 0 | 0 | 0 | NA | NA |
| Ly6c1 | 0 | 0 | 0 | 0 | NA | NA |
| Apol6 | 0 | 0 | 0 | 0 | NA | NA |
| Apol11b | 0 | 0 | 0 | 0 | NA | NA |
| Apol8 | 0 | 0 | 0 | 0 | NA | NA |
| Gm17638 | 0 | 0 | 0 | 0 | NA | NA |
| Cacng2 | 0 | 0 | 0 | 0 | NA | NA |
| Pvalb | 0 | 0 | 0 | 0 | NA | NA |

| | | | | | | |
|---|---|---|---|---|---|---|
| Csf2rb2 | 0 | 0 | 0 | 0 | NA | NA |
| Tex33 | 0 | 0 | 0 | 0 | NA | NA |
| Il2rb | 0 | 0 | 0 | 0 | NA | NA |
| Gm10863 | 0 | 0 | 0 | 0 | NA | NA |
| Gm16059 | 0 | 0 | 0 | 0 | NA | NA |
| Mchr1 | 0 | 0 | 0 | 0 | NA | NA |
| Mpped1 | 0 | 0 | 0 | 0 | NA | NA |
| Mov10l1 | 0 | 0 | 0 | 0 | NA | NA |
| Odf3b | 0 | 0 | 0 | 0 | NA | NA |
| Acr | 0 | 0 | 0 | 0 | NA | NA |
| Syt10 | 0 | 0 | 0 | 0 | NA | NA |
| Lalba | 0 | 0 | 0 | 0 | NA | NA |
| Ddn | 0 | 0 | 0 | 0 | NA | NA |
| B130046B21 | 0 | 0 | 0 | 0 | NA | NA |
| Faim2 | 0 | 0 | 0 | 0 | NA | NA |
| Gm17058 | 0 | 0 | 0 | 0 | NA | NA |
| Gm17057 | 0 | 0 | 0 | 0 | NA | NA |
| Krt4 | 0 | 0 | 0 | 0 | NA | NA |
| Gm9918 | 0 | 0 | 0 | 0 | NA | NA |
| Sp7 | 0 | 0 | 0 | 0 | NA | NA |
| Npff | 0 | 0 | 0 | 0 | NA | NA |
| Hoxc5 | 0 | 0 | 0 | 0 | NA | NA |
| Hoxc8 | 0 | 0 | 0 | 0 | NA | NA |
| Hoxc6 | 0 | 0 | 0 | 0 | NA | NA |
| Gpr84 | 0 | 0 | 0 | 0 | NA | NA |
| Mucl2 | 0 | 0 | 0 | 0 | NA | NA |
| Gm15537 | 0 | 0 | 0 | 0 | NA | NA |
| AU021092 | 0 | 0 | 0 | 0 | NA | NA |
| Grin2a | 0 | 0 | 0 | 0 | NA | NA |
| Rpl39l | 0 | 0 | 0 | 0 | NA | NA |

| | | | | | | |
|---|---|---|---|---|---|---|
| Atf7ip2 | 0 | 0 | 0 | 0 | NA | NA |
| Tnfrsf17 | 0 | 0 | 0 | 0 | NA | NA |
| Pla2g10 | 0 | 0 | 0 | 0 | NA | NA |
| Serpind1 | 0 | 0 | 0 | 0 | NA | NA |
| Gsc2 | 0 | 0 | 0 | 0 | NA | NA |
| Tbx1 | 0 | 0 | 0 | 0 | NA | NA |
| Map6d1 | 0 | 0 | 0 | 0 | NA | NA |
| Teddm3 | 0 | 0 | 0 | 0 | NA | NA |
| Ahsg | 0 | 0 | 0 | 0 | NA | NA |
| Masp1 | 0 | 0 | 0 | 0 | NA | NA |
| A230028O05 | 0 | 0 | 0 | 0 | NA | NA |
| Atp13a4 | 0 | 0 | 0 | 0 | NA | NA |
| Gm26834 | 0 | 0 | 0 | 0 | NA | NA |
| Gm26569 | 0 | 0 | 0 | 0 | NA | NA |
| Pigz | 0 | 0 | 0 | 0 | NA | NA |
| Bex6 | 0 | 0 | 0 | 0 | NA | NA |
| Smco1 | 0 | 0 | 0 | 0 | NA | NA |
| Slc51a | 0 | 0 | 0 | 0 | NA | NA |
| Tnk2os | 0 | 0 | 0 | 0 | NA | NA |
| 1700119H24 | 0 | 0 | 0 | 0 | NA | NA |
| Gm15564 | 0 | 0 | 0 | 0 | NA | NA |
| Csta1 | 0 | 0 | 0 | 0 | NA | NA |
| Gpr156 | 0 | 0 | 0 | 0 | NA | NA |
| Cd80 | 0 | 0 | 0 | 0 | NA | NA |
| Gm17103 | 0 | 0 | 0 | 0 | NA | NA |
| Zdhhc23 | 0 | 0 | 0 | 0 | NA | NA |
| Cd200r4 | 0 | 0 | 0 | 0 | NA | NA |
| Gcsam | 0 | 0 | 0 | 0 | NA | NA |
| Dppa4 | 0 | 0 | 0 | 0 | NA | NA |
| G730013B05 | 0 | 0 | 0 | 0 | NA | NA |

| | | | | | | |
|---|---|---|---|---|---|---|
| Gm16892 | 0 | 0 | 0 | 0 | NA | NA |
| Htr1f | 0 | 0 | 0 | 0 | NA | NA |
| Pou1f1 | 0 | 0 | 0 | 0 | NA | NA |
| Vgll3 | 0 | 0 | 0 | 0 | NA | NA |
| Mir155hg | 0 | 0 | 0 | 0 | NA | NA |
| Map3k7cl | 0 | 0 | 0 | 0 | NA | NA |
| 2810407A14I | 0 | 0 | 0 | 0 | NA | NA |
| Grik1 | 0 | 0 | 0 | 0 | NA | NA |
| Sim2 | 0 | 0 | 0 | 0 | NA | NA |
| Kcnj15 | 0 | 0 | 0 | 0 | NA | NA |
| 2810404F17I | 0 | 0 | 0 | 0 | NA | NA |
| 3300005D01I | 0 | 0 | 0 | 0 | NA | NA |
| Tagap | 0 | 0 | 0 | 0 | NA | NA |
| 1700110C19I | 0 | 0 | 0 | 0 | NA | NA |
| Gm16168 | 0 | 0 | 0 | 0 | NA | NA |
| Gm26873 | 0 | 0 | 0 | 0 | NA | NA |
| Zscan10 | 0 | 0 | 0 | 0 | NA | NA |
| Prss33 | 0 | 0 | 0 | 0 | NA | NA |
| Prss32 | 0 | 0 | 0 | 0 | NA | NA |
| Gm15947 | 0 | 0 | 0 | 0 | NA | NA |
| Dcpp2 | 0 | 0 | 0 | 0 | NA | NA |
| Dcpp3 | 0 | 0 | 0 | 0 | NA | NA |
| Rpl3l | 0 | 0 | 0 | 0 | NA | NA |
| Ptx4 | 0 | 0 | 0 | 0 | NA | NA |
| Ccdc154 | 0 | 0 | 0 | 0 | NA | NA |
| Nhlrc4 | 0 | 0 | 0 | 0 | NA | NA |
| D630044L22I | 0 | 0 | 0 | 0 | NA | NA |
| Nkx2-5 | 0 | 0 | 0 | 0 | NA | NA |
| Gm17382 | 0 | 0 | 0 | 0 | NA | NA |
| Gm10505 | 0 | 0 | 0 | 0 | NA | NA |

| | | | | | | |
|---|---|---|---|---|---|---|
| Slc26a8 | 0 | 0 | 0 | 0 | NA | NA |
| Pnpla1 | 0 | 0 | 0 | 0 | NA | NA |
| Rab44 | 0 | 0 | 0 | 0 | NA | NA |
| Gm26885 | 0 | 0 | 0 | 0 | NA | NA |
| 1700097N02 | 0 | 0 | 0 | 0 | NA | NA |
| Ubash3a | 0 | 0 | 0 | 0 | NA | NA |
| Cryaa | 0 | 0 | 0 | 0 | NA | NA |
| Pglyrp2 | 0 | 0 | 0 | 0 | NA | NA |
| Gm17115 | 0 | 0 | 0 | 0 | NA | NA |
| Myo1f | 0 | 0 | 0 | 0 | NA | NA |
| BC051537 | 0 | 0 | 0 | 0 | NA | NA |
| H2-Ob | 0 | 0 | 0 | 0 | NA | NA |
| Btnl2 | 0 | 0 | 0 | 0 | NA | NA |
| Gm20481 | 0 | 0 | 0 | 0 | NA | NA |
| Ly6g6f | 0 | 0 | 0 | 0 | NA | NA |
| Gm16181 | 0 | 0 | 0 | 0 | NA | NA |
| Cdsn | 0 | 0 | 0 | 0 | NA | NA |
| Trim15 | 0 | 0 | 0 | 0 | NA | NA |
| H2-M5 | 0 | 0 | 0 | 0 | NA | NA |
| H2-M2 | 0 | 0 | 0 | 0 | NA | NA |
| Crisp2 | 0 | 0 | 0 | 0 | NA | NA |
| Ptchd4 | 0 | 0 | 0 | 0 | NA | NA |
| Adgrf4 | 0 | 0 | 0 | 0 | NA | NA |
| Ankrd66 | 0 | 0 | 0 | 0 | NA | NA |
| Runx2os1 | 0 | 0 | 0 | 0 | NA | NA |
| F630040K05I | 0 | 0 | 0 | 0 | NA | NA |
| Crip3 | 0 | 0 | 0 | 0 | NA | NA |
| Gm16494 | 0 | 0 | 0 | 0 | NA | NA |
| Gm5814 | 0 | 0 | 0 | 0 | NA | NA |
| Gm21981 | 0 | 0 | 0 | 0 | NA | NA |

| | | | | | | |
|---|---|---|---|---|---|---|
| Frs3os | 0 | 0 | 0 | 0 | NA | NA |
| Gm15556 | 0 | 0 | 0 | 0 | NA | NA |
| 9830107B12 | 0 | 0 | 0 | 0 | NA | NA |
| A530064D06 | 0 | 0 | 0 | 0 | NA | NA |
| Trem3 | 0 | 0 | 0 | 0 | NA | NA |
| Treml2 | 0 | 0 | 0 | 0 | NA | NA |
| Treml1 | 0 | 0 | 0 | 0 | NA | NA |
| Slc5a7 | 0 | 0 | 0 | 0 | NA | NA |
| Adgre4 | 0 | 0 | 0 | 0 | NA | NA |
| Slc25a41 | 0 | 0 | 0 | 0 | NA | NA |
| Cd70 | 0 | 0 | 0 | 0 | NA | NA |
| Tnfsf14 | 0 | 0 | 0 | 0 | NA | NA |
| Nudt12os | 0 | 0 | 0 | 0 | NA | NA |
| AU016765 | 0 | 0 | 0 | 0 | NA | NA |
| Themis3 | 0 | 0 | 0 | 0 | NA | NA |
| 5031415H12 | 0 | 0 | 0 | 0 | NA | NA |
| Gm20703 | 0 | 0 | 0 | 0 | NA | NA |
| Gm26510 | 0 | 0 | 0 | 0 | NA | NA |
| Cyp1b1 | 0 | 0 | 0 | 0 | NA | NA |
| Haao | 0 | 0 | 0 | 0 | NA | NA |
| Gm15978 | 0 | 0 | 0 | 0 | NA | NA |
| Rpl36-ps4 | 0 | 0 | 0 | 0 | NA | NA |
| Adcyap1 | 0 | 0 | 0 | 0 | NA | NA |
| Lama3 | 0 | 0 | 0 | 0 | NA | NA |
| Gm15328 | 0 | 0 | 0 | 0 | NA | NA |
| Dsc3 | 0 | 0 | 0 | 0 | NA | NA |
| Dsg1c | 0 | 0 | 0 | 0 | NA | NA |
| Gpr17 | 0 | 0 | 0 | 0 | NA | NA |
| Pcdha5 | 0 | 0 | 0 | 0 | NA | NA |
| Pcdha11 | 0 | 0 | 0 | 0 | NA | NA |

| | | | | | | |
|---|---|---|---|---|---|---|
| Pcdhgb2 | 0 | 0 | 0 | 0 | NA | NA |
| Kctd16 | 0 | 0 | 0 | 0 | NA | NA |
| Sh3rf2 | 0 | 0 | 0 | 0 | NA | NA |
| Scgb3a2 | 0 | 0 | 0 | 0 | NA | NA |
| Myot | 0 | 0 | 0 | 0 | NA | NA |
| Gm43425 | 0 | 0 | 0 | 0 | NA | NA |
| Arl14epl | 0 | 0 | 0 | 0 | NA | NA |
| Redrum | 0 | 0 | 0 | 0 | NA | NA |
| Slc27a6 | 0 | 0 | 0 | 0 | NA | NA |
| Adamts19 | 0 | 0 | 0 | 0 | NA | NA |
| Chsy3 | 0 | 0 | 0 | 0 | NA | NA |
| Gm4841 | 0 | 0 | 0 | 0 | NA | NA |
| Iigp1 | 0 | 0 | 0 | 0 | NA | NA |
| Arsi | 0 | 0 | 0 | 0 | NA | NA |
| Gm38165 | 0 | 0 | 0 | 0 | NA | NA |
| 2700046A07 | 0 | 0 | 0 | 0 | NA | NA |
| Apcdd1 | 0 | 0 | 0 | 0 | NA | NA |
| Grp | 0 | 0 | 0 | 0 | NA | NA |
| Mc4r | 0 | 0 | 0 | 0 | NA | NA |
| Dcc | 0 | 0 | 0 | 0 | NA | NA |
| D730045A05 | 0 | 0 | 0 | 0 | NA | NA |
| Slc14a1 | 0 | 0 | 0 | 0 | NA | NA |
| Gm26676 | 0 | 0 | 0 | 0 | NA | NA |
| Sall3 | 0 | 0 | 0 | 0 | NA | NA |
| Cd226 | 0 | 0 | 0 | 0 | NA | NA |
| Dok6 | 0 | 0 | 0 | 0 | NA | NA |
| Gpr152 | 0 | 0 | 0 | 0 | NA | NA |
| Kcnk7 | 0 | 0 | 0 | 0 | NA | NA |
| Slc22a20 | 0 | 0 | 0 | 0 | NA | NA |
| Naaladl1 | 0 | 0 | 0 | 0 | NA | NA |

| | | | | | | |
|---|---|---|---|---|---|---|
| Batf2 | 0 | 0 | 0 | 0 | NA | NA |
| Slc22a30 | 0 | 0 | 0 | 0 | NA | NA |
| Chrm1 | 0 | 0 | 0 | 0 | NA | NA |
| Pga5 | 0 | 0 | 0 | 0 | NA | NA |
| Cd6 | 0 | 0 | 0 | 0 | NA | NA |
| Ptgdr2 | 0 | 0 | 0 | 0 | NA | NA |
| Ms4a15 | 0 | 0 | 0 | 0 | NA | NA |
| Ms4a1 | 0 | 0 | 0 | 0 | NA | NA |
| Ms4a14 | 0 | 0 | 0 | 0 | NA | NA |
| Ms4a4b | 0 | 0 | 0 | 0 | NA | NA |
| Gm19261 | 0 | 0 | 0 | 0 | NA | NA |
| Ms4a4d | 0 | 0 | 0 | 0 | NA | NA |
| Ms4a3 | 0 | 0 | 0 | 0 | NA | NA |
| Glyat | 0 | 0 | 0 | 0 | NA | NA |
| Gm15962 | 0 | 0 | 0 | 0 | NA | NA |
| Tmc1 | 0 | 0 | 0 | 0 | NA | NA |
| Gda | 0 | 0 | 0 | 0 | NA | NA |
| Dmrt1 | 0 | 0 | 0 | 0 | NA | NA |
| Dmrt3 | 0 | 0 | 0 | 0 | NA | NA |
| Dmrt2 | 0 | 0 | 0 | 0 | NA | NA |
| D930032P07 | 0 | 0 | 0 | 0 | NA | NA |
| Lipm | 0 | 0 | 0 | 0 | NA | NA |
| Gm26902 | 0 | 0 | 0 | 0 | NA | NA |
| Ifit3 | 0 | 0 | 0 | 0 | NA | NA |
| Ifit1bl2 | 0 | 0 | 0 | 0 | NA | NA |
| Htr7 | 0 | 0 | 0 | 0 | NA | NA |
| Dntt | 0 | 0 | 0 | 0 | NA | NA |
| 4933411K16I | 0 | 0 | 0 | 0 | NA | NA |
| Loxl4 | 0 | 0 | 0 | 0 | NA | NA |
| Hpse2 | 0 | 0 | 0 | 0 | NA | NA |

| | | | | | | |
|---|---|---|---|---|---|---|
| Gm20467 | 0 | 0 | 0 | 0 | NA | NA |
| Scd4 | 0 | 0 | 0 | 0 | NA | NA |
| Scd1 | 0 | 0 | 0 | 0 | NA | NA |
| Gm26644 | 0 | 0 | 0 | 0 | NA | NA |
| Tlx1 | 0 | 0 | 0 | 0 | NA | NA |
| Lbx1 | 0 | 0 | 0 | 0 | NA | NA |
| Fgf8 | 0 | 0 | 0 | 0 | NA | NA |
| Pitx3 | 0 | 0 | 0 | 0 | NA | NA |
| Sorcs3 | 0 | 0 | 0 | 0 | NA | NA |
| 4833407H14 | 0 | 0 | 0 | 0 | NA | NA |
| Gucy2g | 0 | 0 | 0 | 0 | NA | NA |
| Nrap | 0 | 0 | 0 | 0 | NA | NA |
| Gm17197 | 0 | 0 | 0 | 0 | NA | NA |
| Plekhs1 | 0 | 0 | 0 | 0 | NA | NA |
| Adrb1 | 0 | 0 | 0 | 0 | NA | NA |
| Tdrd1 | 0 | 0 | 0 | 0 | NA | NA |
| 1810007D17 | 0 | 0 | 0 | 0 | NA | NA |
| Emx2os | 0 | 0 | 0 | 0 | NA | NA |
| Emx2 | 0 | 0 | 0 | 0 | NA | NA |
| Il17a | 0 | 0 | 0 | 0 | NA | NA |
| Gm5524 | 0 | 0 | 0 | 0 | NA | NA |
| Gm9898 | 0 | 0 | 0 | 0 | NA | NA |
| Gm3646 | 0 | 0 | 0 | 0 | NA | NA |
| Ctla4 | 0 | 0 | 0 | 0 | NA | NA |
| Ugt1a9 | 0 | 0 | 0 | 0 | NA | NA |
| Serpinb7 | 0 | 0 | 0 | 0 | NA | NA |
| Cd55b | 0 | 0 | 0 | 0 | NA | NA |
| Gm16083 | 0 | 0 | 0 | 0 | NA | NA |
| Gm15848 | 0 | 0 | 0 | 0 | NA | NA |
| Gm29282 | 0 | 0 | 0 | 0 | NA | NA |

| | | | | | | |
|---|---|---|---|---|---|---|
| Gm36945 | 0 | 0 | 0 | 0 | NA | NA |
| Lrrc52 | 0 | 0 | 0 | 0 | NA | NA |
| Fcrla | 0 | 0 | 0 | 0 | NA | NA |
| Gm10521 | 0 | 0 | 0 | 0 | NA | NA |
| Fcer1a | 0 | 0 | 0 | 0 | NA | NA |
| Fut7 | 0 | 0 | 0 | 0 | NA | NA |
| Qrfp | 0 | 0 | 0 | 0 | NA | NA |
| Gm14033 | 0 | 0 | 0 | 0 | NA | NA |
| 4932414N04 | 0 | 0 | 0 | 0 | NA | NA |
| Dhrs9 | 0 | 0 | 0 | 0 | NA | NA |
| Pin1rt1 | 0 | 0 | 0 | 0 | NA | NA |
| Scp2d1 | 0 | 0 | 0 | 0 | NA | NA |
| Rspo4os | 0 | 0 | 0 | 0 | NA | NA |
| Gm14288 | 0 | 0 | 0 | 0 | NA | NA |
| 2210418O10 | 0 | 0 | 0 | 0 | NA | NA |
| Gm14341 | 0 | 0 | 0 | 0 | NA | NA |
| Slc6a14 | 0 | 0 | 0 | 0 | NA | NA |
| Gm9 | 0 | 0 | 0 | 0 | NA | NA |
| Xlr5b | 0 | 0 | 0 | 0 | NA | NA |
| Xlr5c | 0 | 0 | 0 | 0 | NA | NA |
| Avpr2 | 0 | 0 | 0 | 0 | NA | NA |
| Gm14812 | 0 | 0 | 0 | 0 | NA | NA |
| Pof1b | 0 | 0 | 0 | 0 | NA | NA |
| Tcp11x2 | 0 | 0 | 0 | 0 | NA | NA |
| Tmsb15a | 0 | 0 | 0 | 0 | NA | NA |
| Il13ra2 | 0 | 0 | 0 | 0 | NA | NA |
| 1110015O18 | 0 | 0 | 0 | 0 | NA | NA |
| Sirpb1b | 0 | 0 | 0 | 0 | NA | NA |
| Qrfpr | 0 | 0 | 0 | 0 | NA | NA |
| Sh2d2a | 0 | 0 | 0 | 0 | NA | NA |

| | | | | | | |
|---|---|---|---|---|---|---|
| 2210017I01R | 0 | 0 | 0 | 0 | NA | NA |
| Them5 | 0 | 0 | 0 | 0 | NA | NA |
| Gm26594 | 0 | 0 | 0 | 0 | NA | NA |
| Amy2a4 | 0 | 0 | 0 | 0 | NA | NA |
| A730020M07 | 0 | 0 | 0 | 0 | NA | NA |
| Gm30484 | 0 | 0 | 0 | 0 | NA | NA |
| Gm12354 | 0 | 0 | 0 | 0 | NA | NA |
| Gm11264 | 0 | 0 | 0 | 0 | NA | NA |
| Skint5 | 0 | 0 | 0 | 0 | NA | NA |
| Cyp4a12a | 0 | 0 | 0 | 0 | NA | NA |
| Gm12888 | 0 | 0 | 0 | 0 | NA | NA |
| Tssk3 | 0 | 0 | 0 | 0 | NA | NA |
| Gm12974 | 0 | 0 | 0 | 0 | NA | NA |
| Pax7 | 0 | 0 | 0 | 0 | NA | NA |
| Gm13017 | 0 | 0 | 0 | 0 | NA | NA |
| Zfp600 | 0 | 0 | 0 | 0 | NA | NA |
| 1700045H11 | 0 | 0 | 0 | 0 | NA | NA |
| Ankrd65 | 0 | 0 | 0 | 0 | NA | NA |
| Speer4d | 0 | 0 | 0 | 0 | NA | NA |
| Speer4a | 0 | 0 | 0 | 0 | NA | NA |
| En2 | 0 | 0 | 0 | 0 | NA | NA |
| Adgrf3 | 0 | 0 | 0 | 0 | NA | NA |
| Gm7596 | 0 | 0 | 0 | 0 | NA | NA |
| Spata18 | 0 | 0 | 0 | 0 | NA | NA |
| C78283 | 0 | 0 | 0 | 0 | NA | NA |
| Gm15984 | 0 | 0 | 0 | 0 | NA | NA |
| Pdcl2 | 0 | 0 | 0 | 0 | NA | NA |
| Ugt2b1 | 0 | 0 | 0 | 0 | NA | NA |
| Dmp1 | 0 | 0 | 0 | 0 | NA | NA |
| Ibsp | 0 | 0 | 0 | 0 | NA | NA |

| | | | | | | |
|---|---|---|---|---|---|---|
| Gm13822 | 0 | 0 | 0 | 0 | NA | NA |
| Gm43069 | 0 | 0 | 0 | 0 | NA | NA |
| Piwil1 | 0 | 0 | 0 | 0 | NA | NA |
| Gm28616 | 0 | 0 | 0 | 0 | NA | NA |
| Gm43645 | 0 | 0 | 0 | 0 | NA | NA |
| Gm16122 | 0 | 0 | 0 | 0 | NA | NA |
| Cyp3a11 | 0 | 0 | 0 | 0 | NA | NA |
| Gm13855 | 0 | 0 | 0 | 0 | NA | NA |
| 5830405F06I | 0 | 0 | 0 | 0 | NA | NA |
| 2010310C07I | 0 | 0 | 0 | 0 | NA | NA |
| Ghrhr | 0 | 0 | 0 | 0 | NA | NA |
| 6430584L05F | 0 | 0 | 0 | 0 | NA | NA |
| Il23r | 0 | 0 | 0 | 0 | NA | NA |
| Gm1070 | 0 | 0 | 0 | 0 | NA | NA |
| Reg3b | 0 | 0 | 0 | 0 | NA | NA |
| 2310069B03I | 0 | 0 | 0 | 0 | NA | NA |
| Atp6v1b1 | 0 | 0 | 0 | 0 | NA | NA |
| Gm44117 | 0 | 0 | 0 | 0 | NA | NA |
| Cpne9 | 0 | 0 | 0 | 0 | NA | NA |
| 1700030F04I | 0 | 0 | 0 | 0 | NA | NA |
| Clec4e | 0 | 0 | 0 | 0 | NA | NA |
| Vmn2r26 | 0 | 0 | 0 | 0 | NA | NA |
| Klrb1c | 0 | 0 | 0 | 0 | NA | NA |
| Clec2g | 0 | 0 | 0 | 0 | NA | NA |
| Tmem52b | 0 | 0 | 0 | 0 | NA | NA |
| Tarm1 | 0 | 0 | 0 | 0 | NA | NA |
| Gm17067 | 0 | 0 | 0 | 0 | NA | NA |
| Klk1b5 | 0 | 0 | 0 | 0 | NA | NA |
| Gm15413 | 0 | 0 | 0 | 0 | NA | NA |
| Trim34b | 0 | 0 | 0 | 0 | NA | NA |

| | | | | | | |
|---|---|---|---|---|---|---|
| Abca14 | 0 | 0 | 0 | 0 | NA | NA |
| Abca15 | 0 | 0 | 0 | 0 | NA | NA |
| Gm4275 | 0 | 0 | 0 | 0 | NA | NA |
| Taar4 | 0 | 0 | 0 | 0 | NA | NA |
| Tacr2 | 0 | 0 | 0 | 0 | NA | NA |
| 1110002J07F | 0 | 0 | 0 | 0 | NA | NA |
| Kera | 0 | 0 | 0 | 0 | NA | NA |
| Gad1-ps | 0 | 0 | 0 | 0 | NA | NA |
| Myf6 | 0 | 0 | 0 | 0 | NA | NA |
| Glipr1l1 | 0 | 0 | 0 | 0 | NA | NA |
| Rdh16 | 0 | 0 | 0 | 0 | NA | NA |
| Cd209a | 0 | 0 | 0 | 0 | NA | NA |
| Cd209d | 0 | 0 | 0 | 0 | NA | NA |
| 5830468F06I | 0 | 0 | 0 | 0 | NA | NA |
| Gm6040 | 0 | 0 | 0 | 0 | NA | NA |
| Rln3 | 0 | 0 | 0 | 0 | NA | NA |
| Ces1g | 0 | 0 | 0 | 0 | NA | NA |
| Mt4 | 0 | 0 | 0 | 0 | NA | NA |
| Foxl1 | 0 | 0 | 0 | 0 | NA | NA |
| Acox2 | 0 | 0 | 0 | 0 | NA | NA |
| Cysltr2 | 0 | 0 | 0 | 0 | NA | NA |
| Gpr18 | 0 | 0 | 0 | 0 | NA | NA |
| Gsta2 | 0 | 0 | 0 | 0 | NA | NA |
| 1700057G04 | 0 | 0 | 0 | 0 | NA | NA |
| Gm29154 | 0 | 0 | 0 | 0 | NA | NA |
| 2310075C17I | 0 | 0 | 0 | 0 | NA | NA |
| Ccr8 | 0 | 0 | 0 | 0 | NA | NA |
| 4933405E24I | 0 | 0 | 0 | 0 | NA | NA |
| Il3 | 0 | 0 | 0 | 0 | NA | NA |
| Gm12249 | 0 | 0 | 0 | 0 | NA | NA |

| | | | | | | |
|---|---|---|---|---|---|---|
| Gm12298 | 0 | 0 | 0 | 0 | NA | NA |
| Alox12e | 0 | 0 | 0 | 0 | NA | NA |
| Gm12320 | 0 | 0 | 0 | 0 | NA | NA |
| Gm12326 | 0 | 0 | 0 | 0 | NA | NA |
| Wfdc18 | 0 | 0 | 0 | 0 | NA | NA |
| Gm11541 | 0 | 0 | 0 | 0 | NA | NA |
| Gm11534 | 0 | 0 | 0 | 0 | NA | NA |
| Krtap3-3 | 0 | 0 | 0 | 0 | NA | NA |
| Krtap31-2 | 0 | 0 | 0 | 0 | NA | NA |
| Higd1b | 0 | 0 | 0 | 0 | NA | NA |
| Cacng1 | 0 | 0 | 0 | 0 | NA | NA |
| Gm11681 | 0 | 0 | 0 | 0 | NA | NA |
| Akr1c20 | 0 | 0 | 0 | 0 | NA | NA |
| Btn1a1 | 0 | 0 | 0 | 0 | NA | NA |
| Prl6a1 | 0 | 0 | 0 | 0 | NA | NA |
| 1700029N11 | 0 | 0 | 0 | 0 | NA | NA |
| Irx4 | 0 | 0 | 0 | 0 | NA | NA |
| 9330111N05 | 0 | 0 | 0 | 0 | NA | NA |
| Ckmt2 | 0 | 0 | 0 | 0 | NA | NA |
| Gm17190 | 0 | 0 | 0 | 0 | NA | NA |
| 4932411K12I | 0 | 0 | 0 | 0 | NA | NA |
| Mcidas | 0 | 0 | 0 | 0 | NA | NA |
| Gm17024 | 0 | 0 | 0 | 0 | NA | NA |
| Rpl10l | 0 | 0 | 0 | 0 | NA | NA |
| Serpina3n | 0 | 0 | 0 | 0 | NA | NA |
| Anxa13 | 0 | 0 | 0 | 0 | NA | NA |
| Ly6g | 0 | 0 | 0 | 0 | NA | NA |
| Tssk5 | 0 | 0 | 0 | 0 | NA | NA |
| Apol9a | 0 | 0 | 0 | 0 | NA | NA |
| 4930562C15I | 0 | 0 | 0 | 0 | NA | NA |

| | | | | | | |
|---|---|---|---|---|---|---|
| Tnp2 | 0 | 0 | 0 | 0 | NA | NA |
| Stfa1 | 0 | 0 | 0 | 0 | NA | NA |
| Cd200r3 | 0 | 0 | 0 | 0 | NA | NA |
| Btla | 0 | 0 | 0 | 0 | NA | NA |
| 1700029J03F | 0 | 0 | 0 | 0 | NA | NA |
| Itgb2l | 0 | 0 | 0 | 0 | NA | NA |
| Umodl1 | 0 | 0 | 0 | 0 | NA | NA |
| H2-Eb2 | 0 | 0 | 0 | 0 | NA | NA |
| H2-Q1 | 0 | 0 | 0 | 0 | NA | NA |
| Gm16172 | 0 | 0 | 0 | 0 | NA | NA |
| 1700122O11 | 0 | 0 | 0 | 0 | NA | NA |
| Gm19585 | 0 | 0 | 0 | 0 | NA | NA |
| Usp9y | 0 | 0 | 0 | 0 | NA | NA |
| Gm15337 | 0 | 0 | 0 | 0 | NA | NA |
| Plac8l1 | 0 | 0 | 0 | 0 | NA | NA |
| 1700030N03 | 0 | 0 | 0 | 0 | NA | NA |
| Gm28374 | 0 | 0 | 0 | 0 | NA | NA |
| Slc22a27 | 0 | 0 | 0 | 0 | NA | NA |
| Keg1 | 0 | 0 | 0 | 0 | NA | NA |
| Pdcd1lg2 | 0 | 0 | 0 | 0 | NA | NA |
| Cyp2c68 | 0 | 0 | 0 | 0 | NA | NA |
| Gm20538 | 0 | 0 | 0 | 0 | NA | NA |
| Elovl3 | 0 | 0 | 0 | 0 | NA | NA |
| Csprs | 0 | 0 | 0 | 0 | NA | NA |
| Khdc1a | 0 | 0 | 0 | 0 | NA | NA |
| Khdc1c | 0 | 0 | 0 | 0 | NA | NA |
| Gm29156 | 0 | 0 | 0 | 0 | NA | NA |
| Pth2r | 0 | 0 | 0 | 0 | NA | NA |
| Gm29539 | 0 | 0 | 0 | 0 | NA | NA |
| Pax3 | 0 | 0 | 0 | 0 | NA | NA |

| | | | | | | |
|---|---|---|---|---|---|---|
| A530040E14 | 0 | 0 | 0 | 0 | NA | NA |
| Gpr55 | 0 | 0 | 0 | 0 | NA | NA |
| Nmur1 | 0 | 0 | 0 | 0 | NA | NA |
| Gm28187 | 0 | 0 | 0 | 0 | NA | NA |
| Gpr37l1 | 0 | 0 | 0 | 0 | NA | NA |
| Gm37552 | 0 | 0 | 0 | 0 | NA | NA |
| Tex35 | 0 | 0 | 0 | 0 | NA | NA |
| Tnn | 0 | 0 | 0 | 0 | NA | NA |
| Gm31925 | 0 | 0 | 0 | 0 | NA | NA |
| Gm26685 | 0 | 0 | 0 | 0 | NA | NA |
| Gm37756 | 0 | 0 | 0 | 0 | NA | NA |
| Kcnj10 | 0 | 0 | 0 | 0 | NA | NA |
| Slamf8 | 0 | 0 | 0 | 0 | NA | NA |
| Gm5533 | 0 | 0 | 0 | 0 | NA | NA |
| Gm26879 | 0 | 0 | 0 | 0 | NA | NA |
| Gm16897 | 0 | 0 | 0 | 0 | NA | NA |
| Gm10851 | 0 | 0 | 0 | 0 | NA | NA |
| Il2ra | 0 | 0 | 0 | 0 | NA | NA |
| Tmem236 | 0 | 0 | 0 | 0 | NA | NA |
| Malrd1 | 0 | 0 | 0 | 0 | NA | NA |
| Gm13411 | 0 | 0 | 0 | 0 | NA | NA |
| Gm13387 | 0 | 0 | 0 | 0 | NA | NA |
| Lhx3 | 0 | 0 | 0 | 0 | NA | NA |
| Gm13381 | 0 | 0 | 0 | 0 | NA | NA |
| Hc | 0 | 0 | 0 | 0 | NA | NA |
| Gm13561 | 0 | 0 | 0 | 0 | NA | NA |
| Hoxd10 | 0 | 0 | 0 | 0 | NA | NA |
| A930006I01F | 0 | 0 | 0 | 0 | NA | NA |
| Bbox1 | 0 | 0 | 0 | 0 | NA | NA |
| Muc15 | 0 | 0 | 0 | 0 | NA | NA |

| | | | | | | |
|---|---|---|---|---|---|---|
| Duoxa1 | 0 | 0 | 0 | 0 | NA | NA |
| AI847159 | 0 | 0 | 0 | 0 | NA | NA |
| Hao1 | 0 | 0 | 0 | 0 | NA | NA |
| Sstr4 | 0 | 0 | 0 | 0 | NA | NA |
| Gm14198 | 0 | 0 | 0 | 0 | NA | NA |
| Slc32a1 | 0 | 0 | 0 | 0 | NA | NA |
| Wfdc12 | 0 | 0 | 0 | 0 | NA | NA |
| Svs5 | 0 | 0 | 0 | 0 | NA | NA |
| A530013C23 | 0 | 0 | 0 | 0 | NA | NA |
| Bcas1os2 | 0 | 0 | 0 | 0 | NA | NA |
| Pck1 | 0 | 0 | 0 | 0 | NA | NA |
| Srms | 0 | 0 | 0 | 0 | NA | NA |
| Gpr82 | 0 | 0 | 0 | 0 | NA | NA |
| Rhox8 | 0 | 0 | 0 | 0 | NA | NA |
| Sh2d1a | 0 | 0 | 0 | 0 | NA | NA |
| Vgll1 | 0 | 0 | 0 | 0 | NA | NA |
| Gm14809 | 0 | 0 | 0 | 0 | NA | NA |
| Foxr2 | 0 | 0 | 0 | 0 | NA | NA |
| Gm38335 | 0 | 0 | 0 | 0 | NA | NA |
| A930001A20 | 0 | 0 | 0 | 0 | NA | NA |
| Sirpb1a | 0 | 0 | 0 | 0 | NA | NA |
| Dnajc5b | 0 | 0 | 0 | 0 | NA | NA |
| Gm43525 | 0 | 0 | 0 | 0 | NA | NA |
| Ghsr | 0 | 0 | 0 | 0 | NA | NA |
| Gm5103 | 0 | 0 | 0 | 0 | NA | NA |
| Gm20754 | 0 | 0 | 0 | 0 | NA | NA |
| Npy2r | 0 | 0 | 0 | 0 | NA | NA |
| Sprr1a | 0 | 0 | 0 | 0 | NA | NA |
| Gm15444 | 0 | 0 | 0 | 0 | NA | NA |
| Eps8l3 | 0 | 0 | 0 | 0 | NA | NA |

| | | | | | | |
|---|---|---|---|---|---|---|
| Myoz2 | 0 | 0 | 0 | 0 | NA | NA |
| Gm43522 | 0 | 0 | 0 | 0 | NA | NA |
| 4930430E12l | 0 | 0 | 0 | 0 | NA | NA |
| Sit1 | 0 | 0 | 0 | 0 | NA | NA |
| Gm568 | 0 | 0 | 0 | 0 | NA | NA |
| Tyrp1 | 0 | 0 | 0 | 0 | NA | NA |
| Gm12708 | 0 | 0 | 0 | 0 | NA | NA |
| Gm29064 | 0 | 0 | 0 | 0 | NA | NA |
| Gm12963 | 0 | 0 | 0 | 0 | NA | NA |
| Gm13066 | 0 | 0 | 0 | 0 | NA | NA |
| 1700003C15l | 0 | 0 | 0 | 0 | NA | NA |
| Gm42450 | 0 | 0 | 0 | 0 | NA | NA |
| Gm42849 | 0 | 0 | 0 | 0 | NA | NA |
| Gm43042 | 0 | 0 | 0 | 0 | NA | NA |
| AU023070 | 0 | 0 | 0 | 0 | NA | NA |
| Hopxos | 0 | 0 | 0 | 0 | NA | NA |
| Tmprss11g | 0 | 0 | 0 | 0 | NA | NA |
| Cxcl9 | 0 | 0 | 0 | 0 | NA | NA |
| Vmn2r13 | 0 | 0 | 0 | 0 | NA | NA |
| Pitpnm2os2 | 0 | 0 | 0 | 0 | NA | NA |
| Muc3a | 0 | 0 | 0 | 0 | NA | NA |
| Sap25 | 0 | 0 | 0 | 0 | NA | NA |
| 8430423G03 | 0 | 0 | 0 | 0 | NA | NA |
| Gm42529 | 0 | 0 | 0 | 0 | NA | NA |
| Gm5111 | 0 | 0 | 0 | 0 | NA | NA |
| Gm28053 | 0 | 0 | 0 | 0 | NA | NA |
| Gm44226 | 0 | 0 | 0 | 0 | NA | NA |
| Stk31 | 0 | 0 | 0 | 0 | NA | NA |
| Npvf | 0 | 0 | 0 | 0 | NA | NA |
| Hoxa10 | 0 | 0 | 0 | 0 | NA | NA |

| | | | | | | |
|---|---|---|---|---|---|---|
| Tnip3 | 0 | 0 | 0 | 0 | NA | NA |
| Cd8a | 0 | 0 | 0 | 0 | NA | NA |
| Gm44040 | 0 | 0 | 0 | 0 | NA | NA |
| Ssu2 | 0 | 0 | 0 | 0 | NA | NA |
| Clec4a4 | 0 | 0 | 0 | 0 | NA | NA |
| Clec4b2 | 0 | 0 | 0 | 0 | NA | NA |
| Klrb1a | 0 | 0 | 0 | 0 | NA | NA |
| Klrc2 | 0 | 0 | 0 | 0 | NA | NA |
| Klrc1 | 0 | 0 | 0 | 0 | NA | NA |
| Klri1 | 0 | 0 | 0 | 0 | NA | NA |
| Klri2 | 0 | 0 | 0 | 0 | NA | NA |
| Klra5 | 0 | 0 | 0 | 0 | NA | NA |
| A630073D07 | 0 | 0 | 0 | 0 | NA | NA |
| Gm19434 | 0 | 0 | 0 | 0 | NA | NA |
| Nlrp12 | 0 | 0 | 0 | 0 | NA | NA |
| Ceacam16 | 0 | 0 | 0 | 0 | NA | NA |
| Irgc1 | 0 | 0 | 0 | 0 | NA | NA |
| Gm26550 | 0 | 0 | 0 | 0 | NA | NA |
| Igflr1 | 0 | 0 | 0 | 0 | NA | NA |
| Klk1b11 | 0 | 0 | 0 | 0 | NA | NA |
| Klk1b27 | 0 | 0 | 0 | 0 | NA | NA |
| Mrgprb1 | 0 | 0 | 0 | 0 | NA | NA |
| Folh1 | 0 | 0 | 0 | 0 | NA | NA |
| Ccdc83 | 0 | 0 | 0 | 0 | NA | NA |
| Gm15635 | 0 | 0 | 0 | 0 | NA | NA |
| Art2b | 0 | 0 | 0 | 0 | NA | NA |
| Trim30b | 0 | 0 | 0 | 0 | NA | NA |
| Calcb | 0 | 0 | 0 | 0 | NA | NA |
| Umod | 0 | 0 | 0 | 0 | NA | NA |
| Il27 | 0 | 0 | 0 | 0 | NA | NA |

| | | | | | | |
|---|---|---|---|---|---|---|
| Scgb1c1 | 0 | 0 | 0 | 0 | NA | NA |
| Gm16982 | 0 | 0 | 0 | 0 | NA | NA |
| Mrgprd | 0 | 0 | 0 | 0 | NA | NA |
| 4930432B10 | 0 | 0 | 0 | 0 | NA | NA |
| Vnn3 | 0 | 0 | 0 | 0 | NA | NA |
| Samd3 | 0 | 0 | 0 | 0 | NA | NA |
| Prf1 | 0 | 0 | 0 | 0 | NA | NA |
| Ptprq | 0 | 0 | 0 | 0 | NA | NA |
| Cd209b | 0 | 0 | 0 | 0 | NA | NA |
| Cd209g | 0 | 0 | 0 | 0 | NA | NA |
| Myom2 | 0 | 0 | 0 | 0 | NA | NA |
| Spata4 | 0 | 0 | 0 | 0 | NA | NA |
| Slc12a3 | 0 | 0 | 0 | 0 | NA | NA |
| 9330175E14I | 0 | 0 | 0 | 0 | NA | NA |
| Nlrc5 | 0 | 0 | 0 | 0 | NA | NA |
| Ces2e | 0 | 0 | 0 | 0 | NA | NA |
| Kcng4 | 0 | 0 | 0 | 0 | NA | NA |
| 5033426O07 | 0 | 0 | 0 | 0 | NA | NA |
| Gm27011 | 0 | 0 | 0 | 0 | NA | NA |
| Gm8050 | 0 | 0 | 0 | 0 | NA | NA |
| Msmb | 0 | 0 | 0 | 0 | NA | NA |
| A630023A22 | 0 | 0 | 0 | 0 | NA | NA |
| Sh2d4b | 0 | 0 | 0 | 0 | NA | NA |
| Gm16506 | 0 | 0 | 0 | 0 | NA | NA |
| Ear10 | 0 | 0 | 0 | 0 | NA | NA |
| Gm17173 | 0 | 0 | 0 | 0 | NA | NA |
| Tlr11 | 0 | 0 | 0 | 0 | NA | NA |
| Eddm3b | 0 | 0 | 0 | 0 | NA | NA |
| Ang2 | 0 | 0 | 0 | 0 | NA | NA |
| Sgcg | 0 | 0 | 0 | 0 | NA | NA |

| | | | | | | |
|---|---|---|---|---|---|---|
| Gm9922 | 0 | 0 | 0 | 0 | NA | NA |
| Timm8a2 | 0 | 0 | 0 | 0 | NA | NA |
| Mmp12 | 0 | 0 | 0 | 0 | NA | NA |
| Mmp3 | 0 | 0 | 0 | 0 | NA | NA |
| Gm27162 | 0 | 0 | 0 | 0 | NA | NA |
| Crtam | 0 | 0 | 0 | 0 | NA | NA |
| 2900052N01 | 0 | 0 | 0 | 0 | NA | NA |
| Gsta1 | 0 | 0 | 0 | 0 | NA | NA |
| Impg1 | 0 | 0 | 0 | 0 | NA | NA |
| Gm10634 | 0 | 0 | 0 | 0 | NA | NA |
| 4930579C12I | 0 | 0 | 0 | 0 | NA | NA |
| Tdgf1 | 0 | 0 | 0 | 0 | NA | NA |
| Bcl2a1c | 0 | 0 | 0 | 0 | NA | NA |
| C130032M1C | 0 | 0 | 0 | 0 | NA | NA |
| Eomes | 0 | 0 | 0 | 0 | NA | NA |
| Slc22a14 | 0 | 0 | 0 | 0 | NA | NA |
| Scn10a | 0 | 0 | 0 | 0 | NA | NA |
| Abca13 | 0 | 0 | 0 | 0 | NA | NA |
| Gm12022 | 0 | 0 | 0 | 0 | NA | NA |
| Gm12153 | 0 | 0 | 0 | 0 | NA | NA |
| Gm12159 | 0 | 0 | 0 | 0 | NA | NA |
| Tgtp1 | 0 | 0 | 0 | 0 | NA | NA |
| Tgtp2 | 0 | 0 | 0 | 0 | NA | NA |
| Csf2 | 0 | 0 | 0 | 0 | NA | NA |
| Lypd8 | 0 | 0 | 0 | 0 | NA | NA |
| 1810065E05I | 0 | 0 | 0 | 0 | NA | NA |
| Gm12253 | 0 | 0 | 0 | 0 | NA | NA |
| Gm26964 | 0 | 0 | 0 | 0 | NA | NA |
| Slfn4 | 0 | 0 | 0 | 0 | NA | NA |
| Gm53 | 0 | 0 | 0 | 0 | NA | NA |

| | | | | | | |
|---|---|---|---|---|---|---|
| Krt39 | 0 | 0 | 0 | 0 | NA | NA |
| Krt16 | 0 | 0 | 0 | 0 | NA | NA |
| Gm12349 | 0 | 0 | 0 | 0 | NA | NA |
| Dhx58os | 0 | 0 | 0 | 0 | NA | NA |
| Gm11551 | 0 | 0 | 0 | 0 | NA | NA |
| Gm20659 | 0 | 0 | 0 | 0 | NA | NA |
| 9930022D16I | 0 | 0 | 0 | 0 | NA | NA |
| St6galnac1 | 0 | 0 | 0 | 0 | NA | NA |
| Gm12589 | 0 | 0 | 0 | 0 | NA | NA |
| Ptchd3 | 0 | 0 | 0 | 0 | NA | NA |
| Pom121l2 | 0 | 0 | 0 | 0 | NA | NA |
| Hus1b | 0 | 0 | 0 | 0 | NA | NA |
| Stmnd1 | 0 | 0 | 0 | 0 | NA | NA |
| Omd | 0 | 0 | 0 | 0 | NA | NA |
| Hrh2 | 0 | 0 | 0 | 0 | NA | NA |
| 1700066J03F | 0 | 0 | 0 | 0 | NA | NA |
| Gm10037 | 0 | 0 | 0 | 0 | NA | NA |
| Gm15290 | 0 | 0 | 0 | 0 | NA | NA |
| D13Ertd608e | 0 | 0 | 0 | 0 | NA | NA |
| Nkx2-9 | 0 | 0 | 0 | 0 | NA | NA |
| Gm15523 | 0 | 0 | 0 | 0 | NA | NA |
| Bdkrb2 | 0 | 0 | 0 | 0 | NA | NA |
| Rd3l | 0 | 0 | 0 | 0 | NA | NA |
| Gm30948 | 0 | 0 | 0 | 0 | NA | NA |
| Gm19331 | 0 | 0 | 0 | 0 | NA | NA |
| Dcstamp | 0 | 0 | 0 | 0 | NA | NA |
| Gm17473 | 0 | 0 | 0 | 0 | NA | NA |
| Klhl38 | 0 | 0 | 0 | 0 | NA | NA |
| Ly6i | 0 | 0 | 0 | 0 | NA | NA |
| Apol7b | 0 | 0 | 0 | 0 | NA | NA |

| | | | | | | |
|---|---|---|---|---|---|---|
| Apol10a | 0 | 0 | 0 | 0 | NA | NA |
| Apol7e | 0 | 0 | 0 | 0 | NA | NA |
| Wnt10b | 0 | 0 | 0 | 0 | NA | NA |
| Krt6b | 0 | 0 | 0 | 0 | NA | NA |
| Ciita | 0 | 0 | 0 | 0 | NA | NA |
| Gm21859 | 0 | 0 | 0 | 0 | NA | NA |
| Gm9961 | 0 | 0 | 0 | 0 | NA | NA |
| Tssk1 | 0 | 0 | 0 | 0 | NA | NA |
| Tm4sf19 | 0 | 0 | 0 | 0 | NA | NA |
| Gm26838 | 0 | 0 | 0 | 0 | NA | NA |
| Cd200r2 | 0 | 0 | 0 | 0 | NA | NA |
| Trat1 | 0 | 0 | 0 | 0 | NA | NA |
| Adgrg7 | 0 | 0 | 0 | 0 | NA | NA |
| Gpr15 | 0 | 0 | 0 | 0 | NA | NA |
| Speer2 | 0 | 0 | 0 | 0 | NA | NA |
| Olig2 | 0 | 0 | 0 | 0 | NA | NA |
| Fam3b | 0 | 0 | 0 | 0 | NA | NA |
| Has1 | 0 | 0 | 0 | 0 | NA | NA |
| Mmp25 | 0 | 0 | 0 | 0 | NA | NA |
| Dcpp1 | 0 | 0 | 0 | 0 | NA | NA |
| Prss30 | 0 | 0 | 0 | 0 | NA | NA |
| Meiob | 0 | 0 | 0 | 0 | NA | NA |
| Actl9 | 0 | 0 | 0 | 0 | NA | NA |
| Gm20513 | 0 | 0 | 0 | 0 | NA | NA |
| Gm6034 | 0 | 0 | 0 | 0 | NA | NA |
| Gm19684 | 0 | 0 | 0 | 0 | NA | NA |
| H2-T3 | 0 | 0 | 0 | 0 | NA | NA |
| Adgrf1 | 0 | 0 | 0 | 0 | NA | NA |
| Tcte1 | 0 | 0 | 0 | 0 | NA | NA |
| Acer1 | 0 | 0 | 0 | 0 | NA | NA |

| | | | | | | |
|---|---|---|---|---|---|---|
| Vit | 0 | 0 | 0 | 0 | NA | NA |
| Lhcgr | 0 | 0 | 0 | 0 | NA | NA |
| Lvrn | 0 | 0 | 0 | 0 | NA | NA |
| Arhgef37 | 0 | 0 | 0 | 0 | NA | NA |
| Mc2r | 0 | 0 | 0 | 0 | NA | NA |
| Aldh3b3 | 0 | 0 | 0 | 0 | NA | NA |
| Scgb1a1 | 0 | 0 | 0 | 0 | NA | NA |
| Gm28935 | 0 | 0 | 0 | 0 | NA | NA |
| Oosp2 | 0 | 0 | 0 | 0 | NA | NA |
| Oosp1 | 0 | 0 | 0 | 0 | NA | NA |
| Pdzd7 | 0 | 0 | 0 | 0 | NA | NA |
| Gm19557 | 0 | 0 | 0 | 0 | NA | NA |
| Gm37915 | 0 | 0 | 0 | 0 | NA | NA |
| Alppl2 | 0 | 0 | 0 | 0 | NA | NA |
| Prlh | 0 | 0 | 0 | 0 | NA | NA |
| Gal3st2 | 0 | 0 | 0 | 0 | NA | NA |
| Map3k19 | 0 | 0 | 0 | 0 | NA | NA |
| Sele | 0 | 0 | 0 | 0 | NA | NA |
| Gm32391 | 0 | 0 | 0 | 0 | NA | NA |
| Gm32569 | 0 | 0 | 0 | 0 | NA | NA |
| 1700016C15I | 0 | 0 | 0 | 0 | NA | NA |
| Gm33973 | 0 | 0 | 0 | 0 | NA | NA |
| Gm37027 | 0 | 0 | 0 | 0 | NA | NA |
| Gm28035 | 0 | 0 | 0 | 0 | NA | NA |
| Pla2g4f | 0 | 0 | 0 | 0 | NA | NA |
| Astl | 0 | 0 | 0 | 0 | NA | NA |
| Adig | 0 | 0 | 0 | 0 | NA | NA |
| Gm14267 | 0 | 0 | 0 | 0 | NA | NA |
| Gm14321 | 0 | 0 | 0 | 0 | NA | NA |
| Gm14618 | 0 | 0 | 0 | 0 | NA | NA |

| | | | | | | |
|---|---|---|---|---|---|---|
| Gm14565 | 0 | 0 | 0 | 0 | NA | NA |
| Trex2 | 0 | 0 | 0 | 0 | NA | NA |
| Gm614 | 0 | 0 | 0 | 0 | NA | NA |
| Cdx4 | 0 | 0 | 0 | 0 | NA | NA |
| Samd7 | 0 | 0 | 0 | 0 | NA | NA |
| Tshb | 0 | 0 | 0 | 0 | NA | NA |
| 9830132P13I | 0 | 0 | 0 | 0 | NA | NA |
| Gm42706 | 0 | 0 | 0 | 0 | NA | NA |
| Tal2 | 0 | 0 | 0 | 0 | NA | NA |
| Musk | 0 | 0 | 0 | 0 | NA | NA |
| Gm13285 | 0 | 0 | 0 | 0 | NA | NA |
| Gm8439 | 0 | 0 | 0 | 0 | NA | NA |
| Gm16224 | 0 | 0 | 0 | 0 | NA | NA |
| Ctrcos | 0 | 0 | 0 | 0 | NA | NA |
| Perm1 | 0 | 0 | 0 | 0 | NA | NA |
| Gm42435 | 0 | 0 | 0 | 0 | NA | NA |
| Tmprss11d | 0 | 0 | 0 | 0 | NA | NA |
| Sult1e1 | 0 | 0 | 0 | 0 | NA | NA |
| Fam47e | 0 | 0 | 0 | 0 | NA | NA |
| Gm42421 | 0 | 0 | 0 | 0 | NA | NA |
| 1700001J03F | 0 | 0 | 0 | 0 | NA | NA |
| Gm19272 | 0 | 0 | 0 | 0 | NA | NA |
| Asb15 | 0 | 0 | 0 | 0 | NA | NA |
| Gm15624 | 0 | 0 | 0 | 0 | NA | NA |
| Hrh1 | 0 | 0 | 0 | 0 | NA | NA |
| Klra3 | 0 | 0 | 0 | 0 | NA | NA |
| Sbk3 | 0 | 0 | 0 | 0 | NA | NA |
| Gm29638 | 0 | 0 | 0 | 0 | NA | NA |
| Psg29 | 0 | 0 | 0 | 0 | NA | NA |
| Ceacam20 | 0 | 0 | 0 | 0 | NA | NA |

| | | | | | | |
|---|---|---|---|---|---|---|
| A530021J07F | 0 | 0 | 0 | 0 | NA | NA |
| Tmc5 | 0 | 0 | 0 | 0 | NA | NA |
| Pdilt | 0 | 0 | 0 | 0 | NA | NA |
| Acsm1 | 0 | 0 | 0 | 0 | NA | NA |
| Odf3 | 0 | 0 | 0 | 0 | NA | NA |
| Cd209e | 0 | 0 | 0 | 0 | NA | NA |
| Gm4889 | 0 | 0 | 0 | 0 | NA | NA |
| Gm15889 | 0 | 0 | 0 | 0 | NA | NA |
| Gm16156 | 0 | 0 | 0 | 0 | NA | NA |
| Hydin | 0 | 0 | 0 | 0 | NA | NA |
| Gm15395 | 0 | 0 | 0 | 0 | NA | NA |
| Hsd17b2 | 0 | 0 | 0 | 0 | NA | NA |
| D830044D21 | 0 | 0 | 0 | 0 | NA | NA |
| Gm9748 | 0 | 0 | 0 | 0 | NA | NA |
| Acrv1 | 0 | 0 | 0 | 0 | NA | NA |
| Gm26868 | 0 | 0 | 0 | 0 | NA | NA |
| 4930505A04 | 0 | 0 | 0 | 0 | NA | NA |
| Foxi1 | 0 | 0 | 0 | 0 | NA | NA |
| Gm12185 | 0 | 0 | 0 | 0 | NA | NA |
| Slc36a3 | 0 | 0 | 0 | 0 | NA | NA |
| B430319H21 | 0 | 0 | 0 | 0 | NA | NA |
| B130011K05 | 0 | 0 | 0 | 0 | NA | NA |
| 1700086D15 | 0 | 0 | 0 | 0 | NA | NA |
| Ccl8 | 0 | 0 | 0 | 0 | NA | NA |
| Phospho1 | 0 | 0 | 0 | 0 | NA | NA |
| Krt33a | 0 | 0 | 0 | 0 | NA | NA |
| Gm14206 | 0 | 0 | 0 | 0 | NA | NA |
| Gh | 0 | 0 | 0 | 0 | NA | NA |
| Gm11707 | 0 | 0 | 0 | 0 | NA | NA |
| Gm11734 | 0 | 0 | 0 | 0 | NA | NA |

| | | | | | | |
|---|---|---|---|---|---|---|
| Tspan10 | 0 | 0 | 0 | 0 | NA | NA |
| Gm11788 | 0 | 0 | 0 | 0 | NA | NA |
| Calm4 | 0 | 0 | 0 | 0 | NA | NA |
| Gm11373 | 0 | 0 | 0 | 0 | NA | NA |
| Gm15324 | 0 | 0 | 0 | 0 | NA | NA |
| Gzmk | 0 | 0 | 0 | 0 | NA | NA |
| Pax9 | 0 | 0 | 0 | 0 | NA | NA |
| Cpt1b | 0 | 0 | 0 | 0 | NA | NA |
| Glycam1 | 0 | 0 | 0 | 0 | NA | NA |
| Cyp2ab1 | 0 | 0 | 0 | 0 | NA | NA |
| Fbxo40 | 0 | 0 | 0 | 0 | NA | NA |
| Cyp4f40 | 0 | 0 | 0 | 0 | NA | NA |
| Spink5 | 0 | 0 | 0 | 0 | NA | NA |
| Htr4 | 0 | 0 | 0 | 0 | NA | NA |
| Mc5r | 0 | 0 | 0 | 0 | NA | NA |
| Dynap | 0 | 0 | 0 | 0 | NA | NA |
| Kcnk4 | 0 | 0 | 0 | 0 | NA | NA |
| Zp1 | 0 | 0 | 0 | 0 | NA | NA |
| Pou6f2 | 0.00310389 | 0.00280444 | 0.00280458 | -1.35E-07 | 0.5001276 | 0.99999994 |
| F10 | 0.00434561 | 0.00392305 | 0.00392326 | -2.07E-07 | 0.50013985 | 0.99999994 |
| Mutyh | 0.00950327 | 0.00854948 | 0.00854972 | -2.42E-07 | 0.50007513 | 0.99999994 |
| Gm17669 | 0.00254993 | 0.0023045 | 0.0023049 | -4.00E-07 | 0.50046093 | 0.99999994 |
| Gm20089 | 0.00044424 | 0.0004016 | 0.00040212 | -5.27E-07 | 0.50347778 | 0.99999994 |
| Ulbp1 | 0.00044398 | 0.00040113 | 0.00040189 | -7.59E-07 | 0.50501636 | 0.99999994 |
| Laptm5 | 0.01226303 | 0.01101123 | 0.01101202 | -7.89E-07 | 0.50019025 | 0.99999994 |
| Aldoart2 | 0.00044393 | 0.00040105 | 0.00040185 | -7.98E-07 | 0.5052726 | 0.99999994 |
| Gm29543 | 0.00044372 | 0.00040067 | 0.00040166 | -9.91E-07 | 0.50655297 | 0.99999994 |
| Btk | 0.00044359 | 0.00040043 | 0.00040154 | -1.11E-06 | 0.50732053 | 0.99999994 |
| Pdk4 | 0.00044351 | 0.00040028 | 0.00040146 | -1.18E-06 | 0.50783196 | 0.99999994 |
| Wdr27 | 0.00044351 | 0.00040028 | 0.00040146 | -1.18E-06 | 0.50783196 | 0.99999994 |

| | | | | | | |
|---|---|---|---|---|---|---|
| Ugt8a | 0.01882896 | 0.01683235 | 0.01683355 | -1.20E-06 | 0.50018907 | 0.99999994 |
| Sypl2 | 0.00044338 | 0.00040005 | 0.00040135 | -1.30E-06 | 0.50859868 | 0.99999994 |
| Klhl3 | 0.00500509 | 0.00451522 | 0.00451662 | -1.40E-06 | 0.50082219 | 0.99999994 |
| Itm2a | 0.00132849 | 0.00120006 | 0.00120183 | -1.77E-06 | 0.5039083 | 0.99999994 |
| L3mbtl1 | 0.0004424 | 0.00039828 | 0.00040046 | -2.18E-06 | 0.5144599 | 0.99999994 |
| Mcemp1 | 0.00078662 | 0.00070927 | 0.00071188 | -2.62E-06 | 0.50976459 | 0.99999994 |
| Tnfsf13b | 0.00155149 | 0.0014006 | 0.00140336 | -2.76E-06 | 0.50522178 | 0.99999994 |
| Fhl4 | 0.00928053 | 0.00834781 | 0.00835059 | -2.78E-06 | 0.50088339 | 0.99999994 |
| 5730522E02I | 0.00294454 | 0.0026581 | 0.00266089 | -2.78E-06 | 0.50277776 | 0.99999994 |
| Tmem150co: | 0.00128751 | 0.00116182 | 0.0011648 | -2.98E-06 | 0.50679349 | 0.99999994 |
| Popdc2 | 0.00088623 | 0.00079896 | 0.00080198 | -3.01E-06 | 0.50998122 | 0.99999994 |
| Bdh2 | 0.00618673 | 0.00557496 | 0.00557846 | -3.50E-06 | 0.5016684 | 0.99999994 |
| Avpr1b | 0.00165809 | 0.00149613 | 0.00149967 | -3.54E-06 | 0.50626077 | 0.99999994 |
| Tgfb1i1 | 0.0077736 | 0.00699795 | 0.00700179 | -3.84E-06 | 0.50145821 | 0.99999994 |
| Unc13d | 0.00251727 | 0.00227137 | 0.00227543 | -4.06E-06 | 0.50473375 | 0.99999994 |
| Sun3 | 0.00367811 | 0.00331797 | 0.00332213 | -4.17E-06 | 0.50332951 | 0.99999994 |
| Gm10603 | 0.00086703 | 0.00078015 | 0.00078461 | -4.46E-06 | 0.51509063 | 0.99999994 |
| Gm37350 | 0.00499623 | 0.00450401 | 0.00450865 | -4.64E-06 | 0.50273389 | 0.99999994 |
| Fbp1 | 0.01764878 | 0.01578623 | 0.01579096 | -4.73E-06 | 0.50079616 | 0.99999994 |
| Hoxb8 | 0.00043935 | 0.00039282 | 0.0003977 | -4.89E-06 | 0.53260492 | 0.99999994 |
| Zfp85os | 0.00254766 | 0.00229771 | 0.00230285 | -5.14E-06 | 0.50593251 | 0.99999994 |
| Lrat | 0.000439 | 0.00039218 | 0.00039738 | -5.20E-06 | 0.53472579 | 0.99999994 |
| Mageb16 | 0.00043881 | 0.00039184 | 0.00039721 | -5.37E-06 | 0.53584673 | 0.99999994 |
| Krtap31-1 | 0.00130909 | 0.00117892 | 0.0011843 | -5.38E-06 | 0.51205971 | 0.99999994 |
| Dus4l | 0.01759806 | 0.01574069 | 0.01574612 | -5.43E-06 | 0.50091546 | 0.99999994 |
| Ppp1r3b | 0.00043872 | 0.00039169 | 0.00039714 | -5.44E-06 | 0.5363445 | 0.99999994 |
| Tlr1 | 0.00043856 | 0.00039139 | 0.00039698 | -5.59E-06 | 0.53733927 | 0.99999994 |
| Mtmr11 | 0.00391878 | 0.00353325 | 0.00353893 | -5.68E-06 | 0.5042599 | 0.99999994 |
| A530083I20F | 0.00295352 | 0.00266328 | 0.00266899 | -5.71E-06 | 0.50568087 | 0.99999994 |
| Gm2479 | 0.00043835 | 0.00039102 | 0.0003968 | -5.77E-06 | 0.53858127 | 0.99999994 |

| | | | | | | |
|---|---|---|---|---|---|---|
| Nwd1 | 0.00087214 | 0.00078346 | 0.00078923 | -5.78E-06 | 0.51942662 | 0.99999994 |
| 1810012K08I | 0.00043755 | 0.00038961 | 0.00039608 | -6.47E-06 | 0.54328586 | 0.99999994 |
| Gm11527 | 0.00043722 | 0.00038902 | 0.00039578 | -6.76E-06 | 0.54525956 | 0.99999994 |
| Spats1 | 0.0011422 | 0.00102632 | 0.00103343 | -7.12E-06 | 0.51828397 | 0.99999994 |
| Tnfrsf13c | 0.00043668 | 0.00038806 | 0.00039529 | -7.23E-06 | 0.54845762 | 0.99999994 |
| Gm15247 | 0.00679985 | 0.00612142 | 0.00612876 | -7.34E-06 | 0.50317866 | 0.99999994 |
| Kcnj3 | 0.00043631 | 0.0003874 | 0.00039495 | -7.56E-06 | 0.55066492 | 0.99999994 |
| Cd177 | 0.00043631 | 0.0003874 | 0.00039495 | -7.56E-06 | 0.55066492 | 0.99999994 |
| Ifitm5 | 0.00043627 | 0.00038732 | 0.00039491 | -7.59E-06 | 0.55090983 | 0.99999994 |
| Hepacam | 0.00043627 | 0.00038732 | 0.00039491 | -7.59E-06 | 0.55090983 | 0.99999994 |
| Tex14 | 0.00043627 | 0.00038732 | 0.00039491 | -7.59E-06 | 0.55090983 | 0.99999994 |
| Svep1 | 0.00043573 | 0.00038637 | 0.00039443 | -8.06E-06 | 0.5540874 | 0.99999994 |
| 4921507P07I | 0.00359032 | 0.00323494 | 0.00324303 | -8.09E-06 | 0.50662621 | 0.99999994 |
| Klhl41 | 0.00095442 | 0.00085547 | 0.00086364 | -8.17E-06 | 0.52509944 | 0.99999994 |
| Gm13063 | 0.00087716 | 0.0007856 | 0.00079378 | -8.17E-06 | 0.52732187 | 0.99999994 |
| Dkk2 | 0.00043495 | 0.00038499 | 0.00039372 | -8.74E-06 | 0.55871034 | 0.99999994 |
| Cyp2r1 | 0.00043495 | 0.00038499 | 0.00039372 | -8.74E-06 | 0.55871034 | 0.99999994 |
| Abcg8 | 0.00085108 | 0.00076137 | 0.00077019 | -8.82E-06 | 0.53039646 | 0.99999994 |
| Gm11899 | 0.00043479 | 0.00038469 | 0.00039357 | -8.88E-06 | 0.55968034 | 0.99999994 |
| Wdfy4 | 0.0004347 | 0.00038455 | 0.0003935 | -8.95E-06 | 0.56016492 | 0.99999994 |
| Coq3 | 0.04806672 | 0.0421359 | 0.04214503 | -9.13E-06 | 0.50057543 | 0.99999994 |
| Slfn14 | 0.00043429 | 0.00038382 | 0.00039313 | -9.30E-06 | 0.56258352 | 0.99999994 |
| Fgd5 | 0.00043421 | 0.00038368 | 0.00039305 | -9.37E-06 | 0.56306638 | 0.99999994 |
| Sncaip | 0.00115536 | 0.00103594 | 0.00104533 | -9.39E-06 | 0.52383141 | 0.99999994 |
| Cxcl17 | 0.00043409 | 0.00038346 | 0.00039294 | -9.48E-06 | 0.56379013 | 0.99999994 |
| Ermap | 0.00087692 | 0.00078408 | 0.00079356 | -9.48E-06 | 0.53170236 | 0.99999994 |
| Cd69 | 0.0013678 | 0.0012277 | 0.00123736 | -9.66E-06 | 0.52072327 | 0.99999994 |
| Gbp2 | 0.0004336 | 0.0003826 | 0.0003925 | -9.90E-06 | 0.56667863 | 0.99999994 |
| Mall | 0.00086636 | 0.00077368 | 0.00078401 | -1.03E-05 | 0.53493041 | 0.99999994 |
| Klhl30 | 0.00083029 | 0.00074101 | 0.00075139 | -1.04E-05 | 0.53660725 | 0.99999994 |

| | | | | | | |
|---|---|---|---|---|---|---|
| Hsf5 | 0.00084126 | 0.00075079 | 0.00076131 | -1.05E-05 | 0.53665409 | 0.99999994 |
| Nrros | 0.00184072 | 0.00165385 | 0.00166465 | -1.08E-05 | 0.51722646 | 0.99999994 |
| Slc16a7 | 0.00230215 | 0.00207044 | 0.00208129 | -1.08E-05 | 0.51383305 | 0.99999994 |
| Fancd2 | 0.0015001 | 0.00134606 | 0.00135692 | -1.09E-05 | 0.52123742 | 0.99999994 |
| Gm32200 | 0.00063559 | 0.0005642 | 0.00057527 | -1.11E-05 | 0.55093487 | 0.99999994 |
| Gm15222 | 0.00043145 | 0.00037881 | 0.00039055 | -1.17E-05 | 0.5793095 | 0.99999994 |
| Plxnc1 | 0.00253424 | 0.00227888 | 0.00229075 | -1.19E-05 | 0.5137578 | 0.99999994 |
| Kng2 | 0.00077036 | 0.00068519 | 0.00069718 | -1.20E-05 | 0.54555318 | 0.99999994 |
| 9030624G23 | 0.01411505 | 0.01264715 | 0.01265929 | -1.21E-05 | 0.50254672 | 0.99999994 |
| Plin4 | 0.00043092 | 0.00037789 | 0.00039008 | -1.22E-05 | 0.58237549 | 0.99999994 |
| Zfp760 | 0.03887608 | 0.03428211 | 0.03429431 | -1.22E-05 | 0.50094445 | 0.99999994 |
| Ak7 | 0.0014292 | 0.00128065 | 0.00129285 | -1.22E-05 | 0.52504214 | 0.99999994 |
| Lypd3 | 0.00043084 | 0.00037775 | 0.00039 | -1.23E-05 | 0.58284604 | 0.99999994 |
| Scimp | 0.00043064 | 0.0003774 | 0.00038982 | -1.24E-05 | 0.58402108 | 0.99999994 |
| 1700125H03 | 0.00043056 | 0.00037725 | 0.00038975 | -1.25E-05 | 0.58449055 | 0.99999994 |
| Col4a6 | 0.00221688 | 0.00199166 | 0.00200431 | -1.26E-05 | 0.51675002 | 0.99999994 |
| Fbxl13 | 0.00043004 | 0.00037634 | 0.00038928 | -1.29E-05 | 0.58753466 | 0.99999994 |
| Dync1li2 | 0.12875119 | 0.10666702 | 0.10668055 | -1.35E-05 | 0.50033678 | 0.99999994 |
| Ngf | 0.00206421 | 0.00185244 | 0.00186647 | -1.40E-05 | 0.51994904 | 0.99999994 |
| Nr1i2 | 0.00193243 | 0.00173345 | 0.00174748 | -1.40E-05 | 0.52131428 | 0.99999994 |
| Syt1 | 0.00272759 | 0.00245115 | 0.0024652 | -1.41E-05 | 0.51513435 | 0.99999994 |
| Gpr137c | 0.00061235 | 0.00053972 | 0.00055424 | -1.45E-05 | 0.56924269 | 0.99999994 |
| Trim16 | 0.00099963 | 0.00088978 | 0.00090453 | -1.47E-05 | 0.54321522 | 0.99999994 |
| Ttc36 | 0.01393064 | 0.01248064 | 0.01249545 | -1.48E-05 | 0.5031464 | 0.99999994 |
| Ano1 | 0.0053346 | 0.00479798 | 0.00481289 | -1.49E-05 | 0.5082285 | 0.99999994 |
| B230110C06 | 0.00064769 | 0.00057103 | 0.00058621 | -1.52E-05 | 0.56844716 | 0.99999994 |
| Adam28 | 0.0016356 | 0.00146394 | 0.00147935 | -1.54E-05 | 0.52763804 | 0.99999994 |
| Pate2 | 0.00329981 | 0.0029658 | 0.00298121 | -1.54E-05 | 0.51372693 | 0.99999994 |
| Clcnka | 0.00642536 | 0.00577717 | 0.0057927 | -1.55E-05 | 0.50711774 | 0.99999994 |
| Axdnd1 | 0.00058652 | 0.00051503 | 0.00053087 | -1.58E-05 | 0.5786861 | 0.99999994 |

| | | | | | | |
|---|---|---|---|---|---|---|
| Gm43118 | 0.00212784 | 0.0019073 | 0.00192393 | -1.66E-05 | 0.52294694 | 0.99999994 |
| Paqr3 | 0.00666111 | 0.00598761 | 0.00600427 | -1.67E-05 | 0.50736827 | 0.99999994 |
| Ackr3 | 0.00212479 | 0.0019045 | 0.00192118 | -1.67E-05 | 0.52303211 | 0.99999994 |
| Htr1d | 0.00178902 | 0.00160109 | 0.00161795 | -1.69E-05 | 0.52764395 | 0.99999994 |
| Sucnr1 | 0.00042501 | 0.00036759 | 0.00038473 | -1.71E-05 | 0.61657082 | 0.99999994 |
| Scube2 | 0.00191566 | 0.00171519 | 0.00173233 | -1.71E-05 | 0.52625121 | 0.99999994 |
| Platr25 | 0.00160981 | 0.00143873 | 0.00145605 | -1.73E-05 | 0.53155915 | 0.99999994 |
| Anxa8 | 0.00042466 | 0.00036698 | 0.00038441 | -1.74E-05 | 0.61857893 | 0.99999994 |
| 8030423F21I | 0.00042462 | 0.00036691 | 0.00038437 | -1.75E-05 | 0.61880163 | 0.99999994 |
| Kdsr | 0.07136373 | 0.06160847 | 0.061626 | -1.75E-05 | 0.50075539 | 0.99999994 |
| Pdgfc | 0.00359883 | 0.00323288 | 0.0032507 | -1.78E-05 | 0.51456032 | 0.99999994 |
| Pkd2l1 | 0.00042376 | 0.00036543 | 0.0003836 | -1.82E-05 | 0.62368023 | 0.99999994 |
| Mettl7a3 | 0.00042365 | 0.00036523 | 0.00038349 | -1.83E-05 | 0.62434237 | 0.99999994 |
| Cd180 | 0.00067952 | 0.00059664 | 0.000615 | -1.84E-05 | 0.57876722 | 0.99999994 |
| Myh8 | 0.00212636 | 0.00190402 | 0.00192259 | -1.86E-05 | 0.52562058 | 0.99999994 |
| Icam5 | 0.00078774 | 0.00069431 | 0.00071289 | -1.86E-05 | 0.56887699 | 0.99999994 |
| Gpr162 | 0.01350519 | 0.01209867 | 0.01211731 | -1.86E-05 | 0.50408313 | 0.99999994 |
| Sowahd | 0.00042295 | 0.00036403 | 0.00038286 | -1.88E-05 | 0.62829938 | 0.99999994 |
| Klf8 | 0.00383008 | 0.00344016 | 0.00345904 | -1.89E-05 | 0.51448785 | 0.99999994 |
| Rufy4 | 0.00042287 | 0.0003639 | 0.00038279 | -1.89E-05 | 0.62873736 | 0.99999994 |
| Kank2 | 0.04471124 | 0.03927053 | 0.0392898 | -1.93E-05 | 0.50130227 | 0.99999994 |
| Gm43570 | 0.00070745 | 0.00062098 | 0.00064027 | -1.93E-05 | 0.57944744 | 0.99999994 |
| Sema6c | 0.01315823 | 0.01178948 | 0.01180876 | -1.93E-05 | 0.50433696 | 0.99999994 |
| Grm4 | 0.00475281 | 0.00427011 | 0.0042897 | -1.96E-05 | 0.5121254 | 0.99999994 |
| Gm648 | 0.02626279 | 0.02334326 | 0.0233629 | -1.96E-05 | 0.50223275 | 0.99999994 |
| Adgrg2 | 0.00042152 | 0.00036158 | 0.00038157 | -2.00E-05 | 0.63634748 | 0.99999994 |
| Akr1c18 | 0.00075428 | 0.00066262 | 0.00068263 | -2.00E-05 | 0.57736656 | 0.99999994 |
| Gm43062 | 0.00166995 | 0.00149036 | 0.00151039 | -2.00E-05 | 0.53516413 | 0.99999994 |
| Gm26797 | 0.00570824 | 0.0051286 | 0.00514869 | -2.01E-05 | 0.51035697 | 0.99999994 |
| Rras | 0.02272859 | 0.02024668 | 0.02026682 | -2.01E-05 | 0.50263976 | 0.99999994 |

| | | | | | | |
|---|---|---|---|---|---|---|
| Zfp870 | 0.01930032 | 0.01722934 | 0.0172495 | -2.02E-05 | 0.50310324 | 0.99999994 |
| Gm20383 | 0.00042125 | 0.00036112 | 0.00038133 | -2.02E-05 | 0.63785703 | 0.99999994 |
| Prkg1 | 0.00062303 | 0.00054354 | 0.00056391 | -2.04E-05 | 0.59499995 | 0.99999994 |
| Gm43000 | 0.00206135 | 0.00184307 | 0.00186389 | -2.08E-05 | 0.52963144 | 0.99999994 |
| Rel | 0.00409871 | 0.00368008 | 0.00370097 | -2.09E-05 | 0.51498024 | 0.99999994 |
| Gm9970 | 0.0004203 | 0.00035948 | 0.00038046 | -2.10E-05 | 0.64321411 | 0.99999994 |
| Gm30074 | 0.0006926 | 0.00060583 | 0.00062684 | -2.10E-05 | 0.58828568 | 0.99999994 |
| Tppp | 0.00150495 | 0.00134026 | 0.0013613 | -2.10E-05 | 0.54097372 | 0.99999994 |
| Ttc39d | 0.00105332 | 0.00093174 | 0.00095308 | -2.13E-05 | 0.55922417 | 0.99999994 |
| Ceacam1 | 0.00709049 | 0.00636801 | 0.00638945 | -2.14E-05 | 0.50891287 | 0.99999994 |
| Tnf | 0.00115064 | 0.00101955 | 0.00104107 | -2.15E-05 | 0.55471453 | 0.99999994 |
| Gm10814 | 0.00068796 | 0.00060097 | 0.00062264 | -2.17E-05 | 0.59160216 | 0.99999994 |
| Smim3 | 0.00281857 | 0.00252558 | 0.00254727 | -2.17E-05 | 0.52259028 | 0.99999994 |
| Dlx2 | 0.00041934 | 0.00035785 | 0.0003796 | -2.17E-05 | 0.64851756 | 0.99999994 |
| Bglap | 0.00041934 | 0.00035785 | 0.0003796 | -2.17E-05 | 0.64851756 | 0.99999994 |
| Otx1 | 0.00041934 | 0.00035785 | 0.0003796 | -2.17E-05 | 0.64851756 | 0.99999994 |
| Tmem100 | 0.00075064 | 0.00065748 | 0.00067934 | -2.19E-05 | 0.58479468 | 0.99999994 |
| Tnfrsf18 | 0.00304869 | 0.00273287 | 0.0027548 | -2.19E-05 | 0.52113355 | 0.99999994 |
| Cdh24 | 0.00108651 | 0.00096106 | 0.00098309 | -2.20E-05 | 0.55927141 | 0.99999994 |
| Zc3h12b | 0.00152561 | 0.00135794 | 0.00137997 | -2.20E-05 | 0.54232659 | 0.99999994 |
| A230060F14 | 0.00341059 | 0.00305893 | 0.00308106 | -2.21E-05 | 0.51907155 | 0.99999994 |
| Dlgap2 | 0.00130659 | 0.0011599 | 0.00118204 | -2.21E-05 | 0.54960721 | 0.99999994 |
| A630076J17I | 0.00041802 | 0.00035559 | 0.0003784 | -2.28E-05 | 0.65585164 | 0.99999994 |
| Vcan | 0.00287046 | 0.00257118 | 0.00259408 | -2.29E-05 | 0.52342664 | 0.99999994 |
| Cdx1 | 0.00120408 | 0.00106616 | 0.00108937 | -2.32E-05 | 0.55640334 | 0.99999994 |
| Ryr1 | 0.00213732 | 0.00190926 | 0.00193248 | -2.32E-05 | 0.53187286 | 0.99999994 |
| Zbtb8b | 0.00859585 | 0.00771475 | 0.00773809 | -2.33E-05 | 0.50801053 | 0.99999994 |
| Gm26583 | 0.0004173 | 0.00035437 | 0.00037775 | -2.34E-05 | 0.65978842 | 0.99999994 |
| Tac2 | 0.00074926 | 0.00065457 | 0.00067809 | -2.35E-05 | 0.59130657 | 0.99999994 |
| Kif4 | 0.00299644 | 0.00268402 | 0.00270769 | -2.37E-05 | 0.52320187 | 0.99999994 |

| | | | | | | |
|---|---|---|---|---|---|---|
| Aqp9 | 0.00083263 | 0.00072982 | 0.0007535 | -2.37E-05 | 0.58286509 | 0.99999994 |
| 8430436N08 | 0.00041685 | 0.00035361 | 0.00037734 | -2.37E-05 | 0.66225858 | 0.99999994 |
| Ampd1 | 0.00041681 | 0.00035355 | 0.00037731 | -2.38E-05 | 0.66246386 | 0.99999994 |
| 4930519P11I | 0.00041677 | 0.00035348 | 0.00037727 | -2.38E-05 | 0.66266905 | 0.99999994 |
| Tmprss5 | 0.0035196 | 0.00315551 | 0.0031793 | -2.38E-05 | 0.51986706 | 0.99999994 |
| Satb2 | 0.00041655 | 0.0003531 | 0.00037707 | -2.40E-05 | 0.66389836 | 0.99999994 |
| Ccdc150 | 0.00041644 | 0.00035291 | 0.00037697 | -2.41E-05 | 0.66451183 | 0.99999994 |
| Atp12a | 0.00041625 | 0.00035259 | 0.0003768 | -2.42E-05 | 0.66553253 | 0.99999994 |
| Dlk2 | 0.00041625 | 0.00035259 | 0.0003768 | -2.42E-05 | 0.66553253 | 0.99999994 |
| Samd11 | 0.00130503 | 0.00115616 | 0.00118063 | -2.45E-05 | 0.55485064 | 0.99999994 |
| Zfp583 | 0.0224887 | 0.02003163 | 0.02005614 | -2.45E-05 | 0.50324506 | 0.99999994 |
| Chst13 | 0.00212562 | 0.00189727 | 0.00192192 | -2.47E-05 | 0.53402198 | 0.99999994 |
| Fat4 | 0.0136512 | 0.01222224 | 0.01224711 | -2.49E-05 | 0.50539148 | 0.99999994 |
| Tinag | 0.0053987 | 0.00484558 | 0.00487052 | -2.49E-05 | 0.51359271 | 0.99999994 |
| Zfp941 | 0.0087485 | 0.00784975 | 0.0078747 | -2.50E-05 | 0.5084134 | 0.99999994 |
| Lrrc34 | 0.00261164 | 0.0023356 | 0.00236059 | -2.50E-05 | 0.52808295 | 0.99999994 |
| Gm15396 | 0.00041521 | 0.00035082 | 0.00037585 | -2.50E-05 | 0.6712079 | 0.99999994 |
| Ovgp1 | 0.00078812 | 0.00068819 | 0.00071324 | -2.50E-05 | 0.59240096 | 0.99999994 |
| Slurp1 | 0.00134794 | 0.00119426 | 0.00121941 | -2.51E-05 | 0.55458733 | 0.99999994 |
| Wwtr1 | 0.01611018 | 0.01440409 | 0.01442925 | -2.52E-05 | 0.50463026 | 0.99999994 |
| Dyrk4 | 0.00041494 | 0.00035038 | 0.00037562 | -2.52E-05 | 0.67261597 | 0.99999994 |
| Tmem174 | 0.00041494 | 0.00035038 | 0.00037562 | -2.52E-05 | 0.67261597 | 0.99999994 |
| 4930509E16I | 0.00067612 | 0.00058668 | 0.00061193 | -2.53E-05 | 0.60821493 | 0.99999994 |
| Uhrf1 | 0.01185169 | 0.01062025 | 0.0106456 | -2.53E-05 | 0.50632239 | 0.99999994 |
| Gm14391 | 0.00341574 | 0.00306032 | 0.00308571 | -2.54E-05 | 0.5218356 | 0.99999994 |
| Cldn18 | 0.0011527 | 0.00101748 | 0.00104293 | -2.54E-05 | 0.5645111 | 0.99999994 |
| Cybrd1 | 0.00068501 | 0.00059441 | 0.00061998 | -2.56E-05 | 0.60813689 | 0.99999994 |
| C530043A13I | 0.00041446 | 0.00034957 | 0.00037518 | -2.56E-05 | 0.67521951 | 0.99999994 |
| Sox15 | 0.00041446 | 0.00034957 | 0.00037518 | -2.56E-05 | 0.67521951 | 0.99999994 |
| Cep112os1 | 0.00085966 | 0.00075226 | 0.00077795 | -2.57E-05 | 0.58699093 | 0.99999994 |

| | | | | | | |
|---|---|---|---|---|---|---|
| Gm4755 | 0.00071701 | 0.00062306 | 0.00064892 | -2.59E-05 | 0.60458901 | 0.99999994 |
| Gm26631 | 0.00540914 | 0.00485397 | 0.0048799 | -2.59E-05 | 0.51410715 | 0.99999994 |
| Gm9797 | 0.00041402 | 0.00034882 | 0.00037478 | -2.60E-05 | 0.67760955 | 0.99999994 |
| Gm15998 | 0.0004138 | 0.00034845 | 0.00037458 | -2.61E-05 | 0.67879981 | 0.99999994 |
| 1700065D16I | 0.0012194 | 0.00107709 | 0.00110323 | -2.61E-05 | 0.56264826 | 0.99999994 |
| Gm12576 | 0.02149901 | 0.01915965 | 0.01918623 | -2.66E-05 | 0.50367915 | 0.99999994 |
| Hal | 0.00062751 | 0.00054118 | 0.00056795 | -2.68E-05 | 0.62317553 | 0.99999994 |
| Fam151b | 0.00150839 | 0.0013374 | 0.00136442 | -2.70E-05 | 0.55242655 | 0.99999994 |
| Slc1a5 | 0.01172918 | 0.01050934 | 0.01053643 | -2.71E-05 | 0.50682654 | 0.99999994 |
| Gm9725 | 0.00041251 | 0.00034628 | 0.00037341 | -2.71E-05 | 0.68567951 | 0.99999994 |
| 1700039M10 | 0.00300295 | 0.00268639 | 0.00271356 | -2.72E-05 | 0.52656576 | 0.99999994 |
| Dgkz | 0.09456037 | 0.08041929 | 0.08044653 | -2.72E-05 | 0.50089919 | 0.99999994 |
| Mrap | 0.00078563 | 0.00068373 | 0.00071099 | -2.73E-05 | 0.60069817 | 0.99999994 |
| Gm5160 | 0.00399476 | 0.00358007 | 0.00360736 | -2.73E-05 | 0.52008506 | 0.99999994 |
| Fgf13 | 0.00041203 | 0.00034548 | 0.00037298 | -2.75E-05 | 0.68820721 | 0.99999994 |
| Dync1i1 | 0.00705108 | 0.00632654 | 0.00635411 | -2.76E-05 | 0.51152076 | 0.99999994 |
| Gbp3 | 0.00041181 | 0.00034511 | 0.00037278 | -2.77E-05 | 0.68936879 | 0.99999994 |
| Mobp | 0.00041181 | 0.00034511 | 0.00037278 | -2.77E-05 | 0.68936879 | 0.99999994 |
| Arsk | 0.06398791 | 0.05549464 | 0.05552236 | -2.77E-05 | 0.50132583 | 0.99999994 |
| Ppcs | 0.24748086 | 0.18697334 | 0.18700119 | -2.79E-05 | 0.50039554 | 0.99999994 |
| Tmeff2 | 0.00083381 | 0.00072658 | 0.00075456 | -2.80E-05 | 0.59747702 | 0.99999994 |
| Fam161a | 0.01018162 | 0.00912776 | 0.0091558 | -2.80E-05 | 0.50813301 | 0.99999994 |
| 2810442N19 | 0.00041104 | 0.00034383 | 0.00037209 | -2.83E-05 | 0.69340919 | 0.99999994 |
| Crb2 | 0.00079127 | 0.00068773 | 0.00071609 | -2.84E-05 | 0.60395708 | 0.99999994 |
| Tmco4 | 0.00598665 | 0.00537033 | 0.00539878 | -2.85E-05 | 0.51399513 | 0.99999994 |
| Pla2g4c | 0.00095498 | 0.00083559 | 0.00086415 | -2.86E-05 | 0.58707357 | 0.99999994 |
| Soat2 | 0.00083191 | 0.00072428 | 0.00075285 | -2.86E-05 | 0.59973441 | 0.99999994 |
| Sp8 | 0.0004105 | 0.00034292 | 0.00037159 | -2.87E-05 | 0.69627125 | 0.99999994 |
| Gm43140 | 0.00041024 | 0.00034249 | 0.00037136 | -2.89E-05 | 0.69760004 | 0.99999994 |
| A930012L18I | 0.00106381 | 0.00093369 | 0.00096256 | -2.89E-05 | 0.57913756 | 0.99999994 |

| | | | | | | |
|---|---|---|---|---|---|---|
| Cdh5 | 0.00157884 | 0.00139916 | 0.00142807 | -2.89E-05 | 0.55359323 | 0.99999994 |
| Car7 | 0.00110503 | 0.00097086 | 0.00099983 | -2.90E-05 | 0.57646145 | 0.99999994 |
| C1qtnf9 | 0.0004101 | 0.00034225 | 0.00037123 | -2.90E-05 | 0.6983574 | 0.99999994 |
| Slamf9 | 0.00041006 | 0.00034219 | 0.0003712 | -2.90E-05 | 0.69854652 | 0.99999994 |
| Cbln1 | 0.00142421 | 0.00125916 | 0.00128835 | -2.92E-05 | 0.55992765 | 0.99999994 |
| Myoz1 | 0.00040938 | 0.00034104 | 0.00037058 | -2.95E-05 | 0.7021229 | 0.99999994 |
| Gm26634 | 0.00040938 | 0.00034104 | 0.00037058 | -2.95E-05 | 0.7021229 | 0.99999994 |
| Hpcal4 | 0.00040919 | 0.00034074 | 0.00037041 | -2.97E-05 | 0.70305873 | 0.99999994 |
| Unc93a | 0.00040919 | 0.00034074 | 0.00037041 | -2.97E-05 | 0.70305873 | 0.99999994 |
| Mro | 0.00040919 | 0.00034074 | 0.00037041 | -2.97E-05 | 0.70305873 | 0.99999994 |
| Ccdc69 | 0.00079981 | 0.00069413 | 0.00072382 | -2.97E-05 | 0.60757882 | 0.99999994 |
| Mecomos | 0.0004088 | 0.00034008 | 0.00037005 | -3.00E-05 | 0.70510975 | 0.99999994 |
| Tmem221 | 0.0058619 | 0.00525675 | 0.00528673 | -3.00E-05 | 0.515057 | 0.99999994 |
| Prokr1 | 0.00060373 | 0.00051612 | 0.00054644 | -3.03E-05 | 0.64402191 | 0.99999994 |
| Kcng2 | 0.0008516 | 0.00074035 | 0.00077066 | -3.03E-05 | 0.60326994 | 0.99999994 |
| Magix | 0.00040829 | 0.00033924 | 0.0003696 | -3.04E-05 | 0.70770461 | 0.99999994 |
| Xcr1 | 0.00040829 | 0.00033924 | 0.0003696 | -3.04E-05 | 0.70770461 | 0.99999994 |
| Muc13 | 0.00085452 | 0.00074271 | 0.0007733 | -3.06E-05 | 0.60386646 | 0.99999994 |
| Upk2 | 0.0004079 | 0.00033859 | 0.00036924 | -3.07E-05 | 0.70973124 | 0.99999994 |
| Trpc6 | 0.00040765 | 0.00033817 | 0.00036901 | -3.08E-05 | 0.71101533 | 0.99999994 |
| Plce1 | 0.00249272 | 0.00222205 | 0.00225328 | -3.12E-05 | 0.53675857 | 0.99999994 |
| Axl | 0.00205248 | 0.00182464 | 0.00185589 | -3.13E-05 | 0.54462143 | 0.99999994 |
| Rps6ka5 | 0.01732 | 0.01546894 | 0.01550021 | -3.13E-05 | 0.50535684 | 0.99999994 |
| Ackr1 | 0.00074599 | 0.00064334 | 0.00067513 | -3.18E-05 | 0.62304962 | 0.99999994 |
| Gm37933 | 0.00121853 | 0.00107059 | 0.00110244 | -3.18E-05 | 0.57624714 | 0.99999994 |
| Gm15892 | 0.00288902 | 0.00257876 | 0.00261081 | -3.20E-05 | 0.53256095 | 0.99999994 |
| Gm20635 | 0.00119406 | 0.00104824 | 0.00108031 | -3.21E-05 | 0.57832837 | 0.99999994 |
| Atp7b | 0.00237263 | 0.00211253 | 0.0021449 | -3.24E-05 | 0.540014 | 0.99999994 |
| Ly6g6c | 0.00040558 | 0.00033475 | 0.00036715 | -3.24E-05 | 0.7214881 | 0.99999994 |
| Pdzph1 | 0.00040558 | 0.00033475 | 0.00036715 | -3.24E-05 | 0.7214881 | 0.99999994 |

| | | | | | | |
|---|---|---|---|---|---|---|
| Gm11733 | 0.0013591 | 0.00119702 | 0.0012295 | -3.25E-05 | 0.56978498 | 0.99999994 |
| Arntl2 | 0.00170707 | 0.00151127 | 0.00154392 | -3.26E-05 | 0.55596044 | 0.99999994 |
| Cd200r1 | 0.00040505 | 0.00033388 | 0.00036667 | -3.28E-05 | 0.72414819 | 0.99999994 |
| Bnc2 | 0.00107534 | 0.00094019 | 0.00097298 | -3.28E-05 | 0.58874936 | 0.99999994 |
| Srpk3 | 0.00201382 | 0.00178797 | 0.00182097 | -3.30E-05 | 0.54800401 | 0.99999994 |
| 2310047D07 | 0.00176894 | 0.00156678 | 0.00159981 | -3.30E-05 | 0.55465564 | 0.99999994 |
| Gm26692 | 0.00085059 | 0.00073651 | 0.00076974 | -3.32E-05 | 0.61307803 | 0.99999994 |
| Ppp3r2 | 0.00040435 | 0.00033272 | 0.00036603 | -3.33E-05 | 0.72766415 | 0.99999994 |
| 4930563E18I | 0.00040421 | 0.00033249 | 0.0003659 | -3.34E-05 | 0.72836311 | 0.99999994 |
| Slco4c1 | 0.00040417 | 0.00033243 | 0.00036587 | -3.34E-05 | 0.72853763 | 0.99999994 |
| 4732471J01F | 0.00084879 | 0.00073456 | 0.00076811 | -3.35E-05 | 0.61437475 | 0.99999994 |
| Bard1 | 0.0017148 | 0.00151733 | 0.00155091 | -3.36E-05 | 0.55729281 | 0.99999994 |
| Mustn1 | 0.00068063 | 0.00058198 | 0.00061601 | -3.40E-05 | 0.64345639 | 0.99999994 |
| Hmga1 | 0.01053191 | 0.00943407 | 0.00946856 | -3.45E-05 | 0.50967247 | 0.99999994 |
| Rftn2 | 0.00077773 | 0.00066929 | 0.00070384 | -3.46E-05 | 0.6280914 | 0.99999994 |
| Gm42901 | 0.00040242 | 0.00032956 | 0.00036428 | -3.47E-05 | 0.73715182 | 0.99999994 |
| Ryr2 | 0.00072798 | 0.000624 | 0.00065884 | -3.48E-05 | 0.63757824 | 0.99999994 |
| D630003M21 | 0.00619582 | 0.00555172 | 0.00558663 | -3.49E-05 | 0.51658978 | 0.99999994 |
| Ovol2 | 0.19220128 | 0.15149243 | 0.15152745 | -3.50E-05 | 0.50061371 | 0.99999994 |
| Tead4 | 0.00168395 | 0.00148781 | 0.00152304 | -3.52E-05 | 0.56117519 | 0.99999994 |
| Gm15675 | 0.00040173 | 0.00032842 | 0.00036366 | -3.52E-05 | 0.74053626 | 0.99999994 |
| Slc5a5 | 0.00064333 | 0.00054697 | 0.00058227 | -3.53E-05 | 0.65673723 | 0.99999994 |
| Hmcn2 | 0.00513045 | 0.00459376 | 0.00462935 | -3.56E-05 | 0.52040497 | 0.99999994 |
| Txk | 0.00056797 | 0.0004784 | 0.00051409 | -3.57E-05 | 0.67800116 | 0.99999994 |
| 2700069I18R | 0.00040096 | 0.00032717 | 0.00036297 | -3.58E-05 | 0.74421889 | 0.99999994 |
| Apol9b | 0.00080222 | 0.00069014 | 0.000726 | -3.59E-05 | 0.62883206 | 0.99999994 |
| Fgf3 | 0.00040086 | 0.000327 | 0.00036287 | -3.59E-05 | 0.74471781 | 0.99999994 |
| Cebpa | 0.00427557 | 0.00382421 | 0.00386021 | -3.60E-05 | 0.52474904 | 0.99999994 |
| Gpr3 | 0.00084117 | 0.00072511 | 0.00076122 | -3.61E-05 | 0.62391355 | 0.99999994 |
| Slc25a45 | 0.00169718 | 0.00149868 | 0.00153499 | -3.63E-05 | 0.56255148 | 0.99999994 |

| | | | | | | |
|---|---|---|---|---|---|---|
| Hk1 | 0.00127293 | 0.00111507 | 0.00115161 | -3.65E-05 | 0.58364515 | 0.99999994 |
| Tmem232 | 0.00039962 | 0.00032498 | 0.00036175 | -3.68E-05 | 0.75064402 | 0.99999994 |
| Pkia | 0.01491666 | 0.01333407 | 0.013371 | -3.69E-05 | 0.50733466 | 0.99999994 |
| Gm13620 | 0.00204101 | 0.00180853 | 0.00184553 | -3.70E-05 | 0.553076 | 0.99999994 |
| P2rx7 | 0.000399 | 0.00032398 | 0.00036119 | -3.72E-05 | 0.75356518 | 0.99999994 |
| Megf10 | 0.0003988 | 0.00032365 | 0.00036101 | -3.74E-05 | 0.7545327 | 0.99999994 |
| Gm16230 | 0.00488803 | 0.00437398 | 0.00441133 | -3.74E-05 | 0.52247679 | 0.99999994 |
| Il4i1 | 0.00107744 | 0.00093718 | 0.00097488 | -3.77E-05 | 0.60157448 | 0.99999994 |
| Slc6a12 | 0.00039825 | 0.00032276 | 0.00036051 | -3.77E-05 | 0.75709767 | 0.99999994 |
| Tbx21 | 0.00039825 | 0.00032276 | 0.00036051 | -3.77E-05 | 0.75709767 | 0.99999994 |
| Gm37336 | 0.00039805 | 0.00032243 | 0.00036033 | -3.79E-05 | 0.75805388 | 0.99999994 |
| Ugt3a1 | 0.00039801 | 0.00032238 | 0.0003603 | -3.79E-05 | 0.75821295 | 0.99999994 |
| 9430097D07I | 0.0003974 | 0.00032139 | 0.00035974 | -3.84E-05 | 0.76106161 | 0.99999994 |
| A830092H15 | 0.0003974 | 0.00032139 | 0.00035974 | -3.84E-05 | 0.76106161 | 0.99999994 |
| Prss57 | 0.0003974 | 0.00032139 | 0.00035974 | -3.84E-05 | 0.76106161 | 0.99999994 |
| Gm37748 | 0.00125261 | 0.00109475 | 0.00113324 | -3.85E-05 | 0.58942958 | 0.99999994 |
| Mpp4 | 0.0003972 | 0.00032106 | 0.00035956 | -3.85E-05 | 0.76200503 | 0.99999994 |
| Slc6a7 | 0.0003972 | 0.00032106 | 0.00035956 | -3.85E-05 | 0.76200503 | 0.99999994 |
| Uprt | 0.01035885 | 0.00927547 | 0.00931407 | -3.86E-05 | 0.51100314 | 0.99999994 |
| 6430710C18I | 0.00173095 | 0.00152634 | 0.00156549 | -3.92E-05 | 0.56610573 | 0.99999994 |
| Prss35 | 0.002218 | 0.00196603 | 0.00200533 | -3.93E-05 | 0.55189576 | 0.99999994 |
| Gm13344 | 0.00058383 | 0.00048899 | 0.00052843 | -3.94E-05 | 0.69034811 | 0.99999994 |
| Pou2f2 | 0.0054727 | 0.00489759 | 0.00493703 | -3.94E-05 | 0.52120592 | 0.99999994 |
| Ldb2 | 0.00081448 | 0.00069762 | 0.00073708 | -3.95E-05 | 0.63923021 | 0.99999994 |
| Abca8a | 0.00061996 | 0.00052153 | 0.00056112 | -3.96E-05 | 0.68067172 | 0.99999994 |
| Colec12 | 0.00190223 | 0.00168056 | 0.0017202 | -3.96E-05 | 0.56094408 | 0.99999994 |
| Gm13012 | 0.00039558 | 0.00031844 | 0.00035809 | -3.96E-05 | 0.76944257 | 0.99999994 |
| Wfdc17 | 0.00163063 | 0.00143519 | 0.00147487 | -3.97E-05 | 0.57105402 | 0.99999994 |
| Gm16372 | 0.00039531 | 0.00031801 | 0.00035785 | -3.98E-05 | 0.77066328 | 0.99999994 |
| 1700028J19F | 0.01280574 | 0.01145443 | 0.01149516 | -4.07E-05 | 0.50940948 | 0.99999994 |

| | | | | | | |
|---|---|---|---|---|---|---|
| Galnt2 | 0.0011736 | 0.00102104 | 0.00106182 | -4.08E-05 | 0.60088025 | 0.99999994 |
| Inpp5d | 0.00039387 | 0.0003157 | 0.00035655 | -4.08E-05 | 0.77713282 | 0.99999994 |
| Cdh19 | 0.00083357 | 0.00071337 | 0.00075435 | -4.10E-05 | 0.64117881 | 0.99999994 |
| Sphk2 | 0.17971644 | 0.14302397 | 0.14306501 | -4.10E-05 | 0.50076162 | 0.99999994 |
| Gm16958 | 0.00039354 | 0.00031517 | 0.00035625 | -4.11E-05 | 0.77861532 | 0.99999994 |
| Galnt15 | 0.00039354 | 0.00031517 | 0.00035625 | -4.11E-05 | 0.77861532 | 0.99999994 |
| Spata32 | 0.00039354 | 0.00031517 | 0.00035625 | -4.11E-05 | 0.77861532 | 0.99999994 |
| Pnma3 | 0.00039351 | 0.00031512 | 0.00035622 | -4.11E-05 | 0.77876311 | 0.99999994 |
| Gm26822 | 0.00039344 | 0.00031501 | 0.00035616 | -4.11E-05 | 0.77905845 | 0.99999994 |
| Flt4 | 0.00039255 | 0.00031358 | 0.00035535 | -4.18E-05 | 0.78301325 | 0.99999994 |
| Stx17 | 0.09468121 | 0.08050123 | 0.08054311 | -4.19E-05 | 0.50138071 | 0.99999994 |
| Prok2 | 0.00063381 | 0.00053176 | 0.00057366 | -4.19E-05 | 0.68656946 | 0.99999994 |
| 1700056N10 | 0.00154692 | 0.00135731 | 0.00139923 | -4.19E-05 | 0.5790243 | 0.99999994 |
| Tlr5 | 0.00058142 | 0.00048431 | 0.00052625 | -4.19E-05 | 0.7021421 | 0.99999994 |
| Zbp1 | 0.00436649 | 0.0039001 | 0.00394205 | -4.19E-05 | 0.52823412 | 0.99999994 |
| Adcy2 | 0.00084061 | 0.00071871 | 0.00076072 | -4.20E-05 | 0.64339143 | 0.99999994 |
| Cytl1 | 0.00083728 | 0.00071547 | 0.00075771 | -4.22E-05 | 0.64468908 | 0.99999994 |
| Gbx2 | 0.00039169 | 0.00031221 | 0.00035457 | -4.24E-05 | 0.78676502 | 0.99999994 |
| Nol11 | 0.21592611 | 0.16710004 | 0.1671424 | -4.24E-05 | 0.50067303 | 0.99999994 |
| Ccl12 | 0.00136072 | 0.00118835 | 0.00123096 | -4.26E-05 | 0.59111198 | 0.99999994 |
| Acpp | 0.00299535 | 0.00266406 | 0.00270671 | -4.27E-05 | 0.54176726 | 0.99999994 |
| Slc27a2 | 0.00101099 | 0.00087174 | 0.0009148 | -4.31E-05 | 0.62297917 | 0.99999994 |
| Pnldc1 | 0.00039025 | 0.00030991 | 0.00035327 | -4.34E-05 | 0.79298898 | 0.99999994 |
| Gm12951 | 0.00038995 | 0.00030945 | 0.000353 | -4.36E-05 | 0.79424283 | 0.99999994 |
| Twist2 | 0.00038989 | 0.00030934 | 0.00035294 | -4.36E-05 | 0.79452058 | 0.99999994 |
| Mss51 | 0.00794581 | 0.00711236 | 0.00715607 | -4.37E-05 | 0.51621403 | 0.99999994 |
| Grip1os3 | 0.00038969 | 0.00030903 | 0.00035276 | -4.37E-05 | 0.79535191 | 0.99999994 |
| Gm10767 | 0.00947413 | 0.00847987 | 0.00852367 | -4.38E-05 | 0.51364433 | 0.99999994 |
| Gm37359 | 0.00038953 | 0.00030878 | 0.00035262 | -4.38E-05 | 0.79604248 | 0.99999994 |
| Ano9 | 0.01053819 | 0.00943007 | 0.00947417 | -4.41E-05 | 0.5123594 | 0.99999994 |

| | | | | | | |
|---|---|---|---|---|---|---|
| Atp2a1 | 0.00128124 | 0.00111486 | 0.00115913 | -4.43E-05 | 0.60033229 | 0.99999994 |
| Pcdhgc3 | 0.00080202 | 0.00068144 | 0.00072581 | -4.44E-05 | 0.65796999 | 0.99999994 |
| Spef2 | 0.00257452 | 0.00228248 | 0.00232709 | -4.46E-05 | 0.55077361 | 0.99999994 |
| Ankrd26 | 0.08338245 | 0.07140281 | 0.07144753 | -4.47E-05 | 0.5016621 | 0.99999994 |
| Gm15690 | 0.00079992 | 0.00067911 | 0.00072392 | -4.48E-05 | 0.65981772 | 0.99999994 |
| Itgal | 0.00114849 | 0.00099398 | 0.00103912 | -4.51E-05 | 0.61378185 | 0.99999994 |
| Gpc5 | 0.00082441 | 0.00070078 | 0.00074607 | -4.53E-05 | 0.65693018 | 0.99999994 |
| Itga10 | 0.00064534 | 0.00053865 | 0.00058408 | -4.54E-05 | 0.69768604 | 0.99999994 |
| Adra2b | 0.00038704 | 0.00030485 | 0.00035037 | -4.55E-05 | 0.8064274 | 0.99999994 |
| Gm28043 | 0.00038656 | 0.00030409 | 0.00034993 | -4.58E-05 | 0.8083964 | 0.99999994 |
| Nxnl1 | 0.00083558 | 0.0007103 | 0.00075617 | -4.59E-05 | 0.6568004 | 0.99999994 |
| Cd5 | 0.00433066 | 0.00386388 | 0.0039098 | -4.59E-05 | 0.53115475 | 0.99999994 |
| AA467197 | 0.00038621 | 0.00030354 | 0.00034961 | -4.61E-05 | 0.80982926 | 0.99999994 |
| Ccdc187 | 0.00126615 | 0.00109916 | 0.00114548 | -4.63E-05 | 0.60610256 | 0.99999994 |
| Nyx | 0.00064061 | 0.00053339 | 0.00057981 | -4.64E-05 | 0.70297117 | 0.99999994 |
| Tmprss7 | 0.00609366 | 0.00544824 | 0.00549489 | -4.66E-05 | 0.52253146 | 0.99999994 |
| Hao2 | 0.00161611 | 0.00141505 | 0.00146175 | -4.67E-05 | 0.58419667 | 0.99999994 |
| Smim18 | 0.00171254 | 0.00150216 | 0.00154886 | -4.67E-05 | 0.57952754 | 0.99999994 |
| Zfp85 | 0.01574928 | 0.01406237 | 0.01410943 | -4.71E-05 | 0.50885586 | 0.99999994 |
| Dusp15 | 0.00300223 | 0.00266559 | 0.00271291 | -4.73E-05 | 0.54621541 | 0.99999994 |
| Fam124a | 0.0012614 | 0.00109377 | 0.00114119 | -4.74E-05 | 0.60894622 | 0.99999994 |
| Card11 | 0.00074316 | 0.00062513 | 0.00067258 | -4.75E-05 | 0.68068197 | 0.99999994 |
| Hcar2 | 0.00038405 | 0.00030015 | 0.00034766 | -4.75E-05 | 0.81848145 | 0.99999994 |
| Cacng5 | 0.00038405 | 0.00030015 | 0.00034766 | -4.75E-05 | 0.81848145 | 0.99999994 |
| 1700041G16 | 0.00124135 | 0.0010754 | 0.00112307 | -4.77E-05 | 0.61124206 | 0.99999994 |
| Smpd5 | 0.00420708 | 0.00375076 | 0.00379854 | -4.78E-05 | 0.53336593 | 0.99999994 |
| Gm10069 | 0.0101042 | 0.0090385 | 0.00908666 | -4.82E-05 | 0.51406964 | 0.99999994 |
| Runx1 | 0.00262803 | 0.00232701 | 0.00237538 | -4.84E-05 | 0.55391257 | 0.99999994 |
| Mccc1os | 0.00038257 | 0.00029784 | 0.00034632 | -4.85E-05 | 0.82425808 | 0.99999994 |
| Gjc2 | 0.0006028 | 0.00049708 | 0.0005456 | -4.85E-05 | 0.72305021 | 0.99999994 |

| | | | | | | |
|---|---|---|---|---|---|---|
| Hhatl | 0.00038188 | 0.00029677 | 0.0003457 | -4.89E-05 | 0.82690588 | 0.99999994 |
| Casq2 | 0.00080088 | 0.00067576 | 0.00072479 | -4.90E-05 | 0.67373801 | 0.99999994 |
| Slc39a12 | 0.00038154 | 0.00029624 | 0.00034539 | -4.91E-05 | 0.82821648 | 0.99999994 |
| D030055H07 | 0.0008214 | 0.00069408 | 0.00074335 | -4.93E-05 | 0.67045874 | 0.99999994 |
| Tnfrsf8 | 0.0003812 | 0.00029571 | 0.00034508 | -4.94E-05 | 0.82951828 | 0.99999994 |
| Tmprss11a | 0.0003812 | 0.00029571 | 0.00034508 | -4.94E-05 | 0.82951828 | 0.99999994 |
| Impg2 | 0.0018453 | 0.00161933 | 0.00166878 | -4.95E-05 | 0.57818385 | 0.99999994 |
| Dgat2l6 | 0.00038092 | 0.00029528 | 0.00034482 | -4.95E-05 | 0.83057687 | 0.99999994 |
| Clca3b | 0.00038092 | 0.00029528 | 0.00034482 | -4.95E-05 | 0.83057687 | 0.99999994 |
| Insc | 0.00277221 | 0.00245566 | 0.00250545 | -4.98E-05 | 0.55261114 | 0.99999994 |
| Apoo-ps | 0.01120966 | 0.01002344 | 0.01007327 | -4.98E-05 | 0.51313392 | 0.99999994 |
| Il12rb1 | 0.00201508 | 0.00177187 | 0.00182212 | -5.02E-05 | 0.57281688 | 0.99999994 |
| Fbxl21 | 0.00824605 | 0.00737465 | 0.00742496 | -5.03E-05 | 0.51798466 | 0.99999994 |
| Ccdc170 | 0.00037925 | 0.0002927 | 0.00034331 | -5.06E-05 | 0.83680639 | 0.99999994 |
| Thbs2 | 0.00148433 | 0.00129204 | 0.00134267 | -5.06E-05 | 0.59909994 | 0.99999994 |
| Aspm | 0.00201525 | 0.00177156 | 0.00182227 | -5.07E-05 | 0.57347274 | 0.99999994 |
| Gm12860 | 0.00211506 | 0.00186164 | 0.00191239 | -5.08E-05 | 0.57011073 | 0.99999994 |
| Agbl4 | 0.00751149 | 0.00671604 | 0.00676691 | -5.09E-05 | 0.51995367 | 0.99999994 |
| 4933405D12 | 0.0006468 | 0.00053452 | 0.00058541 | -5.09E-05 | 0.71857024 | 0.99999994 |
| Parvg | 0.00037873 | 0.00029189 | 0.00034284 | -5.10E-05 | 0.83872473 | 0.99999994 |
| Cxcl3 | 0.00037873 | 0.00029189 | 0.00034284 | -5.10E-05 | 0.83872473 | 0.99999994 |
| Ly6d | 0.00078887 | 0.00066281 | 0.00071393 | -5.11E-05 | 0.68315462 | 0.99999994 |
| Esrrg | 0.00488749 | 0.00435954 | 0.00441085 | -5.13E-05 | 0.53085841 | 0.99999994 |
| Npc1l1 | 0.00100857 | 0.00086124 | 0.00091261 | -5.14E-05 | 0.64604799 | 0.99999994 |
| Ltf | 0.00080561 | 0.00067766 | 0.00072907 | -5.14E-05 | 0.68058851 | 0.99999994 |
| C1ql2 | 0.00037793 | 0.00029067 | 0.00034212 | -5.15E-05 | 0.84161956 | 0.99999994 |
| Gm44096 | 0.00037793 | 0.00029067 | 0.00034212 | -5.15E-05 | 0.84161956 | 0.99999994 |
| C1qtnf2 | 0.00060877 | 0.00049953 | 0.000551 | -5.15E-05 | 0.73295968 | 0.99999994 |
| Treml4 | 0.00037757 | 0.0002901 | 0.00034179 | -5.17E-05 | 0.84293982 | 0.99999994 |
| Arhgap15 | 0.00037735 | 0.00028978 | 0.0003416 | -5.18E-05 | 0.84370539 | 0.99999994 |

| | | | | | | |
|---|---|---|---|---|---|---|
| Gm14410 | 0.00500335 | 0.00446322 | 0.00451505 | -5.18E-05 | 0.53045246 | 0.99999994 |
| Gbp9 | 0.00037681 | 0.00028894 | 0.0003411 | -5.22E-05 | 0.84565858 | 0.99999994 |
| Slc6a13 | 0.00755042 | 0.00674939 | 0.0068018 | -5.24E-05 | 0.5204523 | 0.99999994 |
| Ush1g | 0.01081428 | 0.00966813 | 0.00972057 | -5.24E-05 | 0.51432168 | 0.99999994 |
| Myom1 | 0.00037617 | 0.00028796 | 0.00034053 | -5.26E-05 | 0.84790942 | 0.99999994 |
| Ccdc3 | 0.00037611 | 0.00028787 | 0.00034047 | -5.26E-05 | 0.84812223 | 0.99999994 |
| Fam186b | 0.00037611 | 0.00028787 | 0.00034047 | -5.26E-05 | 0.84812223 | 0.99999994 |
| Il33 | 0.00037611 | 0.00028787 | 0.00034047 | -5.26E-05 | 0.84812223 | 0.99999994 |
| Cnga1 | 0.00067659 | 0.00055974 | 0.00061236 | -5.26E-05 | 0.71633089 | 0.99999994 |
| Mab21l2 | 0.00037593 | 0.00028759 | 0.00034031 | -5.27E-05 | 0.84875905 | 0.99999994 |
| Scrg1 | 0.00037593 | 0.00028759 | 0.00034031 | -5.27E-05 | 0.84875905 | 0.99999994 |
| Kbtbd6 | 0.00063852 | 0.00052513 | 0.00057792 | -5.28E-05 | 0.72839677 | 0.99999994 |
| Xlr5a | 0.0033622 | 0.00298462 | 0.00303745 | -5.28E-05 | 0.54608863 | 0.99999994 |
| Mst1 | 0.00283961 | 0.00251293 | 0.00256624 | -5.33E-05 | 0.5549942 | 0.99999994 |
| Rnf183 | 0.00037485 | 0.00028594 | 0.00033933 | -5.34E-05 | 0.85252937 | 0.99999994 |
| Prrg4 | 0.00693255 | 0.00619421 | 0.0062478 | -5.36E-05 | 0.52276527 | 0.99999994 |
| Cdh11 | 0.00282288 | 0.00249713 | 0.00255115 | -5.40E-05 | 0.55604689 | 0.99999994 |
| E030037K01I | 0.0007158 | 0.00059347 | 0.00064783 | -5.44E-05 | 0.71176492 | 0.99999994 |
| Rarb | 0.00148388 | 0.0012879 | 0.00134227 | -5.44E-05 | 0.60628252 | 0.99999994 |
| Tepp | 0.00104599 | 0.00089206 | 0.00094645 | -5.44E-05 | 0.64894626 | 0.99999994 |
| Gm42567 | 0.00061003 | 0.00049775 | 0.00055214 | -5.44E-05 | 0.74399705 | 0.99999994 |
| Cd19 | 0.00062687 | 0.00051275 | 0.00056738 | -5.46E-05 | 0.73920222 | 0.99999994 |
| Fgr | 0.00037255 | 0.00028244 | 0.00033725 | -5.48E-05 | 0.86030804 | 0.99999994 |
| Gm26981 | 0.01626675 | 0.01451311 | 0.01456795 | -5.48E-05 | 0.50999525 | 0.99999994 |
| Slc9c1 | 0.0003724 | 0.00028222 | 0.00033712 | -5.49E-05 | 0.86079995 | 0.99999994 |
| Tbx15 | 0.00124336 | 0.00106988 | 0.00112488 | -5.50E-05 | 0.6275905 | 0.99999994 |
| Ttk | 0.00116917 | 0.00100278 | 0.00105781 | -5.50E-05 | 0.63544682 | 0.99999994 |
| Tcf21 | 0.00115267 | 0.00098784 | 0.0010429 | -5.51E-05 | 0.63737493 | 0.99999994 |
| Rgn | 0.01572791 | 0.01403529 | 0.01409049 | -5.52E-05 | 0.51040067 | 0.99999994 |
| Pcdh12 | 0.0006204 | 0.00050626 | 0.00056152 | -5.53E-05 | 0.74378268 | 0.99999994 |

| | | | | | | |
|---|---|---|---|---|---|---|
| Gm3512 | 0.00037178 | 0.00028128 | 0.00033655 | -5.53E-05 | 0.86284863 | 0.99999994 |
| Npm2 | 0.00186964 | 0.00163544 | 0.00169077 | -5.53E-05 | 0.58621541 | 0.99999994 |
| Lpar4 | 0.00177859 | 0.00155294 | 0.00160853 | -5.56E-05 | 0.59096344 | 0.99999994 |
| Pcdhb15 | 0.00449882 | 0.00400554 | 0.00406115 | -5.56E-05 | 0.53631466 | 0.99999994 |
| Gm17134 | 0.00082331 | 0.00068942 | 0.00074508 | -5.57E-05 | 0.69049596 | 0.99999994 |
| Otc | 0.00037078 | 0.00027977 | 0.00033565 | -5.59E-05 | 0.8661069 | 0.99999994 |
| Frmpd1os | 0.00037078 | 0.00027977 | 0.00033565 | -5.59E-05 | 0.8661069 | 0.99999994 |
| Espl1 | 0.00082304 | 0.00068891 | 0.00074483 | -5.59E-05 | 0.69136619 | 0.99999994 |
| A230020J21F | 0.00084755 | 0.00071094 | 0.000767 | -5.61E-05 | 0.68670863 | 0.99999994 |
| 9530052C20I | 0.00037046 | 0.00027928 | 0.00033536 | -5.61E-05 | 0.86714569 | 0.99999994 |
| Slc36a2 | 0.00037046 | 0.00027928 | 0.00033536 | -5.61E-05 | 0.86714569 | 0.99999994 |
| Zap70 | 0.00354775 | 0.00314852 | 0.00320467 | -5.62E-05 | 0.54642633 | 0.99999994 |
| Itgb1bp2 | 0.00109728 | 0.00093666 | 0.00099282 | -5.62E-05 | 0.6467269 | 0.99999994 |
| Ihh | 0.00037011 | 0.00027875 | 0.00033504 | -5.63E-05 | 0.86827044 | 0.99999994 |
| Corin | 0.00037011 | 0.00027875 | 0.00033504 | -5.63E-05 | 0.86827044 | 0.99999994 |
| Gm5145 | 0.00037011 | 0.00027875 | 0.00033504 | -5.63E-05 | 0.86827044 | 0.99999994 |
| C430042M11 | 0.0031336 | 0.00277489 | 0.00283137 | -5.65E-05 | 0.55281369 | 0.99999994 |
| Gkn3 | 0.00036947 | 0.00027779 | 0.00033446 | -5.67E-05 | 0.87030966 | 0.99999994 |
| Car15 | 0.00036947 | 0.00027779 | 0.00033446 | -5.67E-05 | 0.87030966 | 0.99999994 |
| Dpy19l1 | 0.01396895 | 0.01247281 | 0.01252949 | -5.67E-05 | 0.51201127 | 0.99999994 |
| Fam90a1b | 0.00036926 | 0.00027748 | 0.00033428 | -5.68E-05 | 0.87095236 | 0.99999994 |
| Hmmr | 0.0061078 | 0.00545063 | 0.00550759 | -5.70E-05 | 0.52744114 | 0.99999994 |
| Col6a5 | 0.00222917 | 0.00195834 | 0.00201541 | -5.71E-05 | 0.57474916 | 0.99999994 |
| Gm16098 | 0.00036822 | 0.00027592 | 0.00033333 | -5.74E-05 | 0.87421129 | 0.99999994 |
| Defb42 | 0.00081129 | 0.00067659 | 0.0007342 | -5.76E-05 | 0.69925111 | 0.99999994 |
| Gm15612 | 0.00036773 | 0.00027518 | 0.00033289 | -5.77E-05 | 0.87572354 | 0.99999994 |
| Rtp3 | 0.00036773 | 0.00027518 | 0.00033289 | -5.77E-05 | 0.87572354 | 0.99999994 |
| Kif24 | 0.00137836 | 0.00118914 | 0.00124691 | -5.78E-05 | 0.62109798 | 0.99999994 |
| Car5a | 0.00154304 | 0.00133775 | 0.00139573 | -5.80E-05 | 0.60891835 | 0.99999994 |
| Gpat2 | 0.00187599 | 0.00163837 | 0.00169651 | -5.81E-05 | 0.59021459 | 0.99999994 |

| | | | | | | |
|---|---|---|---|---|---|---|
| Gm44317 | 0.00036692 | 0.00027398 | 0.00033216 | -5.82E-05 | 0.87817747 | 0.99999994 |
| Apol10b | 0.00036692 | 0.00027398 | 0.00033216 | -5.82E-05 | 0.87817747 | 0.99999994 |
| Pax2 | 0.00036664 | 0.00027355 | 0.0003319 | -5.83E-05 | 0.87904288 | 0.99999994 |
| Gm15879 | 0.00120026 | 0.00102745 | 0.00108593 | -5.85E-05 | 0.63999589 | 0.99999994 |
| Gabrd | 0.00113322 | 0.00096672 | 0.00102532 | -5.86E-05 | 0.64818975 | 0.99999994 |
| Adtrp | 0.00158193 | 0.00137218 | 0.00143086 | -5.87E-05 | 0.6075753 | 0.99999994 |
| Adgrl2 | 0.00862828 | 0.00770841 | 0.00776712 | -5.87E-05 | 0.52006396 | 0.99999994 |
| Prdm9 | 0.00513753 | 0.00457683 | 0.00463571 | -5.89E-05 | 0.53368843 | 0.99999994 |
| BC034090 | 0.00383137 | 0.00340131 | 0.0034602 | -5.89E-05 | 0.54510347 | 0.99999994 |
| Gmcl1 | 0.20730455 | 0.16147878 | 0.16153768 | -5.89E-05 | 0.50096826 | 0.99999994 |
| Gm10010 | 0.00122363 | 0.00104812 | 0.00110705 | -5.89E-05 | 0.63846048 | 0.99999994 |
| Cmtr1 | 0.01279164 | 0.01142325 | 0.01148261 | -5.94E-05 | 0.51372491 | 0.99999994 |
| Grik5 | 0.05396264 | 0.04707214 | 0.0471315 | -5.94E-05 | 0.50334468 | 0.99999994 |
| Jakmip3 | 0.0008184 | 0.00068125 | 0.00074063 | -5.94E-05 | 0.70320489 | 0.99999994 |
| Gm4767 | 0.00081847 | 0.00068128 | 0.00074069 | -5.94E-05 | 0.70330605 | 0.99999994 |
| Map7d1 | 0.12935076 | 0.10706697 | 0.10712644 | -5.95E-05 | 0.50147399 | 0.99999994 |
| Depdc7 | 0.0044447 | 0.00395288 | 0.00401244 | -5.96E-05 | 0.53935697 | 0.99999994 |
| Lhfpl1 | 0.00256561 | 0.0022594 | 0.00231906 | -5.97E-05 | 0.56797773 | 0.99999994 |
| Gm6878 | 0.00114903 | 0.00097973 | 0.00103961 | -5.99E-05 | 0.64928967 | 0.99999994 |
| Lrrd1 | 0.00108505 | 0.00092184 | 0.00098176 | -5.99E-05 | 0.6577386 | 0.99999994 |
| Zfp114 | 0.00832905 | 0.00743934 | 0.00749927 | -5.99E-05 | 0.52121277 | 0.99999994 |
| Gm13496 | 0.0019345 | 0.00168936 | 0.00174934 | -6.00E-05 | 0.59026875 | 0.99999994 |
| Letm2 | 0.03100634 | 0.02743548 | 0.02749549 | -6.00E-05 | 0.5057961 | 0.99999994 |
| Kcnab3os | 0.00237341 | 0.00208538 | 0.00214561 | -6.02E-05 | 0.57410581 | 0.99999994 |
| Slc23a3 | 0.00036332 | 0.00026862 | 0.00032889 | -6.03E-05 | 0.88874996 | 0.99999994 |
| Tex12 | 0.00036332 | 0.00026862 | 0.00032889 | -6.03E-05 | 0.88874996 | 0.99999994 |
| Dusp27 | 0.00036323 | 0.00026849 | 0.00032882 | -6.03E-05 | 0.88898896 | 0.99999994 |
| Rec8 | 0.00616542 | 0.00549894 | 0.00555933 | -6.04E-05 | 0.52882212 | 0.99999994 |
| Fkbp1b | 0.27425241 | 0.2030103 | 0.20307097 | -6.07E-05 | 0.50079344 | 0.99999994 |
| 4930408O17 | 0.0005421 | 0.00042963 | 0.00049068 | -6.10E-05 | 0.79620339 | 0.99999994 |

| | | | | | | |
|---|---|---|---|---|---|---|
| Hsd11b1 | 0.00075961 | 0.00062639 | 0.00068746 | -6.11E-05 | 0.72285639 | 0.99999994 |
| Gm3411 | 0.00117683 | 0.00100364 | 0.00106474 | -6.11E-05 | 0.64875373 | 0.99999994 |
| Adamts4 | 0.00066841 | 0.00054372 | 0.00060496 | -6.12E-05 | 0.74976937 | 0.99999994 |
| Dysf | 0.00273661 | 0.00241183 | 0.00247333 | -6.15E-05 | 0.56573361 | 0.99999994 |
| Adamts8 | 0.00036097 | 0.00026516 | 0.00032677 | -6.16E-05 | 0.89526189 | 0.99999994 |
| Camkv | 0.00036097 | 0.00026516 | 0.00032677 | -6.16E-05 | 0.89526189 | 0.99999994 |
| Gm17276 | 0.00059844 | 0.00047988 | 0.00054166 | -6.18E-05 | 0.77611158 | 0.99999994 |
| Gm26590 | 0.00114714 | 0.00097602 | 0.0010379 | -6.19E-05 | 0.65427535 | 0.99999994 |
| Cd160 | 0.00036036 | 0.00026427 | 0.00032622 | -6.20E-05 | 0.89690684 | 0.99999994 |
| Gm29629 | 0.00036009 | 0.00026386 | 0.00032597 | -6.21E-05 | 0.89764641 | 0.99999994 |
| Tspan2os | 0.0017316 | 0.00150378 | 0.00156608 | -6.23E-05 | 0.60441443 | 0.99999994 |
| Spag16 | 0.00123118 | 0.00105156 | 0.00111387 | -6.23E-05 | 0.64518592 | 0.99999994 |
| Cep112 | 0.02745232 | 0.02433936 | 0.02440169 | -6.23E-05 | 0.50678266 | 0.99999994 |
| Slc34a1 | 0.00118093 | 0.00100576 | 0.00106844 | -6.27E-05 | 0.65193329 | 0.99999994 |
| Tyro3 | 0.00687504 | 0.00613344 | 0.00619622 | -6.28E-05 | 0.526882 | 0.99999994 |
| Tfr2 | 0.00201189 | 0.00175635 | 0.00181923 | -6.29E-05 | 0.59098331 | 0.99999994 |
| Mcoln2 | 0.0003583 | 0.00026125 | 0.00032436 | -6.31E-05 | 0.90233209 | 0.99999994 |
| Pparg | 0.00073933 | 0.00060574 | 0.00066911 | -6.34E-05 | 0.73578895 | 0.99999994 |
| Zbtb1 | 0.06001829 | 0.05214962 | 0.05221306 | -6.34E-05 | 0.50322641 | 0.99999994 |
| Csgalnact1 | 0.00035759 | 0.00026022 | 0.00032371 | -6.35E-05 | 0.90414852 | 0.99999994 |
| Sbk2 | 0.00096914 | 0.00081327 | 0.00087696 | -6.37E-05 | 0.68559362 | 0.99999994 |
| Sp110 | 0.00065216 | 0.00052649 | 0.00059025 | -6.38E-05 | 0.76395609 | 0.99999994 |
| Grap | 0.00071365 | 0.0005821 | 0.00064588 | -6.38E-05 | 0.74450079 | 0.99999994 |
| Gckr | 0.00151277 | 0.00130455 | 0.00136837 | -6.38E-05 | 0.62188293 | 0.99999994 |
| Mc1r | 0.00081166 | 0.00067063 | 0.00073454 | -6.39E-05 | 0.71874995 | 0.99999994 |
| Col8a1 | 0.0003564 | 0.00025849 | 0.00032264 | -6.41E-05 | 0.90714915 | 0.99999994 |
| Slamf7 | 0.00076227 | 0.00062563 | 0.00068986 | -6.42E-05 | 0.73229273 | 0.99999994 |
| Tnfrsf4 | 0.00196962 | 0.00171679 | 0.00178106 | -6.43E-05 | 0.59490388 | 0.99999994 |
| Steap3 | 0.00136472 | 0.00117027 | 0.00123458 | -6.43E-05 | 0.63559327 | 0.99999994 |
| Ntf5 | 0.00302785 | 0.00267167 | 0.00273602 | -6.44E-05 | 0.5622024 | 0.99999994 |

| | | | | | | |
|---|---|---|---|---|---|---|
| Fam131c | 0.00125285 | 0.00106895 | 0.00113346 | -6.45E-05 | 0.64759051 | 0.99999994 |
| Apbb1ip | 0.00035557 | 0.00025728 | 0.00032188 | -6.46E-05 | 0.90920899 | 0.99999994 |
| Wnt9b | 0.0003553 | 0.00025689 | 0.00032164 | -6.47E-05 | 0.90986404 | 0.99999994 |
| Nsg1 | 0.02211119 | 0.01965966 | 0.01972445 | -6.48E-05 | 0.50872185 | 0.99999994 |
| Gja5 | 0.00035514 | 0.00025666 | 0.00032149 | -6.48E-05 | 0.9102549 | 0.99999994 |
| Cib3 | 0.00631575 | 0.00562935 | 0.0056943 | -6.49E-05 | 0.53025852 | 0.99999994 |
| Adam33 | 0.00035479 | 0.00025616 | 0.00032118 | -6.50E-05 | 0.91109617 | 0.99999994 |
| Cryba4 | 0.0125385 | 0.01119192 | 0.0112573 | -6.54E-05 | 0.51541759 | 0.99999994 |
| Gpr37 | 9.08E-05 | 1.68E-05 | 8.22E-05 | -6.55E-05 | 0.99999994 | 0.99999994 |
| Cacng3 | 9.08E-05 | 1.68E-05 | 8.22E-05 | -6.55E-05 | 0.99999994 | 0.99999994 |
| Tspo2 | 0.00166512 | 0.00144045 | 0.00150603 | -6.56E-05 | 0.61402378 | 0.99999994 |
| Gm44175 | 0.00111001 | 0.00093872 | 0.00100433 | -6.56E-05 | 0.66817634 | 0.99999994 |
| Gm15999 | 0.00035346 | 0.00025424 | 0.00031997 | -6.57E-05 | 0.91426165 | 0.99999994 |
| Xlr | 0.00202435 | 0.00176461 | 0.00183048 | -6.59E-05 | 0.59465252 | 0.99999994 |
| Sycp1 | 9.18E-05 | 1.72E-05 | 8.32E-05 | -6.60E-05 | 0.99999994 | 0.99999994 |
| Gm20649 | 9.18E-05 | 1.72E-05 | 8.32E-05 | -6.60E-05 | 0.99999994 | 0.99999994 |
| Gm20404 | 9.18E-05 | 1.72E-05 | 8.32E-05 | -6.60E-05 | 0.99999994 | 0.99999994 |
| Defb30 | 9.18E-05 | 1.72E-05 | 8.32E-05 | -6.60E-05 | 0.99999994 | 0.99999994 |
| Ctla2b | 0.00080939 | 0.00066627 | 0.00073248 | -6.62E-05 | 0.72631981 | 0.99999994 |
| 5330429C05l | 0.00077926 | 0.00063862 | 0.00070523 | -6.66E-05 | 0.73525085 | 0.99999994 |
| Gm10687 | 0.00035137 | 0.00025125 | 0.00031808 | -6.68E-05 | 0.91904259 | 0.99999994 |
| Bloc1s6os | 0.01077022 | 0.00961432 | 0.00968125 | -6.69E-05 | 0.51835365 | 0.99999994 |
| Ephb6 | 0.00718057 | 0.00640325 | 0.00647024 | -6.70E-05 | 0.52747085 | 0.99999994 |
| Kcnh6 | 0.03552231 | 0.03133856 | 0.03140562 | -6.71E-05 | 0.50567011 | 0.99999994 |
| Col6a2 | 0.00669755 | 0.00596977 | 0.00603697 | -6.72E-05 | 0.52953365 | 0.99999994 |
| Gimap5 | 0.00035064 | 0.0002502 | 0.00031742 | -6.72E-05 | 0.92067434 | 0.99999994 |
| Scnn1a | 0.00517347 | 0.00460071 | 0.00466803 | -6.73E-05 | 0.53823759 | 0.99999994 |
| Gpr65 | 0.00068298 | 0.0005508 | 0.00061813 | -6.73E-05 | 0.76579668 | 0.99999994 |
| Dnm2 | 0.38880715 | 0.26440664 | 0.26447438 | -6.77E-05 | 0.50068019 | 0.99999994 |
| Wscd2 | 0.00574023 | 0.00510963 | 0.00517743 | -6.78E-05 | 0.53472893 | 0.99999994 |

| | | | | | | |
|---|---|---|---|---|---|---|
| Bnip3 | 0.3446078 | 0.2420561 | 0.24212397 | -6.79E-05 | 0.50074438 | 0.99999994 |
| Atg4a | 0.07769834 | 0.06675367 | 0.06682157 | -6.79E-05 | 0.50269824 | 0.99999994 |
| Fgb | 0.001418 | 0.00121482 | 0.00128273 | -6.79E-05 | 0.63772668 | 0.99999994 |
| Gm37986 | 0.00034931 | 0.00024831 | 0.00031622 | -6.79E-05 | 0.9235648 | 0.99999994 |
| Lyzl4 | 0.00078831 | 0.00064541 | 0.00071342 | -6.80E-05 | 0.73713309 | 0.99999994 |
| Sdr42e1 | 0.013773 | 0.01228716 | 0.01235536 | -6.82E-05 | 0.5146557 | 0.99999994 |
| Sult1b1 | 0.00253724 | 0.00222511 | 0.00229346 | -6.84E-05 | 0.57862674 | 0.99999994 |
| Parp3 | 0.00130141 | 0.00110892 | 0.00117735 | -6.84E-05 | 0.650586 | 0.99999994 |
| Pld5 | 0.00034823 | 0.00024677 | 0.00031524 | -6.85E-05 | 0.92586802 | 0.99999994 |
| Gm14661 | 0.00034823 | 0.00024677 | 0.00031524 | -6.85E-05 | 0.92586802 | 0.99999994 |
| Brip1 | 0.00180439 | 0.00156332 | 0.00163184 | -6.85E-05 | 0.61006698 | 0.99999994 |
| Zfp454 | 0.01369408 | 0.01221663 | 0.01228522 | -6.86E-05 | 0.51482137 | 0.99999994 |
| Gm37799 | 0.00034797 | 0.00024641 | 0.00031501 | -6.86E-05 | 0.92640635 | 0.99999994 |
| Gm4208 | 0.00034764 | 0.00024594 | 0.00031471 | -6.88E-05 | 0.92710049 | 0.99999994 |
| Alpk3 | 0.00034759 | 0.00024586 | 0.00031466 | -6.88E-05 | 0.92720671 | 0.99999994 |
| Orm1 | 0.00034759 | 0.00024586 | 0.00031466 | -6.88E-05 | 0.92720671 | 0.99999994 |
| Gm26674 | 0.00034751 | 0.00024576 | 0.00031459 | -6.88E-05 | 0.92736576 | 0.99999994 |
| Gm17430 | 0.00174723 | 0.00151131 | 0.0015802 | -6.89E-05 | 0.61416622 | 0.99999994 |
| Stfa2 | 0.00248322 | 0.00217572 | 0.00224471 | -6.90E-05 | 0.58104894 | 0.99999994 |
| Gm8237 | 0.001236 | 0.00104898 | 0.00111823 | -6.93E-05 | 0.65992191 | 0.99999994 |
| Stk32c | 0.00122166 | 0.00103598 | 0.00110527 | -6.93E-05 | 0.66176729 | 0.99999994 |
| Lyrm7 | 0.00916171 | 0.008175 | 0.00824433 | -6.93E-05 | 0.5223186 | 0.99999994 |
| Krt76 | 0.00034626 | 0.00024399 | 0.00031346 | -6.95E-05 | 0.92991599 | 0.99999994 |
| Lrg1 | 0.00057605 | 0.00045167 | 0.0005214 | -6.97E-05 | 0.81330648 | 0.99999994 |
| Gm37422 | 0.00034512 | 0.00024239 | 0.00031243 | -7.00E-05 | 0.93218071 | 0.99999994 |
| Spata9 | 0.00181413 | 0.00157039 | 0.00164063 | -7.02E-05 | 0.61216864 | 0.99999994 |
| Nlrp3 | 0.00034472 | 0.00024182 | 0.00031206 | -7.02E-05 | 0.93296852 | 0.99999994 |
| Gm16302 | 0.00080161 | 0.00065519 | 0.00072545 | -7.03E-05 | 0.74043625 | 0.99999994 |
| Arhgap30 | 0.00034454 | 0.00024157 | 0.0003119 | -7.03E-05 | 0.93331036 | 0.99999994 |
| Vwa7 | 0.00034437 | 0.00024133 | 0.00031174 | -7.04E-05 | 0.93365049 | 0.99999994 |

| | | | | | | |
|---|---|---|---|---|---|---|
| 2410018L13F | 0.01110206 | 0.00990684 | 0.00997731 | -7.05E-05 | 0.51874849 | 0.99999994 |
| Lrp4 | 0.00079915 | 0.0006527 | 0.00072322 | -7.05E-05 | 0.74184493 | 0.99999994 |
| Tmem132c | 0.00034409 | 0.00024094 | 0.00031149 | -7.06E-05 | 0.93418153 | 0.99999994 |
| BC048679 | 0.00034409 | 0.00024094 | 0.00031149 | -7.06E-05 | 0.93418153 | 0.99999994 |
| Platr19 | 0.00034409 | 0.00024094 | 0.00031149 | -7.06E-05 | 0.93418153 | 0.99999994 |
| Ly75 | 0.00056161 | 0.00043768 | 0.00050833 | -7.07E-05 | 0.82254918 | 0.99999994 |
| Itga7 | 0.00080221 | 0.00065521 | 0.00072599 | -7.08E-05 | 0.74181474 | 0.99999994 |
| Tifab | 0.00034314 | 0.00023961 | 0.00031063 | -7.10E-05 | 0.93598408 | 0.99999994 |
| Col4a3 | 0.00034304 | 0.00023947 | 0.00031054 | -7.11E-05 | 0.93617097 | 0.99999994 |
| A930041C12 | 0.001294 | 0.00109951 | 0.00117066 | -7.12E-05 | 0.65709974 | 0.99999994 |
| Apln | 0.00034271 | 0.00023902 | 0.00031025 | -7.12E-05 | 0.93677467 | 0.99999994 |
| Pebp4 | 0.00034224 | 0.00023836 | 0.00030982 | -7.15E-05 | 0.93764689 | 0.99999994 |
| Snx22 | 0.00437353 | 0.00387661 | 0.00394839 | -7.18E-05 | 0.54815505 | 0.99999994 |
| Gm5616 | 0.00055671 | 0.00043203 | 0.0005039 | -7.19E-05 | 0.82877518 | 0.99999994 |
| D030047H15 | 0.00034115 | 0.00023685 | 0.00030884 | -7.20E-05 | 0.93962145 | 0.99999994 |
| Irx5 | 0.00034115 | 0.00023685 | 0.00030884 | -7.20E-05 | 0.93962145 | 0.99999994 |
| Tunar | 0.00034115 | 0.00023685 | 0.00030884 | -7.20E-05 | 0.93962145 | 0.99999994 |
| Il17rd | 0.00182842 | 0.0015814 | 0.00165354 | -7.21E-05 | 0.61423835 | 0.99999994 |
| Traip | 0.0047748 | 0.00423734 | 0.00430948 | -7.21E-05 | 0.54435966 | 0.99999994 |
| Crygb | 0.00034076 | 0.0002363 | 0.00030848 | -7.22E-05 | 0.94032405 | 0.99999994 |
| Prmt8 | 0.00034061 | 0.00023609 | 0.00030835 | -7.23E-05 | 0.94058543 | 0.99999994 |
| Rab30 | 0.00301553 | 0.00265256 | 0.00272491 | -7.24E-05 | 0.5701426 | 0.99999994 |
| 2310057J18F | 0.00201935 | 0.00175354 | 0.00182597 | -7.24E-05 | 0.60412403 | 0.99999994 |
| Cldn14 | 0.00033993 | 0.00023514 | 0.00030772 | -7.26E-05 | 0.94179034 | 0.99999994 |
| Mybpc3 | 0.00010495 | 2.24E-05 | 9.50E-05 | -7.26E-05 | 0.99999982 | 0.99999994 |
| Ccna1 | 0.00010495 | 2.24E-05 | 9.50E-05 | -7.26E-05 | 0.99999982 | 0.99999994 |
| B230334C09 | 0.00010495 | 2.24E-05 | 9.50E-05 | -7.26E-05 | 0.99999982 | 0.99999994 |
| Gm13010 | 0.00010495 | 2.24E-05 | 9.50E-05 | -7.26E-05 | 0.99999982 | 0.99999994 |
| Reg3g | 0.00010495 | 2.24E-05 | 9.50E-05 | -7.26E-05 | 0.99999982 | 0.99999994 |
| Ptprb | 0.00010495 | 2.24E-05 | 9.50E-05 | -7.26E-05 | 0.99999982 | 0.99999994 |

| | | | | | | |
|---|---|---|---|---|---|---|
| Tmem45b | 0.00010495 | 2.24E-05 | 9.50E-05 | -7.26E-05 | 0.99999982 | 0.99999994 |
| Klk1b4 | 0.00010495 | 2.24E-05 | 9.50E-05 | -7.26E-05 | 0.99999982 | 0.99999994 |
| 1700013H16I | 0.00010495 | 2.24E-05 | 9.50E-05 | -7.26E-05 | 0.99999982 | 0.99999994 |
| Gm29133 | 0.00010495 | 2.24E-05 | 9.50E-05 | -7.26E-05 | 0.99999982 | 0.99999994 |
| 2410004I01R | 0.0011766 | 0.00099189 | 0.00106453 | -7.26E-05 | 0.67517136 | 0.99999994 |
| Susd3 | 0.00069208 | 0.00055363 | 0.00062637 | -7.27E-05 | 0.78025498 | 0.99999994 |
| Ccdc42 | 0.00033905 | 0.00023393 | 0.00030693 | -7.30E-05 | 0.94330401 | 0.99999994 |
| 1600014C23I | 0.00033905 | 0.00023393 | 0.00030693 | -7.30E-05 | 0.94330401 | 0.99999994 |
| Syde1 | 0.00033878 | 0.00023356 | 0.00030669 | -7.31E-05 | 0.94375869 | 0.99999994 |
| Mal | 0.00033851 | 0.00023319 | 0.00030645 | -7.33E-05 | 0.94420975 | 0.99999994 |
| E130114P18I | 0.00033851 | 0.00023319 | 0.00030645 | -7.33E-05 | 0.94420975 | 0.99999994 |
| Plxdc1 | 0.00033851 | 0.00023319 | 0.00030645 | -7.33E-05 | 0.94420975 | 0.99999994 |
| Fstl4 | 0.00196557 | 0.00170406 | 0.0017774 | -7.33E-05 | 0.60821658 | 0.99999994 |
| Xlr3c | 0.00378296 | 0.00334321 | 0.00341659 | -7.34E-05 | 0.55684125 | 0.99999994 |
| Serpinb6c | 0.00010691 | 2.33E-05 | 9.68E-05 | -7.35E-05 | 0.99999979 | 0.99999994 |
| Gm28874 | 0.00242822 | 0.0021212 | 0.00219508 | -7.39E-05 | 0.58862654 | 0.99999994 |
| Gm15965 | 0.0037864 | 0.00334579 | 0.00341969 | -7.39E-05 | 0.55718823 | 0.99999994 |
| Tigd4 | 0.00033706 | 0.0002312 | 0.00030513 | -7.39E-05 | 0.94660757 | 0.99999994 |
| Serpina16 | 0.00033704 | 0.00023117 | 0.00030511 | -7.39E-05 | 0.94664665 | 0.99999994 |
| 9530026P05I | 0.00237986 | 0.00207745 | 0.00215143 | -7.40E-05 | 0.59052294 | 0.99999994 |
| Rex2 | 0.00059583 | 0.00046522 | 0.0005393 | -7.41E-05 | 0.81971256 | 0.99999994 |
| Fbln7 | 0.00158629 | 0.00136056 | 0.0014348 | -7.42E-05 | 0.63472847 | 0.99999994 |
| Hpx | 0.00033608 | 0.00022986 | 0.00030425 | -7.44E-05 | 0.94818685 | 0.99999994 |
| Fcmr | 0.00367159 | 0.00324165 | 0.00331626 | -7.46E-05 | 0.55951627 | 0.99999994 |
| Gm35986 | 0.00160073 | 0.00137321 | 0.00144785 | -7.46E-05 | 0.63425399 | 0.99999994 |
| 2810459M11 | 0.00033551 | 0.00022907 | 0.00030373 | -7.47E-05 | 0.94908961 | 0.99999994 |
| Sox3 | 0.00033551 | 0.00022907 | 0.00030373 | -7.47E-05 | 0.94908961 | 0.99999994 |
| Esr1 | 0.00010978 | 2.45E-05 | 9.94E-05 | -7.49E-05 | 0.99999973 | 0.99999994 |
| Slc9a5 | 0.00010978 | 2.45E-05 | 9.94E-05 | -7.49E-05 | 0.99999973 | 0.99999994 |
| Csf2rb | 0.00010979 | 2.45E-05 | 9.94E-05 | -7.49E-05 | 0.99999973 | 0.99999994 |

| | | | | | | |
|---|---|---|---|---|---|---|
| Gm15489 | 0.00033496 | 0.00022833 | 0.00030323 | -7.49E-05 | 0.94994004 | 0.99999994 |
| Stk10 | 0.00409472 | 0.00362245 | 0.00369738 | -7.49E-05 | 0.55364968 | 0.99999994 |
| Adam18 | 0.00033484 | 0.00022817 | 0.00030312 | -7.50E-05 | 0.95012303 | 0.99999994 |
| Ncf1 | 0.00106304 | 0.00088682 | 0.00096186 | -7.50E-05 | 0.69822017 | 0.99999994 |
| Tmem217 | 0.00654748 | 0.00582709 | 0.0059023 | -7.52E-05 | 0.53379619 | 0.99999994 |
| C2cd4d | 0.00894809 | 0.007978 | 0.00805327 | -7.53E-05 | 0.52480417 | 0.99999994 |
| 9130213A22I | 0.00011136 | 2.52E-05 | 0.00010083 | -7.56E-05 | 0.9999997 | 0.99999994 |
| Ccl11 | 0.00011136 | 2.52E-05 | 0.00010083 | -7.56E-05 | 0.9999997 | 0.99999994 |
| Fsd2 | 0.00083786 | 0.00068239 | 0.00075823 | -7.58E-05 | 0.74724108 | 0.99999994 |
| Spi1 | 0.00220328 | 0.00191618 | 0.00199203 | -7.59E-05 | 0.60004567 | 0.99999994 |
| Ppfia3 | 0.01645293 | 0.01465688 | 0.01473284 | -7.60E-05 | 0.5136891 | 0.99999994 |
| Gm20400 | 0.00033258 | 0.00022509 | 0.00030108 | -7.60E-05 | 0.95350963 | 0.99999994 |
| Ddah1 | 0.00725562 | 0.00646149 | 0.00653753 | -7.60E-05 | 0.53085563 | 0.99999994 |
| Slc1a7 | 0.00033233 | 0.00022475 | 0.00030085 | -7.61E-05 | 0.95388267 | 0.99999994 |
| 4833413E03I | 0.00033233 | 0.00022475 | 0.00030085 | -7.61E-05 | 0.95388267 | 0.99999994 |
| Gm16316 | 0.00011287 | 2.59E-05 | 0.00010219 | -7.63E-05 | 0.99999966 | 0.99999994 |
| Penk | 0.00011287 | 2.59E-05 | 0.00010219 | -7.63E-05 | 0.99999966 | 0.99999994 |
| Lct | 0.00011309 | 2.60E-05 | 0.00010239 | -7.64E-05 | 0.99999966 | 0.99999994 |
| Wfdc13 | 0.00011309 | 2.60E-05 | 0.00010239 | -7.64E-05 | 0.99999966 | 0.99999994 |
| Gramd2 | 0.00226625 | 0.0019725 | 0.00204888 | -7.64E-05 | 0.59798363 | 0.99999994 |
| Sohlh2 | 0.0003317 | 0.0002239 | 0.00030028 | -7.64E-05 | 0.95478566 | 0.99999994 |
| Adarb2 | 0.0003317 | 0.0002239 | 0.00030028 | -7.64E-05 | 0.95478566 | 0.99999994 |
| Pcdh17 | 0.00079398 | 0.00064183 | 0.00071854 | -7.67E-05 | 0.76133513 | 0.99999994 |
| Hoxd4 | 0.00033082 | 0.00022271 | 0.00029948 | -7.68E-05 | 0.95602656 | 0.99999994 |
| Il12rb2 | 0.00011408 | 2.65E-05 | 0.00010328 | -7.68E-05 | 0.99999963 | 0.99999994 |
| Dnase2b | 0.00011422 | 2.66E-05 | 0.00010342 | -7.69E-05 | 0.99999962 | 0.99999994 |
| Cfap52 | 0.00351165 | 0.00309508 | 0.00317214 | -7.71E-05 | 0.56422895 | 0.99999994 |
| Il17f | 0.00033015 | 0.00022181 | 0.00029887 | -7.71E-05 | 0.9569505 | 0.99999994 |
| Klk6 | 0.00032999 | 0.00022159 | 0.00029873 | -7.71E-05 | 0.95717058 | 0.99999994 |
| Dnm3os | 0.00111412 | 0.00093084 | 0.00100805 | -7.72E-05 | 0.69491632 | 0.99999994 |

| | | | | | | |
|---|---|---|---|---|---|---|
| Vav1 | 0.00153603 | 0.00131208 | 0.00138939 | -7.73E-05 | 0.64443933 | 0.99999994 |
| Adm | 0.00127037 | 0.00107194 | 0.0011493 | -7.74E-05 | 0.67292015 | 0.99999994 |
| Ccdc158 | 0.00114214 | 0.00095593 | 0.00103338 | -7.75E-05 | 0.69107068 | 0.99999994 |
| Epha8 | 0.00059702 | 0.0004629 | 0.00054037 | -7.75E-05 | 0.83004524 | 0.99999994 |
| Zp3 | 0.00135452 | 0.00114787 | 0.00122536 | -7.75E-05 | 0.66310043 | 0.99999994 |
| Cetn3 | 0.82983399 | 0.40839169 | 0.40846968 | -7.80E-05 | 0.50050701 | 0.99999994 |
| 4930558K02I | 0.00032793 | 0.00021884 | 0.00029687 | -7.80E-05 | 0.95990132 | 0.99999994 |
| B3gat2 | 0.00011693 | 2.78E-05 | 0.00010587 | -7.80E-05 | 0.99999954 | 0.99999994 |
| Gm12195 | 0.00011693 | 2.78E-05 | 0.00010587 | -7.80E-05 | 0.99999954 | 0.99999994 |
| Gm16192 | 0.00011708 | 2.79E-05 | 0.00010601 | -7.81E-05 | 0.99999953 | 0.99999994 |
| Capn12 | 0.00125052 | 0.00105319 | 0.00113135 | -7.82E-05 | 0.67720147 | 0.99999994 |
| Ptpre | 0.00109383 | 0.00091152 | 0.00098971 | -7.82E-05 | 0.70051123 | 0.99999994 |
| Pla2r1 | 0.00011737 | 2.80E-05 | 0.00010627 | -7.82E-05 | 0.99999952 | 0.99999994 |
| Eme1 | 0.00174907 | 0.00150349 | 0.00158186 | -7.84E-05 | 0.62921906 | 0.99999994 |
| Hmcn1 | 0.00032696 | 0.00021754 | 0.00029599 | -7.84E-05 | 0.96114336 | 0.99999994 |
| Smim17 | 0.00032696 | 0.00021754 | 0.00029599 | -7.84E-05 | 0.96114336 | 0.99999994 |
| Naip5 | 0.00032684 | 0.00021739 | 0.00029588 | -7.85E-05 | 0.96128524 | 0.99999994 |
| Acat3 | 0.00011814 | 2.84E-05 | 0.00010696 | -7.86E-05 | 0.99999949 | 0.99999994 |
| Zfp385c | 0.0032756 | 0.00288073 | 0.00295939 | -7.87E-05 | 0.57021249 | 0.99999994 |
| Gm13861 | 0.00011838 | 2.85E-05 | 0.00010718 | -7.87E-05 | 0.99999948 | 0.99999994 |
| Ctsk | 0.00011863 | 2.86E-05 | 0.00010741 | -7.88E-05 | 0.99999947 | 0.99999994 |
| Epha2 | 0.00114202 | 0.00095448 | 0.00103327 | -7.88E-05 | 0.69411233 | 0.99999994 |
| Il17rb | 0.00076199 | 0.00061081 | 0.00068961 | -7.88E-05 | 0.77653919 | 0.99999994 |
| Itk | 0.0003261 | 0.0002164 | 0.00029521 | -7.88E-05 | 0.9622086 | 0.99999994 |
| Myoc | 0.00121512 | 0.00102025 | 0.00109935 | -7.91E-05 | 0.68400692 | 0.99999994 |
| Chrna5 | 0.00062587 | 0.00048725 | 0.00056647 | -7.92E-05 | 0.8240373 | 0.99999994 |
| Nrl | 0.00011975 | 2.92E-05 | 0.00010843 | -7.92E-05 | 0.99999943 | 0.99999994 |
| Ttl | 0.03487014 | 0.03076312 | 0.03084239 | -7.93E-05 | 0.5068245 | 0.99999994 |
| Lrrc7 | 0.00135678 | 0.00114811 | 0.0012274 | -7.93E-05 | 0.66640459 | 0.99999994 |
| Lin28a | 0.00054955 | 0.000418 | 0.00049742 | -7.94E-05 | 0.85607257 | 0.99999994 |

| | | | | | | |
|---|---|---|---|---|---|---|
| Pkn3 | 0.00121135 | 0.0010164 | 0.00109595 | -7.96E-05 | 0.68550887 | 0.99999994 |
| Polh | 0.00293579 | 0.00257344 | 0.00265299 | -7.96E-05 | 0.5791008 | 0.99999994 |
| Csf3r | 0.00012056 | 2.96E-05 | 0.00010916 | -7.96E-05 | 0.99999939 | 0.99999994 |
| Ms4a2 | 0.00012056 | 2.96E-05 | 0.00010916 | -7.96E-05 | 0.99999939 | 0.99999994 |
| Mfsd2b | 0.00012092 | 2.98E-05 | 0.00010948 | -7.97E-05 | 0.99999937 | 0.99999994 |
| Rsph14 | 0.00245938 | 0.00214335 | 0.0022232 | -7.98E-05 | 0.59447547 | 0.99999994 |
| D930020B18 | 0.00012131 | 2.99E-05 | 0.00010984 | -7.99E-05 | 0.99999936 | 0.99999994 |
| Tmem63c | 0.00012131 | 2.99E-05 | 0.00010984 | -7.99E-05 | 0.99999936 | 0.99999994 |
| Gm13199 | 0.00197964 | 0.0017102 | 0.00179011 | -7.99E-05 | 0.61681759 | 0.99999994 |
| Myom3 | 0.00012177 | 3.02E-05 | 0.00011025 | -8.01E-05 | 0.99999933 | 0.99999994 |
| Lcat | 0.00012177 | 3.02E-05 | 0.00011025 | -8.01E-05 | 0.99999933 | 0.99999994 |
| Gm27019 | 0.00012244 | 3.05E-05 | 0.00011085 | -8.03E-05 | 0.9999993 | 0.99999994 |
| Kazn | 0.04281003 | 0.03758596 | 0.0376664 | -8.04E-05 | 0.50567086 | 0.99999994 |
| Pcdh11x | 0.00160848 | 0.00137435 | 0.00145485 | -8.05E-05 | 0.6436851 | 0.99999994 |
| Gm11464 | 0.00012288 | 3.07E-05 | 0.00011125 | -8.05E-05 | 0.99999928 | 0.99999994 |
| Gbp4 | 0.00012288 | 3.07E-05 | 0.00011125 | -8.05E-05 | 0.99999928 | 0.99999994 |
| Rab7b | 0.00072384 | 0.00057446 | 0.0006551 | -8.06E-05 | 0.79369837 | 0.99999994 |
| Siglec1 | 0.00134322 | 0.00113437 | 0.00121515 | -8.08E-05 | 0.67093355 | 0.99999994 |
| Iqub | 0.00032126 | 0.00021003 | 0.00029083 | -8.08E-05 | 0.96779643 | 0.99999994 |
| Gm15518 | 0.00012357 | 3.11E-05 | 0.00011188 | -8.08E-05 | 0.99999924 | 0.99999994 |
| Gm16638 | 0.01823693 | 0.0162299 | 0.01631075 | -8.09E-05 | 0.51316076 | 0.99999994 |
| Asic3 | 0.00197928 | 0.00170881 | 0.00178978 | -8.10E-05 | 0.61834175 | 0.99999994 |
| Fgg | 0.00120533 | 0.00100948 | 0.00109051 | -8.10E-05 | 0.68954301 | 0.99999994 |
| 2210406O10 | 0.00805414 | 0.00717202 | 0.0072531 | -8.11E-05 | 0.52965757 | 0.99999994 |
| Gm28731 | 0.00105715 | 0.00087542 | 0.00095653 | -8.11E-05 | 0.71377963 | 0.99999994 |
| Gm19461 | 0.00295972 | 0.00259339 | 0.00267457 | -8.12E-05 | 0.58005857 | 0.99999994 |
| Lrrc75b | 0.00032022 | 0.00020867 | 0.00028989 | -8.12E-05 | 0.96890451 | 0.99999994 |
| Fcnb | 0.00012529 | 3.19E-05 | 0.00011344 | -8.15E-05 | 0.99999913 | 0.99999994 |
| Rab6b | 0.00012529 | 3.19E-05 | 0.00011344 | -8.15E-05 | 0.99999913 | 0.99999994 |
| Gm14097 | 0.00012529 | 3.19E-05 | 0.00011344 | -8.15E-05 | 0.99999913 | 0.99999994 |

| | | | | | | |
|---|---|---|---|---|---|---|
| Gm29994 | 0.00031916 | 0.0002073 | 0.00028893 | -8.16E-05 | 0.96999564 | 0.99999994 |
| Gas2l3 | 0.00210897 | 0.00182521 | 0.00190689 | -8.17E-05 | 0.61222436 | 0.99999994 |
| Gm43597 | 0.0025359 | 0.00221041 | 0.00229225 | -8.18E-05 | 0.59391722 | 0.99999994 |
| Klrd1 | 0.0001263 | 3.25E-05 | 0.00011435 | -8.19E-05 | 0.99999906 | 0.99999994 |
| Ribc2 | 0.0001263 | 3.25E-05 | 0.00011435 | -8.19E-05 | 0.99999906 | 0.99999994 |
| Aldh3b2 | 0.0001263 | 3.25E-05 | 0.00011435 | -8.19E-05 | 0.99999906 | 0.99999994 |
| Gm16549 | 0.0001263 | 3.25E-05 | 0.00011435 | -8.19E-05 | 0.99999906 | 0.99999994 |
| 4930500F10I | 0.00250468 | 0.00218208 | 0.00226408 | -8.20E-05 | 0.5952544 | 0.99999994 |
| Slc10a4 | 0.00012681 | 3.27E-05 | 0.00011481 | -8.21E-05 | 0.99999903 | 0.99999994 |
| Lcn4 | 0.00012701 | 3.28E-05 | 0.00011499 | -8.22E-05 | 0.99999901 | 0.99999994 |
| Prss22 | 0.00031775 | 0.00020547 | 0.00028766 | -8.22E-05 | 0.97140387 | 0.99999994 |
| Gm16116 | 0.00012717 | 3.29E-05 | 0.00011514 | -8.22E-05 | 0.999999 | 0.99999994 |
| Gabrq | 0.00012774 | 3.32E-05 | 0.00011565 | -8.24E-05 | 0.99999896 | 0.99999994 |
| Pou2f3 | 0.00012827 | 3.35E-05 | 0.00011614 | -8.27E-05 | 0.99999892 | 0.99999994 |
| Gm15956 | 0.00012827 | 3.35E-05 | 0.00011614 | -8.27E-05 | 0.99999892 | 0.99999994 |
| Cpa5 | 0.00031629 | 0.00020358 | 0.00028633 | -8.28E-05 | 0.97280401 | 0.99999994 |
| C1rb | 0.00031593 | 0.00020312 | 0.00028601 | -8.29E-05 | 0.97313807 | 0.99999994 |
| Tmprss15 | 0.00031593 | 0.00020312 | 0.00028601 | -8.29E-05 | 0.97313807 | 0.99999994 |
| Mas1 | 0.00267498 | 0.00233481 | 0.00241774 | -8.29E-05 | 0.59029457 | 0.99999994 |
| Zfp960 | 0.00305445 | 0.00267683 | 0.00276 | -8.32E-05 | 0.57948628 | 0.99999994 |
| Tmem132e | 0.00031501 | 0.00020194 | 0.00028518 | -8.32E-05 | 0.97398332 | 0.99999994 |
| Vwf | 0.00224958 | 0.00195057 | 0.00203383 | -8.33E-05 | 0.60738853 | 0.99999994 |
| Hspg2 | 0.01572318 | 0.01400297 | 0.0140863 | -8.33E-05 | 0.51570539 | 0.99999994 |
| Gpr150 | 0.00013019 | 3.45E-05 | 0.00011787 | -8.34E-05 | 0.99999875 | 0.99999994 |
| Tnfrsf14 | 0.00106297 | 0.0008783 | 0.0009618 | -8.35E-05 | 0.71833261 | 0.99999994 |
| Apon | 0.00013052 | 3.47E-05 | 0.00011817 | -8.35E-05 | 0.99999872 | 0.99999994 |
| Gm42957 | 0.00013069 | 3.48E-05 | 0.00011832 | -8.36E-05 | 0.99999871 | 0.99999994 |
| Lrrk2 | 0.0001307 | 3.48E-05 | 0.00011834 | -8.36E-05 | 0.9999987 | 0.99999994 |
| Asic5 | 0.0001307 | 3.48E-05 | 0.00011834 | -8.36E-05 | 0.9999987 | 0.99999994 |
| Cyp4a12b | 0.0001307 | 3.48E-05 | 0.00011834 | -8.36E-05 | 0.9999987 | 0.99999994 |

| | | | | | | |
|---|---|---|---|---|---|---|
| Cxcl5 | 0.00066374 | 0.00051712 | 0.00060073 | -8.36E-05 | 0.82288954 | 0.99999994 |
| Cd300ld | 0.00031399 | 0.00020063 | 0.00028425 | -8.36E-05 | 0.97489258 | 0.99999994 |
| Ntrk3 | 0.00013087 | 3.49E-05 | 0.00011849 | -8.36E-05 | 0.99999869 | 0.99999994 |
| Zfp72 | 0.00062856 | 0.00048521 | 0.0005689 | -8.37E-05 | 0.83625776 | 0.99999994 |
| Kcnk2 | 0.00305555 | 0.00267711 | 0.00276099 | -8.39E-05 | 0.58013164 | 0.99999994 |
| Ppp1r32 | 0.00061524 | 0.00047291 | 0.00055685 | -8.39E-05 | 0.8421857 | 0.99999994 |
| 4921507G05 | 0.00179042 | 0.00153522 | 0.00161921 | -8.40E-05 | 0.63506031 | 0.99999994 |
| Cpne1 | 0.00031242 | 0.00019863 | 0.00028283 | -8.42E-05 | 0.97623882 | 0.99999994 |
| Pdgfrb | 0.00220309 | 0.00190756 | 0.00199186 | -8.43E-05 | 0.61092287 | 0.99999994 |
| Slc30a10 | 0.00013293 | 3.60E-05 | 0.00012035 | -8.44E-05 | 0.99999848 | 0.99999994 |
| Sgca | 0.00013309 | 3.60E-05 | 0.0001205 | -8.45E-05 | 0.99999846 | 0.99999994 |
| Gm16152 | 0.00245556 | 0.00213526 | 0.00221975 | -8.45E-05 | 0.60000949 | 0.99999994 |
| Ptger2 | 0.00031146 | 0.00019741 | 0.00028196 | -8.46E-05 | 0.977034 | 0.99999994 |
| Spns3 | 0.00031146 | 0.00019741 | 0.00028196 | -8.46E-05 | 0.977034 | 0.99999994 |
| Cldn19 | 0.00031142 | 0.00019736 | 0.00028192 | -8.46E-05 | 0.97706723 | 0.99999994 |
| Trem1 | 0.00013354 | 3.63E-05 | 0.0001209 | -8.46E-05 | 0.99999841 | 0.99999994 |
| Pcdhgb1 | 0.00013354 | 3.63E-05 | 0.0001209 | -8.46E-05 | 0.99999841 | 0.99999994 |
| Ptafr | 0.00013392 | 3.65E-05 | 0.00012125 | -8.48E-05 | 0.99999836 | 0.99999994 |
| Slc25a2 | 0.00013392 | 3.65E-05 | 0.00012125 | -8.48E-05 | 0.99999836 | 0.99999994 |
| Kdr | 0.00108767 | 0.00089927 | 0.00098413 | -8.49E-05 | 0.71699368 | 0.99999994 |
| Gper1 | 0.01260281 | 0.01122959 | 0.01131455 | -8.50E-05 | 0.51993108 | 0.99999994 |
| Xcl1 | 0.00013465 | 3.69E-05 | 0.00012191 | -8.50E-05 | 0.99999827 | 0.99999994 |
| Scara5 | 0.00031008 | 0.00019566 | 0.00028071 | -8.50E-05 | 0.97813629 | 0.99999994 |
| Gm16519 | 0.00031008 | 0.00019566 | 0.00028071 | -8.50E-05 | 0.97813629 | 0.99999994 |
| Cd3d | 0.00013494 | 3.71E-05 | 0.00012218 | -8.51E-05 | 0.99999824 | 0.99999994 |
| Ston1 | 0.0010327 | 0.0008493 | 0.00093443 | -8.51E-05 | 0.72787751 | 0.99999994 |
| Psors1c2 | 0.00013581 | 3.75E-05 | 0.00012296 | -8.54E-05 | 0.99999812 | 0.99999994 |
| Lrrc2 | 0.00101913 | 0.0008367 | 0.00092216 | -8.55E-05 | 0.73133485 | 0.99999994 |
| Mx1 | 0.00030869 | 0.00019391 | 0.00027945 | -8.55E-05 | 0.97919914 | 0.99999994 |
| 9330175M2C | 0.00013633 | 3.78E-05 | 0.00012343 | -8.56E-05 | 0.99999805 | 0.99999994 |

| | | | | | | |
|---|---|---|---|---|---|---|
| Emc2 | 0.42730875 | 0.28259262 | 0.28267834 | -8.57E-05 | 0.50080525 | 0.99999994 |
| Dram1 | 0.00070247 | 0.00055001 | 0.00063577 | -8.58E-05 | 0.81538207 | 0.99999994 |
| Nmrk1 | 0.04451395 | 0.03903569 | 0.03912153 | -8.58E-05 | 0.50582617 | 0.99999994 |
| Yipf7 | 0.00013706 | 3.82E-05 | 0.0001241 | -8.59E-05 | 0.99999794 | 0.99999994 |
| Rgs13 | 0.00013788 | 3.87E-05 | 0.00012483 | -8.61E-05 | 0.99999782 | 0.99999994 |
| Flt1 | 0.00013797 | 3.87E-05 | 0.00012492 | -8.62E-05 | 0.99999781 | 0.99999994 |
| Myo1a | 0.00013797 | 3.87E-05 | 0.00012492 | -8.62E-05 | 0.99999781 | 0.99999994 |
| Col5a1 | 0.00571239 | 0.00506616 | 0.00515242 | -8.63E-05 | 0.54436302 | 0.99999994 |
| Gm43400 | 0.00013855 | 3.91E-05 | 0.00012544 | -8.64E-05 | 0.99999771 | 0.99999994 |
| Dppa2 | 0.00013855 | 3.91E-05 | 0.00012544 | -8.64E-05 | 0.99999771 | 0.99999994 |
| Slc25a21 | 0.0001387 | 3.91E-05 | 0.00012557 | -8.64E-05 | 0.99999769 | 0.99999994 |
| Plxnb3 | 0.00030611 | 0.00019068 | 0.00027712 | -8.64E-05 | 0.98105904 | 0.99999994 |
| Spta1 | 0.00013879 | 3.92E-05 | 0.00012566 | -8.65E-05 | 0.99999767 | 0.99999994 |
| Srcin1 | 0.00397054 | 0.00349909 | 0.00358555 | -8.65E-05 | 0.56376217 | 0.99999994 |
| 4930447C04I | 0.00030581 | 0.00019032 | 0.00027685 | -8.65E-05 | 0.98126229 | 0.99999994 |
| Lrrc26 | 0.00030573 | 0.00019022 | 0.00027678 | -8.66E-05 | 0.9813161 | 0.99999994 |
| Ttll9 | 0.00094376 | 0.00076742 | 0.00085401 | -8.66E-05 | 0.7501356 | 0.99999994 |
| Dnah6 | 0.00014012 | 4.00E-05 | 0.00012686 | -8.69E-05 | 0.99999744 | 0.99999994 |
| D430041D05 | 0.00070078 | 0.0005472 | 0.00063424 | -8.70E-05 | 0.81950918 | 0.99999994 |
| Tnfrsf26 | 0.00030425 | 0.00018837 | 0.00027543 | -8.71E-05 | 0.98230874 | 0.99999994 |
| Gapdhs | 0.00014108 | 4.05E-05 | 0.00012773 | -8.72E-05 | 0.99999726 | 0.99999994 |
| Ctxn3 | 0.00014108 | 4.05E-05 | 0.00012773 | -8.72E-05 | 0.99999726 | 0.99999994 |
| Casq1 | 0.00014135 | 4.07E-05 | 0.00012798 | -8.73E-05 | 0.99999721 | 0.99999994 |
| Tomt | 0.00014135 | 4.07E-05 | 0.00012798 | -8.73E-05 | 0.99999721 | 0.99999994 |
| D5Ertd615e | 0.00014135 | 4.07E-05 | 0.00012798 | -8.73E-05 | 0.99999721 | 0.99999994 |
| Shcbp1l | 0.00014139 | 4.07E-05 | 0.00012801 | -8.73E-05 | 0.9999972 | 0.99999994 |
| Gm37988 | 0.00014144 | 4.07E-05 | 0.00012806 | -8.73E-05 | 0.99999719 | 0.99999994 |
| Gm10640 | 0.00014144 | 4.07E-05 | 0.00012806 | -8.73E-05 | 0.99999719 | 0.99999994 |
| Gm28777 | 0.00014144 | 4.07E-05 | 0.00012806 | -8.73E-05 | 0.99999719 | 0.99999994 |
| Csmd1 | 0.00137173 | 0.00115355 | 0.00124092 | -8.74E-05 | 0.68034228 | 0.99999994 |

| | | | | | | |
|---|---|---|---|---|---|---|
| Cd300lf | 0.00066813 | 0.00051723 | 0.0006047 | -8.75E-05 | 0.83220068 | 0.99999994 |
| Clgn | 0.00030293 | 0.00018675 | 0.00027424 | -8.75E-05 | 0.98315053 | 0.99999994 |
| Gm20478 | 0.00058251 | 0.00043974 | 0.00052724 | -8.75E-05 | 0.86534633 | 0.99999994 |
| Ear2 | 0.00014289 | 4.15E-05 | 0.00012937 | -8.78E-05 | 0.99999689 | 0.99999994 |
| Phf11d | 0.00014301 | 4.16E-05 | 0.00012948 | -8.79E-05 | 0.99999686 | 0.99999994 |
| Bloodlinc | 0.00014301 | 4.16E-05 | 0.00012948 | -8.79E-05 | 0.99999686 | 0.99999994 |
| Trpa1 | 0.00099328 | 0.00081086 | 0.00089879 | -8.79E-05 | 0.74253282 | 0.99999994 |
| Nxpe2 | 0.00014325 | 4.18E-05 | 0.0001297 | -8.79E-05 | 0.99999681 | 0.99999994 |
| Wdr91 | 0.03859854 | 0.03396773 | 0.03405575 | -8.80E-05 | 0.50686267 | 0.99999994 |
| Osm | 0.00210961 | 0.00181942 | 0.00190747 | -8.81E-05 | 0.62067978 | 0.99999994 |
| Grpr | 0.0001437 | 4.20E-05 | 0.00013011 | -8.81E-05 | 0.9999967 | 0.99999994 |
| Gm44214 | 0.0001437 | 4.20E-05 | 0.00013011 | -8.81E-05 | 0.9999967 | 0.99999994 |
| Gm17087 | 0.0001437 | 4.20E-05 | 0.00013011 | -8.81E-05 | 0.9999967 | 0.99999994 |
| Mos | 0.00014417 | 4.23E-05 | 0.00013053 | -8.82E-05 | 0.99999659 | 0.99999994 |
| Mrgprb2 | 0.00014465 | 4.26E-05 | 0.00013097 | -8.84E-05 | 0.99999648 | 0.99999994 |
| Dhh | 0.00014476 | 4.26E-05 | 0.00013107 | -8.84E-05 | 0.99999645 | 0.99999994 |
| Gm43598 | 0.00014485 | 4.27E-05 | 0.00013114 | -8.84E-05 | 0.99999643 | 0.99999994 |
| G430049J08I | 0.00014485 | 4.27E-05 | 0.00013114 | -8.84E-05 | 0.99999643 | 0.99999994 |
| Ppm1e | 0.0005872 | 0.00044301 | 0.00053149 | -8.85E-05 | 0.86608283 | 0.99999994 |
| Arl9 | 0.00014494 | 4.28E-05 | 0.00013123 | -8.85E-05 | 0.9999964 | 0.99999994 |
| Ggt7 | 0.00599347 | 0.00531642 | 0.00540491 | -8.85E-05 | 0.54339128 | 0.99999994 |
| F420014N23 | 0.00014649 | 4.37E-05 | 0.00013263 | -8.90E-05 | 0.99999599 | 0.99999994 |
| Gm20537 | 0.00029827 | 0.00018104 | 0.00027002 | -8.90E-05 | 0.98586383 | 0.99999994 |
| Cpa4 | 0.0001466 | 4.37E-05 | 0.00013273 | -8.90E-05 | 0.99999596 | 0.99999994 |
| Cd300e | 0.0001466 | 4.37E-05 | 0.00013273 | -8.90E-05 | 0.99999596 | 0.99999994 |
| Clec9a | 0.00014678 | 4.38E-05 | 0.00013289 | -8.90E-05 | 0.99999591 | 0.99999994 |
| Gm8281 | 0.00029791 | 0.00018061 | 0.0002697 | -8.91E-05 | 0.9860535 | 0.99999994 |
| Calr3 | 0.00567721 | 0.00503154 | 0.00512081 | -8.93E-05 | 0.5461872 | 0.99999994 |
| Prdm1 | 0.0001477 | 4.44E-05 | 0.00013373 | -8.93E-05 | 0.99999564 | 0.99999994 |
| Tmem204 | 0.0001477 | 4.44E-05 | 0.00013373 | -8.93E-05 | 0.99999564 | 0.99999994 |

| | | | | | | |
|---|---|---|---|---|---|---|
| Agr3 | 0.00014802 | 4.46E-05 | 0.00013402 | -8.94E-05 | 0.99999554 | 0.99999994 |
| S1pr1 | 0.00014816 | 4.47E-05 | 0.00013415 | -8.95E-05 | 0.99999549 | 0.99999994 |
| Fam78a | 0.00029648 | 0.00017888 | 0.00026841 | -8.95E-05 | 0.98679537 | 0.99999994 |
| Rag2 | 0.00029648 | 0.00017888 | 0.00026841 | -8.95E-05 | 0.98679537 | 0.99999994 |
| Gm15063 | 0.00029648 | 0.00017888 | 0.00026841 | -8.95E-05 | 0.98679537 | 0.99999994 |
| Il12a | 0.00029648 | 0.00017888 | 0.00026841 | -8.95E-05 | 0.98679537 | 0.99999994 |
| Adgrl4 | 0.00029648 | 0.00017888 | 0.00026841 | -8.95E-05 | 0.98679537 | 0.99999994 |
| Pax5 | 0.00029648 | 0.00017888 | 0.00026841 | -8.95E-05 | 0.98679537 | 0.99999994 |
| Hoxa9 | 0.00029648 | 0.00017888 | 0.00026841 | -8.95E-05 | 0.98679537 | 0.99999994 |
| Siglecg | 0.00029648 | 0.00017888 | 0.00026841 | -8.95E-05 | 0.98679537 | 0.99999994 |
| Chst3 | 0.00029648 | 0.00017888 | 0.00026841 | -8.95E-05 | 0.98679537 | 0.99999994 |
| Pou2af1 | 0.00029648 | 0.00017888 | 0.00026841 | -8.95E-05 | 0.98679537 | 0.99999994 |
| Arpp21 | 0.00029648 | 0.00017888 | 0.00026841 | -8.95E-05 | 0.98679537 | 0.99999994 |
| Gm4035 | 0.00029648 | 0.00017888 | 0.00026841 | -8.95E-05 | 0.98679537 | 0.99999994 |
| Erg | 0.00029648 | 0.00017888 | 0.00026841 | -8.95E-05 | 0.98679537 | 0.99999994 |
| Gm8369 | 0.00029648 | 0.00017888 | 0.00026841 | -8.95E-05 | 0.98679537 | 0.99999994 |
| 4930597A21I | 0.00029648 | 0.00017888 | 0.00026841 | -8.95E-05 | 0.98679537 | 0.99999994 |
| Gm11127 | 0.00063744 | 0.00048741 | 0.00057694 | -8.95E-05 | 0.84917884 | 0.99999994 |
| Gm11630 | 0.00393501 | 0.00346398 | 0.00355355 | -8.96E-05 | 0.56661982 | 0.99999994 |
| Phex | 0.00411272 | 0.00362399 | 0.00371359 | -8.96E-05 | 0.56379218 | 0.99999994 |
| Tdrd9 | 0.00029613 | 0.00017846 | 0.00026809 | -8.96E-05 | 0.98697201 | 0.99999994 |
| Slc1a4 | 0.00590319 | 0.00523404 | 0.00532383 | -8.98E-05 | 0.54469168 | 0.99999994 |
| Pou3f2 | 0.00029557 | 0.00017778 | 0.00026757 | -8.98E-05 | 0.98725493 | 0.99999994 |
| Ushbp1 | 0.00014932 | 4.54E-05 | 0.00013519 | -8.98E-05 | 0.99999512 | 0.99999994 |
| Cngb3 | 0.00014947 | 4.55E-05 | 0.00013533 | -8.99E-05 | 0.99999507 | 0.99999994 |
| Efhd1 | 0.00014961 | 4.56E-05 | 0.00013546 | -8.99E-05 | 0.99999502 | 0.99999994 |
| Bpifb4 | 0.00029454 | 0.00017655 | 0.00026665 | -9.01E-05 | 0.98774984 | 0.99999994 |
| Rec114 | 0.00613687 | 0.00544356 | 0.0055337 | -9.01E-05 | 0.54316832 | 0.99999994 |
| 2010110K18I | 0.0002944 | 0.00017637 | 0.00026652 | -9.01E-05 | 0.98781889 | 0.99999994 |
| Cyp26b1 | 0.00015055 | 4.61E-05 | 0.0001363 | -9.02E-05 | 0.99999469 | 0.99999994 |

| | | | | | | |
|---|---|---|---|---|---|---|
| Kndc1 | 0.0002938 | 0.00017566 | 0.00026598 | -9.03E-05 | 0.98809941 | 0.99999994 |
| M5C1000I18 | 0.00319104 | 0.00279281 | 0.00288315 | -9.03E-05 | 0.58261204 | 0.99999994 |
| Rilp | 0.00220774 | 0.0019055 | 0.00199606 | -9.06E-05 | 0.61866815 | 0.99999994 |
| Raet1d | 0.00094551 | 0.00076488 | 0.00085558 | -9.07E-05 | 0.75981126 | 0.99999994 |
| Omp | 0.00029213 | 0.00017366 | 0.00026446 | -9.08E-05 | 0.98885417 | 0.99999994 |
| Rpl7a-ps3 | 0.00505573 | 0.00447136 | 0.00456216 | -9.08E-05 | 0.55270028 | 0.99999994 |
| Slc24a1 | 0.00029173 | 0.0001732 | 0.00026411 | -9.09E-05 | 0.98902526 | 0.99999994 |
| Fam184b | 0.00015342 | 4.79E-05 | 0.0001389 | -9.10E-05 | 0.99999353 | 0.99999994 |
| Shroom4 | 0.00232405 | 0.00200985 | 0.00210105 | -9.12E-05 | 0.61369076 | 0.99999994 |
| Eps8l1 | 0.00668405 | 0.00593362 | 0.00602486 | -9.12E-05 | 0.54014503 | 0.99999994 |
| Mybpc1 | 0.00029001 | 0.00017116 | 0.00026255 | -9.14E-05 | 0.98974741 | 0.99999994 |
| Gypc | 0.00093891 | 0.00075819 | 0.00084962 | -9.14E-05 | 0.76309466 | 0.99999994 |
| Gm33050 | 0.00028979 | 0.00017089 | 0.00026234 | -9.15E-05 | 0.98984033 | 0.99999994 |
| Gm16998 | 0.00015521 | 4.90E-05 | 0.00014053 | -9.15E-05 | 0.99999269 | 0.99999994 |
| 1110002O04 | 0.00015523 | 4.90E-05 | 0.00014054 | -9.15E-05 | 0.99999268 | 0.99999994 |
| Ankrd1 | 0.00015523 | 4.90E-05 | 0.00014054 | -9.15E-05 | 0.99999268 | 0.99999994 |
| Zfp947 | 0.00809865 | 0.00720142 | 0.00729296 | -9.15E-05 | 0.5332946 | 0.99999994 |
| Cacna1i | 0.00317197 | 0.00277428 | 0.00286597 | -9.17E-05 | 0.58431369 | 0.99999994 |
| Gm11646 | 0.00015645 | 4.98E-05 | 0.00014164 | -9.18E-05 | 0.99999205 | 0.99999994 |
| Pkd1l3 | 0.00015645 | 4.98E-05 | 0.00014164 | -9.18E-05 | 0.99999205 | 0.99999994 |
| Shank3 | 0.00028819 | 0.00016902 | 0.0002609 | -9.19E-05 | 0.9904665 | 0.99999994 |
| Apoa2 | 0.01408187 | 0.01253791 | 0.01262982 | -9.19E-05 | 0.51931713 | 0.99999994 |
| Oas1g | 0.00015677 | 5.00E-05 | 0.00014194 | -9.19E-05 | 0.99999187 | 0.99999994 |
| Xdh | 0.00015677 | 5.00E-05 | 0.00014194 | -9.19E-05 | 0.99999187 | 0.99999994 |
| Extl1 | 0.00120213 | 0.00099558 | 0.00108761 | -9.20E-05 | 0.71336371 | 0.99999994 |
| Tnfrsf1b | 0.0001572 | 5.03E-05 | 0.00014232 | -9.20E-05 | 0.99999163 | 0.99999994 |
| Gm37401 | 0.00059328 | 0.00044488 | 0.00053699 | -9.21E-05 | 0.87321896 | 0.99999994 |
| Gm17455 | 0.00015759 | 5.05E-05 | 0.00014268 | -9.21E-05 | 0.99999141 | 0.99999994 |
| Sec16b | 0.00204979 | 0.00176126 | 0.00185345 | -9.22E-05 | 0.62971198 | 0.99999994 |
| BC016548 | 0.00066313 | 0.00050794 | 0.00060018 | -9.22E-05 | 0.84683546 | 0.99999994 |

| | | | | | | |
|---|---|---|---|---|---|---|
| Pgk1-rs7 | 0.00197776 | 0.00169616 | 0.00178841 | -9.23E-05 | 0.63433629 | 0.99999994 |
| Arhgap10 | 0.00724937 | 0.00643961 | 0.00653193 | -9.23E-05 | 0.53747617 | 0.99999994 |
| Nmu | 0.00076497 | 0.0005999 | 0.0006923 | -9.24E-05 | 0.81283904 | 0.99999994 |
| Bin2 | 0.00182589 | 0.00155876 | 0.00165125 | -9.25E-05 | 0.64535462 | 0.99999994 |
| Prom2 | 0.00411857 | 0.00362636 | 0.00371885 | -9.25E-05 | 0.56574773 | 0.99999994 |
| Gm20652 | 0.00015913 | 5.15E-05 | 0.00014407 | -9.25E-05 | 0.99999047 | 0.99999994 |
| Dnase1l3 | 0.00015926 | 5.16E-05 | 0.00014419 | -9.26E-05 | 0.99999038 | 0.99999994 |
| Apol7a | 0.00028537 | 0.00016572 | 0.00025835 | -9.26E-05 | 0.9914946 | 0.99999994 |
| Zan | 0.00015947 | 5.17E-05 | 0.00014438 | -9.26E-05 | 0.99999025 | 0.99999994 |
| Nr6a1os | 0.00016014 | 5.22E-05 | 0.00014499 | -9.28E-05 | 0.9999898 | 0.99999994 |
| Chn1os3 | 0.00028461 | 0.00016484 | 0.00025765 | -9.28E-05 | 0.99175577 | 0.99999994 |
| Zbed5 | 0.02507852 | 0.02223422 | 0.02232708 | -9.29E-05 | 0.51104271 | 0.99999994 |
| BC018473 | 0.00016066 | 5.25E-05 | 0.00014545 | -9.29E-05 | 0.99998944 | 0.99999994 |
| 2610028H24 | 0.00112467 | 0.00092463 | 0.00101759 | -9.30E-05 | 0.72842002 | 0.99999994 |
| Kcng1 | 0.00016078 | 5.26E-05 | 0.00014557 | -9.30E-05 | 0.99998935 | 0.99999994 |
| Gm6377 | 0.00016103 | 5.28E-05 | 0.00014579 | -9.30E-05 | 0.99998918 | 0.99999994 |
| Gm3086 | 0.00028373 | 0.00016382 | 0.00025686 | -9.30E-05 | 0.99204707 | 0.99999994 |
| Lrrc71 | 0.00016134 | 5.30E-05 | 0.00014608 | -9.31E-05 | 0.99998895 | 0.99999994 |
| Nlrp1b | 0.00016134 | 5.30E-05 | 0.00014608 | -9.31E-05 | 0.99998895 | 0.99999994 |
| Gm27022 | 0.00016243 | 5.37E-05 | 0.00014706 | -9.34E-05 | 0.99998812 | 0.99999994 |
| Gm26562 | 0.00016243 | 5.37E-05 | 0.00014706 | -9.34E-05 | 0.99998812 | 0.99999994 |
| Zfp750 | 0.00028224 | 0.0001621 | 0.00025551 | -9.34E-05 | 0.9925207 | 0.99999994 |
| 2310016G11 | 0.00016313 | 5.42E-05 | 0.0001477 | -9.35E-05 | 0.99998755 | 0.99999994 |
| Zpbp | 0.00280288 | 0.00243953 | 0.00253312 | -9.36E-05 | 0.597125 | 0.99999994 |
| Gm26861 | 0.00028139 | 0.00016113 | 0.00025474 | -9.36E-05 | 0.99277859 | 0.99999994 |
| Mndal | 0.00028129 | 0.00016102 | 0.00025465 | -9.36E-05 | 0.99280831 | 0.99999994 |
| Spata20 | 0.00028129 | 0.00016102 | 0.00025465 | -9.36E-05 | 0.99280831 | 0.99999994 |
| Ccdc38 | 0.00028124 | 0.00016096 | 0.00025461 | -9.36E-05 | 0.99282311 | 0.99999994 |
| 0610033M1C | 0.00016359 | 5.45E-05 | 0.00014811 | -9.36E-05 | 0.99998717 | 0.99999994 |
| Gm12216 | 0.00016359 | 5.45E-05 | 0.00014811 | -9.36E-05 | 0.99998717 | 0.99999994 |

| | | | | | | |
|---|---|---|---|---|---|---|
| 3110045C21I | 0.00535672 | 0.00473909 | 0.00483277 | -9.37E-05 | 0.55133256 | 0.99999994 |
| Abcb1b | 0.000164 | 5.47E-05 | 0.00014848 | -9.37E-05 | 0.99998681 | 0.99999994 |
| Mettl7a2 | 0.000164 | 5.47E-05 | 0.00014848 | -9.37E-05 | 0.99998681 | 0.99999994 |
| Il21r | 0.00016414 | 5.48E-05 | 0.00014861 | -9.38E-05 | 0.99998669 | 0.99999994 |
| BC048644 | 0.00016414 | 5.48E-05 | 0.00014861 | -9.38E-05 | 0.99998669 | 0.99999994 |
| Gm2788 | 0.00016417 | 5.48E-05 | 0.00014864 | -9.38E-05 | 0.99998666 | 0.99999994 |
| Phf11b | 0.00016429 | 5.49E-05 | 0.00014875 | -9.38E-05 | 0.99998655 | 0.99999994 |
| Klk1b24 | 0.00016429 | 5.49E-05 | 0.00014875 | -9.38E-05 | 0.99998655 | 0.99999994 |
| Tsnaxip1 | 0.00016435 | 5.50E-05 | 0.0001488 | -9.38E-05 | 0.9999865 | 0.99999994 |
| Wnt2 | 0.00016462 | 5.51E-05 | 0.00014904 | -9.39E-05 | 0.99998626 | 0.99999994 |
| Gm2245 | 0.00016466 | 5.52E-05 | 0.00014908 | -9.39E-05 | 0.99998622 | 0.99999994 |
| Chl1 | 0.00027968 | 0.00015918 | 0.00025319 | -9.40E-05 | 0.99327577 | 0.99999994 |
| 4933433G15 | 0.00016576 | 5.59E-05 | 0.00015007 | -9.42E-05 | 0.99998519 | 0.99999994 |
| Gm15859 | 0.00016576 | 5.59E-05 | 0.00015007 | -9.42E-05 | 0.99998519 | 0.99999994 |
| A730056A06 | 0.00066692 | 0.00050933 | 0.0006036 | -9.43E-05 | 0.85074989 | 0.99999994 |
| Fat2 | 0.00027846 | 0.00015779 | 0.00025209 | -9.43E-05 | 0.99361174 | 0.99999994 |
| Npw | 0.00016653 | 5.64E-05 | 0.00015077 | -9.43E-05 | 0.99998442 | 0.99999994 |
| Vstm2a | 0.00016664 | 5.65E-05 | 0.00015087 | -9.44E-05 | 0.99998431 | 0.99999994 |
| Sox7 | 0.00027715 | 0.00015631 | 0.00025091 | -9.46E-05 | 0.99395469 | 0.99999994 |
| Slfn1 | 0.00027715 | 0.00015631 | 0.00025091 | -9.46E-05 | 0.99395469 | 0.99999994 |
| 4930565N06 | 0.00057444 | 0.0004253 | 0.00051994 | -9.46E-05 | 0.88715859 | 0.99999994 |
| Hr | 0.00016831 | 5.76E-05 | 0.00015238 | -9.47E-05 | 0.9999825 | 0.99999994 |
| Npffr1 | 0.00016833 | 5.77E-05 | 0.0001524 | -9.47E-05 | 0.99998248 | 0.99999994 |
| Gm27184 | 0.00016835 | 5.77E-05 | 0.00015242 | -9.47E-05 | 0.99998246 | 0.99999994 |
| Tmigd1 | 0.00016879 | 5.80E-05 | 0.00015282 | -9.48E-05 | 0.99998194 | 0.99999994 |
| Gm12530 | 0.00016879 | 5.80E-05 | 0.00015282 | -9.48E-05 | 0.99998194 | 0.99999994 |
| Gm15401 | 0.00027589 | 0.00015489 | 0.00024976 | -9.49E-05 | 0.99427041 | 0.99999994 |
| Krt78 | 0.00027563 | 0.00015461 | 0.00024953 | -9.49E-05 | 0.99433219 | 0.99999994 |
| Grifin | 0.00027551 | 0.00015447 | 0.00024942 | -9.50E-05 | 0.99436282 | 0.99999994 |
| Kcna6 | 0.00016948 | 5.85E-05 | 0.00015345 | -9.50E-05 | 0.99998111 | 0.99999994 |

| | | | | | | |
|---|---|---|---|---|---|---|
| Nid2 | 0.00230951 | 0.00199294 | 0.00208793 | -9.50E-05 | 0.61899365 | 0.99999994 |
| Olfml3 | 0.00561147 | 0.00496673 | 0.00506173 | -9.50E-05 | 0.54971082 | 0.99999994 |
| Myh11 | 0.00017088 | 5.94E-05 | 0.00015471 | -9.53E-05 | 0.99997932 | 0.99999994 |
| Abi3bp | 0.00017088 | 5.94E-05 | 0.00015471 | -9.53E-05 | 0.99997932 | 0.99999994 |
| Daam2 | 0.00027384 | 0.0001526 | 0.00024791 | -9.53E-05 | 0.99475242 | 0.99999994 |
| BC043934 | 0.0010783 | 0.00088033 | 0.00097566 | -9.53E-05 | 0.74227026 | 0.99999994 |
| Ttc39aos1 | 0.00017116 | 5.96E-05 | 0.00015496 | -9.53E-05 | 0.99997894 | 0.99999994 |
| 1700095J03F | 0.00027351 | 0.00015223 | 0.00024761 | -9.54E-05 | 0.99482615 | 0.99999994 |
| Tex15 | 0.00027329 | 0.00015199 | 0.00024741 | -9.54E-05 | 0.99487469 | 0.99999994 |
| Iqcf5 | 0.00027329 | 0.00015199 | 0.00024741 | -9.54E-05 | 0.99487469 | 0.99999994 |
| Serpinb11 | 0.00017188 | 6.01E-05 | 0.00015561 | -9.55E-05 | 0.99997794 | 0.99999994 |
| Has3 | 0.00066078 | 0.00050256 | 0.00059805 | -9.55E-05 | 0.85608795 | 0.99999994 |
| Hspa1l | 0.00071888 | 0.0005551 | 0.00065061 | -9.55E-05 | 0.83577228 | 0.99999994 |
| Cped1 | 0.00137149 | 0.00114517 | 0.0012407 | -9.55E-05 | 0.69584897 | 0.99999994 |
| Slamf1 | 0.00777262 | 0.00690531 | 0.00700091 | -9.56E-05 | 0.53621308 | 0.99999994 |
| Cd72 | 0.00017241 | 6.05E-05 | 0.0001561 | -9.56E-05 | 0.99997717 | 0.99999994 |
| Gm15689 | 0.00017257 | 6.06E-05 | 0.00015624 | -9.56E-05 | 0.99997693 | 0.99999994 |
| Gm43409 | 0.00027192 | 0.00015047 | 0.00024618 | -9.57E-05 | 0.99516891 | 0.99999994 |
| Hs3st5 | 0.00017318 | 6.10E-05 | 0.00015679 | -9.58E-05 | 0.99997603 | 0.99999994 |
| Pilra | 0.00017368 | 6.14E-05 | 0.00015724 | -9.59E-05 | 0.99997524 | 0.99999994 |
| Prap1 | 0.00017368 | 6.14E-05 | 0.00015724 | -9.59E-05 | 0.99997524 | 0.99999994 |
| Ccdc33 | 0.00076653 | 0.00059769 | 0.00069371 | -9.60E-05 | 0.82155901 | 0.99999994 |
| Gm26604 | 0.00017452 | 6.20E-05 | 0.000158 | -9.60E-05 | 0.99997387 | 0.99999994 |
| Got1l1 | 0.00017467 | 6.21E-05 | 0.00015814 | -9.61E-05 | 0.99997361 | 0.99999994 |
| Sp100 | 0.00017472 | 6.21E-05 | 0.00015819 | -9.61E-05 | 0.99997353 | 0.99999994 |
| Psg17 | 0.00017474 | 6.21E-05 | 0.0001582 | -9.61E-05 | 0.99997351 | 0.99999994 |
| Gpr68 | 0.00104137 | 0.00084618 | 0.00094227 | -9.61E-05 | 0.75136506 | 0.99999994 |
| Ankrd34a | 0.00017496 | 6.23E-05 | 0.00015841 | -9.61E-05 | 0.99997312 | 0.99999994 |
| Serpina5 | 0.00017499 | 6.23E-05 | 0.00015843 | -9.61E-05 | 0.99997308 | 0.99999994 |
| Lum | 0.00539795 | 0.00477361 | 0.00486983 | -9.62E-05 | 0.55231861 | 0.99999994 |

| | | | | | | |
|---|---|---|---|---|---|---|
| B4galnt3 | 0.00026925 | 0.00014753 | 0.00024376 | -9.62E-05 | 0.99570187 | 0.99999994 |
| Exd1 | 0.00173155 | 0.0014698 | 0.00156604 | -9.62E-05 | 0.65874547 | 0.99999994 |
| Slc16a4 | 0.00017581 | 6.29E-05 | 0.00015917 | -9.63E-05 | 0.99997163 | 0.99999994 |
| Gm2663 | 0.00017609 | 6.31E-05 | 0.00015943 | -9.63E-05 | 0.99997112 | 0.99999994 |
| Zc3h12d | 0.00017613 | 6.31E-05 | 0.00015946 | -9.63E-05 | 0.99997105 | 0.99999994 |
| Adra1a | 0.00026832 | 0.00014651 | 0.00024291 | -9.64E-05 | 0.99587534 | 0.99999994 |
| Gm28321 | 0.00017716 | 6.39E-05 | 0.0001604 | -9.65E-05 | 0.99996909 | 0.99999994 |
| Scarf2 | 0.00237886 | 0.00205394 | 0.00215053 | -9.66E-05 | 0.61751741 | 0.99999994 |
| Tnfrsf19 | 0.00096797 | 0.00077924 | 0.0008759 | -9.67E-05 | 0.76870032 | 0.99999994 |
| Fignl1 | 0.01508079 | 0.01341995 | 0.01351663 | -9.67E-05 | 0.5189868 | 0.99999994 |
| Gpr132 | 0.00017801 | 6.45E-05 | 0.00016117 | -9.67E-05 | 0.99996738 | 0.99999994 |
| Cd33 | 0.00017898 | 6.52E-05 | 0.00016205 | -9.69E-05 | 0.99996532 | 0.99999994 |
| Fes | 0.00228746 | 0.00197114 | 0.00206802 | -9.69E-05 | 0.62241576 | 0.99999994 |
| Clec4d | 0.00026441 | 0.00014227 | 0.00023937 | -9.71E-05 | 0.99653449 | 0.99999994 |
| Ddx43 | 0.00026418 | 0.00014202 | 0.00023916 | -9.71E-05 | 0.99657064 | 0.99999994 |
| Sec14l3 | 0.00018067 | 6.64E-05 | 0.00016357 | -9.71E-05 | 0.99996146 | 0.99999994 |
| A630001G21 | 0.00018075 | 6.65E-05 | 0.00016365 | -9.72E-05 | 0.99996125 | 0.99999994 |
| Cwh43 | 0.00018075 | 6.65E-05 | 0.00016365 | -9.72E-05 | 0.99996125 | 0.99999994 |
| Cfh | 0.00300633 | 0.00261941 | 0.00271661 | -9.72E-05 | 0.59411836 | 0.99999994 |
| Mocos | 0.00018106 | 6.67E-05 | 0.00016392 | -9.72E-05 | 0.99996051 | 0.99999994 |
| Gm26789 | 0.00076406 | 0.00059425 | 0.00069148 | -9.72E-05 | 0.82535228 | 0.99999994 |
| Orc1 | 0.00231175 | 0.00199263 | 0.00208995 | -9.73E-05 | 0.62169887 | 0.99999994 |
| A330032B11 | 0.00018188 | 6.73E-05 | 0.00016467 | -9.74E-05 | 0.99995842 | 0.99999994 |
| Muc16 | 0.00090097 | 0.00071791 | 0.00081531 | -9.74E-05 | 0.78675208 | 0.99999994 |
| Gm16212 | 0.00018255 | 6.78E-05 | 0.00016527 | -9.75E-05 | 0.99995667 | 0.99999994 |
| BC024139 | 0.00167705 | 0.00141933 | 0.0015168 | -9.75E-05 | 0.66558855 | 0.99999994 |
| Htr1b | 0.00018267 | 6.79E-05 | 0.00016538 | -9.75E-05 | 0.99995633 | 0.99999994 |
| Col24a1 | 0.00026207 | 0.00013977 | 0.00023726 | -9.75E-05 | 0.99688243 | 0.99999994 |
| Ggt6 | 0.00026207 | 0.00013977 | 0.00023726 | -9.75E-05 | 0.99688243 | 0.99999994 |
| Dnmbp | 0.00026207 | 0.00013977 | 0.00023726 | -9.75E-05 | 0.99688243 | 0.99999994 |

| | | | | | | |
|---|---|---|---|---|---|---|
| Ifit3b | 0.00148518 | 0.0012459 | 0.00134344 | -9.75E-05 | 0.68555422 | 0.99999994 |
| Sult6b1 | 0.00018355 | 6.86E-05 | 0.00016618 | -9.76E-05 | 0.99995387 | 0.99999994 |
| Fpr1 | 0.00026116 | 0.00013879 | 0.00023643 | -9.76E-05 | 0.9970094 | 0.99999994 |
| Exoc3l4 | 0.00892893 | 0.00793844 | 0.00803613 | -9.77E-05 | 0.53224773 | 0.99999994 |
| Gm14862 | 0.00018423 | 6.91E-05 | 0.00016679 | -9.77E-05 | 0.9999519 | 0.99999994 |
| Inhbb | 0.00463397 | 0.00408493 | 0.00418277 | -9.78E-05 | 0.56186743 | 0.99999994 |
| Barhl1 | 0.00139364 | 0.00116286 | 0.00126072 | -9.79E-05 | 0.69731215 | 0.99999994 |
| Atp6v1c2 | 0.00025969 | 0.00013724 | 0.0002351 | -9.79E-05 | 0.99720404 | 0.99999994 |
| Gm28182 | 0.00018552 | 7.00E-05 | 0.00016797 | -9.79E-05 | 0.9999479 | 0.99999994 |
| Kcnq5 | 0.00018556 | 7.01E-05 | 0.000168 | -9.79E-05 | 0.99994779 | 0.99999994 |
| A730011C13 | 0.00018556 | 7.01E-05 | 0.000168 | -9.79E-05 | 0.99994779 | 0.99999994 |
| Cfb | 0.00018556 | 7.01E-05 | 0.000168 | -9.79E-05 | 0.99994779 | 0.99999994 |
| Snai3 | 0.00025903 | 0.00013654 | 0.00023451 | -9.80E-05 | 0.9972875 | 0.99999994 |
| Adamts12 | 0.00025881 | 0.00013631 | 0.0002343 | -9.80E-05 | 0.9973153 | 0.99999994 |
| Ddx60 | 0.00018607 | 7.05E-05 | 0.00016846 | -9.80E-05 | 0.99994612 | 0.99999994 |
| Gm16630 | 0.00018607 | 7.05E-05 | 0.00016846 | -9.80E-05 | 0.99994612 | 0.99999994 |
| Gm26648 | 0.00025867 | 0.00013616 | 0.00023418 | -9.80E-05 | 0.99733252 | 0.99999994 |
| Clec12a | 0.00018667 | 7.09E-05 | 0.000169 | -9.81E-05 | 0.99994412 | 0.99999994 |
| Gm15494 | 0.00018682 | 7.10E-05 | 0.00016914 | -9.81E-05 | 0.99994358 | 0.99999994 |
| Palb2 | 0.0095944 | 0.00853303 | 0.00863117 | -9.81E-05 | 0.53016688 | 0.99999994 |
| Acap1 | 0.0002573 | 0.00013472 | 0.00023294 | -9.82E-05 | 0.99749672 | 0.99999994 |
| Themis | 0.00018822 | 7.21E-05 | 0.00017041 | -9.83E-05 | 0.99993855 | 0.99999994 |
| Gm29562 | 0.00106245 | 0.00086296 | 0.00096133 | -9.84E-05 | 0.75209833 | 0.99999994 |
| Gzma | 0.0001889 | 7.26E-05 | 0.00017102 | -9.84E-05 | 0.99993597 | 0.99999994 |
| Lhx2 | 0.00025565 | 0.000133 | 0.00023145 | -9.84E-05 | 0.99768132 | 0.99999994 |
| Gm3830 | 0.00025553 | 0.00013288 | 0.00023134 | -9.85E-05 | 0.99769456 | 0.99999994 |
| Wnt6 | 0.00025535 | 0.00013269 | 0.00023118 | -9.85E-05 | 0.99771354 | 0.99999994 |
| Agtr2 | 0.00025535 | 0.00013269 | 0.00023118 | -9.85E-05 | 0.99771354 | 0.99999994 |
| Cd28 | 0.00025526 | 0.00013259 | 0.00023109 | -9.85E-05 | 0.99772369 | 0.99999994 |
| Angptl8 | 0.01163302 | 0.0103522 | 0.01045073 | -9.85E-05 | 0.52502007 | 0.99999994 |

| | | | | | | |
|---|---|---|---|---|---|---|
| Cyp4f14 | 0.00019003 | 7.35E-05 | 0.00017204 | -9.86E-05 | 0.99993142 | 0.99999994 |
| Pde11a | 0.00019033 | 7.37E-05 | 0.00017232 | -9.86E-05 | 0.99993014 | 0.99999994 |
| Gdf9 | 0.00019065 | 7.40E-05 | 0.00017261 | -9.86E-05 | 0.99992877 | 0.99999994 |
| Tlr13 | 0.0001908 | 7.41E-05 | 0.00017274 | -9.87E-05 | 0.99992816 | 0.99999994 |
| C130071C03I | 0.0001908 | 7.41E-05 | 0.00017274 | -9.87E-05 | 0.99992816 | 0.99999994 |
| Ccl20 | 0.00076037 | 0.00058944 | 0.00068814 | -9.87E-05 | 0.8301486 | 0.99999994 |
| Clec4n | 0.00019178 | 7.48E-05 | 0.00017363 | -9.88E-05 | 0.99992378 | 0.99999994 |
| Otog | 0.00025297 | 0.00013023 | 0.00022902 | -9.88E-05 | 0.99795591 | 0.99999994 |
| AA474408 | 0.02207558 | 0.0195943 | 0.01969316 | -9.89E-05 | 0.5133277 | 0.99999994 |
| Vipr2 | 0.00171897 | 0.0014558 | 0.00155467 | -9.89E-05 | 0.66397818 | 0.99999994 |
| Xkrx | 0.00019289 | 7.57E-05 | 0.00017463 | -9.89E-05 | 0.99991851 | 0.99999994 |
| Gm29609 | 0.000193 | 7.58E-05 | 0.00017474 | -9.89E-05 | 0.99991794 | 0.99999994 |
| Gm9968 | 0.000193 | 7.58E-05 | 0.00017474 | -9.89E-05 | 0.99991794 | 0.99999994 |
| Mybl1 | 0.00019357 | 7.63E-05 | 0.00017525 | -9.90E-05 | 0.9999151 | 0.99999994 |
| Chp2 | 0.00019357 | 7.63E-05 | 0.00017525 | -9.90E-05 | 0.9999151 | 0.99999994 |
| Ccr6 | 0.00019451 | 7.70E-05 | 0.0001761 | -9.91E-05 | 0.9999102 | 0.99999994 |
| 2310043M15 | 0.00019451 | 7.70E-05 | 0.0001761 | -9.91E-05 | 0.9999102 | 0.99999994 |
| Mlana | 0.00019542 | 7.77E-05 | 0.00017692 | -9.92E-05 | 0.99990525 | 0.99999994 |
| B430306N03 | 0.00024935 | 0.00012653 | 0.00022575 | -9.92E-05 | 0.99828027 | 0.99999994 |
| Tspear | 0.00538202 | 0.00475625 | 0.00485552 | -9.93E-05 | 0.55412646 | 0.99999994 |
| Tnfsf13os | 0.00024849 | 0.00012566 | 0.00022497 | -9.93E-05 | 0.99835032 | 0.99999994 |
| Sp140 | 0.00151465 | 0.00127068 | 0.00137007 | -9.94E-05 | 0.68539882 | 0.99999994 |
| Ugt2b5 | 0.00024676 | 0.00012391 | 0.0002234 | -9.95E-05 | 0.9984832 | 0.99999994 |
| Cst8 | 0.00024676 | 0.00012391 | 0.0002234 | -9.95E-05 | 0.9984832 | 0.99999994 |
| Mstn | 0.00024614 | 0.00012329 | 0.00022284 | -9.95E-05 | 0.99852821 | 0.99999994 |
| 4921511C10I | 0.00024614 | 0.00012329 | 0.00022284 | -9.95E-05 | 0.99852821 | 0.99999994 |
| Gli1 | 0.00019871 | 8.04E-05 | 0.00017991 | -9.96E-05 | 0.99988486 | 0.99999994 |
| Cav3 | 0.00024601 | 0.00012316 | 0.00022272 | -9.96E-05 | 0.99853721 | 0.99999994 |
| Magel2 | 0.00024601 | 0.00012316 | 0.00022272 | -9.96E-05 | 0.99853721 | 0.99999994 |
| Hsd3b7 | 0.00024601 | 0.00012316 | 0.00022272 | -9.96E-05 | 0.99853721 | 0.99999994 |

| | | | | | | |
|---|---|---|---|---|---|---|
| Gm9951 | 0.00024601 | 0.00012316 | 0.00022272 | -9.96E-05 | 0.99853721 | 0.99999994 |
| Krt33b | 0.00024601 | 0.00012316 | 0.00022272 | -9.96E-05 | 0.99853721 | 0.99999994 |
| Gm867 | 0.00019881 | 8.04E-05 | 0.00018 | -9.96E-05 | 0.99988419 | 0.99999994 |
| Gm26513 | 0.00185917 | 0.00158174 | 0.00168131 | -9.96E-05 | 0.65328813 | 0.99999994 |
| Olfml2a | 0.00024464 | 0.00012179 | 0.00022148 | -9.97E-05 | 0.99863242 | 0.99999994 |
| Slc44a5 | 0.00024464 | 0.00012179 | 0.00022148 | -9.97E-05 | 0.99863242 | 0.99999994 |
| Izumo1r | 0.00024464 | 0.00012179 | 0.00022148 | -9.97E-05 | 0.99863242 | 0.99999994 |
| Gm29431 | 0.00024464 | 0.00012179 | 0.00022148 | -9.97E-05 | 0.99863242 | 0.99999994 |
| Runx3 | 0.00024427 | 0.00012142 | 0.00022114 | -9.97E-05 | 0.99865721 | 0.99999994 |
| Cdhr4 | 0.00024357 | 0.00012073 | 0.00022051 | -9.98E-05 | 0.99870218 | 0.99999994 |
| Olig1 | 0.00024352 | 0.00012068 | 0.00022047 | -9.98E-05 | 0.99870533 | 0.99999994 |
| Gm43915 | 0.00024249 | 0.00011966 | 0.00021954 | -9.99E-05 | 0.99876959 | 0.99999994 |
| G630030J09I | 0.00020241 | 8.34E-05 | 0.00018325 | -9.99E-05 | 0.99985717 | 0.99999994 |
| Gm12905 | 0.00024226 | 0.00011944 | 0.00021933 | -9.99E-05 | 0.99878364 | 0.99999994 |
| Rbakdn | 0.00024209 | 0.00011927 | 0.00021917 | -9.99E-05 | 0.99879388 | 0.99999994 |
| Slc6a4 | 0.00024209 | 0.00011927 | 0.00021917 | -9.99E-05 | 0.99879388 | 0.99999994 |
| Dmc1 | 0.00020315 | 8.40E-05 | 0.00018392 | -9.99E-05 | 0.99985094 | 0.99999994 |
| Cyp26a1 | 0.00020315 | 8.40E-05 | 0.00018392 | -9.99E-05 | 0.99985094 | 0.99999994 |
| Gm28836 | 0.00020315 | 8.40E-05 | 0.00018392 | -9.99E-05 | 0.99985094 | 0.99999994 |
| Zmynd10 | 0.00285566 | 0.00248075 | 0.00258073 | -1.00E-04 | 0.60174595 | 0.99999994 |
| Mdga2 | 0.00020399 | 8.47E-05 | 0.00018469 | -0.0001 | 0.99984347 | 0.99999994 |
| Ch25h | 0.0002045 | 8.51E-05 | 0.00018515 | -0.0001 | 0.99983883 | 0.99999994 |
| Adamts20 | 0.00020451 | 8.51E-05 | 0.00018515 | -0.0001 | 0.99983879 | 0.99999994 |
| Cd247 | 0.00020595 | 8.63E-05 | 0.00018646 | -0.0001001 | 0.99982483 | 0.99999994 |
| Rab38 | 0.0011743 | 0.0009623 | 0.00106245 | -0.0001002 | 0.73482862 | 0.99999994 |
| Il10ra | 0.00020658 | 8.68E-05 | 0.00018703 | -0.0001002 | 0.99981848 | 0.99999994 |
| Gabrp | 0.00020712 | 8.73E-05 | 0.00018751 | -0.0001002 | 0.99981281 | 0.99999994 |
| Comp | 0.0002072 | 8.74E-05 | 0.00018759 | -0.0001002 | 0.99981196 | 0.99999994 |
| H2-Q10 | 0.0010656 | 0.00086385 | 0.00096418 | -0.0001003 | 0.75574626 | 0.99999994 |
| A930006L05I | 0.00023554 | 0.0001129 | 0.00021324 | -0.0001003 | 0.99913263 | 0.99999994 |

| | | | | | | |
|---|---|---|---|---|---|---|
| Cybb | 0.0002343 | 0.00011171 | 0.00021212 | -0.0001004 | 0.99918572 | 0.99999994 |
| E330013P04I | 0.0002343 | 0.00011171 | 0.00021212 | -0.0001004 | 0.99918572 | 0.99999994 |
| Gm19345 | 0.00021053 | 9.02E-05 | 0.0001906 | -0.0001004 | 0.99977292 | 0.99999994 |
| Gm14010 | 0.0002111 | 9.07E-05 | 0.00019112 | -0.0001004 | 0.99976554 | 0.99999994 |
| Gm3510 | 0.00023351 | 0.00011096 | 0.0002114 | -0.0001004 | 0.99921824 | 0.99999994 |
| Gm16045 | 0.00021153 | 9.11E-05 | 0.00019151 | -0.0001005 | 0.99975977 | 0.99999994 |
| Fpr2 | 0.00021195 | 9.14E-05 | 0.00019189 | -0.0001005 | 0.9997541 | 0.99999994 |
| Gjb3 | 0.00021277 | 9.21E-05 | 0.00019263 | -0.0001005 | 0.99974257 | 0.99999994 |
| Clhc1 | 0.00021277 | 9.21E-05 | 0.00019263 | -0.0001005 | 0.99974257 | 0.99999994 |
| L3mbtl4 | 0.00021346 | 9.27E-05 | 0.00019325 | -0.0001005 | 0.99973256 | 0.99999994 |
| Rtl1 | 0.00023131 | 0.00010888 | 0.00020941 | -0.0001005 | 0.99930199 | 0.99999994 |
| Sdr9c7 | 0.00023102 | 0.00010861 | 0.00020915 | -0.0001005 | 0.99931234 | 0.99999994 |
| Fendrr | 0.000231 | 0.00010859 | 0.00020913 | -0.0001005 | 0.99931313 | 0.99999994 |
| Tmem30c | 0.00023098 | 0.00010857 | 0.00020911 | -0.0001005 | 0.99931392 | 0.99999994 |
| Sirpb1c | 0.00023092 | 0.00010852 | 0.00020906 | -0.0001005 | 0.99931589 | 0.99999994 |
| Il25 | 0.00021416 | 9.33E-05 | 0.00019388 | -0.0001006 | 0.99972197 | 0.99999994 |
| Gm5129 | 0.00021475 | 9.39E-05 | 0.00019443 | -0.0001006 | 0.99971259 | 0.99999994 |
| Il13 | 0.00021495 | 9.40E-05 | 0.0001946 | -0.0001006 | 0.99970954 | 0.99999994 |
| Hlx | 0.00021516 | 9.42E-05 | 0.00019479 | -0.0001006 | 0.99970614 | 0.99999994 |
| Itga8 | 0.00021516 | 9.42E-05 | 0.00019479 | -0.0001006 | 0.99970614 | 0.99999994 |
| Fibin | 0.00021516 | 9.42E-05 | 0.00019479 | -0.0001006 | 0.99970614 | 0.99999994 |
| AU015836 | 0.00021516 | 9.42E-05 | 0.00019479 | -0.0001006 | 0.99970614 | 0.99999994 |
| Pcdh18 | 0.00021516 | 9.42E-05 | 0.00019479 | -0.0001006 | 0.99970614 | 0.99999994 |
| Frem1 | 0.00021516 | 9.42E-05 | 0.00019479 | -0.0001006 | 0.99970614 | 0.99999994 |
| Kbtbd12 | 0.00021516 | 9.42E-05 | 0.00019479 | -0.0001006 | 0.99970614 | 0.99999994 |
| Slc7a9 | 0.00021516 | 9.42E-05 | 0.00019479 | -0.0001006 | 0.99970614 | 0.99999994 |
| Tbx2 | 0.00021516 | 9.42E-05 | 0.00019479 | -0.0001006 | 0.99970614 | 0.99999994 |
| Hoxb5os | 0.00021516 | 9.42E-05 | 0.00019479 | -0.0001006 | 0.99970614 | 0.99999994 |
| Tfap2a | 0.00021516 | 9.42E-05 | 0.00019479 | -0.0001006 | 0.99970614 | 0.99999994 |
| F830016B08I | 0.00021548 | 9.45E-05 | 0.00019509 | -0.0001006 | 0.9997008 | 0.99999994 |

| | | | | | | |
|---|---|---|---|---|---|---|
| Slc22a1 | 0.00022902 | 0.00010673 | 0.00020734 | -0.0001006 | 0.99938064 | 0.99999994 |
| Gm28523 | 0.00022686 | 0.00010474 | 0.00020539 | -0.0001007 | 0.9994469 | 0.99999994 |
| Gnb5 | 0.00022686 | 0.00010474 | 0.00020539 | -0.0001007 | 0.9994469 | 0.99999994 |
| Pdzk1 | 0.00022628 | 0.0001042 | 0.00020486 | -0.0001007 | 0.99946373 | 0.99999994 |
| Cend1 | 0.00021947 | 9.80E-05 | 0.0001987 | -0.0001007 | 0.99962771 | 0.99999994 |
| Dsg3 | 0.00022493 | 0.00010296 | 0.00020364 | -0.0001007 | 0.99950078 | 0.99999994 |
| Car4 | 0.00022457 | 0.00010263 | 0.00020332 | -0.0001007 | 0.99951014 | 0.99999994 |
| Dancr | 0.00022178 | 0.00010009 | 0.00020079 | -0.0001007 | 0.99957823 | 0.99999994 |
| Kcne4 | 0.00022232 | 0.00010058 | 0.00020128 | -0.0001007 | 0.99956578 | 0.99999994 |
| Gm42846 | 0.00072573 | 0.00055601 | 0.0006568 | -0.0001008 | 0.8464997 | 0.99999994 |
| Gm17025 | 0.00075868 | 0.00058574 | 0.00068661 | -0.0001009 | 0.835946 | 0.99999994 |
| Gpihbp1 | 0.00053507 | 0.00038315 | 0.00048432 | -0.0001012 | 0.91781141 | 0.99999994 |
| 4933430I17R | 0.00243649 | 0.00210122 | 0.00220254 | -0.0001013 | 0.62027116 | 0.99999994 |
| Cacna2d2 | 0.00233575 | 0.00201013 | 0.00211162 | -0.0001015 | 0.62547634 | 0.99999994 |
| Gm28875 | 0.00064177 | 0.000479 | 0.00058086 | -0.0001019 | 0.87845435 | 0.99999994 |
| Clcnkb | 0.00208558 | 0.00178375 | 0.00188577 | -0.000102 | 0.64061032 | 0.99999994 |
| Gm10655 | 0.00190222 | 0.00161788 | 0.00172019 | -0.0001023 | 0.65390802 | 0.99999994 |
| Sptb | 0.00083279 | 0.00065094 | 0.00075365 | -0.0001027 | 0.81783159 | 0.99999994 |
| Uqcc3 | 0.6533425 | 0.36903837 | 0.36914117 | -0.0001028 | 0.5007395 | 0.99999994 |
| Bco2 | 0.00076743 | 0.00059113 | 0.00069453 | -0.0001034 | 0.83915144 | 0.99999994 |
| Pcdhb18 | 0.00495328 | 0.00436648 | 0.00447002 | -0.0001035 | 0.56126837 | 0.99999994 |
| Gm20692 | 0.00401806 | 0.00352475 | 0.00362835 | -0.0001036 | 0.575365 | 0.99999994 |
| Rmdn2 | 0.01359467 | 0.01209312 | 0.01219686 | -0.0001037 | 0.52257307 | 0.99999994 |
| 2310001K24I | 0.00131992 | 0.00109023 | 0.00119409 | -0.0001039 | 0.71868513 | 0.99999994 |
| Reg3d | 0.00054076 | 0.00038516 | 0.00048947 | -0.0001043 | 0.92198654 | 0.99999994 |
| Foxf1 | 0.00098669 | 0.00078829 | 0.00089282 | -0.0001045 | 0.78211236 | 0.99999994 |
| Ip6k3 | 0.00113 | 0.00091779 | 0.0010224 | -0.0001046 | 0.75208443 | 0.99999994 |
| Khnyn | 0.03328045 | 0.02936268 | 0.02946746 | -0.0001048 | 0.50944077 | 0.99999994 |
| Npas4 | 0.00977831 | 0.00868971 | 0.00879552 | -0.0001058 | 0.53191424 | 0.99999994 |
| Morc4 | 0.02203521 | 0.01955179 | 0.01965767 | -0.0001059 | 0.51430007 | 0.99999994 |

| | | | | | | |
|---|---|---|---|---|---|---|
| Gm13307 | 0.0046451 | 0.00408676 | 0.00419278 | -0.000106 | 0.56683543 | 0.99999994 |
| Nccrp1 | 0.0113042 | 0.01005092 | 0.01015758 | -0.0001067 | 0.52786278 | 0.99999994 |
| Cxcr4 | 0.00300717 | 0.00261059 | 0.00271736 | -0.0001068 | 0.60316453 | 0.99999994 |
| Itgb2 | 0.00061688 | 0.00045143 | 0.00055834 | -0.0001069 | 0.89877334 | 0.99999994 |
| Trim10 | 0.00056277 | 0.00040175 | 0.00050938 | -0.0001076 | 0.92019986 | 0.99999994 |
| C1qtnf7 | 0.00071298 | 0.00053758 | 0.00064527 | -0.0001077 | 0.8667073 | 0.99999994 |
| F2rl3 | 0.00074337 | 0.00056503 | 0.00067276 | -0.0001077 | 0.85676893 | 0.99999994 |
| Gm29502 | 0.00057575 | 0.00041288 | 0.00052113 | -0.0001082 | 0.91661742 | 0.99999994 |
| Abcd2 | 0.00251981 | 0.00216931 | 0.00227772 | -0.0001084 | 0.62430965 | 0.99999994 |
| Igtp | 0.01317715 | 0.01171695 | 0.01182559 | -0.0001086 | 0.52438224 | 0.99999994 |
| Kcnn4 | 0.00288152 | 0.00249524 | 0.00260405 | -0.0001088 | 0.60954542 | 0.99999994 |
| Dusp2 | 0.0026736 | 0.0023073 | 0.00241649 | -0.0001092 | 0.61819782 | 0.99999994 |
| Dars2 | 0.02522713 | 0.02234788 | 0.02245715 | -0.0001093 | 0.51291876 | 0.99999994 |
| Col25a1 | 0.00268276 | 0.0023148 | 0.00242476 | -0.00011 | 0.61861187 | 0.99999994 |
| Cep250 | 0.02030618 | 0.01802541 | 0.01813622 | -0.0001108 | 0.51622085 | 0.99999994 |
| Epsti1 | 0.00169254 | 0.00141989 | 0.0015308 | -0.0001109 | 0.68517956 | 0.99999994 |
| Gm27003 | 0.01563245 | 0.01389488 | 0.01400587 | -0.000111 | 0.5210331 | 0.99999994 |
| Vmn2r29 | 0.00074277 | 0.00056109 | 0.00067222 | -0.0001111 | 0.86442702 | 0.99999994 |
| Lama2 | 0.000518 | 0.00035759 | 0.00046888 | -0.0001113 | 0.9429332 | 0.99999994 |
| 2610206C17I | 0.00097363 | 0.00076962 | 0.00088102 | -0.0001114 | 0.80000959 | 0.99999994 |
| Ppp1r18os | 0.0013265 | 0.00108842 | 0.00120003 | -0.0001116 | 0.73206884 | 0.99999994 |
| Fam229a | 0.00161356 | 0.00134764 | 0.00145944 | -0.0001118 | 0.69494325 | 0.99999994 |
| Ltbr | 0.06850019 | 0.05915117 | 0.05926331 | -0.0001121 | 0.50502489 | 0.99999994 |
| Hvcn1 | 0.00073592 | 0.00055384 | 0.00066603 | -0.0001122 | 0.8689131 | 0.99999994 |
| Rin1 | 0.00092281 | 0.00072257 | 0.00083506 | -0.0001125 | 0.81505136 | 0.99999994 |
| Calml3 | 0.00125767 | 0.00102531 | 0.00113782 | -0.0001125 | 0.74478414 | 0.99999994 |
| Hilpda | 0.13160883 | 0.10868892 | 0.10880169 | -0.0001128 | 0.50275231 | 0.99999994 |
| Lrrtm4 | 0.00203413 | 0.0017265 | 0.00183932 | -0.0001128 | 0.6584607 | 0.99999994 |
| Gm29125 | 0.00134529 | 0.00110406 | 0.00121702 | -0.000113 | 0.73164941 | 0.99999994 |
| Sorbs3 | 0.00165873 | 0.00138696 | 0.00150025 | -0.0001133 | 0.69240477 | 0.99999994 |

| | | | | | | |
|---|---|---|---|---|---|---|
| Frs3 | 0.03927141 | 0.03452052 | 0.03463397 | -0.0001135 | 0.50869814 | 0.99999994 |
| Cks1brt | 0.01395329 | 0.01240203 | 0.01251557 | -0.0001135 | 0.5240775 | 0.99999994 |
| Gm128 | 0.00060719 | 0.00043595 | 0.00054957 | -0.0001136 | 0.91561689 | 0.99999994 |
| Tmem72 | 0.00155305 | 0.00129108 | 0.00140477 | -0.0001137 | 0.70495373 | 0.99999994 |
| Gm16188 | 0.00073562 | 0.00055201 | 0.00066575 | -0.0001137 | 0.87228148 | 0.99999994 |
| 9230111E07l | 0.00054398 | 0.00037857 | 0.00049238 | -0.0001138 | 0.93803857 | 0.99999994 |
| Kel | 0.00073367 | 0.0005501 | 0.00066399 | -0.0001139 | 0.87320478 | 0.99999994 |
| Fam180a | 0.00084769 | 0.00065306 | 0.00076712 | -0.0001141 | 0.83885107 | 0.99999994 |
| Fabp7 | 0.00596141 | 0.00526189 | 0.00537612 | -0.0001142 | 0.55623529 | 0.99999994 |
| Ggta1 | 0.00145074 | 0.00119787 | 0.00131232 | -0.0001144 | 0.71920999 | 0.99999994 |
| Gm20406 | 0.00132252 | 0.00108192 | 0.00119644 | -0.0001145 | 0.73797865 | 0.99999994 |
| Gm16552 | 0.0006512 | 0.00047471 | 0.00058939 | -0.0001147 | 0.90236906 | 0.99999994 |
| Them7 | 0.00071667 | 0.00053361 | 0.00064861 | -0.000115 | 0.88103628 | 0.99999994 |
| BC106179 | 0.00072915 | 0.0005443 | 0.0006599 | -0.0001156 | 0.87820993 | 0.99999994 |
| Pif1 | 0.00057903 | 0.00040845 | 0.00052409 | -0.0001156 | 0.92904404 | 0.99999994 |
| Ablim3 | 0.0008313 | 0.00063656 | 0.0007523 | -0.0001157 | 0.84710128 | 0.99999994 |
| Onecut1 | 0.00767024 | 0.00679336 | 0.00690918 | -0.0001158 | 0.54442209 | 0.99999994 |
| Bglap3 | 0.0007172 | 0.00053325 | 0.0006491 | -0.0001158 | 0.88257058 | 0.99999994 |
| Thrsp | 0.00072598 | 0.00054107 | 0.00065703 | -0.000116 | 0.8799706 | 0.99999994 |
| Lgr5 | 0.00073436 | 0.00054846 | 0.00066462 | -0.0001162 | 0.87765844 | 0.99999994 |
| Mpzl2 | 0.00500251 | 0.00439806 | 0.00451429 | -0.0001162 | 0.56804261 | 0.99999994 |
| Slc8a3 | 0.00053704 | 0.00036977 | 0.0004861 | -0.0001163 | 0.94440883 | 0.99999994 |
| Hsd17b13 | 0.00056683 | 0.00039669 | 0.00051306 | -0.0001164 | 0.93444427 | 0.99999994 |
| Depdc1a | 0.00189347 | 0.00159587 | 0.00171229 | -0.0001164 | 0.67458045 | 0.99999994 |
| E2f3 | 0.02573474 | 0.02278469 | 0.02290124 | -0.0001165 | 0.51351172 | 0.99999994 |
| Traf1 | 0.00056904 | 0.0003981 | 0.00051505 | -0.0001169 | 0.93465436 | 0.99999994 |
| Pdpr | 0.00860583 | 0.00763001 | 0.00774703 | -0.000117 | 0.54004581 | 0.99999994 |
| Lcn5 | 0.00095054 | 0.00074294 | 0.00086014 | -0.0001172 | 0.81777948 | 0.99999994 |
| Batf3 | 0.0006802 | 0.00049743 | 0.00061562 | -0.0001182 | 0.8993607 | 0.99999994 |
| Tacstd2 | 0.0004917 | 0.00032672 | 0.00044508 | -0.0001184 | 0.96163866 | 0.99999994 |

| | | | | | | |
|---|---|---|---|---|---|---|
| Sema3e | 0.00413343 | 0.00361371 | 0.00373224 | -0.0001185 | 0.58370788 | 0.99999994 |
| Rab3il1 | 0.01062922 | 0.00943681 | 0.00955542 | -0.0001186 | 0.53292405 | 0.99999994 |
| 2610035D17I | 0.01780233 | 0.01580785 | 0.01592671 | -0.0001189 | 0.51980892 | 0.99999994 |
| Diaph3 | 0.00303107 | 0.00261993 | 0.00273892 | -0.000119 | 0.61377688 | 0.99999994 |
| Kcnip2 | 0.00337815 | 0.00293275 | 0.00305182 | -0.0001191 | 0.6024594 | 0.99999994 |
| Ccl7 | 0.00052585 | 0.00035666 | 0.00047597 | -0.0001193 | 0.95239927 | 0.99999994 |
| Matn1 | 0.0010065 | 0.0007914 | 0.00091074 | -0.0001193 | 0.80844859 | 0.99999994 |
| Gm26640 | 0.00126515 | 0.00102502 | 0.00114458 | -0.0001196 | 0.75656469 | 0.99999994 |
| Ms4a8a | 0.00053143 | 0.00036144 | 0.00048102 | -0.0001196 | 0.95101648 | 0.99999994 |
| Nckap5los | 0.00064429 | 0.00046338 | 0.00058313 | -0.0001198 | 0.91417513 | 0.99999994 |
| Zdhhc22 | 0.00053699 | 0.0003663 | 0.00048606 | -0.0001198 | 0.94950168 | 0.99999994 |
| Dclk3 | 0.00502498 | 0.00441434 | 0.00453451 | -0.0001202 | 0.57000753 | 0.99999994 |
| Reep1 | 0.00119712 | 0.00096262 | 0.00108309 | -0.0001205 | 0.770454 | 0.99999994 |
| Gm16136 | 0.00562893 | 0.00495667 | 0.00507743 | -0.0001208 | 0.56289555 | 0.99999994 |
| Podxl | 0.01775405 | 0.0157632 | 0.01588403 | -0.0001208 | 0.52019108 | 0.99999994 |
| Cd52 | 0.00798398 | 0.00706935 | 0.00719026 | -0.0001209 | 0.54455809 | 0.99999994 |
| Gm20443 | 0.00055052 | 0.00037739 | 0.0004983 | -0.0001209 | 0.9468644 | 0.99999994 |
| Tns4 | 0.00168112 | 0.00139913 | 0.00152048 | -0.0001214 | 0.70237959 | 0.99999994 |
| Zfp534 | 0.00329386 | 0.00285352 | 0.00297585 | -0.0001223 | 0.60781858 | 0.99999994 |
| Mboat1 | 0.00800523 | 0.00708693 | 0.00720929 | -0.0001224 | 0.54497341 | 0.99999994 |
| Gm13889 | 0.00084824 | 0.00064515 | 0.00076762 | -0.0001225 | 0.85586697 | 0.99999994 |
| Prss34 | 0.00130046 | 0.00105385 | 0.0011765 | -0.0001227 | 0.7561386 | 0.99999994 |
| Gm28809 | 0.00116923 | 0.00093515 | 0.00105787 | -0.0001227 | 0.77999397 | 0.99999994 |
| Anks1b | 0.00096505 | 0.00075044 | 0.00087326 | -0.0001228 | 0.82541406 | 0.99999994 |
| Gm11640 | 0.00050535 | 0.0003344 | 0.00045742 | -0.000123 | 0.96328767 | 0.99999994 |
| Fbxo24 | 0.00114115 | 0.00090922 | 0.00103249 | -0.0001233 | 0.78660897 | 0.99999994 |
| Slitrk1 | 0.00068099 | 0.0004929 | 0.00061634 | -0.0001234 | 0.90875216 | 0.99999994 |
| Lat | 0.00062514 | 0.00044201 | 0.00056581 | -0.0001238 | 0.9273593 | 0.99999994 |
| 1700067K01I | 0.00104329 | 0.00082012 | 0.00094401 | -0.0001239 | 0.80882052 | 0.99999994 |
| D630011A20 | 0.00071778 | 0.00052568 | 0.00064962 | -0.0001239 | 0.8979368 | 0.99999994 |

| | | | | | | |
|---|---|---|---|---|---|---|
| Clec4a2 | 0.0012382 | 0.00099618 | 0.00112022 | -0.000124 | 0.76944524 | 0.99999994 |
| Pkd2l2 | 0.00148558 | 0.00121967 | 0.0013438 | -0.0001241 | 0.73067275 | 0.99999994 |
| Gm26952 | 0.00058384 | 0.00040413 | 0.00052844 | -0.0001243 | 0.94131123 | 0.99999994 |
| Acvr2a | 0.05942913 | 0.05159611 | 0.05172044 | -0.0001243 | 0.50638317 | 0.99999994 |
| Ifitm6 | 0.00121379 | 0.00097327 | 0.00109815 | -0.0001249 | 0.77547101 | 0.99999994 |
| Mmel1 | 0.0080271 | 0.00710388 | 0.00722888 | -0.000125 | 0.54581733 | 0.99999994 |
| Ccdc190 | 0.00506352 | 0.00444371 | 0.00456916 | -0.0001254 | 0.57250449 | 0.99999994 |
| Ppm1n | 0.00158905 | 0.00131181 | 0.0014373 | -0.0001255 | 0.71943267 | 0.99999994 |
| Adcy7 | 0.00050155 | 0.00032843 | 0.00045398 | -0.0001256 | 0.96718273 | 0.99999994 |
| Hnmt | 0.00239083 | 0.00203569 | 0.00216133 | -0.0001256 | 0.65059747 | 0.99999994 |
| Hes7 | 0.00370468 | 0.00322018 | 0.00334607 | -0.0001259 | 0.59887069 | 0.99999994 |
| Aqp5 | 0.00071519 | 0.00052136 | 0.00064727 | -0.0001259 | 0.90230569 | 0.99999994 |
| Syt15 | 0.00088064 | 0.00067093 | 0.00079692 | -0.000126 | 0.85368118 | 0.99999994 |
| Gm11266 | 0.00114053 | 0.00090572 | 0.00103193 | -0.0001262 | 0.79221206 | 0.99999994 |
| Chst15 | 0.01465161 | 0.01300953 | 0.01313576 | -0.0001262 | 0.52550085 | 0.99999994 |
| Unc5cl | 0.00153055 | 0.00125787 | 0.00138444 | -0.0001266 | 0.72857832 | 0.99999994 |
| Pex16 | 0.1706763 | 0.13670282 | 0.13682939 | -0.0001266 | 0.50245637 | 0.99999994 |
| Serpina3a | 0.00104239 | 0.00081635 | 0.0009432 | -0.0001268 | 0.81465853 | 0.99999994 |
| Gm4787 | 0.00184455 | 0.00154077 | 0.0016681 | -0.0001273 | 0.69431986 | 0.99999994 |
| Dnah2os | 0.00049986 | 0.00032484 | 0.00045246 | -0.0001276 | 0.96977279 | 0.99999994 |
| Sp2 | 0.04662892 | 0.04079548 | 0.04092313 | -0.0001276 | 0.5082824 | 0.99999994 |
| Gm6685 | 0.00115074 | 0.00091205 | 0.00104116 | -0.0001291 | 0.79542728 | 0.99999994 |
| Fermt3 | 0.00208955 | 0.0017601 | 0.00188935 | -0.0001293 | 0.67557488 | 0.99999994 |
| Hspb7 | 0.00086386 | 0.00065207 | 0.00078175 | -0.0001297 | 0.86523969 | 0.99999994 |
| Insl5 | 0.00074634 | 0.00054557 | 0.00067545 | -0.0001299 | 0.89971713 | 0.99999994 |
| 1700042O10 | 0.00613985 | 0.00540645 | 0.00553637 | -0.0001299 | 0.56205909 | 0.99999994 |
| Gm10638 | 0.00389443 | 0.00338704 | 0.003517 | -0.00013 | 0.59715101 | 0.99999994 |
| 1700125G22 | 0.00121502 | 0.00096908 | 0.00109927 | -0.0001302 | 0.7847404 | 0.99999994 |
| Ccno | 0.00049657 | 0.00031922 | 0.00044948 | -0.0001303 | 0.97313787 | 0.99999994 |
| Klhl23 | 0.02760163 | 0.02440147 | 0.02453196 | -0.0001305 | 0.51412135 | 0.99999994 |

| | | | | | | |
|---|---|---|---|---|---|---|
| Mgarp | 0.00194503 | 0.00162829 | 0.00175885 | -0.0001306 | 0.68939454 | 0.99999994 |
| Lmod3 | 0.00078076 | 0.00057596 | 0.00070658 | -0.0001306 | 0.89075171 | 0.99999994 |
| A330048O09 | 0.00068968 | 0.0004934 | 0.00062419 | -0.0001308 | 0.91846218 | 0.99999994 |
| Otud7a | 0.00128111 | 0.00102813 | 0.00115901 | -0.0001309 | 0.77387024 | 0.99999994 |
| Mettl15 | 0.05509233 | 0.04795163 | 0.04808253 | -0.0001309 | 0.50722938 | 0.99999994 |
| Sema5b | 0.00067429 | 0.000479 | 0.00061027 | -0.0001313 | 0.92390596 | 0.99999994 |
| Eif1ad | 0.20028471 | 0.1567838 | 0.15691517 | -0.0001314 | 0.50222331 | 0.99999994 |
| Zwilch | 0.00414247 | 0.00360898 | 0.00374037 | -0.0001314 | 0.59243715 | 0.99999994 |
| 6430573P05I | 0.00535501 | 0.00469974 | 0.00483124 | -0.0001315 | 0.57188748 | 0.99999994 |
| 4833413G10 | 0.00227648 | 0.00192623 | 0.00205812 | -0.0001319 | 0.66515072 | 0.99999994 |
| Adam8 | 0.00295643 | 0.00253955 | 0.00267161 | -0.0001321 | 0.62893942 | 0.99999994 |
| Eif1ax | 0.53936589 | 0.3294709 | 0.32960302 | -0.0001321 | 0.50106444 | 0.99999994 |
| Stpg1 | 0.00237202 | 0.00201197 | 0.00214435 | -0.0001324 | 0.65943099 | 0.99999994 |
| Gm44418 | 0.00080277 | 0.00059404 | 0.00072649 | -0.0001325 | 0.88754932 | 0.99999994 |
| Card10 | 0.00831485 | 0.00735365 | 0.00748656 | -0.0001329 | 0.54703516 | 0.99999994 |
| 9530082P21I | 0.0016364 | 0.00134711 | 0.00148007 | -0.000133 | 0.7250678 | 0.99999994 |
| Smg7 | 0.1104859 | 0.09275351 | 0.09288658 | -0.0001331 | 0.50380421 | 0.99999994 |
| Gm14002 | 0.001292 | 0.00103572 | 0.00116885 | -0.0001331 | 0.77582088 | 0.99999994 |
| Nphp4 | 0.02886365 | 0.02549852 | 0.02563202 | -0.0001335 | 0.51382798 | 0.99999994 |
| Tuba8 | 0.00078272 | 0.00057484 | 0.00070836 | -0.0001335 | 0.89520001 | 0.99999994 |
| Gm12279 | 0.00061809 | 0.00042573 | 0.00055943 | -0.0001337 | 0.94417353 | 0.99999994 |
| Ccdc178 | 0.00319987 | 0.00275737 | 0.00289112 | -0.0001337 | 0.62093104 | 0.99999994 |
| Nat1 | 0.0023363 | 0.00197831 | 0.00211211 | -0.0001338 | 0.66336616 | 0.99999994 |
| Adamts13 | 0.00051635 | 0.00033328 | 0.00046737 | -0.0001341 | 0.9719233 | 0.99999994 |
| Epha1 | 0.00937127 | 0.00829755 | 0.00843171 | -0.0001342 | 0.54217479 | 0.99999994 |
| Crygn | 0.00069997 | 0.00049922 | 0.0006335 | -0.0001343 | 0.92087303 | 0.99999994 |
| Dhtkd1 | 0.00501226 | 0.00438807 | 0.00452307 | -0.000135 | 0.57873997 | 0.99999994 |
| Ms4a7 | 0.00182671 | 0.00151693 | 0.00165199 | -0.0001351 | 0.70684367 | 0.99999994 |
| Wipf1 | 0.04695108 | 0.04106144 | 0.04119711 | -0.0001357 | 0.50874436 | 0.99999994 |
| Cdkn2a | 0.00080699 | 0.00059428 | 0.00073031 | -0.000136 | 0.89248009 | 0.99999994 |

| | | | | | | |
|---|---|---|---|---|---|---|
| Nlrc3 | 0.00074518 | 0.0005383 | 0.0006744 | -0.0001361 | 0.9104169 | 0.99999994 |
| Timd4 | 0.00055915 | 0.0003699 | 0.00050611 | -0.0001362 | 0.96338224 | 0.99999994 |
| Icam1 | 0.00175782 | 0.00145335 | 0.00158977 | -0.0001364 | 0.71606502 | 0.99999994 |
| Gm4349 | 0.00054426 | 0.00035603 | 0.00049264 | -0.0001366 | 0.96754343 | 0.99999994 |
| Hcn2 | 0.00432732 | 0.00377011 | 0.00390679 | -0.0001367 | 0.59206808 | 0.99999994 |
| Lrrc10 | 0.00086244 | 0.00064365 | 0.00078046 | -0.0001368 | 0.87836621 | 0.99999994 |
| Ldhal6b | 0.00119025 | 0.00093982 | 0.00107688 | -0.0001371 | 0.80154166 | 0.99999994 |
| Gm26944 | 0.00092055 | 0.0006959 | 0.00083302 | -0.0001371 | 0.86338979 | 0.99999994 |
| Gm5544 | 0.00092965 | 0.00070412 | 0.00084125 | -0.0001371 | 0.86103697 | 0.99999994 |
| Cyp4f15 | 0.00134944 | 0.0010836 | 0.00122077 | -0.0001372 | 0.77274037 | 0.99999994 |
| Far2 | 0.00180139 | 0.00149181 | 0.00162912 | -0.0001373 | 0.71261533 | 0.99999994 |
| Gm9917 | 0.00288767 | 0.00247216 | 0.00260959 | -0.0001374 | 0.63703336 | 0.99999994 |
| Adm2 | 0.00051233 | 0.000326 | 0.00046374 | -0.0001377 | 0.97598282 | 0.99999994 |
| 1700034P13I | 0.00065098 | 0.00045141 | 0.00058919 | -0.0001378 | 0.94021135 | 0.99999994 |
| St8sia4 | 0.00287793 | 0.00246298 | 0.00260081 | -0.0001378 | 0.63786142 | 0.99999994 |
| Phf11c | 0.00163239 | 0.0013386 | 0.00147645 | -0.0001379 | 0.7328631 | 0.99999994 |
| Klhl6 | 0.00093577 | 0.00070891 | 0.00084678 | -0.0001379 | 0.86075875 | 0.99999994 |
| Bach2 | 0.01686852 | 0.01496279 | 0.01510076 | -0.000138 | 0.52424685 | 0.99999994 |
| Sema4f | 0.0015025 | 0.00122108 | 0.00135909 | -0.000138 | 0.75044943 | 0.99999994 |
| Irf5 | 0.00242037 | 0.00204989 | 0.00218798 | -0.0001381 | 0.66279843 | 0.99999994 |
| Gabrr2 | 0.00089658 | 0.00067311 | 0.00081134 | -0.0001382 | 0.87160264 | 0.99999994 |
| Tfdp1 | 0.0100272 | 0.0088796 | 0.00901788 | -0.0001383 | 0.54064917 | 0.99999994 |
| Fanca | 0.00275828 | 0.00235454 | 0.00249288 | -0.0001383 | 0.64408582 | 0.99999994 |
| Gm17477 | 0.00320824 | 0.00276024 | 0.00289866 | -0.0001384 | 0.62470544 | 0.99999994 |
| Aspn | 0.00093558 | 0.00070794 | 0.00084661 | -0.0001387 | 0.86219986 | 0.99999994 |
| Cyp11a1 | 0.00163243 | 0.00133774 | 0.00147649 | -0.0001388 | 0.73418944 | 0.99999994 |
| Slc1a2 | 0.00157805 | 0.00128841 | 0.00142735 | -0.0001389 | 0.74149955 | 0.99999994 |
| Sh3bp2 | 0.00936609 | 0.00828776 | 0.00842708 | -0.0001393 | 0.54381482 | 0.99999994 |
| Gm14019 | 0.00155313 | 0.0012653 | 0.00140484 | -0.0001395 | 0.74574195 | 0.99999994 |
| Rhbdl3 | 0.03709925 | 0.03262568 | 0.03276548 | -0.0001398 | 0.51132849 | 0.99999994 |

| | | | | | | |
|---|---|---|---|---|---|---|
| Lrrc75aos2 | 0.0060622 | 0.00532669 | 0.00546664 | -0.00014 | 0.56765672 | 0.99999994 |
| Gm26793 | 0.00115449 | 0.00090457 | 0.00104455 | -0.00014 | 0.81380699 | 0.99999994 |
| Cacna1s | 0.00050287 | 0.00031518 | 0.00045518 | -0.00014 | 0.97968863 | 0.99999994 |
| Ttc30a2 | 0.00055082 | 0.00035855 | 0.00049857 | -0.00014 | 0.96921496 | 0.99999994 |
| Gpatch1 | 0.0977426 | 0.08284466 | 0.08298497 | -0.0001403 | 0.50448981 | 0.99999994 |
| Eda | 0.00735133 | 0.00648265 | 0.00662334 | -0.0001407 | 0.55622013 | 0.99999994 |
| Pkdrej | 0.0011359 | 0.00088692 | 0.00102773 | -0.0001408 | 0.81912667 | 0.99999994 |
| Zfp804b | 0.00063084 | 0.00043006 | 0.00057096 | -0.0001409 | 0.94977241 | 0.99999994 |
| Nap1l5 | 0.026237 | 0.02319934 | 0.02334037 | -0.000141 | 0.51604086 | 0.99999994 |
| Mab21l3 | 0.00102931 | 0.00079003 | 0.00093137 | -0.0001413 | 0.84379193 | 0.99999994 |
| Prelp | 0.00060882 | 0.00040955 | 0.00055104 | -0.0001415 | 0.95628781 | 0.99999994 |
| Npb | 0.00942643 | 0.00833929 | 0.00848103 | -0.0001417 | 0.54428822 | 0.99999994 |
| Prdm8 | 0.00061792 | 0.00041731 | 0.00055927 | -0.000142 | 0.95445505 | 0.99999994 |
| Oas1a | 0.0009005 | 0.00067287 | 0.00081489 | -0.000142 | 0.87699216 | 0.99999994 |
| Serpina3g | 0.00152181 | 0.00123402 | 0.00137654 | -0.0001425 | 0.75464933 | 0.99999994 |
| Plcb3 | 0.02040427 | 0.01808006 | 0.01822263 | -0.0001426 | 0.52076606 | 0.99999994 |
| Igsf6 | 0.00216672 | 0.00181631 | 0.00195903 | -0.0001427 | 0.68614093 | 0.99999994 |
| Gm28231 | 0.00068461 | 0.00047683 | 0.00061961 | -0.0001428 | 0.93746925 | 0.99999994 |
| E230025N22 | 0.00059158 | 0.00039202 | 0.00053544 | -0.0001434 | 0.96270621 | 0.99999994 |
| Gm14399 | 0.00253584 | 0.00214836 | 0.00229219 | -0.0001438 | 0.66191462 | 0.99999994 |
| Ces5a | 0.00158476 | 0.00128957 | 0.00143342 | -0.0001438 | 0.74791811 | 0.99999994 |
| Rsf1os2 | 0.00100438 | 0.00076452 | 0.00090882 | -0.0001443 | 0.8547234 | 0.99999994 |
| Gm29538 | 0.00235981 | 0.00198893 | 0.00213333 | -0.0001444 | 0.67384317 | 0.99999994 |
| Dach2 | 0.0258044 | 0.0228177 | 0.02296217 | -0.0001445 | 0.51670208 | 0.99999994 |
| Gm15336 | 0.00046156 | 0.00027333 | 0.0004178 | -0.0001445 | 0.98932611 | 0.99999994 |
| Gm43608 | 0.0006337 | 0.00042883 | 0.00057355 | -0.0001447 | 0.95347801 | 0.99999994 |
| Kcnab3 | 0.01457671 | 0.01292443 | 0.01306927 | -0.0001448 | 0.52940171 | 0.99999994 |
| Slc6a20b | 0.00317001 | 0.00271933 | 0.0028642 | -0.0001449 | 0.63181784 | 0.99999994 |
| Ugt2a3 | 0.00096368 | 0.00072706 | 0.00087202 | -0.000145 | 0.86573874 | 0.99999994 |
| Mylk3 | 0.00067239 | 0.00046333 | 0.00060856 | -0.0001452 | 0.94391485 | 0.99999994 |

| | | | | | | |
|---|---|---|---|---|---|---|
| Ttll2 | 0.0006502 | 0.00044308 | 0.00058848 | -0.0001454 | 0.94998297 | 0.99999994 |
| Hmgxb4 | 0.13623288 | 0.11206736 | 0.11221277 | -0.0001454 | 0.50344105 | 0.99999994 |
| Zfp354b | 0.01014453 | 0.00897715 | 0.00912268 | -0.0001455 | 0.54228253 | 0.99999994 |
| Dnaaf1 | 0.00178414 | 0.00146799 | 0.00161354 | -0.0001455 | 0.72588616 | 0.99999994 |
| Awat2 | 0.00232931 | 0.00196024 | 0.0021058 | -0.0001456 | 0.6772846 | 0.99999994 |
| Creb5 | 0.00222513 | 0.00186604 | 0.00201176 | -0.0001457 | 0.68514819 | 0.99999994 |
| Ccl28 | 0.0011108 | 0.0008592 | 0.00100505 | -0.0001458 | 0.83295972 | 0.99999994 |
| Nts | 0.00065327 | 0.00044537 | 0.00059126 | -0.0001459 | 0.94974779 | 0.99999994 |
| Clec2l | 0.00066055 | 0.00045195 | 0.00059784 | -0.0001459 | 0.94785086 | 0.99999994 |
| Gm13205 | 0.01691071 | 0.01499217 | 0.0151381 | -0.0001459 | 0.52558116 | 0.99999994 |
| Pxdc1 | 0.00091013 | 0.00067765 | 0.00082359 | -0.0001459 | 0.88091253 | 0.99999994 |
| Ptch2 | 0.00126412 | 0.00099754 | 0.00114365 | -0.0001461 | 0.80245472 | 0.99999994 |
| Dmrta2 | 0.0010737 | 0.00082495 | 0.0009715 | -0.0001466 | 0.84234654 | 0.99999994 |
| Gm21411 | 0.00051186 | 0.00031662 | 0.00046332 | -0.0001467 | 0.98246562 | 0.99999994 |
| Lypd2 | 0.00095361 | 0.00071618 | 0.00086291 | -0.0001467 | 0.87116233 | 0.99999994 |
| Cyp2j5 | 0.00114388 | 0.00088804 | 0.00103496 | -0.0001469 | 0.82764961 | 0.99999994 |
| Gm17039 | 0.00053932 | 0.00034115 | 0.00048816 | -0.000147 | 0.97749936 | 0.99999994 |
| Slc22a4 | 0.00045817 | 0.0002676 | 0.00041473 | -0.0001471 | 0.99089662 | 0.99999994 |
| Chek2 | 0.00590652 | 0.00517954 | 0.00532682 | -0.0001473 | 0.57300692 | 0.99999994 |
| A730046J19I | 0.00066386 | 0.00045342 | 0.00060084 | -0.0001474 | 0.94878145 | 0.99999994 |
| Slc4a5 | 0.00159556 | 0.00129563 | 0.00144318 | -0.0001475 | 0.7519085 | 0.99999994 |
| Krt79 | 0.00093939 | 0.00070246 | 0.00085005 | -0.0001476 | 0.8761007 | 0.99999994 |
| Pcdhga4 | 0.00118513 | 0.00092448 | 0.00107224 | -0.0001478 | 0.82050123 | 0.99999994 |
| Bag3 | 0.02827154 | 0.02496825 | 0.02511613 | -0.0001479 | 0.51563026 | 0.99999994 |
| Gm16001 | 0.00093276 | 0.00069601 | 0.00084406 | -0.000148 | 0.87849892 | 0.99999994 |
| Zdhhc15 | 0.00445279 | 0.00387128 | 0.00401973 | -0.0001484 | 0.59708799 | 0.99999994 |
| Gm10642 | 0.00370322 | 0.00319623 | 0.00334476 | -0.0001485 | 0.61623014 | 0.99999994 |
| Fads3 | 0.00600663 | 0.00526803 | 0.00541674 | -0.0001487 | 0.57249676 | 0.99999994 |
| Gm16380 | 0.0009184 | 0.00068213 | 0.00083107 | -0.0001489 | 0.88355146 | 0.99999994 |
| Rasl2-9 | 0.00220113 | 0.00184106 | 0.00199009 | -0.000149 | 0.69091725 | 0.99999994 |

| | | | | | | |
|---|---|---|---|---|---|---|
| Atp6v0c | 0.00152788 | 0.00123299 | 0.00138203 | -0.000149 | 0.76356812 | 0.99999994 |
| Col4a4 | 0.00390226 | 0.00337469 | 0.00352406 | -0.0001494 | 0.61107839 | 0.99999994 |
| Eva1c | 0.00061322 | 0.0004054 | 0.00055502 | -0.0001496 | 0.96362211 | 0.99999994 |
| Gm16853 | 0.00251926 | 0.00212755 | 0.00227723 | -0.0001497 | 0.66913176 | 0.99999994 |
| Krt27 | 0.00215545 | 0.00179903 | 0.00194885 | -0.0001498 | 0.69557371 | 0.99999994 |
| 4933415A04I | 0.01232342 | 0.01091594 | 0.0110658 | -0.0001499 | 0.53591444 | 0.99999994 |
| Hsd3b2 | 0.00053446 | 0.0003339 | 0.00048377 | -0.0001499 | 0.98040245 | 0.99999994 |
| 4933423P22I | 0.00066894 | 0.00045551 | 0.00060544 | -0.0001499 | 0.95036381 | 0.99999994 |
| Ssh3 | 0.05339616 | 0.04650397 | 0.04665408 | -0.0001501 | 0.50854332 | 0.99999994 |
| Meikin | 0.0013169 | 0.00104103 | 0.00119135 | -0.0001503 | 0.79952084 | 0.99999994 |
| Ccnd1 | 0.05214323 | 0.04544651 | 0.04559686 | -0.0001504 | 0.50875569 | 0.99999994 |
| Zfp97 | 0.00221908 | 0.00185572 | 0.0020063 | -0.0001506 | 0.69131561 | 0.99999994 |
| Dstyk | 0.04868069 | 0.04251488 | 0.0426661 | -0.0001512 | 0.50941085 | 0.99999994 |
| Abhd17b | 0.10789346 | 0.09074327 | 0.09089514 | -0.0001519 | 0.50443693 | 0.99999994 |
| Efhb | 0.00113216 | 0.00087221 | 0.00102436 | -0.0001521 | 0.83858721 | 0.99999994 |
| Ldb3 | 0.00199971 | 0.00165581 | 0.00180824 | -0.0001524 | 0.71264062 | 0.99999994 |
| Gnat3 | 0.00063447 | 0.00042179 | 0.00057425 | -0.0001525 | 0.96140574 | 0.99999994 |
| Lrrc17 | 0.00183182 | 0.00150412 | 0.00165661 | -0.0001525 | 0.72996756 | 0.99999994 |
| Hoxb1 | 0.00110378 | 0.00084617 | 0.0009987 | -0.0001525 | 0.84533232 | 0.99999994 |
| Sult1a1 | 0.00183017 | 0.00150239 | 0.00165512 | -0.0001527 | 0.73046638 | 0.99999994 |
| Gm13003 | 0.00058043 | 0.00037254 | 0.00052536 | -0.0001528 | 0.97357813 | 0.99999994 |
| Pik3ap1 | 0.00066292 | 0.00044706 | 0.00059999 | -0.0001529 | 0.95511711 | 0.99999994 |
| Icam4 | 0.0006403 | 0.00042617 | 0.00057953 | -0.0001534 | 0.96091346 | 0.99999994 |
| Gm27169 | 0.00054282 | 0.0003379 | 0.00049133 | -0.0001534 | 0.98117724 | 0.99999994 |
| Pde6g | 0.00050206 | 0.00030085 | 0.00045445 | -0.0001536 | 0.98776966 | 0.99999994 |
| Nexn | 0.00137122 | 0.00108673 | 0.00124046 | -0.0001537 | 0.79527975 | 0.99999994 |
| Gm20605 | 0.00063186 | 0.0004179 | 0.00057189 | -0.000154 | 0.96346221 | 0.99999994 |
| Cep55 | 0.00161973 | 0.001311 | 0.00146502 | -0.000154 | 0.75797584 | 0.99999994 |
| Gm11175 | 0.00062795 | 0.00041417 | 0.00056836 | -0.0001542 | 0.96452336 | 0.99999994 |
| Vma21-ps | 0.01220205 | 0.01080345 | 0.01095771 | -0.0001543 | 0.53733002 | 0.99999994 |

| | | | | | | |
|---|---|---|---|---|---|---|
| Lca5l | 0.00753761 | 0.00663603 | 0.00679032 | -0.0001543 | 0.56010719 | 0.99999994 |
| Lair1 | 0.00136276 | 0.00107823 | 0.0012328 | -0.0001546 | 0.79803011 | 0.99999994 |
| Rgs2 | 0.05467543 | 0.04757694 | 0.04773173 | -0.0001548 | 0.50861073 | 0.99999994 |
| Lix1 | 0.00826166 | 0.00728391 | 0.00743894 | -0.000155 | 0.55516544 | 0.99999994 |
| Colec11 | 0.00045772 | 0.00025881 | 0.00041433 | -0.0001555 | 0.99376351 | 0.99999994 |
| Chrdl1 | 0.00038798 | 0.00019557 | 0.00035122 | -0.0001556 | 0.99841009 | 0.99999994 |
| Arhgef28 | 0.06681302 | 0.05771142 | 0.05786712 | -0.0001557 | 0.50714497 | 0.99999994 |
| Tspan32 | 0.00089759 | 0.00065624 | 0.00081225 | -0.000156 | 0.89945558 | 0.99999994 |
| Il2rg | 0.00082332 | 0.00058898 | 0.00074508 | -0.0001561 | 0.918425 | 0.99999994 |
| C030010L15F | 0.00224944 | 0.0018773 | 0.00203371 | -0.0001564 | 0.69564953 | 0.99999994 |
| Gm28535 | 0.00252594 | 0.00212683 | 0.00228326 | -0.0001564 | 0.67581563 | 0.99999994 |
| Itgb3 | 0.00042894 | 0.00023175 | 0.00038828 | -0.0001565 | 0.9963568 | 0.99999994 |
| Gpr171 | 0.00097357 | 0.00072434 | 0.00088097 | -0.0001566 | 0.88167639 | 0.99999994 |
| Sell | 0.00057261 | 0.00036162 | 0.00051829 | -0.0001567 | 0.97789376 | 0.99999994 |
| Tcfl5 | 0.00049186 | 0.00028844 | 0.00044522 | -0.0001568 | 0.99045992 | 0.99999994 |
| Gm12408 | 0.00051501 | 0.00030868 | 0.00046616 | -0.0001575 | 0.98773259 | 0.99999994 |
| Shisa7 | 0.00065162 | 0.00043201 | 0.00058976 | -0.0001578 | 0.96250097 | 0.99999994 |
| Lix1l | 0.00223578 | 0.00186332 | 0.00202137 | -0.000158 | 0.69862385 | 0.99999994 |
| Cyp4f17 | 0.00055689 | 0.000346 | 0.00050406 | -0.0001581 | 0.98157179 | 0.99999994 |
| Myo7b | 0.00770484 | 0.00678184 | 0.00694018 | -0.0001583 | 0.56035277 | 0.99999994 |
| Sspnos | 0.00056328 | 0.00035141 | 0.00050984 | -0.0001584 | 0.98070212 | 0.99999994 |
| Syngr4 | 0.00076479 | 0.0005336 | 0.00069214 | -0.0001585 | 0.93634081 | 0.99999994 |
| Sh3rf3 | 0.00122688 | 0.00095128 | 0.00110999 | -0.0001587 | 0.82938516 | 0.99999994 |
| Gm9949 | 0.00194971 | 0.00160421 | 0.00176308 | -0.0001589 | 0.72568238 | 0.99999994 |
| Matn4 | 0.01309044 | 0.01158956 | 0.01174847 | -0.0001589 | 0.53586958 | 0.99999994 |
| Gm14443 | 0.00212659 | 0.00176366 | 0.0019228 | -0.0001591 | 0.709148 | 0.99999994 |
| 4930445N18 | 0.00025349 | 7.03E-05 | 0.00022949 | -0.0001592 | 0.99999805 | 0.99999994 |
| Ly6c2 | 0.00051025 | 0.0003025 | 0.00046186 | -0.0001594 | 0.98918107 | 0.99999994 |
| Pcdhb3 | 0.00291539 | 0.00247524 | 0.0026346 | -0.0001594 | 0.65638618 | 0.99999994 |
| Nek2 | 0.00123902 | 0.00096143 | 0.00112096 | -0.0001595 | 0.82825383 | 0.99999994 |

| | | | | | | |
|---|---|---|---|---|---|---|
| Osbpl10 | 0.00235737 | 0.00197144 | 0.00213113 | -0.0001597 | 0.69102903 | 0.99999994 |
| Gm26787 | 0.00097311 | 0.00072084 | 0.00088055 | -0.0001597 | 0.88634032 | 0.99999994 |
| Gm9888 | 0.00056292 | 0.0003497 | 0.00050951 | -0.0001598 | 0.98159452 | 0.99999994 |
| Il1rl2 | 0.00108973 | 0.00082607 | 0.000986 | -0.0001599 | 0.85984729 | 0.99999994 |
| Lck | 0.01116306 | 0.00987167 | 0.01003171 | -0.00016 | 0.54228522 | 0.99999994 |
| Hcls1 | 0.00128673 | 0.00100392 | 0.00116409 | -0.0001602 | 0.82013341 | 0.99999994 |
| Sbpl | 0.00100666 | 0.00075063 | 0.00091088 | -0.0001603 | 0.87921312 | 0.99999994 |
| Hsd17b1 | 0.00097076 | 0.00071793 | 0.00087842 | -0.0001605 | 0.88802709 | 0.99999994 |
| Crebrf | 0.08656986 | 0.0738663 | 0.07402679 | -0.0001605 | 0.5057569 | 0.99999994 |
| Zfp825 | 0.05886199 | 0.05108521 | 0.05124586 | -0.0001607 | 0.50832426 | 0.99999994 |
| Gm10451 | 0.00591659 | 0.0051749 | 0.00533586 | -0.000161 | 0.57956711 | 0.99999994 |
| Ly9 | 0.00047355 | 0.00026762 | 0.00042865 | -0.000161 | 0.99379964 | 0.99999994 |
| Amdhd1 | 0.00095536 | 0.00070329 | 0.0008645 | -0.0001612 | 0.89274464 | 0.99999994 |
| Calr4 | 0.00122404 | 0.00094605 | 0.00110742 | -0.0001614 | 0.83395021 | 0.99999994 |
| Myadml2 | 0.0002501 | 6.51E-05 | 0.00022642 | -0.0001614 | 0.99999895 | 0.99999994 |
| Brinp2 | 0.00134947 | 0.00105896 | 0.00122079 | -0.0001618 | 0.81121795 | 0.99999994 |
| Cd38 | 0.00092266 | 0.0006729 | 0.00083492 | -0.000162 | 0.90176668 | 0.99999994 |
| Col17a1 | 0.00095247 | 0.00069964 | 0.00086188 | -0.0001622 | 0.89488949 | 0.99999994 |
| A730098A19 | 0.00214124 | 0.00177362 | 0.00193602 | -0.0001624 | 0.71169842 | 0.99999994 |
| Sema6b | 0.00762944 | 0.0067098 | 0.00687262 | -0.0001628 | 0.56264893 | 0.99999994 |
| Myzap | 0.00112412 | 0.00085403 | 0.00101709 | -0.0001631 | 0.85704383 | 0.99999994 |
| Adap1 | 0.0005035 | 0.00029265 | 0.00045575 | -0.0001631 | 0.99139206 | 0.99999994 |
| Gm14154 | 0.00109617 | 0.00082818 | 0.00099181 | -0.0001636 | 0.86394149 | 0.99999994 |
| Lrrc15 | 0.00025526 | 6.73E-05 | 0.00023109 | -0.0001637 | 0.9999988 | 0.99999994 |
| Gm26760 | 0.003275 | 0.0027949 | 0.00295885 | -0.000164 | 0.64387294 | 0.99999994 |
| Shisa6 | 0.00226112 | 0.00187959 | 0.00204425 | -0.0001647 | 0.70407302 | 0.99999994 |
| AY358078 | 0.00055964 | 0.00034162 | 0.00050655 | -0.0001649 | 0.98489267 | 0.99999994 |
| Mn1 | 0.00111316 | 0.00084219 | 0.00100718 | -0.000165 | 0.86223404 | 0.99999994 |
| Zfp558 | 0.00348511 | 0.00298316 | 0.00314822 | -0.0001651 | 0.63645179 | 0.99999994 |
| 0610039K10I | 0.00187889 | 0.00153402 | 0.00169912 | -0.0001651 | 0.74111116 | 0.99999994 |

| | | | | | | |
|---|---|---|---|---|---|---|
| Adam5 | 0.00170065 | 0.00137264 | 0.00153812 | -0.0001655 | 0.76304633 | 0.99999994 |
| Slco2b1 | 0.00094993 | 0.00069379 | 0.00085958 | -0.0001658 | 0.90040691 | 0.99999994 |
| Uchl4 | 0.00643924 | 0.00563935 | 0.00580515 | -0.0001658 | 0.57538975 | 0.99999994 |
| Mrps5 | 0.38199033 | 0.26096495 | 0.26113077 | -0.0001658 | 0.50168622 | 0.99999994 |
| Sstr1 | 0.00177965 | 0.00144342 | 0.00160948 | -0.0001661 | 0.75389713 | 0.99999994 |
| Lonrf3 | 0.00026127 | 7.04E-05 | 0.00023654 | -0.0001661 | 0.99999852 | 0.99999994 |
| Gal3st3 | 0.00043881 | 0.00023112 | 0.00039721 | -0.0001661 | 0.99730962 | 0.99999994 |
| A230107N01 | 0.00223192 | 0.00185178 | 0.00201789 | -0.0001661 | 0.70813558 | 0.99999994 |
| Bnipl | 0.00104219 | 0.00077669 | 0.00094301 | -0.0001663 | 0.87980718 | 0.99999994 |
| Ntn5 | 0.0003068 | 0.00011115 | 0.00027774 | -0.0001666 | 0.99996731 | 0.99999994 |
| Sgcz | 0.00031871 | 0.00012186 | 0.00028853 | -0.0001667 | 0.99993972 | 0.99999994 |
| Mkrn3 | 0.02368573 | 0.02093961 | 0.02110675 | -0.0001671 | 0.52101893 | 0.99999994 |
| Bpifb3 | 0.00061626 | 0.00039035 | 0.00055778 | -0.0001674 | 0.97714299 | 0.99999994 |
| Exoc3 | 0.27574925 | 0.20378153 | 0.20394901 | -0.0001675 | 0.50218064 | 0.99999994 |
| Fbn1 | 0.00140883 | 0.00110686 | 0.00127444 | -0.0001676 | 0.8092782 | 0.99999994 |
| Hapln4 | 0.00045107 | 0.00024062 | 0.00040831 | -0.0001677 | 0.99686877 | 0.99999994 |
| Iqgap3 | 0.00137781 | 0.00107854 | 0.00124641 | -0.0001679 | 0.81499811 | 0.99999994 |
| Fetub | 0.00026439 | 7.14E-05 | 0.00023936 | -0.0001679 | 0.99999849 | 0.99999994 |
| Dhx58 | 0.00526288 | 0.00458048 | 0.00474842 | -0.0001679 | 0.59305989 | 0.99999994 |
| Plag1 | 0.05368839 | 0.04673229 | 0.04690041 | -0.0001681 | 0.509518 | 0.99999994 |
| Rad54b | 0.00026861 | 7.50E-05 | 0.00024318 | -0.0001681 | 0.99999791 | 0.99999994 |
| Banf2 | 0.0005148 | 0.00029772 | 0.00046598 | -0.0001683 | 0.99188055 | 0.99999994 |
| Prodh | 0.01382822 | 0.01223613 | 0.01240444 | -0.0001683 | 0.53598246 | 0.99999994 |
| Bcl6 | 0.00061307 | 0.00038648 | 0.00055489 | -0.0001684 | 0.97831987 | 0.99999994 |
| Gm26614 | 0.00046523 | 0.00025264 | 0.00042112 | -0.0001685 | 0.99612886 | 0.99999994 |
| a | 0.00049233 | 0.00027692 | 0.00044565 | -0.0001687 | 0.99413477 | 0.99999994 |
| Gm15406 | 0.00084158 | 0.0005927 | 0.0007616 | -0.0001689 | 0.93004201 | 0.99999994 |
| Acss3 | 0.00711493 | 0.00624248 | 0.00641138 | -0.0001689 | 0.56959947 | 0.99999994 |
| Smlr1 | 0.00026936 | 7.48E-05 | 0.00024386 | -0.000169 | 0.99999802 | 0.99999994 |
| Efcab9 | 0.00044525 | 0.00023375 | 0.00040304 | -0.0001693 | 0.99741193 | 0.99999994 |

| | | | | | | |
|---|---|---|---|---|---|---|
| Lbx2 | 0.00036663 | 0.00016171 | 0.0003319 | -0.0001702 | 0.99967885 | 0.99999994 |
| Pstpip2 | 0.00051299 | 0.00029405 | 0.00046434 | -0.0001703 | 0.99267901 | 0.99999994 |
| Pla2g5 | 0.00180942 | 0.00146605 | 0.00163638 | -0.0001703 | 0.75579226 | 0.99999994 |
| Anxa10 | 0.00028914 | 9.14E-05 | 0.00026175 | -0.0001703 | 0.9999926 | 0.99999994 |
| Tnni2 | 0.00027107 | 7.50E-05 | 0.0002454 | -0.0001704 | 0.99999809 | 0.99999994 |
| Oip5 | 0.00212794 | 0.00175357 | 0.00192402 | -0.0001704 | 0.72229964 | 0.99999994 |
| Cldn1 | 0.00049781 | 0.00027994 | 0.00045061 | -0.0001707 | 0.99414989 | 0.99999994 |
| Edn1 | 0.00042496 | 0.0002139 | 0.00038468 | -0.0001708 | 0.99843705 | 0.99999994 |
| Trim6 | 0.00147758 | 0.00116569 | 0.00133657 | -0.0001709 | 0.8026175 | 0.99999994 |
| Catsperg1 | 0.00027332 | 7.62E-05 | 0.00024744 | -0.0001713 | 0.99999796 | 0.99999994 |
| Fam107a | 0.00574714 | 0.0050123 | 0.00518364 | -0.0001713 | 0.58707343 | 0.99999994 |
| Ifnlr1 | 0.0009599 | 0.00069711 | 0.0008686 | -0.0001715 | 0.90561289 | 0.99999994 |
| Gm26717 | 0.0005511 | 0.00032721 | 0.00049882 | -0.0001716 | 0.98899234 | 0.99999994 |
| Gng10 | 0.00564751 | 0.00492249 | 0.00509412 | -0.0001716 | 0.58872297 | 0.99999994 |
| Itga2 | 0.00130626 | 0.00100982 | 0.00118174 | -0.0001719 | 0.83356767 | 0.99999994 |
| Ppara | 0.00136459 | 0.00106253 | 0.00123446 | -0.0001719 | 0.8230468 | 0.99999994 |
| Prok1 | 0.00027735 | 7.87E-05 | 0.00025109 | -0.0001724 | 0.99999756 | 0.99999994 |
| Folr2 | 0.0018118 | 0.00146601 | 0.00163852 | -0.0001725 | 0.75828797 | 0.99999994 |
| Gm26684 | 0.00045563 | 0.00023962 | 0.00041244 | -0.0001728 | 0.99735748 | 0.99999994 |
| Nae1 | 0.36018465 | 0.25001143 | 0.25018443 | -0.000173 | 0.5018362 | 0.99999994 |
| Cby3 | 0.0002809 | 8.12E-05 | 0.0002543 | -0.0001731 | 0.99999707 | 0.99999994 |
| Chad | 0.00198707 | 0.0016236 | 0.00179682 | -0.0001732 | 0.73946233 | 0.99999994 |
| Zfp597 | 0.07180562 | 0.06181651 | 0.06198982 | -0.0001733 | 0.50742377 | 0.99999994 |
| Gcnt1 | 0.01876011 | 0.01659937 | 0.01677278 | -0.0001734 | 0.52743271 | 0.99999994 |
| Nckap1l | 0.0002861 | 8.51E-05 | 0.000259 | -0.0001739 | 0.99999605 | 0.99999994 |
| Rassf1 | 0.17081035 | 0.13674864 | 0.13692253 | -0.0001739 | 0.50337232 | 0.99999994 |
| Plekhg5 | 0.02349121 | 0.02076207 | 0.02093614 | -0.0001741 | 0.52206785 | 0.99999994 |
| Pcdhb13 | 0.00389538 | 0.00334376 | 0.00351786 | -0.0001741 | 0.62908277 | 0.99999994 |
| Nacad | 0.00035913 | 0.000151 | 0.00032511 | -0.0001741 | 0.99981787 | 0.99999994 |
| Hoxb5 | 0.00101301 | 0.0007425 | 0.00091663 | -0.0001741 | 0.89696754 | 0.99999994 |

| | | | | | | |
|---|---|---|---|---|---|---|
| Gtpbp2 | 0.1998328 | 0.15644168 | 0.15661581 | -0.0001741 | 0.50295237 | 0.99999994 |
| Adprhl1 | 0.00051915 | 0.00029565 | 0.00046991 | -0.0001743 | 0.99321484 | 0.99999994 |
| Pld6 | 0.00595973 | 0.00520034 | 0.00537461 | -0.0001743 | 0.58544 | 0.99999994 |
| 4930555F03I | 0.00028959 | 8.78E-05 | 0.00026216 | -0.0001743 | 0.9999952 | 0.99999994 |
| Kirrel | 0.00109722 | 0.0008184 | 0.00099277 | -0.0001744 | 0.87881892 | 0.99999994 |
| Gm15912 | 0.00245002 | 0.00204026 | 0.00221474 | -0.0001745 | 0.70000271 | 0.99999994 |
| Dhrs13os | 0.00052957 | 0.00030481 | 0.00047934 | -0.0001745 | 0.99231691 | 0.99999994 |
| Aldh1l1 | 0.00472745 | 0.00409232 | 0.00426688 | -0.0001746 | 0.6073112 | 0.99999994 |
| Lamp3 | 0.00028247 | 8.11E-05 | 0.00025572 | -0.0001746 | 0.99999724 | 0.99999994 |
| Mrpl19 | 0.23298894 | 0.17783235 | 0.17800694 | -0.0001746 | 0.50260451 | 0.99999994 |
| Alox12 | 0.00087745 | 0.00061907 | 0.00079404 | -0.000175 | 0.92877751 | 0.99999994 |
| Noxo1 | 0.01910819 | 0.01690455 | 0.01707998 | -0.0001754 | 0.52725387 | 0.99999994 |
| Tnnc2 | 0.00042014 | 0.00020466 | 0.00038032 | -0.0001757 | 0.99894533 | 0.99999994 |
| Srpx | 0.00098101 | 0.00071181 | 0.00088769 | -0.0001759 | 0.90638033 | 0.99999994 |
| BC035044 | 0.0006023 | 0.00036902 | 0.00054514 | -0.0001761 | 0.98424432 | 0.99999994 |
| Gm9847 | 0.00081943 | 0.00056513 | 0.00074157 | -0.0001764 | 0.94337082 | 0.99999994 |
| Mbd4 | 0.04221419 | 0.03698016 | 0.03715679 | -0.0001766 | 0.51262105 | 0.99999994 |
| Add2 | 0.00143784 | 0.00112379 | 0.00130066 | -0.0001769 | 0.81730844 | 0.99999994 |
| Gm12580 | 0.000907 | 0.00064367 | 0.00082076 | -0.0001771 | 0.92452519 | 0.99999994 |
| Pram1 | 0.00040463 | 0.00018908 | 0.00036628 | -0.0001772 | 0.99935965 | 0.99999994 |
| Kcnq3 | 0.00146948 | 0.00115202 | 0.00132925 | -0.0001772 | 0.8125997 | 0.99999994 |
| Gbp8 | 0.00165858 | 0.00132275 | 0.00150011 | -0.0001774 | 0.78436283 | 0.99999994 |
| Mrgprf | 0.00075549 | 0.00050636 | 0.00068373 | -0.0001774 | 0.95789323 | 0.99999994 |
| Ccdc24 | 0.00307365 | 0.00259988 | 0.00277731 | -0.0001774 | 0.66467019 | 0.99999994 |
| Rttn | 0.00694457 | 0.00608105 | 0.00625858 | -0.0001775 | 0.57488253 | 0.99999994 |
| Krt6a | 0.000296 | 9.03E-05 | 0.00026797 | -0.0001777 | 0.99999491 | 0.99999994 |
| 4930590J08F | 0.00130327 | 0.00100136 | 0.00117903 | -0.0001777 | 0.84209258 | 0.99999994 |
| Cdh26 | 0.00029938 | 9.33E-05 | 0.00027103 | -0.0001777 | 0.99999365 | 0.99999994 |
| Clec2d | 0.00896582 | 0.00789123 | 0.00806913 | -0.0001779 | 0.55833745 | 0.99999994 |
| Gm16279 | 0.00276622 | 0.00232208 | 0.00250005 | -0.000178 | 0.68219062 | 0.99999994 |

| | | | | | | |
|---|---|---|---|---|---|---|
| Fcgr2b | 0.0013018 | 0.00099967 | 0.00117771 | -0.000178 | 0.84286197 | 0.99999994 |
| Sspo | 0.00107244 | 0.00079232 | 0.00097036 | -0.000178 | 0.88902399 | 0.99999994 |
| 4930589L23F | 0.0002939 | 8.79E-05 | 0.00026607 | -0.0001782 | 0.99999586 | 0.99999994 |
| Grem1 | 0.00029431 | 8.81E-05 | 0.00026644 | -0.0001783 | 0.99999581 | 0.99999994 |
| Gata3 | 0.00075687 | 0.0005066 | 0.00068498 | -0.0001784 | 0.95848492 | 0.99999994 |
| Swap70 | 0.00950822 | 0.00837559 | 0.00855415 | -0.0001786 | 0.55525312 | 0.99999994 |
| Retn | 0.00053135 | 0.00030237 | 0.00048095 | -0.0001786 | 0.99327459 | 0.99999994 |
| 1700104B16I | 0.00029741 | 9.06E-05 | 0.00026924 | -0.0001786 | 0.99999497 | 0.99999994 |
| Mpeg1 | 0.00081427 | 0.00055803 | 0.0007369 | -0.0001789 | 0.94692037 | 0.99999994 |
| Tbkbp1 | 0.00047116 | 0.00024755 | 0.00042649 | -0.0001789 | 0.9973864 | 0.99999994 |
| Gm9945 | 0.0018643 | 0.00150694 | 0.00168594 | -0.000179 | 0.76013903 | 0.99999994 |
| Ces2g | 0.00055158 | 0.00032007 | 0.00049926 | -0.0001792 | 0.99155211 | 0.99999994 |
| Lgi4 | 0.00346663 | 0.00295207 | 0.00313157 | -0.0001795 | 0.64859743 | 0.99999994 |
| Trank1 | 0.00053822 | 0.00030753 | 0.00048717 | -0.0001796 | 0.99294486 | 0.99999994 |
| D330050G23 | 0.00187973 | 0.00151998 | 0.00169988 | -0.0001799 | 0.75942556 | 0.99999994 |
| Arhgap31 | 0.001035 | 0.00075654 | 0.00093651 | -0.00018 | 0.89957871 | 0.99999994 |
| Tnfrsf13b | 0.00030234 | 9.36E-05 | 0.00027371 | -0.0001801 | 0.99999406 | 0.99999994 |
| Krt1 | 0.03555211 | 0.03125123 | 0.03143134 | -0.0001801 | 0.51521301 | 0.99999994 |
| Ceacam10 | 0.00414365 | 0.00356119 | 0.00374144 | -0.0001802 | 0.62576834 | 0.99999994 |
| Gm6871 | 0.00048628 | 0.00025985 | 0.00044017 | -0.0001803 | 0.99680296 | 0.99999994 |
| Abcc2 | 0.00033239 | 0.00012056 | 0.0003009 | -0.0001803 | 0.99996688 | 0.99999994 |
| Psca | 0.00131367 | 0.00100809 | 0.00118843 | -0.0001803 | 0.84377521 | 0.99999994 |
| Ttc9b | 0.00058203 | 0.00034632 | 0.0005268 | -0.0001805 | 0.98870892 | 0.99999994 |
| Gm16006 | 0.00327896 | 0.00278188 | 0.00296242 | -0.0001805 | 0.65750195 | 0.99999994 |
| Rgs7bp | 0.00122599 | 0.00092857 | 0.00110918 | -0.0001806 | 0.86079038 | 0.99999994 |
| 1110028F18I | 0.00128215 | 0.00097925 | 0.00115994 | -0.0001807 | 0.85011146 | 0.99999994 |
| Sbno2 | 0.00243761 | 0.00202278 | 0.00220355 | -0.0001808 | 0.70748425 | 0.99999994 |
| Icos | 0.00030388 | 9.42E-05 | 0.0002751 | -0.0001809 | 0.99999401 | 0.99999994 |
| Gm42902 | 0.00052771 | 0.00029657 | 0.00047766 | -0.0001811 | 0.99419126 | 0.99999994 |
| Dock3 | 0.00031052 | 9.98E-05 | 0.00028111 | -0.0001813 | 0.99999119 | 0.99999994 |

| | | | | | | |
|---|---|---|---|---|---|---|
| Id4 | 0.02581235 | 0.02278773 | 0.02296912 | -0.0001814 | 0.52096073 | 0.99999994 |
| Gpbar1 | 0.00152633 | 0.00119899 | 0.00138062 | -0.0001816 | 0.80939502 | 0.99999994 |
| Lamc2 | 0.01202359 | 0.01061697 | 0.01079876 | -0.0001818 | 0.54460954 | 0.99999994 |
| F730016J06F | 0.00048123 | 0.00025376 | 0.0004356 | -0.0001818 | 0.99727057 | 0.99999994 |
| Cxcl16 | 0.00199976 | 0.00162642 | 0.00180828 | -0.0001819 | 0.74839069 | 0.99999994 |
| Slc2a4 | 0.00055715 | 0.00032236 | 0.0005043 | -0.0001819 | 0.9918343 | 0.99999994 |
| Il17b | 0.00058274 | 0.0003454 | 0.00052745 | -0.0001821 | 0.98920392 | 0.99999994 |
| Prdm14 | 0.00030858 | 9.72E-05 | 0.00027935 | -0.0001821 | 0.99999286 | 0.99999994 |
| Arhgef26 | 0.00304812 | 0.00257203 | 0.00275429 | -0.0001823 | 0.67020341 | 0.99999994 |
| Stat3 | 0.42498538 | 0.28142937 | 0.281612 | -0.0001826 | 0.50172215 | 0.99999994 |
| Clec5a | 0.00051627 | 0.00028461 | 0.00046731 | -0.0001827 | 0.99537045 | 0.99999994 |
| Casp12 | 0.00055838 | 0.0003227 | 0.00050541 | -0.0001827 | 0.99194422 | 0.99999994 |
| Prrg1 | 0.00313689 | 0.00265162 | 0.00283434 | -0.0001827 | 0.66608025 | 0.99999994 |
| Gm17660 | 0.00054824 | 0.00031308 | 0.00049624 | -0.0001832 | 0.99299045 | 0.99999994 |
| Crnde | 0.00031986 | 0.00010639 | 0.00028957 | -0.0001832 | 0.99998728 | 0.99999994 |
| Slc8b1 | 0.01581187 | 0.01398169 | 0.01416491 | -0.0001832 | 0.53430593 | 0.99999994 |
| Gm5091 | 0.00152323 | 0.00119453 | 0.00137783 | -0.0001833 | 0.81206366 | 0.99999994 |
| Mogat1 | 0.00033132 | 0.00011657 | 0.00029993 | -0.0001834 | 0.99997643 | 0.99999994 |
| Tex38 | 0.00031233 | 9.93E-05 | 0.00028275 | -0.0001834 | 0.99999213 | 0.99999994 |
| Tlr6 | 0.00031205 | 9.91E-05 | 0.0002825 | -0.0001834 | 0.99999227 | 0.99999994 |
| Sgcd | 0.00045539 | 0.0002286 | 0.00041222 | -0.0001836 | 0.99848642 | 0.99999994 |
| Gnal | 0.00031634 | 0.0001027 | 0.00028637 | -0.0001837 | 0.99999019 | 0.99999994 |
| Myh1 | 0.0005199 | 0.00028678 | 0.00047059 | -0.0001838 | 0.99533744 | 0.99999994 |
| A530032D15 | 0.00031682 | 0.00010296 | 0.00028681 | -0.0001839 | 0.99999008 | 0.99999994 |
| Hey2 | 0.00031397 | 0.00010034 | 0.00028423 | -0.0001839 | 0.99999171 | 0.99999994 |
| Hhipl2 | 0.00040062 | 0.00017876 | 0.00036265 | -0.0001839 | 0.99963138 | 0.99999994 |
| Ano8 | 0.01301182 | 0.01149462 | 0.01167853 | -0.0001839 | 0.5417414 | 0.99999994 |
| Rinl | 0.00408614 | 0.00350562 | 0.00368965 | -0.000184 | 0.63005605 | 0.99999994 |
| Abcc9 | 0.0003427 | 0.00012617 | 0.00031024 | -0.0001841 | 0.99996079 | 0.99999994 |
| Adam4 | 0.00330945 | 0.00280571 | 0.00298991 | -0.0001842 | 0.65912407 | 0.99999994 |

| | | | | | | |
|---|---|---|---|---|---|---|
| Pcdhgb6 | 0.00323458 | 0.00273774 | 0.00292241 | -0.0001847 | 0.66298631 | 0.99999994 |
| Afap1l1 | 0.00037806 | 0.00015743 | 0.00034224 | -0.0001848 | 0.99983738 | 0.99999994 |
| Lmntd1 | 0.00032124 | 0.00010596 | 0.00029081 | -0.0001848 | 0.99998837 | 0.99999994 |
| Srsf12 | 0.00970076 | 0.00854114 | 0.00872623 | -0.0001851 | 0.55613929 | 0.99999994 |
| Syt12 | 0.00031915 | 0.00010369 | 0.00028892 | -0.0001852 | 0.99999011 | 0.99999994 |
| Gjb5 | 0.0003228 | 0.00010698 | 0.00029222 | -0.0001852 | 0.99998776 | 0.99999994 |
| Smad9 | 0.0003419 | 0.00012421 | 0.00030951 | -0.0001853 | 0.99996628 | 0.99999994 |
| Gm15734 | 0.00033477 | 0.00011772 | 0.00030305 | -0.0001853 | 0.99997657 | 0.99999994 |
| Eppin | 0.00086733 | 0.00059947 | 0.00078488 | -0.0001854 | 0.94207315 | 0.99999994 |
| Gm17552 | 0.00144971 | 0.00112596 | 0.00131139 | -0.0001854 | 0.82669573 | 0.99999994 |
| Ache | 0.00034951 | 0.00013088 | 0.0003164 | -0.0001855 | 0.99995248 | 0.99999994 |
| Tiam2 | 0.000326 | 0.00010941 | 0.00029512 | -0.0001857 | 0.99998597 | 0.99999994 |
| Gm26559 | 0.01033046 | 0.00910277 | 0.00928871 | -0.0001859 | 0.55300155 | 0.99999994 |
| 1700012C14I | 0.00457895 | 0.00394723 | 0.00413326 | -0.000186 | 0.6177536 | 0.99999994 |
| Ros1 | 0.00160973 | 0.00126995 | 0.00145598 | -0.000186 | 0.80247264 | 0.99999994 |
| C1rl | 0.00818962 | 0.00718835 | 0.00737443 | -0.0001861 | 0.56669125 | 0.99999994 |
| Upk1a | 0.00033336 | 0.00011569 | 0.00030178 | -0.0001861 | 0.99997976 | 0.99999994 |
| Slc39a8 | 0.01025099 | 0.00903156 | 0.00921776 | -0.0001862 | 0.55347971 | 0.99999994 |
| Pira2 | 0.00033424 | 0.00011632 | 0.00030258 | -0.0001863 | 0.99997911 | 0.99999994 |
| R3hdml | 0.00082335 | 0.00055845 | 0.00074511 | -0.0001867 | 0.95229294 | 0.99999994 |
| Lamb3 | 0.00043167 | 0.000204 | 0.00039075 | -0.0001867 | 0.99926662 | 0.99999994 |
| Mefv | 0.00035951 | 0.00013855 | 0.00032545 | -0.0001869 | 0.99993396 | 0.99999994 |
| Wfdc21 | 0.00113591 | 0.0008408 | 0.00102775 | -0.0001869 | 0.88701439 | 0.99999994 |
| Nadsyn1 | 0.00813045 | 0.00713447 | 0.00732144 | -0.000187 | 0.56749047 | 0.99999994 |
| Syt6 | 0.00090654 | 0.00063332 | 0.00082034 | -0.000187 | 0.93543873 | 0.99999994 |
| Gm572 | 0.01118096 | 0.00986058 | 0.01004768 | -0.0001871 | 0.54932153 | 0.99999994 |
| Zc3h12a | 0.00311821 | 0.00263035 | 0.00281749 | -0.0001871 | 0.67079933 | 0.99999994 |
| Trim30a | 0.00048004 | 0.00024734 | 0.00043452 | -0.0001872 | 0.99793103 | 0.99999994 |
| Cd86 | 0.00046336 | 0.0002321 | 0.00041943 | -0.0001873 | 0.99852507 | 0.99999994 |
| Gm9830 | 0.00146805 | 0.00114063 | 0.00132796 | -0.0001873 | 0.82613976 | 0.99999994 |

| | | | | | | |
|---|---|---|---|---|---|---|
| Tslp | 0.00032778 | 0.00010934 | 0.00029673 | -0.0001874 | 0.99998686 | 0.99999994 |
| Gm4791 | 0.0004351 | 0.00020627 | 0.00039386 | -0.0001876 | 0.99923866 | 0.99999994 |
| Gm14168 | 0.0013051 | 0.00099295 | 0.00118069 | -0.0001877 | 0.85507136 | 0.99999994 |
| Pmaip1 | 0.00034484 | 0.00012442 | 0.00031217 | -0.0001878 | 0.99996878 | 0.99999994 |
| Gm9923 | 0.00040797 | 0.00018151 | 0.00036931 | -0.0001878 | 0.99964393 | 0.99999994 |
| Mogat2 | 0.0030875 | 0.00260196 | 0.00278981 | -0.0001878 | 0.6729907 | 0.99999994 |
| Fam166b | 0.00408908 | 0.00350441 | 0.0036923 | -0.0001879 | 0.63258843 | 0.99999994 |
| Mlph | 0.00034101 | 0.00012061 | 0.0003087 | -0.0001881 | 0.99997501 | 0.99999994 |
| Ddr2 | 0.00033255 | 0.00011289 | 0.00030105 | -0.0001882 | 0.99998411 | 0.99999994 |
| Filip1l | 0.00435892 | 0.0037469 | 0.00393524 | -0.0001883 | 0.62497291 | 0.99999994 |
| Amy2a3 | 0.00087622 | 0.00060453 | 0.00079292 | -0.0001884 | 0.94312175 | 0.99999994 |
| Lat2 | 0.00312096 | 0.00263152 | 0.00281997 | -0.0001885 | 0.67177827 | 0.99999994 |
| Icam2 | 0.00102295 | 0.00073715 | 0.00092562 | -0.0001885 | 0.91234493 | 0.99999994 |
| Myl10 | 0.00037417 | 0.00015024 | 0.00033872 | -0.0001885 | 0.99989387 | 0.99999994 |
| Gm13398 | 0.00086352 | 0.00059282 | 0.00078144 | -0.0001886 | 0.94593644 | 0.99999994 |
| Cyp27b1 | 0.00039177 | 0.00016602 | 0.00035464 | -0.0001886 | 0.9998001 | 0.99999994 |
| Gm26558 | 0.00241775 | 0.00199697 | 0.00218562 | -0.0001887 | 0.71720352 | 0.99999994 |
| 1700120G07 | 0.0004553 | 0.00022339 | 0.00041213 | -0.0001887 | 0.9988498 | 0.99999994 |
| Morc1 | 0.00036765 | 0.00014396 | 0.00033281 | -0.0001889 | 0.99992069 | 0.99999994 |
| 4933427E11I | 0.00297846 | 0.0025024 | 0.00269148 | -0.0001891 | 0.67996466 | 0.99999994 |
| Gm37584 | 0.00068105 | 0.00042726 | 0.00061639 | -0.0001891 | 0.97944595 | 0.99999994 |
| Ttf2 | 0.01367484 | 0.01207887 | 0.01226812 | -0.0001892 | 0.54089195 | 0.99999994 |
| Klhl40 | 0.00037107 | 0.00014624 | 0.00033591 | -0.0001897 | 0.99991453 | 0.99999994 |
| I830134H01F | 0.00035157 | 0.00012859 | 0.00031827 | -0.0001897 | 0.9999636 | 0.99999994 |
| Ankrd9 | 0.01915916 | 0.01693526 | 0.01712496 | -0.0001897 | 0.52938977 | 0.99999994 |
| Prrx2 | 0.00143851 | 0.00111132 | 0.00130126 | -0.0001899 | 0.83438318 | 0.99999994 |
| Csn3 | 0.00037263 | 0.00014729 | 0.00033732 | -0.00019 | 0.99991158 | 0.99999994 |
| Scel | 0.00034181 | 0.00011937 | 0.00030943 | -0.0001901 | 0.99997829 | 0.99999994 |
| Diras2 | 0.0019958 | 0.00161462 | 0.0018047 | -0.0001901 | 0.75838063 | 0.99999994 |
| 1700003M07 | 0.00159185 | 0.0012496 | 0.00143983 | -0.0001902 | 0.81041723 | 0.99999994 |

| | | | | | | |
|---|---|---|---|---|---|---|
| Gm4961 | 0.00104902 | 0.00075887 | 0.00094918 | -0.0001903 | 0.9090044 | 0.99999994 |
| Gpatch3 | 0.02845976 | 0.02508952 | 0.02528016 | -0.0001906 | 0.52001733 | 0.99999994 |
| Slc25a18 | 0.00036278 | 0.00013773 | 0.0003284 | -0.0001907 | 0.9999444 | 0.99999994 |
| Gas7 | 0.00276841 | 0.00231126 | 0.00250202 | -0.0001908 | 0.69409604 | 0.99999994 |
| Zfp874a | 0.04010605 | 0.03515974 | 0.0353505 | -0.0001908 | 0.51432673 | 0.99999994 |
| A430073D23 | 0.00263728 | 0.00219296 | 0.00238372 | -0.0001908 | 0.70287584 | 0.99999994 |
| Shc3 | 0.00034686 | 0.00012319 | 0.000314 | -0.0001908 | 0.99997383 | 0.99999994 |
| Gjb4 | 0.00051672 | 0.00027683 | 0.00046771 | -0.0001909 | 0.99670273 | 0.99999994 |
| Bves | 0.00110776 | 0.00081137 | 0.0010023 | -0.0001909 | 0.89760143 | 0.99999994 |
| Abca9 | 0.0004252 | 0.00019384 | 0.0003849 | -0.0001911 | 0.99952378 | 0.99999994 |
| Gm10644 | 0.00263766 | 0.00219288 | 0.00238406 | -0.0001912 | 0.70325227 | 0.99999994 |
| Tcap | 0.0009243 | 0.0006451 | 0.00083641 | -0.0001913 | 0.93605275 | 0.99999994 |
| Gcm1 | 0.00043105 | 0.00019869 | 0.00039019 | -0.0001915 | 0.99945615 | 0.99999994 |
| Fbxw18 | 0.00046102 | 0.00022552 | 0.00041731 | -0.0001918 | 0.99889034 | 0.99999994 |
| Gm26771 | 0.00254692 | 0.00211038 | 0.00230219 | -0.0001918 | 0.71040809 | 0.99999994 |
| A930003A15 | 0.00034785 | 0.00012306 | 0.0003149 | -0.0001918 | 0.99997494 | 0.99999994 |
| Papolb | 0.00045879 | 0.00022335 | 0.00041529 | -0.0001919 | 0.99895259 | 0.99999994 |
| Prrx1 | 0.00191785 | 0.00154228 | 0.00173431 | -0.000192 | 0.76944045 | 0.99999994 |
| Gm16105 | 0.00053848 | 0.00029534 | 0.00048741 | -0.0001921 | 0.99564229 | 0.99999994 |
| Zfp934 | 0.00075222 | 0.00048848 | 0.00068077 | -0.0001923 | 0.96995885 | 0.99999994 |
| Bcat1 | 0.00035186 | 0.00012615 | 0.00031853 | -0.0001924 | 0.99997092 | 0.99999994 |
| Kcnmb4os2 | 0.00689155 | 0.00601857 | 0.00621102 | -0.0001924 | 0.58170133 | 0.99999994 |
| Igsf9b | 0.00155071 | 0.00121017 | 0.00140265 | -0.0001925 | 0.81949772 | 0.99999994 |
| 4930426L09F | 0.00036947 | 0.00014192 | 0.00033447 | -0.0001925 | 0.99993643 | 0.99999994 |
| Cers1 | 0.00035438 | 0.00012804 | 0.00032081 | -0.0001928 | 0.99996828 | 0.99999994 |
| Gm4707 | 0.02756783 | 0.02430957 | 0.02450247 | -0.0001929 | 0.52089686 | 0.99999994 |
| Klf12 | 0.00041611 | 0.00018376 | 0.00037667 | -0.0001929 | 0.99967402 | 0.99999994 |
| Lrrc25 | 0.00035534 | 0.00012874 | 0.00032168 | -0.0001929 | 0.99996731 | 0.99999994 |
| Gm15445 | 0.00071822 | 0.00045691 | 0.00065002 | -0.0001931 | 0.97600792 | 0.99999994 |
| Brinp1 | 0.0004473 | 0.0002116 | 0.0004049 | -0.0001933 | 0.99925774 | 0.99999994 |

| | | | | | | |
|---|---|---|---|---|---|---|
| Ifnk | 0.00053212 | 0.00028832 | 0.00048164 | -0.0001933 | 0.9962266 | 0.99999994 |
| Fsip1 | 0.00304743 | 0.00256011 | 0.00275367 | -0.0001936 | 0.68006628 | 0.99999994 |
| Gm16251 | 0.00040778 | 0.0001755 | 0.00036913 | -0.0001936 | 0.99976006 | 0.99999994 |
| Ncapg2 | 0.00511288 | 0.00441978 | 0.00461355 | -0.0001938 | 0.61009589 | 0.99999994 |
| Nrgn | 0.00428624 | 0.00367575 | 0.00386981 | -0.0001941 | 0.63073075 | 0.99999994 |
| Zfp109 | 0.00284486 | 0.00237662 | 0.00257098 | -0.0001944 | 0.69259233 | 0.99999994 |
| Etnppl | 0.00036224 | 0.00013349 | 0.00032792 | -0.0001944 | 0.99996038 | 0.99999994 |
| Plet1os | 0.00136983 | 0.00104465 | 0.0012392 | -0.0001945 | 0.85199435 | 0.99999994 |
| Nkx6-3 | 0.00263571 | 0.00218772 | 0.00238231 | -0.0001946 | 0.70667313 | 0.99999994 |
| 4833412C05I | 0.00062161 | 0.0003677 | 0.00056262 | -0.0001949 | 0.98944649 | 0.99999994 |
| Tmem151b | 0.01884813 | 0.01665549 | 0.01685048 | -0.000195 | 0.53069759 | 0.99999994 |
| Bcl2a1b | 0.00036951 | 0.00013925 | 0.0003345 | -0.0001952 | 0.99994892 | 0.99999994 |
| Egr3 | 0.00039659 | 0.00016351 | 0.00035901 | -0.0001955 | 0.99985541 | 0.99999994 |
| Fli1 | 0.00138557 | 0.00105786 | 0.00125342 | -0.0001956 | 0.85048331 | 0.99999994 |
| Kcnab2 | 0.00117637 | 0.00086865 | 0.00106433 | -0.0001957 | 0.88948156 | 0.99999994 |
| Gm12728 | 0.0021614 | 0.00175854 | 0.00195423 | -0.0001957 | 0.7474704 | 0.99999994 |
| Nr1i3 | 0.000372 | 0.00014083 | 0.00033675 | -0.0001959 | 0.99994618 | 0.99999994 |
| Gm20463 | 0.0004725 | 0.00023167 | 0.0004277 | -0.000196 | 0.9988589 | 0.99999994 |
| 9130204K15I | 0.00042139 | 0.00018535 | 0.00038145 | -0.0001961 | 0.9996891 | 0.99999994 |
| Gm38134 | 0.00072489 | 0.00045992 | 0.00065605 | -0.0001961 | 0.97669871 | 0.99999994 |
| Cxcl1 | 0.00070456 | 0.0004413 | 0.00063766 | -0.0001964 | 0.97980611 | 0.99999994 |
| Gm17196 | 0.0004022 | 0.00016769 | 0.00036409 | -0.0001964 | 0.99983503 | 0.99999994 |
| AI838599 | 0.00184964 | 0.00147629 | 0.0016727 | -0.0001964 | 0.78277734 | 0.99999994 |
| Vstm4 | 0.00037718 | 0.00014473 | 0.00034144 | -0.0001967 | 0.9999372 | 0.99999994 |
| Gm17096 | 0.00040551 | 0.00017004 | 0.00036708 | -0.000197 | 0.9998235 | 0.99999994 |
| 3830408C21I | 0.00102679 | 0.00073198 | 0.00092908 | -0.0001971 | 0.92104778 | 0.99999994 |
| Parvaos | 0.00042603 | 0.00018837 | 0.00038565 | -0.0001973 | 0.99966927 | 0.99999994 |
| Amy2a2 | 0.00038203 | 0.00014843 | 0.00034583 | -0.0001974 | 0.99992749 | 0.99999994 |
| Zfp385a | 0.03151661 | 0.02774107 | 0.02793849 | -0.0001974 | 0.51875754 | 0.99999994 |
| Snx33 | 0.00639891 | 0.00557152 | 0.00576896 | -0.0001974 | 0.59010244 | 0.99999994 |

| | | | | | | |
|---|---|---|---|---|---|---|
| Pld2 | 0.00093806 | 0.00065129 | 0.00084885 | -0.0001976 | 0.93933355 | 0.99999994 |
| Ppm1j | 0.00149304 | 0.00115265 | 0.00135054 | -0.0001979 | 0.83530096 | 0.99999994 |
| Nrn1l | 0.00050289 | 0.0002572 | 0.0004552 | -0.000198 | 0.99810634 | 0.99999994 |
| Slc6a2 | 0.00041369 | 0.00017643 | 0.00037448 | -0.0001981 | 0.9997845 | 0.99999994 |
| Pp2d1 | 0.00042187 | 0.00018371 | 0.00038189 | -0.0001982 | 0.999724 | 0.99999994 |
| Fgf1 | 0.00039132 | 0.00015606 | 0.00035424 | -0.0001982 | 0.99990195 | 0.99999994 |
| Myo3a | 0.00048572 | 0.00024142 | 0.00043966 | -0.0001982 | 0.99865609 | 0.99999994 |
| 1700001L05I | 0.00351027 | 0.00297257 | 0.0031709 | -0.0001983 | 0.661416 | 0.99999994 |
| 1500035N22 | 0.00049761 | 0.0002519 | 0.00045042 | -0.0001985 | 0.99832586 | 0.99999994 |
| 1700093K21I | 0.00045781 | 0.00021585 | 0.00041441 | -0.0001986 | 0.99928672 | 0.99999994 |
| Gm26660 | 0.00224807 | 0.00183358 | 0.00203247 | -0.0001989 | 0.74259057 | 0.99999994 |
| Kif14 | 0.00039785 | 0.00016125 | 0.00036015 | -0.0001989 | 0.99988154 | 0.99999994 |
| Dnah14 | 0.0010146 | 0.00071905 | 0.00091806 | -0.000199 | 0.92547652 | 0.99999994 |
| Nfe2 | 0.0004651 | 0.00022187 | 0.00042101 | -0.0001991 | 0.99917925 | 0.99999994 |
| Gm7854 | 0.00044567 | 0.00020398 | 0.00040342 | -0.0001994 | 0.99950034 | 0.99999994 |
| Ltbp2 | 0.00040962 | 0.00017133 | 0.0003708 | -0.0001995 | 0.99982865 | 0.99999994 |
| Fcgbp | 0.00188083 | 0.00150134 | 0.00170087 | -0.0001995 | 0.78256043 | 0.99999994 |
| Btbd16 | 0.00048683 | 0.00024104 | 0.00044067 | -0.0001996 | 0.99871702 | 0.99999994 |
| Uchl1os | 0.00047627 | 0.0002309 | 0.00043112 | -0.0002002 | 0.99900356 | 0.99999994 |
| Zfp626 | 0.02112404 | 0.01865608 | 0.01885634 | -0.0002003 | 0.52817883 | 0.99999994 |
| Bean1 | 0.00043968 | 0.00019774 | 0.000398 | -0.0002003 | 0.99959493 | 0.99999994 |
| Plek2 | 0.01733886 | 0.01531647 | 0.0155169 | -0.0002004 | 0.53425882 | 0.99999994 |
| Amn | 0.00041534 | 0.00017545 | 0.00037598 | -0.0002005 | 0.99980746 | 0.99999994 |
| Vwa3a | 0.0010186 | 0.00072108 | 0.00092168 | -0.0002006 | 0.92629025 | 0.99999994 |
| Rerg | 0.0007404 | 0.00046947 | 0.00067007 | -0.0002006 | 0.97685473 | 0.99999994 |
| Leap2 | 0.00044766 | 0.00020419 | 0.00040522 | -0.000201 | 0.9995205 | 0.99999994 |
| Cyp17a1 | 0.00046304 | 0.00021805 | 0.00041914 | -0.0002011 | 0.99929723 | 0.99999994 |
| Sctr | 0.00046021 | 0.00021544 | 0.00041658 | -0.0002011 | 0.99934546 | 0.99999994 |
| Gm17249 | 0.00693074 | 0.0060449 | 0.00624618 | -0.0002013 | 0.58492034 | 0.99999994 |
| Snx16 | 0.14570311 | 0.11891827 | 0.1191197 | -0.0002014 | 0.50449031 | 0.99999994 |

| | | | | | | |
|---|---|---|---|---|---|---|
| Pnpla3 | 0.00211508 | 0.00171098 | 0.00191241 | -0.0002014 | 0.75838405 | 0.99999994 |
| Gimap4 | 0.00202675 | 0.00163103 | 0.00183265 | -0.0002016 | 0.76801303 | 0.99999994 |
| Zfp473 | 0.00243081 | 0.00199562 | 0.00219741 | -0.0002018 | 0.72948031 | 0.99999994 |
| Rab8b | 0.01829079 | 0.01615642 | 0.01635833 | -0.0002019 | 0.53273962 | 0.99999994 |
| Liph | 0.00129652 | 0.00097099 | 0.00117293 | -0.0002019 | 0.8741033 | 0.99999994 |
| Gm20878 | 0.00233592 | 0.00190972 | 0.00211177 | -0.0002021 | 0.73789638 | 0.99999994 |
| Rspry1 | 0.15538356 | 0.12586764 | 0.12607002 | -0.0002024 | 0.50426269 | 0.99999994 |
| Gm16596 | 0.00119216 | 0.00087598 | 0.0010786 | -0.0002026 | 0.89443092 | 0.99999994 |
| Gm27010 | 0.00372422 | 0.00316098 | 0.00336368 | -0.0002027 | 0.65582913 | 0.99999994 |
| Dmbt1 | 0.00074016 | 0.00046699 | 0.00066986 | -0.0002029 | 0.97809519 | 0.99999994 |
| Scx | 0.00533616 | 0.00461124 | 0.00481429 | -0.0002031 | 0.61054611 | 0.99999994 |
| Gm26935 | 0.00502117 | 0.00432769 | 0.00453108 | -0.0002034 | 0.61745132 | 0.99999994 |
| Nkg7 | 0.0012353 | 0.00091395 | 0.0011176 | -0.0002036 | 0.88741552 | 0.99999994 |
| 4930540M05 | 0.0013318 | 0.00100109 | 0.00120483 | -0.0002037 | 0.86983555 | 0.99999994 |
| Ptprc | 0.0009724 | 0.00067594 | 0.0008799 | -0.000204 | 0.93857837 | 0.99999994 |
| Spag9 | 0.18372956 | 0.14559935 | 0.14580371 | -0.0002044 | 0.50372198 | 0.99999994 |
| P2ry6 | 0.00300153 | 0.00250756 | 0.00271228 | -0.0002047 | 0.69230956 | 0.99999994 |
| Gm14322 | 0.00456984 | 0.00391986 | 0.00412506 | -0.0002052 | 0.62972294 | 0.99999994 |
| Gm5117 | 0.0018967 | 0.00150961 | 0.0017152 | -0.0002056 | 0.78752179 | 0.99999994 |
| Gm29340 | 0.00131256 | 0.00098166 | 0.00118744 | -0.0002058 | 0.87565213 | 0.99999994 |
| Tube1 | 0.00641656 | 0.00557895 | 0.0057848 | -0.0002059 | 0.59361891 | 0.99999994 |
| Tdrp | 0.02193276 | 0.01936163 | 0.01956762 | -0.000206 | 0.52793173 | 0.99999994 |
| Il22 | 0.00070552 | 0.0004322 | 0.00063853 | -0.0002063 | 0.98425686 | 0.99999994 |
| Astn1 | 0.00136134 | 0.00102504 | 0.00123153 | -0.0002065 | 0.86779999 | 0.99999994 |
| Abcc3 | 0.02392668 | 0.02111134 | 0.02131803 | -0.0002067 | 0.52572844 | 0.99999994 |
| A930004D18 | 0.00441269 | 0.00377694 | 0.00398364 | -0.0002067 | 0.63509548 | 0.99999994 |
| Tnxb | 0.0009246 | 0.00062968 | 0.00083668 | -0.000207 | 0.95020086 | 0.99999994 |
| Pggt1b | 0.10894174 | 0.09149432 | 0.09170141 | -0.0002071 | 0.5059968 | 0.99999994 |
| Rpl21-ps4 | 0.0020595 | 0.00165474 | 0.00186222 | -0.0002075 | 0.77084214 | 0.99999994 |
| Gtpbp10 | 0.05384924 | 0.04682826 | 0.04703596 | -0.0002077 | 0.51172402 | 0.99999994 |

| | | | | | | |
|---|---|---|---|---|---|---|
| Foxs1 | 0.00110802 | 0.00079397 | 0.00100253 | -0.0002086 | 0.91693677 | 0.99999994 |
| Ms4a6b | 0.00220132 | 0.00178145 | 0.00199027 | -0.0002088 | 0.75753221 | 0.99999994 |
| Gm13031 | 0.00087892 | 0.00058637 | 0.00079537 | -0.000209 | 0.95986444 | 0.99999994 |
| Igsf10 | 0.00170484 | 0.00133269 | 0.00154191 | -0.0002092 | 0.81678182 | 0.99999994 |
| Cd1d2 | 0.00146212 | 0.00111321 | 0.0013226 | -0.0002094 | 0.85401827 | 0.99999994 |
| Mmp7 | 0.00193567 | 0.00154088 | 0.00175041 | -0.0002095 | 0.78720564 | 0.99999994 |
| Maml2 | 0.00513859 | 0.0044268 | 0.00463667 | -0.0002099 | 0.61840222 | 0.99999994 |
| Gm29455 | 0.00364674 | 0.0030838 | 0.00329387 | -0.0002101 | 0.66439989 | 0.99999994 |
| Gm28151 | 0.00146421 | 0.00111439 | 0.00132449 | -0.0002101 | 0.85448157 | 0.99999994 |
| Wdr72 | 0.00137592 | 0.00103445 | 0.0012447 | -0.0002103 | 0.86957425 | 0.99999994 |
| Pcsk7 | 0.03348537 | 0.02943458 | 0.02964486 | -0.0002103 | 0.51882866 | 0.99999994 |
| H2-DMb2 | 0.00146045 | 0.00111053 | 0.00132109 | -0.0002106 | 0.85564116 | 0.99999994 |
| Alpk2 | 0.00164532 | 0.00127735 | 0.00148813 | -0.0002108 | 0.82711936 | 0.99999994 |
| 4833422M21 | 0.00100439 | 0.00069781 | 0.00090883 | -0.000211 | 0.93888934 | 0.99999994 |
| Zfp541 | 0.00060434 | 0.00033589 | 0.00054699 | -0.0002111 | 0.99489969 | 0.99999994 |
| Mctp1 | 0.00478186 | 0.00410447 | 0.00431583 | -0.0002114 | 0.62777957 | 0.99999994 |
| Ccl5 | 0.00139115 | 0.00104706 | 0.00125847 | -0.0002114 | 0.86826023 | 0.99999994 |
| Cysltr1 | 0.00099648 | 0.00068997 | 0.00090167 | -0.0002117 | 0.94095421 | 0.99999994 |
| Kif26b | 0.00212958 | 0.00171352 | 0.0019255 | -0.000212 | 0.76816194 | 0.99999994 |
| Dscc1 | 0.01017393 | 0.00893684 | 0.00914894 | -0.0002121 | 0.56131748 | 0.99999994 |
| Tas1r3 | 0.0018345 | 0.00144692 | 0.00165903 | -0.0002121 | 0.80261897 | 0.99999994 |
| Pars2 | 0.01660807 | 0.01465776 | 0.0148702 | -0.0002124 | 0.53788148 | 0.99999994 |
| Rassf4 | 0.05385135 | 0.04682496 | 0.04703774 | -0.0002128 | 0.51201064 | 0.99999994 |
| Pip5kl1 | 0.00171302 | 0.00133642 | 0.0015493 | -0.0002129 | 0.81979557 | 0.99999994 |
| Vegfc | 0.00155889 | 0.00119711 | 0.00141004 | -0.0002129 | 0.84259543 | 0.99999994 |
| Traf3ip3 | 0.00085156 | 0.00055738 | 0.00077062 | -0.0002132 | 0.96725506 | 0.99999994 |
| Rapgef3os2 | 0.0014026 | 0.00105549 | 0.00126882 | -0.0002133 | 0.86846219 | 0.99999994 |
| Ybey | 0.03691958 | 0.03239721 | 0.03261068 | -0.0002135 | 0.51737775 | 0.99999994 |
| Slc6a9 | 0.00474984 | 0.00407353 | 0.00428702 | -0.0002135 | 0.62985892 | 0.99999994 |
| Milr1 | 0.00843075 | 0.0073768 | 0.00759032 | -0.0002135 | 0.57426685 | 0.99999994 |

| | | | | | | |
|---|---|---|---|---|---|---|
| Cdhr2 | 0.01937361 | 0.01710055 | 0.01731415 | -0.0002136 | 0.53272297 | 0.99999994 |
| Ubtf | 0.32610572 | 0.23206721 | 0.2322812 | -0.000214 | 0.50244638 | 0.99999994 |
| Slc25a47 | 0.0018943 | 0.00149893 | 0.00171304 | -0.0002141 | 0.79729329 | 0.99999994 |
| Gm26704 | 0.00093856 | 0.00063486 | 0.00084931 | -0.0002144 | 0.95358658 | 0.99999994 |
| Tmem129 | 0.13361303 | 0.11006858 | 0.11028329 | -0.0002147 | 0.50516984 | 0.99999994 |
| Zfp599 | 0.03473363 | 0.03050888 | 0.03072443 | -0.0002156 | 0.51862362 | 0.99999994 |
| Gm38393 | 0.00357512 | 0.00301339 | 0.00322934 | -0.0002159 | 0.67187932 | 0.99999994 |
| Kif21b | 0.00437306 | 0.00373169 | 0.00394796 | -0.0002163 | 0.64231316 | 0.99999994 |
| Nek10 | 0.00204024 | 0.00162849 | 0.00184483 | -0.0002163 | 0.78247275 | 0.99999994 |
| Hoxb3 | 0.00124872 | 0.00091306 | 0.00112973 | -0.0002167 | 0.89912929 | 0.99999994 |
| Rbm15 | 0.0730992 | 0.06283693 | 0.06305366 | -0.0002167 | 0.50912657 | 0.99999994 |
| Gm5771 | 0.00233832 | 0.00189687 | 0.00211393 | -0.0002171 | 0.75285468 | 0.99999994 |
| Scml2 | 0.00072338 | 0.00043739 | 0.00065468 | -0.0002173 | 0.98642229 | 0.99999994 |
| Etv5 | 0.02438361 | 0.02150111 | 0.0217185 | -0.0002174 | 0.52656061 | 0.99999994 |
| Gm4978 | 0.00183064 | 0.00143806 | 0.00165555 | -0.0002175 | 0.80905562 | 0.99999994 |
| Nat2 | 0.01754847 | 0.01548479 | 0.01570227 | -0.0002175 | 0.53672806 | 0.99999994 |
| Ifi202b | 0.00110585 | 0.00078301 | 0.00100057 | -0.0002176 | 0.92609363 | 0.99999994 |
| Wfdc16 | 0.0087445 | 0.00765353 | 0.00787112 | -0.0002176 | 0.57299504 | 0.99999994 |
| Tgfb2 | 0.02469159 | 0.02177057 | 0.02198829 | -0.0002177 | 0.52627492 | 0.99999994 |
| Gm42918 | 0.00081406 | 0.0005189 | 0.0007367 | -0.0002178 | 0.97545936 | 0.99999994 |
| Cuzd1 | 0.00120549 | 0.00087256 | 0.00109065 | -0.0002181 | 0.90840353 | 0.99999994 |
| Cacna1g | 0.00117184 | 0.00084207 | 0.00106023 | -0.0002182 | 0.91460365 | 0.99999994 |
| Crkl | 0.01570874 | 0.01385519 | 0.0140735 | -0.0002183 | 0.54111877 | 0.99999994 |
| Zscan25 | 0.00066768 | 0.00038584 | 0.0006043 | -0.0002185 | 0.9919415 | 0.99999994 |
| Itih2 | 0.00827768 | 0.00723466 | 0.00745328 | -0.0002186 | 0.57739738 | 0.99999994 |
| Ampd3 | 0.00131587 | 0.00097148 | 0.00119043 | -0.000219 | 0.88957876 | 0.99999994 |
| Prdm10 | 0.01389241 | 0.01224207 | 0.01246148 | -0.0002194 | 0.54664856 | 0.99999994 |
| Zpbp2 | 0.00246314 | 0.00200693 | 0.00222658 | -0.0002197 | 0.74430208 | 0.99999994 |
| Pld4 | 0.00646428 | 0.00560772 | 0.00582763 | -0.0002199 | 0.59916312 | 0.99999994 |
| Clec14a | 0.00080249 | 0.00050543 | 0.00072624 | -0.0002208 | 0.97850645 | 0.99999994 |

| | | | | | | |
|---|---|---|---|---|---|---|
| Slc22a5 | 0.00207518 | 0.00165535 | 0.00187638 | -0.000221 | 0.78350977 | 0.99999994 |
| Cth | 0.00523964 | 0.00450648 | 0.00472752 | -0.000221 | 0.62218753 | 0.99999994 |
| Camk1 | 0.19538365 | 0.15343557 | 0.15365689 | -0.0002213 | 0.50382483 | 0.99999994 |
| Tex11 | 0.00129069 | 0.00094632 | 0.00116767 | -0.0002213 | 0.89648418 | 0.99999994 |
| Cbs | 0.02370629 | 0.02090306 | 0.02112479 | -0.0002217 | 0.52784931 | 0.99999994 |
| Cnn1 | 0.00267026 | 0.00219089 | 0.00241348 | -0.0002226 | 0.73035892 | 0.99999994 |
| Cbln3 | 0.0022047 | 0.0017707 | 0.00199331 | -0.0002226 | 0.77137973 | 0.99999994 |
| Alkbh8 | 0.09262903 | 0.07867808 | 0.07890087 | -0.0002228 | 0.50749782 | 0.99999994 |
| Aspdh | 0.00108533 | 0.00075911 | 0.00098202 | -0.0002229 | 0.93459977 | 0.99999994 |
| Gm6277 | 0.002771 | 0.00228127 | 0.00250436 | -0.0002231 | 0.72339188 | 0.99999994 |
| Phkg1 | 0.0011595 | 0.00082585 | 0.00104907 | -0.0002232 | 0.92165978 | 0.99999994 |
| Nrxn3 | 0.0009163 | 0.00060581 | 0.00082917 | -0.0002234 | 0.96351951 | 0.99999994 |
| Cfap61 | 0.0040084 | 0.0033961 | 0.00361964 | -0.0002235 | 0.65949243 | 0.99999994 |
| Ina | 0.00913004 | 0.00799223 | 0.00821601 | -0.0002238 | 0.5719343 | 0.99999994 |
| Des | 0.00471546 | 0.004032 | 0.00425609 | -0.0002241 | 0.6370062 | 0.99999994 |
| Tbc1d12 | 0.01794445 | 0.01582735 | 0.01605232 | -0.000225 | 0.53716272 | 0.99999994 |
| Gpam | 0.00546209 | 0.00470237 | 0.00492749 | -0.0002251 | 0.61947484 | 0.99999994 |
| Sprtn | 0.00557659 | 0.0048052 | 0.0050304 | -0.0002252 | 0.61714051 | 0.99999994 |
| Tcf7 | 0.00085924 | 0.00055229 | 0.00077757 | -0.0002253 | 0.97310202 | 0.99999994 |
| Zfand4 | 0.00247426 | 0.00201109 | 0.00223662 | -0.0002255 | 0.74895555 | 0.99999994 |
| Ccdc116 | 0.00193329 | 0.00152272 | 0.00174825 | -0.0002255 | 0.80475173 | 0.99999994 |
| Gm13420 | 0.00467606 | 0.00399496 | 0.00422064 | -0.0002257 | 0.63905068 | 0.99999994 |
| Gm5602 | 0.00085143 | 0.00054478 | 0.00077051 | -0.0002257 | 0.97441188 | 0.99999994 |
| Dlec1 | 0.00191332 | 0.00150448 | 0.00173022 | -0.0002257 | 0.80742746 | 0.99999994 |
| Tnfrsf25 | 0.00071114 | 0.00041781 | 0.00064361 | -0.0002258 | 0.99023533 | 0.99999994 |
| Kcnc2 | 0.00362211 | 0.00304565 | 0.00327168 | -0.000226 | 0.67719065 | 0.99999994 |
| Ptpn22 | 0.0011996 | 0.00085858 | 0.00108532 | -0.0002267 | 0.91782626 | 0.99999994 |
| Gm13652 | 0.00081514 | 0.00051086 | 0.00073768 | -0.0002268 | 0.97965399 | 0.99999994 |
| Acacb | 0.00263119 | 0.00215133 | 0.00237823 | -0.0002269 | 0.73730436 | 0.99999994 |
| Pabpc5 | 0.00565956 | 0.00487788 | 0.00510495 | -0.0002271 | 0.61641391 | 0.99999994 |

| | | | | | | |
|---|---|---|---|---|---|---|
| Spag17 | 0.0021014 | 0.00167286 | 0.00190005 | -0.0002272 | 0.78695871 | 0.99999994 |
| Mvb12b | 0.05994634 | 0.05192525 | 0.05215292 | -0.0002277 | 0.51159097 | 0.99999994 |
| Epb41l2 | 0.01452 | 0.01279119 | 0.01301892 | -0.0002277 | 0.54634613 | 0.99999994 |
| Lrrc74b | 0.00665457 | 0.00577061 | 0.0059984 | -0.0002278 | 0.5997809 | 0.99999994 |
| Acot5 | 0.00063462 | 0.00034651 | 0.00057439 | -0.0002279 | 0.99586361 | 0.99999994 |
| Dse | 0.00137373 | 0.00101468 | 0.00124272 | -0.000228 | 0.88904163 | 0.99999994 |
| Aipl1 | 0.00075122 | 0.00045163 | 0.00067987 | -0.0002282 | 0.98727689 | 0.99999994 |
| Slc38a7 | 0.02057694 | 0.0181463 | 0.01837471 | -0.0002284 | 0.53297212 | 0.99999994 |
| Lurap1 | 0.01305899 | 0.01149182 | 0.01172049 | -0.0002287 | 0.55166497 | 0.99999994 |
| Cyb561a3 | 0.0008662 | 0.00055505 | 0.00078387 | -0.0002288 | 0.97399591 | 0.99999994 |
| Fam133b | 0.31614895 | 0.22663244 | 0.2268613 | -0.0002289 | 0.50267896 | 0.99999994 |
| Lancl3 | 0.01009447 | 0.00884897 | 0.00907797 | -0.000229 | 0.56667465 | 0.99999994 |
| Gm5547 | 0.00691473 | 0.0060028 | 0.00623181 | -0.000229 | 0.5966245 | 0.99999994 |
| Mesp2 | 0.0124929 | 0.01098731 | 0.01121671 | -0.0002294 | 0.55414202 | 0.99999994 |
| Nutf2-ps1 | 0.00643168 | 0.0055687 | 0.00579837 | -0.0002297 | 0.60397645 | 0.99999994 |
| 1700030J22F | 0.01288957 | 0.01133972 | 0.01156975 | -0.00023 | 0.55264363 | 0.99999994 |
| AI662270 | 0.00150734 | 0.00113343 | 0.00136347 | -0.00023 | 0.86928821 | 0.99999994 |
| Asb16 | 0.001023 | 0.00069552 | 0.00092566 | -0.0002301 | 0.95102952 | 0.99999994 |
| Ppargc1b | 0.00114157 | 0.00080272 | 0.00103286 | -0.0002301 | 0.93098656 | 0.99999994 |
| Thnsl1 | 0.02700198 | 0.0237779 | 0.02400861 | -0.0002307 | 0.52550014 | 0.99999994 |
| Gm5449 | 0.00077422 | 0.00046989 | 0.00070067 | -0.0002308 | 0.9858262 | 0.99999994 |
| Mfap3l | 0.00114262 | 0.00080271 | 0.00103381 | -0.0002311 | 0.93162444 | 0.99999994 |
| Gm13563 | 0.00146332 | 0.00109248 | 0.00132369 | -0.0002312 | 0.87751816 | 0.99999994 |
| 9330136K24I | 0.00182393 | 0.00141826 | 0.00164948 | -0.0002312 | 0.82460886 | 0.99999994 |
| Gpr21 | 0.00153681 | 0.00115886 | 0.00139009 | -0.0002312 | 0.8659023 | 0.99999994 |
| Zar1 | 0.00283657 | 0.00233167 | 0.00256351 | -0.0002318 | 0.72640269 | 0.99999994 |
| Gm14827 | 0.00124524 | 0.00089441 | 0.00112658 | -0.0002322 | 0.91493278 | 0.99999994 |
| Vsir | 0.00341018 | 0.00284758 | 0.00308069 | -0.0002331 | 0.69275793 | 0.99999994 |
| Mettl2 | 0.04958884 | 0.04320292 | 0.04343607 | -0.0002331 | 0.51425051 | 0.99999994 |
| Pole2 | 0.00870665 | 0.00760407 | 0.00783725 | -0.0002332 | 0.57849452 | 0.99999994 |

| | | | | | | |
|---|---|---|---|---|---|---|
| Fads2 | 0.00805577 | 0.00702108 | 0.00725456 | -0.0002335 | 0.58481306 | 0.99999994 |
| Atg14 | 0.06871538 | 0.05920739 | 0.05944117 | -0.0002338 | 0.51044324 | 0.99999994 |
| Efhc2 | 0.00729514 | 0.00633886 | 0.00657297 | -0.0002341 | 0.59370131 | 0.99999994 |
| Rnf32 | 0.04422659 | 0.03864214 | 0.03887635 | -0.0002342 | 0.51599372 | 0.99999994 |
| Areg | 0.00072138 | 0.00041849 | 0.00065287 | -0.0002344 | 0.99156741 | 0.99999994 |
| Tmed11 | 0.0025197 | 0.00204313 | 0.00227762 | -0.0002345 | 0.75342498 | 0.99999994 |
| Cables1 | 0.01075225 | 0.00943046 | 0.00966522 | -0.0002348 | 0.56421934 | 0.99999994 |
| Gm12764 | 0.0010596 | 0.00072361 | 0.00095875 | -0.0002351 | 0.94871498 | 0.99999994 |
| Wnt5b | 0.01045533 | 0.00916491 | 0.0094002 | -0.0002353 | 0.56616037 | 0.99999994 |
| Gm15743 | 0.0017957 | 0.00138862 | 0.00162399 | -0.0002354 | 0.83265558 | 0.99999994 |
| Senp2 | 0.06759228 | 0.05827691 | 0.05851237 | -0.0002355 | 0.51068455 | 0.99999994 |
| Cxcl10 | 0.00104545 | 0.00071026 | 0.00094596 | -0.0002357 | 0.95139388 | 0.99999994 |
| Ctsh | 0.06502673 | 0.05614971 | 0.05638556 | -0.0002359 | 0.51110607 | 0.99999994 |
| Klhl35 | 0.00182433 | 0.00141377 | 0.00164984 | -0.0002361 | 0.82955976 | 0.99999994 |
| Ngef | 0.00266034 | 0.0021684 | 0.00240452 | -0.0002361 | 0.74332989 | 0.99999994 |
| Fmo4 | 0.00340163 | 0.00283679 | 0.00307298 | -0.0002362 | 0.69554036 | 0.99999994 |
| Nudt15 | 0.00517747 | 0.00443507 | 0.00467162 | -0.0002366 | 0.63196409 | 0.99999994 |
| Bst1 | 0.00171043 | 0.00131001 | 0.00154695 | -0.0002369 | 0.84602823 | 0.99999994 |
| Gm10330 | 0.00256603 | 0.00208213 | 0.00231943 | -0.0002373 | 0.75206788 | 0.99999994 |
| 2810433D01 | 0.00329579 | 0.00273994 | 0.00297759 | -0.0002376 | 0.70237741 | 0.99999994 |
| Gm17529 | 0.00190699 | 0.00148658 | 0.0017245 | -0.0002379 | 0.82077358 | 0.99999994 |
| Ggps1 | 0.13239032 | 0.10914204 | 0.10937997 | -0.0002379 | 0.50577626 | 0.99999994 |
| Tpd52l1 | 0.0256928 | 0.02262589 | 0.02286456 | -0.0002387 | 0.52769788 | 0.99999994 |
| Tigd3 | 0.00059517 | 0.00029967 | 0.0005387 | -0.000239 | 0.99842879 | 0.99999994 |
| Gm8909 | 0.00127297 | 0.00091248 | 0.00115165 | -0.0002392 | 0.91657383 | 0.99999994 |
| Slc1a1 | 0.00055873 | 0.00026654 | 0.00050572 | -0.0002392 | 0.99917864 | 0.99999994 |
| Pzp | 0.00168485 | 0.00128407 | 0.00152385 | -0.0002398 | 0.85253606 | 0.99999994 |
| Ctnna3 | 0.00093546 | 0.00060647 | 0.0008465 | -0.00024 | 0.97045081 | 0.99999994 |
| Erbb2 | 0.01649609 | 0.01453075 | 0.01477106 | -0.0002403 | 0.54311784 | 0.99999994 |
| Dock2 | 0.00075071 | 0.00043893 | 0.00067941 | -0.0002405 | 0.9907635 | 0.99999994 |

| | | | | | | |
|---|---|---|---|---|---|---|
| Chdh | 0.00112654 | 0.00077849 | 0.00101928 | -0.0002408 | 0.94207829 | 0.99999994 |
| Gm13091 | 0.00068558 | 0.00037929 | 0.00062049 | -0.0002412 | 0.99516649 | 0.99999994 |
| Acot12 | 0.00037141 | 9.48E-05 | 0.00033621 | -0.0002414 | 0.99999912 | 0.99999994 |
| Zfp180 | 0.03972202 | 0.03477948 | 0.03502092 | -0.0002414 | 0.5183005 | 0.99999994 |
| Slc6a15 | 0.00054183 | 0.00024846 | 0.00049043 | -0.000242 | 0.99948854 | 0.99999994 |
| Stam | 0.08708868 | 0.07420332 | 0.07444562 | -0.0002423 | 0.50864249 | 0.99999994 |
| Rgs5 | 0.0018662 | 0.00144534 | 0.00168766 | -0.0002423 | 0.83039733 | 0.99999994 |
| 9530052E02I | 0.0018499 | 0.00143056 | 0.00167294 | -0.0002424 | 0.8325759 | 0.99999994 |
| Sostdc1 | 0.00201573 | 0.00158024 | 0.0018227 | -0.0002425 | 0.81204193 | 0.99999994 |
| Rab26 | 0.01421073 | 0.01250136 | 0.01274428 | -0.0002429 | 0.55048244 | 0.99999994 |
| Gm16287 | 0.00155673 | 0.00116486 | 0.00140809 | -0.0002432 | 0.874883 | 0.99999994 |
| Kif5a | 0.00375376 | 0.00314685 | 0.00339029 | -0.0002434 | 0.68365534 | 0.99999994 |
| Luzp2 | 0.00072153 | 0.00040954 | 0.00065301 | -0.0002435 | 0.9934633 | 0.99999994 |
| Clybl | 0.05291259 | 0.04600271 | 0.04624625 | -0.0002435 | 0.51398134 | 0.99999994 |
| Lrrn4 | 0.00150534 | 0.00111804 | 0.00136166 | -0.0002436 | 0.88315108 | 0.99999994 |
| Cpz | 0.00056593 | 0.0002686 | 0.00051224 | -0.0002436 | 0.99922723 | 0.99999994 |
| Foxr1 | 0.00077218 | 0.00045512 | 0.00069882 | -0.0002437 | 0.98986433 | 0.99999994 |
| Lrr1 | 0.00344747 | 0.00287054 | 0.0031143 | -0.0002438 | 0.69881639 | 0.99999994 |
| Pex5l | 0.08584537 | 0.07319763 | 0.07344146 | -0.0002438 | 0.508816 | 0.99999994 |
| 4930570G19 | 0.0011214 | 0.00077071 | 0.00101463 | -0.0002439 | 0.94522263 | 0.99999994 |
| AI837181 | 0.05082225 | 0.04423637 | 0.04448033 | -0.000244 | 0.51456156 | 0.99999994 |
| Myh7b | 0.00196069 | 0.00152865 | 0.001773 | -0.0002443 | 0.8205181 | 0.99999994 |
| B230217O12 | 0.00091747 | 0.00058552 | 0.00083023 | -0.0002447 | 0.97511515 | 0.99999994 |
| Ap5b1 | 0.00487025 | 0.00415052 | 0.00439534 | -0.0002448 | 0.64459039 | 0.99999994 |
| Gm43568 | 0.00160527 | 0.00120671 | 0.00145195 | -0.0002452 | 0.86955652 | 0.99999994 |
| Utp15 | 0.0421101 | 0.03682243 | 0.03706772 | -0.0002453 | 0.51756667 | 0.99999994 |
| Iba57 | 0.01811486 | 0.01595726 | 0.0162029 | -0.0002456 | 0.54018934 | 0.99999994 |
| Fkbp6 | 0.00556401 | 0.00477341 | 0.00501908 | -0.0002457 | 0.62771861 | 0.99999994 |
| Trpv4 | 0.00121949 | 0.00085741 | 0.00110331 | -0.0002459 | 0.93103558 | 0.99999994 |
| Pdss1 | 0.02939483 | 0.02584828 | 0.02609448 | -0.0002462 | 0.52503826 | 0.99999994 |

| | | | | | | |
|---|---|---|---|---|---|---|
| Pdzd9 | 0.00141589 | 0.00103458 | 0.00128083 | -0.0002463 | 0.89968233 | 0.99999994 |
| Efemp1 | 0.00461577 | 0.0039201 | 0.00416639 | -0.0002463 | 0.65301682 | 0.99999994 |
| Nup210l | 0.00101778 | 0.0006745 | 0.00092094 | -0.0002464 | 0.962564 | 0.99999994 |
| Exosc2 | 0.08700969 | 0.07413533 | 0.07438188 | -0.0002466 | 0.50880152 | 0.99999994 |
| 9330179D12I | 0.00078084 | 0.00046009 | 0.00070666 | -0.0002466 | 0.98989697 | 0.99999994 |
| Agrp | 0.00174081 | 0.00132771 | 0.0015744 | -0.0002467 | 0.85151835 | 0.99999994 |
| Ticam2 | 0.00174136 | 0.00132812 | 0.0015749 | -0.0002468 | 0.85152823 | 0.99999994 |
| Epm2a | 0.00919326 | 0.00802541 | 0.00827255 | -0.0002471 | 0.57881411 | 0.99999994 |
| Nuggc | 0.00111225 | 0.0007592 | 0.00100636 | -0.0002472 | 0.94895331 | 0.99999994 |
| Adgrg3 | 0.00123292 | 0.00086818 | 0.00111545 | -0.0002473 | 0.92997355 | 0.99999994 |
| Cdkn3 | 0.0069211 | 0.00599008 | 0.00623753 | -0.0002475 | 0.60413915 | 0.99999994 |
| Bnc1 | 0.00153513 | 0.00114085 | 0.00138857 | -0.0002477 | 0.88248316 | 0.99999994 |
| Stag3 | 0.00081793 | 0.00049248 | 0.0007402 | -0.0002477 | 0.98704822 | 0.99999994 |
| Pot1b | 0.00556514 | 0.00477216 | 0.0050201 | -0.0002479 | 0.62882957 | 0.99999994 |
| Alpk1 | 0.00057666 | 0.0002738 | 0.00052194 | -0.0002481 | 0.99922345 | 0.99999994 |
| Golga7b | 0.01848635 | 0.01628287 | 0.01653106 | -0.0002482 | 0.53980238 | 0.99999994 |
| Arrdc2 | 0.02647548 | 0.02330056 | 0.02354876 | -0.0002482 | 0.52796507 | 0.99999994 |
| 1700037C06I | 0.00207824 | 0.00163086 | 0.00187915 | -0.0002483 | 0.81043413 | 0.99999994 |
| Tgm4 | 0.00080809 | 0.00048258 | 0.0007313 | -0.0002487 | 0.98820899 | 0.99999994 |
| Mep1a | 0.00065609 | 0.00034471 | 0.00059381 | -0.0002491 | 0.99738305 | 0.99999994 |
| Lin52 | 0.03932095 | 0.03442732 | 0.03467653 | -0.0002492 | 0.5190766 | 0.99999994 |
| St6galnac3 | 0.00381018 | 0.00319169 | 0.00344112 | -0.0002494 | 0.68527038 | 0.99999994 |
| Nfatc2ip | 0.01603417 | 0.01411236 | 0.0143619 | -0.0002495 | 0.54603709 | 0.99999994 |
| Akr1b7 | 0.00068743 | 0.00037223 | 0.00062216 | -0.0002499 | 0.99625188 | 0.99999994 |
| Cfap157 | 0.00073421 | 0.00041397 | 0.00066448 | -0.0002505 | 0.99395346 | 0.99999994 |
| E330034G19 | 0.00073425 | 0.00041395 | 0.00066451 | -0.0002506 | 0.99396024 | 0.99999994 |
| Fam178b | 0.00325798 | 0.0026928 | 0.0029435 | -0.0002507 | 0.71461763 | 0.99999994 |
| Krt17 | 0.0010726 | 0.00071928 | 0.00097051 | -0.0002512 | 0.95756053 | 0.99999994 |
| Melk | 0.00243745 | 0.00195174 | 0.0022034 | -0.0002517 | 0.77644309 | 0.99999994 |
| Cyct | 0.00480945 | 0.00408892 | 0.00434065 | -0.0002517 | 0.65026019 | 0.99999994 |

| | | | | | | |
|---|---|---|---|---|---|---|
| Pear1 | 0.00334587 | 0.00277067 | 0.00302273 | -0.0002521 | 0.71057598 | 0.99999994 |
| Sik2 | 0.0319745 | 0.0280836 | 0.02833575 | -0.0002521 | 0.52361636 | 0.99999994 |
| Rimbp3 | 0.00125396 | 0.00088209 | 0.00113446 | -0.0002524 | 0.9306642 | 0.99999994 |
| Mesp1 | 0.00070986 | 0.00039008 | 0.00064245 | -0.0002524 | 0.99553601 | 0.99999994 |
| Gabrg2 | 0.00109082 | 0.00073448 | 0.00098698 | -0.0002525 | 0.95570653 | 0.99999994 |
| Otud1 | 0.00473667 | 0.00402258 | 0.00427517 | -0.0002526 | 0.65293992 | 0.99999994 |
| B430219N15 | 0.00246465 | 0.00197517 | 0.00222795 | -0.0002528 | 0.77493935 | 0.99999994 |
| Gm4076 | 0.00082932 | 0.00049756 | 0.00075051 | -0.0002529 | 0.98756417 | 0.99999994 |
| Ahcy | 0.00694945 | 0.00600988 | 0.00626296 | -0.0002531 | 0.60602789 | 0.99999994 |
| Opcml | 0.00104326 | 0.00069057 | 0.00094398 | -0.0002534 | 0.96302278 | 0.99999994 |
| Tmem212 | 0.00040376 | 0.00011208 | 0.0003655 | -0.0002534 | 0.99999804 | 0.99999994 |
| 1200007C13I | 0.000712 | 0.0003909 | 0.00064439 | -0.0002535 | 0.99558372 | 0.99999994 |
| Ust | 0.00099153 | 0.00064358 | 0.0008972 | -0.0002536 | 0.97005476 | 0.99999994 |
| Gm16894 | 0.00166197 | 0.0012494 | 0.00150317 | -0.0002538 | 0.86944481 | 0.99999994 |
| Lypd6 | 0.00674471 | 0.00582544 | 0.00607929 | -0.0002538 | 0.60947243 | 0.99999994 |
| Plxnd1 | 0.00515393 | 0.00439654 | 0.00465047 | -0.0002539 | 0.6418676 | 0.99999994 |
| Paox | 0.0228686 | 0.02013563 | 0.02038976 | -0.0002541 | 0.53305888 | 0.99999994 |
| Aadat | 0.00064668 | 0.00033106 | 0.0005853 | -0.0002542 | 0.99808214 | 0.99999994 |
| Taf12 | 0.22754918 | 0.1743214 | 0.17457585 | -0.0002544 | 0.50387038 | 0.99999994 |
| Fam135b | 0.00201325 | 0.00156594 | 0.00182046 | -0.0002545 | 0.82397473 | 0.99999994 |
| Gm1123 | 0.00260583 | 0.00210067 | 0.00235534 | -0.0002547 | 0.7641483 | 0.99999994 |
| Asb5 | 0.00063353 | 0.0003186 | 0.0005734 | -0.0002548 | 0.99845113 | 0.99999994 |
| S100a3 | 0.0005903 | 0.00027889 | 0.00053429 | -0.0002554 | 0.99926816 | 0.99999994 |
| Fancb | 0.00399873 | 0.00335546 | 0.00361094 | -0.0002555 | 0.68116233 | 0.99999994 |
| Gm10848 | 0.0009192 | 0.0005763 | 0.0008318 | -0.0002555 | 0.97955267 | 0.99999994 |
| Epas1 | 0.00271729 | 0.00220013 | 0.0024559 | -0.0002558 | 0.75591808 | 0.99999994 |
| Gm6225 | 0.00044234 | 0.00014408 | 0.00040041 | -0.0002563 | 0.99998983 | 0.99999994 |
| Fam13c | 0.00108592 | 0.00072603 | 0.00098255 | -0.0002565 | 0.95887984 | 0.99999994 |
| Nrip2 | 0.00349178 | 0.00289696 | 0.00315424 | -0.0002573 | 0.70640893 | 0.99999994 |
| Ces2b | 0.00068353 | 0.00036129 | 0.00061863 | -0.0002573 | 0.9971885 | 0.99999994 |

| | | | | | | |
|---|---|---|---|---|---|---|
| Zmynd19 | 0.03863557 | 0.0338302 | 0.03408758 | -0.0002574 | 0.52004226 | 0.99999994 |
| Gm15351 | 0.00119807 | 0.00082648 | 0.00108395 | -0.0002575 | 0.9430673 | 0.99999994 |
| 9430037G07 | 0.00160777 | 0.00119672 | 0.00145421 | -0.0002575 | 0.88072036 | 0.99999994 |
| Zfp874b | 0.01529399 | 0.01344748 | 0.01370575 | -0.0002583 | 0.54990923 | 0.99999994 |
| Nrcam | 0.00192025 | 0.00147819 | 0.00173647 | -0.0002583 | 0.83893112 | 0.99999994 |
| Mettl4 | 0.02620956 | 0.02305809 | 0.02331638 | -0.0002583 | 0.52939012 | 0.99999994 |
| Wdr38 | 0.00204807 | 0.00159311 | 0.0018519 | -0.0002588 | 0.82386271 | 0.99999994 |
| C1qc | 0.01250628 | 0.01096962 | 0.01122861 | -0.000259 | 0.56101032 | 0.99999994 |
| Gm26739 | 0.00135177 | 0.0009634 | 0.00122287 | -0.0002595 | 0.92107612 | 0.99999994 |
| 1700028I16R | 0.00199048 | 0.00154018 | 0.0017999 | -0.0002597 | 0.83159662 | 0.99999994 |
| Gm10392 | 0.00106663 | 0.00070481 | 0.00096511 | -0.0002603 | 0.96369441 | 0.99999994 |
| Gm42800 | 0.00129255 | 0.00090859 | 0.00116935 | -0.0002608 | 0.93113697 | 0.99999994 |
| Gm21992 | 0.0029005 | 0.00236023 | 0.00262117 | -0.0002609 | 0.74622414 | 0.99999994 |
| Gm15496 | 0.0036257 | 0.0030139 | 0.00327491 | -0.000261 | 0.70211934 | 0.99999994 |
| Gm15541 | 0.00149828 | 0.00109356 | 0.00135528 | -0.0002617 | 0.90067148 | 0.99999994 |
| Gm6410 | 0.00278022 | 0.00225077 | 0.00251267 | -0.0002619 | 0.75609923 | 0.99999994 |
| Raet1e | 0.00379448 | 0.003165 | 0.00342697 | -0.000262 | 0.69457085 | 0.99999994 |
| Topors | 0.22814412 | 0.17469049 | 0.17495255 | -0.0002621 | 0.50397765 | 0.99999994 |
| Slc22a21 | 0.00156547 | 0.00115364 | 0.00141599 | -0.0002623 | 0.89126169 | 0.99999994 |
| Rassf9 | 0.01070291 | 0.00935827 | 0.00962118 | -0.0002629 | 0.57216735 | 0.99999994 |
| Clrn3 | 0.00043707 | 0.0001325 | 0.00039564 | -0.0002631 | 0.99999523 | 0.99999994 |
| Gm16863 | 0.00051652 | 0.0002041 | 0.00046753 | -0.0002634 | 0.99991175 | 0.99999994 |
| 5530401A14 | 0.00327029 | 0.00269116 | 0.00295461 | -0.0002634 | 0.72358308 | 0.99999994 |
| Sipa1l2 | 0.00573655 | 0.0049102 | 0.00517412 | -0.0002639 | 0.6328956 | 0.99999994 |
| Zbtb7b | 0.06859036 | 0.05907387 | 0.05933784 | -0.000264 | 0.51181173 | 0.99999994 |
| Gja4 | 0.00096663 | 0.00061055 | 0.00087469 | -0.0002641 | 0.97778672 | 0.99999994 |
| Kif13a | 0.01502887 | 0.01320621 | 0.01347057 | -0.0002644 | 0.55196705 | 0.99999994 |
| Lpxn | 0.00206808 | 0.00160551 | 0.00186997 | -0.0002645 | 0.82673928 | 0.99999994 |
| Cox7b2 | 0.00120735 | 0.00082762 | 0.00109233 | -0.0002647 | 0.94663445 | 0.99999994 |
| Gm12927 | 0.00048302 | 0.00017249 | 0.00043722 | -0.0002647 | 0.99997215 | 0.99999994 |

| | | | | | | |
|---|---|---|---|---|---|---|
| Tmprss9 | 0.00052045 | 0.00020594 | 0.00047109 | -0.0002652 | 0.99991035 | 0.99999994 |
| Sipa1 | 0.02630715 | 0.02313646 | 0.02340167 | -0.0002652 | 0.53006588 | 0.99999994 |
| Ogdhl | 0.0019642 | 0.00151094 | 0.00177617 | -0.0002652 | 0.8398766 | 0.99999994 |
| Ncf4 | 0.00139907 | 0.0010002 | 0.00126562 | -0.0002654 | 0.91863621 | 0.99999994 |
| Camkmt | 0.00784689 | 0.00680197 | 0.00706745 | -0.0002655 | 0.5987213 | 0.99999994 |
| Gm44053 | 0.00117414 | 0.00079667 | 0.00106231 | -0.0002656 | 0.95198857 | 0.99999994 |
| Gm16035 | 0.00388713 | 0.00324412 | 0.00351042 | -0.0002663 | 0.69320342 | 0.99999994 |
| Myof | 0.00429385 | 0.00361028 | 0.00387666 | -0.0002664 | 0.67630295 | 0.99999994 |
| Gm10369 | 0.00508832 | 0.00432493 | 0.00459147 | -0.0002665 | 0.65039812 | 0.99999994 |
| Ppp1r3e | 0.00058594 | 0.00026356 | 0.00053034 | -0.0002668 | 0.99959351 | 0.99999994 |
| Gm5239 | 0.00063518 | 0.00030786 | 0.00057489 | -0.000267 | 0.99900541 | 0.99999994 |
| Tnfaip6 | 0.00069392 | 0.00036091 | 0.00062803 | -0.0002671 | 0.99768086 | 0.99999994 |
| Paqr6 | 0.00657172 | 0.00565628 | 0.00592406 | -0.0002678 | 0.61824956 | 0.99999994 |
| Gm38186 | 0.00121319 | 0.00082967 | 0.00109761 | -0.0002679 | 0.94790898 | 0.99999994 |
| Tgfbr3l | 0.00612125 | 0.00525146 | 0.00551967 | -0.0002682 | 0.62681941 | 0.99999994 |
| Klk1b16 | 0.00317979 | 0.00260475 | 0.00287302 | -0.0002683 | 0.73287248 | 0.99999994 |
| Dok2 | 0.00264336 | 0.0021209 | 0.00238921 | -0.0002683 | 0.77262547 | 0.99999994 |
| Gm29427 | 0.0006351 | 0.00030647 | 0.00057482 | -0.0002684 | 0.99905659 | 0.99999994 |
| Hemgn | 0.00051179 | 0.00019456 | 0.00046326 | -0.0002687 | 0.99994349 | 0.99999994 |
| Ctnna2 | 0.00950976 | 0.0082864 | 0.00855552 | -0.0002691 | 0.58292631 | 0.99999994 |
| Asb2 | 0.00064556 | 0.00031514 | 0.00058428 | -0.0002691 | 0.99891576 | 0.99999994 |
| Cyp39a1 | 0.03435425 | 0.03012715 | 0.03039651 | -0.0002694 | 0.52351887 | 0.99999994 |
| Sorcs1 | 0.00063519 | 0.00030541 | 0.0005749 | -0.0002695 | 0.99909659 | 0.99999994 |
| Zfp457 | 0.00152704 | 0.00111176 | 0.00138127 | -0.0002695 | 0.90298595 | 0.99999994 |
| Tmem240 | 0.00799957 | 0.00693471 | 0.00720422 | -0.0002695 | 0.59832416 | 0.99999994 |
| Cadps2 | 0.00265254 | 0.00212784 | 0.00239749 | -0.0002697 | 0.77296757 | 0.99999994 |
| B230303O12 | 0.00115114 | 0.00077173 | 0.00104151 | -0.0002698 | 0.95766178 | 0.99999994 |
| Gm26947 | 0.00278348 | 0.00224578 | 0.00251562 | -0.0002698 | 0.76238441 | 0.99999994 |
| Aass | 0.00076273 | 0.00042008 | 0.00069027 | -0.0002702 | 0.99541307 | 0.99999994 |
| Acadvl | 0.51211079 | 0.31869703 | 0.31896728 | -0.0002703 | 0.50224996 | 0.99999994 |

| | | | | | | |
|---|---|---|---|---|---|---|
| Rusc2 | 0.00404122 | 0.00337887 | 0.0036492 | -0.0002703 | 0.68902964 | 0.99999994 |
| Xylt2 | 0.0369119 | 0.0323332 | 0.03260407 | -0.0002709 | 0.52205017 | 0.99999994 |
| Spata17 | 0.00242152 | 0.00191807 | 0.00218903 | -0.000271 | 0.79501046 | 0.99999994 |
| Cngb1 | 0.00065919 | 0.00032559 | 0.00059662 | -0.000271 | 0.99875176 | 0.99999994 |
| C230037L18F | 0.01079096 | 0.00942833 | 0.00969976 | -0.0002714 | 0.57388169 | 0.99999994 |
| St3gal4 | 0.05665117 | 0.04912107 | 0.04939254 | -0.0002715 | 0.51459171 | 0.99999994 |
| Nek8 | 0.01605603 | 0.01410974 | 0.01438127 | -0.0002715 | 0.55000666 | 0.99999994 |
| Lmbr1l | 0.07892771 | 0.06755328 | 0.06782496 | -0.0002717 | 0.51063586 | 0.99999994 |
| Ccdc15 | 0.0049111 | 0.00416031 | 0.00443208 | -0.0002718 | 0.65842332 | 0.99999994 |
| Gm20939 | 0.00818009 | 0.00709374 | 0.0073659 | -0.0002722 | 0.59713528 | 0.99999994 |
| Slc39a5 | 0.01163728 | 0.01018231 | 0.01045452 | -0.0002722 | 0.56879832 | 0.99999994 |
| Apobec4 | 0.00262007 | 0.00209591 | 0.00236819 | -0.0002723 | 0.77795793 | 0.99999994 |
| Morn1 | 0.01902754 | 0.01673632 | 0.01700882 | -0.0002725 | 0.54246384 | 0.99999994 |
| Atp1a2 | 0.00074824 | 0.00040443 | 0.00067717 | -0.0002727 | 0.99632906 | 0.99999994 |
| Ak4 | 0.00281196 | 0.00226783 | 0.00254131 | -0.0002735 | 0.76309632 | 0.99999994 |
| Mme | 0.00107899 | 0.0007021 | 0.00097628 | -0.0002742 | 0.96921751 | 0.99999994 |
| Sult1c2 | 0.00512252 | 0.00434799 | 0.00462222 | -0.0002742 | 0.65354578 | 0.99999994 |
| 1700037C18I | 0.06054776 | 0.05238114 | 0.05265545 | -0.0002743 | 0.51383095 | 0.99999994 |
| Tle2 | 0.14449738 | 0.11797186 | 0.11824618 | -0.0002743 | 0.50616036 | 0.99999994 |
| Agtr1a | 0.00151796 | 0.00109847 | 0.00137306 | -0.0002746 | 0.90842957 | 0.99999994 |
| 6430562O15 | 0.00126171 | 0.00086678 | 0.00114147 | -0.0002747 | 0.94540293 | 0.99999994 |
| Ift43 | 0.14393721 | 0.11756506 | 0.11783978 | -0.0002747 | 0.50619046 | 0.99999994 |
| Vwc2 | 0.00051935 | 0.00019532 | 0.00047009 | -0.0002748 | 0.99995003 | 0.99999994 |
| 1810062O18 | 0.00852957 | 0.00740387 | 0.00767878 | -0.0002749 | 0.59417542 | 0.99999994 |
| Gm20554 | 0.00784017 | 0.0067863 | 0.00706143 | -0.0002751 | 0.6023181 | 0.99999994 |
| Xaf1 | 0.0021582 | 0.00167617 | 0.00195134 | -0.0002752 | 0.82604311 | 0.99999994 |
| Zbtb39 | 0.01788467 | 0.01572418 | 0.01599948 | -0.0002753 | 0.54559438 | 0.99999994 |
| Col12a1 | 0.00139695 | 0.00098827 | 0.00126371 | -0.0002754 | 0.92658344 | 0.99999994 |
| Gabrb2 | 0.00084323 | 0.00048752 | 0.00076309 | -0.0002756 | 0.99188714 | 0.99999994 |
| Lrrc18 | 0.00216493 | 0.00168178 | 0.00195742 | -0.0002756 | 0.82570403 | 0.99999994 |

| | | | | | | |
|---|---|---|---|---|---|---|
| Rnasek | 0.00054771 | 0.00022006 | 0.00049575 | -0.0002757 | 0.99989288 | 0.99999994 |
| Cdh15 | 0.0012188 | 0.00082691 | 0.00110268 | -0.0002758 | 0.95201669 | 0.99999994 |
| Neu2 | 0.0013622 | 0.00095626 | 0.0012323 | -0.000276 | 0.93202585 | 0.99999994 |
| Adh4 | 0.00047367 | 0.00015257 | 0.00042876 | -0.0002762 | 0.99999097 | 0.99999994 |
| Kcng3 | 0.00049846 | 0.00017498 | 0.00045119 | -0.0002762 | 0.99997698 | 0.99999994 |
| A930007I19F | 0.00460674 | 0.00388157 | 0.00415826 | -0.0002767 | 0.67108839 | 0.99999994 |
| Il20rb | 0.00053781 | 0.00020946 | 0.0004868 | -0.0002773 | 0.99992536 | 0.99999994 |
| Osr1 | 0.00322659 | 0.00263786 | 0.0029152 | -0.0002773 | 0.73672373 | 0.99999994 |
| Clec10a | 0.00057337 | 0.00024155 | 0.00051897 | -0.0002774 | 0.99981326 | 0.99999994 |
| Gm15614 | 0.0166169 | 0.01460033 | 0.01487802 | -0.0002777 | 0.54943645 | 0.99999994 |
| Ctf2 | 0.00120992 | 0.00081672 | 0.00109466 | -0.0002779 | 0.95449273 | 0.99999994 |
| Gemin4 | 0.00108259 | 0.00070141 | 0.00097954 | -0.0002781 | 0.97062114 | 0.99999994 |
| Gjc3 | 0.00105914 | 0.00068014 | 0.00095834 | -0.0002782 | 0.9733403 | 0.99999994 |
| Pcdhb16 | 0.01843451 | 0.01620703 | 0.01648527 | -0.0002782 | 0.54472595 | 0.99999994 |
| Tnfrsf22 | 0.00353148 | 0.00291175 | 0.00319001 | -0.0002783 | 0.71925482 | 0.99999994 |
| Trim36 | 0.00120545 | 0.00081207 | 0.00109061 | -0.0002785 | 0.95543428 | 0.99999994 |
| Ube2ql1 | 0.0058256 | 0.0049754 | 0.00525412 | -0.0002787 | 0.63799811 | 0.99999994 |
| Plekhg2 | 0.0253741 | 0.022307 | 0.02258576 | -0.0002788 | 0.53273808 | 0.99999994 |
| Slc7a7 | 0.009084 | 0.00789544 | 0.00817484 | -0.0002794 | 0.58998246 | 0.99999994 |
| Gm960 | 0.00053801 | 0.00020752 | 0.00048698 | -0.0002795 | 0.99993323 | 0.99999994 |
| 1500015L24F | 0.00172632 | 0.00128161 | 0.00156131 | -0.0002797 | 0.8834589 | 0.99999994 |
| Slc11a1 | 0.00070423 | 0.0003575 | 0.00063736 | -0.0002799 | 0.99826539 | 0.99999994 |
| Tmem255b | 0.00182922 | 0.00137333 | 0.00165426 | -0.0002809 | 0.87084424 | 0.99999994 |
| Wdr17 | 0.00277669 | 0.00222837 | 0.00250949 | -0.0002811 | 0.77206632 | 0.99999994 |
| Catsperd | 0.00148142 | 0.00105886 | 0.00134004 | -0.0002812 | 0.91875388 | 0.99999994 |
| Dgka | 0.02233409 | 0.01963893 | 0.01992032 | -0.0002814 | 0.53745537 | 0.99999994 |
| Clca3a2 | 0.00050703 | 0.00017731 | 0.00045894 | -0.0002816 | 0.99997794 | 0.99999994 |
| 4933431E20I | 0.00607492 | 0.00519632 | 0.00547807 | -0.0002817 | 0.63395386 | 0.99999994 |
| Fsbp | 0.00096294 | 0.00058949 | 0.00087135 | -0.0002819 | 0.98434708 | 0.99999994 |
| Fam76a | 0.21823485 | 0.16834821 | 0.16863019 | -0.000282 | 0.50444041 | 0.99999994 |

| | | | | | | |
|---|---|---|---|---|---|---|
| Adcy1 | 0.00657308 | 0.00564302 | 0.00592528 | -0.0002823 | 0.62441022 | 0.99999994 |
| Pank4 | 0.01157431 | 0.01011573 | 0.01039839 | -0.0002827 | 0.57179262 | 0.99999994 |
| BC048507 | 0.00053656 | 0.00020295 | 0.00048566 | -0.0002827 | 0.99994664 | 0.99999994 |
| Prdm5 | 0.00711818 | 0.00613156 | 0.00641429 | -0.0002827 | 0.61539001 | 0.99999994 |
| Slc17a4 | 0.00166916 | 0.00122649 | 0.00150968 | -0.0002832 | 0.8940945 | 0.99999994 |
| Gm11100 | 0.00211253 | 0.00162681 | 0.0019101 | -0.0002833 | 0.83822999 | 0.99999994 |
| Adamts6 | 0.00536265 | 0.00455473 | 0.00483811 | -0.0002834 | 0.65168549 | 0.99999994 |
| 1810012K16I | 0.00051617 | 0.00018372 | 0.00046722 | -0.0002835 | 0.99997316 | 0.99999994 |
| Gm10714 | 0.00064774 | 0.00030268 | 0.00058625 | -0.0002836 | 0.99935829 | 0.99999994 |
| Krt12 | 0.00105572 | 0.00067165 | 0.00095525 | -0.0002836 | 0.97592986 | 0.99999994 |
| Pdlim2 | 0.0041866 | 0.00349602 | 0.0037801 | -0.0002841 | 0.69154586 | 0.99999994 |
| F8 | 0.00401552 | 0.00334186 | 0.00362606 | -0.0002842 | 0.69906234 | 0.99999994 |
| Ccdc87 | 0.0008231 | 0.00046065 | 0.00074488 | -0.0002842 | 0.9944556 | 0.99999994 |
| Nrg4 | 0.00061695 | 0.00027399 | 0.0005584 | -0.0002844 | 0.99965088 | 0.99999994 |
| Nid1 | 0.00258505 | 0.00205172 | 0.00233659 | -0.0002849 | 0.79147029 | 0.99999994 |
| Il2 | 0.01413355 | 0.01239083 | 0.01267572 | -0.0002849 | 0.55946167 | 0.99999994 |
| Pdgfra | 0.00123469 | 0.00083208 | 0.00111705 | -0.000285 | 0.9552541 | 0.99999994 |
| 9530046B11 | 0.00234245 | 0.00183261 | 0.00211766 | -0.000285 | 0.81486828 | 0.99999994 |
| Adam11 | 0.00274695 | 0.00219746 | 0.00248266 | -0.0002852 | 0.7777636 | 0.99999994 |
| Popdc3 | 0.00054115 | 0.00020434 | 0.00048982 | -0.0002855 | 0.99994765 | 0.99999994 |
| Zscan22 | 0.02510607 | 0.02206554 | 0.02235119 | -0.0002856 | 0.53389633 | 0.99999994 |
| Medag | 0.00113272 | 0.00073908 | 0.00102486 | -0.0002858 | 0.96828184 | 0.99999994 |
| Zfp423 | 0.01141887 | 0.00997404 | 0.01025983 | -0.0002858 | 0.57354716 | 0.99999994 |
| Rspo3 | 0.00063753 | 0.00029122 | 0.00057702 | -0.0002858 | 0.99951123 | 0.99999994 |
| Fbxw10 | 0.00159334 | 0.00115499 | 0.00144117 | -0.0002862 | 0.90687826 | 0.99999994 |
| Disc1 | 0.00288292 | 0.00231893 | 0.00260531 | -0.0002864 | 0.7678228 | 0.99999994 |
| M1ap | 0.00171049 | 0.00126029 | 0.00154701 | -0.0002867 | 0.89133853 | 0.99999994 |
| Cilp2 | 0.00275462 | 0.00220275 | 0.00248958 | -0.0002868 | 0.77842744 | 0.99999994 |
| Gm17435 | 0.00875271 | 0.00759144 | 0.00787847 | -0.000287 | 0.59580476 | 0.99999994 |
| Syk | 0.00425669 | 0.0035561 | 0.00384321 | -0.0002871 | 0.69049698 | 0.99999994 |

| | | | | | | |
|---|---|---|---|---|---|---|
| Chst2 | 0.0049649 | 0.00419314 | 0.00448047 | -0.0002873 | 0.66526429 | 0.99999994 |
| Fmod | 0.0031306 | 0.00254108 | 0.00282866 | -0.0002876 | 0.75071375 | 0.99999994 |
| Mpv17l | 0.03635194 | 0.03183373 | 0.0321214 | -0.0002877 | 0.52376785 | 0.99999994 |
| Prkch | 0.01002298 | 0.00872631 | 0.00901411 | -0.0002878 | 0.58415249 | 0.99999994 |
| Hacd4 | 0.00622405 | 0.00532376 | 0.00561198 | -0.0002882 | 0.63377117 | 0.99999994 |
| 2200002D01 | 0.00506225 | 0.00427979 | 0.00456802 | -0.0002882 | 0.66275344 | 0.99999994 |
| Kdm6bos | 0.00085349 | 0.00048394 | 0.00077237 | -0.0002884 | 0.99353551 | 0.99999994 |
| Lingo3 | 0.00249849 | 0.00196931 | 0.00225849 | -0.0002892 | 0.80297448 | 0.99999994 |
| Ccr9 | 0.00106162 | 0.00067109 | 0.00096058 | -0.0002895 | 0.97757237 | 0.99999994 |
| Spry4 | 0.00623508 | 0.00533236 | 0.00562188 | -0.0002895 | 0.63412153 | 0.99999994 |
| Kif23 | 0.00351894 | 0.00288894 | 0.00317871 | -0.0002898 | 0.72800258 | 0.99999994 |
| Trnp1 | 0.00846229 | 0.00732863 | 0.00761855 | -0.0002899 | 0.59998311 | 0.99999994 |
| C3ar1 | 0.00216666 | 0.00166895 | 0.00195898 | -0.00029 | 0.8378039 | 0.99999994 |
| Gm7535 | 0.00196364 | 0.00148564 | 0.00177566 | -0.00029 | 0.86153037 | 0.99999994 |
| Stfa2l1 | 0.00078012 | 0.00041597 | 0.00070601 | -0.00029 | 0.9968764 | 0.99999994 |
| Ccne1 | 0.02652345 | 0.02330039 | 0.02359067 | -0.0002903 | 0.53263958 | 0.99999994 |
| Ikzf1 | 0.00056548 | 0.00022144 | 0.00051183 | -0.0002904 | 0.99992049 | 0.99999994 |
| Adamtsl1 | 0.00187923 | 0.00140889 | 0.00169943 | -0.0002905 | 0.87244113 | 0.99999994 |
| Spata45 | 0.00141046 | 0.00098534 | 0.00127592 | -0.0002906 | 0.93523193 | 0.99999994 |
| Efcab5 | 0.0010979 | 0.00070281 | 0.00099339 | -0.0002906 | 0.97423816 | 0.99999994 |
| Mpo | 0.00198377 | 0.00150322 | 0.00179384 | -0.0002906 | 0.85957193 | 0.99999994 |
| Eno4 | 0.00416083 | 0.00346617 | 0.00375691 | -0.0002907 | 0.6967624 | 0.99999994 |
| 4932413F04I | 0.00054599 | 0.00020342 | 0.0004942 | -0.0002908 | 0.99995507 | 0.99999994 |
| Ctc1 | 0.05865396 | 0.05078057 | 0.0510717 | -0.0002911 | 0.51513432 | 0.99999994 |
| 1700040D17 | 0.00967774 | 0.00841445 | 0.00870566 | -0.0002912 | 0.58810096 | 0.99999994 |
| 2500002B13 | 0.01069461 | 0.00932255 | 0.00961378 | -0.0002912 | 0.57989975 | 0.99999994 |
| Pcdhga2 | 0.00605999 | 0.00517341 | 0.00546466 | -0.0002912 | 0.6386152 | 0.99999994 |
| Ppfia2 | 0.00363133 | 0.00298872 | 0.00327999 | -0.0002913 | 0.72277047 | 0.99999994 |
| Nat10 | 0.02701767 | 0.02373079 | 0.02402231 | -0.0002915 | 0.53219043 | 0.99999994 |
| 1700012B07 | 0.00058387 | 0.00023689 | 0.00052847 | -0.0002916 | 0.99987994 | 0.99999994 |

| | | | | | | |
|---|---|---|---|---|---|---|
| Lrrc55 | 0.00068755 | 0.00033059 | 0.00062227 | -0.0002917 | 0.99909615 | 0.99999994 |
| Nlrc4 | 0.00251353 | 0.00198019 | 0.00227206 | -0.0002919 | 0.80374596 | 0.99999994 |
| Avil | 0.00214483 | 0.00164737 | 0.00193927 | -0.0002919 | 0.84180344 | 0.99999994 |
| Cxcl13 | 0.00179782 | 0.00133392 | 0.0016259 | -0.000292 | 0.88402621 | 0.99999994 |
| Pole | 0.00127035 | 0.00085701 | 0.00114928 | -0.0002923 | 0.95474704 | 0.99999994 |
| Ccdc120 | 0.02377857 | 0.02089576 | 0.02118818 | -0.0002924 | 0.53659697 | 0.99999994 |
| Gm12278 | 0.00083735 | 0.00046475 | 0.00075777 | -0.000293 | 0.99497238 | 0.99999994 |
| Gm14486 | 0.0007687 | 0.00040264 | 0.00069568 | -0.000293 | 0.99747513 | 0.99999994 |
| Zfp235 | 0.03084794 | 0.0270648 | 0.02735792 | -0.0002931 | 0.52842796 | 0.99999994 |
| Mfsd7a | 0.00180305 | 0.00133736 | 0.00163062 | -0.0002933 | 0.88436649 | 0.99999994 |
| Rpap3 | 0.10716911 | 0.09004389 | 0.09033722 | -0.0002933 | 0.50862182 | 0.99999994 |
| Gm11521 | 0.00236847 | 0.00184737 | 0.00214115 | -0.0002938 | 0.81945446 | 0.99999994 |
| Gsdma | 0.00063202 | 0.0002778 | 0.00057203 | -0.0002942 | 0.99969116 | 0.99999994 |
| Kcnn1 | 0.0067964 | 0.00583133 | 0.00612566 | -0.0002943 | 0.62545262 | 0.99999994 |
| Stat6 | 0.01839538 | 0.01615629 | 0.01645071 | -0.0002944 | 0.54741342 | 0.99999994 |
| 2900072N19 | 0.0006297 | 0.00027532 | 0.00056994 | -0.0002946 | 0.99971005 | 0.99999994 |
| B4galnt1 | 0.0039237 | 0.00324834 | 0.00354337 | -0.000295 | 0.71028214 | 0.99999994 |
| Nos1 | 0.00149718 | 0.00105924 | 0.00135428 | -0.000295 | 0.92649044 | 0.99999994 |
| Slc22a29 | 0.00205399 | 0.00156213 | 0.00185725 | -0.0002951 | 0.85490552 | 0.99999994 |
| Pirt | 0.00118376 | 0.00077581 | 0.001071 | -0.0002952 | 0.96672149 | 0.99999994 |
| Gm43625 | 0.00169328 | 0.00123626 | 0.00153147 | -0.0002952 | 0.90027004 | 0.99999994 |
| Gm20531 | 0.00125286 | 0.00083815 | 0.00113347 | -0.0002953 | 0.95856494 | 0.99999994 |
| Gm17102 | 0.0017498 | 0.0012869 | 0.00158252 | -0.0002956 | 0.89314589 | 0.99999994 |
| Nhlrc1 | 0.00886181 | 0.00768039 | 0.00797609 | -0.0002957 | 0.59745603 | 0.99999994 |
| Gm12454 | 0.0014446 | 0.00101105 | 0.00130677 | -0.0002957 | 0.93400756 | 0.99999994 |
| Gm17655 | 0.00084003 | 0.00046425 | 0.00076019 | -0.0002959 | 0.99521947 | 0.99999994 |
| Esyt3 | 0.00121132 | 0.00079984 | 0.00109592 | -0.0002961 | 0.96393965 | 0.99999994 |
| Purg | 0.0199849 | 0.01755688 | 0.01785312 | -0.0002962 | 0.54397438 | 0.99999994 |
| Dazl | 0.0012826 | 0.00086406 | 0.00116035 | -0.0002963 | 0.95540242 | 0.99999994 |
| Dgat1 | 0.10052456 | 0.08489164 | 0.08518846 | -0.0002968 | 0.50925145 | 0.99999994 |

| | | | | | | |
|---|---|---|---|---|---|---|
| 5730420D15 | 0.00066823 | 0.00030761 | 0.00060479 | -0.0002972 | 0.9994637 | 0.99999994 |
| Sdha | 0.56722245 | 0.33969653 | 0.33999374 | -0.0002972 | 0.50232138 | 0.99999994 |
| Hs3st1 | 0.00299376 | 0.00240769 | 0.00270527 | -0.0002976 | 0.7679789 | 0.99999994 |
| Kcp | 0.00310676 | 0.00250958 | 0.00280717 | -0.0002976 | 0.75979337 | 0.99999994 |
| Ehhadh | 0.00187936 | 0.00140189 | 0.00169954 | -0.0002977 | 0.87814637 | 0.99999994 |
| E030044B06 | 0.00065464 | 0.00029475 | 0.0005925 | -0.0002977 | 0.99958857 | 0.99999994 |
| Gen1 | 0.00203163 | 0.001539 | 0.00183706 | -0.0002981 | 0.8599246 | 0.99999994 |
| Fzd1 | 0.00508597 | 0.00429119 | 0.00458935 | -0.0002982 | 0.66729266 | 0.99999994 |
| Zfp160 | 0.04718997 | 0.04110156 | 0.04140021 | -0.0002986 | 0.51914823 | 0.99999994 |
| Gm2415 | 0.00746077 | 0.00642253 | 0.00672144 | -0.0002989 | 0.61638954 | 0.99999994 |
| Kcnt1 | 0.00198949 | 0.00150007 | 0.00179901 | -0.0002989 | 0.86565005 | 0.99999994 |
| Gm11992 | 0.00805904 | 0.0069581 | 0.00725748 | -0.0002994 | 0.60818189 | 0.99999994 |
| Foxn1 | 0.00089265 | 0.00050831 | 0.00080778 | -0.0002995 | 0.99320194 | 0.99999994 |
| Ercc1 | 0.18691271 | 0.14766401 | 0.1479635 | -0.0002995 | 0.5053748 | 0.99999994 |
| Tyw3 | 0.00742868 | 0.00639315 | 0.00669268 | -0.0002995 | 0.61710892 | 0.99999994 |
| Rasa4 | 0.00487473 | 0.00409973 | 0.00439937 | -0.0002996 | 0.67485479 | 0.99999994 |
| Pf4 | 0.01715868 | 0.01505786 | 0.01535752 | -0.0002997 | 0.55166841 | 0.99999994 |
| Trim14 | 0.00167094 | 0.00121086 | 0.00151128 | -0.0003004 | 0.90710949 | 0.99999994 |
| Aldh1l2 | 0.00068477 | 0.00031892 | 0.00061976 | -0.0003008 | 0.99938332 | 0.99999994 |
| Trpm3 | 0.00069349 | 0.00032678 | 0.00062765 | -0.0003009 | 0.99929042 | 0.99999994 |
| Arl10 | 0.0251991 | 0.02213126 | 0.02243262 | -0.0003014 | 0.53562649 | 0.99999994 |
| Postn | 0.0017952 | 0.00132168 | 0.00162353 | -0.0003019 | 0.89206663 | 0.99999994 |
| Tmem120b | 0.05186419 | 0.04505915 | 0.04536117 | -0.000302 | 0.51767472 | 0.99999994 |
| Grik3 | 0.0015483 | 0.00109843 | 0.00140048 | -0.000302 | 0.92444392 | 0.99999994 |
| Fbxw15 | 0.00298525 | 0.00239505 | 0.0026976 | -0.0003025 | 0.77232933 | 0.99999994 |
| Gm43136 | 0.00111093 | 0.0007026 | 0.00100517 | -0.0003026 | 0.9774463 | 0.99999994 |
| Col11a1 | 0.00112555 | 0.00071518 | 0.00101838 | -0.0003032 | 0.97624738 | 0.99999994 |
| Xkr8 | 0.00281287 | 0.00223891 | 0.00254213 | -0.0003032 | 0.78638596 | 0.99999994 |
| Pagr1a | 0.00601427 | 0.00512035 | 0.00542359 | -0.0003032 | 0.6451154 | 0.99999994 |
| Lox | 0.00062512 | 0.0002622 | 0.00056579 | -0.0003036 | 0.99982263 | 0.99999994 |

| | | | | | | |
|---|---|---|---|---|---|---|
| Spdya | 0.00096002 | 0.00056488 | 0.00086871 | -0.0003038 | 0.99004488 | 0.99999994 |
| Erbb4 | 0.00210437 | 0.00159889 | 0.00190273 | -0.0003038 | 0.85609117 | 0.99999994 |
| Porcn | 0.02430495 | 0.02134558 | 0.02164958 | -0.000304 | 0.53723286 | 0.99999994 |
| Hpdl | 0.00136565 | 0.00093131 | 0.00123542 | -0.0003041 | 0.94934429 | 0.99999994 |
| Zkscan16 | 0.00270038 | 0.00213585 | 0.00244065 | -0.0003048 | 0.79708667 | 0.99999994 |
| Glrb | 0.00291129 | 0.00232593 | 0.0026309 | -0.000305 | 0.77981855 | 0.99999994 |
| Fam227a | 0.02323006 | 0.02040176 | 0.02070702 | -0.0003053 | 0.53908442 | 0.99999994 |
| Pms1 | 0.02137215 | 0.01876887 | 0.01907464 | -0.0003058 | 0.54248769 | 0.99999994 |
| Dnali1 | 0.00119332 | 0.00077378 | 0.00107965 | -0.0003059 | 0.97033523 | 0.99999994 |
| Tchh | 0.00161116 | 0.00115124 | 0.00145727 | -0.000306 | 0.9189233 | 0.99999994 |
| Ect2 | 0.00458507 | 0.00383271 | 0.00413877 | -0.0003061 | 0.68872082 | 0.99999994 |
| Ccr10 | 0.00195039 | 0.00145731 | 0.0017637 | -0.0003064 | 0.87622708 | 0.99999994 |
| Ccdc63 | 0.00252269 | 0.00197389 | 0.00228032 | -0.0003064 | 0.81447165 | 0.99999994 |
| Adhfe1 | 0.00166141 | 0.001196 | 0.00150267 | -0.0003067 | 0.9128423 | 0.99999994 |
| Zscan18 | 0.00967439 | 0.00839572 | 0.00870266 | -0.0003069 | 0.5928059 | 0.99999994 |
| Eno1b | 0.01551778 | 0.01359715 | 0.01390421 | -0.0003071 | 0.55843265 | 0.99999994 |
| Ccdc39 | 0.00563435 | 0.00477519 | 0.0050823 | -0.0003071 | 0.65624126 | 0.99999994 |
| Csrnp2 | 0.01533154 | 0.01343139 | 0.01373905 | -0.0003077 | 0.55924619 | 0.99999994 |
| Npr1 | 0.00280672 | 0.00222871 | 0.00253658 | -0.0003079 | 0.79042427 | 0.99999994 |
| Btg3 | 0.00079065 | 0.00040759 | 0.00071553 | -0.0003079 | 0.9979128 | 0.99999994 |
| Cfap58 | 0.00114312 | 0.00072584 | 0.00103426 | -0.0003084 | 0.9764257 | 0.99999994 |
| Gm12932 | 0.00086376 | 0.00047269 | 0.00078166 | -0.000309 | 0.99574432 | 0.99999994 |
| Shroom1 | 0.01398474 | 0.01223442 | 0.01254352 | -0.0003091 | 0.56514572 | 0.99999994 |
| Faap20 | 0.32526549 | 0.23151754 | 0.23182713 | -0.0003096 | 0.5035461 | 0.99999994 |
| Gm17146 | 0.00239931 | 0.00185935 | 0.00216899 | -0.0003096 | 0.82900417 | 0.99999994 |
| Kif18a | 0.00307015 | 0.00246446 | 0.00277416 | -0.0003097 | 0.77128884 | 0.99999994 |
| Cacna2d4 | 0.00383083 | 0.0031499 | 0.00345971 | -0.0003098 | 0.72443674 | 0.99999994 |
| Pcdhb4 | 0.00119923 | 0.00077507 | 0.00108499 | -0.0003099 | 0.97136961 | 0.99999994 |
| Gm43579 | 0.00641067 | 0.00546944 | 0.00577951 | -0.0003101 | 0.63949944 | 0.99999994 |
| Muc20 | 0.01125223 | 0.00980073 | 0.01011123 | -0.0003105 | 0.58098294 | 0.99999994 |

| | | | | | | |
|---|---|---|---|---|---|---|
| Rorb | 0.00320153 | 0.00258194 | 0.00289262 | -0.0003107 | 0.76267007 | 0.99999994 |
| Cfap69 | 0.0114322 | 0.00996089 | 0.01027171 | -0.0003108 | 0.57981347 | 0.99999994 |
| Slc2a3 | 0.00346588 | 0.00281987 | 0.00313089 | -0.000311 | 0.74576892 | 0.99999994 |
| 2410004P03I | 0.00155038 | 0.001091 | 0.00140236 | -0.0003114 | 0.93027669 | 0.99999994 |
| Nrp2 | 0.00800238 | 0.00689479 | 0.00720674 | -0.000312 | 0.61337552 | 0.99999994 |
| E130018N17 | 0.00215334 | 0.00163499 | 0.00194695 | -0.000312 | 0.8569093 | 0.99999994 |
| Mb21d2 | 0.04103078 | 0.03583104 | 0.03614344 | -0.0003124 | 0.5229402 | 0.99999994 |
| Ppargc1a | 0.00293329 | 0.00233811 | 0.00265073 | -0.0003126 | 0.78378854 | 0.99999994 |
| Bcl2l14 | 0.00523346 | 0.00440888 | 0.00472197 | -0.0003131 | 0.67051819 | 0.99999994 |
| 9530059O14 | 0.00151984 | 0.00106155 | 0.00137476 | -0.0003132 | 0.93530475 | 0.99999994 |
| Gprc6a | 0.00345196 | 0.00280488 | 0.00311835 | -0.0003135 | 0.74828869 | 0.99999994 |
| Map2k6 | 0.02420364 | 0.02124715 | 0.0215608 | -0.0003136 | 0.53856933 | 0.99999994 |
| Gm19705 | 0.00219461 | 0.00167047 | 0.00198421 | -0.0003137 | 0.85371306 | 0.99999994 |
| Ephx3 | 0.00105277 | 0.00063861 | 0.00095257 | -0.000314 | 0.98587971 | 0.99999994 |
| Slc6a8 | 0.02867601 | 0.02515449 | 0.02546858 | -0.0003141 | 0.53271155 | 0.99999994 |
| Tmem106a | 0.02221941 | 0.01950534 | 0.01981955 | -0.0003142 | 0.54202203 | 0.99999994 |
| Lama4 | 0.00145577 | 0.00100244 | 0.00131686 | -0.0003144 | 0.94400539 | 0.99999994 |
| Tbl1xr1 | 0.08759916 | 0.07454258 | 0.07485745 | -0.0003149 | 0.51116857 | 0.99999994 |
| Gm14616 | 0.00088923 | 0.0004897 | 0.00080469 | -0.000315 | 0.99541439 | 0.99999994 |
| Gpr182 | 0.00241612 | 0.00186915 | 0.00218415 | -0.000315 | 0.83146487 | 0.99999994 |
| 4930537H20I | 0.00457579 | 0.00381532 | 0.00413042 | -0.0003151 | 0.69419946 | 0.99999994 |
| Sbspon | 0.00889752 | 0.00769282 | 0.00800803 | -0.0003152 | 0.60334254 | 0.99999994 |
| Neurl1a | 0.01736641 | 0.01522588 | 0.01554126 | -0.0003154 | 0.5537254 | 0.99999994 |
| Hspb1 | 0.04030081 | 0.0352015 | 0.03551759 | -0.0003161 | 0.52361867 | 0.99999994 |
| Cdk5r1 | 0.0560012 | 0.04853022 | 0.04884665 | -0.0003164 | 0.517197 | 0.99999994 |
| Plb1 | 0.00218939 | 0.00166307 | 0.0019795 | -0.0003164 | 0.85634753 | 0.99999994 |
| Gm42851 | 0.00202789 | 0.00151694 | 0.00183368 | -0.0003167 | 0.87488767 | 0.99999994 |
| Gngt2 | 0.00087993 | 0.0004793 | 0.00079628 | -0.000317 | 0.99597213 | 0.99999994 |
| Gm5784 | 0.00628654 | 0.00535077 | 0.00566808 | -0.0003173 | 0.64529102 | 0.99999994 |
| 5430402O13 | 0.00600034 | 0.00509343 | 0.00541108 | -0.0003177 | 0.65201153 | 0.99999994 |

| | | | | | | |
|---|---|---|---|---|---|---|
| Dnajc27 | 0.01284394 | 0.01121145 | 0.01152915 | -0.0003177 | 0.57276691 | 0.99999994 |
| Tmem201 | 0.04161161 | 0.03632324 | 0.036641 | -0.0003178 | 0.52301668 | 0.99999994 |
| Nphs1 | 0.00417291 | 0.00344992 | 0.00376778 | -0.0003179 | 0.71278431 | 0.99999994 |
| Mis18bp1 | 0.00330837 | 0.00267104 | 0.00298893 | -0.0003179 | 0.76050977 | 0.99999994 |
| Hgd | 0.00103215 | 0.00061602 | 0.00093393 | -0.0003179 | 0.98826826 | 0.99999994 |
| Gm10184 | 0.00324507 | 0.00261346 | 0.00293186 | -0.0003184 | 0.76512525 | 0.99999994 |
| A3galt2 | 0.00146124 | 0.00100251 | 0.0013218 | -0.0003193 | 0.94606911 | 0.99999994 |
| Micall2 | 0.00492611 | 0.0041262 | 0.00444558 | -0.0003194 | 0.68375124 | 0.99999994 |
| Ccdc13 | 0.00542863 | 0.00457801 | 0.00489741 | -0.0003194 | 0.66789767 | 0.99999994 |
| Ndnf | 0.00099921 | 0.0005847 | 0.00090415 | -0.0003194 | 0.99065817 | 0.99999994 |
| Hcar1 | 0.00083335 | 0.00043464 | 0.00075415 | -0.0003195 | 0.99759955 | 0.99999994 |
| Zbtb49 | 0.02052964 | 0.01801312 | 0.01833306 | -0.0003199 | 0.54623825 | 0.99999994 |
| Fcrls | 0.00380575 | 0.00311687 | 0.00343712 | -0.0003203 | 0.73243794 | 0.99999994 |
| Mcpt8 | 0.00129739 | 0.00085347 | 0.00117372 | -0.0003203 | 0.96533172 | 0.99999994 |
| Bhlhe40 | 0.00534974 | 0.00450611 | 0.0048265 | -0.0003204 | 0.670707 | 0.99999994 |
| 1810019N24 | 0.00143436 | 0.00097701 | 0.00129752 | -0.0003205 | 0.94993334 | 0.99999994 |
| Pappa | 0.00161079 | 0.00113596 | 0.00145694 | -0.000321 | 0.92874229 | 0.99999994 |
| Batf | 0.00170229 | 0.0012186 | 0.0015396 | -0.000321 | 0.91740626 | 0.99999994 |
| Kif27 | 0.00258435 | 0.00201455 | 0.00233597 | -0.0003214 | 0.8201329 | 0.99999994 |
| Frmd8os | 0.00612556 | 0.00520197 | 0.00552354 | -0.0003216 | 0.65081444 | 0.99999994 |
| Gm13075 | 0.00336983 | 0.0027225 | 0.00304433 | -0.0003218 | 0.75918142 | 0.99999994 |
| Pde1b | 0.00126473 | 0.00082216 | 0.0011442 | -0.000322 | 0.96949899 | 0.99999994 |
| Gm826 | 0.00324978 | 0.00261301 | 0.00293611 | -0.0003231 | 0.76806992 | 0.99999994 |
| Gzmm | 0.00162498 | 0.00114651 | 0.00146976 | -0.0003233 | 0.92839792 | 0.99999994 |
| Hspbap1 | 0.04430668 | 0.03862018 | 0.03894469 | -0.0003245 | 0.52211586 | 0.99999994 |
| Gabra4 | 0.00141906 | 0.00095915 | 0.00128369 | -0.0003245 | 0.95379902 | 0.99999994 |
| Hrh3 | 0.00398977 | 0.00327817 | 0.00360287 | -0.0003247 | 0.72570894 | 0.99999994 |
| Icmt | 0.03317018 | 0.02904675 | 0.02937197 | -0.0003252 | 0.52937642 | 0.99999994 |
| Slc22a2 | 0.00170492 | 0.00121651 | 0.00154198 | -0.0003255 | 0.91998529 | 0.99999994 |
| Lepr | 0.0042781 | 0.00353679 | 0.00386248 | -0.0003257 | 0.71269323 | 0.99999994 |

| | | | | | | |
|---|---|---|---|---|---|---|
| Ppfia4 | 0.00278956 | 0.00219528 | 0.0025211 | -0.0003258 | 0.80517352 | 0.99999994 |
| Fam83b | 0.00483772 | 0.00404024 | 0.00436607 | -0.0003258 | 0.69031979 | 0.99999994 |
| Klhl29 | 0.00152606 | 0.00105427 | 0.00138038 | -0.0003261 | 0.94208716 | 0.99999994 |
| Dclk2 | 0.0025853 | 0.00201067 | 0.00233682 | -0.0003262 | 0.82356262 | 0.99999994 |
| Cort | 0.00217989 | 0.00164466 | 0.00197092 | -0.0003263 | 0.86474187 | 0.99999994 |
| Rnf43 | 0.00725459 | 0.00621007 | 0.00653661 | -0.0003265 | 0.63025032 | 0.99999994 |
| Myo1g | 0.00075407 | 0.0003558 | 0.00068245 | -0.0003266 | 0.9992786 | 0.99999994 |
| Lamb2 | 0.00909779 | 0.00786049 | 0.00818717 | -0.0003267 | 0.6047279 | 0.99999994 |
| Pdp1 | 0.01788668 | 0.01567397 | 0.01600126 | -0.0003273 | 0.55414873 | 0.99999994 |
| Pou6f1 | 0.01433172 | 0.01252384 | 0.01285174 | -0.0003279 | 0.56742821 | 0.99999994 |
| Pcdhga7 | 0.01979569 | 0.01735822 | 0.01768635 | -0.0003281 | 0.54914172 | 0.99999994 |
| Gtf2a1l | 0.00430979 | 0.00356142 | 0.00389101 | -0.0003296 | 0.71356587 | 0.99999994 |
| Aox1 | 0.00051244 | 0.00013386 | 0.00046384 | -0.00033 | 0.99999891 | 0.99999994 |
| Zdhhc18 | 0.00958525 | 0.00829266 | 0.00862299 | -0.0003303 | 0.60063523 | 0.99999994 |
| Nr0b2 | 0.01580144 | 0.01382509 | 0.01415567 | -0.0003306 | 0.56176482 | 0.99999994 |
| Pdzrn4 | 0.0383934 | 0.0335487 | 0.03387936 | -0.0003307 | 0.52589883 | 0.99999994 |
| Gm10112 | 0.00076804 | 0.00036439 | 0.00069508 | -0.0003307 | 0.99922939 | 0.99999994 |
| Gnb1l | 0.02747987 | 0.02409503 | 0.02442572 | -0.0003307 | 0.53590354 | 0.99999994 |
| Gm16099 | 0.00086419 | 0.0004512 | 0.00078205 | -0.0003308 | 0.99756874 | 0.99999994 |
| Ptcd3 | 0.10194539 | 0.08596285 | 0.08629413 | -0.0003313 | 0.51019321 | 0.99999994 |
| Sema3d | 0.00116252 | 0.00072031 | 0.0010518 | -0.0003315 | 0.98204113 | 0.99999994 |
| Elane | 0.00097113 | 0.00054686 | 0.00087876 | -0.0003319 | 0.99403247 | 0.99999994 |
| Ngb | 0.0014479 | 0.00097682 | 0.00130975 | -0.0003329 | 0.95467862 | 0.99999994 |
| Pde6a | 0.00087108 | 0.0004547 | 0.00078828 | -0.0003336 | 0.99757473 | 0.99999994 |
| Gm15821 | 0.00096773 | 0.00054201 | 0.00087568 | -0.0003337 | 0.9943987 | 0.99999994 |
| Aldh3a1 | 0.00139845 | 0.00093127 | 0.00126506 | -0.0003338 | 0.96048138 | 0.99999994 |
| Gm43254 | 0.0037206 | 0.00302652 | 0.00336041 | -0.0003339 | 0.74581725 | 0.99999994 |
| A830018L16I | 0.00174214 | 0.00124107 | 0.0015756 | -0.0003345 | 0.9212119 | 0.99999994 |
| Clip3 | 0.14339689 | 0.1171128 | 0.11744742 | -0.0003346 | 0.50756534 | 0.99999994 |
| Gm7808 | 0.00748849 | 0.00641152 | 0.00674629 | -0.0003348 | 0.62941668 | 0.99999994 |

| | | | | | | |
|---|---|---|---|---|---|---|
| Cenpo | 0.01365639 | 0.0119165 | 0.01225172 | -0.0003352 | 0.57225664 | 0.99999994 |
| C1s1 | 0.00091493 | 0.00049267 | 0.00082793 | -0.0003353 | 0.99648471 | 0.99999994 |
| Eda2r | 0.00089466 | 0.00047417 | 0.0008096 | -0.0003354 | 0.99709129 | 0.99999994 |
| Stab1 | 0.00523728 | 0.00438968 | 0.0047254 | -0.0003357 | 0.68185862 | 0.99999994 |
| Adamts16 | 0.00372994 | 0.00303304 | 0.00336882 | -0.0003358 | 0.74648344 | 0.99999994 |
| Morn5 | 0.00268389 | 0.00208984 | 0.00242577 | -0.0003359 | 0.82168701 | 0.99999994 |
| AU020206 | 0.00829315 | 0.00713113 | 0.00746713 | -0.000336 | 0.61772705 | 0.99999994 |
| Gm14264 | 0.00229392 | 0.00173767 | 0.00207386 | -0.0003362 | 0.8597163 | 0.99999994 |
| Gm15326 | 0.00087194 | 0.00045264 | 0.00078906 | -0.0003364 | 0.99772994 | 0.99999994 |
| Klhl14 | 0.00124197 | 0.0007871 | 0.00112363 | -0.0003365 | 0.97690317 | 0.99999994 |
| Fosl1 | 0.00144852 | 0.00097375 | 0.00131032 | -0.0003366 | 0.95634127 | 0.99999994 |
| Ddx11 | 0.0073388 | 0.00627486 | 0.00661211 | -0.0003372 | 0.63288583 | 0.99999994 |
| Ccr2 | 0.00073248 | 0.00032524 | 0.00066291 | -0.0003377 | 0.99965126 | 0.99999994 |
| Cbr3 | 0.00362266 | 0.00293448 | 0.00327218 | -0.0003377 | 0.75394707 | 0.99999994 |
| Gm3604 | 0.00825574 | 0.00709589 | 0.00743363 | -0.0003377 | 0.61883781 | 0.99999994 |
| Zgrf1 | 0.00385735 | 0.00314568 | 0.0034836 | -0.0003379 | 0.7407588 | 0.99999994 |
| Gp2 | 0.00056384 | 0.00017233 | 0.00051035 | -0.000338 | 0.9999948 | 0.99999994 |
| AI429214 | 0.02745841 | 0.02406869 | 0.024407 | -0.0003383 | 0.53675623 | 0.99999994 |
| 2810032G03 | 0.00219522 | 0.00164642 | 0.00198476 | -0.0003383 | 0.87175406 | 0.99999994 |
| Casd1 | 0.10642482 | 0.08942416 | 0.08976324 | -0.0003391 | 0.51003013 | 0.99999994 |
| Gm20109 | 0.00284269 | 0.00222992 | 0.00256903 | -0.0003391 | 0.81019747 | 0.99999994 |
| Gm20442 | 0.00334775 | 0.00268527 | 0.00302443 | -0.0003392 | 0.7723 | 0.99999994 |
| Aqp8 | 0.00325096 | 0.00259712 | 0.00293718 | -0.0003401 | 0.77954051 | 0.99999994 |
| Gm12204 | 0.00120361 | 0.0007484 | 0.00108895 | -0.0003406 | 0.9813135 | 0.99999994 |
| 9130008F23I | 0.00521682 | 0.00436609 | 0.004707 | -0.0003409 | 0.68513449 | 0.99999994 |
| Chil1 | 0.00682887 | 0.00581386 | 0.00615479 | -0.0003409 | 0.64383054 | 0.99999994 |
| Ank1 | 0.00078061 | 0.00036531 | 0.00070645 | -0.0003411 | 0.99934616 | 0.99999994 |
| Actrt3 | 0.00739173 | 0.00631803 | 0.00665956 | -0.0003415 | 0.63358551 | 0.99999994 |
| Gtf2ird2 | 0.02298139 | 0.02014719 | 0.02048877 | -0.0003416 | 0.54418129 | 0.99999994 |
| Adamts18 | 0.00076785 | 0.00035278 | 0.00069491 | -0.0003421 | 0.9994757 | 0.99999994 |

| | | | | | | |
|---|---|---|---|---|---|---|
| Ly6k | 0.00223644 | 0.0016791 | 0.00202197 | -0.0003429 | 0.87049489 | 0.99999994 |
| Ugt2b36 | 0.00144629 | 0.00096449 | 0.0013083 | -0.0003438 | 0.959872 | 0.99999994 |
| Gm19426 | 0.01212394 | 0.01054378 | 0.01088814 | -0.0003444 | 0.58336992 | 0.99999994 |
| Il18 | 0.00526325 | 0.00440405 | 0.00474875 | -0.0003447 | 0.68551178 | 0.99999994 |
| Dclk1 | 0.00145419 | 0.00097046 | 0.00131544 | -0.000345 | 0.95956191 | 0.99999994 |
| Ankrd45 | 0.00715375 | 0.00610095 | 0.00644618 | -0.0003452 | 0.63926317 | 0.99999994 |
| Rhoj | 0.00189954 | 0.00137222 | 0.00171777 | -0.0003455 | 0.90971504 | 0.99999994 |
| Aldh1a2 | 0.00484455 | 0.0040266 | 0.00437222 | -0.0003456 | 0.70061741 | 0.99999994 |
| Ptgs1 | 0.00281892 | 0.00220181 | 0.00254758 | -0.0003458 | 0.81685622 | 0.99999994 |
| Tceal5 | 0.00544311 | 0.00456448 | 0.00491043 | -0.0003459 | 0.68044568 | 0.99999994 |
| Dtwd2 | 0.0193201 | 0.01692042 | 0.01726694 | -0.0003465 | 0.55313482 | 0.99999994 |
| Gm379 | 0.00056777 | 0.00016729 | 0.00051391 | -0.0003466 | 0.99999643 | 0.99999994 |
| Oasl1 | 0.00323741 | 0.00257824 | 0.00292496 | -0.0003467 | 0.78494936 | 0.99999994 |
| Mbd1 | 0.07936709 | 0.0678364 | 0.06818319 | -0.0003468 | 0.51350401 | 0.99999994 |
| Mxd3 | 0.00253658 | 0.00194573 | 0.00229286 | -0.0003471 | 0.84320975 | 0.99999994 |
| Gm43714 | 0.00106538 | 0.00061676 | 0.00096398 | -0.0003472 | 0.99174796 | 0.99999994 |
| Angptl4 | 0.00481894 | 0.00400192 | 0.00434918 | -0.0003473 | 0.70245527 | 0.99999994 |
| Phrf1 | 0.13064537 | 0.10774044 | 0.10808773 | -0.0003473 | 0.50853165 | 0.99999994 |
| Mkln1 | 0.39437135 | 0.26682871 | 0.26717617 | -0.0003475 | 0.5034534 | 0.99999994 |
| Mfhas1 | 0.04987176 | 0.04332807 | 0.04367575 | -0.0003477 | 0.52112921 | 0.99999994 |
| Eri2 | 0.02354005 | 0.02063082 | 0.02097898 | -0.0003482 | 0.54398071 | 0.99999994 |
| Synm | 0.00178525 | 0.00126637 | 0.00161455 | -0.0003482 | 0.92442165 | 0.99999994 |
| Mynn | 0.11009631 | 0.09223962 | 0.09258785 | -0.0003482 | 0.50998637 | 0.99999994 |
| Btbd19 | 0.00865866 | 0.00744586 | 0.00779431 | -0.0003484 | 0.61698499 | 0.99999994 |
| Loxl3 | 0.00700703 | 0.00596587 | 0.00631461 | -0.0003487 | 0.64341931 | 0.99999994 |
| Eapp | 0.38410426 | 0.26182204 | 0.26217151 | -0.0003495 | 0.50353965 | 0.99999994 |
| Lipk | 0.00058287 | 0.00017767 | 0.00052756 | -0.0003499 | 0.99999494 | 0.99999994 |
| Rab27b | 0.00650306 | 0.00551228 | 0.00586244 | -0.0003502 | 0.65452828 | 0.99999994 |
| Amy2a5 | 0.00250498 | 0.00191402 | 0.00226435 | -0.0003503 | 0.84845527 | 0.99999994 |
| Fap | 0.00183161 | 0.00130585 | 0.00165642 | -0.0003506 | 0.92054664 | 0.99999994 |

| | | | | | | |
|---|---|---|---|---|---|---|
| Jchain | 0.00180353 | 0.00128016 | 0.00163105 | -0.0003509 | 0.92393173 | 0.99999994 |
| Pdgfb | 0.00064894 | 0.00023619 | 0.00058734 | -0.0003512 | 0.99996549 | 0.99999994 |
| Hbq1b | 0.00065097 | 0.00023788 | 0.00058918 | -0.0003513 | 0.9999639 | 0.99999994 |
| Ap5z1 | 0.03649725 | 0.03189506 | 0.03224669 | -0.0003516 | 0.52893079 | 0.99999994 |
| Tmem40 | 0.00095422 | 0.00051172 | 0.00086347 | -0.0003517 | 0.99665175 | 0.99999994 |
| Tmem169 | 0.00446716 | 0.00368056 | 0.00403266 | -0.0003521 | 0.71943926 | 0.99999994 |
| Phldb3 | 0.02516429 | 0.02204986 | 0.02240215 | -0.0003523 | 0.54168323 | 0.99999994 |
| Gm31373 | 0.00129911 | 0.00082284 | 0.00117528 | -0.0003524 | 0.97703738 | 0.99999994 |
| Gm15829 | 0.00234982 | 0.00177171 | 0.00212432 | -0.0003526 | 0.86538922 | 0.99999994 |
| A830009L08I | 0.00116634 | 0.00070251 | 0.00105526 | -0.0003527 | 0.98696152 | 0.99999994 |
| Gm15688 | 0.00103658 | 0.00058515 | 0.00093794 | -0.0003528 | 0.99385414 | 0.99999994 |
| Adh7 | 0.00058701 | 0.00017782 | 0.00053131 | -0.0003535 | 0.99999526 | 0.99999994 |
| Fscn2 | 0.00189067 | 0.00135571 | 0.00170976 | -0.0003541 | 0.91595542 | 0.99999994 |
| 4933424G06 | 0.00070641 | 0.00028495 | 0.00063933 | -0.0003544 | 0.99988771 | 0.99999994 |
| Cdk5 | 0.2475905 | 0.18671381 | 0.18706844 | -0.0003546 | 0.50503396 | 0.99999994 |
| Ska2 | 0.09767786 | 0.08257863 | 0.08293343 | -0.0003548 | 0.51135913 | 0.99999994 |
| Stfa3 | 0.00153033 | 0.00102918 | 0.00138424 | -0.0003551 | 0.9561238 | 0.99999994 |
| Sardh | 0.00375158 | 0.00303286 | 0.00338832 | -0.0003555 | 0.75750739 | 0.99999994 |
| Rpl13-ps3 | 0.27995306 | 0.206048 | 0.20640378 | -0.0003558 | 0.50457726 | 0.99999994 |
| Abcc10 | 0.00897418 | 0.00772076 | 0.00807661 | -0.0003559 | 0.61534415 | 0.99999994 |
| Gm10110 | 0.01006484 | 0.00869511 | 0.00905151 | -0.0003564 | 0.60337559 | 0.99999994 |
| Gm27045 | 0.0009092 | 0.00046614 | 0.00082275 | -0.0003566 | 0.99804341 | 0.99999994 |
| Gpha2 | 0.00275723 | 0.00213509 | 0.00249194 | -0.0003568 | 0.82975194 | 0.99999994 |
| Hs6st3 | 0.00308863 | 0.00243396 | 0.00279082 | -0.0003569 | 0.80266095 | 0.99999994 |
| Pcsk5 | 0.00311663 | 0.00245871 | 0.00281607 | -0.0003574 | 0.80086001 | 0.99999994 |
| B430203G13 | 0.00416415 | 0.00340251 | 0.00375989 | -0.0003574 | 0.73653439 | 0.99999994 |
| Bmp3 | 0.00227783 | 0.00170187 | 0.00205933 | -0.0003575 | 0.8760435 | 0.99999994 |
| Traf3ip1 | 0.0581696 | 0.05030839 | 0.05066601 | -0.0003576 | 0.518737 | 0.99999994 |
| Cldn15 | 0.00283098 | 0.00220083 | 0.00255846 | -0.0003576 | 0.82392977 | 0.99999994 |
| 9430038I01R | 0.07950349 | 0.06793624 | 0.06829436 | -0.0003581 | 0.51392191 | 0.99999994 |

| | | | | | | |
|---|---|---|---|---|---|---|
| A330094K24 | 0.00065103 | 0.00023054 | 0.00058923 | -0.0003587 | 0.99997461 | 0.99999994 |
| Pm20d1 | 0.00213269 | 0.00156953 | 0.00192831 | -0.0003588 | 0.89222921 | 0.99999994 |
| C4bp | 0.00403693 | 0.00328654 | 0.00364534 | -0.0003588 | 0.74382127 | 0.99999994 |
| Apoa1 | 0.00210974 | 0.00154863 | 0.00190759 | -0.000359 | 0.89481399 | 0.99999994 |
| Gm38004 | 0.00082137 | 0.00038435 | 0.00074331 | -0.000359 | 0.99934669 | 0.99999994 |
| C920006O11 | 0.00616745 | 0.00520176 | 0.00556116 | -0.0003594 | 0.66646377 | 0.99999994 |
| A230009B12 | 0.00062367 | 0.00020375 | 0.00056448 | -0.0003607 | 0.99998949 | 0.99999994 |
| Polr3b | 0.03951308 | 0.03448075 | 0.03484152 | -0.0003608 | 0.52747539 | 0.99999994 |
| Gstt3 | 0.00805927 | 0.00689687 | 0.00725769 | -0.0003608 | 0.62964911 | 0.99999994 |
| Mkx | 0.00246882 | 0.0018708 | 0.00223172 | -0.0003609 | 0.85914572 | 0.99999994 |
| Glt8d2 | 0.00064797 | 0.00022536 | 0.00058646 | -0.0003611 | 0.99997921 | 0.99999994 |
| Taf1c | 0.02735277 | 0.02395359 | 0.02431482 | -0.0003612 | 0.53938667 | 0.99999994 |
| Gm44036 | 0.00113834 | 0.0006681 | 0.00102995 | -0.0003618 | 0.99031994 | 0.99999994 |
| Shisa9 | 0.00111202 | 0.00064422 | 0.00100615 | -0.0003619 | 0.99167671 | 0.99999994 |
| Hsd11b2 | 0.00168106 | 0.00115842 | 0.00152043 | -0.000362 | 0.94350175 | 0.99999994 |
| Vtcn1 | 0.00257119 | 0.00196201 | 0.00232409 | -0.0003621 | 0.85013317 | 0.99999994 |
| Ptgis | 0.01028554 | 0.00888643 | 0.00924861 | -0.0003622 | 0.60282399 | 0.99999994 |
| Xkr4 | 0.00069878 | 0.00027018 | 0.00063243 | -0.0003622 | 0.99993127 | 0.99999994 |
| Pla2g7 | 0.00175901 | 0.00122804 | 0.00159084 | -0.0003628 | 0.93549342 | 0.99999994 |
| Hbq1a | 0.00070397 | 0.00027413 | 0.00063713 | -0.000363 | 0.99992541 | 0.99999994 |
| Mr1 | 0.00497208 | 0.0041239 | 0.00448693 | -0.000363 | 0.70490232 | 0.99999994 |
| Nrbp2 | 0.01355007 | 0.0117941 | 0.01215721 | -0.0003631 | 0.57879422 | 0.99999994 |
| Slfn2 | 0.00361316 | 0.00290007 | 0.00326361 | -0.0003635 | 0.77079748 | 0.99999994 |
| Parvb | 0.00301214 | 0.00235814 | 0.00272185 | -0.0003637 | 0.81311933 | 0.99999994 |
| Gm37634 | 0.00146173 | 0.00095838 | 0.00132225 | -0.0003639 | 0.96650573 | 0.99999994 |
| Gm26912 | 0.0030286 | 0.00237258 | 0.00273669 | -0.0003641 | 0.81208735 | 0.99999994 |
| Ptprcap | 0.00364799 | 0.00293071 | 0.003295 | -0.0003643 | 0.76910853 | 0.99999994 |
| Gm32950 | 0.00068898 | 0.00025907 | 0.00062357 | -0.0003645 | 0.99995005 | 0.99999994 |
| Rwdd2a | 0.05668244 | 0.04905414 | 0.04941879 | -0.0003647 | 0.51958664 | 0.99999994 |
| Hand2 | 0.00298305 | 0.00233065 | 0.00269561 | -0.000365 | 0.81626077 | 0.99999994 |

| | | | | | | |
|---|---|---|---|---|---|---|
| Gm27200 | 0.00086053 | 0.00041363 | 0.00077873 | -0.0003651 | 0.99909814 | 0.99999994 |
| 1810041H14I | 0.01171354 | 0.01015727 | 0.01052249 | -0.0003652 | 0.59135382 | 0.99999994 |
| A930016O22 | 0.00093813 | 0.00048311 | 0.00084891 | -0.0003658 | 0.99793643 | 0.99999994 |
| Il23a | 0.00584982 | 0.00490985 | 0.00527589 | -0.000366 | 0.67789328 | 0.99999994 |
| Gm10300 | 0.00089346 | 0.00044236 | 0.00080852 | -0.0003662 | 0.99871288 | 0.99999994 |
| Zmiz1os1 | 0.00548171 | 0.00457863 | 0.00494512 | -0.0003665 | 0.68910426 | 0.99999994 |
| Mob3a | 0.05891166 | 0.05092017 | 0.05128744 | -0.0003673 | 0.51900922 | 0.99999994 |
| Iqcf4 | 0.00064579 | 0.00021713 | 0.00058449 | -0.0003674 | 0.99998565 | 0.99999994 |
| Wdr19 | 0.01682593 | 0.01469523 | 0.01506306 | -0.0003678 | 0.56456219 | 0.99999994 |
| Acoxl | 0.00111153 | 0.00063777 | 0.0010057 | -0.0003679 | 0.99255826 | 0.99999994 |
| Upk1b | 0.00386543 | 0.00312258 | 0.00349088 | -0.0003683 | 0.75874321 | 0.99999994 |
| Sfxn2 | 0.25209295 | 0.18945151 | 0.18981992 | -0.0003684 | 0.50515378 | 0.99999994 |
| 4632411P08I | 0.01833367 | 0.0160275 | 0.01639621 | -0.0003687 | 0.55949326 | 0.99999994 |
| Ccdc171 | 0.00296934 | 0.00231454 | 0.00268325 | -0.0003687 | 0.81981526 | 0.99999994 |
| Fchsd1 | 0.01246652 | 0.0108243 | 0.01119322 | -0.0003689 | 0.58682456 | 0.99999994 |
| Wfdc3 | 0.00460903 | 0.0037913 | 0.00416033 | -0.000369 | 0.72254881 | 0.99999994 |
| Nfkb2 | 0.02553387 | 0.02235622 | 0.02272554 | -0.0003693 | 0.54307176 | 0.99999994 |
| Bora | 0.00934847 | 0.00804173 | 0.00841133 | -0.0003696 | 0.61504259 | 0.99999994 |
| S100a14 | 0.00243814 | 0.00183387 | 0.00220403 | -0.0003702 | 0.86819441 | 0.99999994 |
| Dock4 | 0.00883549 | 0.00758235 | 0.00795253 | -0.0003702 | 0.62166068 | 0.99999994 |
| Qtrt1 | 0.23159563 | 0.17676076 | 0.17713096 | -0.0003702 | 0.50554968 | 0.99999994 |
| Akap12 | 0.00623252 | 0.00524934 | 0.00561957 | -0.0003702 | 0.66950109 | 0.99999994 |
| Slc39a4 | 0.00601248 | 0.00505172 | 0.00542199 | -0.0003703 | 0.67528858 | 0.99999994 |
| Lgi1 | 0.00200877 | 0.00144611 | 0.00181641 | -0.0003703 | 0.91260893 | 0.99999994 |
| Shc4 | 0.00225111 | 0.00166467 | 0.00203521 | -0.0003705 | 0.88722338 | 0.99999994 |
| C130060C02I | 0.00288658 | 0.00223805 | 0.00260861 | -0.0003706 | 0.82781175 | 0.99999994 |
| Pxmp2 | 0.00958833 | 0.00825516 | 0.00862575 | -0.0003706 | 0.61255299 | 0.99999994 |
| Gm16196 | 0.0020417 | 0.00147493 | 0.00184615 | -0.0003712 | 0.90962564 | 0.99999994 |
| Ccdc142 | 0.00098174 | 0.00051688 | 0.00088835 | -0.0003715 | 0.99731069 | 0.99999994 |
| 1700058P15I | 0.00275205 | 0.00211577 | 0.00248726 | -0.0003715 | 0.83992962 | 0.99999994 |

| | | | | | | |
|---|---|---|---|---|---|---|
| Stra8 | 0.00067988 | 0.00024327 | 0.00061533 | -0.0003721 | 0.99997149 | 0.99999994 |
| Synpo2 | 0.0007274 | 0.00028562 | 0.00065832 | -0.0003727 | 0.99991787 | 0.99999994 |
| Xrcc2 | 0.0051518 | 0.00427583 | 0.00464855 | -0.0003727 | 0.70322743 | 0.99999994 |
| Slc16a8 | 0.00318968 | 0.00250917 | 0.00288193 | -0.0003728 | 0.80536847 | 0.99999994 |
| Acad10 | 0.00902659 | 0.00775069 | 0.00812349 | -0.0003728 | 0.61999549 | 0.99999994 |
| Dock5 | 0.00215243 | 0.00157237 | 0.00194613 | -0.0003738 | 0.89944144 | 0.99999994 |
| 4930593A02I | 0.00066489 | 0.00022785 | 0.00060177 | -0.0003739 | 0.99998233 | 0.99999994 |
| Zkscan5 | 0.01626429 | 0.01419154 | 0.01456577 | -0.0003742 | 0.56789538 | 0.99999994 |
| Gnrh1 | 0.00067303 | 0.00023472 | 0.00060913 | -0.0003744 | 0.99997856 | 0.99999994 |
| Ttc21a | 0.00267033 | 0.00203843 | 0.00241354 | -0.0003751 | 0.84955196 | 0.99999994 |
| Gm14412 | 0.00148706 | 0.00096981 | 0.00134514 | -0.0003753 | 0.96836665 | 0.99999994 |
| Car5b | 0.00114508 | 0.00066059 | 0.00103604 | -0.0003754 | 0.99207108 | 0.99999994 |
| Kcnj4 | 0.00141909 | 0.00090763 | 0.00128372 | -0.0003761 | 0.97441814 | 0.99999994 |
| Trmt10a | 0.0857309 | 0.07297275 | 0.07334893 | -0.0003762 | 0.51361656 | 0.99999994 |
| Nes | 0.00206971 | 0.00149437 | 0.00187144 | -0.0003771 | 0.91006961 | 0.99999994 |
| Stmn3 | 0.02593134 | 0.02269551 | 0.02307316 | -0.0003777 | 0.54337816 | 0.99999994 |
| Ice1 | 0.06141506 | 0.05300169 | 0.05337944 | -0.0003778 | 0.51878544 | 0.99999994 |
| 4930553P18I | 0.0008702 | 0.00040849 | 0.00078749 | -0.000379 | 0.9993212 | 0.99999994 |
| Samd12 | 0.00121354 | 0.00071858 | 0.00109793 | -0.0003793 | 0.98927167 | 0.99999994 |
| Star | 0.00285832 | 0.00220378 | 0.00258313 | -0.0003793 | 0.83584645 | 0.99999994 |
| 9430041J12F | 0.00306087 | 0.00238563 | 0.00276579 | -0.0003802 | 0.81988641 | 0.99999994 |
| Phf19 | 0.00156751 | 0.00103735 | 0.00141784 | -0.0003805 | 0.96297085 | 0.99999994 |
| Ncoa4 | 0.00520871 | 0.00431921 | 0.00469971 | -0.0003805 | 0.7050297 | 0.99999994 |
| 1110046J04F | 0.02194379 | 0.0191965 | 0.01957732 | -0.0003808 | 0.55151191 | 0.99999994 |
| Plin5 | 0.01118333 | 0.00966896 | 0.01004979 | -0.0003808 | 0.59957077 | 0.99999994 |
| Ephx1 | 0.01147763 | 0.00993102 | 0.01031221 | -0.0003812 | 0.59717908 | 0.99999994 |
| Pias2 | 0.20790716 | 0.16155064 | 0.16193195 | -0.0003813 | 0.50625282 | 0.99999994 |
| Radil | 0.0045368 | 0.00371357 | 0.00409533 | -0.0003818 | 0.7325356 | 0.99999994 |
| Mybl2 | 0.00375659 | 0.00301102 | 0.00339283 | -0.0003818 | 0.77309281 | 0.99999994 |
| P4ha3 | 0.00699753 | 0.00592367 | 0.00630608 | -0.0003824 | 0.65676723 | 0.99999994 |

| | | | | | | |
|---|---|---|---|---|---|---|
| Wdr4 | 0.05547847 | 0.04802482 | 0.04840729 | -0.0003825 | 0.5209719 | 0.99999994 |
| Mtr | 0.00342836 | 0.00271371 | 0.00309708 | -0.0003834 | 0.79501762 | 0.99999994 |
| Zfp384 | 0.00792542 | 0.00675425 | 0.0071378 | -0.0003835 | 0.6397084 | 0.99999994 |
| Hlf | 0.00139811 | 0.00088058 | 0.00126475 | -0.0003842 | 0.97840652 | 0.99999994 |
| Rab4b | 0.01522227 | 0.01325767 | 0.01364213 | -0.0003845 | 0.57440187 | 0.99999994 |
| Wdr37 | 0.06999621 | 0.06011421 | 0.06049877 | -0.0003846 | 0.51687458 | 0.99999994 |
| Rep15 | 0.00556953 | 0.00463911 | 0.00502405 | -0.0003849 | 0.69497551 | 0.99999994 |
| Abca4 | 0.00106421 | 0.0005779 | 0.00096292 | -0.000385 | 0.99611013 | 0.99999994 |
| Sp6 | 0.00085112 | 0.00038482 | 0.00077022 | -0.0003854 | 0.99956685 | 0.99999994 |
| Aff3 | 0.0023586 | 0.00174607 | 0.00213224 | -0.0003862 | 0.88600151 | 0.99999994 |
| Cfap73 | 0.00177862 | 0.00122228 | 0.00160855 | -0.0003863 | 0.94502688 | 0.99999994 |
| Cntnap3 | 0.00348668 | 0.00276328 | 0.00314964 | -0.0003864 | 0.79290112 | 0.99999994 |
| Cldn34c1 | 0.00676147 | 0.00570777 | 0.00609433 | -0.0003866 | 0.66356175 | 0.99999994 |
| Best3 | 0.00147797 | 0.0009503 | 0.00133692 | -0.0003866 | 0.97288155 | 0.99999994 |
| Ddo | 0.00639938 | 0.00538224 | 0.00576937 | -0.0003871 | 0.67243817 | 0.99999994 |
| Ripk3 | 0.00720057 | 0.00610028 | 0.00648817 | -0.0003879 | 0.65466609 | 0.99999994 |
| Nutf2 | 0.1488893 | 0.1210312 | 0.12141933 | -0.0003881 | 0.50848789 | 0.99999994 |
| Zfp449 | 0.02614293 | 0.02286941 | 0.02325814 | -0.0003887 | 0.54429165 | 0.99999994 |
| Noxred1 | 0.0023339 | 0.00172075 | 0.00210995 | -0.0003892 | 0.89024055 | 0.99999994 |
| Aqp3 | 0.00179474 | 0.00123379 | 0.00162311 | -0.0003893 | 0.94482263 | 0.99999994 |
| Gm14327 | 0.0044501 | 0.0036275 | 0.0040173 | -0.0003898 | 0.74081925 | 0.99999994 |
| Elfn2 | 0.00350741 | 0.00277833 | 0.00316832 | -0.00039 | 0.79370499 | 0.99999994 |
| Ifi27l2a | 0.00422767 | 0.00342707 | 0.00381708 | -0.00039 | 0.75178063 | 0.99999994 |
| Dgat2 | 0.01218143 | 0.01054918 | 0.01093935 | -0.0003902 | 0.59383183 | 0.99999994 |
| Irf2 | 0.29877759 | 0.21680263 | 0.21719313 | -0.0003905 | 0.50477426 | 0.99999994 |
| Apol7d | 0.00295502 | 0.00227962 | 0.00267034 | -0.0003907 | 0.8349559 | 0.99999994 |
| Crygc | 0.00084898 | 0.00037733 | 0.00076829 | -0.000391 | 0.9996469 | 0.99999994 |
| Rp2 | 0.10903164 | 0.0913795 | 0.09177049 | -0.000391 | 0.51131241 | 0.99999994 |
| Fxyd5 | 0.00735986 | 0.00623979 | 0.00663099 | -0.0003912 | 0.6527271 | 0.99999994 |
| Col7a1 | 0.00266931 | 0.00202113 | 0.00241262 | -0.0003915 | 0.85995156 | 0.99999994 |

| | | | | | | |
|---|---|---|---|---|---|---|
| Ankef1 | 0.00285926 | 0.00219237 | 0.00258397 | -0.0003916 | 0.84345983 | 0.99999994 |
| Slc16a5 | 0.01082565 | 0.00933882 | 0.00973071 | -0.0003919 | 0.60567997 | 0.99999994 |
| Dpep1 | 0.00112764 | 0.00062767 | 0.00102027 | -0.0003926 | 0.99478633 | 0.99999994 |
| Cd53 | 0.00276306 | 0.00210411 | 0.0024972 | -0.0003931 | 0.85262992 | 0.99999994 |
| Htra1 | 0.004913 | 0.00404063 | 0.00443379 | -0.0003932 | 0.7224855 | 0.99999994 |
| Ifi27 | 0.0852319 | 0.0725518 | 0.0729454 | -0.0003936 | 0.51432571 | 0.99999994 |
| Slc38a3 | 0.00898912 | 0.00769595 | 0.00808997 | -0.000394 | 0.62710675 | 0.99999994 |
| Zfp808 | 0.00371681 | 0.00296284 | 0.003357 | -0.0003942 | 0.78276536 | 0.99999994 |
| Gm16191 | 0.00388549 | 0.00311414 | 0.00350895 | -0.0003948 | 0.77305262 | 0.99999994 |
| Caprin2 | 0.02182906 | 0.01908158 | 0.01947646 | -0.0003949 | 0.55367565 | 0.99999994 |
| Evi2a | 0.00096121 | 0.0004748 | 0.00086979 | -0.000395 | 0.99874756 | 0.99999994 |
| Ogn | 0.00126183 | 0.00074628 | 0.00114158 | -0.0003953 | 0.989414 | 0.99999994 |
| Arhgap18 | 0.0503698 | 0.04370215 | 0.04409747 | -0.0003953 | 0.52379108 | 0.99999994 |
| D930030I03F | 0.00658341 | 0.00553908 | 0.00593455 | -0.0003955 | 0.67132431 | 0.99999994 |
| Timm23 | 0.01385235 | 0.01203013 | 0.01242588 | -0.0003958 | 0.5839465 | 0.99999994 |
| Fam110c | 0.00721519 | 0.00610535 | 0.00650128 | -0.0003959 | 0.65739895 | 0.99999994 |
| Slfn9 | 0.0022558 | 0.0016429 | 0.00203945 | -0.0003965 | 0.90221106 | 0.99999994 |
| Pcdhgb5 | 0.00358434 | 0.00284051 | 0.00323765 | -0.0003971 | 0.79289008 | 0.99999994 |
| Gm43848 | 0.00561043 | 0.00466351 | 0.00506081 | -0.0003973 | 0.69935661 | 0.99999994 |
| Cacnb4 | 0.00219797 | 0.0015899 | 0.00198724 | -0.0003973 | 0.90839227 | 0.99999994 |
| Hotairm1 | 0.00089873 | 0.00041555 | 0.00081328 | -0.0003977 | 0.99943362 | 0.99999994 |
| Msantd1 | 0.00174342 | 0.00117877 | 0.00157676 | -0.000398 | 0.95353353 | 0.99999994 |
| Ammecr1l | 0.16365468 | 0.13152466 | 0.13192265 | -0.000398 | 0.50801075 | 0.99999994 |
| Ttc30b | 0.08496062 | 0.07232677 | 0.07272592 | -0.0003991 | 0.51457116 | 0.99999994 |
| Cdc25c | 0.00172978 | 0.00116394 | 0.00156444 | -0.0004005 | 0.95581311 | 0.99999994 |
| Pcdhgb4 | 0.0051128 | 0.00421296 | 0.00461348 | -0.0004005 | 0.71832794 | 0.99999994 |
| Nek3 | 0.03964503 | 0.03455425 | 0.03495482 | -0.0004006 | 0.53040193 | 0.99999994 |
| Gm43713 | 0.00150519 | 0.00096007 | 0.00136152 | -0.0004015 | 0.97515751 | 0.99999994 |
| Lrrc4 | 0.00109035 | 0.00058507 | 0.00098656 | -0.0004015 | 0.99662445 | 0.99999994 |
| Rasef | 0.00204806 | 0.00145013 | 0.00185189 | -0.0004018 | 0.92564179 | 0.99999994 |

| | | | | | | |
|---|---|---|---|---|---|---|
| Cd48 | 0.00114951 | 0.0006379 | 0.00104004 | -0.0004021 | 0.99497013 | 0.99999994 |
| Gls2 | 0.00515851 | 0.00425226 | 0.00465458 | -0.0004023 | 0.7174683 | 0.99999994 |
| Grik4 | 0.00140836 | 0.00087154 | 0.00127402 | -0.0004025 | 0.9822589 | 0.99999994 |
| Palm | 0.0819079 | 0.06984804 | 0.07025074 | -0.0004027 | 0.51521871 | 0.99999994 |
| Emilin2 | 0.00126848 | 0.00074456 | 0.00114759 | -0.000403 | 0.9902986 | 0.99999994 |
| Pvt1 | 0.00195009 | 0.00136009 | 0.00176343 | -0.0004033 | 0.93605464 | 0.99999994 |
| Arhgap28 | 0.01335465 | 0.01157962 | 0.01198345 | -0.0004038 | 0.58874377 | 0.99999994 |
| Adamtsl4 | 0.00119705 | 0.00067846 | 0.00108302 | -0.0004046 | 0.99354967 | 0.99999994 |
| Mylk | 0.00340395 | 0.00267047 | 0.00307508 | -0.0004046 | 0.80943972 | 0.99999994 |
| Ints4 | 0.08479056 | 0.07218329 | 0.07258828 | -0.000405 | 0.51481247 | 0.99999994 |
| Degs2 | 0.00208798 | 0.00148211 | 0.00188794 | -0.0004058 | 0.92376233 | 0.99999994 |
| Chst7 | 0.00225872 | 0.00163602 | 0.00204208 | -0.0004061 | 0.90718358 | 0.99999994 |
| Trib2 | 0.00486942 | 0.00398836 | 0.00439459 | -0.0004062 | 0.73082314 | 0.99999994 |
| 2310001H17I | 0.00822411 | 0.00699895 | 0.00740531 | -0.0004064 | 0.64254075 | 0.99999994 |
| Tdh | 0.00260437 | 0.00194752 | 0.00235403 | -0.0004065 | 0.87481522 | 0.99999994 |
| 1810063I02R | 0.00151703 | 0.00096543 | 0.00137222 | -0.0004068 | 0.97576664 | 0.99999994 |
| Bank1 | 0.00146734 | 0.0009203 | 0.00132732 | -0.000407 | 0.97938208 | 0.99999994 |
| Trim2 | 0.00766799 | 0.00650008 | 0.00690716 | -0.0004071 | 0.65258028 | 0.99999994 |
| Fut4 | 0.00997026 | 0.00855979 | 0.00896702 | -0.0004072 | 0.61878609 | 0.99999994 |
| Arg2 | 0.0030686 | 0.00236513 | 0.00277276 | -0.0004076 | 0.83610322 | 0.99999994 |
| Rasa1 | 0.18364669 | 0.14533944 | 0.14574734 | -0.0004079 | 0.50743146 | 0.99999994 |
| Gstm2 | 0.03687582 | 0.0321645 | 0.03257298 | -0.0004085 | 0.53326285 | 0.99999994 |
| Cys1 | 0.02424619 | 0.02118921 | 0.02159809 | -0.0004089 | 0.55013947 | 0.99999994 |
| Zscan12 | 0.03093567 | 0.02702511 | 0.02743412 | -0.000409 | 0.53952572 | 0.99999994 |
| 4921536K21I | 0.00449992 | 0.00365301 | 0.00406214 | -0.0004091 | 0.74870566 | 0.99999994 |
| Sgip1 | 0.00107287 | 0.00056083 | 0.00097075 | -0.0004099 | 0.99752919 | 0.99999994 |
| Gm20390 | 0.00257661 | 0.00191823 | 0.00232898 | -0.0004108 | 0.87979449 | 0.99999994 |
| Fbp2 | 0.00151086 | 0.00095583 | 0.00136665 | -0.0004108 | 0.97729877 | 0.99999994 |
| Slc26a9 | 0.00315213 | 0.00243717 | 0.00284808 | -0.0004109 | 0.83156258 | 0.99999994 |
| Frmd6 | 0.00789264 | 0.00669752 | 0.00710844 | -0.0004109 | 0.64980023 | 0.99999994 |

| | | | | | | |
|---|---|---|---|---|---|---|
| Gm15261 | 0.00169574 | 0.00112268 | 0.00153368 | -0.000411 | 0.96277192 | 0.99999994 |
| Vav3 | 0.00426868 | 0.0034428 | 0.003854 | -0.0004112 | 0.76121221 | 0.99999994 |
| Tdrd5 | 0.00514216 | 0.00422851 | 0.00463988 | -0.0004114 | 0.72245674 | 0.99999994 |
| AK157302 | 0.00438943 | 0.00355129 | 0.0039627 | -0.0004114 | 0.7552363 | 0.99999994 |
| Emp1 | 0.00139486 | 0.00085008 | 0.00126181 | -0.0004117 | 0.9850713 | 0.99999994 |
| Mtpap | 0.11746509 | 0.09779413 | 0.09820601 | -0.0004119 | 0.51113589 | 0.99999994 |
| Ggt1 | 0.02197987 | 0.01919706 | 0.01960904 | -0.000412 | 0.55560931 | 0.99999994 |
| Gm15938 | 0.00112949 | 0.00060962 | 0.00102194 | -0.0004123 | 0.99638077 | 0.99999994 |
| Ism2 | 0.00202932 | 0.00142255 | 0.00183497 | -0.0004124 | 0.93268006 | 0.99999994 |
| Slc2a9 | 0.00181354 | 0.00122649 | 0.0016401 | -0.0004136 | 0.95338881 | 0.99999994 |
| Zfp69 | 0.00455362 | 0.00369652 | 0.00411047 | -0.000414 | 0.74867931 | 0.99999994 |
| Sorbs1 | 0.00685996 | 0.00576822 | 0.00618268 | -0.0004145 | 0.67227704 | 0.99999994 |
| Rasgef1b | 0.007285 | 0.00614888 | 0.00656387 | -0.000415 | 0.66306195 | 0.99999994 |
| Itgae | 0.00270155 | 0.00202609 | 0.0024417 | -0.0004156 | 0.87139636 | 0.99999994 |
| Gm5087 | 0.00698626 | 0.00588026 | 0.00629597 | -0.0004157 | 0.66985388 | 0.99999994 |
| Tgfb3 | 0.00338366 | 0.00264093 | 0.00305679 | -0.0004159 | 0.81741515 | 0.99999994 |
| Tnni1 | 0.00394291 | 0.00314464 | 0.00356067 | -0.000416 | 0.78163706 | 0.99999994 |
| Cdnf | 0.00308518 | 0.00237128 | 0.00278771 | -0.0004164 | 0.8399714 | 0.99999994 |
| Osbp2 | 0.0014543 | 0.00089833 | 0.00131554 | -0.0004172 | 0.98261306 | 0.99999994 |
| Lyl1 | 0.00205619 | 0.00144137 | 0.00185923 | -0.0004179 | 0.9326776 | 0.99999994 |
| Fgf11 | 0.00834918 | 0.00709901 | 0.00751729 | -0.0004183 | 0.64445013 | 0.99999994 |
| Appbp2os | 0.00498859 | 0.0040822 | 0.00450178 | -0.0004196 | 0.73250198 | 0.99999994 |
| Srpx2 | 0.00155064 | 0.00098254 | 0.00140259 | -0.00042 | 0.97689334 | 0.99999994 |
| Dpys | 0.00398681 | 0.00317998 | 0.0036002 | -0.0004202 | 0.78140487 | 0.99999994 |
| Zfp41 | 0.01352553 | 0.0117147 | 0.01213539 | -0.0004207 | 0.59124565 | 0.99999994 |
| Coa4 | 0.14315928 | 0.11685393 | 0.11727477 | -0.0004208 | 0.50952846 | 0.99999994 |
| Tdrd6 | 0.00072605 | 0.00023604 | 0.0006571 | -0.0004211 | 0.99999002 | 0.99999994 |
| Edn2 | 0.00356697 | 0.00280068 | 0.00322199 | -0.0004213 | 0.80795562 | 0.99999994 |
| Inpp5k | 0.0709203 | 0.06083854 | 0.06126071 | -0.0004222 | 0.51829349 | 0.99999994 |
| Csf1r | 0.00091171 | 0.00040262 | 0.00082502 | -0.0004224 | 0.99967276 | 0.99999994 |

| | | | | | | |
|---|---|---|---|---|---|---|
| Gm9903 | 0.00665525 | 0.00557625 | 0.00599902 | -0.0004228 | 0.68049802 | 0.99999994 |
| 2200002J24F | 0.00513438 | 0.00420993 | 0.00463288 | -0.000423 | 0.72830129 | 0.99999994 |
| Trim24 | 0.10820155 | 0.09070878 | 0.09113224 | -0.0004235 | 0.51233736 | 0.99999994 |
| Mapkapk3 | 0.00253486 | 0.00186777 | 0.00229131 | -0.0004235 | 0.89072649 | 0.99999994 |
| Serac1 | 0.00384835 | 0.00305181 | 0.0034755 | -0.0004237 | 0.79144922 | 0.99999994 |
| Gm26612 | 0.00130657 | 0.00075752 | 0.00118202 | -0.0004245 | 0.99158778 | 0.99999994 |
| Cdkn2aipnl | 0.13833038 | 0.11332689 | 0.11375167 | -0.0004248 | 0.50991533 | 0.99999994 |
| Lipo2 | 0.01040503 | 0.00893033 | 0.0093553 | -0.000425 | 0.6188147 | 0.99999994 |
| Prox2 | 0.00516111 | 0.0042317 | 0.00465692 | -0.0004252 | 0.7283376 | 0.99999994 |
| Psd4 | 0.00397935 | 0.00316824 | 0.00359348 | -0.0004252 | 0.78456256 | 0.99999994 |
| Doc2g | 0.00588237 | 0.00487972 | 0.00530513 | -0.0004254 | 0.7032446 | 0.99999994 |
| Irak3 | 0.00336339 | 0.00261277 | 0.00303852 | -0.0004258 | 0.82450883 | 0.99999994 |
| Lrriq1 | 0.00540489 | 0.00445028 | 0.00487608 | -0.0004258 | 0.71946783 | 0.99999994 |
| Acot4 | 0.00191639 | 0.00130677 | 0.00173299 | -0.0004262 | 0.94919735 | 0.99999994 |
| Mtus2 | 0.02493365 | 0.02177378 | 0.02220025 | -0.0004265 | 0.55087361 | 0.99999994 |
| Aifm2 | 0.05987326 | 0.05166533 | 0.05209183 | -0.0004265 | 0.52173122 | 0.99999994 |
| Ugt2b35 | 0.00494588 | 0.00403496 | 0.00446336 | -0.0004284 | 0.73855499 | 0.99999994 |
| Zfp459 | 0.00341539 | 0.00265691 | 0.00308539 | -0.0004285 | 0.82236258 | 0.99999994 |
| Nyap2 | 0.00126959 | 0.0007194 | 0.0011486 | -0.0004292 | 0.9935641 | 0.99999994 |
| Eps8 | 0.18685858 | 0.14749754 | 0.14792686 | -0.0004293 | 0.50770644 | 0.99999994 |
| Lipo4 | 0.01170832 | 0.01008843 | 0.01051784 | -0.0004294 | 0.60709638 | 0.99999994 |
| Gm7598 | 0.00240807 | 0.00174747 | 0.00217688 | -0.0004294 | 0.90541742 | 0.99999994 |
| Cfap70 | 0.00307964 | 0.00235324 | 0.00278272 | -0.0004295 | 0.84786691 | 0.99999994 |
| Plekha2 | 0.00123382 | 0.00068633 | 0.00111626 | -0.0004299 | 0.99482197 | 0.99999994 |
| C230035I16R | 0.00849193 | 0.00721504 | 0.00764509 | -0.00043 | 0.64595668 | 0.99999994 |
| Hdac9 | 0.00247037 | 0.00180304 | 0.00223311 | -0.0004301 | 0.90006641 | 0.99999994 |
| Gm14133 | 0.00147387 | 0.00090255 | 0.00133322 | -0.0004307 | 0.9842301 | 0.99999994 |
| Smarcd3 | 0.00850073 | 0.00722187 | 0.00765296 | -0.0004311 | 0.64615075 | 0.99999994 |
| Ephx4 | 0.02239257 | 0.01954041 | 0.01997169 | -0.0004313 | 0.55714869 | 0.99999994 |
| Gli2 | 0.00167214 | 0.00108101 | 0.00151236 | -0.0004314 | 0.97118559 | 0.99999994 |

| | | | | | | |
|---|---|---|---|---|---|---|
| Lhfp | 0.00339338 | 0.00263375 | 0.00306555 | -0.0004318 | 0.82577344 | 0.99999994 |
| Rsph4a | 0.0021054 | 0.00147162 | 0.00190366 | -0.000432 | 0.9345663 | 0.99999994 |
| Cpa6 | 0.00230557 | 0.00165192 | 0.00208438 | -0.0004325 | 0.91636594 | 0.99999994 |
| Adamts7 | 0.00314502 | 0.00240857 | 0.00284167 | -0.0004331 | 0.84482332 | 0.99999994 |
| Gstm3 | 0.00606116 | 0.00503242 | 0.0054657 | -0.0004333 | 0.7011381 | 0.99999994 |
| Rasgrf2 | 0.00391087 | 0.00309817 | 0.00353181 | -0.0004336 | 0.79311296 | 0.99999994 |
| Gm17231 | 0.00072683 | 0.00022378 | 0.0006578 | -0.000434 | 0.99999438 | 0.99999994 |
| Dzip1 | 0.00490125 | 0.00398758 | 0.00442322 | -0.0004356 | 0.74395684 | 0.99999994 |
| Abca8b | 0.00527079 | 0.00431978 | 0.00475553 | -0.0004358 | 0.72904393 | 0.99999994 |
| Man2a1 | 0.06920812 | 0.05941102 | 0.05984824 | -0.0004372 | 0.51939185 | 0.99999994 |
| Wnt5a | 0.00234551 | 0.00168311 | 0.00212042 | -0.0004373 | 0.91509223 | 0.99999994 |
| Fhod1 | 0.00114612 | 0.00059962 | 0.00103698 | -0.0004374 | 0.99750287 | 0.99999994 |
| Henmt1 | 0.0026928 | 0.00199626 | 0.00243381 | -0.0004376 | 0.88428529 | 0.99999994 |
| Gm12962 | 0.00075028 | 0.0002412 | 0.00067902 | -0.0004378 | 0.99999114 | 0.99999994 |
| Trip10 | 0.02805928 | 0.02449302 | 0.02493109 | -0.0004381 | 0.5465546 | 0.99999994 |
| Ajuba | 0.0048522 | 0.00394078 | 0.0043791 | -0.0004383 | 0.74737481 | 0.99999994 |
| Neurl3 | 0.00118714 | 0.00063567 | 0.00107406 | -0.0004384 | 0.99670486 | 0.99999994 |
| 4930403P22I | 0.00348766 | 0.00271183 | 0.00315052 | -0.0004387 | 0.82300063 | 0.99999994 |
| Pfn3 | 0.00086951 | 0.00034805 | 0.00078686 | -0.0004388 | 0.99989723 | 0.99999994 |
| Acot6 | 0.00163524 | 0.00104014 | 0.00147903 | -0.0004389 | 0.97587911 | 0.99999994 |
| Mcm9 | 0.0076826 | 0.00648043 | 0.00692025 | -0.0004398 | 0.66387082 | 0.99999994 |
| Gm5464 | 0.00169387 | 0.00109209 | 0.00153199 | -0.0004399 | 0.97201853 | 0.99999994 |
| Entpd5 | 0.06941766 | 0.05958092 | 0.06002127 | -0.0004403 | 0.51947424 | 0.99999994 |
| Zeb2 | 0.00884917 | 0.00752424 | 0.00796477 | -0.0004405 | 0.64362576 | 0.99999994 |
| Pygm | 0.00404268 | 0.00320972 | 0.00365052 | -0.0004408 | 0.78922731 | 0.99999994 |
| Nell2 | 0.00260757 | 0.00191601 | 0.00235692 | -0.0004409 | 0.89347113 | 0.99999994 |
| Btc | 0.00678525 | 0.00567472 | 0.00611566 | -0.0004409 | 0.68436068 | 0.99999994 |
| Pde3a | 0.00125566 | 0.00069504 | 0.001136 | -0.000441 | 0.99511366 | 0.99999994 |
| Fmnl3 | 0.00604488 | 0.00500985 | 0.00545108 | -0.0004412 | 0.70498383 | 0.99999994 |
| Mdh1b | 0.00165471 | 0.0010547 | 0.00149662 | -0.0004419 | 0.97532026 | 0.99999994 |

| | | | | | | |
|---|---|---|---|---|---|---|
| Gm29107 | 0.00358896 | 0.0027984 | 0.00324181 | -0.0004434 | 0.81870958 | 0.99999994 |
| Plat | 0.00761947 | 0.00642022 | 0.00686368 | -0.0004435 | 0.66642669 | 0.99999994 |
| Lingo4 | 0.00531325 | 0.0043502 | 0.0047937 | -0.0004435 | 0.73099842 | 0.99999994 |
| Proc | 0.02002733 | 0.01744651 | 0.01789052 | -0.000444 | 0.56560662 | 0.99999994 |
| Stc1 | 0.00146631 | 0.00088211 | 0.00132639 | -0.0004443 | 0.98711164 | 0.99999994 |
| Gm28041 | 0.00101669 | 0.00047566 | 0.00091996 | -0.0004443 | 0.99934716 | 0.99999994 |
| Tnni3k | 0.00113115 | 0.00057897 | 0.00102345 | -0.0004445 | 0.99807889 | 0.99999994 |
| Tgm3 | 0.00074653 | 0.00023096 | 0.00067562 | -0.0004447 | 0.99999409 | 0.99999994 |
| Polk | 0.02370602 | 0.02067963 | 0.02112455 | -0.0004449 | 0.55574733 | 0.99999994 |
| Tat | 0.00252435 | 0.00183684 | 0.00228182 | -0.000445 | 0.90286908 | 0.99999994 |
| Ankrd6 | 0.01060083 | 0.00908487 | 0.00953008 | -0.0004452 | 0.62208326 | 0.99999994 |
| Eng | 0.00230299 | 0.00163677 | 0.00208205 | -0.0004453 | 0.92271378 | 0.99999994 |
| Grhl3 | 0.00176572 | 0.0011514 | 0.0015969 | -0.0004455 | 0.9683442 | 0.99999994 |
| Thbd | 0.00096242 | 0.00042504 | 0.00087088 | -0.0004458 | 0.99967235 | 0.99999994 |
| Timm8a1 | 0.00263272 | 0.00193371 | 0.00237961 | -0.0004459 | 0.89385596 | 0.99999994 |
| Fbxo43 | 0.00161742 | 0.00101687 | 0.00146293 | -0.0004461 | 0.97880016 | 0.99999994 |
| Zc2hc1a | 0.07615879 | 0.06511666 | 0.06556277 | -0.0004461 | 0.5180627 | 0.99999994 |
| Scrt1 | 0.00076027 | 0.00024178 | 0.00068805 | -0.0004463 | 0.99999211 | 0.99999994 |
| Tuba1c | 0.15666532 | 0.12653586 | 0.12698217 | -0.0004463 | 0.50933247 | 0.99999994 |
| Pcdhgc5 | 0.01184661 | 0.0101944 | 0.01064108 | -0.0004467 | 0.61003442 | 0.99999994 |
| Pex13 | 0.22484109 | 0.17240986 | 0.17285659 | -0.0004467 | 0.50686249 | 0.99999994 |
| Itpripl1 | 0.00461082 | 0.00371504 | 0.00416194 | -0.0004469 | 0.7626156 | 0.99999994 |
| Gm15860 | 0.0196211 | 0.01708534 | 0.01753241 | -0.0004471 | 0.56739128 | 0.99999994 |
| Exoc6 | 0.07116437 | 0.06101452 | 0.06146179 | -0.0004473 | 0.51931729 | 0.99999994 |
| 2600014E21I | 0.00252554 | 0.00183546 | 0.0022829 | -0.0004474 | 0.90398798 | 0.99999994 |
| Mmp15 | 0.02961292 | 0.02583629 | 0.02628426 | -0.000448 | 0.54516138 | 0.99999994 |
| Krt81 | 0.00174754 | 0.00113244 | 0.00158048 | -0.000448 | 0.97041617 | 0.99999994 |
| Cep41 | 0.06211288 | 0.05351321 | 0.05396136 | -0.0004482 | 0.52204295 | 0.99999994 |
| L3hypdh | 0.03454544 | 0.03011343 | 0.0305618 | -0.0004484 | 0.53889655 | 0.99999994 |
| Myo16 | 0.00127712 | 0.0007068 | 0.0011554 | -0.0004486 | 0.99512264 | 0.99999994 |

| | | | | | | |
|---|---|---|---|---|---|---|
| 4930442H23I | 0.00083053 | 0.00030294 | 0.0007516 | -0.0004487 | 0.99996458 | 0.99999994 |
| D930048N14 | 0.00283058 | 0.00210881 | 0.0025581 | -0.0004493 | 0.87881497 | 0.99999994 |
| Tsx | 0.00082371 | 0.00029564 | 0.00074543 | -0.0004498 | 0.99997046 | 0.99999994 |
| Mill2 | 0.00155624 | 0.00095755 | 0.00140765 | -0.0004501 | 0.98334757 | 0.99999994 |
| Spns2 | 0.00264438 | 0.00193991 | 0.00239013 | -0.0004502 | 0.89504803 | 0.99999994 |
| Smagp | 0.00767617 | 0.0064642 | 0.00691449 | -0.0004503 | 0.66766627 | 0.99999994 |
| C1qtnf1 | 0.00133685 | 0.00075849 | 0.00120939 | -0.0004509 | 0.99346245 | 0.99999994 |
| Ltc4s | 0.0032081 | 0.00244733 | 0.00289853 | -0.0004512 | 0.84993851 | 0.99999994 |
| Slc25a16 | 0.07241433 | 0.06203916 | 0.06249064 | -0.0004515 | 0.51917782 | 0.99999994 |
| Gm16751 | 0.00615348 | 0.00509709 | 0.00554861 | -0.0004515 | 0.7059764 | 0.99999994 |
| Rbp2 | 0.05438672 | 0.04703706 | 0.04748868 | -0.0004516 | 0.52523722 | 0.99999994 |
| Hp | 0.00182916 | 0.00120249 | 0.00165421 | -0.0004517 | 0.96544097 | 0.99999994 |
| Dok3 | 0.00370726 | 0.00289653 | 0.00334839 | -0.0004519 | 0.81547967 | 0.99999994 |
| Trip12 | 0.31822769 | 0.22754744 | 0.22799994 | -0.0004525 | 0.5052701 | 0.99999994 |
| Dnase1l2 | 0.00120844 | 0.0006407 | 0.00109332 | -0.0004526 | 0.99707106 | 0.99999994 |
| Txnrd3 | 0.01700959 | 0.01477291 | 0.0152256 | -0.0004527 | 0.57844015 | 0.99999994 |
| Kbtbd11 | 0.00227377 | 0.00160262 | 0.00205567 | -0.0004531 | 0.9288139 | 0.99999994 |
| Gm12802 | 0.00613336 | 0.0050772 | 0.00553055 | -0.0004533 | 0.70734074 | 0.99999994 |
| Ribc1 | 0.02801263 | 0.02443694 | 0.02489042 | -0.0004535 | 0.54826185 | 0.99999994 |
| Atad2 | 0.01698062 | 0.01474608 | 0.01519997 | -0.0004539 | 0.57877754 | 0.99999994 |
| Gan | 0.05498026 | 0.04753405 | 0.04798826 | -0.0004542 | 0.52511747 | 0.99999994 |
| A330008L17I | 0.00077009 | 0.00024257 | 0.00069693 | -0.0004544 | 0.99999286 | 0.99999994 |
| Mtbp | 0.01400782 | 0.01210959 | 0.01256402 | -0.0004544 | 0.59512826 | 0.99999994 |
| Gm29650 | 0.00360741 | 0.00280372 | 0.00325843 | -0.0004547 | 0.82352518 | 0.99999994 |
| BC039966 | 0.00210097 | 0.00144484 | 0.00189967 | -0.0004548 | 0.94449783 | 0.99999994 |
| Tmod4 | 0.00342819 | 0.00264209 | 0.00309693 | -0.0004548 | 0.83586618 | 0.99999994 |
| Khdc3 | 0.00121154 | 0.00064121 | 0.00109612 | -0.0004549 | 0.99713208 | 0.99999994 |
| Grin2b | 0.00096166 | 0.00041523 | 0.0008702 | -0.000455 | 0.99974943 | 0.99999994 |
| Ckap2 | 0.00795552 | 0.00670897 | 0.00716477 | -0.0004558 | 0.66401748 | 0.99999994 |
| Vangl1 | 0.01055023 | 0.00902888 | 0.00948492 | -0.000456 | 0.62553229 | 0.99999994 |

| | | | | | | |
|---|---|---|---|---|---|---|
| Gm27017 | 0.00276857 | 0.00204521 | 0.00250216 | -0.0004569 | 0.88792976 | 0.99999994 |
| Tbx6 | 0.00238587 | 0.0016997 | 0.00215685 | -0.0004571 | 0.9208505 | 0.99999994 |
| Dgkeos | 0.00209232 | 0.00143441 | 0.00189186 | -0.0004575 | 0.94624695 | 0.99999994 |
| Abcb4 | 0.00102341 | 0.00046857 | 0.00092603 | -0.0004575 | 0.9994959 | 0.99999994 |
| Nhs | 0.00288799 | 0.00215229 | 0.00260988 | -0.0004576 | 0.87840817 | 0.99999994 |
| Gm15834 | 0.00173398 | 0.00111035 | 0.00156823 | -0.0004579 | 0.97401866 | 0.99999994 |
| Rarres1 | 0.00627229 | 0.00519727 | 0.00565528 | -0.000458 | 0.70508612 | 0.99999994 |
| Gm11963 | 0.00104323 | 0.00048557 | 0.00094395 | -0.0004584 | 0.99938598 | 0.99999994 |
| Gm16062 | 0.00781367 | 0.00657885 | 0.00703769 | -0.0004588 | 0.6678481 | 0.99999994 |
| Csdc2 | 0.0142656 | 0.01233399 | 0.01279301 | -0.000459 | 0.59438319 | 0.99999994 |
| Aph1c | 0.00604743 | 0.00499413 | 0.00545337 | -0.0004592 | 0.7124494 | 0.99999994 |
| Dctd | 0.0070858 | 0.00592596 | 0.00638525 | -0.0004593 | 0.68395794 | 0.99999994 |
| Il17d | 0.00450407 | 0.00360568 | 0.00406588 | -0.0004602 | 0.77439802 | 0.99999994 |
| Parp14 | 0.00085561 | 0.00031408 | 0.00077429 | -0.0004602 | 0.99996193 | 0.99999994 |
| 4933417D19 | 0.00081626 | 0.00027827 | 0.0007387 | -0.0004604 | 0.9999833 | 0.99999994 |
| Katnal2 | 0.02398568 | 0.02090926 | 0.02136976 | -0.0004605 | 0.55702735 | 0.99999994 |
| Fbxo41 | 0.00242537 | 0.00173186 | 0.0021925 | -0.0004606 | 0.91901705 | 0.99999994 |
| Mylip | 0.02545811 | 0.02219797 | 0.02265926 | -0.0004613 | 0.55389565 | 0.99999994 |
| Cdc42ep1 | 0.00715712 | 0.00598791 | 0.00644921 | -0.0004613 | 0.68300392 | 0.99999994 |
| Clcf1 | 0.00957971 | 0.00815673 | 0.00861805 | -0.0004613 | 0.63919472 | 0.99999994 |
| Usp13 | 0.00333679 | 0.00255299 | 0.00301454 | -0.0004616 | 0.84593393 | 0.99999994 |
| Cwf19l1 | 0.02732725 | 0.02383095 | 0.02429254 | -0.0004616 | 0.55032387 | 0.99999994 |
| Dclre1b | 0.02137645 | 0.01861551 | 0.01907842 | -0.0004629 | 0.56415356 | 0.99999994 |
| Cebpb | 0.00670069 | 0.00557683 | 0.00603979 | -0.000463 | 0.6950514 | 0.99999994 |
| Il18r1 | 0.00182791 | 0.00118985 | 0.00165307 | -0.0004632 | 0.96892462 | 0.99999994 |
| Zfp189 | 0.01998343 | 0.01738852 | 0.01785183 | -0.0004633 | 0.56857677 | 0.99999994 |
| Eaf2 | 0.00735653 | 0.00616411 | 0.006628 | -0.0004639 | 0.679348 | 0.99999994 |
| Plppr2 | 0.08805776 | 0.0747632 | 0.07522719 | -0.000464 | 0.5163743 | 0.99999994 |
| Kcnab1 | 0.00620641 | 0.00513132 | 0.00559613 | -0.0004648 | 0.709826 | 0.99999994 |
| Gm27042 | 0.00084005 | 0.00029515 | 0.00076021 | -0.0004651 | 0.9999767 | 0.99999994 |

| | | | | | | |
|---|---|---|---|---|---|---|
| Gna15 | 0.00186713 | 0.0012232 | 0.0016885 | -0.0004653 | 0.9666963 | 0.99999994 |
| Ptar1 | 0.01023059 | 0.00873347 | 0.00919954 | -0.0004661 | 0.63202794 | 0.99999994 |
| Slc2a12 | 0.00636985 | 0.00527667 | 0.00574286 | -0.0004662 | 0.70552073 | 0.99999994 |
| Gm20628 | 0.00169185 | 0.00106349 | 0.00153017 | -0.0004667 | 0.9788267 | 0.99999994 |
| Gm16712 | 0.00194826 | 0.00129462 | 0.00176177 | -0.0004671 | 0.96121717 | 0.99999994 |
| Gp5 | 0.00090515 | 0.00035166 | 0.00081909 | -0.0004674 | 0.99992723 | 0.99999994 |
| 4930539J05F | 0.00936512 | 0.00795845 | 0.00842622 | -0.0004678 | 0.64412847 | 0.99999994 |
| Cd93 | 0.0038141 | 0.0029767 | 0.00344465 | -0.0004679 | 0.81706767 | 0.99999994 |
| Cpsf4 | 0.09492993 | 0.0802726 | 0.08074185 | -0.0004692 | 0.51542909 | 0.99999994 |
| Agbl2 | 0.00121112 | 0.00062589 | 0.00109574 | -0.0004698 | 0.99784271 | 0.99999994 |
| Cmklr1 | 0.00323961 | 0.00245641 | 0.00292694 | -0.0004705 | 0.85770406 | 0.99999994 |
| Gm20517 | 0.00422296 | 0.00334175 | 0.00381284 | -0.0004711 | 0.79458196 | 0.99999994 |
| Kif20a | 0.0054755 | 0.00446826 | 0.00493954 | -0.0004713 | 0.73731264 | 0.99999994 |
| Gm26532 | 0.00701479 | 0.00585027 | 0.00632156 | -0.0004713 | 0.69014167 | 0.99999994 |
| Chek1 | 0.00941062 | 0.00799557 | 0.0084669 | -0.0004713 | 0.64451155 | 0.99999994 |
| Plaur | 0.00340813 | 0.00260748 | 0.00307885 | -0.0004714 | 0.84591774 | 0.99999994 |
| C5ar1 | 0.0015033 | 0.0008884 | 0.00135982 | -0.0004714 | 0.98948977 | 0.99999994 |
| Was | 0.00173388 | 0.00109576 | 0.00156814 | -0.0004724 | 0.97752613 | 0.99999994 |
| Neu3 | 0.00187728 | 0.00122527 | 0.00169766 | -0.0004724 | 0.96800155 | 0.99999994 |
| Gm13594 | 0.00338679 | 0.0025872 | 0.00305961 | -0.0004724 | 0.84796842 | 0.99999994 |
| Sync | 0.00620486 | 0.00512202 | 0.00559474 | -0.0004727 | 0.71308559 | 0.99999994 |
| Dapk2 | 0.00223484 | 0.0015478 | 0.00202053 | -0.0004727 | 0.94030502 | 0.99999994 |
| A830005F24 | 0.00301636 | 0.00225285 | 0.00272566 | -0.0004728 | 0.87587978 | 0.99999994 |
| Sh3kbp1 | 0.10315404 | 0.08675944 | 0.08723267 | -0.0004732 | 0.51440277 | 0.99999994 |
| Plekhh3 | 0.03778938 | 0.03288558 | 0.03335972 | -0.0004741 | 0.53768604 | 0.99999994 |
| Rasa3 | 0.00363429 | 0.00280827 | 0.00328265 | -0.0004744 | 0.83195385 | 0.99999994 |
| Chrna4 | 0.0100018 | 0.00852066 | 0.0089952 | -0.0004745 | 0.63726251 | 0.99999994 |
| Ddb2 | 0.02377595 | 0.02071109 | 0.02118588 | -0.0004748 | 0.5592915 | 0.99999994 |
| 1810014B01 | 0.02556059 | 0.022274 | 0.02274892 | -0.0004749 | 0.55525913 | 0.99999994 |
| Tmem116 | 0.02162293 | 0.01881914 | 0.01929521 | -0.0004761 | 0.56522523 | 0.99999994 |

| | | | | | | |
|---|---|---|---|---|---|---|
| Rbak | 0.03414557 | 0.02973932 | 0.03021606 | -0.0004767 | 0.54182111 | 0.99999994 |
| Sag | 0.00468062 | 0.00374798 | 0.00422475 | -0.0004768 | 0.77372611 | 0.99999994 |
| Vash1 | 0.00389838 | 0.00304364 | 0.00352056 | -0.0004769 | 0.81639863 | 0.99999994 |
| Slc2a8 | 0.09201414 | 0.07793076 | 0.07840797 | -0.0004772 | 0.51615754 | 0.99999994 |
| Phf3 | 0.26838141 | 0.19912938 | 0.19960678 | -0.0004774 | 0.50635097 | 0.99999994 |
| Tmc8 | 0.00643157 | 0.0053204 | 0.00579828 | -0.0004779 | 0.70836024 | 0.99999994 |
| Phka2 | 0.0156562 | 0.01354821 | 0.01402692 | -0.0004787 | 0.58985456 | 0.99999994 |
| Tnnt2 | 0.00154179 | 0.00091515 | 0.0013946 | -0.0004794 | 0.98894181 | 0.99999994 |
| Il7r | 0.00245192 | 0.0017368 | 0.00221646 | -0.0004797 | 0.92513484 | 0.99999994 |
| Vezt | 0.10534607 | 0.08845005 | 0.08893011 | -0.0004801 | 0.51433178 | 0.99999994 |
| Aff2 | 0.01568742 | 0.01357377 | 0.0140546 | -0.0004808 | 0.59006951 | 0.99999994 |
| Apobec1 | 0.00148559 | 0.00086222 | 0.00134381 | -0.0004816 | 0.99147183 | 0.99999994 |
| Otud7b | 0.06707469 | 0.05760214 | 0.05808386 | -0.0004817 | 0.52201247 | 0.99999994 |
| Cacnb1 | 0.01331612 | 0.01146641 | 0.01194919 | -0.0004828 | 0.60600913 | 0.99999994 |
| Elac1 | 0.02169871 | 0.01887902 | 0.01936185 | -0.0004828 | 0.56591752 | 0.99999994 |
| Lamb1 | 0.10307297 | 0.08668688 | 0.08716977 | -0.0004829 | 0.51470721 | 0.99999994 |
| Gm16861 | 0.01549749 | 0.01340186 | 0.01388622 | -0.0004844 | 0.59179893 | 0.99999994 |
| Oxtr | 0.00266245 | 0.0019218 | 0.00240643 | -0.0004846 | 0.90996201 | 0.99999994 |
| Gm11508 | 0.0071255 | 0.00593622 | 0.00642085 | -0.0004846 | 0.69231076 | 0.99999994 |
| Sapcd2 | 0.00334182 | 0.00253435 | 0.00301908 | -0.0004847 | 0.85740069 | 0.99999994 |
| Suv39h1 | 0.03723174 | 0.03239465 | 0.03287961 | -0.000485 | 0.53910468 | 0.99999994 |
| Slc25a15 | 0.01947031 | 0.01691394 | 0.01739944 | -0.0004855 | 0.5736734 | 0.99999994 |
| Faxc | 0.02049891 | 0.01781992 | 0.01830599 | -0.0004861 | 0.5701448 | 0.99999994 |
| 9930004E17I | 0.00184039 | 0.00117721 | 0.00166435 | -0.0004871 | 0.97430728 | 0.99999994 |
| Ryr3 | 0.00595696 | 0.00488429 | 0.00537212 | -0.0004878 | 0.72722957 | 0.99999994 |
| Tbc1d2 | 0.01684029 | 0.01458781 | 0.01507577 | -0.000488 | 0.58529093 | 0.99999994 |
| Caly | 0.00250269 | 0.00177385 | 0.00226227 | -0.0004884 | 0.92465675 | 0.99999994 |
| Zfp182 | 0.01439506 | 0.0124192 | 0.01290798 | -0.0004888 | 0.59949987 | 0.99999994 |
| 1500002F19I | 0.0012106 | 0.00060586 | 0.00109527 | -0.0004894 | 0.9985317 | 0.99999994 |
| Rassf5 | 0.00422264 | 0.00332294 | 0.00381255 | -0.0004896 | 0.80367045 | 0.99999994 |

| | | | | | | |
|---|---|---|---|---|---|---|
| Pik3cg | 0.00170715 | 0.00105409 | 0.00154399 | -0.0004899 | 0.98265762 | 0.99999994 |
| 5330439K02I | 0.00762082 | 0.00637463 | 0.00686489 | -0.0004903 | 0.68273537 | 0.99999994 |
| Rnd2 | 0.00899089 | 0.00760127 | 0.00809155 | -0.0004903 | 0.65664415 | 0.99999994 |
| Necab3 | 0.00944361 | 0.00800591 | 0.00849639 | -0.0004905 | 0.649607 | 0.99999994 |
| Xylb | 0.00354614 | 0.00271256 | 0.00320322 | -0.0004907 | 0.84604063 | 0.99999994 |
| Tmem74b | 0.00280561 | 0.00204477 | 0.00253558 | -0.0004908 | 0.9012067 | 0.99999994 |
| Slc41a1 | 0.02385298 | 0.02076257 | 0.02125341 | -0.0004908 | 0.56108815 | 0.99999994 |
| 1810064F22I | 0.00607797 | 0.00498945 | 0.0054808 | -0.0004914 | 0.72465993 | 0.99999994 |
| Ccdc57 | 0.00720824 | 0.00600291 | 0.00649505 | -0.0004921 | 0.69299568 | 0.99999994 |
| Gm17690 | 0.00981466 | 0.00833584 | 0.00882801 | -0.0004922 | 0.64471711 | 0.99999994 |
| Trmt44 | 0.02637191 | 0.02296596 | 0.02345827 | -0.0004923 | 0.55554902 | 0.99999994 |
| Krt222 | 0.00158365 | 0.00093992 | 0.00143241 | -0.0004925 | 0.98894791 | 0.99999994 |
| D830039M14 | 0.00147316 | 0.00083998 | 0.00133258 | -0.0004926 | 0.993064 | 0.99999994 |
| Cx3cr1 | 0.00255145 | 0.00181342 | 0.00230628 | -0.0004929 | 0.92255606 | 0.99999994 |
| Oasl2 | 0.00141543 | 0.00078736 | 0.00128041 | -0.000493 | 0.99481364 | 0.99999994 |
| Nme3 | 0.02052578 | 0.01783644 | 0.01832966 | -0.0004932 | 0.57107421 | 0.99999994 |
| Prob1 | 0.00304726 | 0.00225954 | 0.00275352 | -0.000494 | 0.88378747 | 0.99999994 |
| Dpyd | 0.01078007 | 0.00919517 | 0.00969004 | -0.0004949 | 0.63304793 | 0.99999994 |
| Pla2g2c | 0.01229002 | 0.01054071 | 0.01103605 | -0.0004953 | 0.61744146 | 0.99999994 |
| Gm42596 | 0.0012067 | 0.00059586 | 0.00109175 | -0.0004959 | 0.99875036 | 0.99999994 |
| Wdr93 | 0.00552724 | 0.00448868 | 0.00498604 | -0.0004974 | 0.74664625 | 0.99999994 |
| Slc2a13 | 0.00237576 | 0.0016496 | 0.00214772 | -0.0004981 | 0.93868378 | 0.99999994 |
| Rcc1 | 0.0621803 | 0.05351857 | 0.05401756 | -0.000499 | 0.52451471 | 0.99999994 |
| Bcap29 | 0.10019214 | 0.0844303 | 0.0849294 | -0.0004991 | 0.51560163 | 0.99999994 |
| 1700034H15I | 0.00198906 | 0.00129951 | 0.00179862 | -0.0004991 | 0.96763515 | 0.99999994 |
| Phtf1os | 0.01255552 | 0.01077333 | 0.01127245 | -0.0004991 | 0.61590038 | 0.99999994 |
| 4933438K21I | 0.00157753 | 0.00092765 | 0.00142689 | -0.0004992 | 0.99006876 | 0.99999994 |
| Wasf3 | 0.00846429 | 0.00712104 | 0.00762034 | -0.0004993 | 0.66862984 | 0.99999994 |
| Ptprd | 0.0121812 | 0.0104397 | 0.01093914 | -0.0004994 | 0.6194009 | 0.99999994 |
| Itga1 | 0.00204147 | 0.00134644 | 0.00184594 | -0.0004995 | 0.96416159 | 0.99999994 |

| | | | | | | |
|---|---|---|---|---|---|---|
| Cdan1 | 0.01360557 | 0.01170699 | 0.01220655 | -0.0004996 | 0.60734917 | 0.99999994 |
| Gmeb2 | 0.02677702 | 0.02331259 | 0.02381216 | -0.0004996 | 0.55553181 | 0.99999994 |
| Impa2 | 0.0042074 | 0.00329899 | 0.00379883 | -0.0004998 | 0.80943805 | 0.99999994 |
| Atad3aos | 0.00485497 | 0.00388145 | 0.00438159 | -0.0005001 | 0.77631395 | 0.99999994 |
| Ccdc9 | 0.08705909 | 0.07392151 | 0.07442174 | -0.0005002 | 0.51784343 | 0.99999994 |
| Gm5424 | 0.00768952 | 0.00642584 | 0.00692645 | -0.0005006 | 0.68477122 | 0.99999994 |
| 4933424M12 | 0.00182741 | 0.00115193 | 0.00165262 | -0.0005007 | 0.97813392 | 0.99999994 |
| Tmem106b | 0.18801513 | 0.1482082 | 0.14870893 | -0.0005007 | 0.50894072 | 0.99999994 |
| Cdc42ep5 | 0.03003177 | 0.02614749 | 0.02664858 | -0.0005011 | 0.54980264 | 0.99999994 |
| Slc38a8 | 0.00490254 | 0.00392252 | 0.00442439 | -0.0005019 | 0.77488584 | 0.99999994 |
| Abcb8 | 0.16333547 | 0.131196 | 0.13169826 | -0.0005023 | 0.51012627 | 0.99999994 |
| B430218F22 | 0.00465645 | 0.00370058 | 0.00420299 | -0.0005024 | 0.7868915 | 0.99999994 |
| Gm1976 | 0.02365356 | 0.02057564 | 0.02107854 | -0.0005029 | 0.5630907 | 0.99999994 |
| P2rx6 | 0.0042738 | 0.00335562 | 0.00385861 | -0.000503 | 0.80721757 | 0.99999994 |
| Alms1 | 0.03477683 | 0.03025871 | 0.03076177 | -0.0005031 | 0.54334095 | 0.99999994 |
| Bvht | 0.00483915 | 0.00386393 | 0.00436736 | -0.0005034 | 0.77854581 | 0.99999994 |
| Arhgap6 | 0.00353185 | 0.00268659 | 0.00319034 | -0.0005038 | 0.85337851 | 0.99999994 |
| Cachd1 | 0.00413277 | 0.00322758 | 0.00373163 | -0.0005041 | 0.81570473 | 0.99999994 |
| Ablim2 | 0.00347716 | 0.00263641 | 0.00314106 | -0.0005046 | 0.85756112 | 0.99999994 |
| Arl16 | 0.23465786 | 0.17854877 | 0.17905364 | -0.0005049 | 0.50748727 | 0.99999994 |
| Gpr183 | 0.00472264 | 0.00375752 | 0.00426255 | -0.000505 | 0.78484176 | 0.99999994 |
| Rnf152 | 0.01248483 | 0.01070443 | 0.01120952 | -0.0005051 | 0.61788536 | 0.99999994 |
| Apc2 | 0.0013473 | 0.00071311 | 0.00121883 | -0.0005057 | 0.99712649 | 0.99999994 |
| C530044C16I | 0.0029911 | 0.00219701 | 0.00270287 | -0.0005059 | 0.89357694 | 0.99999994 |
| Cd68 | 0.0119468 | 0.01022433 | 0.01073034 | -0.000506 | 0.62319888 | 0.99999994 |
| Nek7 | 0.15123334 | 0.12259703 | 0.12310358 | -0.0005065 | 0.51092547 | 0.99999994 |
| Def6 | 0.00725451 | 0.00602995 | 0.00653653 | -0.0005066 | 0.69702631 | 0.99999994 |
| Rsl1 | 0.00744908 | 0.00620406 | 0.00671097 | -0.0005069 | 0.69243932 | 0.99999994 |
| 1700048O20 | 0.02790093 | 0.02428588 | 0.02479301 | -0.0005071 | 0.55415024 | 0.99999994 |
| Grb2 | 0.3080119 | 0.22185871 | 0.22236606 | -0.0005073 | 0.5060585 | 0.99999994 |

| | | | | | | |
|---|---|---|---|---|---|---|
| Pus10 | 0.03982084 | 0.03459818 | 0.03510574 | -0.0005076 | 0.53833379 | 0.99999994 |
| Ccdc122 | 0.00838382 | 0.00704027 | 0.00754831 | -0.000508 | 0.67292315 | 0.99999994 |
| Glt28d2 | 0.01141113 | 0.00974472 | 0.01025292 | -0.0005082 | 0.62927525 | 0.99999994 |
| Zfyve1 | 0.01384292 | 0.01190922 | 0.0124175 | -0.0005083 | 0.60736695 | 0.99999994 |
| Ttc34 | 0.00277321 | 0.00199804 | 0.00250635 | -0.0005083 | 0.91148686 | 0.99999994 |
| Man2c1os | 0.00972512 | 0.00823957 | 0.008748 | -0.0005084 | 0.65057152 | 0.99999994 |
| Srrd | 0.12075044 | 0.10017901 | 0.10068895 | -0.0005099 | 0.51344629 | 0.99999994 |
| Gabra2 | 0.00321688 | 0.00239624 | 0.00290645 | -0.0005102 | 0.87869419 | 0.99999994 |
| Ints8 | 0.09576067 | 0.08089428 | 0.08140521 | -0.0005109 | 0.51666234 | 0.99999994 |
| Gm26533 | 0.00343126 | 0.00258759 | 0.00309969 | -0.0005121 | 0.86427086 | 0.99999994 |
| Taf1b | 0.08133713 | 0.06927471 | 0.06978688 | -0.0005122 | 0.51948099 | 0.99999994 |
| Mypopos | 0.00737176 | 0.0061292 | 0.00664166 | -0.0005125 | 0.69623232 | 0.99999994 |
| Rbl1 | 0.01665158 | 0.01439624 | 0.01490872 | -0.0005125 | 0.59049233 | 0.99999994 |
| A830011K09 | 0.00276369 | 0.00198521 | 0.00249776 | -0.0005126 | 0.91401821 | 0.99999994 |
| Cd84 | 0.00184091 | 0.00115223 | 0.00166482 | -0.0005126 | 0.97979042 | 0.99999994 |
| Slco1a5 | 0.00364491 | 0.00277913 | 0.00329222 | -0.0005131 | 0.85022379 | 0.99999994 |
| Ugt2b34 | 0.02319158 | 0.02015753 | 0.02067326 | -0.0005157 | 0.56594227 | 0.99999994 |
| Chd5 | 0.00159784 | 0.00092905 | 0.00144524 | -0.0005162 | 0.99128348 | 0.99999994 |
| Psme3 | 0.20134939 | 0.15710298 | 0.15761961 | -0.0005166 | 0.50870329 | 0.99999994 |
| Gm14703 | 0.00639165 | 0.0052455 | 0.00576244 | -0.0005169 | 0.72478945 | 0.99999994 |
| A230072E10 | 0.02116134 | 0.01837189 | 0.01888916 | -0.0005173 | 0.57231849 | 0.99999994 |
| Ccdc80 | 0.00799858 | 0.00668562 | 0.00720334 | -0.0005177 | 0.68381795 | 0.99999994 |
| Mppe1 | 0.00289383 | 0.00209738 | 0.00261515 | -0.0005178 | 0.90622766 | 0.99999994 |
| Slc2a10 | 0.00315544 | 0.00233311 | 0.00285106 | -0.000518 | 0.88671882 | 0.99999994 |
| Dna2 | 0.00756049 | 0.00629246 | 0.00681082 | -0.0005184 | 0.69378465 | 0.99999994 |
| Enox1 | 0.01200646 | 0.01026504 | 0.01078349 | -0.0005185 | 0.62552697 | 0.99999994 |
| Camkk1 | 0.00891634 | 0.00750593 | 0.00802486 | -0.0005189 | 0.66655976 | 0.99999994 |
| Lgals9 | 0.00349297 | 0.00263632 | 0.00315531 | -0.000519 | 0.86320644 | 0.99999994 |
| Tmem145 | 0.00657542 | 0.00540839 | 0.00592738 | -0.000519 | 0.71999165 | 0.99999994 |
| Dab2 | 0.00541811 | 0.00436866 | 0.00488796 | -0.0005193 | 0.76027291 | 0.99999994 |

| | | | | | | |
|---|---|---|---|---|---|---|
| Lmcd1 | 0.00405342 | 0.00313919 | 0.00366019 | -0.000521 | 0.82830128 | 0.99999994 |
| Nde1 | 0.04590508 | 0.0397859 | 0.0403071 | -0.0005212 | 0.53429522 | 0.99999994 |
| Plekhf1 | 0.00328465 | 0.00244632 | 0.00296755 | -0.0005212 | 0.87882647 | 0.99999994 |
| Cnnm4 | 0.01319793 | 0.01132233 | 0.01184407 | -0.0005217 | 0.61532215 | 0.99999994 |
| AA986860 | 0.00159622 | 0.00092136 | 0.00144377 | -0.0005224 | 0.99199161 | 0.99999994 |
| Xpo5 | 0.01400492 | 0.01203839 | 0.01256145 | -0.0005231 | 0.60917698 | 0.99999994 |
| C2cd4c | 0.06355941 | 0.05464259 | 0.05516596 | -0.0005234 | 0.52517691 | 0.99999994 |
| Fabp3-ps1 | 0.00721413 | 0.00597663 | 0.00650033 | -0.0005237 | 0.7041131 | 0.99999994 |
| Casp7 | 0.04289154 | 0.03721226 | 0.03773608 | -0.0005238 | 0.53680913 | 0.99999994 |
| Gm12999 | 0.00176255 | 0.00107017 | 0.00159404 | -0.0005239 | 0.9856492 | 0.99999994 |
| Kcna2 | 0.00195166 | 0.00124092 | 0.00176484 | -0.0005239 | 0.97592572 | 0.99999994 |
| Phlda3 | 0.01348166 | 0.01157239 | 0.01209639 | -0.000524 | 0.61345788 | 0.99999994 |
| Gm10382 | 0.0037559 | 0.00286741 | 0.00339222 | -0.0005248 | 0.84844734 | 0.99999994 |
| 4930578M01 | 0.03132582 | 0.02724709 | 0.02777289 | -0.0005258 | 0.5501412 | 0.99999994 |
| Gm5493 | 0.00253726 | 0.0017668 | 0.00229347 | -0.0005267 | 0.93681161 | 0.99999994 |
| Zfp658 | 0.00827881 | 0.00692735 | 0.00745429 | -0.0005269 | 0.68101503 | 0.99999994 |
| Cpne5 | 0.00783294 | 0.00652775 | 0.00705496 | -0.0005272 | 0.69055204 | 0.99999994 |
| Dtx3l | 0.01131214 | 0.00963698 | 0.01016466 | -0.0005277 | 0.63516082 | 0.99999994 |
| Glis2 | 0.07560673 | 0.0645827 | 0.06511077 | -0.0005281 | 0.52152666 | 0.99999994 |
| C8b | 0.0015766 | 0.00089689 | 0.00142605 | -0.0005292 | 0.99324316 | 0.99999994 |
| Dbndd2 | 0.00484673 | 0.00384492 | 0.00437418 | -0.0005293 | 0.78970328 | 0.99999994 |
| Cep192 | 0.03441049 | 0.02991584 | 0.03044513 | -0.0005293 | 0.54606348 | 0.99999994 |
| Ccdc73 | 0.00702273 | 0.00579901 | 0.00632869 | -0.0005297 | 0.71127362 | 0.99999994 |
| Gfod1 | 0.01460971 | 0.01256883 | 0.01309857 | -0.0005297 | 0.60611192 | 0.99999994 |
| Mrap2 | 0.00135732 | 0.00069807 | 0.00122789 | -0.0005298 | 0.99796145 | 0.99999994 |
| Tchp | 0.06102456 | 0.05252347 | 0.05305356 | -0.0005301 | 0.52651336 | 0.99999994 |
| Sowahb | 0.00146404 | 0.00079421 | 0.00132434 | -0.0005301 | 0.99614483 | 0.99999994 |
| Prdm6 | 0.00299987 | 0.00218024 | 0.00271079 | -0.0005305 | 0.90367122 | 0.99999994 |
| Tceal3 | 0.01296864 | 0.01110893 | 0.01164011 | -0.0005312 | 0.61934214 | 0.99999994 |
| Mical1 | 0.01107408 | 0.00942109 | 0.00995235 | -0.0005313 | 0.63882225 | 0.99999994 |

| | | | | | | |
|---|---|---|---|---|---|---|
| Tmem144 | 0.00376356 | 0.00286705 | 0.00339912 | -0.0005321 | 0.85127865 | 0.99999994 |
| Cyp46a1 | 0.00494817 | 0.0039331 | 0.00446543 | -0.0005323 | 0.78625893 | 0.99999994 |
| Klhl5 | 0.00333848 | 0.00248338 | 0.00301607 | -0.0005327 | 0.88012582 | 0.99999994 |
| Crhr1 | 0.00100249 | 0.00037429 | 0.00090712 | -0.0005328 | 0.99995382 | 0.99999994 |
| Cd274 | 0.00261228 | 0.00182758 | 0.00236117 | -0.0005336 | 0.93374098 | 0.99999994 |
| 2610027F03I | 0.0012851 | 0.00062839 | 0.00116261 | -0.0005342 | 0.99888819 | 0.99999994 |
| Cdca7 | 0.01994289 | 0.01728158 | 0.0178161 | -0.0005345 | 0.57914432 | 0.99999994 |
| Usp28 | 0.02281925 | 0.01981137 | 0.02034643 | -0.0005351 | 0.56947769 | 0.99999994 |
| Vamp5 | 0.00594791 | 0.00482858 | 0.00536399 | -0.0005354 | 0.74678708 | 0.99999994 |
| Tmc7 | 0.00437859 | 0.00341686 | 0.00395294 | -0.0005361 | 0.81665719 | 0.99999994 |
| Gm26691 | 0.00657121 | 0.00538719 | 0.0059236 | -0.0005364 | 0.72666715 | 0.99999994 |
| Arhgap11a | 0.00539994 | 0.00433487 | 0.00487163 | -0.0005368 | 0.76834386 | 0.99999994 |
| Ccl9 | 0.00319048 | 0.00234546 | 0.00288265 | -0.0005372 | 0.89259059 | 0.99999994 |
| Gm29666 | 0.00616666 | 0.00502263 | 0.00556044 | -0.0005378 | 0.74014943 | 0.99999994 |
| Aurkb | 0.00493417 | 0.00391495 | 0.00445284 | -0.0005379 | 0.78932036 | 0.99999994 |
| Slc19a3 | 0.00188647 | 0.00116801 | 0.00170597 | -0.000538 | 0.9820927 | 0.99999994 |
| Cfap44 | 0.00487624 | 0.00386243 | 0.00440073 | -0.0005383 | 0.79223607 | 0.99999994 |
| Evc2 | 0.00606418 | 0.00492927 | 0.00546842 | -0.0005391 | 0.74417479 | 0.99999994 |
| Map3k15 | 0.08724915 | 0.07403539 | 0.07457511 | -0.0005397 | 0.51921113 | 0.99999994 |
| B930094E09 | 0.00403427 | 0.00310162 | 0.00364294 | -0.0005413 | 0.83869119 | 0.99999994 |
| Moxd1 | 0.00169802 | 0.00099425 | 0.00153574 | -0.0005415 | 0.99053669 | 0.99999994 |
| H2-Eb1 | 0.00414301 | 0.00319738 | 0.00374086 | -0.0005435 | 0.83324003 | 0.99999994 |
| Cd320 | 0.19290024 | 0.151452 | 0.15199609 | -0.0005441 | 0.50950474 | 0.99999994 |
| Panx2 | 0.00335443 | 0.00248617 | 0.00303045 | -0.0005443 | 0.88405535 | 0.99999994 |
| Gm16344 | 0.00178327 | 0.00106843 | 0.00161276 | -0.0005443 | 0.98766743 | 0.99999994 |
| Zfp605 | 0.05763041 | 0.04966953 | 0.0502141 | -0.0005446 | 0.52877386 | 0.99999994 |
| Gm15013 | 0.01032261 | 0.00873705 | 0.00928171 | -0.0005447 | 0.65195231 | 0.99999994 |
| Cyp2d22 | 0.00855824 | 0.00715908 | 0.00770443 | -0.0005454 | 0.68123874 | 0.99999994 |
| Zfp879 | 0.00250388 | 0.001718 | 0.00226335 | -0.0005454 | 0.94562666 | 0.99999994 |
| Tmem120a | 0.11286429 | 0.09416054 | 0.09470623 | -0.0005457 | 0.51529701 | 0.99999994 |

| | | | | | | |
|---|---|---|---|---|---|---|
| Rgs14 | 0.00671408 | 0.00550576 | 0.00605181 | -0.000546 | 0.7259435 | 0.99999994 |
| Ctla2a | 0.00307582 | 0.00223299 | 0.00277927 | -0.0005463 | 0.90461984 | 0.99999994 |
| Chst10 | 0.00860097 | 0.00719536 | 0.00774267 | -0.0005473 | 0.68101118 | 0.99999994 |
| Slc23a1 | 0.00330258 | 0.00243614 | 0.00298371 | -0.0005476 | 0.88906575 | 0.99999994 |
| Tmem255a | 0.00115008 | 0.00049268 | 0.00104056 | -0.0005479 | 0.99977147 | 0.99999994 |
| Fam217b | 0.00531056 | 0.0042428 | 0.00479128 | -0.0005485 | 0.77696619 | 0.99999994 |
| Mocs1 | 0.02274753 | 0.01973397 | 0.02028345 | -0.0005495 | 0.57155099 | 0.99999994 |
| Hcn1 | 0.00168068 | 0.00096975 | 0.00152008 | -0.0005503 | 0.99202088 | 0.99999994 |
| Gm43653 | 0.00343558 | 0.00255266 | 0.00310358 | -0.0005509 | 0.88131567 | 0.99999994 |
| Fbxo47 | 0.00317596 | 0.00231799 | 0.00286956 | -0.0005516 | 0.89963073 | 0.99999994 |
| Anxa1 | 0.00459982 | 0.00360006 | 0.00415204 | -0.000552 | 0.81189614 | 0.99999994 |
| Ttc16 | 0.01085144 | 0.00920168 | 0.00975373 | -0.000552 | 0.64681632 | 0.99999994 |
| Car9 | 0.00782748 | 0.00649736 | 0.00705006 | -0.0005527 | 0.69910875 | 0.99999994 |
| Mtcp1 | 0.01624123 | 0.0139923 | 0.01454534 | -0.000553 | 0.59989934 | 0.99999994 |
| Glp1r | 0.00137159 | 0.0006876 | 0.00124079 | -0.0005532 | 0.99849956 | 0.99999994 |
| Glyctk | 0.00433729 | 0.00336246 | 0.00391577 | -0.0005533 | 0.82653332 | 0.99999994 |
| Mak | 0.0049482 | 0.00391214 | 0.00446545 | -0.0005533 | 0.79525358 | 0.99999994 |
| Ebf4 | 0.00738437 | 0.0060975 | 0.00665296 | -0.0005555 | 0.71080458 | 0.99999994 |
| Rbm12b2 | 0.01662555 | 0.01433008 | 0.01488568 | -0.0005556 | 0.59810491 | 0.99999994 |
| Tia1 | 0.56767513 | 0.33960307 | 0.34015867 | -0.0005556 | 0.50433731 | 0.99999994 |
| Car8 | 0.03906386 | 0.03389987 | 0.03445567 | -0.0005558 | 0.54275368 | 0.99999994 |
| Krt77 | 0.00106735 | 0.00040979 | 0.00096576 | -0.000556 | 0.99993644 | 0.99999994 |
| 9230105E05I | 0.00474063 | 0.00372248 | 0.00427874 | -0.0005563 | 0.80657926 | 0.99999994 |
| Golga1 | 0.23073465 | 0.17603156 | 0.17658868 | -0.0005571 | 0.50837724 | 0.99999994 |
| Epha4 | 0.0014265 | 0.00073188 | 0.00129041 | -0.0005585 | 0.99801832 | 0.99999994 |
| 2600006K01I | 0.01099773 | 0.00932539 | 0.00988424 | -0.0005588 | 0.64666854 | 0.99999994 |
| Gm20696 | 0.01429965 | 0.01226402 | 0.01282326 | -0.0005592 | 0.61420204 | 0.99999994 |
| Dgkh | 0.0010028 | 0.00034801 | 0.0009074 | -0.0005594 | 0.99997965 | 0.99999994 |
| Adgrd1 | 0.00156093 | 0.0008521 | 0.00141189 | -0.0005598 | 0.99584409 | 0.99999994 |
| Pde8b | 0.002892 | 0.00205288 | 0.0026135 | -0.0005606 | 0.92333332 | 0.99999994 |

| | | | | | | |
|---|---|---|---|---|---|---|
| Nxnl2 | 0.00713813 | 0.00587019 | 0.00643218 | -0.000562 | 0.71957417 | 0.99999994 |
| Ms4a6d | 0.00144045 | 0.00074096 | 0.00130302 | -0.0005621 | 0.99795542 | 0.99999994 |
| Trim34a | 0.00363787 | 0.00272272 | 0.00328588 | -0.0005632 | 0.87302822 | 0.99999994 |
| Cdh7 | 0.0016797 | 0.00095553 | 0.0015192 | -0.0005637 | 0.99323841 | 0.99999994 |
| Helz2 | 0.01323261 | 0.01131122 | 0.01187492 | -0.0005637 | 0.62398951 | 0.99999994 |
| Uri1 | 0.18154335 | 0.14374994 | 0.14431395 | -0.000564 | 0.5103772 | 0.99999994 |
| Serpine1 | 0.0042657 | 0.00328598 | 0.00385132 | -0.0005653 | 0.83573968 | 0.99999994 |
| Isoc2a | 0.00340655 | 0.00251184 | 0.00307743 | -0.0005656 | 0.88939947 | 0.99999994 |
| Gm28424 | 0.00344221 | 0.00254365 | 0.00310957 | -0.0005659 | 0.88713142 | 0.99999994 |
| Sirt1 | 0.02680847 | 0.0232737 | 0.02383963 | -0.0005659 | 0.56277792 | 0.99999994 |
| H2-DMb1 | 0.00793379 | 0.00657926 | 0.0071453 | -0.000566 | 0.70100759 | 0.99999994 |
| Timp1 | 0.01478245 | 0.01268443 | 0.0132519 | -0.0005675 | 0.61219066 | 0.99999994 |
| Krt23 | 0.00270116 | 0.00187365 | 0.00244135 | -0.0005677 | 0.93916847 | 0.99999994 |
| Pcdhb11 | 0.00598243 | 0.00482703 | 0.005395 | -0.000568 | 0.75827498 | 0.99999994 |
| Gm28800 | 0.00268965 | 0.00186227 | 0.00243097 | -0.0005687 | 0.94028841 | 0.99999994 |
| Tm6sf2 | 0.00224815 | 0.00146336 | 0.00203254 | -0.0005692 | 0.96883903 | 0.99999994 |
| Slc12a4 | 0.01370535 | 0.01172568 | 0.01229524 | -0.0005696 | 0.62108991 | 0.99999994 |
| Gm12166 | 0.0113318 | 0.00961253 | 0.01018219 | -0.0005697 | 0.64520237 | 0.99999994 |
| Ccdc78 | 0.00382164 | 0.00288136 | 0.00345144 | -0.0005701 | 0.86421119 | 0.99999994 |
| Pih1d2 | 0.07511094 | 0.06413365 | 0.06470456 | -0.0005709 | 0.52341718 | 0.99999994 |
| Zfp518b | 0.02882366 | 0.02502625 | 0.02559719 | -0.0005709 | 0.55901351 | 0.99999994 |
| Gabpa | 0.06554945 | 0.05624796 | 0.05681946 | -0.0005715 | 0.52668961 | 0.99999994 |
| 1700001C19I | 0.00227379 | 0.00148343 | 0.00205569 | -0.0005723 | 0.96805672 | 0.99999994 |
| 4930525G20 | 0.00441973 | 0.00341757 | 0.00398997 | -0.0005724 | 0.83019021 | 0.99999994 |
| Irf7 | 0.01933064 | 0.01670357 | 0.01727624 | -0.0005727 | 0.58731532 | 0.99999994 |
| Kazald1 | 0.00391113 | 0.00295928 | 0.00353205 | -0.0005728 | 0.85979982 | 0.99999994 |
| Nusap1 | 0.01057948 | 0.00893817 | 0.00951103 | -0.0005729 | 0.65575976 | 0.99999994 |
| Kcnc3 | 0.10934553 | 0.09143709 | 0.09201162 | -0.0005745 | 0.51657625 | 0.99999994 |
| Gm10614 | 0.00848127 | 0.00706063 | 0.00763554 | -0.0005749 | 0.69187966 | 0.99999994 |
| Fxyd4 | 0.005682 | 0.0045486 | 0.00512511 | -0.0005765 | 0.77299825 | 0.99999994 |

| | | | | | | |
|---|---|---|---|---|---|---|
| Zfp3 | 0.03181967 | 0.02762493 | 0.02820145 | -0.0005765 | 0.55411919 | 0.99999994 |
| Klhl1 | 0.01259291 | 0.01072895 | 0.01130573 | -0.0005768 | 0.63291513 | 0.99999994 |
| Igf1 | 0.01629463 | 0.01401566 | 0.01459264 | -0.000577 | 0.60379733 | 0.99999994 |
| Flrt3 | 0.01037311 | 0.00874977 | 0.0093268 | -0.000577 | 0.6597628 | 0.99999994 |
| Rad9a | 0.03570873 | 0.03098939 | 0.03156652 | -0.0005771 | 0.54843147 | 0.99999994 |
| Fam228b | 0.00353625 | 0.00261712 | 0.00319431 | -0.0005772 | 0.88546593 | 0.99999994 |
| Ccdc162 | 0.0092128 | 0.00771272 | 0.00829002 | -0.0005773 | 0.67850825 | 0.99999994 |
| Arnt2 | 0.01755986 | 0.01513491 | 0.01571234 | -0.0005774 | 0.59662598 | 0.99999994 |
| Fxyd7 | 0.00256715 | 0.00174286 | 0.00232044 | -0.0005776 | 0.95122262 | 0.99999994 |
| Gm11696 | 0.00509101 | 0.00401574 | 0.00459388 | -0.0005781 | 0.79890255 | 0.99999994 |
| B3gnt3 | 0.00855465 | 0.0071221 | 0.00770122 | -0.0005791 | 0.69165631 | 0.99999994 |
| Fam162b | 0.00820836 | 0.00681116 | 0.00739121 | -0.0005801 | 0.69929745 | 0.99999994 |
| Ccdc84 | 0.05060973 | 0.04372039 | 0.04430053 | -0.0005801 | 0.53473082 | 0.99999994 |
| Rab39 | 0.00569035 | 0.00455239 | 0.00513261 | -0.0005802 | 0.77411741 | 0.99999994 |
| Gm17484 | 0.03802452 | 0.03298151 | 0.03356206 | -0.0005806 | 0.54583313 | 0.99999994 |
| Cfap46 | 0.00301727 | 0.00214516 | 0.00272647 | -0.0005813 | 0.92207829 | 0.99999994 |
| Nim1k | 0.02581064 | 0.02238622 | 0.02296762 | -0.0005814 | 0.56690467 | 0.99999994 |
| Gm11767 | 0.00260523 | 0.00177292 | 0.00235481 | -0.0005819 | 0.94999696 | 0.99999994 |
| Angptl1 | 0.00357126 | 0.00264379 | 0.00322586 | -0.0005821 | 0.88513743 | 0.99999994 |
| 9430069I07R | 0.00131424 | 0.00060586 | 0.00118896 | -0.0005831 | 0.99945165 | 0.99999994 |
| Ucma | 0.00097851 | 0.00030219 | 0.00088543 | -0.0005832 | 0.99999419 | 0.99999994 |
| Cdk20 | 0.02798306 | 0.02428098 | 0.02486463 | -0.0005836 | 0.5620796 | 0.99999994 |
| Gm17509 | 0.00321069 | 0.00231707 | 0.00290087 | -0.0005838 | 0.90981088 | 0.99999994 |
| Fhl1 | 0.00217152 | 0.00137934 | 0.00196336 | -0.000584 | 0.97614807 | 0.99999994 |
| Adssl1 | 0.01239586 | 0.01054615 | 0.01113031 | -0.0005842 | 0.63658624 | 0.99999994 |
| Insl6 | 0.00688013 | 0.00561591 | 0.00620078 | -0.0005849 | 0.73494639 | 0.99999994 |
| Esam | 0.01022882 | 0.00861298 | 0.00919796 | -0.000585 | 0.66397276 | 0.99999994 |
| Calb1 | 0.01037266 | 0.008741 | 0.00932639 | -0.0005854 | 0.66195391 | 0.99999994 |
| Tfpi2 | 0.00626297 | 0.00506077 | 0.00564692 | -0.0005861 | 0.75519308 | 0.99999994 |
| Cds1 | 0.00590151 | 0.004736 | 0.00532232 | -0.0005863 | 0.76830544 | 0.99999994 |

| | | | | | | |
|---|---|---|---|---|---|---|
| Ccdc22 | 0.06091787 | 0.0523777 | 0.0529645 | -0.0005868 | 0.5293943 | 0.99999994 |
| Cep135 | 0.03018384 | 0.02619392 | 0.02678081 | -0.0005869 | 0.5579886 | 0.99999994 |
| Fbxl20 | 0.07513164 | 0.06413454 | 0.06472153 | -0.000587 | 0.5240694 | 0.99999994 |
| Ddias | 0.00277849 | 0.00192322 | 0.00251112 | -0.0005879 | 0.94042745 | 0.99999994 |
| Ntn1 | 0.00345576 | 0.00253385 | 0.00312178 | -0.0005879 | 0.89500611 | 0.99999994 |
| Herc6 | 0.00655842 | 0.00532381 | 0.00591212 | -0.0005883 | 0.74613521 | 0.99999994 |
| Gm5105 | 0.00775214 | 0.00639409 | 0.00698257 | -0.0005885 | 0.71259658 | 0.99999994 |
| Tgfbr3 | 0.0087695 | 0.00730498 | 0.00789349 | -0.0005885 | 0.69014917 | 0.99999994 |
| Tubgcp2 | 0.09903154 | 0.08342097 | 0.08401022 | -0.0005892 | 0.51861899 | 0.99999994 |
| Zfp287 | 0.01044565 | 0.0088022 | 0.00939156 | -0.0005894 | 0.66192227 | 0.99999994 |
| Krt25 | 0.00156754 | 0.00082821 | 0.00141786 | -0.0005896 | 0.99718159 | 0.99999994 |
| Mcm10 | 0.00320396 | 0.00230515 | 0.0028948 | -0.0005897 | 0.9124287 | 0.99999994 |
| Tm7sf2 | 0.04181221 | 0.03622282 | 0.03681275 | -0.0005899 | 0.54247421 | 0.99999994 |
| Gpr173 | 0.02835094 | 0.02459524 | 0.02518533 | -0.0005901 | 0.5619672 | 0.99999994 |
| Nln | 0.03587412 | 0.03111889 | 0.03170925 | -0.0005904 | 0.54931389 | 0.99999994 |
| Ecm1 | 0.00320716 | 0.00230694 | 0.00289769 | -0.0005908 | 0.91261581 | 0.99999994 |
| Pard6b | 0.02348471 | 0.02033935 | 0.02093044 | -0.0005911 | 0.57455456 | 0.99999994 |
| Slc29a2 | 0.00328474 | 0.00237648 | 0.00296763 | -0.0005911 | 0.90756868 | 0.99999994 |
| Gdpd3 | 0.01277846 | 0.01087909 | 0.01147088 | -0.0005918 | 0.63435326 | 0.99999994 |
| Mknk2 | 0.04023077 | 0.03486553 | 0.0354575 | -0.000592 | 0.54424333 | 0.99999994 |
| Klf1 | 0.00475992 | 0.00370391 | 0.00429609 | -0.0005922 | 0.82056437 | 0.99999994 |
| Taf4b | 0.00352972 | 0.00259592 | 0.00318843 | -0.0005925 | 0.89194676 | 0.99999994 |
| Diras1 | 0.00376099 | 0.0028042 | 0.0033968 | -0.0005926 | 0.87723194 | 0.99999994 |
| Gm20186 | 0.02058424 | 0.01778841 | 0.01838114 | -0.0005927 | 0.58497776 | 0.99999994 |
| Cenpu | 0.00517497 | 0.00407653 | 0.00466938 | -0.0005928 | 0.80097964 | 0.99999994 |
| Spon1 | 0.00579563 | 0.00463413 | 0.0052272 | -0.0005931 | 0.77494282 | 0.99999994 |
| Dach1 | 0.17571408 | 0.13972243 | 0.14031583 | -0.0005934 | 0.51122873 | 0.99999994 |
| Aqp12 | 0.00407718 | 0.00308729 | 0.00368159 | -0.0005943 | 0.85870099 | 0.99999994 |
| Fam13a | 0.00393107 | 0.00295543 | 0.00355 | -0.0005946 | 0.86753759 | 0.99999994 |
| Mettl22 | 0.05343457 | 0.0460919 | 0.04668647 | -0.0005946 | 0.53377834 | 0.99999994 |

| | | | | | | |
|---|---|---|---|---|---|---|
| Fzd8 | 0.00647016 | 0.00523812 | 0.00583291 | -0.0005948 | 0.75135249 | 0.99999994 |
| Lrrc46 | 0.02084798 | 0.01801847 | 0.01861336 | -0.0005949 | 0.58423534 | 0.99999994 |
| 9530053A07 | 0.00907951 | 0.00757581 | 0.00817082 | -0.000595 | 0.68606347 | 0.99999994 |
| Aldh16a1 | 0.03587654 | 0.03111623 | 0.03171133 | -0.0005951 | 0.54970411 | 0.99999994 |
| Fzd6 | 0.01418179 | 0.01212327 | 0.01271857 | -0.0005953 | 0.62230988 | 0.99999994 |
| Gm15559 | 0.00842008 | 0.00698539 | 0.00758077 | -0.0005954 | 0.69943374 | 0.99999994 |
| Asah2 | 0.00329885 | 0.00238429 | 0.00298034 | -0.000596 | 0.90844267 | 0.99999994 |
| Abcb11 | 0.00368014 | 0.00272744 | 0.00332396 | -0.0005965 | 0.88386965 | 0.99999994 |
| Wdr18 | 0.35300647 | 0.24589812 | 0.24649478 | -0.0005967 | 0.5064275 | 0.99999994 |
| Tcf7l1 | 0.01278893 | 0.01088338 | 0.0114802 | -0.0005968 | 0.63534465 | 0.99999994 |
| Trnau1ap | 0.25961131 | 0.19377209 | 0.194369 | -0.0005969 | 0.5081546 | 0.99999994 |
| Mmp16 | 0.00845588 | 0.00701585 | 0.00761282 | -0.000597 | 0.69915318 | 0.99999994 |
| Prrt3 | 0.00331254 | 0.00239569 | 0.00299269 | -0.000597 | 0.90788693 | 0.99999994 |
| Gm16141 | 0.03534916 | 0.03065838 | 0.03125613 | -0.0005978 | 0.55064785 | 0.99999994 |
| Zxdc | 0.05338918 | 0.04605037 | 0.04664819 | -0.0005978 | 0.5339903 | 0.99999994 |
| Cnnm1 | 0.00346158 | 0.00252919 | 0.00312702 | -0.0005978 | 0.89841584 | 0.99999994 |
| Rgcc | 0.00382752 | 0.00285868 | 0.00345673 | -0.000598 | 0.87526018 | 0.99999994 |
| Hif3a | 0.01004597 | 0.00843633 | 0.00903465 | -0.0005983 | 0.67032761 | 0.99999994 |
| 8430432A02 | 0.00199663 | 0.00120604 | 0.00180545 | -0.0005994 | 0.98644472 | 0.99999994 |
| Zfp455 | 0.00990596 | 0.00830996 | 0.00890957 | -0.0005996 | 0.67291224 | 0.99999994 |
| Gm9008 | 0.00262013 | 0.00176839 | 0.00236825 | -0.0005999 | 0.95410258 | 0.99999994 |
| Zfand1 | 0.14378431 | 0.11712835 | 0.11772878 | -0.0006004 | 0.51354074 | 0.99999994 |
| Gm9920 | 0.00872674 | 0.00725417 | 0.00785523 | -0.0006011 | 0.69473903 | 0.99999994 |
| Ephb2 | 0.04708786 | 0.04071109 | 0.04131341 | -0.0006023 | 0.53865471 | 0.99999994 |
| Gm12689 | 0.00401364 | 0.00302171 | 0.00362436 | -0.0006027 | 0.86581033 | 0.99999994 |
| Dhx38 | 0.06697927 | 0.05740175 | 0.05800484 | -0.0006031 | 0.52758755 | 0.99999994 |
| Tusc1 | 0.01935533 | 0.01669411 | 0.01729802 | -0.0006039 | 0.59188097 | 0.99999994 |
| Grn | 0.22494605 | 0.17231935 | 0.17292336 | -0.000604 | 0.50927466 | 0.99999994 |
| Pvr | 0.00333617 | 0.00240883 | 0.00301398 | -0.0006052 | 0.9093021 | 0.99999994 |
| Upp2 | 0.03571952 | 0.03097038 | 0.03157583 | -0.0006055 | 0.55078008 | 0.99999994 |

| | | | | | | |
|---|---|---|---|---|---|---|
| Mki67 | 0.01021844 | 0.00858316 | 0.00918869 | -0.0006055 | 0.66954327 | 0.99999994 |
| Ube2l6 | 0.01142029 | 0.00965481 | 0.01026108 | -0.0006063 | 0.6529462 | 0.99999994 |
| Zfp414 | 0.20764277 | 0.16115268 | 0.16175902 | -0.0006063 | 0.50995295 | 0.99999994 |
| Xrra1 | 0.0034305 | 0.0024923 | 0.00309901 | -0.0006067 | 0.90373773 | 0.99999994 |
| Tmem178 | 0.01171647 | 0.00991831 | 0.0105251 | -0.0006068 | 0.64941873 | 0.99999994 |
| Dnhd1 | 0.00664594 | 0.00538379 | 0.00599066 | -0.0006069 | 0.74994303 | 0.99999994 |
| Rhbdf2 | 0.03057987 | 0.0265181 | 0.02712502 | -0.0006069 | 0.5591983 | 0.99999994 |
| Chrnb1 | 0.01023239 | 0.00859298 | 0.00920115 | -0.0006082 | 0.67001794 | 0.99999994 |
| Aebp1 | 0.00840376 | 0.00695736 | 0.00756616 | -0.0006088 | 0.7038796 | 0.99999994 |
| Ecel1 | 0.00345915 | 0.00251583 | 0.00312483 | -0.000609 | 0.902729 | 0.99999994 |
| Samd4 | 0.0049178 | 0.0038278 | 0.00443811 | -0.0006103 | 0.81999692 | 0.99999994 |
| Trpc7 | 0.00214408 | 0.00132782 | 0.00193859 | -0.0006108 | 0.98200897 | 0.99999994 |
| Zfp526 | 0.00929598 | 0.00775323 | 0.0083644 | -0.0006112 | 0.68664512 | 0.99999994 |
| Tmprss2 | 0.00787702 | 0.00648323 | 0.00709444 | -0.0006112 | 0.71683726 | 0.99999994 |
| Susd1 | 0.00329072 | 0.00236159 | 0.00297302 | -0.0006114 | 0.91449054 | 0.99999994 |
| 1700112J16F | 0.00400643 | 0.00300568 | 0.00361788 | -0.0006122 | 0.86999081 | 0.99999994 |
| Apoc3 | 0.00148217 | 0.00072842 | 0.00134072 | -0.0006123 | 0.99881672 | 0.99999994 |
| Gm16150 | 0.00360214 | 0.00264058 | 0.00325368 | -0.0006131 | 0.8951302 | 0.99999994 |
| Teddm2 | 0.00446994 | 0.00342143 | 0.00403516 | -0.0006137 | 0.84432788 | 0.99999994 |
| Rph3a | 0.0015468 | 0.00078522 | 0.00139912 | -0.0006139 | 0.99825344 | 0.99999994 |
| Zfp52 | 0.00649497 | 0.00524093 | 0.00585518 | -0.0006142 | 0.75750387 | 0.99999994 |
| Fhad1 | 0.08730532 | 0.07400596 | 0.07462043 | -0.0006145 | 0.5218561 | 0.99999994 |
| Chpt1 | 0.04987501 | 0.04306369 | 0.04367851 | -0.0006148 | 0.53732387 | 0.99999994 |
| Glis1 | 0.02350939 | 0.02033637 | 0.02095209 | -0.0006157 | 0.5775425 | 0.99999994 |
| Cryab | 0.01690852 | 0.01452017 | 0.01513615 | -0.000616 | 0.60676257 | 0.99999994 |
| Gm15816 | 0.00197497 | 0.00116986 | 0.00178589 | -0.000616 | 0.9891639 | 0.99999994 |
| Acsf2 | 0.00417201 | 0.00315069 | 0.00376697 | -0.0006163 | 0.8619215 | 0.99999994 |
| Scd3 | 0.02067187 | 0.01784125 | 0.01845831 | -0.0006171 | 0.58804658 | 0.99999994 |
| C78197 | 0.00275451 | 0.00187242 | 0.00248949 | -0.0006171 | 0.95051947 | 0.99999994 |
| Pcolce2 | 0.00313971 | 0.00221974 | 0.00283688 | -0.0006171 | 0.92619619 | 0.99999994 |

| | | | | | | |
|---|---|---|---|---|---|---|
| Pde4c | 0.00163505 | 0.00086101 | 0.00147885 | -0.0006178 | 0.99728982 | 0.99999994 |
| C1d | 0.73973253 | 0.3907278 | 0.39134598 | -0.0006182 | 0.50419462 | 0.99999994 |
| Ano6 | 0.07251504 | 0.06195477 | 0.06257346 | -0.0006187 | 0.5262374 | 0.99999994 |
| Rarg | 0.00576735 | 0.00458271 | 0.00520179 | -0.0006191 | 0.78587811 | 0.99999994 |
| Arhgdib | 0.01133239 | 0.0095636 | 0.01018271 | -0.0006191 | 0.65715503 | 0.99999994 |
| Mrc1 | 0.00397274 | 0.00296795 | 0.00358753 | -0.0006196 | 0.87484166 | 0.99999994 |
| Tec | 0.00786975 | 0.00646829 | 0.00708793 | -0.0006196 | 0.71968815 | 0.99999994 |
| Thbs3 | 0.04070483 | 0.03524364 | 0.03586406 | -0.0006204 | 0.54583711 | 0.99999994 |
| Camp | 0.00393602 | 0.00293397 | 0.00355446 | -0.0006205 | 0.87737804 | 0.99999994 |
| Gm15627 | 0.00444802 | 0.00339473 | 0.00401543 | -0.0006207 | 0.84824491 | 0.99999994 |
| Sap30bp | 0.17623829 | 0.1400557 | 0.140677 | -0.0006213 | 0.51172635 | 0.99999994 |
| Pidd1 | 0.00302121 | 0.00210857 | 0.00273003 | -0.0006215 | 0.9351425 | 0.99999994 |
| Cemip | 0.00405873 | 0.00304331 | 0.00366497 | -0.0006217 | 0.87056712 | 0.99999994 |
| Zfp850 | 0.00580253 | 0.00461141 | 0.0052334 | -0.000622 | 0.78555925 | 0.99999994 |
| Pmel | 0.00455644 | 0.00349056 | 0.00411301 | -0.0006224 | 0.84311705 | 0.99999994 |
| Gm12981 | 0.00384166 | 0.00284642 | 0.00346947 | -0.0006231 | 0.88402589 | 0.99999994 |
| Gp1bb | 0.00365904 | 0.00268171 | 0.00330495 | -0.0006232 | 0.89530514 | 0.99999994 |
| Inpp1 | 0.03420195 | 0.02964136 | 0.03026482 | -0.0006235 | 0.55453257 | 0.99999994 |
| Zcchc17 | 0.48962262 | 0.30920372 | 0.3098278 | -0.0006241 | 0.50534873 | 0.99999994 |
| Ccdc163 | 0.05907625 | 0.05079918 | 0.0514252 | -0.000626 | 0.53229141 | 0.99999994 |
| Rad51c | 0.01702378 | 0.01461197 | 0.01523816 | -0.0006262 | 0.60777774 | 0.99999994 |
| Fam169a | 0.00801867 | 0.00659399 | 0.00722133 | -0.0006273 | 0.71845556 | 0.99999994 |
| Arsg | 0.00689153 | 0.00558304 | 0.006211 | -0.000628 | 0.74952378 | 0.99999994 |
| Lrrc36 | 0.00183859 | 0.00103355 | 0.00166272 | -0.0006292 | 0.99411173 | 0.99999994 |
| Zfp655 | 0.12606565 | 0.10404882 | 0.10467806 | -0.0006292 | 0.51595846 | 0.99999994 |
| Antxr1 | 0.00597789 | 0.00476143 | 0.00539092 | -0.0006295 | 0.78149789 | 0.99999994 |
| Zfp281 | 0.09752371 | 0.08218063 | 0.08281069 | -0.0006301 | 0.52019567 | 0.99999994 |
| Nop14 | 0.1452358 | 0.11815111 | 0.11878135 | -0.0006302 | 0.51408672 | 0.99999994 |
| Junos | 0.01201825 | 0.01016356 | 0.010794 | -0.0006304 | 0.65127913 | 0.99999994 |
| Gm13093 | 0.00111907 | 0.00038091 | 0.00101253 | -0.0006316 | 0.99998354 | 0.99999994 |

| | | | | | | |
|---|---|---|---|---|---|---|
| Gm26901 | 0.00395703 | 0.00294152 | 0.00357339 | -0.0006319 | 0.88040561 | 0.99999994 |
| Tmem154 | 0.00217086 | 0.00133016 | 0.00196277 | -0.0006326 | 0.98403839 | 0.99999994 |
| Rnasel | 0.016312 | 0.01397473 | 0.01460803 | -0.0006333 | 0.61354595 | 0.99999994 |
| Car11 | 0.03168681 | 0.02745244 | 0.02808619 | -0.0006337 | 0.55969462 | 0.99999994 |
| Sufu | 0.01231929 | 0.01042834 | 0.01106212 | -0.0006338 | 0.64853158 | 0.99999994 |
| Fez1 | 0.00589702 | 0.00468398 | 0.00531829 | -0.0006343 | 0.78637308 | 0.99999994 |
| Izumo4 | 0.05994039 | 0.05151347 | 0.05214795 | -0.0006345 | 0.53227366 | 0.99999994 |
| Gm27217 | 0.02695702 | 0.02333467 | 0.02396936 | -0.0006347 | 0.56995343 | 0.99999994 |
| Zfp14 | 0.01815347 | 0.01560154 | 0.01623702 | -0.0006355 | 0.60276626 | 0.99999994 |
| Gm4792 | 0.00171739 | 0.0009171 | 0.00155324 | -0.0006361 | 0.9967963 | 0.99999994 |
| Slc6a20a | 0.00982451 | 0.00820015 | 0.00883681 | -0.0006367 | 0.68423154 | 0.99999994 |
| Gm13613 | 0.00251309 | 0.00163464 | 0.00227166 | -0.000637 | 0.96901971 | 0.99999994 |
| Arhgef38 | 0.00637565 | 0.00511078 | 0.00574808 | -0.0006373 | 0.76974372 | 0.99999994 |
| Ppp1r3c | 0.00468881 | 0.00359442 | 0.00423212 | -0.0006377 | 0.842066 | 0.99999994 |
| Slc12a8 | 0.00326582 | 0.00231283 | 0.00295057 | -0.0006377 | 0.92488348 | 0.99999994 |
| Limch1 | 0.01752068 | 0.01503813 | 0.0156777 | -0.0006396 | 0.6070133 | 0.99999994 |
| Il18bp | 0.00384985 | 0.00283678 | 0.00347685 | -0.0006401 | 0.88978827 | 0.99999994 |
| 3110056K07I | 0.0330099 | 0.02859302 | 0.02923317 | -0.0006401 | 0.55794483 | 0.99999994 |
| Hs3st3b1 | 0.00217441 | 0.00132558 | 0.00196597 | -0.0006404 | 0.98492916 | 0.99999994 |
| Slc41a2 | 0.00120895 | 0.00045259 | 0.00109377 | -0.0006412 | 0.99995231 | 0.99999994 |
| Strada | 0.03743972 | 0.03241715 | 0.03305871 | -0.0006416 | 0.55139167 | 0.99999994 |
| Pwp2 | 0.02155799 | 0.01859626 | 0.0192381 | -0.0006418 | 0.58787162 | 0.99999994 |
| AY074887 | 0.00201794 | 0.0011819 | 0.00182469 | -0.0006428 | 0.99048292 | 0.99999994 |
| 4930509G22 | 0.00237947 | 0.00150756 | 0.00215108 | -0.0006435 | 0.97677689 | 0.99999994 |
| Lrp2bp | 0.00205433 | 0.00121387 | 0.00185756 | -0.0006437 | 0.98946053 | 0.99999994 |
| Arf4os | 0.00219066 | 0.00133659 | 0.00198064 | -0.000644 | 0.98478502 | 0.99999994 |
| Atad5 | 0.02671769 | 0.02311586 | 0.02376035 | -0.0006445 | 0.57163954 | 0.99999994 |
| Rasgrp2 | 0.00813248 | 0.00667797 | 0.00732326 | -0.0006453 | 0.72123969 | 0.99999994 |
| Sult4a1 | 0.04997536 | 0.04311794 | 0.0437635 | -0.0006456 | 0.53910851 | 0.99999994 |
| Ttc39c | 0.01936257 | 0.01665884 | 0.01730441 | -0.0006456 | 0.59805927 | 0.99999994 |

| | | | | | | |
|---|---|---|---|---|---|---|
| Fank1 | 0.01007642 | 0.00841606 | 0.00906185 | -0.0006458 | 0.68237784 | 0.99999994 |
| Zfp963 | 0.03702542 | 0.032056 | 0.03270188 | -0.0006459 | 0.55229704 | 0.99999994 |
| Dnm1 | 0.02071133 | 0.01784615 | 0.01849305 | -0.0006469 | 0.59205924 | 0.99999994 |
| Shisa2 | 0.05393166 | 0.04645805 | 0.04710541 | -0.0006474 | 0.53644268 | 0.99999994 |
| Enpep | 0.00807471 | 0.00662412 | 0.00727152 | -0.0006474 | 0.72328706 | 0.99999994 |
| Zfp951 | 0.0044388 | 0.00335912 | 0.00400713 | -0.000648 | 0.85913249 | 0.99999994 |
| Fstl3 | 0.01532932 | 0.01308898 | 0.01373709 | -0.0006481 | 0.6232548 | 0.99999994 |
| Acsbg1 | 0.0070431 | 0.00569854 | 0.00634696 | -0.0006484 | 0.75175431 | 0.99999994 |
| Gm42912 | 0.0123558 | 0.01044618 | 0.01109463 | -0.0006485 | 0.65138021 | 0.99999994 |
| Usp54 | 0.01480645 | 0.01262417 | 0.01327319 | -0.000649 | 0.62759035 | 0.99999994 |
| Fbn2 | 0.00264376 | 0.00174 | 0.00238956 | -0.0006496 | 0.96480697 | 0.99999994 |
| Pir | 0.01253092 | 0.01060003 | 0.01125055 | -0.0006505 | 0.64983495 | 0.99999994 |
| Cdk19 | 0.02698748 | 0.02334501 | 0.02399595 | -0.0006509 | 0.57164725 | 0.99999994 |
| Cpne2 | 0.0032007 | 0.00224061 | 0.00289187 | -0.0006513 | 0.93306565 | 0.99999994 |
| 1810009A15I | 0.00904677 | 0.00748978 | 0.00814154 | -0.0006518 | 0.70293436 | 0.99999994 |
| Tnfrsf11b | 0.00296797 | 0.0020289 | 0.00268201 | -0.0006531 | 0.94748438 | 0.99999994 |
| Trmt11 | 0.06305233 | 0.05409037 | 0.05474395 | -0.0006536 | 0.53167033 | 0.99999994 |
| Metap1d | 0.08064732 | 0.06857166 | 0.06922581 | -0.0006541 | 0.52507661 | 0.99999994 |
| Akr1e1 | 0.05573009 | 0.04796461 | 0.04861881 | -0.0006542 | 0.53568416 | 0.99999994 |
| Atp6v1g2 | 0.01523707 | 0.01300105 | 0.01365526 | -0.0006542 | 0.62509876 | 0.99999994 |
| Susd2 | 0.00724928 | 0.00587746 | 0.00653184 | -0.0006544 | 0.74756701 | 0.99999994 |
| Ccdc160 | 0.00822391 | 0.00674828 | 0.00740513 | -0.0006569 | 0.72255084 | 0.99999994 |
| Myrip | 0.00180466 | 0.00097517 | 0.00163208 | -0.0006569 | 0.99630958 | 0.99999994 |
| Etaa1os | 0.01302585 | 0.01103255 | 0.01169101 | -0.0006585 | 0.64613121 | 0.99999994 |
| 0610040F04I | 0.00265781 | 0.00174368 | 0.00240225 | -0.0006586 | 0.96598523 | 0.99999994 |
| 2900009J06F | 0.01388737 | 0.01179822 | 0.012457 | -0.0006588 | 0.63758819 | 0.99999994 |
| B3gat3 | 0.18795046 | 0.14800621 | 0.14866524 | -0.000659 | 0.51177002 | 0.99999994 |
| Uckl1os | 0.00494102 | 0.00379967 | 0.00445899 | -0.0006593 | 0.83749945 | 0.99999994 |
| Gm10334 | 0.00374495 | 0.00272297 | 0.00338235 | -0.0006594 | 0.90279474 | 0.99999994 |
| Gm26782 | 0.00974083 | 0.00810256 | 0.00876203 | -0.0006595 | 0.6918095 | 0.99999994 |

| | | | | | | |
|---|---|---|---|---|---|---|
| Rgs1 | 0.00263071 | 0.00171779 | 0.0023778 | -0.00066 | 0.96766947 | 0.99999994 |
| Fbxo5 | 0.01475276 | 0.01256549 | 0.01322555 | -0.0006601 | 0.63013209 | 0.99999994 |
| Caskin1 | 0.0015355 | 0.00072786 | 0.00138891 | -0.000661 | 0.9992327 | 0.99999994 |
| Aurkc | 0.00141514 | 0.00061768 | 0.00128014 | -0.0006625 | 0.99971405 | 0.99999994 |
| Efnb2 | 0.00294517 | 0.00199839 | 0.00266146 | -0.0006631 | 0.95137614 | 0.99999994 |
| Erich5 | 0.03144337 | 0.02721011 | 0.02787493 | -0.0006648 | 0.56306814 | 0.99999994 |
| 6720427I07R | 0.0116726 | 0.0098204 | 0.010486 | -0.0006656 | 0.66367346 | 0.99999994 |
| Coro2a | 0.01098786 | 0.00920951 | 0.00987543 | -0.0006659 | 0.67323123 | 0.99999994 |
| Alox12b | 0.00142112 | 0.00061932 | 0.00128555 | -0.0006662 | 0.99971933 | 0.99999994 |
| Rdh11 | 0.06585847 | 0.05640941 | 0.05707584 | -0.0006664 | 0.53097515 | 0.99999994 |
| Disp1 | 0.01225522 | 0.01033852 | 0.01100507 | -0.0006665 | 0.65658389 | 0.99999994 |
| Strip1 | 0.14008453 | 0.11436789 | 0.11503466 | -0.0006668 | 0.51538803 | 0.99999994 |
| Kirrel3 | 0.0025542 | 0.00164177 | 0.00230876 | -0.000667 | 0.97275989 | 0.99999994 |
| Frrs1l | 0.00184565 | 0.00100185 | 0.0016691 | -0.0006672 | 0.99610423 | 0.99999994 |
| Lrrc8e | 0.00702066 | 0.00565941 | 0.00632683 | -0.0006674 | 0.75871816 | 0.99999994 |
| 6030443J06F | 0.00392575 | 0.00287743 | 0.00354521 | -0.0006678 | 0.89504304 | 0.99999994 |
| Cct6b | 0.00581326 | 0.00457497 | 0.00524304 | -0.0006681 | 0.80182482 | 0.99999994 |
| Adamts2 | 0.00299194 | 0.00203487 | 0.00270364 | -0.0006688 | 0.95016867 | 0.99999994 |
| Kcnj12 | 0.00172582 | 0.00089069 | 0.00156086 | -0.0006702 | 0.99786828 | 0.99999994 |
| Yjefn3 | 0.01484312 | 0.01263534 | 0.01330574 | -0.0006704 | 0.63132936 | 0.99999994 |
| Slc41a3 | 0.0199937 | 0.01718992 | 0.01786088 | -0.000671 | 0.5987267 | 0.99999994 |
| Ctrl | 0.00571252 | 0.00447986 | 0.00515253 | -0.0006727 | 0.80757672 | 0.99999994 |
| Tmem81 | 0.0017141 | 0.0008774 | 0.00155027 | -0.0006729 | 0.99806814 | 0.99999994 |
| 9130023H24 | 0.01348305 | 0.01142437 | 0.01209763 | -0.0006733 | 0.64447341 | 0.99999994 |
| Myo5b | 0.0205144 | 0.01764581 | 0.01831963 | -0.0006738 | 0.5967064 | 0.99999994 |
| Tmem121 | 0.02015112 | 0.01732485 | 0.0179996 | -0.0006748 | 0.59852326 | 0.99999994 |
| Cilp | 0.00207709 | 0.00120304 | 0.0018781 | -0.0006751 | 0.99163399 | 0.99999994 |
| Ppm1f | 0.00701219 | 0.00564405 | 0.00631923 | -0.0006752 | 0.76152079 | 0.99999994 |
| Phgdh | 0.12109713 | 0.10027353 | 0.1009502 | -0.0006767 | 0.51779375 | 0.99999994 |
| Gm12682 | 0.00309148 | 0.00211635 | 0.00279339 | -0.000677 | 0.94666035 | 0.99999994 |

| | | | | | | |
|---|---|---|---|---|---|---|
| Zfp869 | 0.01273911 | 0.01075863 | 0.01143586 | -0.0006772 | 0.65328016 | 0.99999994 |
| Nova1 | 0.00591976 | 0.00466125 | 0.00533871 | -0.0006775 | 0.8008509 | 0.99999994 |
| Arhgap33 | 0.01381059 | 0.01171117 | 0.01238878 | -0.0006776 | 0.64209642 | 0.99999994 |
| Tfcp2l1 | 0.01061707 | 0.00886237 | 0.00954458 | -0.0006822 | 0.68288117 | 0.99999994 |
| Hnf4g | 0.00744336 | 0.00602356 | 0.00670584 | -0.0006823 | 0.75087378 | 0.99999994 |
| Nlrx1 | 0.01040657 | 0.00867345 | 0.00935667 | -0.0006832 | 0.68653268 | 0.99999994 |
| Notch4 | 0.00667704 | 0.00533365 | 0.00601857 | -0.0006849 | 0.77562561 | 0.99999994 |
| Kcnc1 | 0.01956403 | 0.01679704 | 0.01748209 | -0.0006851 | 0.60289095 | 0.99999994 |
| Tenm2 | 0.00564006 | 0.00440223 | 0.00508743 | -0.0006852 | 0.81500886 | 0.99999994 |
| Lonrf1 | 0.02390654 | 0.02061504 | 0.02130038 | -0.0006853 | 0.58479216 | 0.99999994 |
| Gm10516 | 0.00470496 | 0.00356024 | 0.00424664 | -0.0006864 | 0.85901468 | 0.99999994 |
| Slc22a15 | 0.02553384 | 0.02203875 | 0.02272552 | -0.0006868 | 0.57971142 | 0.99999994 |
| Caps2 | 0.00683048 | 0.00546947 | 0.00615624 | -0.0006868 | 0.7711268 | 0.99999994 |
| Ccdc17 | 0.01497847 | 0.01273899 | 0.01342585 | -0.0006869 | 0.63327398 | 0.99999994 |
| Lmln | 0.01363028 | 0.01154127 | 0.01222851 | -0.0006872 | 0.64583124 | 0.99999994 |
| Prr36 | 0.04452303 | 0.03844166 | 0.03912927 | -0.0006876 | 0.54655805 | 0.99999994 |
| 6330403L08F | 0.00317061 | 0.0021761 | 0.00286474 | -0.0006886 | 0.94521052 | 0.99999994 |
| Zscan2 | 0.07004242 | 0.05984755 | 0.06053689 | -0.0006893 | 0.53020962 | 0.99999994 |
| Gm11665 | 0.0012775 | 0.00046565 | 0.00115574 | -0.0006901 | 0.99996474 | 0.99999994 |
| Ube2t | 0.01473116 | 0.01251561 | 0.01320638 | -0.0006908 | 0.63614184 | 0.99999994 |
| Sec14l4 | 0.00741033 | 0.00598506 | 0.00667623 | -0.0006912 | 0.75462418 | 0.99999994 |
| Spred1 | 0.01948811 | 0.01672311 | 0.01741514 | -0.000692 | 0.60430337 | 0.99999994 |
| Mpp1 | 0.01339086 | 0.01132177 | 0.01201565 | -0.0006939 | 0.6496572 | 0.99999994 |
| Gm16794 | 0.00494888 | 0.00377171 | 0.00446606 | -0.0006943 | 0.84963734 | 0.99999994 |
| Nipa2 | 0.26356771 | 0.19604582 | 0.19674095 | -0.0006951 | 0.50938164 | 0.99999994 |
| Pgap3 | 0.02707741 | 0.02337926 | 0.02407446 | -0.0006952 | 0.5762139 | 0.99999994 |
| Cldn8 | 0.0055375 | 0.0042998 | 0.00499527 | -0.0006955 | 0.82296793 | 0.99999994 |
| Casp8 | 0.05253697 | 0.04523375 | 0.04592929 | -0.0006955 | 0.54014609 | 0.99999994 |
| Gm13715 | 0.00174601 | 0.00088339 | 0.00157909 | -0.0006957 | 0.99831937 | 0.99999994 |
| Fyb | 0.01288649 | 0.0108709 | 0.01156701 | -0.0006961 | 0.65563706 | 0.99999994 |

| | | | | | | |
|---|---|---|---|---|---|---|
| Gtdc1 | 0.03860422 | 0.03336437 | 0.03406063 | -0.0006963 | 0.5541158 | 0.99999994 |
| 4930426D05I | 0.00303149 | 0.00204279 | 0.0027393 | -0.0006965 | 0.95472346 | 0.99999994 |
| Sh3tc1 | 0.00347809 | 0.00244537 | 0.00314189 | -0.0006965 | 0.92998005 | 0.99999994 |
| Ltbp4 | 0.00519648 | 0.00399187 | 0.00468872 | -0.0006969 | 0.83873857 | 0.99999994 |
| AW112010 | 0.00186501 | 0.00098942 | 0.00168658 | -0.0006972 | 0.99703368 | 0.99999994 |
| Pbx4 | 0.02739267 | 0.02365219 | 0.02434964 | -0.0006974 | 0.57560283 | 0.99999994 |
| 4930556G22 | 0.00228581 | 0.00136891 | 0.00206654 | -0.0006976 | 0.98768192 | 0.99999994 |
| Wdr12 | 0.18746617 | 0.14764008 | 0.1483379 | -0.0006978 | 0.51249013 | 0.99999994 |
| Amph | 0.00817859 | 0.0066665 | 0.00736455 | -0.0006981 | 0.73595566 | 0.99999994 |
| Ttc23 | 0.01705664 | 0.01456917 | 0.01526724 | -0.0006981 | 0.61956846 | 0.99999994 |
| Slc16a11 | 0.01662047 | 0.01418302 | 0.01488119 | -0.0006982 | 0.6225896 | 0.99999994 |
| Gm42595 | 0.00551679 | 0.00427829 | 0.00497665 | -0.0006984 | 0.824866 | 0.99999994 |
| 1700066M21 | 0.01422573 | 0.01205856 | 0.01275761 | -0.000699 | 0.64234323 | 0.99999994 |
| Cd34 | 0.00256992 | 0.00162342 | 0.00232295 | -0.0006995 | 0.97749089 | 0.99999994 |
| Slc25a40 | 0.01480018 | 0.0125672 | 0.01326763 | -0.0007004 | 0.63735748 | 0.99999994 |
| Tpcn2 | 0.01022044 | 0.00848977 | 0.00919047 | -0.0007007 | 0.69409561 | 0.99999994 |
| Hbegf | 0.02944556 | 0.02543783 | 0.02613862 | -0.0007008 | 0.57081956 | 0.99999994 |
| Dnah8 | 0.00494461 | 0.00376071 | 0.00446222 | -0.0007015 | 0.85232506 | 0.99999994 |
| Atxn3 | 0.04098079 | 0.03539874 | 0.03610061 | -0.0007019 | 0.55148411 | 0.99999994 |
| Zfp619 | 0.00974088 | 0.00806016 | 0.00876208 | -0.0007019 | 0.70306395 | 0.99999994 |
| Snx29 | 0.03813065 | 0.03295135 | 0.03365336 | -0.000702 | 0.55521684 | 0.99999994 |
| Sfxn5 | 0.01363844 | 0.01153331 | 0.01223576 | -0.0007024 | 0.64882023 | 0.99999994 |
| Kcnh1 | 0.00132851 | 0.00049928 | 0.00120185 | -0.0007026 | 0.99995011 | 0.99999994 |
| Gm14419 | 0.00985676 | 0.00816208 | 0.00886562 | -0.0007035 | 0.70132712 | 0.99999994 |
| Tnfsf10 | 0.00193315 | 0.0010434 | 0.00174813 | -0.0007047 | 0.99635614 | 0.99999994 |
| Slc35f1 | 0.00432208 | 0.00319714 | 0.00390208 | -0.0007049 | 0.88541753 | 0.99999994 |
| Sfmbt2 | 0.0098077 | 0.00811625 | 0.00882178 | -0.0007055 | 0.70275835 | 0.99999994 |
| Layn | 0.00788405 | 0.00639511 | 0.00710074 | -0.0007056 | 0.74584463 | 0.99999994 |
| Plcl2 | 0.02478261 | 0.02136225 | 0.022068 | -0.0007057 | 0.58428755 | 0.99999994 |
| Patz1 | 0.11887244 | 0.09856483 | 0.09927125 | -0.0007064 | 0.51888982 | 0.99999994 |

| | | | | | | |
|---|---|---|---|---|---|---|
| Klf4 | 0.05781982 | 0.04966551 | 0.05037289 | -0.0007074 | 0.53723645 | 0.99999994 |
| Lgals7 | 0.00593106 | 0.0046403 | 0.00534886 | -0.0007086 | 0.81104883 | 0.99999994 |
| Rd3 | 0.00634354 | 0.00501054 | 0.00571925 | -0.0007087 | 0.79526756 | 0.99999994 |
| Gm19710 | 0.00477604 | 0.00360185 | 0.00431059 | -0.0007087 | 0.86311707 | 0.99999994 |
| Ntrk1 | 0.00807154 | 0.00655993 | 0.00726869 | -0.0007088 | 0.7418471 | 0.99999994 |
| Wdr31 | 0.04787189 | 0.04127081 | 0.04197959 | -0.0007088 | 0.54474122 | 0.99999994 |
| Napsa | 0.00299162 | 0.00199455 | 0.00270335 | -0.0007088 | 0.95952784 | 0.99999994 |
| B4galt7 | 0.09916389 | 0.0834066 | 0.0841154 | -0.0007088 | 0.52236489 | 0.99999994 |
| Neil3 | 0.00249543 | 0.00154658 | 0.00225572 | -0.0007091 | 0.9818072 | 0.99999994 |
| Rmi2 | 0.00588195 | 0.00459498 | 0.00530475 | -0.0007098 | 0.81343058 | 0.99999994 |
| Fut10 | 0.00447228 | 0.00332747 | 0.00403727 | -0.0007098 | 0.87905295 | 0.99999994 |
| Fastkd5 | 0.02788346 | 0.02406632 | 0.02477778 | -0.0007115 | 0.57578775 | 0.99999994 |
| Klhl26 | 0.0553156 | 0.04755811 | 0.04827033 | -0.0007122 | 0.53911896 | 0.99999994 |
| Haus5 | 0.03237674 | 0.02797217 | 0.02868454 | -0.0007124 | 0.56564882 | 0.99999994 |
| Ric1 | 0.04468821 | 0.03855778 | 0.03927016 | -0.0007124 | 0.54805532 | 0.99999994 |
| Cndp1 | 0.01265092 | 0.01064454 | 0.01135737 | -0.0007128 | 0.66194497 | 0.99999994 |
| Asprv1 | 0.00655193 | 0.00519335 | 0.0059063 | -0.0007129 | 0.78915334 | 0.99999994 |
| Dact1 | 0.02604518 | 0.02245956 | 0.02317269 | -0.0007131 | 0.58115464 | 0.99999994 |
| 4632428C04l | 0.00131076 | 0.00047208 | 0.0011858 | -0.0007137 | 0.99996917 | 0.99999994 |
| Plppr4 | 0.00320324 | 0.00218041 | 0.00289416 | -0.0007138 | 0.94966124 | 0.99999994 |
| Zfp169 | 0.00847725 | 0.00691774 | 0.00763194 | -0.0007142 | 0.73332863 | 0.99999994 |
| Gcnt4 | 0.01225962 | 0.01029443 | 0.01100898 | -0.0007146 | 0.66714428 | 0.99999994 |
| Zfp420 | 0.02079677 | 0.01785302 | 0.01856827 | -0.0007153 | 0.60118044 | 0.99999994 |
| Serpini2 | 0.0106505 | 0.00885879 | 0.00957441 | -0.0007156 | 0.69058789 | 0.99999994 |
| Gng7 | 0.00817222 | 0.00664139 | 0.00735885 | -0.0007175 | 0.74182106 | 0.99999994 |
| Plekhn1 | 0.01055468 | 0.00877093 | 0.00948889 | -0.000718 | 0.69274366 | 0.99999994 |
| Tekt5 | 0.00525016 | 0.00401827 | 0.00473698 | -0.0007187 | 0.843734 | 0.99999994 |
| Fam221b | 0.00834989 | 0.00679703 | 0.00751793 | -0.0007209 | 0.73835523 | 0.99999994 |
| Brca2 | 0.03614266 | 0.03121987 | 0.03194091 | -0.000721 | 0.55972105 | 0.99999994 |
| 4930470P17l | 0.00150019 | 0.00063517 | 0.001357 | -0.0007218 | 0.99980047 | 0.99999994 |

| | | | | | | |
|---|---|---|---|---|---|---|
| Fbxo45 | 0.03655455 | 0.03157408 | 0.03229609 | -0.000722 | 0.55914783 | 0.99999994 |
| Gm15634 | 0.00177124 | 0.00087953 | 0.00160189 | -0.0007224 | 0.99865735 | 0.99999994 |
| Gm27168 | 0.00319849 | 0.00216715 | 0.00288987 | -0.0007227 | 0.95201135 | 0.99999994 |
| Dlgap1 | 0.00627174 | 0.00493131 | 0.0056548 | -0.0007235 | 0.80278932 | 0.99999994 |
| Syne1 | 0.05842083 | 0.05015088 | 0.05087647 | -0.0007256 | 0.53781528 | 0.99999994 |
| Agxt2 | 0.00124194 | 0.000398 | 0.0011236 | -0.0007256 | 0.9999914 | 0.99999994 |
| Ptprm | 0.00449369 | 0.00333079 | 0.00405653 | -0.0007257 | 0.88314748 | 0.99999994 |
| Ube2w | 0.58468862 | 0.34554374 | 0.3462695 | -0.0007258 | 0.50556557 | 0.99999994 |
| Cep95 | 0.04715971 | 0.04064845 | 0.04137449 | -0.000726 | 0.54649268 | 0.99999994 |
| Crb1 | 0.00246934 | 0.00150572 | 0.00223218 | -0.0007265 | 0.98485626 | 0.99999994 |
| Heatr1 | 0.01444661 | 0.01222727 | 0.01295376 | -0.0007265 | 0.64554178 | 0.99999994 |
| 1700025G04 | 0.03299484 | 0.0284934 | 0.02922012 | -0.0007267 | 0.56574339 | 0.99999994 |
| Wrn | 0.04066595 | 0.03510334 | 0.03583072 | -0.0007274 | 0.55374474 | 0.99999994 |
| Sla | 0.00171692 | 0.00082529 | 0.00155282 | -0.0007275 | 0.99909164 | 0.99999994 |
| Gm42669 | 0.00139 | 0.00052956 | 0.00125743 | -0.0007279 | 0.99994167 | 0.99999994 |
| Slc35b1 | 0.74882117 | 0.3926137 | 0.39334159 | -0.0007279 | 0.50491397 | 0.99999994 |
| Rnf157 | 0.00901218 | 0.00738201 | 0.0081106 | -0.0007286 | 0.72506267 | 0.99999994 |
| Snapc1 | 0.07089308 | 0.06050953 | 0.06123827 | -0.0007287 | 0.53156796 | 0.99999994 |
| Gm7628 | 0.01005252 | 0.00831152 | 0.0090405 | -0.000729 | 0.70427488 | 0.99999994 |
| Cbx7 | 0.02422474 | 0.02084946 | 0.02157929 | -0.0007298 | 0.5890582 | 0.99999994 |
| Col28a1 | 0.00221609 | 0.00127362 | 0.0020036 | -0.00073 | 0.99234868 | 0.99999994 |
| Cnbd2 | 0.00833246 | 0.00677078 | 0.00750233 | -0.0007315 | 0.74184886 | 0.99999994 |
| Eya3 | 0.05873927 | 0.05041056 | 0.05114313 | -0.0007326 | 0.53797953 | 0.99999994 |
| Zfp940 | 0.01566849 | 0.01330503 | 0.01403782 | -0.0007328 | 0.63588057 | 0.99999994 |
| 4933427D14 | 0.04093233 | 0.03532586 | 0.03605907 | -0.0007332 | 0.55383206 | 0.99999994 |
| Gm26810 | 0.00382767 | 0.00272294 | 0.00345687 | -0.0007339 | 0.92120223 | 0.99999994 |
| Gm13544 | 0.00430418 | 0.00315169 | 0.00388596 | -0.0007343 | 0.89575687 | 0.99999994 |
| Pcdhga9 | 0.00406481 | 0.00293568 | 0.00367045 | -0.0007348 | 0.90865176 | 0.99999994 |
| Incenp | 0.02107401 | 0.01807745 | 0.01881231 | -0.0007349 | 0.60257356 | 0.99999994 |
| Snx9 | 0.07425756 | 0.06326942 | 0.06400477 | -0.0007354 | 0.53047949 | 0.99999994 |

| | | | | | | |
|---|---|---|---|---|---|---|
| Rab42 | 0.00518299 | 0.00394107 | 0.00467659 | -0.0007355 | 0.85242424 | 0.99999994 |
| Itgb8 | 0.00566203 | 0.00437158 | 0.00510716 | -0.0007356 | 0.8311487 | 0.99999994 |
| Rab11fip4 | 0.026993 | 0.02326504 | 0.02400077 | -0.0007357 | 0.58084091 | 0.99999994 |
| Dynlt1b | 0.00714119 | 0.00569871 | 0.00643493 | -0.0007362 | 0.77683675 | 0.99999994 |
| Gm26812 | 0.00129179 | 0.00043106 | 0.00116866 | -0.0007376 | 0.99998672 | 0.99999994 |
| Kcnh3 | 0.01045907 | 0.00866587 | 0.00940353 | -0.0007377 | 0.69921984 | 0.99999994 |
| Kxd1 | 0.25579934 | 0.19133172 | 0.19206947 | -0.0007378 | 0.51019889 | 0.99999994 |
| Zfp712 | 0.00539716 | 0.00413071 | 0.00486913 | -0.0007384 | 0.84362212 | 0.99999994 |
| Uox | 0.00512921 | 0.00388903 | 0.00462823 | -0.0007392 | 0.85613645 | 0.99999994 |
| Arhgef6 | 0.00364625 | 0.00255396 | 0.00329343 | -0.0007395 | 0.93248364 | 0.99999994 |
| Fcgrt | 0.01332032 | 0.01121291 | 0.01195293 | -0.00074 | 0.65986693 | 0.99999994 |
| Casp2 | 0.08752964 | 0.07406136 | 0.07480138 | -0.00074 | 0.52625219 | 0.99999994 |
| S100b | 0.01042041 | 0.0086283 | 0.00936902 | -0.0007407 | 0.70064408 | 0.99999994 |
| Rapgefl1 | 0.00766391 | 0.00616222 | 0.00690351 | -0.0007413 | 0.76261806 | 0.99999994 |
| Eefsec | 0.12335227 | 0.10190442 | 0.10264594 | -0.0007415 | 0.51917622 | 0.99999994 |
| Fah | 0.04350068 | 0.0375145 | 0.0382566 | -0.0007421 | 0.55136824 | 0.99999994 |
| Ipcef1 | 0.0050649 | 0.00382816 | 0.00457041 | -0.0007422 | 0.86015363 | 0.99999994 |
| Cd59b | 0.01511849 | 0.01280744 | 0.01355008 | -0.0007426 | 0.64237432 | 0.99999994 |
| Zkscan7 | 0.0049813 | 0.00375257 | 0.00449522 | -0.0007427 | 0.8642685 | 0.99999994 |
| Mkln1os | 0.01715666 | 0.01461192 | 0.01535573 | -0.0007438 | 0.626435 | 0.99999994 |
| Hyal2 | 0.16771272 | 0.13402041 | 0.13476519 | -0.0007448 | 0.51467223 | 0.99999994 |
| Rnd1 | 0.00137057 | 0.00049509 | 0.00123987 | -0.0007448 | 0.99996811 | 0.99999994 |
| Nrg2 | 0.00442087 | 0.00324618 | 0.00399099 | -0.0007448 | 0.89292579 | 0.99999994 |
| Plec | 0.0545707 | 0.04689843 | 0.04764358 | -0.0007451 | 0.5414573 | 0.99999994 |
| Gm26542 | 0.00256156 | 0.0015696 | 0.0023154 | -0.0007458 | 0.98398511 | 0.99999994 |
| Paqr9 | 0.01133402 | 0.00943785 | 0.01018417 | -0.0007463 | 0.68715256 | 0.99999994 |
| P2rx1 | 0.0067892 | 0.0053722 | 0.0061192 | -0.000747 | 0.79176793 | 0.99999994 |
| Ccdc88b | 0.00227209 | 0.00130604 | 0.00205415 | -0.0007481 | 0.99232928 | 0.99999994 |
| Foxj1 | 0.00760115 | 0.00609871 | 0.00684726 | -0.0007485 | 0.76659342 | 0.99999994 |
| 1700008O03 | 0.0167936 | 0.01428585 | 0.01503444 | -0.0007486 | 0.62984141 | 0.99999994 |

| | | | | | | |
|---|---|---|---|---|---|---|
| Gm16308 | 0.00158446 | 0.00068378 | 0.00143315 | -0.0007494 | 0.99974975 | 0.99999994 |
| Gm42809 | 0.0049119 | 0.00368297 | 0.0044328 | -0.0007498 | 0.86990967 | 0.99999994 |
| Rap2a | 0.03804782 | 0.032832 | 0.0335821 | -0.0007501 | 0.55909628 | 0.99999994 |
| Atp8b5 | 0.00516855 | 0.0039128 | 0.0046636 | -0.0007508 | 0.85805657 | 0.99999994 |
| Gm26620 | 0.00833138 | 0.00674948 | 0.00750135 | -0.0007519 | 0.74767176 | 0.99999994 |
| Lad1 | 0.0092304 | 0.00755363 | 0.00830576 | -0.0007521 | 0.72666994 | 0.99999994 |
| 9330159F19I | 0.00407686 | 0.00292814 | 0.00368129 | -0.0007532 | 0.91337341 | 0.99999994 |
| Arhgef25 | 0.01239958 | 0.01038034 | 0.01113361 | -0.0007533 | 0.67377299 | 0.99999994 |
| Manea | 0.02831607 | 0.02440143 | 0.02515494 | -0.0007535 | 0.5790204 | 0.99999994 |
| Pbk | 0.0089441 | 0.00729559 | 0.0080497 | -0.0007541 | 0.73354895 | 0.99999994 |
| A930024E05 | 0.00422817 | 0.00306333 | 0.00381753 | -0.0007542 | 0.90575376 | 0.99999994 |
| Dzank1 | 0.00955219 | 0.00783924 | 0.00859345 | -0.0007542 | 0.7204575 | 0.99999994 |
| Pcdhga5 | 0.01119385 | 0.00930491 | 0.01005918 | -0.0007543 | 0.691149 | 0.99999994 |
| 1700084E18I | 0.01313322 | 0.01103189 | 0.01178652 | -0.0007546 | 0.66500893 | 0.99999994 |
| Irgm2 | 0.00373641 | 0.00261841 | 0.00337466 | -0.0007563 | 0.93210796 | 0.99999994 |
| Crtc2 | 0.05925257 | 0.05081637 | 0.05157273 | -0.0007564 | 0.53888398 | 0.99999994 |
| Gm15265 | 0.00298788 | 0.00194225 | 0.00269997 | -0.0007577 | 0.96912309 | 0.99999994 |
| Zfp641 | 0.00336795 | 0.00228442 | 0.00304263 | -0.0007582 | 0.95141509 | 0.99999994 |
| Itga2b | 0.00881611 | 0.0071762 | 0.00793519 | -0.000759 | 0.737833 | 0.99999994 |
| Gnmt | 0.00467809 | 0.00346282 | 0.00422247 | -0.0007597 | 0.8844488 | 0.99999994 |
| Gm6999 | 0.00250391 | 0.00150362 | 0.00226338 | -0.0007598 | 0.98727129 | 0.99999994 |
| Cftr | 0.00210693 | 0.00114513 | 0.00190505 | -0.0007599 | 0.99603677 | 0.99999994 |
| Slc8a1 | 0.01181681 | 0.00985317 | 0.01061452 | -0.0007613 | 0.68347606 | 0.99999994 |
| Abtb2 | 0.0129623 | 0.01087252 | 0.01163446 | -0.0007619 | 0.66855462 | 0.99999994 |
| Zfp354a | 0.01351151 | 0.01135976 | 0.01212293 | -0.0007632 | 0.6624055 | 0.99999994 |
| Zmat4 | 0.00221647 | 0.00124077 | 0.00200394 | -0.0007632 | 0.99437743 | 0.99999994 |
| Mpped2 | 0.16128531 | 0.12949038 | 0.13025435 | -0.000764 | 0.51557113 | 0.99999994 |
| Arl14ep | 0.0716379 | 0.06108645 | 0.06185175 | -0.0007653 | 0.53281983 | 0.99999994 |
| Eya1 | 0.01548991 | 0.01311276 | 0.01387949 | -0.0007667 | 0.64345583 | 0.99999994 |
| 4933406C10I | 0.00519251 | 0.00391638 | 0.00468515 | -0.0007688 | 0.86263046 | 0.99999994 |

| | | | | | | |
|---|---|---|---|---|---|---|
| Azin2 | 0.04134937 | 0.03564719 | 0.0364164 | -0.0007692 | 0.55590745 | 0.99999994 |
| Elf4 | 0.01627798 | 0.01380856 | 0.0145779 | -0.0007693 | 0.63731327 | 0.99999994 |
| Ms4a6c | 0.00544132 | 0.0041387 | 0.00490882 | -0.0007701 | 0.85182394 | 0.99999994 |
| Egfl8 | 0.00748293 | 0.00597115 | 0.00674131 | -0.0007702 | 0.77650898 | 0.99999994 |
| Mex3d | 0.04282164 | 0.03690574 | 0.03767633 | -0.0007706 | 0.55414541 | 0.99999994 |
| Hlcs | 0.01451945 | 0.01224722 | 0.01301844 | -0.0007712 | 0.65332923 | 0.99999994 |
| Tmem151a | 0.01127 | 0.00935555 | 0.01012708 | -0.0007715 | 0.69396372 | 0.99999994 |
| Cd7 | 0.00274212 | 0.00170608 | 0.0024783 | -0.0007722 | 0.98096436 | 0.99999994 |
| Ikzf3 | 0.00196969 | 0.00100866 | 0.00178112 | -0.0007725 | 0.99805415 | 0.99999994 |
| Atg4c | 0.01615738 | 0.01369837 | 0.01447107 | -0.0007727 | 0.63886352 | 0.99999994 |
| A230065N10 | 0.00177533 | 0.00083251 | 0.00160558 | -0.0007731 | 0.99932459 | 0.99999994 |
| Gpr62 | 0.00479722 | 0.00355517 | 0.00432965 | -0.0007745 | 0.88311017 | 0.99999994 |
| Setd4 | 0.01754916 | 0.01492762 | 0.01570288 | -0.0007753 | 0.62878119 | 0.99999994 |
| Pik3cd | 0.00280596 | 0.00175998 | 0.00253589 | -0.0007759 | 0.97915738 | 0.99999994 |
| Col27a1 | 0.0383643 | 0.0330746 | 0.03385433 | -0.0007797 | 0.5609223 | 0.99999994 |
| Lgr4 | 0.03520804 | 0.0303531 | 0.03113427 | -0.0007812 | 0.56631888 | 0.99999994 |
| Ism1 | 0.00379874 | 0.00264936 | 0.0034308 | -0.0007814 | 0.93525856 | 0.99999994 |
| Pdgfd | 0.00490612 | 0.00364535 | 0.00442761 | -0.0007823 | 0.88020869 | 0.99999994 |
| Cdk15 | 0.00392164 | 0.00275799 | 0.00354151 | -0.0007835 | 0.92957628 | 0.99999994 |
| Ldlrap1 | 0.03613978 | 0.0311532 | 0.03193843 | -0.0007852 | 0.56499674 | 0.99999994 |
| Ankrd34b | 0.00398272 | 0.00281027 | 0.00359652 | -0.0007862 | 0.92718805 | 0.99999994 |
| Stom | 0.00522416 | 0.00392632 | 0.00471361 | -0.0007873 | 0.86688181 | 0.99999994 |
| Cyp4f16 | 0.09124574 | 0.07700406 | 0.07779147 | -0.0007874 | 0.52685881 | 0.99999994 |
| Sytl3 | 0.00339544 | 0.0022797 | 0.00306741 | -0.0007877 | 0.95630654 | 0.99999994 |
| B230317F23 | 0.01273964 | 0.0106486 | 0.01143633 | -0.0007877 | 0.67669869 | 0.99999994 |
| Cdcp1 | 0.04601235 | 0.03961003 | 0.04039843 | -0.0007884 | 0.5516783 | 0.99999994 |
| Stoml1 | 0.06284281 | 0.05377974 | 0.0545695 | -0.0007898 | 0.53837237 | 0.99999994 |
| Spic | 0.00261228 | 0.00157138 | 0.00236117 | -0.0007898 | 0.98700908 | 0.99999994 |
| Gm4724 | 0.00298497 | 0.00190745 | 0.00269735 | -0.0007899 | 0.97436752 | 0.99999994 |
| Rnf169 | 0.02841573 | 0.02445126 | 0.02524179 | -0.0007905 | 0.58256798 | 0.99999994 |

| | | | | | | |
|---|---|---|---|---|---|---|
| 1810007C17I | 0.00171696 | 0.00076191 | 0.00155286 | -0.000791 | 0.99965108 | 0.99999994 |
| Gm16104 | 0.00300588 | 0.00192477 | 0.0027162 | -0.0007914 | 0.97377883 | 0.99999994 |
| Eogt | 0.01168145 | 0.00970177 | 0.01049389 | -0.0007921 | 0.69232353 | 0.99999994 |
| Cenpn | 0.01029604 | 0.00846536 | 0.00925799 | -0.0007926 | 0.71562157 | 0.99999994 |
| Sik1 | 0.01889034 | 0.01609443 | 0.01688773 | -0.0007933 | 0.62273925 | 0.99999994 |
| Abcb6 | 0.0567192 | 0.04865602 | 0.04944965 | -0.0007936 | 0.54253794 | 0.99999994 |
| Top1mt | 0.01799862 | 0.01530592 | 0.01610018 | -0.0007943 | 0.62868649 | 0.99999994 |
| Actg2 | 0.00176804 | 0.00080204 | 0.001599 | -0.000797 | 0.99954601 | 0.99999994 |
| Cabp2 | 0.00322282 | 0.00211485 | 0.00291181 | -0.000797 | 0.96576012 | 0.99999994 |
| Xxylt1 | 0.00591441 | 0.00453668 | 0.0053339 | -0.0007972 | 0.84010359 | 0.99999994 |
| Klhdc9 | 0.03806478 | 0.03279939 | 0.03359669 | -0.0007973 | 0.5627586 | 0.99999994 |
| Syngr3 | 0.00317253 | 0.00206858 | 0.00286647 | -0.0007979 | 0.96804576 | 0.99999994 |
| Rnaseh2a | 0.20387109 | 0.15848471 | 0.15928329 | -0.0007986 | 0.5133111 | 0.99999994 |
| Atcayos | 0.00407178 | 0.00287796 | 0.00367672 | -0.0007988 | 0.92592302 | 0.99999994 |
| Myo5c | 0.00425422 | 0.00304218 | 0.00384099 | -0.0007988 | 0.91686907 | 0.99999994 |
| Car6 | 0.00313553 | 0.00203404 | 0.00283311 | -0.0007991 | 0.96976864 | 0.99999994 |
| Ace2 | 0.00692351 | 0.00544052 | 0.0062397 | -0.0007992 | 0.8030442 | 0.99999994 |
| Poglut1 | 0.12444831 | 0.10266851 | 0.10346787 | -0.0007994 | 0.52050665 | 0.99999994 |
| Zfp759 | 0.01343128 | 0.0112522 | 0.01205159 | -0.0007994 | 0.67058153 | 0.99999994 |
| Ahnak | 0.00667655 | 0.00521833 | 0.00601813 | -0.0007998 | 0.81182012 | 0.99999994 |
| Tnfaip3 | 0.00574095 | 0.0043779 | 0.00517808 | -0.0008002 | 0.84816957 | 0.99999994 |
| Rbm3os | 0.00599128 | 0.00460242 | 0.00540295 | -0.0008005 | 0.83799226 | 0.99999994 |
| Chst14 | 0.01290107 | 0.01077889 | 0.01157999 | -0.0008011 | 0.67741596 | 0.99999994 |
| Cabp1 | 0.00731613 | 0.00579058 | 0.00659178 | -0.0008012 | 0.79075639 | 0.99999994 |
| Grid1 | 0.00296532 | 0.00187748 | 0.00267962 | -0.0008021 | 0.97684665 | 0.99999994 |
| Wdr83 | 0.22976013 | 0.17517161 | 0.17597399 | -0.0008024 | 0.51210629 | 0.99999994 |
| Zfp933 | 0.01274344 | 0.01063705 | 0.01143971 | -0.0008027 | 0.67976496 | 0.99999994 |
| Epdr1 | 0.03892985 | 0.03353747 | 0.03434052 | -0.0008031 | 0.56184915 | 0.99999994 |
| Gm10501 | 0.00437888 | 0.0031499 | 0.0039532 | -0.0008033 | 0.91190652 | 0.99999994 |
| 4930518I15R | 0.00808048 | 0.00647297 | 0.00727669 | -0.0008037 | 0.76889417 | 0.99999994 |

| | | | | | | |
|---|---|---|---|---|---|---|
| Ap1ar | 0.15375363 | 0.12410351 | 0.12490736 | -0.0008039 | 0.51708455 | 0.99999994 |
| Slpi | 0.00471367 | 0.00345042 | 0.00425448 | -0.0008041 | 0.8958025 | 0.99999994 |
| Gpc6 | 0.01044031 | 0.00858265 | 0.00938679 | -0.0008041 | 0.71573939 | 0.99999994 |
| Gspt2 | 0.06851979 | 0.05847497 | 0.05927951 | -0.0008045 | 0.53599176 | 0.99999994 |
| Magi2 | 0.01701133 | 0.01442186 | 0.01522714 | -0.0008053 | 0.63758869 | 0.99999994 |
| Lacc1 | 0.00610974 | 0.00470351 | 0.00550933 | -0.0008058 | 0.83486944 | 0.99999994 |
| Rtel1 | 0.01648552 | 0.01395546 | 0.0147617 | -0.0008062 | 0.64190272 | 0.99999994 |
| Skida1 | 0.00919128 | 0.00746443 | 0.00827078 | -0.0008063 | 0.74181517 | 0.99999994 |
| Negr1 | 0.00225163 | 0.00122928 | 0.00203569 | -0.0008064 | 0.99581563 | 0.99999994 |
| Slc26a11 | 0.00426062 | 0.00304014 | 0.00384674 | -0.0008066 | 0.91860397 | 0.99999994 |
| Gm26802 | 0.00409232 | 0.00288823 | 0.00369522 | -0.000807 | 0.92698138 | 0.99999994 |
| Atg7 | 0.05305861 | 0.04556229 | 0.04636942 | -0.0008071 | 0.54612015 | 0.99999994 |
| Spata1 | 0.02462283 | 0.02112075 | 0.02192806 | -0.0008073 | 0.59679658 | 0.99999994 |
| Lrrc61 | 0.04728771 | 0.04067571 | 0.04148328 | -0.0008076 | 0.55155135 | 0.99999994 |
| Hipk2 | 0.18906051 | 0.14860666 | 0.14941457 | -0.0008079 | 0.51435578 | 0.99999994 |
| Tank | 0.09212661 | 0.07768756 | 0.07849816 | -0.0008106 | 0.5273999 | 0.99999994 |
| St14 | 0.07885122 | 0.0669511 | 0.06776258 | -0.0008115 | 0.53176702 | 0.99999994 |
| Pde7b | 0.0021837 | 0.00116263 | 0.00197435 | -0.0008117 | 0.99689675 | 0.99999994 |
| Pcdhb8 | 0.00557556 | 0.00421732 | 0.00502947 | -0.0008122 | 0.85878165 | 0.99999994 |
| Tmem71 | 0.00162651 | 0.0006576 | 0.00147114 | -0.0008135 | 0.99988381 | 0.99999994 |
| Kiss1r | 0.00389279 | 0.00270168 | 0.00351552 | -0.0008138 | 0.93833527 | 0.99999994 |
| Ccl2 | 0.00178877 | 0.00080311 | 0.00161772 | -0.0008146 | 0.99959865 | 0.99999994 |
| 2810405F17l | 0.01229606 | 0.01022679 | 0.01104143 | -0.0008146 | 0.68832568 | 0.99999994 |
| Itga9 | 0.00884239 | 0.00714395 | 0.00795871 | -0.0008148 | 0.75220044 | 0.99999994 |
| Fbxl12 | 0.12277123 | 0.10139469 | 0.10220963 | -0.0008149 | 0.52116299 | 0.99999994 |
| Nxph3 | 0.00360158 | 0.00243791 | 0.00325318 | -0.0008153 | 0.95235466 | 0.99999994 |
| Zfp956 | 0.01534509 | 0.0129357 | 0.01375107 | -0.0008154 | 0.6534628 | 0.99999994 |
| Gss | 0.14623937 | 0.1186922 | 0.11950761 | -0.0008154 | 0.51811247 | 0.99999994 |
| Sord | 0.01269048 | 0.01057676 | 0.01139258 | -0.0008158 | 0.68319877 | 0.99999994 |
| Cela1 | 0.09893648 | 0.08311873 | 0.08393466 | -0.0008159 | 0.52579634 | 0.99999994 |

| | | | | | | |
|---|---|---|---|---|---|---|
| Uba52 | 0.35762398 | 0.24805683 | 0.24887303 | -0.0008162 | 0.50870822 | 0.99999994 |
| Gramd4 | 0.00969655 | 0.00790617 | 0.00872247 | -0.0008163 | 0.73333674 | 0.99999994 |
| Rad50 | 0.06781663 | 0.05788157 | 0.05869801 | -0.0008164 | 0.53688354 | 0.99999994 |
| Lag3 | 0.01688232 | 0.01429482 | 0.01511297 | -0.0008181 | 0.64070569 | 0.99999994 |
| Prpf40a | 0.77982969 | 0.39886765 | 0.39968849 | -0.0008208 | 0.50545343 | 0.99999994 |
| Cnp | 0.16150399 | 0.12958763 | 0.1304086 | -0.000821 | 0.51671239 | 0.99999994 |
| Unc5c | 0.01595887 | 0.0134734 | 0.01429518 | -0.0008218 | 0.64901163 | 0.99999994 |
| Dtx4 | 0.00521031 | 0.00387661 | 0.00470116 | -0.0008245 | 0.87848969 | 0.99999994 |
| Wbscr25 | 0.00674508 | 0.00525466 | 0.00607962 | -0.000825 | 0.81679475 | 0.99999994 |
| Nav3 | 0.00437416 | 0.00312311 | 0.00394896 | -0.0008258 | 0.91804502 | 0.99999994 |
| Ttll13 | 0.00908724 | 0.00735155 | 0.00817774 | -0.0008262 | 0.74936194 | 0.99999994 |
| Simc1 | 0.0318265 | 0.02738106 | 0.02820738 | -0.0008263 | 0.57730078 | 0.99999994 |
| 6330418K02I | 0.06586307 | 0.05625277 | 0.05707965 | -0.0008269 | 0.53840931 | 0.99999994 |
| Zkscan17 | 0.03055924 | 0.02628017 | 0.0271071 | -0.0008269 | 0.58045495 | 0.99999994 |
| Fmo5 | 0.00858077 | 0.00689767 | 0.0077246 | -0.0008269 | 0.76194516 | 0.99999994 |
| Klhl10 | 0.00380518 | 0.00260967 | 0.00343661 | -0.0008269 | 0.94538864 | 0.99999994 |
| Mrps2 | 0.10869395 | 0.09068364 | 0.09151095 | -0.0008273 | 0.52399259 | 0.99999994 |
| Stard8 | 0.00398885 | 0.00277436 | 0.00360204 | -0.0008277 | 0.93693054 | 0.99999994 |
| Fzr1 | 0.28440737 | 0.20815856 | 0.20898682 | -0.0008283 | 0.51052313 | 0.99999994 |
| Nup37 | 0.10609719 | 0.08868207 | 0.08951036 | -0.0008283 | 0.52455746 | 0.99999994 |
| Klhl18 | 0.02037335 | 0.01736523 | 0.0181954 | -0.0008302 | 0.61932063 | 0.99999994 |
| Tmem132a | 0.14329076 | 0.11654008 | 0.11737032 | -0.0008302 | 0.5187771 | 0.99999994 |
| Thumpd2 | 0.02809397 | 0.02413046 | 0.02496133 | -0.0008309 | 0.58767441 | 0.99999994 |
| 1110035H17 | 0.02074807 | 0.01769379 | 0.0185254 | -0.0008316 | 0.61745545 | 0.99999994 |
| Cdca3 | 0.01643918 | 0.01388904 | 0.01472066 | -0.0008316 | 0.6465573 | 0.99999994 |
| Pik3r5 | 0.00333949 | 0.00218514 | 0.00301698 | -0.0008318 | 0.96676871 | 0.99999994 |
| Fam168b | 0.09130401 | 0.07700609 | 0.07783824 | -0.0008322 | 0.52836556 | 0.99999994 |
| Urb2 | 0.00914072 | 0.00739305 | 0.00822556 | -0.0008325 | 0.74974732 | 0.99999994 |
| S100g | 0.00262329 | 0.00153851 | 0.0023711 | -0.0008326 | 0.9902886 | 0.99999994 |
| Lrp1 | 0.01101294 | 0.00906514 | 0.00989781 | -0.0008327 | 0.7122515 | 0.99999994 |

| | | | | | | |
|---|---|---|---|---|---|---|
| Map3k8 | 0.00743993 | 0.00587009 | 0.00670277 | -0.0008327 | 0.79585769 | 0.99999994 |
| Anapc4 | 0.26380412 | 0.19604799 | 0.19688221 | -0.0008342 | 0.51125017 | 0.99999994 |
| Vwa1 | 0.02470955 | 0.02116919 | 0.02200401 | -0.0008348 | 0.59968778 | 0.99999994 |
| Asb11 | 0.00286166 | 0.00175059 | 0.00258614 | -0.0008355 | 0.98424488 | 0.99999994 |
| Snx32 | 0.02245129 | 0.01918741 | 0.02002328 | -0.0008359 | 0.60944284 | 0.99999994 |
| Slc5a10 | 0.00155515 | 0.00056993 | 0.00140667 | -0.0008367 | 0.99996244 | 0.99999994 |
| Pnoc | 0.01202238 | 0.00995964 | 0.01079767 | -0.000838 | 0.69728747 | 0.99999994 |
| Efcab11 | 0.00397918 | 0.00275517 | 0.00359333 | -0.0008382 | 0.93974331 | 0.99999994 |
| Sfrp5 | 0.0054896 | 0.00411395 | 0.00495222 | -0.0008383 | 0.87007255 | 0.99999994 |
| Fabp4 | 0.00385999 | 0.00264767 | 0.00348598 | -0.0008383 | 0.94528112 | 0.99999994 |
| Ppp1r14d | 0.0135392 | 0.01130893 | 0.01214755 | -0.0008386 | 0.67707185 | 0.99999994 |
| Mecom | 0.01456182 | 0.01221656 | 0.01305605 | -0.0008395 | 0.66567288 | 0.99999994 |
| Col6a1 | 0.00234891 | 0.00128395 | 0.00212349 | -0.0008395 | 0.99575154 | 0.99999994 |
| Pfkfb3 | 0.00582743 | 0.00441622 | 0.00525577 | -0.0008395 | 0.85617164 | 0.99999994 |
| Ncmap | 0.00637657 | 0.00490893 | 0.0057489 | -0.00084 | 0.83461271 | 0.99999994 |
| 2310040G07 | 0.00157119 | 0.00058103 | 0.00142116 | -0.0008401 | 0.99995839 | 0.99999994 |
| Ccna2 | 0.01232407 | 0.01022561 | 0.01106638 | -0.0008408 | 0.69347269 | 0.99999994 |
| 3300002I08R | 0.01107484 | 0.00911192 | 0.00995303 | -0.0008411 | 0.71311805 | 0.99999994 |
| D630024D03 | 0.00245298 | 0.00137506 | 0.00221742 | -0.0008424 | 0.99427453 | 0.99999994 |
| Gm24362 | 0.00728532 | 0.00572056 | 0.00656416 | -0.0008436 | 0.80384754 | 0.99999994 |
| Med7 | 0.33710768 | 0.23732565 | 0.23816927 | -0.0008436 | 0.50940511 | 0.99999994 |
| Adora3 | 0.00492111 | 0.00359686 | 0.00444109 | -0.0008442 | 0.89712497 | 0.99999994 |
| Gm42670 | 0.0077478 | 0.00613419 | 0.00697868 | -0.0008445 | 0.78972817 | 0.99999994 |
| Nuf2 | 0.00815375 | 0.00649715 | 0.00734231 | -0.0008452 | 0.77822131 | 0.99999994 |
| Arhgef19 | 0.03014708 | 0.02590359 | 0.02674884 | -0.0008453 | 0.5832983 | 0.99999994 |
| Neb | 0.00226929 | 0.00120593 | 0.00205163 | -0.0008457 | 0.99696353 | 0.99999994 |
| 1110017D15 | 0.08933398 | 0.07540898 | 0.07625499 | -0.000846 | 0.5294347 | 0.99999994 |
| Mfsd8 | 0.03687823 | 0.03172883 | 0.03257506 | -0.0008462 | 0.56864195 | 0.99999994 |
| Vopp1 | 0.04498633 | 0.03867728 | 0.03952435 | -0.0008471 | 0.55671909 | 0.99999994 |
| Syt9 | 0.00859737 | 0.00689223 | 0.00773945 | -0.0008472 | 0.76689584 | 0.99999994 |

| | | | | | | |
|---|---|---|---|---|---|---|
| Slc3a1 | 0.00356356 | 0.00237143 | 0.00321892 | -0.0008475 | 0.96015729 | 0.99999994 |
| Lrrc38 | 0.00274219 | 0.00163076 | 0.00247837 | -0.0008476 | 0.98859134 | 0.99999994 |
| Hpgds | 0.02265206 | 0.01935134 | 0.02019962 | -0.0008483 | 0.61008155 | 0.99999994 |
| Rnft2 | 0.00353004 | 0.00234006 | 0.00318871 | -0.0008486 | 0.96176411 | 0.99999994 |
| Tm4sf1 | 0.00369668 | 0.00249013 | 0.00333886 | -0.0008487 | 0.95468096 | 0.99999994 |
| Prkd1 | 0.00249902 | 0.0014101 | 0.00225896 | -0.0008489 | 0.9938131 | 0.99999994 |
| Ank2 | 0.06324605 | 0.05405615 | 0.05490521 | -0.0008491 | 0.54099228 | 0.99999994 |
| Cdh13 | 0.00193363 | 0.00089903 | 0.00174856 | -0.0008495 | 0.99938961 | 0.99999994 |
| Cdc42ep2 | 0.00553226 | 0.00414073 | 0.00499056 | -0.0008498 | 0.87149553 | 0.99999994 |
| Rragd | 0.03370033 | 0.02898007 | 0.0298309 | -0.0008508 | 0.57528695 | 0.99999994 |
| Ngp | 0.00366175 | 0.00245632 | 0.00330739 | -0.0008511 | 0.95662887 | 0.99999994 |
| Eme2 | 0.01764825 | 0.014939 | 0.01579049 | -0.0008515 | 0.64018006 | 0.99999994 |
| H2-Q7 | 0.00367462 | 0.0024666 | 0.00331899 | -0.0008524 | 0.95632091 | 0.99999994 |
| Gm29394 | 0.00493488 | 0.00360107 | 0.00445347 | -0.0008524 | 0.89867578 | 0.99999994 |
| Cntnap2 | 0.00614488 | 0.00468794 | 0.00554088 | -0.0008529 | 0.84723671 | 0.99999994 |
| Slc16a13 | 0.02564683 | 0.02197075 | 0.02282435 | -0.0008536 | 0.59829587 | 0.99999994 |
| Fn3krp | 0.0106232 | 0.00869641 | 0.00955005 | -0.0008536 | 0.72407388 | 0.99999994 |
| Tmem252 | 0.0061384 | 0.00468128 | 0.00553507 | -0.0008538 | 0.84772922 | 0.99999994 |
| Pola2 | 0.02489586 | 0.02131243 | 0.02216716 | -0.0008547 | 0.60127877 | 0.99999994 |
| Lysmd1 | 0.01655294 | 0.01396563 | 0.0148214 | -0.0008558 | 0.64963254 | 0.99999994 |
| Pcf11 | 0.16123961 | 0.12936589 | 0.13022212 | -0.0008562 | 0.51745459 | 0.99999994 |
| Hadhb | 0.59951436 | 0.35059662 | 0.35145477 | -0.0008581 | 0.50648367 | 0.99999994 |
| Lrig1 | 0.01437023 | 0.01202679 | 0.01288594 | -0.0008591 | 0.67140722 | 0.99999994 |
| Rab17 | 0.01145057 | 0.00942846 | 0.01028808 | -0.0008596 | 0.71095391 | 0.99999994 |
| Tcof1 | 0.08566501 | 0.07243585 | 0.07329566 | -0.0008598 | 0.53111936 | 0.99999994 |
| Cabin1 | 0.14968953 | 0.12113488 | 0.12199504 | -0.0008602 | 0.51871631 | 0.99999994 |
| Extl3 | 0.06575112 | 0.05612588 | 0.05698679 | -0.0008609 | 0.54004947 | 0.99999994 |
| Csrnp1 | 0.02000436 | 0.01700826 | 0.01787028 | -0.000862 | 0.62592758 | 0.99999994 |
| Erf | 0.08246218 | 0.0698382 | 0.07070088 | -0.0008627 | 0.53236654 | 0.99999994 |
| Gna14 | 0.00672282 | 0.00519572 | 0.00605964 | -0.0008639 | 0.82868895 | 0.99999994 |

| | | | | | | |
|---|---|---|---|---|---|---|
| Fbxl19 | 0.04097518 | 0.03523147 | 0.03609579 | -0.0008643 | 0.56331873 | 0.99999994 |
| Gm44109 | 0.00344728 | 0.00224978 | 0.00311413 | -0.0008643 | 0.96766394 | 0.99999994 |
| Entpd6 | 0.02122599 | 0.0180816 | 0.01894604 | -0.0008644 | 0.61932507 | 0.99999994 |
| Inca1 | 0.01395912 | 0.01165539 | 0.01252075 | -0.0008654 | 0.67725978 | 0.99999994 |
| Ubox5 | 0.02104255 | 0.01791868 | 0.01878462 | -0.0008659 | 0.6205211 | 0.99999994 |
| Arhgef10l | 0.02042315 | 0.01737322 | 0.01823926 | -0.000866 | 0.62402126 | 0.99999994 |
| Map3k3 | 0.01170976 | 0.00965218 | 0.01051912 | -0.0008669 | 0.70835638 | 0.99999994 |
| Ano7 | 0.00220871 | 0.0011289 | 0.00199694 | -0.000868 | 0.99809469 | 0.99999994 |
| Tmem161b | 0.04304354 | 0.03699764 | 0.03786601 | -0.0008684 | 0.56066216 | 0.99999994 |
| A430078I02F | 0.00391905 | 0.00267051 | 0.00353918 | -0.0008687 | 0.94884521 | 0.99999994 |
| Prickle1 | 0.01068416 | 0.00873528 | 0.00960446 | -0.0008692 | 0.72654021 | 0.99999994 |
| Zfp748 | 0.02162628 | 0.01842854 | 0.01929816 | -0.0008696 | 0.61789186 | 0.99999994 |
| Smg8 | 0.02168914 | 0.01848292 | 0.01935344 | -0.0008705 | 0.61768409 | 0.99999994 |
| Mgat4a | 0.02073337 | 0.01764187 | 0.01851246 | -0.0008706 | 0.62287117 | 0.99999994 |
| 4932438H23I | 0.00973725 | 0.00788778 | 0.00875883 | -0.000871 | 0.74600219 | 0.99999994 |
| Tmem243 | 0.09099359 | 0.07671754 | 0.07758903 | -0.0008715 | 0.52979926 | 0.99999994 |
| Dock7 | 0.23787182 | 0.18018921 | 0.18106151 | -0.0008723 | 0.51279131 | 0.99999994 |
| Peli3 | 0.00784645 | 0.00619473 | 0.00706705 | -0.0008723 | 0.79435642 | 0.99999994 |
| Dock6 | 0.02814935 | 0.02413624 | 0.02500962 | -0.0008734 | 0.59190571 | 0.99999994 |
| Catsper4 | 0.00204864 | 0.00097875 | 0.00185241 | -0.0008737 | 0.99915345 | 0.99999994 |
| Mavs | 0.05639484 | 0.04830309 | 0.04917731 | -0.0008742 | 0.54709667 | 0.99999994 |
| Gdf3 | 0.00181731 | 0.00076657 | 0.0016435 | -0.0008769 | 0.9998086 | 0.99999994 |
| 1110002E22I | 0.00274701 | 0.00160499 | 0.00248272 | -0.0008777 | 0.99069478 | 0.99999994 |
| Ighmbp2 | 0.02232599 | 0.01903497 | 0.0199132 | -0.0008782 | 0.61545495 | 0.99999994 |
| Xbp1 | 0.68288546 | 0.37686435 | 0.37774284 | -0.0008785 | 0.50617549 | 0.99999994 |
| Ctnnd2 | 0.0045755 | 0.00325027 | 0.00413016 | -0.0008799 | 0.92191437 | 0.99999994 |
| Cpa3 | 0.00714332 | 0.00555509 | 0.00643683 | -0.0008817 | 0.81906681 | 0.99999994 |
| 2310015A10I | 0.03786862 | 0.03254599 | 0.03342791 | -0.0008819 | 0.56970042 | 0.99999994 |
| Reg1 | 0.01063084 | 0.00867494 | 0.00955686 | -0.0008819 | 0.73047919 | 0.99999994 |
| Zfp131 | 0.14027401 | 0.11428939 | 0.11517302 | -0.0008836 | 0.52036476 | 0.99999994 |

| | | | | | | |
|---|---|---|---|---|---|---|
| Spin1 | 0.16049472 | 0.12881248 | 0.12969614 | -0.0008837 | 0.51808659 | 0.99999994 |
| Haus8 | 0.0363385 | 0.03122601 | 0.03210981 | -0.0008838 | 0.57268336 | 0.99999994 |
| Pcgf5 | 0.07981899 | 0.06766735 | 0.06855142 | -0.0008841 | 0.53420441 | 0.99999994 |
| Rimkla | 0.00437193 | 0.0030628 | 0.00394695 | -0.0008842 | 0.93202999 | 0.99999994 |
| Plcg2 | 0.0030599 | 0.00187929 | 0.00276492 | -0.0008856 | 0.98350024 | 0.99999994 |
| Fam172a | 0.13381236 | 0.10954441 | 0.11043038 | -0.000886 | 0.52129475 | 0.99999994 |
| Gm7102 | 0.00598458 | 0.00451072 | 0.00539693 | -0.0008862 | 0.86280637 | 0.99999994 |
| Ror1 | 0.00607578 | 0.0045924 | 0.00547884 | -0.0008864 | 0.85924786 | 0.99999994 |
| Zfp202 | 0.0048873 | 0.00352356 | 0.00441067 | -0.0008871 | 0.90968038 | 0.99999994 |
| Tlr3 | 0.00348645 | 0.00226169 | 0.00314943 | -0.0008877 | 0.96968959 | 0.99999994 |
| Aarsd1 | 0.01187685 | 0.00978015 | 0.01066802 | -0.0008879 | 0.71020727 | 0.99999994 |
| Gm15747 | 0.0038127 | 0.00255539 | 0.00344339 | -0.000888 | 0.95697185 | 0.99999994 |
| Mitf | 0.02164393 | 0.01842548 | 0.01931368 | -0.0008882 | 0.62024057 | 0.99999994 |
| Idi1 | 0.07907526 | 0.06705682 | 0.06794528 | -0.0008885 | 0.53467981 | 0.99999994 |
| Nfkbie | 0.00847228 | 0.00673838 | 0.0076275 | -0.0008891 | 0.78110145 | 0.99999994 |
| Lin37 | 0.24760568 | 0.18618798 | 0.18707775 | -0.0008898 | 0.51262778 | 0.99999994 |
| Zfp846 | 0.02041599 | 0.01734317 | 0.01823296 | -0.0008898 | 0.6273474 | 0.99999994 |
| Natd1 | 0.03927138 | 0.03374397 | 0.03463394 | -0.00089 | 0.56790615 | 0.99999994 |
| Zfp319 | 0.01984627 | 0.01683967 | 0.01773094 | -0.0008913 | 0.6310329 | 0.99999994 |
| Greb1l | 0.00449703 | 0.00316823 | 0.00405954 | -0.0008913 | 0.92805655 | 0.99999994 |
| U2af2 | 0.4458139 | 0.29013793 | 0.29103034 | -0.0008924 | 0.50814222 | 0.99999994 |
| Ccdc71l | 0.00907224 | 0.00727139 | 0.00816432 | -0.0008929 | 0.76669393 | 0.99999994 |
| Kcnj5 | 0.00305399 | 0.00186637 | 0.00275959 | -0.0008932 | 0.98440051 | 0.99999994 |
| Lyn | 0.00740059 | 0.00577333 | 0.0066675 | -0.0008942 | 0.81398254 | 0.99999994 |
| Ppp6r2 | 0.01421798 | 0.01185648 | 0.01275072 | -0.0008942 | 0.67968757 | 0.99999994 |
| Gm11837 | 0.01631104 | 0.01371263 | 0.01460718 | -0.0008946 | 0.65823051 | 0.99999994 |
| Notch2 | 0.00934726 | 0.0075155 | 0.00841025 | -0.0008948 | 0.76057607 | 0.99999994 |
| Stradb | 0.05797184 | 0.04960516 | 0.0505003 | -0.0008951 | 0.546961 | 0.99999994 |
| A830052D11 | 0.00301233 | 0.00182664 | 0.00272202 | -0.0008954 | 0.98572107 | 0.99999994 |
| Rhof | 0.01044296 | 0.00849376 | 0.00938916 | -0.0008954 | 0.73721479 | 0.99999994 |

| | | | | | | |
|---|---|---|---|---|---|---|
| 2410080I02R | 0.00160597 | 0.00055689 | 0.00145258 | -0.0008957 | 0.99997974 | 0.99999994 |
| Cdca2 | 0.00494305 | 0.00356385 | 0.00446082 | -0.000897 | 0.90962399 | 0.99999994 |
| Lcn2 | 0.00434322 | 0.0030225 | 0.0039211 | -0.0008986 | 0.93642561 | 0.99999994 |
| Chka | 0.3606152 | 0.24950571 | 0.25040435 | -0.0008986 | 0.50952907 | 0.99999994 |
| C330002G04 | 0.00348886 | 0.00225183 | 0.00315161 | -0.0008998 | 0.97130776 | 0.99999994 |
| Shkbp1 | 0.15677511 | 0.12616002 | 0.12706022 | -0.0009002 | 0.5188067 | 0.99999994 |
| Vipr1 | 0.00181208 | 0.00073846 | 0.00163877 | -0.0009003 | 0.99987236 | 0.99999994 |
| Cttnbp2nl | 0.05988646 | 0.05120203 | 0.05210286 | -0.0009008 | 0.54581129 | 0.99999994 |
| Abhd3 | 0.01959886 | 0.0166117 | 0.0175128 | -0.0009011 | 0.63401084 | 0.99999994 |
| Scoc | 0.42213306 | 0.27939551 | 0.28029739 | -0.0009019 | 0.50854356 | 0.99999994 |
| Gm15420 | 0.01191602 | 0.00980104 | 0.01070291 | -0.0009019 | 0.71256601 | 0.99999994 |
| Wipf3 | 0.01245853 | 0.01028382 | 0.0111861 | -0.0009023 | 0.70433365 | 0.99999994 |
| Sh3rf1 | 0.0170312 | 0.01434232 | 0.01524472 | -0.0009024 | 0.65321622 | 0.99999994 |
| Gm5860 | 0.00156162 | 0.00051006 | 0.00141251 | -0.0009025 | 0.99998944 | 0.99999994 |
| Gm9958 | 0.00982713 | 0.00793667 | 0.00883915 | -0.0009025 | 0.7516271 | 0.99999994 |
| Tmem69 | 0.03444867 | 0.0295753 | 0.03047815 | -0.0009028 | 0.57815642 | 0.99999994 |
| Hcst | 0.00214526 | 0.00103669 | 0.00193966 | -0.000903 | 0.99902826 | 0.99999994 |
| Sema4g | 0.01062917 | 0.00865208 | 0.00955537 | -0.0009033 | 0.73540392 | 0.99999994 |
| Scn1b | 0.00167575 | 0.00061208 | 0.00151563 | -0.0009036 | 0.9999638 | 0.99999994 |
| Gm11713 | 0.01777449 | 0.01499794 | 0.01590209 | -0.0009042 | 0.64745702 | 0.99999994 |
| Rab11fip2 | 0.17160921 | 0.13657238 | 0.13747715 | -0.0009048 | 0.51747094 | 0.99999994 |
| Nos3 | 0.00618144 | 0.0046672 | 0.00557371 | -0.0009065 | 0.86050513 | 0.99999994 |
| Senp7 | 0.15812363 | 0.12711002 | 0.12801768 | -0.0009077 | 0.51882076 | 0.99999994 |
| Ttc9 | 0.01661991 | 0.01397091 | 0.01488069 | -0.0009098 | 0.65797951 | 0.99999994 |
| Smc2os | 0.0064025 | 0.00486239 | 0.00577218 | -0.0009098 | 0.85294666 | 0.99999994 |
| Slc27a3 | 0.01850742 | 0.01563909 | 0.01654966 | -0.0009106 | 0.64290629 | 0.99999994 |
| Il1r2 | 0.00890719 | 0.00710528 | 0.00801668 | -0.0009114 | 0.77539961 | 0.99999994 |
| Apba1 | 0.00627901 | 0.0047477 | 0.00566132 | -0.0009136 | 0.85863297 | 0.99999994 |
| Dnah2 | 0.00300827 | 0.00180403 | 0.00271836 | -0.0009143 | 0.9874187 | 0.99999994 |
| Kbtbd8 | 0.00376813 | 0.00248886 | 0.00340323 | -0.0009144 | 0.96314489 | 0.99999994 |

| | | | | | | |
|---|---|---|---|---|---|---|
| Tcirg1 | 0.03783461 | 0.03248286 | 0.03339864 | -0.0009158 | 0.57241088 | 0.99999994 |
| Lgals4 | 0.00518304 | 0.00376009 | 0.00467663 | -0.0009165 | 0.90397474 | 0.99999994 |
| Gm15492 | 0.00703517 | 0.00542326 | 0.00633985 | -0.0009166 | 0.83206215 | 0.99999994 |
| Dennd1b | 0.05365146 | 0.04595206 | 0.04686929 | -0.0009172 | 0.55182188 | 0.99999994 |
| Krt86 | 0.00414166 | 0.00282177 | 0.00373964 | -0.0009179 | 0.9488459 | 0.99999994 |
| Plp1 | 0.01938311 | 0.01640431 | 0.01732253 | -0.0009182 | 0.63789306 | 0.99999994 |
| Ddx19b | 0.05052085 | 0.04330701 | 0.0442253 | -0.0009183 | 0.55496398 | 0.99999994 |
| Crym | 0.01310907 | 0.01084649 | 0.01176504 | -0.0009185 | 0.69835971 | 0.99999994 |
| Shpk | 0.01385056 | 0.01150493 | 0.01242429 | -0.0009194 | 0.68883458 | 0.99999994 |
| C4b | 0.00545203 | 0.00399898 | 0.00491845 | -0.0009195 | 0.89331689 | 0.99999994 |
| Iqsec2 | 0.0059541 | 0.00444952 | 0.00536955 | -0.00092 | 0.87296484 | 0.99999994 |
| Brca1 | 0.00242123 | 0.00126824 | 0.00218876 | -0.0009205 | 0.99744054 | 0.99999994 |
| Nedd9 | 0.0069781 | 0.00536753 | 0.00628866 | -0.0009211 | 0.83521675 | 0.99999994 |
| Parp12 | 0.02013143 | 0.01706063 | 0.01798225 | -0.0009216 | 0.63350496 | 0.99999994 |
| Fam122b | 0.01240791 | 0.01021922 | 0.01114103 | -0.0009218 | 0.70909732 | 0.99999994 |
| Dsel | 0.01001752 | 0.00808741 | 0.00900923 | -0.0009218 | 0.75208985 | 0.99999994 |
| Nalcn | 0.00571109 | 0.00422858 | 0.00515125 | -0.0009227 | 0.88341845 | 0.99999994 |
| Polrmt | 0.0306909 | 0.02629875 | 0.0272215 | -0.0009227 | 0.58925754 | 0.99999994 |
| Mir17hg | 0.0117479 | 0.0096295 | 0.01055311 | -0.0009236 | 0.71990742 | 0.99999994 |
| Gm17399 | 0.0036212 | 0.00234677 | 0.00327086 | -0.0009241 | 0.96998423 | 0.99999994 |
| Smad3 | 0.00832933 | 0.00657543 | 0.00749953 | -0.0009241 | 0.79394835 | 0.99999994 |
| Recql5 | 0.01926614 | 0.01629416 | 0.01721935 | -0.0009252 | 0.63969429 | 0.99999994 |
| Ankrd29 | 0.00276697 | 0.00157542 | 0.00250072 | -0.0009253 | 0.99311017 | 0.99999994 |
| Jmjd7 | 0.0464919 | 0.03988115 | 0.04080656 | -0.0009254 | 0.55999397 | 0.99999994 |
| 2810021J22F | 0.03107829 | 0.02663209 | 0.02755797 | -0.0009259 | 0.58848038 | 0.99999994 |
| Gm15952 | 0.0052318 | 0.00379405 | 0.00472048 | -0.0009264 | 0.90428448 | 0.99999994 |
| Vcl | 0.02855831 | 0.02443947 | 0.02536603 | -0.0009266 | 0.59605149 | 0.99999994 |
| Lhx6 | 0.00529476 | 0.00385025 | 0.00477708 | -0.0009268 | 0.90172403 | 0.99999994 |
| Zfp953 | 0.00687291 | 0.00526677 | 0.0061943 | -0.0009275 | 0.84055011 | 0.99999994 |
| Wdr54 | 0.02625949 | 0.02243095 | 0.02336002 | -0.0009291 | 0.60439363 | 0.99999994 |

| | | | | | | |
|---|---|---|---|---|---|---|
| Mtmr9 | 0.02887677 | 0.02471337 | 0.02564344 | -0.0009301 | 0.59538627 | 0.99999994 |
| Vsig10l | 0.00605692 | 0.00453174 | 0.0054619 | -0.0009302 | 0.87151431 | 0.99999994 |
| Ky | 0.00708237 | 0.00545064 | 0.00638217 | -0.0009315 | 0.83436413 | 0.99999994 |
| Phactr2 | 0.01437272 | 0.01195588 | 0.01288815 | -0.0009323 | 0.68491531 | 0.99999994 |
| Grwd1 | 0.07359201 | 0.06252606 | 0.06345847 | -0.0009324 | 0.53895584 | 0.99999994 |
| Ccdc159 | 0.010832 | 0.00880292 | 0.00973638 | -0.0009335 | 0.73831796 | 0.99999994 |
| Coq10a | 0.04332823 | 0.03717515 | 0.03810928 | -0.0009341 | 0.56480393 | 0.99999994 |
| B3galt2 | 0.00487094 | 0.0034617 | 0.00439596 | -0.0009343 | 0.92141367 | 0.99999994 |
| Ccl4 | 0.00950985 | 0.00762128 | 0.0085556 | -0.0009343 | 0.76635962 | 0.99999994 |
| C130080G10 | 0.00503855 | 0.00361223 | 0.00454671 | -0.0009345 | 0.9143563 | 0.99999994 |
| Orai2 | 0.04462417 | 0.03827687 | 0.03921554 | -0.0009387 | 0.56329437 | 0.99999994 |
| Rab11fip3 | 0.01381922 | 0.01145733 | 0.01239644 | -0.0009391 | 0.69296199 | 0.99999994 |
| Zfp397 | 0.15951054 | 0.12806063 | 0.12900023 | -0.0009396 | 0.51933436 | 0.99999994 |
| Scara3 | 0.01280307 | 0.01055309 | 0.01149278 | -0.0009397 | 0.70686487 | 0.99999994 |
| Lmtk3 | 0.01061476 | 0.00860239 | 0.00954252 | -0.0009401 | 0.74401818 | 0.99999994 |
| Col5a2 | 0.00389877 | 0.0025788 | 0.00352091 | -0.0009421 | 0.96255063 | 0.99999994 |
| Apaf1 | 0.02526235 | 0.02154559 | 0.02248797 | -0.0009424 | 0.60985542 | 0.99999994 |
| Malt1 | 0.01208694 | 0.00991249 | 0.01085518 | -0.0009427 | 0.71838603 | 0.99999994 |
| Tmem39b | 0.08064501 | 0.06828123 | 0.06922393 | -0.0009427 | 0.53611341 | 0.99999994 |
| D630033O11 | 0.0040252 | 0.00269186 | 0.00363478 | -0.0009429 | 0.95789443 | 0.99999994 |
| Ss18l1 | 0.01746962 | 0.01468958 | 0.01563255 | -0.000943 | 0.65597912 | 0.99999994 |
| Ccdc113 | 0.01622361 | 0.0135866 | 0.01452974 | -0.0009431 | 0.66715477 | 0.99999994 |
| 1190005I06R | 0.01729304 | 0.01453268 | 0.01547637 | -0.0009437 | 0.65758512 | 0.99999994 |
| Zfp59 | 0.00886111 | 0.0070307 | 0.00797545 | -0.0009448 | 0.78479605 | 0.99999994 |
| Asphd2 | 0.00734234 | 0.00567034 | 0.00661528 | -0.0009449 | 0.82914905 | 0.99999994 |
| Asap1 | 0.18674915 | 0.14690691 | 0.14785279 | -0.0009459 | 0.5169834 | 0.99999994 |
| Zfp689 | 0.02982104 | 0.02551941 | 0.02646531 | -0.0009459 | 0.59402282 | 0.99999994 |
| Gm26881 | 0.02251667 | 0.01913326 | 0.02008071 | -0.0009475 | 0.62326309 | 0.99999994 |
| 5033403F01I | 0.00496868 | 0.00353626 | 0.00448387 | -0.0009476 | 0.92024803 | 0.99999994 |
| L1cam | 0.0055679 | 0.00407433 | 0.00502258 | -0.0009483 | 0.89557109 | 0.99999994 |

| | | | | | | |
|---|---|---|---|---|---|---|
| Guca2a | 0.00571311 | 0.00420452 | 0.00515307 | -0.0009485 | 0.88976205 | 0.99999994 |
| Ptges3l | 0.03515874 | 0.03014286 | 0.03109169 | -0.0009488 | 0.58048462 | 0.99999994 |
| Mul1 | 0.14287597 | 0.11611977 | 0.11706884 | -0.0009491 | 0.52151739 | 0.99999994 |
| Gm15491 | 0.00819579 | 0.0064305 | 0.00737996 | -0.0009495 | 0.8041027 | 0.99999994 |
| Hcn4 | 0.00483551 | 0.00341391 | 0.00436409 | -0.0009502 | 0.92636891 | 0.99999994 |
| Prr15 | 0.00514963 | 0.00369641 | 0.0046466 | -0.0009502 | 0.91327045 | 0.99999994 |
| Pqlc3 | 0.01058047 | 0.00856169 | 0.0095119 | -0.0009502 | 0.74695895 | 0.99999994 |
| Unc5b | 0.01289495 | 0.01062427 | 0.01157454 | -0.0009503 | 0.70763412 | 0.99999994 |
| Stk25 | 0.24346878 | 0.18358114 | 0.18453226 | -0.0009511 | 0.51368427 | 0.99999994 |
| Ccdc138 | 0.01241799 | 0.01019748 | 0.01115001 | -0.0009525 | 0.71520084 | 0.99999994 |
| Myo19 | 0.01001254 | 0.00805219 | 0.00900479 | -0.0009526 | 0.75933329 | 0.99999994 |
| Mrto4 | 0.22765176 | 0.17368594 | 0.17464082 | -0.0009549 | 0.5145163 | 0.99999994 |
| Tesk2 | 0.01146514 | 0.00934609 | 0.01030107 | -0.000955 | 0.73141403 | 0.99999994 |
| 6430550D23I | 0.00674513 | 0.00512359 | 0.00607966 | -0.0009561 | 0.85239472 | 0.99999994 |
| Rrp12 | 0.0161496 | 0.01350797 | 0.01446417 | -0.0009562 | 0.67004606 | 0.99999994 |
| Lgals3 | 0.02119194 | 0.01795959 | 0.01891608 | -0.0009565 | 0.63178345 | 0.99999994 |
| 2310074N15 | 0.00490131 | 0.00346645 | 0.00442328 | -0.0009568 | 0.92504941 | 0.99999994 |
| Pacsin2 | 0.06926691 | 0.05893963 | 0.05989679 | -0.0009572 | 0.54235573 | 0.99999994 |
| Cspg5 | 0.01154815 | 0.00941784 | 0.01037507 | -0.0009572 | 0.73043588 | 0.99999994 |
| D130043K22 | 0.00201272 | 0.0008617 | 0.00181998 | -0.0009583 | 0.99977153 | 0.99999994 |
| H2-Q6 | 0.00518831 | 0.00372253 | 0.00468137 | -0.0009588 | 0.91361553 | 0.99999994 |
| Ccdc40 | 0.01167285 | 0.00952724 | 0.01048622 | -0.000959 | 0.7286503 | 0.99999994 |
| Arl6ip5 | 0.61815268 | 0.35682802 | 0.35778769 | -0.0009597 | 0.5071223 | 0.99999994 |
| Kctd14 | 0.00976614 | 0.00782468 | 0.00878466 | -0.00096 | 0.76650778 | 0.99999994 |
| Rbm41 | 0.01564934 | 0.01306008 | 0.01402084 | -0.0009608 | 0.6758468 | 0.99999994 |
| Trdmt1 | 0.02629659 | 0.02243121 | 0.02339245 | -0.0009612 | 0.60777371 | 0.99999994 |
| Zfp763 | 0.02595261 | 0.02212958 | 0.02309176 | -0.0009622 | 0.60924645 | 0.99999994 |
| Vangl2 | 0.10252876 | 0.08578493 | 0.08674735 | -0.0009624 | 0.52943474 | 0.99999994 |
| Mfsd5 | 0.04957072 | 0.04245737 | 0.04342071 | -0.0009633 | 0.55870217 | 0.99999994 |
| Naa16 | 0.10735615 | 0.08951784 | 0.09048134 | -0.0009635 | 0.52825388 | 0.99999994 |

| | | | | | | |
|---|---|---|---|---|---|---|
| Paupar | 0.00377376 | 0.00244476 | 0.00340831 | -0.0009635 | 0.970067 | 0.99999994 |
| Clic5 | 0.00179062 | 0.00065517 | 0.00161939 | -0.0009642 | 0.99996304 | 0.99999994 |
| Dtx1 | 0.01660095 | 0.01389938 | 0.01486391 | -0.0009645 | 0.66710493 | 0.99999994 |
| Klhl25 | 0.02986776 | 0.02554142 | 0.02650595 | -0.0009645 | 0.59569449 | 0.99999994 |
| Srrm4os | 0.00397629 | 0.00262605 | 0.00359072 | -0.0009647 | 0.96313399 | 0.99999994 |
| 1700010I14R | 0.04135634 | 0.03545755 | 0.03642237 | -0.0009648 | 0.56998138 | 0.99999994 |
| Osbpl7 | 0.01444929 | 0.0119913 | 0.01295614 | -0.0009648 | 0.68994468 | 0.99999994 |
| Itpkc | 0.01826868 | 0.01537354 | 0.0163388 | -0.0009653 | 0.65292931 | 0.99999994 |
| Pnisr | 0.64695959 | 0.36617944 | 0.36714483 | -0.0009654 | 0.50698217 | 0.99999994 |
| Ino80c | 0.06012345 | 0.05133458 | 0.05230095 | -0.0009664 | 0.54894263 | 0.99999994 |
| Golt1a | 0.00296908 | 0.00171603 | 0.00268302 | -0.000967 | 0.99178066 | 0.99999994 |
| Nin | 0.01804429 | 0.0151735 | 0.01614055 | -0.000967 | 0.65498307 | 0.99999994 |
| Cpxm2 | 0.02199575 | 0.01865447 | 0.019623 | -0.0009685 | 0.6287469 | 0.99999994 |
| C030034I22R | 0.02617616 | 0.02231741 | 0.02328719 | -0.0009698 | 0.60918701 | 0.99999994 |
| Nxpe4 | 0.00952134 | 0.00759604 | 0.00856587 | -0.0009698 | 0.77446574 | 0.99999994 |
| Vwa5b1 | 0.00928889 | 0.00738774 | 0.00835806 | -0.0009703 | 0.78016859 | 0.99999994 |
| Rhebl1 | 0.03051305 | 0.02609592 | 0.02706696 | -0.000971 | 0.59436933 | 0.99999994 |
| Scn2b | 0.00699077 | 0.00532872 | 0.00630002 | -0.0009713 | 0.8476095 | 0.99999994 |
| Nop2 | 0.08146849 | 0.06892137 | 0.06989366 | -0.0009723 | 0.53688829 | 0.99999994 |
| Cldn2 | 0.00714719 | 0.00546721 | 0.00644031 | -0.0009731 | 0.84272899 | 0.99999994 |
| E430018J23F | 0.02565813 | 0.02186018 | 0.02283424 | -0.0009741 | 0.61177413 | 0.99999994 |
| Adcy5 | 0.01253467 | 0.0102788 | 0.01125389 | -0.0009751 | 0.71794106 | 0.99999994 |
| Phldb1 | 0.02264941 | 0.01921998 | 0.02019729 | -0.0009773 | 0.62630777 | 0.99999994 |
| Gm15473 | 0.00378559 | 0.00244141 | 0.00341896 | -0.0009775 | 0.97149179 | 0.99999994 |
| Jak3 | 0.00691847 | 0.00525747 | 0.00623517 | -0.0009777 | 0.8516995 | 0.99999994 |
| Cd14 | 0.00940691 | 0.0074852 | 0.00846358 | -0.0009784 | 0.77919329 | 0.99999994 |
| Spata33 | 0.01566351 | 0.01305462 | 0.0140334 | -0.0009788 | 0.67876849 | 0.99999994 |
| Lrrc20 | 0.01170017 | 0.0095308 | 0.01051058 | -0.0009798 | 0.7325339 | 0.99999994 |
| Dpp7 | 0.04718202 | 0.04041312 | 0.04139345 | -0.0009803 | 0.56263156 | 0.99999994 |
| Oxsr1 | 0.06297549 | 0.05369963 | 0.05467998 | -0.0009803 | 0.5474972 | 0.99999994 |

| | | | | | | |
|---|---|---|---|---|---|---|
| Ppp1r3d | 0.0024142 | 0.00120073 | 0.00218242 | -0.0009817 | 0.99862387 | 0.99999994 |
| Tpm2 | 0.00510456 | 0.00362409 | 0.00460607 | -0.000982 | 0.92206241 | 0.99999994 |
| Mzf1 | 0.02360357 | 0.02005266 | 0.0210347 | -0.000982 | 0.62200872 | 0.99999994 |
| Pdlim3 | 0.01047454 | 0.00843407 | 0.00941735 | -0.0009833 | 0.75647151 | 0.99999994 |
| Fzd5 | 0.0034992 | 0.00217759 | 0.00316092 | -0.0009833 | 0.98080748 | 0.99999994 |
| Med30 | 0.33203251 | 0.23448262 | 0.23546638 | -0.0009838 | 0.51109311 | 0.99999994 |
| Dpp10 | 0.00648608 | 0.0048634 | 0.0058472 | -0.0009838 | 0.86862946 | 0.99999994 |
| Hyal3 | 0.01048577 | 0.00844308 | 0.00942737 | -0.0009843 | 0.75646526 | 0.99999994 |
| Hinfp | 0.01478509 | 0.01226934 | 0.01325423 | -0.0009849 | 0.68956569 | 0.99999994 |
| Zfp27 | 0.01481742 | 0.01229742 | 0.01328293 | -0.0009855 | 0.68929794 | 0.99999994 |
| Gpr179 | 0.00703083 | 0.0053491 | 0.00633595 | -0.0009868 | 0.85007463 | 0.99999994 |
| Cdk1 | 0.03018607 | 0.02579569 | 0.02678274 | -0.000987 | 0.59689296 | 0.99999994 |
| Gm14325 | 0.01449633 | 0.01200934 | 0.01299791 | -0.0009886 | 0.69366045 | 0.99999994 |
| Ipo11 | 0.0200312 | 0.01690498 | 0.01789393 | -0.0009889 | 0.643518 | 0.99999994 |
| C2 | 0.0037681 | 0.00241164 | 0.0034032 | -0.0009916 | 0.97377385 | 0.99999994 |
| Tmem79 | 0.0121103 | 0.0098836 | 0.01087599 | -0.0009924 | 0.72819719 | 0.99999994 |
| Gm14418 | 0.0243636 | 0.02070744 | 0.02170097 | -0.0009935 | 0.61971948 | 0.99999994 |
| Card14 | 0.0026267 | 0.00138036 | 0.00237418 | -0.0009938 | 0.99733437 | 0.99999994 |
| Catip | 0.00489426 | 0.00342271 | 0.00441693 | -0.0009942 | 0.9329736 | 0.99999994 |
| Klhl11 | 0.01841003 | 0.01546935 | 0.01646365 | -0.0009943 | 0.65615846 | 0.99999994 |
| Cobl | 0.00892783 | 0.00704015 | 0.00803515 | -0.000995 | 0.79510345 | 0.99999994 |
| 4930532G15 | 0.00793883 | 0.00615441 | 0.00714982 | -0.0009954 | 0.82296004 | 0.99999994 |
| Grk4 | 0.01200159 | 0.00978247 | 0.01077916 | -0.0009967 | 0.73088055 | 0.99999994 |
| Slc5a6 | 0.01378305 | 0.01136697 | 0.0123643 | -0.0009973 | 0.70433763 | 0.99999994 |
| Nek5 | 0.00739991 | 0.00566932 | 0.00666689 | -0.0009976 | 0.84037345 | 0.99999994 |
| Irak4 | 0.01244117 | 0.01017193 | 0.01117064 | -0.0009987 | 0.72411659 | 0.99999994 |
| Rnf215 | 0.16937773 | 0.1349273 | 0.13592612 | -0.0009988 | 0.5195055 | 0.99999994 |
| Ndufa4l2 | 0.00566868 | 0.00411403 | 0.00511315 | -0.0009991 | 0.90331389 | 0.99999994 |
| Agr2 | 0.00269845 | 0.00143882 | 0.00243891 | -0.0010001 | 0.99682823 | 0.99999994 |
| Adcy3 | 0.00883399 | 0.00695088 | 0.0079512 | -0.0010003 | 0.79881975 | 0.99999994 |

| | | | | | | |
|---|---|---|---|---|---|---|
| Nme6 | 0.11674335 | 0.09665831 | 0.09765876 | -0.0010005 | 0.5271829 | 0.99999994 |
| Atp11c | 0.00845099 | 0.00660786 | 0.00760843 | -0.0010006 | 0.80931387 | 0.99999994 |
| Mcm8 | 0.00856618 | 0.00671052 | 0.00771154 | -0.001001 | 0.8062204 | 0.99999994 |
| Gm26876 | 0.00619977 | 0.00458906 | 0.00559017 | -0.0010011 | 0.88337875 | 0.99999994 |
| Zdhhc1 | 0.07535754 | 0.06390549 | 0.06490664 | -0.0010012 | 0.54088804 | 0.99999994 |
| Svopl | 0.00327425 | 0.00195683 | 0.00295817 | -0.0010013 | 0.98787619 | 0.99999994 |
| Mamstr | 0.02436442 | 0.02069893 | 0.02170169 | -0.0010028 | 0.62079401 | 0.99999994 |
| Pold1 | 0.01998966 | 0.01685425 | 0.01785732 | -0.0010031 | 0.64575881 | 0.99999994 |
| Tmem140 | 0.00407097 | 0.0026717 | 0.00367599 | -0.0010043 | 0.96550888 | 0.99999994 |
| Top3a | 0.01493814 | 0.01238549 | 0.01339006 | -0.0010046 | 0.69124439 | 0.99999994 |
| Cx3cl1 | 0.01961196 | 0.01651965 | 0.01752435 | -0.0010047 | 0.64862926 | 0.99999994 |
| Ranbp17 | 0.00788594 | 0.00609718 | 0.00710244 | -0.0010053 | 0.82693435 | 0.99999994 |
| Gm20632 | 0.00557467 | 0.00402273 | 0.00502867 | -0.0010059 | 0.90849346 | 0.99999994 |
| Zfp964 | 0.00539548 | 0.00386138 | 0.00486762 | -0.0010062 | 0.91559003 | 0.99999994 |
| Cntd1 | 0.0046211 | 0.00316489 | 0.00417119 | -0.0010063 | 0.94584591 | 0.99999994 |
| Erc1 | 0.12712734 | 0.1044644 | 0.10547076 | -0.0010064 | 0.52532087 | 0.99999994 |
| Tmem220 | 0.01373954 | 0.01131879 | 0.01232562 | -0.0010068 | 0.70668787 | 0.99999994 |
| Chsy1 | 0.00903092 | 0.00711663 | 0.00812736 | -0.0010107 | 0.79610487 | 0.99999994 |
| Pitpnc1 | 0.1540496 | 0.12410707 | 0.12511871 | -0.0010116 | 0.52146065 | 0.99999994 |
| Gm26826 | 0.00539542 | 0.00385539 | 0.00486757 | -0.0010122 | 0.91684186 | 0.99999994 |
| Amn1 | 0.31740731 | 0.22653854 | 0.22755102 | -0.0010125 | 0.51181389 | 0.99999994 |
| Gm26652 | 0.00479942 | 0.00331846 | 0.00433162 | -0.0010132 | 0.94025471 | 0.99999994 |
| Orai1 | 0.05646337 | 0.0482213 | 0.04923486 | -0.0010136 | 0.55449601 | 0.99999994 |
| Npl | 0.00793079 | 0.00612902 | 0.00714261 | -0.0010136 | 0.82756513 | 0.99999994 |
| Fam120c | 0.01256464 | 0.01026678 | 0.01128057 | -0.0010138 | 0.72515007 | 0.99999994 |
| Kctd1 | 0.03876999 | 0.03318838 | 0.03420313 | -0.0010147 | 0.5782751 | 0.99999994 |
| Evpl | 0.03217997 | 0.02749837 | 0.02851393 | -0.0010156 | 0.59370137 | 0.99999994 |
| Frrs1 | 0.13441191 | 0.10985569 | 0.11087252 | -0.0010168 | 0.52433883 | 0.99999994 |
| Pusl1 | 0.05976419 | 0.05098324 | 0.05200064 | -0.0010174 | 0.55180877 | 0.99999994 |
| Cep57 | 0.20360601 | 0.15809098 | 0.15910871 | -0.0010177 | 0.51698073 | 0.99999994 |

| | | | | | | |
|---|---|---|---|---|---|---|
| Mex3c | 0.04850228 | 0.041496 | 0.04251473 | -0.0010187 | 0.563362 | 0.99999994 |
| Ulk2 | 0.04354412 | 0.03727481 | 0.03829371 | -0.0010189 | 0.5702882 | 0.99999994 |
| Zbtb3 | 0.01735021 | 0.01450744 | 0.01552694 | -0.0010195 | 0.66896523 | 0.99999994 |
| Galnt4 | 0.01169596 | 0.00948713 | 0.01050682 | -0.0010197 | 0.74086115 | 0.99999994 |
| Alox5ap | 0.01822523 | 0.01528009 | 0.01630042 | -0.0010203 | 0.66153493 | 0.99999994 |
| Glb1l | 0.01093614 | 0.00880784 | 0.0098293 | -0.0010215 | 0.75544598 | 0.99999994 |
| Gar1 | 0.30756057 | 0.22109255 | 0.222115 | -0.0010224 | 0.5122219 | 0.99999994 |
| Stk26 | 0.00261177 | 0.00133824 | 0.0023607 | -0.0010225 | 0.99803033 | 0.99999994 |
| Zfp273 | 0.00918698 | 0.00724408 | 0.00826693 | -0.0010229 | 0.79491066 | 0.99999994 |
| Tab3 | 0.031219 | 0.0266572 | 0.02768016 | -0.001023 | 0.5971567 | 0.99999994 |
| Dda1 | 0.53485083 | 0.32685019 | 0.327874 | -0.0010238 | 0.50829142 | 0.99999994 |
| Zfp24 | 0.2049758 | 0.15898569 | 0.16001 | -0.0010243 | 0.51699412 | 0.99999994 |
| Eno2 | 0.01176552 | 0.00954361 | 0.01056881 | -0.0010252 | 0.74075825 | 0.99999994 |
| Rab11fip5 | 0.0073845 | 0.00562759 | 0.00665308 | -0.0010255 | 0.84755394 | 0.99999994 |
| Eln | 0.00759567 | 0.00581674 | 0.00684235 | -0.0010256 | 0.84079618 | 0.99999994 |
| Palmd | 0.00386993 | 0.00246723 | 0.00349493 | -0.0010277 | 0.97484597 | 0.99999994 |
| Dnm1l | 0.22246444 | 0.17031377 | 0.1713416 | -0.0010278 | 0.5159254 | 0.99999994 |
| Tox4 | 0.31047948 | 0.22270743 | 0.22373553 | -0.0010281 | 0.51220056 | 0.99999994 |
| Msh3 | 0.09630969 | 0.08081504 | 0.08184323 | -0.0010282 | 0.53332197 | 0.99999994 |
| Smad2 | 0.14292218 | 0.11607408 | 0.11710243 | -0.0010284 | 0.52330645 | 0.99999994 |
| Rfx1 | 0.01750897 | 0.01463878 | 0.01566734 | -0.0010286 | 0.66894016 | 0.99999994 |
| Atp8b2 | 0.00726755 | 0.00551942 | 0.00654823 | -0.0010288 | 0.85217373 | 0.99999994 |
| Sv2b | 0.00346291 | 0.00209803 | 0.00312821 | -0.0010302 | 0.98581306 | 0.99999994 |
| Gpld1 | 0.004983 | 0.00346342 | 0.00449675 | -0.0010333 | 0.93694216 | 0.99999994 |
| Neurl2 | 0.00767109 | 0.00587545 | 0.00690994 | -0.0010345 | 0.84050172 | 0.99999994 |
| Erich6 | 0.00353476 | 0.00215809 | 0.00319297 | -0.0010349 | 0.98451302 | 0.99999994 |
| Tcp11l1 | 0.12181527 | 0.10045589 | 0.10149087 | -0.001035 | 0.52705942 | 0.99999994 |
| Nr1h3 | 0.00672607 | 0.00502715 | 0.00606256 | -0.0010354 | 0.87219307 | 0.99999994 |
| 1700054A03 | 0.00426515 | 0.00281281 | 0.00385082 | -0.001038 | 0.96361425 | 0.99999994 |
| Slc18b1 | 0.01738987 | 0.0145239 | 0.01556201 | -0.0010381 | 0.67149034 | 0.99999994 |

| | | | | | | |
|---|---|---|---|---|---|---|
| Ifih1 | 0.00731946 | 0.00555646 | 0.00659476 | -0.0010383 | 0.85267985 | 0.99999994 |
| Gmeb1 | 0.08525143 | 0.07192272 | 0.0729612 | -0.0010385 | 0.53774013 | 0.99999994 |
| Miip | 0.01971312 | 0.01657298 | 0.01761355 | -0.0010406 | 0.65293014 | 0.99999994 |
| Nup54 | 0.10364756 | 0.08657417 | 0.08761536 | -0.0010412 | 0.53152415 | 0.99999994 |
| Zfp81 | 0.05233071 | 0.04471243 | 0.04575517 | -0.0010427 | 0.56028662 | 0.99999994 |
| Syde2 | 0.02608761 | 0.02216458 | 0.02320979 | -0.0010452 | 0.61781848 | 0.99999994 |
| Zfp354c | 0.02362482 | 0.0200081 | 0.02105334 | -0.0010452 | 0.6294777 | 0.99999994 |
| Gm9530 | 0.00300692 | 0.00167118 | 0.00271714 | -0.001046 | 0.99480162 | 0.99999994 |
| Dzip1l | 0.01583054 | 0.0131341 | 0.01418146 | -0.0010474 | 0.68849783 | 0.99999994 |
| Zfp217 | 0.02868866 | 0.02443059 | 0.0254796 | -0.001049 | 0.60797581 | 0.99999994 |
| Abcd4 | 0.01758055 | 0.01468134 | 0.01573064 | -0.0010493 | 0.67148108 | 0.99999994 |
| Enkur | 0.00461215 | 0.00311382 | 0.00416314 | -0.0010493 | 0.95330071 | 0.99999994 |
| Rnf180 | 0.00505908 | 0.00351534 | 0.00456517 | -0.0010498 | 0.93708237 | 0.99999994 |
| Irgm1 | 0.00726323 | 0.0054944 | 0.00654435 | -0.00105 | 0.85722226 | 0.99999994 |
| Agbl3 | 0.02446223 | 0.02073615 | 0.02178738 | -0.0010512 | 0.62595751 | 0.99999994 |
| Mterf2 | 0.05015014 | 0.04286021 | 0.0439115 | -0.0010513 | 0.56330764 | 0.99999994 |
| Gm17300 | 0.00669788 | 0.00498419 | 0.00603727 | -0.0010531 | 0.87719203 | 0.99999994 |
| Gm5786 | 0.00805966 | 0.00620468 | 0.00725805 | -0.0010534 | 0.83298811 | 0.99999994 |
| 2700097O09 | 0.16483698 | 0.13169835 | 0.13275273 | -0.0010544 | 0.52108144 | 0.99999994 |
| Tbp | 0.07964635 | 0.06735586 | 0.06841076 | -0.0010549 | 0.5408766 | 0.99999994 |
| Emp3 | 0.01702834 | 0.01418618 | 0.01524219 | -0.001056 | 0.67766042 | 0.99999994 |
| Cd96 | 0.0031756 | 0.00181316 | 0.00286923 | -0.0010561 | 0.99285695 | 0.99999994 |
| 1700017B05 | 0.01678128 | 0.01396714 | 0.01502354 | -0.0010564 | 0.68012645 | 0.99999994 |
| Cnot11 | 0.20155386 | 0.15669657 | 0.15775476 | -0.0010582 | 0.51780659 | 0.99999994 |
| Serp2 | 0.03227142 | 0.02753491 | 0.02859323 | -0.0010583 | 0.59730271 | 0.99999994 |
| Dclre1a | 0.02513699 | 0.02131978 | 0.02237826 | -0.0010585 | 0.62355891 | 0.99999994 |
| Polm | 0.01398521 | 0.01148503 | 0.01254394 | -0.0010589 | 0.71290972 | 0.99999994 |
| Map2k4 | 0.13824901 | 0.11263301 | 0.11369207 | -0.0010591 | 0.52472049 | 0.99999994 |
| Zkscan4 | 0.00912732 | 0.00715349 | 0.00821358 | -0.0010601 | 0.80485751 | 0.99999994 |
| Spata2 | 0.01162026 | 0.00937921 | 0.01043936 | -0.0010601 | 0.75047038 | 0.99999994 |

| | | | | | | |
|---|---|---|---|---|---|---|
| Odf2l | 0.08831482 | 0.07437376 | 0.07543435 | -0.0010606 | 0.53728133 | 0.99999994 |
| Dok7 | 0.02500042 | 0.02119802 | 0.02225871 | -0.0010607 | 0.62445184 | 0.99999994 |
| Arhgef5 | 0.03048889 | 0.02598512 | 0.02704596 | -0.0010608 | 0.60298697 | 0.99999994 |
| Tpra1 | 0.06203145 | 0.05283254 | 0.05389349 | -0.001061 | 0.55212738 | 0.99999994 |
| Ankrd42 | 0.02165351 | 0.01825668 | 0.01932211 | -0.0010654 | 0.64320233 | 0.99999994 |
| Padi1 | 0.01342416 | 0.01097947 | 0.01204526 | -0.0010658 | 0.72205945 | 0.99999994 |
| Mapkapk5 | 0.01198902 | 0.00970214 | 0.01076796 | -0.0010658 | 0.7450043 | 0.99999994 |
| Rnf123 | 0.03626049 | 0.03097648 | 0.03204253 | -0.0010661 | 0.58763159 | 0.99999994 |
| Efr3b | 0.00480887 | 0.0032739 | 0.00434013 | -0.0010662 | 0.94900096 | 0.99999994 |
| Rps2-ps6 | 0.00905012 | 0.0070782 | 0.00814454 | -0.0010663 | 0.80825763 | 0.99999994 |
| Jrk | 0.01926638 | 0.01615287 | 0.01721955 | -0.0010667 | 0.65995297 | 0.99999994 |
| 4732440D04 | 0.02900046 | 0.02468196 | 0.02575116 | -0.0010692 | 0.60886953 | 0.99999994 |
| Noc3l | 0.05578708 | 0.04759641 | 0.04866672 | -0.0010703 | 0.55819327 | 0.99999994 |
| Gm38102 | 0.0076531 | 0.00582241 | 0.00689382 | -0.0010714 | 0.8495508 | 0.99999994 |
| Pnma2 | 0.00963289 | 0.00759305 | 0.00866557 | -0.0010725 | 0.7949849 | 0.99999994 |
| Slc44a4 | 0.00746579 | 0.00565281 | 0.00672594 | -0.0010731 | 0.85588876 | 0.99999994 |
| Fkbp7 | 0.0425919 | 0.03640641 | 0.03747988 | -0.0010735 | 0.57559907 | 0.99999994 |
| Taf8 | 0.05949211 | 0.0506996 | 0.05177311 | -0.0010735 | 0.55488751 | 0.99999994 |
| Bbs7 | 0.05236171 | 0.04470694 | 0.04578133 | -0.0010744 | 0.56206672 | 0.99999994 |
| Rsph3b | 0.00798564 | 0.00611655 | 0.00719174 | -0.0010752 | 0.84016811 | 0.99999994 |
| St8sia3 | 0.10236099 | 0.08554173 | 0.08661705 | -0.0010753 | 0.53292968 | 0.99999994 |
| Sfxn3 | 0.04211005 | 0.03599137 | 0.03706768 | -0.0010763 | 0.57662899 | 0.99999994 |
| Synpo | 0.0083107 | 0.00640582 | 0.00748285 | -0.001077 | 0.83098502 | 0.99999994 |
| AW011738 | 0.00310668 | 0.00172955 | 0.0028071 | -0.0010776 | 0.99469292 | 0.99999994 |
| Zfp553 | 0.15306402 | 0.12333626 | 0.12441453 | -0.0010783 | 0.52300186 | 0.99999994 |
| Umps | 0.30008914 | 0.21685393 | 0.21793259 | -0.0010787 | 0.51314103 | 0.99999994 |
| Pde6c | 0.00734873 | 0.00554191 | 0.00662101 | -0.0010791 | 0.86100854 | 0.99999994 |
| Mycbpap | 0.01643472 | 0.0136376 | 0.01471671 | -0.0010791 | 0.68725324 | 0.99999994 |
| Kifap3 | 0.23335269 | 0.17715615 | 0.17823531 | -0.0010792 | 0.51607398 | 0.99999994 |
| Ccl25 | 0.07835887 | 0.06628095 | 0.06736089 | -0.0010799 | 0.54249263 | 0.99999994 |

| | | | | | | |
|---|---|---|---|---|---|---|
| Fancf | 0.01055103 | 0.00840538 | 0.00948563 | -0.0010803 | 0.77578849 | 0.99999994 |
| Bend4 | 0.00376563 | 0.00231962 | 0.00340098 | -0.0010814 | 0.98284554 | 0.99999994 |
| Arhgef18 | 0.02719416 | 0.02309484 | 0.02417638 | -0.0010815 | 0.61706268 | 0.99999994 |
| Rnf208 | 0.09231995 | 0.07757125 | 0.07865316 | -0.0010819 | 0.53647656 | 0.99999994 |
| Mbtd1 | 0.21280141 | 0.16403773 | 0.16512002 | -0.0010823 | 0.51740012 | 0.99999994 |
| Gab3 | 0.00259215 | 0.00126056 | 0.002343 | -0.0010824 | 0.99894807 | 0.99999994 |
| Mettl16 | 0.23809166 | 0.18011592 | 0.18119843 | -0.0010825 | 0.51586016 | 0.99999994 |
| Sema3b | 0.00473685 | 0.00319193 | 0.00427534 | -0.0010834 | 0.95417762 | 0.99999994 |
| 4732414G09 | 0.00847489 | 0.00654576 | 0.00762983 | -0.0010841 | 0.82786306 | 0.99999994 |
| Zfyve26 | 0.00922316 | 0.00721342 | 0.00829929 | -0.0010859 | 0.80809736 | 0.99999994 |
| Rabep1 | 0.3016 | 0.21769621 | 0.21878236 | -0.0010861 | 0.51318085 | 0.99999994 |
| Crlf3 | 0.03364966 | 0.02870065 | 0.02978705 | -0.0010864 | 0.59590937 | 0.99999994 |
| Cand2 | 0.02778646 | 0.02360609 | 0.02469318 | -0.0010871 | 0.61525347 | 0.99999994 |
| E230013L22I | 0.00288322 | 0.00151771 | 0.00260558 | -0.0010879 | 0.99727475 | 0.99999994 |
| Naa40 | 0.06440741 | 0.05478138 | 0.05587108 | -0.0010897 | 0.55164754 | 0.99999994 |
| Lipt1 | 0.03056787 | 0.02602478 | 0.0271146 | -0.0010898 | 0.60547282 | 0.99999994 |
| 4833445I07R | 0.00353358 | 0.00210194 | 0.0031919 | -0.00109 | 0.98848712 | 0.99999994 |
| Ccl3 | 0.00748336 | 0.00565066 | 0.00674169 | -0.001091 | 0.85930988 | 0.99999994 |
| Ankrd31 | 0.00661388 | 0.00487052 | 0.0059619 | -0.0010914 | 0.88848301 | 0.99999994 |
| Tssk6 | 0.0027137 | 0.00136095 | 0.00245267 | -0.0010917 | 0.99847593 | 0.99999994 |
| Ciao1 | 0.19930471 | 0.15517368 | 0.15626571 | -0.001092 | 0.5185506 | 0.99999994 |
| Gm14326 | 0.0107694 | 0.00858673 | 0.00968052 | -0.0010938 | 0.77400295 | 0.99999994 |
| A530017D24 | 0.00604791 | 0.00435984 | 0.00545381 | -0.001094 | 0.90909149 | 0.99999994 |
| Ankrd55 | 0.00756137 | 0.00571704 | 0.00681161 | -0.0010946 | 0.85760511 | 0.99999994 |
| 9230117E06I | 0.00335925 | 0.00194019 | 0.00303479 | -0.0010946 | 0.99182123 | 0.99999994 |
| Fcho1 | 0.01327992 | 0.01082039 | 0.011917 | -0.0010966 | 0.72990383 | 0.99999994 |
| Pcdhb14 | 0.02198519 | 0.01851692 | 0.01961371 | -0.0010968 | 0.64513563 | 0.99999994 |
| 4930515G01 | 0.00841677 | 0.00648043 | 0.0075778 | -0.0010974 | 0.83246055 | 0.99999994 |
| Gm17354 | 0.00724312 | 0.00542853 | 0.00652632 | -0.0010978 | 0.86857104 | 0.99999994 |
| Hunk | 0.01239561 | 0.01003218 | 0.01113008 | -0.0010979 | 0.74428133 | 0.99999994 |

| | | | | | | |
|---|---|---|---|---|---|---|
| 2810029C07I | 0.01070548 | 0.00852556 | 0.00962348 | -0.0010979 | 0.77619605 | 0.99999994 |
| Nol9 | 0.05503461 | 0.04693576 | 0.04803398 | -0.0010982 | 0.56048055 | 0.99999994 |
| Smim6 | 0.01547737 | 0.01277 | 0.01386837 | -0.0010984 | 0.70096597 | 0.99999994 |
| Hmox1 | 0.02302855 | 0.01943176 | 0.02053017 | -0.0010984 | 0.63912855 | 0.99999994 |
| Zfp493 | 0.00727852 | 0.00545951 | 0.00655806 | -0.0010986 | 0.86757876 | 0.99999994 |
| Pgk1 | 0.02705023 | 0.02294982 | 0.02405073 | -0.0011009 | 0.61970028 | 0.99999994 |
| Ptprr | 0.00777698 | 0.00590226 | 0.00700482 | -0.0011026 | 0.85261553 | 0.99999994 |
| Samd13 | 0.00537156 | 0.00374352 | 0.00484611 | -0.0011026 | 0.93504499 | 0.99999994 |
| Oacyl | 0.00274122 | 0.00137484 | 0.0024775 | -0.0011027 | 0.99847448 | 0.99999994 |
| Hsd17b10 | 0.50558715 | 0.31524741 | 0.31635049 | -0.0011031 | 0.5092586 | 0.99999994 |
| Trmt5 | 0.0312183 | 0.02657631 | 0.02767955 | -0.0011032 | 0.6046132 | 0.99999994 |
| Gm7967 | 0.00253347 | 0.0011859 | 0.00229005 | -0.0011042 | 0.99933488 | 0.99999994 |
| Srrt | 0.38544874 | 0.26172624 | 0.26283161 | -0.0011054 | 0.5111666 | 0.99999994 |
| Nuak2 | 0.01224928 | 0.00989435 | 0.01099977 | -0.0011054 | 0.74823045 | 0.99999994 |
| Aldh1a1 | 0.00718155 | 0.00536385 | 0.00647112 | -0.0011073 | 0.87264124 | 0.99999994 |
| Cxcl2 | 0.00715557 | 0.00534041 | 0.00644782 | -0.0011074 | 0.87352854 | 0.99999994 |
| Mgp | 0.00451709 | 0.00297017 | 0.0040776 | -0.0011074 | 0.96468072 | 0.99999994 |
| Fbxl22 | 0.00714535 | 0.00533093 | 0.00643866 | -0.0011077 | 0.87393239 | 0.99999994 |
| Gm42715 | 0.01188698 | 0.0095679 | 0.01067705 | -0.0011092 | 0.75536634 | 0.99999994 |
| Pfn4 | 0.00773331 | 0.00585641 | 0.00696569 | -0.0011093 | 0.85543151 | 0.99999994 |
| H2-Aa | 0.00937466 | 0.00732526 | 0.00843474 | -0.0011095 | 0.80936541 | 0.99999994 |
| Prpf39 | 0.09752251 | 0.08170006 | 0.08280974 | -0.0011097 | 0.53553753 | 0.99999994 |
| Phf12 | 0.13154974 | 0.10764747 | 0.10875793 | -0.0011105 | 0.52709267 | 0.99999994 |
| Bcas3 | 0.03236419 | 0.02756301 | 0.02867365 | -0.0011106 | 0.60172926 | 0.99999994 |
| Exo5 | 0.05129068 | 0.04376543 | 0.04487649 | -0.0011111 | 0.56544899 | 0.99999994 |
| Prim2 | 0.02009744 | 0.01684019 | 0.0179523 | -0.0011121 | 0.65995848 | 0.99999994 |
| Gca | 0.06545405 | 0.05562766 | 0.05674029 | -0.0011126 | 0.55192477 | 0.99999994 |
| Ano2 | 0.00369045 | 0.00221993 | 0.00333325 | -0.0011133 | 0.98690097 | 0.99999994 |
| Dyrk2 | 0.02048167 | 0.01717688 | 0.01829081 | -0.0011139 | 0.65740013 | 0.99999994 |
| Zbtb32 | 0.00416949 | 0.00265074 | 0.0037647 | -0.001114 | 0.97555785 | 0.99999994 |

| | | | | | | |
|---|---|---|---|---|---|---|
| Mark1 | 0.0077122 | 0.00583268 | 0.00694677 | -0.0011141 | 0.85713145 | 0.99999994 |
| AV099323 | 0.02486489 | 0.02102518 | 0.02214005 | -0.0011149 | 0.63125657 | 0.99999994 |
| Atl1 | 0.01389982 | 0.01135292 | 0.01246806 | -0.0011151 | 0.72419345 | 0.99999994 |
| 4933406I18R | 0.00519056 | 0.00356656 | 0.0046834 | -0.0011168 | 0.94378011 | 0.99999994 |
| Clip4 | 0.01904303 | 0.01590404 | 0.01702249 | -0.0011184 | 0.66907155 | 0.99999994 |
| Armc2 | 0.00774738 | 0.00585766 | 0.0069783 | -0.0011206 | 0.85745045 | 0.99999994 |
| Ubxn2b | 0.08127073 | 0.06861121 | 0.06973289 | -0.0011217 | 0.54263338 | 0.99999994 |
| Tusc2 | 0.26423186 | 0.19601586 | 0.19713765 | -0.0011218 | 0.51510729 | 0.99999994 |
| 1700108N11 | 0.00569051 | 0.00401057 | 0.00513276 | -0.0011222 | 0.92720649 | 0.99999994 |
| Tspan4 | 0.04404053 | 0.03759523 | 0.03871756 | -0.0011223 | 0.57650156 | 0.99999994 |
| Fbxo34 | 0.07156314 | 0.06066786 | 0.06179021 | -0.0011223 | 0.54811691 | 0.99999994 |
| Anp32e | 0.49062568 | 0.30911983 | 0.3102426 | -0.0011228 | 0.50960932 | 0.99999994 |
| Pop1 | 0.03202285 | 0.02725446 | 0.02837769 | -0.0011232 | 0.60390515 | 0.99999994 |
| Aifm3 | 0.01684283 | 0.01395436 | 0.01507802 | -0.0011237 | 0.6900715 | 0.99999994 |
| Rtn4ip1 | 0.02762088 | 0.02342365 | 0.02454875 | -0.0011251 | 0.61984382 | 0.99999994 |
| Bmp2k | 0.0081574 | 0.00621991 | 0.00734558 | -0.0011257 | 0.84614626 | 0.99999994 |
| Bbs12 | 0.01586526 | 0.0130857 | 0.01421223 | -0.0011265 | 0.70111678 | 0.99999994 |
| 2810030D12I | 0.00483733 | 0.00323803 | 0.00436573 | -0.0011277 | 0.95722795 | 0.99999994 |
| Tnfrsf11a | 0.00194026 | 0.00062539 | 0.00175455 | -0.0011292 | 0.99999081 | 0.99999994 |
| Dhdh | 0.01264522 | 0.01022177 | 0.0113523 | -0.0011305 | 0.7462956 | 0.99999994 |
| Atg12 | 0.21868147 | 0.16778519 | 0.16891737 | -0.0011322 | 0.5177927 | 0.99999994 |
| Gm11110 | 0.01635436 | 0.01351321 | 0.01464555 | -0.0011323 | 0.69659979 | 0.99999994 |
| C820005J03F | 0.00357519 | 0.00209692 | 0.0032294 | -0.0011325 | 0.99020865 | 0.99999994 |
| Cela3a | 0.00712215 | 0.00528479 | 0.00641785 | -0.0011331 | 0.8800354 | 0.99999994 |
| Ino80dos | 0.00650975 | 0.00473425 | 0.00586844 | -0.0011342 | 0.90086021 | 0.99999994 |
| Ccdc65 | 0.01748019 | 0.01450591 | 0.01564189 | -0.001136 | 0.68559864 | 0.99999994 |
| Gm12758 | 0.0087876 | 0.00677141 | 0.00790969 | -0.0011383 | 0.83094666 | 0.99999994 |
| Syn2 | 0.04955263 | 0.04226685 | 0.04340538 | -0.0011385 | 0.56930233 | 0.99999994 |
| Tmem86b | 0.01071441 | 0.0084928 | 0.00963145 | -0.0011387 | 0.78433861 | 0.99999994 |
| Casp8ap2 | 0.17059171 | 0.13563157 | 0.13677061 | -0.001139 | 0.52210403 | 0.99999994 |

| | | | | | | |
|---|---|---|---|---|---|---|
| Emsy | 0.16502852 | 0.13174798 | 0.13288707 | -0.0011391 | 0.52275023 | 0.99999994 |
| Apobec2 | 0.00750708 | 0.00562373 | 0.00676295 | -0.0011392 | 0.86891223 | 0.99999994 |
| Sp3 | 0.16891737 | 0.13446419 | 0.13560545 | -0.0011413 | 0.52233702 | 0.99999994 |
| Gm16537 | 0.00521009 | 0.00355797 | 0.00470096 | -0.001143 | 0.94721349 | 0.99999994 |
| Golm1 | 0.06993983 | 0.05930868 | 0.06045225 | -0.0011436 | 0.55010133 | 0.99999994 |
| Sema3f | 0.01478739 | 0.01211184 | 0.01325628 | -0.0011444 | 0.71723817 | 0.99999994 |
| 2610318N02 | 0.0207842 | 0.01741252 | 0.01855721 | -0.0011447 | 0.65931387 | 0.99999994 |
| Sall2 | 0.01327181 | 0.0107626 | 0.01190978 | -0.0011472 | 0.73929052 | 0.99999994 |
| Chm | 0.09639122 | 0.08076104 | 0.08190825 | -0.0011472 | 0.53713929 | 0.99999994 |
| Wtip | 0.01297152 | 0.0104943 | 0.01164267 | -0.0011484 | 0.74426235 | 0.99999994 |
| Foxo6 | 0.00610283 | 0.00435369 | 0.00550313 | -0.0011494 | 0.91778251 | 0.99999994 |
| Taco1os | 0.00398379 | 0.00244792 | 0.00359748 | -0.0011496 | 0.98329045 | 0.99999994 |
| A330102I10F | 0.00548505 | 0.00379847 | 0.00494813 | -0.0011497 | 0.93901377 | 0.99999994 |
| 2210411M09 | 0.00504674 | 0.00340125 | 0.00455407 | -0.0011528 | 0.95400552 | 0.99999994 |
| Zfp438 | 0.02643158 | 0.02235662 | 0.0235104 | -0.0011538 | 0.62803875 | 0.99999994 |
| Pcna-ps2 | 0.01042168 | 0.00821621 | 0.00937016 | -0.0011539 | 0.79381781 | 0.99999994 |
| Naif1 | 0.00913346 | 0.00706502 | 0.00821907 | -0.001154 | 0.82500886 | 0.99999994 |
| Otub1 | 0.89357088 | 0.41616728 | 0.41732176 | -0.0011545 | 0.50734582 | 0.99999994 |
| Gins3 | 0.01664057 | 0.01374394 | 0.01489898 | -0.001155 | 0.69708442 | 0.99999994 |
| Dhx35 | 0.01938362 | 0.0161678 | 0.01732298 | -0.0011552 | 0.67143444 | 0.99999994 |
| F13a1 | 0.0066348 | 0.0048252 | 0.00598067 | -0.0011555 | 0.90078063 | 0.99999994 |
| Dnah7a | 0.01281815 | 0.01035014 | 0.0115062 | -0.0011561 | 0.74818327 | 0.99999994 |
| Ift22 | 0.53770654 | 0.32781181 | 0.32896903 | -0.0011572 | 0.50934049 | 0.99999994 |
| Ado | 0.08019686 | 0.06770042 | 0.06885915 | -0.0011587 | 0.54459203 | 0.99999994 |
| Dennd6b | 0.04574524 | 0.03901183 | 0.04017099 | -0.0011592 | 0.57615739 | 0.99999994 |
| Pecam1 | 0.00876386 | 0.00672756 | 0.00788844 | -0.0011609 | 0.83634903 | 0.99999994 |
| Gm16253 | 0.00331467 | 0.00183295 | 0.00299461 | -0.0011617 | 0.99509013 | 0.99999994 |
| 2210408I21R | 0.01890017 | 0.01573338 | 0.0168964 | -0.001163 | 0.67658634 | 0.99999994 |
| Scnn1g | 0.00241635 | 0.0010195 | 0.00218436 | -0.0011649 | 0.99980713 | 0.99999994 |
| Zfp770 | 0.0420801 | 0.03587714 | 0.03704205 | -0.0011649 | 0.58290548 | 0.99999994 |

| | | | | | | |
|---|---|---|---|---|---|---|
| Aqp1 | 0.00678591 | 0.00495041 | 0.00611626 | -0.0011658 | 0.8977426 | 0.99999994 |
| Mgmt | 0.04796185 | 0.04089001 | 0.04205599 | -0.001166 | 0.57320647 | 0.99999994 |
| Naip2 | 0.00302877 | 0.00157057 | 0.00273684 | -0.0011663 | 0.9977197 | 0.99999994 |
| Ints9 | 0.0577887 | 0.04918051 | 0.0503468 | -0.0011663 | 0.56127188 | 0.99999994 |
| Mbtps2 | 0.09133059 | 0.07669025 | 0.07785957 | -0.0011693 | 0.53981498 | 0.99999994 |
| Gm10699 | 0.00751822 | 0.0056029 | 0.00677294 | -0.00117 | 0.87490832 | 0.99999994 |
| Camta1 | 0.31100047 | 0.22285356 | 0.22402399 | -0.0011704 | 0.51387096 | 0.99999994 |
| AU022252 | 0.15584549 | 0.12522833 | 0.12639896 | -0.0011706 | 0.52457826 | 0.99999994 |
| Nfatc1 | 0.01906967 | 0.01587512 | 0.017046 | -0.0011709 | 0.67624319 | 0.99999994 |
| 7630403G23 | 0.0036326 | 0.00210998 | 0.00328113 | -0.0011712 | 0.99124661 | 0.99999994 |
| Helb | 0.01995646 | 0.01665598 | 0.01782806 | -0.0011721 | 0.66916623 | 0.99999994 |
| Prr5l | 0.00512182 | 0.0034492 | 0.00462159 | -0.0011724 | 0.95434816 | 0.99999994 |
| A930019D19 | 0.0110897 | 0.00879353 | 0.00996628 | -0.0011727 | 0.78326377 | 0.99999994 |
| Myh7 | 0.00640784 | 0.00460408 | 0.00577697 | -0.0011729 | 0.91172083 | 0.99999994 |
| Fdxacb1 | 0.04230163 | 0.0360567 | 0.0372316 | -0.0011749 | 0.5831863 | 0.99999994 |
| Map3k14 | 0.01711723 | 0.01414554 | 0.01532084 | -0.0011753 | 0.69519279 | 0.99999994 |
| Tmem59l | 0.00648559 | 0.00466898 | 0.00584676 | -0.0011778 | 0.91001811 | 0.99999994 |
| Zfp784 | 0.03387768 | 0.02880597 | 0.02998435 | -0.0011784 | 0.60318237 | 0.99999994 |
| Sh2d5 | 0.01092442 | 0.00864021 | 0.00981884 | -0.0011786 | 0.78785899 | 0.99999994 |
| Cript | 0.68797458 | 0.37791412 | 0.37909303 | -0.0011789 | 0.5082575 | 0.99999994 |
| Pus3 | 0.09098679 | 0.07640332 | 0.07758357 | -0.0011802 | 0.54032818 | 0.99999994 |
| Rnf214 | 0.17357607 | 0.13765928 | 0.13883965 | -0.0011804 | 0.52256413 | 0.99999994 |
| Lrrc27 | 0.00518297 | 0.00349594 | 0.00467657 | -0.0011806 | 0.95356472 | 0.99999994 |
| Tcte2 | 0.01590999 | 0.013071 | 0.01425187 | -0.0011809 | 0.70936448 | 0.99999994 |
| Akap6 | 0.00397038 | 0.00240446 | 0.00358541 | -0.001181 | 0.98582658 | 0.99999994 |
| Kctd9 | 0.03234351 | 0.0274744 | 0.02865572 | -0.0011813 | 0.60811349 | 0.99999994 |
| Map7d2 | 0.03329772 | 0.02830073 | 0.02948241 | -0.0011817 | 0.6051853 | 0.99999994 |
| Ddx59 | 0.03894062 | 0.03316655 | 0.03434978 | -0.0011832 | 0.59067698 | 0.99999994 |
| Zfp239 | 0.01740015 | 0.01438718 | 0.01557111 | -0.0011839 | 0.69360979 | 0.99999994 |
| Rbm4 | 0.03208285 | 0.0272456 | 0.02842972 | -0.0011841 | 0.60920265 | 0.99999994 |

| | | | | | | |
|---|---|---|---|---|---|---|
| Gm26982 | 0.00528626 | 0.00358425 | 0.00476943 | -0.0011852 | 0.95094463 | 0.99999994 |
| Pnkp | 0.08666265 | 0.07291455 | 0.07410172 | -0.0011872 | 0.54246307 | 0.99999994 |
| Slc39a14 | 0.01909364 | 0.01587956 | 0.01706715 | -0.0011876 | 0.67837944 | 0.99999994 |
| Cntn4 | 0.0092967 | 0.00717575 | 0.00836505 | -0.0011893 | 0.82801943 | 0.99999994 |
| Tyrobp | 0.01844698 | 0.01530579 | 0.01649628 | -0.0011905 | 0.68451895 | 0.99999994 |
| Myrf | 0.06600939 | 0.05601009 | 0.05720101 | -0.0011909 | 0.55511066 | 0.99999994 |
| Zfp37 | 0.05659904 | 0.04815774 | 0.04934877 | -0.001191 | 0.56381628 | 0.99999994 |
| Me3 | 0.00863889 | 0.00658554 | 0.00777661 | -0.0011911 | 0.84601665 | 0.99999994 |
| Ankrd35 | 0.00351598 | 0.00198368 | 0.00317604 | -0.0011924 | 0.99377194 | 0.99999994 |
| Rdh12 | 0.00275673 | 0.00129901 | 0.00249148 | -0.0011925 | 0.99927837 | 0.99999994 |
| Zfp40 | 0.02772901 | 0.02345039 | 0.02464307 | -0.0011927 | 0.62633267 | 0.99999994 |
| Ubiad1 | 0.03366854 | 0.02860911 | 0.02980339 | -0.0011943 | 0.60516211 | 0.99999994 |
| Gstcd | 0.01756307 | 0.01452018 | 0.01571518 | -0.001195 | 0.6936262 | 0.99999994 |
| Tanc1 | 0.02954197 | 0.02502687 | 0.02622253 | -0.0011957 | 0.61924809 | 0.99999994 |
| Als2 | 0.01074589 | 0.00846344 | 0.00965954 | -0.0011961 | 0.79509636 | 0.99999994 |
| Inpp5j | 0.00467821 | 0.0030262 | 0.00422257 | -0.0011964 | 0.97034752 | 0.99999994 |
| Gm44148 | 0.00413259 | 0.00253467 | 0.00373148 | -0.0011968 | 0.98361585 | 0.99999994 |
| Gpr137b | 0.03505081 | 0.02980159 | 0.03099848 | -0.0011969 | 0.60141379 | 0.99999994 |
| Qpct | 0.02027088 | 0.01690738 | 0.01810512 | -0.0011977 | 0.6701585 | 0.99999994 |
| Wdr5b | 0.03827972 | 0.03258283 | 0.03378159 | -0.0011988 | 0.59336347 | 0.99999994 |
| C1qb | 0.01906129 | 0.0158387 | 0.0170386 | -0.0011999 | 0.68037738 | 0.99999994 |
| Gypa | 0.00473011 | 0.00306837 | 0.00426927 | -0.0012009 | 0.96942169 | 0.99999994 |
| Cracr2b | 0.02330367 | 0.01957071 | 0.02077161 | -0.0012009 | 0.64982042 | 0.99999994 |
| Uhrf2 | 0.43852045 | 0.28656641 | 0.28776899 | -0.0012026 | 0.51109582 | 0.99999994 |
| Agap3 | 0.09244904 | 0.07755322 | 0.07875663 | -0.0012034 | 0.5405063 | 0.99999994 |
| 4933440M02 | 0.00331614 | 0.00179001 | 0.00299593 | -0.0012059 | 0.99631131 | 0.99999994 |
| Gm7160 | 0.01454044 | 0.01183099 | 0.01303707 | -0.0012061 | 0.73098447 | 0.99999994 |
| Sh3d19 | 0.06921515 | 0.05864781 | 0.05985405 | -0.0012062 | 0.55335613 | 0.99999994 |
| Ap3s1 | 0.87321839 | 0.41361519 | 0.4148233 | -0.0012081 | 0.5077333 | 0.99999994 |
| Snhg11 | 0.00664959 | 0.00478566 | 0.00599394 | -0.0012083 | 0.91017108 | 0.99999994 |

| | | | | | | |
|---|---|---|---|---|---|---|
| Pcnt | 0.0534137 | 0.04545556 | 0.04666886 | -0.0012133 | 0.5686949 | 0.99999994 |
| Rbm43 | 0.01631205 | 0.01339461 | 0.01460807 | -0.0012135 | 0.70984459 | 0.99999994 |
| Vps26a | 0.12178737 | 0.10025131 | 0.10146988 | -0.0012186 | 0.53185652 | 0.99999994 |
| Ang | 0.14026388 | 0.11394701 | 0.11516563 | -0.0012186 | 0.52807596 | 0.99999994 |
| Angptl7 | 0.00403451 | 0.00242361 | 0.00364316 | -0.0012196 | 0.98706746 | 0.99999994 |
| Zkscan3 | 0.29443638 | 0.21351122 | 0.21473375 | -0.0012225 | 0.51511481 | 0.99999994 |
| Gpatch2l | 0.0194701 | 0.01617654 | 0.01739925 | -0.0012227 | 0.68002486 | 0.99999994 |
| Slc44a2 | 0.02307378 | 0.01934665 | 0.02056987 | -0.0012232 | 0.65388476 | 0.99999994 |
| Cdh4 | 0.01423992 | 0.01154673 | 0.01277021 | -0.0012235 | 0.73817387 | 0.99999994 |
| Isy1 | 0.10145421 | 0.08468802 | 0.08591218 | -0.0012242 | 0.5377815 | 0.99999994 |
| Cox6a2 | 0.00240708 | 0.00095079 | 0.002176 | -0.0012252 | 0.99991084 | 0.99999994 |
| Nras | 0.33360849 | 0.23508227 | 0.23630803 | -0.0012258 | 0.51377167 | 0.99999994 |
| Tnip2 | 0.03363739 | 0.02855046 | 0.02977643 | -0.001226 | 0.60798028 | 0.99999994 |
| U2af1l4 | 0.01133808 | 0.00896123 | 0.01018779 | -0.0012266 | 0.78854848 | 0.99999994 |
| Mettl5os | 0.00679547 | 0.00489817 | 0.00612483 | -0.0012267 | 0.90875313 | 0.99999994 |
| Lars2 | 0.03247243 | 0.02753943 | 0.02876748 | -0.001228 | 0.61185307 | 0.99999994 |
| Slx4 | 0.01508821 | 0.01229492 | 0.01352321 | -0.0012283 | 0.72727301 | 0.99999994 |
| Zfp2 | 0.03651004 | 0.03102658 | 0.03225772 | -0.0012311 | 0.60026956 | 0.99999994 |
| Pde4b | 0.02000101 | 0.01663585 | 0.01786732 | -0.0012315 | 0.67680263 | 0.99999994 |
| Stk35 | 0.03177485 | 0.02693102 | 0.02816257 | -0.0012315 | 0.61450547 | 0.99999994 |
| Upf2 | 0.30252022 | 0.2180669 | 0.21929893 | -0.001232 | 0.51491517 | 0.99999994 |
| Nkrf | 0.0109484 | 0.00860632 | 0.00984023 | -0.0012339 | 0.79804842 | 0.99999994 |
| Tnfaip8l1 | 0.00757141 | 0.00558609 | 0.00682061 | -0.0012345 | 0.88585781 | 0.99999994 |
| Arntl | 0.02698909 | 0.0227628 | 0.02399736 | -0.0012346 | 0.63398987 | 0.99999994 |
| Nupl2 | 0.03784769 | 0.03217471 | 0.0334099 | -0.0012352 | 0.59719359 | 0.99999994 |
| Pou5f2 | 0.01447075 | 0.01173866 | 0.01297519 | -0.0012365 | 0.73707419 | 0.99999994 |
| Supt3 | 0.0868518 | 0.07301527 | 0.07425443 | -0.0012392 | 0.54422424 | 0.99999994 |
| Matk | 0.00926869 | 0.00710077 | 0.00833999 | -0.0012392 | 0.8386823 | 0.99999994 |
| Dhx34 | 0.04853065 | 0.04129821 | 0.0425388 | -0.0012406 | 0.5769605 | 0.99999994 |
| 2810414N06 | 0.00762691 | 0.00562963 | 0.00687035 | -0.0012407 | 0.88533319 | 0.99999994 |

| | | | | | | |
|---|---|---|---|---|---|---|
| Dock1 | 0.01947616 | 0.01616359 | 0.0174046 | -0.001241 | 0.68247121 | 0.99999994 |
| Epb41l4aos | 0.17147567 | 0.13614178 | 0.13738449 | -0.0012427 | 0.52400576 | 0.99999994 |
| Actn2 | 0.00596082 | 0.00413195 | 0.00537559 | -0.0012436 | 0.93821293 | 0.99999994 |
| Gm12996 | 0.00614803 | 0.00429992 | 0.00554371 | -0.0012438 | 0.93233754 | 0.99999994 |
| Cacna1e | 0.00655268 | 0.0046631 | 0.00590697 | -0.0012439 | 0.9194931 | 0.99999994 |
| 6430571L13F | 0.01310396 | 0.01051566 | 0.01176049 | -0.0012448 | 0.75945947 | 0.99999994 |
| Lrrc28 | 0.04772784 | 0.04061113 | 0.04185725 | -0.0012461 | 0.57854152 | 0.99999994 |
| Cdca4 | 0.0975778 | 0.08160687 | 0.08285376 | -0.0012469 | 0.53989669 | 0.99999994 |
| Zfp28 | 0.01503086 | 0.01222441 | 0.01347233 | -0.0012479 | 0.73124025 | 0.99999994 |
| Pfkp | 0.00710647 | 0.00515566 | 0.00640378 | -0.0012481 | 0.90274521 | 0.99999994 |
| Zfp595 | 0.00718209 | 0.00522281 | 0.0064716 | -0.0012488 | 0.90050624 | 0.99999994 |
| E2f2 | 0.00762074 | 0.00561538 | 0.00686482 | -0.0012494 | 0.88714795 | 0.99999994 |
| Dmtn | 0.00608502 | 0.00423766 | 0.00548713 | -0.0012495 | 0.93520499 | 0.99999994 |
| Lpin3 | 0.01229624 | 0.00979204 | 0.0110416 | -0.0012496 | 0.77436189 | 0.99999994 |
| Relb | 0.01463513 | 0.01187136 | 0.01312113 | -0.0012498 | 0.73696217 | 0.99999994 |
| Impdh2 | 0.48954152 | 0.30854427 | 0.30979423 | -0.00125 | 0.51071321 | 0.99999994 |
| Camsap3 | 0.04714486 | 0.04010976 | 0.04136186 | -0.0012521 | 0.57984634 | 0.99999994 |
| Gm37305 | 0.01201978 | 0.00954211 | 0.01079536 | -0.0012533 | 0.78016426 | 0.99999994 |
| Dzip3 | 0.11579451 | 0.09568495 | 0.09693835 | -0.0012534 | 0.53429303 | 0.99999994 |
| D730045B01 | 0.01388423 | 0.01120017 | 0.01245422 | -0.001254 | 0.74864907 | 0.99999994 |
| Rbm10 | 0.23914141 | 0.18059702 | 0.18185152 | -0.0012545 | 0.51831245 | 0.99999994 |
| Acaa1b | 0.01577126 | 0.01287435 | 0.01412891 | -0.0012546 | 0.72275498 | 0.99999994 |
| Ush2a | 0.00705686 | 0.00510388 | 0.0063593 | -0.0012554 | 0.90558625 | 0.99999994 |
| Vamp1 | 0.0136252 | 0.01096775 | 0.01222399 | -0.0012562 | 0.75303366 | 0.99999994 |
| Hace1 | 0.03796989 | 0.03225859 | 0.03351505 | -0.0012565 | 0.59852963 | 0.99999994 |
| Parp11 | 0.04376885 | 0.03722882 | 0.03848562 | -0.0012568 | 0.58604086 | 0.99999994 |
| Slc8a2 | 0.00806808 | 0.00600874 | 0.00726558 | -0.0012568 | 0.87522708 | 0.99999994 |
| Coro1a | 0.01560044 | 0.01272047 | 0.01397749 | -0.001257 | 0.72528666 | 0.99999994 |
| Dapl1 | 0.00701071 | 0.00506084 | 0.0063179 | -0.0012571 | 0.90731567 | 0.99999994 |
| Apex2 | 0.03799586 | 0.03227923 | 0.03353739 | -0.0012582 | 0.59859624 | 0.99999994 |

| | | | | | | |
|---|---|---|---|---|---|---|
| Tctn1 | 0.03305126 | 0.0280104 | 0.02926899 | -0.0012586 | 0.61264892 | 0.99999994 |
| Aven | 0.02207182 | 0.01842983 | 0.01968985 | -0.00126 | 0.66493266 | 0.99999994 |
| Gm11940 | 0.00825846 | 0.00617535 | 0.00743607 | -0.0012607 | 0.87045247 | 0.99999994 |
| Pafah2 | 0.02459704 | 0.0206446 | 0.02190548 | -0.0012609 | 0.6491918 | 0.99999994 |
| Arhgap23 | 0.01391549 | 0.01121993 | 0.01248199 | -0.0012621 | 0.74953453 | 0.99999994 |
| Ccdc167 | 0.33461995 | 0.23558431 | 0.2368471 | -0.0012628 | 0.51415519 | 0.99999994 |
| Ppp1r1b | 0.01065863 | 0.00831786 | 0.00958167 | -0.0012638 | 0.81001872 | 0.99999994 |
| Zfp937 | 0.03714749 | 0.0315417 | 0.03280703 | -0.0012653 | 0.60130585 | 0.99999994 |
| Car2 | 0.00862209 | 0.00649584 | 0.00776158 | -0.0012657 | 0.86115124 | 0.99999994 |
| Plekha8 | 0.06413481 | 0.05437853 | 0.05564449 | -0.001266 | 0.56018484 | 0.99999994 |
| Hdac7 | 0.03686592 | 0.03129617 | 0.03256445 | -0.0012683 | 0.60227648 | 0.99999994 |
| Crat | 0.06070718 | 0.05151997 | 0.05278859 | -0.0012686 | 0.56354611 | 0.99999994 |
| Rnf25 | 0.17036705 | 0.13534564 | 0.13661445 | -0.0012688 | 0.52464748 | 0.99999994 |
| Lrtm2 | 0.00863832 | 0.00650636 | 0.0077761 | -0.0012697 | 0.86145934 | 0.99999994 |
| Frs2 | 0.03727577 | 0.03164757 | 0.03291753 | -0.00127 | 0.60133402 | 0.99999994 |
| 9330104G04 | 0.0301821 | 0.02550833 | 0.02677929 | -0.001271 | 0.6239668 | 0.99999994 |
| Clstn2 | 0.01304853 | 0.0104395 | 0.01171119 | -0.0012717 | 0.7650975 | 0.99999994 |
| Fdps | 0.45368735 | 0.29323577 | 0.2945089 | -0.0012731 | 0.51147779 | 0.99999994 |
| Zfp955a | 0.02706152 | 0.02278673 | 0.02406059 | -0.0012739 | 0.6377355 | 0.99999994 |
| Gm6576 | 0.01411251 | 0.01138296 | 0.01265703 | -0.0012741 | 0.74858354 | 0.99999994 |
| Tjp3 | 0.02599717 | 0.02185629 | 0.02313072 | -0.0012744 | 0.64309472 | 0.99999994 |
| Slc26a6 | 0.00802184 | 0.00594753 | 0.00722417 | -0.0012766 | 0.88026233 | 0.99999994 |
| Gm17491 | 0.01861906 | 0.01537155 | 0.01664824 | -0.0012767 | 0.69513249 | 0.99999994 |
| Ddhd1 | 0.04409188 | 0.03748446 | 0.03876139 | -0.0012769 | 0.58678494 | 0.99999994 |
| Zfp948 | 0.04937174 | 0.04197468 | 0.04325209 | -0.0012774 | 0.57792566 | 0.99999994 |
| Bahcc1 | 0.04787686 | 0.040705 | 0.04198381 | -0.0012788 | 0.58033487 | 0.99999994 |
| Cnrip1 | 0.01386455 | 0.01115769 | 0.01243673 | -0.001279 | 0.75319049 | 0.99999994 |
| Tspan18 | 0.01428401 | 0.01152974 | 0.01280937 | -0.0012796 | 0.7469599 | 0.99999994 |
| Tgfa | 0.01050126 | 0.00816126 | 0.0094412 | -0.0012799 | 0.81657668 | 0.99999994 |
| Ndst2 | 0.02996801 | 0.02531276 | 0.02659313 | -0.0012804 | 0.62569828 | 0.99999994 |

| | | | | | | |
|---|---|---|---|---|---|---|
| Itln1 | 0.00472563 | 0.00298468 | 0.00426524 | -0.0012806 | 0.97716619 | 0.99999994 |
| Gtf2e1 | 0.06528925 | 0.05532233 | 0.05660351 | -0.0012812 | 0.55987888 | 0.99999994 |
| Krba1 | 0.03106354 | 0.02626346 | 0.02754517 | -0.0012817 | 0.62161572 | 0.99999994 |
| Nxt2 | 0.04355181 | 0.03701841 | 0.03830027 | -0.0012819 | 0.58814641 | 0.99999994 |
| Fan1 | 0.01690766 | 0.01385047 | 0.0151354 | -0.0012849 | 0.71399577 | 0.99999994 |
| Pum1 | 0.4227478 | 0.27929621 | 0.28058131 | -0.0012851 | 0.5121607 | 0.99999994 |
| Top2a | 0.01551036 | 0.01261139 | 0.01389762 | -0.0012862 | 0.73106881 | 0.99999994 |
| Fbxw4 | 0.03593548 | 0.0304757 | 0.03176219 | -0.0012865 | 0.6062687 | 0.99999994 |
| Rsbn1 | 0.20142016 | 0.15637975 | 0.15766639 | -0.0012866 | 0.52165961 | 0.99999994 |
| Dab2ip | 0.09325862 | 0.0781168 | 0.07940515 | -0.0012884 | 0.54300199 | 0.99999994 |
| Trim21 | 0.01263034 | 0.01004909 | 0.01133905 | -0.00129 | 0.77554891 | 0.99999994 |
| Mllt1 | 0.01810057 | 0.01489938 | 0.01619028 | -0.0012909 | 0.70219733 | 0.99999994 |
| Dsg2 | 0.10047013 | 0.08385261 | 0.08514605 | -0.0012934 | 0.54027067 | 0.99999994 |
| Stau1 | 0.25135661 | 0.18807776 | 0.18937129 | -0.0012935 | 0.51813245 | 0.99999994 |
| Tpsab1 | 0.00890551 | 0.00672134 | 0.00801518 | -0.0012938 | 0.85869853 | 0.99999994 |
| Usp6nl | 0.01560038 | 0.0126835 | 0.01397744 | -0.0012939 | 0.73111935 | 0.99999994 |
| Sirt6 | 0.13118266 | 0.10719128 | 0.10848601 | -0.0012947 | 0.53165889 | 0.99999994 |
| Platr26 | 0.00426243 | 0.00255335 | 0.00384837 | -0.001295 | 0.98745261 | 0.99999994 |
| Tph1 | 0.00763974 | 0.00558671 | 0.00688184 | -0.0012951 | 0.8948413 | 0.99999994 |
| St3gal2 | 0.01399967 | 0.01126065 | 0.01255678 | -0.0012961 | 0.75398415 | 0.99999994 |
| Pianp | 0.04350532 | 0.03696436 | 0.03826056 | -0.0012962 | 0.58920685 | 0.99999994 |
| Oxsm | 0.02320884 | 0.0193918 | 0.0206884 | -0.0012966 | 0.66172387 | 0.99999994 |
| Armc7 | 0.01573603 | 0.01279943 | 0.01409768 | -0.0012983 | 0.73005566 | 0.99999994 |
| Angel1 | 0.00812626 | 0.00601868 | 0.00731769 | -0.001299 | 0.88131945 | 0.99999994 |
| Mov10 | 0.1039708 | 0.08656636 | 0.08786585 | -0.0012995 | 0.53921005 | 0.99999994 |
| Nsun4 | 0.05415806 | 0.04599566 | 0.04729612 | -0.0013005 | 0.57261086 | 0.99999994 |
| Zfp316 | 0.01932334 | 0.01596867 | 0.0172698 | -0.0013011 | 0.69198974 | 0.99999994 |
| Lmntd2 | 0.01087632 | 0.00847304 | 0.00977593 | -0.0013029 | 0.81249418 | 0.99999994 |
| Ccnyl1 | 0.01064583 | 0.00826618 | 0.00957024 | -0.0013041 | 0.81779922 | 0.99999994 |
| Cdh8 | 0.00399574 | 0.00230414 | 0.00360824 | -0.0013041 | 0.99193084 | 0.99999994 |

| | | | | | | |
|---|---|---|---|---|---|---|
| R3hcc1l | 0.05760338 | 0.04888707 | 0.05019143 | -0.0013044 | 0.56866778 | 0.99999994 |
| Zfp563 | 0.02018736 | 0.01672419 | 0.01803154 | -0.0013073 | 0.68531327 | 0.99999994 |
| Saxo2 | 0.04134949 | 0.03510825 | 0.03641651 | -0.0013083 | 0.59449748 | 0.99999994 |
| Ftsj3 | 0.14051883 | 0.11404326 | 0.11535175 | -0.0013085 | 0.53009406 | 0.99999994 |
| Zfp839 | 0.02402656 | 0.02009623 | 0.02140559 | -0.0013094 | 0.65805609 | 0.99999994 |
| Tnnt1 | 0.0203986 | 0.01690773 | 0.01821764 | -0.0013099 | 0.6838934 | 0.99999994 |
| Spaca1 | 0.00491639 | 0.00312606 | 0.00443684 | -0.0013108 | 0.97537955 | 0.99999994 |
| Atp11b | 0.16706154 | 0.1329987 | 0.13431031 | -0.0013116 | 0.52591414 | 0.99999994 |
| Tacc3 | 0.02414013 | 0.02019283 | 0.02150514 | -0.0013123 | 0.65769867 | 0.99999994 |
| Mta3 | 0.23503422 | 0.17797601 | 0.1792893 | -0.0013133 | 0.51944371 | 0.99999994 |
| Zp2 | 0.00313094 | 0.00151552 | 0.00282897 | -0.0013135 | 0.99900079 | 0.99999994 |
| Slc12a5 | 0.00419715 | 0.0024759 | 0.0037896 | -0.0013137 | 0.98948641 | 0.99999994 |
| Lats2 | 0.01545834 | 0.01253627 | 0.0138515 | -0.0013152 | 0.73631695 | 0.99999994 |
| D630036H23 | 0.00776511 | 0.00567869 | 0.00699418 | -0.0013155 | 0.89470616 | 0.99999994 |
| Wdr53 | 0.08755152 | 0.07350305 | 0.07481903 | -0.001316 | 0.54660081 | 0.99999994 |
| Gm12524 | 0.01067038 | 0.00827531 | 0.00959215 | -0.0013168 | 0.81958948 | 0.99999994 |
| Zfp943 | 0.06316004 | 0.05351617 | 0.05483361 | -0.0013174 | 0.56353055 | 0.99999994 |
| Jade2 | 0.00549232 | 0.00363699 | 0.00495466 | -0.0013177 | 0.96165574 | 0.99999994 |
| Tango6 | 0.01280995 | 0.01018103 | 0.0114989 | -0.0013179 | 0.77723091 | 0.99999994 |
| Lrguk | 0.00559053 | 0.00372475 | 0.00504292 | -0.0013182 | 0.95906201 | 0.99999994 |
| Ltb | 0.00744875 | 0.00539164 | 0.00671067 | -0.001319 | 0.9046233 | 0.99999994 |
| Thg1l | 0.04146344 | 0.03519398 | 0.03651411 | -0.0013201 | 0.59508878 | 0.99999994 |
| Ophn1 | 0.04048903 | 0.03435843 | 0.03567902 | -0.0013206 | 0.5973031 | 0.99999994 |
| Cpne8 | 0.02363478 | 0.01974079 | 0.02106207 | -0.0013213 | 0.66186917 | 0.99999994 |
| Tcea1 | 0.37550747 | 0.25659482 | 0.25791708 | -0.0013223 | 0.5136113 | 0.99999994 |
| Rnf24 | 0.01059649 | 0.0082039 | 0.00952621 | -0.0013223 | 0.82224455 | 0.99999994 |
| Snx24 | 0.11394461 | 0.09420805 | 0.09553043 | -0.0013224 | 0.53670667 | 0.99999994 |
| Hirip3 | 0.08609864 | 0.07232278 | 0.07364615 | -0.0013234 | 0.54760378 | 0.99999994 |
| Sacs | 0.01268179 | 0.01006131 | 0.01138485 | -0.0013235 | 0.78048538 | 0.99999994 |
| Cep70 | 0.21306833 | 0.1639696 | 0.16529317 | -0.0013236 | 0.52125372 | 0.99999994 |

| | | | | | | |
|---|---|---|---|---|---|---|
| Mrps6 | 0.17922418 | 0.14140293 | 0.14272785 | -0.0013249 | 0.5246348 | 0.99999994 |
| Slc20a2 | 0.02260149 | 0.01882928 | 0.0201552 | -0.0013259 | 0.66926839 | 0.99999994 |
| Prss1 | 0.00568417 | 0.00380076 | 0.00512706 | -0.0013263 | 0.95745708 | 0.99999994 |
| Samd9l | 0.0154916 | 0.01255315 | 0.01388099 | -0.0013278 | 0.73785251 | 0.99999994 |
| Macrod1 | 0.12066941 | 0.09929762 | 0.10062787 | -0.0013303 | 0.53505922 | 0.99999994 |
| Scml4 | 0.00715771 | 0.00511828 | 0.00644974 | -0.0013315 | 0.91529604 | 0.99999994 |
| Slc35e3 | 0.01498905 | 0.01210322 | 0.01343524 | -0.001332 | 0.74535238 | 0.99999994 |
| Nbl1 | 0.01653093 | 0.01346983 | 0.0148019 | -0.0013321 | 0.72542358 | 0.99999994 |
| Birc3 | 0.00819405 | 0.00604597 | 0.0073784 | -0.0013324 | 0.88532531 | 0.99999994 |
| Telo2 | 0.04289665 | 0.03640754 | 0.03774045 | -0.0013329 | 0.59293016 | 0.99999994 |
| Btrc | 0.25082938 | 0.18771589 | 0.18904973 | -0.0013338 | 0.51872904 | 0.99999994 |
| St3gal3 | 0.02712313 | 0.02278034 | 0.02411438 | -0.001334 | 0.64365182 | 0.99999994 |
| Gm10399 | 0.03039735 | 0.02563235 | 0.02696641 | -0.0013341 | 0.62903446 | 0.99999994 |
| Vasn | 0.03758123 | 0.03184634 | 0.03318054 | -0.0013342 | 0.60551768 | 0.99999994 |
| Kcnk3 | 0.0330626 | 0.02794313 | 0.02927881 | -0.0013357 | 0.61930612 | 0.99999994 |
| Aoc2 | 0.00601027 | 0.00408328 | 0.00542001 | -0.0013367 | 0.94966819 | 0.99999994 |
| Trh | 0.0082844 | 0.00612173 | 0.0074593 | -0.0013376 | 0.88368089 | 0.99999994 |
| Arhgap24 | 0.01515524 | 0.01224222 | 0.01358268 | -0.0013405 | 0.74438014 | 0.99999994 |
| Ttc33 | 0.27226365 | 0.20055975 | 0.20190113 | -0.0013414 | 0.5176368 | 0.99999994 |
| Gpbp1l1 | 0.21965086 | 0.16819796 | 0.16953998 | -0.001342 | 0.52101034 | 0.99999994 |
| Scarb1 | 0.11158489 | 0.09238564 | 0.09372826 | -0.0013426 | 0.53798144 | 0.99999994 |
| Masp2 | 0.00835823 | 0.00618186 | 0.0075254 | -0.0013435 | 0.88265866 | 0.99999994 |
| Seh1l | 0.20912769 | 0.16138558 | 0.16272933 | -0.0013438 | 0.52191708 | 0.99999994 |
| Zdhhc9 | 0.01931522 | 0.01591883 | 0.01726264 | -0.0013438 | 0.69782782 | 0.99999994 |
| Zfp551 | 0.01447577 | 0.01163418 | 0.01297965 | -0.0013455 | 0.75490237 | 0.99999994 |
| Pcdhga12 | 0.02223416 | 0.01848686 | 0.01983252 | -0.0013457 | 0.67423562 | 0.99999994 |
| Aldh7a1 | 0.19693463 | 0.15334426 | 0.15469077 | -0.0013465 | 0.52310206 | 0.99999994 |
| Wdr76 | 0.01846598 | 0.01516634 | 0.01651306 | -0.0013467 | 0.70638733 | 0.99999994 |
| Fam81a | 0.00998519 | 0.00763273 | 0.00898036 | -0.0013476 | 0.84107311 | 0.99999994 |
| Tbc1d4 | 0.0092904 | 0.00701177 | 0.00835941 | -0.0013476 | 0.85838449 | 0.99999994 |

| | | | | | | |
|---|---|---|---|---|---|---|
| Gm10762 | 0.04230453 | 0.03588543 | 0.03723408 | -0.0013487 | 0.59526096 | 0.99999994 |
| Eepd1 | 0.01856066 | 0.01524686 | 0.01659667 | -0.0013498 | 0.70587211 | 0.99999994 |
| Il4ra | 0.01131176 | 0.00881403 | 0.01016432 | -0.0013503 | 0.81172582 | 0.99999994 |
| Cep97 | 0.05316048 | 0.0451026 | 0.04645535 | -0.0013527 | 0.57684482 | 0.99999994 |
| Ccdc117 | 0.07209906 | 0.06087796 | 0.0622313 | -0.0013533 | 0.55754785 | 0.99999994 |
| Mknk1 | 0.08977478 | 0.0752561 | 0.0766096 | -0.0013535 | 0.54680819 | 0.99999994 |
| Rab29 | 0.0203403 | 0.01681267 | 0.01816628 | -0.0013536 | 0.69004579 | 0.99999994 |
| Tubgcp6 | 0.01029995 | 0.00790784 | 0.00926147 | -0.0013536 | 0.83469159 | 0.99999994 |
| Zdhhc14 | 0.01431956 | 0.01148688 | 0.01284094 | -0.0013541 | 0.75863002 | 0.99999994 |
| Zfp592 | 0.03345388 | 0.0282625 | 0.0296176 | -0.0013551 | 0.61964491 | 0.99999994 |
| Diaph2 | 0.02897699 | 0.02437495 | 0.02573072 | -0.0013558 | 0.63710445 | 0.99999994 |
| Sh3pxd2b | 0.01251319 | 0.00987863 | 0.01123477 | -0.0013561 | 0.78915253 | 0.99999994 |
| Dph7 | 0.02697152 | 0.02262563 | 0.02398202 | -0.0013564 | 0.6467175 | 0.99999994 |
| Nagpa | 0.03451655 | 0.02917842 | 0.03053682 | -0.0013584 | 0.61642417 | 0.99999994 |
| Sparcl1 | 0.00932029 | 0.00702759 | 0.00838613 | -0.0013585 | 0.85955425 | 0.99999994 |
| Coq5 | 0.22249862 | 0.16999991 | 0.17136343 | -0.0013635 | 0.52111973 | 0.99999994 |
| Zfp575 | 0.01838294 | 0.01507569 | 0.01643972 | -0.001364 | 0.7096258 | 0.99999994 |
| Acox3 | 0.02493813 | 0.02083705 | 0.02220417 | -0.0013671 | 0.65903618 | 0.99999994 |
| Camsap2 | 0.08040662 | 0.06766247 | 0.06902991 | -0.0013674 | 0.55245188 | 0.99999994 |
| Slc29a3 | 0.02687163 | 0.02252572 | 0.02389479 | -0.0013691 | 0.64854032 | 0.99999994 |
| Zkscan2 | 0.01443474 | 0.01157383 | 0.01294322 | -0.0013694 | 0.75935828 | 0.99999994 |
| Sgk3 | 0.02845056 | 0.02390195 | 0.02527215 | -0.0013702 | 0.64090988 | 0.99999994 |
| Serpina9 | 0.01467501 | 0.01178576 | 0.01315653 | -0.0013708 | 0.75600858 | 0.99999994 |
| Slc19a1 | 0.03879644 | 0.03285457 | 0.03422586 | -0.0013713 | 0.60514759 | 0.99999994 |
| Reln | 0.00341881 | 0.00171657 | 0.00308848 | -0.0013719 | 0.99844558 | 0.99999994 |
| 2610044O15 | 0.05028511 | 0.04265383 | 0.04402577 | -0.0013719 | 0.58216161 | 0.99999994 |
| Zfp786 | 0.00868844 | 0.00644841 | 0.00782096 | -0.0013726 | 0.87863066 | 0.99999994 |
| Plekha4 | 0.00657602 | 0.00455478 | 0.00592791 | -0.0013731 | 0.9384477 | 0.99999994 |
| Tmem117 | 0.01251157 | 0.00986002 | 0.01123333 | -0.0013733 | 0.79210813 | 0.99999994 |
| Col19a1 | 0.00479066 | 0.00294981 | 0.00432375 | -0.0013739 | 0.98279181 | 0.99999994 |

| | | | | | | |
|---|---|---|---|---|---|---|
| Stk11ip | 0.03040102 | 0.02559538 | 0.02696959 | -0.0013742 | 0.63275868 | 0.99999994 |
| B3gntl1 | 0.01708192 | 0.01390616 | 0.0152896 | -0.0013834 | 0.72650729 | 0.99999994 |
| Dusp8 | 0.03606587 | 0.03049093 | 0.03187467 | -0.0013837 | 0.613696 | 0.99999994 |
| Itih1 | 0.00862769 | 0.00638237 | 0.0077666 | -0.0013842 | 0.88225797 | 0.99999994 |
| Zfp112 | 0.03140402 | 0.02645465 | 0.02784077 | -0.0013861 | 0.62982965 | 0.99999994 |
| G630016G05 | 0.02525291 | 0.02109348 | 0.02247971 | -0.0013862 | 0.6592686 | 0.99999994 |
| Ankrd13d | 0.04718397 | 0.04000868 | 0.04139511 | -0.0013864 | 0.58820836 | 0.99999994 |
| Rhobtb2 | 0.01645895 | 0.01335071 | 0.01473817 | -0.0013875 | 0.73454952 | 0.99999994 |
| Efnb3 | 0.02284966 | 0.01898459 | 0.02037313 | -0.0013885 | 0.67496481 | 0.99999994 |
| Slc46a1 | 0.02301693 | 0.0191312 | 0.02051997 | -0.0013888 | 0.67382304 | 0.99999994 |
| Ruvbl1 | 0.31763553 | 0.22628708 | 0.22767596 | -0.0013889 | 0.51619474 | 0.99999994 |
| Haghl | 0.3066405 | 0.22021345 | 0.22160263 | -0.0013892 | 0.51664203 | 0.99999994 |
| Tirap | 0.01560286 | 0.01259002 | 0.01397964 | -0.0013896 | 0.74590498 | 0.99999994 |
| Nicn1 | 0.15624425 | 0.12529177 | 0.12668273 | -0.001391 | 0.52913099 | 0.99999994 |
| Arhgef9 | 0.05766801 | 0.04885433 | 0.05024563 | -0.0013913 | 0.57311645 | 0.99999994 |
| Klhl31 | 0.00604105 | 0.00405603 | 0.00544764 | -0.0013916 | 0.95547043 | 0.99999994 |
| 4930412M03 | 0.00671284 | 0.00465867 | 0.0060507 | -0.001392 | 0.93716067 | 0.99999994 |
| Ago4 | 0.02947701 | 0.02477382 | 0.02616599 | -0.0013922 | 0.63838868 | 0.99999994 |
| Spryd4 | 0.0905882 | 0.07587101 | 0.07726343 | -0.0013924 | 0.54774208 | 0.99999994 |
| Btbd7 | 0.06780618 | 0.05729619 | 0.05868937 | -0.0013932 | 0.5627755 | 0.99999994 |
| Agbl5 | 0.04467951 | 0.03786769 | 0.03926273 | -0.001395 | 0.59348006 | 0.99999994 |
| Dok1 | 0.03856654 | 0.03263291 | 0.03402823 | -0.0013953 | 0.60755155 | 0.99999994 |
| Ppp2r2c | 0.00745167 | 0.00531758 | 0.00671329 | -0.0013957 | 0.9168015 | 0.99999994 |
| Cblb | 0.0374559 | 0.03167524 | 0.03307264 | -0.0013974 | 0.61073975 | 0.99999994 |
| A430105J06I | 0.01296512 | 0.01023853 | 0.01163697 | -0.0013984 | 0.78811841 | 0.99999994 |
| Ube2cbp | 0.02961969 | 0.02489136 | 0.02629015 | -0.0013988 | 0.63838965 | 0.99999994 |
| Ptges | 0.00487803 | 0.00300216 | 0.00440233 | -0.0014002 | 0.98287316 | 0.99999994 |
| Ice2 | 0.03396905 | 0.02866259 | 0.03006339 | -0.0014008 | 0.62177761 | 0.99999994 |
| Tph2 | 0.00336516 | 0.00163904 | 0.00304012 | -0.0014011 | 0.99892135 | 0.99999994 |
| Zfp407 | 0.03689417 | 0.03118713 | 0.0325888 | -0.0014017 | 0.61267393 | 0.99999994 |

| | | | | | | |
|---|---|---|---|---|---|---|
| Zbtb42 | 0.01066283 | 0.00818234 | 0.00958542 | -0.0014031 | 0.83505498 | 0.99999994 |
| Gm26549 | 0.01963565 | 0.01614202 | 0.01754525 | -0.0014032 | 0.70276142 | 0.99999994 |
| Ppp1r13l | 0.01165794 | 0.00906912 | 0.01047294 | -0.0014038 | 0.81386643 | 0.99999994 |
| Rbm12b1 | 0.01321154 | 0.01045223 | 0.01185618 | -0.0014039 | 0.78471334 | 0.99999994 |
| Poli | 0.01937552 | 0.01591037 | 0.01731583 | -0.0014055 | 0.7054955 | 0.99999994 |
| Rbm20 | 0.0041688 | 0.00235844 | 0.00376408 | -0.0014056 | 0.99353568 | 0.99999994 |
| Fbxl14 | 0.03162223 | 0.02662415 | 0.02803015 | -0.001406 | 0.63076611 | 0.99999994 |
| Gm1818 | 0.02348024 | 0.01951681 | 0.02092652 | -0.0014097 | 0.67306987 | 0.99999994 |
| Mblac1 | 0.0466544 | 0.03953501 | 0.0409448 | -0.0014098 | 0.59063889 | 0.99999994 |
| Ogg1 | 0.07608103 | 0.06408834 | 0.06549912 | -0.0014108 | 0.55700157 | 0.99999994 |
| Kantr | 0.01562652 | 0.01258949 | 0.01400062 | -0.0014111 | 0.74885639 | 0.99999994 |
| Coro1c | 0.14832586 | 0.11960235 | 0.12101354 | -0.0014112 | 0.53093577 | 0.99999994 |
| Bcl3 | 0.01529022 | 0.01229058 | 0.0137024 | -0.0014118 | 0.75359142 | 0.99999994 |
| Gipr | 0.25416318 | 0.18966634 | 0.19107818 | -0.0014118 | 0.5196131 | 0.99999994 |
| Gtf2a1 | 0.05025685 | 0.04258788 | 0.04400185 | -0.001414 | 0.58468636 | 0.99999994 |
| Rgl2 | 0.14267632 | 0.11550871 | 0.11692365 | -0.0014149 | 0.53210049 | 0.99999994 |
| Bcl7b | 0.35615213 | 0.24670092 | 0.24811684 | -0.0014159 | 0.51515047 | 0.99999994 |
| Dvl2 | 0.08519645 | 0.07150073 | 0.07291673 | -0.001416 | 0.55142494 | 0.99999994 |
| Tnfrsf21 | 0.01133713 | 0.0087693 | 0.01018695 | -0.0014177 | 0.82286064 | 0.99999994 |
| Efcab7 | 0.06358508 | 0.05376885 | 0.05518732 | -0.0014185 | 0.56792245 | 0.99999994 |
| Orc2 | 0.10027287 | 0.0835713 | 0.08499233 | -0.001421 | 0.54430714 | 0.99999994 |
| Gramd1b | 0.01994842 | 0.01639912 | 0.01782097 | -0.0014219 | 0.70231878 | 0.99999994 |
| Trim23 | 0.05914484 | 0.05006061 | 0.05148259 | -0.001422 | 0.5729356 | 0.99999994 |
| Cdk18 | 0.01406465 | 0.01119103 | 0.01261452 | -0.0014235 | 0.77371519 | 0.99999994 |
| Zfp51 | 0.05915308 | 0.05006472 | 0.05148949 | -0.0014248 | 0.57306705 | 0.99999994 |
| Scn8a | 0.02564117 | 0.02139461 | 0.0228194 | -0.0014248 | 0.66115178 | 0.99999994 |
| 1700018L02R | 0.00846136 | 0.00619292 | 0.00761772 | -0.0014248 | 0.89343221 | 0.99999994 |
| BC025920 | 0.01884926 | 0.01542645 | 0.01685147 | -0.001425 | 0.71324376 | 0.99999994 |
| Nap1l3 | 0.0431185 | 0.03650383 | 0.03793007 | -0.0014262 | 0.59881902 | 0.99999994 |
| Ddit4l | 0.01161356 | 0.00900458 | 0.01043338 | -0.0014288 | 0.81899918 | 0.99999994 |

| | | | | | | |
|---|---|---|---|---|---|---|
| Mpst | 0.13822677 | 0.11224584 | 0.11367578 | -0.0014299 | 0.53336481 | 0.99999994 |
| Gas1 | 0.01191526 | 0.00927034 | 0.01070224 | -0.0014319 | 0.81342254 | 0.99999994 |
| Jph3 | 0.00578305 | 0.00378398 | 0.0052159 | -0.0014319 | 0.96617812 | 0.99999994 |
| Tyw5 | 0.08470001 | 0.07108255 | 0.07251499 | -0.0014324 | 0.55230527 | 0.99999994 |
| Ifi27l2b | 0.00596006 | 0.00394234 | 0.0053749 | -0.0014326 | 0.96197773 | 0.99999994 |
| Pgbd1 | 0.01177896 | 0.00914801 | 0.01058079 | -0.0014328 | 0.81630188 | 0.99999994 |
| Lbhd1 | 0.01357741 | 0.01074831 | 0.01218151 | -0.0014332 | 0.78322946 | 0.99999994 |
| Pcsk4 | 0.02086926 | 0.01719706 | 0.01863209 | -0.001435 | 0.69590787 | 0.99999994 |
| Kif19a | 0.04876827 | 0.04130412 | 0.04274039 | -0.0014363 | 0.58849858 | 0.99999994 |
| Gdf11 | 0.0452937 | 0.03834982 | 0.03978632 | -0.0014365 | 0.59496298 | 0.99999994 |
| Mpi | 0.11781888 | 0.09703626 | 0.09847403 | -0.0014378 | 0.5387107 | 0.99999994 |
| Nfkbid | 0.00964739 | 0.0072407 | 0.00867853 | -0.0014378 | 0.86494336 | 0.99999994 |
| Taf5l | 0.07715714 | 0.06494018 | 0.06637935 | -0.0014392 | 0.55737485 | 0.99999994 |
| 1700086O06 | 0.03982336 | 0.0336673 | 0.0351079 | -0.0014406 | 0.60762539 | 0.99999994 |
| Eno3 | 0.02443997 | 0.02032677 | 0.02176788 | -0.0014411 | 0.67027494 | 0.99999994 |
| Ppp3cc | 0.01621546 | 0.01308131 | 0.01452251 | -0.0014412 | 0.74555637 | 0.99999994 |
| Aasdhppt | 0.25748606 | 0.19164606 | 0.19308869 | -0.0014426 | 0.51983206 | 0.99999994 |
| Mon1b | 0.02186188 | 0.01806228 | 0.01950531 | -0.001443 | 0.68879821 | 0.99999994 |
| Vsnl1 | 0.00612065 | 0.0040759 | 0.00551913 | -0.0014432 | 0.959124 | 0.99999994 |
| Fam135a | 0.34318216 | 0.23993262 | 0.24137587 | -0.0014432 | 0.51587371 | 0.99999994 |
| Ikzf5 | 0.11164248 | 0.09232614 | 0.09377233 | -0.0014462 | 0.54088212 | 0.99999994 |
| 9430015G10 | 0.01273605 | 0.00998663 | 0.01143314 | -0.0014465 | 0.80014949 | 0.99999994 |
| Card6 | 0.00694208 | 0.00480903 | 0.00625635 | -0.0014473 | 0.93820305 | 0.99999994 |
| Plpp7 | 0.05520008 | 0.04672455 | 0.04817318 | -0.0014486 | 0.57932398 | 0.99999994 |
| Ap3m2 | 0.03300189 | 0.02777745 | 0.02922623 | -0.0014488 | 0.62928586 | 0.99999994 |
| Casr | 0.0046034 | 0.00270615 | 0.00415526 | -0.0014491 | 0.98986579 | 0.99999994 |
| Fam118a | 0.04764131 | 0.04033436 | 0.04178374 | -0.0014494 | 0.59130147 | 0.99999994 |
| Rras2 | 0.07786053 | 0.0655046 | 0.06695404 | -0.0014494 | 0.55728875 | 0.99999994 |
| Wiz | 0.0649178 | 0.05484561 | 0.0562951 | -0.0014495 | 0.5680406 | 0.99999994 |
| Bcl9l | 0.01449502 | 0.01154667 | 0.01299674 | -0.0014501 | 0.7711577 | 0.99999994 |

| | | | | | | |
|---|---|---|---|---|---|---|
| Six4 | 0.02772432 | 0.02318881 | 0.02463898 | -0.0014502 | 0.65238643 | 0.99999994 |
| Pnpla2 | 0.19888846 | 0.15453764 | 0.15598954 | -0.0014519 | 0.52470072 | 0.99999994 |
| Rell2 | 0.00960171 | 0.00718502 | 0.00863771 | -0.0014527 | 0.868528 | 0.99999994 |
| Lpin2 | 0.03859708 | 0.03260085 | 0.03405449 | -0.0014536 | 0.61184555 | 0.99999994 |
| Hgfac | 0.00899332 | 0.00664007 | 0.00809373 | -0.0014537 | 0.88405446 | 0.99999994 |
| Ehd3 | 0.00959513 | 0.0071775 | 0.00863183 | -0.0014543 | 0.86896067 | 0.99999994 |
| Il1b | 0.00529979 | 0.00332682 | 0.0047816 | -0.0014548 | 0.97857506 | 0.99999994 |
| 9430065F17I | 0.00622023 | 0.00415364 | 0.00560854 | -0.0014549 | 0.95789088 | 0.99999994 |
| Trim67 | 0.01085703 | 0.00830278 | 0.00975872 | -0.0014559 | 0.83966595 | 0.99999994 |
| Mmachc | 0.04686463 | 0.03966728 | 0.0411236 | -0.0014563 | 0.59317662 | 0.99999994 |
| Slc38a6 | 0.02036895 | 0.01673439 | 0.01819152 | -0.0014571 | 0.70304202 | 0.99999994 |
| Cklf | 0.10082298 | 0.0839635 | 0.08542089 | -0.0014574 | 0.54520896 | 0.99999994 |
| Lrrc8b | 0.03691324 | 0.03114784 | 0.03260522 | -0.0014574 | 0.61696734 | 0.99999994 |
| Armt1 | 0.08141147 | 0.06838923 | 0.06984731 | -0.0014581 | 0.55525656 | 0.99999994 |
| Tbcel | 0.04952883 | 0.041927 | 0.04338522 | -0.0014582 | 0.58851504 | 0.99999994 |
| Gucy2c | 0.00834569 | 0.00605522 | 0.00751417 | -0.001459 | 0.90189031 | 0.99999994 |
| Nhsl2 | 0.0252881 | 0.02105102 | 0.0225105 | -0.0014595 | 0.66697282 | 0.99999994 |
| Srek1 | 0.39433941 | 0.26569947 | 0.26716074 | -0.0014613 | 0.51452133 | 0.99999994 |
| Zmym6 | 0.00941668 | 0.00701085 | 0.00847232 | -0.0014615 | 0.87455975 | 0.99999994 |
| Myo1e | 0.12556472 | 0.10284154 | 0.10430357 | -0.001462 | 0.53716842 | 0.99999994 |
| Gm15234 | 0.01293625 | 0.01014922 | 0.01161129 | -0.0014621 | 0.79902815 | 0.99999994 |
| Tcf12 | 0.24714657 | 0.18533341 | 0.18679608 | -0.0014627 | 0.52078392 | 0.99999994 |
| Prkd3 | 0.04836557 | 0.04093374 | 0.04239872 | -0.001465 | 0.59095148 | 0.99999994 |
| Rpgrip1l | 0.04473135 | 0.03784127 | 0.03930695 | -0.0014657 | 0.59801125 | 0.99999994 |
| Serhl | 0.09758277 | 0.08139199 | 0.08285772 | -0.0014657 | 0.54686689 | 0.99999994 |
| Fbxo10 | 0.01267421 | 0.00991046 | 0.0113781 | -0.0014676 | 0.80471695 | 0.99999994 |
| Pabpc1l | 0.03655495 | 0.03082643 | 0.03229643 | -0.00147 | 0.6190438 | 0.99999994 |
| Zfp93 | 0.02561775 | 0.0213287 | 0.02279892 | -0.0014702 | 0.66612786 | 0.99999994 |
| Pbx3 | 0.0667253 | 0.05632163 | 0.05779445 | -0.0014728 | 0.56734888 | 0.99999994 |
| Rnf103 | 0.11024124 | 0.0912257 | 0.092699 | -0.0014733 | 0.54212633 | 0.99999994 |

| | | | | | | |
|---|---|---|---|---|---|---|
| Ercc6 | 0.03631055 | 0.03061211 | 0.03208571 | -0.0014736 | 0.62008618 | 0.99999994 |
| Pglyrp1 | 0.00931869 | 0.00691045 | 0.0083847 | -0.0014743 | 0.87907267 | 0.99999994 |
| Ppm1h | 0.07617864 | 0.06410445 | 0.06557901 | -0.0014746 | 0.55948742 | 0.99999994 |
| Capn5 | 0.00923247 | 0.00683281 | 0.00830761 | -0.0014748 | 0.88132921 | 0.99999994 |
| Bach1 | 0.07702993 | 0.06479901 | 0.06627536 | -0.0014763 | 0.55893774 | 0.99999994 |
| Tomm34 | 0.27897196 | 0.20435582 | 0.20583234 | -0.0014765 | 0.51904176 | 0.99999994 |
| Fam222b | 0.13387272 | 0.10899717 | 0.11047491 | -0.0014777 | 0.53547393 | 0.99999994 |
| Chrna1os | 0.02454973 | 0.02038623 | 0.02186403 | -0.0014778 | 0.67361189 | 0.99999994 |
| Atp8a1 | 0.02121626 | 0.01745931 | 0.01893748 | -0.0014782 | 0.69831677 | 0.99999994 |
| Kptn | 0.10943847 | 0.09060464 | 0.09208298 | -0.0014783 | 0.54255172 | 0.99999994 |
| Aar2 | 0.1323599 | 0.10787907 | 0.10935747 | -0.0014784 | 0.53585129 | 0.99999994 |
| Pde5a | 0.0182067 | 0.01480538 | 0.01628404 | -0.0014787 | 0.72721952 | 0.99999994 |
| Kcnk16 | 0.04567848 | 0.03863532 | 0.04011413 | -0.0014788 | 0.59692148 | 0.99999994 |
| Atmin | 0.05983877 | 0.05058247 | 0.052063 | -0.0014805 | 0.57506654 | 0.99999994 |
| Syngap1 | 0.00479487 | 0.00284691 | 0.00432753 | -0.0014806 | 0.98861877 | 0.99999994 |
| Stx12 | 0.32259884 | 0.22889918 | 0.23038202 | -0.0014828 | 0.51708671 | 0.99999994 |
| Dip2c | 0.04328003 | 0.03658421 | 0.0380681 | -0.0014839 | 0.60236171 | 0.99999994 |
| Psmc3 | 0.94506289 | 0.42100012 | 0.42248579 | -0.0014857 | 0.50933722 | 0.99999994 |
| Plscr1 | 0.12302543 | 0.10091445 | 0.10240056 | -0.0014861 | 0.53847872 | 0.99999994 |
| Dennd4a | 0.01611136 | 0.01294384 | 0.0144303 | -0.0014865 | 0.75353817 | 0.99999994 |
| Elmo2 | 0.02981084 | 0.02496898 | 0.02645644 | -0.0014875 | 0.64588705 | 0.99999994 |
| Dnase1l1 | 0.01090657 | 0.00831496 | 0.00980291 | -0.001488 | 0.84383622 | 0.99999994 |
| Psd3 | 0.04756114 | 0.04022645 | 0.04171563 | -0.0014892 | 0.59391235 | 0.99999994 |
| Foxo4 | 0.03376289 | 0.0283915 | 0.02988503 | -0.0014935 | 0.63030374 | 0.99999994 |
| A130014A01 | 0.0355953 | 0.02997426 | 0.03146863 | -0.0014944 | 0.6240344 | 0.99999994 |
| Sco1 | 0.03832865 | 0.03232821 | 0.03382367 | -0.0014955 | 0.61573567 | 0.99999994 |
| Col14a1 | 0.00617512 | 0.00407097 | 0.00556804 | -0.0014971 | 0.9632481 | 0.99999994 |
| Zfp882 | 0.03914039 | 0.03302296 | 0.03452142 | -0.0014985 | 0.61368291 | 0.99999994 |
| Ly6g6d | 0.02218478 | 0.01829063 | 0.01978913 | -0.0014985 | 0.69288401 | 0.99999994 |
| Homer1 | 0.02514276 | 0.02088351 | 0.02238331 | -0.0014998 | 0.67220553 | 0.99999994 |

| | | | | | | |
|---|---|---|---|---|---|---|
| Spidr | 0.02355833 | 0.01949363 | 0.02099502 | -0.0015014 | 0.68296504 | 0.99999994 |
| Amt | 0.06016991 | 0.05083757 | 0.05233977 | -0.0015022 | 0.5757544 | 0.99999994 |
| Gm16973 | 0.02424117 | 0.02009001 | 0.02159369 | -0.0015037 | 0.67850241 | 0.99999994 |
| Guca1b | 0.00342468 | 0.00158976 | 0.00309377 | -0.001504 | 0.99939374 | 0.99999994 |
| Zw10 | 0.06284489 | 0.05306717 | 0.05457123 | -0.0015041 | 0.57278082 | 0.99999994 |
| Lsp1 | 0.01455126 | 0.01154141 | 0.01304668 | -0.0015053 | 0.77875105 | 0.99999994 |
| Htatip2 | 0.10316023 | 0.08573098 | 0.08723746 | -0.0015065 | 0.54575634 | 0.99999994 |
| Spin4 | 0.00836929 | 0.00602775 | 0.0075353 | -0.0015076 | 0.90852194 | 0.99999994 |
| Gm15972 | 0.01209121 | 0.00935135 | 0.01085899 | -0.0015076 | 0.82229647 | 0.99999994 |
| Ankrd52 | 0.01958878 | 0.01599505 | 0.01750392 | -0.0015089 | 0.71694416 | 0.99999994 |
| Trim65 | 0.02303933 | 0.01903049 | 0.02053963 | -0.0015091 | 0.68760509 | 0.99999994 |
| Stim2 | 0.08215788 | 0.06894402 | 0.0704538 | -0.0015098 | 0.55671321 | 0.99999994 |
| Abi2 | 0.06941857 | 0.0585112 | 0.06002202 | -0.0015108 | 0.56653039 | 0.99999994 |
| Gm3336 | 0.00628145 | 0.00415246 | 0.00566351 | -0.001511 | 0.96212864 | 0.99999994 |
| Ccdc62 | 0.01536984 | 0.01226026 | 0.01377302 | -0.0015128 | 0.7676413 | 0.99999994 |
| 1110002L01R | 0.03169713 | 0.02658234 | 0.02809514 | -0.0015128 | 0.6399842 | 0.99999994 |
| 4931414P19I | 0.04453657 | 0.03762778 | 0.03914082 | -0.001513 | 0.60153027 | 0.99999994 |
| Ccdc157 | 0.01911998 | 0.01557648 | 0.01709039 | -0.0015139 | 0.7222822 | 0.99999994 |
| B230322F03 | 0.01244214 | 0.00965622 | 0.01117151 | -0.0015153 | 0.81670047 | 0.99999994 |
| Bak1 | 0.16494444 | 0.13131196 | 0.1328281 | -0.0015161 | 0.53028159 | 0.99999994 |
| Pgbd5 | 0.01611698 | 0.01291907 | 0.01443527 | -0.0015162 | 0.75776968 | 0.99999994 |
| Gm9726 | 0.03166355 | 0.02654938 | 0.028066 | -0.0015166 | 0.64046136 | 0.99999994 |
| Prickle3 | 0.0247912 | 0.02055818 | 0.02207552 | -0.0015173 | 0.67635152 | 0.99999994 |
| Igsf9 | 0.07739477 | 0.06505614 | 0.06657356 | -0.0015174 | 0.56029567 | 0.99999994 |
| Zfp629 | 0.0468571 | 0.03959926 | 0.0411172 | -0.0015179 | 0.59705609 | 0.99999994 |
| 4930471D02 | 0.00679832 | 0.0046092 | 0.00612739 | -0.0015182 | 0.95045133 | 0.99999994 |
| Magi1 | 0.0307206 | 0.02572881 | 0.0272473 | -0.0015185 | 0.64466542 | 0.99999994 |
| Fastkd1 | 0.01639851 | 0.01316614 | 0.01468464 | -0.0015185 | 0.75437311 | 0.99999994 |
| Sash1 | 0.04688734 | 0.03962311 | 0.04114292 | -0.0015198 | 0.59711412 | 0.99999994 |
| Zfp532 | 0.03603416 | 0.03032702 | 0.03184732 | -0.0015203 | 0.62466505 | 0.99999994 |

| | | | | | | |
|---|---|---|---|---|---|---|
| Msln | 0.04701678 | 0.0397321 | 0.04125297 | -0.0015209 | 0.59692687 | 0.99999994 |
| Cpne3 | 0.13497808 | 0.10976867 | 0.11128965 | -0.001521 | 0.53624241 | 0.99999994 |
| Sh3bp4 | 0.02995503 | 0.02506043 | 0.02658185 | -0.0015214 | 0.64839061 | 0.99999994 |
| Cops8 | 0.46110746 | 0.29622246 | 0.29774554 | -0.0015231 | 0.51358127 | 0.99999994 |
| Sv2c | 0.01489992 | 0.01183253 | 0.01335615 | -0.0015236 | 0.77617292 | 0.99999994 |
| Trmo | 0.04059861 | 0.03424899 | 0.03577298 | -0.001524 | 0.61163128 | 0.99999994 |
| Cmss1 | 0.02772581 | 0.02311495 | 0.02464028 | -0.0015253 | 0.65985027 | 0.99999994 |
| Vrk1 | 0.14977387 | 0.12052885 | 0.12205567 | -0.0015268 | 0.53317984 | 0.99999994 |
| Dhx37 | 0.01796272 | 0.01454076 | 0.01606846 | -0.0015277 | 0.73658472 | 0.99999994 |
| Retnlg | 0.00801753 | 0.00569168 | 0.00722031 | -0.0015286 | 0.9206177 | 0.99999994 |
| Idua | 0.00997863 | 0.00744564 | 0.0089745 | -0.0015289 | 0.87159233 | 0.99999994 |
| Setdb2 | 0.01438991 | 0.01137408 | 0.01290341 | -0.0015293 | 0.78492094 | 0.99999994 |
| Nvl | 0.1308561 | 0.10671423 | 0.10824398 | -0.0015297 | 0.53747341 | 0.99999994 |
| Inpp5e | 0.04736791 | 0.0400199 | 0.04155145 | -0.0015315 | 0.59690629 | 0.99999994 |
| Zfp873 | 0.02698586 | 0.02246229 | 0.02399454 | -0.0015322 | 0.66460421 | 0.99999994 |
| Rgs7 | 0.01726913 | 0.01392264 | 0.01545522 | -0.0015326 | 0.74539343 | 0.99999994 |
| Ly96 | 0.03607062 | 0.03034585 | 0.03187877 | -0.0015329 | 0.62554485 | 0.99999994 |
| Tram2 | 0.00753705 | 0.00525666 | 0.00678981 | -0.0015332 | 0.93358299 | 0.99999994 |
| Haus2 | 0.09192804 | 0.07680571 | 0.07833892 | -0.0015332 | 0.55182573 | 0.99999994 |
| Ovca2 | 0.01675911 | 0.01346877 | 0.01500391 | -0.0015351 | 0.75208052 | 0.99999994 |
| Pole4 | 0.15906718 | 0.12715118 | 0.12868638 | -0.0015352 | 0.53164632 | 0.99999994 |
| Zyg11b | 0.07730496 | 0.06496496 | 0.06650017 | -0.0015352 | 0.56106379 | 0.99999994 |
| Bbs5 | 0.08087215 | 0.06787203 | 0.06940873 | -0.0015367 | 0.55858056 | 0.99999994 |
| C1qa | 0.21283358 | 0.16360282 | 0.1651409 | -0.0015381 | 0.52471678 | 0.99999994 |
| Polr3e | 0.06405494 | 0.05403988 | 0.05557809 | -0.0015382 | 0.57308144 | 0.99999994 |
| Wwox | 0.0861748 | 0.07216944 | 0.07370768 | -0.0015382 | 0.55524142 | 0.99999994 |
| Dcaf4 | 0.053309 | 0.04504225 | 0.04658059 | -0.0015383 | 0.58699793 | 0.99999994 |
| Phlpp1 | 0.0142799 | 0.01126555 | 0.01280571 | -0.0015402 | 0.78831076 | 0.99999994 |
| Ndufb2 | 1.27066354 | 0.42489598 | 0.42643663 | -0.0015406 | 0.50959298 | 0.99999994 |
| Tspyl3 | 0.01461897 | 0.0115642 | 0.01310679 | -0.0015426 | 0.78330689 | 0.99999994 |

| | | | | | | |
|---|---|---|---|---|---|---|
| St7 | 0.01683037 | 0.01352056 | 0.01506699 | -0.0015464 | 0.75275571 | 0.99999994 |
| Rexo4 | 0.1691752 | 0.13423853 | 0.13578507 | -0.0015465 | 0.53021628 | 0.99999994 |
| Zfp653 | 0.02897283 | 0.02417908 | 0.0257271 | -0.001548 | 0.65561281 | 0.99999994 |
| Lbr | 0.0522976 | 0.04417906 | 0.04572721 | -0.0015481 | 0.58914996 | 0.99999994 |
| Wdr45 | 0.13073606 | 0.10660617 | 0.10815498 | -0.0015488 | 0.53797003 | 0.99999994 |
| Clasrp | 0.09498111 | 0.07923393 | 0.08078274 | -0.0015488 | 0.550775 | 0.99999994 |
| Gatb | 0.04828753 | 0.04078046 | 0.04233248 | -0.001552 | 0.59640008 | 0.99999994 |
| Pym1 | 0.18434831 | 0.1446717 | 0.1462244 | -0.0015527 | 0.52817423 | 0.99999994 |
| Med25 | 0.51717834 | 0.31942824 | 0.32098127 | -0.001553 | 0.51284615 | 0.99999994 |
| 9130019O22 | 0.01860878 | 0.01508562 | 0.01663916 | -0.0015535 | 0.73285947 | 0.99999994 |
| Stat5a | 0.02155196 | 0.01767876 | 0.0192328 | -0.001554 | 0.7046568 | 0.99999994 |
| Rab10os | 0.08215721 | 0.06889859 | 0.07045325 | -0.0015547 | 0.55838805 | 0.99999994 |
| Rgs10 | 0.00853016 | 0.0061238 | 0.0076793 | -0.0015555 | 0.91121883 | 0.99999994 |
| Spice1 | 0.04110905 | 0.03465447 | 0.03621051 | -0.001556 | 0.61257552 | 0.99999994 |
| Snca | 0.01672768 | 0.01341884 | 0.01497609 | -0.0015573 | 0.75557676 | 0.99999994 |
| Tvp23a | 0.00703027 | 0.00477813 | 0.00633545 | -0.0015573 | 0.94910194 | 0.99999994 |
| Lmbrd2 | 0.06468278 | 0.05454159 | 0.05609988 | -0.0015583 | 0.57334393 | 0.99999994 |
| Mtif3 | 0.13498031 | 0.10973235 | 0.11129129 | -0.0015589 | 0.53714371 | 0.99999994 |
| 5031425E22I | 0.22837492 | 0.17353914 | 0.1750986 | -0.0015595 | 0.52363643 | 0.99999994 |
| Ppp1r3f | 0.01632021 | 0.01305437 | 0.0146153 | -0.0015609 | 0.76142842 | 0.99999994 |
| Il15ra | 0.00679024 | 0.00455841 | 0.00612014 | -0.0015617 | 0.95529933 | 0.99999994 |
| Fbxl8 | 0.00451683 | 0.0025151 | 0.00407736 | -0.0015623 | 0.99462025 | 0.99999994 |
| Unc119 | 0.26697572 | 0.19720977 | 0.19877217 | -0.0015624 | 0.52086344 | 0.99999994 |
| Pard3b | 0.03187388 | 0.02668496 | 0.02824848 | -0.0015635 | 0.64371982 | 0.99999994 |
| Diablo | 0.5140215 | 0.31816426 | 0.31972858 | -0.0015643 | 0.51299015 | 0.99999994 |
| Slc25a26 | 0.0254891 | 0.02112152 | 0.02268638 | -0.0015649 | 0.67693418 | 0.99999994 |
| Lipe | 0.02112517 | 0.01729222 | 0.01885733 | -0.0015651 | 0.70968323 | 0.99999994 |
| 4933404O12 | 0.01286057 | 0.00997627 | 0.01154395 | -0.0015677 | 0.81698657 | 0.99999994 |
| Zfp367 | 0.00694256 | 0.00468907 | 0.00625678 | -0.0015677 | 0.95232513 | 0.99999994 |
| Acta2 | 0.00869165 | 0.00625445 | 0.00782383 | -0.0015694 | 0.90909391 | 0.99999994 |

| | | | | | | |
|---|---|---|---|---|---|---|
| Tmem199 | 0.40023835 | 0.26842893 | 0.2699984 | -0.0015695 | 0.51543229 | 0.99999994 |
| Nop16 | 0.19846152 | 0.15413653 | 0.15570608 | -0.0015696 | 0.52674807 | 0.99999994 |
| Gm13610 | 0.01016385 | 0.00756981 | 0.00913993 | -0.0015701 | 0.87357971 | 0.99999994 |
| Umad1 | 0.11370739 | 0.09377871 | 0.09534957 | -0.0015709 | 0.54366126 | 0.99999994 |
| Stxbp6 | 0.05955776 | 0.05025501 | 0.05182802 | -0.001573 | 0.58005118 | 0.99999994 |
| Tspan2 | 0.05639955 | 0.04760821 | 0.04918127 | -0.0015731 | 0.5842988 | 0.99999994 |
| Ptk2b | 0.0320715 | 0.02684576 | 0.02841987 | -0.0015741 | 0.64381748 | 0.99999994 |
| Nup107 | 0.03411724 | 0.02861603 | 0.03019156 | -0.0015755 | 0.63583871 | 0.99999994 |
| Casc1 | 0.00953769 | 0.00700438 | 0.00858049 | -0.0015761 | 0.88927339 | 0.99999994 |
| Ndrg4 | 0.01085012 | 0.00817634 | 0.00975255 | -0.0015762 | 0.85899239 | 0.99999994 |
| Casp9 | 0.0445253 | 0.03755447 | 0.03913121 | -0.0015767 | 0.60573069 | 0.99999994 |
| Xpnpep3 | 0.03463745 | 0.02906307 | 0.03064132 | -0.0015782 | 0.63414438 | 0.99999994 |
| Iqcg | 0.02556608 | 0.02117378 | 0.02275372 | -0.0015799 | 0.67802854 | 0.99999994 |
| Arl6 | 0.27345024 | 0.20101888 | 0.20259956 | -0.0015807 | 0.52070882 | 0.99999994 |
| Zfp955b | 0.02685732 | 0.02230001 | 0.02388229 | -0.0015823 | 0.67039555 | 0.99999994 |
| Gstm5 | 0.407408 | 0.27182794 | 0.27341168 | -0.0015837 | 0.5153783 | 0.99999994 |
| Heca | 0.0071436 | 0.00485307 | 0.00643708 | -0.001584 | 0.94928573 | 0.99999994 |
| Paqr4 | 0.02043838 | 0.01666836 | 0.01825268 | -0.0015843 | 0.71828927 | 0.99999994 |
| Ercc8 | 0.03534832 | 0.02967074 | 0.03125541 | -0.0015847 | 0.63212324 | 0.99999994 |
| 9130221H12 | 0.01042219 | 0.00778526 | 0.00937062 | -0.0015854 | 0.8699482 | 0.99999994 |
| F2rl1 | 0.05825765 | 0.04914901 | 0.05073978 | -0.0015908 | 0.58265399 | 0.99999994 |
| B230307C23 | 0.009811 | 0.00723319 | 0.00882474 | -0.0015915 | 0.88502359 | 0.99999994 |
| Slc46a3 | 0.01520364 | 0.01203289 | 0.01362561 | -0.0015927 | 0.78173604 | 0.99999994 |
| Cln3 | 0.09848673 | 0.08198416 | 0.08357707 | -0.0015929 | 0.55047615 | 0.99999994 |
| Caskin2 | 0.01606139 | 0.01279292 | 0.01438602 | -0.0015931 | 0.76947553 | 0.99999994 |
| Gm27030 | 0.01761237 | 0.01416436 | 0.01575877 | -0.0015944 | 0.74967408 | 0.99999994 |
| Gm17324 | 0.00597645 | 0.00379319 | 0.00538963 | -0.0015964 | 0.97567506 | 0.99999994 |
| Lockd | 0.02126304 | 0.01738173 | 0.01897865 | -0.0015969 | 0.71228935 | 0.99999994 |
| Rpusd3 | 0.08743599 | 0.07312874 | 0.07472584 | -0.0015971 | 0.55656468 | 0.99999994 |
| Nck2 | 0.01521859 | 0.01204136 | 0.01363887 | -0.0015975 | 0.78220278 | 0.99999994 |

| | | | | | | |
|---|---|---|---|---|---|---|
| Lrrc3 | 0.00763371 | 0.00527842 | 0.00687644 | -0.001598 | 0.93905333 | 0.99999994 |
| Ttll1 | 0.10856352 | 0.08981238 | 0.09141067 | -0.0015983 | 0.54632586 | 0.99999994 |
| Npas3 | 0.02130881 | 0.01741961 | 0.01901892 | -0.0015993 | 0.71216989 | 0.99999994 |
| Tmem52 | 0.00964567 | 0.00707634 | 0.008677 | -0.0016007 | 0.8902591 | 0.99999994 |
| Snx5 | 0.49252755 | 0.30942469 | 0.31102726 | -0.0016026 | 0.51367983 | 0.99999994 |
| Slc5a3 | 0.01671352 | 0.0133598 | 0.01496356 | -0.0016038 | 0.7622049 | 0.99999994 |
| Urb1 | 0.01657103 | 0.01323243 | 0.01483741 | -0.001605 | 0.76424574 | 0.99999994 |
| Tstd1 | 0.01266952 | 0.00976875 | 0.01137392 | -0.0016052 | 0.82623637 | 0.99999994 |
| Zfp697 | 0.01356229 | 0.01056211 | 0.01216807 | -0.001606 | 0.8101681 | 0.99999994 |
| Spire1 | 0.03034105 | 0.02531072 | 0.02691747 | -0.0016068 | 0.65443818 | 0.99999994 |
| Josd1 | 0.07829915 | 0.06570413 | 0.06731214 | -0.001608 | 0.5631708 | 0.99999994 |
| Cog8 | 0.24369587 | 0.18306421 | 0.18467242 | -0.0016082 | 0.52311235 | 0.99999994 |
| B4gat1 | 0.11543535 | 0.09505665 | 0.09666537 | -0.0016087 | 0.54410302 | 0.99999994 |
| Zfp580 | 0.11227868 | 0.09265008 | 0.09425884 | -0.0016088 | 0.54522533 | 0.99999994 |
| Cetn4 | 0.04857231 | 0.04096157 | 0.04257415 | -0.0016126 | 0.59952722 | 0.99999994 |
| Gm16364 | 0.00447045 | 0.00242206 | 0.00403562 | -0.0016136 | 0.99610924 | 0.99999994 |
| Hjurp | 0.02613924 | 0.02164015 | 0.02325491 | -0.0016148 | 0.67803109 | 0.99999994 |
| Zfp146 | 0.0229757 | 0.01886845 | 0.02048378 | -0.0016153 | 0.70017852 | 0.99999994 |
| Gdpd5 | 0.02476257 | 0.02043324 | 0.02205045 | -0.0016172 | 0.68729071 | 0.99999994 |
| Tbc1d22a | 0.04755083 | 0.04008845 | 0.04170687 | -0.0016184 | 0.60191137 | 0.99999994 |
| Foxred2 | 0.00947291 | 0.00690407 | 0.00852258 | -0.0016185 | 0.89690298 | 0.99999994 |
| Dusp7 | 0.01909575 | 0.01544505 | 0.01706901 | -0.001624 | 0.73672895 | 0.99999994 |
| Zscan20 | 0.01054398 | 0.00785532 | 0.00947934 | -0.001624 | 0.87293597 | 0.99999994 |
| Prss16 | 0.00681444 | 0.00451691 | 0.00614185 | -0.0016249 | 0.96088605 | 0.99999994 |
| Zfr2 | 0.02932231 | 0.02440544 | 0.02603136 | -0.0016259 | 0.661206 | 0.99999994 |
| Dnajc18 | 0.13170726 | 0.10724844 | 0.10887456 | -0.0016261 | 0.5395965 | 0.99999994 |
| 9030622O22 | 0.03872922 | 0.03254174 | 0.03416808 | -0.0016263 | 0.62431423 | 0.99999994 |
| Entpd2 | 0.00482142 | 0.00272498 | 0.00435141 | -0.0016264 | 0.99357594 | 0.99999994 |
| Slc35e2 | 0.04684287 | 0.03947841 | 0.04110509 | -0.0016267 | 0.60388539 | 0.99999994 |
| Parp9 | 0.01459622 | 0.01145984 | 0.01308659 | -0.0016267 | 0.79600262 | 0.99999994 |

| | | | | | | |
|---|---|---|---|---|---|---|
| Spata5 | 0.04476231 | 0.03770598 | 0.03933335 | -0.0016274 | 0.6084947 | 0.99999994 |
| Lhfpl5 | 0.02205044 | 0.01804257 | 0.01967107 | -0.0016285 | 0.70920263 | 0.99999994 |
| Mre11a | 0.04395886 | 0.03701928 | 0.03864784 | -0.0016286 | 0.61044898 | 0.99999994 |
| Cdkl3 | 0.0221794 | 0.01815546 | 0.01978439 | -0.0016289 | 0.70816961 | 0.99999994 |
| Zfp54 | 0.02325431 | 0.01909895 | 0.0207283 | -0.0016293 | 0.69959065 | 0.99999994 |
| A530020G20 | 0.02796904 | 0.02322271 | 0.02485241 | -0.0016297 | 0.66876012 | 0.99999994 |
| Tmem125 | 0.01845302 | 0.01487132 | 0.01650162 | -0.0016303 | 0.74460975 | 0.99999994 |
| G2e3 | 0.08232434 | 0.06895792 | 0.07058897 | -0.001631 | 0.56111764 | 0.99999994 |
| Pik3r3 | 0.05293524 | 0.0446338 | 0.04626536 | -0.0016316 | 0.59279385 | 0.99999994 |
| Rpp40 | 0.03488436 | 0.02922282 | 0.03085467 | -0.0016319 | 0.63759735 | 0.99999994 |
| Cyb561d1 | 0.01401513 | 0.01093834 | 0.01257052 | -0.0016322 | 0.80628202 | 0.99999994 |
| Hs1bp3 | 0.01910311 | 0.01544296 | 0.0170755 | -0.0016325 | 0.73774092 | 0.99999994 |
| Gpn1 | 0.16401667 | 0.1305442 | 0.13217696 | -0.0016328 | 0.53276619 | 0.99999994 |
| Gm43948 | 0.01700624 | 0.01358832 | 0.01522264 | -0.0016343 | 0.76258185 | 0.99999994 |
| Xrcc1 | 0.23766077 | 0.17929508 | 0.18092998 | -0.0016349 | 0.52398097 | 0.99999994 |
| Prdm11 | 0.01771922 | 0.01421818 | 0.01585323 | -0.001635 | 0.75380547 | 0.99999994 |
| Ptpn14 | 0.02704603 | 0.02241107 | 0.02404707 | -0.001636 | 0.6746726 | 0.99999994 |
| Rpp25l | 0.32596891 | 0.23057062 | 0.23220731 | -0.0016367 | 0.5187101 | 0.99999994 |
| Nudcd1 | 0.0729562 | 0.06129936 | 0.06293614 | -0.0016368 | 0.56871844 | 0.99999994 |
| Gm26682 | 0.01237103 | 0.0094678 | 0.0111082 | -0.0016404 | 0.83718894 | 0.99999994 |
| Tmem18 | 0.04440931 | 0.03739157 | 0.03903226 | -0.0016407 | 0.61018255 | 0.99999994 |
| Hdx | 0.00963223 | 0.00702357 | 0.00866499 | -0.0016414 | 0.89633056 | 0.99999994 |
| Hyal1 | 0.0618555 | 0.05210366 | 0.05374679 | -0.0016431 | 0.5806266 | 0.99999994 |
| Acsl1 | 0.01914917 | 0.0154728 | 0.01711614 | -0.0016433 | 0.73861564 | 0.99999994 |
| Garnl3 | 0.01119981 | 0.0084199 | 0.01006448 | -0.0016446 | 0.86163189 | 0.99999994 |
| Dubr | 0.01188028 | 0.00902294 | 0.01067107 | -0.0016481 | 0.84804017 | 0.99999994 |
| Taf1a | 0.05289227 | 0.04458008 | 0.04622911 | -0.001649 | 0.59384116 | 0.99999994 |
| A930033H14 | 0.01446232 | 0.01131494 | 0.01296771 | -0.0016528 | 0.80188339 | 0.99999994 |
| Gm20498 | 0.02299856 | 0.018851 | 0.02050384 | -0.0016528 | 0.70422078 | 0.99999994 |
| Kctd21 | 0.01298827 | 0.01000405 | 0.01165758 | -0.0016535 | 0.82745284 | 0.99999994 |

| | | | | | | |
|---|---|---|---|---|---|---|
| Erbb3 | 0.01820818 | 0.01462965 | 0.01628535 | -0.0016557 | 0.75071404 | 0.99999994 |
| Ptcd1 | 0.05385291 | 0.04538325 | 0.04703906 | -0.0016558 | 0.59262732 | 0.99999994 |
| Tap1 | 0.01097686 | 0.00820963 | 0.00986563 | -0.001656 | 0.86806817 | 0.99999994 |
| Zfp229 | 0.02775724 | 0.02301047 | 0.0246677 | -0.0016572 | 0.67263068 | 0.99999994 |
| Slc25a10 | 0.06262218 | 0.0527282 | 0.05438575 | -0.0016575 | 0.58038161 | 0.99999994 |
| Tdg | 0.36326408 | 0.25009577 | 0.25175364 | -0.0016579 | 0.51748164 | 0.99999994 |
| Gm807 | 0.00586832 | 0.00363419 | 0.00529251 | -0.0016583 | 0.98149533 | 0.99999994 |
| Fam167a | 0.00810896 | 0.00564345 | 0.00730219 | -0.0016587 | 0.93473744 | 0.99999994 |
| Ppp4r4 | 0.01299233 | 0.01000215 | 0.01166118 | -0.001659 | 0.82817916 | 0.99999994 |
| Farp2 | 0.0181038 | 0.01453356 | 0.01619313 | -0.0016596 | 0.7524383 | 0.99999994 |
| Rhbdd1 | 0.02850509 | 0.02365666 | 0.02531966 | -0.001663 | 0.66901338 | 0.99999994 |
| Dcaf11 | 0.34256131 | 0.23938535 | 0.24104955 | -0.0016642 | 0.51832704 | 0.99999994 |
| Borcs5 | 0.08001062 | 0.06704229 | 0.0687075 | -0.0016652 | 0.56408123 | 0.99999994 |
| Vps18 | 0.03314947 | 0.02768842 | 0.02935404 | -0.0016656 | 0.64717237 | 0.99999994 |
| E2f5 | 0.09994657 | 0.08307141 | 0.08473703 | -0.0016656 | 0.55204902 | 0.99999994 |
| Ccnj | 0.02475423 | 0.02037628 | 0.02204315 | -0.0016669 | 0.69263861 | 0.99999994 |
| Ngly1 | 0.10924442 | 0.0902659 | 0.09193396 | -0.0016681 | 0.54806479 | 0.99999994 |
| Tiprl | 0.59951538 | 0.34978564 | 0.35145513 | -0.0016695 | 0.51261211 | 0.99999994 |
| Atf7ip | 0.13772996 | 0.11164045 | 0.11331169 | -0.0016712 | 0.5391032 | 0.99999994 |
| Cog5 | 0.01597356 | 0.01263663 | 0.0143082 | -0.0016716 | 0.78160902 | 0.99999994 |
| Cdk5rap1 | 0.03281533 | 0.02739294 | 0.02906462 | -0.0016717 | 0.64908327 | 0.99999994 |
| Clip2 | 0.02675646 | 0.02212199 | 0.02379421 | -0.0016722 | 0.68003594 | 0.99999994 |
| Suz12 | 0.15055047 | 0.12094084 | 0.12261358 | -0.0016727 | 0.53617755 | 0.99999994 |
| Zfp369 | 0.04059696 | 0.03409733 | 0.03577157 | -0.0016742 | 0.62230578 | 0.99999994 |
| Ptpdc1 | 0.0226733 | 0.01854397 | 0.02021827 | -0.0016743 | 0.70925921 | 0.99999994 |
| Calca | 0.00561983 | 0.00339342 | 0.00506925 | -0.0016758 | 0.98611462 | 0.99999994 |
| Gpkow | 0.04074219 | 0.03421912 | 0.03589609 | -0.001677 | 0.62208704 | 0.99999994 |
| Slain1 | 0.03678312 | 0.03081385 | 0.0324931 | -0.0016792 | 0.63457712 | 0.99999994 |
| Pank2 | 0.13676629 | 0.11092456 | 0.11260463 | -0.0016801 | 0.53955511 | 0.99999994 |
| Gsap | 0.00886206 | 0.0062955 | 0.0079763 | -0.0016808 | 0.91964038 | 0.99999994 |

| | | | | | | |
|---|---|---|---|---|---|---|
| Gm14403 | 0.01872837 | 0.0150632 | 0.01674475 | -0.0016815 | 0.74807556 | 0.99999994 |
| Zfp78 | 0.01317468 | 0.01014183 | 0.0118234 | -0.0016816 | 0.82810145 | 0.99999994 |
| Otoa | 0.00584326 | 0.00358787 | 0.00526999 | -0.0016821 | 0.98319129 | 0.99999994 |
| Hps5 | 0.03866021 | 0.03242642 | 0.03410876 | -0.0016823 | 0.62866162 | 0.99999994 |
| Ccdc141 | 0.00453362 | 0.00240853 | 0.00409247 | -0.0016839 | 0.99691777 | 0.99999994 |
| Vti1a | 0.0448892 | 0.03775652 | 0.03944155 | -0.001685 | 0.61193812 | 0.99999994 |
| Socs1 | 0.0194598 | 0.01570422 | 0.01739017 | -0.001686 | 0.7406413 | 0.99999994 |
| Mrpl39 | 0.28012021 | 0.20481332 | 0.20650105 | -0.0016877 | 0.5216927 | 0.99999994 |
| Snap91 | 0.00871138 | 0.00615368 | 0.00784149 | -0.0016878 | 0.92403065 | 0.99999994 |
| Trib3 | 0.00869001 | 0.00613012 | 0.00782236 | -0.0016922 | 0.92506493 | 0.99999994 |
| Edaradd | 0.00800181 | 0.00551374 | 0.00720623 | -0.0016925 | 0.94101438 | 0.99999994 |
| Pign | 0.04993856 | 0.04203787 | 0.04373233 | -0.0016945 | 0.60176073 | 0.99999994 |
| Adam22 | 0.01949376 | 0.01572296 | 0.01742012 | -0.0016972 | 0.7416679 | 0.99999994 |
| Gpatch4 | 0.07333736 | 0.06155143 | 0.06324933 | -0.0016979 | 0.57090818 | 0.99999994 |
| Ndor1 | 0.0222802 | 0.01817475 | 0.01987297 | -0.0016982 | 0.7152577 | 0.99999994 |
| Tmem241 | 0.0164745 | 0.01305357 | 0.01475194 | -0.0016984 | 0.778262 | 0.99999994 |
| Gm7008 | 0.01236256 | 0.00940217 | 0.01110065 | -0.0016985 | 0.84577359 | 0.99999994 |
| Parg | 0.1416996 | 0.11451371 | 0.11621271 | -0.001699 | 0.53876151 | 0.99999994 |
| Prkd2 | 0.03714741 | 0.031107 | 0.03280697 | -0.0017 | 0.63492042 | 0.99999994 |
| Slc22a18 | 0.01973579 | 0.01593104 | 0.01763354 | -0.0017025 | 0.73977854 | 0.99999994 |
| Flii | 0.12915664 | 0.10527788 | 0.10698213 | -0.0017042 | 0.54222348 | 0.99999994 |
| Lrrc57 | 0.11670576 | 0.09592592 | 0.09763025 | -0.0017043 | 0.54625296 | 0.99999994 |
| Mtm1 | 0.03818325 | 0.03199364 | 0.03369861 | -0.001705 | 0.63185755 | 0.99999994 |
| Fgd1 | 0.02176683 | 0.01771654 | 0.01942175 | -0.0017052 | 0.72053162 | 0.99999994 |
| Ccrl2 | 0.01613714 | 0.01274674 | 0.01445314 | -0.0017064 | 0.78403059 | 0.99999994 |
| Snapc4 | 0.02279913 | 0.01862179 | 0.02032876 | -0.001707 | 0.71189296 | 0.99999994 |
| Tamm41 | 0.04467322 | 0.03755005 | 0.03925737 | -0.0017073 | 0.61389574 | 0.99999994 |
| 1700030K09I | 0.04614 | 0.03879961 | 0.0405071 | -0.0017075 | 0.6104849 | 0.99999994 |
| Spata24 | 0.11076969 | 0.09139532 | 0.09310407 | -0.0017088 | 0.54861592 | 0.99999994 |
| Bsn | 0.0158787 | 0.01251426 | 0.01422413 | -0.0017099 | 0.78818998 | 0.99999994 |

| | | | | | | |
|---|---|---|---|---|---|---|
| Lcorl | 0.24481951 | 0.18365498 | 0.18536523 | -0.0017102 | 0.52448517 | 0.99999994 |
| Fadd | 0.0569997 | 0.04797458 | 0.04968507 | -0.0017105 | 0.5906261 | 0.99999994 |
| Rin2 | 0.00961382 | 0.00693719 | 0.00864853 | -0.0017113 | 0.90610303 | 0.99999994 |
| Cox15 | 0.02100789 | 0.01704195 | 0.01875411 | -0.0017122 | 0.72830438 | 0.99999994 |
| AW554918 | 0.0355818 | 0.02974396 | 0.03145697 | -0.001713 | 0.64150174 | 0.99999994 |
| Fut11 | 0.06191087 | 0.05207995 | 0.05379296 | -0.001713 | 0.58393425 | 0.99999994 |
| Plxna3 | 0.0343342 | 0.02866607 | 0.03037918 | -0.0017131 | 0.64630317 | 0.99999994 |
| A930038B10 | 0.02778541 | 0.02297815 | 0.02469227 | -0.0017141 | 0.67798762 | 0.99999994 |
| Nipa1 | 0.0197471 | 0.01592874 | 0.01764351 | -0.0017148 | 0.74116177 | 0.99999994 |
| Try10 | 0.00960059 | 0.00692034 | 0.0086367 | -0.0017164 | 0.9070509 | 0.99999994 |
| Tmbim4 | 0.83101302 | 0.40693286 | 0.40865113 | -0.0017183 | 0.51116421 | 0.99999994 |
| Tmem256 | 0.9690203 | 0.4226107 | 0.42432905 | -0.0017184 | 0.51075237 | 0.99999994 |
| Stoml2 | 0.48204918 | 0.3049512 | 0.30667344 | -0.0017222 | 0.5149095 | 0.99999994 |
| Decr1 | 0.09314959 | 0.077594 | 0.07931786 | -0.0017239 | 0.55751261 | 0.99999994 |
| Rap2c | 0.03294483 | 0.02745273 | 0.0291768 | -0.0017241 | 0.65296109 | 0.99999994 |
| Zfp790 | 0.02169433 | 0.01763186 | 0.019358 | -0.0017261 | 0.72359123 | 0.99999994 |
| Evi5l | 0.03840944 | 0.03216576 | 0.03389315 | -0.0017274 | 0.63278832 | 0.99999994 |
| Alg10b | 0.00947211 | 0.0067939 | 0.00852187 | -0.001728 | 0.91144376 | 0.99999994 |
| Zfp275 | 0.02871584 | 0.02377334 | 0.02550328 | -0.0017299 | 0.67419014 | 0.99999994 |
| Cic | 0.0848716 | 0.07092332 | 0.07265387 | -0.0017306 | 0.56298798 | 0.99999994 |
| Slc12a1 | 0.00803358 | 0.00550353 | 0.00723469 | -0.0017312 | 0.94439378 | 0.99999994 |
| Nubpl | 0.06412811 | 0.05390697 | 0.05563893 | -0.001732 | 0.58207432 | 0.99999994 |
| Spef1 | 0.01880336 | 0.0150778 | 0.01681095 | -0.0017332 | 0.75372144 | 0.99999994 |
| Gm14508 | 0.00830513 | 0.00574387 | 0.00747786 | -0.001734 | 0.93864481 | 0.99999994 |
| Cmtr2 | 0.02354164 | 0.01924617 | 0.02098038 | -0.0017342 | 0.70891077 | 0.99999994 |
| Perp | 0.03477865 | 0.02902819 | 0.03076334 | -0.0017351 | 0.6463331 | 0.99999994 |
| Mrpl18 | 0.66270457 | 0.37027012 | 0.37200748 | -0.0017374 | 0.51239986 | 0.99999994 |
| Wwp1 | 0.06609216 | 0.0555302 | 0.05726965 | -0.0017394 | 0.58010906 | 0.99999994 |
| 9430060I03R | 0.02382883 | 0.01949246 | 0.02123224 | -0.0017398 | 0.70726933 | 0.99999994 |
| Sh3d21 | 0.02871083 | 0.02375893 | 0.02549891 | -0.00174 | 0.675162 | 0.99999994 |

| | | | | | | |
|---|---|---|---|---|---|---|
| Tpsb2 | 0.01115825 | 0.00828606 | 0.01002742 | -0.0017414 | 0.87614812 | 0.99999994 |
| Adgrf5 | 0.00877497 | 0.00615502 | 0.00789839 | -0.0017434 | 0.92911325 | 0.99999994 |
| Gm9776 | 0.03257312 | 0.02711127 | 0.02885475 | -0.0017435 | 0.65622953 | 0.99999994 |
| Txnl4b | 0.01937072 | 0.01556793 | 0.0173116 | -0.0017437 | 0.74871296 | 0.99999994 |
| Wdhd1 | 0.01233097 | 0.00932743 | 0.01107252 | -0.0017451 | 0.85292918 | 0.99999994 |
| Rrn3 | 0.086041 | 0.07185131 | 0.07359957 | -0.0017483 | 0.56281601 | 0.99999994 |
| Inip | 0.09222541 | 0.07682903 | 0.07857737 | -0.0017483 | 0.55886925 | 0.99999994 |
| S100a4 | 0.00438023 | 0.00220556 | 0.00395442 | -0.0017489 | 0.99837893 | 0.99999994 |
| Traf3 | 0.05536838 | 0.0465649 | 0.04831472 | -0.0017498 | 0.59525093 | 0.99999994 |
| Pop5 | 0.75127935 | 0.39211962 | 0.39387145 | -0.0017518 | 0.51180925 | 0.99999994 |
| Il17ra | 0.02588428 | 0.02127983 | 0.02303202 | -0.0017522 | 0.69370846 | 0.99999994 |
| Pibf1 | 0.07452169 | 0.06246926 | 0.06422145 | -0.0017522 | 0.57205536 | 0.99999994 |
| Mmp24 | 0.0054794 | 0.00319052 | 0.00494305 | -0.0017525 | 0.99086192 | 0.99999994 |
| Tbc1d32 | 0.01828035 | 0.01459463 | 0.01634911 | -0.0017545 | 0.7624837 | 0.99999994 |
| Tns2 | 0.01054179 | 0.00772243 | 0.00947739 | -0.001755 | 0.89113269 | 0.99999994 |
| Sptan1 | 0.230787 | 0.17486667 | 0.17662167 | -0.001755 | 0.52636718 | 0.99999994 |
| Xpo1 | 0.15994006 | 0.12754755 | 0.12930408 | -0.0017565 | 0.53602453 | 0.99999994 |
| Ung | 0.04954269 | 0.041638 | 0.04339696 | -0.001759 | 0.60634106 | 0.99999994 |
| Vps45 | 0.14217645 | 0.11480036 | 0.11655994 | -0.0017596 | 0.54002005 | 0.99999994 |
| AA386476 | 0.01282531 | 0.0097516 | 0.01151257 | -0.001761 | 0.84569969 | 0.99999994 |
| Plpp3 | 0.01690241 | 0.0133692 | 0.01513075 | -0.0017615 | 0.78081391 | 0.99999994 |
| Llgl1 | 0.0301824 | 0.02501755 | 0.02677955 | -0.001762 | 0.66929437 | 0.99999994 |
| Rasgrp1 | 0.00353205 | 0.00142671 | 0.00319052 | -0.0017638 | 0.99988331 | 0.99999994 |
| Arhgap44 | 0.06202407 | 0.05212328 | 0.05388734 | -0.0017641 | 0.58624747 | 0.99999994 |
| 1700086P04I | 0.0068532 | 0.00441167 | 0.00617662 | -0.0017649 | 0.9714164 | 0.99999994 |
| Ccdc68 | 0.02169419 | 0.01759027 | 0.01935788 | -0.0017676 | 0.72834199 | 0.99999994 |
| Kdm3b | 0.07974158 | 0.06672001 | 0.06848835 | -0.0017683 | 0.56822786 | 0.99999994 |
| Bpgm | 0.21419957 | 0.16425775 | 0.16602619 | -0.0017684 | 0.52826158 | 0.99999994 |
| Prps1l3 | 0.01318761 | 0.01006488 | 0.0118349 | -0.00177 | 0.84025732 | 0.99999994 |
| Retsat | 0.01130312 | 0.00838588 | 0.01015662 | -0.0017707 | 0.87707553 | 0.99999994 |

| | | | | | | |
|---|---|---|---|---|---|---|
| Zfp429 | 0.01493111 | 0.01161237 | 0.01338382 | -0.0017715 | 0.81084634 | 0.99999994 |
| Ctrc | 0.01062991 | 0.00778386 | 0.00955603 | -0.0017722 | 0.89147814 | 0.99999994 |
| Chd1l | 0.0355358 | 0.02964499 | 0.03141727 | -0.0017723 | 0.64635269 | 0.99999994 |
| Gm16845 | 0.02518499 | 0.02064787 | 0.02242026 | -0.0017724 | 0.70062751 | 0.99999994 |
| Acadsb | 0.20273529 | 0.15675827 | 0.15853478 | -0.0017765 | 0.52972953 | 0.99999994 |
| Tef | 0.06266368 | 0.05264346 | 0.05442032 | -0.0017769 | 0.58602585 | 0.99999994 |
| Pdzd4 | 0.01235055 | 0.00931286 | 0.01108996 | -0.0017771 | 0.85693282 | 0.99999994 |
| Zfp58 | 0.03060329 | 0.02536748 | 0.02714537 | -0.0017779 | 0.66856655 | 0.99999994 |
| 1700008J07F | 0.06824218 | 0.05727015 | 0.05905 | -0.0017799 | 0.57950693 | 0.99999994 |
| Atr | 0.04764007 | 0.04000154 | 0.04178269 | -0.0017812 | 0.61170027 | 0.99999994 |
| Gm16183 | 0.04459773 | 0.03741178 | 0.03919298 | -0.0017812 | 0.61886879 | 0.99999994 |
| Nol4 | 0.16319721 | 0.12981939 | 0.13160103 | -0.0017816 | 0.53590207 | 0.99999994 |
| Brsk2 | 0.01347681 | 0.01030993 | 0.01209208 | -0.0017821 | 0.83670948 | 0.99999994 |
| Rnf20 | 0.2464068 | 0.18455848 | 0.18634179 | -0.0017833 | 0.5253962 | 0.99999994 |
| Dnajc30 | 0.25673824 | 0.19085224 | 0.19263715 | -0.0017849 | 0.52458942 | 0.99999994 |
| 1700055D18 | 0.01011946 | 0.00731538 | 0.00910029 | -0.0017849 | 0.9041497 | 0.99999994 |
| Rab2b | 0.13758394 | 0.11141959 | 0.11320462 | -0.001785 | 0.54179573 | 0.99999994 |
| Dagla | 0.00711011 | 0.00462192 | 0.00640705 | -0.0017851 | 0.96817283 | 0.99999994 |
| Cma1 | 0.01584407 | 0.01240711 | 0.01419345 | -0.0017863 | 0.79891338 | 0.99999994 |
| Bend3 | 0.01265074 | 0.00957061 | 0.01135721 | -0.0017866 | 0.85247586 | 0.99999994 |
| Top3b | 0.13055531 | 0.10623413 | 0.10802093 | -0.0017868 | 0.54383673 | 0.99999994 |
| Nt5dc1 | 0.02203861 | 0.01787353 | 0.01966066 | -0.0017871 | 0.72742928 | 0.99999994 |
| Hykk | 0.00881077 | 0.00614279 | 0.00793041 | -0.0017876 | 0.93324879 | 0.99999994 |
| Dctn5 | 0.20214985 | 0.15635896 | 0.15814844 | -0.0017895 | 0.53001909 | 0.99999994 |
| Arhgap36 | 0.14620452 | 0.11769125 | 0.11948241 | -0.0017912 | 0.53974275 | 0.99999994 |
| Rnf122 | 0.02845918 | 0.02348773 | 0.02527966 | -0.0017919 | 0.68147646 | 0.99999994 |
| Zfp747 | 0.03263614 | 0.02711689 | 0.02890936 | -0.0017925 | 0.66009219 | 0.99999994 |
| Arl8a | 0.11239628 | 0.09255551 | 0.09434871 | -0.0017932 | 0.55033659 | 0.99999994 |
| Cep170 | 0.2010209 | 0.15560688 | 0.1574024 | -0.0017955 | 0.53026274 | 0.99999994 |
| Zfp90 | 0.08203221 | 0.0685554 | 0.07035173 | -0.0017963 | 0.56747935 | 0.99999994 |

| | | | | | | |
|---|---|---|---|---|---|---|
| Ubash3b | 0.01256582 | 0.00948505 | 0.01128162 | -0.0017966 | 0.85542929 | 0.99999994 |
| Msto1 | 0.09988453 | 0.08289098 | 0.08468778 | -0.0017968 | 0.55615442 | 0.99999994 |
| Cav1 | 0.01041554 | 0.00756728 | 0.00936468 | -0.0017974 | 0.89930075 | 0.99999994 |
| Arfrp1 | 0.27502011 | 0.20172395 | 0.20352156 | -0.0017976 | 0.5234413 | 0.99999994 |
| E130102H24 | 0.00973022 | 0.00695404 | 0.00875256 | -0.0017985 | 0.91430997 | 0.99999994 |
| Pigo | 0.04017063 | 0.03360593 | 0.03540591 | -0.0018 | 0.63246849 | 0.99999994 |
| Slmap | 0.10941477 | 0.09026466 | 0.09206478 | -0.0018001 | 0.55177637 | 0.99999994 |
| Irf9 | 0.03522564 | 0.02934825 | 0.03114947 | -0.0018012 | 0.64984588 | 0.99999994 |
| Trpc1 | 0.06301547 | 0.05291169 | 0.05471326 | -0.0018016 | 0.58674405 | 0.99999994 |
| Map2k3os | 0.00599379 | 0.00360208 | 0.0054052 | -0.0018031 | 0.98680746 | 0.99999994 |
| Echdc1 | 0.14869546 | 0.11947562 | 0.12127977 | -0.0018041 | 0.53943856 | 0.99999994 |
| Lzic | 0.06808401 | 0.0571149 | 0.0589192 | -0.0018043 | 0.58076043 | 0.99999994 |
| C6 | 0.00955309 | 0.0067897 | 0.00859425 | -0.0018045 | 0.91889009 | 0.99999994 |
| Chil6 | 0.00351527 | 0.00137042 | 0.0031754 | -0.001805 | 0.99992273 | 0.99999994 |
| Ccl6 | 0.00606747 | 0.00366534 | 0.00547137 | -0.001806 | 0.98599604 | 0.99999994 |
| Shank1 | 0.01154341 | 0.00856358 | 0.01037085 | -0.0018073 | 0.87696855 | 0.99999994 |
| Arhgap32 | 0.05007593 | 0.04204116 | 0.04384867 | -0.0018075 | 0.60810557 | 0.99999994 |
| Ing5 | 0.04093467 | 0.03425287 | 0.03606108 | -0.0018082 | 0.63072351 | 0.99999994 |
| Car10 | 0.00461074 | 0.00235206 | 0.00416186 | -0.0018098 | 0.99810149 | 0.99999994 |
| Ube3b | 0.0428812 | 0.03591597 | 0.03772725 | -0.0018113 | 0.62535256 | 0.99999994 |
| Rnf115 | 0.30020775 | 0.21618809 | 0.21799938 | -0.0018113 | 0.52205251 | 0.99999994 |
| Mtx3 | 0.02096381 | 0.01690046 | 0.01871532 | -0.0018149 | 0.74069183 | 0.99999994 |
| Farp1 | 0.0360963 | 0.03008512 | 0.03190092 | -0.0018158 | 0.64761193 | 0.99999994 |
| Intu | 0.03458454 | 0.0287753 | 0.03059559 | -0.0018203 | 0.65395488 | 0.99999994 |
| Gm16740 | 0.03057454 | 0.02529961 | 0.02712039 | -0.0018208 | 0.67252232 | 0.99999994 |
| Hmgcs1 | 0.15873499 | 0.12662955 | 0.12845107 | -0.0018215 | 0.53760094 | 0.99999994 |
| Pmp22 | 0.01145053 | 0.00846573 | 0.01028805 | -0.0018223 | 0.88080961 | 0.99999994 |
| Zmym1 | 0.04406195 | 0.03691243 | 0.03873584 | -0.0018234 | 0.62298663 | 0.99999994 |
| Pax6os1 | 0.04411849 | 0.03695853 | 0.0387841 | -0.0018256 | 0.62297937 | 0.99999994 |
| Gadd45b | 0.10915641 | 0.09003908 | 0.09186635 | -0.0018273 | 0.55266561 | 0.99999994 |

| | | | | | | |
|---|---|---|---|---|---|---|
| Zfp174 | 0.09396338 | 0.07814137 | 0.07996917 | -0.0018278 | 0.56046157 | 0.99999994 |
| Csf1 | 0.00639342 | 0.0039361 | 0.00576403 | -0.0018279 | 0.98260972 | 0.99999994 |
| Ssc4d | 0.01691658 | 0.01331483 | 0.01514329 | -0.0018285 | 0.78921934 | 0.99999994 |
| Ttll4 | 0.01521085 | 0.01180139 | 0.01363201 | -0.0018306 | 0.81430471 | 0.99999994 |
| Utp14b | 0.0118797 | 0.00883962 | 0.01067056 | -0.0018309 | 0.87330441 | 0.99999994 |
| Msh6 | 0.03225017 | 0.02674359 | 0.0285748 | -0.0018312 | 0.66515469 | 0.99999994 |
| Shb | 0.04984289 | 0.04181956 | 0.0436513 | -0.0018317 | 0.61000061 | 0.99999994 |
| Upf3a | 0.12518003 | 0.10218306 | 0.10401578 | -0.0018327 | 0.54668201 | 0.99999994 |
| Nol10 | 0.05899188 | 0.04951971 | 0.05135459 | -0.0018349 | 0.5939929 | 0.99999994 |
| Mob2 | 0.22097335 | 0.16855043 | 0.17038783 | -0.0018374 | 0.52861125 | 0.99999994 |
| BC024978 | 0.03788877 | 0.03160755 | 0.03344525 | -0.0018377 | 0.64272126 | 0.99999994 |
| Npc1 | 0.04416578 | 0.03698351 | 0.03882446 | -0.001841 | 0.62385723 | 0.99999994 |
| Oas1c | 0.00667493 | 0.00417529 | 0.00601667 | -0.0018414 | 0.97918279 | 0.99999994 |
| Traf5 | 0.0102499 | 0.00737496 | 0.00921678 | -0.0018418 | 0.90826808 | 0.99999994 |
| Etaa1 | 0.0463284 | 0.03882375 | 0.04066744 | -0.0018437 | 0.61858692 | 0.99999994 |
| Arhgef40 | 0.02324497 | 0.01887534 | 0.02072011 | -0.0018448 | 0.72328635 | 0.99999994 |
| Hacl1 | 0.0519824 | 0.04361597 | 0.04546102 | -0.0018451 | 0.60647799 | 0.99999994 |
| Zfp111 | 0.01851315 | 0.01470876 | 0.01655472 | -0.001846 | 0.77102541 | 0.99999994 |
| Slx4ip | 0.05966218 | 0.05006892 | 0.05191535 | -0.0018464 | 0.59357074 | 0.99999994 |
| Arhgap1 | 0.17847503 | 0.14036731 | 0.14221424 | -0.0018469 | 0.53444398 | 0.99999994 |
| Isg20l2 | 0.06996149 | 0.05862271 | 0.06047012 | -0.0018474 | 0.58057202 | 0.99999994 |
| Pofut2 | 0.51832835 | 0.31958575 | 0.32143577 | -0.00185 | 0.51528 | 0.99999994 |
| Nr1h4 | 0.0224097 | 0.0181365 | 0.01998674 | -0.0018502 | 0.7311201 | 0.99999994 |
| Bivm | 0.02427069 | 0.01976884 | 0.02161956 | -0.0018507 | 0.71559611 | 0.99999994 |
| Nup62 | 0.14325863 | 0.11549412 | 0.11734697 | -0.0018529 | 0.54185204 | 0.99999994 |
| Mta2 | 0.23824828 | 0.17944193 | 0.18129593 | -0.001854 | 0.52713527 | 0.99999994 |
| B3galt5 | 0.01594901 | 0.01243217 | 0.01428645 | -0.0018543 | 0.80619193 | 0.99999994 |
| Rufy1 | 0.10893656 | 0.08984141 | 0.09169743 | -0.001856 | 0.55358701 | 0.99999994 |
| Gm42997 | 0.0227471 | 0.01842697 | 0.02028308 | -0.0018561 | 0.7287787 | 0.99999994 |
| Ankrd61 | 0.01536272 | 0.01191 | 0.01376671 | -0.0018567 | 0.81533819 | 0.99999994 |

| | | | | | | |
|---|---|---|---|---|---|---|
| Ucn3 | 0.10788427 | 0.08903004 | 0.09088807 | -0.001858 | 0.55411945 | 0.99999994 |
| Fbxo27 | 0.013658 | 0.01039408 | 0.01225315 | -0.0018591 | 0.84373181 | 0.99999994 |
| Slc9a9 | 0.01251621 | 0.0093783 | 0.01123746 | -0.0018592 | 0.86460482 | 0.99999994 |
| Slc7a6 | 0.02701261 | 0.02215404 | 0.02401789 | -0.0018638 | 0.69726499 | 0.99999994 |
| Erp27 | 0.01635347 | 0.01278052 | 0.01464475 | -0.0018642 | 0.80159573 | 0.99999994 |
| Rassf7 | 0.20569905 | 0.15862071 | 0.16048508 | -0.0018644 | 0.53081853 | 0.99999994 |
| Atp4a | 0.03213644 | 0.02660969 | 0.02847619 | -0.0018665 | 0.66868993 | 0.99999994 |
| Isoc2b | 0.01343101 | 0.01018474 | 0.01205135 | -0.0018666 | 0.84872805 | 0.99999994 |
| Serpinb9 | 0.01169247 | 0.0086368 | 0.01050371 | -0.0018669 | 0.8816127 | 0.99999994 |
| Pdxk | 0.01981945 | 0.01583964 | 0.01770729 | -0.0018677 | 0.75868238 | 0.99999994 |
| Zfp7 | 0.0304512 | 0.02514518 | 0.02701321 | -0.001868 | 0.67735063 | 0.99999994 |
| Jazf1 | 0.03774485 | 0.03144858 | 0.03332139 | -0.0018728 | 0.64584187 | 0.99999994 |
| Minpp1 | 0.12550979 | 0.10238912 | 0.10426249 | -0.0018734 | 0.54760011 | 0.99999994 |
| Skap2 | 0.10255259 | 0.08489147 | 0.08676586 | -0.0018744 | 0.55716948 | 0.99999994 |
| Styk1 | 0.01189167 | 0.0088067 | 0.01068122 | -0.0018745 | 0.87863071 | 0.99999994 |
| Pxylp1 | 0.02790532 | 0.02292014 | 0.02479684 | -0.0018767 | 0.69278952 | 0.99999994 |
| Mettl13 | 0.04278434 | 0.03576646 | 0.03764443 | -0.001878 | 0.63008085 | 0.99999994 |
| Dnajc6 | 0.01139484 | 0.00835963 | 0.0102384 | -0.0018788 | 0.88904295 | 0.99999994 |
| Mtmr12 | 0.01274478 | 0.00956183 | 0.01144091 | -0.0018791 | 0.86285888 | 0.99999994 |
| Iqce | 0.05218025 | 0.04374899 | 0.04562812 | -0.0018791 | 0.60800874 | 0.99999994 |
| Dnm3 | 0.00803889 | 0.00535982 | 0.00723944 | -0.0018796 | 0.95802696 | 0.99999994 |
| Adora1 | 0.0077936 | 0.00513991 | 0.00701971 | -0.0018798 | 0.96266649 | 0.99999994 |
| Zc3h11a | 0.03244894 | 0.0268671 | 0.02874711 | -0.00188 | 0.66833295 | 0.99999994 |
| Gm16223 | 0.01736493 | 0.01365868 | 0.01553996 | -0.0018813 | 0.78982717 | 0.99999994 |
| Col1a2 | 0.02312648 | 0.01873395 | 0.02061612 | -0.0018822 | 0.72830623 | 0.99999994 |
| Hexdc | 0.03469865 | 0.02881094 | 0.03069421 | -0.0018833 | 0.65851226 | 0.99999994 |
| Rundc3b | 0.01821424 | 0.01440499 | 0.0162907 | -0.0018857 | 0.77949778 | 0.99999994 |
| Znrf3 | 0.05450521 | 0.04570223 | 0.04758844 | -0.0018862 | 0.60404487 | 0.99999994 |
| S100a13 | 0.2679541 | 0.19746515 | 0.19935327 | -0.0018881 | 0.52513419 | 0.99999994 |
| Zswim3 | 0.02029472 | 0.01623692 | 0.01812613 | -0.0018892 | 0.75608521 | 0.99999994 |

| | | | | | | |
|---|---|---|---|---|---|---|
| Irs2 | 0.04819873 | 0.04036715 | 0.0422571 | -0.00189 | 0.61703626 | 0.99999994 |
| Ccdc82 | 0.02432091 | 0.01977084 | 0.02166357 | -0.0018927 | 0.71956833 | 0.99999994 |
| Ptpn3 | 0.01726965 | 0.01356219 | 0.01545568 | -0.0018935 | 0.79260001 | 0.99999994 |
| Rps6kl1 | 0.01220197 | 0.00906347 | 0.01095764 | -0.0018942 | 0.87505958 | 0.99999994 |
| Prkcd | 0.05554368 | 0.04656755 | 0.04846212 | -0.0018946 | 0.60265382 | 0.99999994 |
| Adal | 0.0490515 | 0.04108524 | 0.0429806 | -0.0018954 | 0.61544233 | 0.99999994 |
| Myh9 | 0.16639838 | 0.13194843 | 0.13384657 | -0.0018981 | 0.53760294 | 0.99999994 |
| Atp2b4 | 0.03181288 | 0.02629712 | 0.02819556 | -0.0018984 | 0.6729883 | 0.99999994 |
| Cchcr1 | 0.02098741 | 0.01683619 | 0.01873609 | -0.0018999 | 0.75015492 | 0.99999994 |
| Zfp335 | 0.02129298 | 0.01710429 | 0.01900499 | -0.0019007 | 0.74719998 | 0.99999994 |
| Kcnmb4 | 0.01196438 | 0.00884383 | 0.010746 | -0.0019022 | 0.88065317 | 0.99999994 |
| Magee1 | 0.0398236 | 0.03320523 | 0.03510811 | -0.0019029 | 0.64086707 | 0.99999994 |
| Rgs9 | 0.02067971 | 0.01656197 | 0.01846521 | -0.0019032 | 0.7536699 | 0.99999994 |
| Ube2c | 0.0573472 | 0.04807294 | 0.0499766 | -0.0019037 | 0.60007759 | 0.99999994 |
| Ik | 0.5952372 | 0.34806693 | 0.34997286 | -0.0019059 | 0.51445855 | 0.99999994 |
| Fgfr1 | 0.0364302 | 0.03028135 | 0.03218888 | -0.0019075 | 0.65337852 | 0.99999994 |
| Bcor | 0.10132983 | 0.08390388 | 0.08581542 | -0.0019115 | 0.55893554 | 0.99999994 |
| Gm27029 | 0.02331849 | 0.0188729 | 0.02078461 | -0.0019117 | 0.72980818 | 0.99999994 |
| Cenpm | 0.02212998 | 0.01782911 | 0.01974096 | -0.0019119 | 0.74042228 | 0.99999994 |
| Elk1 | 0.03651987 | 0.0303541 | 0.03226619 | -0.0019121 | 0.65337753 | 0.99999994 |
| Tnfrsf23 | 0.0074016 | 0.00475513 | 0.0066684 | -0.0019133 | 0.97192127 | 0.99999994 |
| Abhd4 | 0.09863595 | 0.08178108 | 0.08369574 | -0.0019147 | 0.56051457 | 0.99999994 |
| Mtrf1l | 0.07385574 | 0.06175995 | 0.063675 | -0.0019151 | 0.57933533 | 0.99999994 |
| Depdc5 | 0.03755833 | 0.0312446 | 0.03316083 | -0.0019162 | 0.64974954 | 0.99999994 |
| Limk1 | 0.02395407 | 0.01942492 | 0.02134205 | -0.0019171 | 0.7250576 | 0.99999994 |
| Adnp2 | 0.02104918 | 0.01687265 | 0.01879046 | -0.0019178 | 0.75154776 | 0.99999994 |
| Misp | 0.10486133 | 0.08663729 | 0.08855526 | -0.001918 | 0.55731549 | 0.99999994 |
| Gm4631 | 0.01259833 | 0.0093914 | 0.01131055 | -0.0019192 | 0.87064348 | 0.99999994 |
| Bcl9 | 0.14413577 | 0.1160625 | 0.11798388 | -0.0019214 | 0.54316056 | 0.99999994 |
| Brms1 | 0.18425193 | 0.14423702 | 0.1461589 | -0.0019219 | 0.53487345 | 0.99999994 |

| | | | | | | |
|---|---|---|---|---|---|---|
| Gpr158 | 0.00354352 | 0.00127869 | 0.00320087 | -0.0019222 | 0.99996796 | 0.99999994 |
| Zbtb44 | 0.06425838 | 0.05382486 | 0.05574722 | -0.0019224 | 0.59077324 | 0.99999994 |
| Cep83os | 0.0250637 | 0.02038981 | 0.0223141 | -0.0019243 | 0.71702392 | 0.99999994 |
| Pnpt1 | 0.08268895 | 0.06896025 | 0.07088495 | -0.0019247 | 0.57171271 | 0.99999994 |
| Srcap | 0.04381728 | 0.03660095 | 0.03852697 | -0.001926 | 0.63034284 | 0.99999994 |
| Hps3 | 0.02753397 | 0.02254682 | 0.02447293 | -0.0019261 | 0.69981904 | 0.99999994 |
| Dnaaf2 | 0.05592547 | 0.04685676 | 0.04878302 | -0.0019263 | 0.6036609 | 0.99999994 |
| Amigo1 | 0.06045521 | 0.05065099 | 0.05257814 | -0.0019271 | 0.59637494 | 0.99999994 |
| Plekha7 | 0.01695464 | 0.01324847 | 0.01517697 | -0.0019285 | 0.80117023 | 0.99999994 |
| Rngtt | 0.11504685 | 0.0944405 | 0.09636987 | -0.0019294 | 0.55300733 | 0.99999994 |
| Tmtc1 | 0.01060129 | 0.00760107 | 0.00953049 | -0.0019294 | 0.91110017 | 0.99999994 |
| Rffl | 0.01810737 | 0.01426657 | 0.01619629 | -0.0019297 | 0.78613333 | 0.99999994 |
| Zfp472 | 0.03640332 | 0.03023401 | 0.0321657 | -0.0019317 | 0.65532737 | 0.99999994 |
| Thap4 | 0.12424221 | 0.10138067 | 0.10331343 | -0.0019328 | 0.54955022 | 0.99999994 |
| Fgd4 | 0.17881687 | 0.14051333 | 0.14244868 | -0.0019353 | 0.53602929 | 0.99999994 |
| Shmt1 | 0.01337607 | 0.01006687 | 0.01200251 | -0.0019356 | 0.85846838 | 0.99999994 |
| Zmat1 | 0.02146531 | 0.01721933 | 0.01915658 | -0.0019373 | 0.74957037 | 0.99999994 |
| Map3k1 | 0.03070749 | 0.02529862 | 0.02723591 | -0.0019373 | 0.68205971 | 0.99999994 |
| Mnd1 | 0.01505233 | 0.01155406 | 0.01349138 | -0.0019373 | 0.83042142 | 0.99999994 |
| Pnliprp2 | 0.00807349 | 0.00533025 | 0.00727043 | -0.0019402 | 0.96215906 | 0.99999994 |
| Gm3294 | 0.00542361 | 0.00295209 | 0.0048929 | -0.0019408 | 0.99585836 | 0.99999994 |
| Zfand2a | 0.03668205 | 0.03046351 | 0.03240599 | -0.0019425 | 0.6550516 | 0.99999994 |
| Ahi1 | 0.22933246 | 0.17376083 | 0.17570393 | -0.0019431 | 0.5293405 | 0.99999994 |
| Map4k4 | 0.19161153 | 0.14918843 | 0.15113159 | -0.0019432 | 0.53410128 | 0.99999994 |
| Agrn | 0.03427537 | 0.02838488 | 0.03032831 | -0.0019434 | 0.66514183 | 0.99999994 |
| Ldhd | 0.00856628 | 0.00576803 | 0.00771163 | -0.0019436 | 0.95329058 | 0.99999994 |
| Gmnn | 0.0850412 | 0.07084673 | 0.07279112 | -0.0019444 | 0.57056173 | 0.99999994 |
| Lipt2 | 0.06407602 | 0.05364904 | 0.05559562 | -0.0019466 | 0.59214279 | 0.99999994 |
| Fez2 | 0.06849519 | 0.05731186 | 0.05925918 | -0.0019473 | 0.58657214 | 0.99999994 |
| Rpl13a-ps1 | 0.04785646 | 0.04001908 | 0.04196648 | -0.0019474 | 0.62129385 | 0.99999994 |

| | | | | | | |
|---|---|---|---|---|---|---|
| Pinx1 | 0.06457563 | 0.05406135 | 0.05601086 | -0.0019495 | 0.59160704 | 0.99999994 |
| Mcur1 | 0.10935183 | 0.09006642 | 0.09201645 | -0.00195 | 0.55608972 | 0.99999994 |
| Brsk1 | 0.02415736 | 0.01956976 | 0.02152024 | -0.0019505 | 0.72684464 | 0.99999994 |
| Cdc45 | 0.02162237 | 0.01734261 | 0.01929472 | -0.0019521 | 0.74966811 | 0.99999994 |
| Ccdc126 | 0.09407041 | 0.07810198 | 0.08005477 | -0.0019528 | 0.56449289 | 0.99999994 |
| Gm10244 | 0.01891232 | 0.01495232 | 0.01690713 | -0.0019548 | 0.77923467 | 0.99999994 |
| Cd82 | 0.18396552 | 0.14400843 | 0.14596419 | -0.0019558 | 0.53553372 | 0.99999994 |
| Zfp579 | 0.1304494 | 0.10598625 | 0.10794235 | -0.0019561 | 0.54800574 | 0.99999994 |
| Ankrd37 | 0.23418724 | 0.17680218 | 0.17875877 | -0.0019566 | 0.52903983 | 0.99999994 |
| Tsacc | 0.03054385 | 0.02513317 | 0.02709372 | -0.0019605 | 0.68497819 | 0.99999994 |
| Nipsnap3b | 0.40820723 | 0.27182771 | 0.27378956 | -0.0019618 | 0.51902089 | 0.99999994 |
| Svop | 0.01623041 | 0.01257255 | 0.01453576 | -0.0019632 | 0.81567744 | 0.99999994 |
| Zdhhc16 | 0.19449067 | 0.15109542 | 0.15306046 | -0.001965 | 0.53405085 | 0.99999994 |
| Ghitm | 0.38166447 | 0.25900481 | 0.26097003 | -0.0019652 | 0.51998878 | 0.99999994 |
| Pitpnm2 | 0.04155094 | 0.0346238 | 0.03658905 | -0.0019652 | 0.63964926 | 0.99999994 |
| Kcnd1 | 0.00897856 | 0.00611384 | 0.00808053 | -0.0019667 | 0.947391 | 0.99999994 |
| Pid1 | 0.00688908 | 0.00424212 | 0.00620881 | -0.0019667 | 0.9825044 | 0.99999994 |
| Atxn7l1 | 0.08087123 | 0.06744124 | 0.06940798 | -0.0019667 | 0.5748024 | 0.99999994 |
| Dnajc1 | 0.3607923 | 0.24852777 | 0.25049474 | -0.001967 | 0.52084243 | 0.99999994 |
| Zfp248 | 0.02377305 | 0.01921513 | 0.02118333 | -0.0019682 | 0.73186333 | 0.99999994 |
| Kctd8 | 0.03663668 | 0.03039227 | 0.03236688 | -0.0019746 | 0.65765989 | 0.99999994 |
| 4933406P04I | 0.02391393 | 0.01933135 | 0.02130685 | -0.0019755 | 0.73143383 | 0.99999994 |
| Ptpn21 | 0.0446994 | 0.03730311 | 0.0392797 | -0.0019766 | 0.6311702 | 0.99999994 |
| Usp21 | 0.06317343 | 0.05286625 | 0.05484476 | -0.0019785 | 0.59488404 | 0.99999994 |
| Try4 | 0.00767123 | 0.0049304 | 0.00691006 | -0.0019797 | 0.97173849 | 0.99999994 |
| Zc3h12c | 0.04045962 | 0.03367233 | 0.0356538 | -0.0019815 | 0.64428252 | 0.99999994 |
| Sat2 | 0.10475445 | 0.08649063 | 0.08847257 | -0.0019819 | 0.55926833 | 0.99999994 |
| Pacsin1 | 0.09201968 | 0.07643007 | 0.07841241 | -0.0019823 | 0.56681825 | 0.99999994 |
| Myo1d | 0.02658568 | 0.0216618 | 0.02364504 | -0.0019832 | 0.7116816 | 0.99999994 |
| Commd8 | 0.33886548 | 0.2371167 | 0.23910035 | -0.0019836 | 0.52201966 | 0.99999994 |

| | | | | | | |
|---|---|---|---|---|---|---|
| Adgrg6 | 0.01060274 | 0.00754701 | 0.00953179 | -0.0019848 | 0.91713146 | 0.99999994 |
| Dcaf12l1 | 0.03132909 | 0.02578944 | 0.02777573 | -0.0019863 | 0.6829657 | 0.99999994 |
| Zfp408 | 0.02201255 | 0.01765126 | 0.01963776 | -0.0019865 | 0.74963335 | 0.99999994 |
| Myh10 | 0.09803141 | 0.08122828 | 0.08321484 | -0.0019866 | 0.5631282 | 0.99999994 |
| Ppa2 | 0.35863059 | 0.24740185 | 0.24938918 | -0.0019873 | 0.52115126 | 0.99999994 |
| Nfkbia | 0.28037638 | 0.20466266 | 0.20665007 | -0.0019874 | 0.52552118 | 0.99999994 |
| C2cd2 | 0.02235492 | 0.01795108 | 0.01993862 | -0.0019875 | 0.74650274 | 0.99999994 |
| Serinc5 | 0.03168978 | 0.02610074 | 0.02808876 | -0.001988 | 0.68122055 | 0.99999994 |
| Aplf | 0.02832014 | 0.02316651 | 0.02515848 | -0.001992 | 0.70091374 | 0.99999994 |
| Ap1s3 | 0.08007204 | 0.06676424 | 0.06875751 | -0.0019933 | 0.57650745 | 0.99999994 |
| Ppif | 0.09160603 | 0.07608714 | 0.07808061 | -0.0019935 | 0.56747232 | 0.99999994 |
| Btbd10 | 0.18270674 | 0.14311297 | 0.14510738 | -0.0019944 | 0.53644726 | 0.99999994 |
| Cacna1b | 0.01809529 | 0.0141899 | 0.01618561 | -0.0019957 | 0.79410253 | 0.99999994 |
| Micall1 | 0.02613481 | 0.02125352 | 0.02325105 | -0.0019975 | 0.7162904 | 0.99999994 |
| Tbc1d31 | 0.01392005 | 0.01048577 | 0.01248604 | -0.0020003 | 0.85686705 | 0.99999994 |
| Rangrf | 0.14774762 | 0.1185961 | 0.1205967 | -0.0020006 | 0.54396321 | 0.99999994 |
| Efnb1 | 0.06725606 | 0.05623217 | 0.05823405 | -0.0020019 | 0.59049625 | 0.99999994 |
| Il13ra1 | 0.04392181 | 0.03661382 | 0.03861622 | -0.0020024 | 0.63501209 | 0.99999994 |
| Napg | 0.16071269 | 0.12784758 | 0.12985012 | -0.0020025 | 0.54088122 | 0.99999994 |
| 5730409E04I | 0.01735153 | 0.01352541 | 0.0155281 | -0.0020027 | 0.80468772 | 0.99999994 |
| Cdr2l | 0.0298654 | 0.02449916 | 0.02650389 | -0.0020047 | 0.69268727 | 0.99999994 |
| Lrwd1 | 0.09384935 | 0.0778725 | 0.07987794 | -0.0020054 | 0.56636074 | 0.99999994 |
| Samd1 | 0.03263069 | 0.0268966 | 0.02890464 | -0.002008 | 0.6781115 | 0.99999994 |
| Atxn7l2 | 0.04296823 | 0.03579347 | 0.03780164 | -0.0020082 | 0.63818491 | 0.99999994 |
| Srgap3 | 0.01059136 | 0.00751341 | 0.00952163 | -0.0020082 | 0.91982625 | 0.99999994 |
| Wdr81 | 0.01395241 | 0.01050512 | 0.01251479 | -0.0020097 | 0.85744302 | 0.99999994 |
| Zbtb17 | 0.04817109 | 0.04022364 | 0.04223364 | -0.00201 | 0.62429935 | 0.99999994 |
| Pias4 | 0.2793735 | 0.20405512 | 0.20606633 | -0.0020112 | 0.52589951 | 0.99999994 |
| Crk | 0.20066529 | 0.15515427 | 0.15716712 | -0.0020129 | 0.53396814 | 0.99999994 |
| Lca5 | 0.05001433 | 0.04178322 | 0.0437965 | -0.0020133 | 0.62019213 | 0.99999994 |

| | | | | | | |
|---|---|---|---|---|---|---|
| Pus7l | 0.02633822 | 0.02141313 | 0.02342882 | -0.0020157 | 0.71656875 | 0.99999994 |
| 2610021A01 | 0.0201936 | 0.0160203 | 0.01803703 | -0.0020167 | 0.77163816 | 0.99999994 |
| Gys1 | 0.03290537 | 0.02712568 | 0.02914262 | -0.0020169 | 0.67748719 | 0.99999994 |
| Rhoq | 0.03765732 | 0.0312289 | 0.03324605 | -0.0020171 | 0.65684324 | 0.99999994 |
| Arrb2 | 0.0991793 | 0.08210886 | 0.08412764 | -0.0020188 | 0.56345341 | 0.99999994 |
| Rgl1 | 0.02297065 | 0.01845823 | 0.02047935 | -0.0020211 | 0.74438366 | 0.99999994 |
| Smim11 | 0.39477188 | 0.26534785 | 0.26736968 | -0.0020218 | 0.5200722 | 0.99999994 |
| Ttc27 | 0.03725644 | 0.03087834 | 0.03290088 | -0.0020225 | 0.65879995 | 0.99999994 |
| Ypel4 | 0.0263285 | 0.02139669 | 0.02342033 | -0.0020236 | 0.71740261 | 0.99999994 |
| Zfp46 | 0.02505256 | 0.02028025 | 0.02230435 | -0.0020241 | 0.72709721 | 0.99999994 |
| Mettl14 | 0.04753531 | 0.03966943 | 0.04169369 | -0.0020243 | 0.62671717 | 0.99999994 |
| Tlr2 | 0.02568318 | 0.02083108 | 0.02285615 | -0.0020251 | 0.72232302 | 0.99999994 |
| Pklr | 0.02821358 | 0.02304017 | 0.02506561 | -0.0020254 | 0.70466648 | 0.99999994 |
| Smcr8 | 0.0217173 | 0.01735129 | 0.0193782 | -0.0020269 | 0.75685997 | 0.99999994 |
| Acbd4 | 0.02006097 | 0.01589227 | 0.01792016 | -0.0020279 | 0.77435031 | 0.99999994 |
| Tcaim | 0.03029704 | 0.02485065 | 0.02687921 | -0.0020286 | 0.69228991 | 0.99999994 |
| Gnb4 | 0.01023076 | 0.00717102 | 0.00919969 | -0.0020287 | 0.9289244 | 0.99999994 |
| Ikbke | 0.01166176 | 0.00844274 | 0.01047634 | -0.0020336 | 0.90183724 | 0.99999994 |
| Plod1 | 0.0581561 | 0.04862073 | 0.0506547 | -0.002034 | 0.60537214 | 0.99999994 |
| Osbpl5 | 0.03137747 | 0.02578291 | 0.02781772 | -0.0020348 | 0.68683514 | 0.99999994 |
| Sema6d | 0.02888619 | 0.02361433 | 0.02565165 | -0.0020373 | 0.70148087 | 0.99999994 |
| Gm4924 | 0.01351838 | 0.01009109 | 0.01212904 | -0.0020379 | 0.86830528 | 0.99999994 |
| Arhgef4 | 0.01626961 | 0.0125299 | 0.01457048 | -0.0020406 | 0.82438823 | 0.99999994 |
| Camk4 | 0.01301142 | 0.00963643 | 0.01167816 | -0.0020417 | 0.87773803 | 0.99999994 |
| Ephx2 | 0.02804495 | 0.02287271 | 0.0249186 | -0.0020459 | 0.70764074 | 0.99999994 |
| Zfp758 | 0.03899785 | 0.03235153 | 0.03439895 | -0.0020474 | 0.6540161 | 0.99999994 |
| Pradc1 | 0.10031547 | 0.0829779 | 0.08502553 | -0.0020476 | 0.56367836 | 0.99999994 |
| Crispld2 | 0.00448867 | 0.00200429 | 0.00405201 | -0.0020477 | 0.99961569 | 0.99999994 |
| Adgra3 | 0.03325257 | 0.02739537 | 0.02944332 | -0.0020479 | 0.67831309 | 0.99999994 |
| Slc22a23 | 0.01255687 | 0.00922492 | 0.01127365 | -0.0020487 | 0.88679219 | 0.99999994 |

| | | | | | | |
|---|---|---|---|---|---|---|
| Usp49 | 0.0178129 | 0.01388527 | 0.01593605 | -0.0020508 | 0.80416524 | 0.99999994 |
| 5330434G04 | 0.01369236 | 0.0102308 | 0.0122837 | -0.0020529 | 0.86702472 | 0.99999994 |
| H2-Ab1 | 0.00724742 | 0.00447672 | 0.00653017 | -0.0020535 | 0.98183138 | 0.99999994 |
| Rfxank | 0.08351541 | 0.06950174 | 0.07155534 | -0.0020536 | 0.57575037 | 0.99999994 |
| 9530034E10I | 0.02648948 | 0.02150728 | 0.023561 | -0.0020537 | 0.71911291 | 0.99999994 |
| Zbtb24 | 0.07395982 | 0.06170632 | 0.06376044 | -0.0020541 | 0.58489882 | 0.99999994 |
| Tbc1d5 | 0.02647231 | 0.02148976 | 0.02354599 | -0.0020562 | 0.71947707 | 0.99999994 |
| Elovl2 | 0.02685132 | 0.02182076 | 0.02387706 | -0.0020563 | 0.71676124 | 0.99999994 |
| Wwc1 | 0.03883092 | 0.03219589 | 0.0342555 | -0.0020596 | 0.65550056 | 0.99999994 |
| Apobec3 | 0.10521302 | 0.0867671 | 0.08882725 | -0.0020602 | 0.56134481 | 0.99999994 |
| Gm2861 | 0.02240248 | 0.01791946 | 0.0199804 | -0.0020609 | 0.75382941 | 0.99999994 |
| Borcs7 | 0.3500661 | 0.24291022 | 0.24497124 | -0.002061 | 0.52232983 | 0.99999994 |
| Cage1 | 0.01472292 | 0.01113802 | 0.01319906 | -0.002061 | 0.85068939 | 0.99999994 |
| Gstt2 | 0.05069449 | 0.04231059 | 0.04437224 | -0.0020616 | 0.62144252 | 0.99999994 |
| Map10 | 0.01209888 | 0.00880185 | 0.01086582 | -0.002064 | 0.89695325 | 0.99999994 |
| Ncapd2 | 0.01757432 | 0.01366112 | 0.01572513 | -0.002064 | 0.80885421 | 0.99999994 |
| Lrrc49 | 0.05000679 | 0.04172572 | 0.04379012 | -0.0020644 | 0.62316383 | 0.99999994 |
| Glg1 | 0.24720493 | 0.18476671 | 0.1868319 | -0.0020652 | 0.5293267 | 0.99999994 |
| Smim24 | 0.15449351 | 0.12336866 | 0.12543551 | -0.0020668 | 0.54366813 | 0.99999994 |
| Phf6 | 0.06050886 | 0.05055601 | 0.05262295 | -0.0020669 | 0.60312784 | 0.99999994 |
| Hspa12b | 0.0151916 | 0.01154786 | 0.01361493 | -0.0020671 | 0.84389449 | 0.99999994 |
| Zfp709 | 0.01517037 | 0.01152854 | 0.0135961 | -0.0020676 | 0.84428646 | 0.99999994 |
| Mettl25 | 0.03867627 | 0.03205374 | 0.03412257 | -0.0020688 | 0.65673563 | 0.99999994 |
| 2010315B03I | 0.0547517 | 0.04572639 | 0.04779593 | -0.0020695 | 0.61340972 | 0.99999994 |
| Rwdd2b | 0.02373638 | 0.01908145 | 0.02115117 | -0.0020697 | 0.7425894 | 0.99999994 |
| Ppme1 | 0.30904104 | 0.22086714 | 0.22293787 | -0.0020707 | 0.52464955 | 0.99999994 |
| Tctn3 | 0.05853199 | 0.0488983 | 0.05096957 | -0.0020713 | 0.60661109 | 0.99999994 |
| Tbc1d10c | 0.01320926 | 0.00978047 | 0.01185415 | -0.0020737 | 0.8778722 | 0.99999994 |
| Sgsh | 0.00394098 | 0.00148502 | 0.00355893 | -0.0020739 | 0.99994753 | 0.99999994 |
| Th | 0.04769982 | 0.03975759 | 0.04183344 | -0.0020759 | 0.62941357 | 0.99999994 |

| | | | | | | |
|---|---|---|---|---|---|---|
| Arhgap29 | 0.05578691 | 0.0465903 | 0.04866657 | -0.0020763 | 0.61178777 | 0.99999994 |
| Gtf2h4 | 0.09875202 | 0.08171131 | 0.08378803 | -0.0020767 | 0.56552027 | 0.99999994 |
| Phc1 | 0.04829613 | 0.04026231 | 0.04233978 | -0.0020775 | 0.6280166 | 0.99999994 |
| 1810055G02 | 0.01694689 | 0.01309012 | 0.01517012 | -0.00208 | 0.81929068 | 0.99999994 |
| Gatm | 0.01772315 | 0.01377477 | 0.01585671 | -0.0020819 | 0.80893046 | 0.99999994 |
| BC049715 | 0.03741301 | 0.03095285 | 0.03303571 | -0.0020829 | 0.66263838 | 0.99999994 |
| Nipbl | 0.44622 | 0.28912706 | 0.29121082 | -0.0020838 | 0.51899431 | 0.99999994 |
| Cpm | 0.03746862 | 0.03099891 | 0.0330836 | -0.0020847 | 0.66255078 | 0.99999994 |
| Ddx1 | 0.43459586 | 0.2839116 | 0.28599745 | -0.0020858 | 0.51935957 | 0.99999994 |
| Ddx51 | 0.03683278 | 0.03044943 | 0.03253589 | -0.0020865 | 0.66525858 | 0.99999994 |
| Ctsb | 0.89365283 | 0.41524149 | 0.41733104 | -0.0020895 | 0.51329345 | 0.99999994 |
| Eid2b | 0.17811908 | 0.13987979 | 0.14196998 | -0.0020902 | 0.53903382 | 0.99999994 |
| Ppp1r21 | 0.07771407 | 0.06474313 | 0.06683443 | -0.0020913 | 0.58249554 | 0.99999994 |
| Fbxw24 | 0.01510672 | 0.01144783 | 0.01353963 | -0.0020918 | 0.84811249 | 0.99999994 |
| Maml1 | 0.04014699 | 0.03329382 | 0.03538562 | -0.0020918 | 0.65302055 | 0.99999994 |
| Cdpf1 | 0.17616716 | 0.13853312 | 0.14062801 | -0.0020949 | 0.5394934 | 0.99999994 |
| Foxp2 | 0.02124941 | 0.01687134 | 0.01896665 | -0.0020953 | 0.76893707 | 0.99999994 |
| Lst1 | 0.01129477 | 0.00805354 | 0.01014917 | -0.0020956 | 0.9153449 | 0.99999994 |
| Syt3 | 0.03070483 | 0.02513699 | 0.0272336 | -0.0020966 | 0.69583142 | 0.99999994 |
| Upk3b | 0.01670086 | 0.01285529 | 0.01495235 | -0.0020971 | 0.82473284 | 0.99999994 |
| Ptk2 | 0.05668807 | 0.0473245 | 0.04942352 | -0.002099 | 0.61129488 | 0.99999994 |
| Nostrin | 0.01166977 | 0.00838285 | 0.01048348 | -0.0021006 | 0.90886009 | 0.99999994 |
| Ncapd3 | 0.08884315 | 0.07375782 | 0.0758599 | -0.0021021 | 0.57316896 | 0.99999994 |
| Cep131 | 0.07627972 | 0.06355862 | 0.06566173 | -0.0021031 | 0.58441447 | 0.99999994 |
| Psmb3 | 1.43904175 | 0.4129598 | 0.41506342 | -0.0021036 | 0.51345601 | 0.99999994 |
| Vav2 | 0.01475835 | 0.0111258 | 0.01323051 | -0.0021047 | 0.85517576 | 0.99999994 |
| 9030407P20I | 0.01393579 | 0.01039506 | 0.01250002 | -0.002105 | 0.86883673 | 0.99999994 |
| Pigw | 0.03528335 | 0.0290941 | 0.03119931 | -0.0021052 | 0.67333639 | 0.99999994 |
| Trpm2 | 0.00936643 | 0.00632206 | 0.00842739 | -0.0021053 | 0.95183226 | 0.99999994 |
| Kctd7 | 0.01024735 | 0.00710826 | 0.0092145 | -0.0021062 | 0.93593386 | 0.99999994 |

| | | | | | | |
|---|---|---|---|---|---|---|
| Tmpo | 0.42394412 | 0.27902499 | 0.28113282 | -0.0021078 | 0.51990171 | 0.99999994 |
| Atp2b2 | 0.01044862 | 0.00728498 | 0.00939421 | -0.0021092 | 0.93247935 | 0.99999994 |
| Fam168a | 0.10817507 | 0.08900082 | 0.09111187 | -0.0021111 | 0.56128471 | 0.99999994 |
| Rabggta | 0.11877074 | 0.09708325 | 0.09919435 | -0.0021111 | 0.55632731 | 0.99999994 |
| Ccdc66 | 0.09915111 | 0.08199375 | 0.08410524 | -0.0021115 | 0.56635836 | 0.99999994 |
| Ccdc60 | 0.01593142 | 0.01215852 | 0.01427086 | -0.0021123 | 0.83775141 | 0.99999994 |
| Nif3l1 | 0.13642972 | 0.11024335 | 0.11235742 | -0.0021141 | 0.54983439 | 0.99999994 |
| Dnajc28 | 0.02039564 | 0.01610059 | 0.01821503 | -0.0021144 | 0.78014571 | 0.99999994 |
| Bms1 | 0.0885138 | 0.07347851 | 0.07559466 | -0.0021161 | 0.57390834 | 0.99999994 |
| Iyd | 0.00890089 | 0.00589452 | 0.00801104 | -0.0021165 | 0.96067815 | 0.99999994 |
| Ptpn2 | 0.20045632 | 0.15491056 | 0.1570288 | -0.0021182 | 0.5357734 | 0.99999994 |
| Gpc1 | 0.04016562 | 0.03328271 | 0.03540161 | -0.0021189 | 0.65483383 | 0.99999994 |
| Fsd1 | 0.03230508 | 0.02650328 | 0.02862241 | -0.0021191 | 0.68893019 | 0.99999994 |
| Rlf | 0.13742645 | 0.11096996 | 0.11308912 | -0.0021192 | 0.54963222 | 0.99999994 |
| Wtap | 0.53561131 | 0.32604405 | 0.3281661 | -0.002122 | 0.51716619 | 0.99999994 |
| Myo9a | 0.04622833 | 0.03845991 | 0.04058228 | -0.0021224 | 0.63612216 | 0.99999994 |
| Phactr4 | 0.13640046 | 0.11021271 | 0.11233592 | -0.0021232 | 0.5500584 | 0.99999994 |
| Tmem9b | 0.43290156 | 0.28310511 | 0.28522916 | -0.002124 | 0.51976688 | 0.99999994 |
| Kif1c | 0.02173218 | 0.01726451 | 0.01939128 | -0.0021268 | 0.76731967 | 0.99999994 |
| Ttc26 | 0.01620826 | 0.01238899 | 0.01451614 | -0.0021272 | 0.83531957 | 0.99999994 |
| Adgb | 0.0146252 | 0.01098293 | 0.01311231 | -0.0021294 | 0.8601437 | 0.99999994 |
| Wdr25 | 0.01680984 | 0.01291848 | 0.01504882 | -0.0021303 | 0.82697737 | 0.99999994 |
| Zcchc4 | 0.03648287 | 0.03010379 | 0.03223429 | -0.0021305 | 0.67001181 | 0.99999994 |
| Zfp280c | 0.02614511 | 0.02112795 | 0.02326004 | -0.0021321 | 0.72911688 | 0.99999994 |
| 1500002C15I | 0.01159682 | 0.00828562 | 0.01041846 | -0.0021328 | 0.91350654 | 0.99999994 |
| Foxn3 | 0.12485053 | 0.10163458 | 0.10376913 | -0.0021346 | 0.5544541 | 0.99999994 |
| Asb9 | 0.00490508 | 0.00229146 | 0.00442666 | -0.0021352 | 0.99933792 | 0.99999994 |
| Hpn | 0.12256749 | 0.09992052 | 0.10205654 | -0.002136 | 0.55539966 | 0.99999994 |
| Gab2 | 0.01682998 | 0.01292716 | 0.01506664 | -0.0021395 | 0.82772484 | 0.99999994 |
| Cyp2j6 | 0.0622099 | 0.05190219 | 0.05404223 | -0.00214 | 0.60395135 | 0.99999994 |

| | | | | | | |
|---|---|---|---|---|---|---|
| Fnta | 0.56099168 | 0.33556846 | 0.33771078 | -0.0021423 | 0.51684061 | 0.99999994 |
| Wdr59 | 0.02587136 | 0.02087705 | 0.02302072 | -0.0021437 | 0.73231616 | 0.99999994 |
| Xndc1 | 0.03419752 | 0.02811578 | 0.03026099 | -0.0021452 | 0.68149033 | 0.99999994 |
| Zfp119b | 0.02388588 | 0.01913477 | 0.02128226 | -0.0021475 | 0.74910054 | 0.99999994 |
| Etv6 | 0.02909467 | 0.0236849 | 0.02583319 | -0.0021483 | 0.71005468 | 0.99999994 |
| Tma16 | 0.09074478 | 0.07523742 | 0.07738921 | -0.0021518 | 0.57341626 | 0.99999994 |
| Eif3l | 0.76296848 | 0.3941719 | 0.39632371 | -0.0021518 | 0.51441468 | 0.99999994 |
| Orc3 | 0.14935249 | 0.11960083 | 0.12175265 | -0.0021518 | 0.54682461 | 0.99999994 |
| Barx2 | 0.01450456 | 0.01085297 | 0.01300521 | -0.0021522 | 0.86467525 | 0.99999994 |
| Vamp8 | 1.01024596 | 0.42480432 | 0.42695794 | -0.0021536 | 0.51339217 | 0.99999994 |
| Agk | 0.08489247 | 0.07051362 | 0.07267077 | -0.0021571 | 0.57831569 | 0.99999994 |
| Poc5 | 0.06921097 | 0.05769223 | 0.0598506 | -0.0021584 | 0.59485227 | 0.99999994 |
| Map3k9 | 0.01877817 | 0.01462975 | 0.01678872 | -0.002159 | 0.80399681 | 0.99999994 |
| Rell1 | 0.05632458 | 0.04695783 | 0.04911831 | -0.0021605 | 0.61515507 | 0.99999994 |
| Sdsl | 0.15767368 | 0.12553705 | 0.12769844 | -0.0021614 | 0.54485142 | 0.99999994 |
| Hexim2 | 0.02324454 | 0.01855797 | 0.02071972 | -0.0021618 | 0.75630893 | 0.99999994 |
| Vars2 | 0.02251719 | 0.01791939 | 0.02008117 | -0.0021618 | 0.76318019 | 0.99999994 |
| Spg11 | 0.0279995 | 0.02271471 | 0.02487897 | -0.0021643 | 0.71872129 | 0.99999994 |
| Eps15l1 | 0.10719579 | 0.08819236 | 0.09035777 | -0.0021654 | 0.56336999 | 0.99999994 |
| Lig4 | 0.06101532 | 0.05087957 | 0.05304585 | -0.0021663 | 0.60712432 | 0.99999994 |
| Sidt2 | 0.05692673 | 0.04745747 | 0.04962384 | -0.0021664 | 0.61431697 | 0.99999994 |
| Rnpepl1 | 0.21601085 | 0.16502956 | 0.1671971 | -0.0021675 | 0.53438321 | 0.99999994 |
| Abhd15 | 0.08651087 | 0.07181134 | 0.07397916 | -0.0021678 | 0.57732392 | 0.99999994 |
| Elk3 | 0.01557024 | 0.01178249 | 0.01395072 | -0.0021682 | 0.84955675 | 0.99999994 |
| Lrrn1 | 0.04730449 | 0.03932884 | 0.04149755 | -0.0021687 | 0.63603049 | 0.99999994 |
| Tfb2m | 0.08206152 | 0.06820506 | 0.07037553 | -0.0021705 | 0.58132737 | 0.99999994 |
| Mipol1 | 0.05314124 | 0.04426806 | 0.04643912 | -0.0021711 | 0.62217144 | 0.99999994 |
| Ccdc191 | 0.11629204 | 0.09514442 | 0.09731625 | -0.0021718 | 0.55904612 | 0.99999994 |
| 4930503L19F | 0.01533326 | 0.01156836 | 0.01374058 | -0.0021722 | 0.85366638 | 0.99999994 |
| Exosc1 | 0.24276469 | 0.18192312 | 0.18409734 | -0.0021742 | 0.53132978 | 0.99999994 |

| | | | | | | |
|---|---|---|---|---|---|---|
| Med12 | 0.06079474 | 0.05068624 | 0.05286171 | -0.0021755 | 0.60793257 | 0.99999994 |
| Ptk7 | 0.102043 | 0.08419359 | 0.08636999 | -0.0021764 | 0.56660251 | 0.99999994 |
| Scrn2 | 0.08204361 | 0.06818444 | 0.07036099 | -0.0021765 | 0.58156857 | 0.99999994 |
| 4930522L14R | 0.02657787 | 0.02145961 | 0.02363822 | -0.0021786 | 0.73022038 | 0.99999994 |
| Bbip1 | 0.64161839 | 0.36327908 | 0.36545784 | -0.0021788 | 0.51582718 | 0.99999994 |
| Zfp119a | 0.03409033 | 0.02798952 | 0.03016829 | -0.0021788 | 0.68464429 | 0.99999994 |
| Metrnl | 0.01548079 | 0.01169229 | 0.01387141 | -0.0021791 | 0.85214488 | 0.99999994 |
| Col1a1 | 0.01888402 | 0.01470277 | 0.01688215 | -0.0021794 | 0.80491039 | 0.99999994 |
| Sergef | 0.06559444 | 0.05467652 | 0.05685679 | -0.0021803 | 0.60073468 | 0.99999994 |
| Zfp84 | 0.06054357 | 0.0504709 | 0.05265194 | -0.002181 | 0.60862286 | 0.99999994 |
| Slc7a1 | 0.05493267 | 0.04576592 | 0.04794821 | -0.0021823 | 0.61903752 | 0.99999994 |
| Plekhb2 | 0.06898291 | 0.05747977 | 0.05966222 | -0.0021825 | 0.59618744 | 0.99999994 |
| Hdac3 | 0.3959585 | 0.26575923 | 0.26794224 | -0.002183 | 0.5216246 | 0.99999994 |
| Atp1b2 | 0.00917 | 0.00606673 | 0.00825175 | -0.002185 | 0.96101357 | 0.99999994 |
| Bcorl1 | 0.02616437 | 0.02109016 | 0.02327688 | -0.0021867 | 0.73412033 | 0.99999994 |
| Zfp677 | 0.01444135 | 0.01076226 | 0.01294909 | -0.0021868 | 0.86951888 | 0.99999994 |
| Micu3 | 0.11743723 | 0.09599727 | 0.09818489 | -0.0021876 | 0.55895002 | 0.99999994 |
| Daxx | 0.12576354 | 0.10226435 | 0.10445225 | -0.0021879 | 0.55544346 | 0.99999994 |
| Arid5a | 0.02021435 | 0.01586735 | 0.01805532 | -0.002188 | 0.79006011 | 0.99999994 |
| Hccs | 0.15842155 | 0.12604039 | 0.12822893 | -0.0021885 | 0.54522533 | 0.99999994 |
| Hook2 | 0.04333888 | 0.03592979 | 0.03811838 | -0.0021886 | 0.64883599 | 0.99999994 |
| Nuak1 | 0.01517096 | 0.01140763 | 0.01359662 | -0.002189 | 0.85805982 | 0.99999994 |
| Ddx56 | 0.23452321 | 0.17677926 | 0.1789693 | -0.00219 | 0.5324594 | 0.99999994 |
| Hcn3 | 0.02048329 | 0.01610127 | 0.01829224 | -0.002191 | 0.78735295 | 0.99999994 |
| Shroom3 | 0.04604617 | 0.03823481 | 0.04042722 | -0.0021924 | 0.64094307 | 0.99999994 |
| Igsf11 | 0.0092707 | 0.00614887 | 0.0083418 | -0.0021929 | 0.95992853 | 0.99999994 |
| Usp5 | 0.21040615 | 0.16136974 | 0.16356289 | -0.0021932 | 0.53555949 | 0.99999994 |
| Adpgk | 0.08818977 | 0.07313965 | 0.07533359 | -0.0021939 | 0.57685447 | 0.99999994 |
| Zfp445 | 0.04192794 | 0.03471731 | 0.03691182 | -0.0021945 | 0.65385155 | 0.99999994 |
| Tbc1d13 | 0.00966015 | 0.0064954 | 0.00868993 | -0.0021945 | 0.95361634 | 0.99999994 |

| | | | | | | |
|---|---|---|---|---|---|---|
| Sc5d | 0.08156314 | 0.06777424 | 0.0699706 | -0.0021964 | 0.58275302 | 0.99999994 |
| Catsper2 | 0.02411218 | 0.01928336 | 0.02148064 | -0.0021973 | 0.7520288 | 0.99999994 |
| Bloc1s4 | 0.09954954 | 0.08222296 | 0.08442177 | -0.0021988 | 0.56881923 | 0.99999994 |
| Clstn3 | 0.02772519 | 0.02244086 | 0.02463974 | -0.0021989 | 0.72375086 | 0.99999994 |
| Cry1 | 0.11704946 | 0.09569024 | 0.09789094 | -0.0022007 | 0.5594767 | 0.99999994 |
| Fech | 0.11851687 | 0.09680143 | 0.09900235 | -0.0022009 | 0.55881951 | 0.99999994 |
| Lrsam1 | 0.01540274 | 0.01159935 | 0.01380219 | -0.0022028 | 0.85596394 | 0.99999994 |
| Tecpr2 | 0.03156376 | 0.02577652 | 0.0279794 | -0.0022029 | 0.69988573 | 0.99999994 |
| Bbs10 | 0.01890682 | 0.01469879 | 0.01690227 | -0.0022035 | 0.80723804 | 0.99999994 |
| Cobll1 | 0.04753213 | 0.03948642 | 0.04169098 | -0.0022046 | 0.63757273 | 0.99999994 |
| Rps6kc1 | 0.02787237 | 0.02256325 | 0.02476811 | -0.0022049 | 0.72325872 | 0.99999994 |
| Ube2v1 | 0.02205698 | 0.01746897 | 0.01967681 | -0.0022078 | 0.77243055 | 0.99999994 |
| Wdr43 | 0.2365195 | 0.17800999 | 0.18021793 | -0.0022079 | 0.53249798 | 0.99999994 |
| Asb3 | 0.11294194 | 0.09255659 | 0.09476551 | -0.0022089 | 0.56165053 | 0.99999994 |
| Dis3 | 0.07366966 | 0.06131306 | 0.06352223 | -0.0022092 | 0.59153466 | 0.99999994 |
| Gas2l1 | 0.05737482 | 0.04778986 | 0.04999977 | -0.0022099 | 0.61569749 | 0.99999994 |
| Cd27 | 0.04081374 | 0.03374674 | 0.03595742 | -0.0022107 | 0.65881674 | 0.99999994 |
| Zbtb45 | 0.09827453 | 0.08119657 | 0.08340828 | -0.0022117 | 0.57005132 | 0.99999994 |
| Ints6 | 0.02720598 | 0.02197494 | 0.0241867 | -0.0022118 | 0.72863519 | 0.99999994 |
| Msh5 | 0.01649154 | 0.01255462 | 0.01476703 | -0.0022124 | 0.84068051 | 0.99999994 |
| Klhl8 | 0.03875122 | 0.03197184 | 0.034187 | -0.0022152 | 0.66687493 | 0.99999994 |
| Mid2 | 0.0150526 | 0.01127584 | 0.01349162 | -0.0022158 | 0.86284654 | 0.99999994 |
| Kifc2 | 0.01346196 | 0.00986287 | 0.01207887 | -0.002216 | 0.88899149 | 0.99999994 |
| Cgref1 | 0.03099467 | 0.02526819 | 0.02748536 | -0.0022172 | 0.70434929 | 0.99999994 |
| Chn1 | 0.05930535 | 0.04939944 | 0.05161689 | -0.0022174 | 0.61254379 | 0.99999994 |
| Zfp110 | 0.06005863 | 0.05002923 | 0.05224678 | -0.0022175 | 0.61122774 | 0.99999994 |
| Ubtd1 | 0.04841243 | 0.0402204 | 0.04243848 | -0.0022181 | 0.63604195 | 0.99999994 |
| Kcnn2 | 0.00733592 | 0.00439073 | 0.00660953 | -0.0022188 | 0.9872755 | 0.99999994 |
| Tuft1 | 0.03598772 | 0.02958809 | 0.03180726 | -0.0022192 | 0.67882224 | 0.99999994 |
| Tdp1 | 0.08352953 | 0.0693475 | 0.07156679 | -0.0022193 | 0.58176607 | 0.99999994 |

| | | | | | | |
|---|---|---|---|---|---|---|
| Ints12 | 0.0855859 | 0.07101221 | 0.0732317 | -0.0022195 | 0.57993958 | 0.99999994 |
| Arv1 | 0.03895254 | 0.03214018 | 0.03436002 | -0.0022198 | 0.66641529 | 0.99999994 |
| Zfp30 | 0.14731619 | 0.11806493 | 0.12028544 | -0.0022205 | 0.54889844 | 0.99999994 |
| Gm16536 | 0.01962596 | 0.01531508 | 0.0175367 | -0.0022216 | 0.80045625 | 0.99999994 |
| Pomt1 | 0.0359904 | 0.02958617 | 0.03180957 | -0.0022234 | 0.67912768 | 0.99999994 |
| Lamc1 | 0.08697755 | 0.07213189 | 0.07435594 | -0.002224 | 0.57890656 | 0.99999994 |
| Nrm | 0.0301384 | 0.02451616 | 0.0267413 | -0.0022251 | 0.71016731 | 0.99999994 |
| Dbr1 | 0.08945998 | 0.07413078 | 0.07635637 | -0.0022256 | 0.57691833 | 0.99999994 |
| Gstm7 | 0.07940417 | 0.06598768 | 0.06821341 | -0.0022257 | 0.58597009 | 0.99999994 |
| 1810024B03 | 0.05187464 | 0.04314395 | 0.04537 | -0.0022261 | 0.62801104 | 0.99999994 |
| 4632404H12 | 0.02052031 | 0.01609855 | 0.01832484 | -0.0022263 | 0.79065059 | 0.99999994 |
| Gmip | 0.04022081 | 0.03322048 | 0.03544896 | -0.0022285 | 0.66219036 | 0.99999994 |
| Stard4 | 0.03010302 | 0.02448099 | 0.02671053 | -0.0022295 | 0.71076154 | 0.99999994 |
| Fbxl4 | 0.05843837 | 0.0486612 | 0.05089116 | -0.00223 | 0.61473044 | 0.99999994 |
| Mif4gd | 0.08602659 | 0.07135788 | 0.07358793 | -0.00223 | 0.57993127 | 0.99999994 |
| Gas8 | 0.21662009 | 0.16535921 | 0.16759019 | -0.002231 | 0.53530418 | 0.99999994 |
| Sugp2 | 0.03030537 | 0.02465542 | 0.02688645 | -0.002231 | 0.70964361 | 0.99999994 |
| Med1 | 0.15446188 | 0.12318149 | 0.12541294 | -0.0022315 | 0.5471388 | 0.99999994 |
| Wdpcp | 0.03386461 | 0.02774049 | 0.02997304 | -0.0022325 | 0.68998235 | 0.99999994 |
| Rabl3 | 0.08420356 | 0.06987705 | 0.07211299 | -0.0022359 | 0.58175592 | 0.99999994 |
| Abcc1 | 0.02687368 | 0.02166007 | 0.02389658 | -0.0022365 | 0.73334965 | 0.99999994 |
| Slc25a42 | 0.01816231 | 0.01400806 | 0.01624482 | -0.0022368 | 0.82030296 | 0.99999994 |
| Klhl22 | 0.1310362 | 0.10613966 | 0.10837748 | -0.0022378 | 0.55465971 | 0.99999994 |
| Gm5113 | 0.02512875 | 0.02013212 | 0.02237104 | -0.0022389 | 0.74735063 | 0.99999994 |
| Gpa33 | 0.01530443 | 0.01147467 | 0.01371501 | -0.0022403 | 0.86155051 | 0.99999994 |
| Cbwd1 | 0.09121674 | 0.07552596 | 0.07776819 | -0.0022422 | 0.57609672 | 0.99999994 |
| Nr2c1 | 0.04647728 | 0.03855112 | 0.04079412 | -0.002243 | 0.64281339 | 0.99999994 |
| Srrm1 | 0.81384156 | 0.40371436 | 0.40595771 | -0.0022433 | 0.51467113 | 0.99999994 |
| Parp4 | 0.04283262 | 0.03544155 | 0.03768571 | -0.0022442 | 0.65408843 | 0.99999994 |
| Mgme1 | 0.03882348 | 0.03200422 | 0.0342491 | -0.0022449 | 0.66868898 | 0.99999994 |

| | | | | | | |
|---|---|---|---|---|---|---|
| Rtca | 0.2068509 | 0.15899403 | 0.16124056 | -0.0022465 | 0.53694554 | 0.99999994 |
| Zfp944 | 0.0575492 | 0.04789738 | 0.05014601 | -0.0022486 | 0.61733209 | 0.99999994 |
| Mkrn2 | 0.11094893 | 0.0909903 | 0.09324138 | -0.0022511 | 0.56383567 | 0.99999994 |
| Gla | 0.02669063 | 0.02148483 | 0.02373671 | -0.0022519 | 0.73613775 | 0.99999994 |
| Naaa | 0.03501673 | 0.02871313 | 0.03096903 | -0.0022559 | 0.68611756 | 0.99999994 |
| Hectd1 | 0.32447441 | 0.22914244 | 0.23139907 | -0.0022566 | 0.52587849 | 0.99999994 |
| Hif1an | 0.1138807 | 0.09322456 | 0.09548171 | -0.0022572 | 0.56251698 | 0.99999994 |
| Ptrh2 | 0.23040905 | 0.17412571 | 0.17638341 | -0.0022577 | 0.53394917 | 0.99999994 |
| 1700105P06I | 0.01198738 | 0.0085088 | 0.01076649 | -0.0022577 | 0.91861487 | 0.99999994 |
| Pmepa1 | 0.06209087 | 0.0516833 | 0.05394302 | -0.0022597 | 0.60981659 | 0.99999994 |
| Marveld1 | 0.00724975 | 0.00427186 | 0.00653227 | -0.0022604 | 0.9893702 | 0.99999994 |
| Wdr48 | 0.08356259 | 0.06932973 | 0.07159359 | -0.0022639 | 0.58335347 | 0.99999994 |
| Zfp868 | 0.03775363 | 0.03106454 | 0.03332895 | -0.0022644 | 0.67445202 | 0.99999994 |
| Zfp710 | 0.04488112 | 0.03716949 | 0.03943466 | -0.0022652 | 0.64889816 | 0.99999994 |
| Cep76 | 0.03628271 | 0.02979539 | 0.0320617 | -0.0022663 | 0.68100458 | 0.99999994 |
| Zbtb34 | 0.02935001 | 0.02378913 | 0.02605547 | -0.0022663 | 0.71869805 | 0.99999994 |
| Esrra | 0.0505243 | 0.04196167 | 0.04422822 | -0.0022666 | 0.63349227 | 0.99999994 |
| 3930402G23 | 0.00926337 | 0.00606851 | 0.00833524 | -0.0022667 | 0.96486501 | 0.99999994 |
| Rdh13 | 0.04605517 | 0.03816808 | 0.04043488 | -0.0022668 | 0.64548483 | 0.99999994 |
| Nup85 | 0.07779617 | 0.06463446 | 0.06690148 | -0.002267 | 0.58922718 | 0.99999994 |
| Blvra | 0.16591403 | 0.13123916 | 0.13350755 | -0.0022684 | 0.5450228 | 0.99999994 |
| Zfp617 | 0.07628138 | 0.06339418 | 0.06566309 | -0.0022689 | 0.59095538 | 0.99999994 |
| Slc9a1 | 0.01886885 | 0.01459972 | 0.01686877 | -0.002269 | 0.814702 | 0.99999994 |
| Ulk4 | 0.02598077 | 0.02084692 | 0.02311638 | -0.0022695 | 0.7432801 | 0.99999994 |
| Clcn7 | 0.01495969 | 0.01113585 | 0.01340918 | -0.0022733 | 0.87044341 | 0.99999994 |
| Wdr1 | 0.36045926 | 0.24805113 | 0.25032474 | -0.0022736 | 0.52410429 | 0.99999994 |
| Slc25a24 | 0.02791684 | 0.02253076 | 0.02480689 | -0.0022761 | 0.72931698 | 0.99999994 |
| Myl6b | 0.06995972 | 0.05819159 | 0.06046866 | -0.0022771 | 0.59896091 | 0.99999994 |
| D130017N08 | 0.01571849 | 0.01180495 | 0.01408214 | -0.0022772 | 0.85912259 | 0.99999994 |
| Oxr1 | 0.11719665 | 0.0957252 | 0.09800255 | -0.0022773 | 0.56146243 | 0.99999994 |

| | | | | | | |
|---|---|---|---|---|---|---|
| Cables2 | 0.03994337 | 0.03293282 | 0.0352109 | -0.0022781 | 0.66663998 | 0.99999994 |
| Frem2 | 0.02059781 | 0.01611098 | 0.01839309 | -0.0022821 | 0.79556431 | 0.99999994 |
| Neurl1b | 0.01645026 | 0.01244742 | 0.01473047 | -0.002283 | 0.84888352 | 0.99999994 |
| Sarm1 | 0.0173812 | 0.0132707 | 0.01555435 | -0.0022837 | 0.83578227 | 0.99999994 |
| Tmem64 | 0.03477718 | 0.02847782 | 0.03076207 | -0.0022842 | 0.68944218 | 0.99999994 |
| Cstf1 | 0.0933103 | 0.07716194 | 0.07944653 | -0.0022846 | 0.57589869 | 0.99999994 |
| Aste1 | 0.02559113 | 0.02048904 | 0.02277564 | -0.0022866 | 0.74802076 | 0.99999994 |
| Fastkd3 | 0.04520049 | 0.03741946 | 0.0397069 | -0.0022874 | 0.64931064 | 0.99999994 |
| Bloc1s3 | 0.05646641 | 0.04694823 | 0.04923741 | -0.0022892 | 0.62151889 | 0.99999994 |
| Gm10643 | 0.01080181 | 0.00741967 | 0.00970944 | -0.0022898 | 0.94176484 | 0.99999994 |
| Pced1a | 0.03351012 | 0.02737597 | 0.02966628 | -0.0022903 | 0.69633509 | 0.99999994 |
| Nudt1 | 0.09435618 | 0.07799246 | 0.0802833 | -0.0022908 | 0.57531999 | 0.99999994 |
| Gtf3c3 | 0.08291726 | 0.06877933 | 0.07107021 | -0.0022909 | 0.58494547 | 0.99999994 |
| Klhl20 | 0.03995668 | 0.03293096 | 0.03522232 | -0.0022914 | 0.6675015 | 0.99999994 |
| Mslnl | 0.01314898 | 0.00950895 | 0.01180054 | -0.0022916 | 0.90192856 | 0.99999994 |
| Ctdp1 | 0.0277868 | 0.02240084 | 0.02469348 | -0.0022926 | 0.73171278 | 0.99999994 |
| Kif16b | 0.07607723 | 0.06320324 | 0.06549601 | -0.0022928 | 0.59212484 | 0.99999994 |
| Mus81 | 0.08563585 | 0.07097725 | 0.07327209 | -0.0022948 | 0.5825703 | 0.99999994 |
| Vps33b | 0.03409085 | 0.02787384 | 0.03016874 | -0.0022949 | 0.69369121 | 0.99999994 |
| 1700037H04I | 0.2058705 | 0.15830075 | 0.16059762 | -0.0022969 | 0.53792177 | 0.99999994 |
| Ccdc142os | 0.01745596 | 0.01332238 | 0.01562046 | -0.0022981 | 0.83627941 | 0.99999994 |
| Dis3l | 0.05147286 | 0.04273199 | 0.04503049 | -0.0022985 | 0.63298186 | 0.99999994 |
| Stard6 | 0.01223802 | 0.0086912 | 0.01098975 | -0.0022985 | 0.91806946 | 0.99999994 |
| Nudt16 | 0.07380841 | 0.06133754 | 0.06363615 | -0.0022986 | 0.59500361 | 0.99999994 |
| Gemin8 | 0.0471651 | 0.03907898 | 0.04137907 | -0.0023001 | 0.64430848 | 0.99999994 |
| Bmi1 | 0.10296132 | 0.08478235 | 0.08708314 | -0.0023008 | 0.5697999 | 0.99999994 |
| Stk38l | 0.04277812 | 0.03533747 | 0.03763912 | -0.0023016 | 0.65801027 | 0.99999994 |
| Ap4e1 | 0.01957995 | 0.01519439 | 0.01749613 | -0.0023017 | 0.80940084 | 0.99999994 |
| Smurf1 | 0.0204394 | 0.01595155 | 0.01825357 | -0.002302 | 0.79939313 | 0.99999994 |
| Dmkn | 0.01164733 | 0.0081599 | 0.01046348 | -0.0023036 | 0.9285976 | 0.99999994 |

| | | | | | | |
|---|---|---|---|---|---|---|
| 4933428C19l | 0.00695581 | 0.00396508 | 0.00626867 | -0.0023036 | 0.99277854 | 0.99999994 |
| Utp20 | 0.0236083 | 0.01873504 | 0.02103885 | -0.0023038 | 0.76696538 | 0.99999994 |
| Cdc16 | 0.32818551 | 0.23109797 | 0.23340255 | -0.0023046 | 0.52620114 | 0.99999994 |
| Gltp | 0.15918991 | 0.12646333 | 0.12877328 | -0.00231 | 0.54752179 | 0.99999994 |
| Nhlrc3 | 0.05044483 | 0.04185081 | 0.04416097 | -0.0023102 | 0.63615877 | 0.99999994 |
| Igfbp6 | 0.01164672 | 0.0081516 | 0.01046294 | -0.0023113 | 0.9292781 | 0.99999994 |
| Gbp7 | 0.01397521 | 0.01022198 | 0.01253505 | -0.0023131 | 0.89033022 | 0.99999994 |
| Aars2 | 0.02540643 | 0.02029994 | 0.02261404 | -0.0023141 | 0.75211134 | 0.99999994 |
| Stard13 | 0.03491278 | 0.02856472 | 0.03087923 | -0.0023145 | 0.69108127 | 0.99999994 |
| Cpeb2 | 0.02065705 | 0.01612862 | 0.01844526 | -0.0023166 | 0.79842488 | 0.99999994 |
| Prelid2 | 0.03973651 | 0.03271598 | 0.03503335 | -0.0023174 | 0.67014071 | 0.99999994 |
| Il17rc | 0.01779512 | 0.01359932 | 0.01592033 | -0.002321 | 0.83408269 | 0.99999994 |
| Nubp1 | 0.18291122 | 0.14292535 | 0.14524668 | -0.0023213 | 0.54236024 | 0.99999994 |
| Fam161b | 0.0300478 | 0.02433942 | 0.02666252 | -0.0023231 | 0.71903178 | 0.99999994 |
| Gchfr | 0.09796934 | 0.08084196 | 0.08316545 | -0.0023235 | 0.57376454 | 0.99999994 |
| Vezf1 | 0.33425435 | 0.23432744 | 0.23665235 | -0.0023249 | 0.52606951 | 0.99999994 |
| Rpp21 | 0.52919646 | 0.32336483 | 0.32569004 | -0.0023252 | 0.5189514 | 0.99999994 |
| Wnk2 | 0.0273666 | 0.0220014 | 0.02432689 | -0.0023255 | 0.73771064 | 0.99999994 |
| Dck | 0.02441551 | 0.01942088 | 0.02174645 | -0.0023256 | 0.76171535 | 0.99999994 |
| Cblc | 0.03311059 | 0.02699163 | 0.02932037 | -0.0023287 | 0.70148389 | 0.99999994 |
| Zfp639 | 0.09643667 | 0.07961438 | 0.08194448 | -0.0023301 | 0.57506116 | 0.99999994 |
| Sumf2 | 0.09027963 | 0.07468443 | 0.07701548 | -0.002331 | 0.57983433 | 0.99999994 |
| Zfp433 | 0.04804065 | 0.03979173 | 0.04212289 | -0.0023312 | 0.64370333 | 0.99999994 |
| Zfp949 | 0.1271135 | 0.10312767 | 0.10546043 | -0.0023328 | 0.55852767 | 0.99999994 |
| Tgds | 0.20972708 | 0.16078756 | 0.16312035 | -0.0023328 | 0.53791911 | 0.99999994 |
| Sytl1 | 0.13370394 | 0.10801585 | 0.11035038 | -0.0023345 | 0.55599368 | 0.99999994 |
| 4833420G17 | 0.18065558 | 0.14136895 | 0.14370749 | -0.0023385 | 0.54312844 | 0.99999994 |
| Tfcp2 | 0.05935845 | 0.04932252 | 0.05166132 | -0.0023388 | 0.6184241 | 0.99999994 |
| Cpsf2 | 0.20725811 | 0.15916613 | 0.16150729 | -0.0023412 | 0.5384336 | 0.99999994 |
| Glmp | 0.48732473 | 0.30653389 | 0.30887505 | -0.0023412 | 0.52011922 | 0.99999994 |

| | | | | | | |
|---|---|---|---|---|---|---|
| Stk4 | 0.10571906 | 0.08687631 | 0.08921834 | -0.002342 | 0.56935519 | 0.99999994 |
| Ppp1r12c | 0.02692092 | 0.02159489 | 0.02393783 | -0.0023429 | 0.74263737 | 0.99999994 |
| Rab40c | 0.10027483 | 0.08264966 | 0.08499386 | -0.0023442 | 0.5728316 | 0.99999994 |
| Tmem51os1 | 0.01316066 | 0.00946639 | 0.01181093 | -0.0023445 | 0.90680376 | 0.99999994 |
| Zfp668 | 0.06851894 | 0.05693393 | 0.05927882 | -0.0023449 | 0.6038421 | 0.99999994 |
| Nipal2 | 0.07327356 | 0.06084948 | 0.06319691 | -0.0023474 | 0.59764191 | 0.99999994 |
| Insr | 0.07484526 | 0.06213927 | 0.06448679 | -0.0023475 | 0.59573088 | 0.99999994 |
| Tnfaip8 | 0.06372451 | 0.05295269 | 0.05530331 | -0.0023506 | 0.61138177 | 0.99999994 |
| Ubap2 | 0.18754175 | 0.1460351 | 0.148389 | -0.0023539 | 0.54204617 | 0.99999994 |
| B4galt6 | 0.23192903 | 0.1749857 | 0.17734075 | -0.002355 | 0.53521863 | 0.99999994 |
| Zdhhc21 | 0.07426521 | 0.06165493 | 0.06401106 | -0.0023561 | 0.59677494 | 0.99999994 |
| Fer1l6 | 0.00439428 | 0.00161028 | 0.00396706 | -0.0023568 | 0.99996165 | 0.99999994 |
| A630072M18 | 0.02981238 | 0.02409896 | 0.02645778 | -0.0023588 | 0.72355766 | 0.99999994 |
| Usp39 | 0.10893272 | 0.08933492 | 0.09169448 | -0.0023596 | 0.56800094 | 0.99999994 |
| Sugp1 | 0.12294494 | 0.09998037 | 0.10234011 | -0.0023597 | 0.56099052 | 0.99999994 |
| Fbf1 | 0.0304252 | 0.02463039 | 0.02699061 | -0.0023602 | 0.71974085 | 0.99999994 |
| Apod | 0.00670165 | 0.00367947 | 0.00604065 | -0.0023612 | 0.99536382 | 0.99999994 |
| Cycs | 1.2848743 | 0.42339512 | 0.42575689 | -0.0023618 | 0.51472733 | 0.99999994 |
| Rad52 | 0.04916208 | 0.0407116 | 0.04307436 | -0.0023628 | 0.64248945 | 0.99999994 |
| Chrnb2 | 0.02138123 | 0.01671943 | 0.01908263 | -0.0023632 | 0.79512214 | 0.99999994 |
| Recql | 0.03534147 | 0.02888559 | 0.03124949 | -0.0023639 | 0.69270323 | 0.99999994 |
| Gm4577 | 0.01194927 | 0.0083676 | 0.01073254 | -0.0023649 | 0.92877747 | 0.99999994 |
| Zfp809 | 0.02074272 | 0.01615361 | 0.01852069 | -0.0023671 | 0.80254079 | 0.99999994 |
| Kntc1 | 0.00521513 | 0.00233759 | 0.00470549 | -0.0023679 | 0.99959539 | 0.99999994 |
| Tnfrsf1a | 0.03373796 | 0.02749495 | 0.02986346 | -0.0023685 | 0.70122382 | 0.99999994 |
| 4930523C07I | 0.01375616 | 0.0099688 | 0.0123404 | -0.0023716 | 0.89959247 | 0.99999994 |
| Cntrob | 0.02860767 | 0.02303694 | 0.02540904 | -0.0023721 | 0.73283645 | 0.99999994 |
| 1110059G10 | 0.20417522 | 0.15711137 | 0.15948348 | -0.0023721 | 0.53943272 | 0.99999994 |
| Actr3b | 0.01366989 | 0.0098894 | 0.01226372 | -0.0023743 | 0.90124994 | 0.99999994 |
| Slc9a6 | 0.05155748 | 0.04272696 | 0.04510201 | -0.002375 | 0.63702591 | 0.99999994 |

| | | | | | | |
|---|---|---|---|---|---|---|
| Syt16 | 0.0179837 | 0.01371192 | 0.01608701 | -0.0023751 | 0.83713336 | 0.99999994 |
| AI661453 | 0.0221863 | 0.01741283 | 0.01979046 | -0.0023776 | 0.7880571 | 0.99999994 |
| Tex9 | 0.09836081 | 0.08109887 | 0.08347692 | -0.0023781 | 0.57519784 | 0.99999994 |
| Itpk1 | 0.03238292 | 0.02631172 | 0.02868989 | -0.0023782 | 0.70944375 | 0.99999994 |
| Usp42 | 0.07102143 | 0.05896454 | 0.06134404 | -0.0023795 | 0.60187201 | 0.99999994 |
| Gm17546 | 0.01937953 | 0.01493806 | 0.01731937 | -0.0023813 | 0.81995907 | 0.99999994 |
| Zfp386 | 0.17717855 | 0.13893992 | 0.14132391 | -0.002384 | 0.54470159 | 0.99999994 |
| Dtnb | 0.11455197 | 0.09360652 | 0.09599317 | -0.0023867 | 0.56572293 | 0.99999994 |
| Spata2l | 0.0196345 | 0.01515654 | 0.01754423 | -0.0023877 | 0.81750561 | 0.99999994 |
| 1010001B22 | 0.05080733 | 0.04207789 | 0.04446771 | -0.0023898 | 0.63972777 | 0.99999994 |
| Galc | 0.0196627 | 0.01517739 | 0.01756909 | -0.0023917 | 0.81756861 | 0.99999994 |
| Ctu1 | 0.04860599 | 0.04020982 | 0.04260272 | -0.0023929 | 0.64575059 | 0.99999994 |
| Rnf213 | 0.02396374 | 0.01895742 | 0.02135052 | -0.0023931 | 0.77219258 | 0.99999994 |
| Mark4 | 0.0836747 | 0.06928579 | 0.07168447 | -0.0023987 | 0.588128 | 0.99999994 |
| Pnpla7 | 0.00769441 | 0.0045305 | 0.00693083 | -0.0024003 | 0.98942428 | 0.99999994 |
| Efna1 | 0.02265924 | 0.01780411 | 0.02020592 | -0.0024018 | 0.78559212 | 0.99999994 |
| Gm26759 | 0.01558825 | 0.01156433 | 0.01396669 | -0.0024024 | 0.87388212 | 0.99999994 |
| Axin2 | 0.02727915 | 0.02184732 | 0.02425057 | -0.0024032 | 0.74526029 | 0.99999994 |
| Plekhm3 | 0.02847851 | 0.02289214 | 0.0252965 | -0.0024044 | 0.73652504 | 0.99999994 |
| Lncpint | 0.00955078 | 0.00618778 | 0.00859219 | -0.0024044 | 0.96874738 | 0.99999994 |
| Zfp141 | 0.02852823 | 0.02293394 | 0.02533982 | -0.0024059 | 0.73630211 | 0.99999994 |
| Zfp39 | 0.02784944 | 0.02234203 | 0.02474811 | -0.0024061 | 0.74123289 | 0.99999994 |
| Zdhhc13 | 0.09707388 | 0.08004527 | 0.0824524 | -0.0024071 | 0.57703979 | 0.99999994 |
| Klhl15 | 0.0275711 | 0.02209791 | 0.02450532 | -0.0024074 | 0.74341934 | 0.99999994 |
| Casp3 | 0.09368878 | 0.07733718 | 0.07974946 | -0.0024123 | 0.57978486 | 0.99999994 |
| Dnmt3b | 0.03209077 | 0.02602426 | 0.02843659 | -0.0024123 | 0.71385088 | 0.99999994 |
| Pan2 | 0.02659582 | 0.02123811 | 0.02365389 | -0.0024158 | 0.75168959 | 0.99999994 |
| Eral1 | 0.06541982 | 0.05429534 | 0.05671188 | -0.0024165 | 0.61165391 | 0.99999994 |
| P2rx4 | 0.09872159 | 0.08134725 | 0.08376383 | -0.0024166 | 0.57614268 | 0.99999994 |
| Numb | 0.05407236 | 0.04480695 | 0.04722393 | -0.002417 | 0.63332801 | 0.99999994 |

| | | | | | | |
|---|---|---|---|---|---|---|
| Syne2 | 0.15846162 | 0.12583791 | 0.12825733 | -0.0024194 | 0.54996151 | 0.99999994 |
| Jak2 | 0.0269081 | 0.02150717 | 0.02392664 | -0.0024195 | 0.74955569 | 0.99999994 |
| Gm37459 | 0.00665523 | 0.00357723 | 0.005999 | -0.0024218 | 0.99639672 | 0.99999994 |
| Tsn | 0.86698879 | 0.41157006 | 0.41399219 | -0.0024221 | 0.5155325 | 0.99999994 |
| Dennd3 | 0.01383783 | 0.00999084 | 0.01241298 | -0.0024221 | 0.90300308 | 0.99999994 |
| Zfp623 | 0.03152211 | 0.02552071 | 0.02794326 | -0.0024226 | 0.71805568 | 0.99999994 |
| Whamm | 0.0481386 | 0.03978284 | 0.04220605 | -0.0024232 | 0.64883124 | 0.99999994 |
| AI597479 | 0.03867391 | 0.03169726 | 0.03412054 | -0.0024233 | 0.68180085 | 0.99999994 |
| Ddr1 | 0.34855437 | 0.24176127 | 0.24418517 | -0.0024239 | 0.52634064 | 0.99999994 |
| Egf | 0.01156519 | 0.00796633 | 0.01039026 | -0.0024239 | 0.93977126 | 0.99999994 |
| Ppil4 | 0.3682027 | 0.25182959 | 0.25425419 | -0.0024246 | 0.52530629 | 0.99999994 |
| Wdr44 | 0.03634817 | 0.02969182 | 0.03211815 | -0.0024263 | 0.69246436 | 0.99999994 |
| Lgals1 | 0.21696848 | 0.16538568 | 0.1678148 | -0.0024291 | 0.53837904 | 0.99999994 |
| Ncoa3 | 0.05255253 | 0.04351283 | 0.04594242 | -0.0024296 | 0.63758459 | 0.99999994 |
| 4732491K20I | 0.02279351 | 0.01789129 | 0.02032382 | -0.0024325 | 0.78718524 | 0.99999994 |
| Ubtd2 | 0.02845243 | 0.0228406 | 0.02527378 | -0.0024332 | 0.73918233 | 0.99999994 |
| Pla1a | 0.02664429 | 0.02126304 | 0.02369623 | -0.0024332 | 0.75285243 | 0.99999994 |
| Zhx1 | 0.15360226 | 0.12236468 | 0.12479924 | -0.0024346 | 0.5516577 | 0.99999994 |
| Osgin1 | 0.02132243 | 0.01659411 | 0.0190309 | -0.0024368 | 0.80297717 | 0.99999994 |
| 9130604C24I | 0.02297453 | 0.01804453 | 0.02048275 | -0.0024382 | 0.78592412 | 0.99999994 |
| Rasgef1c | 0.01305144 | 0.00927439 | 0.01171377 | -0.0024394 | 0.91715444 | 0.99999994 |
| Atg2a | 0.04645732 | 0.03833566 | 0.04077714 | -0.0024415 | 0.65488442 | 0.99999994 |
| Hmgxb3 | 0.04007001 | 0.0328774 | 0.03531957 | -0.0024422 | 0.67733343 | 0.99999994 |
| Nr1d1 | 0.00704499 | 0.00390392 | 0.00634865 | -0.0024447 | 0.99481478 | 0.99999994 |
| Gpr19 | 0.01949135 | 0.01497209 | 0.01741799 | -0.0024459 | 0.82503119 | 0.99999994 |
| Wdr47 | 0.13846172 | 0.11140124 | 0.11384786 | -0.0024466 | 0.5568733 | 0.99999994 |
| Polr1e | 0.0505532 | 0.04180529 | 0.04425268 | -0.0024474 | 0.6436118 | 0.99999994 |
| Taf15 | 0.18767018 | 0.1460266 | 0.14847583 | -0.0024492 | 0.54371676 | 0.99999994 |
| B3galnt2 | 0.03173193 | 0.02567399 | 0.02812533 | -0.0024513 | 0.71909401 | 0.99999994 |
| Ripk1 | 0.06853306 | 0.05683816 | 0.05929049 | -0.0024523 | 0.608463 | 0.99999994 |

| | | | | | | |
|---|---|---|---|---|---|---|
| Lrriq3 | 0.00952876 | 0.00611941 | 0.0085725 | -0.0024531 | 0.97159525 | 0.99999994 |
| Pura | 0.31300998 | 0.22267879 | 0.22513436 | -0.0024556 | 0.52893847 | 0.99999994 |
| Gfod2 | 0.16005398 | 0.12692847 | 0.12938464 | -0.0024562 | 0.55027684 | 0.99999994 |
| Ggct | 0.15864817 | 0.1259292 | 0.12838955 | -0.0024603 | 0.5507501 | 0.99999994 |
| Rhot1 | 0.16299704 | 0.12899841 | 0.13146023 | -0.0024618 | 0.54960036 | 0.99999994 |
| Ipo13 | 0.0512527 | 0.042382 | 0.04484437 | -0.0024624 | 0.64262967 | 0.99999994 |
| Plgrkt | 0.35447485 | 0.24479021 | 0.24725296 | -0.0024628 | 0.52643065 | 0.99999994 |
| Sertad4 | 0.0097815 | 0.00633463 | 0.00879838 | -0.0024637 | 0.96883454 | 0.99999994 |
| Gm14286 | 0.00872988 | 0.00539424 | 0.00785804 | -0.0024638 | 0.98155699 | 0.99999994 |
| Rab23 | 0.07367916 | 0.06106568 | 0.06353003 | -0.0024644 | 0.60187463 | 0.99999994 |
| Cep85 | 0.02751354 | 0.02199065 | 0.0244551 | -0.0024645 | 0.74882344 | 0.99999994 |
| Ap5m1 | 0.04596034 | 0.03788883 | 0.04035415 | -0.0024653 | 0.6578679 | 0.99999994 |
| Wdr20 | 0.12381182 | 0.10052502 | 0.10299074 | -0.0024657 | 0.5633077 | 0.99999994 |
| A430033K04 | 0.04534648 | 0.03736212 | 0.0398313 | -0.0024692 | 0.66006265 | 0.99999994 |
| Sema4a | 0.02307465 | 0.01809698 | 0.02057063 | -0.0024737 | 0.78827135 | 0.99999994 |
| A930029G22 | 0.01732317 | 0.0130262 | 0.01550302 | -0.0024768 | 0.85620986 | 0.99999994 |
| Soga3 | 0.03422422 | 0.02780572 | 0.03028408 | -0.0024784 | 0.70703293 | 0.99999994 |
| Mansc1 | 0.04264771 | 0.03504871 | 0.03752761 | -0.0024789 | 0.66991814 | 0.99999994 |
| B4galt2 | 0.01825528 | 0.01384724 | 0.01632696 | -0.0024797 | 0.84398213 | 0.99999994 |
| Prps2 | 0.03800366 | 0.03106336 | 0.03354411 | -0.0024807 | 0.68873537 | 0.99999994 |
| Fzd7 | 0.01104778 | 0.00744772 | 0.00992889 | -0.0024812 | 0.95188203 | 0.99999994 |
| Borcs6 | 0.12149124 | 0.09876529 | 0.101247 | -0.0024817 | 0.56480234 | 0.99999994 |
| Cpt1a | 0.03764511 | 0.03075358 | 0.03323553 | -0.002482 | 0.69043405 | 0.99999994 |
| Poc1b | 0.04977861 | 0.04111324 | 0.04359685 | -0.0024836 | 0.64772927 | 0.99999994 |
| Fosl2 | 0.02503959 | 0.01980933 | 0.022293 | -0.0024837 | 0.77083039 | 0.99999994 |
| Bhlhe41 | 0.01484931 | 0.01082513 | 0.01331123 | -0.0024861 | 0.89310087 | 0.99999994 |
| Adcy9 | 0.02543286 | 0.02015084 | 0.02263717 | -0.0024863 | 0.76763976 | 0.99999994 |
| Inpp5b | 0.02959694 | 0.02378252 | 0.02627036 | -0.0024878 | 0.73577225 | 0.99999994 |
| Enkd1 | 0.04168825 | 0.03421862 | 0.03670663 | -0.002488 | 0.67406786 | 0.99999994 |
| Lrrfip1 | 0.11448952 | 0.09345639 | 0.09594561 | -0.0024892 | 0.56855371 | 0.99999994 |

| | | | | | | |
|---|---|---|---|---|---|---|
| Kdm4b | 0.06481712 | 0.05372205 | 0.05621147 | -0.0024894 | 0.61592226 | 0.99999994 |
| Tfap4 | 0.05634751 | 0.04664611 | 0.04913756 | -0.0024915 | 0.63213079 | 0.99999994 |
| Egfr | 0.0051393 | 0.00214582 | 0.00463731 | -0.0024915 | 0.99982573 | 0.99999994 |
| Ccdc88a | 0.28615525 | 0.20750133 | 0.20999502 | -0.0024937 | 0.53150117 | 0.99999994 |
| Dip2a | 0.02426777 | 0.01912222 | 0.021617 | -0.0024948 | 0.77881499 | 0.99999994 |
| Rfc2 | 0.34979847 | 0.24233669 | 0.24483222 | -0.0024955 | 0.52704629 | 0.99999994 |
| Tceanc2 | 0.06017829 | 0.0498492 | 0.05234677 | -0.0024976 | 0.62460139 | 0.99999994 |
| Kcnq4 | 0.03495643 | 0.02841806 | 0.03091694 | -0.0024989 | 0.70471269 | 0.99999994 |
| Pcmtd2 | 0.23233843 | 0.17509753 | 0.17759821 | -0.0025007 | 0.53733623 | 0.99999994 |
| Mertk | 0.00911763 | 0.00570323 | 0.00820491 | -0.0025017 | 0.97879791 | 0.99999994 |
| Fam163a | 0.03522362 | 0.02864564 | 0.03114773 | -0.0025021 | 0.70357225 | 0.99999994 |
| Hnf1a | 0.05686745 | 0.0470704 | 0.04957409 | -0.0025037 | 0.63162945 | 0.99999994 |
| Kif3c | 0.01737784 | 0.01304728 | 0.01555138 | -0.0025041 | 0.85809792 | 0.99999994 |
| Xylt1 | 0.01383935 | 0.00990761 | 0.01241433 | -0.0025067 | 0.91053513 | 0.99999994 |
| Asxl2 | 0.28922503 | 0.20924838 | 0.21175893 | -0.0025106 | 0.53145021 | 0.99999994 |
| Cdc26 | 0.75647819 | 0.39245832 | 0.39497052 | -0.0025122 | 0.51688524 | 0.99999994 |
| Mphosph6 | 0.32848579 | 0.23105148 | 0.23356414 | -0.0025127 | 0.52854321 | 0.99999994 |
| Cyth3 | 0.04986037 | 0.04115326 | 0.04366611 | -0.0025128 | 0.64915968 | 0.99999994 |
| Alg8 | 0.05636527 | 0.04663869 | 0.04915248 | -0.0025138 | 0.63323144 | 0.99999994 |
| Abcd1 | 0.0425616 | 0.03494018 | 0.03745397 | -0.0025138 | 0.67247376 | 0.99999994 |
| Lonp2 | 0.19321343 | 0.14969123 | 0.1522059 | -0.0025147 | 0.54378473 | 0.99999994 |
| Fam162a | 0.35851193 | 0.24681091 | 0.24932838 | -0.0025175 | 0.52679241 | 0.99999994 |
| Hip1 | 0.0172858 | 0.01295236 | 0.01546997 | -0.0025176 | 0.86065471 | 0.99999994 |
| 9530077C05I | 0.01656302 | 0.01231255 | 0.01483032 | -0.0025178 | 0.87077454 | 0.99999994 |
| Ccdc93 | 0.03543261 | 0.0288101 | 0.03132818 | -0.0025181 | 0.70368216 | 0.99999994 |
| Atl3 | 0.02905537 | 0.02327761 | 0.02579897 | -0.0025214 | 0.74232669 | 0.99999994 |
| Panx1 | 0.06547459 | 0.05423593 | 0.05675733 | -0.0025214 | 0.61627199 | 0.99999994 |
| C330018D20 | 0.08209155 | 0.06787672 | 0.07039992 | -0.0025232 | 0.59428045 | 0.99999994 |
| Btbd3 | 0.08196164 | 0.06777062 | 0.0702944 | -0.0025238 | 0.59444091 | 0.99999994 |
| Gstk1 | 0.01905033 | 0.01450496 | 0.01702893 | -0.002524 | 0.83807645 | 0.99999994 |

| | | | | | | |
|---|---|---|---|---|---|---|
| Mrpl15 | 0.44999066 | 0.29035624 | 0.29288108 | -0.0025248 | 0.52287976 | 0.99999994 |
| Slc12a2 | 0.02809891 | 0.0224407 | 0.02496564 | -0.0025249 | 0.74959043 | 0.99999994 |
| Asxl1 | 0.03717572 | 0.03030635 | 0.03283135 | -0.002525 | 0.69564952 | 0.99999994 |
| Enpp4 | 0.0918217 | 0.07572848 | 0.07825363 | -0.0025252 | 0.58503562 | 0.99999994 |
| Gfm1 | 0.08066367 | 0.06671335 | 0.06923911 | -0.0025258 | 0.59592588 | 0.99999994 |
| Filip1 | 0.07163874 | 0.05932514 | 0.06185245 | -0.0025273 | 0.6071816 | 0.99999994 |
| Zfp646 | 0.0955011 | 0.07867042 | 0.08119802 | -0.0025276 | 0.58207494 | 0.99999994 |
| Ctbs | 0.0374759 | 0.03056167 | 0.03308986 | -0.0025282 | 0.69447371 | 0.99999994 |
| Lgi2 | 0.01014545 | 0.00659458 | 0.0091235 | -0.0025289 | 0.96748585 | 0.99999994 |
| Zfp335os | 0.02346379 | 0.01838268 | 0.02091209 | -0.0025294 | 0.78962351 | 0.99999994 |
| Sptbn1 | 0.30508586 | 0.21820566 | 0.22073518 | -0.0025295 | 0.53040131 | 0.99999994 |
| Lnpep | 0.02502467 | 0.01975025 | 0.02227993 | -0.0025297 | 0.77510592 | 0.99999994 |
| Tnk1 | 0.07307016 | 0.06049967 | 0.0630298 | -0.0025301 | 0.60534156 | 0.99999994 |
| Zfp729b | 0.02937894 | 0.02354938 | 0.02608065 | -0.0025313 | 0.74087193 | 0.99999994 |
| Tmem30b | 0.07839545 | 0.06485837 | 0.06739074 | -0.0025324 | 0.59875625 | 0.99999994 |
| Zfp251 | 0.02624152 | 0.02081187 | 0.02334432 | -0.0025324 | 0.76488223 | 0.99999994 |
| Tdrd3 | 0.05821858 | 0.04817343 | 0.05070705 | -0.0025336 | 0.63027834 | 0.99999994 |
| Rad51d | 0.06306756 | 0.05222176 | 0.05475663 | -0.0025349 | 0.62101415 | 0.99999994 |
| Slc39a10 | 0.11377427 | 0.09286552 | 0.09540057 | -0.002535 | 0.5701976 | 0.99999994 |
| Slc22a17 | 0.15026252 | 0.11987037 | 0.12240679 | -0.0025364 | 0.55485135 | 0.99999994 |
| Akap10 | 0.04187219 | 0.03432678 | 0.0368641 | -0.0025373 | 0.6765778 | 0.99999994 |
| Lmo2 | 0.0146497 | 0.01059467 | 0.01313407 | -0.0025394 | 0.90094764 | 0.99999994 |
| Ankrd54 | 0.55182214 | 0.33176874 | 0.33430814 | -0.0025394 | 0.52016262 | 0.99999994 |
| Tcf19 | 0.0119642 | 0.00820587 | 0.01074584 | -0.00254 | 0.94218108 | 0.99999994 |
| Aldh3b1 | 0.02075649 | 0.0159924 | 0.01853281 | -0.0025404 | 0.81922879 | 0.99999994 |
| Pkhd1 | 0.01439205 | 0.01036403 | 0.01290531 | -0.0025413 | 0.90502916 | 0.99999994 |
| Cstf2 | 0.07634264 | 0.06317144 | 0.06571323 | -0.0025418 | 0.60159112 | 0.99999994 |
| Zbtb21 | 0.04453863 | 0.03660075 | 0.03914258 | -0.0025418 | 0.66721977 | 0.99999994 |
| Rspo4 | 0.02630021 | 0.02084943 | 0.0233956 | -0.0025462 | 0.76559568 | 0.99999994 |
| Gtf2f1 | 0.23295282 | 0.17543748 | 0.17798426 | -0.0025468 | 0.53794016 | 0.99999994 |

| | | | | | | |
|---|---|---|---|---|---|---|
| Slf1 | 0.07542316 | 0.06241362 | 0.0649604 | -0.0025468 | 0.60293979 | 0.99999994 |
| Irs1 | 0.01168133 | 0.00794381 | 0.01049378 | -0.00255 | 0.94711315 | 0.99999994 |
| Tifa | 0.01758968 | 0.01318866 | 0.0157387 | -0.00255 | 0.85959227 | 0.99999994 |
| Tob2 | 0.13707619 | 0.11028075 | 0.11283213 | -0.0025514 | 0.55982064 | 0.99999994 |
| Dgcr8 | 0.07503123 | 0.06208498 | 0.06463923 | -0.0025543 | 0.60373615 | 0.99999994 |
| Pdcd2l | 0.18764599 | 0.14590349 | 0.14845948 | -0.002556 | 0.54561926 | 0.99999994 |
| Gm13830 | 0.03189754 | 0.02571281 | 0.028269 | -0.0025562 | 0.72637773 | 0.99999994 |
| Ubl7 | 0.4569786 | 0.29339154 | 0.29594925 | -0.0025577 | 0.5229372 | 0.99999994 |
| Zdhhc5 | 0.03496816 | 0.02836926 | 0.03092708 | -0.0025578 | 0.70901601 | 0.99999994 |
| St8sia1 | 0.01293684 | 0.00905316 | 0.01161181 | -0.0025587 | 0.92877435 | 0.99999994 |
| Ptprj | 0.03546328 | 0.02879573 | 0.03135466 | -0.0025589 | 0.70651835 | 0.99999994 |
| Ttll3 | 0.04412756 | 0.03623275 | 0.03879184 | -0.0025591 | 0.66971212 | 0.99999994 |
| Obsl1 | 0.05516455 | 0.04558417 | 0.04814329 | -0.0025591 | 0.6382661 | 0.99999994 |
| Lnx2 | 0.03571356 | 0.02900867 | 0.03157069 | -0.002562 | 0.70546222 | 0.99999994 |
| Gm15728 | 0.03566982 | 0.02897078 | 0.03153295 | -0.0025622 | 0.70569575 | 0.99999994 |
| Cd74 | 0.01104095 | 0.00736051 | 0.0099228 | -0.0025623 | 0.95717637 | 0.99999994 |
| Elp6 | 0.10784473 | 0.08829135 | 0.09085763 | -0.0025663 | 0.5745652 | 0.99999994 |
| Ccdc30 | 0.02261109 | 0.01759683 | 0.02016364 | -0.0025668 | 0.80159937 | 0.99999994 |
| Nup133 | 0.03473193 | 0.02815614 | 0.03072297 | -0.0025668 | 0.71093587 | 0.99999994 |
| Tsix | 0.04764292 | 0.03921826 | 0.04178511 | -0.0025668 | 0.65869289 | 0.99999994 |
| Anapc2 | 0.20302334 | 0.15615746 | 0.15872473 | -0.0025673 | 0.54286833 | 0.99999994 |
| Rxra | 0.02683881 | 0.02129748 | 0.02386613 | -0.0025687 | 0.76313021 | 0.99999994 |
| Gm12500 | 0.00927647 | 0.0057772 | 0.00834696 | -0.0025698 | 0.97978258 | 0.99999994 |
| Clec1b | 0.00664446 | 0.00341853 | 0.00598933 | -0.0025708 | 0.9978624 | 0.99999994 |
| Lrfn1 | 0.03439325 | 0.0278589 | 0.03043023 | -0.0025713 | 0.71309813 | 0.99999994 |
| Qser1 | 0.07692305 | 0.06361532 | 0.06618797 | -0.0025727 | 0.60207652 | 0.99999994 |
| Hat1 | 0.16538301 | 0.13056209 | 0.13313557 | -0.0025735 | 0.55118884 | 0.99999994 |
| Rbl2 | 0.0264733 | 0.02097099 | 0.02354686 | -0.0025759 | 0.76674256 | 0.99999994 |
| C030037D09 | 0.01661226 | 0.0122974 | 0.01487392 | -0.0025765 | 0.87555282 | 0.99999994 |
| 3110082I17R | 0.096542 | 0.07945188 | 0.08202847 | -0.0025766 | 0.58280819 | 0.99999994 |

| | | | | | | |
|---|---|---|---|---|---|---|
| Fancc | 0.0222375 | 0.01725853 | 0.01983545 | -0.0025769 | 0.80641488 | 0.99999994 |
| Foxn2 | 0.03678075 | 0.02991406 | 0.03249105 | -0.002577 | 0.70123039 | 0.99999994 |
| Nup98 | 0.0208565 | 0.01604334 | 0.01862086 | -0.0025775 | 0.82157371 | 0.99999994 |
| Zc3hav1 | 0.00922141 | 0.00571791 | 0.00829772 | -0.0025798 | 0.98075035 | 0.99999994 |
| Paxbp1 | 0.0589543 | 0.04874304 | 0.05132313 | -0.0025801 | 0.63104598 | 0.99999994 |
| Hhat | 0.01185661 | 0.00806886 | 0.01064998 | -0.0025811 | 0.94665292 | 0.99999994 |
| Ccdc85b | 0.02836272 | 0.0226137 | 0.02519559 | -0.0025819 | 0.75241333 | 0.99999994 |
| Tmtc2 | 0.04339933 | 0.03558767 | 0.03817003 | -0.0025824 | 0.67376338 | 0.99999994 |
| Zfp961 | 0.07071074 | 0.0585031 | 0.06108799 | -0.0025849 | 0.61089751 | 0.99999994 |
| Cttnbp2 | 0.01104946 | 0.0073417 | 0.00993039 | -0.0025887 | 0.95864771 | 0.99999994 |
| Sae1 | 0.4007022 | 0.26763122 | 0.2702204 | -0.0025892 | 0.52542712 | 0.99999994 |
| Pacs2 | 0.04640539 | 0.03814122 | 0.04073295 | -0.0025917 | 0.66404452 | 0.99999994 |
| Faf2 | 0.37629316 | 0.25571664 | 0.25830847 | -0.0025918 | 0.52662513 | 0.99999994 |
| Trim12c | 0.03267205 | 0.02634682 | 0.02894047 | -0.0025936 | 0.72459387 | 0.99999994 |
| Cd99l2 | 0.07465723 | 0.06173862 | 0.06433261 | -0.002594 | 0.60580224 | 0.99999994 |
| Sdk1 | 0.04746719 | 0.03904155 | 0.04163581 | -0.0025943 | 0.66083603 | 0.99999994 |
| Sdad1 | 0.0834019 | 0.06886602 | 0.0714633 | -0.0025973 | 0.59557933 | 0.99999994 |
| Gipc2 | 0.0172322 | 0.01281957 | 0.01542255 | -0.002603 | 0.86937565 | 0.99999994 |
| Ddhd2 | 0.16763849 | 0.13210915 | 0.13471336 | -0.0026042 | 0.55119345 | 0.99999994 |
| Acsl4 | 0.02743104 | 0.02177817 | 0.02438312 | -0.0026049 | 0.76149697 | 0.99999994 |
| Ano10 | 0.07035703 | 0.05818989 | 0.06079638 | -0.0026065 | 0.61232148 | 0.99999994 |
| Tagap1 | 0.03436569 | 0.02779738 | 0.0304064 | -0.002609 | 0.71605042 | 0.99999994 |
| Mmgt1 | 0.03690595 | 0.02998909 | 0.03259894 | -0.0026099 | 0.70295016 | 0.99999994 |
| Pde7a | 0.04397682 | 0.03605303 | 0.03866317 | -0.0026101 | 0.67341724 | 0.99999994 |
| Zfp458 | 0.02274157 | 0.01766659 | 0.02027822 | -0.0026116 | 0.80435306 | 0.99999994 |
| Ccdc173 | 0.07071853 | 0.05847856 | 0.06109441 | -0.0026159 | 0.61217893 | 0.99999994 |
| Kctd18 | 0.04123254 | 0.03369979 | 0.03631632 | -0.0026165 | 0.68423678 | 0.99999994 |
| Mdc1 | 0.02621311 | 0.02070197 | 0.02331948 | -0.0026175 | 0.77251322 | 0.99999994 |
| Slc4a3 | 0.0384009 | 0.03126724 | 0.03388581 | -0.0026186 | 0.69650642 | 0.99999994 |
| Arhgap26 | 0.03186204 | 0.02561962 | 0.02823821 | -0.0026186 | 0.73146799 | 0.99999994 |

| | | | | | | |
|---|---|---|---|---|---|---|
| Fam53c | 0.04751415 | 0.03905485 | 0.04167571 | -0.0026209 | 0.66224514 | 0.99999994 |
| Mtrf1 | 0.02743501 | 0.02176473 | 0.02438658 | -0.0026219 | 0.76289273 | 0.99999994 |
| Gpd1l | 0.02758069 | 0.02189178 | 0.02451368 | -0.0026219 | 0.76174971 | 0.99999994 |
| P2ry12 | 0.00677208 | 0.0034816 | 0.00610384 | -0.0026222 | 0.99787913 | 0.99999994 |
| Neurl4 | 0.06117595 | 0.0505575 | 0.05317992 | -0.0026224 | 0.62863478 | 0.99999994 |
| Adam15 | 0.0206874 | 0.01584837 | 0.01847198 | -0.0026236 | 0.82777654 | 0.99999994 |
| Tbc1d19 | 0.15767174 | 0.12507322 | 0.12769707 | -0.0026238 | 0.55439412 | 0.99999994 |
| Itsn1 | 0.03730925 | 0.03032171 | 0.03294636 | -0.0026246 | 0.70203904 | 0.99999994 |
| Trmt2b | 0.05099901 | 0.04200509 | 0.04462984 | -0.0026247 | 0.65227521 | 0.99999994 |
| Rhno1 | 0.15857168 | 0.12570818 | 0.12833534 | -0.0026272 | 0.55419326 | 0.99999994 |
| Net1 | 0.0279288 | 0.02218944 | 0.02481732 | -0.0026279 | 0.75954361 | 0.99999994 |
| Pias1 | 0.15827554 | 0.12549648 | 0.12812541 | -0.0026289 | 0.55431768 | 0.99999994 |
| Rrm2 | 0.03783908 | 0.03077293 | 0.03340249 | -0.0026296 | 0.69986476 | 0.99999994 |
| Tbc1d8b | 0.02179937 | 0.01681999 | 0.01945036 | -0.0026304 | 0.81598543 | 0.99999994 |
| Blvrb | 0.07359919 | 0.06083335 | 0.06346437 | -0.002631 | 0.60870664 | 0.99999994 |
| Cep120 | 0.05745797 | 0.04743797 | 0.0500695 | -0.0026315 | 0.6367712 | 0.99999994 |
| Champ1 | 0.04079697 | 0.03330997 | 0.03594304 | -0.0026331 | 0.68709173 | 0.99999994 |
| Ccdc77 | 0.07588084 | 0.06270174 | 0.06533524 | -0.0026335 | 0.60576601 | 0.99999994 |
| Nbn | 0.03511514 | 0.02841934 | 0.03105404 | -0.0026347 | 0.71387515 | 0.99999994 |
| Mief1 | 0.03432773 | 0.02773771 | 0.03037358 | -0.0026359 | 0.71825001 | 0.99999994 |
| Gxylt1 | 0.07695123 | 0.06357316 | 0.06621101 | -0.0026379 | 0.60456855 | 0.99999994 |
| Osbpl6 | 0.02475168 | 0.01940249 | 0.02204091 | -0.0026384 | 0.78721777 | 0.99999994 |
| Ermp1 | 0.03295984 | 0.02655084 | 0.02918981 | -0.002639 | 0.72634017 | 0.99999994 |
| Actr8 | 0.1517262 | 0.12081755 | 0.1234569 | -0.0026394 | 0.55658029 | 0.99999994 |
| Trappc8 | 0.04397986 | 0.03602571 | 0.03866577 | -0.0026401 | 0.67526198 | 0.99999994 |
| Ndst1 | 0.04092291 | 0.03340645 | 0.036051 | -0.0026445 | 0.68732469 | 0.99999994 |
| Chac1 | 0.00879011 | 0.00526723 | 0.00791193 | -0.0026447 | 0.98695971 | 0.99999994 |
| Ptdss2 | 0.14518261 | 0.11609568 | 0.11874282 | -0.0026471 | 0.55898281 | 0.99999994 |
| Sgk2 | 0.01988635 | 0.01511908 | 0.01776626 | -0.0026472 | 0.83935222 | 0.99999994 |
| Hira | 0.05760068 | 0.04754178 | 0.05018917 | -0.0026474 | 0.63724751 | 0.99999994 |

| | | | | | | |
|---|---|---|---|---|---|---|
| Stat2 | 0.01942109 | 0.01470746 | 0.01735603 | -0.0026486 | 0.84513169 | 0.99999994 |
| Gm16315 | 0.00793513 | 0.00449777 | 0.0071465 | -0.0026487 | 0.99318867 | 0.99999994 |
| Galnt6 | 0.00697782 | 0.00363908 | 0.0062884 | -0.0026493 | 0.99747908 | 0.99999994 |
| Dppa5a | 0.0189641 | 0.01430348 | 0.01695283 | -0.0026494 | 0.85088663 | 0.99999994 |
| Fam131a | 0.07921569 | 0.06541002 | 0.06805978 | -0.0026498 | 0.60224039 | 0.99999994 |
| Cstf2t | 0.09631337 | 0.07919587 | 0.08184616 | -0.0026503 | 0.58532768 | 0.99999994 |
| 4930524O07 | 0.01115805 | 0.00737665 | 0.01002724 | -0.0026506 | 0.96075679 | 0.99999994 |
| Nr6a1 | 0.03344449 | 0.02695843 | 0.02960947 | -0.002651 | 0.72440172 | 0.99999994 |
| Wbp1l | 0.08136146 | 0.06715536 | 0.06980665 | -0.0026513 | 0.59979325 | 0.99999994 |
| Phf20l1 | 0.30758927 | 0.21947806 | 0.22213097 | -0.0026529 | 0.53168129 | 0.99999994 |
| Dus2 | 0.05396914 | 0.04448322 | 0.04713698 | -0.0026538 | 0.64607504 | 0.99999994 |
| Gng13 | 0.05954336 | 0.04916176 | 0.05181598 | -0.0026542 | 0.63343503 | 0.99999994 |
| Tor1aip1 | 0.1884752 | 0.14636314 | 0.14901963 | -0.0026565 | 0.54722734 | 0.99999994 |
| Slc30a1 | 0.02445889 | 0.0191274 | 0.02178446 | -0.0026571 | 0.79156907 | 0.99999994 |
| Mia2 | 0.01767877 | 0.01316031 | 0.01581747 | -0.0026572 | 0.86825856 | 0.99999994 |
| Lrch3 | 0.06628629 | 0.05477341 | 0.0574306 | -0.0026572 | 0.62094971 | 0.99999994 |
| Selenbp1 | 0.03467165 | 0.02801146 | 0.03067087 | -0.0026594 | 0.71808448 | 0.99999994 |
| Dhx40 | 0.22163472 | 0.16815064 | 0.17081116 | -0.0026605 | 0.54128754 | 0.99999994 |
| Slc27a1 | 0.06833774 | 0.05646814 | 0.05912902 | -0.0026609 | 0.61773763 | 0.99999994 |
| Narf | 0.14180039 | 0.11362503 | 0.11628613 | -0.0026611 | 0.56053456 | 0.99999994 |
| Stim1 | 0.02811086 | 0.02231104 | 0.02497606 | -0.002665 | 0.7612267 | 0.99999994 |
| Tanc2 | 0.02264728 | 0.01753015 | 0.02019542 | -0.0026653 | 0.81015424 | 0.99999994 |
| Exoc2 | 0.04866079 | 0.03998386 | 0.04264922 | -0.0026654 | 0.66129059 | 0.99999994 |
| Vps25 | 0.04257213 | 0.03479668 | 0.03746298 | -0.0026663 | 0.68215867 | 0.99999994 |
| Brip1os | 0.05057669 | 0.04160535 | 0.04427256 | -0.0026672 | 0.655795 | 0.99999994 |
| Yod1 | 0.10936289 | 0.08935753 | 0.09202494 | -0.0026674 | 0.57649681 | 0.99999994 |
| Spaca6 | 0.02771113 | 0.02195808 | 0.02462748 | -0.0026694 | 0.76469603 | 0.99999994 |
| Wdr49 | 0.01229669 | 0.00837011 | 0.011042 | -0.0026719 | 0.94637425 | 0.99999994 |
| Zbtb6 | 0.04420539 | 0.03618618 | 0.03885826 | -0.0026721 | 0.67642399 | 0.99999994 |
| Echdc3 | 0.01871012 | 0.01405635 | 0.01672864 | -0.0026723 | 0.85617868 | 0.99999994 |

| | | | | | | |
|---|---|---|---|---|---|---|
| Rab12 | 0.08403498 | 0.06930225 | 0.07197643 | -0.0026742 | 0.59766483 | 0.99999994 |
| Stxbp5 | 0.0313474 | 0.02511688 | 0.02779162 | -0.0026747 | 0.73911684 | 0.99999994 |
| Arhgef11 | 0.03258764 | 0.02618967 | 0.02886733 | -0.0026777 | 0.73152302 | 0.99999994 |
| Zfp185 | 0.02040991 | 0.01554869 | 0.0182276 | -0.0026789 | 0.83603317 | 0.99999994 |
| Zeb1 | 0.03939965 | 0.03206409 | 0.03474412 | -0.00268 | 0.69618002 | 0.99999994 |
| Socs6 | 0.13086193 | 0.10556815 | 0.1082483 | -0.0026801 | 0.56545189 | 0.99999994 |
| Zfp35 | 0.04353275 | 0.0356029 | 0.038284 | -0.0026811 | 0.67944846 | 0.99999994 |
| Mphosph10 | 0.20904356 | 0.15999051 | 0.16267442 | -0.0026839 | 0.54372428 | 0.99999994 |
| Cyp2u1 | 0.01638974 | 0.0119909 | 0.01467688 | -0.002686 | 0.88841893 | 0.99999994 |
| Plxna1 | 0.0178443 | 0.01327703 | 0.0159638 | -0.0026868 | 0.86870501 | 0.99999994 |
| Pde8a | 0.01453798 | 0.01034694 | 0.01303488 | -0.0026879 | 0.91506433 | 0.99999994 |
| Rap1gapos | 0.02277292 | 0.01761688 | 0.02030575 | -0.0026889 | 0.81095726 | 0.99999994 |
| Rfng | 0.07077733 | 0.0584524 | 0.06114288 | -0.0026905 | 0.61520085 | 0.99999994 |
| Herc3 | 0.03825507 | 0.03106872 | 0.03376039 | -0.0026917 | 0.70218749 | 0.99999994 |
| Tmem43 | 0.06885019 | 0.05686085 | 0.05955257 | -0.0026917 | 0.61823968 | 0.99999994 |
| Zg16 | 0.02758609 | 0.02182486 | 0.0245184 | -0.0026935 | 0.76768714 | 0.99999994 |
| Cdh22 | 0.01079774 | 0.00701225 | 0.00970581 | -0.0026936 | 0.96764608 | 0.99999994 |
| Hpse | 0.00852308 | 0.00497494 | 0.00767297 | -0.002698 | 0.99037249 | 0.99999994 |
| Pla2g15 | 0.02759207 | 0.0218219 | 0.02452362 | -0.0027017 | 0.76831713 | 0.99999994 |
| Mbd5 | 0.05150188 | 0.04235279 | 0.04505502 | -0.0027022 | 0.65513549 | 0.99999994 |
| Lrrc56 | 0.03493189 | 0.0281931 | 0.03089574 | -0.0027026 | 0.71980898 | 0.99999994 |
| Foxj2 | 0.01609738 | 0.01171397 | 0.01441791 | -0.0027039 | 0.89404616 | 0.99999994 |
| Tesc | 0.01279088 | 0.00877759 | 0.01148194 | -0.0027043 | 0.94153342 | 0.99999994 |
| Nfu1 | 0.50951862 | 0.31522517 | 0.31793075 | -0.0027056 | 0.52258617 | 0.99999994 |
| Ercc2 | 0.04598169 | 0.03766456 | 0.04037233 | -0.0027078 | 0.67236072 | 0.99999994 |
| Taf2 | 0.06913048 | 0.0570762 | 0.05978412 | -0.0027079 | 0.61848294 | 0.99999994 |
| Nsun3 | 0.02537022 | 0.01987401 | 0.02258236 | -0.0027084 | 0.78765523 | 0.99999994 |
| Ccnt1 | 0.34145194 | 0.237755 | 0.24046566 | -0.0027107 | 0.52990619 | 0.99999994 |
| Cyth1 | 0.08714832 | 0.07178307 | 0.07449375 | -0.0027107 | 0.59569209 | 0.99999994 |
| Zfp777 | 0.0244351 | 0.01905138 | 0.02176362 | -0.0027122 | 0.79659821 | 0.99999994 |

| | | | | | | |
|---|---|---|---|---|---|---|
| Eif2b5 | 0.25208174 | 0.18709995 | 0.1898131 | -0.0027131 | 0.53790001 | 0.99999994 |
| Pfas | 0.04327986 | 0.03535434 | 0.03806796 | -0.0027136 | 0.68242427 | 0.99999994 |
| Zfp212 | 0.09211057 | 0.07577164 | 0.07848529 | -0.0027137 | 0.59101062 | 0.99999994 |
| Six5 | 0.01602564 | 0.01163872 | 0.01435435 | -0.0027156 | 0.8960357 | 0.99999994 |
| Men1 | 0.16087815 | 0.12724949 | 0.12996697 | -0.0027175 | 0.55534503 | 0.99999994 |
| Tprn | 0.02136006 | 0.01634623 | 0.019064 | -0.0027178 | 0.82867229 | 0.99999994 |
| Vipas39 | 0.18414138 | 0.14336582 | 0.14608375 | -0.0027179 | 0.54928053 | 0.99999994 |
| Strn | 0.01766665 | 0.01308821 | 0.01580676 | -0.0027186 | 0.87385512 | 0.99999994 |
| Lrrtm2 | 0.00933441 | 0.00567889 | 0.00839876 | -0.0027199 | 0.98444291 | 0.99999994 |
| Glod4 | 0.39434312 | 0.26444221 | 0.26716253 | -0.0027203 | 0.52701836 | 0.99999994 |
| Pacs1 | 0.02197403 | 0.01688074 | 0.0196039 | -0.0027232 | 0.82242038 | 0.99999994 |
| Ctns | 0.04563839 | 0.03735682 | 0.04007999 | -0.0027232 | 0.67445737 | 0.99999994 |
| Pcgf3 | 0.20107764 | 0.15471657 | 0.15743992 | -0.0027234 | 0.54583292 | 0.99999994 |
| Zfp1 | 0.07670661 | 0.06328644 | 0.06601098 | -0.0027245 | 0.60824018 | 0.99999994 |
| Unkl | 0.01708707 | 0.01256939 | 0.01529415 | -0.0027248 | 0.88216491 | 0.99999994 |
| Anks6 | 0.04345241 | 0.03549015 | 0.03821537 | -0.0027252 | 0.68249175 | 0.99999994 |
| Gm14305 | 0.04916263 | 0.04034703 | 0.04307482 | -0.0027278 | 0.66331456 | 0.99999994 |
| 3830406C13I | 0.22253651 | 0.1686591 | 0.17138763 | -0.0027285 | 0.54219722 | 0.99999994 |
| 2700033N17 | 0.08647675 | 0.0712182 | 0.0739516 | -0.0027334 | 0.59717162 | 0.99999994 |
| Hps6 | 0.0250029 | 0.01952702 | 0.02226087 | -0.0027339 | 0.79316903 | 0.99999994 |
| Gm17259 | 0.04406733 | 0.03600586 | 0.03874043 | -0.0027346 | 0.68076981 | 0.99999994 |
| Tjp2 | 0.03597358 | 0.02905766 | 0.03179506 | -0.0027374 | 0.71670466 | 0.99999994 |
| Zfp365 | 0.00802479 | 0.00448846 | 0.00722681 | -0.0027383 | 0.99416875 | 0.99999994 |
| 9030612E09I | 0.01388424 | 0.00971564 | 0.01245422 | -0.0027386 | 0.92835939 | 0.99999994 |
| Met | 0.02153991 | 0.01648149 | 0.0192222 | -0.0027407 | 0.82870666 | 0.99999994 |
| Usp10 | 0.14842387 | 0.1183426 | 0.12108415 | -0.0027416 | 0.55989828 | 0.99999994 |
| Cdt1 | 0.02796314 | 0.02210546 | 0.02484726 | -0.0027418 | 0.76867949 | 0.99999994 |
| Gdpgp1 | 0.04086121 | 0.03325408 | 0.03599812 | -0.002744 | 0.69406134 | 0.99999994 |
| Lrrc51 | 0.03385518 | 0.02721884 | 0.02996488 | -0.002746 | 0.72906953 | 0.99999994 |
| Mast1 | 0.01957856 | 0.01474867 | 0.01749491 | -0.0027462 | 0.85195917 | 0.99999994 |

| | | | | | | |
|---|---|---|---|---|---|---|
| Zim1 | 0.07536518 | 0.06216634 | 0.0649129 | -0.0027466 | 0.61089043 | 0.99999994 |
| Crnkl1 | 0.21924913 | 0.16653326 | 0.16928208 | -0.0027488 | 0.54303646 | 0.99999994 |
| Src | 0.06169714 | 0.0508657 | 0.05361473 | -0.002749 | 0.63355979 | 0.99999994 |
| Ccnc | 0.16651176 | 0.13117555 | 0.13392589 | -0.0027503 | 0.5543648 | 0.99999994 |
| 1810034E14I | 0.01956898 | 0.01473586 | 0.01748645 | -0.0027506 | 0.85245855 | 0.99999994 |
| Nme7 | 0.08010046 | 0.0660291 | 0.06878066 | -0.0027516 | 0.60499066 | 0.99999994 |
| Sesn1 | 0.1047453 | 0.08571393 | 0.08846549 | -0.0027516 | 0.58200856 | 0.99999994 |
| Rictor | 0.05166142 | 0.04243748 | 0.04518984 | -0.0027524 | 0.65741416 | 0.99999994 |
| Tulp3 | 0.07694797 | 0.06345542 | 0.06620835 | -0.0027529 | 0.60902168 | 0.99999994 |
| Hspb6 | 0.05844109 | 0.0481404 | 0.05089344 | -0.002753 | 0.64060282 | 0.99999994 |
| Lrp12 | 0.0421972 | 0.0343873 | 0.03714225 | -0.0027549 | 0.68924861 | 0.99999994 |
| Hmg20a | 0.06563152 | 0.05413258 | 0.05688756 | -0.002755 | 0.62640917 | 0.99999994 |
| Trmt1l | 0.24436852 | 0.18233122 | 0.18508731 | -0.0027561 | 0.53947774 | 0.99999994 |
| Ing1 | 0.29669431 | 0.21325882 | 0.21601502 | -0.0027562 | 0.53384171 | 0.99999994 |
| Kdm6a | 0.14267688 | 0.1141672 | 0.11692406 | -0.0027569 | 0.56235589 | 0.99999994 |
| Ccdc71 | 0.11979695 | 0.09721284 | 0.0999697 | -0.0027569 | 0.57282155 | 0.99999994 |
| Zbtb33 | 0.05010229 | 0.04111238 | 0.04387099 | -0.0027586 | 0.66222809 | 0.99999994 |
| Ift140 | 0.01776461 | 0.01313438 | 0.01589336 | -0.002759 | 0.87605703 | 0.99999994 |
| Hdac10 | 0.03869131 | 0.03137606 | 0.0341355 | -0.0027594 | 0.70474122 | 0.99999994 |
| Amdhd2 | 0.12399404 | 0.10036552 | 0.10312739 | -0.0027619 | 0.57074248 | 0.99999994 |
| Plekhm1 | 0.01848801 | 0.01377035 | 0.01653252 | -0.0027622 | 0.86695457 | 0.99999994 |
| Ppa1 | 0.40155275 | 0.26786252 | 0.27062707 | -0.0027646 | 0.52710601 | 0.99999994 |
| Txndc16 | 0.03224804 | 0.02580612 | 0.02857296 | -0.0027668 | 0.74039489 | 0.99999994 |
| H2-DMa | 0.03463491 | 0.0278717 | 0.03063912 | -0.0027674 | 0.72615475 | 0.99999994 |
| Engase | 0.02646016 | 0.02076776 | 0.02353538 | -0.0027676 | 0.78311191 | 0.99999994 |
| Prdm15 | 0.03029614 | 0.02411073 | 0.02687843 | -0.0027677 | 0.75345781 | 0.99999994 |
| Ttc38 | 0.0273101 | 0.02150609 | 0.02427758 | -0.0027715 | 0.77633608 | 0.99999994 |
| Wdr6 | 0.32753503 | 0.23027832 | 0.23305223 | -0.0027739 | 0.53157412 | 0.99999994 |
| Nfatc3 | 0.0466008 | 0.03812214 | 0.04089921 | -0.0027771 | 0.67435405 | 0.99999994 |
| Rap1gds1 | 0.03824683 | 0.03097595 | 0.0337533 | -0.0027774 | 0.70805478 | 0.99999994 |

| | | | | | | |
|---|---|---|---|---|---|---|
| 2310068J16F | 0.02488866 | 0.01938275 | 0.02216086 | -0.0027781 | 0.79798549 | 0.99999994 |
| Cers6 | 0.10945296 | 0.08931504 | 0.09209411 | -0.0027791 | 0.57959789 | 0.99999994 |
| Gm12976 | 0.01160011 | 0.0076354 | 0.01042139 | -0.002786 | 0.96241868 | 0.99999994 |
| Tmtc4 | 0.1151212 | 0.09363987 | 0.09642644 | -0.0027866 | 0.5762692 | 0.99999994 |
| Pigg | 0.02449722 | 0.01903046 | 0.02181803 | -0.0027876 | 0.80246148 | 0.99999994 |
| Gm26724 | 0.0829743 | 0.06832861 | 0.0711165 | -0.0027879 | 0.6029308 | 0.99999994 |
| Lsm14b | 0.08091742 | 0.06665696 | 0.06944556 | -0.0027886 | 0.60537605 | 0.99999994 |
| Nup155 | 0.04594106 | 0.03754661 | 0.04033773 | -0.0027911 | 0.67744965 | 0.99999994 |
| Phf20 | 0.25224431 | 0.18712012 | 0.18991207 | -0.0027919 | 0.53897703 | 0.99999994 |
| Avpi1 | 0.09470105 | 0.07776667 | 0.08055897 | -0.0027923 | 0.59123373 | 0.99999994 |
| Gpsm2 | 0.03502493 | 0.02818366 | 0.03097612 | -0.0027925 | 0.72576808 | 0.99999994 |
| Ric3 | 0.05905394 | 0.04861178 | 0.05140653 | -0.0027947 | 0.64127787 | 0.99999994 |
| Gabbr2 | 0.03094326 | 0.02464551 | 0.02744072 | -0.0027952 | 0.75112638 | 0.99999994 |
| Pip4k2b | 0.02579654 | 0.0201593 | 0.02295529 | -0.002796 | 0.79124686 | 0.99999994 |
| Zfp946 | 0.03628275 | 0.02926574 | 0.03206173 | -0.002796 | 0.71920383 | 0.99999994 |
| Eif4ebp2 | 0.06305413 | 0.05194933 | 0.05474545 | -0.0027961 | 0.63306128 | 0.99999994 |
| Cd276 | 0.04313025 | 0.03514075 | 0.03794012 | -0.0027994 | 0.68833344 | 0.99999994 |
| Thada | 0.01513822 | 0.01076628 | 0.01356758 | -0.0028013 | 0.91533212 | 0.99999994 |
| Phactr3 | 0.00575323 | 0.00238538 | 0.00518911 | -0.0028037 | 0.99983874 | 0.99999994 |
| Amer2 | 0.0450137 | 0.03674324 | 0.03954768 | -0.0028044 | 0.68154434 | 0.99999994 |
| Zfp36l2 | 0.03627266 | 0.02924682 | 0.03205304 | -0.0028062 | 0.71997207 | 0.99999994 |
| Med24 | 0.08419218 | 0.0692955 | 0.07210377 | -0.0028083 | 0.60227873 | 0.99999994 |
| Rb1 | 0.06005266 | 0.04943259 | 0.05224179 | -0.0028092 | 0.63980312 | 0.99999994 |
| Abcg1 | 0.08569361 | 0.07050896 | 0.07331879 | -0.0028098 | 0.60067487 | 0.99999994 |
| Cdk17 | 0.03877303 | 0.03139536 | 0.03420575 | -0.0028104 | 0.70777473 | 0.99999994 |
| Trmt12 | 0.08622237 | 0.07093436 | 0.07374612 | -0.0028118 | 0.60017133 | 0.99999994 |
| Slc37a1 | 0.13082301 | 0.10540738 | 0.10821945 | -0.0028121 | 0.56866033 | 0.99999994 |
| Akap8l | 0.32689446 | 0.22989472 | 0.23270689 | -0.0028122 | 0.53205611 | 0.99999994 |
| Zbtb26 | 0.03522706 | 0.02833845 | 0.0311507 | -0.0028122 | 0.72605616 | 0.99999994 |
| Tex26 | 0.01269883 | 0.00858689 | 0.01140001 | -0.0028131 | 0.9497623 | 0.99999994 |

| | | | | | | |
|---|---|---|---|---|---|---|
| Klhdc3 | 0.20128396 | 0.15476251 | 0.15757636 | -0.0028138 | 0.547308 | 0.99999994 |
| Stard3 | 0.12622417 | 0.10198055 | 0.10479651 | -0.002816 | 0.57097657 | 0.99999994 |
| Cars | 0.10803057 | 0.08818365 | 0.09100068 | -0.002817 | 0.58162588 | 0.99999994 |
| Carm1 | 0.0503032 | 0.04122347 | 0.04404109 | -0.0028176 | 0.66489263 | 0.99999994 |
| Usp1 | 0.0964019 | 0.07909756 | 0.08191676 | -0.0028192 | 0.59059724 | 0.99999994 |
| Rnpc3 | 0.15230433 | 0.12104974 | 0.12387099 | -0.0028213 | 0.56025015 | 0.99999994 |
| Dtymk | 0.38650013 | 0.2605254 | 0.26334684 | -0.0028214 | 0.52842636 | 0.99999994 |
| Plcg1 | 0.0675707 | 0.05566853 | 0.05849451 | -0.002826 | 0.62611503 | 0.99999994 |
| Gm42726 | 0.03685885 | 0.02973075 | 0.03255836 | -0.0028276 | 0.71839721 | 0.99999994 |
| Maneal | 0.00731943 | 0.00376574 | 0.00659474 | -0.002829 | 0.9978511 | 0.99999994 |
| Nrde2 | 0.04775427 | 0.03905033 | 0.0418797 | -0.0028294 | 0.67353659 | 0.99999994 |
| Ttll12 | 0.06501765 | 0.05354809 | 0.05637802 | -0.0028299 | 0.63085546 | 0.99999994 |
| Ogfod1 | 0.02980478 | 0.02362077 | 0.02645116 | -0.0028304 | 0.76184843 | 0.99999994 |
| Gm14091 | 0.01237731 | 0.00828165 | 0.01111378 | -0.0028321 | 0.95508042 | 0.99999994 |
| Kpna4 | 0.1154572 | 0.09384904 | 0.09668198 | -0.0028329 | 0.5773202 | 0.99999994 |
| Fchsd2 | 0.0668677 | 0.05507853 | 0.05791242 | -0.0028339 | 0.62768302 | 0.99999994 |
| Pml | 0.02406403 | 0.01860401 | 0.02143843 | -0.0028344 | 0.81058478 | 0.99999994 |
| Plcd1 | 0.01673458 | 0.01214756 | 0.01498221 | -0.0028346 | 0.89605519 | 0.99999994 |
| Pick1 | 0.39103619 | 0.26272489 | 0.2655599 | -0.002835 | 0.52832517 | 0.99999994 |
| Adgrv1 | 0.02672668 | 0.02093041 | 0.0237682 | -0.0028378 | 0.78661516 | 0.99999994 |
| Mapkap1 | 0.12795735 | 0.10325028 | 0.10608952 | -0.0028392 | 0.57069431 | 0.99999994 |
| Unc79 | 0.01949531 | 0.01458171 | 0.01742148 | -0.0028398 | 0.86104211 | 0.99999994 |
| Agps | 0.0559072 | 0.04592625 | 0.04876767 | -0.0028414 | 0.65092818 | 0.99999994 |
| Tmem143 | 0.02938414 | 0.02324357 | 0.02608518 | -0.0028416 | 0.7658091 | 0.99999994 |
| Clmn | 0.01385559 | 0.00958713 | 0.01242876 | -0.0028416 | 0.93597863 | 0.99999994 |
| Cntn3 | 0.01387881 | 0.00960756 | 0.0124494 | -0.0028418 | 0.93567595 | 0.99999994 |
| Gm7334 | 0.03634312 | 0.02927188 | 0.03211379 | -0.0028419 | 0.72208784 | 0.99999994 |
| Brdt | 0.03259772 | 0.02603309 | 0.02887606 | -0.002843 | 0.74387796 | 0.99999994 |
| Gusb | 0.16582513 | 0.13060148 | 0.13344529 | -0.0028438 | 0.55640147 | 0.99999994 |
| Atp11a | 0.05794548 | 0.04763416 | 0.05047821 | -0.0028441 | 0.64618228 | 0.99999994 |

| | | | | | | |
|---|---|---|---|---|---|---|
| Pik3cb | 0.01386832 | 0.00959094 | 0.01244007 | -0.0028491 | 0.93630768 | 0.99999994 |
| Stx1a | 0.15061455 | 0.11981043 | 0.12265958 | -0.0028491 | 0.56143736 | 0.99999994 |
| Nars2 | 0.04117233 | 0.03341395 | 0.03626474 | -0.0028508 | 0.69960305 | 0.99999994 |
| Celsr3 | 0.03767549 | 0.03041084 | 0.03326169 | -0.0028509 | 0.71583562 | 0.99999994 |
| Dnajc17 | 0.15163629 | 0.12053971 | 0.12339247 | -0.0028528 | 0.56115197 | 0.99999994 |
| Slc6a17 | 0.01315934 | 0.00895687 | 0.01180975 | -0.0028529 | 0.94607982 | 0.99999994 |
| Pitpnm3 | 0.00967868 | 0.00585284 | 0.00870649 | -0.0028537 | 0.98543406 | 0.99999994 |
| Pja1 | 0.48174602 | 0.30369083 | 0.30654634 | -0.0028555 | 0.52472044 | 0.99999994 |
| Med22 | 0.09758906 | 0.08000707 | 0.08286273 | -0.0028557 | 0.59071923 | 0.99999994 |
| Rpp30 | 0.2009365 | 0.1544905 | 0.15734657 | -0.0028561 | 0.54808415 | 0.99999994 |
| Bloc1s6 | 0.12313785 | 0.09962883 | 0.10248498 | -0.0028561 | 0.57358379 | 0.99999994 |
| Fbxw17 | 0.05306689 | 0.04351926 | 0.04637641 | -0.0028572 | 0.65912815 | 0.99999994 |
| Plxna2 | 0.01860703 | 0.01378004 | 0.01663762 | -0.0028576 | 0.8735344 | 0.99999994 |
| Uxs1 | 0.07802229 | 0.06422758 | 0.06708613 | -0.0028586 | 0.61165256 | 0.99999994 |
| Hus1 | 0.04779112 | 0.03904999 | 0.04191099 | -0.002861 | 0.67522522 | 0.99999994 |
| Tom1 | 0.05310996 | 0.04355092 | 0.04641274 | -0.0028618 | 0.65925588 | 0.99999994 |
| Ppp2r5e | 0.18924677 | 0.14667833 | 0.14954018 | -0.0028618 | 0.55068293 | 0.99999994 |
| Rcc2 | 0.15075404 | 0.11989096 | 0.1227597 | -0.0028687 | 0.56180658 | 0.99999994 |
| Cers2 | 0.32496734 | 0.22879687 | 0.23166586 | -0.002869 | 0.53284891 | 0.99999994 |
| Mterf1a | 0.02470668 | 0.01913175 | 0.0220015 | -0.0028698 | 0.80736261 | 0.99999994 |
| Akip1 | 0.10519604 | 0.08594408 | 0.08881412 | -0.00287 | 0.58515587 | 0.99999994 |
| Bcl2l1 | 0.17754742 | 0.1387071 | 0.14157741 | -0.0028703 | 0.55367421 | 0.99999994 |
| Tmcc2 | 0.02766784 | 0.02171933 | 0.02458971 | -0.0028704 | 0.78142185 | 0.99999994 |
| Cep57l1 | 0.04408969 | 0.03588781 | 0.03875952 | -0.0028717 | 0.68905448 | 0.99999994 |
| Wdr33 | 0.19139061 | 0.14810618 | 0.15098317 | -0.002877 | 0.55046546 | 0.99999994 |
| Kif13b | 0.03088117 | 0.02450461 | 0.02738678 | -0.0028822 | 0.75819634 | 0.99999994 |
| Dtnbp1 | 0.07743529 | 0.06372406 | 0.06660667 | -0.0028826 | 0.61335507 | 0.99999994 |
| Gpalpp1 | 0.08105959 | 0.06667752 | 0.0695612 | -0.0028837 | 0.60870313 | 0.99999994 |
| Kctd11 | 0.06616354 | 0.05444142 | 0.05732884 | -0.0028874 | 0.63128315 | 0.99999994 |
| Atad3a | 0.15087592 | 0.11995855 | 0.12284717 | -0.0028886 | 0.56218736 | 0.99999994 |

| | | | | | | |
|---|---|---|---|---|---|---|
| Ak5 | 0.01517914 | 0.01071507 | 0.01360388 | -0.0028888 | 0.92124408 | 0.99999994 |
| Anxa11 | 0.11032423 | 0.08987299 | 0.09276263 | -0.0028896 | 0.58213236 | 0.99999994 |
| Arih1 | 0.12840801 | 0.10353373 | 0.10642514 | -0.0028914 | 0.57175467 | 0.99999994 |
| Dhx33 | 0.02027653 | 0.01521795 | 0.0181101 | -0.0028922 | 0.85611098 | 0.99999994 |
| Mlst8 | 0.03446443 | 0.02759934 | 0.03049176 | -0.0028924 | 0.73611559 | 0.99999994 |
| Atp9b | 0.03299201 | 0.02632475 | 0.02921767 | -0.0028929 | 0.74507302 | 0.99999994 |
| Edc3 | 0.09803267 | 0.0803207 | 0.08321585 | -0.0028952 | 0.59156814 | 0.99999994 |
| Plekhb1 | 0.0580847 | 0.04769711 | 0.05059488 | -0.0028978 | 0.64848622 | 0.99999994 |
| Tnk2 | 0.01575988 | 0.01122001 | 0.01411883 | -0.0028988 | 0.91413352 | 0.99999994 |
| Ttc22 | 0.01390772 | 0.00957605 | 0.01247508 | -0.002899 | 0.93904933 | 0.99999994 |
| Dst | 0.4658724 | 0.29690303 | 0.29980396 | -0.0029009 | 0.52567713 | 0.99999994 |
| Usp2 | 0.0331192 | 0.02642522 | 0.02932783 | -0.0029026 | 0.74498401 | 0.99999994 |
| Sntg1 | 0.01062174 | 0.0066442 | 0.00954875 | -0.0029045 | 0.97855292 | 0.99999994 |
| 0610040B10 | 0.07623015 | 0.06271543 | 0.06562117 | -0.0029057 | 0.61590652 | 0.99999994 |
| Rnmt | 0.164162 | 0.12937092 | 0.13227903 | -0.0029081 | 0.55817263 | 0.99999994 |
| Arl13b | 0.10249531 | 0.08381237 | 0.08672138 | -0.002909 | 0.58834217 | 0.99999994 |
| Grin1 | 0.00707948 | 0.00346934 | 0.00637958 | -0.0029102 | 0.9988033 | 0.99999994 |
| Jmjd1c | 0.14041042 | 0.11236044 | 0.11527261 | -0.0029122 | 0.56677196 | 0.99999994 |
| Zfp799 | 0.03305722 | 0.02635847 | 0.02927415 | -0.0029157 | 0.74632471 | 0.99999994 |
| Polr2h | 0.29658236 | 0.21303541 | 0.21595161 | -0.0029162 | 0.53581151 | 0.99999994 |
| Inafm2 | 0.02841221 | 0.02231723 | 0.02523873 | -0.0029215 | 0.77949946 | 0.99999994 |
| Atxn1l | 0.04405481 | 0.03580783 | 0.03872975 | -0.0029219 | 0.69222934 | 0.99999994 |
| Pgpep1 | 0.18969324 | 0.14691691 | 0.14984111 | -0.0029242 | 0.55167763 | 0.99999994 |
| Cmpk2 | 0.0190697 | 0.0141203 | 0.01704603 | -0.0029257 | 0.87337227 | 0.99999994 |
| Yars2 | 0.07182729 | 0.05907933 | 0.06200765 | -0.0029283 | 0.62337158 | 0.99999994 |
| Zfp334 | 0.02971134 | 0.02343893 | 0.02636988 | -0.002931 | 0.77030114 | 0.99999994 |
| Def8 | 0.09221412 | 0.07563632 | 0.07856832 | -0.002932 | 0.59808772 | 0.99999994 |
| AW146154 | 0.05708059 | 0.04682072 | 0.04975294 | -0.0029322 | 0.65258072 | 0.99999994 |
| Clcn6 | 0.02210632 | 0.01678662 | 0.01972017 | -0.0029336 | 0.83896032 | 0.99999994 |
| Stambp | 0.18626869 | 0.14459323 | 0.14752741 | -0.0029342 | 0.55266167 | 0.99999994 |

| | | | | | | |
|---|---|---|---|---|---|---|
| Dhx16 | 0.07095925 | 0.05835679 | 0.0612928 | -0.002936 | 0.62507933 | 0.99999994 |
| Spink13 | 0.01471364 | 0.01025092 | 0.01319082 | -0.0029399 | 0.93103384 | 0.99999994 |
| Tbc1d2b | 0.02254464 | 0.01716479 | 0.02010527 | -0.0029405 | 0.83485268 | 0.99999994 |
| Ak2 | 0.59064496 | 0.3454273 | 0.34836848 | -0.0029412 | 0.52240785 | 0.99999994 |
| Ppp2r5d | 0.07429686 | 0.0610943 | 0.06403702 | -0.0029427 | 0.62015314 | 0.99999994 |
| Grpel2 | 0.17501559 | 0.1368884 | 0.13983412 | -0.0029457 | 0.55575761 | 0.99999994 |
| Cln8 | 0.08896797 | 0.07301421 | 0.0759604 | -0.0029462 | 0.60186292 | 0.99999994 |
| Dcun1d4 | 0.03683706 | 0.02959327 | 0.03253958 | -0.0029463 | 0.72664708 | 0.99999994 |
| Cc2d2b | 0.00919055 | 0.00532377 | 0.00827013 | -0.0029464 | 0.99114025 | 0.99999994 |
| Gnptab | 0.01783451 | 0.01300876 | 0.01595514 | -0.0029464 | 0.89050322 | 0.99999994 |
| Polr3f | 0.09545282 | 0.07821304 | 0.08115947 | -0.0029464 | 0.59547564 | 0.99999994 |
| Susd4 | 0.01586923 | 0.01126796 | 0.01421575 | -0.0029478 | 0.91624758 | 0.99999994 |
| Nckipsd | 0.01984017 | 0.01477692 | 0.01772556 | -0.0029486 | 0.86591402 | 0.99999994 |
| Dtx3 | 0.28331225 | 0.20540488 | 0.20835359 | -0.0029487 | 0.53752631 | 0.99999994 |
| Pomc | 0.11006362 | 0.08961313 | 0.09256277 | -0.0029496 | 0.58399101 | 0.99999994 |
| Dkc1 | 0.127476 | 0.10277746 | 0.10573078 | -0.0029533 | 0.57374975 | 0.99999994 |
| Orc5 | 0.05184111 | 0.04238785 | 0.04534167 | -0.0029538 | 0.66772278 | 0.99999994 |
| Pomk | 0.01867344 | 0.01374213 | 0.01669625 | -0.0029541 | 0.88054788 | 0.99999994 |
| Knstrn | 0.03226215 | 0.0256307 | 0.02858519 | -0.0029545 | 0.75426828 | 0.99999994 |
| Irf3 | 0.24765237 | 0.18415186 | 0.18710638 | -0.0029545 | 0.54185487 | 0.99999994 |
| Zfp780b | 0.04489582 | 0.03649095 | 0.03944719 | -0.0029562 | 0.69105335 | 0.99999994 |
| Bahd1 | 0.03350938 | 0.02670756 | 0.02966564 | -0.0029581 | 0.74656921 | 0.99999994 |
| Fads1 | 0.02752857 | 0.02150961 | 0.02446822 | -0.0029586 | 0.78955023 | 0.99999994 |
| A130010J15F | 0.12427612 | 0.10037546 | 0.10333884 | -0.0029634 | 0.57569034 | 0.99999994 |
| Zfp931 | 0.05559249 | 0.04553973 | 0.04850315 | -0.0029634 | 0.65788087 | 0.99999994 |
| Ddx55 | 0.18400388 | 0.14302675 | 0.14599028 | -0.0029635 | 0.5537417 | 0.99999994 |
| Ino80d | 0.15368612 | 0.12189523 | 0.12485915 | -0.0029639 | 0.56277514 | 0.99999994 |
| Slc5a1 | 0.02214179 | 0.01678689 | 0.01975134 | -0.0029645 | 0.84111328 | 0.99999994 |
| Gk5 | 0.03441839 | 0.02748638 | 0.03045197 | -0.0029656 | 0.74158355 | 0.99999994 |
| Gipc1 | 0.58073243 | 0.34189838 | 0.34486422 | -0.0029658 | 0.52282493 | 0.99999994 |

| | | | | | | |
|---|---|---|---|---|---|---|
| Crebl2 | 0.03374388 | 0.02689842 | 0.02986858 | -0.0029702 | 0.74598738 | 0.99999994 |
| Yme1l1 | 0.37773772 | 0.25605235 | 0.2590265 | -0.0029742 | 0.530461 | 0.99999994 |
| Zfp707 | 0.08256978 | 0.06781227 | 0.07078822 | -0.002976 | 0.61019801 | 0.99999994 |
| Dusp12 | 0.17310048 | 0.13553413 | 0.13851059 | -0.0029765 | 0.55687027 | 0.99999994 |
| Tmem25 | 0.0149173 | 0.01039345 | 0.01337157 | -0.0029781 | 0.93089665 | 0.99999994 |
| Tada1 | 0.14283237 | 0.11405814 | 0.11703713 | -0.002979 | 0.56726955 | 0.99999994 |
| Crybb3 | 0.01374028 | 0.00934686 | 0.01232629 | -0.0029794 | 0.94618425 | 0.99999994 |
| Qrsl1 | 0.0506774 | 0.04137772 | 0.04435779 | -0.0029801 | 0.67263124 | 0.99999994 |
| Vps16 | 0.08116504 | 0.06666638 | 0.06964695 | -0.0029806 | 0.61212512 | 0.99999994 |
| Zfp800 | 0.12263879 | 0.0991287 | 0.10211012 | -0.0029814 | 0.57705013 | 0.99999994 |
| Rasal2 | 0.02449423 | 0.01883245 | 0.02181541 | -0.002983 | 0.81863304 | 0.99999994 |
| Dusp14 | 0.0176052 | 0.01276562 | 0.01575243 | -0.0029868 | 0.8965452 | 0.99999994 |
| Ruvbl2 | 0.24797373 | 0.18431541 | 0.1873034 | -0.002988 | 0.54228301 | 0.99999994 |
| Dpy19l4 | 0.05072085 | 0.04140573 | 0.04439454 | -0.0029888 | 0.67297088 | 0.99999994 |
| Zfp74 | 0.03847486 | 0.0309592 | 0.03394941 | -0.0029902 | 0.7211563 | 0.99999994 |
| Matn2 | 0.00908867 | 0.00518643 | 0.00817901 | -0.0029926 | 0.99256359 | 0.99999994 |
| Ttc7 | 0.03111946 | 0.02460063 | 0.02759373 | -0.0029931 | 0.76485804 | 0.99999994 |
| Gm14295 | 0.02965908 | 0.02333082 | 0.02632442 | -0.0029936 | 0.77546214 | 0.99999994 |
| Szt2 | 0.01964885 | 0.01456286 | 0.01755689 | -0.002994 | 0.87183926 | 0.99999994 |
| Aatf | 0.13171749 | 0.10588677 | 0.10888213 | -0.0029954 | 0.57264712 | 0.99999994 |
| Lrp3 | 0.01777858 | 0.01291015 | 0.01590571 | -0.0029956 | 0.89500806 | 0.99999994 |
| Plagl2 | 0.04945243 | 0.0403244 | 0.04332047 | -0.0029961 | 0.67737025 | 0.99999994 |
| Naprt | 0.04319645 | 0.03500035 | 0.03799668 | -0.0029963 | 0.70017533 | 0.99999994 |
| 1110020A21 | 0.11181995 | 0.09091126 | 0.09390809 | -0.0029968 | 0.58411031 | 0.99999994 |
| Atg9a | 0.07434803 | 0.06108035 | 0.064079 | -0.0029987 | 0.62228664 | 0.99999994 |
| Klk1 | 0.01585454 | 0.01120369 | 0.01420273 | -0.002999 | 0.92007138 | 0.99999994 |
| Pprc1 | 0.01826682 | 0.01333749 | 0.01633715 | -0.0029997 | 0.88917911 | 0.99999994 |
| Slc9a7 | 0.02289591 | 0.01740795 | 0.02041374 | -0.0030058 | 0.83648239 | 0.99999994 |
| Fcho2 | 0.14325639 | 0.1143381 | 0.11734534 | -0.0030072 | 0.5677247 | 0.99999994 |
| Ankrd16 | 0.0961133 | 0.07867812 | 0.08168658 | -0.0030085 | 0.59682942 | 0.99999994 |

| | | | | | | |
|---|---|---|---|---|---|---|
| Hs2st1 | 0.09254831 | 0.07582766 | 0.07883619 | -0.0030085 | 0.60025947 | 0.99999994 |
| Chmp6 | 0.21607136 | 0.16422737 | 0.16723616 | -0.0030088 | 0.54766194 | 0.99999994 |
| Rbm19 | 0.02995954 | 0.02357583 | 0.02658577 | -0.0030099 | 0.77445782 | 0.99999994 |
| Mtmr10 | 0.02388452 | 0.01826892 | 0.02128107 | -0.0030122 | 0.82694196 | 0.99999994 |
| Pla2g1b | 0.03899043 | 0.0313791 | 0.03439258 | -0.0030135 | 0.72013041 | 0.99999994 |
| Sigirr | 0.05458436 | 0.04463896 | 0.04765507 | -0.0030161 | 0.66322595 | 0.99999994 |
| Gm15446 | 0.03520779 | 0.02811761 | 0.03113405 | -0.0030164 | 0.74050621 | 0.99999994 |
| Map2k5 | 0.11043231 | 0.08982901 | 0.0928455 | -0.0030165 | 0.5856078 | 0.99999994 |
| Gzf1 | 0.08767391 | 0.07189838 | 0.07491773 | -0.0030193 | 0.60575255 | 0.99999994 |
| Csnk2a2 | 0.02856207 | 0.02234869 | 0.02536931 | -0.0030206 | 0.78597824 | 0.99999994 |
| Celsr1 | 0.03316988 | 0.02635076 | 0.02937171 | -0.003021 | 0.75320897 | 0.99999994 |
| Zfp92 | 0.03493109 | 0.02787372 | 0.03089505 | -0.0030213 | 0.74246024 | 0.99999994 |
| Zbtb25 | 0.03185523 | 0.02520993 | 0.02823229 | -0.0030224 | 0.76194902 | 0.99999994 |
| Osbpl3 | 0.01209273 | 0.00783795 | 0.01086034 | -0.0030224 | 0.96801877 | 0.99999994 |
| Rnf31 | 0.02959292 | 0.02324348 | 0.02626686 | -0.0030234 | 0.77821052 | 0.99999994 |
| Mapkbp1 | 0.02760506 | 0.02151139 | 0.02453495 | -0.0030236 | 0.79397642 | 0.99999994 |
| Zkscan8 | 0.03917159 | 0.03152385 | 0.03454823 | -0.0030244 | 0.71995299 | 0.99999994 |
| Scrn3 | 0.07542298 | 0.06193381 | 0.06496025 | -0.0030264 | 0.62176259 | 0.99999994 |
| Nbeal2 | 0.04709603 | 0.03829358 | 0.04132035 | -0.0030268 | 0.68707896 | 0.99999994 |
| Slco1a6 | 0.01243448 | 0.00813725 | 0.01116469 | -0.0030274 | 0.96445906 | 0.99999994 |
| Cep164 | 0.15762271 | 0.12463377 | 0.12766226 | -0.0030285 | 0.56273473 | 0.99999994 |
| Atl2 | 0.14470685 | 0.11536858 | 0.11839806 | -0.0030295 | 0.56761987 | 0.99999994 |
| Slc12a6 | 0.05924917 | 0.04853895 | 0.05156989 | -0.0030309 | 0.65218141 | 0.99999994 |
| Rarres2 | 0.01796456 | 0.01303817 | 0.01607008 | -0.0030319 | 0.89541342 | 0.99999994 |
| Napb | 0.02920009 | 0.02288984 | 0.02592497 | -0.0030351 | 0.78209427 | 0.99999994 |
| Cog1 | 0.2183484 | 0.1656662 | 0.16870323 | -0.003037 | 0.54769075 | 0.99999994 |
| Pum3 | 0.20905058 | 0.15963996 | 0.162679 | -0.003039 | 0.54948053 | 0.99999994 |
| Tpk1 | 0.05520526 | 0.04513839 | 0.04817753 | -0.0030391 | 0.66271952 | 0.99999994 |
| Pogk | 0.069873 | 0.05735614 | 0.06039711 | -0.003041 | 0.63124205 | 0.99999994 |
| Zfp566 | 0.03240751 | 0.02566873 | 0.0287112 | -0.0030425 | 0.75970778 | 0.99999994 |

| | | | | | | |
|---|---|---|---|---|---|---|
| Chml | 0.03108372 | 0.02451835 | 0.0275627 | -0.0030443 | 0.768893 | 0.99999994 |
| Slc25a37 | 0.03248338 | 0.02572908 | 0.02877697 | -0.0030479 | 0.75959528 | 0.99999994 |
| Med14 | 0.03786491 | 0.03037566 | 0.03342472 | -0.0030491 | 0.72814264 | 0.99999994 |
| Ubp1 | 0.08533992 | 0.06997932 | 0.07303278 | -0.0030535 | 0.60960848 | 0.99999994 |
| Mrm1 | 0.04486428 | 0.03636674 | 0.03942029 | -0.0030535 | 0.69693667 | 0.99999994 |
| Tubg2 | 0.02116756 | 0.01583924 | 0.01889463 | -0.0030554 | 0.85912115 | 0.99999994 |
| Dcaf15 | 0.0561227 | 0.04589266 | 0.04894873 | -0.0030561 | 0.66114278 | 0.99999994 |
| Grb14 | 0.13089717 | 0.1052153 | 0.10827443 | -0.0030591 | 0.5745872 | 0.99999994 |
| Gorab | 0.08405721 | 0.0689345 | 0.07199443 | -0.0030599 | 0.61137759 | 0.99999994 |
| Papolg | 0.02619004 | 0.0202349 | 0.02329932 | -0.0030644 | 0.80934151 | 0.99999994 |
| Ntm | 0.01504517 | 0.0104202 | 0.01348503 | -0.0030648 | 0.93483901 | 0.99999994 |
| Chadl | 0.04800562 | 0.03902812 | 0.04209315 | -0.003065 | 0.68605 | 0.99999994 |
| Zbtb40 | 0.02275269 | 0.01722284 | 0.02028799 | -0.0030651 | 0.84270933 | 0.99999994 |
| Dnajb14 | 0.13075412 | 0.10510269 | 0.10816838 | -0.0030657 | 0.57481773 | 0.99999994 |
| Nfatc2 | 0.10956262 | 0.0891121 | 0.0921783 | -0.0030662 | 0.58761564 | 0.99999994 |
| Ccdc181 | 0.13592496 | 0.10891999 | 0.11198642 | -0.0030664 | 0.57231284 | 0.99999994 |
| Pnpla6 | 0.03804494 | 0.03051234 | 0.03357962 | -0.0030673 | 0.7284115 | 0.99999994 |
| Pigv | 0.08828143 | 0.07233872 | 0.07540744 | -0.0030687 | 0.60675897 | 0.99999994 |
| Rabgap1l | 0.12709792 | 0.10237859 | 0.10544881 | -0.0030702 | 0.57683569 | 0.99999994 |
| Rnf217 | 0.07121127 | 0.05842998 | 0.06150043 | -0.0030704 | 0.63017732 | 0.99999994 |
| Ufm1 | 0.87417134 | 0.41187745 | 0.41494898 | -0.0030715 | 0.51964854 | 0.99999994 |
| Cep68 | 0.06434175 | 0.05274478 | 0.05581651 | -0.0030717 | 0.64293563 | 0.99999994 |
| C330013E15I | 0.02385429 | 0.01818044 | 0.02125456 | -0.0030741 | 0.83215911 | 0.99999994 |
| Ahcyl2 | 0.21474779 | 0.16330391 | 0.16638095 | -0.003077 | 0.54898689 | 0.99999994 |
| Ankrd50 | 0.03696441 | 0.0295717 | 0.03264931 | -0.0030776 | 0.734817 | 0.99999994 |
| Tbc1d24 | 0.03388661 | 0.02691203 | 0.02999207 | -0.00308 | 0.7528788 | 0.99999994 |
| Dnttip2 | 0.36691719 | 0.25052493 | 0.25360519 | -0.0030803 | 0.53221833 | 0.99999994 |
| Ethe1 | 0.12999844 | 0.10452671 | 0.10760761 | -0.0030809 | 0.57557162 | 0.99999994 |
| Timmdc1 | 0.08945813 | 0.07327374 | 0.07635488 | -0.0030811 | 0.60588256 | 0.99999994 |
| Mcu | 0.05432745 | 0.04435606 | 0.04743878 | -0.0030827 | 0.66732929 | 0.99999994 |

| | | | | | | |
|---|---|---|---|---|---|---|
| Zfyve16 | 0.08182851 | 0.0671029 | 0.07018624 | -0.0030833 | 0.61501703 | 0.99999994 |
| Itgb5 | 0.02832755 | 0.02208132 | 0.02516494 | -0.0030836 | 0.7926493 | 0.99999994 |
| Zfand2b | 0.28118039 | 0.20403274 | 0.20711738 | -0.0030846 | 0.53948427 | 0.99999994 |
| Pofut1 | 0.05497604 | 0.04489978 | 0.04798471 | -0.0030849 | 0.66565084 | 0.99999994 |
| Dio1 | 0.01546772 | 0.01077382 | 0.01385981 | -0.003086 | 0.93084193 | 0.99999994 |
| Stx2 | 0.03506995 | 0.02792883 | 0.031015 | -0.0030862 | 0.74612565 | 0.99999994 |
| Klc2 | 0.17495298 | 0.13670422 | 0.13979091 | -0.0030867 | 0.55842522 | 0.99999994 |
| 2610008E11I | 0.06616376 | 0.05424119 | 0.05732902 | -0.0030878 | 0.64002341 | 0.99999994 |
| Thap1 | 0.05864862 | 0.04797642 | 0.05106723 | -0.0030908 | 0.65647891 | 0.99999994 |
| Senp8 | 0.1055025 | 0.08595918 | 0.08905102 | -0.0030918 | 0.59138457 | 0.99999994 |
| Acvr1b | 0.04076869 | 0.03282592 | 0.03591881 | -0.0030929 | 0.7167329 | 0.99999994 |
| Unc5a | 0.03758983 | 0.03009401 | 0.03318795 | -0.0030939 | 0.7325477 | 0.99999994 |
| 1500015A07I | 0.13207419 | 0.10605099 | 0.10914613 | -0.0030951 | 0.57485925 | 0.99999994 |
| Clasp2 | 0.04081518 | 0.03286271 | 0.03595866 | -0.0030959 | 0.71671004 | 0.99999994 |
| Zfp764 | 0.02768891 | 0.02151212 | 0.02460809 | -0.003096 | 0.79882875 | 0.99999994 |
| Rbm48 | 0.08172052 | 0.06700172 | 0.07009849 | -0.0030968 | 0.61564424 | 0.99999994 |
| Acot11 | 0.00875254 | 0.00478098 | 0.00787831 | -0.0030973 | 0.99556369 | 0.99999994 |
| D3Ertd751e | 0.05945936 | 0.04864684 | 0.05174572 | -0.0030989 | 0.65491609 | 0.99999994 |
| Tmem186 | 0.09188146 | 0.07520182 | 0.07830156 | -0.0030997 | 0.60392011 | 0.99999994 |
| Atf7 | 0.07962363 | 0.06529205 | 0.06839226 | -0.0031002 | 0.61857147 | 0.99999994 |
| Scyl3 | 0.03721958 | 0.02976868 | 0.03286913 | -0.0031005 | 0.73495712 | 0.99999994 |
| Zfp446 | 0.03233018 | 0.02554342 | 0.02864417 | -0.0031007 | 0.76440808 | 0.99999994 |
| Cc2d2a | 0.08539659 | 0.0699773 | 0.07307861 | -0.0031013 | 0.61121321 | 0.99999994 |
| Arhgef7 | 0.11174889 | 0.09075222 | 0.09385373 | -0.0031015 | 0.58705188 | 0.99999994 |
| Myo9b | 0.04025112 | 0.03237234 | 0.03547496 | -0.0031026 | 0.71977112 | 0.99999994 |
| Zfp94 | 0.03281216 | 0.02595704 | 0.02906188 | -0.0031048 | 0.76149946 | 0.99999994 |
| Tm9sf1 | 0.04167646 | 0.03359123 | 0.03669653 | -0.0031053 | 0.71337377 | 0.99999994 |
| Tmem218 | 0.26952488 | 0.19717838 | 0.20028405 | -0.0031057 | 0.54110413 | 0.99999994 |
| Jpx | 0.029533 | 0.02310789 | 0.02621472 | -0.0031068 | 0.78490808 | 0.99999994 |
| Rabep2 | 0.06571749 | 0.05385171 | 0.05695888 | -0.0031072 | 0.64173924 | 0.99999994 |

| | | | | | | |
|---|---|---|---|---|---|---|
| Pelp1 | 0.13676741 | 0.10949336 | 0.11260545 | -0.0031121 | 0.57297858 | 0.99999994 |
| 3110070M22 | 0.01995452 | 0.01471341 | 0.01782635 | -0.0031129 | 0.87745783 | 0.99999994 |
| Dmxl1 | 0.13267095 | 0.10647392 | 0.10958745 | -0.0031135 | 0.57499915 | 0.99999994 |
| Tmem128 | 0.38227018 | 0.25815406 | 0.26126874 | -0.0031147 | 0.53162406 | 0.99999994 |
| Nkiras2 | 0.37826969 | 0.2561753 | 0.25929051 | -0.0031152 | 0.53187019 | 0.99999994 |
| Btd | 0.06388231 | 0.05231852 | 0.05543455 | -0.003116 | 0.64585755 | 0.99999994 |
| D630039A03 | 0.08914848 | 0.07298809 | 0.0761057 | -0.0031176 | 0.6074476 | 0.99999994 |
| Focad | 0.01643492 | 0.01159898 | 0.01471689 | -0.0031179 | 0.92076125 | 0.99999994 |
| Ldlrad4 | 0.03014552 | 0.02362955 | 0.02674749 | -0.0031179 | 0.78110398 | 0.99999994 |
| Zmym4 | 0.14491553 | 0.11543018 | 0.11854931 | -0.0031191 | 0.56951309 | 0.99999994 |
| Apip | 0.25670511 | 0.18949588 | 0.19261713 | -0.0031213 | 0.54294767 | 0.99999994 |
| Dcn | 0.04085413 | 0.03286885 | 0.03599205 | -0.0031232 | 0.7182341 | 0.99999994 |
| Ercc5 | 0.03707111 | 0.0296164 | 0.03274124 | -0.0031248 | 0.73737943 | 0.99999994 |
| Igfbp7 | 0.19653056 | 0.15129249 | 0.15442167 | -0.0031292 | 0.55364791 | 0.99999994 |
| Abcc5 | 0.14217228 | 0.1134256 | 0.1165569 | -0.0031313 | 0.57096177 | 0.99999994 |
| Cnnm3 | 0.05742521 | 0.04691025 | 0.05004202 | -0.0031318 | 0.66150497 | 0.99999994 |
| Ift80 | 0.04720963 | 0.03828513 | 0.04141692 | -0.0031318 | 0.69263343 | 0.99999994 |
| Zfp628 | 0.04661946 | 0.03778316 | 0.04091508 | -0.0031319 | 0.69480697 | 0.99999994 |
| Herc4 | 0.03808196 | 0.03047942 | 0.03361147 | -0.0031321 | 0.73245892 | 0.99999994 |
| Fig4 | 0.02338503 | 0.01771085 | 0.020843 | -0.0031321 | 0.84141087 | 0.99999994 |
| Dis3l2 | 0.07171604 | 0.05878351 | 0.06191608 | -0.0031326 | 0.63185487 | 0.99999994 |
| Tnrc6b | 0.33667517 | 0.23480706 | 0.23793977 | -0.0031327 | 0.53491765 | 0.99999994 |
| Zfp692 | 0.02833466 | 0.02203787 | 0.02517114 | -0.0031333 | 0.79632737 | 0.99999994 |
| Tmem165 | 0.26941945 | 0.1970848 | 0.20022167 | -0.0031369 | 0.54152841 | 0.99999994 |
| Zswim5 | 0.02224867 | 0.01670758 | 0.01984527 | -0.0031377 | 0.85369712 | 0.99999994 |
| Map4k2 | 0.03120218 | 0.0245275 | 0.02766555 | -0.0031381 | 0.77488126 | 0.99999994 |
| Plekha3 | 0.03721738 | 0.02972784 | 0.03286724 | -0.0031394 | 0.73754657 | 0.99999994 |
| Rnf185 | 0.06857972 | 0.05618871 | 0.05932905 | -0.0031403 | 0.63770334 | 0.99999994 |
| Mcoln1 | 0.05368276 | 0.04375511 | 0.04689567 | -0.0031406 | 0.67211814 | 0.99999994 |
| Spink1 | 0.09937582 | 0.08114289 | 0.08428378 | -0.0031409 | 0.59795341 | 0.99999994 |

| | | | | | | |
|---|---|---|---|---|---|---|
| Fam83h | 0.01871713 | 0.01359393 | 0.01673483 | -0.0031409 | 0.89422365 | 0.99999994 |
| Comtd1 | 0.0815244 | 0.06679704 | 0.06993911 | -0.0031421 | 0.61754605 | 0.99999994 |
| Phc2 | 0.12690455 | 0.10216244 | 0.10530453 | -0.0031421 | 0.57871749 | 0.99999994 |
| Guca1a | 0.05099421 | 0.04148363 | 0.04462579 | -0.0031422 | 0.68035203 | 0.99999994 |
| Smu1 | 0.49014652 | 0.30690192 | 0.31004453 | -0.0031426 | 0.52689576 | 0.99999994 |
| Cdkl1 | 0.02244212 | 0.01687048 | 0.02001522 | -0.0031447 | 0.85218056 | 0.99999994 |
| 2810402E24I | 0.1081526 | 0.08794808 | 0.09109458 | -0.0031465 | 0.590922 | 0.99999994 |
| 2810004N23 | 0.39255371 | 0.26314759 | 0.26629648 | -0.0031489 | 0.53136826 | 0.99999994 |
| Gramd1c | 0.02094044 | 0.01554313 | 0.01869475 | -0.0031516 | 0.86910072 | 0.99999994 |
| Ttc21b | 0.02895116 | 0.02255625 | 0.02570823 | -0.003152 | 0.79278118 | 0.99999994 |
| Crebzf | 0.05748371 | 0.04693816 | 0.05009109 | -0.0031529 | 0.66238348 | 0.99999994 |
| Gosr1 | 0.07701164 | 0.06310699 | 0.0662604 | -0.0031534 | 0.62429628 | 0.99999994 |
| Hps1 | 0.05064255 | 0.04117329 | 0.04432829 | -0.003155 | 0.68216351 | 0.99999994 |
| Camta2 | 0.0348493 | 0.02766796 | 0.03082439 | -0.0031564 | 0.75225841 | 0.99999994 |
| Exosc10 | 0.15507419 | 0.122692 | 0.12584958 | -0.0031576 | 0.56631843 | 0.99999994 |
| Sart3 | 0.12844935 | 0.10329699 | 0.10645592 | -0.0031589 | 0.57828892 | 0.99999994 |
| Zfp729a | 0.02305435 | 0.0173935 | 0.02055281 | -0.0031593 | 0.84689223 | 0.99999994 |
| Rbm33 | 0.09756104 | 0.07967888 | 0.08284042 | -0.0031615 | 0.60026543 | 0.99999994 |
| Tapt1 | 0.02950725 | 0.02302975 | 0.02619231 | -0.0031626 | 0.78921934 | 0.99999994 |
| Smyd5 | 0.04391709 | 0.0354491 | 0.03861219 | -0.0031631 | 0.70722104 | 0.99999994 |
| E4f1 | 0.07036758 | 0.05764173 | 0.06080508 | -0.0031633 | 0.63543869 | 0.99999994 |
| Rpp38 | 0.08784945 | 0.07189068 | 0.07505927 | -0.0031686 | 0.61064297 | 0.99999994 |
| Snx25 | 0.01315813 | 0.00863865 | 0.01180868 | -0.00317 | 0.96302265 | 0.99999994 |
| Tyk2 | 0.01904745 | 0.01385621 | 0.01702639 | -0.0031702 | 0.89239423 | 0.99999994 |
| Angpt2 | 0.01826962 | 0.01316915 | 0.01633963 | -0.0031705 | 0.90174746 | 0.99999994 |
| Dhcr7 | 0.14634402 | 0.1164122 | 0.11958326 | -0.0031711 | 0.57005364 | 0.99999994 |
| 5033417F24I | 0.02488641 | 0.01898244 | 0.02215889 | -0.0031764 | 0.83000353 | 0.99999994 |
| Acss2 | 0.02335945 | 0.01764191 | 0.02082055 | -0.0031786 | 0.84523666 | 0.99999994 |
| C1qtnf4 | 0.24037153 | 0.17943473 | 0.18261544 | -0.0031807 | 0.54614894 | 0.99999994 |
| 1700096K18I | 0.13266616 | 0.10640284 | 0.10958391 | -0.0031811 | 0.57660865 | 0.99999994 |

| | | | | | | |
|---|---|---|---|---|---|---|
| Nup93 | 0.1292908 | 0.10389854 | 0.10708187 | -0.0031833 | 0.57843063 | 0.99999994 |
| Unk | 0.05476807 | 0.04462156 | 0.04780971 | -0.0031881 | 0.67143289 | 0.99999994 |
| Rassf6 | 0.03528063 | 0.0280072 | 0.03119696 | -0.0031898 | 0.75193297 | 0.99999994 |
| Pygo2 | 0.05723617 | 0.04669257 | 0.04988347 | -0.0031909 | 0.66486875 | 0.99999994 |
| Tmem209 | 0.03955832 | 0.0316888 | 0.03488037 | -0.0031916 | 0.72875734 | 0.99999994 |
| Meis3 | 0.17370982 | 0.13573881 | 0.13893215 | -0.0031933 | 0.56079917 | 0.99999994 |
| Gria3 | 0.02109445 | 0.01563404 | 0.0188303 | -0.0031963 | 0.87073092 | 0.99999994 |
| Vcpip1 | 0.11280497 | 0.09146446 | 0.09466092 | -0.0031965 | 0.58892046 | 0.99999994 |
| Stxbp4 | 0.09327912 | 0.07622298 | 0.07942157 | -0.0031986 | 0.60567916 | 0.99999994 |
| Elk4 | 0.04721277 | 0.03822031 | 0.04141958 | -0.0031993 | 0.69642353 | 0.99999994 |
| Ttc30a1 | 0.03338676 | 0.02636001 | 0.02955949 | -0.0031995 | 0.76438435 | 0.99999994 |
| Opa3 | 0.14632729 | 0.11637027 | 0.11957117 | -0.0032009 | 0.5707129 | 0.99999994 |
| Ccdc112 | 0.12965938 | 0.10415483 | 0.10735578 | -0.0032009 | 0.57866047 | 0.99999994 |
| Mettl8 | 0.04277335 | 0.03443312 | 0.03763505 | -0.0032019 | 0.71440873 | 0.99999994 |
| Gcc1 | 0.03769918 | 0.03007806 | 0.03328208 | -0.003204 | 0.73917331 | 0.99999994 |
| Hmbox1 | 0.07730797 | 0.06329793 | 0.06650262 | -0.0032047 | 0.6258049 | 0.99999994 |
| Rfx7 | 0.086531 | 0.07079014 | 0.07399541 | -0.0032053 | 0.61345476 | 0.99999994 |
| Sdc4 | 0.03410279 | 0.02697321 | 0.03017907 | -0.0032059 | 0.76024491 | 0.99999994 |
| Smc5 | 0.15438282 | 0.12215037 | 0.12535654 | -0.0032062 | 0.56759142 | 0.99999994 |
| Gm2447 | 0.02465067 | 0.01874569 | 0.02195245 | -0.0032068 | 0.83456018 | 0.99999994 |
| Ift81 | 0.31184795 | 0.22128532 | 0.2244927 | -0.0032074 | 0.53788265 | 0.99999994 |
| Spr | 0.1965639 | 0.15123347 | 0.15444387 | -0.0032104 | 0.55502394 | 0.99999994 |
| Ddx23 | 0.06159755 | 0.05032033 | 0.05353168 | -0.0032113 | 0.65516852 | 0.99999994 |
| Fgl2 | 0.00723476 | 0.00330465 | 0.00651883 | -0.0032142 | 0.99948462 | 0.99999994 |
| Lrig2 | 0.03095739 | 0.02423753 | 0.02745299 | -0.0032155 | 0.78219666 | 0.99999994 |
| Myl4 | 0.01281993 | 0.00829047 | 0.01150778 | -0.0032173 | 0.96862563 | 0.99999994 |
| Agfg2 | 0.05792252 | 0.0472415 | 0.05045897 | -0.0032175 | 0.66437619 | 0.99999994 |
| Gm12258 | 0.02828063 | 0.02190531 | 0.02512405 | -0.0032187 | 0.80310693 | 0.99999994 |
| Kpna2 | 0.05620484 | 0.04579814 | 0.04901773 | -0.0032196 | 0.66901845 | 0.99999994 |
| Gls | 0.09234569 | 0.07545419 | 0.07867379 | -0.0032196 | 0.60734317 | 0.99999994 |

| | | | | | | |
|---|---|---|---|---|---|---|
| mt-Nd6 | 0.0309376 | 0.02421273 | 0.0274358 | -0.0032231 | 0.78288356 | 0.99999994 |
| Smarcal1 | 0.03733193 | 0.02973952 | 0.03296589 | -0.0032264 | 0.74262408 | 0.99999994 |
| Oscp1 | 0.07626571 | 0.06242161 | 0.06565027 | -0.0032287 | 0.62830125 | 0.99999994 |
| Rela | 0.07462148 | 0.06107366 | 0.06430329 | -0.0032296 | 0.63092929 | 0.99999994 |
| Ptbp2 | 0.11777569 | 0.09521092 | 0.09844132 | -0.0032304 | 0.58645355 | 0.99999994 |
| Slc25a22 | 0.09999983 | 0.08154797 | 0.08477931 | -0.0032313 | 0.60013825 | 0.99999994 |
| Adcy6 | 0.02983279 | 0.02324375 | 0.02647553 | -0.0032318 | 0.79175335 | 0.99999994 |
| Enox2 | 0.04485113 | 0.03617166 | 0.03940908 | -0.0032374 | 0.70774682 | 0.99999994 |
| Gm4285 | 0.02400734 | 0.01815102 | 0.02138874 | -0.0032377 | 0.8431778 | 0.99999994 |
| Man1c1 | 0.03821786 | 0.0304885 | 0.03372838 | -0.0032399 | 0.73871777 | 0.99999994 |
| Jade3 | 0.1566094 | 0.12370235 | 0.12694242 | -0.0032401 | 0.56745424 | 0.99999994 |
| Lin7b | 0.01989392 | 0.01453171 | 0.01777294 | -0.0032412 | 0.88760929 | 0.99999994 |
| Nthl1 | 0.05881782 | 0.04796747 | 0.05120889 | -0.0032414 | 0.66324443 | 0.99999994 |
| Nudt6 | 0.02576617 | 0.01968494 | 0.02292873 | -0.0032438 | 0.82682547 | 0.99999994 |
| Cacna1c | 0.02251799 | 0.01683757 | 0.02008187 | -0.0032443 | 0.85889305 | 0.99999994 |
| Otub2 | 0.02980161 | 0.02320304 | 0.02644841 | -0.0032454 | 0.79297094 | 0.99999994 |
| Jag2 | 0.01124732 | 0.00686104 | 0.01010686 | -0.0032458 | 0.98372909 | 0.99999994 |
| Adk | 0.18515173 | 0.14352217 | 0.14677003 | -0.0032479 | 0.55855153 | 0.99999994 |
| Rapgef2 | 0.02218054 | 0.01653367 | 0.0197854 | -0.0032517 | 0.86301598 | 0.99999994 |
| Mxd1 | 0.14733709 | 0.11704833 | 0.12030052 | -0.0032522 | 0.57140307 | 0.99999994 |
| Arhgef16 | 0.16465564 | 0.12937281 | 0.13262552 | -0.0032527 | 0.56483948 | 0.99999994 |
| Cldn10 | 0.03174311 | 0.0248809 | 0.02813503 | -0.0032541 | 0.77931508 | 0.99999994 |
| Rgl3 | 0.06886152 | 0.05630742 | 0.05956193 | -0.0032545 | 0.64196227 | 0.99999994 |
| Ank3 | 0.2007276 | 0.15395259 | 0.15720835 | -0.0032558 | 0.55482138 | 0.99999994 |
| Ppp1r26 | 0.0188099 | 0.01356055 | 0.01681672 | -0.0032562 | 0.90127494 | 0.99999994 |
| Tspan33 | 0.09154507 | 0.07477491 | 0.0780317 | -0.0032568 | 0.60942263 | 0.99999994 |
| Nmi | 0.05031198 | 0.04079096 | 0.04404852 | -0.0032576 | 0.68873346 | 0.99999994 |
| Tmem177 | 0.05388523 | 0.04380858 | 0.04706629 | -0.0032577 | 0.67750078 | 0.99999994 |
| Klf5 | 0.06787943 | 0.05549199 | 0.05874997 | -0.003258 | 0.6439837 | 0.99999994 |
| Nova2 | 0.038992 | 0.03113484 | 0.03439392 | -0.0032591 | 0.73589452 | 0.99999994 |

| | | | | | | |
|---|---|---|---|---|---|---|
| Pcyt1a | 0.05875931 | 0.04789549 | 0.05115991 | -0.0032644 | 0.66448331 | 0.99999994 |
| Eml2 | 0.27354455 | 0.19939038 | 0.20265502 | -0.0032646 | 0.54269648 | 0.99999994 |
| Dlgap5 | 0.04001232 | 0.03200482 | 0.03527007 | -0.0032653 | 0.73113077 | 0.99999994 |
| Usp46 | 0.08658311 | 0.07077157 | 0.0740375 | -0.0032659 | 0.61547747 | 0.99999994 |
| Ino80 | 0.05749338 | 0.04683301 | 0.0500992 | -0.0032662 | 0.66784047 | 0.99999994 |
| 9330151L19I | 0.03910519 | 0.03122496 | 0.03449118 | -0.0032662 | 0.73576314 | 0.99999994 |
| Srsf4 | 0.22867924 | 0.17202442 | 0.17529108 | -0.0032667 | 0.54936035 | 0.99999994 |
| Nsmaf | 0.05469037 | 0.04447518 | 0.04774431 | -0.0032691 | 0.67572256 | 0.99999994 |
| Homez | 0.06183965 | 0.05046253 | 0.05373358 | -0.0032711 | 0.65733773 | 0.99999994 |
| Gm4890 | 0.03364548 | 0.0265112 | 0.02978343 | -0.0032722 | 0.76770696 | 0.99999994 |
| Ppard | 0.03314166 | 0.02606778 | 0.02934728 | -0.0032795 | 0.77151216 | 0.99999994 |
| Grk5 | 0.03094613 | 0.02416196 | 0.02744321 | -0.0032813 | 0.7869458 | 0.99999994 |
| Dffb | 0.04023286 | 0.03217732 | 0.03545929 | -0.003282 | 0.73108124 | 0.99999994 |
| Ttc17 | 0.01828877 | 0.01307105 | 0.01635654 | -0.0032855 | 0.90939561 | 0.99999994 |
| Kif9 | 0.0227555 | 0.01700288 | 0.02029045 | -0.0032876 | 0.8595943 | 0.99999994 |
| Sphkap | 0.30004516 | 0.21461795 | 0.21790782 | -0.0032899 | 0.54002413 | 0.99999994 |
| Hivep2 | 0.01263286 | 0.00805095 | 0.01134129 | -0.0032903 | 0.97326744 | 0.99999994 |
| Rpap2 | 0.08359599 | 0.06833016 | 0.07162066 | -0.0032905 | 0.62012777 | 0.99999994 |
| Zfp672 | 0.10028857 | 0.08171048 | 0.08500457 | -0.0032941 | 0.60177571 | 0.99999994 |
| Osgepl1 | 0.05014923 | 0.04061616 | 0.04391073 | -0.0032946 | 0.69125792 | 0.99999994 |
| Cel | 0.03506978 | 0.0277175 | 0.03101486 | -0.0032974 | 0.76042614 | 0.99999994 |
| Cux2 | 0.0135214 | 0.00883084 | 0.01213172 | -0.0033009 | 0.96493769 | 0.99999994 |
| Stbd1 | 0.08529759 | 0.06969418 | 0.07299854 | -0.0033044 | 0.61840994 | 0.99999994 |
| Tti2 | 0.04638356 | 0.03740989 | 0.04071438 | -0.0033045 | 0.70548672 | 0.99999994 |
| Apeh | 0.1908091 | 0.14728751 | 0.15059225 | -0.0033047 | 0.55806835 | 0.99999994 |
| Sumf1 | 0.18526569 | 0.14354217 | 0.14684736 | -0.0033052 | 0.55954633 | 0.99999994 |
| Ilvbl | 0.09673801 | 0.0788769 | 0.08218473 | -0.0033078 | 0.60561588 | 0.99999994 |
| Kctd12 | 0.12498267 | 0.10055996 | 0.10386807 | -0.0033081 | 0.58394623 | 0.99999994 |
| Prdm4 | 0.03928415 | 0.0313363 | 0.03464492 | -0.0033086 | 0.73751016 | 0.99999994 |
| Nol6 | 0.02090242 | 0.01535223 | 0.01866128 | -0.0033091 | 0.88106297 | 0.99999994 |

| | | | | | | |
|---|---|---|---|---|---|---|
| Kmt2e | 0.61350571 | 0.35291703 | 0.35622883 | -0.0033118 | 0.52467207 | 0.99999994 |
| Sftpd | 0.01877871 | 0.01347695 | 0.01678919 | -0.0033122 | 0.90544118 | 0.99999994 |
| Cwc25 | 0.17491452 | 0.13645005 | 0.13976437 | -0.0033143 | 0.56271104 | 0.99999994 |
| Acy1 | 0.04828945 | 0.03901866 | 0.04233411 | -0.0033154 | 0.69892128 | 0.99999994 |
| Mob3c | 0.10829571 | 0.0878892 | 0.09120468 | -0.0033155 | 0.59559932 | 0.99999994 |
| Cdkal1 | 0.10326424 | 0.08400207 | 0.08731814 | -0.0033161 | 0.59978389 | 0.99999994 |
| Ptch1 | 0.06987152 | 0.05707956 | 0.06039589 | -0.0033163 | 0.64263074 | 0.99999994 |
| Sez6l | 0.03655989 | 0.02898361 | 0.03230069 | -0.0033171 | 0.75287542 | 0.99999994 |
| Add3 | 0.08539028 | 0.06975501 | 0.07307351 | -0.0033185 | 0.6187831 | 0.99999994 |
| Rgp1 | 0.05529097 | 0.04493067 | 0.04824962 | -0.003319 | 0.67647765 | 0.99999994 |
| Acap3 | 0.04002625 | 0.031963 | 0.03528203 | -0.003319 | 0.73439935 | 0.99999994 |
| Uhrf1bp1l | 0.08658095 | 0.0707161 | 0.07403575 | -0.0033197 | 0.61732435 | 0.99999994 |
| 5033430I15R | 0.03345474 | 0.02629804 | 0.02961834 | -0.0033203 | 0.77222579 | 0.99999994 |
| Tubd1 | 0.04464254 | 0.03590329 | 0.03923121 | -0.0033279 | 0.71384325 | 0.99999994 |
| Hspa9 | 0.65324262 | 0.36577924 | 0.3691101 | -0.0033309 | 0.5239489 | 0.99999994 |
| Me2 | 0.10317308 | 0.08391636 | 0.08724744 | -0.0033311 | 0.60030541 | 0.99999994 |
| Nkapl | 0.03687069 | 0.02923727 | 0.03256856 | -0.0033313 | 0.75201616 | 0.99999994 |
| 2310022A10 | 0.07268489 | 0.0593784 | 0.06271312 | -0.0033347 | 0.63831085 | 0.99999994 |
| Polr3g | 0.02279124 | 0.01698703 | 0.02032183 | -0.0033348 | 0.86265071 | 0.99999994 |
| Zfp26 | 0.04285939 | 0.03437315 | 0.0377086 | -0.0033355 | 0.72199483 | 0.99999994 |
| Ppm1a | 0.0753353 | 0.06155249 | 0.06488842 | -0.0033359 | 0.63390186 | 0.99999994 |
| Spsb2 | 0.12564053 | 0.1010232 | 0.10436027 | -0.0033371 | 0.58427681 | 0.99999994 |
| Mfn1 | 0.05723917 | 0.0465473 | 0.04988598 | -0.0033387 | 0.67201531 | 0.99999994 |
| Defb1 | 0.01138918 | 0.00689218 | 0.01023335 | -0.0033412 | 0.98512087 | 0.99999994 |
| Phf7 | 0.02694801 | 0.02061944 | 0.02396148 | -0.003342 | 0.8234001 | 0.99999994 |
| Fam185a | 0.03023781 | 0.02348406 | 0.02682773 | -0.0033437 | 0.79662161 | 0.99999994 |
| Capns1 | 1.03479794 | 0.42479348 | 0.42813821 | -0.0033447 | 0.52073615 | 0.99999994 |
| Pno1 | 0.28633541 | 0.206754 | 0.21009877 | -0.0033448 | 0.54219666 | 0.99999994 |
| Gas2 | 0.02571514 | 0.01953849 | 0.0228841 | -0.0033456 | 0.83475948 | 0.99999994 |
| Zfyve27 | 0.14732754 | 0.11694743 | 0.12029363 | -0.0033462 | 0.573448 | 0.99999994 |

| | | | | | | |
|---|---|---|---|---|---|---|
| Wdr55 | 0.06354236 | 0.05180403 | 0.05515178 | -0.0033478 | 0.65691042 | 0.99999994 |
| Hars2 | 0.07872341 | 0.06430962 | 0.06765833 | -0.0033487 | 0.62909296 | 0.99999994 |
| Celsr2 | 0.15846385 | 0.12491015 | 0.12825891 | -0.0033488 | 0.56898625 | 0.99999994 |
| Pcgf2 | 0.08072343 | 0.06593851 | 0.06928774 | -0.0033492 | 0.62618105 | 0.99999994 |
| Gsn | 0.02836732 | 0.0218497 | 0.0251996 | -0.0033499 | 0.81188439 | 0.99999994 |
| Fgfr2 | 0.03948072 | 0.03146265 | 0.03481373 | -0.0033511 | 0.73914902 | 0.99999994 |
| Zmym2 | 0.14383769 | 0.11441583 | 0.11776754 | -0.0033517 | 0.57512685 | 0.99999994 |
| Edc4 | 0.05623101 | 0.04568502 | 0.04903971 | -0.0033547 | 0.67557004 | 0.99999994 |
| Cab39l | 0.11151297 | 0.09031835 | 0.09367323 | -0.0033549 | 0.5942128 | 0.99999994 |
| Efcab2 | 0.09667367 | 0.07877822 | 0.08213344 | -0.0033552 | 0.60715734 | 0.99999994 |
| Shroom2 | 0.02012022 | 0.01461607 | 0.01797238 | -0.0033563 | 0.89307784 | 0.99999994 |
| Mcm3ap | 0.05351434 | 0.04339738 | 0.04675371 | -0.0033563 | 0.6836193 | 0.99999994 |
| Sdhaf2 | 0.27658547 | 0.20108104 | 0.20443861 | -0.0033576 | 0.54352561 | 0.99999994 |
| Zmym3 | 0.07929422 | 0.0647658 | 0.06812379 | -0.003358 | 0.62858517 | 0.99999994 |
| Gpt | 0.01905295 | 0.0136731 | 0.01703124 | -0.0033581 | 0.90531906 | 0.99999994 |
| Gemin5 | 0.02320292 | 0.01732456 | 0.0206832 | -0.0033586 | 0.8601279 | 0.99999994 |
| Lipo1 | 0.03303529 | 0.0258957 | 0.02925516 | -0.0033595 | 0.77767684 | 0.99999994 |
| Ap3b1 | 0.16308946 | 0.12816538 | 0.13152525 | -0.0033599 | 0.56751056 | 0.99999994 |
| Zfp609 | 0.04044473 | 0.03228111 | 0.03564103 | -0.0033599 | 0.73483495 | 0.99999994 |
| Cdc42bpg | 0.03447975 | 0.02714197 | 0.03050501 | -0.003363 | 0.76847488 | 0.99999994 |
| Vac14 | 0.04826023 | 0.03894468 | 0.04230931 | -0.0033646 | 0.70171649 | 0.99999994 |
| Mafk | 0.02916867 | 0.02253223 | 0.02589762 | -0.0033654 | 0.80647838 | 0.99999994 |
| Nt5dc3 | 0.01701641 | 0.0118647 | 0.01523163 | -0.0033669 | 0.9294057 | 0.99999994 |
| Pfdn4 | 0.89830529 | 0.41448383 | 0.41785306 | -0.0033692 | 0.52140156 | 0.99999994 |
| Dyrk3 | 0.04404347 | 0.0353501 | 0.03872007 | -0.00337 | 0.71881683 | 0.99999994 |
| Daglb | 0.04959475 | 0.04007073 | 0.04344107 | -0.0033703 | 0.69722247 | 0.99999994 |
| Clk2 | 0.05744109 | 0.04668368 | 0.05005534 | -0.0033717 | 0.67305456 | 0.99999994 |
| Amacr | 0.08460641 | 0.0690661 | 0.07243922 | -0.0033731 | 0.62170121 | 0.99999994 |
| Rprd2 | 0.15470687 | 0.1222145 | 0.1255877 | -0.0033732 | 0.57094679 | 0.99999994 |
| Sarnp | 0.93795382 | 0.41849749 | 0.42187148 | -0.003374 | 0.52122777 | 0.99999994 |

| | | | | | | |
|---|---|---|---|---|---|---|
| Manba | 0.02599425 | 0.01975182 | 0.02312817 | -0.0033764 | 0.83440607 | 0.99999994 |
| Nefh | 0.01964768 | 0.01417734 | 0.01755585 | -0.0033785 | 0.89989859 | 0.99999994 |
| Lancl2 | 0.05878472 | 0.04780245 | 0.05118119 | -0.0033787 | 0.66982166 | 0.99999994 |
| Zfp740 | 0.0506026 | 0.04091064 | 0.04429449 | -0.0033839 | 0.69445177 | 0.99999994 |
| Gtf3c5 | 0.06429726 | 0.05239507 | 0.05577954 | -0.0033845 | 0.65684938 | 0.99999994 |
| mt-Nd2 | 1.83323494 | 0.37462665 | 0.37801122 | -0.0033846 | 0.52376233 | 0.99999994 |
| Tmem203 | 0.19725016 | 0.15151599 | 0.15490079 | -0.0033848 | 0.55782245 | 0.99999994 |
| Atxn7l3 | 0.06044281 | 0.0491787 | 0.05256779 | -0.0033891 | 0.66608976 | 0.99999994 |
| Rbbp8 | 0.0830888 | 0.06781913 | 0.07120938 | -0.0033903 | 0.6243428 | 0.99999994 |
| Tacc2 | 0.0186504 | 0.01328562 | 0.01667591 | -0.0033903 | 0.91201546 | 0.99999994 |
| 2310011J03F | 0.3229047 | 0.2271576 | 0.23054808 | -0.0033905 | 0.53899003 | 0.99999994 |
| Oma1 | 0.0657757 | 0.05361664 | 0.05700717 | -0.0033905 | 0.65390703 | 0.99999994 |
| Tmcc1 | 0.1321231 | 0.10579166 | 0.10918232 | -0.0033907 | 0.58188306 | 0.99999994 |
| Mospd2 | 0.09655194 | 0.07864308 | 0.0820364 | -0.0033933 | 0.60846895 | 0.99999994 |
| Ipmk | 0.0567725 | 0.04610032 | 0.04949439 | -0.0033941 | 0.67596989 | 0.99999994 |
| Pik3c3 | 0.06011133 | 0.04889397 | 0.05229082 | -0.0033969 | 0.66727542 | 0.99999994 |
| Cmc2 | 0.19339924 | 0.14893241 | 0.15233033 | -0.0033979 | 0.55901745 | 0.99999994 |
| Kansl1 | 0.13926651 | 0.1110387 | 0.11443681 | -0.0033981 | 0.57834267 | 0.99999994 |
| Zfp410 | 0.21152275 | 0.16088914 | 0.16428952 | -0.0034004 | 0.55478884 | 0.99999994 |
| Abhd12 | 0.15756631 | 0.1242203 | 0.12762223 | -0.0034019 | 0.57041589 | 0.99999994 |
| Nek4 | 0.02065107 | 0.01503736 | 0.01843999 | -0.0034026 | 0.89032582 | 0.99999994 |
| Pigl | 0.03434007 | 0.0269814 | 0.03038426 | -0.0034029 | 0.77200708 | 0.99999994 |
| Rc3h1 | 0.10740959 | 0.08711962 | 0.09052252 | -0.0034029 | 0.59879301 | 0.99999994 |
| Traf6 | 0.04139421 | 0.03305188 | 0.03645481 | -0.0034029 | 0.73281469 | 0.99999994 |
| Eml3 | 0.037725 | 0.0299011 | 0.03330431 | -0.0034032 | 0.75180513 | 0.99999994 |
| Rfx5 | 0.01837659 | 0.0130305 | 0.01643412 | -0.0034036 | 0.91598743 | 0.99999994 |
| Nme4 | 0.1503411 | 0.11905901 | 0.12246324 | -0.0034042 | 0.57339842 | 0.99999994 |
| Fut8 | 0.06539348 | 0.05328396 | 0.05669002 | -0.0034061 | 0.65539555 | 0.99999994 |
| 5930430L01F | 0.04783739 | 0.03854292 | 0.04195029 | -0.0034074 | 0.70562735 | 0.99999994 |
| Cgn | 0.05351316 | 0.04333574 | 0.04675271 | -0.003417 | 0.68668923 | 0.99999994 |

| | | | | | | |
|---|---|---|---|---|---|---|
| Zfp746 | 0.03983387 | 0.03169865 | 0.03511693 | -0.0034183 | 0.74148421 | 0.99999994 |
| Mks1 | 0.06227374 | 0.050677 | 0.05409544 | -0.0034184 | 0.66298869 | 0.99999994 |
| Gpat4 | 0.23992951 | 0.17892148 | 0.18234112 | -0.0034196 | 0.54967246 | 0.99999994 |
| Armc10 | 0.32273607 | 0.22703659 | 0.23045653 | -0.0034199 | 0.5393433 | 0.99999994 |
| Ppl | 0.0322369 | 0.02513992 | 0.0285633 | -0.0034234 | 0.78750119 | 0.99999994 |
| Cyp4f13 | 0.06931917 | 0.05651534 | 0.05993995 | -0.0034246 | 0.64814036 | 0.99999994 |
| Fnbp1 | 0.05393827 | 0.04368581 | 0.04711097 | -0.0034252 | 0.68578768 | 0.99999994 |
| Cenpw | 0.05945316 | 0.04831491 | 0.05174054 | -0.0034256 | 0.67028602 | 0.99999994 |
| 3110040N11 | 0.13911349 | 0.1108991 | 0.11432489 | -0.0034258 | 0.57904868 | 0.99999994 |
| Zfc3h1 | 0.09205047 | 0.07500978 | 0.0784371 | -0.0034273 | 0.61441723 | 0.99999994 |
| Exog | 0.0756487 | 0.06171644 | 0.06514514 | -0.0034287 | 0.63695648 | 0.99999994 |
| Lsm10 | 0.24566914 | 0.18245704 | 0.18588826 | -0.0034312 | 0.54889359 | 0.99999994 |
| Heatr5a | 0.01361926 | 0.00878562 | 0.01221872 | -0.0034331 | 0.96927464 | 0.99999994 |
| Usp29 | 0.20886054 | 0.15911622 | 0.16255493 | -0.0034387 | 0.55598988 | 0.99999994 |
| Baiap2 | 0.10402484 | 0.08446762 | 0.08790771 | -0.0034401 | 0.60275459 | 0.99999994 |
| Usp38 | 0.05238287 | 0.04235904 | 0.0457992 | -0.0034402 | 0.69145748 | 0.99999994 |
| Slc35b3 | 0.08182045 | 0.06673931 | 0.07017969 | -0.0034404 | 0.62790547 | 0.99999994 |
| Mfsd3 | 0.03310921 | 0.02587822 | 0.02931918 | -0.003441 | 0.78265859 | 0.99999994 |
| Doc2b | 0.05684408 | 0.04611291 | 0.04955447 | -0.0034416 | 0.67806151 | 0.99999994 |
| Mtmr1 | 0.03115249 | 0.02418033 | 0.02762241 | -0.0034421 | 0.79657819 | 0.99999994 |
| Pex6 | 0.11979253 | 0.09652316 | 0.09996637 | -0.0034432 | 0.5906703 | 0.99999994 |
| Hectd3 | 0.06090544 | 0.04950847 | 0.05295413 | -0.0034457 | 0.66753655 | 0.99999994 |
| Tmem8b | 0.02437302 | 0.01826209 | 0.02170922 | -0.0034471 | 0.8547307 | 0.99999994 |
| Tnrc6c | 0.12244517 | 0.09851718 | 0.1019646 | -0.0034474 | 0.5890302 | 0.99999994 |
| Ulk3 | 0.02725598 | 0.02078236 | 0.02423034 | -0.003448 | 0.82823153 | 0.99999994 |
| Dennd5b | 0.03400176 | 0.02664207 | 0.03009169 | -0.0034496 | 0.77728752 | 0.99999994 |
| Prune2 | 0.02267259 | 0.01676727 | 0.02021764 | -0.0034504 | 0.87201825 | 0.99999994 |
| Gm20522 | 0.02795388 | 0.02138864 | 0.02483918 | -0.0034505 | 0.82243147 | 0.99999994 |
| Uba6 | 0.08534302 | 0.06958474 | 0.07303529 | -0.0034506 | 0.62342155 | 0.99999994 |
| Acp2 | 0.10916926 | 0.08842431 | 0.09187623 | -0.0034519 | 0.59874042 | 0.99999994 |

| | | | | | | |
|---|---|---|---|---|---|---|
| Sco2 | 0.06129446 | 0.04982285 | 0.05327882 | -0.003456 | 0.66704534 | 0.99999994 |
| Trim41 | 0.13982743 | 0.11139079 | 0.11484684 | -0.0034561 | 0.5793802 | 0.99999994 |
| Zfp574 | 0.06134266 | 0.04986153 | 0.05331904 | -0.0034575 | 0.66699695 | 0.99999994 |
| Polr1b | 0.02033422 | 0.01470233 | 0.01816093 | -0.0034586 | 0.89753844 | 0.99999994 |
| Zdhhc7 | 0.06524151 | 0.05310122 | 0.05656388 | -0.0034627 | 0.65817268 | 0.99999994 |
| H2-T22 | 0.0290737 | 0.0223501 | 0.02581493 | -0.0034648 | 0.81417938 | 0.99999994 |
| Kremen1 | 0.03828559 | 0.03031912 | 0.03378664 | -0.0034675 | 0.75274093 | 0.99999994 |
| Zhx2 | 0.02086077 | 0.01515583 | 0.01862462 | -0.0034688 | 0.89246251 | 0.99999994 |
| Sap130 | 0.02980391 | 0.02298099 | 0.02645041 | -0.0034694 | 0.80869424 | 0.99999994 |
| Samd8 | 0.08546949 | 0.06966789 | 0.07313757 | -0.0034697 | 0.62391555 | 0.99999994 |
| Kctd2 | 0.02248005 | 0.01657856 | 0.02004854 | -0.00347 | 0.87535536 | 0.99999994 |
| Kl | 0.01216437 | 0.00745333 | 0.01092415 | -0.0034708 | 0.98277888 | 0.99999994 |
| Crls1 | 0.04459855 | 0.03572132 | 0.03919368 | -0.0034724 | 0.72231042 | 0.99999994 |
| 2700038G22 | 0.03691424 | 0.0291312 | 0.03260608 | -0.0034749 | 0.7609554 | 0.99999994 |
| Mrps18a | 0.53995522 | 0.32635134 | 0.32982779 | -0.0034765 | 0.5279666 | 0.99999994 |
| Pds5a | 0.1750394 | 0.13637327 | 0.13985055 | -0.0034773 | 0.56572689 | 0.99999994 |
| Napepld | 0.0362622 | 0.02856613 | 0.03204402 | -0.0034779 | 0.76498959 | 0.99999994 |
| Per2 | 0.02877988 | 0.02207946 | 0.02555906 | -0.0034796 | 0.81758358 | 0.99999994 |
| Nfya | 0.05309285 | 0.0429158 | 0.04639831 | -0.0034825 | 0.69132351 | 0.99999994 |
| Lemd3 | 0.01918788 | 0.01366703 | 0.0171503 | -0.0034833 | 0.91180005 | 0.99999994 |
| Eid2 | 0.02419981 | 0.0180729 | 0.02155744 | -0.0034845 | 0.8590213 | 0.99999994 |
| Rab40b | 0.04730125 | 0.03800916 | 0.04149479 | -0.0034856 | 0.71196855 | 0.99999994 |
| Nom1 | 0.11960307 | 0.09633638 | 0.09982332 | -0.0034869 | 0.59193051 | 0.99999994 |
| Lsg1 | 0.18326866 | 0.14200267 | 0.14549007 | -0.0034874 | 0.56338341 | 0.99999994 |
| Nedd1 | 0.05765414 | 0.0467465 | 0.05023399 | -0.0034875 | 0.67800159 | 0.99999994 |
| Calu | 0.08778938 | 0.0715211 | 0.07501084 | -0.0034897 | 0.62159596 | 0.99999994 |
| Nhlrc2 | 0.0687575 | 0.05598452 | 0.05947598 | -0.0034915 | 0.6520099 | 0.99999994 |
| Pstk | 0.09716488 | 0.07903246 | 0.08252489 | -0.0034924 | 0.61091139 | 0.99999994 |
| Cetn2 | 0.58848862 | 0.34411745 | 0.34761092 | -0.0034935 | 0.52666772 | 0.99999994 |
| Zbtb37 | 0.04623902 | 0.03709522 | 0.04059137 | -0.0034962 | 0.71678454 | 0.99999994 |

| | | | | | | |
|---|---|---|---|---|---|---|
| Zfp959 | 0.04445711 | 0.03557686 | 0.03907304 | -0.0034962 | 0.72427776 | 0.99999994 |
| Zfp236 | 0.03716421 | 0.02932512 | 0.03282144 | -0.0034963 | 0.76085991 | 0.99999994 |
| Ccdc97 | 0.09885333 | 0.08036942 | 0.08386857 | -0.0034991 | 0.60938442 | 0.99999994 |
| Brpf3 | 0.03622918 | 0.0285159 | 0.03201553 | -0.0034996 | 0.76657404 | 0.99999994 |
| Lztfl1 | 0.39734228 | 0.26510804 | 0.26860857 | -0.0035005 | 0.53456342 | 0.99999994 |
| 3222401L13F | 0.02042563 | 0.0147409 | 0.01824145 | -0.0035005 | 0.89926245 | 0.99999994 |
| Dcp1a | 0.03214345 | 0.02498045 | 0.02848227 | -0.0035018 | 0.79343088 | 0.99999994 |
| Rbm14 | 0.17457579 | 0.13602521 | 0.13953053 | -0.0035053 | 0.56640256 | 0.99999994 |
| Aasdh | 0.02962216 | 0.02278548 | 0.02629229 | -0.0035068 | 0.81267929 | 0.99999994 |
| Uspl1 | 0.0871112 | 0.07095572 | 0.0744638 | -0.0035081 | 0.62308345 | 0.99999994 |
| 1700007L15F | 0.04862374 | 0.03910836 | 0.04261779 | -0.0035094 | 0.70819792 | 0.99999994 |
| Racgap1 | 0.03078445 | 0.02379317 | 0.02730277 | -0.0035096 | 0.80389961 | 0.99999994 |
| Gatad2a | 0.11290126 | 0.09122323 | 0.09473446 | -0.0035112 | 0.59743395 | 0.99999994 |
| Crocc | 0.02256959 | 0.01661551 | 0.02012719 | -0.0035117 | 0.87725323 | 0.99999994 |
| Bmp1 | 0.14264771 | 0.11339109 | 0.11690284 | -0.0035118 | 0.57924276 | 0.99999994 |
| Mdn1 | 0.03753817 | 0.0296308 | 0.03314347 | -0.0035127 | 0.75974023 | 0.99999994 |
| Dhcr24 | 0.0171568 | 0.01184252 | 0.01535585 | -0.0035133 | 0.9361124 | 0.99999994 |
| Prpf38b | 0.44086633 | 0.28530609 | 0.2888221 | -0.003516 | 0.53229174 | 0.99999994 |
| Dvl1 | 0.03857684 | 0.0305195 | 0.03403709 | -0.0035176 | 0.75424431 | 0.99999994 |
| Coa7 | 0.17996711 | 0.13971804 | 0.14323659 | -0.0035185 | 0.56494182 | 0.99999994 |
| Nme2 | 0.04058391 | 0.0322405 | 0.03576038 | -0.0035199 | 0.74382505 | 0.99999994 |
| Tmem259 | 0.06776126 | 0.05513223 | 0.0586522 | -0.00352 | 0.65522882 | 0.99999994 |
| Dhrs11 | 0.04745189 | 0.03810066 | 0.04162281 | -0.0035221 | 0.71337253 | 0.99999994 |
| Trim32 | 0.07848772 | 0.06394332 | 0.06746604 | -0.0035227 | 0.63591433 | 0.99999994 |
| Rala | 0.87104768 | 0.41101085 | 0.41453559 | -0.0035247 | 0.52256732 | 0.99999994 |
| Smim15 | 0.16124399 | 0.1266995 | 0.1302252 | -0.0035257 | 0.57150759 | 0.99999994 |
| Mtrr | 0.04014623 | 0.03185868 | 0.03538498 | -0.0035263 | 0.74644342 | 0.99999994 |
| Kpna1 | 0.10897726 | 0.0882016 | 0.0917287 | -0.0035271 | 0.60100409 | 0.99999994 |
| AU041133 | 0.05238048 | 0.04226902 | 0.04579719 | -0.0035282 | 0.69595382 | 0.99999994 |
| Plch1 | 0.07106439 | 0.05784886 | 0.06137943 | -0.0035306 | 0.64909444 | 0.99999994 |

| | | | | | | |
|---|---|---|---|---|---|---|
| Padi2 | 0.03980084 | 0.03155508 | 0.03508857 | -0.0035335 | 0.74866955 | 0.99999994 |
| Map3k10 | 0.05916778 | 0.0479668 | 0.05150179 | -0.003535 | 0.67612143 | 0.99999994 |
| Cdc25b | 0.02741055 | 0.02082941 | 0.02436524 | -0.0035358 | 0.83296725 | 0.99999994 |
| Ppp4r1 | 0.06690374 | 0.05440569 | 0.05794227 | -0.0035366 | 0.65773141 | 0.99999994 |
| Lamtor3 | 0.11577015 | 0.093382 | 0.09691985 | -0.0035378 | 0.59598792 | 0.99999994 |
| 1810010H24 | 0.02748089 | 0.02088854 | 0.02442662 | -0.0035381 | 0.83251263 | 0.99999994 |
| Ube2u | 0.01331664 | 0.0084104 | 0.01194965 | -0.0035393 | 0.97566577 | 0.99999994 |
| Mettl21a | 0.08694591 | 0.07079067 | 0.0743304 | -0.0035397 | 0.62437287 | 0.99999994 |
| Zfp266 | 0.19707065 | 0.15124121 | 0.15478132 | -0.0035401 | 0.56050206 | 0.99999994 |
| Epg5 | 0.01802849 | 0.01258543 | 0.01612659 | -0.0035412 | 0.92807981 | 0.99999994 |
| Mfsd2a | 0.03222293 | 0.02500887 | 0.02855119 | -0.0035423 | 0.79555502 | 0.99999994 |
| Zfp263 | 0.09081276 | 0.07390031 | 0.07744381 | -0.0035435 | 0.61965171 | 0.99999994 |
| Slc4a2 | 0.06876248 | 0.05593631 | 0.05948009 | -0.0035438 | 0.65416175 | 0.99999994 |
| Zc2hc1c | 0.0391545 | 0.03098945 | 0.03453354 | -0.0035441 | 0.75273597 | 0.99999994 |
| Chpf2 | 0.04333906 | 0.0345721 | 0.03811854 | -0.0035464 | 0.73213694 | 0.99999994 |
| Atrn | 0.05846231 | 0.04736455 | 0.05091121 | -0.0035467 | 0.67857116 | 0.99999994 |
| Mtfmt | 0.08204652 | 0.06681434 | 0.07036335 | -0.003549 | 0.63146495 | 0.99999994 |
| Tep1 | 0.02388636 | 0.01773333 | 0.02128268 | -0.0035494 | 0.86652062 | 0.99999994 |
| Col3a1 | 0.02355461 | 0.01744029 | 0.02099176 | -0.0035515 | 0.86994909 | 0.99999994 |
| Snx7 | 0.09064777 | 0.07375861 | 0.07731128 | -0.0035527 | 0.62015155 | 0.99999994 |
| Rbks | 0.05445202 | 0.04399091 | 0.04754367 | -0.0035528 | 0.69054782 | 0.99999994 |
| Ccdc51 | 0.05395273 | 0.04356976 | 0.04712315 | -0.0035534 | 0.69214244 | 0.99999994 |
| Gtf2ird1 | 0.06774734 | 0.05508482 | 0.05864068 | -0.0035559 | 0.65675718 | 0.99999994 |
| Csnk1g2 | 0.13700843 | 0.10922563 | 0.11278239 | -0.0035568 | 0.58313427 | 0.99999994 |
| Msra | 0.06136202 | 0.04977766 | 0.05333519 | -0.0035575 | 0.6714743 | 0.99999994 |
| Arrdc1 | 0.16889425 | 0.13203041 | 0.13558933 | -0.0035589 | 0.56934823 | 0.99999994 |
| Cep162 | 0.09668183 | 0.07857857 | 0.08213995 | -0.0035614 | 0.61355723 | 0.99999994 |
| Polr3a | 0.03090133 | 0.02384221 | 0.02740429 | -0.0035621 | 0.80653835 | 0.99999994 |
| Tnks1bp1 | 0.02125056 | 0.01540139 | 0.01896767 | -0.0035663 | 0.89462723 | 0.99999994 |
| Rhpn1 | 0.03450071 | 0.02695626 | 0.03052313 | -0.0035669 | 0.78167064 | 0.99999994 |

| | | | | | | |
|---|---|---|---|---|---|---|
| Col6a6 | 0.01875425 | 0.01320003 | 0.0167676 | -0.0035676 | 0.92164826 | 0.99999994 |
| Slc45a4 | 0.02993428 | 0.02299499 | 0.0265638 | -0.0035688 | 0.81441214 | 0.99999994 |
| Ptgr1 | 0.0374112 | 0.02946428 | 0.03303416 | -0.0035699 | 0.76403031 | 0.99999994 |
| Tmem170b | 0.03520809 | 0.02756414 | 0.03113431 | -0.0035702 | 0.77735348 | 0.99999994 |
| Pigs | 0.3171969 | 0.22386543 | 0.22743579 | -0.0035704 | 0.54161112 | 0.99999994 |
| Prkci | 0.02937136 | 0.02250304 | 0.02607405 | -0.003571 | 0.81901854 | 0.99999994 |
| Gskip | 0.1930741 | 0.14853994 | 0.15211257 | -0.0035726 | 0.562116 | 0.99999994 |
| 9930021J03F | 0.11155843 | 0.09013368 | 0.09370801 | -0.0035743 | 0.60021199 | 0.99999994 |
| Rassf8 | 0.02560507 | 0.01921247 | 0.02278783 | -0.0035754 | 0.8518422 | 0.99999994 |
| Xpot | 0.08563807 | 0.06969689 | 0.07327389 | -0.003577 | 0.62738742 | 0.99999994 |
| Setd7 | 0.04950934 | 0.03978888 | 0.04336871 | -0.0035798 | 0.70864913 | 0.99999994 |
| Mier1 | 0.24046348 | 0.17909026 | 0.18267249 | -0.0035822 | 0.55192717 | 0.99999994 |
| Ankrd13b | 0.01931617 | 0.01367965 | 0.01726348 | -0.0035838 | 0.91648838 | 0.99999994 |
| Susd6 | 0.05028321 | 0.04043915 | 0.04402417 | -0.003585 | 0.70610678 | 0.99999994 |
| Rhot2 | 0.08513472 | 0.0692811 | 0.07286679 | -0.0035857 | 0.62837445 | 0.99999994 |
| Chmp2b | 0.2539771 | 0.1873781 | 0.19096528 | -0.0035872 | 0.54975249 | 0.99999994 |
| Mlh1 | 0.07993064 | 0.06505505 | 0.06864236 | -0.0035873 | 0.63603002 | 0.99999994 |
| Trim3 | 0.04541827 | 0.03629927 | 0.03989247 | -0.0035932 | 0.72559793 | 0.99999994 |
| Cebpz | 0.2685842 | 0.19613234 | 0.19972701 | -0.0035947 | 0.54767947 | 0.99999994 |
| Vwa8 | 0.02359049 | 0.01742842 | 0.02102323 | -0.0035948 | 0.87247833 | 0.99999994 |
| Lonrf2 | 0.01393589 | 0.00890193 | 0.01250012 | -0.0035982 | 0.97232053 | 0.99999994 |
| Pfkfb4 | 0.02110658 | 0.01524175 | 0.01884097 | -0.0035992 | 0.89823762 | 0.99999994 |
| Cutc | 0.05635055 | 0.04554089 | 0.04914011 | -0.0035992 | 0.687062 | 0.99999994 |
| Supv3l1 | 0.03600062 | 0.02821697 | 0.03181839 | -0.0036014 | 0.77439841 | 0.99999994 |
| Ccdc92 | 0.11771095 | 0.09478881 | 0.09839228 | -0.0036035 | 0.59629899 | 0.99999994 |
| Map3k4 | 0.08964528 | 0.07290183 | 0.07650544 | -0.0036036 | 0.62306184 | 0.99999994 |
| Spsb1 | 0.06152788 | 0.04986946 | 0.05347356 | -0.0036041 | 0.67315281 | 0.99999994 |
| Rbm34 | 0.16241272 | 0.1274442 | 0.13104896 | -0.0036048 | 0.57263885 | 0.99999994 |
| Rbm12 | 0.07165239 | 0.05825794 | 0.06186368 | -0.0036057 | 0.65098015 | 0.99999994 |
| Kank3 | 0.03283112 | 0.02547163 | 0.0290783 | -0.0036067 | 0.79548678 | 0.99999994 |

| | | | | | | |
|---|---|---|---|---|---|---|
| Patl1 | 0.05328626 | 0.04295416 | 0.04656141 | -0.0036073 | 0.696964 | 0.99999994 |
| Tfb1m | 0.02601159 | 0.01953291 | 0.02314332 | -0.0036104 | 0.85045998 | 0.99999994 |
| Nsun5 | 0.06386606 | 0.05181036 | 0.05542103 | -0.0036107 | 0.66773276 | 0.99999994 |
| Ell | 0.0780873 | 0.06352597 | 0.06713921 | -0.0036132 | 0.63991264 | 0.99999994 |
| Nfkbiz | 0.03598017 | 0.02818651 | 0.03180075 | -0.0036142 | 0.77532882 | 0.99999994 |
| Dok5 | 0.01189752 | 0.00706764 | 0.01068644 | -0.0036188 | 0.9879042 | 0.99999994 |
| Tmem175 | 0.08097053 | 0.06586802 | 0.06948876 | -0.0036207 | 0.63564138 | 0.99999994 |
| Tmem104 | 0.01885467 | 0.01323319 | 0.01685625 | -0.0036231 | 0.92374314 | 0.99999994 |
| Urod | 0.38001489 | 0.25653196 | 0.26015505 | -0.0036231 | 0.5369294 | 0.99999994 |
| Gcfc2 | 0.0459426 | 0.03671556 | 0.04033904 | -0.0036235 | 0.72505055 | 0.99999994 |
| Ppp6r1 | 0.05949245 | 0.04814928 | 0.0517734 | -0.0036241 | 0.67937179 | 0.99999994 |
| Ugp2 | 0.26207559 | 0.19222247 | 0.19584814 | -0.0036257 | 0.54903636 | 0.99999994 |
| Eif2ak3 | 0.02829948 | 0.0215136 | 0.02514048 | -0.0036269 | 0.83154195 | 0.99999994 |
| Dph6 | 0.15351038 | 0.12110402 | 0.12473359 | -0.0036296 | 0.57679101 | 0.99999994 |
| Cars2 | 0.06380051 | 0.05173352 | 0.05536652 | -0.003633 | 0.6688618 | 0.99999994 |
| Nudcd2 | 0.51431105 | 0.31621027 | 0.31984375 | -0.0036335 | 0.53013824 | 0.99999994 |
| B4galt1 | 0.06800647 | 0.05522097 | 0.05885506 | -0.0036341 | 0.65946627 | 0.99999994 |
| Aph1b | 0.02625144 | 0.01971863 | 0.02335298 | -0.0036343 | 0.84987725 | 0.99999994 |
| Tmem51 | 0.16949069 | 0.13236922 | 0.13600477 | -0.0036355 | 0.57061147 | 0.99999994 |
| Efemp2 | 0.11376557 | 0.09175591 | 0.09539393 | -0.003638 | 0.60019514 | 0.99999994 |
| Zfp606 | 0.087668 | 0.07127397 | 0.07491297 | -0.003639 | 0.62678161 | 0.99999994 |
| Pex1 | 0.03949995 | 0.03119027 | 0.03483025 | -0.00364 | 0.75667042 | 0.99999994 |
| Gm16124 | 0.02846594 | 0.02164303 | 0.02528555 | -0.0036425 | 0.83119153 | 0.99999994 |
| Fam174b | 0.47761945 | 0.30116739 | 0.30481014 | -0.0036427 | 0.53170212 | 0.99999994 |
| Mllt6 | 0.03811927 | 0.02999685 | 0.03364357 | -0.0036467 | 0.76469959 | 0.99999994 |
| Ptpn13 | 0.0283865 | 0.02156748 | 0.02521632 | -0.0036488 | 0.83227228 | 0.99999994 |
| Rft1 | 0.05553028 | 0.04480158 | 0.04845085 | -0.0036493 | 0.69193764 | 0.99999994 |
| Zrsr1 | 0.05950819 | 0.04813407 | 0.05178656 | -0.0036525 | 0.68063329 | 0.99999994 |
| Traf2 | 0.11764744 | 0.0946867 | 0.09834417 | -0.0036575 | 0.59776001 | 0.99999994 |
| Zbed4 | 0.05726525 | 0.0462472 | 0.04990786 | -0.0036607 | 0.68730748 | 0.99999994 |

| | | | | | | |
|---|---|---|---|---|---|---|
| Trim45 | 0.05605428 | 0.045229 | 0.04889125 | -0.0036622 | 0.69096988 | 0.99999994 |
| Kalrn | 0.05187524 | 0.04170771 | 0.0453705 | -0.0036628 | 0.70449463 | 0.99999994 |
| Dpy19l3 | 0.0242295 | 0.01792004 | 0.02158346 | -0.0036634 | 0.87071994 | 0.99999994 |
| Bace1 | 0.23023983 | 0.17261218 | 0.17627668 | -0.0036645 | 0.55502794 | 0.99999994 |
| Ttll5 | 0.02840733 | 0.02156707 | 0.02523447 | -0.0036674 | 0.83332527 | 0.99999994 |
| Slc23a2 | 0.02123872 | 0.01528762 | 0.01895725 | -0.0036696 | 0.90121334 | 0.99999994 |
| Hdc | 0.0123868 | 0.0074473 | 0.01112224 | -0.0036749 | 0.9860735 | 0.99999994 |
| Spred2 | 0.03314883 | 0.0256772 | 0.02935348 | -0.0036763 | 0.79776188 | 0.99999994 |
| Arsa | 0.03460498 | 0.02693597 | 0.03061326 | -0.0036773 | 0.78801873 | 0.99999994 |
| Fggy | 0.06106839 | 0.04941196 | 0.05309015 | -0.0036782 | 0.67766024 | 0.99999994 |
| Carf | 0.05066531 | 0.04066699 | 0.04434755 | -0.0036806 | 0.70967457 | 0.99999994 |
| Fancm | 0.04692905 | 0.03749755 | 0.04117838 | -0.0036808 | 0.7240768 | 0.99999994 |
| Cep89 | 0.08510068 | 0.06915786 | 0.07283925 | -0.0036814 | 0.63172322 | 0.99999994 |
| Tardbp | 0.53875244 | 0.32568716 | 0.32936884 | -0.0036817 | 0.52965575 | 0.99999994 |
| Apof | 0.00981499 | 0.00514545 | 0.0088283 | -0.0036829 | 0.9972551 | 0.99999994 |
| Phc3 | 0.04532368 | 0.03612821 | 0.03981187 | -0.0036837 | 0.73101654 | 0.99999994 |
| Polg | 0.04197673 | 0.03326883 | 0.03695358 | -0.0036848 | 0.74656712 | 0.99999994 |
| Vps50 | 0.09979651 | 0.08093221 | 0.0846179 | -0.0036857 | 0.61406518 | 0.99999994 |
| Cd59a | 0.09436836 | 0.07660714 | 0.08029304 | -0.0036859 | 0.62003157 | 0.99999994 |
| Pcsk6 | 0.03296168 | 0.0255051 | 0.0291914 | -0.0036863 | 0.79970496 | 0.99999994 |
| Fbxo31 | 0.07449014 | 0.06050731 | 0.06419557 | -0.0036883 | 0.64892912 | 0.99999994 |
| Fuz | 0.11396573 | 0.09185815 | 0.09554653 | -0.0036884 | 0.60139334 | 0.99999994 |
| Anks4b | 0.01054118 | 0.00578606 | 0.00947684 | -0.0036908 | 0.99523369 | 0.99999994 |
| Gtf2h3 | 0.20291317 | 0.15495999 | 0.15865209 | -0.0036921 | 0.56155186 | 0.99999994 |
| D630045J12F | 0.027696 | 0.02091922 | 0.02461428 | -0.0036951 | 0.84116113 | 0.99999994 |
| Tatdn2 | 0.02362034 | 0.01735057 | 0.02104941 | -0.0036988 | 0.87893225 | 0.99999994 |
| Slc10a3 | 0.04362489 | 0.03466205 | 0.03836269 | -0.0037006 | 0.73959678 | 0.99999994 |
| Ddx52 | 0.17558494 | 0.13652048 | 0.14022681 | -0.0037063 | 0.56982722 | 0.99999994 |
| Cul4b | 0.08009649 | 0.06507005 | 0.06877742 | -0.0037074 | 0.64012904 | 0.99999994 |
| Snrnp27 | 0.91445505 | 0.41587843 | 0.4195863 | -0.0037079 | 0.523453 | 0.99999994 |

| | | | | | | |
|---|---|---|---|---|---|---|
| Pde12 | 0.09526914 | 0.07730333 | 0.0810128 | -0.0037095 | 0.6197353 | 0.99999994 |
| Cdc14a | 0.02015066 | 0.01428896 | 0.0179992 | -0.0037102 | 0.91498386 | 0.99999994 |
| Slc25a38 | 0.12776707 | 0.10223575 | 0.10594775 | -0.003712 | 0.59220183 | 0.99999994 |
| Dhrs13 | 0.05050749 | 0.04050174 | 0.044214 | -0.0037123 | 0.71187694 | 0.99999994 |
| Xrcc3 | 0.03499609 | 0.0272377 | 0.0309512 | -0.0037135 | 0.78774619 | 0.99999994 |
| Usp30 | 0.05310919 | 0.0426954 | 0.04641209 | -0.0037167 | 0.70299787 | 0.99999994 |
| Slc20a1 | 0.02234734 | 0.01621524 | 0.01993196 | -0.0037167 | 0.89273619 | 0.99999994 |
| Srrm3 | 0.06268882 | 0.05072426 | 0.05444126 | -0.003717 | 0.67525309 | 0.99999994 |
| Klhdc1 | 0.03672263 | 0.02872377 | 0.03244097 | -0.0037172 | 0.77718143 | 0.99999994 |
| Myo15b | 0.01715914 | 0.0116403 | 0.01535792 | -0.0037176 | 0.94644007 | 0.99999994 |
| Pias3 | 0.08428634 | 0.06846164 | 0.07218004 | -0.0037184 | 0.63416571 | 0.99999994 |
| Epb41l1 | 0.03635436 | 0.02840421 | 0.03212348 | -0.0037193 | 0.77955066 | 0.99999994 |
| Cpt1c | 0.04149929 | 0.03282542 | 0.03654482 | -0.0037194 | 0.75094268 | 0.99999994 |
| Yeats2 | 0.06021348 | 0.04865639 | 0.05237618 | -0.0037198 | 0.68180014 | 0.99999994 |
| Lyst | 0.02986709 | 0.02278433 | 0.02650536 | -0.003721 | 0.82496987 | 0.99999994 |
| Ppox | 0.0627763 | 0.05079264 | 0.05451411 | -0.0037215 | 0.6752311 | 0.99999994 |
| Fam199x | 0.04041027 | 0.03188929 | 0.03561147 | -0.0037222 | 0.75670294 | 0.99999994 |
| Zc3h18 | 0.13887778 | 0.11042984 | 0.11415243 | -0.0037226 | 0.58592248 | 0.99999994 |
| Snph | 0.01597727 | 0.01058876 | 0.01431149 | -0.0037227 | 0.95831524 | 0.99999994 |
| Ugdh | 0.1240041 | 0.09941139 | 0.10313493 | -0.0037235 | 0.59495835 | 0.99999994 |
| Hivep1 | 0.06168593 | 0.04988126 | 0.05360539 | -0.0037241 | 0.67811597 | 0.99999994 |
| Ormdl1 | 0.14356324 | 0.11383945 | 0.11756825 | -0.0037288 | 0.58360072 | 0.99999994 |
| Acbd7 | 0.05674092 | 0.04573891 | 0.04946788 | -0.003729 | 0.69208465 | 0.99999994 |
| 1700052K11l | 0.13485925 | 0.10747184 | 0.11120214 | -0.0037303 | 0.58834469 | 0.99999994 |
| Ttpal | 0.04223659 | 0.03344312 | 0.03717596 | -0.0037328 | 0.7480497 | 0.99999994 |
| Zfp346 | 0.0275359 | 0.02074056 | 0.02447461 | -0.0037341 | 0.84507825 | 0.99999994 |
| Ankrd49 | 0.04009121 | 0.03160325 | 0.03533777 | -0.0037345 | 0.75911193 | 0.99999994 |
| Pkmyt1 | 0.04930565 | 0.03946043 | 0.04319607 | -0.0037356 | 0.71757336 | 0.99999994 |
| Cox11 | 0.13690701 | 0.10897153 | 0.11270794 | -0.0037364 | 0.58732401 | 0.99999994 |
| Kcnq1 | 0.01967042 | 0.01383923 | 0.0175759 | -0.0037367 | 0.92148818 | 0.99999994 |

| | | | | | | |
|---|---|---|---|---|---|---|
| Mthfr | 0.02724903 | 0.02048629 | 0.02422427 | -0.003738 | 0.84781889 | 0.99999994 |
| Plekhm2 | 0.03012669 | 0.02299297 | 0.02673112 | -0.0037381 | 0.82403241 | 0.99999994 |
| Pyroxd2 | 0.01618946 | 0.01075918 | 0.01449948 | -0.0037403 | 0.95701798 | 0.99999994 |
| Stag1 | 0.20160702 | 0.15404806 | 0.15778989 | -0.0037418 | 0.56271203 | 0.99999994 |
| Ubac1 | 0.20022827 | 0.15313528 | 0.1568778 | -0.0037425 | 0.56308519 | 0.99999994 |
| Gm21988 | 0.01956951 | 0.01374392 | 0.01748692 | -0.003743 | 0.92288745 | 0.99999994 |
| Dennd1a | 0.06231085 | 0.05038217 | 0.05412636 | -0.0037442 | 0.67740464 | 0.99999994 |
| St6gal1 | 0.03789009 | 0.02970206 | 0.03344639 | -0.0037443 | 0.77191857 | 0.99999994 |
| Ripk4 | 0.07307515 | 0.05928777 | 0.06303389 | -0.0037461 | 0.6537959 | 0.99999994 |
| Ublcp1 | 0.09074922 | 0.07364501 | 0.07739277 | -0.0037478 | 0.62640066 | 0.99999994 |
| Dlg3 | 0.03825827 | 0.03001405 | 0.03376314 | -0.0037491 | 0.77008529 | 0.99999994 |
| Fpgt | 0.03421855 | 0.0265299 | 0.03027918 | -0.0037493 | 0.79509119 | 0.99999994 |
| Gprin3 | 0.06245957 | 0.05050093 | 0.05425028 | -0.0037493 | 0.67725416 | 0.99999994 |
| Dnajc11 | 0.15669341 | 0.12324856 | 0.12700215 | -0.0037536 | 0.57798092 | 0.99999994 |
| Txlng | 0.07845152 | 0.06368073 | 0.0674365 | -0.0037558 | 0.6445752 | 0.99999994 |
| Lgals12 | 0.01774862 | 0.01212227 | 0.01587923 | -0.003757 | 0.94235623 | 0.99999994 |
| Snrnp40 | 0.44198447 | 0.28556435 | 0.28932268 | -0.0037583 | 0.53445227 | 0.99999994 |
| Adat1 | 0.02510521 | 0.01859169 | 0.02235043 | -0.0037587 | 0.86851997 | 0.99999994 |
| Man2b1 | 0.08090208 | 0.0656743 | 0.06943308 | -0.0037588 | 0.64070503 | 0.99999994 |
| Irf1 | 0.04863046 | 0.03886173 | 0.04262348 | -0.0037618 | 0.72155253 | 0.99999994 |
| Scai | 0.06802469 | 0.05510722 | 0.05887013 | -0.0037629 | 0.66475185 | 0.99999994 |
| Cdkl5 | 0.02787264 | 0.02100433 | 0.02476834 | -0.003764 | 0.84412545 | 0.99999994 |
| Gtf2b | 0.53364966 | 0.32364714 | 0.32741192 | -0.0037648 | 0.53050475 | 0.99999994 |
| 2310057M21 | 0.11632813 | 0.09357791 | 0.09734364 | -0.0037657 | 0.60160387 | 0.99999994 |
| Dnajc16 | 0.02785692 | 0.0209884 | 0.02475464 | -0.0037662 | 0.84440197 | 0.99999994 |
| Nubp2 | 0.18591872 | 0.14352177 | 0.14729025 | -0.0037685 | 0.56761525 | 0.99999994 |
| Raf1 | 0.21697439 | 0.16404827 | 0.16781861 | -0.0037703 | 0.55943894 | 0.99999994 |
| Vps39 | 0.07012664 | 0.05683475 | 0.06060637 | -0.0037716 | 0.66064886 | 0.99999994 |
| Hspa12a | 0.01002564 | 0.00524107 | 0.00901649 | -0.0037754 | 0.99734152 | 0.99999994 |
| Ppp1r12b | 0.03731644 | 0.02917607 | 0.03295255 | -0.0037765 | 0.7772214 | 0.99999994 |

| | | | | | | |
|---|---|---|---|---|---|---|
| Cep78 | 0.07411692 | 0.06011014 | 0.06388937 | -0.0037792 | 0.65311417 | 0.99999994 |
| Ttc4 | 0.2475489 | 0.18326348 | 0.18704292 | -0.0037794 | 0.55349653 | 0.99999994 |
| Ebi3 | 0.01536278 | 0.00998727 | 0.01376675 | -0.0037795 | 0.96618155 | 0.99999994 |
| Klhl21 | 0.02647248 | 0.01976432 | 0.02354614 | -0.0037818 | 0.85748751 | 0.99999994 |
| Foxk1 | 0.03203318 | 0.02460473 | 0.02838664 | -0.0037819 | 0.81241171 | 0.99999994 |
| Camsap1 | 0.11563447 | 0.09303431 | 0.09681674 | -0.0037824 | 0.60258725 | 0.99999994 |
| Slc39a11 | 0.07156514 | 0.05800913 | 0.06179185 | -0.0037827 | 0.65817325 | 0.99999994 |
| Socs4 | 0.13016938 | 0.10395096 | 0.10773453 | -0.0037836 | 0.59241726 | 0.99999994 |
| Vrk3 | 0.21874575 | 0.16517381 | 0.16895868 | -0.0037849 | 0.55926667 | 0.99999994 |
| Gmps | 0.47920109 | 0.30169108 | 0.30547694 | -0.0037859 | 0.53287308 | 0.99999994 |
| Rnf19a | 0.10590701 | 0.08557648 | 0.08936351 | -0.003787 | 0.61105963 | 0.99999994 |
| Ankrd27 | 0.031943 | 0.02451818 | 0.02830843 | -0.0037902 | 0.81359647 | 0.99999994 |
| Phykpl | 0.07572803 | 0.06141874 | 0.06521011 | -0.0037914 | 0.65062432 | 0.99999994 |
| Pomgnt1 | 0.12029017 | 0.09654983 | 0.1003419 | -0.0037921 | 0.59930697 | 0.99999994 |
| Ralgapa2 | 0.05443423 | 0.04373587 | 0.04752869 | -0.0037928 | 0.70235328 | 0.99999994 |
| C8g | 0.02234359 | 0.01613491 | 0.01992867 | -0.0037938 | 0.89744892 | 0.99999994 |
| Lclat1 | 0.02547442 | 0.01887852 | 0.02267353 | -0.003795 | 0.86738544 | 0.99999994 |
| Cldn7 | 0.88908613 | 0.41299571 | 0.41679341 | -0.0037977 | 0.52418133 | 0.99999994 |
| Zfp691 | 0.07685141 | 0.06233164 | 0.0661294 | -0.0037978 | 0.6488692 | 0.99999994 |
| Lemd2 | 0.0555504 | 0.04466866 | 0.04846777 | -0.0037991 | 0.69907905 | 0.99999994 |
| Fam83g | 0.02352078 | 0.01716246 | 0.02096208 | -0.0037996 | 0.88619567 | 0.99999994 |
| Rad51ap2 | 0.01799373 | 0.01229597 | 0.01609586 | -0.0037999 | 0.94195817 | 0.99999994 |
| Nop9 | 0.11674108 | 0.09385448 | 0.09765704 | -0.0038026 | 0.60225367 | 0.99999994 |
| Poll | 0.0340557 | 0.02633508 | 0.03013834 | -0.0038033 | 0.79954249 | 0.99999994 |
| Rbbp5 | 0.0475509 | 0.03789936 | 0.04170693 | -0.0038076 | 0.72830065 | 0.99999994 |
| Lrba | 0.08614146 | 0.06986731 | 0.07368074 | -0.0038134 | 0.6347677 | 0.99999994 |
| Rnf113a2 | 0.1807771 | 0.13997665 | 0.14379056 | -0.0038139 | 0.57007053 | 0.99999994 |
| Ccdc91 | 0.16288343 | 0.12756614 | 0.1313803 | -0.0038142 | 0.57661598 | 0.99999994 |
| Dlc1 | 0.0344765 | 0.02668802 | 0.0305022 | -0.0038142 | 0.79739447 | 0.99999994 |
| Nufip1 | 0.11861632 | 0.09526084 | 0.09907757 | -0.0038167 | 0.60118683 | 0.99999994 |

| | | | | | | |
|---|---|---|---|---|---|---|
| Ttbk2 | 0.05282245 | 0.04235098 | 0.0461702 | -0.0038192 | 0.70905173 | 0.99999994 |
| Wrap53 | 0.07271698 | 0.05891987 | 0.0627395 | -0.0038196 | 0.65735131 | 0.99999994 |
| Pgm1 | 0.08264942 | 0.06703319 | 0.07085287 | -0.0038197 | 0.64014416 | 0.99999994 |
| B4galt4 | 0.04815756 | 0.03840215 | 0.04222215 | -0.00382 | 0.7264881 | 0.99999994 |
| Arl15 | 0.03419004 | 0.02643387 | 0.03025453 | -0.0038207 | 0.79971103 | 0.99999994 |
| Zbtb22 | 0.06035756 | 0.04867558 | 0.05249656 | -0.003821 | 0.68597945 | 0.99999994 |
| Kri1 | 0.06555358 | 0.05300169 | 0.05682289 | -0.0038212 | 0.67278763 | 0.99999994 |
| Hs6st1 | 0.08418083 | 0.0682709 | 0.07209458 | -0.0038237 | 0.63796825 | 0.99999994 |
| Copg2 | 0.11477363 | 0.09233722 | 0.09616193 | -0.0038247 | 0.60439802 | 0.99999994 |
| Ttf1 | 0.11349003 | 0.09135714 | 0.0951838 | -0.0038267 | 0.60549795 | 0.99999994 |
| Toe1 | 0.07772185 | 0.06301396 | 0.06684078 | -0.0038268 | 0.64843361 | 0.99999994 |
| Phka1 | 0.01999058 | 0.01402963 | 0.01785813 | -0.0038285 | 0.92321252 | 0.99999994 |
| Rnf141 | 0.1387113 | 0.11020181 | 0.11403059 | -0.0038288 | 0.58842629 | 0.99999994 |
| Clp1 | 0.07026409 | 0.05688975 | 0.06071973 | -0.00383 | 0.6627059 | 0.99999994 |
| Proscos | 0.02886767 | 0.02179984 | 0.02563552 | -0.0038357 | 0.8403622 | 0.99999994 |
| Tmem130 | 0.02198466 | 0.01577677 | 0.01961324 | -0.0038365 | 0.90354595 | 0.99999994 |
| G6pdx | 0.06277537 | 0.05067581 | 0.05451334 | -0.0038375 | 0.68031574 | 0.99999994 |
| Scaf4 | 0.05028784 | 0.04018691 | 0.04402808 | -0.0038412 | 0.7192873 | 0.99999994 |
| Gga3 | 0.04139239 | 0.03261126 | 0.03645326 | -0.003842 | 0.75851961 | 0.99999994 |
| Tha1 | 0.0272868 | 0.02041407 | 0.02425724 | -0.0038432 | 0.85419449 | 0.99999994 |
| Amigo3 | 0.0131552 | 0.00796191 | 0.01180607 | -0.0038442 | 0.98489677 | 0.99999994 |
| Kif2a | 0.08747385 | 0.07091201 | 0.07475639 | -0.0038444 | 0.63393965 | 0.99999994 |
| Ubxn8 | 0.11660691 | 0.09370981 | 0.09755524 | -0.0038454 | 0.60348575 | 0.99999994 |
| Gba2 | 0.03904485 | 0.0305936 | 0.03443933 | -0.0038457 | 0.77134869 | 0.99999994 |
| Pkp2 | 0.05517569 | 0.04430589 | 0.04815266 | -0.0038468 | 0.70255235 | 0.99999994 |
| Ncdn | 0.04696156 | 0.03735723 | 0.04120603 | -0.0038488 | 0.73293968 | 0.99999994 |
| Tars2 | 0.0718754 | 0.05819795 | 0.06204725 | -0.0038493 | 0.660178 | 0.99999994 |
| Arid3b | 0.04961135 | 0.03960582 | 0.04345514 | -0.0038493 | 0.72228082 | 0.99999994 |
| Usp24 | 0.03257337 | 0.02500546 | 0.02885496 | -0.0038495 | 0.81273351 | 0.99999994 |
| Mast2 | 0.03467227 | 0.02682165 | 0.03067141 | -0.0038498 | 0.79827611 | 0.99999994 |

| | | | | | | |
|---|---|---|---|---|---|---|
| Zdhhc3 | 0.25147881 | 0.18559557 | 0.18944577 | -0.0038502 | 0.55380494 | 0.99999994 |
| Trpt1 | 0.06213757 | 0.05013126 | 0.05398195 | -0.0038507 | 0.68254137 | 0.99999994 |
| Rprd1a | 0.14029405 | 0.11133631 | 0.11518766 | -0.0038514 | 0.5880604 | 0.99999994 |
| Reep6 | 0.13085798 | 0.10439071 | 0.10824537 | -0.0038547 | 0.5936856 | 0.99999994 |
| Dph2 | 0.04368393 | 0.03455681 | 0.03841311 | -0.0038563 | 0.74800724 | 0.99999994 |
| Avl9 | 0.2087649 | 0.15863373 | 0.16249248 | -0.0038588 | 0.56279927 | 0.99999994 |
| Hltf | 0.06236905 | 0.05031552 | 0.05417486 | -0.0038593 | 0.68231761 | 0.99999994 |
| Lyz2 | 0.0375164 | 0.0292653 | 0.03312474 | -0.0038594 | 0.78099165 | 0.99999994 |
| Zxdb | 0.03354587 | 0.02583756 | 0.02969722 | -0.0038597 | 0.80651017 | 0.99999994 |
| Adh1 | 0.01300952 | 0.00781676 | 0.01167648 | -0.0038597 | 0.98610697 | 0.99999994 |
| Vps54 | 0.04845715 | 0.03860971 | 0.04247644 | -0.0038667 | 0.72772374 | 0.99999994 |
| Mad1l1 | 0.05747393 | 0.04621057 | 0.05008288 | -0.0038723 | 0.69660369 | 0.99999994 |
| Mios | 0.0569115 | 0.04573803 | 0.04961105 | -0.003873 | 0.69834537 | 0.99999994 |
| Asrgl1 | 0.0566438 | 0.04551126 | 0.04938635 | -0.0038751 | 0.69926563 | 0.99999994 |
| Pigq | 0.11479781 | 0.09230374 | 0.09618034 | -0.0038766 | 0.60576135 | 0.99999994 |
| Atxn1 | 0.05100702 | 0.04075707 | 0.04463662 | -0.0038795 | 0.71854699 | 0.99999994 |
| Fbxw8 | 0.05844567 | 0.04701609 | 0.05089728 | -0.0038812 | 0.69412686 | 0.99999994 |
| Srr | 0.04371379 | 0.03455729 | 0.03843861 | -0.0038813 | 0.74924733 | 0.99999994 |
| Usp51 | 0.13048414 | 0.10408575 | 0.10796813 | -0.0038824 | 0.59458403 | 0.99999994 |
| Dph5 | 0.06588326 | 0.05321107 | 0.0570964 | -0.0038853 | 0.6747098 | 0.99999994 |
| Kbtbd7 | 0.03154402 | 0.02407432 | 0.02796228 | -0.003888 | 0.82265234 | 0.99999994 |
| Sec23a | 0.0405354 | 0.03182966 | 0.03571878 | -0.0038891 | 0.76569889 | 0.99999994 |
| Kdelr3 | 0.02035882 | 0.01429134 | 0.01818259 | -0.0038913 | 0.92285273 | 0.99999994 |
| Tjap1 | 0.08780398 | 0.07113081 | 0.07502261 | -0.0038918 | 0.6350653 | 0.99999994 |
| Gtf2h1 | 0.12759185 | 0.10192091 | 0.10581715 | -0.0038962 | 0.59680691 | 0.99999994 |
| Cyp4v3 | 0.04136738 | 0.03253522 | 0.03643184 | -0.0038966 | 0.76174922 | 0.99999994 |
| Mycbp | 0.46109574 | 0.2938431 | 0.29774045 | -0.0038974 | 0.53471602 | 0.99999994 |
| Tysnd1 | 0.12585171 | 0.10061363 | 0.10451816 | -0.0039045 | 0.59818995 | 0.99999994 |
| Mpdz | 0.04643349 | 0.03684995 | 0.04075687 | -0.0039069 | 0.73828702 | 0.99999994 |
| Rcor3 | 0.15303179 | 0.12048411 | 0.12439149 | -0.0039074 | 0.58281054 | 0.99999994 |

| | | | | | | |
|---|---|---|---|---|---|---|
| Krr1 | 0.12599988 | 0.10071781 | 0.10462891 | -0.0039111 | 0.59824977 | 0.99999994 |
| Maob | 0.28285552 | 0.204177 | 0.2080891 | -0.0039121 | 0.54979378 | 0.99999994 |
| Sptbn4 | 0.01686483 | 0.01118403 | 0.01509749 | -0.0039135 | 0.95777349 | 0.99999994 |
| Hdac8 | 0.06421608 | 0.05179813 | 0.05571206 | -0.0039139 | 0.6799745 | 0.99999994 |
| Tada2b | 0.03404353 | 0.02621097 | 0.03012782 | -0.0039168 | 0.80658396 | 0.99999994 |
| Ddx47 | 0.29287922 | 0.20993014 | 0.21384745 | -0.0039173 | 0.54852377 | 0.99999994 |
| Blm | 0.03054077 | 0.02317304 | 0.02709105 | -0.003918 | 0.83214251 | 0.99999994 |
| Vps51 | 0.12852929 | 0.1025972 | 0.10651542 | -0.0039182 | 0.59671663 | 0.99999994 |
| Anapc7 | 0.1598673 | 0.12533419 | 0.12925263 | -0.0039184 | 0.57996139 | 0.99999994 |
| Atp5k | 1.18679445 | 0.42537996 | 0.42929997 | -0.00392 | 0.52423288 | 0.99999994 |
| Mapk10 | 0.01011363 | 0.00517194 | 0.00909508 | -0.0039231 | 0.99795546 | 0.99999994 |
| Cxxc1 | 0.20127724 | 0.15364833 | 0.15757191 | -0.0039236 | 0.56582124 | 0.99999994 |
| Gm4353 | 0.02612801 | 0.01932126 | 0.0232451 | -0.0039238 | 0.86941136 | 0.99999994 |
| Bscl2 | 0.20643852 | 0.15704033 | 0.16097025 | -0.0039299 | 0.56454717 | 0.99999994 |
| Vcpkmt | 0.13111453 | 0.10450284 | 0.10843553 | -0.0039327 | 0.59538116 | 0.99999994 |
| Adam32 | 0.0156948 | 0.01012554 | 0.01406114 | -0.0039356 | 0.96877309 | 0.99999994 |
| Mfsd6 | 0.0727897 | 0.05886351 | 0.06279929 | -0.0039358 | 0.66172298 | 0.99999994 |
| Sirt2 | 0.31551482 | 0.22257708 | 0.22651321 | -0.0039361 | 0.54604212 | 0.99999994 |
| Usp31 | 0.02232585 | 0.01597688 | 0.01991308 | -0.0039362 | 0.90587196 | 0.99999994 |
| Csk | 0.171573 | 0.13351375 | 0.13745203 | -0.0039383 | 0.57562679 | 0.99999994 |
| Tubgcp3 | 0.05518552 | 0.04422128 | 0.04816092 | -0.0039396 | 0.70695162 | 0.99999994 |
| Ctdspl2 | 0.15539858 | 0.12213928 | 0.12608071 | -0.0039414 | 0.58241891 | 0.99999994 |
| Zfp827 | 0.04890031 | 0.03891007 | 0.04285239 | -0.0039423 | 0.72985362 | 0.99999994 |
| Per3 | 0.03247608 | 0.02482636 | 0.02877064 | -0.0039443 | 0.81925931 | 0.99999994 |
| Ankzf1 | 0.08276576 | 0.06700233 | 0.07094728 | -0.003945 | 0.64435361 | 0.99999994 |
| Galns | 0.12221552 | 0.09784671 | 0.10179194 | -0.0039452 | 0.60179099 | 0.99999994 |
| Dolpp1 | 0.06486693 | 0.05230676 | 0.05625285 | -0.0039461 | 0.67972794 | 0.99999994 |
| Ascc2 | 0.07301503 | 0.05903722 | 0.06298449 | -0.0039473 | 0.6617182 | 0.99999994 |
| Pikfyve | 0.05491494 | 0.04398224 | 0.04793329 | -0.0039511 | 0.70838444 | 0.99999994 |
| Mipep | 0.05814088 | 0.04668985 | 0.05064196 | -0.0039521 | 0.69828118 | 0.99999994 |

| | | | | | | |
|---|---|---|---|---|---|---|
| Csad | 0.07223303 | 0.05838826 | 0.06234152 | -0.0039533 | 0.66352386 | 0.99999994 |
| Opa1 | 0.11289776 | 0.09077306 | 0.09473178 | -0.0039587 | 0.6095556 | 0.99999994 |
| Stk32b | 0.01361872 | 0.00825459 | 0.01221824 | -0.0039637 | 0.98458859 | 0.99999994 |
| S100a9 | 0.01883796 | 0.01287262 | 0.01684149 | -0.0039689 | 0.94163432 | 0.99999994 |
| Cd47 | 0.38688592 | 0.25956631 | 0.26353568 | -0.0039694 | 0.53993026 | 0.99999994 |
| Gm10053 | 0.07219309 | 0.05833586 | 0.06230866 | -0.0039728 | 0.66436662 | 0.99999994 |
| G730003C15 | 0.04598411 | 0.03640123 | 0.04037439 | -0.0039732 | 0.74377865 | 0.99999994 |
| Tatdn1 | 0.12376825 | 0.09898457 | 0.10295806 | -0.0039735 | 0.60136844 | 0.99999994 |
| Cpped1 | 0.05704019 | 0.04574483 | 0.04971904 | -0.0039742 | 0.70265966 | 0.99999994 |
| Zfp13 | 0.03120537 | 0.02369309 | 0.02766832 | -0.0039752 | 0.83055224 | 0.99999994 |
| Sirt4 | 0.07465249 | 0.06035258 | 0.06432872 | -0.0039761 | 0.65962067 | 0.99999994 |
| Ptges2 | 0.09963665 | 0.08051435 | 0.08449095 | -0.0039766 | 0.62296766 | 0.99999994 |
| Abcc4 | 0.04960733 | 0.03947387 | 0.04345173 | -0.0039779 | 0.72885963 | 0.99999994 |
| Gsto2 | 0.01289581 | 0.00758805 | 0.01157531 | -0.0039873 | 0.98907208 | 0.99999994 |
| Rcan1 | 0.09851807 | 0.07961471 | 0.083602 | -0.0039873 | 0.62455516 | 0.99999994 |
| Armc8 | 0.18890732 | 0.14532149 | 0.14931124 | -0.0039898 | 0.5705854 | 0.99999994 |
| Zhx3 | 0.06165748 | 0.04959114 | 0.05358166 | -0.0039905 | 0.6899596 | 0.99999994 |
| Zfp362 | 0.0474016 | 0.03758912 | 0.04158007 | -0.003991 | 0.73855359 | 0.99999994 |
| Snx19 | 0.03239927 | 0.024713 | 0.02870406 | -0.0039911 | 0.82264982 | 0.99999994 |
| Lhfpl4 | 0.01934115 | 0.0132913 | 0.01728552 | -0.0039942 | 0.93798828 | 0.99999994 |
| Bace2 | 0.14370673 | 0.11367726 | 0.11767245 | -0.0039952 | 0.58939744 | 0.99999994 |
| Znfx1 | 0.05403062 | 0.04318994 | 0.04718877 | -0.0039988 | 0.71364582 | 0.99999994 |
| Rrp15 | 0.13957639 | 0.1106621 | 0.11466338 | -0.0040013 | 0.59183943 | 0.99999994 |
| Tex10 | 0.08965567 | 0.07251028 | 0.0765138 | -0.0040035 | 0.63618803 | 0.99999994 |
| Arnt | 0.06590685 | 0.05311223 | 0.05711597 | -0.0040037 | 0.67960718 | 0.99999994 |
| Acad9 | 0.04753291 | 0.03768619 | 0.04169165 | -0.0040055 | 0.738751 | 0.99999994 |
| Med20 | 0.11817476 | 0.09473583 | 0.09874347 | -0.0040076 | 0.60648033 | 0.99999994 |
| Gm14296 | 0.04555036 | 0.03599432 | 0.040005 | -0.0040107 | 0.7477193 | 0.99999994 |
| Adamts17 | 0.01711719 | 0.01130962 | 0.01532081 | -0.0040112 | 0.95930884 | 0.99999994 |
| Arl4d | 0.02868844 | 0.02146791 | 0.0254794 | -0.0040115 | 0.85267637 | 0.99999994 |

| | | | | | | |
|---|---|---|---|---|---|---|
| Tceanc | 0.04380166 | 0.03450125 | 0.03851364 | -0.0040124 | 0.75599118 | 0.99999994 |
| Cdkn2b | 0.03982613 | 0.03109738 | 0.03511028 | -0.0040129 | 0.77660683 | 0.99999994 |
| Rfxap | 0.11911921 | 0.09544442 | 0.09945778 | -0.0040134 | 0.60588128 | 0.99999994 |
| Map2k7 | 0.17278664 | 0.13427901 | 0.13829331 | -0.0040143 | 0.57660541 | 0.99999994 |
| Slc35a5 | 0.06487639 | 0.05224487 | 0.0562607 | -0.0040158 | 0.68265078 | 0.99999994 |
| Bmpr2 | 0.07072409 | 0.0570819 | 0.061099 | -0.0040171 | 0.66917589 | 0.99999994 |
| Tstd2 | 0.05019517 | 0.03993005 | 0.04394963 | -0.0040196 | 0.72866556 | 0.99999994 |
| Mrgpre | 0.02742338 | 0.02035646 | 0.02437643 | -0.00402 | 0.86383525 | 0.99999994 |
| Ccdc137 | 0.07290831 | 0.05887526 | 0.06289678 | -0.0040215 | 0.66479948 | 0.99999994 |
| Dnmt3bos | 0.0128919 | 0.00754995 | 0.01157183 | -0.0040219 | 0.98965156 | 0.99999994 |
| Soga1 | 0.02036041 | 0.01416155 | 0.01818399 | -0.0040224 | 0.92954884 | 0.99999994 |
| Cog2 | 0.08342806 | 0.06746144 | 0.07148452 | -0.0040231 | 0.64602604 | 0.99999994 |
| Fry | 0.03473975 | 0.026704 | 0.03072972 | -0.0040257 | 0.80839883 | 0.99999994 |
| Ythdc2 | 0.06405023 | 0.05154824 | 0.05557418 | -0.0040259 | 0.68516934 | 0.99999994 |
| Ccnk | 0.07281159 | 0.05879063 | 0.06281728 | -0.0040266 | 0.66519347 | 0.99999994 |
| Acad12 | 0.03084165 | 0.02332538 | 0.02735245 | -0.0040271 | 0.83645901 | 0.99999994 |
| Snap29 | 0.09906351 | 0.08000532 | 0.08403563 | -0.0040303 | 0.62522604 | 0.99999994 |
| 4833418N02 | 0.03554877 | 0.0273977 | 0.03142846 | -0.0040308 | 0.80336198 | 0.99999994 |
| Zfp280d | 0.20098696 | 0.15334801 | 0.15737995 | -0.0040319 | 0.56770339 | 0.99999994 |
| Flrt1 | 0.06485436 | 0.05220739 | 0.05624241 | -0.004035 | 0.68351442 | 0.99999994 |
| Commd2 | 0.3858144 | 0.25897568 | 0.2630109 | -0.0040352 | 0.54067109 | 0.99999994 |
| Cebpg | 0.20460192 | 0.1557288 | 0.15976419 | -0.0040354 | 0.56675925 | 0.99999994 |
| Clec16a | 0.0349575 | 0.02688192 | 0.03091787 | -0.0040359 | 0.80755075 | 0.99999994 |
| Primpol | 0.03903682 | 0.03039636 | 0.03443244 | -0.0040361 | 0.78237565 | 0.99999994 |
| Tsga10 | 0.03433879 | 0.02634694 | 0.03038315 | -0.0040362 | 0.81172107 | 0.99999994 |
| Saal1 | 0.08489685 | 0.06863527 | 0.07267431 | -0.004039 | 0.64428731 | 0.99999994 |
| Nup205 | 0.04533264 | 0.0357796 | 0.03981951 | -0.0040399 | 0.75026507 | 0.99999994 |
| Gm9755 | 0.07082891 | 0.05714516 | 0.06118539 | -0.0040402 | 0.66986366 | 0.99999994 |
| Chfr | 0.18207534 | 0.14063667 | 0.14467695 | -0.0040403 | 0.57373338 | 0.99999994 |
| Slc7a4 | 0.17709775 | 0.1372272 | 0.14126836 | -0.0040412 | 0.57550749 | 0.99999994 |

| | | | | | | |
|---|---|---|---|---|---|---|
| Nphp3 | 0.01695372 | 0.01113399 | 0.01517616 | -0.0040422 | 0.96187901 | 0.99999994 |
| Pex7 | 0.4046991 | 0.2680829 | 0.27212667 | -0.0040438 | 0.53939613 | 0.99999994 |
| Pan3 | 0.12768779 | 0.10184444 | 0.10588867 | -0.0040442 | 0.60034012 | 0.99999994 |
| Coro7 | 0.05249521 | 0.04184931 | 0.04589404 | -0.0040447 | 0.72127637 | 0.99999994 |
| Swsap1 | 0.08484767 | 0.06858921 | 0.07263451 | -0.0040453 | 0.64457648 | 0.99999994 |
| Map3k11 | 0.05153537 | 0.04103777 | 0.04508332 | -0.0040456 | 0.72484912 | 0.99999994 |
| Fn3k | 0.01931867 | 0.01321976 | 0.01726568 | -0.0040459 | 0.94059695 | 0.99999994 |
| Fam193b | 0.06550278 | 0.05273332 | 0.05678073 | -0.0040474 | 0.68241915 | 0.99999994 |
| Rad17 | 0.24041572 | 0.17859154 | 0.18264286 | -0.0040513 | 0.55869015 | 0.99999994 |
| Gm13269 | 0.03220273 | 0.02447882 | 0.02853367 | -0.0040549 | 0.82790532 | 0.99999994 |
| Harbi1 | 0.15761818 | 0.12360408 | 0.12765905 | -0.004055 | 0.5837252 | 0.99999994 |
| Srf | 0.07580088 | 0.06121279 | 0.06526977 | -0.004057 | 0.66046756 | 0.99999994 |
| Parp16 | 0.04743999 | 0.03755559 | 0.04161269 | -0.0040571 | 0.74182237 | 0.99999994 |
| Ulk1 | 0.08089201 | 0.0653669 | 0.06942488 | -0.004058 | 0.65138919 | 0.99999994 |
| Cbarp | 0.16396186 | 0.12808028 | 0.13213847 | -0.0040582 | 0.58099024 | 0.99999994 |
| Lrp6 | 0.12819929 | 0.10221154 | 0.10626973 | -0.0040582 | 0.60032603 | 0.99999994 |
| Fam13b | 0.0526212 | 0.04193998 | 0.04600038 | -0.0040604 | 0.72158254 | 0.99999994 |
| Rpusd4 | 0.06677581 | 0.05377428 | 0.0578363 | -0.004062 | 0.67992756 | 0.99999994 |
| E430024P14I | 0.04128178 | 0.03229624 | 0.03635851 | -0.0040623 | 0.77147184 | 0.99999994 |
| Fam210a | 0.04938223 | 0.03919832 | 0.04326097 | -0.0040627 | 0.73404772 | 0.99999994 |
| C920021L13I | 0.02815718 | 0.02095276 | 0.02501644 | -0.0040637 | 0.86020941 | 0.99999994 |
| Rev1 | 0.06447687 | 0.05186396 | 0.0559288 | -0.0040648 | 0.68572785 | 0.99999994 |
| Eml6 | 0.01285187 | 0.00747088 | 0.01153621 | -0.0040653 | 0.99050391 | 0.99999994 |
| Tcerg1 | 0.22752892 | 0.17049603 | 0.17456301 | -0.004067 | 0.56162101 | 0.99999994 |
| Rybp | 0.14175209 | 0.1121832 | 0.11625094 | -0.0040677 | 0.59208553 | 0.99999994 |
| Pi4k2a | 0.03703874 | 0.02864314 | 0.03271335 | -0.0040702 | 0.79621766 | 0.99999994 |
| Alg9 | 0.06418931 | 0.05161892 | 0.0556898 | -0.0040709 | 0.68672011 | 0.99999994 |
| Gde1 | 0.43754416 | 0.28325698 | 0.28732947 | -0.0040725 | 0.53758218 | 0.99999994 |
| Ubac2 | 0.10862756 | 0.0873871 | 0.09145991 | -0.0040728 | 0.61654327 | 0.99999994 |
| D130020L05I | 0.03452745 | 0.02647252 | 0.03054624 | -0.0040737 | 0.81265123 | 0.99999994 |

| | | | | | | |
|---|---|---|---|---|---|---|
| 4933434E20I | 0.42595728 | 0.27798458 | 0.28205854 | -0.004074 | 0.53829618 | 0.99999994 |
| Cep63 | 0.13513302 | 0.10732906 | 0.11140374 | -0.0040747 | 0.59617597 | 0.99999994 |
| Mak16 | 0.23192314 | 0.17326063 | 0.17733704 | -0.0040764 | 0.56080411 | 0.99999994 |
| Rnf166 | 0.12050139 | 0.09642251 | 0.10050119 | -0.0040787 | 0.60647295 | 0.99999994 |
| Ap4b1 | 0.05431489 | 0.04334822 | 0.0474282 | -0.00408 | 0.71654521 | 0.99999994 |
| Cluap1 | 0.18321189 | 0.14137027 | 0.14545143 | -0.0040812 | 0.57407861 | 0.99999994 |
| Fbxo42 | 0.03622153 | 0.02792765 | 0.03200894 | -0.0040813 | 0.80198076 | 0.99999994 |
| Ccdc134 | 0.04875083 | 0.03864411 | 0.0427256 | -0.0040815 | 0.73756898 | 0.99999994 |
| Slc10a7 | 0.03235897 | 0.02458457 | 0.02866913 | -0.0040846 | 0.82852337 | 0.99999994 |
| Stat1 | 0.03951355 | 0.03075713 | 0.03484193 | -0.0040848 | 0.78241616 | 0.99999994 |
| Slc12a7 | 0.07363594 | 0.05940501 | 0.06349454 | -0.0040895 | 0.66593667 | 0.99999994 |
| Zfp598 | 0.04078431 | 0.03183922 | 0.0359322 | -0.004093 | 0.77583871 | 0.99999994 |
| Nell1 | 0.01041605 | 0.00527032 | 0.00936513 | -0.0040948 | 0.99819533 | 0.99999994 |
| Katnal1 | 0.05910287 | 0.04735023 | 0.05144747 | -0.0040972 | 0.70198172 | 0.99999994 |
| Leo1 | 0.26715326 | 0.19477888 | 0.19887769 | -0.0040988 | 0.55455793 | 0.99999994 |
| Tmem132b | 0.01456442 | 0.00895802 | 0.01305836 | -0.0041003 | 0.98169586 | 0.99999994 |
| Pold3 | 0.08667175 | 0.0700085 | 0.07410907 | -0.0041006 | 0.64367777 | 0.99999994 |
| Erlin1 | 0.03913902 | 0.03041742 | 0.03452024 | -0.0041028 | 0.78556493 | 0.99999994 |
| Ifrd2 | 0.07576722 | 0.06113934 | 0.0652422 | -0.0041029 | 0.6622443 | 0.99999994 |
| Mfsd12 | 0.02395924 | 0.01723393 | 0.02134657 | -0.0041126 | 0.9001521 | 0.99999994 |
| Grcc10 | 0.39045032 | 0.26115819 | 0.26527496 | -0.0041168 | 0.54113716 | 0.99999994 |
| Tcaf1 | 0.24451451 | 0.1810572 | 0.18517729 | -0.0041201 | 0.55886828 | 0.99999994 |
| Ptprn | 0.19182259 | 0.14715305 | 0.15127334 | -0.0041203 | 0.57193467 | 0.99999994 |
| Gigyf1 | 0.05941094 | 0.04758457 | 0.05170522 | -0.0041206 | 0.70210992 | 0.99999994 |
| Mvk | 0.08976351 | 0.07247904 | 0.07660053 | -0.0041215 | 0.63988278 | 0.99999994 |
| Zfp715 | 0.02643186 | 0.0193881 | 0.02351065 | -0.0041225 | 0.8784308 | 0.99999994 |
| Dnmt3a | 0.72813932 | 0.38465702 | 0.38878184 | -0.0041248 | 0.52815046 | 0.99999994 |
| Ppip5k2 | 0.10077336 | 0.08125699 | 0.08538225 | -0.0041253 | 0.62612345 | 0.99999994 |
| Usp45 | 0.04047212 | 0.03153711 | 0.03566452 | -0.0041274 | 0.77944853 | 0.99999994 |
| Zkscan6 | 0.05018906 | 0.03981689 | 0.04394446 | -0.0041276 | 0.73408389 | 0.99999994 |

| | | | | | | |
|---|---|---|---|---|---|---|
| Slc25a46 | 0.17461553 | 0.13542974 | 0.13955798 | -0.0041282 | 0.57804733 | 0.99999994 |
| Ostm1 | 0.08242546 | 0.06654175 | 0.07067107 | -0.0041293 | 0.65133675 | 0.99999994 |
| Sars2 | 0.05310715 | 0.04227825 | 0.04641037 | -0.0041321 | 0.72329033 | 0.99999994 |
| Kank1 | 0.03013179 | 0.02260309 | 0.02673555 | -0.0041325 | 0.84822741 | 0.99999994 |
| Sipa1l1 | 0.05927288 | 0.04745723 | 0.05158973 | -0.0041325 | 0.70305058 | 0.99999994 |
| Cyb5rl | 0.06306367 | 0.0506192 | 0.05475339 | -0.0041342 | 0.69237486 | 0.99999994 |
| Etf1 | 0.38991159 | 0.26087626 | 0.26501271 | -0.0041364 | 0.5413739 | 0.99999994 |
| Zfp12 | 0.03216023 | 0.02436013 | 0.02849682 | -0.0041367 | 0.83304262 | 0.99999994 |
| Slc25a28 | 0.03226579 | 0.02444855 | 0.02858835 | -0.0041398 | 0.83244913 | 0.99999994 |
| Dnajb12 | 0.09661264 | 0.07794421 | 0.08208479 | -0.0041406 | 0.63147597 | 0.99999994 |
| Itch | 0.20947887 | 0.15881397 | 0.16295848 | -0.0041445 | 0.56721587 | 0.99999994 |
| Sos2 | 0.06152176 | 0.04932335 | 0.05346845 | -0.0041451 | 0.69708283 | 0.99999994 |
| 5730455P16I | 0.11164421 | 0.08962756 | 0.09377365 | -0.0041461 | 0.61573681 | 0.99999994 |
| Tdo2 | 0.01280218 | 0.00734391 | 0.01149199 | -0.0041481 | 0.99186138 | 0.99999994 |
| Rufy3 | 0.24097421 | 0.17884078 | 0.18298917 | -0.0041484 | 0.55997288 | 0.99999994 |
| Actr1a | 0.2253148 | 0.16900787 | 0.17315786 | -0.00415 | 0.56337429 | 0.99999994 |
| Abt1 | 0.15199479 | 0.11949774 | 0.12364932 | -0.0041516 | 0.58842313 | 0.99999994 |
| Lrch1 | 0.03979809 | 0.0309346 | 0.03508622 | -0.0041516 | 0.78454098 | 0.99999994 |
| Ylpm1 | 0.2067433 | 0.15701838 | 0.16117005 | -0.0041517 | 0.5680709 | 0.99999994 |
| Acap2 | 0.10412044 | 0.0838292 | 0.08798176 | -0.0041526 | 0.62330286 | 0.99999994 |
| Dffa | 0.08868963 | 0.0715825 | 0.07573628 | -0.0041538 | 0.64247017 | 0.99999994 |
| Zfp120 | 0.10712315 | 0.08614735 | 0.09030179 | -0.0041544 | 0.62028649 | 0.99999994 |
| Zfp865 | 0.11807653 | 0.09451203 | 0.09866912 | -0.0041571 | 0.61043125 | 0.99999994 |
| Tsr2 | 0.10962162 | 0.08806599 | 0.09222359 | -0.0041576 | 0.61794213 | 0.99999994 |
| Hnf4a | 0.04181004 | 0.03265199 | 0.0368109 | -0.0041589 | 0.77398582 | 0.99999994 |
| Uckl1 | 0.21713418 | 0.16376201 | 0.16792159 | -0.0041596 | 0.56548248 | 0.99999994 |
| Dnajb3 | 0.05981688 | 0.04788291 | 0.05204469 | -0.0041618 | 0.70273346 | 0.99999994 |
| Prss2 | 0.0487353 | 0.03854885 | 0.04271242 | -0.0041636 | 0.74178455 | 0.99999994 |
| Gnl3 | 0.15253102 | 0.11986926 | 0.12403325 | -0.004164 | 0.588413 | 0.99999994 |
| Zfp142 | 0.02607463 | 0.01903434 | 0.02319844 | -0.0041641 | 0.8839191 | 0.99999994 |

| | | | | | | |
|---|---|---|---|---|---|---|
| Cwc22 | 0.1743366 | 0.13519985 | 0.13936534 | -0.0041655 | 0.57884983 | 0.99999994 |
| Tcf4 | 0.23667375 | 0.17614847 | 0.18031424 | -0.0041658 | 0.5611087 | 0.99999994 |
| Nrxn1 | 0.01598598 | 0.01015248 | 0.0143192 | -0.0041667 | 0.97362249 | 0.99999994 |
| Ddx19a | 0.14915834 | 0.1174462 | 0.12161297 | -0.0041668 | 0.59020155 | 0.99999994 |
| Notch1 | 0.04842236 | 0.03827798 | 0.04244691 | -0.0041689 | 0.74336522 | 0.99999994 |
| Mgat5 | 0.06557465 | 0.05266983 | 0.05684037 | -0.0041705 | 0.68736566 | 0.99999994 |
| Aldh4a1 | 0.04123916 | 0.03215016 | 0.036322 | -0.0041718 | 0.77772429 | 0.99999994 |
| Wnt4 | 0.04258961 | 0.0333052 | 0.03747793 | -0.0041727 | 0.77068585 | 0.99999994 |
| Polr1a | 0.03803158 | 0.02939515 | 0.03356813 | -0.004173 | 0.79601662 | 0.99999994 |
| Zfp866 | 0.07565455 | 0.0609763 | 0.06514994 | -0.0041736 | 0.66509892 | 0.99999994 |
| Tmem178b | 0.03864625 | 0.02991857 | 0.03409676 | -0.0041782 | 0.79265423 | 0.99999994 |
| Prmt7 | 0.14326611 | 0.11317337 | 0.1173524 | -0.004179 | 0.59368699 | 0.99999994 |
| S100a6 | 0.04809951 | 0.03798819 | 0.04217286 | -0.0041847 | 0.74552986 | 0.99999994 |
| Med13l | 0.0624114 | 0.05002445 | 0.05421015 | -0.0041857 | 0.69635789 | 0.99999994 |
| Fcna | 0.01032599 | 0.0050975 | 0.00928473 | -0.0041872 | 0.99865842 | 0.99999994 |
| Tmppe | 0.02626176 | 0.01917461 | 0.023362 | -0.0041874 | 0.8835816 | 0.99999994 |
| Alkbh3 | 0.09836413 | 0.07929194 | 0.08347956 | -0.0041876 | 0.63077463 | 0.99999994 |
| Arsb | 0.0433623 | 0.03394968 | 0.03813839 | -0.0041887 | 0.7676297 | 0.99999994 |
| Klhl28 | 0.05572302 | 0.04442326 | 0.04861287 | -0.0041896 | 0.71690329 | 0.99999994 |
| Cyb5r1 | 0.0351901 | 0.02692757 | 0.03111878 | -0.0041912 | 0.81500892 | 0.99999994 |
| Lipa | 0.04203198 | 0.03280466 | 0.03700087 | -0.0041962 | 0.77484152 | 0.99999994 |
| Mapre2 | 0.28733459 | 0.20647414 | 0.21067369 | -0.0041995 | 0.55277956 | 0.99999994 |
| Rsph3a | 0.02822824 | 0.0208769 | 0.02507838 | -0.0042015 | 0.86761029 | 0.99999994 |
| Pigm | 0.06252687 | 0.05010446 | 0.05430635 | -0.0042019 | 0.69673332 | 0.99999994 |
| Rnft1 | 0.06164017 | 0.04936472 | 0.05356722 | -0.0042025 | 0.69923814 | 0.99999994 |
| Myh14 | 0.05089325 | 0.04033359 | 0.04454039 | -0.0042068 | 0.73522302 | 0.99999994 |
| Cdc25a | 0.10426948 | 0.08388858 | 0.08809719 | -0.0042086 | 0.62475355 | 0.99999994 |
| Slc15a2 | 0.05780542 | 0.04615146 | 0.05036081 | -0.0042094 | 0.71101899 | 0.99999994 |
| Ide | 0.07640688 | 0.06155507 | 0.06576579 | -0.0042107 | 0.66501165 | 0.99999994 |
| Got2 | 0.36875494 | 0.25032187 | 0.2545326 | -0.0042107 | 0.5438413 | 0.99999994 |

| | | | | | | |
|---|---|---|---|---|---|---|
| Dcaf6 | 0.08008964 | 0.06456081 | 0.06877185 | -0.004211 | 0.65820975 | 0.99999994 |
| Med17 | 0.06601602 | 0.05299505 | 0.05720651 | -0.0042115 | 0.68794405 | 0.99999994 |
| Tnfsf12 | 0.01675864 | 0.01079199 | 0.01500351 | -0.0042115 | 0.96915066 | 0.99999994 |
| Eml1 | 0.03731039 | 0.02873099 | 0.03294734 | -0.0042164 | 0.80284583 | 0.99999994 |
| Rims2 | 0.06925215 | 0.05566822 | 0.05988461 | -0.0042164 | 0.68034519 | 0.99999994 |
| Polg2 | 0.04383355 | 0.03432381 | 0.03854087 | -0.0042171 | 0.76679234 | 0.99999994 |
| Mapk8ip1 | 0.17227562 | 0.13372078 | 0.13793928 | -0.0042185 | 0.58065393 | 0.99999994 |
| Zbtb38 | 0.08694326 | 0.0701052 | 0.07432826 | -0.0042231 | 0.64735545 | 0.99999994 |
| Polr3c | 0.12304363 | 0.0981872 | 0.10241423 | -0.004227 | 0.60823946 | 0.99999994 |
| Ccdc148 | 0.04223445 | 0.03294683 | 0.03717412 | -0.0042273 | 0.77545526 | 0.99999994 |
| Zfp637 | 0.27946359 | 0.20189055 | 0.2061188 | -0.0042282 | 0.55430519 | 0.99999994 |
| Foxred1 | 0.07314798 | 0.05886302 | 0.06309374 | -0.0042307 | 0.67232343 | 0.99999994 |
| Aup1 | 0.21858293 | 0.16462098 | 0.16885402 | -0.004233 | 0.56626384 | 0.99999994 |
| Adora2b | 0.01441047 | 0.00868679 | 0.01292167 | -0.0042349 | 0.98542738 | 0.99999994 |
| Tnik | 0.06965662 | 0.05598363 | 0.06021853 | -0.0042349 | 0.68014767 | 0.99999994 |
| Sclt1 | 0.05066845 | 0.0401148 | 0.04435021 | -0.0042354 | 0.73750673 | 0.99999994 |
| Slc12a9 | 0.05594888 | 0.04456632 | 0.04880269 | -0.0042364 | 0.71830425 | 0.99999994 |
| 1810062G17 | 0.00950747 | 0.00431682 | 0.00855347 | -0.0042367 | 0.99951136 | 0.99999994 |
| Rmi1 | 0.09223905 | 0.07434665 | 0.0785883 | -0.0042417 | 0.64030139 | 0.99999994 |
| Endod1 | 0.01942311 | 0.01311604 | 0.01735781 | -0.0042418 | 0.94809196 | 0.99999994 |
| Terf2 | 0.08439947 | 0.06802866 | 0.07227166 | -0.004243 | 0.6520232 | 0.99999994 |
| Tufm | 0.12740547 | 0.10143207 | 0.1056782 | -0.0042461 | 0.60543911 | 0.99999994 |
| Prr3 | 0.15563957 | 0.12200574 | 0.12625236 | -0.0042466 | 0.58857975 | 0.99999994 |
| Dclre1c | 0.02645458 | 0.01928302 | 0.0235305 | -0.0042475 | 0.8852282 | 0.99999994 |
| Isca1 | 0.04664675 | 0.03669073 | 0.0409383 | -0.0042476 | 0.75510172 | 0.99999994 |
| Utp23 | 0.10606681 | 0.08523884 | 0.0894869 | -0.0042481 | 0.62399346 | 0.99999994 |
| Mrps30 | 0.227944 | 0.17057716 | 0.17482588 | -0.0042487 | 0.56425526 | 0.99999994 |
| Mrpl49 | 0.45831295 | 0.29228229 | 0.29653105 | -0.0042488 | 0.53799102 | 0.99999994 |
| Wdfy3 | 0.07946156 | 0.06400991 | 0.06826018 | -0.0042503 | 0.66073244 | 0.99999994 |
| Foxo1 | 0.19202863 | 0.1471611 | 0.15141161 | -0.0042505 | 0.57411529 | 0.99999994 |

| | | | | | | |
|---|---|---|---|---|---|---|
| Timm10 | 0.21078962 | 0.1595566 | 0.16381258 | -0.004256 | 0.56864967 | 0.99999994 |
| Gal3st1 | 0.03641682 | 0.02792088 | 0.03217735 | -0.0042565 | 0.81070904 | 0.99999994 |
| Gtf2h2 | 0.12796819 | 0.10183973 | 0.1060976 | -0.0042579 | 0.60531562 | 0.99999994 |
| Paxip1 | 0.1040836 | 0.08369502 | 0.08795323 | -0.0042582 | 0.62637361 | 0.99999994 |
| Ugcg | 0.06551282 | 0.05252887 | 0.05678906 | -0.0042602 | 0.6912321 | 0.99999994 |
| Lypla1 | 0.2438689 | 0.18051866 | 0.18477919 | -0.0042605 | 0.56098945 | 0.99999994 |
| Crtap | 0.03211515 | 0.02419699 | 0.02845773 | -0.0042607 | 0.84052087 | 0.99999994 |
| Rab31 | 0.02787817 | 0.02051197 | 0.02477317 | -0.0042612 | 0.87388539 | 0.99999994 |
| Nsdhl | 0.11890375 | 0.09503147 | 0.09929492 | -0.0042635 | 0.61248668 | 0.99999994 |
| Fer | 0.14758674 | 0.11621527 | 0.12048065 | -0.0042654 | 0.59315004 | 0.99999994 |
| Snhg20 | 0.14814503 | 0.11661755 | 0.12088323 | -0.0042657 | 0.59285163 | 0.99999994 |
| Nup160 | 0.0287022 | 0.02122463 | 0.02549139 | -0.0042668 | 0.8673915 | 0.99999994 |
| Sart1 | 0.22531349 | 0.16888976 | 0.17315703 | -0.0042673 | 0.56514981 | 0.99999994 |
| Mettl3 | 0.10901771 | 0.08749151 | 0.09175979 | -0.0042683 | 0.62157715 | 0.99999994 |
| Snhg12 | 0.1042983 | 0.08384982 | 0.0881195 | -0.0042697 | 0.62647163 | 0.99999994 |
| Pyroxd1 | 0.13043659 | 0.10366017 | 0.10793285 | -0.0042727 | 0.60391798 | 0.99999994 |
| Setd1a | 0.04106143 | 0.03189613 | 0.03616971 | -0.0042736 | 0.78420474 | 0.99999994 |
| Zfp958 | 0.04369864 | 0.0341515 | 0.03842567 | -0.0042742 | 0.7704714 | 0.99999994 |
| Tmem260 | 0.03257611 | 0.02458247 | 0.02885733 | -0.0042749 | 0.83794774 | 0.99999994 |
| Dtwd1 | 0.08950314 | 0.07211584 | 0.07639109 | -0.0042752 | 0.6452489 | 0.99999994 |
| Lsm1 | 0.48414945 | 0.30327513 | 0.30755232 | -0.0042772 | 0.53687789 | 0.99999994 |
| Zfp213 | 0.05058225 | 0.03999654 | 0.04427727 | -0.0042807 | 0.74006255 | 0.99999994 |
| Setd6 | 0.07170831 | 0.05762759 | 0.06190972 | -0.0042821 | 0.67738617 | 0.99999994 |
| Mgat3 | 0.03873644 | 0.02989179 | 0.03417429 | -0.0042825 | 0.7978969 | 0.99999994 |
| Smim10l2a | 0.03527702 | 0.0269092 | 0.03119384 | -0.0042846 | 0.81971683 | 0.99999994 |
| Noct | 0.15019906 | 0.11807456 | 0.12236121 | -0.0042867 | 0.59219341 | 0.99999994 |
| Dusp22 | 0.0863324 | 0.06954725 | 0.073835 | -0.0042878 | 0.65045478 | 0.99999994 |
| Slc18a1 | 0.02441551 | 0.0174569 | 0.02174645 | -0.0042896 | 0.90540643 | 0.99999994 |
| Pycard | 0.09308687 | 0.07497741 | 0.07926763 | -0.0042902 | 0.64067497 | 0.99999994 |
| Klhl42 | 0.04794995 | 0.03775563 | 0.04204588 | -0.0042902 | 0.75149349 | 0.99999994 |

| | | | | | | |
|---|---|---|---|---|---|---|
| Lactb2 | 0.06552002 | 0.05250359 | 0.05679504 | -0.0042914 | 0.69250239 | 0.99999994 |
| Sp4 | 0.05917211 | 0.04721255 | 0.05150541 | -0.0042929 | 0.71048166 | 0.99999994 |
| Ift172 | 0.02418079 | 0.01724721 | 0.02154077 | -0.0042936 | 0.90770639 | 0.99999994 |
| Mgat4b | 0.04448441 | 0.03480234 | 0.03909633 | -0.004294 | 0.7676332 | 0.99999994 |
| Pou2f1 | 0.10650258 | 0.08552748 | 0.08982324 | -0.0042958 | 0.62488541 | 0.99999994 |
| Mtor | 0.06337839 | 0.05071475 | 0.05501534 | -0.0043006 | 0.6985841 | 0.99999994 |
| Mob1b | 0.15306519 | 0.12011275 | 0.12441537 | -0.0043026 | 0.59102978 | 0.99999994 |
| Rnf130 | 0.54983509 | 0.32926142 | 0.33356418 | -0.0043028 | 0.53421221 | 0.99999994 |
| Acsf3 | 0.09109993 | 0.07337021 | 0.07767441 | -0.0043042 | 0.64388049 | 0.99999994 |
| Cdc123 | 0.52536829 | 0.31989002 | 0.32419993 | -0.0043099 | 0.53525623 | 0.99999994 |
| Zadh2 | 0.04325588 | 0.03373713 | 0.03804747 | -0.0043103 | 0.77460158 | 0.99999994 |
| Abl2 | 0.04459067 | 0.03487575 | 0.03918697 | -0.0043112 | 0.7680101 | 0.99999994 |
| Hmgcl | 0.22852333 | 0.17088073 | 0.17519248 | -0.0043117 | 0.56506465 | 0.99999994 |
| Csgalnact2 | 0.02673945 | 0.01946726 | 0.02377935 | -0.0043121 | 0.88629347 | 0.99999994 |
| Med16 | 0.11641469 | 0.09309687 | 0.09740936 | -0.0043125 | 0.61588418 | 0.99999994 |
| Dnmt1 | 0.09186071 | 0.07397207 | 0.07828492 | -0.0043128 | 0.6430823 | 0.99999994 |
| 1700019D03I | 0.03396741 | 0.02574826 | 0.03006197 | -0.0043137 | 0.83024819 | 0.99999994 |
| Dcdc2a | 0.03144232 | 0.02355455 | 0.02787402 | -0.0043195 | 0.84884582 | 0.99999994 |
| Slitrk6 | 0.13138727 | 0.10431733 | 0.1086376 | -0.0043203 | 0.60438278 | 0.99999994 |
| Gm14966 | 0.0508349 | 0.04016967 | 0.04449104 | -0.0043214 | 0.74103039 | 0.99999994 |
| Tbc1d8 | 0.07929212 | 0.06379902 | 0.06812208 | -0.0043231 | 0.66363927 | 0.99999994 |
| Itga3 | 0.02276486 | 0.01597547 | 0.02029867 | -0.0043232 | 0.92185619 | 0.99999994 |
| Zfx | 0.13196491 | 0.10473936 | 0.10906526 | -0.0043259 | 0.60411552 | 0.99999994 |
| Ap1s1 | 0.68109662 | 0.37293868 | 0.37726494 | -0.0043263 | 0.53042222 | 0.99999994 |
| Gatc | 0.16867627 | 0.13110972 | 0.13543741 | -0.0043277 | 0.58421684 | 0.99999994 |
| Rpp25 | 0.01541084 | 0.00948033 | 0.01380938 | -0.004329 | 0.9815413 | 0.99999994 |
| Tsen54 | 0.08885357 | 0.07153814 | 0.07586829 | -0.0043302 | 0.64799353 | 0.99999994 |
| Zfp426 | 0.05683269 | 0.04521209 | 0.04954491 | -0.0043328 | 0.7197551 | 0.99999994 |
| Erich3 | 0.01175568 | 0.00622416 | 0.01056004 | -0.0043359 | 0.9968625 | 0.99999994 |
| Leng8 | 0.08566574 | 0.068958 | 0.07329625 | -0.0043383 | 0.65320058 | 0.99999994 |

| | | | | | | |
|---|---|---|---|---|---|---|
| Ptpn9 | 0.08971053 | 0.07221782 | 0.07655792 | -0.0043401 | 0.64704388 | 0.99999994 |
| Card19 | 0.2726432 | 0.197784 | 0.20212468 | -0.0043407 | 0.55683395 | 0.99999994 |
| Map3k12 | 0.05023109 | 0.03963894 | 0.04398004 | -0.0043411 | 0.74441688 | 0.99999994 |
| Rcn3 | 0.11879071 | 0.09486824 | 0.09920946 | -0.0043412 | 0.61457743 | 0.99999994 |
| Adam17 | 0.04613875 | 0.03616346 | 0.04050603 | -0.0043426 | 0.76226753 | 0.99999994 |
| Smug1 | 0.05545169 | 0.04404123 | 0.04838477 | -0.0043435 | 0.72492907 | 0.99999994 |
| Eef2k | 0.02867837 | 0.02112353 | 0.02547063 | -0.0043471 | 0.87203179 | 0.99999994 |
| Abl1 | 0.05542055 | 0.04401118 | 0.04835858 | -0.0043474 | 0.72521421 | 0.99999994 |
| Cbx8 | 0.0620559 | 0.04956564 | 0.05391387 | -0.0043482 | 0.70431264 | 0.99999994 |
| Dnlz | 0.34408226 | 0.2374994 | 0.24184839 | -0.004349 | 0.54763822 | 0.99999994 |
| Dnajc25 | 0.08868446 | 0.07138306 | 0.07573211 | -0.004349 | 0.64886315 | 0.99999994 |
| Phb | 0.1859396 | 0.14295296 | 0.14730441 | -0.0043514 | 0.57794232 | 0.99999994 |
| Tenm3 | 0.04096628 | 0.03173663 | 0.03608816 | -0.0043515 | 0.78890529 | 0.99999994 |
| Fkbpl | 0.11411554 | 0.09130783 | 0.09566071 | -0.0043529 | 0.61901182 | 0.99999994 |
| Nr1d2 | 0.01943036 | 0.01300979 | 0.0173642 | -0.0043544 | 0.95246233 | 0.99999994 |
| Rapgef6 | 0.08152832 | 0.06558762 | 0.06994229 | -0.0043547 | 0.66072077 | 0.99999994 |
| Tbc1d10b | 0.06994664 | 0.05610189 | 0.06045787 | -0.004356 | 0.68424039 | 0.99999994 |
| Asns | 0.1666586 | 0.12967039 | 0.1340286 | -0.0043582 | 0.58567947 | 0.99999994 |
| Fnbp4 | 0.14332514 | 0.11303698 | 0.1173953 | -0.0043583 | 0.59759174 | 0.99999994 |
| Ifngr2 | 0.11486886 | 0.09187544 | 0.09623442 | -0.004359 | 0.61848445 | 0.99999994 |
| Agpat2 | 0.16644475 | 0.12951739 | 0.13387901 | -0.0043616 | 0.58583972 | 0.99999994 |
| Mccc1 | 0.05420437 | 0.04297338 | 0.04733512 | -0.0043617 | 0.73017972 | 0.99999994 |
| Mtap | 0.06775129 | 0.05428184 | 0.05864395 | -0.0043621 | 0.68974238 | 0.99999994 |
| Ttll7 | 0.39678595 | 0.26397729 | 0.26834086 | -0.0043636 | 0.54309772 | 0.99999994 |
| 5430416N02 | 0.07265894 | 0.05832523 | 0.06269179 | -0.0043666 | 0.678541 | 0.99999994 |
| Gm14167 | 0.03554819 | 0.02705856 | 0.03142796 | -0.0043694 | 0.82262825 | 0.99999994 |
| Ppat | 0.10586404 | 0.08495984 | 0.08933033 | -0.0043705 | 0.62766004 | 0.99999994 |
| Ptpn23 | 0.0491203 | 0.03866665 | 0.04303893 | -0.0043723 | 0.75054713 | 0.99999994 |
| Arhgap21 | 0.05981243 | 0.0476686 | 0.05204097 | -0.0043724 | 0.71200543 | 0.99999994 |
| Trak2 | 0.04380581 | 0.03414435 | 0.03851718 | -0.0043728 | 0.77508284 | 0.99999994 |

| | | | | | | |
|---|---|---|---|---|---|---|
| Zfp651 | 0.02496182 | 0.01785011 | 0.02222491 | -0.0043748 | 0.90494483 | 0.99999994 |
| Prpf31 | 0.27050145 | 0.19648543 | 0.20086147 | -0.004376 | 0.55765162 | 0.99999994 |
| Mtmr2 | 0.15917886 | 0.12438416 | 0.12876545 | -0.0043813 | 0.58958796 | 0.99999994 |
| Adgrl3 | 0.01681117 | 0.01066795 | 0.01505 | -0.004382 | 0.97369477 | 0.99999994 |
| Gclc | 0.01771412 | 0.01146641 | 0.01584872 | -0.0043823 | 0.96715568 | 0.99999994 |
| Nup214 | 0.10788247 | 0.08650064 | 0.09088668 | -0.004386 | 0.6259798 | 0.99999994 |
| Fxr2 | 0.14837029 | 0.11665643 | 0.12104555 | -0.0043891 | 0.59536207 | 0.99999994 |
| Zfp324 | 0.02549428 | 0.01830167 | 0.02269091 | -0.0043892 | 0.90105395 | 0.99999994 |
| Adck1 | 0.04392365 | 0.0342258 | 0.03861779 | -0.004392 | 0.77548255 | 0.99999994 |
| Smim13 | 0.10086315 | 0.08106001 | 0.08545218 | -0.0043922 | 0.63387391 | 0.99999994 |
| Ino80e | 0.23946759 | 0.1776607 | 0.18205423 | -0.0043935 | 0.56381097 | 0.99999994 |
| Dbt | 0.05219794 | 0.04124682 | 0.04564306 | -0.0043962 | 0.73927922 | 0.99999994 |
| Adar | 0.03052541 | 0.02268143 | 0.0270777 | -0.0043963 | 0.86008683 | 0.99999994 |
| Chp1 | 0.20238448 | 0.15390473 | 0.15830332 | -0.0043986 | 0.5733668 | 0.99999994 |
| Mthfsl | 0.23634365 | 0.17570824 | 0.1801081 | -0.0043999 | 0.56458653 | 0.99999994 |
| Med23 | 0.04428515 | 0.03452636 | 0.03892632 | -0.0044 | 0.77409137 | 0.99999994 |
| Lcmt2 | 0.0983806 | 0.07909194 | 0.08349266 | -0.0044007 | 0.63714678 | 0.99999994 |
| Tmem150a | 0.06436982 | 0.05143855 | 0.05583984 | -0.0044013 | 0.70008802 | 0.99999994 |
| Sdc2 | 0.09316688 | 0.07492762 | 0.0793317 | -0.0044041 | 0.64413214 | 0.99999994 |
| Mtus1 | 0.01911647 | 0.01268286 | 0.01708729 | -0.0044044 | 0.95689568 | 0.99999994 |
| Rcan2 | 0.11243348 | 0.08997034 | 0.09437714 | -0.0044068 | 0.62202691 | 0.99999994 |
| Brd8 | 0.37202807 | 0.25176894 | 0.25617767 | -0.0044087 | 0.54560054 | 0.99999994 |
| Ptdss1 | 0.14037395 | 0.11083498 | 0.11524599 | -0.004411 | 0.60054984 | 0.99999994 |
| Sntg2 | 0.01573083 | 0.00968204 | 0.01409308 | -0.004411 | 0.98139247 | 0.99999994 |
| Cst6 | 0.04047811 | 0.0312575 | 0.03566966 | -0.0044122 | 0.79484765 | 0.99999994 |
| Zranb3 | 0.02172173 | 0.01496972 | 0.01938209 | -0.0044124 | 0.93515604 | 0.99999994 |
| Otud4 | 0.07252423 | 0.05816814 | 0.06258102 | -0.0044129 | 0.68059778 | 0.99999994 |
| Fam98b | 0.18065615 | 0.13929297 | 0.14370788 | -0.0044149 | 0.58101581 | 0.99999994 |
| Plekhj1 | 0.72638624 | 0.38397413 | 0.3883929 | -0.0044188 | 0.53018292 | 0.99999994 |
| Sspn | 0.0407248 | 0.03146196 | 0.03588118 | -0.0044192 | 0.79383905 | 0.99999994 |

| | | | | | | |
|---|---|---|---|---|---|---|
| Timm21 | 0.15547515 | 0.12171395 | 0.12613526 | -0.0044213 | 0.59224335 | 0.99999994 |
| Mpp2 | 0.0329646 | 0.02477183 | 0.02919393 | -0.0044221 | 0.84333504 | 0.99999994 |
| Cipc | 0.03749155 | 0.02868058 | 0.03310333 | -0.0044228 | 0.81308449 | 0.99999994 |
| Lims1 | 0.21207542 | 0.1602238 | 0.16464869 | -0.0044249 | 0.57098722 | 0.99999994 |
| Rbsn | 0.05294958 | 0.04185086 | 0.04627746 | -0.0044266 | 0.73784111 | 0.99999994 |
| Mrpl3 | 0.20550949 | 0.15593055 | 0.16036061 | -0.0044301 | 0.57294865 | 0.99999994 |
| Tyms | 0.13023092 | 0.10335014 | 0.10778021 | -0.0044301 | 0.607802 | 0.99999994 |
| Eif2ak1 | 0.08460278 | 0.06800596 | 0.07243628 | -0.0044303 | 0.65803757 | 0.99999994 |
| Zfp687 | 0.07521955 | 0.0603609 | 0.06479357 | -0.0044327 | 0.67558078 | 0.99999994 |
| Vps9d1 | 0.09876332 | 0.07936357 | 0.08379701 | -0.0044334 | 0.63764359 | 0.99999994 |
| Pgap2 | 0.11594919 | 0.09262174 | 0.09705587 | -0.0044341 | 0.61947618 | 0.99999994 |
| Tfip11 | 0.07118025 | 0.05703948 | 0.06147487 | -0.0044354 | 0.68447673 | 0.99999994 |
| Ddi2 | 0.07010257 | 0.05614954 | 0.06058651 | -0.004437 | 0.68703697 | 0.99999994 |
| Zfp719 | 0.05824214 | 0.04628703 | 0.05072679 | -0.0044398 | 0.71991363 | 0.99999994 |
| Uhrf1bp1 | 0.01997695 | 0.01340563 | 0.01784612 | -0.0044405 | 0.95116358 | 0.99999994 |
| Cog3 | 0.029663 | 0.02188355 | 0.02632783 | -0.0044443 | 0.86941473 | 0.99999994 |
| Chst11 | 0.01955264 | 0.01302671 | 0.01747204 | -0.0044453 | 0.95482431 | 0.99999994 |
| Gm11423 | 0.04286937 | 0.03327027 | 0.03771713 | -0.0044469 | 0.78371073 | 0.99999994 |
| Cask | 0.04149002 | 0.03208937 | 0.03653688 | -0.0044475 | 0.79110301 | 0.99999994 |
| Ankle2 | 0.10227712 | 0.08210244 | 0.08655191 | -0.0044495 | 0.63389632 | 0.99999994 |
| 1810030O07 | 0.10861208 | 0.08699658 | 0.091448 | -0.0044514 | 0.62703552 | 0.99999994 |
| Supt20 | 0.22872809 | 0.17087054 | 0.17532197 | -0.0044514 | 0.56710372 | 0.99999994 |
| Ints2 | 0.06684047 | 0.05343718 | 0.05788986 | -0.0044527 | 0.69566873 | 0.99999994 |
| Cradd | 0.07092531 | 0.05681186 | 0.06126483 | -0.004453 | 0.68573982 | 0.99999994 |
| Pdzd8 | 0.12701357 | 0.10093252 | 0.10538588 | -0.0044534 | 0.61075313 | 0.99999994 |
| Map4k3 | 0.04057335 | 0.03129719 | 0.03575132 | -0.0044541 | 0.79653055 | 0.99999994 |
| Usf3 | 0.04820118 | 0.03780453 | 0.04225918 | -0.0044547 | 0.75855592 | 0.99999994 |
| Ntmt1 | 0.3604496 | 0.24586415 | 0.25031981 | -0.0044557 | 0.54715728 | 0.99999994 |
| Nup188 | 0.03297849 | 0.02474883 | 0.02920596 | -0.0044571 | 0.84514477 | 0.99999994 |
| Cacnb3 | 0.07248498 | 0.05809062 | 0.06254875 | -0.0044581 | 0.68240278 | 0.99999994 |

| | | | | | | |
|---|---|---|---|---|---|---|
| Nprl3 | 0.05260249 | 0.04152625 | 0.04598459 | -0.0044583 | 0.74065086 | 0.99999994 |
| Rtn2 | 0.09483966 | 0.07621024 | 0.08066973 | -0.0044595 | 0.64355194 | 0.99999994 |
| Them6 | 0.03685005 | 0.02809069 | 0.03255078 | -0.0044601 | 0.8191291 | 0.99999994 |
| Slc35d1 | 0.02841764 | 0.02078291 | 0.02524346 | -0.0044605 | 0.88023959 | 0.99999994 |
| Zc3h6 | 0.05758689 | 0.04571546 | 0.0501776 | -0.0044621 | 0.72305112 | 0.99999994 |
| Pcgf1 | 0.08798359 | 0.07070303 | 0.07516741 | -0.0044644 | 0.6537032 | 0.99999994 |
| Cdip1 | 0.18332433 | 0.14106224 | 0.14552797 | -0.0044657 | 0.58092472 | 0.99999994 |
| Ecd | 0.15481295 | 0.1211945 | 0.12566334 | -0.0044688 | 0.59356048 | 0.99999994 |
| Tfdp2 | 0.13378875 | 0.1059417 | 0.11041296 | -0.0044713 | 0.60624859 | 0.99999994 |
| Zfp871 | 0.10129742 | 0.08131867 | 0.0857902 | -0.0044715 | 0.63568184 | 0.99999994 |
| Gtpbp8 | 0.09607936 | 0.07718453 | 0.0816595 | -0.004475 | 0.64235886 | 0.99999994 |
| Acaca | 0.05172285 | 0.04076667 | 0.04524176 | -0.0044751 | 0.74486348 | 0.99999994 |
| Mpz | 0.02335382 | 0.01633861 | 0.02081562 | -0.004477 | 0.92387727 | 0.99999994 |
| Eif5 | 1.21948403 | 0.42395338 | 0.42843219 | -0.0044788 | 0.52773811 | 0.99999994 |
| Acvr1 | 0.04982051 | 0.03915254 | 0.04363234 | -0.0044798 | 0.75282868 | 0.99999994 |
| Bspry | 0.09670948 | 0.07768013 | 0.08216199 | -0.0044819 | 0.64173431 | 0.99999994 |
| Pik3ca | 0.10886656 | 0.08716055 | 0.09164364 | -0.0044831 | 0.6276441 | 0.99999994 |
| Git1 | 0.05425025 | 0.04288802 | 0.04737376 | -0.0044857 | 0.73574394 | 0.99999994 |
| Lztr1 | 0.1320842 | 0.10466705 | 0.10915354 | -0.0044865 | 0.60780152 | 0.99999994 |
| C1galt1c1 | 0.329084 | 0.22939802 | 0.23388584 | -0.0044878 | 0.55081579 | 0.99999994 |
| Unc13a | 0.11896008 | 0.09484874 | 0.0993375 | -0.0044888 | 0.61820928 | 0.99999994 |
| Zfp830 | 0.2589336 | 0.1894705 | 0.1939612 | -0.0044907 | 0.56123931 | 0.99999994 |
| Nfrkb | 0.1045768 | 0.08384376 | 0.0883351 | -0.0044913 | 0.63248338 | 0.99999994 |
| Sec22c | 0.03079636 | 0.02281807 | 0.02731312 | -0.0044951 | 0.86334431 | 0.99999994 |
| Zc3h14 | 0.40195189 | 0.26631954 | 0.27081772 | -0.0044982 | 0.54401727 | 0.99999994 |
| Cdk13 | 0.1014438 | 0.08140417 | 0.08590409 | -0.0044999 | 0.6363352 | 0.99999994 |
| Per1 | 0.04173946 | 0.03224235 | 0.03675047 | -0.0045081 | 0.79289966 | 0.99999994 |
| Naa60 | 0.08194943 | 0.06577609 | 0.07028448 | -0.0045084 | 0.66529997 | 0.99999994 |
| Klhdc4 | 0.09284568 | 0.07456533 | 0.07907444 | -0.0045091 | 0.64786769 | 0.99999994 |
| Dcbld2 | 0.03158387 | 0.02348499 | 0.02799685 | -0.0045119 | 0.85829996 | 0.99999994 |

| | | | | | | |
|---|---|---|---|---|---|---|
| Heatr5b | 0.04186002 | 0.03234054 | 0.03685369 | -0.0045131 | 0.79250562 | 0.99999994 |
| Btbd6 | 0.06502971 | 0.05187357 | 0.05638803 | -0.0045145 | 0.70295271 | 0.99999994 |
| Brpf1 | 0.06905975 | 0.05521071 | 0.0597257 | -0.004515 | 0.69258475 | 0.99999994 |
| Kdf1 | 0.27564491 | 0.19936976 | 0.20388787 | -0.0045181 | 0.55863268 | 0.99999994 |
| Cep83 | 0.14835883 | 0.11651847 | 0.12103729 | -0.0045188 | 0.59812992 | 0.99999994 |
| Ccdc149 | 0.02307712 | 0.0160535 | 0.02057279 | -0.0045193 | 0.9281602 | 0.99999994 |
| Tmem68 | 0.10926092 | 0.08742604 | 0.09194663 | -0.0045206 | 0.62826538 | 0.99999994 |
| Ubxn2a | 0.22769971 | 0.17014642 | 0.1746712 | -0.0045248 | 0.56845017 | 0.99999994 |
| Med12l | 0.02397241 | 0.01683298 | 0.02135812 | -0.0045251 | 0.92077145 | 0.99999994 |
| B3gnt9 | 0.01317522 | 0.00729727 | 0.01182388 | -0.0045266 | 0.99458745 | 0.99999994 |
| Ikzf4 | 0.04955554 | 0.03887962 | 0.04340786 | -0.0045282 | 0.75627788 | 0.99999994 |
| Mt3 | 0.04324665 | 0.03351107 | 0.03803958 | -0.0045285 | 0.78594275 | 0.99999994 |
| Fam184a | 0.02103452 | 0.01424803 | 0.01877756 | -0.0045295 | 0.94582444 | 0.99999994 |
| Plagl1 | 0.04825209 | 0.03777087 | 0.0423024 | -0.0045315 | 0.76209076 | 0.99999994 |
| Neil2 | 0.03365536 | 0.0252583 | 0.02979198 | -0.0045337 | 0.844458 | 0.99999994 |
| Trpm7 | 0.10231583 | 0.08204636 | 0.08658197 | -0.0045356 | 0.63634068 | 0.99999994 |
| Nol4l | 0.08722035 | 0.0700157 | 0.07455187 | -0.0045362 | 0.65726564 | 0.99999994 |
| Ctif | 0.03802253 | 0.02902325 | 0.03356034 | -0.0045371 | 0.81590956 | 0.99999994 |
| Scamp1 | 0.14922134 | 0.11710957 | 0.1216583 | -0.0045487 | 0.59827209 | 0.99999994 |
| Dlg5 | 0.05480889 | 0.04329332 | 0.04784405 | -0.0045507 | 0.73667438 | 0.99999994 |
| Cda | 0.12331546 | 0.0980662 | 0.10261832 | -0.0045521 | 0.61610768 | 0.99999994 |
| Edrf1 | 0.05339119 | 0.04209719 | 0.04664989 | -0.0045527 | 0.74202885 | 0.99999994 |
| Mfn2 | 0.05272332 | 0.04153136 | 0.04608655 | -0.0045552 | 0.74470353 | 0.99999994 |
| Bud13 | 0.06349849 | 0.05055453 | 0.05511528 | -0.0045607 | 0.7091176 | 0.99999994 |
| Ppwd1 | 0.08839954 | 0.07093938 | 0.07550261 | -0.0045632 | 0.65626728 | 0.99999994 |
| Slc39a3 | 0.06488524 | 0.05170361 | 0.05626805 | -0.0045644 | 0.70538797 | 0.99999994 |
| Mrps25 | 0.1921279 | 0.14691028 | 0.15147822 | -0.0045679 | 0.57954428 | 0.99999994 |
| Slc35g1 | 0.02919015 | 0.02134833 | 0.02591631 | -0.004568 | 0.87965108 | 0.99999994 |
| Rpl13 | 3.30248278 | 0.27714609 | 0.28171771 | -0.0045716 | 0.54300895 | 0.99999994 |
| Surf6 | 0.07100782 | 0.0567604 | 0.06133282 | -0.0045724 | 0.69013663 | 0.99999994 |

| | | | | | | |
|---|---|---|---|---|---|---|
| Slc52a2 | 0.09244718 | 0.07417632 | 0.07875514 | -0.0045788 | 0.65062198 | 0.99999994 |
| Npat | 0.08956878 | 0.07186503 | 0.0764439 | -0.0045789 | 0.65494504 | 0.99999994 |
| Sall1 | 0.02769095 | 0.02002953 | 0.02460988 | -0.0045803 | 0.89230316 | 0.99999994 |
| Dram2 | 0.21900455 | 0.1645437 | 0.16912498 | -0.0045813 | 0.5715447 | 0.99999994 |
| Atg16l1 | 0.07210322 | 0.05765218 | 0.06223472 | -0.0045825 | 0.6879786 | 0.99999994 |
| Nlgn2 | 0.04614933 | 0.03593 | 0.04051503 | -0.004585 | 0.7743627 | 0.99999994 |
| Tom1l1 | 0.13212738 | 0.10459943 | 0.10918548 | -0.0045861 | 0.61010125 | 0.99999994 |
| 2410022M11 | 0.04869078 | 0.03808708 | 0.04267466 | -0.0045876 | 0.76287051 | 0.99999994 |
| Tdrkh | 0.20249464 | 0.15378766 | 0.15837602 | -0.0045884 | 0.57645911 | 0.99999994 |
| Phf10 | 0.07166077 | 0.05728208 | 0.06187059 | -0.0045885 | 0.68922514 | 0.99999994 |
| Rpl29 | 0.16158842 | 0.12587659 | 0.13046813 | -0.0045915 | 0.5926069 | 0.99999994 |
| Slc2a6 | 0.01354923 | 0.00756488 | 0.01215646 | -0.0045916 | 0.99403408 | 0.99999994 |
| Gpr27 | 0.03222359 | 0.02395573 | 0.02855176 | -0.004596 | 0.85802442 | 0.99999994 |
| Ccdc59 | 0.60853834 | 0.34995136 | 0.35454799 | -0.0045966 | 0.53438525 | 0.99999994 |
| Parp2 | 0.18773373 | 0.14392004 | 0.14851878 | -0.0045987 | 0.58164652 | 0.99999994 |
| Mrps10 | 0.50353798 | 0.31092157 | 0.31552288 | -0.0046013 | 0.53866487 | 0.99999994 |
| Plekhg1 | 0.0282687 | 0.0205073 | 0.02511366 | -0.0046064 | 0.88893999 | 0.99999994 |
| C1qbp | 0.43765308 | 0.28277128 | 0.28737856 | -0.0046073 | 0.5424926 | 0.99999994 |
| Cdk7 | 0.16943361 | 0.13135571 | 0.13596503 | -0.0046093 | 0.58926545 | 0.99999994 |
| Sav1 | 0.03435176 | 0.0257843 | 0.03039436 | -0.0046101 | 0.84365703 | 0.99999994 |
| Wdr41 | 0.08190907 | 0.06564114 | 0.07025169 | -0.0046106 | 0.66889021 | 0.99999994 |
| Prkx | 0.06246344 | 0.04964156 | 0.0542535 | -0.0046119 | 0.71424819 | 0.99999994 |
| Ltbp3 | 0.04570786 | 0.03552665 | 0.04013915 | -0.0046125 | 0.77783484 | 0.99999994 |
| Dmwd | 0.07301969 | 0.05837157 | 0.06298832 | -0.0046167 | 0.68718252 | 0.99999994 |
| Nap1l1 | 0.95324662 | 0.4185132 | 0.42313324 | -0.00462 | 0.52896901 | 0.99999994 |
| Gucd1 | 0.07041941 | 0.0562254 | 0.06084782 | -0.0046224 | 0.69345345 | 0.99999994 |
| Rif1 | 0.18344834 | 0.14098774 | 0.14561237 | -0.0046246 | 0.58371449 | 0.99999994 |
| Fancl | 0.10618352 | 0.08495236 | 0.08957701 | -0.0046246 | 0.63444581 | 0.99999994 |
| Slc25a29 | 0.05253235 | 0.04129751 | 0.04592539 | -0.0046279 | 0.74881265 | 0.99999994 |
| Pcid2 | 0.09176484 | 0.07357668 | 0.07820802 | -0.0046313 | 0.65327497 | 0.99999994 |

| | | | | | | |
|---|---|---|---|---|---|---|
| Tm9sf3 | 0.73807879 | 0.3863441 | 0.39097836 | -0.0046343 | 0.531443 | 0.99999994 |
| Ccser1 | 0.01754504 | 0.01106348 | 0.01569924 | -0.0046358 | 0.97532338 | 0.99999994 |
| Commd9 | 0.48055333 | 0.30140699 | 0.3060457 | -0.0046387 | 0.54018104 | 0.99999994 |
| Ror2 | 0.07103668 | 0.05671021 | 0.0613566 | -0.0046464 | 0.69289387 | 0.99999994 |
| Slc4a8 | 0.05839025 | 0.04620419 | 0.05085086 | -0.0046467 | 0.72848823 | 0.99999994 |
| Srsf5 | 0.92565308 | 0.41606372 | 0.42071209 | -0.0046484 | 0.52931374 | 0.99999994 |
| Mrpl9 | 0.21379701 | 0.16111639 | 0.16576549 | -0.0046491 | 0.57404661 | 0.99999994 |
| Cxxc4 | 0.32608105 | 0.22761581 | 0.23226787 | -0.0046521 | 0.55302961 | 0.99999994 |
| Plaa | 0.24070129 | 0.17816769 | 0.18281998 | -0.0046523 | 0.56725385 | 0.99999994 |
| Mtss1 | 0.32493076 | 0.22699199 | 0.23164606 | -0.0046541 | 0.55319389 | 0.99999994 |
| Itgb1bp1 | 0.27630437 | 0.199619 | 0.2042741 | -0.0046551 | 0.56028375 | 0.99999994 |
| Myo1c | 0.01893761 | 0.01227415 | 0.01692945 | -0.0046553 | 0.96640341 | 0.99999994 |
| Mtf1 | 0.12938466 | 0.10249607 | 0.10715163 | -0.0046556 | 0.61378892 | 0.99999994 |
| Fnip2 | 0.05085277 | 0.03984976 | 0.04450615 | -0.0046564 | 0.75691416 | 0.99999994 |
| Pex12 | 0.0837761 | 0.06710794 | 0.07176665 | -0.0046587 | 0.66716395 | 0.99999994 |
| Usb1 | 0.13228373 | 0.10464155 | 0.10930114 | -0.0046596 | 0.61170494 | 0.99999994 |
| Abcb7 | 0.03733117 | 0.02830564 | 0.03296524 | -0.0046596 | 0.82661392 | 0.99999994 |
| Gm7467 | 0.03183221 | 0.02355168 | 0.02821233 | -0.0046606 | 0.86425184 | 0.99999994 |
| Ddx10 | 0.11403605 | 0.09093776 | 0.09560013 | -0.0046624 | 0.62726837 | 0.99999994 |
| Ldah | 0.15141511 | 0.11856849 | 0.12323392 | -0.0046654 | 0.59947899 | 0.99999994 |
| Ippk | 0.05930492 | 0.04694911 | 0.05161653 | -0.0046674 | 0.72637985 | 0.99999994 |
| Slc40a1 | 0.05808534 | 0.04592571 | 0.05059541 | -0.0046697 | 0.73050734 | 0.99999994 |
| Fhod3 | 0.03264763 | 0.02424659 | 0.02891931 | -0.0046727 | 0.85892904 | 0.99999994 |
| Fbxl3 | 0.07531814 | 0.06019976 | 0.06487436 | -0.0046746 | 0.68425532 | 0.99999994 |
| Samd10 | 0.20838197 | 0.15756746 | 0.16224233 | -0.0046749 | 0.57604957 | 0.99999994 |
| Dgkg | 0.01525065 | 0.00899218 | 0.01366731 | -0.0046751 | 0.98860436 | 0.99999994 |
| Mrps27 | 0.11289085 | 0.09004695 | 0.09472651 | -0.0046796 | 0.62885736 | 0.99999994 |
| Mxra8 | 0.04173555 | 0.03206742 | 0.03674713 | -0.0046797 | 0.80169193 | 0.99999994 |
| Dus3l | 0.13545081 | 0.10695757 | 0.11163765 | -0.0046801 | 0.60989613 | 0.99999994 |
| Myadm | 0.0585754 | 0.04632477 | 0.05100593 | -0.0046812 | 0.72936644 | 0.99999994 |

| | | | | | | |
|---|---|---|---|---|---|---|
| Rnf41 | 0.15520694 | 0.12126251 | 0.12594418 | -0.0046817 | 0.59771386 | 0.99999994 |
| Cdon | 0.03675162 | 0.02778205 | 0.03246595 | -0.0046839 | 0.83155325 | 0.99999994 |
| Rmnd1 | 0.05925908 | 0.04689191 | 0.05157818 | -0.0046863 | 0.72733763 | 0.99999994 |
| Nynrin | 0.03897532 | 0.0296932 | 0.0343796 | -0.0046864 | 0.81788892 | 0.99999994 |
| Pcif1 | 0.43761046 | 0.28267267 | 0.28735935 | -0.0046867 | 0.54322492 | 0.99999994 |
| Fbxl18 | 0.06109933 | 0.04842708 | 0.05311597 | -0.0046889 | 0.72159979 | 0.99999994 |
| Dock8 | 0.02615576 | 0.01857919 | 0.02326936 | -0.0046902 | 0.91014591 | 0.99999994 |
| Tmem80 | 0.19791757 | 0.15065276 | 0.15534467 | -0.0046919 | 0.57966759 | 0.99999994 |
| Zbtb8a | 0.05504849 | 0.04335364 | 0.04804566 | -0.004692 | 0.74216779 | 0.99999994 |
| Trmt61a | 0.03467403 | 0.02597935 | 0.03067293 | -0.0046936 | 0.84579323 | 0.99999994 |
| Plekhh2 | 0.09674569 | 0.07749666 | 0.08219086 | -0.0046942 | 0.64808939 | 0.99999994 |
| Lrrc45 | 0.08691743 | 0.06961256 | 0.07430741 | -0.0046949 | 0.66296137 | 0.99999994 |
| Trim46 | 0.03448542 | 0.0258126 | 0.03050992 | -0.0046973 | 0.84727484 | 0.99999994 |
| Ercc6l2 | 0.04943168 | 0.03860258 | 0.04330289 | -0.0047003 | 0.76501073 | 0.99999994 |
| Plekha5 | 0.13279009 | 0.10497463 | 0.10967551 | -0.0047009 | 0.61229395 | 0.99999994 |
| Cnep1r1 | 0.20105858 | 0.15272604 | 0.15742732 | -0.0047013 | 0.57878258 | 0.99999994 |
| Cdk8 | 0.22852411 | 0.17048993 | 0.17519297 | -0.004703 | 0.57090925 | 0.99999994 |
| Xpr1 | 0.10920627 | 0.08719891 | 0.09190465 | -0.0047057 | 0.63338108 | 0.99999994 |
| Ikbkg | 0.05608173 | 0.04420712 | 0.04891431 | -0.0047072 | 0.73909727 | 0.99999994 |
| Tti1 | 0.03296622 | 0.02448742 | 0.02919533 | -0.0047079 | 0.85844854 | 0.99999994 |
| Zfp738 | 0.04719723 | 0.03669692 | 0.04140638 | -0.0047095 | 0.77549842 | 0.99999994 |
| Mllt3 | 0.07583203 | 0.06058322 | 0.06529527 | -0.0047121 | 0.68451362 | 0.99999994 |
| Pxk | 0.07807426 | 0.06241377 | 0.06712856 | -0.0047148 | 0.67993298 | 0.99999994 |
| Pptc7 | 0.04913067 | 0.03833157 | 0.04304772 | -0.0047162 | 0.76707467 | 0.99999994 |
| Dcxr | 0.05477726 | 0.04310058 | 0.04781743 | -0.0047169 | 0.74428205 | 0.99999994 |
| Elf1 | 0.1930834 | 0.14740053 | 0.1521188 | -0.0047183 | 0.58178436 | 0.99999994 |
| Rnf4 | 0.28536464 | 0.20482024 | 0.20953933 | -0.0047191 | 0.55958203 | 0.99999994 |
| Gm15545 | 0.04127577 | 0.03163421 | 0.03635335 | -0.0047191 | 0.80622905 | 0.99999994 |
| Apmap | 0.18628405 | 0.14281783 | 0.14753782 | -0.00472 | 0.58431648 | 0.99999994 |
| Zfp280b | 0.04058789 | 0.03104174 | 0.03576379 | -0.0047221 | 0.81026393 | 0.99999994 |

| | | | | | | |
|---|---|---|---|---|---|---|
| Inpp5f | 0.06403748 | 0.05083944 | 0.05556357 | -0.0047241 | 0.71428201 | 0.99999994 |
| Arel1 | 0.03316696 | 0.02464288 | 0.02936918 | -0.0047263 | 0.8579582 | 0.99999994 |
| Myo5a | 0.02210817 | 0.01499098 | 0.0197218 | -0.0047308 | 0.94483508 | 0.99999994 |
| Aldh3a2 | 0.05431402 | 0.04269409 | 0.04742747 | -0.0047334 | 0.74675776 | 0.99999994 |
| Uap1l1 | 0.03783438 | 0.02866419 | 0.03339844 | -0.0047343 | 0.82729849 | 0.99999994 |
| Flcn | 0.06130048 | 0.04854876 | 0.05328384 | -0.0047351 | 0.72291356 | 0.99999994 |
| Abcf3 | 0.2640915 | 0.19231865 | 0.19705385 | -0.0047352 | 0.56354053 | 0.99999994 |
| Mri1 | 0.09009161 | 0.07212892 | 0.07686434 | -0.0047354 | 0.6591271 | 0.99999994 |
| Ssh2 | 0.07241279 | 0.05775387 | 0.06248937 | -0.0047355 | 0.69301876 | 0.99999994 |
| Ftx | 0.02090468 | 0.01392662 | 0.01866327 | -0.0047366 | 0.95442383 | 0.99999994 |
| Asb7 | 0.09768443 | 0.07820073 | 0.08293866 | -0.0047379 | 0.64811977 | 0.99999994 |
| Sppl2b | 0.07418509 | 0.05920737 | 0.06394531 | -0.0047379 | 0.68906216 | 0.99999994 |
| Kiz | 0.17693452 | 0.13641765 | 0.14115612 | -0.0047385 | 0.58840631 | 0.99999994 |
| Nsun6 | 0.03827812 | 0.02903939 | 0.03378021 | -0.0047408 | 0.82489309 | 0.99999994 |
| Acad11 | 0.04849895 | 0.03776592 | 0.0425119 | -0.004746 | 0.77129212 | 0.99999994 |
| Elac2 | 0.07890019 | 0.06305337 | 0.06780252 | -0.0047492 | 0.67947719 | 0.99999994 |
| 6030458C11I | 0.03972262 | 0.03027172 | 0.03502143 | -0.0047497 | 0.81667227 | 0.99999994 |
| Nub1 | 0.18610762 | 0.14266776 | 0.14741829 | -0.0047505 | 0.58492137 | 0.99999994 |
| Fam234b | 0.04857478 | 0.03782525 | 0.04257624 | -0.004751 | 0.77118941 | 0.99999994 |
| AI480526 | 0.05314643 | 0.04169018 | 0.04644349 | -0.0047533 | 0.75214406 | 0.99999994 |
| Gtf3c4 | 0.08986034 | 0.07192332 | 0.0766784 | -0.0047551 | 0.66011714 | 0.99999994 |
| Med11 | 0.09826638 | 0.07864351 | 0.0834018 | -0.0047583 | 0.64793903 | 0.99999994 |
| Ncbp3 | 0.18025594 | 0.1386727 | 0.14343421 | -0.0047615 | 0.58744121 | 0.99999994 |
| Fbxw7 | 0.09287146 | 0.07433238 | 0.07909509 | -0.0047627 | 0.65572097 | 0.99999994 |
| Dgke | 0.03205501 | 0.02364284 | 0.02840558 | -0.0047627 | 0.86780285 | 0.99999994 |
| Cerkl | 0.06341826 | 0.05028531 | 0.05504852 | -0.0047632 | 0.71767825 | 0.99999994 |
| Ctdsp1 | 0.11305797 | 0.09008885 | 0.0948541 | -0.0047652 | 0.63096087 | 0.99999994 |
| Tmem135 | 0.08271581 | 0.06613749 | 0.07090675 | -0.0047693 | 0.67282001 | 0.99999994 |
| Tmtc3 | 0.06573014 | 0.0521993 | 0.05696938 | -0.0047701 | 0.71135252 | 0.99999994 |
| Ipo8 | 0.09829429 | 0.07865269 | 0.083424 | -0.0047713 | 0.64828736 | 0.99999994 |

| | | | | | | |
|---|---|---|---|---|---|---|
| Myc | 0.02301702 | 0.01574844 | 0.02052005 | -0.0047716 | 0.93916824 | 0.99999994 |
| Cox10 | 0.08586095 | 0.06868091 | 0.07345406 | -0.0047732 | 0.66732545 | 0.99999994 |
| Usp32 | 0.04798276 | 0.03729763 | 0.04207374 | -0.0047761 | 0.77505968 | 0.99999994 |
| Nup153 | 0.07159959 | 0.05704361 | 0.06182021 | -0.0047766 | 0.69648253 | 0.99999994 |
| Gm17767 | 0.0115503 | 0.00559802 | 0.01037699 | -0.004779 | 0.99891342 | 0.99999994 |
| Impa1 | 0.37754384 | 0.25415051 | 0.25893023 | -0.0047797 | 0.54889616 | 0.99999994 |
| Rnf182 | 0.01438283 | 0.00811655 | 0.01289712 | -0.0047806 | 0.99319338 | 0.99999994 |
| Mef2d | 0.07324903 | 0.058396 | 0.06317676 | -0.0047808 | 0.69276555 | 0.99999994 |
| 1700113A16I | 0.05520648 | 0.0433972 | 0.04817855 | -0.0047814 | 0.74556336 | 0.99999994 |
| Det1 | 0.0471156 | 0.03655175 | 0.04133699 | -0.0047852 | 0.77951356 | 0.99999994 |
| Pus7 | 0.06508798 | 0.05165037 | 0.05643641 | -0.004786 | 0.71378854 | 0.99999994 |
| Fbxo6 | 0.15502681 | 0.12102674 | 0.1258158 | -0.0047891 | 0.60000823 | 0.99999994 |
| Mto1 | 0.07829657 | 0.06251802 | 0.06731004 | -0.004792 | 0.6822099 | 0.99999994 |
| Fbxo17 | 0.04982421 | 0.03884104 | 0.04363548 | -0.0047944 | 0.76772207 | 0.99999994 |
| Fktn | 0.08195913 | 0.06549782 | 0.07029236 | -0.0047945 | 0.6750929 | 0.99999994 |
| Mat1a | 0.01327292 | 0.00711294 | 0.01191077 | -0.0047978 | 0.99633292 | 0.99999994 |
| Ext1 | 0.02110303 | 0.01403991 | 0.01883785 | -0.0047979 | 0.95499582 | 0.99999994 |
| Mcph1 | 0.04385246 | 0.03375888 | 0.03855701 | -0.0047981 | 0.79625935 | 0.99999994 |
| C730034F03I | 0.03836223 | 0.02905146 | 0.03385255 | -0.0048011 | 0.82742242 | 0.99999994 |
| Arl5a | 0.09285408 | 0.07427997 | 0.07908118 | -0.0048012 | 0.65693914 | 0.99999994 |
| Fam219b | 0.05953157 | 0.04700443 | 0.05180612 | -0.0048017 | 0.73136611 | 0.99999994 |
| Lnx1 | 0.03661198 | 0.02754313 | 0.03234559 | -0.0048025 | 0.83849527 | 0.99999994 |
| Zfp395 | 0.08044777 | 0.06425465 | 0.06906341 | -0.0048088 | 0.6784861 | 0.99999994 |
| Slc36a4 | 0.05060175 | 0.0394843 | 0.04429377 | -0.0048095 | 0.76508493 | 0.99999994 |
| Slc4a1ap | 0.13494215 | 0.10645354 | 0.11126319 | -0.0048097 | 0.61322688 | 0.99999994 |
| Gfm2 | 0.09256428 | 0.07403764 | 0.07884898 | -0.0048113 | 0.65769073 | 0.99999994 |
| Tgfbrap1 | 0.03048958 | 0.0222336 | 0.02704656 | -0.004813 | 0.88189338 | 0.99999994 |
| Cep44 | 0.11354563 | 0.09041238 | 0.09522621 | -0.0048138 | 0.63174882 | 0.99999994 |
| Crtc3 | 0.11791549 | 0.09373017 | 0.09854719 | -0.004817 | 0.62754766 | 0.99999994 |
| Snx13 | 0.05322401 | 0.0416883 | 0.04650892 | -0.0048206 | 0.75488026 | 0.99999994 |

| | | | | | | |
|---|---|---|---|---|---|---|
| Tspan9 | 0.06064433 | 0.04791499 | 0.0527361 | -0.0048211 | 0.72857927 | 0.99999994 |
| Spink10 | 0.01710704 | 0.01049057 | 0.01531182 | -0.0048212 | 0.98195421 | 0.99999994 |
| Faap100 | 0.10964576 | 0.08742022 | 0.09224213 | -0.0048219 | 0.63606392 | 0.99999994 |
| Ttyh3 | 0.11917328 | 0.0946767 | 0.09949865 | -0.0048219 | 0.62649465 | 0.99999994 |
| Mxra7 | 0.05351746 | 0.04193371 | 0.04675634 | -0.0048226 | 0.75382101 | 0.99999994 |
| Mapkapk2 | 0.07369564 | 0.05872037 | 0.06354357 | -0.0048232 | 0.69330596 | 0.99999994 |
| Ankrd46 | 0.12547286 | 0.09941075 | 0.10423487 | -0.0048241 | 0.62098325 | 0.99999994 |
| Bbs9 | 0.0385771 | 0.02921312 | 0.03403732 | -0.0048242 | 0.82726744 | 0.99999994 |
| Dyrk1a | 0.07446351 | 0.05934788 | 0.06417373 | -0.0048259 | 0.69165851 | 0.99999994 |
| Rab5b | 0.13549306 | 0.10684271 | 0.11166873 | -0.004826 | 0.61320021 | 0.99999994 |
| Pa2g4 | 0.62370085 | 0.35480505 | 0.35963353 | -0.0048285 | 0.53560563 | 0.99999994 |
| Clk4 | 0.26705749 | 0.19399117 | 0.19882077 | -0.0048296 | 0.56422547 | 0.99999994 |
| Uba3 | 0.32201404 | 0.22523385 | 0.23006428 | -0.0048304 | 0.55557398 | 0.99999994 |
| Sema4d | 0.03348265 | 0.02481112 | 0.0296425 | -0.0048314 | 0.86101895 | 0.99999994 |
| St8sia5 | 0.01762438 | 0.01093597 | 0.01576939 | -0.0048334 | 0.97933554 | 0.99999994 |
| Cwc27 | 0.14595512 | 0.11446787 | 0.11930205 | -0.0048342 | 0.60631186 | 0.99999994 |
| Rapgef1 | 0.04760853 | 0.0369148 | 0.04175589 | -0.0048411 | 0.77986121 | 0.99999994 |
| Itpr3 | 0.1797959 | 0.13827714 | 0.1431194 | -0.0048423 | 0.58909179 | 0.99999994 |
| Gphn | 0.09417477 | 0.07529593 | 0.08013823 | -0.0048423 | 0.65623378 | 0.99999994 |
| Atat1 | 0.30099503 | 0.21359955 | 0.21844234 | -0.0048428 | 0.55865856 | 0.99999994 |
| Rae1 | 0.334478 | 0.23192752 | 0.23677149 | -0.004844 | 0.55415997 | 0.99999994 |
| Nkap | 0.18423465 | 0.14130148 | 0.14614716 | -0.0048457 | 0.58733665 | 0.99999994 |
| Ppil3 | 0.140659 | 0.11060785 | 0.11545404 | -0.0048462 | 0.61003147 | 0.99999994 |
| Mib2 | 0.04571679 | 0.03530023 | 0.04014676 | -0.0048465 | 0.78917379 | 0.99999994 |
| Clasp1 | 0.11309545 | 0.09003534 | 0.09488271 | -0.0048474 | 0.63309439 | 0.99999994 |
| Kpna6 | 0.04728416 | 0.03663006 | 0.04148026 | -0.0048502 | 0.78180809 | 0.99999994 |
| Git2 | 0.13032429 | 0.10299141 | 0.10784951 | -0.0048581 | 0.6178493 | 0.99999994 |
| Zfp654 | 0.03902632 | 0.02956494 | 0.03442342 | -0.0048585 | 0.82625509 | 0.99999994 |
| Ppip5k1 | 0.03517322 | 0.02624136 | 0.0311042 | -0.0048628 | 0.85098227 | 0.99999994 |
| Slc15a4 | 0.06366743 | 0.05039277 | 0.05525582 | -0.004863 | 0.72100318 | 0.99999994 |

| | | | | | | |
|---|---|---|---|---|---|---|
| Ccser2 | 0.10496507 | 0.08376925 | 0.08863551 | -0.0048663 | 0.64261091 | 0.99999994 |
| Trnt1 | 0.19764302 | 0.15028992 | 0.15516213 | -0.0048722 | 0.58278159 | 0.99999994 |
| Morn2 | 0.28144347 | 0.20239744 | 0.20727016 | -0.0048727 | 0.5621745 | 0.99999994 |
| Pigb | 0.03438266 | 0.02554792 | 0.03042107 | -0.0048732 | 0.85685194 | 0.99999994 |
| Pdss2 | 0.07247852 | 0.05766713 | 0.06254343 | -0.0048763 | 0.69811045 | 0.99999994 |
| Rnf113a1 | 0.02993177 | 0.02168181 | 0.02656162 | -0.0048798 | 0.88931322 | 0.99999994 |
| Ipp | 0.08322889 | 0.06644253 | 0.07132301 | -0.0048805 | 0.67561803 | 0.99999994 |
| Atrnl1 | 0.02676563 | 0.01892154 | 0.02380222 | -0.0048807 | 0.91385722 | 0.99999994 |
| P4ha2 | 0.05164538 | 0.04029543 | 0.04517629 | -0.0048809 | 0.76397736 | 0.99999994 |
| Rev3l | 0.22290059 | 0.16673805 | 0.17162005 | -0.004882 | 0.57509094 | 0.99999994 |
| Prkab1 | 0.16317177 | 0.12669943 | 0.13158314 | -0.0048837 | 0.59756557 | 0.99999994 |
| Rpl36-ps3 | 0.14837042 | 0.11616166 | 0.12104565 | -0.004884 | 0.60587086 | 0.99999994 |
| Mterf3 | 0.20156455 | 0.15287697 | 0.15776182 | -0.0048849 | 0.58164512 | 0.99999994 |
| Fads6 | 0.03056818 | 0.0222288 | 0.02711486 | -0.0048861 | 0.88482671 | 0.99999994 |
| Zfp62 | 0.11725652 | 0.09316073 | 0.09804793 | -0.0048872 | 0.62997479 | 0.99999994 |
| Naa30 | 0.0711382 | 0.05655043 | 0.06144024 | -0.0048898 | 0.70185935 | 0.99999994 |
| Dennd5a | 0.06412052 | 0.05074136 | 0.05563262 | -0.0048913 | 0.72080393 | 0.99999994 |
| Faah | 0.17764651 | 0.13675332 | 0.14164549 | -0.0048922 | 0.59091458 | 0.99999994 |
| Med9os | 0.03387576 | 0.02508948 | 0.02998269 | -0.0048932 | 0.86133246 | 0.99999994 |
| Crybg3 | 0.03058939 | 0.02223969 | 0.0271333 | -0.0048936 | 0.88502812 | 0.99999994 |
| Ilf3 | 0.2141835 | 0.16111179 | 0.16601579 | -0.004904 | 0.57793923 | 0.99999994 |
| Hnrnph2 | 0.42843157 | 0.27828682 | 0.28319226 | -0.0049054 | 0.54589668 | 0.99999994 |
| Zmat3 | 0.08062145 | 0.06429734 | 0.06920476 | -0.0049074 | 0.6815411 | 0.99999994 |
| Ap2m1 | 1.34098261 | 0.41771466 | 0.42262374 | -0.0049091 | 0.53081487 | 0.99999994 |
| Pyurf | 0.30803901 | 0.21746704 | 0.22238113 | -0.0049141 | 0.55846936 | 0.99999994 |
| Specc1 | 0.03059002 | 0.02221961 | 0.02713384 | -0.0049142 | 0.88600216 | 0.99999994 |
| Terf2ip | 0.18385507 | 0.1409748 | 0.14588908 | -0.0049143 | 0.58870689 | 0.99999994 |
| Plekha1 | 0.08688214 | 0.06936116 | 0.07427892 | -0.0049178 | 0.67028331 | 0.99999994 |
| Tmem70 | 0.36743146 | 0.24894717 | 0.25386499 | -0.0049178 | 0.55129958 | 0.99999994 |
| Fbxl5 | 0.11716525 | 0.09305851 | 0.09797874 | -0.0049202 | 0.63091018 | 0.99999994 |

| | | | | | | |
|---|---|---|---|---|---|---|
| Etv3 | 0.05747969 | 0.04516546 | 0.05008772 | -0.0049223 | 0.74348865 | 0.99999994 |
| Zfp932 | 0.05110879 | 0.03979783 | 0.04472269 | -0.0049249 | 0.76822029 | 0.99999994 |
| Rps27a | 3.25597984 | 0.28459871 | 0.28952437 | -0.0049257 | 0.5450815 | 0.99999994 |
| Rfx3 | 0.19297364 | 0.14711892 | 0.15204527 | -0.0049263 | 0.58537733 | 0.99999994 |
| Ubl4a | 0.40931168 | 0.26938438 | 0.27431096 | -0.0049266 | 0.5475791 | 0.99999994 |
| Kcnb1 | 0.02239614 | 0.01504407 | 0.01997483 | -0.0049308 | 0.94982041 | 0.99999994 |
| Fn1 | 0.0629458 | 0.04972289 | 0.05465526 | -0.0049324 | 0.72598121 | 0.99999994 |
| Cerk | 0.02338932 | 0.01591262 | 0.02084676 | -0.0049341 | 0.94242578 | 0.99999994 |
| Tspan5 | 0.16276159 | 0.12635839 | 0.13129456 | -0.0049362 | 0.59880445 | 0.99999994 |
| Pcna | 0.07168669 | 0.05695221 | 0.06189192 | -0.0049397 | 0.70237949 | 0.99999994 |
| Slc35e4 | 0.02046987 | 0.01333849 | 0.01828042 | -0.0049419 | 0.96402846 | 0.99999994 |
| Zswim1 | 0.06071948 | 0.04785443 | 0.05279886 | -0.0049444 | 0.73347078 | 0.99999994 |
| Cherp | 0.09120595 | 0.07281497 | 0.07775953 | -0.0049446 | 0.66394843 | 0.99999994 |
| Eps15 | 0.077122 | 0.06140561 | 0.06635062 | -0.004945 | 0.69008396 | 0.99999994 |
| Slc44a1 | 0.06464868 | 0.05112469 | 0.05607156 | -0.0049469 | 0.72148257 | 0.99999994 |
| Cpsf7 | 0.09629666 | 0.07688024 | 0.08183284 | -0.0049526 | 0.65647036 | 0.99999994 |
| Tmem231 | 0.04455441 | 0.03420229 | 0.03915603 | -0.0049537 | 0.80013848 | 0.99999994 |
| Trp53bp2 | 0.09545496 | 0.07620699 | 0.08116118 | -0.0049542 | 0.65774385 | 0.99999994 |
| Rrnad1 | 0.13047793 | 0.10300757 | 0.10796352 | -0.0049559 | 0.62002743 | 0.99999994 |
| Dolk | 0.1158795 | 0.09204438 | 0.09700293 | -0.0049585 | 0.63316826 | 0.99999994 |
| Mrps11 | 0.58570578 | 0.34166989 | 0.34662937 | -0.0049595 | 0.5379368 | 0.99999994 |
| Zfp954 | 0.07957286 | 0.0633899 | 0.06835088 | -0.004961 | 0.68549665 | 0.99999994 |
| Fam229b | 0.08196252 | 0.06533212 | 0.07029512 | -0.004963 | 0.68080607 | 0.99999994 |
| Dcun1d3 | 0.06870939 | 0.054467 | 0.05943623 | -0.0049692 | 0.71106945 | 0.99999994 |
| Dcaf13 | 0.18208791 | 0.13971584 | 0.14468552 | -0.0049697 | 0.59042307 | 0.99999994 |
| Nphp1 | 0.12904574 | 0.10192829 | 0.10689965 | -0.0049714 | 0.62154987 | 0.99999994 |
| Snupn | 0.08452724 | 0.06740065 | 0.07237512 | -0.0049745 | 0.67634545 | 0.99999994 |
| 1700029J07F | 0.03428826 | 0.02536437 | 0.03033946 | -0.0049751 | 0.86247675 | 0.99999994 |
| Ppp2r1b | 0.08756619 | 0.06985126 | 0.07483086 | -0.0049796 | 0.67109734 | 0.99999994 |
| Lpin1 | 0.05303785 | 0.04137224 | 0.04635191 | -0.0049797 | 0.76272804 | 0.99999994 |

| | | | | | | |
|---|---|---|---|---|---|---|
| Usp53 | 0.08937646 | 0.07130786 | 0.07628917 | -0.0049813 | 0.66808381 | 0.99999994 |
| Cela2a | 0.1063786 | 0.08474202 | 0.08972758 | -0.0049856 | 0.64425261 | 0.99999994 |
| Ube2q2 | 0.10616674 | 0.08457669 | 0.08956405 | -0.0049874 | 0.64455392 | 0.99999994 |
| 1110006O24 | 0.05487529 | 0.0429112 | 0.04789993 | -0.0049887 | 0.75592444 | 0.99999994 |
| Sh2b1 | 0.12045832 | 0.09547846 | 0.10046872 | -0.0049903 | 0.62953402 | 0.99999994 |
| Ttc7b | 0.04385842 | 0.03357121 | 0.0385621 | -0.0049909 | 0.80551783 | 0.99999994 |
| Rrp1b | 0.08580679 | 0.06841014 | 0.07341028 | -0.0050001 | 0.67486236 | 0.99999994 |
| Ptrhd1 | 0.22146847 | 0.16570037 | 0.17070478 | -0.0050044 | 0.57735798 | 0.99999994 |
| Kansl2 | 0.28611738 | 0.20496754 | 0.2099732 | -0.0050057 | 0.56304121 | 0.99999994 |
| Utp14a | 0.09660504 | 0.07707095 | 0.08207873 | -0.0050078 | 0.65767158 | 0.99999994 |
| Uqcc1 | 0.13089039 | 0.10326158 | 0.1082694 | -0.0050078 | 0.62091265 | 0.99999994 |
| Nek9 | 0.07132032 | 0.05657798 | 0.06159025 | -0.0050123 | 0.70598745 | 0.99999994 |
| Fam98a | 0.1641975 | 0.12729082 | 0.13230395 | -0.0050131 | 0.59956301 | 0.99999994 |
| Gcat | 0.26044168 | 0.18985349 | 0.19486807 | -0.0050146 | 0.56800187 | 0.99999994 |
| Acly | 0.9345079 | 0.41653779 | 0.42155265 | -0.0050149 | 0.53155729 | 0.99999994 |
| Fam117b | 0.08024885 | 0.06388519 | 0.06890148 | -0.0050163 | 0.6860228 | 0.99999994 |
| Apba3 | 0.0831338 | 0.06622873 | 0.07124588 | -0.0050172 | 0.68037302 | 0.99999994 |
| Srsf1 | 0.60551597 | 0.34849949 | 0.35351699 | -0.0050175 | 0.53763383 | 0.99999994 |
| Efhc1 | 0.02984869 | 0.02147141 | 0.02648936 | -0.0050179 | 0.89633312 | 0.99999994 |
| Cpsf1 | 0.05858509 | 0.04599553 | 0.05101403 | -0.0050185 | 0.74370778 | 0.99999994 |
| Ctps | 0.12947351 | 0.10219623 | 0.10721767 | -0.0050214 | 0.62238257 | 0.99999994 |
| Rreb1 | 0.05020674 | 0.03893758 | 0.04395943 | -0.0050218 | 0.77649517 | 0.99999994 |
| Zfhx3 | 0.1871384 | 0.14309083 | 0.14811622 | -0.0050254 | 0.58933765 | 0.99999994 |
| Pskh1 | 0.08416678 | 0.06705758 | 0.0720832 | -0.0050256 | 0.67870462 | 0.99999994 |
| Tango2 | 0.09515674 | 0.07589705 | 0.08092303 | -0.005026 | 0.66034859 | 0.99999994 |
| Man1a | 0.02339362 | 0.0158221 | 0.02085053 | -0.0050284 | 0.94578456 | 0.99999994 |
| Sgf29 | 0.11714133 | 0.09293088 | 0.0979606 | -0.0050297 | 0.63373675 | 0.99999994 |
| Bap1 | 0.08897251 | 0.07093123 | 0.07596406 | -0.0050328 | 0.67039421 | 0.99999994 |
| Acbd5 | 0.23078467 | 0.17158571 | 0.17662021 | -0.0050345 | 0.57524237 | 0.99999994 |
| Tubgcp5 | 0.08116729 | 0.06461389 | 0.06964878 | -0.0050349 | 0.68480784 | 0.99999994 |

| | | | | | | |
|---|---|---|---|---|---|---|
| Slc17a9 | 0.01773257 | 0.0108245 | 0.01586504 | -0.0050405 | 0.98277725 | 0.99999994 |
| Dnajb2 | 0.30610235 | 0.21626139 | 0.2213026 | -0.0050412 | 0.56026065 | 0.99999994 |
| Cabp7 | 0.02097293 | 0.01368172 | 0.01872334 | -0.0050416 | 0.96346172 | 0.99999994 |
| Scpep1 | 0.54853969 | 0.32803458 | 0.33307791 | -0.0050433 | 0.54014056 | 0.99999994 |
| Cpox | 0.22875664 | 0.17029153 | 0.17534002 | -0.0050485 | 0.57599326 | 0.99999994 |
| Fam216a | 0.59473833 | 0.34475023 | 0.34979936 | -0.0050491 | 0.53827154 | 0.99999994 |
| Uhmk1 | 0.06588896 | 0.05205155 | 0.05710113 | -0.0050496 | 0.72194764 | 0.99999994 |
| Arhgef10 | 0.0241609 | 0.01647268 | 0.02152334 | -0.0050507 | 0.94085361 | 0.99999994 |
| Tmem62 | 0.0557521 | 0.04358608 | 0.04863732 | -0.0050512 | 0.7553104 | 0.99999994 |
| Zbtb7c | 0.0503892 | 0.03906184 | 0.04411388 | -0.005052 | 0.77706057 | 0.99999994 |
| Fbrs | 0.06394433 | 0.05043306 | 0.05548613 | -0.0050531 | 0.72780488 | 0.99999994 |
| Ezh1 | 0.0637548 | 0.0502752 | 0.0553285 | -0.0050533 | 0.72838727 | 0.99999994 |
| Syn1 | 0.03124923 | 0.02265166 | 0.0277064 | -0.0050547 | 0.88769924 | 0.99999994 |
| Mcl1 | 0.20285079 | 0.15355487 | 0.15861096 | -0.0050561 | 0.58401938 | 0.99999994 |
| Parp10 | 0.03422547 | 0.02522776 | 0.03028516 | -0.0050574 | 0.86683709 | 0.99999994 |
| Arhgap39 | 0.06348407 | 0.05004433 | 0.05510328 | -0.005059 | 0.72943679 | 0.99999994 |
| Rnaset2b | 0.1506886 | 0.11765307 | 0.12271274 | -0.0050597 | 0.60813216 | 0.99999994 |
| Sema3c | 0.04550249 | 0.03490022 | 0.03996422 | -0.005064 | 0.80051153 | 0.99999994 |
| Lmtk2 | 0.02997387 | 0.02153407 | 0.02659823 | -0.0050642 | 0.89748205 | 0.99999994 |
| Snx1 | 0.26796449 | 0.19429365 | 0.19935944 | -0.0050658 | 0.5671568 | 0.99999994 |
| Plxnb1 | 0.07039382 | 0.05576022 | 0.06082672 | -0.0050665 | 0.71035945 | 0.99999994 |
| Gpr119 | 0.07982156 | 0.0634851 | 0.06855351 | -0.0050684 | 0.68868564 | 0.99999994 |
| 1700123O20 | 0.43709011 | 0.28205458 | 0.28712474 | -0.0050702 | 0.54678411 | 0.99999994 |
| Frmd4b | 0.05816757 | 0.04559265 | 0.05066431 | -0.0050717 | 0.74739753 | 0.99999994 |
| Mmaa | 0.08784385 | 0.06997951 | 0.07505475 | -0.0050752 | 0.67368337 | 0.99999994 |
| Kcnma1 | 0.08702048 | 0.06931452 | 0.07439059 | -0.0050761 | 0.67515715 | 0.99999994 |
| Ercc3 | 0.05854465 | 0.04590198 | 0.05098017 | -0.0050782 | 0.74634987 | 0.99999994 |
| Asb6 | 0.08883115 | 0.07077189 | 0.07585024 | -0.0050784 | 0.67207816 | 0.99999994 |
| C030014I23R | 0.03543796 | 0.02625402 | 0.0313328 | -0.0050788 | 0.85969278 | 0.99999994 |
| Zgpat | 0.12334164 | 0.09755746 | 0.10263796 | -0.0050805 | 0.62910529 | 0.99999994 |

| | | | | | | |
|---|---|---|---|---|---|---|
| Ccdc184 | 0.06596748 | 0.05208375 | 0.05716625 | -0.0050825 | 0.72300757 | 0.99999994 |
| Rnf40 | 0.06573344 | 0.05188419 | 0.05697212 | -0.0050879 | 0.7238944 | 0.99999994 |
| Ncoa5 | 0.10175051 | 0.08105437 | 0.08614262 | -0.0050883 | 0.65290527 | 0.99999994 |
| Naa25 | 0.0877877 | 0.06991967 | 0.07500948 | -0.0050898 | 0.67424709 | 0.99999994 |
| Cdkl2 | 0.07713827 | 0.06127392 | 0.06636393 | -0.00509 | 0.6951608 | 0.99999994 |
| Gpatch8 | 0.28391218 | 0.20361013 | 0.20870068 | -0.0050906 | 0.56448859 | 0.99999994 |
| Fbxo33 | 0.06994891 | 0.05536792 | 0.06045975 | -0.0050918 | 0.71246095 | 0.99999994 |
| Usp12 | 0.0754342 | 0.05987438 | 0.06496945 | -0.0050951 | 0.69916776 | 0.99999994 |
| Cwf19l2 | 0.10112353 | 0.08055818 | 0.08565488 | -0.0050967 | 0.65397375 | 0.99999994 |
| Tnrc18 | 0.08781227 | 0.06993052 | 0.0750293 | -0.0050988 | 0.67449075 | 0.99999994 |
| Fbxo4 | 0.05065761 | 0.03923624 | 0.04434104 | -0.0051048 | 0.77825666 | 0.99999994 |
| Parp1 | 0.28530649 | 0.20439643 | 0.2095058 | -0.0051094 | 0.56447826 | 0.99999994 |
| Rnf135 | 0.08364439 | 0.06654612 | 0.0716599 | -0.0051138 | 0.68261008 | 0.99999994 |
| Spata7 | 0.08881278 | 0.07072102 | 0.07583545 | -0.0051144 | 0.67325238 | 0.99999994 |
| Gm20342 | 0.0573846 | 0.0448925 | 0.05000797 | -0.0051155 | 0.75203254 | 0.99999994 |
| Lin7c | 0.30055496 | 0.21307902 | 0.21819481 | -0.0051158 | 0.56200894 | 0.99999994 |
| Tmed5 | 0.19078531 | 0.14545669 | 0.15057626 | -0.0051196 | 0.58952184 | 0.99999994 |
| Chtf8 | 0.21232955 | 0.15969411 | 0.16481375 | -0.0051196 | 0.58190428 | 0.99999994 |
| Wipf2 | 0.05443611 | 0.04241029 | 0.04753027 | -0.00512 | 0.76332076 | 0.99999994 |
| Map4 | 0.07361159 | 0.05835332 | 0.06347455 | -0.0051212 | 0.70438206 | 0.99999994 |
| Gpc4 | 0.20980896 | 0.15805039 | 0.16317373 | -0.0051233 | 0.58277461 | 0.99999994 |
| Map3k5 | 0.02365497 | 0.0159531 | 0.02107978 | -0.0051267 | 0.94726045 | 0.99999994 |
| Vps37c | 0.03315967 | 0.02422711 | 0.02936287 | -0.0051358 | 0.87783692 | 0.99999994 |
| Heg1 | 0.02594856 | 0.01795044 | 0.02308822 | -0.0051378 | 0.93072693 | 0.99999994 |
| Gm9828 | 0.02274678 | 0.01514369 | 0.0202828 | -0.0051391 | 0.95415417 | 0.99999994 |
| Nup43 | 0.14125402 | 0.11074839 | 0.115888 | -0.0051396 | 0.61608264 | 0.99999994 |
| Cmc1 | 0.25483037 | 0.18634111 | 0.19148272 | -0.0051416 | 0.57092655 | 0.99999994 |
| Inha | 0.03841862 | 0.02875854 | 0.03390104 | -0.0051425 | 0.84368363 | 0.99999994 |
| Elmod3 | 0.05457956 | 0.04250488 | 0.04765103 | -0.0051462 | 0.76388776 | 0.99999994 |
| Trmt1 | 0.11853431 | 0.09386862 | 0.09901554 | -0.0051469 | 0.63533005 | 0.99999994 |

| | | | | | | |
|---|---|---|---|---|---|---|
| Zbtb4 | 0.04669108 | 0.0358248 | 0.040976 | -0.0051512 | 0.79864393 | 0.99999994 |
| Slc26a2 | 0.02768055 | 0.01944867 | 0.0246008 | -0.0051521 | 0.91834712 | 0.99999994 |
| Pkig | 0.58176751 | 0.34007518 | 0.34523276 | -0.0051576 | 0.53960639 | 0.99999994 |
| Aurkaip1 | 0.45758373 | 0.29105544 | 0.29621325 | -0.0051578 | 0.54613558 | 0.99999994 |
| Tigd2 | 0.12783488 | 0.10084039 | 0.10599828 | -0.0051579 | 0.6269928 | 0.99999994 |
| Megf8 | 0.03391067 | 0.02485235 | 0.03001289 | -0.0051605 | 0.87379611 | 0.99999994 |
| Rbbp6 | 0.57206882 | 0.33659257 | 0.34175319 | -0.0051606 | 0.54003181 | 0.99999994 |
| Rabepk | 0.13711924 | 0.10770087 | 0.11286372 | -0.0051629 | 0.6196224 | 0.99999994 |
| Dcun1d2 | 0.04407011 | 0.03357985 | 0.0387428 | -0.005163 | 0.81247168 | 0.99999994 |
| Phkb | 0.06874176 | 0.05429236 | 0.05946297 | -0.0051706 | 0.71863929 | 0.99999994 |
| Yae1d1 | 0.23730014 | 0.1755341 | 0.18070512 | -0.005171 | 0.57553235 | 0.99999994 |
| Smc4 | 0.13255166 | 0.10432698 | 0.10949927 | -0.0051723 | 0.6233984 | 0.99999994 |
| Ciz1 | 0.09806434 | 0.07806694 | 0.08324105 | -0.0051741 | 0.66047036 | 0.99999994 |
| Arhgap17 | 0.06446142 | 0.05074011 | 0.05591597 | -0.0051759 | 0.73110056 | 0.99999994 |
| Slc48a1 | 0.29657572 | 0.21077099 | 0.21594784 | -0.0051769 | 0.5633904 | 0.99999994 |
| Noc4l | 0.14166914 | 0.11100919 | 0.11619052 | -0.0051813 | 0.61670172 | 0.99999994 |
| Atm | 0.07648665 | 0.06064521 | 0.06583105 | -0.0051858 | 0.69998412 | 0.99999994 |
| Nle1 | 0.0571866 | 0.04465421 | 0.04984188 | -0.0051877 | 0.75578614 | 0.99999994 |
| Tmem242 | 0.41520063 | 0.27188684 | 0.27707569 | -0.0051889 | 0.54960164 | 0.99999994 |
| Pgam5 | 0.14768055 | 0.11535865 | 0.12054833 | -0.0051897 | 0.61277551 | 0.99999994 |
| Pcsk2os1 | 0.26889627 | 0.19471947 | 0.1999119 | -0.0051924 | 0.56863083 | 0.99999994 |
| Luc7l3 | 0.68853501 | 0.37404491 | 0.37923786 | -0.005193 | 0.53631184 | 0.99999994 |
| Plod3 | 0.11632251 | 0.09214487 | 0.09733938 | -0.0051945 | 0.63878627 | 0.99999994 |
| E230001N04 | 0.0299386 | 0.0213718 | 0.02656756 | -0.0051958 | 0.90350228 | 0.99999994 |
| Ash1l | 0.43204285 | 0.27963968 | 0.28483897 | -0.0051993 | 0.54835313 | 0.99999994 |
| Fpgs | 0.05611726 | 0.04374459 | 0.04894416 | -0.0051996 | 0.76025895 | 0.99999994 |
| Fam50a | 0.30432435 | 0.21510898 | 0.2203095 | -0.0052005 | 0.56242747 | 0.99999994 |
| Cops7b | 0.14223886 | 0.11140056 | 0.11660537 | -0.0052048 | 0.61681026 | 0.99999994 |
| Nek1 | 0.09881854 | 0.07863564 | 0.08384091 | -0.0052053 | 0.66029203 | 0.99999994 |
| Lrp5 | 0.05129089 | 0.03967056 | 0.04487666 | -0.0052061 | 0.78000153 | 0.99999994 |

| | | | | | | |
|---|---|---|---|---|---|---|
| Pigh | 0.09761426 | 0.07767115 | 0.0828828 | -0.0052116 | 0.66222754 | 0.99999994 |
| Casc3 | 0.1349834 | 0.10608062 | 0.11129357 | -0.0052129 | 0.62239659 | 0.99999994 |
| Arpc5l | 1.10133912 | 0.42455554 | 0.42977193 | -0.0052164 | 0.53219637 | 0.99999994 |
| Tcp1 | 1.13816819 | 0.42469171 | 0.42990952 | -0.0052178 | 0.53219477 | 0.99999994 |
| Dtd2 | 0.22864504 | 0.17005108 | 0.17526945 | -0.0052184 | 0.57854884 | 0.99999994 |
| Mphosph9 | 0.04055355 | 0.0305135 | 0.03573435 | -0.0052208 | 0.83460033 | 0.99999994 |
| Rprml | 0.02969918 | 0.02113813 | 0.0263593 | -0.0052212 | 0.90632684 | 0.99999994 |
| Armc5 | 0.07195395 | 0.05688952 | 0.0621119 | -0.0052224 | 0.71214672 | 0.99999994 |
| Csnk2a1 | 0.36944692 | 0.24965307 | 0.25488109 | -0.005228 | 0.55429985 | 0.99999994 |
| Gtf3a | 0.41232726 | 0.27050157 | 0.27572994 | -0.0052284 | 0.55022005 | 0.99999994 |
| Aip | 0.34946139 | 0.23942445 | 0.24465703 | -0.0052326 | 0.55660303 | 0.99999994 |
| Zfp593 | 0.09757362 | 0.07761756 | 0.08285044 | -0.0052329 | 0.66291045 | 0.99999994 |
| Heatr3 | 0.06706973 | 0.05284687 | 0.05807976 | -0.0052329 | 0.7256557 | 0.99999994 |
| Baiap2l1 | 0.14909941 | 0.11633723 | 0.12157056 | -0.0052333 | 0.61276794 | 0.99999994 |
| Fam171a1 | 0.07233994 | 0.05719527 | 0.06242946 | -0.0052342 | 0.71160504 | 0.99999994 |
| Tial1 | 0.29138008 | 0.20775719 | 0.21299209 | -0.0052349 | 0.56497668 | 0.99999994 |
| Wnk4 | 0.04702199 | 0.03601786 | 0.04125741 | -0.0052395 | 0.80103772 | 0.99999994 |
| Shoc2 | 0.24854056 | 0.18240691 | 0.18765067 | -0.0052438 | 0.5737804 | 0.99999994 |
| 0610009L18F | 0.10881816 | 0.08635792 | 0.09160643 | -0.0052485 | 0.64853575 | 0.99999994 |
| Thtpa | 0.10474957 | 0.08321973 | 0.0884688 | -0.0052491 | 0.65355556 | 0.99999994 |
| Rpap1 | 0.05194477 | 0.04017539 | 0.04542924 | -0.0052539 | 0.7792914 | 0.99999994 |
| Zfp952 | 0.05248998 | 0.04063522 | 0.04588962 | -0.0052544 | 0.77701763 | 0.99999994 |
| Ralgds | 0.12417832 | 0.09801006 | 0.10326554 | -0.0052555 | 0.63260528 | 0.99999994 |
| Caml | 0.30709142 | 0.21659521 | 0.22185383 | -0.0052586 | 0.5626834 | 0.99999994 |
| Sbf1 | 0.0567767 | 0.04423923 | 0.04949791 | -0.0052587 | 0.76027106 | 0.99999994 |
| Gna12 | 0.04933407 | 0.03796027 | 0.04322015 | -0.0052599 | 0.79105234 | 0.99999994 |
| Exoc8 | 0.05043053 | 0.03888841 | 0.04414887 | -0.0052605 | 0.78614694 | 0.99999994 |
| H2-M3 | 0.02358683 | 0.0157588 | 0.02102002 | -0.0052612 | 0.95214901 | 0.99999994 |
| Pkd2 | 0.13766412 | 0.10800174 | 0.11326342 | -0.0052617 | 0.621424 | 0.99999994 |
| Xpo7 | 0.11011629 | 0.08733759 | 0.09260317 | -0.0052656 | 0.64746755 | 0.99999994 |

| | | | | | | |
|---|---|---|---|---|---|---|
| Mien1 | 1.19913087 | 0.42375275 | 0.42902064 | -0.0052679 | 0.53257035 | 0.99999994 |
| Vps13d | 0.03755449 | 0.02788819 | 0.03315753 | -0.0052693 | 0.85493037 | 0.99999994 |
| Smim12 | 0.11497596 | 0.0910457 | 0.09631593 | -0.0052702 | 0.64215458 | 0.99999994 |
| Sdhaf3 | 0.11469002 | 0.09082552 | 0.09609828 | -0.0052728 | 0.642528 | 0.99999994 |
| Mnt | 0.05467436 | 0.04245644 | 0.04773084 | -0.0052744 | 0.76899654 | 0.99999994 |
| Zfp788 | 0.07288062 | 0.05759891 | 0.06287402 | -0.0052751 | 0.71173712 | 0.99999994 |
| Nampt | 0.06209393 | 0.04866884 | 0.05394557 | -0.0052767 | 0.74250799 | 0.99999994 |
| Bsdc1 | 0.18415189 | 0.14081276 | 0.1460909 | -0.0052781 | 0.59502424 | 0.99999994 |
| Tmem185b | 0.07723683 | 0.06116562 | 0.06644448 | -0.0052789 | 0.70153919 | 0.99999994 |
| Slc7a6os | 0.15885745 | 0.12325622 | 0.12853782 | -0.0052816 | 0.607769 | 0.99999994 |
| Tet1 | 0.05359814 | 0.04154065 | 0.04682434 | -0.0052837 | 0.77370635 | 0.99999994 |
| Ctdsp2 | 0.15172813 | 0.11816979 | 0.12345828 | -0.0052885 | 0.61222897 | 0.99999994 |
| Crlf2 | 0.07476083 | 0.05912882 | 0.06441756 | -0.0052887 | 0.70763656 | 0.99999994 |
| Phf21a | 0.22396936 | 0.16701201 | 0.17230158 | -0.0052896 | 0.58095874 | 0.99999994 |
| Pphln1 | 0.11326921 | 0.08971995 | 0.09501532 | -0.0052954 | 0.6446701 | 0.99999994 |
| Mrrf | 0.15660991 | 0.12164471 | 0.12694278 | -0.0052981 | 0.60942059 | 0.99999994 |
| Pi4kb | 0.07288816 | 0.05758102 | 0.06288022 | -0.0052992 | 0.71258789 | 0.99999994 |
| Manbal | 0.39538746 | 0.26236602 | 0.26766684 | -0.0053008 | 0.5524371 | 0.99999994 |
| Samhd1 | 0.04804276 | 0.03681965 | 0.04212468 | -0.005305 | 0.79906262 | 0.99999994 |
| Cul2 | 0.15189443 | 0.11827143 | 0.12357743 | -0.005306 | 0.6124851 | 0.99999994 |
| Xpo4 | 0.07882675 | 0.06243636 | 0.06774262 | -0.0053063 | 0.69895267 | 0.99999994 |
| Dlgap3 | 0.01966181 | 0.0122613 | 0.01756831 | -0.005307 | 0.97782376 | 0.99999994 |
| Cdk12 | 0.28976617 | 0.20676081 | 0.21206897 | -0.0053082 | 0.56616198 | 0.99999994 |
| Senp1 | 0.09462434 | 0.07518909 | 0.08049766 | -0.0053086 | 0.66965687 | 0.99999994 |
| Mief2 | 0.06711021 | 0.05279917 | 0.05811328 | -0.0053141 | 0.72863282 | 0.99999994 |
| Osbpl8 | 0.05854668 | 0.04566612 | 0.05098187 | -0.0053158 | 0.75617197 | 0.99999994 |
| Mms19 | 0.05426164 | 0.04206722 | 0.04738336 | -0.0053161 | 0.77240933 | 0.99999994 |
| Rnf186 | 0.05123169 | 0.03951002 | 0.04482661 | -0.0053166 | 0.78508044 | 0.99999994 |
| 2510009E07I | 0.05318062 | 0.0411471 | 0.04647233 | -0.0053252 | 0.77719304 | 0.99999994 |
| Cnksr1 | 0.01820315 | 0.01095208 | 0.01628091 | -0.0053288 | 0.98532157 | 0.99999994 |

| | | | | | | |
|---|---|---|---|---|---|---|
| Smim8 | 0.22981695 | 0.17066135 | 0.17600985 | -0.0053485 | 0.58014727 | 0.99999994 |
| Utp6 | 0.08045779 | 0.06372189 | 0.06907157 | -0.0053497 | 0.69691326 | 0.99999994 |
| Cyb5r4 | 0.29071731 | 0.20726327 | 0.21261329 | -0.00535 | 0.56650965 | 0.99999994 |
| Eif4enif1 | 0.09743592 | 0.07738946 | 0.08274079 | -0.0053513 | 0.66658586 | 0.99999994 |
| Abhd6 | 0.05430792 | 0.04206952 | 0.04742233 | -0.0053528 | 0.77377553 | 0.99999994 |
| Tnpo2 | 0.07955109 | 0.06298009 | 0.06833314 | -0.0053531 | 0.69897093 | 0.99999994 |
| Pnpla8 | 0.22467148 | 0.16739486 | 0.17274867 | -0.0053538 | 0.58171905 | 0.99999994 |
| Igfals | 0.05000467 | 0.03843114 | 0.04378832 | -0.0053572 | 0.79227974 | 0.99999994 |
| Lmf2 | 0.06222312 | 0.04869512 | 0.05405325 | -0.0053581 | 0.74531648 | 0.99999994 |
| Lrp8 | 0.15972119 | 0.12378951 | 0.12914928 | -0.0053598 | 0.60881915 | 0.99999994 |
| Kdm4a | 0.08380723 | 0.0664315 | 0.07179188 | -0.0053604 | 0.69040561 | 0.99999994 |
| Cdadc1 | 0.10231627 | 0.08122142 | 0.08658232 | -0.0053609 | 0.65988167 | 0.99999994 |
| Abhd5 | 0.0895791 | 0.07108716 | 0.0764522 | -0.005365 | 0.67978957 | 0.99999994 |
| Spg21 | 0.48231126 | 0.30141793 | 0.30678327 | -0.0053653 | 0.54633746 | 0.99999994 |
| Zfp523 | 0.11118282 | 0.08805462 | 0.0934205 | -0.0053659 | 0.64889064 | 0.99999994 |
| Taf1d | 0.2048662 | 0.15457206 | 0.15993796 | -0.0053659 | 0.58835663 | 0.99999994 |
| Nudt9 | 0.11938196 | 0.09428699 | 0.09965631 | -0.0053693 | 0.64006564 | 0.99999994 |
| Kansl3 | 0.11375899 | 0.09001916 | 0.09538892 | -0.0053698 | 0.64606132 | 0.99999994 |
| Prkdc | 0.03283289 | 0.02370975 | 0.02907983 | -0.0053701 | 0.89050343 | 0.99999994 |
| Fyco1 | 0.0670119 | 0.0526576 | 0.05803186 | -0.0053743 | 0.73119705 | 0.99999994 |
| Setd2 | 0.21471199 | 0.16098088 | 0.16635779 | -0.0053769 | 0.58517203 | 0.99999994 |
| Hgh1 | 0.08316003 | 0.06589 | 0.07126716 | -0.0053772 | 0.69224592 | 0.99999994 |
| Hgs | 0.20182137 | 0.15255119 | 0.15793151 | -0.0053803 | 0.58969667 | 0.99999994 |
| Psmb8 | 0.01852809 | 0.01118676 | 0.01656792 | -0.0053812 | 0.98468879 | 0.99999994 |
| Pcbd1 | 1.11757894 | 0.42453512 | 0.42991727 | -0.0053821 | 0.53320588 | 0.99999994 |
| Ermard | 0.07930674 | 0.06275138 | 0.068134 | -0.0053826 | 0.7005057 | 0.99999994 |
| S100z | 0.02507357 | 0.0169384 | 0.02232274 | -0.0053843 | 0.94581277 | 0.99999994 |
| Morc2a | 0.07120006 | 0.0561064 | 0.06149119 | -0.0053848 | 0.72001917 | 0.99999994 |
| Map1s | 0.06110138 | 0.04773208 | 0.05311768 | -0.0053856 | 0.75012647 | 0.99999994 |
| Sgtb | 0.05067804 | 0.03897256 | 0.04435832 | -0.0053858 | 0.79050676 | 0.99999994 |

| | | | | | | |
|---|---|---|---|---|---|---|
| Pi4k2b | 0.1957589 | 0.14852034 | 0.15390727 | -0.0053869 | 0.5921116 | 0.99999994 |
| Slc35a3 | 0.04137804 | 0.03105313 | 0.03644096 | -0.0053878 | 0.83747886 | 0.99999994 |
| Nmral1 | 0.23736271 | 0.17535514 | 0.18074414 | -0.005389 | 0.57865884 | 0.99999994 |
| Galnt12 | 0.19340712 | 0.14694503 | 0.1523356 | -0.0053906 | 0.59310634 | 0.99999994 |
| Smim1 | 0.2633738 | 0.19123179 | 0.19662504 | -0.0053933 | 0.57243546 | 0.99999994 |
| Appl2 | 0.13949768 | 0.10921177 | 0.11460584 | -0.0053941 | 0.62296987 | 0.99999994 |
| Cptp | 0.07376046 | 0.05820201 | 0.06359678 | -0.0053948 | 0.71384197 | 0.99999994 |
| Trim33 | 0.09974975 | 0.07918532 | 0.08458077 | -0.0053954 | 0.66444035 | 0.99999994 |
| Inpp4b | 0.01900343 | 0.0115909 | 0.01698754 | -0.0053966 | 0.98276823 | 0.99999994 |
| Phf13 | 0.08049819 | 0.06370681 | 0.06910445 | -0.0053976 | 0.69843926 | 0.99999994 |
| Atg16l2 | 0.08437667 | 0.06685283 | 0.0722532 | -0.0054004 | 0.69058577 | 0.99999994 |
| S100pbp | 0.1015771 | 0.08060716 | 0.08600777 | -0.0054006 | 0.66201441 | 0.99999994 |
| Wdr36 | 0.06393123 | 0.05007265 | 0.05547523 | -0.0054026 | 0.74158643 | 0.99999994 |
| Slc16a6 | 0.02905492 | 0.02039505 | 0.02579858 | -0.0054035 | 0.91836816 | 0.99999994 |
| Plcb1 | 0.03397152 | 0.02465996 | 0.03006553 | -0.0054056 | 0.884301 | 0.99999994 |
| Tmem198 | 0.07023971 | 0.05529144 | 0.06069963 | -0.0054082 | 0.72343121 | 0.99999994 |
| Bdp1 | 0.189977 | 0.14462299 | 0.15003224 | -0.0054093 | 0.59483022 | 0.99999994 |
| Reps2 | 0.03410003 | 0.02476728 | 0.03017668 | -0.0054094 | 0.88360529 | 0.99999994 |
| Cpeb1 | 0.01459134 | 0.00767243 | 0.01308226 | -0.0054098 | 0.99704345 | 0.99999994 |
| Gon4l | 0.31246559 | 0.21942166 | 0.2248339 | -0.0054122 | 0.56365132 | 0.99999994 |
| Rexo2 | 0.45184681 | 0.28827851 | 0.29369963 | -0.0054211 | 0.54889227 | 0.99999994 |
| Cep19 | 0.14839066 | 0.11563793 | 0.12106023 | -0.0054223 | 0.61720144 | 0.99999994 |
| Shq1 | 0.05380932 | 0.04157969 | 0.04700232 | -0.0054226 | 0.77873863 | 0.99999994 |
| Ranbp10 | 0.05213752 | 0.04016349 | 0.04559203 | -0.0054285 | 0.78598232 | 0.99999994 |
| Lmbr1 | 0.05269929 | 0.04063173 | 0.04606628 | -0.0054346 | 0.78385147 | 0.99999994 |
| Trim39 | 0.07396968 | 0.0583332 | 0.06376853 | -0.0054353 | 0.71476394 | 0.99999994 |
| Eftud2 | 0.17388522 | 0.13361637 | 0.13905341 | -0.005437 | 0.6026709 | 0.99999994 |
| Hsd17b4 | 0.18873859 | 0.14375903 | 0.1491974 | -0.0054384 | 0.59585402 | 0.99999994 |
| Tnpo3 | 0.24589289 | 0.18058736 | 0.18602588 | -0.0054385 | 0.57714716 | 0.99999994 |
| Asb8 | 0.14162246 | 0.11071703 | 0.11615651 | -0.0054395 | 0.62236949 | 0.99999994 |

| | | | | | | |
|---|---|---|---|---|---|---|
| Slc11a2 | 0.03824719 | 0.02831351 | 0.03375361 | -0.0054401 | 0.85832203 | 0.99999994 |
| Hfe | 0.01794741 | 0.01061443 | 0.01605493 | -0.0054405 | 0.98795296 | 0.99999994 |
| Cd2bp2 | 0.19473894 | 0.14778582 | 0.15322637 | -0.0054405 | 0.59341736 | 0.99999994 |
| Bysl | 0.11969821 | 0.09445258 | 0.09989516 | -0.0054426 | 0.64156978 | 0.99999994 |
| Rrp8 | 0.10571556 | 0.08377296 | 0.08921564 | -0.0054427 | 0.65765703 | 0.99999994 |
| Rrp9 | 0.10636482 | 0.08427263 | 0.08971694 | -0.0054443 | 0.65686778 | 0.99999994 |
| Tmx3 | 0.0974175 | 0.07728133 | 0.08272611 | -0.0054448 | 0.66934352 | 0.99999994 |
| 5031439G07 | 0.0476464 | 0.03633852 | 0.04178807 | -0.0054496 | 0.80731668 | 0.99999994 |
| Elmod2 | 0.0736276 | 0.05803715 | 0.0634877 | -0.0054506 | 0.71615683 | 0.99999994 |
| Tbc1d30 | 0.02531926 | 0.01708608 | 0.02253777 | -0.0054517 | 0.9463136 | 0.99999994 |
| Tbce | 0.14454789 | 0.11283047 | 0.11828281 | -0.0054523 | 0.62051362 | 0.99999994 |
| Coro1b | 0.32013929 | 0.22358902 | 0.22904371 | -0.0054547 | 0.56297701 | 0.99999994 |
| N4bp2l1 | 0.09256012 | 0.07338696 | 0.07884565 | -0.0054587 | 0.67754233 | 0.99999994 |
| Rpe | 0.11591329 | 0.09156982 | 0.0970286 | -0.0054588 | 0.64597768 | 0.99999994 |
| Tmem150c | 0.05889713 | 0.04581307 | 0.05127528 | -0.0054622 | 0.76086368 | 0.99999994 |
| Bcl2l11 | 0.09123725 | 0.0723209 | 0.07778465 | -0.0054637 | 0.67994875 | 0.99999994 |
| Lsr | 0.63211721 | 0.35692876 | 0.36239284 | -0.0054641 | 0.53997199 | 0.99999994 |
| Pced1b | 0.03386812 | 0.02451198 | 0.02997608 | -0.0054641 | 0.88749902 | 0.99999994 |
| Lmnb2 | 0.07732742 | 0.06105194 | 0.06651852 | -0.0054666 | 0.70782048 | 0.99999994 |
| Spata13 | 0.04926221 | 0.03769256 | 0.04315924 | -0.0054667 | 0.800418 | 0.99999994 |
| Traf3ip2 | 0.05821363 | 0.04523335 | 0.05070291 | -0.0054696 | 0.76363733 | 0.99999994 |
| Zfp524 | 0.08887681 | 0.07041564 | 0.07588701 | -0.0054714 | 0.68435633 | 0.99999994 |
| Trmu | 0.08616312 | 0.06821975 | 0.07369825 | -0.0054785 | 0.68963385 | 0.99999994 |
| Ptpru | 0.0480791 | 0.03667435 | 0.04215554 | -0.0054812 | 0.80661211 | 0.99999994 |
| Rnls | 0.05629981 | 0.04361493 | 0.0490975 | -0.0054826 | 0.77134868 | 0.99999994 |
| Smarcb1 | 0.64199542 | 0.36009316 | 0.36557741 | -0.0054843 | 0.53977075 | 0.99999994 |
| Tarsl2 | 0.03557886 | 0.02596957 | 0.03145443 | -0.0054849 | 0.8771183 | 0.99999994 |
| Ccni | 0.41527581 | 0.27162466 | 0.27711082 | -0.0054862 | 0.55242105 | 0.99999994 |
| Ptpn4 | 0.05841751 | 0.04538513 | 0.05087369 | -0.0054886 | 0.76366089 | 0.99999994 |
| Cdkn2d | 0.3097584 | 0.21784632 | 0.2233359 | -0.0054896 | 0.56498193 | 0.99999994 |

| | | | | | | |
|---|---|---|---|---|---|---|
| Snx12 | 0.18775879 | 0.14303926 | 0.14853572 | -0.0054965 | 0.59728033 | 0.99999994 |
| Polr2d | 0.33217574 | 0.23004092 | 0.23554296 | -0.005502 | 0.5617808 | 0.99999994 |
| Smyd4 | 0.04581751 | 0.03472805 | 0.04023254 | -0.0055045 | 0.81877381 | 0.99999994 |
| Rars2 | 0.29421148 | 0.20910138 | 0.21460588 | -0.0055045 | 0.56778233 | 0.99999994 |
| Ier5 | 0.09982482 | 0.07913403 | 0.08464037 | -0.0055063 | 0.66750519 | 0.99999994 |
| Krt10 | 0.2864685 | 0.20466456 | 0.21017541 | -0.0055108 | 0.56927602 | 0.99999994 |
| Scfd2 | 0.0431356 | 0.03242985 | 0.03794468 | -0.0055148 | 0.83333449 | 0.99999994 |
| Reps1 | 0.04939206 | 0.03775301 | 0.04326931 | -0.0055163 | 0.80194867 | 0.99999994 |
| Kifc3 | 0.0672409 | 0.05270419 | 0.0582215 | -0.0055173 | 0.7359092 | 0.99999994 |
| Klhl32 | 0.22061167 | 0.16463821 | 0.17015613 | -0.0055179 | 0.5854501 | 0.99999994 |
| Brf1 | 0.1162112 | 0.09173352 | 0.09725487 | -0.0055213 | 0.64724544 | 0.99999994 |
| Sgsm1 | 0.04862632 | 0.03709525 | 0.04261997 | -0.0055247 | 0.8058893 | 0.99999994 |
| Rab3ip | 0.29380814 | 0.20884926 | 0.21437643 | -0.0055272 | 0.5681311 | 0.99999994 |
| Tsnax | 0.33638364 | 0.23225672 | 0.23778499 | -0.0055283 | 0.56149232 | 0.99999994 |
| Mylpf | 0.06554642 | 0.0512877 | 0.05681695 | -0.0055292 | 0.74143502 | 0.99999994 |
| Trim11 | 0.12514512 | 0.09845438 | 0.10398966 | -0.0055353 | 0.63844754 | 0.99999994 |
| Ttc13 | 0.13034933 | 0.10232647 | 0.10786809 | -0.0055416 | 0.63381101 | 0.99999994 |
| 2900026A02 | 0.12074288 | 0.09514151 | 0.10068325 | -0.0055417 | 0.64295734 | 0.99999994 |
| Cluh | 0.04661688 | 0.03537088 | 0.04091289 | -0.005542 | 0.81638167 | 0.99999994 |
| Sh3yl1 | 0.07777233 | 0.06133791 | 0.06688201 | -0.0055441 | 0.70944703 | 0.99999994 |
| Chmp4c | 0.04857941 | 0.03703586 | 0.04258017 | -0.0055443 | 0.80695124 | 0.99999994 |
| Slk | 0.06891558 | 0.05406109 | 0.0596066 | -0.0055455 | 0.73213241 | 0.99999994 |
| Idh3b | 0.49930787 | 0.30825988 | 0.31380541 | -0.0055455 | 0.54681972 | 0.99999994 |
| Med15 | 0.07505736 | 0.05911197 | 0.06466065 | -0.0055487 | 0.71606696 | 0.99999994 |
| D6Wsu163e | 0.06567965 | 0.05137735 | 0.0569275 | -0.0055501 | 0.7418184 | 0.99999994 |
| Dctn4 | 0.13891877 | 0.10863197 | 0.11418243 | -0.0055505 | 0.62686754 | 0.99999994 |
| Prpf3 | 0.08065234 | 0.06367668 | 0.0692299 | -0.0055532 | 0.70330629 | 0.99999994 |
| Folr1 | 0.026847 | 0.01831929 | 0.02387329 | -0.005554 | 0.93925726 | 0.99999994 |
| Galnt1 | 0.14045991 | 0.10975351 | 0.11530874 | -0.0055552 | 0.62577476 | 0.99999994 |
| C2cd3 | 0.09523405 | 0.07542895 | 0.08098478 | -0.0055558 | 0.67603814 | 0.99999994 |

| | | | | | | |
|---|---|---|---|---|---|---|
| Mfap1a | 0.12759321 | 0.10025807 | 0.10581817 | -0.0055601 | 0.63673719 | 0.99999994 |
| Ppm1k | 0.05428031 | 0.04183867 | 0.04739908 | -0.0055604 | 0.78255767 | 0.99999994 |
| Zscan26 | 0.08574234 | 0.06779686 | 0.07335818 | -0.0055613 | 0.69308771 | 0.99999994 |
| Ndufaf3 | 0.47706979 | 0.29901578 | 0.30457801 | -0.0055622 | 0.54837584 | 0.99999994 |
| Zfp322a | 0.19231237 | 0.14603914 | 0.15160197 | -0.0055628 | 0.5964799 | 0.99999994 |
| Tsen2 | 0.08491551 | 0.06712553 | 0.07268941 | -0.0055639 | 0.69479866 | 0.99999994 |
| Frmd4a | 0.05582533 | 0.04313399 | 0.04869887 | -0.0055649 | 0.77656102 | 0.99999994 |
| Noa1 | 0.08882866 | 0.07027969 | 0.07584823 | -0.0055685 | 0.68746904 | 0.99999994 |
| Unc119b | 0.08189989 | 0.06467233 | 0.07024424 | -0.0055719 | 0.70124862 | 0.99999994 |
| Gdap1 | 0.10843131 | 0.08573658 | 0.09130899 | -0.0055724 | 0.65771127 | 0.99999994 |
| Colgalt1 | 0.08593954 | 0.06794424 | 0.07351757 | -0.0055733 | 0.69308569 | 0.99999994 |
| Nup50 | 0.21418551 | 0.16044242 | 0.16601709 | -0.0055747 | 0.5884319 | 0.99999994 |
| Gtpbp1 | 0.04672952 | 0.03543199 | 0.04100869 | -0.0055767 | 0.81731663 | 0.99999994 |
| Dcp2 | 0.0731515 | 0.05751863 | 0.06309663 | -0.005578 | 0.72188469 | 0.99999994 |
| Mfap3 | 0.17641422 | 0.13521923 | 0.14079814 | -0.0055789 | 0.60401296 | 0.99999994 |
| Alg11 | 0.08609117 | 0.0680586 | 0.07364012 | -0.0055815 | 0.6930503 | 0.99999994 |
| Cebpzos | 0.42931378 | 0.27801249 | 0.28359541 | -0.0055829 | 0.55212764 | 0.99999994 |
| Slc25a19 | 0.07137568 | 0.05605112 | 0.06163584 | -0.0055847 | 0.72678598 | 0.99999994 |
| Ccdc130 | 0.07859632 | 0.06196646 | 0.06755464 | -0.0055882 | 0.70905278 | 0.99999994 |
| Zfp612 | 0.1142022 | 0.0901358 | 0.09572674 | -0.0055909 | 0.65127524 | 0.99999994 |
| Amz2 | 0.27335424 | 0.19694991 | 0.20254311 | -0.0055932 | 0.57292039 | 0.99999994 |
| Ppfibp1 | 0.07801172 | 0.06148299 | 0.0670775 | -0.0055945 | 0.71060905 | 0.99999994 |
| Nob1 | 0.25322401 | 0.18491137 | 0.190508 | -0.0055966 | 0.57751738 | 0.99999994 |
| Tle3 | 0.18903276 | 0.14379872 | 0.14939586 | -0.0055971 | 0.59846717 | 0.99999994 |
| Maml3 | 0.19520455 | 0.14793912 | 0.15353734 | -0.0055982 | 0.59588163 | 0.99999994 |
| Ykt6 | 0.19358811 | 0.14685661 | 0.15245677 | -0.0056002 | 0.59658062 | 0.99999994 |
| Aff1 | 0.03563443 | 0.02590016 | 0.0315024 | -0.0056022 | 0.88174124 | 0.99999994 |
| Man1b1 | 0.08113371 | 0.06401245 | 0.06962147 | -0.005609 | 0.70411254 | 0.99999994 |
| Gtpbp6 | 0.16716081 | 0.12877049 | 0.13437968 | -0.0056092 | 0.60943452 | 0.99999994 |
| Wdr45b | 0.30009678 | 0.21232636 | 0.21793689 | -0.0056105 | 0.56802968 | 0.99999994 |

| | | | | | | |
|---|---|---|---|---|---|---|
| Smg9 | 0.10455409 | 0.08270669 | 0.08831752 | -0.0056108 | 0.6638087 | 0.99999994 |
| Vps41 | 0.15593614 | 0.12085003 | 0.12646349 | -0.0056135 | 0.6161796 | 0.99999994 |
| Ptprt | 0.03564171 | 0.02589257 | 0.03150868 | -0.0056161 | 0.8822736 | 0.99999994 |
| Gpr155 | 0.0226305 | 0.01456301 | 0.02018068 | -0.0056177 | 0.9680532 | 0.99999994 |
| Zdhhc24 | 0.0476267 | 0.03615212 | 0.04177133 | -0.0056192 | 0.81472266 | 0.99999994 |
| Mmd | 0.08083253 | 0.06375713 | 0.0693765 | -0.0056194 | 0.70510932 | 0.99999994 |
| Rrm2b | 0.09844069 | 0.0779208 | 0.08354045 | -0.0056197 | 0.67283822 | 0.99999994 |
| Serinc1 | 0.83446936 | 0.40357514 | 0.40919713 | -0.005622 | 0.53643335 | 0.99999994 |
| Tasp1 | 0.07574414 | 0.05960048 | 0.0652233 | -0.0056228 | 0.71696078 | 0.99999994 |
| Tdrd7 | 0.04360606 | 0.0327194 | 0.03834661 | -0.0056272 | 0.83566416 | 0.99999994 |
| Pms2 | 0.05849364 | 0.04530814 | 0.05093745 | -0.0056293 | 0.76901953 | 0.99999994 |
| Ndfip2 | 0.60330057 | 0.34712236 | 0.35275808 | -0.0056357 | 0.54234488 | 0.99999994 |
| Ndufaf7 | 0.10094661 | 0.07987704 | 0.08551716 | -0.0056401 | 0.66967189 | 0.99999994 |
| Galnt10 | 0.03741586 | 0.02739723 | 0.03303816 | -0.0056409 | 0.87212732 | 0.99999994 |
| Hspa1b | 0.01995711 | 0.01218436 | 0.01782863 | -0.0056443 | 0.98245426 | 0.99999994 |
| Prr14 | 0.0871581 | 0.06885457 | 0.07450165 | -0.0056471 | 0.69305863 | 0.99999994 |
| Rhd | 0.02352822 | 0.01532036 | 0.02096861 | -0.0056482 | 0.96351362 | 0.99999994 |
| Wdr7 | 0.03829782 | 0.02814683 | 0.03379715 | -0.0056503 | 0.86710982 | 0.99999994 |
| Usp11 | 0.07445729 | 0.05851316 | 0.06416863 | -0.0056555 | 0.72128211 | 0.99999994 |
| Ralgps1 | 0.0627137 | 0.04880573 | 0.05446198 | -0.0056562 | 0.75531391 | 0.99999994 |
| Scamp5 | 0.04808589 | 0.03650148 | 0.04216131 | -0.0056598 | 0.81422238 | 0.99999994 |
| Cln5 | 0.07696644 | 0.0605572 | 0.06622345 | -0.0056663 | 0.71550376 | 0.99999994 |
| Rhbdl2 | 0.02214496 | 0.01408526 | 0.01975414 | -0.0056689 | 0.97194543 | 0.99999994 |
| Ggnbp1 | 0.08713832 | 0.06881391 | 0.07448568 | -0.0056718 | 0.69387116 | 0.99999994 |
| Lypla2 | 0.27747843 | 0.19928721 | 0.20496073 | -0.0056735 | 0.57309313 | 0.99999994 |
| 4930481A15 | 0.02147375 | 0.01348861 | 0.019164 | -0.0056754 | 0.97565355 | 0.99999994 |
| Mier2 | 0.06837237 | 0.0534821 | 0.05915765 | -0.0056755 | 0.7384607 | 0.99999994 |
| Ttc19 | 0.10882579 | 0.08593601 | 0.09161229 | -0.0056763 | 0.65998668 | 0.99999994 |
| Lgals8 | 0.08553715 | 0.06751382 | 0.07319228 | -0.0056785 | 0.69721896 | 0.99999994 |
| Zbtb48 | 0.03591164 | 0.02606313 | 0.03174162 | -0.0056785 | 0.88313476 | 0.99999994 |

| | | | | | | |
|---|---|---|---|---|---|---|
| Ipo9 | 0.08333188 | 0.06572798 | 0.07140653 | -0.0056785 | 0.7017157 | 0.99999994 |
| Alg6 | 0.044627 | 0.03353653 | 0.03921795 | -0.0056814 | 0.83255035 | 0.99999994 |
| Nudt13 | 0.08558139 | 0.06754604 | 0.07322805 | -0.005682 | 0.69724384 | 0.99999994 |
| A930015D03 | 0.08120359 | 0.06399517 | 0.0696783 | -0.0056831 | 0.70640359 | 0.99999994 |
| Slc30a4 | 0.05070596 | 0.03869618 | 0.04438195 | -0.0056858 | 0.80309967 | 0.99999994 |
| 2810001G20 | 0.09872381 | 0.07807928 | 0.0837656 | -0.0056863 | 0.67431438 | 0.99999994 |
| Psmb9 | 0.02042243 | 0.01254613 | 0.01823863 | -0.0056925 | 0.98112368 | 0.99999994 |
| Uck1 | 0.24530094 | 0.17996882 | 0.18566168 | -0.0056929 | 0.58086266 | 0.99999994 |
| Cacul1 | 0.11461007 | 0.09034365 | 0.09603741 | -0.0056938 | 0.65344372 | 0.99999994 |
| Sestd1 | 0.05754339 | 0.04444733 | 0.05014114 | -0.0056938 | 0.77513564 | 0.99999994 |
| Riok3 | 0.40126902 | 0.26479763 | 0.27049147 | -0.0056938 | 0.55571599 | 0.99999994 |
| Sgpp1 | 0.08267494 | 0.06517591 | 0.07087358 | -0.0056977 | 0.7037158 | 0.99999994 |
| Ptprs | 0.16619618 | 0.12800349 | 0.13370507 | -0.0057016 | 0.61173607 | 0.99999994 |
| Enah | 0.24114986 | 0.17739476 | 0.18309803 | -0.0057033 | 0.5821267 | 0.99999994 |
| Ssh1 | 0.05005256 | 0.03812328 | 0.04382887 | -0.0057056 | 0.80689648 | 0.99999994 |
| Syap1 | 0.22862673 | 0.16955193 | 0.17525788 | -0.0057059 | 0.58578437 | 0.99999994 |
| Coq6 | 0.14331167 | 0.11167742 | 0.11738551 | -0.0057081 | 0.62690887 | 0.99999994 |
| Rbm6 | 0.29466699 | 0.20915611 | 0.21486482 | -0.0057087 | 0.57018756 | 0.99999994 |
| Pigu | 0.13510012 | 0.10567035 | 0.11137951 | -0.0057092 | 0.63352195 | 0.99999994 |
| Dock9 | 0.04951957 | 0.03766766 | 0.04337737 | -0.0057097 | 0.80953054 | 0.99999994 |
| Crebbp | 0.28292269 | 0.20241679 | 0.208128 | -0.0057112 | 0.57246615 | 0.99999994 |
| Stat5b | 0.04116852 | 0.03055013 | 0.03626147 | -0.0057113 | 0.85277396 | 0.99999994 |
| Ebag9 | 0.12929931 | 0.1013731 | 0.1070882 | -0.0057151 | 0.63879412 | 0.99999994 |
| Cdc37l1 | 0.14268076 | 0.11120317 | 0.11692688 | -0.0057237 | 0.62772616 | 0.99999994 |
| Gpatch2 | 0.17487943 | 0.13401301 | 0.13974016 | -0.0057272 | 0.60749886 | 0.99999994 |
| Tab1 | 0.1006106 | 0.07952789 | 0.08525548 | -0.0057276 | 0.67262893 | 0.99999994 |
| Ndufaf2 | 0.15652781 | 0.1211517 | 0.1268844 | -0.0057327 | 0.61819283 | 0.99999994 |
| Clic4 | 0.08090474 | 0.06370201 | 0.06943525 | -0.0057332 | 0.70870735 | 0.99999994 |
| 1810044D09 | 0.07913202 | 0.06225799 | 0.06799157 | -0.0057336 | 0.7127089 | 0.99999994 |
| Mcat | 0.1472709 | 0.11451625 | 0.12025275 | -0.0057365 | 0.62458045 | 0.99999994 |

| | | | | | | |
|---|---|---|---|---|---|---|
| Wdr35 | 0.08565382 | 0.06754535 | 0.07328662 | -0.0057413 | 0.69897625 | 0.99999994 |
| Ccz1 | 0.31313773 | 0.21945553 | 0.22520483 | -0.0057493 | 0.56746799 | 0.99999994 |
| Rabgef1 | 0.07678967 | 0.06032869 | 0.0660789 | -0.0057502 | 0.71878614 | 0.99999994 |
| Tap2 | 0.03133241 | 0.0220279 | 0.02777861 | -0.0057507 | 0.91589762 | 0.99999994 |
| Ikzf2 | 0.04874589 | 0.03696969 | 0.04272141 | -0.0057517 | 0.81491808 | 0.99999994 |
| Ppcdc | 0.05787544 | 0.04466705 | 0.05041951 | -0.0057525 | 0.77620447 | 0.99999994 |
| Nprl2 | 0.16182767 | 0.12488197 | 0.1306368 | -0.0057548 | 0.61532561 | 0.99999994 |
| Ahdc1 | 0.10032762 | 0.07927952 | 0.085035 | -0.0057555 | 0.67383461 | 0.99999994 |
| Rnase1 | 0.10175814 | 0.08038479 | 0.08614855 | -0.0057638 | 0.67196421 | 0.99999994 |
| 2810454H06I | 0.05453264 | 0.04184651 | 0.04761154 | -0.005765 | 0.78987366 | 0.99999994 |
| Plekhh1 | 0.03156858 | 0.02221826 | 0.02798359 | -0.0057653 | 0.91487169 | 0.99999994 |
| Dnal1 | 0.0587838 | 0.04541053 | 0.05118041 | -0.0057699 | 0.77349645 | 0.99999994 |
| Itpr1 | 0.17901521 | 0.13681333 | 0.14258465 | -0.0057713 | 0.60619894 | 0.99999994 |
| Kat7 | 0.11272276 | 0.08882594 | 0.09459814 | -0.0057722 | 0.65768691 | 0.99999994 |
| Airn | 0.06189336 | 0.0480033 | 0.05377836 | -0.0057751 | 0.76263431 | 0.99999994 |
| Gna13 | 0.11650637 | 0.09170202 | 0.09747895 | -0.0057769 | 0.65338593 | 0.99999994 |
| Agtpbp1 | 0.05269285 | 0.04027108 | 0.04606084 | -0.0057898 | 0.79861715 | 0.99999994 |
| Akap8 | 0.18756824 | 0.14261684 | 0.14840691 | -0.0057901 | 0.60245113 | 0.99999994 |
| Mpv17l2 | 0.21091503 | 0.15810282 | 0.16389421 | -0.0057914 | 0.59297722 | 0.99999994 |
| Slc3a2 | 0.58654846 | 0.34113513 | 0.34692706 | -0.0057919 | 0.5442424 | 0.99999994 |
| Tmx4 | 0.19643637 | 0.14856539 | 0.15435891 | -0.0057935 | 0.59864123 | 0.99999994 |
| Dcaf17 | 0.14541208 | 0.11310821 | 0.11890901 | -0.0058008 | 0.62730365 | 0.99999994 |
| Zfp113 | 0.0909638 | 0.07175314 | 0.07756511 | -0.005812 | 0.69102646 | 0.99999994 |
| Serinc3 | 0.1625244 | 0.12531263 | 0.1311276 | -0.005815 | 0.61607258 | 0.99999994 |
| Atg4b | 0.29709998 | 0.21042823 | 0.21624474 | -0.0058165 | 0.57104737 | 0.99999994 |
| Rnf11 | 0.20078854 | 0.15143185 | 0.15724868 | -0.0058168 | 0.59724671 | 0.99999994 |
| Etfdh | 0.17099824 | 0.13123328 | 0.13705304 | -0.0058198 | 0.61127839 | 0.99999994 |
| Rhog | 0.0784193 | 0.0615865 | 0.0674102 | -0.0058237 | 0.71737347 | 0.99999994 |
| Dynlt1c | 0.18844128 | 0.14317066 | 0.14899673 | -0.0058261 | 0.60267489 | 0.99999994 |
| Rcl1 | 0.08544826 | 0.06729362 | 0.0731204 | -0.0058268 | 0.70209234 | 0.99999994 |

| | | | | | | |
|---|---|---|---|---|---|---|
| Ddx25 | 0.04760933 | 0.03592788 | 0.04175657 | -0.0058287 | 0.82359257 | 0.99999994 |
| Ttc9c | 0.11771315 | 0.0925645 | 0.09839394 | -0.0058294 | 0.6533433 | 0.99999994 |
| Nadk | 0.17723131 | 0.13553039 | 0.14136018 | -0.0058298 | 0.60815463 | 0.99999994 |
| Lsm8 | 0.53152289 | 0.32075677 | 0.32659117 | -0.0058344 | 0.54732774 | 0.99999994 |
| Rps6kb1 | 0.29562895 | 0.20957348 | 0.21541105 | -0.0058376 | 0.57157492 | 0.99999994 |
| Wdr73 | 0.12822128 | 0.10044801 | 0.10628611 | -0.0058381 | 0.64267635 | 0.99999994 |
| Tsc2 | 0.05312974 | 0.04058974 | 0.04642942 | -0.0058397 | 0.79876138 | 0.99999994 |
| Erlin2 | 0.0937667 | 0.07397098 | 0.07981181 | -0.0058408 | 0.68691661 | 0.99999994 |
| Fbxo7 | 0.08121393 | 0.06384516 | 0.0696867 | -0.0058415 | 0.71156918 | 0.99999994 |
| Mlxip | 0.04381934 | 0.03268592 | 0.03852873 | -0.0058428 | 0.84361434 | 0.99999994 |
| Rab19 | 0.02466293 | 0.01611927 | 0.02196318 | -0.0058439 | 0.96172879 | 0.99999994 |
| Sppl3 | 0.08881414 | 0.06998895 | 0.07583655 | -0.0058476 | 0.69611449 | 0.99999994 |
| Crem | 0.14631144 | 0.11371081 | 0.1195597 | -0.0058489 | 0.6276482 | 0.99999994 |
| Kbtbd3 | 0.12631155 | 0.09900638 | 0.10486178 | -0.0058554 | 0.64493662 | 0.99999994 |
| Nt5c3 | 0.41433911 | 0.27081682 | 0.27667283 | -0.005856 | 0.55602059 | 0.99999994 |
| Tor3a | 0.05319399 | 0.04062759 | 0.04648361 | -0.005856 | 0.79914486 | 0.99999994 |
| Tmem38b | 0.06827195 | 0.05321329 | 0.05907462 | -0.0058613 | 0.74551356 | 0.99999994 |
| Ehd4 | 0.05668476 | 0.04355854 | 0.04942074 | -0.0058622 | 0.78510906 | 0.99999994 |
| Msl3 | 0.10027591 | 0.07913213 | 0.08499469 | -0.0058626 | 0.67692837 | 0.99999994 |
| Gba | 0.12338261 | 0.09680598 | 0.10266872 | -0.0058627 | 0.64806485 | 0.99999994 |
| Zcchc10 | 0.09351238 | 0.07374474 | 0.07960829 | -0.0058635 | 0.68802991 | 0.99999994 |
| Get4 | 0.3436383 | 0.23575066 | 0.24161541 | -0.0058647 | 0.56417781 | 0.99999994 |
| Helq | 0.03336649 | 0.02366924 | 0.02954195 | -0.0058727 | 0.90712081 | 0.99999994 |
| Mprip | 0.1434591 | 0.11161889 | 0.11749262 | -0.0058737 | 0.6303449 | 0.99999994 |
| Tmc6 | 0.06654758 | 0.05177121 | 0.05764718 | -0.005876 | 0.75126697 | 0.99999994 |
| Rbpms | 0.44804489 | 0.28614377 | 0.29202043 | -0.0058767 | 0.5532803 | 0.99999994 |
| Rmdn3 | 0.17139703 | 0.13145289 | 0.13732991 | -0.005877 | 0.61212367 | 0.99999994 |
| Rnf168 | 0.07639176 | 0.05987468 | 0.06575342 | -0.0058787 | 0.72411736 | 0.99999994 |
| H6pd | 0.01764822 | 0.00990817 | 0.01579046 | -0.0058823 | 0.99342421 | 0.99999994 |
| Rabif | 0.15348706 | 0.11883457 | 0.12471693 | -0.0058824 | 0.62322118 | 0.99999994 |

| | | | | | | |
|---|---|---|---|---|---|---|
| Myd88 | 0.03306709 | 0.0233911 | 0.0292827 | -0.0058916 | 0.90975396 | 0.99999994 |
| Spopl | 0.09135196 | 0.07198291 | 0.07787673 | -0.0058938 | 0.69278565 | 0.99999994 |
| Tomm40l | 0.09113297 | 0.07180623 | 0.07770094 | -0.0058947 | 0.69321287 | 0.99999994 |
| Fam222a | 0.11883049 | 0.09333923 | 0.09923953 | -0.0059003 | 0.65385707 | 0.99999994 |
| Ring1 | 0.16024369 | 0.12361681 | 0.12951875 | -0.0059019 | 0.61917642 | 0.99999994 |
| Eif2b1 | 0.33105401 | 0.22904028 | 0.23494276 | -0.0059025 | 0.56640465 | 0.99999994 |
| Map4k5 | 0.08807189 | 0.069335 | 0.07523858 | -0.0059036 | 0.69926583 | 0.99999994 |
| Trp53rka | 0.15522624 | 0.12005174 | 0.12595793 | -0.0059062 | 0.62252455 | 0.99999994 |
| Sfi1 | 0.08163276 | 0.0641209 | 0.07002717 | -0.0059063 | 0.7127425 | 0.99999994 |
| Rtkn2 | 0.01790017 | 0.01010625 | 0.01601318 | -0.0059069 | 0.9929635 | 0.99999994 |
| Rxrb | 0.09901511 | 0.07808924 | 0.08399716 | -0.0059079 | 0.68016989 | 0.99999994 |
| Csnk1g1 | 0.14464114 | 0.11244108 | 0.11835042 | -0.0059093 | 0.63019043 | 0.99999994 |
| Tefm | 0.13501909 | 0.10540909 | 0.11131985 | -0.0059108 | 0.63811833 | 0.99999994 |
| Stam2 | 0.17912425 | 0.13674841 | 0.14265937 | -0.005911 | 0.60864922 | 0.99999994 |
| Rnaseh2b | 0.12012125 | 0.09429911 | 0.10021446 | -0.0059153 | 0.65280476 | 0.99999994 |
| Cbx5 | 0.33308503 | 0.23011266 | 0.23602871 | -0.0059161 | 0.56625259 | 0.99999994 |
| Uimc1 | 0.12564277 | 0.0984376 | 0.10436195 | -0.0059243 | 0.64723398 | 0.99999994 |
| Lrrc24 | 0.12014024 | 0.09430321 | 0.10022878 | -0.0059256 | 0.65303485 | 0.99999994 |
| Rad21 | 0.53043763 | 0.32024333 | 0.32617122 | -0.0059279 | 0.54814403 | 0.99999994 |
| Pcca | 0.05716577 | 0.04389468 | 0.04982441 | -0.0059297 | 0.78587508 | 0.99999994 |
| Phlpp2 | 0.03374056 | 0.02393456 | 0.02986571 | -0.0059311 | 0.90690111 | 0.99999994 |
| Glrx5 | 0.75185368 | 0.38806018 | 0.39399353 | -0.0059334 | 0.53992375 | 0.99999994 |
| Usp35 | 0.03280961 | 0.02312574 | 0.02905967 | -0.0059339 | 0.91296074 | 0.99999994 |
| Sbf2 | 0.09663491 | 0.07616591 | 0.08210254 | -0.0059366 | 0.68485015 | 0.99999994 |
| Dlg1 | 0.07926134 | 0.06215806 | 0.06809699 | -0.0059389 | 0.71921775 | 0.99999994 |
| Tmem131 | 0.10668707 | 0.08402428 | 0.08996556 | -0.0059413 | 0.66988037 | 0.99999994 |
| Bicc1 | 0.10484574 | 0.08259888 | 0.0885432 | -0.0059443 | 0.67251641 | 0.99999994 |
| Sfmbt1 | 0.04050086 | 0.02974328 | 0.03568916 | -0.0059459 | 0.86627649 | 0.99999994 |
| Rbmx2 | 0.21976385 | 0.16366063 | 0.16961249 | -0.0059519 | 0.59234357 | 0.99999994 |
| Cmc4 | 0.08404684 | 0.06602688 | 0.07198604 | -0.0059592 | 0.70919271 | 0.99999994 |

| | | | | | | |
|---|---|---|---|---|---|---|
| Foxj3 | 0.0771723 | 0.06043149 | 0.06639174 | -0.0059602 | 0.72493577 | 0.99999994 |
| Ctu2 | 0.1273966 | 0.09971089 | 0.10567158 | -0.0059607 | 0.64634442 | 0.99999994 |
| Kmt2d | 0.11691561 | 0.09182853 | 0.09778943 | -0.0059609 | 0.65753613 | 0.99999994 |
| Mrpl50 | 0.37560783 | 0.25200524 | 0.25796711 | -0.0059619 | 0.56113007 | 0.99999994 |
| Mfsd6l | 0.03498353 | 0.02497723 | 0.03094036 | -0.0059631 | 0.90023255 | 0.99999994 |
| Rufy2 | 0.11629874 | 0.09135462 | 0.09732133 | -0.0059667 | 0.6583987 | 0.99999994 |
| Abhd14a | 0.15930788 | 0.12288917 | 0.1288568 | -0.0059676 | 0.62106149 | 0.99999994 |
| Decr2 | 0.11904373 | 0.09343212 | 0.09940074 | -0.0059686 | 0.65530681 | 0.99999994 |
| S100a8 | 0.02964129 | 0.02034006 | 0.02630894 | -0.0059689 | 0.93450007 | 0.99999994 |
| Capn9 | 0.02632925 | 0.01745042 | 0.02342098 | -0.0059706 | 0.95513887 | 0.99999994 |
| C77080 | 0.04202654 | 0.03102384 | 0.03699621 | -0.0059724 | 0.85871135 | 0.99999994 |
| Dhodh | 0.0779041 | 0.06101026 | 0.06698962 | -0.0059794 | 0.72378857 | 0.99999994 |
| Fntb | 0.07318679 | 0.05714055 | 0.06312562 | -0.0059851 | 0.73601355 | 0.99999994 |
| Prmt1 | 1.58676975 | 0.39392711 | 0.399914 | -0.0059869 | 0.53968835 | 0.99999994 |
| Dhx57 | 0.05043258 | 0.03815944 | 0.0441506 | -0.0059912 | 0.8168051 | 0.99999994 |
| Creb1 | 0.13849526 | 0.10787983 | 0.11387242 | -0.0059926 | 0.63694104 | 0.99999994 |
| Zfp61 | 0.04302555 | 0.03185755 | 0.03785063 | -0.0059931 | 0.85404355 | 0.99999994 |
| Ppp1r8 | 0.19034805 | 0.14428563 | 0.15028206 | -0.0059964 | 0.60472496 | 0.99999994 |
| Rwdd1 | 0.86619828 | 0.40788587 | 0.41388553 | -0.0059997 | 0.53843433 | 0.99999994 |
| Prdm2 | 0.12497156 | 0.09785798 | 0.10385975 | -0.0060018 | 0.64975257 | 0.99999994 |
| Tfpt | 0.20075127 | 0.15122029 | 0.15722402 | -0.0060037 | 0.60032108 | 0.99999994 |
| Apold1 | 0.02494312 | 0.0162033 | 0.02220854 | -0.0060052 | 0.96406152 | 0.99999994 |
| Asb1 | 0.0390877 | 0.0284709 | 0.03447616 | -0.0060053 | 0.87686205 | 0.99999994 |
| Tbl3 | 0.1343789 | 0.10484285 | 0.11084819 | -0.0060053 | 0.64080683 | 0.99999994 |
| Dusp28 | 0.14546469 | 0.11294021 | 0.11894711 | -0.0060069 | 0.63162111 | 0.99999994 |
| Bag4 | 0.21701941 | 0.1618357 | 0.16784762 | -0.0060119 | 0.59422065 | 0.99999994 |
| Pitpnm1 | 0.02244681 | 0.01400168 | 0.02001935 | -0.0060177 | 0.9772958 | 0.99999994 |
| Lgalsl | 0.04832784 | 0.03634863 | 0.04236669 | -0.0060181 | 0.8278023 | 0.99999994 |
| Usp40 | 0.07603208 | 0.05944074 | 0.06545905 | -0.0060183 | 0.72972664 | 0.99999994 |
| Fam166a | 0.02283795 | 0.01434284 | 0.02036284 | -0.00602 | 0.9754581 | 0.99999994 |

| | | | | | | |
|---|---|---|---|---|---|---|
| Stx3 | 0.21686874 | 0.16173048 | 0.16775051 | -0.00602 | 0.59439883 | 0.99999994 |
| Secisbp2l | 0.09001189 | 0.07077784 | 0.07680025 | -0.0060224 | 0.69915414 | 0.99999994 |
| Cyb5d2 | 0.0501482 | 0.03788514 | 0.04390986 | -0.0060247 | 0.81945625 | 0.99999994 |
| Eif3j2 | 0.18278017 | 0.13913156 | 0.14515741 | -0.0060259 | 0.60884971 | 0.99999994 |
| Cstf3 | 0.37955498 | 0.25389634 | 0.25992745 | -0.0060311 | 0.56137168 | 0.99999994 |
| Snrnp35 | 0.10818966 | 0.08508594 | 0.0911231 | -0.0060372 | 0.67039446 | 0.99999994 |
| Znhit6 | 0.14778641 | 0.11458698 | 0.12062468 | -0.0060377 | 0.63049875 | 0.99999994 |
| Flot1 | 0.26039649 | 0.18879515 | 0.19484092 | -0.0060458 | 0.58181621 | 0.99999994 |
| Tmco6 | 0.04210848 | 0.03101971 | 0.03706633 | -0.0060466 | 0.86122728 | 0.99999994 |
| Aamdc | 0.08984473 | 0.07061749 | 0.07666584 | -0.0060484 | 0.70025575 | 0.99999994 |
| Zfp511 | 0.15314468 | 0.11842048 | 0.12447221 | -0.0060517 | 0.62688949 | 0.99999994 |
| Trappc10 | 0.06491342 | 0.05023873 | 0.05629146 | -0.0060527 | 0.76291917 | 0.99999994 |
| Zpr1 | 0.17517014 | 0.13388754 | 0.13994075 | -0.0060532 | 0.61329759 | 0.99999994 |
| Fbxo16 | 0.13359066 | 0.10421136 | 0.11026677 | -0.0060554 | 0.64264608 | 0.99999994 |
| Asl | 0.21496449 | 0.16046405 | 0.16652109 | -0.006057 | 0.59565591 | 0.99999994 |
| Tmem94 | 0.0603825 | 0.04645915 | 0.0525174 | -0.0060583 | 0.77871413 | 0.99999994 |
| Gtpbp3 | 0.06400923 | 0.04947865 | 0.05554009 | -0.0060614 | 0.76621837 | 0.99999994 |
| B3galt4 | 0.05813964 | 0.04457933 | 0.05064091 | -0.0060616 | 0.78720131 | 0.99999994 |
| Arih2 | 0.10743205 | 0.08447623 | 0.09053982 | -0.0060636 | 0.67212417 | 0.99999994 |
| Rpl14 | 3.2518593 | 0.28413606 | 0.29020193 | -0.0060659 | 0.55532737 | 0.99999994 |
| Isca2 | 0.34466882 | 0.23608607 | 0.24215595 | -0.0060699 | 0.56625534 | 0.99999994 |
| Dync2li1 | 0.33187079 | 0.22930991 | 0.2353799 | -0.00607 | 0.5681454 | 0.99999994 |
| Pcyt2 | 0.16176862 | 0.12451884 | 0.13059518 | -0.0060763 | 0.62160824 | 0.99999994 |
| Tesk1 | 0.05150953 | 0.03898473 | 0.04506148 | -0.0060768 | 0.81531148 | 0.99999994 |
| Ppil1 | 0.19748846 | 0.14898011 | 0.15505933 | -0.0060792 | 0.60294127 | 0.99999994 |
| Zfat | 0.03428572 | 0.0242555 | 0.03033726 | -0.0060818 | 0.90896544 | 0.99999994 |
| Zbtb11 | 0.07516873 | 0.05866944 | 0.06475193 | -0.0060825 | 0.73410194 | 0.99999994 |
| Stard3nl | 0.29725874 | 0.21025169 | 0.2163346 | -0.0060829 | 0.57423396 | 0.99999994 |
| Deptor | 0.28929773 | 0.20571266 | 0.2118006 | -0.0060879 | 0.57586587 | 0.99999994 |
| Opn3 | 0.04832038 | 0.03627019 | 0.04236036 | -0.0060902 | 0.83071339 | 0.99999994 |

| | | | | | | |
|---|---|---|---|---|---|---|
| Pcnp | 0.6202225 | 0.35238687 | 0.35847825 | -0.0060914 | 0.54502715 | 0.99999994 |
| Eml5 | 0.14455041 | 0.11219267 | 0.11828463 | -0.006092 | 0.63413288 | 0.99999994 |
| Zfp157 | 0.12622824 | 0.0987067 | 0.10479955 | -0.0060929 | 0.65061754 | 0.99999994 |
| Ddx28 | 0.07970616 | 0.0623658 | 0.06845949 | -0.0060937 | 0.72324004 | 0.99999994 |
| Ncaph2 | 0.19425371 | 0.14680739 | 0.15290205 | -0.0060947 | 0.60461918 | 0.99999994 |
| Vegfb | 0.0511597 | 0.03867038 | 0.04476573 | -0.0060954 | 0.81762144 | 0.99999994 |
| Dnttip1 | 0.19215495 | 0.14539734 | 0.15149637 | -0.006099 | 0.6056409 | 0.99999994 |
| Acp6 | 0.13411793 | 0.1045554 | 0.11065578 | -0.0061004 | 0.64317579 | 0.99999994 |
| Memo1 | 0.17093992 | 0.13090968 | 0.13701253 | -0.0061029 | 0.61657196 | 0.99999994 |
| Rnf139 | 0.07588785 | 0.05923541 | 0.06534098 | -0.0061056 | 0.7330229 | 0.99999994 |
| Snhg9 | 0.03819701 | 0.02760476 | 0.03371045 | -0.0061057 | 0.88601601 | 0.99999994 |
| Tcf20 | 0.068886 | 0.05347638 | 0.05958216 | -0.0061058 | 0.75241877 | 0.99999994 |
| Ambra1 | 0.04328172 | 0.03196339 | 0.03806954 | -0.0061062 | 0.85715945 | 0.99999994 |
| Cpeb3 | 0.02797491 | 0.01874713 | 0.02485753 | -0.0061104 | 0.94910574 | 0.99999994 |
| Dmap1 | 0.16266063 | 0.12511043 | 0.13122351 | -0.0061131 | 0.62175293 | 0.99999994 |
| Wdr90 | 0.03187718 | 0.02213817 | 0.02825134 | -0.0061132 | 0.92511391 | 0.99999994 |
| Tmem216 | 0.16557447 | 0.12715458 | 0.13326972 | -0.0061151 | 0.61998071 | 0.99999994 |
| Scnm1 | 0.19607638 | 0.14800307 | 0.15411898 | -0.0061159 | 0.6041656 | 0.99999994 |
| Ankib1 | 0.06130236 | 0.04716501 | 0.05328541 | -0.0061204 | 0.77773127 | 0.99999994 |
| Azi2 | 0.23436578 | 0.17274364 | 0.17887066 | -0.006127 | 0.59017927 | 0.99999994 |
| Mras | 0.08801476 | 0.06906535 | 0.07519254 | -0.0061272 | 0.70622991 | 0.99999994 |
| Scd2 | 0.18510102 | 0.14060643 | 0.14673561 | -0.0061292 | 0.60950839 | 0.99999994 |
| Ubxn10 | 0.07648219 | 0.05969758 | 0.0658274 | -0.0061298 | 0.73231781 | 0.99999994 |
| Bdh1 | 0.03436199 | 0.02427033 | 0.03040321 | -0.0061329 | 0.91031676 | 0.99999994 |
| Supt7l | 0.09750264 | 0.07666044 | 0.08279391 | -0.0061335 | 0.6890329 | 0.99999994 |
| Sgsm3 | 0.10024969 | 0.07883838 | 0.08497426 | -0.0061359 | 0.68461624 | 0.99999994 |
| Rab21 | 0.04302869 | 0.03171728 | 0.03785332 | -0.006136 | 0.85970566 | 0.99999994 |
| Stk40 | 0.07796659 | 0.06090422 | 0.06704065 | -0.0061364 | 0.72882921 | 0.99999994 |
| Gsk3a | 0.09128756 | 0.07168537 | 0.07782503 | -0.0061397 | 0.70025144 | 0.99999994 |
| Sncb | 0.02500561 | 0.01611936 | 0.02226325 | -0.0061439 | 0.96688957 | 0.99999994 |

| | | | | | | |
|---|---|---|---|---|---|---|
| 4932422M17 | 0.05820207 | 0.04454623 | 0.05069322 | -0.006147 | 0.79020727 | 0.99999994 |
| Ift52 | 0.20676669 | 0.15503798 | 0.16118537 | -0.0061474 | 0.60019991 | 0.99999994 |
| Rps6ka3 | 0.09340274 | 0.07336405 | 0.07952053 | -0.0061565 | 0.69684055 | 0.99999994 |
| Msl2 | 0.08582949 | 0.06727207 | 0.07342863 | -0.0061566 | 0.71161081 | 0.99999994 |
| Cdk5r2 | 0.03159945 | 0.02185228 | 0.02801038 | -0.0061581 | 0.92832169 | 0.99999994 |
| Ikbkb | 0.11315908 | 0.08876698 | 0.09493127 | -0.0061643 | 0.66721119 | 0.99999994 |
| Cpsf3 | 0.25793769 | 0.18719517 | 0.19336113 | -0.006166 | 0.58404798 | 0.99999994 |
| AW209491 | 0.17783703 | 0.13560902 | 0.14177634 | -0.0061673 | 0.6139212 | 0.99999994 |
| Nudt16l1 | 0.48841571 | 0.30315689 | 0.30932783 | -0.0061709 | 0.55282065 | 0.99999994 |
| Mink1 | 0.04467904 | 0.0330911 | 0.03926233 | -0.0061712 | 0.85227556 | 0.99999994 |
| Khsrp | 0.04877285 | 0.0365726 | 0.04274428 | -0.0061717 | 0.83174521 | 0.99999994 |
| Nacc1 | 0.07335663 | 0.05709091 | 0.06326515 | -0.0061742 | 0.74202946 | 0.99999994 |
| Ptpn6 | 0.12800675 | 0.0999514 | 0.10612633 | -0.0061749 | 0.65073232 | 0.99999994 |
| Igf1r | 0.23251229 | 0.17152833 | 0.17770749 | -0.0061792 | 0.59151798 | 0.99999994 |
| 2310022B05 | 0.46339873 | 0.29255478 | 0.2987373 | -0.0061825 | 0.55478364 | 0.99999994 |
| Mepce | 0.07886707 | 0.06159205 | 0.06777551 | -0.0061835 | 0.72817028 | 0.99999994 |
| Dsp | 0.08278236 | 0.06477449 | 0.07096076 | -0.0061863 | 0.71914125 | 0.99999994 |
| Trappc5 | 0.30176404 | 0.21268229 | 0.2188745 | -0.0061922 | 0.57468553 | 0.99999994 |
| Snx27 | 0.13396847 | 0.10435309 | 0.11054555 | -0.0061925 | 0.64537764 | 0.99999994 |
| Mapk8 | 0.12988138 | 0.10132662 | 0.10752069 | -0.0061941 | 0.64931075 | 0.99999994 |
| Trim25 | 0.06474803 | 0.04995884 | 0.05615409 | -0.0061952 | 0.76863793 | 0.99999994 |
| Ddx39b | 1.20245365 | 0.42273813 | 0.42893762 | -0.0061995 | 0.53832122 | 0.99999994 |
| Cttn | 0.33951674 | 0.23324504 | 0.23944466 | -0.0061996 | 0.56841652 | 0.99999994 |
| Pcyox1 | 0.09245368 | 0.07255367 | 0.07876035 | -0.0062067 | 0.70005258 | 0.99999994 |
| Tkfc | 0.08620367 | 0.06752433 | 0.07373101 | -0.0062067 | 0.71237379 | 0.99999994 |
| Serf1 | 0.74120275 | 0.38546173 | 0.39167163 | -0.0062099 | 0.54202464 | 0.99999994 |
| Ssx2ip | 0.06467406 | 0.04987994 | 0.05609264 | -0.0062127 | 0.76951284 | 0.99999994 |
| Arhgef12 | 0.07689091 | 0.05993873 | 0.06616169 | -0.006223 | 0.73436741 | 0.99999994 |
| Vamp3 | 0.39583583 | 0.2616598 | 0.2678831 | -0.0062233 | 0.56144606 | 0.99999994 |
| Add1 | 0.23300369 | 0.17178924 | 0.1780162 | -0.006227 | 0.59205636 | 0.99999994 |

| | | | | | | |
|---|---|---|---|---|---|---|
| Mon1a | 0.06068464 | 0.04654226 | 0.05276976 | -0.0062275 | 0.78392986 | 0.99999994 |
| Rita1 | 0.10265449 | 0.08061738 | 0.08684498 | -0.0062276 | 0.68343411 | 0.99999994 |
| Sypl | 0.39356824 | 0.26056019 | 0.2667878 | -0.0062276 | 0.56173871 | 0.99999994 |
| Thsd4 | 0.02436465 | 0.01546984 | 0.02170189 | -0.0062321 | 0.97202866 | 0.99999994 |
| Myo18a | 0.08437317 | 0.06601511 | 0.07225037 | -0.0062353 | 0.71716781 | 0.99999994 |
| Zfp810 | 0.06106291 | 0.04685029 | 0.05308558 | -0.0062353 | 0.78284504 | 0.99999994 |
| Gp9 | 0.02817166 | 0.0187871 | 0.02502907 | -0.006242 | 0.95154494 | 0.99999994 |
| Atp1b3 | 0.20524025 | 0.15394154 | 0.16018377 | -0.0062422 | 0.60233395 | 0.99999994 |
| Inpp5a | 0.03267811 | 0.02269401 | 0.02894573 | -0.0062517 | 0.92473279 | 0.99999994 |
| Gripap1 | 0.25863074 | 0.18752599 | 0.19377882 | -0.0062528 | 0.58503318 | 0.99999994 |
| Hibch | 0.16806732 | 0.12875377 | 0.1350127 | -0.0062589 | 0.62117788 | 0.99999994 |
| Pgs1 | 0.16199432 | 0.12449012 | 0.13075424 | -0.0062641 | 0.62509517 | 0.99999994 |
| Igf2bp3 | 0.27403171 | 0.19667576 | 0.20294133 | -0.0062656 | 0.58141196 | 0.99999994 |
| Maea | 0.33203588 | 0.22920142 | 0.23546818 | -0.0062668 | 0.57030581 | 0.99999994 |
| Zfp444 | 0.24581428 | 0.17971074 | 0.18597753 | -0.0062668 | 0.58873651 | 0.99999994 |
| Lrrc41 | 0.27338425 | 0.19629336 | 0.20256075 | -0.0062674 | 0.5815863 | 0.99999994 |
| Myo10 | 0.13209316 | 0.10289274 | 0.10916017 | -0.0062674 | 0.64883399 | 0.99999994 |
| Leng9 | 0.04867001 | 0.03638873 | 0.04265704 | -0.0062683 | 0.83599491 | 0.99999994 |
| Rere | 0.28887528 | 0.20528583 | 0.21155839 | -0.0062726 | 0.57822573 | 0.99999994 |
| Trmt13 | 0.14408721 | 0.11167484 | 0.11794864 | -0.0062738 | 0.63835223 | 0.99999994 |
| Gprin1 | 0.03870533 | 0.02786575 | 0.03414755 | -0.0062818 | 0.88961937 | 0.99999994 |
| Ift57 | 0.31123238 | 0.21787019 | 0.22415231 | -0.0062821 | 0.57399401 | 0.99999994 |
| Arhgef3 | 0.02731558 | 0.01799961 | 0.02428236 | -0.0062828 | 0.95748769 | 0.99999994 |
| Btaf1 | 0.06043814 | 0.04627894 | 0.05256388 | -0.0062849 | 0.78695076 | 0.99999994 |
| Tsfm | 0.32961567 | 0.227885 | 0.23417149 | -0.0062865 | 0.5709112 | 0.99999994 |
| Naa50 | 0.26678063 | 0.19236754 | 0.19865619 | -0.0062887 | 0.58344455 | 0.99999994 |
| Ddx31 | 0.04303641 | 0.03156578 | 0.03785991 | -0.0062941 | 0.86576798 | 0.99999994 |
| Eif4e3 | 0.24614552 | 0.17988679 | 0.18618121 | -0.0062944 | 0.58902525 | 0.99999994 |
| Sin3a | 0.10947163 | 0.0858138 | 0.09210845 | -0.0062946 | 0.67540654 | 0.99999994 |
| Ofd1 | 0.10119445 | 0.0794154 | 0.08571007 | -0.0062947 | 0.68752696 | 0.99999994 |

| | | | | | | |
|---|---|---|---|---|---|---|
| Zc3h10 | 0.08187858 | 0.06392959 | 0.07022692 | -0.0062973 | 0.72470732 | 0.99999994 |
| Gtf3c2 | 0.16208126 | 0.12451249 | 0.1308155 | -0.006303 | 0.62578857 | 0.99999994 |
| Pdik1l | 0.0852032 | 0.06661428 | 0.07292218 | -0.0063079 | 0.7176204 | 0.99999994 |
| 2700062C07I | 0.13554758 | 0.10539782 | 0.11170885 | -0.006311 | 0.64656146 | 0.99999994 |
| Gabbr1 | 0.07259788 | 0.05632912 | 0.06264158 | -0.0063125 | 0.74881611 | 0.99999994 |
| Lrrc14 | 0.09248393 | 0.07247186 | 0.07878459 | -0.0063127 | 0.70310471 | 0.99999994 |
| Ddx18 | 0.31530068 | 0.22008203 | 0.22639558 | -0.0063135 | 0.5736315 | 0.99999994 |
| Fam177a | 0.18521812 | 0.14049834 | 0.14681509 | -0.0063167 | 0.61271059 | 0.99999994 |
| Osbpl9 | 0.11454846 | 0.08967249 | 0.0959905 | -0.006318 | 0.66934863 | 0.99999994 |
| Limk2 | 0.16049747 | 0.12337756 | 0.12969808 | -0.0063205 | 0.62717531 | 0.99999994 |
| Kbtbd4 | 0.0870542 | 0.06809634 | 0.0744178 | -0.0063215 | 0.71410848 | 0.99999994 |
| Tmem168 | 0.06705435 | 0.05174425 | 0.05806701 | -0.0063228 | 0.76571059 | 0.99999994 |
| Tada3 | 0.21522065 | 0.16036345 | 0.16668669 | -0.0063232 | 0.59967597 | 0.99999994 |
| Ube3c | 0.17142507 | 0.13102479 | 0.13734937 | -0.0063246 | 0.62039105 | 0.99999994 |
| Akr1b10 | 0.04156195 | 0.0302719 | 0.03659848 | -0.0063266 | 0.87506123 | 0.99999994 |
| Sgpl1 | 0.08738672 | 0.06835896 | 0.0746861 | -0.0063271 | 0.71359002 | 0.99999994 |
| Fbxo38 | 0.07374217 | 0.05725378 | 0.06358177 | -0.006328 | 0.74616689 | 0.99999994 |
| Ubr7 | 0.1142543 | 0.08943728 | 0.09576644 | -0.0063292 | 0.67000091 | 0.99999994 |
| Cdc73 | 0.19430815 | 0.14660554 | 0.15293845 | -0.0063329 | 0.60858307 | 0.99999994 |
| Ier3 | 0.05225491 | 0.03935801 | 0.04569116 | -0.0063332 | 0.82189816 | 0.99999994 |
| Vhl | 0.12380974 | 0.09665583 | 0.10298918 | -0.0063334 | 0.658853 | 0.99999994 |
| Pogz | 0.09320141 | 0.07302314 | 0.07935934 | -0.0063362 | 0.70244866 | 0.99999994 |
| Arfip1 | 0.18619026 | 0.14113586 | 0.14747428 | -0.0063384 | 0.61259512 | 0.99999994 |
| Trmt10b | 0.09845702 | 0.07721063 | 0.08355344 | -0.0063428 | 0.69332787 | 0.99999994 |
| Pcdh1 | 0.0643029 | 0.04944055 | 0.05578422 | -0.0063437 | 0.7754594 | 0.99999994 |
| Itga6 | 0.08765935 | 0.0685616 | 0.074906 | -0.0063444 | 0.71354832 | 0.99999994 |
| Ppt1 | 0.14172995 | 0.1098871 | 0.11623482 | -0.0063477 | 0.64188863 | 0.99999994 |
| Shmt2 | 0.1276266 | 0.09949512 | 0.10584306 | -0.0063479 | 0.65513255 | 0.99999994 |
| Tigar | 0.05958168 | 0.04549905 | 0.05184803 | -0.006349 | 0.79249118 | 0.99999994 |
| Mcrs1 | 0.30230322 | 0.21282055 | 0.21917718 | -0.0063566 | 0.57653988 | 0.99999994 |

| | | | | | | |
|---|---|---|---|---|---|---|
| Maz | 0.22625743 | 0.16739681 | 0.1737567 | -0.0063599 | 0.59624432 | 0.99999994 |
| Nupr1l | 0.06709873 | 0.05174069 | 0.05810377 | -0.0063631 | 0.7669849 | 0.99999994 |
| Commd7 | 0.42306258 | 0.27436294 | 0.28072653 | -0.0063636 | 0.55996803 | 0.99999994 |
| Rad23b | 0.63731527 | 0.35770775 | 0.3640756 | -0.0063678 | 0.54634125 | 0.99999994 |
| Surf1 | 0.21878112 | 0.16261255 | 0.16898141 | -0.0063689 | 0.5990452 | 0.99999994 |
| A430005L14I | 0.32989596 | 0.227953 | 0.23432199 | -0.006369 | 0.57178607 | 0.99999994 |
| Zkscan1 | 0.2588442 | 0.18753641 | 0.19390738 | -0.006371 | 0.58655786 | 0.99999994 |
| Pag1 | 0.01866443 | 0.01031525 | 0.0166883 | -0.006373 | 0.99448863 | 0.99999994 |
| Ncoa6 | 0.08616429 | 0.06732423 | 0.07369919 | -0.006375 | 0.71761572 | 0.99999994 |
| Kcnip1 | 0.02031919 | 0.01176895 | 0.01814768 | -0.0063787 | 0.99034896 | 0.99999994 |
| Map3k7 | 0.07528526 | 0.05846588 | 0.06484741 | -0.0063815 | 0.74377966 | 0.99999994 |
| Ankra2 | 0.30026499 | 0.21164759 | 0.2180316 | -0.006384 | 0.57726426 | 0.99999994 |
| Pak4 | 0.18951077 | 0.14333216 | 0.14971814 | -0.006386 | 0.61176253 | 0.99999994 |
| Srbd1 | 0.05276097 | 0.03972899 | 0.04611833 | -0.0063893 | 0.82178434 | 0.99999994 |
| Cep350 | 0.09271859 | 0.07257743 | 0.07897263 | -0.0063952 | 0.70506741 | 0.99999994 |
| Tmem14a | 0.06781123 | 0.05229742 | 0.05869354 | -0.0063961 | 0.76588929 | 0.99999994 |
| Tyw1 | 0.08767532 | 0.06851482 | 0.07491887 | -0.0064041 | 0.71531656 | 0.99999994 |
| Sox12 | 0.0603874 | 0.04611675 | 0.05252149 | -0.0064047 | 0.79152128 | 0.99999994 |
| Mark3 | 0.14141648 | 0.10960111 | 0.11600642 | -0.0064053 | 0.64338872 | 0.99999994 |
| Abca2 | 0.03608466 | 0.02548275 | 0.03189089 | -0.0064081 | 0.90947438 | 0.99999994 |
| Chrm3 | 0.02392597 | 0.01490883 | 0.02131741 | -0.0064086 | 0.97730797 | 0.99999994 |
| Brd1 | 0.09088732 | 0.07108992 | 0.07750369 | -0.0064138 | 0.70912932 | 0.99999994 |
| Erc2 | 0.02252599 | 0.01367062 | 0.0200889 | -0.0064183 | 0.98327605 | 0.99999994 |
| Klhl12 | 0.07107736 | 0.05497164 | 0.06139012 | -0.0064185 | 0.7567661 | 0.99999994 |
| Kif5c | 0.07573942 | 0.05880031 | 0.06521944 | -0.0064191 | 0.74381199 | 0.99999994 |
| Zdhhc8 | 0.04471388 | 0.03287169 | 0.03929205 | -0.0064204 | 0.86162624 | 0.99999994 |
| Pik3r4 | 0.05453254 | 0.04119103 | 0.04761145 | -0.0064204 | 0.81530387 | 0.99999994 |
| Urm1 | 0.30943293 | 0.21673282 | 0.22315537 | -0.0064225 | 0.57596209 | 0.99999994 |
| Qpctl | 0.13387548 | 0.10405158 | 0.11047695 | -0.0064254 | 0.65067222 | 0.99999994 |
| Scmh1 | 0.11521247 | 0.09007006 | 0.09649588 | -0.0064258 | 0.67121044 | 0.99999994 |

| | | | | | | |
|---|---|---|---|---|---|---|
| Egln2 | 0.4814369 | 0.29998269 | 0.30641668 | -0.006434 | 0.55557789 | 0.99999994 |
| Chchd10 | 0.4800469 | 0.2993853 | 0.30583284 | -0.0064475 | 0.5557998 | 0.99999994 |
| Rac3 | 0.09736159 | 0.07623088 | 0.08268159 | -0.0064507 | 0.69823126 | 0.99999994 |
| Alkbh7 | 0.27920242 | 0.19951528 | 0.20596665 | -0.0064514 | 0.58257799 | 0.99999994 |
| H2-Q4 | 0.02933349 | 0.0195884 | 0.02604109 | -0.0064527 | 0.9504642 | 0.99999994 |
| Trip4 | 0.13871802 | 0.10757867 | 0.11403551 | -0.0064568 | 0.64687279 | 0.99999994 |
| Abhd18 | 0.11463264 | 0.08959646 | 0.09605459 | -0.0064581 | 0.67275501 | 0.99999994 |
| Arid4a | 0.18494215 | 0.14016897 | 0.14662776 | -0.0064588 | 0.61531745 | 0.99999994 |
| Fance | 0.06878298 | 0.0530345 | 0.05949704 | -0.0064625 | 0.7651632 | 0.99999994 |
| Aebp2 | 0.07015036 | 0.0541623 | 0.06062593 | -0.0064636 | 0.76104468 | 0.99999994 |
| Rtcb | 0.5901027 | 0.34171435 | 0.34817822 | -0.0064639 | 0.54917374 | 0.99999994 |
| Xkr6 | 0.05014215 | 0.0374402 | 0.04390473 | -0.0064645 | 0.83647717 | 0.99999994 |
| Acadm | 0.24888521 | 0.18139232 | 0.18786167 | -0.0064693 | 0.59065233 | 0.99999994 |
| Cbll1 | 0.1518111 | 0.1170413 | 0.12351773 | -0.0064764 | 0.63646072 | 0.99999994 |
| Plin3 | 0.20668684 | 0.15465584 | 0.16113304 | -0.0064772 | 0.60548523 | 0.99999994 |
| Mrpl16 | 0.21584369 | 0.16060482 | 0.16708919 | -0.0064844 | 0.60191949 | 0.99999994 |
| Wdr5 | 0.10574745 | 0.08275512 | 0.08924027 | -0.0064852 | 0.68570845 | 0.99999994 |
| Fam149b | 0.05946932 | 0.04526607 | 0.05175405 | -0.006488 | 0.79798731 | 0.99999994 |
| Arid2 | 0.15698797 | 0.12071581 | 0.12721149 | -0.0064957 | 0.63302925 | 0.99999994 |
| Diaph1 | 0.14709609 | 0.11362611 | 0.12012656 | -0.0065005 | 0.64065041 | 0.99999994 |
| Tor1a | 0.36951752 | 0.24841235 | 0.25491663 | -0.0065043 | 0.56743166 | 0.99999994 |
| Armcx4 | 0.04689736 | 0.03464652 | 0.04115143 | -0.0065049 | 0.8536433 | 0.99999994 |
| Nol8 | 0.10514694 | 0.08227042 | 0.08877616 | -0.0065057 | 0.68715246 | 0.99999994 |
| 0610030E20I | 0.09843706 | 0.07703052 | 0.08353757 | -0.0065071 | 0.69794121 | 0.99999994 |
| Emc9 | 0.20787996 | 0.15540705 | 0.16191416 | -0.0065071 | 0.60546186 | 0.99999994 |
| Ralgapa1 | 0.10200188 | 0.0798291 | 0.08633803 | -0.0065089 | 0.69210147 | 0.99999994 |
| Xiap | 0.16153216 | 0.12391916 | 0.13042846 | -0.0065093 | 0.63013056 | 0.99999994 |
| 5830454E08I | 0.03324987 | 0.02292296 | 0.02944098 | -0.006518 | 0.9297145 | 0.99999994 |
| Abr | 0.11148085 | 0.08712625 | 0.09364864 | -0.0065224 | 0.67853237 | 0.99999994 |
| Snip1 | 0.11638518 | 0.09086241 | 0.09738695 | -0.0065245 | 0.67218326 | 0.99999994 |

| | | | | | | |
|---|---|---|---|---|---|---|
| Reep2 | 0.05345668 | 0.04017391 | 0.0467051 | -0.0065312 | 0.82402606 | 0.99999994 |
| Smg6 | 0.16697777 | 0.12771881 | 0.13425175 | -0.0065329 | 0.62699732 | 0.99999994 |
| Usp15 | 0.18728374 | 0.14168101 | 0.14821453 | -0.0065335 | 0.61540033 | 0.99999994 |
| Mbd6 | 0.09713181 | 0.07596405 | 0.08249854 | -0.0065345 | 0.70098399 | 0.99999994 |
| Aff4 | 0.29986776 | 0.21127273 | 0.21780791 | -0.0065352 | 0.57914994 | 0.99999994 |
| Fhdc1 | 0.03803947 | 0.02703972 | 0.03357492 | -0.0065352 | 0.90244835 | 0.99999994 |
| Nap1l4 | 0.65529616 | 0.36321177 | 0.36974795 | -0.0065362 | 0.54683429 | 0.99999994 |
| Htt | 0.07418932 | 0.05741021 | 0.06394879 | -0.0065386 | 0.75193559 | 0.99999994 |
| Hscb | 0.07722237 | 0.05989123 | 0.06643267 | -0.0065414 | 0.74390551 | 0.99999994 |
| Rrp36 | 0.09717611 | 0.07599152 | 0.08253384 | -0.0065423 | 0.70112472 | 0.99999994 |
| Cul5 | 0.35255697 | 0.23971889 | 0.24626233 | -0.0065434 | 0.57019324 | 0.99999994 |
| Tradd | 0.12157266 | 0.09476285 | 0.10130829 | -0.0065454 | 0.66642448 | 0.99999994 |
| Srgap1 | 0.05116697 | 0.03822445 | 0.04477189 | -0.0065474 | 0.83482909 | 0.99999994 |
| Epb41l4a | 0.07337595 | 0.05673262 | 0.06328102 | -0.0065484 | 0.7545282 | 0.99999994 |
| Fsd1l | 0.06160683 | 0.0469895 | 0.05353942 | -0.0065499 | 0.79227135 | 0.99999994 |
| Arpin | 0.10661973 | 0.08336339 | 0.08991361 | -0.0065502 | 0.68613229 | 0.99999994 |
| Yars | 0.18442196 | 0.13972312 | 0.14627445 | -0.0065513 | 0.61719587 | 0.99999994 |
| Phf21b | 0.0496471 | 0.03693333 | 0.04348543 | -0.0065521 | 0.84205435 | 0.99999994 |
| Nqo2 | 0.07888494 | 0.06123468 | 0.06779008 | -0.0065554 | 0.74010841 | 0.99999994 |
| Prss36 | 0.04684288 | 0.03454814 | 0.04110511 | -0.006557 | 0.85583736 | 0.99999994 |
| Sh3glb2 | 0.24858706 | 0.18111958 | 0.18767915 | -0.0065596 | 0.59198191 | 0.99999994 |
| Hddc3 | 0.23442791 | 0.17234832 | 0.17890959 | -0.0065613 | 0.59642828 | 0.99999994 |
| Rnf19b | 0.03337401 | 0.02298647 | 0.02954846 | -0.006562 | 0.9303247 | 0.99999994 |
| Map6 | 0.03968085 | 0.02841884 | 0.03498557 | -0.0065667 | 0.89423829 | 0.99999994 |
| Mfap1b | 0.16538507 | 0.12656503 | 0.13313701 | -0.006572 | 0.62876133 | 0.99999994 |
| Fahd2a | 0.17767655 | 0.13509289 | 0.14166612 | -0.0065732 | 0.62128274 | 0.99999994 |
| Parp8 | 0.04035773 | 0.02899045 | 0.03556641 | -0.006576 | 0.89078409 | 0.99999994 |
| Cuta | 0.79632906 | 0.39625136 | 0.40282734 | -0.006576 | 0.5432648 | 0.99999994 |
| Zcchc8 | 0.10111561 | 0.07907257 | 0.08564871 | -0.0065761 | 0.69535166 | 0.99999994 |
| Tbc1d14 | 0.05082123 | 0.03789679 | 0.04447947 | -0.0065827 | 0.83771227 | 0.99999994 |

| | | | | | | |
|---|---|---|---|---|---|---|
| Cdk14 | 0.10442071 | 0.081627 | 0.08821428 | -0.0065873 | 0.69042462 | 0.99999994 |
| Pip4k2c | 0.12710637 | 0.09886691 | 0.10545512 | -0.0065882 | 0.66124002 | 0.99999994 |
| Zfpl1 | 0.22603476 | 0.16702601 | 0.17361532 | -0.0065893 | 0.59972378 | 0.99999994 |
| Gstm4 | 0.08983987 | 0.07007163 | 0.07666194 | -0.0065903 | 0.71641327 | 0.99999994 |
| Atxn7 | 0.07025609 | 0.05412258 | 0.06071314 | -0.0065906 | 0.76502608 | 0.99999994 |
| Htatsf1 | 0.24108147 | 0.17646484 | 0.18305564 | -0.0065908 | 0.59470236 | 0.99999994 |
| Ppp2r5a | 0.12055794 | 0.09394886 | 0.10054383 | -0.006595 | 0.66880304 | 0.99999994 |
| Asah1 | 0.41522071 | 0.27048935 | 0.27708507 | -0.0065957 | 0.56294793 | 0.99999994 |
| Vamp4 | 0.31430438 | 0.21925179 | 0.22584772 | -0.0065959 | 0.57706874 | 0.99999994 |
| Kat2b | 0.06448325 | 0.04933638 | 0.0559341 | -0.0065977 | 0.78381795 | 0.99999994 |
| Ei24 | 0.51753869 | 0.31452173 | 0.32112378 | -0.006602 | 0.55442506 | 0.99999994 |
| Pknox1 | 0.07971912 | 0.06186774 | 0.06847006 | -0.0066023 | 0.73951433 | 0.99999994 |
| Map3k2 | 0.11946612 | 0.0931167 | 0.09971989 | -0.0066032 | 0.67030908 | 0.99999994 |
| Pitrm1 | 0.05259607 | 0.03937047 | 0.04597917 | -0.0066087 | 0.83064941 | 0.99999994 |
| Mccc2 | 0.0972228 | 0.07596062 | 0.08257104 | -0.0066104 | 0.70294528 | 0.99999994 |
| Tmem102 | 0.05241792 | 0.03921835 | 0.0458288 | -0.0066104 | 0.83150461 | 0.99999994 |
| Sdcbp2 | 0.04647791 | 0.03417917 | 0.04079466 | -0.0066155 | 0.85980253 | 0.99999994 |
| Arhgef17 | 0.09483231 | 0.07404824 | 0.08066386 | -0.0066156 | 0.70743905 | 0.99999994 |
| Nfkbil1 | 0.14417809 | 0.11139848 | 0.11801458 | -0.0066161 | 0.64548795 | 0.99999994 |
| Glce | 0.05848819 | 0.04431542 | 0.05093289 | -0.0066175 | 0.80643383 | 0.99999994 |
| Slc25a25 | 0.06864253 | 0.05275592 | 0.05938097 | -0.006625 | 0.77115003 | 0.99999994 |
| Fuca2 | 0.06394127 | 0.04885597 | 0.05548358 | -0.0066276 | 0.78672613 | 0.99999994 |
| Nf1 | 0.10143321 | 0.07926737 | 0.08589585 | -0.0066285 | 0.69625641 | 0.99999994 |
| Nup88 | 0.27616965 | 0.19756382 | 0.20419523 | -0.0066314 | 0.58557266 | 0.99999994 |
| Wdr77 | 0.22182706 | 0.16429997 | 0.17093418 | -0.0066342 | 0.60192902 | 0.99999994 |
| Bbs2 | 0.05642587 | 0.04256574 | 0.04920337 | -0.0066376 | 0.81539562 | 0.99999994 |
| Zfr | 0.66181775 | 0.36510021 | 0.37173964 | -0.0066394 | 0.54731694 | 0.99999994 |
| Csde1 | 0.61514651 | 0.35014094 | 0.35678059 | -0.0066397 | 0.54929256 | 0.99999994 |
| Ears2 | 0.08152953 | 0.06330058 | 0.06994328 | -0.0066427 | 0.73636281 | 0.99999994 |
| Wdr26 | 0.28772544 | 0.20425343 | 0.21089833 | -0.0066449 | 0.5830591 | 0.99999994 |

| | | | | | | |
|---|---|---|---|---|---|---|
| Zc3h4 | 0.04025502 | 0.02883294 | 0.0354783 | -0.0066454 | 0.89376224 | 0.99999994 |
| Hcfc1 | 0.15861406 | 0.1217119 | 0.12836537 | -0.0066535 | 0.63495834 | 0.99999994 |
| Dusp18 | 0.18678094 | 0.14121891 | 0.14787431 | -0.0066554 | 0.61775273 | 0.99999994 |
| Sfswap | 0.15725965 | 0.12074874 | 0.12740449 | -0.0066558 | 0.63598058 | 0.99999994 |
| Thrap3 | 0.61283535 | 0.34934605 | 0.35600299 | -0.0066569 | 0.54952758 | 0.99999994 |
| Pwp1 | 0.10236696 | 0.07996357 | 0.08662169 | -0.0066581 | 0.69554737 | 0.99999994 |
| Cenpv | 0.05822628 | 0.04405517 | 0.0507135 | -0.0066583 | 0.8089228 | 0.99999994 |
| Arfgef2 | 0.03953471 | 0.02820117 | 0.03486009 | -0.0066589 | 0.89822246 | 0.99999994 |
| Zc3h7b | 0.10560884 | 0.08247332 | 0.08913318 | -0.0066599 | 0.69052901 | 0.99999994 |
| Fam117a | 0.06261767 | 0.04771371 | 0.05438199 | -0.0066683 | 0.79280648 | 0.99999994 |
| Phf8 | 0.09001851 | 0.07013392 | 0.07680558 | -0.0066717 | 0.7184352 | 0.99999994 |
| Nsf | 0.16660557 | 0.12731883 | 0.1339915 | -0.0066727 | 0.62985951 | 0.99999994 |
| Svil | 0.18857729 | 0.14241565 | 0.14908854 | -0.0066729 | 0.61711968 | 0.99999994 |
| Pkd1 | 0.04876634 | 0.03606521 | 0.04273876 | -0.0066736 | 0.85068465 | 0.99999994 |
| Uvrag | 0.05780638 | 0.0436879 | 0.05036162 | -0.0066737 | 0.81113149 | 0.99999994 |
| Tmem141 | 0.13250409 | 0.10279 | 0.10946409 | -0.0066741 | 0.65757086 | 0.99999994 |
| Gmpr2 | 0.15087158 | 0.11616511 | 0.12284406 | -0.006679 | 0.64128729 | 0.99999994 |
| Pex26 | 0.06056589 | 0.04599042 | 0.05267059 | -0.0066802 | 0.80072708 | 0.99999994 |
| Mapk6 | 0.11266574 | 0.08787332 | 0.09455458 | -0.0066813 | 0.6809442 | 0.99999994 |
| Gid4 | 0.08194697 | 0.06359943 | 0.07028249 | -0.0066831 | 0.73661465 | 0.99999994 |
| Pes1 | 0.15787974 | 0.12115919 | 0.12784467 | -0.0066855 | 0.63611228 | 0.99999994 |
| Shprh | 0.08429466 | 0.06550089 | 0.07218678 | -0.0066859 | 0.73122019 | 0.99999994 |
| Stxbp3 | 0.13420383 | 0.10403199 | 0.11071912 | -0.0066871 | 0.65616739 | 0.99999994 |
| Cnnm2 | 0.04709905 | 0.03463406 | 0.04132291 | -0.0066889 | 0.85936049 | 0.99999994 |
| 2210016L21F | 0.46273209 | 0.29175912 | 0.29844912 | -0.00669 | 0.55930524 | 0.99999994 |
| Mast3 | 0.09063584 | 0.07061096 | 0.0773017 | -0.0066907 | 0.71773672 | 0.99999994 |
| Arrdc3 | 0.14652767 | 0.11301218 | 0.11971599 | -0.0067038 | 0.64532903 | 0.99999994 |
| Lig3 | 0.06386124 | 0.04870702 | 0.05541703 | -0.00671 | 0.78986853 | 0.99999994 |
| Oard1 | 0.36758845 | 0.24723 | 0.25394425 | -0.0067142 | 0.56985033 | 0.99999994 |
| Dym | 0.05452873 | 0.04089305 | 0.04760824 | -0.0067152 | 0.82610538 | 0.99999994 |

| | | | | | | |
|---|---|---|---|---|---|---|
| Trpm4 | 0.04114518 | 0.02952175 | 0.03624147 | -0.0067197 | 0.89143411 | 0.99999994 |
| Sgsm2 | 0.04339818 | 0.03144518 | 0.03816904 | -0.0067239 | 0.87951877 | 0.99999994 |
| 1600020E01I | 0.16192371 | 0.1239805 | 0.13070448 | -0.006724 | 0.63398671 | 0.99999994 |
| Pdia2 | 0.03159026 | 0.02127543 | 0.02800241 | -0.006727 | 0.94509503 | 0.99999994 |
| Tacc1 | 0.08034601 | 0.06225266 | 0.06898058 | -0.0067279 | 0.74190139 | 0.99999994 |
| Thap2 | 0.11441029 | 0.089157 | 0.09588527 | -0.0067283 | 0.6797774 | 0.99999994 |
| Axin1 | 0.04501489 | 0.0328202 | 0.03954869 | -0.0067285 | 0.8712791 | 0.99999994 |
| Cmtm4 | 0.19618408 | 0.14746171 | 0.15419077 | -0.0067291 | 0.61427973 | 0.99999994 |
| Man2c1 | 0.10206512 | 0.07965595 | 0.08638718 | -0.0067312 | 0.69800067 | 0.99999994 |
| Wdr13 | 0.09578426 | 0.07468068 | 0.08142404 | -0.0067434 | 0.709274 | 0.99999994 |
| Gatad2b | 0.37418113 | 0.25051081 | 0.25725515 | -0.0067443 | 0.56926637 | 0.99999994 |
| Brox | 0.14778071 | 0.11387065 | 0.12062056 | -0.0067499 | 0.64523569 | 0.99999994 |
| Galnt14 | 0.02008677 | 0.01119208 | 0.0179429 | -0.0067508 | 0.99386673 | 0.99999994 |
| Asphd1 | 0.03956733 | 0.028134 | 0.03488811 | -0.0067541 | 0.90123611 | 0.99999994 |
| Mboat7 | 0.1692701 | 0.12909687 | 0.13585117 | -0.0067543 | 0.6296563 | 0.99999994 |
| Invs | 0.07014674 | 0.05386819 | 0.06062294 | -0.0067548 | 0.77085559 | 0.99999994 |
| Adipor2 | 0.14968297 | 0.11523553 | 0.12199032 | -0.0067548 | 0.64377678 | 0.99999994 |
| Ap4s1 | 0.12768802 | 0.09913236 | 0.10588883 | -0.0067565 | 0.66448092 | 0.99999994 |
| Hdac5 | 0.10676168 | 0.08326651 | 0.0900231 | -0.0067566 | 0.69131672 | 0.99999994 |
| R3hdm4 | 0.45951145 | 0.29028869 | 0.29705258 | -0.0067639 | 0.56023506 | 0.99999994 |
| Gpr107 | 0.15781426 | 0.12103375 | 0.12779821 | -0.0067645 | 0.63770321 | 0.99999994 |
| Scyl2 | 0.08302056 | 0.0643881 | 0.07115403 | -0.0067659 | 0.73661436 | 0.99999994 |
| Smurf2 | 0.06260575 | 0.04760514 | 0.05437206 | -0.0067669 | 0.79628448 | 0.99999994 |
| Zbtb11os1 | 0.05257291 | 0.0391907 | 0.04595962 | -0.0067689 | 0.8365423 | 0.99999994 |
| Ube2h | 0.10669543 | 0.08319992 | 0.08997201 | -0.0067721 | 0.69182002 | 0.99999994 |
| Fuom | 0.26926816 | 0.19335438 | 0.20013211 | -0.0067777 | 0.58917611 | 0.99999994 |
| Mrps7 | 0.4499144 | 0.28606851 | 0.2928474 | -0.0067789 | 0.56122771 | 0.99999994 |
| Gapvd1 | 0.23116183 | 0.17007762 | 0.17685783 | -0.0067802 | 0.60070998 | 0.99999994 |
| Ahcyl1 | 0.27246079 | 0.19523582 | 0.20201726 | -0.0067814 | 0.58840539 | 0.99999994 |
| Cactin | 0.10526702 | 0.08208744 | 0.088869 | -0.0067816 | 0.6942539 | 0.99999994 |

| | | | | | | |
|---|---|---|---|---|---|---|
| Dcaf5 | 0.07959242 | 0.06158304 | 0.06836683 | -0.0067838 | 0.74552881 | 0.99999994 |
| Itsn2 | 0.10126227 | 0.07897901 | 0.08576285 | -0.0067838 | 0.70073573 | 0.99999994 |
| B9d1 | 0.70988629 | 0.37779457 | 0.38457892 | -0.0067844 | 0.54673827 | 0.99999994 |
| Epc1 | 0.1364933 | 0.10561378 | 0.11240413 | -0.0067903 | 0.6561989 | 0.99999994 |
| Eri1 | 0.12512297 | 0.0971806 | 0.10397308 | -0.0067925 | 0.66816611 | 0.99999994 |
| Esyt2 | 0.06088535 | 0.04614177 | 0.05293735 | -0.0067956 | 0.8035802 | 0.99999994 |
| Senp5 | 0.08931942 | 0.06944526 | 0.07624327 | -0.006798 | 0.7235748 | 0.99999994 |
| Rbm27 | 0.23416542 | 0.1719457 | 0.17874509 | -0.0067994 | 0.59994515 | 0.99999994 |
| Wdr11 | 0.05021878 | 0.03716923 | 0.04396962 | -0.0068004 | 0.84837258 | 0.99999994 |
| Psd | 0.04555562 | 0.03320843 | 0.04000948 | -0.0068011 | 0.87107524 | 0.99999994 |
| Usp34 | 0.41581712 | 0.27056055 | 0.27736362 | -0.0068031 | 0.56484498 | 0.99999994 |
| Letm1 | 0.24965702 | 0.18152412 | 0.18833375 | -0.0068096 | 0.59509677 | 0.99999994 |
| C1qtnf6 | 0.23582421 | 0.17296186 | 0.17978349 | -0.0068216 | 0.59969825 | 0.99999994 |
| Ltn1 | 0.08091671 | 0.06262316 | 0.06944498 | -0.0068218 | 0.74340459 | 0.99999994 |
| Api5 | 0.1703926 | 0.12980994 | 0.13663222 | -0.0068223 | 0.63019255 | 0.99999994 |
| Lmo7 | 0.0598671 | 0.04526144 | 0.05208668 | -0.0068252 | 0.8084567 | 0.99999994 |
| Akap13 | 0.15345997 | 0.11787211 | 0.12469756 | -0.0068255 | 0.64219902 | 0.99999994 |
| Wls | 0.29576727 | 0.20866124 | 0.21548953 | -0.0068283 | 0.58352603 | 0.99999994 |
| B230354K17 | 0.06750002 | 0.05160575 | 0.058436 | -0.0068303 | 0.78172267 | 0.99999994 |
| Prpf18 | 0.10182206 | 0.07936519 | 0.08619826 | -0.0068331 | 0.7011322 | 0.99999994 |
| Sp1 | 0.06024355 | 0.04556357 | 0.05240131 | -0.0068377 | 0.80745998 | 0.99999994 |
| Slc2a1 | 0.05744348 | 0.0432171 | 0.05005735 | -0.0068403 | 0.81847795 | 0.99999994 |
| Cd2ap | 0.23119947 | 0.17004086 | 0.17688153 | -0.0068407 | 0.60157511 | 0.99999994 |
| Setdb1 | 0.09146664 | 0.07112308 | 0.07796876 | -0.0068457 | 0.72053454 | 0.99999994 |
| Gabpb2 | 0.0982629 | 0.07655233 | 0.08339903 | -0.0068467 | 0.70762469 | 0.99999994 |
| Rai1 | 0.07100774 | 0.05447844 | 0.06133276 | -0.0068543 | 0.77152834 | 0.99999994 |
| Msl1 | 0.30509659 | 0.21388318 | 0.22074118 | -0.006858 | 0.58191697 | 0.99999994 |
| Clcc1 | 0.11464612 | 0.08920339 | 0.09606486 | -0.0068615 | 0.68276064 | 0.99999994 |
| Tram1 | 0.81383718 | 0.39909504 | 0.40595694 | -0.0068619 | 0.5447915 | 0.99999994 |
| Dr1 | 0.0998751 | 0.07781366 | 0.08468029 | -0.0068666 | 0.70531776 | 0.99999994 |

| | | | | | | |
|---|---|---|---|---|---|---|
| Thumpd3 | 0.20834348 | 0.15534906 | 0.16221718 | -0.0068681 | 0.61096064 | 0.99999994 |
| Adsl | 0.16904609 | 0.12882611 | 0.13569513 | -0.006869 | 0.63192315 | 0.99999994 |
| Slc27a4 | 0.09971389 | 0.07768192 | 0.08455229 | -0.0068704 | 0.70570086 | 0.99999994 |
| Ndufaf5 | 0.07057298 | 0.05410153 | 0.06097444 | -0.0068729 | 0.77346013 | 0.99999994 |
| Ppp1r37 | 0.31879024 | 0.22143249 | 0.22830743 | -0.0068749 | 0.57943203 | 0.99999994 |
| Optn | 0.05989247 | 0.04523234 | 0.05210789 | -0.0068756 | 0.81010834 | 0.99999994 |
| Ccdc25 | 0.29991518 | 0.21095822 | 0.21783461 | -0.0068764 | 0.58321361 | 0.99999994 |
| Gak | 0.0736819 | 0.05665543 | 0.06353229 | -0.0068769 | 0.76439053 | 0.99999994 |
| Dpcd | 0.49637752 | 0.30573112 | 0.31260842 | -0.0068773 | 0.55821222 | 0.99999994 |
| Fcer1g | 0.04450207 | 0.03222817 | 0.03911139 | -0.0068832 | 0.87929181 | 0.99999994 |
| Epb41l4b | 0.04396962 | 0.03177214 | 0.03865703 | -0.0068849 | 0.88209194 | 0.99999994 |
| Aftph | 0.13940222 | 0.10764776 | 0.11453605 | -0.0068883 | 0.65553917 | 0.99999994 |
| Hcfc2 | 0.11284025 | 0.08779657 | 0.09468787 | -0.0068913 | 0.68596464 | 0.99999994 |
| Nlk | 0.12174051 | 0.09454077 | 0.10143462 | -0.0068939 | 0.67450573 | 0.99999994 |
| Mfsd1 | 0.1126884 | 0.08767572 | 0.0945719 | -0.0068962 | 0.68629706 | 0.99999994 |
| Ddt | 0.31520902 | 0.21944654 | 0.22634522 | -0.0068987 | 0.58038412 | 0.99999994 |
| Mrpl46 | 0.3504357 | 0.23826296 | 0.24516313 | -0.0069002 | 0.57430484 | 0.99999994 |
| Eaf1 | 0.06724207 | 0.05131894 | 0.05822246 | -0.0069035 | 0.78501835 | 0.99999994 |
| Sos1 | 0.07396753 | 0.05686313 | 0.06376677 | -0.0069036 | 0.76443526 | 0.99999994 |
| Tlk2 | 0.16566218 | 0.12642361 | 0.13333117 | -0.0069076 | 0.63489616 | 0.99999994 |
| Kat6a | 0.09026284 | 0.07008892 | 0.07700198 | -0.0069131 | 0.72494412 | 0.99999994 |
| Lpcat1 | 0.07668125 | 0.05907707 | 0.06599024 | -0.0069132 | 0.7571997 | 0.99999994 |
| Gm26670 | 0.04202078 | 0.03007467 | 0.03699128 | -0.0069166 | 0.89336105 | 0.99999994 |
| Dpp8 | 0.05869489 | 0.0441886 | 0.05110597 | -0.0069174 | 0.81619553 | 0.99999994 |
| Tbc1d7 | 0.17209182 | 0.13089157 | 0.13781187 | -0.0069203 | 0.63090558 | 0.99999994 |
| Cab39 | 0.14812414 | 0.11394117 | 0.12086817 | -0.006927 | 0.64857583 | 0.99999994 |
| Tmem63a | 0.0839014 | 0.06493978 | 0.07186819 | -0.0069284 | 0.73946657 | 0.99999994 |
| Rubcn | 0.0496932 | 0.03659551 | 0.04352449 | -0.006929 | 0.85535352 | 0.99999994 |
| Fbxo8 | 0.15412551 | 0.11824074 | 0.1251729 | -0.0069322 | 0.64379948 | 0.99999994 |
| Bst2 | 0.04851174 | 0.03559011 | 0.04252276 | -0.0069326 | 0.86108591 | 0.99999994 |

| | | | | | | |
|---|---|---|---|---|---|---|
| Rsph1 | 0.07292059 | 0.05597324 | 0.06290687 | -0.0069336 | 0.76842427 | 0.99999994 |
| Slc9a8 | 0.06605101 | 0.05030104 | 0.05723552 | -0.0069345 | 0.79001248 | 0.99999994 |
| C3 | 0.01728175 | 0.00853171 | 0.01546638 | -0.0069347 | 0.99857987 | 0.99999994 |
| Mettl17 | 0.07930154 | 0.06119109 | 0.06812976 | -0.0069387 | 0.75108769 | 0.99999994 |
| Msl3l2 | 0.06223781 | 0.04711876 | 0.05406549 | -0.0069467 | 0.80379576 | 0.99999994 |
| Tmem161a | 0.20561116 | 0.15347916 | 0.16042737 | -0.0069482 | 0.61343807 | 0.99999994 |
| Cux1 | 0.22444745 | 0.16565732 | 0.17260606 | -0.0069487 | 0.60563886 | 0.99999994 |
| Thap7 | 0.22053925 | 0.16315975 | 0.17010972 | -0.00695 | 0.60717015 | 0.99999994 |
| Ppfia1 | 0.1102843 | 0.08578044 | 0.09273201 | -0.0069516 | 0.69110608 | 0.99999994 |
| Zfp148 | 0.28419452 | 0.20191173 | 0.20886385 | -0.0069521 | 0.58767174 | 0.99999994 |
| Commd4 | 0.38137781 | 0.25387402 | 0.26082856 | -0.0069545 | 0.57043435 | 0.99999994 |
| Cfap43 | 0.02161636 | 0.01233342 | 0.01928944 | -0.006956 | 0.99181072 | 0.99999994 |
| Anks1 | 0.04184296 | 0.02988088 | 0.03683908 | -0.0069582 | 0.89566994 | 0.99999994 |
| Pih1d1 | 0.19692931 | 0.14772605 | 0.15468723 | -0.0069612 | 0.61773864 | 0.99999994 |
| Plekhg3 | 0.04553458 | 0.03303006 | 0.03999156 | -0.0069615 | 0.87671057 | 0.99999994 |
| 3110009E18I | 0.0632513 | 0.0479383 | 0.05490957 | -0.0069713 | 0.80096843 | 0.99999994 |
| Mlh3 | 0.05465283 | 0.04073674 | 0.04771271 | -0.006976 | 0.83477586 | 0.99999994 |
| Lss | 0.04017704 | 0.02843393 | 0.03541141 | -0.0069775 | 0.9051677 | 0.99999994 |
| Wdr24 | 0.04362855 | 0.03138795 | 0.03836581 | -0.0069779 | 0.88698309 | 0.99999994 |
| Ankrd13a | 0.08177002 | 0.06315623 | 0.07013872 | -0.0069825 | 0.74622487 | 0.99999994 |
| Slc30a6 | 0.07983968 | 0.06158265 | 0.06856827 | -0.0069856 | 0.75115795 | 0.99999994 |
| Fnip1 | 0.07198729 | 0.0551517 | 0.06213933 | -0.0069876 | 0.77292492 | 0.99999994 |
| Trim8 | 0.10734956 | 0.08348784 | 0.09047627 | -0.0069884 | 0.69642301 | 0.99999994 |
| Ap1m1 | 0.28728205 | 0.20364993 | 0.21064348 | -0.0069936 | 0.58745266 | 0.99999994 |
| Tmem19 | 0.1006401 | 0.07828237 | 0.08527846 | -0.0069961 | 0.70749303 | 0.99999994 |
| Mdm1 | 0.158416 | 0.12122854 | 0.12822499 | -0.0069965 | 0.64177108 | 0.99999994 |
| Glb1l2 | 0.08430061 | 0.06519004 | 0.0721916 | -0.0070016 | 0.74072169 | 0.99999994 |
| Gpn2 | 0.07430971 | 0.05704569 | 0.06404756 | -0.0070019 | 0.76659925 | 0.99999994 |
| Pdha1 | 0.2441291 | 0.1779352 | 0.18493968 | -0.0070045 | 0.59952031 | 0.99999994 |
| Osbpl11 | 0.051955 | 0.03843297 | 0.04543788 | -0.0070049 | 0.84759512 | 0.99999994 |

| | | | | | | |
|---|---|---|---|---|---|---|
| Osbpl2 | 0.17462558 | 0.13255839 | 0.13956491 | -0.0070065 | 0.63087306 | 0.99999994 |
| Ucp2 | 0.40252745 | 0.26408278 | 0.27109243 | -0.0070097 | 0.56832597 | 0.99999994 |
| Fbxw11 | 0.10255975 | 0.07976171 | 0.08677142 | -0.0070097 | 0.70461644 | 0.99999994 |
| Spock2 | 0.2941402 | 0.2075548 | 0.21456534 | -0.0070105 | 0.5860846 | 0.99999994 |
| Gm2a | 0.07672523 | 0.05900959 | 0.06602621 | -0.0070166 | 0.76033249 | 0.99999994 |
| Cnot10 | 0.12047862 | 0.09346726 | 0.10048403 | -0.0070168 | 0.67896666 | 0.99999994 |
| Trub2 | 0.09401455 | 0.07299256 | 0.08001009 | -0.0070175 | 0.7203281 | 0.99999994 |
| Fam193a | 0.13409525 | 0.10361481 | 0.11063906 | -0.0070242 | 0.66370595 | 0.99999994 |
| Phf1 | 0.09421613 | 0.07314429 | 0.08017131 | -0.007027 | 0.72019824 | 0.99999994 |
| Zbtb14 | 0.08155861 | 0.06293398 | 0.06996691 | -0.0070329 | 0.74827891 | 0.99999994 |
| Rbck1 | 0.21039412 | 0.15652036 | 0.16355506 | -0.0070347 | 0.61267494 | 0.99999994 |
| Zfp512b | 0.06656632 | 0.05061276 | 0.05766271 | -0.00705 | 0.79212563 | 0.99999994 |
| Glb1 | 0.1231492 | 0.09544317 | 0.1024935 | -0.0070503 | 0.67647999 | 0.99999994 |
| Gnat2 | 0.01638546 | 0.00762105 | 0.01467309 | -0.007052 | 0.99931075 | 0.99999994 |
| Ppih | 0.22979777 | 0.16894566 | 0.17599775 | -0.0070521 | 0.60515562 | 0.99999994 |
| Tmf1 | 0.17945066 | 0.13582842 | 0.142883 | -0.0070546 | 0.6287881 | 0.99999994 |
| Mtfr1l | 0.38732202 | 0.25669334 | 0.26374901 | -0.0070557 | 0.5706649 | 0.99999994 |
| Anapc1 | 0.08590088 | 0.06641891 | 0.07348633 | -0.0070674 | 0.73896701 | 0.99999994 |
| Tm7sf3 | 0.14732558 | 0.11322351 | 0.12029221 | -0.0070687 | 0.65215539 | 0.99999994 |
| Vwa5a | 0.0570804 | 0.04266309 | 0.04975279 | -0.0070897 | 0.82856793 | 0.99999994 |
| Zswim8 | 0.11384913 | 0.08836665 | 0.09545765 | -0.007091 | 0.68951021 | 0.99999994 |
| Baz2b | 0.43186129 | 0.27765168 | 0.28475641 | -0.0071047 | 0.56595204 | 0.99999994 |
| Gmpr | 0.10531454 | 0.0817992 | 0.08890574 | -0.0071065 | 0.70265951 | 0.99999994 |
| Tom1l2 | 0.04424157 | 0.03176769 | 0.03888913 | -0.0071214 | 0.8885633 | 0.99999994 |
| Slc7a2 | 0.25266603 | 0.18304494 | 0.19016872 | -0.0071238 | 0.59845468 | 0.99999994 |
| Pitpna | 0.25884586 | 0.18678284 | 0.19390838 | -0.0071255 | 0.59661727 | 0.99999994 |
| Tnip1 | 0.10778783 | 0.08368346 | 0.09081382 | -0.0071304 | 0.69938484 | 0.99999994 |
| Zfp64 | 0.12144277 | 0.09407526 | 0.1012105 | -0.0071352 | 0.68056046 | 0.99999994 |
| Arhgap12 | 0.08505578 | 0.06566709 | 0.07280292 | -0.0071358 | 0.74293719 | 0.99999994 |
| Rptor | 0.0560908 | 0.04178576 | 0.04892193 | -0.0071362 | 0.83421448 | 0.99999994 |

| | | | | | | |
|---|---|---|---|---|---|---|
| Emc1 | 0.0668988 | 0.05080144 | 0.05793818 | -0.0071367 | 0.79386926 | 0.99999994 |
| Dnal4 | 0.22207119 | 0.16395186 | 0.17109029 | -0.0071384 | 0.60938807 | 0.99999994 |
| Chtop | 0.49141147 | 0.30342709 | 0.31056708 | -0.00714 | 0.56081301 | 0.99999994 |
| Zdhhc17 | 0.05751998 | 0.04297954 | 0.0501215 | -0.007142 | 0.8285584 | 0.99999994 |
| Mob1a | 0.0892186 | 0.0690183 | 0.07616214 | -0.0071438 | 0.73380176 | 0.99999994 |
| Mdm4 | 0.15442015 | 0.11823907 | 0.12538317 | -0.0071441 | 0.64775509 | 0.99999994 |
| Paf1 | 0.16724013 | 0.12728682 | 0.13443511 | -0.0071483 | 0.63830684 | 0.99999994 |
| Bag5 | 0.21509658 | 0.15945655 | 0.1666065 | -0.0071499 | 0.61243019 | 0.99999994 |
| Zfp207 | 0.81521824 | 0.39904556 | 0.40619621 | -0.0071507 | 0.54664054 | 0.99999994 |
| Senp3 | 0.13367056 | 0.1031731 | 0.11032575 | -0.0071526 | 0.66696405 | 0.99999994 |
| Cs | 0.2084637 | 0.15513951 | 0.16229574 | -0.0071562 | 0.61543151 | 0.99999994 |
| 1500026H17 | 0.07613508 | 0.05838603 | 0.06554337 | -0.0071573 | 0.76634844 | 0.99999994 |
| Brd2 | 0.42705067 | 0.27540195 | 0.28256008 | -0.0071581 | 0.56695476 | 0.99999994 |
| Ubr2 | 0.13995087 | 0.10776962 | 0.11493703 | -0.0071674 | 0.66096084 | 0.99999994 |
| Tspyl5 | 0.0642197 | 0.04854693 | 0.05571507 | -0.0071681 | 0.80410824 | 0.99999994 |
| Il3ra | 0.08665121 | 0.06692346 | 0.07409249 | -0.007169 | 0.74023008 | 0.99999994 |
| Ak1 | 0.27726374 | 0.1976645 | 0.20483527 | -0.0071708 | 0.59212792 | 0.99999994 |
| Btbd9 | 0.17639459 | 0.13361132 | 0.14078463 | -0.0071733 | 0.6327542 | 0.99999994 |
| Phax | 0.50418845 | 0.30861213 | 0.31578588 | -0.0071738 | 0.56009633 | 0.99999994 |
| Dimt1 | 0.12336506 | 0.09548079 | 0.10265555 | -0.0071748 | 0.67911433 | 0.99999994 |
| Tmem87a | 0.12096868 | 0.0936784 | 0.10085342 | -0.007175 | 0.68209081 | 0.99999994 |
| Tmem127 | 0.07872561 | 0.06048271 | 0.06766012 | -0.0071774 | 0.75993984 | 0.99999994 |
| Ppp1r9b | 0.06846221 | 0.05205047 | 0.05923192 | -0.0071815 | 0.79017722 | 0.99999994 |
| Wdr70 | 0.0967111 | 0.07498108 | 0.08216328 | -0.0071822 | 0.71966858 | 0.99999994 |
| Odf2 | 0.14940181 | 0.11460523 | 0.12178813 | -0.0071829 | 0.6526861 | 0.99999994 |
| Insig2 | 0.41419161 | 0.2694207 | 0.27660381 | -0.0071831 | 0.56861873 | 0.99999994 |
| Gga1 | 0.18955125 | 0.14256129 | 0.14974542 | -0.0071841 | 0.62526686 | 0.99999994 |
| Furin | 0.0869625 | 0.06715894 | 0.07434379 | -0.0071848 | 0.73998325 | 0.99999994 |
| Bin3 | 0.17833996 | 0.13493658 | 0.14212157 | -0.007185 | 0.63175829 | 0.99999994 |
| Dlg4 | 0.06077858 | 0.04565849 | 0.05284821 | -0.0071897 | 0.81741628 | 0.99999994 |

| | | | | | | |
|---|---|---|---|---|---|---|
| Cox16 | 0.06704293 | 0.05086432 | 0.05805756 | -0.0071932 | 0.79523258 | 0.99999994 |
| Mthfsd | 0.18579763 | 0.1400134 | 0.14720817 | -0.0071948 | 0.62753364 | 0.99999994 |
| Arpc2 | 1.42660264 | 0.40897278 | 0.41616765 | -0.0071949 | 0.54580824 | 0.99999994 |
| Mtmr3 | 0.08054327 | 0.06194563 | 0.06914114 | -0.0071955 | 0.7557591 | 0.99999994 |
| Trim37 | 0.13327932 | 0.10283542 | 0.11003693 | -0.0072015 | 0.66844874 | 0.99999994 |
| C2cd4a | 0.06211978 | 0.04676363 | 0.05396712 | -0.0072035 | 0.81286032 | 0.99999994 |
| Fam8a1 | 0.07497608 | 0.05738657 | 0.06459403 | -0.0072075 | 0.77117497 | 0.99999994 |
| Mpg | 0.1814832 | 0.1370636 | 0.14427288 | -0.0072093 | 0.63028826 | 0.99999994 |
| Xk | 0.02570474 | 0.01566539 | 0.02287501 | -0.0072096 | 0.98204363 | 0.99999994 |
| Tk2 | 0.06719597 | 0.05097099 | 0.05818429 | -0.0072133 | 0.79537396 | 0.99999994 |
| Ranbp9 | 0.08977659 | 0.06939364 | 0.07661105 | -0.0072174 | 0.73469821 | 0.99999994 |
| Xrn1 | 0.16679828 | 0.12690636 | 0.13412627 | -0.0072199 | 0.63994292 | 0.99999994 |
| Mon2 | 0.05626612 | 0.04184921 | 0.04906921 | -0.00722 | 0.83631131 | 0.99999994 |
| Ubr1 | 0.13534626 | 0.10433669 | 0.11156071 | -0.007224 | 0.66677556 | 0.99999994 |
| Snx11 | 0.06503498 | 0.04916704 | 0.05639241 | -0.0072254 | 0.80313075 | 0.99999994 |
| Fubp3 | 0.18221661 | 0.13754226 | 0.14477329 | -0.007231 | 0.63023162 | 0.99999994 |
| Ganc | 0.06402808 | 0.04831701 | 0.05555577 | -0.0072388 | 0.80711223 | 0.99999994 |
| Vps13b | 0.06125326 | 0.04600515 | 0.05324444 | -0.0072393 | 0.81727261 | 0.99999994 |
| Cdr2 | 0.0308111 | 0.02008521 | 0.02732592 | -0.0072407 | 0.96111501 | 0.99999994 |
| Tsr1 | 0.08927119 | 0.06896204 | 0.07620446 | -0.0072424 | 0.73650769 | 0.99999994 |
| Kat6b | 0.08854713 | 0.06837062 | 0.0756215 | -0.0072509 | 0.73834022 | 0.99999994 |
| Cc2d1a | 0.07997322 | 0.06141805 | 0.06867704 | -0.007259 | 0.75914661 | 0.99999994 |
| Adck2 | 0.07074954 | 0.05386093 | 0.06111998 | -0.007259 | 0.78539867 | 0.99999994 |
| Bcdin3d | 0.1092615 | 0.08466878 | 0.09194707 | -0.0072783 | 0.70086841 | 0.99999994 |
| Ksr2 | 0.0475201 | 0.03440062 | 0.04168077 | -0.0072801 | 0.87750976 | 0.99999994 |
| Nceh1 | 0.0301762 | 0.01949261 | 0.02677416 | -0.0072816 | 0.96487347 | 0.99999994 |
| Bri3bp | 0.19377896 | 0.14529823 | 0.1525845 | -0.0072863 | 0.62470316 | 0.99999994 |
| Mroh7 | 0.01870124 | 0.00943104 | 0.0167208 | -0.0072898 | 0.99814587 | 0.99999994 |
| Ndufaf4 | 0.23594483 | 0.17256664 | 0.17985889 | -0.0072923 | 0.60637269 | 0.99999994 |
| Gck | 0.50308749 | 0.30804707 | 0.31534058 | -0.0072935 | 0.56117733 | 0.99999994 |

| | | | | | | |
|---|---|---|---|---|---|---|
| Snx4 | 0.44470965 | 0.28324302 | 0.29053898 | -0.007296 | 0.56637522 | 0.99999994 |
| Gnaz | 0.25005011 | 0.18127609 | 0.18857397 | -0.0072979 | 0.60164347 | 0.99999994 |
| Nf2 | 0.0814452 | 0.0625761 | 0.06987473 | -0.0072986 | 0.75655924 | 0.99999994 |
| Pycrl | 0.37299431 | 0.24936295 | 0.25666167 | -0.0072987 | 0.57506628 | 0.99999994 |
| Angptl6 | 0.05032233 | 0.03675194 | 0.04405728 | -0.0073053 | 0.86514284 | 0.99999994 |
| St6galnac5 | 0.03926821 | 0.02732516 | 0.03463122 | -0.0073061 | 0.9198816 | 0.99999994 |
| Xrcc6 | 0.1065152 | 0.08252689 | 0.08983297 | -0.0073061 | 0.70586991 | 0.99999994 |
| Plekha6 | 0.08716127 | 0.06719629 | 0.07450421 | -0.0073079 | 0.74309085 | 0.99999994 |
| Zcchc2 | 0.04961201 | 0.03614714 | 0.0434557 | -0.0073086 | 0.86853539 | 0.99999994 |
| Ccar2 | 0.08681485 | 0.06690993 | 0.07422461 | -0.0073147 | 0.7440781 | 0.99999994 |
| Trak1 | 0.06849682 | 0.05193907 | 0.05926053 | -0.0073215 | 0.79456755 | 0.99999994 |
| Slc25a12 | 0.05140821 | 0.03765019 | 0.04497584 | -0.0073257 | 0.86085873 | 0.99999994 |
| Aco1 | 0.06961155 | 0.05285419 | 0.06018134 | -0.0073272 | 0.79115053 | 0.99999994 |
| Kirrel3os | 0.02613809 | 0.01592215 | 0.02325391 | -0.0073318 | 0.98207794 | 0.99999994 |
| Cpb1 | 0.08316938 | 0.06394249 | 0.07127475 | -0.0073323 | 0.75325252 | 0.99999994 |
| Pik3r2 | 0.13146702 | 0.1013641 | 0.10869667 | -0.0073326 | 0.67329546 | 0.99999994 |
| Bcs1l | 0.07209845 | 0.05489718 | 0.0622308 | -0.0073336 | 0.78360373 | 0.99999994 |
| Herpud2 | 0.08571972 | 0.06600263 | 0.07333989 | -0.0073373 | 0.74727287 | 0.99999994 |
| Ufsp1 | 0.06629152 | 0.05009303 | 0.05743494 | -0.0073419 | 0.80258251 | 0.99999994 |
| Cyb561d2 | 0.14550942 | 0.11163727 | 0.11897949 | -0.0073422 | 0.65937709 | 0.99999994 |
| Ankfy1 | 0.07672638 | 0.05868256 | 0.06602715 | -0.0073446 | 0.77047921 | 0.99999994 |
| Dgcr2 | 0.12776844 | 0.09860108 | 0.10594877 | -0.0073477 | 0.67782572 | 0.99999994 |
| Abhd16a | 0.41789633 | 0.27098243 | 0.27833267 | -0.0073502 | 0.56976732 | 0.99999994 |
| Aptx | 0.20298111 | 0.15134557 | 0.1586969 | -0.0073513 | 0.62108923 | 0.99999994 |
| Abcd3 | 0.40729112 | 0.26600349 | 0.27335638 | -0.0073529 | 0.57104935 | 0.99999994 |
| Csf2ra | 0.0907592 | 0.07004499 | 0.0774008 | -0.0073558 | 0.736499 | 0.99999994 |
| Csnk1d | 0.33772285 | 0.23113947 | 0.23849541 | -0.0073559 | 0.58133215 | 0.99999994 |
| Lifr | 0.04785906 | 0.03461103 | 0.04196869 | -0.0073577 | 0.87838416 | 0.99999994 |
| Sh3gl2 | 0.06046877 | 0.04523066 | 0.05258947 | -0.0073588 | 0.82418181 | 0.99999994 |
| Crcp | 0.19890868 | 0.14863975 | 0.15600296 | -0.0073632 | 0.62330571 | 0.99999994 |

| | | | | | | |
|---|---|---|---|---|---|---|
| Kmt2b | 0.08348618 | 0.06416318 | 0.07153165 | -0.0073685 | 0.75353954 | 0.99999994 |
| Nudt19 | 0.14411133 | 0.11058969 | 0.11796614 | -0.0073764 | 0.66137669 | 0.99999994 |
| Slc16a9 | 0.06842567 | 0.05180636 | 0.05920171 | -0.0073954 | 0.79715381 | 0.99999994 |
| Iqgap2 | 0.16922386 | 0.12842204 | 0.13581896 | -0.0073969 | 0.64151675 | 0.99999994 |
| Tsc22d2 | 0.1022335 | 0.07911831 | 0.08651802 | -0.0073997 | 0.71542535 | 0.99999994 |
| Rassf10 | 0.04420707 | 0.03145914 | 0.0388597 | -0.0074006 | 0.89753958 | 0.99999994 |
| Golga7 | 0.48878601 | 0.30207631 | 0.30948133 | -0.007405 | 0.56327114 | 0.99999994 |
| Ubr5 | 0.15613704 | 0.11920003 | 0.12660646 | -0.0074064 | 0.65150763 | 0.99999994 |
| Pmm2 | 0.1809686 | 0.13651432 | 0.14392143 | -0.0074071 | 0.63404206 | 0.99999994 |
| Ncstn | 0.30194472 | 0.21156817 | 0.21897596 | -0.0074078 | 0.58908022 | 0.99999994 |
| Pfkm | 0.06630066 | 0.05003397 | 0.05744251 | -0.0074085 | 0.80468949 | 0.99999994 |
| Pip5k1c | 0.05212116 | 0.03816774 | 0.04557822 | -0.0074105 | 0.86042806 | 0.99999994 |
| Cc2d1b | 0.07631574 | 0.05827862 | 0.06569121 | -0.0074126 | 0.77370449 | 0.99999994 |
| BC005624 | 0.4160292 | 0.27004957 | 0.27746261 | -0.007413 | 0.57057556 | 0.99999994 |
| Oxld1 | 0.06672465 | 0.0503806 | 0.05779391 | -0.0074133 | 0.80339783 | 0.99999994 |
| Ckmt1 | 0.13617595 | 0.10475563 | 0.11217094 | -0.0074153 | 0.67004341 | 0.99999994 |
| Tex2 | 0.04084896 | 0.02857053 | 0.03598762 | -0.0074171 | 0.91495006 | 0.99999994 |
| Fam171b | 0.05024896 | 0.03657509 | 0.04399517 | -0.0074201 | 0.86920459 | 0.99999994 |
| Impact | 0.07405703 | 0.05641839 | 0.06384022 | -0.0074218 | 0.78048837 | 0.99999994 |
| Arap1 | 0.06305704 | 0.04732511 | 0.05474787 | -0.0074228 | 0.81659777 | 0.99999994 |
| Atic | 0.1463546 | 0.11216788 | 0.1195909 | -0.007423 | 0.66025577 | 0.99999994 |
| Ints5 | 0.06578789 | 0.04959425 | 0.05701729 | -0.007423 | 0.80691571 | 0.99999994 |
| Ggcx | 0.06172037 | 0.0462063 | 0.05363411 | -0.0074278 | 0.82169447 | 0.99999994 |
| Ralgapb | 0.06341543 | 0.0476183 | 0.05504616 | -0.0074279 | 0.81546071 | 0.99999994 |
| Rnaseh1 | 0.08147515 | 0.0624638 | 0.06989907 | -0.0074353 | 0.76054214 | 0.99999994 |
| Abhd11 | 0.11104007 | 0.0858697 | 0.09331119 | -0.0074415 | 0.70223476 | 0.99999994 |
| Tmem181a | 0.10398141 | 0.08042715 | 0.08787407 | -0.0074469 | 0.71365522 | 0.99999994 |
| Spg20 | 0.06933495 | 0.05249953 | 0.05995298 | -0.0074535 | 0.79603142 | 0.99999994 |
| Ccdc32 | 0.10555698 | 0.08163632 | 0.08909312 | -0.0074568 | 0.71127547 | 0.99999994 |
| Klhdc10 | 0.09691905 | 0.07487156 | 0.08232902 | -0.0074575 | 0.72672604 | 0.99999994 |

| | | | | | | |
|---|---|---|---|---|---|---|
| Cnot3 | 0.29414598 | 0.20710844 | 0.21456862 | -0.0074602 | 0.59150968 | 0.99999994 |
| Crhr2 | 0.052835 | 0.03871923 | 0.04618079 | -0.0074616 | 0.8589217 | 0.99999994 |
| Mbnl1 | 0.05516639 | 0.04068107 | 0.04814483 | -0.0074638 | 0.84894469 | 0.99999994 |
| Fam114a1 | 0.02514007 | 0.01491274 | 0.02238095 | -0.0074682 | 0.98682903 | 0.99999994 |
| Zer1 | 0.09025136 | 0.06951755 | 0.07699275 | -0.0074752 | 0.74094574 | 0.99999994 |
| Zfp87 | 0.06910769 | 0.05228917 | 0.05976529 | -0.0074761 | 0.79747967 | 0.99999994 |
| Kat8 | 0.14634174 | 0.11210368 | 0.11958161 | -0.0074779 | 0.66138629 | 0.99999994 |
| Afg3l1 | 0.26185654 | 0.18823645 | 0.1957169 | -0.0074804 | 0.60041101 | 0.99999994 |
| Cinp | 0.10526671 | 0.08138786 | 0.08886876 | -0.0074809 | 0.71237185 | 0.99999994 |
| Uggt2 | 0.06987947 | 0.05291976 | 0.06040245 | -0.0074827 | 0.79519868 | 0.99999994 |
| Mtg1 | 0.1233138 | 0.0951311 | 0.10261707 | -0.007486 | 0.68636934 | 0.99999994 |
| Zbtb5 | 0.07303505 | 0.0555141 | 0.06300094 | -0.0074868 | 0.78553416 | 0.99999994 |
| 9330160F10I | 0.08026551 | 0.06142725 | 0.06891504 | -0.0074878 | 0.76523061 | 0.99999994 |
| Svip | 0.22356112 | 0.16455149 | 0.17204139 | -0.0074899 | 0.61401062 | 0.99999994 |
| Aatk | 0.05436225 | 0.0399777 | 0.04746808 | -0.0074904 | 0.85322405 | 0.99999994 |
| Qars | 0.27235065 | 0.19445727 | 0.20195239 | -0.0074951 | 0.59756114 | 0.99999994 |
| Abhd13 | 0.11870845 | 0.09165189 | 0.09914725 | -0.0074954 | 0.69259151 | 0.99999994 |
| Rc3h2 | 0.21132538 | 0.15666535 | 0.16416118 | -0.0074958 | 0.61941263 | 0.99999994 |
| Gpr137 | 0.10374781 | 0.08019559 | 0.08769307 | -0.0074975 | 0.71535479 | 0.99999994 |
| Mgat1 | 0.10360832 | 0.08008065 | 0.08758494 | -0.0075043 | 0.71576893 | 0.99999994 |
| Zdhhc6 | 0.17853759 | 0.13474941 | 0.14225715 | -0.0075077 | 0.63731637 | 0.99999994 |
| Rad1 | 0.12301512 | 0.09487866 | 0.10239282 | -0.0075142 | 0.6873954 | 0.99999994 |
| Golim4 | 0.42640647 | 0.27474826 | 0.28226469 | -0.0075164 | 0.5703449 | 0.99999994 |
| Akap9 | 0.74928717 | 0.38592139 | 0.39344279 | -0.0075214 | 0.55062826 | 0.99999994 |
| Ankrd39 | 0.15551373 | 0.11863673 | 0.12616274 | -0.007526 | 0.65434322 | 0.99999994 |
| Pex14 | 0.25117052 | 0.18173052 | 0.18925782 | -0.0075273 | 0.60439581 | 0.99999994 |
| Cant1 | 0.10852177 | 0.0838512 | 0.09137856 | -0.0075274 | 0.70826432 | 0.99999994 |
| Snapc2 | 0.15571425 | 0.11876975 | 0.12630553 | -0.0075358 | 0.6543676 | 0.99999994 |
| Ddx6 | 0.73410365 | 0.38255572 | 0.39009303 | -0.0075373 | 0.55116803 | 0.99999994 |
| Cacnb2 | 0.05716398 | 0.04228488 | 0.04982291 | -0.007538 | 0.84305256 | 0.99999994 |

| | | | | | | |
|---|---|---|---|---|---|---|
| Vma21 | 0.48309334 | 0.29957227 | 0.30711075 | -0.0075385 | 0.56489424 | 0.99999994 |
| Pomt2 | 0.08801328 | 0.06765262 | 0.07519134 | -0.0075387 | 0.74773206 | 0.99999994 |
| Mpnd | 0.41971574 | 0.27163915 | 0.27917801 | -0.0075389 | 0.5713243 | 0.99999994 |
| Emg1 | 0.67563644 | 0.36825623 | 0.37579544 | -0.0075392 | 0.55311675 | 0.99999994 |
| Gne | 0.04056287 | 0.02819447 | 0.03574234 | -0.0075479 | 0.92008645 | 0.99999994 |
| Dennd2d | 0.03093811 | 0.01988615 | 0.02743624 | -0.0075501 | 0.96650499 | 0.99999994 |
| Sema4b | 0.08181609 | 0.06261011 | 0.07017615 | -0.007566 | 0.76351505 | 0.99999994 |
| Rsbn1l | 0.24839784 | 0.17999608 | 0.18756326 | -0.0075672 | 0.60586152 | 0.99999994 |
| Nmd3 | 0.26204884 | 0.18826426 | 0.19583212 | -0.0075679 | 0.60150034 | 0.99999994 |
| Senp6 | 0.49231087 | 0.30336552 | 0.31093798 | -0.0075725 | 0.56438898 | 0.99999994 |
| Golga2 | 0.08251243 | 0.06316802 | 0.07074167 | -0.0075737 | 0.76196393 | 0.99999994 |
| Wasl | 0.21999704 | 0.16218683 | 0.16976209 | -0.0075753 | 0.61677635 | 0.99999994 |
| Thoc5 | 0.14379786 | 0.11015918 | 0.11773862 | -0.0075794 | 0.66585775 | 0.99999994 |
| Cnst | 0.05654719 | 0.04172298 | 0.04930525 | -0.0075823 | 0.84699482 | 0.99999994 |
| Kat2a | 0.10615698 | 0.08197405 | 0.08955652 | -0.0075825 | 0.71347768 | 0.99999994 |
| Tmem38a | 0.14476906 | 0.11085899 | 0.11844315 | -0.0075842 | 0.66502654 | 0.99999994 |
| Lrrc1 | 0.0465146 | 0.03323658 | 0.04082587 | -0.0075893 | 0.89202679 | 0.99999994 |
| Akt3 | 0.06552435 | 0.04920379 | 0.05679863 | -0.0075948 | 0.81328212 | 0.99999994 |
| Pds5b | 0.11040045 | 0.08522172 | 0.09282107 | -0.0075994 | 0.70711046 | 0.99999994 |
| Actr1b | 0.33417519 | 0.2290086 | 0.23661017 | -0.0076016 | 0.58466739 | 0.99999994 |
| Dnajc14 | 0.17167033 | 0.12991608 | 0.13751956 | -0.0076035 | 0.64357492 | 0.99999994 |
| Socs7 | 0.09088938 | 0.06989687 | 0.07750534 | -0.0076085 | 0.7432622 | 0.99999994 |
| Arhgef1 | 0.1816445 | 0.1367728 | 0.14438299 | -0.0076102 | 0.63714791 | 0.99999994 |
| Mau2 | 0.2058309 | 0.15295514 | 0.16057163 | -0.0076165 | 0.6238982 | 0.99999994 |
| Syt4 | 0.0208438 | 0.01099288 | 0.01860968 | -0.0076168 | 0.99677905 | 0.99999994 |
| Ticam1 | 0.06195497 | 0.04620974 | 0.05382973 | -0.00762 | 0.82696927 | 0.99999994 |
| Ncbp1 | 0.14353504 | 0.10992662 | 0.11754777 | -0.0076212 | 0.66697008 | 0.99999994 |
| Zfp282 | 0.09234739 | 0.07105303 | 0.07867515 | -0.0076221 | 0.74049567 | 0.99999994 |
| Prr14l | 0.13100743 | 0.10073361 | 0.10835615 | -0.0076225 | 0.68019912 | 0.99999994 |
| Appbp2 | 0.37694415 | 0.25100842 | 0.25863226 | -0.0076238 | 0.57777829 | 0.99999994 |

| | | | | | | |
|---|---|---|---|---|---|---|
| Jade1 | 0.16740896 | 0.12692437 | 0.13455306 | -0.0076287 | 0.64705894 | 0.99999994 |
| Sec24b | 0.0828269 | 0.06336778 | 0.0709969 | -0.0076291 | 0.76277982 | 0.99999994 |
| Tmem33 | 0.54456783 | 0.3239438 | 0.33157968 | -0.0076359 | 0.56091438 | 0.99999994 |
| Ctnnd1 | 0.11759934 | 0.09067182 | 0.09830773 | -0.0076359 | 0.69742983 | 0.99999994 |
| Tmed6 | 0.11982189 | 0.09235243 | 0.09998853 | -0.0076361 | 0.69439416 | 0.99999994 |
| Unc93b1 | 0.05960237 | 0.04422829 | 0.05186533 | -0.007637 | 0.83648973 | 0.99999994 |
| Rbm28 | 0.15791343 | 0.12022449 | 0.12786857 | -0.0076441 | 0.65465626 | 0.99999994 |
| Ppp2cb | 0.10511246 | 0.08110335 | 0.0887495 | -0.0076461 | 0.71683799 | 0.99999994 |
| D730003I15F | 0.06973141 | 0.05263277 | 0.06028026 | -0.0076475 | 0.80079449 | 0.99999994 |
| Nfic | 0.18023838 | 0.13576915 | 0.1434222 | -0.007653 | 0.63877448 | 0.99999994 |
| Fam120b | 0.088927 | 0.06826994 | 0.07592741 | -0.0076575 | 0.74898863 | 0.99999994 |
| Nolc1 | 0.30050239 | 0.21050772 | 0.21816522 | -0.0076575 | 0.59236589 | 0.99999994 |
| Mrgbp | 0.13399598 | 0.10290418 | 0.11056584 | -0.0076617 | 0.67769121 | 0.99999994 |
| Adam9 | 0.03463508 | 0.02297382 | 0.03063927 | -0.0076654 | 0.95207476 | 0.99999994 |
| Eif2s1 | 0.68819555 | 0.37147727 | 0.37915015 | -0.0076729 | 0.55357749 | 0.99999994 |
| Ppp1r16a | 0.15036633 | 0.1147927 | 0.12248135 | -0.0076887 | 0.66197036 | 0.99999994 |
| Ptp4a2 | 0.32414984 | 0.2235345 | 0.23122328 | -0.0076888 | 0.58758662 | 0.99999994 |
| Syt14 | 0.0683876 | 0.05146956 | 0.05917024 | -0.0077007 | 0.80683395 | 0.99999994 |
| Rasa2 | 0.10293945 | 0.07936371 | 0.08706617 | -0.0077025 | 0.72202454 | 0.99999994 |
| Eea1 | 0.17053805 | 0.12902184 | 0.13673332 | -0.0077115 | 0.64632027 | 0.99999994 |
| Coro2b | 0.03385769 | 0.0222543 | 0.02996705 | -0.0077127 | 0.95665987 | 0.99999994 |
| Fbxl17 | 0.07754371 | 0.05898197 | 0.06669525 | -0.0077133 | 0.77929227 | 0.99999994 |
| Irgq | 0.09996766 | 0.07704025 | 0.08475377 | -0.0077135 | 0.72767673 | 0.99999994 |
| Dhx15 | 0.32571113 | 0.22435187 | 0.23206803 | -0.0077162 | 0.58757879 | 0.99999994 |
| Kpna3 | 0.16177009 | 0.12288003 | 0.13059621 | -0.0077162 | 0.65294588 | 0.99999994 |
| Mvd | 0.18281031 | 0.13745117 | 0.14517794 | -0.0077268 | 0.63843169 | 0.99999994 |
| Zcchc7 | 0.27466689 | 0.19558261 | 0.20331431 | -0.0077317 | 0.59991611 | 0.99999994 |
| Mtx2 | 0.4939788 | 0.30388898 | 0.31162442 | -0.0077354 | 0.56561868 | 0.99999994 |
| Inafm1 | 0.11968524 | 0.09214912 | 0.09988536 | -0.0077362 | 0.69691328 | 0.99999994 |
| Gnai2 | 0.94965395 | 0.41511516 | 0.42285216 | -0.007737 | 0.54846834 | 0.99999994 |

| | | | | | | |
|---|---|---|---|---|---|---|
| Pemt | 0.05556002 | 0.04073785 | 0.04847585 | -0.007738 | 0.85600168 | 0.99999994 |
| Nfil3 | 0.05513045 | 0.04036577 | 0.0481146 | -0.0077488 | 0.85814059 | 0.99999994 |
| Lias | 0.28212144 | 0.19990847 | 0.20766359 | -0.0077551 | 0.59815678 | 0.99999994 |
| Armc6 | 0.06943634 | 0.05228098 | 0.06003669 | -0.0077557 | 0.80507433 | 0.99999994 |
| Cyp51 | 0.13109398 | 0.10066235 | 0.1084203 | -0.007758 | 0.68306677 | 0.99999994 |
| Cpn1 | 0.16297982 | 0.12368722 | 0.13144812 | -0.0077609 | 0.65284087 | 0.99999994 |
| Arfgap2 | 0.31140155 | 0.21648268 | 0.22424589 | -0.0077632 | 0.59112412 | 0.99999994 |
| Tes | 0.45365551 | 0.28672562 | 0.29449493 | -0.0077693 | 0.56969853 | 0.99999994 |
| Fam122a | 0.48115534 | 0.29852863 | 0.30629852 | -0.0077699 | 0.56704352 | 0.99999994 |
| Gopc | 0.12000887 | 0.09235845 | 0.10012966 | -0.0077712 | 0.69728575 | 0.99999994 |
| Egln1 | 0.07022779 | 0.0529113 | 0.0606898 | -0.0077785 | 0.80320717 | 0.99999994 |
| Fbxw9 | 0.21449843 | 0.15843981 | 0.16621963 | -0.0077798 | 0.62230757 | 0.99999994 |
| Sowahc | 0.06875219 | 0.05168939 | 0.05947159 | -0.0077822 | 0.80812917 | 0.99999994 |
| Cdc42bpb | 0.07354388 | 0.05563578 | 0.06341895 | -0.0077832 | 0.79301 | 0.99999994 |
| Gstp1 | 0.0664214 | 0.04975813 | 0.0575426 | -0.0077845 | 0.81606888 | 0.99999994 |
| AU040320 | 0.05151406 | 0.03727844 | 0.04506531 | -0.0077869 | 0.87496036 | 0.99999994 |
| C1galt1 | 0.1246068 | 0.09579895 | 0.1035866 | -0.0077877 | 0.69161242 | 0.99999994 |
| 2310039H08 | 0.53937319 | 0.32181511 | 0.32960581 | -0.0077907 | 0.56250858 | 0.99999994 |
| Pus1 | 0.09685073 | 0.07448247 | 0.08227458 | -0.0077921 | 0.73578827 | 0.99999994 |
| Sipa1l3 | 0.07386781 | 0.05588831 | 0.06368492 | -0.0077966 | 0.79243609 | 0.99999994 |
| Vmp1 | 0.40811522 | 0.26594584 | 0.27374609 | -0.0078003 | 0.57521591 | 0.99999994 |
| Epn2 | 0.11183659 | 0.08611936 | 0.09392082 | -0.0078015 | 0.7098359 | 0.99999994 |
| Ubap2l | 0.70102961 | 0.37461938 | 0.38242297 | -0.0078036 | 0.55402111 | 0.99999994 |
| Spen | 0.08937613 | 0.06848489 | 0.07628891 | -0.007804 | 0.75203758 | 0.99999994 |
| Sh3bp5l | 0.12903795 | 0.09908288 | 0.10689386 | -0.007811 | 0.68665661 | 0.99999994 |
| Arxes2 | 0.18497876 | 0.1388399 | 0.14665262 | -0.0078127 | 0.63855854 | 0.99999994 |
| Oxnad1 | 0.09118502 | 0.06992833 | 0.07774272 | -0.0078144 | 0.74827474 | 0.99999994 |
| Smyd3 | 0.09398965 | 0.07217536 | 0.07999018 | -0.0078148 | 0.74225334 | 0.99999994 |
| Trappc13 | 0.24902093 | 0.18012895 | 0.18794472 | -0.0078158 | 0.60903618 | 0.99999994 |
| Rce1 | 0.23311721 | 0.17026703 | 0.17808749 | -0.0078205 | 0.61497305 | 0.99999994 |

| | | | | | | |
|---|---|---|---|---|---|---|
| Tbck | 0.09664672 | 0.07428918 | 0.08211196 | -0.0078228 | 0.73700819 | 0.99999994 |
| Rilpl1 | 0.0867546 | 0.06635082 | 0.07417596 | -0.0078251 | 0.7587251 | 0.99999994 |
| Alkbh2 | 0.0963606 | 0.07405859 | 0.08188383 | -0.0078252 | 0.73764943 | 0.99999994 |
| Chd8 | 0.23695958 | 0.17266401 | 0.18049266 | -0.0078286 | 0.61359887 | 0.99999994 |
| Mtg2 | 0.13321986 | 0.10215721 | 0.10999302 | -0.0078358 | 0.68231904 | 0.99999994 |
| Cops5 | 0.83223794 | 0.40100769 | 0.40884525 | -0.0078376 | 0.55076814 | 0.99999994 |
| Cul9 | 0.04717959 | 0.03354575 | 0.04139139 | -0.0078456 | 0.89647104 | 0.99999994 |
| Ggnbp2 | 0.47210162 | 0.29462257 | 0.30247071 | -0.0078481 | 0.56856067 | 0.99999994 |
| Satb1 | 0.07436649 | 0.05623649 | 0.06409414 | -0.0078577 | 0.79276124 | 0.99999994 |
| Mapk9 | 0.08046466 | 0.06121612 | 0.06907716 | -0.007861 | 0.77562453 | 0.99999994 |
| Sgcb | 0.16788816 | 0.1270255 | 0.13488766 | -0.0078622 | 0.65098348 | 0.99999994 |
| Ccdc86 | 0.18046841 | 0.13571736 | 0.14357953 | -0.0078622 | 0.64225411 | 0.99999994 |
| Flot2 | 0.13579085 | 0.10402115 | 0.1118878 | -0.0078666 | 0.68010688 | 0.99999994 |
| Nudcd3 | 0.29571046 | 0.20759049 | 0.2154573 | -0.0078668 | 0.596012 | 0.99999994 |
| Pten | 0.16167976 | 0.12266289 | 0.13053253 | -0.0078696 | 0.65590166 | 0.99999994 |
| Fbxo11 | 0.14738185 | 0.11246028 | 0.12033282 | -0.0078725 | 0.66839279 | 0.99999994 |
| Pgd | 0.28662511 | 0.20239262 | 0.21026556 | -0.0078729 | 0.59840967 | 0.99999994 |
| 4933417E11I | 0.02968157 | 0.01846926 | 0.02634399 | -0.0078747 | 0.9766888 | 0.99999994 |
| Use1 | 0.65908303 | 0.3630356 | 0.37091111 | -0.0078755 | 0.55619682 | 0.99999994 |
| Exosc3 | 0.21026328 | 0.15559359 | 0.16346983 | -0.0078762 | 0.62578646 | 0.99999994 |
| Neo1 | 0.11536029 | 0.08873034 | 0.0966083 | -0.007878 | 0.70636457 | 0.99999994 |
| Tmem237 | 0.19831396 | 0.14771827 | 0.15560807 | -0.0078898 | 0.63212906 | 0.99999994 |
| Plrg1 | 0.16084782 | 0.12205332 | 0.12994555 | -0.0078922 | 0.65699403 | 0.99999994 |
| Ogfod3 | 0.13387127 | 0.10258116 | 0.11047385 | -0.0078927 | 0.68280467 | 0.99999994 |
| Slc38a2 | 0.19043734 | 0.14244487 | 0.15034215 | -0.0078973 | 0.6366993 | 0.99999994 |
| Cacna1d | 0.05466918 | 0.03981925 | 0.04772647 | -0.0079072 | 0.86494532 | 0.99999994 |
| Elmo3 | 0.06137324 | 0.04542152 | 0.05334455 | -0.007923 | 0.83858015 | 0.99999994 |
| Trio | 0.0858398 | 0.06550715 | 0.07343696 | -0.0079298 | 0.76385604 | 0.99999994 |
| Spryd7 | 0.13082082 | 0.10028717 | 0.10821783 | -0.0079307 | 0.6871563 | 0.99999994 |
| Bcl2l13 | 0.08605238 | 0.06567252 | 0.07360877 | -0.0079362 | 0.76351832 | 0.99999994 |

| | | | | | | |
|---|---|---|---|---|---|---|
| Arxes1 | 0.13609255 | 0.10417193 | 0.11210963 | -0.0079377 | 0.68128327 | 0.99999994 |
| Atp6v0a2 | 0.09650761 | 0.07406194 | 0.08200105 | -0.0079391 | 0.74035811 | 0.99999994 |
| Dcaf10 | 0.13876445 | 0.10612794 | 0.11406949 | -0.0079416 | 0.67846745 | 0.99999994 |
| Ap5s1 | 0.11260126 | 0.08656311 | 0.09450534 | -0.0079422 | 0.71205577 | 0.99999994 |
| Coq4 | 0.15636297 | 0.11882025 | 0.12676718 | -0.0079469 | 0.66176378 | 0.99999994 |
| Emc3 | 0.30569969 | 0.21312736 | 0.22107794 | -0.0079506 | 0.59459499 | 0.99999994 |
| Map2k3 | 0.09576739 | 0.07345823 | 0.08141058 | -0.0079523 | 0.74222046 | 0.99999994 |
| Sqle | 0.2150644 | 0.15863194 | 0.16658569 | -0.0079538 | 0.6246864 | 0.99999994 |
| Nrep | 0.32164244 | 0.22190824 | 0.22986224 | -0.007954 | 0.5910831 | 0.99999994 |
| Slx1b | 0.06858778 | 0.05138163 | 0.05933572 | -0.0079541 | 0.81388382 | 0.99999994 |
| Rpa1 | 0.11644369 | 0.08947677 | 0.09743137 | -0.0079546 | 0.70658846 | 0.99999994 |
| 2410002F23I | 0.27559762 | 0.19590366 | 0.20386016 | -0.0079565 | 0.60248803 | 0.99999994 |
| Spast | 0.09727917 | 0.07465649 | 0.08261594 | -0.0079595 | 0.73933364 | 0.99999994 |
| Lmbrd1 | 0.17056849 | 0.12879002 | 0.13675447 | -0.0079645 | 0.65086616 | 0.99999994 |
| Ckap5 | 0.16953498 | 0.12807096 | 0.1360356 | -0.0079646 | 0.65162747 | 0.99999994 |
| Gad1 | 0.09378444 | 0.07186105 | 0.079826 | -0.007965 | 0.74670734 | 0.99999994 |
| Tjp1 | 0.16277353 | 0.12333213 | 0.13130297 | -0.0079708 | 0.65692272 | 0.99999994 |
| Morc3 | 0.10835153 | 0.08327646 | 0.09124763 | -0.0079712 | 0.71954253 | 0.99999994 |
| Pex10 | 0.06521853 | 0.0485633 | 0.0565448 | -0.0079815 | 0.82628044 | 0.99999994 |
| Rab26os | 0.05818523 | 0.04269758 | 0.05067911 | -0.0079815 | 0.85275441 | 0.99999994 |
| Atf1 | 0.17009872 | 0.12844532 | 0.13642787 | -0.0079825 | 0.65153603 | 0.99999994 |
| Ppp1r13b | 0.06273341 | 0.04648876 | 0.05447839 | -0.0079896 | 0.83551748 | 0.99999994 |
| Crot | 0.2117854 | 0.15646978 | 0.16446026 | -0.0079905 | 0.62680626 | 0.99999994 |
| 2310061I04R | 0.19150645 | 0.14306422 | 0.15106101 | -0.0079968 | 0.6377198 | 0.99999994 |
| Tfam | 0.23140518 | 0.16900636 | 0.17701107 | -0.0080047 | 0.61829672 | 0.99999994 |
| Osbpl1a | 0.10824179 | 0.08315641 | 0.0911632 | -0.0080068 | 0.72059924 | 0.99999994 |
| Tmem60 | 0.3263291 | 0.22439032 | 0.23240182 | -0.0080115 | 0.59074507 | 0.99999994 |
| Med13 | 0.13180303 | 0.10093262 | 0.10894545 | -0.0080128 | 0.68778053 | 0.99999994 |
| Taok1 | 0.22007606 | 0.16179698 | 0.16981277 | -0.0080158 | 0.62331718 | 0.99999994 |
| Lyrm2 | 0.34900918 | 0.23640452 | 0.24442186 | -0.0080173 | 0.58641664 | 0.99999994 |

| | | | | | | |
|---|---|---|---|---|---|---|
| Slc35e1 | 0.05199471 | 0.03745073 | 0.04547142 | -0.0080207 | 0.87982519 | 0.99999994 |
| Sf1 | 0.34281418 | 0.23316099 | 0.24118249 | -0.0080215 | 0.58760333 | 0.99999994 |
| Degs1 | 0.48671131 | 0.30059814 | 0.30862012 | -0.008022 | 0.56868179 | 0.99999994 |
| Megf9 | 0.04897358 | 0.03489223 | 0.04291453 | -0.0080223 | 0.89330848 | 0.99999994 |
| Ip6k2 | 0.64021997 | 0.35698724 | 0.36500971 | -0.0080225 | 0.55815723 | 0.99999994 |
| Srp9 | 1.21347723 | 0.42060647 | 0.42863069 | -0.0080242 | 0.54958422 | 0.99999994 |
| Ivns1abp | 0.21967383 | 0.16152978 | 0.16955472 | -0.0080249 | 0.62363517 | 0.99999994 |
| Zfp318 | 0.10879444 | 0.0835576 | 0.0915882 | -0.0080306 | 0.72026881 | 0.99999994 |
| Cep104 | 0.14393396 | 0.10980395 | 0.11783742 | -0.0080335 | 0.67501061 | 0.99999994 |
| Aktip | 0.26437196 | 0.18918761 | 0.19722129 | -0.0080337 | 0.60685922 | 0.99999994 |
| Ppid | 0.60235455 | 0.34439574 | 0.35243318 | -0.0080374 | 0.56032847 | 0.99999994 |
| Aqr | 0.13514825 | 0.10337272 | 0.11141495 | -0.0080422 | 0.68455483 | 0.99999994 |
| Eri3 | 0.38440552 | 0.25427092 | 0.26231954 | -0.0080486 | 0.58091462 | 0.99999994 |
| Twf2 | 0.26950255 | 0.19221892 | 0.20027084 | -0.0080519 | 0.60550408 | 0.99999994 |
| Taf6 | 0.20891464 | 0.15453241 | 0.16259026 | -0.0080579 | 0.62925624 | 0.99999994 |
| Iars2 | 0.10636227 | 0.081655 | 0.08971497 | -0.00806 | 0.72508197 | 0.99999994 |
| Ints1 | 0.03907303 | 0.02639865 | 0.03446355 | -0.0080649 | 0.94034434 | 0.99999994 |
| Gart | 0.19612755 | 0.14608504 | 0.15415309 | -0.0080681 | 0.63622255 | 0.99999994 |
| Actl6b | 0.06423528 | 0.04765893 | 0.05572801 | -0.0080691 | 0.83242719 | 0.99999994 |
| Secisbp2 | 0.11012203 | 0.08453796 | 0.09260757 | -0.0080696 | 0.7190492 | 0.99999994 |
| Abce1 | 0.2023665 | 0.15022019 | 0.15829145 | -0.0080713 | 0.63284841 | 0.99999994 |
| Rsph9 | 0.22516187 | 0.16498928 | 0.17306063 | -0.0080713 | 0.62188866 | 0.99999994 |
| Tbc1d23 | 0.07510601 | 0.05661762 | 0.06470053 | -0.0080829 | 0.79717289 | 0.99999994 |
| Fam120a | 0.29827309 | 0.20882358 | 0.21690819 | -0.0080846 | 0.59796997 | 0.99999994 |
| Edem3 | 0.06139238 | 0.04526748 | 0.05336052 | -0.008093 | 0.84364489 | 0.99999994 |
| Sik3 | 0.05850036 | 0.04284917 | 0.05094308 | -0.0080939 | 0.85487464 | 0.99999994 |
| Gdap2 | 0.09837615 | 0.07539307 | 0.08348912 | -0.0080961 | 0.7406907 | 0.99999994 |
| Mrps15 | 0.39654717 | 0.26012347 | 0.26822588 | -0.0081024 | 0.57967858 | 0.99999994 |
| Nup35 | 0.13894029 | 0.10609243 | 0.11419817 | -0.0081057 | 0.68170276 | 0.99999994 |
| Dicer1 | 0.21067508 | 0.15562197 | 0.16373802 | -0.0081161 | 0.62927648 | 0.99999994 |

| | | | | | | |
|---|---|---|---|---|---|---|
| Cdc42se2 | 0.21825331 | 0.16052569 | 0.16864206 | -0.0081164 | 0.62565077 | 0.99999994 |
| Gm15417 | 0.07370728 | 0.05543232 | 0.06355312 | -0.0081208 | 0.80248978 | 0.99999994 |
| Ntpcr | 0.10252789 | 0.07862583 | 0.08674668 | -0.0081209 | 0.73340359 | 0.99999994 |
| Trim56 | 0.05686616 | 0.04144904 | 0.049573 | -0.008124 | 0.86232732 | 0.99999994 |
| Pbx2 | 0.1092445 | 0.08380413 | 0.09193402 | -0.0081299 | 0.7219464 | 0.99999994 |
| Wdtc1 | 0.07352711 | 0.05527353 | 0.06340518 | -0.0081316 | 0.80335618 | 0.99999994 |
| Usp19 | 0.17149038 | 0.12925516 | 0.1373947 | -0.0081395 | 0.65333358 | 0.99999994 |
| Sympk | 0.1373474 | 0.10488851 | 0.11303113 | -0.0081426 | 0.68421396 | 0.99999994 |
| Gkap1 | 0.15868007 | 0.12026676 | 0.12841215 | -0.0081454 | 0.6635691 | 0.99999994 |
| Atg2b | 0.05376189 | 0.03881656 | 0.04696235 | -0.0081458 | 0.87586656 | 0.99999994 |
| Gm21092 | 0.36847333 | 0.24623765 | 0.25439066 | -0.008153 | 0.58446551 | 0.99999994 |
| Bckdk | 0.28820843 | 0.2030219 | 0.21117574 | -0.0081538 | 0.60141559 | 0.99999994 |
| Ncald | 0.08642565 | 0.06575233 | 0.07391033 | -0.008158 | 0.76874142 | 0.99999994 |
| Fgfr1op2 | 0.39897746 | 0.26123347 | 0.26939408 | -0.0081606 | 0.5798999 | 0.99999994 |
| Bphl | 0.33224766 | 0.22741869 | 0.23558141 | -0.0081627 | 0.59120193 | 0.99999994 |
| Adarb1 | 0.05752022 | 0.04195463 | 0.05012171 | -0.0081671 | 0.86095498 | 0.99999994 |
| Smchd1 | 0.21656092 | 0.15938265 | 0.16755203 | -0.0081694 | 0.62723757 | 0.99999994 |
| Synj1 | 0.06688939 | 0.04976035 | 0.05793039 | -0.00817 | 0.82607294 | 0.99999994 |
| Drosha | 0.08211278 | 0.06224399 | 0.07041717 | -0.0081732 | 0.78011746 | 0.99999994 |
| Dbnl | 0.48340135 | 0.29905778 | 0.30723961 | -0.0081818 | 0.57034771 | 0.99999994 |
| Myo6 | 0.24694414 | 0.17848726 | 0.18667182 | -0.0081846 | 0.61479769 | 0.99999994 |
| Nfyc | 0.26368274 | 0.18862385 | 0.19680968 | -0.0081858 | 0.60905451 | 0.99999994 |
| Vamp2 | 0.46480903 | 0.29115628 | 0.29934595 | -0.0081897 | 0.57225112 | 0.99999994 |
| Fkbp14 | 0.07486629 | 0.05631204 | 0.06450403 | -0.008192 | 0.80104357 | 0.99999994 |
| Spsb3 | 0.13692803 | 0.1045313 | 0.11272337 | -0.0081921 | 0.6857165 | 0.99999994 |
| Pmpcb | 0.3976432 | 0.26055285 | 0.26875328 | -0.0082004 | 0.58047328 | 0.99999994 |
| Ramp2 | 0.14325921 | 0.10913839 | 0.11734739 | -0.008209 | 0.67926475 | 0.99999994 |
| Hspb8 | 0.15967456 | 0.12090422 | 0.12911629 | -0.0082121 | 0.66398322 | 0.99999994 |
| Micu2 | 0.2532508 | 0.18231011 | 0.19052428 | -0.0082142 | 0.61293572 | 0.99999994 |
| Apc | 0.13283733 | 0.10149458 | 0.10971043 | -0.0082158 | 0.6909245 | 0.99999994 |

| | | | | | | |
|---|---|---|---|---|---|---|
| Poldip2 | 0.3520642 | 0.23779005 | 0.24600732 | -0.0082173 | 0.58797507 | 0.99999994 |
| Wrnip1 | 0.14567546 | 0.11088111 | 0.11909969 | -0.0082186 | 0.67700062 | 0.99999994 |
| Nptxr | 0.03426756 | 0.02210035 | 0.03032156 | -0.0082212 | 0.96444436 | 0.99999994 |
| Alkbh1 | 0.25464796 | 0.18314995 | 0.19137216 | -0.0082222 | 0.61255598 | 0.99999994 |
| Wdsub1 | 0.10185508 | 0.07800125 | 0.08622392 | -0.0082227 | 0.73721329 | 0.99999994 |
| Pnkd | 0.68065948 | 0.36891684 | 0.37714788 | -0.008231 | 0.55775168 | 0.99999994 |
| Mecp2 | 0.1538349 | 0.11673381 | 0.12496541 | -0.0082316 | 0.66947395 | 0.99999994 |
| Jph4 | 0.05395635 | 0.03889349 | 0.04712621 | -0.0082327 | 0.87755143 | 0.99999994 |
| Sgpp2 | 0.02782236 | 0.01649169 | 0.02472449 | -0.0082328 | 0.98666946 | 0.99999994 |
| Dedd | 0.15299106 | 0.1161285 | 0.12436236 | -0.0082339 | 0.67028785 | 0.99999994 |
| Lap3 | 0.31655347 | 0.21884658 | 0.22708317 | -0.0082366 | 0.59539087 | 0.99999994 |
| Kidins220 | 0.16060448 | 0.12153662 | 0.12977369 | -0.0082371 | 0.66366852 | 0.99999994 |
| Ppp3r1 | 0.27601302 | 0.19586575 | 0.20410352 | -0.0082378 | 0.60590257 | 0.99999994 |
| Nploc4 | 0.07874464 | 0.05943652 | 0.06767565 | -0.0082391 | 0.79113727 | 0.99999994 |
| Rpl30 | 2.08813827 | 0.36025987 | 0.36850209 | -0.0082422 | 0.55917647 | 0.99999994 |
| Micu1 | 0.14762707 | 0.11226208 | 0.12050975 | -0.0082477 | 0.67564681 | 0.99999994 |
| Tceal1 | 0.23509445 | 0.1710777 | 0.179327 | -0.0082493 | 0.62027453 | 0.99999994 |
| C87436 | 0.12537908 | 0.09590848 | 0.10416472 | -0.0082562 | 0.7011088 | 0.99999994 |
| Med9 | 0.32860722 | 0.22536591 | 0.23362947 | -0.0082636 | 0.59306553 | 0.99999994 |
| Dennd6a | 0.12952597 | 0.09899236 | 0.10725666 | -0.0082643 | 0.69598335 | 0.99999994 |
| Piga | 0.05986722 | 0.04382065 | 0.05208678 | -0.0082661 | 0.8545977 | 0.99999994 |
| Ern1 | 0.06388458 | 0.04716601 | 0.05543644 | -0.0082704 | 0.83965623 | 0.99999994 |
| Mcc | 0.02696047 | 0.01570175 | 0.02397237 | -0.0082706 | 0.98917595 | 0.99999994 |
| Sike1 | 0.20384123 | 0.15099214 | 0.15926363 | -0.0082715 | 0.63521722 | 0.99999994 |
| Pgm2 | 0.16055432 | 0.12146424 | 0.12973825 | -0.008274 | 0.66440185 | 0.99999994 |
| Pcyox1l | 0.15284334 | 0.11597862 | 0.12425671 | -0.0082781 | 0.67128071 | 0.99999994 |
| 2410021H03I | 0.0449629 | 0.03122428 | 0.03950438 | -0.0082801 | 0.91851768 | 0.99999994 |
| Ret | 0.08300254 | 0.06285913 | 0.07113941 | -0.0082803 | 0.78076142 | 0.99999994 |
| Hdhd2 | 0.2530321 | 0.18211048 | 0.19039136 | -0.0082809 | 0.61390501 | 0.99999994 |
| Crtc1 | 0.09461687 | 0.07221076 | 0.08049169 | -0.0082809 | 0.7532655 | 0.99999994 |

| | | | | | | |
|---|---|---|---|---|---|---|
| Ncoa2 | 0.12878726 | 0.09842124 | 0.10670739 | -0.0082861 | 0.6973822 | 0.99999994 |
| Sbno1 | 0.45902049 | 0.28854976 | 0.29683906 | -0.0082893 | 0.57373066 | 0.99999994 |
| Aspscr1 | 0.34826025 | 0.23573996 | 0.24403201 | -0.008292 | 0.5894688 | 0.99999994 |
| Ctdspl | 0.07170018 | 0.05360693 | 0.06190303 | -0.0082961 | 0.81382198 | 0.99999994 |
| Slc25a14 | 0.16469503 | 0.12435668 | 0.13265315 | -0.0082965 | 0.66140692 | 0.99999994 |
| Gramd3 | 0.19034617 | 0.14197052 | 0.15028079 | -0.0083103 | 0.64359522 | 0.99999994 |
| Hsd17b12 | 0.39843225 | 0.26082041 | 0.2691324 | -0.008312 | 0.58143951 | 0.99999994 |
| Emp2 | 0.06243434 | 0.04591246 | 0.05422926 | -0.0083168 | 0.84633575 | 0.99999994 |
| Hsdl1 | 0.10754735 | 0.08230447 | 0.09062864 | -0.0083242 | 0.72952625 | 0.99999994 |
| Trrap | 0.11403482 | 0.0872748 | 0.09559919 | -0.0083244 | 0.71891073 | 0.99999994 |
| Dhrs1 | 0.2232945 | 0.16354196 | 0.17187138 | -0.0083294 | 0.62649625 | 0.99999994 |
| Ep400 | 0.15204603 | 0.11535387 | 0.12368603 | -0.0083322 | 0.67307073 | 0.99999994 |
| Cbl | 0.1102257 | 0.08435071 | 0.09268708 | -0.0083364 | 0.72530673 | 0.99999994 |
| Ensa | 0.16665212 | 0.12568584 | 0.13402406 | -0.0083382 | 0.66060725 | 0.99999994 |
| Mtif2 | 0.10885391 | 0.0832847 | 0.09163391 | -0.0083492 | 0.72790636 | 0.99999994 |
| Ube2q1 | 0.17749239 | 0.1331876 | 0.14153961 | -0.008352 | 0.65275908 | 0.99999994 |
| Gpn3 | 0.127624 | 0.09748622 | 0.10584112 | -0.0083549 | 0.70036143 | 0.99999994 |
| Zfp219 | 0.0966541 | 0.07376051 | 0.08211784 | -0.0083573 | 0.75093454 | 0.99999994 |
| Rnf170 | 0.10121731 | 0.07736993 | 0.08572787 | -0.0083579 | 0.74181595 | 0.99999994 |
| Cenpc1 | 0.12387064 | 0.09467474 | 0.10303485 | -0.0083601 | 0.70543192 | 0.99999994 |
| Grip1 | 0.03974785 | 0.02668083 | 0.03504309 | -0.0083623 | 0.94390124 | 0.99999994 |
| Ankmy2 | 0.09549319 | 0.07282852 | 0.0811917 | -0.0083632 | 0.75353003 | 0.99999994 |
| Apbb2 | 0.11873216 | 0.09079687 | 0.09916518 | -0.0083683 | 0.71284446 | 0.99999994 |
| 2510039O18 | 0.10138438 | 0.07748916 | 0.08585786 | -0.0083687 | 0.74176796 | 0.99999994 |
| Mttp | 0.10179346 | 0.07779068 | 0.08617602 | -0.0083853 | 0.74140889 | 0.99999994 |
| Nfs1 | 0.26515218 | 0.18929963 | 0.19768668 | -0.0083871 | 0.6111826 | 0.99999994 |
| Slc7a5 | 0.1032947 | 0.07895376 | 0.08734177 | -0.008388 | 0.73867119 | 0.99999994 |
| Celf1 | 0.40876087 | 0.26566164 | 0.27405104 | -0.0083894 | 0.58073262 | 0.99999994 |
| Agap1 | 0.10213149 | 0.07804336 | 0.08643876 | -0.0083954 | 0.74102263 | 0.99999994 |
| Abi1 | 0.16634156 | 0.12540751 | 0.13380681 | -0.0083993 | 0.66196503 | 0.99999994 |

| | | | | | | |
|---|---|---|---|---|---|---|
| Rnf216 | 0.14467627 | 0.1099743 | 0.11837589 | -0.0084016 | 0.68168907 | 0.99999994 |
| Cpeb4 | 0.09243755 | 0.07033961 | 0.07874742 | -0.0084078 | 0.76136332 | 0.99999994 |
| Zranb1 | 0.25084058 | 0.18063876 | 0.18905656 | -0.0084178 | 0.6165287 | 0.99999994 |
| Noc2l | 0.08677361 | 0.06577138 | 0.07419131 | -0.0084199 | 0.77500346 | 0.99999994 |
| Hspbp1 | 0.2615443 | 0.18709708 | 0.19552974 | -0.0084327 | 0.61297072 | 0.99999994 |
| Pex3 | 0.14015531 | 0.10665085 | 0.11508635 | -0.0084355 | 0.68718689 | 0.99999994 |
| Irf6 | 0.12282872 | 0.09381346 | 0.10225282 | -0.0084394 | 0.70862674 | 0.99999994 |
| Mettl7a1 | 0.07911459 | 0.0595377 | 0.06797735 | -0.0084397 | 0.795715 | 0.99999994 |
| Ube2z | 0.15122158 | 0.11465262 | 0.12309514 | -0.0084425 | 0.67599508 | 0.99999994 |
| Lonp1 | 0.18753621 | 0.13993862 | 0.14838525 | -0.0084466 | 0.64762065 | 0.99999994 |
| Scaf1 | 0.13138975 | 0.10019229 | 0.10863943 | -0.0084471 | 0.69761671 | 0.99999994 |
| Armc9 | 0.11515299 | 0.08799831 | 0.09645062 | -0.0084523 | 0.72016117 | 0.99999994 |
| Osgep | 0.24031359 | 0.1741203 | 0.1825795 | -0.0084592 | 0.62111086 | 0.99999994 |
| Abhd14b | 0.17284792 | 0.12987564 | 0.13833575 | -0.0084601 | 0.65802483 | 0.99999994 |
| Malsu1 | 0.30480479 | 0.21211201 | 0.22057812 | -0.0084661 | 0.60082411 | 0.99999994 |
| Gale | 0.2557479 | 0.18357215 | 0.19203835 | -0.0084662 | 0.61541167 | 0.99999994 |
| Mbp | 0.16088692 | 0.1215009 | 0.12997316 | -0.0084723 | 0.66781683 | 0.99999994 |
| Rragc | 0.15304731 | 0.11592516 | 0.12440259 | -0.0084774 | 0.67494087 | 0.99999994 |
| Drg2 | 0.27520904 | 0.19514242 | 0.20363236 | -0.0084899 | 0.60930959 | 0.99999994 |
| Mkrn2os | 0.05389744 | 0.03858396 | 0.04707657 | -0.0084926 | 0.88508697 | 0.99999994 |
| Arhgap35 | 0.09092183 | 0.06903267 | 0.0775314 | -0.0084987 | 0.76719716 | 0.99999994 |
| Zbtb41 | 0.07141924 | 0.05316678 | 0.06167171 | -0.0085049 | 0.82067898 | 0.99999994 |
| Mrps22 | 0.1698699 | 0.12776295 | 0.13626869 | -0.0085057 | 0.66110566 | 0.99999994 |
| Sec62 | 1.2117584 | 0.42017475 | 0.42868597 | -0.0085112 | 0.55256979 | 0.99999994 |
| Sgms1 | 0.07765783 | 0.05827495 | 0.06678848 | -0.0085135 | 0.80191653 | 0.99999994 |
| Afg3l2 | 0.12174236 | 0.09292056 | 0.10143601 | -0.0085154 | 0.71184894 | 0.99999994 |
| Cse1l | 0.11390202 | 0.08698204 | 0.09549796 | -0.0085159 | 0.72360126 | 0.99999994 |
| Nsun2 | 0.24577822 | 0.17743771 | 0.18595536 | -0.0085177 | 0.61977628 | 0.99999994 |
| Cyld | 0.09750165 | 0.07427439 | 0.08279313 | -0.0085187 | 0.75329185 | 0.99999994 |
| Braf | 0.14735796 | 0.11179155 | 0.12031558 | -0.008524 | 0.68138847 | 0.99999994 |

| | | | | | | |
|---|---|---|---|---|---|---|
| Iars | 0.07202329 | 0.05364445 | 0.06216895 | -0.0085245 | 0.8193021 | 0.99999994 |
| Gabarapl1 | 0.3238614 | 0.22253747 | 0.23106699 | -0.0085295 | 0.59704674 | 0.99999994 |
| Adprm | 0.10285436 | 0.0784684 | 0.08700013 | -0.0085317 | 0.74304469 | 0.99999994 |
| Prr12 | 0.08003714 | 0.06019726 | 0.0687291 | -0.0085318 | 0.79568344 | 0.99999994 |
| Dot1l | 0.13175137 | 0.1003712 | 0.10890722 | -0.008536 | 0.69906647 | 0.99999994 |
| Nr3c1 | 0.22215298 | 0.16260612 | 0.17114258 | -0.0085365 | 0.63006067 | 0.99999994 |
| Zfp942 | 0.07856942 | 0.0589933 | 0.06753269 | -0.0085394 | 0.80001789 | 0.99999994 |
| Chmp1a | 0.37609131 | 0.24966723 | 0.25820801 | -0.0085408 | 0.5871319 | 0.99999994 |
| Rock2 | 0.11546587 | 0.08814418 | 0.09668857 | -0.0085444 | 0.72180899 | 0.99999994 |
| Sdk2 | 0.0353306 | 0.02269324 | 0.03124011 | -0.0085469 | 0.96570331 | 0.99999994 |
| Utp18 | 0.24101923 | 0.17446847 | 0.18301708 | -0.0085486 | 0.62206701 | 0.99999994 |
| Fam3c | 0.20247599 | 0.149814 | 0.1583637 | -0.0085497 | 0.64033783 | 0.99999994 |
| Ppp1r27 | 0.02639984 | 0.01493272 | 0.02348267 | -0.0085499 | 0.99231525 | 0.99999994 |
| Klf9 | 0.04238169 | 0.02874982 | 0.03730009 | -0.0085503 | 0.93647404 | 0.99999994 |
| Szrd1 | 0.25927295 | 0.18561428 | 0.19416546 | -0.0085512 | 0.61529641 | 0.99999994 |
| Akt2 | 0.18268929 | 0.13654036 | 0.14509548 | -0.0085551 | 0.65264536 | 0.99999994 |
| Mzt2 | 0.20074806 | 0.14866039 | 0.1572219 | -0.0085615 | 0.64150012 | 0.99999994 |
| Ift46 | 0.24819391 | 0.17887618 | 0.18743833 | -0.0085621 | 0.61945937 | 0.99999994 |
| Ube4a | 0.09380482 | 0.07127721 | 0.07984231 | -0.0085651 | 0.76240575 | 0.99999994 |
| Scaf8 | 0.0907199 | 0.06880388 | 0.07736923 | -0.0085653 | 0.76941086 | 0.99999994 |
| Slc39a9 | 0.05183472 | 0.03676964 | 0.04533627 | -0.0085666 | 0.89576227 | 0.99999994 |
| Ralgps2 | 0.0697854 | 0.05175651 | 0.06032482 | -0.0085683 | 0.82778385 | 0.99999994 |
| Ankrd10 | 0.21693474 | 0.15922141 | 0.16779306 | -0.0085716 | 0.63308591 | 0.99999994 |
| Zcchc14 | 0.077763 | 0.05829982 | 0.06687438 | -0.0085746 | 0.80330097 | 0.99999994 |
| Plp2 | 0.26516183 | 0.18911436 | 0.19769244 | -0.0085781 | 0.61364279 | 0.99999994 |
| Bmpr1a | 0.16986995 | 0.12768809 | 0.13626872 | -0.0085806 | 0.66244329 | 0.99999994 |
| Smad5 | 0.11738418 | 0.08956107 | 0.09814469 | -0.0085836 | 0.71976978 | 0.99999994 |
| Aifm1 | 0.17974031 | 0.13449231 | 0.14308135 | -0.008589 | 0.65526514 | 0.99999994 |
| Bfar | 0.22747836 | 0.16593843 | 0.17453098 | -0.0085926 | 0.6284336 | 0.99999994 |
| Otud6b | 0.11458877 | 0.08742818 | 0.09602119 | -0.008593 | 0.72430624 | 0.99999994 |

| | | | | | | |
|---|---|---|---|---|---|---|
| 1700056E22I | 0.12943512 | 0.09859559 | 0.10718914 | -0.0085936 | 0.70320899 | 0.99999994 |
| Tceal8 | 0.83637172 | 0.40089745 | 0.40949535 | -0.0085979 | 0.5555748 | 0.99999994 |
| Thap11 | 0.32460849 | 0.22286053 | 0.23147166 | -0.0086111 | 0.59778868 | 0.99999994 |
| Rapgef4 | 0.09956811 | 0.07582109 | 0.08443652 | -0.0086154 | 0.75148747 | 0.99999994 |
| Pabpn1 | 0.31491754 | 0.21756697 | 0.22618502 | -0.0086181 | 0.60010509 | 0.99999994 |
| Smg5 | 0.10480026 | 0.07988334 | 0.08850802 | -0.0086247 | 0.74171104 | 0.99999994 |
| Hexim1 | 0.1389591 | 0.10558255 | 0.11421194 | -0.0086294 | 0.6924922 | 0.99999994 |
| Daam1 | 0.23984126 | 0.17365669 | 0.18228632 | -0.0086296 | 0.62366441 | 0.99999994 |
| Ncoa7 | 0.13433174 | 0.10216935 | 0.11081342 | -0.0086441 | 0.69820109 | 0.99999994 |
| Dhx29 | 0.10685509 | 0.08144865 | 0.09009513 | -0.0086465 | 0.73852817 | 0.99999994 |
| Slc25a44 | 0.14374663 | 0.10905351 | 0.11770142 | -0.0086479 | 0.68760199 | 0.99999994 |
| Rab20 | 0.04730691 | 0.03284828 | 0.0414996 | -0.0086513 | 0.91737586 | 0.99999994 |
| Mical3 | 0.07661039 | 0.05728092 | 0.06593228 | -0.0086514 | 0.80878272 | 0.99999994 |
| Rprd1b | 0.14351963 | 0.10888161 | 0.11753658 | -0.008655 | 0.68798631 | 0.99999994 |
| Rhobtb1 | 0.03779773 | 0.02470327 | 0.03336691 | -0.0086636 | 0.95803346 | 0.99999994 |
| Rnf26 | 0.09560853 | 0.07261856 | 0.08128378 | -0.0086652 | 0.76102328 | 0.99999994 |
| Gpi1 | 0.59097233 | 0.33981796 | 0.34848326 | -0.0086653 | 0.56573088 | 0.99999994 |
| Mettl5 | 0.16534105 | 0.12443895 | 0.13310616 | -0.0086672 | 0.66764831 | 0.99999994 |
| Ccm2 | 0.20474888 | 0.15119201 | 0.15986083 | -0.0086688 | 0.64093402 | 0.99999994 |
| Fkrp | 0.11012727 | 0.0839362 | 0.09261159 | -0.0086754 | 0.73353333 | 0.99999994 |
| Atp7a | 0.07240085 | 0.0538041 | 0.06247955 | -0.0086755 | 0.8223228 | 0.99999994 |
| Oxct1 | 0.14144782 | 0.10734335 | 0.11602926 | -0.0086859 | 0.69086116 | 0.99999994 |
| Psmf1 | 0.22583766 | 0.16480356 | 0.17349014 | -0.0086866 | 0.63053896 | 0.99999994 |
| Atp13a2 | 0.09537295 | 0.0724078 | 0.0810957 | -0.0086879 | 0.76210966 | 0.99999994 |
| Mrpl12 | 0.51099176 | 0.30982822 | 0.31852034 | -0.0086921 | 0.57206965 | 0.99999994 |
| Suox | 0.10228235 | 0.07785603 | 0.08655596 | -0.0086999 | 0.74826679 | 0.99999994 |
| Zfp512 | 0.17414836 | 0.13053429 | 0.13923528 | -0.008701 | 0.66128205 | 0.99999994 |
| Erap1 | 0.08802089 | 0.06649613 | 0.07519748 | -0.0087013 | 0.77941792 | 0.99999994 |
| Slc35g2 | 0.21184785 | 0.15579728 | 0.16450084 | -0.0087036 | 0.63764861 | 0.99999994 |
| Zfp507 | 0.09710911 | 0.07377343 | 0.08248046 | -0.008707 | 0.75887157 | 0.99999994 |

| | | | | | | |
|---|---|---|---|---|---|---|
| Polr3d | 0.21032078 | 0.15479891 | 0.16350728 | -0.0087084 | 0.63852418 | 0.99999994 |
| Gigyf2 | 0.2585164 | 0.18500027 | 0.19370994 | -0.0087097 | 0.61763916 | 0.99999994 |
| Atf6 | 0.14596678 | 0.11059872 | 0.11931049 | -0.0087118 | 0.68652755 | 0.99999994 |
| Coa5 | 0.08276325 | 0.06222863 | 0.07094525 | -0.0087166 | 0.79327205 | 0.99999994 |
| Hnrnpul1 | 0.33638687 | 0.22906757 | 0.23778671 | -0.0087191 | 0.59641348 | 0.99999994 |
| Tbc1d20 | 0.09991265 | 0.07598334 | 0.0847101 | -0.0087268 | 0.75355897 | 0.99999994 |
| Tspo | 0.19752072 | 0.14635336 | 0.15508078 | -0.0087274 | 0.64602007 | 0.99999994 |
| Map7 | 0.22797952 | 0.16611799 | 0.17484837 | -0.0087304 | 0.63019107 | 0.99999994 |
| Thap3 | 0.13004615 | 0.09891108 | 0.10764304 | -0.008732 | 0.70538814 | 0.99999994 |
| Acox1 | 0.12198675 | 0.09288784 | 0.10161988 | -0.008732 | 0.71632907 | 0.99999994 |
| Csrp1 | 0.19635839 | 0.14557405 | 0.15430695 | -0.0087329 | 0.64680616 | 0.99999994 |
| Uaca | 0.08916407 | 0.06737746 | 0.07611825 | -0.0087408 | 0.77767396 | 0.99999994 |
| Ttc32 | 0.12358322 | 0.09407799 | 0.10281925 | -0.0087413 | 0.71426776 | 0.99999994 |
| Ppp1r15b | 0.09036194 | 0.06833956 | 0.07708163 | -0.0087421 | 0.7748509 | 0.99999994 |
| Znhit2 | 0.26038155 | 0.18608429 | 0.19483195 | -0.0087477 | 0.61747681 | 0.99999994 |
| Hps4 | 0.16080127 | 0.12115907 | 0.12991268 | -0.0087536 | 0.67310735 | 0.99999994 |
| Usp22 | 0.17705344 | 0.13248335 | 0.14123789 | -0.0087545 | 0.66004666 | 0.99999994 |
| Numa1 | 0.17979828 | 0.13435896 | 0.14312103 | -0.0087621 | 0.658184 | 0.99999994 |
| Zzz3 | 0.15316528 | 0.11572427 | 0.12448693 | -0.0087627 | 0.68030161 | 0.99999994 |
| Mtmr4 | 0.08856356 | 0.06687132 | 0.07563474 | -0.0087634 | 0.77971726 | 0.99999994 |
| Pafah1b2 | 0.6741511 | 0.366626 | 0.37539287 | -0.0087669 | 0.56176739 | 0.99999994 |
| Scap | 0.06367648 | 0.04649621 | 0.05526335 | -0.0087671 | 0.8545123 | 0.99999994 |
| Tmigd3 | 0.05044435 | 0.03538021 | 0.04416057 | -0.0087804 | 0.90716055 | 0.99999994 |
| Scaf11 | 0.5912324 | 0.33978809 | 0.3485744 | -0.0087863 | 0.56662297 | 0.99999994 |
| Rimbp2 | 0.2404837 | 0.17389203 | 0.18268503 | -0.008793 | 0.62566137 | 0.99999994 |
| Tmem184b | 0.10355912 | 0.07874301 | 0.0875468 | -0.0088038 | 0.7483703 | 0.99999994 |
| Srpk2 | 0.48744109 | 0.300118 | 0.30892338 | -0.0088054 | 0.57523931 | 0.99999994 |
| Cntn1 | 0.02927926 | 0.01718674 | 0.02599389 | -0.0088071 | 0.98794215 | 0.99999994 |
| Mrpl24 | 0.72010025 | 0.37817702 | 0.38698487 | -0.0088079 | 0.56020935 | 0.99999994 |
| Atp13a3 | 0.0817459 | 0.0613094 | 0.07011912 | -0.0088097 | 0.79850718 | 0.99999994 |

| | | | | | | |
|---|---|---|---|---|---|---|
| Hexb | 0.07853185 | 0.05869183 | 0.06750204 | -0.0088102 | 0.80751425 | 0.99999994 |
| Cadps | 0.18593578 | 0.13848532 | 0.14730182 | -0.0088165 | 0.6548342 | 0.99999994 |
| Gjd2 | 0.28664486 | 0.20145842 | 0.21027692 | -0.0088185 | 0.60993567 | 0.99999994 |
| Nipal1 | 0.19699941 | 0.14591085 | 0.1547339 | -0.008823 | 0.64786048 | 0.99999994 |
| Psen1 | 0.15323596 | 0.1157115 | 0.12453747 | -0.008826 | 0.68144236 | 0.99999994 |
| Slco3a1 | 0.06935677 | 0.05114403 | 0.05997099 | -0.008827 | 0.83639055 | 0.99999994 |
| Poldip3 | 0.19971619 | 0.14771109 | 0.15653852 | -0.0088274 | 0.6463043 | 0.99999994 |
| Cfap97 | 0.20744339 | 0.1528004 | 0.16162856 | -0.0088282 | 0.64191059 | 0.99999994 |
| Strn4 | 0.26117803 | 0.18648083 | 0.19531007 | -0.0088292 | 0.61825812 | 0.99999994 |
| Bcr | 0.0549686 | 0.03914572 | 0.04797845 | -0.0088327 | 0.88979176 | 0.99999994 |
| Naa15 | 0.33899907 | 0.23033737 | 0.239171 | -0.0088336 | 0.59710025 | 0.99999994 |
| Mvp | 0.13367162 | 0.10149232 | 0.11032653 | -0.0088342 | 0.70298203 | 0.99999994 |
| Plcb4 | 0.09613907 | 0.0728721 | 0.08170713 | -0.008835 | 0.76416219 | 0.99999994 |
| Nelfa | 0.23903935 | 0.17295029 | 0.18178808 | -0.0088378 | 0.6268813 | 0.99999994 |
| Hemk1 | 0.0782986 | 0.05846386 | 0.06731169 | -0.0088478 | 0.80919939 | 0.99999994 |
| Sirpa | 0.03881949 | 0.02539775 | 0.03424567 | -0.0088479 | 0.95726366 | 0.99999994 |
| Fkbp15 | 0.06638293 | 0.04865977 | 0.05751072 | -0.0088509 | 0.84718119 | 0.99999994 |
| Ddx21 | 0.157869 | 0.1189861 | 0.12783705 | -0.0088509 | 0.67755071 | 0.99999994 |
| Ago3 | 0.06563485 | 0.04803272 | 0.05689032 | -0.0088576 | 0.84998509 | 0.99999994 |
| Zmiz2 | 0.12017653 | 0.09139631 | 0.10025617 | -0.0088599 | 0.72181331 | 0.99999994 |
| Man2b2 | 0.05138253 | 0.03609399 | 0.04495413 | -0.0088601 | 0.90522656 | 0.99999994 |
| Mrpl38 | 0.43358986 | 0.27667921 | 0.28554152 | -0.0088623 | 0.58183588 | 0.99999994 |
| Zswim7 | 0.18108076 | 0.13513285 | 0.14399805 | -0.0088652 | 0.65902265 | 0.99999994 |
| Igf2r | 0.15231551 | 0.1150133 | 0.12387899 | -0.0088657 | 0.68309744 | 0.99999994 |
| Acbd3 | 0.28496527 | 0.20043685 | 0.20930893 | -0.0088721 | 0.61108436 | 0.99999994 |
| Sh3gl1 | 0.24916107 | 0.17915505 | 0.18803047 | -0.0088754 | 0.62331297 | 0.99999994 |
| Ptp4a3 | 0.20666543 | 0.15224338 | 0.16111901 | -0.0088756 | 0.64307172 | 0.99999994 |
| Actr2 | 0.24253591 | 0.17507976 | 0.18395592 | -0.0088762 | 0.62596292 | 0.99999994 |
| Adck5 | 0.20112991 | 0.14859035 | 0.15747449 | -0.0088841 | 0.64636585 | 0.99999994 |
| Tfe3 | 0.12012474 | 0.09133101 | 0.10021709 | -0.0088861 | 0.7224746 | 0.99999994 |

| | | | | | | |
|---|---|---|---|---|---|---|
| Fra10ac1 | 0.15974554 | 0.12027696 | 0.1291665 | -0.0088895 | 0.67656193 | 0.99999994 |
| Mtmr14 | 0.10287507 | 0.07812395 | 0.0870162 | -0.0088923 | 0.7518169 | 0.99999994 |
| Nit1 | 0.24731109 | 0.17800285 | 0.18689704 | -0.0088942 | 0.62428991 | 0.99999994 |
| Gorasp1 | 0.06550345 | 0.04788691 | 0.05678129 | -0.0088944 | 0.85144936 | 0.99999994 |
| Vsig10 | 0.08581176 | 0.06451688 | 0.0734143 | -0.0088974 | 0.79008464 | 0.99999994 |
| Trappc11 | 0.09345013 | 0.07065949 | 0.07955847 | -0.008899 | 0.77172617 | 0.99999994 |
| Agpat3 | 0.29289236 | 0.20495062 | 0.21385493 | -0.0089043 | 0.60916822 | 0.99999994 |
| Slc25a36 | 0.17830888 | 0.13318655 | 0.14210024 | -0.0089137 | 0.66185874 | 0.99999994 |
| Phf2 | 0.07497272 | 0.0556768 | 0.06459127 | -0.0089145 | 0.82086522 | 0.99999994 |
| Zfp771 | 0.17399555 | 0.13021453 | 0.13912968 | -0.0089151 | 0.66513768 | 0.99999994 |
| Zfp938 | 0.16791544 | 0.12598608 | 0.1349067 | -0.0089206 | 0.67008489 | 0.99999994 |
| Shtn1 | 0.09921605 | 0.07523117 | 0.08415684 | -0.0089257 | 0.75989773 | 0.99999994 |
| Trappc12 | 0.0987415 | 0.0748512 | 0.08377966 | -0.0089285 | 0.76095351 | 0.99999994 |
| Polr2b | 0.1677451 | 0.12585702 | 0.13478779 | -0.0089308 | 0.67040699 | 0.99999994 |
| Rmnd5a | 0.0973721 | 0.07375908 | 0.08268995 | -0.0089309 | 0.76390362 | 0.99999994 |
| Nabp1 | 0.03904573 | 0.02550751 | 0.0344401 | -0.0089326 | 0.95786518 | 0.99999994 |
| Mmadhc | 0.50027709 | 0.30526634 | 0.31420009 | -0.0089338 | 0.5750564 | 0.99999994 |
| Ifnar1 | 0.10743584 | 0.08160209 | 0.09054274 | -0.0089407 | 0.7445004 | 0.99999994 |
| Mpp7 | 0.07211609 | 0.05330231 | 0.06224531 | -0.008943 | 0.83055048 | 0.99999994 |
| 1300002E11I | 0.26424712 | 0.18819797 | 0.19714676 | -0.0089488 | 0.61872812 | 0.99999994 |
| Slc35a4 | 0.21103368 | 0.15502243 | 0.16397142 | -0.008949 | 0.64180246 | 0.99999994 |
| Ldlr | 0.09272038 | 0.07002423 | 0.07897407 | -0.0089498 | 0.77467763 | 0.99999994 |
| Clstn1 | 0.05976161 | 0.04304858 | 0.05199849 | -0.0089499 | 0.87403687 | 0.99999994 |
| Sars | 0.64699827 | 0.35820499 | 0.36715699 | -0.008952 | 0.56446321 | 0.99999994 |
| Elp2 | 0.1923471 | 0.14265823 | 0.15162527 | -0.008967 | 0.65308153 | 0.99999994 |
| Hnrnpul2 | 0.37615415 | 0.24927029 | 0.25823929 | -0.008969 | 0.59141417 | 0.99999994 |
| Ccdc166 | 0.09078262 | 0.06844854 | 0.0774196 | -0.0089711 | 0.77973879 | 0.99999994 |
| Exph5 | 0.04312926 | 0.0289556 | 0.03793926 | -0.0089837 | 0.94250592 | 0.99999994 |
| Snx2 | 0.60311295 | 0.34370905 | 0.35269368 | -0.0089846 | 0.56732433 | 0.99999994 |
| Vps52 | 0.14240417 | 0.10774102 | 0.11672567 | -0.0089846 | 0.6957989 | 0.99999994 |

| | | | | | | |
|---|---|---|---|---|---|---|
| Pik3c2a | 0.16273977 | 0.12229306 | 0.13127921 | -0.0089861 | 0.67567269 | 0.99999994 |
| Dynlt1a | 0.13506996 | 0.10236747 | 0.11135731 | -0.0089898 | 0.70449252 | 0.99999994 |
| Nip7 | 0.25540937 | 0.18284034 | 0.19183344 | -0.0089931 | 0.62249802 | 0.99999994 |
| Prkaa1 | 0.08585503 | 0.0644545 | 0.07344927 | -0.0089948 | 0.79250761 | 0.99999994 |
| Rhpn2 | 0.11158021 | 0.08471786 | 0.09372468 | -0.0090068 | 0.73880559 | 0.99999994 |
| Nkiras1 | 0.23352666 | 0.16933161 | 0.17834449 | -0.0090129 | 0.63171089 | 0.99999994 |
| Idh1 | 0.18090393 | 0.134863 | 0.14387723 | -0.0090142 | 0.66167388 | 0.99999994 |
| Pck2 | 0.13448521 | 0.10191213 | 0.11092655 | -0.0090144 | 0.7057212 | 0.99999994 |
| Hs3st6 | 0.05639069 | 0.04015753 | 0.04917383 | -0.0090163 | 0.88885869 | 0.99999994 |
| Fam76b | 0.12758294 | 0.09679389 | 0.10581051 | -0.0090166 | 0.71471761 | 0.99999994 |
| Parn | 0.13409187 | 0.10161868 | 0.11063656 | -0.0090179 | 0.7062814 | 0.99999994 |
| Mnat1 | 0.2659466 | 0.18914016 | 0.19815996 | -0.0090198 | 0.61904844 | 0.99999994 |
| Mybbp1a | 0.14470888 | 0.1093747 | 0.11839953 | -0.0090248 | 0.69405274 | 0.99999994 |
| Ahctf1 | 0.11767775 | 0.08934177 | 0.09836713 | -0.0090254 | 0.72931174 | 0.99999994 |
| Nr2c2ap | 0.18057628 | 0.13462697 | 0.14365328 | -0.0090263 | 0.66211624 | 0.99999994 |
| Lta4h | 0.10491085 | 0.07956341 | 0.08859357 | -0.0090302 | 0.75126262 | 0.99999994 |
| Slc35a2 | 0.17820677 | 0.1329929 | 0.14203017 | -0.0090373 | 0.66404894 | 0.99999994 |
| Srebf2 | 0.08157153 | 0.06093934 | 0.06997742 | -0.0090381 | 0.80502698 | 0.99999994 |
| Prkab2 | 0.05991286 | 0.04308439 | 0.05212493 | -0.0090405 | 0.87584776 | 0.99999994 |
| 9330020H09 | 0.06676522 | 0.0487867 | 0.05782752 | -0.0090408 | 0.85098385 | 0.99999994 |
| Vps4a | 0.23820067 | 0.17222148 | 0.18126629 | -0.0090448 | 0.63010764 | 0.99999994 |
| Ezr | 0.28302536 | 0.1991387 | 0.20818747 | -0.0090488 | 0.61383009 | 0.99999994 |
| Cmtm6 | 0.0924129 | 0.06967665 | 0.07872766 | -0.009051 | 0.77794026 | 0.99999994 |
| Slc44a3 | 0.27940087 | 0.19702937 | 0.20608227 | -0.0090529 | 0.61501123 | 0.99999994 |
| Nrf1 | 0.11311154 | 0.0858382 | 0.09489499 | -0.0090568 | 0.73737821 | 0.99999994 |
| Sptlc2 | 0.14075005 | 0.10646116 | 0.11552047 | -0.0090593 | 0.69916585 | 0.99999994 |
| Polr2a | 0.18287947 | 0.13616517 | 0.14522505 | -0.0090599 | 0.66102332 | 0.99999994 |
| Vps35 | 0.48723439 | 0.29977334 | 0.30883752 | -0.0090642 | 0.57744432 | 0.99999994 |
| Dync2h1 | 0.1701664 | 0.1274056 | 0.13647494 | -0.0090693 | 0.67087957 | 0.99999994 |
| Pex11b | 0.16222781 | 0.1218454 | 0.13091874 | -0.0090733 | 0.67771527 | 0.99999994 |

| | | | | | | |
|---|---|---|---|---|---|---|
| Wdr82 | 0.10500497 | 0.07959279 | 0.08866637 | -0.0090736 | 0.75211841 | 0.99999994 |
| Mrpl4 | 0.38891685 | 0.2554468 | 0.26452789 | -0.0090811 | 0.59037475 | 0.99999994 |
| Col4a1 | 0.07469531 | 0.05528203 | 0.06436383 | -0.0090818 | 0.82619091 | 0.99999994 |
| Gnpat | 0.12732572 | 0.09653674 | 0.10561872 | -0.009082 | 0.71646426 | 0.99999994 |
| Clec12b | 0.02398062 | 0.01227592 | 0.02136532 | -0.0090894 | 0.99768566 | 0.99999994 |
| Foxp4 | 0.18579209 | 0.13811141 | 0.14720441 | -0.009093 | 0.65952666 | 0.99999994 |
| Fbxl6 | 0.16064878 | 0.12070741 | 0.12980498 | -0.0090976 | 0.67957846 | 0.99999994 |
| BC023719 | 0.04299502 | 0.02872591 | 0.03782454 | -0.0090986 | 0.94533104 | 0.99999994 |
| 1110038F14I | 0.35892976 | 0.24044064 | 0.24954242 | -0.0091018 | 0.59591333 | 0.99999994 |
| Exoc5 | 0.16382139 | 0.12292922 | 0.13203979 | -0.0091106 | 0.67698328 | 0.99999994 |
| Acad8 | 0.18067506 | 0.13460927 | 0.14372081 | -0.0091115 | 0.66348614 | 0.99999994 |
| Ablim1 | 0.03868274 | 0.02501527 | 0.03412813 | -0.0091129 | 0.96224667 | 0.99999994 |
| Armcx2 | 0.20567161 | 0.15135038 | 0.16046707 | -0.0091167 | 0.64734813 | 0.99999994 |
| Ythdf3 | 0.16684014 | 0.12503262 | 0.13415554 | -0.0091229 | 0.67459937 | 0.99999994 |
| Alg3 | 0.1283709 | 0.09727394 | 0.10639752 | -0.0091236 | 0.71592622 | 0.99999994 |
| Trp53bp1 | 0.17807761 | 0.13281686 | 0.14194151 | -0.0091247 | 0.66563857 | 0.99999994 |
| Bag6 | 0.30268596 | 0.21025686 | 0.21939189 | -0.009135 | 0.60917028 | 0.99999994 |
| Synrg | 0.25481422 | 0.18232691 | 0.19147293 | -0.009146 | 0.62473936 | 0.99999994 |
| Ryk | 0.13911933 | 0.10518147 | 0.11432916 | -0.0091477 | 0.7028406 | 0.99999994 |
| Rnf13 | 0.12849365 | 0.0973385 | 0.10648889 | -0.0091504 | 0.71632834 | 0.99999994 |
| Arf2 | 0.17330834 | 0.12950156 | 0.13865444 | -0.0091529 | 0.66981449 | 0.99999994 |
| C2cd5 | 0.11702309 | 0.08871109 | 0.09787095 | -0.0091599 | 0.73334991 | 0.99999994 |
| Stk19 | 0.13715028 | 0.1037204 | 0.1128865 | -0.0091661 | 0.70556673 | 0.99999994 |
| Ppan | 0.12798513 | 0.09693812 | 0.10611023 | -0.0091721 | 0.71747993 | 0.99999994 |
| Capn7 | 0.11883638 | 0.09006614 | 0.09924399 | -0.0091778 | 0.73090883 | 0.99999994 |
| Myg1 | 0.27708352 | 0.19555123 | 0.20472992 | -0.0091787 | 0.61731079 | 0.99999994 |
| Kdm2a | 0.17064342 | 0.12762094 | 0.13680655 | -0.0091856 | 0.67253645 | 0.99999994 |
| Mrps18b | 0.2628823 | 0.18714448 | 0.1963311 | -0.0091866 | 0.62227485 | 0.99999994 |
| Thra | 0.16456843 | 0.12336511 | 0.13256432 | -0.0091992 | 0.67792836 | 0.99999994 |
| Cited4 | 0.05399729 | 0.03796077 | 0.04716069 | -0.0091999 | 0.90294158 | 0.99999994 |

| | | | | | | |
|---|---|---|---|---|---|---|
| Ankhd1 | 0.177596 | 0.13240863 | 0.14161079 | -0.0092022 | 0.66732691 | 0.99999994 |
| Ak3 | 0.12230431 | 0.09265111 | 0.1018587 | -0.0092076 | 0.72631526 | 0.99999994 |
| Emilin1 | 0.04556856 | 0.03080681 | 0.04002051 | -0.0092137 | 0.93729497 | 0.99999994 |
| Ankrd12 | 0.39945447 | 0.26040861 | 0.26962284 | -0.0092142 | 0.58997393 | 0.99999994 |
| Mical2 | 0.14627125 | 0.11030954 | 0.11953065 | -0.0092211 | 0.6961993 | 0.99999994 |
| Bop1 | 0.13940389 | 0.10531449 | 0.11453727 | -0.0092228 | 0.70401519 | 0.99999994 |
| Dnajb4 | 0.09533152 | 0.07183914 | 0.08106262 | -0.0092235 | 0.77558734 | 0.99999994 |
| Dmtf1 | 0.16423451 | 0.12310544 | 0.13232994 | -0.0092245 | 0.67867751 | 0.99999994 |
| Kctd3 | 0.08253548 | 0.06153237 | 0.07076039 | -0.009228 | 0.80732195 | 0.99999994 |
| Rp9 | 0.33557352 | 0.2281253 | 0.23735451 | -0.0092292 | 0.60211432 | 0.99999994 |
| Golga5 | 0.11858487 | 0.08982025 | 0.09905379 | -0.0092335 | 0.73253182 | 0.99999994 |
| Podxl2 | 0.21862348 | 0.15964035 | 0.16888009 | -0.0092397 | 0.64213878 | 0.99999994 |
| Frmd5 | 0.0446603 | 0.03000294 | 0.03924635 | -0.0092434 | 0.94152715 | 0.99999994 |
| Mrpl41 | 0.64381696 | 0.3569057 | 0.36615407 | -0.0092484 | 0.56675839 | 0.99999994 |
| Clec11a | 0.06776725 | 0.04940546 | 0.05865715 | -0.0092517 | 0.85311185 | 0.99999994 |
| Exoc1 | 0.08565125 | 0.06403085 | 0.07328453 | -0.0092537 | 0.79968673 | 0.99999994 |
| Brwd1 | 0.27087396 | 0.19182255 | 0.20108148 | -0.0092589 | 0.62038672 | 0.99999994 |
| Kcnu1 | 0.03175882 | 0.01888429 | 0.02814866 | -0.0092644 | 0.98576541 | 0.99999994 |
| Uvssa | 0.10001136 | 0.07552169 | 0.08478846 | -0.0092668 | 0.76653788 | 0.99999994 |
| Tubg1 | 0.35578795 | 0.23865682 | 0.24792947 | -0.0092726 | 0.59830028 | 0.99999994 |
| Pdgfrl | 0.06269919 | 0.04517556 | 0.05444989 | -0.0092743 | 0.87155286 | 0.99999994 |
| Prpsap2 | 0.1025899 | 0.07750667 | 0.08679483 | -0.0092882 | 0.76186529 | 0.99999994 |
| Srxn1 | 0.08811545 | 0.06598525 | 0.07527369 | -0.0092884 | 0.79428087 | 0.99999994 |
| Pfkfb2 | 0.0398068 | 0.02580288 | 0.03509369 | -0.0092908 | 0.96098423 | 0.99999994 |
| Mrpl22 | 0.36591164 | 0.24378438 | 0.25309661 | -0.0093122 | 0.59673638 | 0.99999994 |
| Ppp1r11 | 0.76154929 | 0.38671421 | 0.3960296 | -0.0093154 | 0.56220832 | 0.99999994 |
| Enoph1 | 0.08367277 | 0.06236742 | 0.0716829 | -0.0093155 | 0.8064862 | 0.99999994 |
| 4930453N24 | 0.1820254 | 0.13532667 | 0.14464288 | -0.0093162 | 0.66593842 | 0.99999994 |
| Rlim | 0.19507824 | 0.14412995 | 0.153453 | -0.009323 | 0.65704338 | 0.99999994 |
| Ccl27a | 0.13312658 | 0.10059214 | 0.10992412 | -0.009332 | 0.71399373 | 0.99999994 |

| | | | | | | |
|---|---|---|---|---|---|---|
| Pappa2 | 0.05149015 | 0.03570497 | 0.0450451 | -0.0093401 | 0.91623766 | 0.99999994 |
| Rpf1 | 0.29962221 | 0.20832079 | 0.21766955 | -0.0093488 | 0.61251734 | 0.99999994 |
| B830012L14I | 0.06462638 | 0.0467028 | 0.05605303 | -0.0093502 | 0.86657586 | 0.99999994 |
| Immt | 0.48378515 | 0.29803923 | 0.30740008 | -0.0093609 | 0.58031424 | 0.99999994 |
| Grhl1 | 0.03188667 | 0.01889852 | 0.02825957 | -0.009361 | 0.98627064 | 0.99999994 |
| Tmub1 | 0.14855127 | 0.11181329 | 0.12117593 | -0.0093626 | 0.69647973 | 0.99999994 |
| Zfyve19 | 0.19986979 | 0.14727608 | 0.15664032 | -0.0093642 | 0.65465881 | 0.99999994 |
| Nufip2 | 0.18295761 | 0.13591396 | 0.14527828 | -0.0093643 | 0.66605808 | 0.99999994 |
| Mrpl55 | 0.34630329 | 0.23364557 | 0.24301116 | -0.0093656 | 0.60123087 | 0.99999994 |
| Rhbdd3 | 0.08020357 | 0.05949746 | 0.06886461 | -0.0093672 | 0.81737753 | 0.99999994 |
| Tmem245 | 0.10640074 | 0.08037708 | 0.08974466 | -0.0093676 | 0.75640742 | 0.99999994 |
| Vat1l | 0.11775233 | 0.08905369 | 0.09842362 | -0.0093699 | 0.73685706 | 0.99999994 |
| Anks3 | 0.21076941 | 0.154418 | 0.16379943 | -0.0093814 | 0.64848572 | 0.99999994 |
| Mapk1ip1 | 0.12129394 | 0.09171585 | 0.10109844 | -0.0093826 | 0.73163096 | 0.99999994 |
| Gcnt2 | 0.05306631 | 0.03699133 | 0.04637592 | -0.0093846 | 0.91100581 | 0.99999994 |
| Sycn | 0.07250602 | 0.05318069 | 0.06256604 | -0.0093854 | 0.84098176 | 0.99999994 |
| Pcbp2 | 1.40323022 | 0.40873817 | 0.41813008 | -0.0093919 | 0.55942561 | 0.99999994 |
| Cdipt | 0.34150515 | 0.23110048 | 0.24049369 | -0.0093932 | 0.60256203 | 0.99999994 |
| Sdhaf4 | 0.62680944 | 0.35126599 | 0.36065969 | -0.0093937 | 0.56881996 | 0.99999994 |
| Mcts1 | 0.50950982 | 0.30853352 | 0.31792723 | -0.0093937 | 0.57795851 | 0.99999994 |
| Smg1 | 0.10483617 | 0.0791411 | 0.0885358 | -0.0093947 | 0.76000484 | 0.99999994 |
| Srm | 0.35087318 | 0.23599242 | 0.24539013 | -0.0093977 | 0.60060702 | 0.99999994 |
| Blmh | 0.58604633 | 0.33735186 | 0.34674972 | -0.0093979 | 0.57158285 | 0.99999994 |
| Vkorc1l1 | 0.10383241 | 0.07835961 | 0.08775862 | -0.009399 | 0.76204649 | 0.99999994 |
| Gpaa1 | 0.24590932 | 0.17663459 | 0.18603598 | -0.0094014 | 0.63170828 | 0.99999994 |
| Mrps26 | 0.52067853 | 0.31294535 | 0.322362 | -0.0094166 | 0.5770849 | 0.99999994 |
| Prrc2b | 0.28893581 | 0.2021748 | 0.21159311 | -0.0094183 | 0.61649334 | 0.99999994 |
| Cmip | 0.41331375 | 0.2667683 | 0.27619265 | -0.0094243 | 0.58983895 | 0.99999994 |
| Sumo3 | 0.77906174 | 0.39010572 | 0.39953903 | -0.0094333 | 0.56244056 | 0.99999994 |
| Gm15706 | 0.05371946 | 0.03749315 | 0.0469266 | -0.0094334 | 0.90956771 | 0.99999994 |

| | | | | | | |
|---|---|---|---|---|---|---|
| Prkar1b | 0.20985383 | 0.1537626 | 0.16320299 | -0.0094404 | 0.64989366 | 0.99999994 |
| Zfp945 | 0.13966668 | 0.10528307 | 0.11472937 | -0.0094463 | 0.70817295 | 0.99999994 |
| Kbtbd2 | 0.1582204 | 0.11863983 | 0.12808631 | -0.0094465 | 0.68825573 | 0.99999994 |
| Pip5k1b | 0.06097036 | 0.04356053 | 0.05300832 | -0.0094478 | 0.88226323 | 0.99999994 |
| Epb41 | 0.10871438 | 0.08207846 | 0.09152666 | -0.0094482 | 0.75399903 | 0.99999994 |
| Pigf | 0.24503759 | 0.17604929 | 0.18549955 | -0.0094503 | 0.6327356 | 0.99999994 |
| St7l | 0.22100691 | 0.16095071 | 0.17040932 | -0.0094586 | 0.64410853 | 0.99999994 |
| Krcc1 | 0.54498188 | 0.32227369 | 0.33173631 | -0.0094626 | 0.57529428 | 0.99999994 |
| Ffar3 | 0.02509507 | 0.01287659 | 0.02234156 | -0.009465 | 0.99759851 | 0.99999994 |
| Tfrc | 0.06818091 | 0.04953003 | 0.05899934 | -0.0094693 | 0.85731534 | 0.99999994 |
| Zfp622 | 0.27364363 | 0.19323967 | 0.20271327 | -0.0094736 | 0.62212931 | 0.99999994 |
| Cenpb | 0.08371882 | 0.06223948 | 0.07172023 | -0.0094808 | 0.8105457 | 0.99999994 |
| Znhit3 | 0.19850626 | 0.14624192 | 0.1557358 | -0.0094939 | 0.65754819 | 0.99999994 |
| Fmnl2 | 0.09129222 | 0.06833483 | 0.07782878 | -0.0094939 | 0.79159718 | 0.99999994 |
| BC051226 | 0.06262585 | 0.04486111 | 0.0543888 | -0.0095277 | 0.87820151 | 0.99999994 |
| Dtx2 | 0.09999024 | 0.07524228 | 0.08477169 | -0.0095294 | 0.77284813 | 0.99999994 |
| Suclg2 | 0.19094385 | 0.14114981 | 0.15068287 | -0.0095331 | 0.66316602 | 0.99999994 |
| Cul3 | 0.25698279 | 0.18324441 | 0.19278487 | -0.0095405 | 0.62907603 | 0.99999994 |
| Stag2 | 0.21375013 | 0.15619376 | 0.16573513 | -0.0095414 | 0.64921658 | 0.99999994 |
| Trappc2l | 1.03625006 | 0.41866115 | 0.4282053 | -0.0095442 | 0.55897139 | 0.99999994 |
| Camkk2 | 0.06660478 | 0.04814668 | 0.05769458 | -0.0095479 | 0.86467416 | 0.99999994 |
| Urgcp | 0.07233654 | 0.0528783 | 0.06242666 | -0.0095484 | 0.84568759 | 0.99999994 |
| Rpl7l1 | 0.37163111 | 0.24642843 | 0.25597861 | -0.0095502 | 0.59806385 | 0.99999994 |
| Scly | 0.33314809 | 0.22650424 | 0.23606237 | -0.0095581 | 0.60623326 | 0.99999994 |
| Mllt10 | 0.28198302 | 0.19802502 | 0.20758329 | -0.0095583 | 0.62038627 | 0.99999994 |
| Mgrn1 | 0.16185487 | 0.12109024 | 0.13065597 | -0.0095657 | 0.68698863 | 0.99999994 |
| Mypop | 0.06101962 | 0.04348064 | 0.05304944 | -0.0095688 | 0.8850549 | 0.99999994 |
| Elovl5 | 0.06013812 | 0.04274284 | 0.0523132 | -0.0095704 | 0.88833813 | 0.99999994 |
| Slc36a1 | 0.08997889 | 0.0671993 | 0.07677373 | -0.0095744 | 0.79676232 | 0.99999994 |
| Vstm2l | 0.02264682 | 0.01061769 | 0.02019502 | -0.0095773 | 0.99920225 | 0.99999994 |

| | | | | | | |
|---|---|---|---|---|---|---|
| Ifnar2 | 0.06109264 | 0.04352937 | 0.05311039 | -0.009581 | 0.88508476 | 0.99999994 |
| Pacrg | 0.11097417 | 0.08367939 | 0.09326071 | -0.0095813 | 0.75296634 | 0.99999994 |
| Mblac2 | 0.05857302 | 0.04142001 | 0.05100393 | -0.0095839 | 0.89449006 | 0.99999994 |
| B4galt3 | 0.33246383 | 0.22611012 | 0.23569694 | -0.0095868 | 0.6067056 | 0.99999994 |
| Tmem101 | 0.32515903 | 0.22217901 | 0.23176955 | -0.0095905 | 0.60851002 | 0.99999994 |
| Itgav | 0.05623121 | 0.03944312 | 0.04903988 | -0.0095968 | 0.90364474 | 0.99999994 |
| Mtdh | 0.71165265 | 0.37540124 | 0.38500026 | -0.009599 | 0.56590566 | 0.99999994 |
| Asb13 | 0.13572113 | 0.10223473 | 0.11183652 | -0.0096018 | 0.71616476 | 0.99999994 |
| Mecr | 0.14082892 | 0.10597243 | 0.11557801 | -0.0096056 | 0.70993808 | 0.99999994 |
| Kif3b | 0.09484833 | 0.07106744 | 0.08067666 | -0.0096092 | 0.7860598 | 0.99999994 |
| Robo1 | 0.05576918 | 0.03904108 | 0.04865167 | -0.0096106 | 0.90572629 | 0.99999994 |
| mt-Atp8 | 0.1800566 | 0.13368626 | 0.14329783 | -0.0096116 | 0.67237068 | 0.99999994 |
| Cds2 | 0.15017216 | 0.1127299 | 0.12234189 | -0.009612 | 0.69950152 | 0.99999994 |
| Mpv17 | 0.26583172 | 0.18847955 | 0.19809156 | -0.009612 | 0.62664733 | 0.99999994 |
| Loxl2 | 0.03208853 | 0.01881853 | 0.02843465 | -0.0096161 | 0.98780934 | 0.99999994 |
| Relt | 0.07504676 | 0.05503451 | 0.06465197 | -0.0096175 | 0.8389574 | 0.99999994 |
| Tspyl4 | 0.14866834 | 0.11164276 | 0.12126024 | -0.0096175 | 0.70122742 | 0.99999994 |
| Slc16a12 | 0.13864487 | 0.10436158 | 0.11398196 | -0.0096204 | 0.71287859 | 0.99999994 |
| Tmcc3 | 0.07821457 | 0.05761525 | 0.0672431 | -0.0096279 | 0.82971921 | 0.99999994 |
| Hcfc1r1 | 0.79810428 | 0.39352724 | 0.40315726 | -0.00963 | 0.56316444 | 0.99999994 |
| Fbxo22 | 0.24110418 | 0.17343162 | 0.18306972 | -0.0096381 | 0.63699508 | 0.99999994 |
| Usp37 | 0.08847335 | 0.06591893 | 0.07556207 | -0.0096431 | 0.80219175 | 0.99999994 |
| Tsc1 | 0.08865195 | 0.06605461 | 0.07570593 | -0.0096513 | 0.80194284 | 0.99999994 |
| Rab39b | 0.09021991 | 0.06729362 | 0.07696748 | -0.0096739 | 0.79859744 | 0.99999994 |
| Thoc2 | 0.4112343 | 0.26554056 | 0.27521643 | -0.0096759 | 0.59251514 | 0.99999994 |
| Ubap1 | 0.16089543 | 0.12030142 | 0.12997917 | -0.0096778 | 0.68991411 | 0.99999994 |
| Ttc39b | 0.0533037 | 0.03689778 | 0.04657612 | -0.0096783 | 0.91669288 | 0.99999994 |
| Dapp1 | 0.14033574 | 0.10553671 | 0.1152181 | -0.0096814 | 0.71202529 | 0.99999994 |
| Pak2 | 0.35752954 | 0.23914276 | 0.24882457 | -0.0096818 | 0.60218234 | 0.99999994 |
| Cdh2 | 0.06315743 | 0.04512811 | 0.05483144 | -0.0097033 | 0.88059222 | 0.99999994 |

| | | | | | | |
|---|---|---|---|---|---|---|
| Gm8186 | 0.18998347 | 0.140331 | 0.1500366 | -0.0097056 | 0.66661598 | 0.99999994 |
| Trappc6a | 0.59180791 | 0.33906539 | 0.34877596 | -0.0097106 | 0.57351304 | 0.99999994 |
| Gm16174 | 0.02844155 | 0.01555252 | 0.0252643 | -0.0097118 | 0.99474757 | 0.99999994 |
| Mrpl44 | 0.28066966 | 0.19710707 | 0.2068206 | -0.0097135 | 0.62271613 | 0.99999994 |
| Parp6 | 0.08618606 | 0.06399252 | 0.07371678 | -0.0097243 | 0.81004582 | 0.99999994 |
| Smad4 | 0.11391706 | 0.08577597 | 0.09550943 | -0.0097335 | 0.75122701 | 0.99999994 |
| Rab34 | 0.25861253 | 0.18402952 | 0.19376785 | -0.0097383 | 0.63101149 | 0.99999994 |
| Krit1 | 0.35464805 | 0.23759729 | 0.24734227 | -0.009745 | 0.60343656 | 0.99999994 |
| Sharpin | 0.15786669 | 0.11808418 | 0.12783541 | -0.0097512 | 0.69418358 | 0.99999994 |
| Guf1 | 0.15864143 | 0.11862649 | 0.12838477 | -0.0097583 | 0.69354957 | 0.99999994 |
| Alg1 | 0.09806276 | 0.07347958 | 0.08323979 | -0.0097602 | 0.78244732 | 0.99999994 |
| Entpd3 | 0.34244519 | 0.2312267 | 0.24098848 | -0.0097618 | 0.60627778 | 0.99999994 |
| Foxk2 | 0.06602235 | 0.04744977 | 0.05721176 | -0.009762 | 0.87197305 | 0.99999994 |
| Txnl4a | 0.60207898 | 0.3425747 | 0.35233845 | -0.0097637 | 0.57317216 | 0.99999994 |
| Cndp2 | 0.16777846 | 0.12504508 | 0.13481109 | -0.009766 | 0.68516844 | 0.99999994 |
| Alkbh5 | 0.18893317 | 0.13956112 | 0.14932868 | -0.0097676 | 0.66836071 | 0.99999994 |
| Ube2d3 | 1.58151565 | 0.39065275 | 0.40043434 | -0.0097816 | 0.56458248 | 0.99999994 |
| Mapk1ip1l | 0.16649362 | 0.12411648 | 0.1339132 | -0.0097967 | 0.68685687 | 0.99999994 |
| Ncor2 | 0.15304767 | 0.11460577 | 0.12440284 | -0.0097971 | 0.69993266 | 0.99999994 |
| Cox7a1 | 0.066872 | 0.04811747 | 0.05791598 | -0.0097985 | 0.86995013 | 0.99999994 |
| Tmprss4 | 0.07025058 | 0.0509018 | 0.06070859 | -0.0098068 | 0.85887074 | 0.99999994 |
| Cdkn1b | 0.21634827 | 0.15760577 | 0.16741486 | -0.0098091 | 0.65173391 | 0.99999994 |
| Trmt2a | 0.17301562 | 0.12863168 | 0.13845185 | -0.0098202 | 0.68158124 | 0.99999994 |
| Rnpep | 0.29594652 | 0.20576791 | 0.21559119 | -0.0098233 | 0.61916666 | 0.99999994 |
| Atg13 | 0.10572351 | 0.07939175 | 0.08922177 | -0.00983 | 0.76833215 | 0.99999994 |
| Las1l | 0.19457035 | 0.14327822 | 0.15311372 | -0.0098355 | 0.66552208 | 0.99999994 |
| Ndufaf6 | 0.08880943 | 0.06598778 | 0.07583275 | -0.009845 | 0.80624998 | 0.99999994 |
| Tbccd1 | 0.0620123 | 0.04403235 | 0.05387752 | -0.0098452 | 0.88807099 | 0.99999994 |
| Lrrc8d | 0.10386885 | 0.0779389 | 0.08778686 | -0.009848 | 0.77238128 | 0.99999994 |
| Ythdf1 | 0.1130135 | 0.08496986 | 0.09482015 | -0.0098503 | 0.75537078 | 0.99999994 |

| | | | | | | |
|---|---|---|---|---|---|---|
| Rbm3 | 1.38704492 | 0.40956005 | 0.4194114 | -0.0098514 | 0.56212066 | 0.99999994 |
| Atp6v1h | 0.35148371 | 0.23584993 | 0.24570666 | -0.0098567 | 0.60527519 | 0.99999994 |
| Rbm5 | 0.62861812 | 0.35139277 | 0.36125197 | -0.0098592 | 0.57207669 | 0.99999994 |
| Dct | 0.08291269 | 0.0612011 | 0.07106651 | -0.0098654 | 0.8222649 | 0.99999994 |
| Myt1l | 0.0427199 | 0.02772386 | 0.03758934 | -0.0098655 | 0.95968042 | 0.99999994 |
| Iah1 | 0.13685231 | 0.10279624 | 0.11266778 | -0.0098715 | 0.72012267 | 0.99999994 |
| Slc39a13 | 0.07070787 | 0.05121371 | 0.06108563 | -0.0098719 | 0.85897381 | 0.99999994 |
| Nr1h2 | 0.11571587 | 0.08699777 | 0.0968786 | -0.0098808 | 0.75139761 | 0.99999994 |
| Dctn1 | 0.14429184 | 0.10821555 | 0.1180971 | -0.0098816 | 0.71122204 | 0.99999994 |
| Med8 | 0.35281815 | 0.23651345 | 0.24639741 | -0.009884 | 0.60527018 | 0.99999994 |
| Eif4e2 | 0.58852822 | 0.33773358 | 0.34762485 | -0.0098913 | 0.57510999 | 0.99999994 |
| C2cd2l | 0.06584149 | 0.04717022 | 0.05706175 | -0.0098915 | 0.87572234 | 0.99999994 |
| Rab5a | 0.49898266 | 0.30378073 | 0.31367285 | -0.0098921 | 0.58313401 | 0.99999994 |
| Hars | 0.27611134 | 0.19426786 | 0.20416109 | -0.0098932 | 0.62648325 | 0.99999994 |
| Cuedc1 | 0.21124309 | 0.15421339 | 0.16410766 | -0.0098943 | 0.65590787 | 0.99999994 |
| Prrc2a | 0.21309789 | 0.15541212 | 0.16531234 | -0.0099002 | 0.65491925 | 0.99999994 |
| Tnpo1 | 0.20789299 | 0.15202 | 0.16192269 | -0.0099027 | 0.65802614 | 0.99999994 |
| Dip2b | 0.13619475 | 0.10226144 | 0.11218475 | -0.0099233 | 0.72199847 | 0.99999994 |
| Isoc1 | 0.18800401 | 0.13877448 | 0.14870142 | -0.0099269 | 0.67160952 | 0.99999994 |
| P4htm | 0.09224722 | 0.06866344 | 0.07859485 | -0.0099314 | 0.79985733 | 0.99999994 |
| Ski | 0.08442895 | 0.06236187 | 0.07229553 | -0.0099337 | 0.81980106 | 0.99999994 |
| Dhx9 | 0.32697247 | 0.22281501 | 0.23274897 | -0.009934 | 0.61183318 | 0.99999994 |
| Nagk | 0.4333311 | 0.27548903 | 0.28542413 | -0.0099351 | 0.59161286 | 0.99999994 |
| Acyp1 | 0.38722484 | 0.25375367 | 0.26370148 | -0.0099478 | 0.5991327 | 0.99999994 |
| Slc24a2 | 0.03242426 | 0.01877581 | 0.02872573 | -0.0099499 | 0.98943369 | 0.99999994 |
| Scrn1 | 0.12126932 | 0.09112953 | 0.10107989 | -0.0099504 | 0.74384856 | 0.99999994 |
| Tars | 0.23347798 | 0.16835703 | 0.17831394 | -0.0099569 | 0.64493666 | 0.99999994 |
| Mkks | 0.22140308 | 0.1607049 | 0.17066294 | -0.009958 | 0.65113741 | 0.99999994 |
| Lrfn4 | 0.08855843 | 0.06567019 | 0.0756306 | -0.0099604 | 0.8096562 | 0.99999994 |
| Pigc | 0.17437777 | 0.12943202 | 0.13939378 | -0.0099618 | 0.68285436 | 0.99999994 |

| | | | | | | |
|---|---|---|---|---|---|---|
| Tet2 | 0.10528821 | 0.0789229 | 0.08888538 | -0.0099625 | 0.77218442 | 0.99999994 |
| Nudt5 | 0.37754184 | 0.24896268 | 0.25892924 | -0.0099666 | 0.60110675 | 0.99999994 |
| Sv2a | 0.05207671 | 0.03556831 | 0.04554069 | -0.0099724 | 0.92752327 | 0.99999994 |
| Nudt18 | 0.12905043 | 0.096929 | 0.10690315 | -0.0099741 | 0.73271439 | 0.99999994 |
| As3mt | 0.23706736 | 0.17058484 | 0.18055991 | -0.0099751 | 0.64346343 | 0.99999994 |
| Cdc27 | 0.13100959 | 0.09838101 | 0.10835776 | -0.0099768 | 0.73001763 | 0.99999994 |
| Atxn2 | 0.11927453 | 0.08959807 | 0.09957516 | -0.0099771 | 0.74759603 | 0.99999994 |
| Rab9 | 0.20661268 | 0.15109819 | 0.16108443 | -0.0099862 | 0.66006961 | 0.99999994 |
| Samm50 | 0.42505534 | 0.27165375 | 0.28164416 | -0.0099904 | 0.59332738 | 0.99999994 |
| Tbc1d22b | 0.06969034 | 0.05024363 | 0.06024637 | -0.0100027 | 0.86545464 | 0.99999994 |
| Marveld3 | 0.20638535 | 0.15093103 | 0.16093538 | -0.0100044 | 0.660484 | 0.99999994 |
| Rbfa | 0.22682701 | 0.16411058 | 0.1741181 | -0.0100075 | 0.64898333 | 0.99999994 |
| Tmem230 | 0.22389754 | 0.1622347 | 0.17225582 | -0.0100211 | 0.65071024 | 0.99999994 |
| Ip6k1 | 0.13316521 | 0.09992621 | 0.10995265 | -0.0100264 | 0.72806955 | 0.99999994 |
| Hdgf | 0.54595255 | 0.32207607 | 0.33210309 | -0.010027 | 0.57963984 | 0.99999994 |
| Spock1 | 0.10619118 | 0.07954945 | 0.08958292 | -0.0100335 | 0.772023 | 0.99999994 |
| Supt6 | 0.23972055 | 0.1721773 | 0.18221136 | -0.0100341 | 0.64302421 | 0.99999994 |
| Prkag1 | 0.26142067 | 0.18540886 | 0.19545561 | -0.0100467 | 0.63388025 | 0.99999994 |
| Snrnp200 | 0.15614918 | 0.11656767 | 0.12661509 | -0.0100474 | 0.70132285 | 0.99999994 |
| Atox1 | 0.88677839 | 0.40646597 | 0.41651781 | -0.0100518 | 0.56381083 | 0.99999994 |
| Nedd4 | 1.07829173 | 0.41938556 | 0.42944122 | -0.0100557 | 0.56192958 | 0.99999994 |
| Il1r1 | 0.19900341 | 0.14600827 | 0.15606582 | -0.0100576 | 0.66602544 | 0.99999994 |
| Man2a2 | 0.06311784 | 0.04473992 | 0.05479849 | -0.0100586 | 0.88911065 | 0.99999994 |
| Pef1 | 0.25720774 | 0.18285923 | 0.1929207 | -0.0100615 | 0.63576141 | 0.99999994 |
| Tead1 | 0.09411559 | 0.0700287 | 0.08009091 | -0.0100622 | 0.79849848 | 0.99999994 |
| Rbm15b | 0.09577639 | 0.07135431 | 0.08141776 | -0.0100634 | 0.79467296 | 0.99999994 |
| Ltv1 | 0.19232489 | 0.14154513 | 0.15161037 | -0.0100652 | 0.67072217 | 0.99999994 |
| Mthfd1l | 0.06299256 | 0.04462395 | 0.05469419 | -0.0100702 | 0.88981829 | 0.99999994 |
| Fbxl2 | 0.0562207 | 0.0389532 | 0.04903105 | -0.0100779 | 0.91436736 | 0.99999994 |
| Arhgdig | 0.08856648 | 0.06555638 | 0.07563709 | -0.0100807 | 0.8124988 | 0.99999994 |

| | | | | | | |
|---|---|---|---|---|---|---|
| Wdr46 | 0.12336261 | 0.09255795 | 0.10265371 | -0.0100958 | 0.7436494 | 0.99999994 |
| Metap1 | 0.17206826 | 0.12769495 | 0.13779554 | -0.0101006 | 0.68719662 | 0.99999994 |
| Osbp | 0.04920326 | 0.0330072 | 0.04310927 | -0.0101021 | 0.9405996 | 0.99999994 |
| Stox2 | 0.09980347 | 0.07452109 | 0.08462342 | -0.0101023 | 0.78658719 | 0.99999994 |
| Prep | 0.09343265 | 0.06943576 | 0.07954448 | -0.0101087 | 0.80119763 | 0.99999994 |
| Ndrg3 | 0.22178306 | 0.16079316 | 0.17090604 | -0.0101129 | 0.65316194 | 0.99999994 |
| Pycr2 | 0.88157557 | 0.40577253 | 0.41588826 | -0.0101157 | 0.56430921 | 0.99999994 |
| Ext2 | 0.23625308 | 0.16993307 | 0.18005152 | -0.0101184 | 0.64582416 | 0.99999994 |
| Txnrd2 | 0.09669646 | 0.07203223 | 0.08215161 | -0.0101194 | 0.79387072 | 0.99999994 |
| Pkdcc | 0.15101514 | 0.11281768 | 0.12294706 | -0.0101294 | 0.70829314 | 0.99999994 |
| Mdp1 | 0.31478659 | 0.21598037 | 0.22611303 | -0.0101327 | 0.61725799 | 0.99999994 |
| Ep300 | 0.14014311 | 0.10494046 | 0.11507744 | -0.010137 | 0.72117907 | 0.99999994 |
| Nr2c2 | 0.10436126 | 0.07802961 | 0.08816825 | -0.0101386 | 0.77799151 | 0.99999994 |
| Fam234a | 0.1605304 | 0.11957619 | 0.12972135 | -0.0101452 | 0.69866892 | 0.99999994 |
| Vars | 0.12089657 | 0.09064977 | 0.10079909 | -0.0101493 | 0.74864085 | 0.99999994 |
| Gin1 | 0.09071273 | 0.0672128 | 0.07736346 | -0.0101507 | 0.8087675 | 0.99999994 |
| Rcbtb1 | 0.10677242 | 0.07987821 | 0.09003139 | -0.0101532 | 0.77357213 | 0.99999994 |
| Rom1 | 0.07410832 | 0.05371472 | 0.06388232 | -0.0101676 | 0.85529907 | 0.99999994 |
| Kcmf1 | 0.18463661 | 0.13624822 | 0.14642028 | -0.0101721 | 0.67811282 | 0.99999994 |
| Dxo | 0.17159405 | 0.1272936 | 0.13746663 | -0.010173 | 0.68885052 | 0.99999994 |
| Psmb10 | 0.14490361 | 0.10836375 | 0.11854067 | -0.0101769 | 0.71615607 | 0.99999994 |
| Abat | 0.085398 | 0.06289988 | 0.07307976 | -0.0101799 | 0.82309333 | 0.99999994 |
| Nipal3 | 0.10394302 | 0.07764553 | 0.08784433 | -0.0101988 | 0.78018215 | 0.99999994 |
| Aldh2 | 0.21329913 | 0.15523638 | 0.16544283 | -0.0102065 | 0.65933254 | 0.99999994 |
| Col6a3 | 0.06256758 | 0.04413119 | 0.05434026 | -0.0102091 | 0.89445073 | 0.99999994 |
| Stxbp1 | 0.21998293 | 0.15953932 | 0.16975304 | -0.0102137 | 0.65560599 | 0.99999994 |
| Samd4b | 0.14178699 | 0.10606197 | 0.11627637 | -0.0102144 | 0.72063652 | 0.99999994 |
| Mns1 | 0.09673993 | 0.07196785 | 0.08218626 | -0.0102184 | 0.79605177 | 0.99999994 |
| Tvp23b | 0.68594371 | 0.36833562 | 0.37855589 | -0.0102203 | 0.57130935 | 0.99999994 |
| Raver1 | 0.15939478 | 0.11869746 | 0.12891831 | -0.0102208 | 0.70115541 | 0.99999994 |

| | | | | | | |
|---|---|---|---|---|---|---|
| Snx30 | 0.12036739 | 0.09017126 | 0.10040014 | -0.0102289 | 0.75116387 | 0.99999994 |
| 4930550C14I | 0.04349884 | 0.0280244 | 0.03825503 | -0.0102306 | 0.96247222 | 0.99999994 |
| Sltm | 0.52675429 | 0.31450961 | 0.32474048 | -0.0102309 | 0.58305182 | 0.99999994 |
| Gm11789 | 0.03676198 | 0.02224174 | 0.03247488 | -0.0102331 | 0.98202453 | 0.99999994 |
| Llph | 0.60858682 | 0.34432314 | 0.35456448 | -0.0102413 | 0.57623234 | 0.99999994 |
| Eif2a | 0.36055587 | 0.24013262 | 0.25037407 | -0.0102414 | 0.60729494 | 0.99999994 |
| Psmc4 | 0.99130985 | 0.41562093 | 0.42586735 | -0.0102464 | 0.56361966 | 0.99999994 |
| Socs2 | 0.05746436 | 0.03982831 | 0.05007486 | -0.0102466 | 0.91340971 | 0.99999994 |
| Aagab | 0.14717163 | 0.10992757 | 0.12018109 | -0.0102535 | 0.71494932 | 0.99999994 |
| Snx10 | 0.11264589 | 0.08428518 | 0.09453943 | -0.0102543 | 0.76484792 | 0.99999994 |
| Unc13b | 0.0778473 | 0.0566887 | 0.06694324 | -0.0102545 | 0.84605 | 0.99999994 |
| Fbxo3 | 0.27396173 | 0.19263934 | 0.20290021 | -0.0102609 | 0.63179795 | 0.99999994 |
| Snx21 | 0.11748859 | 0.08795681 | 0.09822381 | -0.010267 | 0.75671246 | 0.99999994 |
| Smpd1 | 0.23105538 | 0.16651773 | 0.17679077 | -0.010273 | 0.65054453 | 0.99999994 |
| 2310009B15I | 0.22510092 | 0.16274813 | 0.17302187 | -0.0102737 | 0.65366815 | 0.99999994 |
| Ric8b | 0.0843511 | 0.061958 | 0.07223249 | -0.0102745 | 0.82813297 | 0.99999994 |
| Cpd | 0.09684683 | 0.07198621 | 0.08227146 | -0.0102853 | 0.79733832 | 0.99999994 |
| Ssr1 | 0.60502098 | 0.34306083 | 0.35334766 | -0.0102868 | 0.57682719 | 0.99999994 |
| Tmem164 | 0.06584689 | 0.04677584 | 0.05706623 | -0.0102904 | 0.88498826 | 0.99999994 |
| Eif2b4 | 0.24600187 | 0.17579791 | 0.1860929 | -0.010295 | 0.6436534 | 0.99999994 |
| Atp8b1 | 0.41423604 | 0.26632349 | 0.2766246 | -0.0103011 | 0.59788384 | 0.99999994 |
| Mtx1 | 0.31413584 | 0.215448 | 0.22575492 | -0.0103069 | 0.61939738 | 0.99999994 |
| Nfxl1 | 0.10030813 | 0.07471001 | 0.08501981 | -0.0103098 | 0.79021674 | 0.99999994 |
| Bola2 | 0.596584 | 0.34013052 | 0.35044059 | -0.0103101 | 0.57762926 | 0.99999994 |
| Pepd | 0.29861083 | 0.20678871 | 0.21709897 | -0.0103103 | 0.62404338 | 0.99999994 |
| Znrf2 | 0.12433193 | 0.09306949 | 0.10338067 | -0.0103112 | 0.74662432 | 0.99999994 |
| Slc25a23 | 0.12695595 | 0.09502985 | 0.10534288 | -0.010313 | 0.74268819 | 0.99999994 |
| 2510046G10 | 0.11255712 | 0.08414298 | 0.09447161 | -0.0103286 | 0.7666146 | 0.99999994 |
| Dpm3 | 0.69742957 | 0.37118454 | 0.38151384 | -0.0103293 | 0.57150909 | 0.99999994 |
| Ap1g1 | 0.15987221 | 0.11889913 | 0.1292561 | -0.010357 | 0.70310677 | 0.99999994 |

| | | | | | | |
|---|---|---|---|---|---|---|
| Dido1 | 0.13019877 | 0.09739919 | 0.10775634 | -0.0103572 | 0.73884389 | 0.99999994 |
| Macf1 | 0.2404074 | 0.17227877 | 0.1826377 | -0.0103589 | 0.64711325 | 0.99999994 |
| Slc6a6 | 0.45374116 | 0.28417176 | 0.29453251 | -0.0103607 | 0.59256577 | 0.99999994 |
| Mrpl13 | 0.35603871 | 0.23769761 | 0.2480585 | -0.0103609 | 0.6095024 | 0.99999994 |
| Lrfn3 | 0.1152444 | 0.08615911 | 0.09652016 | -0.0103611 | 0.76255122 | 0.99999994 |
| Zfp260 | 0.20164736 | 0.1474543 | 0.15781655 | -0.0103622 | 0.66896494 | 0.99999994 |
| Extl2 | 0.07821234 | 0.05687894 | 0.06724129 | -0.0103623 | 0.84750485 | 0.99999994 |
| Gsto1 | 0.23128094 | 0.16656923 | 0.17693284 | -0.0103636 | 0.6516901 | 0.99999994 |
| Fhit | 0.06462547 | 0.04568793 | 0.05605227 | -0.0103643 | 0.89079996 | 0.99999994 |
| Dnajc21 | 0.16081986 | 0.11955644 | 0.12992581 | -0.0103694 | 0.70237492 | 0.99999994 |
| Sorl1 | 0.11007941 | 0.08220089 | 0.09257488 | -0.010374 | 0.77214063 | 0.99999994 |
| Ets2 | 0.13626941 | 0.10186227 | 0.11223962 | -0.0103773 | 0.73086376 | 0.99999994 |
| Ttc8 | 0.21059057 | 0.15330053 | 0.163683 | -0.0103825 | 0.66356601 | 0.99999994 |
| Ccdc107 | 0.16177535 | 0.12021672 | 0.13059992 | -0.0103832 | 0.70166879 | 0.99999994 |
| Sgta | 0.54149664 | 0.32001969 | 0.33041466 | -0.010395 | 0.58293655 | 0.99999994 |
| Dpf3 | 0.03723241 | 0.02248316 | 0.03288018 | -0.010397 | 0.9823457 | 0.99999994 |
| 1700088E04I | 0.17664833 | 0.13056155 | 0.14095926 | -0.0103977 | 0.68828654 | 0.99999994 |
| Abca7 | 0.06881191 | 0.04911816 | 0.05952094 | -0.0104028 | 0.87765929 | 0.99999994 |
| Nsd1 | 0.47264576 | 0.29228714 | 0.30270232 | -0.0104152 | 0.59057626 | 0.99999994 |
| Alg14 | 0.12365908 | 0.0924517 | 0.10287617 | -0.0104245 | 0.75000032 | 0.99999994 |
| Mrps9 | 0.30467642 | 0.21007923 | 0.22050637 | -0.0104271 | 0.62352837 | 0.99999994 |
| Uchl5 | 0.2467946 | 0.1761504 | 0.18658 | -0.0104296 | 0.64508426 | 0.99999994 |
| Igsf5 | 0.08560862 | 0.06282008 | 0.07325007 | -0.01043 | 0.82838202 | 0.99999994 |
| Aida | 0.15670541 | 0.11657287 | 0.12701067 | -0.0104378 | 0.70781861 | 0.99999994 |
| Psma4 | 1.37595265 | 0.40979853 | 0.42023804 | -0.0104395 | 0.56566835 | 0.99999994 |
| Eif4ebp1 | 0.15388732 | 0.11456308 | 0.12500284 | -0.0104398 | 0.71086513 | 0.99999994 |
| Fahd1 | 0.16154673 | 0.11999695 | 0.13043873 | -0.0104418 | 0.70293123 | 0.99999994 |
| Usp20 | 0.09823954 | 0.07293429 | 0.08338044 | -0.0104462 | 0.79783937 | 0.99999994 |
| Zmynd11 | 0.34278507 | 0.23071866 | 0.24116719 | -0.0104485 | 0.61347419 | 0.99999994 |
| Uchl3 | 0.46200151 | 0.28768122 | 0.29813295 | -0.0104517 | 0.59225663 | 0.99999994 |

| | | | | | | |
|---|---|---|---|---|---|---|
| Ppp2r3a | 0.07999493 | 0.05823969 | 0.06869472 | -0.010455 | 0.84448606 | 0.99999994 |
| Atg10 | 0.17023578 | 0.12606502 | 0.13652318 | -0.0104582 | 0.69493949 | 0.99999994 |
| Iffo1 | 0.06270521 | 0.04399378 | 0.05445491 | -0.0104611 | 0.89950366 | 0.99999994 |
| Zbtb8os | 0.24090907 | 0.17248658 | 0.18294879 | -0.0104622 | 0.64827007 | 0.99999994 |
| Zfp704 | 0.10124868 | 0.07528698 | 0.08575228 | -0.0104653 | 0.79170429 | 0.99999994 |
| Slc30a9 | 0.26438939 | 0.18676138 | 0.19723169 | -0.0104703 | 0.63809136 | 0.99999994 |
| Stx7 | 0.50267261 | 0.30470129 | 0.31517257 | -0.0104713 | 0.58751238 | 0.99999994 |
| Fip1l1 | 0.62693764 | 0.35022742 | 0.36070173 | -0.0104743 | 0.57663483 | 0.99999994 |
| Esco1 | 0.18913349 | 0.13897741 | 0.14946379 | -0.0104864 | 0.67975373 | 0.99999994 |
| Mrps35 | 0.28480929 | 0.19872494 | 0.2092189 | -0.010494 | 0.63076051 | 0.99999994 |
| Snx8 | 0.08878253 | 0.06530454 | 0.07581109 | -0.0105066 | 0.82186486 | 0.99999994 |
| Taf1 | 0.12252172 | 0.09151279 | 0.10202214 | -0.0105093 | 0.75354079 | 0.99999994 |
| Capg | 0.1051641 | 0.07827678 | 0.08878943 | -0.0105126 | 0.78468414 | 0.99999994 |
| Pom121 | 0.09070888 | 0.06684331 | 0.07736037 | -0.0105171 | 0.81724739 | 0.99999994 |
| Kti12 | 0.16441378 | 0.12193336 | 0.13245579 | -0.0105224 | 0.70152378 | 0.99999994 |
| Pdk1 | 0.07781835 | 0.05639431 | 0.06691959 | -0.0105253 | 0.85243383 | 0.99999994 |
| Uros | 0.10716755 | 0.07979764 | 0.09033601 | -0.0105384 | 0.78127722 | 0.99999994 |
| Setd1b | 0.15130626 | 0.11261227 | 0.12315587 | -0.0105436 | 0.71561466 | 0.99999994 |
| Rpp14 | 0.16812076 | 0.12450036 | 0.13504999 | -0.0105496 | 0.69845813 | 0.99999994 |
| Chchd6 | 0.30643972 | 0.21093708 | 0.22149072 | -0.0105536 | 0.62444029 | 0.99999994 |
| Cers5 | 0.2219653 | 0.16046675 | 0.17102259 | -0.0105558 | 0.6594062 | 0.99999994 |
| Flnb | 0.05434186 | 0.03688677 | 0.04745091 | -0.0105641 | 0.93081991 | 0.99999994 |
| Arid1b | 0.08978097 | 0.06604919 | 0.07661457 | -0.0105654 | 0.82067284 | 0.99999994 |
| Rora | 0.0853375 | 0.0624645 | 0.07303082 | -0.0105663 | 0.8322419 | 0.99999994 |
| Fryl | 0.23229301 | 0.16699883 | 0.17756965 | -0.0105708 | 0.65404223 | 0.99999994 |
| Tbc1d9b | 0.07205979 | 0.05162096 | 0.06219899 | -0.010578 | 0.871189 | 0.99999994 |
| N4bp2 | 0.2070294 | 0.15077931 | 0.16135751 | -0.0105782 | 0.66871746 | 0.99999994 |
| Tshz1 | 0.29243735 | 0.20301421 | 0.21359554 | -0.0105813 | 0.62920561 | 0.99999994 |
| Nfe2l1 | 0.30749382 | 0.21149528 | 0.22207785 | -0.0105826 | 0.62445111 | 0.99999994 |
| Acyp2 | 0.12253321 | 0.09144441 | 0.10203077 | -0.0105864 | 0.75510442 | 0.99999994 |

| | | | | | | |
|---|---|---|---|---|---|---|
| Slc31a1 | 0.33574338 | 0.22684997 | 0.23744482 | -0.0105948 | 0.6167699 | 0.99999994 |
| Wfikkn2 | 0.04173866 | 0.02613591 | 0.03674979 | -0.0106139 | 0.97272564 | 0.99999994 |
| Ift122 | 0.11255973 | 0.08385386 | 0.0944736 | -0.0106197 | 0.77284189 | 0.99999994 |
| Cib2 | 0.33268046 | 0.22519271 | 0.23581267 | -0.01062 | 0.61782489 | 0.99999994 |
| Cby1 | 0.31687992 | 0.21664168 | 0.22726212 | -0.0106204 | 0.62212465 | 0.99999994 |
| Usp8 | 0.19579822 | 0.14330511 | 0.1539335 | -0.0106284 | 0.6770949 | 0.99999994 |
| Aen | 0.14470627 | 0.10776492 | 0.11839764 | -0.0106327 | 0.72500503 | 0.99999994 |
| Lcor | 0.20258972 | 0.14780218 | 0.15843875 | -0.0106366 | 0.67251402 | 0.99999994 |
| Inpp4a | 0.07582974 | 0.05464947 | 0.0652934 | -0.0106439 | 0.8610612 | 0.99999994 |
| Xpnpep1 | 0.25098119 | 0.17848553 | 0.18914235 | -0.0106568 | 0.64618398 | 0.99999994 |
| Fam110b | 0.1208026 | 0.09005788 | 0.10072827 | -0.0106704 | 0.75963346 | 0.99999994 |
| Herc1 | 0.23179573 | 0.16658619 | 0.17725688 | -0.0106707 | 0.65568181 | 0.99999994 |
| Ipo4 | 0.10104556 | 0.07492015 | 0.08559419 | -0.010674 | 0.79675393 | 0.99999994 |
| Oat | 0.18100502 | 0.13327063 | 0.14394631 | -0.0106757 | 0.68922827 | 0.99999994 |
| Smarcc2 | 0.22583352 | 0.16280437 | 0.17348751 | -0.0106831 | 0.65905669 | 0.99999994 |
| Fgf12 | 0.07395298 | 0.05307046 | 0.06375482 | -0.0106844 | 0.86768332 | 0.99999994 |
| Dgkd | 0.10849222 | 0.0806688 | 0.09135584 | -0.010687 | 0.78191512 | 0.99999994 |
| Psmg3 | 0.2671761 | 0.18819859 | 0.19889126 | -0.0106927 | 0.63977403 | 0.99999994 |
| Nifk | 0.2994944 | 0.20690319 | 0.21759752 | -0.0106943 | 0.62821972 | 0.99999994 |
| Txndc15 | 0.47275673 | 0.29205316 | 0.30274953 | -0.0106964 | 0.59296323 | 0.99999994 |
| Cbx4 | 0.07379626 | 0.05292661 | 0.06362618 | -0.0106996 | 0.86850531 | 0.99999994 |
| Hipk1 | 0.12694687 | 0.09463125 | 0.10533611 | -0.0107049 | 0.75062492 | 0.99999994 |
| Mdm2 | 0.25774441 | 0.18253915 | 0.19324456 | -0.0107054 | 0.64384323 | 0.99999994 |
| Snap47 | 0.33616773 | 0.22696458 | 0.23767032 | -0.0107057 | 0.61784741 | 0.99999994 |
| Mrpl32 | 0.63067286 | 0.35121361 | 0.36192271 | -0.0107091 | 0.57806979 | 0.99999994 |
| Rab6a | 0.48593566 | 0.29758535 | 0.30829742 | -0.0107121 | 0.59145176 | 0.99999994 |
| Ankrd13c | 0.16636886 | 0.12311152 | 0.13382591 | -0.0107144 | 0.70295624 | 0.99999994 |
| Mark2 | 0.25184817 | 0.17894628 | 0.18967084 | -0.0107246 | 0.64667927 | 0.99999994 |
| Gfer | 0.20188763 | 0.14725043 | 0.15797527 | -0.0107248 | 0.67432801 | 0.99999994 |
| Rab14 | 1.16783051 | 0.4188915 | 0.42961783 | -0.0107263 | 0.56599638 | 0.99999994 |

| | | | | | | |
|---|---|---|---|---|---|---|
| Lars | 0.32491971 | 0.22091198 | 0.23164009 | -0.0107281 | 0.62106559 | 0.99999994 |
| Galk2 | 0.3503389 | 0.23438053 | 0.24511289 | -0.0107324 | 0.61464684 | 0.99999994 |
| Birc2 | 0.11785207 | 0.08776087 | 0.09849916 | -0.0107383 | 0.76597949 | 0.99999994 |
| Slc25a39 | 0.38152569 | 0.25013661 | 0.26090155 | -0.0107649 | 0.60820542 | 0.99999994 |
| Gna11 | 0.36911879 | 0.2439508 | 0.25471588 | -0.0107651 | 0.61076687 | 0.99999994 |
| Sirt3 | 0.14764234 | 0.10975376 | 0.12052076 | -0.010767 | 0.72396492 | 0.99999994 |
| Tor1b | 0.25315293 | 0.17969691 | 0.19046481 | -0.0107679 | 0.64665909 | 0.99999994 |
| Gpr89 | 0.23761802 | 0.17013087 | 0.18090334 | -0.0107725 | 0.65408542 | 0.99999994 |
| Atg4d | 0.18967438 | 0.13905288 | 0.1498284 | -0.0107755 | 0.68393081 | 0.99999994 |
| Akr7a5 | 0.42051344 | 0.26877034 | 0.27954787 | -0.0107775 | 0.601266 | 0.99999994 |
| Lats1 | 0.1268509 | 0.09448397 | 0.10526449 | -0.0107805 | 0.75228631 | 0.99999994 |
| Irak1bp1 | 0.18917679 | 0.1387067 | 0.14949299 | -0.0107863 | 0.68448332 | 0.99999994 |
| Usp47 | 0.17286179 | 0.12755873 | 0.13834535 | -0.0107866 | 0.69811552 | 0.99999994 |
| Ggact | 0.04337054 | 0.02735022 | 0.03814543 | -0.0107952 | 0.9702012 | 0.99999994 |
| Ogfr | 0.35590936 | 0.23718843 | 0.24799195 | -0.0108035 | 0.61408328 | 0.99999994 |
| Igsf1 | 0.14055042 | 0.10454761 | 0.1153748 | -0.0108272 | 0.73390175 | 0.99999994 |
| Dcaf12 | 0.11207787 | 0.08327774 | 0.09410533 | -0.0108276 | 0.77812287 | 0.99999994 |
| Eif3c | 0.91287328 | 0.40859252 | 0.41942313 | -0.0108306 | 0.56823594 | 0.99999994 |
| Dgcr6 | 0.43701714 | 0.27626009 | 0.28709182 | -0.0108317 | 0.59914661 | 0.99999994 |
| Stx18 | 0.25284939 | 0.17943436 | 0.19028025 | -0.0108459 | 0.64780815 | 0.99999994 |
| Smarca2 | 0.28519865 | 0.19859727 | 0.20944359 | -0.0108463 | 0.63484325 | 0.99999994 |
| Taf7 | 0.41519737 | 0.26622762 | 0.27707416 | -0.0108465 | 0.60278968 | 0.99999994 |
| Nudt3 | 0.18654996 | 0.13686157 | 0.14771793 | -0.0108564 | 0.68765037 | 0.99999994 |
| Nrbp1 | 0.26190589 | 0.18488673 | 0.19574647 | -0.0108597 | 0.64404246 | 0.99999994 |
| Traf7 | 0.14794173 | 0.10986915 | 0.12073668 | -0.0108675 | 0.72546013 | 0.99999994 |
| Gm11808 | 0.551435 | 0.32329472 | 0.33416339 | -0.0108687 | 0.58570022 | 0.99999994 |
| St6galnac6 | 0.0708041 | 0.05029502 | 0.06116495 | -0.0108699 | 0.88158251 | 0.99999994 |
| Prkag2 | 0.04466271 | 0.02837643 | 0.0392484 | -0.010872 | 0.96739673 | 0.99999994 |
| Zdhhc12 | 0.36381106 | 0.24115857 | 0.25203156 | -0.010873 | 0.61300683 | 0.99999994 |
| Mitd1 | 0.12740965 | 0.09480632 | 0.10568132 | -0.010875 | 0.75331666 | 0.99999994 |

| | | | | | | |
|---|---|---|---|---|---|---|
| Cad | 0.04324802 | 0.02716464 | 0.03804075 | -0.0108761 | 0.97148609 | 0.99999994 |
| Ap3m1 | 0.22527243 | 0.16225233 | 0.17313093 | -0.0108786 | 0.6621174 | 0.99999994 |
| Atp13a1 | 0.10141938 | 0.07500385 | 0.08588509 | -0.0108812 | 0.80047793 | 0.99999994 |
| Esyt1 | 0.07911173 | 0.05709271 | 0.06797502 | -0.0108823 | 0.85670565 | 0.99999994 |
| Cyfip1 | 0.21751202 | 0.15727787 | 0.16816498 | -0.0108871 | 0.66674288 | 0.99999994 |
| Gm30173 | 0.03986802 | 0.02425783 | 0.03514624 | -0.0108884 | 0.98040372 | 0.99999994 |
| Aldh18a1 | 0.10143908 | 0.07501032 | 0.08590042 | -0.0108901 | 0.80062763 | 0.99999994 |
| Atp6v0a1 | 0.10779131 | 0.07992493 | 0.0908165 | -0.0108916 | 0.78764983 | 0.99999994 |
| Pdcd11 | 0.09722094 | 0.07167762 | 0.08256955 | -0.0108919 | 0.81004296 | 0.99999994 |
| Fam187b | 0.11747039 | 0.08731402 | 0.09821003 | -0.010896 | 0.76989339 | 0.99999994 |
| Ctdnep1 | 0.14358654 | 0.10667854 | 0.11758518 | -0.0109066 | 0.73151856 | 0.99999994 |
| Sacm1l | 0.14029156 | 0.10427424 | 0.11518584 | -0.0109116 | 0.73583582 | 0.99999994 |
| Zfp688 | 0.10802492 | 0.08008466 | 0.09099633 | -0.0109117 | 0.78761864 | 0.99999994 |
| Glis3 | 0.1020263 | 0.07544336 | 0.08635701 | -0.0109137 | 0.79988738 | 0.99999994 |
| Nudt7 | 0.22994324 | 0.16517358 | 0.17608956 | -0.010916 | 0.66006285 | 0.99999994 |
| Dhps | 0.21782927 | 0.1574342 | 0.16836923 | -0.010935 | 0.66724137 | 0.99999994 |
| Lancl1 | 0.24186556 | 0.17260619 | 0.18354125 | -0.0109351 | 0.6541591 | 0.99999994 |
| Ap2a1 | 0.18492498 | 0.13567724 | 0.1466161 | -0.0109389 | 0.69027267 | 0.99999994 |
| Usp50 | 0.22055618 | 0.15917983 | 0.17012057 | -0.0109407 | 0.66570417 | 0.99999994 |
| Nmt2 | 0.29302537 | 0.20297915 | 0.21393072 | -0.0109516 | 0.63335536 | 0.99999994 |
| Cisd3 | 0.27069698 | 0.19002149 | 0.20097697 | -0.0109555 | 0.64163986 | 0.99999994 |
| Eif2d | 0.15193137 | 0.11264841 | 0.1236039 | -0.0109555 | 0.72239748 | 0.99999994 |
| Mapk8ip2 | 0.14545328 | 0.1079628 | 0.11893884 | -0.010976 | 0.73048126 | 0.99999994 |
| Stard7 | 0.24303811 | 0.17328338 | 0.18426629 | -0.0109829 | 0.65422113 | 0.99999994 |
| Rgs11 | 0.04597571 | 0.02938263 | 0.04036723 | -0.0109846 | 0.96495333 | 0.99999994 |
| Slc37a3 | 0.09621612 | 0.07078234 | 0.08176859 | -0.0109863 | 0.81442837 | 0.99999994 |
| Gnl2 | 0.26191191 | 0.18474249 | 0.19575008 | -0.0110076 | 0.64591167 | 0.99999994 |
| Dnaja2 | 0.64231871 | 0.35467069 | 0.36567989 | -0.0110092 | 0.57941497 | 0.99999994 |
| Armc1 | 0.44976218 | 0.28176826 | 0.29278016 | -0.0110119 | 0.59884379 | 0.99999994 |
| Ghdc | 0.09512437 | 0.06988179 | 0.08089718 | -0.0110154 | 0.81762873 | 0.99999994 |

| | | | | | | |
|---|---|---|---|---|---|---|
| Lpgat1 | 0.12233864 | 0.09086391 | 0.10188452 | -0.0110206 | 0.76423826 | 0.99999994 |
| Zfp652 | 0.09756188 | 0.07181725 | 0.08284109 | -0.0110238 | 0.81213082 | 0.99999994 |
| Ccdc47 | 0.41379865 | 0.26539537 | 0.27641983 | -0.0110245 | 0.60467847 | 0.99999994 |
| P4ha1 | 0.16589115 | 0.12246241 | 0.13349153 | -0.0110291 | 0.70882346 | 0.99999994 |
| Chd9 | 0.1689011 | 0.12455975 | 0.1355941 | -0.0110344 | 0.70598053 | 0.99999994 |
| Nfx1 | 0.19215907 | 0.14045838 | 0.15149914 | -0.0110408 | 0.68619492 | 0.99999994 |
| Son | 1.39789586 | 0.40752286 | 0.4185641 | -0.0110412 | 0.56969082 | 0.99999994 |
| Zzef1 | 0.09193226 | 0.0673004 | 0.0783423 | -0.0110419 | 0.82592307 | 0.99999994 |
| Gm10138 | 0.07036487 | 0.04975256 | 0.06080285 | -0.0110503 | 0.88680768 | 0.99999994 |
| Necap1 | 0.28005411 | 0.19541138 | 0.20646259 | -0.0110512 | 0.63918868 | 0.99999994 |
| Mettl1 | 0.30434929 | 0.20927212 | 0.22032345 | -0.0110513 | 0.63076496 | 0.99999994 |
| Sobp | 0.09701956 | 0.07134855 | 0.08240911 | -0.0110606 | 0.81417452 | 0.99999994 |
| Serpinb6a | 0.2722727 | 0.19083804 | 0.20190647 | -0.0110684 | 0.64240647 | 0.99999994 |
| Magt1 | 0.51540474 | 0.30920922 | 0.32027827 | -0.0110691 | 0.59097301 | 0.99999994 |
| Grtp1 | 0.12098976 | 0.08980018 | 0.1008693 | -0.0110691 | 0.7674426 | 0.99999994 |
| Gm5617 | 0.41524235 | 0.26602413 | 0.27709518 | -0.011071 | 0.60486031 | 0.99999994 |
| Pla2g6 | 0.21498771 | 0.15546021 | 0.16653611 | -0.0110759 | 0.67100777 | 0.99999994 |
| Clint1 | 0.24692336 | 0.17558192 | 0.18665906 | -0.0110771 | 0.65358473 | 0.99999994 |
| Stk32a | 0.04084407 | 0.0249023 | 0.03598342 | -0.0110811 | 0.97980953 | 0.99999994 |
| Ddx54 | 0.22210566 | 0.16002946 | 0.17111232 | -0.0110829 | 0.66681277 | 0.99999994 |
| Sdc3 | 0.05974173 | 0.04089573 | 0.05198186 | -0.0110861 | 0.92213734 | 0.99999994 |
| Ngdn | 0.52726275 | 0.31385207 | 0.32493849 | -0.0110864 | 0.58982898 | 0.99999994 |
| Pccb | 0.19388661 | 0.14156378 | 0.15265652 | -0.0110927 | 0.68569434 | 0.99999994 |
| Nbea | 0.21075636 | 0.15269493 | 0.16379093 | -0.011096 | 0.67398161 | 0.99999994 |
| Bcl2l2 | 0.04786047 | 0.03086033 | 0.04196989 | -0.0111096 | 0.9609619 | 0.99999994 |
| Cul7 | 0.12779966 | 0.09485661 | 0.10597204 | -0.0111154 | 0.75746969 | 0.99999994 |
| Uxt | 0.2243358 | 0.16141411 | 0.17253497 | -0.0111209 | 0.66605302 | 0.99999994 |
| Fem1a | 0.10798417 | 0.07984357 | 0.09096496 | -0.0111214 | 0.79212238 | 0.99999994 |
| Agpat5 | 0.11834348 | 0.0877424 | 0.09887116 | -0.0111288 | 0.7731399 | 0.99999994 |
| Ifrd1 | 0.17525427 | 0.12886842 | 0.13999878 | -0.0111304 | 0.70166751 | 0.99999994 |

| | | | | | | |
|---|---|---|---|---|---|---|
| 9530068E07I | 0.64317145 | 0.35481927 | 0.36594991 | -0.0111306 | 0.58022074 | 0.99999994 |
| Ocel1 | 0.09323719 | 0.06825529 | 0.07938799 | -0.0111327 | 0.8247013 | 0.99999994 |
| Pde4dip | 0.07206666 | 0.05107093 | 0.06220464 | -0.0111337 | 0.88324974 | 0.99999994 |
| Got1 | 0.09433211 | 0.06912683 | 0.08026405 | -0.0111372 | 0.8221574 | 0.99999994 |
| Dtnbos | 0.16700949 | 0.12313616 | 0.13427393 | -0.0111378 | 0.70957116 | 0.99999994 |
| Cdk11b | 0.3652771 | 0.24163616 | 0.25277525 | -0.0111391 | 0.61536385 | 0.99999994 |
| B4galt5 | 0.07996341 | 0.05752518 | 0.06866905 | -0.0111439 | 0.85997749 | 0.99999994 |
| Wdfy1 | 0.26880277 | 0.1887082 | 0.19985652 | -0.0111483 | 0.64479377 | 0.99999994 |
| Xrcc5 | 0.20695114 | 0.15014778 | 0.16130624 | -0.0111585 | 0.67740621 | 0.99999994 |
| Wbp11 | 0.52469188 | 0.31277691 | 0.32393569 | -0.0111588 | 0.59068021 | 0.99999994 |
| Prkcz | 0.11658754 | 0.08638043 | 0.09754054 | -0.0111601 | 0.77684689 | 0.99999994 |
| Ybx3 | 0.08904003 | 0.06484781 | 0.0760184 | -0.0111706 | 0.8359936 | 0.99999994 |
| Gtf3c1 | 0.19983186 | 0.14544449 | 0.15661519 | -0.0111707 | 0.68252441 | 0.99999994 |
| Sephs2 | 0.38641179 | 0.25213081 | 0.26330358 | -0.0111728 | 0.61120061 | 0.99999994 |
| Mmgt2 | 0.12494698 | 0.09266595 | 0.10384135 | -0.0111754 | 0.76311449 | 0.99999994 |
| Rnf14 | 0.21248297 | 0.15373756 | 0.16491335 | -0.0111758 | 0.67403611 | 0.99999994 |
| Faim | 0.29704475 | 0.20503091 | 0.21621347 | -0.0111826 | 0.63467726 | 0.99999994 |
| Fbxl15 | 0.26380391 | 0.18568688 | 0.19688208 | -0.0111952 | 0.64746927 | 0.99999994 |
| Il6ra | 0.08259462 | 0.05960847 | 0.07080839 | -0.0111999 | 0.85379523 | 0.99999994 |
| Pld3 | 0.5441832 | 0.32021795 | 0.33143409 | -0.0112161 | 0.58911131 | 0.99999994 |
| Rab10 | 0.55556092 | 0.32448135 | 0.33570156 | -0.0112202 | 0.58802857 | 0.99999994 |
| Ngrn | 0.39453295 | 0.25603007 | 0.26725426 | -0.0112242 | 0.61009044 | 0.99999994 |
| Ift88 | 0.15104983 | 0.11174672 | 0.12297195 | -0.0112252 | 0.72827742 | 0.99999994 |
| Dpm1 | 0.25428393 | 0.17992459 | 0.19115143 | -0.0112268 | 0.6520813 | 0.99999994 |
| Anxa7 | 0.21741343 | 0.15686346 | 0.16810148 | -0.011238 | 0.67183831 | 0.99999994 |
| Rab3gap1 | 0.13886161 | 0.10290198 | 0.1141406 | -0.0112386 | 0.74389644 | 0.99999994 |
| Rb1cc1 | 0.29366107 | 0.20305135 | 0.21429273 | -0.0112414 | 0.63652277 | 0.99999994 |
| Tmem115 | 0.20542959 | 0.14906264 | 0.16030814 | -0.0112455 | 0.6797282 | 0.99999994 |
| Med21 | 0.50946507 | 0.30665832 | 0.3179093 | -0.011251 | 0.59311759 | 0.99999994 |
| Aprt | 0.60452915 | 0.34192818 | 0.35317928 | -0.0112511 | 0.58396563 | 0.99999994 |

| | | | | | | |
|---|---|---|---|---|---|---|
| Ube2o | 0.08632626 | 0.06257799 | 0.07383004 | -0.0112521 | 0.84481861 | 0.99999994 |
| Ncln | 0.1699086 | 0.12504021 | 0.13629562 | -0.0112554 | 0.70873312 | 0.99999994 |
| Tspan17 | 0.13016309 | 0.09647415 | 0.10772986 | -0.0112557 | 0.75661706 | 0.99999994 |
| Kmt2c | 0.23638592 | 0.16887799 | 0.1801345 | -0.0112565 | 0.6612776 | 0.99999994 |
| Tnfrsf9 | 0.03430785 | 0.019098 | 0.0303564 | -0.0112584 | 0.9932192 | 0.99999994 |
| Cct7 | 1.59470937 | 0.38786016 | 0.39912292 | -0.0112628 | 0.57449673 | 0.99999994 |
| Rab33b | 0.14316099 | 0.1059973 | 0.11727602 | -0.0112787 | 0.73896511 | 0.99999994 |
| Nnt | 0.10362093 | 0.0763118 | 0.08759472 | -0.0112829 | 0.8043864 | 0.99999994 |
| Tmco3 | 0.12917212 | 0.09570609 | 0.10699363 | -0.0112875 | 0.75873155 | 0.99999994 |
| Mettl23 | 0.20910396 | 0.15142532 | 0.16271384 | -0.0112885 | 0.67788578 | 0.99999994 |
| Rpusd1 | 0.15492379 | 0.11445104 | 0.12574236 | -0.0112913 | 0.7249876 | 0.99999994 |
| Ivd | 0.33909177 | 0.22792433 | 0.23922002 | -0.0112957 | 0.62335554 | 0.99999994 |
| Ccny | 0.12535637 | 0.09284544 | 0.10414772 | -0.0113023 | 0.76496311 | 0.99999994 |
| Tor1aip2 | 0.44283078 | 0.278393 | 0.28970097 | -0.011308 | 0.60249841 | 0.99999994 |
| Llgl2 | 0.22466122 | 0.16143256 | 0.17274213 | -0.0113096 | 0.66850811 | 0.99999994 |
| Cap1 | 0.58728366 | 0.33587684 | 0.34718645 | -0.0113096 | 0.58582996 | 0.99999994 |
| Npnt | 0.07739094 | 0.05525659 | 0.06657043 | -0.0113138 | 0.87102883 | 0.99999994 |
| Cr1l | 0.26196594 | 0.18446599 | 0.19578245 | -0.0113165 | 0.64978699 | 0.99999994 |
| Ireb2 | 0.17109269 | 0.12579775 | 0.13711863 | -0.0113209 | 0.70869172 | 0.99999994 |
| Lsm5 | 0.57745836 | 0.33237152 | 0.34369386 | -0.0113223 | 0.5867844 | 0.99999994 |
| Lsm4 | 0.79487297 | 0.39122391 | 0.40255563 | -0.0113317 | 0.57431587 | 0.99999994 |
| Nfat5 | 0.25271631 | 0.1788658 | 0.1901993 | -0.0113335 | 0.65418198 | 0.99999994 |
| Luc7l2 | 0.72844101 | 0.37751485 | 0.38884861 | -0.0113338 | 0.57691708 | 0.99999994 |
| Abca5 | 0.06502503 | 0.04504889 | 0.05638415 | -0.0113353 | 0.90957927 | 0.99999994 |
| Mrpl36 | 0.36965978 | 0.24364189 | 0.25498822 | -0.0113463 | 0.61645788 | 0.99999994 |
| Spns1 | 0.1660317 | 0.12223933 | 0.13358994 | -0.0113506 | 0.71415502 | 0.99999994 |
| Setx | 0.09111574 | 0.06632932 | 0.0776871 | -0.0113578 | 0.83476145 | 0.99999994 |
| Ddit3 | 0.20874085 | 0.15110347 | 0.16247678 | -0.0113733 | 0.67937167 | 0.99999994 |
| Hnrnph1 | 0.70718147 | 0.37255558 | 0.3839305 | -0.0113749 | 0.57816905 | 0.99999994 |
| Coprs | 0.30109942 | 0.20712233 | 0.21850104 | -0.0113787 | 0.6355684 | 0.99999994 |

| | | | | | | |
|---|---|---|---|---|---|---|
| Clpx | 0.21682918 | 0.15633556 | 0.167725 | -0.0113894 | 0.67436664 | 0.99999994 |
| Khdrbs3 | 0.08850198 | 0.06418345 | 0.07558513 | -0.0114017 | 0.84233052 | 0.99999994 |
| Wbp2 | 0.56041918 | 0.32609503 | 0.33749979 | -0.0114048 | 0.58898358 | 0.99999994 |
| Elovl6 | 0.06561446 | 0.04546653 | 0.05687341 | -0.0114069 | 0.90906833 | 0.99999994 |
| Ube3a | 0.68882141 | 0.36790365 | 0.37931181 | -0.0114082 | 0.57933646 | 0.99999994 |
| 2010320M18 | 0.26393219 | 0.18554923 | 0.19695871 | -0.0114095 | 0.65010027 | 0.99999994 |
| Ilkap | 0.51296676 | 0.30789905 | 0.31930862 | -0.0114096 | 0.5939996 | 0.99999994 |
| Arid1a | 0.39958671 | 0.25826875 | 0.26968623 | -0.0114175 | 0.61095337 | 0.99999994 |
| Ech1 | 0.41925283 | 0.26754011 | 0.27896316 | -0.0114231 | 0.60740666 | 0.99999994 |
| Coq10b | 0.23965803 | 0.17074345 | 0.18217253 | -0.0114291 | 0.66188188 | 0.99999994 |
| Ranbp3 | 0.31544506 | 0.2150447 | 0.22647489 | -0.0114302 | 0.63154435 | 0.99999994 |
| Phyh | 0.50031176 | 0.30277014 | 0.3142142 | -0.0114441 | 0.59577752 | 0.99999994 |
| Golph3 | 0.24614411 | 0.17473574 | 0.18618034 | -0.0114446 | 0.65879245 | 0.99999994 |
| Tmed8 | 0.19597763 | 0.14260353 | 0.15405314 | -0.0114496 | 0.68960004 | 0.99999994 |
| Cry2 | 0.09419411 | 0.06870073 | 0.08015371 | -0.011453 | 0.82922602 | 0.99999994 |
| Scaper | 0.1404276 | 0.10383163 | 0.11528516 | -0.0114535 | 0.7457919 | 0.99999994 |
| Clns1a | 0.47643041 | 0.29284712 | 0.30430775 | -0.0114606 | 0.59897204 | 0.99999994 |
| Ccdc6 | 0.07418346 | 0.05248126 | 0.06394397 | -0.0114627 | 0.88361046 | 0.99999994 |
| Rab1b | 0.27103757 | 0.18971134 | 0.20117808 | -0.0114667 | 0.64780407 | 0.99999994 |
| Gm20604 | 0.1907798 | 0.1390935 | 0.15057255 | -0.0114791 | 0.69408198 | 0.99999994 |
| Apoo | 0.33731001 | 0.22678842 | 0.23827657 | -0.0114881 | 0.6258669 | 0.99999994 |
| Sigmar1 | 0.3118213 | 0.21298863 | 0.22447797 | -0.0114893 | 0.63333118 | 0.99999994 |
| Lamtor4 | 0.82437324 | 0.39613465 | 0.40763188 | -0.0114972 | 0.57446057 | 0.99999994 |
| Vldlr | 0.09559126 | 0.06976677 | 0.08126999 | -0.0115032 | 0.82694548 | 0.99999994 |
| Vta1 | 0.48008836 | 0.29434101 | 0.30585027 | -0.0115093 | 0.59889235 | 0.99999994 |
| Pnpo | 0.10419971 | 0.0765234 | 0.08804315 | -0.0115198 | 0.80810434 | 0.99999994 |
| Tpd52l2 | 0.46526281 | 0.28801746 | 0.29954149 | -0.011524 | 0.60105738 | 0.99999994 |
| Dalrd3 | 0.24835063 | 0.1760032 | 0.18753435 | -0.0115311 | 0.65883562 | 0.99999994 |
| Npepps | 0.43264836 | 0.27358177 | 0.28511416 | -0.0115324 | 0.60612691 | 0.99999994 |
| Morf4l2 | 1.55923295 | 0.39130111 | 0.40283866 | -0.0115375 | 0.57559704 | 0.99999994 |

| | | | | | | |
|---|---|---|---|---|---|---|
| Srebf1 | 0.06503785 | 0.04484913 | 0.05639479 | -0.0115457 | 0.9135195 | 0.99999994 |
| Atg5 | 0.26849347 | 0.1881222 | 0.19967324 | -0.011551 | 0.64990658 | 0.99999994 |
| Ppp3cb | 0.22206798 | 0.15952237 | 0.17108824 | -0.0115659 | 0.67363816 | 0.99999994 |
| Mapk14 | 0.09279447 | 0.06745935 | 0.07903342 | -0.0115741 | 0.83516835 | 0.99999994 |
| Gm9843 | 0.66267154 | 0.3604199 | 0.37199752 | -0.0115776 | 0.58205938 | 0.99999994 |
| Clip1 | 0.1529564 | 0.11275168 | 0.12433758 | -0.0115859 | 0.73245208 | 0.99999994 |
| Abca3 | 0.10380693 | 0.07614097 | 0.08773887 | -0.0115979 | 0.81053826 | 0.99999994 |
| Pcx | 0.05556319 | 0.0368805 | 0.04847852 | -0.011598 | 0.94436039 | 0.99999994 |
| Zfp398 | 0.11009241 | 0.08098438 | 0.09258485 | -0.0116005 | 0.79786293 | 0.99999994 |
| Prpf4 | 0.13130778 | 0.09697664 | 0.10857872 | -0.0116021 | 0.76153817 | 0.99999994 |
| Nebl | 0.04517406 | 0.02806769 | 0.03968437 | -0.0116167 | 0.97432212 | 0.99999994 |
| Rangap1 | 0.5562292 | 0.32432889 | 0.33594975 | -0.0116209 | 0.59105144 | 0.99999994 |
| U2af1 | 1.09754106 | 0.41808787 | 0.42971687 | -0.011629 | 0.57147646 | 0.99999994 |
| Stk16 | 0.43794701 | 0.27588103 | 0.28751096 | -0.0116299 | 0.60613223 | 0.99999994 |
| Exoc4 | 0.15301589 | 0.11274394 | 0.12438012 | -0.0116362 | 0.73326727 | 0.99999994 |
| Eif4g3 | 0.37678276 | 0.24691038 | 0.25855202 | -0.0116416 | 0.6178013 | 0.99999994 |
| Ubr3 | 0.12974681 | 0.095768 | 0.10742074 | -0.0116527 | 0.76487289 | 0.99999994 |
| Srd5a3 | 0.15271381 | 0.11249954 | 0.12416404 | -0.0116645 | 0.73412175 | 0.99999994 |
| 6430548M08 | 0.03554459 | 0.01974754 | 0.03142485 | -0.0116773 | 0.99330955 | 0.99999994 |
| Becn1 | 0.41890026 | 0.26711214 | 0.27879942 | -0.0116873 | 0.60989088 | 0.99999994 |
| Cltb | 0.79920956 | 0.39166823 | 0.40336192 | -0.0116937 | 0.57650936 | 0.99999994 |
| Blzf1 | 0.14326055 | 0.10563473 | 0.11734836 | -0.0117136 | 0.74679334 | 0.99999994 |
| Ankrd17 | 0.29686442 | 0.20439542 | 0.21611137 | -0.011716 | 0.64089727 | 0.99999994 |
| Vgf | 0.06001084 | 0.0404854 | 0.05220683 | -0.0117214 | 0.93247476 | 0.99999994 |
| Nudt22 | 0.09772319 | 0.07124701 | 0.08296952 | -0.0117225 | 0.82650835 | 0.99999994 |
| Gm19412 | 0.10199714 | 0.0746104 | 0.08633435 | -0.0117239 | 0.81697935 | 0.99999994 |
| Clcn4 | 0.19165864 | 0.13942471 | 0.15116324 | -0.0117385 | 0.69738598 | 0.99999994 |
| Hivep3 | 0.06038952 | 0.04078232 | 0.05252327 | -0.0117409 | 0.93161781 | 0.99999994 |
| Dnajc13 | 0.15167207 | 0.11167374 | 0.12341811 | -0.0117444 | 0.73676822 | 0.99999994 |
| Trim44 | 0.18567639 | 0.13537632 | 0.14712596 | -0.0117496 | 0.702493 | 0.99999994 |

| | | | | | | |
|---|---|---|---|---|---|---|
| Sf3b3 | 0.14020654 | 0.10337286 | 0.11512376 | -0.0117509 | 0.75156631 | 0.99999994 |
| Abcb10 | 0.10830703 | 0.07946104 | 0.0912134 | -0.0117524 | 0.80445322 | 0.99999994 |
| Atp2c1 | 0.27153889 | 0.18971153 | 0.20147389 | -0.0117624 | 0.65121719 | 0.99999994 |
| Anapc11 | 0.74869717 | 0.38154406 | 0.39331464 | -0.0117706 | 0.57894777 | 0.99999994 |
| Lrif1 | 0.18408924 | 0.13427733 | 0.14604831 | -0.011771 | 0.70418621 | 0.99999994 |
| Cyp20a1 | 0.1181786 | 0.08697307 | 0.09874638 | -0.0117733 | 0.78629239 | 0.99999994 |
| Pold2 | 0.15918761 | 0.11699151 | 0.12877165 | -0.0117801 | 0.72871422 | 0.99999994 |
| Rabl6 | 0.32276026 | 0.21868591 | 0.23046967 | -0.0117838 | 0.63319791 | 0.99999994 |
| Acot8 | 0.21090016 | 0.15209972 | 0.16388453 | -0.0117848 | 0.68390753 | 0.99999994 |
| Gmfb | 0.18819837 | 0.13703878 | 0.1488327 | -0.0117939 | 0.70106594 | 0.99999994 |
| Amfr | 0.18252441 | 0.13318912 | 0.14498312 | -0.011794 | 0.70591071 | 0.99999994 |
| Prkcb | 0.10099406 | 0.07375886 | 0.08555409 | -0.0117952 | 0.82061288 | 0.99999994 |
| Il6st | 0.06076214 | 0.04103656 | 0.05283448 | -0.0117979 | 0.93140831 | 0.99999994 |
| Lysmd3 | 0.13754961 | 0.10136257 | 0.11317945 | -0.0118169 | 0.75646583 | 0.99999994 |
| Sppl2a | 0.10264828 | 0.07501618 | 0.08684016 | -0.011824 | 0.81761575 | 0.99999994 |
| Paip2b | 0.08668294 | 0.06229081 | 0.07411811 | -0.0118273 | 0.85592297 | 0.99999994 |
| H2-K1 | 0.35801209 | 0.23723273 | 0.24907214 | -0.0118394 | 0.62415191 | 0.99999994 |
| Hsdl2 | 0.17139276 | 0.12548173 | 0.13732695 | -0.0118452 | 0.71706612 | 0.99999994 |
| Lgals3bp | 0.29502124 | 0.20321086 | 0.21506607 | -0.0118552 | 0.64316151 | 0.99999994 |
| Mrpl23 | 0.87783844 | 0.40357201 | 0.41542898 | -0.011857 | 0.57533862 | 0.99999994 |
| Mrpl11 | 0.52891865 | 0.31372325 | 0.32558221 | -0.011859 | 0.59578435 | 0.99999994 |
| Rpl38 | 2.25235901 | 0.35792402 | 0.36978344 | -0.0118594 | 0.58452286 | 0.99999994 |
| Psen2 | 0.1327038 | 0.09774623 | 0.10961173 | -0.0118655 | 0.76440763 | 0.99999994 |
| Mtmr6 | 0.10343615 | 0.07558581 | 0.08745146 | -0.0118657 | 0.81677759 | 0.99999994 |
| Nsfl1c | 0.59459537 | 0.33787032 | 0.34974954 | -0.0118792 | 0.58943144 | 0.99999994 |
| Polr3h | 0.19118034 | 0.13895817 | 0.15084186 | -0.0118837 | 0.70000129 | 0.99999994 |
| Pi4ka | 0.09209554 | 0.066584 | 0.07847324 | -0.0118892 | 0.84338904 | 0.99999994 |
| Dmxl2 | 0.08434927 | 0.06033451 | 0.07223101 | -0.0118965 | 0.86352783 | 0.99999994 |
| Matr3 | 0.56792307 | 0.3283453 | 0.34024895 | -0.0119036 | 0.59207016 | 0.99999994 |
| Ctnnbip1 | 0.64890835 | 0.35584177 | 0.36775659 | -0.0119148 | 0.58537244 | 0.99999994 |

| | | | | | | |
|---|---|---|---|---|---|---|
| Mroh1 | 0.08738073 | 0.06275593 | 0.07468127 | -0.0119253 | 0.85608822 | 0.99999994 |
| Fam20a | 0.03872486 | 0.02223741 | 0.03416434 | -0.0119269 | 0.98993095 | 0.99999994 |
| Ehmt2 | 0.26150958 | 0.18357883 | 0.19550892 | -0.0119301 | 0.65769318 | 0.99999994 |
| Sec24a | 0.10115579 | 0.07374698 | 0.08567999 | -0.011933 | 0.82305191 | 0.99999994 |
| R3hdm2 | 0.15209774 | 0.11178783 | 0.12372306 | -0.0119352 | 0.73960267 | 0.99999994 |
| Ctso | 0.18407588 | 0.13409799 | 0.14603923 | -0.0119412 | 0.70687305 | 0.99999994 |
| Thop1 | 0.16667064 | 0.12209316 | 0.13403702 | -0.0119439 | 0.72345586 | 0.99999994 |
| Gle1 | 0.15725741 | 0.11545876 | 0.1274029 | -0.0119441 | 0.73369911 | 0.99999994 |
| Uap1 | 0.198747 | 0.14394918 | 0.15589564 | -0.0119465 | 0.69500387 | 0.99999994 |
| Tlk1 | 0.15719732 | 0.11541286 | 0.12736021 | -0.0119474 | 0.73382306 | 0.99999994 |
| Ncor1 | 0.50931538 | 0.30589673 | 0.3178493 | -0.0119526 | 0.59882586 | 0.99999994 |
| Tm2d3 | 0.45848128 | 0.28465134 | 0.29660436 | -0.011953 | 0.60574582 | 0.99999994 |
| Mrps17 | 0.68953828 | 0.36754142 | 0.37949672 | -0.0119553 | 0.58304736 | 0.99999994 |
| Qrich1 | 0.27894031 | 0.1938486 | 0.20581389 | -0.0119653 | 0.65061344 | 0.99999994 |
| Mib1 | 0.15946685 | 0.11700093 | 0.12896932 | -0.0119684 | 0.73161727 | 0.99999994 |
| Alg12 | 0.06553354 | 0.04481322 | 0.05680626 | -0.011993 | 0.92003434 | 0.99999994 |
| Ptcd2 | 0.16868237 | 0.12344606 | 0.13544166 | -0.0119956 | 0.72224722 | 0.99999994 |
| Fto | 0.1173697 | 0.08612636 | 0.09813372 | -0.0120074 | 0.79230727 | 0.99999994 |
| Bcat2 | 0.08723684 | 0.06255309 | 0.07456518 | -0.0120121 | 0.85820905 | 0.99999994 |
| Usp7 | 0.13445338 | 0.098889 | 0.11090309 | -0.0120141 | 0.76457002 | 0.99999994 |
| Max | 0.30046264 | 0.20609788 | 0.21814285 | -0.012045 | 0.64339084 | 0.99999994 |
| Galnt11 | 0.19872379 | 0.14383258 | 0.15588023 | -0.0120476 | 0.69653325 | 0.99999994 |
| Tpcn1 | 0.14097891 | 0.10363462 | 0.11568741 | -0.0120528 | 0.75599799 | 0.99999994 |
| Abhd17c | 0.08861829 | 0.06361905 | 0.07567882 | -0.0120598 | 0.85559169 | 0.99999994 |
| Ddx41 | 0.40372286 | 0.25959613 | 0.27166218 | -0.012066 | 0.61624994 | 0.99999994 |
| Clptm1 | 0.51991388 | 0.30999471 | 0.32206102 | -0.0120663 | 0.59846916 | 0.99999994 |
| Naglu | 0.09322904 | 0.06731206 | 0.07938147 | -0.0120694 | 0.84424376 | 0.99999994 |
| Slc35c1 | 0.05200225 | 0.03340372 | 0.04547779 | -0.0120741 | 0.96140446 | 0.99999994 |
| Kdm5c | 0.1968738 | 0.14257496 | 0.15465027 | -0.0120753 | 0.69837675 | 0.99999994 |
| Emc8 | 0.39507512 | 0.25543905 | 0.2675161 | -0.012077 | 0.61810285 | 0.99999994 |

| | | | | | | |
|---|---|---|---|---|---|---|
| Tollip | 0.12956584 | 0.09520204 | 0.10728628 | -0.0120842 | 0.77329498 | 0.99999994 |
| Polr1c | 0.3904709 | 0.25319349 | 0.26528497 | -0.0120915 | 0.61920429 | 0.99999994 |
| Disp2 | 0.27425312 | 0.19097116 | 0.20307139 | -0.0121002 | 0.65417846 | 0.99999994 |
| Ormdl3 | 0.26131934 | 0.18326774 | 0.19539484 | -0.0121271 | 0.66024213 | 0.99999994 |
| Fkbp5 | 0.11484772 | 0.08408964 | 0.09621833 | -0.0121287 | 0.79928202 | 0.99999994 |
| Fdx1 | 0.39940939 | 0.25746187 | 0.26960123 | -0.0121394 | 0.6178033 | 0.99999994 |
| Gnaq | 0.25049769 | 0.1767073 | 0.1888473 | -0.01214 | 0.66563884 | 0.99999994 |
| Mapk1 | 0.42916878 | 0.27137752 | 0.28352919 | -0.0121517 | 0.61228608 | 0.99999994 |
| Fam207a | 0.24913349 | 0.17585817 | 0.18801359 | -0.0121554 | 0.66652794 | 0.99999994 |
| Nemf | 0.34582807 | 0.23059996 | 0.24276278 | -0.0121628 | 0.63061955 | 0.99999994 |
| Mga | 0.24488355 | 0.17323513 | 0.18540467 | -0.0121695 | 0.66891105 | 0.99999994 |
| Pak1ip1 | 0.43992424 | 0.27621898 | 0.28839965 | -0.0121807 | 0.61069611 | 0.99999994 |
| Supt5 | 0.35978692 | 0.23779481 | 0.24998109 | -0.0121863 | 0.62721624 | 0.99999994 |
| Mrpl10 | 0.29679078 | 0.20386876 | 0.21606966 | -0.0122009 | 0.64649235 | 0.99999994 |
| Ccpg1 | 0.19162476 | 0.13893146 | 0.15114048 | -0.012209 | 0.70460648 | 0.99999994 |
| Fkbp1a | 1.70261368 | 0.37656724 | 0.38878341 | -0.0122162 | 0.58283559 | 0.99999994 |
| Lsm2 | 0.36588217 | 0.24085765 | 0.25308169 | -0.012224 | 0.62608696 | 0.99999994 |
| Larp4 | 0.21338651 | 0.15327393 | 0.16549948 | -0.0122255 | 0.68853755 | 0.99999994 |
| Tbc1d15 | 0.21548482 | 0.15462919 | 0.1668574 | -0.0122282 | 0.68715911 | 0.99999994 |
| Aacs | 0.35743773 | 0.23654349 | 0.24877744 | -0.0122339 | 0.6282927 | 0.99999994 |
| Tpd52 | 1.24212666 | 0.41539882 | 0.42763725 | -0.0122384 | 0.57553991 | 0.99999994 |
| Fam219a | 0.14419828 | 0.10578647 | 0.11802923 | -0.0122428 | 0.75504141 | 0.99999994 |
| Prkar2a | 0.12082152 | 0.08849699 | 0.10074253 | -0.0122455 | 0.79077099 | 0.99999994 |
| Slc25a1 | 0.1708151 | 0.12467882 | 0.13692583 | -0.012247 | 0.72419932 | 0.99999994 |
| Hip1r | 0.19068865 | 0.13825374 | 0.15051124 | -0.0122575 | 0.70612094 | 0.99999994 |
| Vps26b | 0.12602896 | 0.09238906 | 0.10465064 | -0.0122616 | 0.78227649 | 0.99999994 |
| Tmem87b | 0.18867864 | 0.13689505 | 0.14915695 | -0.0122619 | 0.70788143 | 0.99999994 |
| Smpd2 | 0.33881206 | 0.22680319 | 0.23907209 | -0.0122689 | 0.63367277 | 0.99999994 |
| Mrpl43 | 0.54814466 | 0.32065717 | 0.33292932 | -0.0122722 | 0.59691152 | 0.99999994 |
| Dcaf7 | 0.25300501 | 0.17809714 | 0.19037489 | -0.0122777 | 0.66614208 | 0.99999994 |

| | | | | | | |
|---|---|---|---|---|---|---|
| Gm6483 | 0.28345809 | 0.19615095 | 0.208438 | -0.0122871 | 0.65261146 | 0.99999994 |
| Slc35b4 | 0.06085758 | 0.04062632 | 0.05291417 | -0.0122878 | 0.93891807 | 0.99999994 |
| Med31 | 0.4757023 | 0.29171088 | 0.30399964 | -0.0122888 | 0.60606357 | 0.99999994 |
| Ubxn1 | 0.89302218 | 0.40496826 | 0.41725832 | -0.0122901 | 0.5777178 | 0.99999994 |
| Tbl2 | 0.06452353 | 0.04367551 | 0.05596757 | -0.0122921 | 0.92811906 | 0.99999994 |
| Asnsd1 | 0.15367542 | 0.11255606 | 0.1248515 | -0.0122954 | 0.74393397 | 0.99999994 |
| Golph3l | 0.11497925 | 0.08402087 | 0.09631843 | -0.0122976 | 0.80229802 | 0.99999994 |
| Mysm1 | 0.23722722 | 0.16835934 | 0.18065964 | -0.0123003 | 0.67479808 | 0.99999994 |
| Pold4 | 0.25058369 | 0.1765993 | 0.1888998 | -0.0123005 | 0.66765201 | 0.99999994 |
| Usp4 | 0.19306453 | 0.13980533 | 0.15210616 | -0.0123008 | 0.70481507 | 0.99999994 |
| Jagn1 | 0.7584343 | 0.38307761 | 0.39538056 | -0.012303 | 0.58204355 | 0.99999994 |
| H2-D1 | 0.32973675 | 0.22193301 | 0.23423652 | -0.0123035 | 0.63669244 | 0.99999994 |
| Jmjd6 | 0.31366612 | 0.21318659 | 0.22549615 | -0.0123096 | 0.641833 | 0.99999994 |
| Xpo6 | 0.12561473 | 0.09201636 | 0.10434098 | -0.0123246 | 0.78413653 | 0.99999994 |
| Ap1b1 | 0.19161674 | 0.13880083 | 0.15113509 | -0.0123343 | 0.70651461 | 0.99999994 |
| Rpl6 | 2.72904417 | 0.34831171 | 0.36064656 | -0.0123348 | 0.59004512 | 0.99999994 |
| Nop56 | 0.40162523 | 0.25832592 | 0.2706617 | -0.0123358 | 0.61919712 | 0.99999994 |
| Kif1a | 0.14831451 | 0.10866922 | 0.12100536 | -0.0123361 | 0.75130304 | 0.99999994 |
| Gpd1 | 0.07650109 | 0.05350614 | 0.06584286 | -0.0123367 | 0.89383354 | 0.99999994 |
| Tbc1d10a | 0.16140014 | 0.11797559 | 0.13033536 | -0.0123598 | 0.73605352 | 0.99999994 |
| Rbm47 | 0.18735056 | 0.1358807 | 0.14825972 | -0.012379 | 0.71081804 | 0.99999994 |
| Zmpste24 | 0.28755779 | 0.19842266 | 0.21080199 | -0.0123793 | 0.6520622 | 0.99999994 |
| Ydjc | 0.07592066 | 0.05298628 | 0.06536784 | -0.0123816 | 0.89631042 | 0.99999994 |
| Pdcl | 0.15206566 | 0.1113148 | 0.12370009 | -0.0123853 | 0.74744076 | 0.99999994 |
| Psph | 0.14941294 | 0.10940977 | 0.12179613 | -0.0123864 | 0.75077632 | 0.99999994 |
| Leprotl1 | 0.38123091 | 0.24835679 | 0.26075603 | -0.0123992 | 0.62419505 | 0.99999994 |
| Tcn2 | 0.21527578 | 0.15432289 | 0.16672232 | -0.0123994 | 0.6897158 | 0.99999994 |
| Usp36 | 0.07395225 | 0.05135106 | 0.06375423 | -0.0124032 | 0.90233506 | 0.99999994 |
| Tmem251 | 0.19963514 | 0.14406684 | 0.1564848 | -0.012418 | 0.70132665 | 0.99999994 |
| Pcnx | 0.06592113 | 0.04470885 | 0.05712781 | -0.012419 | 0.92605418 | 0.99999994 |

| | | | | | | |
|---|---|---|---|---|---|---|
| Hnrnpa0 | 0.19941868 | 0.14390575 | 0.15634129 | -0.0124355 | 0.70175442 | 0.99999994 |
| Rcor1 | 0.10321728 | 0.07484375 | 0.08728172 | -0.012438 | 0.82857675 | 0.99999994 |
| Babam1 | 0.37659604 | 0.24601806 | 0.25845916 | -0.0124411 | 0.62567052 | 0.99999994 |
| Rps13 | 2.83954115 | 0.34111894 | 0.35356725 | -0.0124483 | 0.59264545 | 0.99999994 |
| Ndufs1 | 0.36326087 | 0.23930155 | 0.25175201 | -0.0124505 | 0.62899471 | 0.99999994 |
| Pomgnt2 | 0.21969144 | 0.15711333 | 0.16956602 | -0.0124527 | 0.68752032 | 0.99999994 |
| Bola3 | 0.42256803 | 0.26804534 | 0.28049834 | -0.012453 | 0.61619999 | 0.99999994 |
| Pycr1 | 0.04389554 | 0.02613393 | 0.03859379 | -0.0124599 | 0.98418204 | 0.99999994 |
| Orai3 | 0.08780169 | 0.0625544 | 0.07502076 | -0.0124664 | 0.86565865 | 0.99999994 |
| Cgrrf1 | 0.31053134 | 0.21128598 | 0.22376426 | -0.0124783 | 0.64475332 | 0.99999994 |
| Atp1a1 | 0.65196171 | 0.35618156 | 0.36871112 | -0.0125296 | 0.58947572 | 0.99999994 |
| B3galt6 | 0.19080272 | 0.13805501 | 0.15058797 | -0.012533 | 0.71020628 | 0.99999994 |
| Sdr39u1 | 0.11158744 | 0.08119546 | 0.09373021 | -0.0125347 | 0.81331257 | 0.99999994 |
| Hacd3 | 0.23147473 | 0.16451543 | 0.17705486 | -0.0125394 | 0.68132903 | 0.99999994 |
| Psmg2 | 0.32889598 | 0.22123833 | 0.23378476 | -0.0125464 | 0.63953792 | 0.99999994 |
| Slc35a1 | 0.32589302 | 0.21961112 | 0.23216631 | -0.0125552 | 0.64056348 | 0.99999994 |
| Ffar1 | 0.2308009 | 0.1640504 | 0.17663043 | -0.01258 | 0.68227914 | 0.99999994 |
| Tnfaip1 | 0.26567695 | 0.18541646 | 0.19799939 | -0.0125829 | 0.66385743 | 0.99999994 |
| Nfasc | 0.06267444 | 0.04184292 | 0.05442928 | -0.0125864 | 0.93812623 | 0.99999994 |
| Acsl5 | 0.10196045 | 0.07371662 | 0.08630583 | -0.0125892 | 0.83421231 | 0.99999994 |
| Papola | 0.78962363 | 0.38897642 | 0.40156777 | -0.0125913 | 0.58266393 | 0.99999994 |
| 1600012H06 | 0.20374662 | 0.14659125 | 0.15920132 | -0.0126101 | 0.70098617 | 0.99999994 |
| Camk2b | 0.24293532 | 0.17158973 | 0.18420278 | -0.0126131 | 0.67572797 | 0.99999994 |
| Rps29 | 2.86819026 | 0.33865188 | 0.3512707 | -0.0126188 | 0.59449321 | 0.99999994 |
| Cspp1 | 0.49111876 | 0.29782556 | 0.31044626 | -0.0126207 | 0.60665206 | 0.99999994 |
| Lgmn | 0.24361855 | 0.17199973 | 0.1846247 | -0.012625 | 0.67550797 | 0.99999994 |
| Pitpnb | 0.3387358 | 0.22640582 | 0.23903175 | -0.0126259 | 0.63742952 | 0.99999994 |
| Keap1 | 0.32231718 | 0.21760111 | 0.23022902 | -0.0126279 | 0.64248053 | 0.99999994 |
| Wipi1 | 0.22600195 | 0.16096563 | 0.17359448 | -0.0126289 | 0.68589385 | 0.99999994 |
| Slc4a7 | 0.20946851 | 0.15031591 | 0.16295171 | -0.0126358 | 0.69712712 | 0.99999994 |

| | | | | | | |
|---|---|---|---|---|---|---|
| Hdac4 | 0.08141599 | 0.05721356 | 0.06985098 | -0.0126374 | 0.8857567 | 0.99999994 |
| Pde3b | 0.13172122 | 0.09623049 | 0.10888489 | -0.0126544 | 0.78041321 | 0.99999994 |
| Nxt1 | 0.19959068 | 0.14377987 | 0.15645533 | -0.0126755 | 0.70515132 | 0.99999994 |
| Ankrd40 | 0.08921544 | 0.06346706 | 0.0761596 | -0.0126925 | 0.86635581 | 0.99999994 |
| Vmac | 0.17273855 | 0.12556724 | 0.13826001 | -0.0126928 | 0.72942585 | 0.99999994 |
| Epb41l3 | 0.28802511 | 0.1983739 | 0.21107047 | -0.0126966 | 0.65556954 | 0.99999994 |
| Zmynd8 | 0.30807169 | 0.20969036 | 0.22239931 | -0.0127089 | 0.64816672 | 0.99999994 |
| Mis12 | 0.13065612 | 0.09538025 | 0.1080957 | -0.0127154 | 0.78318545 | 0.99999994 |
| Rps24 | 2.64567431 | 0.35184285 | 0.36455975 | -0.0127169 | 0.59180591 | 0.99999994 |
| Aqp4 | 0.0552115 | 0.03545509 | 0.04818278 | -0.0127277 | 0.96065221 | 0.99999994 |
| Phkg2 | 0.19717894 | 0.14210932 | 0.15485339 | -0.0127441 | 0.70808631 | 0.99999994 |
| Mospd3 | 0.19690857 | 0.14191951 | 0.15467342 | -0.0127539 | 0.70845047 | 0.99999994 |
| Scfd1 | 0.52903379 | 0.31286805 | 0.32562691 | -0.0127589 | 0.6028814 | 0.99999994 |
| Twf1 | 0.30354138 | 0.2071105 | 0.21987128 | -0.0127608 | 0.65036953 | 0.99999994 |
| Saysd1 | 0.27420535 | 0.19027975 | 0.20304333 | -0.0127636 | 0.66218277 | 0.99999994 |
| Ints3 | 0.09046451 | 0.06439455 | 0.07716405 | -0.0127695 | 0.86466652 | 0.99999994 |
| Cntfr | 0.14682317 | 0.10714533 | 0.11992948 | -0.0127841 | 0.76100807 | 0.99999994 |
| Efr3a | 0.1394922 | 0.10181508 | 0.11460184 | -0.0127868 | 0.77116431 | 0.99999994 |
| Ubn2 | 0.37234238 | 0.24354207 | 0.25633519 | -0.0127931 | 0.63013237 | 0.99999994 |
| Atf2 | 0.22011946 | 0.15704746 | 0.1698406 | -0.0127931 | 0.69195027 | 0.99999994 |
| Tns3 | 0.0866163 | 0.06126662 | 0.0740643 | -0.0127977 | 0.87496137 | 0.99999994 |
| Cdh1 | 0.31324363 | 0.21244993 | 0.22526324 | -0.0128133 | 0.64751626 | 0.99999994 |
| Unc45a | 0.23167657 | 0.16436686 | 0.1771819 | -0.012815 | 0.6848962 | 0.99999994 |
| Meaf6 | 0.26198064 | 0.18296583 | 0.19579126 | -0.0128254 | 0.66859216 | 0.99999994 |
| Stx6 | 0.30409547 | 0.20734935 | 0.22018146 | -0.0128321 | 0.65096623 | 0.99999994 |
| Lasp1 | 0.25805521 | 0.18059724 | 0.19343199 | -0.0128347 | 0.6706352 | 0.99999994 |
| Ecsit | 0.15427576 | 0.11244191 | 0.12528014 | -0.0128382 | 0.75241784 | 0.99999994 |
| Nbeal1 | 0.18561904 | 0.13424214 | 0.14708707 | -0.0128449 | 0.719478 | 0.99999994 |
| Tomm6 | 0.16609081 | 0.1207494 | 0.13363131 | -0.0128819 | 0.73945488 | 0.99999994 |
| Rdh14 | 0.27235902 | 0.18907345 | 0.20195732 | -0.0128839 | 0.66445136 | 0.99999994 |

| | | | | | | |
|---|---|---|---|---|---|---|
| Syp | 0.1002757 | 0.07210208 | 0.08499453 | -0.0128924 | 0.84367414 | 0.99999994 |
| Nudt14 | 0.36965716 | 0.24208312 | 0.2549869 | -0.0129038 | 0.63188479 | 0.99999994 |
| Atxn2l | 0.13338113 | 0.0971848 | 0.1101121 | -0.0129273 | 0.78273554 | 0.99999994 |
| Cdc42bpa | 0.1246193 | 0.09066583 | 0.10359597 | -0.0129301 | 0.79695699 | 0.99999994 |
| Slc7a8 | 0.06995463 | 0.04753195 | 0.06046446 | -0.0129325 | 0.92273376 | 0.99999994 |
| Nt5m | 0.25048939 | 0.17590812 | 0.18884224 | -0.0129341 | 0.67577081 | 0.99999994 |
| Yipf1 | 0.28887899 | 0.19862085 | 0.21156052 | -0.0129397 | 0.65804101 | 0.99999994 |
| Nav2 | 0.34851437 | 0.23120924 | 0.24416435 | -0.0129551 | 0.638023 | 0.99999994 |
| Ube2d2a | 1.09839942 | 0.41677288 | 0.42972968 | -0.0129568 | 0.57953196 | 0.99999994 |
| Agpat1 | 0.12482657 | 0.09079375 | 0.1037512 | -0.0129574 | 0.79710079 | 0.99999994 |
| Oaz2 | 0.71145544 | 0.37199514 | 0.3849533 | -0.0129582 | 0.58864626 | 0.99999994 |
| Huwe1 | 0.43217699 | 0.27193487 | 0.28489996 | -0.0129651 | 0.61902242 | 0.99999994 |
| Nomo1 | 0.10407086 | 0.07497674 | 0.08794336 | -0.0129666 | 0.83681014 | 0.99999994 |
| Itfg1 | 0.45469586 | 0.28198356 | 0.29495105 | -0.0129675 | 0.61510367 | 0.99999994 |
| Sec14l1 | 0.0702728 | 0.04775263 | 0.06072692 | -0.0129743 | 0.9225057 | 0.99999994 |
| Sec24c | 0.16529119 | 0.12009549 | 0.13307121 | -0.0129757 | 0.74184982 | 0.99999994 |
| Rars | 0.40541269 | 0.25948916 | 0.27246573 | -0.0129766 | 0.62438528 | 0.99999994 |
| Larp1 | 0.11486548 | 0.08323657 | 0.09623184 | -0.0129953 | 0.81564251 | 0.99999994 |
| Alyref2 | 0.15443825 | 0.11239385 | 0.12539609 | -0.0130022 | 0.75496372 | 0.99999994 |
| Ccdc186 | 0.16734028 | 0.12149009 | 0.13450508 | -0.013015 | 0.74023928 | 0.99999994 |
| Ttc3 | 0.91929833 | 0.40703888 | 0.42007965 | -0.0130408 | 0.5818533 | 0.99999994 |
| Wsb2 | 0.17685729 | 0.12802797 | 0.14110301 | -0.013075 | 0.73131614 | 0.99999994 |
| Sez6l2 | 0.34855036 | 0.23110052 | 0.24418308 | -0.0130826 | 0.63931435 | 0.99999994 |
| Ifi35 | 0.13529937 | 0.09844267 | 0.1115262 | -0.0130835 | 0.78256362 | 0.99999994 |
| Gpx1 | 0.53452262 | 0.31466342 | 0.32774783 | -0.0130844 | 0.60477891 | 0.99999994 |
| Chmp7 | 0.23340952 | 0.16518377 | 0.17827098 | -0.0130872 | 0.68745737 | 0.99999994 |
| Acbd6 | 0.43151892 | 0.27150222 | 0.28460065 | -0.0130984 | 0.62033149 | 0.99999994 |
| Ogdh | 0.21534737 | 0.15366528 | 0.16676859 | -0.0131033 | 0.69951351 | 0.99999994 |
| Chmp2a | 1.20019653 | 0.41589077 | 0.4289943 | -0.0131035 | 0.58055631 | 0.99999994 |
| Enpp5 | 0.10289994 | 0.07393133 | 0.0870355 | -0.0131042 | 0.84187386 | 0.99999994 |

| | | | | | | |
|---|---|---|---|---|---|---|
| Ginm1 | 0.59074486 | 0.3352992 | 0.34840351 | -0.0131043 | 0.5988463 | 0.99999994 |
| Mogs | 0.07754406 | 0.05357551 | 0.06669554 | -0.01312 | 0.90480176 | 0.99999994 |
| Dhrs7b | 0.1241045 | 0.09008976 | 0.10321021 | -0.0131204 | 0.80127376 | 0.99999994 |
| Rab22a | 0.38223749 | 0.24812935 | 0.26125263 | -0.0131233 | 0.63094729 | 0.99999994 |
| B230217C12 | 0.09256272 | 0.06570894 | 0.07884773 | -0.0131388 | 0.86632418 | 0.99999994 |
| Gcgr | 0.06932279 | 0.04679703 | 0.05994294 | -0.0131459 | 0.9278256 | 0.99999994 |
| Araf | 0.47438141 | 0.29028 | 0.30343977 | -0.0131598 | 0.61358582 | 0.99999994 |
| Arhgap5 | 0.35274161 | 0.2331925 | 0.24635784 | -0.0131653 | 0.63897296 | 0.99999994 |
| Ascc1 | 0.32590151 | 0.21900367 | 0.2321709 | -0.0131672 | 0.64710098 | 0.99999994 |
| Hipk3 | 0.0984861 | 0.07040279 | 0.08357657 | -0.0131738 | 0.85295936 | 0.99999994 |
| Sh3glb1 | 0.60129934 | 0.33888501 | 0.3520702 | -0.0131852 | 0.59842926 | 0.99999994 |
| Adprh | 0.4135004 | 0.26308775 | 0.27628012 | -0.0131924 | 0.62469723 | 0.99999994 |
| Rpl6l | 0.5070618 | 0.30373769 | 0.31694439 | -0.0132067 | 0.60924914 | 0.99999994 |
| P3h1 | 0.08390513 | 0.05865786 | 0.07187121 | -0.0132133 | 0.88947771 | 0.99999994 |
| Dpagt1 | 0.13777005 | 0.10012424 | 0.11334108 | -0.0132168 | 0.78118629 | 0.99999994 |
| Mea1 | 0.52928824 | 0.31248234 | 0.32572566 | -0.0132433 | 0.60666354 | 0.99999994 |
| Rps6ka2 | 0.05812949 | 0.03738868 | 0.05063241 | -0.0132437 | 0.95916512 | 0.99999994 |
| Mta1 | 0.10390351 | 0.07455276 | 0.08781371 | -0.013261 | 0.84259786 | 0.99999994 |
| Entpd7 | 0.07963883 | 0.05514162 | 0.06840464 | -0.013263 | 0.90157971 | 0.99999994 |
| Armcx1 | 0.2172529 | 0.15473286 | 0.16799808 | -0.0132652 | 0.70041338 | 0.99999994 |
| Rpia | 0.22095984 | 0.157113 | 0.17037917 | -0.0132662 | 0.69786906 | 0.99999994 |
| Fundc1 | 0.2122776 | 0.15150929 | 0.16478001 | -0.0132707 | 0.70404707 | 0.99999994 |
| Slc39a6 | 0.18988708 | 0.13669543 | 0.14997168 | -0.0132762 | 0.72215464 | 0.99999994 |
| N4bp1 | 0.10370254 | 0.07437692 | 0.08765797 | -0.0132811 | 0.84339381 | 0.99999994 |
| Dag1 | 0.18557959 | 0.13377722 | 0.14706031 | -0.0132831 | 0.72615648 | 0.99999994 |
| Tmem167 | 0.94781614 | 0.40941651 | 0.42270647 | -0.01329 | 0.58288401 | 0.99999994 |
| Gpr45 | 0.04692737 | 0.02787919 | 0.04117695 | -0.0132978 | 0.9842072 | 0.99999994 |
| Tmem42 | 0.15698815 | 0.11390596 | 0.12721162 | -0.0133057 | 0.75685436 | 0.99999994 |
| Dusp3 | 0.09210265 | 0.06517318 | 0.07847894 | -0.0133058 | 0.87045751 | 0.99999994 |
| Ropn1l | 0.09927395 | 0.07089654 | 0.08420285 | -0.0133063 | 0.85357337 | 0.99999994 |

| | | | | | | |
|---|---|---|---|---|---|---|
| Zfp560 | 0.22997647 | 0.16277913 | 0.17611053 | -0.0133314 | 0.69282712 | 0.99999994 |
| Wwp2 | 0.104021 | 0.07456662 | 0.08790474 | -0.0133381 | 0.84375112 | 0.99999994 |
| Specc1l | 0.15457269 | 0.11215048 | 0.12549199 | -0.0133415 | 0.76042226 | 0.99999994 |
| Suco | 0.19140475 | 0.13764925 | 0.15099267 | -0.0133434 | 0.72181135 | 0.99999994 |
| Dlst | 0.19172069 | 0.1378459 | 0.15120491 | -0.013359 | 0.72176457 | 0.99999994 |
| Ipo5 | 0.11524817 | 0.08314858 | 0.09652303 | -0.0133744 | 0.82181909 | 0.99999994 |
| Slain2 | 0.17880343 | 0.12906243 | 0.14243946 | -0.013377 | 0.73405517 | 0.99999994 |
| Mrpl21 | 0.58031102 | 0.33133627 | 0.34471401 | -0.0133777 | 0.60192055 | 0.99999994 |
| Gid8 | 0.26013066 | 0.18128873 | 0.19468122 | -0.0133925 | 0.67648909 | 0.99999994 |
| Pabpc4 | 0.14711147 | 0.10673622 | 0.12013766 | -0.0134014 | 0.77111432 | 0.99999994 |
| Nptn | 0.15007559 | 0.10887097 | 0.1222725 | -0.0134015 | 0.76717108 | 0.99999994 |
| Srpyd3 | 0.13607459 | 0.09868612 | 0.11209643 | -0.0134103 | 0.78707495 | 0.99999994 |
| Cfap126 | 0.10463424 | 0.07496768 | 0.08837955 | -0.0134119 | 0.84378189 | 0.99999994 |
| Tnrc6a | 0.29262478 | 0.20028582 | 0.21370242 | -0.0134166 | 0.6619889 | 0.99999994 |
| Gm37035 | 0.03533306 | 0.01782539 | 0.03124223 | -0.0134168 | 0.99787205 | 0.99999994 |
| Adra2a | 0.16541343 | 0.11973067 | 0.13315688 | -0.0134262 | 0.74894085 | 0.99999994 |
| Eci1 | 0.50441461 | 0.30244892 | 0.31587726 | -0.0134283 | 0.61140012 | 0.99999994 |
| Prkce | 0.11961145 | 0.08639824 | 0.09982964 | -0.0134314 | 0.81475825 | 0.99999994 |
| Mrpl40 | 0.42650572 | 0.26887398 | 0.28231022 | -0.0134362 | 0.624303 | 0.99999994 |
| Arfgef3 | 0.05810831 | 0.03717506 | 0.05061466 | -0.0134396 | 0.9614234 | 0.99999994 |
| Plppr1 | 0.08220652 | 0.05705053 | 0.0704933 | -0.0134428 | 0.89783991 | 0.99999994 |
| Cep170b | 0.07553724 | 0.0516025 | 0.06505385 | -0.0134514 | 0.91564368 | 0.99999994 |
| Prrc2c | 0.84286339 | 0.39704432 | 0.4105008 | -0.0134565 | 0.58636327 | 0.99999994 |
| Rnf145 | 0.18289404 | 0.13177826 | 0.14523498 | -0.0134567 | 0.73129806 | 0.99999994 |
| Gcsh | 0.37855781 | 0.24597579 | 0.25943341 | -0.0134576 | 0.6350603 | 0.99999994 |
| Commd5 | 0.1680287 | 0.12152133 | 0.13498574 | -0.0134644 | 0.74663925 | 0.99999994 |
| Slc43a2 | 0.1814117 | 0.13074119 | 0.14422407 | -0.0134829 | 0.73311913 | 0.99999994 |
| Spa17 | 0.15054869 | 0.10912487 | 0.1226123 | -0.0134874 | 0.76797774 | 0.99999994 |
| Nrbf2 | 0.24377578 | 0.17120587 | 0.18472174 | -0.0135159 | 0.68688421 | 0.99999994 |
| Plcxd2 | 0.06663694 | 0.04419835 | 0.05772123 | -0.0135229 | 0.94055453 | 0.99999994 |

| | | | | | | |
|---|---|---|---|---|---|---|
| Nelfb | 0.27509787 | 0.19004421 | 0.20356717 | -0.013523 | 0.67082104 | 0.99999994 |
| Dcps | 0.49402946 | 0.29811729 | 0.31164524 | -0.0135279 | 0.61368645 | 0.99999994 |
| Tmem11 | 0.5062543 | 0.30308587 | 0.31661935 | -0.0135335 | 0.61199301 | 0.99999994 |
| Nfkb1 | 0.07914542 | 0.05445621 | 0.06800249 | -0.0135463 | 0.90757427 | 0.99999994 |
| Pcbp3 | 0.09468507 | 0.06699973 | 0.0805462 | -0.0135465 | 0.86853295 | 0.99999994 |
| Chkb | 0.24661039 | 0.17291192 | 0.18646686 | -0.0135549 | 0.68576254 | 0.99999994 |
| Taok2 | 0.16700586 | 0.12070005 | 0.13427139 | -0.0135713 | 0.74945777 | 0.99999994 |
| B3galnt1 | 0.07237996 | 0.04887903 | 0.06246238 | -0.0135833 | 0.92612446 | 0.99999994 |
| mt-Cytb | 3.52426012 | 0.24585452 | 0.25943912 | -0.0135846 | 0.63628132 | 0.99999994 |
| Gm5914 | 0.07989165 | 0.05500345 | 0.0686106 | -0.0136071 | 0.90660212 | 0.99999994 |
| Slc30a7 | 0.08449071 | 0.05873719 | 0.07234554 | -0.0136083 | 0.89472868 | 0.99999994 |
| Smap2 | 0.25329939 | 0.17693694 | 0.1905538 | -0.0136169 | 0.68284151 | 0.99999994 |
| Pafah1b3 | 0.64450861 | 0.35275534 | 0.36637257 | -0.0136172 | 0.59770126 | 0.99999994 |
| Fitm2 | 0.14344809 | 0.10386569 | 0.11748462 | -0.0136189 | 0.77982759 | 0.99999994 |
| Ebp | 0.26422653 | 0.18350672 | 0.19713447 | -0.0136278 | 0.67729536 | 0.99999994 |
| Itgb1 | 0.46783394 | 0.28701817 | 0.30064682 | -0.0136287 | 0.61857491 | 0.99999994 |
| Lcmt1 | 0.22983769 | 0.16238719 | 0.17602295 | -0.0136358 | 0.69694705 | 0.99999994 |
| Ten1 | 0.4798653 | 0.29211116 | 0.30575646 | -0.0136453 | 0.61678962 | 0.99999994 |
| Cyb561 | 0.25516949 | 0.17804259 | 0.19168818 | -0.0136456 | 0.68219372 | 0.99999994 |
| Dap3 | 0.34516547 | 0.2287651 | 0.24241615 | -0.0136511 | 0.64610878 | 0.99999994 |
| Trappc4 | 0.70617653 | 0.37003242 | 0.38368859 | -0.0136562 | 0.59363744 | 0.99999994 |
| Itgb3bp | 0.24042082 | 0.16898613 | 0.18264602 | -0.0136599 | 0.69069509 | 0.99999994 |
| Ctnnbl1 | 0.35020329 | 0.23138084 | 0.24504248 | -0.0136616 | 0.64472078 | 0.99999994 |
| Rnaset2a | 0.18858093 | 0.13542603 | 0.149091 | -0.013665 | 0.72909897 | 0.99999994 |
| Nfkbib | 0.168367 | 0.12153531 | 0.13522176 | -0.0136864 | 0.74975605 | 0.99999994 |
| Synj2bp | 0.29372198 | 0.20062593 | 0.2143274 | -0.0137015 | 0.66477338 | 0.99999994 |
| Tln1 | 0.3134119 | 0.21164439 | 0.22535602 | -0.0137116 | 0.65726182 | 0.99999994 |
| Scyl1 | 0.20676922 | 0.14747128 | 0.16118703 | -0.0137158 | 0.71444038 | 0.99999994 |
| Cln6 | 0.20633144 | 0.1471775 | 0.16090003 | -0.0137225 | 0.71487872 | 0.99999994 |
| Mfsd11 | 0.12348867 | 0.08901683 | 0.10274831 | -0.0137315 | 0.81315119 | 0.99999994 |

| | | | | | | |
|---|---|---|---|---|---|---|
| Hexa | 0.4026958 | 0.25743863 | 0.27117273 | -0.0137341 | 0.63198866 | 0.99999994 |
| Shisa5 | 0.39078761 | 0.25170431 | 0.26543903 | -0.0137347 | 0.63473576 | 0.99999994 |
| Emd | 0.41827279 | 0.26475789 | 0.27850778 | -0.0137499 | 0.62877981 | 0.99999994 |
| Gnl3l | 0.14424265 | 0.10430871 | 0.11806142 | -0.0137527 | 0.78094371 | 0.99999994 |
| Yipf4 | 0.82899081 | 0.39458217 | 0.40834121 | -0.013759 | 0.58873251 | 0.99999994 |
| Fbxw2 | 0.32020932 | 0.21530676 | 0.22908188 | -0.0137751 | 0.65551727 | 0.99999994 |
| Ccng1 | 0.10204416 | 0.07259547 | 0.08637089 | -0.0137754 | 0.8557988 | 0.99999994 |
| Rps19bp1 | 0.42913874 | 0.26973664 | 0.28351547 | -0.0137788 | 0.62684113 | 0.99999994 |
| Kctd13 | 0.3186547 | 0.2144297 | 0.22823337 | -0.0138037 | 0.65637133 | 0.99999994 |
| Nbr1 | 0.21448067 | 0.15238983 | 0.16620813 | -0.0138183 | 0.7100052 | 0.99999994 |
| 2310030G06 | 0.24793257 | 0.17345577 | 0.18727817 | -0.0138224 | 0.68838739 | 0.99999994 |
| Tnr | 0.03965829 | 0.0211362 | 0.03496621 | -0.01383 | 0.99576524 | 0.99999994 |
| Gm10941 | 0.16821903 | 0.12128357 | 0.13511854 | -0.013835 | 0.75223937 | 0.99999994 |
| Tmem167b | 0.13868604 | 0.10017702 | 0.1140121 | -0.0138351 | 0.79037757 | 0.99999994 |
| Ehd1 | 0.22512996 | 0.15919754 | 0.17304033 | -0.0138428 | 0.70280733 | 0.99999994 |
| Bhlha15 | 0.05067469 | 0.03050111 | 0.04435549 | -0.0138544 | 0.98119618 | 0.99999994 |
| Fh1 | 0.35997597 | 0.23621997 | 0.25007775 | -0.0138578 | 0.64388193 | 0.99999994 |
| Dhx32 | 0.11576418 | 0.0830514 | 0.09691531 | -0.0138639 | 0.82950325 | 0.99999994 |
| Usp33 | 0.14031321 | 0.10132667 | 0.11520165 | -0.013875 | 0.78863592 | 0.99999994 |
| Atp9a | 0.1775248 | 0.12768678 | 0.14156187 | -0.0138751 | 0.74293441 | 0.99999994 |
| Dpm2 | 0.38769965 | 0.25004263 | 0.26393361 | -0.013891 | 0.6369537 | 0.99999994 |
| Ppp6c | 0.38550925 | 0.24894921 | 0.26286129 | -0.0139121 | 0.63769021 | 0.99999994 |
| Fbxl16 | 0.1424393 | 0.10283622 | 0.11675123 | -0.013915 | 0.78620974 | 0.99999994 |
| Alg2 | 0.46361866 | 0.28491611 | 0.2988323 | -0.0139162 | 0.62171081 | 0.99999994 |
| Smc3 | 0.68752929 | 0.36504996 | 0.37897544 | -0.0139255 | 0.59661243 | 0.99999994 |
| Comt | 0.53117141 | 0.31252668 | 0.32645524 | -0.0139286 | 0.61179501 | 0.99999994 |
| Cuedc2 | 0.47401372 | 0.28935493 | 0.30328372 | -0.0139288 | 0.62008494 | 0.99999994 |
| Gnl1 | 0.20623054 | 0.14687955 | 0.16083385 | -0.0139543 | 0.71820598 | 0.99999994 |
| Ampd2 | 0.11055485 | 0.07898057 | 0.09293943 | -0.0139589 | 0.84128148 | 0.99999994 |
| Mtmr7 | 0.20455035 | 0.14576309 | 0.15973028 | -0.0139672 | 0.71973266 | 0.99999994 |

| | | | | | | |
|---|---|---|---|---|---|---|
| Slc16a10 | 0.14444967 | 0.10423998 | 0.11821158 | -0.0139716 | 0.78427857 | 0.99999994 |
| Sucla2 | 0.55098106 | 0.32000694 | 0.33399353 | -0.0139866 | 0.6097806 | 0.99999994 |
| Naa10 | 0.31300873 | 0.21114096 | 0.22513367 | -0.0139927 | 0.66045746 | 0.99999994 |
| Fasn | 0.17147721 | 0.12338794 | 0.13738556 | -0.0139976 | 0.75117361 | 0.99999994 |
| Dpp9 | 0.13760025 | 0.0992137 | 0.11321658 | -0.0140029 | 0.7948237 | 0.99999994 |
| Meox1 | 0.05807899 | 0.036571 | 0.05059009 | -0.0140191 | 0.96744871 | 0.99999994 |
| Ift27 | 0.28166752 | 0.19337624 | 0.20740022 | -0.014024 | 0.67367696 | 0.99999994 |
| Psmd1 | 0.5769149 | 0.32947232 | 0.34349893 | -0.0140266 | 0.60711555 | 0.99999994 |
| Skap1 | 0.05802664 | 0.03651887 | 0.05054622 | -0.0140274 | 0.9676439 | 0.99999994 |
| Elp5 | 0.31745531 | 0.21354099 | 0.2275773 | -0.0140363 | 0.65929691 | 0.99999994 |
| Ywhag | 0.41312837 | 0.26206636 | 0.27610575 | -0.0140394 | 0.63249236 | 0.99999994 |
| Tpr | 0.87086162 | 0.40046867 | 0.41451083 | -0.0140422 | 0.58920246 | 0.99999994 |
| Abcb9 | 0.20296427 | 0.14462507 | 0.15868579 | -0.0140607 | 0.72233836 | 0.99999994 |
| Dusp19 | 0.1459318 | 0.10521555 | 0.11928519 | -0.0140696 | 0.78380688 | 0.99999994 |
| Dld | 0.35508305 | 0.2334932 | 0.24756648 | -0.0140733 | 0.64742967 | 0.99999994 |
| Tmem30a | 0.50346174 | 0.30141868 | 0.31549204 | -0.0140734 | 0.61673795 | 0.99999994 |
| Lrp10 | 0.23991448 | 0.1682584 | 0.18233179 | -0.0140734 | 0.69629536 | 0.99999994 |
| Msrb1 | 0.21297931 | 0.1511595 | 0.16523543 | -0.0140759 | 0.71465517 | 0.99999994 |
| Mtcl1 | 0.10162315 | 0.07196003 | 0.08604359 | -0.0140836 | 0.86203504 | 0.99999994 |
| Cstb | 1.25600024 | 0.41299653 | 0.42708195 | -0.0140854 | 0.58688171 | 0.99999994 |
| Pdcd6ip | 0.28955521 | 0.19785917 | 0.21194814 | -0.014089 | 0.67092635 | 0.99999994 |
| Rap1gap | 0.09469196 | 0.06645789 | 0.0805517 | -0.0140938 | 0.87791693 | 0.99999994 |
| Tmem126a | 0.47182496 | 0.28824887 | 0.30235287 | -0.014104 | 0.62191047 | 0.99999994 |
| Ccnh | 0.29586449 | 0.20142639 | 0.21554467 | -0.0141183 | 0.66857978 | 0.99999994 |
| Clptm1l | 0.28460479 | 0.19496418 | 0.20910084 | -0.0141367 | 0.67365043 | 0.99999994 |
| Zc3h3 | 0.19469677 | 0.13904575 | 0.15319819 | -0.0141524 | 0.73069417 | 0.99999994 |
| Cct8 | 1.24580597 | 0.41334197 | 0.42750054 | -0.0141586 | 0.58724152 | 0.99999994 |
| Aak1 | 0.125552 | 0.09012724 | 0.10429406 | -0.0141668 | 0.8170471 | 0.99999994 |
| E2f4 | 0.22413229 | 0.15823005 | 0.17240537 | -0.0141753 | 0.70791082 | 0.99999994 |
| Rab37 | 0.11070242 | 0.0788756 | 0.09305253 | -0.0141769 | 0.844738 | 0.99999994 |

| | | | | | | |
|---|---|---|---|---|---|---|
| Lrrfip2 | 0.17992746 | 0.12902377 | 0.14320946 | -0.0141857 | 0.74516408 | 0.99999994 |
| Gtf2a2 | 1.11947691 | 0.41573415 | 0.4299229 | -0.0141887 | 0.58693975 | 0.99999994 |
| Ank | 0.25325798 | 0.17633857 | 0.19052864 | -0.0141901 | 0.68996388 | 0.99999994 |
| Mlx | 0.28142211 | 0.1930654 | 0.20725776 | -0.0141924 | 0.67573541 | 0.99999994 |
| Ube2a | 0.49007546 | 0.29581593 | 0.31001514 | -0.0141992 | 0.61976833 | 0.99999994 |
| Rilpl2 | 0.15014629 | 0.10811372 | 0.1223233 | -0.0142096 | 0.78030696 | 0.99999994 |
| Ergic2 | 0.74464846 | 0.37821 | 0.39243056 | -0.0142206 | 0.59530282 | 0.99999994 |
| Unc80 | 0.11300457 | 0.08057679 | 0.09481333 | -0.0142365 | 0.84121758 | 0.99999994 |
| Ikbip | 0.09015755 | 0.0626719 | 0.07691735 | -0.0142455 | 0.89117264 | 0.99999994 |
| 4931406C07I | 0.11794752 | 0.08431804 | 0.09857144 | -0.0142534 | 0.83210257 | 0.99999994 |
| Sptlc1 | 0.15830786 | 0.1138902 | 0.12814832 | -0.0142581 | 0.7705333 | 0.99999994 |
| Agt | 0.11243968 | 0.08011998 | 0.09438188 | -0.0142619 | 0.84275182 | 0.99999994 |
| Vapa | 1.09735865 | 0.41544656 | 0.42971412 | -0.0142676 | 0.58745665 | 0.99999994 |
| Parva | 0.18360751 | 0.1314467 | 0.14572068 | -0.014274 | 0.74280571 | 0.99999994 |
| Cnot8 | 0.23168425 | 0.16290334 | 0.17718673 | -0.0142834 | 0.70422212 | 0.99999994 |
| Pank3 | 0.11488054 | 0.08195916 | 0.09624331 | -0.0142842 | 0.83840339 | 0.99999994 |
| Copz2 | 0.09075289 | 0.06310944 | 0.07739573 | -0.0142863 | 0.89040419 | 0.99999994 |
| Rab8a | 0.49666721 | 0.2984153 | 0.312727 | -0.0143117 | 0.6196733 | 0.99999994 |
| Tgoln1 | 0.35325111 | 0.23227242 | 0.24662122 | -0.0143488 | 0.65072443 | 0.99999994 |
| Cish | 0.07790565 | 0.05263098 | 0.06699089 | -0.0143599 | 0.92318574 | 0.99999994 |
| Dnah9 | 0.09404055 | 0.06566214 | 0.08003089 | -0.0143687 | 0.88397264 | 0.99999994 |
| Tmem14c | 0.74629534 | 0.37842104 | 0.39279117 | -0.0143701 | 0.59619896 | 0.99999994 |
| Mrps21 | 1.13206284 | 0.41555523 | 0.42993497 | -0.0143797 | 0.58808847 | 0.99999994 |
| Rusc1 | 0.08029349 | 0.05455593 | 0.06893782 | -0.0143819 | 0.91753161 | 0.99999994 |
| 2510002D24I | 0.477497 | 0.29037594 | 0.30475844 | -0.0143825 | 0.62329048 | 0.99999994 |
| Ildr1 | 0.19319998 | 0.13780701 | 0.15219689 | -0.0143899 | 0.73543807 | 0.99999994 |
| Nudt2 | 0.14809836 | 0.10645358 | 0.12084958 | -0.014396 | 0.78608995 | 0.99999994 |
| Ankrd33b | 0.04544716 | 0.02551721 | 0.03991708 | -0.0143999 | 0.9918303 | 0.99999994 |
| Gfpt1 | 0.44737816 | 0.27732331 | 0.29172491 | -0.0144016 | 0.62877272 | 0.99999994 |
| Akap11 | 0.15255034 | 0.10964527 | 0.12404708 | -0.0144018 | 0.7801796 | 0.99999994 |

| | | | | | | |
|---|---|---|---|---|---|---|
| Tmem205 | 0.31971083 | 0.21440687 | 0.22881004 | -0.0144032 | 0.66239389 | 0.99999994 |
| Lyve1 | 0.10993823 | 0.07806107 | 0.09246657 | -0.0144055 | 0.85013294 | 0.99999994 |
| Nars | 0.38894509 | 0.25011328 | 0.26454167 | -0.0144284 | 0.64171436 | 0.99999994 |
| Pum2 | 0.31701388 | 0.21287413 | 0.22733553 | -0.0144614 | 0.66400856 | 0.99999994 |
| Ppp1r2 | 0.58691669 | 0.33259522 | 0.34705701 | -0.0144618 | 0.60925165 | 0.99999994 |
| Pdpk1 | 0.28506985 | 0.1949049 | 0.20936928 | -0.0144644 | 0.67719082 | 0.99999994 |
| Helz | 0.27306575 | 0.18790492 | 0.20237339 | -0.0144685 | 0.68292336 | 0.99999994 |
| Sstr3 | 0.05070196 | 0.0298973 | 0.04437856 | -0.0144813 | 0.98507403 | 0.99999994 |
| Plxnb2 | 0.20461805 | 0.14528922 | 0.1597748 | -0.0144856 | 0.72690596 | 0.99999994 |
| Aimp2 | 0.27775245 | 0.19062539 | 0.2051208 | -0.0144954 | 0.68096159 | 0.99999994 |
| Eif3m | 0.95358195 | 0.40865589 | 0.42315923 | -0.0145033 | 0.59023117 | 0.99999994 |
| Uck2 | 0.22270004 | 0.15698424 | 0.17149204 | -0.0145078 | 0.71332052 | 0.99999994 |
| Irf2bp2 | 0.1797906 | 0.12860339 | 0.14311578 | -0.0145124 | 0.75015429 | 0.99999994 |
| Thoc7 | 0.87262426 | 0.40022953 | 0.41474475 | -0.0145152 | 0.59210336 | 0.99999994 |
| 2610528J11F | 0.19511178 | 0.13895348 | 0.15347539 | -0.0145219 | 0.73559638 | 0.99999994 |
| Zfp787 | 0.25941125 | 0.17971206 | 0.19424867 | -0.0145366 | 0.69080337 | 0.99999994 |
| Alg5 | 0.22583098 | 0.15894239 | 0.17348589 | -0.0145435 | 0.71158101 | 0.99999994 |
| Zcchc9 | 0.30277367 | 0.20489406 | 0.21944107 | -0.014547 | 0.67048688 | 0.99999994 |
| Eef1d | 1.13534711 | 0.41537344 | 0.42993098 | -0.0145575 | 0.58916035 | 0.99999994 |
| Slc39a7 | 0.55234277 | 0.31993192 | 0.33450268 | -0.0145708 | 0.61407178 | 0.99999994 |
| Nck1 | 0.32463669 | 0.21689648 | 0.23148692 | -0.0145904 | 0.66259104 | 0.99999994 |
| Chuk | 0.17155244 | 0.12284398 | 0.13743777 | -0.0145938 | 0.76015557 | 0.99999994 |
| Mocs2 | 0.51145613 | 0.30410752 | 0.31870591 | -0.0145984 | 0.61977725 | 0.99999994 |
| Vdac1 | 0.78025771 | 0.38516414 | 0.39977167 | -0.0146075 | 0.59608298 | 0.99999994 |
| Ddx3x | 0.46686287 | 0.28559908 | 0.30022989 | -0.0146308 | 0.62717398 | 0.99999994 |
| Nbas | 0.09072103 | 0.06273289 | 0.07737013 | -0.0146372 | 0.89603547 | 0.99999994 |
| Wnk1 | 0.35670061 | 0.23376049 | 0.24839884 | -0.0146383 | 0.65256867 | 0.99999994 |
| B230219D22 | 0.58313864 | 0.33106905 | 0.34572 | -0.014651 | 0.61106034 | 0.99999994 |
| Gtf2i | 0.39871366 | 0.25461136 | 0.26926749 | -0.0146561 | 0.6414372 | 0.99999994 |
| Tomm70a | 0.2493335 | 0.17346284 | 0.18813594 | -0.0146731 | 0.69816943 | 0.99999994 |

| | | | | | | |
|---|---|---|---|---|---|---|
| Clta | 1.52892051 | 0.39134972 | 0.4060301 | -0.0146804 | 0.59509322 | 0.99999994 |
| Fndc3a | 0.75838858 | 0.38067951 | 0.395371 | -0.0146915 | 0.59767815 | 0.99999994 |
| Hspa4 | 0.50021745 | 0.29948051 | 0.31417582 | -0.0146953 | 0.62223106 | 0.99999994 |
| Utrn | 0.08033963 | 0.05426141 | 0.06897539 | -0.014714 | 0.92218872 | 0.99999994 |
| Fam53a | 0.17443521 | 0.12471201 | 0.13943345 | -0.0147214 | 0.75890387 | 0.99999994 |
| Slc35c2 | 0.17741687 | 0.12675699 | 0.14148771 | -0.0147307 | 0.75584962 | 0.99999994 |
| Qsox2 | 0.15153843 | 0.10857637 | 0.12332233 | -0.014746 | 0.78695991 | 0.99999994 |
| Tmod2 | 0.11351046 | 0.08045173 | 0.09519938 | -0.0147476 | 0.84876671 | 0.99999994 |
| Safb2 | 0.3107036 | 0.20909615 | 0.22385964 | -0.0147635 | 0.66966388 | 0.99999994 |
| Syt7 | 0.33374266 | 0.22156488 | 0.23637959 | -0.0148147 | 0.66172535 | 0.99999994 |
| Stt3b | 0.18513864 | 0.13193042 | 0.14676114 | -0.0148307 | 0.74941442 | 0.99999994 |
| Sumo2 | 1.9996977 | 0.35590564 | 0.37075555 | -0.0148499 | 0.60511443 | 0.99999994 |
| Wdr89 | 0.82118226 | 0.39228461 | 0.40713592 | -0.0148513 | 0.59592272 | 0.99999994 |
| Chmp3 | 0.28162556 | 0.1925245 | 0.20737587 | -0.0148514 | 0.68321194 | 0.99999994 |
| Ncoa1 | 0.11024846 | 0.07785254 | 0.09270454 | -0.014852 | 0.85687869 | 0.99999994 |
| 2210016F16l | 0.28851437 | 0.19648394 | 0.21135134 | -0.0148674 | 0.68019284 | 0.99999994 |
| Gnpnat1 | 0.28754895 | 0.19592732 | 0.21079691 | -0.0148696 | 0.68065766 | 0.99999994 |
| Endog | 0.1569771 | 0.11233265 | 0.12720376 | -0.0148711 | 0.78176865 | 0.99999994 |
| Azin1 | 0.2509644 | 0.17425485 | 0.18913211 | -0.0148773 | 0.69971936 | 0.99999994 |
| Banp | 0.37794199 | 0.24424583 | 0.25912791 | -0.0148821 | 0.6488746 | 0.99999994 |
| Mrps23 | 0.32522061 | 0.21691012 | 0.23180286 | -0.0148927 | 0.66554848 | 0.99999994 |
| Herc2 | 0.19661646 | 0.13957346 | 0.15447889 | -0.0149054 | 0.73964844 | 0.99999994 |
| Thyn1 | 0.33061868 | 0.21980279 | 0.23470953 | -0.0149067 | 0.66376298 | 0.99999994 |
| Mrps18c | 0.64494525 | 0.35160145 | 0.36651038 | -0.0149089 | 0.60671511 | 0.99999994 |
| Klhl9 | 0.2512918 | 0.17442179 | 0.18933177 | -0.01491 | 0.69992746 | 0.99999994 |
| Esrp1 | 0.19694064 | 0.13978168 | 0.15469477 | -0.0149131 | 0.73946438 | 0.99999994 |
| Trub1 | 0.10909131 | 0.07688868 | 0.09181634 | -0.0149277 | 0.86041873 | 0.99999994 |
| Tspyl2 | 0.14014264 | 0.10014897 | 0.1150771 | -0.0149281 | 0.80606362 | 0.99999994 |
| Cmbl | 0.07401804 | 0.04886923 | 0.06380823 | -0.014939 | 0.9404318 | 0.99999994 |
| Chic1 | 0.27775197 | 0.19017348 | 0.20512052 | -0.014947 | 0.6861781 | 0.99999994 |

| | | | | | | |
|---|---|---|---|---|---|---|
| Ola1 | 0.41294706 | 0.26107195 | 0.27602074 | -0.0149488 | 0.64076202 | 0.99999994 |
| Ddx24 | 0.61960139 | 0.34330714 | 0.35827127 | -0.0149641 | 0.60950149 | 0.99999994 |
| Fbxo9 | 0.25687415 | 0.17774528 | 0.19271926 | -0.014974 | 0.69748765 | 0.99999994 |
| Snap25 | 0.26263047 | 0.18120594 | 0.1961804 | -0.0149745 | 0.69430149 | 0.99999994 |
| Sun1 | 0.13784876 | 0.09841935 | 0.11339878 | -0.0149794 | 0.81037336 | 0.99999994 |
| Rfesd | 0.16463187 | 0.11761236 | 0.13260884 | -0.0149965 | 0.77420133 | 0.99999994 |
| Wdr74 | 0.20717272 | 0.14644118 | 0.16145138 | -0.0150102 | 0.73199054 | 0.99999994 |
| Rrp7a | 0.28434671 | 0.19394037 | 0.20895178 | -0.0150114 | 0.68374709 | 0.99999994 |
| Slc50a1 | 0.31989559 | 0.21389915 | 0.22891082 | -0.0150117 | 0.66876855 | 0.99999994 |
| Adgrl1 | 0.11726243 | 0.08303085 | 0.09805241 | -0.0150216 | 0.84607628 | 0.99999994 |
| Abracl | 0.6574224 | 0.35537981 | 0.37040609 | -0.0150263 | 0.6064309 | 0.99999994 |
| Pop7 | 0.39804887 | 0.25392085 | 0.26894825 | -0.0150274 | 0.64502455 | 0.99999994 |
| Sepsecs | 0.10504956 | 0.0736727 | 0.08870086 | -0.0150282 | 0.87028823 | 0.99999994 |
| Efcab1 | 0.54355122 | 0.31616616 | 0.33119468 | -0.0150285 | 0.61869052 | 0.99999994 |
| Rims3 | 0.05871311 | 0.03607091 | 0.05112123 | -0.0150503 | 0.97498236 | 0.99999994 |
| Psmc1 | 0.82337564 | 0.39240168 | 0.40747734 | -0.0150757 | 0.59726295 | 0.99999994 |
| Eif1a | 0.42020465 | 0.26432704 | 0.27940475 | -0.0150777 | 0.64027792 | 0.99999994 |
| Rbm42 | 0.66165342 | 0.35659133 | 0.37168997 | -0.0150986 | 0.60657058 | 0.99999994 |
| Snx14 | 0.19835927 | 0.14053599 | 0.15563817 | -0.0151022 | 0.74082546 | 0.99999994 |
| Bub3 | 0.58907955 | 0.33270505 | 0.34781878 | -0.0151137 | 0.61380041 | 0.99999994 |
| Ppp6r3 | 0.13352084 | 0.09509082 | 0.11021524 | -0.0151244 | 0.81949087 | 0.99999994 |
| Zmiz1 | 0.1774545 | 0.12638629 | 0.14151357 | -0.0151273 | 0.76162522 | 0.99999994 |
| Spg7 | 0.21619548 | 0.15218772 | 0.16731627 | -0.0151285 | 0.72636644 | 0.99999994 |
| Ddx49 | 0.17982822 | 0.12800061 | 0.14314153 | -0.0151409 | 0.75930915 | 0.99999994 |
| Ranbp2 | 0.16752758 | 0.11949307 | 0.13463591 | -0.0151428 | 0.77296806 | 0.99999994 |
| Cul4a | 0.2433754 | 0.16932696 | 0.1844746 | -0.0151476 | 0.70766268 | 0.99999994 |
| Rmnd5b | 0.132788 | 0.09452601 | 0.10967397 | -0.015148 | 0.82104728 | 0.99999994 |
| Bnip1 | 0.17294644 | 0.12324075 | 0.13840395 | -0.0151632 | 0.76707697 | 0.99999994 |
| BC029722 | 0.14579947 | 0.10402287 | 0.11918944 | -0.0151666 | 0.8015039 | 0.99999994 |
| Mthfd2 | 0.1877241 | 0.13333834 | 0.14851227 | -0.0151739 | 0.75177924 | 0.99999994 |

| | | | | | | |
|---|---|---|---|---|---|---|
| Efcab14 | 0.21707813 | 0.15269296 | 0.16788547 | -0.0151925 | 0.72653125 | 0.99999994 |
| Rbm25 | 0.98825095 | 0.41047586 | 0.42567017 | -0.0151943 | 0.59390387 | 0.99999994 |
| Tmod1 | 0.06591882 | 0.04192262 | 0.05712589 | -0.0152033 | 0.96176106 | 0.99999994 |
| Gsk3b | 0.58846652 | 0.332395 | 0.34760314 | -0.0152081 | 0.61456051 | 0.99999994 |
| Mbtps1 | 0.15701788 | 0.1120143 | 0.12723274 | -0.0152184 | 0.78703448 | 0.99999994 |
| Rdx | 0.59954051 | 0.3362341 | 0.3514638 | -0.0152297 | 0.61349299 | 0.99999994 |
| Prkca | 0.09837118 | 0.06825287 | 0.08348517 | -0.0152323 | 0.88771899 | 0.99999994 |
| Sub1 | 2.12893316 | 0.35321341 | 0.36845027 | -0.0152369 | 0.60844417 | 0.99999994 |
| Vegfa | 0.2053955 | 0.14502722 | 0.16028576 | -0.0152585 | 0.73684785 | 0.99999994 |
| Bckdhb | 0.11499861 | 0.08107458 | 0.09633316 | -0.0152586 | 0.85413317 | 0.99999994 |
| Dhdds | 0.17176135 | 0.12230141 | 0.1375827 | -0.0152813 | 0.77014374 | 0.99999994 |
| Cdk16 | 0.24202054 | 0.16833868 | 0.18363716 | -0.0152985 | 0.71039159 | 0.99999994 |
| Gaa | 0.27429491 | 0.18779269 | 0.20309593 | -0.0153032 | 0.69200985 | 0.99999994 |
| Bet1 | 0.53343303 | 0.31201747 | 0.32732849 | -0.015311 | 0.62223493 | 0.99999994 |
| Gnai3 | 0.46732279 | 0.28511277 | 0.30042744 | -0.0153147 | 0.63281422 | 0.99999994 |
| Ap2a2 | 0.23433051 | 0.16351629 | 0.17884856 | -0.0153323 | 0.71587609 | 0.99999994 |
| Hbs1l | 0.25313885 | 0.17511399 | 0.19045626 | -0.0153423 | 0.7040916 | 0.99999994 |
| Dusp11 | 0.25912244 | 0.17873022 | 0.19407488 | -0.0153447 | 0.7006565 | 0.99999994 |
| Fam20b | 0.12506622 | 0.08858388 | 0.1039306 | -0.0153467 | 0.83717142 | 0.99999994 |
| Ptgr2 | 0.21027852 | 0.14812617 | 0.16347975 | -0.0153536 | 0.7340629 | 0.99999994 |
| Acp1 | 0.63613218 | 0.3483321 | 0.36369386 | -0.0153618 | 0.61070325 | 0.99999994 |
| Tmed7 | 0.26484674 | 0.18213838 | 0.19750456 | -0.0153662 | 0.69772743 | 0.99999994 |
| Nat9 | 0.1949467 | 0.13798534 | 0.15336515 | -0.0153798 | 0.74777834 | 0.99999994 |
| Sfxn1 | 0.31097472 | 0.2086299 | 0.22400973 | -0.0153798 | 0.67616644 | 0.99999994 |
| Atf5 | 0.18695942 | 0.13261413 | 0.14799511 | -0.015381 | 0.75546487 | 0.99999994 |
| Coa6 | 0.26928224 | 0.18472912 | 0.20014045 | -0.0154113 | 0.69586927 | 0.99999994 |
| Tpp2 | 0.20974135 | 0.14768165 | 0.16312966 | -0.015448 | 0.73576456 | 0.99999994 |
| Yipf6 | 0.1789514 | 0.1270885 | 0.14254091 | -0.0154524 | 0.76472809 | 0.99999994 |
| 1110065P20I | 0.52434131 | 0.30833226 | 0.32379863 | -0.0154664 | 0.62473615 | 0.99999994 |
| 0610040J01F | 0.35292133 | 0.23098294 | 0.24645077 | -0.0154678 | 0.66194186 | 0.99999994 |

| | | | | | | |
|---|---|---|---|---|---|---|
| Far1 | 0.13070635 | 0.09265175 | 0.10813295 | -0.0154812 | 0.82969961 | 0.99999994 |
| Actn3 | 0.18051796 | 0.12811456 | 0.1436134 | -0.0154988 | 0.76373181 | 0.99999994 |
| Dnaja3 | 0.21177477 | 0.14895406 | 0.16445334 | -0.0154993 | 0.73478311 | 0.99999994 |
| Ralbp1 | 0.75086283 | 0.37825497 | 0.39378281 | -0.0155278 | 0.60352307 | 0.99999994 |
| Meig1 | 0.06592264 | 0.04159198 | 0.05712906 | -0.0155371 | 0.96487491 | 0.99999994 |
| Coasy | 0.21956346 | 0.15390719 | 0.16948388 | -0.0155767 | 0.7296528 | 0.99999994 |
| Gnptg | 0.54163387 | 0.31488567 | 0.33046684 | -0.0155812 | 0.62317893 | 0.99999994 |
| Ktn1 | 0.4659313 | 0.28424222 | 0.29982931 | -0.0155871 | 0.63534412 | 0.99999994 |
| Vps13c | 0.09948398 | 0.06877047 | 0.0843697 | -0.0155992 | 0.89078309 | 0.99999994 |
| Kpnb1 | 0.23761619 | 0.16528872 | 0.1809022 | -0.0156135 | 0.71718634 | 0.99999994 |
| Yipf2 | 0.25265736 | 0.17454829 | 0.19016345 | -0.0156152 | 0.70766729 | 0.99999994 |
| Syne4 | 0.15668455 | 0.11137646 | 0.12699585 | -0.0156194 | 0.79351382 | 0.99999994 |
| Mthfd1 | 0.14468899 | 0.10274622 | 0.11838511 | -0.0156389 | 0.81038528 | 0.99999994 |
| Fastk | 0.25786654 | 0.17765013 | 0.19331823 | -0.0156681 | 0.70522422 | 0.99999994 |
| Sec13 | 0.7053778 | 0.36782466 | 0.38349593 | -0.0156713 | 0.60719173 | 0.99999994 |
| Atp6v1c1 | 0.51819101 | 0.30570943 | 0.32138154 | -0.0156721 | 0.62725653 | 0.99999994 |
| Trpc4ap | 0.23934625 | 0.16630571 | 0.18197883 | -0.0156731 | 0.71677452 | 0.99999994 |
| Ppm1g | 0.43396422 | 0.27003773 | 0.28571127 | -0.0156735 | 0.64250189 | 0.99999994 |
| Purb | 0.25099466 | 0.17346923 | 0.18915057 | -0.0156813 | 0.70947052 | 0.99999994 |
| Dnajc4 | 0.1926224 | 0.13612231 | 0.15180986 | -0.0156876 | 0.75422549 | 0.99999994 |
| Rnps1 | 0.56617892 | 0.32389688 | 0.3396131 | -0.0157162 | 0.62097222 | 0.99999994 |
| Dap | 0.65145457 | 0.35283071 | 0.36855291 | -0.0157222 | 0.61177774 | 0.99999994 |
| Pbdc1 | 0.51454871 | 0.30420897 | 0.31993824 | -0.0157293 | 0.62826039 | 0.99999994 |
| 2900093K20I | 0.13217658 | 0.09348394 | 0.10922188 | -0.0157379 | 0.83125187 | 0.99999994 |
| Prkar2b | 0.10514729 | 0.07303182 | 0.08877643 | -0.0157446 | 0.88110187 | 0.99999994 |
| Hibadh | 0.41762518 | 0.26244924 | 0.27820648 | -0.0157572 | 0.64691578 | 0.99999994 |
| Vps29 | 0.81677803 | 0.39068105 | 0.40644359 | -0.0157625 | 0.60185223 | 0.99999994 |
| Arfgef1 | 0.19929626 | 0.14049648 | 0.1562601 | -0.0157636 | 0.74904819 | 0.99999994 |
| Yipf3 | 0.48811065 | 0.2934275 | 0.3092013 | -0.0157738 | 0.63291047 | 0.99999994 |
| R3hdm1 | 0.41534762 | 0.2613682 | 0.27714437 | -0.0157762 | 0.64762138 | 0.99999994 |

| | | | | | | |
|---|---|---|---|---|---|---|
| Ndfip1 | 0.26182674 | 0.17992104 | 0.19569904 | -0.015778 | 0.70424905 | 0.99999994 |
| Aig1 | 0.43140795 | 0.2687676 | 0.28455015 | -0.0157825 | 0.64400852 | 0.99999994 |
| Cpsf6 | 0.40103656 | 0.25459098 | 0.27038034 | -0.0157894 | 0.65125474 | 0.99999994 |
| Prepl | 0.106487 | 0.07401668 | 0.08981122 | -0.0157945 | 0.87912262 | 0.99999994 |
| Mrpl42 | 0.71773438 | 0.37061918 | 0.3864383 | -0.0158191 | 0.60737448 | 0.99999994 |
| Mpp3 | 0.1291117 | 0.09112283 | 0.10694871 | -0.0158259 | 0.83768388 | 0.99999994 |
| Mirg | 0.09724662 | 0.06675229 | 0.08259001 | -0.0158377 | 0.89910078 | 0.99999994 |
| Khk | 0.12665758 | 0.08927536 | 0.10512019 | -0.0158448 | 0.84214605 | 0.99999994 |
| Ppp2r5c | 0.14210879 | 0.10062717 | 0.11651069 | -0.0158835 | 0.81791276 | 0.99999994 |
| Gpx4 | 1.61976239 | 0.38084129 | 0.39674575 | -0.0159045 | 0.60520205 | 0.99999994 |
| Nav1 | 0.1703318 | 0.12068021 | 0.13658995 | -0.0159097 | 0.7809246 | 0.99999994 |
| Agl | 0.12342586 | 0.08679138 | 0.10270117 | -0.0159098 | 0.84876699 | 0.99999994 |
| Dus1l | 0.1498298 | 0.10616445 | 0.12209587 | -0.0159314 | 0.80745073 | 0.99999994 |
| Sqstm1 | 0.66072905 | 0.35547286 | 0.37141027 | -0.0159374 | 0.61241842 | 0.99999994 |
| Dohh | 0.5570126 | 0.32030152 | 0.33624035 | -0.0159388 | 0.6238211 | 0.99999994 |
| Chchd4 | 0.33218992 | 0.21960793 | 0.23555054 | -0.0159426 | 0.67383141 | 0.99999994 |
| Farsa | 0.46375767 | 0.28294827 | 0.29889233 | -0.0159441 | 0.63873355 | 0.99999994 |
| Aqp11 | 0.1108835 | 0.0772437 | 0.09319126 | -0.0159476 | 0.87266601 | 0.99999994 |
| Pithd1 | 0.3035676 | 0.20392128 | 0.21988596 | -0.0159647 | 0.68555175 | 0.99999994 |
| Rsrp1 | 1.38332131 | 0.40372091 | 0.41969121 | -0.0159703 | 0.59997831 | 0.99999994 |
| Strap | 0.60788268 | 0.3383535 | 0.35432475 | -0.0159713 | 0.6179254 | 0.99999994 |
| Laptm4b | 0.38534264 | 0.2467891 | 0.26277958 | -0.0159905 | 0.65727889 | 0.99999994 |
| Mrps28 | 0.46378519 | 0.28290103 | 0.29890422 | -0.0160032 | 0.6392213 | 0.99999994 |
| Iqgap1 | 0.42094052 | 0.26370035 | 0.2797457 | -0.0160453 | 0.64869073 | 0.99999994 |
| Rab3gap2 | 0.09355794 | 0.06358787 | 0.07964476 | -0.0160569 | 0.91019622 | 0.99999994 |
| Lrrc73 | 0.10492814 | 0.07254006 | 0.08860694 | -0.0160669 | 0.88628039 | 0.99999994 |
| Frzb | 0.15599739 | 0.11043934 | 0.12650708 | -0.0160677 | 0.80106399 | 0.99999994 |
| Elp3 | 0.15584962 | 0.11033309 | 0.1264019 | -0.0160688 | 0.8012761 | 0.99999994 |
| Fbl | 0.54075902 | 0.31406318 | 0.330134 | -0.0160708 | 0.62704219 | 0.99999994 |
| Ppil6 | 0.09569322 | 0.06526779 | 0.08135138 | -0.0160836 | 0.90591236 | 0.99999994 |

| | | | | | | |
|---|---|---|---|---|---|---|
| Gm4419 | 0.07670251 | 0.0499196 | 0.06600763 | -0.016088 | 0.94763607 | 0.99999994 |
| Fem1b | 0.19991913 | 0.14052168 | 0.15667302 | -0.0161513 | 0.75370666 | 0.99999994 |
| Alkbh6 | 0.391733 | 0.24974664 | 0.26589845 | -0.0161518 | 0.65701707 | 0.99999994 |
| Hint3 | 0.19068737 | 0.13434599 | 0.15051038 | -0.0161644 | 0.76265648 | 0.99999994 |
| Ndufa9 | 0.58835029 | 0.33134136 | 0.34756223 | -0.0162209 | 0.62196853 | 0.99999994 |
| Lpp | 0.27106036 | 0.18496086 | 0.20119153 | -0.0162307 | 0.70436163 | 0.99999994 |
| Slirp | 0.73255758 | 0.37352243 | 0.38975493 | -0.0162325 | 0.60919586 | 0.99999994 |
| Snrpf | 1.08669584 | 0.41328805 | 0.42953644 | -0.0162484 | 0.59940022 | 0.99999994 |
| Sdccag8 | 0.18086048 | 0.12758888 | 0.14384754 | -0.0162587 | 0.77407916 | 0.99999994 |
| Tmem107 | 0.2826998 | 0.19172054 | 0.20799887 | -0.0162783 | 0.69879484 | 0.99999994 |
| Cmas | 0.36922839 | 0.23847681 | 0.25477108 | -0.0162943 | 0.66484307 | 0.99999994 |
| Cnot1 | 0.20967067 | 0.14677404 | 0.16308357 | -0.0163095 | 0.74719333 | 0.99999994 |
| Por | 0.49871569 | 0.2972236 | 0.31356397 | -0.0163404 | 0.63565717 | 0.99999994 |
| Fbxw5 | 0.22702811 | 0.15789919 | 0.17424562 | -0.0163464 | 0.73383396 | 0.99999994 |
| Mboat2 | 0.14628332 | 0.10319256 | 0.11953938 | -0.0163468 | 0.81865369 | 0.99999994 |
| Arfip2 | 0.32314741 | 0.21432502 | 0.2306798 | -0.0163548 | 0.68150931 | 0.99999994 |
| Glt8d1 | 0.20658239 | 0.14470605 | 0.16106458 | -0.0163585 | 0.75049653 | 0.99999994 |
| Lamtor1 | 0.63484293 | 0.34688832 | 0.36327702 | -0.0163887 | 0.61802265 | 0.99999994 |
| Spty2d1 | 0.18881157 | 0.13285517 | 0.14924664 | -0.0163915 | 0.76762732 | 0.99999994 |
| Gpr108 | 0.23399664 | 0.1622462 | 0.17863928 | -0.0163931 | 0.72934387 | 0.99999994 |
| Tnks2 | 0.34402711 | 0.22542035 | 0.24181946 | -0.0163991 | 0.67415083 | 0.99999994 |
| Tmem248 | 0.20830847 | 0.1457951 | 0.1621943 | -0.0163992 | 0.74953068 | 0.99999994 |
| Pkp4 | 0.19870138 | 0.13945001 | 0.15586536 | -0.0164153 | 0.75835813 | 0.99999994 |
| Nucks1 | 0.48577445 | 0.29180797 | 0.3082303 | -0.0164223 | 0.63857288 | 0.99999994 |
| Pard6a | 0.13179835 | 0.09251647 | 0.10894199 | -0.0164255 | 0.84221416 | 0.99999994 |
| Tm9sf4 | 0.16776823 | 0.11835769 | 0.13480394 | -0.0164463 | 0.79162833 | 0.99999994 |
| Fam174a | 0.31981729 | 0.21241098 | 0.22886811 | -0.0164571 | 0.68390129 | 0.99999994 |
| Golgb1 | 0.8040645 | 0.3877629 | 0.40423429 | -0.0164714 | 0.60689199 | 0.99999994 |
| Snrpa | 0.6097962 | 0.33848372 | 0.35497559 | -0.0164919 | 0.62143391 | 0.99999994 |
| Zfp950 | 0.15338302 | 0.10814448 | 0.12464258 | -0.0164981 | 0.81085655 | 0.99999994 |

| | | | | | | |
|---|---|---|---|---|---|---|
| Nsmce1 | 0.54442826 | 0.3150272 | 0.33152686 | -0.0164997 | 0.62978286 | 0.99999994 |
| Vps8 | 0.16745997 | 0.11807924 | 0.13458869 | -0.0165094 | 0.79289527 | 0.99999994 |
| 0610010K14I | 0.20406206 | 0.14288314 | 0.15940901 | -0.0165259 | 0.75492239 | 0.99999994 |
| Zfp9 | 0.07564248 | 0.04860282 | 0.06514005 | -0.0165372 | 0.95447283 | 0.99999994 |
| Lmf1 | 0.13670926 | 0.0960038 | 0.11256274 | -0.0165589 | 0.83626117 | 0.99999994 |
| Chst12 | 0.08224719 | 0.05396368 | 0.07052632 | -0.0165626 | 0.94099746 | 0.99999994 |
| Ankrd24 | 0.12685629 | 0.0886994 | 0.10526852 | -0.0165691 | 0.85261197 | 0.99999994 |
| Deaf1 | 0.16993177 | 0.11974193 | 0.13631174 | -0.0165698 | 0.79078114 | 0.99999994 |
| Anxa6 | 0.60951237 | 0.33828937 | 0.35487918 | -0.0165898 | 0.62216424 | 0.99999994 |
| Ebna1bp2 | 0.54543114 | 0.31528812 | 0.33190614 | -0.016618 | 0.63053605 | 0.99999994 |
| Slc25a51 | 0.37042668 | 0.23874651 | 0.25537388 | -0.0166274 | 0.66763601 | 0.99999994 |
| Limd1 | 0.18826464 | 0.13224168 | 0.14887746 | -0.0166358 | 0.77149856 | 0.99999994 |
| Knop1 | 0.551684 | 0.31761932 | 0.3342565 | -0.0166372 | 0.62979544 | 0.99999994 |
| Ube2g2 | 0.33273934 | 0.21919091 | 0.23584413 | -0.0166532 | 0.68082708 | 0.99999994 |
| Tomm5 | 0.55466533 | 0.31871098 | 0.33536854 | -0.0166576 | 0.62953323 | 0.99999994 |
| Nelfe | 0.26193704 | 0.17910592 | 0.19576514 | -0.0166592 | 0.71445352 | 0.99999994 |
| Rasgrf1 | 0.17799433 | 0.12522241 | 0.14188435 | -0.0166619 | 0.78279788 | 0.99999994 |
| Tcta | 0.27565631 | 0.18718874 | 0.20389455 | -0.0167058 | 0.70725361 | 0.99999994 |
| Sort1 | 0.12624772 | 0.08809609 | 0.1048141 | -0.016718 | 0.8558139 | 0.99999994 |
| Ddrgk1 | 0.44068265 | 0.27201626 | 0.28873978 | -0.0167235 | 0.65007749 | 0.99999994 |
| Syngr2 | 0.34865192 | 0.22749935 | 0.24423595 | -0.0167366 | 0.67585386 | 0.99999994 |
| mt-Nd4l | 0.31972132 | 0.21205305 | 0.22881576 | -0.0167627 | 0.6870958 | 0.99999994 |
| Coq9 | 0.38686109 | 0.24675283 | 0.26352353 | -0.0167707 | 0.66407571 | 0.99999994 |
| Pex11g | 0.10647821 | 0.07303224 | 0.08980443 | -0.0167722 | 0.89309648 | 0.99999994 |
| Ipo7 | 0.16677728 | 0.11732941 | 0.13411159 | -0.0167822 | 0.79756341 | 0.99999994 |
| Hid1 | 0.29235559 | 0.19676217 | 0.21354891 | -0.0167867 | 0.6995984 | 0.99999994 |
| Rab4a | 0.21785253 | 0.15159663 | 0.1683842 | -0.0167876 | 0.7465351 | 0.99999994 |
| Etnk1 | 0.1993843 | 0.13952798 | 0.15631849 | -0.0167905 | 0.76268748 | 0.99999994 |
| Tmem215 | 0.20763362 | 0.14496161 | 0.16175303 | -0.0167914 | 0.75521407 | 0.99999994 |
| Scnn1b | 0.05962318 | 0.03507765 | 0.05188273 | -0.0168051 | 0.98445927 | 0.99999994 |

| | | | | | | |
|---|---|---|---|---|---|---|
| Morf4l1 | 2.12667171 | 0.3516461 | 0.36845335 | -0.0168073 | 0.61929677 | 0.99999994 |
| Nktr | 0.71229028 | 0.36833546 | 0.38515195 | -0.0168165 | 0.6143327 | 0.99999994 |
| Atad1 | 0.28711356 | 0.19371469 | 0.21054659 | -0.0168319 | 0.70268365 | 0.99999994 |
| Adh5 | 0.64544761 | 0.34982313 | 0.3666688 | -0.0168457 | 0.62012519 | 0.99999994 |
| Trappc3 | 0.56301149 | 0.32160134 | 0.33845361 | -0.0168523 | 0.62984127 | 0.99999994 |
| Rpl4 | 2.7779568 | 0.34065461 | 0.3575339 | -0.0168793 | 0.623334 | 0.99999994 |
| Rheb | 0.90294543 | 0.40147982 | 0.41836478 | -0.016885 | 0.60589923 | 0.99999994 |
| Katna1 | 0.25603527 | 0.17532549 | 0.1922122 | -0.0168867 | 0.7206559 | 0.99999994 |
| Pja2 | 0.50930425 | 0.30095573 | 0.31784484 | -0.0168891 | 0.63821607 | 0.99999994 |
| Pfkl | 0.45678465 | 0.27895724 | 0.29586458 | -0.0169073 | 0.64816875 | 0.99999994 |
| Siva1 | 0.44562305 | 0.27403012 | 0.29094548 | -0.0169154 | 0.65062037 | 0.99999994 |
| Cdc5l | 0.52050476 | 0.30537242 | 0.32229363 | -0.0169212 | 0.63663371 | 0.99999994 |
| Golga3 | 0.12875792 | 0.08976041 | 0.10668556 | -0.0169252 | 0.85451577 | 0.99999994 |
| Rnf181 | 0.38528816 | 0.24581955 | 0.26275285 | -0.0169333 | 0.66602885 | 0.99999994 |
| Eif3b | 0.50284168 | 0.29830487 | 0.31524105 | -0.0169362 | 0.63968237 | 0.99999994 |
| Jak1 | 0.23164562 | 0.16022567 | 0.17716242 | -0.0169368 | 0.73772544 | 0.99999994 |
| Gnb1 | 0.64370681 | 0.34913579 | 0.36611925 | -0.0169835 | 0.6212531 | 0.99999994 |
| Chpf | 0.23716072 | 0.16363326 | 0.18061816 | -0.0169849 | 0.73432544 | 0.99999994 |
| Ssr3 | 0.66912054 | 0.3569049 | 0.3739331 | -0.0170282 | 0.61909995 | 0.99999994 |
| Pcmt1 | 0.68393269 | 0.36099187 | 0.37802182 | -0.01703 | 0.6178619 | 0.99999994 |
| Ttc39a | 0.14175218 | 0.09922098 | 0.11625101 | -0.01703 | 0.83524809 | 0.99999994 |
| Ube2j1 | 0.23985615 | 0.16526036 | 0.18229557 | -0.0170352 | 0.73303741 | 0.99999994 |
| Farsb | 0.31933405 | 0.21153698 | 0.22860442 | -0.0170674 | 0.69039088 | 0.99999994 |
| Banf1 | 0.52491594 | 0.30694745 | 0.32402325 | -0.0170758 | 0.63712534 | 0.99999994 |
| Ptbp1 | 0.40320007 | 0.25432707 | 0.27141314 | -0.0170861 | 0.6624039 | 0.99999994 |
| Hacd2 | 0.24022388 | 0.16543534 | 0.18252383 | -0.0170885 | 0.73342037 | 0.99999994 |
| Ube4b | 0.17716532 | 0.12422201 | 0.14131482 | -0.0170928 | 0.78962573 | 0.99999994 |
| Ccdc12 | 0.83085298 | 0.39153037 | 0.40862572 | -0.0170953 | 0.60967678 | 0.99999994 |
| Pgap1 | 0.18626428 | 0.13039657 | 0.14752443 | -0.0171279 | 0.78018413 | 0.99999994 |
| Gm10076 | 0.89960896 | 0.40086644 | 0.41799773 | -0.0171313 | 0.60749856 | 0.99999994 |

| | | | | | | |
|---|---|---|---|---|---|---|
| Exosc7 | 0.36975882 | 0.23790172 | 0.25503805 | -0.0171363 | 0.6726524 | 0.99999994 |
| Isg20 | 0.28648281 | 0.193009 | 0.21018364 | -0.0171746 | 0.70674764 | 0.99999994 |
| Ap3s2 | 0.27717115 | 0.18757571 | 0.20478115 | -0.0172054 | 0.71200728 | 0.99999994 |
| Psmg4 | 0.26316715 | 0.17929087 | 0.19650148 | -0.0172106 | 0.72005275 | 0.99999994 |
| Lyplal1 | 0.3568889 | 0.23125326 | 0.24849559 | -0.0172423 | 0.67791053 | 0.99999994 |
| Vps4b | 0.37167009 | 0.23873096 | 0.25599816 | -0.0172672 | 0.67327498 | 0.99999994 |
| Tbc1d1 | 0.08286907 | 0.05370864 | 0.07103111 | -0.0173225 | 0.94773707 | 0.99999994 |
| Ergic1 | 0.18949934 | 0.13236931 | 0.14971044 | -0.0173411 | 0.77965048 | 0.99999994 |
| Cog7 | 0.18624029 | 0.1301505 | 0.14750818 | -0.0173577 | 0.78326601 | 0.99999994 |
| Tapbp | 0.24667801 | 0.16913847 | 0.1865084 | -0.0173699 | 0.73234514 | 0.99999994 |
| Pex2 | 0.32192078 | 0.21263449 | 0.23001359 | -0.0173791 | 0.69249442 | 0.99999994 |
| F8a | 0.20826279 | 0.14477358 | 0.16216445 | -0.0173909 | 0.76233767 | 0.99999994 |
| Cops3 | 0.67521846 | 0.35829039 | 0.37568228 | -0.0173919 | 0.62101601 | 0.99999994 |
| Jmjd8 | 0.20972465 | 0.14569367 | 0.16311877 | -0.0174251 | 0.76147733 | 0.99999994 |
| Napa | 1.09531879 | 0.41221645 | 0.42968271 | -0.0174663 | 0.60664129 | 0.99999994 |
| Epm2aip1 | 0.20438014 | 0.14209449 | 0.15961832 | -0.0175238 | 0.76754127 | 0.99999994 |
| Cdhr1 | 0.09232396 | 0.06112606 | 0.07865637 | -0.0175303 | 0.93103135 | 0.99999994 |
| Tulp4 | 0.30150122 | 0.20118696 | 0.21872686 | -0.0175399 | 0.70325286 | 0.99999994 |
| Rundc3a | 0.4087686 | 0.25651239 | 0.2740547 | -0.0175423 | 0.66497344 | 0.99999994 |
| Mrpl2 | 0.4102768 | 0.25722196 | 0.27476583 | -0.0175439 | 0.66458543 | 0.99999994 |
| Borcs8 | 0.33121626 | 0.21747047 | 0.23502964 | -0.0175592 | 0.69051102 | 0.99999994 |
| Fmn2 | 0.09697919 | 0.06481706 | 0.08237695 | -0.0175599 | 0.92203636 | 0.99999994 |
| Jakmip1 | 0.08458486 | 0.05483529 | 0.07242177 | -0.0175865 | 0.94699513 | 0.99999994 |
| Timm9 | 0.18984364 | 0.13233167 | 0.14994242 | -0.0176108 | 0.782831 | 0.99999994 |
| Bcar1 | 0.20036839 | 0.13934509 | 0.15697059 | -0.0176255 | 0.77259241 | 0.99999994 |
| Zfp667 | 0.15400059 | 0.10740337 | 0.12508372 | -0.0176804 | 0.8266127 | 0.99999994 |
| Creg1 | 0.24307899 | 0.16658773 | 0.18429155 | -0.0177038 | 0.73873081 | 0.99999994 |
| Creld1 | 0.24961025 | 0.17058551 | 0.18830516 | -0.0177197 | 0.73446328 | 0.99999994 |
| Prorsd1 | 0.23480721 | 0.16142718 | 0.17914717 | -0.01772 | 0.74485966 | 0.99999994 |
| Tmem106c | 0.25526935 | 0.17402042 | 0.19174866 | -0.0177282 | 0.73086043 | 0.99999994 |

| | | | | | | |
|---|---|---|---|---|---|---|
| Trim27 | 0.38557811 | 0.24516588 | 0.26289506 | -0.0177292 | 0.6732456 | 0.99999994 |
| Trabd | 0.37484659 | 0.23985697 | 0.25758743 | -0.0177305 | 0.67658688 | 0.99999994 |
| Cog6 | 0.16824662 | 0.11739119 | 0.13513779 | -0.0177466 | 0.80897571 | 0.99999994 |
| Tm2d1 | 0.71914618 | 0.36898823 | 0.38676933 | -0.0177811 | 0.62020354 | 0.99999994 |
| Prkcsh | 0.64304601 | 0.348095 | 0.36591022 | -0.0178152 | 0.62706157 | 0.99999994 |
| Dennd4c | 0.10600224 | 0.07161595 | 0.08943705 | -0.0178211 | 0.90763591 | 0.99999994 |
| Auh | 0.39193592 | 0.24817539 | 0.26599691 | -0.0178215 | 0.67219022 | 0.99999994 |
| Polr2l | 0.35338079 | 0.22884036 | 0.24668822 | -0.0178479 | 0.68495019 | 0.99999994 |
| Trappc9 | 0.12787438 | 0.08817146 | 0.10602771 | -0.0178563 | 0.86885718 | 0.99999994 |
| Ppp1r10 | 0.36498988 | 0.23477277 | 0.25262969 | -0.0178569 | 0.68100078 | 0.99999994 |
| Tbcd | 0.10524452 | 0.0709806 | 0.0888516 | -0.017871 | 0.90968444 | 0.99999994 |
| Ibtk | 0.1320984 | 0.09129012 | 0.10916404 | -0.0178739 | 0.86211606 | 0.99999994 |
| Commd1 | 0.82052901 | 0.38914225 | 0.40703381 | -0.0178916 | 0.61508087 | 0.99999994 |
| Vti1b | 0.43352873 | 0.26760688 | 0.28551379 | -0.0179069 | 0.66183444 | 0.99999994 |
| Txndc5 | 0.40874758 | 0.25613646 | 0.27404477 | -0.0179083 | 0.66821167 | 0.99999994 |
| Hdlbp | 0.58764613 | 0.32938368 | 0.34731422 | -0.0179305 | 0.63444261 | 0.99999994 |
| Ccdc127 | 0.30192577 | 0.20101075 | 0.21896532 | -0.0179546 | 0.70739949 | 0.99999994 |
| Txn2 | 0.49590738 | 0.29445199 | 0.31241585 | -0.0179639 | 0.64904353 | 0.99999994 |
| Tsen34 | 0.84914183 | 0.39344598 | 0.41144705 | -0.0180011 | 0.61455843 | 0.99999994 |
| Mafg | 0.27283885 | 0.18423446 | 0.20223986 | -0.0180054 | 0.72328061 | 0.99999994 |
| Ndufa13 | 1.62602172 | 0.3781062 | 0.39611631 | -0.0180101 | 0.61891891 | 0.99999994 |
| Slit1 | 0.10482203 | 0.07051007 | 0.08852486 | -0.0180148 | 0.91222233 | 0.99999994 |
| G6pc3 | 0.23146009 | 0.15902637 | 0.17704564 | -0.0180193 | 0.75094535 | 0.99999994 |
| Tmem229b | 0.07028752 | 0.04271933 | 0.06073906 | -0.0180197 | 0.97585297 | 0.99999994 |
| Sec22a | 0.22026504 | 0.15190889 | 0.16993395 | -0.0180251 | 0.7599203 | 0.99999994 |
| Zc3h15 | 0.738349 | 0.37300704 | 0.39103852 | -0.0180315 | 0.62055416 | 0.99999994 |
| Tmem263 | 0.22241032 | 0.15326379 | 0.17130704 | -0.0180433 | 0.75837784 | 0.99999994 |
| mt-Nd5 | 1.01304837 | 0.40904162 | 0.42709355 | -0.0180519 | 0.61077663 | 0.99999994 |
| Hspe1 | 1.38945051 | 0.40117317 | 0.41922951 | -0.0180563 | 0.61282566 | 0.99999994 |
| Scarb2 | 0.22800563 | 0.1567992 | 0.1748649 | -0.0180657 | 0.75417402 | 0.99999994 |

| | | | | | | |
|---|---|---|---|---|---|---|
| Slc35f5 | 0.16720921 | 0.1163374 | 0.13441351 | -0.0180761 | 0.81464808 | 0.99999994 |
| Blcap | 0.29045963 | 0.19438674 | 0.21246591 | -0.0180792 | 0.71444015 | 0.99999994 |
| Zfp516 | 0.27395367 | 0.18479627 | 0.20289548 | -0.0180992 | 0.72366959 | 0.99999994 |
| Tex261 | 0.27681126 | 0.18645113 | 0.2045707 | -0.0181196 | 0.72226322 | 0.99999994 |
| Cyb5b | 0.30632788 | 0.20330862 | 0.22142837 | -0.0181197 | 0.70701925 | 0.99999994 |
| Mrpl48 | 0.48167019 | 0.2883625 | 0.30651454 | -0.018152 | 0.6532814 | 0.99999994 |
| Dynlrb1 | 1.37687518 | 0.40201189 | 0.42017052 | -0.0181586 | 0.61320023 | 0.99999994 |
| Rab18 | 0.65870551 | 0.3526272 | 0.37079643 | -0.0181692 | 0.62785032 | 0.99999994 |
| Pin4 | 0.63037724 | 0.34365123 | 0.36182635 | -0.0181751 | 0.63094589 | 0.99999994 |
| Amd1 | 0.21514402 | 0.14845799 | 0.16663716 | -0.0181792 | 0.76613301 | 0.99999994 |
| Mfsd10 | 0.1764927 | 0.12267108 | 0.14085217 | -0.0181811 | 0.80488423 | 0.99999994 |
| Cand1 | 0.1684338 | 0.11708259 | 0.13526834 | -0.0181858 | 0.81457826 | 0.99999994 |
| Ube2k | 0.84039157 | 0.39192172 | 0.41012017 | -0.0181985 | 0.61614275 | 0.99999994 |
| Metap2 | 1.0441017 | 0.41029729 | 0.42849945 | -0.0182022 | 0.61131745 | 0.99999994 |
| Eif3a | 1.02212383 | 0.40929482 | 0.42752318 | -0.0182284 | 0.61172121 | 0.99999994 |
| Fam220a | 0.33441459 | 0.21850459 | 0.23673772 | -0.0182331 | 0.69590586 | 0.99999994 |
| Pdzd11 | 0.43924873 | 0.26985412 | 0.28809635 | -0.0182422 | 0.66329766 | 0.99999994 |
| Rab27a | 0.26178747 | 0.17742724 | 0.19567551 | -0.0182483 | 0.7326203 | 0.99999994 |
| Iscu | 0.68058422 | 0.35886417 | 0.37712772 | -0.0182636 | 0.62640784 | 0.99999994 |
| Mapk8ip3 | 0.14822794 | 0.10267184 | 0.12094298 | -0.0182711 | 0.84269394 | 0.99999994 |
| Gmppb | 0.28953616 | 0.19364731 | 0.21193722 | -0.0182899 | 0.71716335 | 0.99999994 |
| Lysmd2 | 0.3744128 | 0.23907596 | 0.25737087 | -0.0182949 | 0.6819503 | 0.99999994 |
| Ntrk2 | 0.07005276 | 0.04224965 | 0.06054542 | -0.0182958 | 0.97786028 | 0.99999994 |
| Arf3 | 0.27563607 | 0.18558053 | 0.20388269 | -0.0183022 | 0.72492268 | 0.99999994 |
| Sgms2 | 0.10879424 | 0.07325191 | 0.09158805 | -0.0183361 | 0.90867147 | 0.99999994 |
| Tmem39a | 0.18231867 | 0.12649745 | 0.14484288 | -0.0183454 | 0.80040734 | 0.99999994 |
| Psme4 | 0.23498116 | 0.16090304 | 0.17925609 | -0.0183531 | 0.75222499 | 0.99999994 |
| Mbd2 | 0.1042495 | 0.06971074 | 0.08808171 | -0.018371 | 0.91747696 | 0.99999994 |
| Gcdh | 0.17744654 | 0.1231248 | 0.1415081 | -0.0183833 | 0.8064055 | 0.99999994 |
| Rwdd4a | 0.4163613 | 0.25923395 | 0.27761755 | -0.0183836 | 0.67031329 | 0.99999994 |

| | | | | | | |
|---|---|---|---|---|---|---|
| Hax1 | 0.53088901 | 0.30795756 | 0.32634597 | -0.0183884 | 0.64619269 | 0.99999994 |
| H3f3a | 2.76562069 | 0.33995697 | 0.35836999 | -0.018413 | 0.63382576 | 0.99999994 |
| Preb | 0.35910923 | 0.23119479 | 0.24963432 | -0.0184395 | 0.68852707 | 0.99999994 |
| Rbm8a | 1.13194146 | 0.41148177 | 0.42993506 | -0.0184533 | 0.61244535 | 0.99999994 |
| Prrc1 | 0.18018314 | 0.12492872 | 0.14338441 | -0.0184557 | 0.80421439 | 0.99999994 |
| Sil1 | 0.23089163 | 0.15822317 | 0.1766876 | -0.0184644 | 0.75666381 | 0.99999994 |
| Bpnt1 | 0.37885615 | 0.24110152 | 0.2595813 | -0.0184798 | 0.68220404 | 0.99999994 |
| Ube2s | 1.20127892 | 0.41048344 | 0.42896727 | -0.0184838 | 0.6128737 | 0.99999994 |
| Nus1 | 0.25343623 | 0.17212785 | 0.19063692 | -0.0185091 | 0.74094786 | 0.99999994 |
| Mapre1 | 0.70947974 | 0.36595775 | 0.38448171 | -0.018524 | 0.62577996 | 0.99999994 |
| Snrpg | 0.94611505 | 0.40403181 | 0.42257047 | -0.0185387 | 0.61486514 | 0.99999994 |
| Trip11 | 0.33951175 | 0.22087658 | 0.23944202 | -0.0185654 | 0.69711092 | 0.99999994 |
| Bola1 | 0.38259921 | 0.24285138 | 0.26143089 | -0.0185795 | 0.68191398 | 0.99999994 |
| Spcs3 | 0.29563668 | 0.1968228 | 0.21541544 | -0.0185926 | 0.71719012 | 0.99999994 |
| Cox18 | 0.212151 | 0.14607678 | 0.16469779 | -0.018621 | 0.77414852 | 0.99999994 |
| Fndc3b | 0.11547936 | 0.07806015 | 0.09669883 | -0.0186387 | 0.90025604 | 0.99999994 |
| Tmem50b | 0.12683639 | 0.08659318 | 0.10525366 | -0.0186605 | 0.88102161 | 0.99999994 |
| Tsg101 | 0.93863747 | 0.40325946 | 0.42193418 | -0.0186747 | 0.61585374 | 0.99999994 |
| Rabggtb | 0.37060396 | 0.23677233 | 0.25546296 | -0.0186906 | 0.68687229 | 0.99999994 |
| Abhd8 | 0.17958094 | 0.12427557 | 0.14297222 | -0.0186966 | 0.80797545 | 0.99999994 |
| Mrps12 | 0.66738108 | 0.35469521 | 0.37341314 | -0.0187179 | 0.63068214 | 0.99999994 |
| Pkm | 1.91290783 | 0.35473148 | 0.37345069 | -0.0187192 | 0.63067809 | 0.99999994 |
| Tab2 | 0.26876277 | 0.18110516 | 0.19983282 | -0.0187277 | 0.73362421 | 0.99999994 |
| St3gal5 | 0.16117177 | 0.11144166 | 0.13017424 | -0.0187326 | 0.83093733 | 0.99999994 |
| Gpsm1 | 0.23325132 | 0.1593793 | 0.17817169 | -0.0187924 | 0.758682 | 0.99999994 |
| Pelo | 0.17411881 | 0.12040428 | 0.13921487 | -0.0188106 | 0.81578042 | 0.99999994 |
| Shc1 | 0.1604609 | 0.11084666 | 0.12967224 | -0.0188256 | 0.83306745 | 0.99999994 |
| Ncbp2 | 0.47113207 | 0.28322575 | 0.30205751 | -0.0188318 | 0.66092551 | 0.99999994 |
| Cbr1 | 0.43536781 | 0.26750126 | 0.28634681 | -0.0188456 | 0.66933578 | 0.99999994 |
| Kif1b | 0.39970594 | 0.25088954 | 0.26974337 | -0.0188538 | 0.67912345 | 0.99999994 |

| | | | | | | |
|---|---|---|---|---|---|---|
| Abhd17a | 0.2378384 | 0.16215589 | 0.1810407 | -0.0188848 | 0.75626586 | 0.99999994 |
| Mpc2 | 0.8215758 | 0.38828622 | 0.40719734 | -0.0189111 | 0.62139084 | 0.99999994 |
| Tbl1x | 0.38772449 | 0.24501273 | 0.26394575 | -0.018933 | 0.68348464 | 0.99999994 |
| Nrd1 | 0.44511753 | 0.27178388 | 0.29072058 | -0.0189367 | 0.66770177 | 0.99999994 |
| Syvn1 | 0.12084673 | 0.08181234 | 0.10076153 | -0.0189492 | 0.89467732 | 0.99999994 |
| Scamp3 | 0.44811288 | 0.27308548 | 0.29205055 | -0.0189651 | 0.66721945 | 0.99999994 |
| Pgrmc2 | 0.31438339 | 0.20691175 | 0.22589122 | -0.0189795 | 0.71201122 | 0.99999994 |
| Zdhhc2 | 0.12853888 | 0.08753064 | 0.10652256 | -0.0189919 | 0.88233945 | 0.99999994 |
| Ndel1 | 0.24825033 | 0.16847895 | 0.1874729 | -0.0189939 | 0.74996943 | 0.99999994 |
| Usp48 | 0.18706138 | 0.12902673 | 0.1480641 | -0.0190374 | 0.80395755 | 0.99999994 |
| Cnpy3 | 0.20821589 | 0.14309338 | 0.1621338 | -0.0190404 | 0.7828043 | 0.99999994 |
| Ttc14 | 0.4168419 | 0.25878375 | 0.27784164 | -0.0190579 | 0.67601247 | 0.99999994 |
| Vps53 | 0.21391242 | 0.14677565 | 0.16584025 | -0.0190646 | 0.77792357 | 0.99999994 |
| Cnih1 | 0.55962841 | 0.31814327 | 0.33720805 | -0.0190648 | 0.64666479 | 0.99999994 |
| Pigk | 0.28104038 | 0.18795866 | 0.20703604 | -0.0190774 | 0.73017808 | 0.99999994 |
| Map2 | 0.18328973 | 0.12640501 | 0.14550442 | -0.0190994 | 0.80886905 | 0.99999994 |
| Kars | 0.29648261 | 0.19679417 | 0.21589509 | -0.0191009 | 0.72203889 | 0.99999994 |
| Impdh1 | 0.19292875 | 0.13291227 | 0.15201519 | -0.0191029 | 0.79855414 | 0.99999994 |
| Nmt1 | 0.37139205 | 0.23674979 | 0.25585867 | -0.0191089 | 0.69045201 | 0.99999994 |
| Ap3d1 | 0.21472915 | 0.14725671 | 0.16636889 | -0.0191122 | 0.77776636 | 0.99999994 |
| Zfp330 | 0.37789455 | 0.23999014 | 0.25910436 | -0.0191142 | 0.68830157 | 0.99999994 |
| Eef2kmt | 0.22762644 | 0.15550899 | 0.17462479 | -0.0191158 | 0.76689426 | 0.99999994 |
| Fam3a | 0.22027665 | 0.15079269 | 0.16994139 | -0.0191487 | 0.77338072 | 0.99999994 |
| Upb1 | 0.17530967 | 0.12086399 | 0.140037 | -0.019173 | 0.81894391 | 0.99999994 |
| Triap1 | 0.33083776 | 0.21563078 | 0.23482692 | -0.0191961 | 0.70682407 | 0.99999994 |
| Tbc1d16 | 0.23442194 | 0.15970297 | 0.17890585 | -0.0192029 | 0.76252889 | 0.99999994 |
| Timm10b | 0.67433036 | 0.3562374 | 0.37544152 | -0.0192041 | 0.6332516 | 0.99999994 |
| Ap2b1 | 0.29647787 | 0.19666685 | 0.2158924 | -0.0192256 | 0.72332876 | 0.99999994 |
| Madd | 0.13888153 | 0.09490942 | 0.11415518 | -0.0192458 | 0.86911453 | 0.99999994 |
| Cops7a | 0.45891558 | 0.2775048 | 0.29679341 | -0.0192886 | 0.66734614 | 0.99999994 |

| | | | | | | |
|---|---|---|---|---|---|---|
| Robo2 | 0.08298537 | 0.05180303 | 0.07112548 | -0.0193224 | 0.96472108 | 0.99999994 |
| Zfyve21 | 0.32010386 | 0.20968815 | 0.22902439 | -0.0193362 | 0.71293787 | 0.99999994 |
| Yif1a | 0.54971942 | 0.3141773 | 0.33352081 | -0.0193435 | 0.65027198 | 0.99999994 |
| Calm1 | 2.81382263 | 0.33593186 | 0.35529627 | -0.0193644 | 0.6416175 | 0.99999994 |
| Slc37a4 | 0.283386 | 0.18902695 | 0.20839628 | -0.0193693 | 0.7319342 | 0.99999994 |
| Tra2b | 0.92286058 | 0.40104757 | 0.42043652 | -0.019389 | 0.62056546 | 0.99999994 |
| 2300009A05I | 0.36114281 | 0.2312722 | 0.25067356 | -0.0194014 | 0.69678694 | 0.99999994 |
| Ascc3 | 0.23277594 | 0.15846641 | 0.17787315 | -0.0194067 | 0.76615352 | 0.99999994 |
| Creb3 | 0.39371159 | 0.24743719 | 0.26685716 | -0.01942 | 0.68594989 | 0.99999994 |
| Prpf8 | 0.18251133 | 0.12552804 | 0.14497421 | -0.0194462 | 0.81403218 | 0.99999994 |
| Anapc16 | 0.68273354 | 0.3582436 | 0.37770232 | -0.0194587 | 0.63417321 | 0.99999994 |
| Zcrb1 | 0.89532261 | 0.3980603 | 0.41751943 | -0.0194591 | 0.62180686 | 0.99999994 |
| Hopx | 0.14352539 | 0.0980493 | 0.11754076 | -0.0194915 | 0.86516121 | 0.99999994 |
| Znhit1 | 0.73025485 | 0.36975407 | 0.38924905 | -0.019495 | 0.63057406 | 0.99999994 |
| Stub1 | 1.04572062 | 0.40904399 | 0.42855762 | -0.0195136 | 0.61908772 | 0.99999994 |
| Acot13 | 0.49393601 | 0.29204296 | 0.31160683 | -0.0195639 | 0.66199417 | 0.99999994 |
| Tonsl | 0.9784012 | 0.40541104 | 0.42500016 | -0.0195891 | 0.62050411 | 0.99999994 |
| Gns | 0.1305686 | 0.08842807 | 0.10803078 | -0.0196027 | 0.88644343 | 0.99999994 |
| Fbxo21 | 0.20765227 | 0.14215771 | 0.16176523 | -0.0196075 | 0.79011359 | 0.99999994 |
| Cfap20 | 0.82854325 | 0.38865203 | 0.40827289 | -0.0196209 | 0.62547511 | 0.99999994 |
| Pdxdc1 | 0.1883906 | 0.12933074 | 0.14896251 | -0.0196318 | 0.80982235 | 0.99999994 |
| Mtch2 | 0.9975633 | 0.40661716 | 0.42625325 | -0.0196361 | 0.62043997 | 0.99999994 |
| G3bp2 | 0.65168763 | 0.34895298 | 0.36862563 | -0.0196726 | 0.63879269 | 0.99999994 |
| Ubr4 | 0.20806046 | 0.14234879 | 0.16203219 | -0.0196834 | 0.7906283 | 0.99999994 |
| Sec23ip | 0.1887907 | 0.12954429 | 0.14923256 | -0.0196883 | 0.81007543 | 0.99999994 |
| 9130401M01 | 0.19493425 | 0.13363937 | 0.15335684 | -0.0197175 | 0.80395257 | 0.99999994 |
| Dnajc24 | 0.43162895 | 0.26491366 | 0.28465072 | -0.0197371 | 0.67779344 | 0.99999994 |
| Sorbs2 | 0.1966376 | 0.13474948 | 0.15449297 | -0.0197435 | 0.8025187 | 0.99999994 |
| M6pr | 0.31336829 | 0.2055791 | 0.22533198 | -0.0197529 | 0.72022318 | 0.99999994 |
| Ppp1cc | 0.85253082 | 0.39216437 | 0.41194564 | -0.0197813 | 0.62537653 | 0.99999994 |

| | | | | | | |
|---|---|---|---|---|---|---|
| Atp6v0e2 | 0.25486442 | 0.17172134 | 0.19150336 | -0.019782 | 0.75414512 | 0.99999994 |
| Ufsp2 | 0.36551185 | 0.23310977 | 0.25289418 | -0.0197844 | 0.69872604 | 0.99999994 |
| Cpq | 0.29830079 | 0.1970886 | 0.21692384 | -0.0198352 | 0.72862143 | 0.99999994 |
| Dync1h1 | 0.31382921 | 0.20570891 | 0.22558602 | -0.0198771 | 0.72123459 | 0.99999994 |
| Psmd11 | 0.84214421 | 0.39051201 | 0.41039034 | -0.0198783 | 0.62643203 | 0.99999994 |
| Slc2a5 | 0.27434435 | 0.18324473 | 0.20312496 | -0.0198802 | 0.74262863 | 0.99999994 |
| Cdk5rap3 | 0.42158009 | 0.26014282 | 0.2800417 | -0.0198989 | 0.68188623 | 0.99999994 |
| Ganab | 0.23128457 | 0.15702658 | 0.17693513 | -0.0199085 | 0.77306122 | 0.99999994 |
| Aga | 0.27962784 | 0.18628689 | 0.20621446 | -0.0199276 | 0.73996522 | 0.99999994 |
| Mcf2l | 0.22062683 | 0.15023285 | 0.17016585 | -0.019933 | 0.78222115 | 0.99999994 |
| Tex264 | 0.25230772 | 0.17001688 | 0.18995068 | -0.0199338 | 0.75757748 | 0.99999994 |
| Tra2a | 0.53508069 | 0.30800934 | 0.32796233 | -0.019953 | 0.65724948 | 0.99999994 |
| Ormdl2 | 0.41435664 | 0.25669684 | 0.27668104 | -0.0199842 | 0.68466258 | 0.99999994 |
| Txlna | 0.29759704 | 0.19653237 | 0.216526 | -0.0199936 | 0.73060257 | 0.99999994 |
| Cox14 | 1.17692569 | 0.40948863 | 0.42949316 | -0.0200045 | 0.62173235 | 0.99999994 |
| Mgat2 | 0.34598232 | 0.22283198 | 0.24284342 | -0.0200114 | 0.70833088 | 0.99999994 |
| Cisd2 | 0.4560857 | 0.27553165 | 0.29555923 | -0.0200276 | 0.67402174 | 0.99999994 |
| Bicd1 | 0.2056115 | 0.14038856 | 0.1604276 | -0.020039 | 0.79713941 | 0.99999994 |
| Larp4b | 0.30450433 | 0.20035282 | 0.22041016 | -0.0200573 | 0.7276526 | 0.99999994 |
| Atp6ap1 | 0.64265691 | 0.34571742 | 0.36578703 | -0.0200696 | 0.6425241 | 0.99999994 |
| Pecr | 0.11098854 | 0.07320033 | 0.09327173 | -0.0200714 | 0.92398527 | 0.99999994 |
| Tor2a | 0.5527593 | 0.3145781 | 0.3346582 | -0.0200801 | 0.65519839 | 0.99999994 |
| 2900076A07 | 0.19855976 | 0.13569036 | 0.15577133 | -0.020081 | 0.8045791 | 0.99999994 |
| Lrrc59 | 0.40393593 | 0.25167312 | 0.27176362 | -0.0200905 | 0.68866869 | 0.99999994 |
| Gmds | 0.22944511 | 0.15563672 | 0.17577508 | -0.0201384 | 0.77715247 | 0.99999994 |
| Clock | 0.26995564 | 0.18039649 | 0.20053886 | -0.0201424 | 0.74811594 | 0.99999994 |
| Akr1b3 | 0.38035768 | 0.24017874 | 0.26032458 | -0.0201458 | 0.69676482 | 0.99999994 |
| Mrpl57 | 0.9052803 | 0.39847149 | 0.41861893 | -0.0201474 | 0.62565212 | 0.99999994 |
| Cnpy2 | 1.08063035 | 0.40930144 | 0.42946985 | -0.0201684 | 0.62270409 | 0.99999994 |
| Spag7 | 0.80978452 | 0.38507161 | 0.40524795 | -0.0201763 | 0.62983049 | 0.99999994 |

| | | | | | | |
|---|---|---|---|---|---|---|
| Nedd8 | 1.41529516 | 0.39695086 | 0.41713117 | -0.0201803 | 0.62628372 | 0.99999994 |
| Psmg1 | 0.2386001 | 0.16133171 | 0.18151488 | -0.0201832 | 0.77039253 | 0.99999994 |
| Sec16a | 0.13368204 | 0.09014665 | 0.11033421 | -0.0201876 | 0.88836835 | 0.99999994 |
| Pts | 0.30471196 | 0.2003358 | 0.22052624 | -0.0201904 | 0.72887951 | 0.99999994 |
| Atf6b | 0.36943483 | 0.23468229 | 0.25487501 | -0.0201927 | 0.70102696 | 0.99999994 |
| Med28 | 1.18252338 | 0.40918878 | 0.42938651 | -0.0201977 | 0.62289981 | 0.99999994 |
| Setd3 | 0.27423452 | 0.18283447 | 0.20306046 | -0.020226 | 0.74633847 | 0.99999994 |
| Arfgap1 | 0.17109024 | 0.11685315 | 0.13711693 | -0.0202638 | 0.83737038 | 0.99999994 |
| Trpm5 | 0.13360327 | 0.09000462 | 0.11027609 | -0.0202715 | 0.88944895 | 0.99999994 |
| Slc35b2 | 0.29829652 | 0.19660079 | 0.21692143 | -0.0203206 | 0.73353858 | 0.99999994 |
| Scamp2 | 0.35337213 | 0.2263487 | 0.24668375 | -0.020335 | 0.7083971 | 0.99999994 |
| Rpl36al | 2.12556874 | 0.34810283 | 0.36845476 | -0.0203519 | 0.64344029 | 0.99999994 |
| Ppm1l | 0.17801796 | 0.12152994 | 0.14190057 | -0.0203706 | 0.83035213 | 0.99999994 |
| Neu1 | 0.22603091 | 0.15323523 | 0.17361287 | -0.0203776 | 0.78268211 | 0.99999994 |
| Lrp11 | 0.35419307 | 0.22670052 | 0.2471076 | -0.0204071 | 0.70873992 | 0.99999994 |
| Gbf1 | 0.23786963 | 0.16064055 | 0.18106015 | -0.0204196 | 0.77358051 | 0.99999994 |
| Zfhx2 | 0.17799137 | 0.12144988 | 0.14188231 | -0.0204324 | 0.83111485 | 0.99999994 |
| Fkbp9 | 0.27641738 | 0.18388805 | 0.20434025 | -0.0204522 | 0.74736724 | 0.99999994 |
| Nans | 0.52482884 | 0.30352998 | 0.32398922 | -0.0204592 | 0.66287961 | 0.99999994 |
| Sptssa | 0.4885047 | 0.28890384 | 0.30936472 | -0.0204609 | 0.67011865 | 0.99999994 |
| Phb2 | 1.02561834 | 0.40717542 | 0.42769807 | -0.0205226 | 0.62528781 | 0.99999994 |
| Timm17b | 0.58102522 | 0.32441708 | 0.34496853 | -0.0205515 | 0.65415093 | 0.99999994 |
| Dhrs7 | 0.1975169 | 0.13451808 | 0.15507825 | -0.0205602 | 0.81124556 | 0.99999994 |
| Rps14 | 3.49245561 | 0.23951845 | 0.26012048 | -0.020602 | 0.70097123 | 0.99999994 |
| Mcee | 0.4051245 | 0.25169497 | 0.27232884 | -0.0206339 | 0.69298887 | 0.99999994 |
| Tkt | 0.36888605 | 0.23394615 | 0.25459865 | -0.0206525 | 0.70538415 | 0.99999994 |
| Bet1l | 0.34299581 | 0.22061462 | 0.24127795 | -0.0206633 | 0.71568098 | 0.99999994 |
| Mrpl14 | 0.80540538 | 0.38380582 | 0.40447328 | -0.0206675 | 0.63311746 | 0.99999994 |
| Ctnnb1 | 0.67118349 | 0.3538603 | 0.37454773 | -0.0206874 | 0.6434334 | 0.99999994 |
| Prdx6 | 0.89470385 | 0.3967424 | 0.41744975 | -0.0207074 | 0.62936983 | 0.99999994 |

| | | | | | | |
|---|---|---|---|---|---|---|
| Slc7a14 | 0.20556873 | 0.13963729 | 0.16039951 | -0.0207622 | 0.8055472 | 0.99999994 |
| Mlec | 0.35069051 | 0.22451694 | 0.24529537 | -0.0207784 | 0.71357029 | 0.99999994 |
| Prdx3 | 0.53760887 | 0.30814201 | 0.32893166 | -0.0207897 | 0.66301492 | 0.99999994 |
| Dnajc10 | 0.38958754 | 0.24406097 | 0.26485486 | -0.0207939 | 0.69937157 | 0.99999994 |
| Ptpn11 | 0.23968094 | 0.16138892 | 0.18218676 | -0.0207978 | 0.77633329 | 0.99999994 |
| Rint1 | 0.18824704 | 0.12806151 | 0.14886557 | -0.0208041 | 0.82387411 | 0.99999994 |
| Polr2g | 1.0936963 | 0.40882954 | 0.42965686 | -0.0208273 | 0.62652455 | 0.99999994 |
| Birc6 | 0.33803984 | 0.2178174 | 0.23866334 | -0.0208459 | 0.71951341 | 0.99999994 |
| Cltc | 0.47105135 | 0.28111662 | 0.30202308 | -0.0209065 | 0.67751546 | 0.99999994 |
| Dcaf8 | 0.55863365 | 0.31590198 | 0.33684052 | -0.0209385 | 0.66047867 | 0.99999994 |
| Pttg1ip | 0.28488667 | 0.1883243 | 0.20926357 | -0.0209393 | 0.74730929 | 0.99999994 |
| Cflar | 0.16499458 | 0.11182727 | 0.13286326 | -0.021036 | 0.85403315 | 0.99999994 |
| Gga2 | 0.25223442 | 0.1688409 | 0.18990606 | -0.0210652 | 0.76984922 | 0.99999994 |
| Exosc4 | 0.58558686 | 0.32549956 | 0.34658732 | -0.0210878 | 0.65726106 | 0.99999994 |
| Polr2i | 0.67155337 | 0.35356989 | 0.3746577 | -0.0210878 | 0.6460421 | 0.99999994 |
| Hmox2 | 0.43041429 | 0.26298888 | 0.28409753 | -0.0211087 | 0.68954899 | 0.99999994 |
| Usp16 | 0.2950847 | 0.19397774 | 0.21510211 | -0.0211244 | 0.74334613 | 0.99999994 |
| Tomm40 | 0.59524334 | 0.32876324 | 0.349975 | -0.0212118 | 0.65668682 | 0.99999994 |
| Golt1b | 0.24617278 | 0.16498169 | 0.18619796 | -0.0212163 | 0.77591116 | 0.99999994 |
| P2ry1 | 0.16358293 | 0.11064045 | 0.13187222 | -0.0212318 | 0.85807054 | 0.99999994 |
| Ccdc124 | 0.53388477 | 0.30626601 | 0.32750243 | -0.0212364 | 0.66699188 | 0.99999994 |
| Ficd | 0.10188901 | 0.0649902 | 0.0862503 | -0.0212601 | 0.94957495 | 0.99999994 |
| D5Ertd579e | 0.18332394 | 0.12425651 | 0.1455277 | -0.0212712 | 0.83470665 | 0.99999994 |
| Gpr180 | 0.18628245 | 0.1262623 | 0.14753674 | -0.0212744 | 0.83142974 | 0.99999994 |
| Rnf5 | 0.79019265 | 0.38037946 | 0.40167548 | -0.021296 | 0.63792081 | 0.99999994 |
| Pigyl | 0.52457187 | 0.30256641 | 0.32388878 | -0.0213224 | 0.66938074 | 0.99999994 |
| Sec61a1 | 0.50111593 | 0.2932112 | 0.31454119 | -0.02133 | 0.67414495 | 0.99999994 |
| Gng4 | 0.19648559 | 0.13304232 | 0.15439171 | -0.0213494 | 0.82132996 | 0.99999994 |
| Pigt | 0.55069719 | 0.31251476 | 0.33388725 | -0.0213725 | 0.6649755 | 0.99999994 |
| Akt1 | 0.2066296 | 0.13971032 | 0.16109552 | -0.0213852 | 0.81154989 | 0.99999994 |

| | | | | | | |
|---|---|---|---|---|---|---|
| Ywhab | 1.00672006 | 0.40539622 | 0.4267835 | -0.0213873 | 0.63064723 | 0.99999994 |
| Syndig1l | 0.11841043 | 0.07751247 | 0.09892182 | -0.0214093 | 0.92515329 | 0.99999994 |
| Trappc6b | 0.58239043 | 0.32403322 | 0.34545425 | -0.021421 | 0.6601049 | 0.99999994 |
| Eif3j1 | 0.3244371 | 0.20993923 | 0.23137886 | -0.0214396 | 0.73131006 | 0.99999994 |
| Aph1a | 0.27206271 | 0.18029468 | 0.20178273 | -0.0214881 | 0.76078893 | 0.99999994 |
| Txndc11 | 0.14012087 | 0.09356761 | 0.1150612 | -0.0214936 | 0.89314282 | 0.99999994 |
| Dnpep | 0.6561631 | 0.34848443 | 0.37001658 | -0.0215321 | 0.65075122 | 0.99999994 |
| Ddx17 | 0.42971121 | 0.26222333 | 0.28377684 | -0.0215535 | 0.69341928 | 0.99999994 |
| Ddb1 | 0.41256487 | 0.25427544 | 0.27584145 | -0.021566 | 0.69861161 | 0.99999994 |
| Hmbs | 0.29264035 | 0.19212985 | 0.21371129 | -0.0215814 | 0.74926747 | 0.99999994 |
| Nol12 | 0.2361969 | 0.15843303 | 0.18001642 | -0.0215834 | 0.78758678 | 0.99999994 |
| Npdc1 | 0.75160397 | 0.37214742 | 0.39394048 | -0.0217931 | 0.6436507 | 0.99999994 |
| Trp53inp2 | 0.16142175 | 0.10848612 | 0.13035059 | -0.0218645 | 0.86789814 | 0.99999994 |
| Irak1 | 0.19633626 | 0.13241639 | 0.1542922 | -0.0218758 | 0.82735119 | 0.99999994 |
| Cacfd1 | 0.33478278 | 0.21505029 | 0.2369338 | -0.0218835 | 0.73065323 | 0.99999994 |
| Rpl24 | 2.87894133 | 0.32845854 | 0.35041669 | -0.0219582 | 0.66170001 | 0.99999994 |
| Arl8b | 0.24213059 | 0.16174696 | 0.18370525 | -0.0219583 | 0.7868777 | 0.99999994 |
| Pgm3 | 0.15181999 | 0.10154671 | 0.1235241 | -0.0219774 | 0.88185251 | 0.99999994 |
| Ube2n | 1.21254246 | 0.40667837 | 0.42866083 | -0.0219825 | 0.63357975 | 0.99999994 |
| Gemin7 | 0.48698197 | 0.28674331 | 0.30873263 | -0.0219893 | 0.68228417 | 0.99999994 |
| Stx8 | 0.28097621 | 0.18499033 | 0.20699876 | -0.0220084 | 0.76043873 | 0.99999994 |
| Sdhb | 1.03275606 | 0.40603396 | 0.4280427 | -0.0220087 | 0.63391901 | 0.99999994 |
| Tmem41b | 0.27049953 | 0.17882429 | 0.20086033 | -0.022036 | 0.76738365 | 0.99999994 |
| Smyd2 | 0.18562709 | 0.12502629 | 0.14709252 | -0.0220662 | 0.84099613 | 0.99999994 |
| Psmd2 | 0.85887655 | 0.39073451 | 0.41286571 | -0.0221312 | 0.63938165 | 0.99999994 |
| Gramd1a | 0.23805872 | 0.15900873 | 0.18117791 | -0.0221692 | 0.79231534 | 0.99999994 |
| Slc30a5 | 0.16496577 | 0.11065932 | 0.13284306 | -0.0221837 | 0.86683605 | 0.99999994 |
| Galnt18 | 0.14182497 | 0.09408643 | 0.11630403 | -0.0222176 | 0.8982364 | 0.99999994 |
| Ogfod2 | 0.26056389 | 0.17271339 | 0.19494146 | -0.0222281 | 0.77606982 | 0.99999994 |
| Cyb5a | 0.46436646 | 0.27691808 | 0.29915509 | -0.022237 | 0.68962406 | 0.99999994 |

| | | | | | | |
|---|---|---|---|---|---|---|
| Arl6ip1 | 0.88688329 | 0.39428998 | 0.41653039 | -0.0222404 | 0.63885976 | 0.99999994 |
| Sec63 | 0.24004068 | 0.16009286 | 0.18241013 | -0.0223173 | 0.79228728 | 0.99999994 |
| Ap3b2 | 0.18882699 | 0.12692137 | 0.14925705 | -0.0223357 | 0.84039808 | 0.99999994 |
| Setd5 | 0.45542868 | 0.27292975 | 0.29527189 | -0.0223421 | 0.6927506 | 0.99999994 |
| Dctn3 | 1.45103734 | 0.39160337 | 0.41395389 | -0.0223505 | 0.64035027 | 0.99999994 |
| Chchd3 | 0.48898467 | 0.28720946 | 0.30956364 | -0.0223542 | 0.68462387 | 0.99999994 |
| Ufl1 | 0.27664765 | 0.18211473 | 0.204475 | -0.0223603 | 0.76666353 | 0.99999994 |
| Vkorc1 | 0.42924135 | 0.26118037 | 0.28356233 | -0.022382 | 0.70034586 | 0.99999994 |
| Rab11b | 0.82274889 | 0.38499106 | 0.40738001 | -0.0223889 | 0.64275128 | 0.99999994 |
| Ssu72 | 0.95310809 | 0.40070924 | 0.42312249 | -0.0224133 | 0.63780378 | 0.99999994 |
| Rfk | 0.35335254 | 0.22424273 | 0.24667362 | -0.0224309 | 0.72750547 | 0.99999994 |
| Smim20 | 0.41061041 | 0.25248236 | 0.27492291 | -0.0224405 | 0.70654676 | 0.99999994 |
| Zc3h7a | 0.17812328 | 0.11952604 | 0.14197287 | -0.0224468 | 0.85369503 | 0.99999994 |
| Slc38a4 | 0.17466871 | 0.11712285 | 0.13959469 | -0.0224718 | 0.85803555 | 0.99999994 |
| Snf8 | 0.60169101 | 0.32970542 | 0.352205 | -0.0224996 | 0.66466254 | 0.99999994 |
| Rab5c | 0.50347589 | 0.29297859 | 0.31549777 | -0.0225192 | 0.68264522 | 0.99999994 |
| Erp44 | 0.66329394 | 0.3496652 | 0.37218526 | -0.0225201 | 0.65643786 | 0.99999994 |
| Hspa13 | 0.26250779 | 0.17358043 | 0.19610696 | -0.0225265 | 0.77774713 | 0.99999994 |
| Cyfip2 | 0.13244658 | 0.0868314 | 0.10942158 | -0.0225902 | 0.91531184 | 0.99999994 |
| Pdhb | 0.67870296 | 0.35399851 | 0.37662273 | -0.0226242 | 0.65536874 | 0.99999994 |
| Pafah1b1 | 0.90628216 | 0.39608502 | 0.41872729 | -0.0226423 | 0.64055253 | 0.99999994 |
| Tmem65 | 0.13669962 | 0.08991255 | 0.11255566 | -0.0226431 | 0.90972717 | 0.99999994 |
| Actn4 | 0.24835652 | 0.16483717 | 0.18753795 | -0.0227008 | 0.78980004 | 0.99999994 |
| Emc4 | 0.52789931 | 0.30248126 | 0.32518615 | -0.0227049 | 0.67894655 | 0.99999994 |
| Ifi30 | 0.2400611 | 0.15971508 | 0.18242281 | -0.0227077 | 0.79632717 | 0.99999994 |
| Sh3pxd2a | 0.08527851 | 0.05027086 | 0.07298311 | -0.0227122 | 0.9808416 | 0.99999994 |
| Lamp2 | 0.45978224 | 0.27444059 | 0.29717027 | -0.0227297 | 0.6946678 | 0.99999994 |
| Ptma | 3.09938022 | 0.29653802 | 0.31944013 | -0.0229021 | 0.68339882 | 0.99999994 |
| Twsg1 | 0.23804093 | 0.15823568 | 0.18116684 | -0.0229312 | 0.80025358 | 0.99999994 |
| Tomm22 | 0.65299942 | 0.34607977 | 0.36903441 | -0.0229546 | 0.66057689 | 0.99999994 |

| | | | | | | |
|---|---|---|---|---|---|---|
| Cfap36 | 0.56844154 | 0.31747904 | 0.34043761 | -0.0229586 | 0.67324603 | 0.99999994 |
| Orc6 | 0.38392226 | 0.2390936 | 0.26208204 | -0.0229884 | 0.72034281 | 0.99999994 |
| Tapbpl | 0.14119702 | 0.09285141 | 0.11584645 | -0.022995 | 0.90679258 | 0.99999994 |
| Dnajc15 | 0.50048821 | 0.29126825 | 0.31428599 | -0.0230177 | 0.68704783 | 0.99999994 |
| Chmp4b | 0.8657609 | 0.39079356 | 0.4138264 | -0.0230328 | 0.64448795 | 0.99999994 |
| Nucb1 | 0.57423663 | 0.31948625 | 0.34253579 | -0.0230495 | 0.67289176 | 0.99999994 |
| Yeats4 | 0.45195077 | 0.27068561 | 0.2937454 | -0.0230598 | 0.69935336 | 0.99999994 |
| Slc33a1 | 0.1725609 | 0.11507696 | 0.13813695 | -0.02306 | 0.8667554 | 0.99999994 |
| Psmc5 | 1.0858589 | 0.406466 | 0.42952791 | -0.0230619 | 0.63959951 | 0.99999994 |
| Gosr2 | 0.24023431 | 0.15934034 | 0.18253031 | -0.02319 | 0.801132 | 0.99999994 |
| Wbp1 | 0.47495044 | 0.28048164 | 0.3036811 | -0.0231995 | 0.69445105 | 0.99999994 |
| Gorasp2 | 0.55607624 | 0.31261976 | 0.33589296 | -0.0232732 | 0.67767342 | 0.99999994 |
| Srp19 | 0.70999053 | 0.36127557 | 0.38460384 | -0.0233283 | 0.6567993 | 0.99999994 |
| Chid1 | 0.3111911 | 0.20077808 | 0.22412947 | -0.0233514 | 0.75600409 | 0.99999994 |
| Desi1 | 0.34830999 | 0.22068449 | 0.24405792 | -0.0233734 | 0.73809375 | 0.99999994 |
| BC004004 | 0.43607448 | 0.26325926 | 0.28666626 | -0.023407 | 0.70660987 | 0.99999994 |
| Hook1 | 0.2345375 | 0.15556331 | 0.17897825 | -0.0234149 | 0.80807434 | 0.99999994 |
| Psmd3 | 0.58357773 | 0.32241873 | 0.34587581 | -0.0234571 | 0.67416014 | 0.99999994 |
| Eif6 | 0.75594878 | 0.37137934 | 0.39485922 | -0.0234799 | 0.65387858 | 0.99999994 |
| Eny2 | 0.90440865 | 0.39501611 | 0.41852431 | -0.0235082 | 0.64575422 | 0.99999994 |
| Emc6 | 0.62807127 | 0.33755789 | 0.36107308 | -0.0235152 | 0.66767459 | 0.99999994 |
| Eif4g1 | 0.59475892 | 0.32618101 | 0.34980652 | -0.0236255 | 0.67348699 | 0.99999994 |
| Zfp568 | 0.14458385 | 0.09466832 | 0.11830888 | -0.0236406 | 0.90825373 | 0.99999994 |
| Cyhr1 | 0.34874635 | 0.22064346 | 0.2442851 | -0.0236416 | 0.74027544 | 0.99999994 |
| Snx3 | 0.82649116 | 0.38431031 | 0.40795845 | -0.0236481 | 0.65019583 | 0.99999994 |
| Larp1b | 0.09406839 | 0.05628602 | 0.08005316 | -0.0237671 | 0.97593504 | 0.99999994 |
| Csnk1a1 | 1.31126531 | 0.40060931 | 0.42437768 | -0.0237684 | 0.6453538 | 0.99999994 |
| Tmem176b | 1.32938043 | 0.39953685 | 0.42333795 | -0.0238011 | 0.64588591 | 0.99999994 |
| Txndc9 | 0.83241117 | 0.38505975 | 0.40887265 | -0.0238129 | 0.65086941 | 0.99999994 |
| Snx17 | 0.61491785 | 0.33278024 | 0.35670379 | -0.0239235 | 0.67235709 | 0.99999994 |

| | | | | | | |
|---|---|---|---|---|---|---|
| Snrpb | 1.27927673 | 0.40208456 | 0.42603352 | -0.023949 | 0.64586448 | 0.99999994 |
| Adi1 | 0.20737841 | 0.13761298 | 0.16158603 | -0.0239731 | 0.83831185 | 0.99999994 |
| Ssb | 1.24590977 | 0.4035183 | 0.42749639 | -0.0239781 | 0.64555686 | 0.99999994 |
| Gtf2h5 | 1.07226607 | 0.40537019 | 0.4293598 | -0.0239896 | 0.64502027 | 0.99999994 |
| Atp6v0e | 1.33285317 | 0.39911315 | 0.42312664 | -0.0240135 | 0.64719742 | 0.99999994 |
| Uba5 | 0.40720871 | 0.24930316 | 0.27331739 | -0.0240142 | 0.72067276 | 0.99999994 |
| Ninj1 | 0.43773618 | 0.26339085 | 0.28741601 | -0.0240252 | 0.7110334 | 0.99999994 |
| Tmem160 | 0.4951966 | 0.28806377 | 0.31212445 | -0.0240607 | 0.69606545 | 0.99999994 |
| BC031181 | 1.03351858 | 0.40398829 | 0.42807853 | -0.0240902 | 0.6460167 | 0.99999994 |
| Naga | 0.19388453 | 0.12855768 | 0.15265513 | -0.0240974 | 0.85331159 | 0.99999994 |
| BC005537 | 0.29563554 | 0.19131702 | 0.21541479 | -0.0240978 | 0.77175099 | 0.99999994 |
| Itpkb | 0.10277979 | 0.06280012 | 0.08694225 | -0.0241421 | 0.96772326 | 0.99999994 |
| Mrpl34 | 0.81235544 | 0.38152611 | 0.4056986 | -0.0241725 | 0.65416939 | 0.99999994 |
| Idh3a | 0.42622159 | 0.2580022 | 0.28217986 | -0.0241777 | 0.71577251 | 0.99999994 |
| Rab25 | 0.47111309 | 0.2778545 | 0.30204942 | -0.0241949 | 0.7030493 | 0.99999994 |
| Gm10073 | 0.34602802 | 0.21859756 | 0.2428673 | -0.0242697 | 0.74702916 | 0.99999994 |
| Tmem234 | 0.97918883 | 0.40075283 | 0.42505502 | -0.0243022 | 0.64823961 | 0.99999994 |
| Atp2a3 | 0.26672825 | 0.17421637 | 0.19862505 | -0.0244087 | 0.79331797 | 0.99999994 |
| Snd1 | 0.51700986 | 0.29649334 | 0.32091462 | -0.0244213 | 0.69376193 | 0.99999994 |
| Snrnp70 | 0.81091362 | 0.38100637 | 0.40544624 | -0.0244399 | 0.65587768 | 0.99999994 |
| Ndufs4 | 1.11096769 | 0.40536217 | 0.4298682 | -0.024506 | 0.64782952 | 0.99999994 |
| Dguok | 0.36746082 | 0.22928066 | 0.25387981 | -0.0245992 | 0.7405215 | 0.99999994 |
| Chchd1 | 0.71278458 | 0.36063244 | 0.38526939 | -0.024637 | 0.66482087 | 0.99999994 |
| Rcan3 | 0.17813112 | 0.11730498 | 0.14197825 | -0.0246733 | 0.87631379 | 0.99999994 |
| Taf6l | 0.55937444 | 0.31243222 | 0.33711428 | -0.0246821 | 0.68699479 | 0.99999994 |
| Sdhaf1 | 0.37734469 | 0.23412479 | 0.25883131 | -0.0247065 | 0.73740872 | 0.99999994 |
| Dnajc5 | 0.31522468 | 0.20155872 | 0.22635382 | -0.0247951 | 0.76704393 | 0.99999994 |
| Snrpd1 | 0.94415955 | 0.39757773 | 0.42241275 | -0.024835 | 0.65223033 | 0.99999994 |
| Tmem214 | 0.29287078 | 0.1889964 | 0.21384263 | -0.0248462 | 0.78035646 | 0.99999994 |
| Bccip | 0.53121936 | 0.30160015 | 0.32647379 | -0.0248736 | 0.69397165 | 0.99999994 |

| | | | | | | |
|---|---|---|---|---|---|---|
| Ndufa4 | 2.14124261 | 0.34363978 | 0.3685241 | -0.0248843 | 0.67345213 | 0.99999994 |
| Ubl3 | 0.55162681 | 0.30930427 | 0.33423512 | -0.0249308 | 0.69023012 | 0.99999994 |
| Dnajc19 | 0.56927594 | 0.31580757 | 0.34074092 | -0.0249334 | 0.68689634 | 0.99999994 |
| Ubfd1 | 0.40001721 | 0.2449216 | 0.2698925 | -0.0249709 | 0.73100558 | 0.99999994 |
| Ctsd | 0.6635454 | 0.34723256 | 0.37226106 | -0.0250285 | 0.67275439 | 0.99999994 |
| Naa38 | 0.6872291 | 0.35376468 | 0.37889611 | -0.0251314 | 0.67057557 | 0.99999994 |
| Phf5a | 0.6060339 | 0.32851315 | 0.35369403 | -0.0251809 | 0.68221173 | 0.99999994 |
| Papss1 | 0.43724815 | 0.26198582 | 0.28719601 | -0.0252102 | 0.72049007 | 0.99999994 |
| Sod1 | 0.36148048 | 0.22562714 | 0.25084573 | -0.0252186 | 0.7483135 | 0.99999994 |
| Plpp5 | 0.24279611 | 0.15889242 | 0.18411676 | -0.0252243 | 0.81909815 | 0.99999994 |
| Copg1 | 0.55067737 | 0.30861196 | 0.33387983 | -0.0252679 | 0.69278152 | 0.99999994 |
| Atp6v0d1 | 0.86804307 | 0.38885066 | 0.41413401 | -0.0252834 | 0.65776766 | 0.99999994 |
| Rer1 | 0.63649022 | 0.33849874 | 0.36380947 | -0.0253107 | 0.6783508 | 0.99999994 |
| Ubxn6 | 0.48015775 | 0.28051677 | 0.30587944 | -0.0253627 | 0.70950069 | 0.99999994 |
| Pdia5 | 0.25457146 | 0.16594573 | 0.19132579 | -0.0253801 | 0.81137918 | 0.99999994 |
| Polr2c | 0.87528046 | 0.3895195 | 0.41509476 | -0.0255753 | 0.65914049 | 0.99999994 |
| Nabp2 | 0.45498899 | 0.26948382 | 0.29507943 | -0.0255956 | 0.71815931 | 0.99999994 |
| Ube2e1 | 0.71796196 | 0.3608488 | 0.38649173 | -0.0256429 | 0.67062355 | 0.99999994 |
| Map2k2 | 0.99543259 | 0.40042231 | 0.42612331 | -0.025701 | 0.65596237 | 0.99999994 |
| Saraf | 1.04137294 | 0.40269703 | 0.42839949 | -0.0257025 | 0.6551853 | 0.99999994 |
| Sec22b | 0.59382453 | 0.32375753 | 0.34948074 | -0.0257232 | 0.68791012 | 0.99999994 |
| Tmem219 | 0.20509257 | 0.1343604 | 0.16008674 | -0.0257263 | 0.85762128 | 0.99999994 |
| Tmed2 | 1.57655225 | 0.37522044 | 0.40097135 | -0.0257509 | 0.66548519 | 0.99999994 |
| Sod2 | 0.90128281 | 0.3924115 | 0.41818244 | -0.0257709 | 0.6591723 | 0.99999994 |
| Gmppa | 0.34911905 | 0.21870014 | 0.24447901 | -0.0257789 | 0.75862109 | 0.99999994 |
| Ndufa6 | 1.64530549 | 0.36848207 | 0.39426178 | -0.0257797 | 0.66830633 | 0.99999994 |
| Trmt112 | 0.80539284 | 0.37868695 | 0.40447105 | -0.0257841 | 0.66433651 | 0.99999994 |
| Rps28 | 2.3170336 | 0.34453969 | 0.37037812 | -0.0258384 | 0.67880609 | 0.99999994 |
| Tmed4 | 1.02061895 | 0.40150836 | 0.42744661 | -0.0259382 | 0.65686426 | 0.99999994 |
| Zfp106 | 0.35217735 | 0.22007117 | 0.24606589 | -0.0259947 | 0.75903021 | 0.99999994 |

| | | | | | | |
|---|---|---|---|---|---|---|
| Mrpl54 | 1.02054563 | 0.40142352 | 0.42744319 | -0.0260197 | 0.65733154 | 0.99999994 |
| Arfgap3 | 0.29263029 | 0.18767423 | 0.21370555 | -0.0260313 | 0.79126152 | 0.99999994 |
| Denr | 0.74208136 | 0.36582457 | 0.39186572 | -0.0260411 | 0.67087964 | 0.99999994 |
| Phactr1 | 0.1637803 | 0.10591752 | 0.13201092 | -0.0260934 | 0.90586294 | 0.99999994 |
| Aco2 | 0.84111563 | 0.38413434 | 0.41023195 | -0.0260976 | 0.66401859 | 0.99999994 |
| Rab3d | 0.46502484 | 0.27329795 | 0.29943896 | -0.026141 | 0.7194134 | 0.99999994 |
| Wfs1 | 0.19248661 | 0.12554175 | 0.15171882 | -0.0261771 | 0.87461192 | 0.99999994 |
| Aldh9a1 | 0.20814423 | 0.13573541 | 0.16208696 | -0.0263515 | 0.8604111 | 0.99999994 |
| Nol7 | 0.7179633 | 0.36007742 | 0.38649204 | -0.0264146 | 0.6754187 | 0.99999994 |
| Atp6v1e1 | 1.250245 | 0.40087998 | 0.42732097 | -0.026441 | 0.65978265 | 0.99999994 |
| Prkacb | 0.23517075 | 0.15290446 | 0.17937476 | -0.0264703 | 0.83701839 | 0.99999994 |
| Bclaf1 | 0.69269289 | 0.35380952 | 0.3803071 | -0.0264976 | 0.67859593 | 0.99999994 |
| Edem1 | 0.14470448 | 0.09189727 | 0.11839634 | -0.0264991 | 0.93186206 | 0.99999994 |
| Vbp1 | 0.74726715 | 0.36646925 | 0.39300332 | -0.0265341 | 0.67343266 | 0.99999994 |
| Eif3d | 0.67719711 | 0.34966068 | 0.37621712 | -0.0265564 | 0.68077229 | 0.99999994 |
| Derl1 | 0.36369584 | 0.22539481 | 0.25197304 | -0.0265782 | 0.75869643 | 0.99999994 |
| Sesn3 | 0.28912356 | 0.18511108 | 0.21170076 | -0.0265897 | 0.79843397 | 0.99999994 |
| Ilk | 0.39035711 | 0.23863874 | 0.2652296 | -0.0265909 | 0.74772244 | 0.99999994 |
| H13 | 0.76701317 | 0.37051773 | 0.39715647 | -0.0266387 | 0.67236982 | 0.99999994 |
| Eif4a3 | 0.68720783 | 0.35218999 | 0.37889049 | -0.0267005 | 0.68049203 | 0.99999994 |
| Ppig | 0.79999672 | 0.37678016 | 0.40350412 | -0.026724 | 0.6703387 | 0.99999994 |
| Psmd6 | 0.9696757 | 0.39756361 | 0.42437702 | -0.0268134 | 0.66296494 | 0.99999994 |
| Vapb | 0.65546597 | 0.34294591 | 0.3698006 | -0.0268547 | 0.68558727 | 0.99999994 |
| Mia3 | 0.3488944 | 0.21746162 | 0.24436214 | -0.0269005 | 0.76814846 | 0.99999994 |
| Uso1 | 0.38925554 | 0.23778351 | 0.26469305 | -0.0269095 | 0.7507039 | 0.99999994 |
| Lamtor2 | 0.98811815 | 0.39875049 | 0.42566136 | -0.0269109 | 0.66305812 | 0.99999994 |
| Mvb12a | 0.3265896 | 0.20561375 | 0.23254242 | -0.0269287 | 0.77959132 | 0.99999994 |
| Copa | 0.35182611 | 0.21893925 | 0.24588403 | -0.0269448 | 0.76712934 | 0.99999994 |
| Smim10l1 | 0.45045713 | 0.26610975 | 0.29308702 | -0.0269773 | 0.72995808 | 0.99999994 |
| Srprb | 0.39682117 | 0.24126837 | 0.26835781 | -0.0270894 | 0.74918509 | 0.99999994 |

| | | | | | | |
|---|---|---|---|---|---|---|
| Ndufs7 | 0.99842335 | 0.39917954 | 0.42630525 | -0.0271257 | 0.66405024 | 0.99999994 |
| Tbrg1 | 0.86306064 | 0.3863315 | 0.41345953 | -0.027128 | 0.66884961 | 0.99999994 |
| Nudc | 0.84943356 | 0.38434218 | 0.41149017 | -0.027148 | 0.66972432 | 0.99999994 |
| Suclg1 | 0.91586535 | 0.39249654 | 0.41973092 | -0.0272344 | 0.66709298 | 0.99999994 |
| Ndufv1 | 0.59789488 | 0.32364371 | 0.35089487 | -0.0272512 | 0.69740286 | 0.99999994 |
| Upf3b | 0.44459908 | 0.2631942 | 0.29048973 | -0.0272955 | 0.73416443 | 0.99999994 |
| Chchd7 | 0.47184359 | 0.27504687 | 0.3023608 | -0.0273139 | 0.72618185 | 0.99999994 |
| Nckap1 | 0.25302079 | 0.16301122 | 0.19038449 | -0.0273733 | 0.83072737 | 0.99999994 |
| Sec23b | 0.38548705 | 0.23546525 | 0.2628504 | -0.0273852 | 0.75600044 | 0.99999994 |
| Sdhd | 0.83068063 | 0.3812057 | 0.40859834 | -0.0273926 | 0.67229006 | 0.99999994 |
| Emc7 | 0.80188445 | 0.37643753 | 0.40384393 | -0.0274064 | 0.67426681 | 0.99999994 |
| Sec31a | 0.30835033 | 0.19509298 | 0.2225542 | -0.0274612 | 0.79427245 | 0.99999994 |
| Sec24d | 0.14360763 | 0.0899814 | 0.11760049 | -0.0276191 | 0.94100666 | 0.99999994 |
| Atp5l | 2.89695603 | 0.32132005 | 0.34895556 | -0.0276355 | 0.70095523 | 0.99999994 |
| Tm9sf2 | 0.40943474 | 0.24673165 | 0.274369 | -0.0276374 | 0.7487301 | 0.99999994 |
| Bcap31 | 0.71503403 | 0.35814339 | 0.38580221 | -0.0276588 | 0.68339219 | 0.99999994 |
| Ywhaz | 0.67533105 | 0.34803548 | 0.37571277 | -0.0276773 | 0.68805607 | 0.99999994 |
| Ptges3 | 0.88438586 | 0.38852127 | 0.41622973 | -0.0277085 | 0.67115855 | 0.99999994 |
| Grpel1 | 0.74720929 | 0.36516201 | 0.3929907 | -0.0278287 | 0.681306 | 0.99999994 |
| Rnf10 | 0.51357269 | 0.29166948 | 0.31954997 | -0.0278805 | 0.71929959 | 0.99999994 |
| Unc50 | 0.53135844 | 0.29849888 | 0.32652758 | -0.0280287 | 0.71612658 | 0.99999994 |
| Ddx50 | 0.3558081 | 0.21988667 | 0.24793983 | -0.0280532 | 0.77431528 | 0.99999994 |
| D8Ertd738e | 1.27315132 | 0.39825213 | 0.42632173 | -0.0280696 | 0.66940239 | 0.99999994 |
| Arl3 | 0.74388672 | 0.36406755 | 0.3922633 | -0.0281957 | 0.6838374 | 0.99999994 |
| Clpp | 0.55353635 | 0.30656771 | 0.33494805 | -0.0283803 | 0.71362193 | 0.99999994 |
| Higd2a | 0.94371533 | 0.39398856 | 0.42237671 | -0.0283882 | 0.67269862 | 0.99999994 |
| Ciapin1 | 0.8376951 | 0.38119791 | 0.40970185 | -0.0285039 | 0.67835321 | 0.99999994 |
| Acaa1a | 0.51352898 | 0.29097716 | 0.31953256 | -0.0285554 | 0.72402919 | 0.99999994 |
| Mrpl52 | 0.83635676 | 0.38087604 | 0.40949301 | -0.028617 | 0.67909596 | 0.99999994 |
| Mrps16 | 0.61175721 | 0.32702219 | 0.35563924 | -0.028617 | 0.70388728 | 0.99999994 |

| | | | | | | |
|---|---|---|---|---|---|---|
| Creb3l2 | 0.17824927 | 0.11341366 | 0.14205934 | -0.0286457 | 0.9102388 | 0.99999994 |
| Polr2e | 0.65985119 | 0.3424271 | 0.37114424 | -0.0287171 | 0.69673503 | 0.99999994 |
| Uggt1 | 0.2828954 | 0.17934585 | 0.20811219 | -0.0287663 | 0.82123356 | 0.99999994 |
| Ssbp1 | 0.73239671 | 0.36090775 | 0.38971968 | -0.0288119 | 0.688677 | 0.99999994 |
| Sap18 | 0.63290104 | 0.33371086 | 0.36264751 | -0.0289366 | 0.70233535 | 0.99999994 |
| Arcn1 | 0.33357526 | 0.20733799 | 0.23629031 | -0.0289523 | 0.79263386 | 0.99999994 |
| Pin1 | 1.03929564 | 0.39932352 | 0.42832178 | -0.0289983 | 0.67387872 | 0.99999994 |
| Gps1 | 0.59003755 | 0.31909491 | 0.34815535 | -0.0290604 | 0.71075942 | 0.99999994 |
| Wdr83os | 0.43606957 | 0.25755911 | 0.28666404 | -0.0291049 | 0.75042202 | 0.99999994 |
| Ndufs8 | 0.91231993 | 0.39019066 | 0.41936581 | -0.0291752 | 0.67834678 | 0.99999994 |
| Pebp1 | 1.9561714 | 0.342229 | 0.3714464 | -0.0292174 | 0.69971428 | 0.99999994 |
| Actr10 | 0.68481275 | 0.34901713 | 0.37825581 | -0.0292387 | 0.69655845 | 0.99999994 |
| Mrps36 | 0.68041496 | 0.34782414 | 0.37708236 | -0.0292582 | 0.6972383 | 0.99999994 |
| Ndufa11 | 1.6298398 | 0.36642895 | 0.39573076 | -0.0293018 | 0.68894731 | 0.99999994 |
| Dph3 | 0.50349707 | 0.28618201 | 0.31550633 | -0.0293243 | 0.73193194 | 0.99999994 |
| Smim7 | 0.65556523 | 0.34042183 | 0.36983137 | -0.0294095 | 0.70170946 | 0.99999994 |
| Lamtor5 | 0.74995627 | 0.36405178 | 0.3935879 | -0.0295361 | 0.69129163 | 0.99999994 |
| Lman2 | 0.76063472 | 0.36621344 | 0.39583954 | -0.0296261 | 0.69082372 | 0.99999994 |
| Yipf5 | 0.43166535 | 0.25501699 | 0.28466728 | -0.0296503 | 0.7559428 | 0.99999994 |
| Ndufa3 | 0.98819399 | 0.3960068 | 0.42566639 | -0.0296596 | 0.67860517 | 0.99999994 |
| Epn1 | 0.45872311 | 0.26703007 | 0.29670964 | -0.0296796 | 0.7472398 | 0.99999994 |
| Csnk2b | 0.72516339 | 0.35839332 | 0.38812063 | -0.0297273 | 0.69491478 | 0.99999994 |
| Rab11a | 1.11447803 | 0.40005838 | 0.42989578 | -0.0298374 | 0.67795651 | 0.99999994 |
| Cops4 | 0.69088754 | 0.34998315 | 0.37984399 | -0.0298608 | 0.69960999 | 0.99999994 |
| Prdx2 | 2.33383321 | 0.34060685 | 0.37049416 | -0.0298873 | 0.70435247 | 0.99999994 |
| Srpr | 0.69399529 | 0.35064376 | 0.38064011 | -0.0299963 | 0.70005348 | 0.99999994 |
| Golga4 | 0.4441736 | 0.26026914 | 0.29030015 | -0.030031 | 0.75445863 | 0.99999994 |
| Adipor1 | 0.60568697 | 0.32347079 | 0.35357546 | -0.0301047 | 0.7145566 | 0.99999994 |
| Copb1 | 0.64402891 | 0.33604055 | 0.36622105 | -0.0301805 | 0.70834445 | 0.99999994 |
| Slc38a10 | 0.24878413 | 0.15759041 | 0.1877998 | -0.0302094 | 0.85785649 | 0.99999994 |

| | | | | | | |
|---|---|---|---|---|---|---|
| Erlec1 | 0.41333727 | 0.24598159 | 0.27620368 | -0.0302221 | 0.76679599 | 0.99999994 |
| Pdcd6 | 0.76034758 | 0.36555473 | 0.39577978 | -0.030225 | 0.6943908 | 0.99999994 |
| Stx5a | 0.45962902 | 0.26686759 | 0.29710368 | -0.0302361 | 0.75092838 | 0.99999994 |
| Uba1 | 0.51892165 | 0.29139915 | 0.32166992 | -0.0302708 | 0.73447271 | 0.99999994 |
| Hagh | 0.75618145 | 0.36463243 | 0.39490815 | -0.0302757 | 0.69508324 | 0.99999994 |
| Psme2 | 1.02538432 | 0.39725058 | 0.42768648 | -0.0304359 | 0.68214069 | 0.99999994 |
| Ndufa5 | 1.07997532 | 0.39895939 | 0.429462 | -0.0305026 | 0.68181096 | 0.99999994 |
| 0610009B22 | 0.6756402 | 0.34512034 | 0.37579645 | -0.0306761 | 0.70655761 | 0.99999994 |
| Jtb | 0.62124973 | 0.32810608 | 0.3588201 | -0.030714 | 0.7155818 | 0.99999994 |
| Cdc37 | 0.9709165 | 0.39369932 | 0.42446739 | -0.0307681 | 0.68527275 | 0.99999994 |
| Tmem208 | 0.7229497 | 0.35678996 | 0.38762573 | -0.0308358 | 0.70177617 | 0.99999994 |
| Sdhc | 0.56867848 | 0.30967046 | 0.34052379 | -0.0308533 | 0.72677978 | 0.99999994 |
| Trappc1 | 0.767756 | 0.36643978 | 0.39730543 | -0.0308656 | 0.69746224 | 0.99999994 |
| Arhgdia | 0.82259676 | 0.3764675 | 0.40735635 | -0.0308889 | 0.69312772 | 0.99999994 |
| Dpp3 | 0.4355803 | 0.25552318 | 0.2864429 | -0.0309197 | 0.76377924 | 0.99999994 |
| Os9 | 0.56445535 | 0.30804833 | 0.33898302 | -0.0309347 | 0.72822105 | 0.99999994 |
| Yif1b | 0.35512001 | 0.21664434 | 0.24758552 | -0.0309412 | 0.79725501 | 0.99999994 |
| Sys1 | 0.65835144 | 0.33974654 | 0.3706888 | -0.0309423 | 0.71076527 | 0.99999994 |
| Psenen | 1.2931178 | 0.39438604 | 0.42533834 | -0.0309523 | 0.68594542 | 0.99999994 |
| Tpt1 | 2.83196368 | 0.32311743 | 0.35411114 | -0.0309937 | 0.71991876 | 0.99999994 |
| Akr1a1 | 1.51549986 | 0.3763681 | 0.40740164 | -0.0310335 | 0.69393784 | 0.99999994 |
| Pdcd5 | 1.00822557 | 0.39576691 | 0.42685851 | -0.0310916 | 0.68610401 | 0.99999994 |
| Sar1a | 0.61572321 | 0.32583474 | 0.35697417 | -0.0311394 | 0.71926006 | 0.99999994 |
| Prpf19 | 0.64429115 | 0.33507247 | 0.3663039 | -0.0312314 | 0.7148221 | 0.99999994 |
| Tmem50a | 0.99655259 | 0.39489849 | 0.42619181 | -0.0312933 | 0.68748924 | 0.99999994 |
| Cyc1 | 0.9598283 | 0.39232566 | 0.42363962 | -0.031314 | 0.68864548 | 0.99999994 |
| Ociad1 | 0.76400323 | 0.36522148 | 0.39653752 | -0.031316 | 0.70044378 | 0.99999994 |
| Eid1 | 0.65906167 | 0.33956836 | 0.37090463 | -0.0313363 | 0.71306784 | 0.99999994 |
| Ndufb10 | 1.21087587 | 0.3973458 | 0.42871409 | -0.0313683 | 0.68688309 | 0.99999994 |
| Eif3g | 1.01158014 | 0.39560881 | 0.42702284 | -0.031414 | 0.68781857 | 0.99999994 |

| | | | | | | |
|---|---|---|---|---|---|---|
| Bcas2 | 0.818522 | 0.37527352 | 0.40671882 | -0.0314453 | 0.69659549 | 0.99999994 |
| Psmb2 | 1.58400588 | 0.36868961 | 0.40018804 | -0.0314984 | 0.69983143 | 0.99999994 |
| Polr2f | 0.86852194 | 0.38265339 | 0.41419827 | -0.0315449 | 0.69390183 | 0.99999994 |
| Txnl1 | 0.90178592 | 0.38664425 | 0.41823774 | -0.0315935 | 0.69245395 | 0.99999994 |
| Cd63 | 2.39501286 | 0.33886171 | 0.37046082 | -0.0315991 | 0.71490301 | 0.99999994 |
| Cisd1 | 0.88774166 | 0.38496756 | 0.41663312 | -0.0316656 | 0.69353876 | 0.99999994 |
| Kif5b | 1.29374938 | 0.39350912 | 0.42530573 | -0.0317966 | 0.69063148 | 0.99999994 |
| Churc1 | 0.60846159 | 0.32267409 | 0.35452187 | -0.0318478 | 0.72506698 | 0.99999994 |
| Mrps14 | 0.92525884 | 0.38869805 | 0.42067395 | -0.0319759 | 0.69355667 | 0.99999994 |
| Eif2s2 | 1.29625782 | 0.39315796 | 0.42517607 | -0.0320181 | 0.69190613 | 0.99999994 |
| Cmpk1 | 0.70842777 | 0.35221149 | 0.38422976 | -0.0320183 | 0.71044367 | 0.99999994 |
| Ndufb11 | 1.82740793 | 0.34647084 | 0.37850337 | -0.0320325 | 0.71339286 | 0.99999994 |
| Mff | 0.67797459 | 0.34436198 | 0.37642669 | -0.0320647 | 0.71464314 | 0.99999994 |
| Cct5 | 1.61482309 | 0.36495628 | 0.39721098 | -0.0322547 | 0.70557933 | 0.99999994 |
| Mrpl17 | 0.75293581 | 0.36194813 | 0.39422311 | -0.032275 | 0.70710772 | 0.99999994 |
| Srp68 | 0.50213118 | 0.28266138 | 0.31495315 | -0.0322918 | 0.75252859 | 0.99999994 |
| Hnrnpa3 | 1.41574827 | 0.38478442 | 0.4170929 | -0.0323085 | 0.69693618 | 0.99999994 |
| Mdh1 | 1.43206095 | 0.38313569 | 0.41569284 | -0.0325572 | 0.69893063 | 0.99999994 |
| 4833439L19F | 0.63525088 | 0.33076102 | 0.36340902 | -0.032648 | 0.72507837 | 0.99999994 |
| Ppp2r1a | 0.83955042 | 0.37722734 | 0.40999003 | -0.0327627 | 0.70260787 | 0.99999994 |
| Aimp1 | 0.66288201 | 0.33929032 | 0.37206103 | -0.0327707 | 0.72115775 | 0.99999994 |
| Pigp | 0.723982 | 0.35499406 | 0.38785684 | -0.0328628 | 0.71361867 | 0.99999994 |
| Arf4 | 0.95301898 | 0.39006394 | 0.42311558 | -0.0330516 | 0.69845459 | 0.99999994 |
| Eif4g2 | 1.25626224 | 0.39394564 | 0.42707091 | -0.0331253 | 0.69717502 | 0.99999994 |
| Tm2d2 | 0.75666672 | 0.36187457 | 0.39501012 | -0.0331356 | 0.71170439 | 0.99999994 |
| Rnh1 | 0.71644567 | 0.35294529 | 0.38613521 | -0.0331899 | 0.71638694 | 0.99999994 |
| Commd6 | 0.62036391 | 0.32529666 | 0.35852534 | -0.0332287 | 0.73135679 | 0.99999994 |
| Timm17a | 0.96498591 | 0.39060845 | 0.42403032 | -0.0334219 | 0.70008708 | 0.99999994 |
| Mrpl51 | 0.65747152 | 0.33695766 | 0.37042105 | -0.0334634 | 0.72618902 | 0.99999994 |
| Ube2l3 | 1.0885653 | 0.39604387 | 0.42957002 | -0.0335261 | 0.69829428 | 0.99999994 |

| | | | | | | |
|---|---|---|---|---|---|---|
| Ndufb7 | 1.4749863 | 0.37802583 | 0.41157412 | -0.0335483 | 0.70628846 | 0.99999994 |
| Sdf4 | 0.86941264 | 0.3806903 | 0.41431754 | -0.0336272 | 0.70548759 | 0.99999994 |
| Cfdp1 | 0.71751541 | 0.35261675 | 0.38638686 | -0.0337701 | 0.71963712 | 0.99999994 |
| Tfg | 0.82432375 | 0.37376234 | 0.40762422 | -0.0338619 | 0.70985204 | 0.99999994 |
| Mettl9 | 0.64148415 | 0.33154847 | 0.36541486 | -0.0338664 | 0.7313518 | 0.99999994 |
| Mrps34 | 0.68396004 | 0.34415831 | 0.3780291 | -0.0338708 | 0.72454558 | 0.99999994 |
| Tbcb | 1.07280985 | 0.39544813 | 0.4293676 | -0.0339195 | 0.70050149 | 0.99999994 |
| Bud31 | 0.9467575 | 0.38859389 | 0.42262196 | -0.0340281 | 0.70400244 | 0.99999994 |
| Copz1 | 0.83659129 | 0.37500152 | 0.40952966 | -0.0345281 | 0.71267239 | 0.99999994 |
| Rraga | 0.78639374 | 0.36631992 | 0.40095343 | -0.0346335 | 0.71734154 | 0.99999994 |
| Cdc42 | 1.55255108 | 0.36859817 | 0.4035538 | -0.0349556 | 0.71788516 | 0.99999994 |
| Tmem147 | 0.84573623 | 0.37596819 | 0.41093978 | -0.0349716 | 0.7144615 | 0.99999994 |
| Rabac1 | 0.87837323 | 0.38047159 | 0.41549507 | -0.0350235 | 0.71263082 | 0.99999994 |
| Itpa | 0.53317321 | 0.29213537 | 0.32722838 | -0.035093 | 0.76233972 | 0.99999994 |
| Pam16 | 0.7023294 | 0.34735194 | 0.38274951 | -0.0353976 | 0.73091929 | 0.99999994 |
| Dctn2 | 0.85376768 | 0.37670191 | 0.41212635 | -0.0354244 | 0.71638957 | 0.99999994 |
| Fam104a | 0.96259116 | 0.38841062 | 0.42385014 | -0.0354395 | 0.71108605 | 0.99999994 |
| Rbx1 | 1.86475062 | 0.34056167 | 0.37624429 | -0.0356826 | 0.73607194 | 0.99999994 |
| Rps25 | 2.43914536 | 0.33446833 | 0.37031299 | -0.0358447 | 0.7403115 | 0.99999994 |
| Scp2 | 0.97336206 | 0.38874364 | 0.42464386 | -0.0359002 | 0.71319456 | 0.99999994 |
| Itm2b | 1.98628509 | 0.3347499 | 0.37070794 | -0.035958 | 0.7407487 | 0.99999994 |
| Tma7 | 2.01469085 | 0.33439269 | 0.37039935 | -0.0360067 | 0.74120573 | 0.99999994 |
| Anapc5 | 0.81517592 | 0.37012929 | 0.40618949 | -0.0360602 | 0.72271637 | 0.99999994 |
| Med29 | 0.57617029 | 0.30715162 | 0.34323157 | -0.03608 | 0.75794443 | 0.99999994 |
| Pfdn6 | 0.64628926 | 0.33082941 | 0.36693393 | -0.0361045 | 0.74375115 | 0.99999994 |
| Ndufs6 | 1.19373099 | 0.39288484 | 0.42914998 | -0.0362651 | 0.71310981 | 0.99999994 |
| Naca | 2.47315473 | 0.3334686 | 0.36987203 | -0.0364034 | 0.74380798 | 0.99999994 |
| Cwc15 | 0.71301686 | 0.34845819 | 0.38532454 | -0.0368663 | 0.73789011 | 0.99999994 |
| Rpl18 | 2.83662644 | 0.31689896 | 0.35380055 | -0.0369016 | 0.75624094 | 0.99999994 |
| Erh | 2.01692251 | 0.33332241 | 0.37033503 | -0.0370126 | 0.74705803 | 0.99999994 |

| | | | | | | |
|---|---|---|---|---|---|---|
| Tmed9 | 1.11760115 | 0.39288572 | 0.42991734 | -0.0370316 | 0.71679821 | 0.99999994 |
| Snrpc | 0.84506368 | 0.37353744 | 0.41083734 | -0.0372999 | 0.727188 | 0.99999994 |
| Atp6ap2 | 0.57009637 | 0.30369589 | 0.34103875 | -0.0373429 | 0.76695451 | 0.99999994 |
| Timm8b | 1.17849494 | 0.39202988 | 0.42946405 | -0.0374342 | 0.71911025 | 0.99999994 |
| Pomp | 1.3860672 | 0.38192942 | 0.41948508 | -0.0375557 | 0.72438911 | 0.99999994 |
| Sra1 | 0.79049078 | 0.36411135 | 0.40173186 | -0.0376205 | 0.73346947 | 0.99999994 |
| Polr2j | 0.80915536 | 0.36749081 | 0.4051372 | -0.0376464 | 0.73188565 | 0.99999994 |
| Eprs | 0.52390222 | 0.28597444 | 0.32362685 | -0.0376524 | 0.78066318 | 0.99999994 |
| Tcf25 | 0.81171776 | 0.36793122 | 0.40558711 | -0.0376559 | 0.7317109 | 0.99999994 |
| Taldo1 | 1.27005108 | 0.38874243 | 0.42646472 | -0.0377223 | 0.72199493 | 0.99999994 |
| G3bp1 | 1.02671768 | 0.38997819 | 0.42775224 | -0.0377741 | 0.72167053 | 0.99999994 |
| Imp3 | 0.61631179 | 0.31936962 | 0.35717156 | -0.0378019 | 0.75943564 | 0.99999994 |
| Eif4h | 0.81332716 | 0.36796804 | 0.40586814 | -0.0379001 | 0.7328886 | 0.99999994 |
| Rpl27a | 2.76122333 | 0.32038254 | 0.35865474 | -0.0382722 | 0.7612418 | 0.99999994 |
| Ndufb6 | 0.98373554 | 0.38706952 | 0.42536733 | -0.0382978 | 0.72551459 | 0.99999994 |
| Park7 | 1.39339979 | 0.38043482 | 0.41892664 | -0.0384918 | 0.72959863 | 0.99999994 |
| Prelid1 | 0.71665907 | 0.34750237 | 0.38618546 | -0.0386831 | 0.74753354 | 0.99999994 |
| Aamp | 0.96746631 | 0.38535498 | 0.42421469 | -0.0388597 | 0.7289857 | 0.99999994 |
| Vcp | 1.28110392 | 0.38693123 | 0.4259439 | -0.0390127 | 0.72895749 | 0.99999994 |
| Mrpl28 | 0.7143134 | 0.34661293 | 0.38563181 | -0.0390189 | 0.74968585 | 0.99999994 |
| Mrps24 | 0.64202898 | 0.32652587 | 0.36558805 | -0.0390622 | 0.76152464 | 0.99999994 |
| Mydgf | 0.90281415 | 0.37918384 | 0.41835043 | -0.0391666 | 0.73341623 | 0.99999994 |
| Mrpl20 | 0.92546475 | 0.38148223 | 0.42069388 | -0.0392116 | 0.73250924 | 0.99999994 |
| Psmd8 | 1.07908472 | 0.39006421 | 0.42945111 | -0.0393869 | 0.7292296 | 0.99999994 |
| Sf3b5 | 0.6244832 | 0.32036352 | 0.35989248 | -0.039529 | 0.76764222 | 0.99999994 |
| Ppp1ca | 1.77241275 | 0.34275962 | 0.38244495 | -0.0396853 | 0.75512709 | 0.99999994 |
| Psmd13 | 0.82217763 | 0.36735178 | 0.40729113 | -0.0399394 | 0.74303442 | 0.99999994 |
| Psma5 | 1.15680984 | 0.38931981 | 0.4297145 | -0.0403947 | 0.73425113 | 0.99999994 |
| Txn1 | 1.36150515 | 0.38067074 | 0.42127781 | -0.0406071 | 0.73943614 | 0.99999994 |
| Copb2 | 0.62434196 | 0.31919637 | 0.35984575 | -0.0406494 | 0.77395174 | 0.99999994 |

| | | | | | | |
|---|---|---|---|---|---|---|
| Cox5a | 1.3774341 | 0.37942875 | 0.42012922 | -0.0407005 | 0.74048516 | 0.99999994 |
| Rpl27 | 1.78690475 | 0.34067589 | 0.38143476 | -0.0407589 | 0.76154357 | 0.99999994 |
| Pfdn1 | 1.02060303 | 0.38657252 | 0.4274458 | -0.0408733 | 0.73777314 | 0.99999994 |
| Ywhaq | 1.89162509 | 0.33340103 | 0.37477975 | -0.0413787 | 0.76880462 | 0.99999994 |
| Idh3g | 0.78363852 | 0.35903708 | 0.40042543 | -0.0413883 | 0.75427594 | 0.99999994 |
| Hsbp1 | 1.52864973 | 0.36429388 | 0.40605791 | -0.041764 | 0.75320887 | 0.99999994 |
| Rhoa | 1.14425122 | 0.38806753 | 0.42985597 | -0.0417884 | 0.74121561 | 0.99999994 |
| Srp72 | 0.67861862 | 0.33478067 | 0.37660004 | -0.0418194 | 0.77009268 | 0.99999994 |
| Psmb6 | 1.78802552 | 0.33948972 | 0.38135601 | -0.0418663 | 0.76753495 | 0.99999994 |
| Pfdn2 | 1.33311069 | 0.38123011 | 0.42311089 | -0.0418808 | 0.74500914 | 0.99999994 |
| Uqcrb | 2.27428903 | 0.32816083 | 0.37007356 | -0.0419127 | 0.77453534 | 0.99999994 |
| Uqcrc1 | 1.25154754 | 0.3852492 | 0.42726749 | -0.0420183 | 0.74363694 | 0.99999994 |
| Arpp19 | 1.0464407 | 0.38656042 | 0.42858322 | -0.0420228 | 0.74301201 | 0.99999994 |
| Arl1 | 0.88683653 | 0.37448529 | 0.41652479 | -0.0420395 | 0.74915106 | 0.99999994 |
| Rtn3 | 0.85866675 | 0.37064252 | 0.41283557 | -0.042193 | 0.75184273 | 0.99999994 |
| Uqcrfs1 | 1.26396156 | 0.38447289 | 0.42673992 | -0.042267 | 0.74514456 | 0.99999994 |
| Ndufc2 | 1.36592482 | 0.37866577 | 0.42096329 | -0.0422975 | 0.74819389 | 0.99999994 |
| Gdi2 | 1.00168505 | 0.38406749 | 0.42650011 | -0.0424326 | 0.74609225 | 0.99999994 |
| Tmbim6 | 1.40214207 | 0.37571669 | 0.41821895 | -0.0425023 | 0.75062712 | 0.99999994 |
| Uqcr10 | 1.77611508 | 0.3395415 | 0.38218831 | -0.0426468 | 0.7711851 | 0.99999994 |
| Ergic3 | 0.99015222 | 0.3831377 | 0.42579441 | -0.0426567 | 0.74756399 | 0.99999994 |
| Rps15 | 2.64663324 | 0.32179719 | 0.3645232 | -0.042726 | 0.78236229 | 0.99999994 |
| Kdelr1 | 0.86277126 | 0.37064066 | 0.41342002 | -0.0427794 | 0.75451862 | 0.99999994 |
| Ubxn4 | 1.09398305 | 0.38685874 | 0.42966149 | -0.0428027 | 0.74636813 | 0.99999994 |
| Atp5g3 | 1.21356562 | 0.38536896 | 0.42862782 | -0.0432589 | 0.74914024 | 0.99999994 |
| Ufc1 | 0.83627504 | 0.3662174 | 0.40948023 | -0.0432628 | 0.75905453 | 0.99999994 |
| Prkar1a | 1.45151358 | 0.37058715 | 0.41390686 | -0.0433197 | 0.75699158 | 0.99999994 |
| Rap1a | 0.80941653 | 0.3617787 | 0.4051832 | -0.0434045 | 0.76209256 | 0.99999994 |
| Psma1 | 1.25215948 | 0.38356257 | 0.42724224 | -0.0436797 | 0.75191284 | 0.99999994 |
| Tspan3 | 1.27486668 | 0.38245182 | 0.42624177 | -0.0437899 | 0.75296219 | 0.99999994 |

| | | | | | | |
|---|---|---|---|---|---|---|
| Eif3i | 0.94712405 | 0.37864517 | 0.42265127 | -0.0440061 | 0.75586231 | 0.99999994 |
| Atraid | 0.84074245 | 0.36609551 | 0.41017437 | -0.0440789 | 0.76279339 | 0.99999994 |
| Psma2 | 1.62351767 | 0.35220502 | 0.39636851 | -0.0441635 | 0.77085129 | 0.99999994 |
| Tmco1 | 0.72600639 | 0.34411984 | 0.38830841 | -0.0441886 | 0.77561809 | 0.99999994 |
| Fkbp8 | 0.90905418 | 0.37482912 | 0.419025 | -0.0441959 | 0.75868133 | 0.99999994 |
| Ran | 1.5424443 | 0.35976683 | 0.40462309 | -0.0448563 | 0.76971836 | 0.99999994 |
| Pgls | 1.06242728 | 0.38412174 | 0.42909988 | -0.0449781 | 0.75732315 | 0.99999994 |
| Mrpl30 | 0.81923322 | 0.36158041 | 0.40683066 | -0.0452502 | 0.77045973 | 0.99999994 |
| Ubl5 | 1.70817585 | 0.34252301 | 0.38828448 | -0.0457615 | 0.78362324 | 0.99999994 |
| Polr1d | 1.00879473 | 0.38088293 | 0.42688666 | -0.0460037 | 0.76341383 | 0.99999994 |
| Fis1 | 1.11610644 | 0.38361883 | 0.4299074 | -0.0462886 | 0.76321944 | 0.99999994 |
| Sec11a | 1.35775821 | 0.37469479 | 0.42153183 | -0.046837 | 0.77022799 | 0.99999994 |
| Ndufs3 | 0.99042972 | 0.37896704 | 0.42581194 | -0.0468449 | 0.76800319 | 0.99999994 |
| Arpc3 | 1.11627976 | 0.38287578 | 0.4299086 | -0.0470328 | 0.76676001 | 0.99999994 |
| Ost4 | 0.85102234 | 0.36466666 | 0.41172434 | -0.0470577 | 0.77660657 | 0.99999994 |
| Gabarapl2 | 1.58124955 | 0.35335269 | 0.40046063 | -0.0471079 | 0.78319128 | 0.99999994 |
| Fau | 2.81172544 | 0.30822333 | 0.35542783 | -0.0472045 | 0.81166091 | 0.99999994 |
| Psma6 | 1.16019619 | 0.38244378 | 0.42968811 | -0.0472443 | 0.76787558 | 0.99999994 |
| Sdf2 | 0.69468784 | 0.33343714 | 0.38081681 | -0.0473797 | 0.79620917 | 0.99999994 |
| Slc25a4 | 1.74466863 | 0.33754617 | 0.38495571 | -0.0474095 | 0.79382618 | 0.99999994 |
| Slc25a3 | 1.58490731 | 0.35247124 | 0.40009874 | -0.0476275 | 0.78592709 | 0.99999994 |
| Emc10 | 0.86890509 | 0.36650933 | 0.41424962 | -0.0477403 | 0.77849299 | 0.99999994 |
| Uqcrc2 | 0.93938715 | 0.3742254 | 0.42200274 | -0.0477773 | 0.77445656 | 0.99999994 |
| Psmb4 | 1.30583436 | 0.37676301 | 0.42467224 | -0.0479092 | 0.77365392 | 0.99999994 |
| Esd | 1.1718745 | 0.38160061 | 0.42957461 | -0.047974 | 0.77137063 | 0.99999994 |
| Atp5e | 2.02182431 | 0.32219535 | 0.37018031 | -0.047985 | 0.80588632 | 0.99999994 |
| Psmb7 | 1.25986616 | 0.37880777 | 0.42691751 | -0.0481097 | 0.77340616 | 0.99999994 |
| Rpl37a | 3.44112782 | 0.21175484 | 0.26012646 | -0.0483716 | 0.89210084 | 0.99999994 |
| Gnb2 | 2.02093727 | 0.32138225 | 0.37021858 | -0.0488363 | 0.81004293 | 0.99999994 |
| Cox6a1 | 2.01705509 | 0.3214667 | 0.3703312 | -0.0488645 | 0.81010784 | 0.99999994 |

| | | | | | | |
|---|---|---|---|---|---|---|
| Myl6 | 1.89140548 | 0.32582979 | 0.37479137 | -0.0489616 | 0.80773116 | 0.99999994 |
| Capzb | 1.16810467 | 0.3798253 | 0.42961503 | -0.0497897 | 0.77977419 | 0.99999994 |
| Pgam1 | 1.11510617 | 0.37998217 | 0.42990036 | -0.0499182 | 0.7802115 | 0.99999994 |
| Ndufa7 | 1.50417531 | 0.35844377 | 0.40862148 | -0.0501777 | 0.79314558 | 0.99999994 |
| Ndufa8 | 1.02498521 | 0.377311 | 0.42766668 | -0.0503557 | 0.78340776 | 0.99999994 |
| Ndufab1 | 1.45273871 | 0.3632936 | 0.41378575 | -0.0504922 | 0.79167337 | 0.99999994 |
| Sf3b6 | 1.27144958 | 0.37568236 | 0.42640046 | -0.0507181 | 0.78574351 | 0.99999994 |
| Vdac2 | 1.24671911 | 0.37660059 | 0.42746395 | -0.0508634 | 0.7858287 | 0.99999994 |
| Eif3k | 1.3554412 | 0.37065752 | 0.4216862 | -0.0510287 | 0.78973228 | 0.99999994 |
| Cd164 | 1.07196703 | 0.37814213 | 0.42935547 | -0.0512133 | 0.78639284 | 0.99999994 |
| Ndufb3 | 0.96635699 | 0.37287956 | 0.42413252 | -0.051253 | 0.78940736 | 0.99999994 |
| Tbca | 1.62042657 | 0.34507453 | 0.39667912 | -0.0516046 | 0.80673729 | 0.99999994 |
| Chmp5 | 1.2201062 | 0.37649571 | 0.42841116 | -0.0519155 | 0.79005919 | 0.99999994 |
| Scand1 | 1.21433875 | 0.37573907 | 0.42860272 | -0.0528637 | 0.79417457 | 0.99999994 |
| Psap | 0.95279828 | 0.36898088 | 0.42309843 | -0.0541176 | 0.80272628 | 0.99999994 |
| Psma3 | 1.64036415 | 0.34038912 | 0.39470822 | -0.0543191 | 0.82017629 | 0.99999994 |
| Timm13 | 1.39607178 | 0.36416021 | 0.41871154 | -0.0545513 | 0.80708807 | 0.99999994 |
| Mlf2 | 1.07401389 | 0.3739808 | 0.42938455 | -0.0554038 | 0.80479499 | 0.99999994 |
| Txndc17 | 0.84974923 | 0.35571556 | 0.41153679 | -0.0558212 | 0.81670374 | 0.99999994 |
| Rplp1 | 2.50924739 | 0.31346678 | 0.36956034 | -0.0560936 | 0.84383198 | 0.99999994 |
| Brk1 | 1.03353796 | 0.37141321 | 0.42807944 | -0.0566662 | 0.81087441 | 0.99999994 |
| Rps27 | 2.77443068 | 0.30037979 | 0.3577731 | -0.0573933 | 0.85719484 | 0.99999994 |
| Arpc1a | 1.14797211 | 0.37124464 | 0.42981837 | -0.0585737 | 0.81782271 | 0.99999994 |
| Rbm39 | 2.04740846 | 0.31052049 | 0.36923179 | -0.0587113 | 0.8550687 | 0.99999994 |
| Sar1b | 0.81785034 | 0.34778162 | 0.40661299 | -0.0588314 | 0.83224745 | 0.99999994 |
| Ndufv2 | 1.11180628 | 0.37009555 | 0.4298751 | -0.0597796 | 0.8226862 | 0.99999994 |
| Ftl1 | 3.57287497 | 0.19688427 | 0.25697832 | -0.0600941 | 0.94021498 | 0.99999994 |
| Rps16 | 2.95638301 | 0.28241666 | 0.34254644 | -0.0601298 | 0.87868557 | 0.99999994 |
| Rpl11 | 2.90590363 | 0.2873535 | 0.34814065 | -0.0607872 | 0.87742988 | 0.99999994 |
| Srp14 | 1.69501126 | 0.32860638 | 0.3894616 | -0.0608552 | 0.85085036 | 0.99999994 |

| | | | | | | |
|---|---|---|---|---|---|---|
| Laptm4a | 1.44347962 | 0.35380227 | 0.41465994 | -0.0608577 | 0.83569612 | 0.99999994 |
| Atp6v1g1 | 1.92482039 | 0.31128279 | 0.37264293 | -0.0613601 | 0.86347141 | 0.99999994 |
| Snrpd2 | 1.49104117 | 0.34830124 | 0.40997046 | -0.0616692 | 0.84165261 | 0.99999994 |
| Gng5 | 1.79438415 | 0.31917525 | 0.38085511 | -0.0616799 | 0.85948395 | 0.99999994 |
| Cox7b | 1.43695874 | 0.35289562 | 0.41525191 | -0.0623563 | 0.84123627 | 0.99999994 |
| Rab2a | 1.35075626 | 0.35945991 | 0.4219956 | -0.0625357 | 0.83803358 | 0.99999994 |
| Ap2s1 | 1.01147621 | 0.36447101 | 0.42701781 | -0.0625468 | 0.83521492 | 0.99999994 |
| Rab1a | 1.12980115 | 0.36656031 | 0.429936 | -0.0633757 | 0.83675183 | 0.99999994 |
| Ndufb5 | 1.44924031 | 0.34954791 | 0.41413109 | -0.0645832 | 0.85037661 | 0.99999994 |
| Swi5 | 1.82021845 | 0.31391278 | 0.37895008 | -0.0650373 | 0.87335556 | 0.99999994 |
| Tomm20 | 1.48617258 | 0.3453294 | 0.4104582 | -0.0651288 | 0.85455859 | 0.99999994 |
| Rpl10 | 3.04884001 | 0.26123341 | 0.32742457 | -0.0661912 | 0.91078894 | 0.99999994 |
| Atp5a1 | 1.7587424 | 0.31564583 | 0.38362419 | -0.0679784 | 0.88090229 | 0.99999994 |
| Atp5c1 | 1.62411491 | 0.32740124 | 0.3963084 | -0.0689072 | 0.87643697 | 0.99999994 |
| Ndufs5 | 1.53108191 | 0.33663719 | 0.4058078 | -0.0691706 | 0.87172585 | 0.99999994 |
| Edf1 | 1.64030528 | 0.32550679 | 0.39471353 | -0.0692067 | 0.87841227 | 0.99999994 |
| Psma7 | 1.97285154 | 0.30155776 | 0.37085526 | -0.0692975 | 0.89321301 | 0.99999994 |
| Spint2 | 2.08752551 | 0.29900639 | 0.36850245 | -0.0694961 | 0.89531975 | 0.99999994 |
| Atp5o | 1.71797897 | 0.31784481 | 0.38741869 | -0.0695739 | 0.88402873 | 0.99999994 |
| Rps6 | 3.01902715 | 0.26363939 | 0.33321356 | -0.0695742 | 0.91770991 | 0.99999994 |
| Rpl19 | 2.8750881 | 0.28083677 | 0.35071691 | -0.0698801 | 0.90762577 | 0.99999994 |
| Rps18 | 2.93826868 | 0.27295236 | 0.34450399 | -0.0715516 | 0.91658932 | 0.99999994 |
| Psmb5 | 1.47544429 | 0.33993079 | 0.41152863 | -0.0715978 | 0.87658322 | 0.99999994 |
| Mdh2 | 1.61874816 | 0.32325853 | 0.39684743 | -0.0735889 | 0.89145691 | 0.99999994 |
| Cfl1 | 2.28977847 | 0.29647565 | 0.37022024 | -0.0737446 | 0.90756152 | 0.99999994 |
| Ndufb9 | 1.51579629 | 0.33279903 | 0.40736953 | -0.0745705 | 0.88847746 | 0.99999994 |
| Uqcrh | 1.84613617 | 0.30221021 | 0.3769849 | -0.0747747 | 0.90662988 | 0.99999994 |
| Ndufa2 | 1.61657253 | 0.32181282 | 0.39704911 | -0.0752363 | 0.89640037 | 0.99999994 |
| Eif5a | 1.96087419 | 0.29576316 | 0.37126186 | -0.0754987 | 0.91207069 | 0.99999994 |
| mt-Co1 | 3.75299143 | 0.16607187 | 0.24371648 | -0.0776446 | 0.98302315 | 0.99999994 |

| | | | | | | |
|---|---|---|---|---|---|---|
| Atp6v1f | 1.68316322 | 0.3118132 | 0.39063609 | -0.0788229 | 0.91038434 | 0.99999994 |
| Rpl35a | 2.78645702 | 0.27429251 | 0.35695658 | -0.0826641 | 0.93840008 | 0.99999994 |
| Atp5d | 1.57678929 | 0.31686175 | 0.40094563 | -0.0840839 | 0.91863234 | 0.99999994 |
| Atp5g2 | 1.84461463 | 0.2914092 | 0.37704434 | -0.0856351 | 0.93470847 | 0.99999994 |
| Ywhae | 2.22975511 | 0.28290464 | 0.36946158 | -0.0865569 | 0.94055413 | 0.99999994 |
| Atp5j | 1.94639394 | 0.28379074 | 0.37182574 | -0.088035 | 0.94248558 | 0.99999994 |
| Serf2 | 2.02771153 | 0.27899088 | 0.3699252 | -0.0909343 | 0.94910671 | 0.99999994 |
| Cox7a2 | 1.97579247 | 0.27850627 | 0.37082399 | -0.0923177 | 0.95125237 | 0.99999994 |
| Arf1 | 1.62300961 | 0.30355168 | 0.39641962 | -0.0928679 | 0.9405448 | 0.99999994 |
| Atp5h | 1.97050179 | 0.27781323 | 0.37087986 | -0.0930666 | 0.95257111 | 0.99999994 |
| Gabarap | 1.91225985 | 0.27986536 | 0.37349877 | -0.0936334 | 0.9524116 | 0.99999994 |
| Atp5j2 | 1.9278487 | 0.27511494 | 0.37253001 | -0.0974151 | 0.95912405 | 0.99999994 |
| Rps2 | 2.41617895 | 0.27280278 | 0.37046291 | -0.0976601 | 0.96034688 | 0.99999994 |
| Hint1 | 1.90349183 | 0.27611477 | 0.37414702 | -0.0980323 | 0.95942637 | 0.99999994 |
| mt-Nd1 | 2.9593508 | 0.2441154 | 0.34216874 | -0.0980533 | 0.97177041 | 0.99999994 |
| Pfdn5 | 1.90438392 | 0.27529738 | 0.37408126 | -0.0987839 | 0.96060487 | 0.99999994 |
| Cox5b | 1.88192947 | 0.27601341 | 0.37529002 | -0.0992766 | 0.96086613 | 0.99999994 |
| Cox6c | 2.00238297 | 0.27036529 | 0.37074884 | -0.1003836 | 0.96424839 | 0.99999994 |
| Eif1 | 3.00690747 | 0.23486366 | 0.33546149 | -0.1005978 | 0.97703856 | 0.99999994 |
| Rps27l | 1.75944903 | 0.28128826 | 0.38356354 | -0.1022753 | 0.96204149 | 0.99999994 |
| Rpl22l1 | 2.13655889 | 0.26608842 | 0.36846354 | -0.1023751 | 0.96780172 | 0.99999994 |
| mt-Co2 | 3.20797544 | 0.19439573 | 0.299056 | -0.1046603 | 0.99008417 | 0.99999994 |
| Rplp2 | 2.826018 | 0.24464626 | 0.3545068 | -0.1098605 | 0.98043344 | 0.99999994 |
| Atp5b | 2.39373698 | 0.25725843 | 0.37046 | -0.1132016 | 0.97902346 | 0.99999994 |
| Psmb1 | 2.12660155 | 0.25025141 | 0.36845344 | -0.118202 | 0.98363578 | 0.99999994 |
| Rpl36 | 2.68934773 | 0.24062198 | 0.36269406 | -0.1220721 | 0.98746534 | 0.99999994 |
| Rpl37 | 2.80243056 | 0.23365668 | 0.35601018 | -0.1223535 | 0.98892121 | 0.99999994 |
| Rpl31 | 2.56163765 | 0.24292598 | 0.36739606 | -0.1244701 | 0.98793669 | 0.99999994 |
| Atpif1 | 2.34535911 | 0.24512026 | 0.37066398 | -0.1255437 | 0.98791781 | 0.99999994 |
| Rpl34 | 2.80375141 | 0.2225426 | 0.3559275 | -0.1333849 | 0.99369309 | 0.99999994 |

| | | | | | | |
|---|---|---|---|---|---|---|
| Ppia | 3.02254733 | 0.1961096 | 0.33252564 | -0.136416 | 0.99684018 | 0.99999994 |
| Cox6b1 | 2.06844212 | 0.23205978 | 0.36890471 | -0.1368449 | 0.99322868 | 0.99999994 |
| Cox8a | 2.23681894 | 0.22499904 | 0.3695647 | -0.1445657 | 0.99539047 | 0.99999994 |
| Fth1 | 2.43255285 | 0.22482375 | 0.37044053 | -0.1456168 | 0.99555888 | 0.99999994 |
| Bsg | 2.29478746 | 0.22198785 | 0.37025209 | -0.1482642 | 0.99615608 | 0.99999994 |
| Rpl41 | 3.71694303 | 0.09742959 | 0.24645206 | -0.1490225 | 0.9999715 | 0.99999994 |
| Oaz1 | 2.55201582 | 0.21876523 | 0.367845 | -0.1490798 | 0.99650878 | 0.99999994 |
| Cox4i1 | 2.51009963 | 0.21889758 | 0.36953151 | -0.1506339 | 0.9966698 | 0.99999994 |
| Cox7c | 2.35134145 | 0.21816485 | 0.37067021 | -0.1525054 | 0.99691526 | 0.99999994 |
| Chchd2 | 2.62986376 | 0.20552329 | 0.36520425 | -0.159681 | 0.99819522 | 0.99999994 |
| Mrfap1 | 2.50944644 | 0.20475892 | 0.36955361 | -0.1647947 | 0.99850256 | 0.99999994 |
| mt-Nd4 | 2.77105345 | 0.19055522 | 0.35800204 | -0.1674468 | 0.99907512 | 0.99999994 |

ꞔ

ꟼMs for all cellular genes.
of log-CPMs for all cellular genes.
ological component of the variance for all genes.
echnical component of the variance for all genes.
