## Supplemental Table 2 for "Differential CpG methylation at *Nnat* in the early establishment of beta cell heterogeneity"

| | P value | log2FC | pct.1 | pct.2 | P adj |
|---|---|---|---|---|---|
| Nnat | 1.65E-34 | 6.58273925 | 1 | 0 | 3.04E-30 |
| Ins2 | 4.37E-32 | 4.87033507 | 1 | 1 | 8.04E-28 |
| Npy | 1.43E-12 | 4.73843395 | 0.575 | 0.056 | 2.64E-08 |
| Ins1 | 5.54E-30 | 4.59606362 | 0.999 | 0.944 | 1.02E-25 |
| Iapp | 7.70E-31 | 3.32260997 | 1 | 0.741 | 1.42E-26 |
| Sdf2l1 | 5.26E-28 | 2.45602443 | 0.981 | 0.444 | 9.69E-24 |
| Ppp1r1a | 8.24E-29 | 2.43902473 | 0.962 | 0.204 | 1.52E-24 |
| Dlk1 | 2.76E-25 | 2.30025687 | 0.994 | 0.537 | 5.08E-21 |
| Gng12 | 6.82E-29 | 2.07206851 | 0.988 | 0.37 | 1.26E-24 |
| Slc2a2 | 8.78E-26 | 1.935078 | 0.871 | 0.056 | 1.62E-21 |
| Scg2 | 3.39E-21 | 1.84958002 | 0.98 | 0.593 | 6.23E-17 |
| Manf | 1.79E-25 | 1.83872939 | 0.999 | 0.833 | 3.30E-21 |
| G6pc2 | 8.07E-19 | 1.81121706 | 0.733 | 0.056 | 1.49E-14 |
| Slc30a8 | 4.72E-23 | 1.79411126 | 0.967 | 0.352 | 8.70E-19 |
| Prss53 | 3.23E-22 | 1.7321276 | 0.822 | 0.056 | 5.95E-18 |
| Creld2 | 3.46E-21 | 1.66691549 | 0.938 | 0.444 | 6.37E-17 |
| Hadh | 8.59E-23 | 1.6616633 | 0.988 | 0.426 | 1.58E-18 |
| Ero1lb | 8.35E-28 | 1.65798497 | 0.945 | 0.204 | 1.54E-23 |
| Sytl4 | 7.68E-27 | 1.6474081 | 0.948 | 0.148 | 1.41E-22 |
| Pdia6 | 6.24E-24 | 1.6321682 | 0.999 | 0.815 | 1.15E-19 |
| Hspa5 | 1.45E-23 | 1.61554141 | 0.999 | 0.907 | 2.68E-19 |
| Pcsk1 | 1.22E-22 | 1.57931895 | 0.946 | 0.315 | 2.24E-18 |
| Fkbp11 | 8.89E-20 | 1.54602207 | 0.804 | 0.111 | 1.64E-15 |
| Sec61b | 4.68E-24 | 1.54570683 | 0.997 | 0.907 | 8.61E-20 |
| Atp2a2 | 7.84E-23 | 1.53276312 | 0.971 | 0.444 | 1.44E-18 |
| Nucb2 | 1.71E-22 | 1.47186531 | 0.916 | 0.278 | 3.16E-18 |
| Scg3 | 4.60E-23 | 1.43402875 | 0.999 | 0.759 | 8.46E-19 |
| Hsp90b1 | 2.29E-22 | 1.38357314 | 1 | 0.87 | 4.22E-18 |
| Fam151a | 7.56E-19 | 1.36105172 | 0.78 | 0.13 | 1.39E-14 |
| Krtcap2 | 3.84E-24 | 1.35516339 | 0.993 | 0.704 | 7.08E-20 |
| Dnajc3 | 1.44E-20 | 1.34077898 | 0.929 | 0.519 | 2.65E-16 |
| Serp1 | 1.94E-23 | 1.32100054 | 0.959 | 0.5 | 3.57E-19 |
| Gm10320 | 2.12E-07 | 1.32011898 | 0.704 | 0.556 | 0.00390229 |
| Ssr4 | 2.57E-24 | 1.31585906 | 1 | 0.926 | 4.74E-20 |
| Fkbp2 | 4.61E-24 | 1.310648 | 0.996 | 0.704 | 8.49E-20 |
| Ostc | 5.52E-23 | 1.30445031 | 0.997 | 0.796 | 1.02E-18 |
| Chgb | 8.32E-17 | 1.2741408 | 1 | 0.63 | 1.53E-12 |
| Papss2 | 4.62E-22 | 1.27106466 | 0.919 | 0.241 | 8.50E-18 |
| Calr | 3.01E-18 | 1.26765849 | 1 | 0.907 | 5.54E-14 |
| Ccnd2 | 1.42E-07 | 1.2582746 | 0.654 | 0.296 | 0.00260814 |
| Pdia4 | 4.98E-17 | 1.23218918 | 0.972 | 0.741 | 9.17E-13 |
| Mlxipl | 4.65E-18 | 1.21013132 | 0.881 | 0.315 | 8.56E-14 |
| Pdia3 | 6.92E-21 | 1.20242377 | 1 | 0.944 | 1.27E-16 |

| | | | | | |
|---|---|---|---|---|---|
| Rpn2 | 6.12E-21 | 1.19660934 | 0.991 | 0.741 | 1.13E-16 |
| S100a10 | 2.34E-14 | 1.19433134 | 0.784 | 0.204 | 4.31E-10 |
| Sec61g | 1.27E-23 | 1.18264894 | 1 | 0.981 | 2.34E-19 |
| Chga | 1.72E-13 | 1.18178741 | 0.999 | 0.87 | 3.17E-09 |
| Tmem215 | 8.66E-21 | 1.15332303 | 0.765 | 0.037 | 1.59E-16 |
| Spcs1 | 5.53E-21 | 1.13571024 | 0.999 | 0.796 | 1.02E-16 |
| Sec11c | 3.04E-18 | 1.13518312 | 0.932 | 0.389 | 5.60E-14 |
| Dnajb9 | 4.56E-18 | 1.09915547 | 0.919 | 0.463 | 8.39E-14 |
| Surf4 | 2.32E-19 | 1.09387264 | 0.943 | 0.537 | 4.27E-15 |
| Scg5 | 2.94E-18 | 1.09363029 | 1 | 0.87 | 5.41E-14 |
| Vimp | 4.83E-18 | 1.08956164 | 0.943 | 0.519 | 8.89E-14 |
| Spcs2 | 9.02E-20 | 1.08623772 | 0.997 | 0.833 | 1.66E-15 |
| Selk | 1.82E-21 | 1.08178994 | 0.996 | 0.907 | 3.35E-17 |
| Ddost | 5.63E-19 | 1.08060019 | 0.997 | 0.741 | 1.04E-14 |
| Rpl36al | 3.15E-22 | 1.07811099 | 0.997 | 0.963 | 5.81E-18 |
| Tmed3 | 1.28E-18 | 1.07501991 | 0.997 | 0.778 | 2.36E-14 |
| P4hb | 1.34E-17 | 1.07422459 | 0.949 | 0.463 | 2.46E-13 |
| Lman1 | 6.21E-18 | 1.07159974 | 0.945 | 0.574 | 1.14E-13 |
| Bace2 | 9.61E-13 | 1.06074463 | 0.557 | 0.019 | 1.77E-08 |
| Derl3 | 6.64E-13 | 1.05855031 | 0.586 | 0.037 | 1.22E-08 |
| Slc35b1 | 1.65E-17 | 1.04211137 | 0.928 | 0.519 | 3.05E-13 |
| Ufm1 | 1.71E-17 | 1.02434611 | 0.948 | 0.593 | 3.14E-13 |
| Fkbp1b | 1.05E-14 | 1.02214315 | 0.687 | 0.111 | 1.93E-10 |
| Cryba2 | 2.87E-10 | 1.01563548 | 0.962 | 0.722 | 5.28E-06 |
| Fmo1 | 5.12E-09 | 0.98965041 | 0.507 | 0.093 | 9.43E-05 |
| Cope | 1.16E-17 | 0.97430775 | 0.993 | 0.852 | 2.13E-13 |
| Tmed2 | 1.07E-17 | 0.9728583 | 0.991 | 0.815 | 1.96E-13 |
| Pdx1 | 7.80E-14 | 0.96427109 | 0.942 | 0.37 | 1.44E-09 |
| Ssr2 | 4.37E-16 | 0.95895328 | 0.999 | 0.926 | 8.04E-12 |
| Atp5g1 | 1.78E-16 | 0.95759786 | 0.999 | 0.907 | 3.28E-12 |
| Ffar1 | 1.77E-15 | 0.95357405 | 0.77 | 0.185 | 3.27E-11 |
| Sec11a | 2.15E-16 | 0.95176913 | 0.977 | 0.704 | 3.95E-12 |
| Dnajb11 | 1.17E-13 | 0.94892582 | 0.925 | 0.519 | 2.15E-09 |
| Guk1 | 5.91E-15 | 0.94563856 | 0.906 | 0.5 | 1.09E-10 |
| P2ry1 | 5.77E-15 | 0.93914479 | 0.643 | 0.037 | 1.06E-10 |
| Derl2 | 2.18E-15 | 0.93914479 | 0.893 | 0.519 | 4.01E-11 |
| Trpm5 | 3.65E-14 | 0.9347763 | 0.572 | 0 | 6.72E-10 |
| Nkx6-1 | 2.61E-14 | 0.92987815 | 0.912 | 0.333 | 4.81E-10 |
| Hyou1 | 1.71E-14 | 0.92701486 | 0.83 | 0.259 | 3.14E-10 |
| Tspan13 | 2.48E-15 | 0.92529683 | 0.916 | 0.481 | 4.57E-11 |
| Rasgrf1 | 5.47E-14 | 0.91676084 | 0.617 | 0.056 | 1.01E-09 |
| Tmem258 | 1.32E-15 | 0.91469175 | 0.997 | 0.815 | 2.44E-11 |
| Ckb | 4.96E-09 | 0.90783525 | 0.696 | 0.315 | 9.13E-05 |
| Ndufa1 | 8.50E-15 | 0.90667795 | 0.991 | 0.722 | 1.56E-10 |

| | | | | | |
|---|---|---|---|---|---|
| Zc3h3 | 3.85E-12 | 0.88885667 | 0.552 | 0.037 | 7.09E-08 |
| Anxa5 | 1.52E-13 | 0.88369861 | 0.933 | 0.537 | 2.80E-09 |
| Rwdd1 | 4.69E-14 | 0.8784218 | 0.919 | 0.537 | 8.64E-10 |
| Vps8 | 5.33E-13 | 0.87265896 | 0.607 | 0.074 | 9.81E-09 |
| Rab3a | 1.41E-13 | 0.87153791 | 0.936 | 0.519 | 2.59E-09 |
| Stt3a | 4.31E-14 | 0.86954794 | 0.829 | 0.333 | 7.93E-10 |
| Smim14 | 1.18E-14 | 0.86764527 | 0.967 | 0.685 | 2.18E-10 |
| Tssc4 | 1.40E-12 | 0.85805558 | 0.843 | 0.444 | 2.58E-08 |
| Edem2 | 6.34E-13 | 0.85713014 | 0.862 | 0.389 | 1.17E-08 |
| Ociad2 | 2.05E-13 | 0.85629941 | 0.939 | 0.5 | 3.77E-09 |
| Dad1 | 5.22E-16 | 0.84516275 | 0.997 | 0.944 | 9.61E-12 |
| Gnas | 2.90E-11 | 0.84300757 | 1 | 0.944 | 5.33E-07 |
| Uqcrq | 2.25E-16 | 0.83361421 | 0.993 | 0.926 | 4.14E-12 |
| Dpm3 | 1.37E-13 | 0.83189224 | 0.913 | 0.463 | 2.51E-09 |
| Srp9 | 6.53E-15 | 0.82853288 | 0.988 | 0.722 | 1.20E-10 |
| 1110008P14F | 1.08E-12 | 0.8258741 | 0.897 | 0.481 | 1.98E-08 |
| Kdelr2 | 2.09E-11 | 0.82086078 | 0.878 | 0.407 | 3.85E-07 |
| Ucn3 | 2.73E-08 | 0.81803571 | 0.409 | 0.019 | 0.00050242 |
| Syndig1l | 1.10E-12 | 0.81272143 | 0.529 | 0 | 2.03E-08 |
| Mien1 | 9.86E-13 | 0.81128303 | 0.984 | 0.778 | 1.81E-08 |
| Ift20 | 8.25E-12 | 0.80312545 | 0.917 | 0.648 | 1.52E-07 |
| Tram1 | 4.41E-13 | 0.79780576 | 0.906 | 0.556 | 8.13E-09 |
| Cntfr | 4.65E-11 | 0.79760663 | 0.538 | 0.056 | 8.55E-07 |
| Gpx3 | 6.22E-10 | 0.79749602 | 0.972 | 0.778 | 1.15E-05 |
| Frzb | 1.81E-12 | 0.79304113 | 0.57 | 0.037 | 3.32E-08 |
| Canx | 2.74E-11 | 0.79072525 | 0.949 | 0.704 | 5.04E-07 |
| Exoc7 | 1.15E-10 | 0.78735978 | 0.722 | 0.278 | 2.11E-06 |
| Uso1 | 1.19E-12 | 0.78478989 | 0.755 | 0.278 | 2.18E-08 |
| Meg3 | 8.61E-10 | 0.78162013 | 1 | 1 | 1.58E-05 |
| Slc2a5 | 5.75E-12 | 0.77954583 | 0.774 | 0.296 | 1.06E-07 |
| Arhgap36 | 1.33E-11 | 0.77799956 | 0.528 | 0.019 | 2.46E-07 |
| Cycs | 2.88E-14 | 0.77544819 | 0.951 | 0.648 | 5.29E-10 |
| Uqcc2 | 7.87E-14 | 0.77279252 | 0.994 | 0.852 | 1.45E-09 |
| Minos1 | 2.61E-13 | 0.77262682 | 0.999 | 0.907 | 4.81E-09 |
| Ssr3 | 7.24E-12 | 0.77208374 | 0.867 | 0.463 | 1.33E-07 |
| Magt1 | 1.68E-12 | 0.7715636 | 0.874 | 0.426 | 3.08E-08 |
| Nt5dc2 | 1.34E-09 | 0.77043415 | 0.852 | 0.537 | 2.48E-05 |
| Bet1 | 5.10E-11 | 0.76673222 | 0.765 | 0.315 | 9.39E-07 |
| Cd164 | 1.94E-13 | 0.76508471 | 0.972 | 0.722 | 3.58E-09 |
| Srm | 5.75E-11 | 0.76415731 | 0.662 | 0.185 | 1.06E-06 |
| Rpn1 | 4.66E-12 | 0.76366112 | 0.959 | 0.759 | 8.58E-08 |
| Ufc1 | 2.13E-11 | 0.76138255 | 0.881 | 0.5 | 3.91E-07 |
| Phactr1 | 6.17E-13 | 0.75783227 | 0.645 | 0.111 | 1.14E-08 |
| Acly | 3.34E-12 | 0.75694146 | 0.951 | 0.704 | 6.15E-08 |

| | | | | | |
|---|---|---|---|---|---|
| Tspan7 | 1.64E-10 | 0.75539245 | 0.984 | 0.741 | 3.02E-06 |
| Entpd3 | 3.05E-13 | 0.7550693 | 0.754 | 0.222 | 5.62E-09 |
| Spock2 | 2.91E-12 | 0.75344717 | 0.613 | 0.074 | 5.36E-08 |
| Gfpt1 | 2.85E-12 | 0.75321384 | 0.803 | 0.352 | 5.25E-08 |
| Cdk2ap2 | 2.59E-10 | 0.75169593 | 0.867 | 0.444 | 4.77E-06 |
| Gadd45g | 4.69E-08 | 0.74496008 | 0.871 | 0.611 | 0.00086436 |
| Ssr1 | 2.08E-13 | 0.74412881 | 0.862 | 0.426 | 3.84E-09 |
| C1qa | 1.72E-07 | 0.74316865 | 0.538 | 0.148 | 0.0031659 |
| Hopx | 3.38E-10 | 0.74284981 | 0.549 | 0.074 | 6.21E-06 |
| Cst3 | 1.60E-13 | 0.74281311 | 1 | 0.926 | 2.94E-09 |
| Igsf1 | 2.75E-09 | 0.7383449 | 0.528 | 0.093 | 5.07E-05 |
| Cks1b | 8.32E-08 | 0.73794853 | 0.642 | 0.278 | 0.00153169 |
| Fam174b | 6.64E-11 | 0.72760289 | 0.807 | 0.389 | 1.22E-06 |
| H13 | 4.37E-11 | 0.72514016 | 0.87 | 0.463 | 8.05E-07 |
| Pcsk2 | 2.85E-10 | 0.72427543 | 0.999 | 0.759 | 5.24E-06 |
| Gmppb | 1.21E-10 | 0.7241319 | 0.629 | 0.148 | 2.24E-06 |
| Cib2 | 1.09E-10 | 0.72097387 | 0.635 | 0.167 | 2.00E-06 |
| Mif | 2.00E-10 | 0.71843727 | 0.971 | 0.741 | 3.68E-06 |
| Gch1 | 4.62E-10 | 0.71751976 | 0.98 | 0.741 | 8.51E-06 |
| Maob | 6.51E-10 | 0.70920392 | 0.713 | 0.259 | 1.20E-05 |
| Scgn | 8.72E-10 | 0.70696073 | 0.97 | 0.704 | 1.61E-05 |
| Uggt1 | 8.04E-11 | 0.70614427 | 0.652 | 0.204 | 1.48E-06 |
| Adra2a | 1.92E-10 | 0.70366392 | 0.536 | 0.074 | 3.53E-06 |
| Sar1b | 6.16E-11 | 0.69562695 | 0.87 | 0.574 | 1.13E-06 |
| Ece1 | 1.54E-10 | 0.69291794 | 0.872 | 0.481 | 2.83E-06 |
| Cpe | 6.82E-10 | 0.69231263 | 0.999 | 0.907 | 1.26E-05 |
| Wls | 6.88E-11 | 0.69063044 | 0.654 | 0.148 | 1.27E-06 |
| Derl1 | 2.47E-10 | 0.6896638 | 0.622 | 0.167 | 4.55E-06 |
| Txn1 | 7.43E-11 | 0.68493263 | 0.986 | 0.815 | 1.37E-06 |
| Pdia5 | 7.83E-11 | 0.68461051 | 0.668 | 0.204 | 1.44E-06 |
| Abracl | 7.67E-10 | 0.6779828 | 0.793 | 0.37 | 1.41E-05 |
| Srpr | 5.19E-10 | 0.67784128 | 0.867 | 0.593 | 9.56E-06 |
| Rab1a | 1.15E-10 | 0.67761398 | 0.958 | 0.722 | 2.11E-06 |
| Nans | 1.84E-09 | 0.67428175 | 0.806 | 0.426 | 3.40E-05 |
| Rrbp1 | 4.73E-09 | 0.66928563 | 0.922 | 0.593 | 8.71E-05 |
| Acvr1c | 2.17E-08 | 0.66784277 | 0.499 | 0.093 | 0.00040008 |
| Usmg5 | 1.56E-10 | 0.66599532 | 0.994 | 0.926 | 2.88E-06 |
| Hdlbp | 1.15E-08 | 0.66562495 | 0.817 | 0.463 | 0.0002115 |
| Lpl | 4.83E-06 | 0.66343656 | 0.59 | 0.241 | 0.08890266 |
| Atp2a3 | 6.36E-10 | 0.66205685 | 0.617 | 0.185 | 1.17E-05 |
| Tmed9 | 2.39E-10 | 0.65329269 | 0.952 | 0.759 | 4.41E-06 |
| Ndufv3 | 6.02E-10 | 0.64942935 | 0.991 | 0.778 | 1.11E-05 |
| Xbp1 | 5.92E-09 | 0.64566254 | 0.872 | 0.556 | 0.00010904 |
| Prnp | 1.40E-09 | 0.64534671 | 0.959 | 0.778 | 2.59E-05 |

| | | | | | |
|---|---|---|---|---|---|
| Yif1b | 2.90E-09 | 0.64531724 | 0.645 | 0.222 | 5.34E-05 |
| Ppib | 3.25E-09 | 0.64509697 | 0.991 | 0.87 | 5.98E-05 |
| Wfs1 | 9.30E-11 | 0.63654876 | 0.548 | 0.056 | 1.71E-06 |
| Ppa1 | 1.19E-08 | 0.6334596 | 0.675 | 0.278 | 0.00021941 |
| Txnl1 | 2.94E-09 | 0.63148977 | 0.922 | 0.667 | 5.41E-05 |
| Erp29 | 2.07E-10 | 0.63127157 | 0.981 | 0.852 | 3.82E-06 |
| Serf2 | 5.44E-10 | 0.63053282 | 0.999 | 0.926 | 1.00E-05 |
| Golga4 | 4.70E-10 | 0.62940105 | 0.743 | 0.296 | 8.66E-06 |
| Enho | 2.15E-09 | 0.62415177 | 0.674 | 0.222 | 3.95E-05 |
| Lyve1 | 6.50E-08 | 0.61989083 | 0.393 | 0.019 | 0.00119725 |
| Sec61a1 | 8.36E-09 | 0.61191389 | 0.807 | 0.407 | 0.00015388 |
| Atp5j2 | 3.47E-11 | 0.60932684 | 0.999 | 0.963 | 6.40E-07 |
| 1500009L16R | 3.82E-08 | 0.59848949 | 0.642 | 0.222 | 0.00070271 |
| Arl1 | 2.33E-09 | 0.58723267 | 0.936 | 0.722 | 4.28E-05 |
| Mafb | 4.73E-06 | 0.58707896 | 0.97 | 0.759 | 0.08701757 |
| Tmem14c | 2.77E-07 | 0.5849625 | 0.81 | 0.481 | 0.00509862 |
| Clptm1l | 4.81E-08 | 0.58212693 | 0.532 | 0.148 | 0.00088507 |
| Aldh9a1 | 7.13E-08 | 0.58104969 | 0.5 | 0.111 | 0.00131327 |
| Gjd2 | 1.64E-09 | 0.57722166 | 0.701 | 0.241 | 3.02E-05 |
| Wipi1 | 9.30E-07 | 0.57675758 | 0.555 | 0.185 | 0.01711522 |
| Slc38a4 | 2.98E-08 | 0.5698292 | 0.496 | 0.093 | 0.00054862 |
| Ndufb10 | 1.81E-09 | 0.56979493 | 0.958 | 0.778 | 3.32E-05 |
| Dpp3 | 3.33E-08 | 0.56928088 | 0.68 | 0.315 | 0.00061245 |
| Mrps33 | 5.15E-08 | 0.56892645 | 0.983 | 0.796 | 0.00094888 |
| S100a11 | 3.78E-07 | 0.56879879 | 0.996 | 0.796 | 0.0069602 |
| Plpp5 | 3.77E-08 | 0.56801699 | 0.645 | 0.278 | 0.00069493 |
| Zfp568 | 1.09E-08 | 0.56591321 | 0.5 | 0.093 | 0.00019986 |
| Dap | 5.86E-08 | 0.56550554 | 0.891 | 0.574 | 0.00107953 |
| Gng4 | 2.51E-08 | 0.56522628 | 0.568 | 0.167 | 0.0004629 |
| Actn3 | 8.07E-08 | 0.563764 | 0.5 | 0.093 | 0.00148543 |
| Ndufb11 | 3.94E-09 | 0.56130269 | 0.997 | 0.87 | 7.25E-05 |
| Tmem167 | 1.63E-08 | 0.55855666 | 0.888 | 0.537 | 0.00030037 |
| Gorasp2 | 1.34E-07 | 0.55517595 | 0.723 | 0.352 | 0.00247369 |
| Srp54b | 1.32E-07 | 0.55220403 | 0.859 | 0.667 | 0.00243652 |
| Sec23b | 2.88E-07 | 0.55140792 | 0.639 | 0.278 | 0.0052944 |
| Bet1l | 4.76E-07 | 0.55018378 | 0.594 | 0.241 | 0.00875696 |
| Nme1 | 4.10E-09 | 0.54761465 | 0.99 | 0.944 | 7.55E-05 |
| Gm10941 | 5.73E-08 | 0.54113971 | 0.467 | 0.074 | 0.00105584 |
| Kcnmb2 | 3.68E-07 | 0.53795634 | 0.571 | 0.204 | 0.00676951 |
| Tbcb | 2.58E-07 | 0.53565958 | 0.906 | 0.722 | 0.00474758 |
| Tmem65 | 9.40E-08 | 0.53523223 | 0.449 | 0.074 | 0.00173131 |
| Sez6l2 | 3.63E-07 | 0.53432636 | 0.791 | 0.407 | 0.00668613 |
| Morf4l2 | 2.89E-08 | 0.53208757 | 0.987 | 0.926 | 0.00053234 |
| Prdx3 | 1.53E-06 | 0.53164567 | 0.735 | 0.463 | 0.02816534 |

| | | | | | |
|---|---|---|---|---|---|
| Sh3pxd2a | 1.38E-08 | 0.53099714 | 0.39 | 0 | 0.0002542 |
| Cox5b | 2.32E-08 | 0.5289863 | 0.996 | 0.926 | 0.00042677 |
| Cyb5a | 2.56E-06 | 0.52586775 | 0.661 | 0.333 | 0.04719801 |
| Nt5c3 | 4.64E-06 | 0.52486204 | 0.664 | 0.389 | 0.08540539 |
| Atp6v1e1 | 1.35E-07 | 0.52482206 | 0.959 | 0.722 | 0.00249075 |
| Dnajc10 | 4.53E-07 | 0.52262768 | 0.641 | 0.259 | 0.00834855 |
| Spcs3 | 1.26E-06 | 0.51850604 | 0.694 | 0.37 | 0.02315999 |
| Ndufb8 | 1.03E-08 | 0.51780701 | 0.996 | 0.852 | 0.00018921 |
| Erp44 | 1.97E-06 | 0.51605502 | 0.799 | 0.519 | 0.03624444 |
| Swi5 | 1.85E-08 | 0.5144081 | 0.996 | 0.926 | 0.00034116 |
| Edem1 | 2.27E-07 | 0.51425951 | 0.393 | 0.037 | 0.00418739 |
| Sec24d | 6.24E-08 | 0.51362185 | 0.514 | 0.111 | 0.00114973 |
| Uqcr11 | 2.56E-08 | 0.51331811 | 0.997 | 0.944 | 0.00047224 |
| Ivd | 2.29E-06 | 0.51126201 | 0.526 | 0.185 | 0.04222398 |
| Itpkb | 7.37E-08 | 0.51084807 | 0.442 | 0.056 | 0.00135619 |
| Ndufs5 | 2.84E-08 | 0.50900673 | 0.987 | 0.87 | 0.00052348 |
| 2310039H08I | 3.24E-06 | 0.50885952 | 0.754 | 0.444 | 0.05966556 |
| Syt13 | 4.28E-06 | 0.50846361 | 0.964 | 0.741 | 0.07882877 |
| Galnt18 | 3.13E-07 | 0.50807033 | 0.526 | 0.148 | 0.00576797 |
| Ndufb6 | 4.31E-06 | 0.50658321 | 0.864 | 0.667 | 0.07930347 |
| Tma7 | 4.45E-07 | 0.50563802 | 0.996 | 0.926 | 0.00820123 |
| Mia3 | 3.66E-07 | 0.50533866 | 0.696 | 0.296 | 0.00673011 |
| Romo1 | 2.90E-06 | 0.50486761 | 0.988 | 0.796 | 0.05340112 |
| Mrps36 | 2.53E-06 | 0.50131271 | 0.775 | 0.5 | 0.04649454 |
| Yipf5 | 3.16E-06 | 0.49668126 | 0.578 | 0.278 | 0.05823468 |
| Atp5o | 1.36E-07 | 0.49641361 | 0.99 | 0.833 | 0.00251051 |
| Sub1 | 2.30E-09 | 0.49589555 | 0.997 | 0.926 | 4.24E-05 |
| Slc37a4 | 3.98E-07 | 0.49547271 | 0.609 | 0.222 | 0.00731971 |
| Uba5 | 2.17E-06 | 0.49435995 | 0.686 | 0.37 | 0.03999682 |
| Ubl5 | 1.36E-06 | 0.49404437 | 0.997 | 0.907 | 0.02498464 |
| 4833439L19R | 4.83E-06 | 0.49350704 | 0.772 | 0.519 | 0.08901326 |
| Ndufa2 | 9.68E-08 | 0.49327644 | 0.994 | 0.852 | 0.00178261 |
| Timm10b | 1.40E-06 | 0.49251625 | 0.755 | 0.444 | 0.02585369 |
| Slc33a1 | 1.76E-07 | 0.48900672 | 0.47 | 0.093 | 0.00323453 |
| Ntrk2 | 1.24E-06 | 0.48872442 | 0.372 | 0.037 | 0.0227429 |
| Tmod2 | 2.50E-06 | 0.48872442 | 0.374 | 0.056 | 0.04597909 |
| Cox7a2 | 1.95E-07 | 0.48707885 | 0.997 | 0.963 | 0.0035836 |
| Naga | 1.67E-06 | 0.48705515 | 0.533 | 0.185 | 0.03073007 |
| Tmed10 | 1.17E-07 | 0.48488033 | 0.996 | 0.833 | 0.00215907 |
| Tvp23b | 1.90E-06 | 0.48080593 | 0.829 | 0.63 | 0.03500112 |
| Fam183b | 4.03E-06 | 0.48022962 | 0.986 | 0.815 | 0.07414891 |
| Gabarap | 1.77E-08 | 0.47976082 | 0.996 | 0.926 | 0.00032581 |
| Gtf2a2 | 2.49E-06 | 0.47959049 | 0.897 | 0.667 | 0.04587321 |
| Smdt1 | 2.97E-06 | 0.47662687 | 0.984 | 0.778 | 0.05469024 |

| | | | | | |
|---|---|---|---|---|---|
| Camk2n1 | 3.64E-06 | 0.47127634 | 0.819 | 0.5 | 0.06709597 |
| Mrps12 | 4.82E-06 | 0.47019643 | 0.801 | 0.537 | 0.08881683 |
| Slit1 | 1.21E-06 | 0.46786215 | 0.396 | 0.056 | 0.02235723 |
| Pclo | 2.45E-06 | 0.46628143 | 0.803 | 0.444 | 0.04516566 |
| Arf1 | 1.21E-06 | 0.46193847 | 0.996 | 0.944 | 0.02228789 |
| Ost4 | 1.53E-06 | 0.46138263 | 0.899 | 0.667 | 0.02822726 |
| Zfyve21 | 2.87E-06 | 0.46038683 | 0.604 | 0.241 | 0.05276708 |
| Scnn1b | 6.68E-07 | 0.4582907 | 0.325 | 0 | 0.01230717 |
| Cox6a1 | 1.88E-06 | 0.45385311 | 0.999 | 1 | 0.03465386 |
| G6pc3 | 2.28E-06 | 0.45234332 | 0.428 | 0.093 | 0.04204452 |
| BC031181 | 1.83E-06 | 0.43685237 | 0.89 | 0.685 | 0.03360356 |
| Sep-15 | 1.56E-06 | 0.43635084 | 0.99 | 0.963 | 0.02872937 |
| Mpp3 | 2.39E-06 | 0.43614534 | 0.406 | 0.074 | 0.04404063 |
| Atp5e | 6.25E-07 | 0.43551738 | 0.999 | 0.981 | 0.01151379 |
| Hspa13 | 3.59E-06 | 0.43381841 | 0.61 | 0.259 | 0.06605131 |
| Thyn1 | 4.00E-06 | 0.43089652 | 0.558 | 0.204 | 0.07356453 |
| Cox8a | 1.25E-07 | 0.42833837 | 0.999 | 1 | 0.00229315 |
| Ghr | 4.91E-06 | 0.42178306 | 0.628 | 0.278 | 0.09045805 |
| Larp1b | 4.80E-06 | 0.41635744 | 0.362 | 0.056 | 0.08834163 |
| Mrfap1 | 1.20E-06 | 0.4131196 | 1 | 0.981 | 0.02209465 |
| Srprb | 5.30E-06 | 0.41079524 | 0.641 | 0.333 | 0.09753858 |
| Uqcrb | 8.07E-07 | 0.40614904 | 0.997 | 0.981 | 0.01486059 |
| Wbp5 | 1.69E-07 | 0.40612262 | 0.999 | 0.981 | 0.00311691 |
| Atpif1 | 4.49E-06 | 0.35961054 | 0.997 | 0.963 | 0.08268626 |
| Prdm10 | 2.49E-06 | -0.1312445 | 0.014 | 0.111 | 0.04583032 |
| Schip1.1 | 2.49E-06 | -0.1312445 | 0.014 | 0.111 | 0.04583032 |
| Gm10069 | 4.89E-07 | -0.1509752 | 0.017 | 0.13 | 0.00900878 |
| Gm20554 | 1.33E-08 | -0.1571537 | 0.013 | 0.13 | 0.00024443 |
| Hbq1a | 3.57E-08 | -0.1682819 | 0.019 | 0.148 | 0.00065641 |
| Sstr2 | 2.47E-16 | -0.1716742 | 0.003 | 0.13 | 4.55E-12 |
| Klb | 4.74E-08 | -0.1785502 | 0.014 | 0.13 | 0.00087353 |
| Ace2 | 2.71E-09 | -0.1826781 | 0.012 | 0.13 | 4.98E-05 |
| Igfbp2 | 4.47E-09 | -0.199127 | 0.038 | 0.222 | 8.23E-05 |
| Galnt13 | 2.23E-21 | -0.1993088 | 0 | 0.13 | 4.10E-17 |
| Mospd1 | 4.42E-08 | -0.2016339 | 0.014 | 0.13 | 0.00081347 |
| Ptprk | 6.66E-08 | -0.2141171 | 0.038 | 0.204 | 0.00122622 |
| Nudt11 | 1.64E-06 | -0.2241041 | 0.046 | 0.204 | 0.03026484 |
| Edn3 | 5.10E-07 | -0.2262576 | 0.023 | 0.148 | 0.00938376 |
| Mpp1 | 3.45E-07 | -0.2301112 | 0.035 | 0.185 | 0.00635098 |
| Tspan12 | 9.49E-12 | -0.2323665 | 0.023 | 0.204 | 1.75E-07 |
| H2-T23 | 3.74E-06 | -0.2398171 | 0.049 | 0.204 | 0.06882596 |
| Mboat4 | 4.00E-11 | -0.2426056 | 0.016 | 0.167 | 7.36E-07 |
| Acot1 | 5.34E-06 | -0.253033 | 0.094 | 0.296 | 0.09836462 |
| Slc25a33 | 1.41E-06 | -0.263896 | 0.032 | 0.167 | 0.02594331 |

| | | | | | |
|---|---|---|---|---|---|
| Ndst4 | 2.26E-12 | -0.2646319 | 0.017 | 0.185 | 4.17E-08 |
| Amer1 | 2.94E-07 | -0.2664201 | 0.057 | 0.241 | 0.0054219 |
| Azin2 | 2.29E-06 | -0.2695727 | 0.072 | 0.259 | 0.04216241 |
| Efnb1 | 9.17E-07 | -0.2754175 | 0.07 | 0.259 | 0.01689125 |
| Drc1 | 1.63E-09 | -0.2801748 | 0.02 | 0.167 | 3.01E-05 |
| Wfdc15b | 7.64E-09 | -0.2832476 | 0.028 | 0.185 | 0.00014063 |
| Lynx1 | 3.21E-17 | -0.2832476 | 0.004 | 0.148 | 5.92E-13 |
| Bcl11a | 3.11E-07 | -0.2880938 | 0.074 | 0.278 | 0.00571915 |
| Cer1 | 6.62E-10 | -0.2904431 | 0.014 | 0.148 | 1.22E-05 |
| Ppp1r14a | 2.58E-11 | -0.2925056 | 0.012 | 0.148 | 4.75E-07 |
| P3h4 | 3.76E-07 | -0.3041683 | 0.216 | 0.537 | 0.00692039 |
| Apom | 4.34E-12 | -0.3049427 | 0.004 | 0.111 | 7.99E-08 |
| Cited1 | 3.69E-11 | -0.305647 | 0.016 | 0.167 | 6.80E-07 |
| Ube2l6 | 4.43E-09 | -0.3141086 | 0.059 | 0.278 | 8.15E-05 |
| Cbfa2t2 | 1.82E-06 | -0.3153088 | 0.155 | 0.407 | 0.03349785 |
| Celf2 | 1.71E-06 | -0.3211343 | 0.141 | 0.389 | 0.03141402 |
| Ppp1r14c | 9.58E-09 | -0.3230486 | 0.035 | 0.204 | 0.00017639 |
| Fam43a | 7.47E-07 | -0.3350213 | 0.117 | 0.352 | 0.01375771 |
| Irx2 | 3.83E-19 | -0.3390747 | 0.01 | 0.204 | 7.05E-15 |
| Ccdc34 | 4.88E-07 | -0.3441559 | 0.232 | 0.556 | 0.00899225 |
| Cdc42ep3 | 9.38E-16 | -0.3474672 | 0.019 | 0.222 | 1.73E-11 |
| Pou6f2 | 7.48E-22 | -0.3494613 | 0.003 | 0.167 | 1.38E-17 |
| Litaf | 5.34E-07 | -0.3497949 | 0.075 | 0.278 | 0.00982884 |
| Nxph1 | 4.74E-27 | -0.3515476 | 0.001 | 0.185 | 8.72E-23 |
| Bcl2l1 | 1.51E-06 | -0.3554192 | 0.155 | 0.407 | 0.02778832 |
| Syne1 | 4.96E-08 | -0.3689037 | 0.087 | 0.315 | 0.00091371 |
| Igsf21 | 1.90E-19 | -0.3699849 | 0.016 | 0.241 | 3.51E-15 |
| Gm609 | 1.96E-20 | -0.3741011 | 0.014 | 0.241 | 3.60E-16 |
| Plk2 | 1.29E-06 | -0.3746707 | 0.072 | 0.259 | 0.0237565 |
| Col9a3 | 1.71E-15 | -0.3799236 | 0.019 | 0.222 | 3.15E-11 |
| Tox | 1.82E-06 | -0.3807306 | 0.196 | 0.463 | 0.03350525 |
| Plscr3 | 3.98E-07 | -0.3817799 | 0.222 | 0.519 | 0.00732433 |
| Tekt2 | 2.17E-06 | -0.3878036 | 0.278 | 0.574 | 0.03993357 |
| Muc1 | 3.46E-11 | -0.3926311 | 0.048 | 0.278 | 6.37E-07 |
| Cdc25b | 3.50E-07 | -0.3935297 | 0.077 | 0.278 | 0.00644689 |
| Wsb1 | 1.85E-09 | -0.3960156 | 0.067 | 0.296 | 3.42E-05 |
| Gypa | 7.65E-07 | -0.3968178 | 0.036 | 0.185 | 0.01408311 |
| Jup | 2.68E-07 | -0.4009928 | 0.274 | 0.611 | 0.00493293 |
| Cbfb | 2.74E-06 | -0.4036776 | 0.216 | 0.481 | 0.0504696 |
| St3gal6 | 8.36E-10 | -0.4050361 | 0.106 | 0.389 | 1.54E-05 |
| Tcp11 | 1.15E-06 | -0.408999 | 0.172 | 0.426 | 0.02113834 |
| Setd8 | 9.56E-08 | -0.4184072 | 0.132 | 0.389 | 0.00176096 |
| Serping1 | 7.70E-08 | -0.4196201 | 0.126 | 0.389 | 0.0014184 |
| Arg1 | 5.37E-10 | -0.421494 | 0.019 | 0.167 | 9.89E-06 |

| | | | | | |
|---|---|---|---|---|---|
| Miat | 1.02E-08 | -0.4246035 | 0.035 | 0.204 | 0.00018864 |
| Rnf146 | 1.13E-06 | -0.4284542 | 0.246 | 0.537 | 0.02087876 |
| Tmed1 | 4.49E-06 | -0.432008 | 0.174 | 0.407 | 0.082632 |
| Ppp3ca | 2.77E-06 | -0.4432023 | 0.31 | 0.593 | 0.05091119 |
| Serpine2 | 4.97E-09 | -0.4459103 | 0.162 | 0.481 | 9.15E-05 |
| Eef2 | 1.62E-06 | -0.4617499 | 0.997 | 0.963 | 0.02985169 |
| Cyba | 5.34E-07 | -0.4669784 | 0.136 | 0.389 | 0.0098271 |
| Npc2 | 2.21E-07 | -0.4857405 | 0.823 | 0.926 | 0.00406021 |
| Irx1 | 4.44E-33 | -0.4888643 | 0.003 | 0.241 | 8.18E-29 |
| Fth1 | 2.57E-06 | -0.4956086 | 0.999 | 1 | 0.04727794 |
| Akap17b | 6.59E-09 | -0.4981162 | 0.109 | 0.37 | 0.00012141 |
| Ccndbp1 | 1.78E-08 | -0.50564 | 0.317 | 0.648 | 0.00032765 |
| Dpp4 | 1.39E-09 | -0.5096035 | 0.052 | 0.259 | 2.55E-05 |
| Blvrb | 1.56E-06 | -0.5199597 | 0.125 | 0.352 | 0.02875606 |
| Tshz2 | 4.55E-06 | -0.520988 | 0.536 | 0.815 | 0.08381693 |
| Dpysl4 | 2.63E-18 | -0.5260752 | 0.014 | 0.222 | 4.84E-14 |
| Sms | 3.90E-08 | -0.5334322 | 0.228 | 0.537 | 0.00071713 |
| App | 1.47E-06 | -0.5369115 | 0.638 | 0.852 | 0.02703956 |
| Cystm1 | 3.62E-07 | -0.5389514 | 0.91 | 0.963 | 0.00665928 |
| Fam210b | 1.15E-06 | -0.5594977 | 0.164 | 0.407 | 0.02120667 |
| Gria2 | 6.11E-18 | -0.5627644 | 0.036 | 0.315 | 1.13E-13 |
| Marcks | 2.49E-06 | -0.5639643 | 0.42 | 0.685 | 0.04587927 |
| Trf | 7.91E-12 | -0.5823276 | 0.039 | 0.259 | 1.46E-07 |
| Arx | 1.15E-26 | -0.5860338 | 0.01 | 0.259 | 2.11E-22 |
| Gltscr2 | 1.01E-07 | -0.5870801 | 0.642 | 0.852 | 0.00186102 |
| Gm43861 | 9.12E-15 | -0.5881022 | 0.097 | 0.444 | 1.68E-10 |
| Nefm | 7.17E-09 | -0.5917967 | 0.08 | 0.315 | 0.00013205 |
| Slc48a1 | 8.25E-11 | -0.5977276 | 0.332 | 0.722 | 1.52E-06 |
| Casp6 | 2.37E-10 | -0.5997622 | 0.148 | 0.463 | 4.37E-06 |
| Sgce | 1.33E-27 | -0.6002769 | 0.025 | 0.352 | 2.46E-23 |
| Rpl18a | 7.47E-10 | -0.6072102 | 1 | 1 | 1.38E-05 |
| Rpl9 | 7.97E-09 | -0.6121415 | 1 | 1 | 0.00014666 |
| Hsp90ab1 | 1.34E-08 | -0.6139207 | 1 | 0.981 | 0.00024651 |
| Prox1 | 3.74E-09 | -0.6315579 | 0.291 | 0.63 | 6.88E-05 |
| Ly6e | 2.54E-06 | -0.6426332 | 0.517 | 0.741 | 0.04667658 |
| Alcam | 3.06E-08 | -0.6705644 | 0.201 | 0.5 | 0.0005635 |
| Pbx1 | 3.00E-09 | -0.6795781 | 0.419 | 0.759 | 5.52E-05 |
| Lpar6 | 6.13E-15 | -0.6963781 | 0.091 | 0.426 | 1.13E-10 |
| Hspa8 | 1.17E-09 | -0.7089191 | 1 | 1 | 2.16E-05 |
| Ctxn2 | 2.75E-30 | -0.7100373 | 0.017 | 0.333 | 5.07E-26 |
| Pax4 | 2.02E-18 | -0.7717149 | 0.026 | 0.278 | 3.72E-14 |
| Mid1ip1 | 1.78E-08 | -0.7762771 | 0.63 | 0.889 | 0.0003278 |
| Marcksl1 | 9.46E-07 | -0.7850325 | 0.645 | 0.852 | 0.01741458 |
| Auts2 | 3.19E-13 | -0.7935713 | 0.191 | 0.574 | 5.87E-09 |

| | | | | | |
|---|---|---|---|---|---|
| Neurod2 | 1.36E-38 | -0.8149681 | 0 | 0.241 | 2.51E-34 |
| Cck | 5.12E-12 | -0.8345886 | 0.052 | 0.296 | 9.43E-08 |
| Slc25a37 | 1.11E-20 | -0.8664034 | 0.103 | 0.537 | 2.05E-16 |
| Mar-02 | 1.18E-15 | -0.8682101 | 0.252 | 0.685 | 2.17E-11 |
| Ctsz | 2.37E-07 | -0.8732457 | 0.425 | 0.685 | 0.00436659 |
| Bnip3l | 3.10E-11 | -0.8749993 | 0.652 | 0.87 | 5.71E-07 |
| Pou3f4 | 1.72E-29 | -0.8861568 | 0.033 | 0.407 | 3.17E-25 |
| Glud1 | 5.15E-10 | -0.886751 | 0.523 | 0.778 | 9.48E-06 |
| Cldn6 | 6.55E-09 | -0.9455972 | 0.481 | 0.759 | 0.00012064 |
| Slc25a5 | 4.91E-08 | -0.9474792 | 0.99 | 0.981 | 0.00090329 |
| Fam220a | 2.63E-13 | -1.0164661 | 0.448 | 0.796 | 4.85E-09 |
| Hn1 | 2.27E-07 | -1.0232271 | 0.619 | 0.926 | 0.00417202 |
| Pgrmc1 | 4.20E-11 | -1.0655116 | 0.565 | 0.815 | 7.73E-07 |
| Runx1t1 | 2.44E-11 | -1.1170142 | 0.209 | 0.556 | 4.49E-07 |
| Etv1 | 9.81E-23 | -1.1344684 | 0.062 | 0.444 | 1.81E-18 |
| Smarca1 | 3.30E-09 | -1.2169585 | 0.229 | 0.537 | 6.08E-05 |
| Peg10 | 2.24E-27 | -1.3256614 | 0.033 | 0.389 | 4.13E-23 |
| Snca | 2.57E-22 | -1.4111494 | 0.061 | 0.444 | 4.73E-18 |
| Fech | 2.51E-29 | -1.5516169 | 0.155 | 0.741 | 4.62E-25 |
| Cd24a | 4.73E-09 | -1.6202088 | 0.312 | 0.611 | 8.71E-05 |
| Mdk | 8.35E-08 | -1.6568836 | 0.13 | 0.37 | 0.00153698 |
| Mkrn1 | 6.04E-14 | -1.7601903 | 0.528 | 0.833 | 1.11E-09 |
| Cdkn1a | 1.28E-13 | -1.957444 | 0.159 | 0.537 | 2.37E-09 |
| Alas2 | 2.37E-48 | -2.6731505 | 0.101 | 0.815 | 4.36E-44 |
| Hba-a2 | 3.03E-29 | -2.8079657 | 0.517 | 0.963 | 5.57E-25 |
| Neurog3 | 5.61E-21 | -3.5590884 | 0.03 | 0.315 | 1.03E-16 |
| Hbb-bt | 6.41E-27 | -3.5743731 | 0.916 | 1 | 1.18E-22 |
| Hbb-bs | 2.04E-25 | -3.6729115 | 1 | 1 | 3.75E-21 |
| Hba-a1 | 5.95E-27 | -3.8048485 | 1 | 1 | 1.10E-22 |
| Ghrl | 4.58E-07 | -6.0877854 | 0.196 | 0.444 | 0.00843844 |
| Gcg | 1.70E-06 | -7.3723647 | 0.329 | 0.519 | 0.03134401 |
