## Supplemental Table 3 for "Differential CpG methylation at *Nnat* in the early establishment of beta cell heterogeneity"

| | mean outside cluster | mean inside cluster | fold change | P value | P adj |
|---|---|---|---|---|---|
| Cel | 0.233640324 | 5.963338639 | 25.5235849 | 0 | 0 |
| Nnat | 19.20454751 | 0.441698532 | 0.02299968 | 0 | 0 |
| Cpb1 | 0.562672546 | 12.46056486 | 22.1453222 | 0 | 0 |
| Cela3b | 3.12855273 | 68.74864244 | 21.9745833 | 0 | 0 |
| Cela2a | 1.111916612 | 29.78551575 | 26.7875445 | 0 | 0 |
| Cpa2 | 0.726918902 | 11.03608894 | 15.1820085 | 0 | 0 |
| Cpa1 | 1.836326842 | 31.09021415 | 16.9306539 | 0 | 0 |
| Try5 | 0.250140275 | 5.726410198 | 22.8927956 | 0 | 0 |
| Prss2 | 0.324514964 | 6.581466182 | 20.2809328 | 0 | 0 |
| Iapp | 91.66209605 | 1.899939259 | 0.02072764 | 0 | 0 |
| Sycn | 0.585674735 | 9.557963026 | 16.3195755 | 0 | 0 |
| Nupr1 | 0.361819147 | 4.79540221 | 13.2535888 | 0 | 0 |
| Zg16 | 0.15703751 | 3.383869576 | 21.5481611 | 0 | 0 |
| Ctrb1 | 20.00913738 | 441.0237366 | 22.0411169 | 0 | 0 |
| Rnase1 | 1.520848532 | 25.49161381 | 16.7614416 | 0 | 0 |
| Serpina6 | 0.489935632 | 8.325823114 | 16.9937081 | 0 | 0 |
| Cela1 | 0.336682883 | 8.435592687 | 25.055009 | 0 | 0 |
| Pdia2 | 0.177429957 | 3.264596644 | 18.3993543 | 0 | 0 |
| Clps | 11.24404441 | 183.6833176 | 16.3360541 | 0 | 0 |
| Reep5 | 1.274027349 | 14.48823196 | 11.3719945 | 0 | 0 |
| Spink1 | 0.595179716 | 10.1271368 | 17.0152586 | 0 | 0 |
| Pnliprp1 | 8.200619212 | 150.8281915 | 18.392293 | 0 | 0 |
| Chga | 9.478339052 | 0.295046166 | 0.03112847 | 5.48E-278 | 4.09E-275 |
| Npy | 4.82089598 | 0.058446729 | 0.01212362 | 6.29E-269 | 4.51E-266 |
| Ins1 | 364.5818638 | 6.25696234 | 0.01716202 | 9.24E-250 | 6.35E-247 |
| Mt1 | 3.079565106 | 22.01346431 | 7.148238 | 1.60E-242 | 1.06E-239 |
| Tmem97 | 0.361515588 | 3.651175641 | 10.0996354 | 6.88E-224 | 4.38E-221 |
| Try10 | 0.107766145 | 2.347919329 | 21.7871701 | 9.46E-220 | 5.81E-217 |
| Pcsk2 | 5.449683042 | 0.30278701 | 0.05556048 | 5.31E-215 | 3.15E-212 |

| | | | | | |
|---|---|---|---|---|---|
| Ins2 | 620.9450669 | 9.381406066 | 0.01510827 | 2.31E-211 | 1.32E-208 |
| Ctrc | 0.066935511 | 2.083693749 | 31.1298699 | 2.76E-210 | 1.53E-207 |
| Arhgdig | 0.180388082 | 2.532888983 | 14.0413322 | 7.82E-209 | 4.20E-206 |
| Fkbp11 | 0.872131518 | 6.196077829 | 7.10452232 | 8.60E-209 | 4.48E-206 |
| Gcg | 21.73144496 | 1.949976526 | 0.08973064 | 3.05E-205 | 1.54E-202 |
| Cpe | 4.617605439 | 0.269479353 | 0.05835911 | 3.50E-205 | 1.72E-202 |
| Erp27 | 0.155214424 | 2.403216809 | 15.4832054 | 5.61E-205 | 2.68E-202 |
| Scg2 | 2.784519721 | 0.076783259 | 0.02757505 | 5.17E-203 | 2.40E-200 |
| H19 | 0.623975279 | 4.626417007 | 7.41442355 | 6.62E-200 | 2.99E-197 |
| Hbb-bt | 29.5257054 | 2.941538457 | 0.09962636 | 1.90E-191 | 8.39E-189 |
| Scg5 | 2.706891532 | 0.099820411 | 0.0368764 | 1.14E-188 | 4.89E-186 |
| Pcsk1n | 4.888500711 | 0.387687076 | 0.07930593 | 6.98E-187 | 2.93E-184 |
| Gm42418 | 5.14962236 | 28.65164872 | 5.56383492 | 1.78E-186 | 7.28E-184 |
| Gpx3 | 2.826883991 | 0.157243123 | 0.05562419 | 1.07E-182 | 4.26E-180 |
| Klk1 | 0.102468138 | 1.973282706 | 19.2575248 | 7.25E-179 | 2.83E-176 |
| Chgb | 5.873318811 | 0.565534307 | 0.09628871 | 3.98E-177 | 1.52E-174 |
| Scd2 | 0.388716687 | 2.958284617 | 7.6103875 | 7.86E-177 | 2.94E-174 |
| Gstm1 | 0.371869889 | 2.76995957 | 7.44873315 | 1.57E-170 | 5.73E-168 |
| Rbp4 | 6.678807246 | 0.721257368 | 0.10799194 | 3.12E-170 | 1.12E-167 |
| Pyy | 16.97982291 | 2.098694354 | 0.12359931 | 1.36E-168 | 4.78E-166 |
| Mt2 | 1.46797597 | 7.66602692 | 5.22217466 | 1.54E-164 | 5.29E-162 |
| Scg3 | 2.232526696 | 0.060719362 | 0.0271976 | 1.79E-162 | 6.03E-160 |
| Gamt | 0.157313786 | 1.992161016 | 12.6636137 | 4.93E-159 | 1.63E-156 |
| Reg1 | 0.167201118 | 2.007037722 | 12.0037339 | 9.39E-157 | 3.04E-154 |
| Prss1 | 0.039528454 | 1.437264496 | 36.3602507 | 2.56E-150 | 8.14E-148 |
| Fabp5 | 0.892492911 | 4.507751682 | 5.05074228 | 6.44E-144 | 2.01E-141 |
| Hba-a1 | 135.758941 | 15.30401976 | 0.11272937 | 2.46E-136 | 7.54E-134 |
| Try4 | 0.074886124 | 1.459501975 | 19.4896183 | 4.54E-134 | 1.37E-131 |
| Tmed6 | 0.215660823 | 1.877987309 | 8.70805963 | 1.48E-128 | 4.40E-126 |
| Aqp12 | 0.066627749 | 1.372606508 | 20.6011238 | 5.33E-128 | 1.55E-125 |

| | | | | | |
|---|---|---|---|---|---|
| Cckar | 0.112955175 | 1.553881428 | 13.7566201 | 6.30E-128 | 1.80E-125 |
| P4hb | 1.225090113 | 5.303989949 | 4.32946923 | 5.35E-126 | 1.51E-123 |
| Gcat | 0.270690844 | 1.974224887 | 7.29328284 | 3.14E-122 | 8.69E-120 |
| Ppy | 1.822329446 | 0.089334729 | 0.04902227 | 6.02E-122 | 1.64E-119 |
| Tmem27 | 1.885736244 | 0.108101487 | 0.05732588 | 3.99E-121 | 1.07E-118 |
| Ppp1r1a | 1.49232816 | 0.02315593 | 0.01551665 | 1.50E-115 | 3.96E-113 |
| Ifitm3 | 0.675699058 | 3.007063703 | 4.45030028 | 1.55E-114 | 4.03E-112 |
| Tuba1a | 2.563601505 | 0.401319189 | 0.15654507 | 1.73E-114 | 4.43E-112 |
| Ppib | 2.323394976 | 8.755072022 | 3.7682237 | 4.45E-113 | 1.12E-110 |
| Slc30a8 | 1.380270244 | 0.018938657 | 0.01372098 | 5.64E-108 | 1.40E-105 |
| Cela3a | 0.065128215 | 1.188117895 | 18.2427522 | 1.02E-107 | 2.51E-105 |
| Slc38a5 | 2.20896331 | 0.3326959 | 0.15061178 | 3.87E-101 | 9.37E-99 |
| Rps27l | 2.456580769 | 8.554578111 | 3.48231095 | 1.27E-100 | 3.03E-98 |
| Pnliprp2 | 0.0638332 | 1.116782653 | 17.495326 | 2.85E-99 | 6.71E-97 |
| Gng12 | 1.591382237 | 0.111132185 | 0.06983375 | 1.73E-98 | 4.01E-96 |
| Gnas | 10.38838883 | 2.413216037 | 0.23229936 | 8.72E-98 | 2.00E-95 |
| Cryba2 | 1.808286472 | 0.192899216 | 0.10667514 | 5.41E-97 | 1.22E-94 |
| Rrbp1 | 1.270828816 | 4.528254859 | 3.5632296 | 6.82E-96 | 1.52E-93 |
| Hes1 | 0.544165459 | 2.312850518 | 4.25027072 | 1.07E-95 | 2.35E-93 |
| Hbb-bs | 168.419001 | 27.44735295 | 0.16297064 | 1.94E-94 | 4.22E-92 |
| 2210010C04l | 0.02590946 | 0.903555159 | 34.8735624 | 2.11E-94 | 4.52E-92 |
| Meg3 | 16.3336588 | 3.917887158 | 0.23986586 | 5.21E-94 | 1.11E-91 |
| Amy2a5 | 0.060126197 | 1.032252587 | 17.1681004 | 2.39E-91 | 5.00E-89 |
| Serf2 | 3.102586844 | 10.07245182 | 3.24646894 | 3.27E-91 | 6.76E-89 |
| Rplp1 | 8.108701615 | 26.55299236 | 3.27462936 | 1.81E-90 | 3.69E-88 |
| Fam183b | 1.256713817 | 0.045675678 | 0.03634533 | 4.85E-89 | 9.81E-87 |
| Rpl36a | 4.187881153 | 12.89499439 | 3.07912138 | 5.17E-84 | 1.03E-81 |
| Rps12 | 4.714259629 | 14.44668783 | 3.06446589 | 2.65E-83 | 5.24E-81 |
| Insig1 | 0.114472305 | 1.133756651 | 9.9042004 | 2.32E-82 | 4.54E-80 |
| Serp1 | 1.182179878 | 3.83920428 | 3.24756355 | 2.70E-81 | 5.21E-79 |

| | | | | | |
|---|---|---|---|---|---|
| Serpini2 | 0.048836925 | 0.890087617 | 18.2257096 | 1.09E-80 | 2.09E-78 |
| Aplp1 | 1.309799904 | 0.09612009 | 0.07338532 | 1.07E-79 | 2.01E-77 |
| Rpl35 | 4.21876409 | 12.55623346 | 2.97628244 | 3.57E-79 | 6.67E-77 |
| Dbi | 1.738733209 | 5.300738747 | 3.0486211 | 1.36E-77 | 2.51E-75 |
| Rbpjl | 0.07677937 | 0.951881219 | 12.397617 | 1.42E-76 | 2.59E-74 |
| Prss3 | 0.01439082 | 0.694630497 | 48.2690021 | 2.18E-75 | 3.95E-73 |
| Aqp8 | 0.025174859 | 0.743097838 | 29.5174578 | 4.33E-75 | 7.76E-73 |
| Gm10334 | 0.012531545 | 0.664103018 | 52.9945032 | 1.79E-74 | 3.17E-72 |
| Dlk1 | 5.43166273 | 1.559993713 | 0.28720371 | 7.58E-74 | 1.33E-71 |
| Mafb | 1.145442911 | 0.071350855 | 0.06229106 | 3.72E-73 | 6.45E-71 |
| S100a11 | 2.869942761 | 0.789560354 | 0.27511362 | 3.89E-73 | 6.69E-71 |
| Ubb | 9.905746072 | 2.925533749 | 0.29533704 | 4.92E-73 | 8.37E-71 |
| Psmb10 | 0.236059242 | 1.353439485 | 5.73347384 | 4.45E-72 | 7.49E-70 |
| Scgn | 1.040711861 | 0.04409348 | 0.04236858 | 7.75E-72 | 1.29E-69 |
| Hadh | 1.799616101 | 0.349821179 | 0.19438656 | 1.20E-71 | 1.98E-69 |
| Tspan7 | 1.41302132 | 0.172974794 | 0.12241485 | 1.58E-71 | 2.58E-69 |
| Igf1 | 0.102154531 | 0.974443385 | 9.538915 | 9.12E-71 | 1.48E-68 |
| Rps24 | 8.130750906 | 22.91260358 | 2.81801814 | 7.60E-70 | 1.22E-67 |
| Pcsk1 | 0.993511299 | 0.038955082 | 0.0392095 | 2.78E-69 | 4.42E-67 |
| Pax6 | 1.226489707 | 0.12031187 | 0.09809448 | 1.01E-68 | 1.59E-66 |
| Mif | 2.341838619 | 6.549896869 | 2.7969036 | 1.39E-68 | 2.16E-66 |
| Ass1 | 0.128214312 | 1.037199197 | 8.08957423 | 1.49E-68 | 2.31E-66 |
| Ldha | 0.872156432 | 2.625248231 | 3.01006578 | 3.47E-67 | 5.32E-65 |
| Rpl12 | 3.376408547 | 9.175084651 | 2.7174095 | 1.80E-66 | 2.73E-64 |
| Rpl38 | 5.914929886 | 16.03145349 | 2.71033703 | 4.22E-66 | 6.36E-64 |
| Gatm | 0.094235602 | 0.90957591 | 9.65214729 | 1.20E-65 | 1.79E-63 |
| Gch1 | 1.112131265 | 0.091679856 | 0.08243618 | 1.25E-65 | 1.85E-63 |
| Map1b | 1.080800659 | 0.08370754 | 0.07744956 | 2.69E-65 | 3.95E-63 |
| Rps23 | 9.234592349 | 25.19543039 | 2.72837494 | 5.63E-65 | 8.19E-63 |
| Rpl22 | 2.445846801 | 6.62917195 | 2.71037906 | 6.45E-65 | 9.31E-63 |

| | | | | | |
|---|---|---|---|---|---|
| Rps8 | 10.28970136 | 28.01109026 | 2.72224521 | 5.12E-64 | 7.33E-62 |
| Dhx34 | 0.079732771 | 0.84223961 | 10.5632804 | 1.05E-63 | 1.49E-61 |
| Rpl5 | 2.65263897 | 7.045057338 | 2.65586739 | 1.32E-62 | 1.85E-60 |
| 1700086L19F | 1.082517678 | 0.102192804 | 0.0944029 | 1.07E-61 | 1.49E-59 |
| Rpl18a | 9.781993262 | 26.03614216 | 2.66163976 | 1.31E-61 | 1.81E-59 |
| Cldn10 | 0.326258536 | 1.432135098 | 4.38957128 | 1.81E-61 | 2.48E-59 |
| Phgdh | 0.317786952 | 1.410402444 | 4.43820124 | 2.88E-61 | 3.93E-59 |
| Rpl36 | 6.833679873 | 17.76270409 | 2.59928829 | 1.84E-60 | 2.49E-58 |
| Edem1 | 0.205053854 | 1.130040315 | 5.51094405 | 2.47E-59 | 3.32E-57 |
| Ang | 0.157606946 | 1.012408865 | 6.42363102 | 4.49E-59 | 5.98E-57 |
| Rpsa | 6.014614415 | 15.39676 | 2.55989145 | 6.76E-59 | 8.94E-57 |
| Rpl32 | 11.62786863 | 29.81549696 | 2.56414119 | 3.75E-56 | 4.91E-54 |
| Syt13 | 0.824434359 | 0.037964165 | 0.04604874 | 8.53E-56 | 1.11E-53 |
| Hmgn3 | 1.1405438 | 0.155752706 | 0.13656004 | 2.51E-55 | 3.24E-53 |
| Ngfrap1 | 2.358938553 | 0.789045943 | 0.33449194 | 2.69E-55 | 3.45E-53 |
| Ggh | 0.142190436 | 0.930092486 | 6.54117472 | 6.69E-55 | 8.52E-53 |
| Rps14 | 12.41105566 | 31.30042321 | 2.52197912 | 7.84E-54 | 9.91E-52 |
| Cbs | 0.119635528 | 0.857267603 | 7.16566074 | 2.99E-53 | 3.76E-51 |
| Tff2 | 0.139437974 | 0.900774227 | 6.46003527 | 3.04E-53 | 3.78E-51 |
| Rnase4 | 0.671203787 | 1.960604157 | 2.92102666 | 4.55E-53 | 5.62E-51 |
| Rps20 | 5.65038623 | 13.73881861 | 2.4314831 | 5.54E-53 | 6.79E-51 |
| Tm4sf4 | 1.131803309 | 0.177967255 | 0.15724221 | 1.04E-51 | 1.27E-49 |
| Rpl37 | 7.738251276 | 18.6291496 | 2.40741079 | 4.46E-51 | 5.39E-49 |
| Ptf1a | 0.068650335 | 0.686290626 | 9.99690133 | 1.60E-50 | 1.92E-48 |
| Calm2 | 2.973546801 | 1.077149625 | 0.36224405 | 3.77E-50 | 4.50E-48 |
| Emb | 0.900819438 | 0.090378038 | 0.10032869 | 7.41E-50 | 8.78E-48 |
| Msrb1 | 0.192966218 | 0.99360874 | 5.14913309 | 7.94E-50 | 9.34E-48 |
| Krtcap2 | 1.935661394 | 4.681447824 | 2.41852622 | 8.87E-50 | 1.04E-47 |
| Neurod1 | 0.789260234 | 0.056067507 | 0.07103805 | 9.68E-49 | 1.12E-46 |
| Slc25a4 | 1.858780333 | 0.580626722 | 0.31236974 | 1.20E-48 | 1.39E-46 |

| | | | | | |
|---|---|---|---|---|---|
| Rpl28 | 7.002099746 | 16.42411279 | 2.34559823 | 1.73E-48 | 1.98E-46 |
| Isl1 | 0.878620703 | 0.089266499 | 0.10159845 | 4.05E-48 | 4.61E-46 |
| Rpl13 | 12.25149556 | 29.27434516 | 2.38945074 | 4.31E-48 | 4.87E-46 |
| Rpl10a | 5.997327794 | 13.84097353 | 2.30785677 | 4.72E-47 | 5.30E-45 |
| Rpl39 | 6.70772315 | 15.50089055 | 2.31090196 | 5.85E-47 | 6.52E-45 |
| Rps28 | 4.639003013 | 10.6717809 | 2.30044707 | 8.43E-47 | 9.35E-45 |
| Hba-a2 | 2.693766012 | 1.013311933 | 0.37616925 | 9.33E-47 | 1.03E-44 |
| Ggt1 | 0.065762898 | 0.63373927 | 9.6367296 | 3.80E-46 | 4.16E-44 |
| Rps21 | 3.735349379 | 8.543393129 | 2.28717377 | 5.82E-46 | 6.33E-44 |
| Slc2a2 | 0.711546639 | 0.044138786 | 0.06203218 | 7.81E-46 | 8.44E-44 |
| Alas2 | 0.701011997 | 0.043568255 | 0.06215051 | 4.56E-45 | 4.89E-43 |
| Rpl41 | 21.87292119 | 52.78939127 | 2.41345867 | 6.37E-45 | 6.80E-43 |
| Rps12-ps3 | 0.52351751 | 1.573571553 | 3.00576681 | 7.04E-45 | 7.47E-43 |
| Rps10 | 5.684622862 | 12.84653614 | 2.25987483 | 8.17E-45 | 8.61E-43 |
| Krt7 | 1.090788702 | 0.206126449 | 0.1889701 | 1.35E-44 | 1.42E-42 |
| Prnp | 0.758498314 | 0.066173292 | 0.0872425 | 1.53E-44 | 1.60E-42 |
| Rplp2 | 8.905602918 | 20.18546741 | 2.26660313 | 4.46E-44 | 4.61E-42 |
| Pdia3 | 3.712833679 | 1.483954307 | 0.39968241 | 3.36E-43 | 3.46E-41 |
| Copz2 | 0.151562009 | 0.825537116 | 5.44686045 | 4.56E-43 | 4.66E-41 |
| Rpl23 | 8.201610992 | 18.3210801 | 2.2338392 | 7.45E-43 | 7.57E-41 |
| P2rx1 | 0.043595995 | 0.525177962 | 12.0464726 | 2.24E-42 | 2.27E-40 |
| Ywhaq | 1.787661984 | 0.613984915 | 0.34345694 | 4.39E-42 | 4.41E-40 |
| Gm15915 | 0.087716799 | 0.656173308 | 7.48058886 | 4.57E-42 | 4.56E-40 |
| Mrfap1 | 3.460804465 | 1.409789464 | 0.40735889 | 1.52E-41 | 1.51E-39 |
| Eef1b2 | 2.434588597 | 5.371465243 | 2.20631332 | 3.16E-41 | 3.12E-39 |
| Ssr1 | 0.597701756 | 1.633147896 | 2.73237929 | 1.48E-40 | 1.45E-38 |
| Prss53 | 0.553511892 | 0.015959565 | 0.02883328 | 2.48E-40 | 2.42E-38 |
| Rps15 | 6.6917147 | 14.45713959 | 2.16045367 | 5.92E-40 | 5.75E-38 |
| G6pc2 | 0.530972431 | 0.012919929 | 0.02433258 | 6.72E-40 | 6.49E-38 |
| Rap1gap | 0.121375238 | 0.716492662 | 5.90312056 | 9.86E-40 | 9.46E-38 |

| | | | | | |
|---|---|---|---|---|---|
| Ndufs6 | 1.125087003 | 2.559362411 | 2.27481288 | 1.05E-39 | 1.00E-37 |
| Bcat2 | 0.118929678 | 0.706637806 | 5.94164398 | 1.11E-39 | 1.05E-37 |
| Gnb2l1 | 3.085218097 | 6.645413142 | 2.15395247 | 1.48E-39 | 1.39E-37 |
| Ttc23 | 0.04046603 | 0.484118478 | 11.9635773 | 7.15E-39 | 6.72E-37 |
| Gnmt | 0.023719672 | 0.41920256 | 17.6732024 | 9.81E-39 | 9.16E-37 |
| Sytl4 | 0.564016511 | 0.023530448 | 0.04171943 | 1.11E-38 | 1.03E-36 |
| Gm10076 | 1.029216603 | 2.327506766 | 2.26143531 | 1.16E-38 | 1.07E-36 |
| Gpc3 | 0.212878672 | 0.905049086 | 4.25147845 | 1.96E-38 | 1.80E-36 |
| Rps26 | 4.576852991 | 9.707642737 | 2.12103005 | 2.23E-38 | 2.04E-36 |
| Mlxipl | 0.543389083 | 0.021060118 | 0.03875698 | 2.33E-38 | 2.12E-36 |
| Chst2 | 0.022886154 | 0.420119413 | 18.356925 | 3.08E-38 | 2.78E-36 |
| Rps27a | 11.63368963 | 24.97803859 | 2.14704358 | 1.01E-37 | 9.10E-36 |
| Meis2 | 1.362845063 | 0.407770156 | 0.29920507 | 1.35E-37 | 1.21E-35 |
| Resp18 | 0.597682991 | 0.042861913 | 0.07171346 | 4.41E-37 | 3.93E-35 |
| Cst3 | 3.33709725 | 1.442509432 | 0.43226473 | 5.31E-37 | 4.70E-35 |
| Rpl17 | 9.181530064 | 19.37260832 | 2.10995424 | 6.33E-37 | 5.58E-35 |
| Rpl37a | 12.4618624 | 26.53852272 | 2.12957918 | 1.09E-36 | 9.56E-35 |
| Atp2a2 | 1.057973554 | 0.249553275 | 0.23587856 | 1.18E-36 | 1.03E-34 |
| Rps4x | 8.513403719 | 17.77762544 | 2.08819246 | 3.64E-36 | 3.16E-34 |
| Rps19 | 11.02751865 | 23.09266542 | 2.09409443 | 1.29E-35 | 1.12E-33 |
| Rpl30 | 3.850298687 | 7.945239017 | 2.06353835 | 1.38E-35 | 1.19E-33 |
| Rpl34 | 6.911547278 | 14.28272486 | 2.0665018 | 1.40E-35 | 1.20E-33 |
| Rps25 | 5.381266261 | 11.08777071 | 2.06043897 | 1.45E-35 | 1.23E-33 |
| Pak3 | 0.600962591 | 0.049954081 | 0.08312345 | 1.86E-35 | 1.58E-33 |
| Pdx1 | 0.783621472 | 0.124841706 | 0.15931379 | 2.13E-35 | 1.79E-33 |
| Aldh1a1 | 0.044745514 | 0.455326785 | 10.1759204 | 6.24E-35 | 5.23E-33 |
| Rpl31 | 6.397079317 | 13.06988227 | 2.04310149 | 1.25E-34 | 1.04E-32 |
| Hist3h2ba | 0.546556024 | 0.038704917 | 0.07081601 | 3.03E-34 | 2.51E-32 |
| Nkx6-1 | 0.570708714 | 0.044882341 | 0.07864317 | 5.65E-34 | 4.66E-32 |
| Bex2 | 1.390150086 | 0.46500595 | 0.33450054 | 7.53E-34 | 6.19E-32 |

| | | | | | |
|---|---|---|---|---|---|
| Ndufa1 | 1.491308604 | 0.523635157 | 0.35112461 | 7.95E-34 | 6.50E-32 |
| Ddx5 | 2.103421377 | 0.920401106 | 0.43757333 | 8.39E-34 | 6.83E-32 |
| Krt12 | 0.416119655 | 0.005513633 | 0.01325011 | 1.33E-33 | 1.07E-31 |
| Zcchc18 | 0.498559728 | 0.030044853 | 0.0602633 | 1.34E-33 | 1.07E-31 |
| Papss2 | 0.498498116 | 0.029833789 | 0.05984735 | 1.34E-33 | 1.07E-31 |
| Rab3a | 0.632553884 | 0.074939762 | 0.11847174 | 1.34E-33 | 1.07E-31 |
| Rpl26 | 8.810546479 | 17.88183265 | 2.02959404 | 1.87E-33 | 1.49E-31 |
| Rpl21 | 6.87900421 | 13.86512005 | 2.0155708 | 2.45E-33 | 1.94E-31 |
| Rps17 | 7.757977848 | 15.6751245 | 2.02051679 | 2.56E-33 | 2.01E-31 |
| Prkar1a | 1.166976062 | 0.343830725 | 0.29463391 | 3.06E-33 | 2.40E-31 |
| Asns | 0.18503587 | 0.775657796 | 4.19193207 | 4.27E-33 | 3.33E-31 |
| Sst | 5.382418708 | 2.494661568 | 0.46348337 | 7.38E-33 | 5.73E-31 |
| Rpl27a | 7.530374446 | 15.12054216 | 2.00794028 | 7.95E-33 | 6.15E-31 |
| Rps11 | 6.885580811 | 13.74693279 | 1.99648122 | 1.81E-32 | 1.39E-30 |
| Qsox1 | 0.143443267 | 0.686372971 | 4.78497865 | 1.90E-32 | 1.46E-30 |
| Rbm39 | 2.229373396 | 1.012784621 | 0.45429116 | 2.44E-32 | 1.86E-30 |
| Sel1l | 0.283949846 | 0.960218677 | 3.38164887 | 2.64E-32 | 2.01E-30 |
| Fam151a | 0.402368809 | 0.005319957 | 0.01322159 | 2.68E-32 | 2.03E-30 |
| Rps6 | 8.182767895 | 16.3516259 | 1.9983001 | 3.21E-32 | 2.42E-30 |
| Selm | 1.078951558 | 0.303074681 | 0.28089739 | 3.46E-32 | 2.59E-30 |
| Tmsb15b2 | 0.49652439 | 0.029173072 | 0.05875456 | 3.81E-32 | 2.85E-30 |
| Gm10073 | 0.361653193 | 1.095851887 | 3.0301181 | 6.65E-32 | 4.94E-30 |
| Snhg18 | 0.155037935 | 0.704714675 | 4.54543384 | 6.67E-32 | 4.94E-30 |
| Sepp1 | 0.895631625 | 1.935431384 | 2.16096811 | 9.73E-32 | 7.17E-30 |
| Rap1b | 0.820814562 | 0.17179313 | 0.20929591 | 1.12E-31 | 8.22E-30 |
| Abcc8 | 0.571313583 | 0.064019487 | 0.11205665 | 1.29E-31 | 9.44E-30 |
| Tmem59 | 1.621325213 | 0.635038836 | 0.39167888 | 1.36E-31 | 9.88E-30 |
| Itm2b | 3.207226237 | 1.489071404 | 0.46428636 | 1.39E-31 | 1.01E-29 |
| Prdx2 | 2.802525769 | 1.302551157 | 0.46477758 | 3.38E-31 | 2.44E-29 |
| Hepacam2 | 0.466293479 | 0.030069542 | 0.0644863 | 3.54E-31 | 2.54E-29 |

| | | | | | |
|---|---|---|---|---|---|
| Kctd14 | 0.040879124 | 0.405300594 | 9.91461053 | 5.49E-31 | 3.92E-29 |
| Ifitm2 | 1.027934735 | 2.13387301 | 2.07588375 | 5.50E-31 | 3.92E-29 |
| Tsga10ip | 0.023106393 | 0.352527851 | 15.2567238 | 7.24E-31 | 5.14E-29 |
| mt-Atp8 | 0.239509205 | 0.856337444 | 3.57538426 | 1.76E-30 | 1.25E-28 |
| Eif5b | 0.819397496 | 1.799027983 | 2.19554977 | 2.81E-30 | 1.98E-28 |
| Ttr | 9.107921766 | 4.401648182 | 0.4832769 | 4.55E-30 | 3.19E-28 |
| Gm5771 | 0.010753046 | 0.286835071 | 26.6747745 | 5.08E-30 | 3.55E-28 |
| Lbh | 0.084426157 | 0.517219464 | 6.12629403 | 5.12E-30 | 3.56E-28 |
| Gstm7 | 0.074190477 | 0.49384756 | 6.65648183 | 6.17E-30 | 4.27E-28 |
| Prdx4 | 0.422117991 | 1.164163271 | 2.75790963 | 7.35E-30 | 5.07E-28 |
| Maged1 | 1.659518701 | 0.689203301 | 0.41530312 | 1.31E-29 | 9.03E-28 |
| Tram1 | 0.731862493 | 1.650770334 | 2.25557444 | 2.72E-29 | 1.86E-27 |
| Rpl14 | 9.500126414 | 18.3614587 | 1.93275941 | 5.14E-29 | 3.50E-27 |
| Egr1 | 0.690243413 | 1.577154718 | 2.28492542 | 5.64E-29 | 3.83E-27 |
| Rpl23a | 6.590690757 | 12.64065959 | 1.91795672 | 5.71E-29 | 3.86E-27 |
| Krt18 | 1.482404622 | 2.921088603 | 1.97050695 | 8.09E-29 | 5.45E-27 |
| Car9 | 0.079042752 | 0.49195008 | 6.22384809 | 9.50E-29 | 6.38E-27 |
| H3f3a | 5.215851515 | 2.564465816 | 0.49166772 | 1.07E-28 | 7.12E-27 |
| Slc39a5 | 0.019242993 | 0.310845357 | 16.1536907 | 1.53E-28 | 1.02E-26 |
| Fkbp1a | 1.607725497 | 0.671038529 | 0.41738377 | 1.83E-28 | 1.22E-26 |
| Rpl13a | 13.16499397 | 25.53332974 | 1.93948663 | 2.16E-28 | 1.43E-26 |
| Mageh1 | 0.670632687 | 0.122113316 | 0.18208673 | 2.21E-28 | 1.45E-26 |
| Nenf | 0.978915436 | 0.285388792 | 0.2915357 | 3.03E-28 | 1.99E-26 |
| Tspan3 | 0.825138243 | 0.203144153 | 0.24619408 | 3.27E-28 | 2.13E-26 |
| Dctpp1 | 0.269705271 | 0.87236403 | 3.23450864 | 4.18E-28 | 2.72E-26 |
| Dnajb9 | 0.679453449 | 0.133050289 | 0.19581958 | 7.99E-28 | 5.18E-26 |
| Edem2 | 0.64672486 | 1.484531185 | 2.29546021 | 1.17E-27 | 7.55E-26 |
| Selk | 2.381489341 | 1.160916018 | 0.48747479 | 1.65E-27 | 1.06E-25 |
| Zbtb20 | 0.797696813 | 0.189945882 | 0.23811789 | 1.71E-27 | 1.09E-25 |
| Rpl35a | 7.73902911 | 14.58142521 | 1.88414141 | 2.72E-27 | 1.74E-25 |

| | | | | | |
|---|---|---|---|---|---|
| Uqcrq | 2.162977335 | 4.102973524 | 1.89691009 | 4.02E-27 | 2.56E-25 |
| Rps16 | 8.993633713 | 16.96069482 | 1.88585564 | 4.03E-27 | 2.56E-25 |
| Serpinf2 | 0.036438594 | 0.361357316 | 9.91688426 | 4.74E-27 | 2.99E-25 |
| Rps15a | 8.247153956 | 15.49251781 | 1.87852899 | 5.91E-27 | 3.72E-25 |
| Rpl3 | 6.601455667 | 12.34555866 | 1.87012672 | 6.51E-27 | 4.08E-25 |
| Gabarapl2 | 1.321486358 | 0.50974955 | 0.38573955 | 9.17E-27 | 5.73E-25 |
| Mkrn1 | 0.749916642 | 0.172642883 | 0.2302161 | 1.55E-26 | 9.66E-25 |
| Ypel3 | 0.650948086 | 0.128941287 | 0.19808229 | 1.58E-26 | 9.83E-25 |
| Enpp2 | 0.482319792 | 0.052321676 | 0.10847922 | 1.61E-26 | 9.95E-25 |
| Atp5k | 1.260931259 | 2.43413305 | 1.93042486 | 2.07E-26 | 1.27E-24 |
| Aldh7a1 | 0.131990727 | 0.581199371 | 4.40333486 | 2.51E-26 | 1.54E-24 |
| Cd24a | 0.99657909 | 1.969449457 | 1.97620989 | 2.84E-26 | 1.74E-24 |
| Pkm | 1.978787179 | 0.956319658 | 0.48328576 | 3.30E-26 | 2.01E-24 |
| Rps13 | 7.489945816 | 13.90057021 | 1.8558973 | 4.07E-26 | 2.47E-24 |
| Sumo2 | 1.914006042 | 0.916989068 | 0.47909413 | 4.21E-26 | 2.55E-24 |
| Lcmt1 | 0.150515459 | 0.61676909 | 4.09771258 | 4.64E-26 | 2.80E-24 |
| Itga6 | 0.108172152 | 0.52751358 | 4.87661168 | 7.26E-26 | 4.36E-24 |
| Dnaja1 | 1.348357039 | 0.539557067 | 0.4001589 | 9.32E-26 | 5.58E-24 |
| Anxa5 | 0.95397973 | 0.29780473 | 0.31217092 | 1.23E-25 | 7.34E-24 |
| Pdia4 | 1.63532203 | 0.73117339 | 0.44711279 | 1.65E-25 | 9.80E-24 |
| Rpl10 | 11.55853015 | 21.47229077 | 1.85770081 | 2.13E-25 | 1.26E-23 |
| Ap1s2 | 0.430202237 | 0.039379277 | 0.09153666 | 2.24E-25 | 1.32E-23 |
| Mgll | 0.066777085 | 0.416874445 | 6.2427769 | 2.27E-25 | 1.34E-23 |
| Rpl22l1 | 3.985079229 | 7.298348086 | 1.83141857 | 2.88E-25 | 1.69E-23 |
| Ffar1 | 0.294003522 | 0 | 0 | 3.39E-25 | 1.98E-23 |
| Chmp5 | 0.861100709 | 0.249998975 | 0.2903249 | 5.49E-25 | 3.20E-23 |
| Rps3 | 7.354608484 | 13.42658598 | 1.82560173 | 7.57E-25 | 4.39E-23 |
| Agr2 | 0.026304129 | 0.310536827 | 11.8056307 | 7.64E-25 | 4.42E-23 |
| Ndn | 0.456280763 | 0.053568673 | 0.11740288 | 8.03E-25 | 4.63E-23 |
| Glul | 0.498454764 | 0.074173709 | 0.1488073 | 8.62E-25 | 4.96E-23 |

| | | | | | |
|---|---|---|---|---|---|
| Tdh | 0.013212224 | 0.257978744 | 19.5257617 | 8.67E-25 | 4.97E-23 |
| Rps29 | 9.774659824 | 17.81651654 | 1.82272497 | 2.72E-24 | 1.55E-22 |
| Rbp2 | 0.074221845 | 0.431159301 | 5.80906202 | 2.84E-24 | 1.62E-22 |
| Junb | 1.088608592 | 0.391773454 | 0.35988459 | 3.36E-24 | 1.90E-22 |
| Gm2000 | 0.196787888 | 0.682828295 | 3.46986952 | 3.41E-24 | 1.93E-22 |
| Peg3 | 1.902897566 | 0.950325049 | 0.49940946 | 5.74E-24 | 3.23E-22 |
| Cotl1 | 0.465361471 | 0.060671184 | 0.13037432 | 7.54E-24 | 4.23E-22 |
| Atp6v1e1 | 0.964809331 | 0.323396946 | 0.3351926 | 9.79E-24 | 5.48E-22 |
| Ffar4 | 0.345221042 | 0.017818151 | 0.05161375 | 1.03E-23 | 5.75E-22 |
| Ece1 | 0.453259933 | 0.056075473 | 0.12371593 | 1.19E-23 | 6.64E-22 |
| Dbpht2 | 0.428079655 | 0.048924058 | 0.11428728 | 1.60E-23 | 8.87E-22 |
| Sez6l2 | 0.34173799 | 0.015791071 | 0.04620812 | 1.61E-23 | 8.92E-22 |
| Rpl6 | 7.455146744 | 13.37406614 | 1.79393734 | 1.78E-23 | 9.80E-22 |
| Rpl11 | 7.914642654 | 14.20226516 | 1.79442911 | 1.95E-23 | 1.07E-21 |
| Gnb2 | 2.048689408 | 1.063260738 | 0.51899558 | 2.63E-23 | 1.44E-21 |
| Atp5o | 1.787233027 | 3.249796183 | 1.81833937 | 2.77E-23 | 1.51E-21 |
| Morf4l1 | 2.335889532 | 1.222066763 | 0.52316976 | 3.10E-23 | 1.68E-21 |
| 5330417C22I | 0.180580997 | 0.643107324 | 3.56132337 | 3.15E-23 | 1.71E-21 |
| Dmbt1 | 0.018236021 | 0.264172934 | 14.4863257 | 3.63E-23 | 1.96E-21 |
| Rpl18 | 6.410297715 | 11.41672343 | 1.78099738 | 4.37E-23 | 2.35E-21 |
| Eef1d | 1.237420179 | 2.285077805 | 1.84664663 | 4.43E-23 | 2.38E-21 |
| Malat1 | 68.98728469 | 33.63969061 | 0.48762161 | 5.05E-23 | 2.70E-21 |
| Sypl | 0.259176215 | 0.774308231 | 2.98757442 | 9.75E-23 | 5.21E-21 |
| S100a10 | 1.138023142 | 0.448177471 | 0.39382105 | 1.45E-22 | 7.74E-21 |
| Mlec | 0.346436268 | 0.921709305 | 2.66054507 | 1.74E-22 | 9.23E-21 |
| Serpinb1a | 0.145748472 | 0.56018422 | 3.84349978 | 1.82E-22 | 9.61E-21 |
| Mrps33 | 1.177555305 | 0.474267899 | 0.40275637 | 1.82E-22 | 9.61E-21 |
| Arpc2 | 1.162367044 | 0.464286735 | 0.39943212 | 2.07E-22 | 1.09E-20 |
| Tubb3 | 0.379054407 | 0.032884761 | 0.08675473 | 2.49E-22 | 1.31E-20 |
| Clu | 2.006984053 | 3.58652952 | 1.78702442 | 2.61E-22 | 1.36E-20 |

| | | | | | |
|---|---|---|---|---|---|
| Cirbp | 0.539695916 | 0.107174509 | 0.19858314 | 2.62E-22 | 1.36E-20 |
| Cd74 | 0.480488763 | 0.079959271 | 0.16641236 | 4.16E-22 | 2.16E-20 |
| Odc1 | 0.162542017 | 0.589930275 | 3.62940171 | 4.23E-22 | 2.19E-20 |
| Gnao1 | 0.322968421 | 0.019422229 | 0.06013662 | 4.30E-22 | 2.22E-20 |
| 1500011B03I | 0.450378185 | 0.063582201 | 0.14117513 | 5.30E-22 | 2.73E-20 |
| Cd81 | 1.792718886 | 0.899237999 | 0.50160569 | 5.66E-22 | 2.90E-20 |
| Eef1g | 3.238358647 | 5.697048465 | 1.75923951 | 5.83E-22 | 2.98E-20 |
| Eif1 | 5.651842389 | 3.076261094 | 0.5442935 | 7.19E-22 | 3.66E-20 |
| Ssr3 | 0.646154181 | 1.368863684 | 2.11847841 | 7.51E-22 | 3.82E-20 |
| Camk2n1 | 0.359589329 | 0.031597752 | 0.08787177 | 7.83E-22 | 3.97E-20 |
| Jund | 0.168506802 | 0.601179332 | 3.56768584 | 8.00E-22 | 4.04E-20 |
| Rpl9 | 6.690266453 | 11.71599769 | 1.75120046 | 8.36E-22 | 4.21E-20 |
| Lyz2 | 0.625534053 | 0.152755979 | 0.2442009 | 1.03E-21 | 5.18E-20 |
| Hspa8 | 5.739389898 | 3.134219733 | 0.54608936 | 1.06E-21 | 5.29E-20 |
| Efcab1 | 0.320459789 | 0.015718705 | 0.04905048 | 1.13E-21 | 5.64E-20 |
| Cuzd1 | 0.008931893 | 0.211272679 | 23.6537416 | 1.29E-21 | 6.42E-20 |
| Sub1 | 2.435065387 | 1.30650089 | 0.53653627 | 1.38E-21 | 6.83E-20 |
| Lpl | 0.34336273 | 0.025718414 | 0.07490159 | 1.45E-21 | 7.18E-20 |
| Fxyd6 | 0.354872534 | 0.032489746 | 0.09155328 | 1.95E-21 | 9.62E-20 |
| Gnai2 | 0.904423072 | 0.31963464 | 0.35341274 | 1.97E-21 | 9.69E-20 |
| Pim2 | 0.342108884 | 0.026389124 | 0.07713662 | 2.37E-21 | 1.16E-19 |
| Rpl24 | 6.644736978 | 11.55958097 | 1.73965968 | 2.43E-21 | 1.19E-19 |
| Actg1 | 7.466889392 | 4.130719301 | 0.55320483 | 5.90E-21 | 2.88E-19 |
| Nisch | 0.852191574 | 0.289592896 | 0.33982136 | 5.98E-21 | 2.91E-19 |
| Maob | 0.259886525 | 0.008248384 | 0.0317384 | 9.02E-21 | 4.38E-19 |
| Espn | 0.043296441 | 0.317952234 | 7.3436114 | 9.47E-21 | 4.58E-19 |
| Tspan1 | 0.026651767 | 0.270810967 | 10.1610887 | 1.09E-20 | 5.26E-19 |
| Tubb2a | 0.507565529 | 0.1017654 | 0.20049707 | 1.18E-20 | 5.69E-19 |
| Immp1l | 0.803472633 | 0.265964257 | 0.33101844 | 1.25E-20 | 6.01E-19 |
| Parm1 | 0.081208908 | 0.402014072 | 4.95036914 | 1.39E-20 | 6.65E-19 |

| | | | | | |
|---|---|---|---|---|---|
| Vwa5b2 | 0.290696341 | 0.013022499 | 0.0447976 | 1.87E-20 | 8.93E-19 |
| Rps5 | 8.934519135 | 15.38274133 | 1.72172012 | 2.62E-20 | 1.25E-18 |
| Pam | 0.490555766 | 0.096189911 | 0.19608354 | 3.06E-20 | 1.45E-18 |
| Rps3a1 | 8.235088505 | 14.14226746 | 1.71731821 | 3.24E-20 | 1.53E-18 |
| Rps2 | 6.245247786 | 10.68692739 | 1.71120951 | 3.29E-20 | 1.55E-18 |
| Gm8730 | 6.012650686 | 10.26930026 | 1.70794892 | 4.25E-20 | 2.00E-18 |
| Mid1ip1 | 0.604975895 | 0.157874616 | 0.26096018 | 6.90E-20 | 3.24E-18 |
| Ap2m1 | 0.968599676 | 0.372819782 | 0.38490595 | 7.25E-20 | 3.39E-18 |
| Atp5j2 | 2.796822688 | 4.785978418 | 1.71121982 | 8.17E-20 | 3.81E-18 |
| Rps7 | 7.75171894 | 13.21201915 | 1.70439863 | 8.70E-20 | 4.05E-18 |
| Srp19 | 0.504617719 | 1.118619699 | 2.21676659 | 8.86E-20 | 4.11E-18 |
| Atp6v0e | 1.112673066 | 0.471734929 | 0.42396544 | 9.77E-20 | 4.53E-18 |
| Hspa1a | 0.376948607 | 0.050273003 | 0.13336832 | 1.03E-19 | 4.76E-18 |
| Tmem215 | 0.247814536 | 0.001237225 | 0.00499255 | 1.77E-19 | 8.16E-18 |
| Npdc1 | 0.49655163 | 0.104491385 | 0.21043408 | 2.22E-19 | 1.02E-17 |
| Sdsl | 0.118023513 | 0.469387027 | 3.97706367 | 2.22E-19 | 1.02E-17 |
| Gm1673 | 0.456991025 | 0.085810315 | 0.18777243 | 3.27E-19 | 1.49E-17 |
| Marcksl1 | 0.946506396 | 0.368696069 | 0.38953363 | 3.49E-19 | 1.59E-17 |
| Rogdi | 0.463613371 | 0.092340694 | 0.19917608 | 4.09E-19 | 1.86E-17 |
| Hdlbp | 0.560474505 | 1.182845164 | 2.11043527 | 5.63E-19 | 2.55E-17 |
| Plk2 | 0.112834414 | 0.45258747 | 4.01107652 | 5.78E-19 | 2.62E-17 |
| Sec11a | 1.235369168 | 0.564279406 | 0.45676986 | 7.22E-19 | 3.25E-17 |
| Hspe1 | 1.689663057 | 2.895993819 | 1.71394753 | 7.22E-19 | 3.25E-17 |
| Gp2 | 0.004799477 | 0.172385294 | 35.9175167 | 9.75E-19 | 4.37E-17 |
| S100a8 | 0.043263645 | 0.293737721 | 6.78948156 | 1.18E-18 | 5.28E-17 |
| Rpl8 | 7.127943756 | 11.93394939 | 1.67424854 | 1.32E-18 | 5.87E-17 |
| Creld2 | 1.385583138 | 0.674566377 | 0.48684656 | 1.43E-18 | 6.37E-17 |
| Sf3b6 | 0.894763631 | 0.348450947 | 0.38943352 | 2.15E-18 | 9.56E-17 |
| Apoe | 1.254519126 | 0.584567966 | 0.46596975 | 2.23E-18 | 9.89E-17 |
| Atraid | 0.632286105 | 0.190904454 | 0.30192733 | 2.54E-18 | 1.12E-16 |

| | | | | | |
|---|---|---|---|---|---|
| Rundc3a | 0.246569197 | 0.010560949 | 0.04283158 | 2.61E-18 | 1.15E-16 |
| Slc2a5 | 0.289875341 | 0.02018811 | 0.06964411 | 2.64E-18 | 1.16E-16 |
| Tob1 | 0.137548656 | 0.488629573 | 3.55241255 | 2.86E-18 | 1.25E-16 |
| Myo15b | 0.036848288 | 0.270905944 | 7.35192753 | 3.06E-18 | 1.34E-16 |
| Marcks | 0.380076044 | 0.899990909 | 2.36792327 | 3.16E-18 | 1.38E-16 |
| Rsrp1 | 1.257818152 | 0.588594872 | 0.4679491 | 3.48E-18 | 1.51E-16 |
| Wdr89 | 0.629221523 | 1.267724338 | 2.01475044 | 3.64E-18 | 1.58E-16 |
| Xist | 2.96344981 | 1.70026683 | 0.57374578 | 4.43E-18 | 1.92E-16 |
| Rpl7a | 3.389988909 | 5.628183229 | 1.66023647 | 4.98E-18 | 2.15E-16 |
| Gstm2 | 0.071622921 | 0.35023459 | 4.88997913 | 5.12E-18 | 2.20E-16 |
| Rpl23a-ps3 | 0.712211499 | 1.378040009 | 1.9348747 | 5.69E-18 | 2.44E-16 |
| Clic1 | 0.971900527 | 0.402397007 | 0.41403106 | 7.41E-18 | 3.18E-16 |
| Cplx2 | 0.338139865 | 0.042645423 | 0.1261177 | 7.49E-18 | 3.20E-16 |
| Larp1 | 0.123446238 | 0.457656352 | 3.70733332 | 9.28E-18 | 3.96E-16 |
| Bhlha15 | 0.086369376 | 0.380663894 | 4.40739429 | 9.32E-18 | 3.96E-16 |
| Rnf130 | 0.306929711 | 0.030451883 | 0.09921452 | 1.02E-17 | 4.31E-16 |
| Eif4a2 | 1.274352701 | 0.61149249 | 0.47984556 | 1.32E-17 | 5.58E-16 |
| Aup1 | 0.132329214 | 0.469316593 | 3.5465834 | 1.49E-17 | 6.28E-16 |
| Fam46c | 0.157035172 | 0.519437071 | 3.30777534 | 1.56E-17 | 6.57E-16 |
| Tusc3 | 0.489865373 | 0.119622436 | 0.24419451 | 1.62E-17 | 6.82E-16 |
| Homer2 | 0.047096321 | 0.288909729 | 6.13444373 | 1.87E-17 | 7.83E-16 |
| Atp6v1g1 | 1.637982346 | 0.880974012 | 0.53784097 | 1.91E-17 | 7.99E-16 |
| Gjd2 | 0.219518119 | 0.007211021 | 0.03284932 | 2.03E-17 | 8.46E-16 |
| Sostdc1 | 0.035205281 | 0.257607822 | 7.3173063 | 2.12E-17 | 8.80E-16 |
| Mapk3 | 0.504686803 | 0.126060046 | 0.24977876 | 2.93E-17 | 1.22E-15 |
| Eva1b | 0.478102667 | 0.115193348 | 0.24093852 | 3.09E-17 | 1.28E-15 |
| Gdi1 | 0.435053338 | 0.092670855 | 0.21301033 | 3.18E-17 | 1.31E-15 |
| Sec61g | 5.651291815 | 9.247109317 | 1.6362824 | 3.30E-17 | 1.36E-15 |
| Prmt1 | 1.11461853 | 0.515844468 | 0.46279911 | 4.46E-17 | 1.83E-15 |
| C1qa | 0.418582926 | 0.934538999 | 2.2326257 | 4.55E-17 | 1.87E-15 |

| | | | | | |
|---|---|---|---|---|---|
| Car2 | 0.36874297 | 0.059987145 | 0.1626801 | 4.88E-17 | 2.00E-15 |
| Exoc7 | 0.359012133 | 0.056528035 | 0.15745439 | 5.52E-17 | 2.25E-15 |
| Ift81 | 0.270892586 | 0.023357421 | 0.08622392 | 6.28E-17 | 2.56E-15 |
| Fmo1 | 0.315663466 | 0.040520933 | 0.12836751 | 6.51E-17 | 2.65E-15 |
| Fkbp1b | 0.256776677 | 0.016444979 | 0.0640439 | 7.42E-17 | 3.01E-15 |
| Tmem147 | 0.578313407 | 1.163794588 | 2.01239427 | 8.24E-17 | 3.33E-15 |
| Slc26a9 | 0.015125778 | 0.192288761 | 12.7126524 | 9.34E-17 | 3.77E-15 |
| Acta2 | 0.303558053 | 0.033683153 | 0.11096116 | 9.59E-17 | 3.86E-15 |
| Ctrl | 0.050788865 | 0.290631282 | 5.72234254 | 1.03E-16 | 4.15E-15 |
| Atp1b1 | 0.825070844 | 0.326374599 | 0.39557161 | 1.15E-16 | 4.61E-15 |
| Neurog3 | 0.226524745 | 0.011334404 | 0.05003605 | 1.16E-16 | 4.65E-15 |
| Prps1 | 0.381448964 | 0.070294953 | 0.18428403 | 1.24E-16 | 4.93E-15 |
| Cfap20 | 0.534467531 | 0.149014122 | 0.27880856 | 1.31E-16 | 5.22E-15 |
| Enho | 0.209732437 | 0.002689251 | 0.01282229 | 1.39E-16 | 5.53E-15 |
| Ybx1 | 0.736528325 | 1.376209773 | 1.86850896 | 1.42E-16 | 5.62E-15 |
| Upk3a | 0.031575617 | 0.241051186 | 7.63409266 | 1.43E-16 | 5.64E-15 |
| Gm10263 | 0.03806945 | 0.254317161 | 6.68034771 | 1.49E-16 | 5.86E-15 |
| Trappc2l | 0.683917706 | 0.234164982 | 0.34238766 | 1.53E-16 | 6.02E-15 |
| Bex4 | 0.527256548 | 1.08371261 | 2.05538009 | 2.14E-16 | 8.40E-15 |
| Rexo2 | 0.362982522 | 0.83810373 | 2.30893687 | 2.18E-16 | 8.54E-15 |
| Bax | 0.812250137 | 0.319732356 | 0.3936378 | 2.22E-16 | 8.67E-15 |
| Sphkap | 0.261935918 | 0.022840933 | 0.08720046 | 2.43E-16 | 9.47E-15 |
| Cabp2 | 0.015437159 | 0.187219866 | 12.127871 | 2.79E-16 | 1.09E-14 |
| Rab3b | 0.220981256 | 0.009225272 | 0.04174685 | 3.00E-16 | 1.16E-14 |
| Il1r1 | 0.250118773 | 0.020108398 | 0.0803954 | 3.16E-16 | 1.22E-14 |
| Pcp4 | 0.251016881 | 0.016243122 | 0.06470928 | 3.16E-16 | 1.22E-14 |
| Bace2 | 0.234652702 | 0.015745994 | 0.0671034 | 3.18E-16 | 1.22E-14 |
| Cd164 | 0.829636859 | 0.330620517 | 0.39851233 | 3.37E-16 | 1.29E-14 |
| Rps9 | 8.297333415 | 13.40111932 | 1.61511158 | 3.56E-16 | 1.36E-14 |
| Atp5g1 | 3.669527069 | 5.920102947 | 1.61331497 | 3.59E-16 | 1.37E-14 |

| | | | | | |
|---|---|---|---|---|---|
| Ctsf | 0.282797691 | 0.031025463 | 0.10970904 | 5.43E-16 | 2.07E-14 |
| Derl2 | 0.5534238 | 0.166916261 | 0.30160658 | 6.20E-16 | 2.36E-14 |
| Cox7a2l | 1.824132151 | 1.05467166 | 0.57817722 | 7.94E-16 | 3.02E-14 |
| Zfp706 | 0.600923711 | 1.16942351 | 1.94604321 | 8.83E-16 | 3.35E-14 |
| Maged2 | 0.558578673 | 0.169953069 | 0.30425986 | 9.07E-16 | 3.43E-14 |
| Ndrg1 | 0.220988993 | 0.600759906 | 2.71850602 | 1.01E-15 | 3.80E-14 |
| Insm1 | 0.242323648 | 0.018159986 | 0.07494104 | 1.19E-15 | 4.48E-14 |
| Npepl1 | 0.443244708 | 0.111347434 | 0.25120985 | 1.21E-15 | 4.56E-14 |
| Ptprn | 0.21552025 | 0.007475895 | 0.03468767 | 1.31E-15 | 4.90E-14 |
| H2afy | 0.841405931 | 0.347996567 | 0.41358939 | 1.48E-15 | 5.54E-14 |
| Nop10 | 0.704253141 | 1.305588924 | 1.85386312 | 1.75E-15 | 6.53E-14 |
| Spcs3 | 0.320085401 | 0.755211538 | 2.35940638 | 1.89E-15 | 7.03E-14 |
| Tmed9 | 1.007899593 | 0.468358512 | 0.46468767 | 1.96E-15 | 7.30E-14 |
| 1110008P14I | 0.572208055 | 0.182293835 | 0.31857964 | 2.55E-15 | 9.47E-14 |
| Cym | 0.010780241 | 0.158900551 | 14.7399811 | 2.59E-15 | 9.61E-14 |
| Gm10260 | 2.313963187 | 3.705943196 | 1.60155668 | 2.70E-15 | 9.97E-14 |
| Eif1ax | 0.36927325 | 0.822914882 | 2.22847141 | 3.51E-15 | 1.30E-13 |
| Srp14 | 1.428185401 | 0.771752307 | 0.54037263 | 3.53E-15 | 1.30E-13 |
| Bckdk | 0.206065567 | 0.564988442 | 2.74178967 | 3.81E-15 | 1.40E-13 |
| Uqcr11 | 2.33803871 | 3.731613212 | 1.59604424 | 3.95E-15 | 1.45E-13 |
| Ube2n | 0.696302981 | 0.266724495 | 0.3830581 | 4.03E-15 | 1.47E-13 |
| Manf | 4.558765216 | 2.785968558 | 0.6111235 | 4.18E-15 | 1.52E-13 |
| Gm8797 | 0.791183145 | 0.325373711 | 0.41124955 | 4.18E-15 | 1.52E-13 |
| Gck | 0.257866993 | 0.024321299 | 0.09431722 | 4.29E-15 | 1.56E-13 |
| Entpd3 | 0.233348949 | 0.018568397 | 0.07957352 | 4.59E-15 | 1.66E-13 |
| Gstt2 | 0.10180987 | 0.374866009 | 3.68202031 | 5.22E-15 | 1.89E-13 |
| Rps18 | 7.713444244 | 12.20566973 | 1.58238905 | 5.36E-15 | 1.93E-13 |
| Tbcb | 0.739691209 | 0.292037432 | 0.39480993 | 6.70E-15 | 2.41E-13 |
| Mapre3 | 0.255630857 | 0.026933849 | 0.10536228 | 6.76E-15 | 2.43E-13 |
| Rasd1 | 0.255919102 | 0.026459919 | 0.10339173 | 6.76E-15 | 2.43E-13 |

| | | | | | |
|---|---|---|---|---|---|
| Muc1 | 0.100074328 | 0.370992915 | 3.70717366 | 6.86E-15 | 2.46E-13 |
| Itln1 | 0.033568277 | 0.228892649 | 6.81871896 | 7.07E-15 | 2.53E-13 |
| Lysmd2 | 0.265742387 | 0.031176783 | 0.11731957 | 7.26E-15 | 2.59E-13 |
| Uqcc2 | 1.832549479 | 1.086561509 | 0.59292342 | 7.85E-15 | 2.79E-13 |
| Amy1 | 0.010077503 | 0.157079297 | 15.5871248 | 8.03E-15 | 2.85E-13 |
| Dnmt3a | 0.488113595 | 0.139720935 | 0.28624676 | 8.86E-15 | 3.14E-13 |
| Pclo | 0.333027104 | 0.063628512 | 0.19106106 | 1.00E-14 | 3.54E-13 |
| 1110008F13I | 0.362913579 | 0.79971805 | 2.20360465 | 1.11E-14 | 3.91E-13 |
| Gjb1 | 0.070778662 | 0.311885033 | 4.4064839 | 1.43E-14 | 5.05E-13 |
| Gm10709 | 2.897563301 | 4.559465997 | 1.57355182 | 1.44E-14 | 5.05E-13 |
| Trpm5 | 0.168049062 | 0.000763657 | 0.00454425 | 1.52E-14 | 5.32E-13 |
| Atp2a3 | 0.184758398 | 0.001747287 | 0.00945714 | 1.72E-14 | 6.02E-13 |
| Anpep | 0.053012334 | 0.267448142 | 5.04501726 | 1.98E-14 | 6.93E-13 |
| Epcam | 1.752031755 | 1.03446921 | 0.59043976 | 2.21E-14 | 7.69E-13 |
| Phactr1 | 0.163503097 | 0.003252912 | 0.01989511 | 2.46E-14 | 8.54E-13 |
| 1500009L16F | 0.248706184 | 0.027303268 | 0.10978122 | 2.75E-14 | 9.55E-13 |
| Atp6v1f | 1.312678606 | 0.707309861 | 0.53882943 | 3.21E-14 | 1.11E-12 |
| Rnaseh2c | 0.263321781 | 0.648330595 | 2.46212293 | 3.48E-14 | 1.20E-12 |
| Naa20 | 0.37736243 | 0.088612981 | 0.23482195 | 4.45E-14 | 1.54E-12 |
| Gfra3 | 0.220965299 | 0.018660986 | 0.08445211 | 4.76E-14 | 1.64E-12 |
| Gatsl2 | 0.306546474 | 0.050034911 | 0.16322129 | 4.87E-14 | 1.67E-12 |
| Amy2a2 | 0.0090257 | 0.144711442 | 16.0332658 | 4.91E-14 | 1.68E-12 |
| Lrpprc | 0.494512576 | 0.153251757 | 0.30990467 | 5.32E-14 | 1.82E-12 |
| Rasgrf1 | 0.193157165 | 0.006769365 | 0.03504589 | 5.43E-14 | 1.86E-12 |
| Tmem206 | 0.20765695 | 0.014959386 | 0.07203894 | 5.60E-14 | 1.91E-12 |
| Cyb5r3 | 1.141776717 | 0.587164659 | 0.51425524 | 6.34E-14 | 2.16E-12 |
| Lyve1 | 0.176550145 | 0.001725742 | 0.0097748 | 7.34E-14 | 2.49E-12 |
| Bola2 | 0.440425404 | 0.897648602 | 2.03813993 | 7.45E-14 | 2.53E-12 |
| Kif5b | 0.987557252 | 0.477936155 | 0.48395792 | 7.73E-14 | 2.62E-12 |
| Ndufa13 | 1.762481209 | 2.773868595 | 1.57384293 | 8.27E-14 | 2.79E-12 |

| | | | | | |
|---|---|---|---|---|---|
| Snap25 | 0.205329504 | 0.013598824 | 0.06622927 | 8.48E-14 | 2.86E-12 |
| Tmsb15l | 0.190671559 | 0.012003601 | 0.06295433 | 8.83E-14 | 2.97E-12 |
| Rplp0 | 5.289481736 | 8.179980715 | 1.54646166 | 8.92E-14 | 2.99E-12 |
| Btbd17 | 0.156690758 | 0.001123555 | 0.00717052 | 1.08E-13 | 3.63E-12 |
| Slc38a3 | 0.012393956 | 0.15089018 | 12.1744969 | 1.09E-13 | 3.63E-12 |
| Ubc | 2.855358965 | 1.784498451 | 0.62496466 | 1.16E-13 | 3.88E-12 |
| Serpini1 | 0.061998885 | 0.282447882 | 4.55569295 | 1.18E-13 | 3.93E-12 |
| Tox3 | 0.216176445 | 0.019179516 | 0.08872158 | 1.20E-13 | 3.98E-12 |
| Psma3 | 1.441026018 | 0.818905603 | 0.56827954 | 1.23E-13 | 4.09E-12 |
| Arl6ip1 | 0.918682558 | 0.436463228 | 0.47509689 | 1.26E-13 | 4.17E-12 |
| Mrps24 | 0.532312744 | 1.027631982 | 1.93050419 | 1.33E-13 | 4.38E-12 |
| Rgs17 | 0.201637387 | 0.011595292 | 0.05750566 | 1.34E-13 | 4.41E-12 |
| Gpd2 | 0.279176801 | 0.041966277 | 0.15032151 | 1.36E-13 | 4.47E-12 |
| Fam129a | 0.012936655 | 0.160846819 | 12.4334161 | 1.44E-13 | 4.74E-12 |
| Nes | 0.012860122 | 0.158803367 | 12.3485114 | 1.44E-13 | 4.74E-12 |
| Syt7 | 0.188805929 | 0.011814987 | 0.06257742 | 1.52E-13 | 4.97E-12 |
| Tst | 0.154872771 | 0.458192503 | 2.95850911 | 1.54E-13 | 5.04E-12 |
| Atp6v0b | 1.084960548 | 0.55186312 | 0.5086481 | 1.56E-13 | 5.08E-12 |
| Rph3al | 0.35044527 | 0.080304224 | 0.22914912 | 1.57E-13 | 5.11E-12 |
| Lypd2 | 0.155095677 | 0 | 0 | 1.91E-13 | 6.21E-12 |
| Rpn1 | 0.970397063 | 1.608914371 | 1.65799592 | 1.98E-13 | 6.41E-12 |
| Ndufa2 | 1.982054464 | 3.082878446 | 1.55539543 | 2.01E-13 | 6.52E-12 |
| 2610524H06 | 0.407352825 | 0.109790649 | 0.26952225 | 2.28E-13 | 7.36E-12 |
| Dctn3 | 0.886276316 | 0.418532185 | 0.47223668 | 2.45E-13 | 7.90E-12 |
| Smim6 | 0.020417807 | 0.172976635 | 8.47185178 | 2.83E-13 | 9.11E-12 |
| Ttc3 | 0.617594192 | 0.238378719 | 0.38597953 | 3.21E-13 | 1.03E-11 |
| Med28 | 0.704119685 | 0.292403648 | 0.41527549 | 3.24E-13 | 1.04E-11 |
| Spock2 | 0.198087014 | 0.012137184 | 0.06127198 | 3.63E-13 | 1.16E-11 |
| Arpc5 | 0.837152914 | 0.387643131 | 0.46304937 | 3.72E-13 | 1.19E-11 |
| Pnmal2 | 0.326849322 | 0.069333792 | 0.21212769 | 3.74E-13 | 1.19E-11 |

| | | | | | |
|---|---|---|---|---|---|
| Guk1 | 0.648452887 | 0.256832862 | 0.3960702 | 4.28E-13 | 1.36E-11 |
| Atp6v0d1 | 0.563789491 | 0.20723046 | 0.36756709 | 4.44E-13 | 1.41E-11 |
| Spp1 | 3.698223549 | 2.349230058 | 0.63523203 | 4.47E-13 | 1.42E-11 |
| Ctnnbip1 | 0.299136941 | 0.058616538 | 0.19595219 | 4.59E-13 | 1.45E-11 |
| Ucn3 | 0.148091909 | 0.001777353 | 0.01200169 | 4.67E-13 | 1.48E-11 |
| Fam216a | 0.318668236 | 0.065171409 | 0.20451178 | 4.69E-13 | 1.48E-11 |
| Nkx2-2 | 0.318488964 | 0.064425276 | 0.20228417 | 4.69E-13 | 1.48E-11 |
| Atp2c2 | 0.008332363 | 0.136216996 | 16.3479433 | 4.77E-13 | 1.50E-11 |
| Gars | 0.713170738 | 0.301736059 | 0.42309091 | 4.96E-13 | 1.55E-11 |
| Mapk15 | 0.179498758 | 0.010221338 | 0.05694378 | 5.85E-13 | 1.83E-11 |
| Zdbf2 | 0.204206829 | 0.019922633 | 0.09756105 | 6.90E-13 | 2.16E-11 |
| Extl3 | 0.050311548 | 0.248775692 | 4.9447036 | 7.17E-13 | 2.24E-11 |
| Sumo1 | 0.842854742 | 0.391988053 | 0.4650719 | 9.04E-13 | 2.81E-11 |
| Pea15a | 0.248521845 | 0.034236922 | 0.13776222 | 9.07E-13 | 2.82E-11 |
| Zc3h3 | 0.177353495 | 0.011283533 | 0.06362171 | 9.14E-13 | 2.84E-11 |
| Ube2d3 | 1.180211986 | 0.644586548 | 0.54616167 | 9.23E-13 | 2.86E-11 |
| Spc24 | 0.153824048 | 0.438358229 | 2.84973797 | 1.04E-12 | 3.22E-11 |
| Arf5 | 1.656421199 | 1.005951674 | 0.60730427 | 1.06E-12 | 3.28E-11 |
| Dcxr | 0.066431977 | 0.275046931 | 4.14027919 | 1.09E-12 | 3.36E-11 |
| mt-Co1 | 16.41347547 | 25.28448025 | 1.54047083 | 1.17E-12 | 3.58E-11 |
| Hnrnpk | 1.794281208 | 1.110831671 | 0.61909564 | 1.21E-12 | 3.71E-11 |
| Tmem234 | 0.624327961 | 1.127774813 | 1.80638203 | 1.26E-12 | 3.87E-11 |
| Galnt18 | 0.144812007 | 0.003244599 | 0.02240559 | 1.37E-12 | 4.19E-11 |
| Pcsk2os1 | 0.16231339 | 0.005079751 | 0.03129595 | 1.38E-12 | 4.22E-11 |
| Pycr1 | 0.064455576 | 0.272759378 | 4.23174216 | 1.39E-12 | 4.24E-11 |
| Atp6v1b2 | 0.407700235 | 0.117173671 | 0.28740153 | 1.47E-12 | 4.48E-11 |
| Gdap1l1 | 0.17373484 | 0.010664336 | 0.06138283 | 1.49E-12 | 4.52E-11 |
| Slc7a14 | 0.176006422 | 0.010780443 | 0.06125028 | 1.51E-12 | 4.56E-11 |
| Bnip3l | 0.655363678 | 0.268964245 | 0.41040456 | 1.54E-12 | 4.66E-11 |
| Ip6k2 | 0.33490106 | 0.077935946 | 0.23271335 | 1.59E-12 | 4.79E-11 |

| | | | | | |
|---|---|---|---|---|---|
| Higd1a | 0.554877857 | 1.029530807 | 1.8554188 | 1.60E-12 | 4.83E-11 |
| Psma1 | 0.903215508 | 0.445984635 | 0.49377433 | 1.83E-12 | 5.49E-11 |
| 1810064F22I | 0.018661954 | 0.164317454 | 8.80494389 | 1.99E-12 | 5.97E-11 |
| Bok | 0.034294203 | 0.200687572 | 5.85193872 | 2.02E-12 | 6.07E-11 |
| Cct5 | 1.539023455 | 0.924736028 | 0.60085896 | 2.08E-12 | 6.21E-11 |
| Chic1 | 0.243936891 | 0.03462014 | 0.14192253 | 2.16E-12 | 6.46E-11 |
| Paip2 | 0.806819107 | 0.378124146 | 0.46866038 | 2.29E-12 | 6.83E-11 |
| Prdx3 | 0.450718007 | 0.14399839 | 0.31948666 | 2.31E-12 | 6.88E-11 |
| Slc41a1 | 0.023331662 | 0.174901838 | 7.49632992 | 2.37E-12 | 7.03E-11 |
| Mgst2 | 0.025424918 | 0.17959735 | 7.06383212 | 2.40E-12 | 7.11E-11 |
| Akr1c13 | 0.259796838 | 0.045631924 | 0.17564465 | 2.52E-12 | 7.47E-11 |
| Tmsb10 | 4.107349673 | 6.188518881 | 1.50669394 | 2.56E-12 | 7.57E-11 |
| Golgb1 | 0.723049447 | 0.317330537 | 0.43887806 | 2.60E-12 | 7.69E-11 |
| Cnn3 | 0.555124148 | 0.209217989 | 0.37688504 | 2.81E-12 | 8.27E-11 |
| Tmem14c | 0.661945866 | 0.279940032 | 0.42290472 | 3.16E-12 | 9.30E-11 |
| Fech | 0.303910101 | 0.069341422 | 0.22816426 | 3.34E-12 | 9.82E-11 |
| Fgd2 | 0.183990611 | 0.016468338 | 0.0895064 | 3.47E-12 | 1.02E-10 |
| P2ry1 | 0.185023563 | 0.011266589 | 0.06089273 | 3.47E-12 | 1.02E-10 |
| Igsf1 | 0.158082558 | 0.003388911 | 0.0214376 | 3.63E-12 | 1.06E-10 |
| Isg20 | 0.311061555 | 0.071884353 | 0.23109366 | 3.85E-12 | 1.12E-10 |
| Hopx | 0.171617351 | 0.004822713 | 0.02810154 | 3.87E-12 | 1.13E-10 |
| Hmgn2 | 1.183078438 | 0.652448583 | 0.55148379 | 4.09E-12 | 1.19E-10 |
| Lrp11 | 0.1945284 | 0.017559224 | 0.09026561 | 4.23E-12 | 1.23E-10 |
| Zwint | 0.400365443 | 0.119278414 | 0.29792385 | 4.42E-12 | 1.28E-10 |
| Celf3 | 0.205363243 | 0.020593064 | 0.10027629 | 4.78E-12 | 1.38E-10 |
| Taldo1 | 1.040695243 | 1.652849644 | 1.58821678 | 5.07E-12 | 1.46E-10 |
| Sepw1 | 1.131938958 | 0.618678235 | 0.54656502 | 5.26E-12 | 1.52E-10 |
| Rps18-ps3 | 1.557844361 | 2.376942367 | 1.52578937 | 5.77E-12 | 1.66E-10 |
| Chd7 | 0.256028389 | 0.042738919 | 0.16693039 | 5.80E-12 | 1.67E-10 |
| Prdx5 | 0.976161328 | 0.504006651 | 0.51631491 | 6.26E-12 | 1.80E-10 |

| | | | | | |
|---|---|---|---|---|---|
| Dap | 0.655676703 | 1.154549923 | 1.76085244 | 6.44E-12 | 1.84E-10 |
| Gadd45g | 0.670458107 | 0.287499244 | 0.42881015 | 6.51E-12 | 1.86E-10 |
| Nr5a2 | 0.059993079 | 0.251912716 | 4.1990296 | 6.61E-12 | 1.89E-10 |
| Slc25a5 | 2.842224181 | 1.841622428 | 0.64795115 | 7.02E-12 | 2.00E-10 |
| Tead2 | 0.071198974 | 0.275257961 | 3.86603831 | 7.10E-12 | 2.02E-10 |
| Ybx3 | 0.140153931 | 0.402071293 | 2.86878355 | 7.37E-12 | 2.09E-10 |
| Wdtc1 | 0.062091817 | 0.260920329 | 4.20216935 | 7.61E-12 | 2.16E-10 |
| Ankrd44 | 0.222932577 | 0.030116905 | 0.13509423 | 7.64E-12 | 2.16E-10 |
| Actg2 | 0.151420687 | 0.000871267 | 0.00575395 | 8.83E-12 | 2.50E-10 |
| Uqcr10 | 1.837859544 | 2.77742285 | 1.51122694 | 9.29E-12 | 2.62E-10 |
| Zcrb1 | 0.578895799 | 0.231841805 | 0.4004897 | 9.66E-12 | 2.72E-10 |
| F3 | 0.189443642 | 0.016249702 | 0.08577592 | 1.05E-11 | 2.94E-10 |
| Fam220a | 0.29638786 | 0.063031305 | 0.21266493 | 1.07E-11 | 3.01E-10 |
| Ube2b | 0.810203281 | 0.393251409 | 0.48537376 | 1.09E-11 | 3.06E-10 |
| Por | 0.32893387 | 0.082850823 | 0.25187684 | 1.17E-11 | 3.27E-10 |
| Slc37a4 | 0.211247447 | 0.025758841 | 0.12193682 | 1.24E-11 | 3.46E-10 |
| Tinag | 0.015333419 | 0.146546279 | 9.55731255 | 1.24E-11 | 3.46E-10 |
| Txn1 | 1.865941157 | 2.813745807 | 1.50794991 | 1.26E-11 | 3.51E-10 |
| Oaz2 | 0.406032189 | 0.126085522 | 0.31053085 | 1.26E-11 | 3.51E-10 |
| Galnt7 | 0.055366943 | 0.244192219 | 4.41043348 | 1.34E-11 | 3.72E-10 |
| Mecom | 0.031609091 | 0.188690803 | 5.96951063 | 1.37E-11 | 3.80E-10 |
| Dbt | 0.048403752 | 0.230423543 | 4.76044794 | 1.39E-11 | 3.85E-10 |
| Dynll1 | 3.070209314 | 2.001366946 | 0.65186661 | 1.41E-11 | 3.88E-10 |
| Ndrg2 | 0.071162655 | 0.271361001 | 3.81325012 | 1.45E-11 | 3.99E-10 |
| Lrmp | 0.018876078 | 0.15131526 | 8.01624483 | 1.52E-11 | 4.20E-10 |
| Mllt11 | 0.162086537 | 0.009974556 | 0.06153846 | 1.53E-11 | 4.21E-10 |
| Irs2 | 0.053693458 | 0.236582433 | 4.40616871 | 1.60E-11 | 4.40E-10 |
| Rpl27 | 1.869611026 | 2.809096834 | 1.50250335 | 1.76E-11 | 4.83E-10 |
| Slc7a2 | 0.258585481 | 0.050561752 | 0.19553206 | 1.77E-11 | 4.85E-10 |
| Acly | 0.753058117 | 0.354341664 | 0.47053694 | 1.88E-11 | 5.15E-10 |

| | | | | | |
|---|---|---|---|---|---|
| Itm2c | 0.640360267 | 0.27647337 | 0.4317466 | 1.98E-11 | 5.39E-10 |
| Mcfd2 | 0.263364679 | 0.59235379 | 2.24917704 | 1.99E-11 | 5.42E-10 |
| Trim35 | 0.31038837 | 0.070767096 | 0.22799532 | 2.10E-11 | 5.72E-10 |
| Prpsap1 | 0.230519924 | 0.540173076 | 2.34328152 | 2.13E-11 | 5.78E-10 |
| Hagh | 0.469126879 | 0.167535091 | 0.35712107 | 2.18E-11 | 5.91E-10 |
| Atp5e | 3.327019846 | 4.933155957 | 1.48275519 | 2.20E-11 | 5.96E-10 |
| Tshz2 | 0.42917565 | 0.147166548 | 0.34290517 | 2.46E-11 | 6.64E-10 |
| Cldn18 | 0.011185786 | 0.131114466 | 11.7215246 | 2.51E-11 | 6.77E-10 |
| Kif12 | 0.254270636 | 0.049129501 | 0.19321736 | 2.54E-11 | 6.84E-10 |
| Rpl7 | 6.125494437 | 9.048860421 | 1.47724572 | 2.78E-11 | 7.48E-10 |
| Aldh1l1 | 0.006924255 | 0.112920385 | 16.3079477 | 2.87E-11 | 7.68E-10 |
| Gata4 | 0.007153458 | 0.111966098 | 15.6520249 | 2.87E-11 | 7.68E-10 |
| Psap | 0.895648432 | 0.458953143 | 0.51242555 | 3.10E-11 | 8.30E-10 |
| Ak3 | 0.104043433 | 0.325428583 | 3.12781475 | 3.28E-11 | 8.75E-10 |
| Ywhae | 2.402858022 | 1.572477513 | 0.65441965 | 3.63E-11 | 9.68E-10 |
| Cyth2 | 0.296940656 | 0.069993645 | 0.23571594 | 3.93E-11 | 1.05E-09 |
| Gip | 0.126902065 | 0.00173379 | 0.01366242 | 3.96E-11 | 1.05E-09 |
| Actn3 | 0.126076997 | 0.002501503 | 0.01984107 | 3.96E-11 | 1.05E-09 |
| Sh3bgrl3 | 0.734687075 | 0.347471175 | 0.47295126 | 3.97E-11 | 1.05E-09 |
| Etfb | 0.845631169 | 1.378353029 | 1.62996952 | 4.02E-11 | 1.07E-09 |
| H13 | 0.660895356 | 1.13781588 | 1.72162789 | 4.08E-11 | 1.08E-09 |
| Rap1gapos | 0.024555871 | 0.167583997 | 6.82459998 | 4.32E-11 | 1.14E-09 |
| Lgals2 | 0.022181297 | 0.156350871 | 7.04877036 | 4.44E-11 | 1.17E-09 |
| Ankrd54 | 0.259689203 | 0.051150348 | 0.19696756 | 4.66E-11 | 1.23E-09 |
| Matn4 | 0.040715613 | 0.203362257 | 4.99469957 | 4.92E-11 | 1.29E-09 |
| Adora1 | 0.041392376 | 0.200276615 | 4.83849044 | 4.92E-11 | 1.29E-09 |
| Iah1 | 0.112816387 | 0.345336565 | 3.06104968 | 5.10E-11 | 1.34E-09 |
| Arl3 | 0.508730098 | 0.200387797 | 0.39389806 | 5.51E-11 | 1.44E-09 |
| Lin37 | 0.139963071 | 0.389905941 | 2.78577726 | 5.90E-11 | 1.54E-09 |
| Fam92b | 0.14126937 | 0.003509497 | 0.02484259 | 5.98E-11 | 1.56E-09 |

| | | | | | |
|---|---|---|---|---|---|
| Ndufs2 | 0.887549779 | 0.460458312 | 0.51879717 | 6.50E-11 | 1.69E-09 |
| Adra2a | 0.141388452 | 0.007427787 | 0.05253461 | 6.54E-11 | 1.70E-09 |
| Gadd45a | 0.211687326 | 0.032093343 | 0.15160729 | 6.71E-11 | 1.74E-09 |
| Arhgap36 | 0.124700671 | 0.002085763 | 0.01672615 | 7.05E-11 | 1.83E-09 |
| Laptm4a | 1.365300861 | 0.824933785 | 0.60421392 | 8.10E-11 | 2.10E-09 |
| Cdc42 | 1.266691713 | 0.753572474 | 0.59491387 | 8.12E-11 | 2.10E-09 |
| Lgals3bp | 0.291505301 | 0.069971878 | 0.24003638 | 8.51E-11 | 2.19E-09 |
| Dnajc24 | 0.291859713 | 0.067112491 | 0.22994777 | 8.51E-11 | 2.19E-09 |
| Eif3i | 0.937090941 | 1.485395235 | 1.58511322 | 8.54E-11 | 2.20E-09 |
| Cldn11 | 0.138626948 | 0.006210053 | 0.04479687 | 9.83E-11 | 2.52E-09 |
| Prlr | 0.138843249 | 0.002241512 | 0.01614419 | 9.83E-11 | 2.52E-09 |
| Mknk1 | 0.04652284 | 0.208892101 | 4.49009779 | 1.00E-10 | 2.56E-09 |
| PISD | 0.384697001 | 0.121118646 | 0.31484167 | 1.03E-10 | 2.62E-09 |
| BC031181 | 0.603145896 | 0.261463193 | 0.43349908 | 1.05E-10 | 2.67E-09 |
| Camk2b | 0.197584672 | 0.025508774 | 0.129103 | 1.07E-10 | 2.72E-09 |
| Smc6 | 0.45369824 | 0.166362744 | 0.36668148 | 1.07E-10 | 2.73E-09 |
| Slc6a6 | 0.313668206 | 0.083389485 | 0.26585253 | 1.09E-10 | 2.77E-09 |
| Pafah1b3 | 0.460554827 | 0.170431408 | 0.37005672 | 1.46E-10 | 3.69E-09 |
| mt-Co2 | 9.602178488 | 14.02592267 | 1.46070214 | 1.50E-10 | 3.79E-09 |
| Plcxd3 | 0.149575071 | 0.009160414 | 0.06124292 | 1.58E-10 | 4.01E-09 |
| Nucb2 | 0.809468901 | 1.309745421 | 1.61803056 | 1.60E-10 | 4.03E-09 |
| Ywhah | 0.565580576 | 0.242081348 | 0.42802274 | 1.60E-10 | 4.03E-09 |
| Kcnmb2 | 0.195310448 | 0.025872001 | 0.13246604 | 1.62E-10 | 4.08E-09 |
| Vtn | 0.010880926 | 0.116616335 | 10.7175012 | 1.72E-10 | 4.34E-09 |
| Fau | 7.052371437 | 10.25544482 | 1.45418388 | 1.85E-10 | 4.64E-09 |
| Pcmtd1 | 0.112432652 | 0.330435861 | 2.93896707 | 1.94E-10 | 4.87E-09 |
| Arl1 | 0.803960137 | 0.405661653 | 0.50457931 | 1.97E-10 | 4.94E-09 |
| 2310036O22 | 0.760097703 | 0.377358969 | 0.49646114 | 1.97E-10 | 4.94E-09 |
| Csrp2 | 0.460066797 | 0.853300727 | 1.85473225 | 2.03E-10 | 5.07E-09 |
| Nav2 | 0.300155001 | 0.079270822 | 0.26409962 | 2.12E-10 | 5.28E-09 |

| | | | | | |
|---|---|---|---|---|---|
| Xbp1 | 0.717518934 | 1.18989345 | 1.65834432 | 2.30E-10 | 5.74E-09 |
| Gnaz | 0.133547731 | 0.005765341 | 0.04317064 | 2.57E-10 | 6.38E-09 |
| Trappc6b | 0.330472919 | 0.0931434 | 0.28184881 | 2.60E-10 | 6.46E-09 |
| Rbpms | 0.256981225 | 0.057487956 | 0.22370489 | 2.68E-10 | 6.65E-09 |
| Ntn1 | 0.015044505 | 0.130451721 | 8.67105413 | 2.76E-10 | 6.84E-09 |
| Clptm1l | 0.180555396 | 0.445261995 | 2.46606862 | 2.85E-10 | 7.04E-09 |
| Ppp1ca | 1.716759106 | 1.126023515 | 0.65590071 | 2.86E-10 | 7.05E-09 |
| Mpv17l | 0.024100738 | 0.1524572 | 6.32583111 | 3.01E-10 | 7.41E-09 |
| Lamtor2 | 0.795602562 | 1.284405604 | 1.61438093 | 3.15E-10 | 7.74E-09 |
| Chd3 | 0.208998613 | 0.035372025 | 0.16924526 | 3.17E-10 | 7.80E-09 |
| Fxyd3 | 0.36968212 | 0.717487226 | 1.94082209 | 3.27E-10 | 8.02E-09 |
| Atp5l | 5.52727846 | 3.747190833 | 0.67794501 | 3.29E-10 | 8.06E-09 |
| Arpc5l | 0.692249911 | 0.333947904 | 0.48240946 | 3.42E-10 | 8.36E-09 |
| Arpc3 | 0.880277585 | 0.469006628 | 0.53279401 | 3.51E-10 | 8.58E-09 |
| Frzb | 0.158760749 | 0.012801991 | 0.080637 | 3.53E-10 | 8.62E-09 |
| Hmgn5 | 0.139345812 | 0.370476942 | 2.6586873 | 3.55E-10 | 8.65E-09 |
| Rhou | 0.047360503 | 0.20715934 | 4.37409498 | 3.69E-10 | 8.97E-09 |
| Galk1 | 0.148980367 | 0.389263053 | 2.61284798 | 3.79E-10 | 9.22E-09 |
| Lrpap1 | 0.557712558 | 0.243142284 | 0.43596344 | 3.86E-10 | 9.37E-09 |
| Atp6v0e2 | 0.178722201 | 0.024736098 | 0.13840529 | 3.95E-10 | 9.58E-09 |
| Stard10 | 0.515054191 | 0.913729123 | 1.77404463 | 4.03E-10 | 9.76E-09 |
| Echdc2 | 0.110585581 | 0.323549695 | 2.92578554 | 4.49E-10 | 1.09E-08 |
| Aldh3a1 | 0.114089118 | 0 | 0 | 4.54E-10 | 1.10E-08 |
| Tpt1 | 8.701822241 | 12.56264154 | 1.4436794 | 4.95E-10 | 1.19E-08 |
| Srpk2 | 0.268295768 | 0.069265798 | 0.25816955 | 5.01E-10 | 1.20E-08 |
| Syndig1l | 0.114633531 | 0 | 0 | 5.07E-10 | 1.22E-08 |
| Baiap3 | 0.156079713 | 0.010634594 | 0.06813566 | 5.27E-10 | 1.27E-08 |
| Gabra4 | 0.004982878 | 0.096923716 | 19.4513514 | 5.51E-10 | 1.32E-08 |
| Kif21a | 0.346388782 | 0.110893091 | 0.32014054 | 5.51E-10 | 1.32E-08 |
| Hsd17b10 | 0.38289129 | 0.732991148 | 1.91435838 | 5.89E-10 | 1.41E-08 |

| | | | | | |
|---|---|---|---|---|---|
| Epb41l3 | 0.186331887 | 0.025598425 | 0.13738081 | 5.93E-10 | 1.41E-08 |
| Sec61a1 | 0.481660933 | 0.871874738 | 1.81014211 | 6.02E-10 | 1.43E-08 |
| Cd164l2 | 0.18727379 | 0.027641728 | 0.14760062 | 6.06E-10 | 1.44E-08 |
| Aacs | 0.17623303 | 0.019698793 | 0.11177696 | 6.08E-10 | 1.45E-08 |
| Fam20c | 0.214669938 | 0.038448859 | 0.17910686 | 6.21E-10 | 1.47E-08 |
| Lxn | 0.078450786 | 0.26179361 | 3.33704253 | 6.25E-10 | 1.48E-08 |
| Snca | 0.216498096 | 0.041236474 | 0.19047038 | 6.43E-10 | 1.52E-08 |
| Psme2 | 0.67969732 | 0.329014346 | 0.48406009 | 6.50E-10 | 1.54E-08 |
| Gpx4 | 1.831042367 | 2.660814125 | 1.45316907 | 6.80E-10 | 1.61E-08 |
| Rnh1 | 0.550738835 | 0.956155174 | 1.73613174 | 6.92E-10 | 1.63E-08 |
| 0610009B22 | 0.386762023 | 0.139904644 | 0.36173315 | 7.08E-10 | 1.67E-08 |
| Fubp1 | 0.543555949 | 0.23446568 | 0.43135519 | 7.36E-10 | 1.73E-08 |
| Sesn3 | 0.222874177 | 0.043924012 | 0.19707986 | 7.75E-10 | 1.82E-08 |
| Syne4 | 0.123092052 | 0.344940873 | 2.80230014 | 8.00E-10 | 1.88E-08 |
| Noxa1 | 0.006986156 | 0.099828609 | 14.2894901 | 8.57E-10 | 2.01E-08 |
| Gtf2e2 | 0.409097038 | 0.150032083 | 0.3667396 | 9.00E-10 | 2.10E-08 |
| Cystm1 | 0.882042368 | 0.476218202 | 0.539904 | 9.09E-10 | 2.12E-08 |
| Spc25 | 0.288541018 | 0.081343633 | 0.28191359 | 9.27E-10 | 2.16E-08 |
| Ubl4a | 0.250143045 | 0.541300282 | 2.16396295 | 9.64E-10 | 2.24E-08 |
| Lgals9 | 0.04001284 | 0.189336871 | 4.73190279 | 9.64E-10 | 2.24E-08 |
| Klk1b3 | 0.003958808 | 0.087653983 | 22.1415062 | 9.76E-10 | 2.27E-08 |
| Glrx | 0.297473156 | 0.085873959 | 0.28867801 | 1.00E-09 | 2.33E-08 |
| Krcc1 | 0.376534444 | 0.133022809 | 0.35328191 | 1.01E-09 | 2.34E-08 |
| Klhl32 | 0.124958887 | 0.003459082 | 0.02768176 | 1.05E-09 | 2.43E-08 |
| Gng4 | 0.140920009 | 0.008568904 | 0.06080686 | 1.10E-09 | 2.55E-08 |
| Hspa5 | 7.081077885 | 4.859890288 | 0.68632069 | 1.15E-09 | 2.66E-08 |
| Gc | 0.043322551 | 0.192807716 | 4.45051625 | 1.21E-09 | 2.78E-08 |
| Arpc1a | 0.812359597 | 0.427246416 | 0.52593263 | 1.28E-09 | 2.94E-08 |
| Cldn4 | 0.285823215 | 0.079882185 | 0.27948109 | 1.33E-09 | 3.05E-08 |
| Tshz1 | 0.229057733 | 0.044748781 | 0.19536027 | 1.39E-09 | 3.19E-08 |

| | | | | | |
|---|---|---|---|---|---|
| Eif3j1 | 0.284910453 | 0.581384995 | 2.04058851 | 1.47E-09 | 3.37E-08 |
| Cgrrf1 | 0.211621332 | 0.039946474 | 0.18876393 | 1.47E-09 | 3.37E-08 |
| Myl9 | 0.172708815 | 0.021802785 | 0.12624014 | 1.56E-09 | 3.56E-08 |
| Kcnq1ot1 | 0.581745266 | 0.26758183 | 0.45996391 | 1.56E-09 | 3.56E-08 |
| Nt5c3 | 0.325708955 | 0.09823782 | 0.30161228 | 1.58E-09 | 3.60E-08 |
| Fdps | 0.352285209 | 0.678146818 | 1.92499373 | 1.62E-09 | 3.69E-08 |
| Hexa | 0.269256992 | 0.067748169 | 0.25161155 | 1.70E-09 | 3.86E-08 |
| Stx7 | 0.269736244 | 0.067149919 | 0.24894659 | 1.70E-09 | 3.86E-08 |
| Atp5g3 | 1.294820871 | 1.880989318 | 1.45270235 | 1.79E-09 | 4.05E-08 |
| 2210013O21 | 0.429108902 | 0.16774987 | 0.3909261 | 1.83E-09 | 4.14E-08 |
| Nagk | 0.284093557 | 0.081579835 | 0.28715834 | 1.94E-09 | 4.38E-08 |
| Dusp1 | 0.423589225 | 0.161734505 | 0.38181922 | 1.97E-09 | 4.44E-08 |
| Pbld2 | 0.036404212 | 0.179585115 | 4.93308612 | 2.01E-09 | 4.53E-08 |
| Eny2 | 0.608212719 | 1.015059776 | 1.66892231 | 2.03E-09 | 4.57E-08 |
| Tubb5 | 2.131057334 | 1.44924091 | 0.68005721 | 2.06E-09 | 4.63E-08 |
| Htra2 | 0.092607607 | 0.288438104 | 3.11462646 | 2.09E-09 | 4.69E-08 |
| Tpst2 | 0.305543478 | 0.094311832 | 0.30866911 | 2.13E-09 | 4.77E-08 |
| Wipi1 | 0.20568142 | 0.040052659 | 0.19473154 | 2.13E-09 | 4.78E-08 |
| Cd200 | 0.180182824 | 0.024434964 | 0.13561206 | 2.14E-09 | 4.79E-08 |
| Syce2 | 0.313874027 | 0.092492193 | 0.29467935 | 2.15E-09 | 4.81E-08 |
| Mgat4a | 0.020730267 | 0.133920615 | 6.46014917 | 2.17E-09 | 4.83E-08 |
| Camk1d | 0.019968784 | 0.13466877 | 6.74396435 | 2.17E-09 | 4.83E-08 |
| Zfr | 0.337017919 | 0.1087277 | 0.32261697 | 2.20E-09 | 4.90E-08 |
| Cfl1 | 2.852191869 | 1.958689678 | 0.68673139 | 2.23E-09 | 4.96E-08 |
| Cisd1 | 0.623255672 | 0.296901063 | 0.47637122 | 2.27E-09 | 5.03E-08 |
| 1810043G02 | 0.169322303 | 0.021151446 | 0.12491825 | 2.29E-09 | 5.08E-08 |
| Hap1 | 0.195729143 | 0.034685374 | 0.17721109 | 2.53E-09 | 5.61E-08 |
| Mdm1 | 0.119579871 | 0.006842252 | 0.0572191 | 2.73E-09 | 6.04E-08 |
| Lrrc59 | 0.3332453 | 0.648280174 | 1.94535429 | 2.75E-09 | 6.06E-08 |
| Wnk3 | 0.134873692 | 0.009075765 | 0.06729085 | 2.82E-09 | 6.21E-08 |

| | | | | | |
|---|---|---|---|---|---|
| Rnf138rt1 | 0.135823149 | 0.006784927 | 0.04995413 | 2.82E-09 | 6.21E-08 |
| Lyplal1 | 0.295780114 | 0.087667772 | 0.29639508 | 2.96E-09 | 6.52E-08 |
| Adam10 | 0.116300945 | 0.320112949 | 2.75245355 | 3.08E-09 | 6.77E-08 |
| Slc25a3 | 1.780754118 | 1.20209775 | 0.67504982 | 3.12E-09 | 6.85E-08 |
| Cyp2c70 | 0.003197886 | 0.083613235 | 26.14641 | 3.19E-09 | 7.00E-08 |
| Pla2g2f | 0.143862198 | 0.012036165 | 0.08366454 | 3.22E-09 | 7.04E-08 |
| Pnmal1 | 0.145442254 | 0.014628744 | 0.10058112 | 3.28E-09 | 7.17E-08 |
| Ppp1r2 | 0.39117458 | 0.143141237 | 0.36592674 | 3.35E-09 | 7.32E-08 |
| Banp | 0.257459811 | 0.066645438 | 0.25885763 | 3.45E-09 | 7.53E-08 |
| Uchl1 | 0.155607676 | 0.01629595 | 0.10472459 | 3.49E-09 | 7.60E-08 |
| Fam122a | 0.263089649 | 0.066232496 | 0.25174877 | 3.57E-09 | 7.76E-08 |
| Pf4 | 0.212687081 | 0.045143046 | 0.212251 | 3.90E-09 | 8.47E-08 |
| Man2a1 | 0.071890353 | 0.236959836 | 3.29612843 | 3.96E-09 | 8.59E-08 |
| Rhov | 0.041432047 | 0.177887456 | 4.29347497 | 4.05E-09 | 8.77E-08 |
| 1700023F06I | 0.130750331 | 0.00927768 | 0.07095722 | 4.17E-09 | 9.01E-08 |
| Orc6 | 0.29318891 | 0.086339378 | 0.29448378 | 4.20E-09 | 9.09E-08 |
| Tmem238 | 0.026902683 | 0.148138635 | 5.50646317 | 4.23E-09 | 9.12E-08 |
| Slc39a11 | 0.048868778 | 0.198219481 | 4.05615787 | 4.23E-09 | 9.12E-08 |
| Ubqln2 | 0.393419566 | 0.149832507 | 0.38084661 | 4.27E-09 | 9.19E-08 |
| Snd1 | 0.321754509 | 0.624909181 | 1.94219246 | 4.42E-09 | 9.51E-08 |
| Elof1 | 0.674539018 | 0.345626454 | 0.51238912 | 4.63E-09 | 9.94E-08 |
| Fam81a | 0.011672401 | 0.108928927 | 9.33217823 | 4.99E-09 | 1.07E-07 |
| Crip1 | 1.159717719 | 0.709387211 | 0.61168955 | 5.01E-09 | 1.07E-07 |
| Stk39 | 0.039695403 | 0.173867493 | 4.380041 | 5.20E-09 | 1.11E-07 |
| Mt3 | 0.05257551 | 0.200243772 | 3.80868914 | 5.31E-09 | 1.13E-07 |
| Ush1c | 0.152668068 | 0.018173028 | 0.11903621 | 5.40E-09 | 1.15E-07 |
| Ripply3 | 0.101049935 | 0 | 0 | 5.47E-09 | 1.17E-07 |
| Yipf4 | 0.350348635 | 0.121321617 | 0.34628825 | 5.57E-09 | 1.19E-07 |
| Phyh | 0.275100368 | 0.07968455 | 0.28965628 | 5.69E-09 | 1.21E-07 |
| Pigp | 0.424552144 | 0.171871952 | 0.40483119 | 5.81E-09 | 1.23E-07 |

| | | | | | |
|---|---|---|---|---|---|
| Mar-02 | 0.25614145 | 0.529132679 | 2.0657831 | 5.83E-09 | 1.24E-07 |
| Mpzl1 | 0.334734954 | 0.113414898 | 0.33882 | 5.85E-09 | 1.24E-07 |
| Gstz1 | 0.327755638 | 0.109706046 | 0.33471902 | 6.02E-09 | 1.27E-07 |
| Lyar | 0.223669049 | 0.4812394 | 2.15156904 | 6.04E-09 | 1.28E-07 |
| Naa10 | 0.211183071 | 0.464505315 | 2.1995386 | 6.29E-09 | 1.33E-07 |
| Smco4 | 0.176525404 | 0.412112456 | 2.33457874 | 6.35E-09 | 1.34E-07 |
| Eif1b | 0.541183913 | 0.248960592 | 0.46002955 | 6.89E-09 | 1.45E-07 |
| Arf1 | 1.728512459 | 1.179378812 | 0.68230854 | 6.97E-09 | 1.47E-07 |
| Saraf | 0.854052144 | 0.480018729 | 0.5620485 | 6.98E-09 | 1.47E-07 |
| Sar1b | 0.520742485 | 0.238590762 | 0.45817418 | 6.99E-09 | 1.47E-07 |
| Dbn1 | 0.183027846 | 0.030290943 | 0.16549909 | 7.03E-09 | 1.47E-07 |
| Rem2 | 0.115391816 | 0.002325517 | 0.02015322 | 7.09E-09 | 1.48E-07 |
| Gtf2a2 | 0.702918954 | 0.364074533 | 0.51794667 | 7.18E-09 | 1.50E-07 |
| 1810058I24R | 0.646079651 | 0.325807465 | 0.50428374 | 7.22E-09 | 1.51E-07 |
| Mrpl12 | 0.341776369 | 0.648001685 | 1.89598153 | 7.60E-09 | 1.59E-07 |
| Disp2 | 0.161458088 | 0.018161361 | 0.11248344 | 8.25E-09 | 1.72E-07 |
| Uck2 | 0.141985645 | 0.357612103 | 2.51864971 | 8.54E-09 | 1.78E-07 |
| Ociad2 | 0.687431872 | 0.353543174 | 0.51429558 | 8.95E-09 | 1.86E-07 |
| Rpl19 | 7.480812113 | 10.48955433 | 1.4021946 | 8.95E-09 | 1.86E-07 |
| Nme1 | 1.799581185 | 2.555978996 | 1.4203188 | 8.96E-09 | 1.86E-07 |
| Gcnt1 | 0.029302994 | 0.14947774 | 5.10110803 | 9.27E-09 | 1.92E-07 |
| Cfap36 | 0.360406696 | 0.131822782 | 0.36576119 | 1.00E-08 | 2.08E-07 |
| Eif3m | 0.72173092 | 1.14730017 | 1.58965085 | 1.04E-08 | 2.14E-07 |
| Dnajb6 | 0.593322153 | 0.291278818 | 0.49092861 | 1.04E-08 | 2.14E-07 |
| Tsc22d1 | 0.785449054 | 0.4274237 | 0.5441775 | 1.04E-08 | 2.15E-07 |
| Tmed11 | 0.008428943 | 0.097222409 | 11.5343532 | 1.04E-08 | 2.15E-07 |
| Mapt | 0.126573776 | 0.011413905 | 0.0901759 | 1.07E-08 | 2.19E-07 |
| Tmem176b | 1.221352924 | 0.767077532 | 0.62805559 | 1.07E-08 | 2.20E-07 |
| Ift20 | 0.831733539 | 1.28265605 | 1.5421478 | 1.09E-08 | 2.23E-07 |
| Dst | 0.262212498 | 0.072679928 | 0.2771795 | 1.12E-08 | 2.29E-07 |

| | | | | | |
|---|---|---|---|---|---|
| Stbd1 | 0.08914864 | 0.267685432 | 3.00268667 | 1.15E-08 | 2.36E-07 |
| Abcb9 | 0.111811814 | 0.005015588 | 0.0448574 | 1.15E-08 | 2.36E-07 |
| Abcf1 | 0.345501346 | 0.64790126 | 1.8752496 | 1.21E-08 | 2.46E-07 |
| Tmc4 | 0.305619556 | 0.100692461 | 0.32946995 | 1.21E-08 | 2.47E-07 |
| Prr15l | 0.122177999 | 0.320549193 | 2.62362451 | 1.30E-08 | 2.64E-07 |
| Slc16a10 | 0.149061826 | 0.015337579 | 0.10289408 | 1.40E-08 | 2.84E-07 |
| Eif4ebp1 | 0.200007246 | 0.439295303 | 2.19639693 | 1.47E-08 | 2.99E-07 |
| Atp2b1 | 0.503306116 | 0.225874328 | 0.44878121 | 1.48E-08 | 3.01E-07 |
| Spdef | 0.012797384 | 0.111483646 | 8.71144008 | 1.51E-08 | 3.06E-07 |
| Fabp4 | 0.013606785 | 0.109538951 | 8.05031838 | 1.51E-08 | 3.06E-07 |
| Crlf1 | 0.015014486 | 0.114529569 | 7.62793794 | 1.56E-08 | 3.15E-07 |
| Lgi2 | 0.016614223 | 0.118344077 | 7.12305812 | 1.56E-08 | 3.15E-07 |
| Nipal1 | 0.178122788 | 0.027381685 | 0.15372365 | 1.64E-08 | 3.30E-07 |
| Ube2s | 1.043105077 | 0.62931021 | 0.60330471 | 1.64E-08 | 3.30E-07 |
| Mylk | 0.124263661 | 0.007068884 | 0.05688617 | 1.64E-08 | 3.30E-07 |
| Tmed3 | 1.977092313 | 2.777311344 | 1.40474541 | 1.65E-08 | 3.31E-07 |
| Snx2 | 0.296808021 | 0.09572187 | 0.32250432 | 1.72E-08 | 3.45E-07 |
| Nup62cl | 0.005184117 | 0.082374103 | 15.8897072 | 1.72E-08 | 3.45E-07 |
| Gm5424 | 0.005058453 | 0.081506941 | 16.1130171 | 1.72E-08 | 3.45E-07 |
| Rangrf | 0.084511364 | 0.254160067 | 3.007407 | 1.75E-08 | 3.50E-07 |
| Kctd13 | 0.165283661 | 0.028638635 | 0.17326961 | 1.77E-08 | 3.53E-07 |
| Gpx2 | 0.166537996 | 0.025654367 | 0.15404513 | 1.77E-08 | 3.53E-07 |
| Bmp7 | 0.031290186 | 0.146445759 | 4.68024566 | 1.96E-08 | 3.90E-07 |
| Fus | 0.943252759 | 0.55527104 | 0.58867683 | 2.04E-08 | 4.06E-07 |
| Abhd16a | 0.210253051 | 0.048410255 | 0.23024758 | 2.16E-08 | 4.30E-07 |
| Taf7 | 0.248491285 | 0.070500199 | 0.28371296 | 2.22E-08 | 4.41E-07 |
| Rcn2 | 0.249097998 | 0.068090675 | 0.27334895 | 2.22E-08 | 4.41E-07 |
| Scpep1 | 0.293956687 | 0.096501066 | 0.32828328 | 2.41E-08 | 4.77E-07 |
| Fos | 2.476689541 | 1.741650903 | 0.70321729 | 2.42E-08 | 4.79E-07 |
| Jun | 1.402580316 | 1.984615285 | 1.41497443 | 2.55E-08 | 5.04E-07 |

| | | | | | |
|---|---|---|---|---|---|
| Akap9 | 0.457759082 | 0.202607811 | 0.44260795 | 2.60E-08 | 5.13E-07 |
| Hcfc1r1 | 0.559207016 | 0.268164153 | 0.47954361 | 2.67E-08 | 5.26E-07 |
| Ddit4 | 0.151386404 | 0.357921871 | 2.36429338 | 2.75E-08 | 5.41E-07 |
| Rpl4 | 5.939650869 | 8.228204141 | 1.38530098 | 2.79E-08 | 5.50E-07 |
| Fam159b | 0.15221284 | 0.02032638 | 0.13353919 | 2.99E-08 | 5.88E-07 |
| Tfg | 0.593366188 | 0.301065754 | 0.5073861 | 3.00E-08 | 5.89E-07 |
| Morf4l2 | 1.441540224 | 0.962270907 | 0.6675297 | 3.09E-08 | 6.05E-07 |
| Slc35g2 | 0.107467912 | 0.006480334 | 0.06030018 | 3.09E-08 | 6.05E-07 |
| Pde10a | 0.107596291 | 0.004256231 | 0.03955741 | 3.09E-08 | 6.05E-07 |
| 1810011O10 | 0.022890359 | 0.135084212 | 5.90135844 | 3.15E-08 | 6.15E-07 |
| Amy2a3 | 0.003310655 | 0.076039501 | 22.9681148 | 3.37E-08 | 6.58E-07 |
| Vps29 | 0.446815856 | 0.198071078 | 0.44329465 | 3.74E-08 | 7.30E-07 |
| Crls1 | 0.036302467 | 0.156026072 | 4.29794685 | 3.76E-08 | 7.33E-07 |
| Pafah1b2 | 0.324431695 | 0.118143922 | 0.36415654 | 4.07E-08 | 7.93E-07 |
| Rab26 | 0.014212762 | 0.102767577 | 7.23065501 | 4.13E-08 | 8.03E-07 |
| 0610040J01F | 0.236324666 | 0.068366147 | 0.28928909 | 4.38E-08 | 8.49E-07 |
| Vamp2 | 0.237601316 | 0.064872203 | 0.27302964 | 4.38E-08 | 8.49E-07 |
| Paip2b | 0.060774057 | 0.206367575 | 3.39565245 | 4.60E-08 | 8.92E-07 |
| Pabpc4 | 0.125942517 | 0.314840818 | 2.49987713 | 4.64E-08 | 8.97E-07 |
| Desi1 | 0.252694442 | 0.072942246 | 0.28865789 | 4.72E-08 | 9.12E-07 |
| Pa2g4 | 0.446076533 | 0.773350579 | 1.73367241 | 4.74E-08 | 9.14E-07 |
| Map9 | 0.128790723 | 0.012846508 | 0.09974716 | 4.75E-08 | 9.16E-07 |
| Gypa | 0.105199129 | 0.002550057 | 0.02424029 | 4.76E-08 | 9.18E-07 |
| Akr1c12 | 0.267392273 | 0.083777349 | 0.31331253 | 4.89E-08 | 9.40E-07 |
| Mrpl30 | 0.550253106 | 0.904352404 | 1.64352076 | 5.05E-08 | 9.71E-07 |
| Tmem50a | 0.623115093 | 0.324230657 | 0.52033831 | 5.29E-08 | 1.02E-06 |
| Cacybp | 0.669347511 | 0.363635468 | 0.54326857 | 5.46E-08 | 1.05E-06 |
| Cpox | 0.13240403 | 0.324865777 | 2.45359433 | 5.48E-08 | 1.05E-06 |
| 1810037I17R | 1.086664226 | 0.679236477 | 0.62506565 | 5.55E-08 | 1.06E-06 |
| Fcer1g | 0.220278961 | 0.056622434 | 0.25704876 | 5.71E-08 | 1.09E-06 |

| | | | | | |
|---|---|---|---|---|---|
| Slc25a13 | 0.036385613 | 0.158011044 | 4.34267921 | 6.24E-08 | 1.19E-06 |
| Ptgr1 | 0.086302168 | 0.248715226 | 2.88191168 | 6.29E-08 | 1.20E-06 |
| Ppp2r5b | 0.115612994 | 0.008938417 | 0.07731326 | 6.57E-08 | 1.25E-06 |
| Zfp207 | 0.442286533 | 0.197419369 | 0.4463608 | 6.58E-08 | 1.25E-06 |
| Rhoa | 1.037742821 | 0.643669924 | 0.62025958 | 6.74E-08 | 1.28E-06 |
| Calr | 5.294156692 | 3.796574552 | 0.71712546 | 6.76E-08 | 1.28E-06 |
| Lgals1 | 1.75268115 | 2.434699068 | 1.38912835 | 6.87E-08 | 1.30E-06 |
| Mff | 0.369336674 | 0.149112453 | 0.40373043 | 6.87E-08 | 1.30E-06 |
| Eef1a1 | 13.61883648 | 18.83865192 | 1.3832791 | 6.89E-08 | 1.30E-06 |
| Ddt | 0.26285655 | 0.517567544 | 1.9690114 | 7.02E-08 | 1.33E-06 |
| Tm2d2 | 0.414045426 | 0.179328055 | 0.43311203 | 7.11E-08 | 1.34E-06 |
| Ndufb9 | 1.419569552 | 1.983401031 | 1.39718482 | 7.18E-08 | 1.35E-06 |
| Exosc2 | 0.055509504 | 0.192253666 | 3.46343695 | 7.31E-08 | 1.38E-06 |
| Actr10 | 0.36200678 | 0.143165543 | 0.39547752 | 7.38E-08 | 1.39E-06 |
| Rimbp2 | 0.137250058 | 0.014947614 | 0.10890789 | 7.53E-08 | 1.41E-06 |
| Hk2 | 0.036260464 | 0.151343694 | 4.17379365 | 7.84E-08 | 1.47E-06 |
| Wls | 0.353943926 | 0.137430637 | 0.38828364 | 7.91E-08 | 1.48E-06 |
| Atp6ap2 | 0.35455927 | 0.143082124 | 0.40354924 | 7.96E-08 | 1.49E-06 |
| Cntfr | 0.102794462 | 0.006707294 | 0.06524957 | 8.05E-08 | 1.51E-06 |
| Krt5 | 0.103693364 | 0.001067982 | 0.01029942 | 8.05E-08 | 1.51E-06 |
| Lbp | 0.215845649 | 0.057194556 | 0.26497896 | 8.15E-08 | 1.52E-06 |
| Cpq | 0.216685861 | 0.055779185 | 0.25741959 | 8.15E-08 | 1.52E-06 |
| Tonsl | 0.593243231 | 0.309043411 | 0.52093879 | 8.35E-08 | 1.56E-06 |
| Dynlt1f | 0.305419902 | 0.111904967 | 0.3663971 | 8.48E-08 | 1.58E-06 |
| Ptprn2 | 0.191106636 | 0.04419501 | 0.23125837 | 8.95E-08 | 1.66E-06 |
| Cap1 | 0.339263945 | 0.131743967 | 0.38832292 | 8.99E-08 | 1.67E-06 |
| Hdac1 | 0.291148322 | 0.101228673 | 0.34768764 | 9.04E-08 | 1.68E-06 |
| Zfp503 | 0.019927609 | 0.118041685 | 5.92352478 | 9.42E-08 | 1.75E-06 |
| Uqcrh | 2.251145895 | 3.097585514 | 1.37600389 | 9.49E-08 | 1.76E-06 |
| Ube2w | 0.253094634 | 0.080378898 | 0.31758436 | 9.65E-08 | 1.79E-06 |

| | | | | | |
|---|---|---|---|---|---|
| Acaa2 | 0.186568333 | 0.402105188 | 2.15527031 | 9.70E-08 | 1.79E-06 |
| Klhdc8b | 0.173119868 | 0.035668413 | 0.20603304 | 9.79E-08 | 1.81E-06 |
| Snx15 | 0.198775834 | 0.046992474 | 0.23640939 | 1.00E-07 | 1.85E-06 |
| Ccbe1 | 0.010867804 | 0.090539285 | 8.33096443 | 1.01E-07 | 1.87E-06 |
| Slc38a4 | 0.199102263 | 0.050416102 | 0.25321712 | 1.03E-07 | 1.89E-06 |
| Tm2d1 | 0.32672035 | 0.123891167 | 0.37919636 | 1.04E-07 | 1.90E-06 |
| Stxbp5l | 0.113116137 | 0.008787092 | 0.07768204 | 1.04E-07 | 1.91E-06 |
| Slit1 | 0.083992142 | 0.001959732 | 0.02333232 | 1.05E-07 | 1.92E-06 |
| Igfbpl1 | 0.085657862 | 0.001527314 | 0.01783041 | 1.06E-07 | 1.94E-06 |
| Psmc2 | 0.696823523 | 0.385955281 | 0.55387809 | 1.08E-07 | 1.98E-06 |
| Zp3 | 0.002599622 | 0.072239286 | 27.7883794 | 1.09E-07 | 1.98E-06 |
| Cdkn2d | 0.206033175 | 0.054447569 | 0.26426603 | 1.10E-07 | 2.01E-06 |
| Rarres2 | 0.473401825 | 0.224792381 | 0.47484477 | 1.10E-07 | 2.01E-06 |
| Fam104a | 0.640312976 | 0.342331426 | 0.53463141 | 1.11E-07 | 2.02E-06 |
| Sdf2l1 | 2.735714063 | 1.963447618 | 0.71770937 | 1.15E-07 | 2.10E-06 |
| Ddr1 | 0.180807401 | 0.040183464 | 0.22224458 | 1.16E-07 | 2.10E-06 |
| Celf4 | 0.12338702 | 0.014255841 | 0.11553761 | 1.16E-07 | 2.10E-06 |
| Ap3b2 | 0.123282704 | 0.012305557 | 0.09981576 | 1.16E-07 | 2.10E-06 |
| Mcf2l | 0.123941996 | 0.010969604 | 0.08850595 | 1.16E-07 | 2.10E-06 |
| Cltb | 0.561061748 | 0.286502393 | 0.51064325 | 1.16E-07 | 2.10E-06 |
| Epb41l4b | 0.045695704 | 0.172212297 | 3.76867588 | 1.20E-07 | 2.16E-06 |
| Blcap | 0.161959791 | 0.029975444 | 0.18507954 | 1.20E-07 | 2.17E-06 |
| Dtnbos | 0.099747276 | 0.003685484 | 0.03694821 | 1.23E-07 | 2.23E-06 |
| Plac8 | 0.125411271 | 0.011686762 | 0.0931875 | 1.25E-07 | 2.25E-06 |
| Tmem107 | 0.230944944 | 0.06207521 | 0.26878791 | 1.35E-07 | 2.43E-06 |
| Hmgcr | 0.249878866 | 0.081190091 | 0.3249178 | 1.37E-07 | 2.46E-06 |
| Polr2c | 0.441139434 | 0.204738324 | 0.4641125 | 1.39E-07 | 2.50E-06 |
| Prss8 | 0.110922512 | 0.286066487 | 2.57897591 | 1.40E-07 | 2.51E-06 |
| Ank | 0.170935036 | 0.036732481 | 0.21489147 | 1.47E-07 | 2.63E-06 |
| Ddx39b | 0.723402181 | 0.410226253 | 0.56707909 | 1.47E-07 | 2.63E-06 |

| | | | | | |
|---|---|---|---|---|---|
| 9530068E07I | 0.351075671 | 0.1389562 | 0.39580128 | 1.48E-07 | 2.65E-06 |
| Shmt2 | 0.103988048 | 0.269631001 | 2.59290376 | 1.53E-07 | 2.74E-06 |
| Rps26-ps1 | 0.161801182 | 0.365581094 | 2.25944637 | 1.54E-07 | 2.75E-06 |
| Slc35b1 | 0.676360394 | 0.373147934 | 0.55169986 | 1.55E-07 | 2.77E-06 |
| Sep-07 | 0.526835199 | 0.261834369 | 0.49699483 | 1.56E-07 | 2.78E-06 |
| Ero1lb | 0.651315214 | 0.3538853 | 0.54333953 | 1.57E-07 | 2.80E-06 |
| Prrc2c | 0.641579752 | 0.347838356 | 0.54215919 | 1.58E-07 | 2.82E-06 |
| Ncam1 | 0.141787851 | 0.023369797 | 0.16482228 | 1.65E-07 | 2.93E-06 |
| Gjb2 | 0.005242453 | 0.073839519 | 14.0849166 | 1.66E-07 | 2.94E-06 |
| Fam162a | 0.340390864 | 0.612662442 | 1.79987922 | 1.66E-07 | 2.94E-06 |
| Rab14 | 0.768704395 | 0.43964229 | 0.57192634 | 1.67E-07 | 2.95E-06 |
| Aqp5 | 0.08242176 | 0.000705613 | 0.008561 | 1.67E-07 | 2.96E-06 |
| Mafg | 0.177545075 | 0.040924267 | 0.23050072 | 1.71E-07 | 3.02E-06 |
| Rbm5 | 0.362513102 | 0.147202161 | 0.40606025 | 1.72E-07 | 3.04E-06 |
| Necab2 | 0.111790346 | 0.00656856 | 0.05875784 | 1.73E-07 | 3.06E-06 |
| Cldn3 | 0.617774191 | 0.973783648 | 1.57627765 | 1.74E-07 | 3.06E-06 |
| Idi1 | 0.081237964 | 0.233204163 | 2.87063033 | 1.80E-07 | 3.17E-06 |
| Tmem199 | 0.160247298 | 0.033232536 | 0.20738282 | 1.83E-07 | 3.21E-06 |
| Ctso | 0.120272124 | 0.013793984 | 0.11468978 | 1.84E-07 | 3.23E-06 |
| Nudt7 | 0.120251593 | 0.012575733 | 0.10457851 | 1.84E-07 | 3.23E-06 |
| Rai2 | 0.122152393 | 0.01419594 | 0.11621499 | 1.85E-07 | 3.24E-06 |
| Atp6v1c1 | 0.249310031 | 0.078615749 | 0.31533328 | 1.94E-07 | 3.40E-06 |
| Pin1 | 0.578089984 | 0.302555197 | 0.52337042 | 1.96E-07 | 3.42E-06 |
| Ddx41 | 0.241208777 | 0.077969962 | 0.32324679 | 1.97E-07 | 3.45E-06 |
| Rab37 | 0.097812365 | 0.004119163 | 0.0421129 | 2.09E-07 | 3.65E-06 |
| Amigo2 | 0.098979098 | 0.001869963 | 0.01889251 | 2.09E-07 | 3.65E-06 |
| Atpif1 | 3.607400418 | 2.614394613 | 0.72473092 | 2.12E-07 | 3.69E-06 |
| Rab7 | 0.546827953 | 0.280810416 | 0.51352608 | 2.13E-07 | 3.70E-06 |
| Psmb1 | 2.279543393 | 1.638978158 | 0.71899406 | 2.14E-07 | 3.73E-06 |
| Grb10 | 0.55765358 | 0.292024563 | 0.52366662 | 2.17E-07 | 3.76E-06 |

| | | | | | |
|---|---|---|---|---|---|
| B9d1 | 0.306202646 | 0.115625431 | 0.37761082 | 2.17E-07 | 3.77E-06 |
| Mapre1 | 0.423367504 | 0.193311754 | 0.45660508 | 2.21E-07 | 3.84E-06 |
| Sod2 | 0.582816624 | 0.305003192 | 0.52332617 | 2.22E-07 | 3.85E-06 |
| Sfxn1 | 0.265188446 | 0.510043735 | 1.92332563 | 2.25E-07 | 3.89E-06 |
| Fam195b | 0.631030642 | 0.341538193 | 0.54123868 | 2.25E-07 | 3.89E-06 |
| Lsm4 | 0.596628436 | 0.94378723 | 1.58186766 | 2.35E-07 | 4.06E-06 |
| Ppm1b | 0.291313503 | 0.105789126 | 0.36314529 | 2.36E-07 | 4.07E-06 |
| Pla2g12a | 0.079511031 | 0.227512321 | 2.86139316 | 2.36E-07 | 4.07E-06 |
| Dcn | 0.853093747 | 0.511886894 | 0.60003592 | 2.38E-07 | 4.09E-06 |
| Prr13 | 0.37761754 | 0.159797436 | 0.42317271 | 2.39E-07 | 4.11E-06 |
| Rab11a | 0.695094665 | 0.389800606 | 0.5607878 | 2.40E-07 | 4.12E-06 |
| Magt1 | 0.586552405 | 0.932747588 | 1.59022038 | 2.42E-07 | 4.16E-06 |
| Ginm1 | 0.274478735 | 0.095384081 | 0.34750991 | 2.53E-07 | 4.33E-06 |
| Cib2 | 0.215623865 | 0.058326033 | 0.27049897 | 2.59E-07 | 4.43E-06 |
| Dtx3 | 0.140023084 | 0.023360832 | 0.16683558 | 2.59E-07 | 4.43E-06 |
| Ncl | 1.512503554 | 2.080310578 | 1.37540872 | 2.67E-07 | 4.57E-06 |
| Tsta3 | 0.170877871 | 0.372101549 | 2.1775877 | 2.68E-07 | 4.58E-06 |
| Nrp1 | 0.040614942 | 0.15642055 | 3.85130549 | 2.69E-07 | 4.59E-06 |
| Padi2 | 0.032250592 | 0.140762378 | 4.36464476 | 2.72E-07 | 4.64E-06 |
| Cxxc4 | 0.232436486 | 0.068631862 | 0.29527146 | 2.77E-07 | 4.72E-06 |
| Npm3 | 0.30300278 | 0.555981797 | 1.83490659 | 2.79E-07 | 4.74E-06 |
| Slitrk6 | 0.1177813 | 0.016410297 | 0.13932854 | 2.83E-07 | 4.81E-06 |
| Slbp | 0.334965993 | 0.136539376 | 0.40762161 | 2.88E-07 | 4.89E-06 |
| Tcea3 | 0.015455565 | 0.098528257 | 6.37493723 | 2.92E-07 | 4.96E-06 |
| Txndc9 | 0.41900115 | 0.195984553 | 0.46774228 | 2.93E-07 | 4.96E-06 |
| Fip1l1 | 0.355851817 | 0.148473785 | 0.41723486 | 3.07E-07 | 5.20E-06 |
| Fermt1 | 0.054649017 | 0.180899015 | 3.31019706 | 3.08E-07 | 5.21E-06 |
| Mydgf | 0.846328412 | 1.243654941 | 1.46947086 | 3.18E-07 | 5.37E-06 |
| Polr2g | 0.582981737 | 0.312858231 | 0.53665186 | 3.20E-07 | 5.39E-06 |
| Pofut2 | 0.374395627 | 0.159550575 | 0.42615502 | 3.20E-07 | 5.39E-06 |

| | | | | | |
|---|---|---|---|---|---|
| Tsg101 | 0.463485387 | 0.223469694 | 0.48215046 | 3.20E-07 | 5.39E-06 |
| H2-Aa | 0.165010517 | 0.039069664 | 0.23677075 | 3.28E-07 | 5.51E-06 |
| C1qtnf4 | 0.166482618 | 0.03141199 | 0.1886803 | 3.28E-07 | 5.51E-06 |
| Aco2 | 0.517491274 | 0.262982037 | 0.50818642 | 3.34E-07 | 5.60E-06 |
| Zfp395 | 0.096383729 | 0.254160101 | 2.63696066 | 3.36E-07 | 5.63E-06 |
| Klhl7 | 0.197876762 | 0.052693886 | 0.26629649 | 3.36E-07 | 5.64E-06 |
| Cdc34 | 0.066170739 | 0.199563961 | 3.01589441 | 3.56E-07 | 5.96E-06 |
| Dhcr24 | 0.036358394 | 0.141047145 | 3.87935576 | 3.61E-07 | 6.03E-06 |
| Ift57 | 0.171818381 | 0.041970485 | 0.24427238 | 3.73E-07 | 6.23E-06 |
| Rbbp4 | 0.437712659 | 0.209540035 | 0.47871596 | 3.78E-07 | 6.31E-06 |
| Vps37a | 0.219339429 | 0.068085869 | 0.31041327 | 3.80E-07 | 6.33E-06 |
| Ppp2r1a | 0.479921877 | 0.240273344 | 0.50065095 | 3.92E-07 | 6.54E-06 |
| Ubxn1 | 0.54682618 | 0.284440748 | 0.52016666 | 3.97E-07 | 6.60E-06 |
| Cnot6l | 0.32424903 | 0.132022603 | 0.40716422 | 4.19E-07 | 6.97E-06 |
| Rhob | 0.327124267 | 0.1317312 | 0.40269468 | 4.19E-07 | 6.97E-06 |
| Cxx1a | 0.179387221 | 0.043123455 | 0.24039313 | 4.20E-07 | 6.98E-06 |
| Ube2e1 | 0.411013139 | 0.188529996 | 0.45869579 | 4.28E-07 | 7.09E-06 |
| 1110004F10Il | 0.669215347 | 0.380090447 | 0.56796433 | 4.28E-07 | 7.09E-06 |
| Zfhx2 | 0.127106104 | 0.015401144 | 0.12116762 | 4.31E-07 | 7.14E-06 |
| Arpp19 | 0.731459442 | 1.097559232 | 1.50050593 | 4.34E-07 | 7.17E-06 |
| Ly6h | 0.106161379 | 0.010010743 | 0.09429741 | 4.38E-07 | 7.23E-06 |
| Tmod2 | 0.105833806 | 0.009450568 | 0.08929631 | 4.38E-07 | 7.23E-06 |
| Mrps36 | 0.525595606 | 0.271619347 | 0.5167839 | 4.39E-07 | 7.24E-06 |
| Cdhr1 | 0.077086341 | 0.003883051 | 0.05037276 | 4.49E-07 | 7.40E-06 |
| B230219D22 | 0.320146324 | 0.125035653 | 0.39055783 | 4.55E-07 | 7.50E-06 |
| Pglyrp1 | 0.023011434 | 0.121740202 | 5.29042228 | 4.57E-07 | 7.51E-06 |
| Dag1 | 0.172889646 | 0.368457503 | 2.13117159 | 4.58E-07 | 7.53E-06 |
| Fam204a | 0.277780117 | 0.098514804 | 0.35465031 | 4.65E-07 | 7.64E-06 |
| Ndufc1 | 1.291762146 | 1.772198706 | 1.37192339 | 4.66E-07 | 7.65E-06 |
| Rpl10-ps3 | 0.221555233 | 0.438085 | 1.97731732 | 4.83E-07 | 7.92E-06 |

| | | | | | |
|---|---|---|---|---|---|
| Rraga | 0.477780434 | 0.233198036 | 0.4880862 | 5.06E-07 | 8.28E-06 |
| Oaz1 | 4.231495698 | 3.102025749 | 0.73308021 | 5.08E-07 | 8.29E-06 |
| Gls2 | 0.005540832 | 0.068089391 | 12.2886577 | 5.08E-07 | 8.29E-06 |
| Slc29a1 | 0.145495252 | 0.024593015 | 0.16902968 | 5.10E-07 | 8.32E-06 |
| Atxn7l3b | 0.79890384 | 0.475388158 | 0.59505054 | 5.18E-07 | 8.45E-06 |
| Cenpm | 0.069548949 | 0.206044664 | 2.96258484 | 5.22E-07 | 8.50E-06 |
| Eif2s2 | 1.037975565 | 1.468465202 | 1.41473966 | 5.36E-07 | 8.72E-06 |
| Jagn1 | 0.425860407 | 0.201130837 | 0.47229288 | 5.41E-07 | 8.79E-06 |
| 2300009A05 | 0.218027449 | 0.065618766 | 0.30096562 | 5.45E-07 | 8.85E-06 |
| Chpf | 0.170832396 | 0.038963869 | 0.22808244 | 5.60E-07 | 9.09E-06 |
| Rufy3 | 0.132882617 | 0.02225154 | 0.1674526 | 5.86E-07 | 9.49E-06 |
| Zfp568 | 0.133506403 | 0.02054183 | 0.153864 | 5.86E-07 | 9.49E-06 |
| Cox14 | 0.950148095 | 0.6047125 | 0.63644026 | 5.89E-07 | 9.52E-06 |
| Nsa2 | 0.409172783 | 0.693271949 | 1.69432567 | 5.89E-07 | 9.52E-06 |
| Pfdn4 | 0.4505024 | 0.216764582 | 0.48116188 | 6.03E-07 | 9.74E-06 |
| Vimp | 0.942798622 | 1.348526098 | 1.43034373 | 6.35E-07 | 1.02E-05 |
| Rps27 | 7.717177107 | 10.35093058 | 1.34128457 | 6.37E-07 | 1.03E-05 |
| Aldoa | 1.689895432 | 1.216965599 | 0.72014255 | 6.54E-07 | 1.05E-05 |
| Ube2m | 0.080315073 | 0.223566911 | 2.78362335 | 6.54E-07 | 1.05E-05 |
| Slpi | 0.124047814 | 0.018135594 | 0.14619842 | 6.56E-07 | 1.05E-05 |
| Lrp8 | 0.103815927 | 0.005268271 | 0.05074627 | 6.56E-07 | 1.05E-05 |
| Npc2 | 0.883552966 | 0.547815565 | 0.6200144 | 6.59E-07 | 1.06E-05 |
| Oser1 | 0.267002398 | 0.098549864 | 0.3690973 | 6.73E-07 | 1.08E-05 |
| Slmo2 | 0.26757344 | 0.095173304 | 0.3556904 | 6.73E-07 | 1.08E-05 |
| Twsg1 | 0.185665076 | 0.048769555 | 0.2626749 | 6.77E-07 | 1.08E-05 |
| Klc3 | 0.18681489 | 0.046751913 | 0.25025796 | 6.77E-07 | 1.08E-05 |
| Anxa4 | 0.799074493 | 0.48024 | 0.60099528 | 6.85E-07 | 1.09E-05 |
| Arid4b | 0.394458303 | 0.180478203 | 0.4575343 | 6.93E-07 | 1.11E-05 |
| Atp13a2 | 0.071585822 | 0.206981862 | 2.89138065 | 7.03E-07 | 1.12E-05 |
| Myo5c | 0.01680784 | 0.099526061 | 5.92140702 | 7.09E-07 | 1.13E-05 |

| | | | | | |
|---|---|---|---|---|---|
| Cops5 | 0.425114567 | 0.198571084 | 0.46710017 | 7.11E-07 | 1.13E-05 |
| Syp | 0.074054498 | 0.002201717 | 0.02973104 | 7.50E-07 | 1.19E-05 |
| Cpn1 | 0.15549145 | 0.34369019 | 2.21034783 | 7.50E-07 | 1.19E-05 |
| Pdgfa | 0.156334952 | 0.340499793 | 2.17801451 | 7.50E-07 | 1.19E-05 |
| Trp53i13 | 0.14188665 | 0.028476159 | 0.20069654 | 7.51E-07 | 1.19E-05 |
| Apbb1 | 0.142241888 | 0.027183645 | 0.19110858 | 7.51E-07 | 1.19E-05 |
| Sdpr | 0.089838794 | 0.238437224 | 2.65405637 | 7.58E-07 | 1.20E-05 |
| Rab27a | 0.208466817 | 0.058767352 | 0.28190267 | 7.64E-07 | 1.21E-05 |
| Aldh1b1 | 0.045536628 | 0.162256106 | 3.56319984 | 7.68E-07 | 1.21E-05 |
| Prkacb | 0.230365005 | 0.074180389 | 0.3220124 | 7.72E-07 | 1.22E-05 |
| Esd | 1.001243375 | 0.647838265 | 0.64703376 | 7.82E-07 | 1.23E-05 |
| Ccdc107 | 0.14983251 | 0.337430453 | 2.25205099 | 7.96E-07 | 1.25E-05 |
| Rpl6l | 0.361269202 | 0.6291789 | 1.74157912 | 8.08E-07 | 1.27E-05 |
| Alg5 | 0.138673687 | 0.317867354 | 2.2921966 | 8.12E-07 | 1.28E-05 |
| Robo2 | 0.089035515 | 0.007724357 | 0.08675591 | 8.26E-07 | 1.29E-05 |
| Ntrk2 | 0.090399007 | 0.003967875 | 0.04389291 | 8.26E-07 | 1.29E-05 |
| Gpr119 | 0.089165199 | 0.003675001 | 0.04121565 | 8.26E-07 | 1.29E-05 |
| Slc25a11 | 0.376624659 | 0.173536729 | 0.46076837 | 8.31E-07 | 1.30E-05 |
| Cd47 | 0.378530938 | 0.174237312 | 0.46029873 | 8.34E-07 | 1.31E-05 |
| Brk1 | 0.741692583 | 0.435054389 | 0.58656969 | 8.35E-07 | 1.31E-05 |
| Tvp23b | 0.471172767 | 0.232940082 | 0.49438358 | 8.46E-07 | 1.32E-05 |
| Ei24 | 0.271635112 | 0.101305598 | 0.37294736 | 8.92E-07 | 1.39E-05 |
| Ahi1 | 0.132319211 | 0.021440892 | 0.16203915 | 8.98E-07 | 1.40E-05 |
| Psmd13 | 0.4536948 | 0.223382642 | 0.49236324 | 9.21E-07 | 1.44E-05 |
| Mrpl13 | 0.220901363 | 0.435570442 | 1.97178703 | 9.34E-07 | 1.45E-05 |
| H2afy2 | 0.150093791 | 0.032771806 | 0.21834219 | 9.36E-07 | 1.46E-05 |
| 2010300C02I | 0.024646499 | 0.118835672 | 4.8216046 | 9.38E-07 | 1.46E-05 |
| Rab27b | 0.026007525 | 0.116075755 | 4.46316034 | 9.38E-07 | 1.46E-05 |
| Rpp21 | 0.349240959 | 0.602076234 | 1.72395654 | 9.51E-07 | 1.47E-05 |
| 2310022B05I | 0.182395578 | 0.048677155 | 0.26687684 | 9.67E-07 | 1.50E-05 |

| | | | | | |
|---|---|---|---|---|---|
| Rsrc2 | 0.642858352 | 0.365077469 | 0.56789722 | 9.67E-07 | 1.50E-05 |
| Vapa | 0.680734867 | 0.393110543 | 0.57747967 | 9.77E-07 | 1.51E-05 |
| Mtus1 | 0.028134579 | 0.130515257 | 4.63896247 | 9.85E-07 | 1.52E-05 |
| Eif5a | 2.444956509 | 3.278182103 | 1.34079363 | 9.92E-07 | 1.53E-05 |
| Slc43a2 | 0.099377527 | 0.008684842 | 0.08739242 | 1.02E-06 | 1.57E-05 |
| Arl6ip5 | 0.413326059 | 0.197628053 | 0.4781408 | 1.03E-06 | 1.59E-05 |
| Sorbs2 | 0.191090956 | 0.052238069 | 0.27336756 | 1.05E-06 | 1.62E-05 |
| Mrpl41 | 0.394455975 | 0.185394748 | 0.47000112 | 1.07E-06 | 1.65E-05 |
| Spg21 | 0.234108164 | 0.078453017 | 0.3351144 | 1.07E-06 | 1.65E-05 |
| Dguok | 0.235667426 | 0.076344801 | 0.32395144 | 1.07E-06 | 1.65E-05 |
| Mtmr7 | 0.109235408 | 0.013752834 | 0.12590088 | 1.07E-06 | 1.65E-05 |
| Cox5a | 1.227276109 | 1.67858394 | 1.36773129 | 1.12E-06 | 1.71E-05 |
| Ftl1 | 11.99456251 | 8.86709826 | 0.73925983 | 1.15E-06 | 1.77E-05 |
| Wfdc15b | 0.131997039 | 0.301208572 | 2.2819343 | 1.16E-06 | 1.77E-05 |
| Rab11b | 0.449639982 | 0.225668319 | 0.50188668 | 1.18E-06 | 1.81E-05 |
| Bmyc | 0.388026107 | 0.179847234 | 0.46349261 | 1.18E-06 | 1.81E-05 |
| Etv1 | 0.164398739 | 0.037269791 | 0.22670363 | 1.19E-06 | 1.82E-05 |
| Vdac2 | 0.890145566 | 0.561707738 | 0.63102908 | 1.20E-06 | 1.83E-05 |
| Nol4 | 0.072436672 | 0.002154638 | 0.02974512 | 1.21E-06 | 1.84E-05 |
| H2afx | 0.461658317 | 0.231193642 | 0.50078951 | 1.22E-06 | 1.86E-05 |
| Nudt22 | 0.067673258 | 0.197105067 | 2.91259907 | 1.24E-06 | 1.89E-05 |
| Cdk4 | 0.955583318 | 0.614396338 | 0.64295423 | 1.26E-06 | 1.91E-05 |
| Sqstm1 | 0.399755164 | 0.188466618 | 0.47145512 | 1.27E-06 | 1.92E-05 |
| Vat1l | 0.086488927 | 0.001969098 | 0.02276705 | 1.31E-06 | 1.98E-05 |
| Tmem176a | 0.508351701 | 0.264820814 | 0.52094016 | 1.31E-06 | 1.99E-05 |
| Tmem108 | 0.128386493 | 0.024651787 | 0.19201231 | 1.35E-06 | 2.05E-05 |
| Tmed5 | 0.152540361 | 0.329630291 | 2.16093818 | 1.38E-06 | 2.08E-05 |
| Eif4e3 | 0.087360655 | 0.007690659 | 0.08803344 | 1.41E-06 | 2.13E-05 |
| Atat1 | 0.130217379 | 0.023941763 | 0.18385997 | 1.43E-06 | 2.16E-05 |
| B4galt3 | 0.180998322 | 0.047252307 | 0.26106489 | 1.45E-06 | 2.18E-05 |

| | | | | | |
|---|---|---|---|---|---|
| Capza2 | 0.478261902 | 0.24765544 | 0.51782389 | 1.45E-06 | 2.18E-05 |
| Selenbp1 | 0.123133869 | 0.288167175 | 2.34027548 | 1.50E-06 | 2.26E-05 |
| Fads1 | 0.031108639 | 0.123551196 | 3.97160404 | 1.52E-06 | 2.29E-05 |
| Tmed2 | 1.516641161 | 2.038215244 | 1.34390078 | 1.54E-06 | 2.32E-05 |
| Map1lc3a | 0.384890792 | 0.179601379 | 0.46662945 | 1.56E-06 | 2.34E-05 |
| Stx6 | 0.154062981 | 0.03703018 | 0.24035742 | 1.57E-06 | 2.35E-05 |
| C1qtnf6 | 0.15407882 | 0.03379487 | 0.21933495 | 1.57E-06 | 2.35E-05 |
| Rbmx2 | 0.155408524 | 0.034368235 | 0.22114768 | 1.62E-06 | 2.43E-05 |
| Fam195a | 0.057570384 | 0.180198098 | 3.13004855 | 1.67E-06 | 2.50E-05 |
| Adamts9 | 0.120652454 | 0.278068589 | 2.30470729 | 1.70E-06 | 2.55E-05 |
| Fam46a | 0.162126058 | 0.03811285 | 0.23508158 | 1.76E-06 | 2.63E-05 |
| Cdk18 | 0.016982388 | 0.094635464 | 5.5725653 | 1.79E-06 | 2.67E-05 |
| Srm | 0.333738522 | 0.576788376 | 1.72826431 | 1.86E-06 | 2.78E-05 |
| Pgrmc1 | 0.486250854 | 0.252562499 | 0.51940783 | 1.87E-06 | 2.78E-05 |
| Chn2 | 0.006382663 | 0.071053254 | 11.1322274 | 1.88E-06 | 2.79E-05 |
| Napa | 0.70013841 | 0.418560828 | 0.59782583 | 1.89E-06 | 2.81E-05 |
| Kars | 0.199739707 | 0.394650808 | 1.9758255 | 1.92E-06 | 2.86E-05 |
| Cnot7 | 0.331935846 | 0.142847802 | 0.43034762 | 1.93E-06 | 2.87E-05 |
| Pfkfb3 | 0.014710813 | 0.094685684 | 6.43646842 | 1.95E-06 | 2.90E-05 |
| Slc1a5 | 0.02888802 | 0.121763099 | 4.21500333 | 1.98E-06 | 2.93E-05 |
| Rbm47 | 0.158438554 | 0.337306944 | 2.12894485 | 1.99E-06 | 2.95E-05 |
| Peg10 | 0.245238287 | 0.08624796 | 0.35169044 | 1.99E-06 | 2.95E-05 |
| Baz2b | 0.237206833 | 0.08393316 | 0.35383955 | 2.02E-06 | 2.99E-05 |
| Rock1 | 0.380827511 | 0.180904211 | 0.47502926 | 2.05E-06 | 3.03E-05 |
| Psmd4 | 0.705682354 | 0.420523055 | 0.59590984 | 2.06E-06 | 3.04E-05 |
| Rpl13-ps3 | 0.171777651 | 0.35770955 | 2.08239866 | 2.06E-06 | 3.04E-05 |
| 9230105E05I | 0.008468715 | 0.075572404 | 8.9237161 | 2.08E-06 | 3.07E-05 |
| Amy2a4 | 0.001356325 | 0.052838084 | 38.9568002 | 2.09E-06 | 3.08E-05 |
| Tpm2 | 0.183920427 | 0.05501 | 0.29909674 | 2.11E-06 | 3.10E-05 |
| Vapb | 0.322824875 | 0.141331185 | 0.43779521 | 2.11E-06 | 3.11E-05 |

| | | | | | |
|---|---|---|---|---|---|
| Ppp1r15a | 0.230159999 | 0.078250962 | 0.33998506 | 2.12E-06 | 3.11E-05 |
| Tagln | 0.085305032 | 0.004659938 | 0.05462677 | 2.13E-06 | 3.13E-05 |
| Cnih2 | 0.084847494 | 0.003453706 | 0.04070487 | 2.13E-06 | 3.13E-05 |
| Zfp36l1 | 0.256436186 | 0.47386739 | 1.84789595 | 2.14E-06 | 3.14E-05 |
| Calm1 | 4.406797188 | 3.286600865 | 0.74580261 | 2.14E-06 | 3.14E-05 |
| Stip1 | 0.391833531 | 0.190141006 | 0.48525966 | 2.17E-06 | 3.18E-05 |
| 1810041H14I | 0.009735539 | 0.082177516 | 8.4409829 | 2.17E-06 | 3.18E-05 |
| Klf15 | 0.010688201 | 0.077580254 | 7.25849489 | 2.17E-06 | 3.18E-05 |
| Lgmn | 0.212667874 | 0.071635857 | 0.33684381 | 2.19E-06 | 3.20E-05 |
| Bambi | 0.214241433 | 0.069876667 | 0.32615851 | 2.20E-06 | 3.20E-05 |
| Hist3h2a | 0.192659058 | 0.05935999 | 0.308109 | 2.20E-06 | 3.21E-05 |
| Fdft1 | 0.174725931 | 0.356383133 | 2.0396694 | 2.22E-06 | 3.24E-05 |
| Hmbs | 0.262106451 | 0.103280659 | 0.39404089 | 2.30E-06 | 3.34E-05 |
| H2-D1 | 0.471262639 | 0.248765675 | 0.52787056 | 2.35E-06 | 3.41E-05 |
| Eef2 | 2.724948498 | 3.607807129 | 1.32399094 | 2.37E-06 | 3.44E-05 |
| Trp53i11 | 0.11544296 | 0.018300147 | 0.15852111 | 2.39E-06 | 3.47E-05 |
| Tcaf1 | 0.116864596 | 0.018059909 | 0.15453704 | 2.44E-06 | 3.54E-05 |
| 2010107E04I | 1.616485729 | 2.157065977 | 1.33441696 | 2.49E-06 | 3.61E-05 |
| Smim24 | 0.09511124 | 0.007843837 | 0.08247014 | 2.53E-06 | 3.66E-05 |
| Sdccag8 | 0.133541352 | 0.025433333 | 0.19045287 | 2.55E-06 | 3.68E-05 |
| Tapbpl | 0.105452147 | 0.012999339 | 0.1232724 | 2.58E-06 | 3.73E-05 |
| Gipr | 0.105645654 | 0.012682969 | 0.12005197 | 2.58E-06 | 3.73E-05 |
| Hnrnpr | 0.451674463 | 0.233323723 | 0.51657497 | 2.60E-06 | 3.75E-05 |
| Smc3 | 0.465574799 | 0.241216151 | 0.51810397 | 2.65E-06 | 3.81E-05 |
| Usp9x | 0.48939168 | 0.256623501 | 0.52437242 | 2.70E-06 | 3.89E-05 |
| Ppp2r2b | 0.105783526 | 0.017225093 | 0.16283342 | 2.71E-06 | 3.89E-05 |
| Eid1 | 0.501183969 | 0.264602665 | 0.52795516 | 2.71E-06 | 3.90E-05 |
| Pld3 | 0.30145406 | 0.126804527 | 0.42064296 | 2.73E-06 | 3.92E-05 |
| Gemin7 | 0.302400812 | 0.128852686 | 0.42609901 | 2.75E-06 | 3.95E-05 |
| Cd63 | 2.888580962 | 2.153822748 | 0.74563351 | 2.76E-06 | 3.95E-05 |

| | | | | | |
|---|---|---|---|---|---|
| Ugt2b34 | 0.041643835 | 0.152446073 | 3.66071169 | 2.78E-06 | 3.97E-05 |
| Ube2v2 | 0.322129979 | 0.137076541 | 0.42553177 | 2.82E-06 | 4.03E-05 |
| Hist1h1e | 0.179245156 | 0.362368848 | 2.02163816 | 2.88E-06 | 4.12E-05 |
| Snrpg | 0.721673663 | 1.061492635 | 1.47087623 | 2.97E-06 | 4.25E-05 |
| Stub1 | 0.712103797 | 0.43108894 | 0.60537374 | 3.00E-06 | 4.28E-05 |
| Pkig | 0.295689515 | 0.122551036 | 0.41445851 | 3.03E-06 | 4.32E-05 |
| Pdk1 | 0.080335354 | 0.209423721 | 2.60686872 | 3.04E-06 | 4.33E-05 |
| Wfs1 | 0.142326092 | 0.031233888 | 0.219453 | 3.04E-06 | 4.33E-05 |
| Blvrb | 0.219890261 | 0.074518974 | 0.33889165 | 3.05E-06 | 4.33E-05 |
| Timm8b | 0.756189393 | 0.466903636 | 0.61744272 | 3.07E-06 | 4.36E-05 |
| Fam213b | 0.122813647 | 0.022510074 | 0.18328642 | 3.11E-06 | 4.42E-05 |
| Zfp516 | 0.204502454 | 0.064700918 | 0.31638211 | 3.13E-06 | 4.44E-05 |
| 4833439L19F | 0.364419168 | 0.167291802 | 0.45906422 | 3.29E-06 | 4.67E-05 |
| Rwdd1 | 0.761777805 | 0.469305496 | 0.61606612 | 3.32E-06 | 4.71E-05 |
| Fxyd1 | 0.075961443 | 0.207935663 | 2.73738433 | 3.36E-06 | 4.74E-05 |
| Kctd12b | 0.080416972 | 0.00653469 | 0.08126008 | 3.36E-06 | 4.74E-05 |
| BC049730 | 0.080731731 | 0.006018631 | 0.074551 | 3.36E-06 | 4.74E-05 |
| Cox6b1 | 2.773755044 | 3.658085037 | 1.31882051 | 3.37E-06 | 4.76E-05 |
| Ak1 | 0.149238455 | 0.034924077 | 0.23401526 | 3.41E-06 | 4.81E-05 |
| Atg9b | 0.003952321 | 0.054833031 | 13.8736286 | 3.49E-06 | 4.91E-05 |
| mt-Nd3 | 0.857007963 | 1.216768897 | 1.41978715 | 3.59E-06 | 5.06E-05 |
| Pfn2 | 0.113002945 | 0.020578534 | 0.18210618 | 3.65E-06 | 5.12E-05 |
| Slc16a12 | 0.112691582 | 0.020738434 | 0.18402825 | 3.65E-06 | 5.12E-05 |
| F13a1 | 0.113048746 | 0.019759174 | 0.17478455 | 3.65E-06 | 5.12E-05 |
| Cacna2d1 | 0.112952142 | 0.017721988 | 0.1568982 | 3.65E-06 | 5.12E-05 |
| Psmc5 | 0.639049946 | 0.376949373 | 0.58985902 | 3.75E-06 | 5.26E-05 |
| Ubl3 | 0.336265027 | 0.153978084 | 0.45790692 | 3.75E-06 | 5.26E-05 |
| Fdx1 | 0.249255459 | 0.455922574 | 1.82913777 | 3.97E-06 | 5.57E-05 |
| Ttc28 | 0.163630929 | 0.046386431 | 0.28348205 | 4.02E-06 | 5.62E-05 |
| Mns1 | 0.091668771 | 0.007395206 | 0.08067312 | 4.03E-06 | 5.63E-05 |

| | | | | | |
|---|---|---|---|---|---|
| Ptp4a2 | 0.284715743 | 0.504178935 | 1.7708151 | 4.05E-06 | 5.66E-05 |
| Spock1 | 0.092968015 | 0.007182107 | 0.07725353 | 4.10E-06 | 5.73E-05 |
| BC005624 | 0.224265761 | 0.082594218 | 0.36828724 | 4.16E-06 | 5.81E-05 |
| Yeats4 | 0.258217056 | 0.104040972 | 0.4029206 | 4.34E-06 | 6.05E-05 |
| Mar-02 | 0.200758063 | 0.066802693 | 0.33275223 | 4.38E-06 | 6.10E-05 |
| Elp4 | 0.138458521 | 0.032582562 | 0.23532363 | 4.46E-06 | 6.20E-05 |
| Pcm1 | 0.342129339 | 0.155165393 | 0.45352846 | 4.47E-06 | 6.22E-05 |
| Sdhaf4 | 0.343191762 | 0.157846 | 0.45993528 | 4.54E-06 | 6.30E-05 |
| Rab3d | 0.279413343 | 0.496245044 | 1.77602486 | 4.62E-06 | 6.41E-05 |
| Aldh1a7 | 0.005372322 | 0.059333624 | 11.0443173 | 4.64E-06 | 6.43E-05 |
| Psmb2 | 1.357635514 | 0.962268759 | 0.70878284 | 4.65E-06 | 6.44E-05 |
| Pdcd6 | 0.452614324 | 0.234104469 | 0.51722726 | 4.85E-06 | 6.71E-05 |
| D5Ertd579e | 0.146215561 | 0.033709041 | 0.23054346 | 5.05E-06 | 6.98E-05 |
| BC023829 | 0.147697474 | 0.033397825 | 0.2261232 | 5.10E-06 | 7.05E-05 |
| Pdzk1ip1 | 0.271572063 | 0.108386747 | 0.3991086 | 5.11E-06 | 7.06E-05 |
| Ube2d2a | 0.631186006 | 0.372510826 | 0.59017599 | 5.25E-06 | 7.25E-05 |
| Ubxn6 | 0.288570541 | 0.12589881 | 0.43628435 | 5.36E-06 | 7.40E-05 |
| Meig1 | 0.064256908 | 0.00164637 | 0.02562168 | 5.38E-06 | 7.42E-05 |
| Vdac3 | 0.43147536 | 0.224550889 | 0.52042575 | 5.39E-06 | 7.42E-05 |
| Hprt | 0.431874689 | 0.220956783 | 0.51162244 | 5.39E-06 | 7.42E-05 |
| Pecr | 0.118691966 | 0.266822203 | 2.24802245 | 5.44E-06 | 7.48E-05 |
| Rpl36-ps3 | 0.075352761 | 0.205531741 | 2.72759404 | 5.50E-06 | 7.55E-05 |
| Pkib | 0.079724527 | 0.001861851 | 0.02335355 | 5.50E-06 | 7.55E-05 |
| Cmtm4 | 0.101437035 | 0.241069993 | 2.3765481 | 5.56E-06 | 7.62E-05 |
| Cep70 | 0.109317468 | 0.019092349 | 0.17465048 | 5.58E-06 | 7.64E-05 |
| Cbarp | 0.11026257 | 0.015830392 | 0.14356995 | 5.58E-06 | 7.64E-05 |
| Clca3a1 | 0.011536757 | 0.077920702 | 6.75412509 | 5.73E-06 | 7.84E-05 |
| Stxbp1 | 0.111205771 | 0.020878713 | 0.18774846 | 5.78E-06 | 7.89E-05 |
| Col9a2 | 0.111528737 | 0.018900509 | 0.16946762 | 5.78E-06 | 7.89E-05 |
| Rsph9 | 0.111962516 | 0.015007982 | 0.1340447 | 5.78E-06 | 7.89E-05 |

| | | | | | |
|---|---|---|---|---|---|
| Suclg2 | 0.163450741 | 0.333591177 | 2.04092789 | 5.81E-06 | 7.92E-05 |
| Hkdc1 | 0.008857205 | 0.070758872 | 7.98884899 | 5.82E-06 | 7.92E-05 |
| Stra6 | 0.008320384 | 0.070678802 | 8.49465591 | 5.82E-06 | 7.92E-05 |
| Bche | 0.0086263 | 0.067782896 | 7.85770257 | 5.82E-06 | 7.92E-05 |
| Pafah1b1 | 0.531658484 | 0.296939896 | 0.55851624 | 5.92E-06 | 8.05E-05 |
| Phlda1 | 0.178639538 | 0.357531257 | 2.00141168 | 5.97E-06 | 8.11E-05 |
| Kras | 0.214574887 | 0.077049642 | 0.35908043 | 6.01E-06 | 8.16E-05 |
| Prkar1b | 0.099287995 | 0.013498638 | 0.13595438 | 6.19E-06 | 8.40E-05 |
| Usp51 | 0.089566372 | 0.007374377 | 0.08233422 | 6.28E-06 | 8.51E-05 |
| Smap2 | 0.182151714 | 0.058527785 | 0.32131339 | 6.32E-06 | 8.56E-05 |
| Ttc5 | 0.184248453 | 0.05726028 | 0.31077754 | 6.33E-06 | 8.56E-05 |
| Pcbp4 | 0.17760955 | 0.053151357 | 0.29925957 | 6.40E-06 | 8.66E-05 |
| Cdv3 | 0.115069309 | 0.263063219 | 2.28612844 | 6.52E-06 | 8.81E-05 |
| Gm26917 | 0.093846495 | 0.226710943 | 2.41576357 | 6.57E-06 | 8.87E-05 |
| Cdkn2aipnl | 0.05981505 | 0.168778838 | 2.82167845 | 6.61E-06 | 8.91E-05 |
| Asna1 | 0.363173444 | 0.176158114 | 0.4850523 | 6.67E-06 | 8.98E-05 |
| Mrpl52 | 0.725491836 | 1.053490185 | 1.45210481 | 6.83E-06 | 9.20E-05 |
| Trappc3 | 0.267535876 | 0.111023362 | 0.41498495 | 6.86E-06 | 9.23E-05 |
| Asl | 0.153865858 | 0.320765584 | 2.08470929 | 6.87E-06 | 9.23E-05 |
| Prom1 | 0.030574387 | 0.119737241 | 3.91625972 | 6.87E-06 | 9.23E-05 |
| Lrrc7 | 0.002272336 | 0.047301976 | 20.8164519 | 6.96E-06 | 9.34E-05 |
| Smim13 | 0.040236982 | 0.14191518 | 3.52698368 | 6.96E-06 | 9.34E-05 |
| Sdhd | 0.552544362 | 0.31674169 | 0.5732421 | 7.14E-06 | 9.57E-05 |
| Pls1 | 0.025564327 | 0.110750591 | 4.33223186 | 7.18E-06 | 9.61E-05 |
| Nt5dc2 | 0.48772106 | 0.756484924 | 1.55106061 | 7.22E-06 | 9.66E-05 |
| Ndufs7 | 0.677363623 | 0.988090768 | 1.45873019 | 7.73E-06 | 0.00010333 |
| Trappc1 | 0.470830911 | 0.251439443 | 0.53403342 | 7.92E-06 | 0.0001058 |
| Abcd3 | 0.252707462 | 0.101087164 | 0.40001654 | 7.94E-06 | 0.00010602 |
| Mrpl21 | 0.337485314 | 0.563612104 | 1.67003446 | 8.14E-06 | 0.00010863 |
| Serinc1 | 0.380468931 | 0.190295851 | 0.50016134 | 8.15E-06 | 0.00010863 |

| | | | | | |
|---|---|---|---|---|---|
| Fam134b | 0.053043524 | 0.155072424 | 2.92349399 | 8.17E-06 | 0.00010876 |
| Becn1 | 0.209420092 | 0.076505903 | 0.36532265 | 8.18E-06 | 0.00010876 |
| Dpysl2 | 0.210376153 | 0.071066546 | 0.33780704 | 8.18E-06 | 0.00010876 |
| Rsf1 | 0.245953113 | 0.093935458 | 0.38192425 | 8.55E-06 | 0.00011351 |
| H2-Eb1 | 0.107956317 | 0.02053564 | 0.19022176 | 8.56E-06 | 0.00011351 |
| Rfesd | 0.107749158 | 0.019273446 | 0.17887328 | 8.56E-06 | 0.00011351 |
| Cers5 | 0.124892465 | 0.028157904 | 0.22545719 | 8.56E-06 | 0.00011351 |
| Gm5617 | 0.272254255 | 0.112796245 | 0.4143048 | 8.59E-06 | 0.00011382 |
| Rfx6 | 0.075886932 | 0.007148421 | 0.09419832 | 8.61E-06 | 0.00011387 |
| BC048546 | 0.076570838 | 0.001527314 | 0.01994642 | 8.61E-06 | 0.00011387 |
| Fbxl16 | 0.062056143 | 0.0032466 | 0.05231715 | 8.71E-06 | 0.00011514 |
| Ildr1 | 0.126426315 | 0.028874363 | 0.22838887 | 8.77E-06 | 0.0001157 |
| Chmp7 | 0.127341866 | 0.024509218 | 0.19246787 | 8.77E-06 | 0.0001157 |
| Nmt2 | 0.187482114 | 0.063145926 | 0.3368104 | 8.85E-06 | 0.00011672 |
| Dctn2 | 0.449897813 | 0.241226262 | 0.53618012 | 8.90E-06 | 0.0001173 |
| Pts | 0.180576119 | 0.05977298 | 0.33101265 | 8.91E-06 | 0.00011731 |
| Plpp3 | 0.042399038 | 0.140898486 | 3.32315294 | 8.93E-06 | 0.00011745 |
| Cisd3 | 0.1726562 | 0.050422259 | 0.29203851 | 8.94E-06 | 0.00011748 |
| Pex19 | 0.173502487 | 0.056173818 | 0.32376376 | 8.96E-06 | 0.00011748 |
| Dync2li1 | 0.174812461 | 0.053029988 | 0.30335359 | 8.96E-06 | 0.00011748 |
| Rab28 | 0.174532645 | 0.052533028 | 0.30099256 | 8.96E-06 | 0.00011748 |
| Phip | 0.341407511 | 0.162088897 | 0.47476664 | 9.19E-06 | 0.00012042 |
| Pdzd11 | 0.238010129 | 0.095420673 | 0.40091014 | 9.24E-06 | 0.00012088 |
| Hhex | 0.239268117 | 0.090993926 | 0.38030109 | 9.24E-06 | 0.00012088 |
| Ndufb7 | 1.238386412 | 1.647441613 | 1.33031306 | 9.38E-06 | 0.00012259 |
| Mien1 | 1.050739483 | 0.719520515 | 0.68477537 | 9.44E-06 | 0.00012336 |
| Hif3a | 0.032943883 | 0.121007385 | 3.67313664 | 9.49E-06 | 0.00012387 |
| Mvb12a | 0.229828573 | 0.419670838 | 1.82601681 | 9.60E-06 | 0.00012519 |
| Tkt | 0.489893993 | 0.759296624 | 1.54992026 | 9.66E-06 | 0.00012591 |
| Cxx1b | 0.133600511 | 0.029856435 | 0.22347545 | 9.80E-06 | 0.0001277 |

| | | | | | |
|---|---|---|---|---|---|
| Pgam2 | 0.000536268 | 0.038762744 | 72.2824219 | 1.00E-05 | 0.00013076 |
| Cnpy2 | 0.851757475 | 1.192122103 | 1.39960275 | 1.01E-05 | 0.00013106 |
| Rbx1 | 1.607654825 | 1.190346259 | 0.74042403 | 1.03E-05 | 0.00013396 |
| Mrpl1 | 0.222744306 | 0.086196389 | 0.3869746 | 1.03E-05 | 0.00013404 |
| Ramp1 | 0.115790969 | 0.021238256 | 0.18341894 | 1.07E-05 | 0.00013923 |
| Gm10941 | 0.115108963 | 0.020740598 | 0.1801823 | 1.07E-05 | 0.00013923 |
| Atf6b | 0.214681745 | 0.080628426 | 0.37557188 | 1.08E-05 | 0.00014003 |
| 1110032A03I | 0.140997467 | 0.034944255 | 0.24783605 | 1.08E-05 | 0.00014003 |
| Sod1 | 0.390804097 | 0.630573828 | 1.61352922 | 1.08E-05 | 0.00014005 |
| Psmb5 | 1.214806374 | 0.858960899 | 0.70707638 | 1.09E-05 | 0.00014081 |
| Tsc22d4 | 0.250927788 | 0.101587727 | 0.40484845 | 1.09E-05 | 0.00014097 |
| Hsp90aa1 | 2.749586623 | 2.087324277 | 0.75914112 | 1.10E-05 | 0.00014189 |
| Upb1 | 0.117095046 | 0.019344365 | 0.16520225 | 1.11E-05 | 0.00014333 |
| Vbp1 | 0.412897116 | 0.213190859 | 0.51632925 | 1.12E-05 | 0.00014402 |
| Prrg2 | 0.207456344 | 0.074690066 | 0.36002787 | 1.13E-05 | 0.00014591 |
| Qsox2 | 0.082429595 | 0.206298877 | 2.50272828 | 1.13E-05 | 0.00014591 |
| Eif4b | 0.244427363 | 0.436233625 | 1.78471682 | 1.14E-05 | 0.00014602 |
| Tpr | 0.630357798 | 0.383321242 | 0.60810106 | 1.16E-05 | 0.00014858 |
| Cdh1 | 0.310257924 | 0.526352147 | 1.69649864 | 1.16E-05 | 0.00014886 |
| Cldnd1 | 0.242031528 | 0.094645924 | 0.39104791 | 1.16E-05 | 0.00014891 |
| S100a1 | 0.287239439 | 0.129872089 | 0.45213878 | 1.17E-05 | 0.00015018 |
| Ptk7 | 0.055855962 | 0.160512935 | 2.87369387 | 1.18E-05 | 0.00015071 |
| Timm17b | 0.372768613 | 0.604865812 | 1.62263075 | 1.18E-05 | 0.00015071 |
| Ghitm | 0.253455808 | 0.453096727 | 1.78767545 | 1.19E-05 | 0.0001517 |
| Nfat5 | 0.201430924 | 0.07208296 | 0.35785449 | 1.19E-05 | 0.00015211 |
| Sephs2 | 0.24607549 | 0.438001567 | 1.77994796 | 1.22E-05 | 0.0001562 |
| Eif3d | 0.45525294 | 0.710794798 | 1.56131841 | 1.24E-05 | 0.00015817 |
| 1700019D03I | 0.033885692 | 0.120690134 | 3.56168423 | 1.24E-05 | 0.0001584 |
| Cldn7 | 0.709136694 | 0.443216794 | 0.62500897 | 1.25E-05 | 0.0001587 |
| Zdhhc12 | 0.186756406 | 0.060425763 | 0.3235539 | 1.25E-05 | 0.00015917 |

| | | | | | |
|---|---|---|---|---|---|
| Tspan13 | 0.68279773 | 0.995396402 | 1.45782032 | 1.25E-05 | 0.0001593 |
| Tssc4 | 0.476680185 | 0.26509062 | 0.5561184 | 1.26E-05 | 0.00016036 |
| Ocrl | 0.164930338 | 0.045790357 | 0.27763453 | 1.26E-05 | 0.00016042 |
| Cox4i1 | 4.044797473 | 5.231066034 | 1.29328256 | 1.26E-05 | 0.00016053 |
| Man1a | 0.060650545 | 0.170147502 | 2.80537465 | 1.28E-05 | 0.00016255 |
| Pdcd5 | 0.740693336 | 0.471708705 | 0.63684751 | 1.28E-05 | 0.00016255 |
| Ckb | 0.567126724 | 0.33196633 | 0.58534771 | 1.28E-05 | 0.00016264 |
| H2-Ab1 | 0.179124341 | 0.056144844 | 0.31344062 | 1.28E-05 | 0.00016264 |
| Gm11808 | 0.365568934 | 0.595305834 | 1.62843661 | 1.29E-05 | 0.00016264 |
| Grhpr | 0.285087115 | 0.490432793 | 1.72029098 | 1.29E-05 | 0.00016317 |
| Itpkb | 0.105559125 | 0.017579946 | 0.16654122 | 1.30E-05 | 0.00016356 |
| C1qc | 0.104543378 | 0.018050795 | 0.17266321 | 1.30E-05 | 0.00016356 |
| Set | 0.561764013 | 0.844110254 | 1.50260649 | 1.35E-05 | 0.00016975 |
| Mrpl15 | 0.248560067 | 0.442207035 | 1.77907513 | 1.35E-05 | 0.00017046 |
| Ak2 | 0.367911847 | 0.592948103 | 1.61165808 | 1.36E-05 | 0.0001709 |
| Ids | 0.106162142 | 0.017515026 | 0.16498373 | 1.36E-05 | 0.00017131 |
| Tmsb15b1 | 0.107354747 | 0.014874979 | 0.13855911 | 1.36E-05 | 0.00017131 |
| Madcam1 | 0.004635228 | 0.056886538 | 12.2726503 | 1.38E-05 | 0.00017333 |
| Rims4 | 0.004993211 | 0.056453238 | 11.3059986 | 1.38E-05 | 0.00017333 |
| Ier2 | 1.022747307 | 1.387326217 | 1.35647017 | 1.39E-05 | 0.00017388 |
| Slc29a4 | 0.072172394 | 0.006813908 | 0.09441156 | 1.40E-05 | 0.00017533 |
| Gast | 0.073128525 | 0.001548463 | 0.02117454 | 1.40E-05 | 0.00017533 |
| Med31 | 0.220571527 | 0.081301606 | 0.3685952 | 1.41E-05 | 0.0001768 |
| Fyn | 0.036392106 | 0.13117708 | 3.60454759 | 1.41E-05 | 0.00017683 |
| Tmem229b | 0.074736414 | 0.005157168 | 0.06900475 | 1.42E-05 | 0.00017697 |
| Meis3 | 0.073892607 | 0.005680778 | 0.07687884 | 1.42E-05 | 0.00017697 |
| Efcab10 | 0.074719473 | 0.003595985 | 0.04812648 | 1.42E-05 | 0.00017697 |
| 2410015M2( | 1.2871889 | 0.924983863 | 0.71860771 | 1.44E-05 | 0.00018005 |
| Scnn1b | 0.059127636 | 0 | 0 | 1.44E-05 | 0.00018005 |
| Ctss | 0.094489928 | 0.015533507 | 0.16439326 | 1.48E-05 | 0.00018413 |

| | | | | | |
|---|---|---|---|---|---|
| Unc80 | 0.094644011 | 0.013348591 | 0.14104 | 1.48E-05 | 0.00018413 |
| Erh | 2.002713869 | 1.524659056 | 0.7612965 | 1.53E-05 | 0.00019083 |
| Pipox | 0.006350679 | 0.06322234 | 9.95520926 | 1.55E-05 | 0.00019216 |
| Slc25a47 | 0.006615142 | 0.061208677 | 9.25281385 | 1.55E-05 | 0.00019216 |
| Slc12a8 | 0.0065471 | 0.060659679 | 9.26512242 | 1.55E-05 | 0.00019216 |
| Cspp1 | 0.267850311 | 0.113620521 | 0.4241941 | 1.57E-05 | 0.00019426 |
| Emc7 | 0.538517628 | 0.30851258 | 0.57289226 | 1.57E-05 | 0.00019444 |
| Map2k3 | 0.098245953 | 0.228920233 | 2.33007291 | 1.57E-05 | 0.00019455 |
| Gstm3 | 0.009870519 | 0.071877773 | 7.28206629 | 1.58E-05 | 0.00019541 |
| Fam13a | 0.009594876 | 0.07142791 | 7.44438077 | 1.58E-05 | 0.00019541 |
| Gpsm1 | 0.138091646 | 0.035237425 | 0.2551742 | 1.59E-05 | 0.0001966 |
| Tinagl1 | 0.239692244 | 0.099959408 | 0.4170323 | 1.60E-05 | 0.00019709 |
| Cstf3 | 0.205979279 | 0.074193104 | 0.36019693 | 1.60E-05 | 0.00019768 |
| Ghrl | 1.762322928 | 1.337922485 | 0.75918123 | 1.62E-05 | 0.00020001 |
| Ubald2 | 0.347002722 | 0.171555547 | 0.49439251 | 1.63E-05 | 0.00020128 |
| Papss1 | 0.346160846 | 0.171106356 | 0.49429725 | 1.63E-05 | 0.00020128 |
| Echs1 | 0.291946815 | 0.500112704 | 1.71302675 | 1.65E-05 | 0.00020327 |
| Clta | 1.32474165 | 0.961878854 | 0.72608788 | 1.67E-05 | 0.00020492 |
| Agtpbp1 | 0.04311353 | 0.131908393 | 3.05955912 | 1.67E-05 | 0.00020511 |
| Gse1 | 0.145056745 | 0.040422847 | 0.2786692 | 1.68E-05 | 0.00020641 |
| Trappc2 | 0.145990092 | 0.038006258 | 0.26033451 | 1.68E-05 | 0.00020641 |
| Dpf2 | 0.189322161 | 0.067989124 | 0.35911868 | 1.71E-05 | 0.00020967 |
| Serinc3 | 0.123080977 | 0.270457664 | 2.19739614 | 1.71E-05 | 0.0002098 |
| Prmt2 | 0.147420834 | 0.036980089 | 0.2508471 | 1.75E-05 | 0.00021386 |
| Tubg1 | 0.183537661 | 0.057748831 | 0.31464295 | 1.75E-05 | 0.00021422 |
| Hdgf | 0.401004308 | 0.634549797 | 1.58240145 | 1.75E-05 | 0.00021439 |
| Hsbp1 | 1.168781819 | 0.83092372 | 0.71093142 | 1.75E-05 | 0.00021439 |
| Srsf2 | 0.196660263 | 0.369082957 | 1.87675411 | 1.77E-05 | 0.0002156 |
| St3gal5 | 0.153649588 | 0.046230176 | 0.30088057 | 1.77E-05 | 0.00021585 |
| Capsl | 0.153414105 | 0.04398239 | 0.28669065 | 1.77E-05 | 0.00021585 |

| | | | | | |
|---|---|---|---|---|---|
| Adk | 0.221306529 | 0.405314989 | 1.83146422 | 1.78E-05 | 0.00021736 |
| H2-Ke6 | 0.394137578 | 0.208604472 | 0.52926816 | 1.80E-05 | 0.00021947 |
| Nktr | 0.613113632 | 0.369203952 | 0.60217867 | 1.81E-05 | 0.00022046 |
| Ptprj | 0.035892709 | 0.120045716 | 3.3445711 | 1.83E-05 | 0.00022204 |
| Ctnnbl1 | 0.168983427 | 0.054616417 | 0.32320576 | 1.83E-05 | 0.00022241 |
| Cby1 | 0.169704466 | 0.050128593 | 0.29538759 | 1.83E-05 | 0.00022241 |
| Aes | 0.859231943 | 1.193786751 | 1.38936496 | 1.85E-05 | 0.00022408 |
| Cetn3 | 0.505301087 | 0.289088901 | 0.57211217 | 1.85E-05 | 0.00022482 |
| Gng5 | 1.880896073 | 2.44382777 | 1.2992891 | 1.88E-05 | 0.0002274 |
| Psma5 | 0.814860069 | 0.535448416 | 0.65710474 | 1.88E-05 | 0.00022774 |
| Araf | 0.269935125 | 0.120056247 | 0.44475963 | 1.88E-05 | 0.00022778 |
| Plscr3 | 0.120139444 | 0.027630999 | 0.22999106 | 1.89E-05 | 0.00022907 |
| Iqgap1 | 0.364383984 | 0.183232592 | 0.50285578 | 1.95E-05 | 0.00023583 |
| Srsf3 | 1.223461816 | 0.875114405 | 0.71527725 | 1.99E-05 | 0.0002408 |
| Trp53inp2 | 0.103547362 | 0.018530035 | 0.17895226 | 2.00E-05 | 0.00024094 |
| Hdac6 | 0.102706763 | 0.018739434 | 0.1824557 | 2.00E-05 | 0.00024094 |
| Pja2 | 0.301519078 | 0.139971056 | 0.46421957 | 2.03E-05 | 0.00024419 |
| Ghr | 0.243889851 | 0.4299761 | 1.76299299 | 2.13E-05 | 0.00025584 |
| Ube2e3 | 0.344516274 | 0.172028178 | 0.49933252 | 2.13E-05 | 0.00025584 |
| Shisa5 | 0.345647247 | 0.170185142 | 0.49236655 | 2.13E-05 | 0.00025584 |
| Runx1t1 | 0.128391513 | 0.033309104 | 0.25943384 | 2.14E-05 | 0.00025747 |
| C1qb | 0.128080771 | 0.032530583 | 0.25398491 | 2.14E-05 | 0.00025747 |
| Stt3b | 0.178031259 | 0.341013944 | 1.91547229 | 2.17E-05 | 0.00026014 |
| Dnm2 | 0.20288633 | 0.073956161 | 0.36452018 | 2.19E-05 | 0.00026208 |
| Sec61b | 5.84204592 | 7.501565574 | 1.28406481 | 2.19E-05 | 0.00026282 |
| Mtcl1 | 0.070568169 | 0.005589515 | 0.07920731 | 2.20E-05 | 0.00026365 |
| Cfap126 | 0.070364296 | 0.005142303 | 0.07308115 | 2.20E-05 | 0.00026365 |
| Stra13 | 0.426299787 | 0.668008728 | 1.56699287 | 2.27E-05 | 0.00027168 |
| Slc35c2 | 0.114561898 | 0.249350956 | 2.17656097 | 2.28E-05 | 0.00027237 |
| Dazap1 | 0.439313888 | 0.242836823 | 0.55276382 | 2.30E-05 | 0.00027439 |

| | | | | | |
|---|---|---|---|---|---|
| BC022687 | 0.090962307 | 0.016346186 | 0.17970285 | 2.30E-05 | 0.00027439 |
| Gm26699 | 0.091257518 | 0.013183823 | 0.14446835 | 2.30E-05 | 0.00027439 |
| Coro1a | 0.135126817 | 0.03578722 | 0.26484173 | 2.31E-05 | 0.00027486 |
| Maf1 | 0.230961936 | 0.0905407 | 0.39201568 | 2.33E-05 | 0.00027777 |
| Gem | 0.092989044 | 0.015456915 | 0.16622297 | 2.34E-05 | 0.00027777 |
| Kif1a | 0.093615376 | 0.013888153 | 0.14835333 | 2.34E-05 | 0.00027777 |
| Cdc14b | 0.092369607 | 0.013141555 | 0.14227142 | 2.34E-05 | 0.00027777 |
| Bpnt1 | 0.187265813 | 0.066345505 | 0.35428519 | 2.38E-05 | 0.00028249 |
| Mfng | 0.081570595 | 0.007168637 | 0.08788262 | 2.43E-05 | 0.00028783 |
| Cacna1a | 0.143376254 | 0.040753969 | 0.2842449 | 2.47E-05 | 0.00029248 |
| Hnrnpa3 | 1.230065129 | 0.888009456 | 0.72192068 | 2.49E-05 | 0.00029471 |
| Tbl1x | 0.285503246 | 0.131423765 | 0.46032319 | 2.49E-05 | 0.00029532 |
| Tcta | 0.172539079 | 0.055186743 | 0.31985069 | 2.52E-05 | 0.00029754 |
| BC005561 | 0.173412469 | 0.053547107 | 0.30878464 | 2.52E-05 | 0.00029754 |
| Tmem55b | 0.150765345 | 0.043793225 | 0.29047275 | 2.52E-05 | 0.00029754 |
| Commd9 | 0.223158089 | 0.090265013 | 0.40448909 | 2.55E-05 | 0.00030072 |
| Ankrd37 | 0.16580855 | 0.05266104 | 0.31760148 | 2.56E-05 | 0.00030222 |
| Sec11c | 0.760461248 | 1.069486347 | 1.40636535 | 2.60E-05 | 0.00030689 |
| Ap1s1 | 0.409854885 | 0.219319493 | 0.53511499 | 2.64E-05 | 0.00031103 |
| Mbnl1 | 0.08810741 | 0.209056889 | 2.37275036 | 2.64E-05 | 0.00031127 |
| Zfyve21 | 0.299833448 | 0.140289707 | 0.46789212 | 2.68E-05 | 0.0003151 |
| Fam213a | 0.240396564 | 0.102628753 | 0.42691439 | 2.68E-05 | 0.00031524 |
| Npepps | 0.261162111 | 0.112700073 | 0.43153302 | 2.76E-05 | 0.00032401 |
| Dhrs7 | 0.199653778 | 0.368133812 | 1.84386098 | 2.78E-05 | 0.00032701 |
| S100a6 | 0.828294027 | 0.555871321 | 0.67110386 | 2.81E-05 | 0.00032956 |
| Tubb4b | 1.14600421 | 0.814724012 | 0.71092584 | 2.85E-05 | 0.00033441 |
| Tbca | 1.288961536 | 0.937508182 | 0.72733604 | 2.86E-05 | 0.00033475 |
| Hsp90b1 | 4.804556645 | 3.711893339 | 0.7725777 | 2.88E-05 | 0.00033786 |
| Trappc4 | 0.366554071 | 0.191926507 | 0.52359672 | 2.93E-05 | 0.00034338 |
| Mtfr1l | 0.234643878 | 0.098636332 | 0.4203661 | 2.96E-05 | 0.00034611 |

| | | | | | |
|---|---|---|---|---|---|
| Actl6a | 0.234611129 | 0.097609213 | 0.41604682 | 2.96E-05 | 0.00034611 |
| Nudc | 0.583615873 | 0.356219585 | 0.61036651 | 2.97E-05 | 0.00034697 |
| Smarcb1 | 0.323230643 | 0.157472936 | 0.48718443 | 2.97E-05 | 0.00034705 |
| Raly | 0.585136281 | 0.355011924 | 0.60671665 | 3.00E-05 | 0.00034955 |
| Ogfr | 0.198980406 | 0.077832289 | 0.39115555 | 3.00E-05 | 0.00034955 |
| Tyrobp | 0.198891622 | 0.074682621 | 0.37549405 | 3.00E-05 | 0.00034955 |
| AU020206 | 0.015402144 | 0.078531561 | 5.09874208 | 3.05E-05 | 0.00035477 |
| Nomo1 | 0.083866738 | 0.202898753 | 2.41929945 | 3.13E-05 | 0.0003644 |
| Ap2s1 | 0.82313381 | 0.546448631 | 0.66386367 | 3.16E-05 | 0.00036754 |
| Eif4e | 0.429816233 | 0.237578574 | 0.55274453 | 3.28E-05 | 0.00038088 |
| Thop1 | 0.097216253 | 0.216467874 | 2.2266634 | 3.30E-05 | 0.00038262 |
| Agpat5 | 0.080012841 | 0.193644002 | 2.42016156 | 3.30E-05 | 0.00038262 |
| Ostc | 2.178848832 | 1.677034851 | 0.76968848 | 3.36E-05 | 0.00038908 |
| Pja1 | 0.265810879 | 0.11789248 | 0.44352014 | 3.36E-05 | 0.00038908 |
| Chchd2 | 4.336819363 | 3.359126202 | 0.77455986 | 3.37E-05 | 0.00039001 |
| Hsd17b13 | 0.004106879 | 0.046979141 | 11.4391331 | 3.37E-05 | 0.00039008 |
| Nccrp1 | 0.012721769 | 0.076072228 | 5.97968954 | 3.45E-05 | 0.00039945 |
| Gnptg | 0.296649056 | 0.138076242 | 0.46545317 | 3.50E-05 | 0.00040472 |
| Dynlt1a | 0.088967602 | 0.01323918 | 0.148809 | 3.52E-05 | 0.00040569 |
| Rpusd1 | 0.090244061 | 0.011923268 | 0.13212247 | 3.52E-05 | 0.00040569 |
| Ttc8 | 0.090337093 | 0.010310456 | 0.11413314 | 3.52E-05 | 0.00040569 |
| Ube2c | 0.450691628 | 0.688099963 | 1.52676446 | 3.53E-05 | 0.00040674 |
| Nkiras2 | 0.169596832 | 0.058089537 | 0.34251546 | 3.56E-05 | 0.00040997 |
| Tmem163 | 0.069134985 | 0.00479724 | 0.06938947 | 3.65E-05 | 0.00041926 |
| Lrrc73 | 0.069954178 | 0.002422611 | 0.03463139 | 3.65E-05 | 0.00041926 |
| Tceal1 | 0.106039284 | 0.023921722 | 0.22559302 | 3.67E-05 | 0.00042201 |
| Fbxo21 | 0.148675528 | 0.044701951 | 0.30066785 | 3.68E-05 | 0.00042323 |
| Pdlim1 | 0.081672547 | 0.196563075 | 2.40672151 | 3.71E-05 | 0.00042585 |
| Adprh | 0.213106942 | 0.085214304 | 0.39986639 | 3.73E-05 | 0.00042734 |
| Cmtm8 | 0.070405196 | 0.183188353 | 2.60191523 | 3.74E-05 | 0.00042889 |

| | | | | | |
|---|---|---|---|---|---|
| Snx17 | 0.386163882 | 0.20454409 | 0.52968208 | 3.75E-05 | 0.00042936 |
| Spag7 | 0.513083544 | 0.300715956 | 0.5860955 | 3.78E-05 | 0.00043272 |
| Pcmt1 | 0.388300187 | 0.206075658 | 0.53071223 | 3.78E-05 | 0.00043272 |
| Cadps | 0.079994972 | 0.008578677 | 0.1072402 | 3.82E-05 | 0.00043663 |
| Vdr | 0.080126874 | 0.00605846 | 0.07561084 | 3.82E-05 | 0.00043663 |
| Taf10 | 0.043902716 | 0.13494812 | 3.07379888 | 3.84E-05 | 0.00043827 |
| Edaradd | 0.012259683 | 0.070895649 | 5.78282883 | 3.84E-05 | 0.00043827 |
| Vars | 0.093005437 | 0.213408282 | 2.29457856 | 3.87E-05 | 0.00044173 |
| Myh11 | 0.054223047 | 0 | 0 | 3.88E-05 | 0.00044188 |
| Sumo3 | 0.529240277 | 0.315713754 | 0.59654143 | 3.98E-05 | 0.00045273 |
| Comt | 0.33183151 | 0.167509665 | 0.50480337 | 3.98E-05 | 0.00045298 |
| Cdc5l | 0.33109721 | 0.167600707 | 0.50619788 | 3.98E-05 | 0.00045298 |
| Ppia | 8.838015617 | 11.26642225 | 1.27476831 | 4.01E-05 | 0.00045586 |
| Krt6a | 0.056126099 | 0 | 0 | 4.04E-05 | 0.0004593 |
| Gm13344 | 0.004616801 | 0.053941564 | 11.6837521 | 4.05E-05 | 0.00045946 |
| Ctla2a | 0.005334775 | 0.051279256 | 9.61226168 | 4.05E-05 | 0.00045946 |
| Hbb-y | 0.004723312 | 0.051381383 | 10.8782522 | 4.05E-05 | 0.00045946 |
| Psmc1 | 0.426086084 | 0.239191597 | 0.56136919 | 4.12E-05 | 0.00046641 |
| Procr | 0.040387595 | 0.128765931 | 3.18825451 | 4.17E-05 | 0.00047222 |
| Dcps | 0.356977862 | 0.182840264 | 0.51218936 | 4.23E-05 | 0.00047879 |
| Fuom | 0.211318778 | 0.383515095 | 1.81486519 | 4.27E-05 | 0.00048246 |
| Tia1 | 0.281512651 | 0.129190928 | 0.45891695 | 4.32E-05 | 0.0004883 |
| Bri3 | 0.203138968 | 0.368307801 | 1.81308296 | 4.33E-05 | 0.0004883 |
| Cth | 0.006238592 | 0.056358205 | 9.03380196 | 4.36E-05 | 0.00049127 |
| Nphs1 | 0.007435398 | 0.053537245 | 7.20032051 | 4.36E-05 | 0.00049127 |
| Pde4a | 0.009127851 | 0.061079422 | 6.69154425 | 4.36E-05 | 0.00049163 |
| Dnajc8 | 0.494386181 | 0.287336893 | 0.58119928 | 4.43E-05 | 0.00049876 |
| Dnajc25 | 0.030214372 | 0.109659108 | 3.62936912 | 4.44E-05 | 0.00049942 |
| Carnmt1 | 0.030301572 | 0.10871651 | 3.5878175 | 4.44E-05 | 0.00049942 |
| Scoc | 0.190886773 | 0.069952681 | 0.36646165 | 4.52E-05 | 0.00050756 |

| | | | | | |
|---|---|---|---|---|---|
| Pde6d | 0.191751896 | 0.067157714 | 0.35023233 | 4.52E-05 | 0.00050756 |
| Serpinb6a | 0.240079327 | 0.415651918 | 1.73131074 | 4.59E-05 | 0.00051549 |
| Nono | 0.65593714 | 0.418433311 | 0.63791678 | 4.61E-05 | 0.00051736 |
| Pvrl3 | 0.097434567 | 0.017946697 | 0.1841923 | 4.64E-05 | 0.00052049 |
| Pop5 | 0.53962499 | 0.324170173 | 0.60073232 | 4.74E-05 | 0.00053075 |
| Cldn6 | 0.371847428 | 0.198064187 | 0.53264907 | 4.78E-05 | 0.00053488 |
| Cox7b | 1.555136731 | 1.999468328 | 1.28571867 | 4.79E-05 | 0.00053622 |
| Rpl9-ps6 | 0.251560995 | 0.435614744 | 1.73164661 | 4.82E-05 | 0.00053861 |
| Ssb | 0.894292953 | 0.617135127 | 0.69008162 | 4.85E-05 | 0.00054129 |
| Atp5j | 2.182902208 | 2.790024164 | 1.27812604 | 4.91E-05 | 0.00054797 |
| Creb3l1 | 0.054099258 | 0.151007715 | 2.79130846 | 4.93E-05 | 0.0005502 |
| Daam1 | 0.168353091 | 0.055505218 | 0.32969527 | 4.98E-05 | 0.00055529 |
| Ost4 | 0.841646444 | 1.15093477 | 1.36748011 | 5.01E-05 | 0.00055857 |
| Tsen34 | 0.630466322 | 0.399624956 | 0.63385615 | 5.10E-05 | 0.00056775 |
| Gm9493 | 0.471277266 | 0.711317286 | 1.50933927 | 5.11E-05 | 0.00056911 |
| Dsg2 | 0.069936511 | 0.172359684 | 2.46451647 | 5.14E-05 | 0.00057143 |
| Acvr1c | 0.146039059 | 0.041847211 | 0.28654807 | 5.19E-05 | 0.00057649 |
| Klf7 | 0.139213205 | 0.037791398 | 0.27146418 | 5.19E-05 | 0.00057649 |
| Pabpc1 | 1.420317094 | 1.824628146 | 1.28466253 | 5.22E-05 | 0.00057981 |
| Zcchc11 | 0.229128811 | 0.093820666 | 0.40946691 | 5.29E-05 | 0.00058672 |
| Scfd1 | 0.352484333 | 0.181743208 | 0.51560649 | 5.33E-05 | 0.00059078 |
| Ccndbp1 | 0.153143246 | 0.050951265 | 0.33270331 | 5.33E-05 | 0.00059095 |
| Usmg5 | 2.387994697 | 1.851589768 | 0.77537432 | 5.39E-05 | 0.00059686 |
| Cdh2 | 0.086580367 | 0.013274396 | 0.15331878 | 5.42E-05 | 0.00059947 |
| Ndufb6 | 0.746565217 | 1.039913298 | 1.39293028 | 5.44E-05 | 0.00060066 |
| Med27 | 0.10462649 | 0.02389153 | 0.22835067 | 5.44E-05 | 0.00060066 |
| Tmed8 | 0.10453851 | 0.023150739 | 0.22145656 | 5.44E-05 | 0.00060066 |
| St18 | 0.105439673 | 0.019441375 | 0.18438387 | 5.44E-05 | 0.00060066 |
| Itpa | 0.299741036 | 0.142500484 | 0.47541199 | 5.49E-05 | 0.00060555 |
| Maml3 | 0.087533049 | 0.015554057 | 0.17769353 | 5.59E-05 | 0.00061559 |

| | | | | | |
|---|---|---|---|---|---|
| Adrbk2 | 0.088613859 | 0.012104696 | 0.13660048 | 5.59E-05 | 0.00061559 |
| Sct | 0.065887583 | 0.006312689 | 0.09580999 | 5.63E-05 | 0.00061851 |
| Myt1 | 0.065683514 | 0.004676099 | 0.07119136 | 5.63E-05 | 0.00061851 |
| Syt5 | 0.065720396 | 0.003988082 | 0.06068256 | 5.63E-05 | 0.00061851 |
| 1700056E22I | 0.066521861 | 0.002089599 | 0.03141221 | 5.63E-05 | 0.00061851 |
| Bpgm | 0.21940661 | 0.092487354 | 0.42153404 | 5.77E-05 | 0.00063336 |
| Sra1 | 0.506794013 | 0.305098672 | 0.60201712 | 5.77E-05 | 0.00063336 |
| Psmd2 | 0.467800137 | 0.269418147 | 0.57592575 | 5.87E-05 | 0.00064292 |
| Fis1 | 0.822078818 | 0.556531245 | 0.6769804 | 5.87E-05 | 0.00064292 |
| Nars | 0.25953856 | 0.436539302 | 1.68198245 | 5.92E-05 | 0.00064805 |
| Tspyl4 | 0.075553305 | 0.00865073 | 0.11449837 | 5.92E-05 | 0.00064805 |
| Cyhr1 | 0.240006585 | 0.10776213 | 0.44899656 | 5.93E-05 | 0.0006481 |
| Sep-15 | 1.631780791 | 1.249236651 | 0.76556646 | 6.00E-05 | 0.0006553 |
| Prox1 | 0.18561716 | 0.073144049 | 0.39405866 | 6.05E-05 | 0.00066097 |
| Hspb11 | 0.187961603 | 0.069581328 | 0.37018905 | 6.12E-05 | 0.00066856 |
| Rab2a | 1.093508093 | 0.785694171 | 0.71850787 | 6.17E-05 | 0.00067264 |
| Gstt3 | 0.024924534 | 0.097845423 | 3.92566701 | 6.19E-05 | 0.00067488 |
| Scn9a | 0.077484428 | 0.011308963 | 0.14595143 | 6.29E-05 | 0.00068488 |
| Nadk2 | 0.016757016 | 0.080837043 | 4.82407139 | 6.31E-05 | 0.00068659 |
| Usp18 | 0.051714093 | 0.0032466 | 0.0627798 | 6.33E-05 | 0.00068794 |
| Ankrd33b | 0.0519754 | 0.001067982 | 0.02054783 | 6.33E-05 | 0.00068794 |
| Aamp | 0.600038948 | 0.37999914 | 0.63329079 | 6.43E-05 | 0.00069826 |
| Rac1 | 0.214315577 | 0.384698507 | 1.79500955 | 6.48E-05 | 0.00070362 |
| Commd7 | 0.233109988 | 0.100485181 | 0.43106339 | 6.53E-05 | 0.00070919 |
| Amn1 | 0.120039776 | 0.030574026 | 0.25469912 | 6.73E-05 | 0.00072981 |
| Slirp | 0.520158338 | 0.76173509 | 1.46442926 | 6.96E-05 | 0.00075459 |
| Fkbp4 | 0.987599547 | 0.69798016 | 0.70674411 | 6.97E-05 | 0.00075459 |
| Ide | 0.067751703 | 0.170373004 | 2.51466749 | 7.15E-05 | 0.00077408 |
| Txndc15 | 0.243006565 | 0.112182482 | 0.46164383 | 7.21E-05 | 0.0007797 |
| Ttc14 | 0.24464332 | 0.110876062 | 0.45321516 | 7.21E-05 | 0.00077973 |

| | | | | | |
|---|---|---|---|---|---|
| Lgals3 | 0.199512488 | 0.078076765 | 0.39133773 | 7.34E-05 | 0.00079314 |
| Rgs11 | 0.049145279 | 0.141181211 | 2.87273192 | 7.45E-05 | 0.0008044 |
| 4930415O20 | 0.001787009 | 0.038433605 | 21.5072276 | 7.53E-05 | 0.00081254 |
| Pgap1 | 0.143519614 | 0.043422907 | 0.3025573 | 7.62E-05 | 0.00082198 |
| Cyb5a | 0.526118053 | 0.774069934 | 1.47128563 | 7.64E-05 | 0.00082378 |
| Son | 1.12049104 | 0.811629171 | 0.72435133 | 7.67E-05 | 0.00082672 |
| Kmt2e | 0.378880926 | 0.206036746 | 0.54380343 | 7.74E-05 | 0.00083358 |
| D8Ertd738e | 0.927736873 | 1.242390301 | 1.33916236 | 7.75E-05 | 0.00083371 |
| Stat3 | 0.218765937 | 0.088667596 | 0.40530805 | 7.77E-05 | 0.00083498 |
| Wdr6 | 0.191078098 | 0.075461776 | 0.39492635 | 7.79E-05 | 0.0008364 |
| Krit1 | 0.191238254 | 0.072964179 | 0.38153548 | 7.79E-05 | 0.0008364 |
| Map2k1 | 0.192028131 | 0.070376945 | 0.36649289 | 7.79E-05 | 0.0008364 |
| Trappc6a | 0.369891227 | 0.578869219 | 1.56497148 | 7.93E-05 | 0.00085087 |
| Nab1 | 0.024179339 | 0.09154289 | 3.78599638 | 7.97E-05 | 0.00085447 |
| Rab6a | 0.238913382 | 0.10584711 | 0.4430355 | 8.02E-05 | 0.00085888 |
| Spcs2 | 1.949700581 | 1.515466086 | 0.77728145 | 8.10E-05 | 0.00086747 |
| Irak1bp1 | 0.100874515 | 0.021451652 | 0.2126568 | 8.15E-05 | 0.00087189 |
| Hmox1 | 0.101730823 | 0.019755204 | 0.19419093 | 8.15E-05 | 0.00087189 |
| Aldh6a1 | 0.030570619 | 0.107101473 | 3.50341199 | 8.32E-05 | 0.00088853 |
| Efemp2 | 0.102745083 | 0.024693555 | 0.24033807 | 8.33E-05 | 0.00088853 |
| Plppr1 | 0.083825616 | 0.013517611 | 0.16125871 | 8.33E-05 | 0.00088853 |
| Vgf | 0.084627918 | 0.011008587 | 0.13008222 | 8.33E-05 | 0.00088853 |
| Rpl15 | 1.313704642 | 1.681771215 | 1.28017452 | 8.34E-05 | 0.00088859 |
| Ctbp2 | 0.065041711 | 0.161638134 | 2.4851458 | 8.43E-05 | 0.00089725 |
| Tenm4 | 0.063864158 | 0.162720305 | 2.54791279 | 8.43E-05 | 0.00089725 |
| Gaa | 0.210965792 | 0.090415757 | 0.42858018 | 8.51E-05 | 0.000905 |
| Rheb | 0.573156632 | 0.359553328 | 0.62732124 | 8.66E-05 | 0.00092115 |
| Nfu1 | 0.249602672 | 0.116020786 | 0.46482189 | 8.73E-05 | 0.0009274 |
| Mycbp | 0.284616434 | 0.466304782 | 1.63836212 | 8.74E-05 | 0.00092786 |
| Gipc1 | 0.280821279 | 0.135050799 | 0.4809137 | 8.78E-05 | 0.00093218 |

| | | | | | |
|---|---|---|---|---|---|
| Akr1c19 | 0.062009258 | 0.007971508 | 0.12855352 | 9.02E-05 | 0.00095558 |
| Rapgef4 | 0.061881197 | 0.005084571 | 0.08216665 | 9.02E-05 | 0.00095558 |
| Ddost | 2.146912937 | 1.674069382 | 0.77975653 | 9.04E-05 | 0.00095766 |
| Tmed4 | 0.518302472 | 0.315042471 | 0.60783517 | 9.08E-05 | 0.00096126 |
| Tprgl | 0.203770844 | 0.083173105 | 0.4081698 | 9.16E-05 | 0.00096922 |
| Sgta | 0.261213147 | 0.119963603 | 0.45925561 | 9.25E-05 | 0.00097785 |
| Mtch2 | 0.579502439 | 0.360120933 | 0.62143126 | 9.30E-05 | 0.00098311 |
| Nipsnap3b | 0.204954551 | 0.084611183 | 0.412829 | 9.34E-05 | 0.00098595 |
| Golim4 | 0.261441108 | 0.126952349 | 0.48558679 | 9.35E-05 | 0.00098595 |
| Jakmip1 | 0.064339893 | 0.004356355 | 0.06770846 | 9.36E-05 | 0.00098595 |
| Sstr3 | 0.064781896 | 0.003259658 | 0.05031743 | 9.36E-05 | 0.00098595 |
| Pde1c | 0.06392657 | 0.004111494 | 0.06431588 | 9.36E-05 | 0.00098595 |
| Metap2 | 0.750773185 | 1.038530769 | 1.38328165 | 9.59E-05 | 0.00100932 |
| Card19 | 0.171452495 | 0.061440391 | 0.35835227 | 9.62E-05 | 0.0010117 |
| Rab18 | 0.31569795 | 0.163716307 | 0.51858527 | 9.64E-05 | 0.00101336 |
| Eif2s3x | 0.126836387 | 0.255787834 | 2.0166755 | 9.67E-05 | 0.00101591 |
| Psat1 | 0.100554175 | 0.214765611 | 2.13581994 | 9.73E-05 | 0.00102185 |
| Kdelr3 | 0.061377343 | 0.156633312 | 2.55197282 | 9.77E-05 | 0.00102504 |
| Dcaf8 | 0.272011315 | 0.130924265 | 0.48131919 | 9.78E-05 | 0.00102565 |
| Myef2 | 0.162635629 | 0.057527859 | 0.35372237 | 9.80E-05 | 0.00102687 |
| Coq10b | 0.117526793 | 0.030545046 | 0.25989858 | 9.82E-05 | 0.00102767 |
| Gprasp1 | 0.117415038 | 0.030634488 | 0.26090771 | 9.82E-05 | 0.00102767 |
| Spink4 | 0.116849236 | 0.030288506 | 0.25921014 | 9.82E-05 | 0.00102767 |
| Psmc4 | 0.522108515 | 0.323838402 | 0.62025114 | 9.83E-05 | 0.00102818 |
| Bbx | 0.196123799 | 0.079481735 | 0.40526308 | 9.92E-05 | 0.00103694 |
| Psmb7 | 0.928700806 | 0.654854194 | 0.70512935 | 9.95E-05 | 0.00103959 |
| Nit1 | 0.117625323 | 0.031457685 | 0.26743973 | 0.0001008 | 0.00105218 |
| 2010320M18 | 0.153475812 | 0.051941271 | 0.33843294 | 0.00010107 | 0.00105374 |
| Gm10093 | 0.153702157 | 0.051353603 | 0.33411114 | 0.00010107 | 0.00105374 |
| Rbm25 | 0.696971716 | 0.461473308 | 0.66211196 | 0.00010174 | 0.00106014 |

| | | | | | |
|---|---|---|---|---|---|
| Vat1 | 0.127923261 | 0.260483371 | 2.03624711 | 0.00010292 | 0.00107179 |
| C1rl | 0.004132803 | 0.043006263 | 10.4060752 | 0.00010307 | 0.0010726 |
| Dusp18 | 0.091011485 | 0.017223033 | 0.18924021 | 0.00010408 | 0.00108252 |
| Fmnl2 | 0.049384759 | 0.001130544 | 0.02289256 | 0.00010429 | 0.00108337 |
| Pappa2 | 0.048671975 | 0.00063061 | 0.01295633 | 0.00010429 | 0.00108337 |
| Morc4 | 0.021874663 | 0.08792888 | 4.01966787 | 0.00010489 | 0.00108898 |
| Rrp1 | 0.570126283 | 0.359433878 | 0.63044608 | 0.00010529 | 0.00109249 |
| As3mt | 0.140002783 | 0.045107885 | 0.32219277 | 0.00010622 | 0.00110144 |
| Unc50 | 0.297768812 | 0.147821239 | 0.49642956 | 0.00010647 | 0.00110333 |
| Cep170 | 0.092372604 | 0.021072997 | 0.22813037 | 0.00010731 | 0.00111078 |
| Agt | 0.093121758 | 0.017827984 | 0.19144811 | 0.00010732 | 0.00111078 |
| Chkb | 0.134253905 | 0.036764904 | 0.27384607 | 0.00010847 | 0.00112206 |
| Banf1 | 0.387891479 | 0.596619455 | 1.53810921 | 0.00010914 | 0.00112833 |
| Hdac2 | 0.38119468 | 0.210413607 | 0.55198464 | 0.00010937 | 0.00113 |
| Relt | 0.050307457 | 0.003319007 | 0.06597445 | 0.00010948 | 0.00113044 |
| Atxn10 | 0.688788036 | 0.455172911 | 0.66083162 | 0.00011445 | 0.00118103 |
| Hnrnpab | 0.587986312 | 0.835276847 | 1.42057192 | 0.00011452 | 0.00118103 |
| Acacb | 0.007969474 | 0.059688603 | 7.48965432 | 0.00011662 | 0.00120059 |
| Scml4 | 0.007974411 | 0.059438471 | 7.4536501 | 0.00011662 | 0.00120059 |
| Thbs1 | 0.008747722 | 0.054201283 | 6.19604543 | 0.00011662 | 0.00120059 |
| Mab21l3 | 0.004536813 | 0.050114766 | 11.0462495 | 0.00011737 | 0.0012076 |
| Cat | 0.138770654 | 0.269679639 | 1.94334775 | 0.00011775 | 0.00121073 |
| Serbp1 | 1.633981913 | 2.071690265 | 1.26787833 | 0.00011816 | 0.00121427 |
| Romo1 | 1.456698169 | 1.850167492 | 1.2701104 | 0.00011835 | 0.0012155 |
| Gale | 0.171507247 | 0.31595746 | 1.84223971 | 0.00011856 | 0.00121689 |
| Cct3 | 0.817586609 | 0.561757257 | 0.68709205 | 0.00012048 | 0.00123587 |
| Arhgef38 | 0.007221589 | 0.053280851 | 7.37799594 | 0.00012075 | 0.00123724 |
| Ssfa2 | 0.032886142 | 0.105309029 | 3.20223115 | 0.00012076 | 0.00123724 |
| Slc39a6 | 0.099612048 | 0.023195911 | 0.2328625 | 0.00012153 | 0.00124437 |
| Puf60 | 0.492220546 | 0.296732406 | 0.60284441 | 0.00012228 | 0.0012514 |

| | | | | | |
|---|---|---|---|---|---|
| Fkbp3 | 0.748605025 | 0.500535494 | 0.66862428 | 0.00012379 | 0.00126606 |
| Pnisr | 0.359673752 | 0.201313545 | 0.55971153 | 0.0001241 | 0.00126845 |
| Eif4g3 | 0.198910652 | 0.087167953 | 0.43822667 | 0.00012458 | 0.00127263 |
| Fam32a | 0.50735266 | 0.31087807 | 0.61274552 | 0.00012598 | 0.00128611 |
| Abcc5 | 0.081065855 | 0.015222789 | 0.187783 | 0.00012696 | 0.00129456 |
| Mapk8ip2 | 0.080800761 | 0.012652962 | 0.15659458 | 0.00012696 | 0.00129456 |
| Mogs | 0.073950606 | 0.176593437 | 2.38799176 | 0.00012703 | 0.00129456 |
| Polr2e | 0.422960694 | 0.636137849 | 1.50401174 | 0.00012916 | 0.00131481 |
| Atp1a1 | 0.6811218 | 0.453256371 | 0.66545568 | 0.00012917 | 0.00131481 |
| Rabep1 | 0.165742473 | 0.059655061 | 0.35992621 | 0.00012934 | 0.00131573 |
| Mettl21b | 0.025418714 | 0.093458419 | 3.67675647 | 0.00012961 | 0.00131774 |
| Mdh2 | 1.472324685 | 1.127992847 | 0.7661305 | 0.0001308 | 0.00132904 |
| Otub1 | 0.400120305 | 0.227746317 | 0.5691946 | 0.00013097 | 0.00132993 |
| Actr3 | 0.495046961 | 0.303211136 | 0.61248964 | 0.00013198 | 0.00133946 |
| Scrn1 | 0.083345968 | 0.013444053 | 0.16130417 | 0.00013293 | 0.00134674 |
| Tmem38a | 0.082172501 | 0.014295647 | 0.17397118 | 0.00013293 | 0.00134674 |
| Spa17 | 0.083006786 | 0.011506643 | 0.13862292 | 0.00013293 | 0.00134674 |
| Armcx1 | 0.107289946 | 0.025136843 | 0.23428889 | 0.00013414 | 0.00135739 |
| Ovol2 | 0.106186873 | 0.025756609 | 0.24255926 | 0.00013414 | 0.00135739 |
| S100a13 | 0.192918733 | 0.077941095 | 0.40400999 | 0.00013465 | 0.00136173 |
| Eif3b | 0.314732513 | 0.501689703 | 1.59401931 | 0.00013697 | 0.00138435 |
| Zdhhc2 | 0.107446823 | 0.0265808 | 0.24738563 | 0.00013827 | 0.00139666 |
| Psmd6 | 0.591213572 | 0.376324316 | 0.63652855 | 0.00013901 | 0.00140333 |
| Fam103a1 | 0.341911862 | 0.182942295 | 0.53505688 | 0.00014161 | 0.00142877 |
| Hmga2 | 0.081336909 | 0.1843412 | 2.26639052 | 0.000142 | 0.00143185 |
| Rdh14 | 0.152252741 | 0.052567967 | 0.34526779 | 0.0001424 | 0.00143503 |
| Rhd | 0.060072974 | 0.004953994 | 0.08246626 | 0.00014356 | 0.00144414 |
| Ppil6 | 0.061192608 | 0.003300147 | 0.05393048 | 0.00014356 | 0.00144414 |
| B230217C12 | 0.061152591 | 0.002809794 | 0.04594725 | 0.00014356 | 0.00144414 |
| Stx1a | 0.071519273 | 0.009401582 | 0.13145523 | 0.00014399 | 0.00144429 |

| | | | | | |
|---|---|---|---|---|---|
| Fam219a | 0.071603086 | 0.009113351 | 0.12727595 | 0.00014399 | 0.00144429 |
| Dcx | 0.070516404 | 0.009618395 | 0.1363994 | 0.00014399 | 0.00144429 |
| Cd79a | 0.071665287 | 0.007952495 | 0.11096719 | 0.00014399 | 0.00144429 |
| Phyhipl | 0.070814438 | 0.007983876 | 0.11274361 | 0.00014399 | 0.00144429 |
| Dnajb2 | 0.145954735 | 0.048829704 | 0.33455375 | 0.00014928 | 0.0014956 |
| Fam135a | 0.144779752 | 0.049660047 | 0.34300409 | 0.00014928 | 0.0014956 |
| Uchl5 | 0.186086152 | 0.3331749 | 1.79043361 | 0.00015026 | 0.00150422 |
| Trabd | 0.307419392 | 0.492189748 | 1.60103676 | 0.00015037 | 0.00150422 |
| 2810428I15R | 0.460780805 | 0.67903016 | 1.47365114 | 0.0001504 | 0.00150422 |
| Ift74 | 0.128959386 | 0.036394076 | 0.28221347 | 0.00015073 | 0.00150661 |
| Unc119 | 0.122847499 | 0.036124275 | 0.29405788 | 0.00015364 | 0.00153387 |
| Phldb2 | 0.124177849 | 0.033863447 | 0.27270119 | 0.00015364 | 0.00153387 |
| Dnah9 | 0.072933362 | 0.006341825 | 0.08695369 | 0.0001549 | 0.00154558 |
| Tapbp | 0.206294176 | 0.089937556 | 0.4359675 | 0.000156 | 0.00155567 |
| Tecr | 1.106067358 | 0.812312003 | 0.73441459 | 0.00015654 | 0.0015601 |
| Podxl2 | 0.089338239 | 0.017512868 | 0.19602881 | 0.00015755 | 0.00156841 |
| Dusp10 | 0.090357684 | 0.013879705 | 0.15360846 | 0.00015755 | 0.00156841 |
| Erp29 | 1.557135708 | 1.205546592 | 0.77420779 | 0.000158 | 0.00157193 |
| Aga | 0.180104563 | 0.066571278 | 0.36962571 | 0.0001581 | 0.00157202 |
| Ndfip2 | 0.335061888 | 0.1780596 | 0.53142302 | 0.00016085 | 0.00159843 |
| Rbbp6 | 0.257930011 | 0.121854099 | 0.47243087 | 0.00016265 | 0.00161541 |
| Myeov2 | 1.068937812 | 0.783491954 | 0.73296308 | 0.00016394 | 0.00162724 |
| Eif2ak3 | 0.023741244 | 0.086790002 | 3.65566361 | 0.00016547 | 0.00164153 |
| Commd1 | 0.602715107 | 0.388287556 | 0.64423067 | 0.00016683 | 0.00165402 |
| Smpd2 | 0.171178966 | 0.063911633 | 0.37336149 | 0.00016767 | 0.00166141 |
| Wtap | 0.269758424 | 0.130112922 | 0.48233126 | 0.00016976 | 0.00168021 |
| Nudt16l1 | 0.269566474 | 0.128305338 | 0.4759692 | 0.00016976 | 0.00168021 |
| Trf | 0.216777347 | 0.096395512 | 0.44467521 | 0.00017064 | 0.00168794 |
| Skap1 | 0.046992283 | 0.001525869 | 0.03247063 | 0.00017088 | 0.00168845 |
| Calml3 | 0.046533124 | 0 | 0 | 0.00017089 | 0.00168845 |

| | | | | | |
|---|---|---|---|---|---|
| Itfg1 | 0.218864913 | 0.096319775 | 0.44008779 | 0.00017288 | 0.00170711 |
| Rbm3 | 1.279134687 | 0.964147076 | 0.75374946 | 0.0001751 | 0.00172767 |
| Mecr | 0.102942308 | 0.215554235 | 2.09393242 | 0.00017523 | 0.00172767 |
| Med21 | 0.281617176 | 0.137158265 | 0.487038 | 0.00017526 | 0.00172767 |
| Nap1l4 | 0.350309956 | 0.190883606 | 0.54489918 | 0.0001775 | 0.00174872 |
| Ssbp4 | 0.248742545 | 0.119114971 | 0.47886851 | 0.00017833 | 0.00175595 |
| Nhp2l1 | 0.709137576 | 0.476866058 | 0.67245916 | 0.00017877 | 0.0017592 |
| Agpat4 | 0.095826566 | 0.023902776 | 0.24943789 | 0.00018 | 0.00177028 |
| Pnn | 0.402212516 | 0.233411101 | 0.58031785 | 0.00018094 | 0.00177857 |
| Acadvl | 0.229415039 | 0.106844109 | 0.46572408 | 0.00018765 | 0.00184342 |
| N6amt1 | 0.158140371 | 0.053022153 | 0.33528537 | 0.0001913 | 0.00187822 |
| Isyna1 | 0.473729229 | 0.290943466 | 0.61415562 | 0.00019201 | 0.00188413 |
| Mapre2 | 0.149181291 | 0.0526248 | 0.35275737 | 0.00019796 | 0.00193919 |
| Mast4 | 0.149557971 | 0.052066947 | 0.3481389 | 0.00019796 | 0.00193919 |
| Stx12 | 0.148900936 | 0.050739629 | 0.34076098 | 0.00019796 | 0.00193919 |
| Taf6l | 0.34373805 | 0.185974067 | 0.54103428 | 0.00020138 | 0.00197155 |
| Xpot | 0.044327556 | 0.121469735 | 2.74027591 | 0.00020165 | 0.00197309 |
| Cryzl1 | 0.183609691 | 0.078741785 | 0.42885419 | 0.0002019 | 0.00197438 |
| Csnk1a1 | 0.869530694 | 0.615884884 | 0.70829574 | 0.00020302 | 0.00198428 |
| Mocs2 | 0.319519383 | 0.170744425 | 0.53437893 | 0.00020404 | 0.00199303 |
| Gpx8 | 0.086294996 | 0.18663874 | 2.16279911 | 0.00020511 | 0.00200238 |
| Fign | 0.014210338 | 0.066205175 | 4.65894453 | 0.00020696 | 0.00201931 |
| Nhp2 | 0.214996446 | 0.368544621 | 1.71418937 | 0.00020724 | 0.00202091 |
| Agtrap | 0.110923898 | 0.033611558 | 0.30301458 | 0.00020761 | 0.00202217 |
| Arl6 | 0.112327246 | 0.030954366 | 0.27557308 | 0.00020761 | 0.00202217 |
| 2210016L21F | 0.251558235 | 0.123721573 | 0.4918208 | 0.00020857 | 0.00202925 |
| Hmox2 | 0.252463246 | 0.122231803 | 0.48415682 | 0.00020857 | 0.00202925 |
| Sec63 | 0.160909321 | 0.295377336 | 1.83567574 | 0.00021326 | 0.00207374 |
| Metrn | 0.028407859 | 0.096576623 | 3.39964458 | 0.00021415 | 0.00207749 |
| 2310033P09I | 0.12727046 | 0.039654015 | 0.31157281 | 0.00021427 | 0.00207749 |

| | | | | | |
|---|---|---|---|---|---|
| Ctxn1 | 0.127644297 | 0.036102035 | 0.28283312 | 0.00021427 | 0.00207749 |
| Acsl1 | 0.022741603 | 0.082582508 | 3.6313406 | 0.00021429 | 0.00207749 |
| Slc39a14 | 0.021946803 | 0.082733107 | 3.76971111 | 0.00021429 | 0.00207749 |
| Ttll7 | 0.176644226 | 0.066754321 | 0.37790265 | 0.00021437 | 0.00207749 |
| Ifi35 | 0.11397234 | 0.027114088 | 0.2379006 | 0.00021619 | 0.00209389 |
| Lsm5 | 0.389310695 | 0.588085254 | 1.51058078 | 0.00021831 | 0.0021132 |
| St5 | 0.040197051 | 0.117387725 | 2.92030687 | 0.00021938 | 0.0021224 |
| Golga4 | 0.358966709 | 0.203116821 | 0.56583749 | 0.00022006 | 0.00212777 |
| Sf3b4 | 0.277303347 | 0.140925803 | 0.50820087 | 0.0002246 | 0.00217049 |
| Map3k15 | 0.066948978 | 0.012257743 | 0.18309081 | 0.0002266 | 0.00218371 |
| Snx24 | 0.067137085 | 0.009718938 | 0.14476259 | 0.0002266 | 0.00218371 |
| Il6ra | 0.067087933 | 0.00959341 | 0.14299754 | 0.0002266 | 0.00218371 |
| Upk3bl | 0.067595788 | 0.007826461 | 0.11578326 | 0.00022661 | 0.00218371 |
| Tmod1 | 0.068193087 | 0.006493409 | 0.09522093 | 0.00022661 | 0.00218371 |
| Nsmce1 | 0.287338951 | 0.148723163 | 0.5175879 | 0.00022683 | 0.00218461 |
| Dnaja4 | 0.069063144 | 0.009628438 | 0.139415 | 0.0002293 | 0.0022072 |
| Pycr2 | 0.679021492 | 0.456085752 | 0.67168088 | 0.00023068 | 0.00221925 |
| Mphosph6 | 0.16861332 | 0.069255419 | 0.41073516 | 0.00023233 | 0.00223262 |
| Atp2c1 | 0.168660179 | 0.065446435 | 0.38803726 | 0.00023233 | 0.00223262 |
| Birc5 | 0.261368969 | 0.424397714 | 1.62374943 | 0.00023323 | 0.00223996 |
| Slc25a53 | 0.086035982 | 0.016300583 | 0.1894624 | 0.00023536 | 0.00225921 |
| Fam43a | 0.059208983 | 0.005116876 | 0.0864206 | 0.00023971 | 0.00229844 |
| Flywch2 | 0.058927609 | 0.004589904 | 0.07789055 | 0.00023972 | 0.00229844 |
| mt-Co3 | 20.40569841 | 25.59844627 | 1.25447538 | 0.00024061 | 0.00230573 |
| Gm13010 | 0.001602506 | 0.035021793 | 21.8543895 | 0.00024439 | 0.00233937 |
| B230334C09 | 0.001799817 | 0.034242594 | 19.0255978 | 0.00024439 | 0.00233937 |
| Tnfaip8 | 0.109803331 | 0.22568066 | 2.05531707 | 0.00024539 | 0.00234759 |
| Bbip1 | 0.403085989 | 0.237839638 | 0.5900469 | 0.00024596 | 0.00234952 |
| Scaf11 | 0.403181243 | 0.234010186 | 0.58040941 | 0.00024596 | 0.00234952 |
| Usf2 | 0.225928471 | 0.10947088 | 0.48453778 | 0.00024614 | 0.00234952 |

| | | | | | |
|---|---|---|---|---|---|
| Limd2 | 0.227198358 | 0.105626183 | 0.46490733 | 0.00024614 | 0.00234952 |
| Prkca | 0.087755002 | 0.016000424 | 0.18233062 | 0.00024635 | 0.00235025 |
| Naga | 0.188461673 | 0.078281497 | 0.41537091 | 0.00024717 | 0.00235674 |
| Vasp | 0.269341627 | 0.136718378 | 0.50760211 | 0.00025023 | 0.00238328 |
| Ncbp2 | 0.26948412 | 0.134644802 | 0.4996391 | 0.00025023 | 0.00238328 |
| Ppp1r15b | 0.061534107 | 0.149450666 | 2.42874517 | 0.0002517 | 0.00239466 |
| Fbxo16 | 0.060651096 | 0.147879395 | 2.43819823 | 0.0002517 | 0.00239466 |
| Thap11 | 0.161624958 | 0.063761289 | 0.3945015 | 0.00025223 | 0.00239834 |
| Lztfl1 | 0.208275429 | 0.090020488 | 0.43221847 | 0.00025421 | 0.00241581 |
| Ergic1 | 0.157395654 | 0.290638803 | 1.84654911 | 0.00025464 | 0.0024186 |
| Snrpd2 | 1.400495076 | 1.760973233 | 1.25739338 | 0.00025485 | 0.00241926 |
| Hpcal1 | 0.184571563 | 0.323850579 | 1.75460712 | 0.00025994 | 0.00246619 |
| Vkorc1 | 0.471226551 | 0.68409425 | 1.45173112 | 0.00026013 | 0.0024666 |
| Hip1r | 0.153415843 | 0.056356921 | 0.36734747 | 0.00026242 | 0.00248651 |
| Ninj1 | 0.378354732 | 0.220373948 | 0.58245326 | 0.00026252 | 0.00248651 |
| Tekt2 | 0.094798456 | 0.023799069 | 0.25104912 | 0.00026951 | 0.00254994 |
| Upp1 | 0.095419898 | 0.019374767 | 0.20304745 | 0.00026951 | 0.00254994 |
| Lrrc42 | 0.180384865 | 0.075005236 | 0.4158067 | 0.00027075 | 0.00256027 |
| Snap47 | 0.217910945 | 0.100383287 | 0.46066198 | 0.00027252 | 0.00257559 |
| Rps6ka6 | 0.01026612 | 0.056201734 | 5.47448632 | 0.00027475 | 0.00259525 |
| Akirin1 | 0.200372372 | 0.085701185 | 0.42770959 | 0.00027657 | 0.00261098 |
| Ccdc88a | 0.147510668 | 0.052808708 | 0.35799925 | 0.00027976 | 0.00263825 |
| Ankra2 | 0.14656059 | 0.052765773 | 0.36002702 | 0.00027976 | 0.00263825 |
| Csf1r | 0.04330341 | 0.003862608 | 0.08919871 | 0.00028087 | 0.00264444 |
| Gcgr | 0.043768875 | 0.002154638 | 0.04922762 | 0.00028087 | 0.00264444 |
| Ppp1r17 | 0.044241653 | 0 | 0 | 0.00028088 | 0.00264444 |
| R3hdm4 | 0.261500318 | 0.131586089 | 0.50319667 | 0.00028189 | 0.00265244 |
| Srsf7 | 0.67846936 | 0.461846002 | 0.68071755 | 0.00028694 | 0.00269852 |
| Fuca1 | 0.273999499 | 0.140951655 | 0.51442304 | 0.00028823 | 0.00270913 |
| Gm10036 | 0.151462015 | 0.280920277 | 1.85472428 | 0.00028861 | 0.00271124 |

| | | | | | |
|---|---|---|---|---|---|
| B4galnt1 | 0.027622068 | 0.090374736 | 3.27183095 | 0.000295 | 0.00276978 |
| Hmgn1 | 2.008379864 | 1.598778016 | 0.7960536 | 0.00029556 | 0.00277353 |
| Gmpr | 0.076234392 | 0.013796927 | 0.18098035 | 0.00029583 | 0.00277455 |
| Rom1 | 0.044919396 | 0.002555411 | 0.0568888 | 0.00029695 | 0.00278057 |
| 1110017D15I | 0.04501367 | 0.002054019 | 0.04563102 | 0.00029695 | 0.00278057 |
| Slurp1 | 0.04552492 | 0 | 0 | 0.00029696 | 0.00278057 |
| Tomm20 | 1.248894571 | 0.944963912 | 0.75664026 | 0.00029886 | 0.00279659 |
| Cdo1 | 0.092493261 | 0.20027004 | 2.16523925 | 0.000299 | 0.00279659 |
| 1700088E04I | 0.103337417 | 0.025200469 | 0.24386587 | 0.00030148 | 0.00281829 |
| Itpr1 | 0.10802106 | 0.032013331 | 0.29636194 | 0.00030252 | 0.00282647 |
| Wars | 0.137307268 | 0.259974743 | 1.89337934 | 0.00030309 | 0.00283028 |
| Pomp | 1.065961901 | 0.789062887 | 0.74023554 | 0.00030504 | 0.00284695 |
| Sltm | 0.360039692 | 0.206490811 | 0.57352235 | 0.00030567 | 0.0028513 |
| Bcl11b | 0.007726566 | 0.053149319 | 6.87877627 | 0.00030596 | 0.00285245 |
| Ramp2 | 0.115972138 | 0.038236293 | 0.3297024 | 0.00030631 | 0.00285416 |
| Taf1d | 0.189862916 | 0.328025293 | 1.72769543 | 0.00030674 | 0.0028566 |
| Kcnq1 | 0.030913629 | 0.095571311 | 3.0915591 | 0.00030802 | 0.00286634 |
| Rap1a | 0.535170436 | 0.342225142 | 0.63946945 | 0.00030812 | 0.00286634 |
| Cbx6 | 0.213031488 | 0.095899487 | 0.45016579 | 0.00030926 | 0.00287539 |
| Reg3d | 0.002618009 | 0.041705227 | 15.9301309 | 0.00031066 | 0.00288222 |
| Ldb3 | 0.002596862 | 0.04121373 | 15.8705887 | 0.00031066 | 0.00288222 |
| Lrrc39 | 0.003951122 | 0.039201007 | 9.92148664 | 0.00031066 | 0.00288222 |
| Pm20d1 | 0.003369007 | 0.037810827 | 11.2231363 | 0.00031067 | 0.00288222 |
| Dynlt1c | 0.076995479 | 0.014622326 | 0.18991149 | 0.0003142 | 0.00291192 |
| Slc24a5 | 0.078049362 | 0.011912463 | 0.15262729 | 0.00031421 | 0.00291192 |
| Trp53 | 0.254171035 | 0.127586129 | 0.50196958 | 0.00031583 | 0.00292538 |
| Gabarapl1 | 0.223945353 | 0.103753796 | 0.46329962 | 0.00032352 | 0.00299501 |
| Cct8 | 0.803476233 | 0.567146464 | 0.70586589 | 0.00032667 | 0.00302249 |
| Pdhb | 0.326679815 | 0.177636802 | 0.54376424 | 0.00032784 | 0.00303175 |
| Vps35 | 0.276903354 | 0.147403965 | 0.53233001 | 0.00032858 | 0.00303603 |

| | | | | | |
|---|---|---|---|---|---|
| Gfra1 | 0.006026178 | 0.050457566 | 8.37306245 | 0.00032866 | 0.00303603 |
| Uhrf2 | 0.225423878 | 0.103085175 | 0.45729483 | 0.00033048 | 0.00305123 |
| Tiprl | 0.278590007 | 0.147519422 | 0.52952159 | 0.00033121 | 0.00305633 |
| Ppid | 0.303412858 | 0.162722484 | 0.53630714 | 0.00033358 | 0.00307651 |
| Liph | 0.005781054 | 0.042606586 | 7.37003785 | 0.00033423 | 0.00307759 |
| Npw | 0.005919991 | 0.041914776 | 7.0802095 | 0.00033423 | 0.00307759 |
| 2200002J24F | 0.005410306 | 0.041595108 | 7.68812504 | 0.00033423 | 0.00307759 |
| Gm5900 | 0.18544275 | 0.081171266 | 0.43771604 | 0.00033821 | 0.00311085 |
| Fam174a | 0.186187719 | 0.07703745 | 0.41376225 | 0.00033821 | 0.00311085 |
| H2afj | 0.803300687 | 1.071790607 | 1.3342334 | 0.0003407 | 0.00313211 |
| Sac3d1 | 0.158515695 | 0.060703021 | 0.38294644 | 0.00034174 | 0.00314 |
| Mrps34 | 0.377798509 | 0.562817185 | 1.48972845 | 0.00034487 | 0.00316706 |
| Lamtor5 | 0.421685594 | 0.251668303 | 0.59681504 | 0.00034725 | 0.00318724 |
| Hspa2 | 0.065694363 | 0.010226409 | 0.15566646 | 0.00034854 | 0.00319735 |
| Sf3b2 | 0.697804067 | 0.477041559 | 0.68363253 | 0.00034975 | 0.00320674 |
| Btg1 | 0.464682852 | 0.289498688 | 0.62300274 | 0.0003513 | 0.00321919 |
| Tma7 | 2.219618604 | 1.771160966 | 0.79795734 | 0.0003525 | 0.00322846 |
| Cdkn1a | 0.245024932 | 0.398450892 | 1.62616469 | 0.00035671 | 0.00326532 |
| Tox | 0.084037777 | 0.020231331 | 0.24074091 | 0.00035794 | 0.00327483 |
| Auh | 0.248011663 | 0.126272071 | 0.50913763 | 0.00036088 | 0.0033 |
| Snx4 | 0.216903651 | 0.100635038 | 0.46396194 | 0.00036136 | 0.00330256 |
| Arl2 | 0.258969995 | 0.131118723 | 0.50630855 | 0.00036396 | 0.00332281 |
| Sucla2 | 0.258883539 | 0.130938948 | 0.50578321 | 0.00036396 | 0.00332281 |
| Clip3 | 0.05609366 | 0.005727034 | 0.10209771 | 0.00036515 | 0.0033302 |
| Meox1 | 0.055307519 | 0.004158307 | 0.07518521 | 0.00036516 | 0.0033302 |
| Creld1 | 0.153014709 | 0.056735315 | 0.3707834 | 0.00037058 | 0.00337605 |
| Phf23 | 0.152758328 | 0.054405028 | 0.35615098 | 0.00037058 | 0.00337605 |
| Trappc10 | 0.04067964 | 0.11538955 | 2.83654303 | 0.000372 | 0.00338726 |
| Pex7 | 0.197500091 | 0.090117611 | 0.45629149 | 0.0003742 | 0.00340543 |
| Fam3a | 0.144100866 | 0.051447138 | 0.35702171 | 0.00038483 | 0.00349849 |

| | | | | | |
|---|---|---|---|---|---|
| Myo6 | 0.144048506 | 0.051130057 | 0.35495028 | 0.00038483 | 0.00349849 |
| Yap1 | 0.018607905 | 0.075911116 | 4.07950893 | 0.0003858 | 0.00350362 |
| Fgfrl1 | 0.018316037 | 0.074137881 | 4.04770321 | 0.0003858 | 0.00350362 |
| Dnajb11 | 0.862655147 | 0.620628437 | 0.71943979 | 0.00038799 | 0.00352165 |
| Cd52 | 0.091201131 | 0.023848046 | 0.26148849 | 0.00039456 | 0.00357749 |
| Tmx4 | 0.091322936 | 0.023566657 | 0.25805847 | 0.00039456 | 0.00357749 |
| Fpgs | 0.045336879 | 0.125661512 | 2.77172835 | 0.0003964 | 0.00359041 |
| Prps2 | 0.045286853 | 0.122571865 | 2.7065662 | 0.0003964 | 0.00359041 |
| Acadsb | 0.091530612 | 0.190841739 | 2.0850045 | 0.00040243 | 0.00364309 |
| Larp4 | 0.119413965 | 0.235290024 | 1.97037275 | 0.00040359 | 0.00365161 |
| Ndufv1 | 0.360121466 | 0.542858142 | 1.50743067 | 0.00040413 | 0.00365462 |
| Slc48a1 | 0.189696779 | 0.080723408 | 0.42553916 | 0.00040757 | 0.00368374 |
| Galnt11 | 0.092632206 | 0.024501929 | 0.26450767 | 0.00041511 | 0.00374795 |
| Npnt | 0.093241595 | 0.023623643 | 0.25335949 | 0.00041511 | 0.00374795 |
| Tyms | 0.189869318 | 0.325806559 | 1.71595159 | 0.0004169 | 0.00376213 |
| Maoa | 0.02530173 | 0.087842941 | 3.47181565 | 0.0004182 | 0.00376993 |
| Ric1 | 0.025773907 | 0.086486982 | 3.35560225 | 0.0004182 | 0.00376993 |
| Rtcb | 0.33667013 | 0.515144248 | 1.53011569 | 0.00042027 | 0.00378536 |
| Usp29 | 0.128142706 | 0.047179988 | 0.36818317 | 0.00042035 | 0.00378536 |
| Tmem30a | 0.299045007 | 0.162128467 | 0.54215407 | 0.00042207 | 0.00379882 |
| Rora | 0.097537696 | 0.026108098 | 0.26767187 | 0.00042333 | 0.00380823 |
| Tes | 0.219516207 | 0.108104764 | 0.49246826 | 0.0004259 | 0.00382532 |
| Scamp2 | 0.220116532 | 0.105729492 | 0.48033417 | 0.0004259 | 0.00382532 |
| Grina | 0.21943939 | 0.104752606 | 0.47736464 | 0.0004259 | 0.00382532 |
| Pcolce | 0.099383614 | 0.02672851 | 0.26894282 | 0.00042804 | 0.00384255 |
| Abhd8 | 0.130500416 | 0.043210158 | 0.33111127 | 0.00043486 | 0.00390034 |
| 1810008I18R | 0.122501246 | 0.038762932 | 0.31642888 | 0.00043493 | 0.00390034 |
| Spcs1 | 2.287005516 | 1.83327489 | 0.80160493 | 0.00044145 | 0.00395674 |
| Slc4a7 | 0.115148251 | 0.033857776 | 0.29403639 | 0.00044815 | 0.00401331 |
| Creg1 | 0.233612087 | 0.381106718 | 1.63136558 | 0.00044823 | 0.00401331 |

| | | | | | |
|---|---|---|---|---|---|
| Pcbp3 | 0.072970108 | 0.014749745 | 0.20213407 | 0.00045427 | 0.00405898 |
| Prkar2b | 0.072146302 | 0.014892293 | 0.20641796 | 0.00045428 | 0.00405898 |
| Slc25a14 | 0.072499248 | 0.014448238 | 0.19928811 | 0.00045428 | 0.00405898 |
| B3galnt1 | 0.073226073 | 0.012133509 | 0.1656993 | 0.00045428 | 0.00405898 |
| Alg2 | 0.203859755 | 0.088152571 | 0.43241772 | 0.0004548 | 0.00406154 |
| AI467606 | 0.073666856 | 0.014804136 | 0.2009606 | 0.00045938 | 0.00409822 |
| Aqp11 | 0.073799779 | 0.013520158 | 0.18320052 | 0.00045939 | 0.00409822 |
| Uba5 | 0.316483633 | 0.485444756 | 1.53387002 | 0.00045994 | 0.00410106 |
| Cacnb2 | 0.042143788 | 0.001937944 | 0.04598409 | 0.00046258 | 0.00411615 |
| Lingo1 | 0.042156247 | 0.001766158 | 0.04189552 | 0.00046258 | 0.00411615 |
| Tcp11 | 0.042321264 | 0 | 0 | 0.00046259 | 0.00411615 |
| Fbxl2 | 0.041711974 | 0.000593961 | 0.01423958 | 0.00046259 | 0.00411615 |
| Pigx | 0.156269529 | 0.061754082 | 0.39517673 | 0.00046511 | 0.00413216 |
| Ddit3 | 0.154911096 | 0.062019671 | 0.40035654 | 0.00046511 | 0.00413216 |
| Twf2 | 0.156015149 | 0.060912724 | 0.39042827 | 0.00046511 | 0.00413216 |
| Dtnb | 0.057990632 | 0.140383619 | 2.42079822 | 0.00046886 | 0.00416327 |
| Smim20 | 0.255474573 | 0.131206346 | 0.51357888 | 0.00047033 | 0.00417199 |
| Dbnl | 0.255401897 | 0.130664173 | 0.5116022 | 0.00047033 | 0.00417199 |
| Vta1 | 0.212575398 | 0.100687854 | 0.47365714 | 0.00047491 | 0.00421044 |
| Gsk3b | 0.355634178 | 0.205392643 | 0.5775391 | 0.00047753 | 0.0042315 |
| Cyba | 0.400104139 | 0.587255912 | 1.46775765 | 0.00048284 | 0.00427634 |
| Zfp771 | 0.087380923 | 0.184032126 | 2.10609045 | 0.00049295 | 0.00436363 |
| Bcl7b | 0.175178477 | 0.076639643 | 0.43749463 | 0.00049889 | 0.00441247 |
| Vps72 | 0.224582482 | 0.111588145 | 0.49686932 | 0.00049898 | 0.00441247 |
| Snapc5 | 0.147393476 | 0.05558591 | 0.37712599 | 0.00050075 | 0.00442362 |
| Tcn2 | 0.147475395 | 0.054870631 | 0.37206634 | 0.00050075 | 0.00442362 |
| Wdfy1 | 0.149733093 | 0.056272516 | 0.37581883 | 0.00050318 | 0.00443826 |
| B4galt6 | 0.148657896 | 0.056097795 | 0.37736169 | 0.00050318 | 0.00443826 |
| Slc25a37 | 0.148931934 | 0.055573409 | 0.37314636 | 0.00050318 | 0.00443826 |
| Tra2a | 0.372881258 | 0.21917667 | 0.58779213 | 0.00050741 | 0.00447321 |

| | | | | | |
|---|---|---|---|---|---|
| Ubl5 | 1.945656976 | 1.560566481 | 0.80207688 | 0.00050999 | 0.00449366 |
| Psmd8 | 0.642688799 | 0.437759347 | 0.68113735 | 0.00052017 | 0.00458101 |
| Ciapin1 | 0.416435567 | 0.254403929 | 0.61090826 | 0.00052581 | 0.00462835 |
| Yipf6 | 0.118726321 | 0.22745525 | 1.91579465 | 0.00052753 | 0.00464108 |
| Pde3b | 0.080454968 | 0.01980837 | 0.24620443 | 0.00052859 | 0.00464799 |
| Swi5 | 2.078731268 | 1.673174348 | 0.80490171 | 0.00053304 | 0.00468476 |
| Tmem33 | 0.260027164 | 0.137300327 | 0.52802301 | 0.00053686 | 0.00471593 |
| Hmg20b | 0.250078772 | 0.127978592 | 0.51175312 | 0.0005387 | 0.00472968 |
| Arhgef17 | 0.06254233 | 0.011052952 | 0.17672754 | 0.0005426 | 0.00475667 |
| Sobp | 0.062114431 | 0.01025503 | 0.16509899 | 0.00054261 | 0.00475667 |
| Sh2d3c | 0.062414893 | 0.009431845 | 0.15111529 | 0.00054261 | 0.00475667 |
| Mad2l2 | 0.141848656 | 0.051732823 | 0.36470436 | 0.0005449 | 0.00477436 |
| Wwc2 | 0.026647186 | 0.08958444 | 3.36187244 | 0.00054848 | 0.00480085 |
| Rhoq | 0.026546941 | 0.087989873 | 3.31450132 | 0.00054849 | 0.00480085 |
| Rnaseh2a | 0.134647846 | 0.251370497 | 1.86687351 | 0.0005508 | 0.00481731 |
| Sgpp1 | 0.059046224 | 0.142074881 | 2.4061637 | 0.00055093 | 0.00481731 |
| Glis3 | 0.064806179 | 0.007968323 | 0.12295623 | 0.00055754 | 0.00487023 |
| Bend7 | 0.064110327 | 0.007984659 | 0.1245456 | 0.00055754 | 0.00487023 |
| Naca | 3.996201779 | 4.89862627 | 1.22582055 | 0.00056959 | 0.00497288 |
| Hes6 | 0.324917216 | 0.182776378 | 0.56253214 | 0.00057498 | 0.00500912 |
| Nefm | 0.052082396 | 0.006106366 | 0.11724435 | 0.00057518 | 0.00500912 |
| Lhx1 | 0.052421901 | 0.00484432 | 0.09241023 | 0.00057518 | 0.00500912 |
| Slc16a9 | 0.052651101 | 0.002201717 | 0.04181712 | 0.00057519 | 0.00500912 |
| Rims3 | 0.052651029 | 0.002201717 | 0.04181718 | 0.00057519 | 0.00500912 |
| Sh3bgrl2 | 0.01150807 | 0.056519334 | 4.91127839 | 0.00058447 | 0.0050814 |
| Mpp3 | 0.087814293 | 0.023134963 | 0.26345327 | 0.00058467 | 0.0050814 |
| Fam177a | 0.088208438 | 0.022382328 | 0.25374362 | 0.00058467 | 0.0050814 |
| Atg10 | 0.088042334 | 0.020466755 | 0.23246493 | 0.00058468 | 0.0050814 |
| Btf3l4 | 0.451613385 | 0.287317479 | 0.63620231 | 0.00059064 | 0.00513061 |
| Arxes1 | 0.089710215 | 0.020570527 | 0.22929972 | 0.00059298 | 0.00514831 |

| | | | | | |
|---|---|---|---|---|---|
| Cops3 | 0.351183169 | 0.207755675 | 0.59158779 | 0.00059407 | 0.00515518 |
| Bclaf1 | 0.407499639 | 0.249913592 | 0.61328543 | 0.00059489 | 0.00515782 |
| Hnrnpa0 | 0.17175229 | 0.299525785 | 1.74394056 | 0.00059497 | 0.00515782 |
| Lpp | 0.241430177 | 0.123081566 | 0.50980191 | 0.00059769 | 0.00517876 |
| Ppp1cc | 0.468455457 | 0.29758694 | 0.63525131 | 0.000598 | 0.00517884 |
| Dhps | 0.127247115 | 0.043856991 | 0.34466 | 0.00059841 | 0.00517982 |
| Igsf8 | 0.178615708 | 0.081155224 | 0.45435659 | 0.00060136 | 0.00520272 |
| Ahsa1 | 0.534048412 | 0.353554976 | 0.66202795 | 0.00060523 | 0.0052327 |
| Ddx3x | 0.344530663 | 0.519576966 | 1.50807177 | 0.00060544 | 0.0052327 |
| Gpc4 | 0.117792485 | 0.043232354 | 0.36702132 | 0.00061173 | 0.0052725 |
| Gt(ROSA)26 | 0.117989949 | 0.040144145 | 0.3402336 | 0.00061173 | 0.0052725 |
| Sccpdh | 0.11909941 | 0.037452834 | 0.314467 | 0.00061173 | 0.0052725 |
| Lrrc16b | 0.054109813 | 0.004528845 | 0.0836973 | 0.00061218 | 0.0052725 |
| Gpd1 | 0.054453107 | 0.004064224 | 0.07463713 | 0.00061219 | 0.0052725 |
| Lrfn3 | 0.054604049 | 0.003644001 | 0.066735 | 0.00061219 | 0.0052725 |
| Gdap1 | 0.054269598 | 0.003611631 | 0.06654981 | 0.00061219 | 0.0052725 |
| Psme1 | 0.676820979 | 0.47104569 | 0.69596792 | 0.00061371 | 0.00528297 |
| Kctd5 | 0.096371826 | 0.026659664 | 0.27663338 | 0.0006147 | 0.00528358 |
| Rhbdd2 | 0.097202523 | 0.024557787 | 0.25264557 | 0.00061471 | 0.00528358 |
| Ap2a1 | 0.09691097 | 0.024568978 | 0.25352113 | 0.00061471 | 0.00528358 |
| Fbxw2 | 0.180546354 | 0.080571632 | 0.44626563 | 0.00061687 | 0.00529956 |
| Klhdc10 | 0.065371574 | 0.148534448 | 2.27215651 | 0.00062241 | 0.00534448 |
| Alkbh6 | 0.210522361 | 0.099341247 | 0.47187979 | 0.00062501 | 0.00536412 |
| Rassf7 | 0.103185559 | 0.031398624 | 0.30429282 | 0.00062706 | 0.00537899 |
| Sparc | 1.946372621 | 1.563840722 | 0.8034642 | 0.00062862 | 0.0053827 |
| Uckl1 | 0.111246099 | 0.035956035 | 0.32321165 | 0.00062874 | 0.0053827 |
| Kctd12 | 0.111162493 | 0.035007355 | 0.31492057 | 0.00062874 | 0.0053827 |
| Rnf114 | 0.111451654 | 0.033729566 | 0.30263854 | 0.00062874 | 0.0053827 |
| Unc45a | 0.120678987 | 0.037469037 | 0.31048518 | 0.00063336 | 0.00541955 |
| Pdlim7 | 0.104131921 | 0.031249609 | 0.30009635 | 0.00064224 | 0.00548969 |

| | | | | | |
|---|---|---|---|---|---|
| Uck1 | 0.104466861 | 0.030471915 | 0.29168978 | 0.00064224 | 0.00548969 |
| Prelid2 | 0.040704135 | 0.107477136 | 2.64044761 | 0.00064252 | 0.00548969 |
| Yipf5 | 0.27258669 | 0.424027374 | 1.55556889 | 0.00064742 | 0.00552878 |
| Slc7a1 | 0.04485635 | 0.119643181 | 2.66725182 | 0.00065245 | 0.005569 |
| Lyrm9 | 0.114291482 | 0.219632545 | 1.92168778 | 0.00066382 | 0.00566042 |
| Rnf187 | 0.114798826 | 0.218205263 | 1.90076216 | 0.00066382 | 0.00566042 |
| Tex264 | 0.172903534 | 0.072483199 | 0.41921179 | 0.00066493 | 0.0056671 |
| Vamp4 | 0.192456412 | 0.088452735 | 0.45959879 | 0.00066956 | 0.00570368 |
| Nucb1 | 0.447039058 | 0.280298821 | 0.62701193 | 0.00067531 | 0.00574986 |
| Camk2g | 0.070681589 | 0.014682032 | 0.20772075 | 0.0006849 | 0.00582857 |
| Bet1 | 0.424632981 | 0.612296079 | 1.44194188 | 0.000687 | 0.00584359 |
| Hnrnph3 | 0.270027033 | 0.14643438 | 0.54229526 | 0.00069494 | 0.00590819 |
| Gstp1 | 0.062057321 | 0.14806518 | 2.38594217 | 0.00070576 | 0.00599426 |
| Comtd1 | 0.062628281 | 0.144598751 | 2.30884115 | 0.00070576 | 0.00599426 |
| Sf3b1 | 0.637320919 | 0.440511633 | 0.6911928 | 0.00072781 | 0.00617849 |
| Slc25a17 | 0.184284632 | 0.084635448 | 0.45926482 | 0.00072893 | 0.00618493 |
| Egln3 | 0.037731023 | 0.102432947 | 2.71482032 | 0.00073699 | 0.00625025 |
| Gpatch8 | 0.137211703 | 0.0539281 | 0.39302843 | 0.00073845 | 0.00625955 |
| Nt5m | 0.139503236 | 0.052365476 | 0.37537105 | 0.00074171 | 0.00628093 |
| Armc10 | 0.13849018 | 0.051109964 | 0.36905118 | 0.00074171 | 0.00628093 |
| Hbp1 | 0.216040571 | 0.106227573 | 0.49170196 | 0.00074476 | 0.0063037 |
| Ndufa7 | 1.394883951 | 1.096431984 | 0.78603814 | 0.00074562 | 0.00630788 |
| Tle6 | 0.144257236 | 0.257216706 | 1.78304198 | 0.00075111 | 0.0063512 |
| Atp6v1d | 0.297285455 | 0.169507687 | 0.57018493 | 0.00075386 | 0.00637129 |
| Tm9sf3 | 0.479352059 | 0.672165551 | 1.40223775 | 0.00075437 | 0.00637245 |
| Slc25a1 | 0.126564603 | 0.238870135 | 1.8873376 | 0.00075532 | 0.00637733 |
| Nod1 | 0.038350897 | 0.002410027 | 0.06284148 | 0.00075798 | 0.00637815 |
| Sdc3 | 0.039185344 | 0.001451973 | 0.03705398 | 0.00075798 | 0.00637815 |
| Hs3st6 | 0.038704136 | 0.000866895 | 0.02239799 | 0.000758 | 0.00637815 |
| Gm15706 | 0.039365381 | 0 | 0 | 0.000758 | 0.00637815 |

| | | | | | |
|---|---|---|---|---|---|
| Cnn1 | 0.03810764 | 0.001073839 | 0.0281791 | 0.000758 | 0.00637815 |
| 4930550C14I | 0.038626485 | 0 | 0 | 0.00075801 | 0.00637815 |
| Tmigd3 | 0.038578851 | 0 | 0 | 0.00075801 | 0.00637815 |
| Fndc3a | 0.471064802 | 0.299312016 | 0.63539457 | 0.00077278 | 0.00649921 |
| Nsd1 | 0.274870811 | 0.148930068 | 0.54181842 | 0.00077576 | 0.00652111 |
| Notch1 | 0.053157036 | 0.127132323 | 2.39163677 | 0.00077892 | 0.00654449 |
| Cyfip2 | 0.130427084 | 0.046869745 | 0.35935592 | 0.000783 | 0.00657555 |
| Serpina16 | 0.00220077 | 0.029840652 | 13.5591882 | 0.00078434 | 0.00658045 |
| Hba-x | 0.001651696 | 0.029226549 | 17.6948719 | 0.00078435 | 0.00658045 |
| Rnf113a2 | 0.077055933 | 0.020036538 | 0.26002589 | 0.00079172 | 0.00662936 |
| Fmn2 | 0.078216768 | 0.017355764 | 0.22189314 | 0.00079172 | 0.00662936 |
| Wdsub1 | 0.07772159 | 0.017465591 | 0.22471994 | 0.00079172 | 0.00662936 |
| Sumf1 | 0.078421445 | 0.015534083 | 0.19808463 | 0.00079173 | 0.00662936 |
| Fth1 | 6.822097978 | 5.556545874 | 0.81449224 | 0.00080474 | 0.00673506 |
| Hpn | 0.076269066 | 0.163498951 | 2.14371252 | 0.00080564 | 0.00673928 |
| Gad1 | 0.041337991 | 0 | 0 | 0.00080738 | 0.00674733 |
| Dapl1 | 0.041047927 | 0 | 0 | 0.00080739 | 0.00674733 |
| Vim | 0.745089556 | 0.528175234 | 0.70887483 | 0.00080857 | 0.00675394 |
| Pdha1 | 0.146190487 | 0.257714009 | 1.76286443 | 0.00081654 | 0.00681724 |
| Rnf5 | 0.459241191 | 0.289988745 | 0.63145195 | 0.00082007 | 0.00684335 |
| S100a9 | 0.033174383 | 0.100209151 | 3.02067871 | 0.00083588 | 0.00696858 |
| Slc12a2 | 0.034172549 | 0.094117193 | 2.75417537 | 0.00083589 | 0.00696858 |
| Pou3f4 | 0.060226116 | 0.010765014 | 0.17874329 | 0.00083914 | 0.006989 |
| Fgf12 | 0.06039191 | 0.0087051 | 0.14414348 | 0.00083915 | 0.006989 |
| Polr3k | 0.313370158 | 0.472765357 | 1.5086483 | 0.00085214 | 0.00709374 |
| Tuba1b | 1.597513783 | 1.281126803 | 0.80195039 | 0.00085605 | 0.00712285 |
| Sf3b5 | 0.474564669 | 0.666683699 | 1.40483214 | 0.00085776 | 0.00712859 |
| Cit | 0.086062658 | 0.02276284 | 0.26449149 | 0.00085799 | 0.00712859 |
| Samd10 | 0.085502606 | 0.022640672 | 0.26479511 | 0.00085799 | 0.00712859 |
| Dnajc19 | 0.35320952 | 0.521042408 | 1.47516525 | 0.0008596 | 0.00713855 |

| | | | | | |
|---|---|---|---|---|---|
| Trappc5 | 0.166374026 | 0.284670656 | 1.71102823 | 0.00086649 | 0.00719232 |
| Atp6ap1 | 0.33052443 | 0.192172943 | 0.58141827 | 0.00086905 | 0.00721008 |
| Tra2b | 0.580498634 | 0.39514832 | 0.680705 | 0.0008695 | 0.00721031 |
| Ube2d1 | 0.116151081 | 0.041496866 | 0.3572663 | 0.00087489 | 0.00724797 |
| Avl9 | 0.116997883 | 0.040080451 | 0.34257416 | 0.00087489 | 0.00724797 |
| Atoh8 | 0.007308343 | 0.042790411 | 5.85500867 | 0.00087635 | 0.00725657 |
| Mib1 | 0.106391503 | 0.207959297 | 1.95466076 | 0.00087949 | 0.00727909 |
| Cebpzos | 0.243399237 | 0.12599372 | 0.51764221 | 0.0008814 | 0.00729139 |
| Lman2l | 0.108505831 | 0.036475151 | 0.33615844 | 0.00088882 | 0.00734924 |
| Snx14 | 0.100800546 | 0.032913317 | 0.32651923 | 0.0009059 | 0.00748329 |
| Trp53bp1 | 0.101184993 | 0.029962645 | 0.29611748 | 0.0009059 | 0.00748329 |
| Arhgap26 | 0.021697435 | 0.074098772 | 3.41509359 | 0.00090764 | 0.00749403 |
| Myl12b | 0.658585788 | 0.459297741 | 0.69740002 | 0.00090874 | 0.00749953 |
| Plekhj1 | 0.41756335 | 0.262755342 | 0.62925863 | 0.00090955 | 0.00750262 |
| Alad | 0.530587165 | 0.357510181 | 0.67380104 | 0.00091142 | 0.00751443 |
| 2900093K20I | 0.095163092 | 0.025934578 | 0.27252769 | 0.00091504 | 0.00754067 |
| Gps1 | 0.346387019 | 0.209393216 | 0.60450653 | 0.00091685 | 0.00755196 |
| Col7a1 | 0.004062039 | 0.035236891 | 8.67468007 | 0.00092077 | 0.00756315 |
| Tnfsf10 | 0.00407461 | 0.033994153 | 8.34292221 | 0.00092078 | 0.00756315 |
| Apol7a | 0.003413484 | 0.033739223 | 9.88410269 | 0.00092079 | 0.00756315 |
| Cebpa | 0.003968065 | 0.032088705 | 8.08673887 | 0.0009208 | 0.00756315 |
| Arxes2 | 0.109184308 | 0.038067502 | 0.3486536 | 0.00092129 | 0.00756315 |
| Snx21 | 0.11078554 | 0.03386601 | 0.3056898 | 0.00092129 | 0.00756315 |
| Tle3 | 0.110239808 | 0.033419867 | 0.30315607 | 0.00092129 | 0.00756315 |
| Cracr2b | 0.029691824 | 0.093993878 | 3.16564847 | 0.00092453 | 0.00758612 |
| Gm5914 | 0.049983225 | 0.006145837 | 0.12295799 | 0.00092605 | 0.00759155 |
| Pdgfrl | 0.051171733 | 0.002178701 | 0.04257626 | 0.00092608 | 0.00759155 |
| Srsf5 | 0.568229846 | 0.386060465 | 0.67940899 | 0.00093176 | 0.0076337 |
| Eif5 | 0.822439262 | 0.598591364 | 0.7278244 | 0.0009329 | 0.0076337 |
| Lipc | 0.004384781 | 0.041795118 | 9.53186029 | 0.00093341 | 0.0076337 |

| | | | | | |
|---|---|---|---|---|---|
| Adhfe1 | 0.005586926 | 0.038973519 | 6.97584263 | 0.00093342 | 0.0076337 |
| Hes7 | 0.00525157 | 0.036950444 | 7.03607558 | 0.00093344 | 0.0076337 |
| Adrm1 | 0.335275933 | 0.19480081 | 0.58101638 | 0.00095302 | 0.00779013 |
| Commd4 | 0.180980762 | 0.083933274 | 0.46376904 | 0.00096544 | 0.00788038 |
| Emp3 | 0.180341364 | 0.083926097 | 0.46537353 | 0.00096544 | 0.00788038 |
| Setd3 | 0.180825693 | 0.082151398 | 0.45431264 | 0.00096544 | 0.00788038 |
| Dtymk | 0.321937367 | 0.483409569 | 1.50156402 | 0.00098031 | 0.00799792 |
| Frg1 | 0.259354875 | 0.137130343 | 0.52873632 | 0.00099499 | 0.0081139 |
| Mrpl37 | 0.101937809 | 0.197792501 | 1.94032521 | 0.00099629 | 0.0081206 |
| Spr | 0.152419929 | 0.265107159 | 1.73932084 | 0.0009997 | 0.00814215 |
| Psma2 | 1.21532943 | 0.94270811 | 0.77568113 | 0.00100059 | 0.00814215 |
| Farsa | 0.237454884 | 0.122795226 | 0.51713076 | 0.00100083 | 0.00814215 |
| Lsm1 | 0.23629311 | 0.12356355 | 0.5229249 | 0.00100083 | 0.00814215 |
| Nipbl | 0.247950866 | 0.133706636 | 0.5392465 | 0.00100569 | 0.00817784 |
| Dynlrb1 | 0.857345255 | 0.634418495 | 0.73998018 | 0.00102353 | 0.00831894 |
| Chd3os | 0.067436285 | 0.015426814 | 0.22876133 | 0.00103893 | 0.0084394 |
| Mettl9 | 0.413591868 | 0.258318277 | 0.62457291 | 0.00103933 | 0.0084394 |
| Ilf2 | 0.368454387 | 0.22400143 | 0.60794888 | 0.00104715 | 0.00849818 |
| Mat1a | 0.016761058 | 0.064095981 | 3.82410126 | 0.00104756 | 0.00849818 |
| Anxa6 | 0.443140319 | 0.2859814 | 0.64535179 | 0.00105212 | 0.00853114 |
| Atp5d | 1.583957765 | 1.938755213 | 1.22399426 | 0.00106029 | 0.00859335 |
| Pgls | 0.841601958 | 1.084790418 | 1.288959 | 0.00106859 | 0.00865356 |
| Akap7 | 0.023290681 | 0.082074775 | 3.52393192 | 0.00106873 | 0.00865356 |
| Meaf6 | 0.126879789 | 0.049997754 | 0.3940561 | 0.00109027 | 0.00881611 |
| Ccdc90b | 0.126769294 | 0.047629582 | 0.3757186 | 0.00109027 | 0.00881611 |
| Dctn6 | 0.15489281 | 0.06946045 | 0.44844205 | 0.00109034 | 0.00881611 |
| Pfdn2 | 0.826327949 | 0.602089148 | 0.7286322 | 0.0011147 | 0.00900887 |
| Dda1 | 0.240578681 | 0.124584188 | 0.51785216 | 0.00112852 | 0.00911626 |
| Eif4a1 | 3.240927459 | 2.647756097 | 0.81697481 | 0.00113586 | 0.0091712 |
| Gys1 | 0.031604154 | 0.09558399 | 3.02441221 | 0.00113778 | 0.00918244 |

| | | | | | |
|---|---|---|---|---|---|
| Grasp | 0.012801944 | 0.057056614 | 4.45687113 | 0.0011432 | 0.00921751 |
| Icmt | 0.014112065 | 0.054847845 | 3.88659254 | 0.0011432 | 0.00921751 |
| Habp2 | 0.082538001 | 0.172139888 | 2.08558344 | 0.00115699 | 0.00932428 |
| Degs1 | 0.305064726 | 0.174786927 | 0.5729503 | 0.00116255 | 0.00936475 |
| 2610301B20I | 0.075637953 | 0.019423218 | 0.25679195 | 0.00117764 | 0.00948179 |
| Smpd1 | 0.121148688 | 0.044986997 | 0.37133705 | 0.00118122 | 0.00950621 |
| Dync1i2 | 0.436289555 | 0.281285796 | 0.64472274 | 0.00119231 | 0.00959096 |
| Ifrd1 | 0.121101381 | 0.22672182 | 1.87216544 | 0.00119558 | 0.00961273 |
| Gde1 | 0.292494659 | 0.168035127 | 0.57448956 | 0.00119654 | 0.00961599 |
| Ptpn11 | 0.166682751 | 0.07805437 | 0.46828103 | 0.00120254 | 0.0096552 |
| Nras | 0.166864116 | 0.076285213 | 0.45716967 | 0.00120255 | 0.0096552 |
| Txnip | 0.347855828 | 0.211573197 | 0.60822094 | 0.00123355 | 0.00989952 |
| Ccdc28b | 0.269832236 | 0.147203951 | 0.54553879 | 0.00125092 | 0.01003422 |
| Pla2g2d | 0.036472613 | 0.003893585 | 0.10675366 | 0.00125552 | 0.01003902 |
| Slc24a2 | 0.036980324 | 0.002392764 | 0.0647037 | 0.00125554 | 0.01003902 |
| Gm30173 | 0.036654759 | 0.002642602 | 0.07209438 | 0.00125554 | 0.01003902 |
| Gm37035 | 0.037634741 | 0 | 0 | 0.00125558 | 0.01003902 |
| H2-Q2 | 0.037042312 | 0 | 0 | 0.00125559 | 0.01003902 |
| Rims2 | 0.036743426 | 0 | 0 | 0.0012556 | 0.01003902 |
| Muc4 | 0.036552038 | 0 | 0 | 0.00125561 | 0.01003902 |
| B9d2 | 0.105302261 | 0.031998572 | 0.30387355 | 0.00126434 | 0.01010415 |
| Slc3a2 | 0.423948459 | 0.272145112 | 0.64192971 | 0.00126561 | 0.01010959 |
| Idh2 | 0.962667101 | 1.224163544 | 1.27163746 | 0.00127477 | 0.01017798 |
| Mrpl17 | 0.546698277 | 0.745633606 | 1.36388505 | 0.00127878 | 0.01020528 |
| Dus1l | 0.084428696 | 0.170851466 | 2.02361844 | 0.00128231 | 0.01022869 |
| Sft2d2 | 0.052587473 | 0.121489907 | 2.31024426 | 0.00128676 | 0.01024348 |
| Asap1 | 0.09742306 | 0.034617339 | 0.35533003 | 0.00128825 | 0.01024348 |
| Dync2h1 | 0.098190234 | 0.029258322 | 0.29797589 | 0.00128826 | 0.01024348 |
| Pfkl | 0.336484429 | 0.202933925 | 0.60310049 | 0.00128897 | 0.01024348 |
| Tmem101 | 0.140555304 | 0.05562995 | 0.39578691 | 0.0012902 | 0.01024348 |

| | | | | | |
|---|---|---|---|---|---|
| Ahdc1 | 0.057302625 | 0.011259665 | 0.19649476 | 0.00129164 | 0.01024348 |
| Rrm2b | 0.057812444 | 0.008607716 | 0.14889036 | 0.00129166 | 0.01024348 |
| Rab39b | 0.0571635 | 0.007566768 | 0.13237062 | 0.00129167 | 0.01024348 |
| Tdrkh | 0.085247943 | 0.021589877 | 0.2532598 | 0.0012925 | 0.01024348 |
| Vps13c | 0.084478608 | 0.022305269 | 0.26403452 | 0.0012925 | 0.01024348 |
| Dab2 | 0.084495935 | 0.021638149 | 0.25608508 | 0.0012925 | 0.01024348 |
| Fam110b | 0.084618423 | 0.021251475 | 0.25114478 | 0.0012925 | 0.01024348 |
| D430042O09 | 0.085318592 | 0.018415076 | 0.21583896 | 0.00129251 | 0.01024348 |
| Arc | 0.085246236 | 0.01822209 | 0.2137583 | 0.00129251 | 0.01024348 |
| Tagln2 | 0.847273351 | 0.62885302 | 0.74220795 | 0.00131064 | 0.01038241 |
| Dnase2a | 0.158945167 | 0.06911085 | 0.43480938 | 0.00131492 | 0.01041151 |
| Tmem51 | 0.09968771 | 0.031949118 | 0.32049205 | 0.00133739 | 0.01057484 |
| Madd | 0.099336106 | 0.031155154 | 0.31363374 | 0.00133739 | 0.01057484 |
| Gm27033 | 0.100625793 | 0.029127239 | 0.28946096 | 0.0013374 | 0.01057484 |
| Pcbp1 | 0.551221703 | 0.752097165 | 1.36441864 | 0.00134397 | 0.01062193 |
| P4htm | 0.058936563 | 0.011233317 | 0.19060014 | 0.00134698 | 0.01063039 |
| Pacrg | 0.059074367 | 0.010527029 | 0.17819961 | 0.00134699 | 0.01063039 |
| Mrc1 | 0.059251398 | 0.008309117 | 0.14023495 | 0.001347 | 0.01063039 |
| Pold4 | 0.160742577 | 0.072131384 | 0.44873851 | 0.00134751 | 0.01063039 |
| Sh3bgrl | 0.124993151 | 0.226967275 | 1.8158377 | 0.00135025 | 0.0106471 |
| Churc1 | 0.445730514 | 0.286990805 | 0.64386618 | 0.00135282 | 0.01066244 |
| Jkamp | 0.191800585 | 0.091028611 | 0.47460028 | 0.00136051 | 0.01071816 |
| Fads2 | 0.01140109 | 0.054369894 | 4.76883319 | 0.00138728 | 0.01092402 |
| Etv5 | 0.024929473 | 0.079305545 | 3.18119617 | 0.00139576 | 0.01098581 |
| Drg1 | 0.313603542 | 0.18341221 | 0.58485376 | 0.00139695 | 0.01099014 |
| Vti1b | 0.273804076 | 0.155162458 | 0.56669156 | 0.00139922 | 0.01100291 |
| Hnrnpa1 | 1.072714375 | 0.823381079 | 0.76756786 | 0.00140188 | 0.0110188 |
| Prkra | 0.134022479 | 0.052683849 | 0.39309711 | 0.0014155 | 0.0111208 |
| Erdr1 | 0.134995911 | 0.244323252 | 1.80985668 | 0.00141644 | 0.01112309 |
| Creb3 | 0.225055636 | 0.118075497 | 0.52465025 | 0.0014438 | 0.01130852 |

| | | | | | |
|---|---|---|---|---|---|
| Nfia | 0.080008463 | 0.162424194 | 2.03008768 | 0.00144463 | 0.01130852 |
| Clns1a | 0.23746701 | 0.12648014 | 0.53262194 | 0.00144517 | 0.01130852 |
| Rtn2 | 0.047528008 | 0.004400142 | 0.09257999 | 0.00144593 | 0.01130852 |
| Atg16l2 | 0.046601259 | 0.004994935 | 0.10718456 | 0.00144593 | 0.01130852 |
| Stk32a | 0.047797847 | 0.003489933 | 0.07301444 | 0.00144594 | 0.01130852 |
| Stab1 | 0.047917646 | 0.002899624 | 0.06051265 | 0.00144595 | 0.01130852 |
| Fam222a | 0.047608684 | 0.002239473 | 0.04703917 | 0.00144596 | 0.01130852 |
| Rsph1 | 0.048107411 | 0.00063061 | 0.01310838 | 0.00144598 | 0.01130852 |
| Faim | 0.152160267 | 0.064421394 | 0.42337855 | 0.00145621 | 0.01138336 |
| Bub3 | 0.374217234 | 0.230893132 | 0.61700293 | 0.00148393 | 0.01159475 |
| St7 | 0.015764191 | 0.060527115 | 3.83953201 | 0.00148593 | 0.01160513 |
| Rab24 | 0.183310393 | 0.08566754 | 0.46733597 | 0.00151042 | 0.01179104 |
| Ctage5 | 0.220186687 | 0.350280667 | 1.5908349 | 0.0015188 | 0.01185108 |
| C1d | 0.421197047 | 0.270583126 | 0.64241458 | 0.00153073 | 0.0119387 |
| 4921524J17F | 0.18496176 | 0.087390686 | 0.47247975 | 0.00153998 | 0.01199999 |
| Rhoc | 0.18432192 | 0.087851847 | 0.47662181 | 0.00153998 | 0.01199999 |
| Zfpm1 | 0.01904069 | 0.063124286 | 3.31523095 | 0.00154344 | 0.01202152 |
| Elavl4 | 0.049007008 | 0.005889203 | 0.12017062 | 0.00155744 | 0.01211992 |
| Glb1l2 | 0.04954671 | 0.002446347 | 0.04937456 | 0.00155749 | 0.01211992 |
| Perp | 0.067169208 | 0.147532472 | 2.19643012 | 0.00156043 | 0.01213731 |
| Zfand6 | 0.507083571 | 0.341693115 | 0.67383984 | 0.00156351 | 0.01215574 |
| Rpl27-ps3 | 0.221760106 | 0.3485397 | 1.57169703 | 0.00157326 | 0.01221176 |
| Sbk1 | 0.065870769 | 0.01447219 | 0.2197058 | 0.00157354 | 0.01221176 |
| 2510046G10 | 0.065530182 | 0.012608634 | 0.19240956 | 0.00157355 | 0.01221176 |
| 1700096K18I | 0.066682551 | 0.010868489 | 0.16298849 | 0.00157356 | 0.01221176 |
| Timm17a | 0.491239516 | 0.326180813 | 0.66399547 | 0.00158562 | 0.01229982 |
| Usp14 | 0.164061343 | 0.073678724 | 0.44909253 | 0.00159093 | 0.01232989 |
| Camta1 | 0.164596976 | 0.071433836 | 0.4339924 | 0.00159093 | 0.01232989 |
| Hadhb | 0.306667036 | 0.179549155 | 0.58548567 | 0.00159241 | 0.01233577 |
| Cdc42se2 | 0.115915291 | 0.045936526 | 0.39629393 | 0.00161173 | 0.01247422 |

| | | | | | |
|---|---|---|---|---|---|
| Taok2 | 0.116993878 | 0.043746743 | 0.37392336 | 0.00161173 | 0.01247422 |
| Msln | 0.118292468 | 0.044982303 | 0.38026346 | 0.00161739 | 0.01251238 |
| Ndufaf3 | 0.219025923 | 0.113897188 | 0.52001693 | 0.00163144 | 0.01261539 |
| Stt3a | 0.519750849 | 0.710434414 | 1.36687495 | 0.00164061 | 0.01268062 |
| Upf3b | 0.349166186 | 0.214167674 | 0.61336888 | 0.00164411 | 0.01270193 |
| Lpcat3 | 0.220618904 | 0.11271465 | 0.51090205 | 0.00165811 | 0.01280435 |
| Nans | 0.5032296 | 0.684987654 | 1.36118315 | 0.00165946 | 0.01280904 |
| Tmem54 | 0.009483747 | 0.050924451 | 5.36965513 | 0.00166619 | 0.01284385 |
| Arhgef26 | 0.009977697 | 0.049248343 | 4.93584288 | 0.00166619 | 0.01284385 |
| Rgmb | 0.01096986 | 0.045335834 | 4.1327633 | 0.00166622 | 0.01284385 |
| Klk1b4 | 0.000427254 | 0.020330896 | 47.5850678 | 0.00167256 | 0.01288698 |
| Ubap2l | 0.364887391 | 0.22769238 | 0.62400726 | 0.00168021 | 0.01294012 |
| Spint2 | 3.018373661 | 3.64096057 | 1.20626568 | 0.00168647 | 0.01298252 |
| Snrpc | 0.569225446 | 0.391185529 | 0.68722425 | 0.00169539 | 0.01304534 |
| Lrif1 | 0.109289308 | 0.041930713 | 0.38366711 | 0.00170008 | 0.01307552 |
| Rnf149 | 0.076462168 | 0.159223369 | 2.08238104 | 0.00171043 | 0.01314927 |
| Spint1 | 0.338955016 | 0.203233885 | 0.59958955 | 0.00172661 | 0.01326184 |
| 1810022K09l | 0.275898082 | 0.41626048 | 1.50874728 | 0.00172662 | 0.01326184 |
| Gchfr | 0.072797596 | 0.016311888 | 0.2240718 | 0.00174139 | 0.01336936 |
| Pomgnt2 | 0.136820182 | 0.056592625 | 0.41362776 | 0.00174677 | 0.01340467 |
| Fnta | 0.269710295 | 0.153749368 | 0.57005376 | 0.00175101 | 0.0134312 |
| Tmem60 | 0.155560985 | 0.070333289 | 0.45212679 | 0.00176164 | 0.01350667 |
| Ndufs1 | 0.187635297 | 0.094680802 | 0.50460017 | 0.00177078 | 0.01357068 |
| Ergic3 | 0.71392481 | 0.518655717 | 0.72648507 | 0.00177523 | 0.01359872 |
| Ric8 | 0.102417599 | 0.039100543 | 0.38177562 | 0.0017854 | 0.01366452 |
| Mgp | 0.102958995 | 0.035454584 | 0.34435635 | 0.00178541 | 0.01366452 |
| Ms4a6c | 0.074939578 | 0.015222501 | 0.20313032 | 0.0018015 | 0.01378155 |
| Zmynd8 | 0.199069546 | 0.103284607 | 0.51883681 | 0.00182088 | 0.01392363 |
| Tmem45a | 0.247362579 | 0.137199788 | 0.55465054 | 0.00182722 | 0.01396584 |
| Lsp1 | 0.094956409 | 0.029545658 | 0.3111497 | 0.0018442 | 0.0140894 |

| | | | | | |
|---|---|---|---|---|---|
| Gprasp2 | 0.080520863 | 0.025493897 | 0.31661232 | 0.00184815 | 0.01409459 |
| Cmbl | 0.080695255 | 0.024735852 | 0.30653415 | 0.00184815 | 0.01409459 |
| Gm20604 | 0.081390406 | 0.021970903 | 0.26994463 | 0.00184816 | 0.01409459 |
| Xlr3a | 0.081109784 | 0.021562208 | 0.26583979 | 0.00184816 | 0.01409459 |
| Gm26825 | 0.087896523 | 0.02548035 | 0.2898903 | 0.00186006 | 0.01417902 |
| Pole4 | 0.124898366 | 0.222340292 | 1.78016974 | 0.00186276 | 0.01418492 |
| Tbc1d1 | 0.096541335 | 0.031166034 | 0.3228258 | 0.00186496 | 0.01418492 |
| Ncoa7 | 0.09688261 | 0.030180333 | 0.31151445 | 0.00186496 | 0.01418492 |
| Cryl1 | 0.09695634 | 0.029306123 | 0.30226103 | 0.00186496 | 0.01418492 |
| Commd10 | 0.096574942 | 0.029559062 | 0.30607383 | 0.00186496 | 0.01418492 |
| Id1 | 0.0714989 | 0.146230845 | 2.04521812 | 0.00188383 | 0.01432214 |
| Gps2 | 0.16720117 | 0.080070207 | 0.47888545 | 0.00189086 | 0.01436282 |
| Pdpk1 | 0.167676925 | 0.078270254 | 0.46679204 | 0.00189086 | 0.01436282 |
| Ndufb5 | 1.201632385 | 1.47843046 | 1.23035171 | 0.00190899 | 0.01449413 |
| Lancl1 | 0.130350011 | 0.05341688 | 0.40979574 | 0.00191426 | 0.01452771 |
| Lfng | 0.024108137 | 0.075571795 | 3.13470079 | 0.00192334 | 0.01459021 |
| Mtx2 | 0.274092832 | 0.158653909 | 0.57883276 | 0.00193043 | 0.01463753 |
| Gas8 | 0.089473779 | 0.025544224 | 0.28549397 | 0.00193518 | 0.01466711 |
| Flrt2 | 0.026369951 | 0.079629136 | 3.01969224 | 0.00195723 | 0.01482117 |
| Sapcd2 | 0.027660536 | 0.076722897 | 2.7737314 | 0.00195724 | 0.01482117 |
| Fam174b | 0.407326789 | 0.263993529 | 0.64811237 | 0.0019697 | 0.01490431 |
| Galnt3 | 0.007708828 | 0.045529293 | 5.90612358 | 0.00197166 | 0.01490431 |
| Rnd2 | 0.008384547 | 0.043196788 | 5.15195227 | 0.00197167 | 0.01490431 |
| Afp | 0.008565664 | 0.04202252 | 4.90592692 | 0.00197169 | 0.01490431 |
| Lsm8 | 0.303264357 | 0.176824872 | 0.58307172 | 0.00197277 | 0.01490596 |
| Ywhab | 0.623555505 | 0.444551384 | 0.71292993 | 0.00198111 | 0.01496242 |
| Gzf1 | 0.046768478 | 0.114346452 | 2.44494703 | 0.00199148 | 0.01502855 |
| Foxa3 | 0.117866966 | 0.218488923 | 1.8536909 | 0.00199162 | 0.01502855 |
| Ten1 | 0.263378619 | 0.151521517 | 0.57529923 | 0.0020016 | 0.0150906 |
| Diablo | 0.264193339 | 0.150274628 | 0.56880551 | 0.0020016 | 0.0150906 |

| | | | | | |
|---|---|---|---|---|---|
| Nol4l | 0.055427125 | 0.011650365 | 0.21019248 | 0.00200715 | 0.01512582 |
| Acsl4 | 0.031444104 | 0.090419711 | 2.87556964 | 0.00201894 | 0.01520799 |
| Grhl1 | 0.033240432 | 0.003703456 | 0.11141421 | 0.00205083 | 0.01542866 |
| Cpb2 | 0.03306846 | 0.002600684 | 0.07864546 | 0.00205088 | 0.01542866 |
| Crybb1 | 0.034453773 | 0 | 0 | 0.00205093 | 0.01542866 |
| Fbl | 0.437444906 | 0.613861532 | 1.40328879 | 0.00206227 | 0.01550307 |
| Smarca1 | 0.192422318 | 0.097375501 | 0.50605097 | 0.00206429 | 0.01550307 |
| Pla2g6 | 0.121093889 | 0.050945934 | 0.42071433 | 0.00206505 | 0.01550307 |
| Eea1 | 0.121056214 | 0.048980405 | 0.40460876 | 0.00206505 | 0.01550307 |
| Arf4 | 0.791534521 | 1.018612578 | 1.28688333 | 0.00206533 | 0.01550307 |
| Pkp4 | 0.238053324 | 0.131098183 | 0.55070932 | 0.0020688 | 0.01552236 |
| Gatad2b | 0.204414419 | 0.105208909 | 0.51468438 | 0.00210046 | 0.015753 |
| Hacd1 | 0.375420345 | 0.537371165 | 1.43138531 | 0.00210939 | 0.01581308 |
| Lmo4 | 0.094651108 | 0.179862164 | 1.90026475 | 0.00214954 | 0.016107 |
| mt-Nd4 | 5.584150777 | 4.626145653 | 0.82844211 | 0.00215267 | 0.01612343 |
| Sptan1 | 0.142761966 | 0.060947956 | 0.42692012 | 0.00216086 | 0.01617775 |
| Acin1 | 0.367731852 | 0.230290081 | 0.62624458 | 0.00220066 | 0.01641176 |
| Tmx1 | 0.367719639 | 0.230032678 | 0.62556539 | 0.00220066 | 0.01641176 |
| Slc36a4 | 0.034896675 | 0.002371574 | 0.06795987 | 0.00220157 | 0.01641176 |
| Nudt11 | 0.035280745 | 0.001692458 | 0.04797116 | 0.00220158 | 0.01641176 |
| Edn3 | 0.035817649 | 0.000763657 | 0.0213207 | 0.0022016 | 0.01641176 |
| Nqo1 | 0.0352381 | 0.000976969 | 0.0277248 | 0.00220162 | 0.01641176 |
| Kcnma1 | 0.036035535 | 0 | 0 | 0.00220163 | 0.01641176 |
| E530001K10I | 0.035786946 | 0 | 0 | 0.00220164 | 0.01641176 |
| Gm11789 | 0.035204532 | 0 | 0 | 0.00220167 | 0.01641176 |
| 2410021H03I | 0.035104084 | 0 | 0 | 0.00220167 | 0.01641176 |
| 1700011H14I | 0.11442856 | 0.047293844 | 0.41330455 | 0.00222471 | 0.01656196 |
| Rars2 | 0.114866572 | 0.046478495 | 0.40463029 | 0.00222471 | 0.01656196 |
| Crot | 0.114861868 | 0.043458533 | 0.37835475 | 0.00222471 | 0.01656196 |
| Fam149a | 0.028565774 | 0.084771041 | 2.9675737 | 0.00223322 | 0.01661812 |

| | | | | | |
|---|---|---|---|---|---|
| Fkbp9 | 0.296342212 | 0.176671 | 0.59617224 | 0.00225011 | 0.0167365 |
| Entpd1 | 0.007075224 | 0.0404005 | 5.71013726 | 0.00228154 | 0.01695595 |
| Ak4 | 0.006854749 | 0.037742857 | 5.50608922 | 0.00228158 | 0.01695595 |
| Rnf44 | 0.04920922 | 0.115815194 | 2.35352629 | 0.00229996 | 0.01708512 |
| Ptn | 0.253561807 | 0.386305089 | 1.5235145 | 0.00231091 | 0.01715903 |
| Dnajc3 | 0.955410412 | 1.205408015 | 1.26166514 | 0.00232052 | 0.01720335 |
| Fbxo32 | 0.04352553 | 0.00652243 | 0.14985297 | 0.00232179 | 0.01720335 |
| 6330403K07I | 0.043163316 | 0.006823216 | 0.15807906 | 0.00232179 | 0.01720335 |
| Clec10a | 0.044106224 | 0.003502733 | 0.07941584 | 0.00232185 | 0.01720335 |
| Nebl | 0.044491518 | 0.001807018 | 0.04061489 | 0.00232188 | 0.01720335 |
| RP23-60P18. | 0.045425885 | 0.007096029 | 0.15621114 | 0.00233975 | 0.01728422 |
| Micalcl | 0.04529004 | 0.006761906 | 0.14930228 | 0.00233976 | 0.01728422 |
| 9330182L06F | 0.045567243 | 0.005498888 | 0.12067635 | 0.00233979 | 0.01728422 |
| Sh3bgr | 0.046074305 | 0.00482991 | 0.10482872 | 0.00233979 | 0.01728422 |
| Aatk | 0.046052021 | 0.004095303 | 0.08892775 | 0.00233981 | 0.01728422 |
| Fam171b | 0.045139342 | 0.004989739 | 0.11054081 | 0.00233981 | 0.01728422 |
| Gpr45 | 0.045373743 | 0.003604349 | 0.07943689 | 0.00233984 | 0.01728422 |
| Klhdc2 | 0.195514031 | 0.101386956 | 0.51856614 | 0.00235456 | 0.01738552 |
| Suox | 0.063120957 | 0.009998687 | 0.15840518 | 0.00235694 | 0.0173956 |
| Ap1m1 | 0.134239924 | 0.057432357 | 0.42783365 | 0.00237044 | 0.01746948 |
| Rgs2 | 0.133814167 | 0.055069249 | 0.41153527 | 0.00237044 | 0.01746948 |
| Armc1 | 0.220432306 | 0.118199448 | 0.53621654 | 0.00237102 | 0.01746948 |
| Ppp5c | 0.22076233 | 0.114666542 | 0.51941172 | 0.00237102 | 0.01746948 |
| Apoa1bp | 0.404430405 | 0.261659984 | 0.64698396 | 0.00237334 | 0.01747911 |
| Rilpl2 | 0.107922024 | 0.040931822 | 0.37927219 | 0.00239314 | 0.01761734 |
| Kin | 0.134986593 | 0.059152356 | 0.43820912 | 0.00239998 | 0.01764508 |
| Haghl | 0.135488699 | 0.056905213 | 0.4199997 | 0.00239998 | 0.01764508 |
| 2900076A07I | 0.135708635 | 0.054957604 | 0.40496763 | 0.00239999 | 0.01764508 |
| Srp54b | 0.626376414 | 0.445171192 | 0.71070874 | 0.00240574 | 0.0176798 |
| Rab1a | 0.944964866 | 0.720236824 | 0.76218371 | 0.00243245 | 0.01786844 |

| | | | | | |
|---|---|---|---|---|---|
| Ewsr1 | 0.51179556 | 0.347218193 | 0.67843143 | 0.00243424 | 0.01787398 |
| Setd5 | 0.299967091 | 0.179394012 | 0.59804564 | 0.00244268 | 0.01792826 |
| Galns | 0.063959693 | 0.015040931 | 0.23516266 | 0.00245729 | 0.01800489 |
| Abhd18 | 0.063773196 | 0.014684894 | 0.2302675 | 0.0024573 | 0.01800489 |
| Ift122 | 0.063805759 | 0.013917178 | 0.21811789 | 0.0024573 | 0.01800489 |
| Tspan17 | 0.064492498 | 0.01273629 | 0.19748482 | 0.00245731 | 0.01800489 |
| Mob4 | 0.34367052 | 0.215398887 | 0.62675986 | 0.00246197 | 0.01803134 |
| Asb16 | 0.000855358 | 0.029841855 | 34.8881314 | 0.00248132 | 0.01815864 |
| Gm11963 | 0.002259952 | 0.024363509 | 10.7805416 | 0.00248146 | 0.01815864 |
| Nr2f6 | 0.082546172 | 0.164263679 | 1.98996119 | 0.00248914 | 0.01820705 |
| Ptpn1 | 0.166009888 | 0.08027181 | 0.48353632 | 0.0025019 | 0.01828488 |
| Max | 0.165299407 | 0.078153019 | 0.47279673 | 0.00250191 | 0.01828488 |
| Zfp511 | 0.099000282 | 0.039342325 | 0.39739609 | 0.00250768 | 0.01831928 |
| Chac2 | 0.090574087 | 0.179155432 | 1.97799877 | 0.0025335 | 0.01850007 |
| 9530052E02I | 0.004755877 | 0.036167713 | 7.60484619 | 0.00254766 | 0.01857997 |
| Clec14a | 0.005318347 | 0.03541352 | 6.65874513 | 0.00254767 | 0.01857997 |
| Smpdl3b | 0.004769065 | 0.034842663 | 7.30597316 | 0.00254769 | 0.01857997 |
| Dnajc1 | 0.197596903 | 0.315726354 | 1.59783048 | 0.00255808 | 0.01864784 |
| Akap11 | 0.101867058 | 0.033479525 | 0.328659 | 0.00256933 | 0.01872192 |
| Rpa3 | 0.24853565 | 0.141039749 | 0.56748297 | 0.00257225 | 0.01873524 |
| Nfic | 0.12643091 | 0.055126342 | 0.4360195 | 0.00259673 | 0.01887483 |
| Hspb8 | 0.126573571 | 0.054053714 | 0.42705372 | 0.00259673 | 0.01887483 |
| Tsen15 | 0.127291934 | 0.052411299 | 0.41174092 | 0.00259674 | 0.01887483 |
| 1110020A21I | 0.070282568 | 0.018154709 | 0.25831027 | 0.00259864 | 0.01887483 |
| Ckmt1 | 0.070567781 | 0.0177522 | 0.25156239 | 0.00259864 | 0.01887483 |
| Alox5ap | 0.070707674 | 0.01679548 | 0.23753405 | 0.00259865 | 0.01887483 |
| mt-Nd5 | 1.085239205 | 1.339209366 | 1.23402229 | 0.00259911 | 0.01887483 |
| Calu | 0.108599546 | 0.198531761 | 1.82810856 | 0.00260977 | 0.01894428 |
| Trnau1ap | 0.145948367 | 0.065977895 | 0.45206326 | 0.00261329 | 0.0189618 |
| Lap3 | 0.187869668 | 0.301419572 | 1.60440786 | 0.0026296 | 0.01907212 |

| | | | | | |
|---|---|---|---|---|---|
| Polr2m | 0.430440527 | 0.282314117 | 0.65587253 | 0.00265682 | 0.01926141 |
| Ift88 | 0.077430574 | 0.024277182 | 0.31353483 | 0.00266929 | 0.01932741 |
| Acyp2 | 0.078136936 | 0.02329786 | 0.29816705 | 0.00266929 | 0.01932741 |
| Ptpn6 | 0.07770468 | 0.021501157 | 0.27670351 | 0.0026693 | 0.01932741 |
| Gm5091 | 0.002545636 | 0.031135092 | 12.2307717 | 0.00267405 | 0.0193536 |
| Cox20 | 0.380949771 | 0.243277298 | 0.63860728 | 0.00268921 | 0.01945513 |
| Snrpa | 0.381639158 | 0.246195482 | 0.64510016 | 0.00271202 | 0.01960749 |
| Tbc1d7 | 0.086044554 | 0.027095341 | 0.31489896 | 0.00271482 | 0.01960749 |
| Anks3 | 0.086418537 | 0.024989028 | 0.28916282 | 0.00271483 | 0.01960749 |
| Stard5 | 0.085908173 | 0.025129138 | 0.29251161 | 0.00271483 | 0.01960749 |
| Itpr2 | 0.031192652 | 0.082848832 | 2.65603677 | 0.0027437 | 0.01980766 |
| Serf1 | 0.365906632 | 0.517175879 | 1.41340942 | 0.0027465 | 0.01981957 |
| Bud31 | 0.560269412 | 0.393487918 | 0.70231912 | 0.00275136 | 0.01983826 |
| Arl5a | 0.057007841 | 0.126411816 | 2.21744613 | 0.0027514 | 0.01983826 |
| Pdcd10 | 0.265637975 | 0.15292181 | 0.57567752 | 0.00277153 | 0.01997501 |
| Bckdhb | 0.093676336 | 0.173761599 | 1.85491456 | 0.00277792 | 0.02001265 |
| Dennd4c | 0.079704751 | 0.022524171 | 0.28259509 | 0.00278575 | 0.02005226 |
| RP23-218K1[illegible] | 0.078955985 | 0.022904189 | 0.29008806 | 0.00278575 | 0.02005226 |
| Sdf4 | 0.643706889 | 0.460903148 | 0.71601401 | 0.00279127 | 0.02008357 |
| Fmo2 | 0.157390192 | 0.076536041 | 0.48628215 | 0.00279619 | 0.02011059 |
| Kmt2a | 0.1588486 | 0.076489491 | 0.48152449 | 0.00281977 | 0.02026323 |
| Clock | 0.15976454 | 0.074814788 | 0.46828156 | 0.00281977 | 0.02026323 |
| Aldh18a1 | 0.077905595 | 0.154088036 | 1.97788152 | 0.00284067 | 0.02040487 |
| Lars | 0.203758195 | 0.317802425 | 1.55970377 | 0.00285015 | 0.02045712 |
| Dynll2 | 0.545186091 | 0.727512183 | 1.3344291 | 0.00285033 | 0.02045712 |
| Psmb3 | 0.97453995 | 1.213140859 | 1.2448344 | 0.00287403 | 0.02061862 |
| Ppp1r12a | 0.13870068 | 0.061991122 | 0.44694173 | 0.00289541 | 0.02076333 |
| Zfp36 | 0.17003191 | 0.08016814 | 0.47148879 | 0.00292461 | 0.02096395 |
| Chmp4c | 0.051726569 | 0.122356887 | 2.36545529 | 0.00293683 | 0.02104278 |
| Irf2bpl | 0.141239903 | 0.060919449 | 0.43131896 | 0.00296422 | 0.02123022 |

| | | | | | |
|---|---|---|---|---|---|
| Birc6 | 0.229199393 | 0.124690865 | 0.5440279 | 0.00298072 | 0.02133947 |
| Tax1bp3 | 0.182169597 | 0.092591049 | 0.5082684 | 0.00302665 | 0.02165927 |
| Acp1 | 0.388806624 | 0.25546968 | 0.65706103 | 0.00304812 | 0.02178712 |
| Abca1 | 0.051741821 | 0.011824928 | 0.22853715 | 0.00305329 | 0.02178712 |
| Mras | 0.052082293 | 0.009376683 | 0.1800359 | 0.00305333 | 0.02178712 |
| Wnt4 | 0.052188191 | 0.008326076 | 0.15953946 | 0.00305335 | 0.02178712 |
| Nfasc | 0.053023406 | 0.006323109 | 0.11925127 | 0.00305337 | 0.02178712 |
| Wdr47 | 0.052286695 | 0.0070585 | 0.13499611 | 0.00305337 | 0.02178712 |
| Olfm1 | 0.052957822 | 0.005312504 | 0.10031576 | 0.00305339 | 0.02178712 |
| Ythdf1 | 0.059278699 | 0.12504032 | 2.10936344 | 0.00306205 | 0.02183986 |
| Ift22 | 0.206253031 | 0.110926318 | 0.53781667 | 0.00307131 | 0.02189676 |
| Cnih1 | 0.313857638 | 0.190220923 | 0.6060739 | 0.00311294 | 0.02218436 |
| St3gal3 | 0.018609078 | 0.061737 | 3.31757431 | 0.00311564 | 0.02218535 |
| Celsr1 | 0.01839845 | 0.059819273 | 3.25132132 | 0.00311566 | 0.02218535 |
| Ndrg3 | 0.11364886 | 0.044232409 | 0.3892024 | 0.00312117 | 0.02219816 |
| Gtf3c1 | 0.114206726 | 0.042641344 | 0.37336981 | 0.00312118 | 0.02219816 |
| Fam114a1 | 0.048159244 | 0.107189584 | 2.22573228 | 0.00312133 | 0.02219816 |
| Lamtor1 | 0.330317145 | 0.206377419 | 0.62478567 | 0.00318634 | 0.02265111 |
| Arg1 | 0.054722923 | 0.008505215 | 0.15542326 | 0.00322821 | 0.02289944 |
| Sh3kbp1 | 0.053320036 | 0.009881778 | 0.18532954 | 0.00322822 | 0.02289944 |
| Btbd10 | 0.054753632 | 0.007757265 | 0.14167581 | 0.00322823 | 0.02289944 |
| Gm19412 | 0.053551994 | 0.008426842 | 0.15735814 | 0.00322824 | 0.02289944 |
| Ttc39b | 0.054838292 | 0.006504288 | 0.1186085 | 0.00322825 | 0.02289944 |
| Tmem256 | 0.848301985 | 1.066495651 | 1.25721225 | 0.00322927 | 0.02289944 |
| Brd3 | 0.301106144 | 0.182515614 | 0.60615041 | 0.00327427 | 0.02320895 |
| Scarb2 | 0.1619987 | 0.081842283 | 0.50520333 | 0.0032844 | 0.0232616 |
| Tctex1d2 | 0.162084993 | 0.078067532 | 0.48164565 | 0.0032844 | 0.0232616 |
| Tubb2b | 0.104432956 | 0.039288755 | 0.37621031 | 0.00330393 | 0.02338066 |
| Kif1bp | 0.105505538 | 0.035748421 | 0.33882981 | 0.00330394 | 0.02338066 |
| Hnrnpa2b1 | 1.808312288 | 1.500586973 | 0.82982734 | 0.003308 | 0.0233998 |

| | | | | | |
|---|---|---|---|---|---|
| Kdm5b | 0.233246463 | 0.130795641 | 0.56076152 | 0.00331674 | 0.02345197 |
| Hsp90ab1 | 5.843485375 | 4.879856901 | 0.83509354 | 0.00334053 | 0.02361048 |
| Commd8 | 0.163666214 | 0.080525905 | 0.492013 | 0.00334541 | 0.02362551 |
| 1110004E09I | 0.164492607 | 0.079402882 | 0.48271399 | 0.00334541 | 0.02362551 |
| Chchd7 | 0.451845402 | 0.61948232 | 1.37100503 | 0.00335578 | 0.02368903 |
| Clptm1 | 0.222120096 | 0.12340145 | 0.55556184 | 0.00338717 | 0.02390079 |
| Kcnu1 | 0.031420903 | 0.002770105 | 0.08816121 | 0.00341955 | 0.02403167 |
| Cerkl | 0.03147145 | 0.002201717 | 0.0699592 | 0.00341958 | 0.02403167 |
| Sdcbp2 | 0.032341902 | 0.001260604 | 0.03897743 | 0.00341959 | 0.02403167 |
| Cntn1 | 0.031588292 | 0.001663642 | 0.05266641 | 0.00341961 | 0.02403167 |
| Rnls | 0.03237959 | 0.000817239 | 0.02523932 | 0.00341962 | 0.02403167 |
| Flrt1 | 0.032169456 | 0.000880784 | 0.0273795 | 0.00341963 | 0.02403167 |
| Slc4a1 | 0.032421384 | 0.000446987 | 0.0137868 | 0.00341964 | 0.02403167 |
| Ccdc136 | 0.031419551 | 0.001394267 | 0.04437579 | 0.00341965 | 0.02403167 |
| Ppp1r27 | 0.031919012 | 0.000595808 | 0.01866625 | 0.00341967 | 0.02403167 |
| Dmrtc1a | 0.031330152 | 0.000705613 | 0.02252184 | 0.00341971 | 0.02403167 |
| Abhd17c | 0.067223033 | 0.139981626 | 2.08234617 | 0.00342363 | 0.0240494 |
| Tmem9b | 0.209176058 | 0.115373277 | 0.55156062 | 0.00343566 | 0.02412007 |
| Ormdl2 | 0.289278094 | 0.174698198 | 0.60391091 | 0.0034365 | 0.02412007 |
| Rcn1 | 0.262695373 | 0.153293929 | 0.58354255 | 0.00344524 | 0.02415244 |
| Ankrd12 | 0.262621883 | 0.151060572 | 0.57520177 | 0.00344524 | 0.02415244 |
| Pop7 | 0.175661298 | 0.090485518 | 0.51511357 | 0.00344673 | 0.02415244 |
| Ppil4 | 0.176037049 | 0.085767418 | 0.48721232 | 0.00344673 | 0.02415244 |
| Rangap1 | 0.35104963 | 0.221138353 | 0.62993473 | 0.00345082 | 0.02417123 |
| D15Ertd621e | 0.116877243 | 0.204334309 | 1.74828138 | 0.00351617 | 0.02461895 |
| Mfap3 | 0.096762098 | 0.036202129 | 0.37413543 | 0.00353658 | 0.02474179 |
| Kctd20 | 0.097112886 | 0.032572437 | 0.33540798 | 0.0035366 | 0.02474179 |
| Tmem237 | 0.09756237 | 0.037930324 | 0.38878026 | 0.00354596 | 0.02479722 |
| Scyl1 | 0.125152563 | 0.053411062 | 0.42676762 | 0.00357242 | 0.02493573 |
| Srsf11 | 0.675825101 | 0.492524679 | 0.72877536 | 0.00357505 | 0.02493573 |

| | | | | | |
|---|---|---|---|---|---|
| Tmco3 | 0.058905226 | 0.015915756 | 0.2701926 | 0.00357685 | 0.02493573 |
| Igf2bp1 | 0.059177619 | 0.014136724 | 0.23888633 | 0.00357687 | 0.02493573 |
| St6galnac6 | 0.058865535 | 0.013715563 | 0.23299819 | 0.00357688 | 0.02493573 |
| Fam105a | 0.060002888 | 0.011662808 | 0.19437078 | 0.00357689 | 0.02493573 |
| Jade3 | 0.059637834 | 0.010637259 | 0.17836428 | 0.00357692 | 0.02493573 |
| Hsf2 | 0.061595639 | 0.013530313 | 0.21966349 | 0.00358171 | 0.02493573 |
| Klhl22 | 0.060520706 | 0.01414428 | 0.23370977 | 0.00358171 | 0.02493573 |
| Khdrbs3 | 0.060804198 | 0.013483249 | 0.22174865 | 0.00358172 | 0.02493573 |
| Gm4419 | 0.060941136 | 0.012255392 | 0.20110213 | 0.00358174 | 0.02493573 |
| 1700021F05I | 0.099809637 | 0.182943445 | 1.83292366 | 0.00361296 | 0.02506142 |
| Gal | 0.042358204 | 0.005549695 | 0.13101818 | 0.00362008 | 0.02506142 |
| Hpgds | 0.04214427 | 0.005411856 | 0.12841262 | 0.0036201 | 0.02506142 |
| Pacsin1 | 0.04148745 | 0.005815652 | 0.14017858 | 0.00362011 | 0.02506142 |
| Kcnb2 | 0.04174919 | 0.005311447 | 0.12722275 | 0.00362011 | 0.02506142 |
| Rgs4 | 0.041963774 | 0.005012758 | 0.11945441 | 0.00362012 | 0.02506142 |
| 1500026H17I | 0.043022416 | 0.003885324 | 0.0903093 | 0.00362012 | 0.02506142 |
| Tff3 | 0.04188339 | 0.004817895 | 0.11503117 | 0.00362013 | 0.02506142 |
| Tnr | 0.042869777 | 0.003718354 | 0.08673603 | 0.00362013 | 0.02506142 |
| Fam229b | 0.04230717 | 0.003678754 | 0.08695345 | 0.00362016 | 0.02506142 |
| Unc13a | 0.04175091 | 0.004026496 | 0.09644092 | 0.00362016 | 0.02506142 |
| Fam20a | 0.042918423 | 0.002215078 | 0.05161136 | 0.00362019 | 0.02506142 |
| Rgs16 | 0.042215276 | 0.002658307 | 0.06297026 | 0.0036202 | 0.02506142 |
| Insrr | 0.042987313 | 0.001731896 | 0.04028853 | 0.00362021 | 0.02506142 |
| Hint2 | 0.323520319 | 0.200035018 | 0.61830743 | 0.00363481 | 0.02515233 |
| Larp1b | 0.10862203 | 0.191944736 | 1.76708846 | 0.0036676 | 0.02536903 |
| Tns3 | 0.089004413 | 0.034134921 | 0.38351942 | 0.00370058 | 0.02555612 |
| Srd5a3 | 0.08927683 | 0.032534696 | 0.36442485 | 0.00370059 | 0.02555612 |
| Hsdl2 | 0.08910351 | 0.031815856 | 0.35706624 | 0.00370059 | 0.02555612 |
| Zfp667 | 0.089416535 | 0.030828874 | 0.34477822 | 0.00370059 | 0.02555612 |
| Idh3b | 0.294139898 | 0.181819673 | 0.61814012 | 0.00373048 | 0.02575218 |

| | | | | | |
|---|---|---|---|---|---|
| Cdc42ep5 | 0.062528182 | 0.132817482 | 2.12412192 | 0.0037559 | 0.02590684 |
| Pah | 0.063244931 | 0.131331377 | 2.07655184 | 0.0037559 | 0.02590684 |
| Cfl2 | 0.226766631 | 0.125918053 | 0.55527593 | 0.00378172 | 0.02607031 |
| Stard13 | 0.027966543 | 0.08225428 | 2.94116724 | 0.00379227 | 0.02607031 |
| Sowahc | 0.067321273 | 0.019698497 | 0.29260434 | 0.00379236 | 0.02607031 |
| 2210019I11R | 0.066947357 | 0.019108463 | 0.28542521 | 0.00379237 | 0.02607031 |
| Cramp1l | 0.067966058 | 0.017397498 | 0.25597333 | 0.00379238 | 0.02607031 |
| Eml5 | 0.068400182 | 0.01458642 | 0.21325119 | 0.00379241 | 0.02607031 |
| Mcts2 | 0.068191864 | 0.014441852 | 0.21178263 | 0.00379241 | 0.02607031 |
| Fitm2 | 0.092199078 | 0.030450343 | 0.33026732 | 0.00379326 | 0.02607031 |
| Scaper | 0.09163876 | 0.030764218 | 0.33571185 | 0.00379326 | 0.02607031 |
| Arcn1 | 0.271047258 | 0.400308145 | 1.47689428 | 0.00380144 | 0.02611608 |
| Gon4l | 0.166770249 | 0.085886886 | 0.51500124 | 0.00383124 | 0.02629979 |
| Ehd1 | 0.167496954 | 0.08143781 | 0.48620473 | 0.00383124 | 0.02629979 |
| Fbxo6 | 0.082950746 | 0.027093425 | 0.32662063 | 0.00383846 | 0.02631798 |
| Odf2 | 0.082409484 | 0.025929421 | 0.31464123 | 0.00383848 | 0.02631798 |
| St3gal6 | 0.08280346 | 0.024412436 | 0.29482386 | 0.00383849 | 0.02631798 |
| Cnbp | 1.032579094 | 0.808404254 | 0.78289814 | 0.00384707 | 0.02636631 |
| Abce1 | 0.100701305 | 0.188085382 | 1.86775516 | 0.00386557 | 0.02647841 |
| Leo1 | 0.116128639 | 0.051829572 | 0.44631172 | 0.00386958 | 0.02647841 |
| Supt20 | 0.116142153 | 0.050674619 | 0.43631548 | 0.00386958 | 0.02647841 |
| Vps4a | 0.11707258 | 0.047382047 | 0.40472369 | 0.00386959 | 0.02647841 |
| Efcab14 | 0.134702747 | 0.064243996 | 0.4769316 | 0.00388458 | 0.02652027 |
| E2f4 | 0.13554112 | 0.06180031 | 0.45595248 | 0.00388458 | 0.02652027 |
| Mysm1 | 0.13532076 | 0.059956063 | 0.44306626 | 0.00388458 | 0.02652027 |
| Rbfox2 | 0.136065836 | 0.057896075 | 0.42550046 | 0.00388459 | 0.02652027 |
| Kiz | 0.073677344 | 0.024853329 | 0.33732662 | 0.00388649 | 0.02652027 |
| Abhd10 | 0.074661819 | 0.021796804 | 0.29194044 | 0.0038865 | 0.02652027 |
| Cep19 | 0.074959024 | 0.020826467 | 0.27783802 | 0.00388651 | 0.02652027 |
| Gtf2b | 0.25301657 | 0.152370442 | 0.60221527 | 0.00390837 | 0.02665881 |

| | | | | | |
|---|---|---|---|---|---|
| Mea1 | 0.282171943 | 0.17262327 | 0.61176624 | 0.00391579 | 0.02669883 |
| Sh3gl1 | 0.137805247 | 0.059828784 | 0.43415462 | 0.0039214 | 0.02672647 |
| Sfr1 | 0.733430938 | 0.545716926 | 0.74406041 | 0.0039265 | 0.02674241 |
| Arhgdia | 0.599095171 | 0.431827037 | 0.72079873 | 0.00392685 | 0.02674241 |
| Dapp1 | 0.075792315 | 0.023161263 | 0.30558854 | 0.00393629 | 0.02677504 |
| 1600014C10I | 0.076649585 | 0.020273882 | 0.26450087 | 0.00393631 | 0.02677504 |
| Rfx3 | 0.075616636 | 0.021177509 | 0.28006415 | 0.00393631 | 0.02677504 |
| Mbnl3 | 0.010854625 | 0.041052047 | 3.78198667 | 0.00394766 | 0.02683142 |
| Ly6e | 0.547180259 | 0.72317587 | 1.32164101 | 0.00394773 | 0.02683142 |
| Mvp | 0.06875341 | 0.020508023 | 0.29828372 | 0.00397622 | 0.02700383 |
| Ces1d | 0.068573205 | 0.01925217 | 0.28075354 | 0.00397624 | 0.02700383 |
| Ccpg1os | 0.118527681 | 0.047491959 | 0.40068243 | 0.00397862 | 0.02700936 |
| Edem3 | 0.050647236 | 0.116261229 | 2.29550987 | 0.00404013 | 0.02740527 |
| Cox16 | 0.0510088 | 0.115260003 | 2.25961014 | 0.00404013 | 0.02740527 |
| Nfkbia | 0.298915205 | 0.187725965 | 0.62802414 | 0.00404741 | 0.02744378 |
| Wtip | 0.013261312 | 0.053213792 | 4.01270948 | 0.00405738 | 0.02750059 |
| Ube4a | 0.058165468 | 0.11959904 | 2.05618633 | 0.00411955 | 0.02791091 |
| Usp4 | 0.086365658 | 0.161944965 | 1.87510835 | 0.00413128 | 0.02797937 |
| Gramd1a | 0.149502569 | 0.06971563 | 0.46631727 | 0.00417503 | 0.02825345 |
| Scly | 0.149692229 | 0.067288659 | 0.44951337 | 0.00417504 | 0.02825345 |
| Nckap1 | 0.230631492 | 0.130001414 | 0.5636759 | 0.00418084 | 0.02827275 |
| Adipor2 | 0.065027597 | 0.131158908 | 2.01697301 | 0.00418118 | 0.02827275 |
| Tspan15 | 0.10983192 | 0.044135298 | 0.401844 | 0.00424237 | 0.02866398 |
| Tada3 | 0.110447065 | 0.041750321 | 0.37801204 | 0.00424238 | 0.02866398 |
| Smarce1 | 0.287395085 | 0.175088467 | 0.60922569 | 0.00425887 | 0.02876414 |
| Wbp2 | 0.258709609 | 0.154617327 | 0.59764818 | 0.00428012 | 0.02888496 |
| 2610001J05F | 0.259277727 | 0.152911333 | 0.58975885 | 0.00428012 | 0.02888496 |
| Sarnp | 0.489836981 | 0.338842759 | 0.69174597 | 0.0042956 | 0.02897802 |
| Lsm10 | 0.12864098 | 0.056053424 | 0.43573536 | 0.00431046 | 0.02906684 |
| Atp6v1h | 0.15824964 | 0.079353512 | 0.50144514 | 0.00431277 | 0.02907104 |

| | | | | | |
|---|---|---|---|---|---|
| Tor1a | 0.160629026 | 0.080828553 | 0.50320017 | 0.00434417 | 0.02925972 |
| Yipf1 | 0.160204553 | 0.080023319 | 0.49950714 | 0.00434417 | 0.02925972 |
| mt-Nd4l | 0.294230642 | 0.42425216 | 1.44190339 | 0.00434849 | 0.02927734 |
| Pddc1 | 0.207199926 | 0.11299124 | 0.54532471 | 0.00437807 | 0.02946492 |
| Rab22a | 0.170703627 | 0.090275265 | 0.52884211 | 0.00441033 | 0.02964939 |
| Fam134a | 0.171430731 | 0.085978794 | 0.50153665 | 0.00441034 | 0.02964939 |
| Coprs | 0.129846415 | 0.059136179 | 0.45543174 | 0.00441065 | 0.02964939 |
| C1galt1 | 0.072682862 | 0.144606593 | 1.98955557 | 0.00443872 | 0.02981726 |
| Mllt10 | 0.173002529 | 0.087995666 | 0.50863803 | 0.00444083 | 0.02981726 |
| Tulp4 | 0.173409971 | 0.08404928 | 0.48468539 | 0.00444083 | 0.02981726 |
| Plxnb2 | 0.183175601 | 0.094882056 | 0.51798414 | 0.00444653 | 0.02984384 |
| Vps8 | 0.196384732 | 0.103583118 | 0.52744995 | 0.00446295 | 0.0299424 |
| Cyp2j6 | 0.040223385 | 0.095508184 | 2.37444423 | 0.00449831 | 0.03015608 |
| Gm10269 | 0.041151247 | 0.093213425 | 2.26514214 | 0.00449831 | 0.03015608 |
| Slc44a1 | 0.051824814 | 0.117305805 | 2.26350652 | 0.00451542 | 0.03025893 |
| Prpf40a | 0.42075754 | 0.287257694 | 0.6827155 | 0.004525 | 0.03031133 |
| Cuta | 0.684911829 | 0.509654756 | 0.74411732 | 0.00452798 | 0.03031949 |
| Bace1 | 0.101327049 | 0.039381395 | 0.38865629 | 0.00457681 | 0.03063452 |
| Tpm3 | 0.557091259 | 0.400037622 | 0.71808275 | 0.00458086 | 0.03064965 |
| Cenpw | 0.125147235 | 0.216851896 | 1.73277417 | 0.00459951 | 0.03076247 |
| Hacd2 | 0.14083955 | 0.066641756 | 0.47317501 | 0.0046338 | 0.03097977 |
| Tmem50b | 0.102619817 | 0.04310638 | 0.42005903 | 0.00465072 | 0.03106871 |
| Dnal4 | 0.102539543 | 0.04170425 | 0.40671383 | 0.00465072 | 0.03106871 |
| Nolc1 | 0.191404987 | 0.295992124 | 1.54641804 | 0.00467224 | 0.03120034 |
| 9330020H09 | 0.048625814 | 0.011914442 | 0.24502298 | 0.00472656 | 0.03152698 |
| Ccdc112 | 0.049047948 | 0.00844931 | 0.17226632 | 0.00472665 | 0.03152698 |
| Filip1 | 0.048413278 | 0.008563445 | 0.17688215 | 0.00472666 | 0.03152698 |
| Pstk | 0.081665616 | 0.150446865 | 1.84223021 | 0.00475383 | 0.0316959 |
| Sepn1 | 0.05039731 | 0.011287556 | 0.2239714 | 0.00476208 | 0.03170239 |
| Ccdc184 | 0.051365268 | 0.008042507 | 0.1565748 | 0.00476214 | 0.03170239 |

| | | | | | |
|---|---|---|---|---|---|
| Ccl4 | 0.050062342 | 0.008895593 | 0.1776903 | 0.00476215 | 0.03170239 |
| Mttp | 0.051295605 | 0.006684192 | 0.1303073 | 0.00476219 | 0.03170239 |
| Tmem218 | 0.121279744 | 0.053629457 | 0.44219632 | 0.00479031 | 0.03185262 |
| Nmd3 | 0.121923981 | 0.052083124 | 0.42717703 | 0.00479031 | 0.03185262 |
| Cox18 | 0.121249099 | 0.052357329 | 0.43181623 | 0.00479031 | 0.03185262 |
| Aurkaip1 | 0.272589795 | 0.400273843 | 1.46841096 | 0.00480837 | 0.03196031 |
| Dram1 | 0.008730102 | 0.040661613 | 4.65763293 | 0.00481082 | 0.03196421 |
| Mzt2 | 0.117408378 | 0.198511901 | 1.6907814 | 0.00482161 | 0.03202354 |
| Kdelr1 | 0.707458056 | 0.901671621 | 1.27452308 | 0.00484952 | 0.03219645 |
| Frk | 0.021864066 | 0.06672141 | 3.051647 | 0.00485766 | 0.03223803 |
| Ifngr1 | 0.094780012 | 0.0349529 | 0.36877923 | 0.00491685 | 0.03260578 |
| Rac2 | 0.09549958 | 0.031253772 | 0.32726607 | 0.00491686 | 0.03260578 |
| Znhit1 | 0.545486725 | 0.38711918 | 0.70967663 | 0.00494229 | 0.03276172 |
| Hdac11 | 0.068178579 | 0.13290213 | 1.94932385 | 0.00496084 | 0.03287204 |
| BC003331 | 0.083307127 | 0.15633865 | 1.87665395 | 0.0049737 | 0.03294454 |
| Pgam1 | 1.149231148 | 0.921146752 | 0.80153305 | 0.00497989 | 0.03297283 |
| Hnrnph1 | 0.465838655 | 0.325141257 | 0.69796968 | 0.0049832 | 0.032982 |
| Sdf2 | 0.466980376 | 0.324881262 | 0.69570646 | 0.00498667 | 0.03299092 |
| Galnt1 | 0.099109213 | 0.18264368 | 1.84285269 | 0.00498838 | 0.03299092 |
| Rab5a | 0.188514412 | 0.100750579 | 0.53444497 | 0.00503164 | 0.03326418 |
| Rrp7a | 0.175516892 | 0.091801361 | 0.52303434 | 0.0050375 | 0.03329015 |
| Emc2 | 0.200659594 | 0.112540171 | 0.56085118 | 0.00505096 | 0.03336622 |
| Ctsz | 0.487014067 | 0.342779761 | 0.70383955 | 0.00505371 | 0.03337159 |
| Dnajc12 | 0.165051019 | 0.085193054 | 0.51616194 | 0.00506571 | 0.03343794 |
| Ctnnb1 | 0.539793811 | 0.71258706 | 1.32010973 | 0.00509438 | 0.03361428 |
| Exosc5 | 0.160705898 | 0.257675646 | 1.60339882 | 0.00513871 | 0.03389381 |
| Vdac1 | 0.529057689 | 0.377038574 | 0.71266061 | 0.00514768 | 0.03393994 |
| Ell2 | 0.091114307 | 0.167611207 | 1.83957067 | 0.0051497 | 0.03394021 |
| Snx6 | 0.285095627 | 0.174097186 | 0.61066242 | 0.00522393 | 0.03441625 |
| Fkbp5 | 0.086073857 | 0.029980835 | 0.34831523 | 0.00524994 | 0.03457432 |

| | | | | | |
|---|---|---|---|---|---|
| Eps8l2 | 0.087516885 | 0.033086485 | 0.3780583 | 0.00526172 | 0.03461214 |
| Eif2s3y | 0.088290895 | 0.031812539 | 0.36031506 | 0.00526172 | 0.03461214 |
| Arf2 | 0.087551665 | 0.032252954 | 0.36838767 | 0.00526172 | 0.03461214 |
| 0610009L18F | 0.058016573 | 0.010109998 | 0.17426052 | 0.00529527 | 0.0348195 |
| Pebp1 | 2.981586842 | 2.508622946 | 0.84137175 | 0.0053046 | 0.03486747 |
| Sae1 | 0.256339542 | 0.151208155 | 0.58987449 | 0.00531263 | 0.03490695 |
| Tubb6 | 0.114377074 | 0.050768699 | 0.44387129 | 0.00533175 | 0.03501914 |
| Mcts1 | 0.256736107 | 0.154672256 | 0.60245619 | 0.005359 | 0.0351847 |
| Ppp3ca | 0.174126663 | 0.278529365 | 1.59957907 | 0.00538225 | 0.03532385 |
| Cpne8 | 0.041765692 | 0.097466896 | 2.33365929 | 0.00539946 | 0.03542328 |
| Mrpl23 | 0.612319922 | 0.79461735 | 1.29771598 | 0.00542779 | 0.03559553 |
| Asah1 | 0.216961426 | 0.119853949 | 0.55242054 | 0.00543421 | 0.03562405 |
| Ficd | 0.079592157 | 0.027652538 | 0.34742792 | 0.00546105 | 0.03577271 |
| Rmnd5b | 0.079120879 | 0.027400062 | 0.34630635 | 0.00546105 | 0.03577271 |
| App | 0.579256628 | 0.41930454 | 0.72386662 | 0.00550713 | 0.03606078 |
| Anapc11 | 0.537660582 | 0.38639589 | 0.71866137 | 0.00555359 | 0.03635118 |
| Pabpn1l | 0.028349419 | 0.003244599 | 0.11445027 | 0.00556858 | 0.03639544 |
| Hemgn | 0.028348552 | 0.003210479 | 0.11325018 | 0.00556859 | 0.03639544 |
| Cpt1c | 0.028273853 | 0.002370973 | 0.08385745 | 0.00556869 | 0.03639544 |
| Jph4 | 0.029562786 | 0 | 0 | 0.00556883 | 0.03639544 |
| Nrp2 | 0.014750352 | 0.056018562 | 3.79777789 | 0.00558026 | 0.03644266 |
| Slc1a3 | 0.015615892 | 0.052428024 | 3.35735055 | 0.0055803 | 0.03644266 |
| Vps51 | 0.06360029 | 0.019172896 | 0.30145926 | 0.00558672 | 0.03645725 |
| Tmem53 | 0.06480278 | 0.014938641 | 0.23052469 | 0.00558677 | 0.03645725 |
| Wdyhv1 | 0.031815315 | 0.085426731 | 2.68508202 | 0.00559254 | 0.03646726 |
| Elovl7 | 0.03132331 | 0.084292229 | 2.69103833 | 0.00559255 | 0.03646726 |
| Slc22a17 | 0.081824746 | 0.026084584 | 0.31878601 | 0.00560188 | 0.03650044 |
| Pard6a | 0.081527283 | 0.025086225 | 0.30770344 | 0.00560189 | 0.03650044 |
| Samd14 | 0.073264609 | 0.020195233 | 0.27564786 | 0.0056107 | 0.03653015 |
| Wfdc10 | 0.073203815 | 0.020190009 | 0.27580543 | 0.0056107 | 0.03653015 |

| | | | | | |
|---|---|---|---|---|---|
| Atp13a1 | 0.077342946 | 0.151873054 | 1.96363161 | 0.00562376 | 0.03660131 |
| Gtf2f2 | 0.157922075 | 0.079268207 | 0.50194507 | 0.00564422 | 0.03672057 |
| Pogz | 0.066211052 | 0.01906916 | 0.2880057 | 0.00566788 | 0.0368328 |
| Tpst1 | 0.066538613 | 0.017294039 | 0.25990982 | 0.0056679 | 0.0368328 |
| Eid2b | 0.065783741 | 0.017925074 | 0.27248487 | 0.0056679 | 0.0368328 |
| Hmgb2 | 1.027177216 | 1.256264049 | 1.22302562 | 0.00572296 | 0.03717649 |
| Ccnd2 | 1.154205954 | 0.927888412 | 0.80391927 | 0.00573398 | 0.03723402 |
| Clk1 | 0.320328816 | 0.206732013 | 0.64537439 | 0.00574427 | 0.03725023 |
| Nrros | 0.038192785 | 0.008190866 | 0.21446109 | 0.00574915 | 0.03725023 |
| Osbpl8 | 0.038418465 | 0.006993403 | 0.18203234 | 0.0057492 | 0.03725023 |
| Wfdc17 | 0.038726574 | 0.005373514 | 0.13875522 | 0.00574927 | 0.03725023 |
| Crhr2 | 0.039274811 | 0.002014001 | 0.05127971 | 0.00574946 | 0.03725023 |
| Syt14 | 0.039124715 | 0.001845176 | 0.04716139 | 0.00574948 | 0.03725023 |
| Fzd4 | 0.007290113 | 0.031851684 | 4.36916166 | 0.00577616 | 0.03738562 |
| Uba2 | 0.232468156 | 0.135975053 | 0.58491905 | 0.0057766 | 0.03738562 |
| Rnf145 | 0.105756454 | 0.045876243 | 0.43379143 | 0.00578249 | 0.03738562 |
| Aida | 0.125013929 | 0.05960161 | 0.47675976 | 0.00578778 | 0.03738562 |
| Bhlhb9 | 0.124999728 | 0.059563694 | 0.47651059 | 0.00578778 | 0.03738562 |
| Serping1 | 0.12486833 | 0.058833236 | 0.47116219 | 0.00578778 | 0.03738562 |
| Flot1 | 0.124903316 | 0.057781806 | 0.46261227 | 0.00578779 | 0.03738562 |
| Akap13 | 0.124730305 | 0.057822337 | 0.4635789 | 0.00578779 | 0.03738562 |
| Tor1aip2 | 0.233698717 | 0.136555605 | 0.5843233 | 0.0058225 | 0.03759574 |
| Iffo1 | 0.040246713 | 0.006603778 | 0.16408243 | 0.00588284 | 0.03792942 |
| Adarb1 | 0.040298869 | 0.006450046 | 0.16005525 | 0.00588285 | 0.03792942 |
| Optn | 0.040670017 | 0.004706881 | 0.11573345 | 0.00588293 | 0.03792942 |
| Plekha6 | 0.040509874 | 0.003540841 | 0.08740687 | 0.00588301 | 0.03792942 |
| Rbm4b | 0.10864308 | 0.041695205 | 0.3837815 | 0.00594341 | 0.03830442 |
| Zyg11a | 0 | 0.018827625 | Inf | 0.00600754 | 0.03858752 |
| Adprhl1 | 0.000461189 | 0.018285857 | 39.6493601 | 0.00600755 | 0.03858752 |
| Klk1b22 | 0.000538905 | 0.017770229 | 32.9747 | 0.00600762 | 0.03858752 |

| | | | | | |
|---|---|---|---|---|---|
| Rfx8 | 0.000235512 | 0.014937702 | 63.4264156 | 0.00600824 | 0.03858752 |
| Dpf3 | 0.029754913 | 0.003415858 | 0.1147998 | 0.00602507 | 0.03858752 |
| Akt3 | 0.030261556 | 0.002215078 | 0.07319776 | 0.00602515 | 0.03858752 |
| Abhd15 | 0.030266051 | 0.002201717 | 0.07274545 | 0.00602515 | 0.03858752 |
| Fam155a | 0.030662183 | 0.000866895 | 0.02827244 | 0.00602527 | 0.03858752 |
| Sgtb | 0.031294121 | 0 | 0 | 0.00602531 | 0.03858752 |
| Acbd7 | 0.030388388 | 0.00081648 | 0.02686815 | 0.00602532 | 0.03858752 |
| Th | 0.031137574 | 0 | 0 | 0.00602533 | 0.03858752 |
| Tnfrsf9 | 0.030920489 | 0 | 0 | 0.00602536 | 0.03858752 |
| Tnnt2 | 0.030059688 | 0.000754043 | 0.02508487 | 0.00602537 | 0.03858752 |
| Ptprt | 0.030647128 | 0 | 0 | 0.0060254 | 0.03858752 |
| Aif1 | 0.030465477 | 0 | 0 | 0.00602542 | 0.03858752 |
| Fbxo2 | 0.030043064 | 0 | 0 | 0.00602549 | 0.03858752 |
| Nap1l5 | 0.029897667 | 0 | 0 | 0.00602551 | 0.03858752 |
| Bad | 0.248893324 | 0.150485654 | 0.60461909 | 0.00605871 | 0.03878566 |
| Eif2b4 | 0.145797536 | 0.234817178 | 1.61057028 | 0.00607264 | 0.03886037 |
| Ptbp3 | 0.325520274 | 0.209194092 | 0.64264536 | 0.00612232 | 0.0391637 |
| Eif3a | 0.825111894 | 1.032937801 | 1.25187603 | 0.00612781 | 0.03918426 |
| Cyb561 | 0.137495569 | 0.064666876 | 0.47031971 | 0.00614573 | 0.03928422 |
| Pbx3 | 0.039076751 | 0.09222258 | 2.36003705 | 0.0062044 | 0.03962985 |
| Ppt2 | 0.039627198 | 0.089146261 | 2.2496231 | 0.00620442 | 0.03962985 |
| Ndufc2 | 1.334471465 | 1.098989739 | 0.82353933 | 0.00624153 | 0.03985208 |
| Nsmce4a | 0.064351203 | 0.127880413 | 1.98722646 | 0.00631047 | 0.04027727 |
| A430005L14I | 0.196460684 | 0.109977567 | 0.55979428 | 0.00632957 | 0.04038419 |
| Tmem42 | 0.098991462 | 0.043821185 | 0.44267641 | 0.00634138 | 0.04042971 |
| Nav1 | 0.100610116 | 0.036892866 | 0.36669141 | 0.00634141 | 0.04042971 |
| D17Wsu92e | 0.07303783 | 0.140904099 | 1.92919338 | 0.00637621 | 0.04063648 |
| Abcb6 | 0.033945676 | 0.083445406 | 2.45820428 | 0.00641817 | 0.04085853 |
| Gna12 | 0.033131147 | 0.083659467 | 2.52510022 | 0.00641818 | 0.04085853 |
| Rnf185 | 0.034520565 | 0.081646104 | 2.3651439 | 0.00641818 | 0.04085853 |

| | | | | | |
|---|---|---|---|---|---|
| Klc1 | 0.117704526 | 0.052462172 | 0.44571074 | 0.00643376 | 0.0409274 |
| Pex14 | 0.118582529 | 0.051434023 | 0.43374031 | 0.00643376 | 0.0409274 |
| Ick | 0.024098671 | 0.064800363 | 2.68895996 | 0.00648055 | 0.04120976 |
| Mia3 | 0.312396328 | 0.204591326 | 0.65490951 | 0.00652264 | 0.04146206 |
| Coq3 | 0.028989334 | 0.074855604 | 2.58217746 | 0.00652603 | 0.04146828 |
| Hnrnpl | 0.399758466 | 0.271209959 | 0.67843456 | 0.00653493 | 0.04150949 |
| Zdhhc20 | 0.174814919 | 0.092010335 | 0.52632999 | 0.00653866 | 0.04151781 |
| Psmd7 | 0.647039113 | 0.482620263 | 0.7458904 | 0.00654391 | 0.04152972 |
| Ndufb3 | 0.765923246 | 0.589645773 | 0.76984969 | 0.00654537 | 0.04152972 |
| Stk16 | 0.251424956 | 0.154134985 | 0.61304569 | 0.00658655 | 0.04177558 |
| Tmbim4 | 0.579575316 | 0.427399693 | 0.73743598 | 0.00660397 | 0.04187063 |
| Cdc25a | 0.066010695 | 0.132854016 | 2.0126135 | 0.00663463 | 0.04204954 |
| Ipo5 | 0.107838505 | 0.188933405 | 1.7520032 | 0.00665381 | 0.04215552 |
| Esf1 | 0.197875396 | 0.301817253 | 1.52528945 | 0.00674156 | 0.04269574 |
| Adm2 | 0.004556546 | 0.031777943 | 6.97412965 | 0.00676472 | 0.04277998 |
| Slc6a9 | 0.005602252 | 0.030135741 | 5.37921889 | 0.00676475 | 0.04277998 |
| Slit3 | 0.004631174 | 0.030003533 | 6.47860239 | 0.00676481 | 0.04277998 |
| Creb3l4 | 0.005075315 | 0.029412288 | 5.79516465 | 0.00676482 | 0.04277998 |
| Tspyl2 | 0.090717965 | 0.038831342 | 0.42804467 | 0.00684042 | 0.04321053 |
| Capn2 | 0.091370927 | 0.036391884 | 0.39828734 | 0.00684043 | 0.04321053 |
| Msantd4 | 0.090701041 | 0.034751109 | 0.38313903 | 0.00684045 | 0.04321053 |
| Gbe1 | 0.020362881 | 0.060164812 | 2.95463155 | 0.00685294 | 0.04327348 |
| Herc2 | 0.130467518 | 0.062256224 | 0.47717796 | 0.00689068 | 0.0434799 |
| Strn4 | 0.129765689 | 0.062372676 | 0.48065615 | 0.00689069 | 0.0434799 |
| Arpc4 | 0.285153885 | 0.182918319 | 0.6414723 | 0.00689388 | 0.0434841 |
| Smim10l1 | 0.242611657 | 0.144617218 | 0.5960852 | 0.00692504 | 0.04366459 |
| Snapin | 0.093771497 | 0.031584887 | 0.33682823 | 0.00694913 | 0.04380045 |
| Prosc | 0.131561166 | 0.063627075 | 0.48363112 | 0.00702548 | 0.04426539 |
| Vamp3 | 0.201782101 | 0.113667082 | 0.56331598 | 0.00703592 | 0.04431497 |
| Top1 | 0.70618764 | 0.534028176 | 0.75621286 | 0.00705739 | 0.04443391 |

| | | | | | |
|---|---|---|---|---|---|
| Mut | 0.073815002 | 0.141075823 | 1.91120802 | 0.00709855 | 0.04467667 |
| Ash1l | 0.256527908 | 0.160921722 | 0.62730688 | 0.00713126 | 0.04486609 |
| Exph5 | 0.046842933 | 0.01180091 | 0.25192509 | 0.00715614 | 0.04495818 |
| Unc93b1 | 0.046994354 | 0.01077882 | 0.22936416 | 0.00715617 | 0.04495818 |
| Sult1d1 | 0.047850099 | 0.005652422 | 0.1181277 | 0.00715637 | 0.04495818 |
| Arhgef1 | 0.111280482 | 0.04998057 | 0.44914049 | 0.00716157 | 0.04495818 |
| Mef2a | 0.11184073 | 0.047701623 | 0.42651388 | 0.00716158 | 0.04495818 |
| Pdlim4 | 0.112488725 | 0.044226114 | 0.39316041 | 0.0071616 | 0.04495818 |
| Zfp91 | 0.120107022 | 0.200774209 | 1.67162756 | 0.00716484 | 0.0449621 |
| Ostf1 | 0.259409717 | 0.15479885 | 0.59673497 | 0.0071963 | 0.04514306 |
| Acyp1 | 0.228640749 | 0.135292049 | 0.59172326 | 0.00722198 | 0.04528758 |
| Hsbp1l1 | 0.039833648 | 0.093341453 | 2.3432816 | 0.00723984 | 0.04538305 |
| Capzb | 0.864021549 | 0.678935203 | 0.78578503 | 0.00725572 | 0.04546602 |
| Psmd9 | 0.191750862 | 0.102856779 | 0.53640843 | 0.0073115 | 0.04579882 |
| Pigk | 0.17705362 | 0.096863175 | 0.54708384 | 0.00732451 | 0.0458469 |
| Ppil2 | 0.177806847 | 0.094039295 | 0.52888455 | 0.00732451 | 0.0458469 |
| Zfp414 | 0.084767629 | 0.030386312 | 0.35846599 | 0.00733404 | 0.04588981 |
| Csrp1 | 0.216408407 | 0.324657987 | 1.50020968 | 0.00740893 | 0.04634155 |
| 1190007I07R | 0.06007277 | 0.126367181 | 2.10356839 | 0.00741647 | 0.04637183 |
| Tmem106c | 0.178520003 | 0.097025971 | 0.54350196 | 0.00743341 | 0.04644397 |
| Ddah2 | 0.180193703 | 0.093160549 | 0.51700225 | 0.00743341 | 0.04644397 |
| Hnrnpu | 0.470972676 | 0.329768562 | 0.70018619 | 0.00751861 | 0.04695919 |
| Pnlip | 0.002827172 | 0.028915159 | 10.2275919 | 0.00757213 | 0.04721003 |
| Pik3c2b | 0.003179634 | 0.027528381 | 8.65772012 | 0.00757221 | 0.04721003 |
| Myo16 | 0.003222487 | 0.02686712 | 8.33738674 | 0.00757225 | 0.04721003 |
| Cxcl15 | 0.00325261 | 0.026477389 | 8.14035193 | 0.00757228 | 0.04721003 |
| Gm5093 | 0.00377687 | 0.023561384 | 6.23833567 | 0.00757251 | 0.04721003 |
| Prrc1 | 0.143993178 | 0.232611717 | 1.61543568 | 0.00758513 | 0.04727154 |
| Snx3 | 0.69199933 | 0.525733072 | 0.75973061 | 0.00761168 | 0.0474198 |
| Plxna2 | 0.025841966 | 0.070549647 | 2.7300418 | 0.00765628 | 0.0476804 |

| | | | | | |
|---|---|---|---|---|---|
| Actrt3 | 0.001650116 | 0.023668049 | 14.343261 | 0.00770629 | 0.0479083 |
| Klk1b8 | 0.001628748 | 0.023023795 | 14.1358854 | 0.00770636 | 0.0479083 |
| Myl10 | 0.000907806 | 0.02360408 | 26.0012383 | 0.00770638 | 0.0479083 |
| Tnc | 0.002523533 | 0.020233343 | 8.01786292 | 0.0077066 | 0.0479083 |
| Banf2 | 0.002396154 | 0.018952913 | 7.909722 | 0.00770682 | 0.0479083 |
| Spop | 0.262291534 | 0.160372646 | 0.61142898 | 0.00771519 | 0.047943 |
| Bin1 | 0.113069144 | 0.193654374 | 1.71270753 | 0.00778545 | 0.04836212 |
| Dync1h1 | 0.204655663 | 0.116884202 | 0.57112616 | 0.00779789 | 0.04842188 |
| Ddb1 | 0.326975675 | 0.455170666 | 1.39206278 | 0.00781086 | 0.04848488 |
| Ddhd2 | 0.07744943 | 0.028856113 | 0.37258006 | 0.00781833 | 0.04849649 |
| Fam98c | 0.077365899 | 0.025744459 | 0.33276235 | 0.00781837 | 0.04849649 |
| Col5a2 | 0.036171734 | 0.082304721 | 2.27538774 | 0.0078353 | 0.04858393 |
| Tsn | 0.496034658 | 0.354754944 | 0.71518177 | 0.00785526 | 0.04869014 |
| Rnf14 | 0.10329221 | 0.045373151 | 0.43926982 | 0.00789489 | 0.04891811 |
| Lrfn4 | 0.053638249 | 0.014763169 | 0.27523586 | 0.00791956 | 0.04896562 |
| N4bp2l1 | 0.053561103 | 0.014292029 | 0.26683596 | 0.00791958 | 0.04896562 |
| Dach1 | 0.053425389 | 0.014229024 | 0.2663345 | 0.00791958 | 0.04896562 |
| Armc9 | 0.054148032 | 0.012087611 | 0.2232327 | 0.00791963 | 0.04896562 |
| Srxn1 | 0.054705165 | 0.011055833 | 0.20209853 | 0.00791965 | 0.04896562 |
| Socs3 | 0.054753167 | 0.010850131 | 0.19816445 | 0.00791965 | 0.04896562 |
| Uso1 | 0.310810197 | 0.20352402 | 0.65481771 | 0.00792709 | 0.04899395 |
| Mboat7 | 0.104324262 | 0.046078948 | 0.44168966 | 0.00798224 | 0.04926404 |
| Fam101b | 0.104405505 | 0.045235612 | 0.43326845 | 0.00798224 | 0.04926404 |
| Arrdc1 | 0.105747945 | 0.042522235 | 0.40210933 | 0.00798225 | 0.04926404 |
| Grb7 | 0.105679333 | 0.041974376 | 0.39718623 | 0.00798225 | 0.04926404 |
| Eif2b2 | 0.169746347 | 0.26144185 | 1.54019132 | 0.00799472 | 0.04932329 |
| Ttc13 | 0.07838117 | 0.142174302 | 1.81388339 | 0.00800978 | 0.04939845 |
| Kat8 | 0.069208018 | 0.023733148 | 0.34292483 | 0.0080524 | 0.04956075 |
| Mettl6 | 0.069012266 | 0.023283169 | 0.33737726 | 0.00805241 | 0.04956075 |
| Rnf41 | 0.069508685 | 0.020866867 | 0.30020518 | 0.00805244 | 0.04956075 |

| | | | | | |
|---|---|---|---|---|---|
| Slc40a1 | 0.068918737 | 0.021083986 | 0.30592531 | 0.00805245 | 0.04956075 |
| Trappc12 | 0.055669329 | 0.014271119 | 0.25635515 | 0.0080591 | 0.04956075 |
| Pip4k2c | 0.056086845 | 0.013217696 | 0.23566482 | 0.00805912 | 0.04956075 |
| Cdc25b | 0.056212591 | 0.012268893 | 0.21825881 | 0.00805915 | 0.04956075 |
| Phf1 | 0.055629852 | 0.012006889 | 0.21583536 | 0.00805917 | 0.04956075 |
| Map2 | 0.135546723 | 0.063997324 | 0.47214217 | 0.0081191 | 0.0499114 |
| Nfatc1 | 0.016163577 | 0.056749909 | 3.51097476 | 0.00821192 | 0.0504102 |
| Ptpn3 | 0.016950743 | 0.053992566 | 3.1852625 | 0.00821196 | 0.0504102 |
| Gstk1 | 0.016238989 | 0.054330564 | 3.34568636 | 0.00821197 | 0.0504102 |
| Dtx4 | 0.017496507 | 0.052632056 | 3.00814651 | 0.00821197 | 0.0504102 |
| Fam114a2 | 0.221268746 | 0.134111568 | 0.60610263 | 0.0082249 | 0.0504715 |
| Pspc1 | 0.071319562 | 0.022346363 | 0.31332726 | 0.00826957 | 0.0507275 |
| Phf10 | 0.055059601 | 0.117795367 | 2.13941554 | 0.0083001 | 0.0508617 |
| Pradc1 | 0.056640377 | 0.109862344 | 1.93964714 | 0.00830013 | 0.0508617 |
| Ilk | 0.265249468 | 0.16600788 | 0.62585566 | 0.00830303 | 0.0508617 |
| Osbp | 0.04875422 | 0.105769447 | 2.16944188 | 0.00830604 | 0.0508617 |
| Zfos1 | 0.524899352 | 0.684651535 | 1.30434822 | 0.00830625 | 0.0508617 |
| Gadd45b | 0.124431064 | 0.207361547 | 1.66647733 | 0.00831617 | 0.05088622 |
| Pih1d1 | 0.124642686 | 0.207055141 | 1.66118966 | 0.00831617 | 0.05088622 |
| Ssbp2 | 0.146558749 | 0.074588838 | 0.50893474 | 0.00832484 | 0.05090302 |
| Arl6ip6 | 0.146861746 | 0.073968769 | 0.5036626 | 0.00832485 | 0.05090302 |
| Wbp11 | 0.267999647 | 0.164749713 | 0.61473854 | 0.00834029 | 0.05097934 |
| Irf2 | 0.158487723 | 0.082893499 | 0.52302789 | 0.00839312 | 0.05126575 |
| Gid8 | 0.159373439 | 0.081579391 | 0.5118757 | 0.00839312 | 0.05126575 |
| Txndc11 | 0.106057612 | 0.185571349 | 1.7497221 | 0.00854318 | 0.05215115 |
| Srrm2 | 1.202084958 | 0.980466568 | 0.81563833 | 0.00854415 | 0.05215115 |
| Nfib | 0.240858265 | 0.350287597 | 1.45433081 | 0.00858345 | 0.05237243 |
| Ing4 | 0.211020595 | 0.12438217 | 0.58943143 | 0.00862901 | 0.05263172 |
| Fam175b | 0.097215162 | 0.037191033 | 0.38256412 | 0.00866952 | 0.05282371 |
| Bcl2l1 | 0.09670116 | 0.037258703 | 0.38529737 | 0.00866953 | 0.05282371 |

| | | | | | |
|---|---|---|---|---|---|
| Gsto1 | 0.303152311 | 0.42512183 | 1.40233742 | 0.0086697 | 0.05282371 |
| Pcf11 | 0.116499294 | 0.054758734 | 0.4700349 | 0.00872096 | 0.05309836 |
| Nkiras1 | 0.115930691 | 0.054151914 | 0.46710593 | 0.00872096 | 0.05309836 |
| Ndufa10 | 0.409324928 | 0.549815946 | 1.34322615 | 0.00875467 | 0.05328471 |
| Mrpl20 | 0.573540275 | 0.426115358 | 0.7429563 | 0.00878897 | 0.05347456 |
| Fam76a | 0.098054581 | 0.041586133 | 0.42411208 | 0.00890786 | 0.05410226 |
| Polr3h | 0.097545654 | 0.041372703 | 0.42413681 | 0.00890786 | 0.05410226 |
| Fbxw9 | 0.097832572 | 0.038622708 | 0.39478373 | 0.00890788 | 0.05410226 |
| Tmem161a | 0.098941194 | 0.037479673 | 0.37880757 | 0.00890788 | 0.05410226 |
| St3gal4 | 0.098493468 | 0.037633333 | 0.38208963 | 0.00890789 | 0.05410226 |
| Nipal2 | 0.032094563 | 0.07761358 | 2.41827816 | 0.00891558 | 0.05412984 |
| Pcnp | 0.31906582 | 0.213428772 | 0.66891769 | 0.00893703 | 0.05423714 |
| Ascc1 | 0.198879819 | 0.11551205 | 0.58081333 | 0.00893956 | 0.05423714 |
| Pdxdc1 | 0.128512395 | 0.211050075 | 1.64225462 | 0.00895323 | 0.05430088 |
| Rtf1 | 0.302757396 | 0.195507276 | 0.64575557 | 0.00898038 | 0.05442119 |
| Gm21092 | 0.227471089 | 0.135050352 | 0.59370337 | 0.00898109 | 0.05442119 |
| Prkrir | 0.026882666 | 0.072375236 | 2.69226408 | 0.00899046 | 0.05442119 |
| Fgfr1 | 0.026574969 | 0.070778005 | 2.66333355 | 0.00899048 | 0.05442119 |
| Ddx20 | 0.027878667 | 0.067392968 | 2.41736692 | 0.00899051 | 0.05442119 |
| Sh3gl2 | 0.037097689 | 0.006532876 | 0.17609928 | 0.00901696 | 0.05442119 |
| Nme5 | 0.037359341 | 0.005962041 | 0.15958636 | 0.00901699 | 0.05442119 |
| Fam101a | 0.036967579 | 0.006322299 | 0.17102281 | 0.00901699 | 0.05442119 |
| Evpl | 0.037251426 | 0.00399361 | 0.1072069 | 0.00901719 | 0.05442119 |
| Klf1 | 0.036593207 | 0.004325844 | 0.11821441 | 0.00901722 | 0.05442119 |
| Zbtb7c | 0.036976579 | 0.003871164 | 0.10469234 | 0.00901723 | 0.05442119 |
| Msl3l2 | 0.037894654 | 0.002501402 | 0.06600936 | 0.00901728 | 0.05442119 |
| Fbxo44 | 0.03788241 | 0.002392764 | 0.06316292 | 0.00901729 | 0.05442119 |
| St8sia3 | 0.037631632 | 0.00163296 | 0.04339328 | 0.00901741 | 0.05442119 |
| Nrep | 0.304672245 | 0.197278693 | 0.64751121 | 0.00904152 | 0.05452411 |
| Dgkz | 0.044212601 | 0.096688389 | 2.18689663 | 0.00904398 | 0.05452411 |

| | | | | | |
|---|---|---|---|---|---|
| Pcca | 0.044838239 | 0.095793058 | 2.13641437 | 0.00904398 | 0.05452411 |
| Gdi2 | 0.673149131 | 0.515517158 | 0.76582905 | 0.00913023 | 0.05502479 |
| Bex1 | 0.472153258 | 0.339373501 | 0.71877827 | 0.00915879 | 0.05517751 |
| Cln6 | 0.129418169 | 0.061112656 | 0.47221079 | 0.00920509 | 0.05541767 |
| Rcan3 | 0.128698703 | 0.059801436 | 0.46466231 | 0.0092051 | 0.05541767 |
| Ubl7 | 0.187048079 | 0.108845793 | 0.58191345 | 0.00920862 | 0.05541942 |
| Reep3 | 0.072788755 | 0.137422037 | 1.88795696 | 0.0092138 | 0.0554312 |
| Dhrs4 | 0.394867169 | 0.269547108 | 0.6826273 | 0.00922438 | 0.0554754 |
| Slc50a1 | 0.241361589 | 0.15053339 | 0.62368412 | 0.00927398 | 0.0557347 |
| Rwdd4a | 0.242134923 | 0.146864944 | 0.60654177 | 0.00927398 | 0.0557347 |
| Pla2g1b | 0.05139483 | 0.105745681 | 2.05751593 | 0.00929729 | 0.05583588 |
| Wwc1 | 0.051549651 | 0.101530063 | 1.9695587 | 0.00929731 | 0.05583588 |
| Fkbp2 | 1.565404483 | 1.837598179 | 1.17388074 | 0.00931043 | 0.05587881 |
| Clpp | 0.292970833 | 0.410096785 | 1.39978707 | 0.00931097 | 0.05587881 |
| Fhdc1 | 0.026562902 | 0.00273192 | 0.10284719 | 0.00935591 | 0.05591854 |
| 1700113A16I | 0.026620866 | 0.0025903 | 0.09730337 | 0.00935593 | 0.05591854 |
| S100z | 0.027825214 | 0 | 0 | 0.00935624 | 0.05591854 |
| Ddx25 | 0.026436957 | 0.001224407 | 0.04631422 | 0.00935627 | 0.05591854 |
| Rltpr | 0.027572172 | 0 | 0 | 0.0093563 | 0.05591854 |
| Tmem198 | 0.026781351 | 0.000763657 | 0.02851451 | 0.0093563 | 0.05591854 |
| Cldn13 | 0.027412841 | 0 | 0 | 0.00935634 | 0.05591854 |
| Cdk5r2 | 0.027105258 | 0 | 0 | 0.00935641 | 0.05591854 |
| 4933417E11I | 0.02706887 | 0 | 0 | 0.00935642 | 0.05591854 |
| Map6 | 0.026735526 | 0 | 0 | 0.00935651 | 0.05591854 |
| Kcnb1 | 0.026483456 | 0 | 0 | 0.00935658 | 0.05591854 |
| Mar-04 | 0.026263154 | 0 | 0 | 0.00935664 | 0.05591854 |
| Dnlz | 0.177436325 | 0.271339102 | 1.52921958 | 0.00940015 | 0.05615904 |
| Ext2 | 0.138917017 | 0.071295735 | 0.51322535 | 0.00941534 | 0.05621074 |
| Adgrg1 | 0.137934321 | 0.070356765 | 0.51007439 | 0.00941535 | 0.05621074 |
| Rbm18 | 0.140394985 | 0.070320223 | 0.50087418 | 0.00947852 | 0.05656823 |

| | | | | | |
|---|---|---|---|---|---|
| Auts2 | 0.162654018 | 0.08604722 | 0.52901995 | 0.00949109 | 0.05660394 |
| Rb1cc1 | 0.16264803 | 0.08509945 | 0.5232123 | 0.00949109 | 0.05660394 |
| Btrc | 0.090099154 | 0.035143138 | 0.39004959 | 0.00951087 | 0.05666231 |
| Smpd3 | 0.089367919 | 0.034472935 | 0.38574172 | 0.00951088 | 0.05666231 |
| Specc1l | 0.090511065 | 0.031181134 | 0.3445008 | 0.00951091 | 0.05666231 |
| Akap8l | 0.151126796 | 0.078682293 | 0.52063761 | 0.00952066 | 0.05666231 |
| 0610010K14I | 0.150409852 | 0.077956748 | 0.5182955 | 0.00952066 | 0.05666231 |
| Arl2bp | 0.1512831 | 0.075544902 | 0.49936115 | 0.00952067 | 0.05666231 |
| Leprot | 0.164228938 | 0.087676193 | 0.53386567 | 0.00955272 | 0.05680869 |
| Proser2 | 0.065902254 | 0.128515954 | 1.95009953 | 0.00955531 | 0.05680869 |
| Rprm | 0.066005781 | 0.125618959 | 1.90315088 | 0.00955532 | 0.05680869 |
| Nsfl1c | 0.307278504 | 0.201794757 | 0.65671615 | 0.00955849 | 0.05680869 |
| Psmd14 | 0.273995145 | 0.175721509 | 0.64133074 | 0.00957278 | 0.05687395 |
| Hmmr | 0.091844639 | 0.161254108 | 1.75572695 | 0.00957842 | 0.05688783 |
| 2310011J03F | 0.152450139 | 0.077775312 | 0.51016885 | 0.0095824 | 0.0568918 |
| Snrpn | 0.108667707 | 0.048440569 | 0.44576784 | 0.0096321 | 0.05714748 |
| Kdf1 | 0.108422824 | 0.04622305 | 0.42632214 | 0.00963212 | 0.05714748 |
| Rab5c | 0.293442897 | 0.18814656 | 0.64116924 | 0.00964537 | 0.05720631 |
| Kif1b | 0.229448987 | 0.142495022 | 0.62103139 | 0.00976647 | 0.05790458 |
| Gcsh | 0.438574477 | 0.31043479 | 0.70782685 | 0.00983084 | 0.05826611 |
| Mthfsl | 0.109527263 | 0.0511956 | 0.46742335 | 0.00984568 | 0.05829385 |
| Ttc33 | 0.109921734 | 0.049825662 | 0.45328308 | 0.00984569 | 0.05829385 |
| Kifap3 | 0.109875418 | 0.048877718 | 0.44484671 | 0.0098457 | 0.05829385 |
| Itgb1 | 0.380517686 | 0.261759076 | 0.68790252 | 0.00989267 | 0.05855182 |
| Zbtb18 | 0.018510149 | 0.056134479 | 3.03263255 | 0.00990849 | 0.05862526 |

| | mean outside cluster | mean inside cluster | fold change | P value | P adj |
|---|---|---|---|---|---|
| Acta2 | 0.146239883 | 3.35157816 | 22.9183591 | 0 | 0 |
| Iapp | 68.41381283 | 0.536298928 | 0.00783904 | 1.39E-41 | 1.20E-37 |
| Nnat | 14.34608312 | 0.115262423 | 0.00803442 | 4.88E-36 | 2.79E-32 |
| Myl9 | 0.085044811 | 1.865532667 | 21.9358789 | 3.85E-34 | 1.66E-30 |
| Chga | 7.102986561 | 0.037410969 | 0.00526694 | 4.64E-32 | 1.59E-28 |
| Pyy | 13.16681127 | 0.307726384 | 0.02337137 | 1.99E-31 | 5.69E-28 |
| Npy | 3.586643828 | 0.015885894 | 0.00442918 | 7.12E-29 | 1.75E-25 |
| Sst | 4.697531177 | 0.055353062 | 0.01178344 | 1.04E-28 | 2.24E-25 |
| Chgb | 4.510457895 | 0.034658975 | 0.00768414 | 2.12E-28 | 4.05E-25 |
| Ins1 | 271.689853 | 4.093629557 | 0.01506729 | 9.40E-28 | 1.62E-24 |
| Actg2 | 0.074352711 | 1.445467237 | 19.4406797 | 1.41E-26 | 2.20E-23 |
| Rbp4 | 5.149344777 | 0.116410947 | 0.02260694 | 1.55E-26 | 2.22E-23 |
| Pcsk2 | 4.120828779 | 0.065559566 | 0.01590932 | 2.80E-26 | 3.70E-23 |
| Reg1 | 0.606470603 | 3.450329504 | 5.68919496 | 4.96E-25 | 6.09E-22 |
| Ins2 | 462.3981436 | 5.894122246 | 0.01274686 | 5.17E-24 | 5.92E-21 |
| Pcsk1n | 3.726351728 | 0.184185445 | 0.04942782 | 8.91E-24 | 9.57E-21 |
| Hbb-bt | 22.65979417 | 1.802233954 | 0.07953444 | 3.88E-23 | 3.92E-20 |
| Gcg | 16.61604678 | 1.346346978 | 0.08102691 | 8.12E-23 | 7.44E-20 |
| Cd74 | 0.319057683 | 2.26626389 | 7.10299112 | 8.22E-23 | 7.44E-20 |
| Mylk | 0.062908848 | 1.183346092 | 18.8104873 | 2.53E-21 | 2.17E-18 |
| Hba-a1 | 104.7289571 | 7.102039856 | 0.06781353 | 1.46E-20 | 1.20E-17 |
| Cpe | 3.488700787 | 0.307756131 | 0.08821511 | 1.45E-19 | 1.13E-16 |
| Scg2 | 2.083972748 | 0.007023125 | 0.00337007 | 3.65E-19 | 2.72E-16 |
| Scg5 | 2.032878419 | 0.01404625 | 0.00690954 | 3.55E-17 | 2.54E-14 |
| Ctrl | 0.091559809 | 1.105496383 | 12.0740355 | 3.56E-16 | 2.45E-13 |
| Ifitm3 | 1.245961032 | 4.317703027 | 3.4653596 | 1.86E-15 | 1.23E-12 |
| Rnase1 | 7.763015919 | 24.57251078 | 3.16533046 | 2.04E-15 | 1.30E-12 |
| Scg3 | 1.670601021 | 0.006079878 | 0.00363934 | 2.31E-15 | 1.42E-12 |
| Tagln | 0.042067195 | 0.852767539 | 20.2715571 | 5.75E-15 | 3.41E-12 |

| | | | | | |
|---|---|---|---|---|---|
| Gpx3 | 2.134508245 | 0.152490047 | 0.07144036 | 1.67E-14 | 9.57E-12 |
| Gstm1 | 0.979979731 | 3.300098144 | 3.36751673 | 1.26E-13 | 6.97E-11 |
| Hbb-bs | 132.1359408 | 16.62596645 | 0.12582471 | 1.63E-13 | 8.74E-11 |
| Ube2c | 0.47697498 | 2.029500471 | 4.25494115 | 1.72E-13 | 8.96E-11 |
| Cpa2 | 3.428879241 | 9.980557898 | 2.91073473 | 3.05E-13 | 1.51E-10 |
| Tmem97 | 1.209086596 | 3.870126439 | 3.20086787 | 3.07E-13 | 1.51E-10 |
| Slc38a5 | 1.727727666 | 0.125014444 | 0.07235773 | 3.71E-13 | 1.77E-10 |
| Ldha | 1.289066728 | 4.063133266 | 3.15199607 | 4.48E-13 | 2.08E-10 |
| Cryba2 | 1.394146411 | 0.007118569 | 0.00510604 | 6.57E-13 | 2.97E-10 |
| Hes1 | 0.982523607 | 3.089422175 | 3.1443745 | 2.63E-12 | 1.16E-09 |
| Fabp5 | 1.809224324 | 5.308080165 | 2.9338983 | 3.08E-12 | 1.32E-09 |
| Gnas | 8.334683569 | 1.841598389 | 0.22095601 | 7.28E-12 | 3.04E-09 |
| Tmem27 | 1.426123092 | 0.051001518 | 0.03576235 | 7.43E-12 | 3.04E-09 |
| Ghrl | 1.684807258 | 0.10014214 | 0.05943834 | 1.03E-11 | 4.13E-09 |
| Myh11 | 0.025238456 | 0.566930146 | 22.462949 | 1.13E-11 | 4.41E-09 |
| Ppy | 1.37388121 | 0.047991191 | 0.03493111 | 2.27E-11 | 8.67E-09 |
| Ptf1a | 0.215164385 | 1.206925327 | 5.6093174 | 7.01E-11 | 2.62E-08 |
| Sycn | 2.960496955 | 7.755846105 | 2.61977844 | 1.11E-10 | 4.07E-08 |
| Ppp1r1a | 1.111303164 | 0.020265226 | 0.01823555 | 1.80E-10 | 6.43E-08 |
| Cckar | 0.482204936 | 1.725806437 | 3.57898957 | 5.63E-10 | 1.97E-07 |
| Slc30a8 | 1.027641263 | 0 | 0 | 1.26E-09 | 4.33E-07 |
| Tpm2 | 0.128570689 | 0.887611571 | 6.90368528 | 2.35E-09 | 7.92E-07 |
| Cnn1 | 0.017339976 | 0.425197609 | 24.5212341 | 2.83E-09 | 9.33E-07 |
| Itln1 | 0.072414705 | 0.678068642 | 9.36368716 | 3.91E-09 | 1.27E-06 |
| Fam183b | 0.943404698 | 0.01404625 | 0.01488889 | 5.41E-09 | 1.72E-06 |
| Dbi | 2.633667739 | 6.404424061 | 2.43175096 | 6.27E-09 | 1.96E-06 |
| Cpa1 | 9.643063723 | 22.79395898 | 2.36376733 | 1.04E-08 | 3.19E-06 |
| H2-Ab1 | 0.127844501 | 0.792552313 | 6.19934615 | 1.21E-08 | 3.65E-06 |
| Hmgb2 | 1.049093289 | 2.681594937 | 2.55610723 | 2.40E-08 | 7.10E-06 |
| Ang | 0.370677446 | 1.341634497 | 3.61941227 | 3.42E-08 | 9.97E-06 |

| | | | | | |
|---|---|---|---|---|---|
| Mt2 | 3.090693875 | 7.097820199 | 2.2965135 | 4.33E-08 | 1.24E-05 |
| Tpm1 | 0.40195812 | 1.405266249 | 3.4960514 | 4.93E-08 | 1.39E-05 |
| Ttr | 7.93512575 | 2.577485745 | 0.32481977 | 5.15E-08 | 1.43E-05 |
| 1700086L19R | 0.830912164 | 0 | 0 | 5.27E-08 | 1.44E-05 |
| Aplp1 | 0.995864112 | 0.062264877 | 0.06252347 | 5.69E-08 | 1.53E-05 |
| Rpl35 | 6.337645763 | 14.2223566 | 2.24410722 | 6.49E-08 | 1.71E-05 |
| Ggh | 0.338470938 | 1.23789649 | 3.65731988 | 7.48E-08 | 1.95E-05 |
| Phgdh | 0.592184018 | 1.753380924 | 2.96087174 | 7.62E-08 | 1.95E-05 |
| H2-Aa | 0.114034514 | 0.734740803 | 6.44314408 | 9.75E-08 | 2.46E-05 |
| Cldn10 | 0.605338425 | 1.727882569 | 2.85440755 | 1.23E-07 | 3.05E-05 |
| Nupr1 | 1.544435555 | 3.558440336 | 2.30403938 | 1.52E-07 | 3.73E-05 |
| Clps | 57.01223875 | 144.2504833 | 2.53016697 | 1.57E-07 | 3.79E-05 |
| Mt1 | 8.101435009 | 17.81655725 | 2.19918536 | 1.65E-07 | 3.93E-05 |
| Scgn | 0.782981268 | 0.01404625 | 0.01793945 | 1.68E-07 | 3.95E-05 |
| Pax6 | 0.941421208 | 0.049161874 | 0.05222091 | 1.72E-07 | 4.00E-05 |
| Mif | 3.40668993 | 7.56497838 | 2.22062428 | 1.78E-07 | 4.09E-05 |
| Hspe1 | 1.962018861 | 4.437447941 | 2.26167446 | 1.90E-07 | 4.29E-05 |
| Erp27 | 0.751084707 | 1.919026699 | 2.55500702 | 2.08E-07 | 4.65E-05 |
| Reep5 | 4.80783321 | 10.45791184 | 2.17518191 | 2.67E-07 | 5.89E-05 |
| Cldn3 | 0.688498487 | 1.795496461 | 2.60784373 | 3.17E-07 | 6.90E-05 |
| Csrp1 | 0.227806042 | 0.958579432 | 4.20787536 | 3.43E-07 | 7.36E-05 |
| Pcsk1 | 0.746925986 | 0 | 0 | 3.60E-07 | 7.63E-05 |
| Serpina6 | 2.59219431 | 5.679455686 | 2.19098378 | 4.07E-07 | 8.53E-05 |
| Mafb | 0.867593787 | 0.042138749 | 0.04856968 | 4.72E-07 | 9.77E-05 |
| Gng12 | 1.205700394 | 0.176005981 | 0.14597821 | 6.27E-07 | 0.00012816 |
| Sepp1 | 1.147508346 | 2.613570783 | 2.27760503 | 6.43E-07 | 0.00012993 |
| Bex4 | 0.655843144 | 1.682488854 | 2.565383 | 7.18E-07 | 0.00014338 |
| Ncl | 1.619738576 | 3.602334681 | 2.22402228 | 8.43E-07 | 0.00016656 |
| Psmb10 | 0.52279661 | 1.471596176 | 2.81485409 | 9.22E-07 | 0.00018008 |
| Gcat | 0.716248444 | 1.835014488 | 2.56198042 | 9.39E-07 | 0.00018132 |

| | | | | | |
|---|---|---|---|---|---|
| Gch1 | 0.848312826 | 0.058045205 | 0.06842429 | 1.08E-06 | 0.00020612 |
| Krt18 | 1.8282564 | 3.960034558 | 2.16601706 | 1.20E-06 | 0.00022697 |
| Rpl22 | 3.512234842 | 7.34206002 | 2.090424 | 1.41E-06 | 0.00026409 |
| Dlk1 | 4.424219009 | 1.679260488 | 0.37956089 | 1.64E-06 | 0.00030264 |
| Tmed6 | 0.653340445 | 1.631951347 | 2.49785753 | 2.22E-06 | 0.00040273 |
| Pnliprp1 | 46.40424862 | 104.9906292 | 2.26252191 | 2.23E-06 | 0.00040273 |
| Rpl5 | 3.773501668 | 7.748970173 | 2.05352239 | 2.26E-06 | 0.00040395 |
| Ppp1r14a | 0.006106504 | 0.222917491 | 36.5049311 | 2.51E-06 | 0.00044417 |
| Rpl36a | 6.441290481 | 13.09269895 | 2.03262048 | 2.70E-06 | 0.00047276 |
| Hba-a2 | 2.261717148 | 0.86669284 | 0.38320125 | 3.03E-06 | 0.00052646 |
| Xbp1 | 0.821398624 | 1.898728288 | 2.31157958 | 3.24E-06 | 0.00055588 |
| Rps27l | 4.047727823 | 8.199820997 | 2.02578369 | 3.64E-06 | 0.00061942 |
| Rpl12 | 4.863216395 | 9.834765165 | 2.02227587 | 3.76E-06 | 0.00062839 |
| Gamt | 0.642803603 | 1.629899238 | 2.53560999 | 3.77E-06 | 0.00062839 |
| Rnase4 | 0.999544007 | 2.193479428 | 2.1944801 | 3.92E-06 | 0.00064513 |
| Tm4sf4 | 0.885677097 | 0.128665073 | 0.14527312 | 3.94E-06 | 0.00064513 |
| Dhx34 | 0.274072712 | 0.973460674 | 3.55183362 | 4.69E-06 | 0.00075579 |
| Tspan7 | 1.091448259 | 0.169435344 | 0.15523901 | 4.71E-06 | 0.00075579 |
| Cd24a | 1.23228204 | 2.602459559 | 2.11190253 | 4.91E-06 | 0.00078116 |
| Syt13 | 0.621054592 | 0.01404625 | 0.02261677 | 5.29E-06 | 0.00083374 |
| Lyar | 0.277623252 | 0.969887416 | 3.49353812 | 5.66E-06 | 0.00088367 |
| Neurod1 | 0.600360309 | 0.007079223 | 0.01179162 | 7.60E-06 | 0.0011769 |
| Ass1 | 0.362711022 | 1.086318107 | 2.99499613 | 7.93E-06 | 0.00121637 |
| Rps12 | 7.243940937 | 14.25273324 | 1.96753858 | 8.43E-06 | 0.00128135 |
| Rps18-ps3 | 1.740955618 | 3.492680683 | 2.00618594 | 9.77E-06 | 0.00147308 |
| Serf2 | 4.921931061 | 9.640326302 | 1.95864716 | 1.08E-05 | 0.00160764 |
| Prnp | 0.58011569 | 0.020366356 | 0.0351074 | 1.10E-05 | 0.00163664 |
| H19 | 1.694699635 | 3.391531382 | 2.00125811 | 1.14E-05 | 0.00167465 |
| Meg3 | 13.07279567 | 5.44665118 | 0.41664012 | 1.46E-05 | 0.00212119 |
| H2afz | 3.138168137 | 6.067548656 | 1.93346831 | 1.49E-05 | 0.00215029 |

| | | | | | |
|---|---|---|---|---|---|
| Gapdh | 3.312750073 | 6.440252394 | 1.94408037 | 1.59E-05 | 0.00226968 |
| Rps8 | 14.85827304 | 29.0876284 | 1.95767222 | 1.63E-05 | 0.00231448 |
| Car9 | 0.180848124 | 0.693432448 | 3.83433586 | 1.71E-05 | 0.00240806 |
| Ifitm2 | 1.300162564 | 2.690607099 | 2.06943899 | 1.72E-05 | 0.00240866 |
| Rbpjl | 0.303661148 | 0.956377574 | 3.14948942 | 1.81E-05 | 0.00251429 |
| Hmgn3 | 0.886262255 | 0.111224928 | 0.12549889 | 2.38E-05 | 0.00327092 |
| Rps20 | 7.723775993 | 14.67884104 | 1.90047472 | 2.51E-05 | 0.00342134 |
| Isl1 | 0.675881274 | 0.012587657 | 0.01862407 | 2.76E-05 | 0.00373059 |
| Rps11 | 8.598973403 | 16.27134095 | 1.89224227 | 2.78E-05 | 0.00373059 |
| Rps4x | 10.8421796 | 20.60289904 | 1.90025436 | 2.88E-05 | 0.00383216 |
| Cdk1 | 0.238519526 | 0.793706002 | 3.32763534 | 2.98E-05 | 0.00393359 |
| Aldh1a1 | 0.146589236 | 0.632423563 | 4.31425648 | 3.31E-05 | 0.00433717 |
| Tuba1b | 1.472068637 | 2.929370703 | 1.98996883 | 3.43E-05 | 0.00445848 |
| Alas2 | 0.531255346 | 0.013804121 | 0.02598397 | 3.62E-05 | 0.00467702 |
| Eno1 | 1.13934353 | 2.269794587 | 1.99219509 | 3.69E-05 | 0.00468103 |
| Rpl17 | 11.75037484 | 22.21142108 | 1.89027341 | 3.70E-05 | 0.00468103 |
| Rpl10a | 8.001526789 | 14.99734829 | 1.87431083 | 3.71E-05 | 0.00468103 |
| Rps23 | 13.37282467 | 25.27086274 | 1.88971764 | 4.25E-05 | 0.00533483 |
| Rpl32 | 16.31421238 | 31.01248533 | 1.90094898 | 4.30E-05 | 0.00534963 |
| Rpsa | 8.438485733 | 15.77079923 | 1.86891342 | 4.36E-05 | 0.00539055 |
| Tmsb4x | 6.367419033 | 11.83221922 | 1.85824416 | 4.44E-05 | 0.0054529 |
| Csrp2 | 0.548709848 | 1.36102781 | 2.4804144 | 4.48E-05 | 0.00546294 |
| Rps2 | 7.320926343 | 13.59341258 | 1.8567886 | 4.66E-05 | 0.00564065 |
| Igf1 | 0.329523764 | 0.932688376 | 2.83041309 | 5.39E-05 | 0.00645134 |
| Tst | 0.225144255 | 0.777742031 | 3.45441651 | 5.41E-05 | 0.00645134 |
| Rpl13 | 16.61422395 | 31.28732682 | 1.88316511 | 5.67E-05 | 0.00672231 |
| Rpl21 | 8.636895554 | 15.92839082 | 1.8442264 | 5.95E-05 | 0.00700497 |
| Bhlha15 | 0.156446607 | 0.618632217 | 3.95427059 | 6.17E-05 | 0.00721234 |
| Rps15a | 10.04527156 | 18.58263553 | 1.84988882 | 6.21E-05 | 0.00721234 |
| Apoe | 1.0887228 | 0.281024065 | 0.2581227 | 6.44E-05 | 0.00742358 |

| | | | | | |
|---|---|---|---|---|---|
| Dctpp1 | 0.419337803 | 1.126821309 | 2.68714459 | 6.81E-05 | 0.00779518 |
| Rpl3 | 8.0294384 | 14.70265863 | 1.83109427 | 7.14E-05 | 0.00812827 |
| Rpl23a | 8.101420746 | 14.86832714 | 1.83527404 | 7.23E-05 | 0.00817243 |
| Rplp1 | 12.96124488 | 23.96435699 | 1.84892402 | 7.39E-05 | 0.00828278 |
| Ran | 1.708018603 | 3.252853702 | 1.90446035 | 7.42E-05 | 0.00828278 |
| Rpl18a | 14.00125287 | 25.9237448 | 1.85153036 | 7.81E-05 | 0.0086555 |
| Rpl39 | 8.965123696 | 16.39517699 | 1.82877309 | 8.46E-05 | 0.00931966 |
| Rpl7a | 3.930776938 | 7.140439288 | 1.81654655 | 0.00010348 | 0.01132512 |
| Rps24 | 11.99262201 | 21.8683351 | 1.82348239 | 0.00010452 | 0.01136663 |
| Pa2g4 | 0.51939552 | 1.213216082 | 2.33582315 | 0.00010564 | 0.01141581 |
| Npm1 | 2.952733704 | 5.380416437 | 1.8221814 | 0.00010904 | 0.01170973 |
| Rpl23 | 10.79317141 | 19.59083699 | 1.81511404 | 0.00011039 | 0.0117807 |
| Ptma | 7.987512428 | 14.36487057 | 1.79841605 | 0.00011876 | 0.01259541 |
| Ppia | 9.32148261 | 16.83182758 | 1.80570284 | 0.00012188 | 0.01284699 |
| Gnb2l1 | 3.994482073 | 7.186733214 | 1.79916522 | 0.00012908 | 0.01352383 |
| Mrpl12 | 0.410803962 | 1.043459846 | 2.54004329 | 0.000132 | 0.01374553 |
| Rpl37 | 10.55389388 | 18.98787137 | 1.7991342 | 0.00013331 | 0.01379861 |
| Top2a | 0.272010718 | 0.817622058 | 3.00584501 | 0.00013549 | 0.01393951 |
| Selk | 2.069935013 | 0.968614848 | 0.46794457 | 0.00013682 | 0.01393951 |
| Prss2 | 2.022959124 | 3.716250101 | 1.83703667 | 0.00013711 | 0.01393951 |
| Rps17 | 9.76476987 | 17.45747321 | 1.78780181 | 0.00014857 | 0.01501611 |
| Npm3 | 0.358213074 | 0.951640918 | 2.65663368 | 0.00015557 | 0.01563189 |
| Resp18 | 0.454336725 | 0.021069375 | 0.04637392 | 0.0001641 | 0.01639272 |
| Rps10 | 7.528839202 | 13.36231749 | 1.77481775 | 0.00016608 | 0.01644255 |
| C1qbp | 0.331480467 | 0.912985234 | 2.75426556 | 0.00016651 | 0.01644255 |
| Tpi1 | 1.410622336 | 2.631295376 | 1.86534362 | 0.00017981 | 0.01765449 |
| Rps6 | 10.25238439 | 18.22779937 | 1.77790831 | 0.00018123 | 0.01769282 |
| Ppib | 4.033386855 | 7.17334009 | 1.77849047 | 0.00019029 | 0.01847228 |
| Eif5a | 2.622537198 | 4.743339635 | 1.80868345 | 0.000196 | 0.01891988 |
| Try4 | 0.443256409 | 1.109787409 | 2.5037143 | 0.00020198 | 0.01938796 |

| | | | | | |
|---|---|---|---|---|---|
| Rps3 | 8.870331244 | 15.68147067 | 1.76785627 | 0.00020365 | 0.01943931 |
| Flna | 0.06861143 | 0.391017388 | 5.69901237 | 0.00020583 | 0.01944761 |
| Des | 0.034568621 | 0.292721648 | 8.46784272 | 0.000206 | 0.01944761 |
| Mgst1 | 0.567643671 | 1.285764038 | 2.26509006 | 0.00020988 | 0.01968579 |
| Edem1 | 0.446451465 | 1.074654794 | 2.4071033 | 0.00021081 | 0.01968579 |
| Gm10260 | 2.649188787 | 4.688316199 | 1.76971767 | 0.00021766 | 0.02021542 |
| Gm10076 | 1.361073265 | 2.514385221 | 1.84735479 | 0.00022039 | 0.02035896 |
| Kctd14 | 0.133004688 | 0.496683652 | 3.73433193 | 0.00022524 | 0.020679 |
| Atp2a2 | 0.84905467 | 0.219756696 | 0.25882514 | 0.00022626 | 0.020679 |
| Rps26 | 5.890357323 | 10.36940561 | 1.76040349 | 0.00023446 | 0.02131435 |
| Abcc8 | 0.441237937 | 0.006423671 | 0.01455829 | 0.00025004 | 0.02261135 |
| Rps25 | 6.838545645 | 11.96098367 | 1.74905372 | 0.00025709 | 0.02304618 |
| Rrbp1 | 2.135765464 | 3.769576094 | 1.76497661 | 0.00025753 | 0.02304618 |
| Rpl7 | 6.808746188 | 11.90132999 | 1.74794737 | 0.00026084 | 0.02322152 |
| Rpl9 | 7.937867623 | 13.84650398 | 1.74436066 | 0.00026501 | 0.02347097 |
| Id3 | 0.160888814 | 0.537954872 | 3.34364372 | 0.00028338 | 0.02496912 |
| Snrpf | 0.945523289 | 1.818067731 | 1.92281645 | 0.00028969 | 0.02529173 |
| Tkt | 0.547701507 | 1.218151304 | 2.22411531 | 0.00028998 | 0.02529173 |
| Rpl28 | 9.44465633 | 16.48109115 | 1.74501756 | 0.00029402 | 0.02551438 |
| Rpl14 | 11.74080452 | 20.56310906 | 1.75142249 | 0.00029605 | 0.0255611 |
| Map1b | 0.81983749 | 0.171818628 | 0.20957645 | 0.00030599 | 0.02628787 |
| Cldn18 | 0.039146503 | 0.250761337 | 6.40571484 | 0.00033345 | 0.02850376 |
| Rpl30 | 4.898012751 | 8.496516419 | 1.73468646 | 0.00034314 | 0.02918703 |
| Prss53 | 0.414491615 | 0 | 0 | 0.00034751 | 0.02941346 |
| Eef1d | 1.497249402 | 2.738478222 | 1.82900606 | 0.00035014 | 0.02949073 |
| Rplp2 | 11.82044709 | 20.60862454 | 1.74347251 | 0.00035282 | 0.02957173 |
| Serpinf2 | 0.117429526 | 0.486472145 | 4.14267315 | 0.0003574 | 0.02966606 |
| Ccnb1 | 0.118388727 | 0.465029165 | 3.92798516 | 0.0003574 | 0.02966606 |
| Eef1g | 3.85114687 | 6.64675275 | 1.72591516 | 0.00036214 | 0.02991474 |
| Cela3b | 20.98330285 | 37.09473651 | 1.76782162 | 0.00037436 | 0.03077605 |

| | | | | | |
|---|---|---|---|---|---|
| Dstn | 1.152248467 | 2.069008301 | 1.79562686 | 0.00038318 | 0.03125755 |
| Eif3m | 0.820029468 | 1.606852257 | 1.95950551 | 0.00038385 | 0.03125755 |
| Txn1 | 2.089569833 | 3.658501968 | 1.75083977 | 0.00039041 | 0.03164159 |
| Rps14 | 17.33551848 | 30.36603067 | 1.75166556 | 0.00039512 | 0.03187264 |
| Rps15 | 8.693626714 | 14.92938553 | 1.71727934 | 0.00045289 | 0.03633154 |
| Mrpl30 | 0.63034705 | 1.351189891 | 2.14356503 | 0.00045785 | 0.03633154 |
| Naca | 4.171594186 | 7.12904192 | 1.70894905 | 0.00045863 | 0.03633154 |
| Rpl26 | 11.13014268 | 19.15414062 | 1.72092498 | 0.00045885 | 0.03633154 |
| Rps19 | 14.12337668 | 24.37793537 | 1.7260699 | 0.0004746 | 0.03740621 |
| Hadh | 1.417857513 | 0.56581187 | 0.39906116 | 0.00048146 | 0.03777354 |
| Rps12-ps3 | 0.796020036 | 1.569008272 | 1.97106631 | 0.00049654 | 0.03877976 |
| G6pc2 | 0.39674557 | 0.007023125 | 0.01770184 | 0.00051268 | 0.03985951 |
| Ddit4 | 0.198399914 | 0.607258071 | 3.06077789 | 0.00051688 | 0.04000366 |
| Hnrnpab | 0.639427776 | 1.318079346 | 2.06134202 | 0.0005192 | 0.04000366 |
| Eef1b2 | 3.19415952 | 5.456982825 | 1.70842527 | 0.00053074 | 0.04071045 |
| Rpl36 | 9.701429083 | 16.51718714 | 1.70255196 | 0.00053436 | 0.04080645 |
| Cald1 | 0.174170924 | 0.569771136 | 3.27133326 | 0.00056313 | 0.04281262 |
| Id2 | 0.458968948 | 1.030345337 | 2.24491295 | 0.00058499 | 0.04420333 |
| Rpl19 | 8.17295685 | 13.84565625 | 1.69408165 | 0.00058656 | 0.04420333 |
| Serp1 | 1.885923757 | 3.287605979 | 1.74323377 | 0.00062354 | 0.04663452 |
| Ndufa1 | 1.241873062 | 0.463690345 | 0.37337982 | 0.00062425 | 0.04663452 |
| Arhgdig | 0.815891156 | 1.572709627 | 1.92759735 | 0.00063182 | 0.04674743 |
| Rpl18 | 7.668238436 | 12.9640295 | 1.69061377 | 0.00063354 | 0.04674743 |
| Ranbp1 | 1.074052711 | 1.942798365 | 1.80884825 | 0.00063393 | 0.04674743 |
| Rpl11 | 9.497698742 | 16.02412709 | 1.68715891 | 0.00064211 | 0.04701462 |
| Serbp1 | 1.723227192 | 2.994923416 | 1.73797363 | 0.00064302 | 0.04701462 |
| Rpl34 | 8.807933871 | 14.87878473 | 1.68924801 | 0.00065986 | 0.047759 |
| Scd2 | 1.081383461 | 1.965545392 | 1.8176211 | 0.00066127 | 0.047759 |
| H13 | 0.774959613 | 1.50444786 | 1.94132421 | 0.00066175 | 0.047759 |
| Krt7 | 0.861041917 | 0.216262057 | 0.25116322 | 0.00066432 | 0.047759 |

| | | | | | |
|---|---|---|---|---|---|
| Tubb5 | 1.906264568 | 3.270604114 | 1.71571364 | 0.00068616 | 0.04912346 |
| Prdx6 | 0.78021903 | 1.536726374 | 1.96960894 | 0.00070338 | 0.05000605 |
| Rplp0 | 5.990129155 | 10.04744015 | 1.67733281 | 0.00070431 | 0.05000605 |
| Galk1 | 0.205346834 | 0.615838075 | 2.99901422 | 0.00071446 | 0.05051777 |
| Lyz2 | 0.506386162 | 0.020134819 | 0.03976179 | 0.00071838 | 0.05058674 |
| Tmsb15b2 | 0.375362458 | 0.026577809 | 0.07080572 | 0.0007447 | 0.05203756 |
| Papss2 | 0.377281211 | 0.016384968 | 0.04342906 | 0.00074806 | 0.05203756 |
| Zcchc18 | 0.377632033 | 0.007079223 | 0.01874635 | 0.00074807 | 0.05203756 |
| Cel | 1.795742503 | 3.081910659 | 1.71623195 | 0.00075628 | 0.052397 |
| Mgll | 0.15519006 | 0.508141085 | 3.27431464 | 0.00076482 | 0.05272125 |
| Ckb | 0.485595153 | 1.112156419 | 2.29029555 | 0.0007671 | 0.05272125 |
| Nop10 | 0.852264755 | 1.608600299 | 1.88744201 | 0.00077697 | 0.0531869 |
| Cox5a | 1.326873867 | 2.342474906 | 1.76540888 | 0.00080009 | 0.0545521 |
| Acaa2 | 0.23525517 | 0.676582942 | 2.87595355 | 0.00080582 | 0.05472584 |
| Atp5o | 2.155963827 | 3.65471976 | 1.69516748 | 0.00081352 | 0.05503129 |
| Rps13 | 9.118110612 | 15.22074847 | 1.66928754 | 0.00084438 | 0.05674032 |
| Rpl27a | 9.4801928 | 15.84481218 | 1.6713597 | 0.00084539 | 0.05674032 |
| Rps18 | 8.813577271 | 14.68112308 | 1.66573942 | 0.00086422 | 0.05777801 |
| Rps9 | 9.553821337 | 15.9633251 | 1.67088378 | 0.00088147 | 0.05870313 |
| Rps3a1 | 9.712468973 | 16.22971655 | 1.67101863 | 0.00092373 | 0.0612801 |
| Rpl10 | 14.07437681 | 23.59147237 | 1.67620014 | 0.00093436 | 0.06174699 |
| Rps27a | 15.08585838 | 25.33278006 | 1.67924021 | 0.00094106 | 0.06195105 |
| Rpl35a | 9.481835439 | 15.80126059 | 1.66647699 | 0.00098366 | 0.06450856 |
| Tnnt2 | 0.017391595 | 0.193271641 | 11.1129336 | 0.00099126 | 0.0647601 |
| Itga6 | 0.213900001 | 0.643363656 | 3.00777771 | 0.0010217 | 0.06649595 |
| Rpl24 | 7.881509093 | 13.00855995 | 1.65051639 | 0.00103536 | 0.06713054 |
| Rab3a | 0.488454968 | 0.054216507 | 0.11099592 | 0.0010395 | 0.0671455 |
| Hsp90aa1 | 2.521079557 | 4.242578236 | 1.68284187 | 0.00104609 | 0.06731827 |
| Idh2 | 1.015271762 | 1.802794222 | 1.77567651 | 0.00105071 | 0.0673631 |
| Rps16 | 11.02493466 | 18.30371829 | 1.66021105 | 0.00107726 | 0.06880853 |

| | | | | | |
|---|---|---|---|---|---|
| Hepacam2 | 0.353188197 | 0.028148597 | 0.07969858 | 0.00108646 | 0.06913912 |
| Rpl6 | 8.953605395 | 14.77627149 | 1.65031525 | 0.00114647 | 0.07268875 |
| Prdx4 | 0.612234668 | 1.25417586 | 2.04852147 | 0.00118383 | 0.0747814 |
| Tacc3 | 0.071697692 | 0.303748882 | 4.23652244 | 0.00125983 | 0.07929069 |
| Eif1ax | 0.480910589 | 1.052306396 | 2.18815393 | 0.00128718 | 0.08042291 |
| Rexo2 | 0.48134146 | 1.023801697 | 2.12697592 | 0.00128718 | 0.08042291 |
| Cyb5a | 0.578858979 | 1.214011875 | 2.0972498 | 0.00130642 | 0.0813291 |
| Peg3 | 1.659124047 | 0.824075386 | 0.49669305 | 0.00133543 | 0.08283528 |
| Ptn | 0.279321026 | 0.715924245 | 2.56308755 | 0.0013979 | 0.08639837 |
| Rps7 | 9.123765111 | 14.89722379 | 1.63279344 | 0.00140438 | 0.08645998 |
| Pak3 | 0.458220595 | 0.04278034 | 0.09336189 | 0.00140896 | 0.08645998 |
| Spc24 | 0.222764112 | 0.623329869 | 2.79816109 | 0.00143229 | 0.08756483 |
| Rpl13a | 16.334386 | 27.01288512 | 1.65374353 | 0.00143716 | 0.08756483 |
| P2rx1 | 0.169090817 | 0.503396973 | 2.97708049 | 0.00144964 | 0.08801335 |
| Krtcap2 | 2.655104725 | 4.408210328 | 1.66027738 | 0.00146184 | 0.08844134 |
| Snrpe | 1.089461359 | 1.908307054 | 1.751606 | 0.00147348 | 0.08883285 |
| 2700094K13I | 0.491755965 | 1.034785891 | 2.10426709 | 0.00150952 | 0.09045603 |
| Bok | 0.074556746 | 0.310874116 | 4.16963094 | 0.00151093 | 0.09045603 |
| Cela3a | 0.365113861 | 0.858133059 | 2.3503163 | 0.00154802 | 0.09235436 |
| Cks2 | 0.400705279 | 0.932824147 | 2.32795572 | 0.00157187 | 0.09345269 |
| Rps5 | 10.56050307 | 17.15639922 | 1.62458162 | 0.00161122 | 0.09546172 |
| Ybx1 | 0.895857676 | 1.627125852 | 1.81627718 | 0.00162976 | 0.09622872 |
| Srsf3 | 1.106885153 | 1.871317417 | 1.6906157 | 0.00163621 | 0.09627836 |
| Fam162a | 0.403807429 | 0.886652805 | 2.19573178 | 0.00166282 | 0.09751032 |
| Agr2 | 0.097163709 | 0.419595071 | 4.31843405 | 0.00167884 | 0.09811488 |
| Ndufs6 | 1.497830989 | 2.532948543 | 1.69107767 | 0.00173995 | 0.10126317 |
| Hmgb1 | 2.10269801 | 3.501587211 | 1.66528298 | 0.00174576 | 0.10126317 |
| Tmed9 | 0.871781563 | 0.32251014 | 0.36994375 | 0.00175039 | 0.10126317 |
| Rgs5 | 0.009037941 | 0.110706756 | 12.2491121 | 0.00176687 | 0.10138605 |
| Selm | 0.878213064 | 0.283208233 | 0.32248237 | 0.00177087 | 0.10138605 |

| | | | | | |
|---|---|---|---|---|---|
| Rps28 | 6.22109009 | 10.01842547 | 1.61039711 | 0.00177385 | 0.10138605 |
| Rpl38 | 8.594989105 | 13.90922158 | 1.61829427 | 0.00177611 | 0.10138605 |
| H2-Eb1 | 0.077029681 | 0.334029719 | 4.33637678 | 0.00179938 | 0.10237412 |
| Asns | 0.337584813 | 0.800642663 | 2.37167856 | 0.00191851 | 0.10879131 |
| Smco4 | 0.231681077 | 0.638412109 | 2.75556431 | 0.00197129 | 0.11134315 |
| P4hb | 2.322088578 | 3.824346674 | 1.64694263 | 0.00197647 | 0.11134315 |
| Slc2a2 | 0.537831697 | 0.066572032 | 0.12377856 | 0.00198688 | 0.11156381 |
| Pfn1 | 3.501420591 | 5.642744833 | 1.61155871 | 0.00200367 | 0.11189549 |
| Nkx6-1 | 0.435303927 | 0.007118569 | 0.0163531 | 0.00200581 | 0.11189549 |
| Cd164 | 0.7038005 | 0.193526813 | 0.27497396 | 0.00202611 | 0.11266201 |
| Lgals1 | 1.910905753 | 3.144912214 | 1.64577045 | 0.00207934 | 0.1152493 |
| Fbl | 0.472435883 | 1.023168312 | 2.16572947 | 0.00210842 | 0.11627053 |
| Cdc20 | 0.205028768 | 0.592082927 | 2.88780415 | 0.0021113 | 0.11627053 |
| Prss8 | 0.152007598 | 0.451252388 | 2.96861732 | 0.00212379 | 0.11658469 |
| Pdia2 | 1.017857735 | 1.758984121 | 1.72812374 | 0.00217622 | 0.11908216 |
| Dut | 0.430926438 | 0.951729416 | 2.20856585 | 0.00220184 | 0.12010176 |
| Gm8730 | 7.089095355 | 11.32280578 | 1.59721448 | 0.00221725 | 0.1205596 |
| Anp32b | 0.811431804 | 1.504832463 | 1.85453966 | 0.00223718 | 0.12073261 |
| Pes1 | 0.103372374 | 0.347188906 | 3.35862372 | 0.00224216 | 0.12073261 |
| Clu | 2.409024745 | 3.878788549 | 1.61010739 | 0.00224745 | 0.12073261 |
| Gm15915 | 0.235016152 | 0.662154893 | 2.81748674 | 0.00224854 | 0.12073261 |
| Rpl8 | 8.337722775 | 13.33583078 | 1.59945721 | 0.0023237 | 0.12399483 |
| Procr | 0.058702108 | 0.303972317 | 5.17821807 | 0.00232373 | 0.12399483 |
| Pdx1 | 0.612180942 | 0.145895591 | 0.23832103 | 0.00243771 | 0.12967426 |
| Ap1s2 | 0.329256776 | 0.022909019 | 0.06957797 | 0.00250452 | 0.13281695 |
| Dbpht2 | 0.331014738 | 0 | 0 | 0.00253329 | 0.13392931 |
| Maged1 | 1.408698289 | 0.655841418 | 0.46556557 | 0.00255543 | 0.13468551 |
| Srm | 0.390010991 | 0.834211685 | 2.13894404 | 0.00261345 | 0.13732212 |
| Adamts9 | 0.157151648 | 0.442772539 | 2.81748582 | 0.00264639 | 0.13862897 |
| Try10 | 0.715200037 | 1.346941378 | 1.8833072 | 0.00265727 | 0.13877564 |

| | | | | | |
|---|---|---|---|---|---|
| Try5 | 1.796322565 | 0.951858129 | 0.52989265 | 0.00268753 | 0.1399307 |
| Ypel3 | 0.515666487 | 0.124146463 | 0.24074953 | 0.00277436 | 0.14401506 |
| Eln | 0.061352172 | 0.306288562 | 4.99230188 | 0.00280983 | 0.14541719 |
| Odc1 | 0.272009398 | 0.642996932 | 2.36387764 | 0.00282897 | 0.14596781 |
| Gpc3 | 0.392996493 | 0.883394542 | 2.24784332 | 0.00284363 | 0.14628495 |
| Sytl4 | 0.424126953 | 0.011685333 | 0.0275515 | 0.00290905 | 0.14903845 |
| Mageh1 | 0.528619922 | 0.111764589 | 0.21142712 | 0.0029145 | 0.14903845 |
| Homer2 | 0.107967183 | 0.359406278 | 3.32884742 | 0.00294397 | 0.15009891 |
| S100a11 | 2.317409222 | 1.279208241 | 0.55199929 | 0.00297264 | 0.15111192 |
| Cnn2 | 0.062686494 | 0.302688523 | 4.82860824 | 0.0030796 | 0.15608754 |
| Dcn | 0.772548875 | 0.209267349 | 0.27087911 | 0.00309651 | 0.15615085 |
| Eef1a1 | 14.84673618 | 23.63092963 | 1.59165822 | 0.00309902 | 0.15615085 |
| Pdlim3 | 0.011603743 | 0.141732536 | 12.2143811 | 0.00311705 | 0.15659981 |
| Myl6 | 2.854355776 | 4.531291476 | 1.58750059 | 0.00320019 | 0.16030813 |
| Echdc2 | 0.161948065 | 0.470992999 | 2.90829655 | 0.00326706 | 0.16318228 |
| Uchl5 | 0.217845506 | 0.576194249 | 2.64496734 | 0.00331907 | 0.16529952 |
| Set | 0.624709287 | 1.235317068 | 1.97742709 | 0.00332928 | 0.1653288 |
| Bex1 | 0.422166484 | 0.930041896 | 2.20302163 | 0.0034602 | 0.1713203 |
| Fam151a | 0.299514933 | 0 | 0 | 0.00346988 | 0.1713203 |
| Rps29 | 11.83942238 | 18.62526986 | 1.57315697 | 0.00352321 | 0.17345524 |
| Ubb | 8.051932535 | 4.565498125 | 0.5670065 | 0.00355496 | 0.17451815 |
| Zyx | 0.111189477 | 0.380071545 | 3.41823303 | 0.00357418 | 0.17496193 |
| Krt12 | 0.309754166 | 0 | 0 | 0.00364156 | 0.17775351 |
| Tspan1 | 0.087331334 | 0.371699536 | 4.256199 | 0.00365933 | 0.17811482 |
| Rpl22l1 | 4.837454482 | 7.567452908 | 1.56434607 | 0.00368417 | 0.17881751 |
| Hsd17b10 | 0.467756454 | 0.959099242 | 2.05042439 | 0.00372559 | 0.17981907 |
| Rpl41 | 30.00785589 | 48.40993501 | 1.61324205 | 0.00372574 | 0.17981907 |
| Gm10709 | 3.311656322 | 5.205543502 | 1.57188518 | 0.00376145 | 0.18103431 |
| Hsp90ab1 | 5.496683424 | 8.559707506 | 1.5572495 | 0.00382821 | 0.18373285 |
| Serpinb1a | 0.25202076 | 0.60698964 | 2.40849063 | 0.00384454 | 0.18400236 |

| | | | | | |
|---|---|---|---|---|---|
| Pcbp1 | 0.592403884 | 1.167232409 | 1.97033213 | 0.00390627 | 0.18643744 |
| Rpl4 | 6.483529632 | 10.11940767 | 1.56078683 | 0.00392658 | 0.18688781 |
| Cntnap2 | 0.013093654 | 0.123290387 | 9.41604149 | 0.00397519 | 0.1886787 |
| Meis2 | 1.112463724 | 0.507951217 | 0.45660025 | 0.00404296 | 0.19136684 |
| Mlxipl | 0.407847604 | 0.022965116 | 0.05630808 | 0.00420599 | 0.19853658 |
| Lrpprc | 0.408739105 | 0.048787001 | 0.11935976 | 0.00423293 | 0.19926076 |
| Cdo1 | 0.115371263 | 0.393277085 | 3.40879588 | 0.004304 | 0.20162291 |
| Lbh | 0.197048973 | 0.503657916 | 2.55600375 | 0.00430658 | 0.20162291 |
| Gstm2 | 0.141487493 | 0.441695885 | 3.12180162 | 0.00435119 | 0.20315797 |
| Uqcrq | 2.664532582 | 4.166693726 | 1.5637616 | 0.00436586 | 0.2032907 |
| Hist3h2ba | 0.414681202 | 0.04398723 | 0.10607481 | 0.00441434 | 0.20499244 |
| Rpl23a-ps3 | 0.880396315 | 1.550183742 | 1.76077945 | 0.00466446 | 0.2160237 |
| Tuba1a | 1.983230051 | 1.141508764 | 0.57558061 | 0.00469841 | 0.21614737 |
| Cpn1 | 0.199585534 | 0.523216889 | 2.6215171 | 0.00470408 | 0.21614737 |
| Gatm | 0.31123619 | 0.700479449 | 2.25063624 | 0.00470487 | 0.21614737 |
| Tpx2 | 0.06873696 | 0.274746011 | 3.99706376 | 0.00472704 | 0.21658668 |
| Rfc4 | 0.092871665 | 0.30939459 | 3.33142075 | 0.00484541 | 0.2211746 |
| Bex2 | 1.148782763 | 0.517686332 | 0.45063901 | 0.00485292 | 0.2211746 |
| Impdh2 | 0.403467154 | 0.878041847 | 2.17624121 | 0.00487199 | 0.22145625 |
| Ndufa9 | 0.349842991 | 0.731570793 | 2.09114035 | 0.00489253 | 0.22146981 |
| Rps21 | 5.003190914 | 7.758988885 | 1.55080808 | 0.00489806 | 0.22146981 |
| Pdia3 | 3.12701053 | 1.774851584 | 0.56758734 | 0.00492917 | 0.22229147 |
| Zg16 | 1.037328385 | 1.74016101 | 1.67754111 | 0.00494785 | 0.22254973 |
| Rph3al | 0.28148259 | 0.038067194 | 0.13523818 | 0.00512417 | 0.22987848 |
| Rap1b | 0.651762646 | 0.198268971 | 0.30420426 | 0.00515638 | 0.23072128 |
| Acly | 0.653916479 | 0.191518867 | 0.29287971 | 0.00519765 | 0.23196361 |
| Hn1 | 0.753775902 | 1.343217249 | 1.78198487 | 0.00526241 | 0.23424551 |
| Tubb3 | 0.29012391 | 0 | 0 | 0.00535561 | 0.23777815 |
| Eif3i | 1.071331669 | 1.788961332 | 1.66984827 | 0.00548865 | 0.24272623 |
| Hspa5 | 6.532057928 | 3.828085194 | 0.58604581 | 0.00549532 | 0.24272623 |

| | | | | | |
|---|---|---|---|---|---|
| Mki67 | 0.175915872 | 0.441400841 | 2.5091587 | 0.00558299 | 0.24578841 |
| Dtymk | 0.355481381 | 0.800391613 | 2.25157113 | 0.00559325 | 0.24578841 |
| Mest | 0.147625871 | 0.442221468 | 2.99555536 | 0.0056484 | 0.24757878 |
| Tuba1c | 0.278972567 | 0.704349197 | 2.52479734 | 0.00567699 | 0.24819872 |
| Hspd1 | 0.849280098 | 1.473711259 | 1.73524761 | 0.00574351 | 0.25038258 |
| Mrpl37 | 0.121482256 | 0.399956895 | 3.2923071 | 0.00576858 | 0.25038258 |
| Jun | 1.541481547 | 2.443570348 | 1.58520895 | 0.00577066 | 0.25038258 |
| Aqp12 | 0.419136426 | 0.850495988 | 2.02916267 | 0.00597611 | 0.25828974 |
| 1190007I07R | 0.072697477 | 0.300101793 | 4.12809086 | 0.00598297 | 0.25828974 |
| Glul | 0.388686744 | 0.063159509 | 0.16249463 | 0.00601355 | 0.2589596 |
| Rps27 | 8.337720039 | 12.73158341 | 1.5269862 | 0.00603582 | 0.25926845 |
| Pam | 0.389492692 | 0.049266155 | 0.12648801 | 0.0060525 | 0.2593367 |
| Nsa2 | 0.475698597 | 0.965697098 | 2.03006085 | 0.00607102 | 0.25948306 |
| Melk | 0.03134208 | 0.180518427 | 5.75961868 | 0.00614982 | 0.26219917 |
| Edem2 | 0.864256572 | 1.476702755 | 1.70863931 | 0.00623294 | 0.26425447 |
| Synpo2 | 0.004326929 | 0.058919555 | 13.616944 | 0.00624416 | 0.26425447 |
| Tmem213 | 0.004358493 | 0.057934687 | 13.2923677 | 0.00624417 | 0.26425447 |
| Sdsl | 0.208087949 | 0.510378813 | 2.45270721 | 0.00632907 | 0.26705121 |
| AU040320 | 0.051725241 | 0.205416363 | 3.97129832 | 0.00634134 | 0.26705121 |
| Apex1 | 0.209652093 | 0.530963373 | 2.53259275 | 0.00659324 | 0.27698056 |
| Dek | 0.382431911 | 0.808623372 | 2.11442442 | 0.00668135 | 0.27999754 |
| Spink1 | 3.206111088 | 4.869607342 | 1.51885172 | 0.00678866 | 0.28274951 |
| Rangrf | 0.124913752 | 0.391120833 | 3.1311271 | 0.00681283 | 0.28274951 |
| Cdca3 | 0.125518054 | 0.387980968 | 3.09103716 | 0.00681283 | 0.28274951 |
| Ptgr1 | 0.125839914 | 0.347209601 | 2.75913731 | 0.00681284 | 0.28274951 |
| Mrpl15 | 0.293629791 | 0.638363377 | 2.17404159 | 0.0068772 | 0.2847327 |
| Polr1d | 0.805845446 | 1.357049957 | 1.68400773 | 0.00693732 | 0.28583555 |
| Hif3a | 0.052761868 | 0.235978476 | 4.47251933 | 0.00694884 | 0.28583555 |
| Pex11a | 0.033201536 | 0.172194808 | 5.18635069 | 0.00695375 | 0.28583555 |
| Insig1 | 0.387815994 | 0.793927896 | 2.04717678 | 0.00705941 | 0.28948647 |

| | | | | | |
|---|---|---|---|---|---|
| Lap3 | 0.21143954 | 0.525042073 | 2.48317828 | 0.00714741 | 0.29239726 |
| Gpx4 | 2.033662651 | 3.140422107 | 1.54421979 | 0.00717913 | 0.29268927 |
| Gm10073 | 0.554175741 | 1.017096466 | 1.83533199 | 0.00720365 | 0.29268927 |
| Zbtb20 | 0.640168318 | 0.185340948 | 0.28951909 | 0.00721764 | 0.29268927 |
| Tmsb10 | 4.625406921 | 7.016725657 | 1.5169964 | 0.00722269 | 0.29268927 |
| Etfb | 0.977078884 | 1.634455352 | 1.67279774 | 0.00724506 | 0.29290509 |
| Birc5 | 0.2969851 | 0.677982356 | 2.2828834 | 0.00733043 | 0.2956605 |
| Cdc42ep5 | 0.076826028 | 0.282350988 | 3.67519959 | 0.00746577 | 0.30041413 |
| Ffar4 | 0.260156743 | 0.023004463 | 0.08842539 | 0.00752519 | 0.3013952 |
| Pim2 | 0.260721285 | 0.007023125 | 0.02693729 | 0.00752523 | 0.3013952 |
| Pclo | 0.264487659 | 0.012627003 | 0.04774137 | 0.0075791 | 0.30211093 |
| Galnt7 | 0.102018276 | 0.332719279 | 3.26136934 | 0.00758671 | 0.30211093 |
| Plk1 | 0.053785671 | 0.221040988 | 4.10966316 | 0.00759585 | 0.30211093 |
| Ube2s | 0.911886482 | 1.536781713 | 1.68527744 | 0.00773216 | 0.30615377 |
| Malat1 | 59.8399988 | 32.81026988 | 0.54829998 | 0.00773314 | 0.30615377 |
| Stmn1 | 0.699783705 | 1.238239298 | 1.76946003 | 0.00803995 | 0.31707866 |
| Rpl13-ps3 | 0.215401361 | 0.532765851 | 2.47336344 | 0.008046 | 0.31707866 |
| Rpl31 | 8.150141388 | 12.22829556 | 1.50037833 | 0.00817574 | 0.32145425 |
| Fau | 7.843184026 | 11.7777184 | 1.50165014 | 0.00822871 | 0.32194304 |
| 2210010C04I | 0.276385493 | 0.036544927 | 0.13222447 | 0.0082379 | 0.32194304 |
| Hdgf | 0.454903936 | 0.888455175 | 1.95306109 | 0.00824438 | 0.32194304 |
| Hmga2 | 0.104096093 | 0.334820232 | 3.21645342 | 0.00854633 | 0.33232831 |
| Mrpl42 | 0.504730862 | 0.972882374 | 1.92752702 | 0.00854901 | 0.33232831 |
| Alyref | 0.07852591 | 0.318671626 | 4.05817173 | 0.00859081 | 0.33271363 |
| Eny2 | 0.707493261 | 1.25274873 | 1.77068645 | 0.00859765 | 0.33271363 |
| H2afv | 0.709065522 | 1.224855766 | 1.72742254 | 0.00868917 | 0.33533427 |
| Cmc2 | 0.159240797 | 0.404342268 | 2.53918767 | 0.00871279 | 0.33533427 |
| Psmb3 | 1.025336031 | 1.634813007 | 1.59441681 | 0.00875323 | 0.33533427 |
| Tmem54 | 0.017799074 | 0.143428978 | 8.05822708 | 0.00875564 | 0.33533427 |
| Postn | 0.035544842 | 0.173378561 | 4.87774177 | 0.00876295 | 0.33533427 |

| | | | | | |
|---|---|---|---|---|---|
| Snrpd2 | 1.479576107 | 2.30917875 | 1.56070292 | 0.00889633 | 0.33968151 |
| Rnaseh2c | 0.359510085 | 0.788261922 | 2.1926003 | 0.00911006 | 0.347071 |
| Matn4 | 0.080480524 | 0.295526858 | 3.6720295 | 0.00919664 | 0.34922509 |
| Prrx1 | 0.005600816 | 0.051664709 | 9.22449642 | 0.00920725 | 0.34922509 |
| Stbd1 | 0.132871498 | 0.366079495 | 2.75513937 | 0.00932208 | 0.35280162 |
| Hacd1 | 0.409978505 | 0.820533457 | 2.00140604 | 0.00974173 | 0.36777986 |
| Emb | 0.687310455 | 0.215082721 | 0.31293387 | 0.00976066 | 0.36777986 |
| Rpl15 | 1.397779052 | 2.115028422 | 1.51313501 | 0.00983719 | 0.36921691 |
| Snrpb | 0.88923688 | 1.474078709 | 1.65768958 | 0.00985737 | 0.36921691 |
| Serpini2 | 0.274726137 | 0.598625826 | 2.17899117 | 0.00986326 | 0.36921691 |
| 1110008F13I | 0.473182301 | 0.915150693 | 1.93403407 | 0.00993831 | 0.37121762 |

| | mean outside cluster | mean inside | fold change | P value | P adj |
|---|---|---|---|---|---|
| pTimer | 1.697318694 | 103.819879 | 61.166992 | 0 | 0 |
| tdTomato | 2.019322728 | 129.826237 | 64.2919705 | 0 | 0 |
| eGFP | 0.280184359 | 14.3989159 | 51.3908627 | 0 | 0 |
| Ins2 | 0.55146718 | 23.1357663 | 41.9531156 | 0 | 0 |
| Ins1 | 0.290633684 | 9.35986078 | 32.2050103 | 0 | 0 |
| Sparc | 3.233562179 | 0.17714077 | 0.05478193 | 8.04E-117 | 1.27E-113 |
| Lgals1 | 2.796090868 | 0.15066406 | 0.05388382 | 3.71E-101 | 5.04E-98 |
| Itm2a | 2.80392957 | 0.20076338 | 0.07160072 | 3.74E-95 | 4.44E-92 |
| Col1a1 | 3.090143773 | 0.33765111 | 0.10926712 | 2.01E-90 | 2.12E-87 |
| Cdkn1c | 2.881537031 | 0.3246935 | 0.11268066 | 1.20E-83 | 1.14E-80 |
| Dlk1 | 5.089391931 | 0.96793995 | 0.19018774 | 3.25E-82 | 2.81E-79 |
| Dcn | 2.147697214 | 0.11127149 | 0.05180967 | 1.04E-78 | 8.22E-76 |
| Col3a1 | 2.26559324 | 0.19647079 | 0.08671936 | 2.52E-73 | 1.84E-70 |
| Ptn | 2.115844126 | 0.14270494 | 0.06744587 | 4.87E-73 | 3.31E-70 |
| Tmsb4x | 3.481598021 | 0.74034389 | 0.21264485 | 1.89E-71 | 1.20E-68 |
| Iapp | 3.101241822 | 10.7118985 | 3.45406748 | 3.72E-69 | 2.21E-66 |
| Nnat | 0.292736352 | 2.01451743 | 6.8816784 | 1.23E-67 | 6.86E-65 |
| Hbb-bs | 1.781034151 | 0.09677236 | 0.05433493 | 5.24E-65 | 2.76E-62 |
| Col1a2 | 2.073138735 | 0.20083366 | 0.0968742 | 1.83E-64 | 9.16E-62 |
| H19 | 1.912811649 | 0.16056536 | 0.08394207 | 1.97E-62 | 9.36E-60 |
| Hba-a1 | 1.480544595 | 0.03866694 | 0.0261167 | 1.80E-60 | 8.12E-58 |
| Gcg | 5.682334612 | 1.52468566 | 0.26832029 | 5.75E-58 | 2.48E-55 |
| Chga | 1.05096559 | 3.16805148 | 3.0144198 | 1.31E-49 | 5.40E-47 |
| Sst | 6.124291736 | 1.94874512 | 0.31819926 | 9.36E-47 | 3.71E-44 |
| Serpinh1 | 1.398775208 | 0.1537784 | 0.1099379 | 4.04E-42 | 1.53E-39 |
| Pcsk2 | 0.742501356 | 2.3998946 | 3.23217538 | 2.24E-41 | 8.18E-39 |
| Mest | 1.273569198 | 0.13887885 | 0.10904696 | 9.64E-38 | 3.39E-35 |
| Crip1 | 1.141230816 | 0.0930605 | 0.08154398 | 1.28E-37 | 4.35E-35 |
| Vim | 1.243735825 | 0.12870062 | 0.10347907 | 1.98E-37 | 6.50E-35 |

| | | | | | |
|---|---|---|---|---|---|
| Rpl39 | 3.623638005 | 1.35579638 | 0.37415337 | 7.04E-37 | 2.23E-34 |
| Apoe | 0.910237047 | 0.03285111 | 0.03609072 | 1.97E-36 | 6.02E-34 |
| S100a6 | 0.960932766 | 0.04812417 | 0.05008068 | 4.80E-36 | 1.42E-33 |
| Rps8 | 4.32930233 | 1.74991963 | 0.40420361 | 7.97E-32 | 2.29E-29 |
| Lum | 0.849514773 | 0.04762814 | 0.05606511 | 2.17E-31 | 6.07E-29 |
| Rps19 | 4.563215717 | 1.89294375 | 0.41482671 | 1.50E-30 | 4.06E-28 |
| Igf2 | 0.859697908 | 0.06039049 | 0.07024617 | 1.22E-29 | 3.21E-27 |
| Hba-a2 | 0.679206518 | 0.013277 | 0.0195478 | 1.09E-28 | 2.79E-26 |
| Rps4x | 3.114400896 | 1.34171044 | 0.43080852 | 1.12E-28 | 2.79E-26 |
| Ifitm2 | 1.08143595 | 0.17155857 | 0.1586396 | 5.01E-28 | 1.22E-25 |
| Rpl18a | 3.858464197 | 1.67978602 | 0.43535094 | 1.13E-27 | 2.69E-25 |
| Rpl36a | 2.061492528 | 0.70535371 | 0.34215681 | 3.74E-27 | 8.67E-25 |
| Pyy | 7.283585737 | 3.1757946 | 0.43602076 | 8.31E-27 | 1.86E-24 |
| Ghrl | 1.434885649 | 0.36253771 | 0.25265965 | 8.40E-27 | 1.86E-24 |
| Rps5 | 3.6430424 | 1.63850438 | 0.44976264 | 3.83E-26 | 8.27E-24 |
| Gsn | 0.725745643 | 0.04854291 | 0.06688695 | 6.12E-26 | 1.29E-23 |
| Tmsb10 | 4.215320946 | 1.91964956 | 0.4553982 | 3.64E-25 | 7.51E-23 |
| Igfbp4 | 0.682959875 | 0.05281999 | 0.07733981 | 3.75E-24 | 7.57E-22 |
| Gpc3 | 0.715968574 | 0.06241448 | 0.08717488 | 1.65E-23 | 3.26E-21 |
| Rps11 | 2.724579724 | 1.22627999 | 0.45008042 | 2.99E-23 | 5.80E-21 |
| Rplp0 | 4.13878564 | 1.96424579 | 0.47459471 | 7.12E-23 | 1.35E-20 |
| Rps2 | 3.251471814 | 1.55396163 | 0.4779256 | 9.25E-23 | 1.72E-20 |
| Anxa2 | 0.666696567 | 0.05130689 | 0.07695689 | 1.92E-22 | 3.50E-20 |
| Rpl23 | 2.971847958 | 1.4475912 | 0.48710136 | 5.58E-22 | 1.00E-19 |
| Rps26 | 2.618094742 | 1.19383523 | 0.4559939 | 6.59E-22 | 1.16E-19 |
| Sparcl1 | 0.601155956 | 0.04187772 | 0.06966199 | 1.02E-21 | 1.76E-19 |
| Col6a1 | 0.648124473 | 0.04972149 | 0.07671595 | 1.18E-21 | 2.00E-19 |
| Rpl9 | 2.797249583 | 1.33659583 | 0.47782502 | 1.39E-21 | 2.31E-19 |
| Rpl23a | 3.763974122 | 1.83752869 | 0.48818845 | 1.63E-21 | 2.67E-19 |
| Rpl35 | 2.927595608 | 1.42697492 | 0.48742214 | 1.92E-21 | 3.10E-19 |

| | | | | | |
|---|---|---|---|---|---|
| Rcn3 | 0.569338361 | 0.0337827 | 0.05933677 | 2.94E-21 | 4.65E-19 |
| Rpl13 | 4.938729143 | 2.41817435 | 0.48963494 | 3.50E-21 | 5.45E-19 |
| Mafb | 0.200630002 | 0.83350158 | 4.15442144 | 3.73E-20 | 5.71E-18 |
| Ppy | 0.876970743 | 0.17901613 | 0.20413011 | 6.71E-20 | 1.01E-17 |
| Rpl37 | 3.752575595 | 1.90602107 | 0.50792343 | 1.41E-19 | 2.10E-17 |
| Hbb-bt | 0.474358408 | 0.01735658 | 0.03658959 | 1.47E-19 | 2.15E-17 |
| Gng12 | 0.193972636 | 0.79908787 | 4.11959074 | 2.56E-19 | 3.69E-17 |
| Gm8730 | 2.440449449 | 1.16583019 | 0.47771126 | 3.93E-19 | 5.51E-17 |
| Rpl17 | 3.291790968 | 1.69257095 | 0.51417935 | 3.94E-19 | 5.51E-17 |
| Mfap2 | 0.604694741 | 0.06548369 | 0.10829214 | 4.68E-19 | 6.44E-17 |
| Rps20 | 2.147376418 | 0.95842639 | 0.44632435 | 4.90E-19 | 6.65E-17 |
| Rpl32 | 3.939931682 | 2.02752381 | 0.51460888 | 8.69E-19 | 1.16E-16 |
| Rps15a | 2.944242405 | 1.53218954 | 0.52040197 | 1.16E-18 | 1.53E-16 |
| S100a10 | 0.688871309 | 0.10946634 | 0.15890681 | 1.19E-18 | 1.54E-16 |
| Mdk | 0.72399273 | 0.12233903 | 0.16897826 | 1.51E-18 | 1.94E-16 |
| Fstl1 | 0.571861502 | 0.05199742 | 0.0909266 | 1.56E-18 | 1.98E-16 |
| Rps23 | 3.484815175 | 1.82101919 | 0.52255833 | 2.56E-18 | 3.20E-16 |
| Sec61b | 0.731237925 | 1.6850435 | 2.30437105 | 3.23E-18 | 3.98E-16 |
| Laptm4a | 1.127013606 | 0.32654035 | 0.28973949 | 3.47E-18 | 4.22E-16 |
| Rps29 | 5.408881913 | 2.85416268 | 0.52768072 | 2.07E-17 | 2.49E-15 |
| Col6a2 | 0.517766793 | 0.04564429 | 0.08815608 | 3.06E-17 | 3.63E-15 |
| Rplp1 | 2.745822959 | 1.45291056 | 0.52913483 | 7.45E-17 | 8.74E-15 |
| Cxcl12 | 0.45612918 | 0.03020406 | 0.06621822 | 1.87E-16 | 2.17E-14 |
| Rpl10a | 2.280266231 | 1.12988596 | 0.49550616 | 2.64E-16 | 3.02E-14 |
| Spp1 | 0.673114359 | 0.12296277 | 0.18267738 | 6.28E-16 | 7.10E-14 |
| Cryba2 | 0.199039481 | 0.73844829 | 3.71005937 | 6.74E-16 | 7.54E-14 |
| Mfap4 | 0.46170366 | 0.04261586 | 0.09230132 | 9.78E-16 | 1.08E-13 |
| Rps18 | 4.139926761 | 2.2893984 | 0.55300457 | 1.37E-15 | 1.50E-13 |
| Rps3 | 2.571220388 | 1.37166481 | 0.53346839 | 1.53E-15 | 1.65E-13 |
| Stmn1 | 0.589859059 | 0.09012367 | 0.15278848 | 1.66E-15 | 1.77E-13 |

| | | | | | |
|---|---|---|---|---|---|
| Rps7 | 2.765246617 | 1.5168152 | 0.548528 | 2.10E-15 | 2.22E-13 |
| Pcsk1n | 0.53615854 | 1.2872677 | 2.4009087 | 4.16E-15 | 4.34E-13 |
| Dbpht2 | 0.224500935 | 0.7602597 | 3.38644335 | 4.29E-15 | 4.42E-13 |
| Rps28 | 4.929433598 | 2.74901234 | 0.55767306 | 4.34E-15 | 4.42E-13 |
| Rpsa | 2.088048775 | 1.0452644 | 0.50059386 | 4.37E-15 | 4.42E-13 |
| Ifitm3 | 0.392563366 | 0.02648462 | 0.06746586 | 8.17E-15 | 8.17E-13 |
| Eef1a1 | 4.864331011 | 2.72810405 | 0.56083849 | 9.08E-15 | 8.98E-13 |
| Ifitm1 | 0.337310878 | 0.01046152 | 0.03101449 | 1.05E-14 | 1.03E-12 |
| Rps27a | 3.758187203 | 2.12168003 | 0.56454879 | 1.60E-14 | 1.55E-12 |
| Pdx1 | 0.097997784 | 0.49375052 | 5.03838451 | 1.88E-14 | 1.81E-12 |
| Rpl3 | 2.759258929 | 1.57050735 | 0.56917723 | 2.65E-14 | 2.52E-12 |
| Rps6 | 2.830642355 | 1.62260561 | 0.57322876 | 2.95E-14 | 2.77E-12 |
| Rpl6 | 3.15602617 | 1.81094193 | 0.57380447 | 3.89E-14 | 3.62E-12 |
| Eln | 0.418065814 | 0.03360008 | 0.08037032 | 5.48E-14 | 5.05E-12 |
| Rps9 | 3.020676711 | 1.74364103 | 0.57723524 | 5.83E-14 | 5.33E-12 |
| Rps3a1 | 3.077028145 | 1.77992418 | 0.57845561 | 9.21E-14 | 8.34E-12 |
| Rps24 | 3.042723984 | 1.76033905 | 0.5785405 | 9.79E-14 | 8.78E-12 |
| Rps27 | 5.129396296 | 2.95378506 | 0.57585433 | 1.30E-13 | 1.16E-11 |
| Cd81 | 1.291231561 | 0.52884108 | 0.40956332 | 1.35E-13 | 1.19E-11 |
| Csrp2 | 0.410601378 | 0.04002915 | 0.09748907 | 1.38E-13 | 1.20E-11 |
| Bgn | 0.399472704 | 0.03684843 | 0.09224267 | 3.27E-13 | 2.82E-11 |
| Cd63 | 1.280938915 | 0.54512019 | 0.42556298 | 3.96E-13 | 3.39E-11 |
| Rpl24 | 2.515845958 | 1.42986235 | 0.56834257 | 4.81E-13 | 4.08E-11 |
| Pcolce | 0.350715699 | 0.02344608 | 0.06685211 | 5.36E-13 | 4.42E-11 |
| B2m | 0.349167414 | 0.02449072 | 0.07014033 | 5.36E-13 | 4.42E-11 |
| Cald1 | 0.350673958 | 0.02203316 | 0.0628309 | 5.36E-13 | 4.42E-11 |
| Ldha | 0.352321058 | 0.02682852 | 0.07614794 | 5.43E-13 | 4.45E-11 |
| Rarres2 | 0.326005471 | 0.01236443 | 0.03792706 | 5.71E-13 | 4.64E-11 |
| Rpl21 | 2.394555467 | 1.3460629 | 0.56213478 | 5.94E-13 | 4.78E-11 |
| Serf2 | 1.261817963 | 0.52508868 | 0.41613664 | 7.78E-13 | 6.21E-11 |

| | | | | | |
|---|---|---|---|---|---|
| Rpl28 | 2.270992885 | 1.24974176 | 0.55030633 | 8.87E-13 | 7.02E-11 |
| Atp5l | 1.156168146 | 2.08351672 | 1.80208798 | 1.14E-12 | 8.94E-11 |
| Ccl21a | 0.260727746 | 0.00369456 | 0.01417019 | 1.23E-12 | 9.58E-11 |
| Akap12 | 0.339887018 | 0.02563691 | 0.07542775 | 1.34E-12 | 1.04E-10 |
| Fbln1 | 0.315343362 | 0.01847798 | 0.05859639 | 1.46E-12 | 1.12E-10 |
| Rps12 | 1.525109537 | 0.72126934 | 0.47292953 | 1.81E-12 | 1.38E-10 |
| Chgb | 1.309981335 | 2.2658007 | 1.7296435 | 2.21E-12 | 1.66E-10 |
| Gnas | 2.580948625 | 4.19267298 | 1.62446975 | 3.32E-12 | 2.48E-10 |
| Emp3 | 0.307755584 | 0.01655405 | 0.05378959 | 3.72E-12 | 2.76E-10 |
| Ndn | 0.623183927 | 0.16022073 | 0.25710023 | 4.11E-12 | 3.03E-10 |
| Postn | 0.370807712 | 0.03735633 | 0.10074313 | 4.72E-12 | 3.45E-10 |
| Rps14 | 3.968093647 | 2.38887896 | 0.60202182 | 5.62E-12 | 4.08E-10 |
| Rps15 | 2.354420157 | 1.36329122 | 0.57903481 | 6.37E-12 | 4.58E-10 |
| Rps17 | 2.432579676 | 1.41223381 | 0.58054987 | 6.53E-12 | 4.66E-10 |
| Rpl4 | 2.509561179 | 1.47657965 | 0.58838161 | 7.06E-12 | 5.00E-10 |
| Scgn | 0.160764088 | 0.55406124 | 3.44642417 | 9.71E-12 | 6.83E-10 |
| Ccdc80 | 0.294556918 | 0.01534355 | 0.05209026 | 1.52E-11 | 1.06E-09 |
| 1700086L19F | 0.256431549 | 0.7291618 | 2.84349488 | 2.67E-11 | 1.85E-09 |
| Ppic | 0.374058824 | 0.04575361 | 0.12231661 | 3.02E-11 | 2.08E-09 |
| Rpl35a | 3.150749989 | 1.93380088 | 0.61375891 | 3.16E-11 | 2.16E-09 |
| Rpl31 | 2.416012741 | 1.43735449 | 0.59492836 | 3.63E-11 | 2.46E-09 |
| Hmgb1 | 1.796547774 | 0.97160328 | 0.54081684 | 4.82E-11 | 3.24E-09 |
| Rps16 | 2.957512242 | 1.82834939 | 0.61820518 | 5.70E-11 | 3.81E-09 |
| Fth1 | 1.489627982 | 0.76887065 | 0.51614944 | 1.38E-10 | 9.14E-09 |
| Cd34 | 0.271486209 | 0.01781835 | 0.06563261 | 1.70E-10 | 1.12E-08 |
| Igf1 | 0.241875935 | 0.01123696 | 0.04645754 | 2.57E-10 | 1.68E-08 |
| Hmgb2 | 0.552194245 | 0.14880564 | 0.2694806 | 3.25E-10 | 2.11E-08 |
| Rpl11 | 2.699709606 | 1.7110935 | 0.6338065 | 5.05E-10 | 3.27E-08 |
| Pcsk1 | 0.073282079 | 0.35234592 | 4.80807751 | 5.12E-10 | 3.29E-08 |
| Nrp1 | 0.281398979 | 0.02729041 | 0.09698121 | 5.27E-10 | 3.34E-08 |

| | | | | | |
|---|---|---|---|---|---|
| Prkcdbp | 0.281884711 | 0.02625869 | 0.093154 | 5.27E-10 | 3.34E-08 |
| Ccnd3 | 0.356326838 | 0.0568624 | 0.15957934 | 5.41E-10 | 3.40E-08 |
| Rpl14 | 3.233384886 | 2.05270183 | 0.63484611 | 5.80E-10 | 3.62E-08 |
| Rpl37a | 5.214802713 | 3.31427784 | 0.63555191 | 9.79E-10 | 6.08E-08 |
| Maged2 | 0.588440548 | 0.18599173 | 0.31607564 | 1.08E-09 | 6.67E-08 |
| Fam183b | 0.236349091 | 0.63101349 | 2.66983677 | 1.09E-09 | 6.69E-08 |
| Pmp22 | 0.271231288 | 0.02156753 | 0.0795171 | 1.29E-09 | 7.83E-08 |
| Scg3 | 0.163646326 | 0.50249793 | 3.07063372 | 1.30E-09 | 7.85E-08 |
| Rpl34 | 2.56959836 | 1.63792691 | 0.63742526 | 1.39E-09 | 8.34E-08 |
| Rpl7 | 2.065171091 | 1.23359725 | 0.59733417 | 1.69E-09 | 1.01E-07 |
| Ebf1 | 0.267944513 | 0.01762234 | 0.06576862 | 2.01E-09 | 1.19E-07 |
| Ppp1r1a | 0.024986311 | 0.21321798 | 8.53339148 | 2.03E-09 | 1.20E-07 |
| Rpl12 | 1.048514654 | 0.48935501 | 0.46671261 | 2.04E-09 | 1.20E-07 |
| Rplp2 | 2.732757581 | 1.76722613 | 0.64668236 | 2.13E-09 | 1.24E-07 |
| Rpl8 | 2.339700276 | 1.46664743 | 0.6268527 | 3.17E-09 | 1.83E-07 |
| Rpl13a | 4.155700221 | 2.70247753 | 0.65030618 | 4.02E-09 | 2.31E-07 |
| Rpl27a | 2.401472483 | 1.52258549 | 0.63402163 | 4.23E-09 | 2.42E-07 |
| Ppia | 3.135090709 | 2.05371445 | 0.65507338 | 6.94E-09 | 3.95E-07 |
| Gpx8 | 0.2088292 | 0.01121513 | 0.05370481 | 7.04E-09 | 3.98E-07 |
| Rpl18 | 2.228594069 | 1.40216704 | 0.62917113 | 7.81E-09 | 4.39E-07 |
| Fabp5 | 0.39104061 | 0.0797264 | 0.20388267 | 9.87E-09 | 5.51E-07 |
| Acta2 | 0.227907578 | 0.01235169 | 0.05419603 | 1.03E-08 | 5.73E-07 |
| Zeb2 | 0.201279557 | 0.01446373 | 0.07185892 | 1.14E-08 | 6.29E-07 |
| Dbi | 0.433102091 | 0.10734917 | 0.24786111 | 1.41E-08 | 7.75E-07 |
| Eef1g | 1.314970894 | 0.71148745 | 0.54106707 | 1.48E-08 | 8.10E-07 |
| Rpl23a-ps3 | 1.042929396 | 0.51097101 | 0.48993825 | 1.65E-08 | 8.95E-07 |
| Mlxipl | 0.06555632 | 0.28956558 | 4.41705063 | 1.90E-08 | 1.03E-06 |
| Tspan7 | 0.148105246 | 0.44637117 | 3.01387817 | 1.95E-08 | 1.04E-06 |
| Eef2 | 1.637991336 | 0.95464481 | 0.58281432 | 2.05E-08 | 1.10E-06 |
| Plagl1 | 0.314036394 | 0.0518701 | 0.16517227 | 2.32E-08 | 1.23E-06 |

| | | | | | |
|---|---|---|---|---|---|
| mt-Nd1 | 2.760686043 | 1.84044683 | 0.66666285 | 2.49E-08 | 1.32E-06 |
| Cthrc1 | 0.21638126 | 0.02284201 | 0.1055637 | 2.59E-08 | 1.36E-06 |
| Fos | 0.50460365 | 0.99281626 | 1.96751699 | 2.86E-08 | 1.49E-06 |
| Nfib | 0.328402853 | 0.06620954 | 0.20161075 | 2.90E-08 | 1.50E-06 |
| Islr | 0.19273281 | 0.01117456 | 0.05797954 | 2.94E-08 | 1.52E-06 |
| Npm1 | 1.237961386 | 0.66338762 | 0.53587101 | 3.01E-08 | 1.54E-06 |
| Myl12a | 0.578911371 | 0.21157453 | 0.36546964 | 3.04E-08 | 1.55E-06 |
| S100a16 | 0.238410031 | 0.02716409 | 0.11393855 | 3.05E-08 | 1.55E-06 |
| Rbp1 | 0.25909318 | 0.02513215 | 0.09700044 | 3.12E-08 | 1.57E-06 |
| Ahnak | 0.237282648 | 0.02403358 | 0.10128672 | 5.07E-08 | 2.54E-06 |
| Itih5 | 0.236384744 | 0.01960353 | 0.08293062 | 5.07E-08 | 2.54E-06 |
| Pax6 | 0.27793254 | 0.65726436 | 2.36483415 | 5.64E-08 | 2.80E-06 |
| Rpl38 | 2.625516202 | 1.76895659 | 0.67375573 | 6.27E-08 | 3.10E-06 |
| Igfbp7 | 0.319557922 | 0.06169828 | 0.19307386 | 6.53E-08 | 3.21E-06 |
| Cd24a | 0.477905416 | 0.15454692 | 0.32338391 | 6.82E-08 | 3.34E-06 |
| Sfrp1 | 0.185704278 | 0.01006761 | 0.05421314 | 8.08E-08 | 3.94E-06 |
| Ctrb1 | 0.288805246 | 0.65995992 | 2.28513828 | 8.26E-08 | 4.00E-06 |
| Rpl41 | 9.081129352 | 6.08264599 | 0.66981162 | 1.02E-07 | 4.94E-06 |
| Sytl4 | 0.017172862 | 0.1666736 | 9.70563909 | 1.04E-07 | 4.99E-06 |
| Rps25 | 1.856114376 | 1.16091714 | 0.6254556 | 1.11E-07 | 5.30E-06 |
| Stmn2 | 0.149093557 | 0.00387064 | 0.02596113 | 1.66E-07 | 7.90E-06 |
| Sepw1 | 0.738917106 | 0.32645031 | 0.44179558 | 1.70E-07 | 8.04E-06 |
| Il11ra1 | 0.219895827 | 0.02585922 | 0.11759761 | 1.85E-07 | 8.70E-06 |
| Tmsb15b2 | 0.269155986 | 0.61827659 | 2.29709394 | 2.10E-07 | 9.82E-06 |
| Rpl26 | 2.749908748 | 1.88673364 | 0.68610773 | 2.15E-07 | 1.00E-05 |
| Tuba1b | 0.880160719 | 0.43000679 | 0.48855486 | 2.32E-07 | 1.08E-05 |
| Rps10 | 1.786478974 | 1.12280461 | 0.62850144 | 3.19E-07 | 1.47E-05 |
| Col4a1 | 0.267132468 | 0.04616912 | 0.1728323 | 3.27E-07 | 1.50E-05 |
| Rpl7a | 1.316880869 | 0.76031478 | 0.57736034 | 3.76E-07 | 1.72E-05 |
| mt-Co3 | 7.173713207 | 4.9202711 | 0.68587508 | 3.90E-07 | 1.77E-05 |

| | | | | | |
|---|---|---|---|---|---|
| Ppib | 0.81214888 | 0.38736584 | 0.47696407 | 3.93E-07 | 1.78E-05 |
| Ociad2 | 0.045194397 | 0.22425901 | 4.96209746 | 4.10E-07 | 1.85E-05 |
| Meis2 | 0.478523226 | 0.90844965 | 1.8984442 | 4.21E-07 | 1.88E-05 |
| Snhg18 | 0.208989671 | 0.02028181 | 0.09704697 | 4.41E-07 | 1.97E-05 |
| Hhex | 0.211121756 | 0.02706729 | 0.12820701 | 4.54E-07 | 2.02E-05 |
| Igfbp6 | 0.168502545 | 0.00451481 | 0.02679373 | 5.47E-07 | 2.42E-05 |
| Ptma | 3.112278105 | 2.16691606 | 0.69624757 | 6.13E-07 | 2.70E-05 |
| Col14a1 | 0.184128099 | 0.01400489 | 0.07606058 | 6.54E-07 | 2.85E-05 |
| Myl9 | 0.1853819 | 0.01045521 | 0.05639825 | 6.54E-07 | 2.85E-05 |
| Tppp3 | 0.185778978 | 0.01680238 | 0.09044284 | 6.56E-07 | 2.85E-05 |
| Mrfap1 | 0.79508442 | 1.31743834 | 1.65697919 | 7.42E-07 | 3.20E-05 |
| Gas6 | 0.161396265 | 0.0122328 | 0.0757936 | 8.25E-07 | 3.54E-05 |
| Rrbp1 | 0.544207968 | 0.20761038 | 0.38149089 | 9.71E-07 | 4.15E-05 |
| Dek | 0.216830683 | 0.03691846 | 0.17026402 | 1.01E-06 | 4.30E-05 |
| Igfbp5 | 0.180130085 | 0.02120093 | 0.11769787 | 1.01E-06 | 4.30E-05 |
| Rpl19 | 2.61376005 | 1.83303107 | 0.70130044 | 1.02E-06 | 4.31E-05 |
| Mmp14 | 0.252760326 | 0.04121221 | 0.16304856 | 1.12E-06 | 4.71E-05 |
| Rpl36 | 2.766866675 | 1.9415534 | 0.70171556 | 1.18E-06 | 4.95E-05 |
| Spats2l | 0.130494015 | 0.00702004 | 0.05379586 | 1.20E-06 | 5.00E-05 |
| Cpe | 0.835622564 | 1.35247508 | 1.61852389 | 1.26E-06 | 5.22E-05 |
| Arf4 | 0.520457651 | 0.20922654 | 0.40200493 | 1.37E-06 | 5.67E-05 |
| Slc30a8 | 0.04890166 | 0.22564367 | 4.6142334 | 1.40E-06 | 5.74E-05 |
| Tpm2 | 0.158906179 | 0.0113294 | 0.07129616 | 1.42E-06 | 5.79E-05 |
| Matn2 | 0.158753185 | 0.01043022 | 0.06570086 | 1.42E-06 | 5.79E-05 |
| Lmna | 0.336912398 | 0.10021129 | 0.2974402 | 1.46E-06 | 5.94E-05 |
| Prdx4 | 0.277685016 | 0.06771649 | 0.24386081 | 1.55E-06 | 6.28E-05 |
| Nid1 | 0.173834785 | 0.01955947 | 0.11251761 | 1.62E-06 | 6.51E-05 |
| Cav1 | 0.195927831 | 0.02466321 | 0.12587906 | 1.67E-06 | 6.68E-05 |
| Ftl1 | 3.207043911 | 2.25933575 | 0.70449168 | 1.68E-06 | 6.70E-05 |
| mt-Co2 | 2.929466869 | 2.06792447 | 0.70590471 | 1.77E-06 | 7.05E-05 |

| | | | | | |
|---|---|---|---|---|---|
| Gm10260 | 0.946923535 | 0.50790524 | 0.53637408 | 1.91E-06 | 7.56E-05 |
| Gm42418 | 0.749960409 | 0.36223941 | 0.48301138 | 2.04E-06 | 8.05E-05 |
| Rps27rt | 0.868629667 | 0.45341484 | 0.52198866 | 2.08E-06 | 8.17E-05 |
| Ptrf | 0.149487003 | 0.01316359 | 0.08805843 | 2.13E-06 | 8.26E-05 |
| Tubb6 | 0.149522863 | 0.01145158 | 0.07658746 | 2.13E-06 | 8.26E-05 |
| Olfml3 | 0.153183493 | 0.00783408 | 0.05114179 | 2.13E-06 | 8.26E-05 |
| Rps13 | 2.312723939 | 1.60697445 | 0.69484058 | 2.19E-06 | 8.45E-05 |
| Ddah2 | 0.276654153 | 0.05892014 | 0.212974 | 2.33E-06 | 8.97E-05 |
| Gnb2l1 | 0.915818982 | 0.49627211 | 0.54188887 | 2.39E-06 | 9.15E-05 |
| Rpl15 | 1.702338312 | 1.10612632 | 0.6497688 | 2.44E-06 | 9.30E-05 |
| Calu | 0.212732706 | 0.03193165 | 0.15010223 | 2.45E-06 | 9.30E-05 |
| Hadh | 0.226780725 | 0.51982011 | 2.29217061 | 2.52E-06 | 9.50E-05 |
| Agtr2 | 0.171511243 | 0.01415396 | 0.08252495 | 2.52E-06 | 9.50E-05 |
| Clu | 0.300807759 | 0.08211106 | 0.27296855 | 2.76E-06 | 0.0001036 |
| Tshz2 | 0.383176857 | 0.13157835 | 0.34338803 | 3.06E-06 | 0.00011427 |
| Cyba | 0.22332528 | 0.03837446 | 0.17183213 | 3.10E-06 | 0.00011541 |
| Cnn2 | 0.121682188 | 0.00603567 | 0.04960189 | 3.23E-06 | 0.00011875 |
| Ndufa4l2 | 0.121476291 | 0.00322586 | 0.02655547 | 3.23E-06 | 0.00011875 |
| Mmp2 | 0.121875358 | 0.00267012 | 0.02190862 | 3.23E-06 | 0.00011875 |
| Kdelr3 | 0.146010036 | 0.01097115 | 0.07513967 | 3.34E-06 | 0.00012245 |
| Arpc1b | 0.365888746 | 0.11936381 | 0.32622979 | 4.37E-06 | 0.00015965 |
| Syt13 | 0.189911287 | 0.45213205 | 2.38075394 | 4.67E-06 | 0.00017004 |
| Tubb3 | 0.135088456 | 0.36505126 | 2.7023128 | 4.76E-06 | 0.00017244 |
| Lyz2 | 0.115400754 | 0.00094864 | 0.00822039 | 5.34E-06 | 0.00019271 |
| Fau | 2.355883704 | 1.66670734 | 0.70746588 | 5.69E-06 | 0.00020466 |
| Rpl22l1 | 1.647211154 | 1.08274967 | 0.65732294 | 5.99E-06 | 0.00021488 |
| Id3 | 0.180328888 | 0.02589622 | 0.14360552 | 6.16E-06 | 0.00021988 |
| Copz2 | 0.162308327 | 0.01795294 | 0.11061007 | 6.26E-06 | 0.00022173 |
| Fbn2 | 0.162313521 | 0.01607348 | 0.09902736 | 6.26E-06 | 0.00022173 |
| Atp2a2 | 0.081011038 | 0.25725144 | 3.17551093 | 6.78E-06 | 0.00023944 |

| | | | | | |
|---|---|---|---|---|---|
| Timp2 | 0.136926076 | 0.01426582 | 0.10418631 | 8.57E-06 | 0.00030145 |
| Serpinf1 | 0.112754519 | 0.00517187 | 0.0458684 | 8.68E-06 | 0.00030425 |
| Mt1 | 0.342692793 | 0.11536362 | 0.33663859 | 9.02E-06 | 0.00031504 |
| Rpl10 | 3.384143444 | 2.45280356 | 0.72479302 | 9.22E-06 | 0.00032082 |
| Col5a1 | 0.175094511 | 0.02380502 | 0.13595526 | 9.64E-06 | 0.00033171 |
| Col5a2 | 0.175359074 | 0.01984432 | 0.11316391 | 9.64E-06 | 0.00033171 |
| Fscn1 | 0.175868803 | 0.01926477 | 0.1095406 | 9.64E-06 | 0.00033171 |
| Rhoc | 0.24132904 | 0.05691445 | 0.23583754 | 9.78E-06 | 0.00033549 |
| Auts2 | 0.207657613 | 0.04393616 | 0.21157984 | 1.07E-05 | 0.00036314 |
| Crip2 | 0.209078843 | 0.0379492 | 0.18150662 | 1.07E-05 | 0.00036314 |
| Tuba4a | 0.100032496 | 0.29640581 | 2.96309521 | 1.20E-05 | 0.00040647 |
| Ccnd2 | 0.295113098 | 0.08347675 | 0.28286357 | 1.25E-05 | 0.00042172 |
| Cplx2 | 0.134046514 | 0.34940616 | 2.60660386 | 1.26E-05 | 0.00042392 |
| Papss2 | 0.026320106 | 0.15223035 | 5.78380455 | 1.26E-05 | 0.00042424 |
| Actb | 4.100663602 | 2.98466698 | 0.72784975 | 1.30E-05 | 0.00043526 |
| Nbl1 | 0.131465616 | 0.01098555 | 0.08356214 | 1.39E-05 | 0.00045588 |
| Osr1 | 0.132282174 | 0.00873766 | 0.06605317 | 1.39E-05 | 0.00045588 |
| Aebp1 | 0.13224564 | 0.00848535 | 0.06416353 | 1.39E-05 | 0.00045588 |
| Hes1 | 0.132438246 | 0.00740044 | 0.05587841 | 1.39E-05 | 0.00045588 |
| Nkain4 | 0.133075689 | 0.00561857 | 0.04222086 | 1.39E-05 | 0.00045588 |
| Cmtm3 | 0.135939505 | 0.00893541 | 0.06573075 | 1.42E-05 | 0.00046465 |
| Hmgn3 | 0.403560283 | 0.74118969 | 1.83662695 | 1.43E-05 | 0.00046605 |
| C1qb | 0.109049152 | 0.00163312 | 0.014976 | 1.50E-05 | 0.00048769 |
| Ero1lb | 0.022862433 | 0.13396839 | 5.85976105 | 1.52E-05 | 0.00049425 |
| Acly | 0.169531269 | 0.39197304 | 2.31209878 | 1.71E-05 | 0.00055334 |
| Tubb5 | 1.157599223 | 0.7125361 | 0.61552918 | 2.17E-05 | 0.00069981 |
| Fxyd1 | 0.128675831 | 0.00530003 | 0.04118903 | 2.20E-05 | 0.00070455 |
| Emp1 | 0.169247208 | 0.01973111 | 0.11658161 | 2.34E-05 | 0.00074876 |
| Vcan | 0.148012033 | 0.01323525 | 0.08942011 | 2.41E-05 | 0.0007684 |
| Nrk | 0.150539242 | 0.01717533 | 0.11409203 | 2.55E-05 | 0.00080153 |

| | | | | | |
|---|---|---|---|---|---|
| Fbn1 | 0.149359368 | 0.01810326 | 0.12120604 | 2.55E-05 | 0.00080153 |
| Mgp | 0.148980845 | 0.01586533 | 0.1064924 | 2.55E-05 | 0.00080153 |
| Gpx7 | 0.15118396 | 0.01359284 | 0.08990928 | 2.55E-05 | 0.00080153 |
| Tpm1 | 0.28930954 | 0.08147596 | 0.2816221 | 2.61E-05 | 0.00081857 |
| Clic1 | 0.51410111 | 0.23929986 | 0.46547236 | 3.28E-05 | 0.00102354 |
| Hnrnpa1 | 0.692019308 | 0.36604537 | 0.52895254 | 3.33E-05 | 0.00103724 |
| Gm13305 | 0.122000301 | 0.01122628 | 0.09201844 | 3.53E-05 | 0.00109556 |
| Pak3 | 0.086853471 | 0.24907068 | 2.86771133 | 3.55E-05 | 0.00109848 |
| Htra3 | 0.125489615 | 0.00584375 | 0.04656758 | 3.65E-05 | 0.00112598 |
| Dpysl3 | 0.145223389 | 0.01191849 | 0.08207001 | 3.82E-05 | 0.00117363 |
| Cxcl13 | 0.100141336 | 0.00330297 | 0.03298309 | 4.05E-05 | 0.00124154 |
| Rpl5 | 1.037397781 | 0.63426038 | 0.61139554 | 4.10E-05 | 0.00125273 |
| Scg2 | 0.180185084 | 0.40727008 | 2.26028745 | 4.15E-05 | 0.00126363 |
| Eef1b2 | 0.730824758 | 0.39571635 | 0.54146544 | 4.20E-05 | 0.00127605 |
| Marcks | 0.405025585 | 0.16579619 | 0.40934745 | 4.23E-05 | 0.00128091 |
| Rpl29 | 1.018226729 | 0.61716462 | 0.60611709 | 4.30E-05 | 0.00129725 |
| Manf | 0.421143653 | 0.73805401 | 1.75249942 | 4.66E-05 | 0.00140013 |
| Sdf2l1 | 0.081795319 | 0.24491873 | 2.99428785 | 4.79E-05 | 0.0014362 |
| Txndc5 | 0.274552416 | 0.09411684 | 0.34280098 | 5.15E-05 | 0.00153707 |
| Rbp4 | 2.225440459 | 2.94670529 | 1.32409981 | 5.27E-05 | 0.0015678 |
| Des | 0.120374869 | 0.00575617 | 0.04781869 | 5.61E-05 | 0.00166674 |
| Csrp1 | 0.136892711 | 0.02030868 | 0.14835472 | 5.89E-05 | 0.00173841 |
| Mxra8 | 0.139134841 | 0.01157374 | 0.08318361 | 5.89E-05 | 0.00173841 |
| Tsc22d1 | 0.523163409 | 0.25742153 | 0.49204803 | 5.98E-05 | 0.00175941 |
| Nedd4 | 0.528845354 | 0.24969116 | 0.472144 | 6.03E-05 | 0.00176786 |
| Rpl22 | 0.827678427 | 0.47658478 | 0.57580911 | 6.10E-05 | 0.001782 |
| Plod2 | 0.093889134 | 0.00616751 | 0.06568933 | 6.31E-05 | 0.0018274 |
| Pdlim2 | 0.09366433 | 0.00475586 | 0.05077552 | 6.31E-05 | 0.0018274 |
| Htra1 | 0.093774498 | 0.00456696 | 0.04870154 | 6.31E-05 | 0.0018274 |
| Tubb4b | 0.313837806 | 0.58125758 | 1.85209546 | 6.72E-05 | 0.00193977 |

| | | | | | |
|---|---|---|---|---|---|
| Immp1l | 0.170902953 | 0.385099 | 2.25331977 | 7.00E-05 | 0.00201561 |
| H2afz | 0.893261094 | 0.5303004 | 0.59366786 | 7.38E-05 | 0.00211748 |
| Ttr | 1.272556567 | 1.78211069 | 1.40041766 | 8.31E-05 | 0.00237775 |
| Sfr1 | 0.48702562 | 0.22922658 | 0.47066636 | 8.35E-05 | 0.00238103 |
| Abcc8 | 0.116064958 | 0.29180701 | 2.51416981 | 8.58E-05 | 0.00243889 |
| Phlda3 | 0.112211091 | 0.01076084 | 0.09589817 | 8.96E-05 | 0.00251819 |
| Cryab | 0.114107256 | 0.00747448 | 0.06550403 | 8.96E-05 | 0.00251819 |
| Igfbp2 | 0.112756631 | 0.00678927 | 0.06021173 | 8.96E-05 | 0.00251819 |
| Smoc2 | 0.114111373 | 0.00374301 | 0.03280133 | 8.96E-05 | 0.00251819 |
| Tm4sf1 | 0.130811028 | 0.02036816 | 0.15570677 | 9.34E-05 | 0.00260226 |
| P4ha2 | 0.132652754 | 0.01345804 | 0.10145313 | 9.34E-05 | 0.00260226 |
| Fkbp7 | 0.133330582 | 0.00920684 | 0.0690527 | 9.34E-05 | 0.00260226 |
| Svbp | 0.133830248 | 0.01869297 | 0.13967674 | 9.38E-05 | 0.0026055 |
| Rps12-ps3 | 0.580531207 | 0.29853635 | 0.51424686 | 0.0001093 | 0.00302694 |
| mt-Atp6 | 6.34314287 | 4.77220579 | 0.7523409 | 0.00011183 | 0.00308789 |
| Peg3 | 0.860210704 | 0.51580649 | 0.59962808 | 0.00011677 | 0.00321511 |
| Naca | 1.281397354 | 0.8619289 | 0.67264764 | 0.00011955 | 0.00328207 |
| Sep-11 | 0.146544928 | 0.0267587 | 0.18259726 | 0.00012636 | 0.00344927 |
| Mfap5 | 0.146491418 | 0.0221052 | 0.15089757 | 0.00012637 | 0.00344927 |
| Rcn1 | 0.272391046 | 0.09648641 | 0.35422022 | 0.00014173 | 0.00385757 |
| Nid2 | 0.110896298 | 0.00979489 | 0.08832474 | 0.00014329 | 0.00386095 |
| Fkbp10 | 0.109990167 | 0.00865109 | 0.07865328 | 0.0001433 | 0.00386095 |
| 6330403K07I | 0.109514853 | 0.00739495 | 0.06752466 | 0.0001433 | 0.00386095 |
| Bst2 | 0.129513093 | 0.01042301 | 0.08047842 | 0.00014348 | 0.00386095 |
| Tagln2 | 0.338599319 | 0.13570789 | 0.40079198 | 0.0001456 | 0.00390701 |
| Tpm4 | 0.249302318 | 0.07772536 | 0.31177152 | 0.00014903 | 0.00398763 |
| Nkx6-1 | 0.061080776 | 0.19692162 | 3.22395406 | 0.00015556 | 0.00415072 |
| Aplp1 | 0.374132778 | 0.64842311 | 1.73313632 | 0.00016233 | 0.00431936 |
| Gadd45g | 0.048543416 | 0.17555125 | 3.61637607 | 0.00016399 | 0.00435111 |
| Cdc42ep2 | 0.085474688 | 0.00423857 | 0.04958863 | 0.00017054 | 0.00451235 |

| | | | | | |
|---|---|---|---|---|---|
| Ap1s2 | 0.132133844 | 0.30696657 | 2.32314875 | 0.00017353 | 0.00457885 |
| Ubqln2 | 0.154796385 | 0.33869146 | 2.18798041 | 0.00017681 | 0.00465251 |
| Gjd2 | 0.027715855 | 0.12887002 | 4.64968604 | 0.00017866 | 0.00467525 |
| Fam159b | 0.028494632 | 0.12366753 | 4.34002893 | 0.00017866 | 0.00467525 |
| Eif3e | 0.492709251 | 0.24755815 | 0.50244267 | 0.00019377 | 0.00505655 |
| Grn | 0.143097909 | 0.02603522 | 0.18193989 | 0.0001962 | 0.00510607 |
| 1110008P14I | 0.079781702 | 0.22321247 | 2.7977903 | 0.00020497 | 0.00531972 |
| Rpl27 | 1.618297454 | 1.15571844 | 0.71415699 | 0.00021238 | 0.00549701 |
| Ramp2 | 0.217982276 | 0.06665866 | 0.30579851 | 0.00021615 | 0.00557942 |
| Ece1 | 0.074467469 | 0.21167855 | 2.84256408 | 0.00022101 | 0.00568937 |
| Col18a1 | 0.122732779 | 0.01746975 | 0.1423397 | 0.00022516 | 0.00574957 |
| 2810417H13 | 0.122859766 | 0.01213674 | 0.09878533 | 0.00022516 | 0.00574957 |
| Fcgrt | 0.123971608 | 0.01009443 | 0.08142532 | 0.00022516 | 0.00574957 |
| Ltbp4 | 0.104622015 | 0.00757358 | 0.07238994 | 0.00022691 | 0.00576328 |
| Msn | 0.103660499 | 0.0080991 | 0.07813103 | 0.00022692 | 0.00576328 |
| Capns1 | 0.389912272 | 0.17564703 | 0.45047833 | 0.00022838 | 0.00578505 |
| Lmo4 | 0.124668032 | 0.01986562 | 0.15934814 | 0.00022948 | 0.00579739 |
| Tpt1 | 2.313493308 | 1.74940776 | 0.75617585 | 0.00023181 | 0.00584085 |
| Emilin1 | 0.107922764 | 0.00903152 | 0.08368503 | 0.00024029 | 0.00602241 |
| Ifi27 | 0.106506192 | 0.0088968 | 0.0835332 | 0.00024029 | 0.00602241 |
| Rpl30 | 1.233454963 | 0.83409128 | 0.67622354 | 0.00024119 | 0.00602909 |
| Hist3h2ba | 0.109482883 | 0.27620609 | 2.52282438 | 0.00025523 | 0.00636321 |
| Hist1h2al | 0.067552905 | 0.19499926 | 2.88661541 | 0.0002597 | 0.00645772 |
| Lhfp | 0.078856712 | 0.00514627 | 0.065261 | 0.00027945 | 0.00691271 |
| Gas1 | 0.078913221 | 0.00249514 | 0.03161876 | 0.00027945 | 0.00691271 |
| Colec12 | 0.081822917 | 0.00618567 | 0.07559823 | 0.00029751 | 0.00732151 |
| Dpt | 0.081722059 | 0.00401917 | 0.049181 | 0.00029752 | 0.00732151 |
| Cd9 | 0.196177457 | 0.05895737 | 0.30053083 | 0.00030074 | 0.00738164 |
| Gm27033 | 0.016446948 | 0.09180117 | 5.58165353 | 0.00031018 | 0.00759391 |
| Sep-07 | 0.425994535 | 0.20882209 | 0.49019899 | 0.00034018 | 0.00830686 |

| | | | | | |
|---|---|---|---|---|---|
| Zfp36l1 | 0.151784447 | 0.02474237 | 0.16300994 | 0.00035467 | 0.00863837 |
| Gng11 | 0.154819334 | 0.02757574 | 0.17811561 | 0.00035808 | 0.00869915 |
| Maf | 0.102349689 | 0.00785479 | 0.07674464 | 0.00036478 | 0.0087502 |
| Cdh11 | 0.101667387 | 0.00752348 | 0.07400094 | 0.00036478 | 0.0087502 |
| Gap43 | 0.101491974 | 0.0076243 | 0.07512215 | 0.00036478 | 0.0087502 |
| Clec3b | 0.102748306 | 0.00497447 | 0.04841418 | 0.00036478 | 0.0087502 |
| Penk | 0.100573586 | 0.00688672 | 0.06847445 | 0.00036478 | 0.0087502 |
| Myl6 | 1.278762462 | 0.87920788 | 0.6875459 | 0.00038429 | 0.00919485 |
| Thra | 0.164172099 | 0.04055476 | 0.24702589 | 0.00039347 | 0.00939083 |
| Rps26-ps1 | 0.306114405 | 0.12714632 | 0.41535555 | 0.00039602 | 0.00942813 |
| Neurod1 | 0.229151381 | 0.43951153 | 1.91799644 | 0.00039957 | 0.00948872 |
| Zfos1 | 0.232309572 | 0.08470979 | 0.36464184 | 0.00041924 | 0.00993096 |
| Mt2 | 0.178066243 | 0.04542668 | 0.25511113 | 0.00042099 | 0.00994767 |
| Peg10 | 0.204666376 | 0.06920686 | 0.33814474 | 0.00043634 | 0.01028485 |
| Mfge8 | 0.131584614 | 0.02494959 | 0.18960869 | 0.00045167 | 0.01059356 |
| Sdpr | 0.132652711 | 0.02099258 | 0.15825217 | 0.00045167 | 0.01059356 |
| Col6a3 | 0.135577941 | 0.02110436 | 0.15566219 | 0.00045976 | 0.01075335 |
| Lama4 | 0.075638497 | 0.00514385 | 0.06800573 | 0.00046187 | 0.01075335 |
| Sfrp2 | 0.075768228 | 0.00066363 | 0.0087587 | 0.00046188 | 0.01075335 |
| Ctsl | 0.385092225 | 0.1847209 | 0.47967964 | 0.00047396 | 0.01100771 |
| Eif3f | 0.696261905 | 0.41178821 | 0.59142718 | 0.00048984 | 0.01134885 |
| Itgb1 | 0.348418293 | 0.16389307 | 0.47039168 | 0.00049196 | 0.01137016 |
| Rps21 | 1.467078519 | 1.04641226 | 0.71326261 | 0.00049754 | 0.01147119 |
| Bex2 | 0.341522068 | 0.57516139 | 1.68411193 | 0.0005309 | 0.01221069 |
| Bloc1s1 | 0.113339418 | 0.01621882 | 0.14309955 | 0.00053992 | 0.01235833 |
| Loxl1 | 0.113992506 | 0.01098739 | 0.09638695 | 0.00053992 | 0.01235833 |
| Top2a | 0.117313889 | 0.01674901 | 0.14277092 | 0.00055838 | 0.01271952 |
| Prss23 | 0.116494268 | 0.01659972 | 0.14249391 | 0.00055838 | 0.01271952 |
| Ogn | 0.095102373 | 0.00538878 | 0.05666294 | 0.00057298 | 0.01295898 |
| Ptgis | 0.095275141 | 0.00450724 | 0.04730763 | 0.00057298 | 0.01295898 |

| | | | | | |
|---|---|---|---|---|---|
| Lox | 0.095381483 | 0.00394624 | 0.04137319 | 0.00057298 | 0.01295898 |
| Efemp2 | 0.175890313 | 0.04147583 | 0.23580506 | 0.00061379 | 0.01365525 |
| Cenpa | 0.09729563 | 0.01258432 | 0.12934105 | 0.00061382 | 0.01365525 |
| Plxdc2 | 0.097395175 | 0.01233152 | 0.12661323 | 0.00061382 | 0.01365525 |
| Ehd2 | 0.097267047 | 0.0122064 | 0.12549372 | 0.00061382 | 0.01365525 |
| Prrx1 | 0.097426755 | 0.00900656 | 0.09244441 | 0.00061383 | 0.01365525 |
| Cldn5 | 0.099635429 | 0.00397352 | 0.03988057 | 0.00061383 | 0.01365525 |
| Lrrc17 | 0.097617014 | 0.00566735 | 0.05805696 | 0.00061383 | 0.01365525 |
| Egr1 | 0.183689743 | 0.36676598 | 1.99666011 | 0.00069569 | 0.01544012 |
| Kcnj8 | 0.069895788 | 0.00301023 | 0.04306734 | 0.00075477 | 0.01671229 |
| Rap1b | 0.382383911 | 0.62784339 | 1.64191894 | 0.00076384 | 0.0168737 |
| Gch1 | 0.175400793 | 0.34897965 | 1.98961272 | 0.00078086 | 0.01720972 |
| Mapt | 0.015091026 | 0.08739236 | 5.79101556 | 0.00079586 | 0.01749964 |
| Abi3bp | 0.072371681 | 0.00582773 | 0.08052499 | 0.00080889 | 0.01762348 |
| Cdc42ep5 | 0.072533876 | 0.00471215 | 0.06496482 | 0.00080889 | 0.01762348 |
| Glipr2 | 0.073679008 | 0.00281871 | 0.03825667 | 0.0008089 | 0.01762348 |
| Lgals7 | 0.072340193 | 0.00144828 | 0.02002036 | 0.00080891 | 0.01762348 |
| Lpl | 0.110365735 | 0.01603274 | 0.14526922 | 0.00083803 | 0.01817472 |
| Cdo1 | 0.111721591 | 0.01159164 | 0.10375469 | 0.00083804 | 0.01817472 |
| Tcf4 | 0.182052557 | 0.05328704 | 0.29270138 | 0.00090387 | 0.01955781 |
| Gpd2 | 0.039734671 | 0.13736212 | 3.4569838 | 0.00092105 | 0.01988425 |
| Eng | 0.092156598 | 0.01011699 | 0.10978047 | 0.00092576 | 0.01989575 |
| Osr2 | 0.092418159 | 0.00462966 | 0.05009475 | 0.00092577 | 0.01989575 |
| Fam151a | 0.006191236 | 0.06536329 | 10.5573885 | 0.00092979 | 0.0199369 |
| Usmg5 | 0.626374511 | 0.93059157 | 1.48567918 | 0.00093452 | 0.01999321 |
| Pabpc1 | 0.657970007 | 0.39492706 | 0.60022045 | 0.00095197 | 0.02032071 |
| Hnrnpf | 0.515188564 | 0.28363669 | 0.55054927 | 0.00098023 | 0.02087724 |
| Anxa5 | 0.407372786 | 0.20535687 | 0.50410062 | 0.00098895 | 0.02101578 |
| Gm9493 | 0.473474597 | 0.25288863 | 0.53411235 | 0.00100524 | 0.02131414 |
| Mrps33 | 0.244138572 | 0.4490437 | 1.83929848 | 0.00105092 | 0.02221344 |

| | | | | | |
|---|---|---|---|---|---|
| Pfn1 | 0.831927108 | 0.53929868 | 0.64825232 | 0.00105233 | 0.02221344 |
| Rab27a | 0.024892783 | 0.10465958 | 4.20441466 | 0.00115894 | 0.02440969 |
| Mif | 0.552755897 | 0.31474487 | 0.56941025 | 0.00116428 | 0.02446798 |
| Lamb1 | 0.13766351 | 0.03491133 | 0.25359901 | 0.00117633 | 0.0246666 |
| Uqcc2 | 0.339580656 | 0.56502061 | 1.66387749 | 0.00120335 | 0.0251775 |
| Rbms1 | 0.151008078 | 0.04171688 | 0.27625595 | 0.00125092 | 0.02595539 |
| Stbd1 | 0.065965374 | 0.00748584 | 0.11348135 | 0.00125415 | 0.02595539 |
| Slc43a3 | 0.067105226 | 0.00376393 | 0.05608999 | 0.00125417 | 0.02595539 |
| Id1 | 0.066841306 | 0.00319799 | 0.04784458 | 0.00125417 | 0.02595539 |
| Tyrobp | 0.066942738 | 0.00130008 | 0.01942071 | 0.00125419 | 0.02595539 |
| Plac8 | 0.103304123 | 0.01586588 | 0.15358422 | 0.00128693 | 0.0265208 |
| Scg5 | 0.140879634 | 0.29252808 | 2.07643981 | 0.00128709 | 0.0265208 |
| Snrpg | 0.688797361 | 0.4286002 | 0.62224425 | 0.00130398 | 0.02681067 |
| Cd248 | 0.107903198 | 0.01370327 | 0.12699597 | 0.0013498 | 0.02769281 |
| Ost4 | 0.369473939 | 0.18266789 | 0.49439993 | 0.00141167 | 0.02889963 |
| Actg1 | 3.276928302 | 2.59914512 | 0.79316509 | 0.00142707 | 0.02915215 |
| Pls3 | 0.085133298 | 0.00960708 | 0.11284756 | 0.00144185 | 0.02926541 |
| Hic1 | 0.087388449 | 0.00550853 | 0.06303501 | 0.00144186 | 0.02926541 |
| Figf | 0.085733697 | 0.00684675 | 0.07986061 | 0.00144186 | 0.02926541 |
| Itm2b | 1.174088229 | 0.83269286 | 0.70922512 | 0.00145534 | 0.02947602 |
| Xist | 1.041742414 | 0.72534645 | 0.69628196 | 0.0015023 | 0.03036243 |
| Slc25a4 | 0.734425467 | 0.46644608 | 0.63511697 | 0.0015179 | 0.03058422 |
| Ppm1b | 0.070084496 | 0.18187515 | 2.59508398 | 0.00151971 | 0.03058422 |
| Pecam1 | 0.088631005 | 0.00915484 | 0.1032916 | 0.00156192 | 0.03103973 |
| Ndrg2 | 0.089295992 | 0.00746233 | 0.08356845 | 0.00156193 | 0.03103973 |
| Filip1l | 0.089342686 | 0.00732565 | 0.08199492 | 0.00156193 | 0.03103973 |
| Rhoj | 0.090020167 | 0.00653923 | 0.07264189 | 0.00156193 | 0.03103973 |
| Axl | 0.088789292 | 0.00624397 | 0.07032345 | 0.00156194 | 0.03103973 |
| Upk3b | 0.089642483 | 0.00243244 | 0.02713485 | 0.00156195 | 0.03103973 |
| Col4a2 | 0.119079453 | 0.02491913 | 0.20926471 | 0.00157224 | 0.031179 |

| | | | | | |
|---|---|---|---|---|---|
| Reep5 | 0.259833585 | 0.11353441 | 0.43695048 | 0.00158978 | 0.03146108 |
| Gm10073 | 0.372890481 | 0.19640113 | 0.52669922 | 0.00160795 | 0.03175443 |
| Eno1 | 0.395081318 | 0.20557473 | 0.52033524 | 0.00162873 | 0.03209814 |
| Nfia | 0.132994467 | 0.03112169 | 0.23400741 | 0.00173575 | 0.03413635 |
| Rsu1 | 0.13398153 | 0.03280365 | 0.24483707 | 0.00180554 | 0.03543567 |
| Serping1 | 0.147188873 | 0.03962214 | 0.26919253 | 0.00184802 | 0.03612018 |
| Aldh2 | 0.148516528 | 0.0365333 | 0.24598809 | 0.00184803 | 0.03612018 |
| Tpi1 | 0.405374747 | 0.2144333 | 0.52897548 | 0.00185733 | 0.03622755 |
| Emb | 0.12776761 | 0.27101615 | 2.12116473 | 0.00189882 | 0.03696079 |
| Tshz1 | 0.04953032 | 0.142545 | 2.87793422 | 0.00198296 | 0.03844121 |
| Gck | 0.049249165 | 0.14014837 | 2.84570044 | 0.00198297 | 0.03844121 |
| Ffar1 | 0.015546637 | 0.07961662 | 5.12114717 | 0.00199365 | 0.03856961 |
| Rpl13-ps3 | 0.328409811 | 0.1602713 | 0.48802228 | 0.00200476 | 0.03865809 |
| Galk1 | 0.10241529 | 0.00907343 | 0.08859444 | 0.00200636 | 0.03865809 |
| Insm1 | 0.059209451 | 0.15459712 | 2.61102095 | 0.0020422 | 0.03898501 |
| Rest | 0.06047889 | 0.0056394 | 0.09324583 | 0.00204382 | 0.03898501 |
| Angptl2 | 0.060698418 | 0.00520757 | 0.08579416 | 0.00204383 | 0.03898501 |
| Gstm2 | 0.061051339 | 0.00403025 | 0.06601404 | 0.00204383 | 0.03898501 |
| Spon1 | 0.06086182 | 0.00291662 | 0.04792205 | 0.00204385 | 0.03898501 |
| Btf3 | 0.996538208 | 0.69281779 | 0.69522452 | 0.0020503 | 0.03902962 |
| Tpd52 | 0.132520983 | 0.2682044 | 2.02386367 | 0.00216354 | 0.04110295 |
| Clec14a | 0.06329274 | 0.00594822 | 0.09397947 | 0.00220559 | 0.0416523 |
| Col12a1 | 0.063798247 | 0.00502224 | 0.07872066 | 0.0022056 | 0.0416523 |
| Trf | 0.062991991 | 0.0046667 | 0.07408405 | 0.00220561 | 0.0416523 |
| Vkorc1 | 0.208556402 | 0.07842691 | 0.37604653 | 0.0022217 | 0.04187284 |
| Kdelr2 | 0.333142822 | 0.17311296 | 0.51963586 | 0.00230246 | 0.0430421 |
| Acvr1c | 0.010866738 | 0.07573127 | 6.96908957 | 0.00230496 | 0.0430421 |
| Slc2a2 | 0.012002078 | 0.07089836 | 5.9071737 | 0.00230498 | 0.0430421 |
| Cdk1 | 0.079223729 | 0.00993584 | 0.1254149 | 0.0023109 | 0.0430421 |
| Pros1 | 0.078592821 | 0.00853267 | 0.10856812 | 0.00231091 | 0.0430421 |

| | | | | | |
|---|---|---|---|---|---|
| Selenbp1 | 0.078785635 | 0.00672652 | 0.08537749 | 0.00231092 | 0.0430421 |
| Ndufa12 | 0.274061353 | 0.45503692 | 1.6603469 | 0.00233795 | 0.04325263 |
| Tinagl1 | 0.083385483 | 0.00932366 | 0.1118139 | 0.00234041 | 0.04325263 |
| Tfpi | 0.084327912 | 0.00519062 | 0.06155283 | 0.00234044 | 0.04325263 |
| Trip6 | 0.081788747 | 0.00714491 | 0.08735811 | 0.00234044 | 0.04325263 |
| Grb10 | 0.399248647 | 0.21413784 | 0.53635208 | 0.00236758 | 0.04366916 |
| G6pc2 | 0.004547339 | 0.04252254 | 9.35108144 | 0.00250017 | 0.04602542 |
| Pcbd1 | 0.138662248 | 0.28000297 | 2.01931656 | 0.00252908 | 0.04646761 |
| Adra2a | 0.008902011 | 0.06378542 | 7.16528221 | 0.00257155 | 0.04711339 |
| Cdk4 | 0.450632334 | 0.2573898 | 0.57117473 | 0.00257415 | 0.04711339 |
| Fbln2 | 0.126570085 | 0.03195408 | 0.25246152 | 0.00258965 | 0.04719778 |
| Tpst1 | 0.126222425 | 0.03032146 | 0.24022242 | 0.00258965 | 0.04719778 |
| Eif4ebp1 | 0.127698119 | 0.03473605 | 0.2720169 | 0.00259863 | 0.04719778 |
| Myadm | 0.129847595 | 0.0318947 | 0.24563178 | 0.00259864 | 0.04719778 |
| Oat | 0.167809218 | 0.05590999 | 0.33317589 | 0.00261709 | 0.04744222 |
| Anxa6 | 0.266524007 | 0.11735856 | 0.44033016 | 0.00264361 | 0.04783179 |
| Egfl7 | 0.140561883 | 0.04026228 | 0.28643811 | 0.00266374 | 0.04799308 |
| Pgrmc1 | 0.345029977 | 0.17276996 | 0.50073898 | 0.00266572 | 0.04799308 |
| Krt19 | 0.153248244 | 0.04532732 | 0.29577706 | 0.00267239 | 0.04799308 |
| Psap | 0.407686199 | 0.22491699 | 0.55169146 | 0.00267274 | 0.04799308 |
| Prss53 | 0.00689858 | 0.05023026 | 7.28124628 | 0.00269671 | 0.04833224 |
| Gm11808 | 0.479549109 | 0.27533865 | 0.57416153 | 0.00274219 | 0.04905475 |
| Gm10076 | 0.853059895 | 0.57980349 | 0.679675 | 0.00286081 | 0.0510806 |
| Basp1 | 0.094057796 | 0.01949439 | 0.20725967 | 0.00304572 | 0.05417894 |
| Adamts5 | 0.095758443 | 0.01248186 | 0.13034738 | 0.00304575 | 0.05417894 |
| Kcnq1ot1 | 0.394345161 | 0.21580856 | 0.54725804 | 0.00305449 | 0.05423298 |
| Cyb5a | 0.205512095 | 0.08036554 | 0.39105017 | 0.00312707 | 0.05541793 |
| Gfra3 | 0.034099185 | 0.11316958 | 3.3188352 | 0.00315083 | 0.05563167 |
| Rab3b | 0.032870198 | 0.11042551 | 3.35944161 | 0.00315084 | 0.05563167 |
| Rpl9-ps6 | 0.236081106 | 0.10448288 | 0.44257197 | 0.00320565 | 0.05649434 |

| | | | | | |
|---|---|---|---|---|---|
| Capn6 | 0.098677345 | 0.0117065 | 0.11863413 | 0.00323628 | 0.0569285 |
| Fus | 0.49710119 | 0.29409479 | 0.59161956 | 0.00331691 | 0.058239 |
| Hsp90ab1 | 2.272367669 | 1.8308775 | 0.80571359 | 0.00332836 | 0.05833228 |
| Rgs5 | 0.057006825 | 0.00686793 | 0.12047557 | 0.00341712 | 0.05934177 |
| Ccnb2 | 0.057654858 | 0.00478051 | 0.08291597 | 0.00341715 | 0.05934177 |
| Thbd | 0.057325954 | 0.00480625 | 0.08384076 | 0.00341715 | 0.05934177 |
| Lockd | 0.058507252 | 0.00278787 | 0.04765 | 0.00341717 | 0.05934177 |
| Tmem119 | 0.059117638 | 0.00104191 | 0.01762437 | 0.0034172 | 0.05934177 |
| Rogdi | 0.069014298 | 0.16152735 | 2.34049113 | 0.00349815 | 0.06063676 |
| Plvap | 0.108591369 | 0.01801472 | 0.16589454 | 0.00356071 | 0.06160879 |
| Sptssa | 0.227983294 | 0.0906991 | 0.39783224 | 0.00357819 | 0.06165483 |
| Tubb2b | 0.11013213 | 0.0256952 | 0.2333125 | 0.00358215 | 0.06165483 |
| mt-Cytb | 3.665129775 | 2.96727292 | 0.80959559 | 0.00358285 | 0.06165483 |
| Plat | 0.077872394 | 0.00880301 | 0.11304401 | 0.00361321 | 0.06173109 |
| Trim47 | 0.07637308 | 0.00842295 | 0.11028687 | 0.00361323 | 0.06173109 |
| Plpp1 | 0.076547426 | 0.00609024 | 0.07956165 | 0.00361326 | 0.06173109 |
| G0s2 | 0.076409431 | 0.0050942 | 0.06666972 | 0.00361327 | 0.06173109 |
| mt-Nd4 | 1.965736106 | 1.56085737 | 0.794032 | 0.00381718 | 0.06501274 |
| Atp8b1 | 0.051105879 | 0.14336348 | 2.80522478 | 0.00381904 | 0.06501274 |
| Gltscr2 | 0.304641503 | 0.14505915 | 0.47616344 | 0.00383084 | 0.06509688 |
| S100a13 | 0.134052905 | 0.04066183 | 0.30332674 | 0.00387201 | 0.06567899 |
| Cks2 | 0.138395465 | 0.03876582 | 0.28010904 | 0.00394004 | 0.06671388 |
| Spock2 | 0.037553695 | 0.11690324 | 3.11296239 | 0.00395354 | 0.06682319 |
| Tmem163 | 0.019619849 | 0.07847522 | 3.99978705 | 0.00398994 | 0.06731867 |
| Rpl6l | 0.437813179 | 0.25228445 | 0.57623767 | 0.00404203 | 0.0680766 |
| Cd164 | 0.155928335 | 0.29207643 | 1.87314529 | 0.00417497 | 0.07019122 |
| Creld2 | 0.082029476 | 0.19222785 | 2.34339973 | 0.004303 | 0.07221596 |
| Vdac1 | 0.290417667 | 0.14252401 | 0.49075529 | 0.00435035 | 0.07288177 |
| Prps1 | 0.072352821 | 0.17734581 | 2.45112497 | 0.004531 | 0.07577462 |
| Pdap1 | 0.436601544 | 0.64512161 | 1.4775981 | 0.0045402 | 0.07579509 |

| | | | | | |
|---|---|---|---|---|---|
| Spint2 | 0.409622706 | 0.62232085 | 1.51925379 | 0.00455848 | 0.07596674 |
| Tead2 | 0.090452877 | 0.01522868 | 0.16836033 | 0.00470737 | 0.07803758 |
| Adamts1 | 0.087825336 | 0.01579006 | 0.17978938 | 0.00470739 | 0.07803758 |
| Slit2 | 0.087996612 | 0.01478383 | 0.16800449 | 0.00470739 | 0.07803758 |
| Selm | 0.40427924 | 0.22843203 | 0.56503527 | 0.0047559 | 0.07815305 |
| Birc5 | 0.092412993 | 0.01774472 | 0.19201547 | 0.00476369 | 0.07815305 |
| Gstm1 | 0.092253115 | 0.01690846 | 0.1832833 | 0.00476369 | 0.07815305 |
| Meis1 | 0.090931264 | 0.0173189 | 0.19046143 | 0.0047637 | 0.07815305 |
| Acaa2 | 0.090726031 | 0.01622839 | 0.17887243 | 0.00476371 | 0.07815305 |
| Sgce | 0.092049867 | 0.01387196 | 0.15070044 | 0.00476372 | 0.07815305 |
| Eef1d | 0.453680168 | 0.27070262 | 0.59668163 | 0.00486074 | 0.07960725 |
| Pclo | 0.038435312 | 0.12362394 | 3.21641564 | 0.00489224 | 0.07998511 |
| Ptp4a2 | 0.155538529 | 0.05569049 | 0.3580495 | 0.0052286 | 0.08533752 |
| Zbtb20 | 0.218948287 | 0.37307892 | 1.70395906 | 0.00533696 | 0.08695668 |
| Celf2 | 0.143796826 | 0.04936909 | 0.34332528 | 0.00544747 | 0.08860534 |
| Mylk | 0.051571015 | 0.00486051 | 0.09424892 | 0.00555385 | 0.08978816 |
| Arhgef25 | 0.051766351 | 0.00454259 | 0.08775188 | 0.00555386 | 0.08978816 |
| Abcg2 | 0.051432734 | 0.00334083 | 0.06495537 | 0.00555392 | 0.08978816 |
| Snhg6 | 0.115875792 | 0.03618933 | 0.31231141 | 0.00556745 | 0.08978816 |
| P3h4 | 0.116162292 | 0.03473689 | 0.2990376 | 0.00556745 | 0.08978816 |
| Tcf7l2 | 0.132244507 | 0.03637508 | 0.27505932 | 0.00560596 | 0.09010337 |
| Rbms3 | 0.132932448 | 0.03431656 | 0.25815033 | 0.00560597 | 0.09010337 |
| Sdc1 | 0.069789022 | 0.0113972 | 0.16330935 | 0.00572893 | 0.09177085 |
| Igfbp3 | 0.07202061 | 0.00389994 | 0.0541503 | 0.00572904 | 0.09177085 |
| Phgdh | 0.072967114 | 0.01146243 | 0.15709036 | 0.00588714 | 0.09258803 |
| Tspan4 | 0.072346774 | 0.01043268 | 0.14420385 | 0.00588717 | 0.09258803 |
| Plpp3 | 0.073619704 | 0.00901158 | 0.12240714 | 0.00588717 | 0.09258803 |
| Klf2 | 0.074085031 | 0.00795094 | 0.10732176 | 0.00588718 | 0.09258803 |
| Lrp1 | 0.074544283 | 0.00669927 | 0.08986966 | 0.0058872 | 0.09258803 |
| Smc2 | 0.072418846 | 0.00845707 | 0.11677993 | 0.00588721 | 0.09258803 |

| | | | | | |
|---|---|---|---|---|---|
| F2r | 0.072938747 | 0.00654914 | 0.08978955 | 0.00588723 | 0.09258803 |
| Lpar4 | 0.073324265 | 0.00521056 | 0.0710619 | 0.00588725 | 0.09258803 |
| Mef2c | 0.073639293 | 0.00476811 | 0.06474954 | 0.00588726 | 0.09258803 |
| Mmp23 | 0.074523203 | 0.00378507 | 0.0507905 | 0.00588726 | 0.09258803 |
| Tgfb1i1 | 0.074809861 | 0.00306452 | 0.0409641 | 0.00588727 | 0.09258803 |
| Srpx | 0.054784842 | 0.00417551 | 0.07621654 | 0.00603556 | 0.09445141 |
| Fam212a | 0.054229556 | 0.00457846 | 0.08442735 | 0.00603556 | 0.09445141 |
| Tcf21 | 0.055047147 | 0.00320722 | 0.05826324 | 0.00603558 | 0.09445141 |
| Cdc42 | 0.558576519 | 0.35305209 | 0.6320568 | 0.00609212 | 0.09517941 |
| Ly6e | 0.247136228 | 0.11858246 | 0.47982628 | 0.006127 | 0.09556715 |
| Tbrg1 | 0.251701622 | 0.12067718 | 0.47944537 | 0.00620065 | 0.09655726 |
| Rab3a | 0.070997859 | 0.16659175 | 2.34643337 | 0.00623667 | 0.09695931 |
| Tmed9 | 0.3582698 | 0.19816486 | 0.55311629 | 0.00638455 | 0.09909611 |
| Gm6133 | 0.291338019 | 0.15142315 | 0.51975075 | 0.00645847 | 0.10007997 |
| Rps18-ps3 | 0.293157982 | 0.15327024 | 0.52282472 | 0.00652939 | 0.10101411 |
| Minos1 | 0.508458834 | 0.73269541 | 1.44101225 | 0.00665817 | 0.10274547 |
| Vwa5b2 | 0.120998764 | 0.23575293 | 1.94839118 | 0.00666293 | 0.10274547 |
| Anxa4 | 0.165386797 | 0.3042148 | 1.83941404 | 0.00704365 | 0.1084402 |
| Flna | 0.084502355 | 0.02249286 | 0.26618025 | 0.00714571 | 0.10952767 |
| Timp3 | 0.086499041 | 0.01111592 | 0.12850921 | 0.00714584 | 0.10952767 |
| 2700094K13I | 0.199020986 | 0.08618544 | 0.43304699 | 0.00715532 | 0.10952767 |
| Aplp2 | 0.2019554 | 0.08728107 | 0.43217995 | 0.0071604 | 0.10952767 |
| Chchd2 | 1.026131238 | 1.3354742 | 1.3014653 | 0.00717508 | 0.10957563 |
| Eif3h | 0.525024207 | 0.33459859 | 0.63730127 | 0.00736494 | 0.1122947 |
| Scn9a | 0.007632542 | 0.04056382 | 5.31458824 | 0.00762659 | 0.11609768 |
| Gip | 0.004369765 | 0.03744676 | 8.56951372 | 0.00775445 | 0.11777962 |
| C1qbp | 0.137013879 | 0.05355709 | 0.39088804 | 0.00777426 | 0.11777962 |
| Cpa1 | 0.136735051 | 0.0514144 | 0.37601479 | 0.00777427 | 0.11777962 |
| Ddx5 | 0.897731061 | 1.18224078 | 1.31692088 | 0.0079049 | 0.11956798 |
| Gabarapl2 | 0.307316896 | 0.47759585 | 1.55408263 | 0.00806413 | 0.12161897 |

| | | | | | |
|---|---|---|---|---|---|
| Vat1 | 0.102159909 | 0.02324018 | 0.2274883 | 0.00806611 | 0.12161897 |
| Ctsd | 0.245652348 | 0.12178949 | 0.49577986 | 0.00811737 | 0.12219798 |
| C1qa | 0.113556468 | 0.03058565 | 0.26934309 | 0.00815446 | 0.12256206 |
| Entpd3 | 0.031509158 | 0.0969449 | 3.0767215 | 0.00828966 | 0.12420162 |
| Slc7a2 | 0.031005481 | 0.09092575 | 2.93257031 | 0.0082897 | 0.12420162 |
| Rph3al | 0.089909553 | 0.18788116 | 2.08966852 | 0.0083417 | 0.12458777 |
| Smim22 | 0.089328217 | 0.18765417 | 2.10072667 | 0.0083417 | 0.12458777 |
| Tmem256 | 0.35794126 | 0.21027695 | 0.58746217 | 0.00885597 | 0.13167737 |
| Cnih2 | 0.040531803 | 0.11087035 | 2.73539147 | 0.00885795 | 0.13167737 |
| Kcnmb2 | 0.040841151 | 0.10739284 | 2.62952526 | 0.00885797 | 0.13167737 |
| Chd7 | 0.059788206 | 0.1397089 | 2.33673006 | 0.00895867 | 0.13265072 |
| Apoc1 | 0.067406929 | 0.01167712 | 0.17323328 | 0.00900707 | 0.13265072 |
| Ntrk2 | 0.067092554 | 0.01196531 | 0.1783404 | 0.00900707 | 0.13265072 |
| Phactr2 | 0.068034174 | 0.0075281 | 0.11065173 | 0.00900717 | 0.13265072 |
| Tagln | 0.067538619 | 0.00606247 | 0.08976303 | 0.00900723 | 0.13265072 |
| Cpxm1 | 0.068727069 | 0.00486369 | 0.07076818 | 0.00900723 | 0.13265072 |
| Ubl5 | 0.44069184 | 0.64656122 | 1.46715043 | 0.00906521 | 0.13329785 |
| Klhl32 | 0.017606365 | 0.07137128 | 4.0537206 | 0.00915494 | 0.13440921 |
| Rnd3 | 0.048186837 | 0.00596571 | 0.12380378 | 0.00935252 | 0.13563772 |
| Fam111a | 0.048439261 | 0.0051827 | 0.10699372 | 0.00935256 | 0.13563772 |
| Fhl1 | 0.048651649 | 0.00444441 | 0.0913516 | 0.00935259 | 0.13563772 |
| Cnrip1 | 0.049002783 | 0.00288865 | 0.05894873 | 0.00935266 | 0.13563772 |
| Pygb | 0.048440519 | 0.00294807 | 0.06085965 | 0.0093527 | 0.13563772 |
| F13a1 | 0.049486683 | 0.0007565 | 0.01528687 | 0.00935278 | 0.13563772 |
| Anxa1 | 0.047581044 | 0.0026172 | 0.05500516 | 0.00935278 | 0.13563772 |
| Gpm6b | 0.047819702 | 0.001513 | 0.03163958 | 0.00935285 | 0.13563772 |
| Cd47 | 0.25236023 | 0.12441392 | 0.4930013 | 0.00938923 | 0.13595772 |
| Ctsb | 0.293238446 | 0.1639759 | 0.55918962 | 0.00946646 | 0.1366594 |
| Arhgdia | 0.295159612 | 0.15972247 | 0.54113933 | 0.00946646 | 0.1366594 |
| Raly | 0.274847968 | 0.14744459 | 0.53645872 | 0.00959384 | 0.13828817 |

| | | | | | |
|---|---|---|---|---|---|
| Pam | 0.27858307 | 0.43836337 | 1.57354634 | 0.00972034 | 0.13989919 |

| | mean outside clu: | mean inside clu: | fold change | P value | P adj |
|---|---|---|---|---|---|
| Gm42418__chr17 | 49.67008029 | 262.3576568 | 5.28200589 | 3.94E-75 | 2.31E-71 |
| Nnat__chr2 | 0.773582648 | 6.607925 | 8.54197676 | 3.28E-19 | 9.60E-16 |
| Gm10800__chr2 | 63.58162519 | 18.43505228 | 0.28994308 | 1.16E-14 | 2.27E-11 |
| Gm10801__chr2 | 7.204485334 | 1.247061154 | 0.17309511 | 3.19E-10 | 4.66E-07 |
| Gm26870__chr9 | 28.20495913 | 11.1051816 | 0.39373153 | 9.72E-08 | 0.00011383 |
| Iapp__chr6 | 208.5153844 | 111.079066 | 0.53271401 | 2.90E-07 | 0.00025671 |
| Gm10717__chr9 | 9.054032755 | 2.72492766 | 0.30096287 | 3.07E-07 | 0.00025671 |
| Gm43359__chr5 | 1.239994427 | 4.213725228 | 3.39818078 | 5.29E-07 | 0.00036151 |
| Gm11168__chr9 | 4.941120132 | 1.093614541 | 0.22132928 | 5.56E-07 | 0.00036151 |
| Gcg__chr2 | 14.16092835 | 26.14863372 | 1.84653386 | 1.42E-05 | 0.00833563 |
| Fos__chr12 | 9.82825758 | 18.36758344 | 1.8688545 | 4.06E-05 | 0.0205652 |
| Chga__chr12 | 82.26401711 | 46.90746249 | 0.57020632 | 4.22E-05 | 0.0205652 |
| Bclaf1__chr10 | 1.763138089 | 4.497327764 | 2.55075186 | 5.26E-05 | 0.02368907 |
| Resp18__chr1 | 7.154827509 | 2.849153229 | 0.3982141 | 0.00010753 | 0.04495425 |
| Pclo__chr5 | 2.19249644 | 5.075875993 | 2.31511254 | 0.00015545 | 0.06065644 |
| Atrnl1__chr19 | 0.350450383 | 1.701260096 | 4.85449633 | 0.00022077 | 0.07993657 |
| Angpt2__chr8 | 1.231826163 | 3.442114026 | 2.794318 | 0.00023218 | 0.07993657 |
| Gm42566__chr5 | 0.387060505 | 1.758290624 | 4.54267641 | 0.0004593 | 0.14934973 |
| Chd7__chr4 | 2.540579159 | 5.400184882 | 2.12557238 | 0.00049659 | 0.15297592 |
| Ccnd1__chr7 | 0.748274698 | 2.280242927 | 3.047334 | 0.0005857 | 0.17140517 |
| Gimap1__chr6 | 0.965887948 | 2.666125321 | 2.76028428 | 0.00080217 | 0.22357688 |
| Hsp90ab1__chr17 | 11.90073758 | 19.60350879 | 1.64725158 | 0.00085381 | 0.22715128 |
| Gm37728__chr6 | 0.985232904 | 2.701047652 | 2.74153212 | 0.00091359 | 0.2324876 |
| Lbh__chr17 | 0.848579848 | 2.372048575 | 2.7953157 | 0.00160367 | 0.39109426 |
| Gria2__chr3 | 0.266285835 | 1.138543981 | 4.27564606 | 0.00283207 | 0.50275719 |
| Snx30__chr4 | 0.497133264 | 1.752353016 | 3.52491604 | 0.0022488 | 0.50275719 |
| Ccpg1__chr9 | 0.710420358 | 2.016724028 | 2.83877567 | 0.00281969 | 0.50275719 |
| BC005537__chr13 | 0.812244542 | 2.072328585 | 2.55136043 | 0.00276828 | 0.50275719 |

| | | | | | |
|---|---|---|---|---|---|
| Srek1__chr13 | 1.189549162 | 2.943686471 | 2.47462363 | 0.00247155 | 0.50275719 |
| Chd4__chr6 | 1.379922724 | 3.17980426 | 2.30433502 | 0.0025544 | 0.50275719 |
| Cox6a2__chr7 | 1.536959393 | 3.449193572 | 2.24416701 | 0.00283461 | 0.50275719 |
| Slc2a2__chr3 | 3.816029669 | 1.582399599 | 0.41467172 | 0.00242998 | 0.50275719 |
| Itm2b__chr14 | 3.326502836 | 1.269087186 | 0.38150792 | 0.00253341 | 0.50275719 |
| Snhg11__chr2 | 0.530557538 | 1.590712409 | 2.9981902 | 0.00313432 | 0.53956437 |
| Ppargc1a__chr5 | 0.282490498 | 1.031027256 | 3.64977676 | 0.00357221 | 0.58078244 |
| Chic1__chrX | 3.57990472 | 6.339934847 | 1.77097866 | 0.00353257 | 0.58078244 |
| Tnpo3__chr6 | 0.463134697 | 1.506066446 | 3.25189725 | 0.00407305 | 0.63516424 |
| Meg3__chr12 | 19.60042121 | 28.99461438 | 1.47928527 | 0.00412374 | 0.63516424 |
| Bdp1__chr13 | 0.47581212 | 1.491707514 | 3.13507675 | 0.00480518 | 0.68744686 |
| Pim1__chr17 | 0.57238567 | 1.660365008 | 2.90078018 | 0.00495971 | 0.68744686 |
| Trim28__chr7 | 0.573910899 | 1.600168836 | 2.78818339 | 0.00495971 | 0.68744686 |
| Gm43322__chr3 | 0.671458449 | 1.803946963 | 2.68660997 | 0.00492441 | 0.68744686 |
| Gm10719__chr9 | 1.99847851 | 0.501044741 | 0.2507131 | 0.00505044 | 0.68744686 |
| Trim27__chr13 | 0.312073568 | 1.041570609 | 3.33758035 | 0.00548261 | 0.72874373 |
| Filip1__chr9 | 0.481338294 | 1.598288441 | 3.32050963 | 0.00563547 | 0.72874373 |
| RP23-343H23.2__c | 1.292458682 | 2.873283436 | 2.22311435 | 0.00597637 | 0.72874373 |
| Akap9__chr5 | 1.633355398 | 3.463385213 | 2.12041128 | 0.00583911 | 0.72874373 |
| Jun__chr4 | 8.3121578 | 12.98288177 | 1.56191474 | 0.00596915 | 0.72874373 |
| Elmsan1__chr12 | 0.408812468 | 1.242681975 | 3.03973599 | 0.00625623 | 0.74730035 |
| Scyl2__chr10 | 0.242506342 | 0.933644886 | 3.84998131 | 0.00665873 | 0.76637474 |
| Brd2__chr17 | 1.317807402 | 2.823327509 | 2.1424432 | 0.00667779 | 0.76637474 |
| Srsf11__chr3 | 1.900357162 | 3.736710863 | 1.9663203 | 0.00681097 | 0.76662749 |
| Tubb4b__chr2 | 1.961396896 | 0.543766731 | 0.27723442 | 0.00709646 | 0.78369074 |
| Nol3__chr8 | 0.338614714 | 1.082747949 | 3.19758092 | 0.00806961 | 0.8746559 |
| 2900060B14Rik__c | 0.436958284 | 1.271284919 | 2.90939654 | 0.00869362 | 0.92515936 |
| Rps19__chr7 | 1.845343587 | 3.542559743 | 1.91972908 | 0.00953071 | 0.99612961 |
| Slc35e1__chr8 | 0.270611645 | 0.918626794 | 3.39463143 | 0.01023353 | 0.99828112 |

| | | | | | |
|---|---|---|---|---|---|
| Polr3gl__chr3 | 0.267301253 | 0.905462623 | 3.38742378 | 0.01023353 | 0.99828112 |
| Arfip1__chr3 | 0.637948525 | 1.628797206 | 2.55317967 | 0.00974183 | 0.99828112 |
| Tm4sf4__chr3 | 3.200190965 | 1.272410996 | 0.39760471 | 0.00996548 | 0.99828112 |
| Ffar3__chr7 | 0.194530925 | 0.763389218 | 3.92425634 | 0.01254793 | 1 |
| Gm37733__chr1 | 0.292312178 | 1.053759569 | 3.60491163 | 0.0149916 | 1 |
| Sema4b__chr7 | 0.241508102 | 0.807386957 | 3.34310506 | 0.02344998 | 1 |
| Zfp37__chr4 | 0.252800254 | 0.840327494 | 3.32407693 | 0.02811421 | 1 |
| RP23-256J14.2__ch | 0.223634074 | 0.743044362 | 3.32259011 | 0.01931578 | 1 |
| Esco1__chr18 | 0.263707708 | 0.873763548 | 3.31337887 | 0.03332607 | 1 |
| Fbxl2__chr9 | 0.250340131 | 0.796380698 | 3.1811947 | 0.02811422 | 1 |
| Ift74__chr4 | 0.379005544 | 1.200059338 | 3.16633716 | 0.01349136 | 1 |
| Slc9b2__chr3 | 0.312495508 | 0.960742714 | 3.07442088 | 0.01786642 | 1 |
| Rab3c__chr13 | 0.319082736 | 0.960713749 | 3.01086095 | 0.02109698 | 1 |
| Ikzf5__chr7 | 0.250873196 | 0.743262825 | 2.96270322 | 0.02811423 | 1 |
| Scn9a__chr2 | 0.287932229 | 0.841189865 | 2.92148561 | 0.0454433 | 1 |
| Cnot8__chr11 | 0.274695401 | 0.79155807 | 2.88158471 | 0.03909921 | 1 |
| Gatad2a__chr8 | 0.387728236 | 1.110738053 | 2.86473346 | 0.0157704 | 1 |
| Vps13c__chr9 | 0.292274742 | 0.836601958 | 2.86238199 | 0.0454433 | 1 |
| Gm37437__chr2 | 0.36115254 | 1.023339922 | 2.83353932 | 0.0330845 | 1 |
| Lrrc16b__chr14 | 0.556091486 | 1.557587105 | 2.80095478 | 0.01155419 | 1 |
| Gm43238__chr5 | 0.429083332 | 1.191994293 | 2.77800186 | 0.02422028 | 1 |
| Hdac4__chr1 | 0.359609996 | 0.996446451 | 2.77090866 | 0.0330845 | 1 |
| Exph5__chr9 | 0.269617783 | 0.745085014 | 2.76348617 | 0.03332608 | 1 |
| Cdk9__chr2 | 0.358298396 | 0.980223865 | 2.7357752 | 0.03308451 | 1 |
| 6330419E04Rik__c | 0.383276641 | 1.042594895 | 2.72021507 | 0.04311871 | 1 |
| Cttnbp2__chr6 | 0.349064052 | 0.949407912 | 2.71986733 | 0.02869165 | 1 |
| Prkx__chrX | 0.358206574 | 0.968572147 | 2.70394855 | 0.03308451 | 1 |
| Zc3hav1__chr6 | 0.364472219 | 0.980162386 | 2.68926501 | 0.0330845 | 1 |
| Gm26809__chr6 | 0.34979153 | 0.929333163 | 2.65682009 | 0.02869165 | 1 |
| Far2os1__chr6 | 0.489756953 | 1.296407002 | 2.64704155 | 0.01559897 | 1 |

| | | | | | |
|---|---|---|---|---|---|
| Dsg2__chr18 | 0.298078538 | 0.787498641 | 2.64191661 | 0.04544331 | 1 |
| Fam219a__chr4 | 0.561594841 | 1.482564117 | 2.63991762 | 0.01155419 | 1 |
| Gm42467__chr5 | 0.296141439 | 0.781075392 | 2.63750792 | 0.04544331 | 1 |
| Spire1__chr18 | 0.291362961 | 0.760907868 | 2.61154632 | 0.04544331 | 1 |
| Akap7__chr10 | 0.297608878 | 0.76304533 | 2.56391992 | 0.04544331 | 1 |
| Pirt__chr11 | 0.448246655 | 1.142923402 | 2.54976449 | 0.02761674 | 1 |
| Vps51__chr19 | 0.296847303 | 0.756440604 | 2.54824819 | 0.04544331 | 1 |
| Trps1__chr15 | 0.280980057 | 0.71379482 | 2.54037538 | 0.03909922 | 1 |
| Clmn__chr12 | 0.372830205 | 0.943116085 | 2.52961287 | 0.03789037 | 1 |
| Ypel4__chr2 | 0.458393573 | 1.146857089 | 2.50190482 | 0.03132196 | 1 |
| Tmem2__chr19 | 0.284047874 | 0.709317354 | 2.49717537 | 0.03909922 | 1 |
| Gm38142__chr1 | 0.376436848 | 0.936249125 | 2.48713464 | 0.03789037 | 1 |
| 2010111I01Rik__ch | 0.456091097 | 1.131035529 | 2.47984566 | 0.03132196 | 1 |
| Gm42631__chr5 | 0.455503991 | 1.12923992 | 2.47909994 | 0.03132196 | 1 |
| Cxxc1__chr18 | 0.423228333 | 1.042431007 | 2.46304636 | 0.02112177 | 1 |
| N4bp2l2__chr5 | 0.833709629 | 2.043543188 | 2.451145 | 0.01169862 | 1 |
| Gm42851__chr5 | 0.531350378 | 1.301063664 | 2.44859836 | 0.02290031 | 1 |
| Ythdc1__chr5 | 0.760522045 | 1.841394693 | 2.42122461 | 0.01661882 | 1 |
| Amer1__chrX | 0.447848616 | 1.080881121 | 2.41349662 | 0.02761674 | 1 |
| Gm43874__chr6 | 0.520093284 | 1.246362236 | 2.39642056 | 0.02024354 | 1 |
| 0610011F06Rik__c | 0.630139232 | 1.506871225 | 2.3913306 | 0.03447401 | 1 |
| Cdc42bpa__chr1 | 0.448559903 | 1.069403332 | 2.38408142 | 0.02761674 | 1 |
| Gadd45b__chr10 | 0.49424774 | 1.175404422 | 2.37816853 | 0.04438238 | 1 |
| Per1__chr11 | 0.555852025 | 1.312028208 | 2.36039116 | 0.02893023 | 1 |
| Htatsf1__chrX | 0.588161717 | 1.386488977 | 2.35732612 | 0.01490703 | 1 |
| Zrsr1__chr11 | 0.725698144 | 1.701530641 | 2.34468099 | 0.02637639 | 1 |
| Rab11fip2__chr19 | 0.391178239 | 0.910427163 | 2.32739727 | 0.04311873 | 1 |
| Lsm4__chr8 | 0.403699793 | 0.931534789 | 2.30749385 | 0.048777 | 1 |
| Gm10687__chr9 | 0.7328602 | 1.673451876 | 2.28345307 | 0.02795373 | 1 |
| Kif12__chr4 | 0.690281969 | 1.538866012 | 2.22932958 | 0.04449738 | 1 |

| | | | | | |
|---|---|---|---|---|---|
| Cwc25__chr11 | 0.405932069 | 0.89820791 | 2.21270498 | 0.048777 | 1 |
| Srrt__chr5 | 0.693355153 | 1.495085408 | 2.15630532 | 0.04449739 | 1 |
| Huwe1__chrX | 0.823837097 | 1.752749025 | 2.12754321 | 0.02125297 | 1 |
| Lpl__chr8 | 1.176507835 | 2.494218397 | 2.12001852 | 0.01263377 | 1 |
| Sdk2__chr11 | 0.710098907 | 1.495703084 | 2.10633064 | 0.0473411 | 1 |
| Atp5k__chr5 | 0.701145494 | 1.471286469 | 2.09840394 | 0.04449739 | 1 |
| Otud7b__chr3 | 0.504213356 | 1.047535826 | 2.07756461 | 0.04438239 | 1 |
| Gm37899__chr12 | 0.961572732 | 1.971596797 | 2.05038759 | 0.03271401 | 1 |
| Gm44080__chr6 | 0.506377035 | 1.037764182 | 2.0493903 | 0.04438239 | 1 |
| Apc__chr18 | 0.506342977 | 1.03124482 | 2.03665276 | 0.04438239 | 1 |
| Rufy3__chr5 | 0.711284526 | 1.426905393 | 2.00609649 | 0.04734111 | 1 |
| Gabpb2__chr3 | 0.851504326 | 1.703354331 | 2.00040596 | 0.02447685 | 1 |
| Rasgrf1__chr9 | 1.041453361 | 2.055675013 | 1.9738522 | 0.0421671 | 1 |
| Sf1__chr19 | 1.52933925 | 3.018431056 | 1.97368312 | 0.01181032 | 1 |
| Gck__chr11 | 1.486370001 | 2.907688515 | 1.95623466 | 0.01755884 | 1 |
| Arid4b__chr13 | 1.417300354 | 2.770568509 | 1.95482101 | 0.01412912 | 1 |
| Klf4__chr4 | 1.308849142 | 2.556482866 | 1.95322959 | 0.01164126 | 1 |
| Rundc3a__chr11 | 1.072436111 | 2.037635918 | 1.90000681 | 0.04678116 | 1 |
| Gm42428__chr5 | 1.781338086 | 3.334360557 | 1.87182915 | 0.01273505 | 1 |
| Arglu1__chr8 | 1.309925711 | 2.39414878 | 1.82769814 | 0.02828599 | 1 |
| PISD__chrJH58430[illegible] | 1.308483019 | 2.353156218 | 1.79838499 | 0.04651862 | 1 |
| Nisch__chr14 | 1.958320572 | 3.435804191 | 1.75446464 | 0.01744111 | 1 |
| Rpl26__chr11 | 2.918525298 | 4.585231908 | 1.57107835 | 0.02734153 | 1 |
| Srrm2__chr17 | 4.95045424 | 7.532329072 | 1.52154302 | 0.01814024 | 1 |
| Rpsa__chr9 | 4.949497821 | 7.528764015 | 1.52111674 | 0.01813482 | 1 |
| Fosb__chr7 | 10.21614493 | 14.50096037 | 1.41941608 | 0.02616176 | 1 |
| Ubb__chr11 | 7.212495649 | 4.317982874 | 0.59868083 | 0.02321447 | 1 |
| G6pc2__chr2 | 7.288269023 | 4.362032992 | 0.59850055 | 0.02030188 | 1 |
| Nkx6-1__chr5 | 5.16105756 | 2.870096581 | 0.55610629 | 0.01480458 | 1 |
| Chchd2__chr5 | 3.721470603 | 1.967752677 | 0.52875674 | 0.03224038 | 1 |

| | | | | | |
|---|---|---|---|---|---|
| Maged1__chrX | 4.40529352 | 2.253554844 | 0.51155612 | 0.01704116 | 1 |
| Cd164__chr10 | 3.965570513 | 2.023056474 | 0.51015521 | 0.01610976 | 1 |
| Srpr__chr9 | 2.835330112 | 1.217867944 | 0.42953303 | 0.01346936 | 1 |
| Mbnl2__chr14 | 2.084791083 | 0.849844139 | 0.40763995 | 0.03383727 | 1 |
| Oaz1__chr10 | 2.346969909 | 0.956301826 | 0.40746233 | 0.03223383 | 1 |
| Pdyn__chr2 | 1.567022023 | 0.583611143 | 0.37243327 | 0.03524366 | 1 |
| Gm21738__chr14 | 2.168461643 | 0.636767112 | 0.29364924 | 0.01061515 | 1 |
| Eif4a1__chr11 | 1.397526078 | 0.392786434 | 0.28105839 | 0.0244639 | 1 |
| Rheb__chr5 | 0.979699977 | 0.268360028 | 0.27392062 | 0.04332568 | 1 |
| Prpf8__chr11 | 0.999211646 | 0.221975444 | 0.22215058 | 0.04396851 | 1 |
| Pgrmc2__chr3 | 0.955945167 | 0.193000382 | 0.20189482 | 0.0427597 | 1 |
| Gm10721__chr9 | 1.013083637 | 0.175359646 | 0.17309494 | 0.04484994 | 1 |
| Srebf1__chr11 | 0.61881668 | 0.1 | 0.16159875 | 0.04751523 | 1 |
